# Cell-type-resolved somatic mosaicism reveals clonal dynamics of the human forebrain

**DOI:** 10.1101/2023.10.24.563814

**Authors:** Changuk Chung, Xiaoxu Yang, Robert F. Hevner, Katie Kennedy, Keng Ioi Vong, Yang Liu, Arzoo Patel, Rahul Nedunuri, Scott T. Barton, Chelsea Barrows, Valentina Stanley, Swapnil Mittal, Martin W. Breuss, Johannes C. M. Schlachetzki, Joseph G. Gleeson

**Affiliations:** Department of Neurosciences, University of California San Diego, La Jolla, CA, 92037, USA; Rady Children’s Institute for Genomic Medicine, San Diego, CA, 92123, USA; Department of Human Genetics, University of Utah, Salt Lake City, UT, 84112, USA; Sanford Consortium for Regenerative Medicine, La Jolla, CA, 92037, USA; Department of Pathology, UC San Diego School of Medicine, University of California, San Diego, La Jolla, CA, 92037, USA; BioSkryb Genomics, Inc., Durham, NC; Division of Medical Education, School of Medicine, University of California, San Diego, La Jolla, CA, 92037, USA; Department of Pediatrics, Section of Clinical Genetics and Metabolism, University of Colorado Aurora, CO, 80045, USA; Department of Cellular and Molecular Medicine, University of California, San Diego, La Jolla, CA, USA

**Keywords:** brain mosaicism, clonal dynamics, whole-genome sequencing, cell type, lineage, inhibitory neurons, somatic, migration

## Abstract

Debate remains around anatomic origins of specific brain cell subtypes and lineage relationships within the human forebrain. Thus, direct observation in the mature human brain is critical for a complete understanding of the structural organization and cellular origins. Here, we utilize brain mosaic variation within specific cell types as distinct indicators for clonal dynamics, denoted as cell-type-specific Mosaic Variant Barcode Analysis. From four hemispheres from two different human neurotypical donors, we identified 287 and 780 mosaic variants (MVs), respectively that were used to deconvolve clonal dynamics. Clonal spread and allelic fractions within the brain reveal that local hippocampal excitatory neurons are more lineage-restricted compared with resident neocortical excitatory neurons or resident basal ganglia GABAergic inhibitory neurons. Furthermore, simultaneous genome-transcriptome analysis at both a cell-type-specific and single-cell level suggests a dorsal neocortical origin for a subgroup of DLX1^+^ inhibitory neurons that disperse radially from an origin shared with excitatory neurons. Finally, the distribution of MVs across 17 locations within one parietal lobe reveals restrictions of clonal spread in the anterior-posterior axis precedes that of the dorsal-ventral axis for both excitatory and inhibitory neurons. Thus cell-type resolved somatic mosaicism can uncover lineage relationships governing the development of the human forebrain.

## Main text

Forebrain development is under the control of morphogens and transcription factors, mediating patterning of three-dimensional structures including the hippocampus, cortex, and basal ganglia ^1–5^. Although clonal dynamics in the mouse forebrain were investigated using single-cell viral barcoding ^6,7^, studies in humans are limited to clonal relationships at broader spatial levels ^8–11^. Given that the human brain comprises diverse cell types originating from various sources that intermingle, and eventually reside in proximal locations, direct mapping of clonal dynamics within distinct human forebrain cell types is essential.

While most forebrain cells are thought to originate from radial glia that line the telencephalic lateral ventricles^12–14^, observations in rodents instead suggest ventral telencephalic progenitors as a source of GABAergic cortical inhibitory neurons ^15^, supported by subsequent investigations in non-human primate, and human fetal tissue ^6,16–20^. However, conflicting findings suggest a recently evolved, potentially primate-specific dorsal telencephalic source of inhibitory neurons based on marker staining or single-cell lineage tracing in cultured human fetal brain tissues^21–26^. Yet none of these studies directly observed lineage relationships of inhibitory neurons within the fully developed human brain, leaving this longstanding debate unresolved.

Postzygotic mutations transmit faithfully to daughter cells that distribute in mosaic patterns, referred to as mosaic variants (MVs). The human brain, like other tissues, acquires MVs in part due to rapid expansions of initial founder cell pools^9,10^, resulting in clonal lineages sharing MVs that can vary in bulk allele fraction (AF) depending upon cell mixing. As neurogenesis predominantly occurs during brain development, the distribution of MVs in adult using Mosaic Variant Barcode Analysis (MVBA) can reveal clonal dynamics and lineage relationships that likely originated during embryogenesis ^9–11^.

Such approaches face challenges when dealing with small cell populations, such as cortical inhibitory neurons, due to technical limitations in obtaining adequate sample quantity from postmortem human tissues, and in deriving high-quality MV call sets. To overcome these challenges, we developed a methanol-fixed nuclei sorting (MFNS) protocol, which was added to prior protocols for cell-type-specific mosaic variant barcode analysis in bulk, sorted nuclei, and single nuclei, allowing DNA isolation from high-quality intact nuclei suitable for library preparation. Additionally, we utilized a recently developed single-cell multi-omics approach generating DNA genotypes with RNA transcriptomes from the same cell ^27^, deconvolving lineages and mapping cell-type-specific clonal dynamics within the human brain.

### Identification of brain MVs

Deep sequencing (300X) of a single biopsy detects dozens of clonal MVs including single nucleotide variants (SNVs) and small insertions or deletions (indels)^28,29^. To examine genomic relationships across distinct cell types in human brains, we further optimized a prior protocol for lower cell number input (100-fold improvement) and greater cell type diversity, termed cell-type-specific mosaic variant barcode analysis (cMVBA), comprising three phases (Fig. 1a) ^10^. In the ‘Tissue collection’ phase, we biopsied (8 mm punch) each lobe of the neocortex (CTX), basal ganglia (BG), hippocampus (HIP), thalamus (TH), and cerebellum (CB) from a previously published donor (ID01)^10^ and from an ascertained new donor (ID05) (Extended Data Fig. 1, Supplementary Data 1). We additionally collected non-brain tissue including heart, liver, both kidneys, adrenal, and skin to define MV distribution across the body. In the ‘MV discovery’ phase, a subset of bulk tissues (35 and 32 tissues from ID01 and ID05, respectively) underwent 300X whole genome sequencing (WGS), followed by state-of-the-art MV calling and filtering based on established methods (Methods). In the ‘Validation and quantification’ phase, we first prepared DNA samples extracted from bulk tissue and sorted nuclear populations, or amplified DNA from single nuclei (Supplementary Data 1). NEUN^+^, DLX1^+^, TBR1^+^, NEUN^-^/LHX2^+^, OLIG2^+^, NEUN^+^/DARPP32^+^, PU.1^+^ nuclei pools represented pan-neurons, GABAergic inhibitory neurons, excitatory neurons, astrocytes, oligodendrocytes, medium spiny neurons, and microglia, respectively^10,30^. To isolate intact DNA from underrepresented cell types such as cortical inhibitory neurons, we implemented MFNS by screening several antibodies targeting DLX1 (pan-inhibitory neuronal marker), TBR1 (excitatory neuronal marker), and COUPTFII (CGE-derived inhibitory neuronal marker; encoded by *NR2F2*) (Methods), and confirmed that each antibody labels a particular cell type within the biopsy (Extended Data Fig. 2a-c).

**Figure 1.**
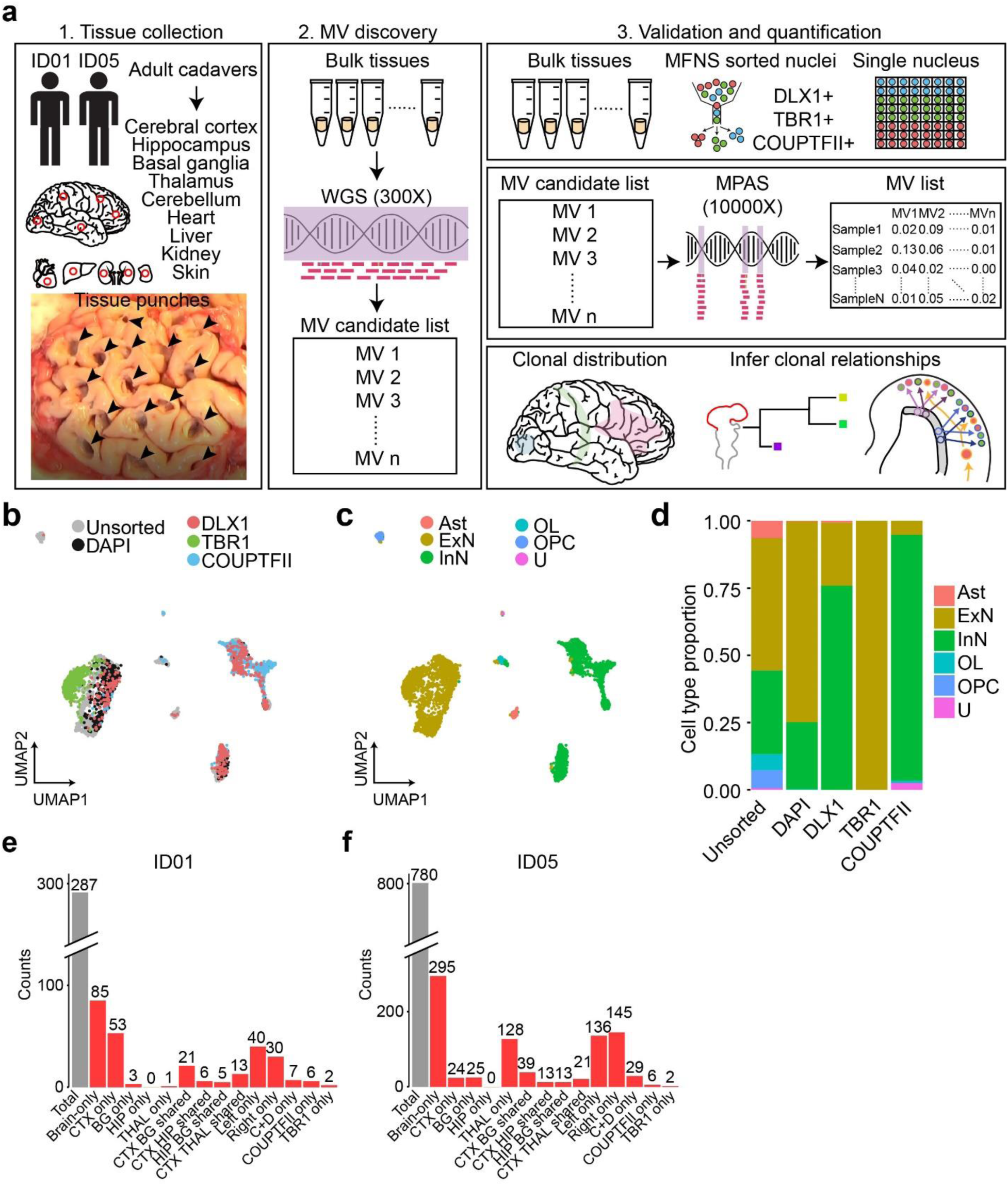
Comprehensive cMVBA identifies cell-type-resolved and region-specific MVs. (a) cMVBA workflow overview consists of three phases: 1. Tissue collection: Cadaveric organs accessed for tissue punches from organs listed (red circles and black arrows: punch locations in organs and frontal lobe) for MV detection. 2. A subset of the bulk tissue punches undergo 300x WGS followed by best-practice MV calling pipelines to generate a list of MV candidates. 3. DNA from each punch bulk tissue, methanol fixed nuclear sorted (MFNS) samples, or individual nuclei are subjected to validation and quantification of MV candidates via massive parallel amplicon sequencing (MPAS). AFs of the validated MVs from MPAS are used to determine clonal dynamics of different neuronal cell types and reconstruct features of brain development. (b) UMAP plot from snRNA-seq with MFNS sorted or unsorted cortical nuclear pools (*n* = 3322). Sorted nuclear groups labeled with distinct colors. (c) Cortical cell types based on marker expression and differentially expressed genes in each cluster. (d) Cell type proportion within each sorted cortical nuclear population. (e-f) Number of MVs categorized by location detected in each donor ID01 or ID05 (Supplementary Data 4). ‘Brain-only’ MVs (i.e., detected only in brain tissue, but not other organs) including subtypes in red. ‘C+D only’: Brain-only MVs exclusively detected in both COUPTFII^+^ and DLX1^+^ populations but not the other cell types. Ast, Astrocyte; ExN, Excitatory neurons; InN, inhibitory neurons; OL, oligodendrocytes; OPC, oligodendrocyte precursor cells; U, undefined.

We further confirmed the cell-type composition of DLX1^+^, TBR1^+^, and COUPTFII^+^ nuclear pools compared with unsorted and DAPI^+^ nuclear pools with single-nucleus RNA-seq (Fig. 1b). Cell types of each cluster were identified based on marker expression patterns (Fig. 1c, Extended Data Fig. 2d and e). DLX1^+^ and COUPTFII^+^ pools contained mostly inhibitory neurons (>75%), and almost 100% of TBR1^+^ nuclear pools were confirmed as excitatory neurons (Fig. 1d). Of note, COUPTFII^+^ nuclei were mostly in a subcluster of inhibitory neuronal clusters, whereas DLX1^+^ nuclei covered all inhibitory neuronal clusters, suggesting that nuclei sorted by COUPTFII antibody reflect a subset (∼38%) of DLX1^+^ nuclei highly expressing *NR2F2* (Extended Data Fig. 2f). Cell type identities of the DLX1^+^ and TBR1^+^ nuclei were further confirmed as inhibitory and excitatory neurons, respectively, by comparing DNA methylation patterns from reference methylomes of inhibitory and excitatory neurons across marker gene regions ^31^ (Extended Data Fig. 2g-j).

Using a total of 321 samples from ID01 and 147 samples (bulk or sorted nuclei) from ID05, we conducted ultra-deep massive parallel amplicon sequencing (MPAS) of each candidate MV (ave. coverage ∼10,000X). This step served two functions: 1] providing orthogonal validation for the MV within the sample. 2] providing accurate assessment of AF of each detected variant, allowing for downstream analyses (Extended Data Fig. 3a,b, Supplementary Data 2 and 3). A total of 287 and 780 MV candidates detected in WGS were thus validated and quantified in ID01 (Extended Data Fig. 3c) and ID05 (Extended Data Fig. 3d), respectively (Methods), subsequently used to annotate MVs according to brain region and cell type (Fig. 1e and f, Extended Data Fig. 4, Supplementary Data 4). The proportion of ’Brain-only’ MVs showed a similar distribution in ID01 and ID05, accounting for 29.6% (85/287) and 37.8% (295/780) of total MVs, respectively. A total of 7 and 29 MVs were exclusively found in DLX1^+^ or COUPTFII^+^ but not in TBR1^+^ neurons (C+D only) in ID01 and ID05, respectively. We observed similar trends of MV hemispheric restriction and microglia distribution as we and others recently reported ^10,32,33^ (Extended Data Fig. 5). This suggests that the cMVBA pipeline reports anatomic- and cell-type-specific MVs that can be used to profile clonal dynamics and reconstruct lineage relationships of specific cell types.

### Genetic similarity of major telencephalic-derived structures

The telencephalon derives from the most rostral part of the neural tube, subsequently committing to CTX, BG, and HIP, composing major structures of the adult forebrain. Since we observed significantly fewer Brain-only MVs shared between HIP and other telencephalic structures such as CTX and BG (Fig. 1e, f), we hypothesized the HIP founder cells are restricted in lineage compared to other brain regions (Fig. 2a). We thus performed hierarchical clustering using allelic fractions (AFs, portion of alternative allele among total alleles) in bulk samples from ID01. We found HIP samples strongly clustered away from CTX and BG samples, suggesting that HIP progenitors are clonally more distinct from CTX or BG cells (Fig. 2b).

**Figure 2.**
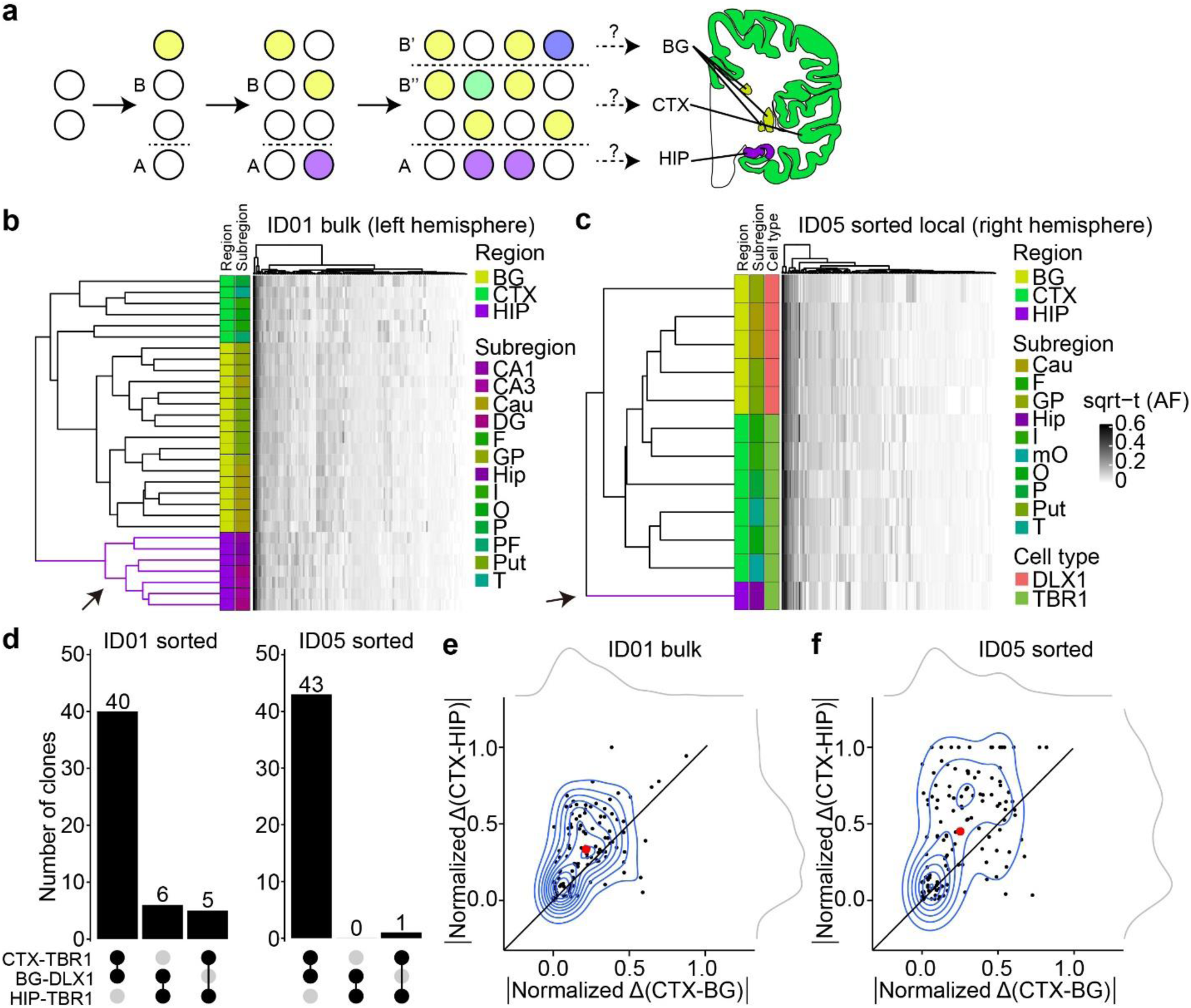
Human hippocampal lineage diverges from the cortex and basal ganglia. (a) Model of clonal dynamics in forebrain anlage. Cells restricted to anlage A, which acquire a new MV (purple) are rarely present in anlage B. Later, B diverges into B’ and B’’, which share more clones (yellow) than with anlage A. This analysis was applied to the geographies of the basal ganglia (BG), cortex (CTX) and hippocampus (HIP). (b-c) Heatmaps with 30 bulk samples from left hemispheric CTX, BG, and HIP (y-axis, b) or 12 selected sorted cell types (y-axis, c), compared with MVs identified in at least two samples (146 MVs, x-axis in (b) or 131 MVs, x-axis in (c)), depicted in Fig. 2d. Dendrograms at right shows greater HIP lineage separation (purple, arrow) compared with CTX or BG (green and yellow) either using bulk tissue (b) sorted nuclei (c), suggesting HIP earlier lineage restriction. (d) Counts of shared MVs across CTX, BG or HIP within sorted nuclear pools showing many more shared MVs between CTX and BG compared with HIP in both donors ID01 and ID05. (permutation *p*-value <0.001) (e-f) Contour plots of informative 113 and 131 MVs from (b) and (c) (blue) and two kernel density estimation plots (grey). Axes show absolute normalized AF difference for each MV averaged across all samples from the respective tissues (CTX, HIP, BG). Black line: identity line, black dots: individual MVs, large red dot: averaged across all MVs, suggesting AF differences are smaller between CTX and BG than between CTX and HIP. Abbreviations: Cau, caudate; DG, dentate gyrus; HIP and Hip, hippocampal tissue, where lowercase Hip refers to hippocampal subregion not specified; I, insular cortex; O, occipital cortex; P, parietal cortex; PF, prefrontal cortex; Put, putamen; T, temporal cortex; mO, medial occipital cortex; GP, globus pallidus; sqrt-t (AF), square-root transformed allele fraction.

Forebrain structures contain heterogeneous cell types, not only derived from local progenitors but also cells migrating from distant brain regions^15,34,35^. To exclude the possibility that migrating cells contributed to these findings, we repeated hierarchical clustering, this time restricting analysis to only locally originating cell types, i.e. excitatory (TBR1^+^ nuclei) in HIP or CTX and inhibitory neurons (DLX1^+^ nuclei) in BG. Hierarchical clustering with Manhattan distances of AFs in sorted nuclei samples from ID05 (Fig. 2c) and ID01 (Extended Data Fig. 6a) replicated the genomic similarity of HIP-TBR1^+^ clones from CTX-TBR1^+^ and BG-DLX1^+^ clones, confirming that cortical excitatory neurons show AF patterns more like inhibitory neurons in BG than excitatory neurons in HIP. Furthermore, CTX-TBR1^+^ nuclei shared many more MVs with BG-DLX1^+^ (40 and 43 for ID01 and ID05, respectively) than with HIP-TBR1^+^ nuclei (5 and 1 for ID01 and ID05, respectively) (Fig. 2d). The AF variation was greater between CTX and HIP than between CTX and BG, exhibiting average vector of data points above the identity line (Fig. 2e, f and Extended Data Fig. 6b). Taken together, this analysis suggests that the clonality of HIP progenitors is unlikely a result of migration of cells from other forebrain structures, and instead the results of locally proliferative cells within the HIP anlage.

### Clonal dynamics of cortical inhibitory neurons

While in vitro analysis of neuronal progenitors from dorsal human brain cortical tissue shows the potential to develop into inhibitory neurons ^25^, direct evidence for a dorsal origin of cortical inhibitory neurons in the mature human brain is lacking. cMVBA allowed for comparison of genomic similarity between different classes of cortical inhibitory neurons and other cell types. Hierarchical clustering was carried out for AFs measured in DLX1^+^, TBR1^+,^ and COUPTFII^+^ nuclear pools isolated from widespread sampling from cortical areas in both ID01 and ID05 (Fig. 3a). Intriguingly, most of COUPTFII^+^ nuclei (i.e. caudal ganglionic eminence (CGE)-derived inhibitory neurons that distribute across cortical areas ^25^) were exclusively clustered together, whereas most DLX1^+^ and TBR1^+^ nuclear populations in the same punch were clustered together (Fig. 3b, c). Bootstrap analysis further statistically validated that many of these dendrogram clusters appeared very unlikely to have arisen by chance (Extended Data Fig. 7).

**Figure 3.**
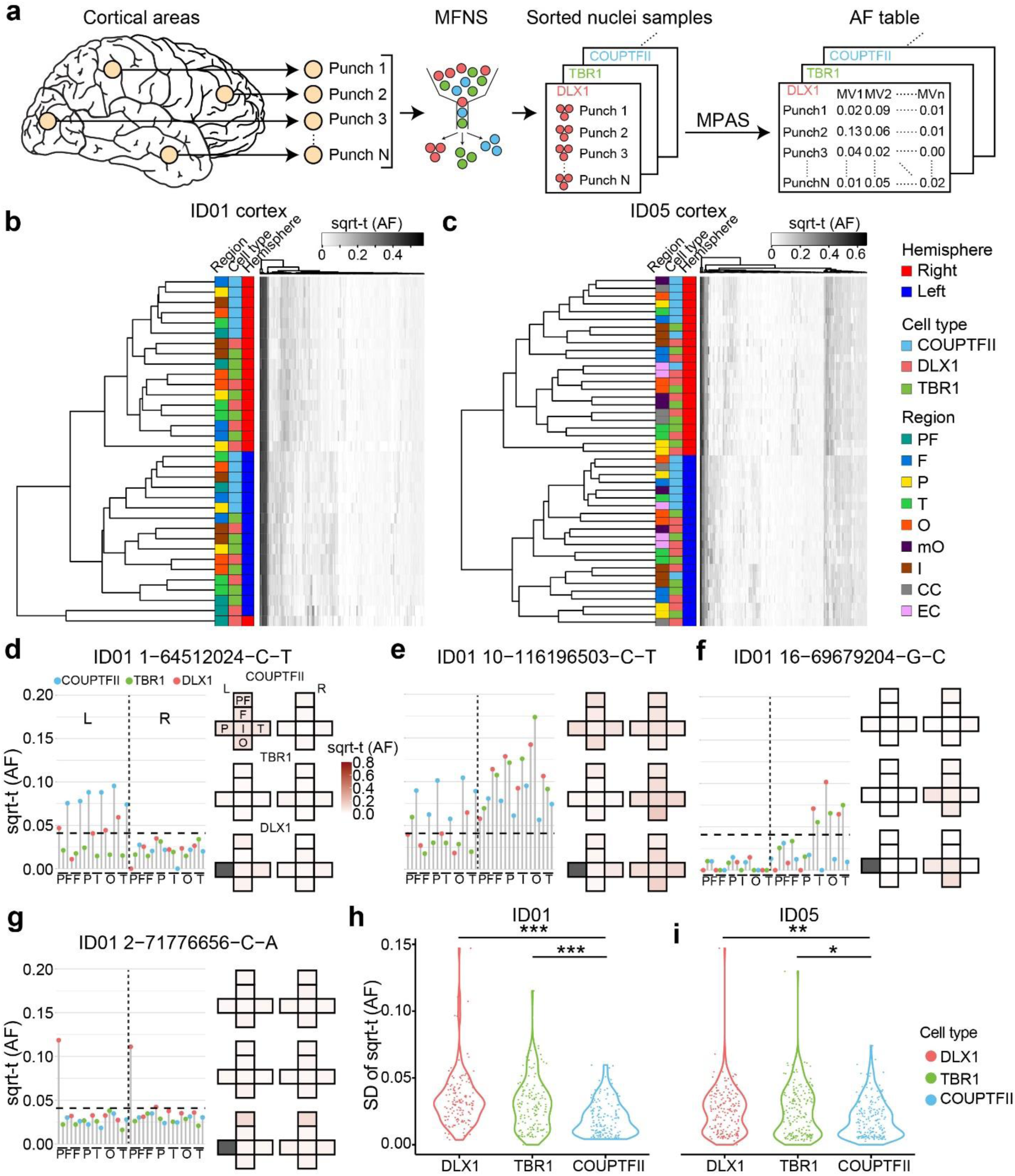
Cell-type-resolved clonal dynamics of cortical excitatory and inhibitory neurons. (a) cMVBA workflow uses MFNS nuclei for MPAS assessment of AFs in cortical punches. (b-c) Heatmaps of sorted nuclei based on AFs of 146 informative shared MVs from ID01 (b) and 186 from ID05 (c) (y-axis) compared with color-coded hemisphere, cell type or region. PF, prefrontal cortex; F, frontal cortex; P, parietal cortex; T, temporal cortex; O, occipital cortex; mO, medial occipital cortex; I, insular cortex; CC, cingulate cortex; EC, Entorhinal cortex. sqrt-t (AF), square-root transformed allelic fraction. Dendrograms indicate that subcortically derived COUTFII samples cluster together (teal), whereas DLX1 and TBR1 samples derived from the same region cluster together (notice interspersed orange and green boxes) suggesting shared lineage relationships. (d-g) Lolliplots comparing regions (x-axis) with sqrt-t AF (y-axis) for representative MVs. Height of individual lollipop: AF; color: cell type; dashed line: threshold. Next to each lolliplots is the ‘geoclone’ representation of sqrt-t AF shaded intensity (pink) from tissue where detected. Gray boxes: not sampled. (d) MV at chr. 1 position 64512024 C-T is found in COUPTFII^+^ cells in the left hemisphere at AFs different from either DLX1^+^ or TBR1^+^ cells, consistent with COUPTFII^+^ cells having different lineage than DLX^+^ or TBR1^+^. MVs in (e-f) also show more AF similarities between DLX^+^ or TBR1^+^ than COUPTFII^+^ cells. MF in (g) shows a representation of DLX1^+^ cells in both prefrontal lobes, consistent with a potential ventrally-derived DLX1^+^ clone distributed across the two hemispheres. (h-i) Standard deviation (SD) of sqrt-t AFs for 146 and 186 MVs in the three different cell types in donor ID01 (h) and ID05 (i), respectively, showing lower variation in COUPTFII^+^ cells compared with DLX1^+^ or TBR1^+^, consistent with their clonal distribution across cortical areas. Each dot: single MV measured in 34 vs. 45 punches across the neocortex in ID01 and ID05, respectively. One-way ANOVA with Tukey’s multiple comparison test. *:P<0.05, **:P<0.001, ***P<0.0001. PF, prefrontal cortex; F, frontal cortex; P, parietal cortex; O, occipital cortex; T, temporal cortex; I, insular cortex; CC, cingulate cortex; mOC, medial occipital cortex; L, left; R, right.

Next, we measured AF correlations between MVs within each cell type (Supplementary Data 5). As expected, MV clustering confirmed hemispheric restrictions of all three cell types (DLX1^+^, TBR1^+^ and COUTPFII^+^ nuclei). Additionally, MVs in TBR1^+^ nuclei showed many small subclusters enriched in particular lobes, in contrast to widespread distributions of COUPTFII^+^ nuclei. This result was replicated in four independent hemispheres from both donors. A similar pattern was also observed in the global DLX1^+^ nuclear pools, although with less intensity. The data suggests wide distribution of COUPTFII^+^ ventrally-derived cortical inhibitory neuronal clones through tangential migration to the dorsal telencephalon as reported in mice^36^, while focal distribution of TBR1^+^ and at least some DLX1^+^ neurons distribute radially from a dorsal telencephalic, likely radial glial, source. Collectively, the result suggests a significant portion of dorsal clones have the potential to differentiate into both cortical excitatory and inhibitory neurons.

Next, we investigated distributions of individual MVs across cell types and cortical areas by mapping each MV AF onto a ‘Lolliplot’ and geographic map, called a ‘Geoclone’ (Supplementary Data 6). As an example, MV 1-64512024-C-T was inhibitory neuron-specific, showing notably uniform enrichment across cortical areas in COUPTFII ^+^ and DLX1^+^ but not TBR1^+^ nuclear pools (Fig. 3d). This suggests a subset of nuclei in DLX1^+^ pools distribute in patterns similar to COUPTFII^+^ nuclei. MV 10-116196503-C-T was present in all three cell types, but with less AF variation in COUPTFII^+^ nuclei than the other two cell types (Fig. 3e), suggesting a wide distribution across cortical regions and tangential migration from a ventral origin. Furthermore, MV 16-69679204-G-C was locally enriched in both DLX1 ^+^ and TBR1^+^ nuclei of the same cortical lobes but absent in COUPTFII^+^ nuclei, implying a subset of DLX1^+^ cells share locally proliferating dorsal telencephalic origins with TBR1^+^ cells (Fig. 3f). Interestingly, we found some MVs (i.e. MV 2-71776656-C-A) that were distinctively enriched in DLX1^+^ nuclei distributed to both prefrontal lobes of ID01 (Fig. 3g), suggesting some inhibitory neurons or their progenitors can populate both hemispheres. However, prefrontal samples were not available in ID05, and more data will be required to confirm these results.

To examine whether COUPTFII^+^ clones are more uniformly distributed than DLX1 or TBR1^+^ clones, standard deviations of AFs of shared MVs in different cortical areas were calculated for the three neuronal populations (Fig. 3h, i). COUPTFII^+^ nuclei standard deviations were significantly lower than the other two cell types in both ID01 and ID05, suggesting that the distribution of CGE-derived inhibitory neuronal clones is wider than excitatory neurons, whereas cortical pan-inhibitory neuronal clones showed a patchy distribution in a fashion similar to excitatory neuronal clones.

### Dual origins of cortical inhibitory neurons

The prior data could not exclude the possibility that the observed genomic similarity between TBR1^+^ and DLX1^+^ sorted nuclear pools derived in part from a rare cell type or from pool contaminants from the sorting protocol. We thus conducted single-cell simultaneous DNA + RNA sequencing (ResolveOME, see Methods), incorporating primary template-directed amplification (PTA) coupled with single-nuclear MPAS (snMPAS) with full-transcript snRNA-seq, in individual NEUN^+^ nuclei from right frontal and temporal cortex in ID05. This allowed for both *a priori* identification of both cell types and MVs in the same cell ^37^ (Fig. 4a, Supplementary Data 7). UMAP clustering with a reference dataset distinguished between cortical excitatory and inhibitory neurons, along with a few minor non-neuronal cell types as expected (Extended Data Fig. 8a-c). The detection frequency of MVs in single-cell genotyping was positively correlated with AFs in sorted populations, as expected (Extended Data Fig. 8d-f). Informative MVs were detected from a total of 85 excitatory and 33 inhibitory neurons, allowing direct observation of single-cell level MV distribution (Fig. 4b).

**Figure 4.**
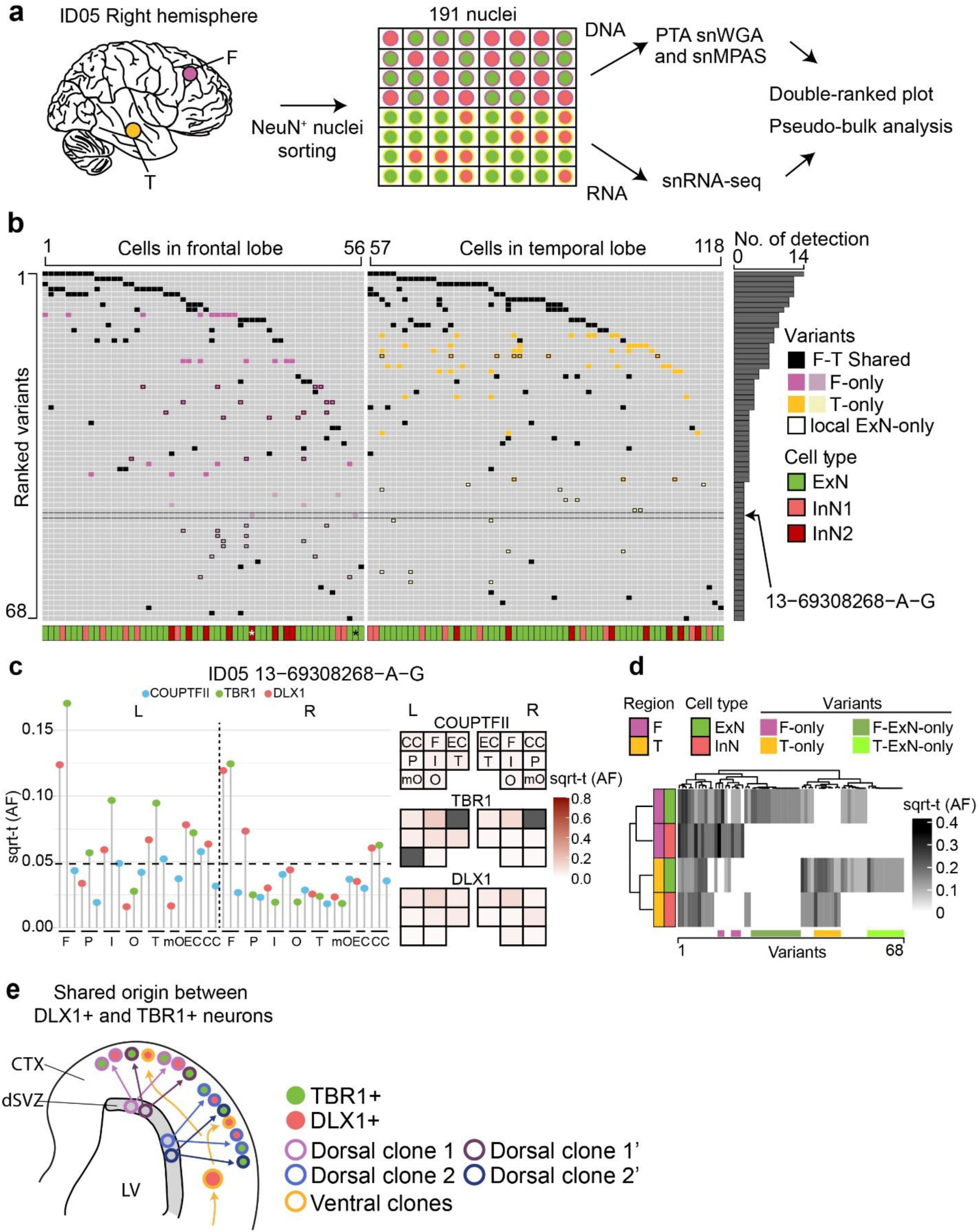
snMPAS incorporating snRNA-seq supports the existence of dorsally derived cortical inhibitory neurons in humans. (a) Frontal and temporal small punches of the right hemisphere of ID05 NEUN^+^ 191 single-nuclei were subjected to primary template-directed amplification (PTA) single nuclear whole genome amplification (snWGA) followed by single nuclear MPAS (snMPAS) genotyping with concurrent snRNA-seq, termed ResolveOME, for analysis. (b) A double-ranked plot divided in half based on brain region (frontal: left, temporal: right). 68 MVs identified in anywhere between 2 to 14 cells (No. of detection). A total of 118 excitatory and inhibitory neurons plotted. MVs were further classified according to whether they were detected in both frontal and temporal lobes (F-T Shared, black) vs. a single lobe F-only (purple) or T-only (yellow). Dark purple or yellow: detected in 3 or more nuclei, light purple or yellow: detected in only 2 nuclei. x-axis: cell type, i.e. ExN: green; InN1 (orange): All inhibitory neurons except for InN2. InN2 (dark red): Inhibitory neurons carrying at least one F- or T-only MV detected in 3 or more nuclei, or the nucleus marked by asterisk. MV 13-69308268-A-G (arrow) was detected in 2 nuclei (*), an ExN and an InN, consistent with a shared lineage. (c) Distribution MV 13-69308268-A-G across cortical areas in ID05. Left: lolliplot and Right: geoclone showing sqrt-t AFs of each sorted nuclear pool in different cortical locations, reproducing single-nucleus data. Note that DLX^+^ and TBR1^+^ nuclei show similar sqrt-t AF (geoclone) and are more similar to each other than COUPTF2+ AF (lolliplots). Dashed line: threshold. (d) Pseudo-bulk analysis after aggregation based on individual MVs (x-axis) and cell types and regions (y-axis). Note that the dendrogram cluster pseudo-bulk cell populations of ExN and InN in F (purple) or T (orange) lobes based upon shared MV, although some MVs are present in a single cell type. (e) Model for the shared origin of local ExN and InN cortical neurons. Dorsal clones can produce both TBR1^+^ excitatory neurons and DLX1^+^ inhibitory neurons (purple and blue arrows), evidenced both by similar spatial clonality from sqrt-t AFs, as well as shared MVs at the single cell level. Ventrally derived DLX1^+^ inhibitory neurons (yellow arrow) tangentially migrate and are more likely to disperse across the cortex.

These genotypes allowed for the assessment of shared MVs in individual excitatory or inhibitory neurons in individual nuclei across various brain regions. As an example, MV 13-69308268-A-G, detected in both excitatory and inhibitory neurons of the right frontal but not temporal lobe (Fig. 4b), was intriguingly enriched in TBR1^+^ and DLX1^+^ nuclear pools with high AFs but below the level of detection in the COUPTFII^+^ nuclear pool (Fig. 4c). Furthermore, overall distribution patterns of this MV in TBR1^+^ and DLX1^+^ nuclei across cortical lobes were very similar to each other, but distinct from that of COUPTFII^+^, supporting this MV is shared between locally born and cortical resident excitatory and inhibitory neurons but underrepresented within cortical inhibitory neurons derived from ventral telencephalic sources (Fig. 4c).

We also calculated the probability of observing ventrally originated cortical inhibitory neurons carrying seemingly locally enriched MVs by chance using MVs shared in more than two cells in one lobe but none in the other lobe. Among the 33 inhibitory neurons analyzed, we identified 15 with distinct local MVs. These MVs were found exclusively in one lobe and were shared with at least two other local cells, including one excitatory neuron within the same lobe. This observation was highly unlikely to occur by chance (one-tailed permutation test *p*<0.0001, Extended Data Fig. 8g, Methods), suggesting the existence of locally derived inhibitory neurons from progenitor cells that also produce local excitatory neurons.

Inhibitory neurons carrying locally enriched MVs (InN2; Fig. 4b) showed comparable inhibitory neuronal marker expression (Extended Data Fig. 8h) but a decreased tendency for RNA expression of CGE markers (Extended Data Fig. 8i), or a RELN^+^ neuronal marker (Extended Data Fig. 8j) compared to those not carrying locally enriched MVs (InN1). Instead, InN2 displayed an increased tendency for a parvalbumin-positive (PV^+^) inhibitory neuronal marker (Extended Data Fig. 8k) and some unique transcription patterns (Extended Data Fig. 8l-m). Thus, a significant portion of dorsally-derived inhibitory neurons may contribute to PV^+^ neurons, whereas COUPTFII^+^ neurons contribute to ventrally-derived inhibitory neurons of other classes.

We further observed 21 MVs specific to only local excitatory neurons but not inhibitory neurons in the same lobe (Fig 4b), implying a subset of fate-restricted dorsal telencephalic neural progenitors generate mostly excitatory neurons. This was also supported by pseudo-bulk analysis, by aggregating cells based on each cell type and region, demonstrating local excitatory neuron-specific MVs (Fig. 4d). Interestingly, we could not find evidence of MVs specific to cortical inhibitory neurons within anatomically defined regions, which may be due to their sparse population from total cortical cells, or a limited number of inhibitory neuron-specific MVs. Taken together, single-cell level genotyping incorporating transcriptomics supports the concept that dorsal telencephalic neural progenitor cells may have the potential to generate both excitatory and inhibitory neurons, even amongst progenitor pool predominantly generating excitatory neurons (Fig. 4e).

### Anterior-posterior restriction within a lobe

We used this same approach to study clonal dynamics within a single human cerebral lobe. Prior data suggests that clonality between lobes is more restricted than within a lobe ^9^. Our prior data suggests a restriction of clonal spread (RCS) along the anterior-posterior axis follows the establishment of the midline RCS, but did not consider cell-type-specific effects ^10^. We thus assessed clonal dynamics of DLX1^+^ and TBR1^+^ cells, selecting the parietal lobe for analysis. We performed high-density biopsies from a total of 17 small punches offset by 1cm distances followed by FANS and MPAS genotyping, capable of distinguishing between A-P and D-V RCSs with samples clustered based on AFs of informative MVs (Fig. 5a-b, Supplementary Data 8). Notably, the main clusters (C1 and C2) were formed along the A-P rather than the D-V axis in both cell types (Fig. 5c). The RCS dominated along the A-P over the D-V axis in both cell types, represented by MVs 18-33999883-C-T, 4-10646818-G-A, and 3-172635725-G-A (Fig. 5d). Absolute values of normalized AF difference through the A-P axis were larger than those through the D-V axis (Fig. 5e, f). These results suggest that the A-P axis RCS is established prior to D-V in both cell types.

**Figure 5.**
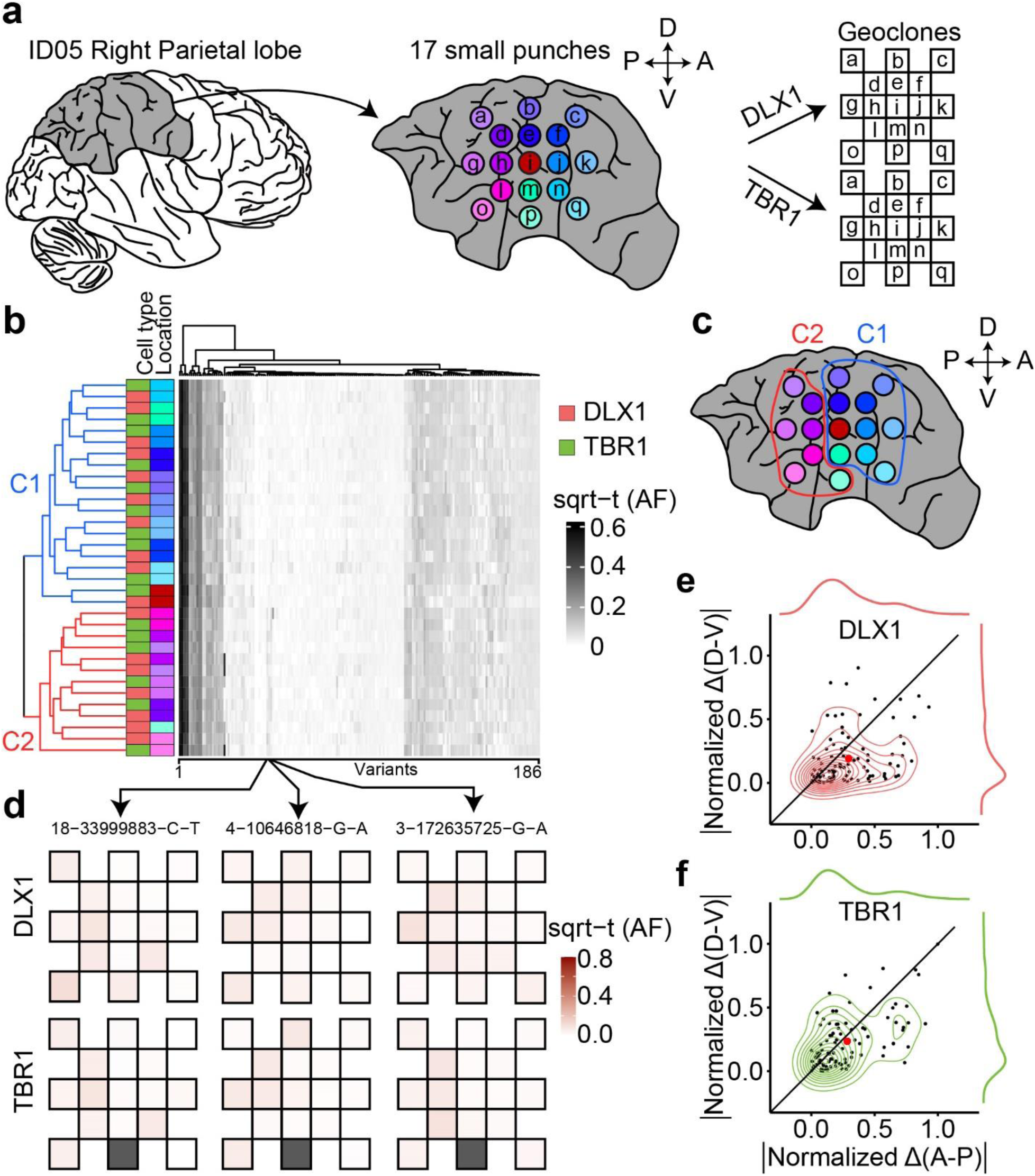
Earlier establishment of the A-P axis compared to D-V restricted clonal spread (RCS) within a cortical lobe. (a) Workflow for the observation of clonal dynamics in a lobe. A total of 17 punches were radially sampled and subject to MFNS to assess MVs. The AF of MVs in different sites were mapped onto the geoclones (checkerboard). (b) Heatmap and dendrogram hierarchical cluster of sorted nuclear pools based on sqrt-t AFs of 186 informative MVs from ID05 detected in the right parietal lobe. Sidebars left of the heatmap: cell type and sample location information, according to the convention in Fig. 3. Colors for lobar location taken from (a). Dendrogram highlights two main clusters (C1:blue; C2:red) that when mapped back onto the sampled spatial coordinates (c) are separated in the AP dimension (red and blue circle). (d) Geoclones of three individual MVs from (b, box, arrows), showing shades of pink are more different in the A-P axis than in the D-V axis. Gray box: not available. (e-f) Cell-type-resolved contour plots of 94 shared MVs from (b) (in the center) with kernel density estimation plots (in the periphery) for DLX1 (top) or TBR1 (bottom), showing MVs from both cell types with a greater normalized difference of sqrt-t AFs in A-P than in D-V. Black line: identity line, dots: individual MVs, large red dot: average across all MVs.

## Discussion

Here using clonal dynamics, we interrogate cellular origins across the neurotypical mature human forebrain. MVs originating in locally born cellular populations demonstrate stronger lineage restriction within the hippocampus than restriction to either the neocortex or basal ganglia. This is consistent with prior viral barcode tracing in mice showing hippocampal lineage restriction prior to neural tube closure at E9.5, at a time even prior to Prox1 expression ^7,38^. We hypothesize that hippocampal lineage restriction may be complete by the time anterior neural plate boundary (aNPB) expresses WNT3A that defines the cortical hem anlage at post-conception week 6 in humans ^39,40^. It is also possible that the pallial and subpallial lineage commitment to future subcortical and cortical structures occurs after aNPB definition at the time of dorsal and ventral axis differentiation under the control of BMP signaling ^41,42^. Future studies might assess if the early hippocampal lineage commitment we observed here occurs in other vertebrate species, where WNT and BMP-mediated neurulation is well-conserved ^43–45^.

While prior studies in mammals suggested most or all cortical inhibitory neurons derive from the ventral telencephalon in mammals ^6,16–20^, subsequent studies in non-human primates, human fetal brain, or stem cell culture suggested a potential primate-specific dorsal pallial contribution ^25,26^. Our cMVBA data provides evidence for such a dorsal source of cortical inhibitory neurons evidenced in the mature human brain. Furthermore, both cortical excitatory and inhibitory neurons showed similar clonal MV dynamics within the same lobe, whereas some excitatory neurons had additional local MVs absent in inhibitory neurons at the single-cell level. This implies that early dorsal progenitors can produce both cell types. However, there may be further fate-restriction occurring in some cell populations or at certain times in development that produce exclusively either excitatory or inhibitory neurons within the same location. Future studies could better define timing of this lineage restriction.

In non-mammalian vertebrates, telencephalic inhibitory neurons are thought to derive exclusively from subpallial structures, suggesting the dorsal origin of inhibitory neurons may have been a relatively late evolutionary adaptation ^46–48^. Could dorsally derived inhibitory neurons have arisen to serve specific functions? For instance, the double bouquet cell (DBC) is a type of cortical inhibitory neuron characterized by vertical cell body and axonal bundling, akin to minicolumns, the signature of radial migration ^49–51^. Thus, it would be appealing to consider whether DBCs might be one type of dorsally derived radially migrating cortical inhibitory neuron. Interestingly, DBCs were described in gyrated mammals, not observed in rodents, lagomorphs, or artiodactyls ^50^.

Despite our findings, there remain notable disparities when compared to *in vitro* human inhibitory neuron studies where cortical inhibitory neurons, derived from human dorsal forebrain organoid outer radial glia, exhibited elevated *NR2F2* expression ^25,26^. In contrast, our study suggests a substantial number of COUPTFII^+^ cortical clones displayed dispersed clonal dynamics, arguing against a purely radial distribution. This discrepancy could reflect differences between in vitro and in vivo experimental conditions and underscores the need for further investigation of cortical inhibitory neuron diversity.

## Figure legends

**Extended Data Fig. 1.**
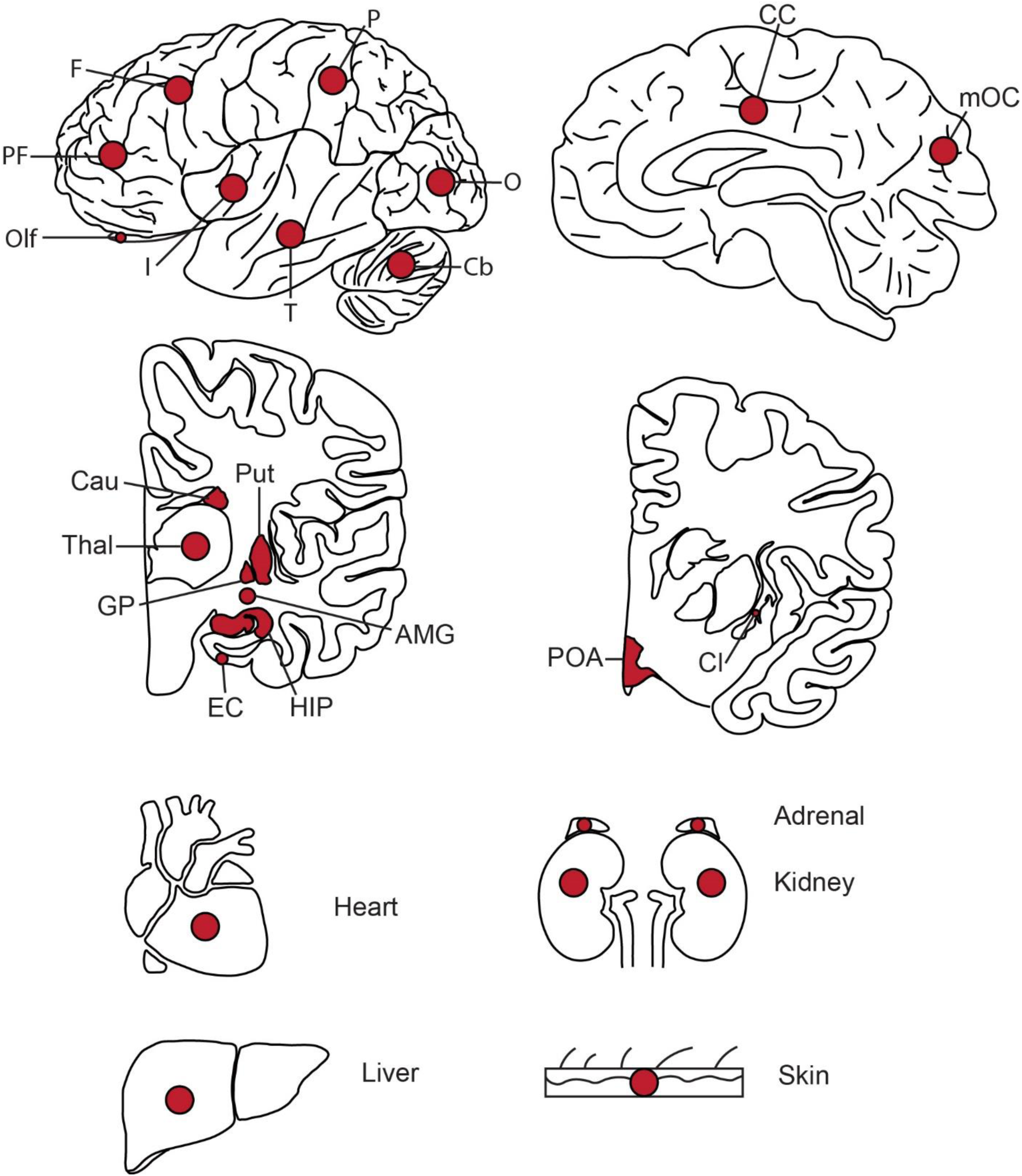
Tissues collected from ID01 and ID05. Red dots indicate approximate sites of punch biopsies. Abbreviations: PF, prefrontal cortex; F, frontal cortex; P, parietal cortex; O, occipital cortex; T, temporal cortex; I, insular cortex; Cb, Cerebellum; CC, cingulate cortex; mOC, medial occipital cortex; Cau, Caudate; Put, Putamen; Thal, Thalamus; GP, globus pallidus; EC, entorhinal cortex; HIP, hippocampus; AMG, amygdala; POA, preoptic area; Cl, Claustrum.

**Extended Data Fig. 2.**
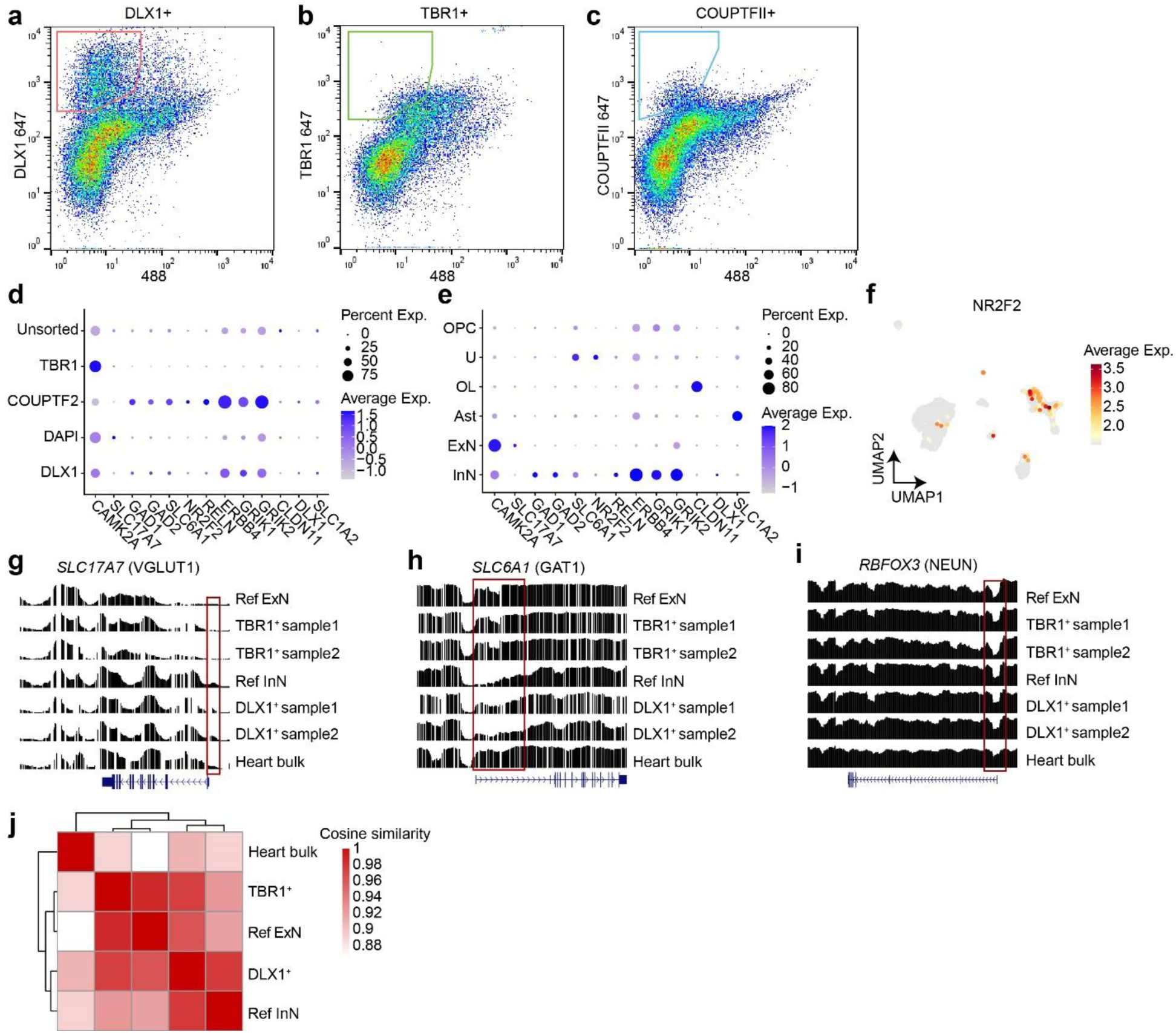
Bisulfite sequencing in sorted TBR1^+^ and DLX1^+^ nuclear pools correlate with excitatory and inhibitory neuron methylome signatures. (a-c) MFNS gating strategy on 30,000 single brain nuclei using DLX1, TBR1, and COUPTFII antibodies. (d) snRNA-seq of post-MFNS nuclei confirming enrichment of targeted nuclear types. (e) Marker expression in assigned nuclear types correlating with targeted nuclear types. (f) UMAP plot *NR2F2* expression pattern (encoding COUPTFII) highlighting a subpopulation of inhibitory neurons (compare with Fig. 1c). (g-i) Reference excitatory and inhibitory neuronal methylome signatures (aggregated from an available public single-nuclei methylome dataset^31^) compared to methylome signatures of sorted nuclei and a bulk heart sample. Normalized relative methylation levels (y-axes) and genomic positions (x-axes) of genes listed at top. (g) Methylome signature of *SLC17A7* encoding VGLUT1, an excitatory neuronal marker in the brain, showing reduced methylation (i.e. representing activation) across the gene body and especially near the transcription start site (TSS, red box) in TBR1^+^ excitatory neuron samples. (h) Methylome signature of *SLC6A1* encoding VGAT1, an inhibitory neuronal marker in the brain, showing reduced methylation across the gene body and especially near the TSS (red box). (i) Methylome signature of RBFOX3 encoding NEUN, a mature neuronal marker in the brain, showing reduced methylation at the TSS in neurons compared with bulk heart. Ref ExN, reference excitatory neurons; Ref InN, reference inhibitory neurons. (j) Heatmap and dendrograms based on cosine similarities of global methylation patterns between groups. Two different TBR1^+^ or DLX1^+^ samples were aggregated. The TBR1^+^ nuclear pool was clustered with Ref ExN while the DLX1^+^ clustered near the pool with Ref InN. The control heart bulk sample was distant from either group.

**Extended Data Fig. 3.**
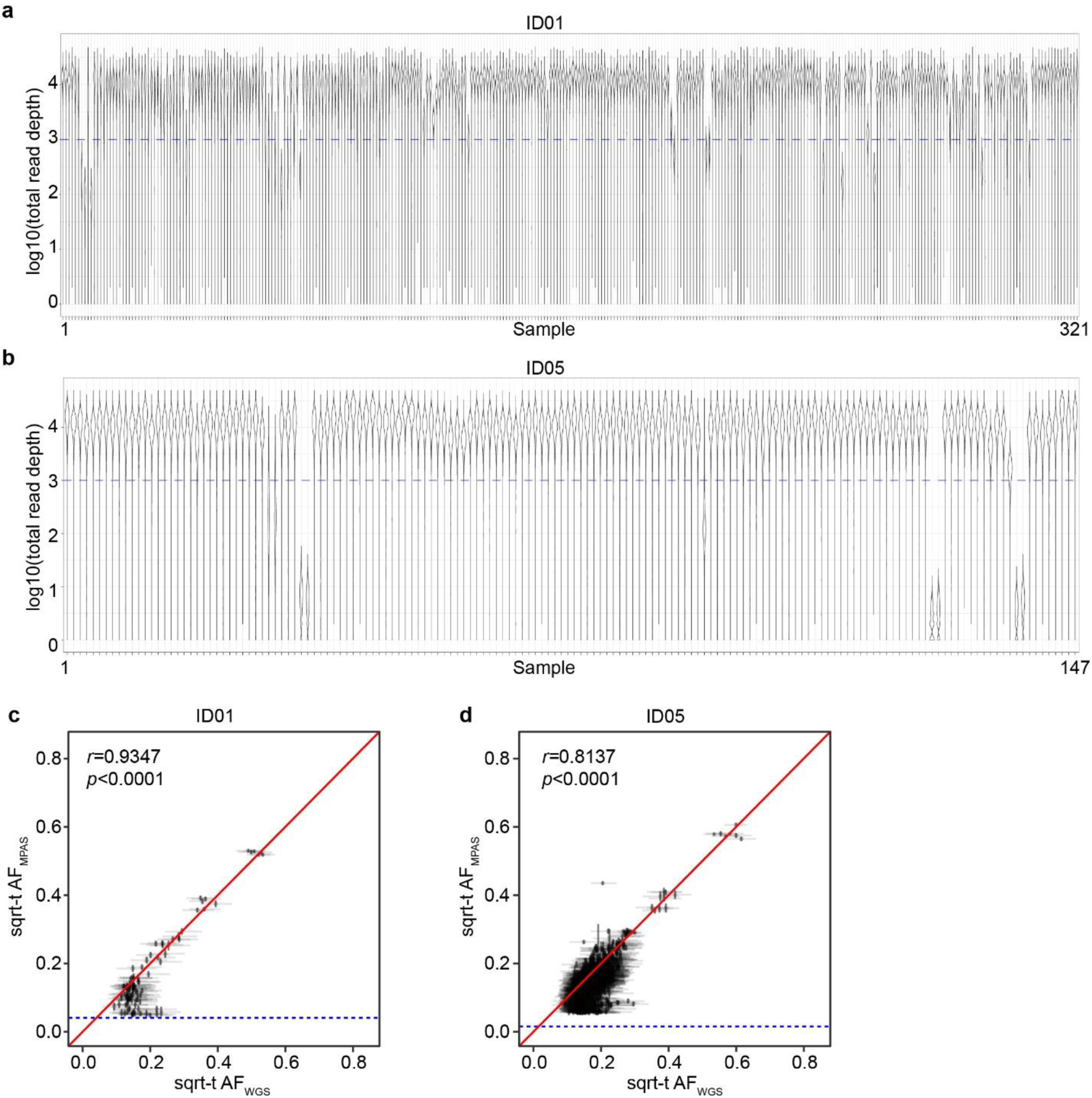
Quality controls of MPAS results. (a-b) Violin plot distribution of log-transformed total read depths (y-axes) of individual variant positions in 321 or 147 samples from ID01 or ID05 (x-axes), respectively. (c-d) Correlation between sqrt-t AF of individual variants from WGS and MPAS. Blue horizontal dashed lines: Lower bound for binomial distribution detection threshold. *r* and *p-*values from Pearson’s Product-Moment correlation. Identity lines (red).

**Extended Data Fig. 4.**
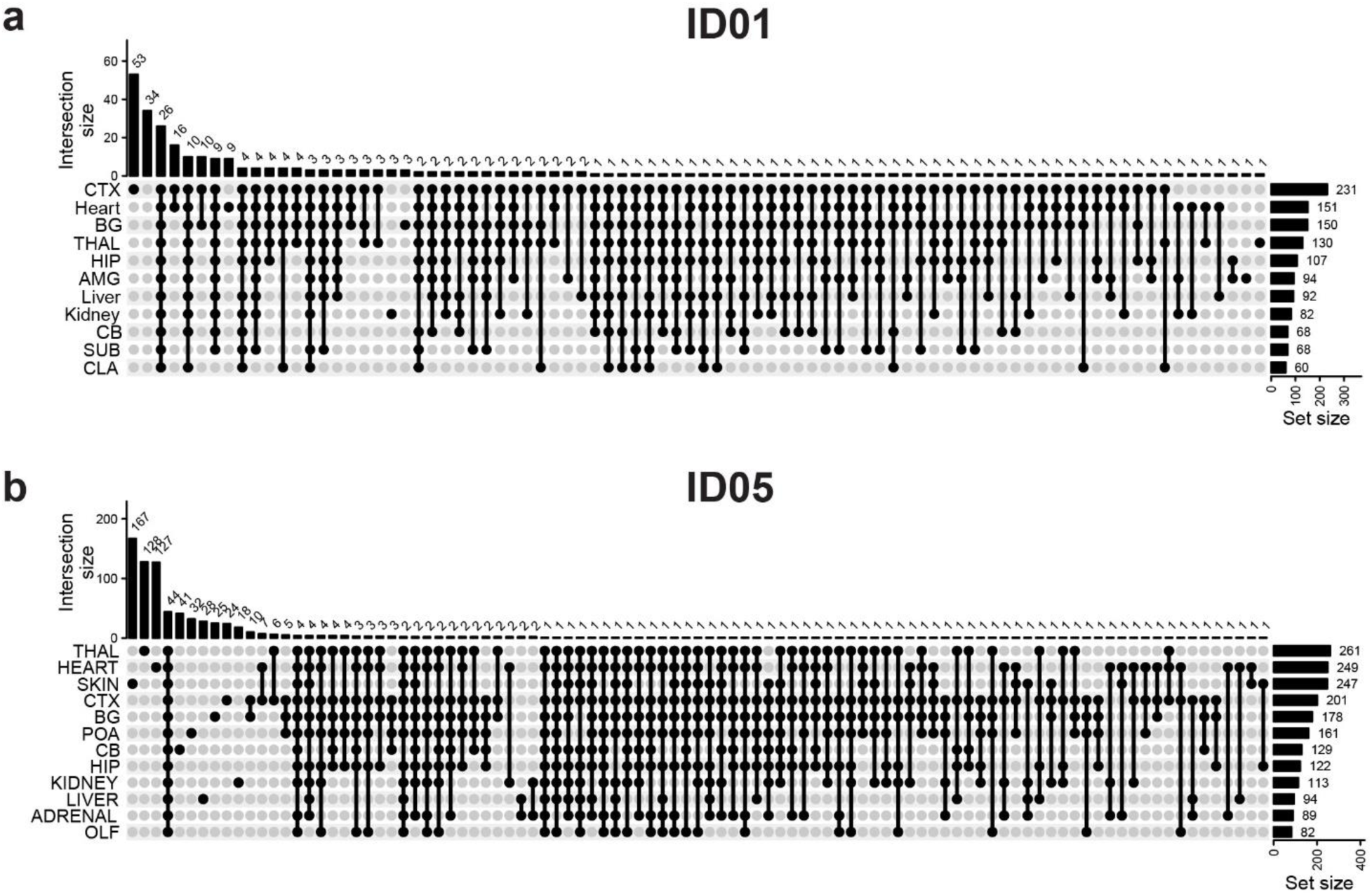
Upset plots showing the intersection of validated MVs detected at different tissue sites. (a) ID01. (b) ID05. CTX, cortex; BG, Basal ganglia; THAL, thalamus; HIP, hippocampus; AMG, amygdala; CB, cerebellum; SUB, subiculum; CLA, Claustrum; POA, preoptic area; OLF, olfactory bulb.

**Extended Data Fig. 5.**
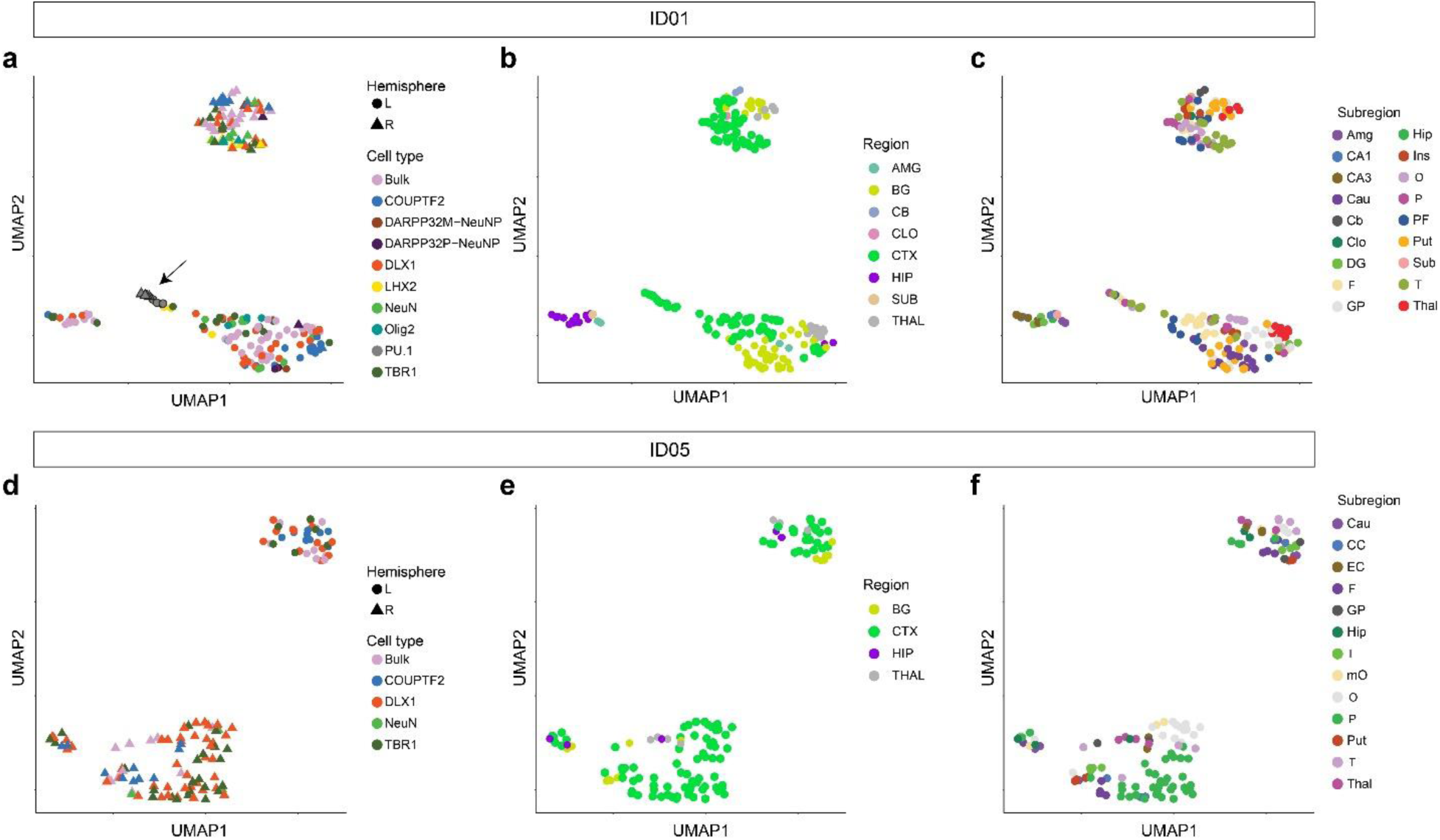
UMAP relationships between samples from the brain based on AFs of validated MVs. Clustering by the same hemisphere validates lateralization of brain-derived cell clones except for the independent origin of microglia (marked by PU.1, arrow). (a-c) UMAP clustering in ID01 samples labeled by (a) cell type, (b) gross region, or (c) subregion, respectively. Clustered samples tend to show similar AF patterns. (d-f) UMAP clustering in ID05 samples labeled by (d) cell type, (e) region, or (f) subregion, respectively. Although PU.1 cells were not sorted in ID05, other findings are similar between ID01 and ID05.

**Extended Data Fig. 6.**
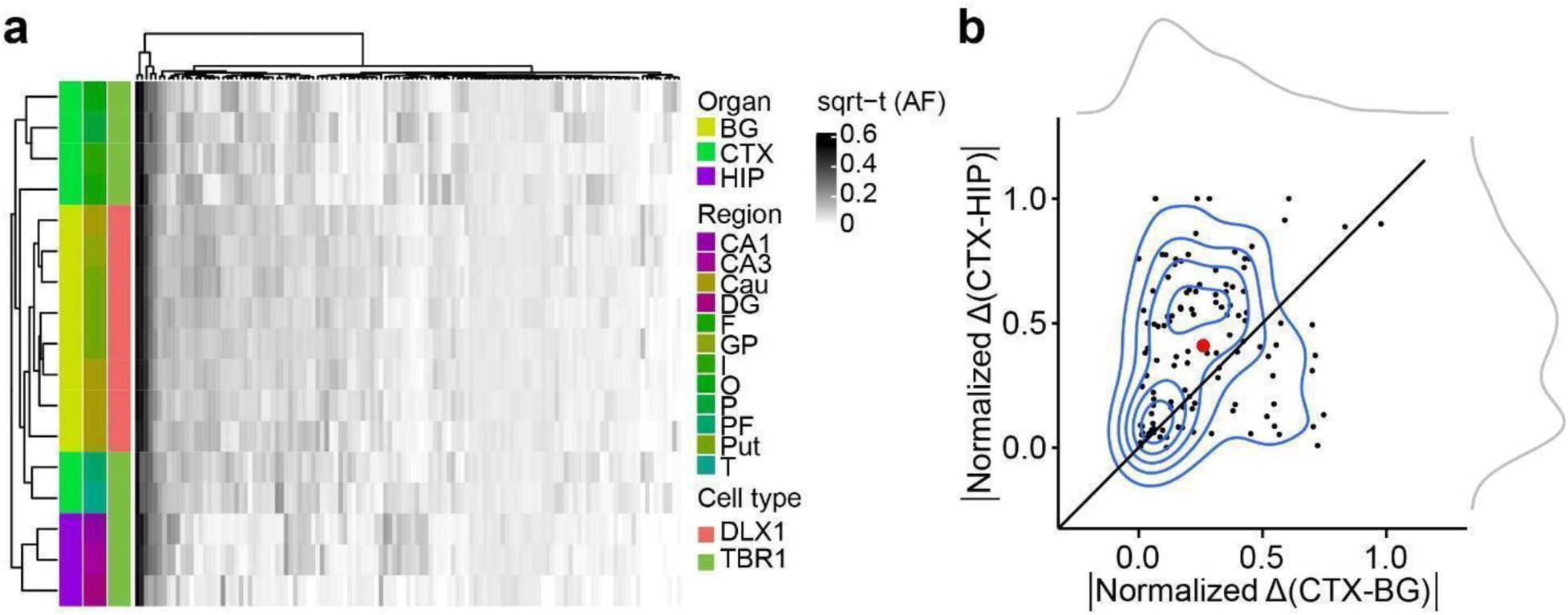
Evidence for HIP lineage restriction occurring prior to CTX or BG in ID01 sorted nuclear pools. (a) Heatmap with 17 sorted nuclear samples based on sqrt-t AFs of 121 informative MVs from ID01, similar to Fig. 2c, showing greater HIP lineage separation compared with CTX or BG (purple compared with green or yellow). (b) Contour plot (at center) with 121 informative MVs derived from (a) and two kernel density estimation plots (at periphery). Axes show the absolute normalized difference value for each MV between the average AF of CTX and BG (CTX-BG) or CTX and HIP regions (CTX-HIP). Solid line: identity. Red dot: averaged x and y values of individual data points. sqrt-t AF, square-root transformed allele fraction; CTX, cortex; BG, basal ganglia; HIP, hippocampus; Cau, caudate; DG, dentate gyrus; HIP and Hip, hippocampal tissue; I, insular cortex; O, occipital cortex; P, parietal cortex; PF, prefrontal cortex; Put, putamen; T, temporal cortex; GP, globus pallidus.

**Extended Data Fig. 7.**
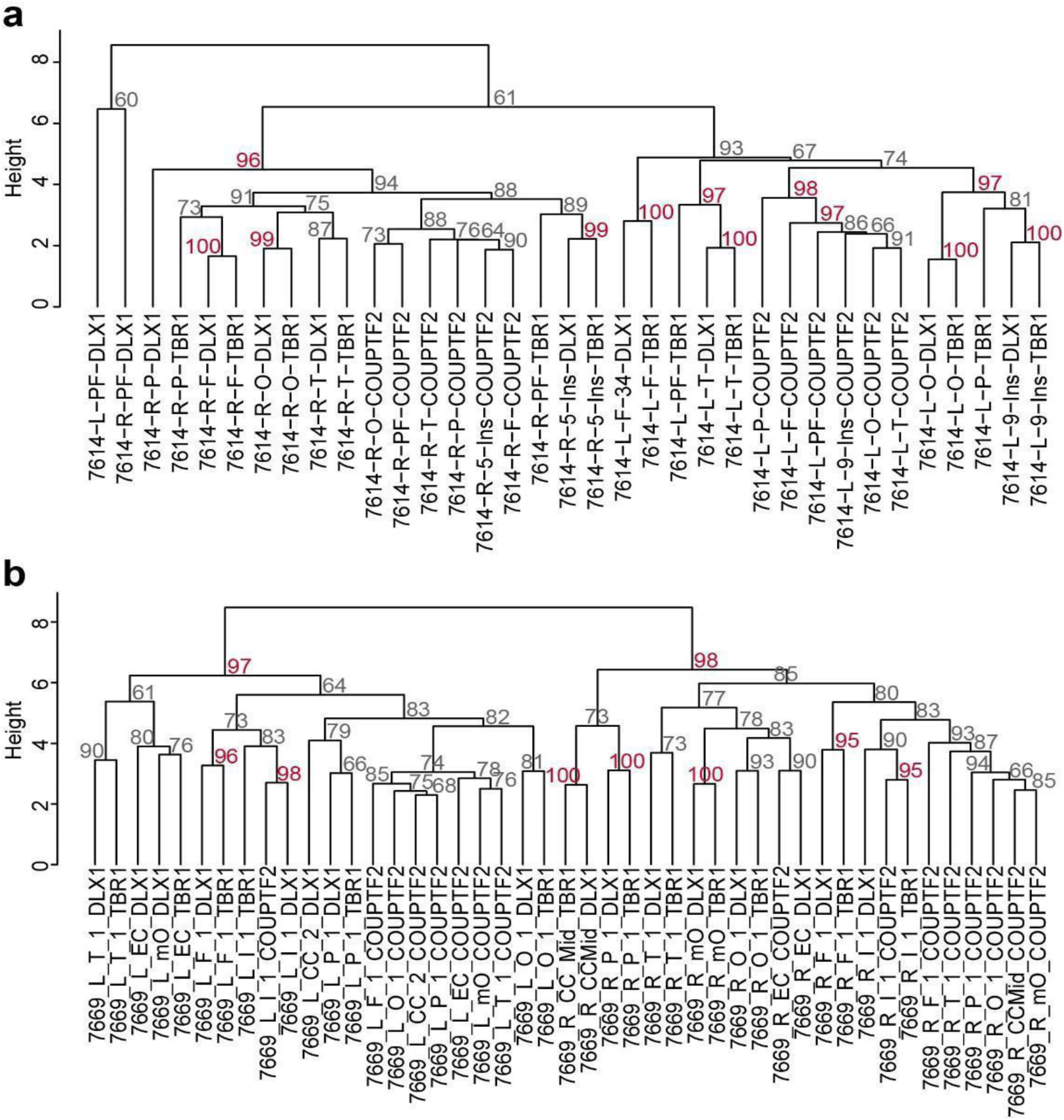
Bootstrapping results for dendrograms with sorted nuclei from cortical areas. (a) Bootstrapping results of ID01. (b) Bootstrapping results of ID05. The percentage of 10,000 replicates showing relationships between sqrt-t AFs for TBR1^+^ and DLX1^+^ nuclei in the same geographic region were more similar than TBR1^+^ nuclei from two different geographic regions (arrow for example). COUPTFII nuclei clustered among themselves, outside of the DLX1 and TBR1 clusters. Approximated unbiased *p*-value > 95% (red): the hypothesis “the cluster does not exist” rejected with a significance level (< 5%).

**Extended Data Fig. 8.**
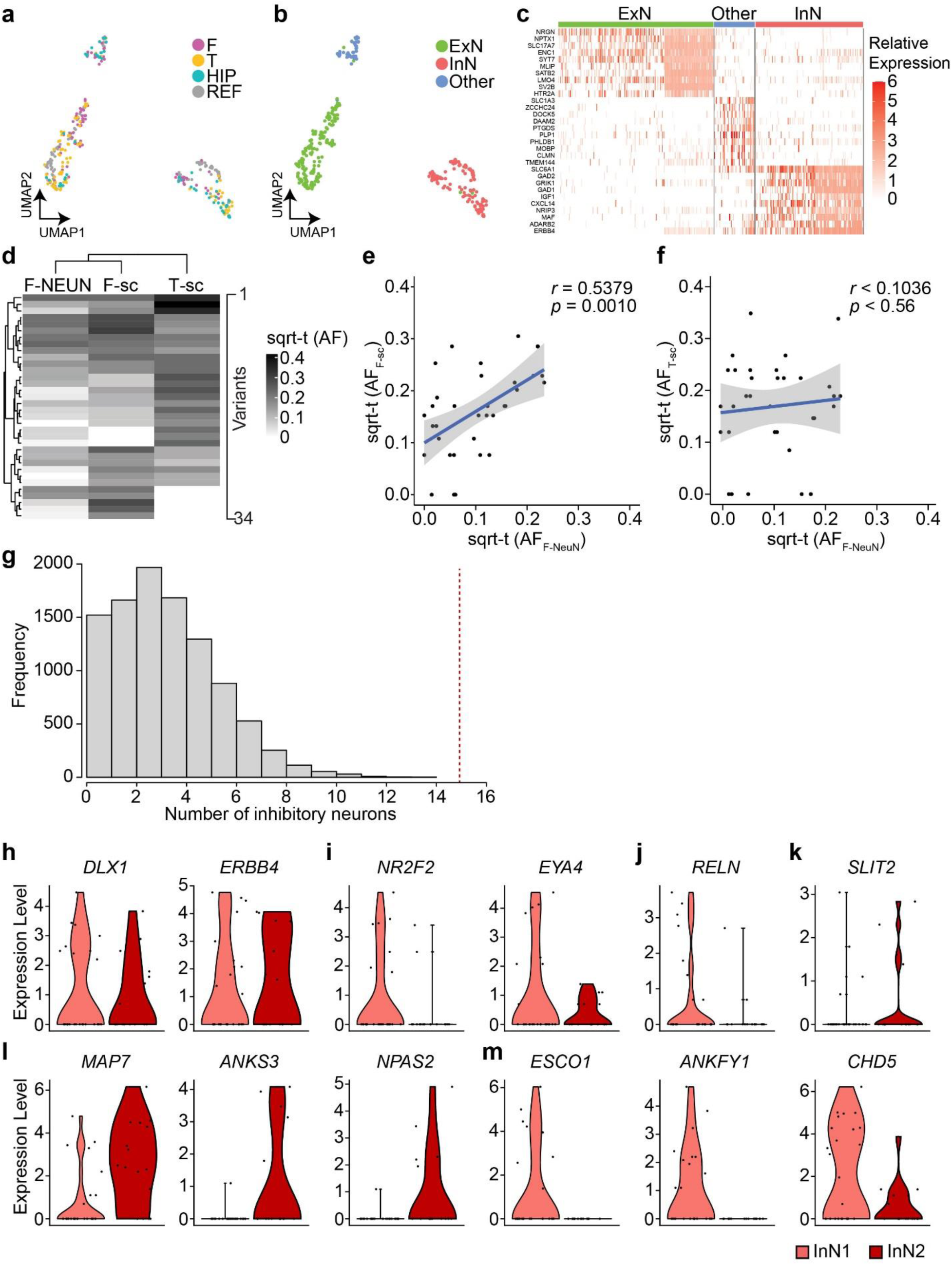
Quality controls of the ResolveOME dataset in ID05. (a) A UMAP plot of snRNA-seq using 225 NEUN ^+^ nuclei and 121 aggregated reference cell types^52,53^. F, frontal; T, temporal; HIP, hippocampus; REF, reference dataset. (b) UMAP labeled by cell types. Note that UMAP clusters separate by cell type (ExN, InN or Other) more than by location. (c) Relative expression of cell type markers within clusters, confirming cell identity. (d) Hierarchical clustering based on sqrt-t AFs of 34 informative MVs shared in 5 to 29 cells in single-nuclear data. F-NEUN, sorted frontal NEUN^+^ nuclei pool; F-sc, pseudo-bulk snMPAS data from a frontal lobe punch; T-sc, snMPAS data from a frontal (F) lobe punch. (e) Correlation between sqrt-t AFs of MVs between F-NEUN and F-sc. (f) Correlation between sqrt-t AFs of MVs between F-NEUN and T-sc. Linear regression with upper and lower 95% prediction intervals displayed by blue solid lines and gray surrounding area; sqrt-t (AF), sqrt-t AF. Pearson’s Product-Moment correlation in e and f. (g) Null distribution of the frequency of the number of inhibitory neurons carrying MVs exclusively detected in one lobe and shared with at least two other local cells, including one excitatory neuron within the same lobe. 10,000 permutations. Portion to the right of the red dashed line, compared to the entire distribution, represents the probability (*p* < 0.0001, one-tailed permutation test) of having 15 or more InNs. (h-m) RNA expression levels of informative genes between InN1 (*n* = 17) and InN2 (*n* = 16) (Fig 4b) in snRNA-seq. (h) Comparable expression levels of inhibitory neuronal markers between both groups. (i) Decreased tendency for the expression of CGE-derived cell markers in InN2 compared to InN1, implying COUPTFII+ inhibitory neurons are unlikely InN1, consistent with previous observations in sorted nuclear populations. (j) RELN^+^ inhibitory neuronal marker showed decreased expression tendency in InN2 compared to InN1. (k) Increased expression tendency for parvalbumin-positive (PV^+^) inhibitory neuronal marker in InN2 compared to InN1, implying dorsally derived inhibitory neurons include PV^+^ neurons. (l, m) top 3 genes increased (l) or decreased (m) in InN2 compared to InN1 among the most variable 3000 protein-coding genes.

---

**Supplementary Data 1. Sample information.**

**Supplementary Data 2. MPAS sequencing results with annotations.**

**Supplementary Data 3. Visualization of AFs of all validated MVs for bulk and sorted nuclei. Supplementary Data 4. Validated variant list with annotations.**

**Supplementary Data 5. Visualization of AFs of all validated MVs for three cell types in the cortex.**

**Supplementary Data 6. Pearson correlation between MVs based on regional AF patterns within the three sorted nuclei groups.** The validated MVs observed in the according cell types among the ‘Sort_main’ group labeled in Supplementary Data 1. Sidebars indicate the locations where the MVs were detected in MPAS.

**Supplementary Data 7. Visualization of AFs of all MVs for 191 NEUN^+^ nuclei. Supplementary Data 8. Visualization of AFs of informative 186 MVs for TBR1^+^ and DLX1^+^ nuclei in 17 positions of the ID05 right parietal cortex.**

**Supplementary Data 9. Targeted genomic regions for MPAS and snMPAS.**

## Material and Methods

### Subject recruitment

The entire cortex, cerebellum, basal ganglia, as well as heart, liver, and kidneys of ID01 and ID05, were collected from the UC San Diego Anatomical Material Program (Case Number UCSD-19-110 and UCSD-21-160, respectively). Organs of ID01 were donated from a 70-year-old female, cause of death was ‘global geriatric decline’ with a contributing cause of ‘post-surgical malabsorption’ as documented^10^. Organs of ID05 were donated from a 73-year-old female, medical history indicated ‘knee replacement, fractured pelvis, hernia, fractured fibula, hypothyroidism, empyema, pulmonary arterial hypertension, and scleroderma’. Both donors were documented to be of European ancestry. Organs were collected within a 26-hour postmortem interval for both donors (ID01: 24 hrs, ID05: 26 hrs). Prior medical history showed no signs of neurological, psychiatric or cancer diseases for either, and tested negative for infection with HIV, Hepatitis B, or COVID-19.

According to 45 CFR 46.102(e)(1), The use of human anatomical cadaver specimens of ID01 and ID05 are exempt from oversight of the University of California, San Diego Human Research Protections Program (IRB) but are subject to oversight by the University of California, San Diego Anatomical Materials Review Committee (AMRC). This study was overseen and approved by the AMRC. The approval number is 106135. Donors met AMRC qualifications: (i) Obtain information or biospecimens through intervention or interaction with the individual, and uses, studies, or analyzes the information or biospecimens; or (ii) Obtains, uses, studies, analyzes, or generates identifiable private information or identifiable biospecimens.

### Tissue dissection

For ID01, after the removal of the meninges, diencephalon regions, and brain stem the cerebral cortical regions including lobes from prefrontal, frontal, parietal, occipital, and temporal cortex, and cerebellum were dissected by a pathologist. Punches were further collected from each of the non-neocortical tissues: from the cerebellum (4 punches from each hemisphere, 2 of which were pooled for WGS analysis), heart, liver, and each kidney, with special attention to avoiding sample cross-contamination, as documented^10^. Frozen brain tissue was further sliced at 1 cm thickness to dissect the I (or Ins), Cau, Put, AMG, GP, HIP (CA1-CA3, and DG wherever distinguishable), thalamus, and other subcortical structures. For ID05, apart from the organ samples collected in ID01, the adrenal gland and leg skin were further sampled. Cau, Put, GP, and HIP samples were collected before frozen. Extensive 13-17 sublobar punch biopsies were collected from all 10 cortical lobes from ID01 as well as the right occipital lobe and the right parietal lobe from ID05 with an 8mm skin punch. Sample information summarized in Extended Data Fig. 1 and Supplementary Data 1. The dissection procedure was conducted on ice at room temperature for ID01 and in a cold room for ID05. During dissection, subsamples and the remnants of the large pieces were immediately labeled and snap-frozen on dry ice, and stored at -80°C.

### Lobar tissue homogenization and nuclei extraction

Frozen brain lobar samples (after the removal of the 8mm biopsies, i.e. Lrg) of ID01 were pulverized in liquid nitrogen and then homogenized in 1% formaldehyde in Dulbecco’s phosphate-buffered saline (DPBS, Corning) using a motorized homogenizer (Fisherbrand PowerGen 125), then incubated on a rocker at room temperature for 10 min, quenched with 0.125 M glycine at room temperature on a rocker for 5 min, then centrifuged at 1,100×g in a swinging bucket centrifuge. The following steps were all performed on ice except where indicated. Homogenates were washed twice with NF1 buffer (10 mM Tris-HCl pH 8.0, 1 mM EDTA, 5mM MgCl_2_, 0.1M sucrose, 0.5% Triton X-100 in UltraPure water) and centrifuged at 1,100×g for 5 min at 4°C in a swinging bucket centrifuge. Next, pellets were resuspended in 5 ml NF1 buffer and Dounce homogenized 5x in a 7 ml Wheaton Dounce Tissue Grinder (DWK Life Sciences) using a ‘loose’ pestle. After 30 minutes of incubation on ice, homogenates were Dounce homogenized 20x with a ‘tight’ pestle and filtered through a 70 μm strainer. To remove myelin debris, homogenates were overlaid on a sucrose cushion (1.2M sucrose, 1 M Tris-HCl pH 8.0, 1 mM MgCl_2_, 0.1 M DTT) and centrifuged at 3,200×g for 30 min with acceleration and brakes on ‘low’. Pellets of nuclei were washed with NF1 buffer and centrifuged at 1,600×g for 5 min and stored at -80°C, same as documented^10,54^.

### DNA extraction of bulk tissue and lobar nuclear fractions

Small cortical biopsies were first cut in half on dry ice. Half of the biopsy was stored as backup and partly used for single nuclei fluorescence-activated nuclei sorting (FANS) for ID01 and ID05. The other half of the cortical biopsy was homogenized with a Pellet Pestle Motor (Kimble, 749540-0000) and resuspended with 450 µL RLT buffer (Qiagen, 40724) in a 1.5 ml microcentrifuge tube (USA Scientific, 1615-5500). The same experimental procedure was carried out on both punches from the cerebellum, heart, liver, and both kidneys. Nuclear preparations were pelleted at 1,000 ×g for 5 min and resuspended with 450 μL RLT buffer in a 1.5 ml microcentrifuge tube. Both homogenates and nuclear preps were then treated with the same protocol: following 1 min vortex, samples were incubated at 70°C for 30 min. 50 μl Bond-Breaker TCEP solution (Thermo Scientific, 77720) and 120 mg stainless steel beads (0.2 mm diameter, Next Advance, SSB02) were added, and cellular/nuclear disruption was performed for 5 min on a DisruptorGenie (Scientific Industries), supernatant was transferred to a DNA Mini Column from an AllPrep DNA/RNA Mini Kit (Qiagen, 80204) and centrifuged at 8500×g for 30 sec, washed with Buffer AW1 (Qiagen, 80204), centrifuged at 8500×g for 30 sec and washed again with Buffer AW2 (Qiagen, 80204), and then centrifuged at full speed for 2 min. DNA was eluted two times with 50 μl of pre-heated (70°C) EB (Qiagen, 80204) through centrifugation at 8,500×g for 1 min as documented previously^10^. For low-input DNA extraction from sorted nuclei, we further developed an on-bead DNA extraction method derived from a published protocol^55^: sorted nuclei were centrifuged down for 1 minute at 1,000g (4°C) in a 200 μl PCR tube and resuspended in 20 μl lysis buffer that consists of 30mM Tris-HCl (pH 8.0), 0.5%(v/v) Tween-20 (Sigma-Aldrich P1379), 0.5%(v/v) IGEPAL CA-630 (Sigma I8896), 1.25 µg/ml protease (Qiagen, 19155) as final concentration in nuclease-free water (Ambion, AM9937). The mixture was lightly vortexed for 10 sec and centrifuged at 1,500g for 1 min (4°C). The tube was then subjected to 50°C for 12 min and 75°C for 30 min on a thermocycler. To each lysate, 20 μl of Agencourt AMPure XP beads (Beckman Coulter, A63881) was added. The final AMPure beads to sample ratio was 1:1. After pipetting to achieve mixing, the mixture was left at room temperature for 5 min, placed on a magnet for 5 min, and the supernatant removed. The remaining material was washed twice with 150 μl of 80% (v/v) ethanol for 1 min each. The DNA was then suspended in Low TE solution from the AmpliSeq for Illumina kit (Illumina, 20019103) and kept on beads at -20℃ until MPAS analyses.

### Whole-genome library preparation and deep sequencing

A total of 1.0 µg of extracted DNA was used for PCR-free library construction using the KAPA HyperPrep PCR-Free Library Prep kit (Roche, KK8505). Mechanical shearing using the Covaris microtube system (Covaris, SKU 520053) was performed to generate fragments with a peak size of approximately 400 base pairs (bp). Each fragmented DNA sample went through multiple enzymatic reactions to generate a library in which an Illumina dual index adapter would be ligated to the DNA fragments. Beads-based double size selection was performed to ensure the fragment size of each sample was between 300-600 bp as measured by an Agilent DNA High Sensitivity NGS Fragment Analysis Kit (Agilent, DNF-474-0500). The concentration of ligated fragments in each library was quantified with the KAPA Library Quantification Kits for Illumina platforms (Roche/KAPA Biosystems, KK4824) on a Roche LightCycler 480 Instrument (Roche). Libraries with concentrations of more than 3 nM and fragments with peak size of 400 bp were sequenced on an Illumina NovaSeq 6000 S4 and/or S2 Flow Cell (FC). Each library was sequenced in 6-8 independent pools. For each sequencing run, 24 WGS libraries were normalized to obtain a final concentration of 2 nM using 10 mM Tris-HCl (pH 8 or 8.5; Fisher Scientific, 50-190-8153). 0.5 to 1% PhiX library was spiked into the library pool as a positive control. The normalized libraries in a pool with a total of 311 µl libraries were incubated with 77 µl of 0.2 N Sodium Hydroxyl (NaOH) (VWR, 82023-092) at room temperature for 8 minutes to denature double-stranded DNA. 78 µl of 400 mM Tris-HCl was used to terminate the denaturing process. The denatured library with a final loading concentration of 400 pM in a pool was loaded on the S4 FC using Illumina SBS kits (Illumina, 20012866) with the following setting on the NovaSeq 6000: PE150:S4 FC, dual Index, Read 1:151, Index_Read2:8; Index_Read3:8; Read 4:151. The target for whole genome sequencing with high-quality sequencing raw data was 120 GB or greater with a Q30 >90% per library per sequencing run. In case the first sequencing run generated less than that, additional sequencing was performed by sequencing the same library on a NovaSeq 6000 S2 FC with a 2×101 read length for ID01 as documented^10^ and all data were generated at 2×151 read length for ID05. FASTQ files generated with Picard’s (v 2.20.7) *SamToFastq* command from the DRAGEN platform were used as input for the bioinformatic pipeline for ID01 and *bcl2fastq2* (v 2.20) generated FASTQ files from raw sequence files were used for ID05.

### Whole-genome sequencing (WGS) data processing

FASTQ files were then aligned to the human_g1k_v37_decoy genome by BWA’s (v 0.7.17) *mem* with *-K 100000000 -Y* parameters. SAM files were compressed to BAM files via SAMtools’s (v 1.7) *view* command. BAM files were subsequently sorted by SAMBAMBA’s (v 0.7.0) *sort* command and duplicated reads marked by its *markdup* command. Reads aligned to the INDEL regions were realigned with GATK’s (v 3.8-1) *RealignerTargetCreator* and *IndelRealigner* following the best practice guideline. Base quality scores were recalibrated using GATK’s (v 3.8.1) *BaseRecalibrator* and *PrintReads*. Germline heterozygous variants were called by GATK’s (v 3.8.1) *HaplotypeCaller*. The distribution of library DNA insertion sizes for each sample was summarized by Picard’s (v 2.20.7) *CollectInsertSizeMetrics*. The depth of coverage of each sample was calculated by BEDTools’s (v2.27.1) *coverage* command. The code and Snakemake wrapper of the pipeline are freely accessible on GitHub (https://github.com/shishenyxx/Human_Inhibitory_Neurons).

### Mosaic SNV/INDEL detection in WGS data

Mosaic single nucleotide variants/mosaic small (typically below 20 bp) INDELs were called by using a combination of four different computational methods: MosaicHunter (single-mode, v 1.0)^56^ with a posterior mosaic probability >0.05 ^10,57^, or Single-mode of GATK’s (v 4.0.4) Mutect2^58^ with “PASS” followed by DeepMosaic (v 1.0.1)^59^ and MosaicForecast (v 8-13-2019)^60^, were implemented for sample-specific or tissue-shared variants; the intersection of variants from the paired-mode of Mutect2 and Strelka2 (v 2.9.2) (set on “pass” for all variant filter criteria)^61^ were collected for sample-specific variants. For the panel of normal samples required for the pipeline of DeepMosaic and MosaicForecast, we employed an in-house panel of similarly (300×) sequenced normal tissues (*n* = 15 sperm and 11 blood samples from 11 individuals)^10^. For ‘tumor’-‘normal’ comparisons, required by Mutect2 and Strelka2 pipelines, we employed left-right combined heart tissues as ‘normal’. Variants were excluded if: 1) residing in segmental duplication regions as annotated in the UCSC genome browser (UCSC SegDup) or RepeatMasker regions, 2) residing within a homopolymer or dinucleotide repeat with more than 3 units, or 3) overlapped with annotated germline INDELs. We further removed any variants with a population allele frequency higher than 0.001 in gnomAD (v 2.1.1)^62^. Finally, variants with a lower CI of AF <0.001 were considered noises from reference homozygous and removed. Fractions of mutant alleles (i.e., AF) for variants called in one sample were calculated in all the other samples together with the exact binomial confidence intervals using scripts described below for MPAS analysis. This bioinformatic pipeline yielded a total of 898 candidate MVs for ID01 and 2195 candidate MVs for ID05 (for skin samples because of the clonal nature only 10% of the total calls were randomly selected) that were interrogated with MPAS. Scripts for variant filtering are provided on GitHub (https://github.com/shishenyxx/Human_Inhibitory_Neurons).

### Nuclear preparation for MFNS or unfixed nuclei sorting

For the MFNS protocol, approximately 200 mg of freshly frozen tissue stored at -80°C was prepared, and subsequent procedures were conducted using solutions maintained at 4°C. The prepared tissue was homogenized in 300 µl of lysis buffer (composed of 10 mM Tris-HCl, pH 7.4, 10 mM NaCl, 3 mM MgCl2, 0.1% NP-40, and 1 mM Dithiothreitol in nuclease-free water) while kept on ice. Following this, an additional 9.7 ml of lysis buffer was added to the homogenate, and the mixture was incubated on ice for 5 minutes. The homogenate was then passed through a 70 µm cell strainer (FALCON, 352350) and centrifuged at 1100g for 5 minutes at 4°C. The supernatant was discarded, and the remaining pellet was gently resuspended and washed with 10 ml of sorting buffer (containing 1% bovine serum albumin, 1 mM EDTA, and 10 mM HEPES in 1x HBSS solution). For the density gradient centrifugation step, the resuspended pellet in 25% Iodixanol solution (OptiPrep™, Millipore D1556) was layered onto a 29% Iodixanol cushion. The centrifugation was carried out at 10,000g with a swinging bucket rotor, employing low acceleration and braking, at 4°C for 40 min, the pellet resuspended in 80% methanol prechilled at -20°C and stored at -20°C for at least 30 minutes before further use. For single-nucleus transcriptome and whole-genome amplification with ResolveOME, brain tissue samples from the right F, T, and HIP of participant ID05 were homogenized in 1 ml of ice-cold NIB (composed of 0.25 M sucrose, 25 mM KCl, 5 mM MgCl2, 10 mM Tris pH 7.5, 100 mM DTT, and 0.1% Triton X-100) and subjected to homogenization, incubated on a rocker for 5 min at 4 °C and then centrifuged at 1,000g employing low acceleration and braking in a swinging-bucket centrifuge, and the pellet was reconstituted in 0.5 ml of sorting buffer and filtered through a 70-um strainer. Nuclei in the flow-through were immediately subjected to staining.

### Fluorescence-activated nuclear sorting

For pellets of defrosted and homogenized brain nuclei were washed twice in sorting buffer and then re-suspended in 0.2 ml sorting buffer and incubated overnight at 4°C. The following antibodies were used: NEUN Alexa Fluor 488 (1:2,500; Millipore Sigma, MAB377), TBR1 unconjugated (1:1,000; Abcam, ab31940), OLIG2 unconjugated (1:1,000; Abcam, ab1091986), LHX2 unconjugated (1:500; Abcam, ab2199883), DLX1 (1:200; Atlas Antibodies, HPA045884), COUPTFII (1:400; Novus Biologicals, PP-H7147-00), PU.1 Alexa Fluor 647 (1:100; BioLegend, 658004). The following day, nuclei were washed with staining buffer and in case an unconjugated antibody was used, nuclei were stained subsequently for 30 minutes with goat anti-rabbit Alexa 647 (1:4,000; ThermoFisher Scientific, A21244) for TBR1, DLX1, or LHX2, goat anti-rabbit Alexa 555 (1:4,000; ThermoFisher Scientific, A32732) for OLIG2, and donkey anti-mouse Alexa 647 (1:4000; ThermoFisher Scientific, A32787) for COUPTFII. Stained nuclei were washed one more time with staining buffer and passed through a 70 μm strainer. Immediately before the sort, nuclei were stained with 0.5 μg/ml DAPI. For single-nucleus transcriptome and whole-genome amplification with ResolveOME, nuclei were stained with Propidium Iodide (1:20; Invitrogen, BMS500PI). Nuclei for the cell type of origin were sorted either on a MoFlo Astrio EQ sorter (Beckman Coulter), BD FACSAria II (Becton-Dickinson), or BD InFlux Cytometer (Becton-Dickinson) similar to previous documentation^10^. For methanol fixed nuclei, at least 200 sorted nuclei were pooled in each tube. Sorted nuclei were pelleted in staining buffer at 1,600×g for 10 minutes. Nuclei for DNA extraction and bisulfite sequencing were stored at -80℃. FANS data was visualized using FlowJo v10 software (Ashland, Oregon). Following MPAS (see below) sorted populations were deemed to be of sufficient overall quality if at least 95% variants were sequenced above >1,000×.

### Bisulfite sequencing of sorted nuclei for cell type of origin

Sorted nuclei DNA on beads were processed by Pico Methyl-Seq Library Prep Kit (Zymo Research, D5455) to generate bisulfite sequencing libraries. Samples were sequenced at PE150 on the Illumina NovaSeq 6000 platform.

### Bisulfite sequencing data processing and data visualization

FASTQ files were analyzed with the Bismark bisulfite mapper and methylation marker (v 0.23.1)^63^, and read pairs were treated as singletons according to the developers’ suggestions. The pipeline was run with Snakemake (v 6.12.3), and bedgraph were generated with BEDTools (v 2.30.0). Python (v 3.10) with argparse, textwrap, and numpy packages and R (4.1.3) with lsa, pheatmap, ggfortify packages were used for visualization. Human single-cell methylation data from published literature^31^ was downloaded, and reads from excitatory neurons and inhibitory neurons were pooled separately and used as positive controls according to the original authors’ cell-type labels. Codes for plotting the cosine similarity and hierarchical clustering of methylation patterns of gold-standard inhibitory neurons, sorted DLX1 positive cells, excitatory neurons, and sorted TBR1 positive cells. Control samples from the heart, cortical microglia, and cortical oligodendrocytes were available on GitHub (https://github.com/shishenyxx/Human_Inhibitory_Neurons).

### Single nucleus transcriptome and whole-genome amplification

After sorting, a total of 192 single nuclei from each of 2 samples (right temporal and frontal cortex of ID05) were snap-frozen on dry ice. Nuclei underwent the ResolveOME™ workflow (BioSkryb Genomics Inc.). Briefly, Biotin-dT-primed first strand cDNA was generated. After termination of the reaction and nuclear lysis, whole genome amplification with primary template-directed amplification was performed ^64^. The mRNA-derived cDNA was affinity purified with streptavidin beads from the combined pool of cDNA and amplified genome. Remaining cDNA were pre-amplified on beads. Independently, amplified cDNA and single-cell genomic DNA from each cell underwent SPRI (Beckman Coulter, B23319) cleanup prior to library preparation. Illumina libraries were prepared using the ResolveOME library preparation kit (BioSkryb Genomics Inc.) with NEXTFLEX Unique Dual Index Barcodes (PerkinElmer Applied Genomics, NOVA-534100). Libraries were sequenced at low-pass (2x 50bp paired end) targeting 2 million reads on a NextSeq (Illumina) instrument. Libraries of interest were identified based on QC sequencing, and were subsequently sequenced at paired-end 150 bp (DNA libraries) and paired-end 100 bp (RNA derived libraries) on a NovaSeq X Plus (Illumina) platform.

### Massive parallel amplicon sequencing (MPAS) and single nucleus MPAS (snMPAS)

Two customized AmpliSeq Custom DNA Panel for Illumina (20020495, Illumina, San Diego, CA, USA) were used for MPAS for ID01 (#203019), as well as MPAS for ID05 (#201745), respectively. Designed genomic regions are provided in Supplementary Data 9. A list of 898 candidate MVs from the MV detection pipeline in ID01 described above was subjected to the AmpliSeq design system. For the first panel, we randomly selected 120 high-confidence heterozygous variants as positive controls. These heterozygous variants presented with estimated AFs between 48-52% for all the 25 sequenced bulk tissues, and with read depths between 270-330×. Of the 120 variants, 45 were private variants and 75 were present in gnomAD at different population allele frequencies. We also randomly selected 30 reference homozygous variants as negative controls. These reference homozygous variants presented with ∼0% AF across all sequenced samples, with average depth 270-330×, and gnomAD (v 2.1.1) allele frequency >0.5 to exclude any potential contamination or amplification bias. For the second panel, 2195 candidate MVs detected from ID05 as well as 152 randomly chosen variants detected as heterozygous and 59 as alternative homozygous in ID05 were subjected to the AmpliSeq design system. The AmpliSeq design software determined 1048 pairs of primers suitable for multiplex PCR reaction in a single pool after optimization for the ID01 panel and 1811 pairs for the ID05 panel. DNA from extracted tissue, nuclei, amplified single nuclei, and a duplicate unrelated control sample was diluted to 5 ng/µl in low TE provided in AmpliSeq Library PLUS (384 Reactions) kit (Illumina, 20019103). AmpliSeq was carried out following the manufacturer’s protocol (document 1000000036408v07). For amplification in bulk samples, 14 cycles each with 8 minutes were used, for amplification in low-input sorted nuclei, additional cycles and time is added accordingly based on the recommended table from the manufacturer. After amplification and FUPA treatment, libraries were barcoded with AmpliSeq CD Indexes (Illumina, 20031676) and pooled with similar molecular numbers based on measurements made with a Qubit dsDNA High Sensitivity kit (Thermo Fisher Scientific, Q32854) and a plate reader (Eppendorf, PlateReader AF2200). To avoid index hopping, the two MPAS library pools for ID01 and ID05 and the snMPAS library pool for ID05) were sequenced on separate lanes on different NovaSeq 6000 runs. 192 GB of FASTQ data were obtained from the ID01 MPAS libraries, 339 GB of FASTQ data were obtained from the ID05 MPAS libraries, 193 GB of FASTQ data were obtained from the ID05 snMPAS libraries, all aiming for an average of 10000×for each variant.

### Data analysis for MPAS and snMPAS

Raw reads from MPAS and snMPAS were mapped to the human_g1k_v37_decoy genome with BWA’s (v 0.7.17) *mem* command. BAM files were processed without removing PCR duplicates. Reads near insertion/deletions were re-aligned with GATK’s (v 3.8.1) *IndelRealigner* and base qualities scores were recalibrated with GATK’s (v 3.8.1) *BaseRecalibrator*. The final BAM files were parsed by SAMtools’s (v 1.7) *mpileup* and the 95% confidence intervals (CIs) of the measured AF of all the candidate MVs, together with the homozygous (negative control) and heterozygous (positive control) variants were estimated based on an exact binomial estimation (https://github.com/shishenyxx/Human_Inhibitory_Neurons). Following depth calculation, regions of 639 mosaic candidates, 120 heterozygous variants (positive controls), and 30 homozygous variants (negative controls) were detected and subjected to the next genotyping steps with 259 previously validated MVs ^10^ for ID01, and 2195 mosaic candidates, 152 heterozygous variants (positive controls), and 59 homozygous variants for ID05. The genotypes of candidate MVs from MPAS were determined by comparing them to the AF distribution of the reference homozygous and heterozygous variants, and by comparing the AF distribution of MVs in the two individuals, in which the variant was not originally detected. The exact binomial lower bounds of all reference homozygous variants with >30 read depth were estimated and the 95% single-tail confidence threshold for the lower bound was calculated to be 0.001677998 (ID01) and 0.002360687 (ID05). The distribution of the exact binomial upper bond of all heterozygous variants was calculated and 0.3923302 (ID01) and 0.4562841 (ID05) were the threshold for the upper bond based on ∼5% false discovery rate (FDR) and manual inspection. Mosaic candidates from WGS were considered positive if variants met the following criteria at the same DNA samples: 1) the 95% exact binomial lower bound was >0.001677998 (ID01) or >0.002360687 (ID05), respectively, 2) the lower CI of the unrelated control sample was <0.001677998 (ID01) or <0.002360687 (ID05), 3) the 95% exact binomial upper bound was <0.3923302 (ID01) or <0.4562841 (ID05), respectively, 4) the sequencing depth was >30, and 5) the assessed alternative allele was supported by ≥3 reads. 6) for ID01, variants are detected in GP, Cau, Put, and Thal in WGS. These criteria ensured the FDR for each variant was under 5%. After the MPAS quantification, 287 MVs (28 newly validated variants and original 259 variants^10^) from ID01 and 780 MVs from ID05 were considered positively validated in the sample where the variant was originally detected and used for all the analysis presented throughout the manuscript. In snMPAS, mosaic candidates from WGS were considered positive if the lower CI of AF was larger than the upper CI of AF in the normal control sample.

### Permutation test for the significance of the snMPAS result

Regarding MVs presented more than 2 nuclei and a maximum of 14 nuclei from Fig. 4b, detection events of each MV were randomly re-assigned within a total of 118 nuclei (16 and 17 InNs in F and T respectively; 40 and 45 ExNs in F and T, respectively), maintaining the original detection frequency. The number of inhibitory neurons sharing MVs exclusively in one lobe and shared with at least two other local cells, including one excitatory neuron, were used as the outcome and a null distribution was generated from the 10,000 permutations. The probability of having more than or equal to 15 inhibitory neurons sharing MVs exclusively in one lobe and shared with at least two other local cells, including one excitatory neuron was calculated and used as the one-tailed permutation *p-* value.

### Single-nucleus RNA sequencing with Chromium platform

Nuclei that underwent MFNS (16000 nuclei) were resuspended in a sorting buffer to make the desired concentration (800∼1000 nuclei/ul) targeting 10000 nuclei per reaction. Gel beads emulsion (GEM) generation, cDNA, and sequencing library constructions were performed in accordance with instructions in the Chromium Single Cell 3’ Reagent Kits User Guide (v3.1). Each library pool was sequenced with 200 million read pairs using NovaSeq 6000.

### Single-nucleus RNA-seq bioinformatics pipeline

For the snRNA-seq data made with Chromium platform, fastq files from single-nucleus libraries were processed through Cell Ranger (v6.0.2) analysis pipeline with –include-introns option and hg19 reference genome. Seurat (v4.0.5) package was used to handle single nuclei data objects. Nuclei passed a control filter (nCount > 400, nFeature_RNA < 2000, percentage of mitochondrial gene < 10%) was used for downstream analysis. A total of 15,896 protein-coding genes were used for further downstream analysis. To balance the nucleus number for each group, a total of 500 random nuclei for each DAPI^+^, DLX1^+^, TBR1^+^ and COUPTFII^+^ group were selected. Data were normalized and scaled with the most variable 1000 features using the ‘*ScaleData*’ functions. Dimensionality reduction by PCA and UMAP embedding was performed using *runPCA* and *runUMAP* functions.

Clustering was performed by *FindNeighbors* and *FindClusters* functions. For the full-length transcript snRNA-seq through ResolveOME platform, quality control were carried out for raw FASTQ from single nucleus RNA sequencing from ResolveOME files using fastqc (v0.11.8). Preprocessing was carried out with cutadapt (v1.16). Cleaned FASTQ files were aligned to GRCh38 human genome and gencode (v27) gtf annotation using STAR (v2.6.0c). Aligned BAM files were indexed with SAMtools (v1.7). PCR duplicates were marked with Picard (v2.20.7) MarkDuplicates. Post alignment quality control was carried out with Picard (v2.20.7) CollectRnaSeqMetrics, CollectInsertSizeMetrics, CollectGcBiasMetrics as well as qualimap (v2.2.2-dev). Raw read counts were collected with featureCounts (v2.0.0). Transcripts were collected with rsem (v1.3.1) with seed 12345 using the same gtf file. We also employed trimmed-means data from Human Multiple Cortical Areas SMART-seq dataset (https://portal.brain-map.org/atlases-and-data/rnaseq/human-multiple-cortical-areas-smart-seq) for reference group. Seurat (v4.0.5) package was used for further analysis. Nuclei passed a control filter (nCount > 1000, nFeature_RNA >500, percentage of mitochondrial gene < 30%) was used for downstream analysis. *SCTransform* function was used for normalizing and scaling data. For both snRNA-seq data made with Chromium platform and ResolveOME, cell type identification was performed using known cell type markers expressed in the brain including excitatory (RORB, CUX2, SATB2), inhibitory neuron (*GAD1*, *GAD2*), astrocyte (*SLC1A2*, *SLC1A3*), oligodendrocyte (*MOBP*, *PLP1*), oligodendrocyte precursor cell (*PDGFRA*), microglia (*PTPRC*), and endothelial cell markers (*CLDN5*, *ID1*) as well as using positive markers found by *FindAllMarkers* function with 1000 most variable features in scaled data. The final visualization of various snRNA-seq data was performed by r-ggplot2 (v3.3.5).

### Statistical tests and packages for customized plots

Hierarchical clustering with *p*-value via bootstrap resampling was performed using r-pvclust (v2.2.0) package. Pearson’s product-moment correlation is calculated using *cor.test()* in R. One-way ANOVA with Tukey multiple comparisons of means was performed with *aov()* and *TukeyHSD()* function. Various heatmaps with dendrograms and sidebars were generated by ComplexHeatmap (v2.16.0) package. Various plots including violine plots, scatter plots, contour plots, bar plots, upset plots lolliplots were generated using r-ggplot2 (v3.4.3). The oncoplot was generated using maftools (v2.16.0) in R. UMAP analysis was performed with r-umap (v0.2.10.0) package. Single-nucleus RNA-seq data was analyzed and plotted using Seurat4 (4.2.0) package in R.

## Data availability statement

Raw whole genome sequencing and massive parallel amplicon sequencing data (MPAS) and single nucleus MPAS (snMPAS) are available through SRA (accession number: PRJNA799597) and NDA (accession number: study 919) for ID01 and ID05. The 300× WGS panel of normal is available on SRA (accession number: PRJNA660493).

## Code availability statement

Details and codes for the data processing and annotation are provided on GitHub (https://github.com/shishenyxx/Human_Inhibitory_Neurons).

## Acknowledgments

We thank the individuals who donate their bodies and tissues for the advancement of research. T. Komiyama for feedback. This work was supported by the National Institute of Mental Health (NIMH) (grants U01MH108898 and R01MH124890 to J.G.G. and R21MH134401 to X.Y. and J.C.M.S.), Eunice Kennedy Shriver National Institute of Child Health and Human Development (NICHD) (grant K99HD111686 to X.Y.), 2021 NARSAD Young Investigator Grant from the Brain & Behavior Research Foundation (30598 to C.C.) and Rady Children’s Institute for Genomic Medicine. We thank the San Diego Supercomputer Center (grant no. TG-IBN190021 to X.Y. and J.G.G.) for computational help. This publication includes data generated at the UC San Diego IGM Genomics Center utilizing an Illumina NovaSeq 6000 that was purchased with funding from a National Institutes of Health SIG grant (no. S10OD026929 C.C., X.Y., and J.G.G.). We are grateful to C. Fine, M. Espinoza, and M. Banihassan (UCSD) for technical assistance with flow cytometry experiments. This work was additionally made possible by the UC San Diego Stem Cell Program and a CIRM Major Facilities grant (FA1-00607) to the Sanford Consortium for Regenerative Medicine. This publication includes data generated at the UCSD Human Embryonic Stem Cell Core Facility, using the BD Biosciences Influx, FACS Aria Fusion, and FACS Aria II Flow Cytometry Sorters.

## Author contributions

C.C., X.Y., and J.G.G. designed the study. C.C., X.Y., R.F.H., K.I.V., C.B., V.S., S.M., M.W.B., J.C.M.S. and S.T.B. organized, handled and sequenced human samples. C.C., X.Y., and K.K. performed ResolveOME experiment. C.C. and Y.L. optimized and performed MFNS experiment. C.C., X.Y., A.P., and R.N. performed the bioinformatics and data analyses. C.C., X.Y. and J.G.G. wrote the manuscript. All authors reviewed the manuscript. C.C. and X.Y. contributed equally to this work.

