## Supplementary Data 6 for "Cell-type-resolved somatic mosaicism reveals clonal dynamics of the human forebrain"

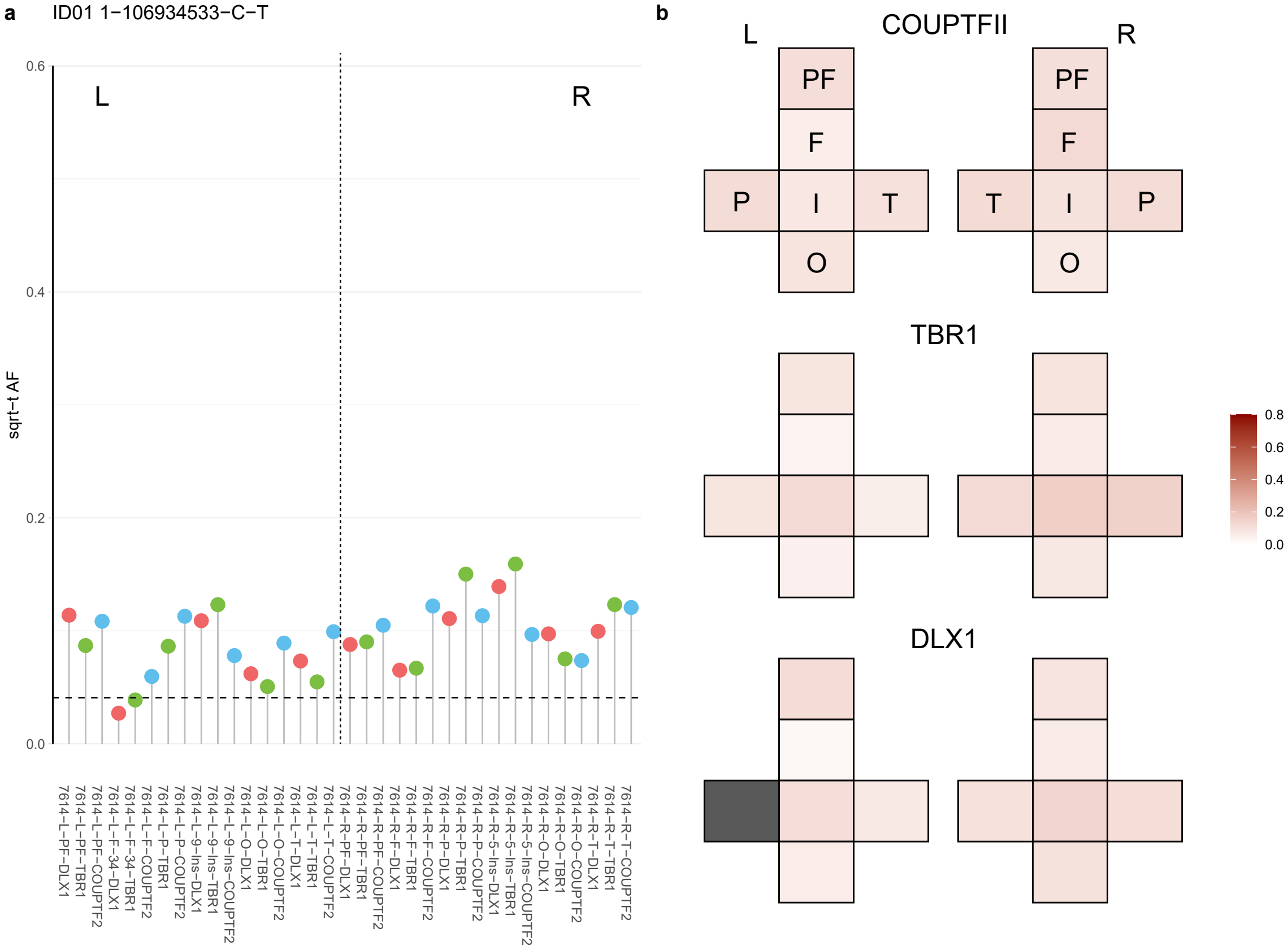

**a** ID01 1-168467950-G-A

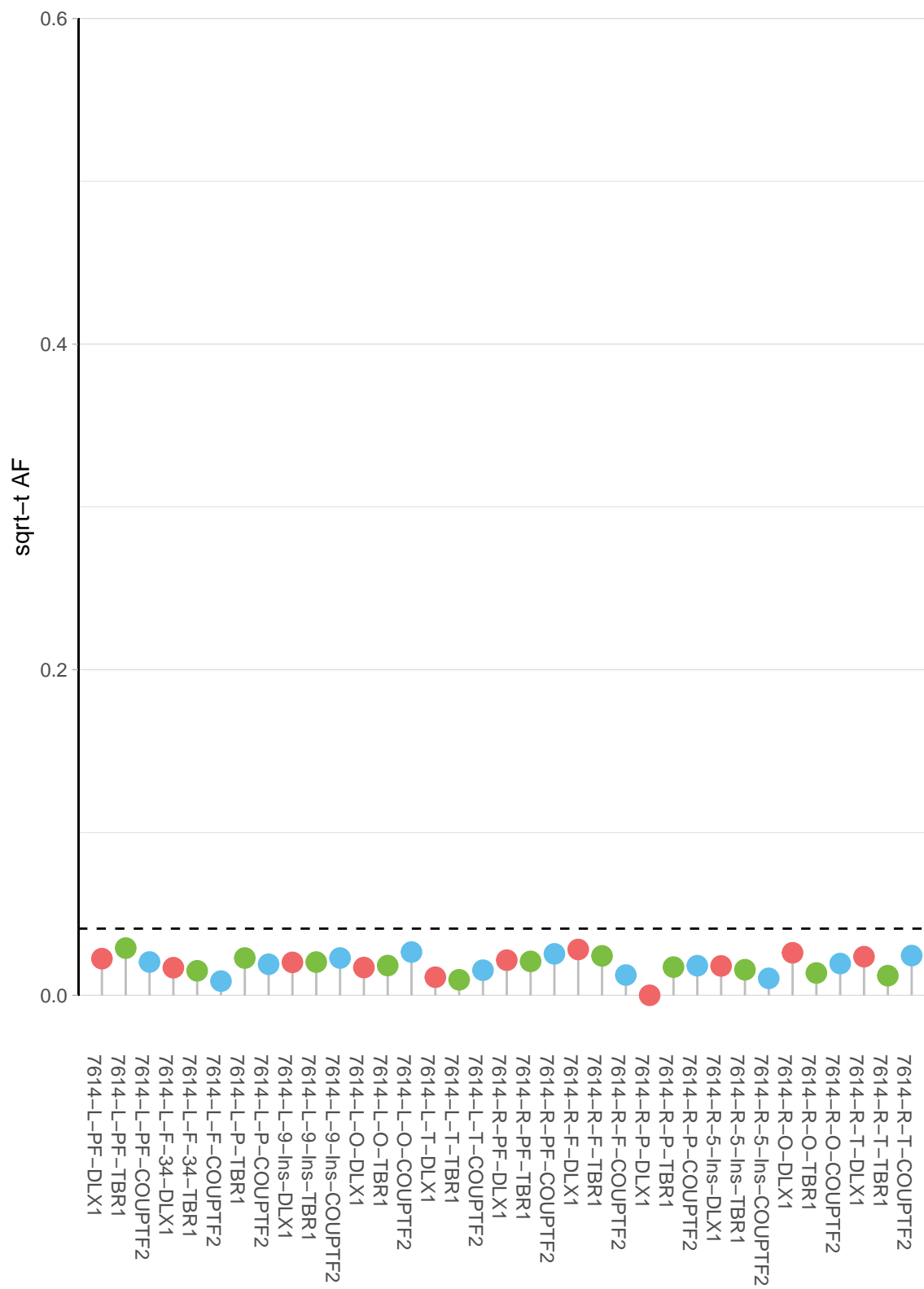

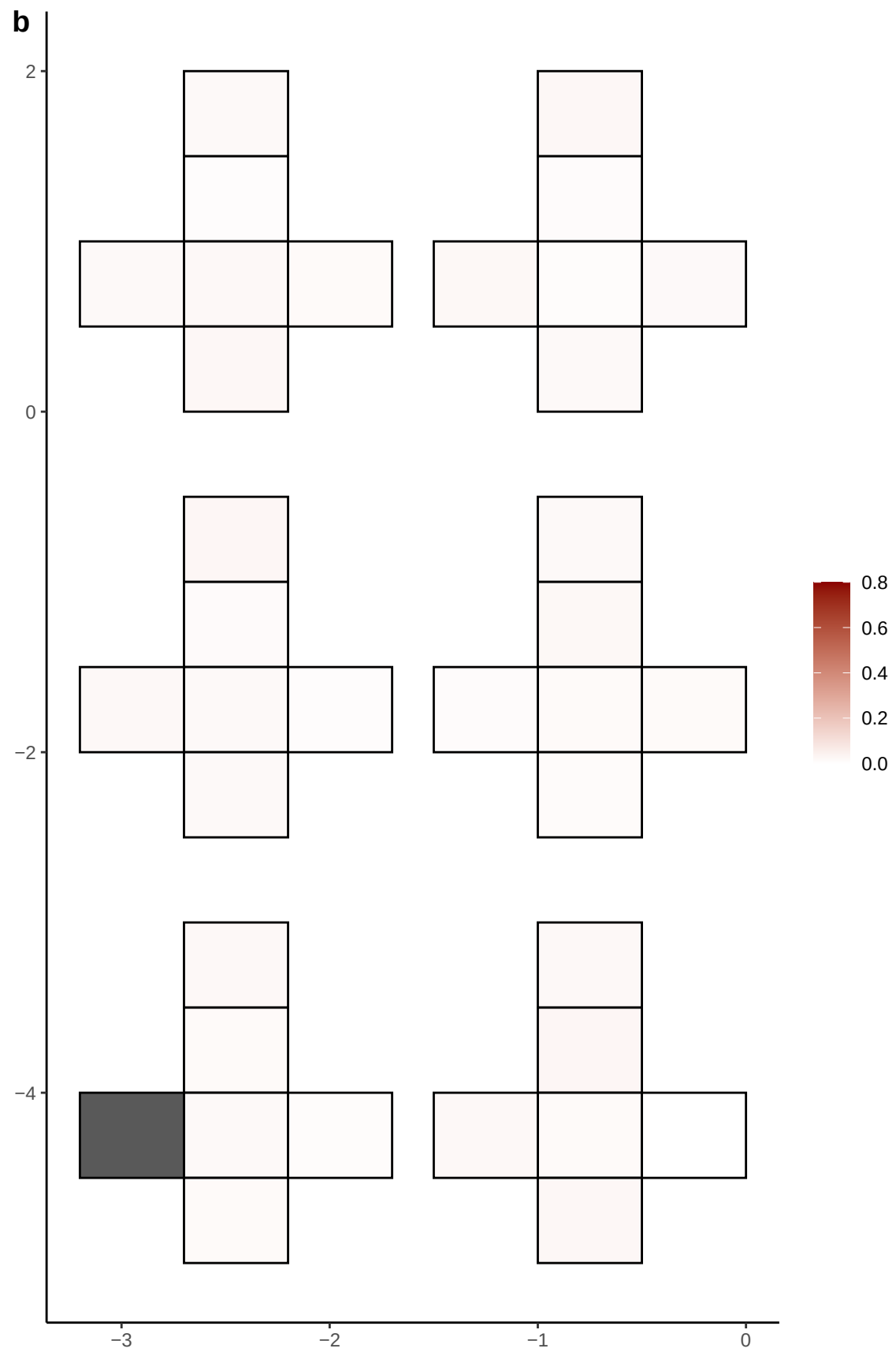

**a** ID01 1-169329191-G-A

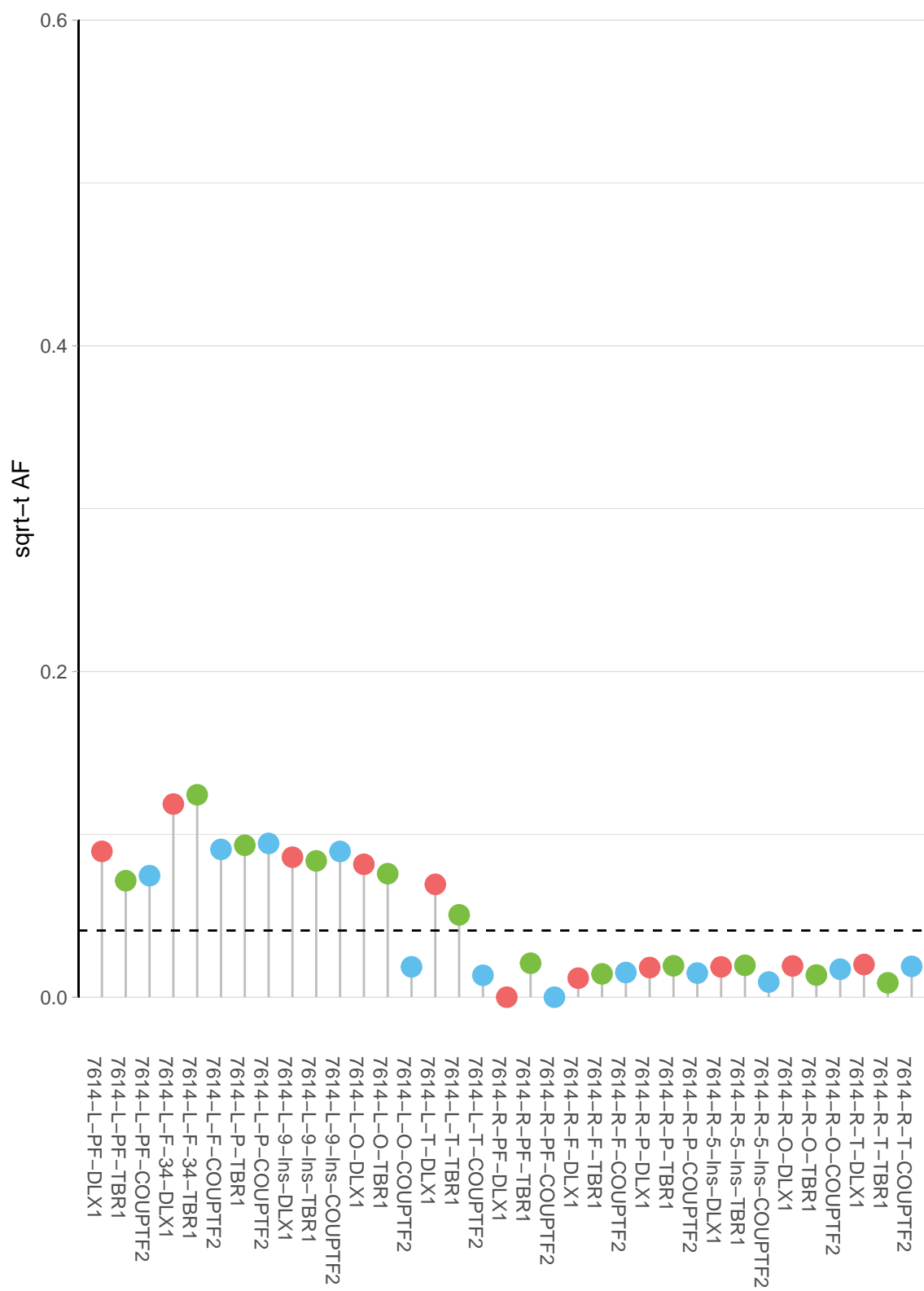

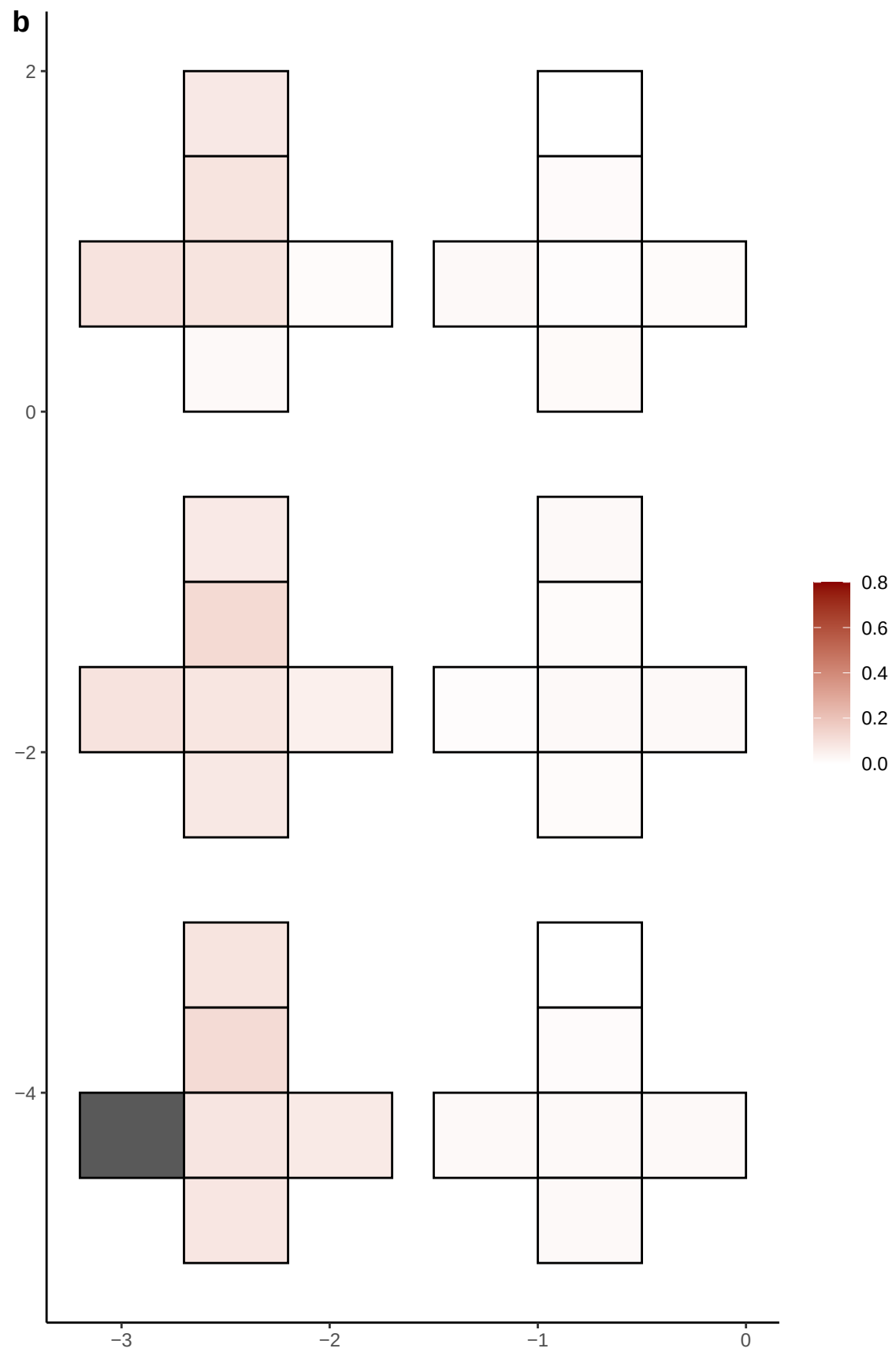

**a** ID01 1-180856518-T-G

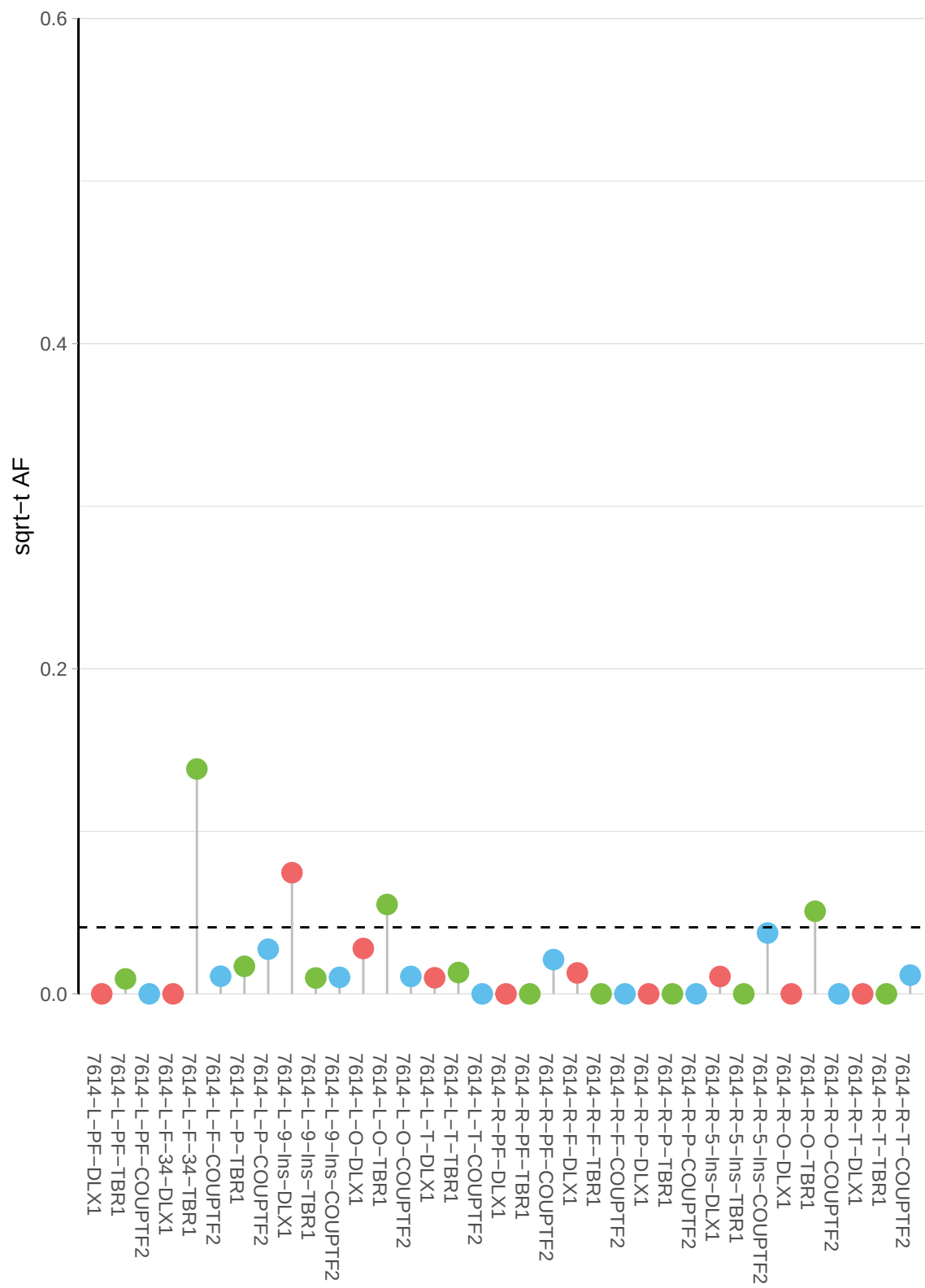

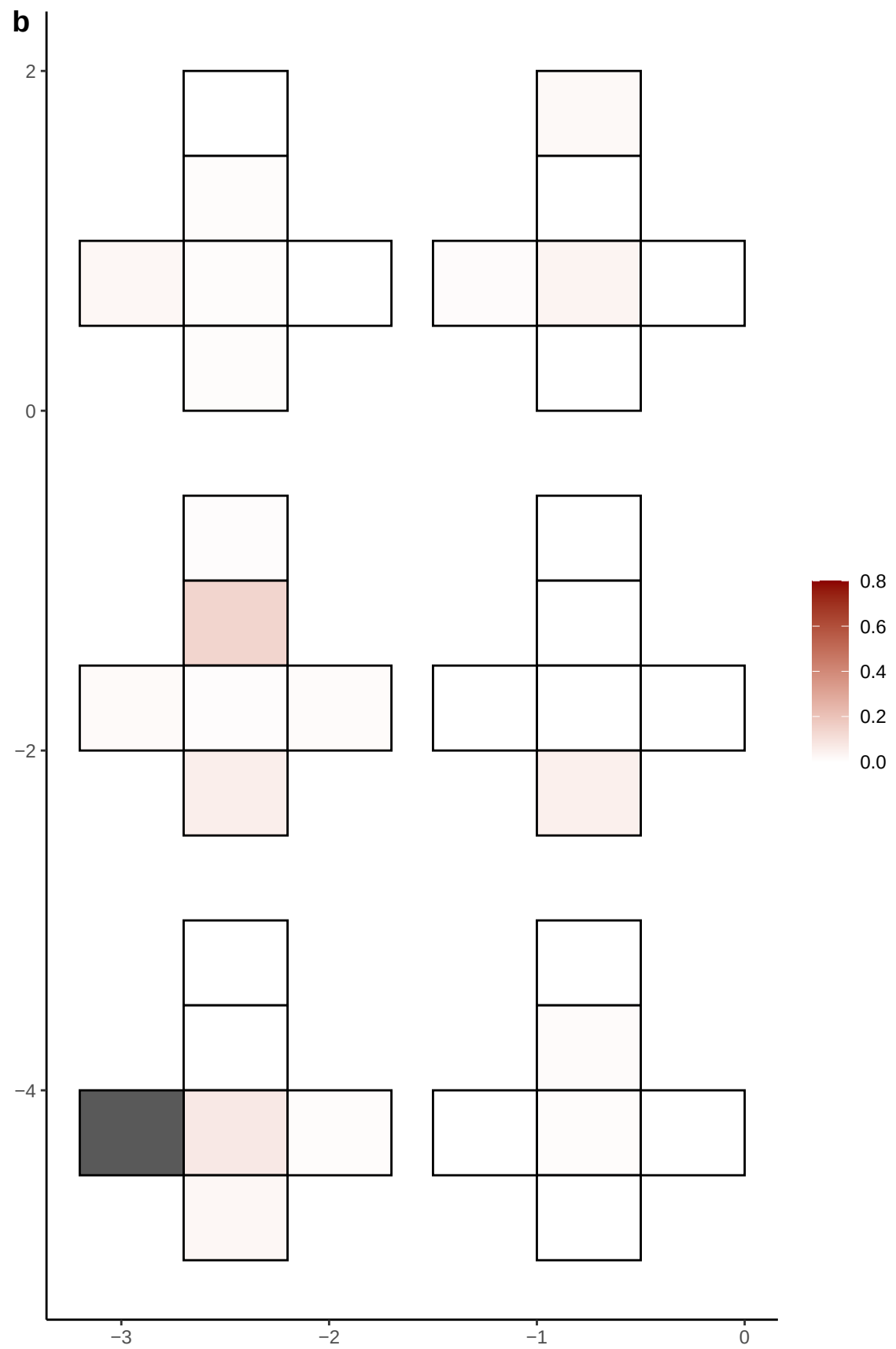

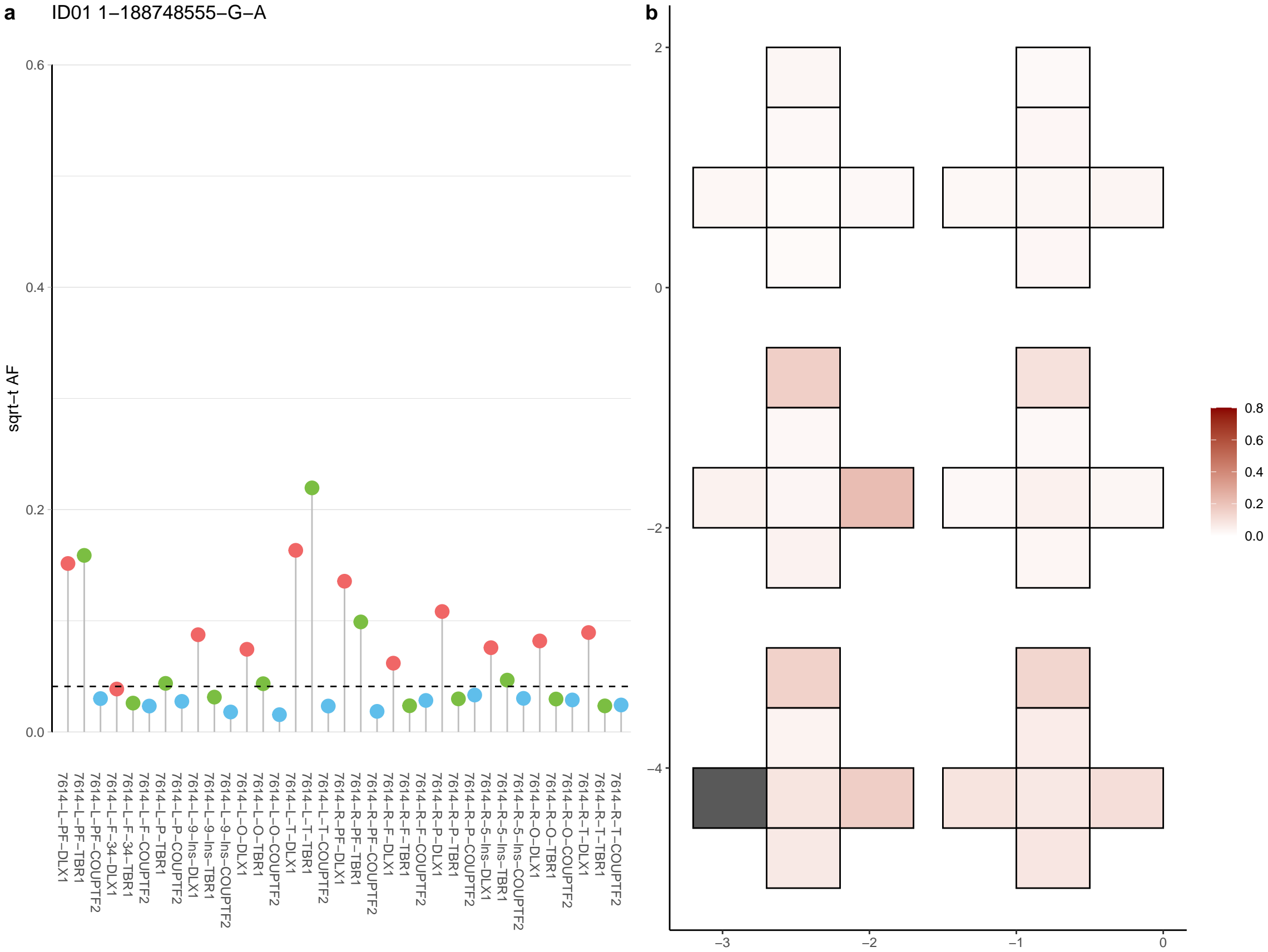

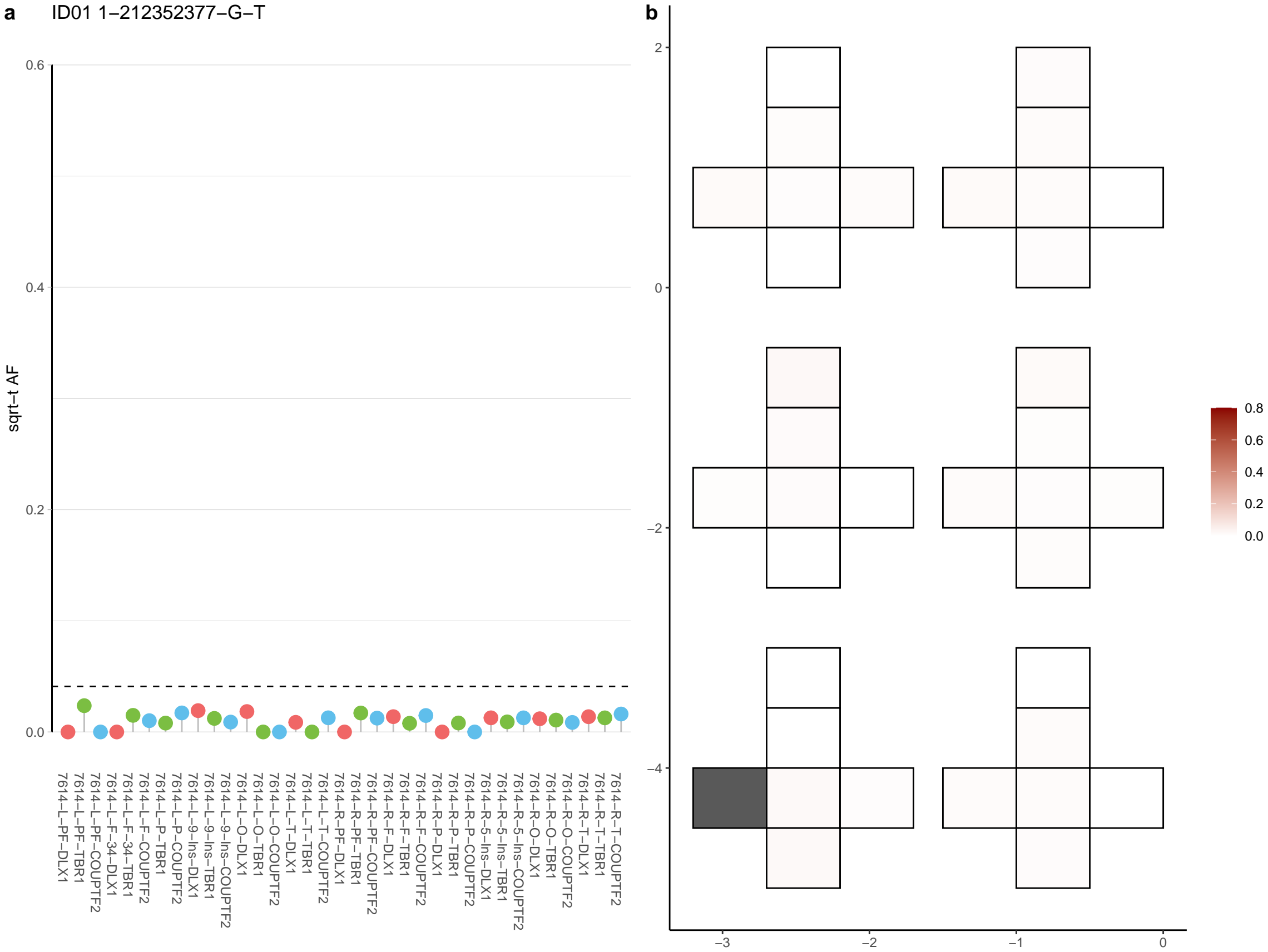

**a** ID01 1-221888654-T-C

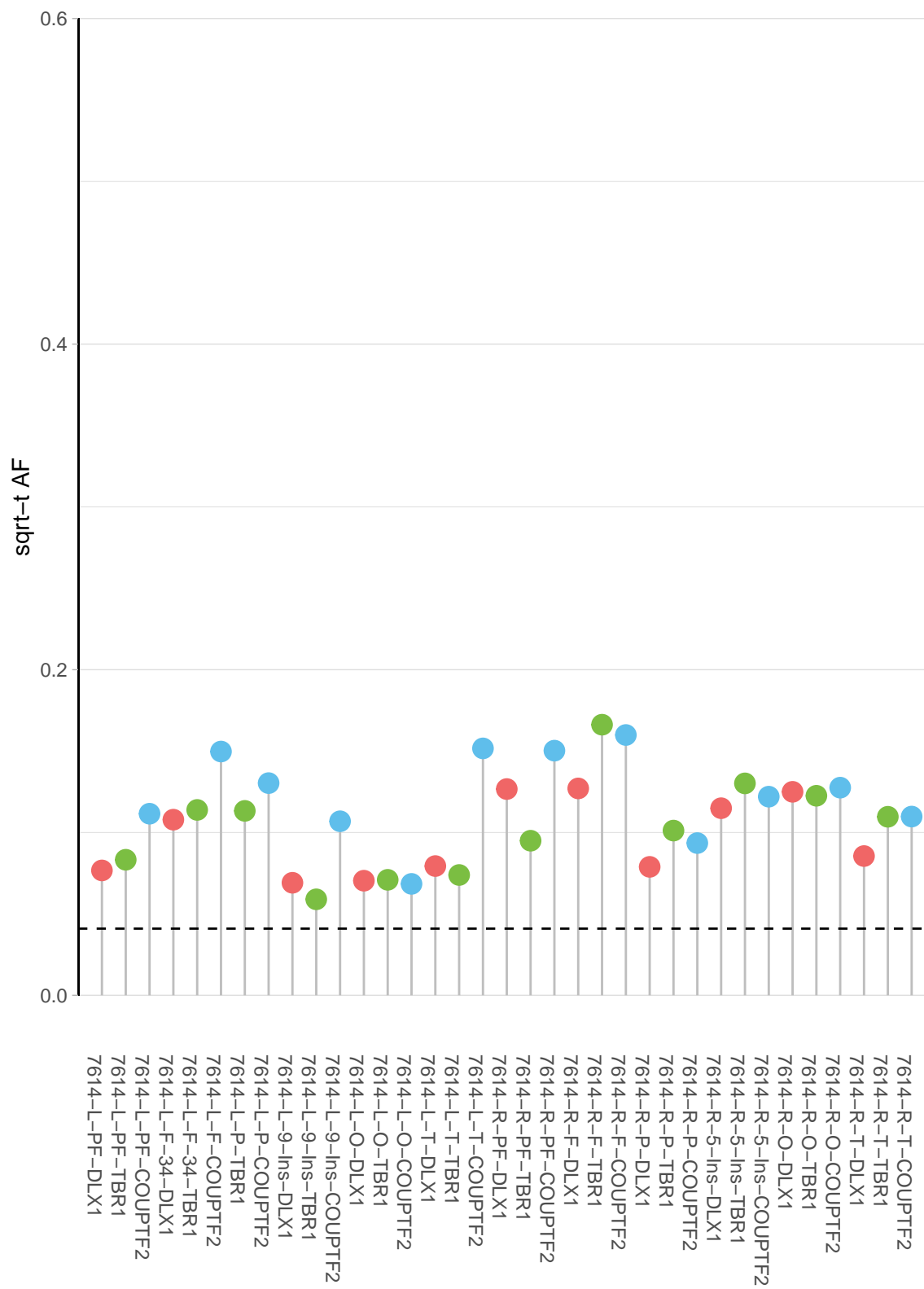

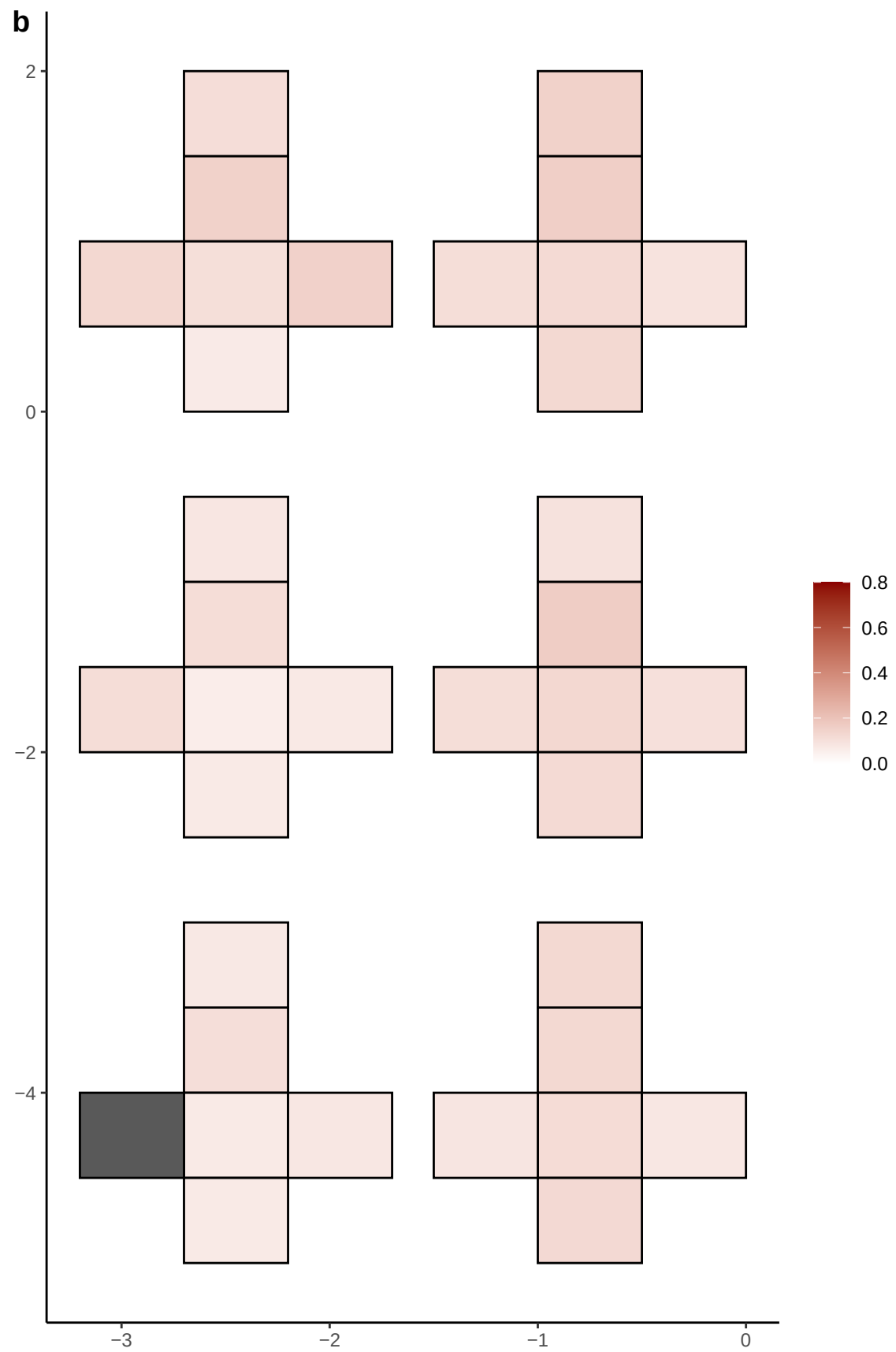

**a** ID01 1-239397797-T-A

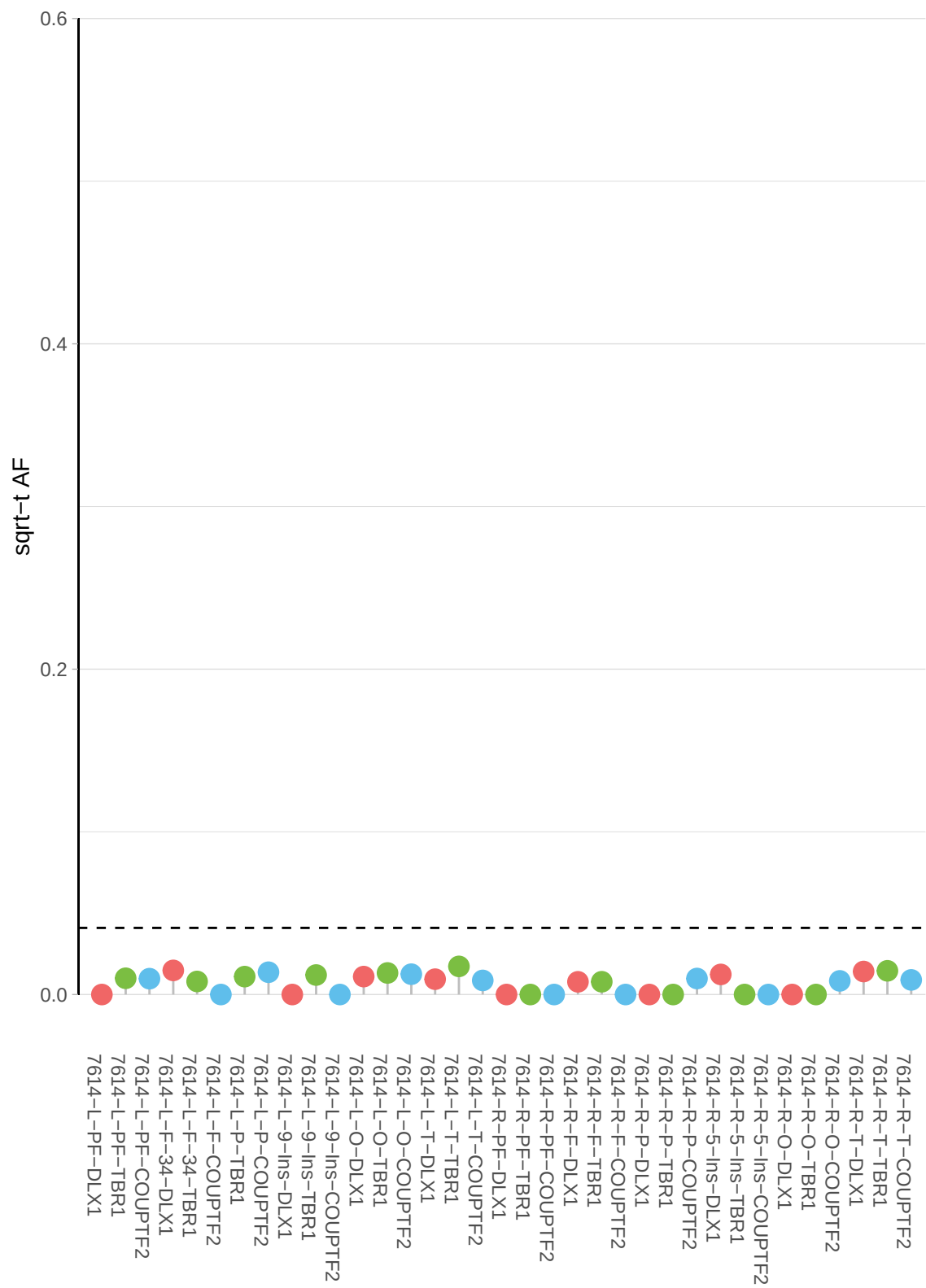

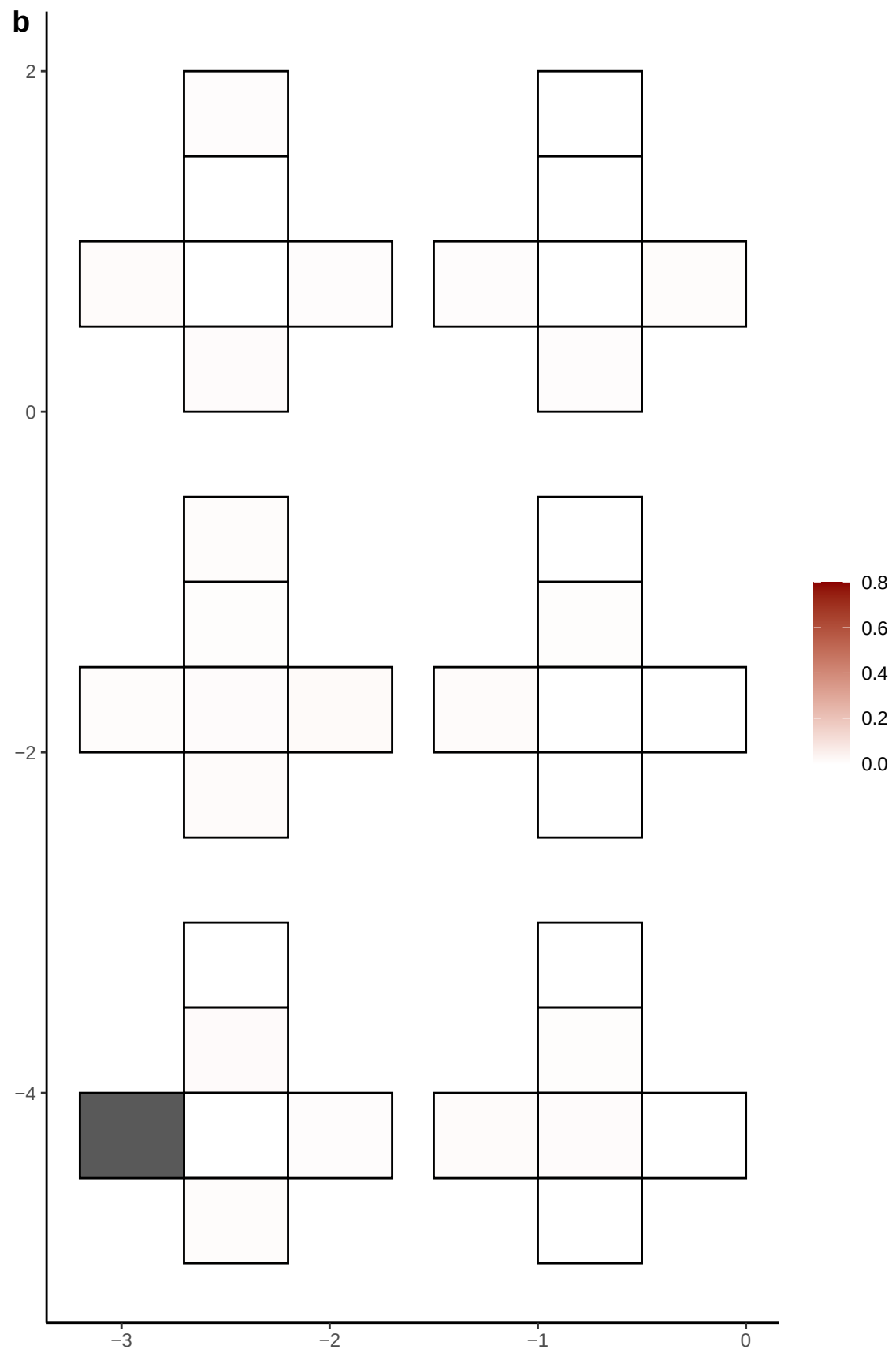

**a** ID01 1-240507547-A-T

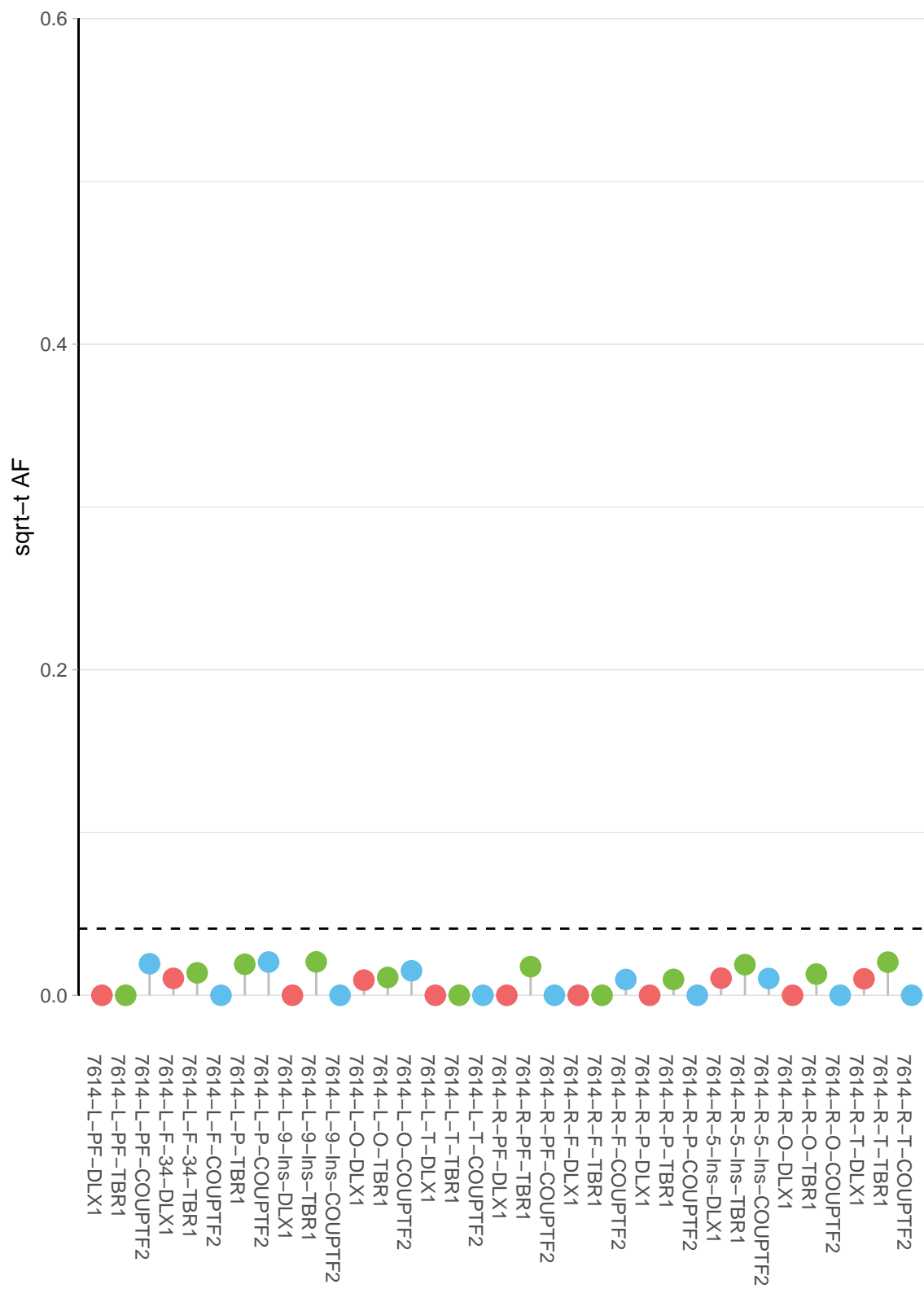

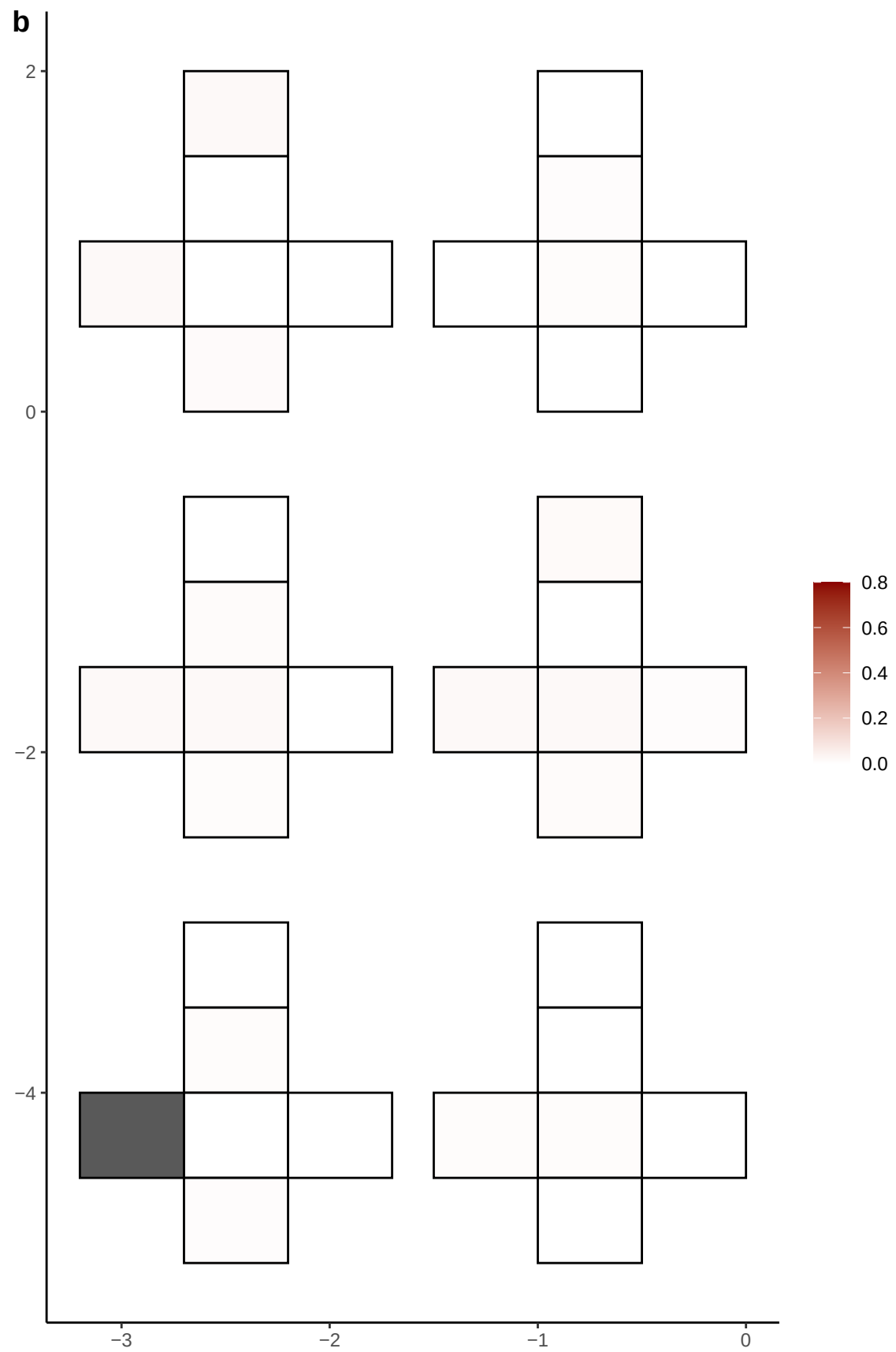

**a** ID01 1-241434408-C-T

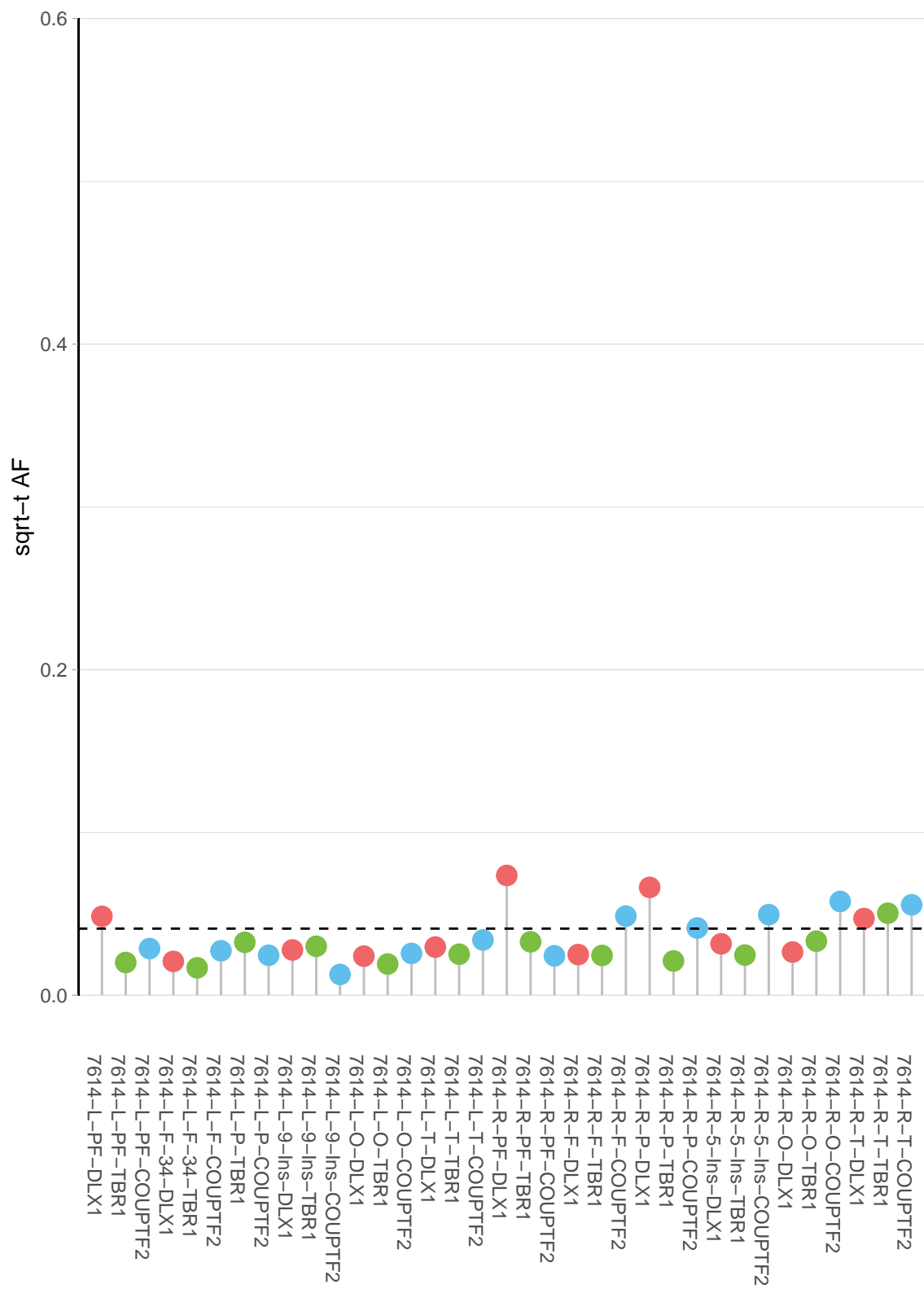

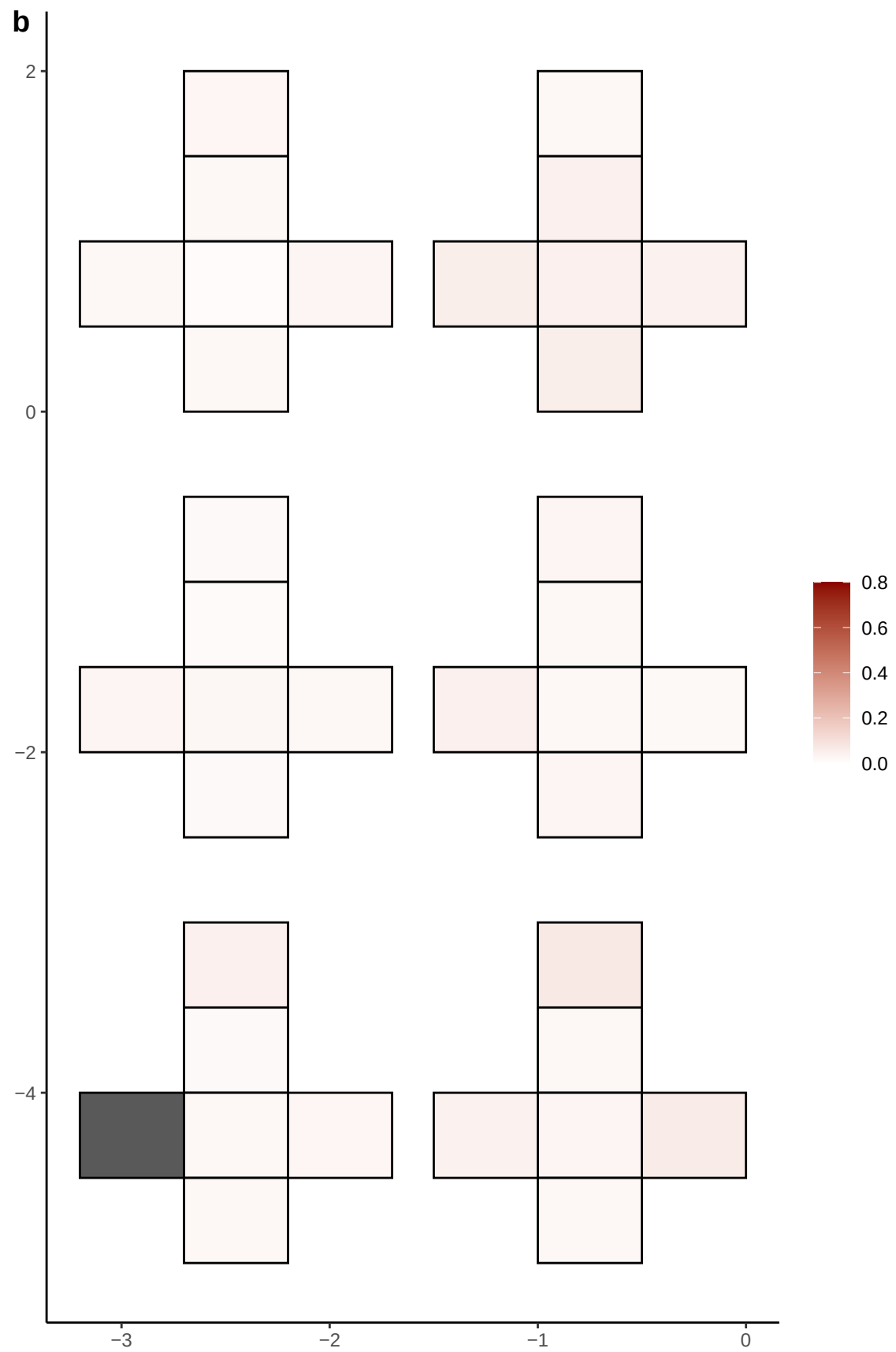

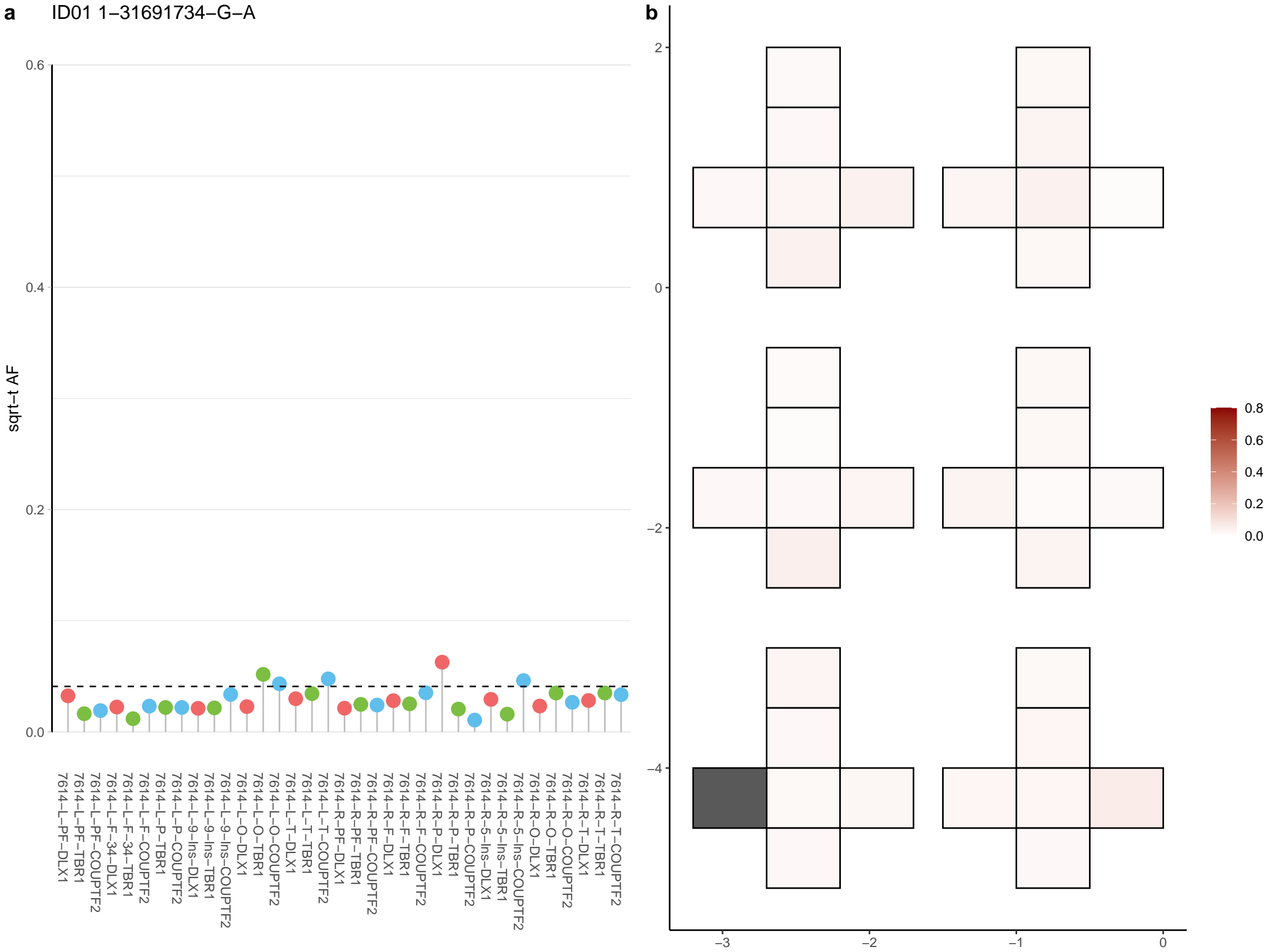

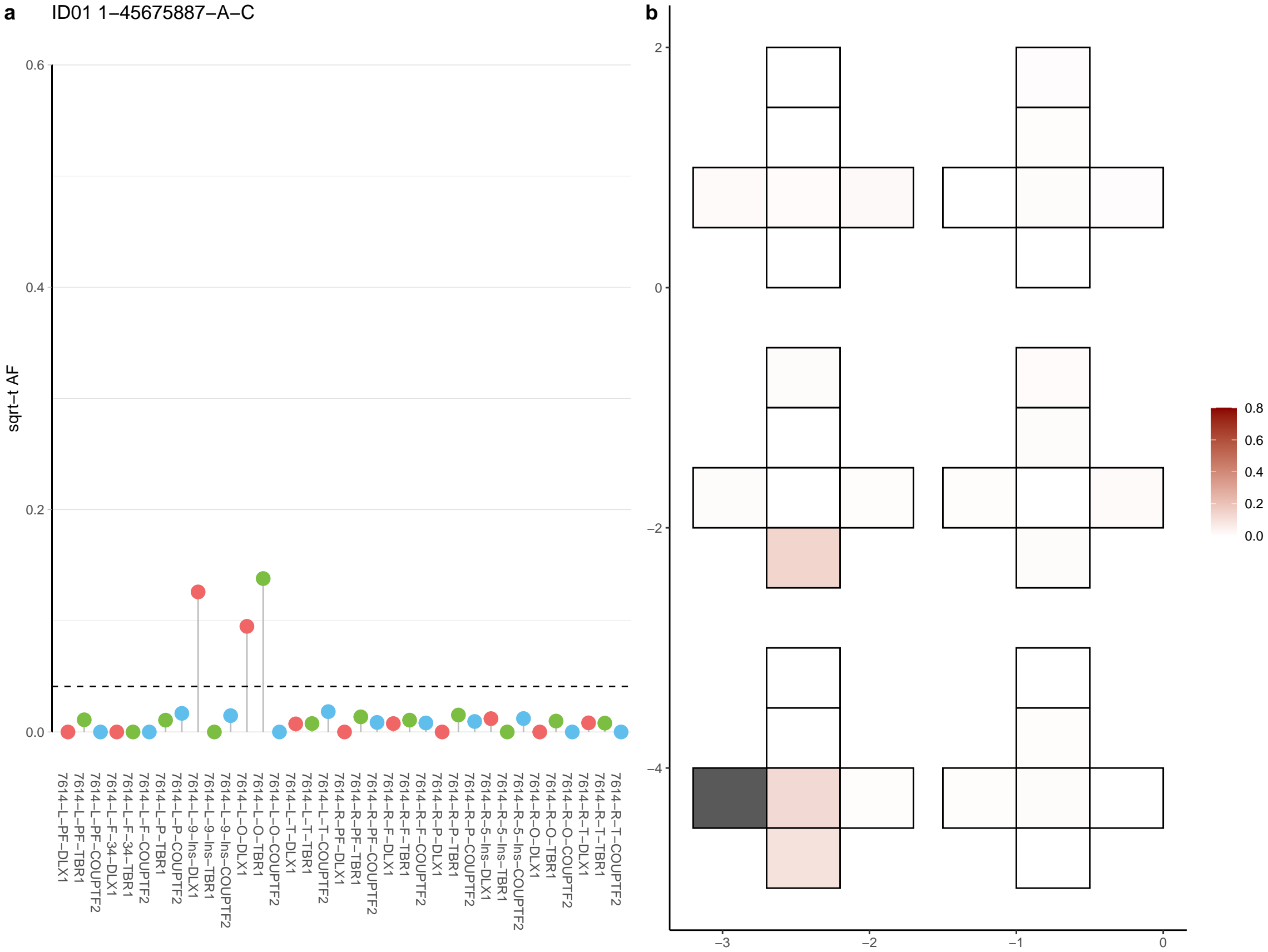

**a** ID01 1-64512024-C-T

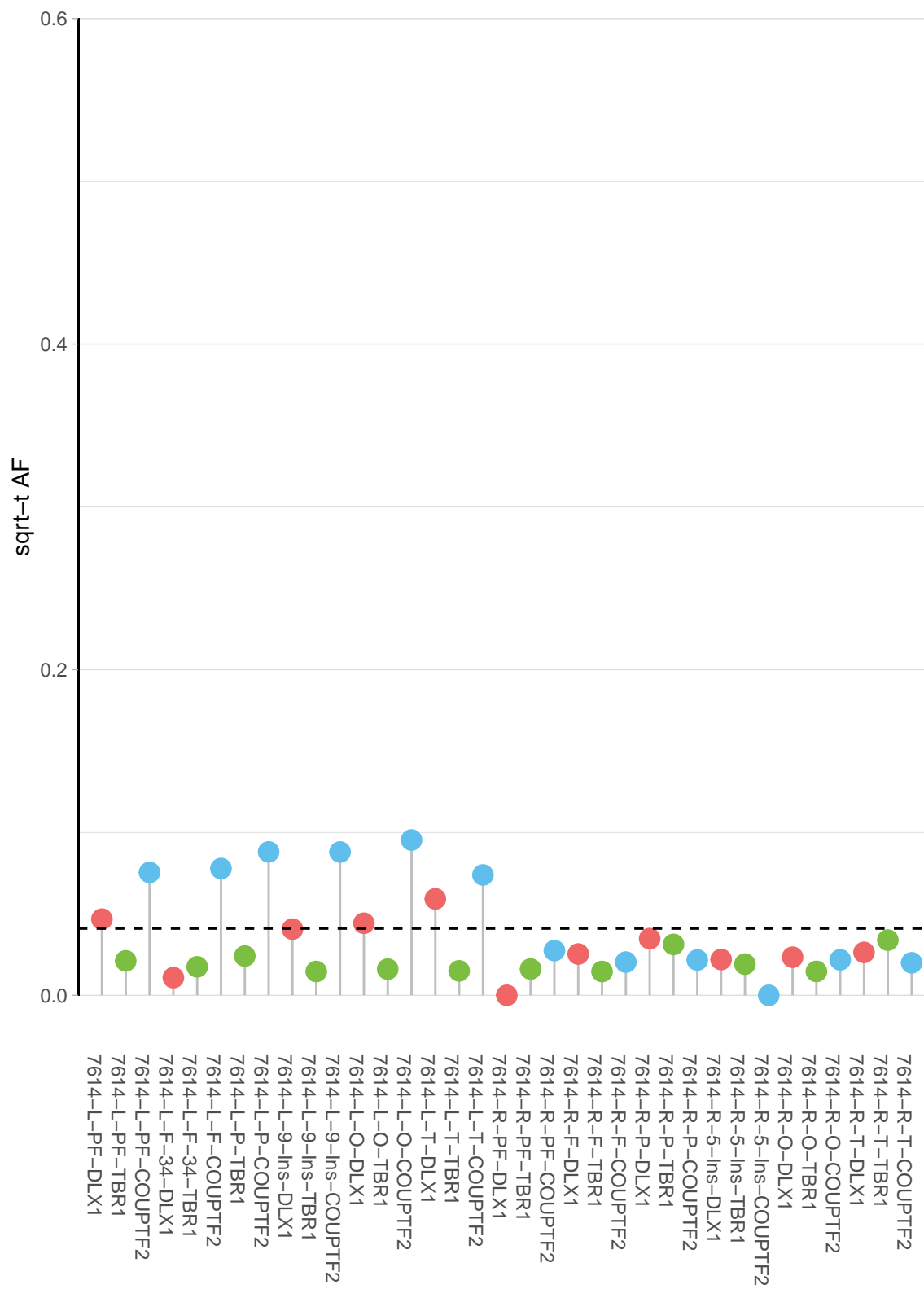

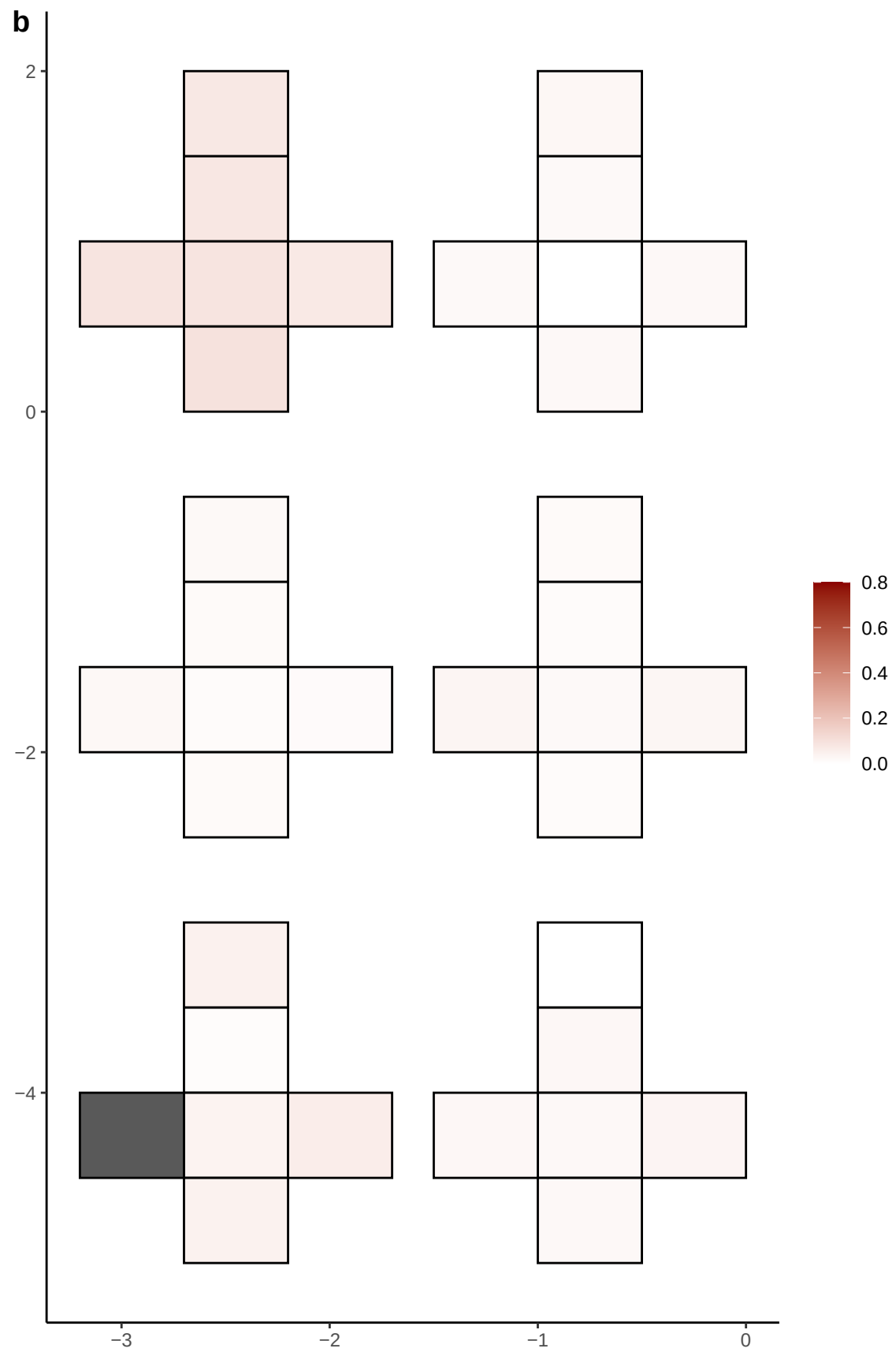

**a** ID01 1-98426749-G-A

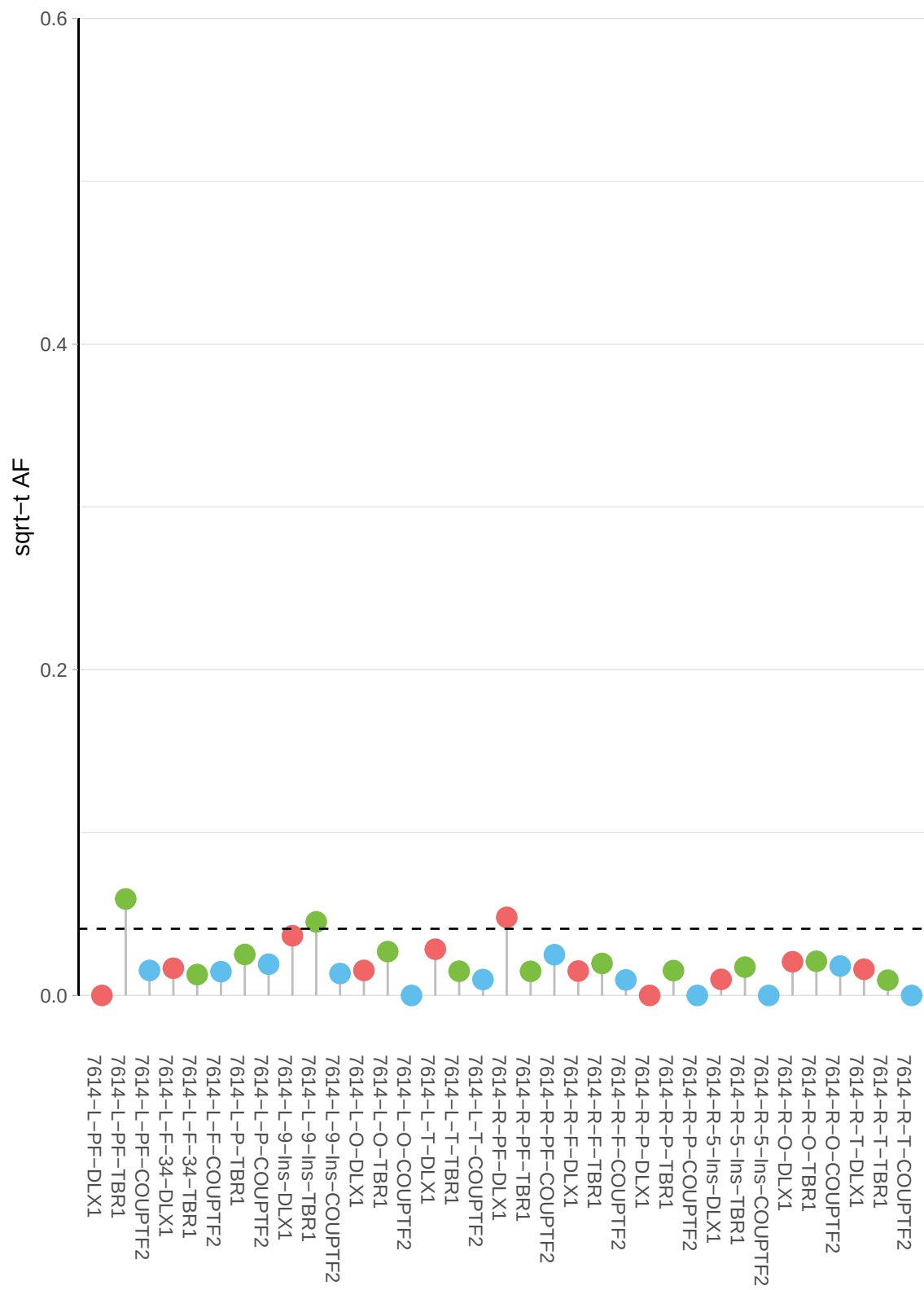

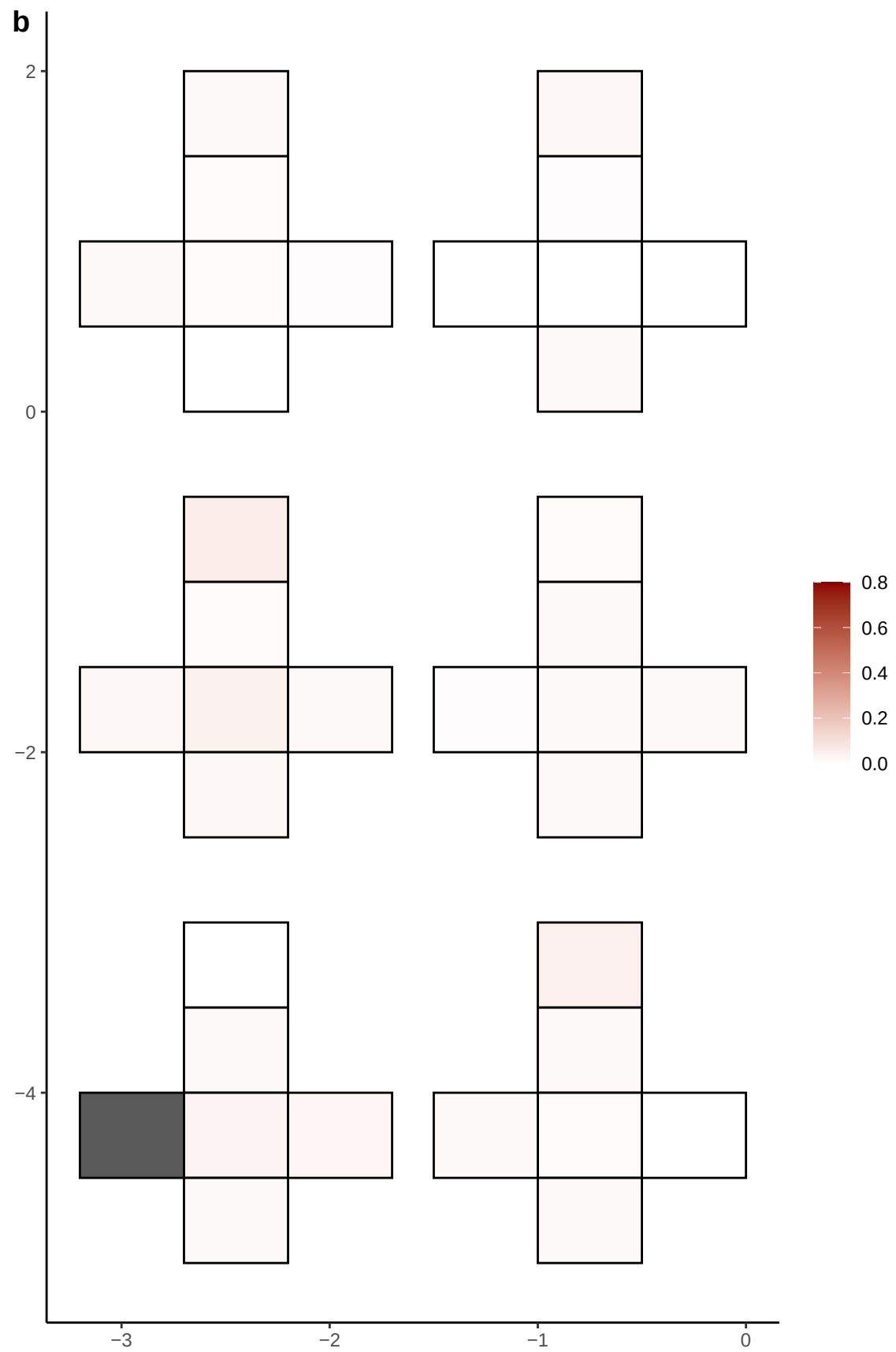

**a** ID01 1-99581604-C-G

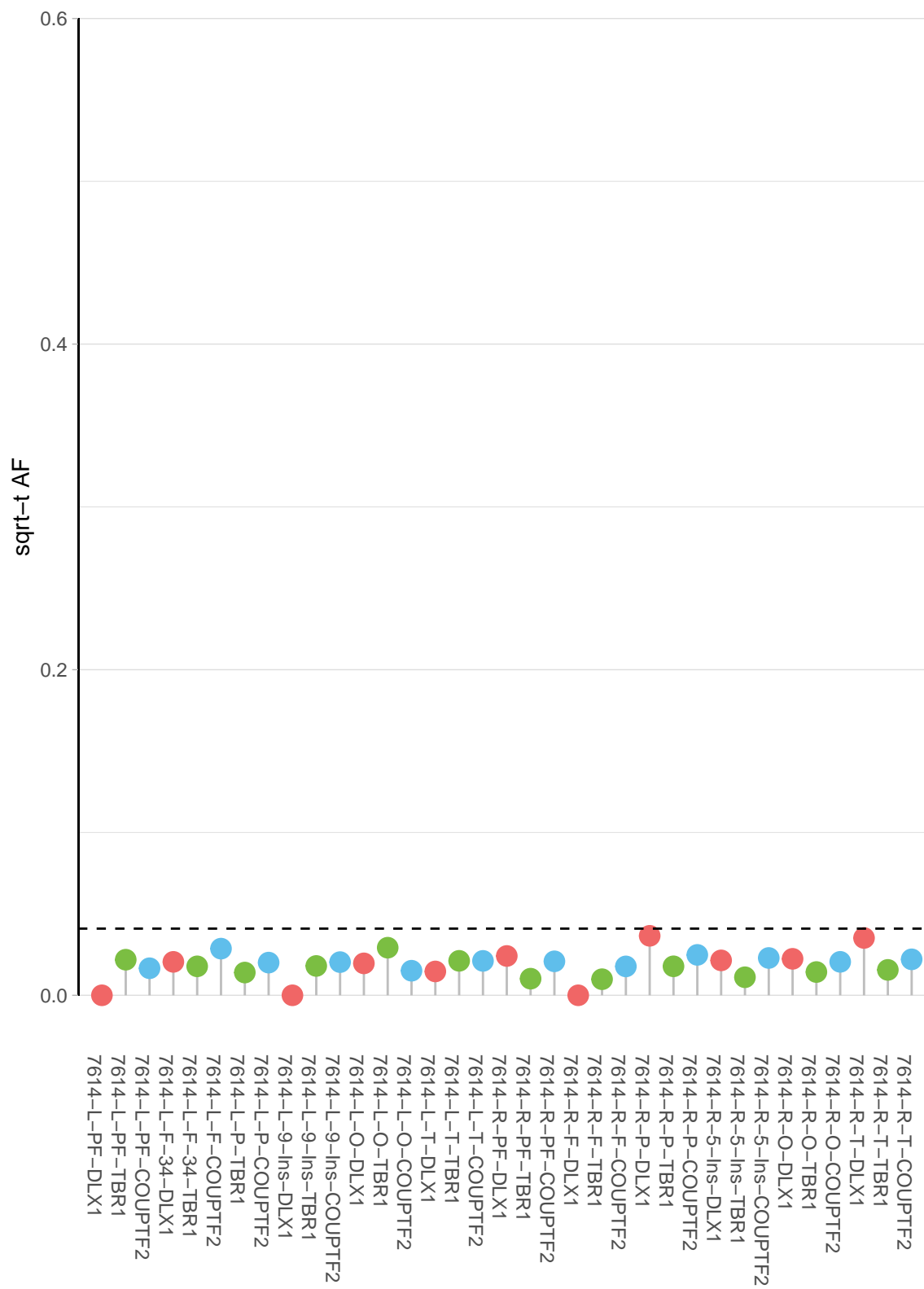

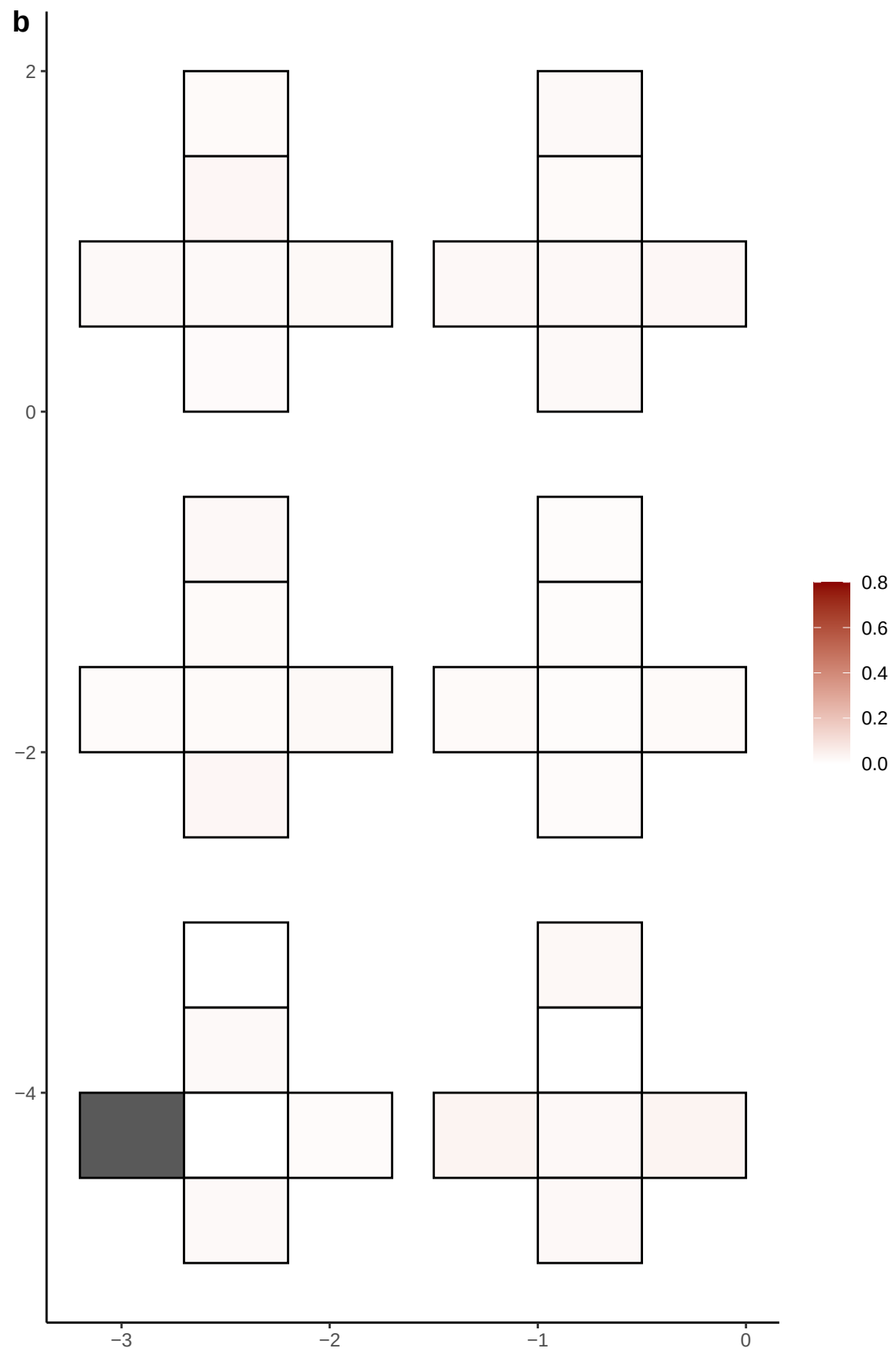

**a** ID01 10-116196503-C-T

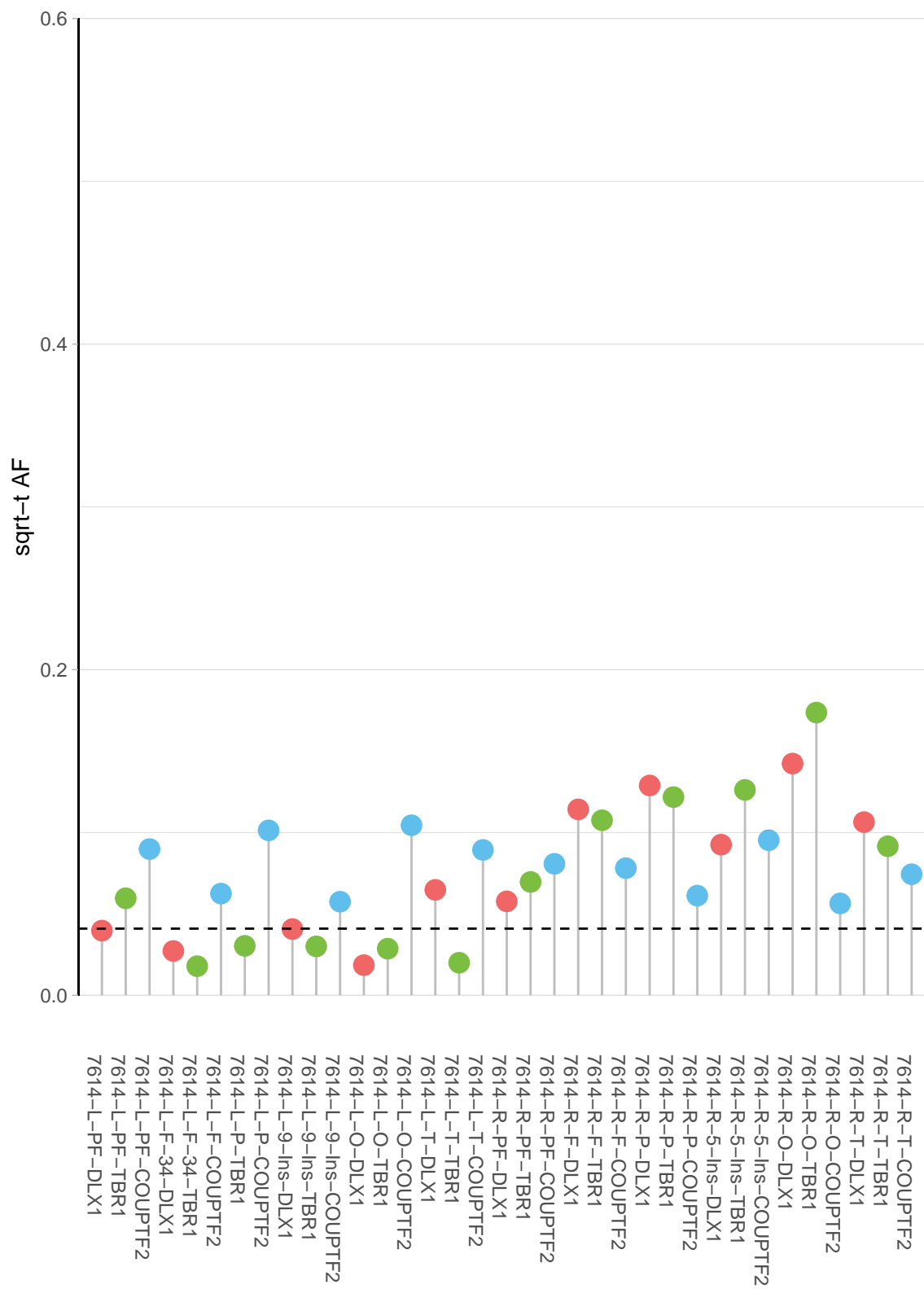

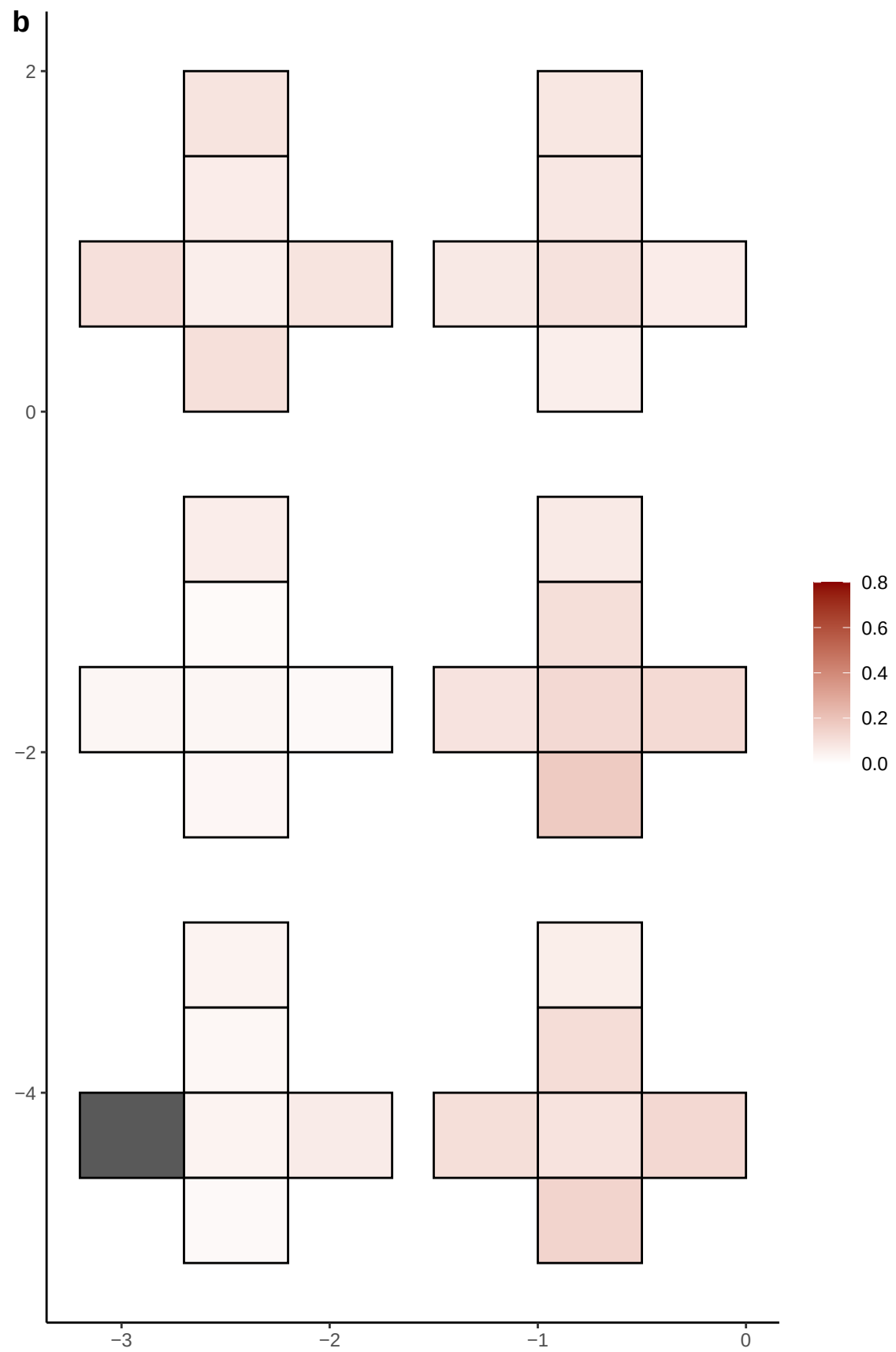

**a** ID01 10-127139902-G-A

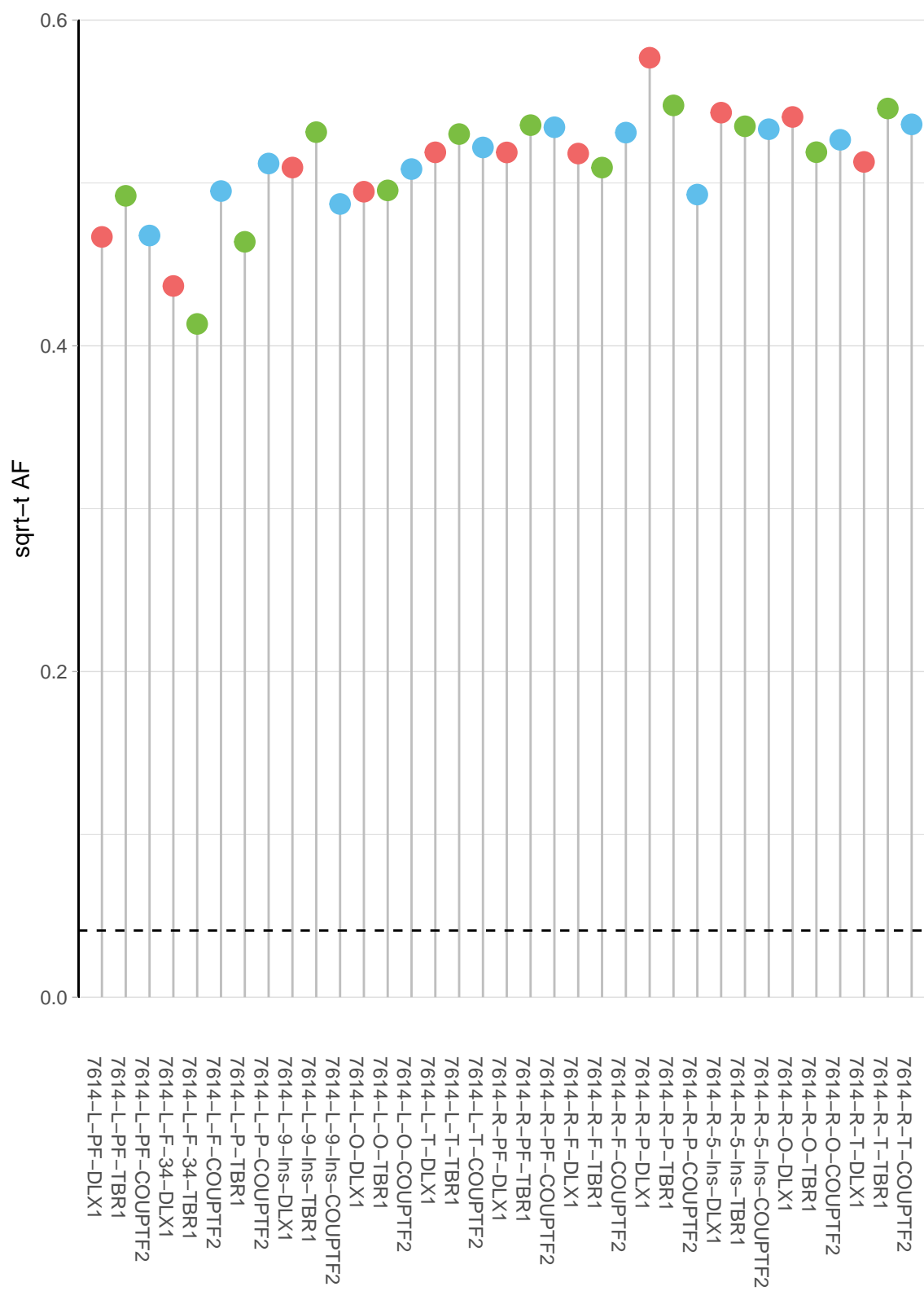

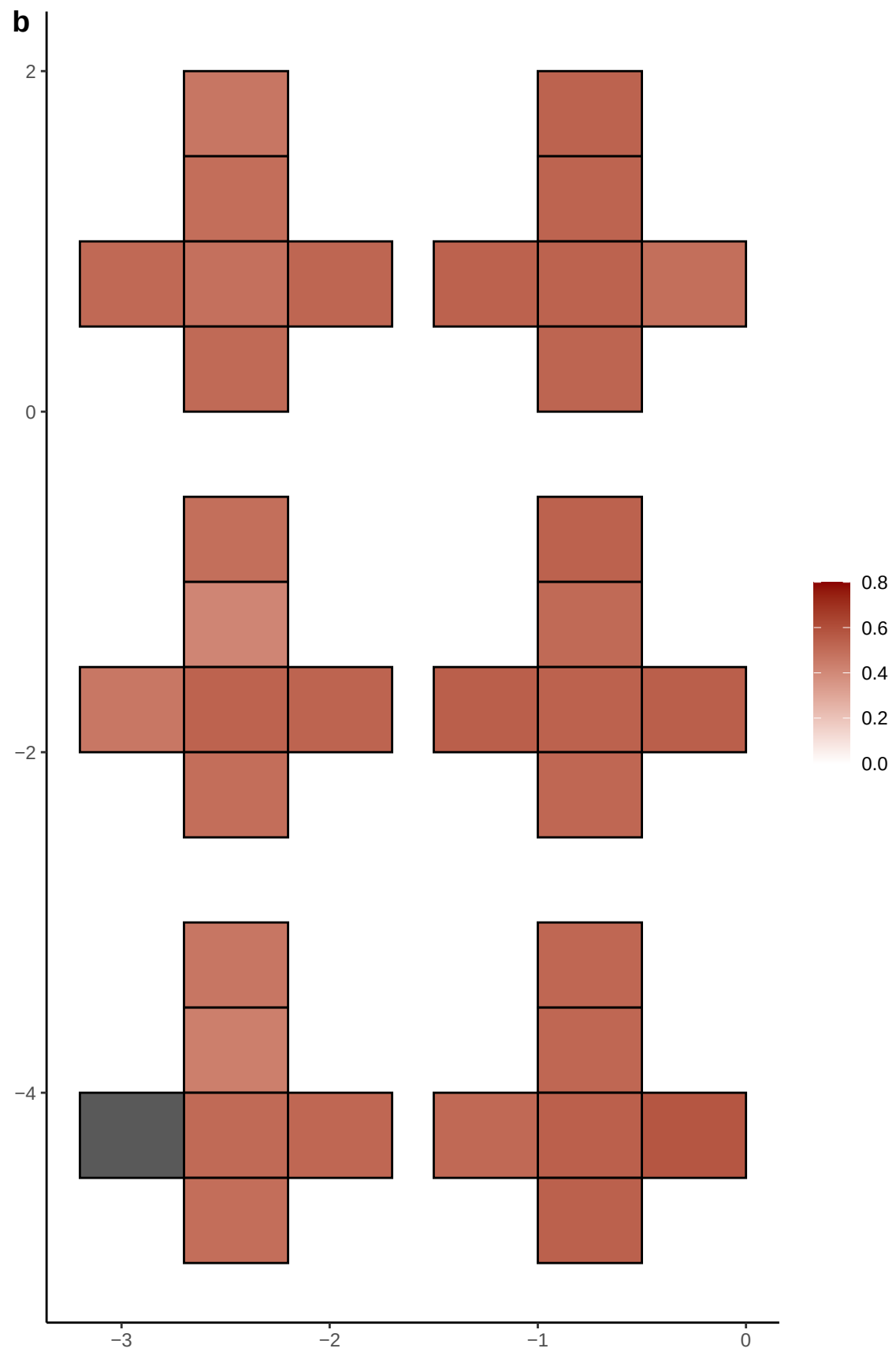

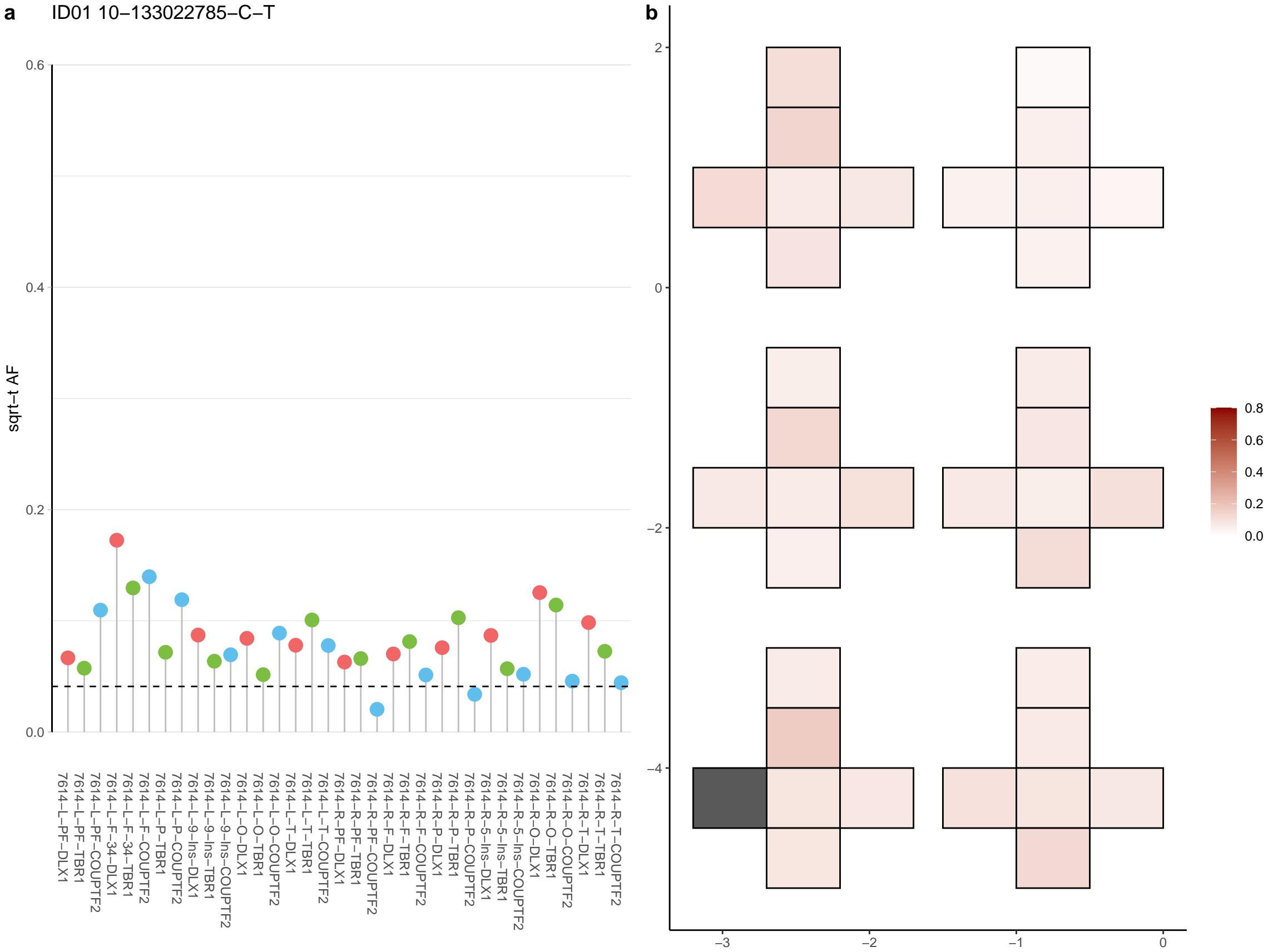

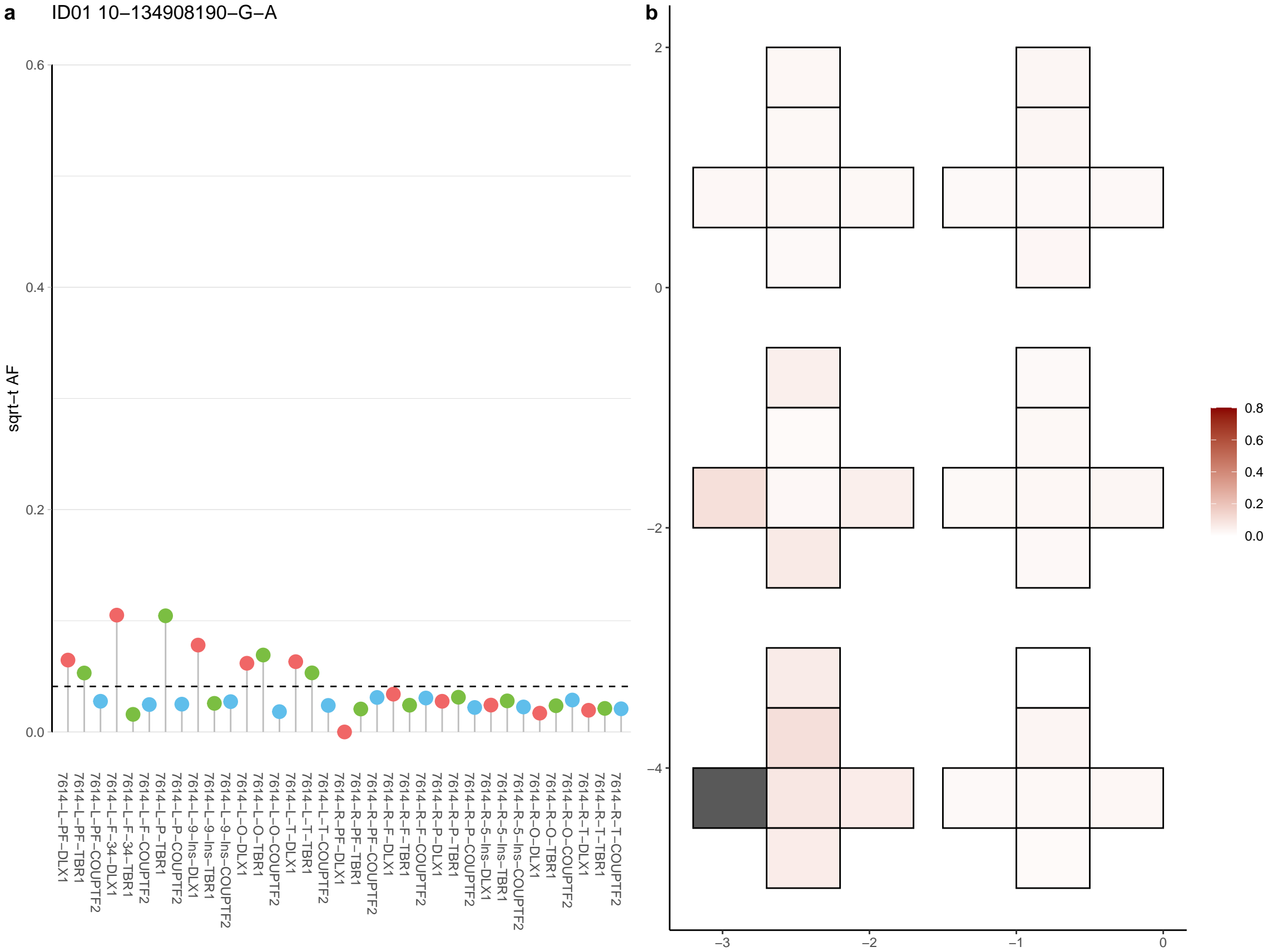

**a** ID01 10-23042694-GAGTT-G

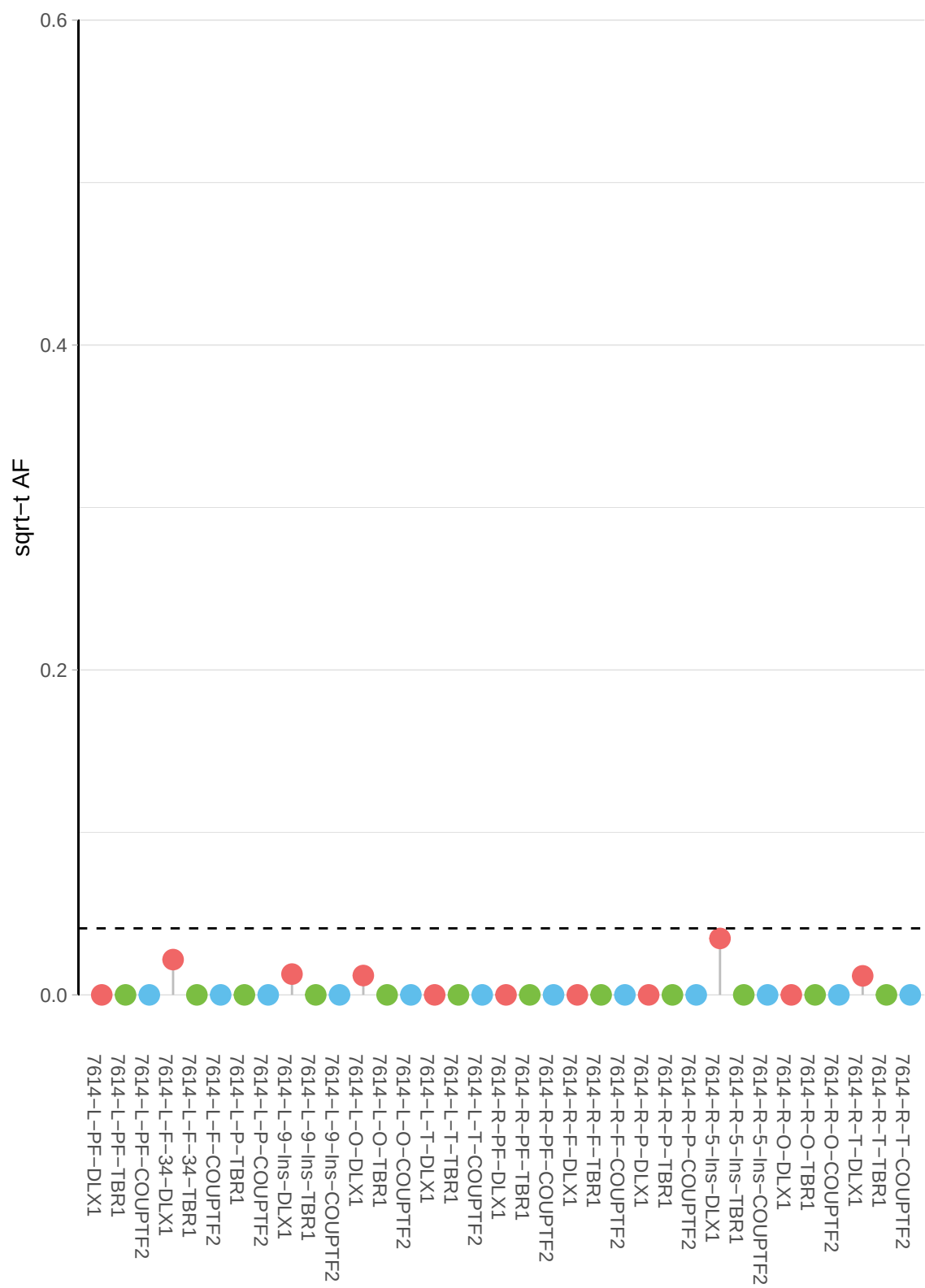

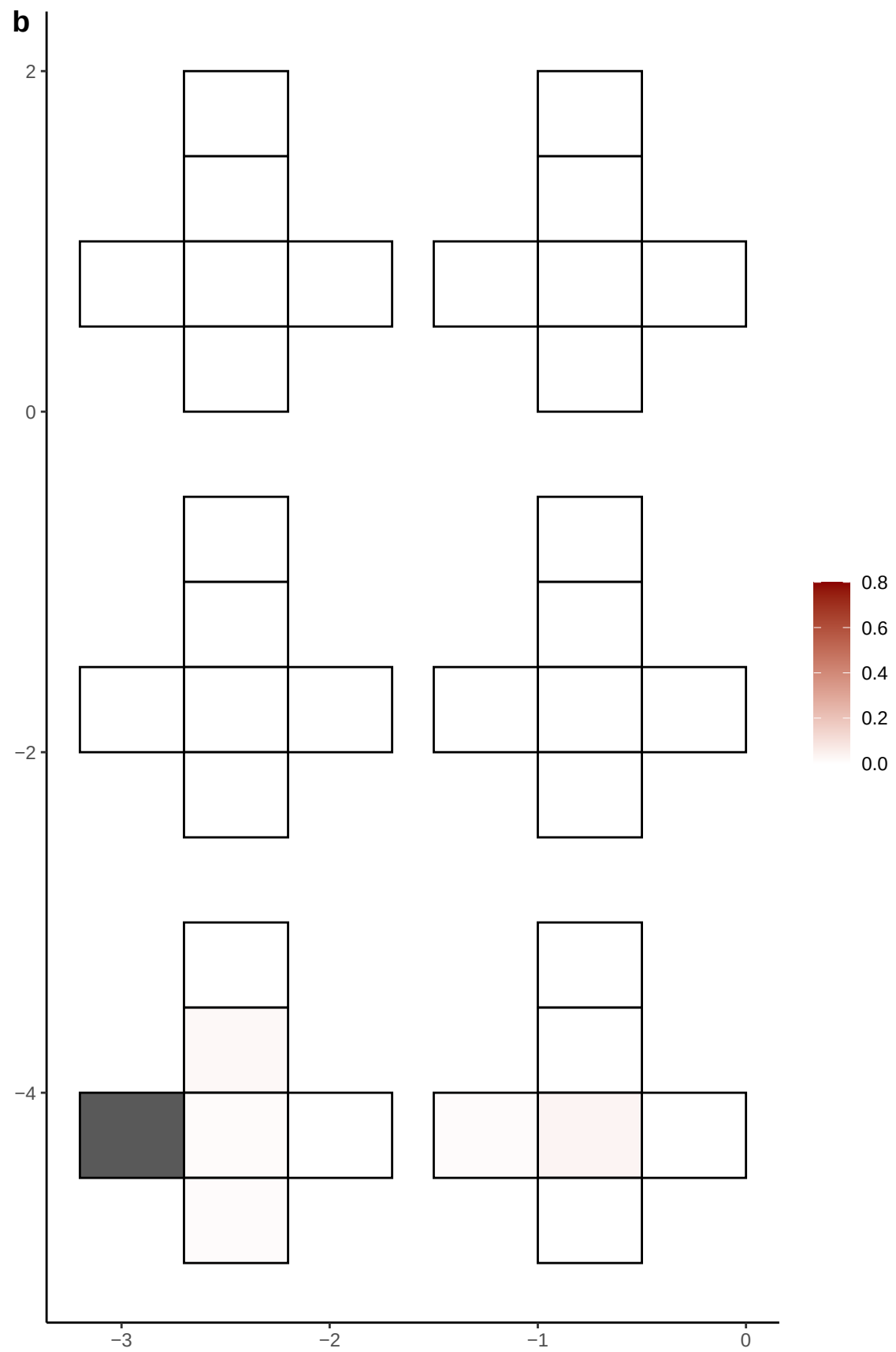

**a** ID01 10-71631216-G-A

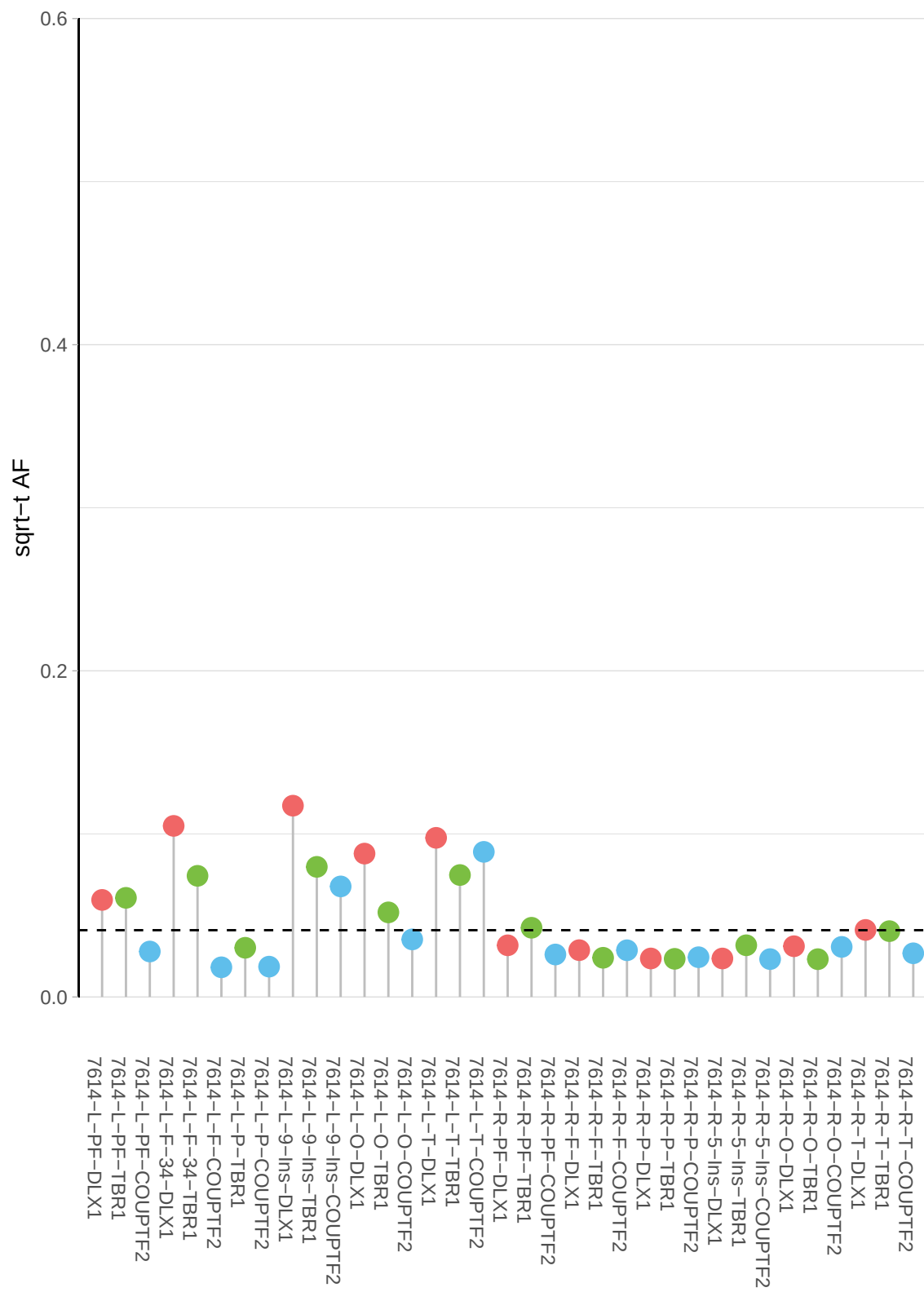

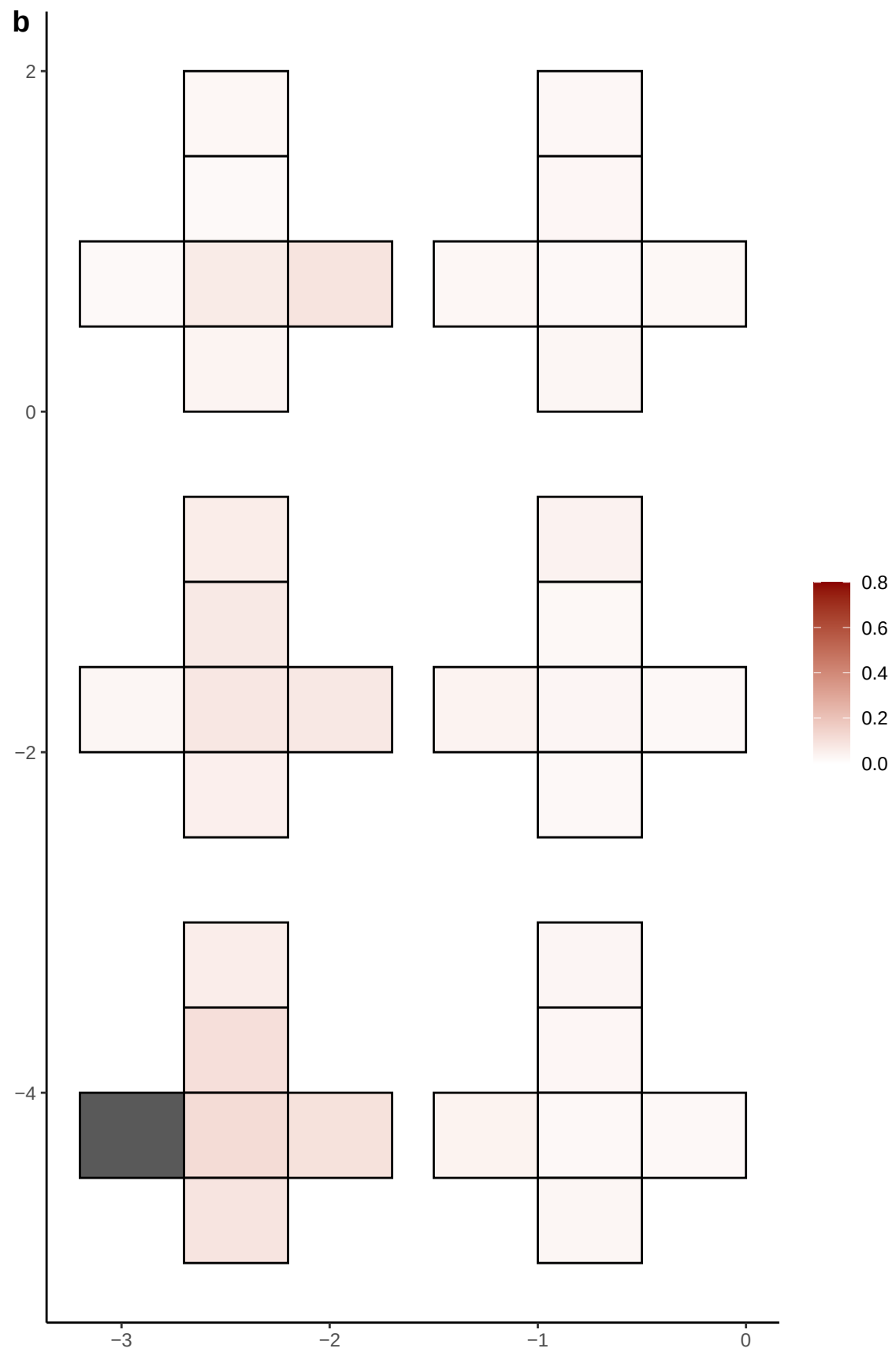

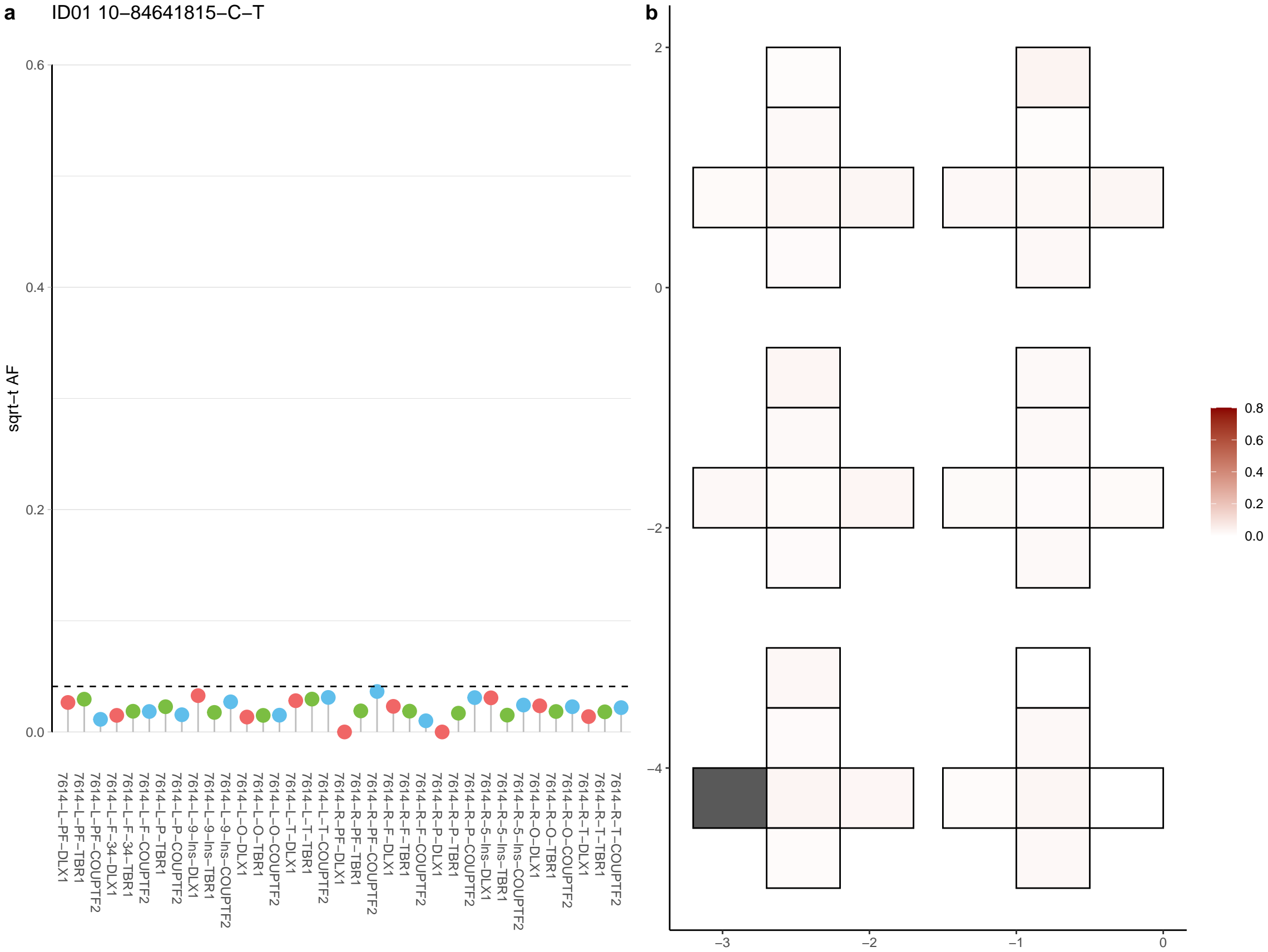

**a** ID01 10-97951160-C-T

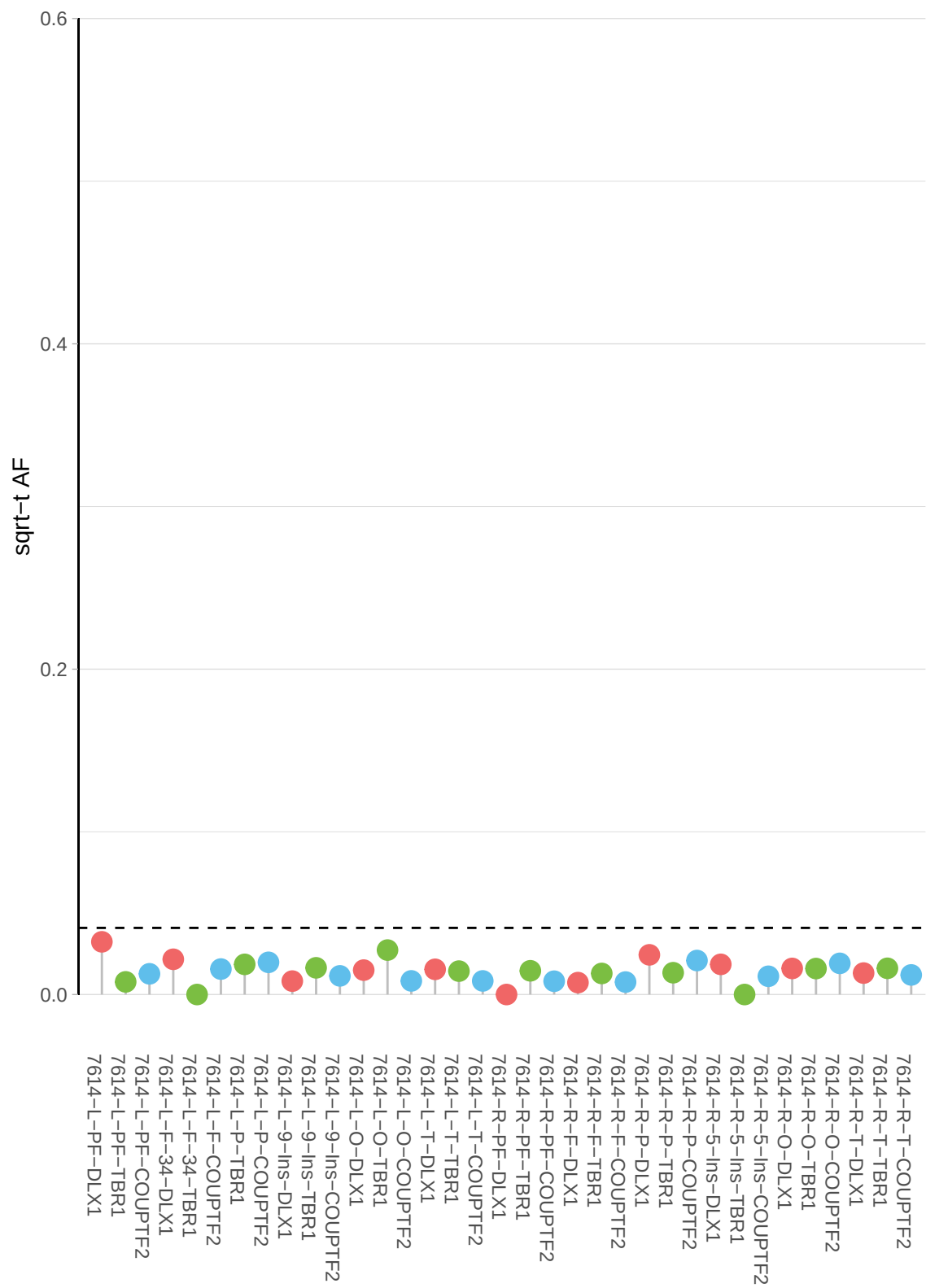

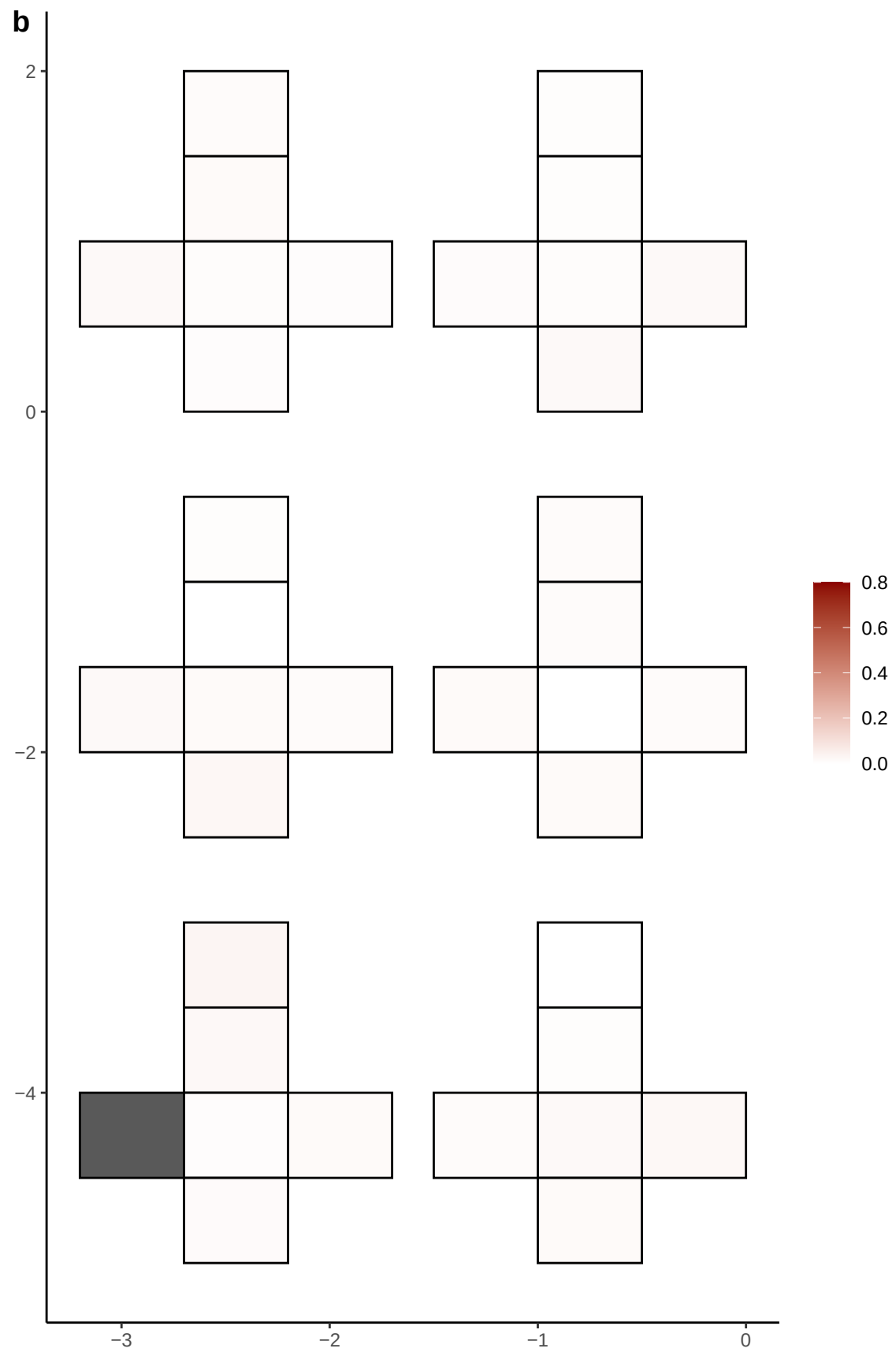

**a** ID01 10-98982494-C-CT

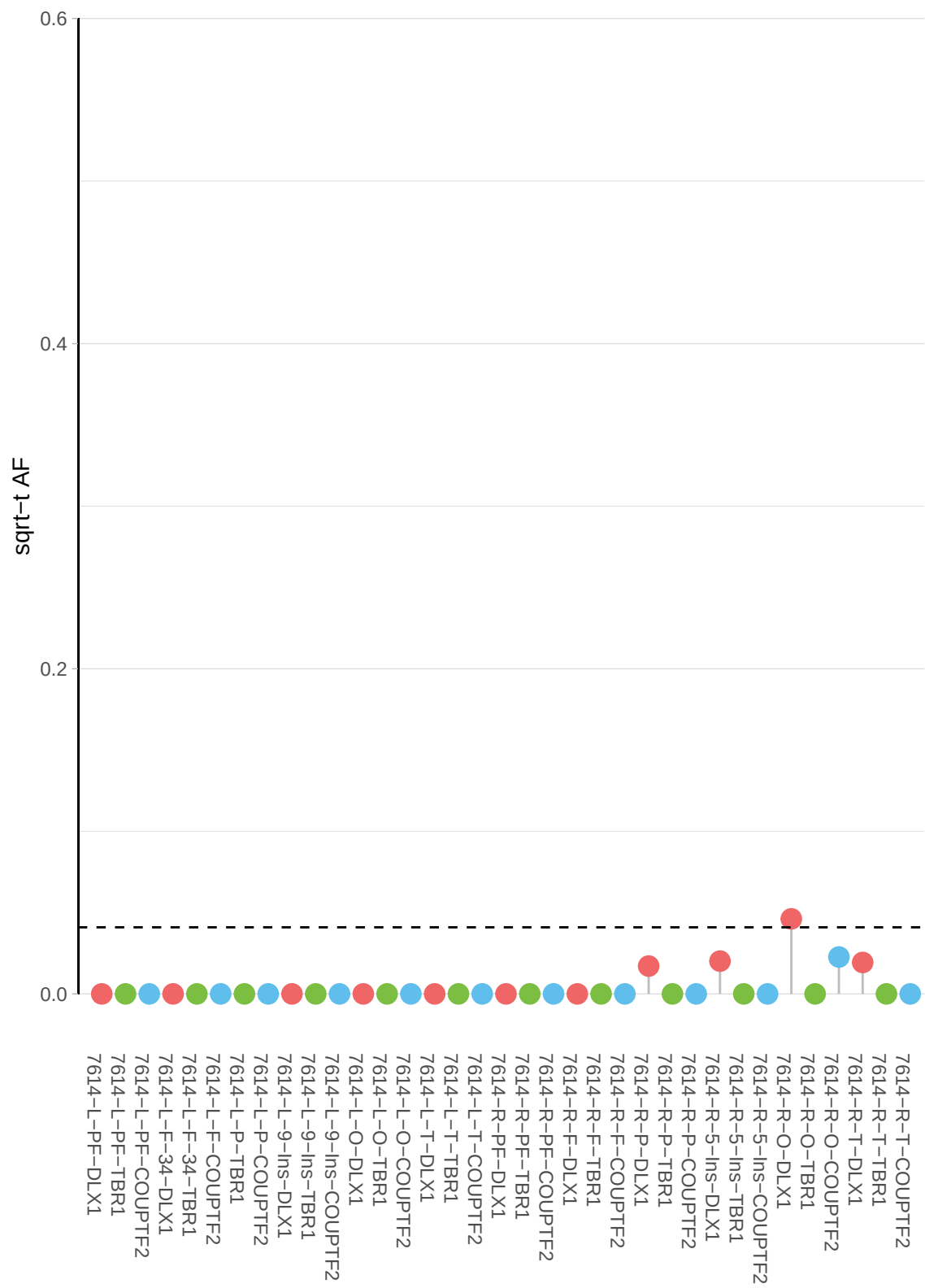

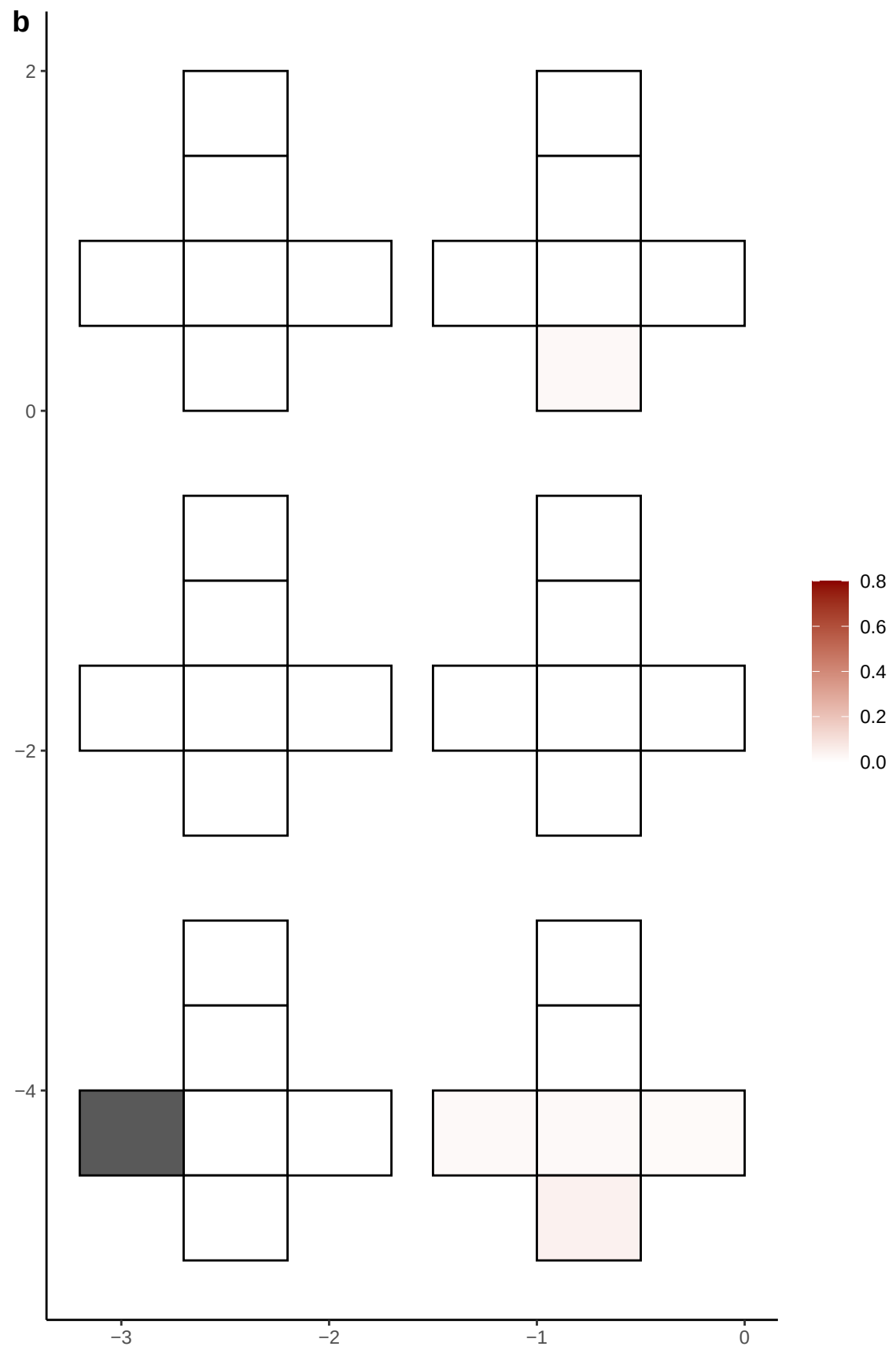

**a** ID01 11-110591774-A-C

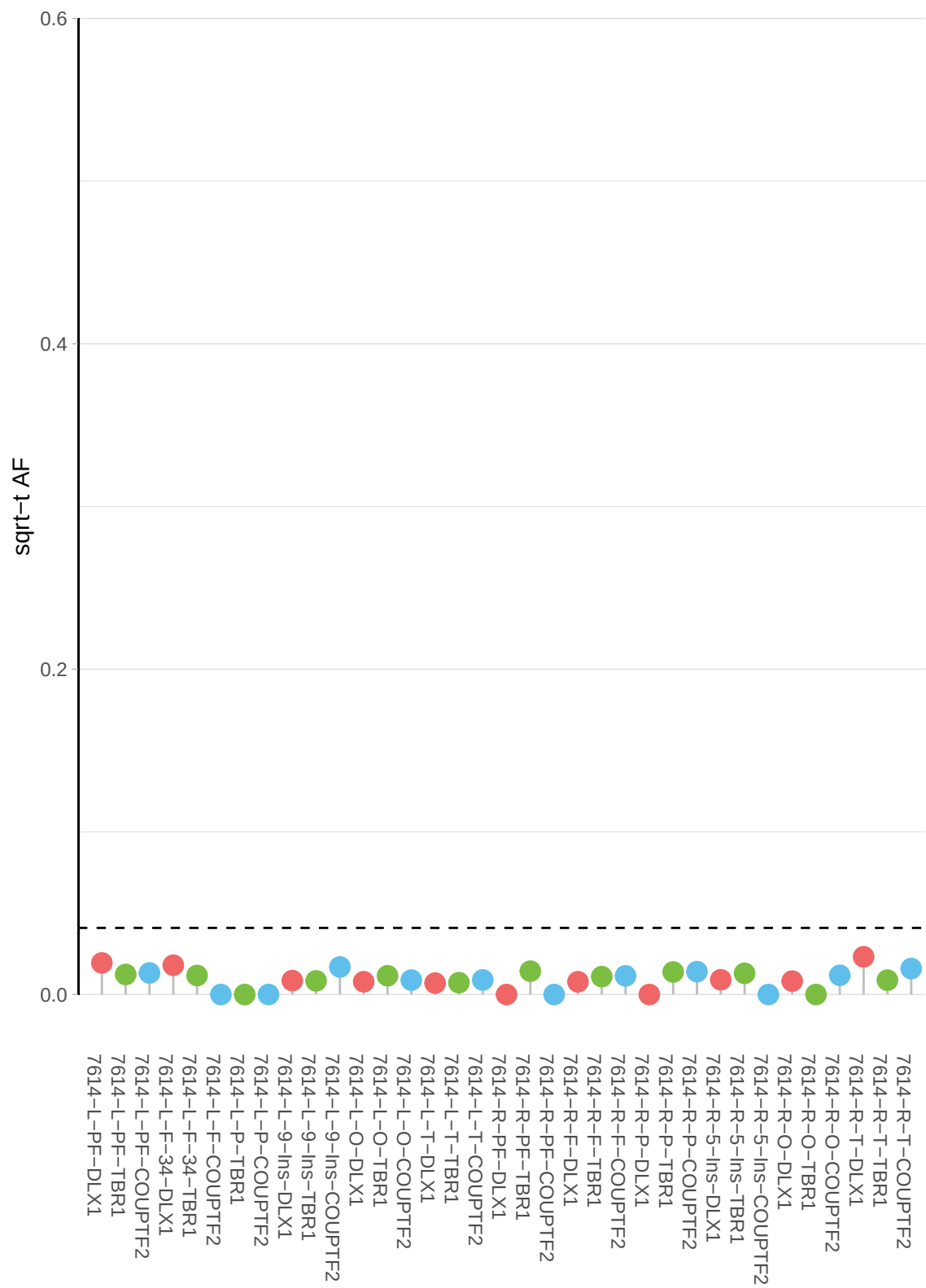

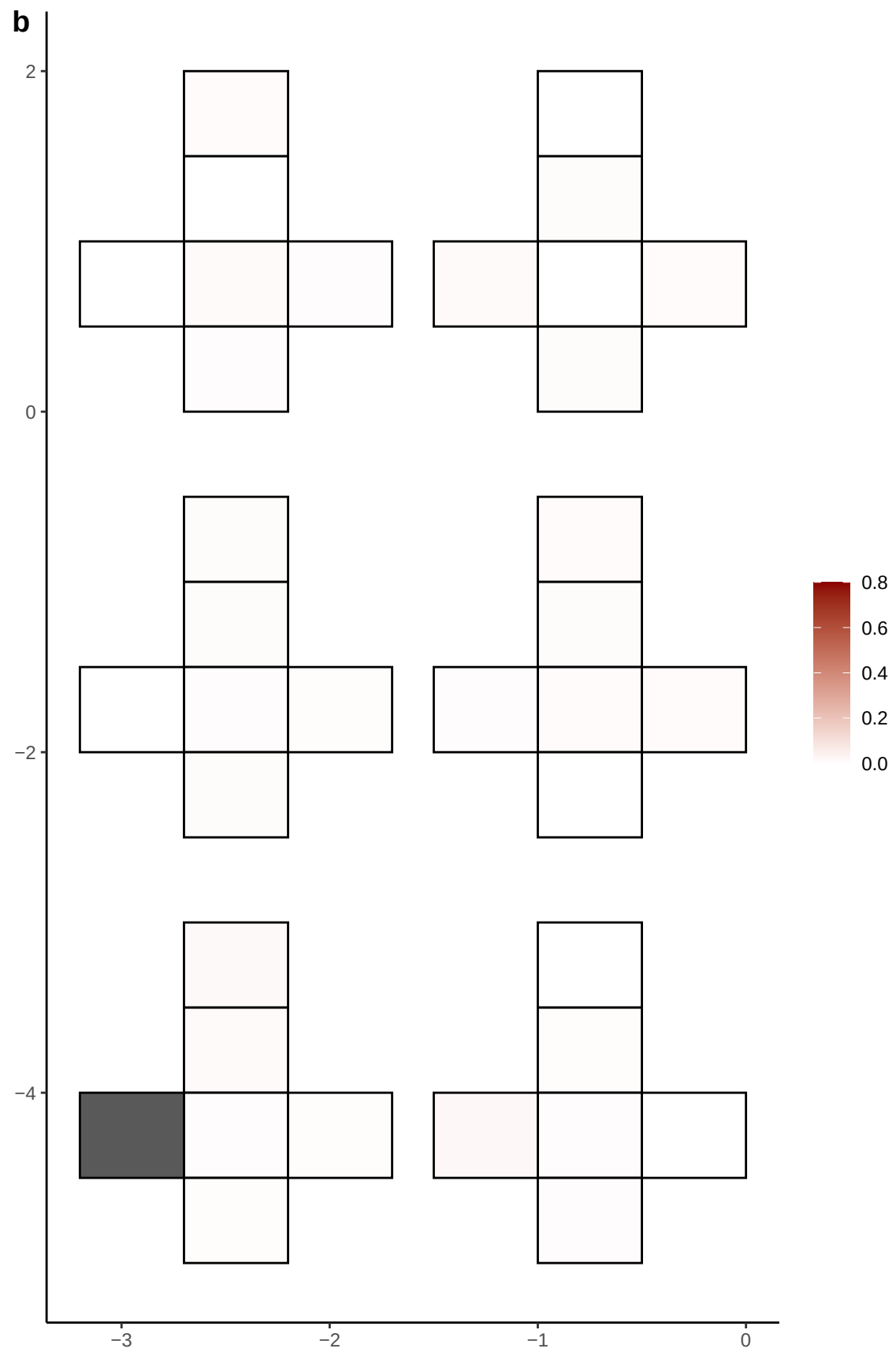

**a** ID01 11-114745430-C-A

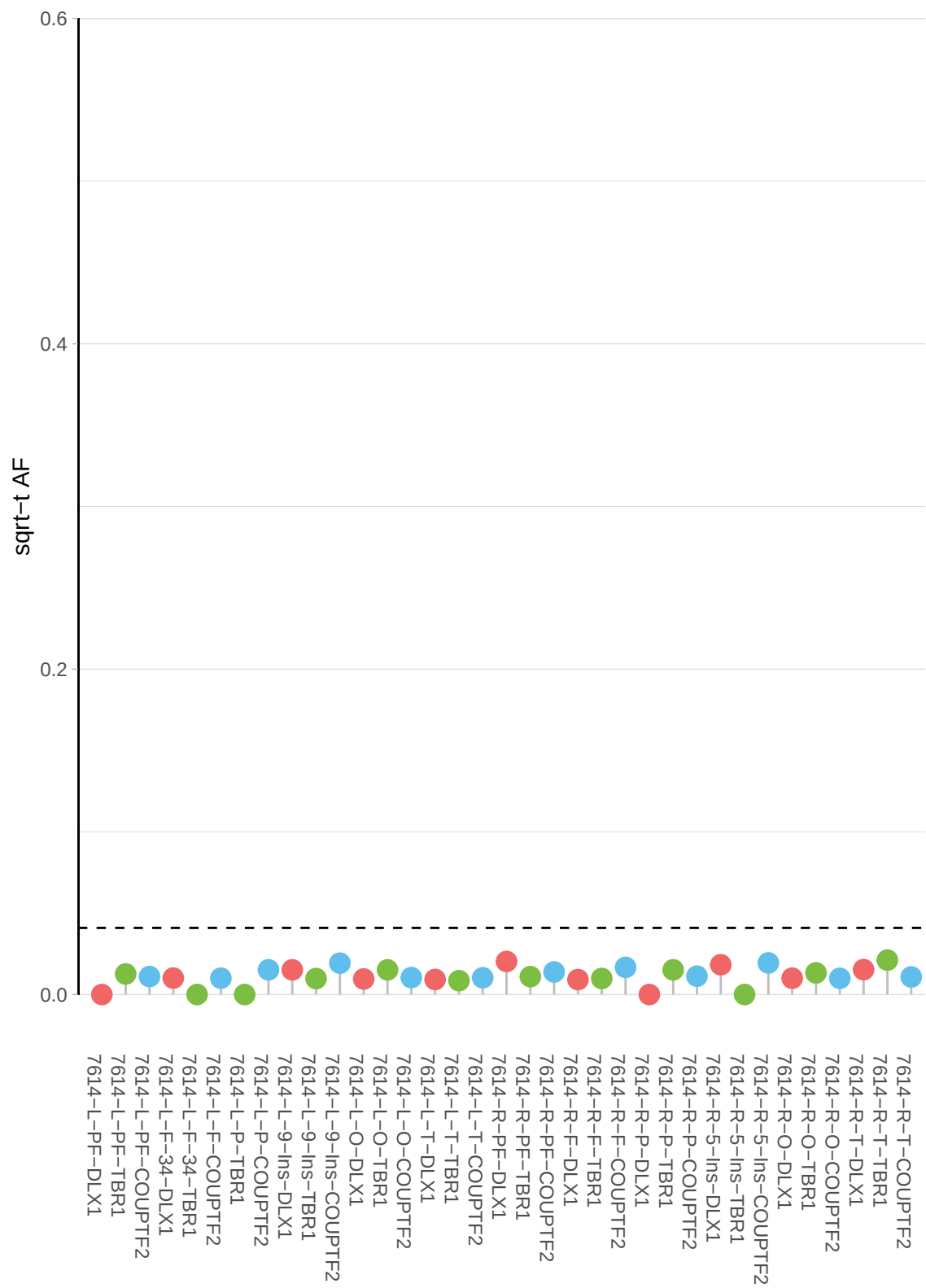

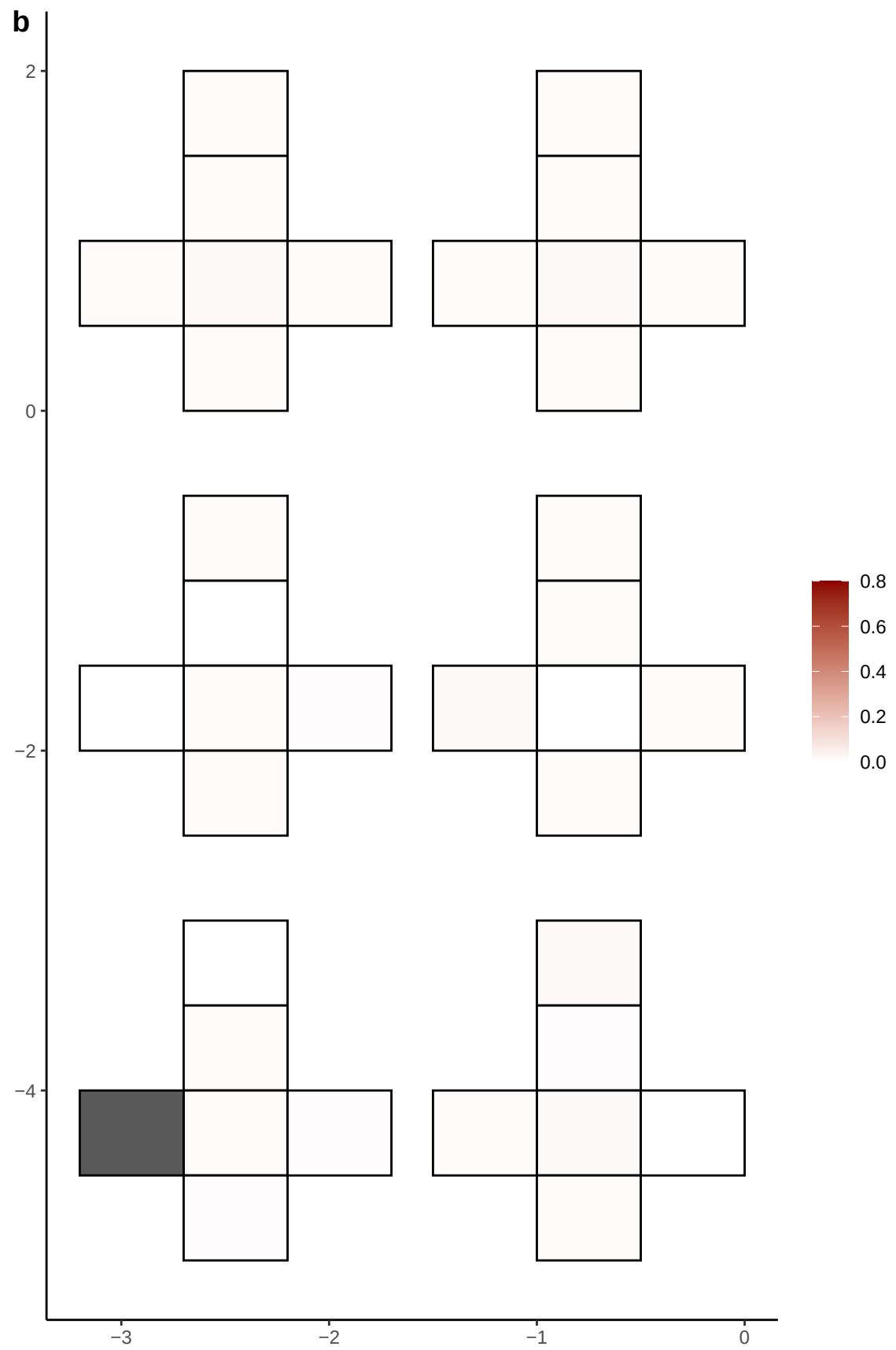

**a** ID01 11-115043033-C-T

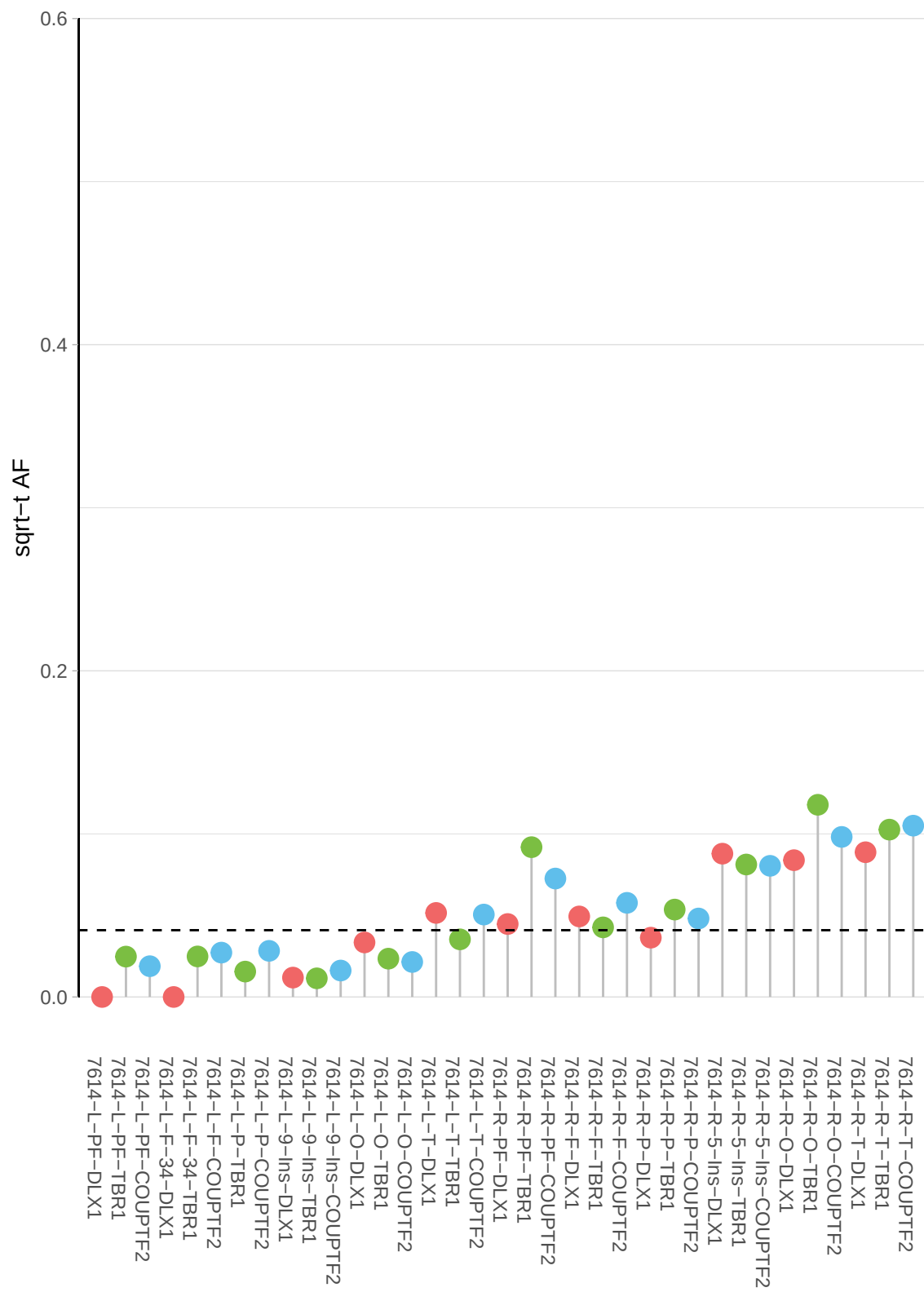

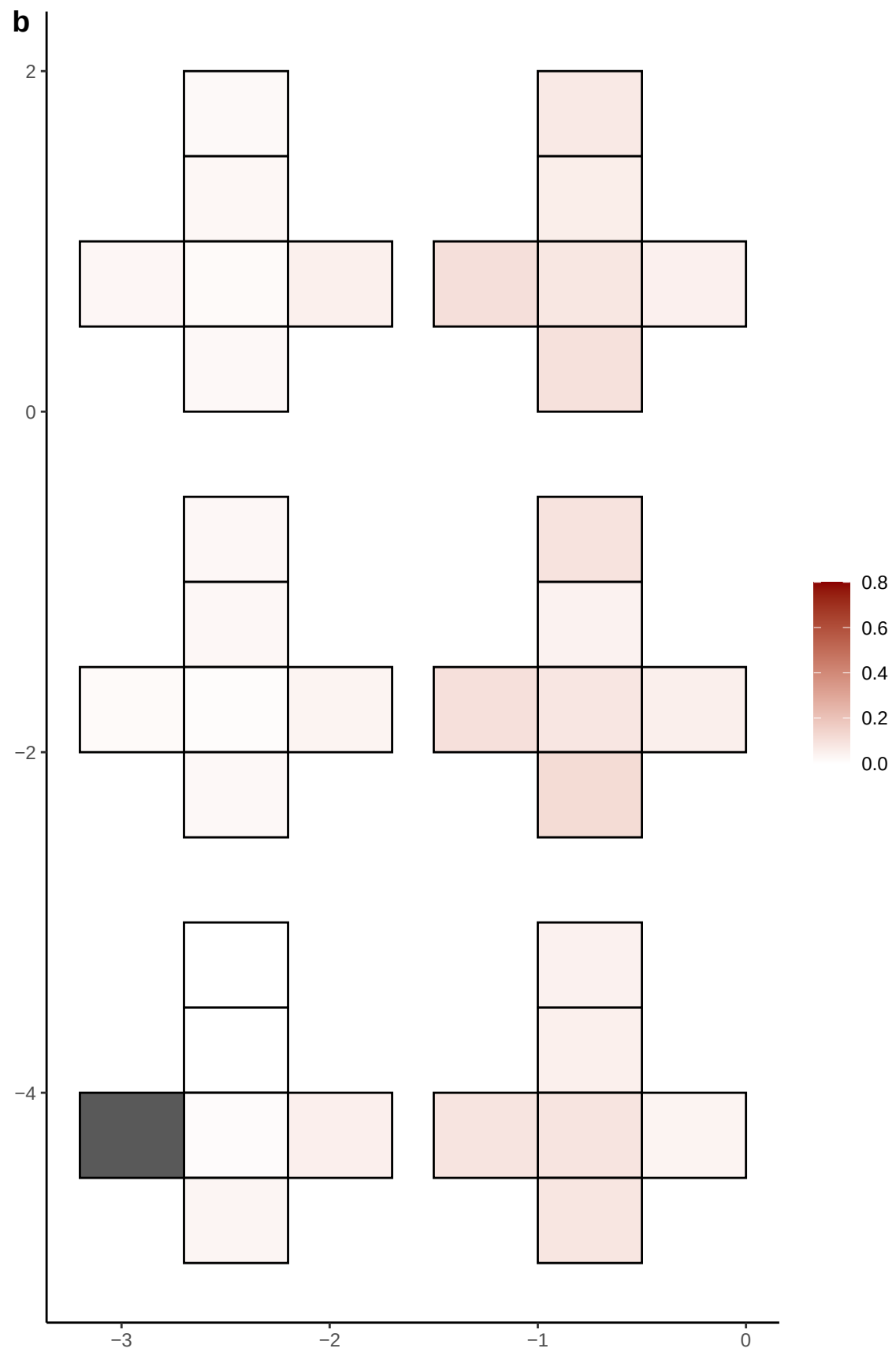

**a** ID01 11-117506570-A-C

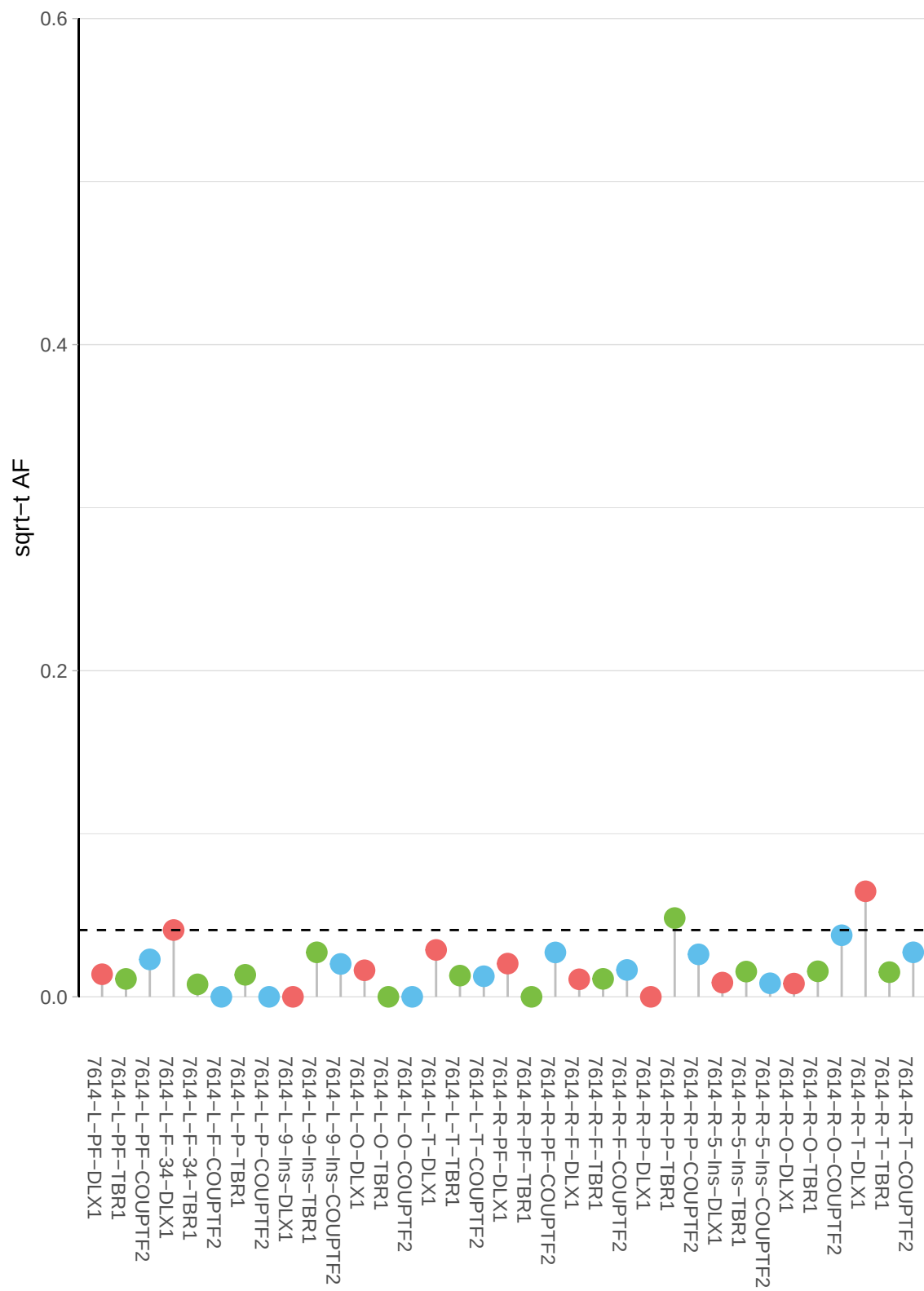

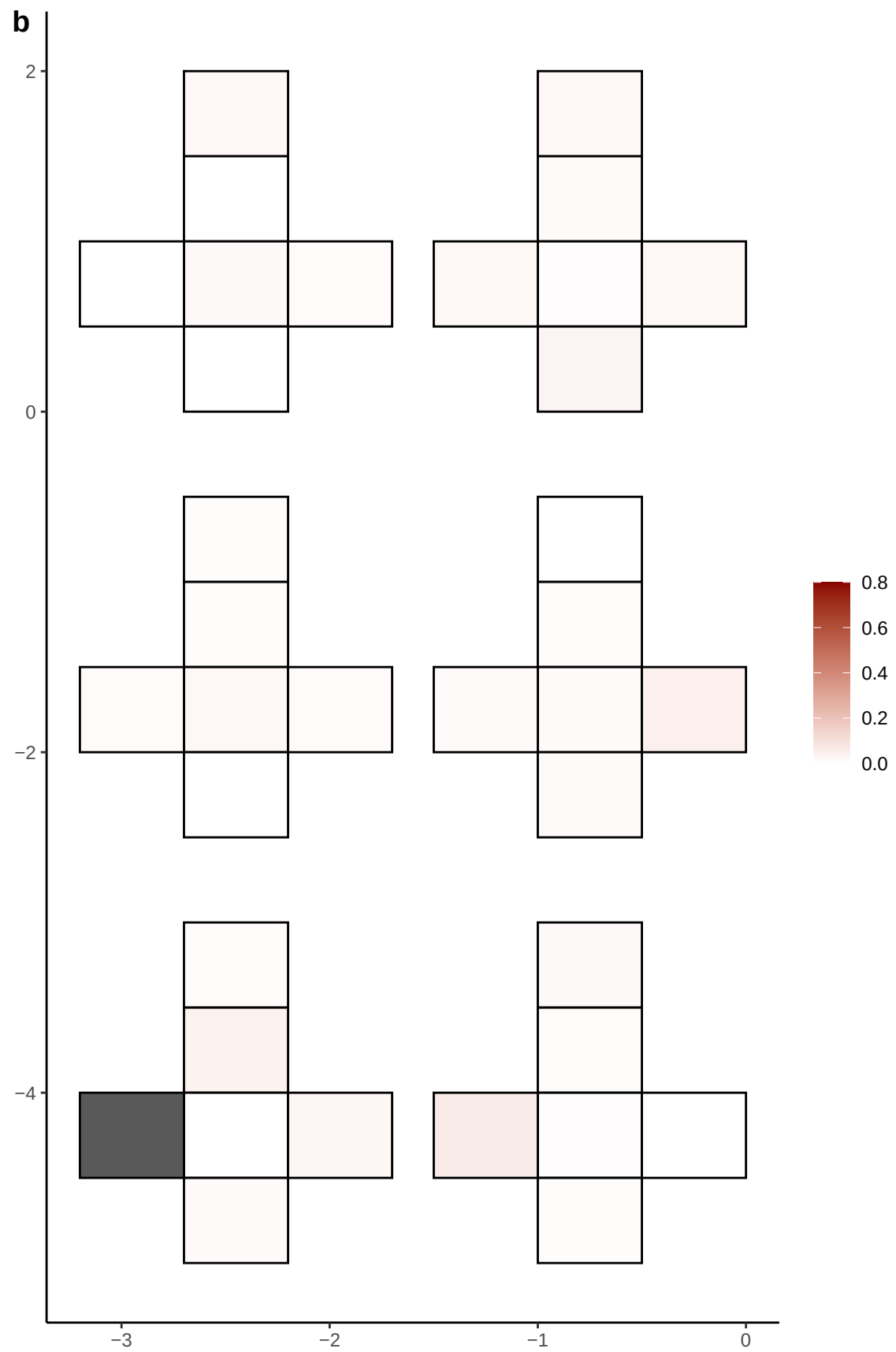

**a** ID01 11-123516528-G-A

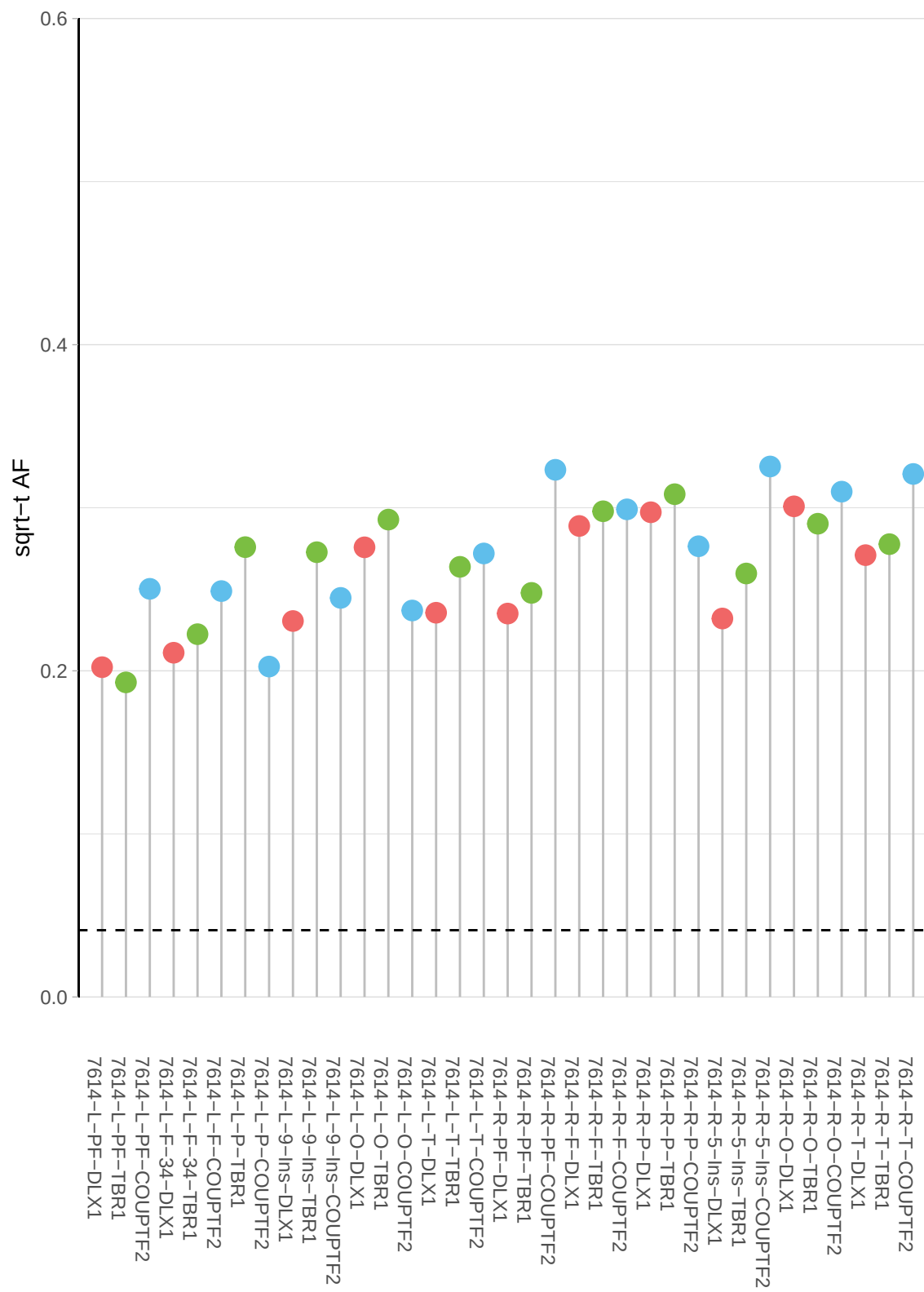

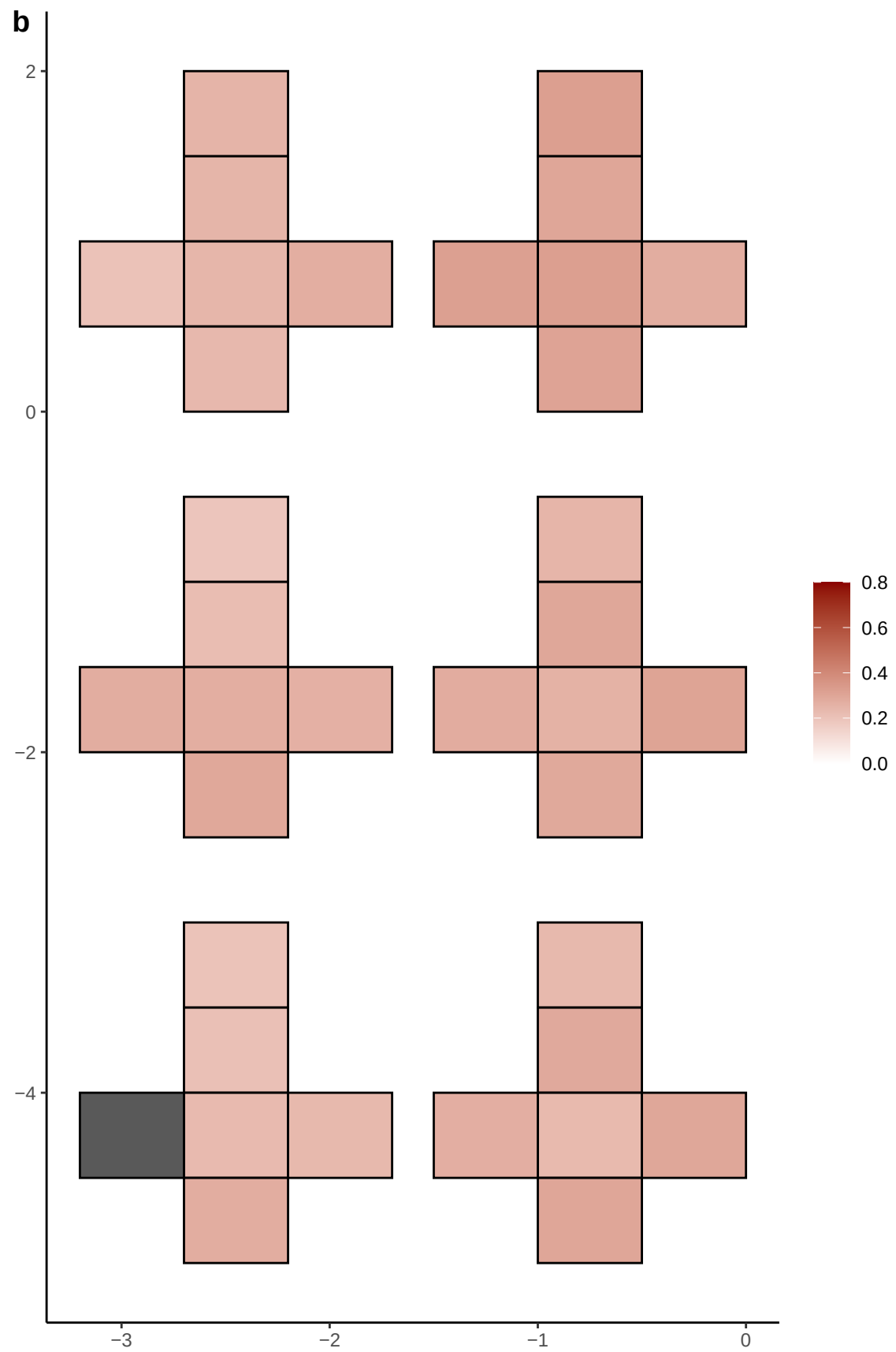

**a** ID01 11-129560658-C-T

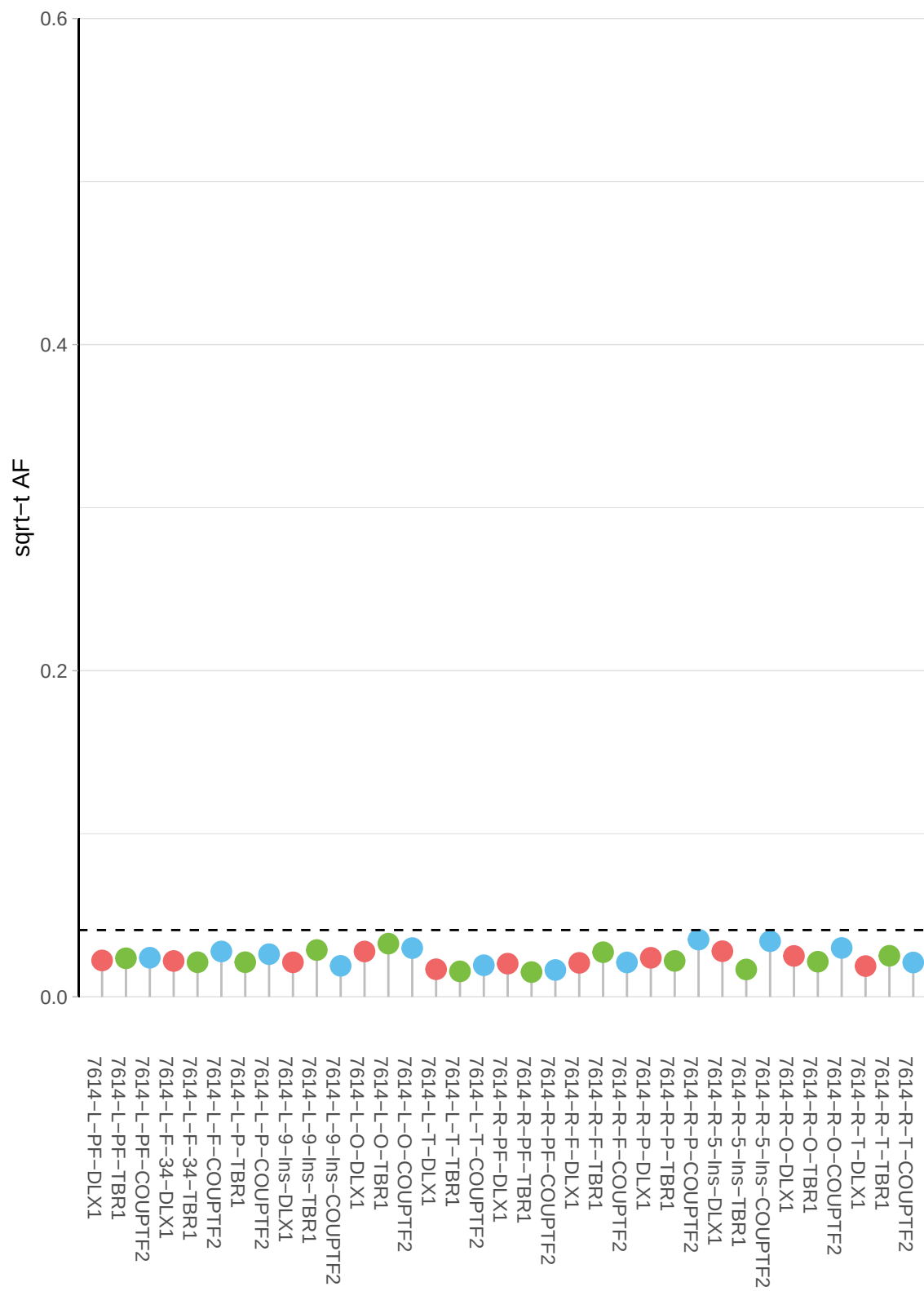

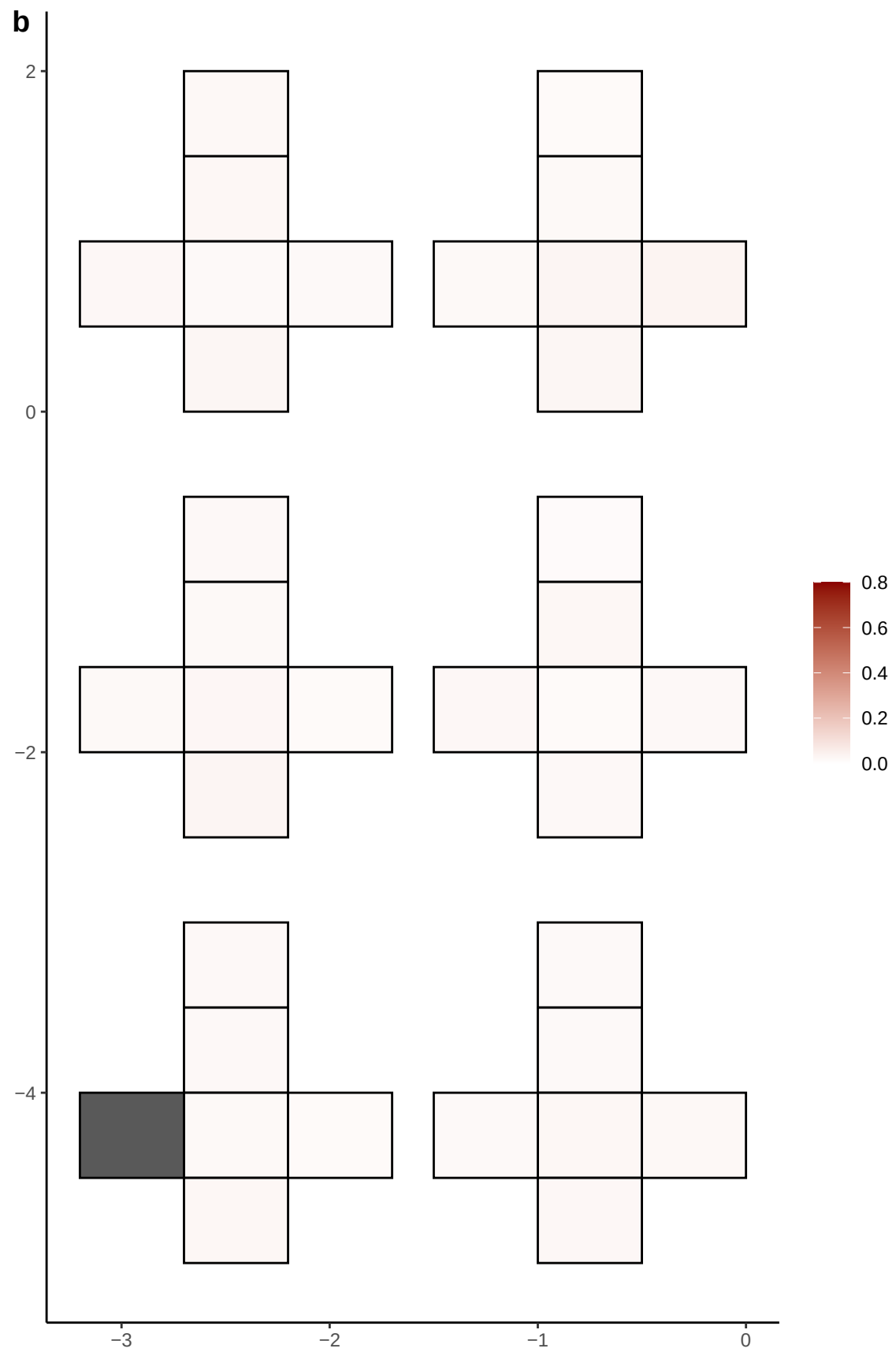

**a** ID01 11-34161341-C-T

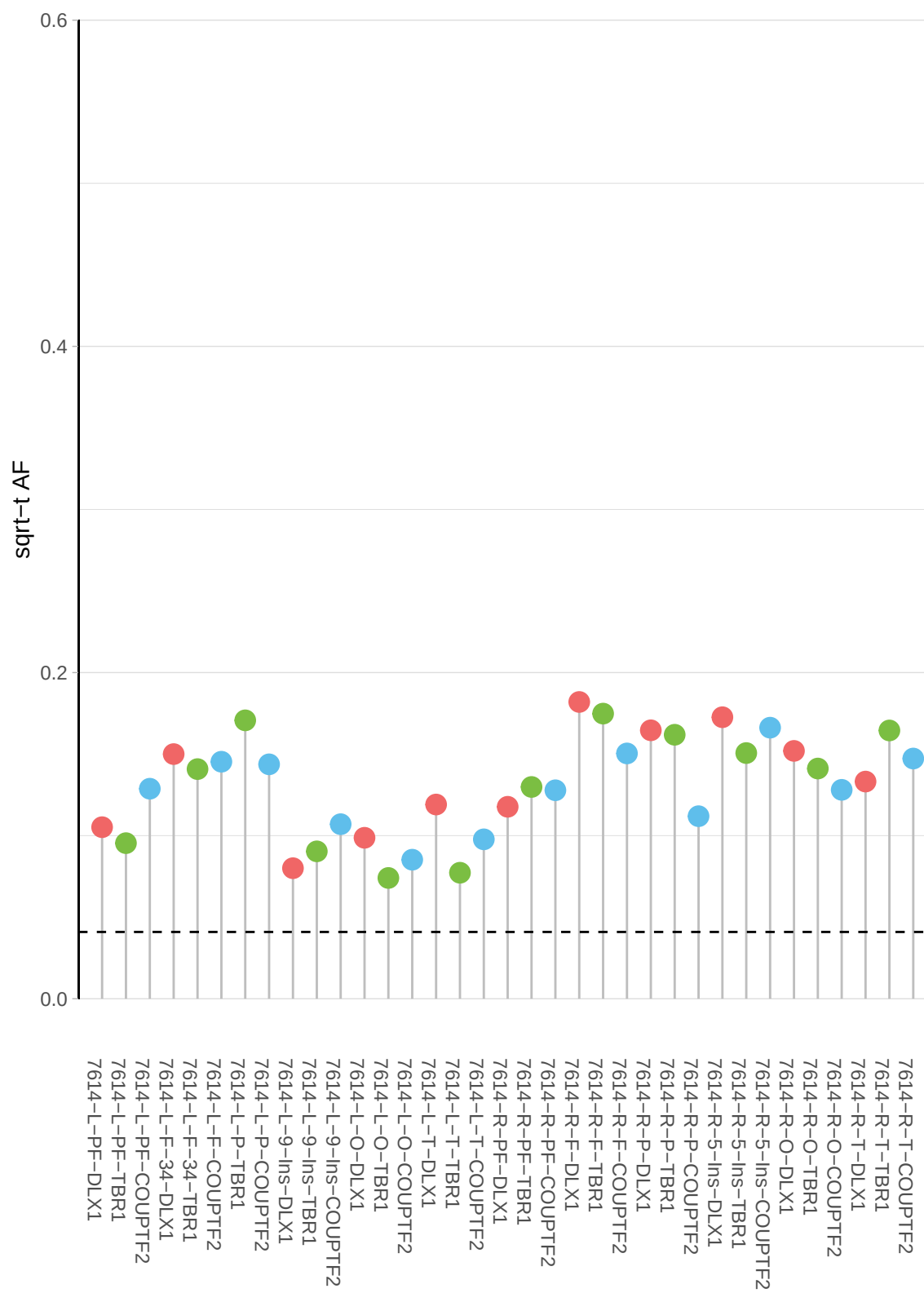

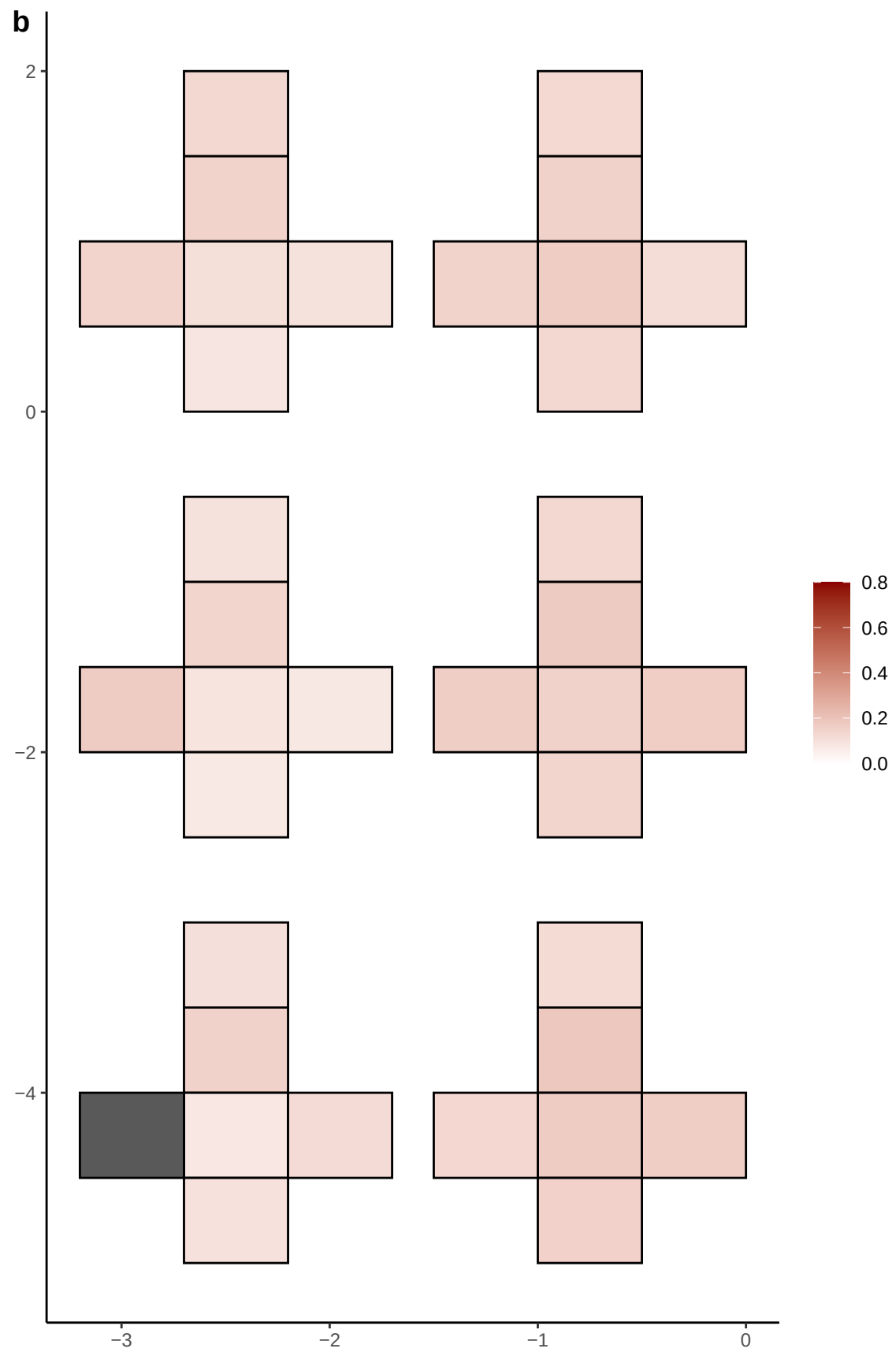

**a** ID01 11-43099503-A-C

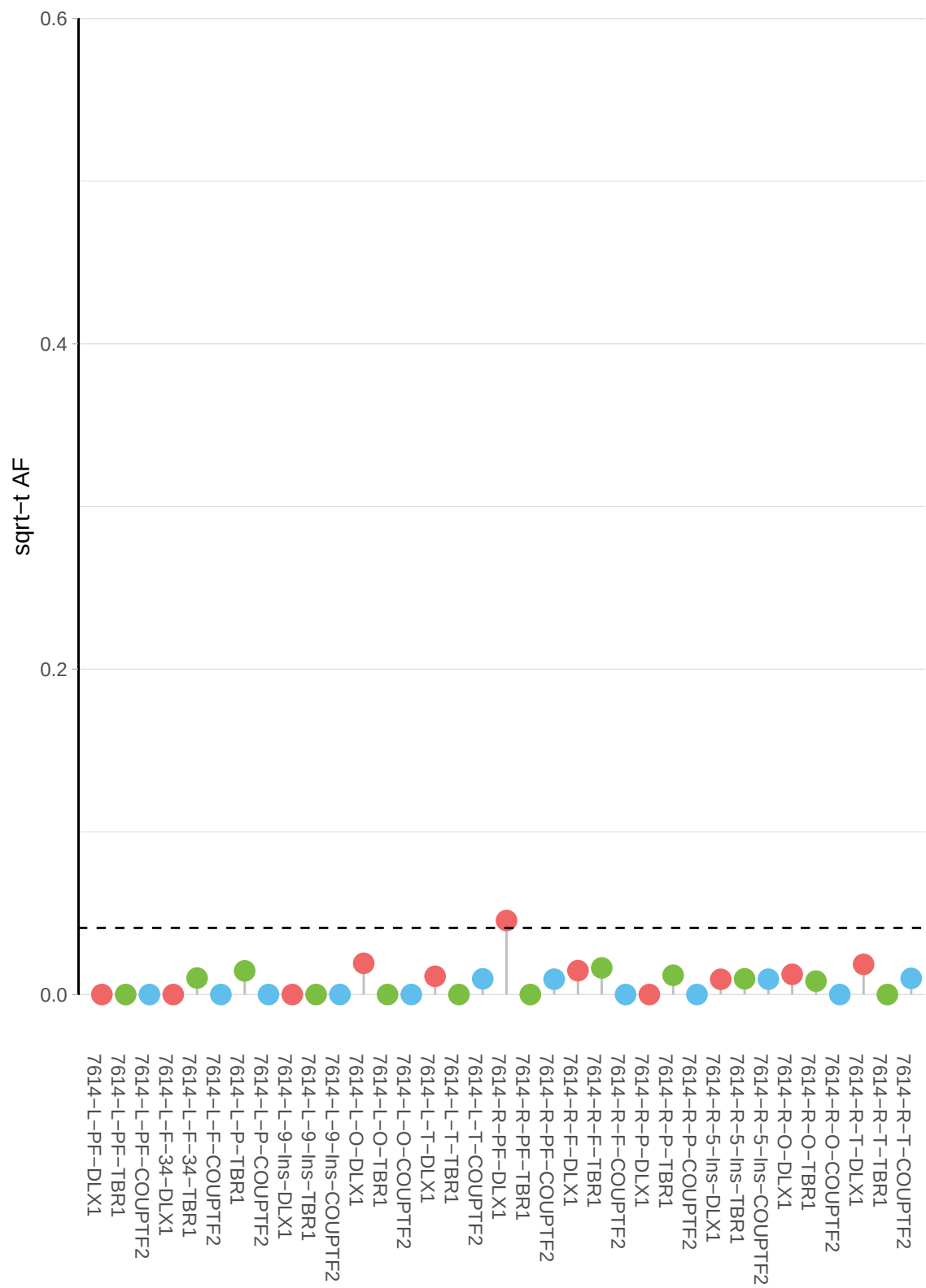

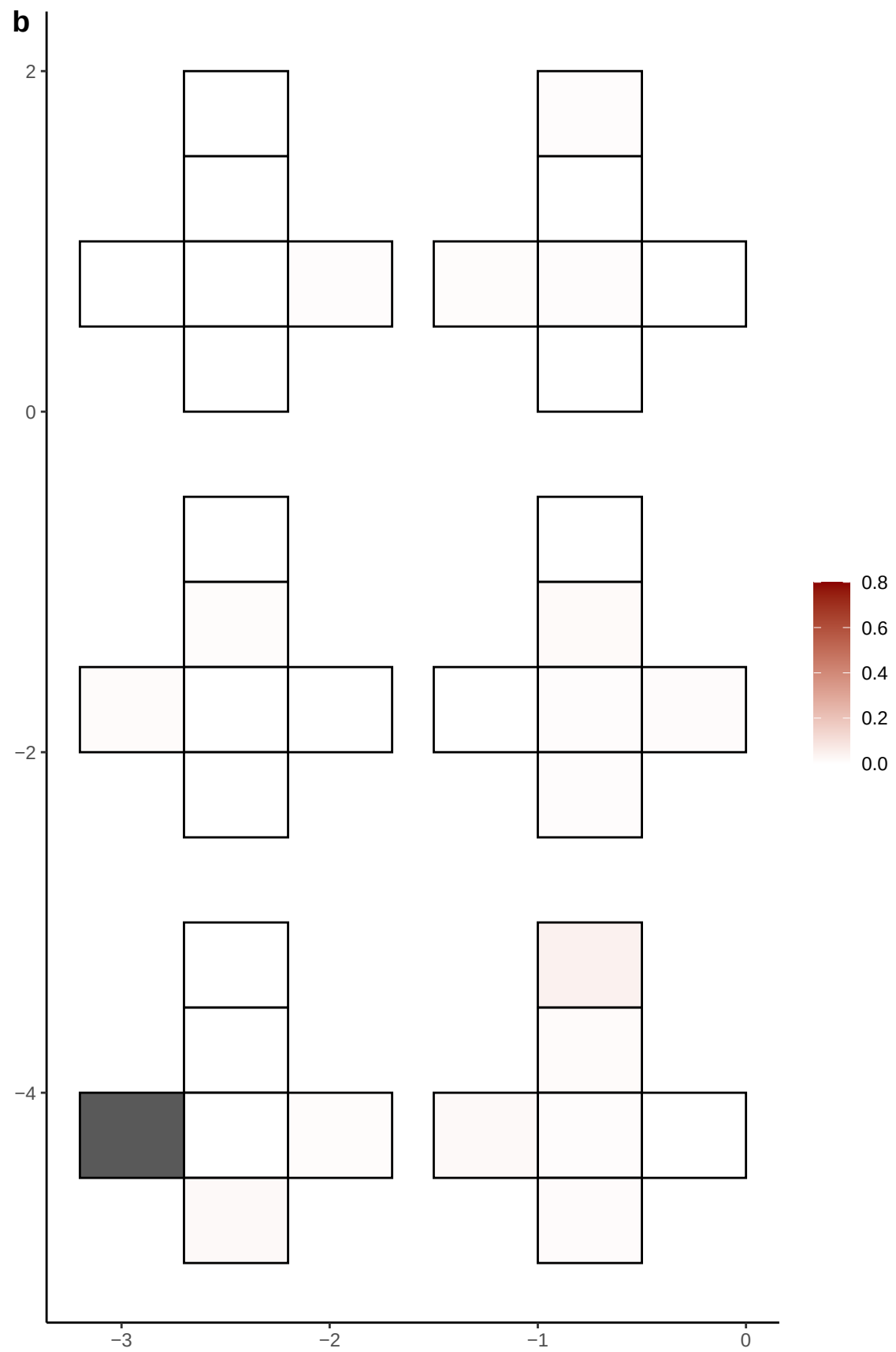

**a** ID01 11-5470408-T-C

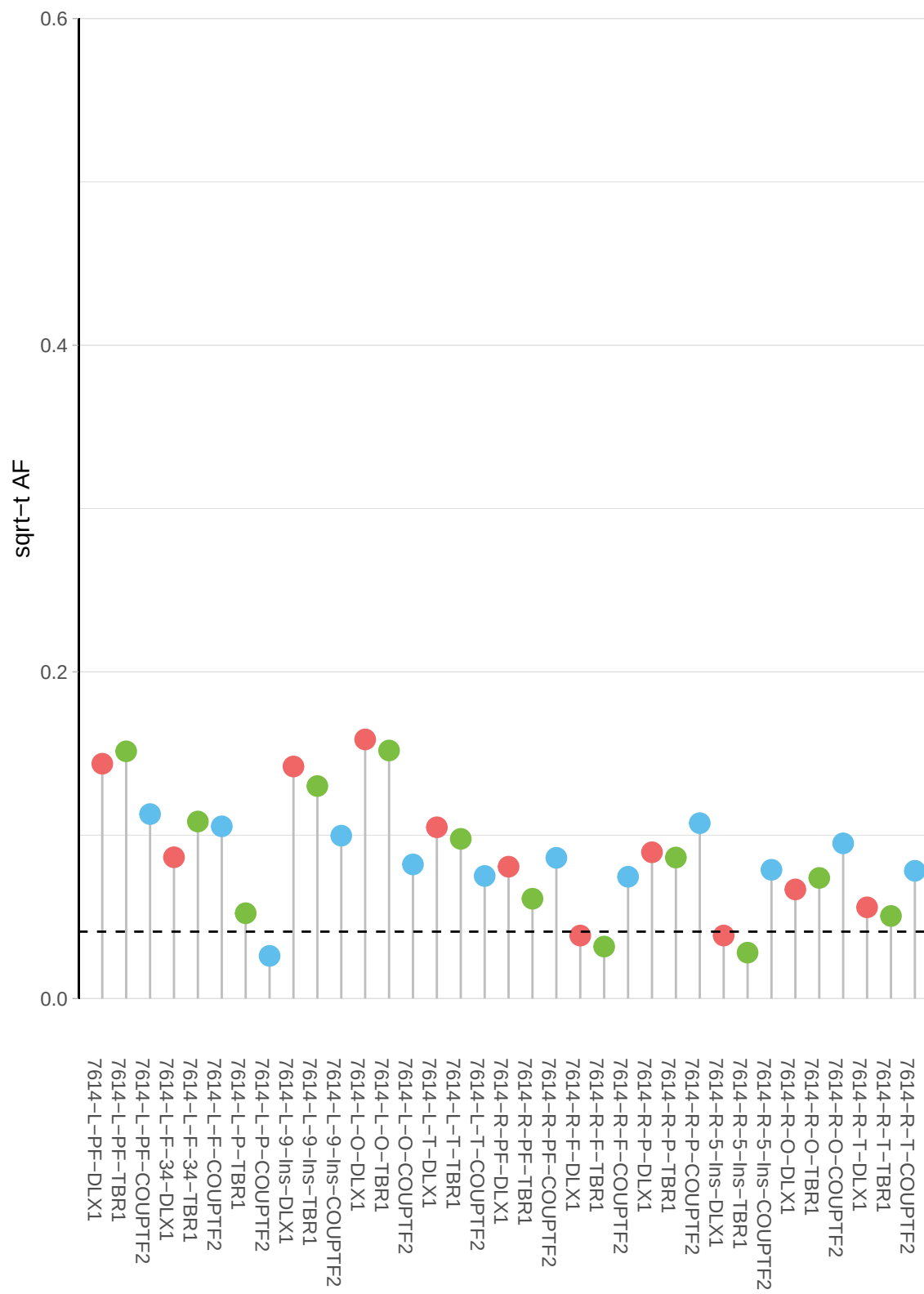

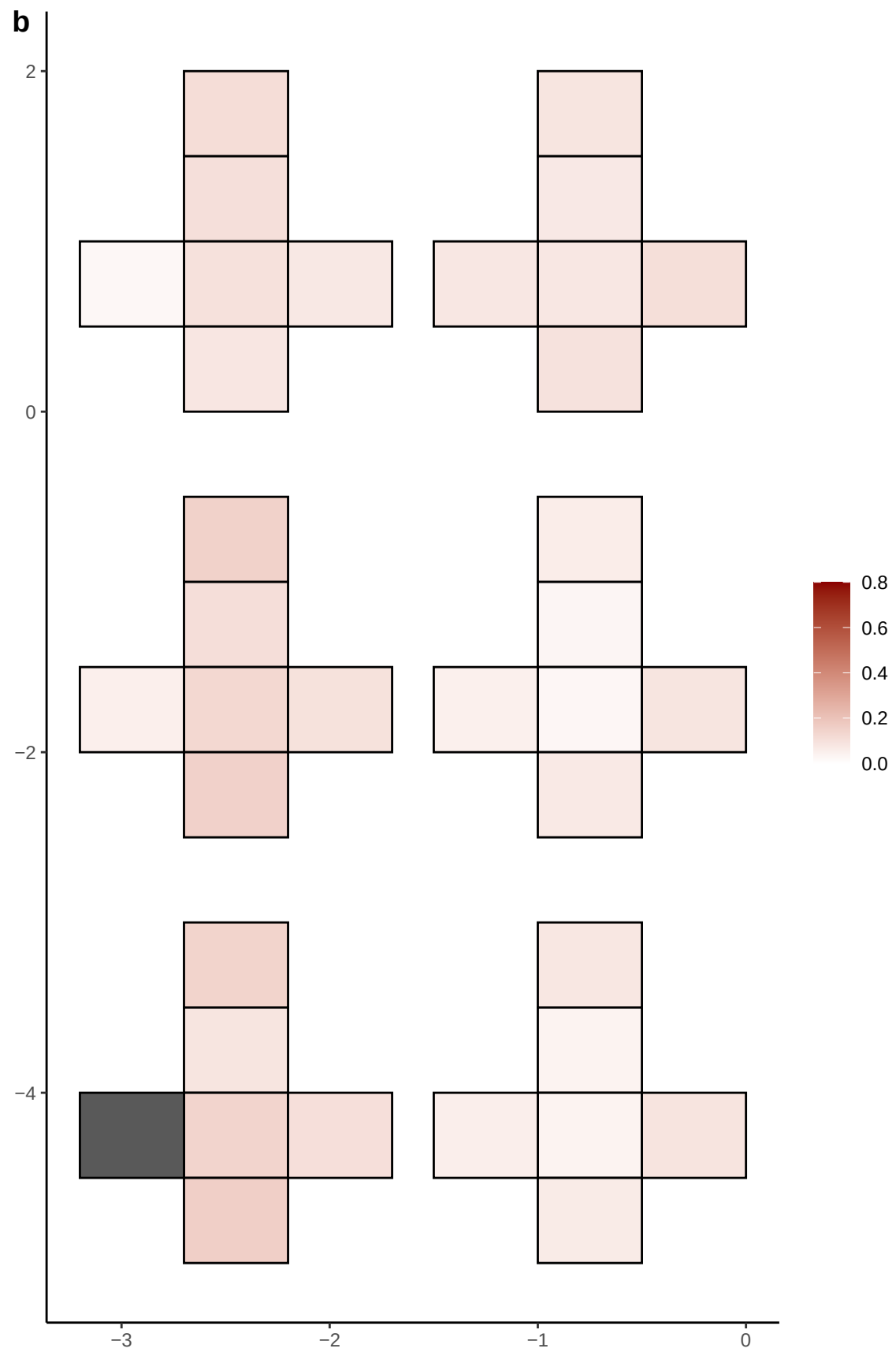

**a** ID01 11-8350555-G-A

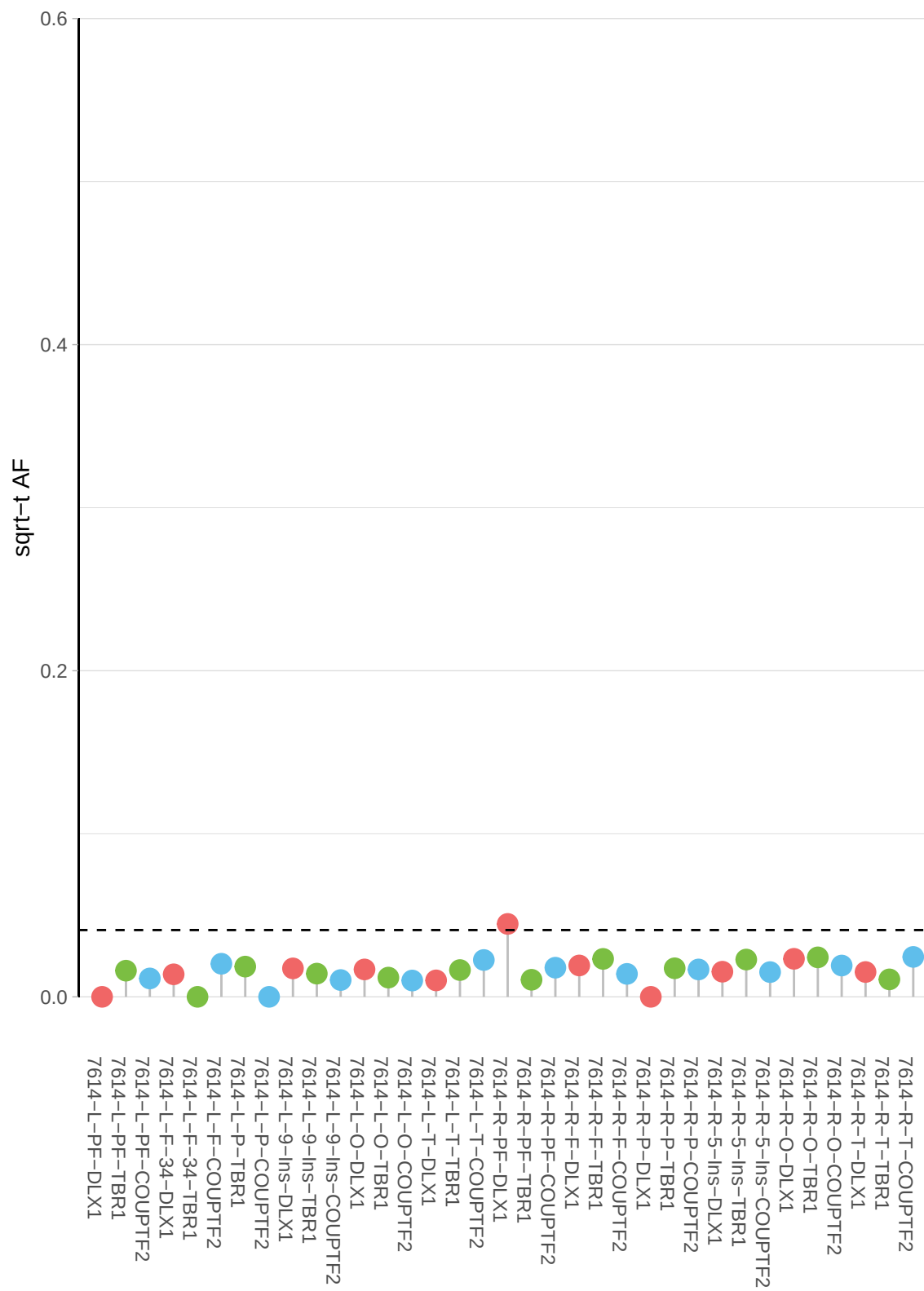

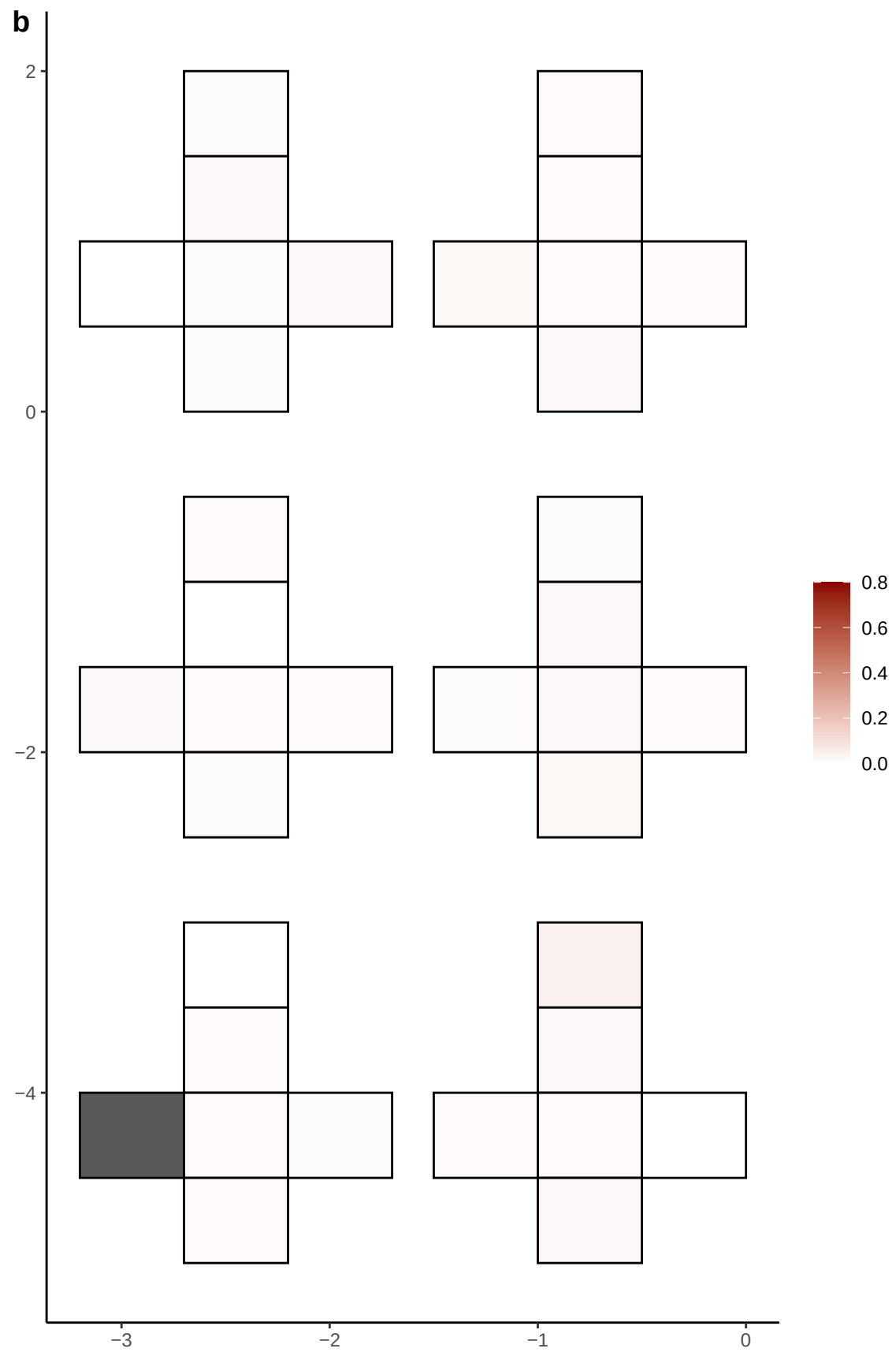

**a** ID01 11-84980734-G-A

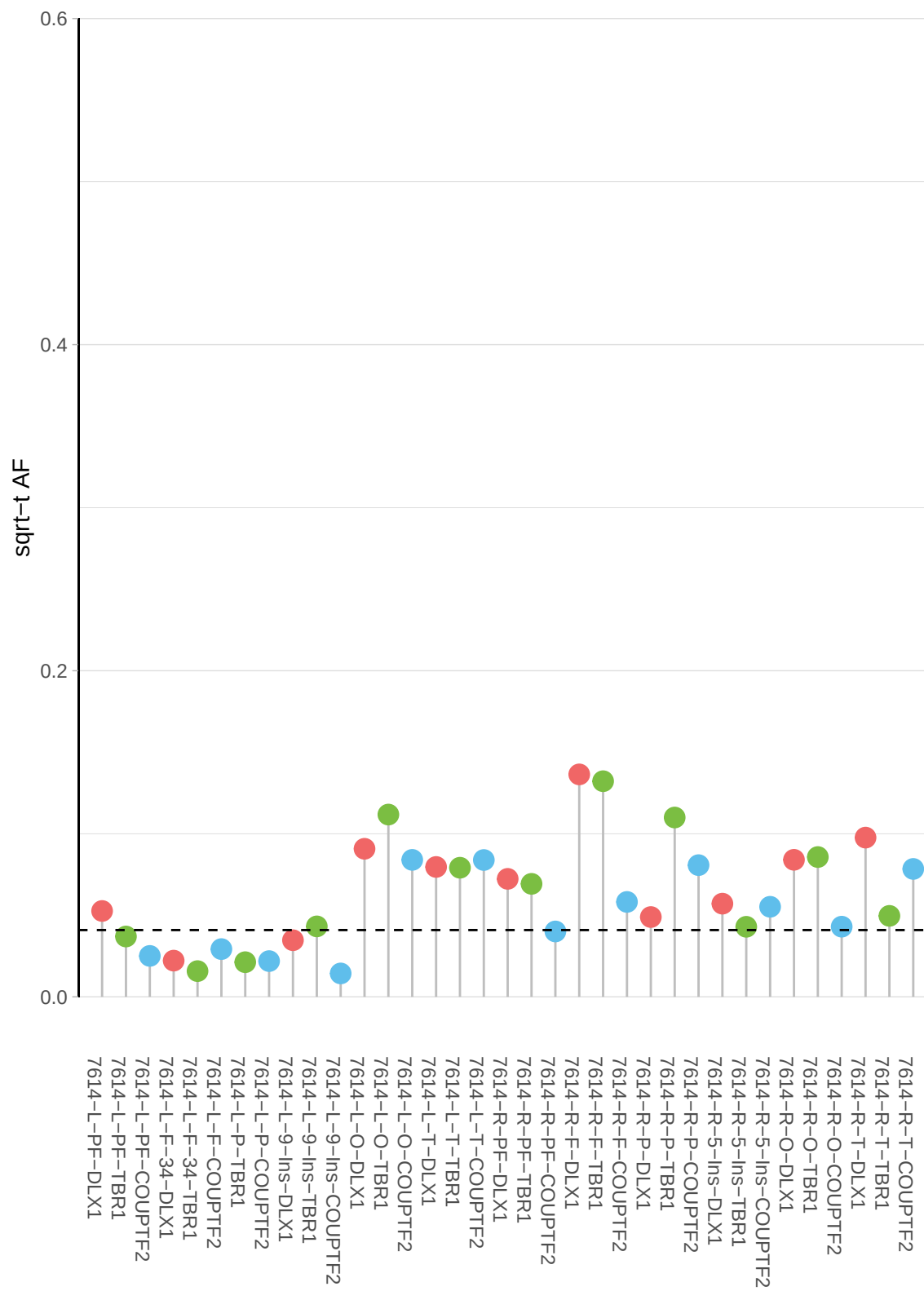

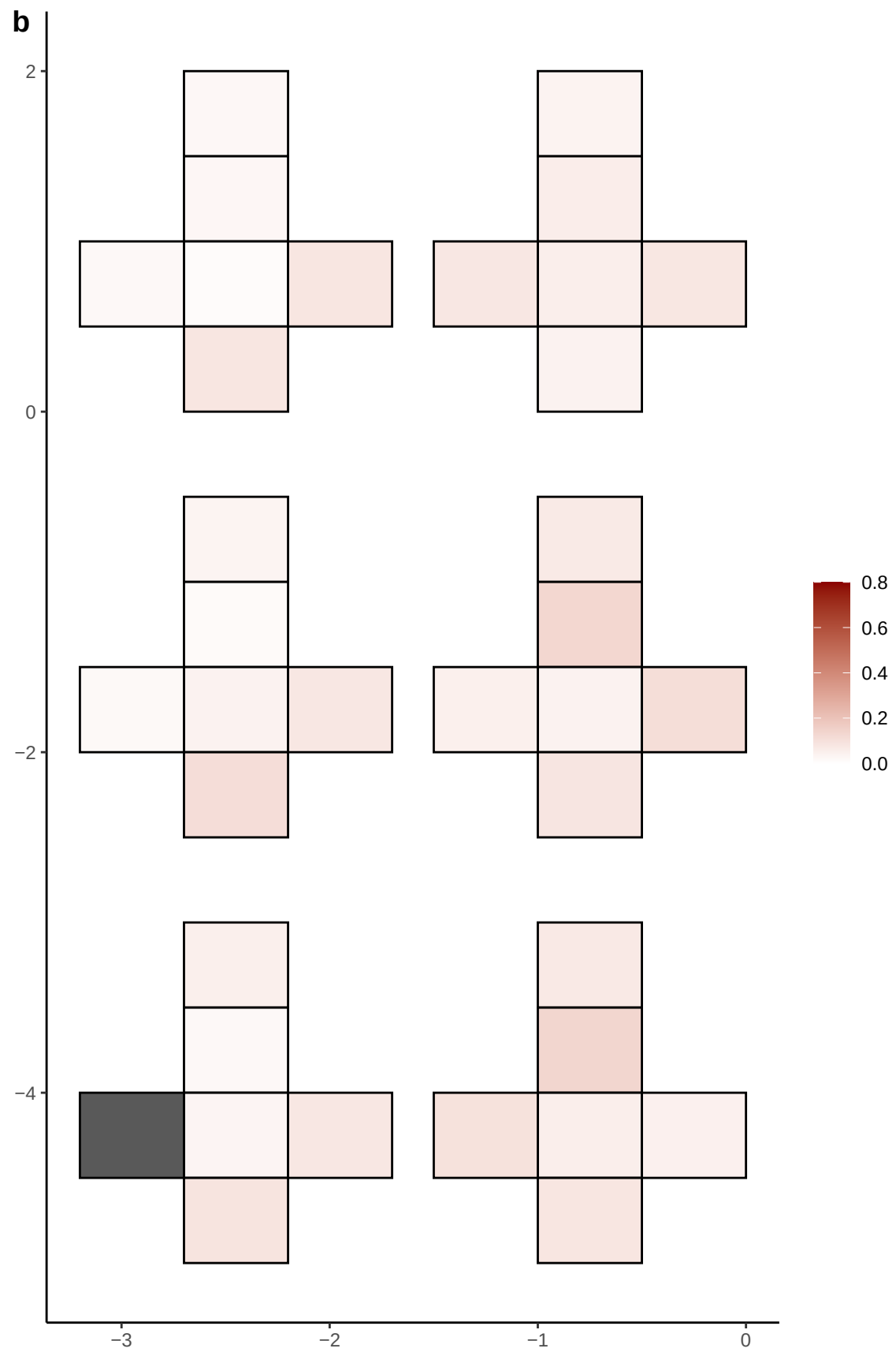

**a** ID01 11-88087255-A-C

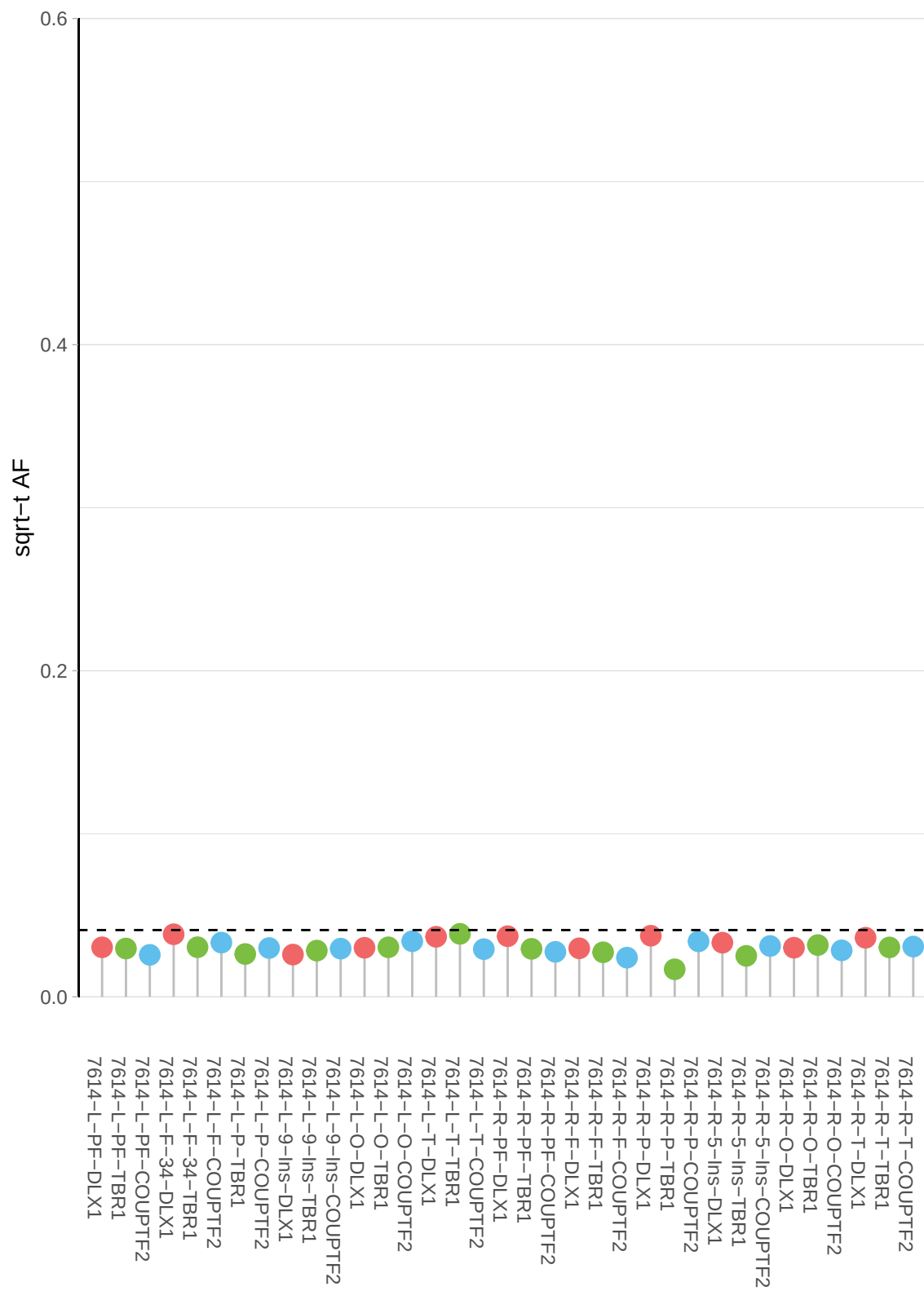

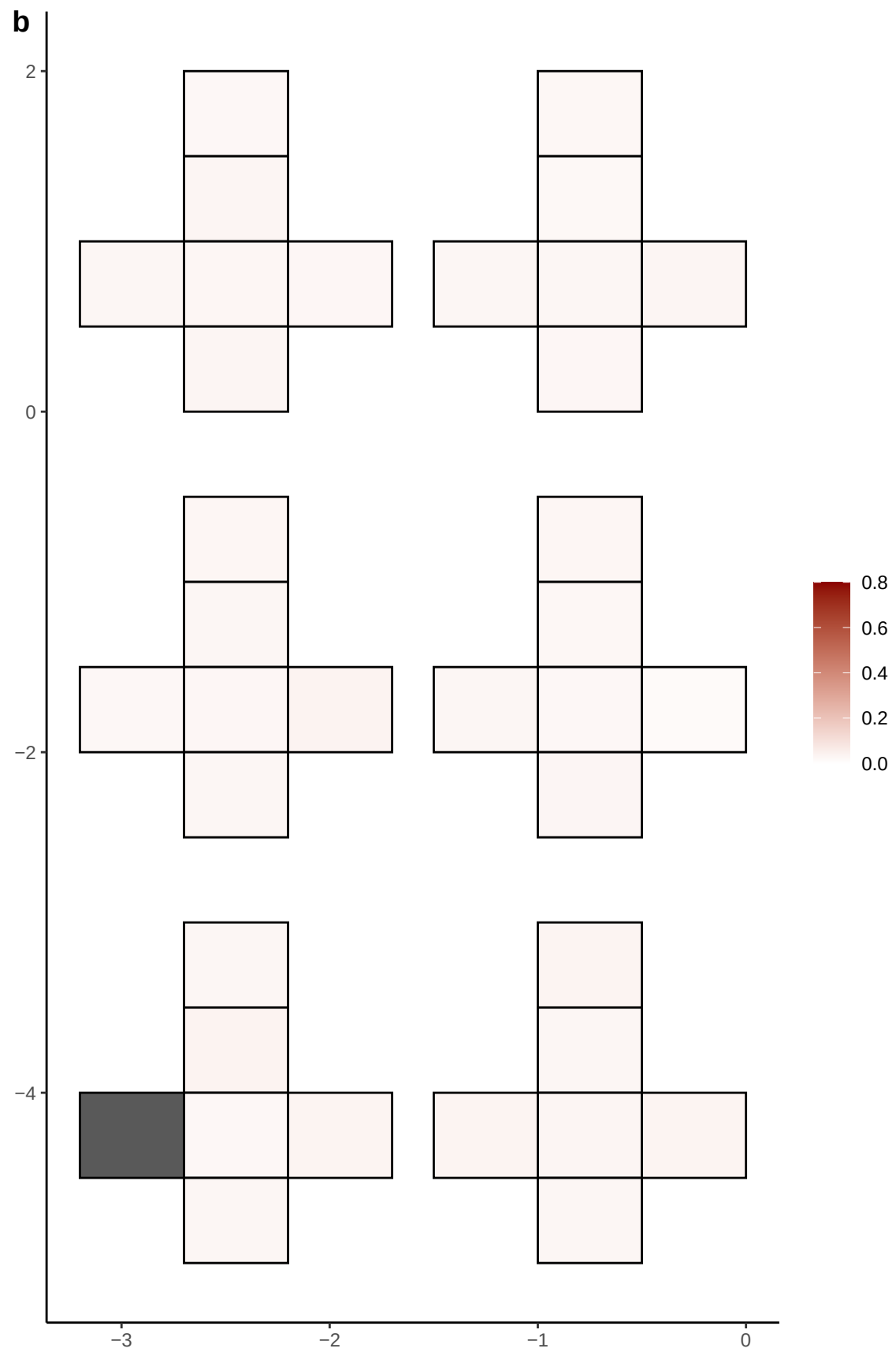

**a** ID01 11-92861647-G-A

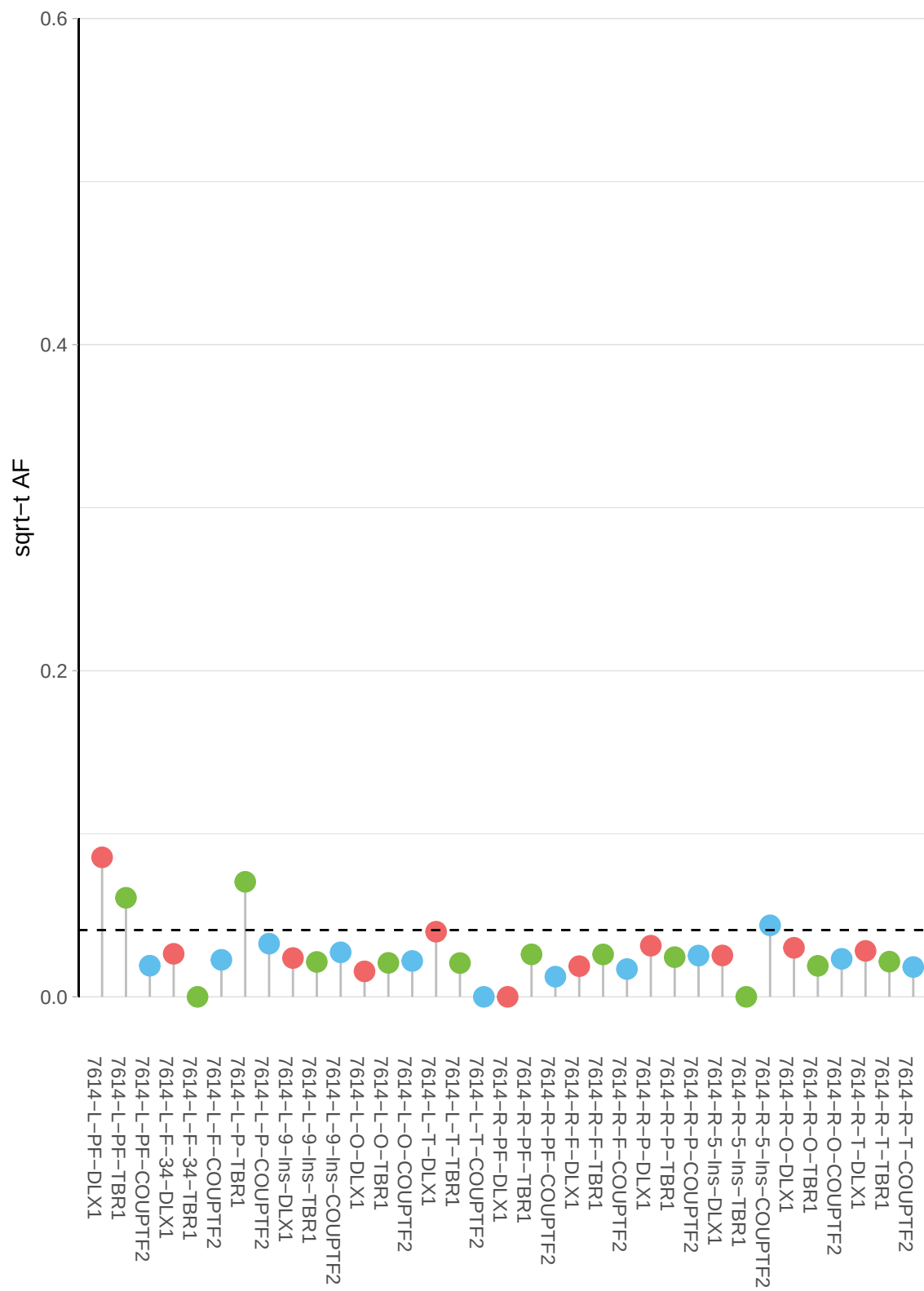

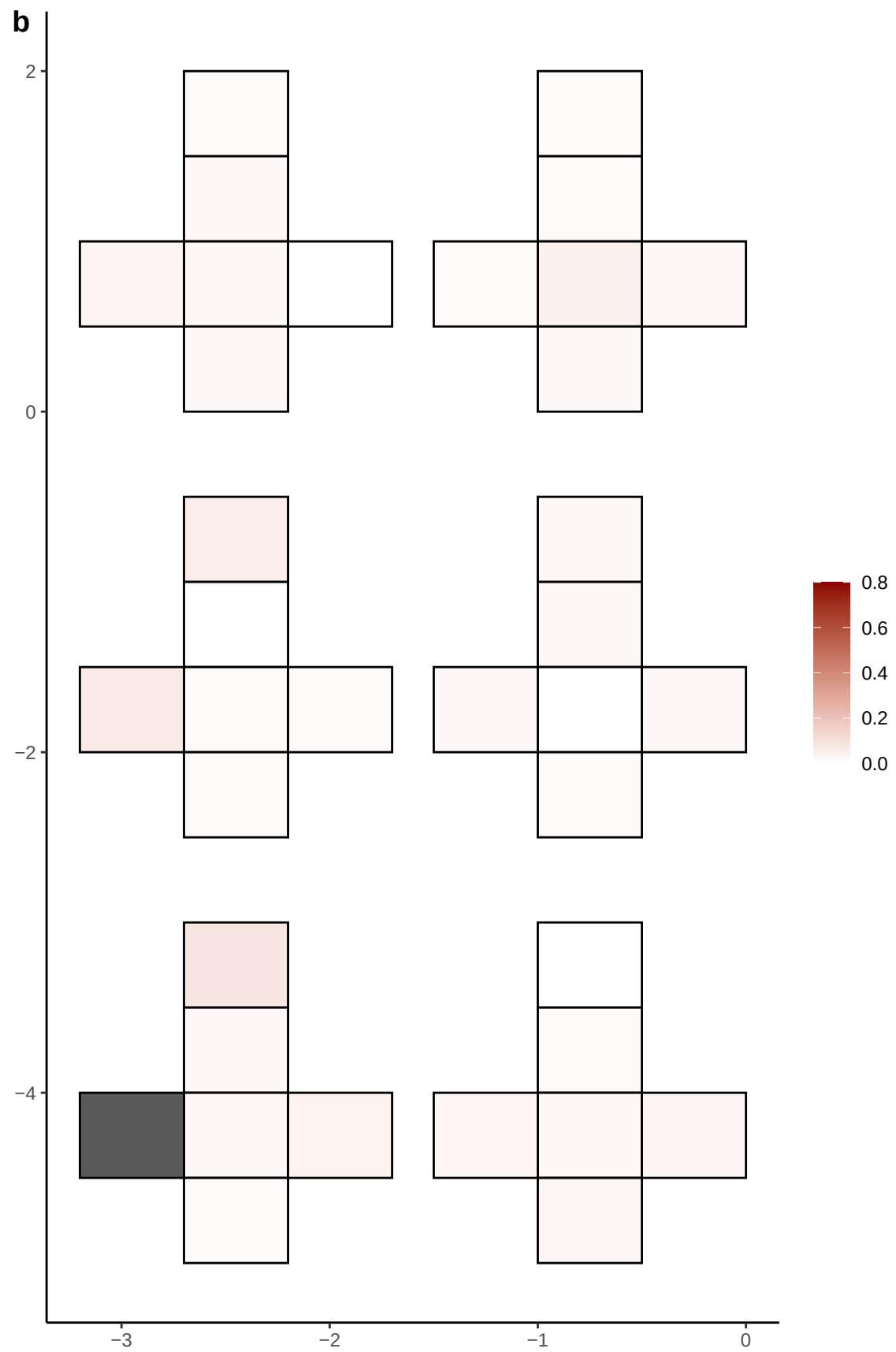

**a** ID01 11-94045731-T-C

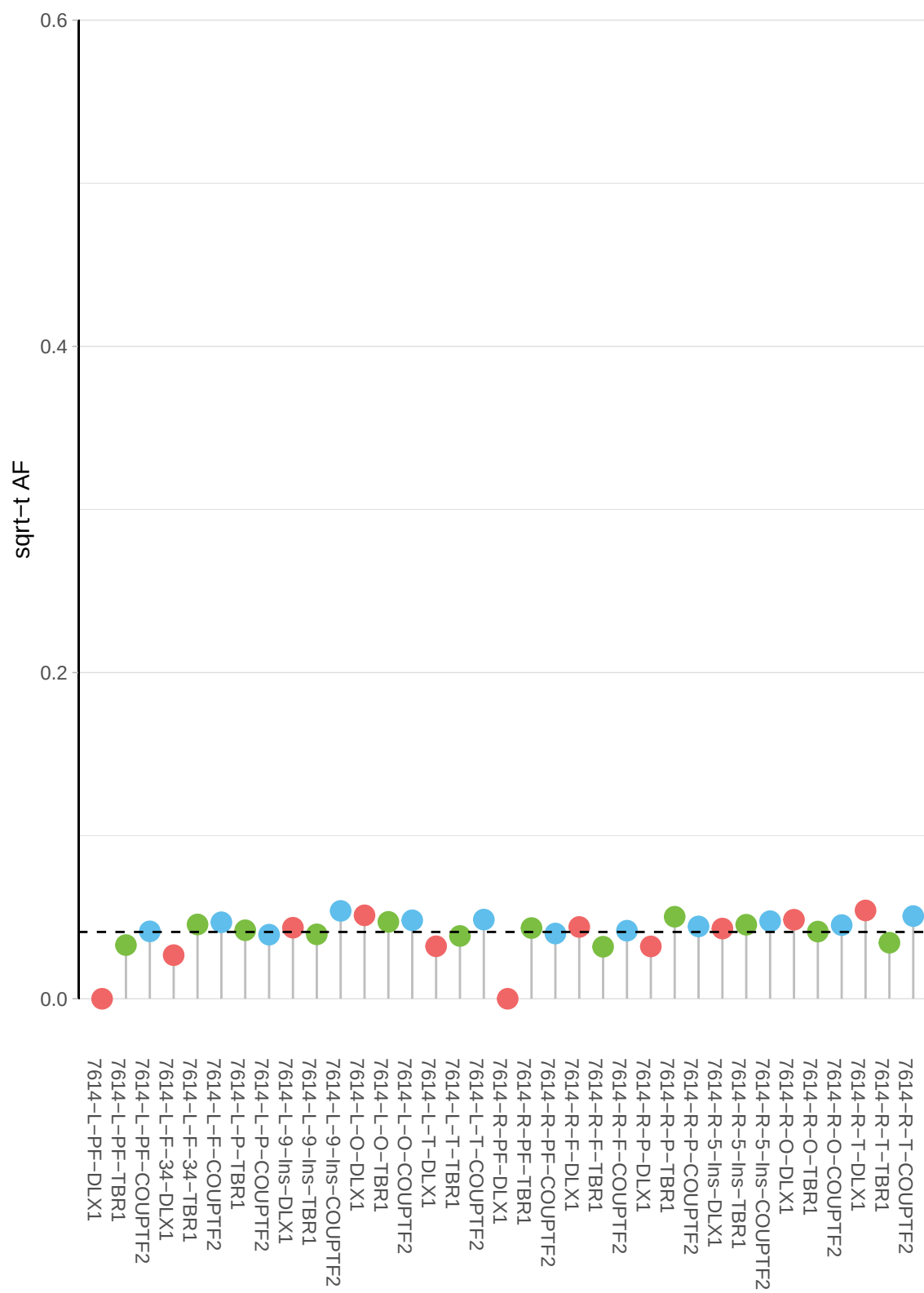

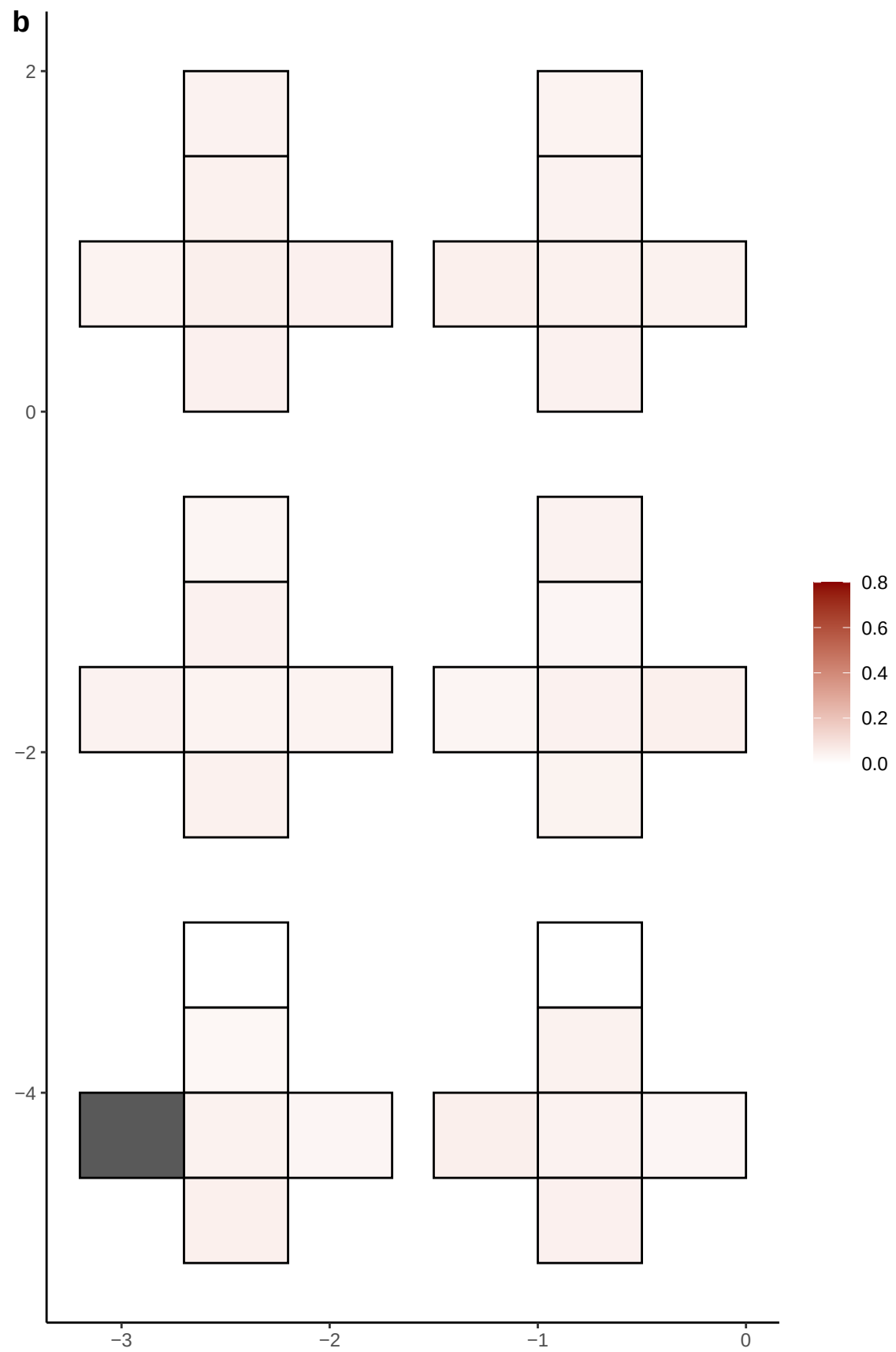

**a** ID01 11-94048896-G-A

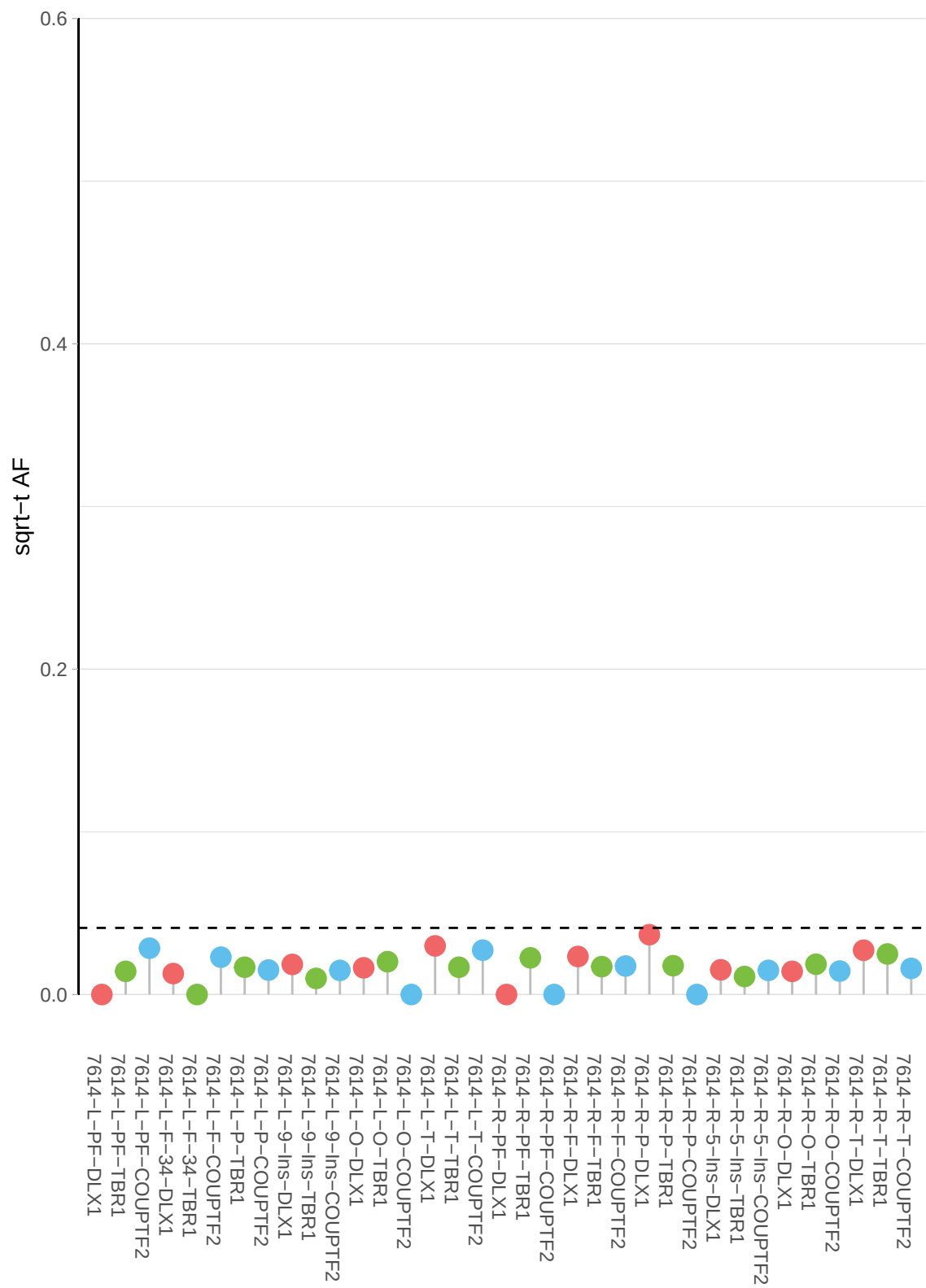

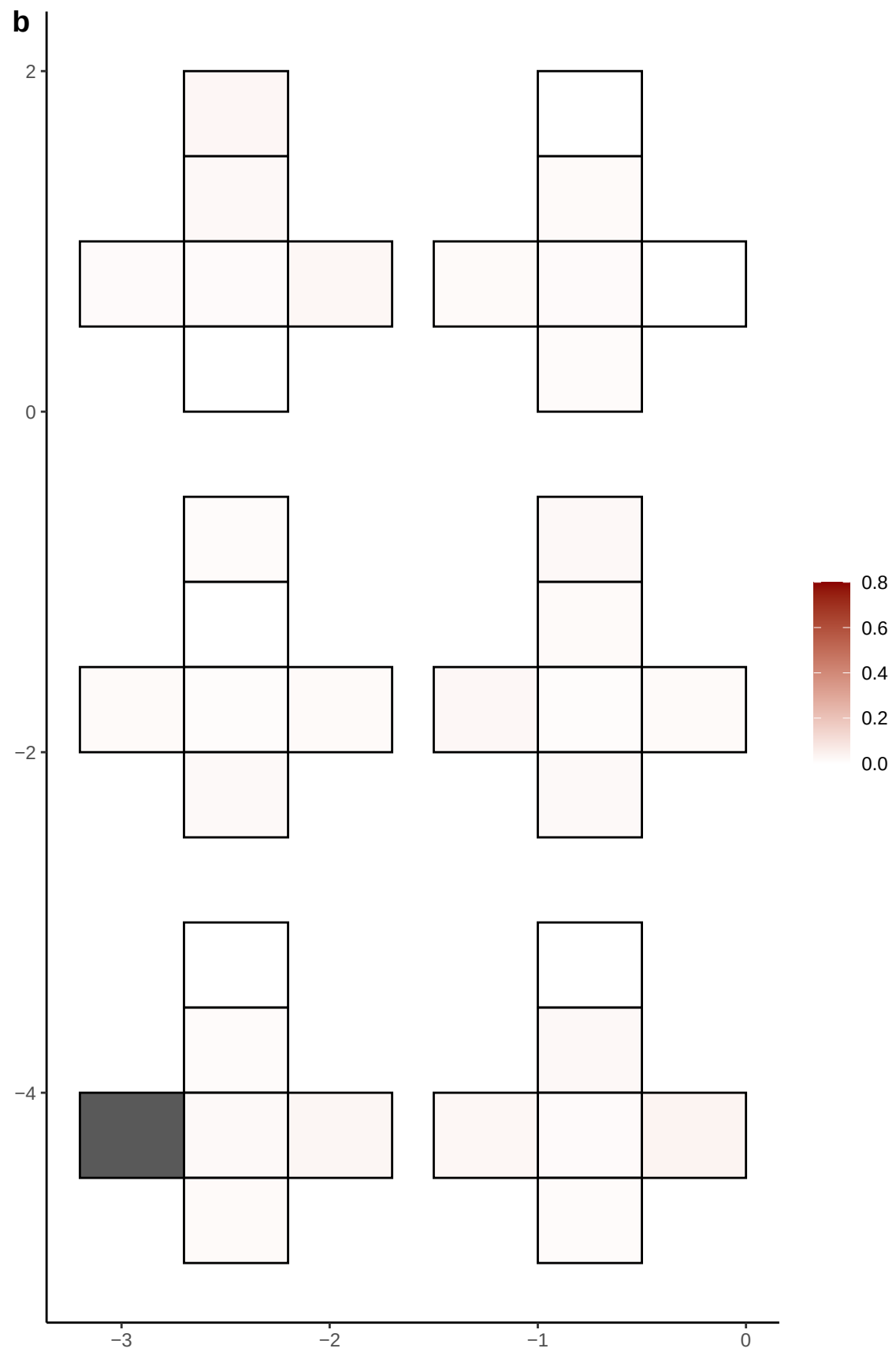

**a** ID01 11-97326785-G-A

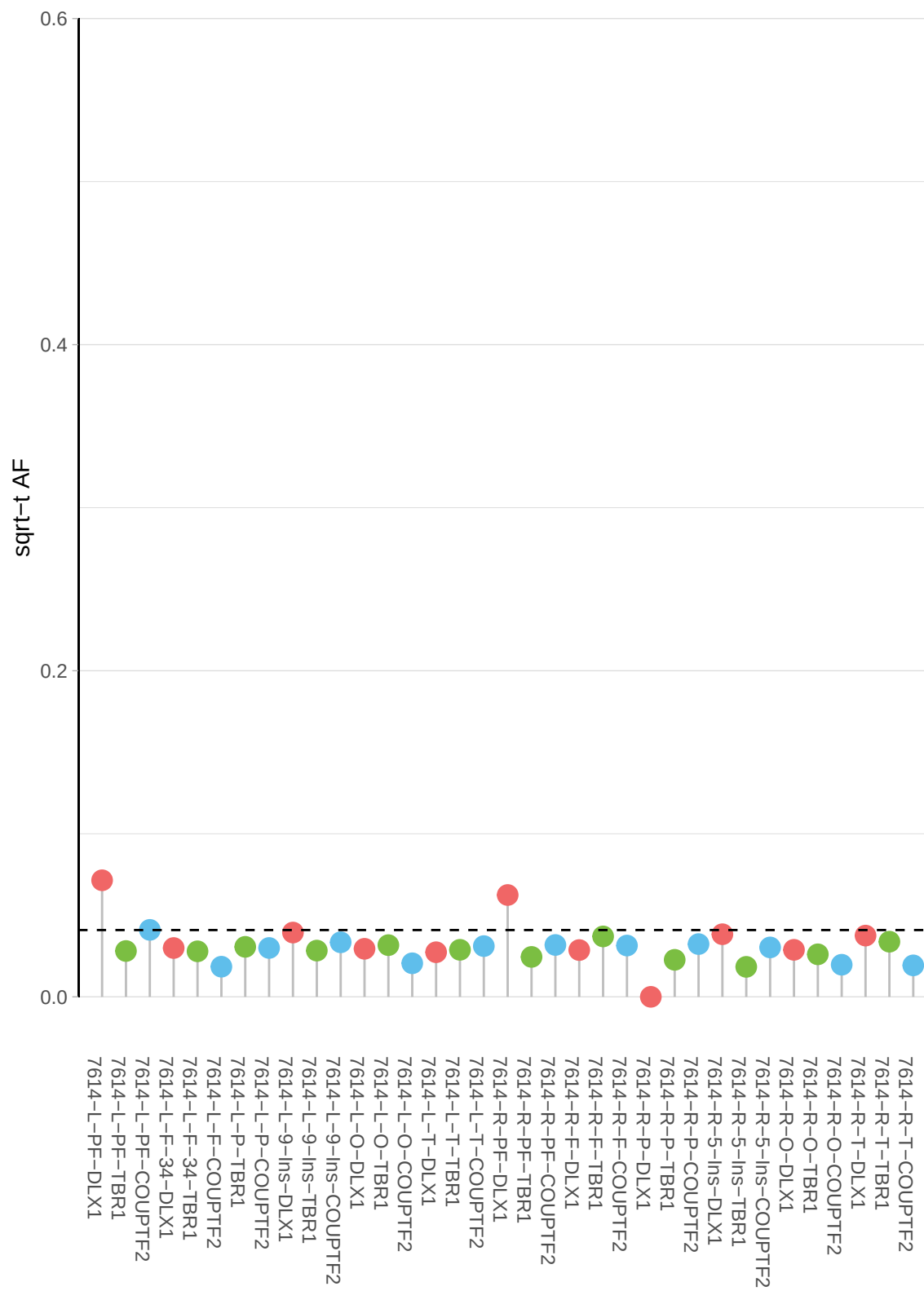

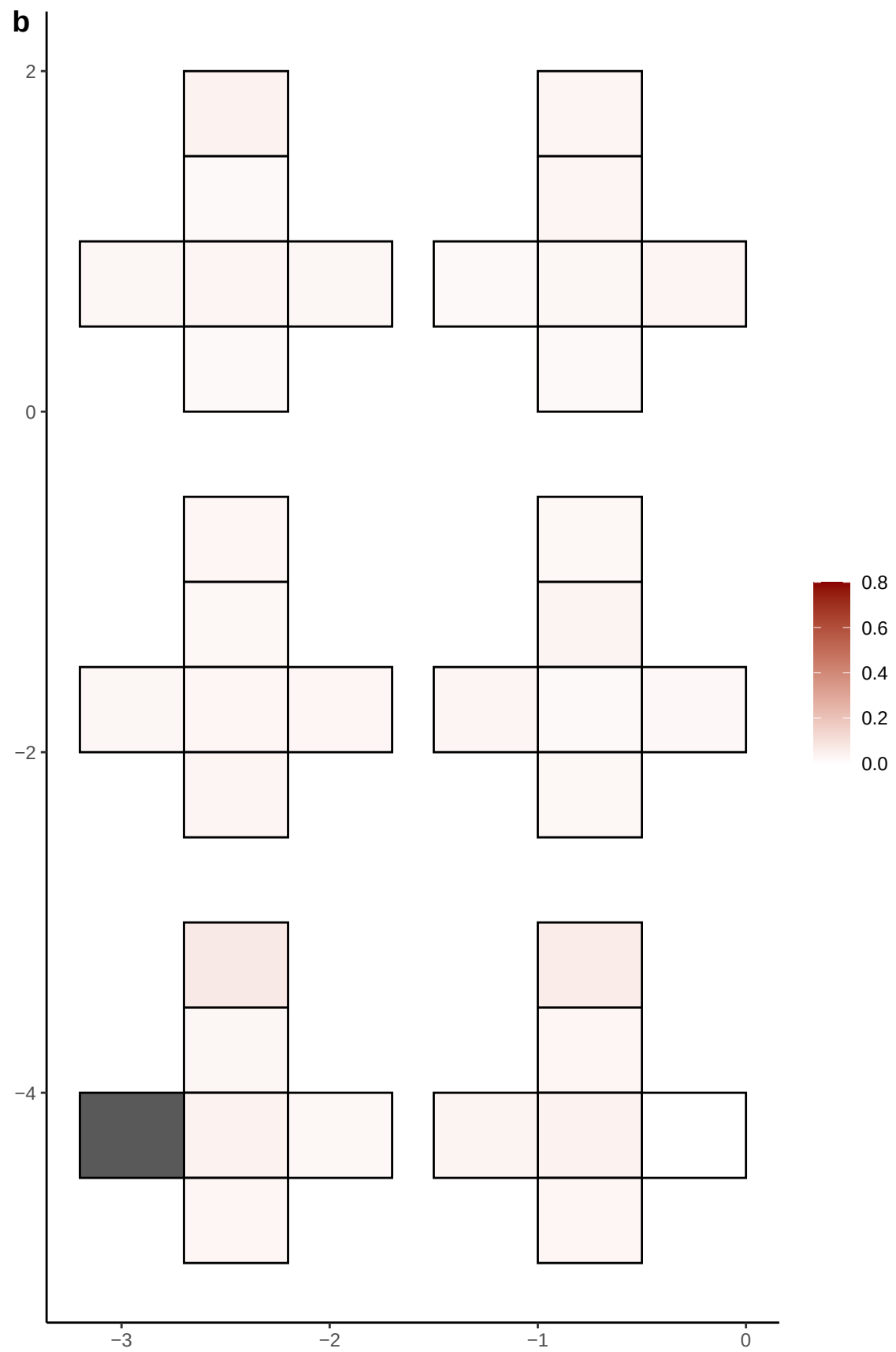

**a** ID01 11-99258149-G-A

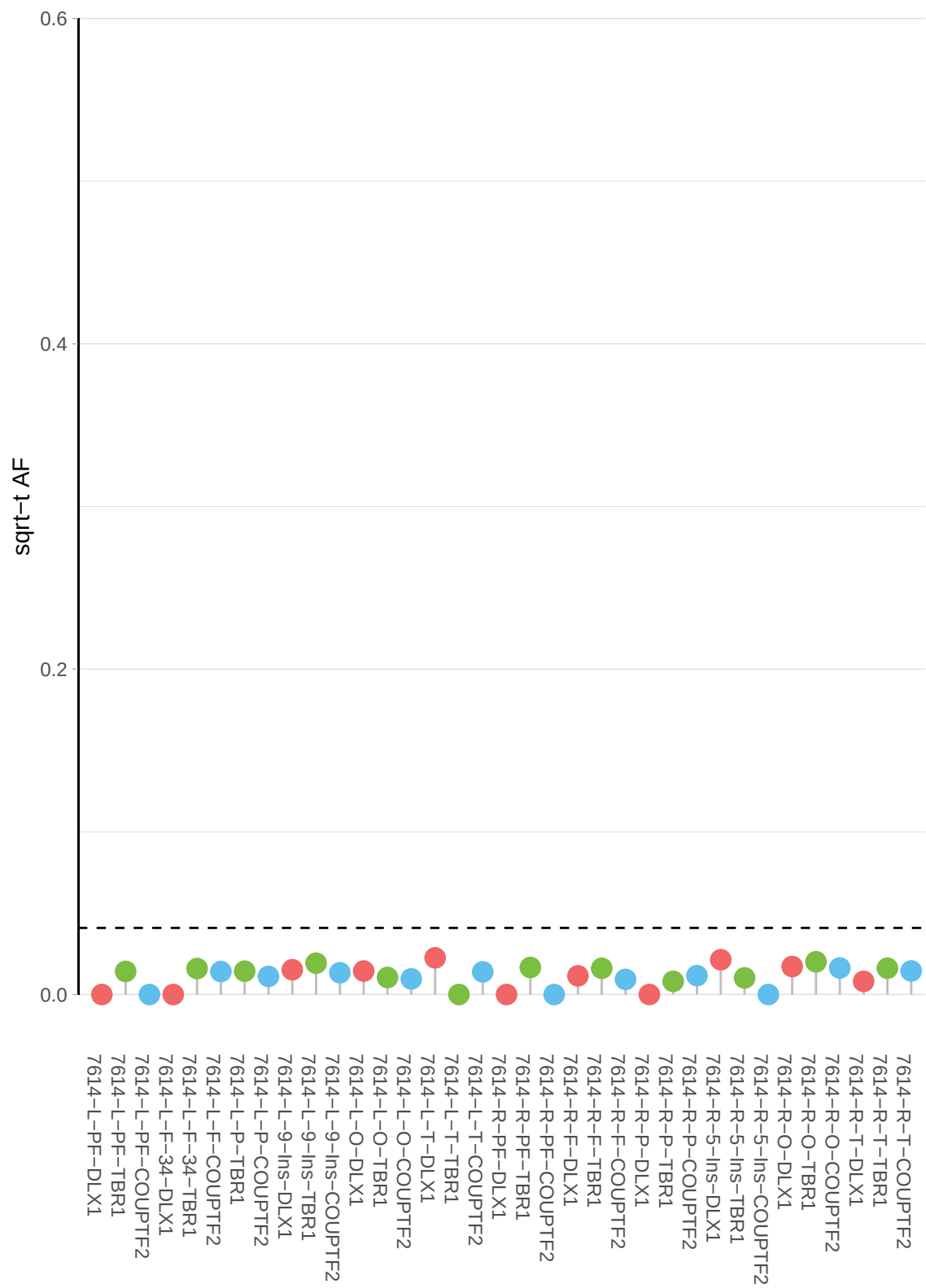

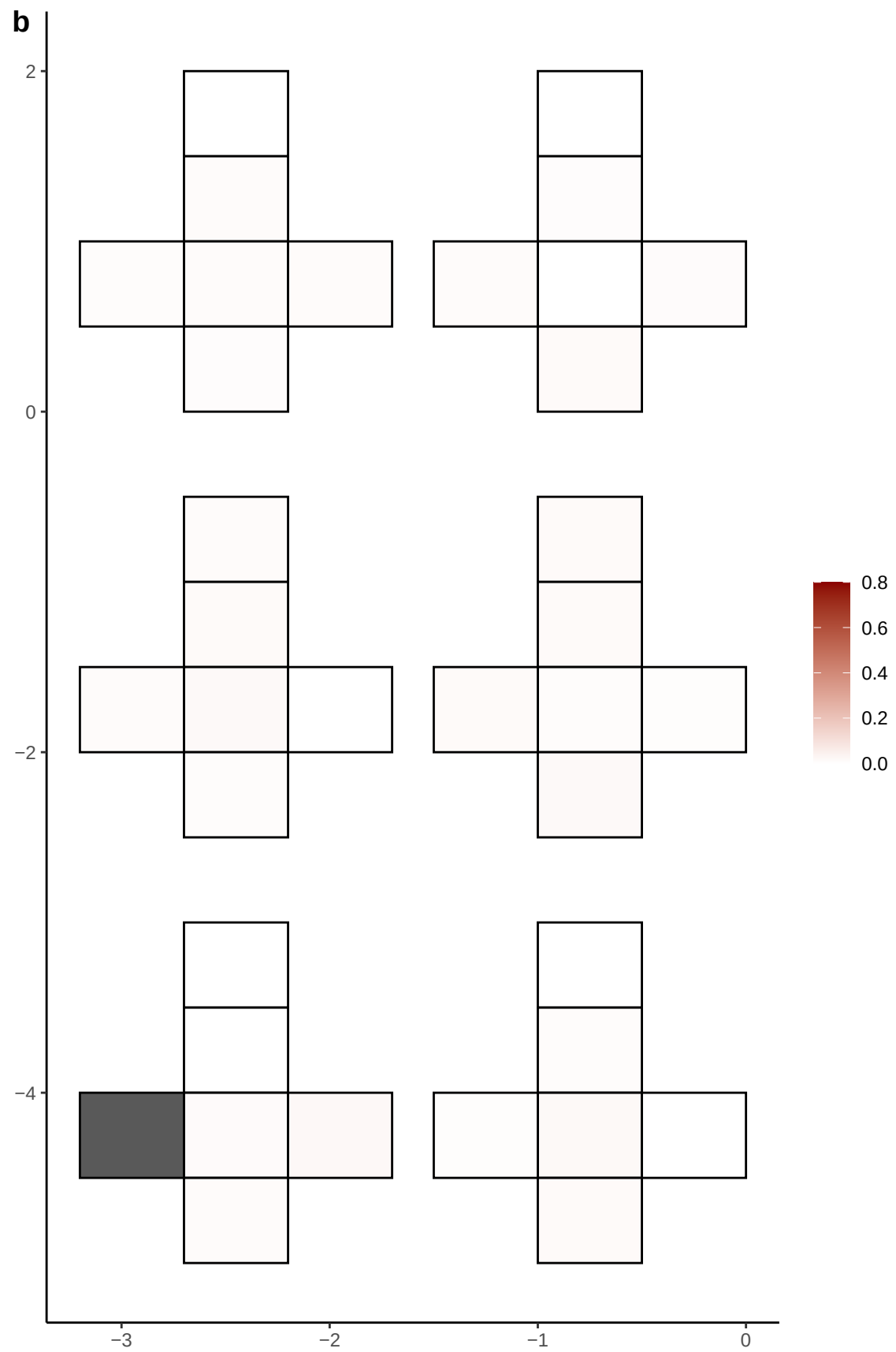

**a** ID01 12-107509134-G-A

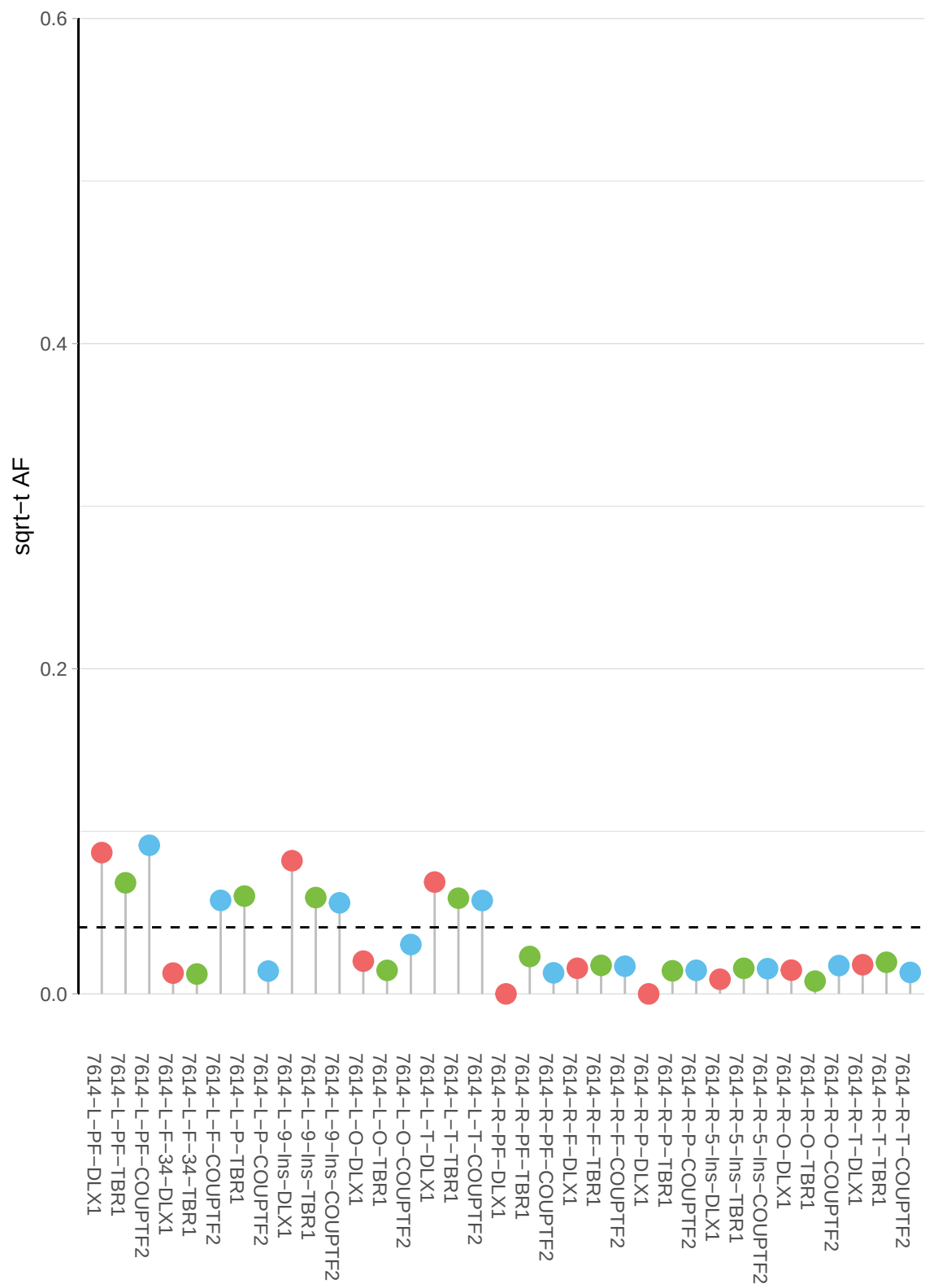

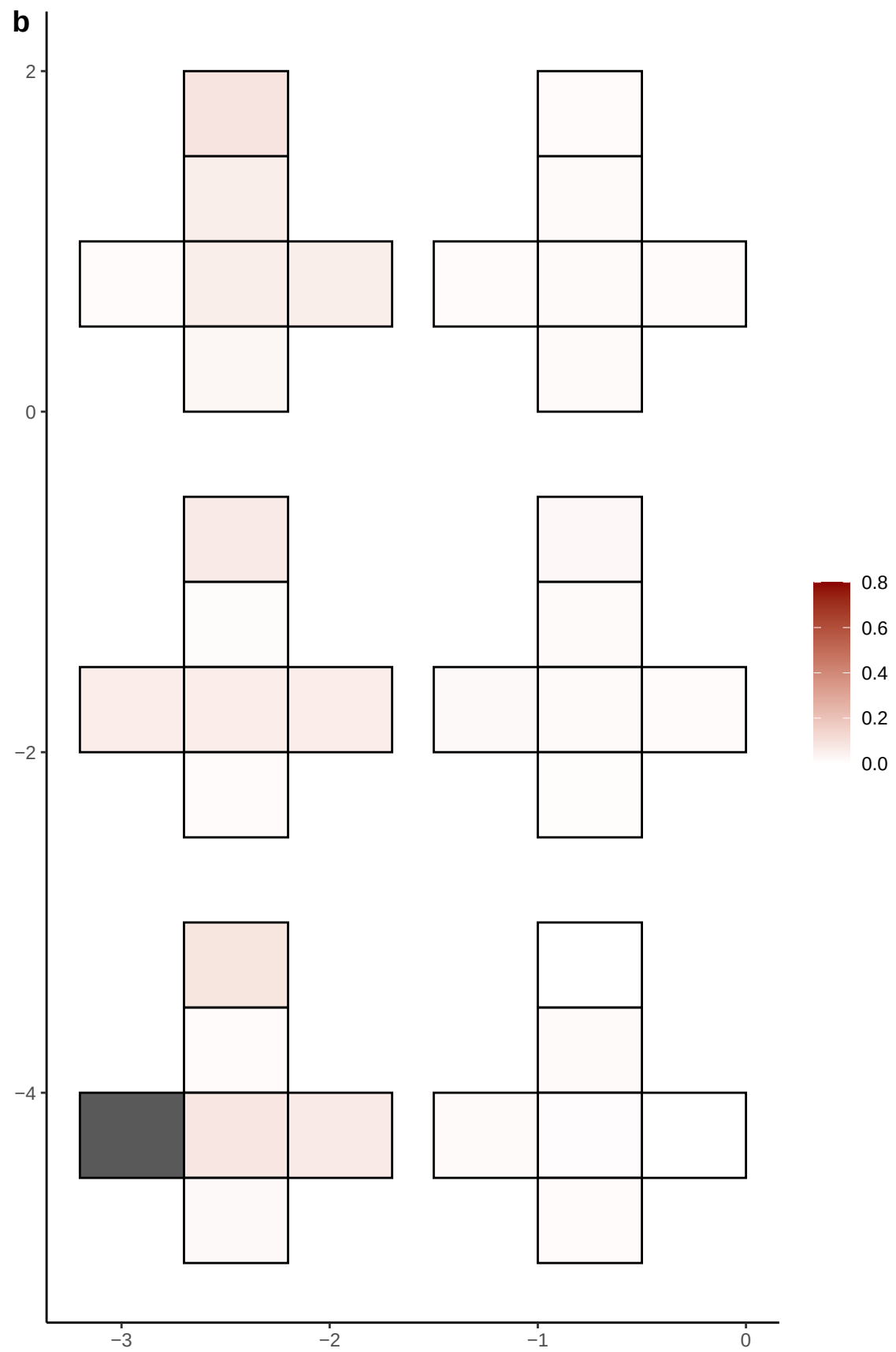

**a** ID01 12-116308531-A-G

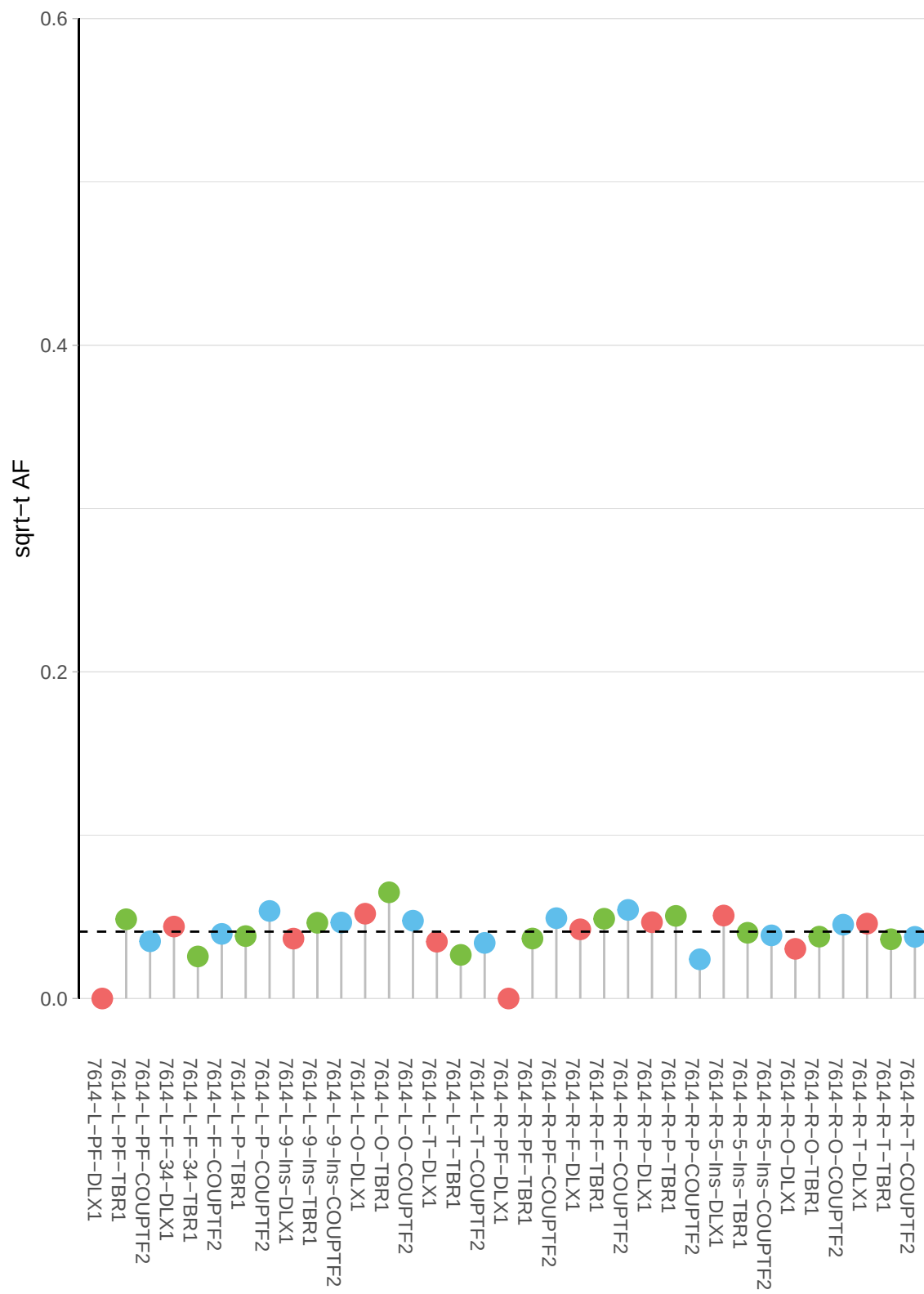

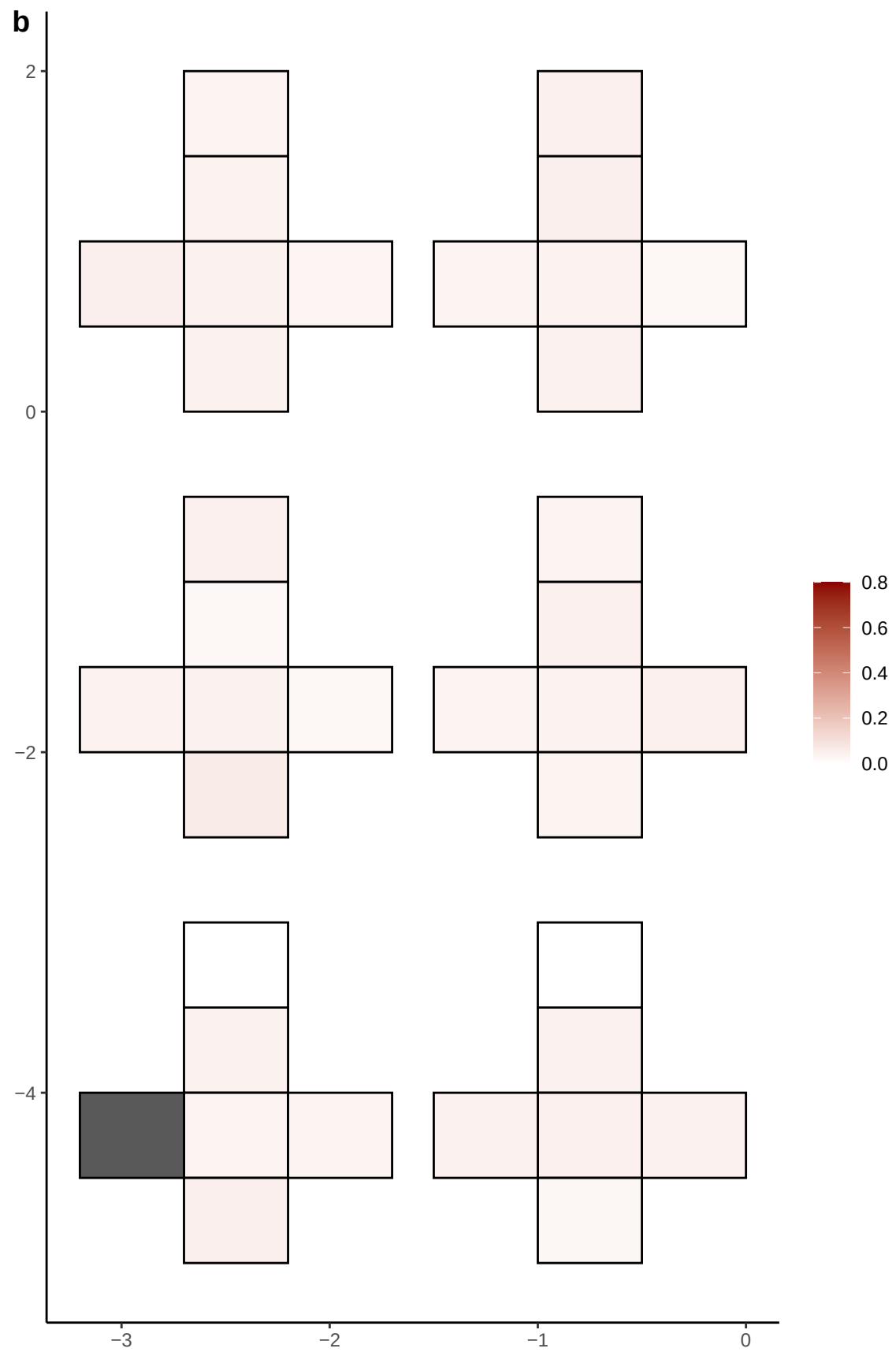

**a** ID01 12-30637099-C-G

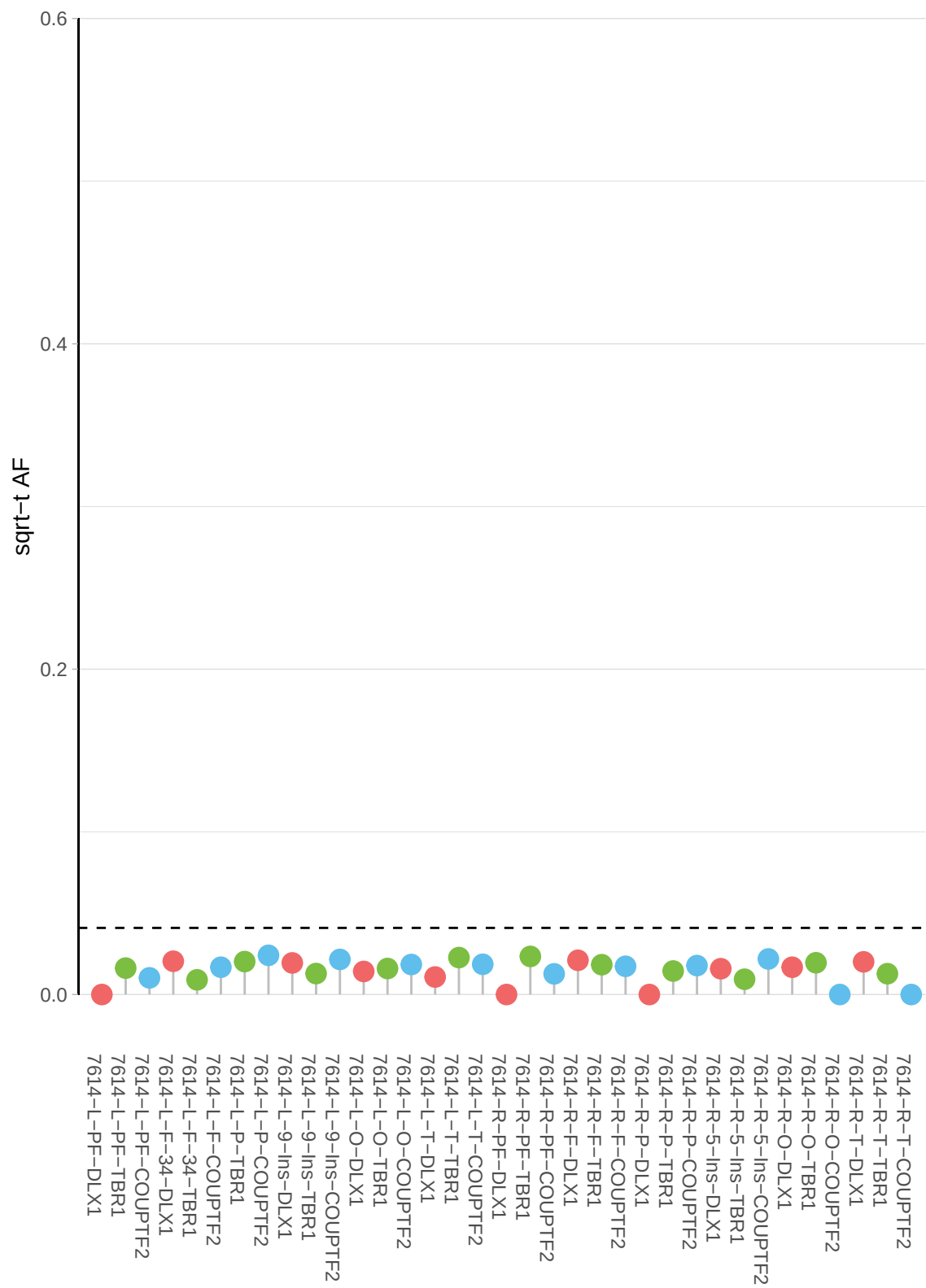

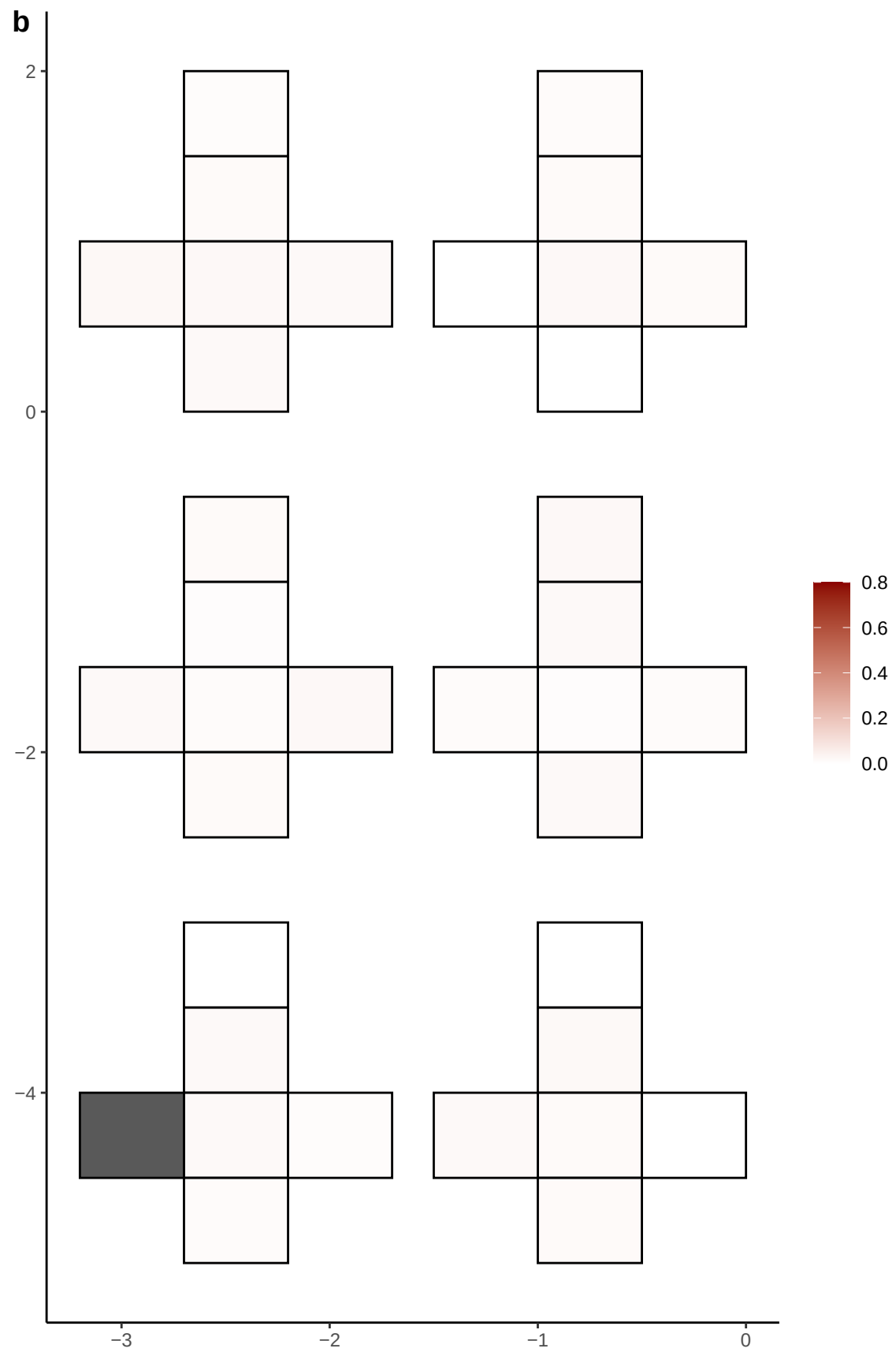

**a** ID01 12-30749782-G-T

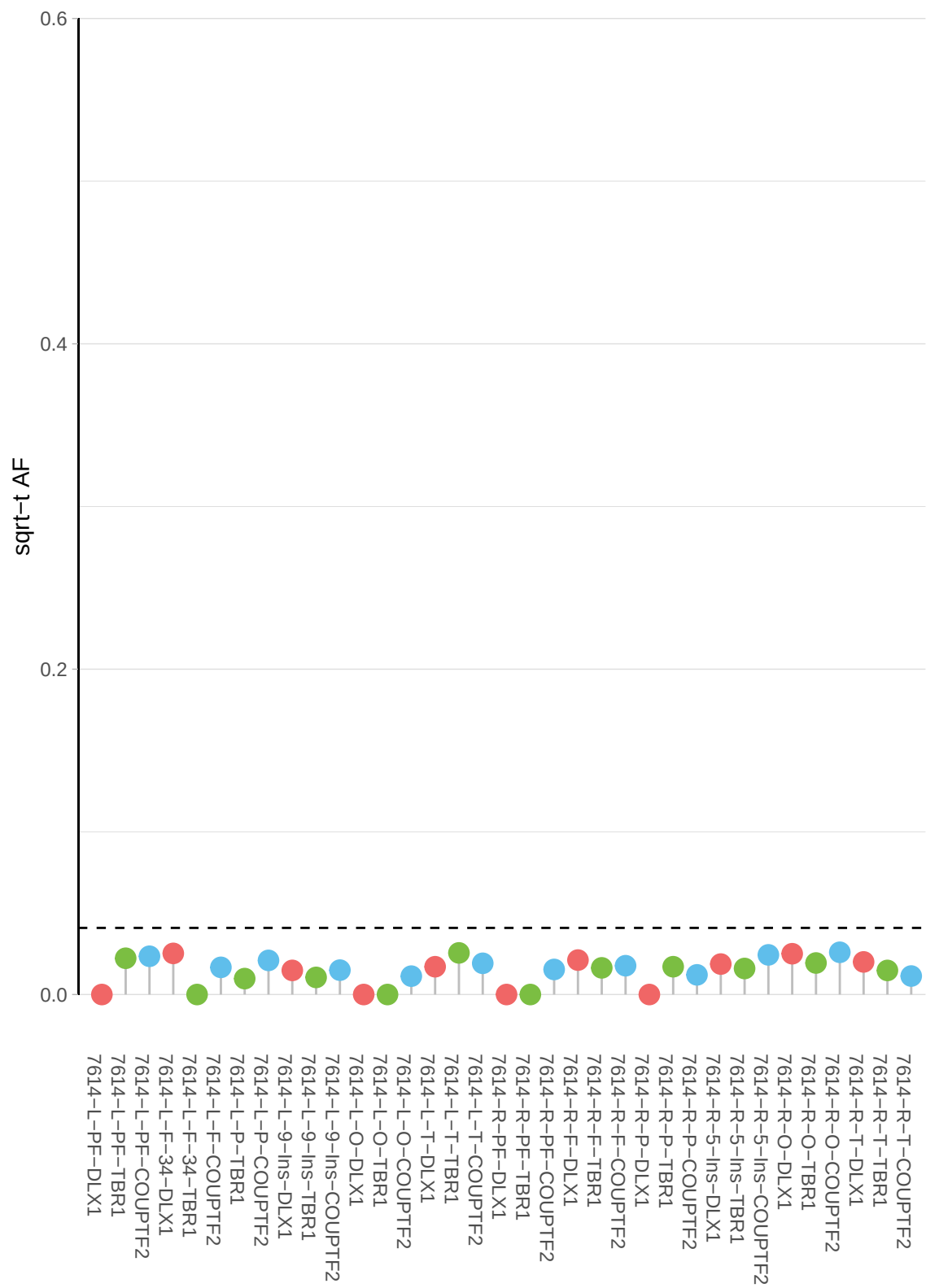

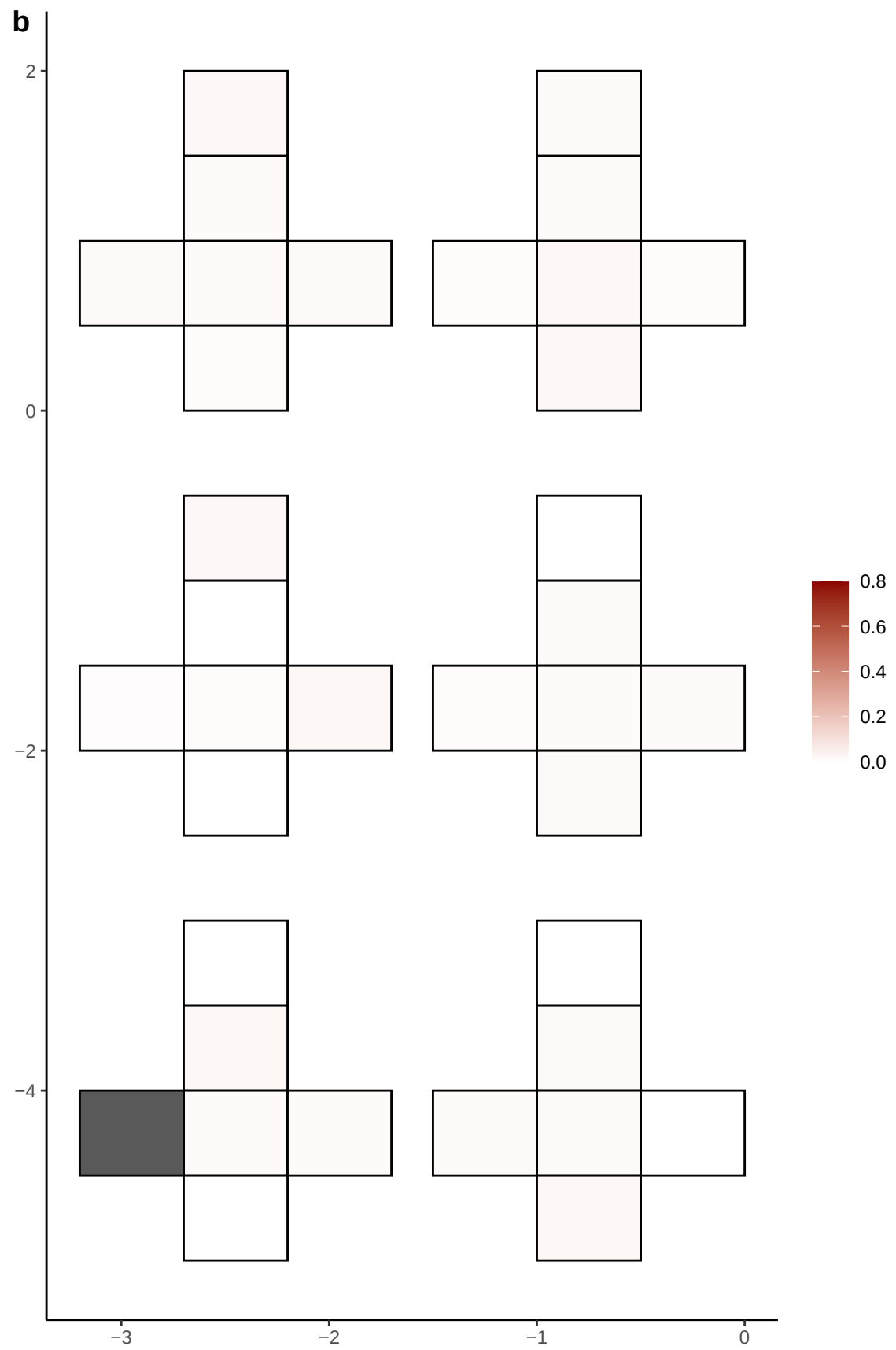

**a** ID01 12-40717911-G-A

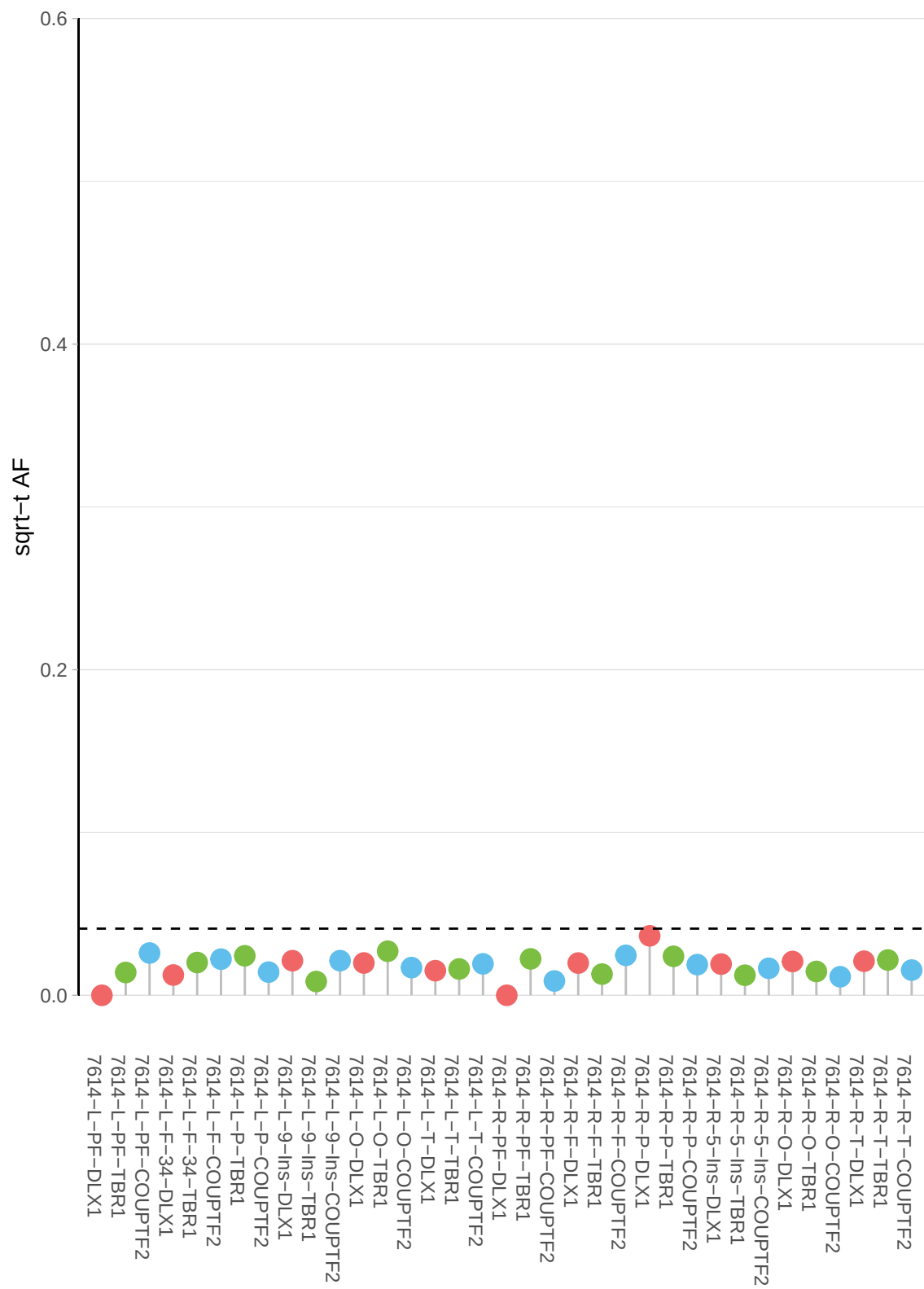

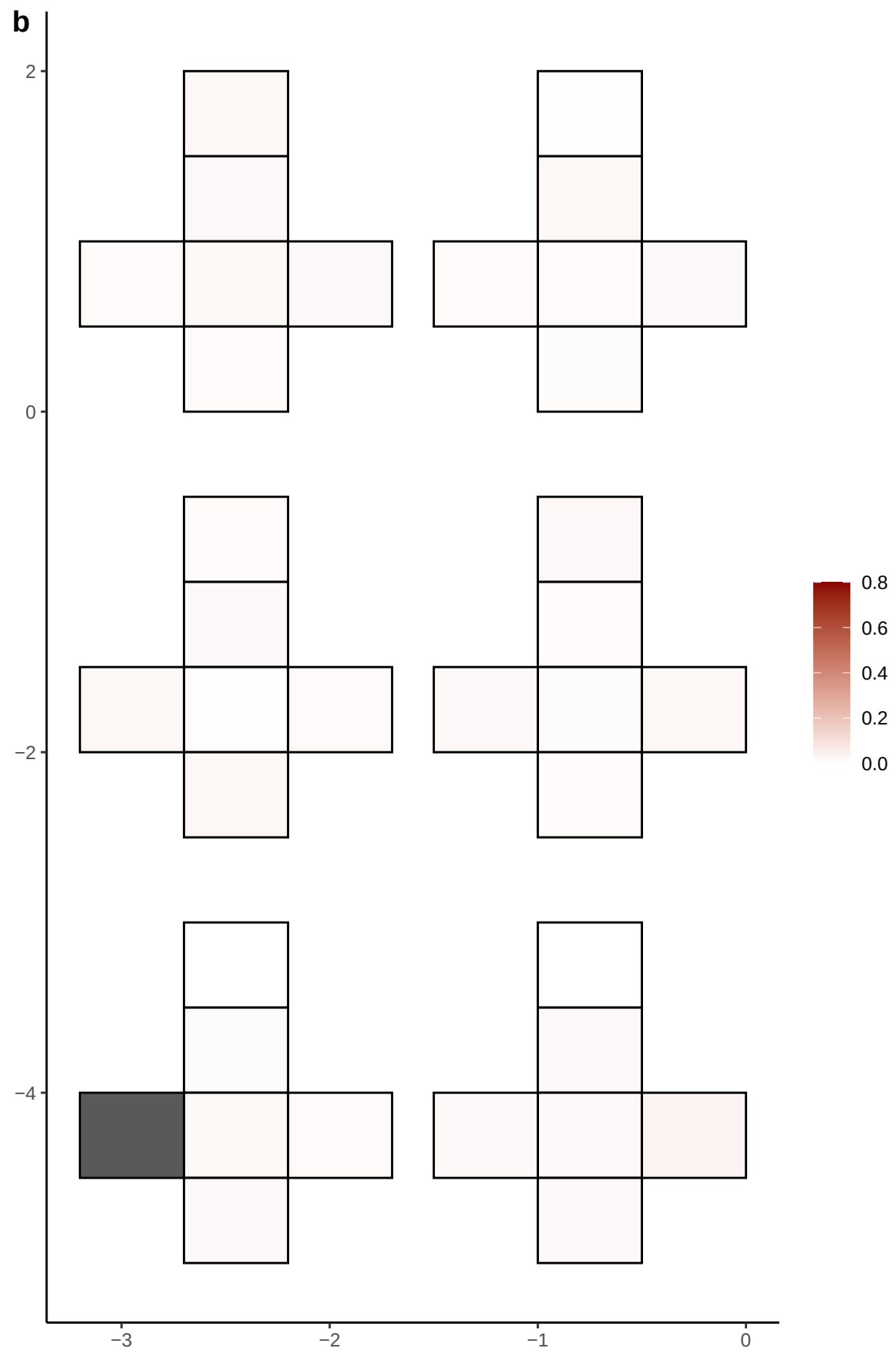

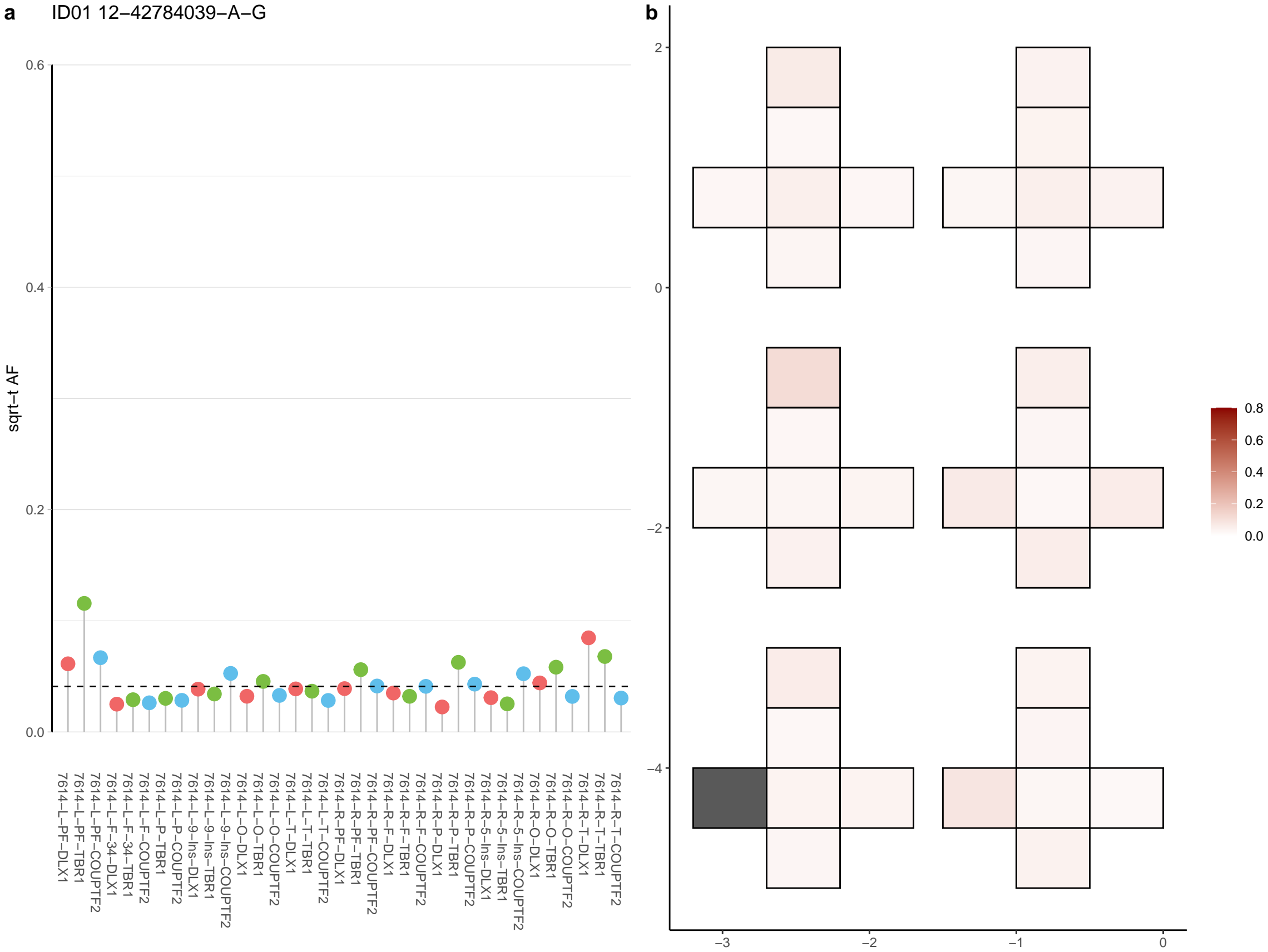

**a** ID01 12-99984882-G-A

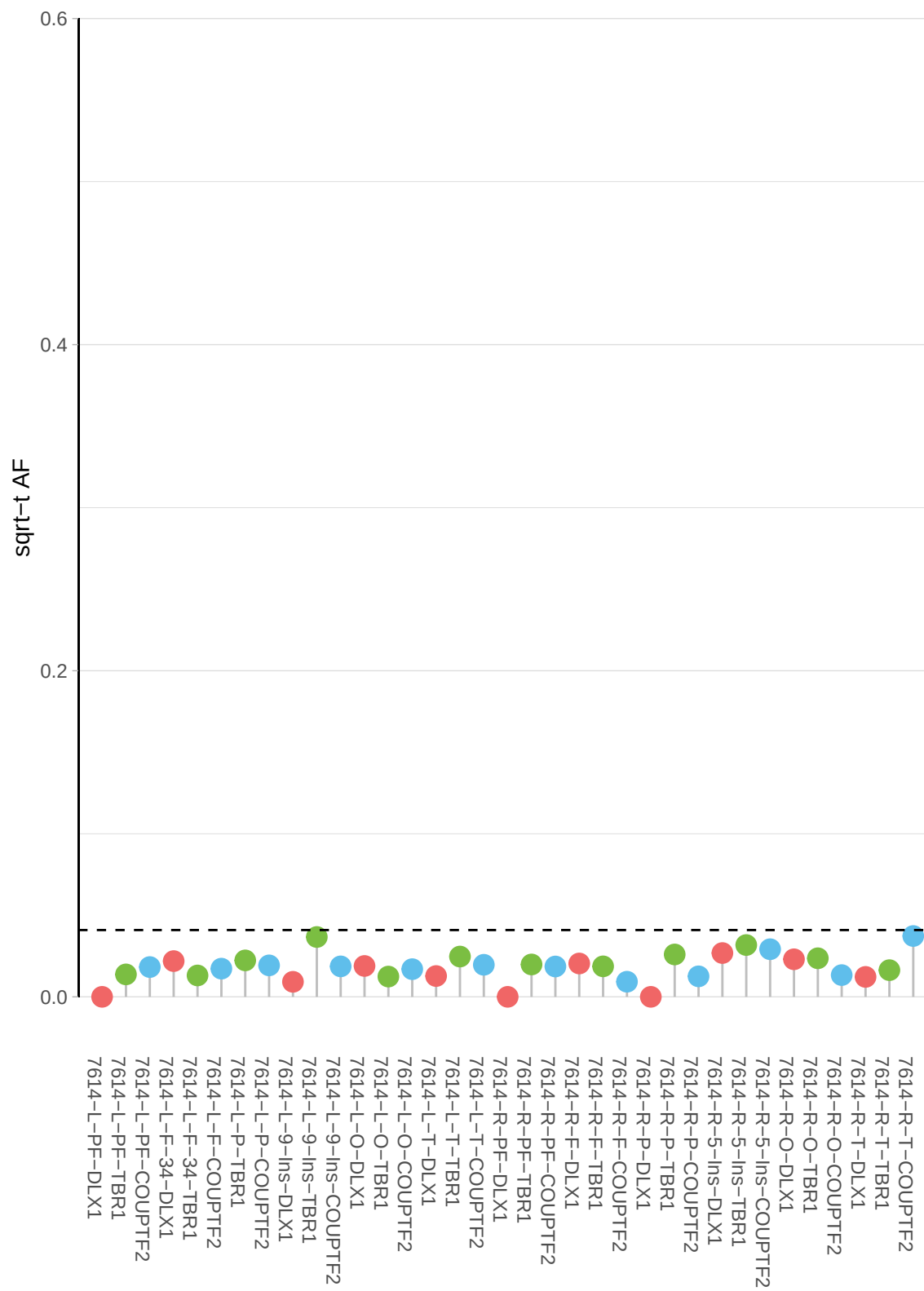

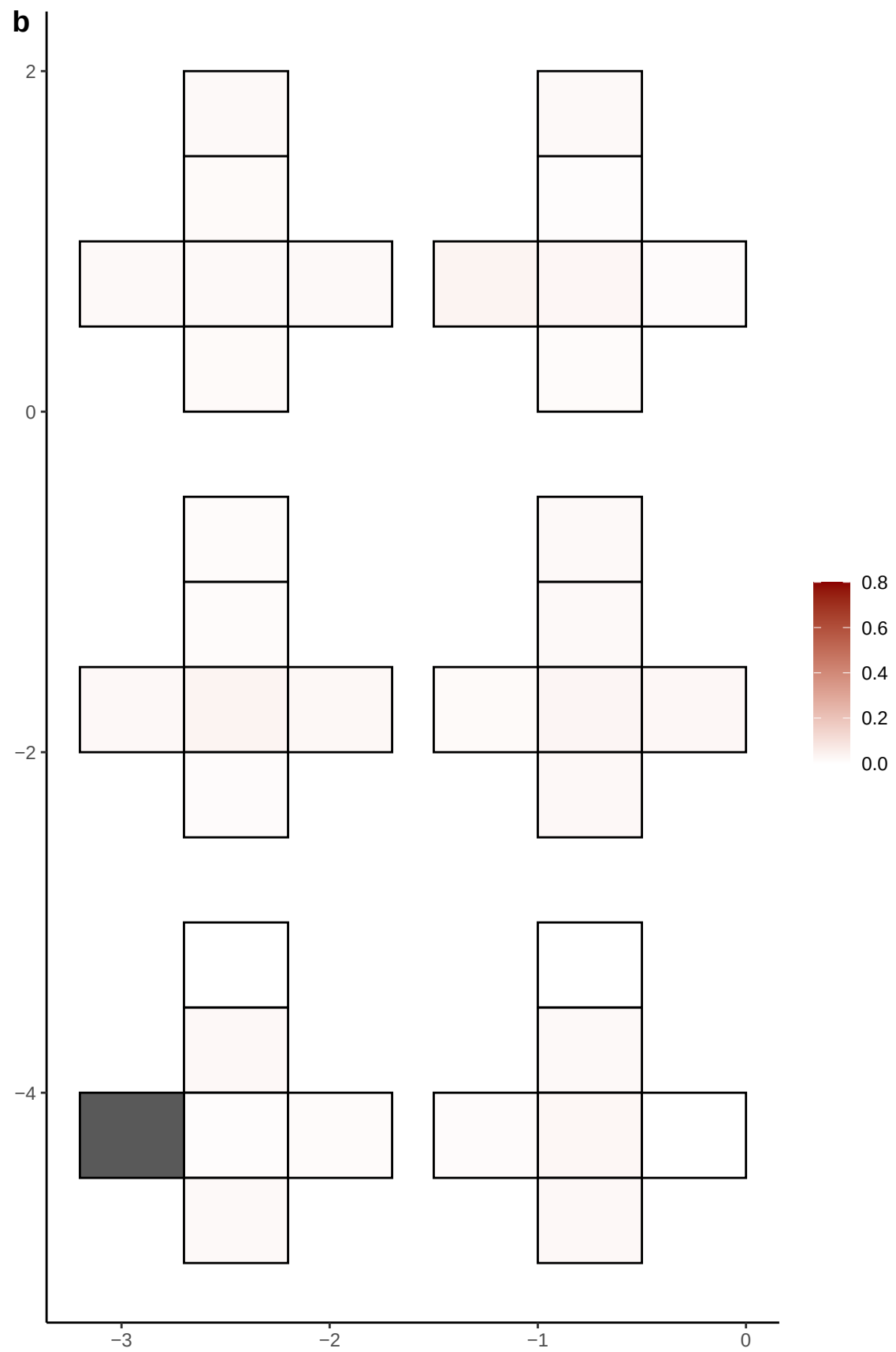

**a** ID01 13-104312064-G-A

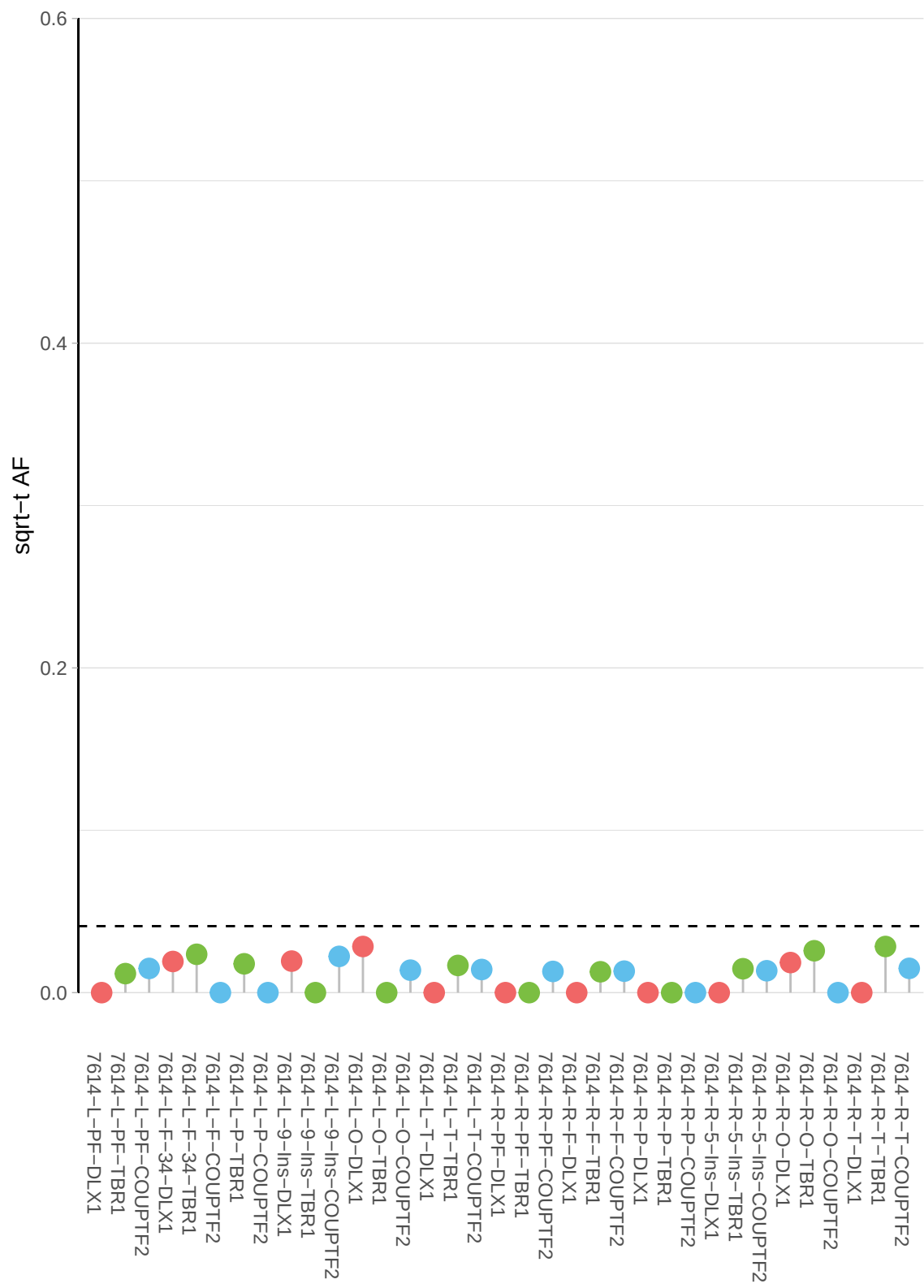

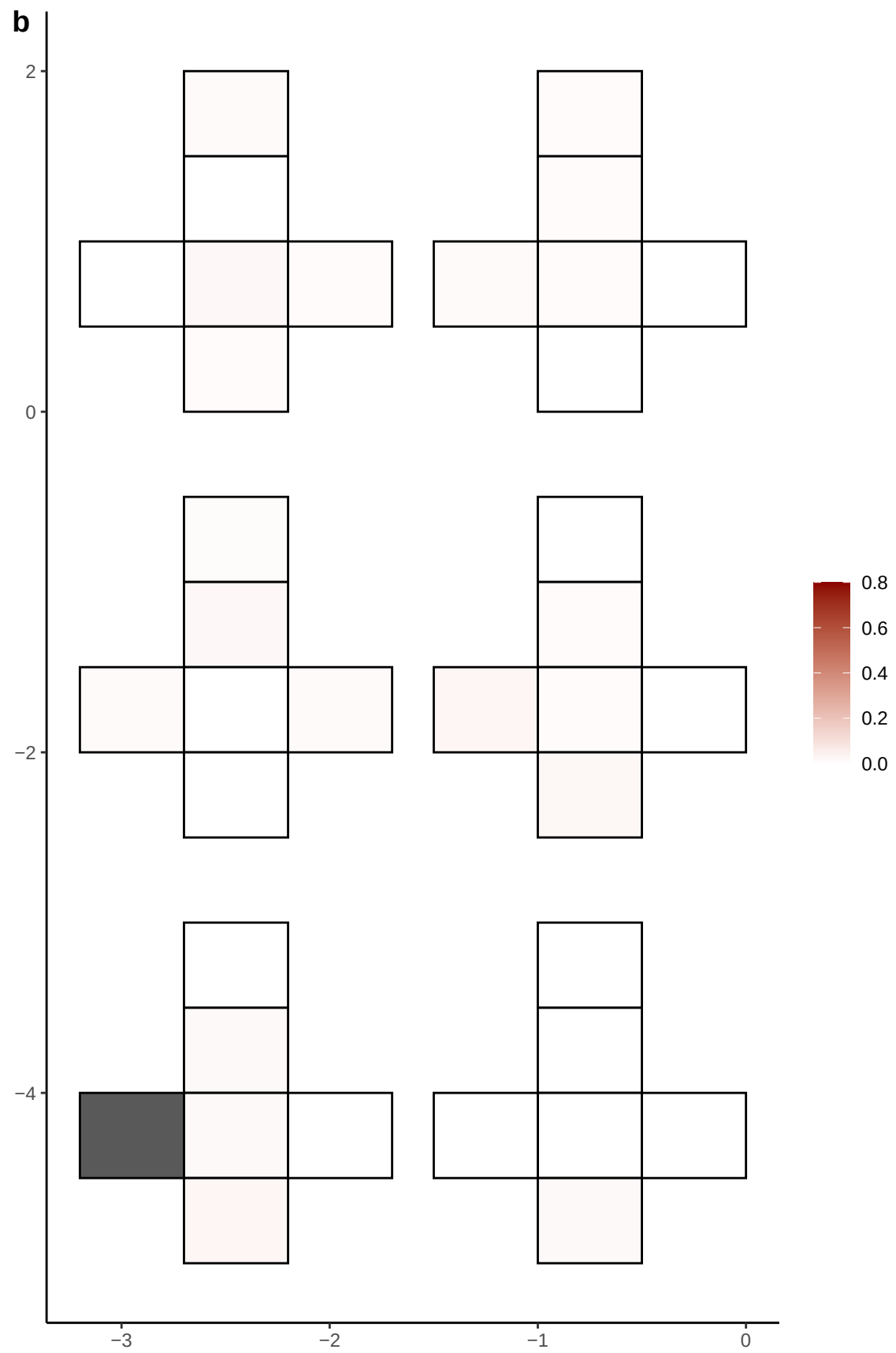

**a** ID01 13-104815490-G-A

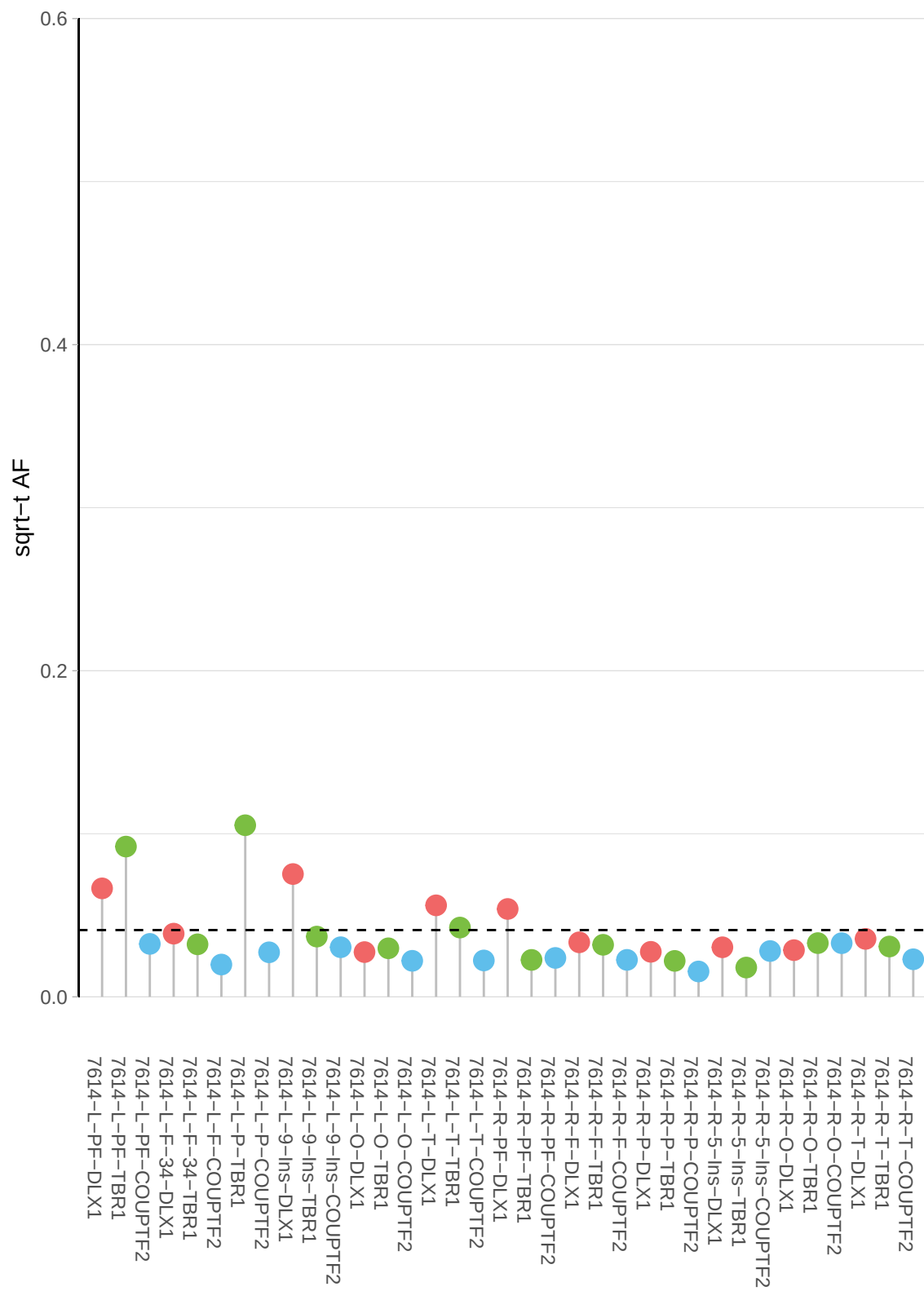

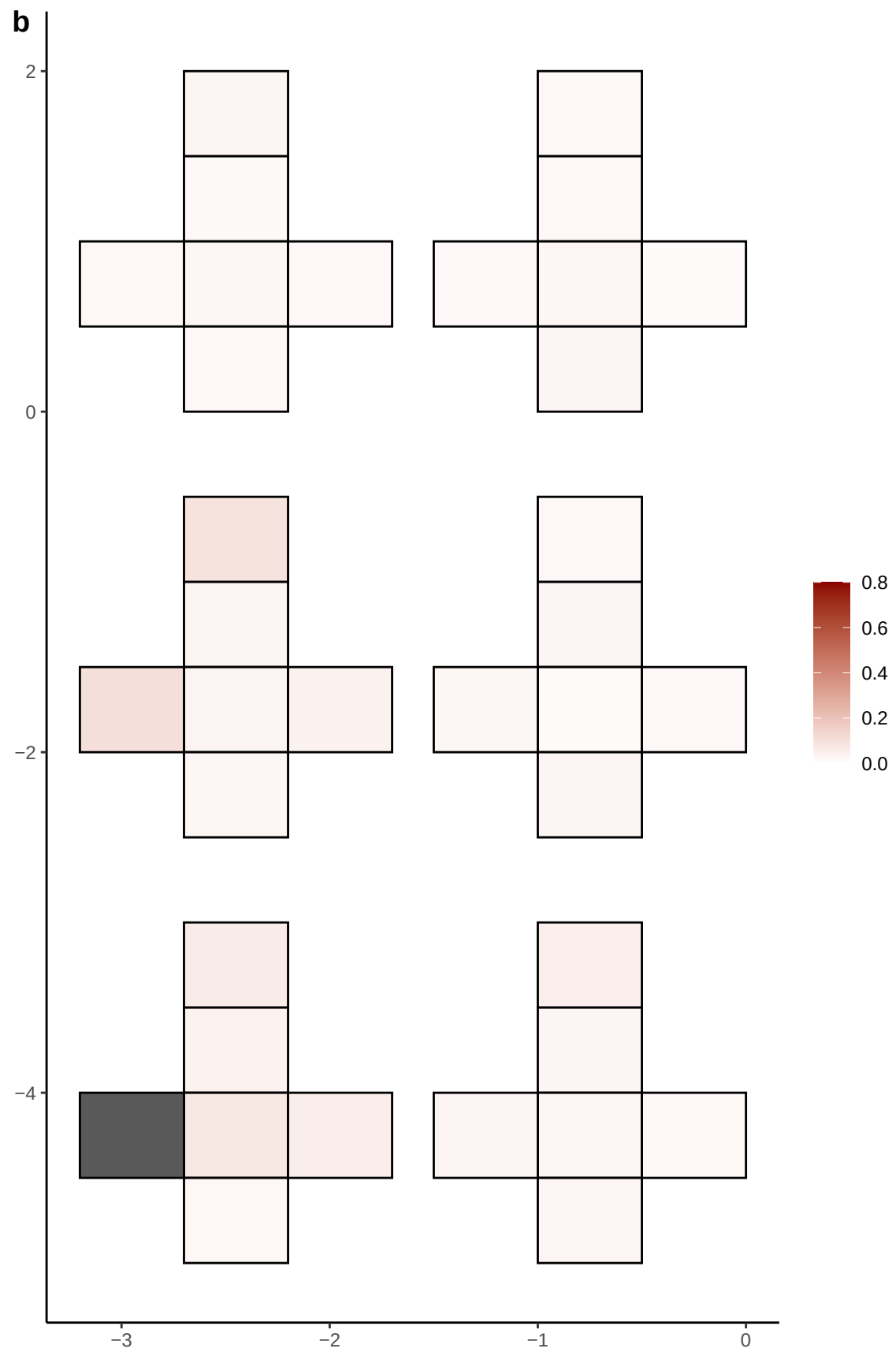

**a** ID01 13-113887705-CCAG-C

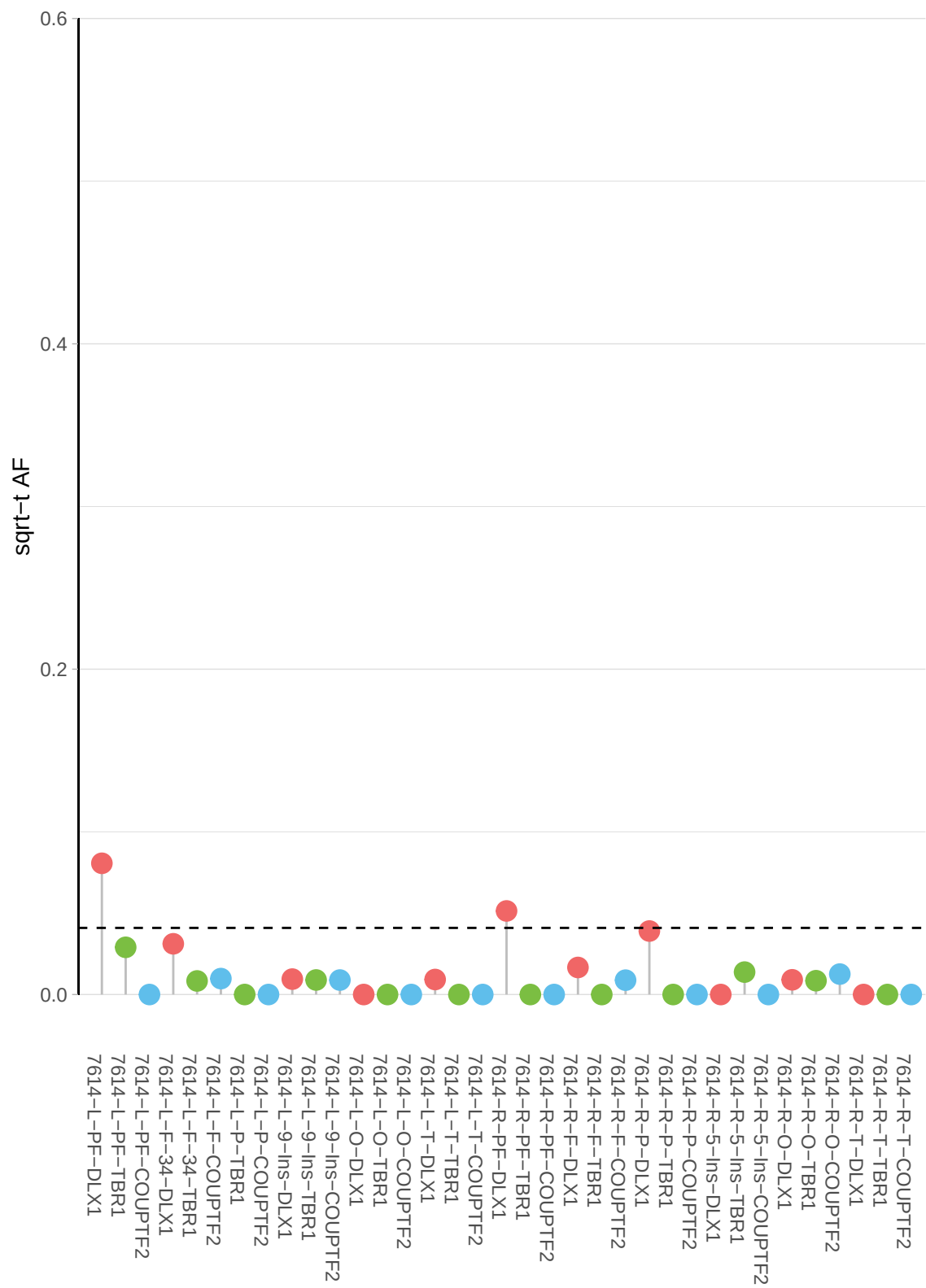

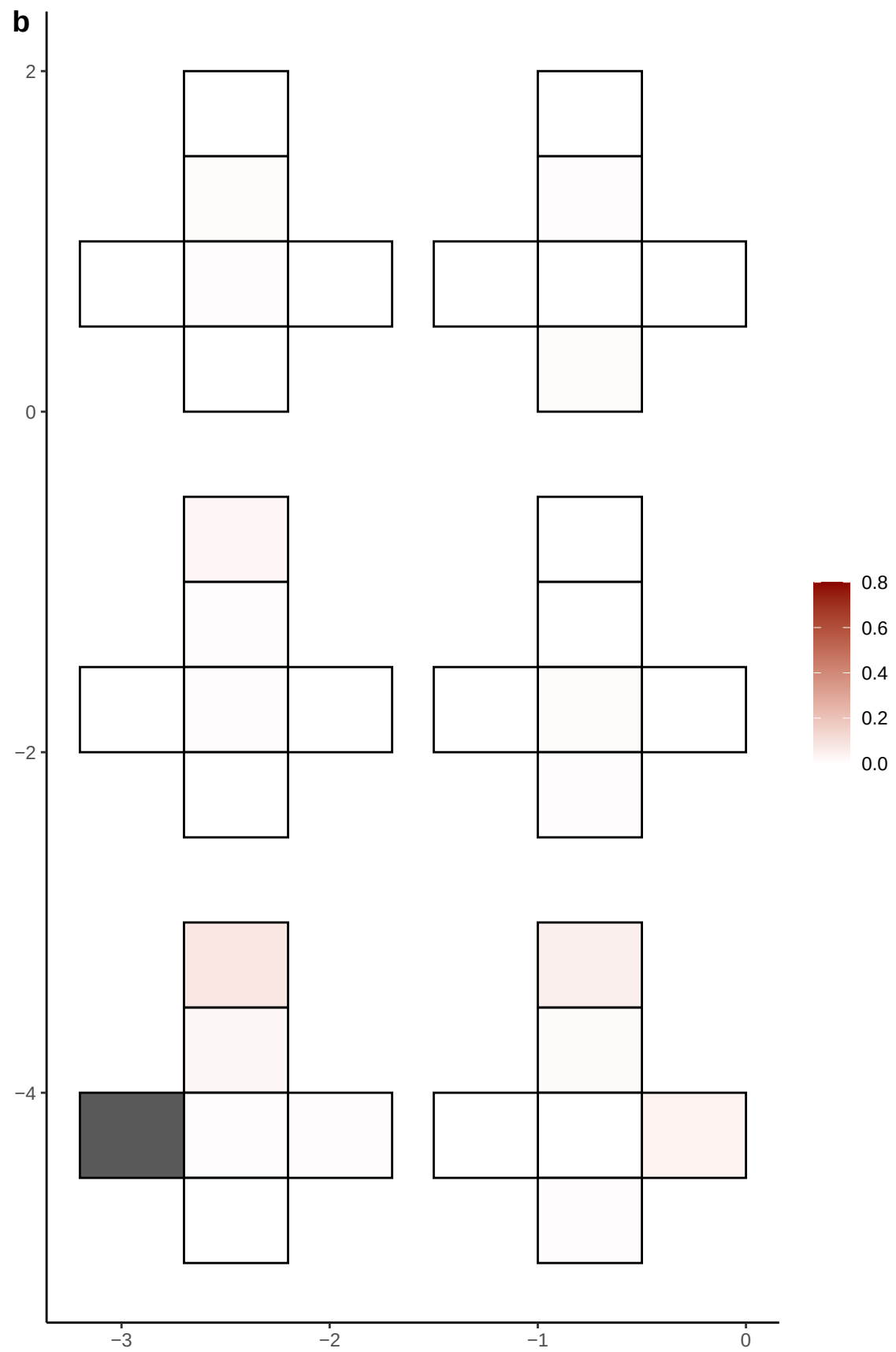

**a** ID01 13-29160961-C-A

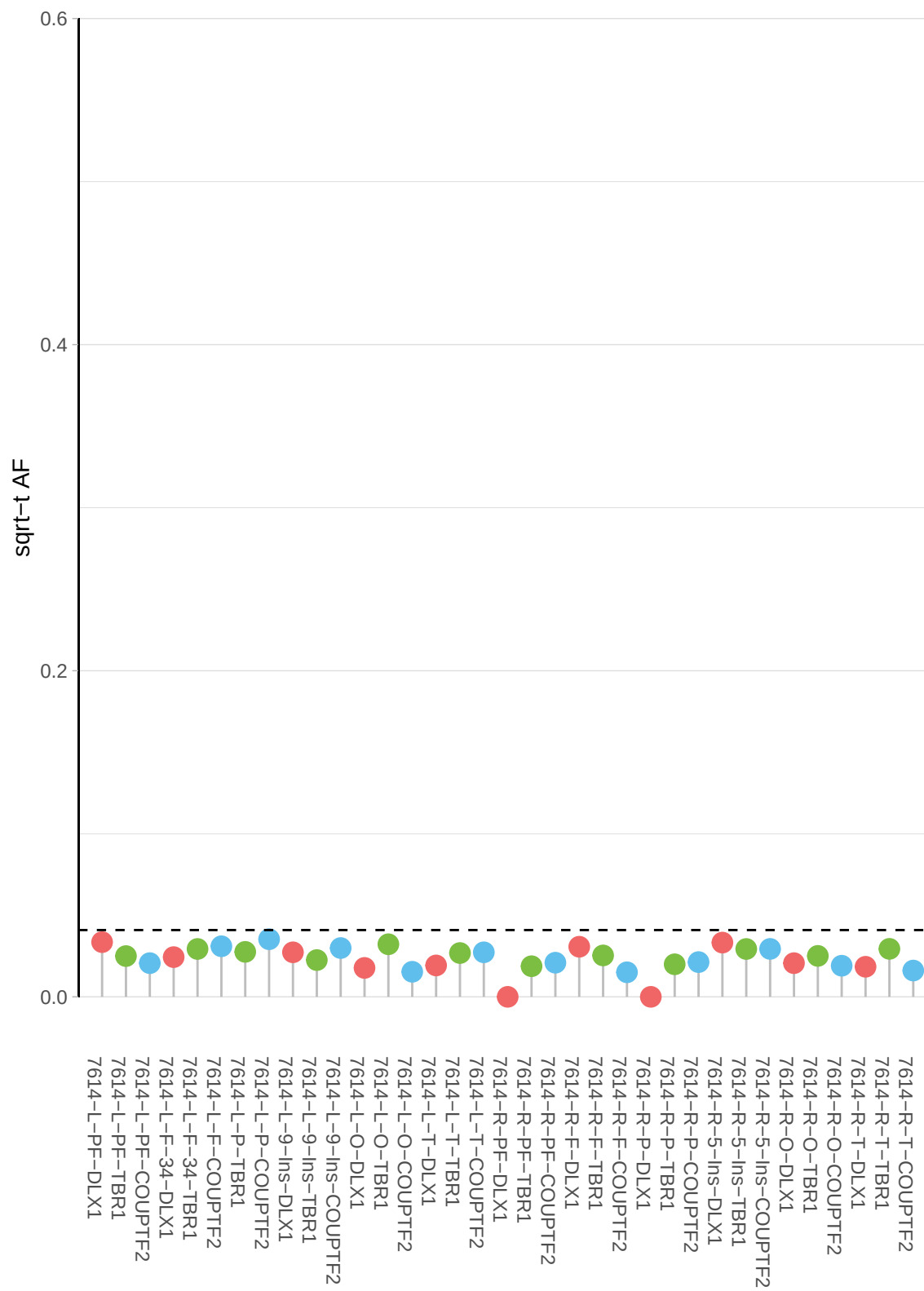

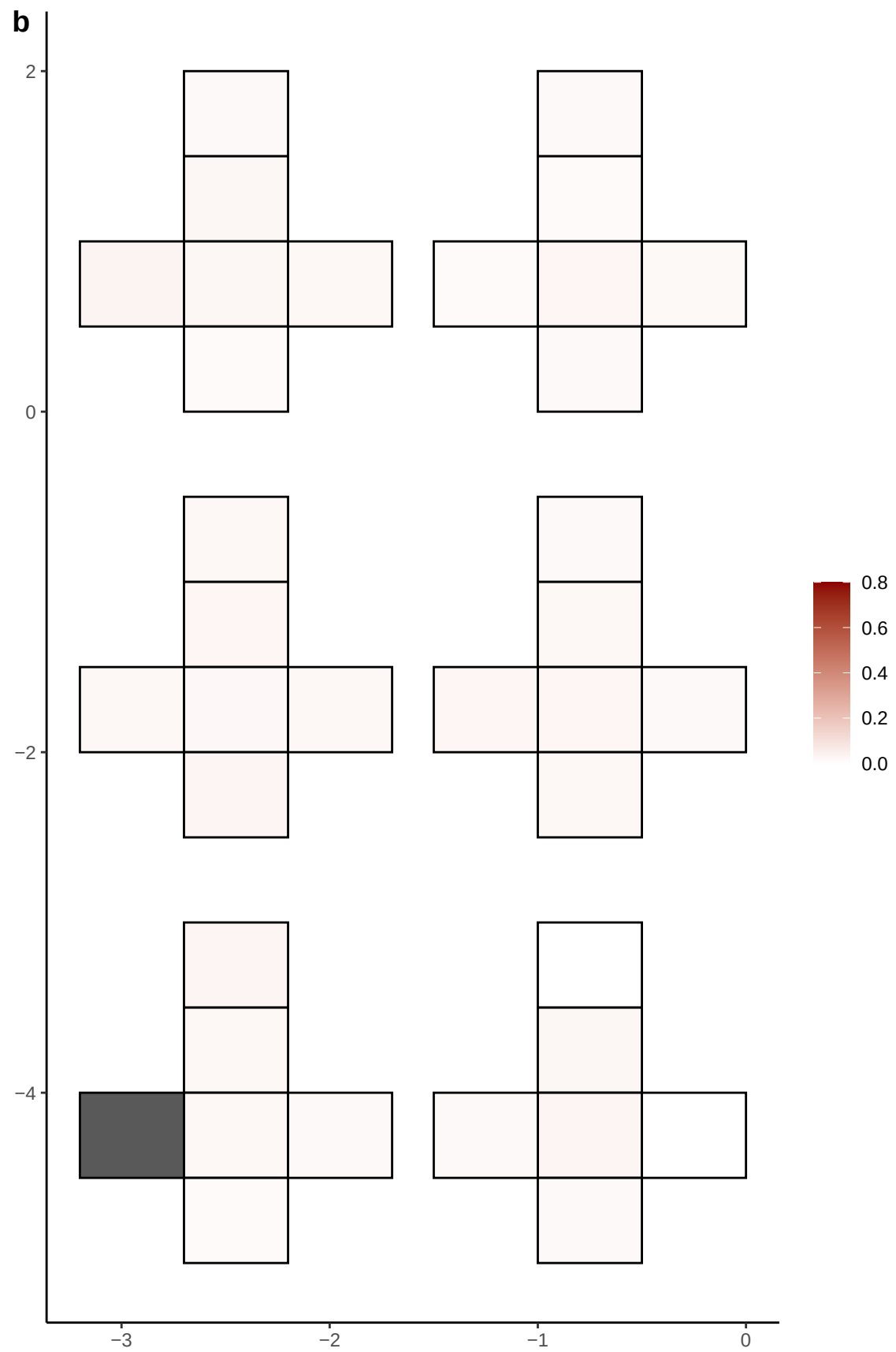

**a** ID01 13-40455905-T-C

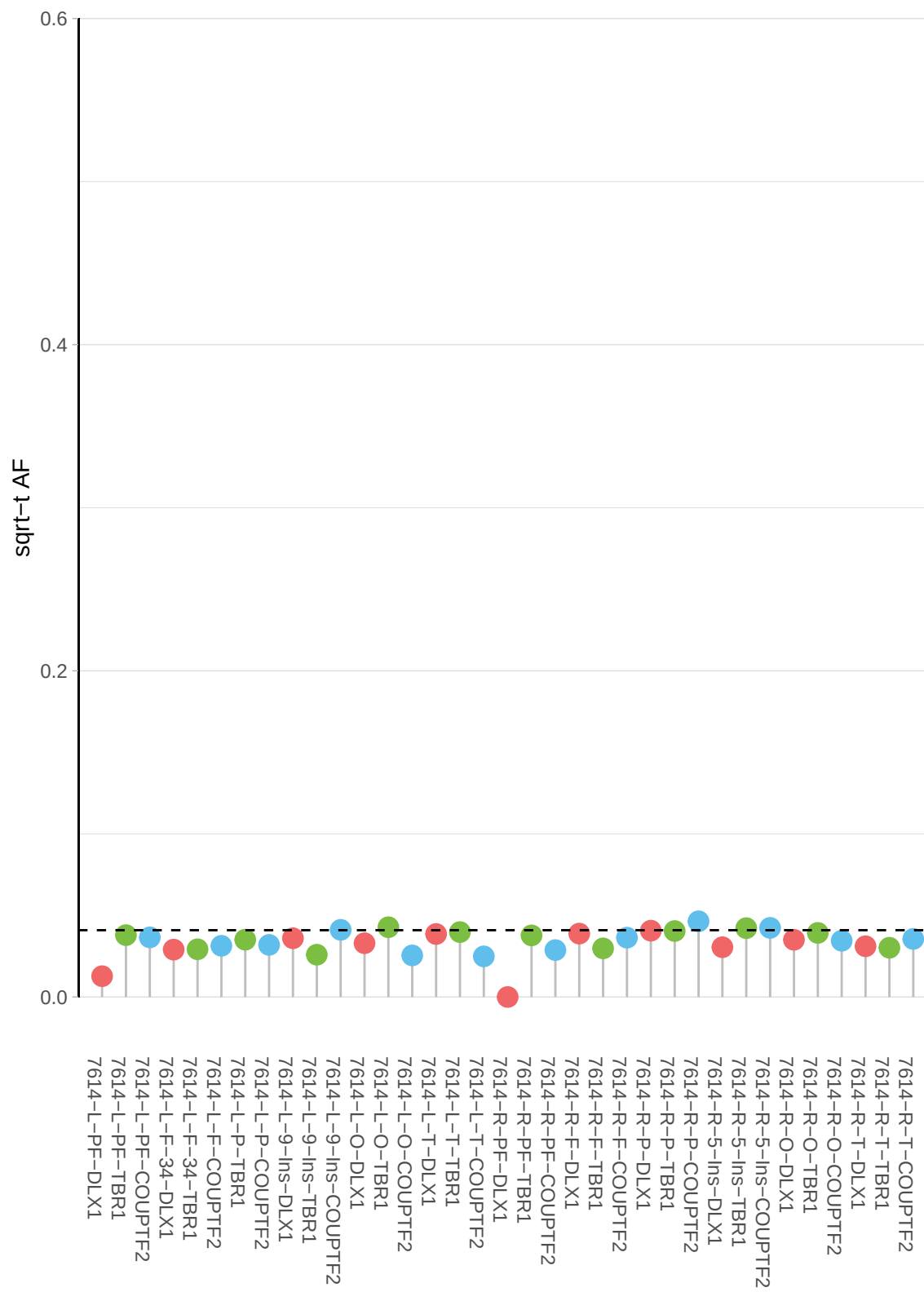

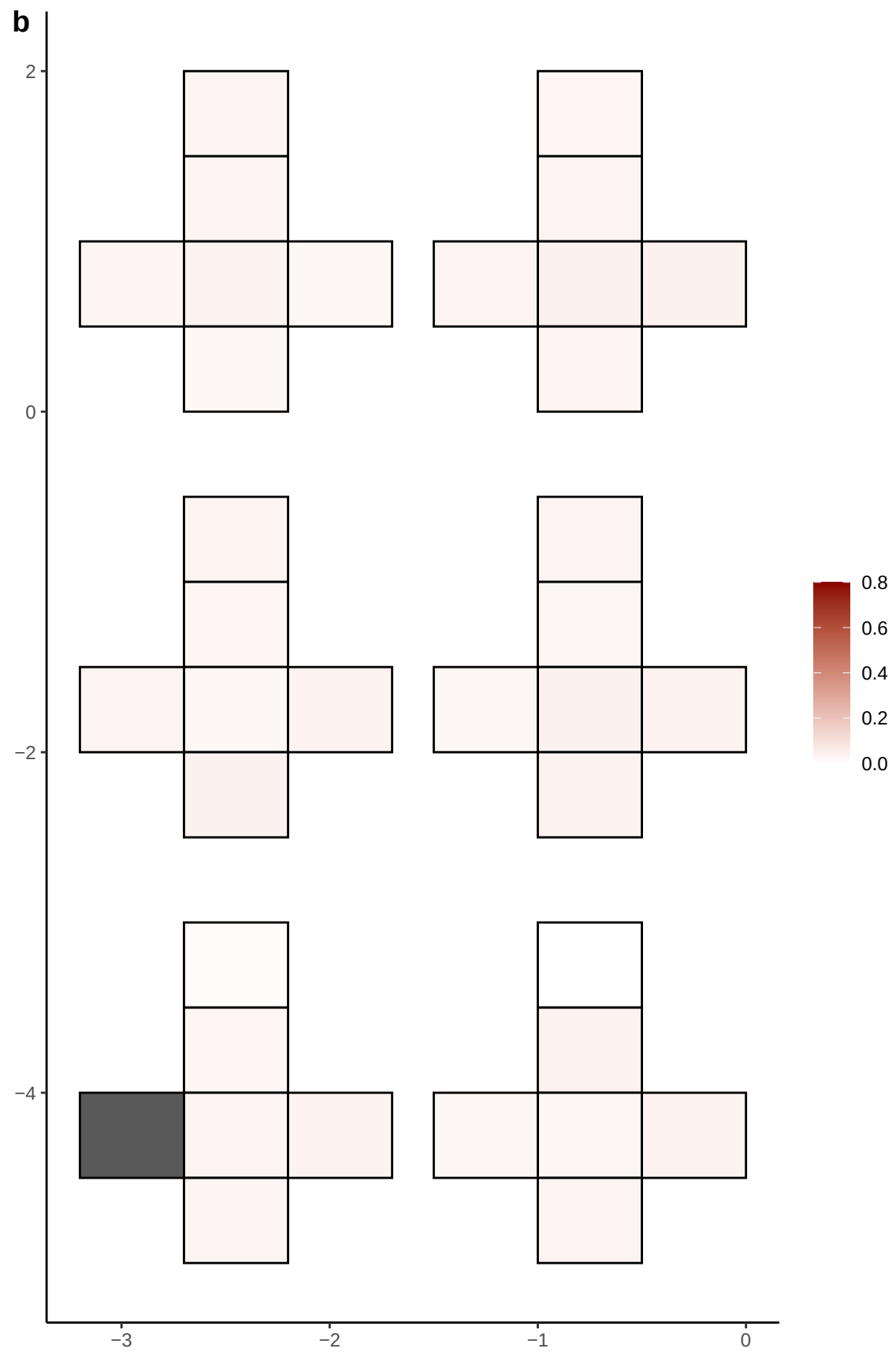

**a** ID01 13-47430361-C-T

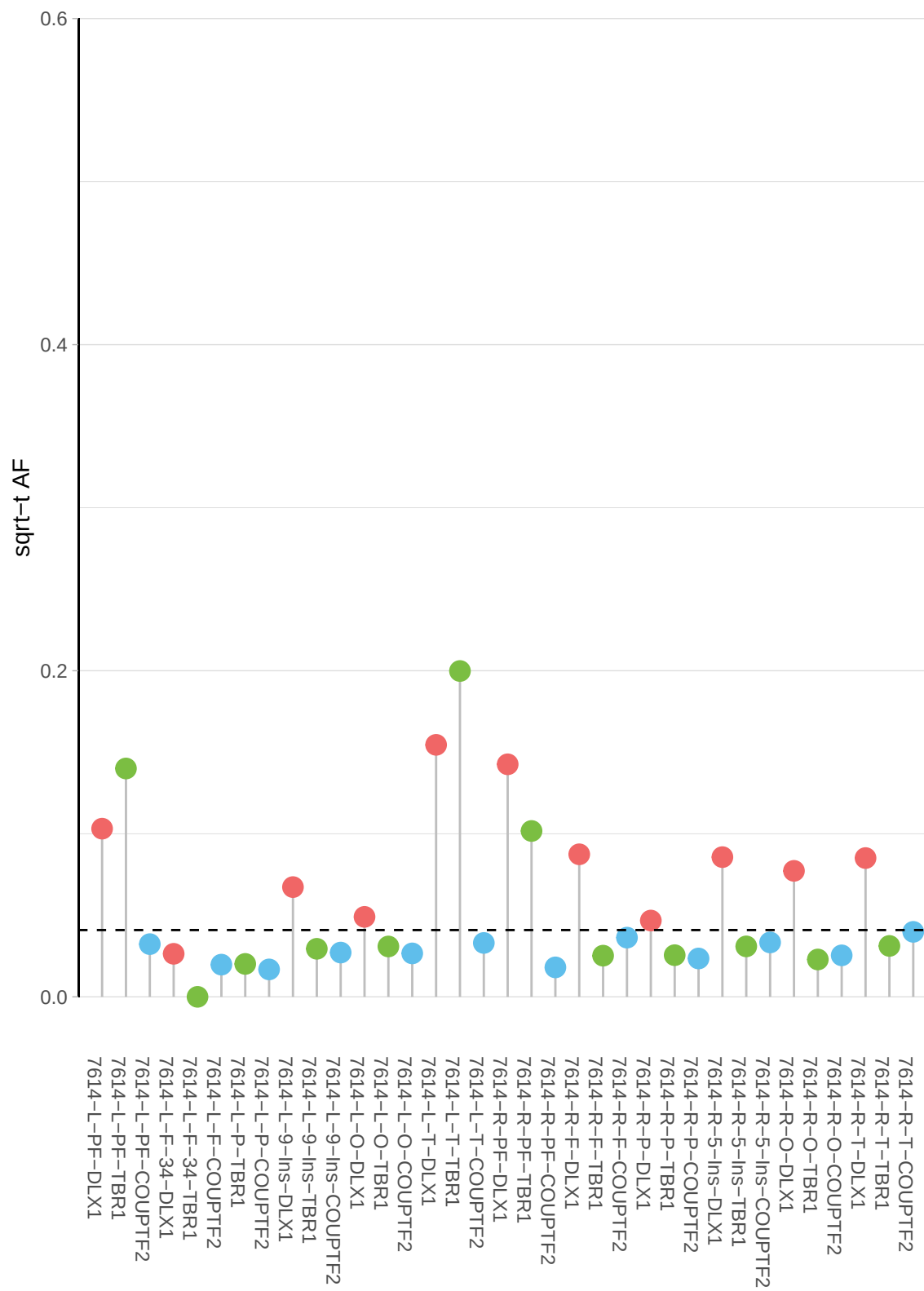

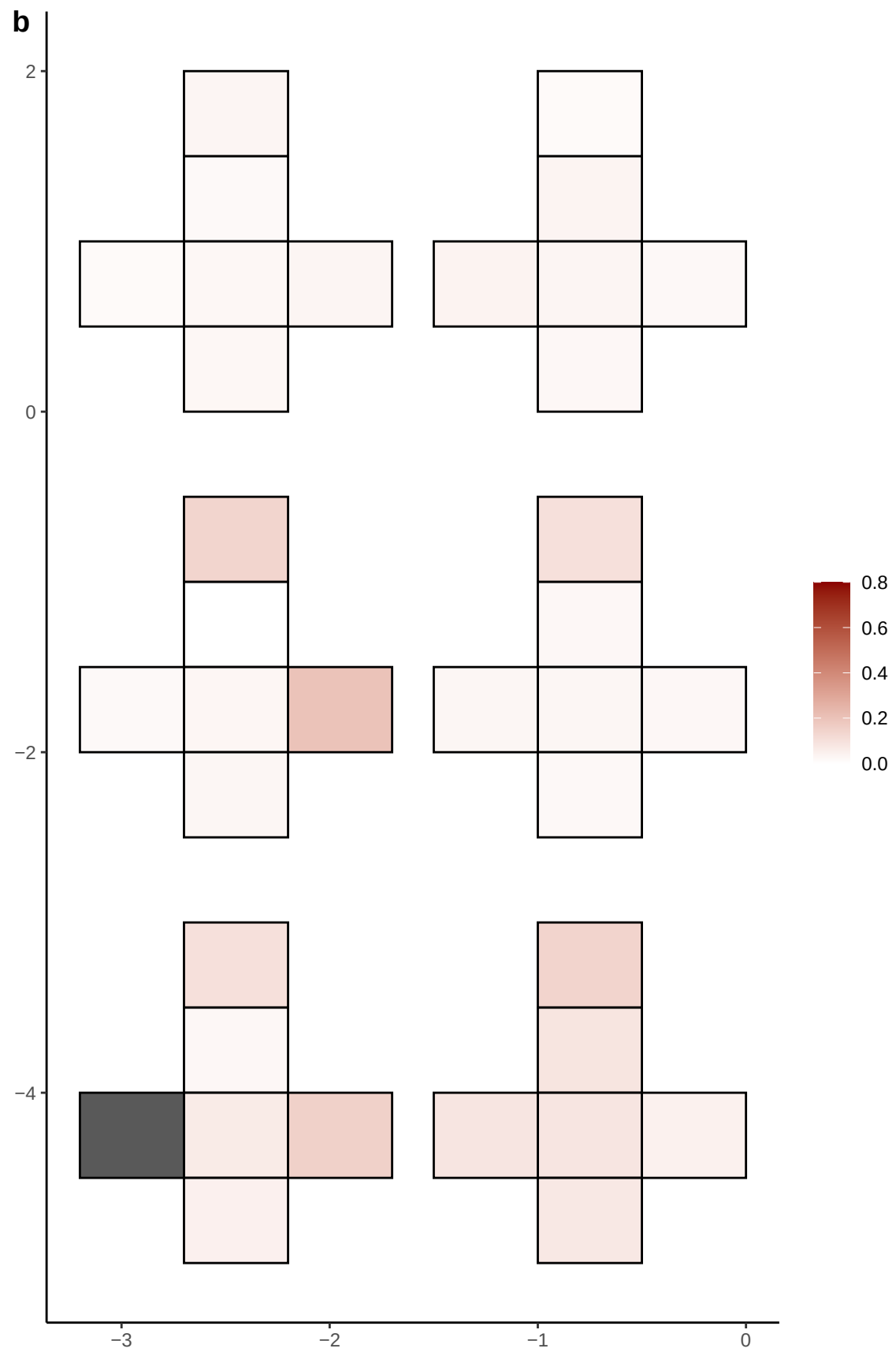

**a** ID01 13-66931942-G-A

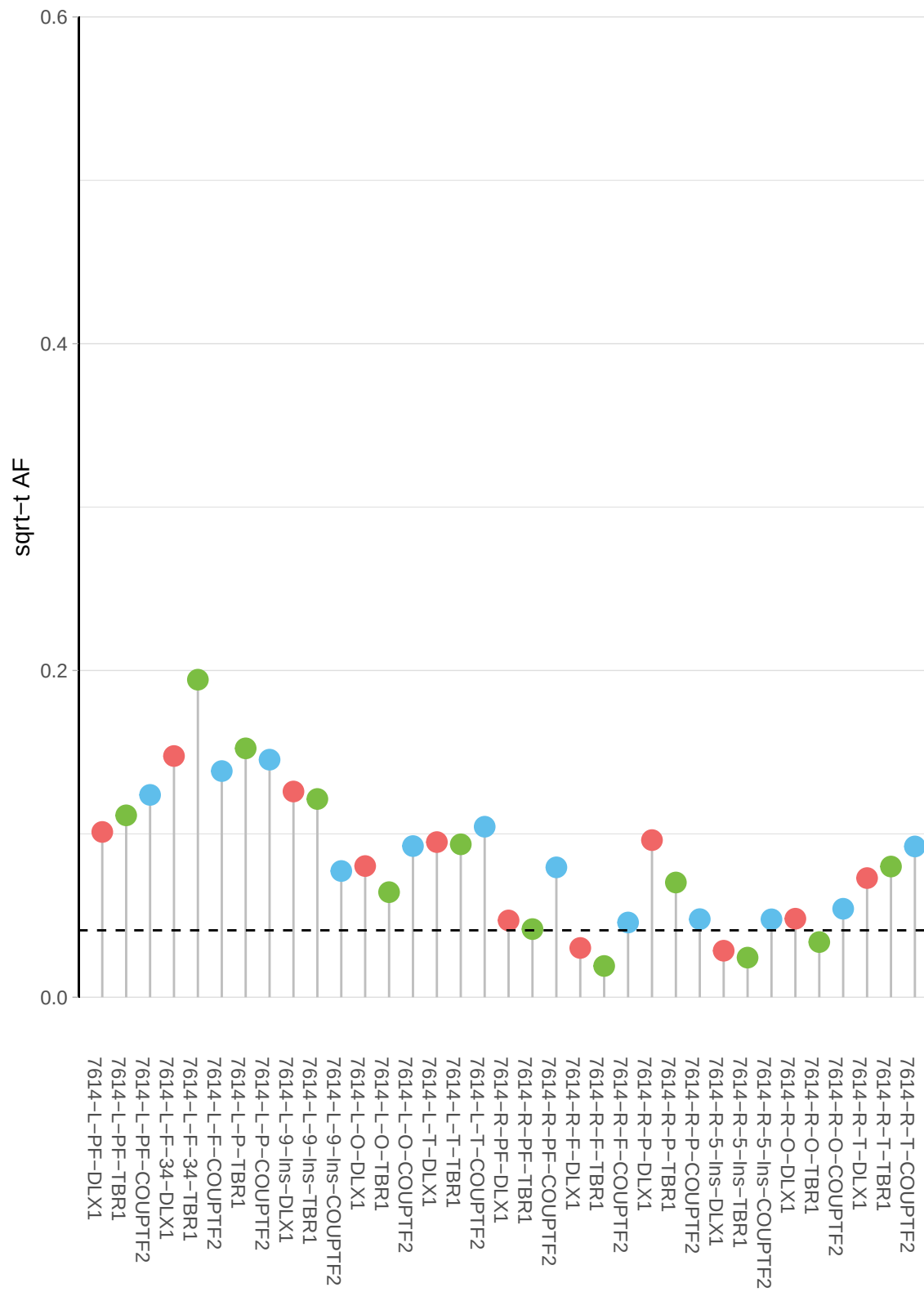

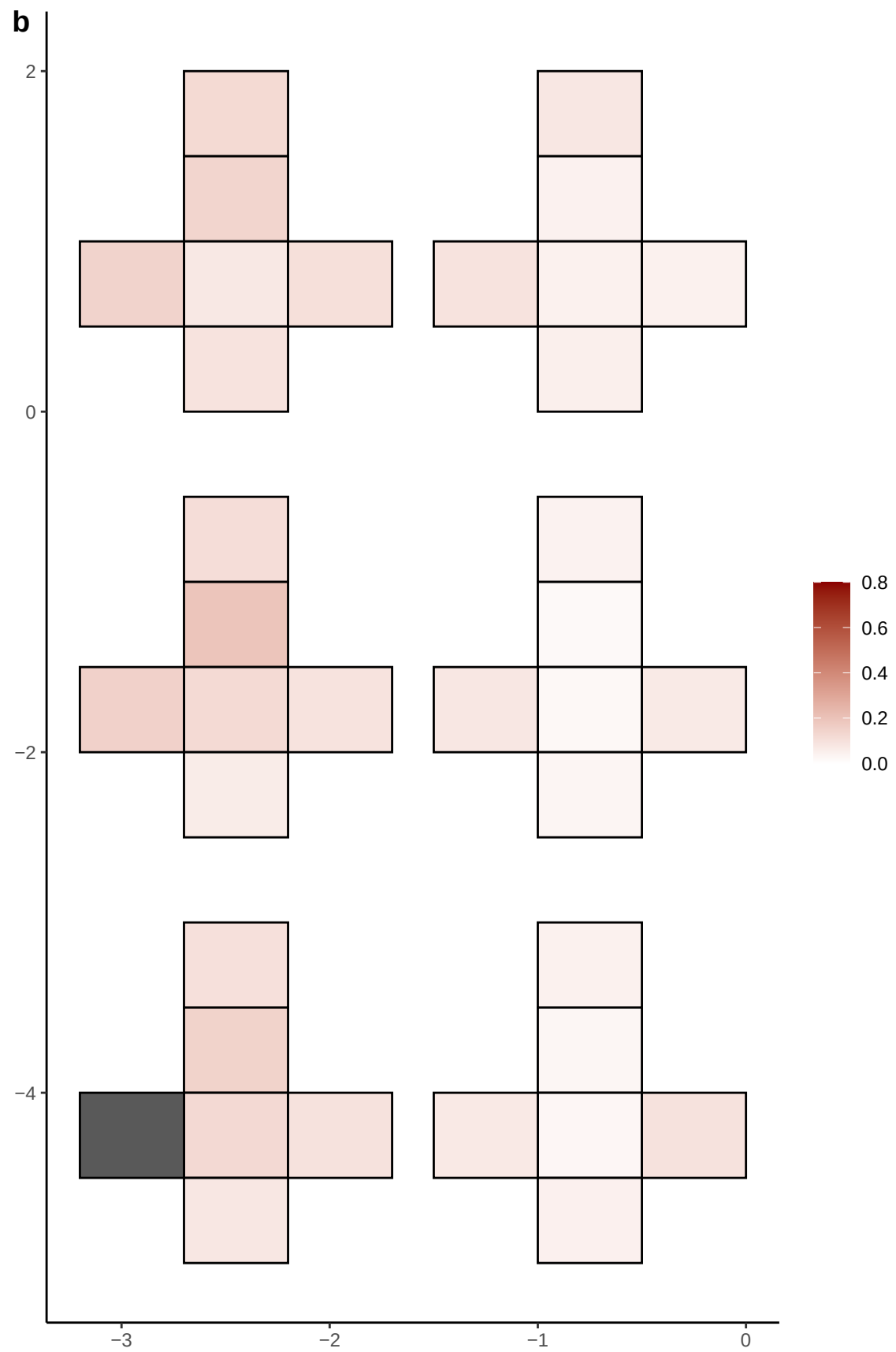

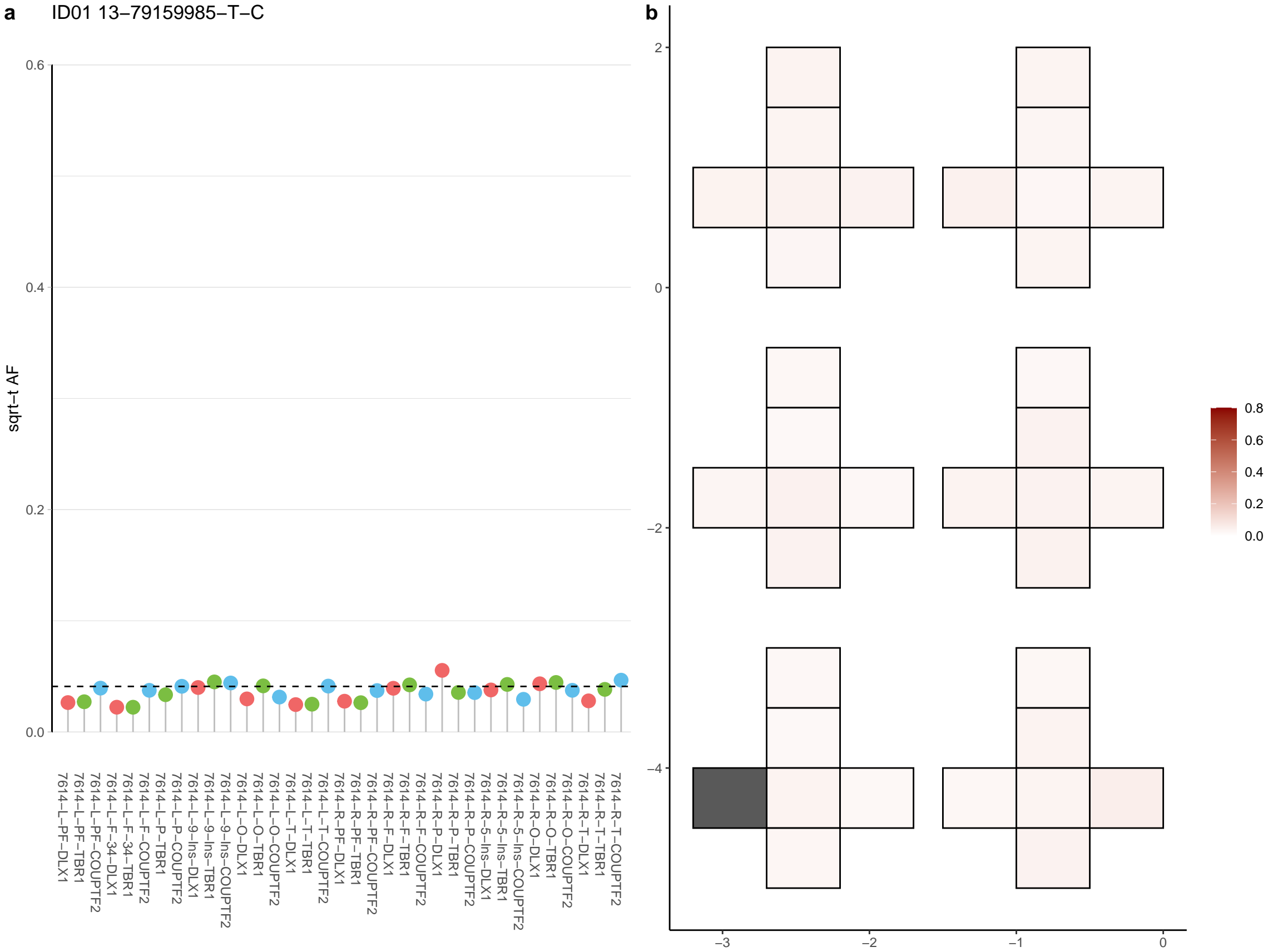

**a** ID01 13-84595018-A-G

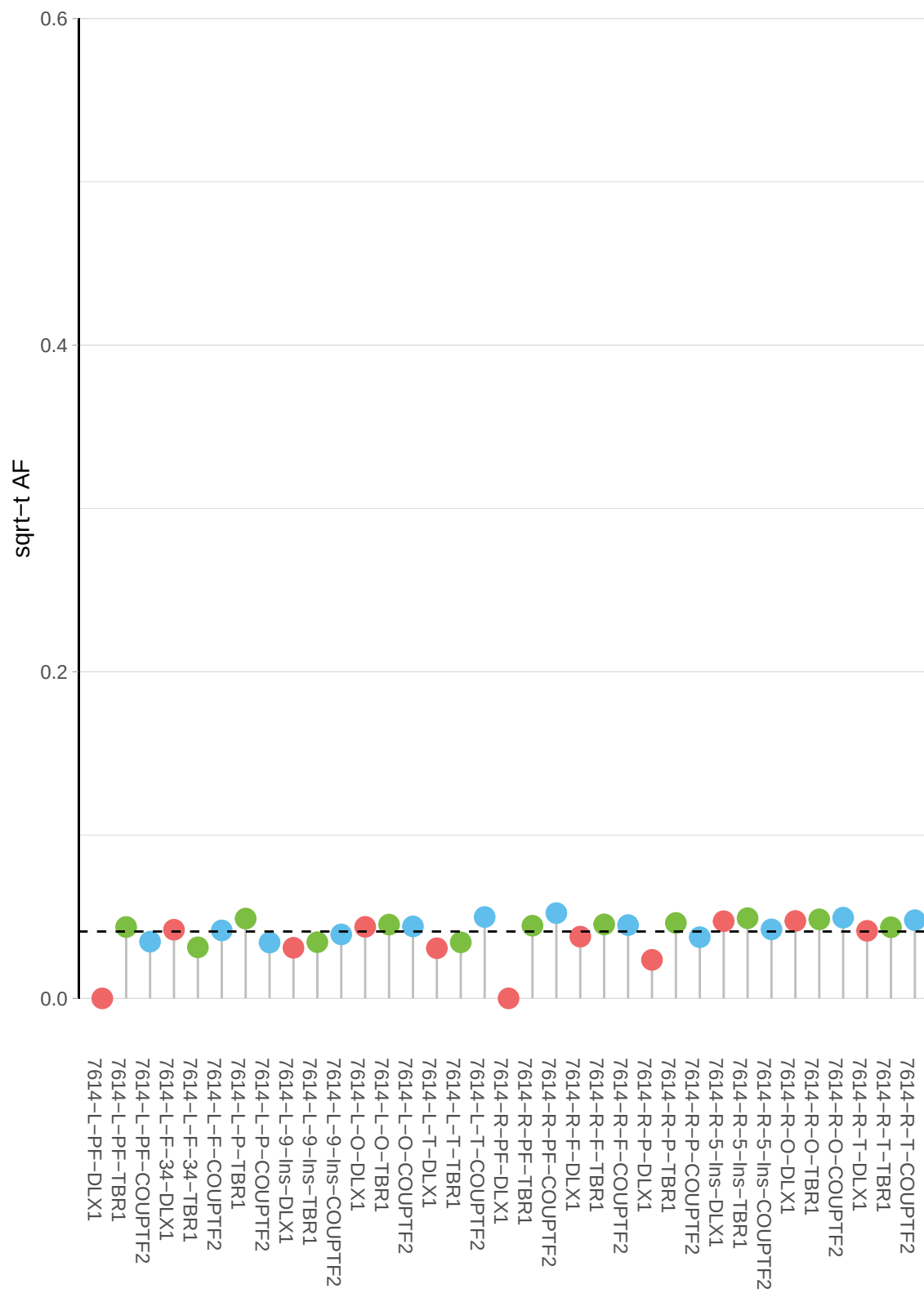

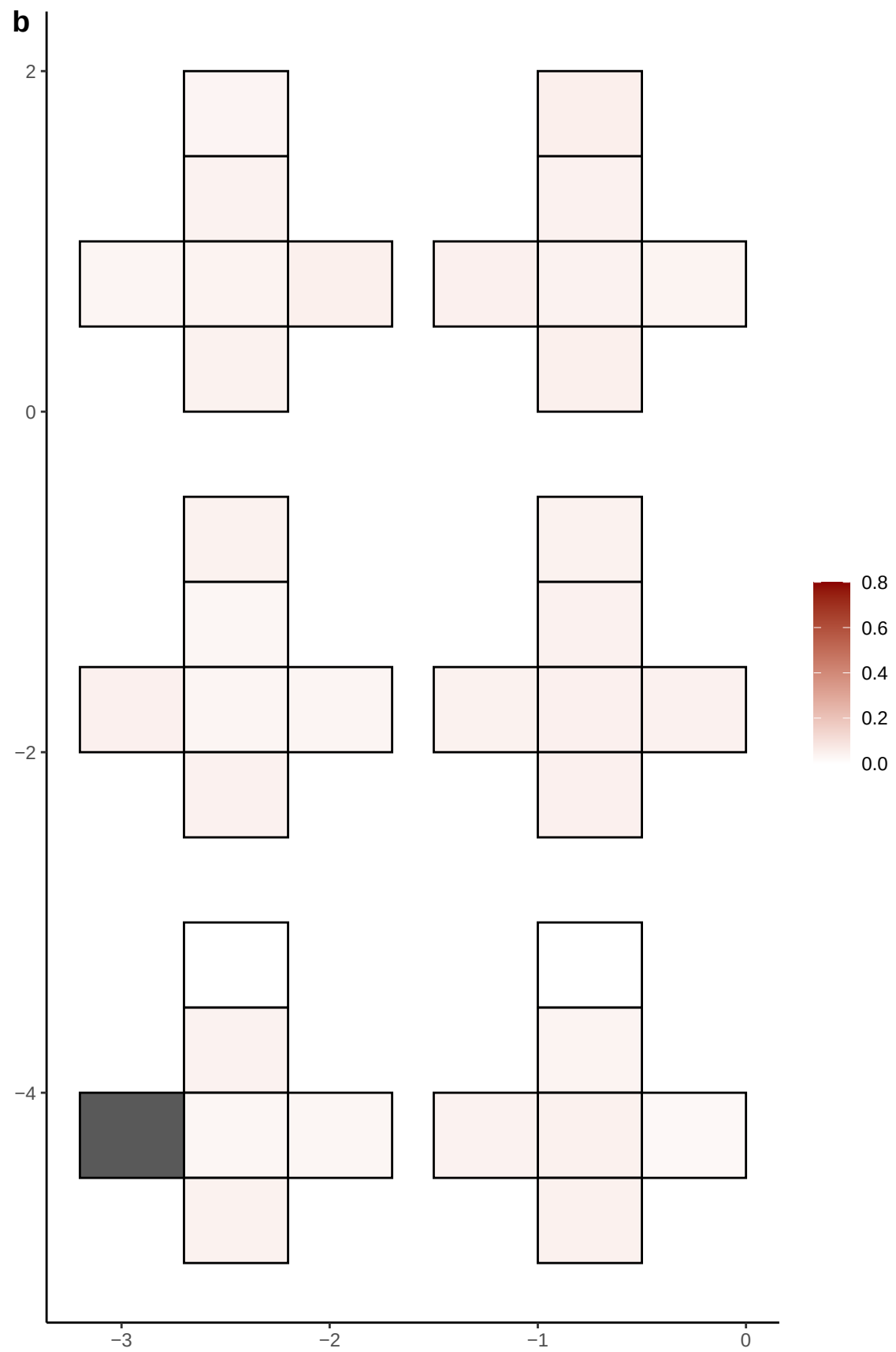

**a** ID01 13-92654445-A-T

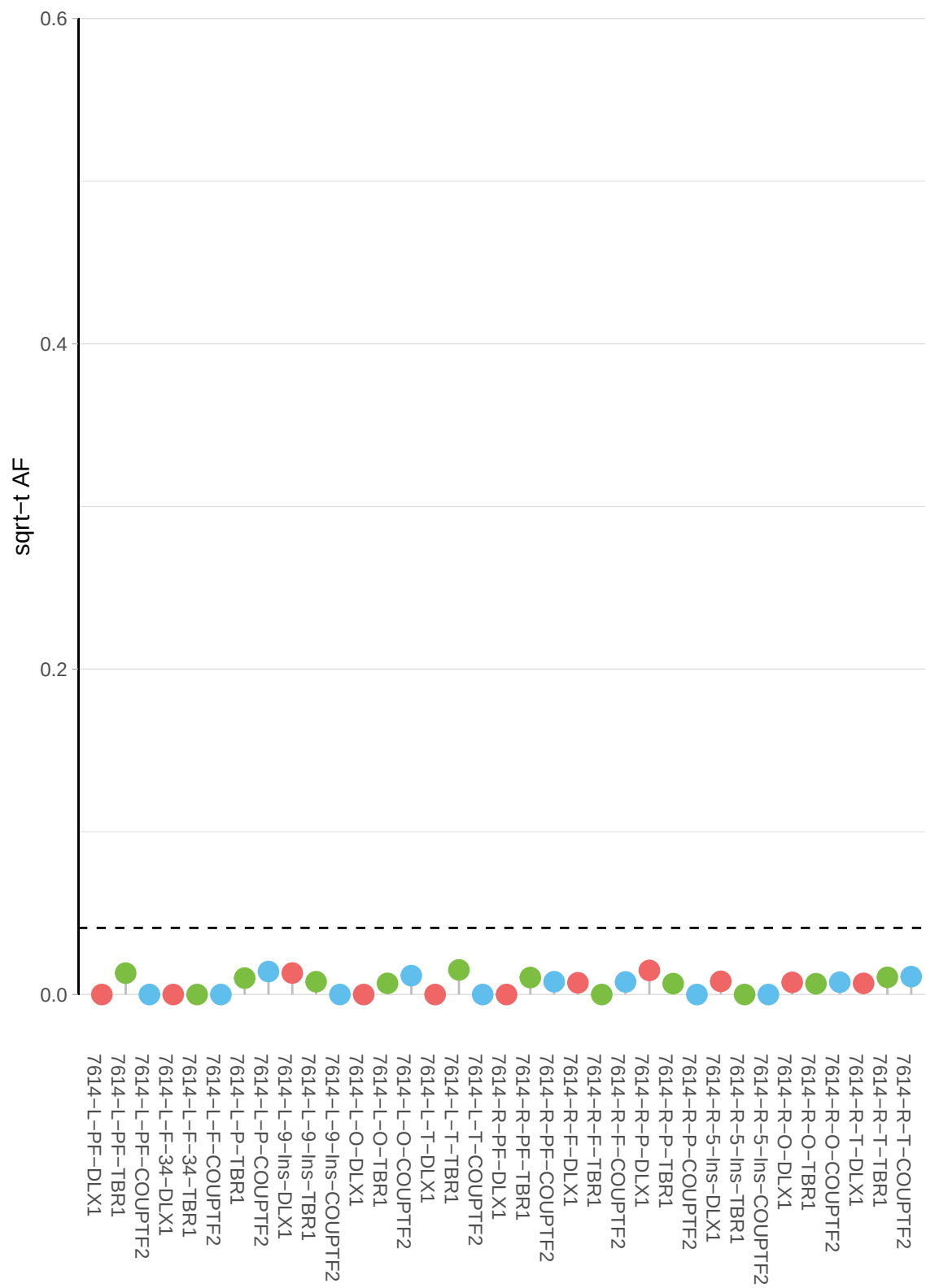

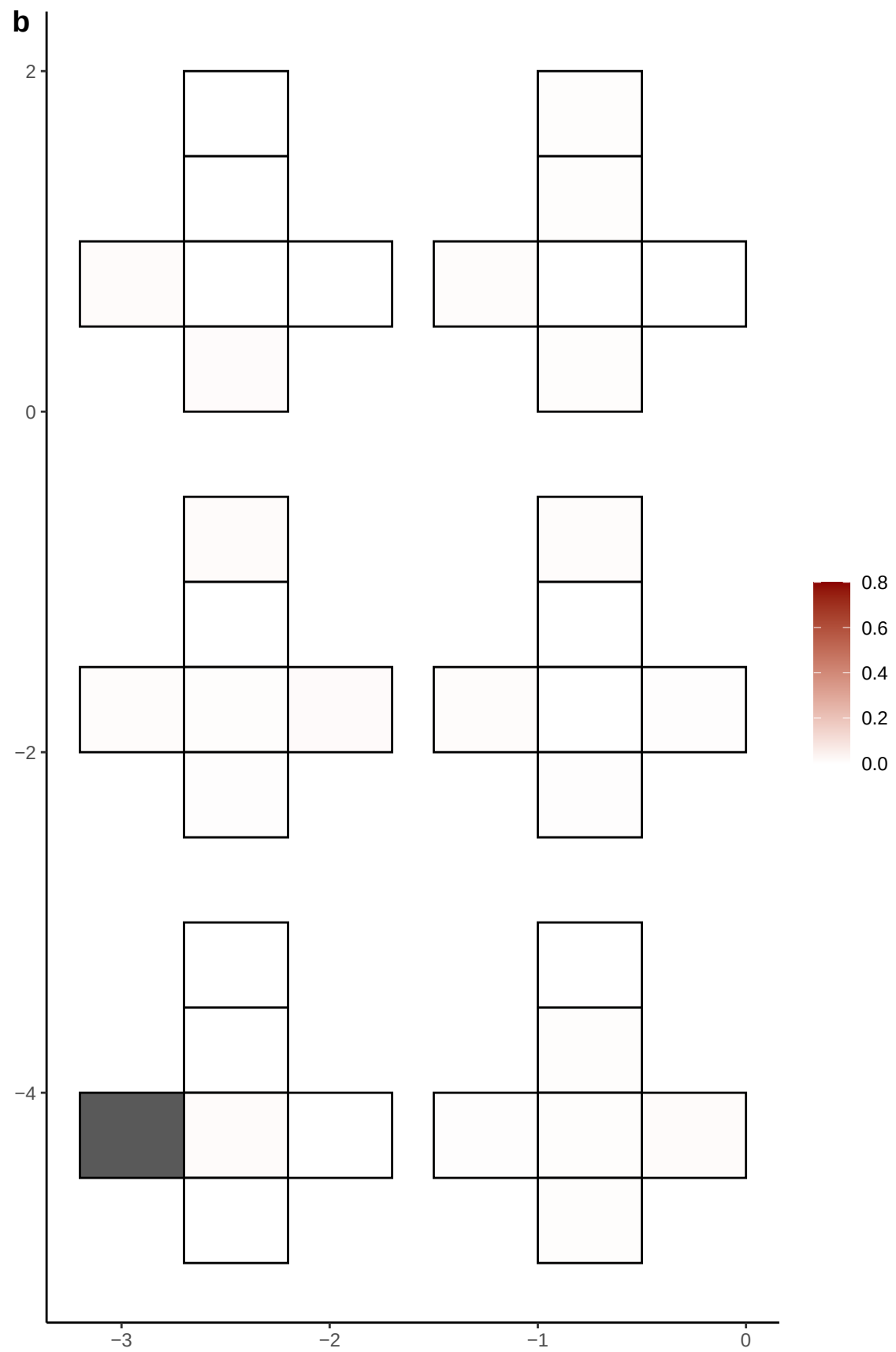

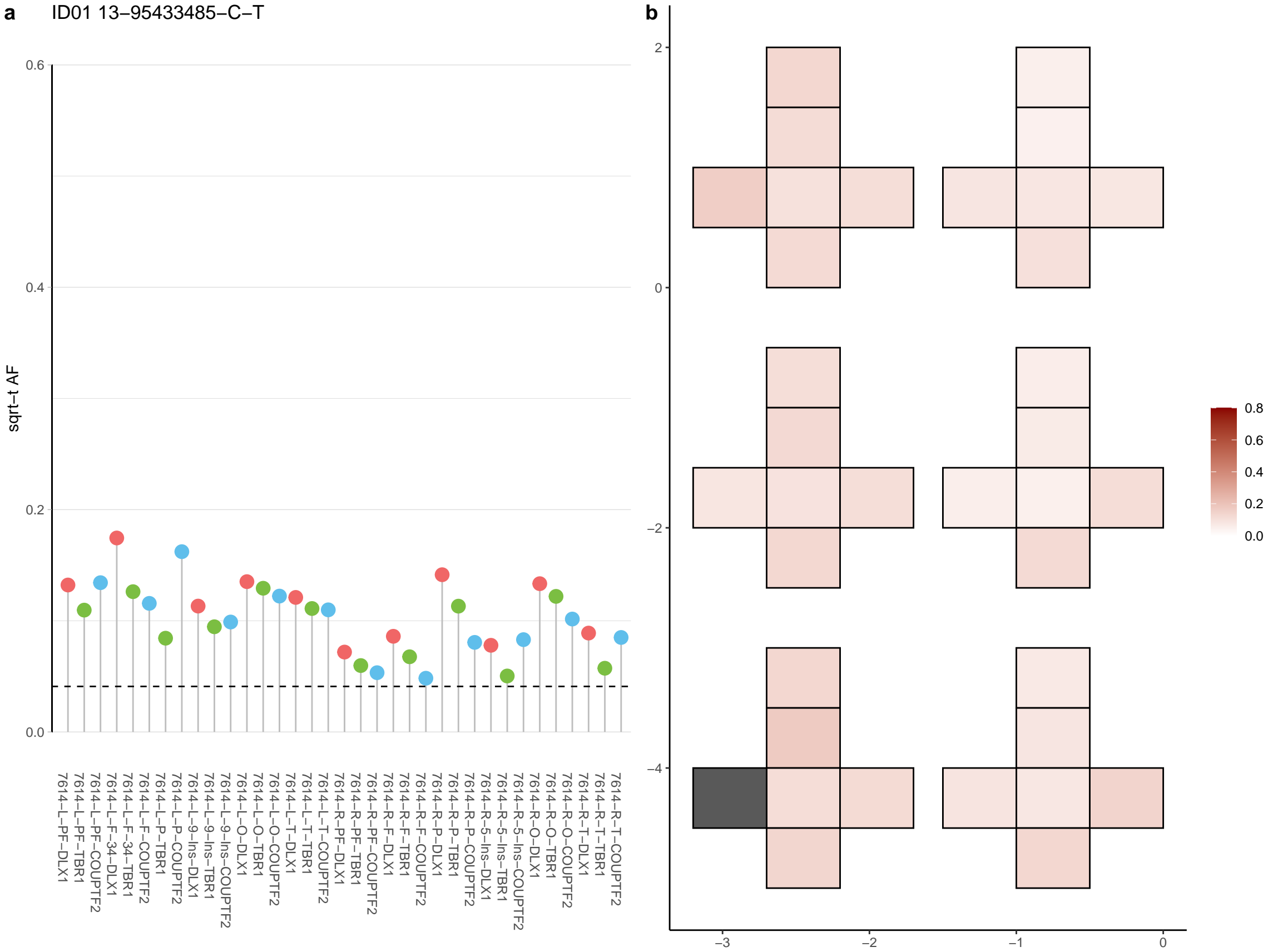

**a** ID01 13-97971976-C-T

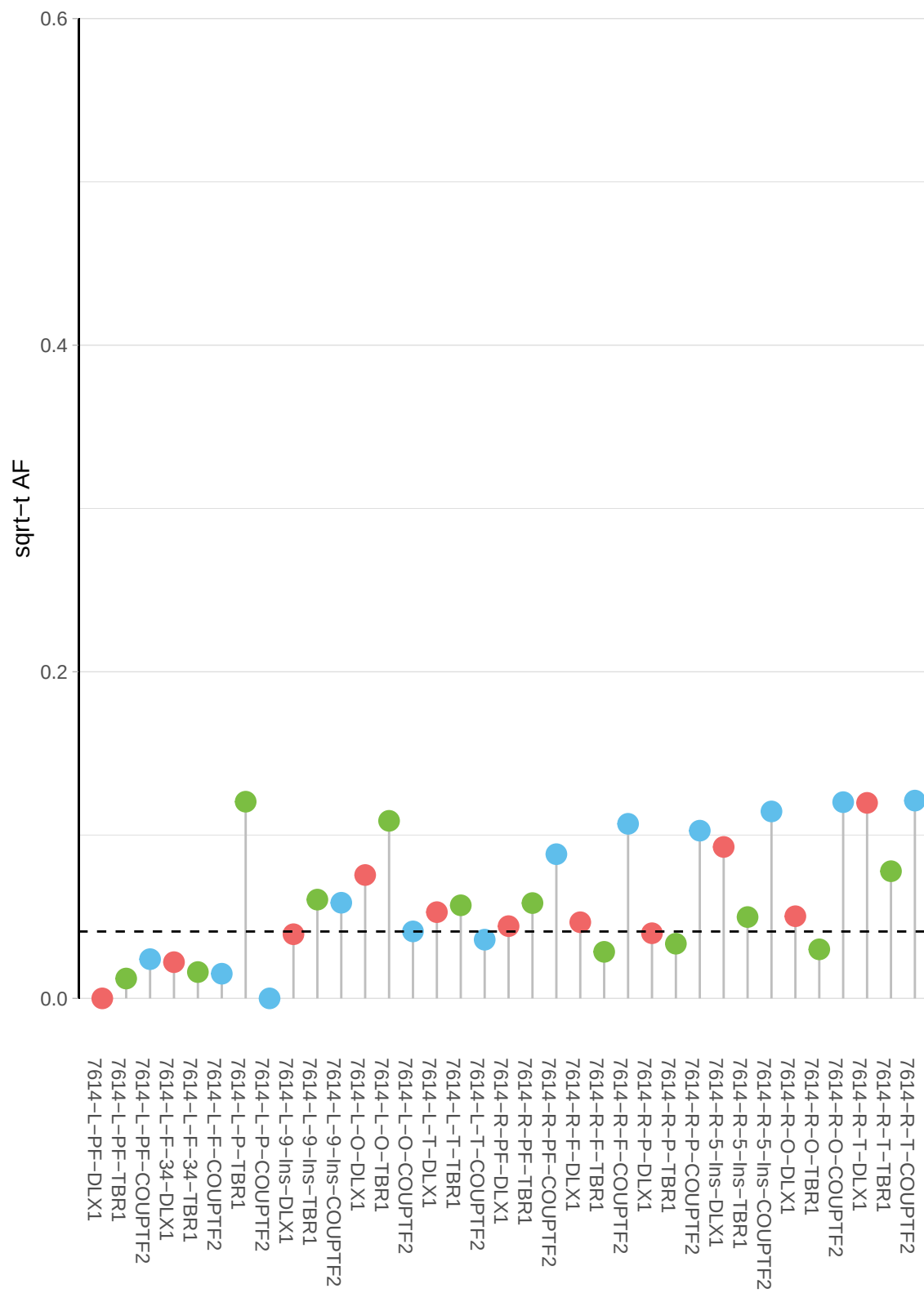

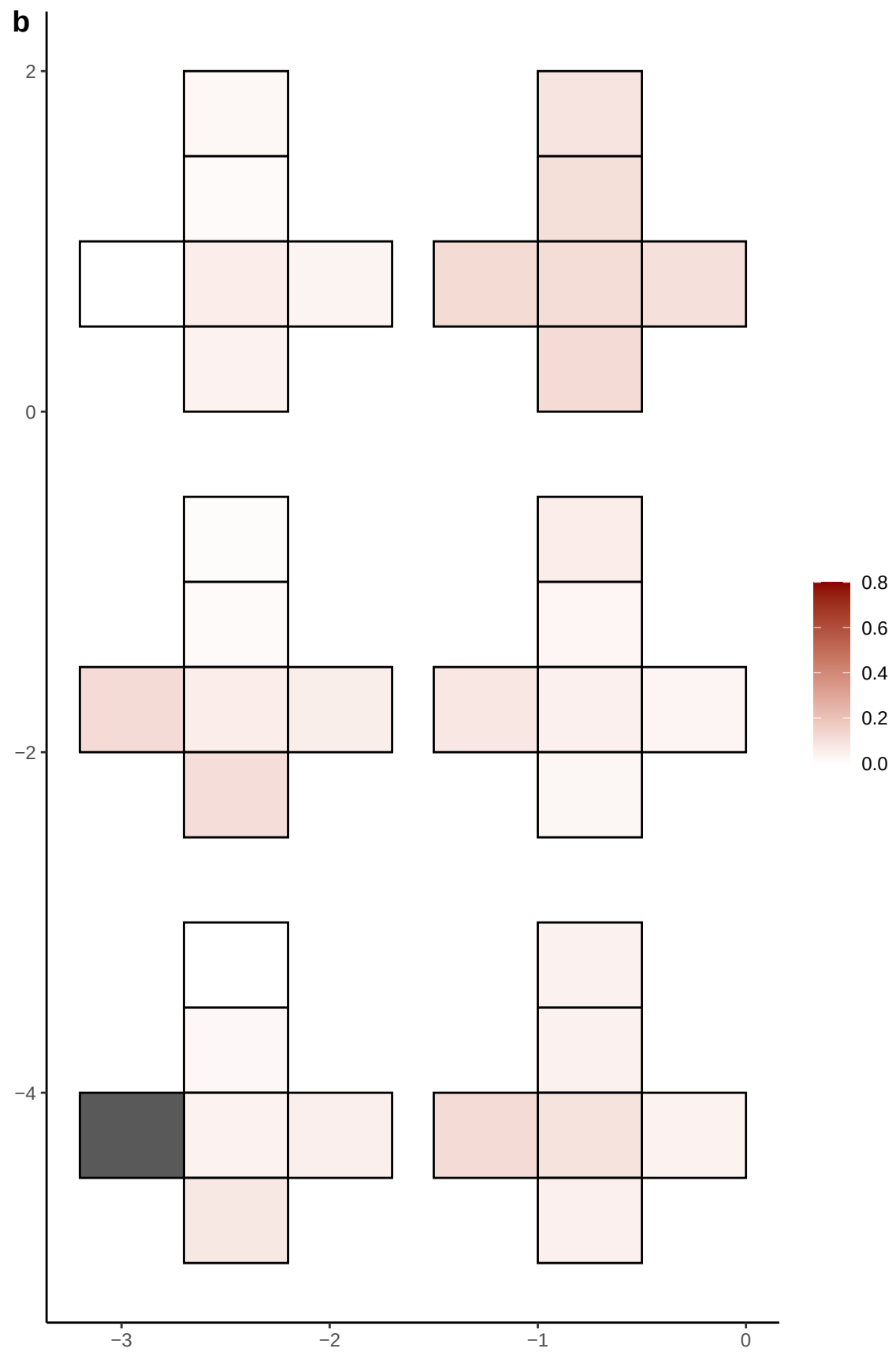

**a** ID01 14-38033902-G-T

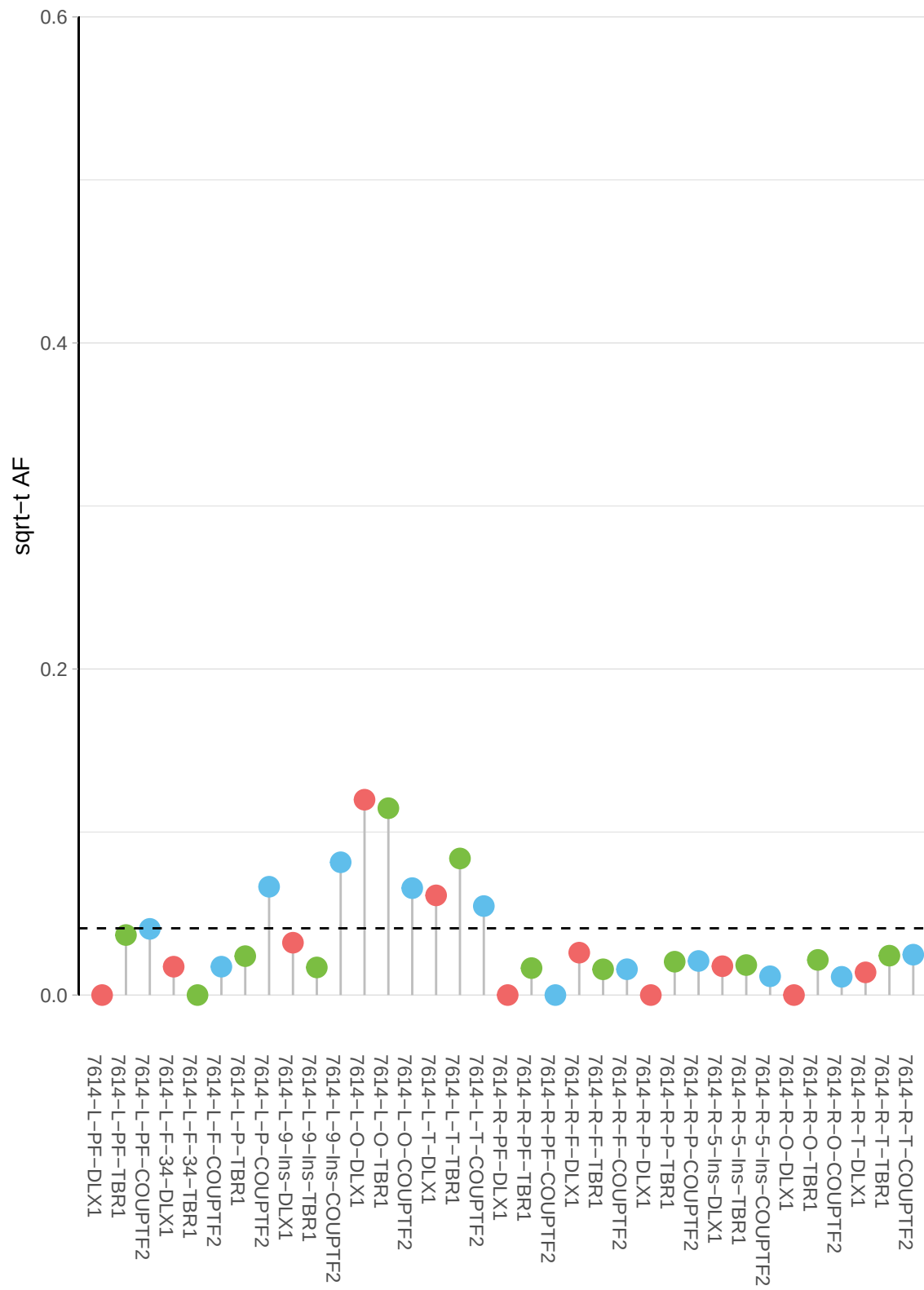

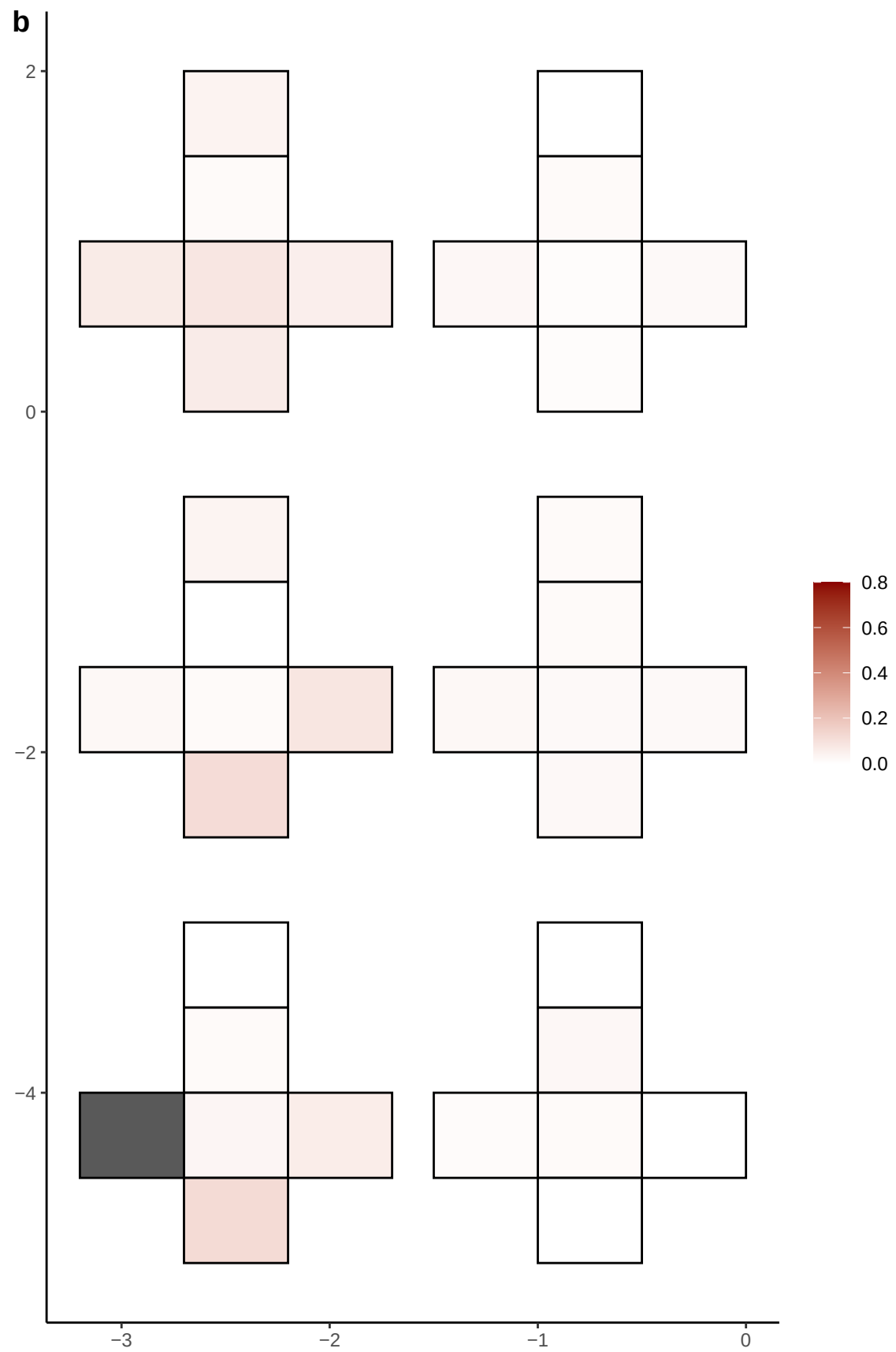

**a** ID01 14-44933484-T-C

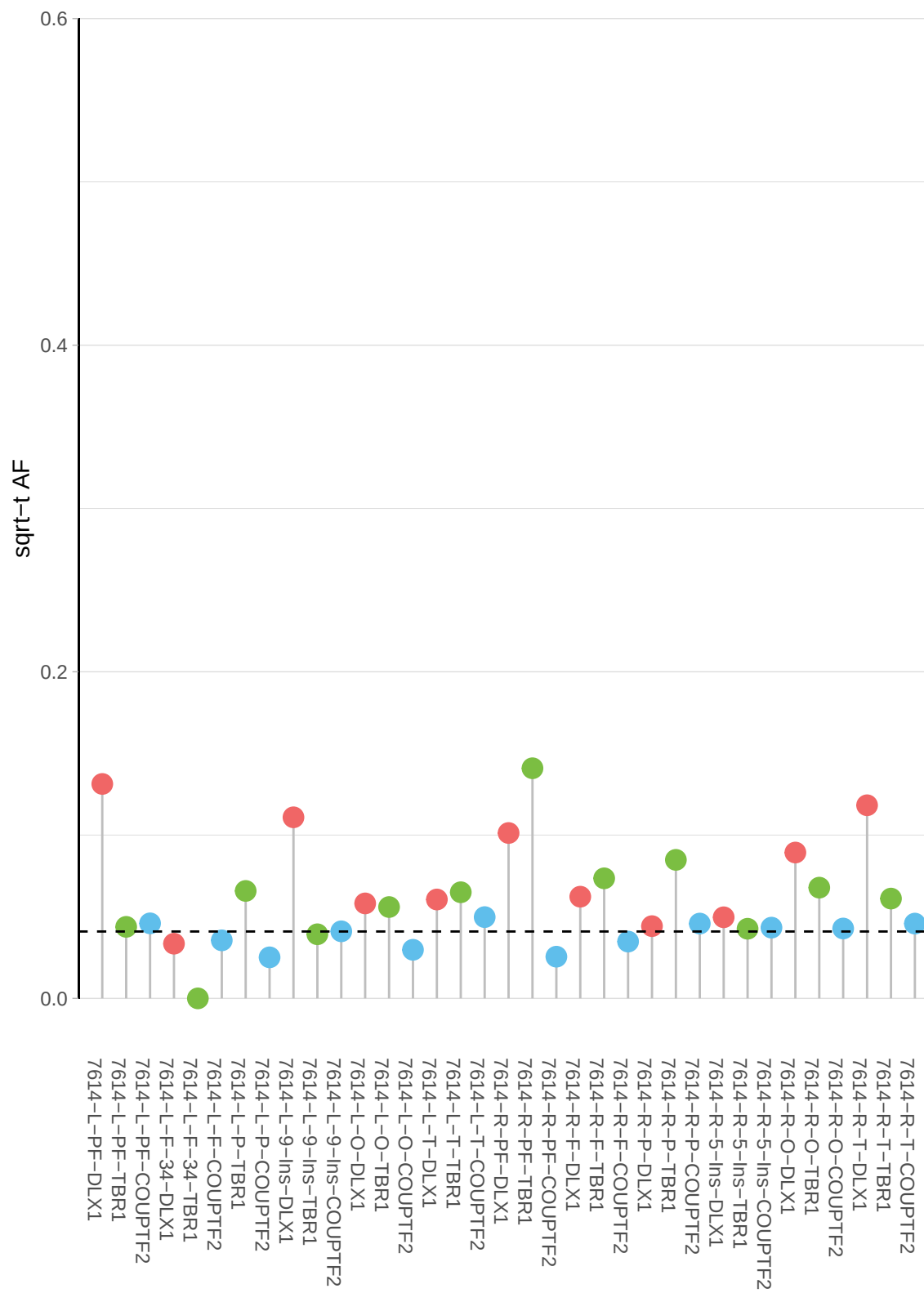

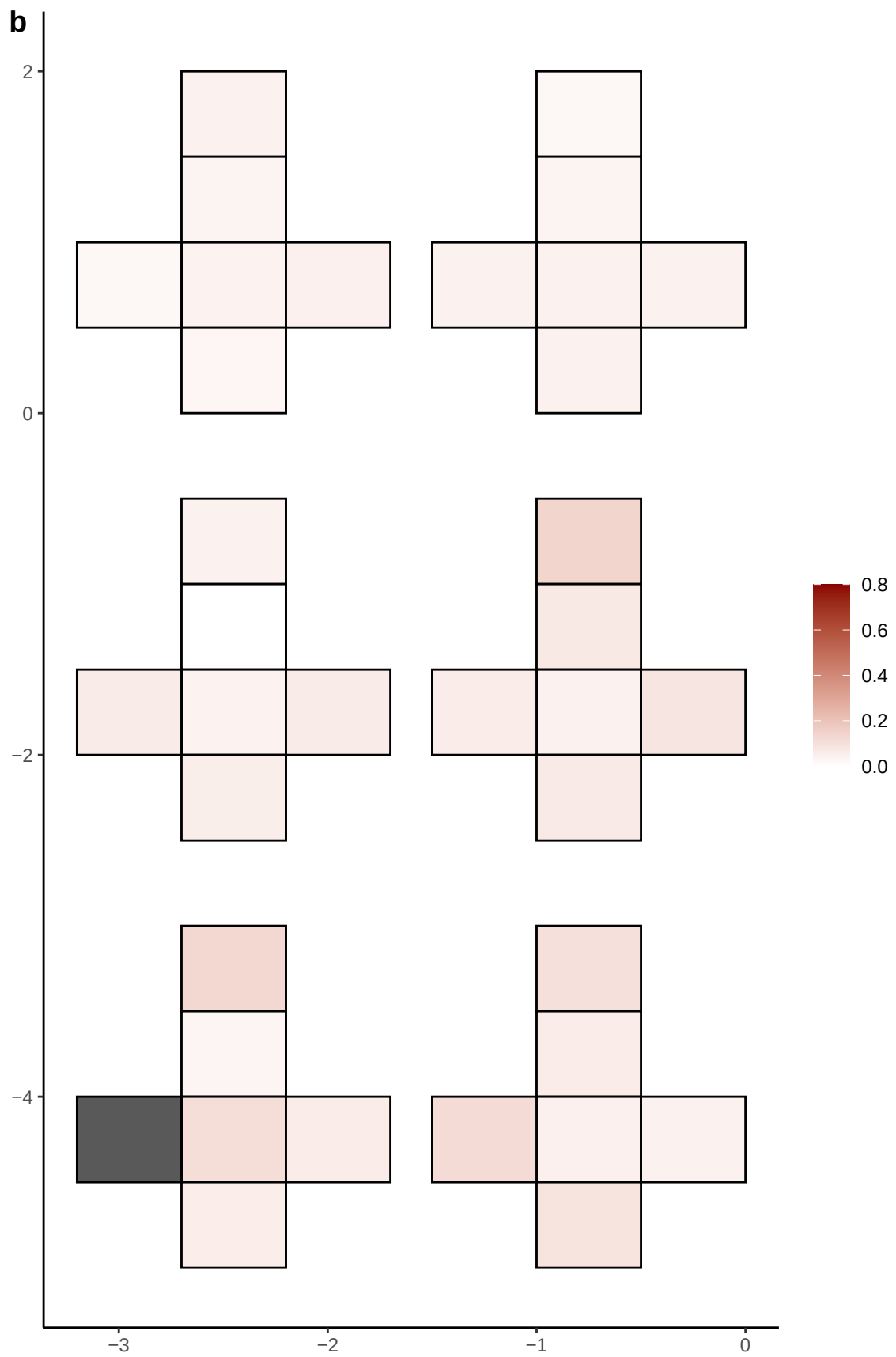

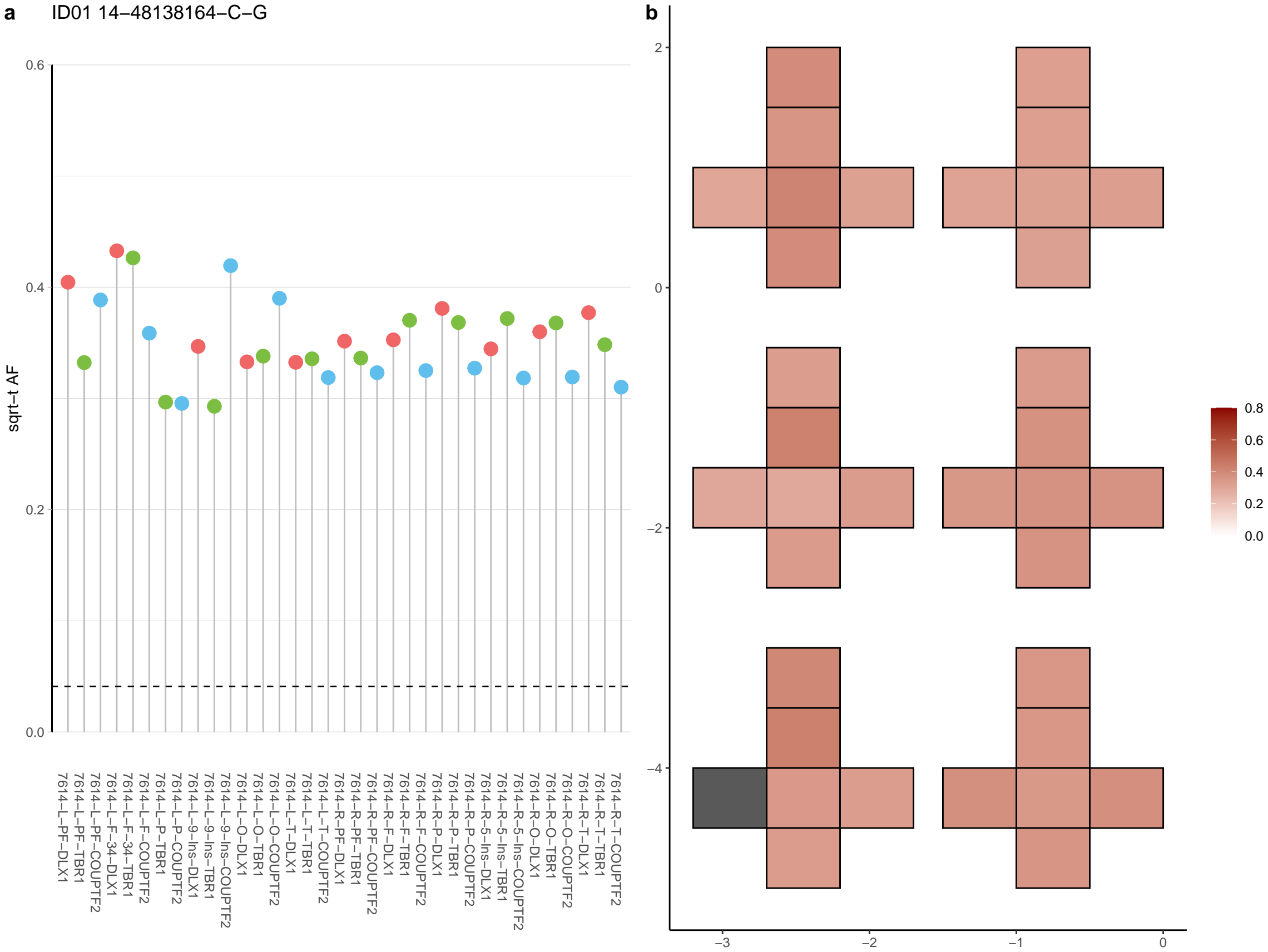

**a** ID01 14-48579727-C-T

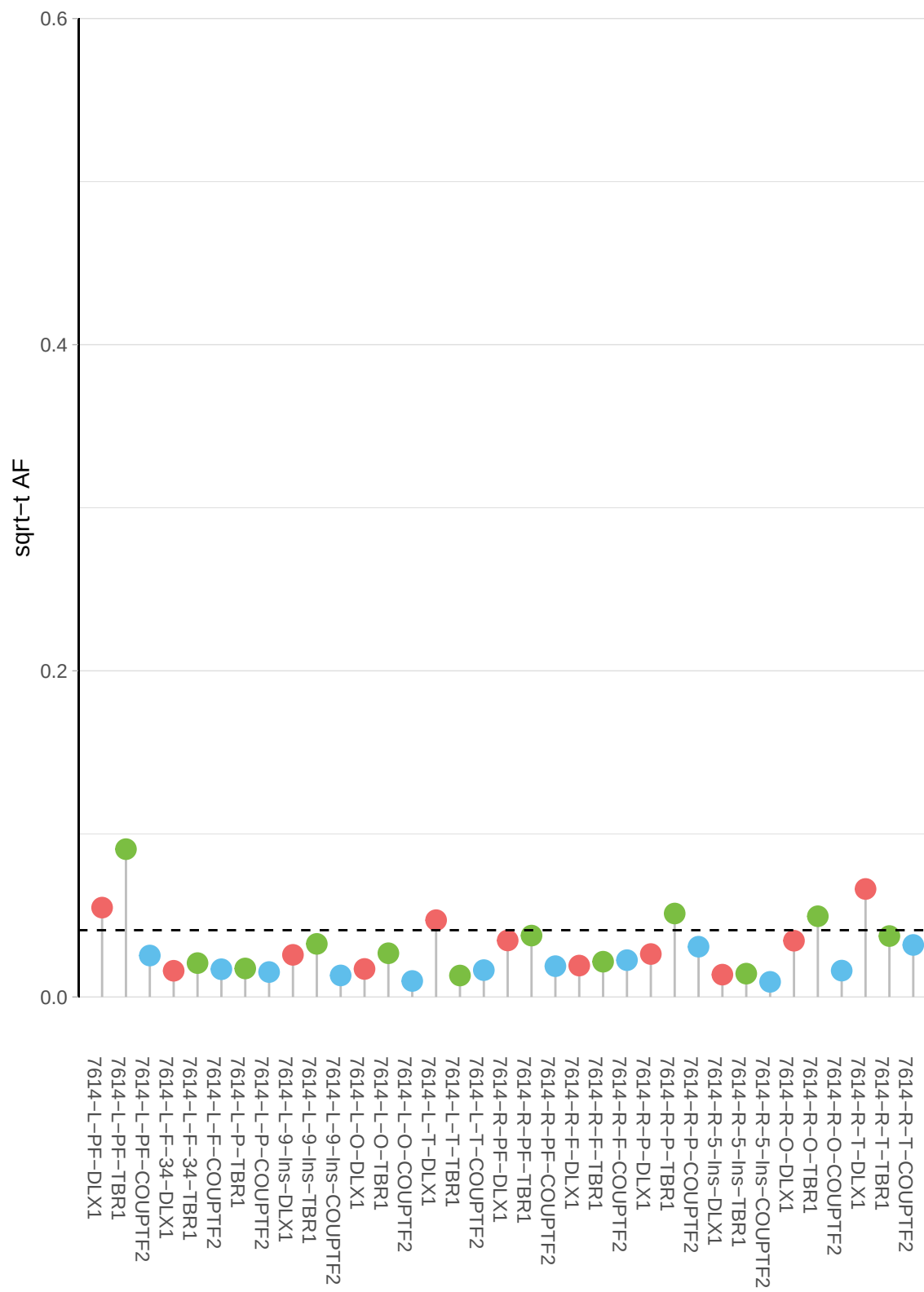

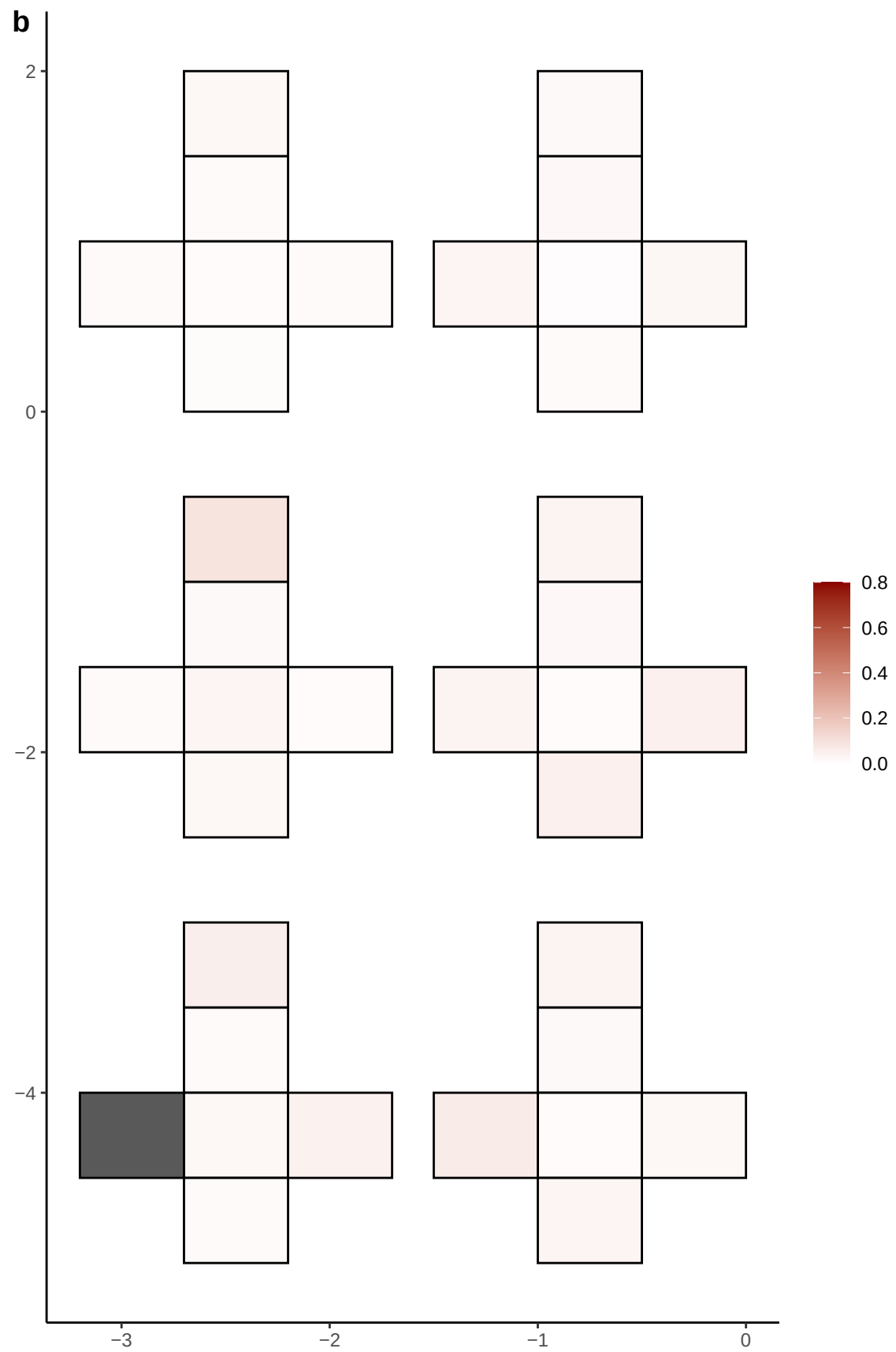

**a** ID01 14-54432218-C-T

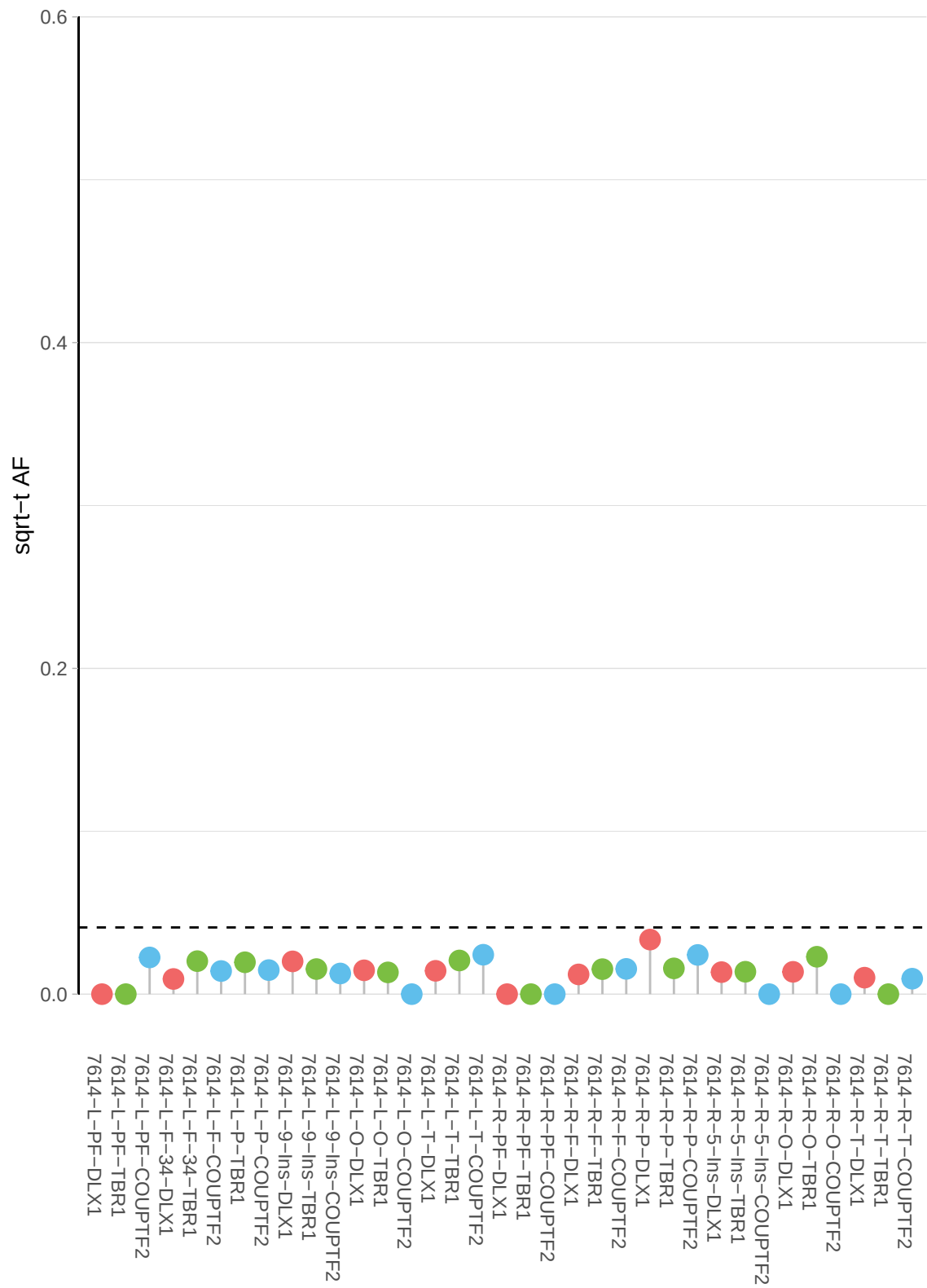

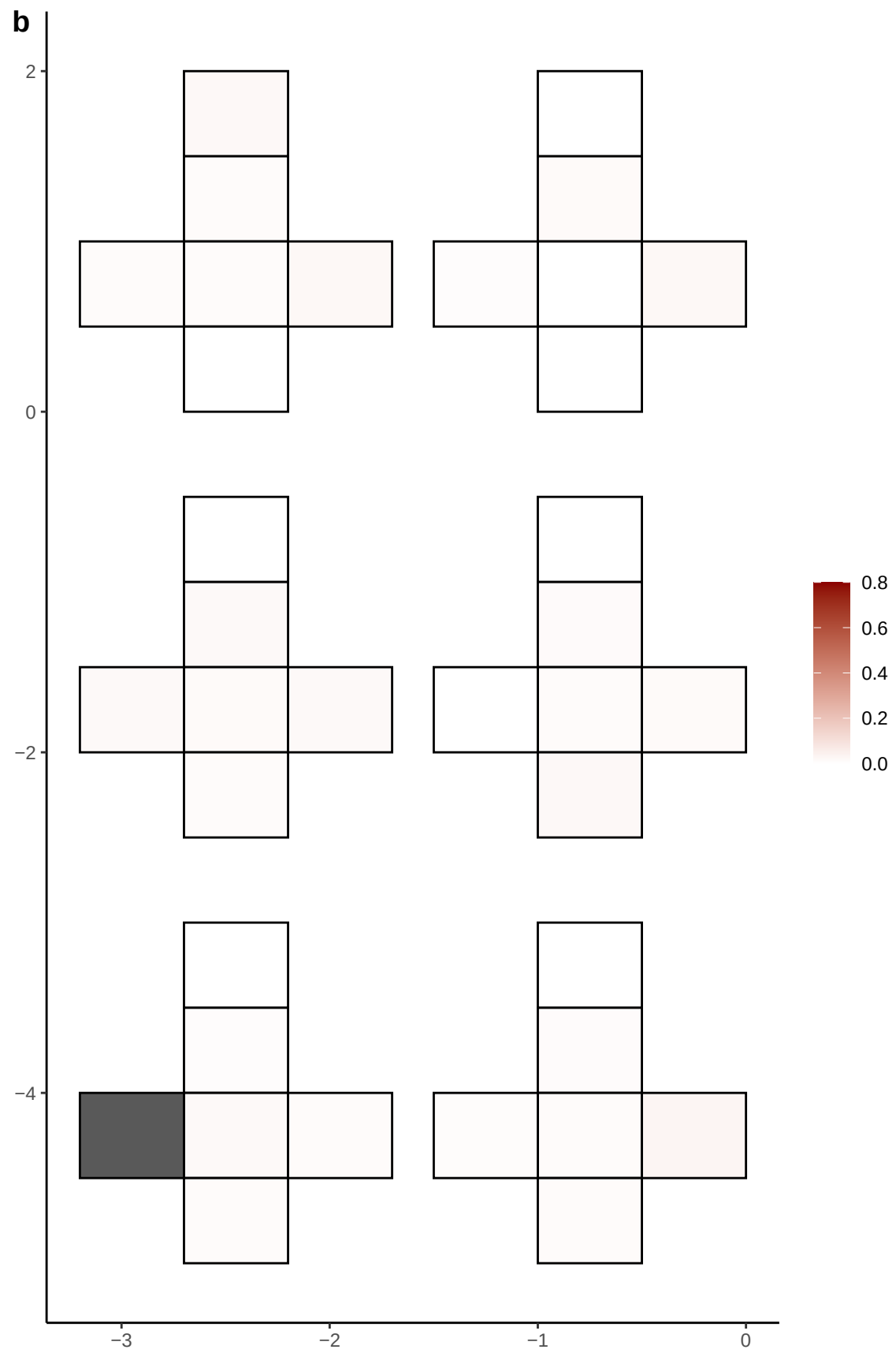

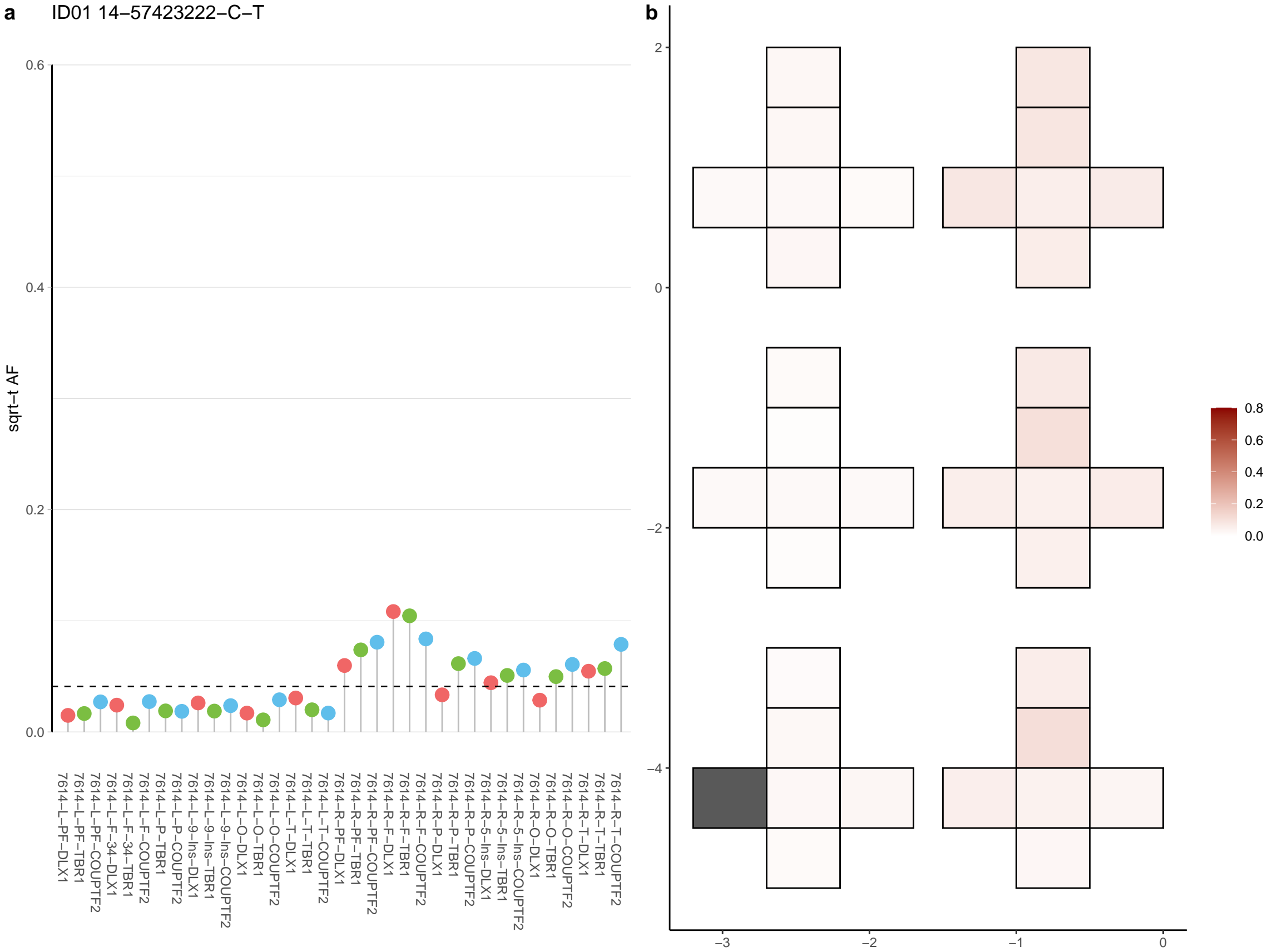

**a** ID01 14-59303124-G-A

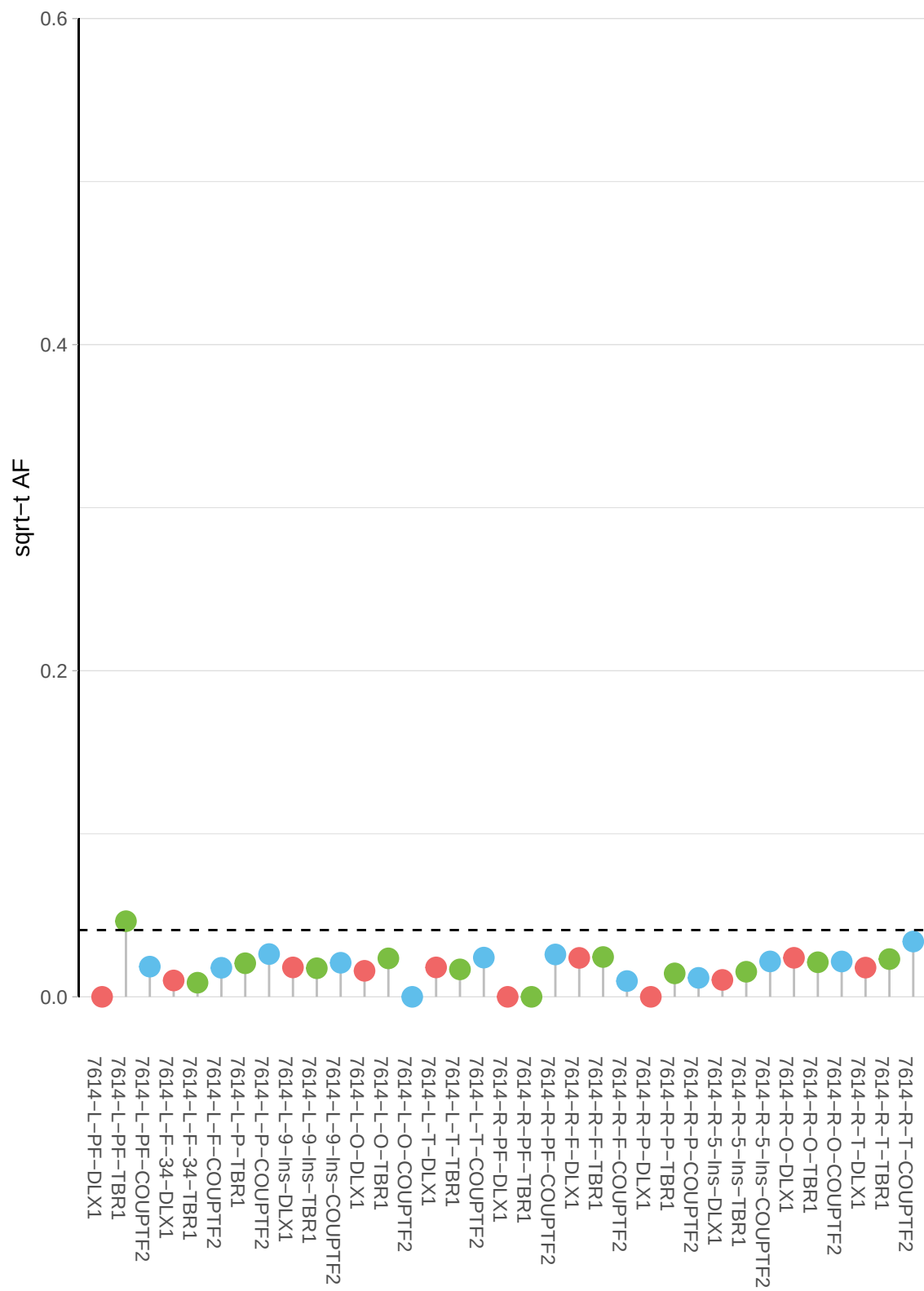

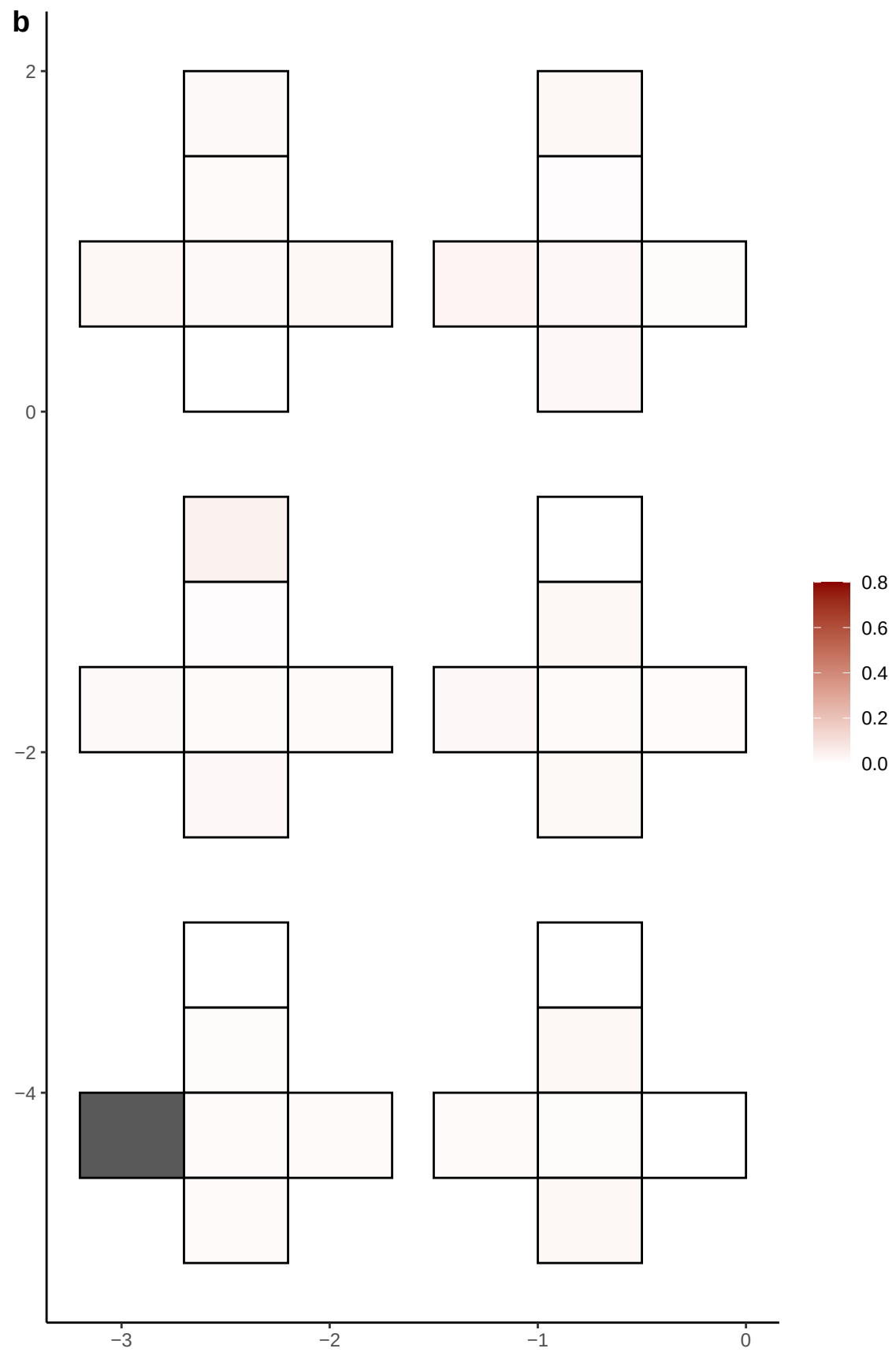

**a** ID01 14-59583360-A-G

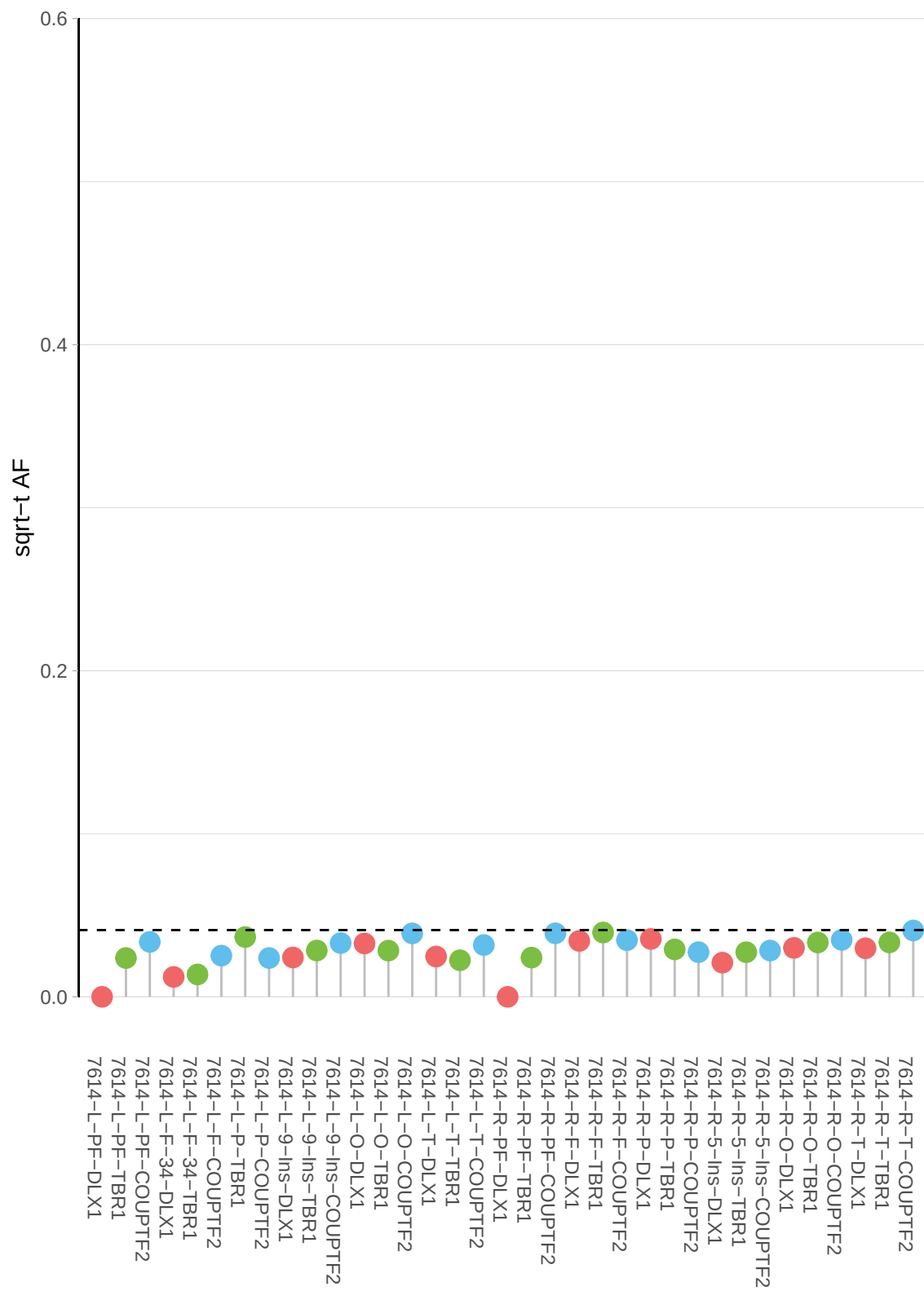

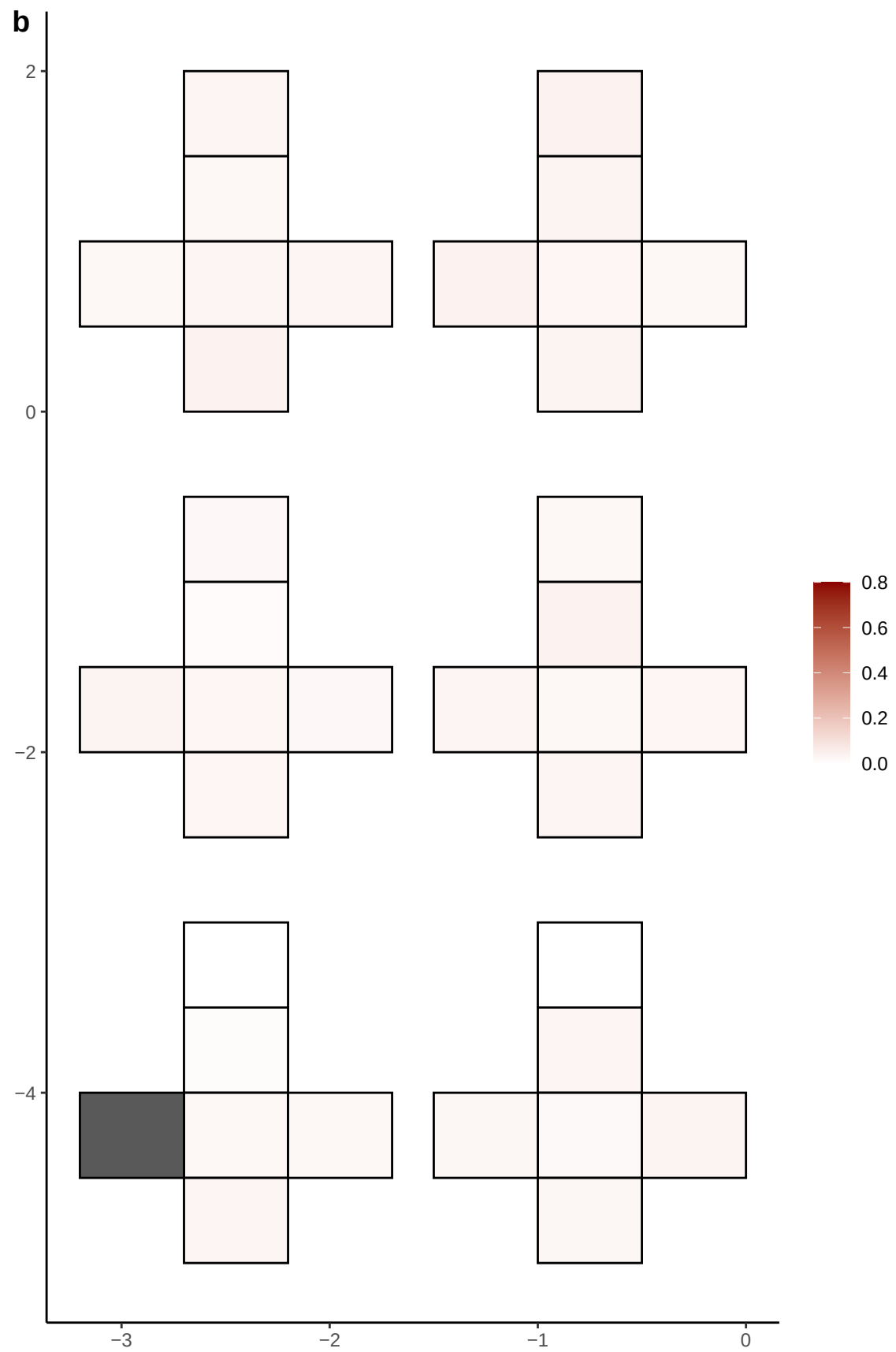

**a** ID01 14-65962309-A-C

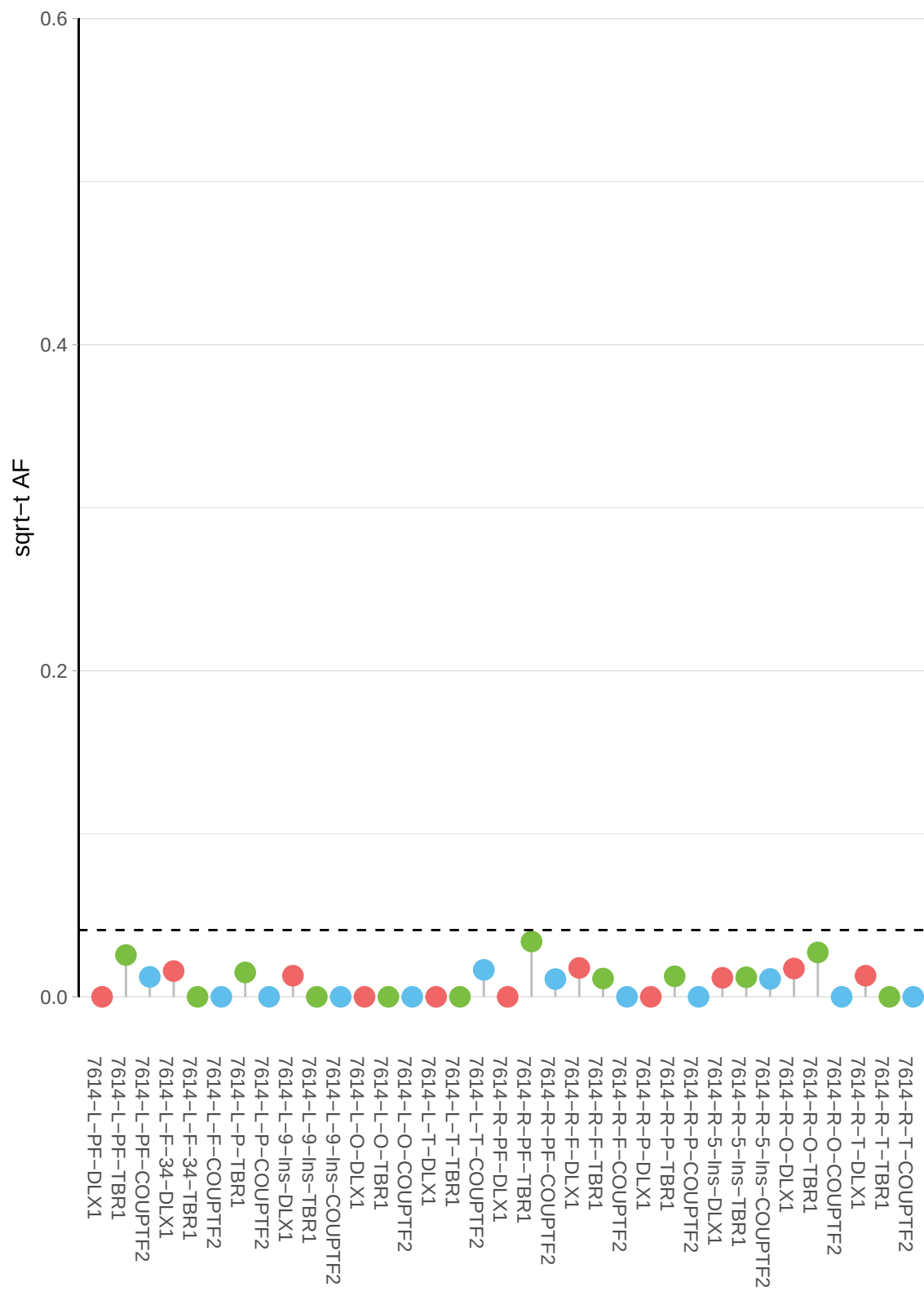

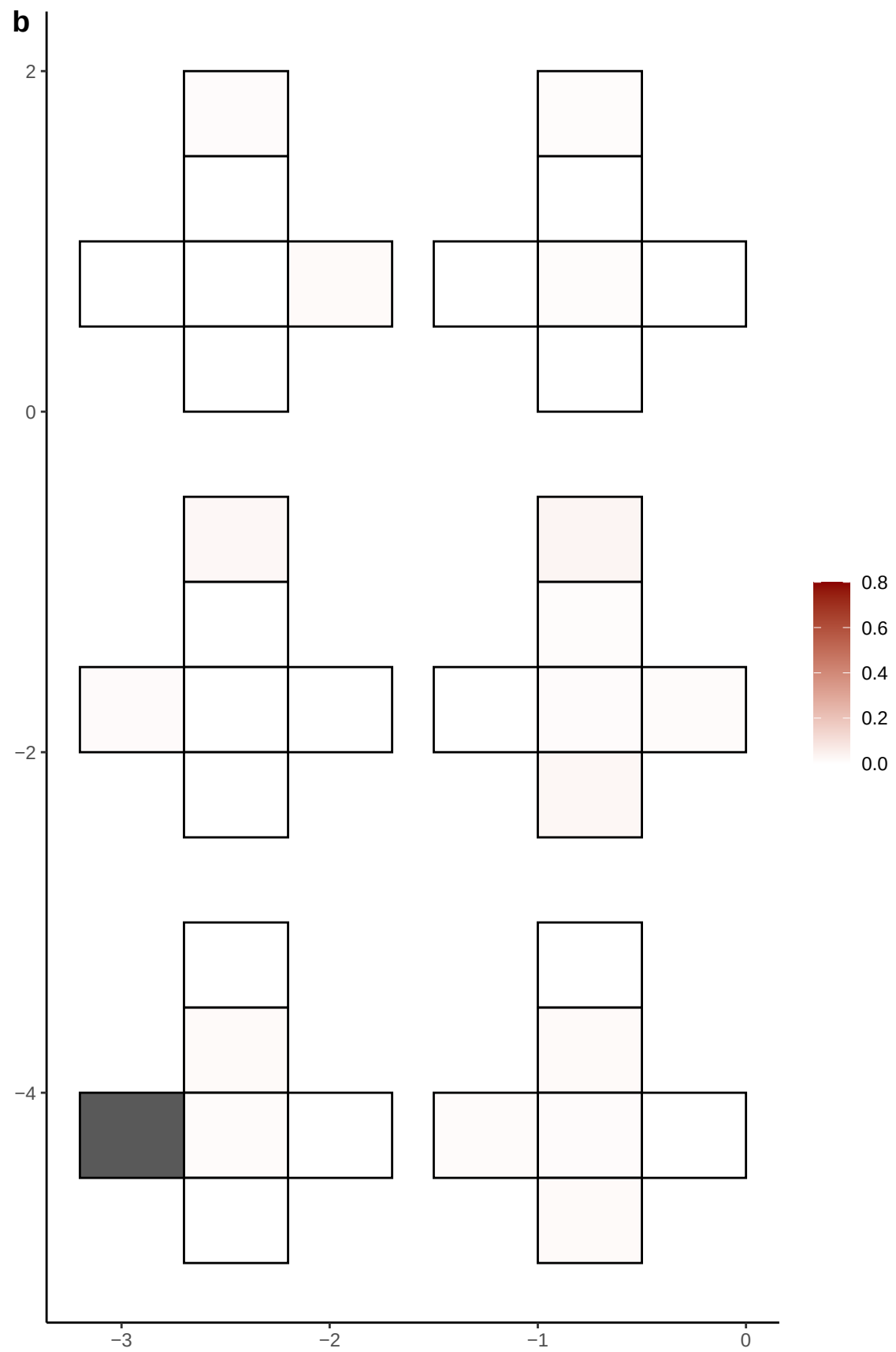

**a** ID01 14-78677839-C-T

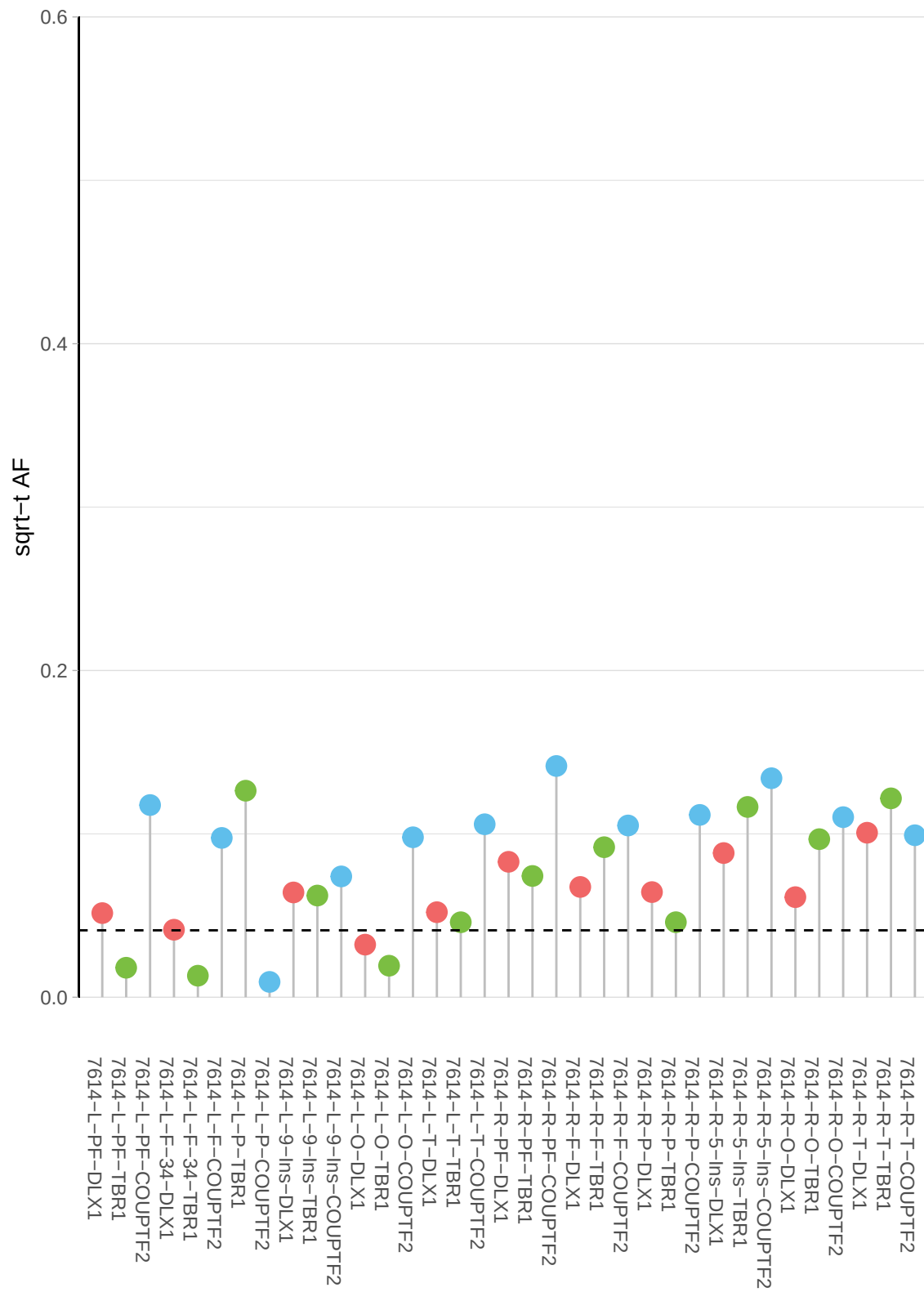

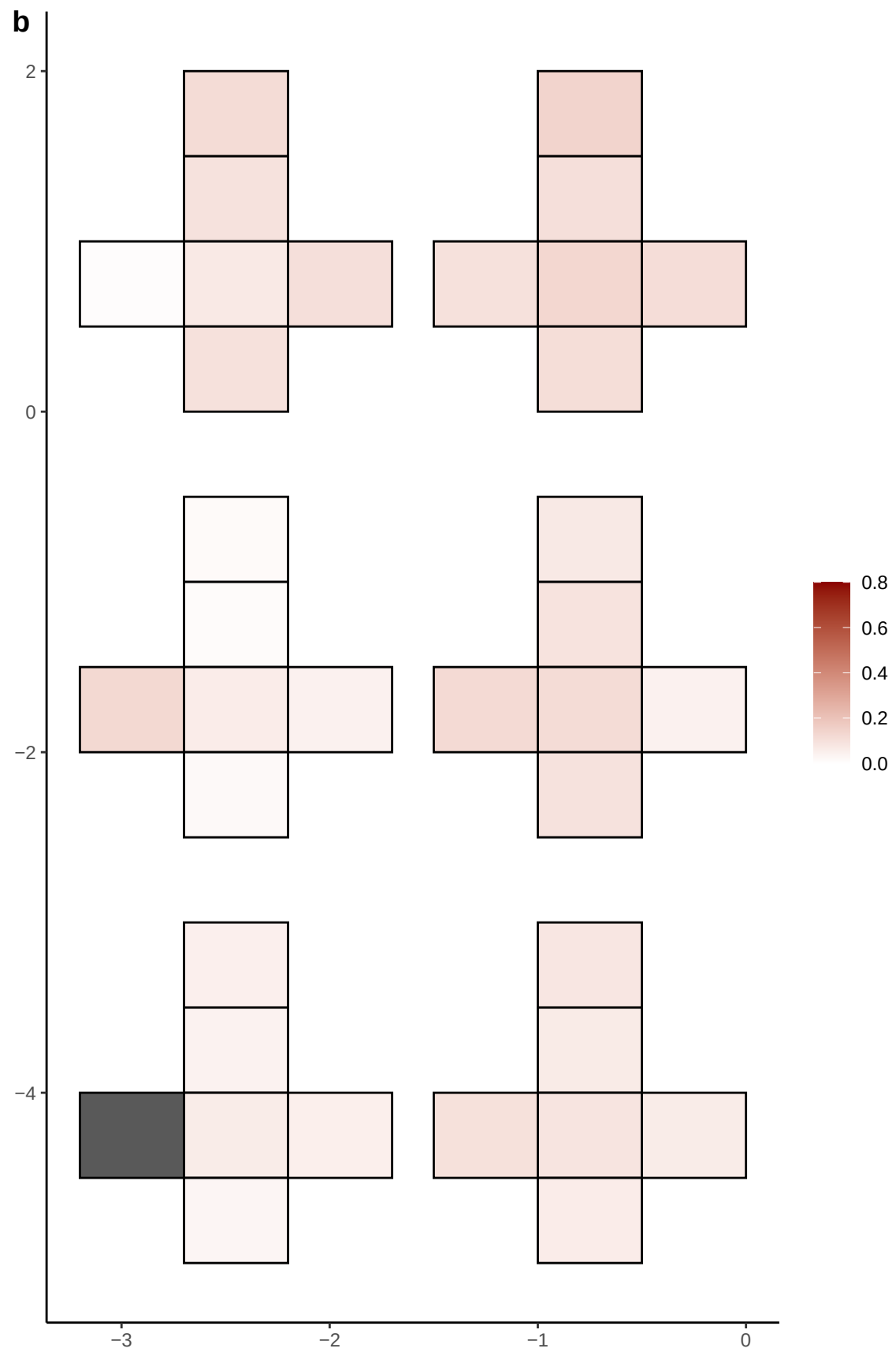

**a** ID01 14-82666170-G-A

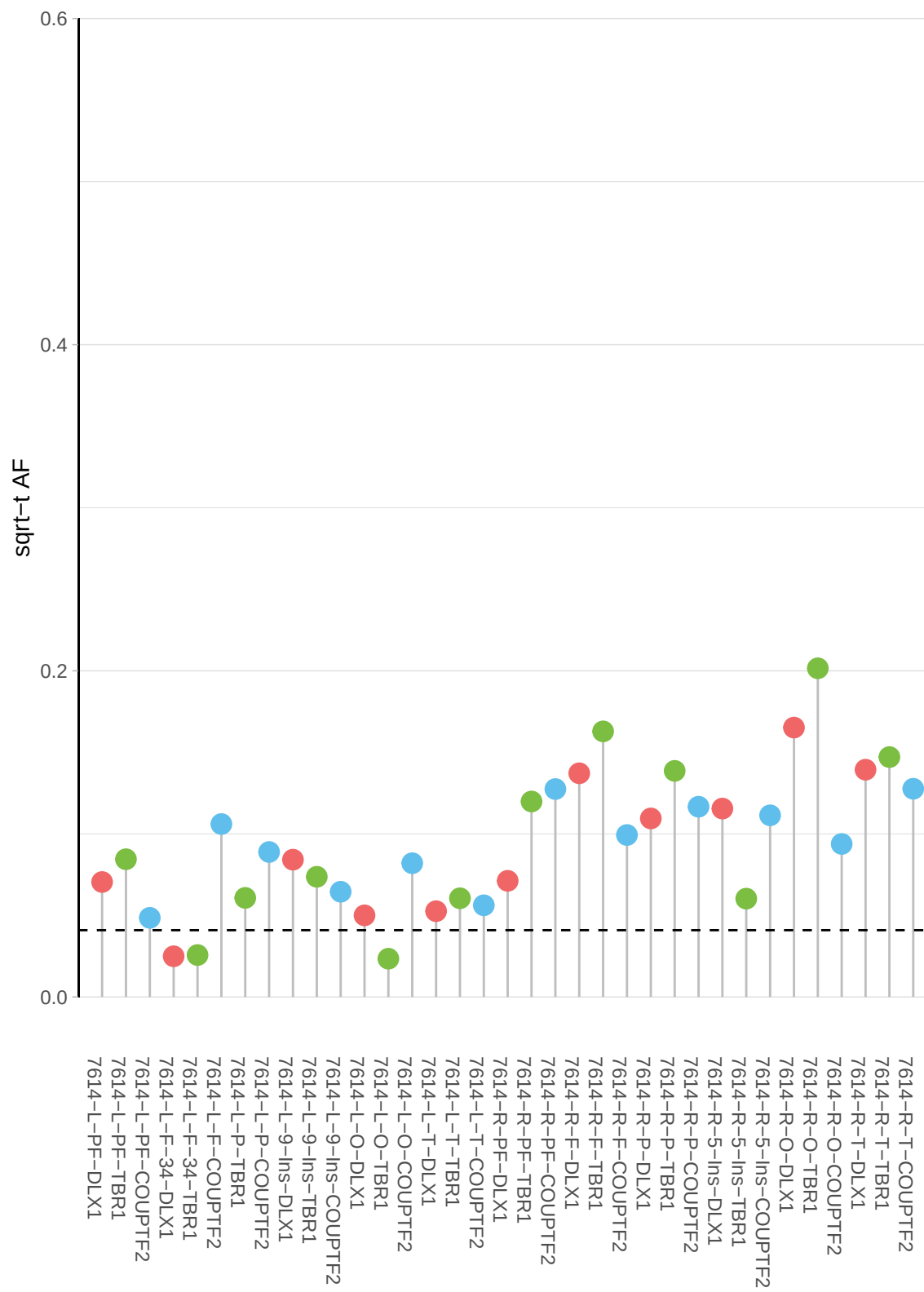

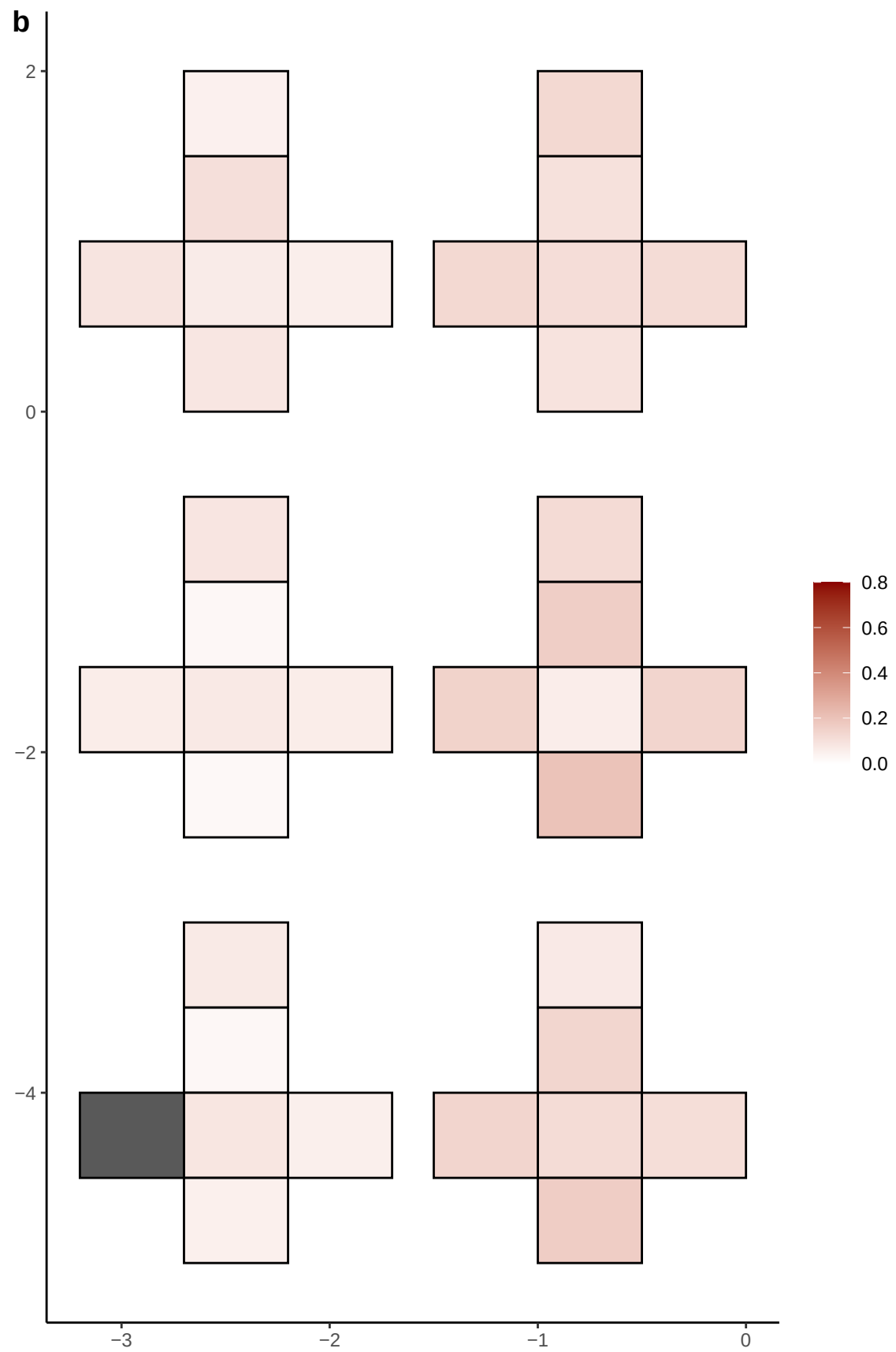

**a** ID01 15-42431069-T-C

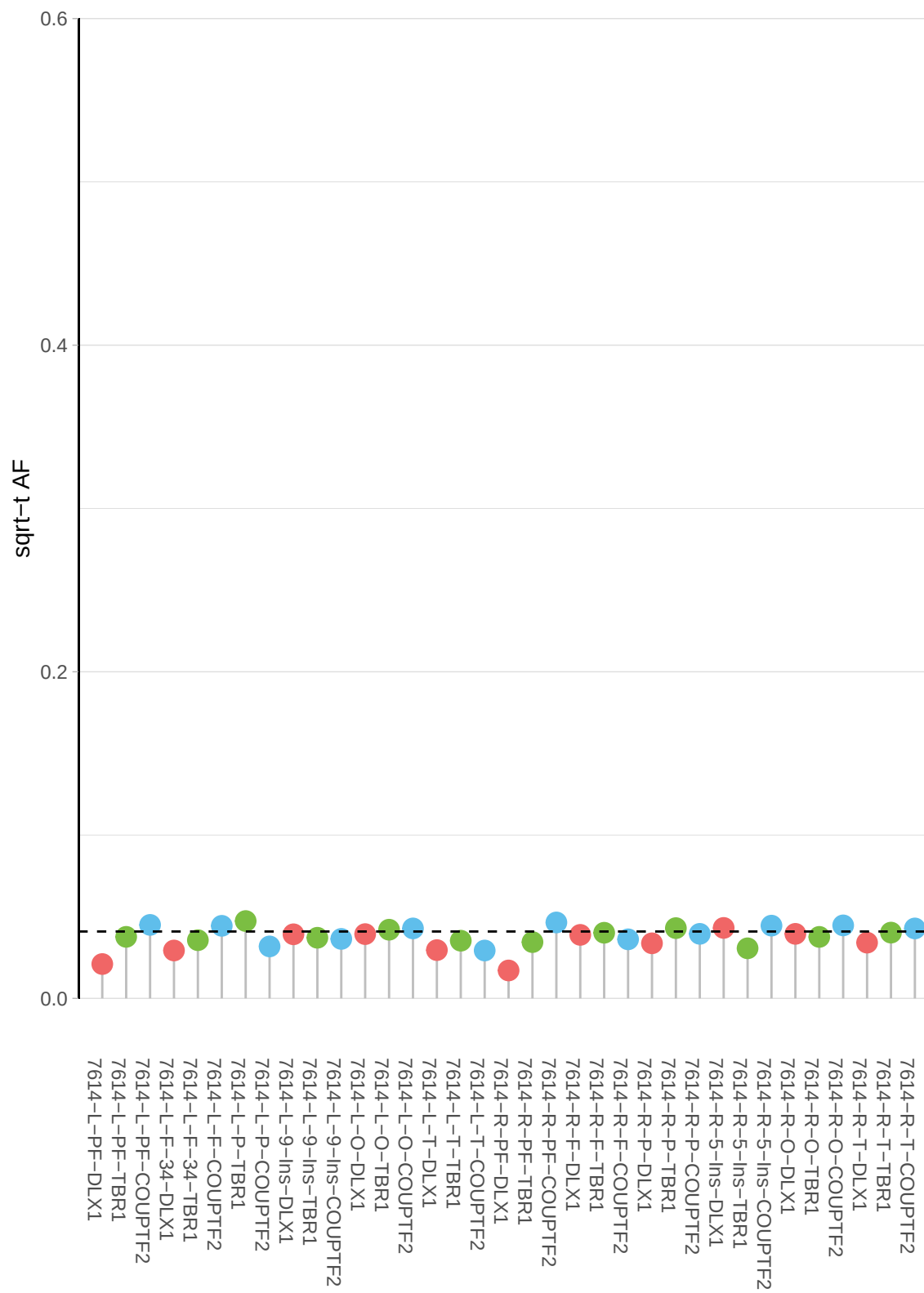

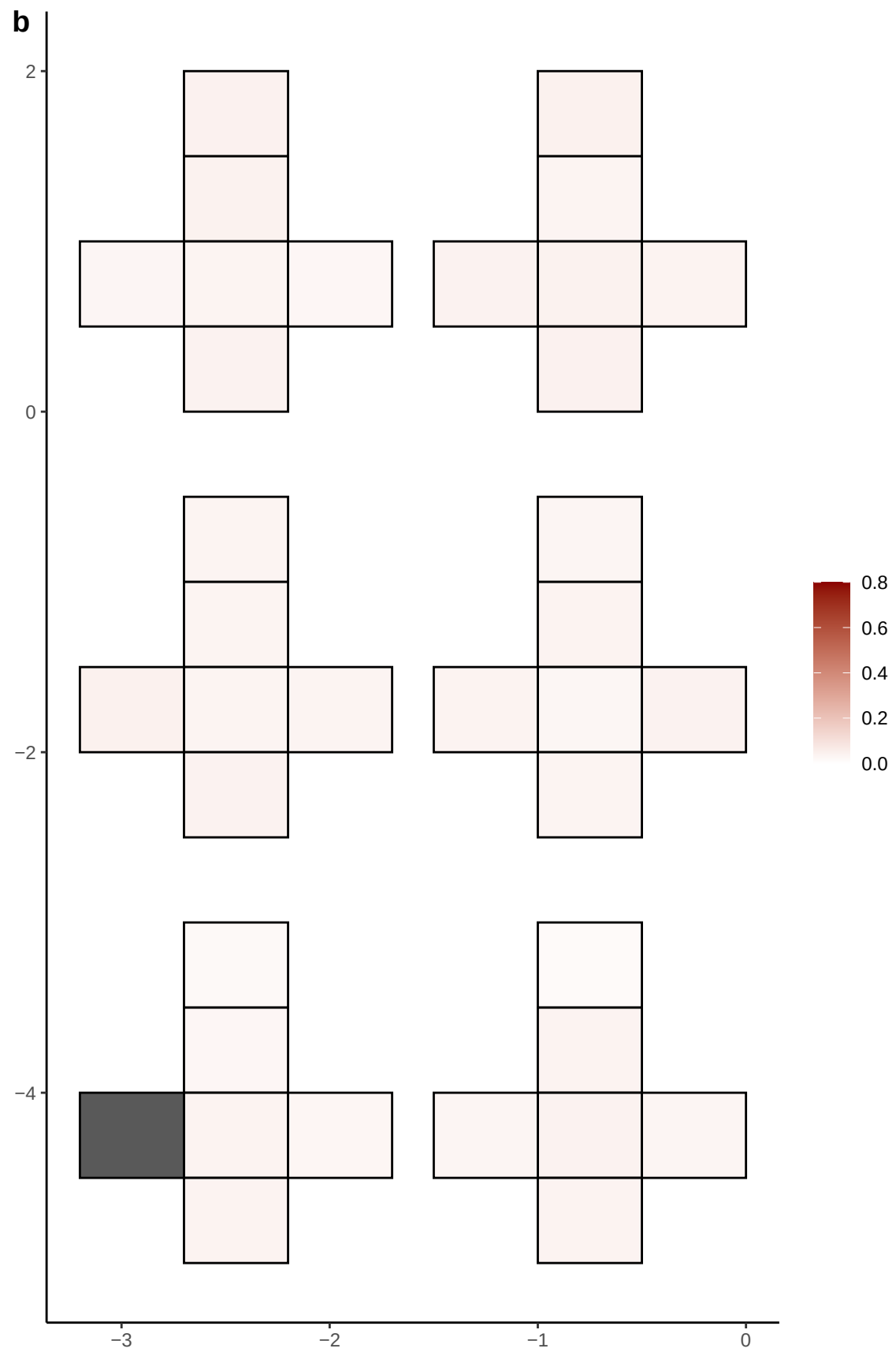

**a** ID01 15-65325135-G-A

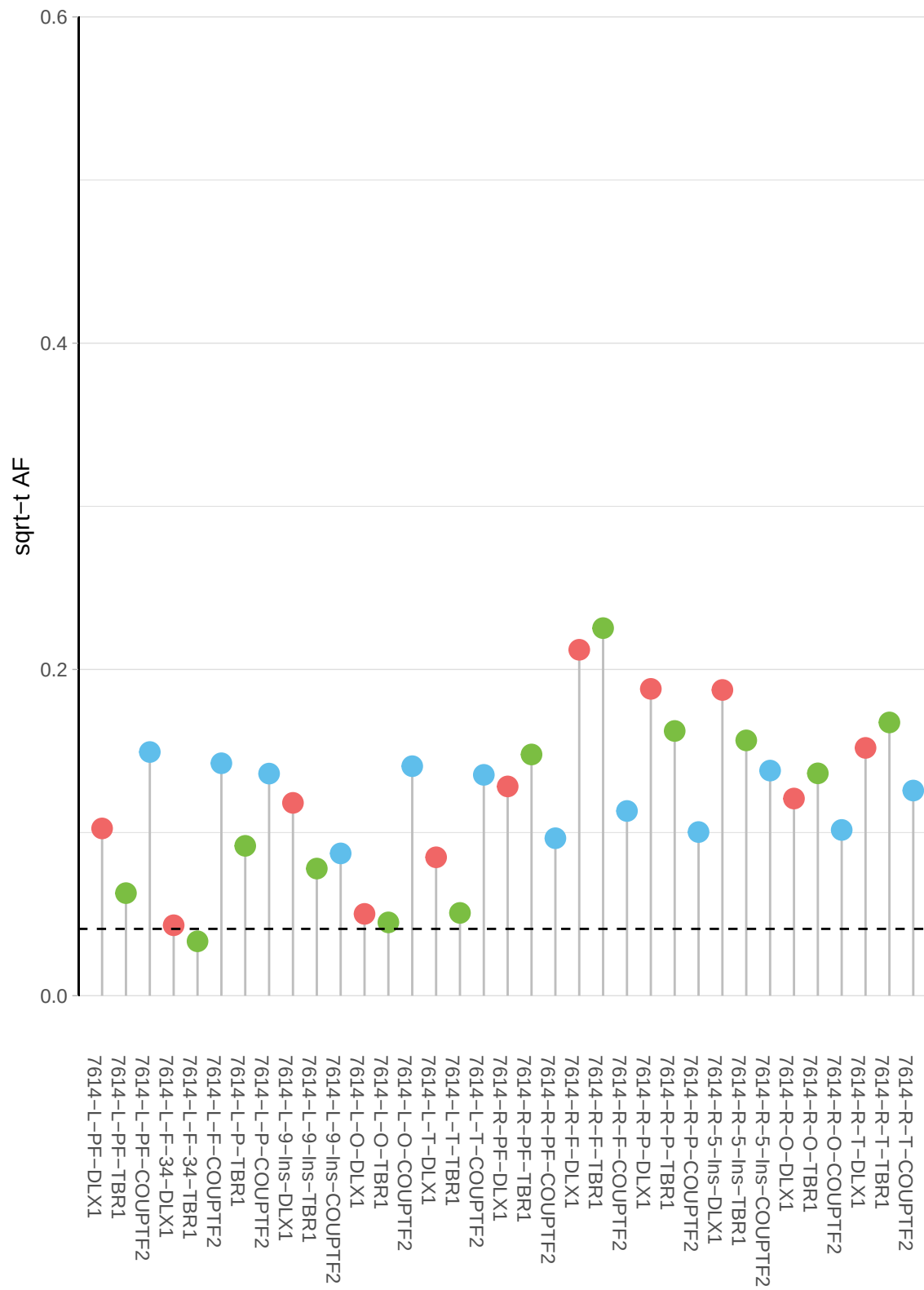

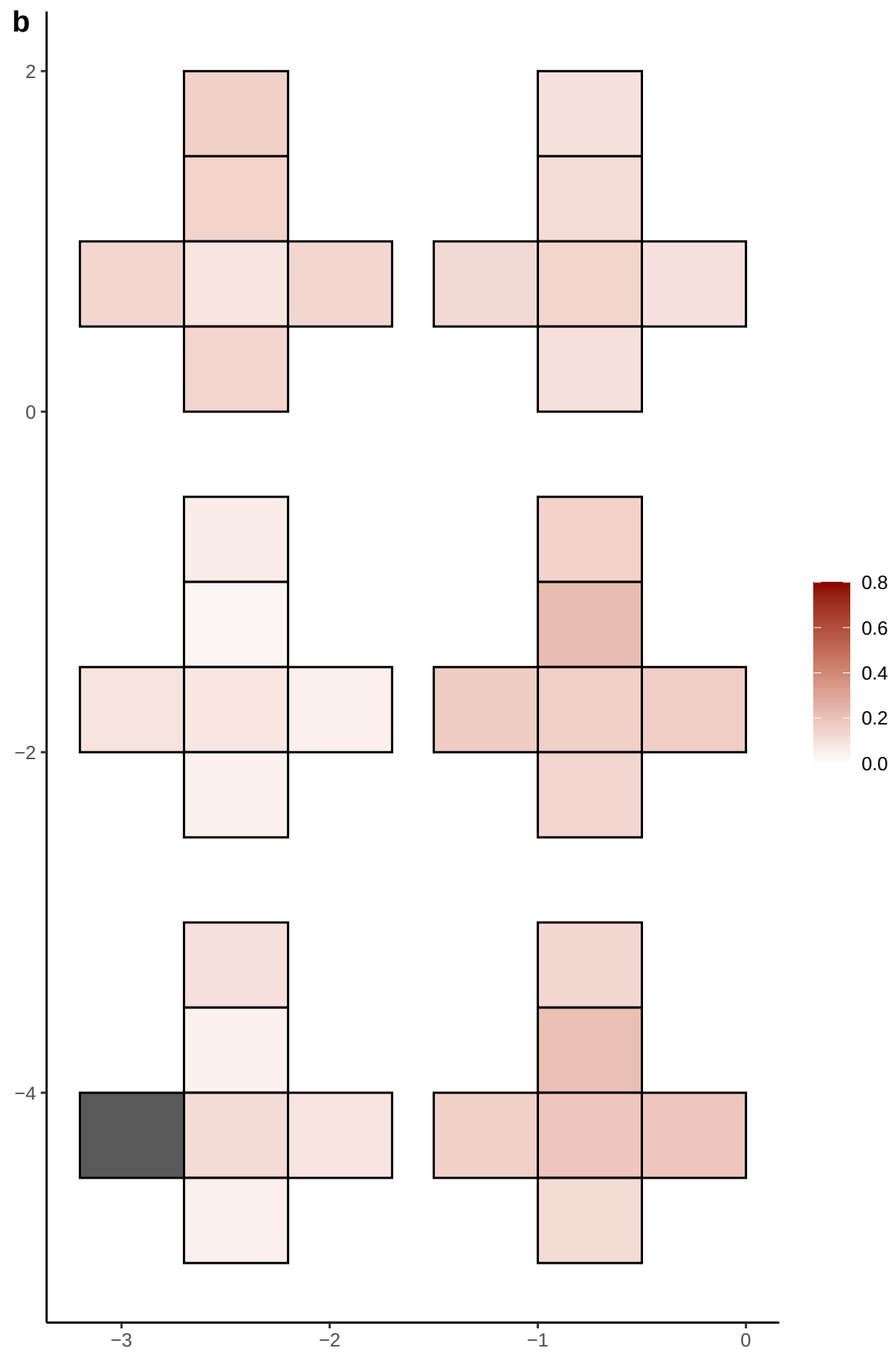

**a** ID01 15-89933068-T-G

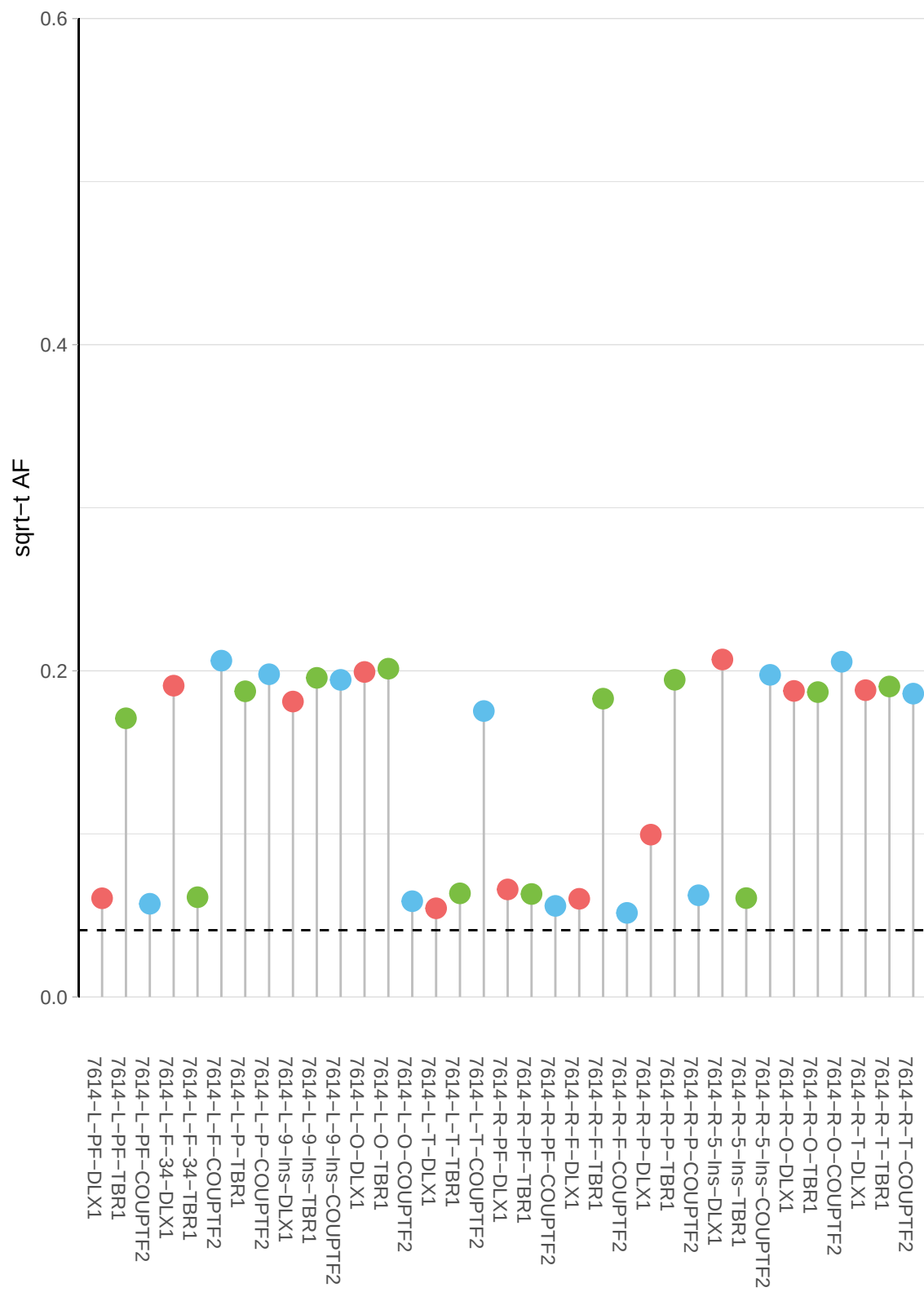

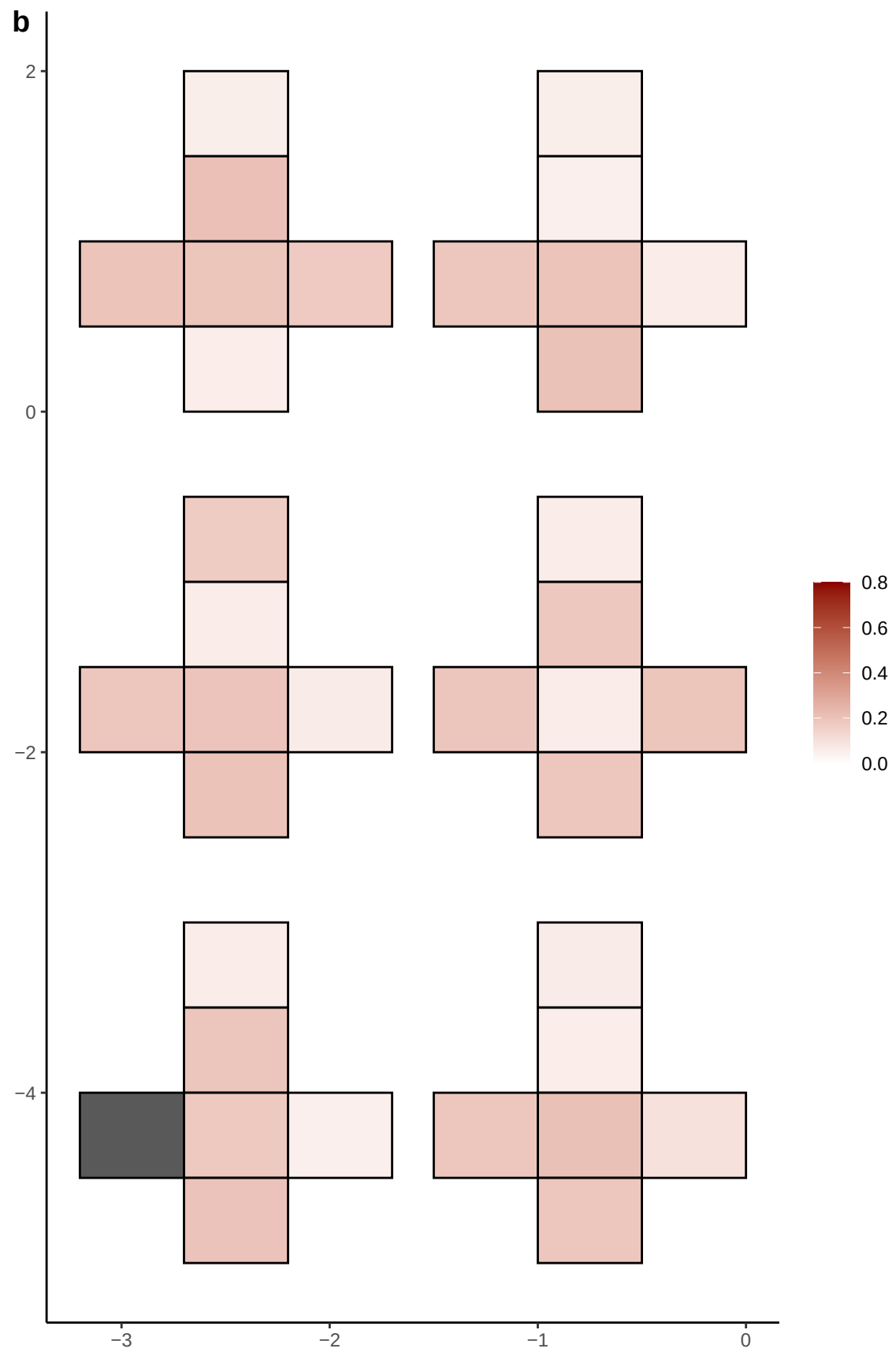

**a** ID01 15-93474236-T-G

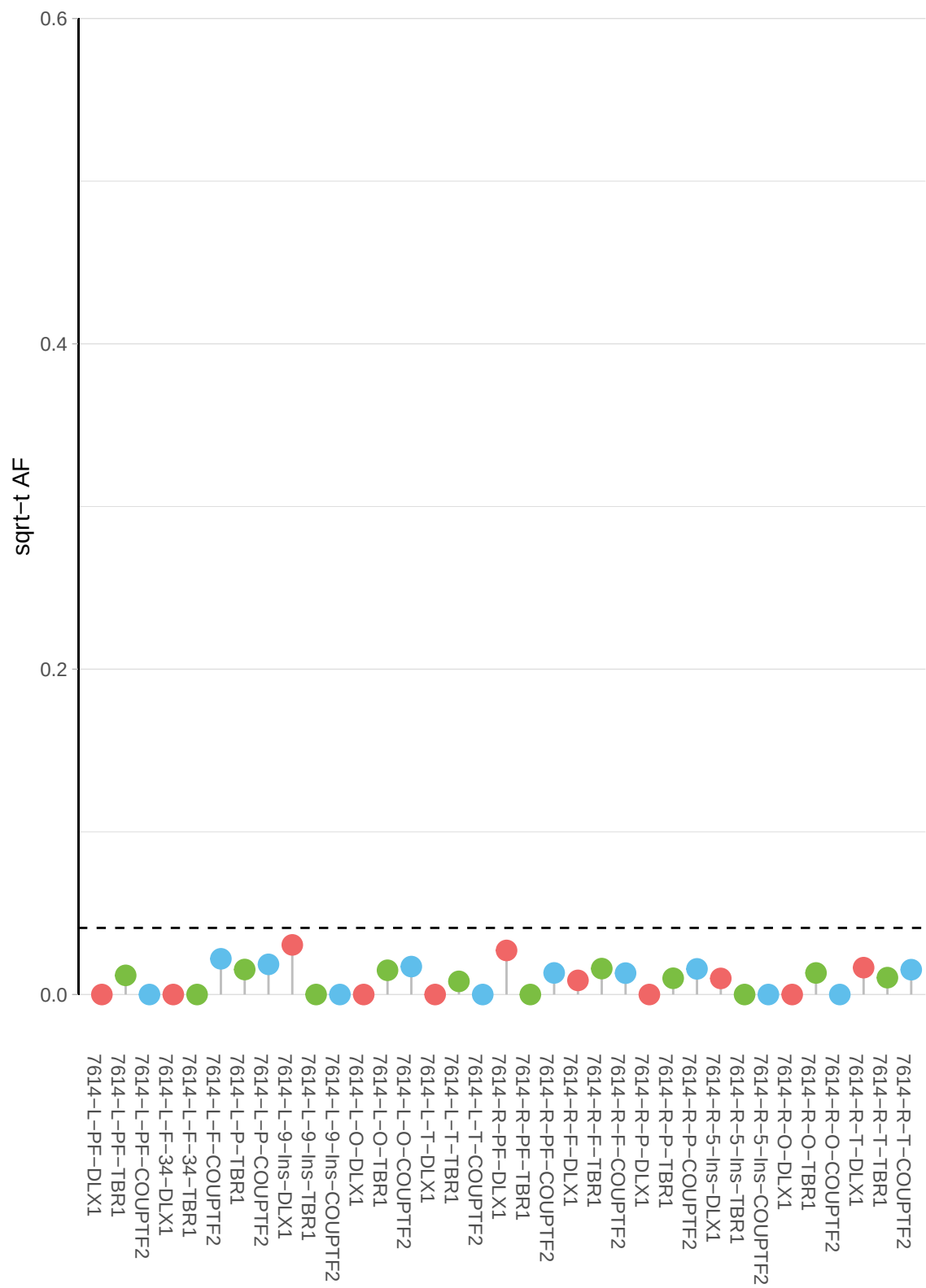

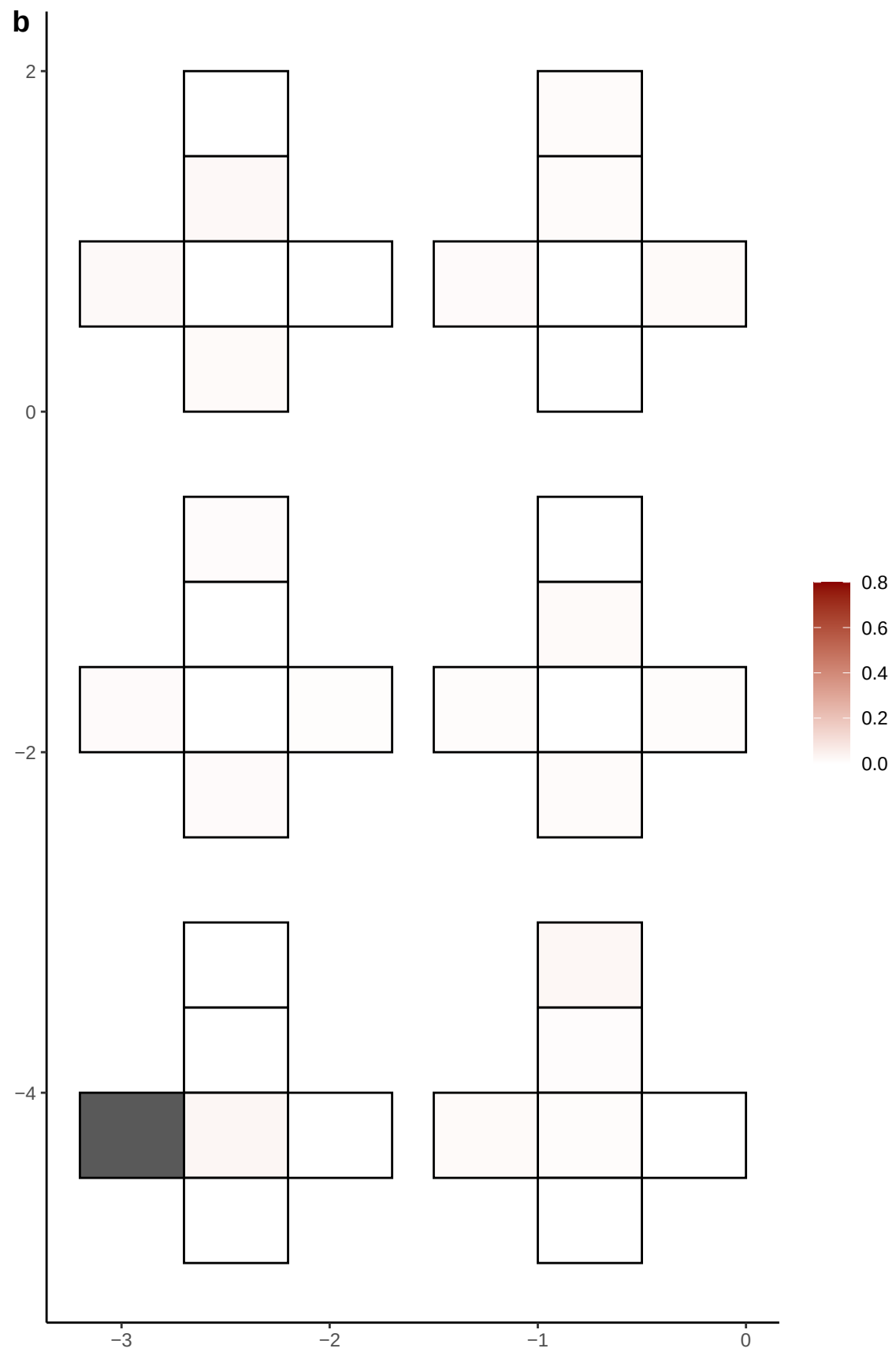

**a** ID01 16-27872227-G-A

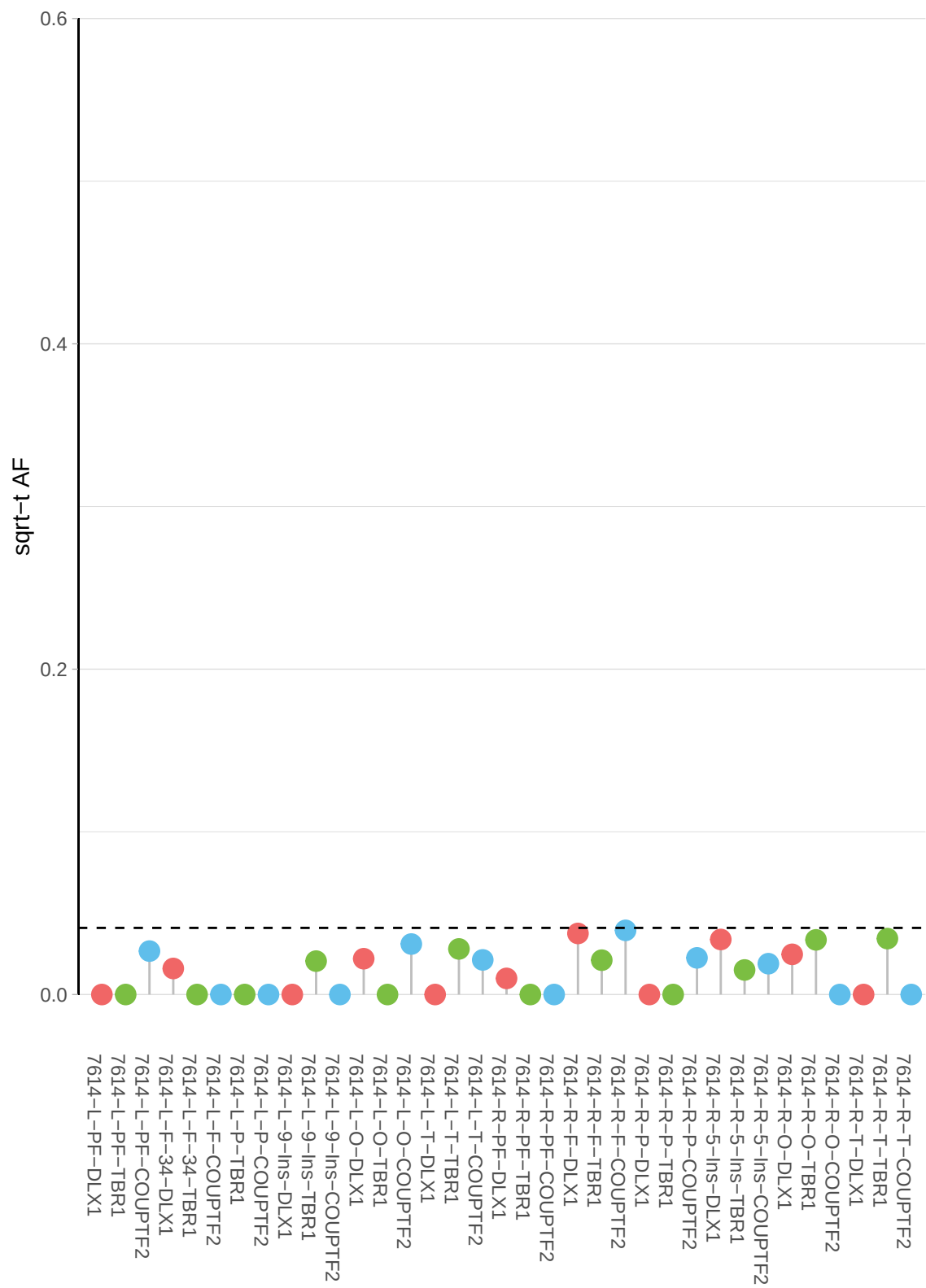

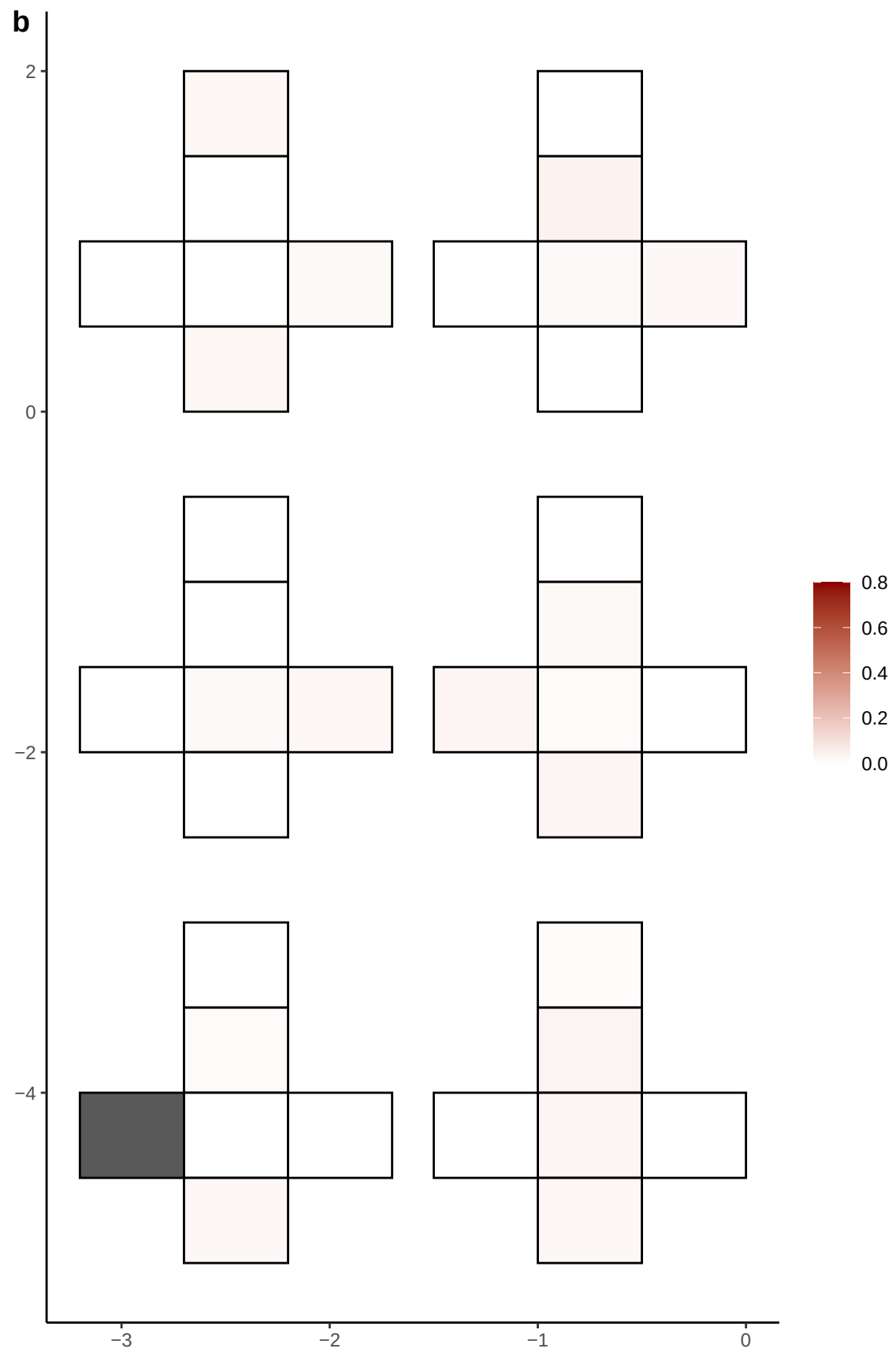

**a** ID01 16-27878370-C-T

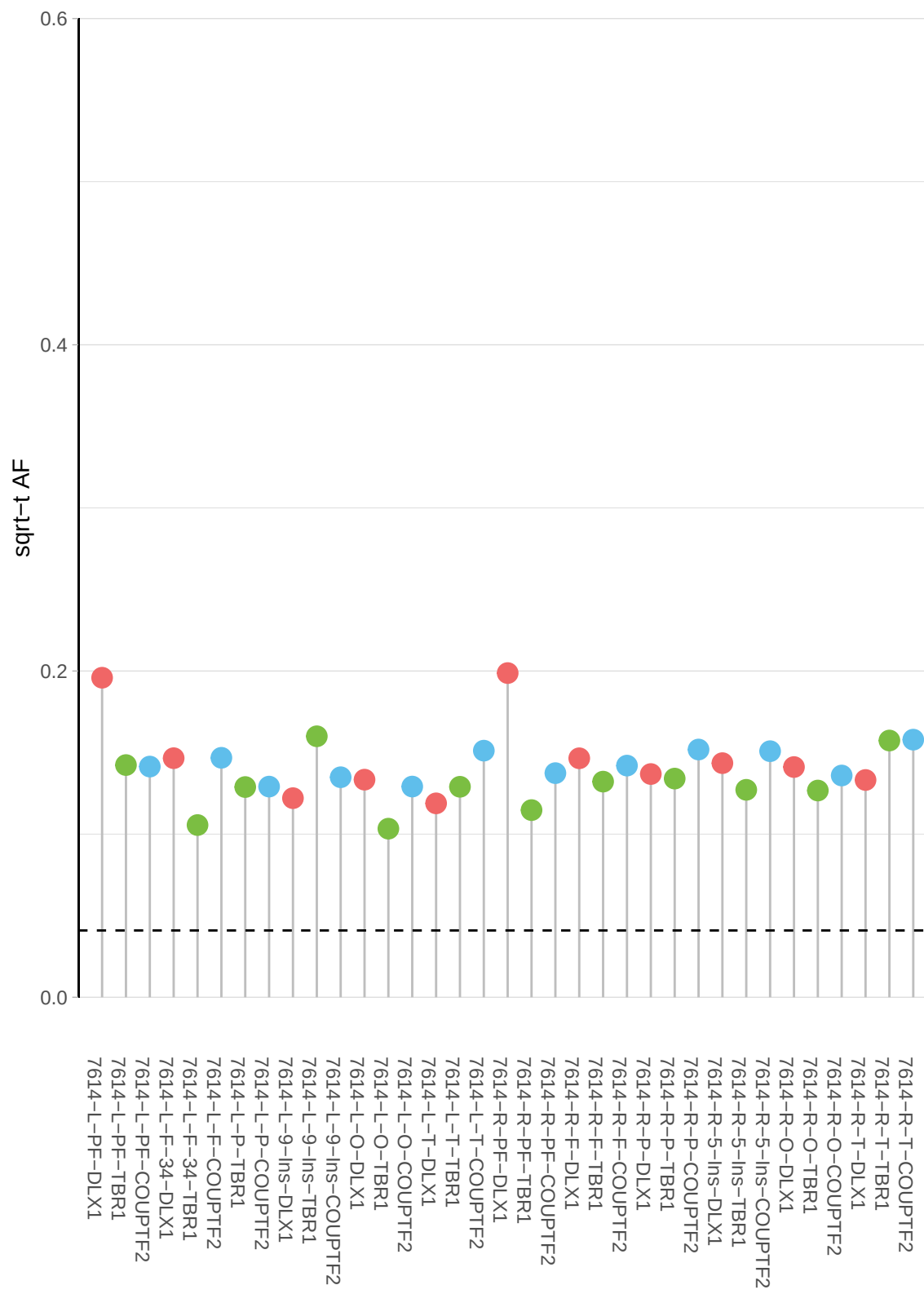

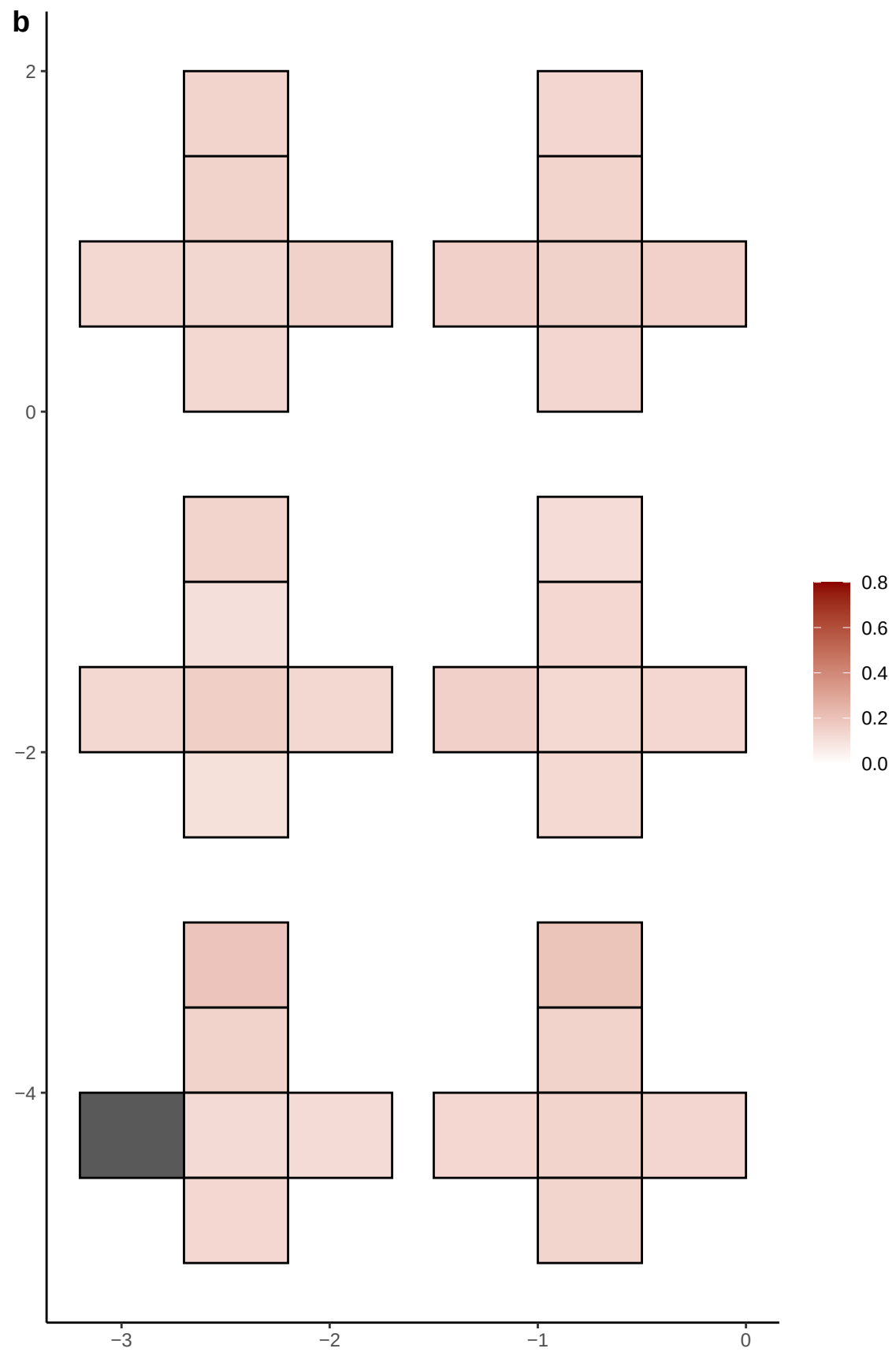

**a** ID01 16-27878380-T-C

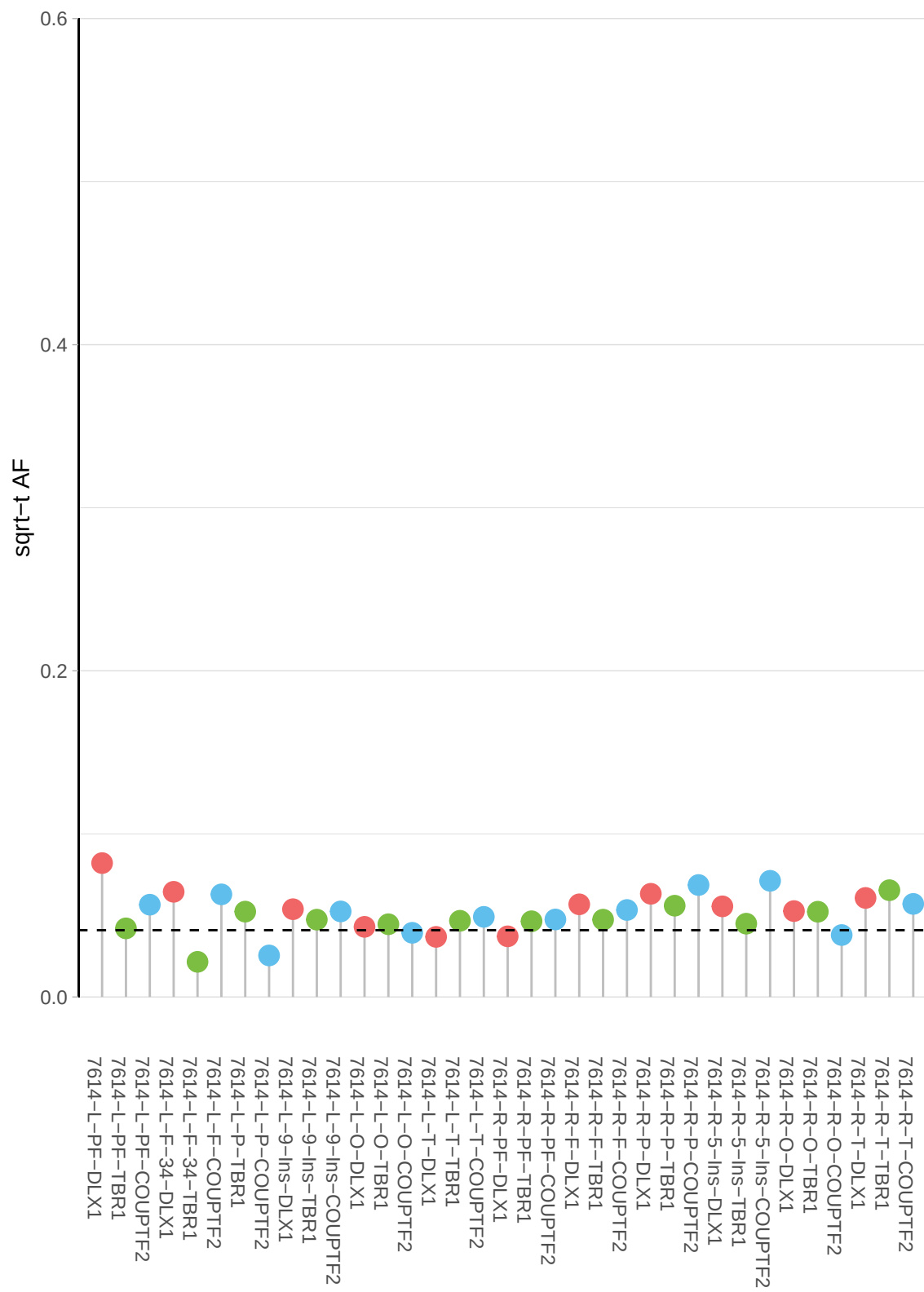

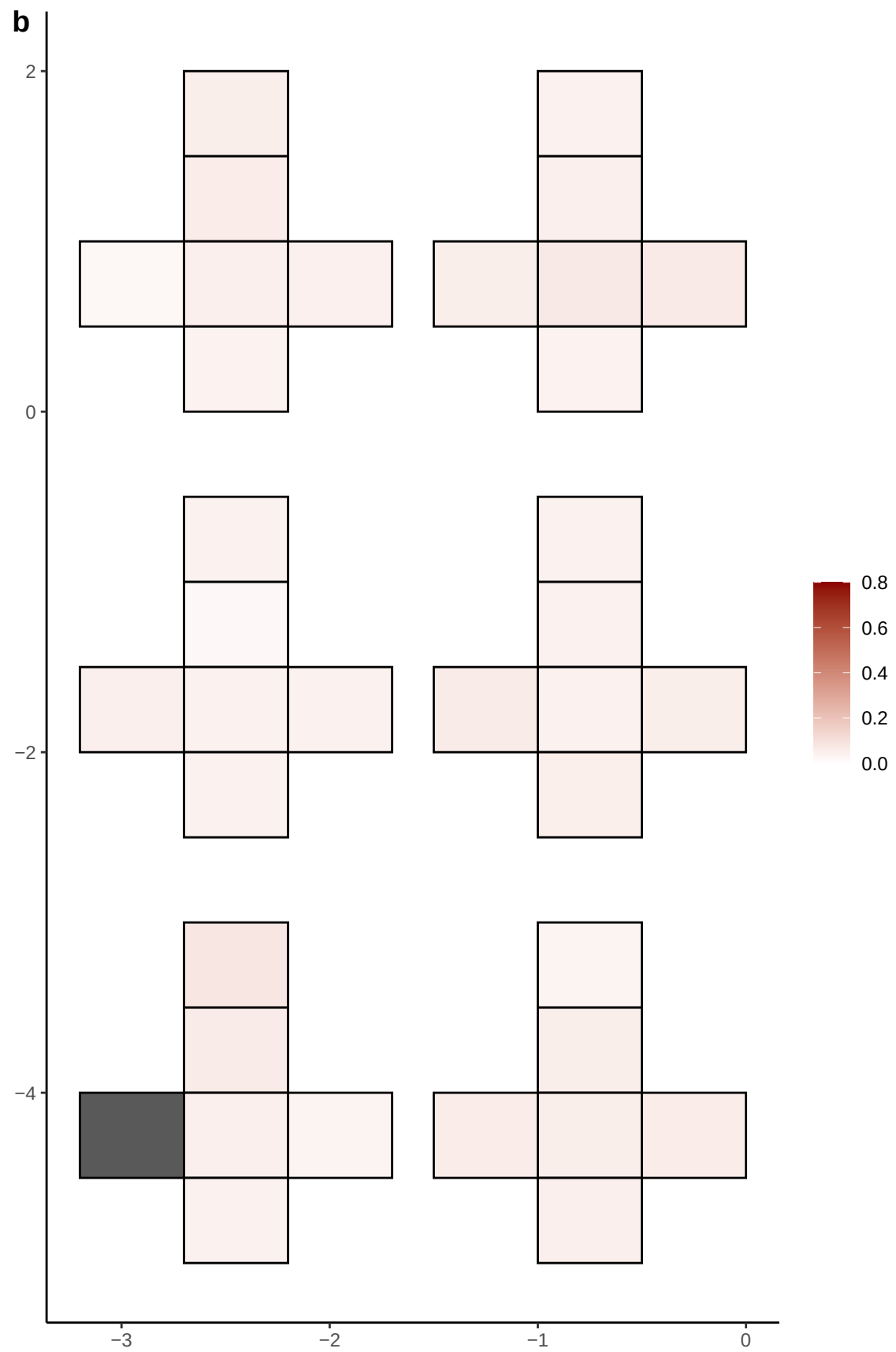

**a** ID01 16-28153814-G-A

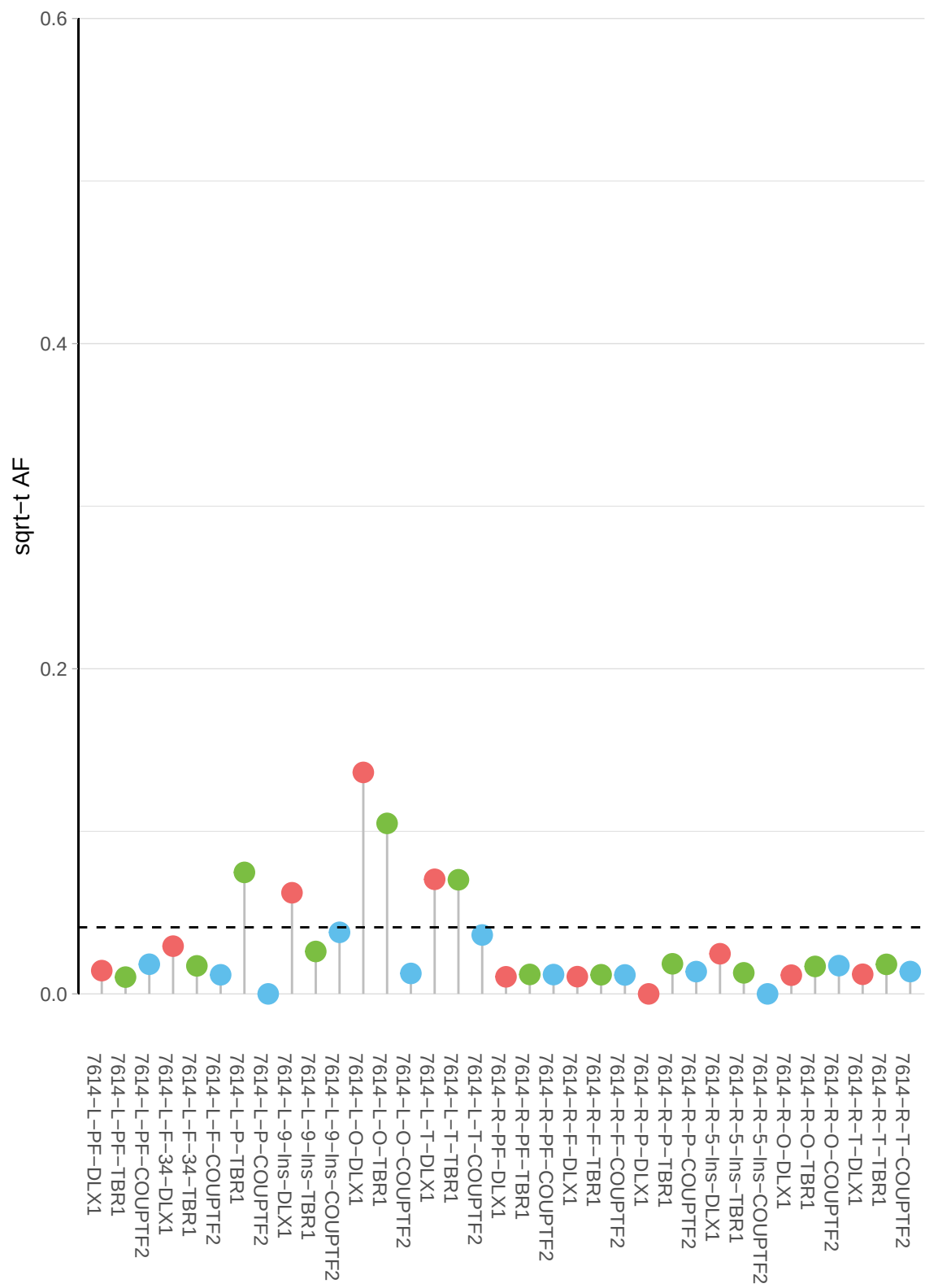

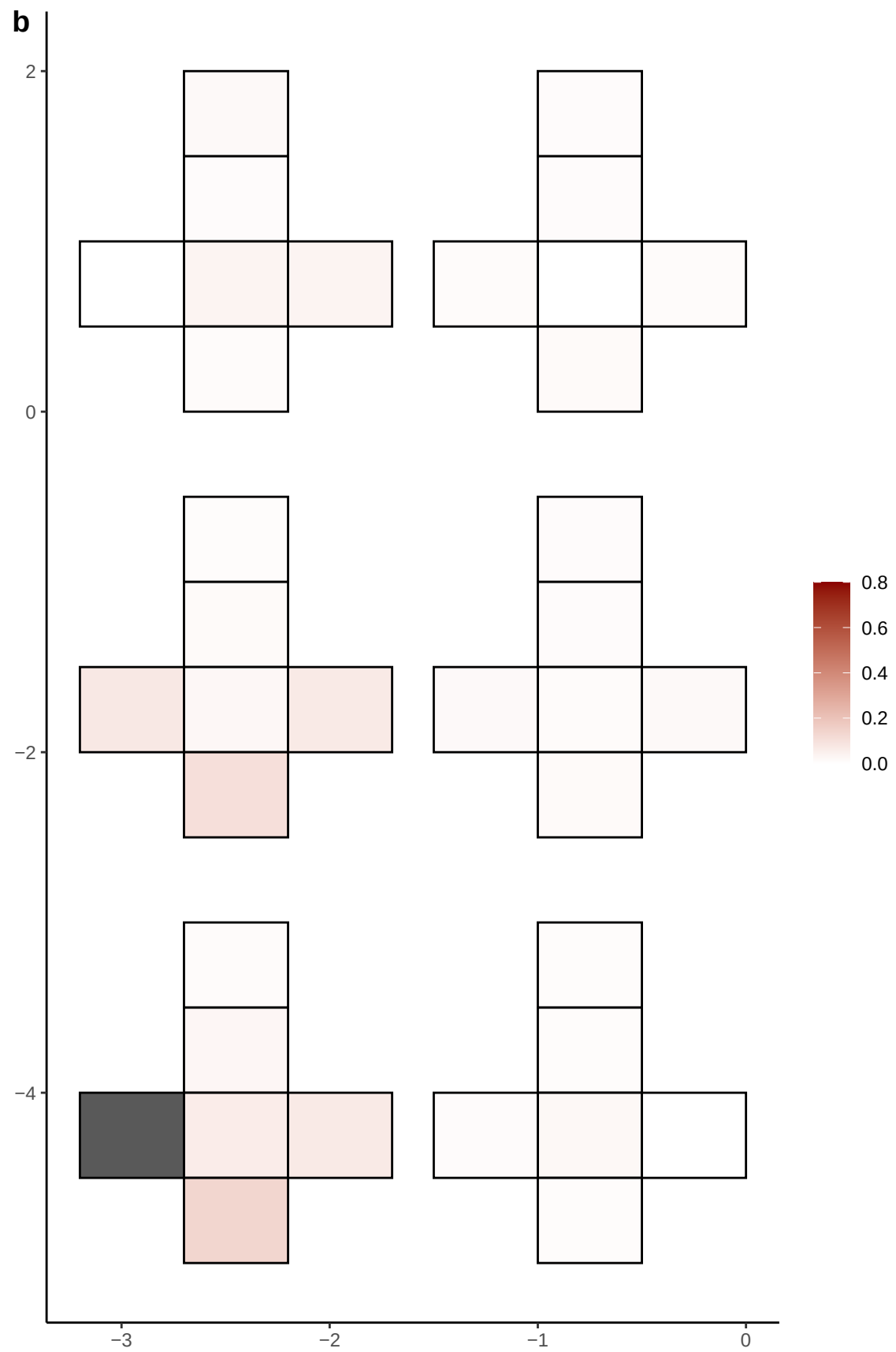

**a** ID01 16-58426889-C-A

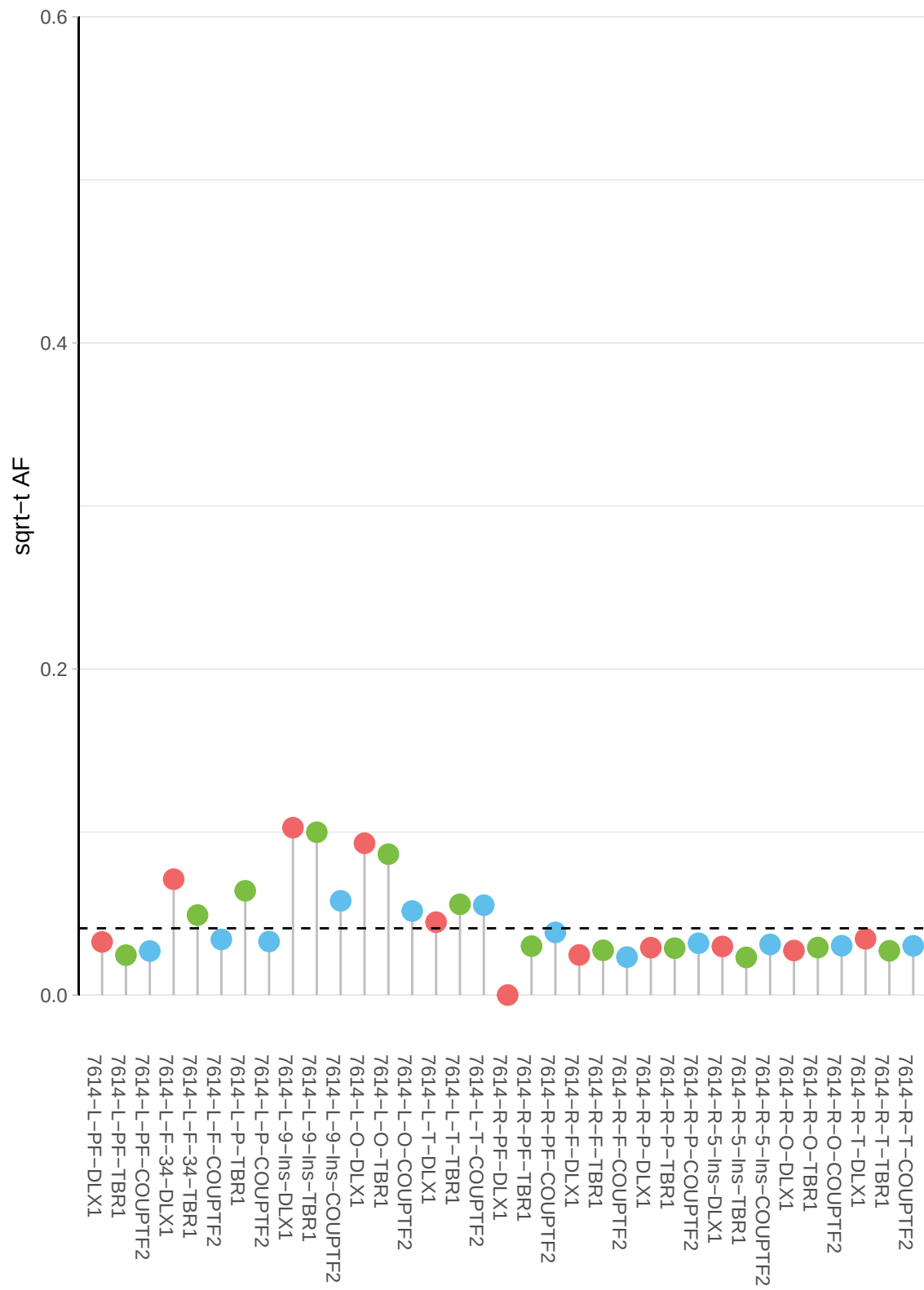

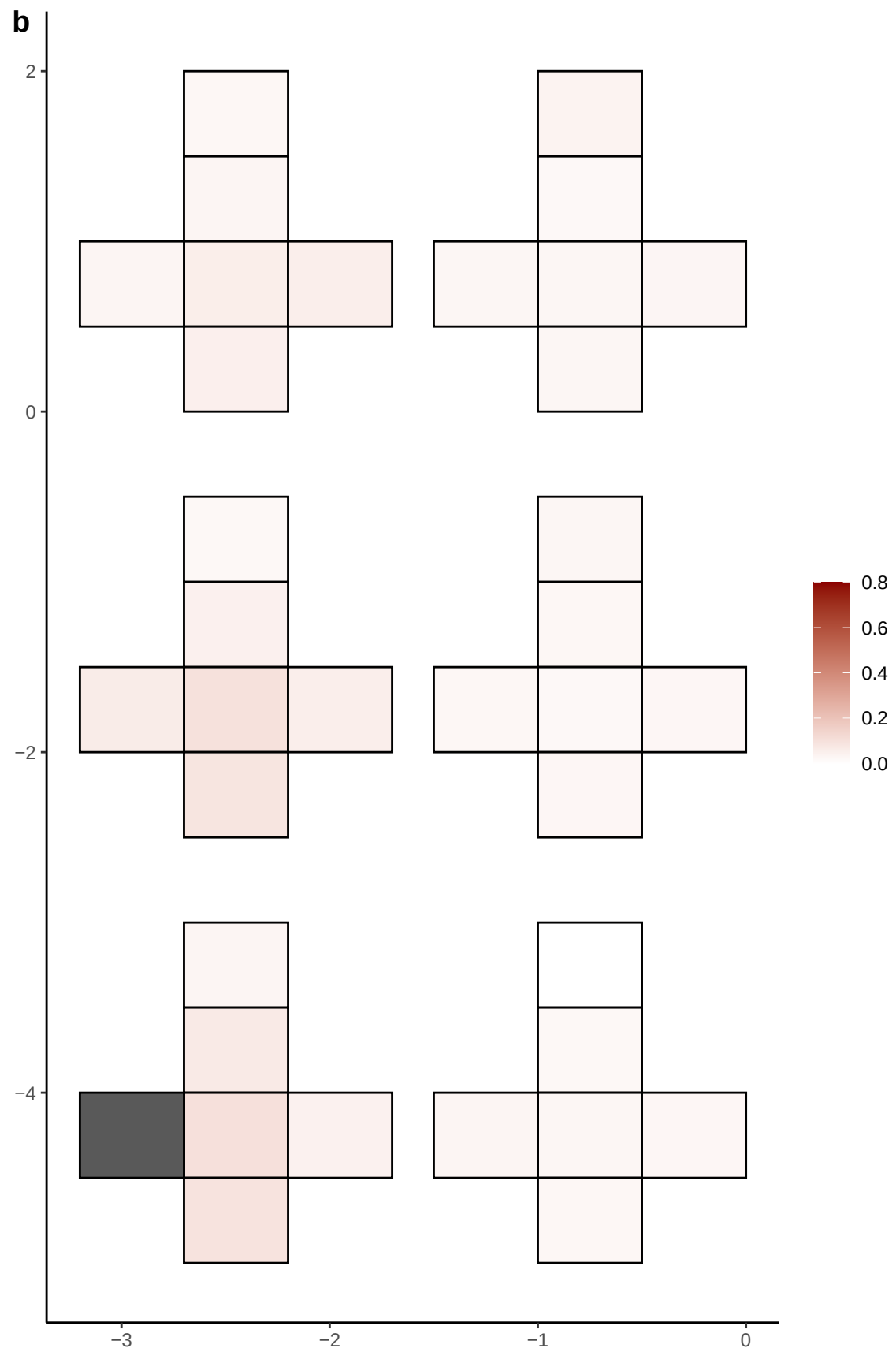

**a** ID01 16-69679204-G-C

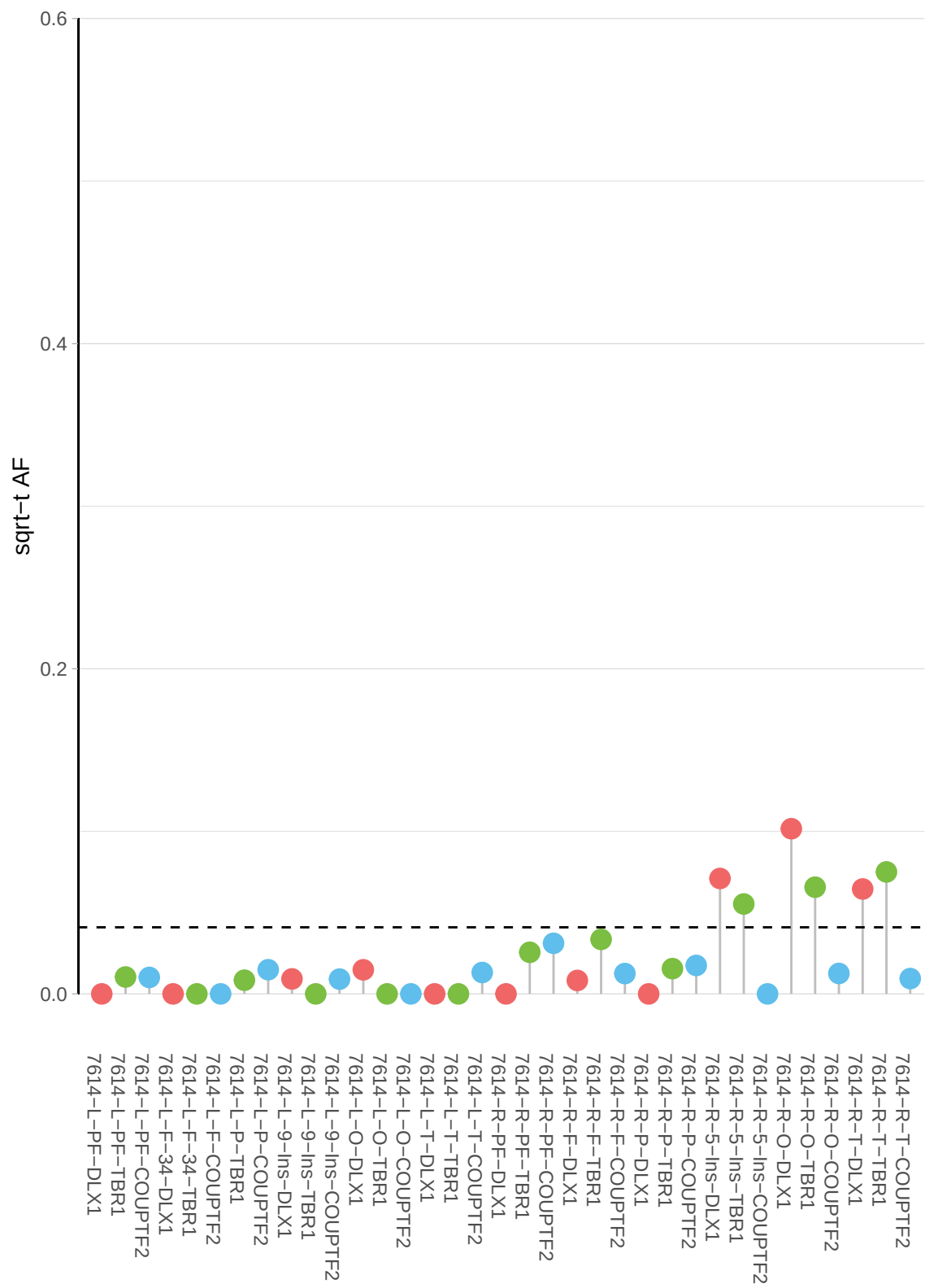

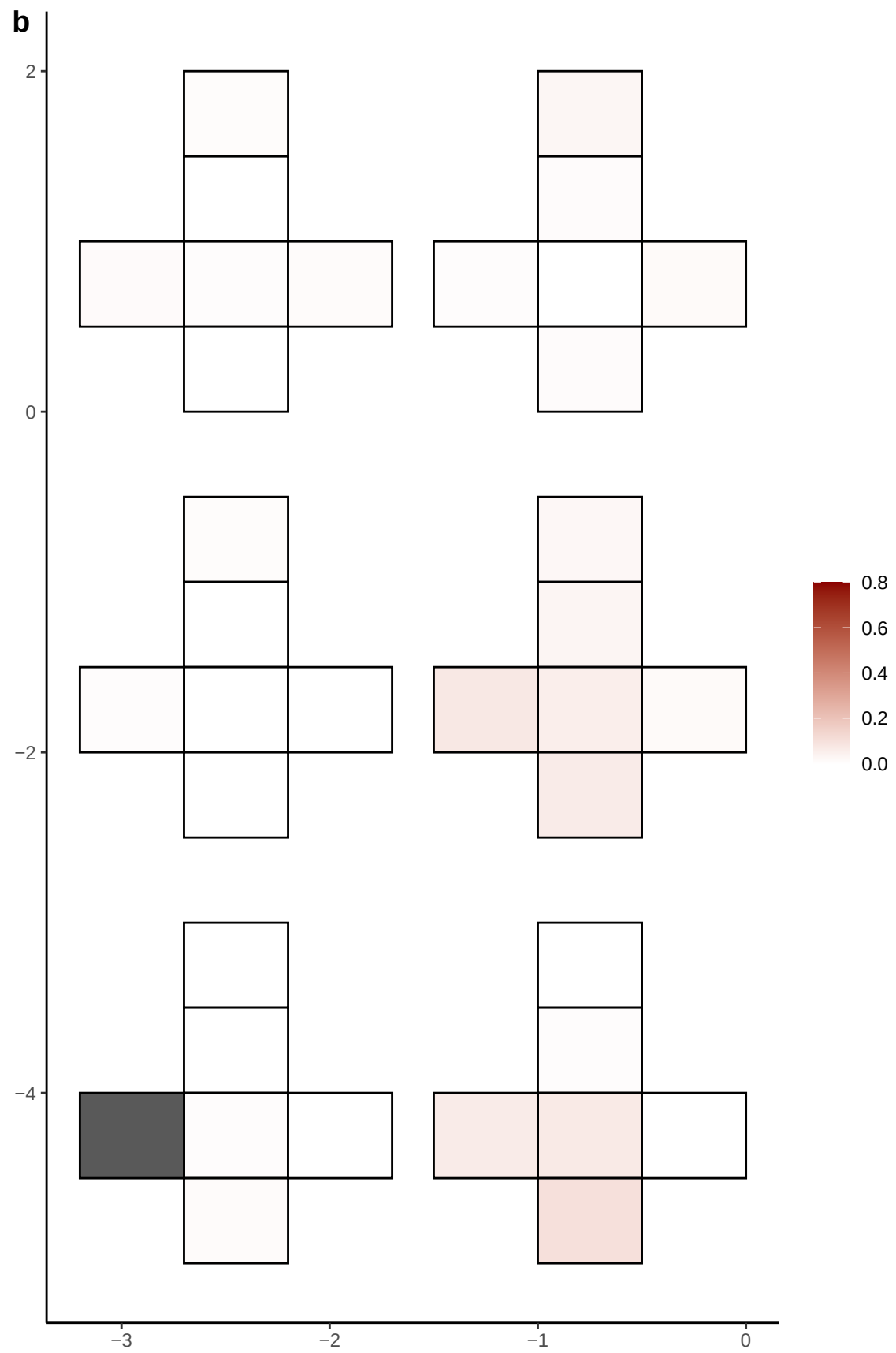

**a** ID01 16-73458900-G-A

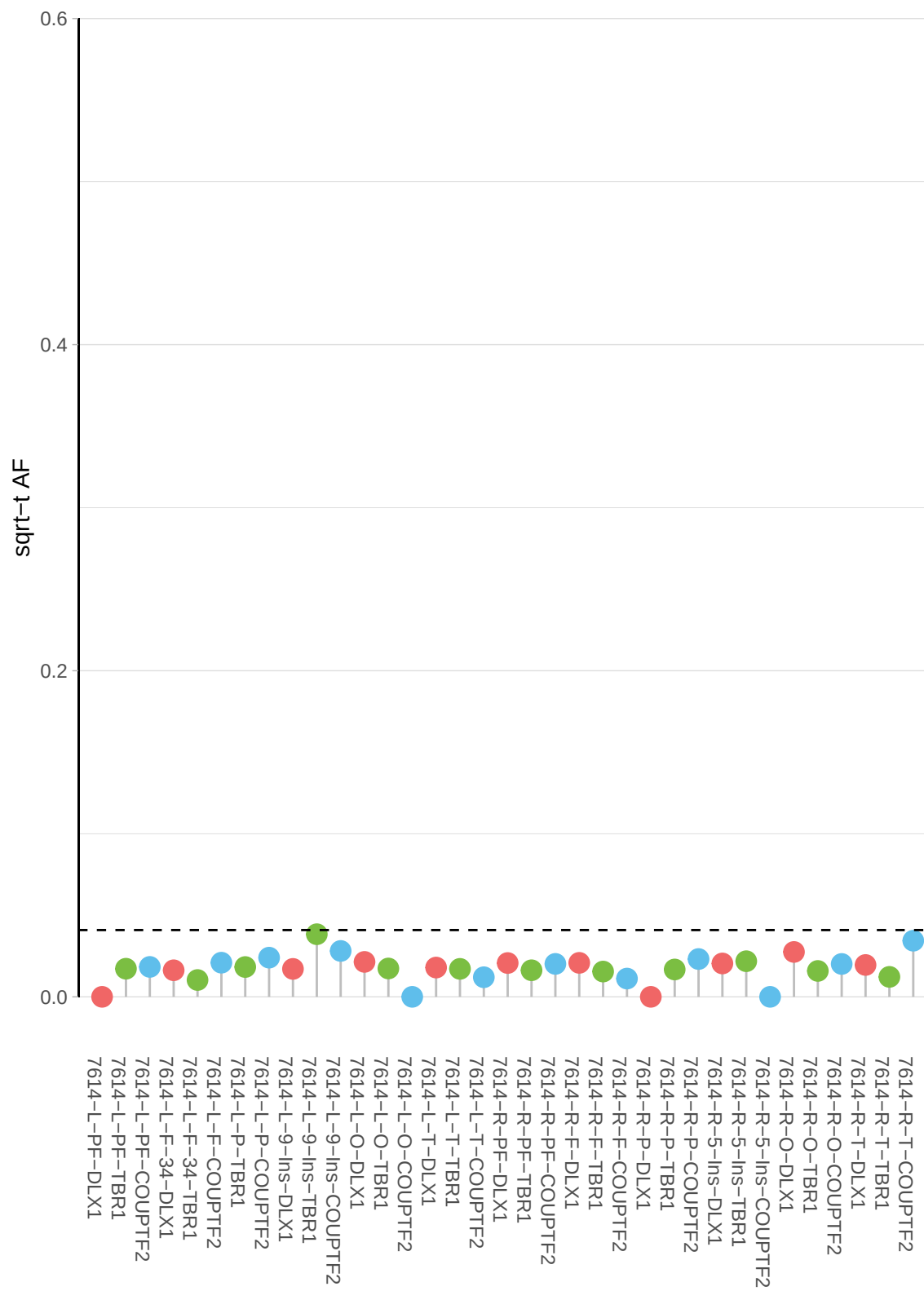

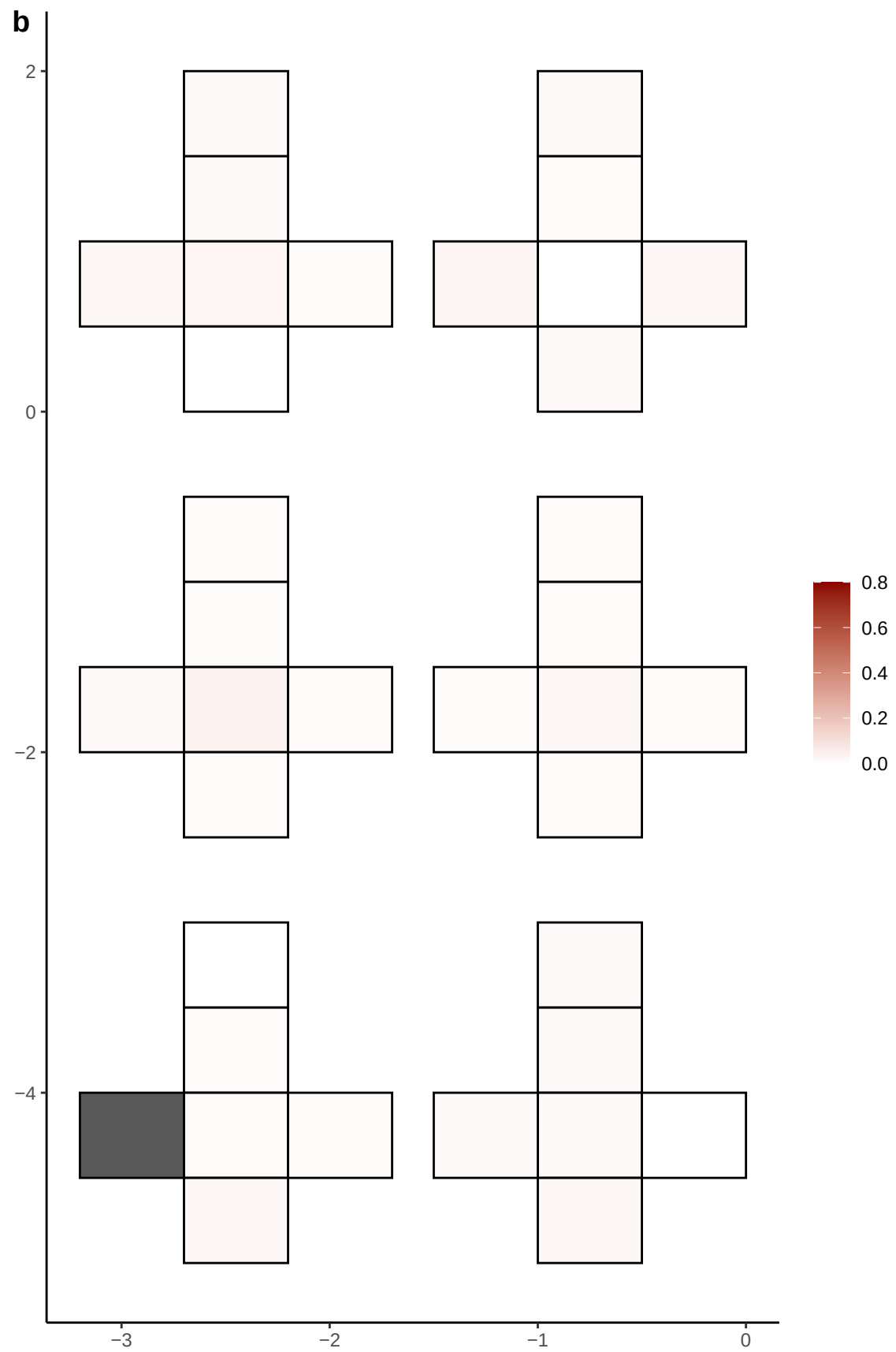

**a** ID01 16-73751747-A-G

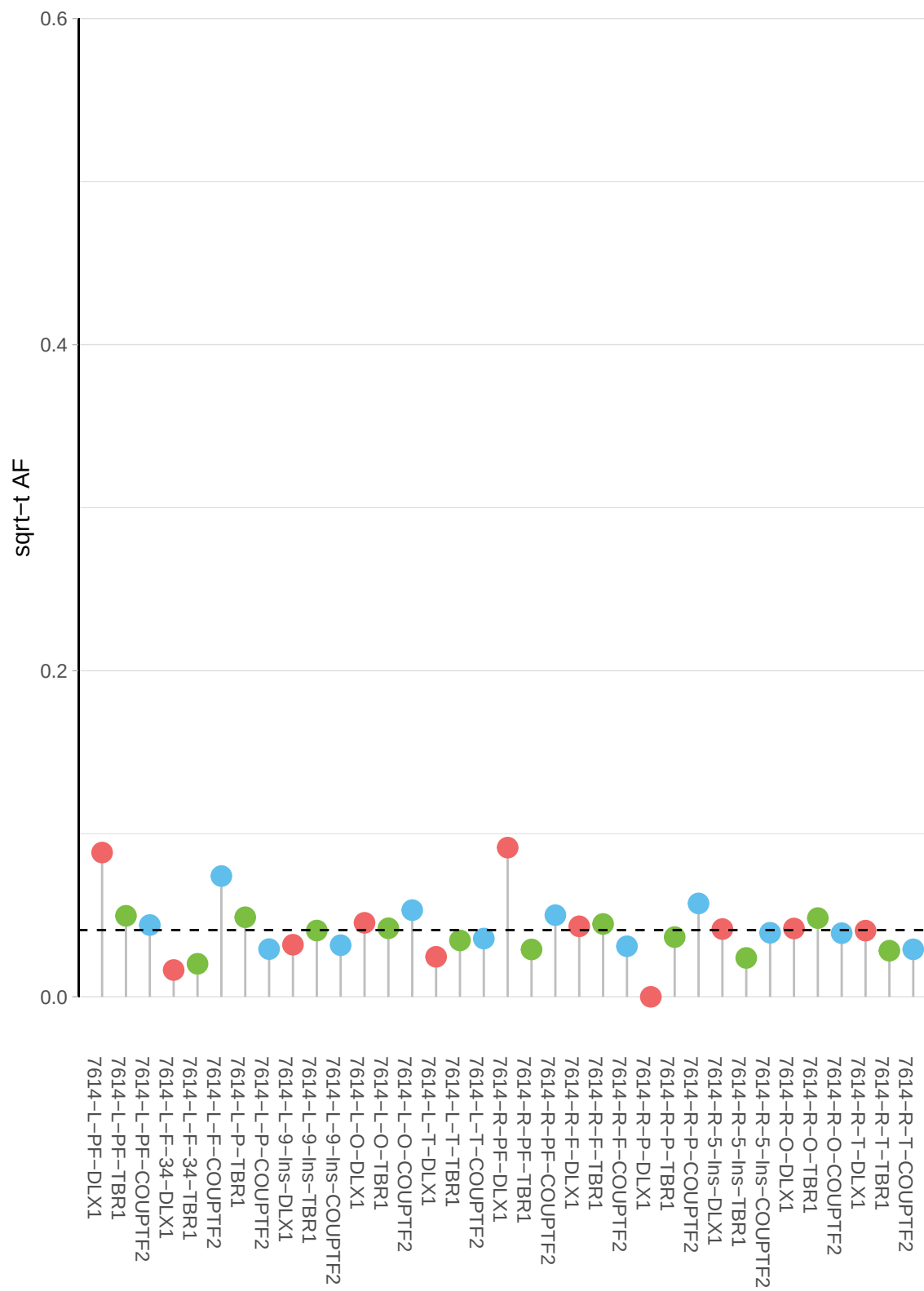

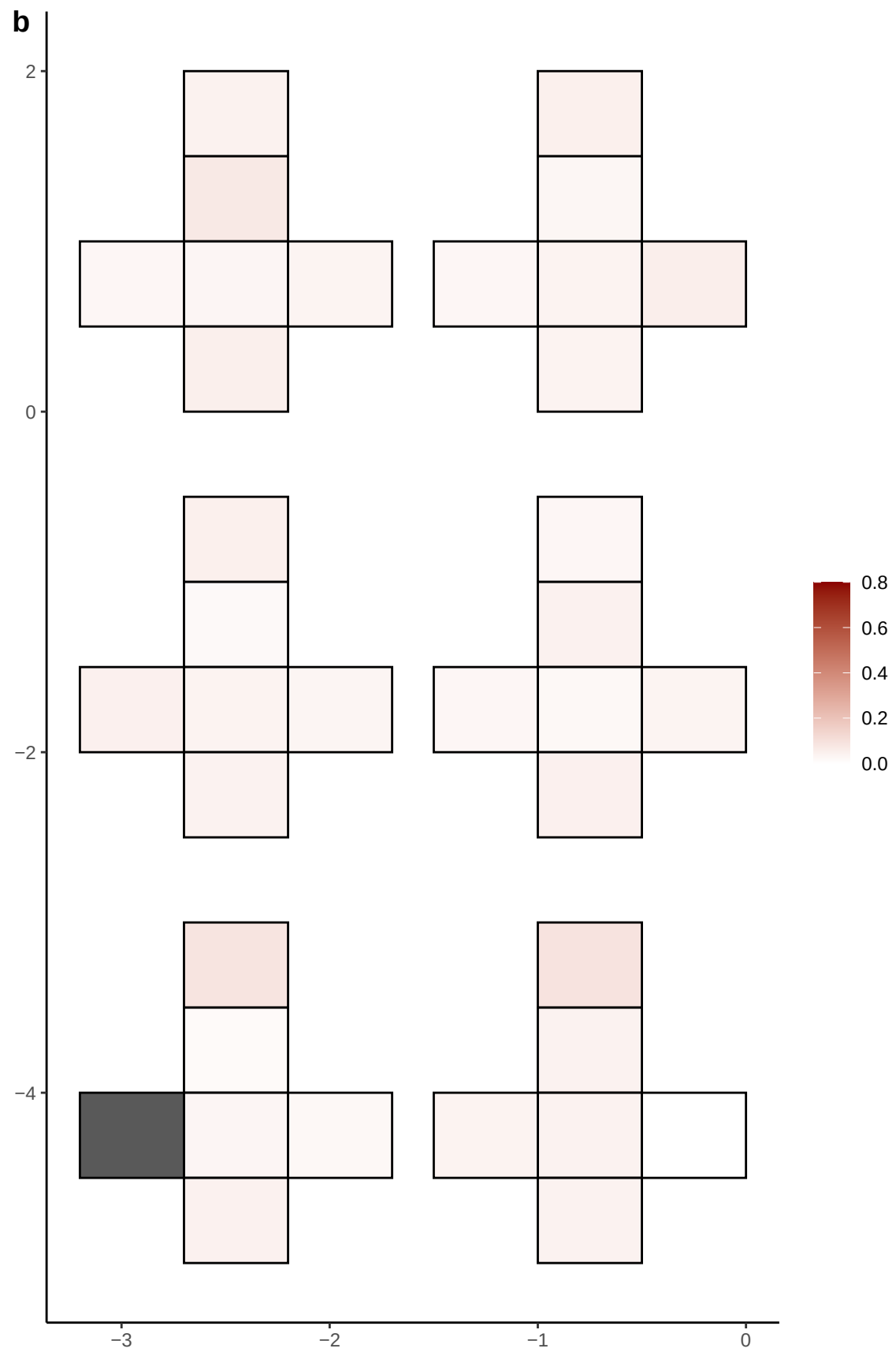

**a** ID01 16-78187280-G-A

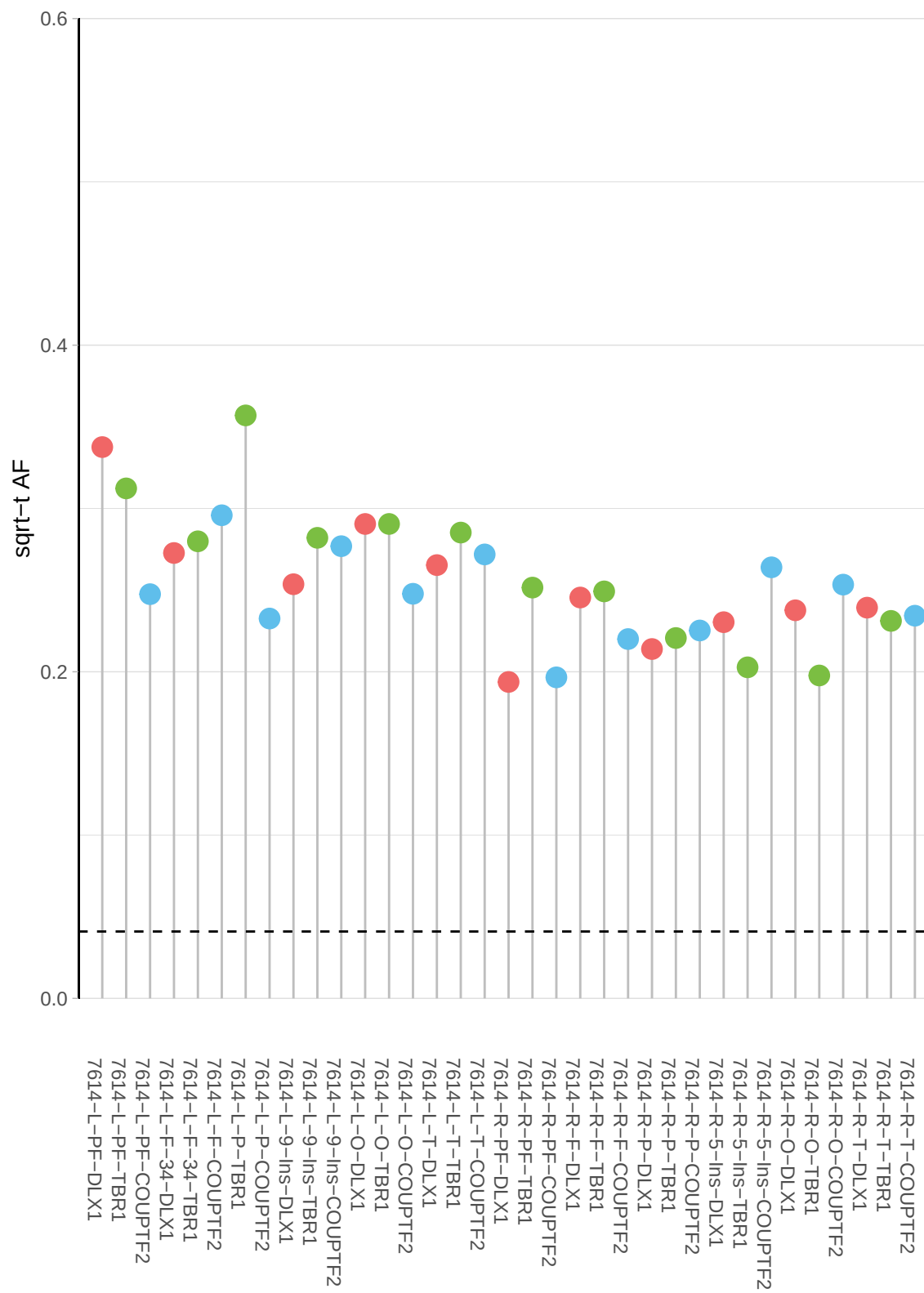

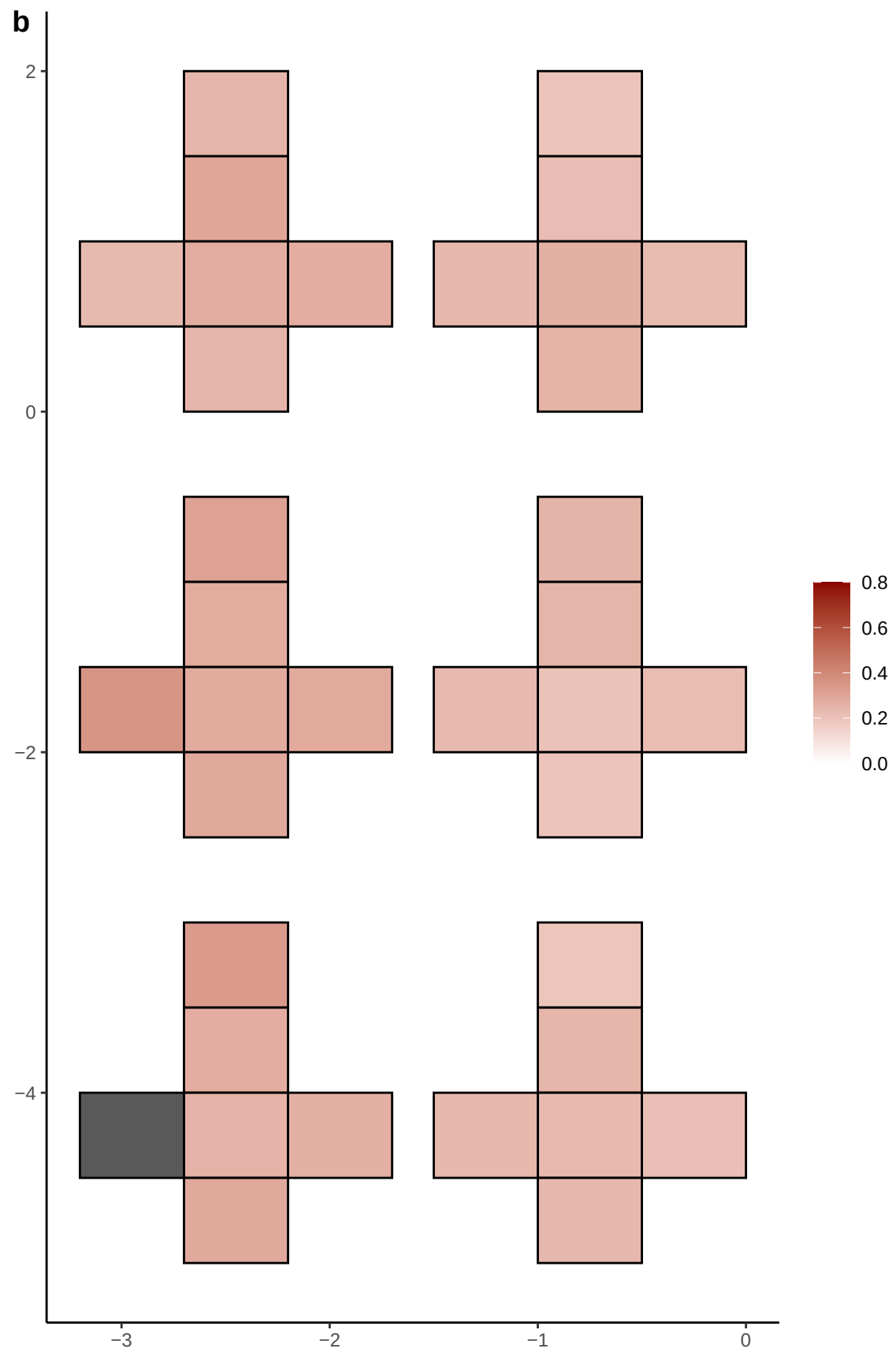

**a** ID01 16-78774167-C-T

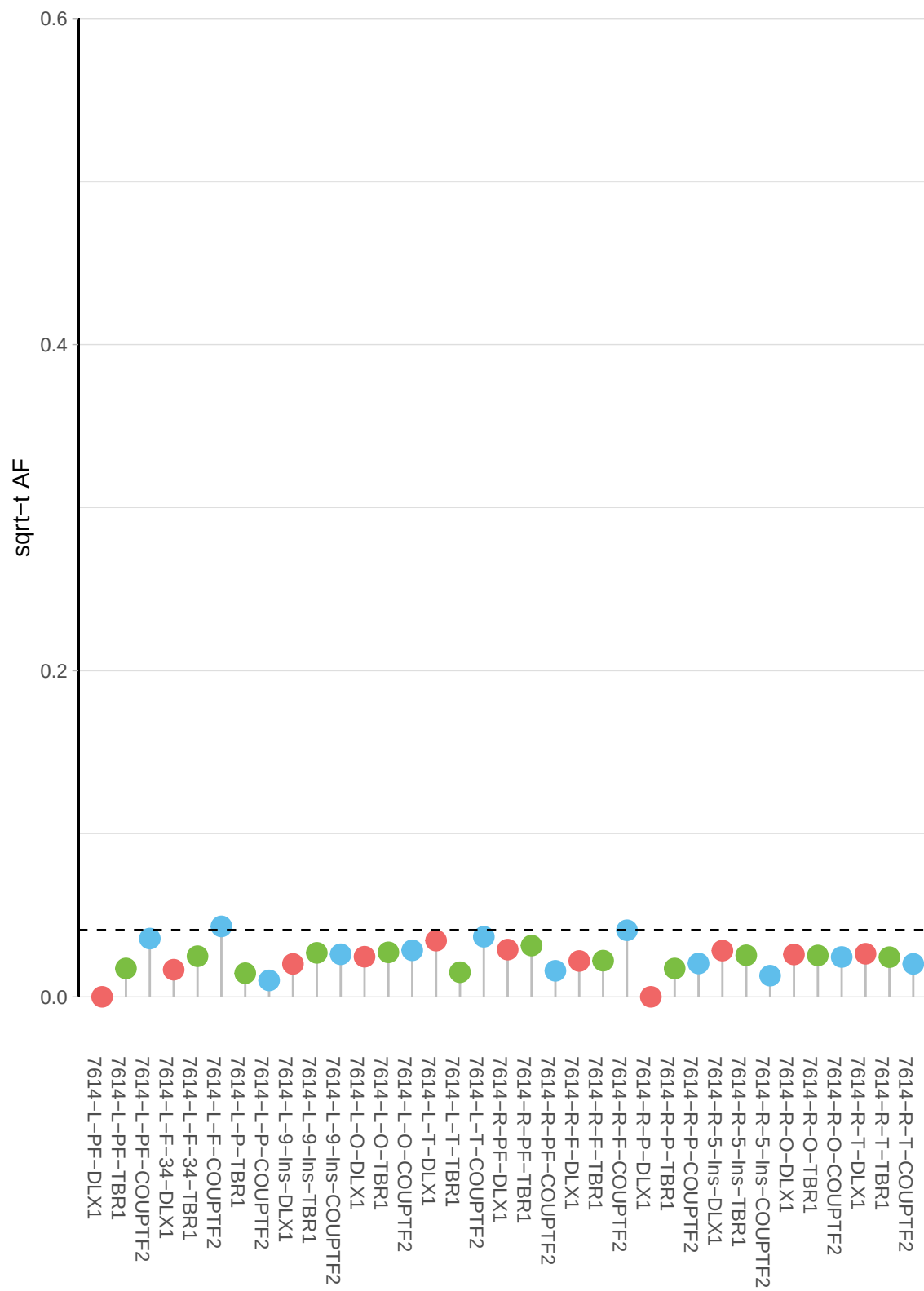

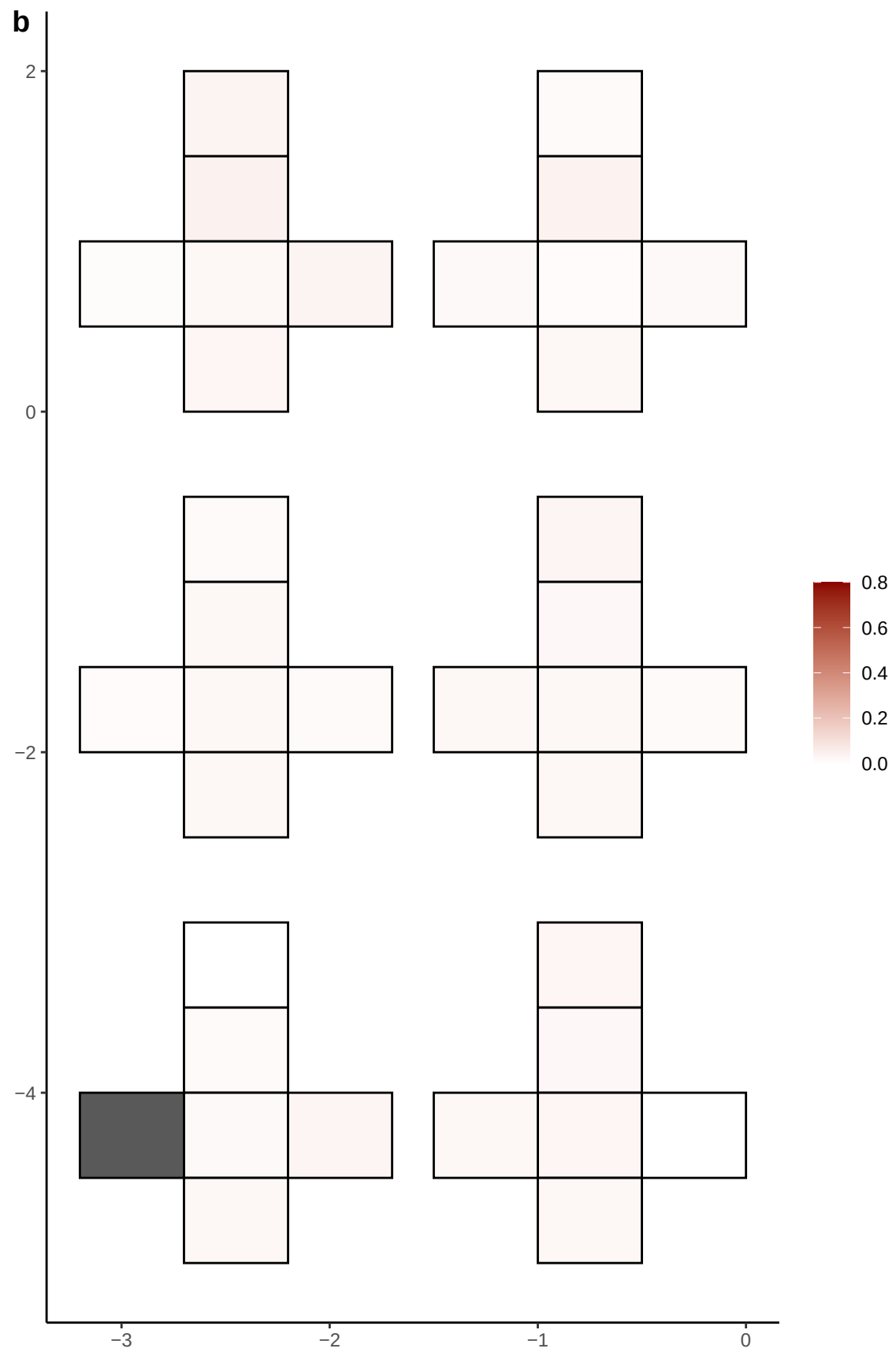

**a** ID01 16-80604095-C-T

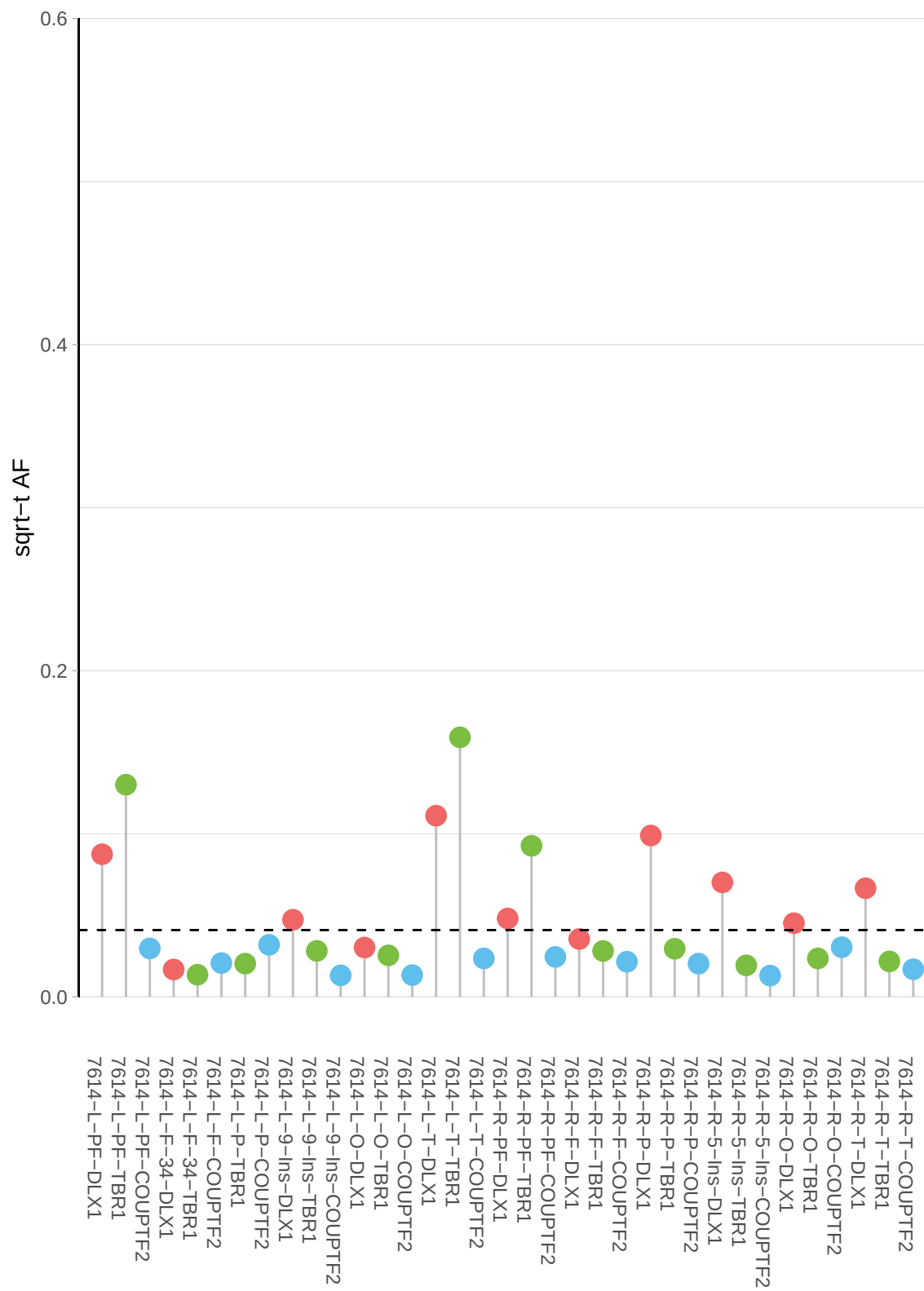

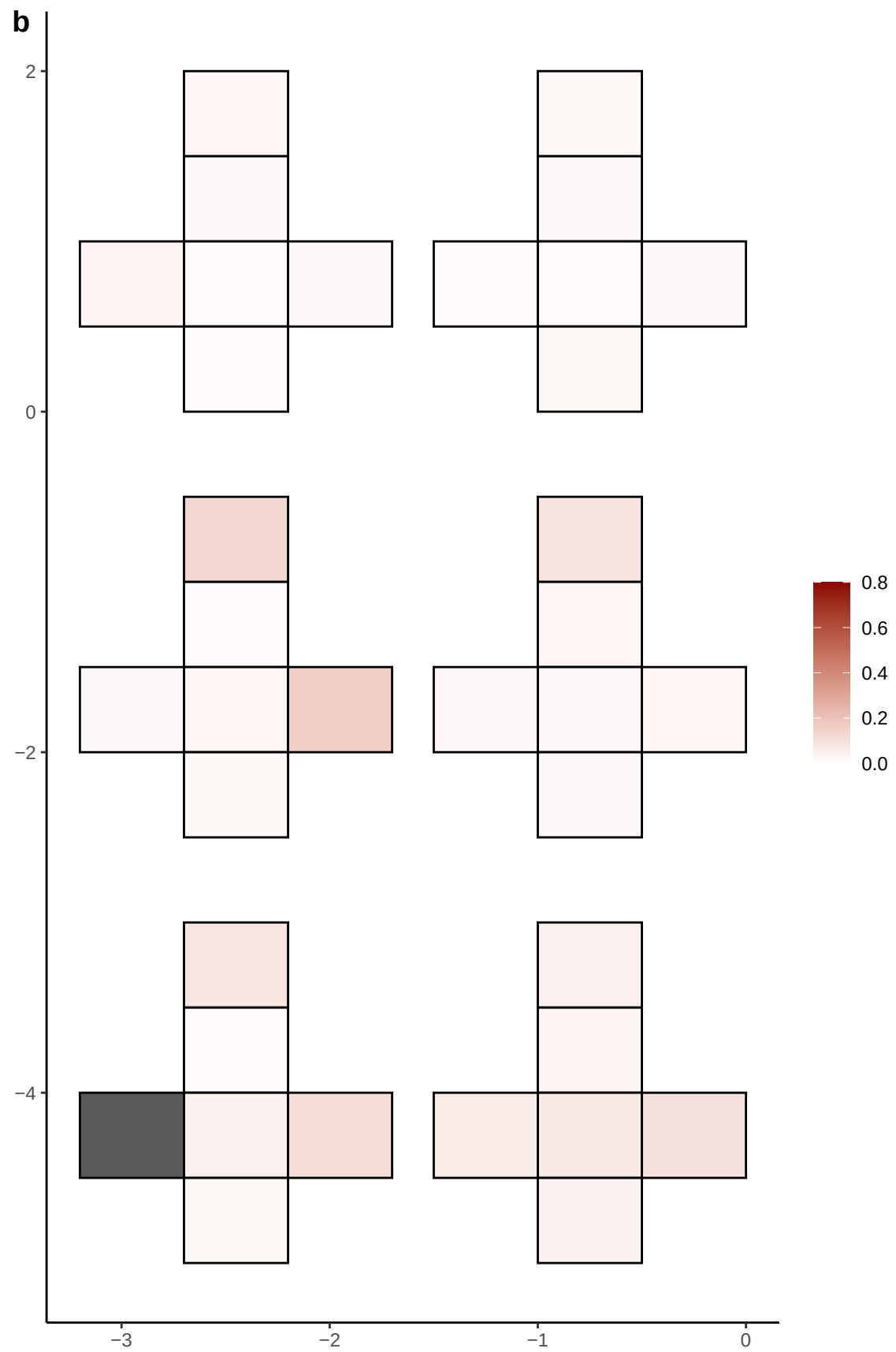

**a** ID01 16-83174915-T-C

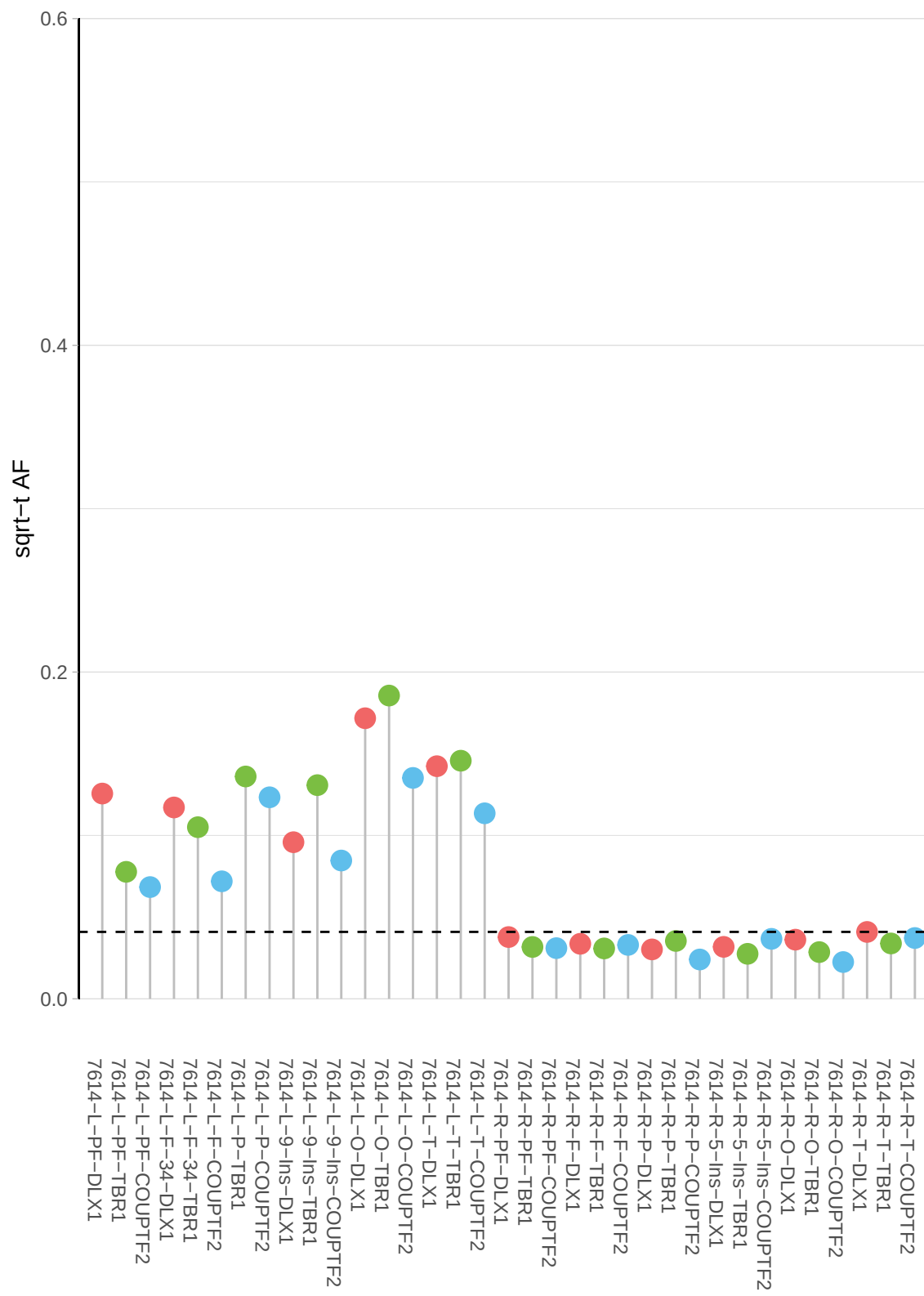

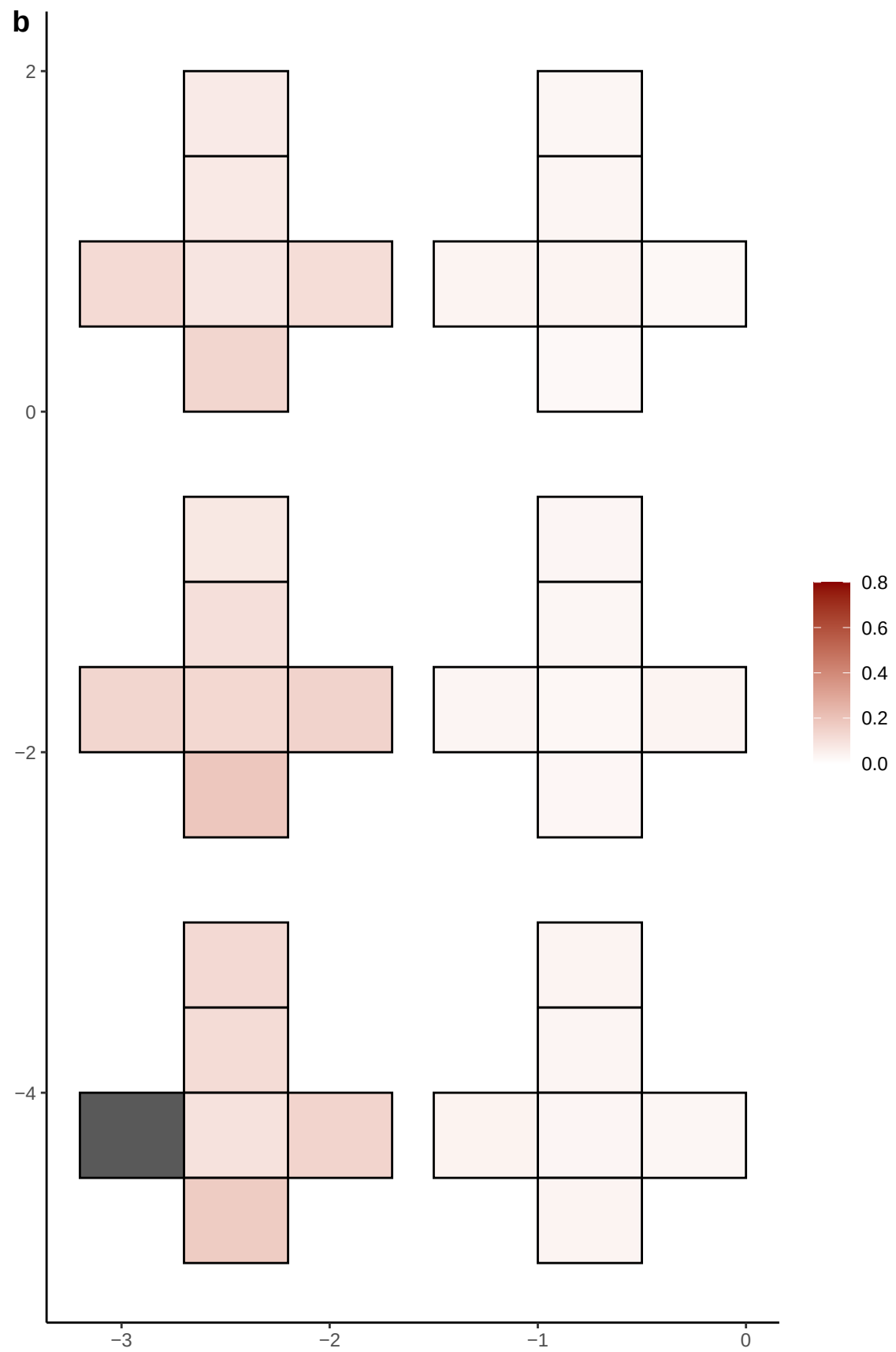

**a** ID01 17-30813389-C-G

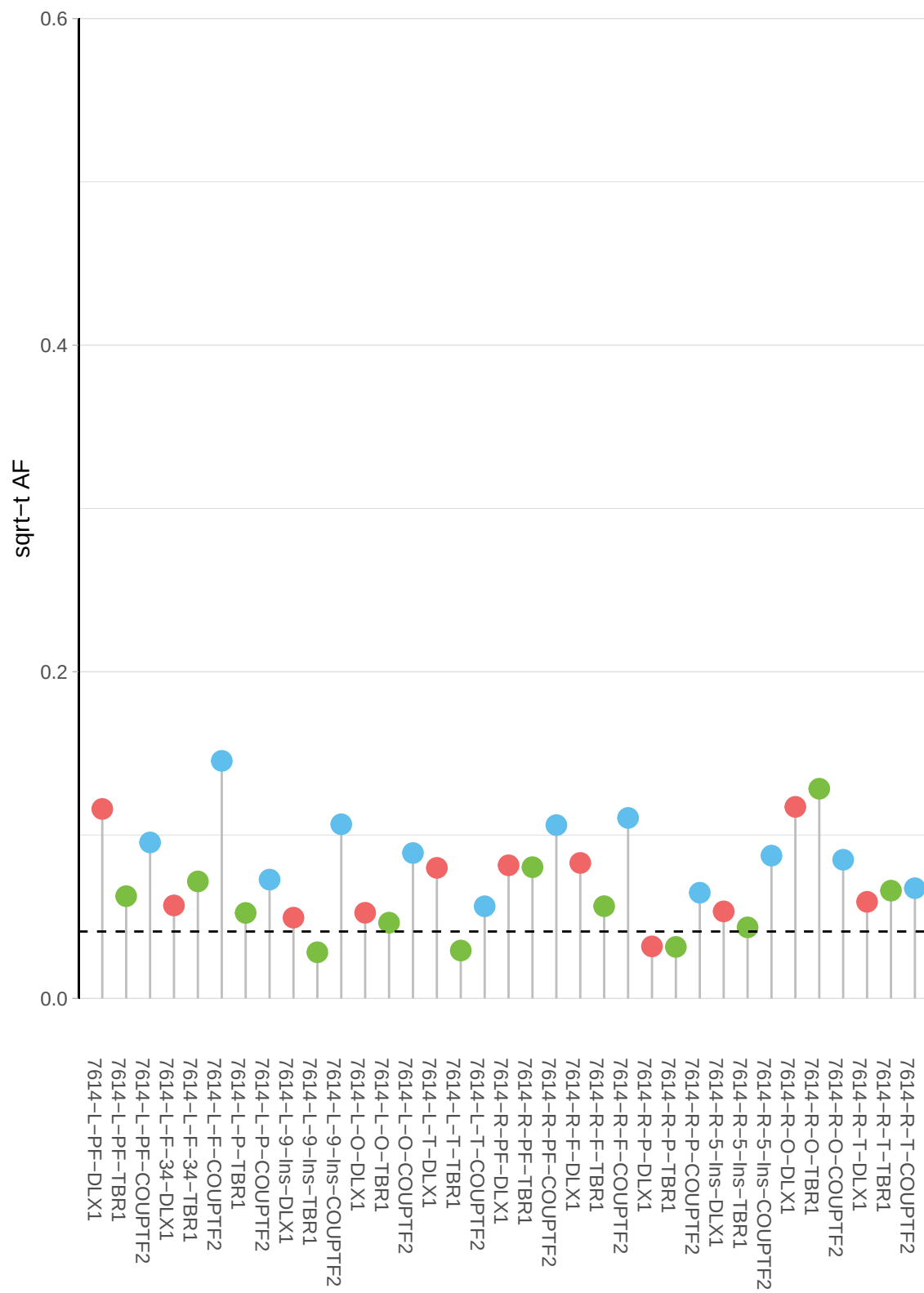

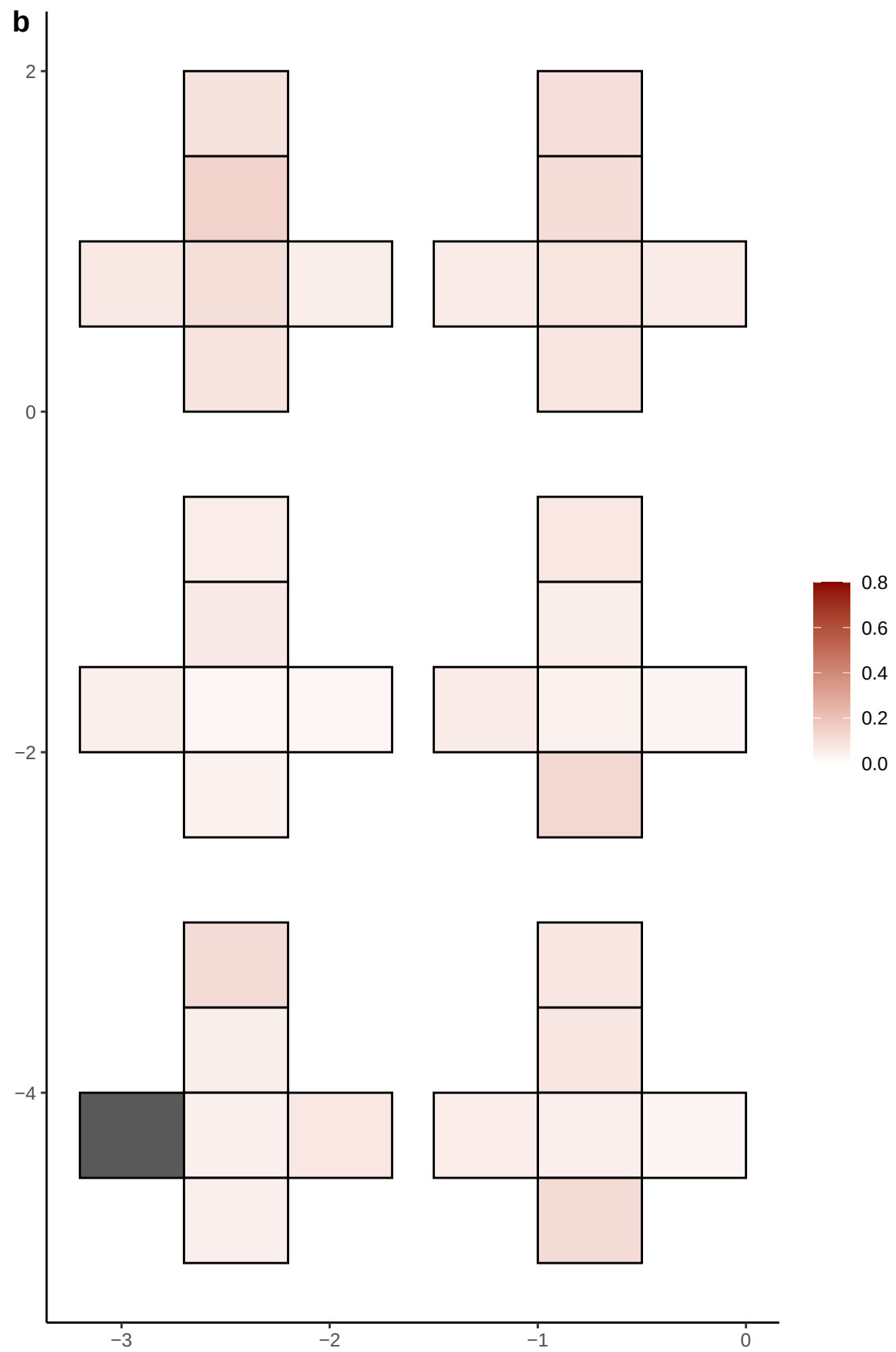

**a** ID01 17-31854088-G-A

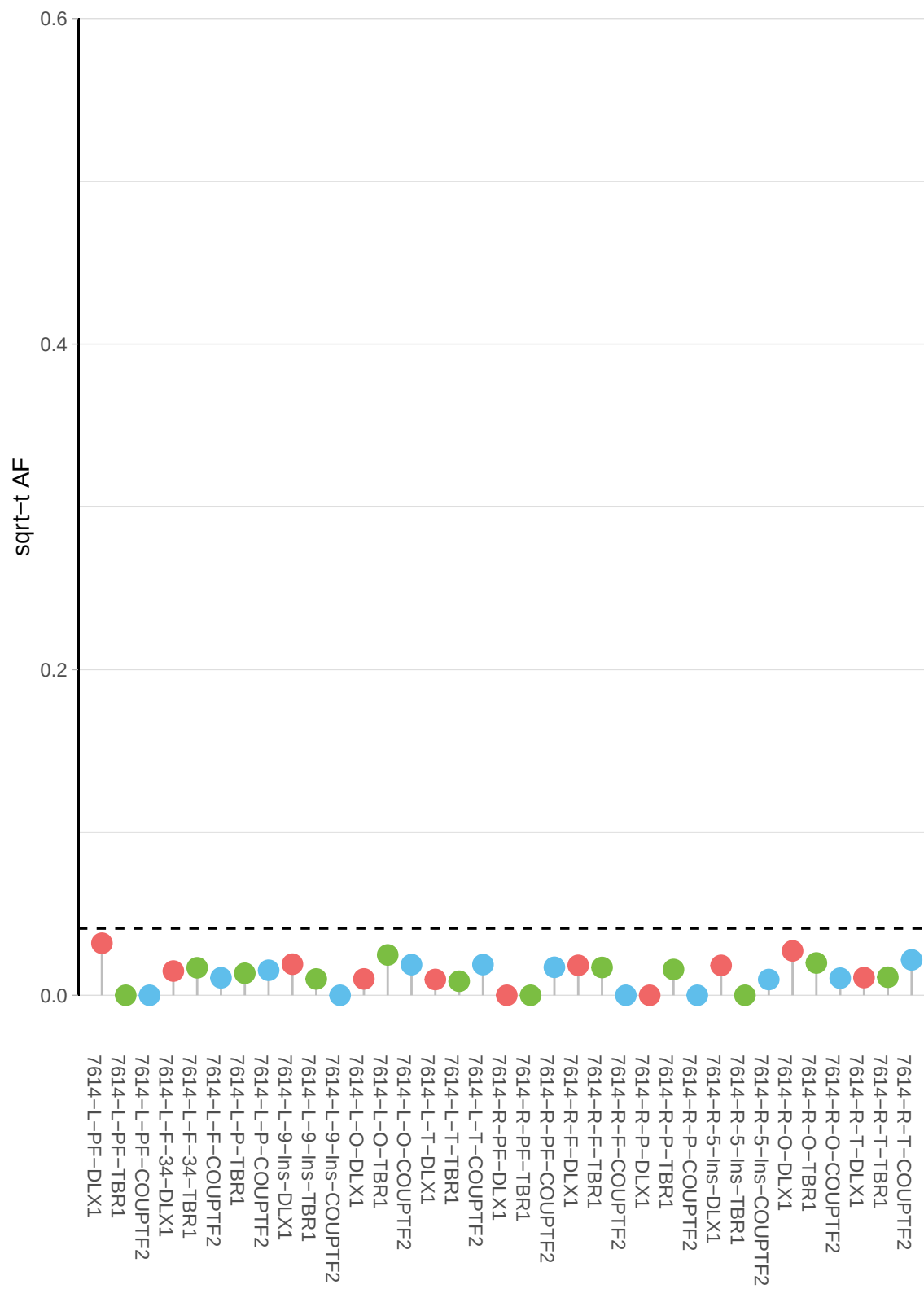

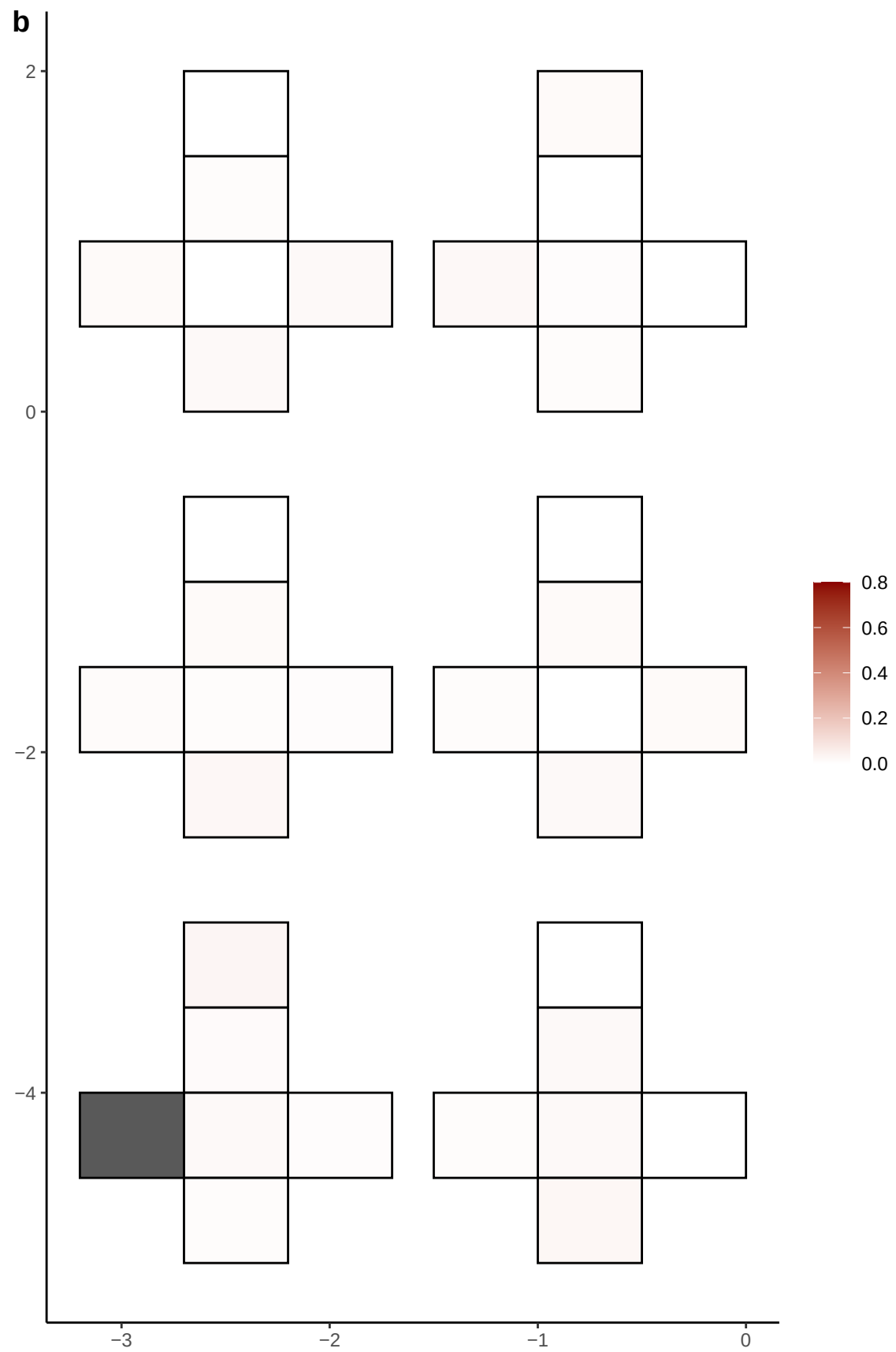

**a** ID01 17-32373696-G-A

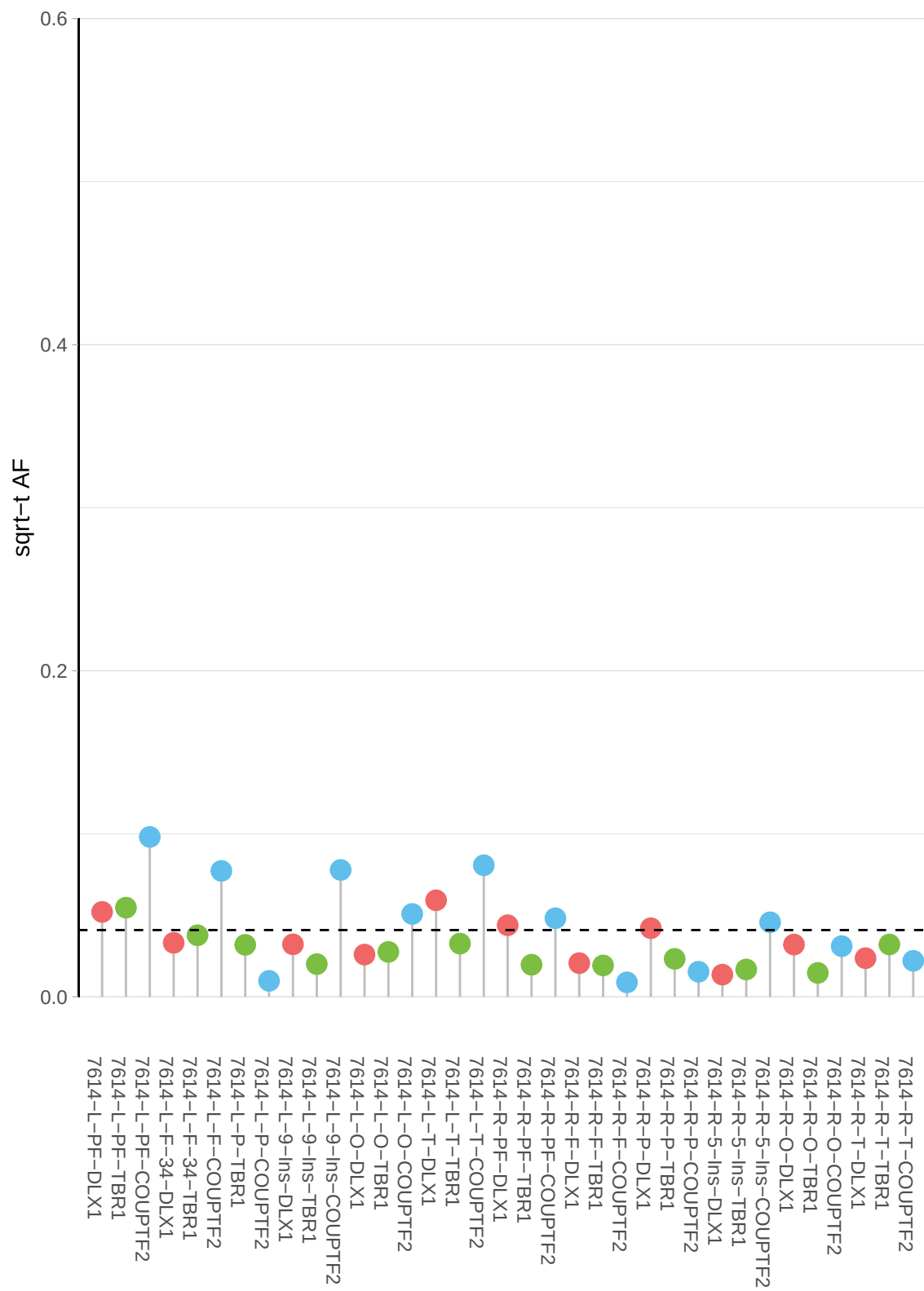

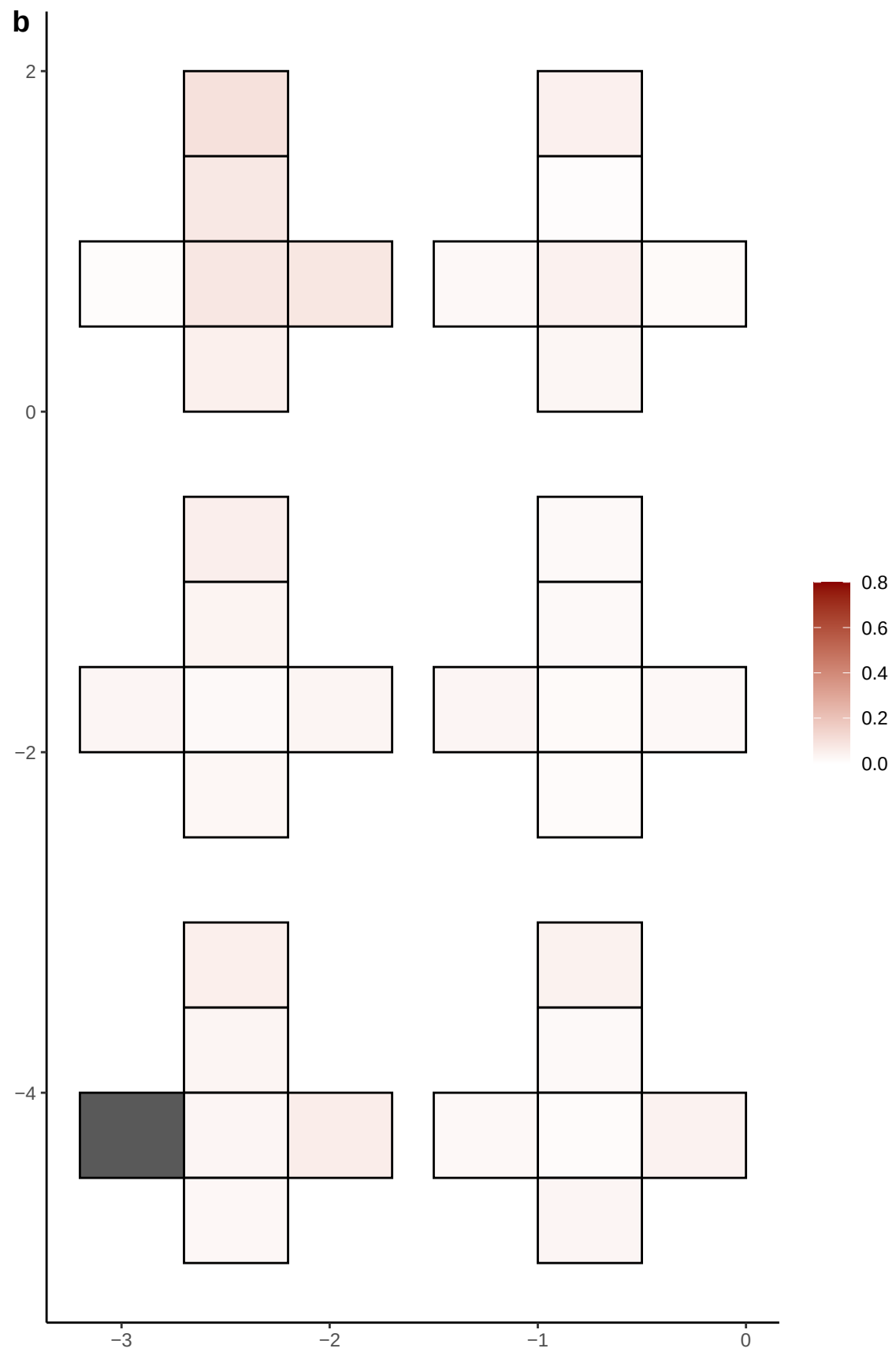

**a** ID01 17-5657235-A-C

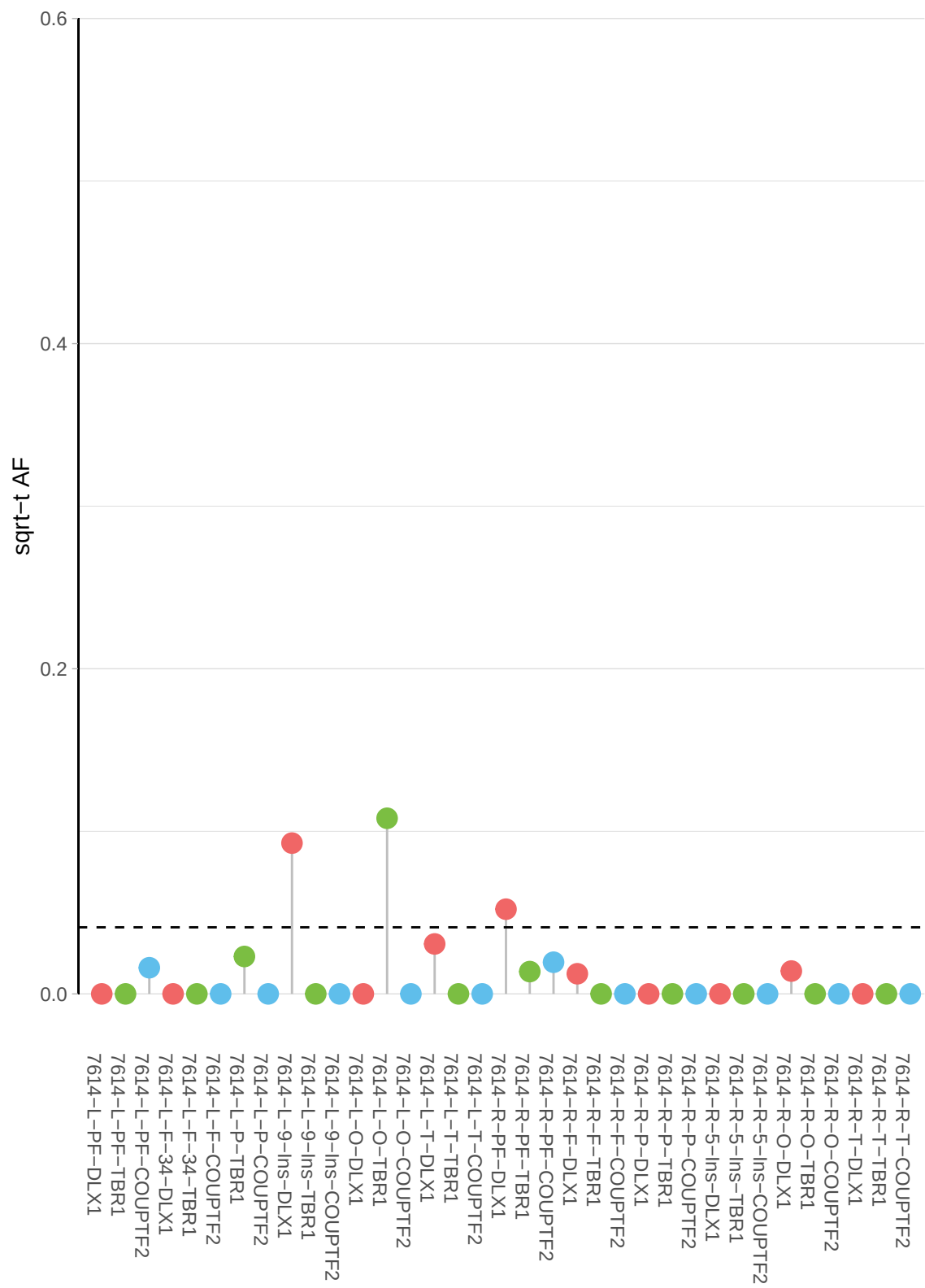

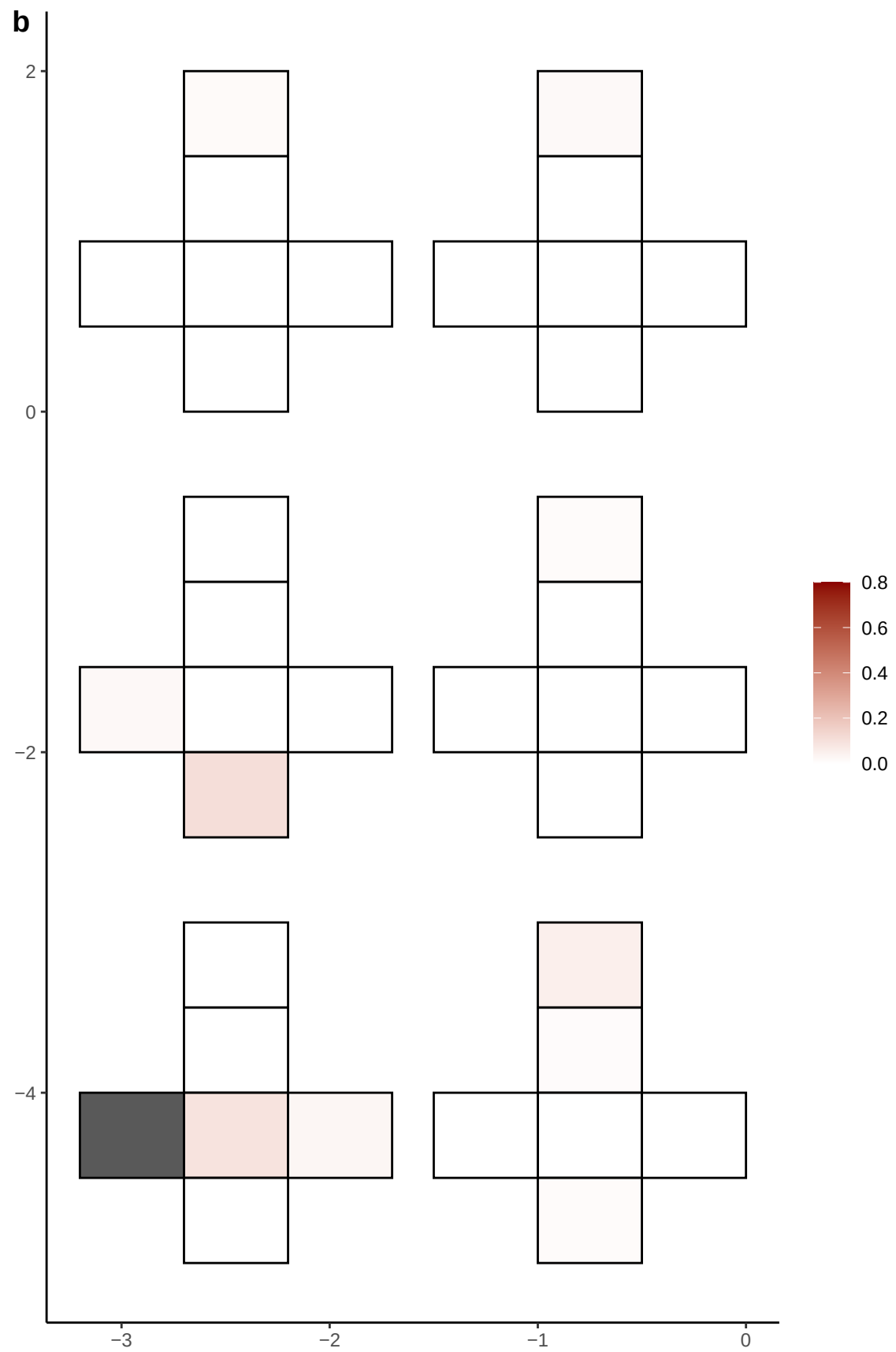

**a** ID01 17-56786985-T-C

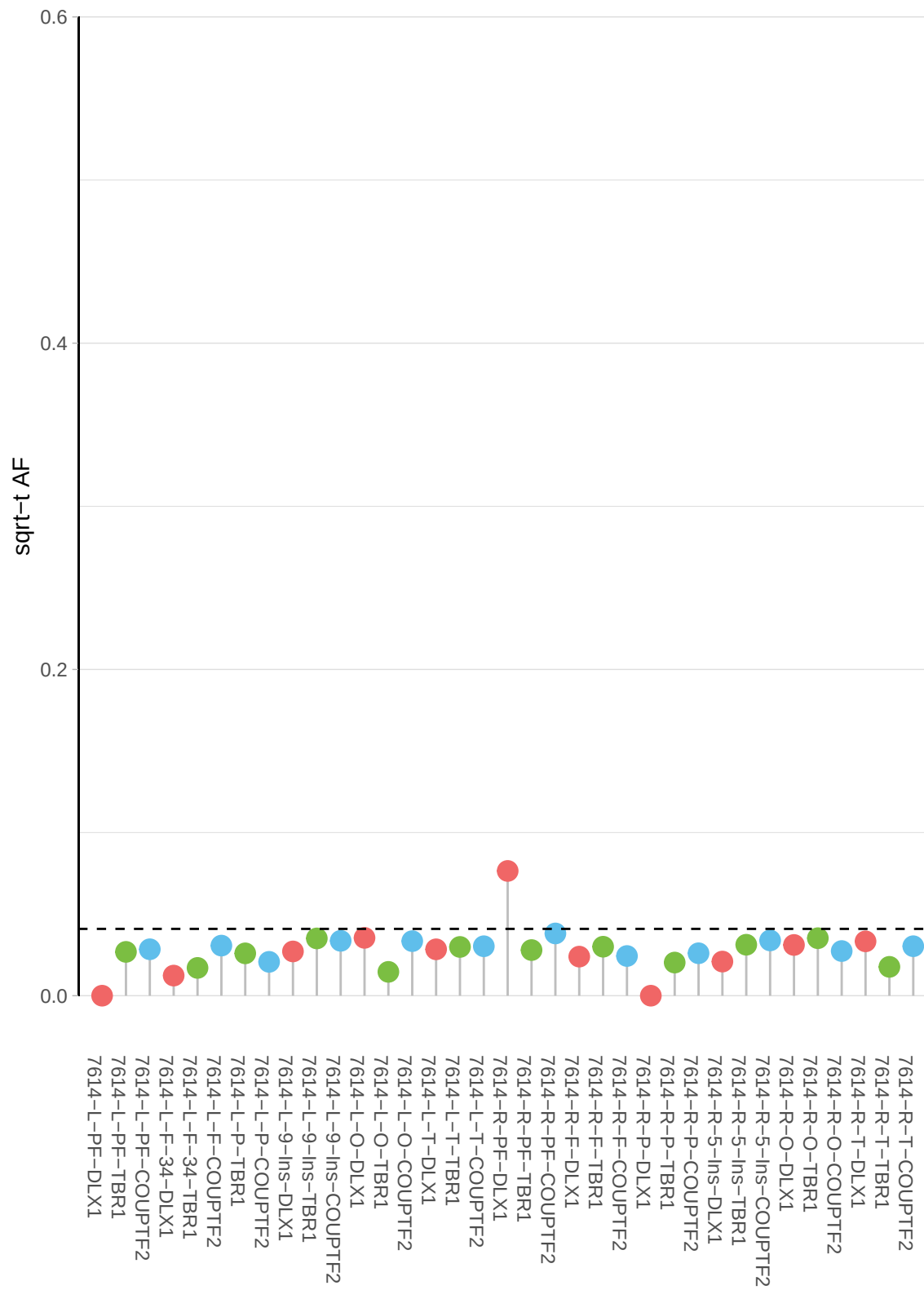

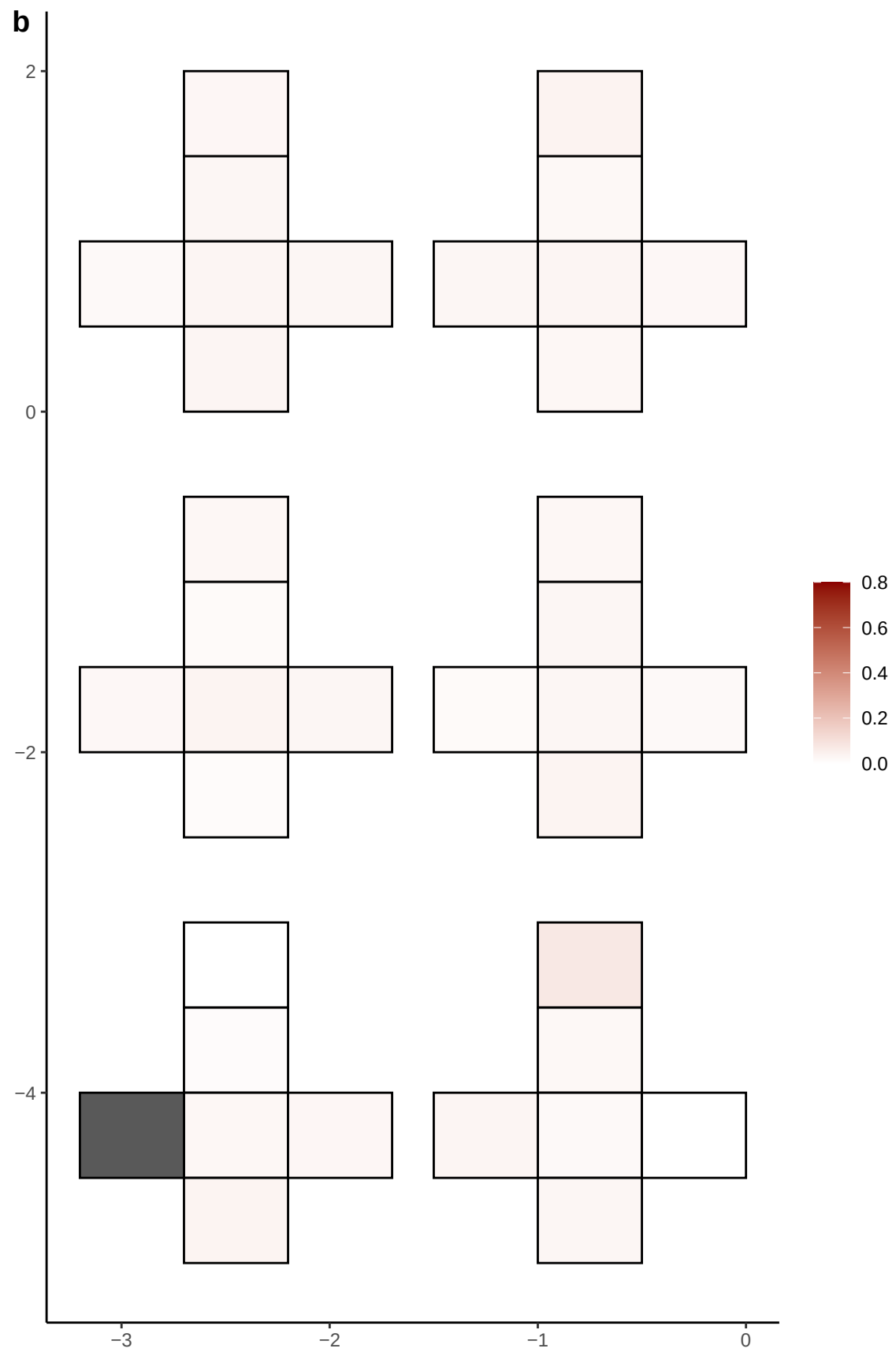

**a** ID01 18-12016169-T-C

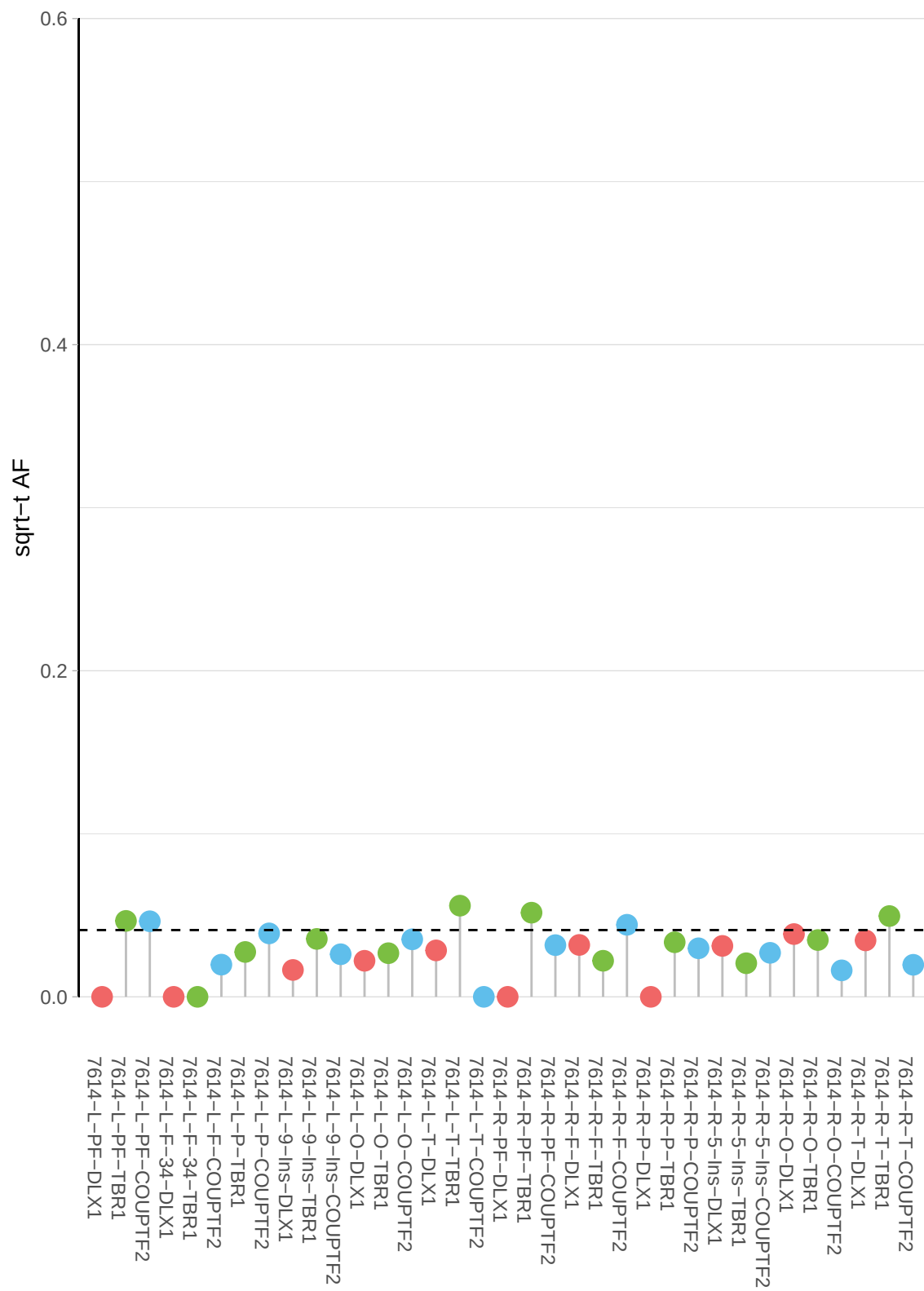

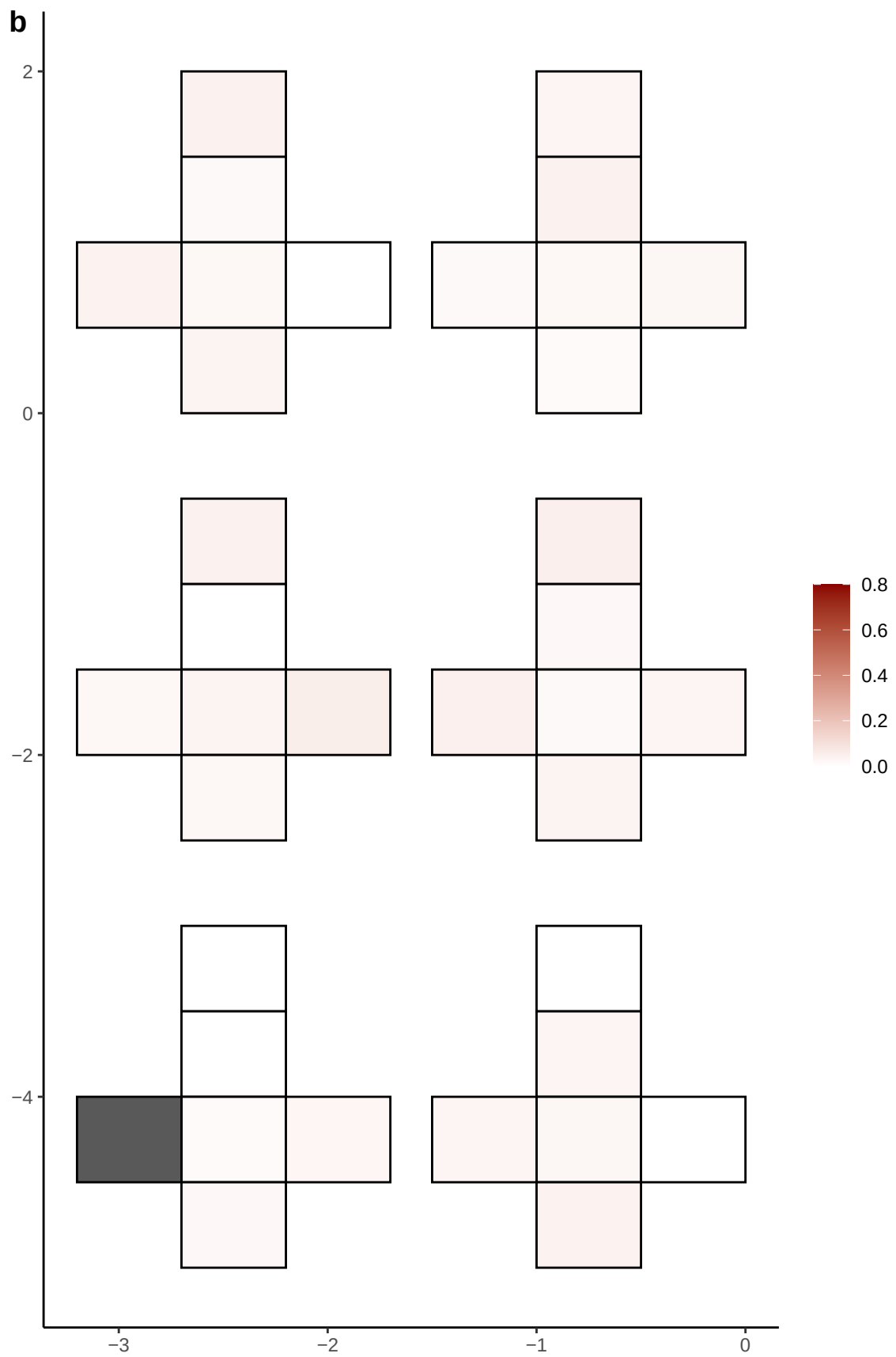

**a** ID01 18-27189441-TA-T

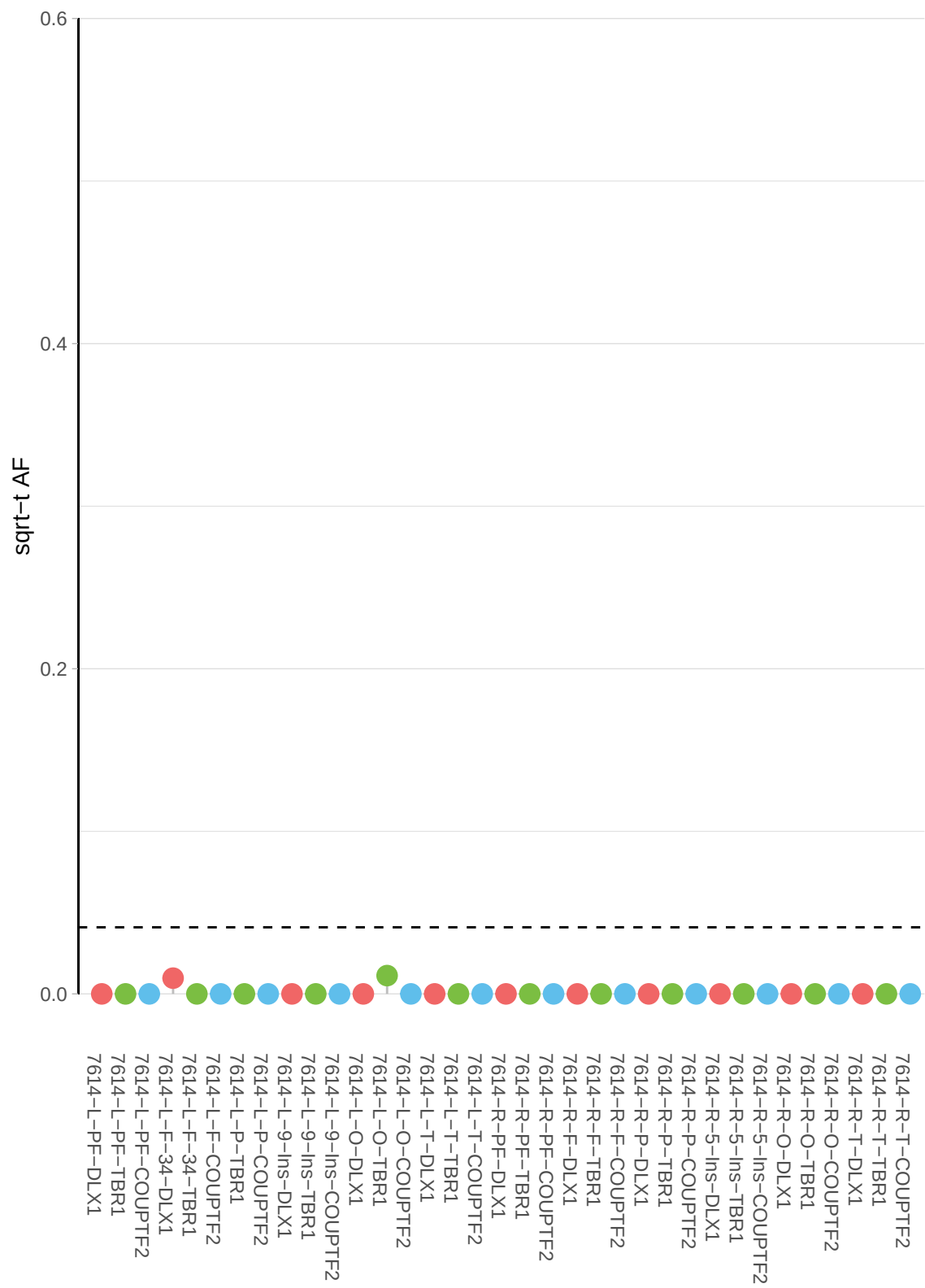

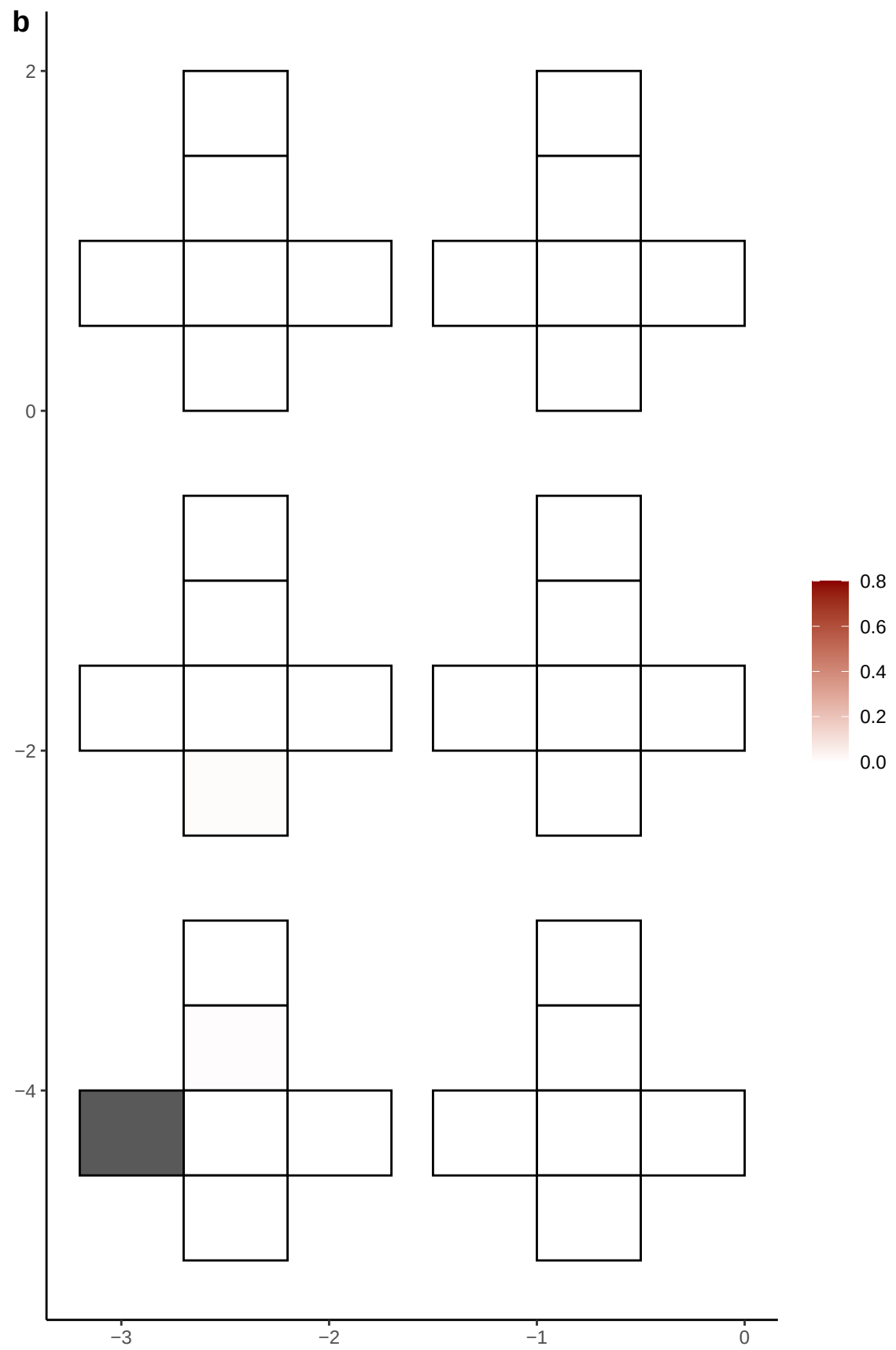

**a** ID01 18-39171826-C-G

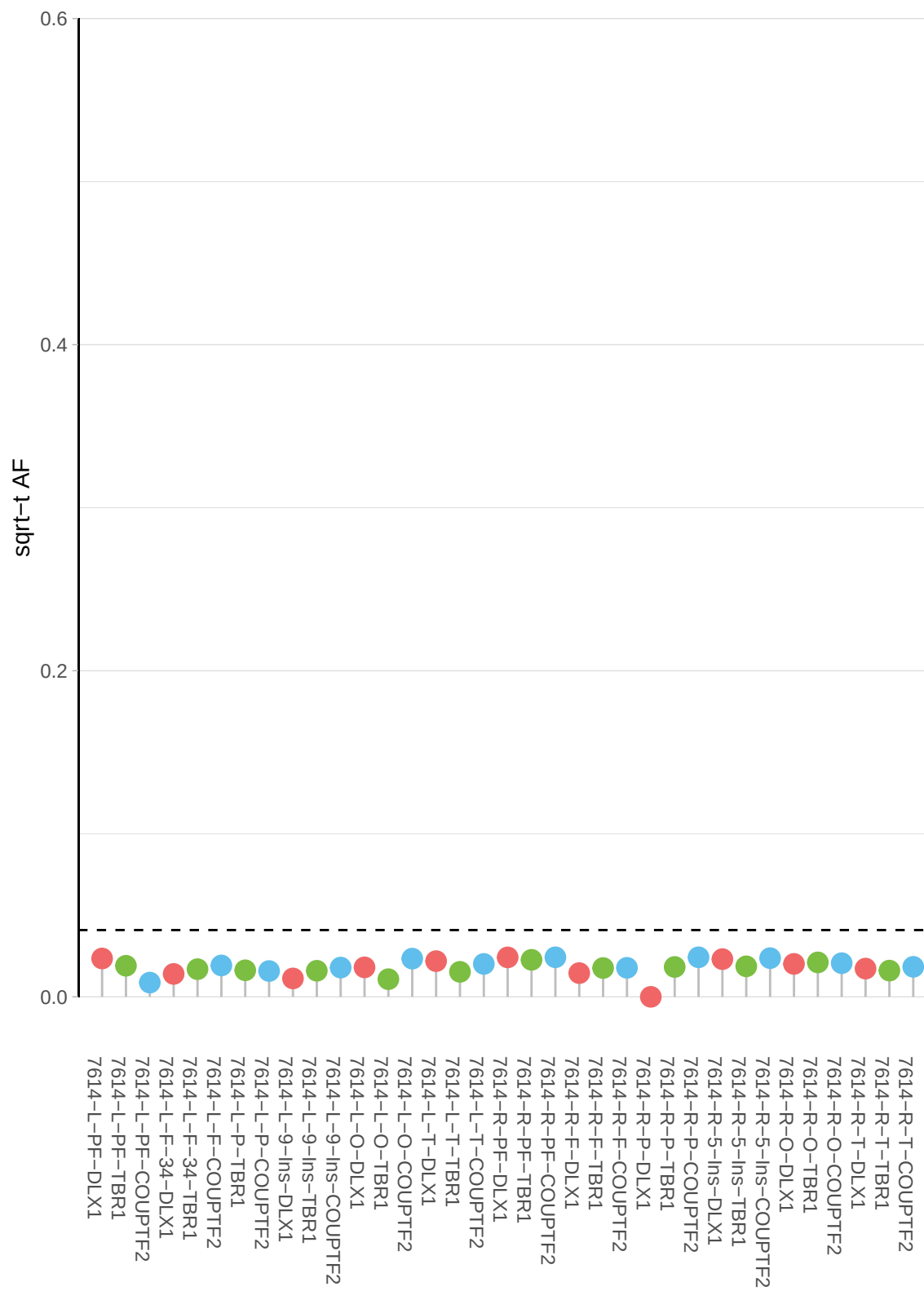

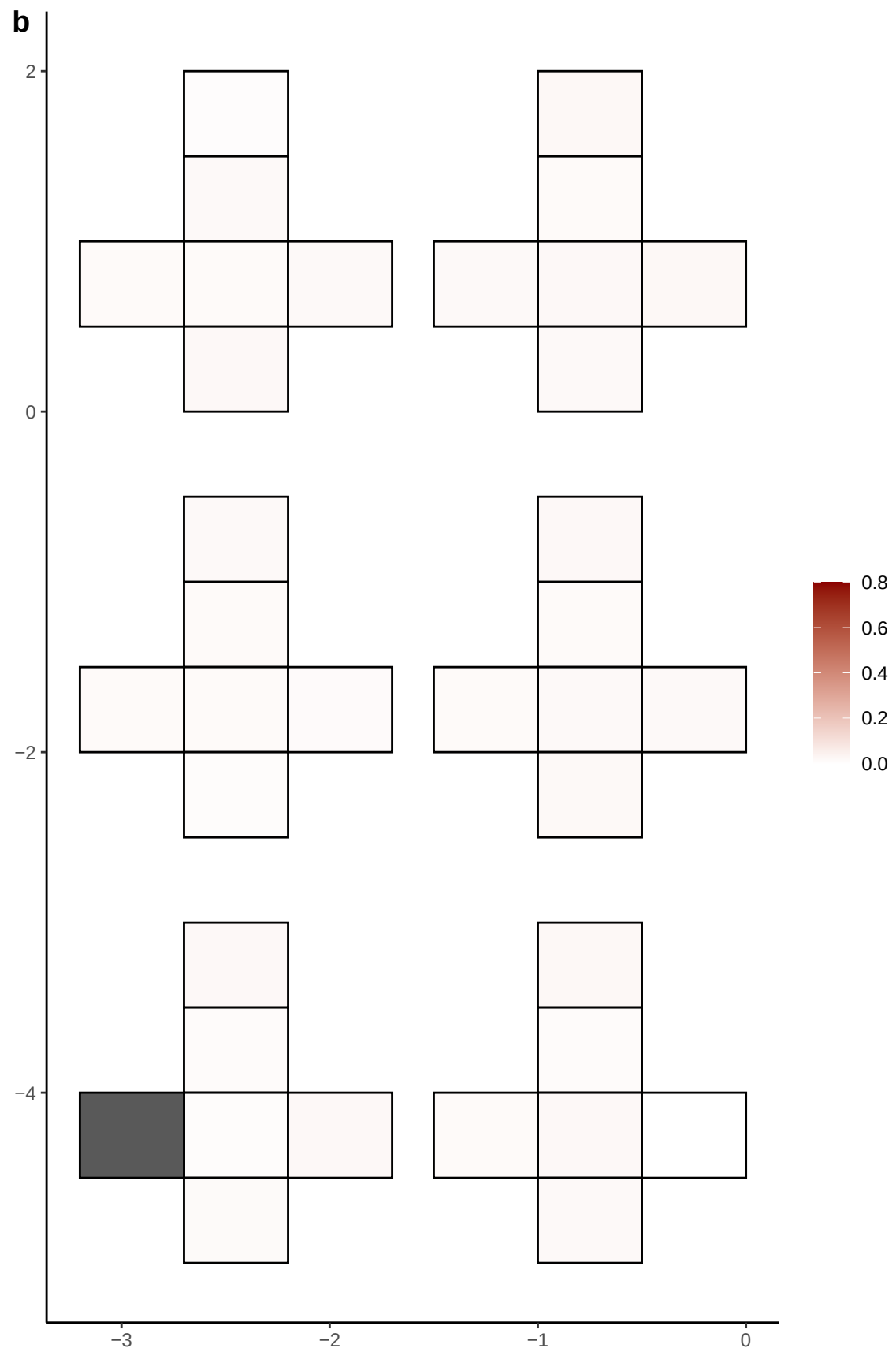

**a** ID01 18-40074413-T-C

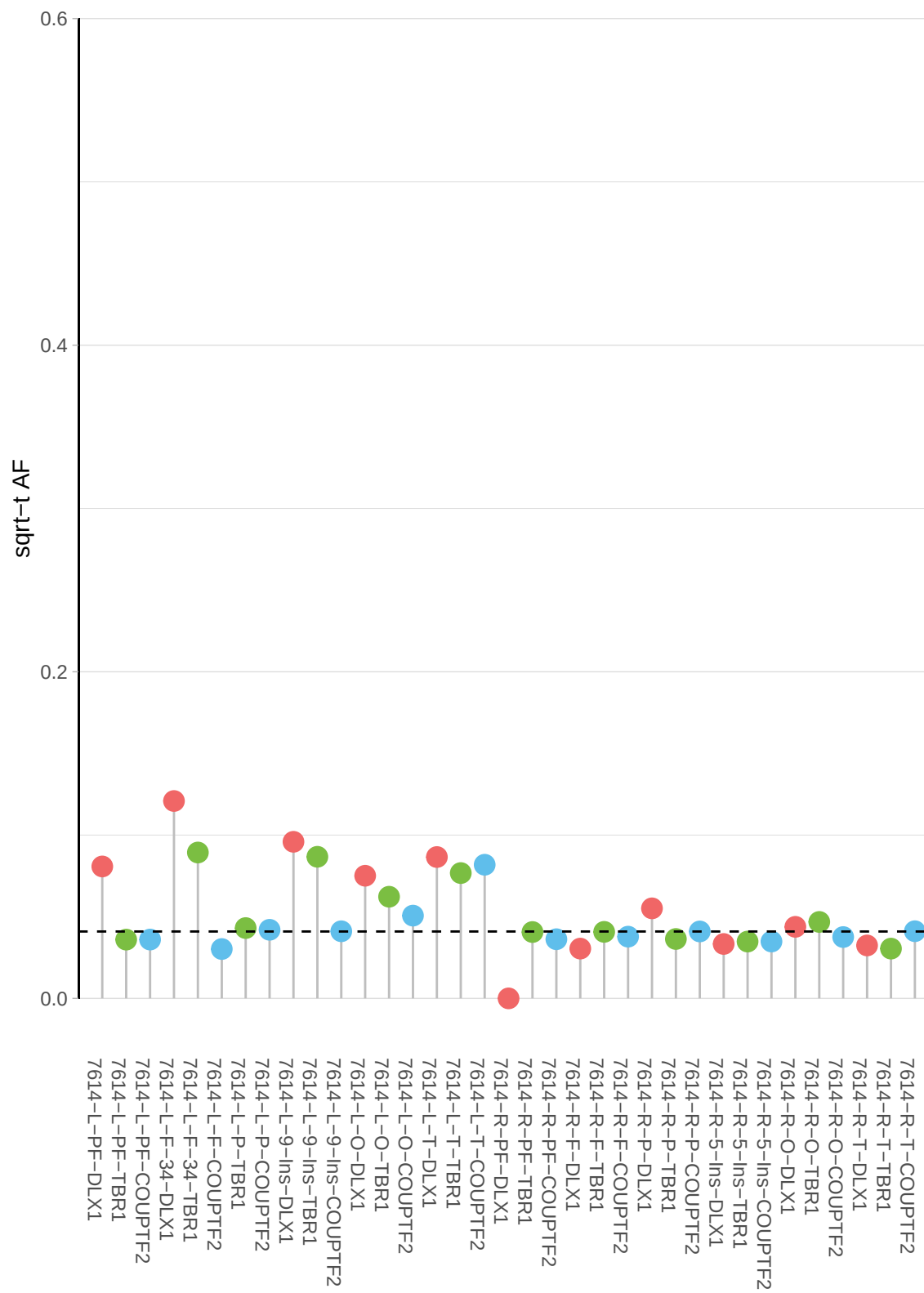

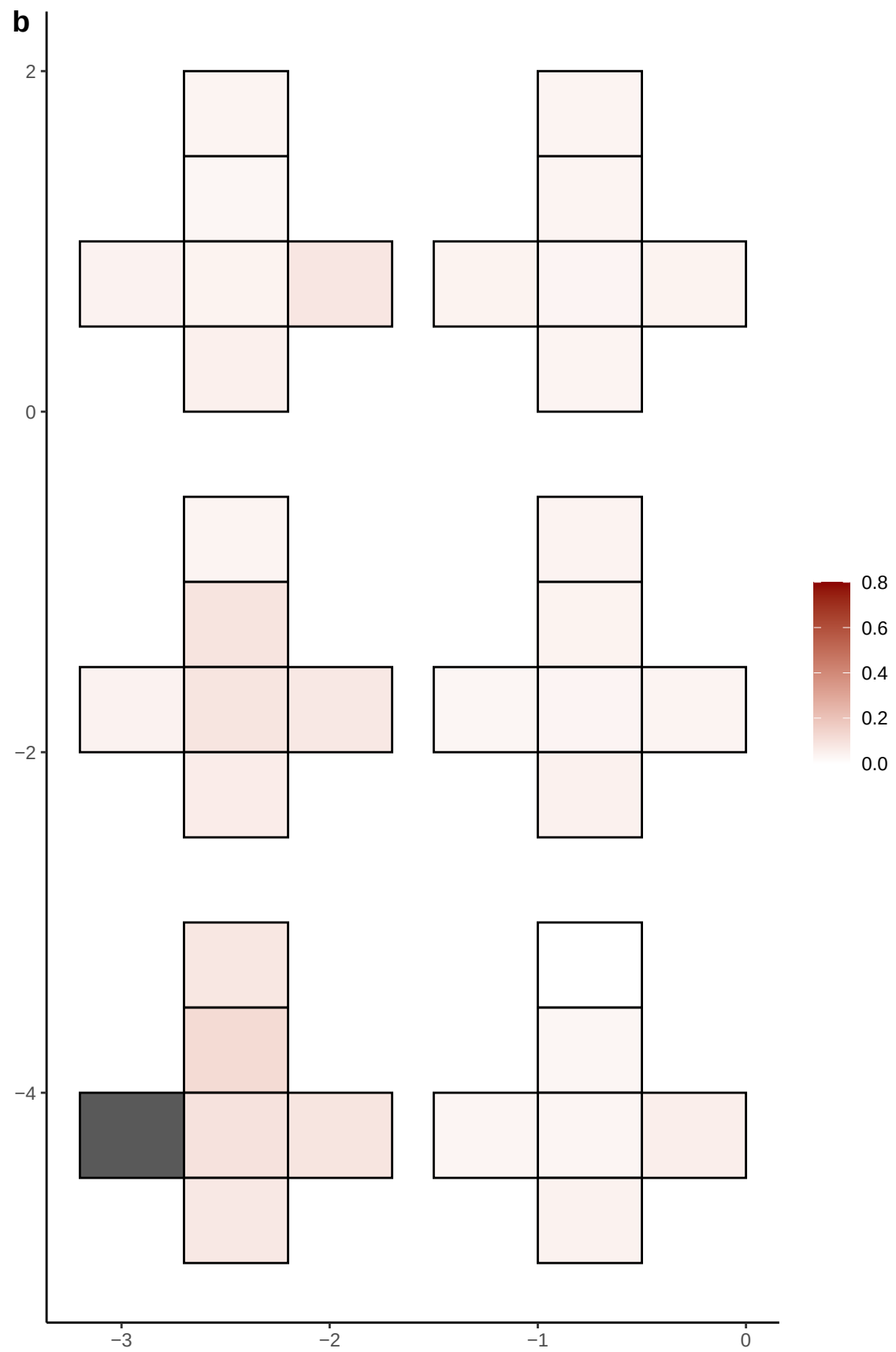

**a** ID01 18-56480516-G-A

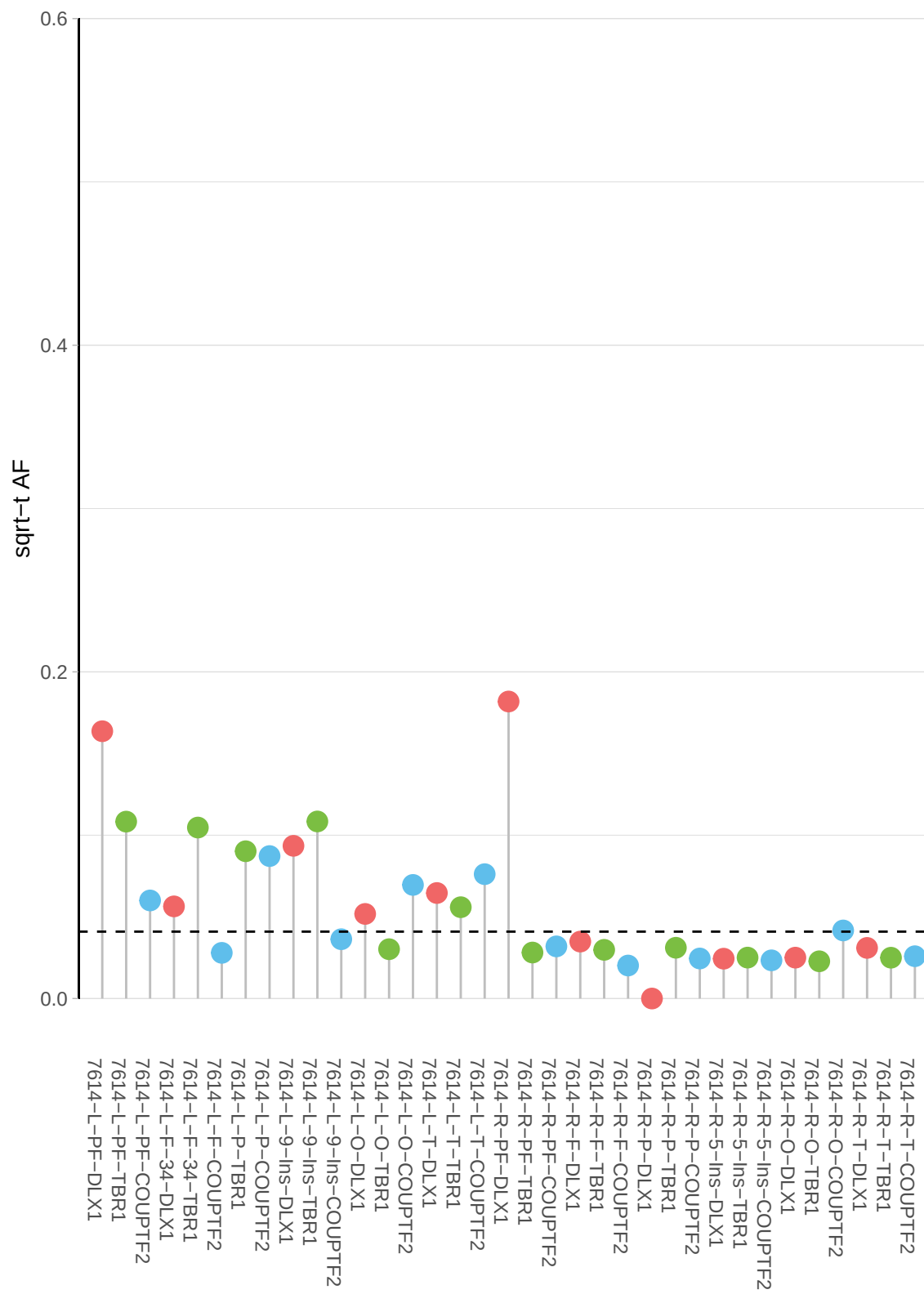

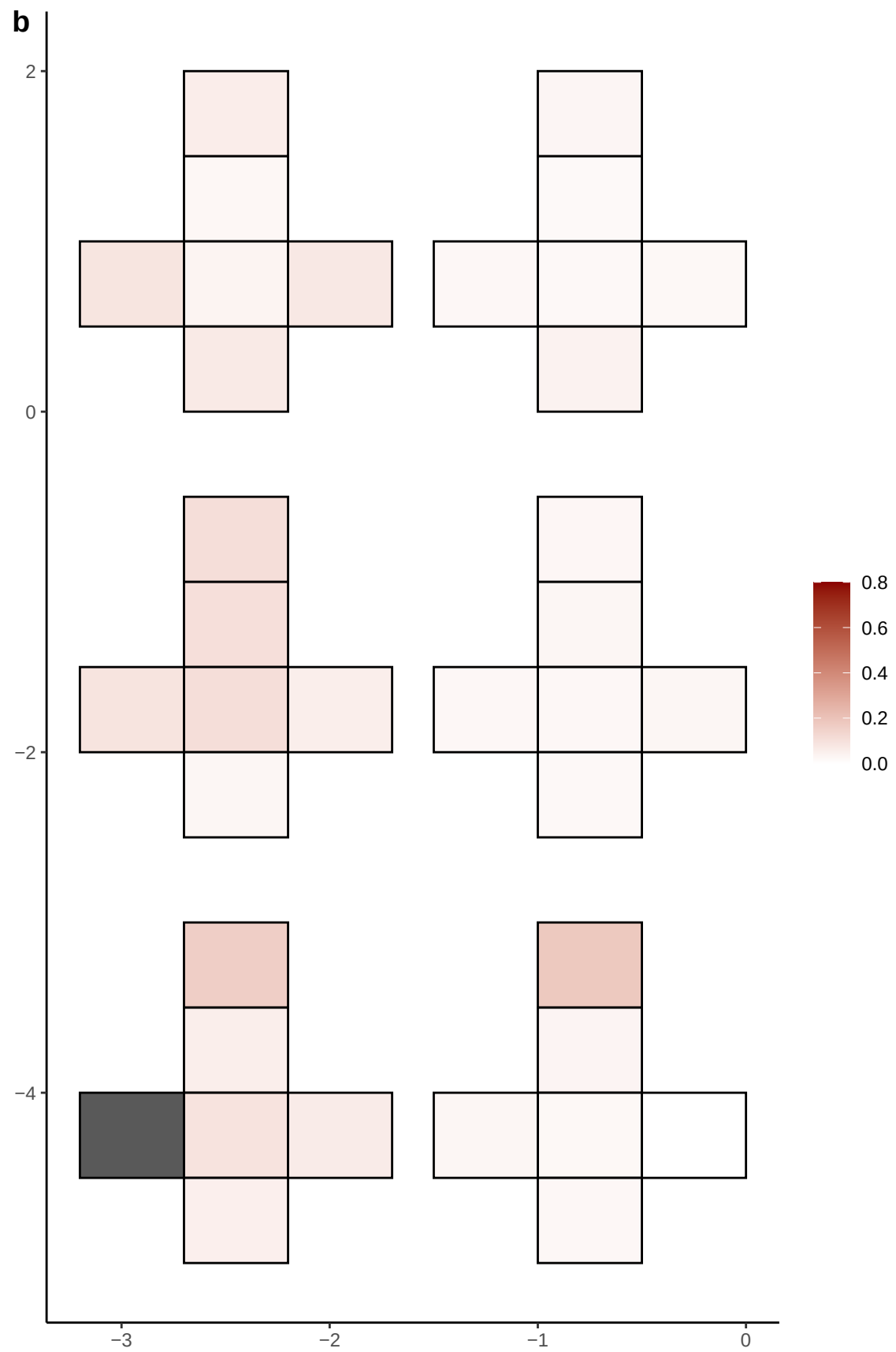

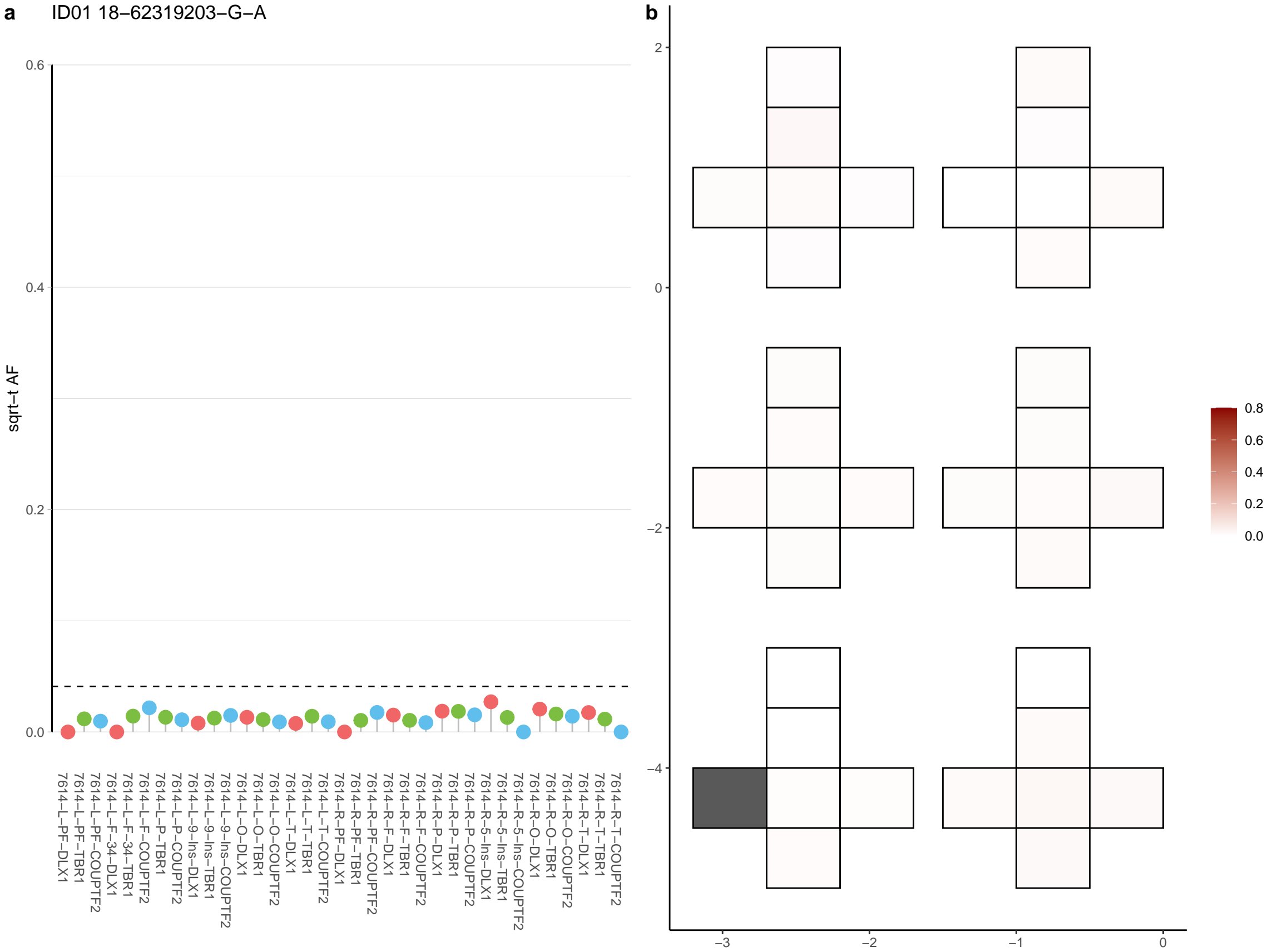

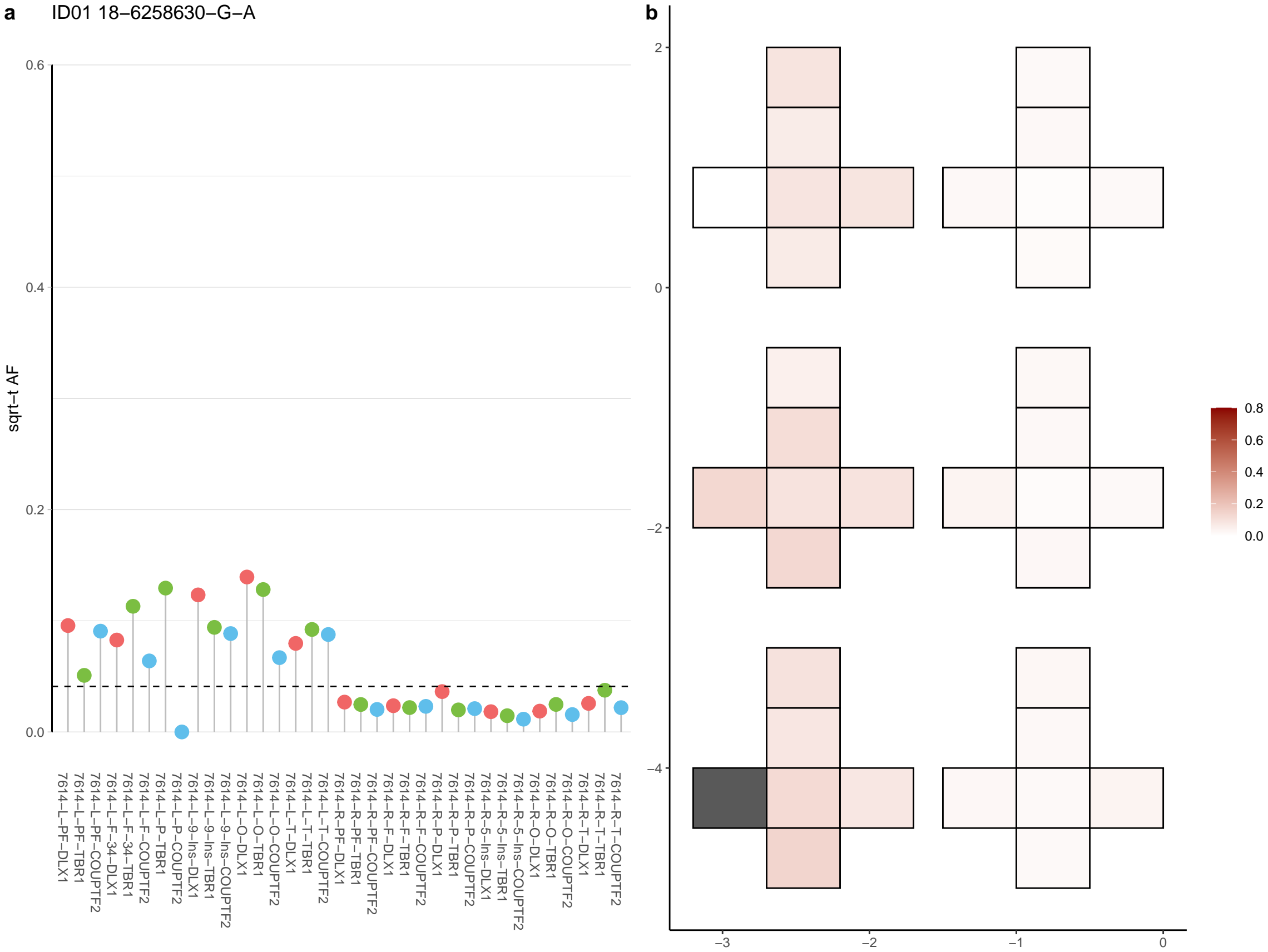

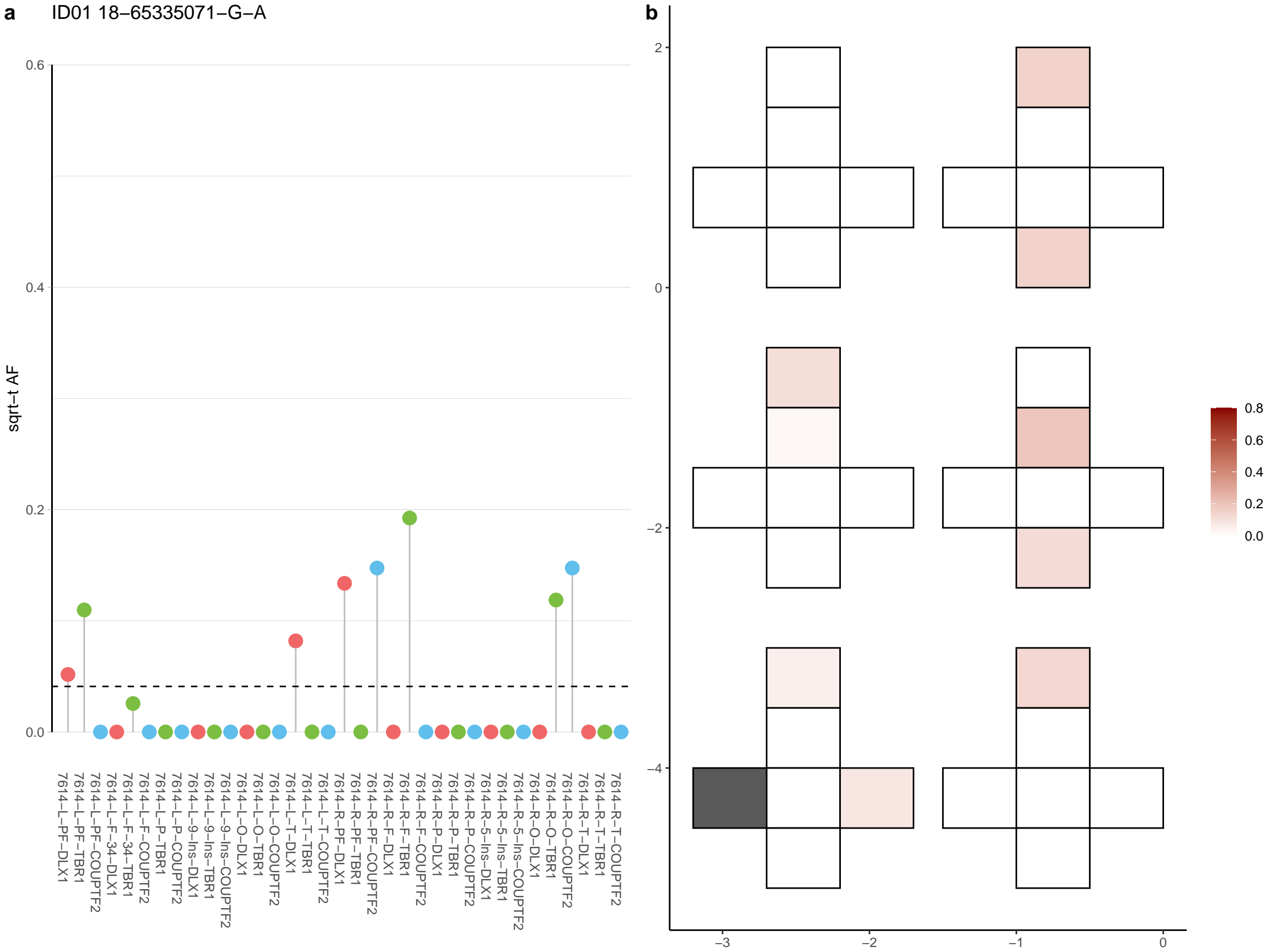

**a** ID01 18-72031228-C-T

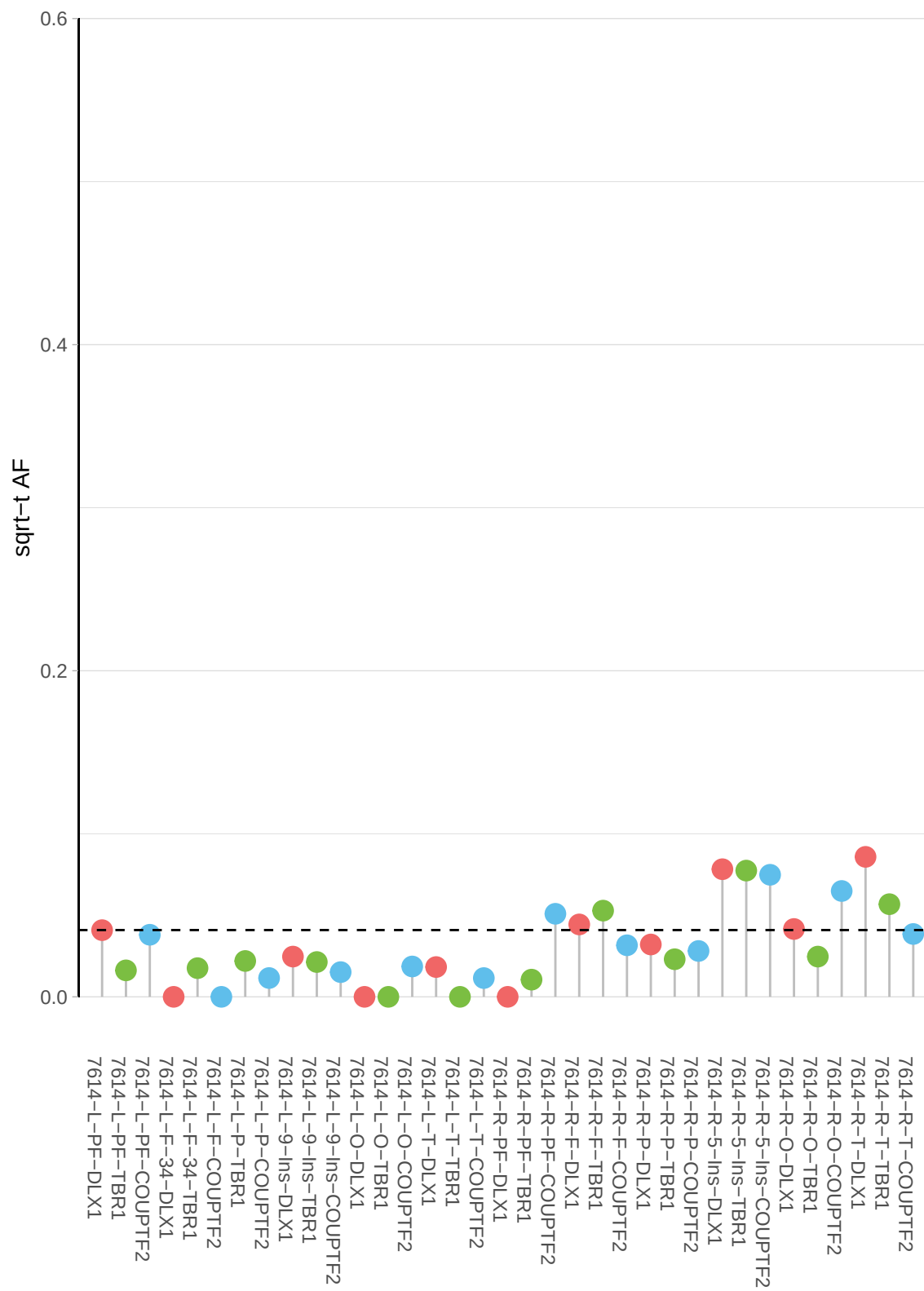

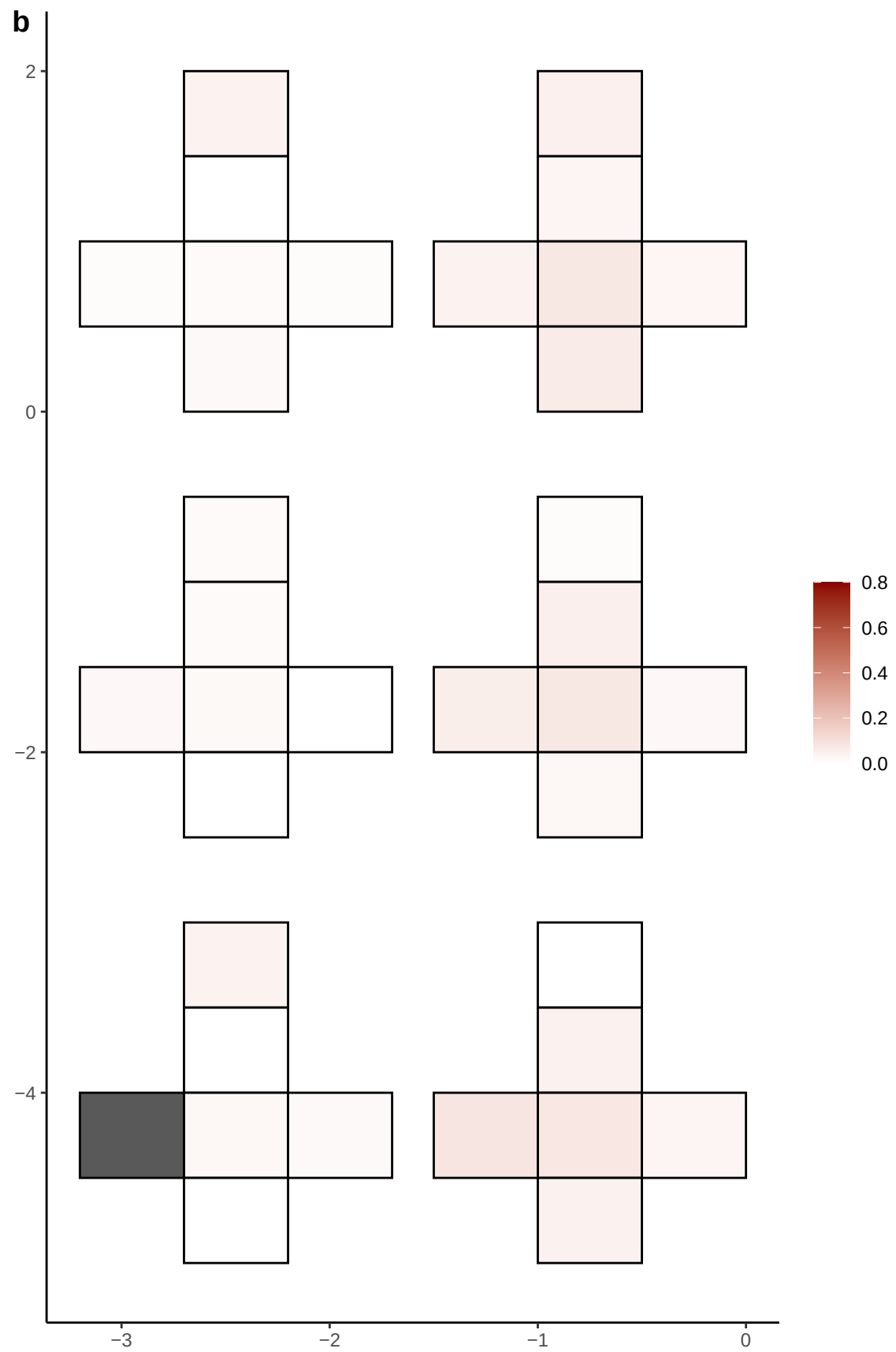

**a** ID01 18-73702872-A-C

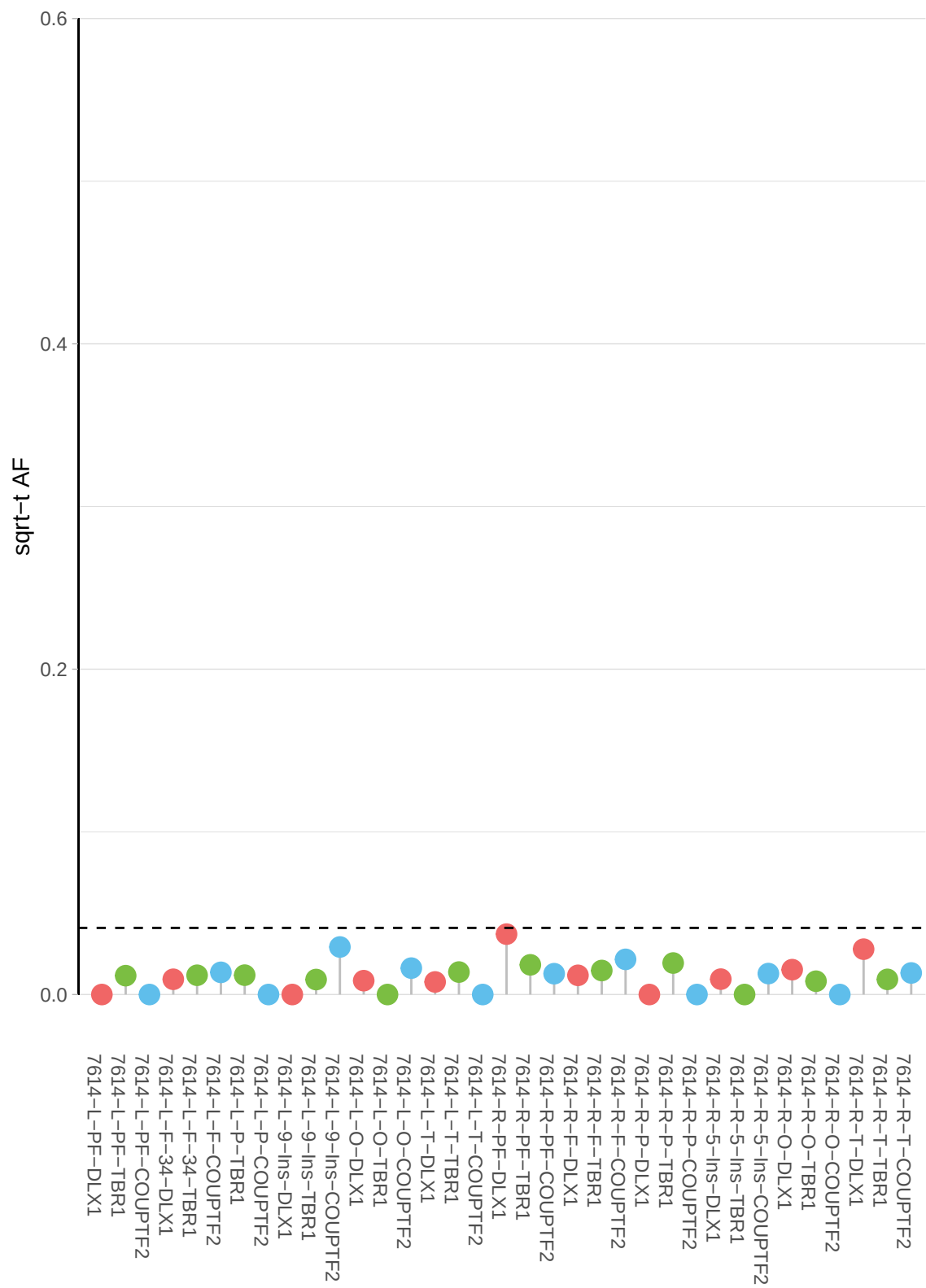

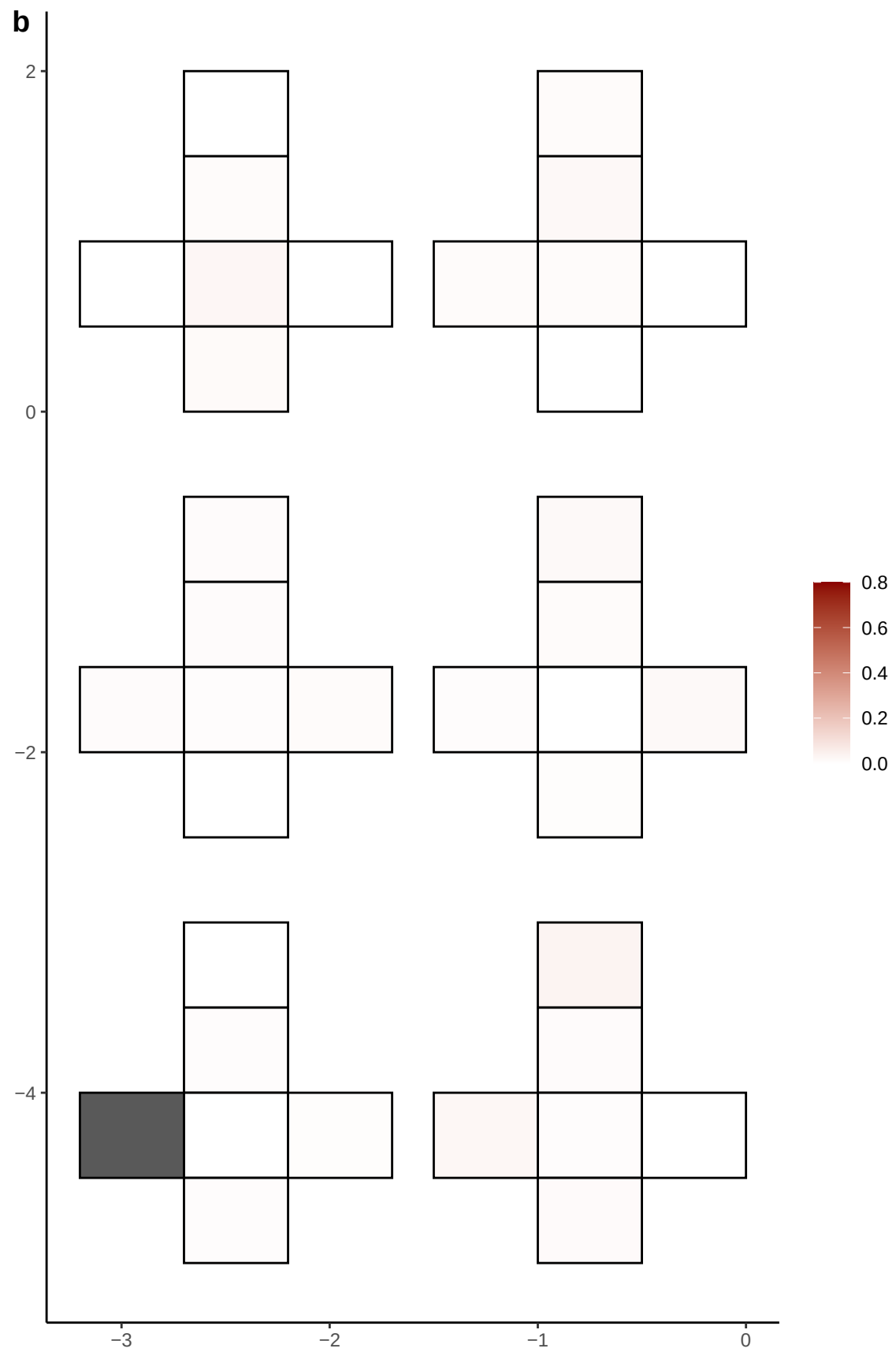

**a** ID01 19-23463606-T-C

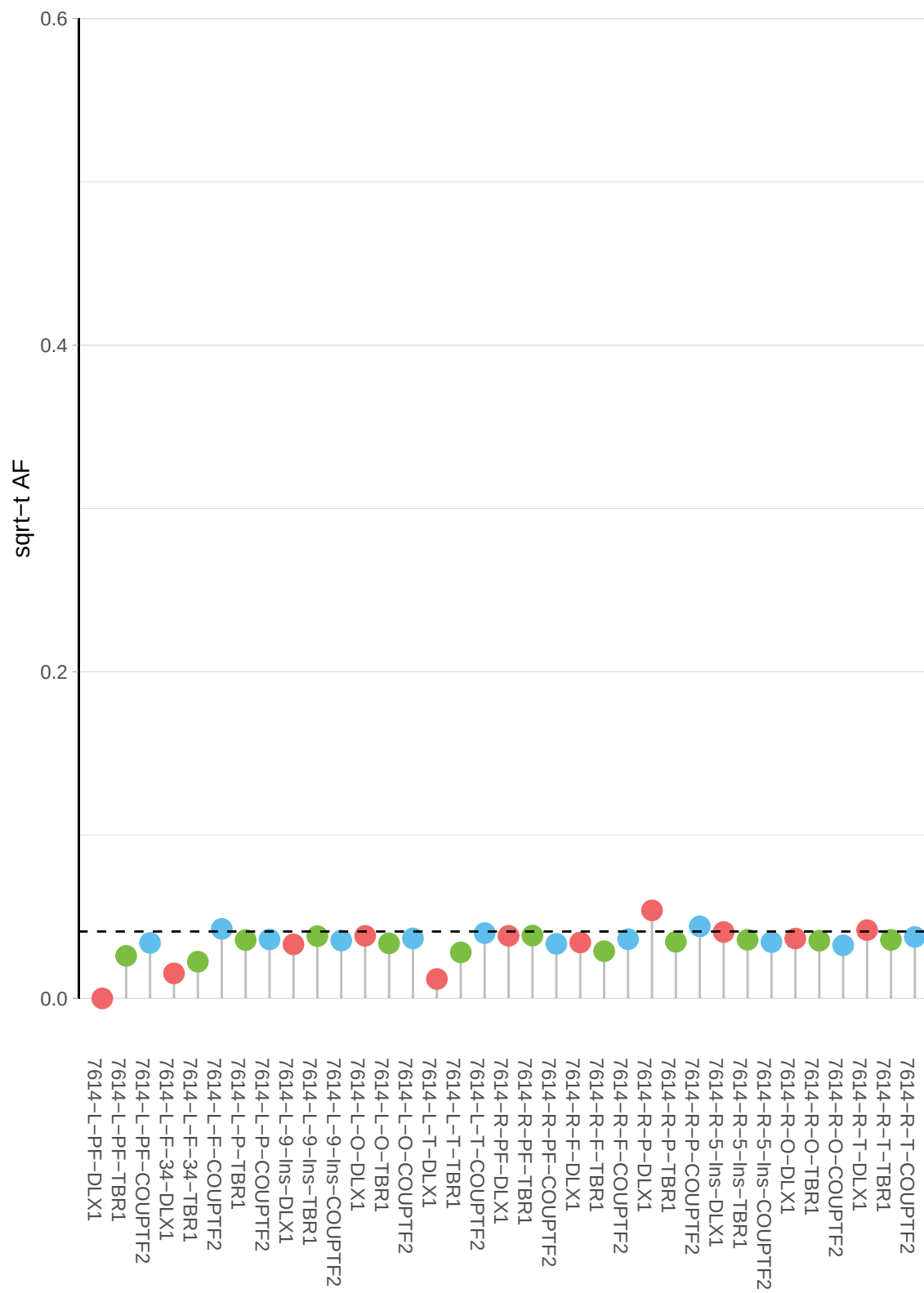

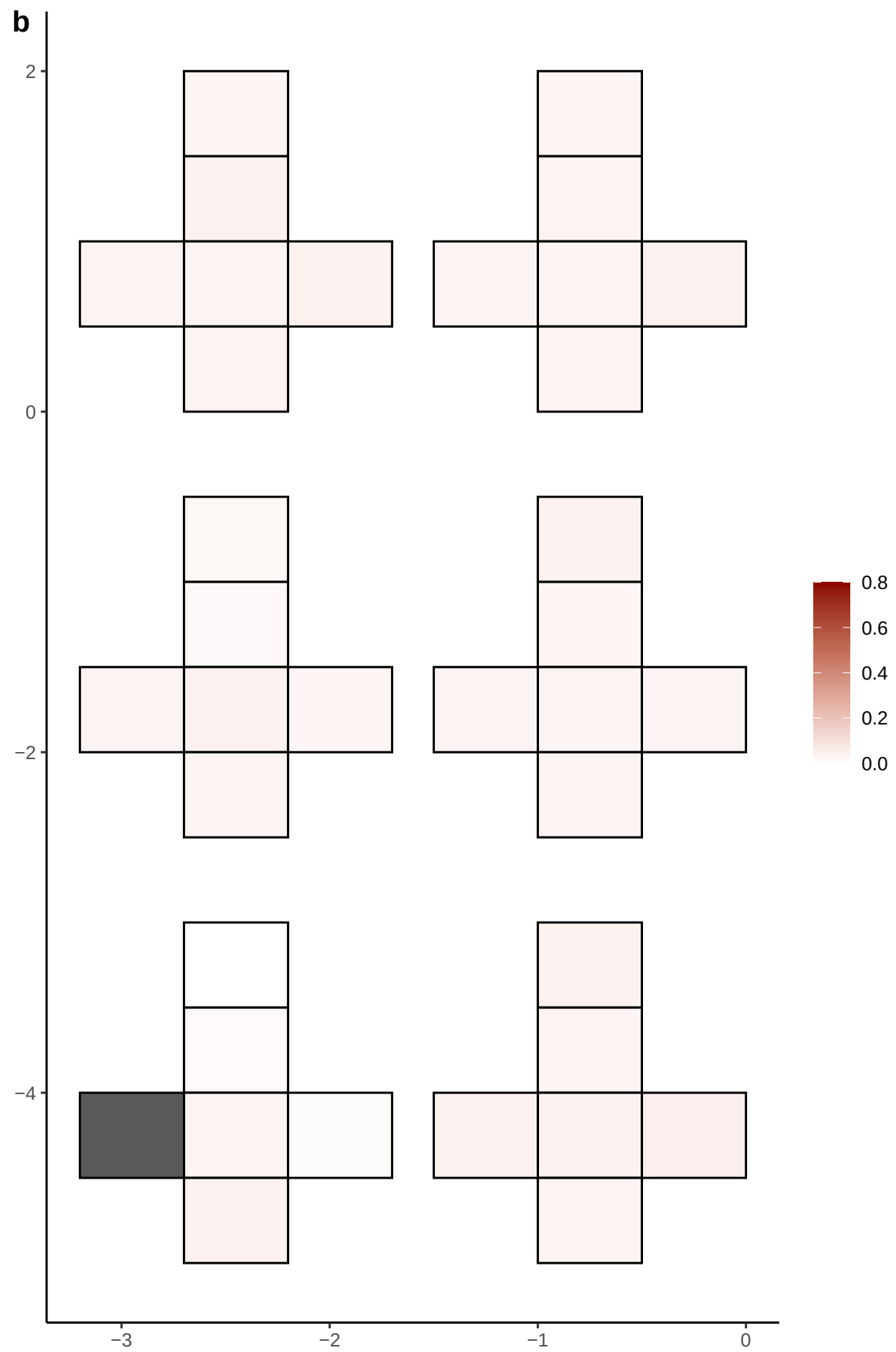

**a** ID01 19-29424630-G-T

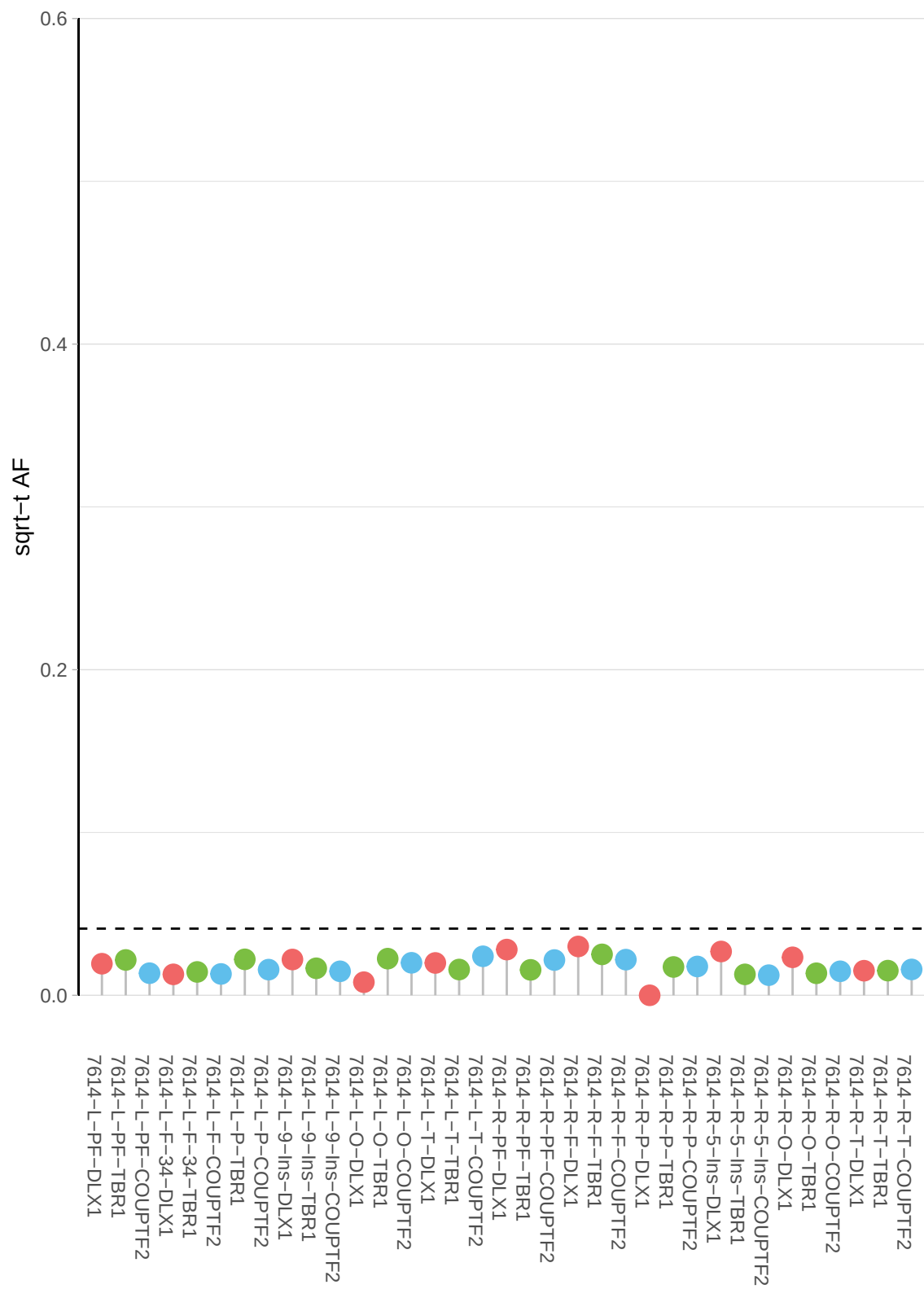

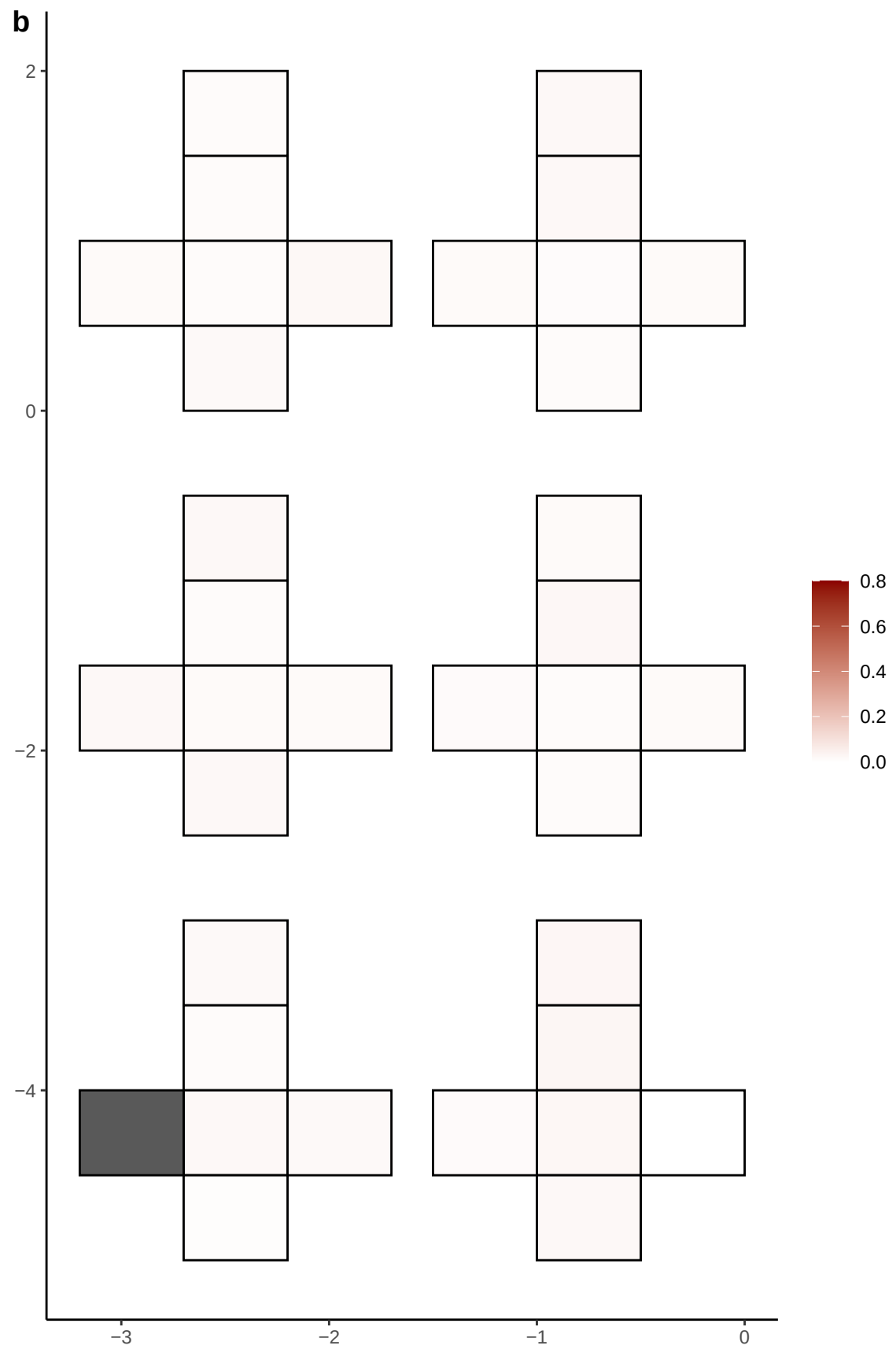

**a** ID01 19-31142255-G-A

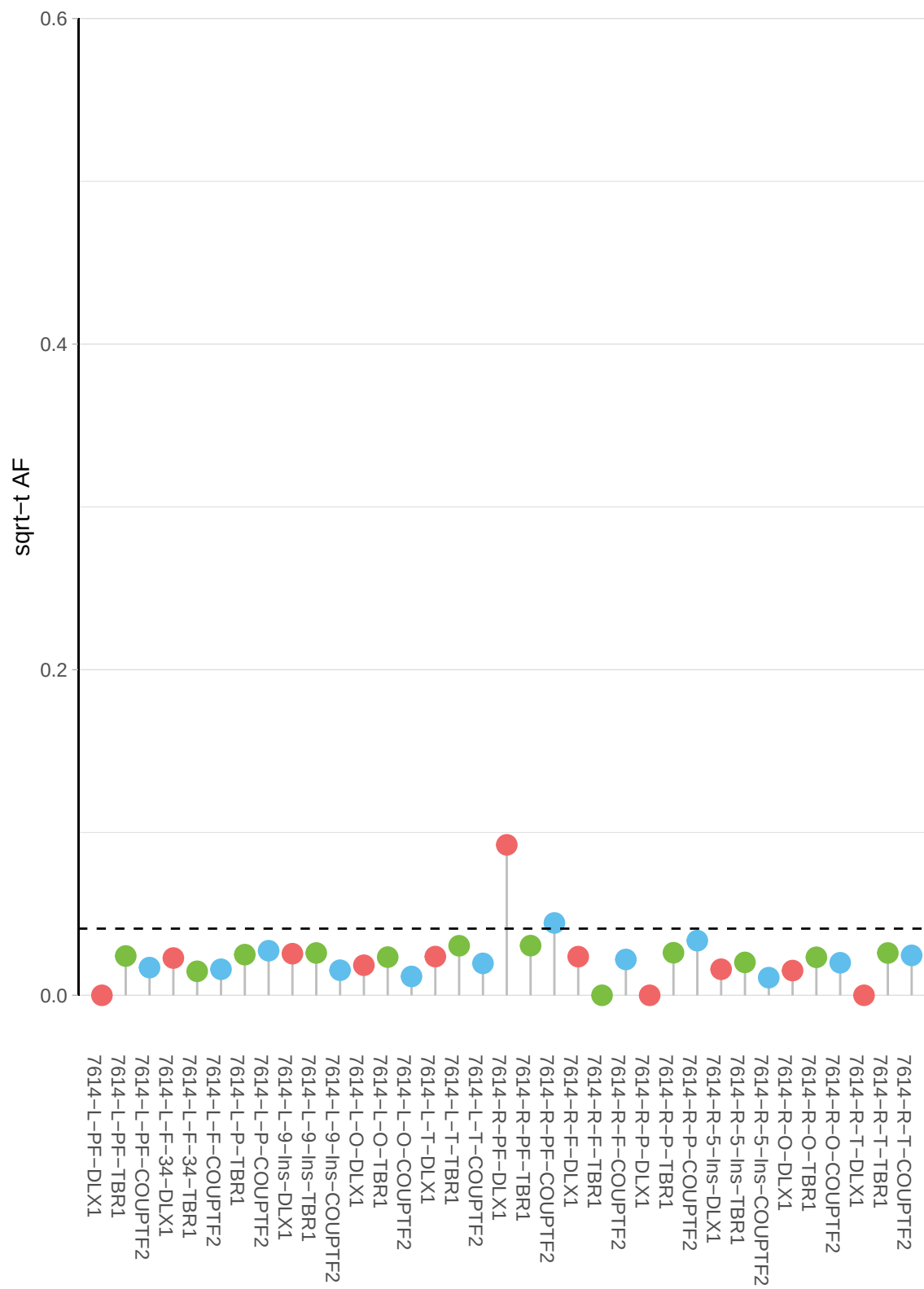

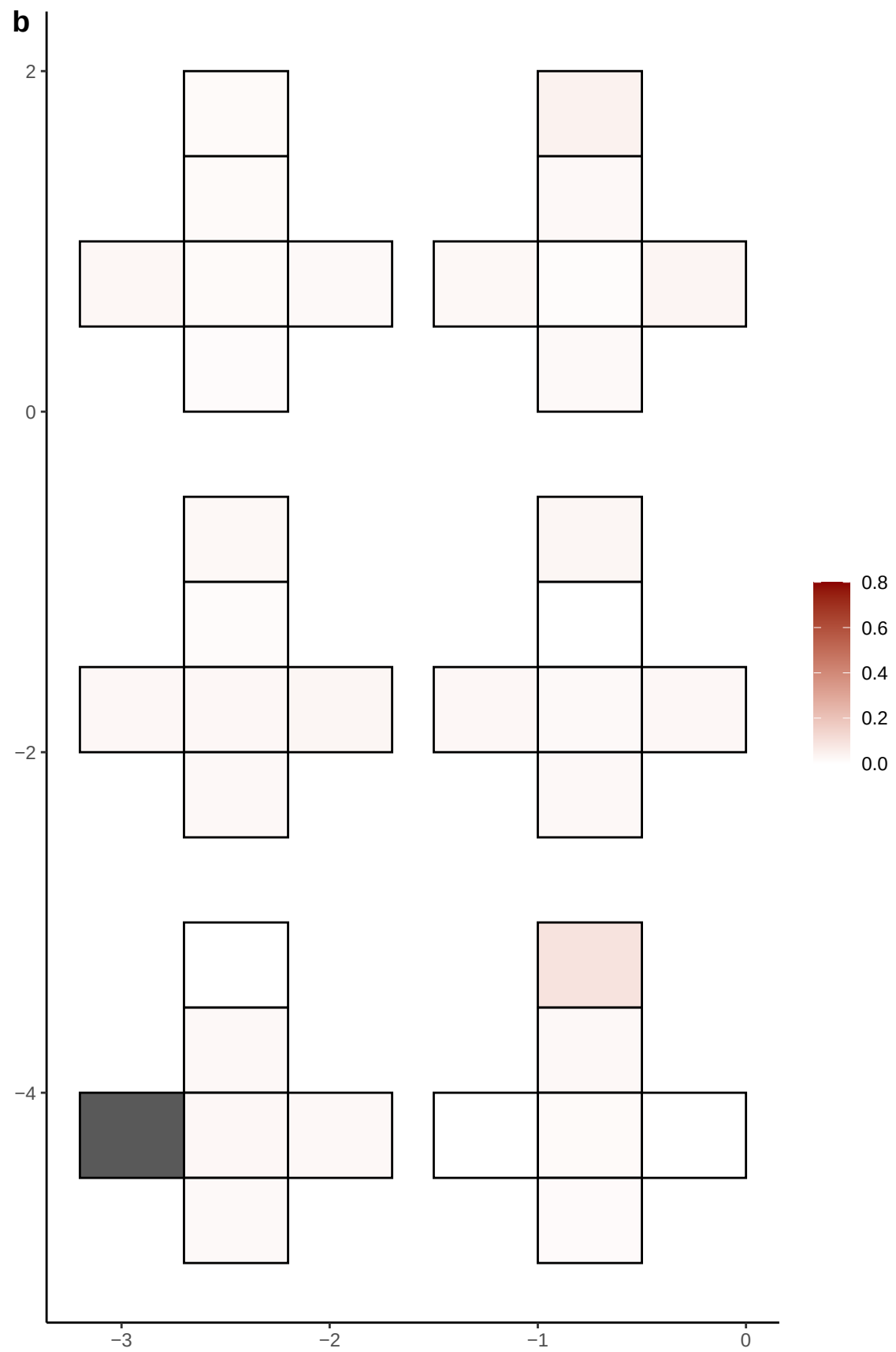

**a** ID01 19-39211568-A-G

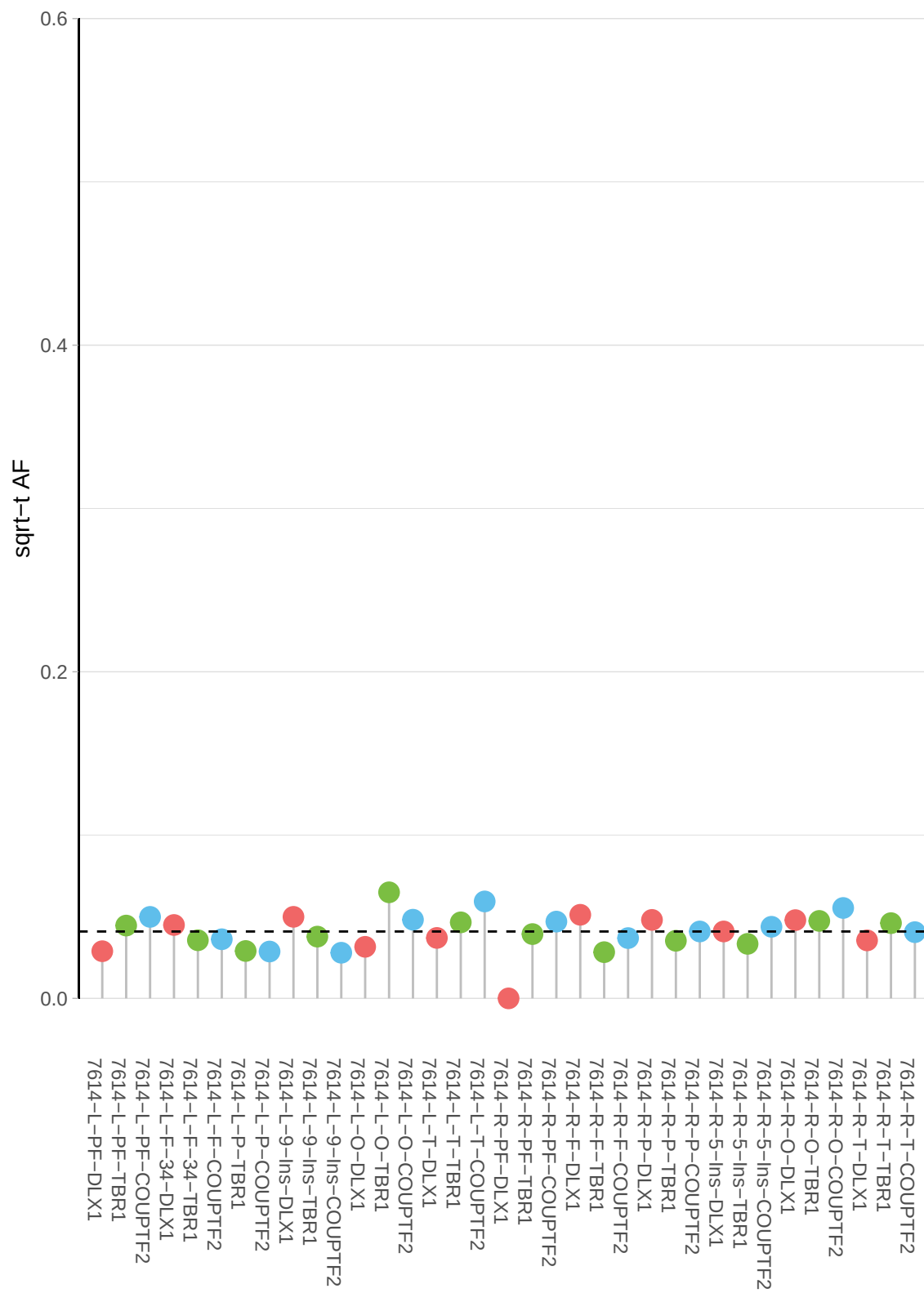

**b**

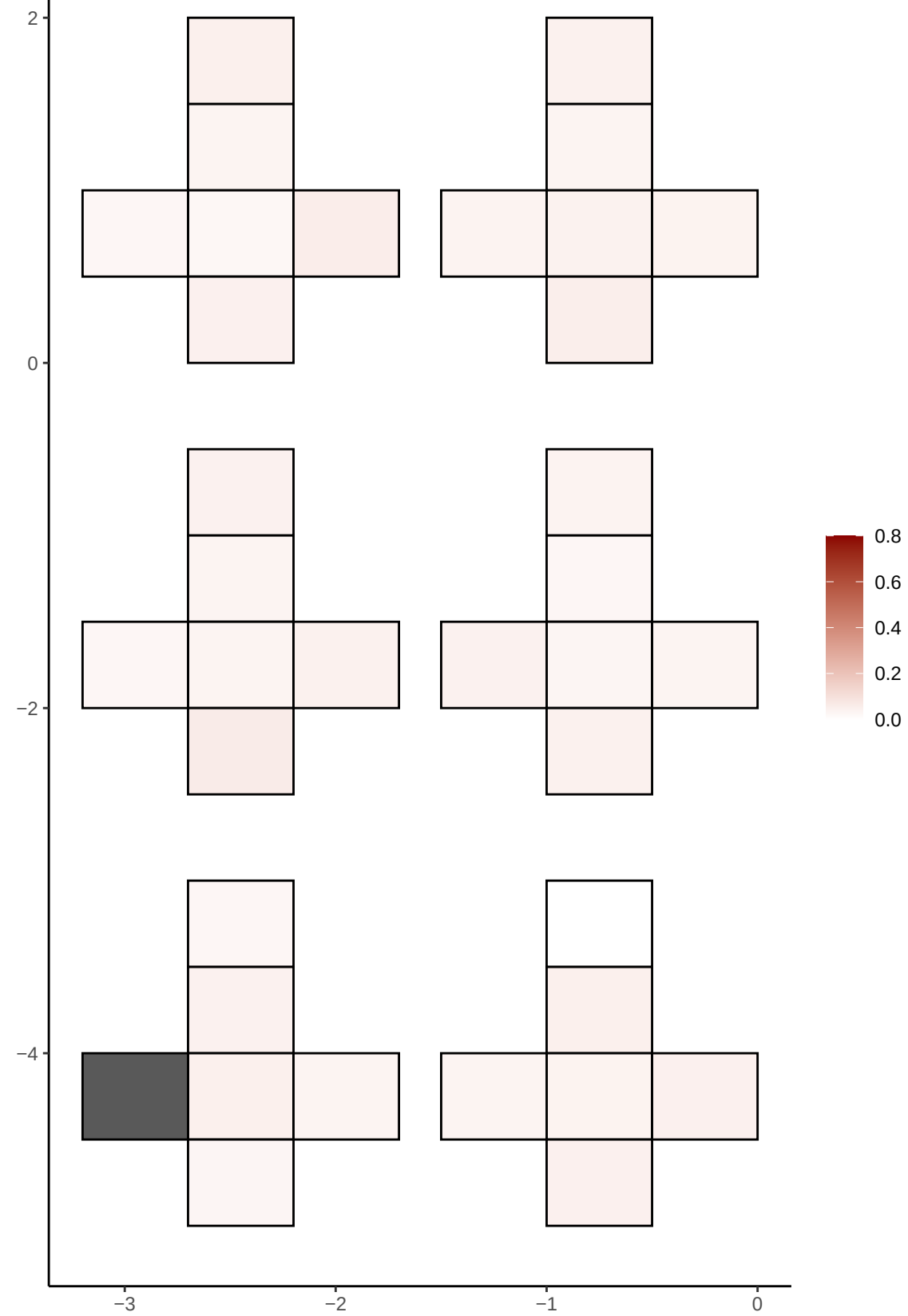

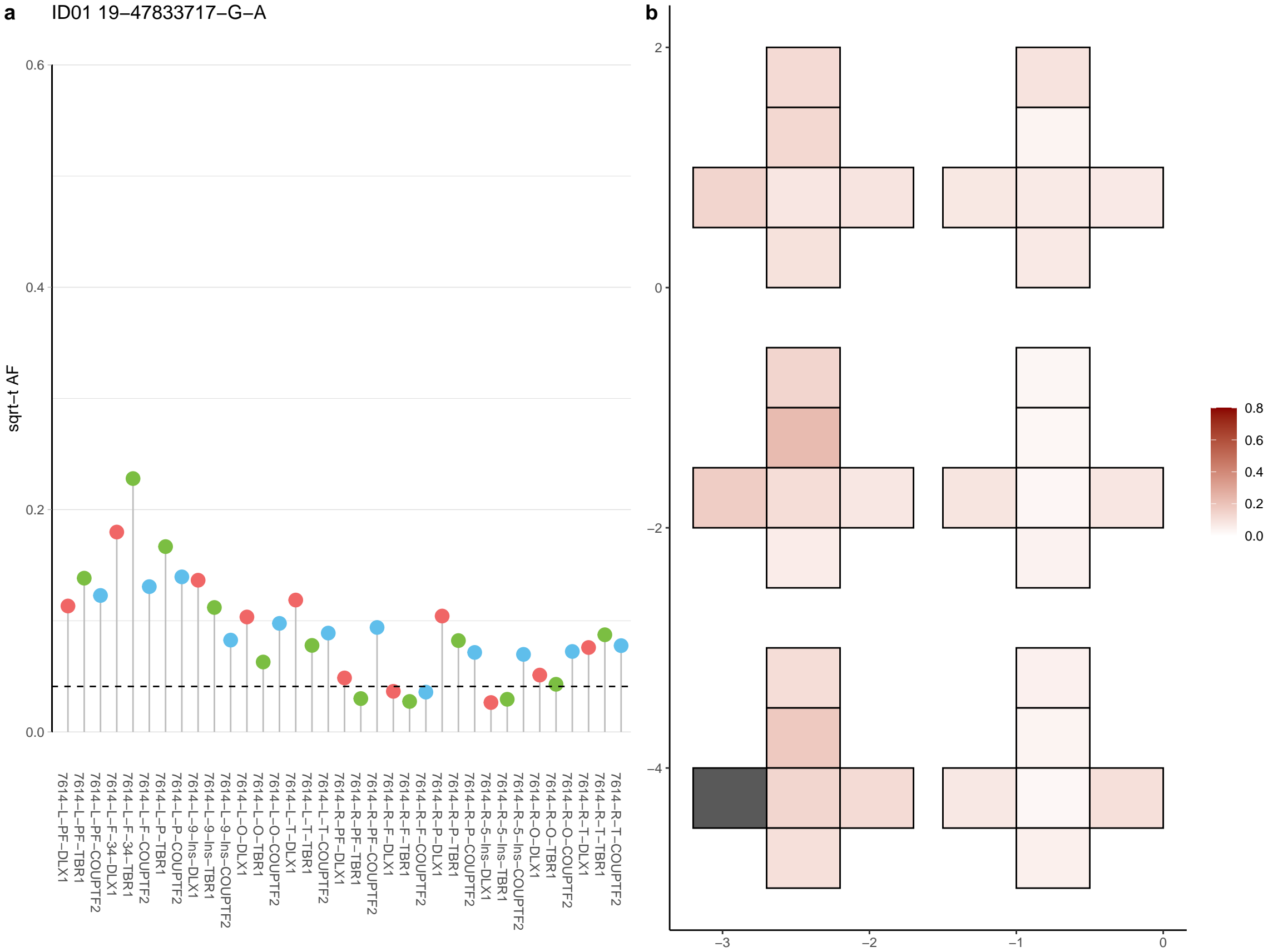

**a** ID01 19-58435948-C-A

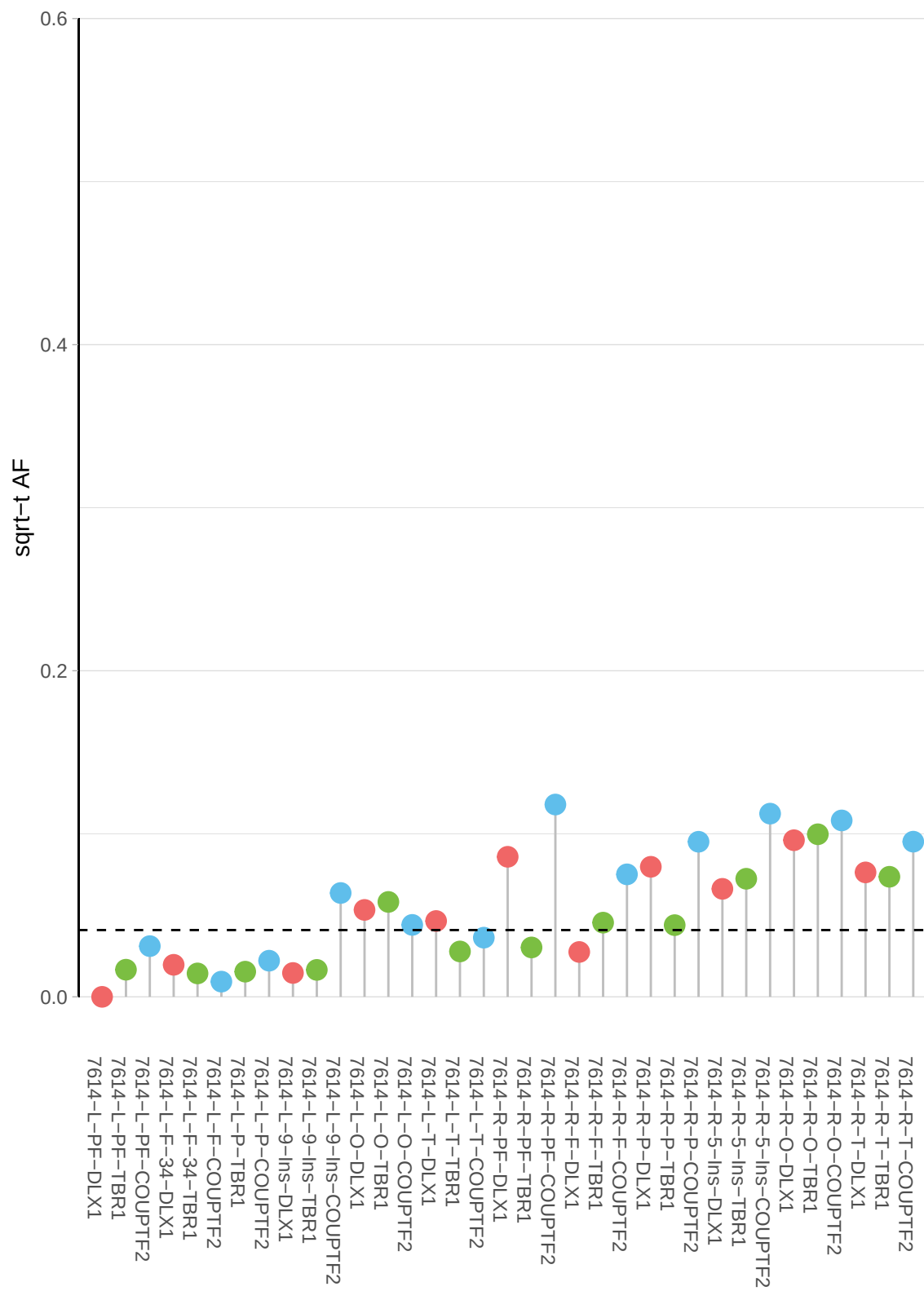

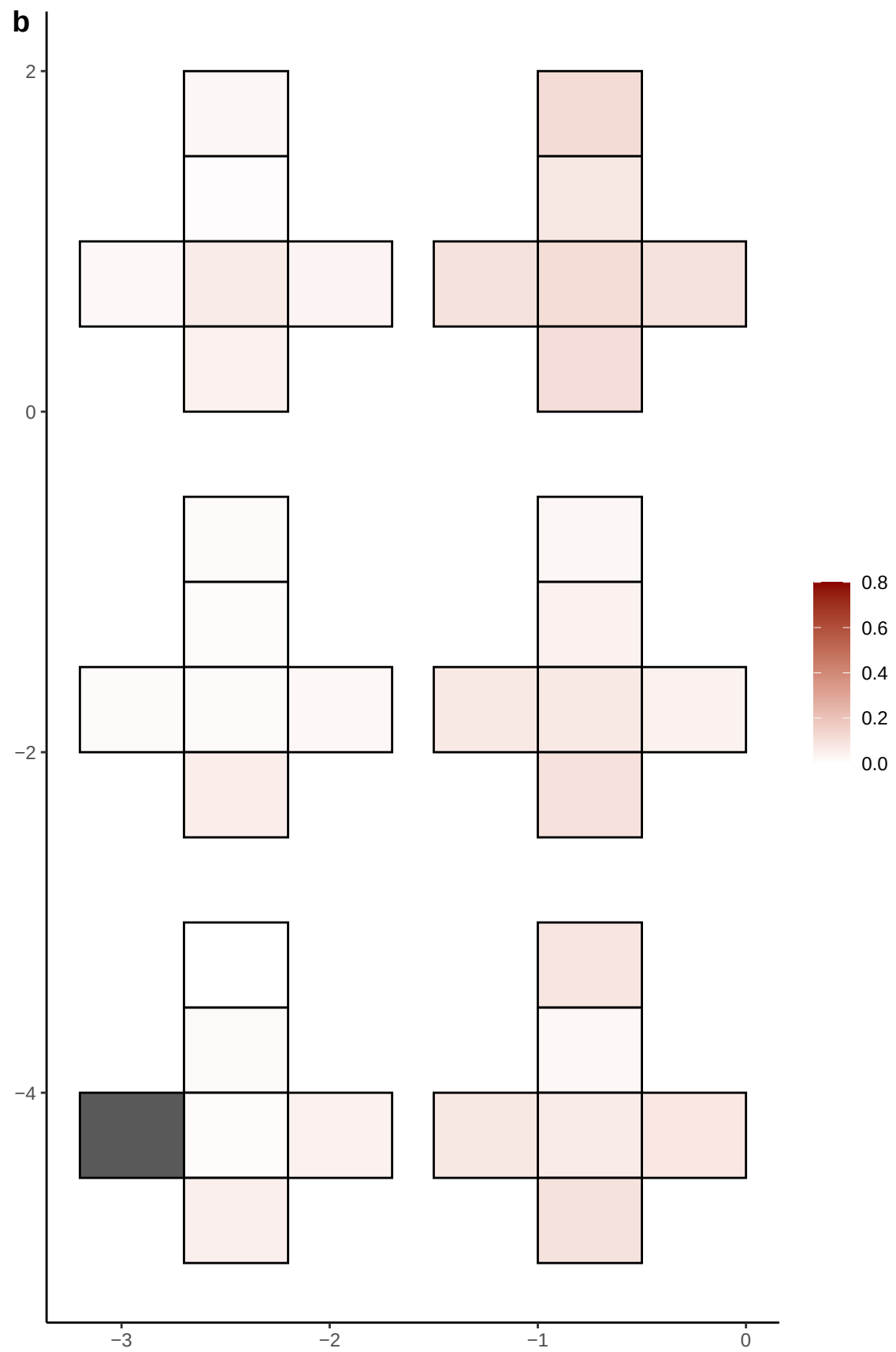

**a** ID01 2-101097017-C-T

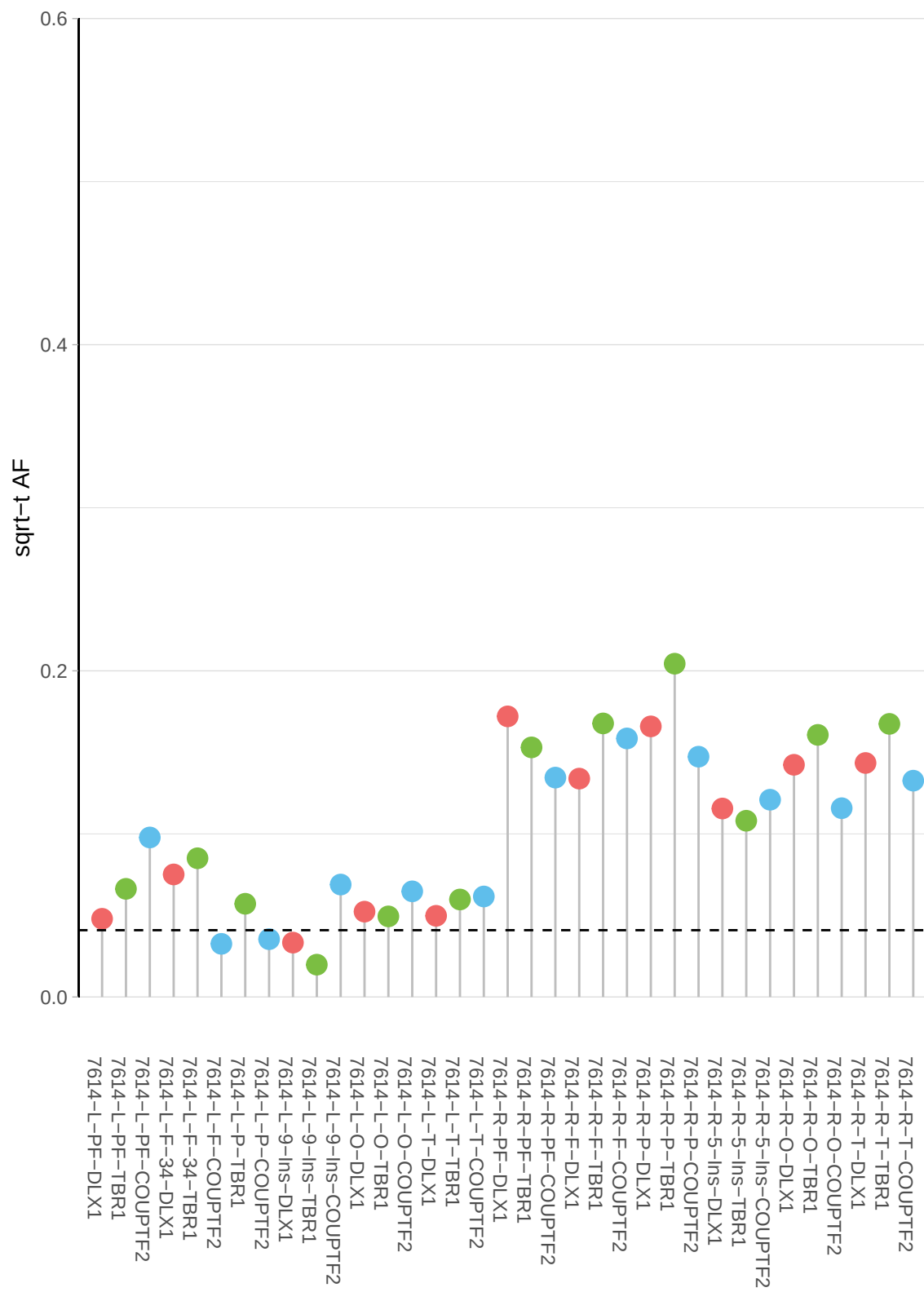

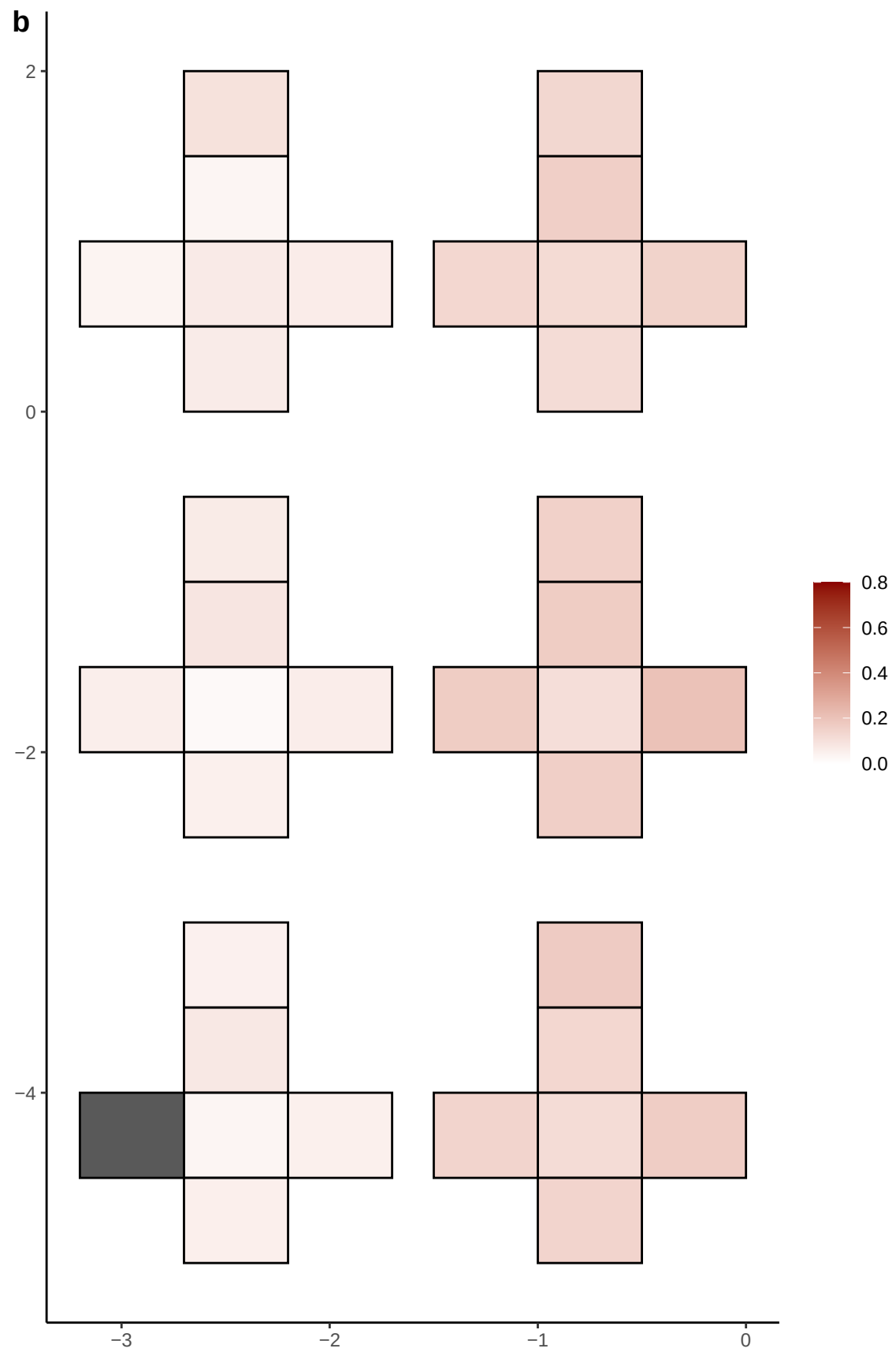

**a** ID01 2-119890951-G-A

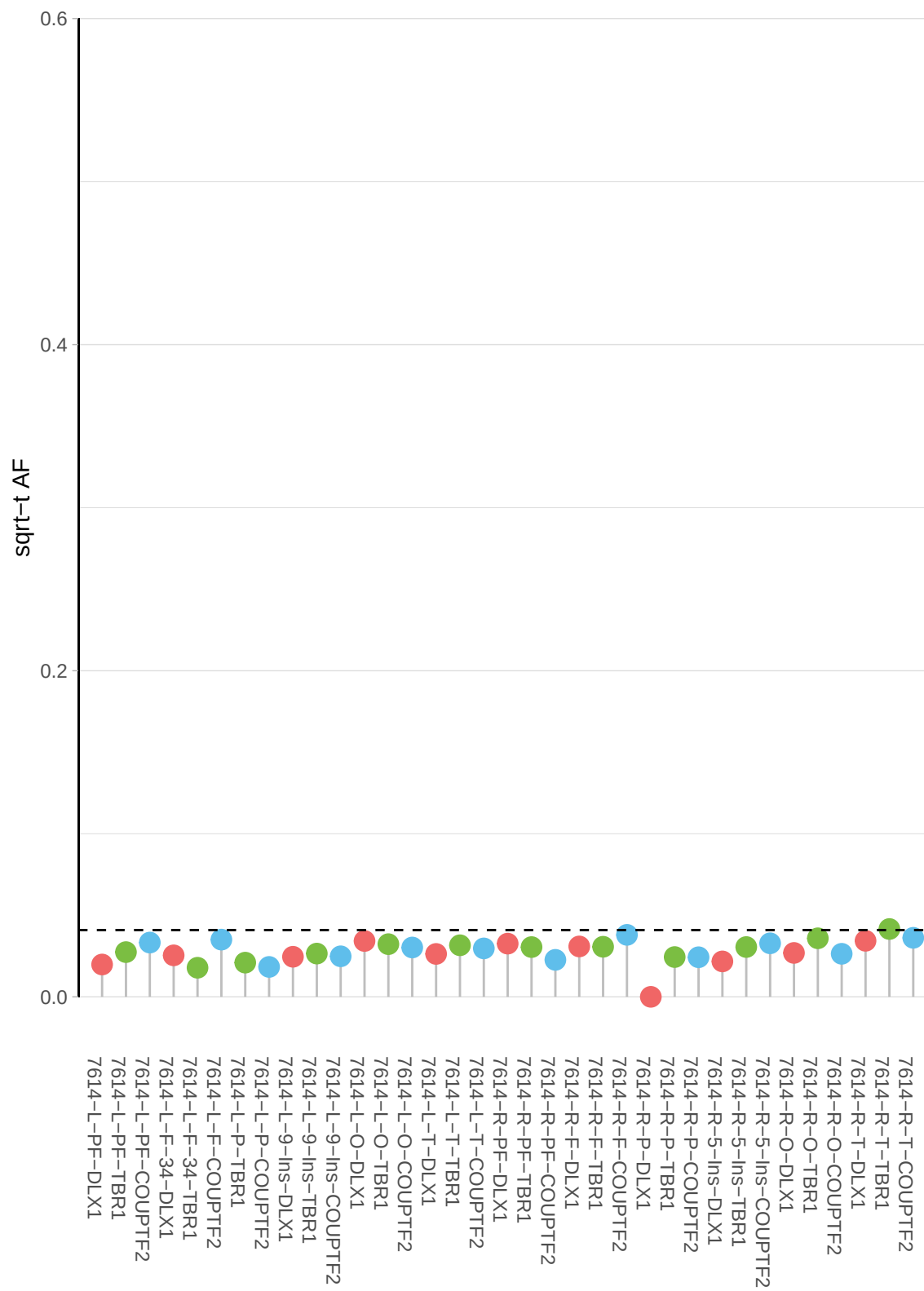

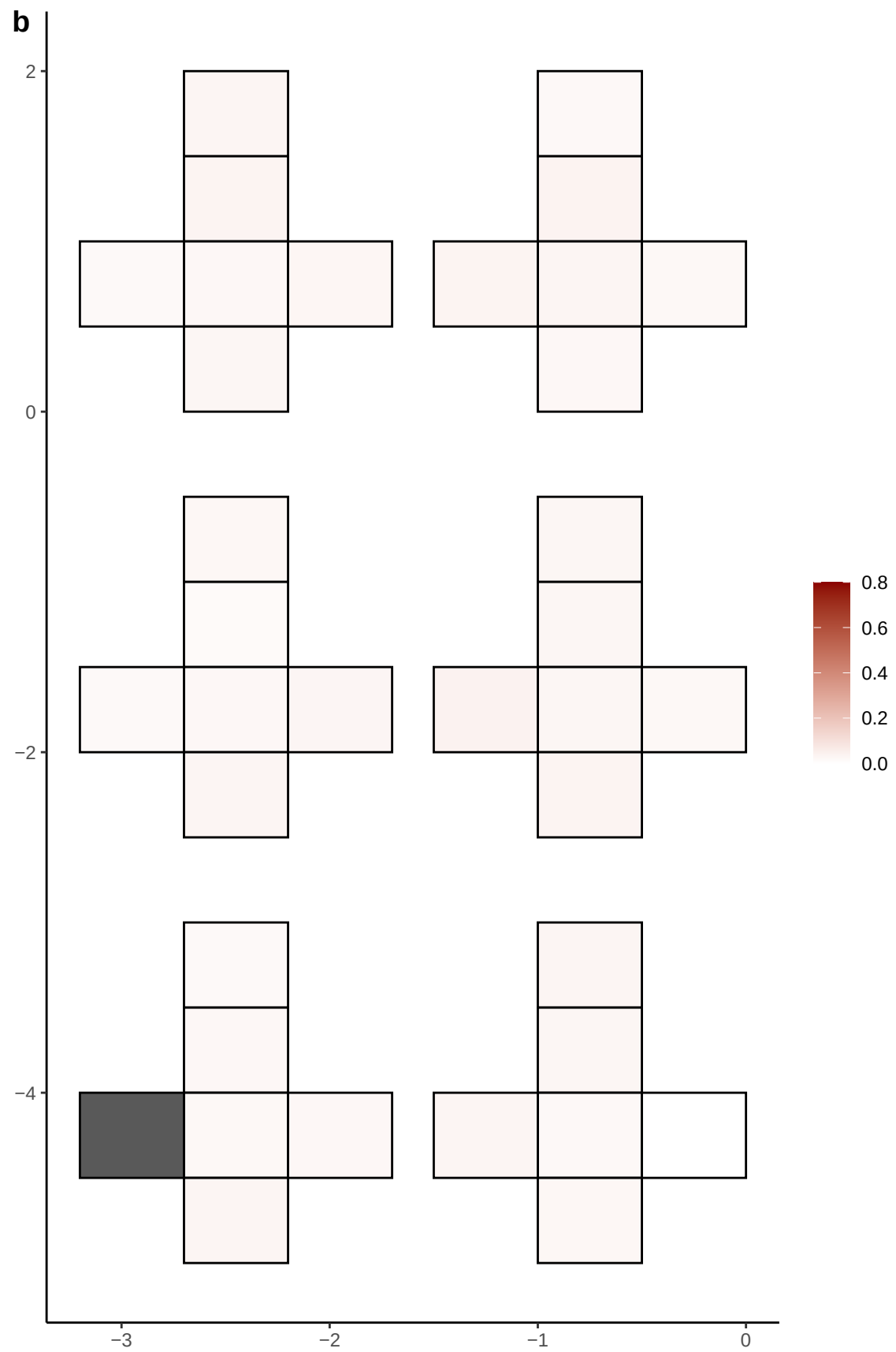

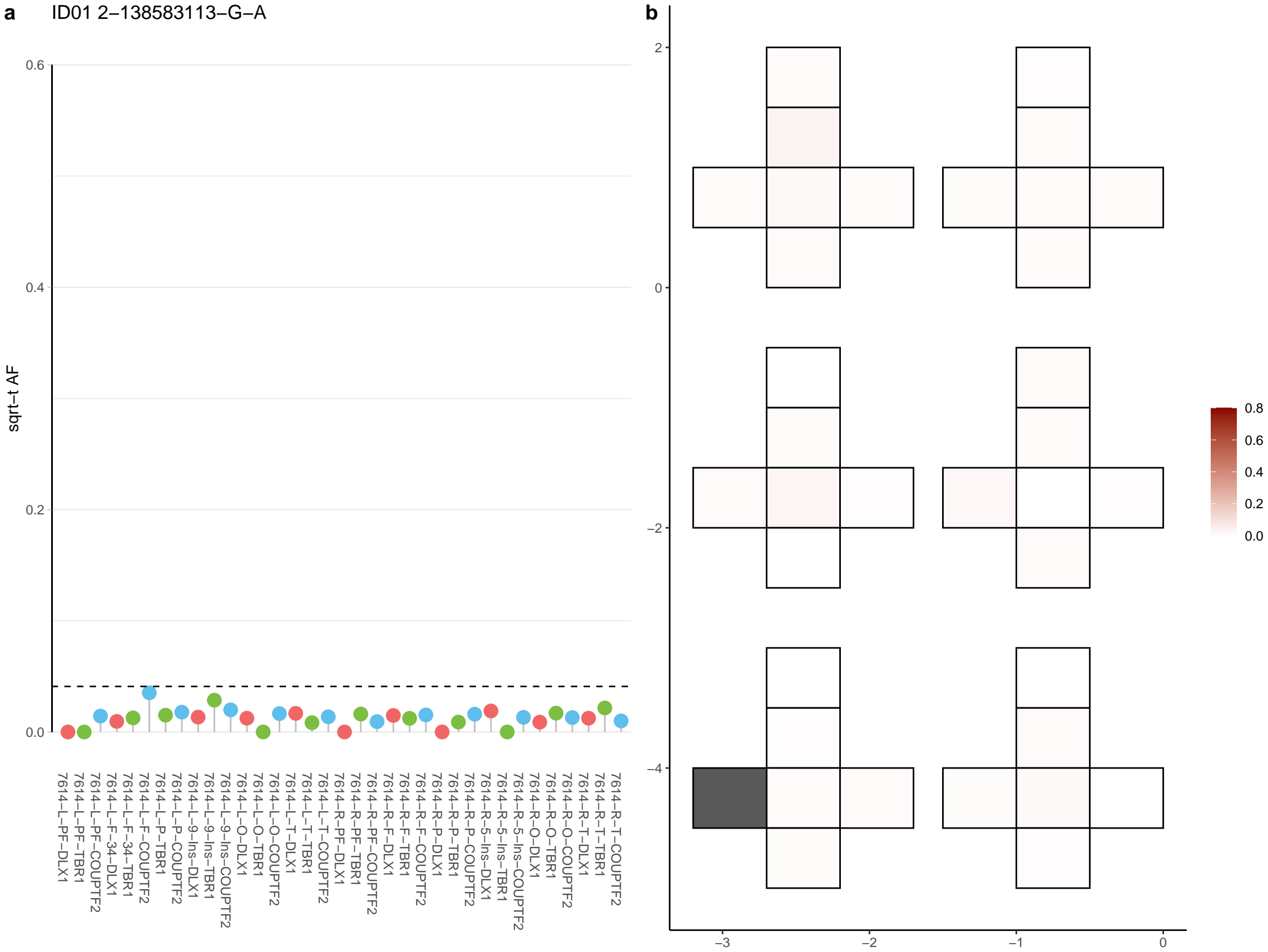

**a** ID01 2-139753954-C-T

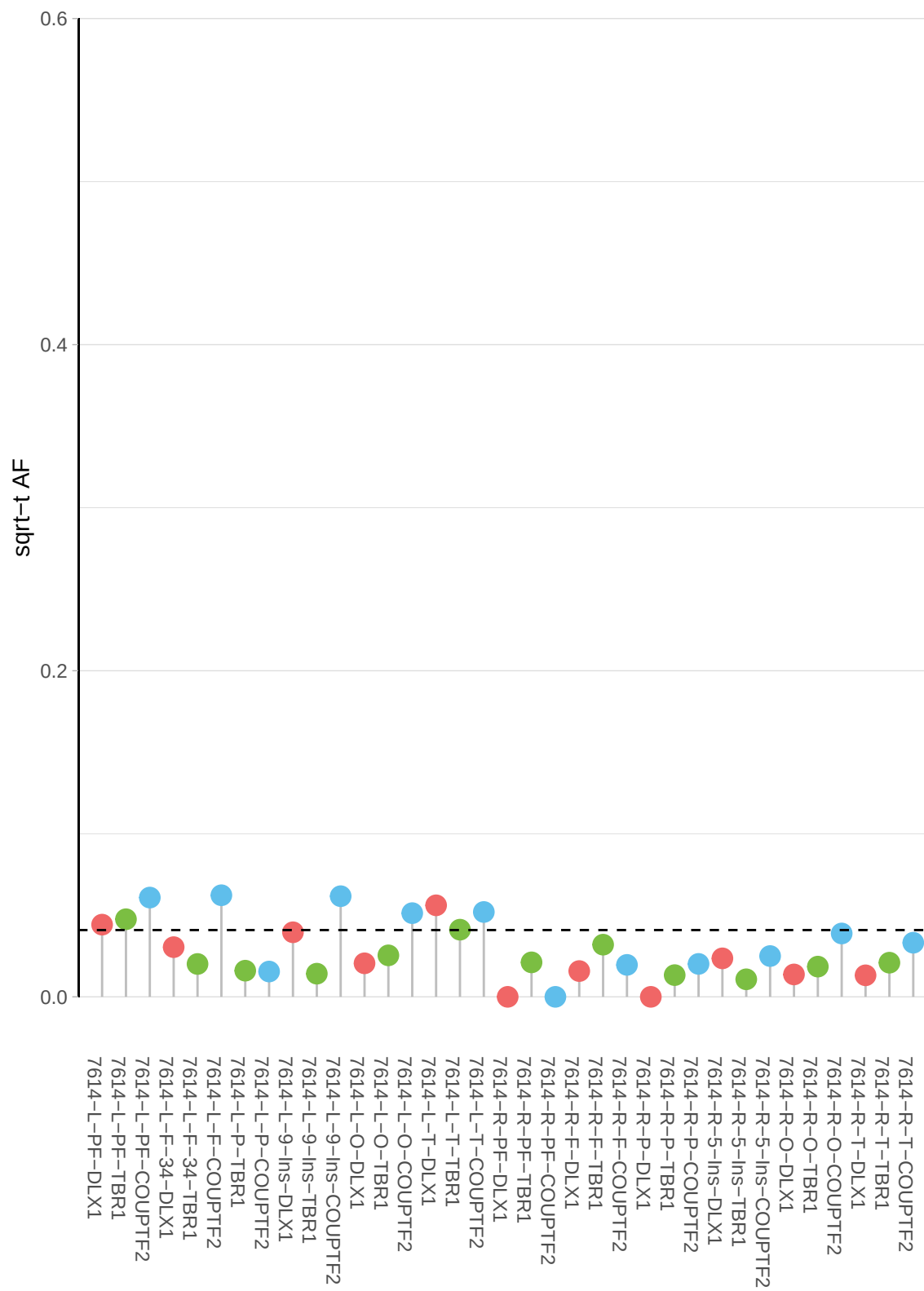

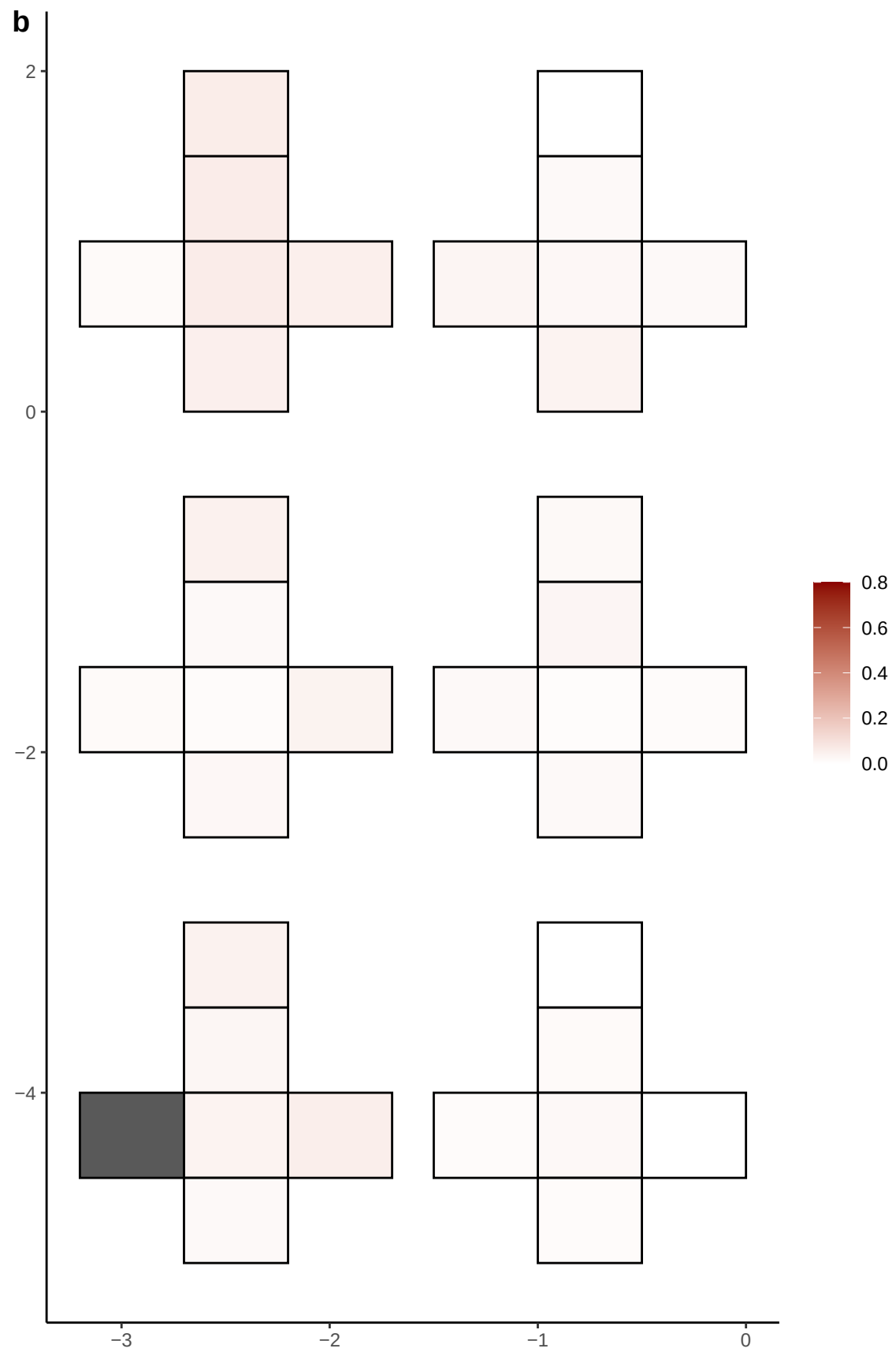

**a** ID01 2-142968696-G-T

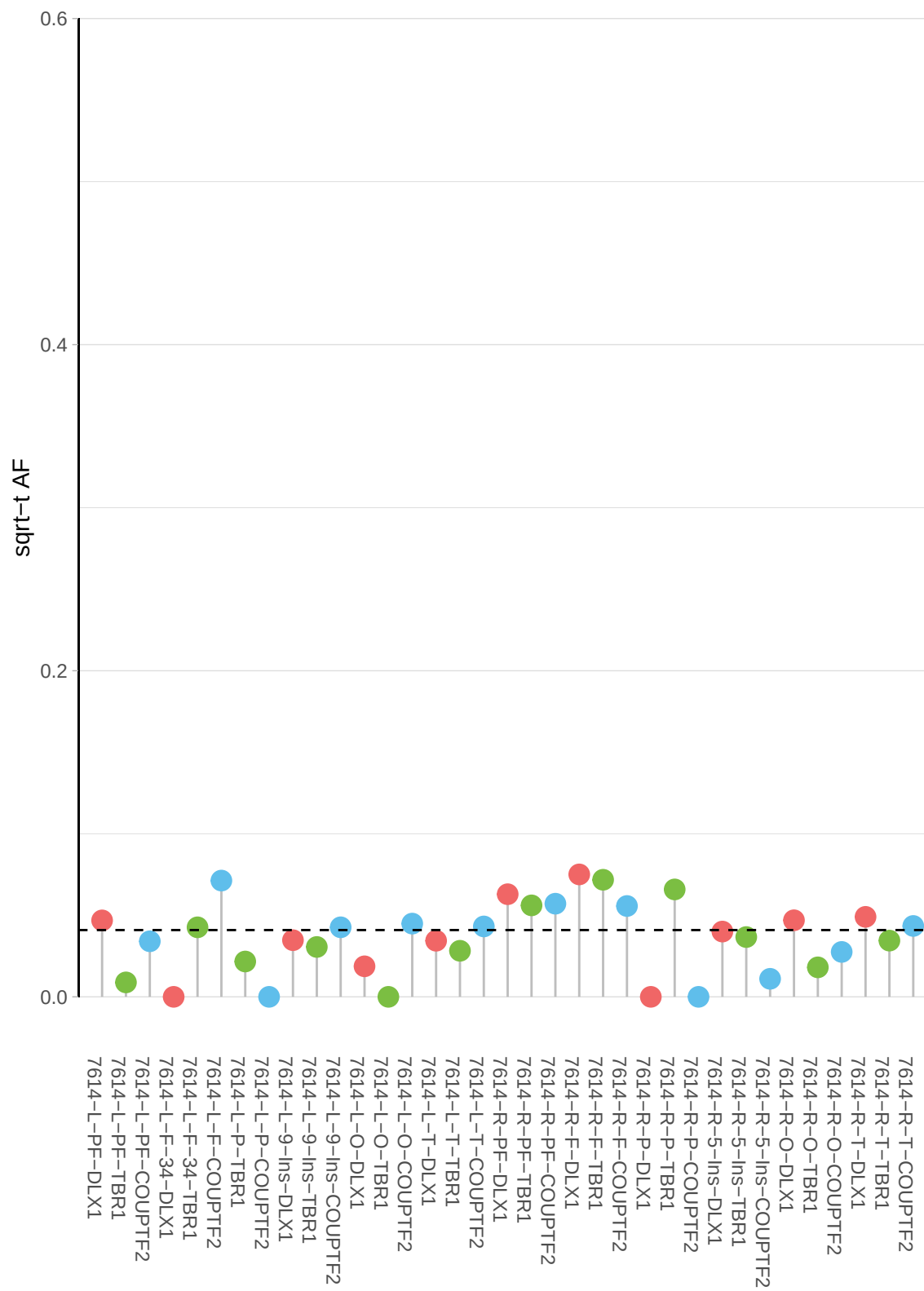

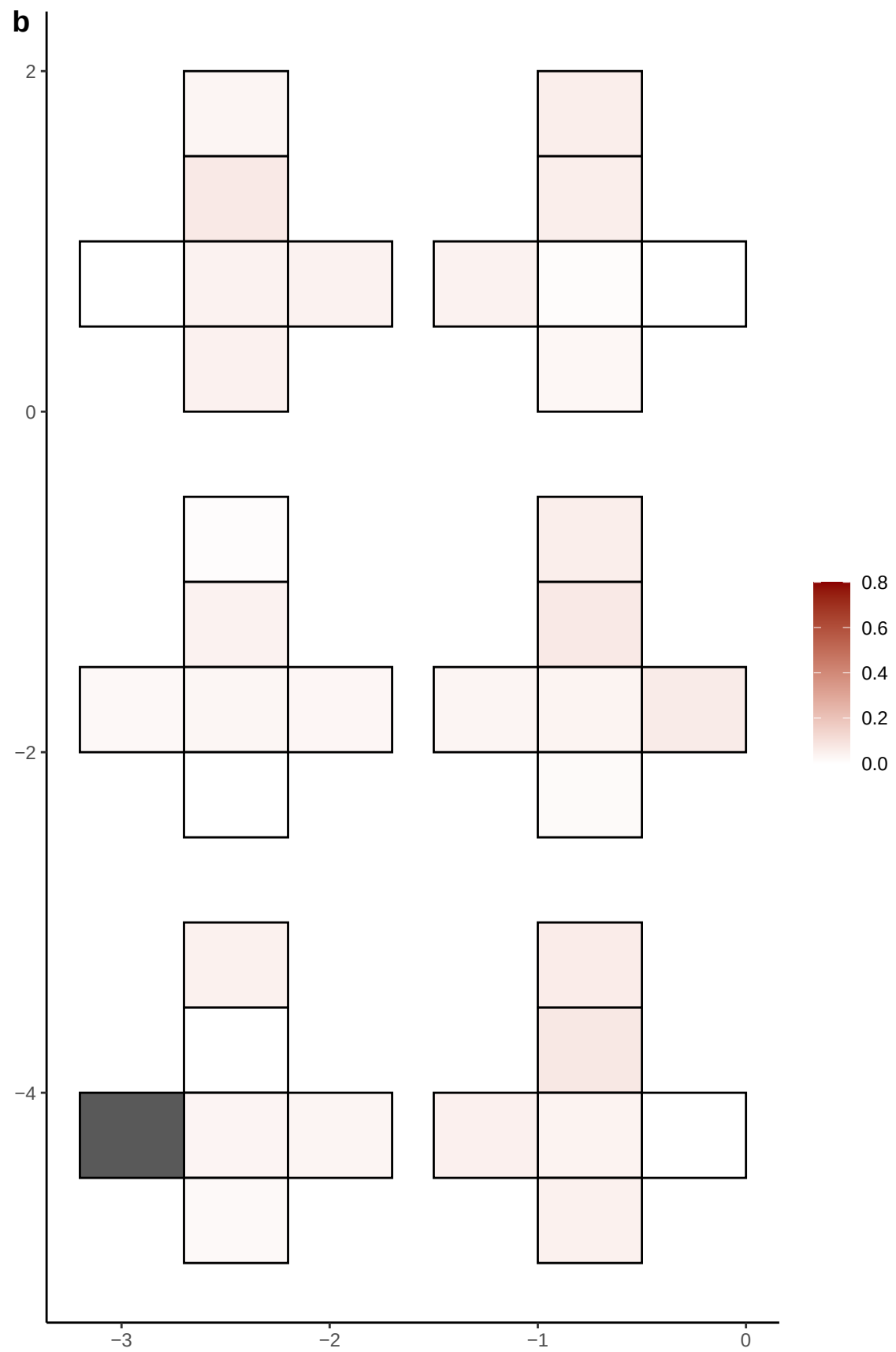

**a** ID01 2-143727379-C-T

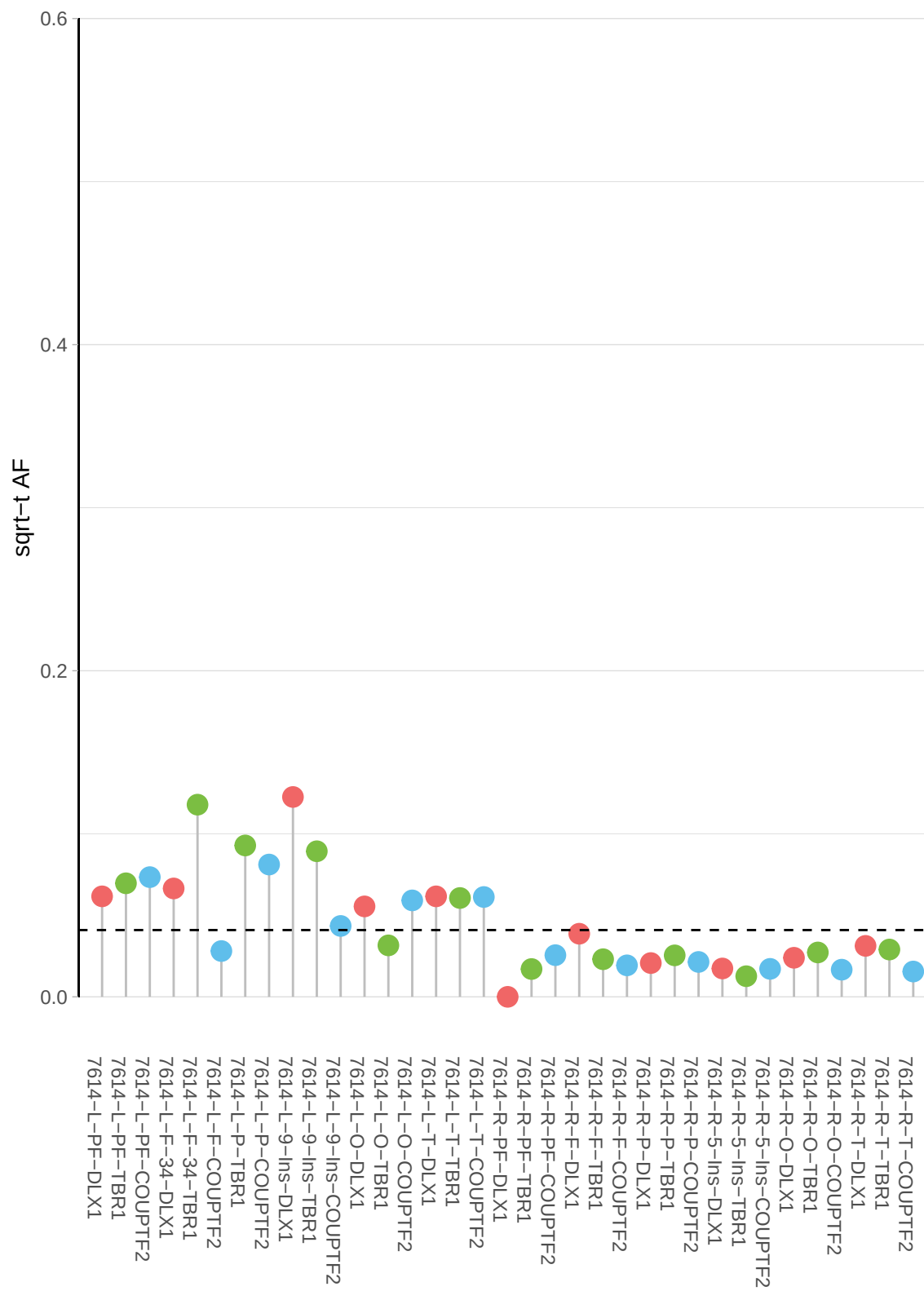

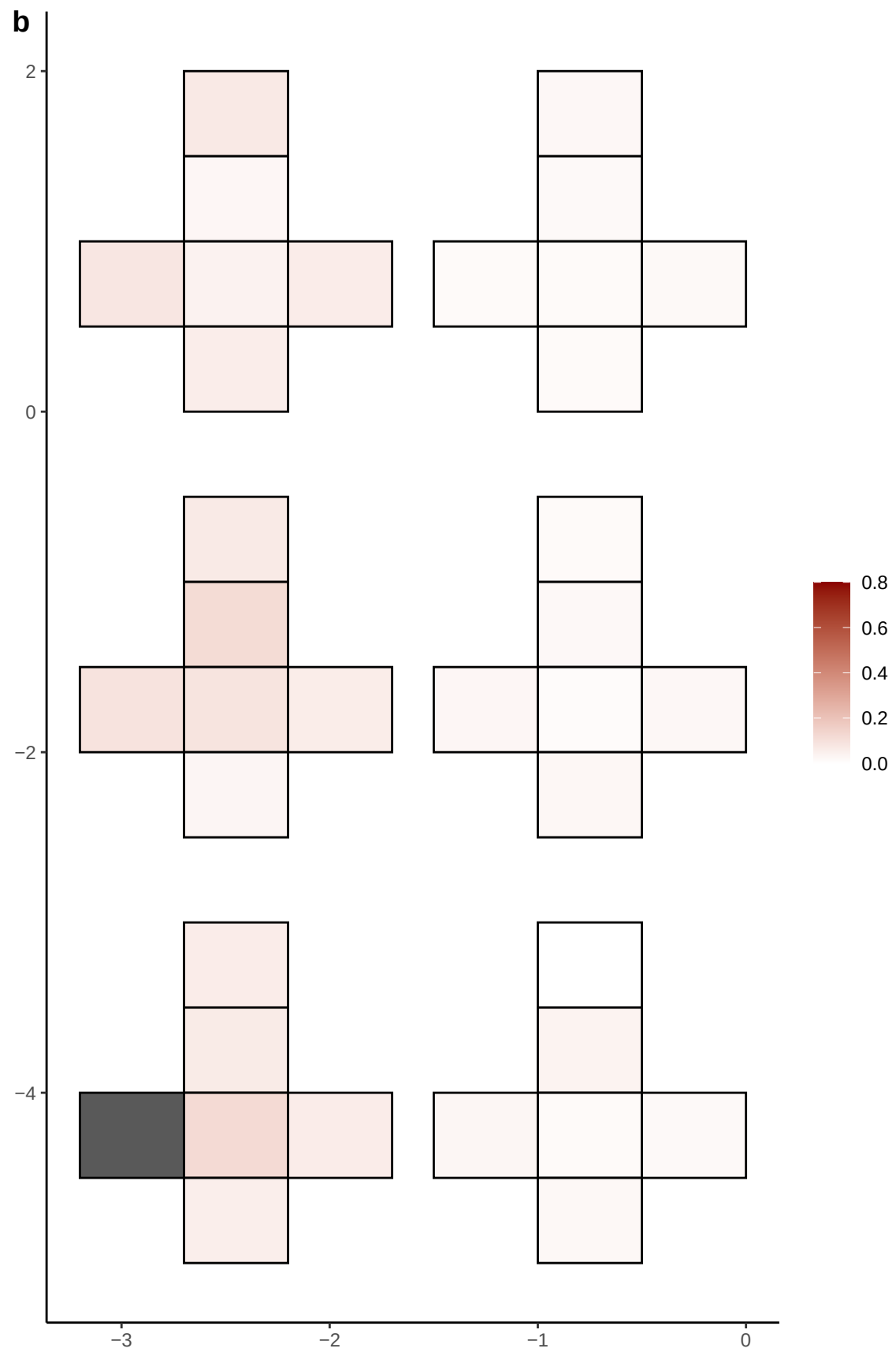

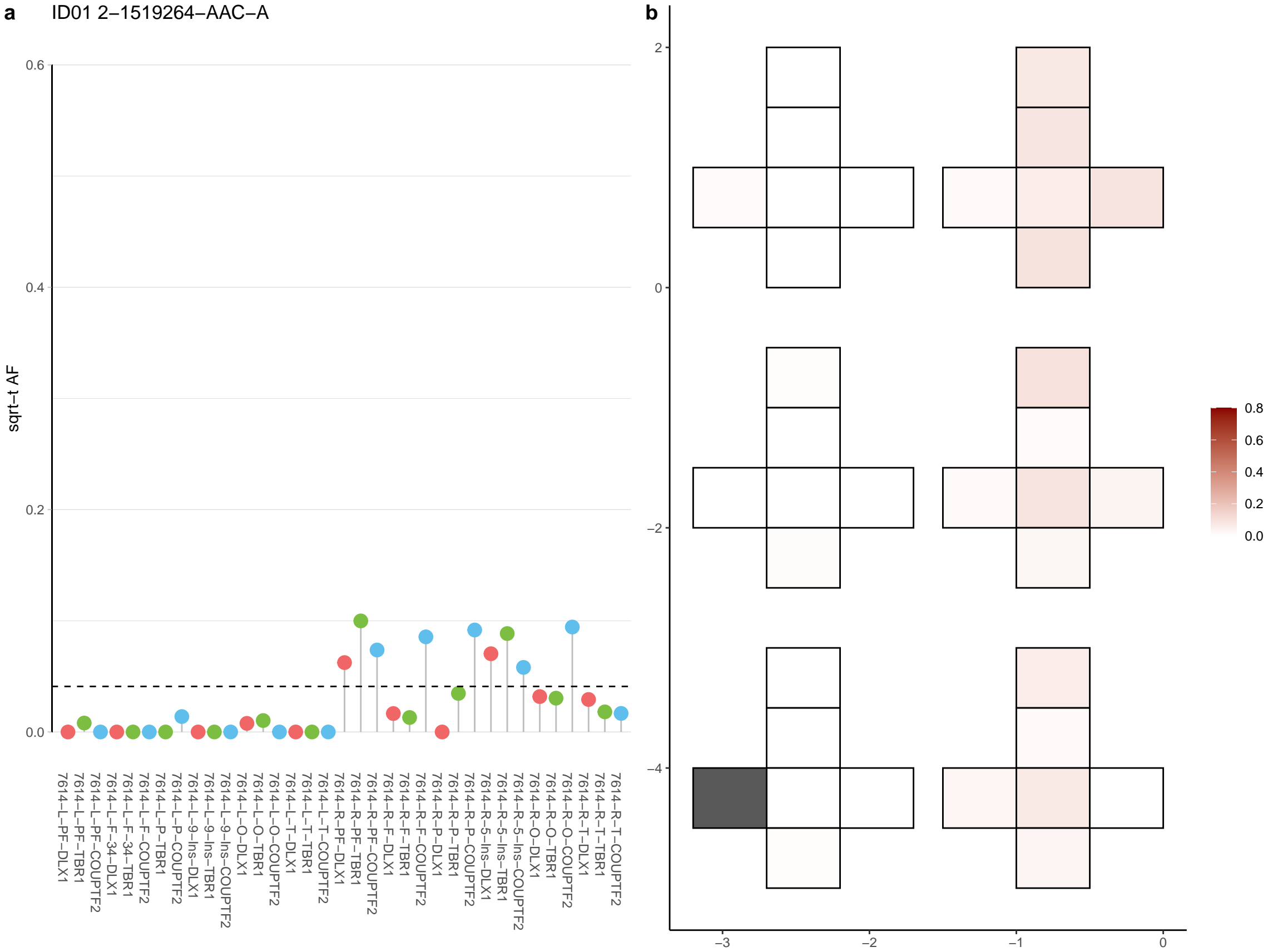

**a** ID01 2-170746611-A-G

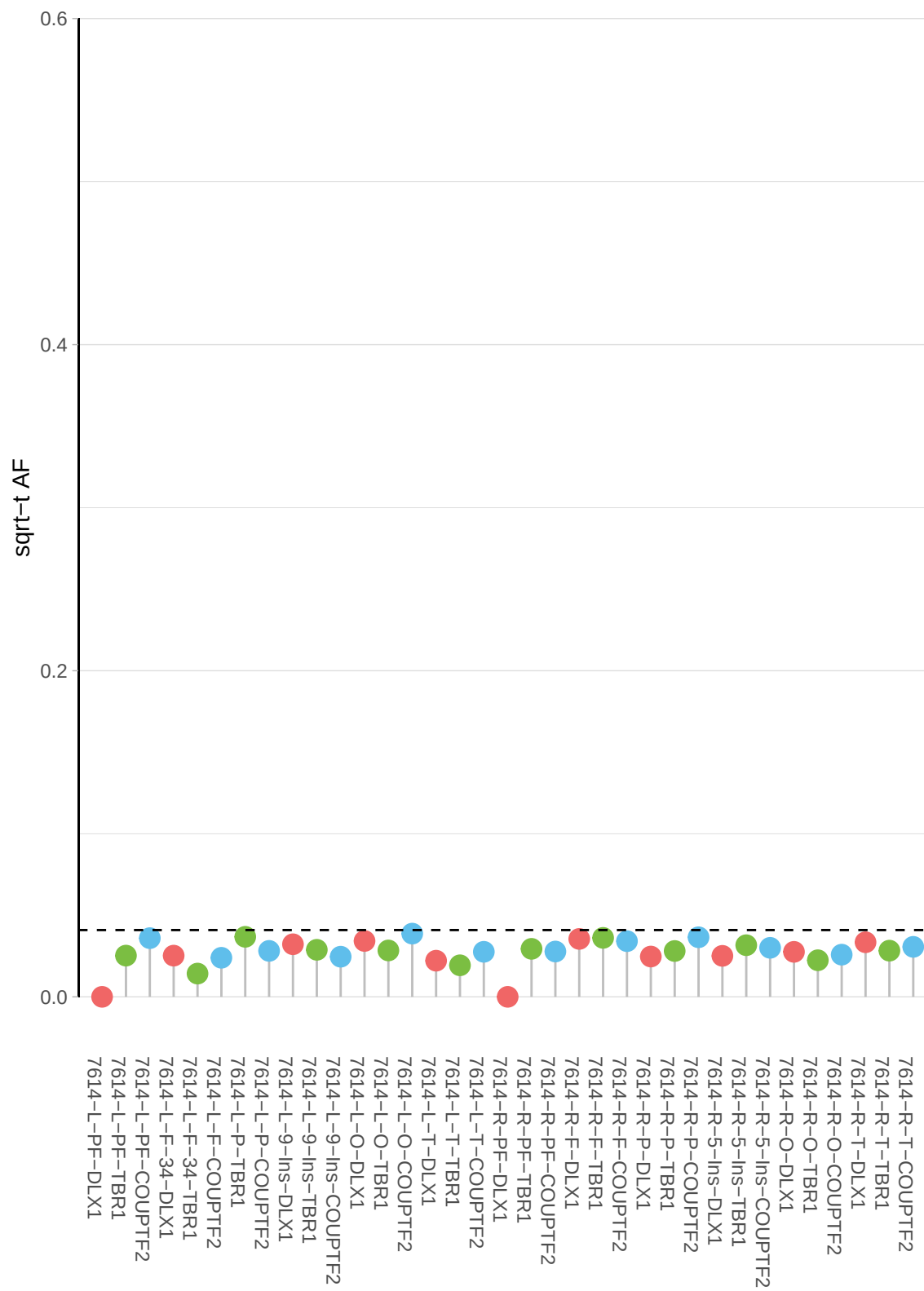

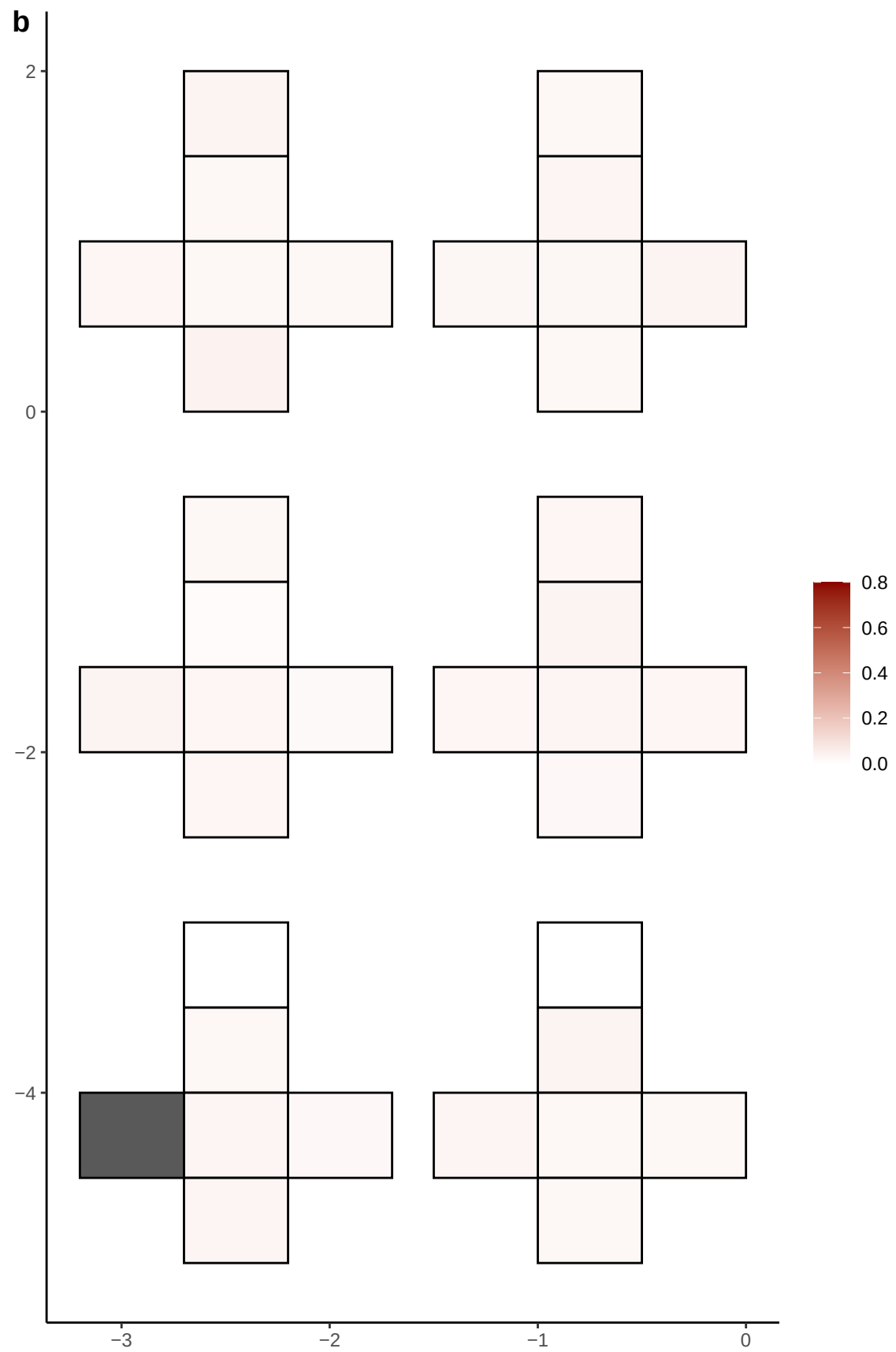

**a** ID01 2-189956910-G-A

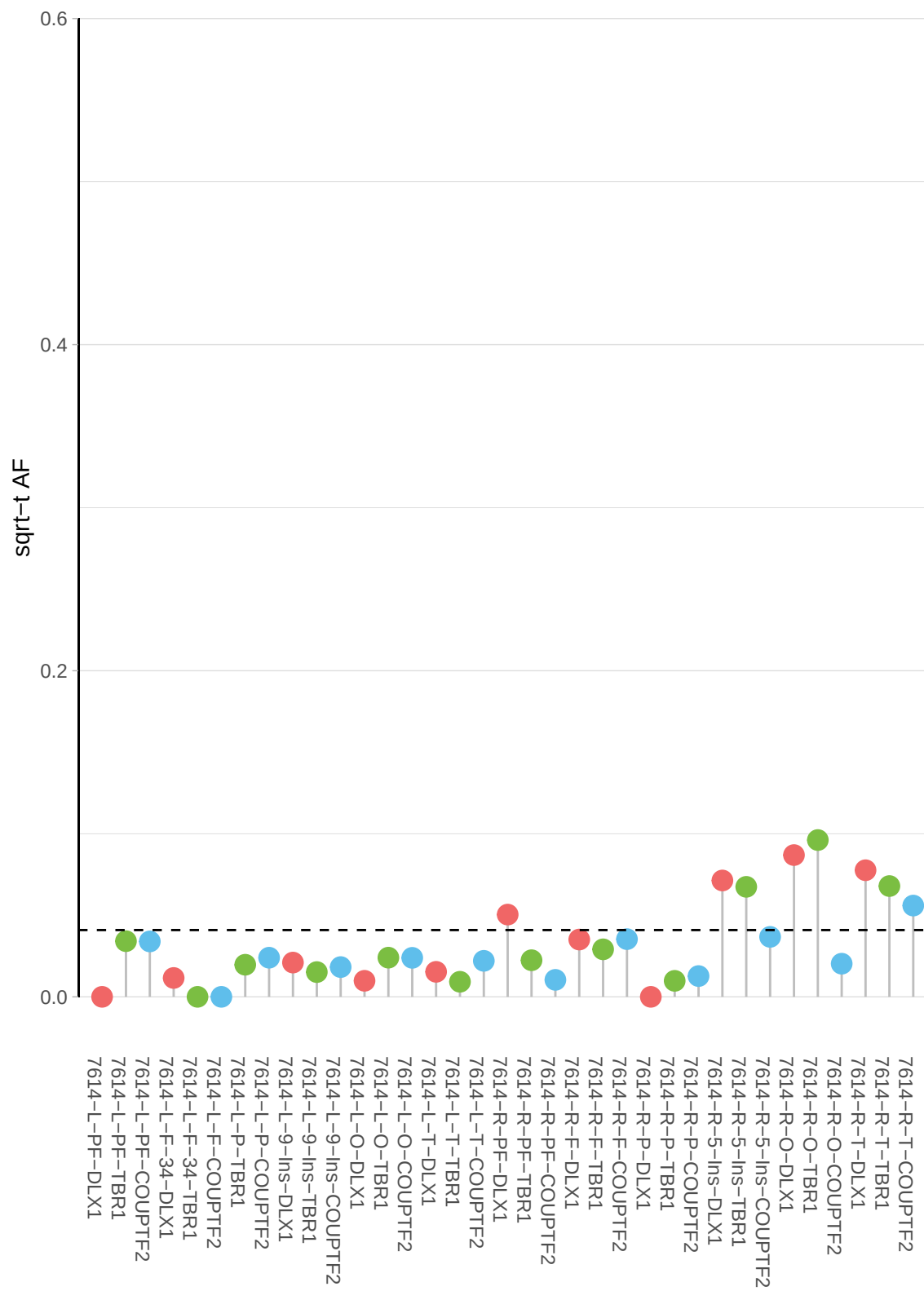

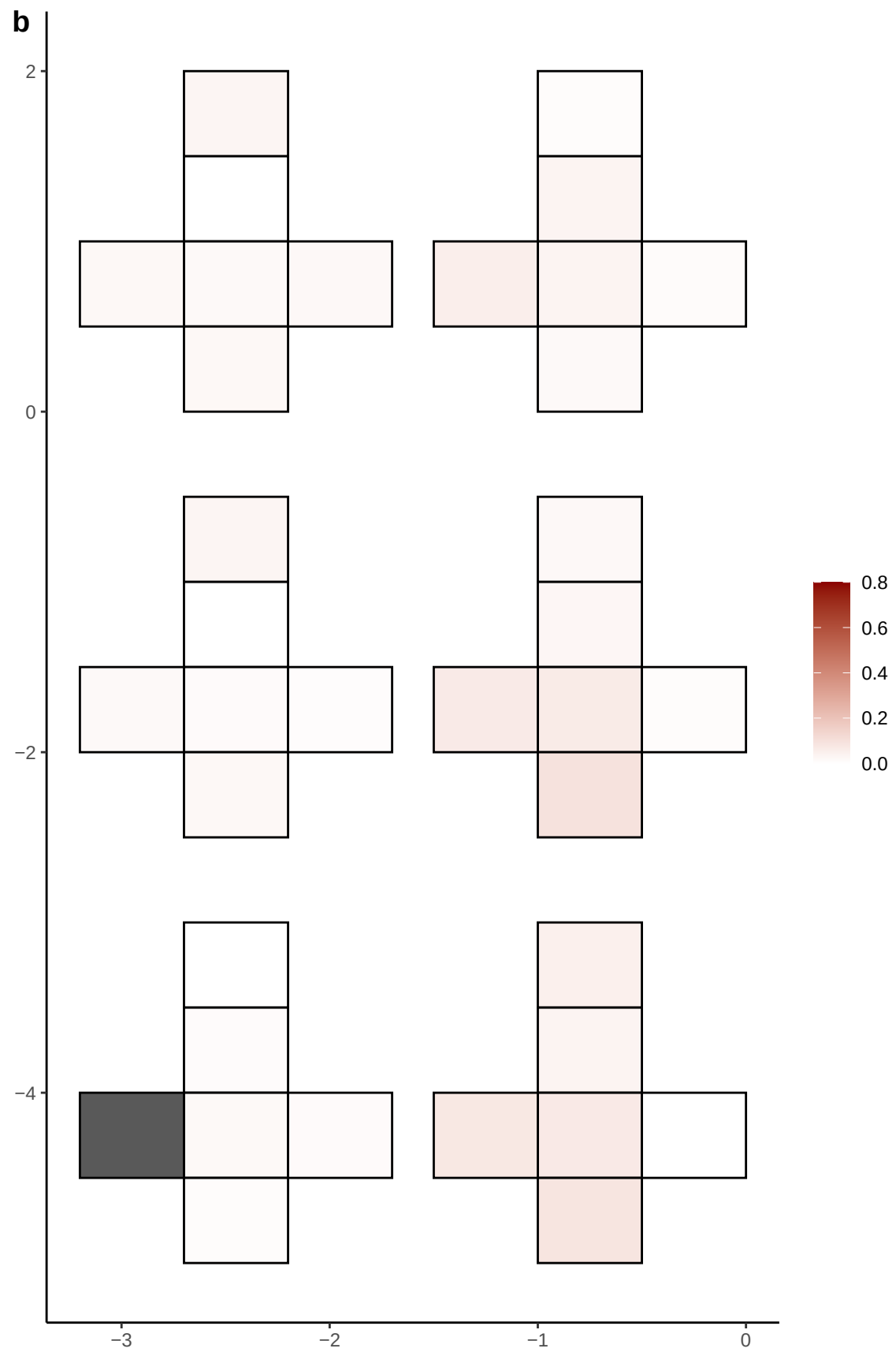

**a** ID01 2-194793292-CTT-C

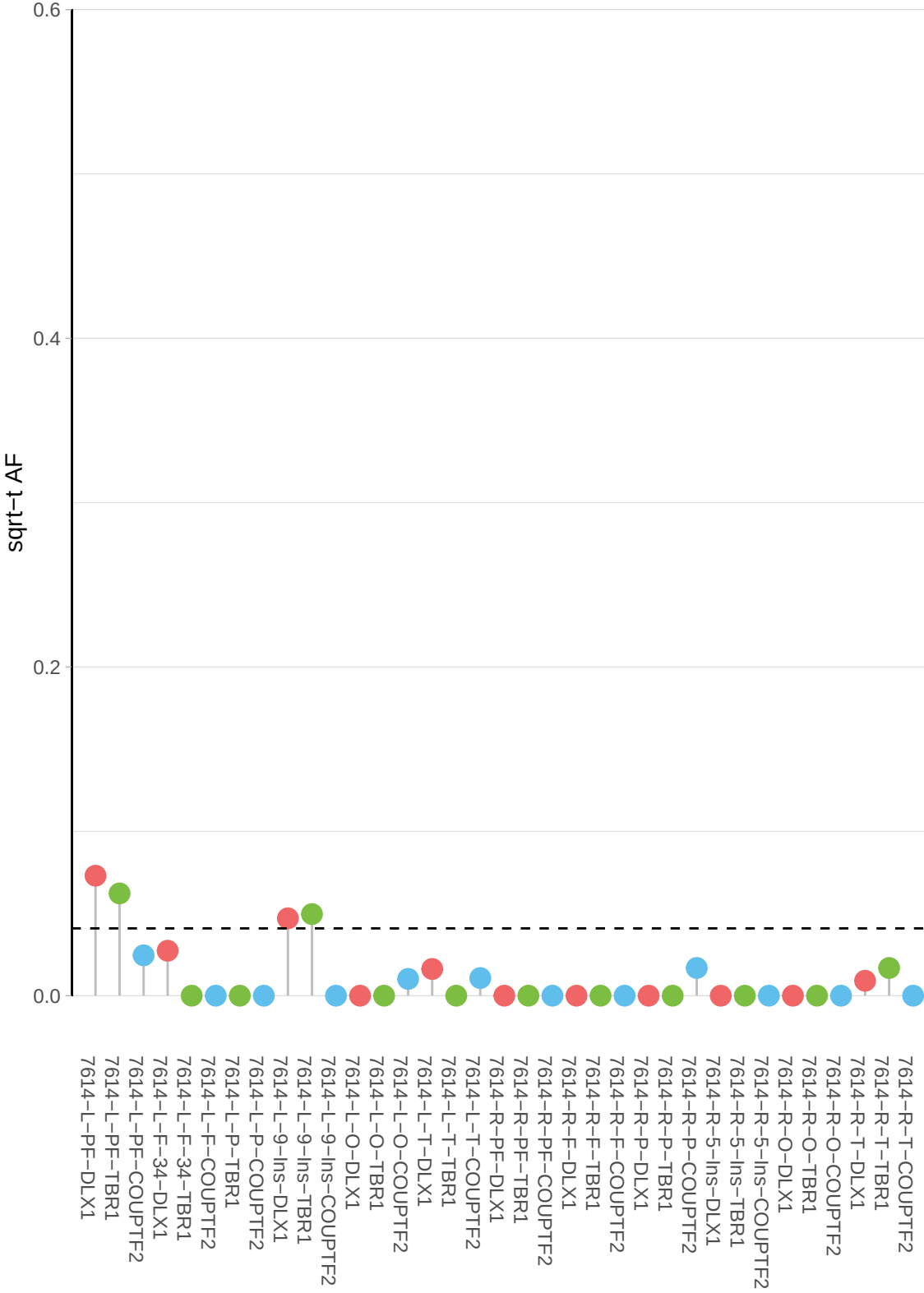

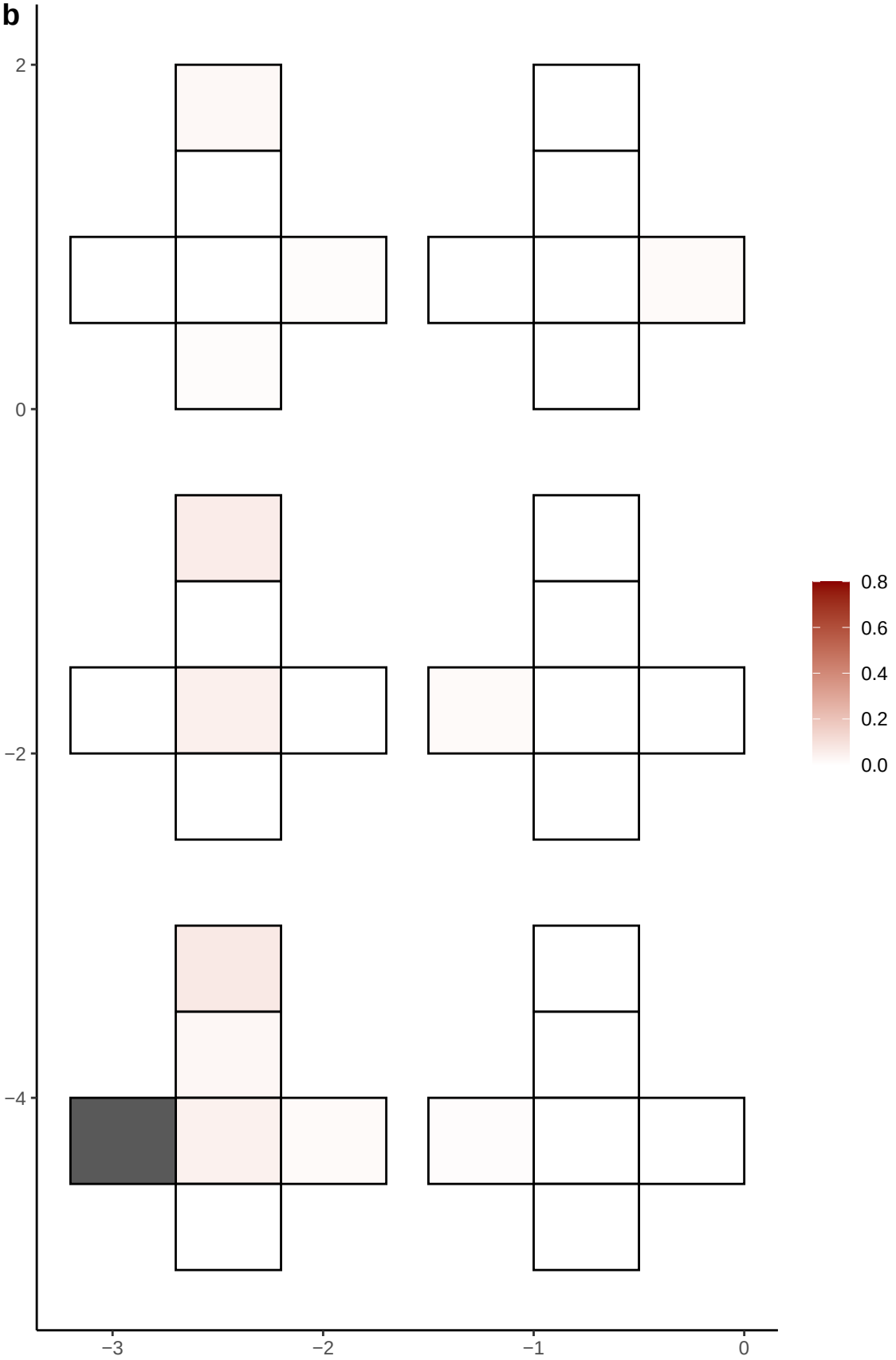

**a** ID01 2-195046625-G-A

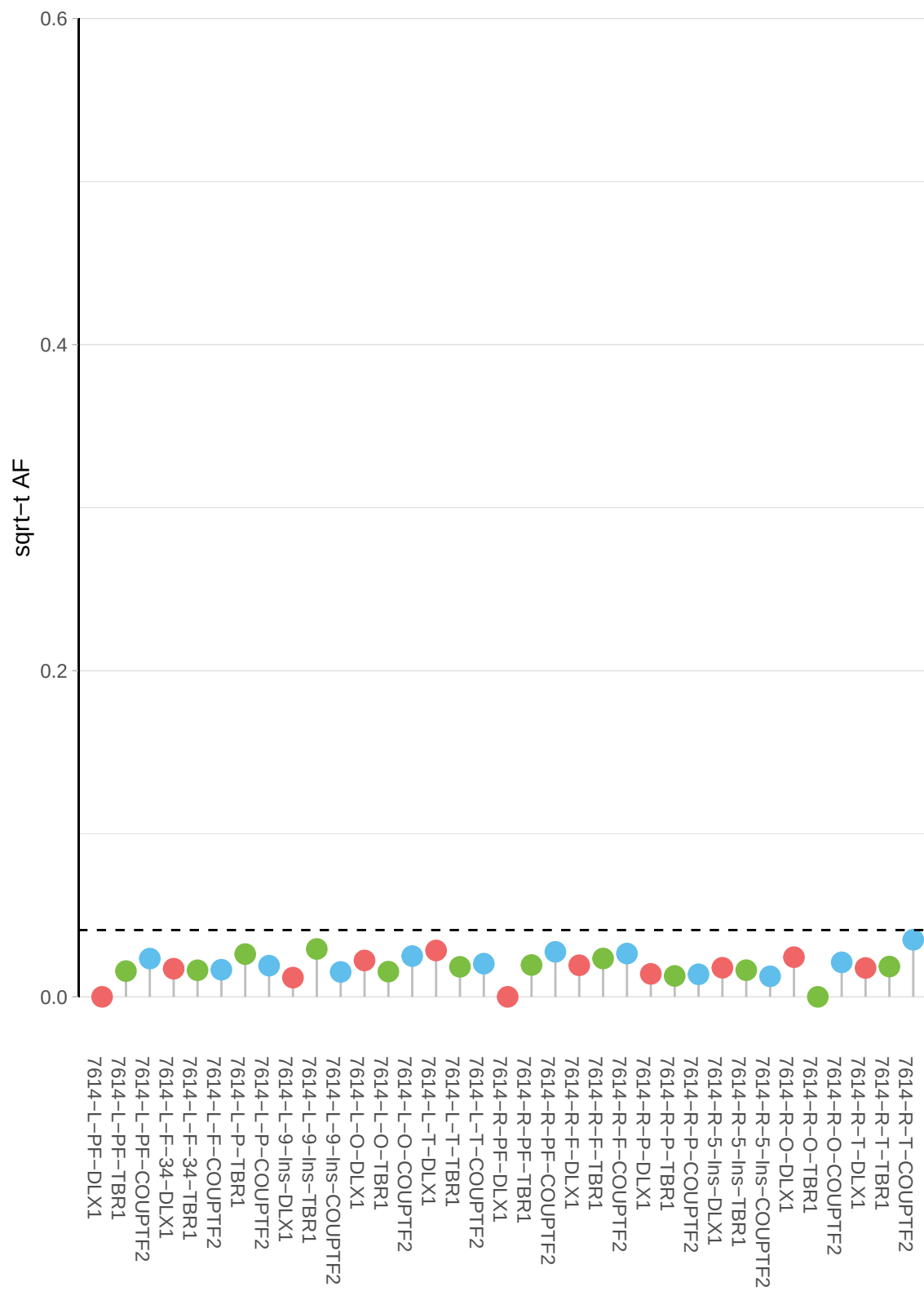

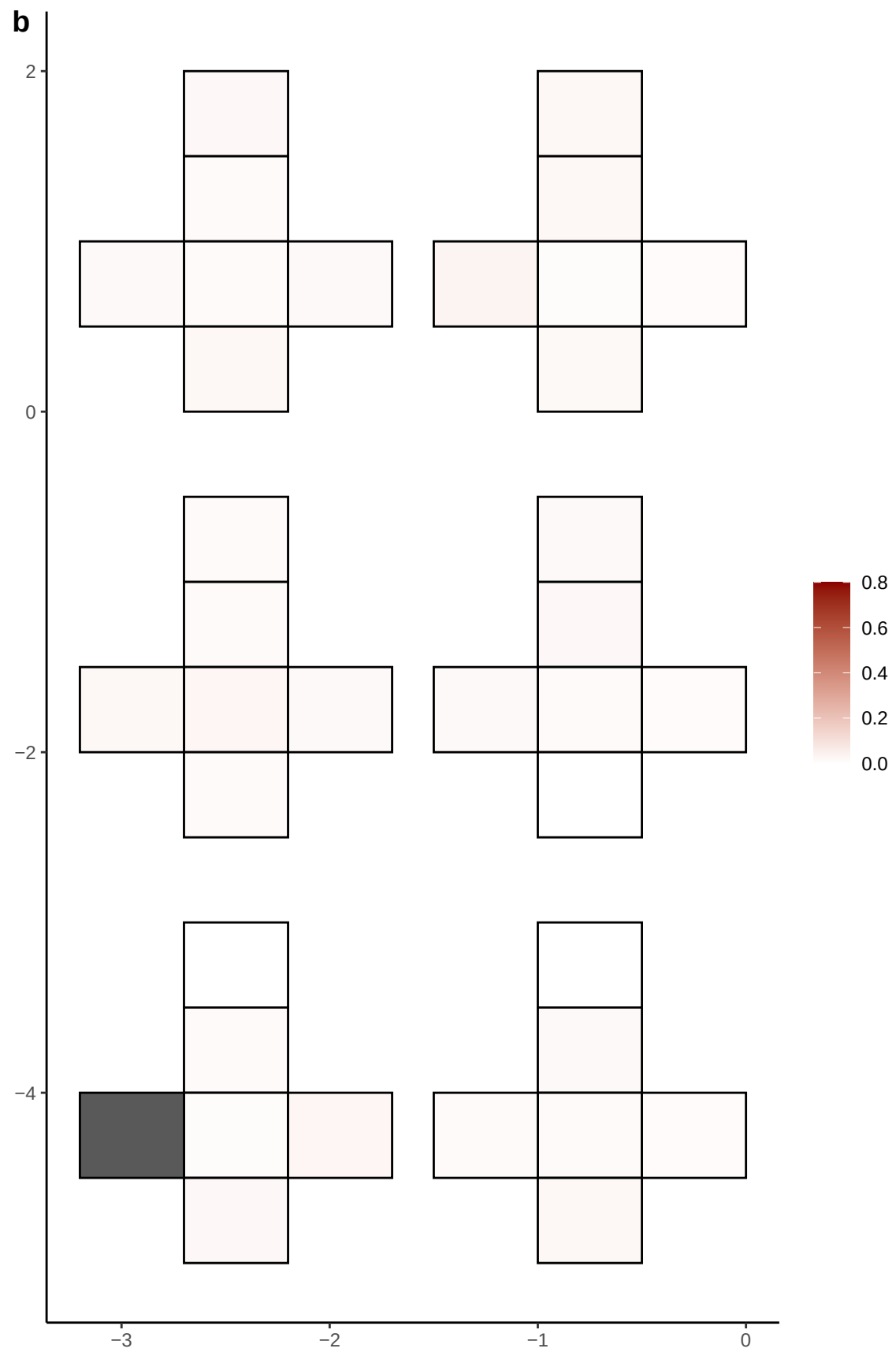

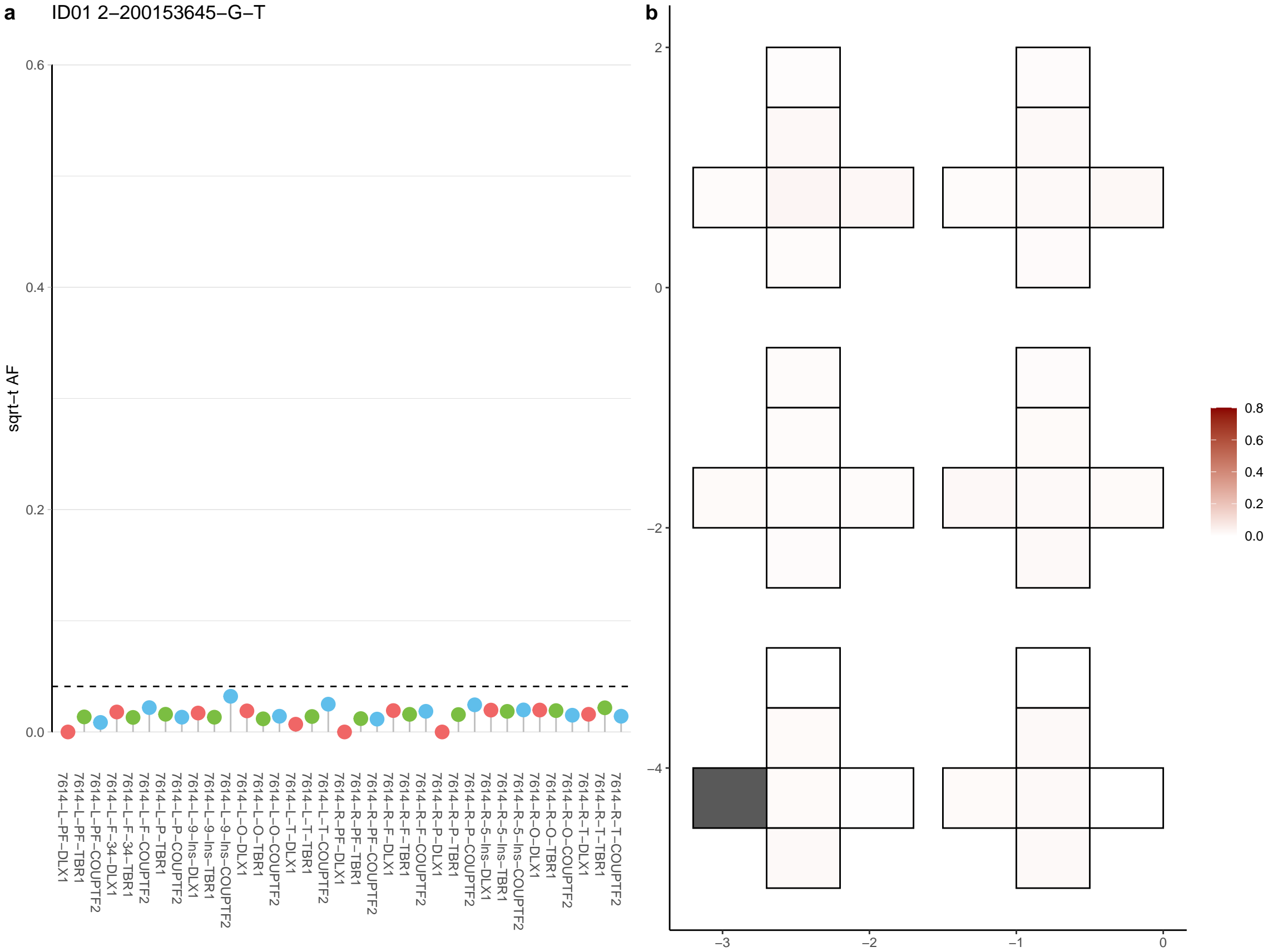

**a** ID01 2-209589692-G-A

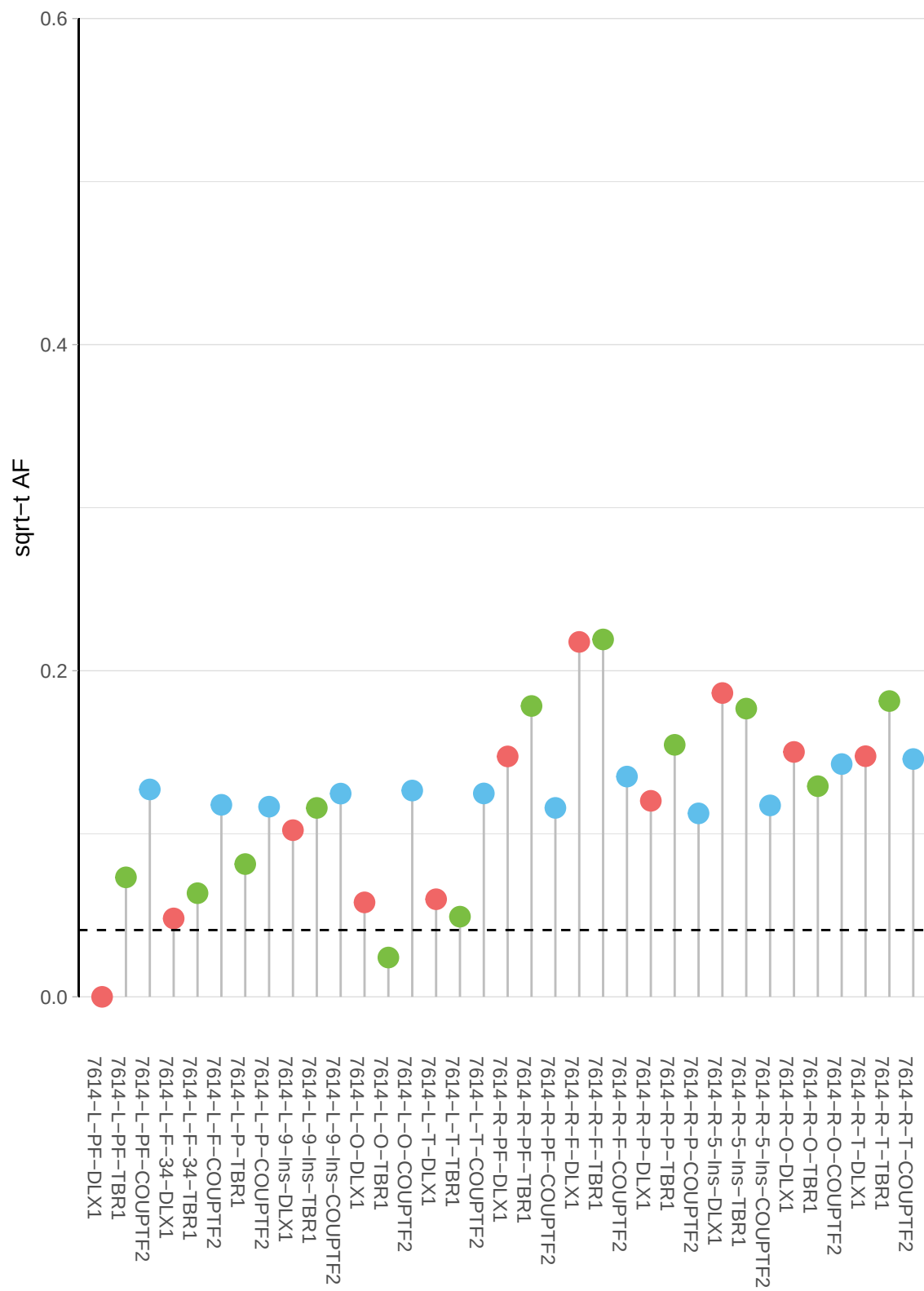

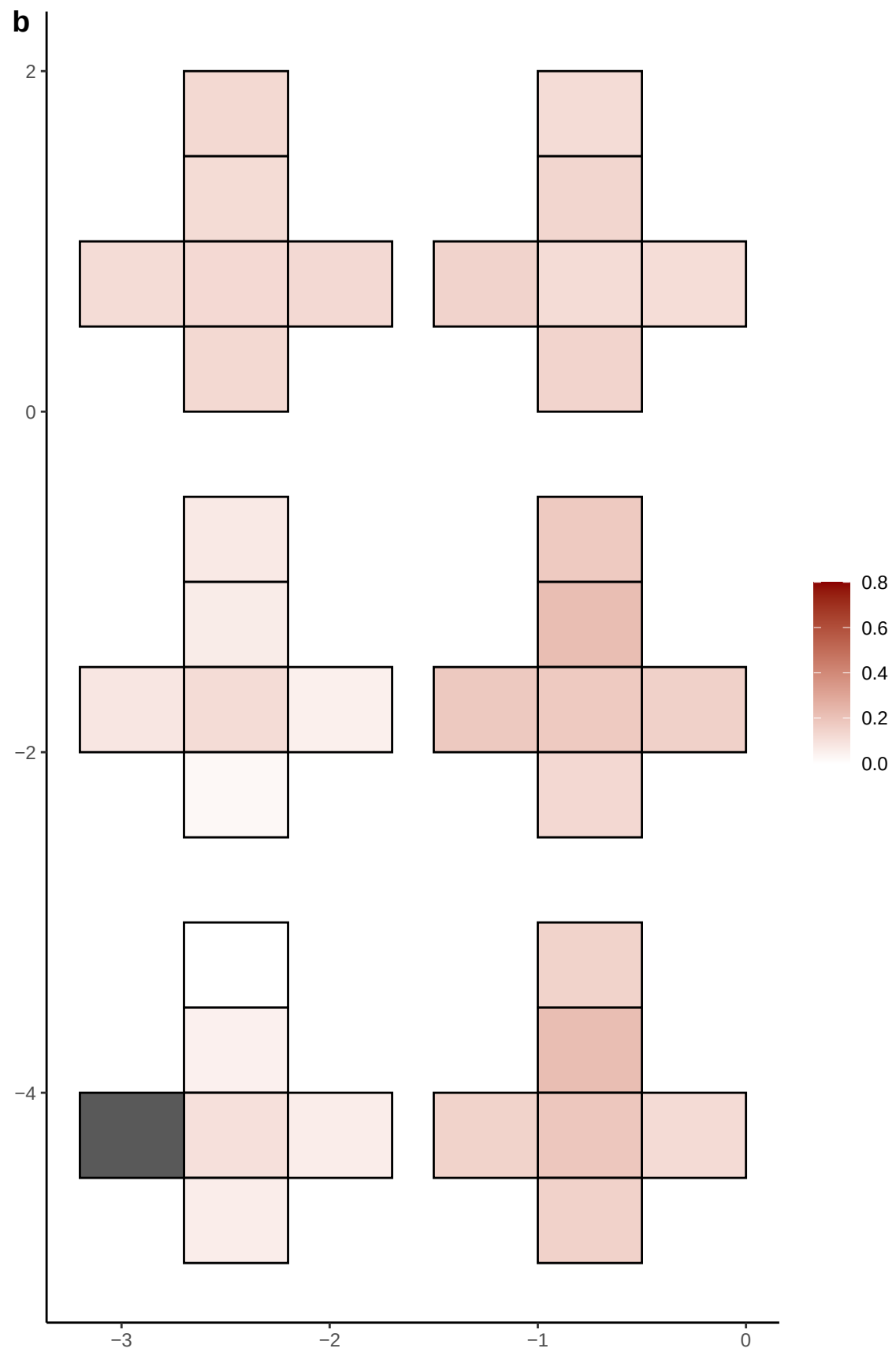

**a** ID01 2-231841419-C-T

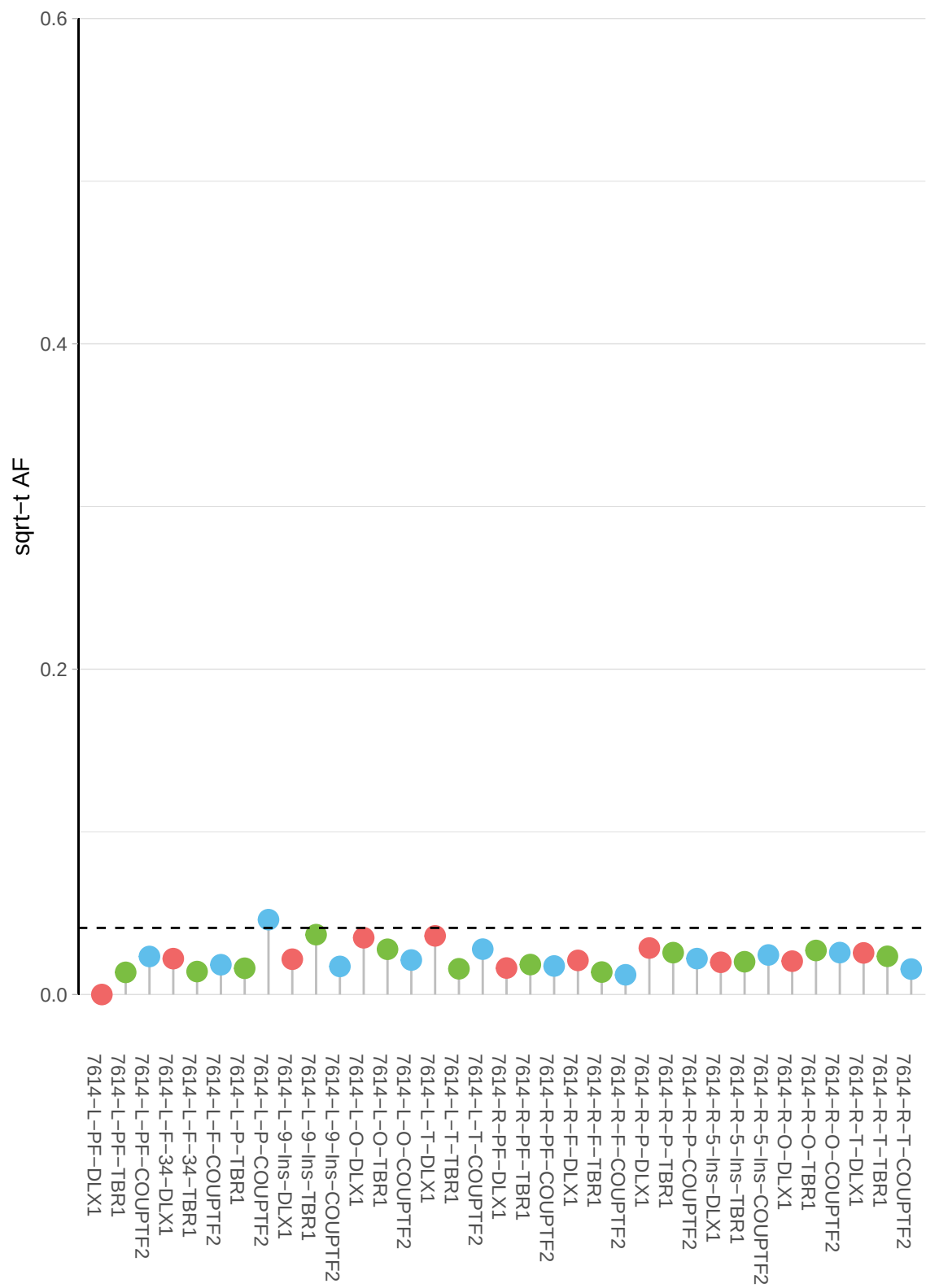

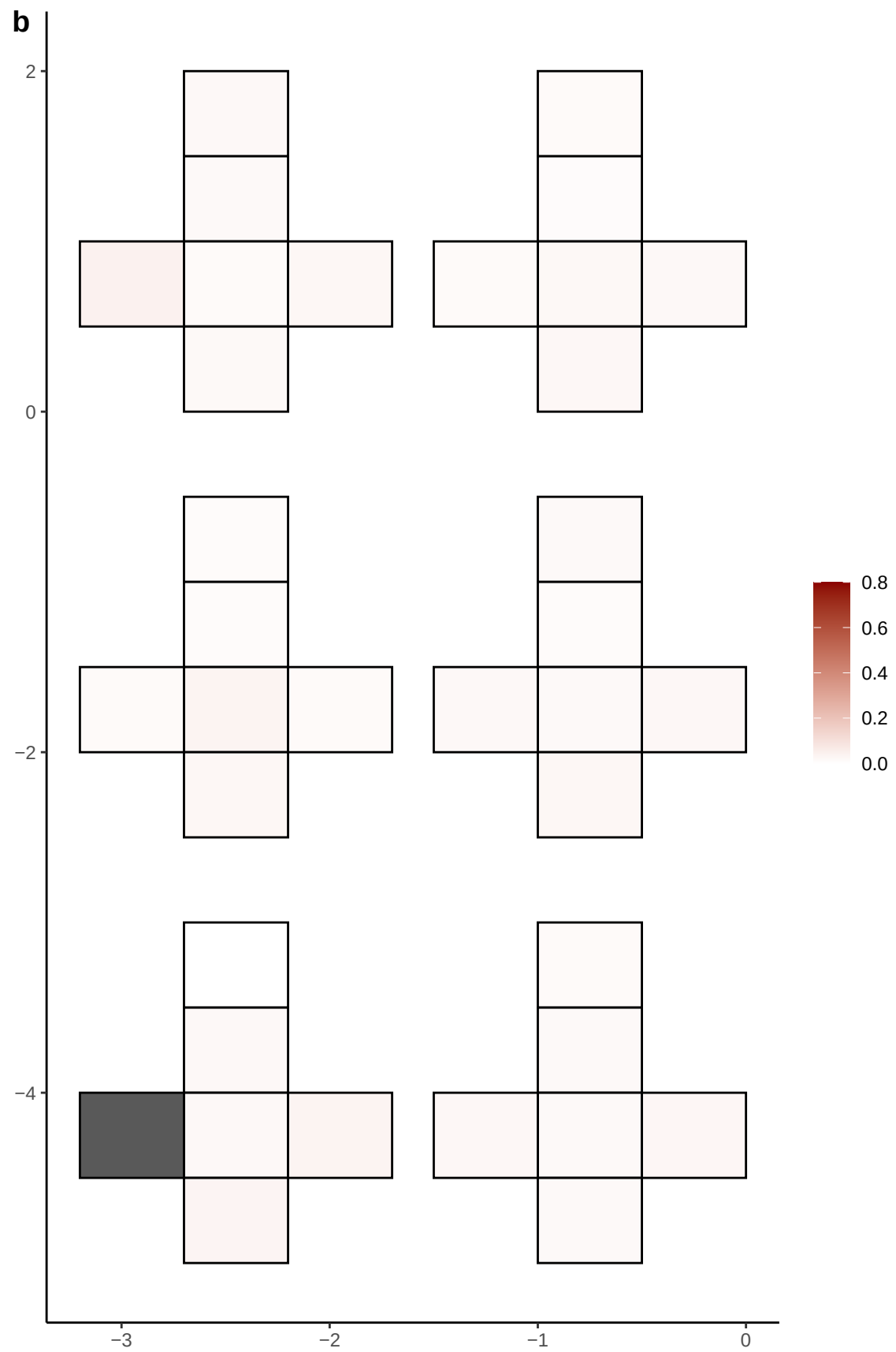

**a** ID01 2-235210131-C-T

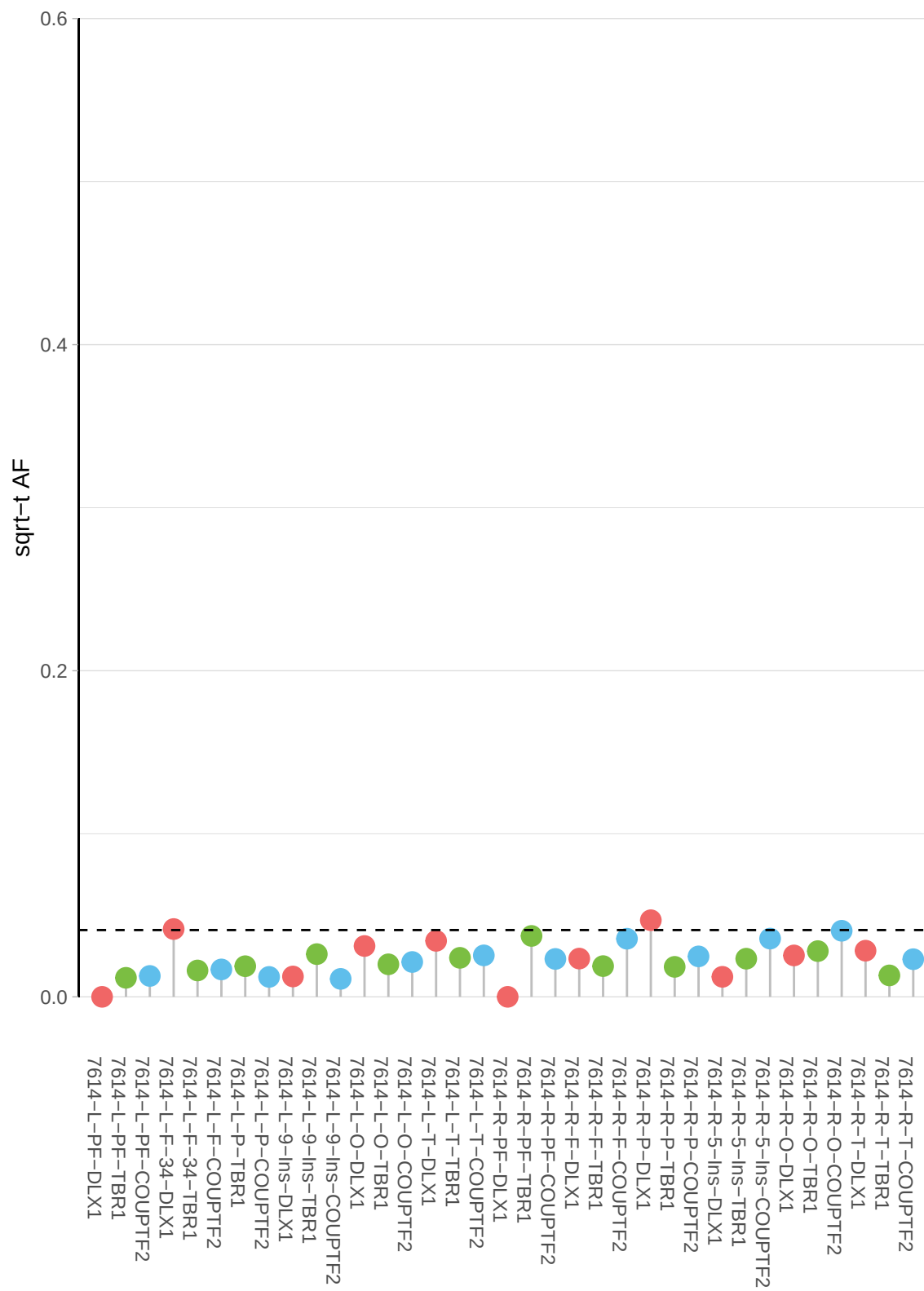

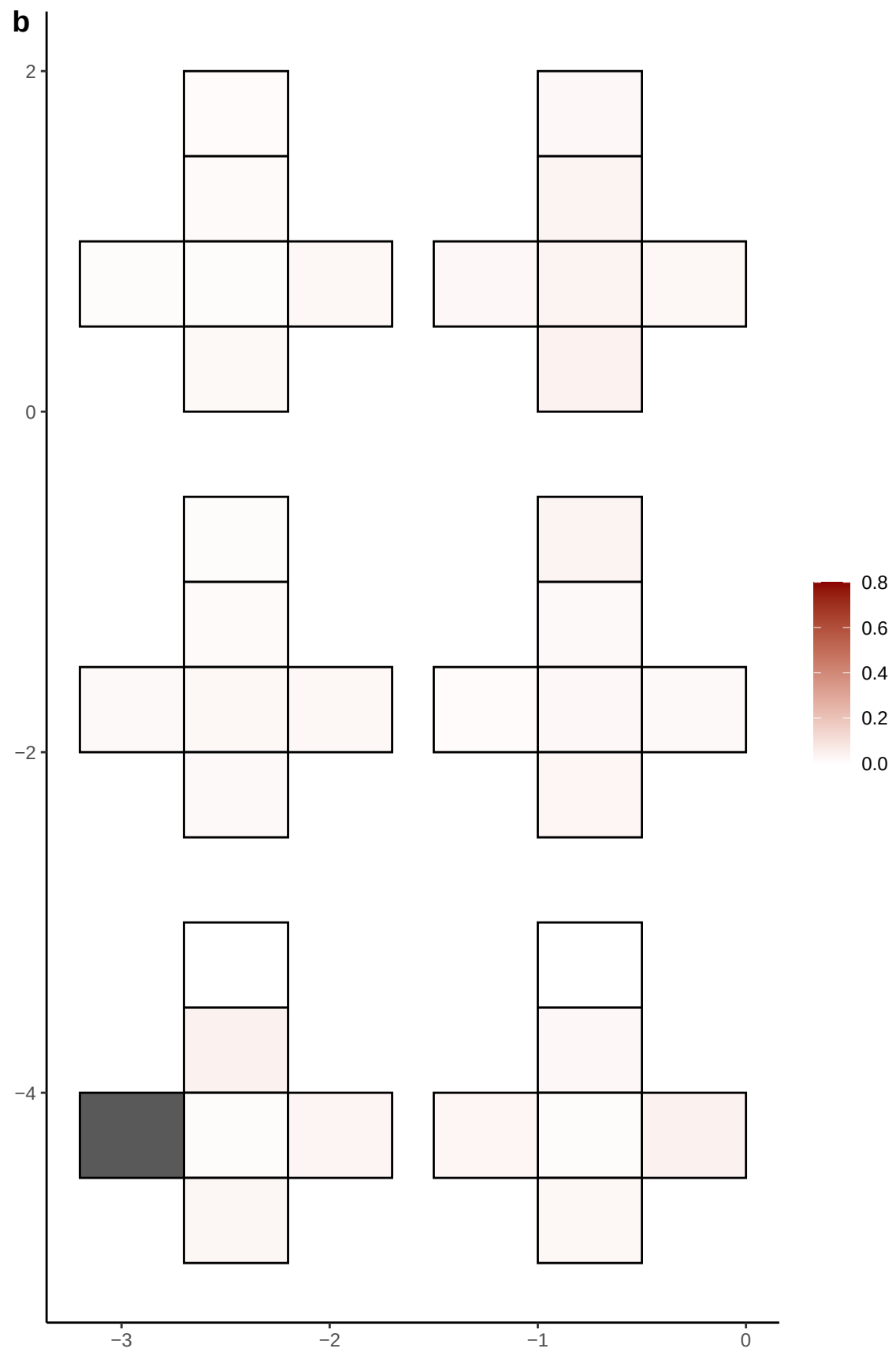

**a** ID01 2-241281331-C-A

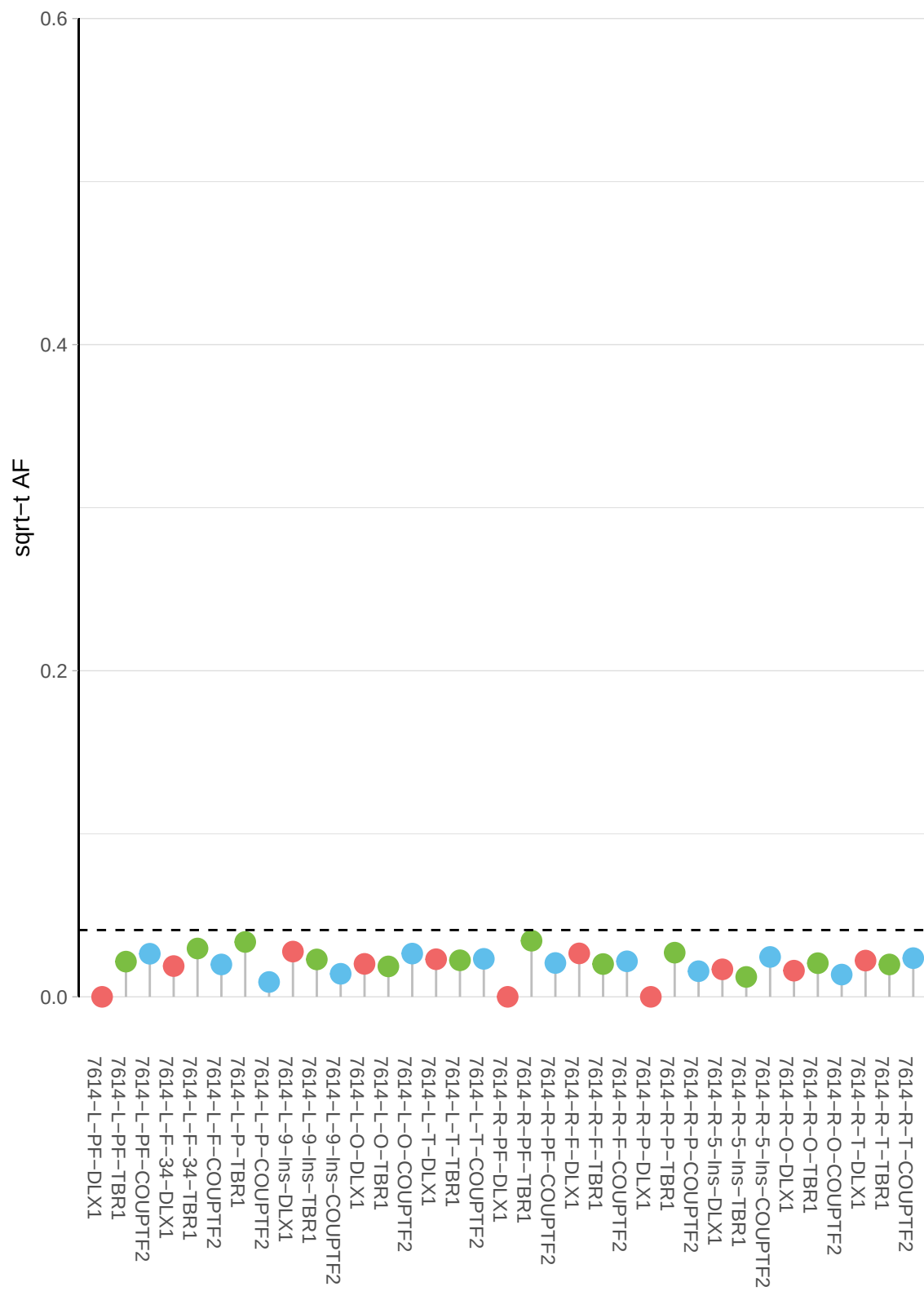

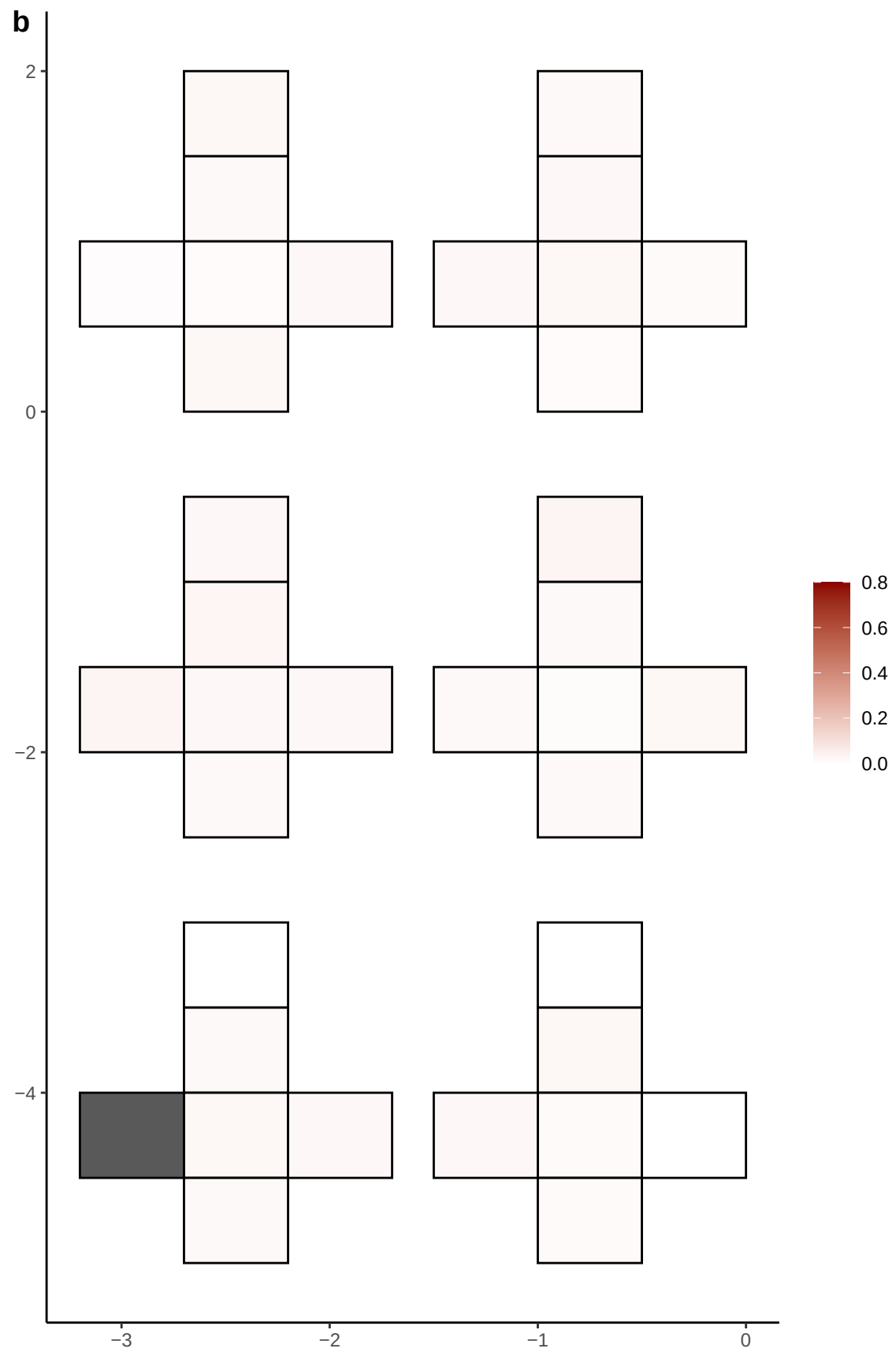

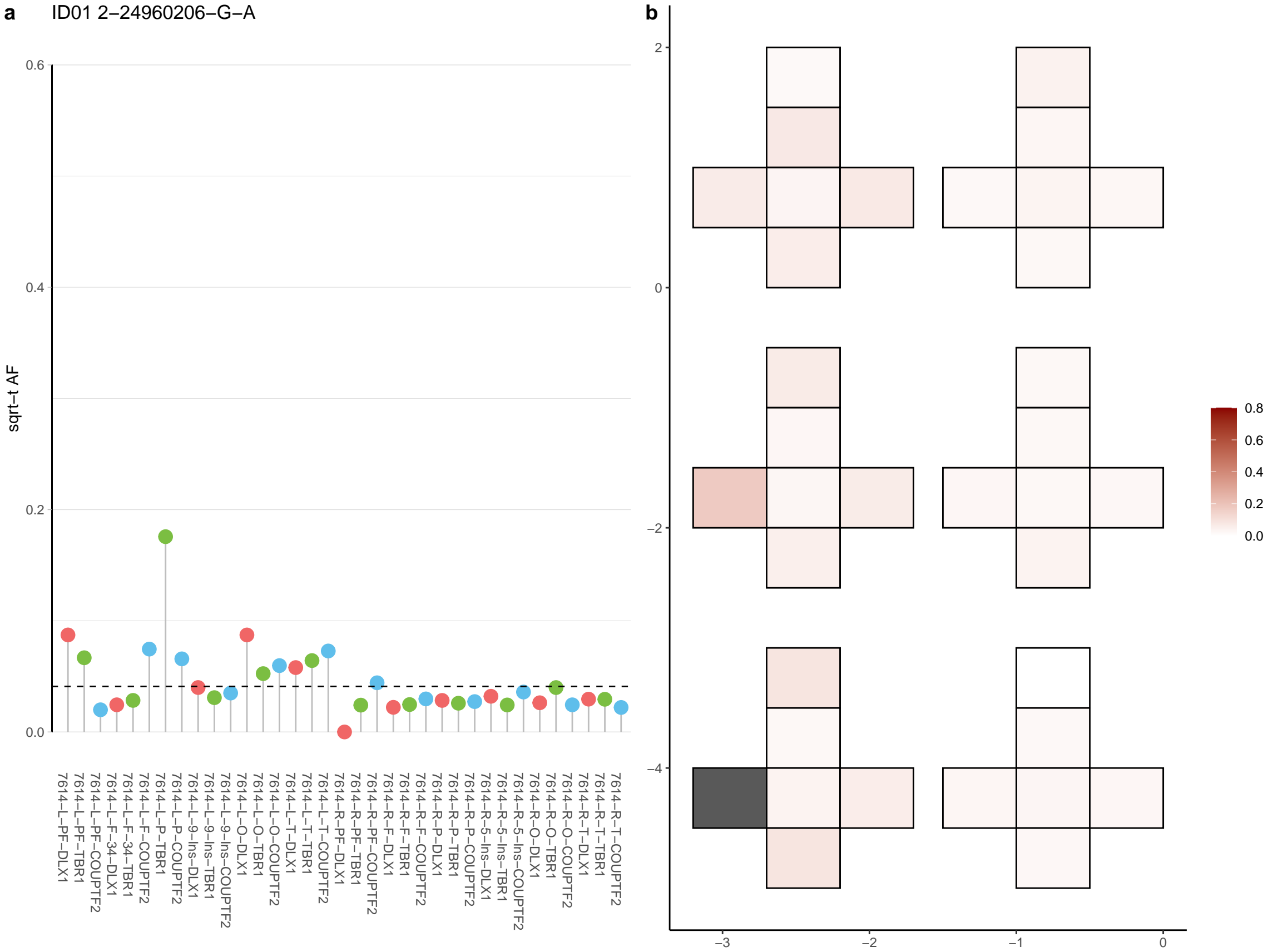

**a** ID01 2-34606921-T-A

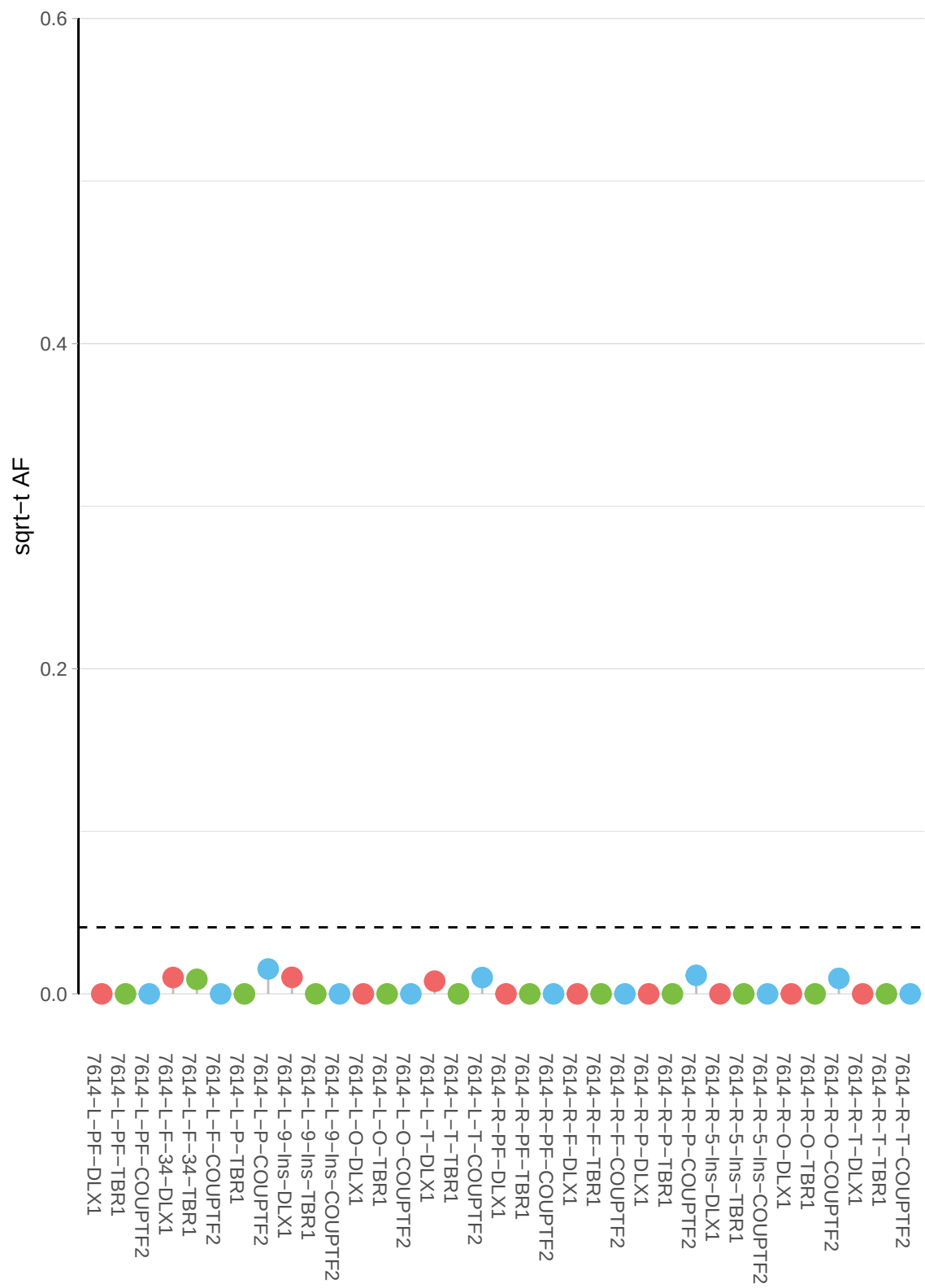

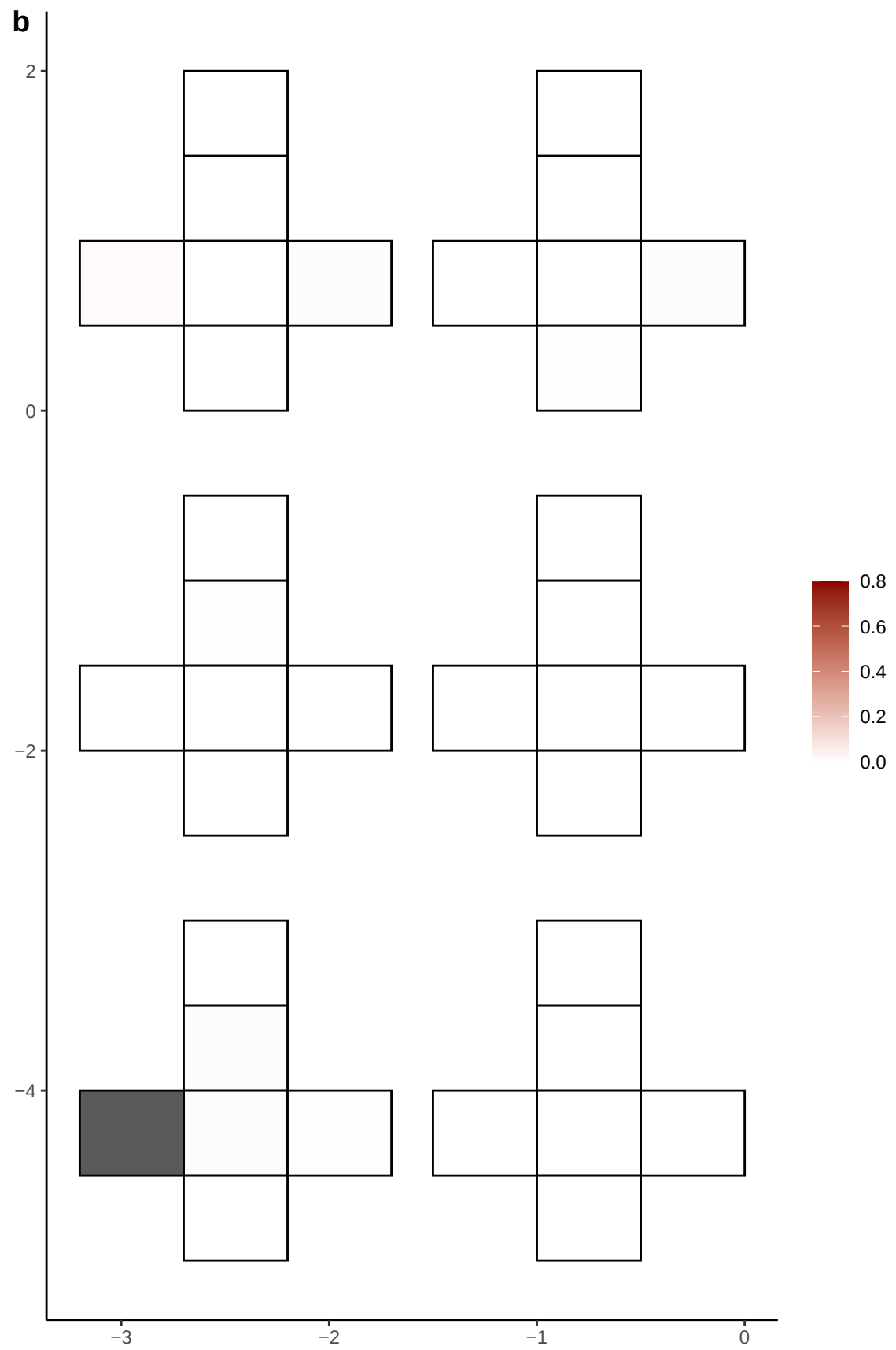

**a** ID01 2-49268814-A-C

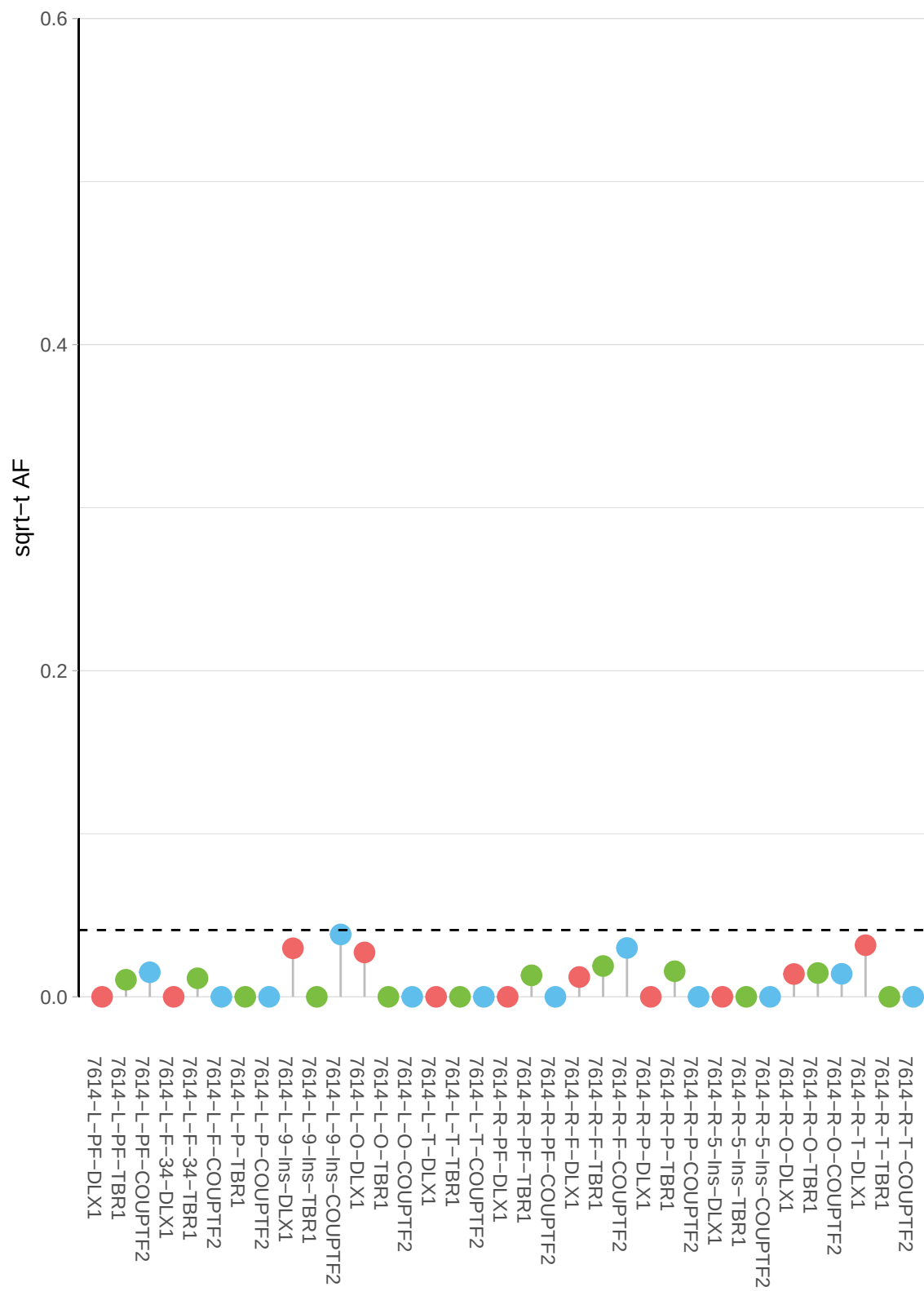

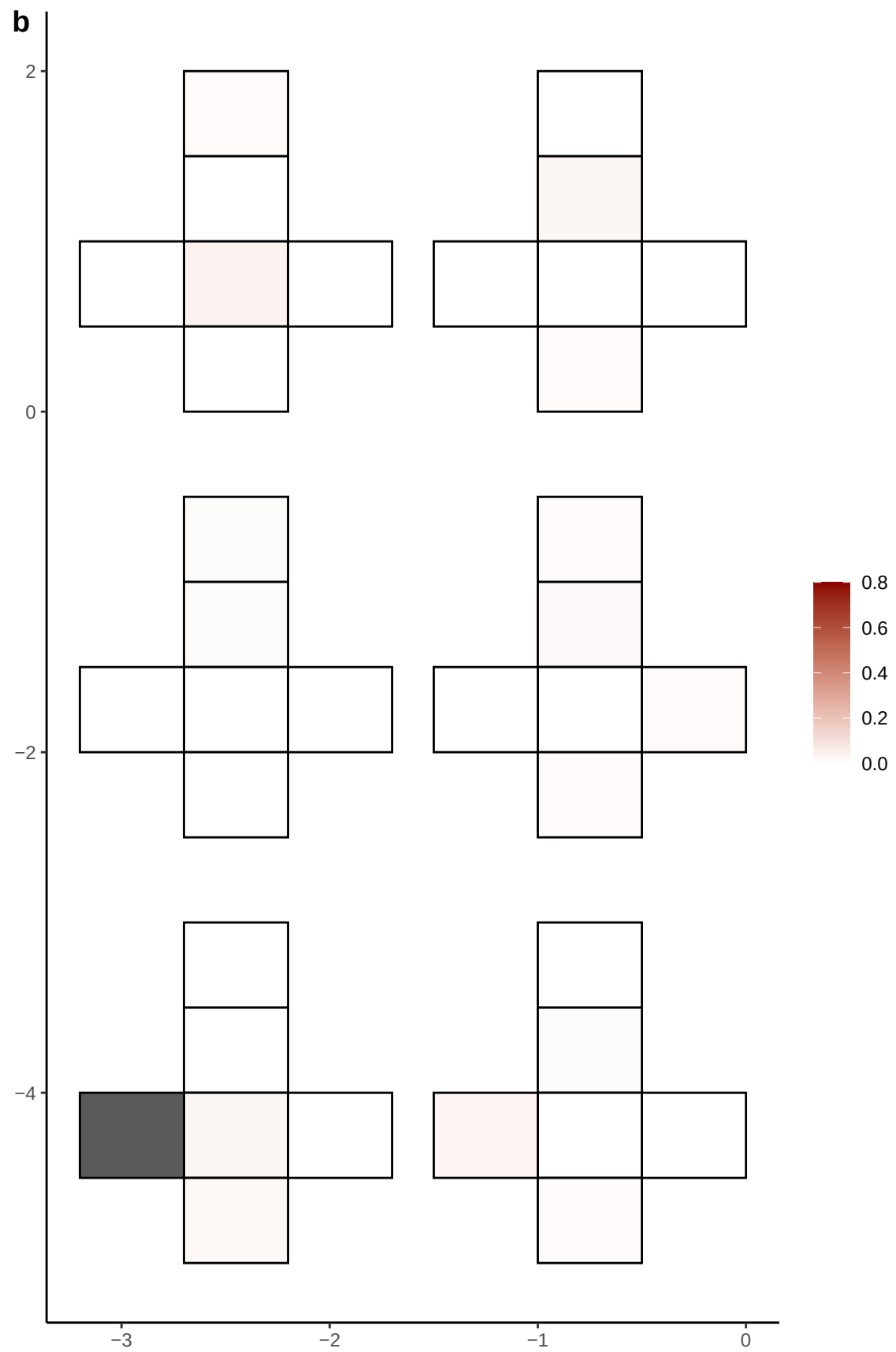

**a** ID01 2-53799113-A-T

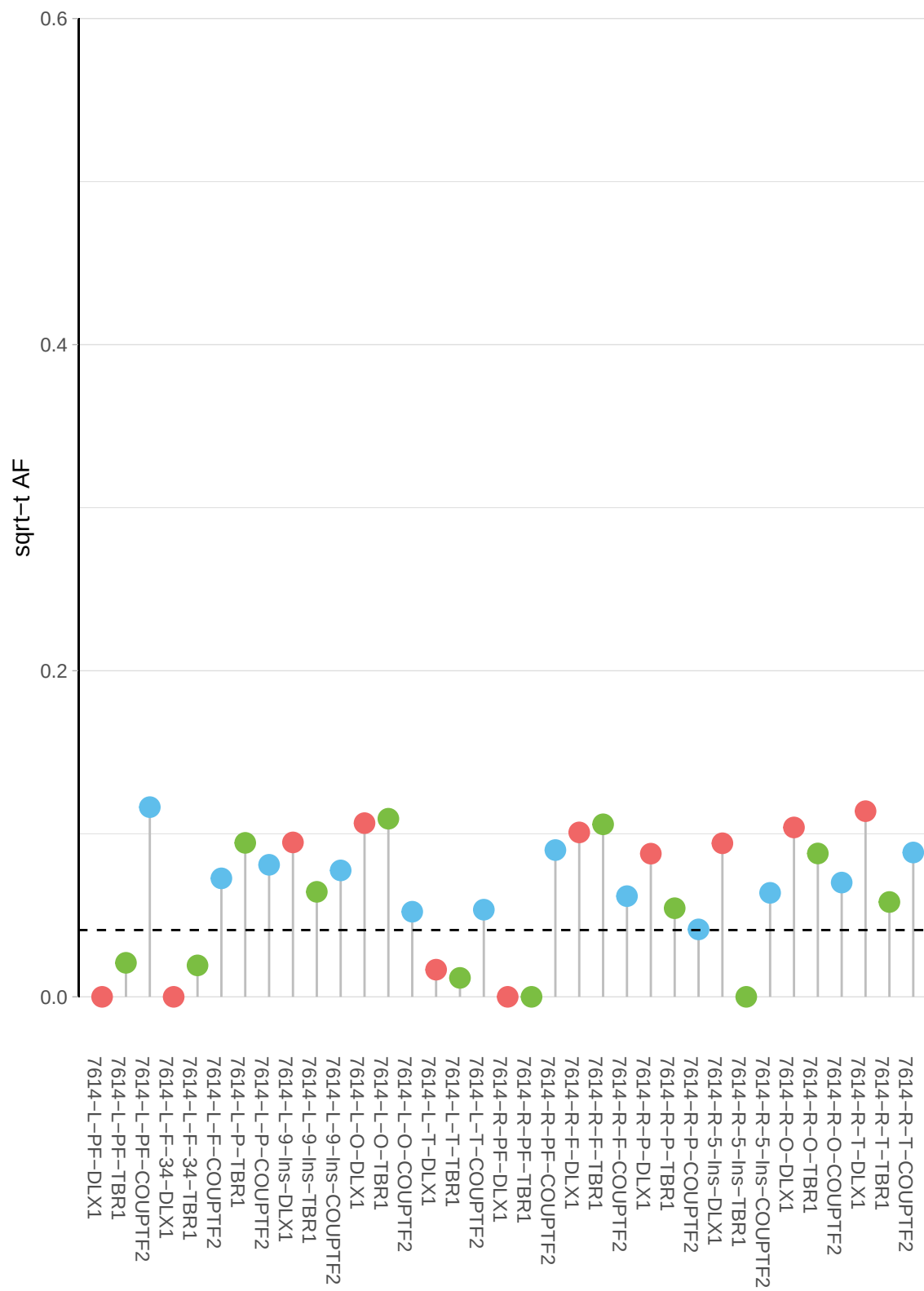

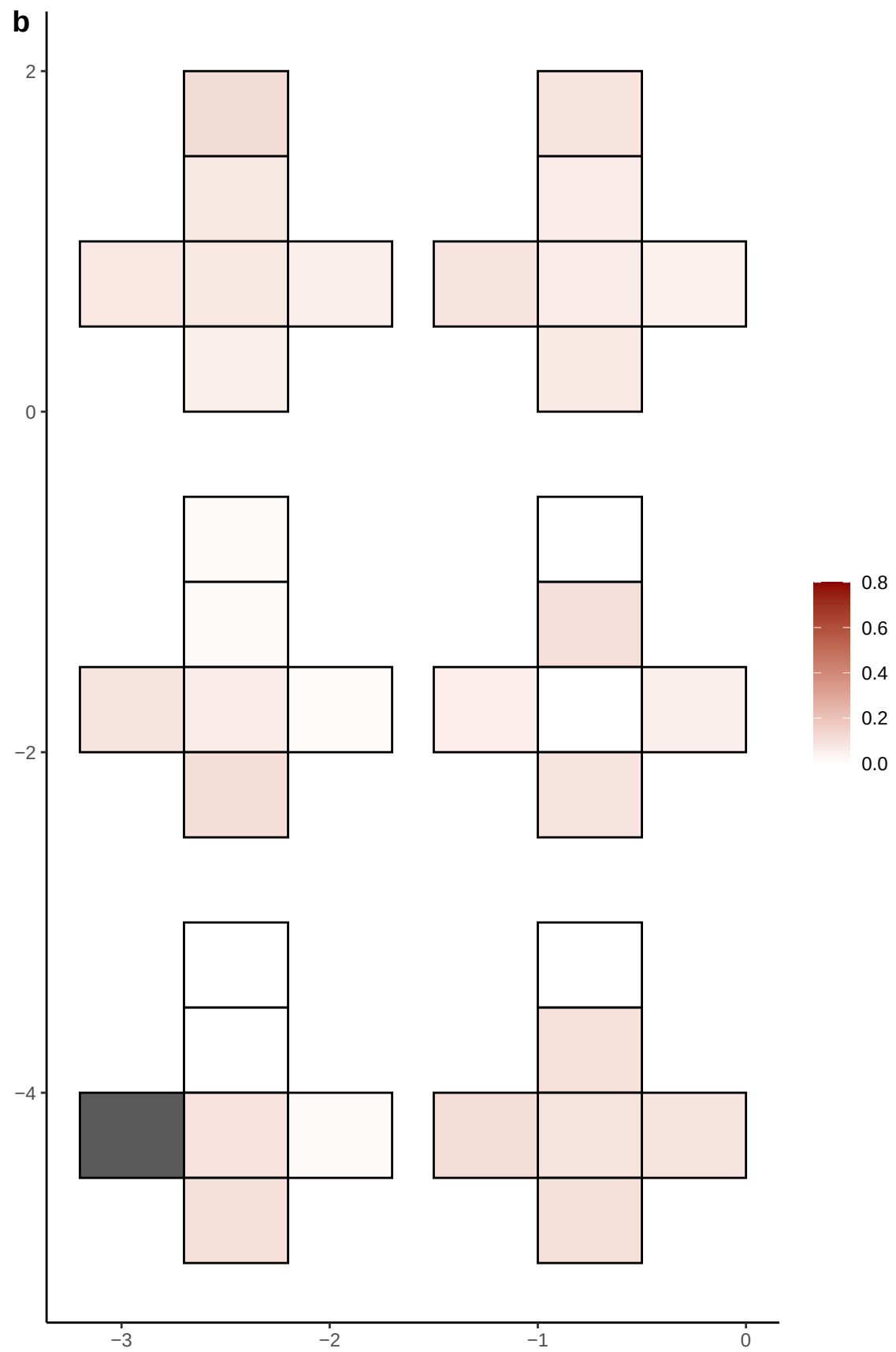

**a** ID01 2-56448245-G-A

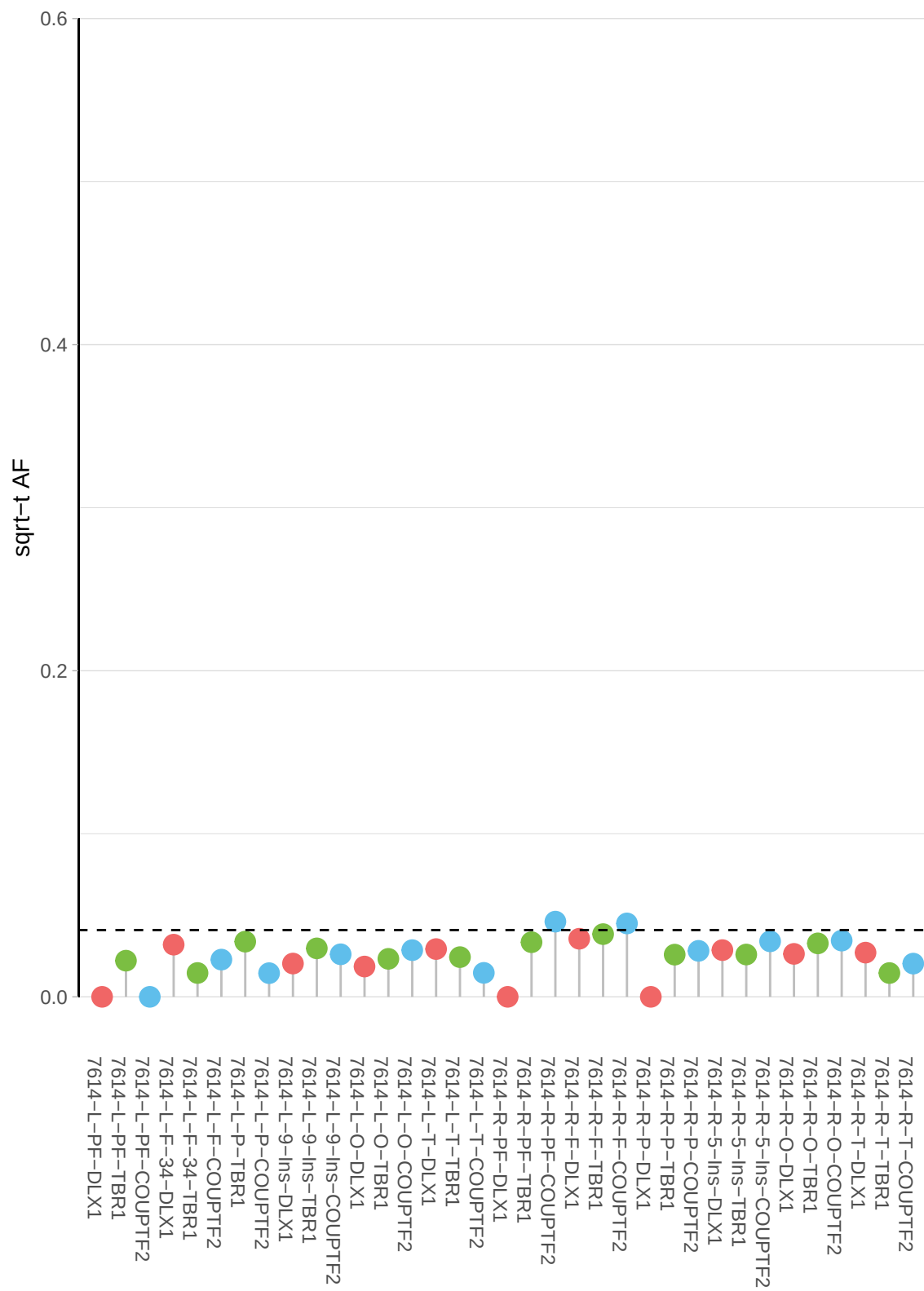

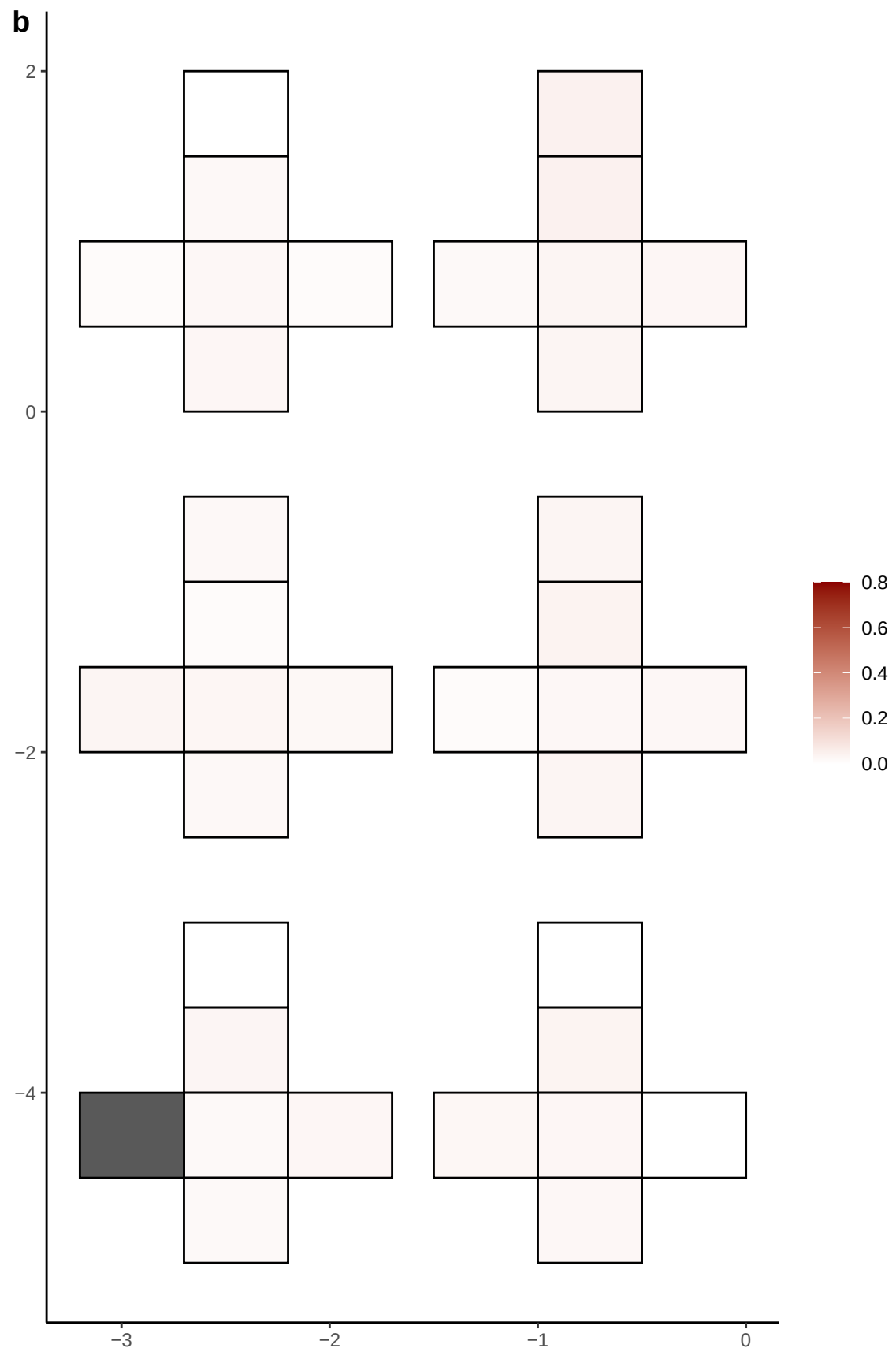

**a** ID01 2-5965706-G-A

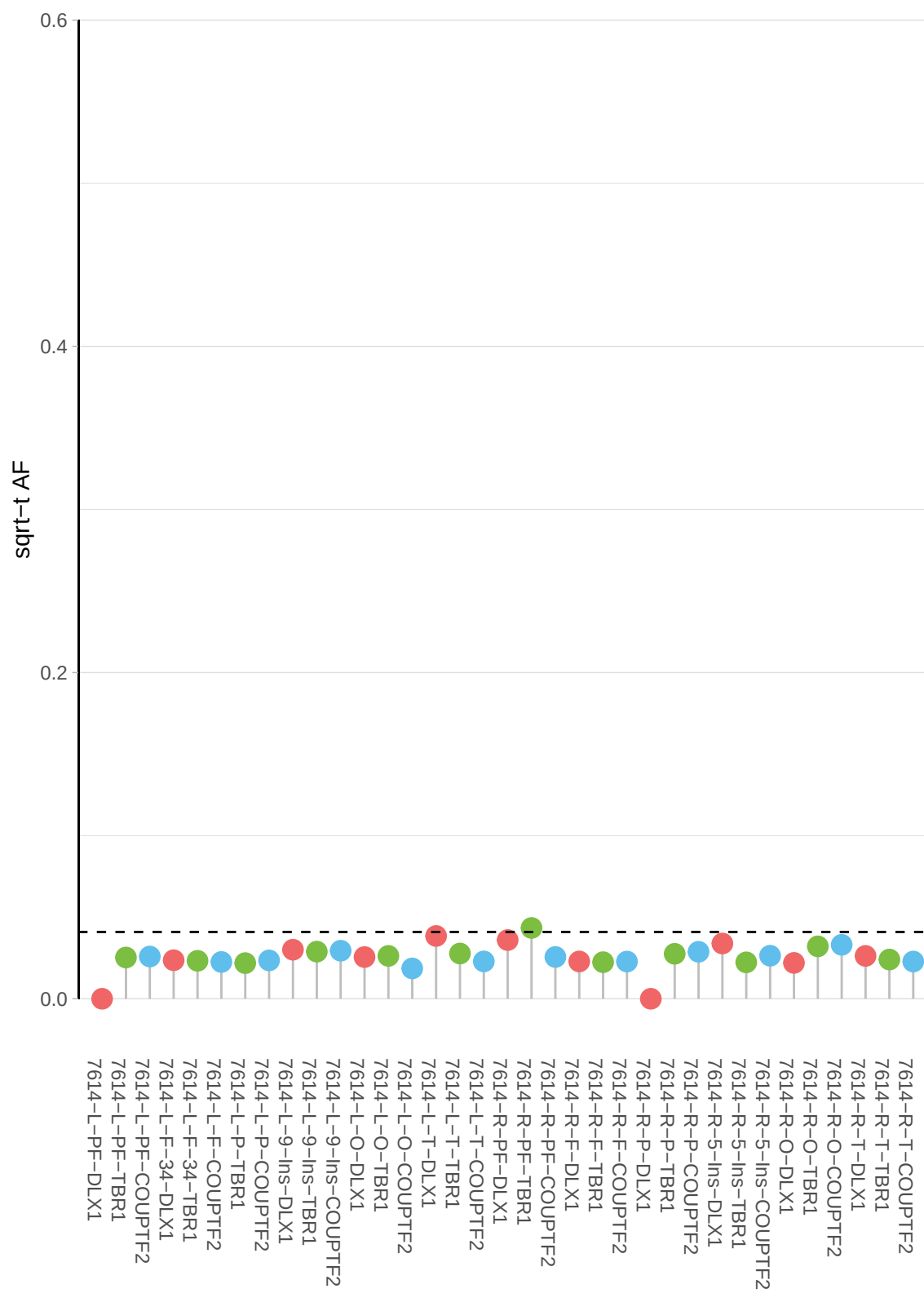

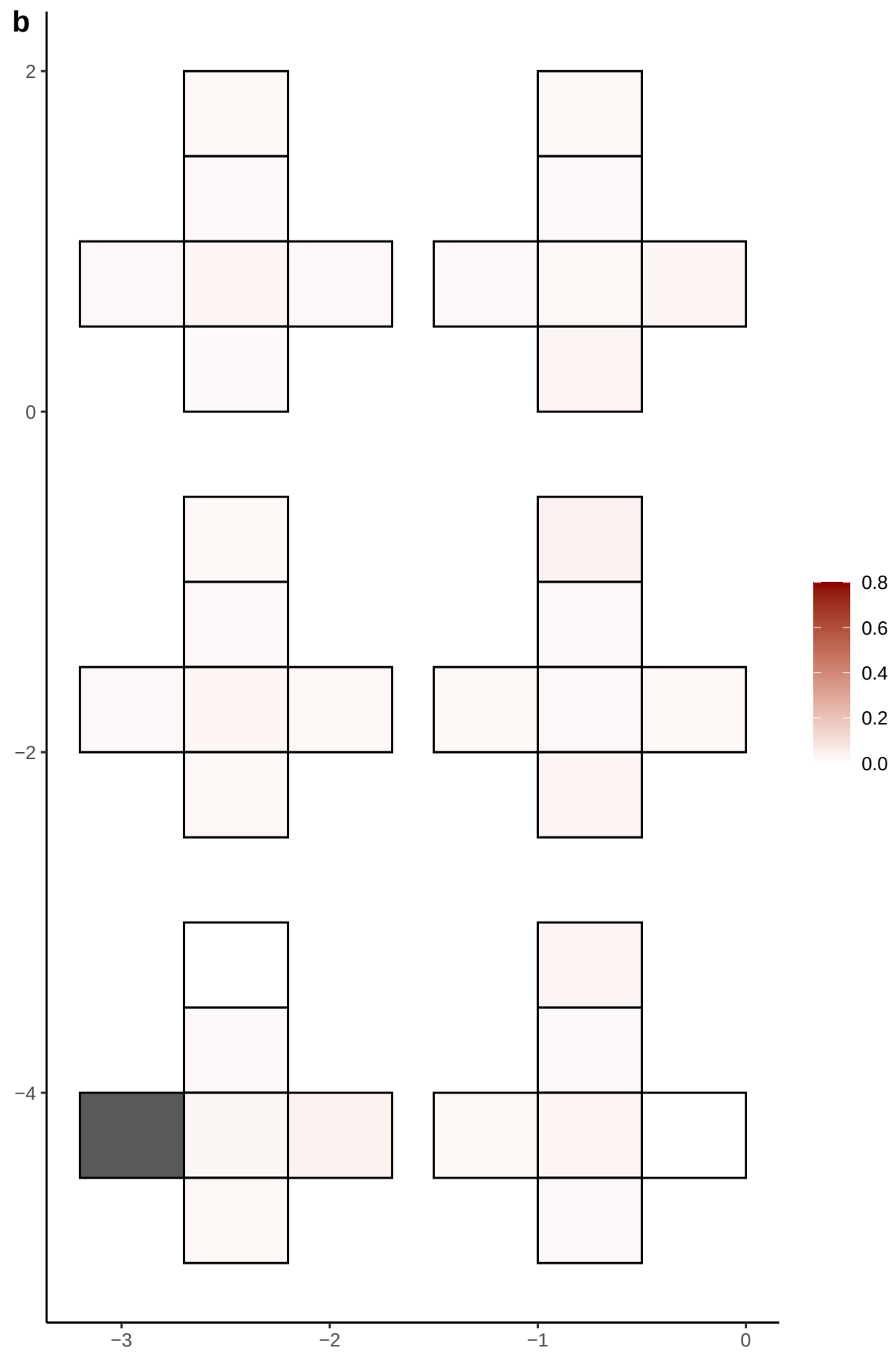

**a** ID01 2-71776656-C-A

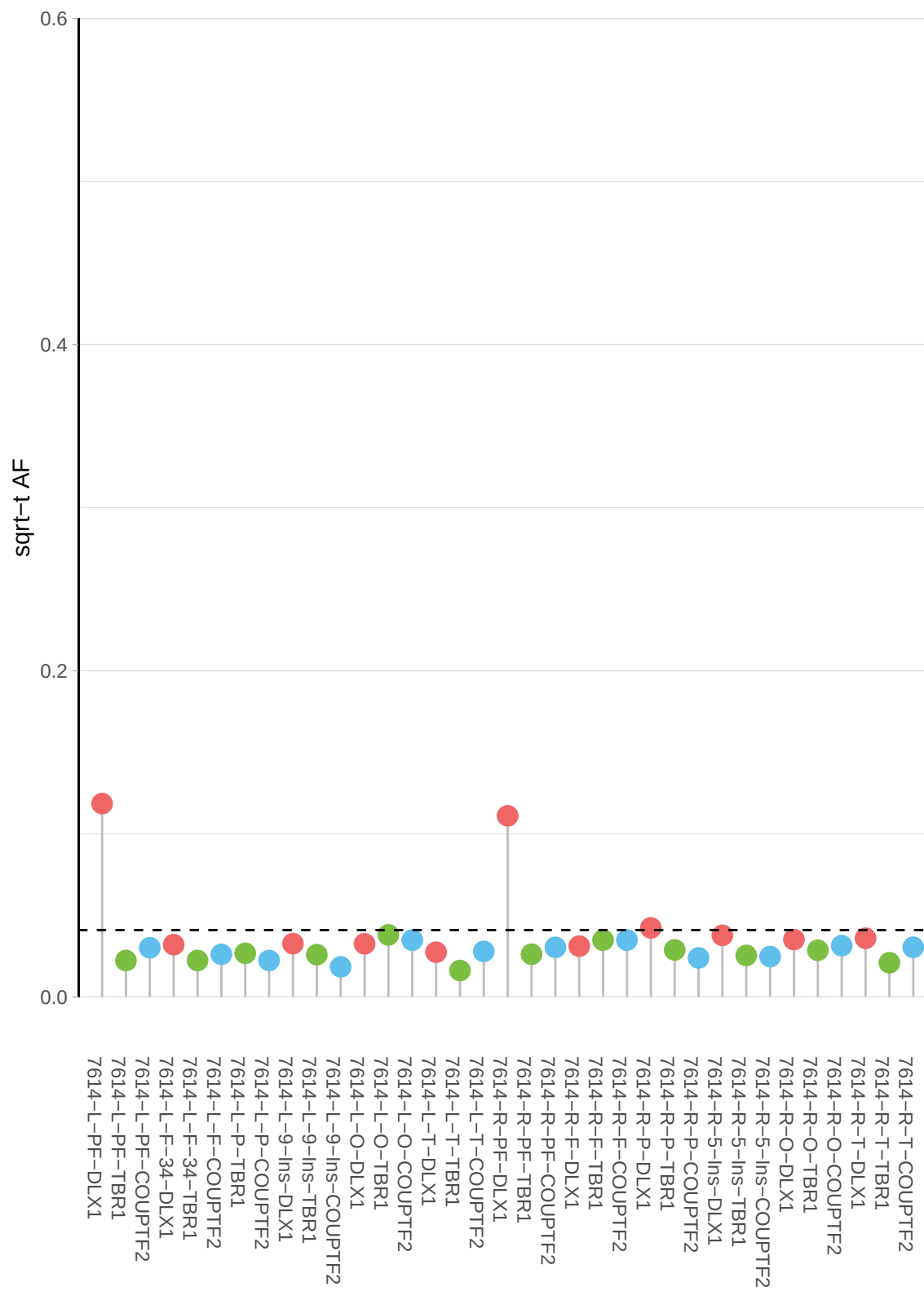

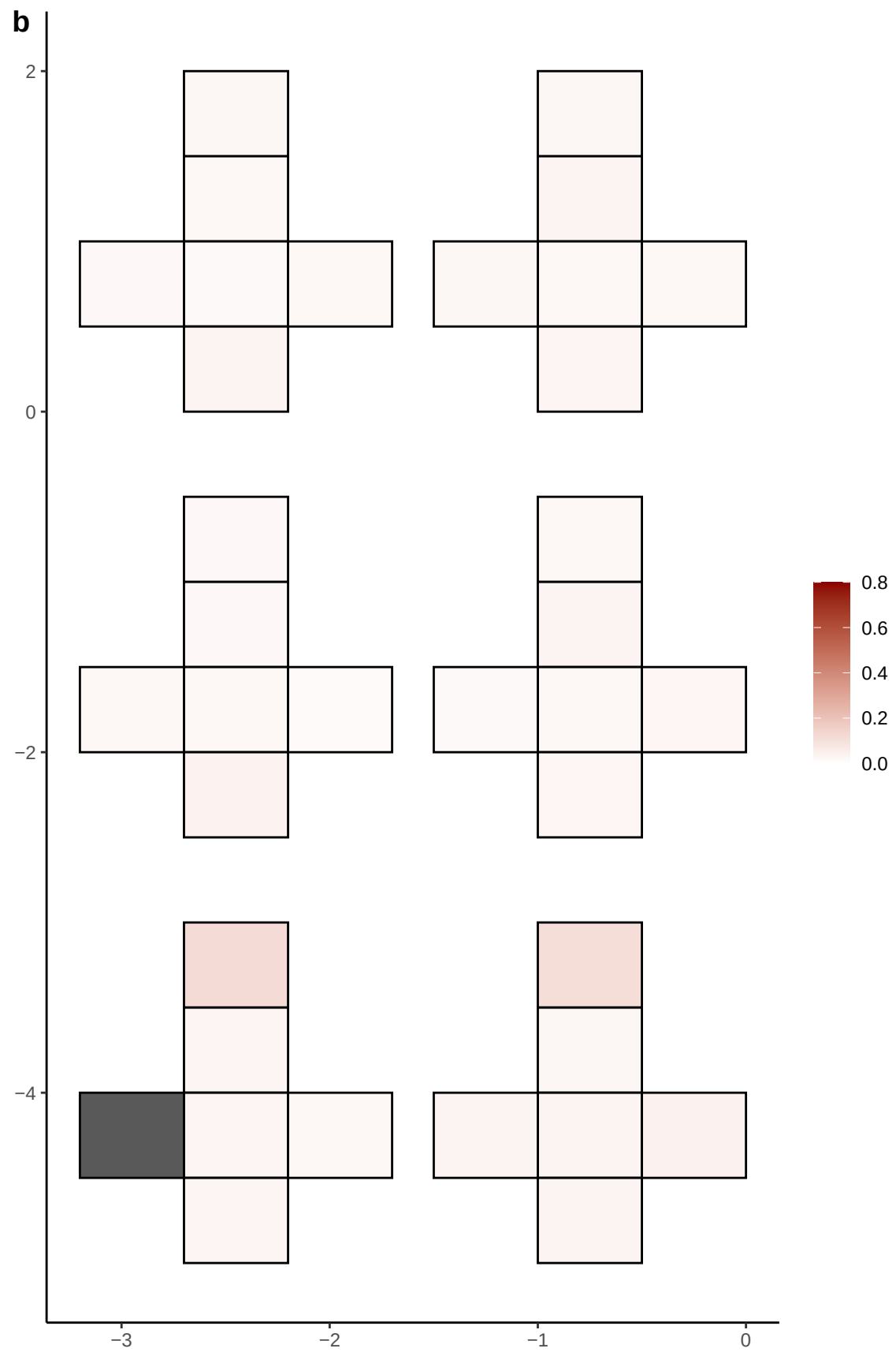

**a** ID01 2-8566334-G-T

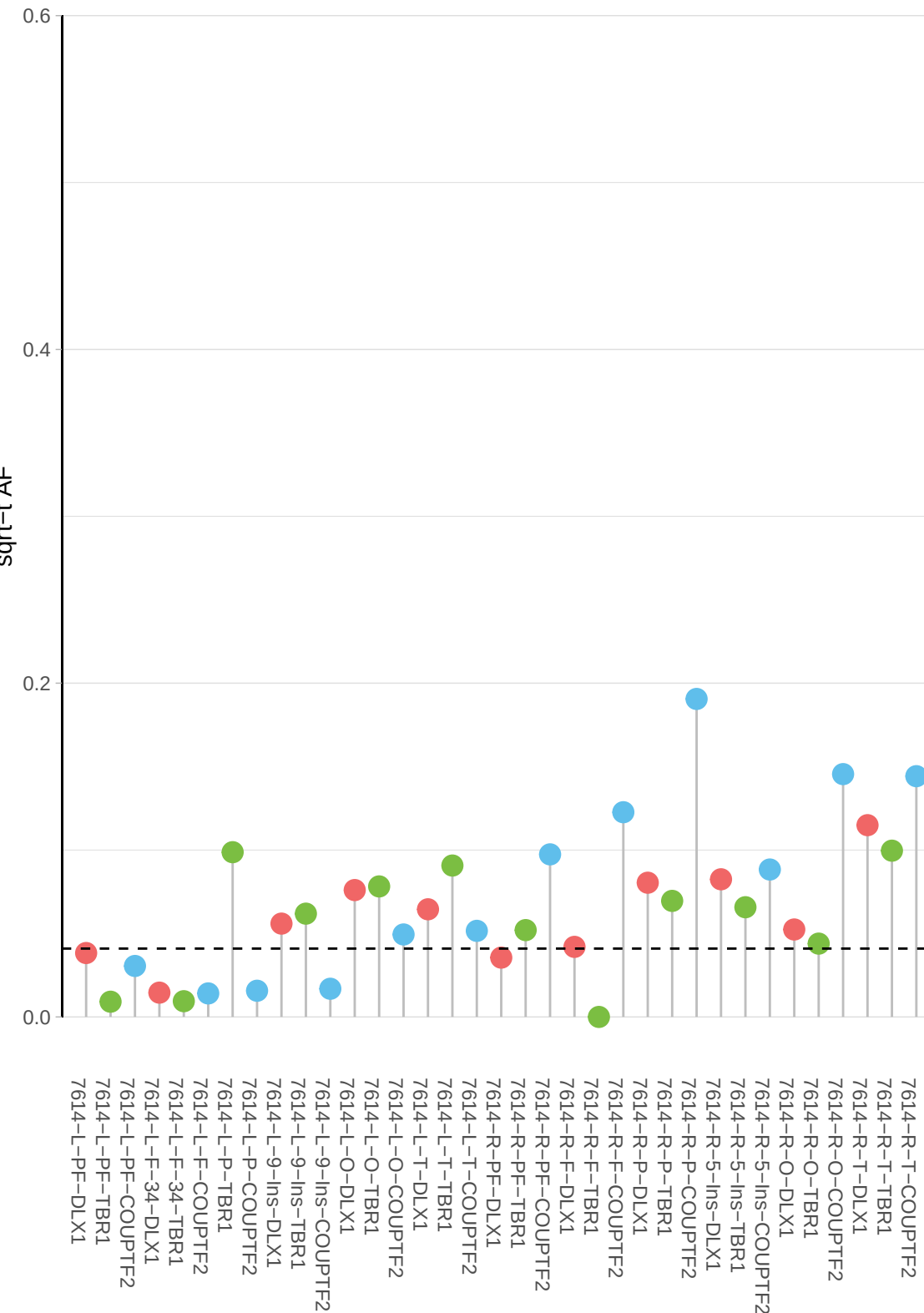

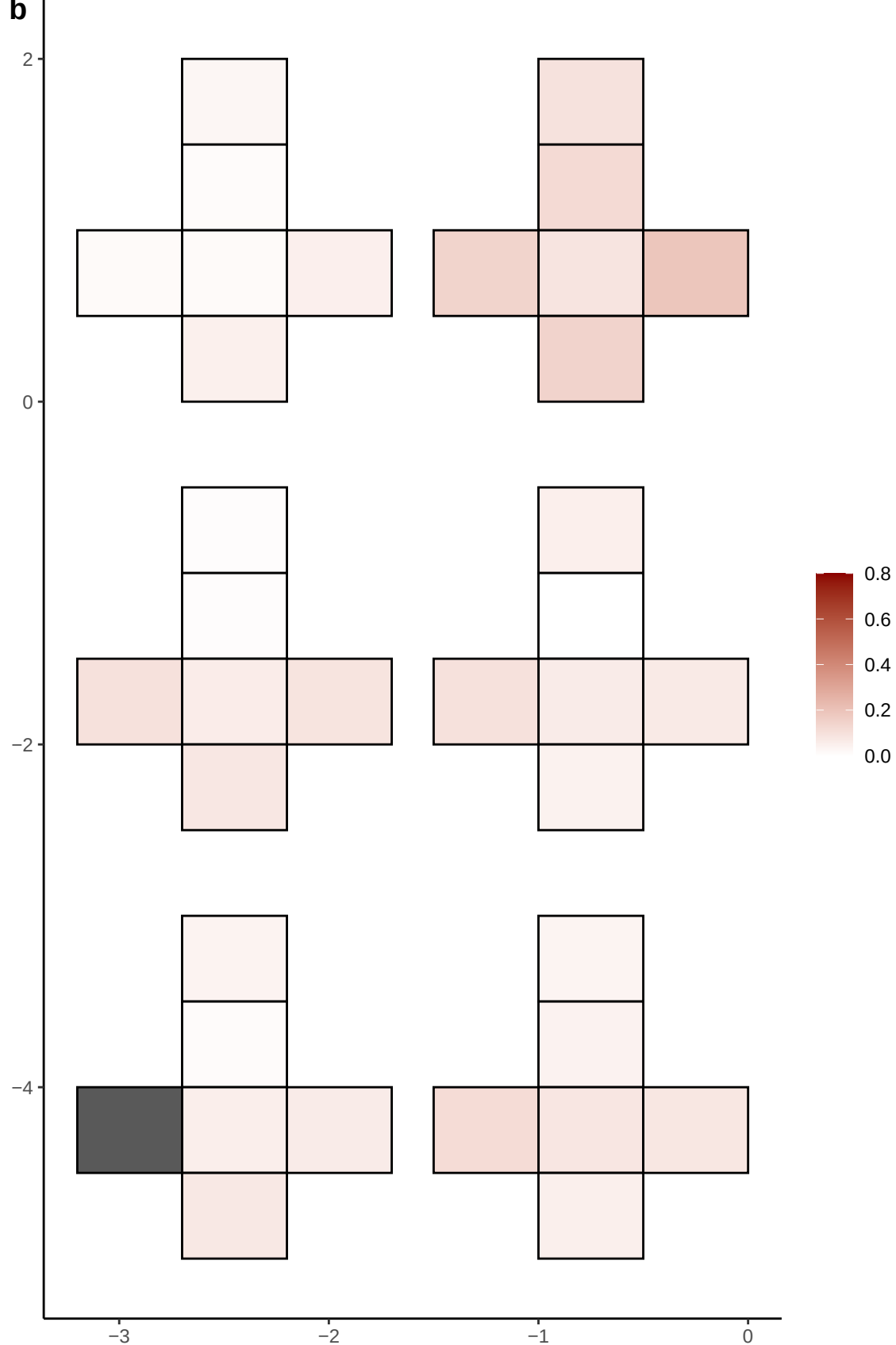

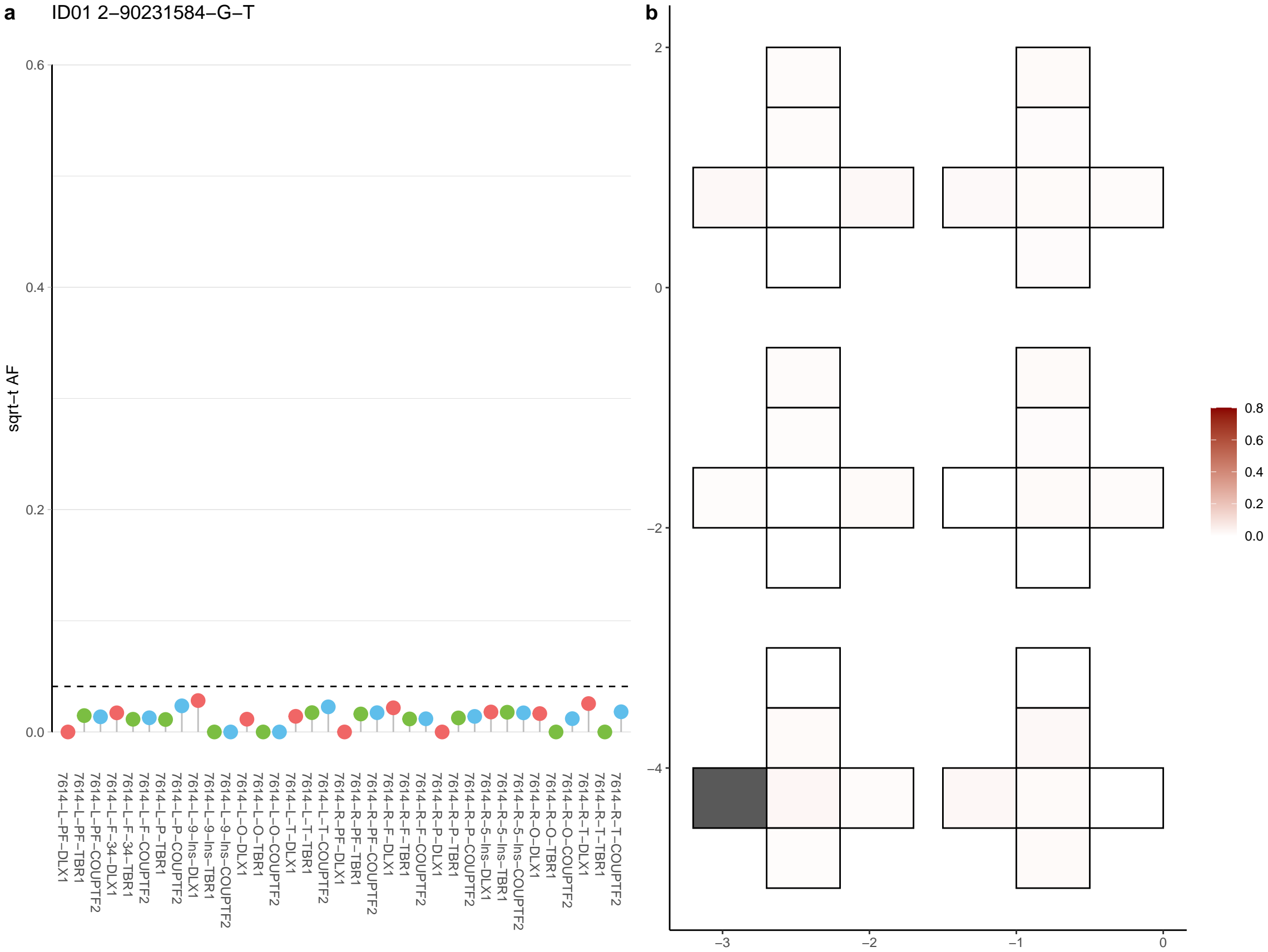

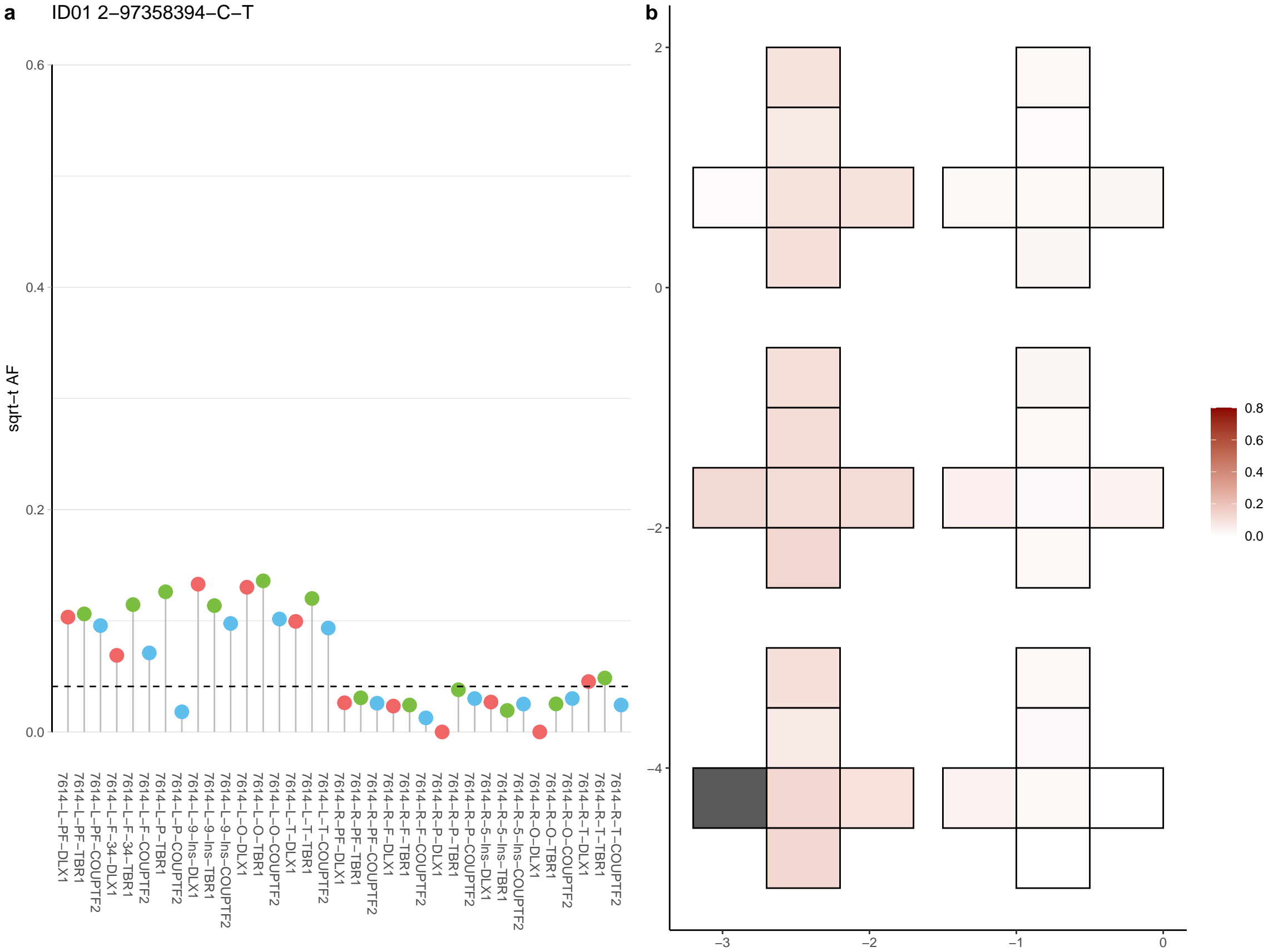

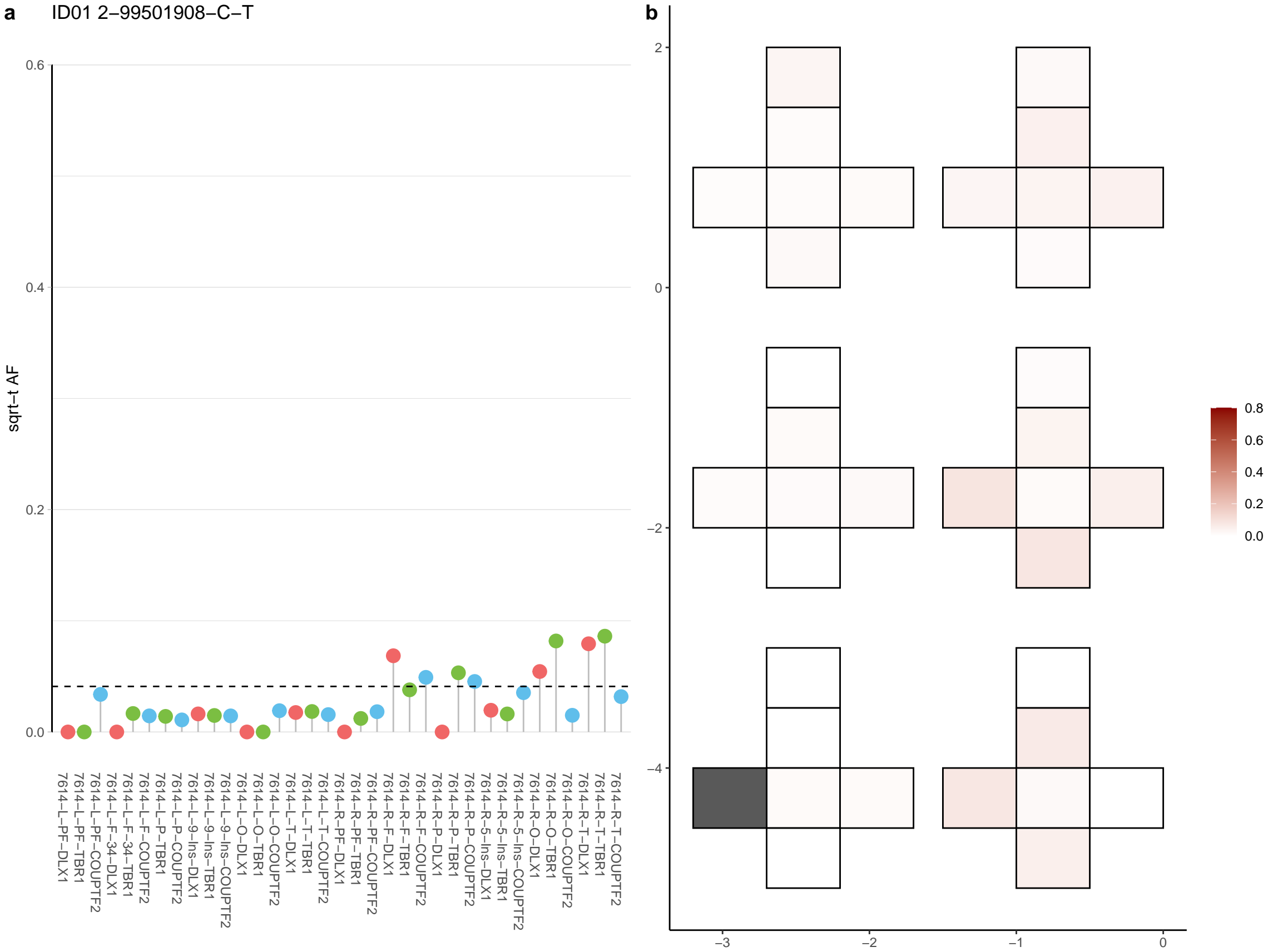

**a** ID01 20-13255185-C-A

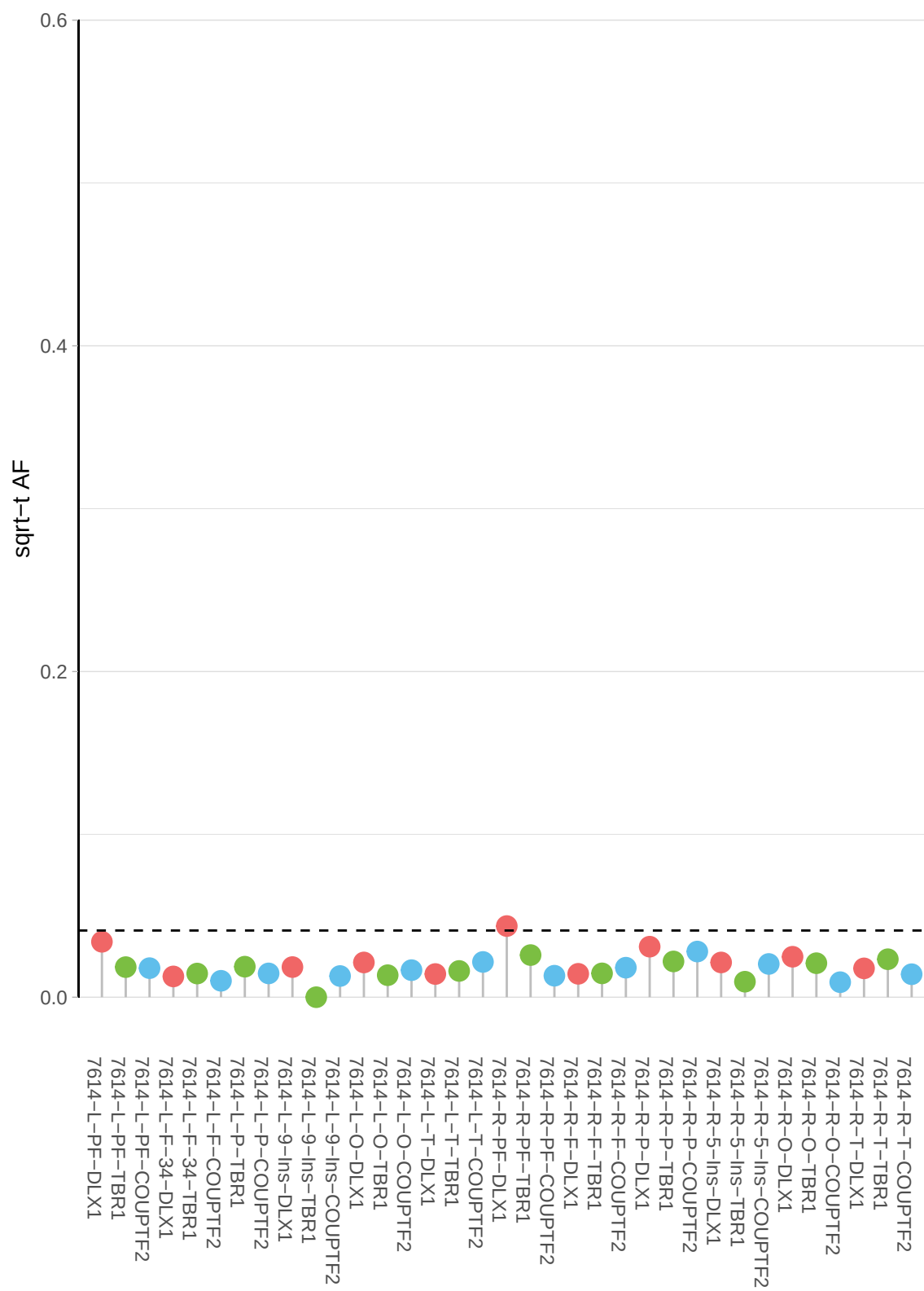

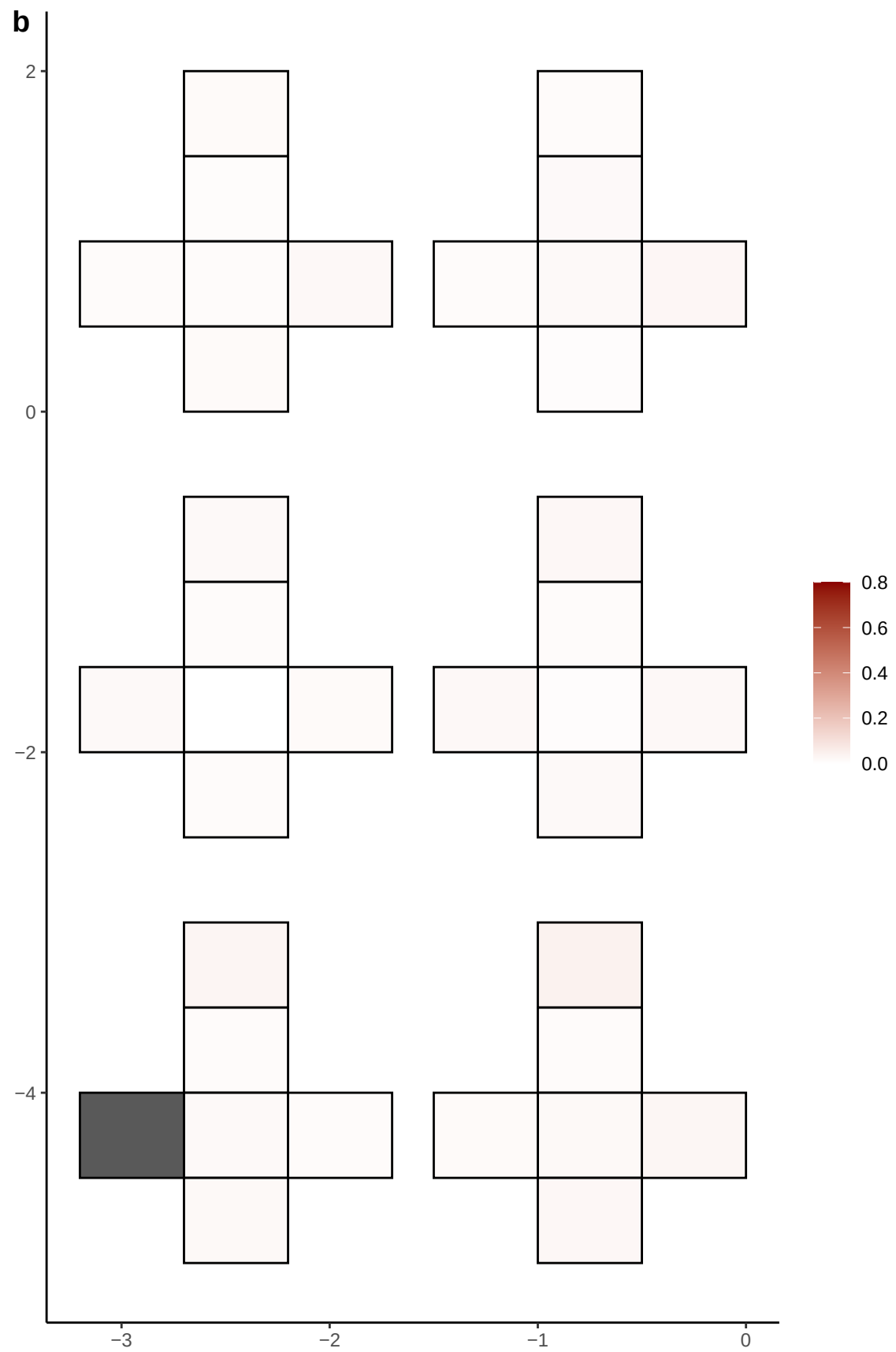

**a** ID01 20-13515886-C-T

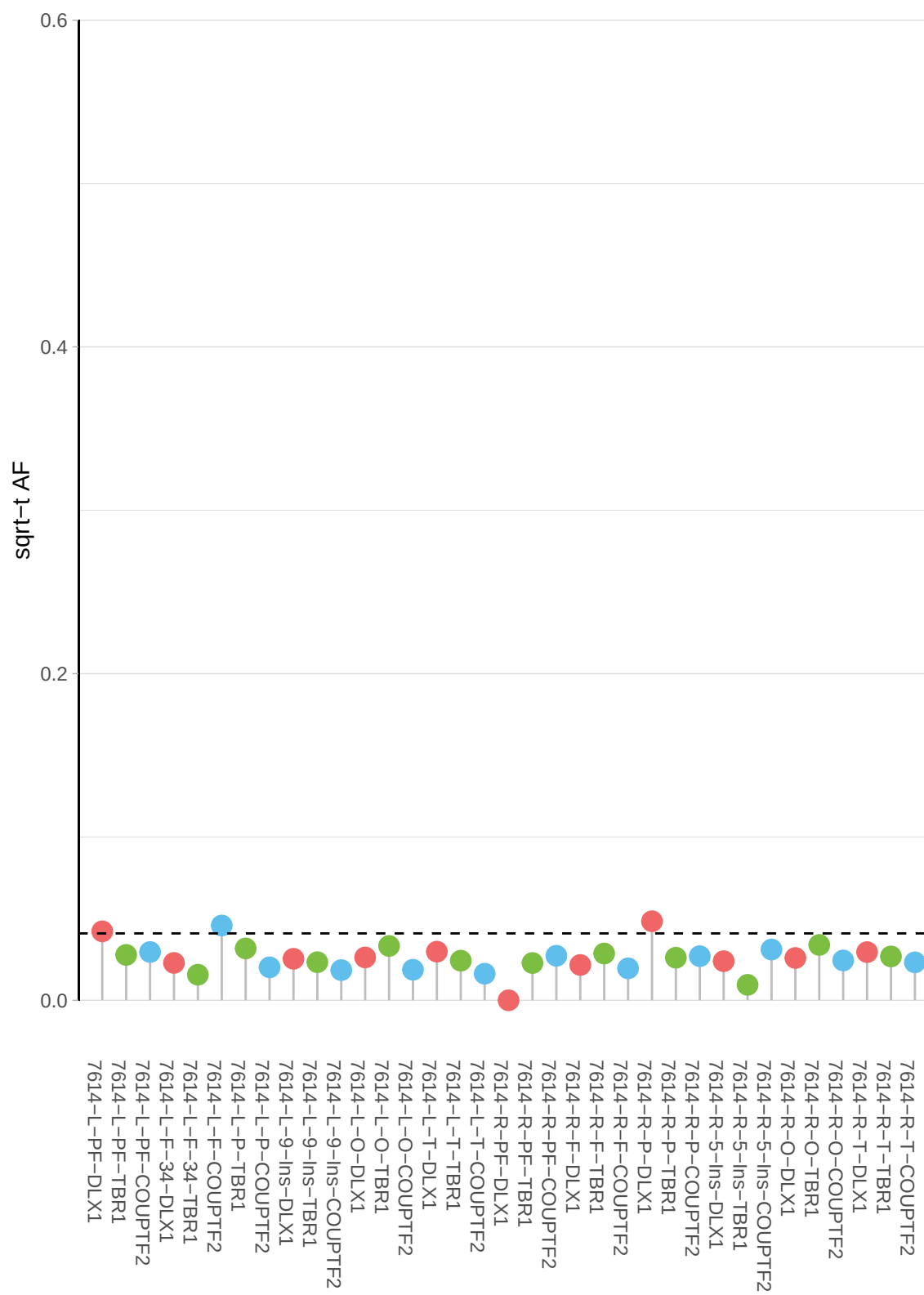

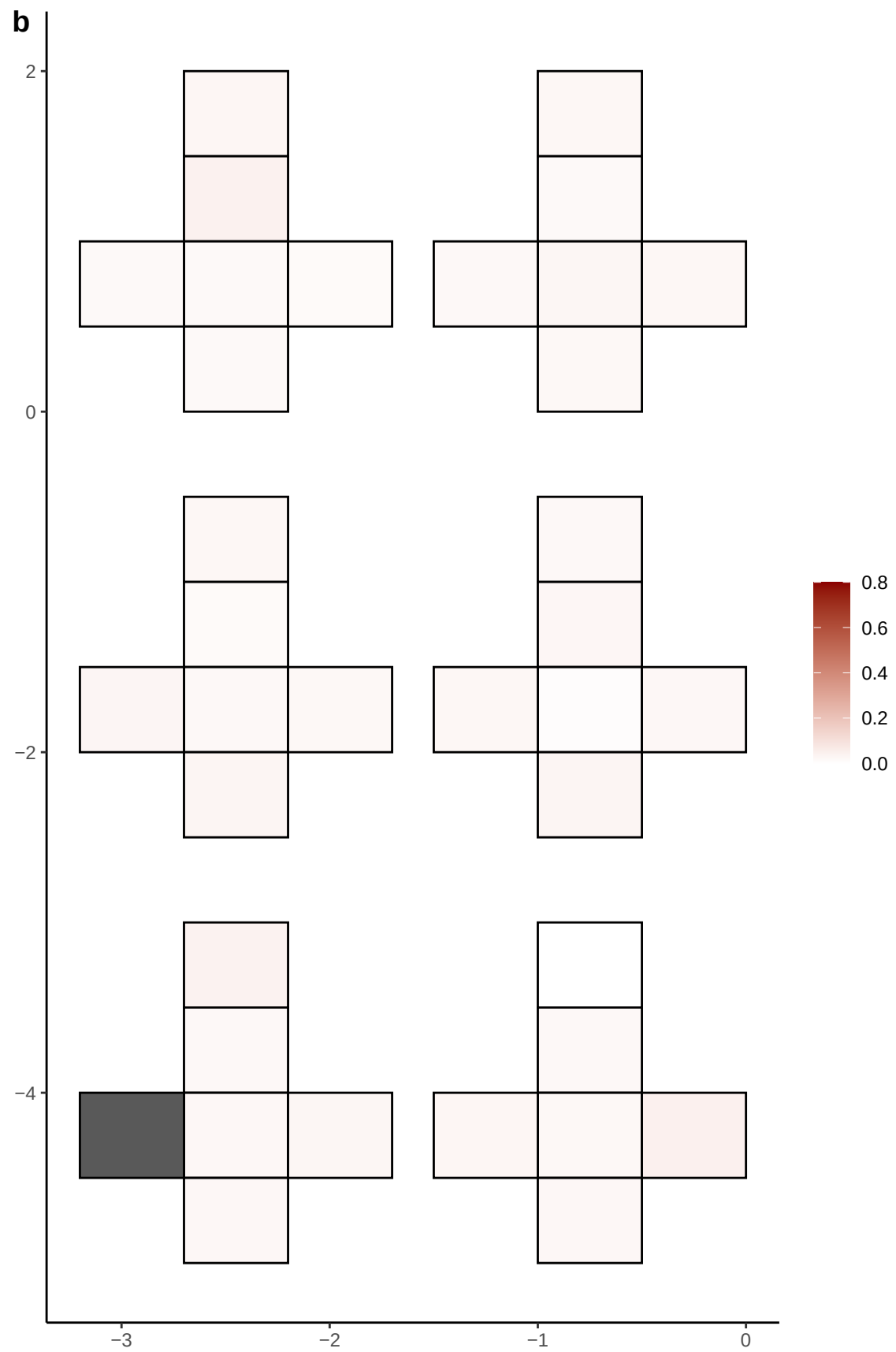

**a** ID01 20-1547810-C-T

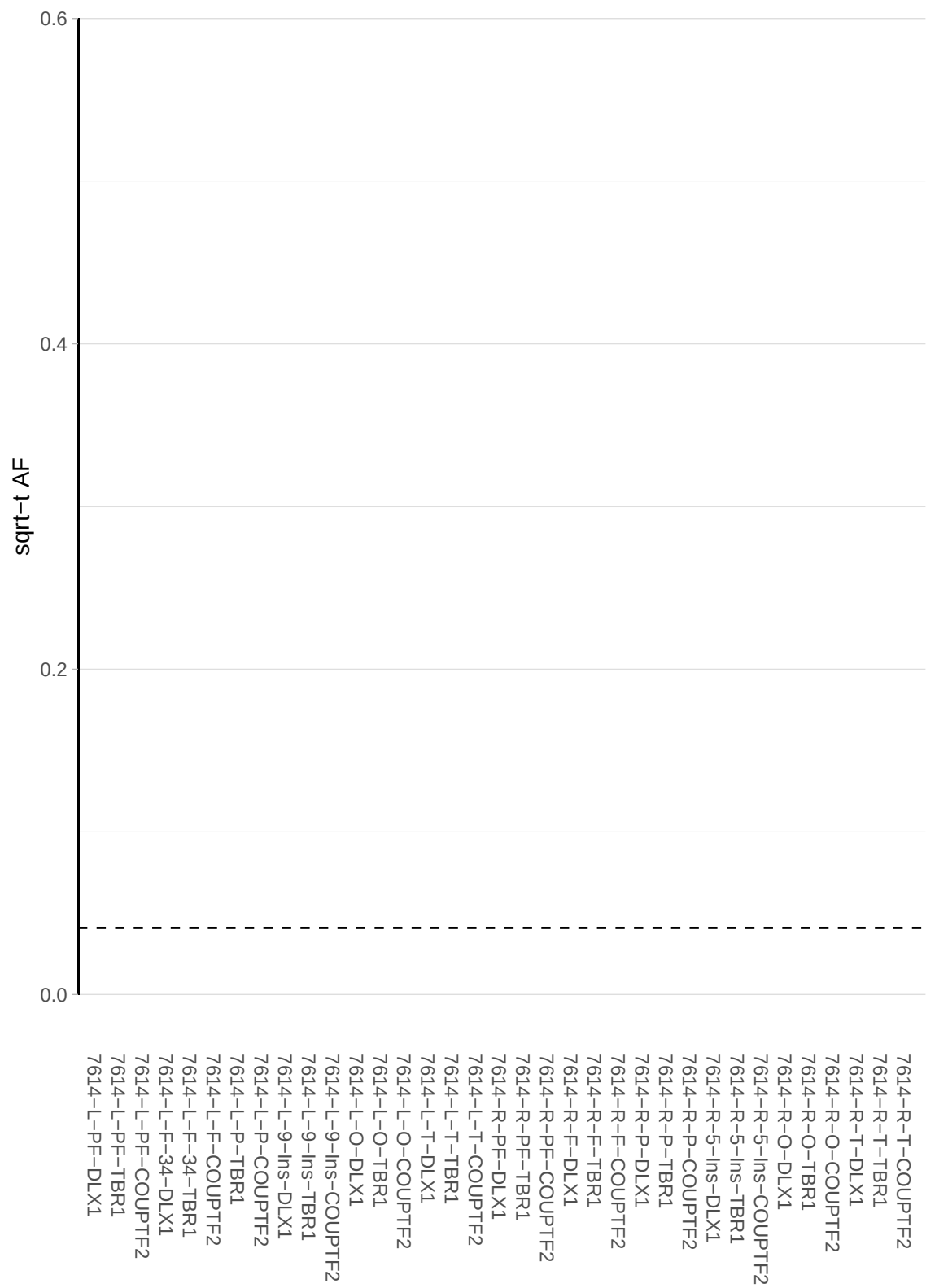

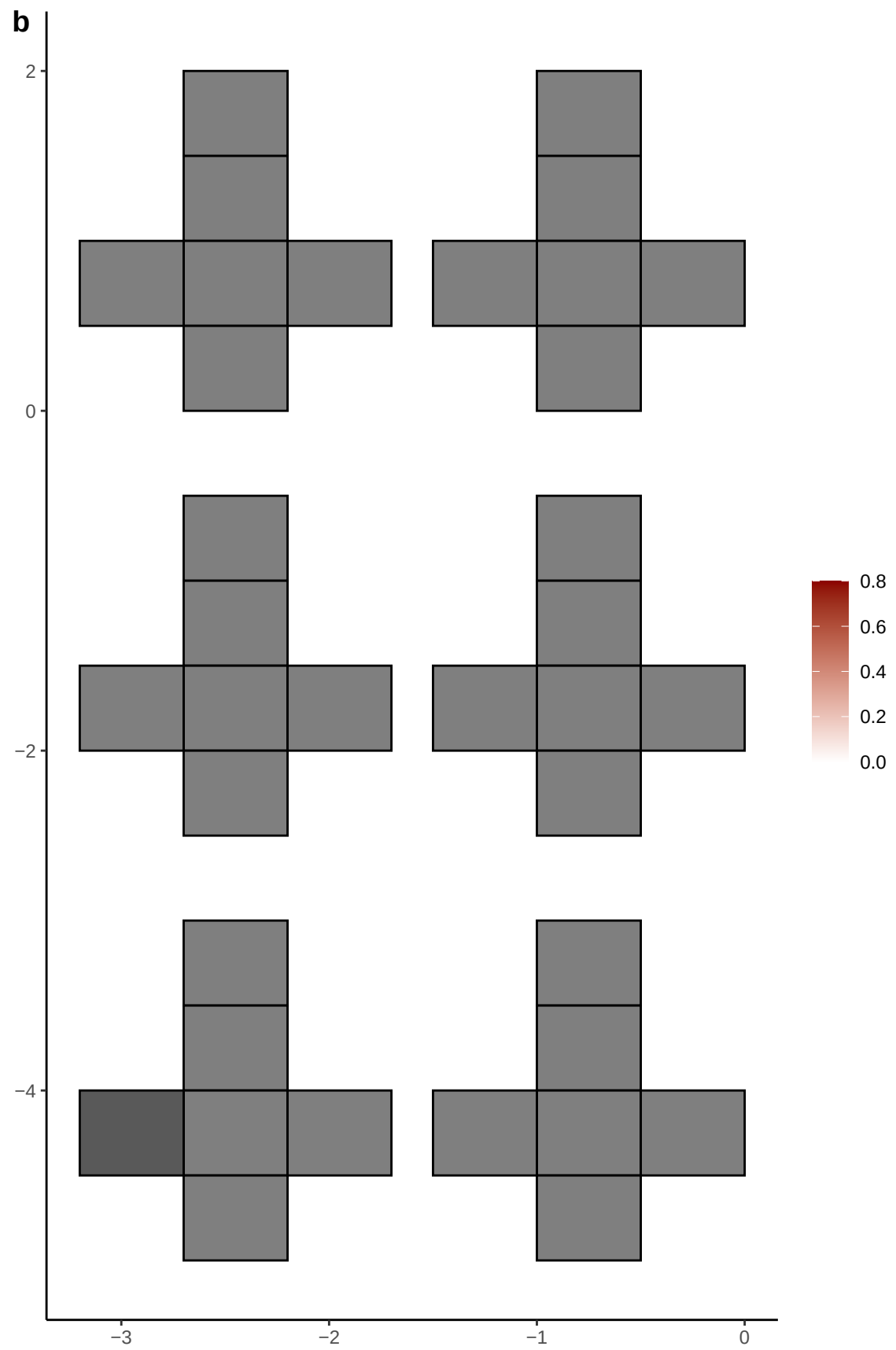

**a** ID01 20-29913114-C-T

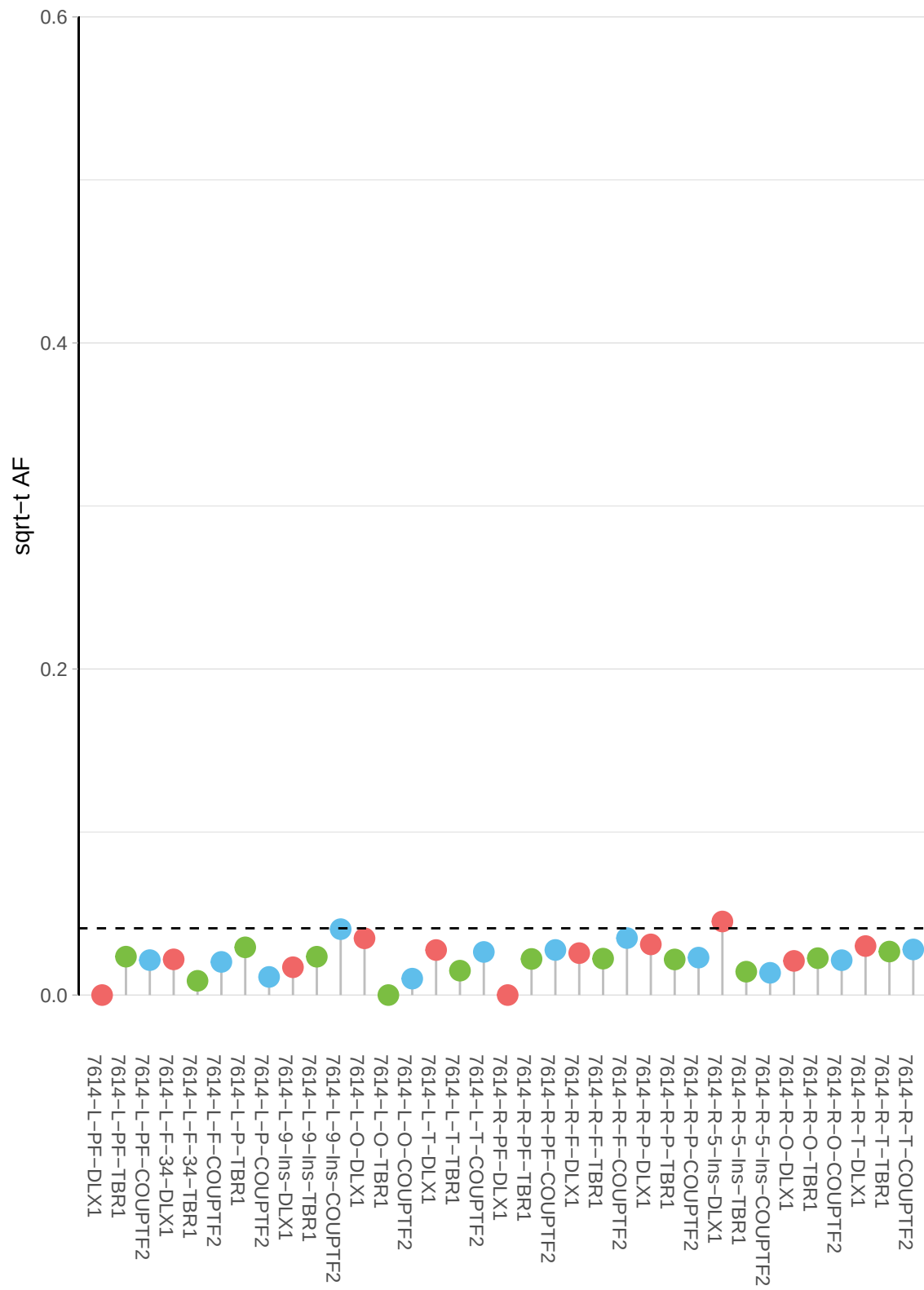

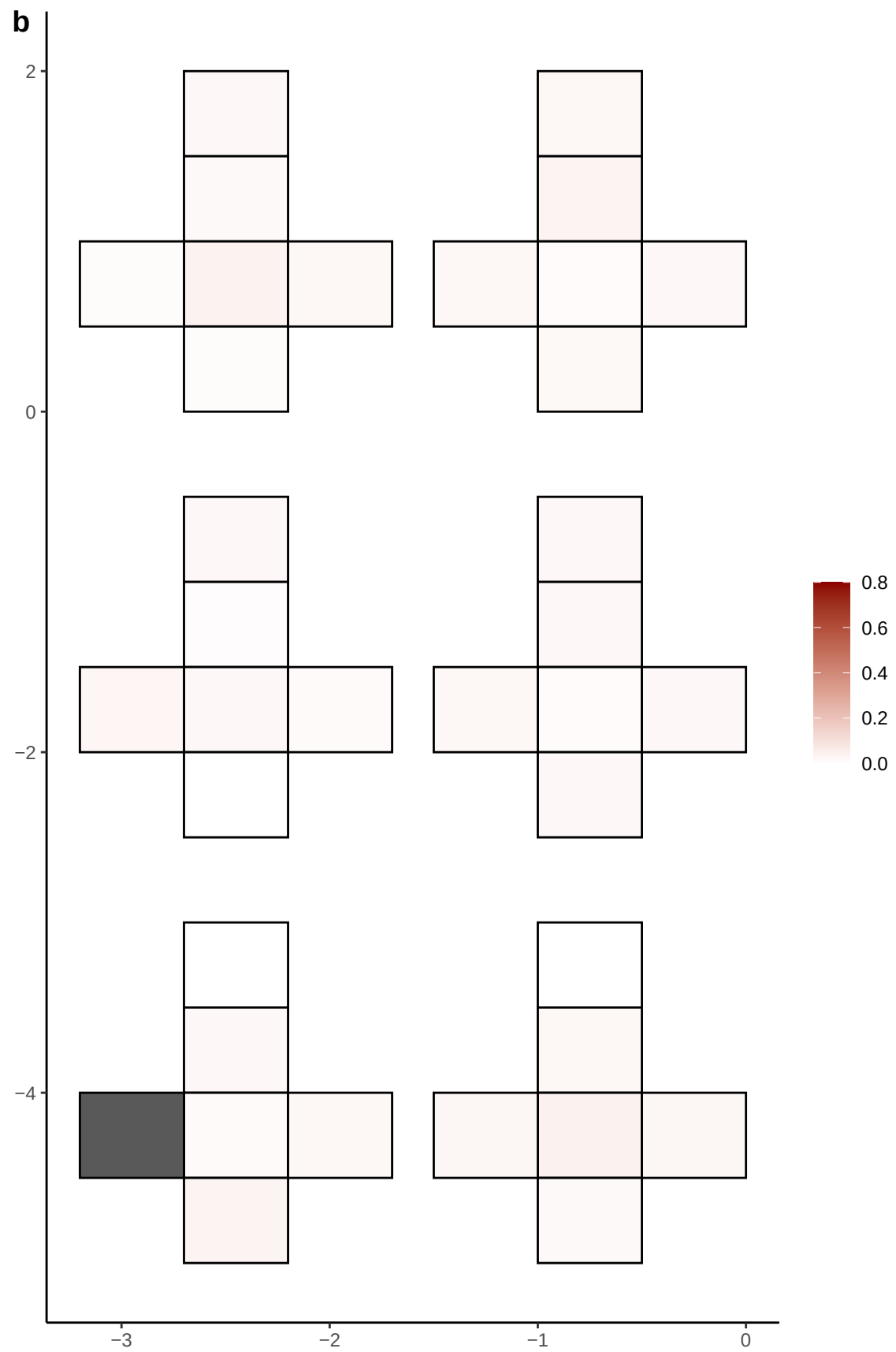

**a** ID01 20-52915779-G-T

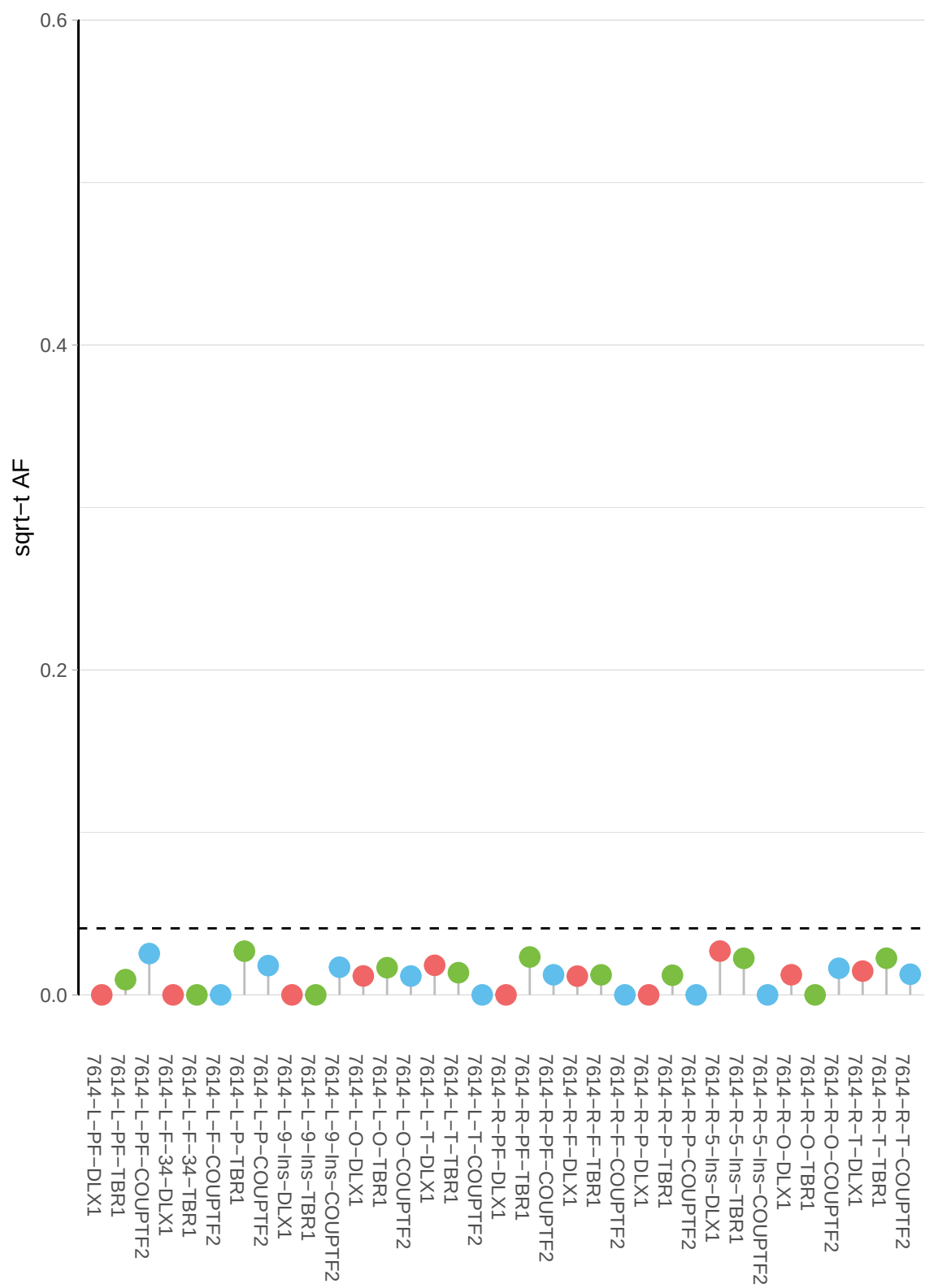

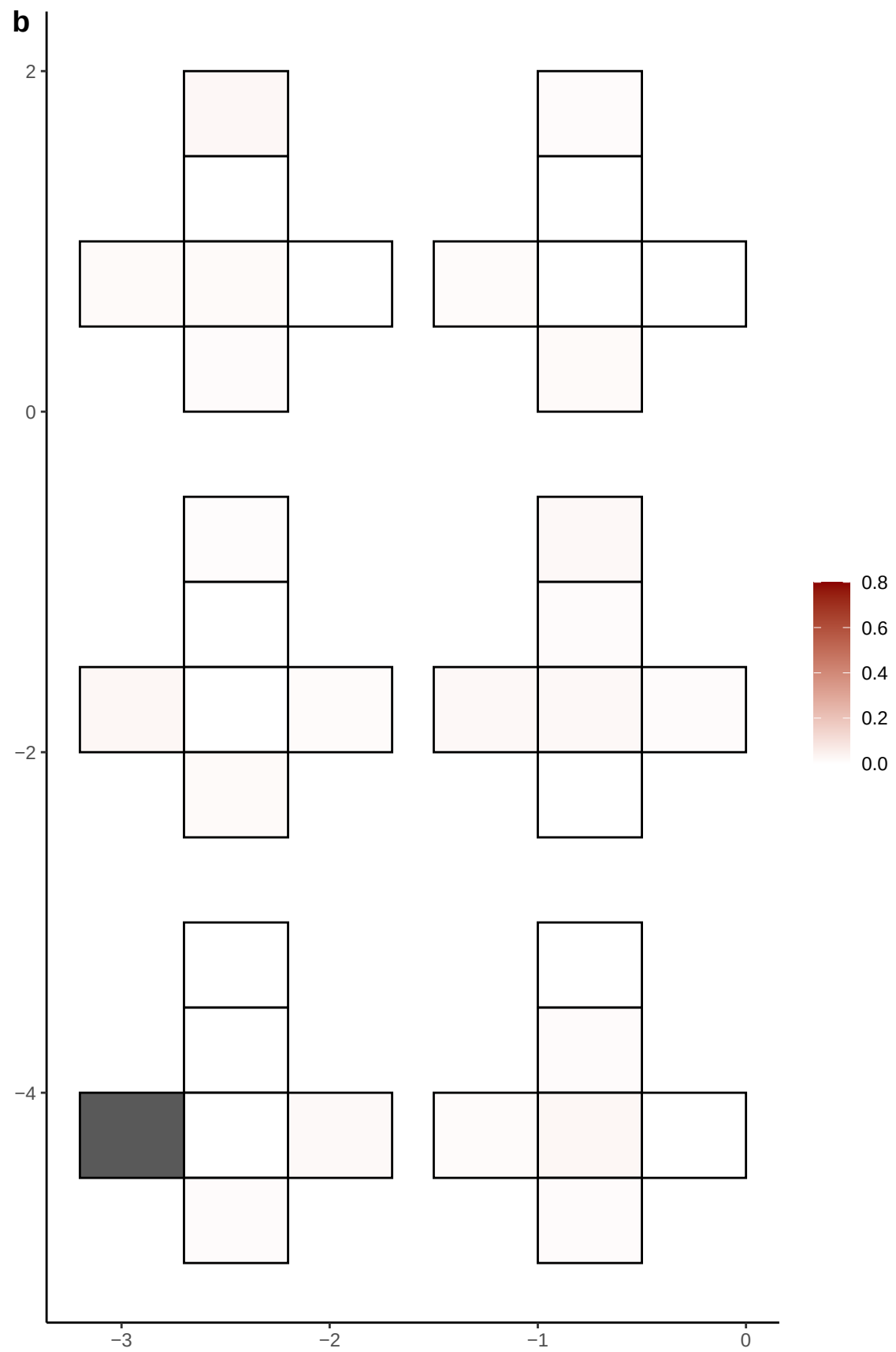

**a** ID01 20-52916352-A-C

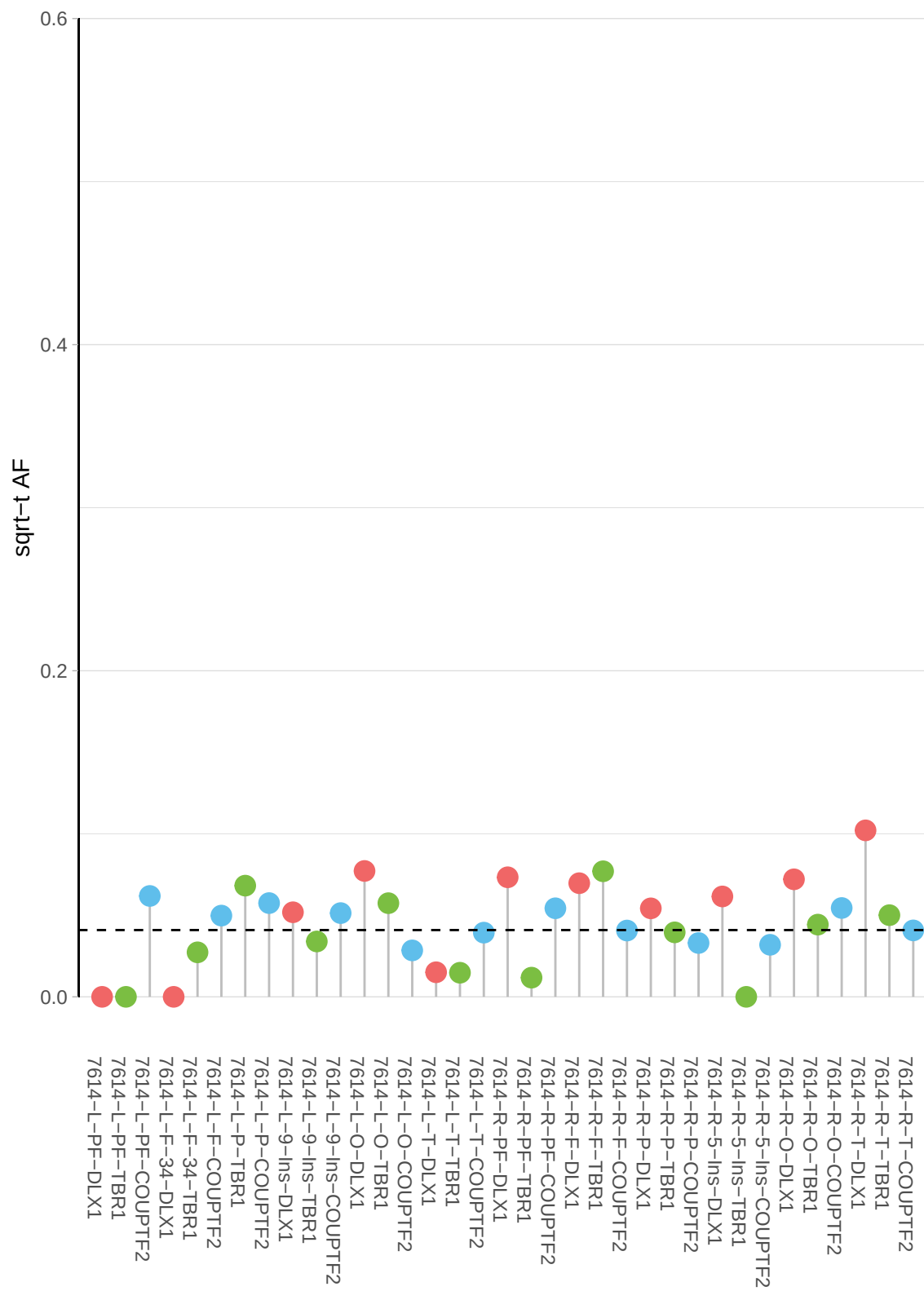

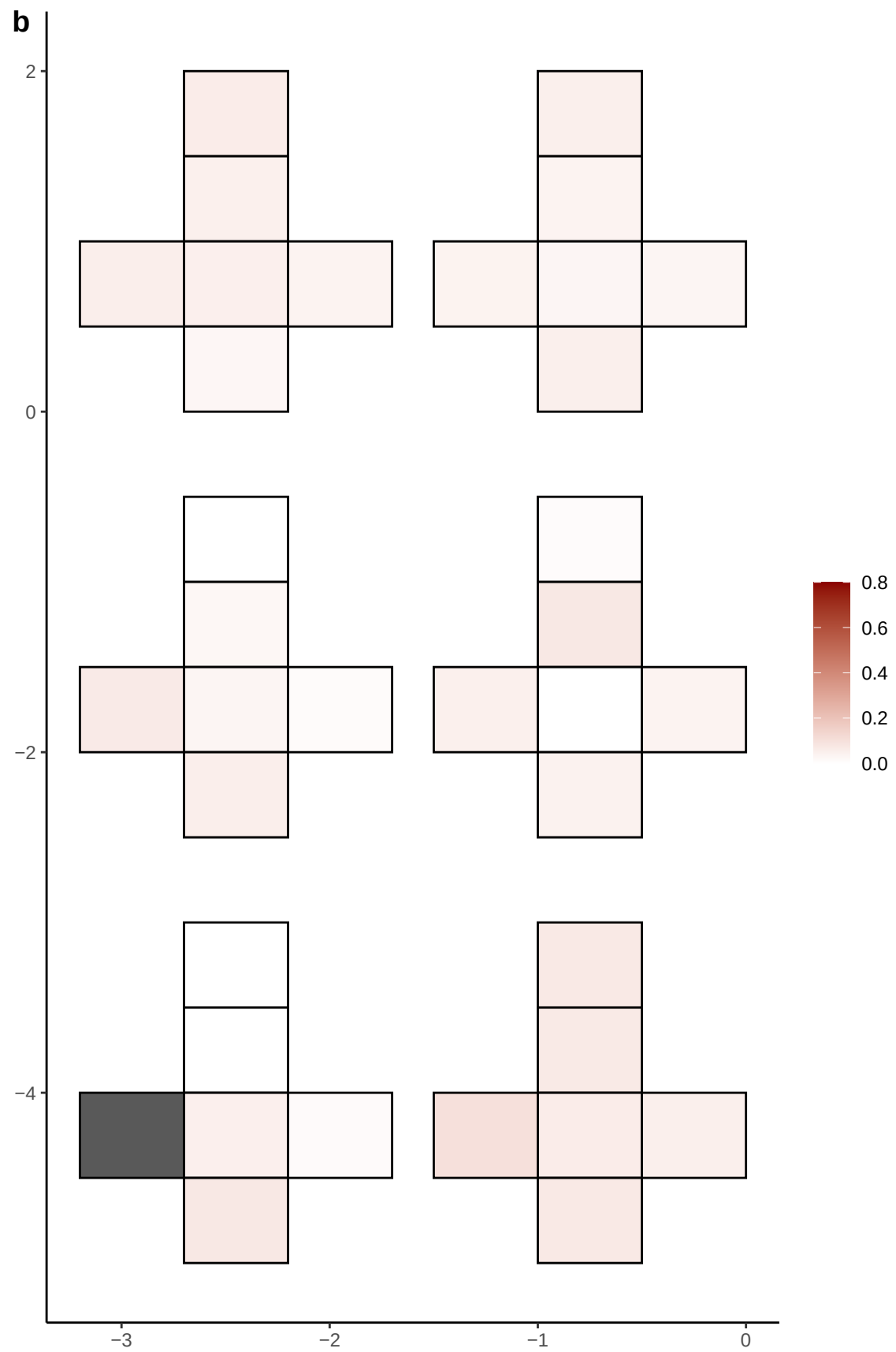

**a** ID01 20-53672093-T-A

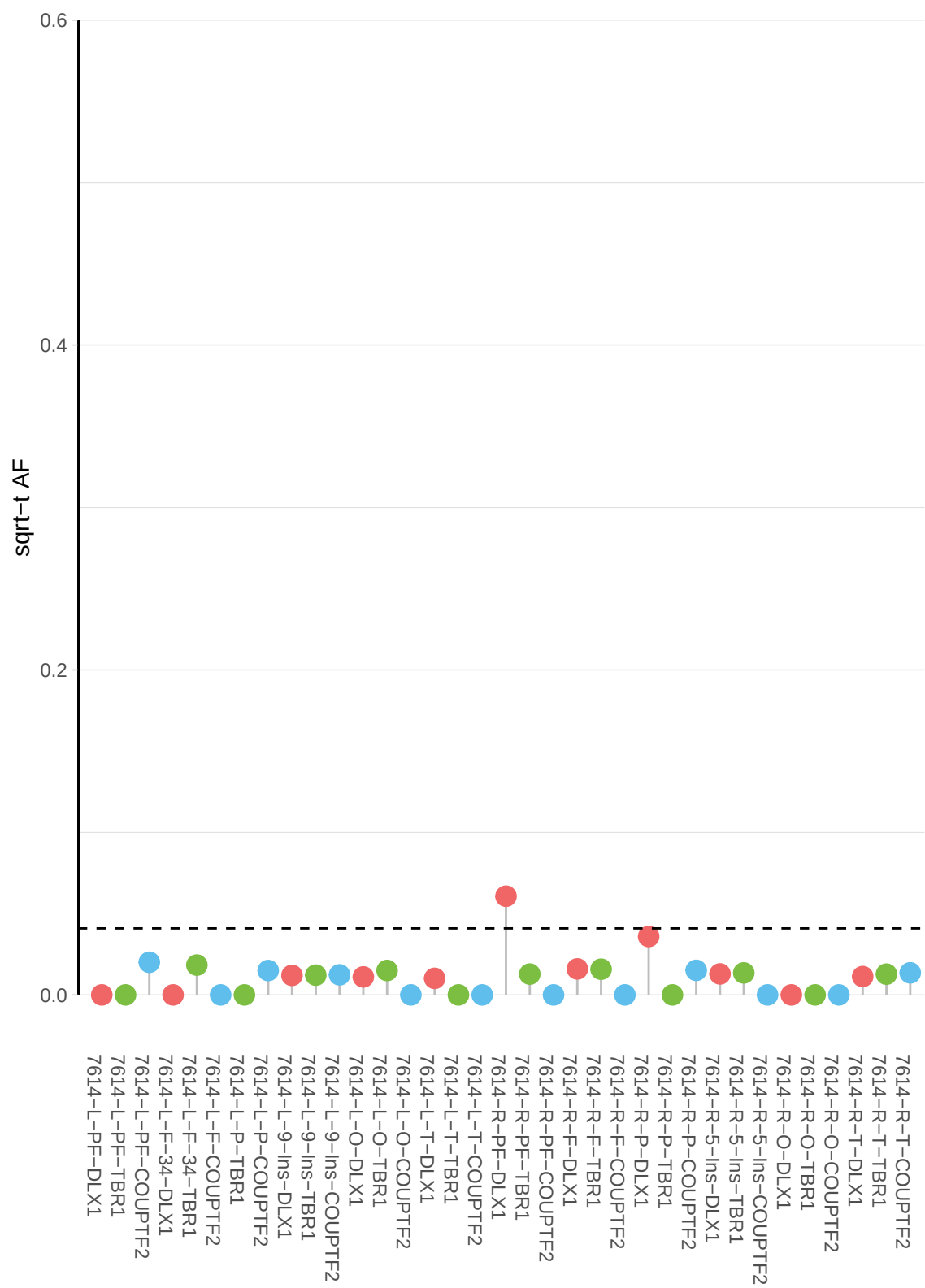

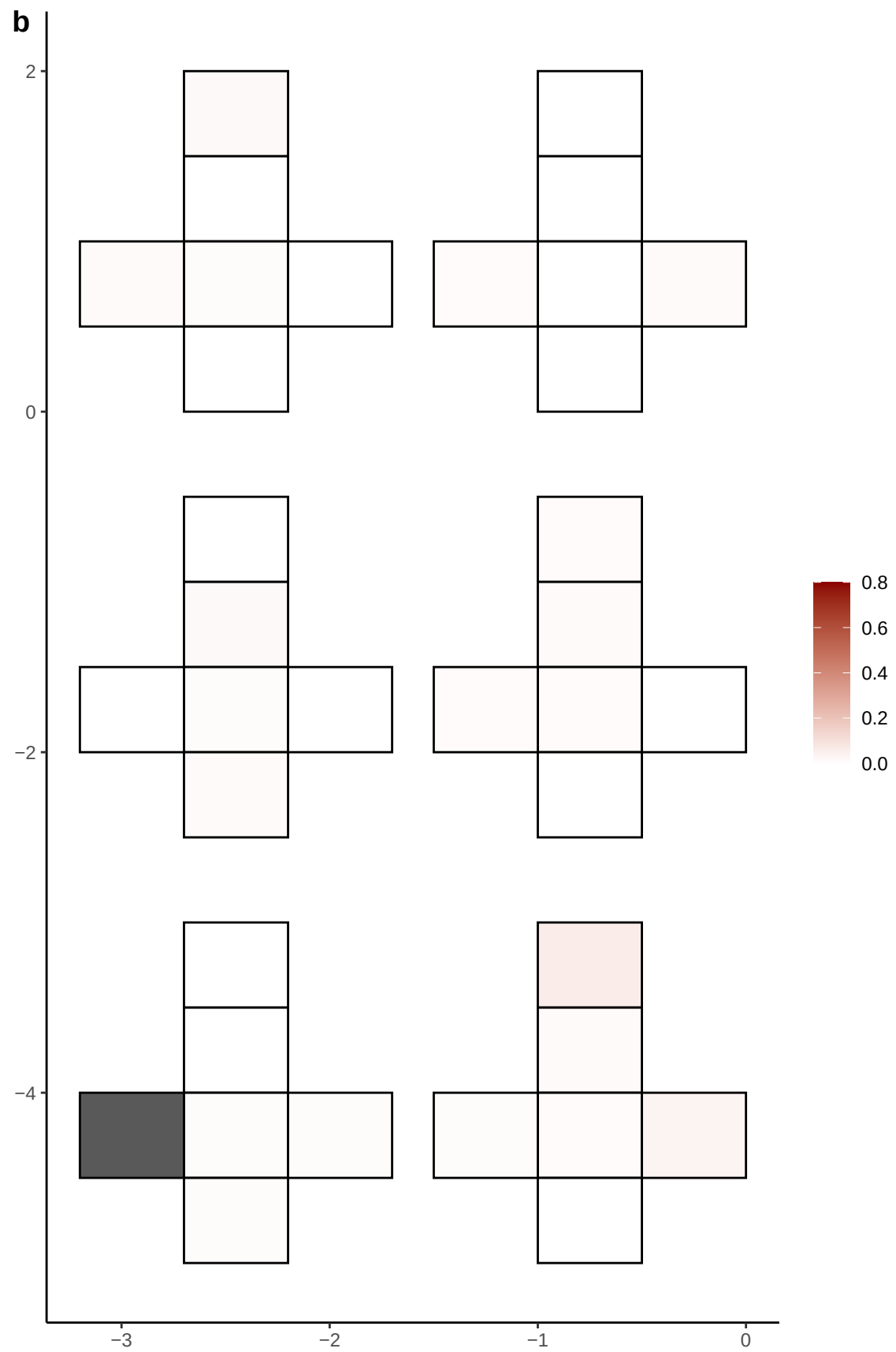

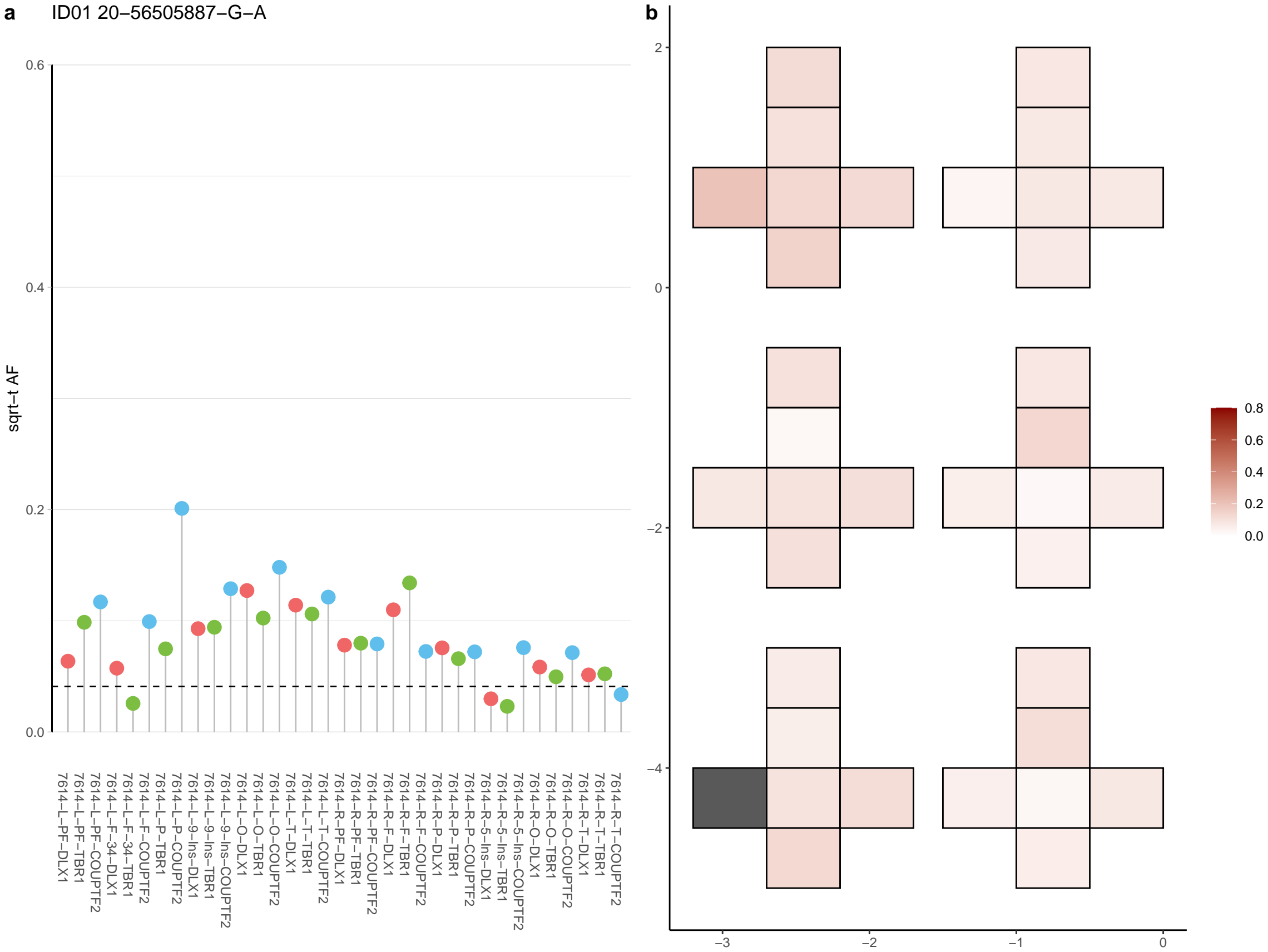

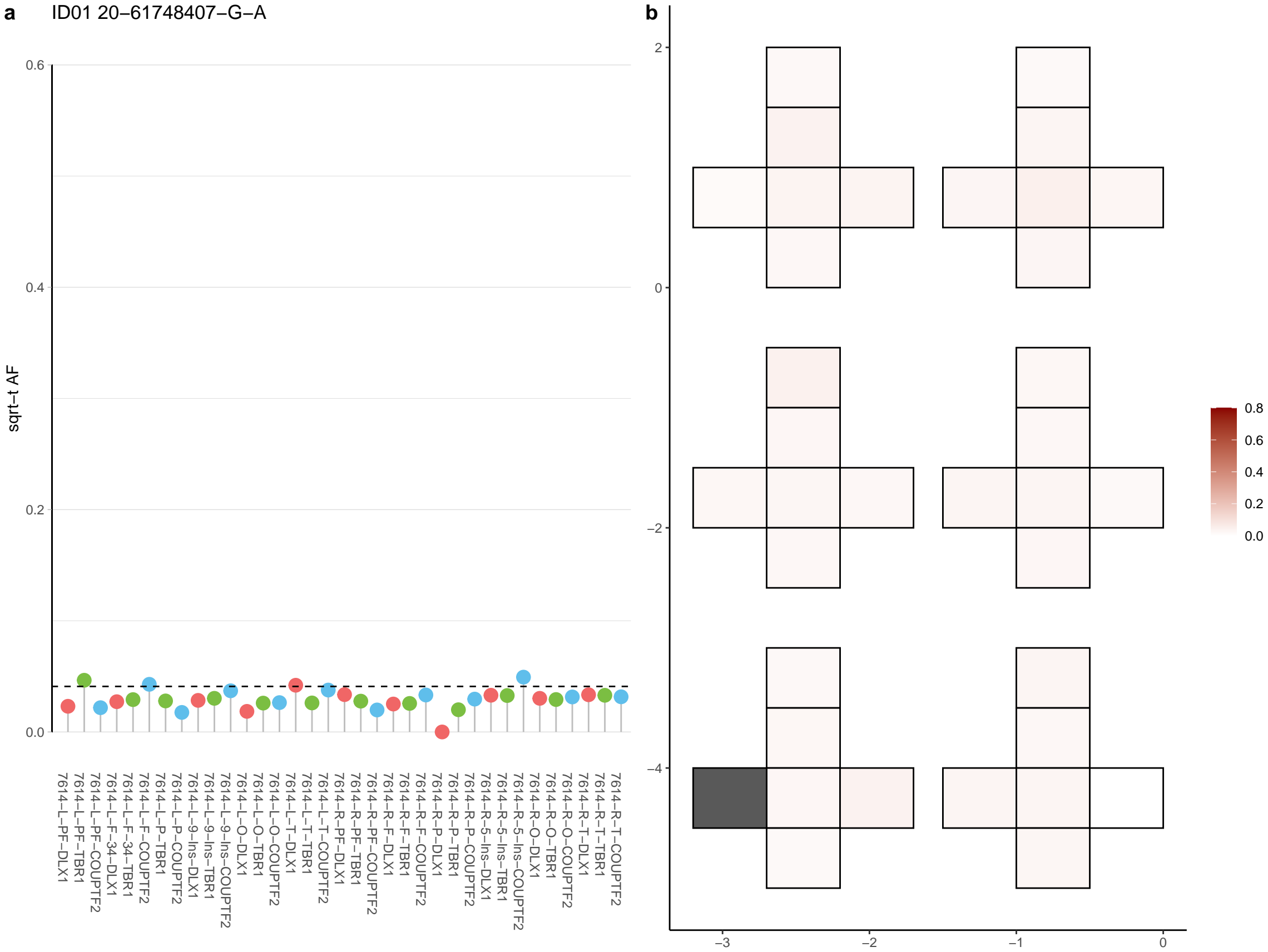

**a** ID01 21-17683648-TTA-T

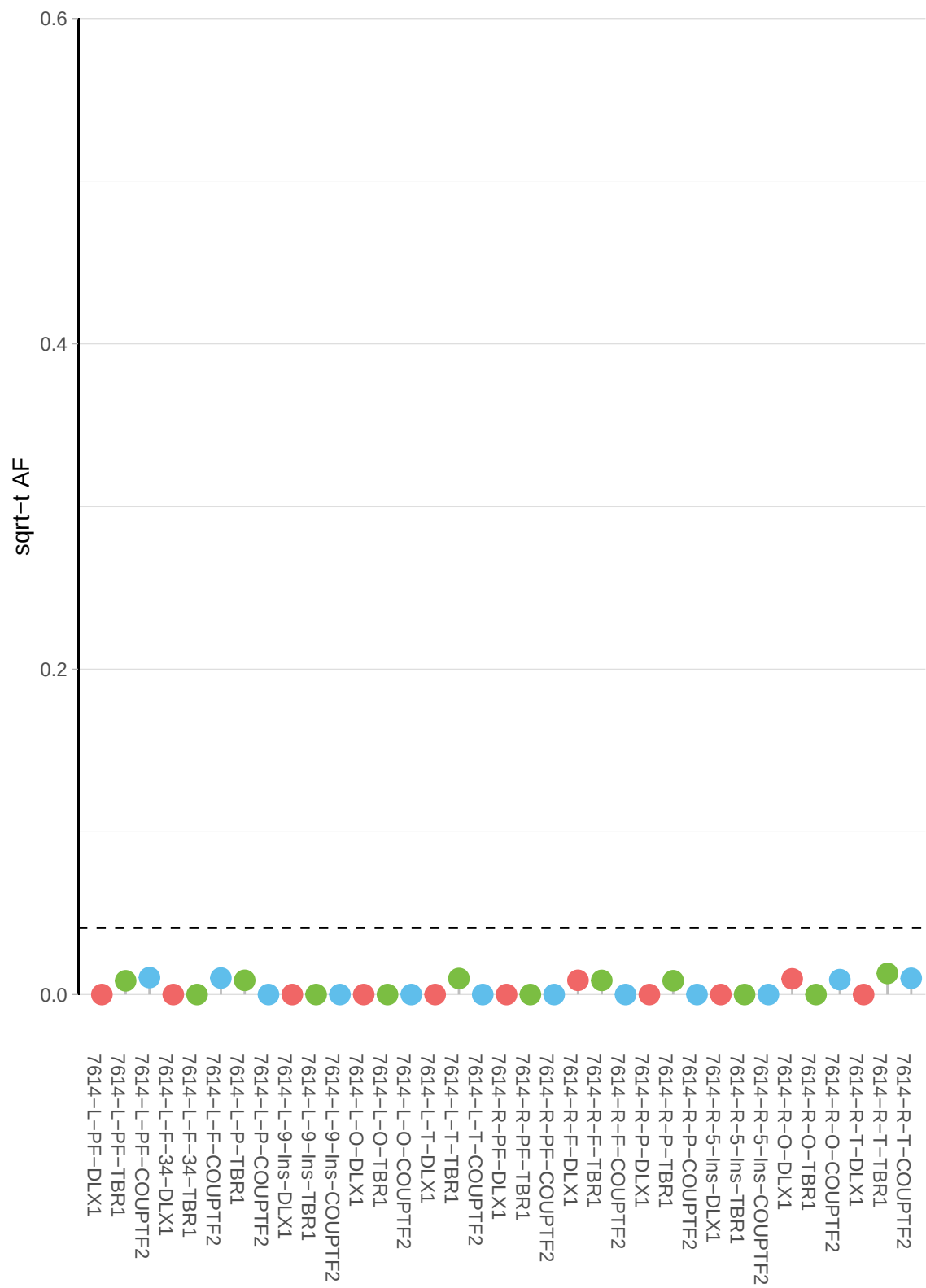

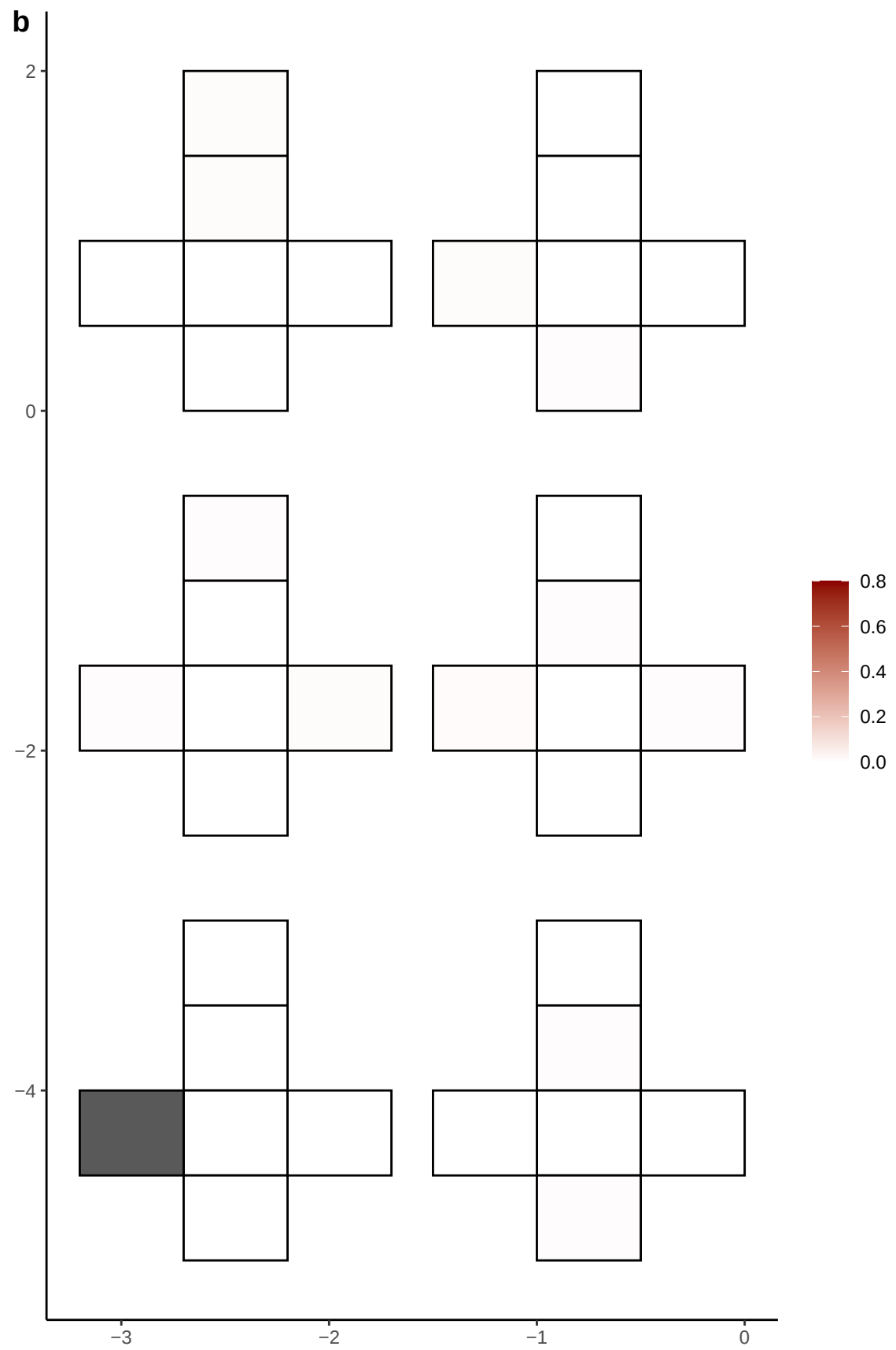

**a** ID01 21-35693543-G-A

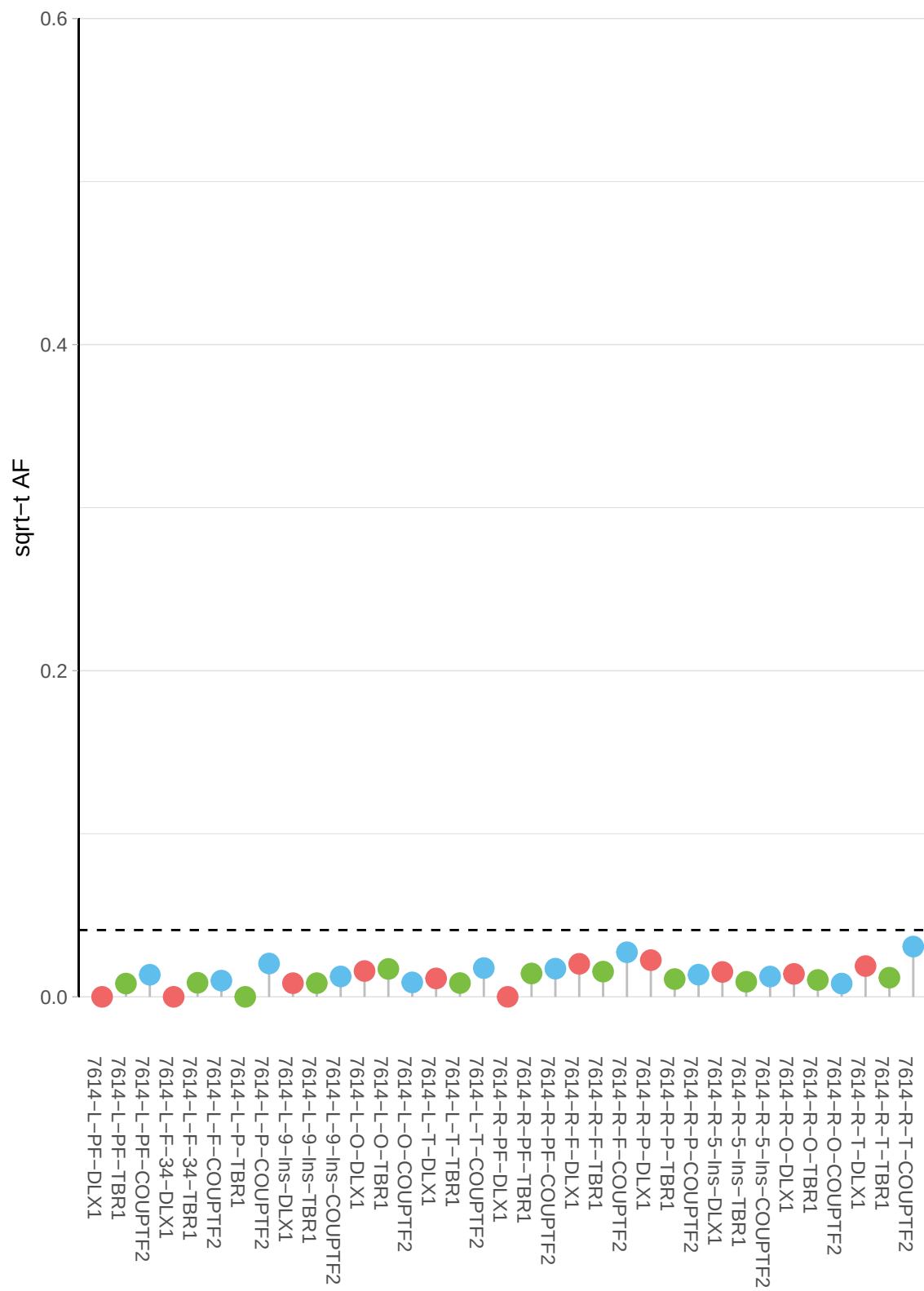

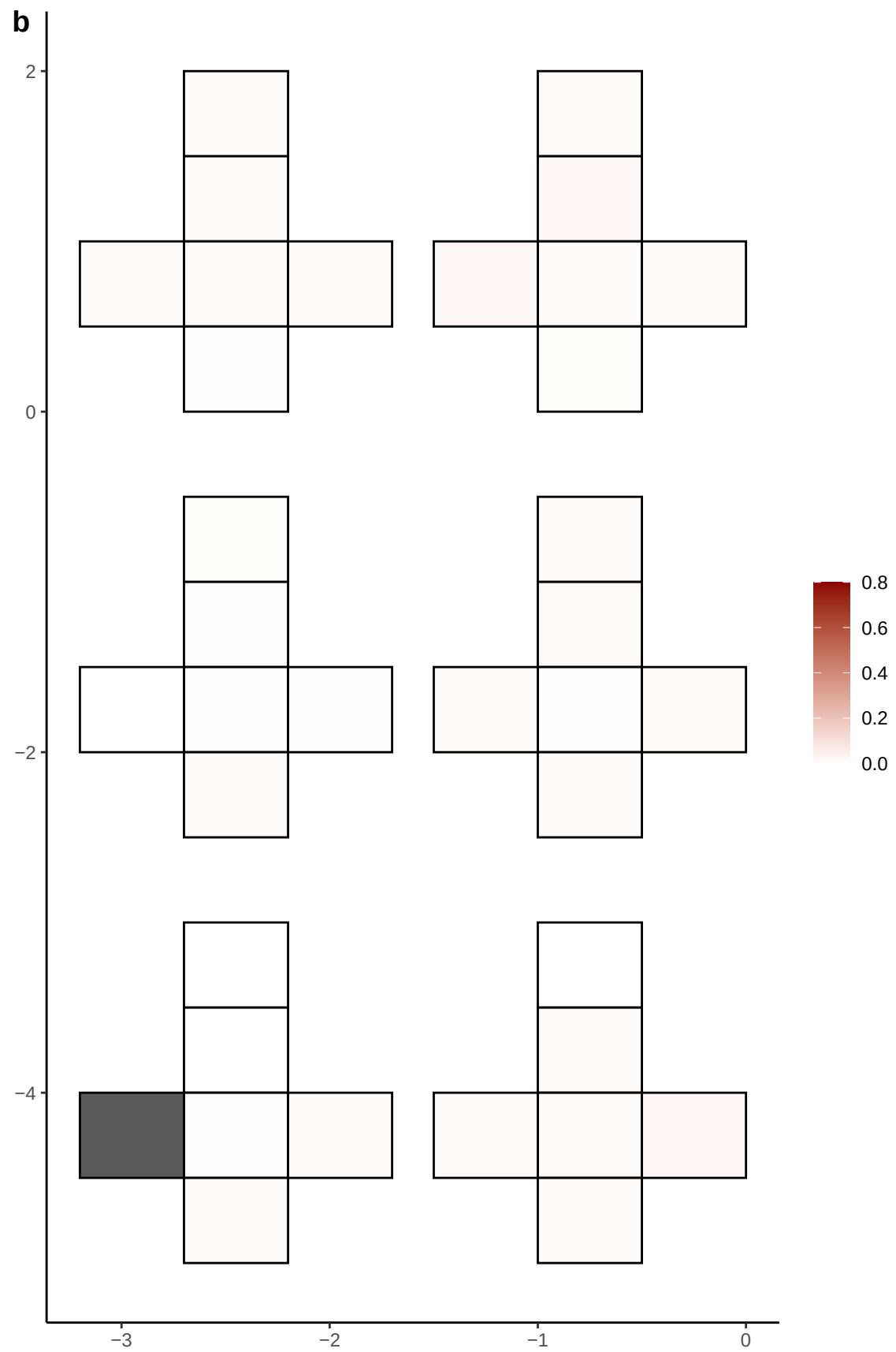

**a** ID01 21-42801460-G-A

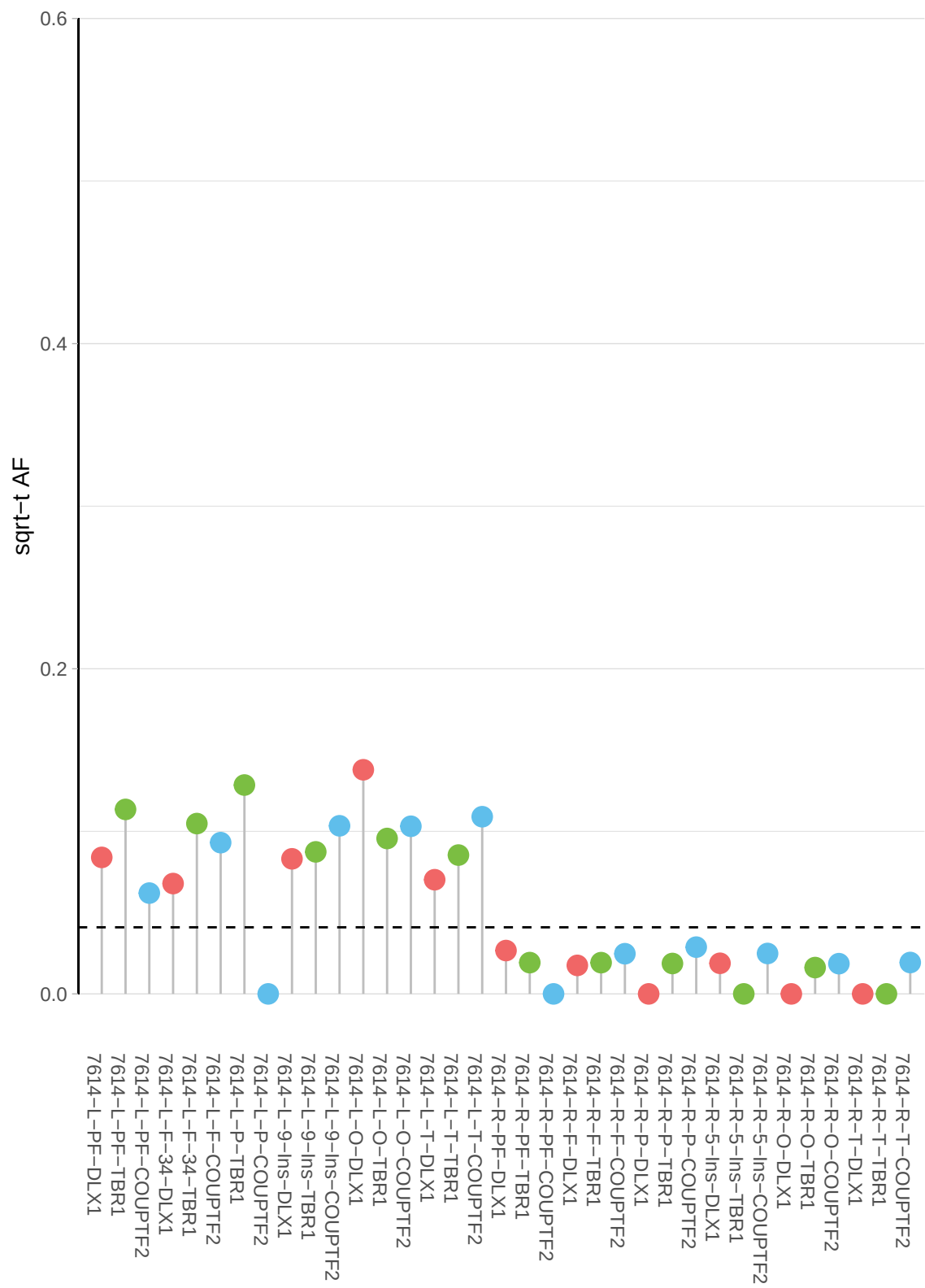

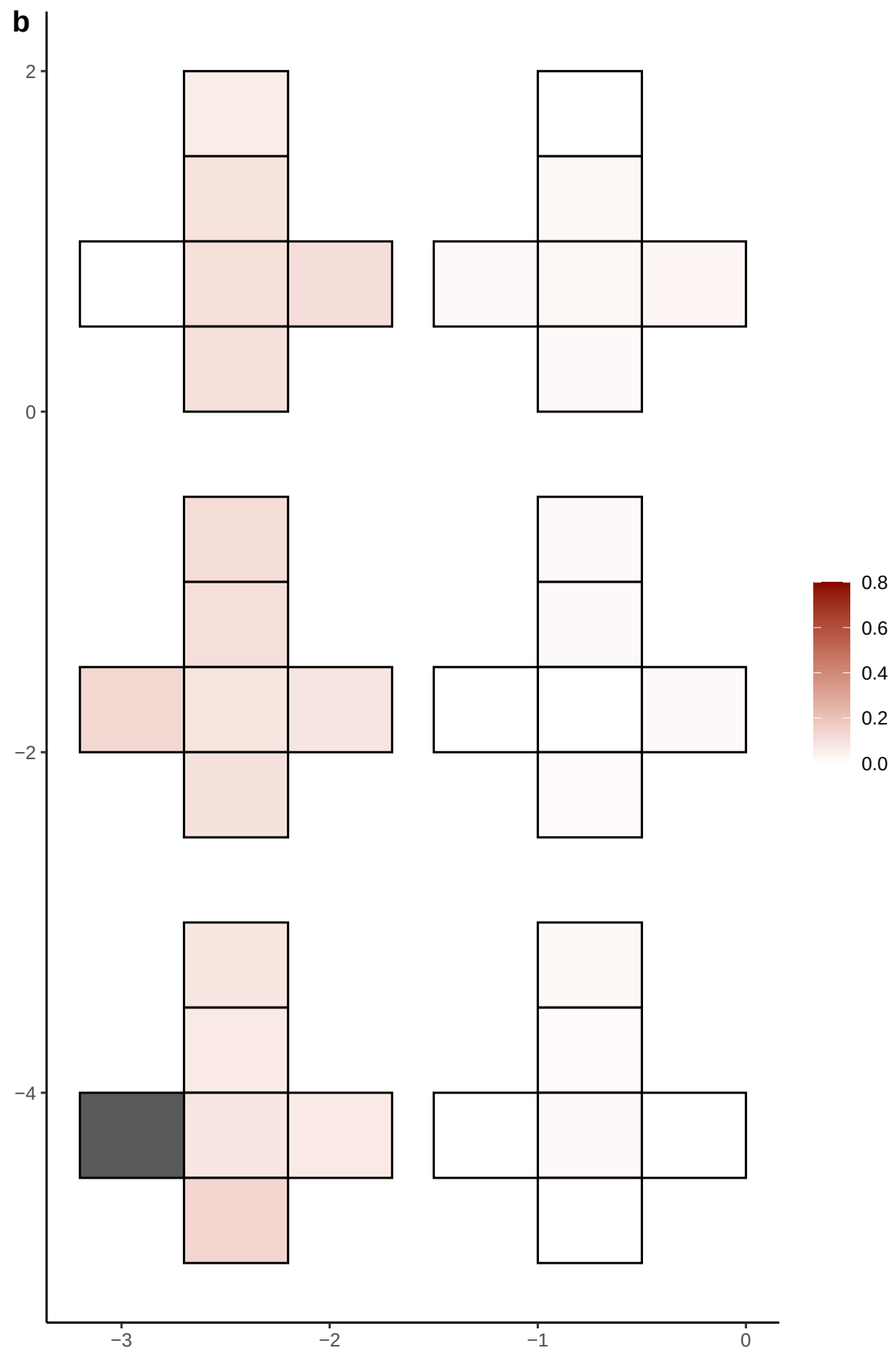

**a** ID01 21-43101791-G-A

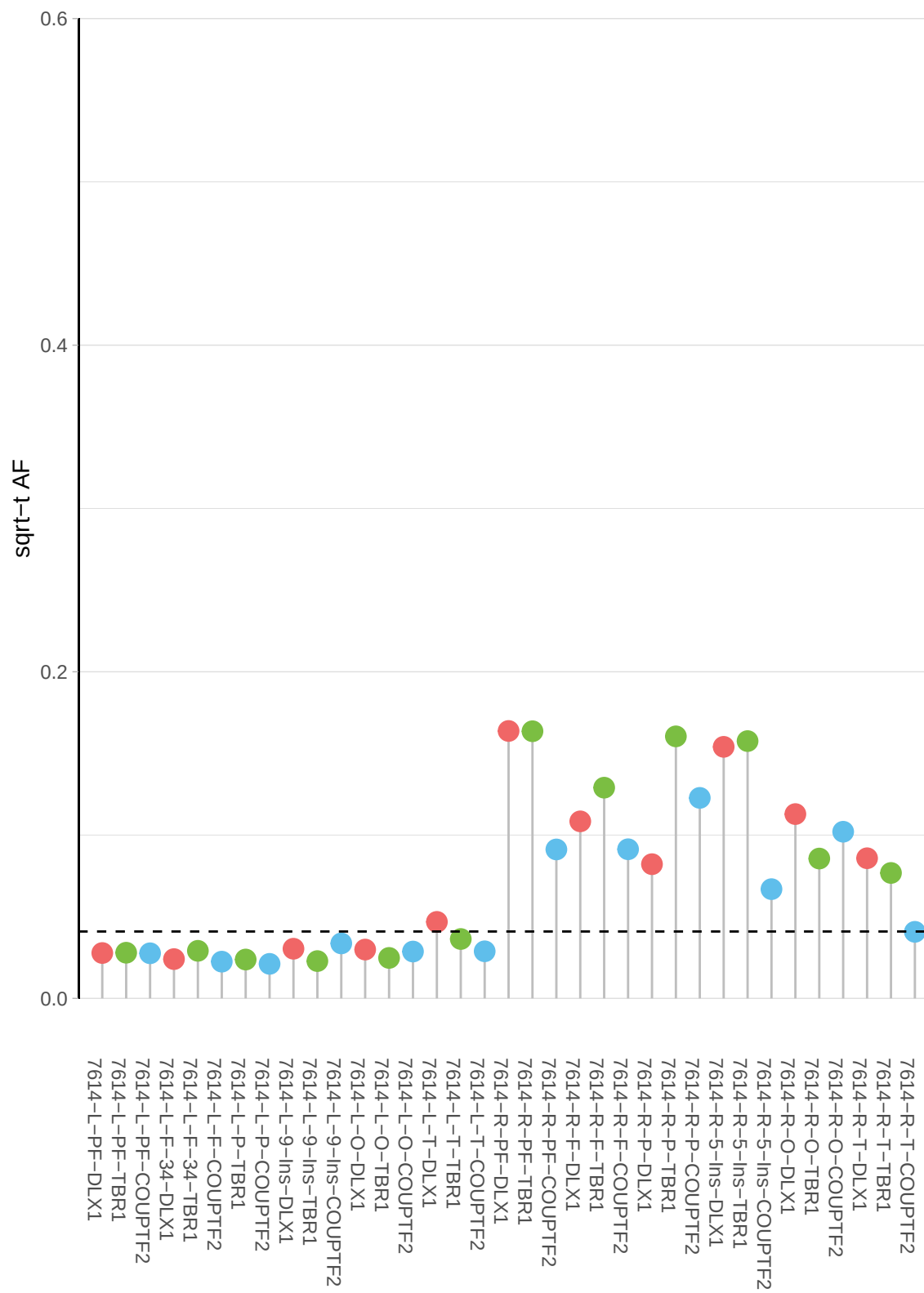

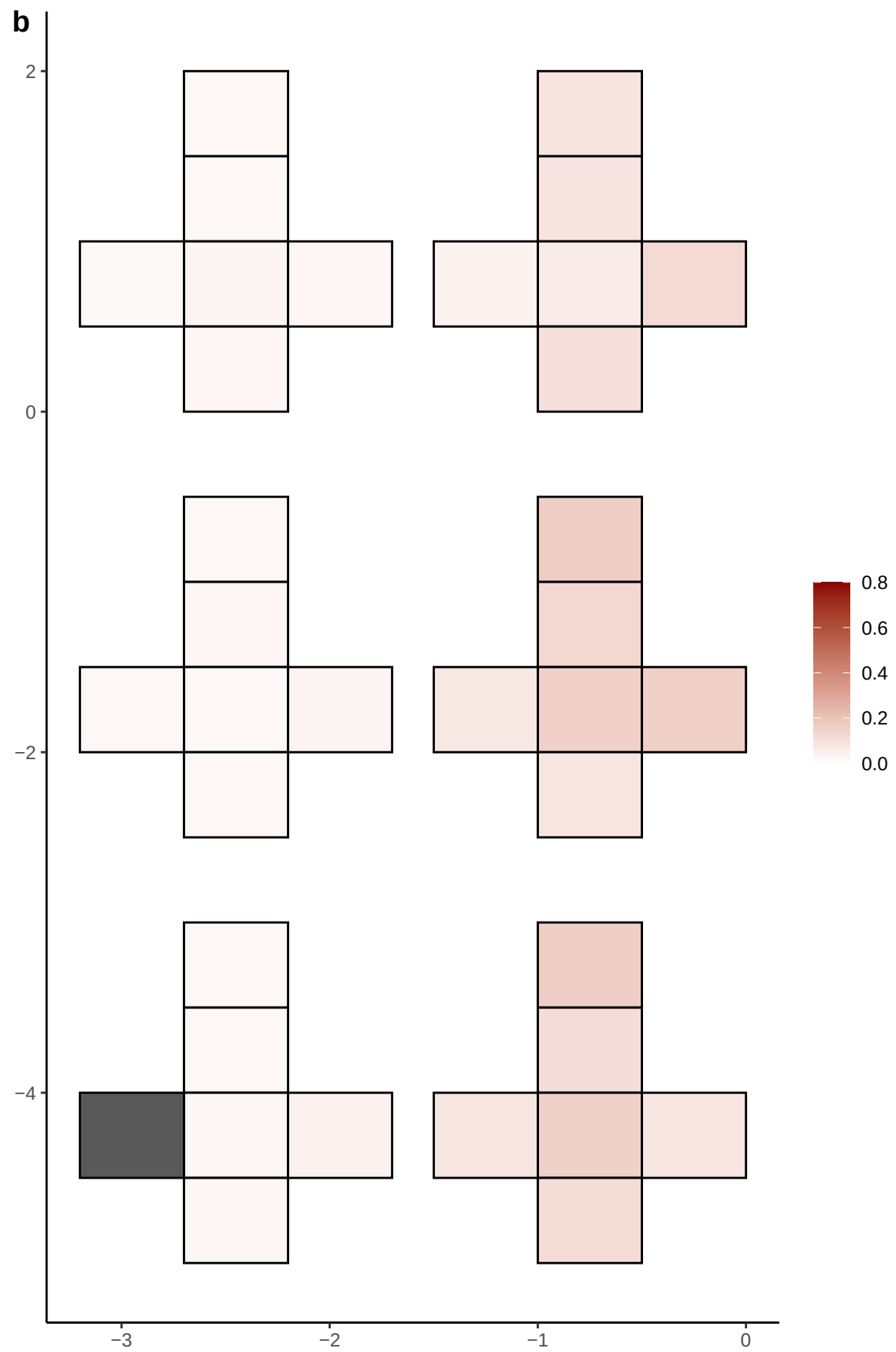

**a** ID01 22-18444369-T-C

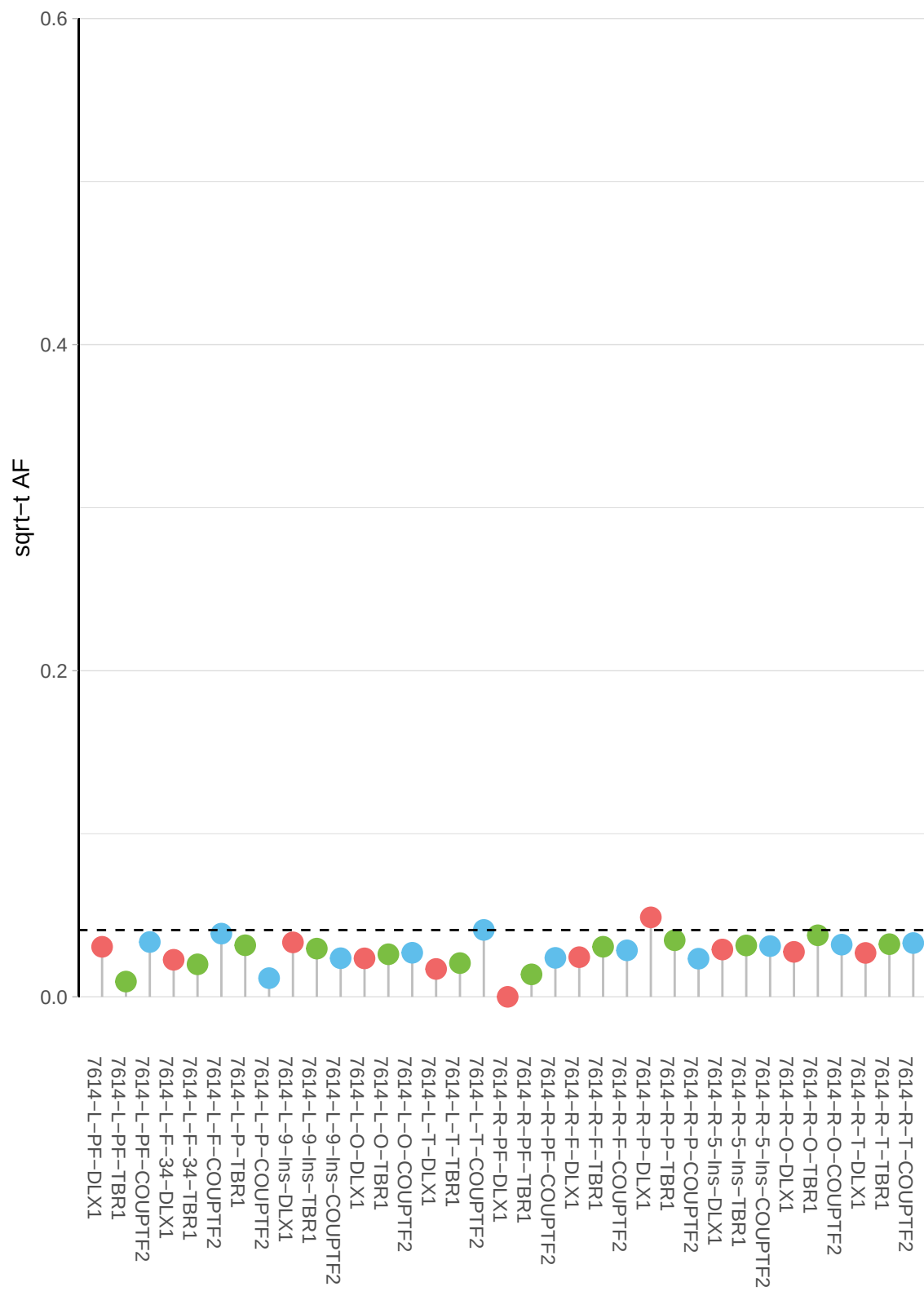

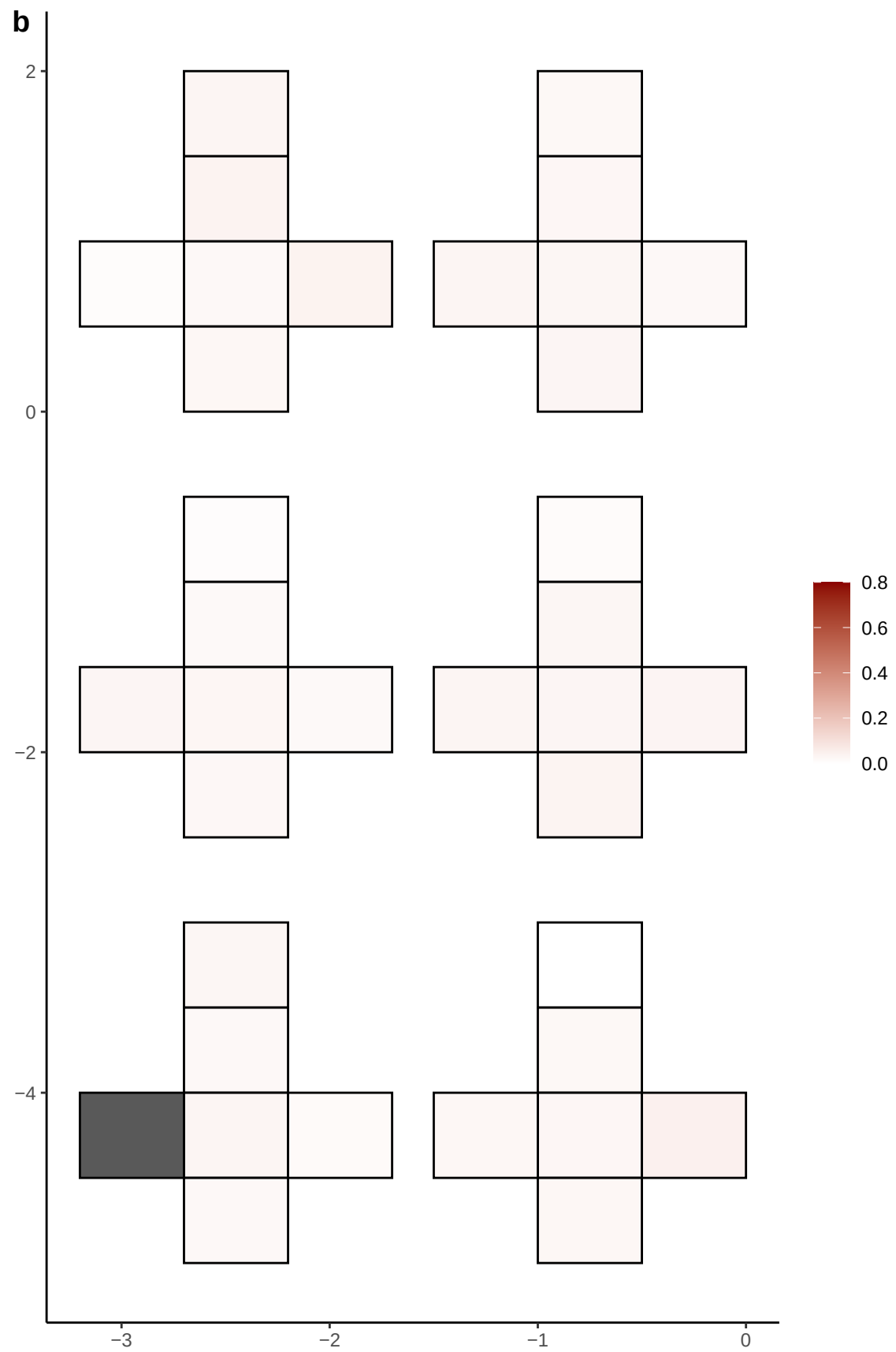

**a** ID01 22-19049947-T-TCCTC

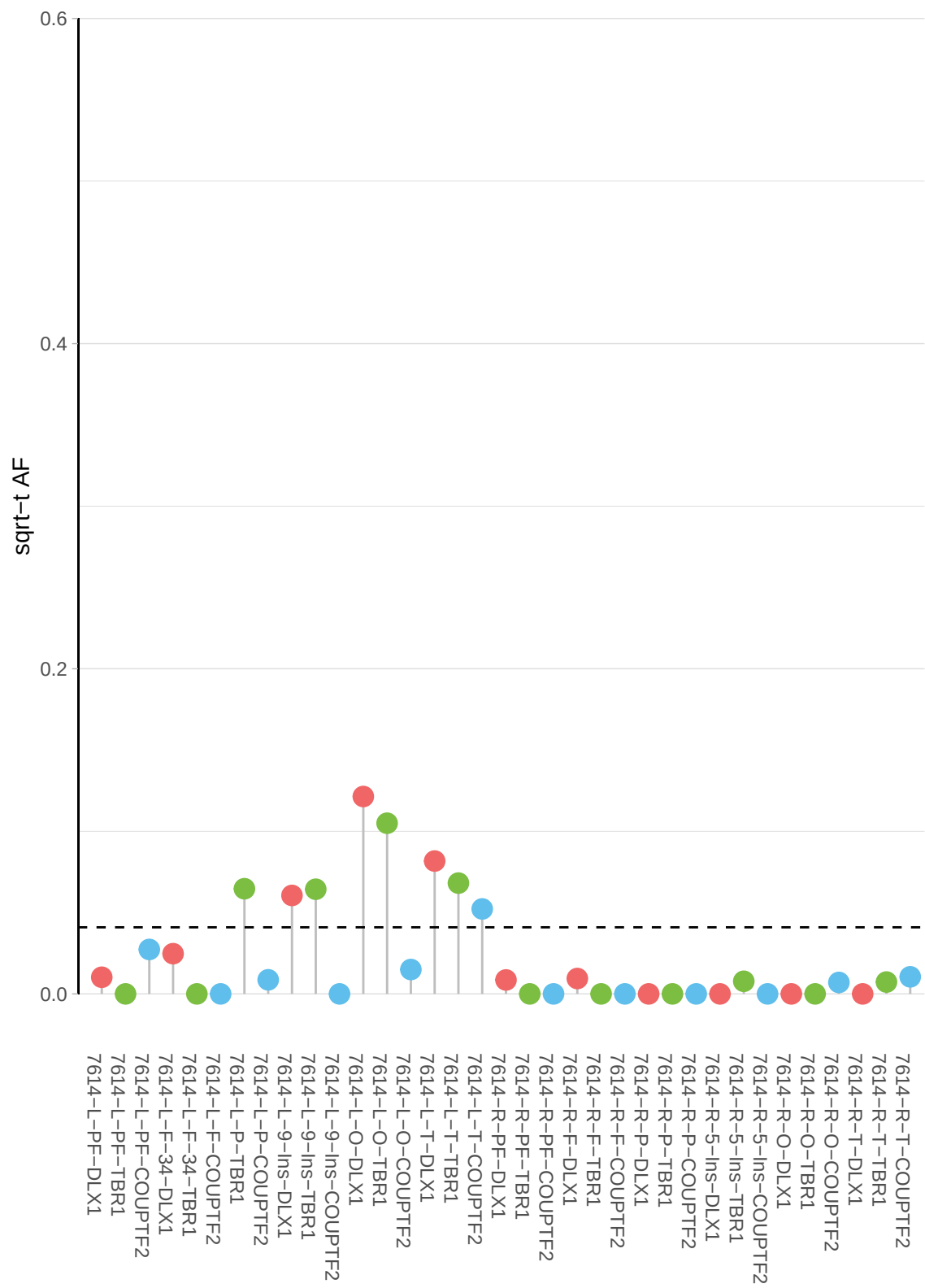

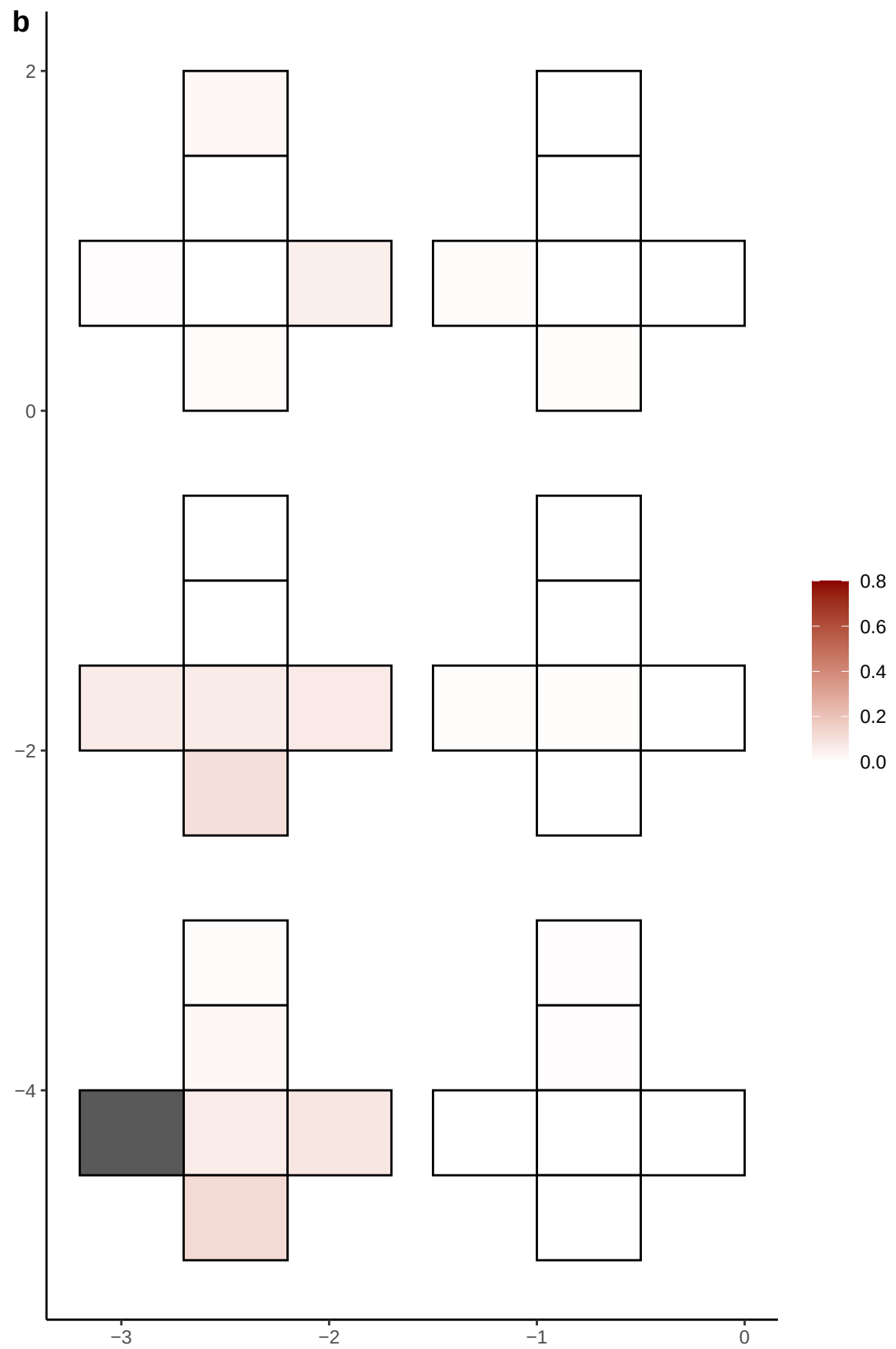

**a** ID01 22-35279008-G-A

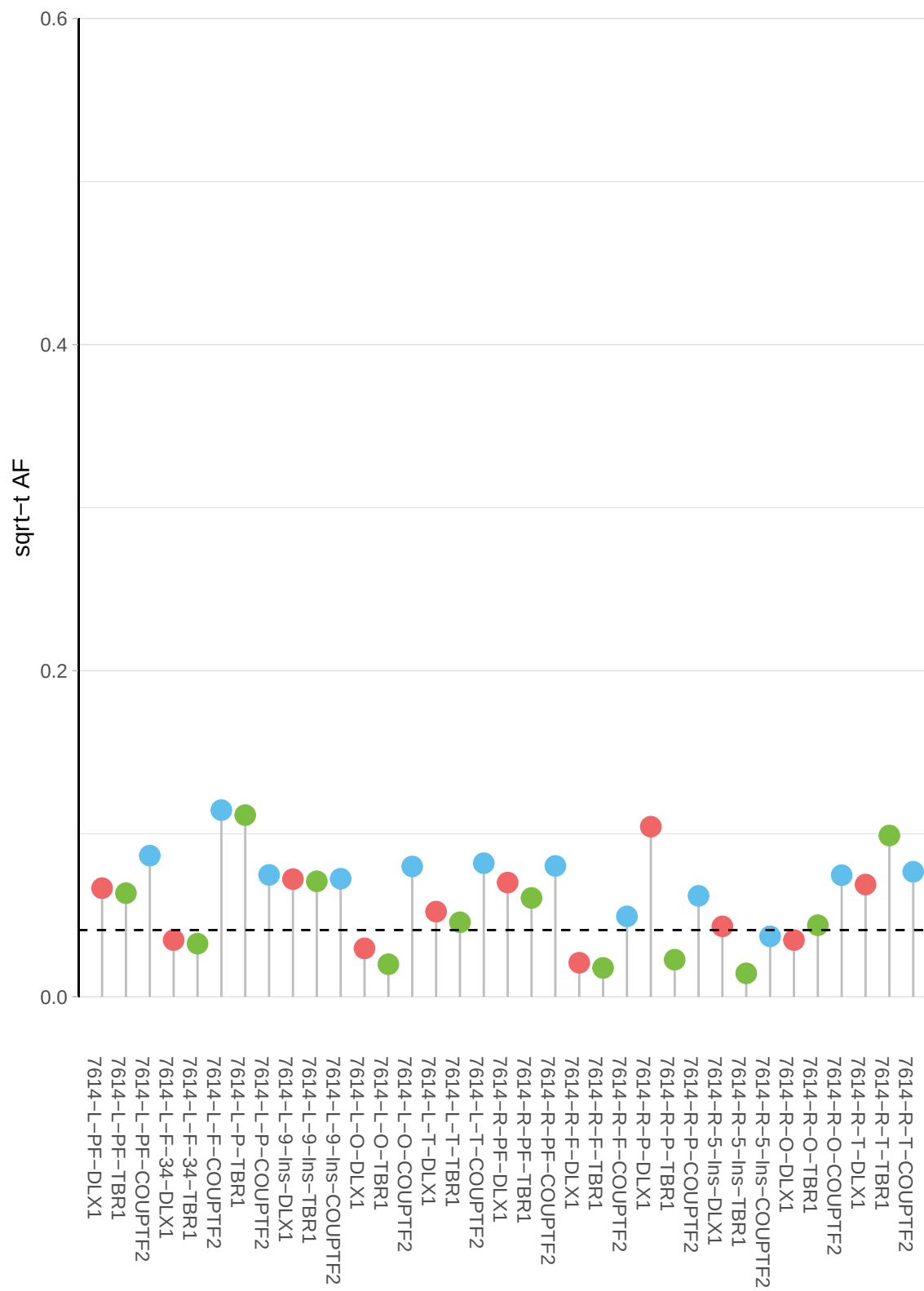

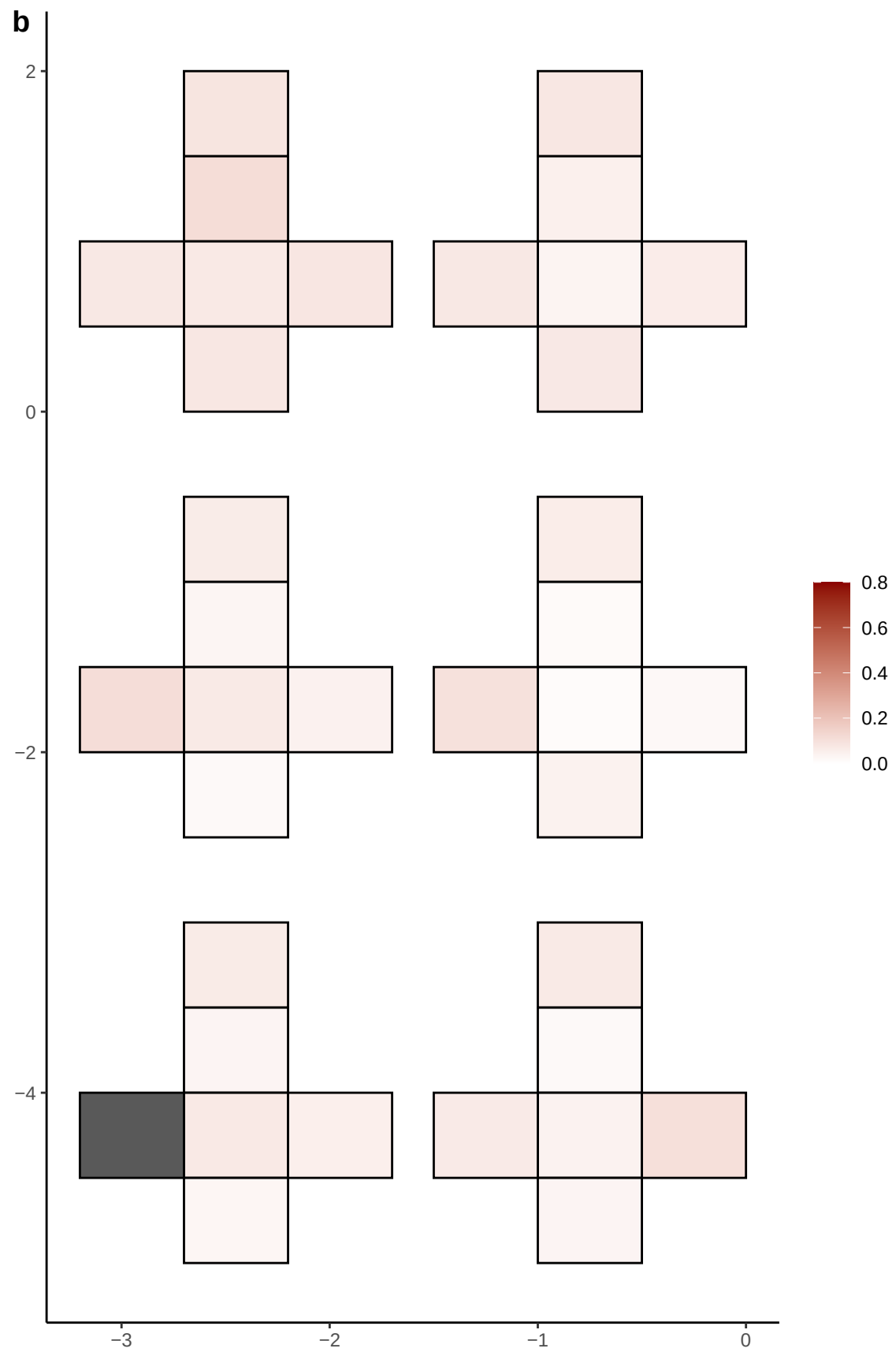

**a** ID01 22-46947808-G-A

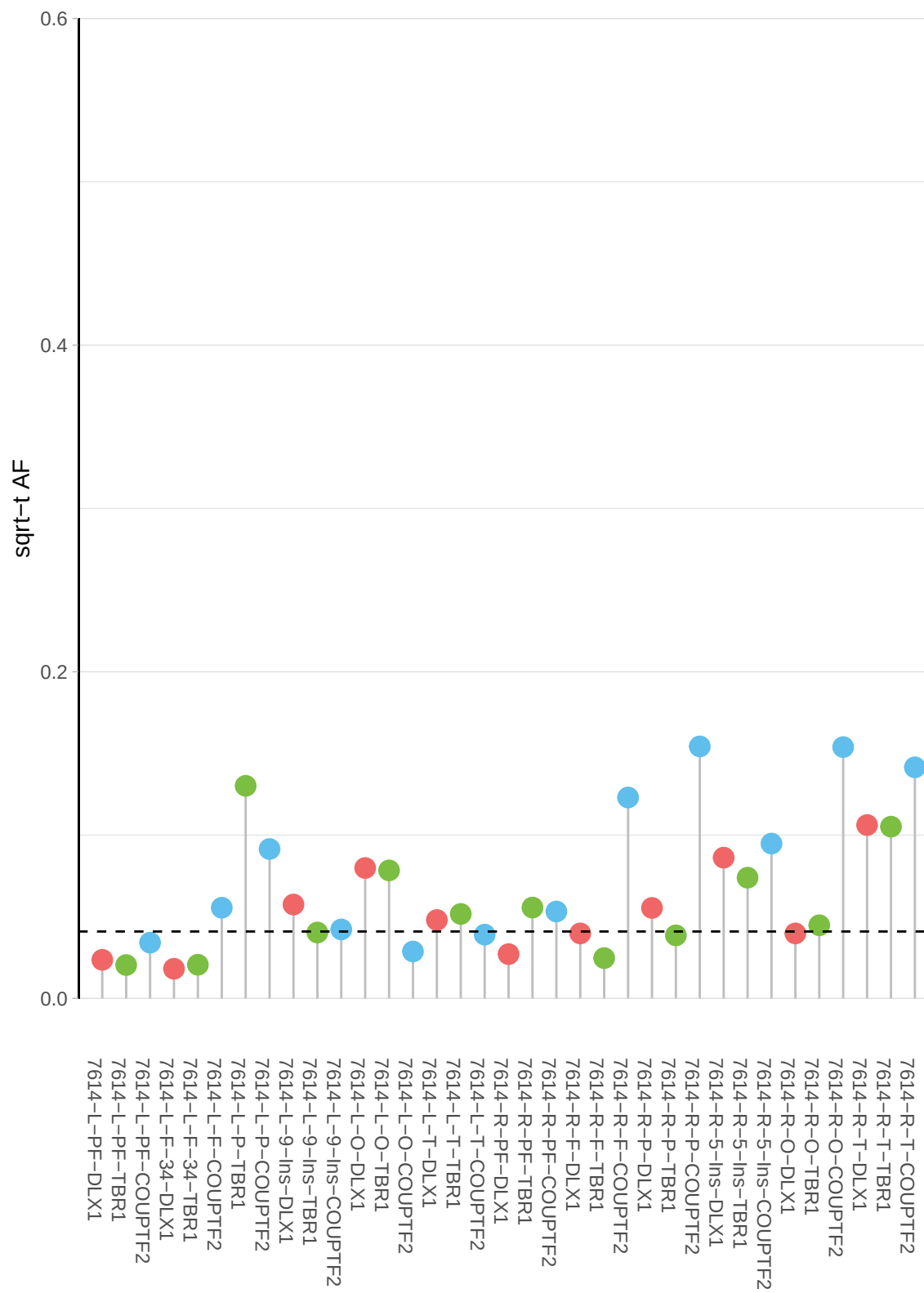

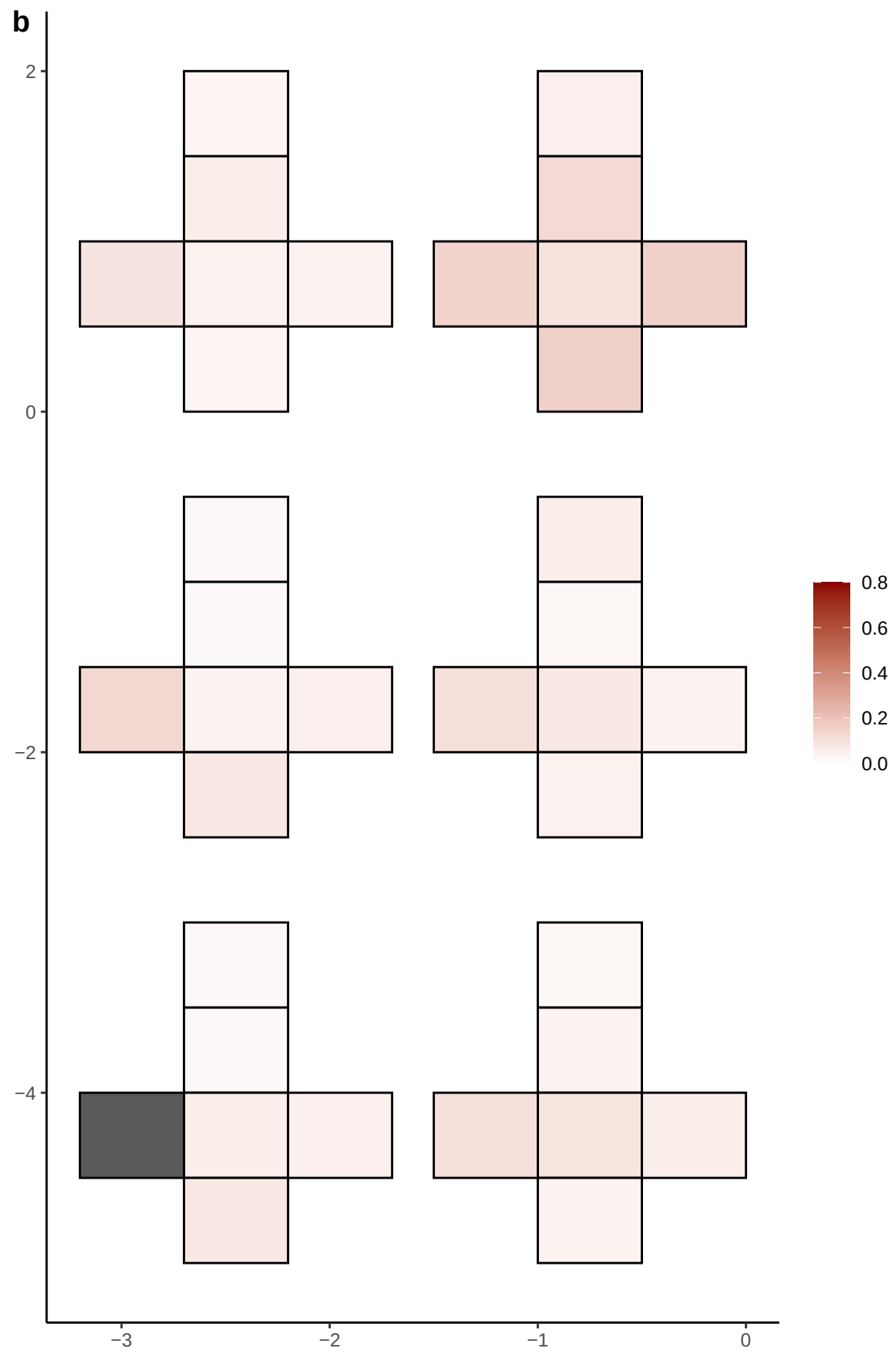

**a** ID01 3-107731469-C-T

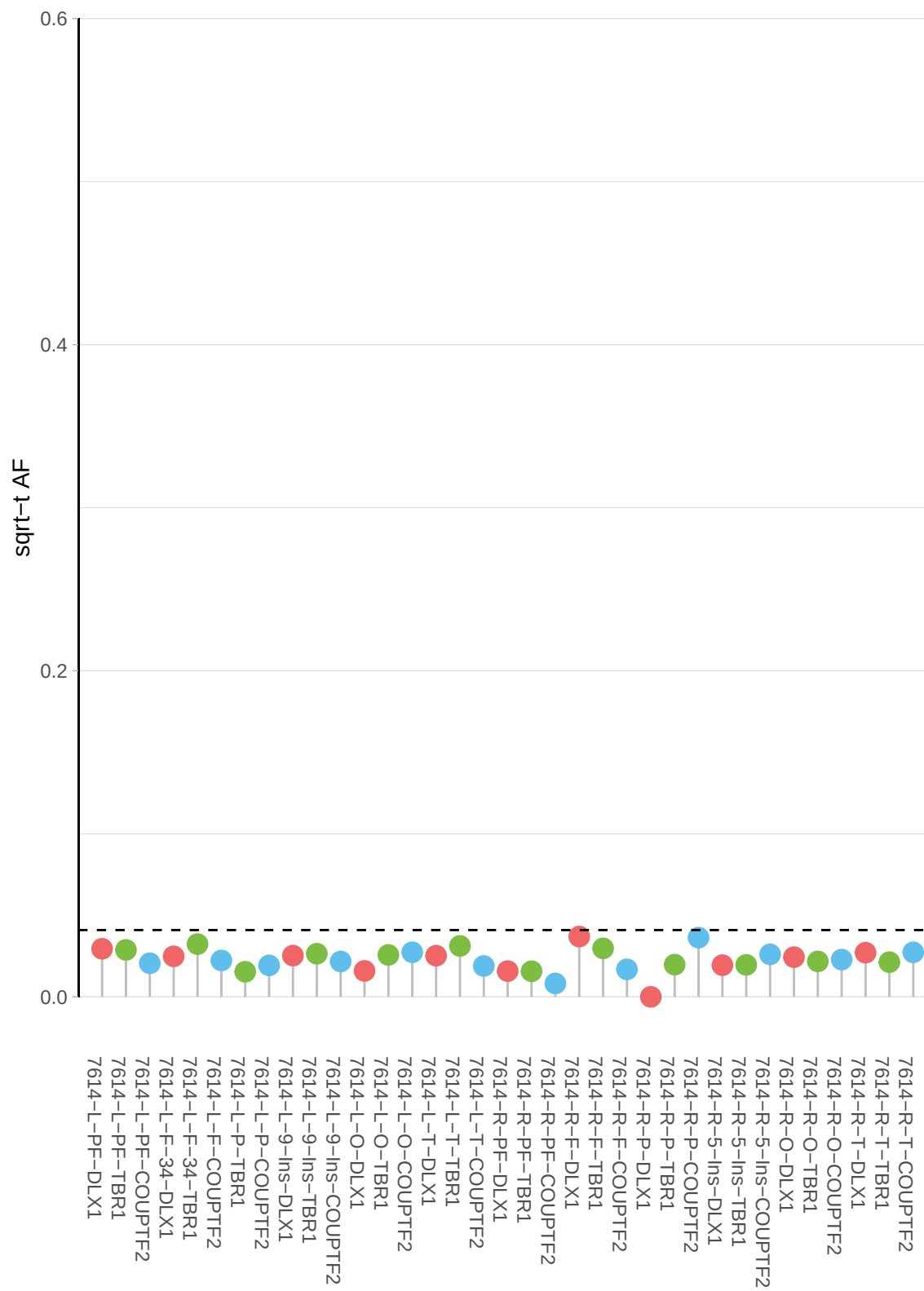

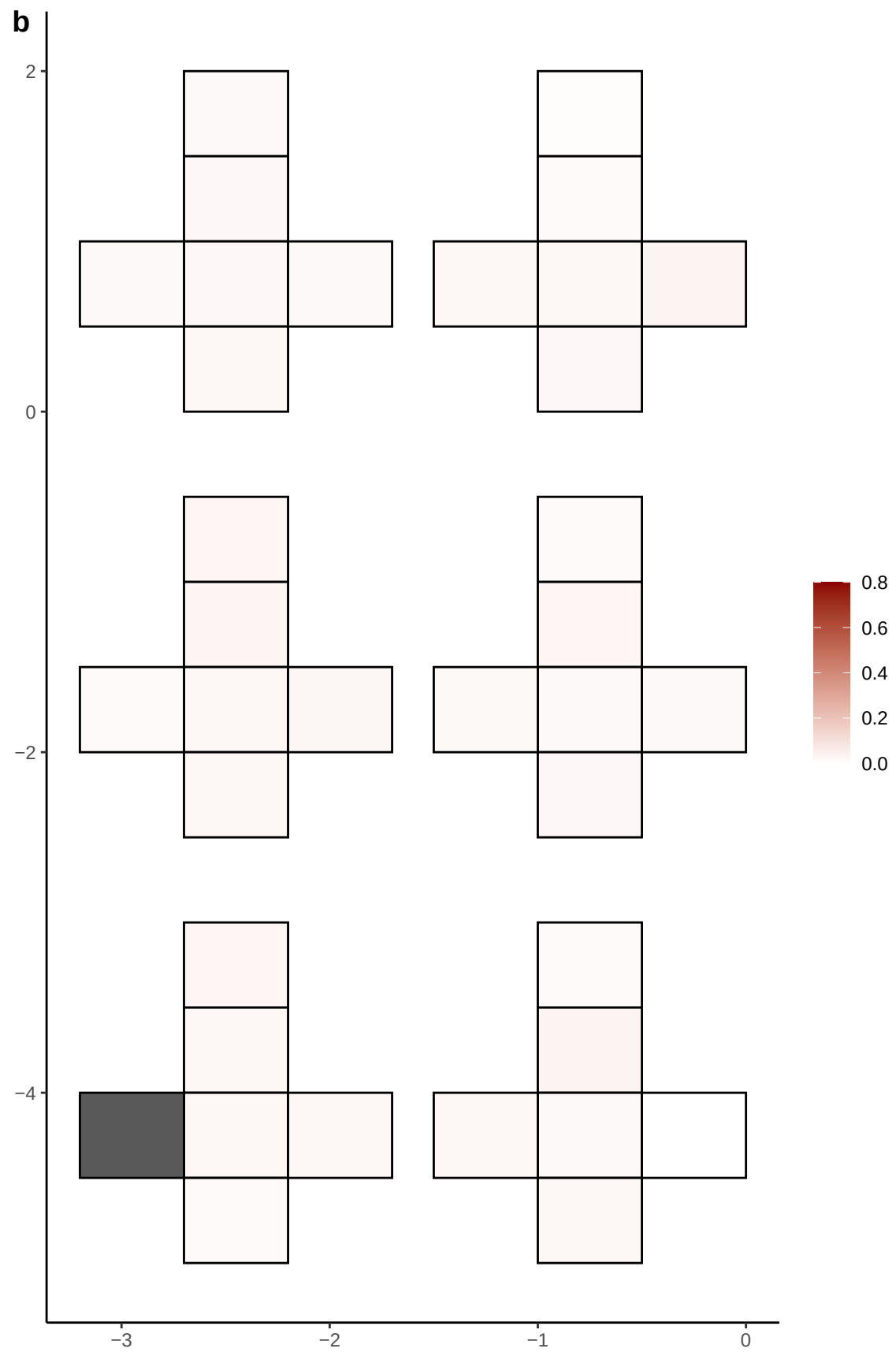

**a** ID01 3-111155080-C-T

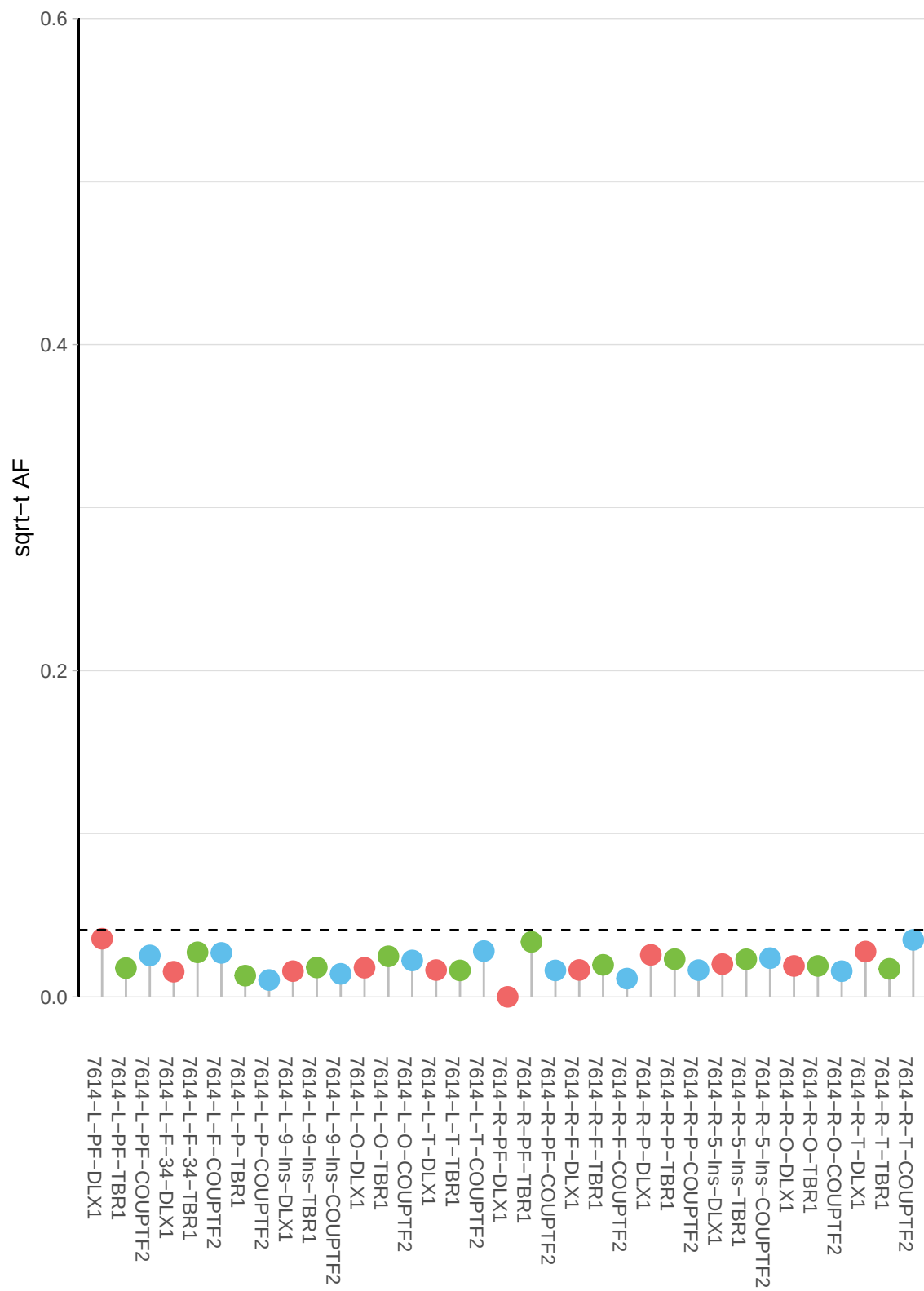

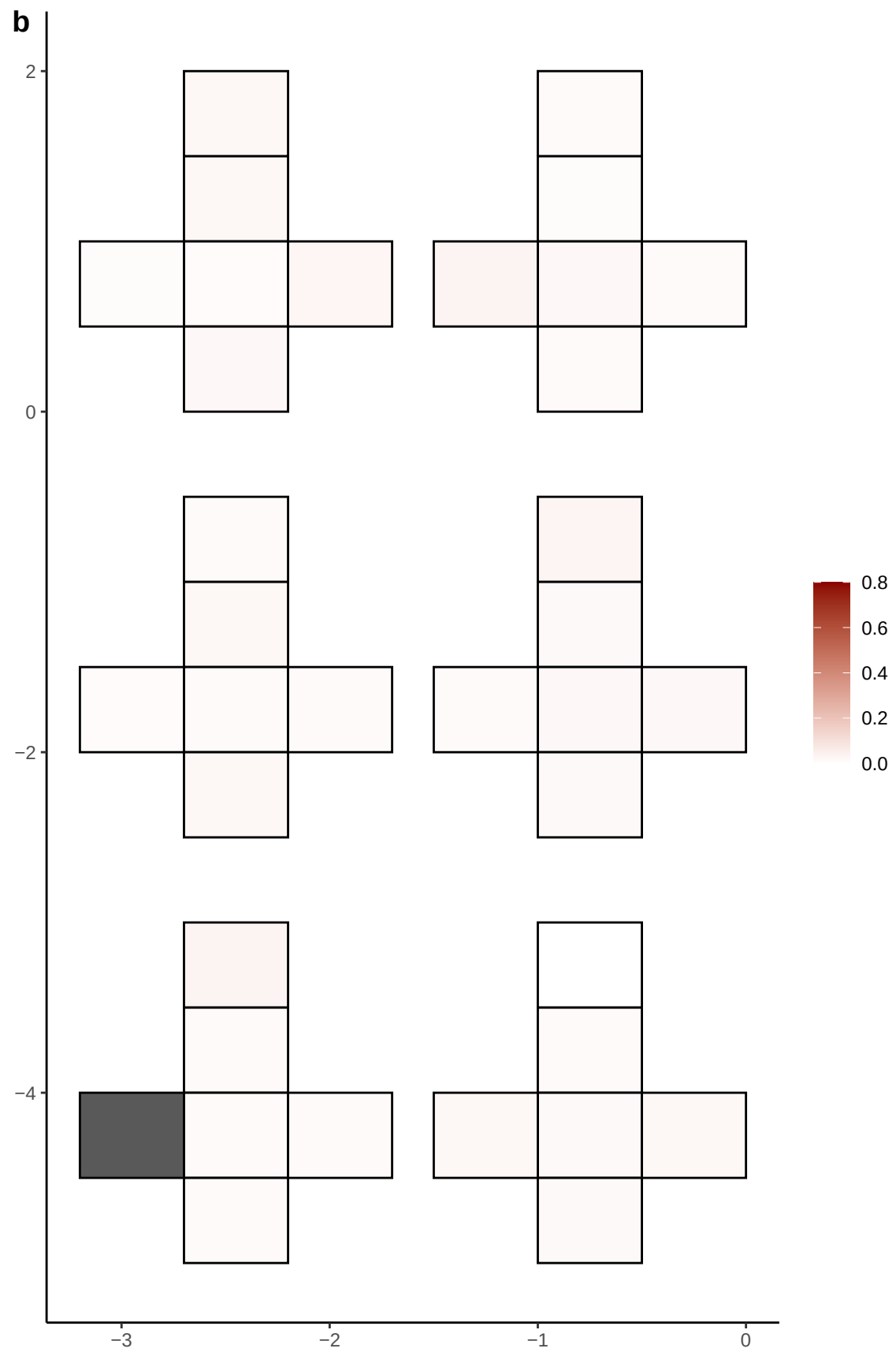

**a** ID01 3-112163628-C-T

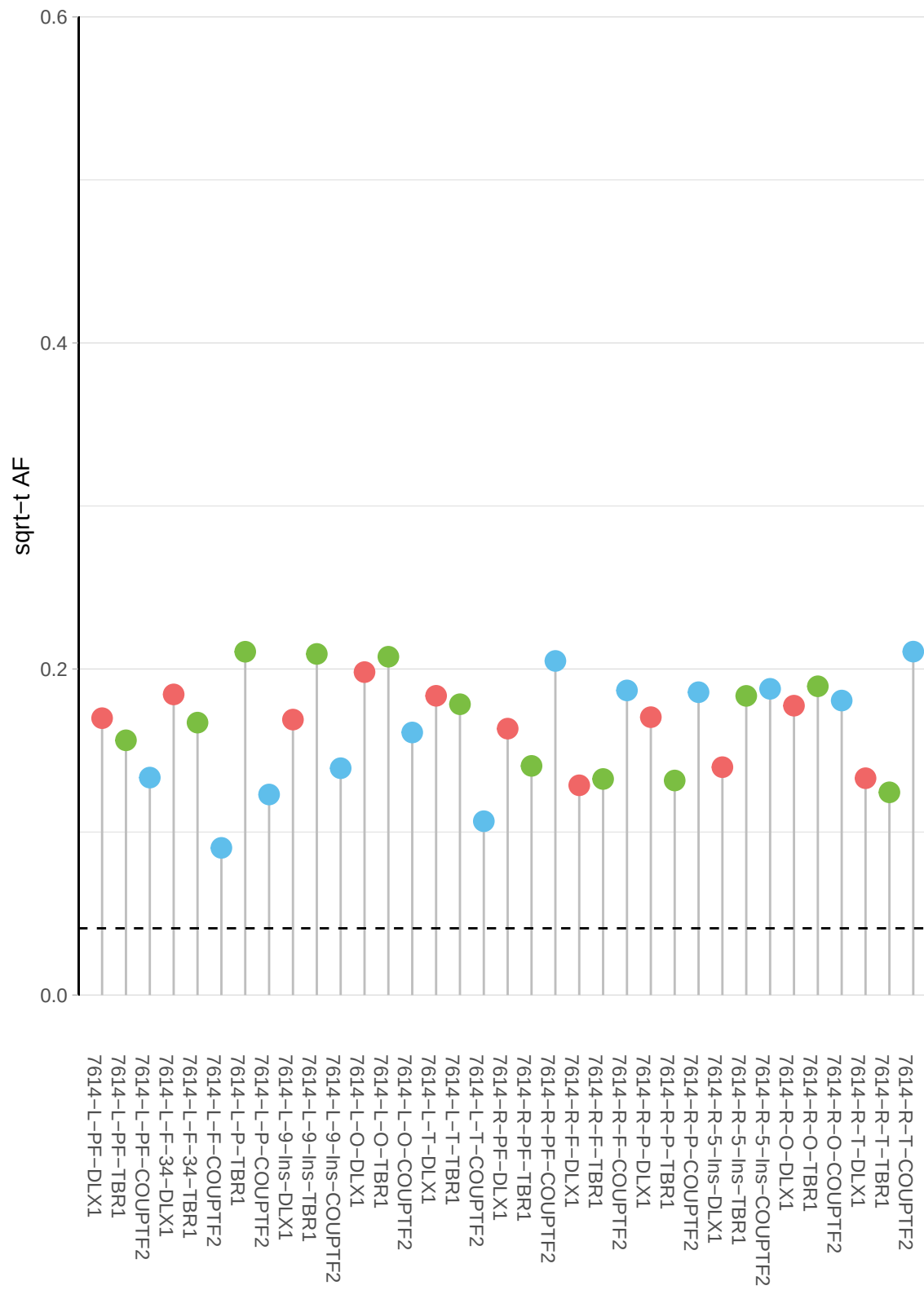

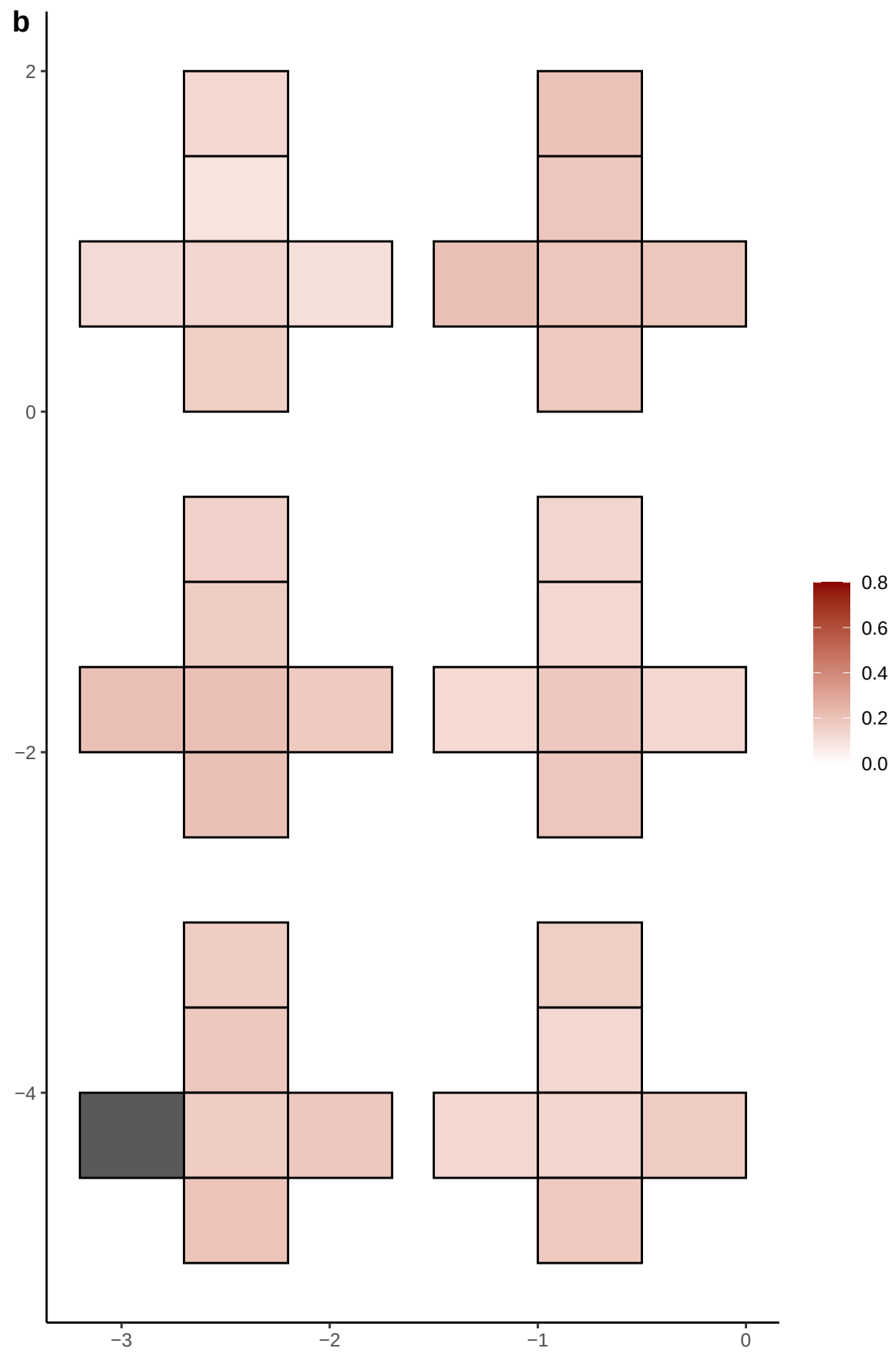

**a** ID01 3-152697890-C-T

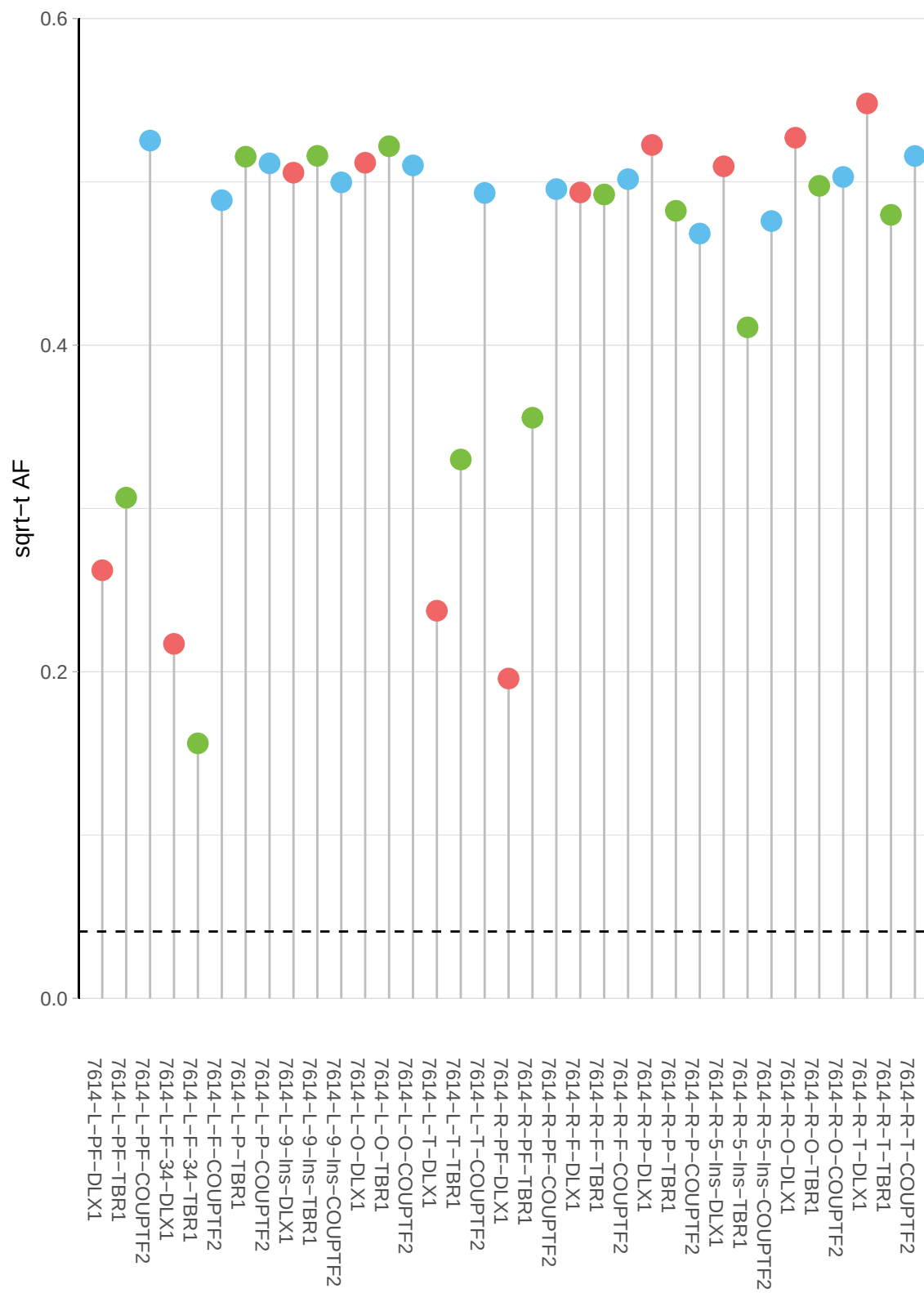

**b**

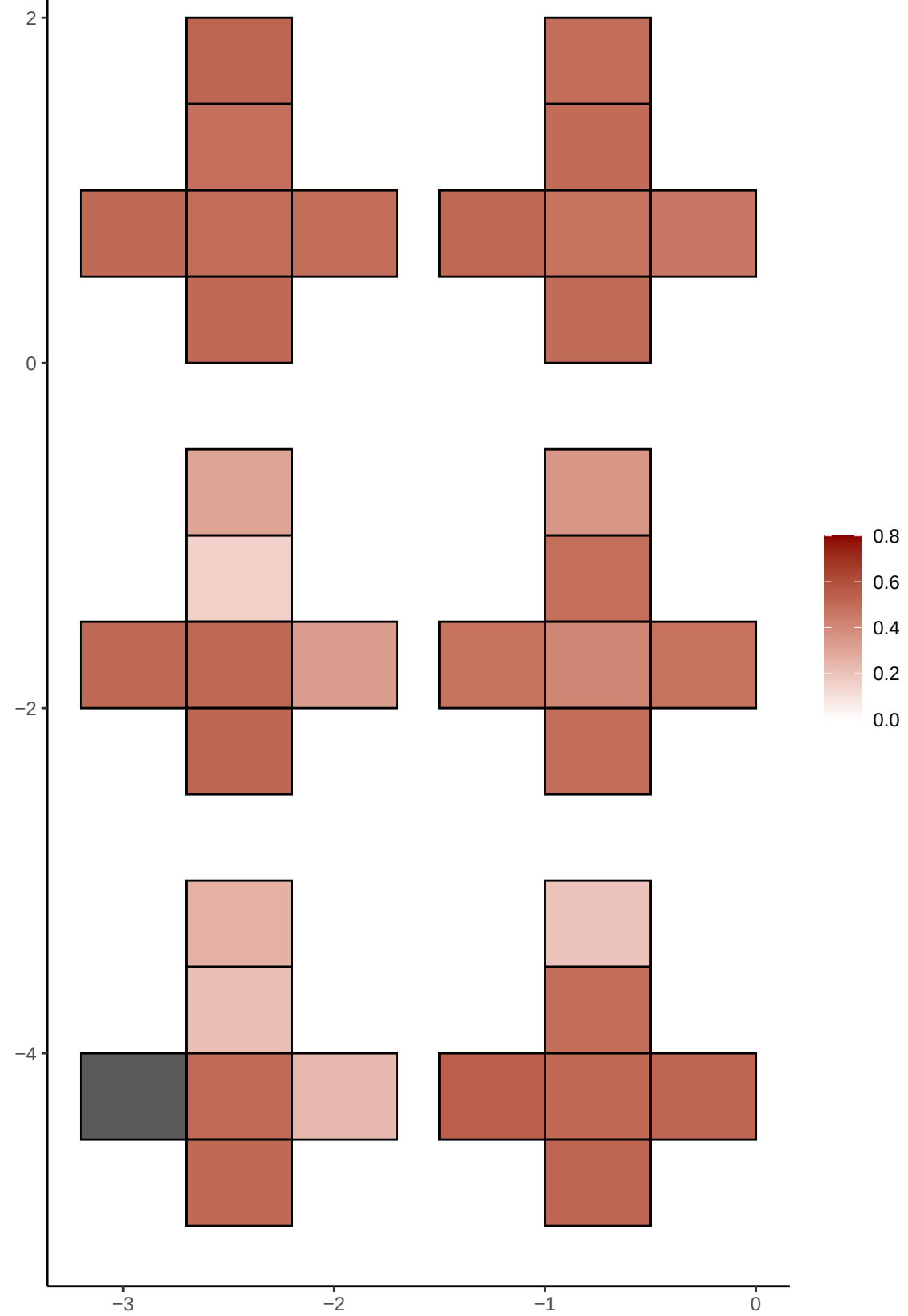

**a** ID01 3-162456631-G-A

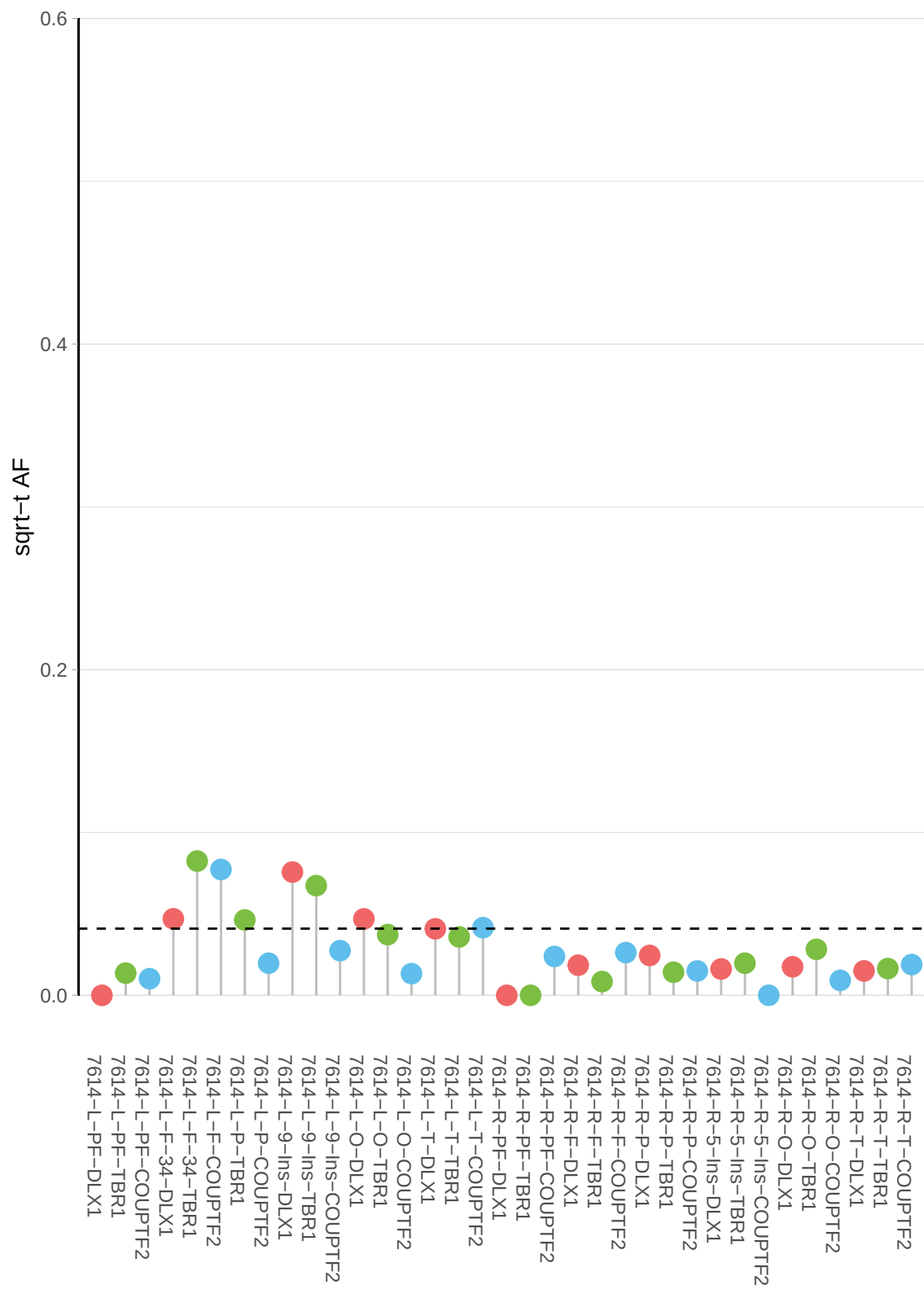

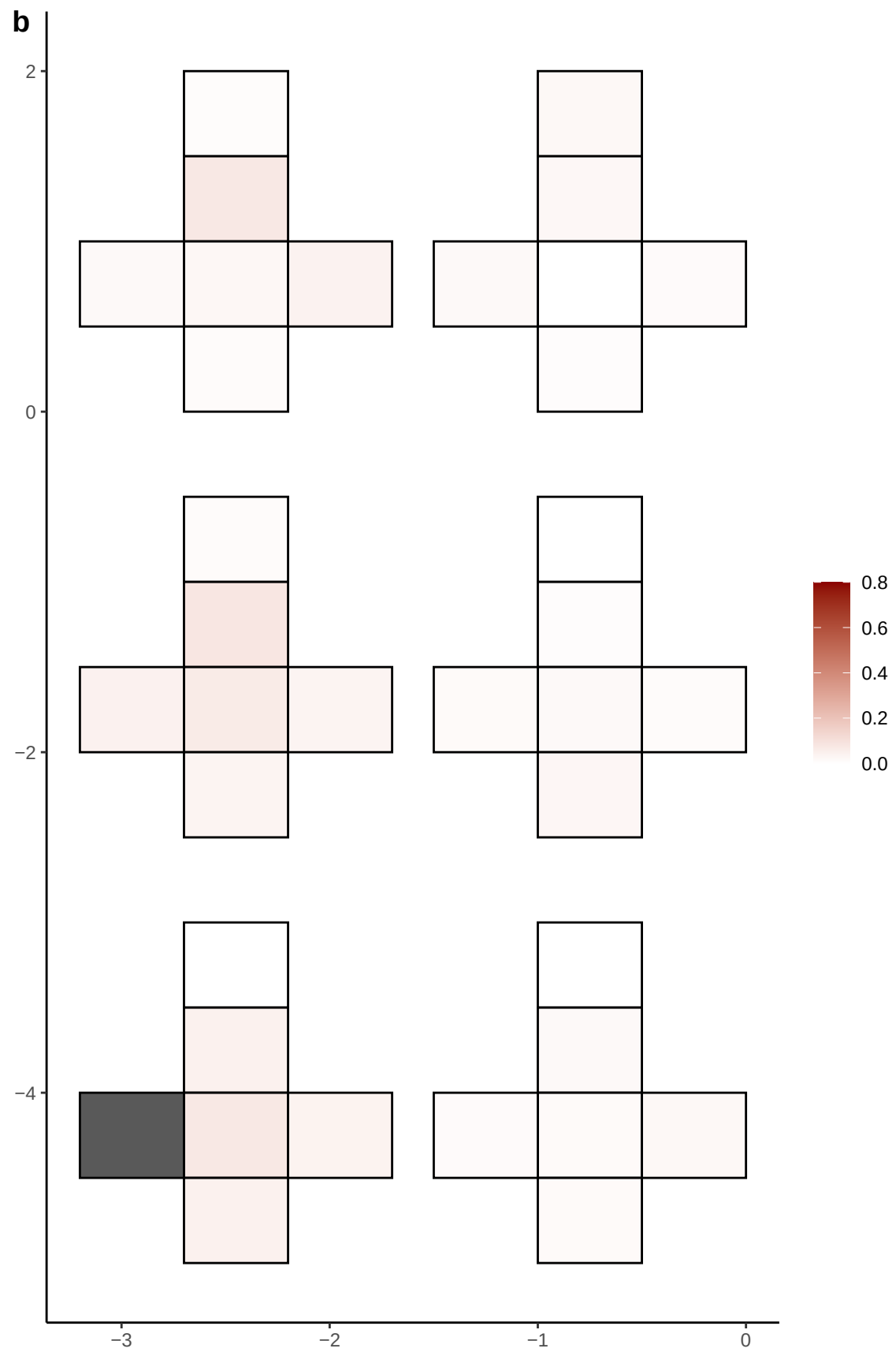

**a** ID01 3-184022071-G-A

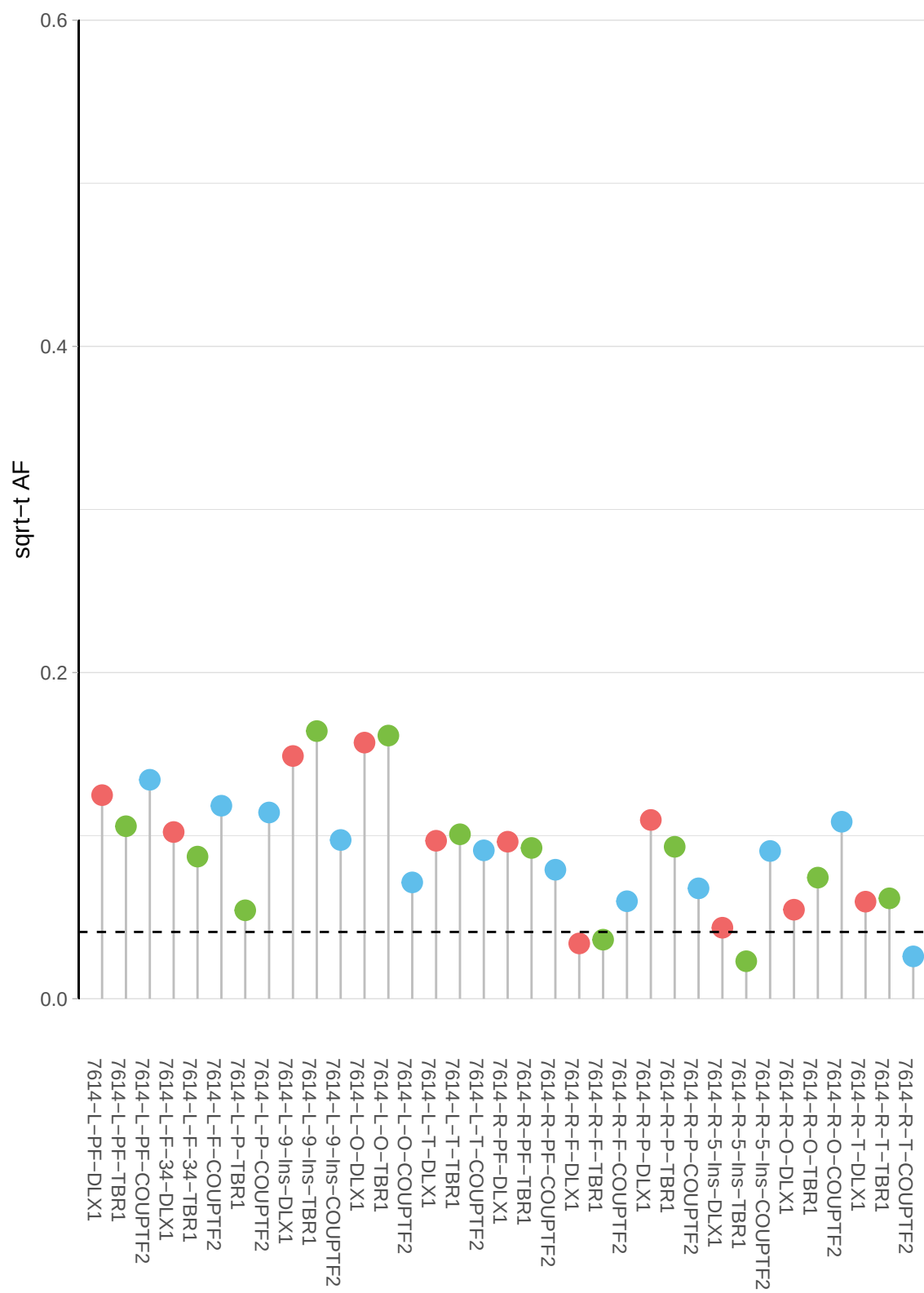

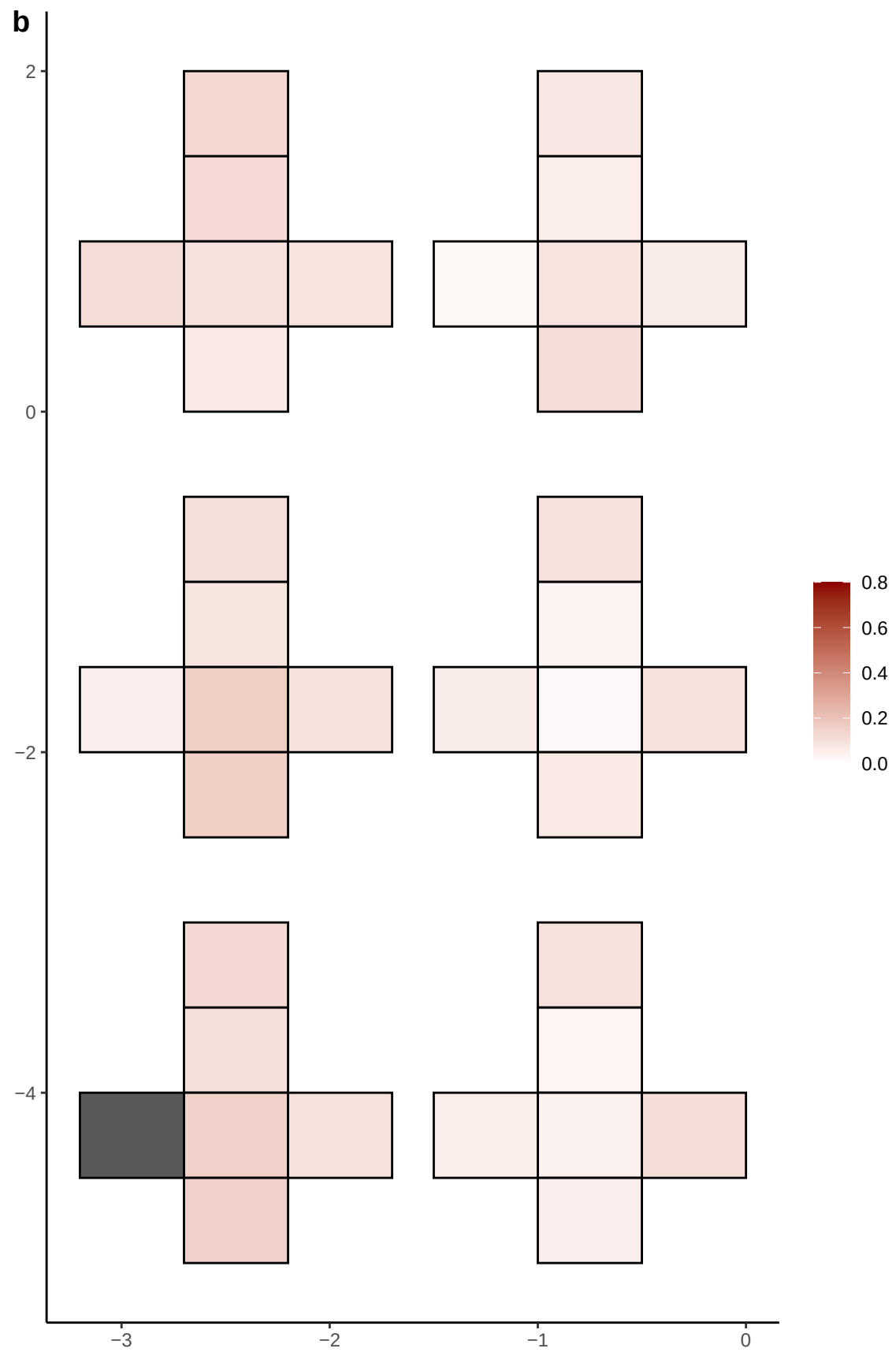

**a** ID01 3-196886773-C-T

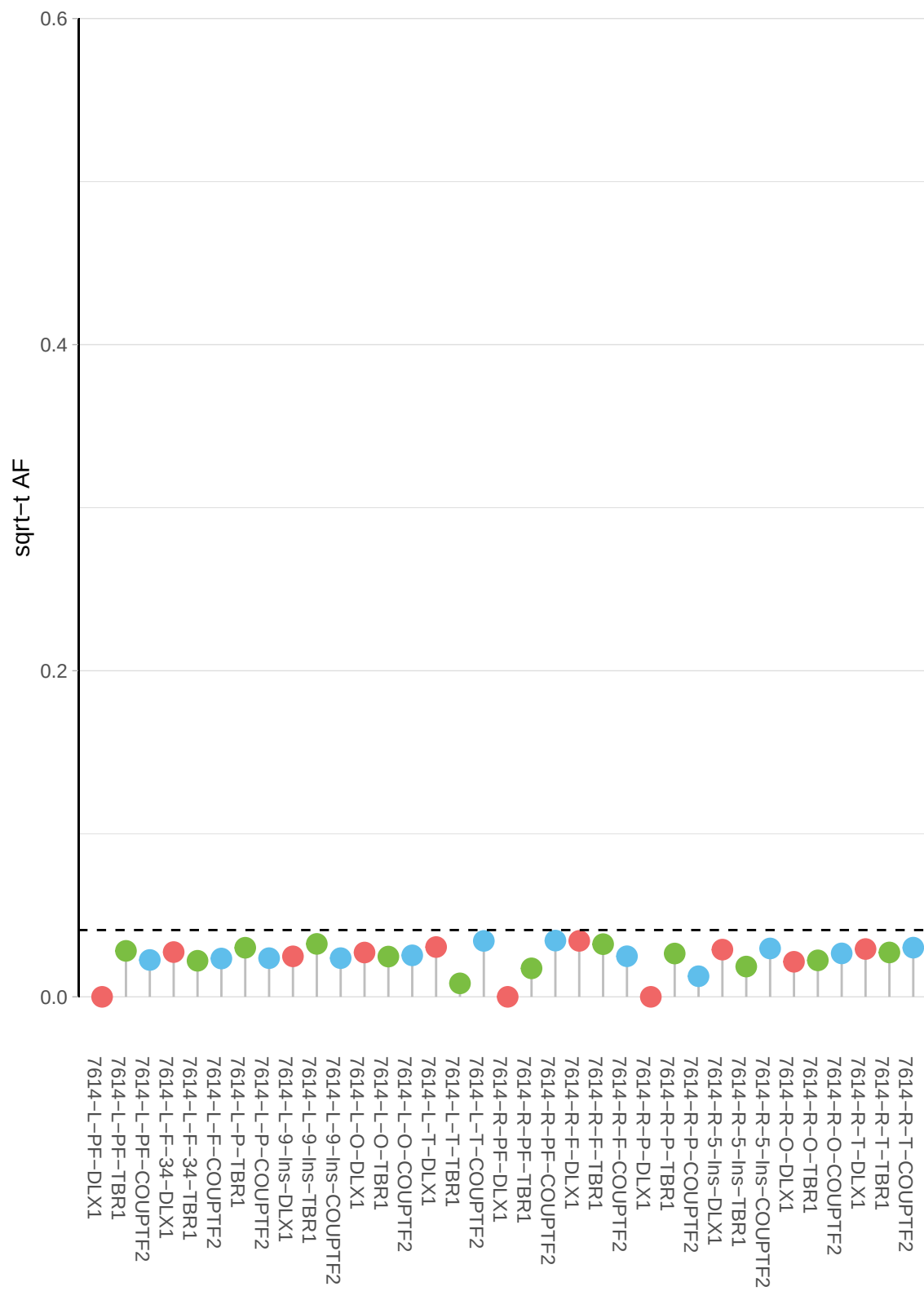

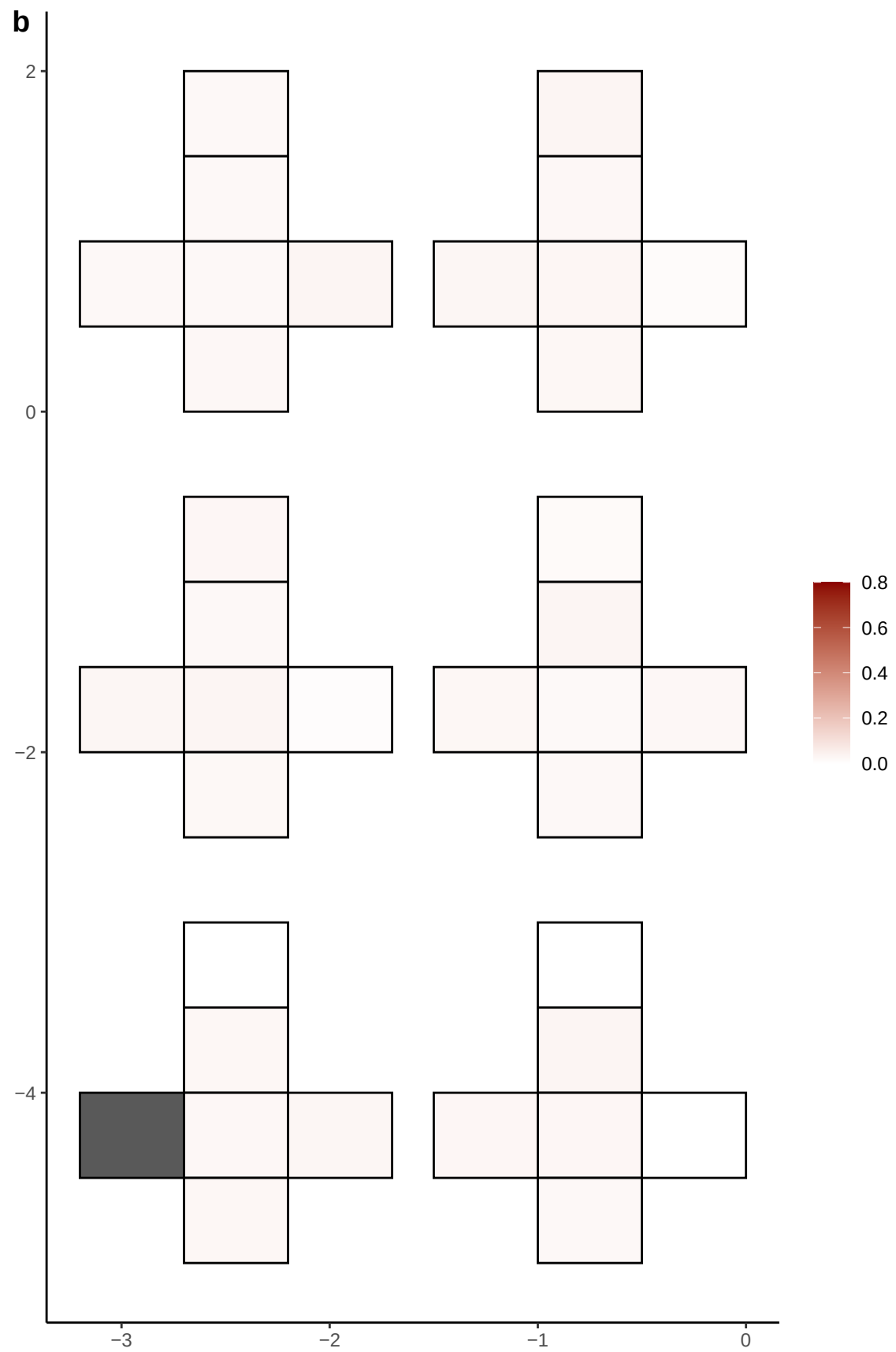

**a** ID01 3-23502134-AGAAAG-A

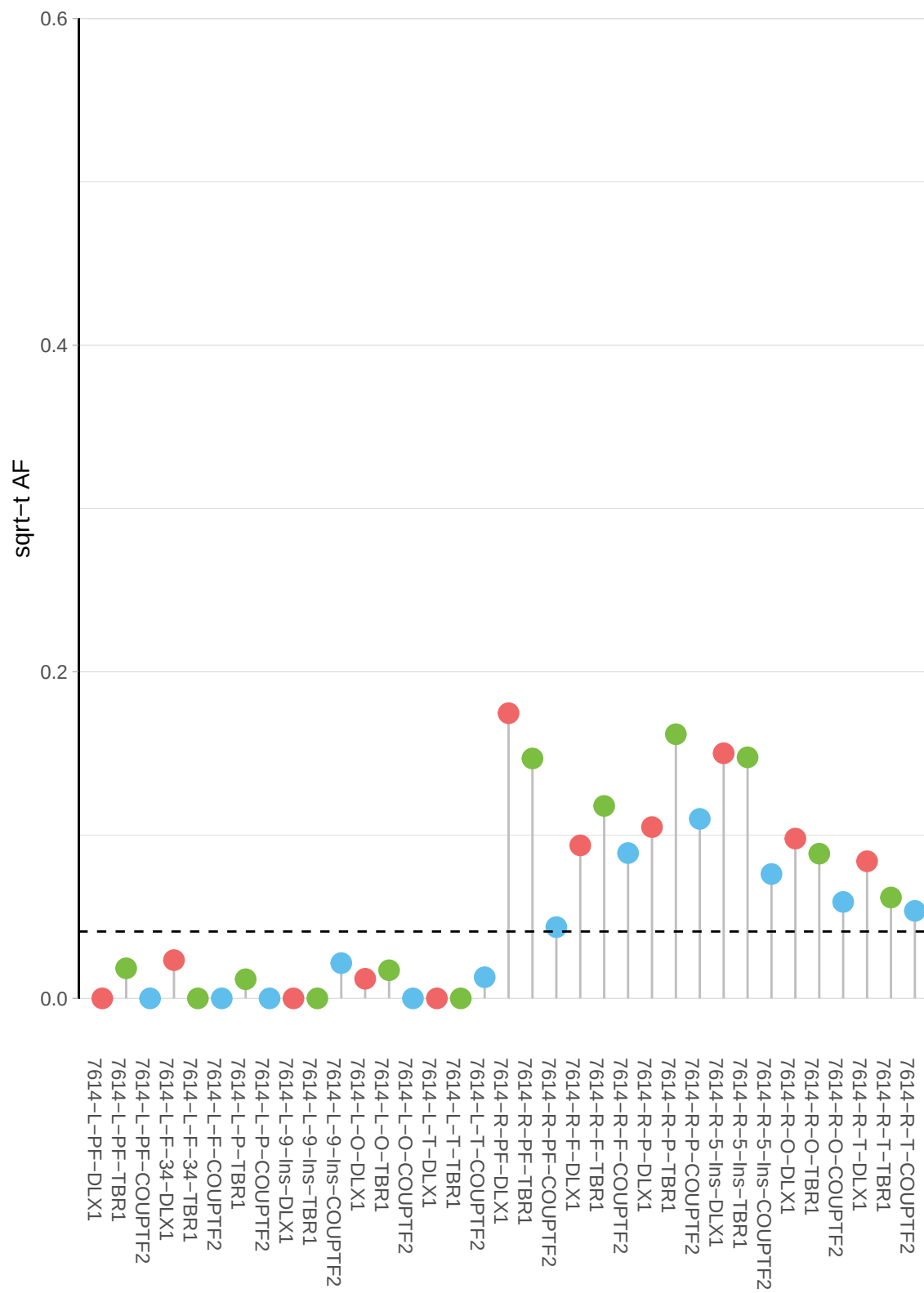

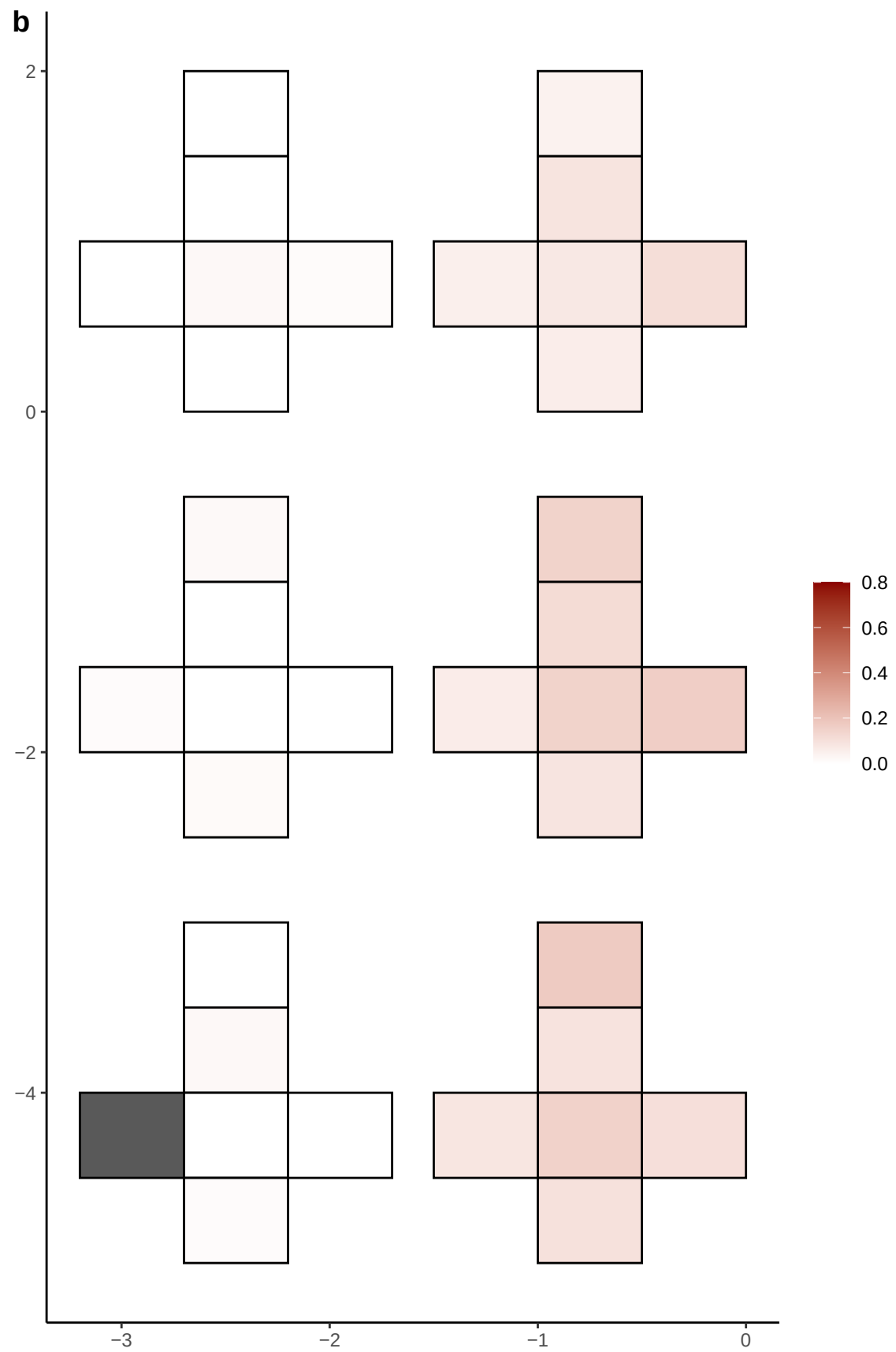

**a** ID01 3-25576791-C-T

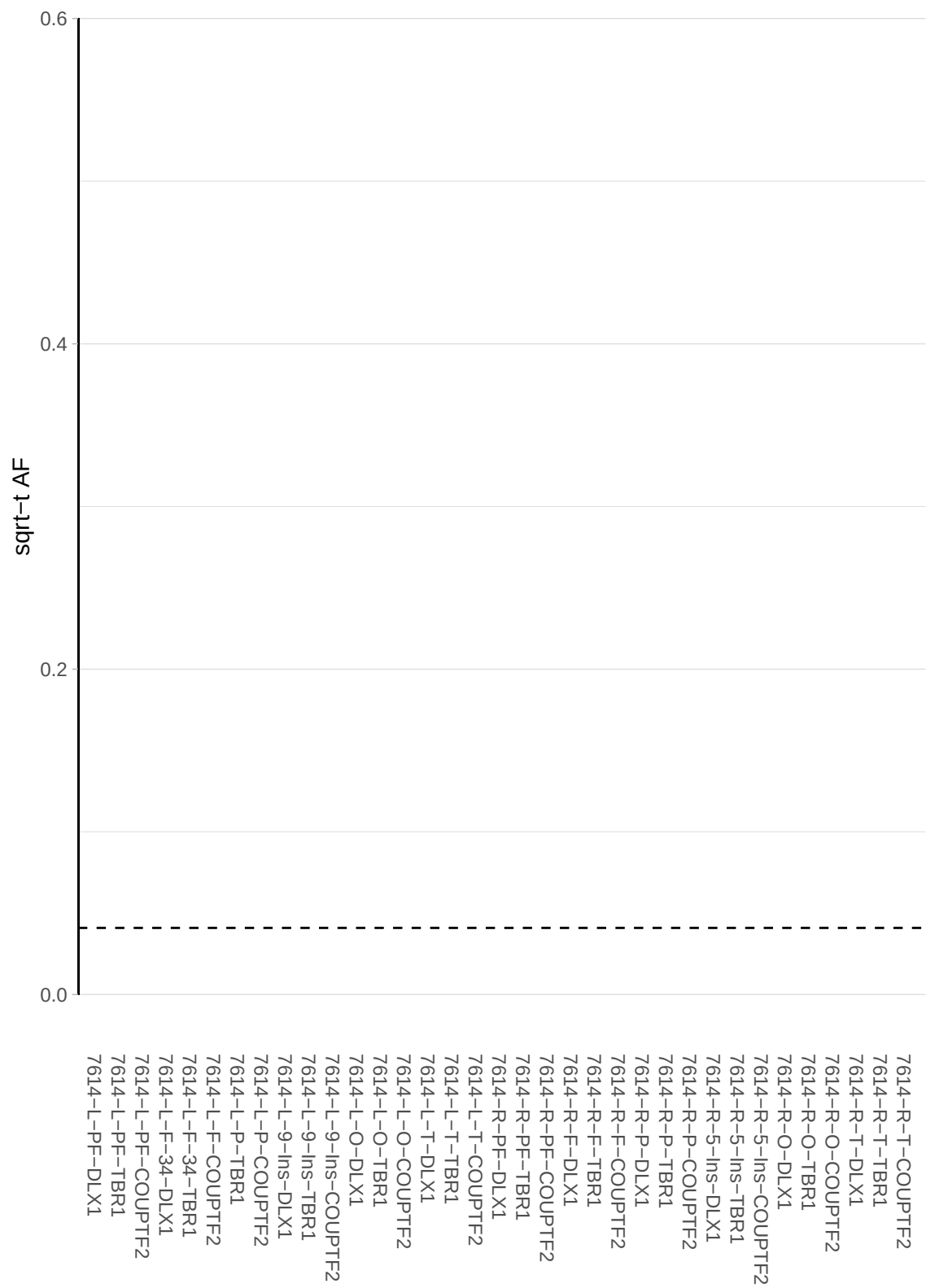

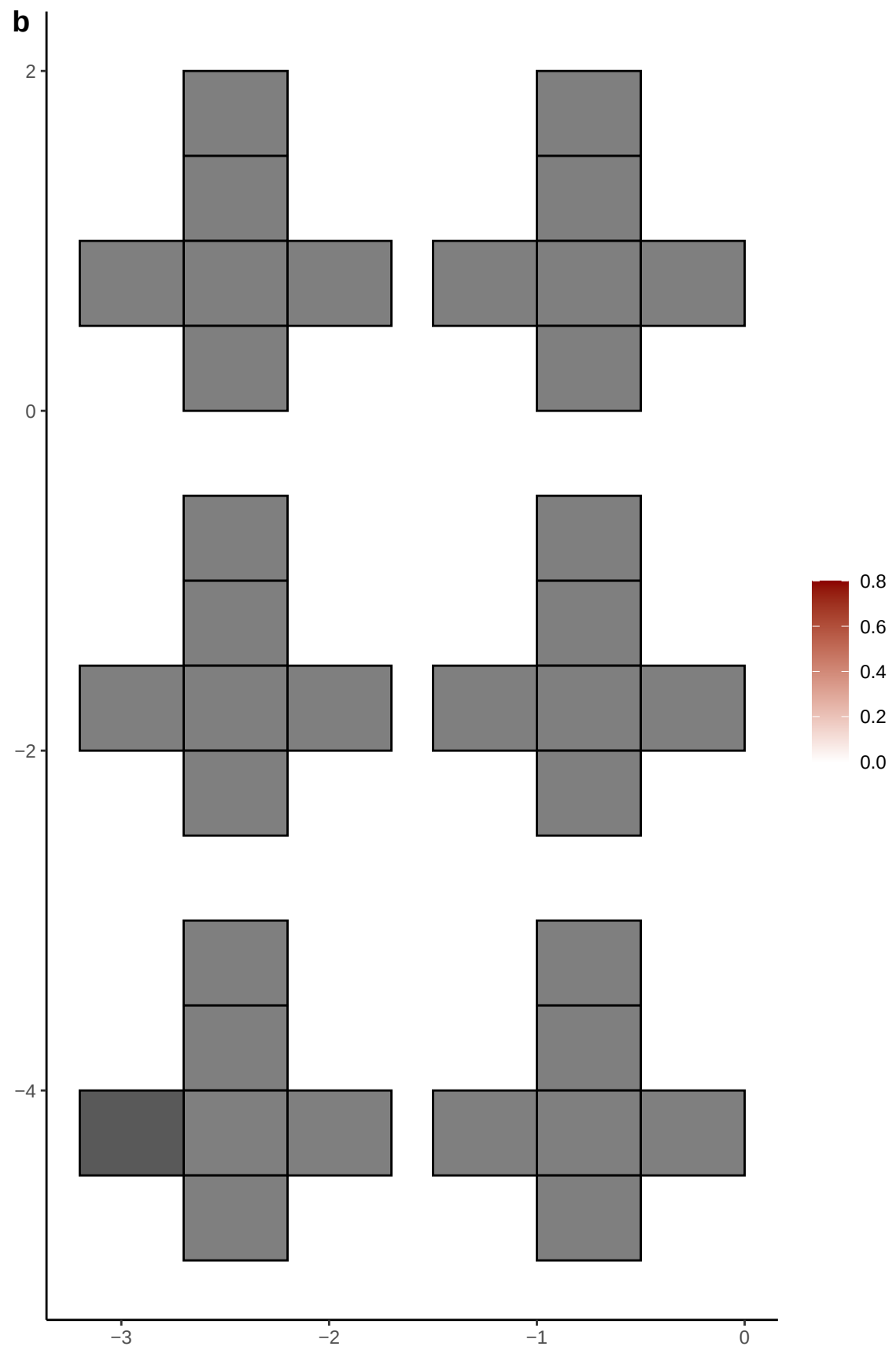

**a** ID01 3-31854915-A-C

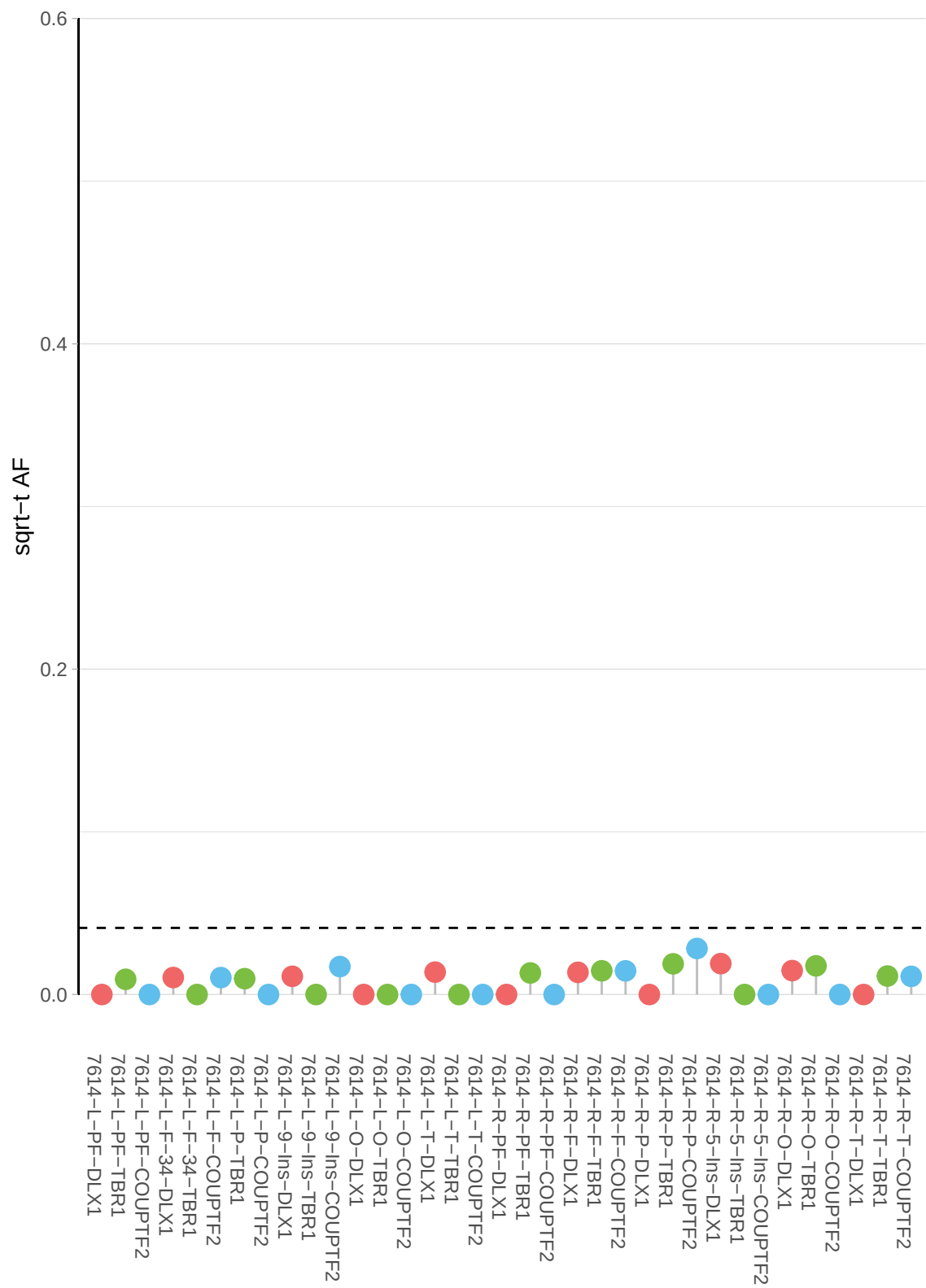

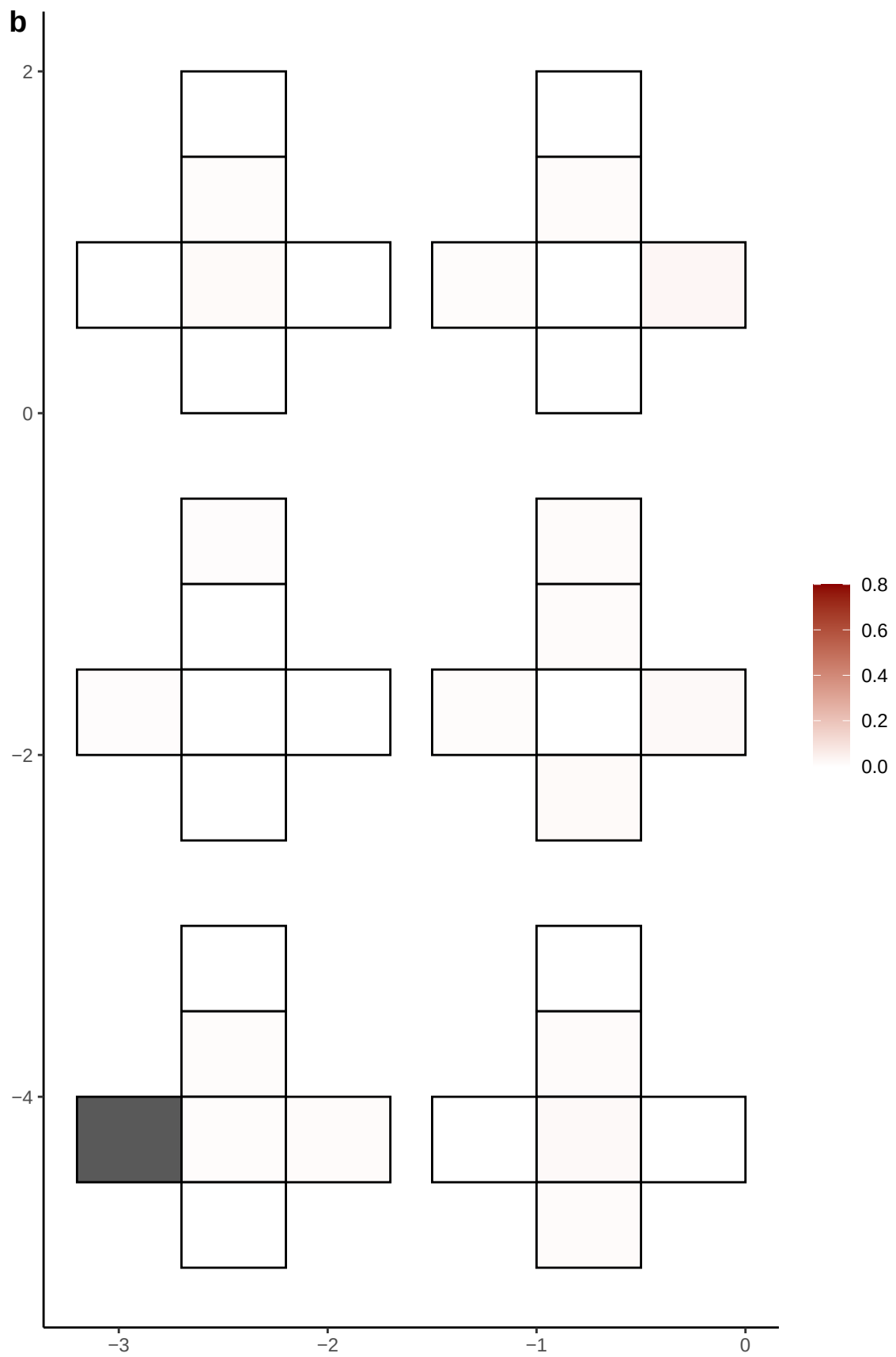

**a** ID01 3-34583283-C-T

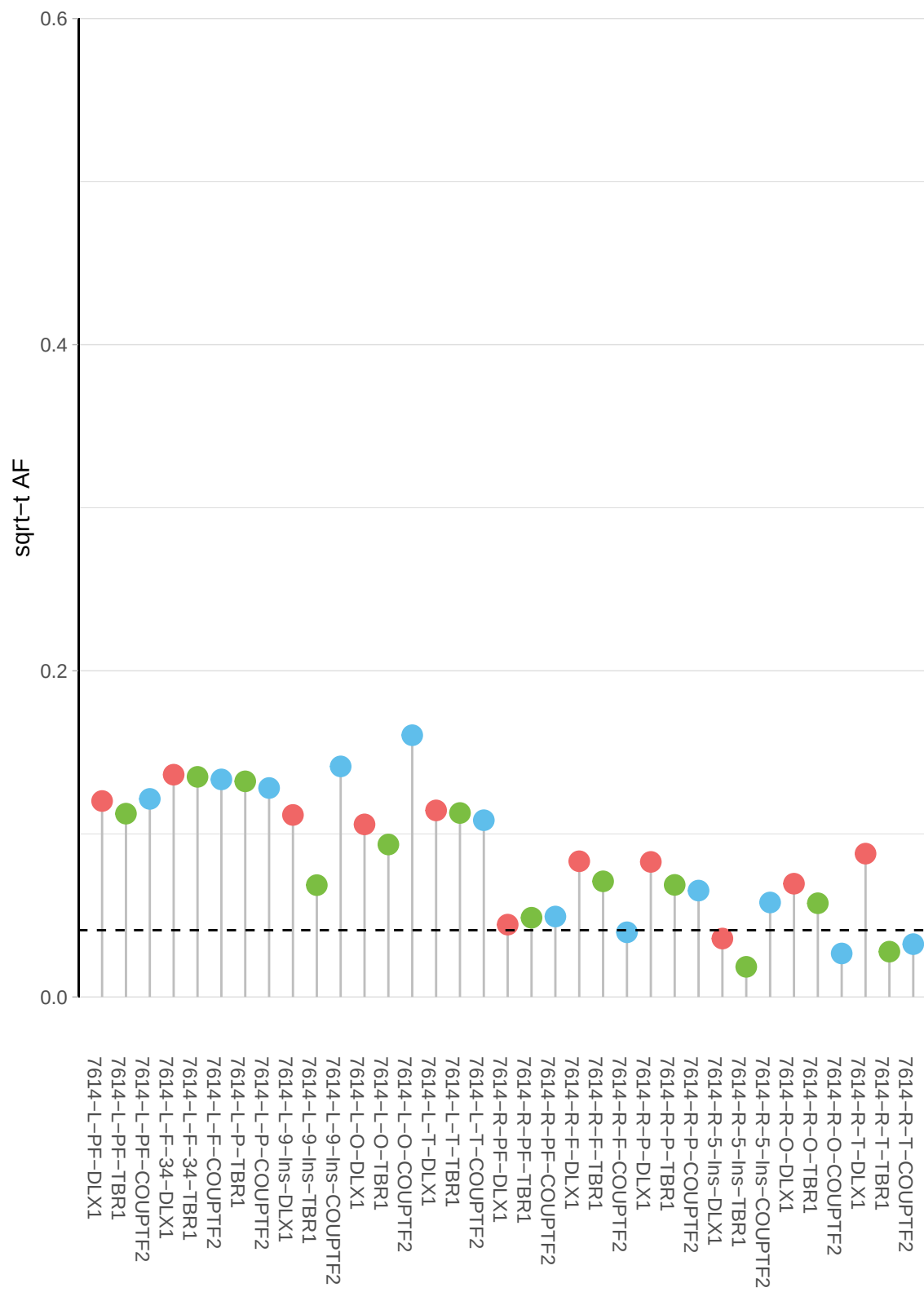

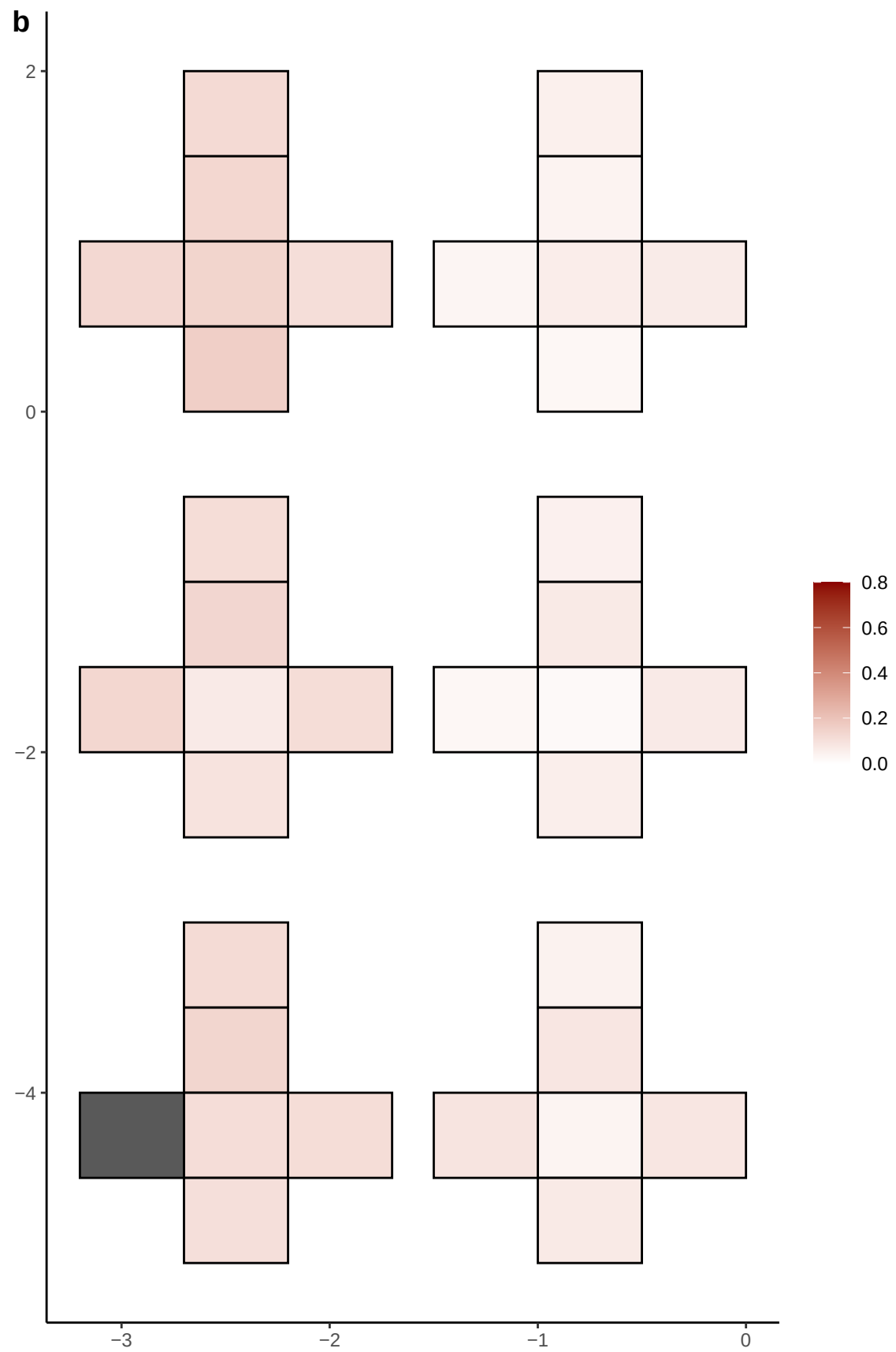

**a** ID01 3-36132231-T-G

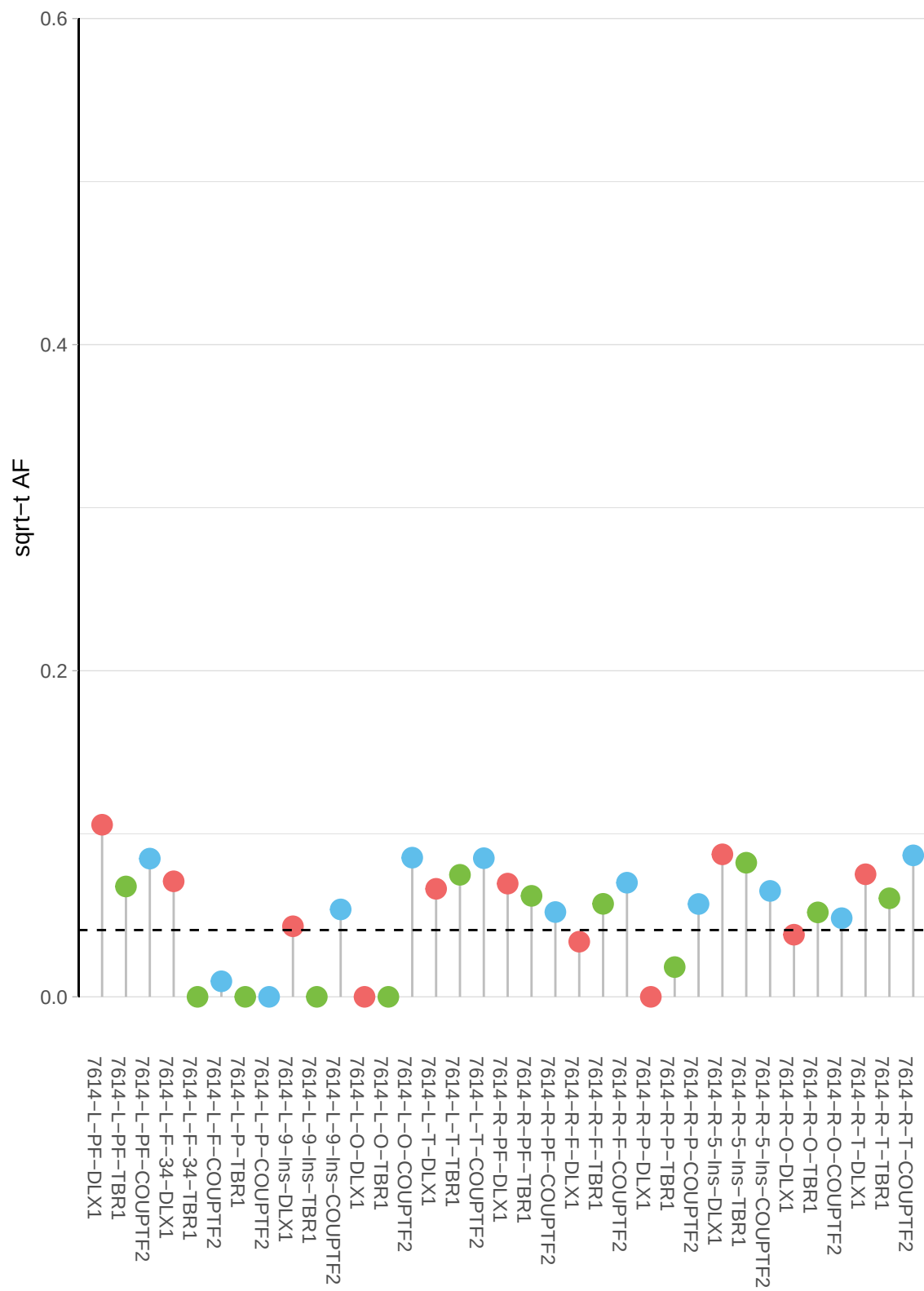

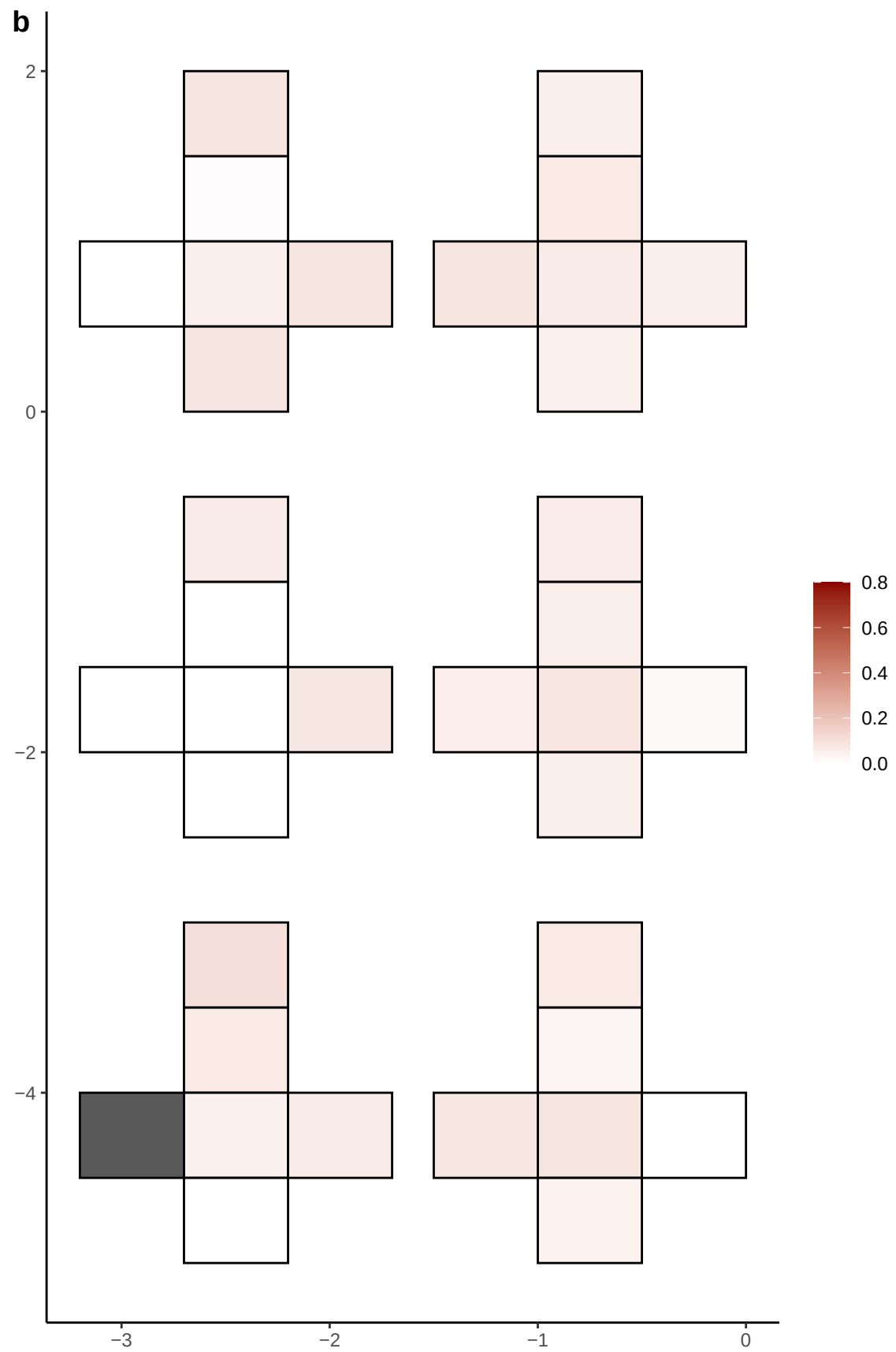

**a** ID01 3-49012324-G-T

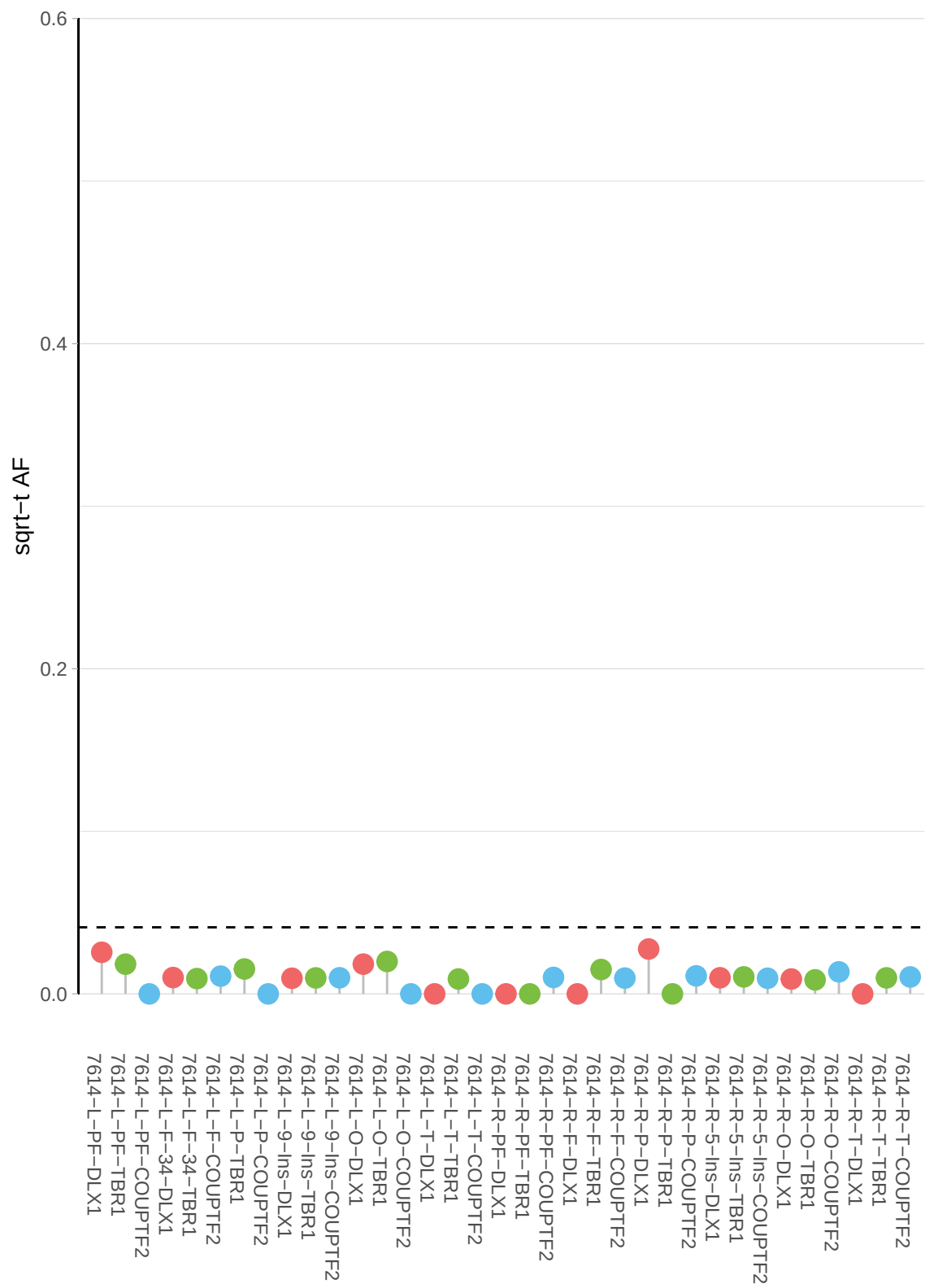

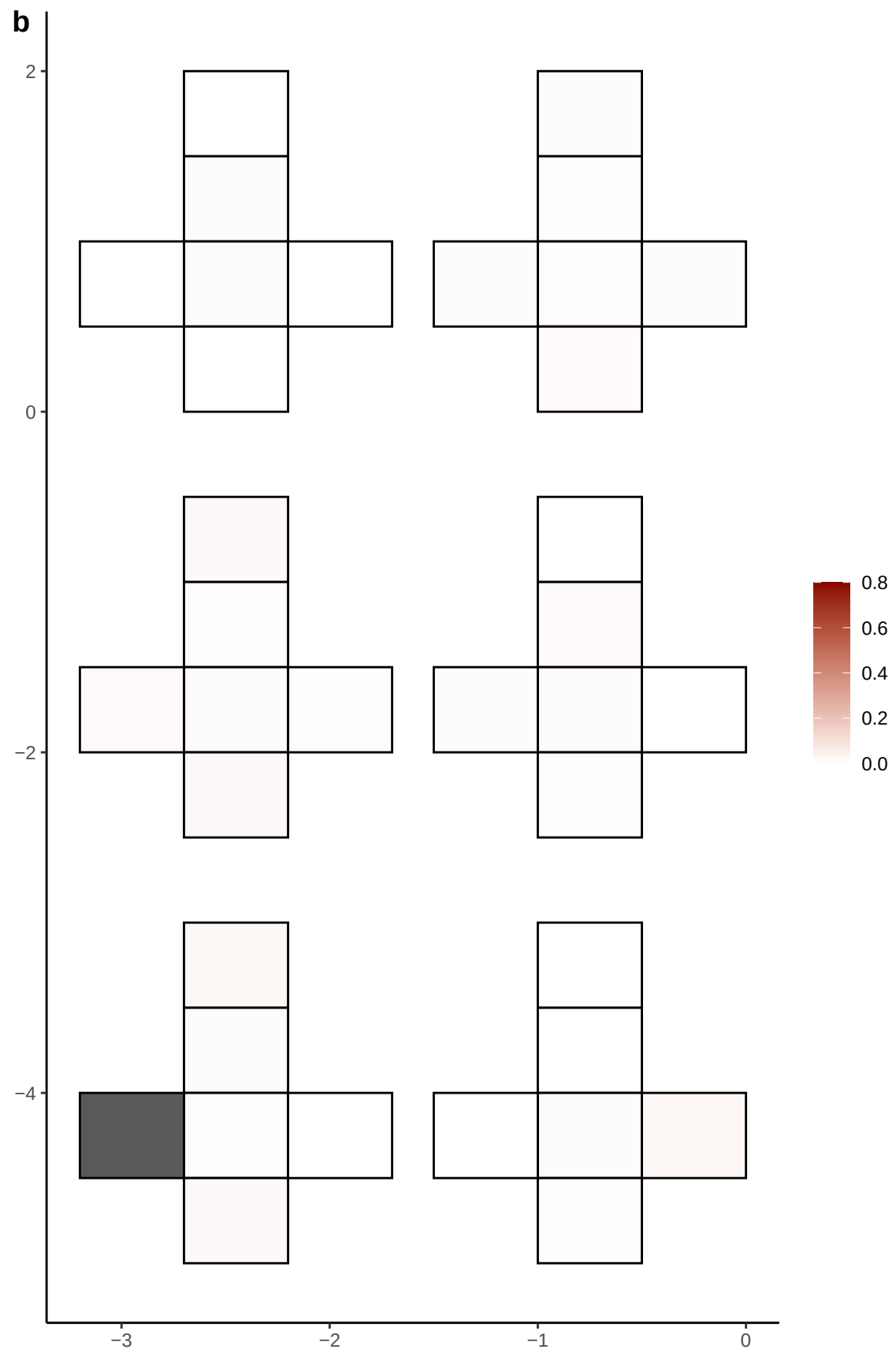

**a** ID01 3-53696469-T-C

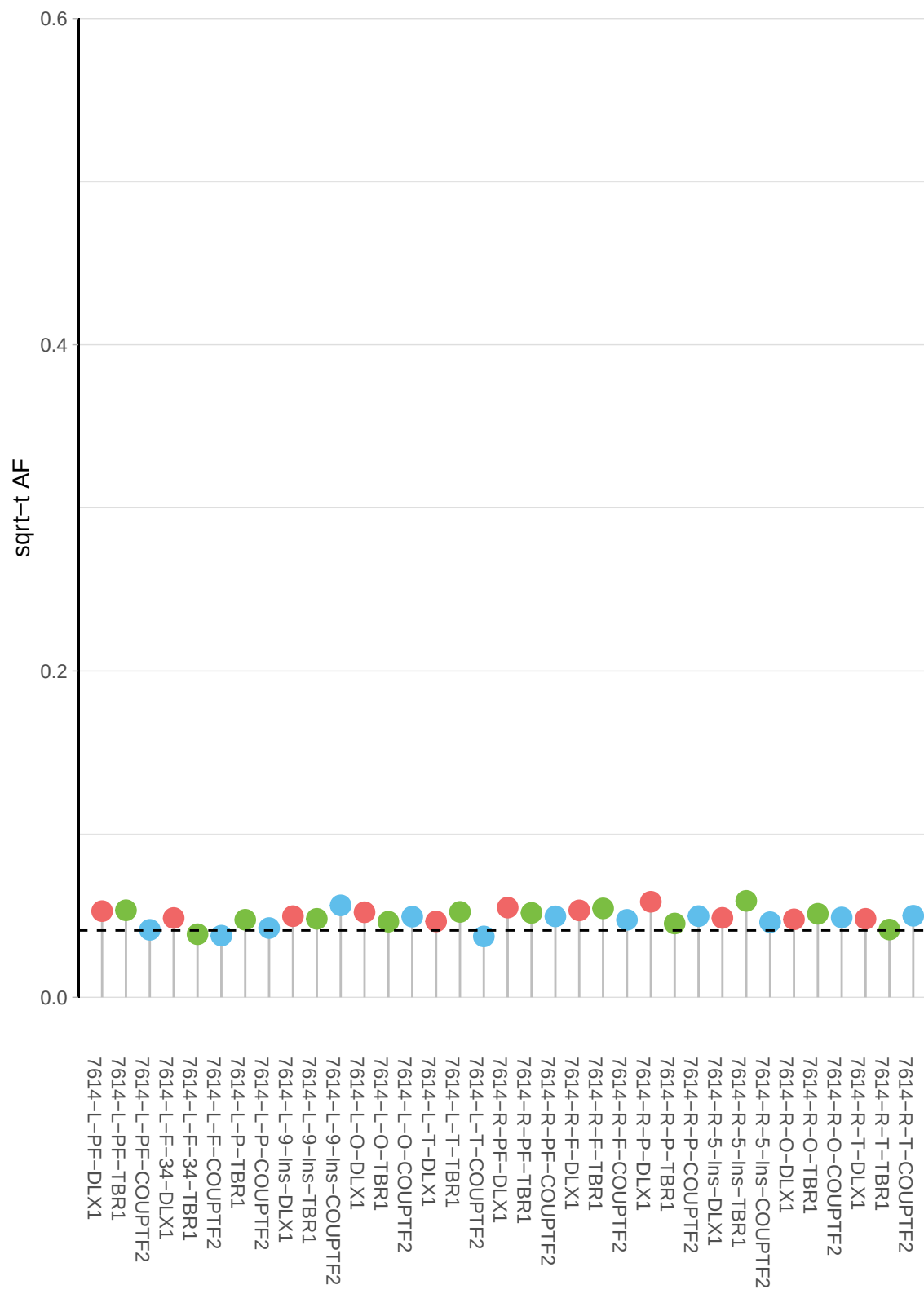

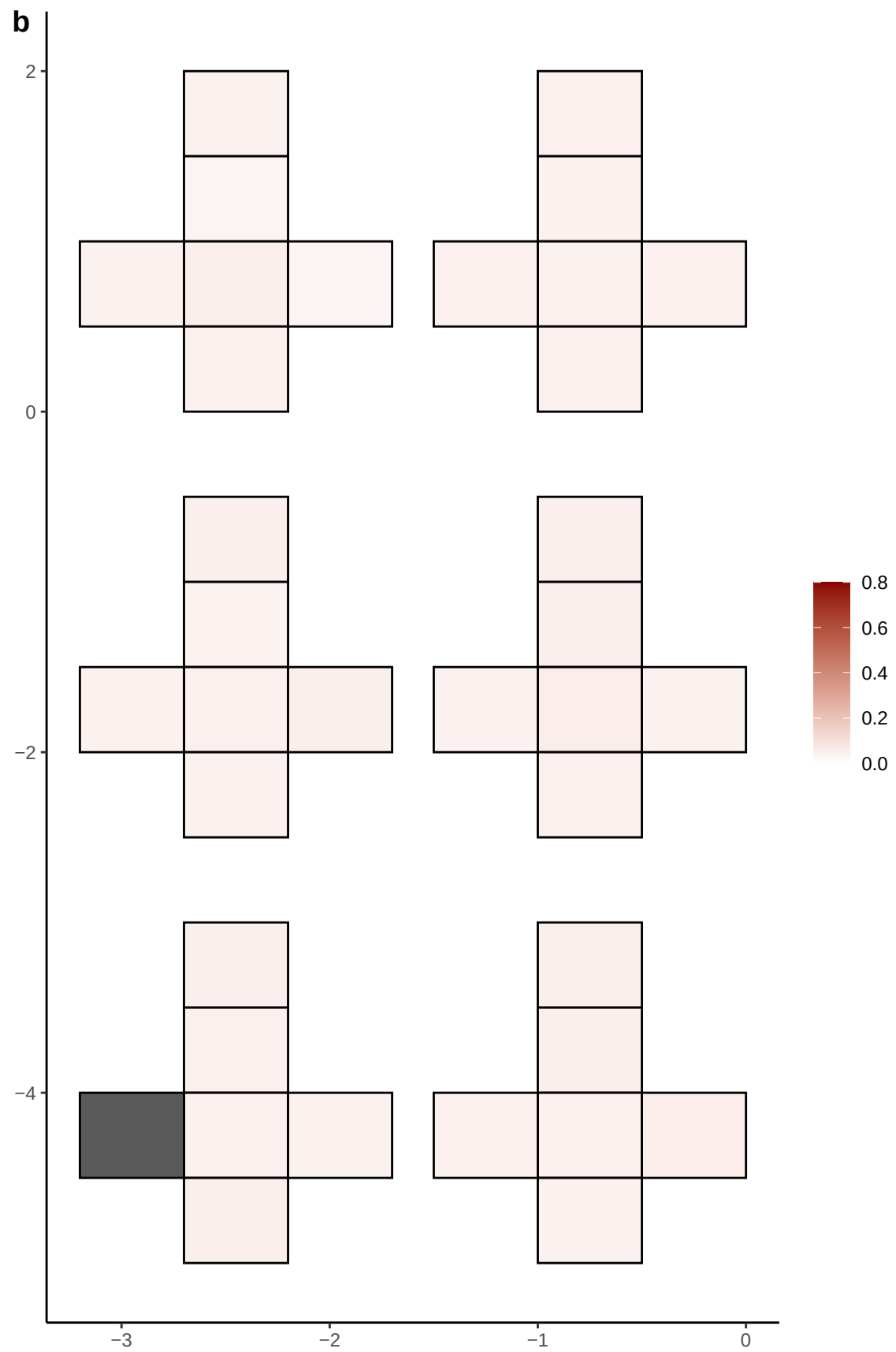

**a** ID01 3-6341933-TATC-T

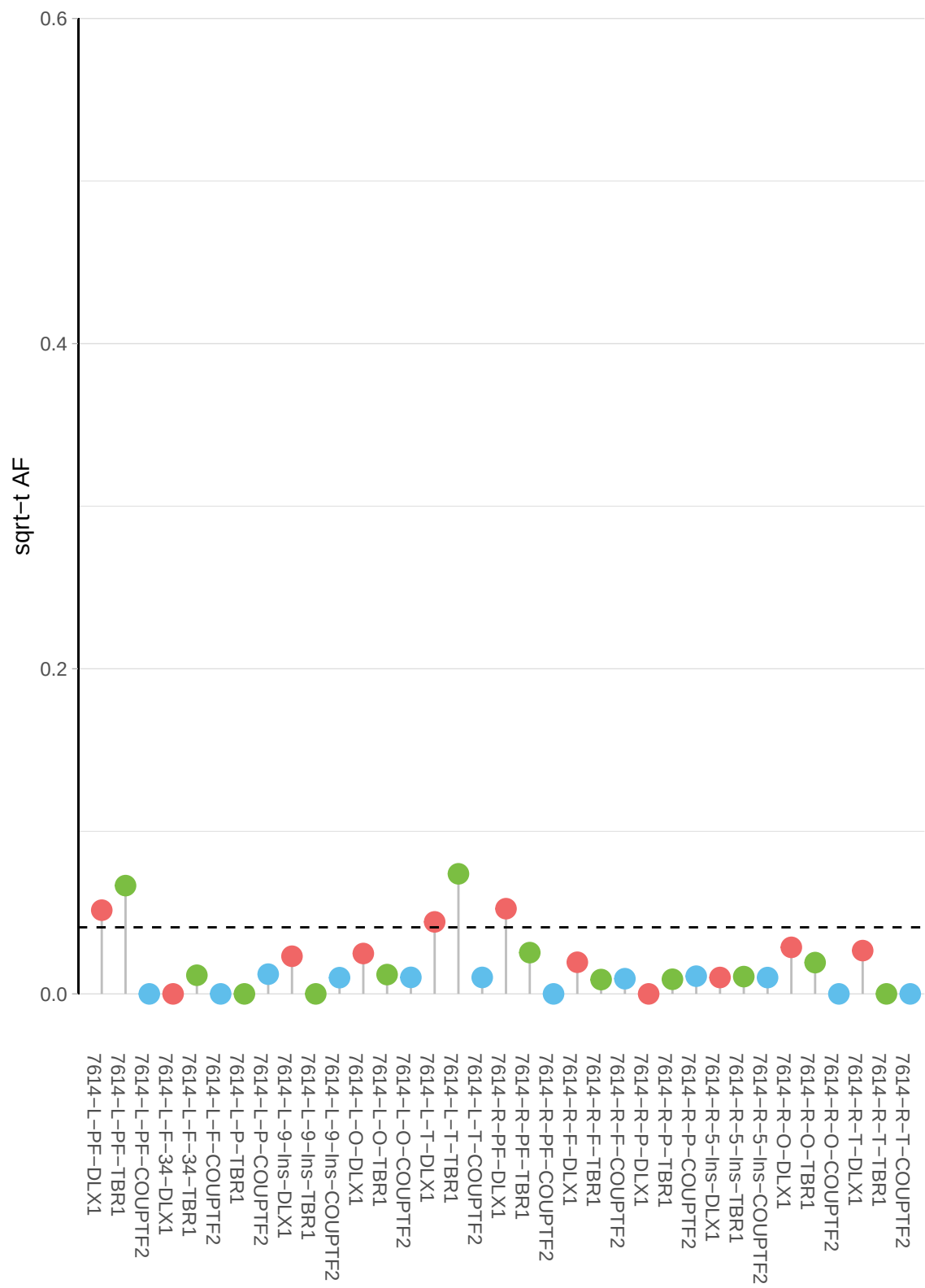

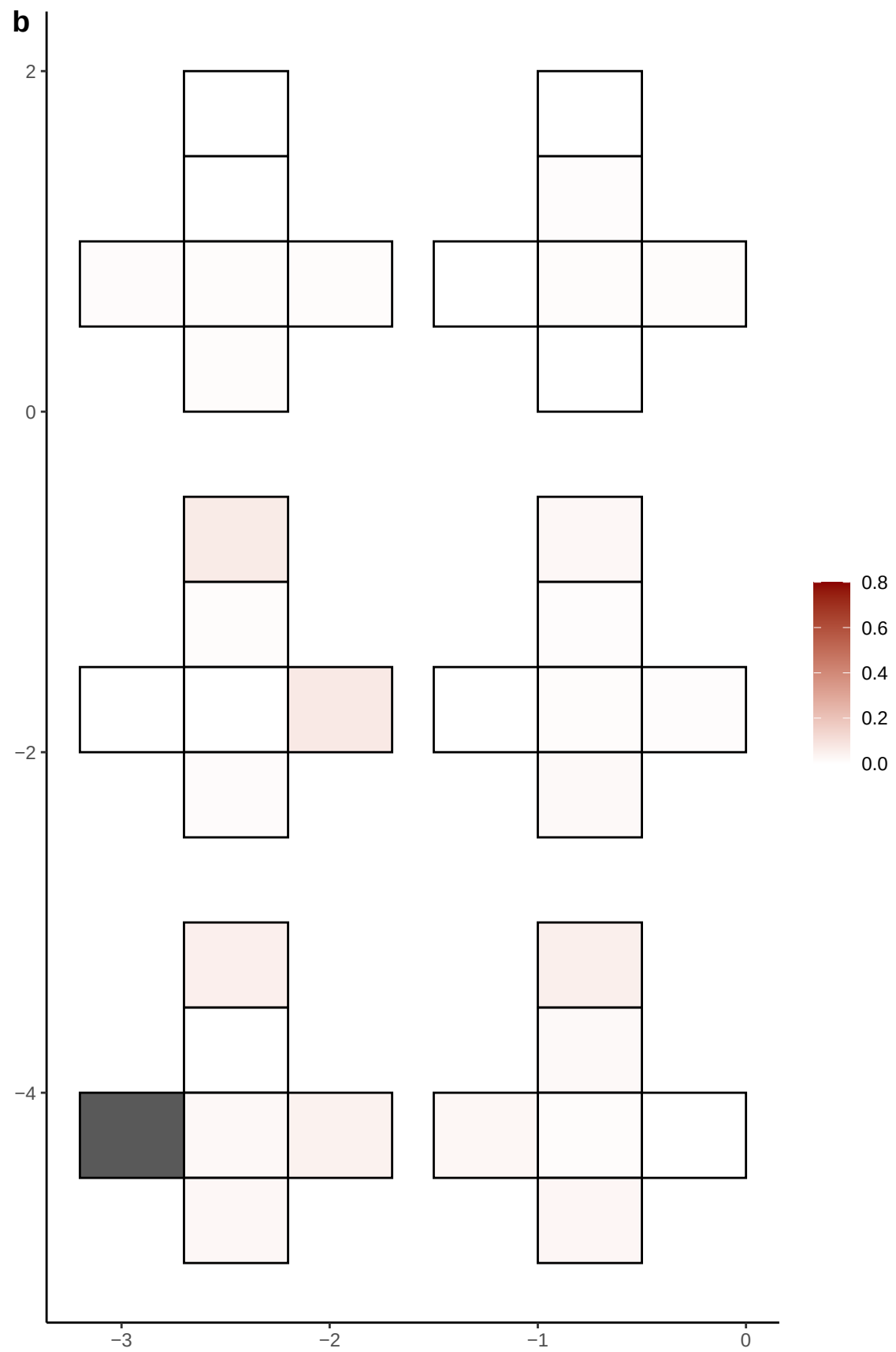

**a** ID01 3-63769495-C-T

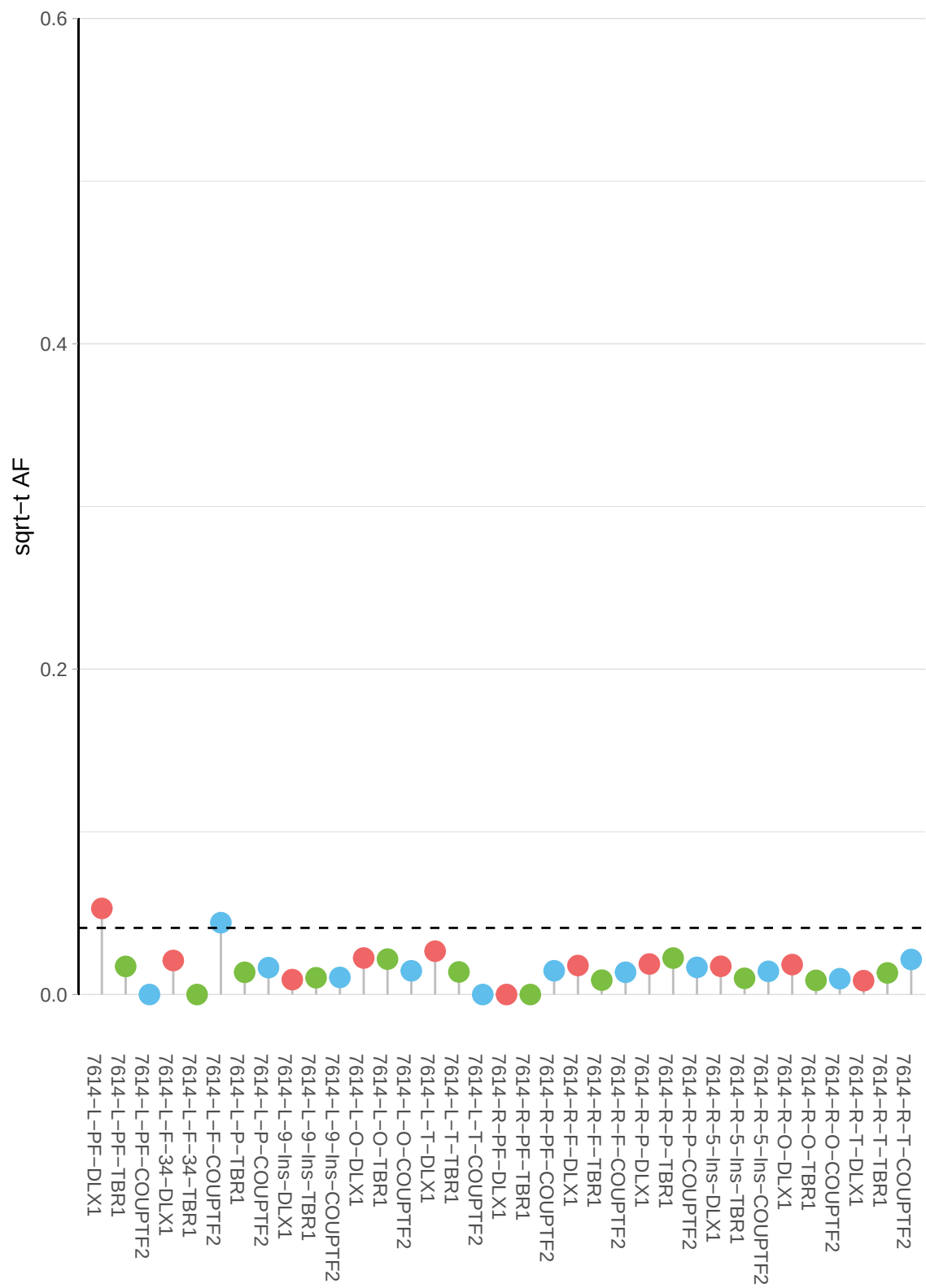

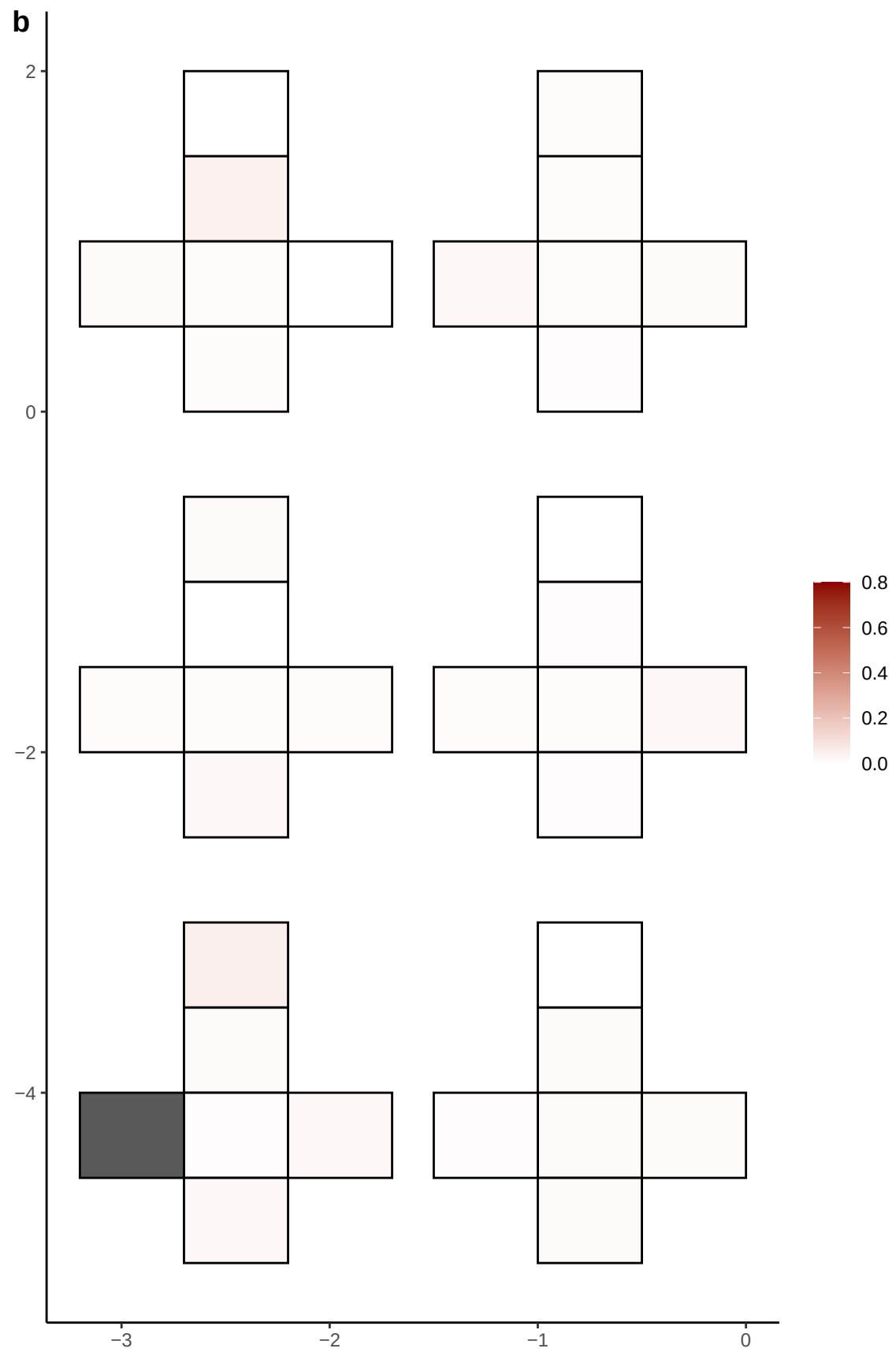

**a** ID01 3-70199639-G-C

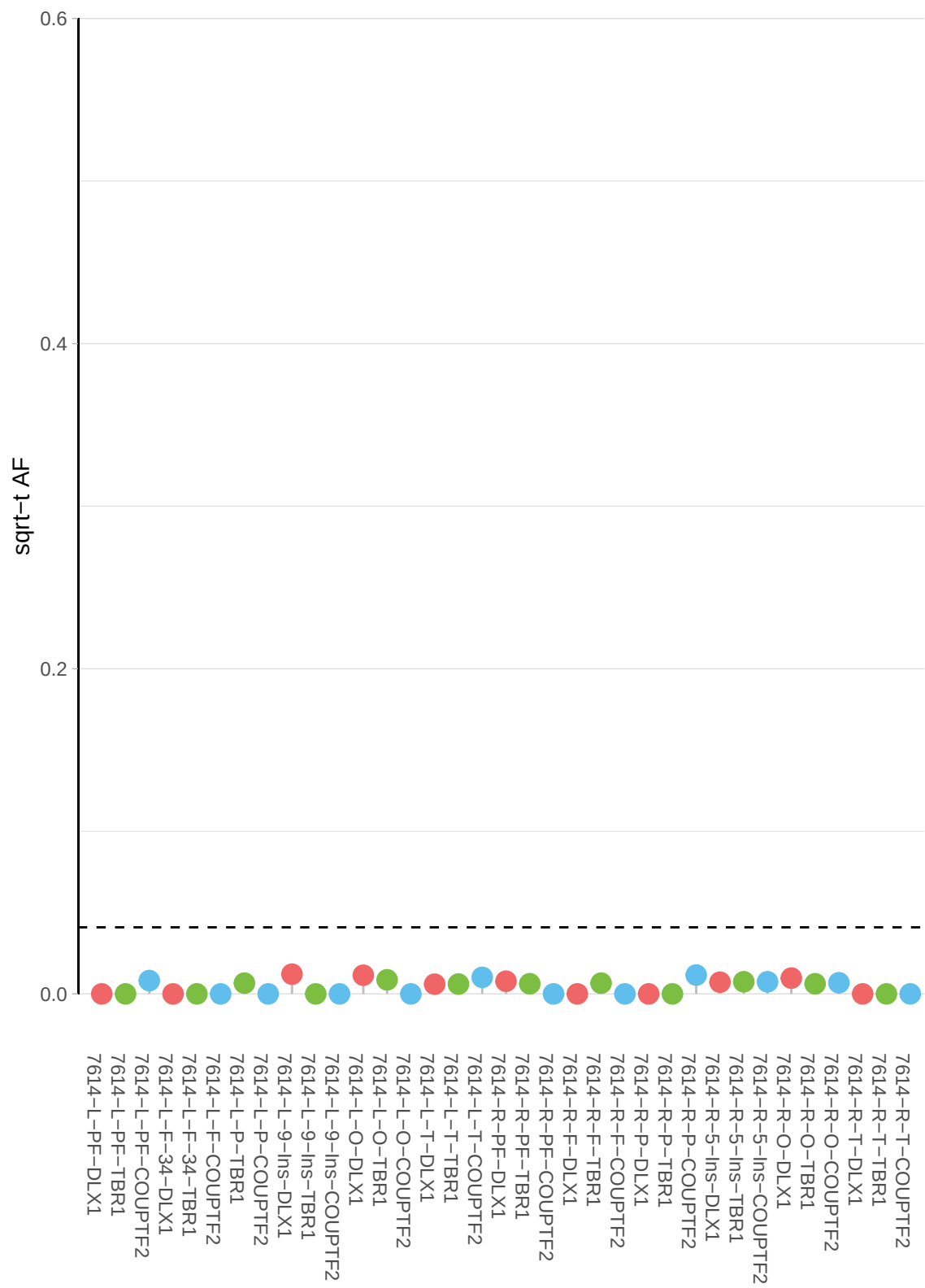

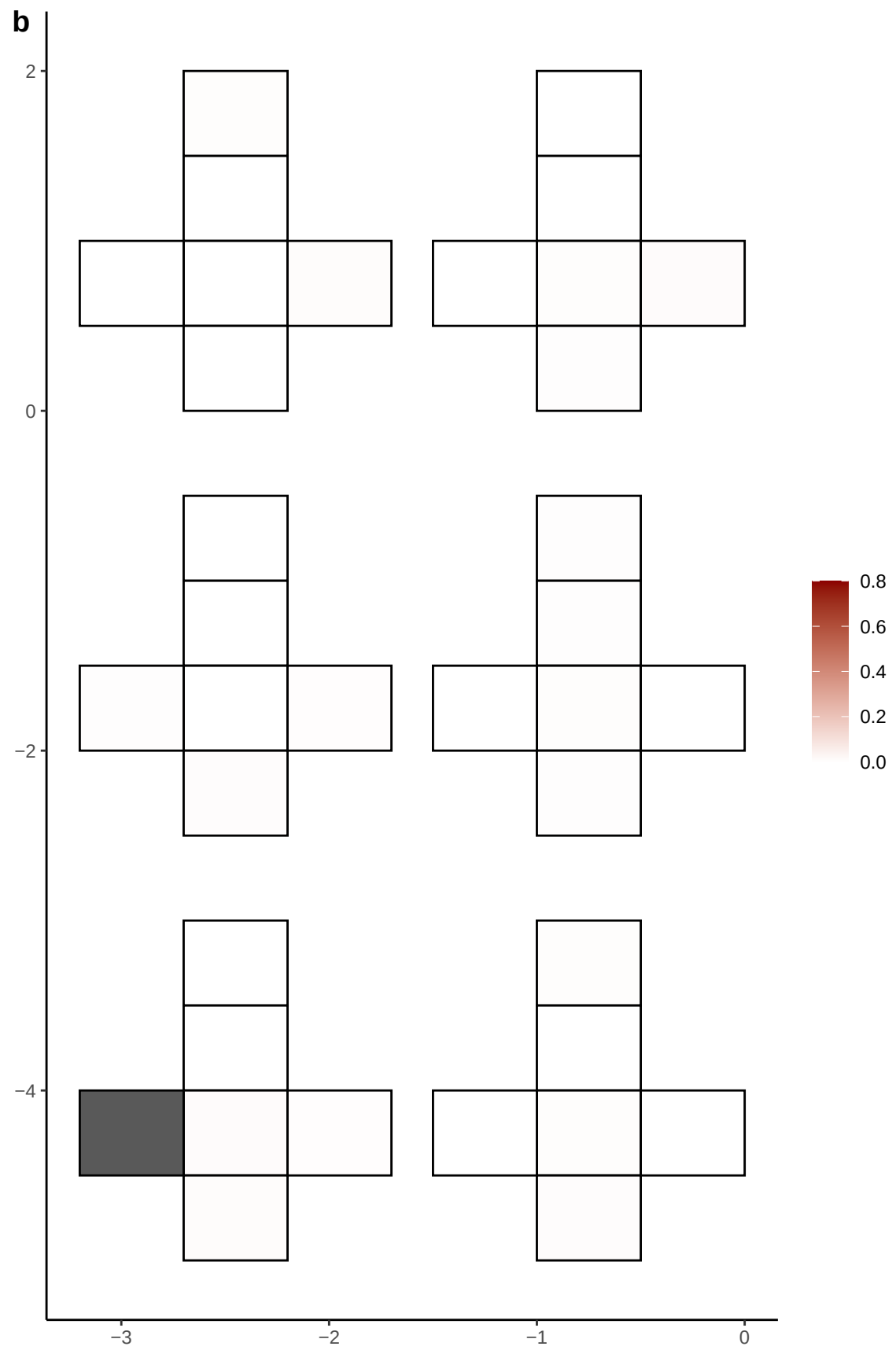

**a** ID01 3-83951534-A-G

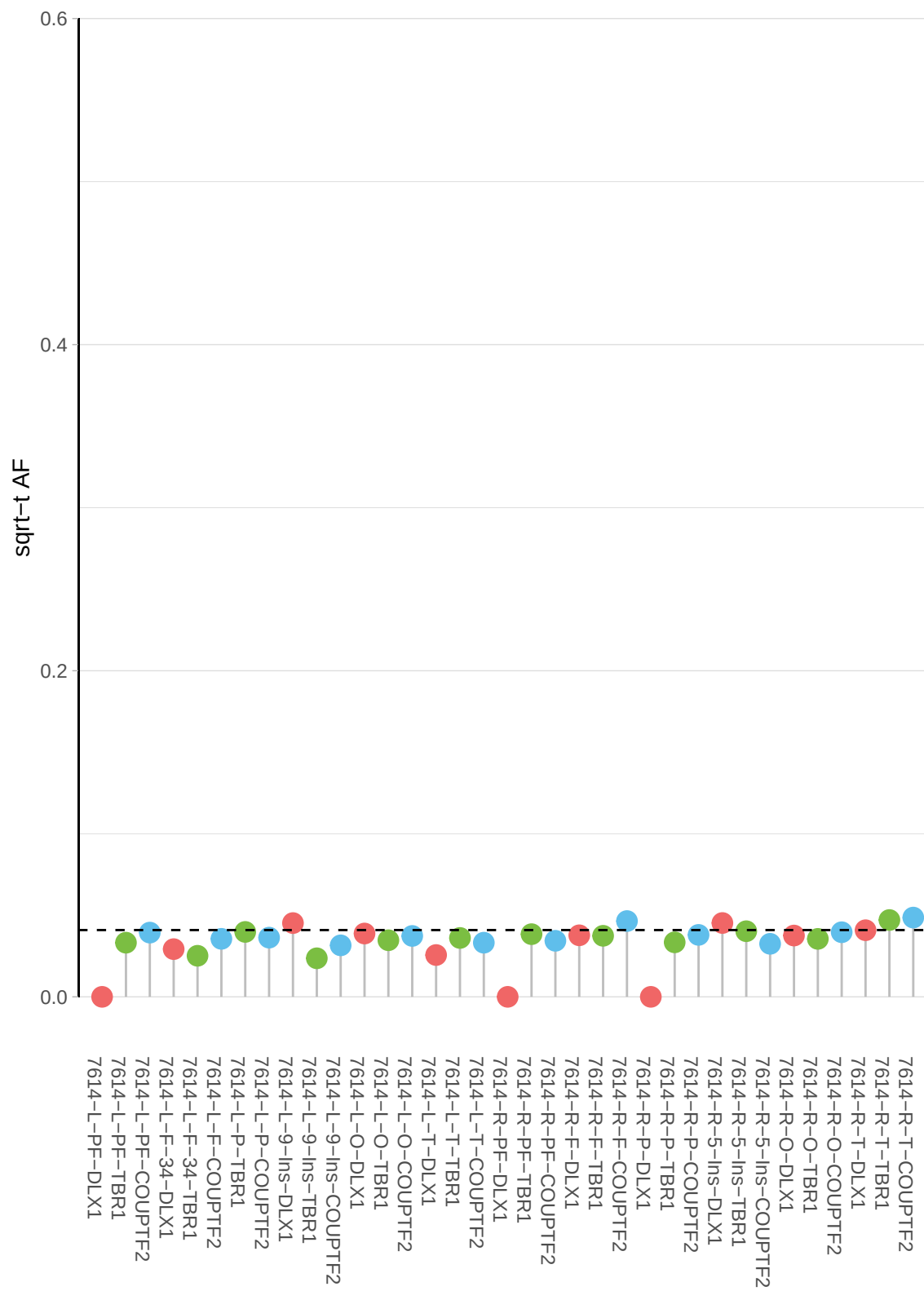

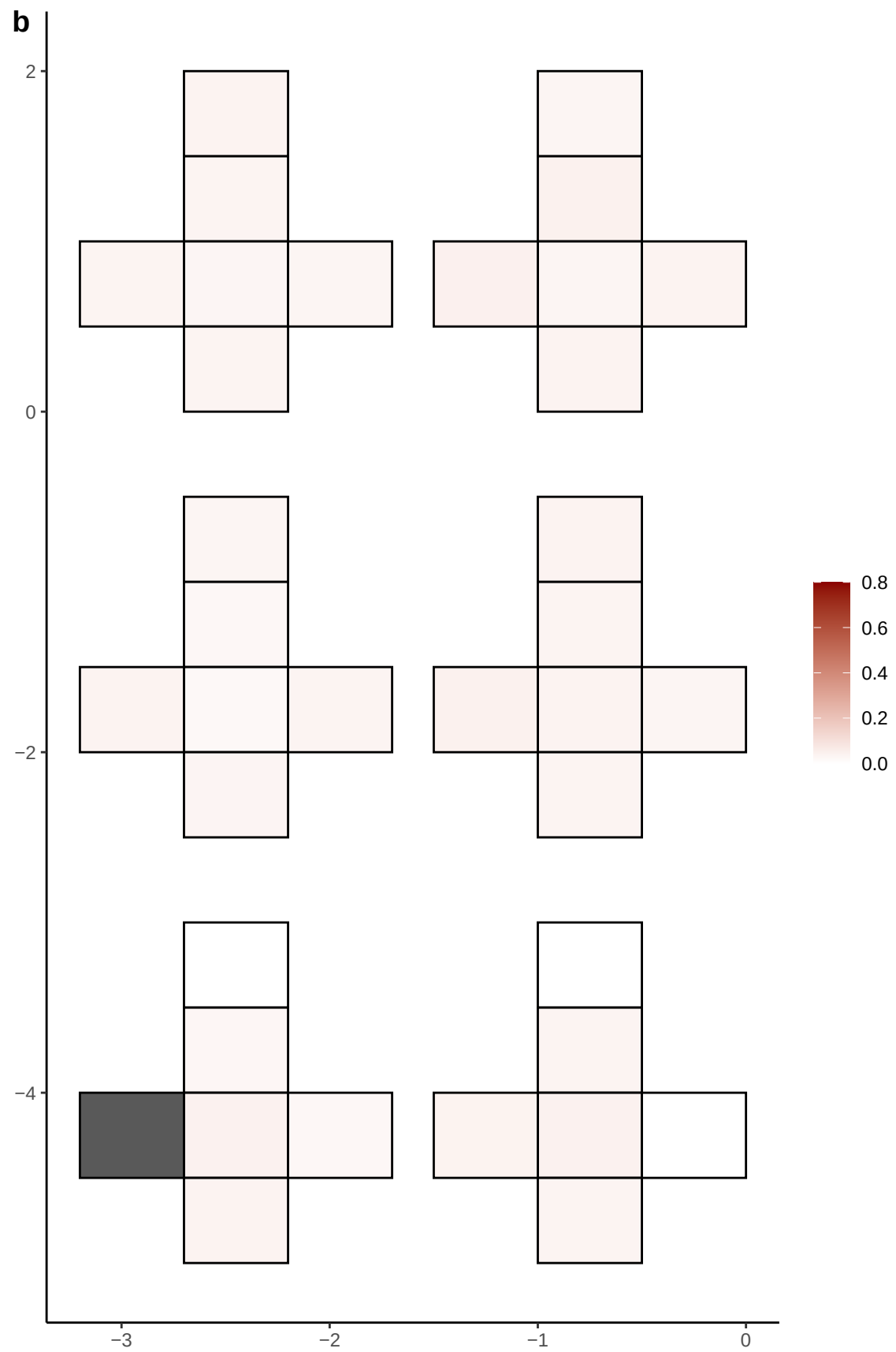

**a** ID01 3-84719043-C-T

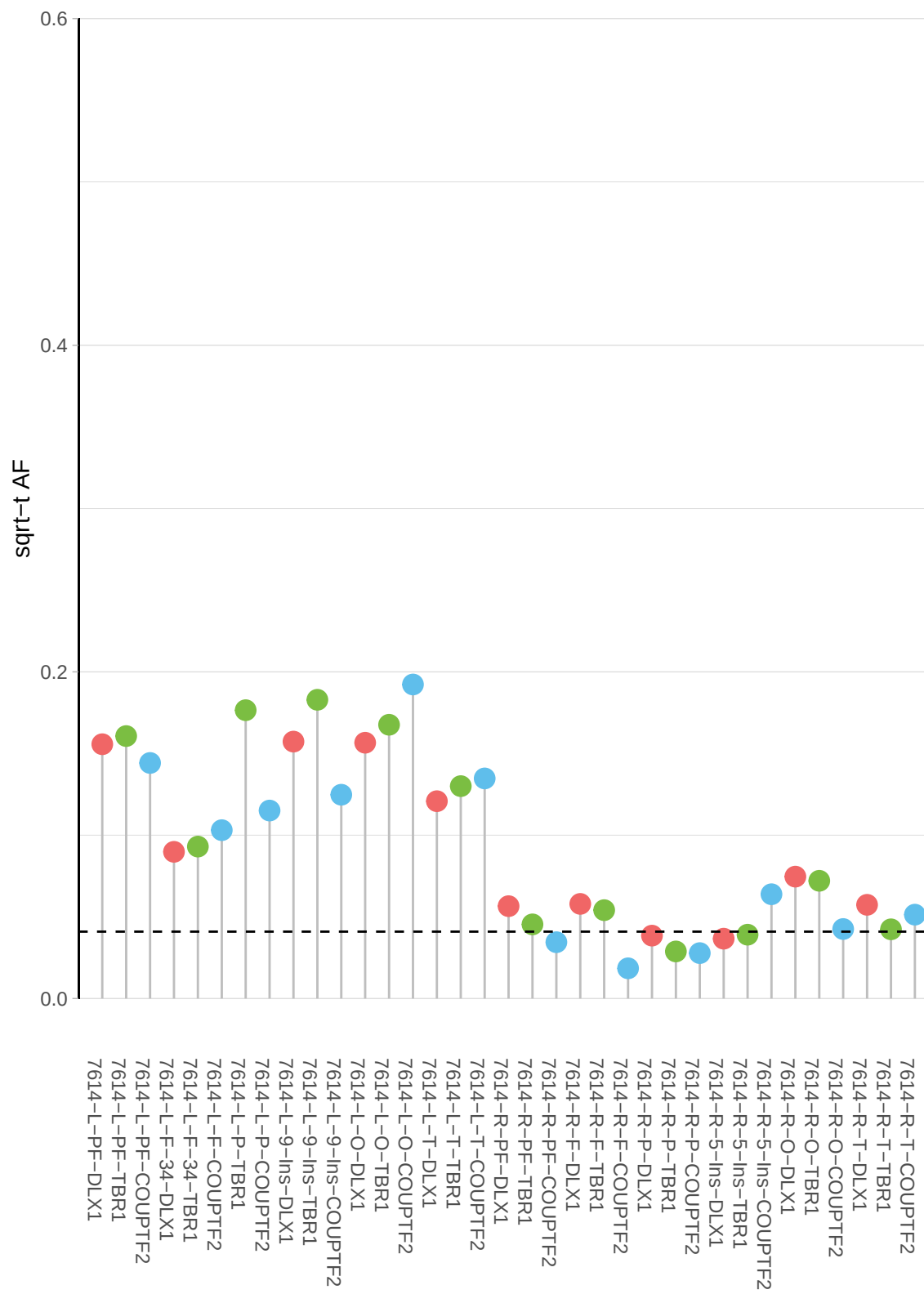

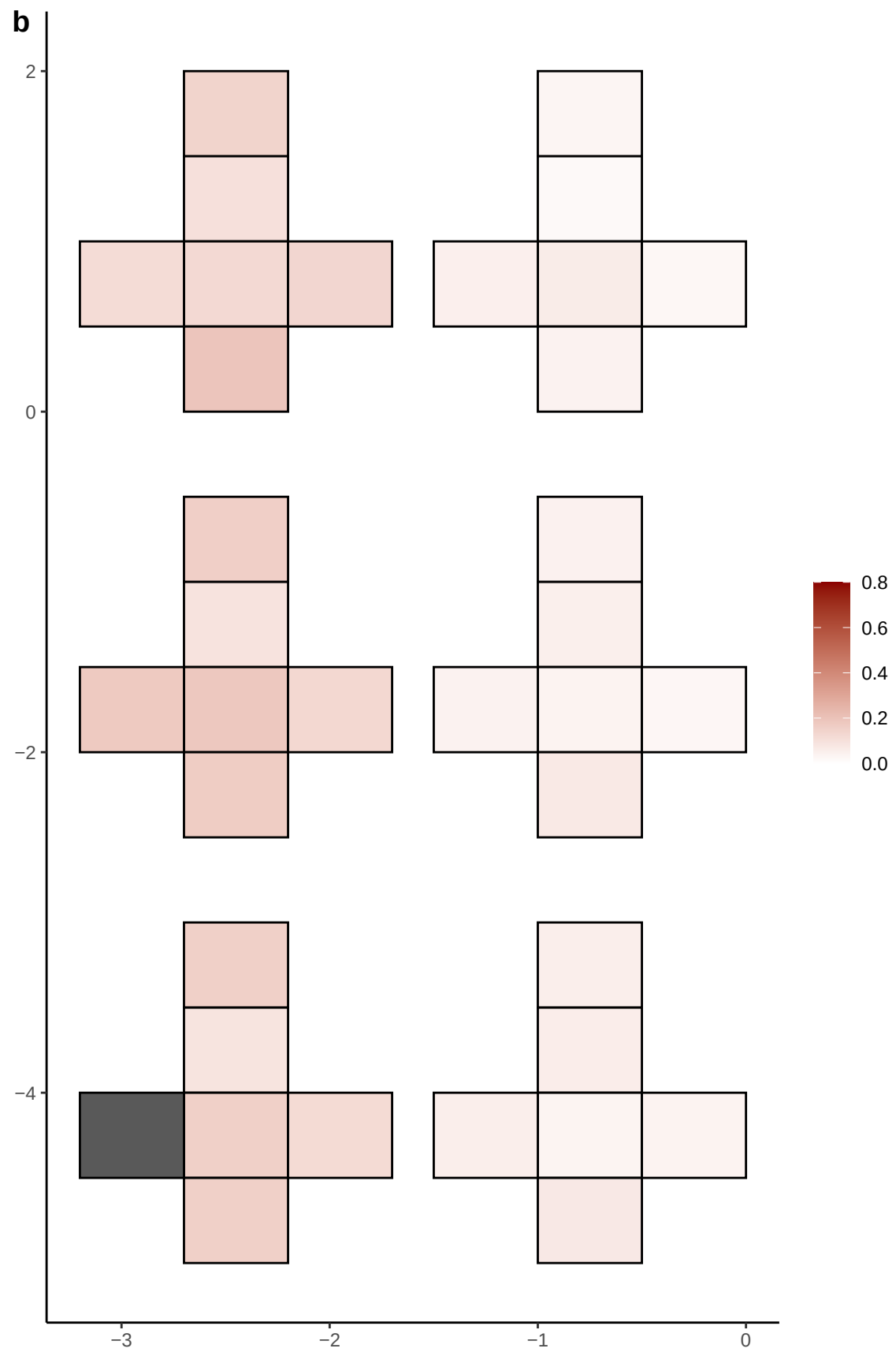

**a** ID01 3-84856980-T-A

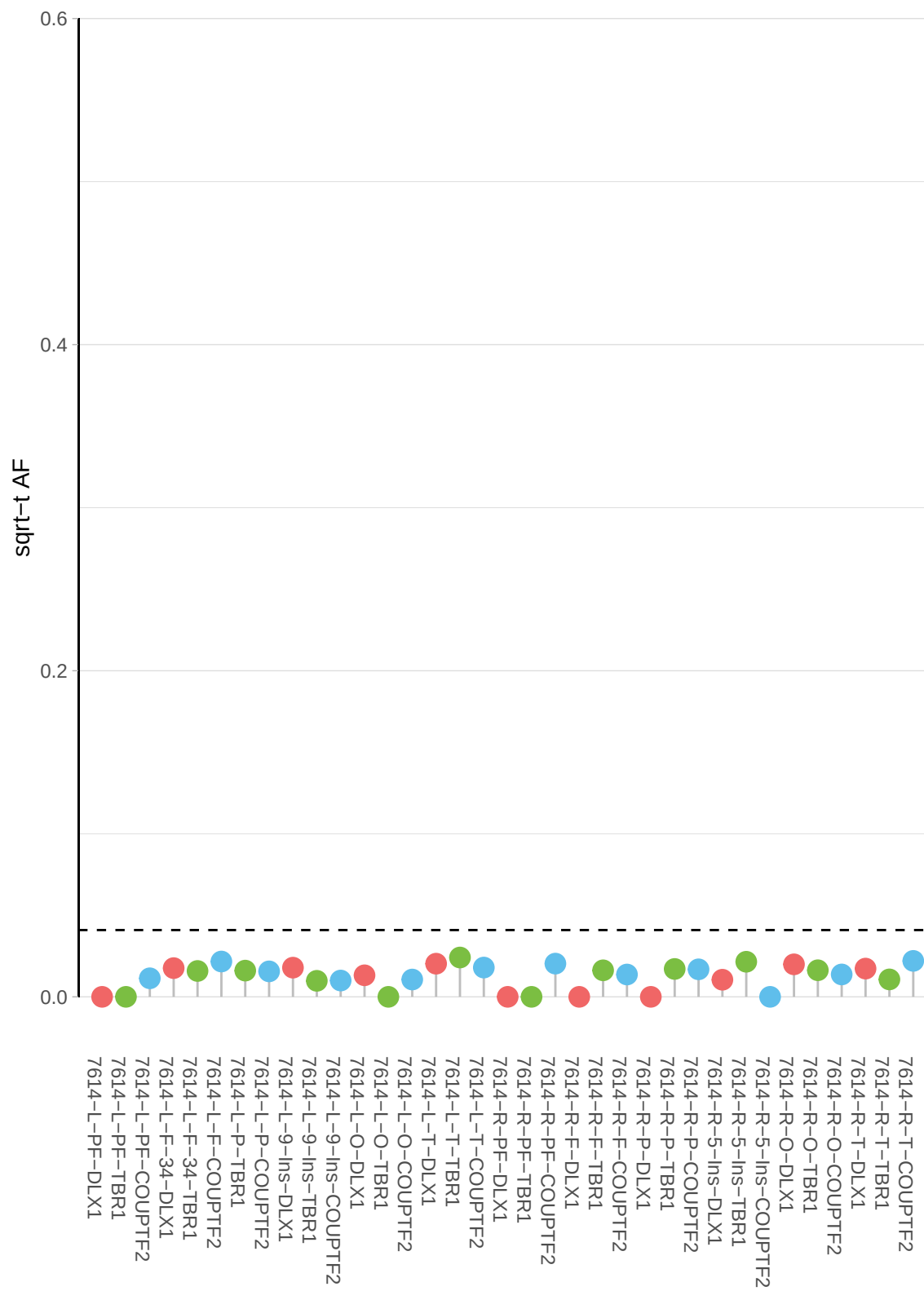

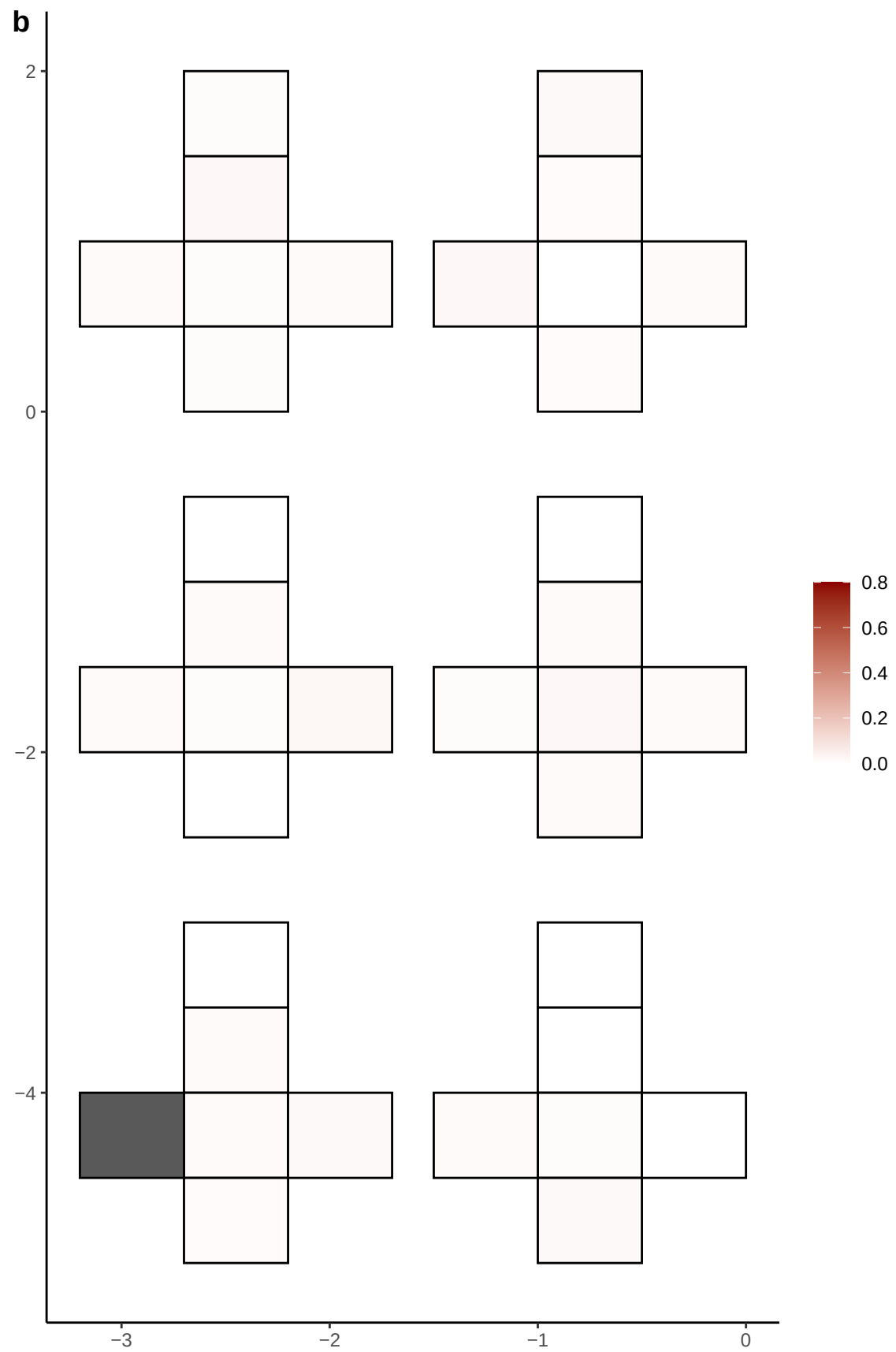

**a** ID01 3-85794290-A-C

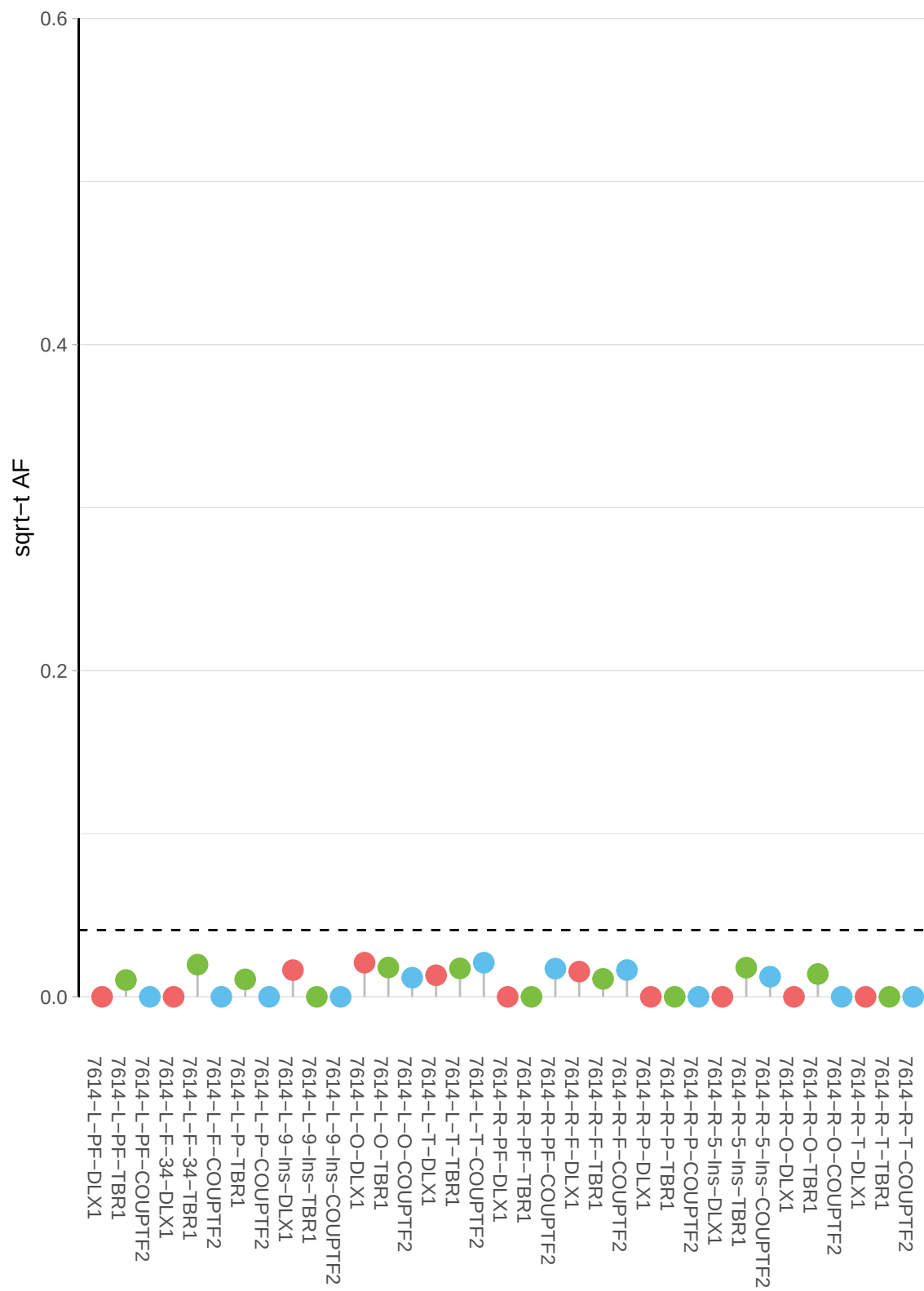

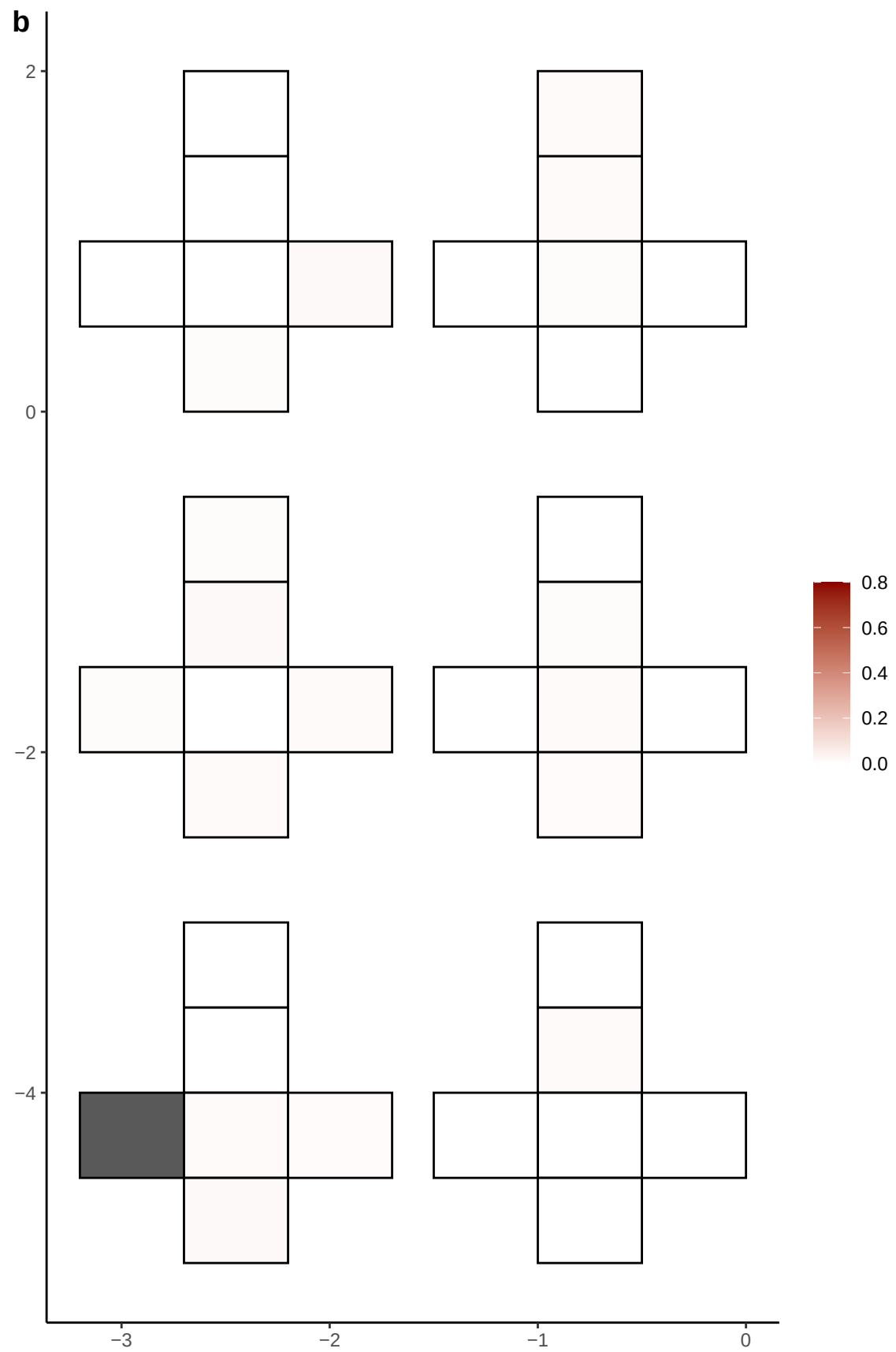

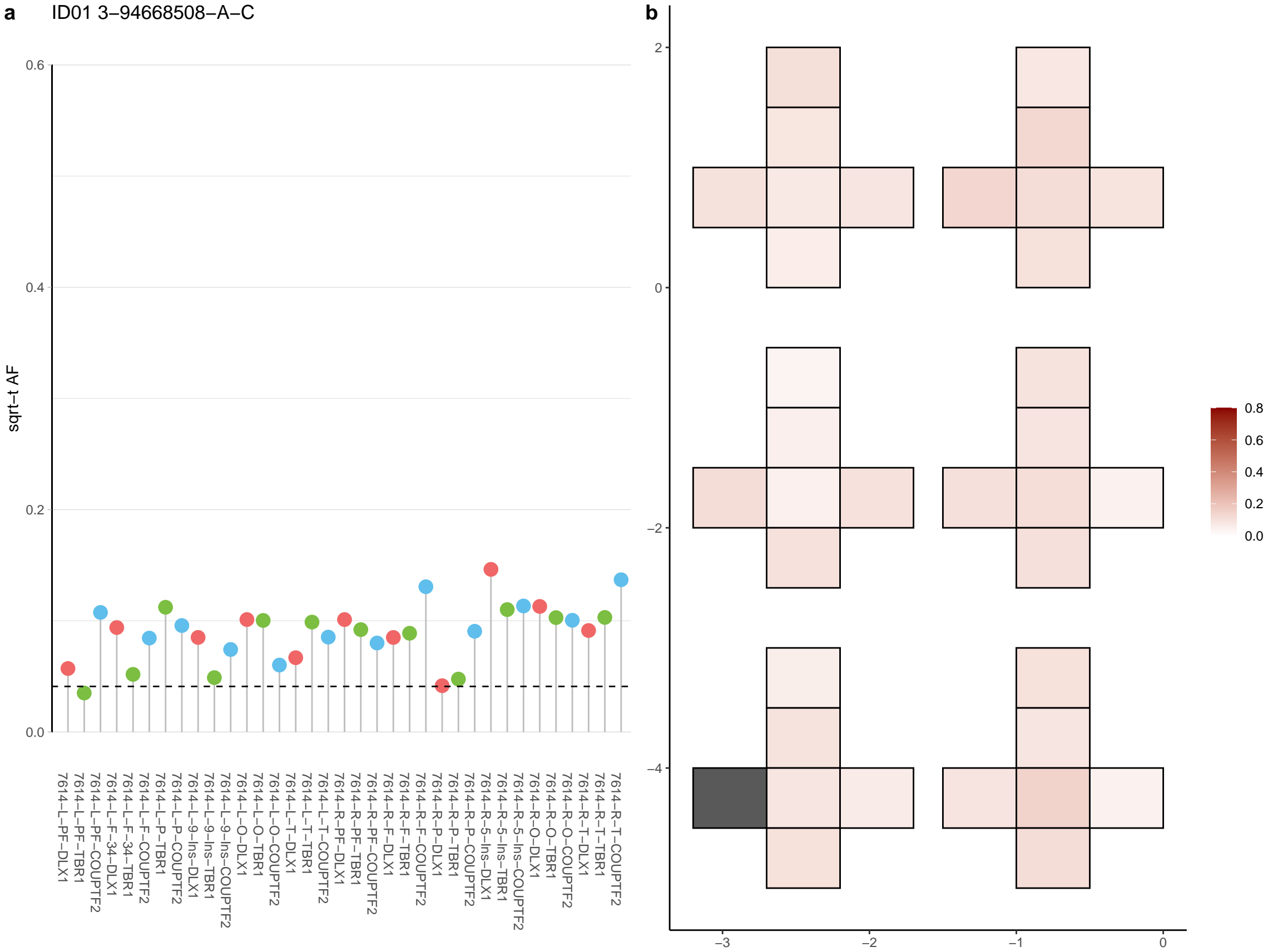

**a** ID01 3-96090255-C-T

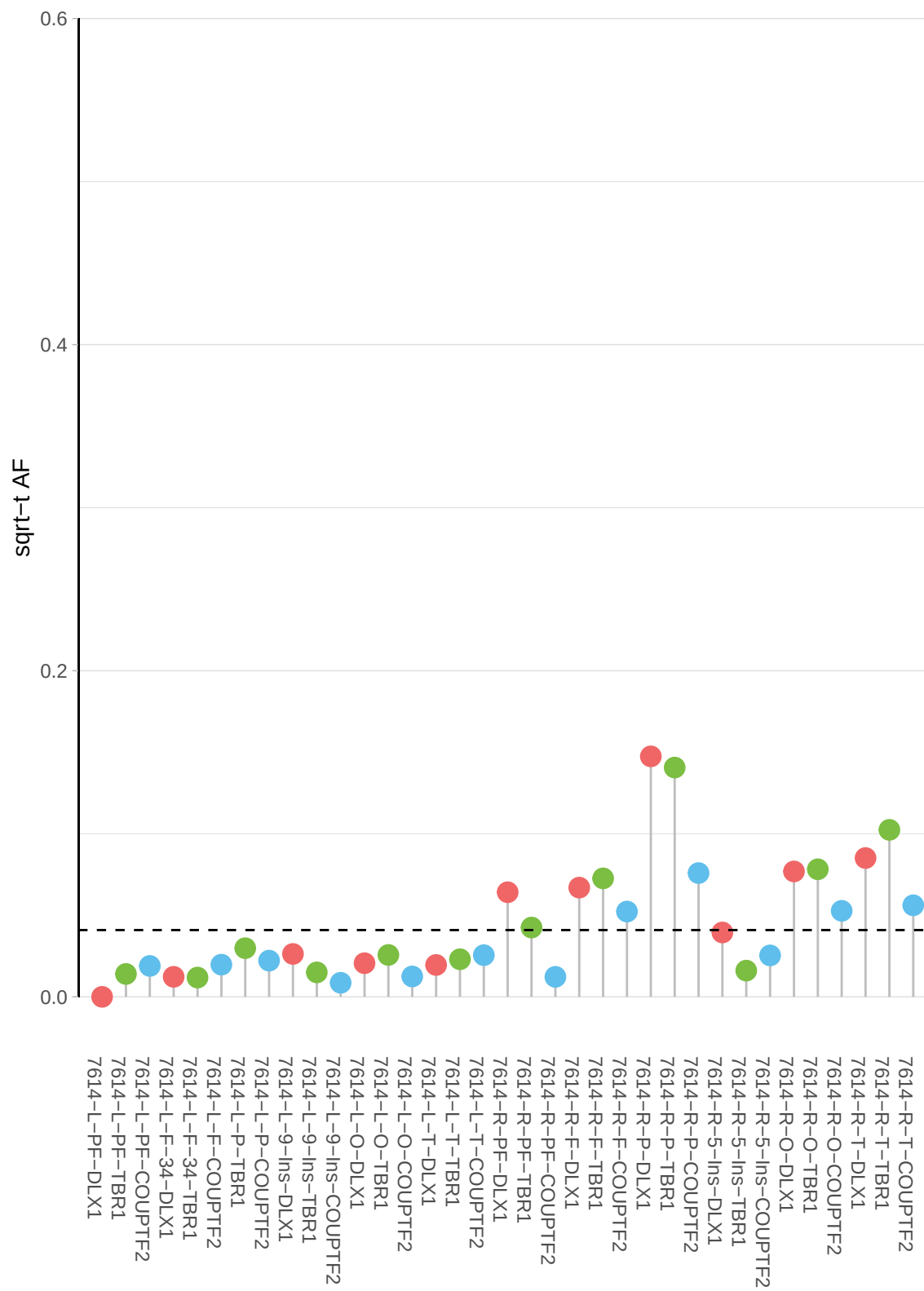

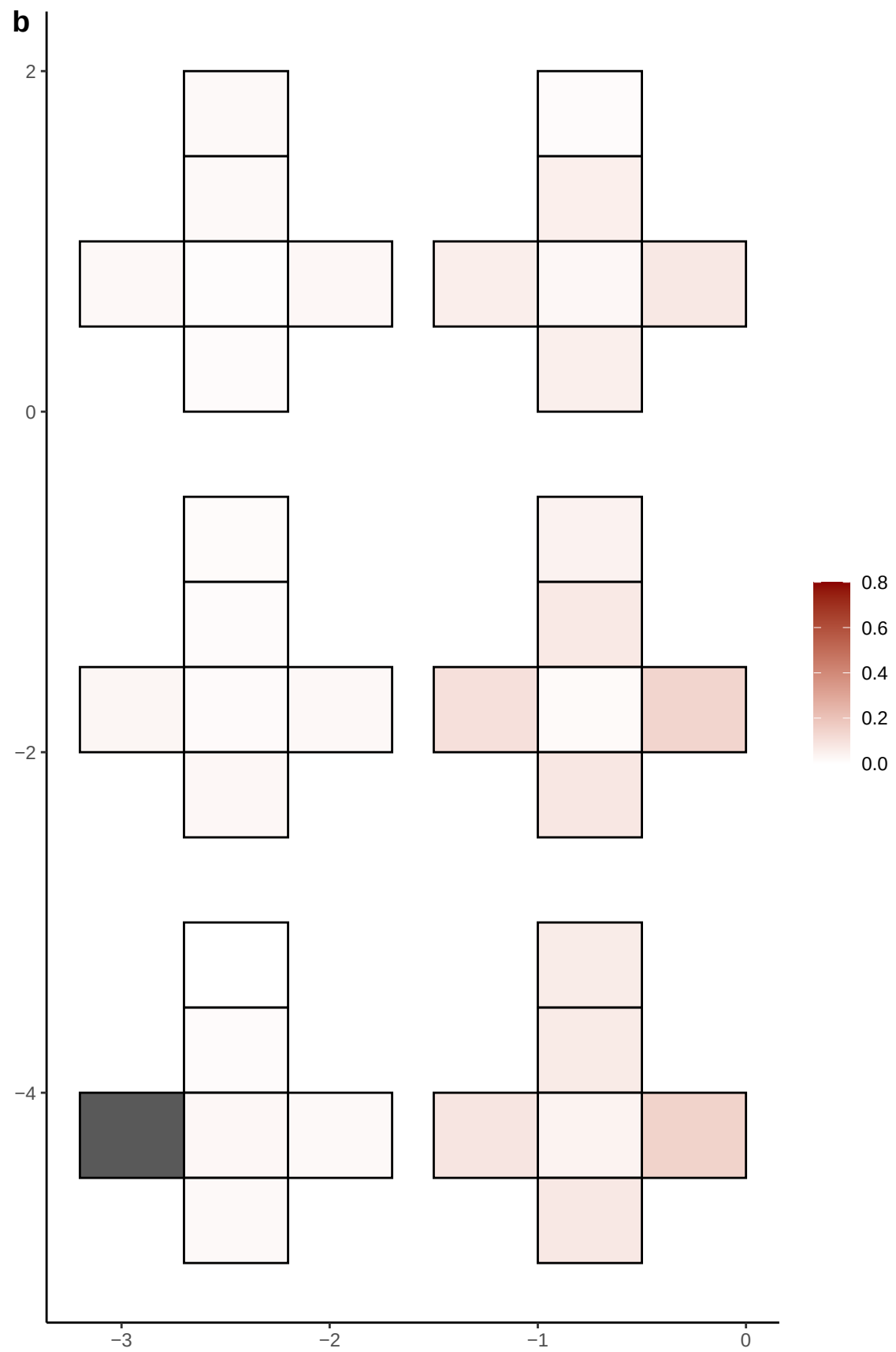

**a** ID01 4-130387276-A-G

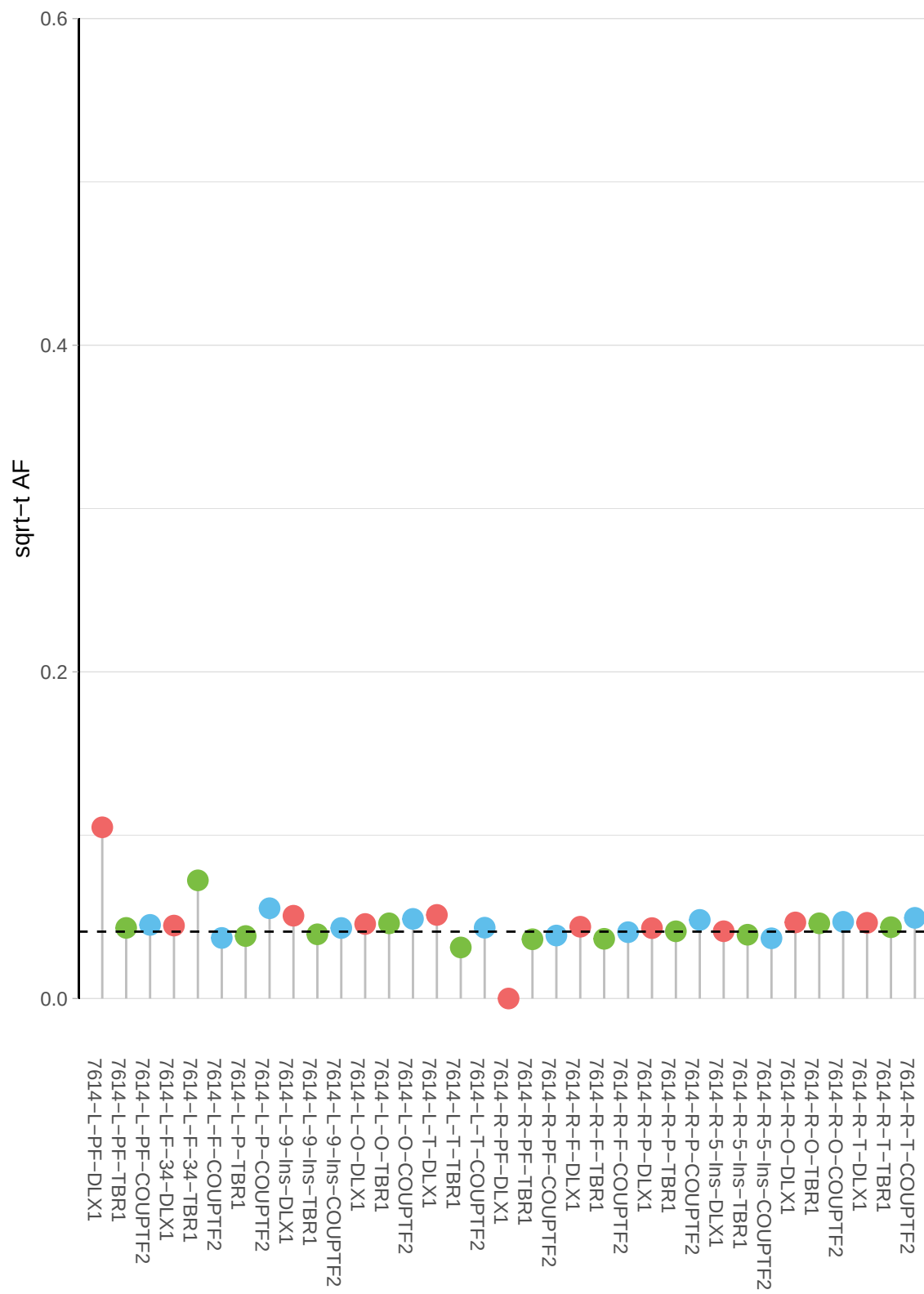

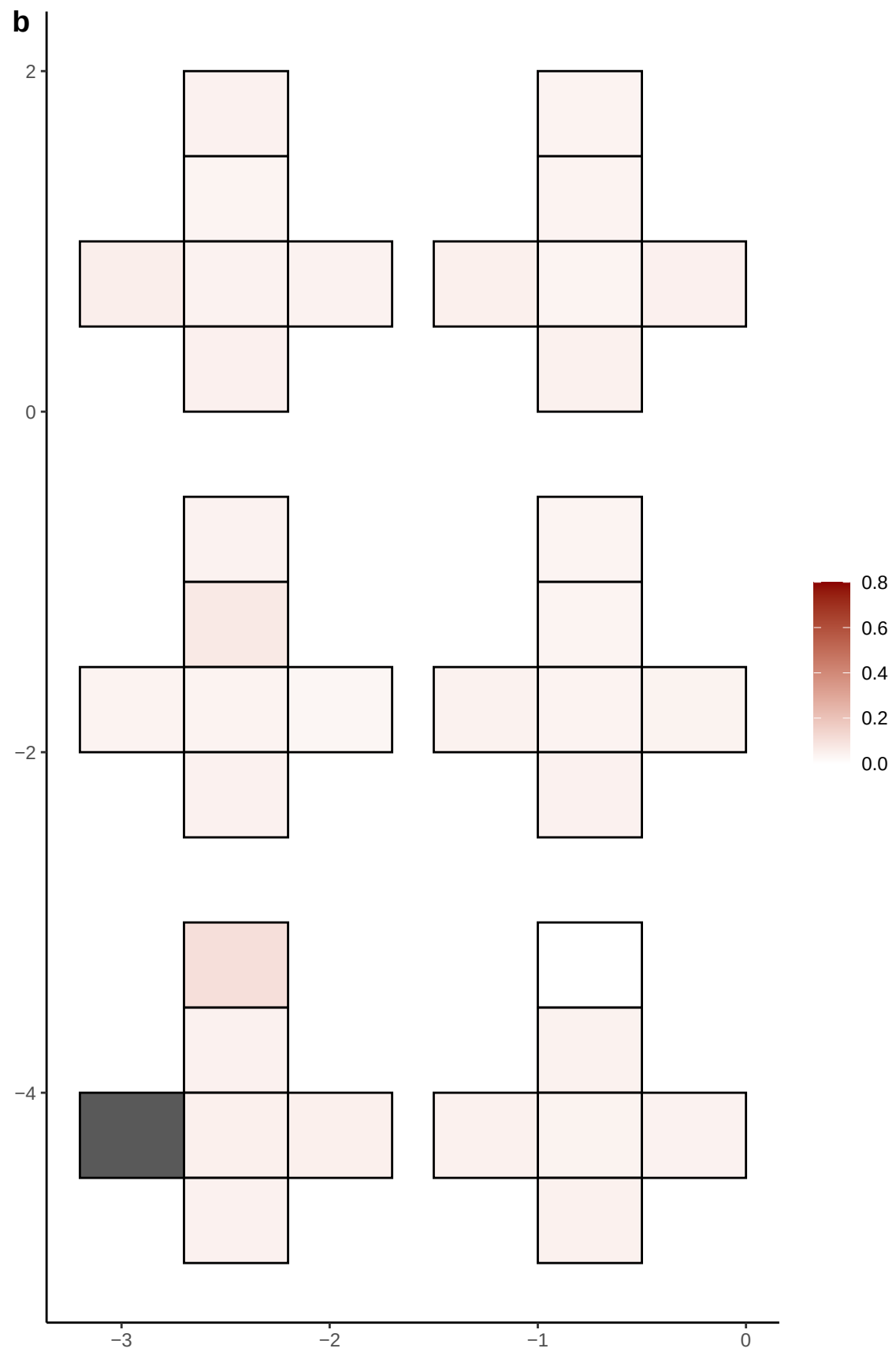

**a** ID01 4-149096018-G-A

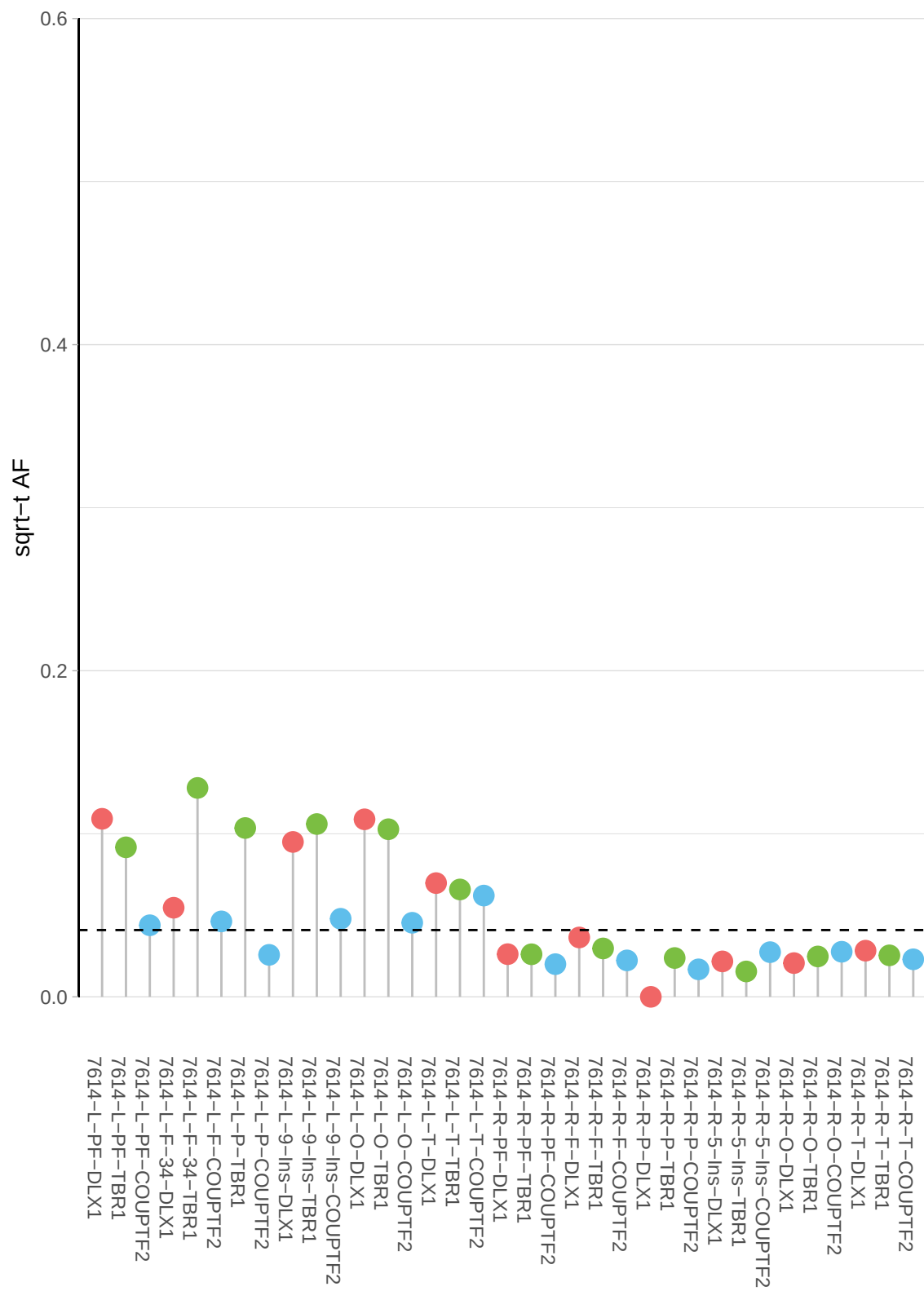

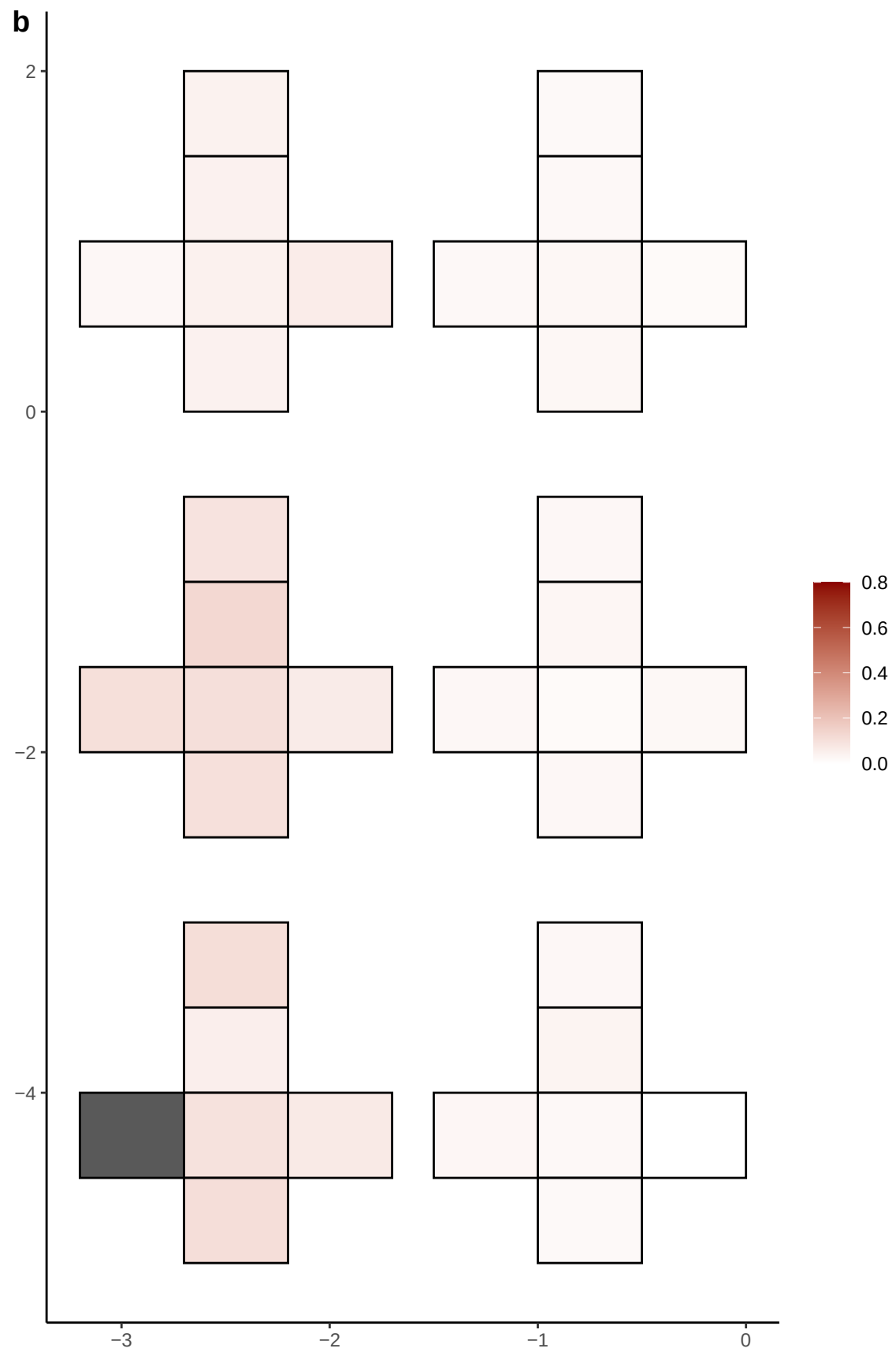

**a** ID01 4-149368759-C-T

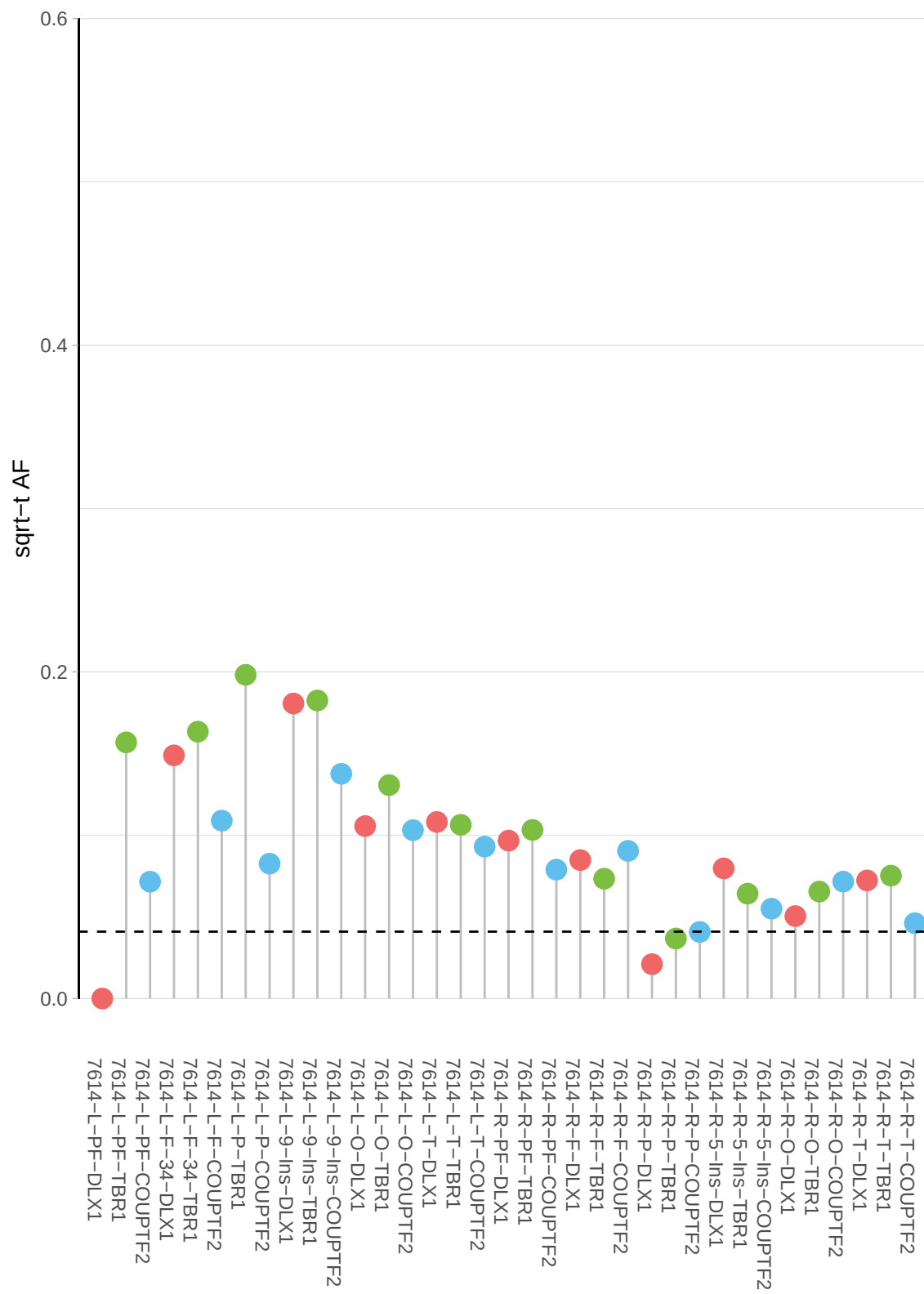

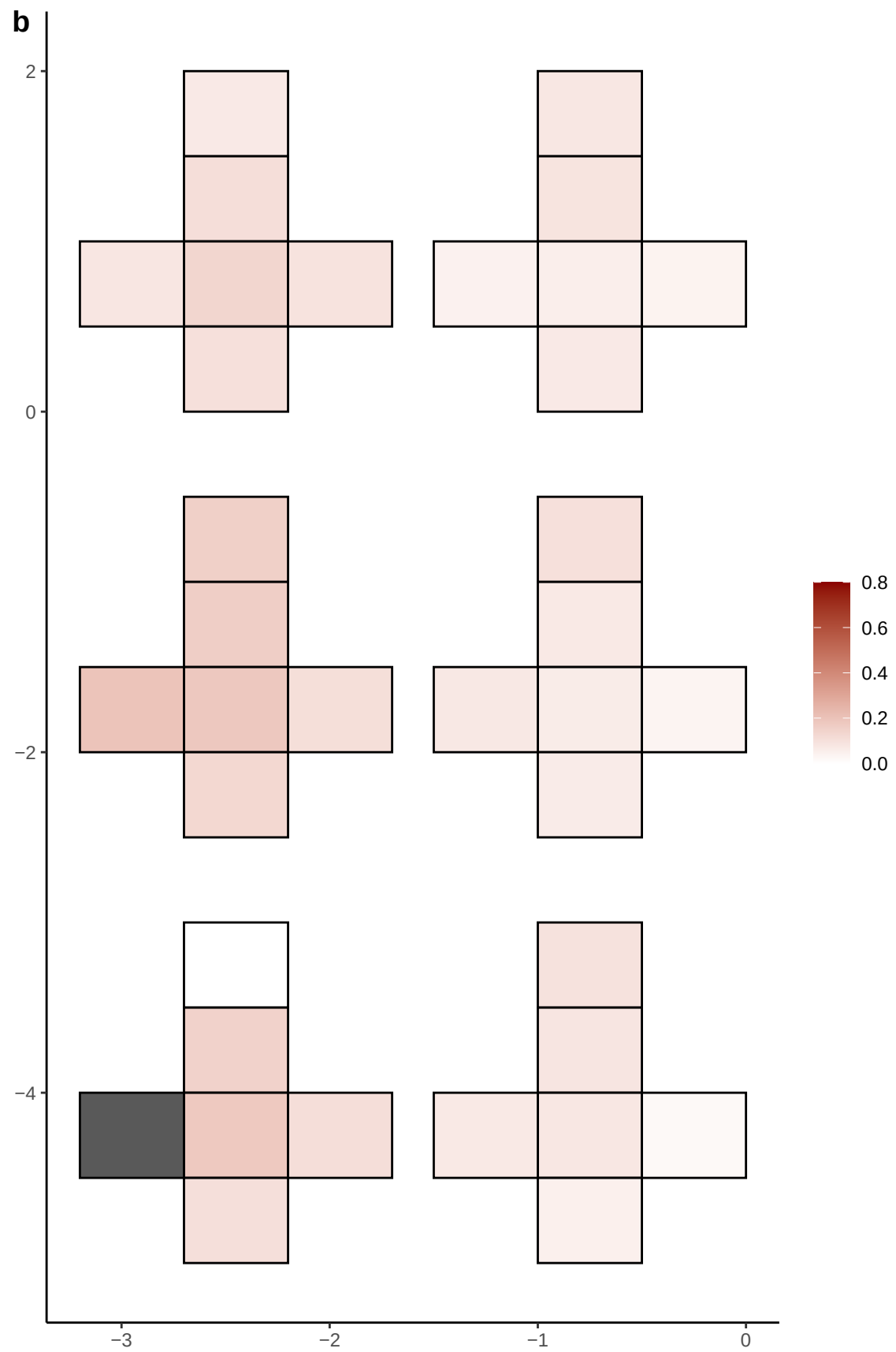

**a** ID01 4-159172058-G-A

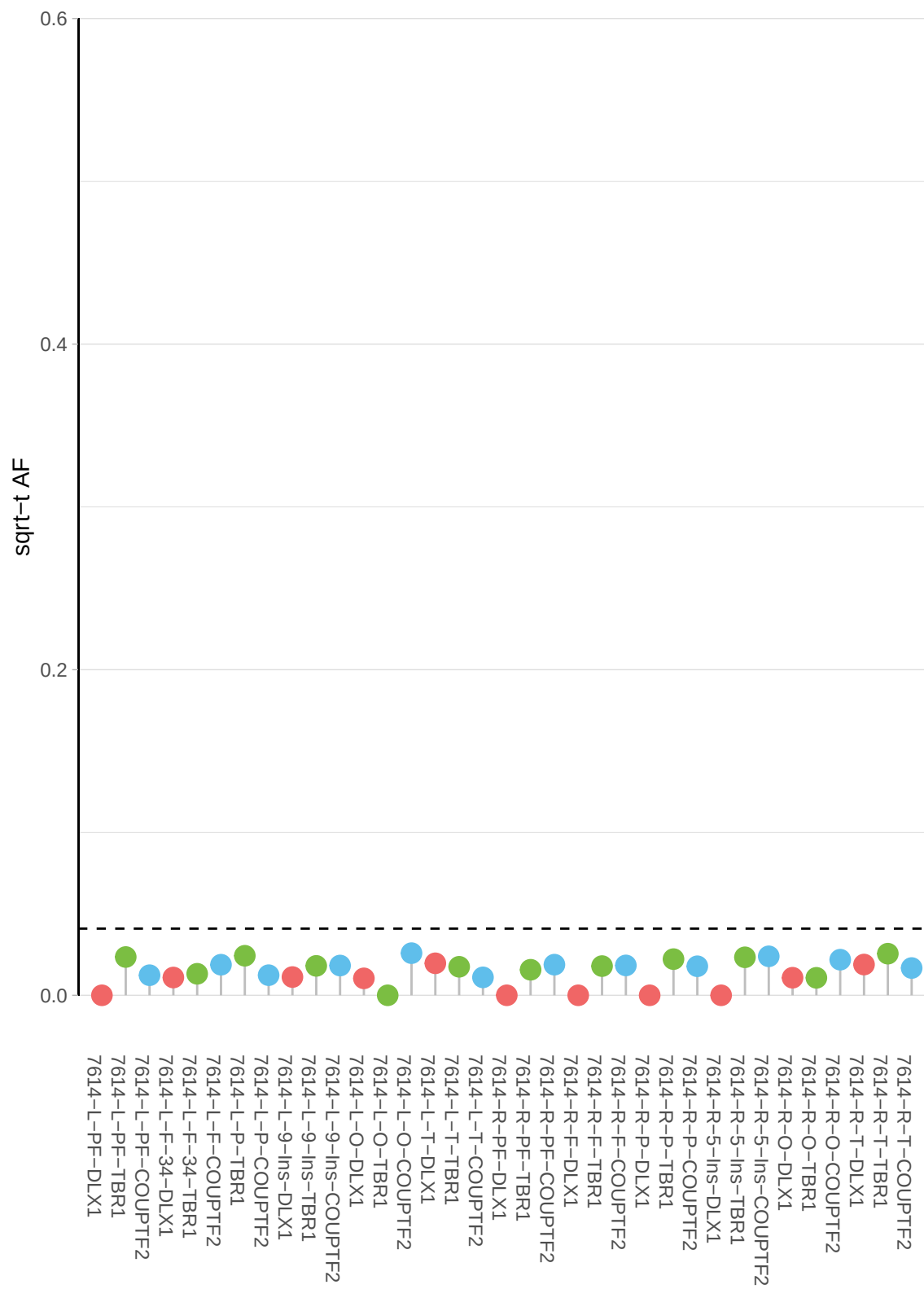

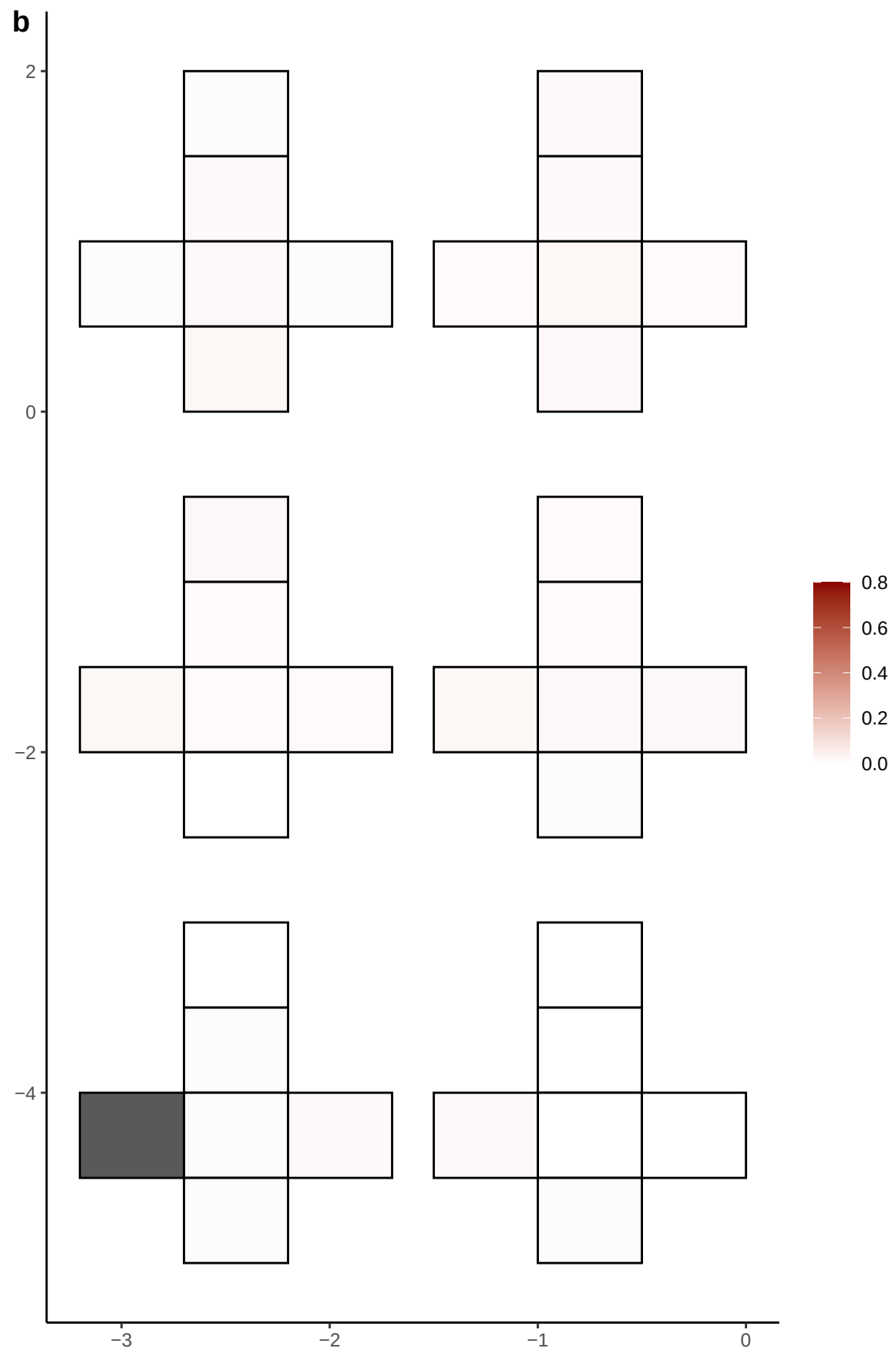

**a** ID01 4-163244475-A-T

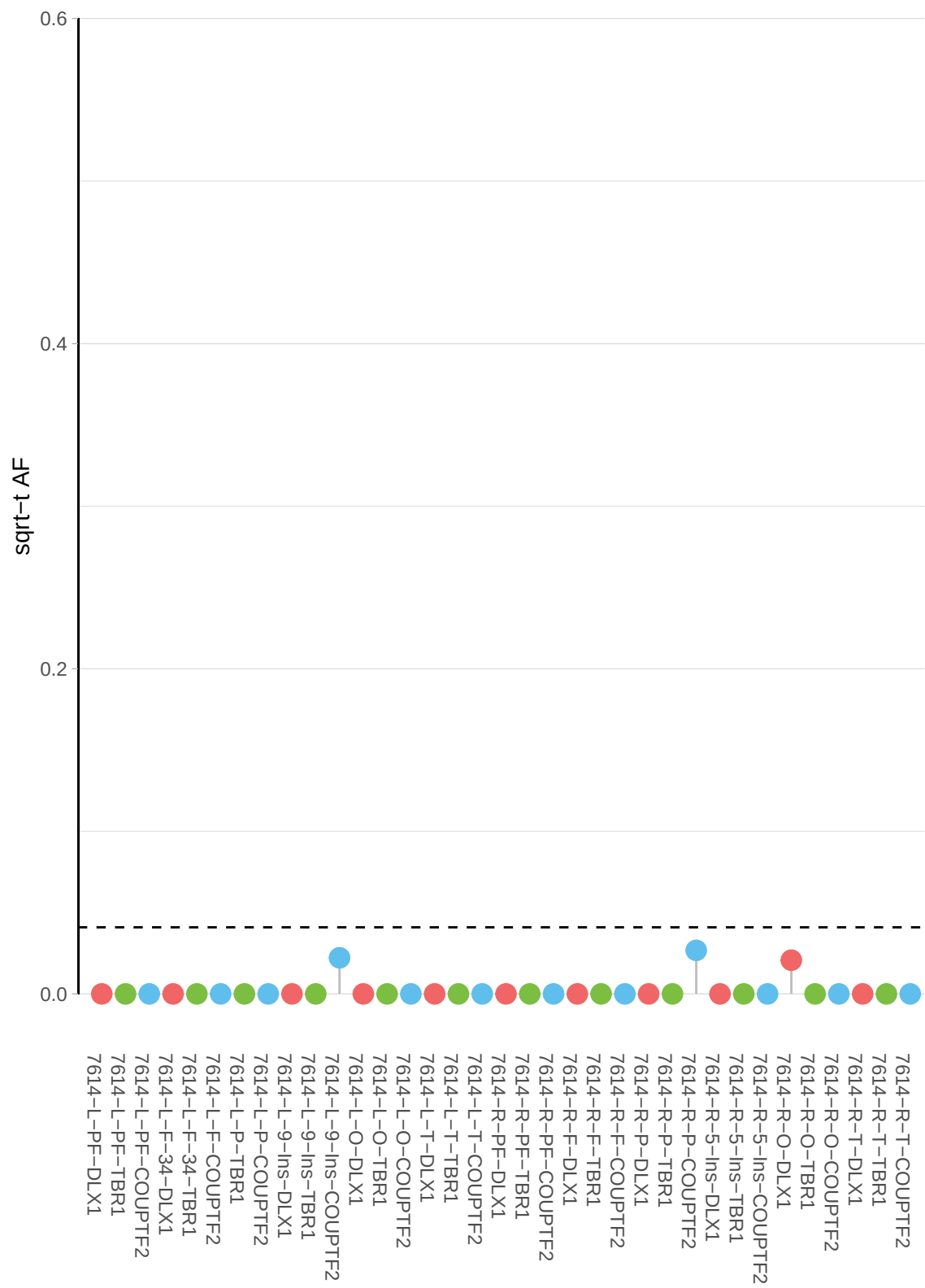

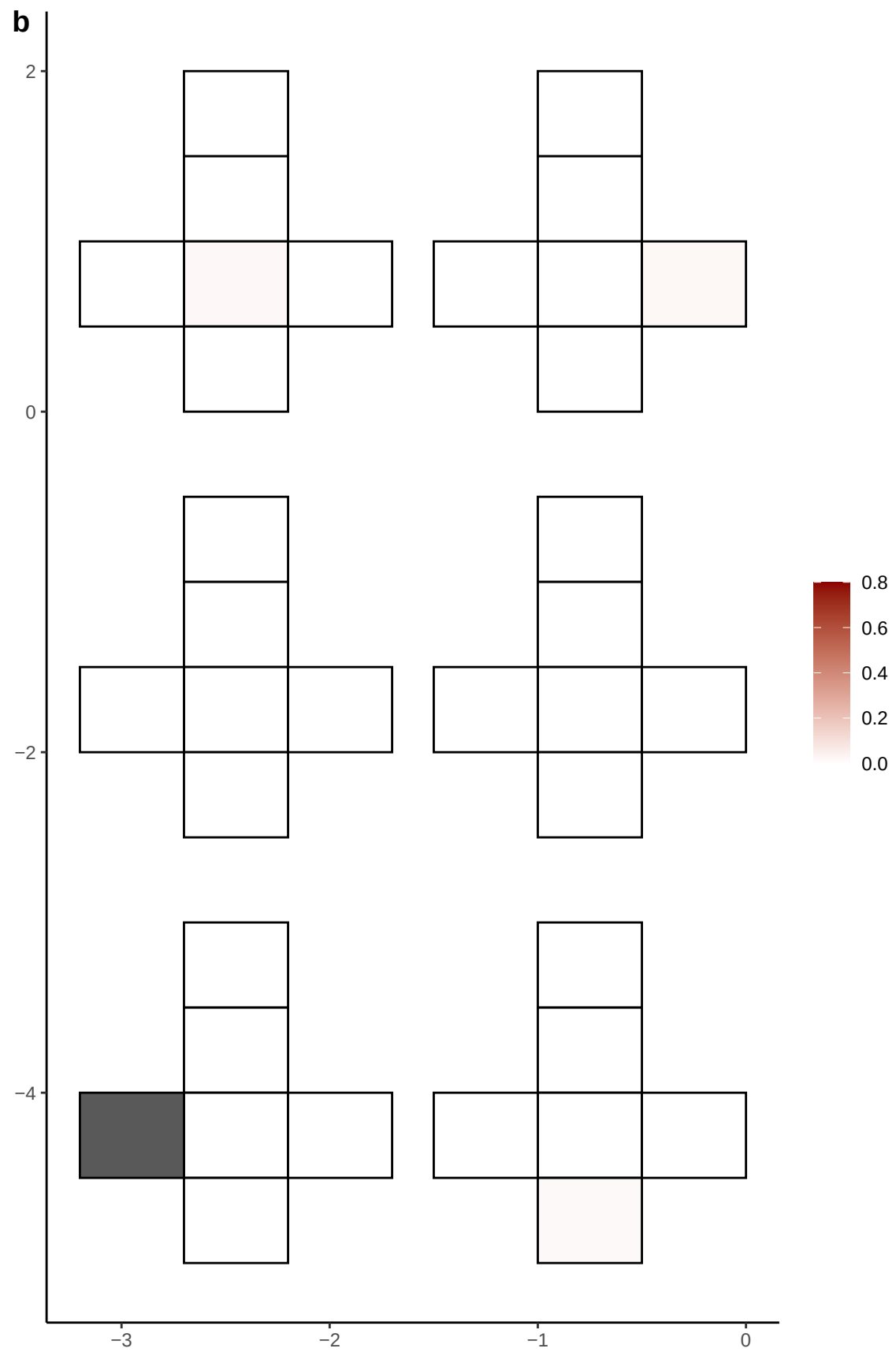

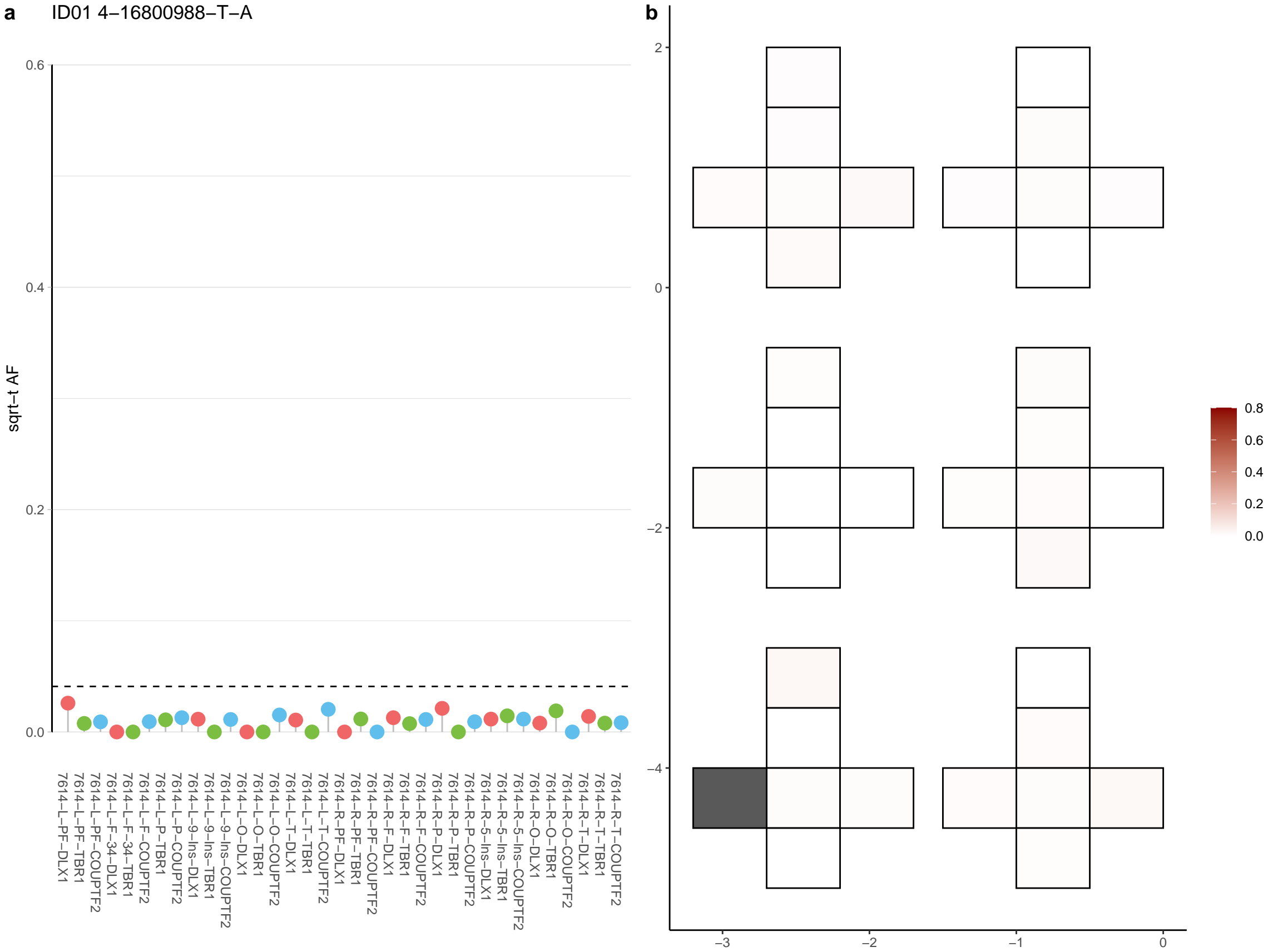

**a** ID01 4-176659497-G-A

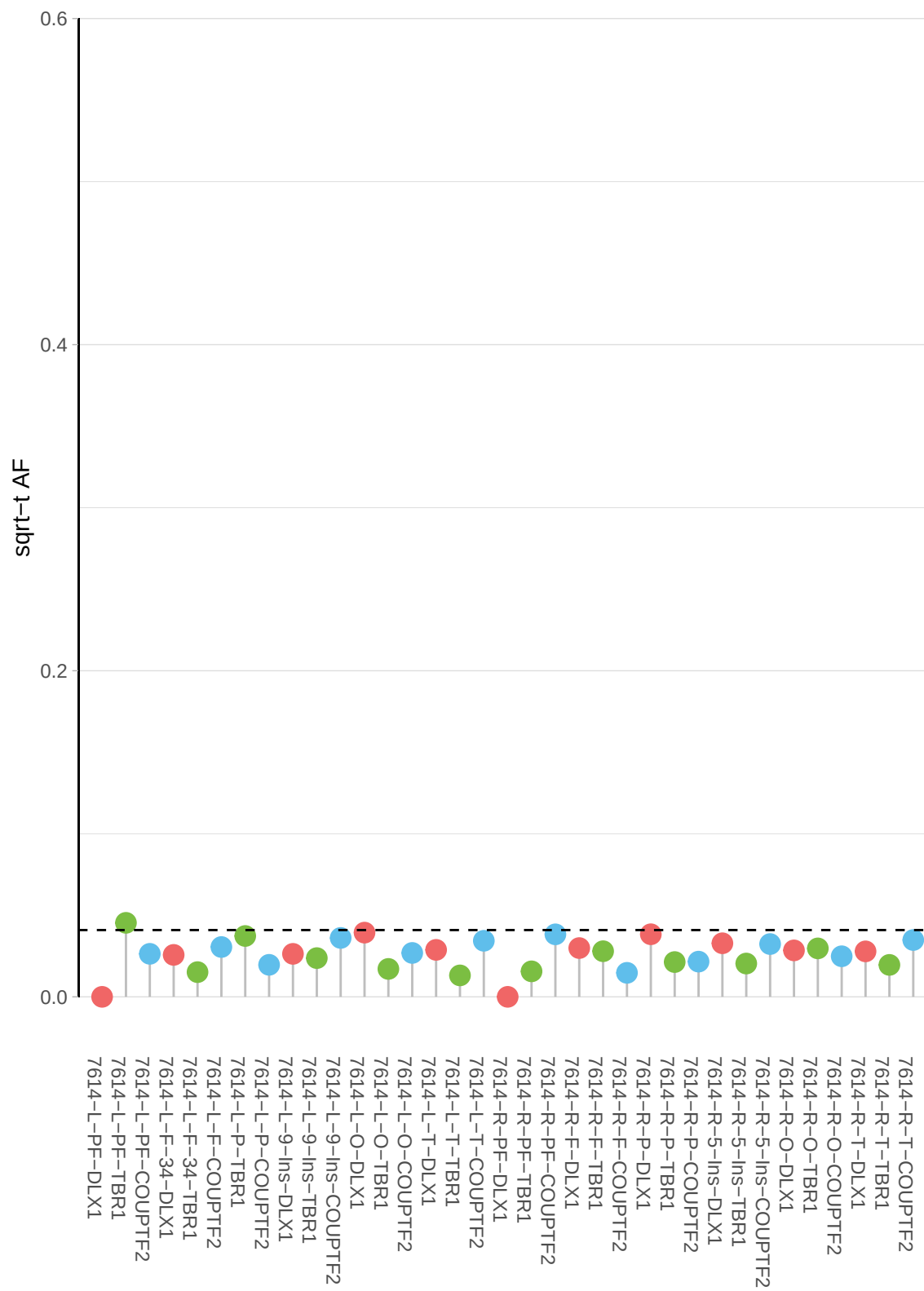

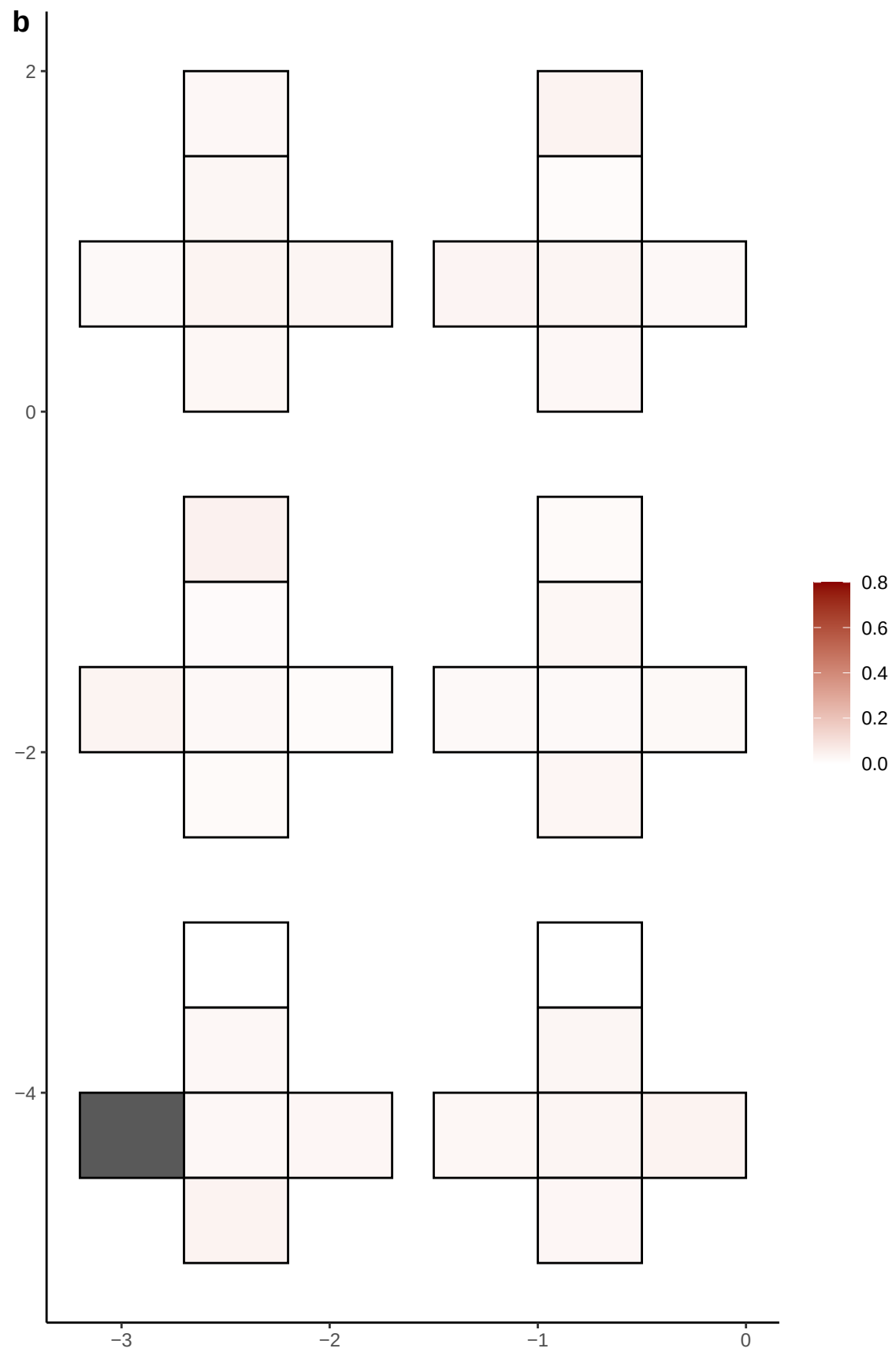

**a** ID01 4-181698552-T-C

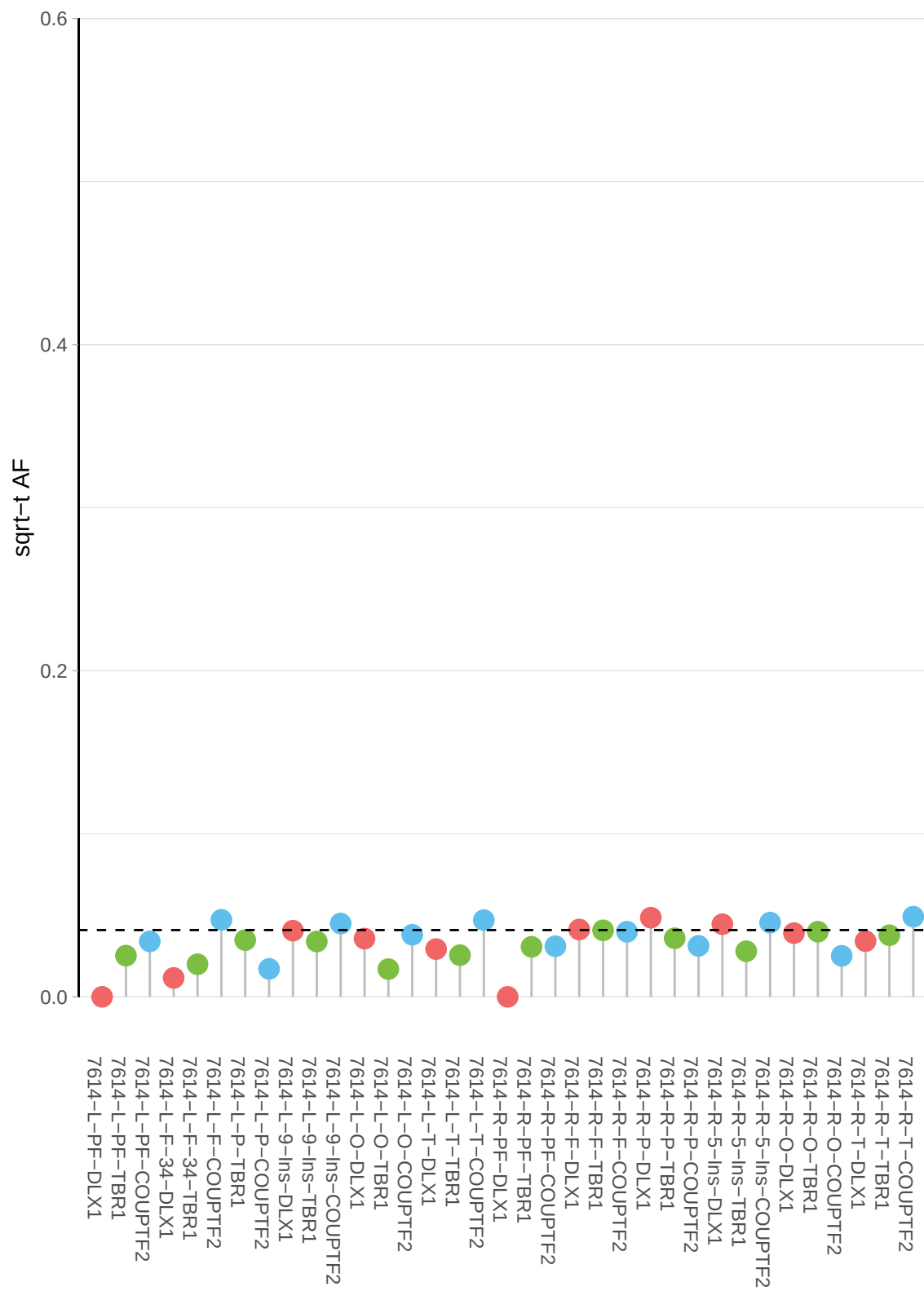

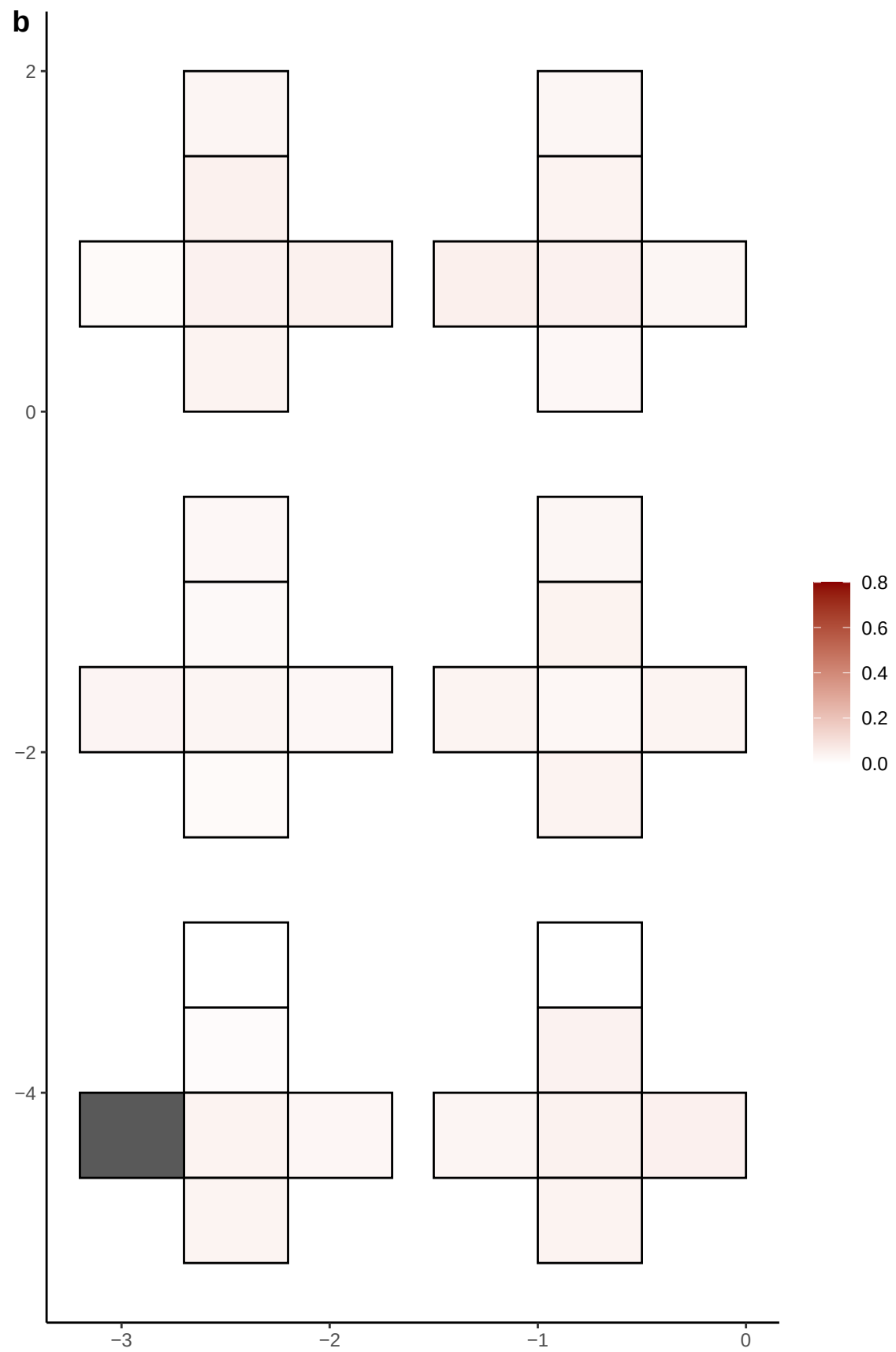

**a** ID01 4-184808022-A-G

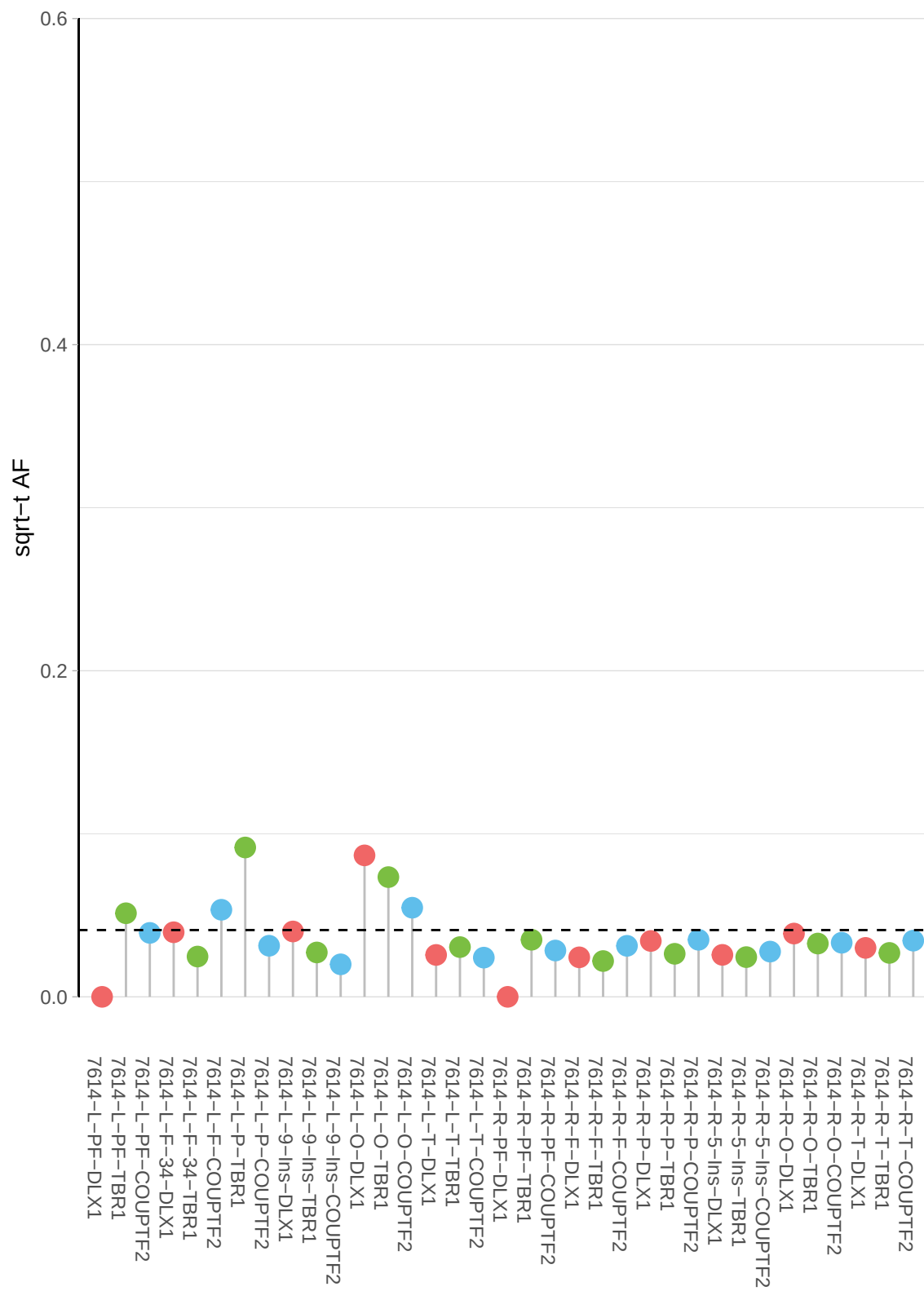

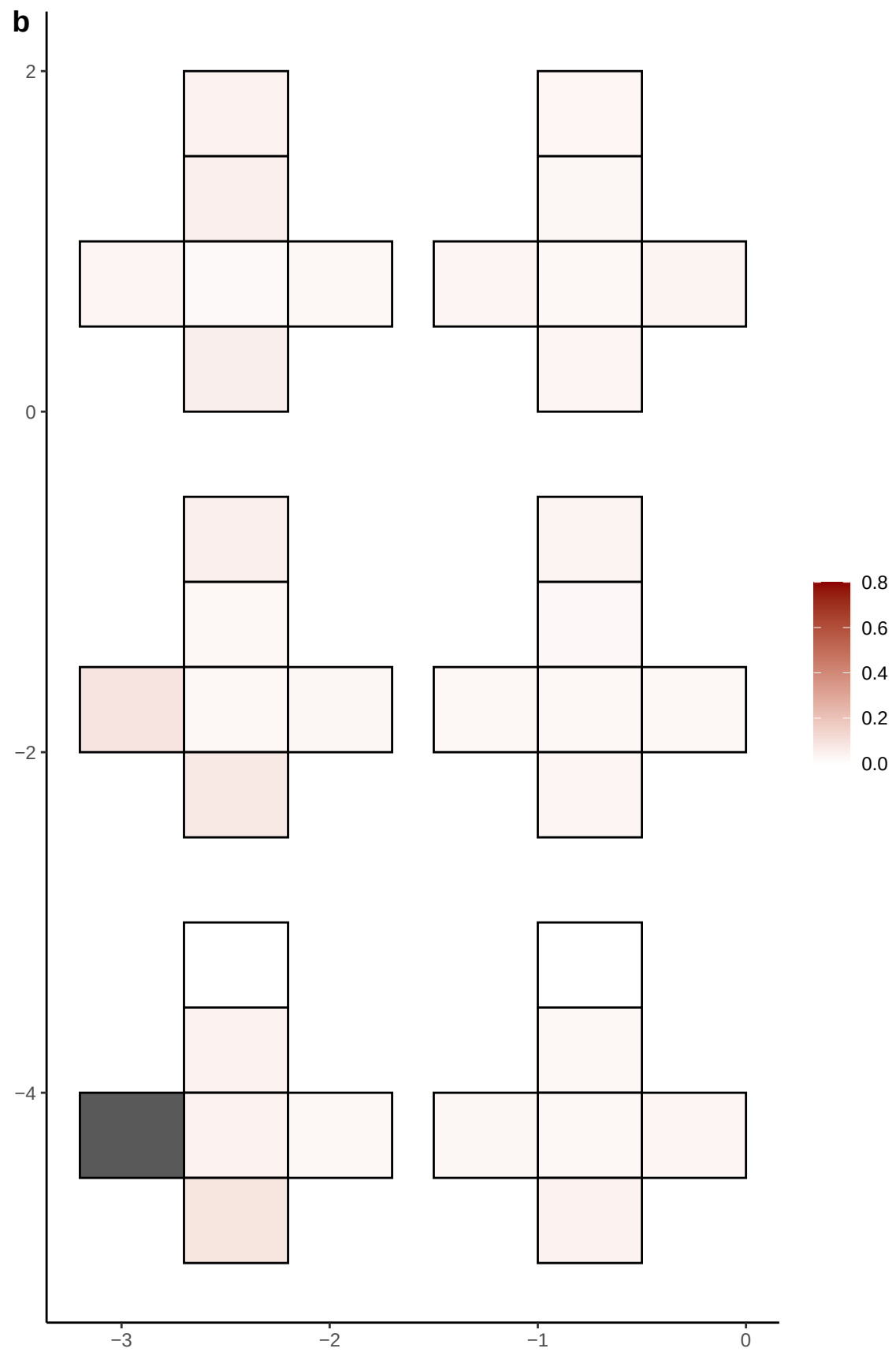

**a** ID01 4-185375793-C-T

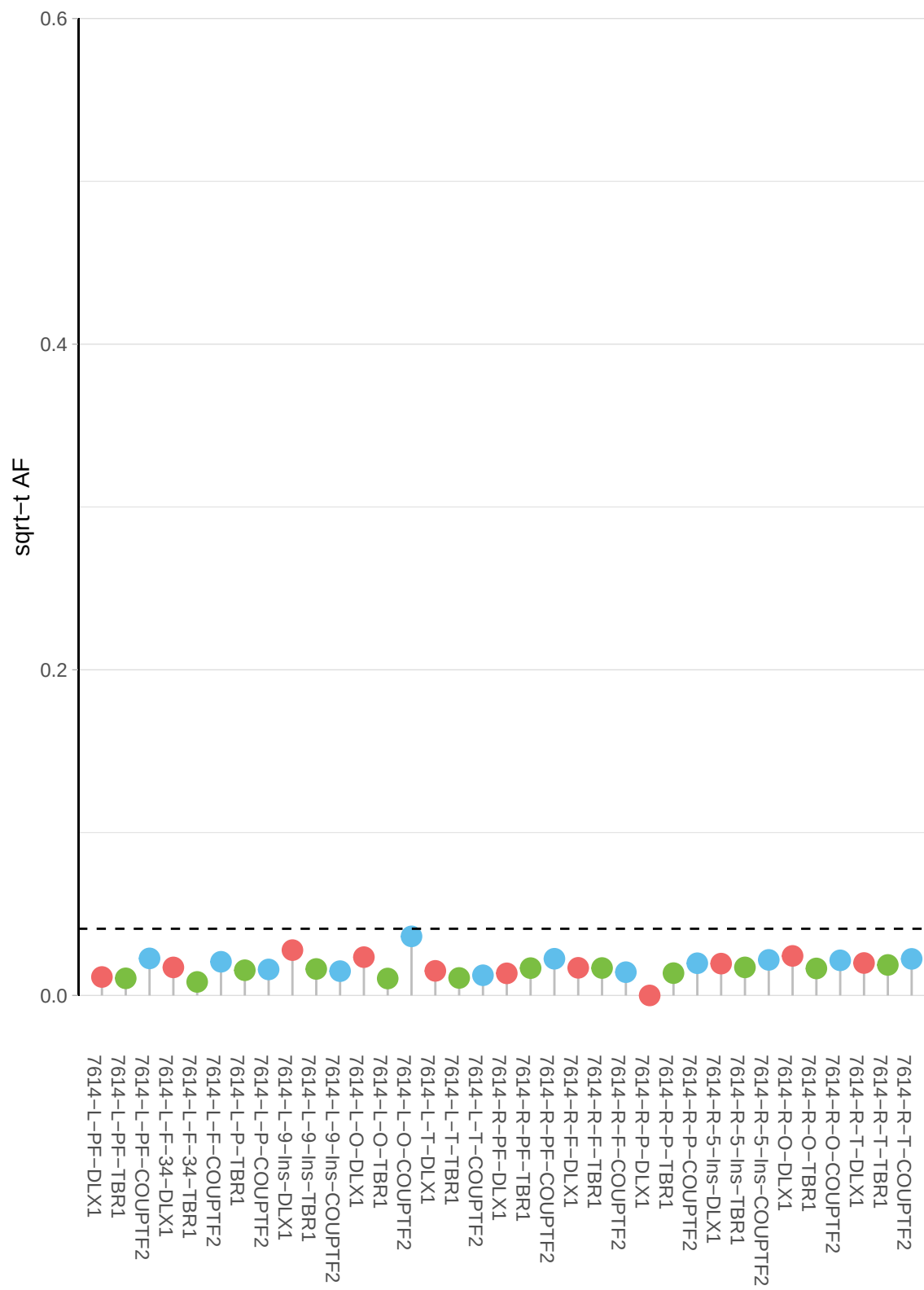

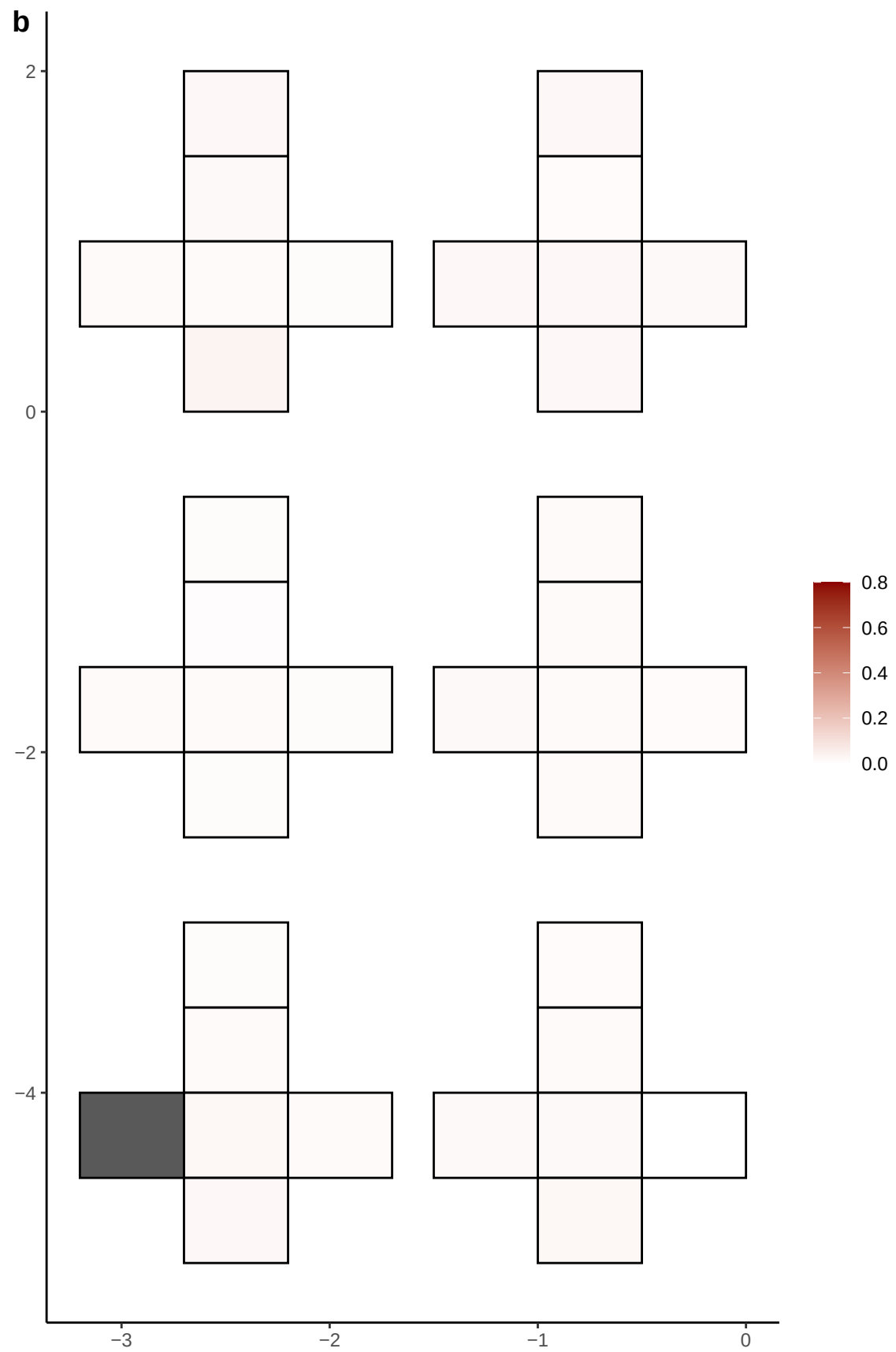

**a** ID01 4-189162568-G-A

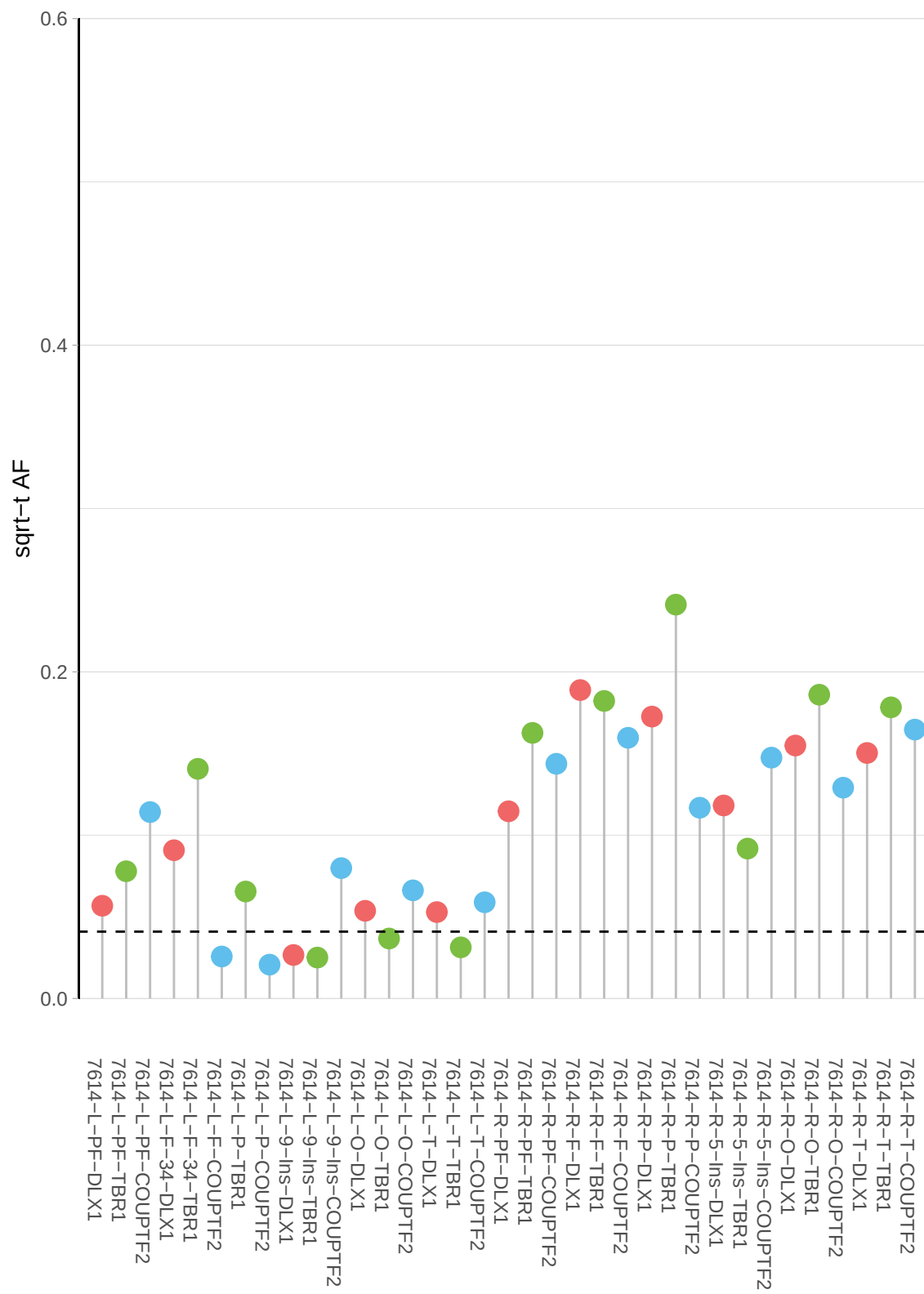

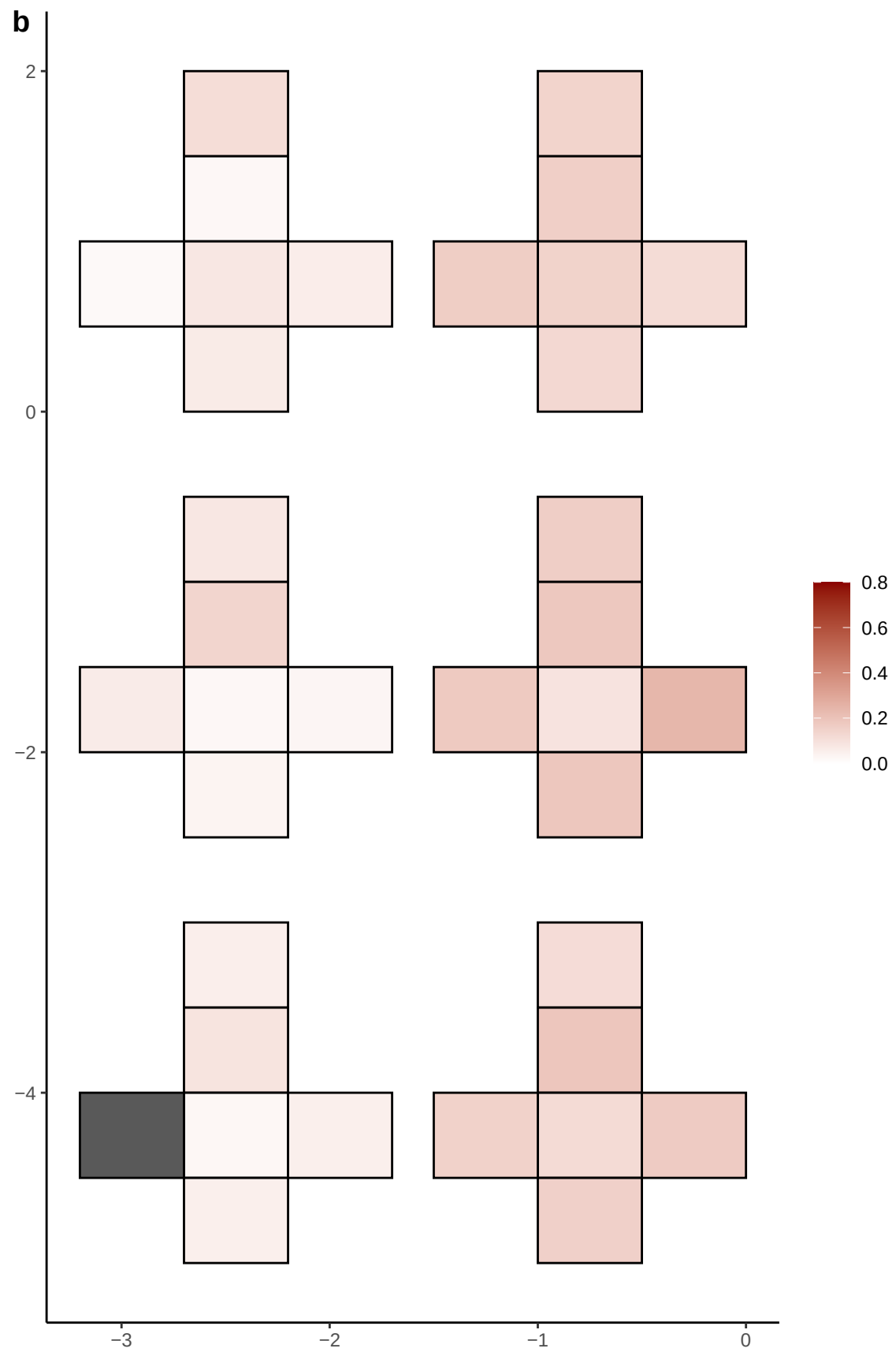

**a** ID01 4-28629401-C-T

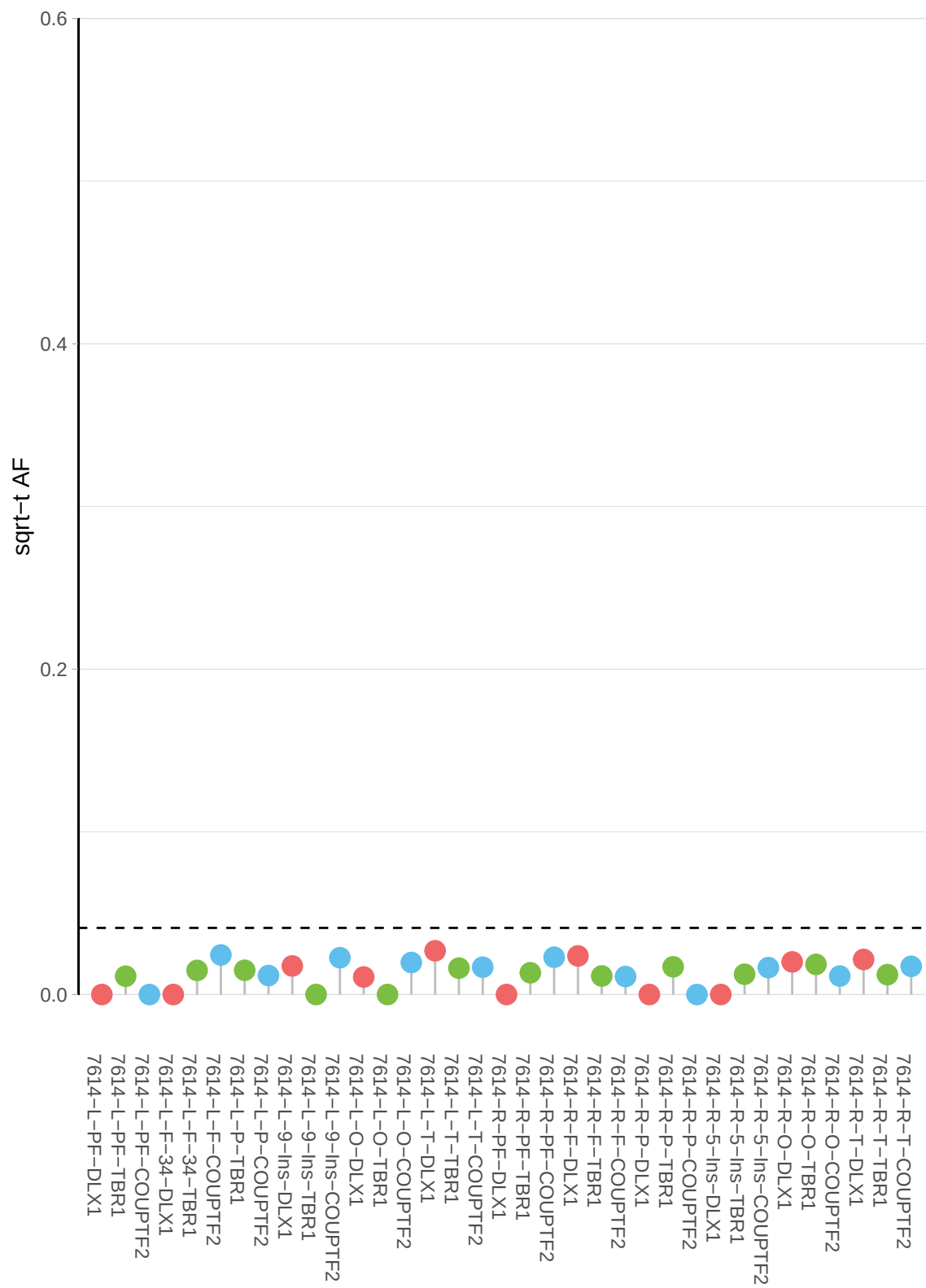

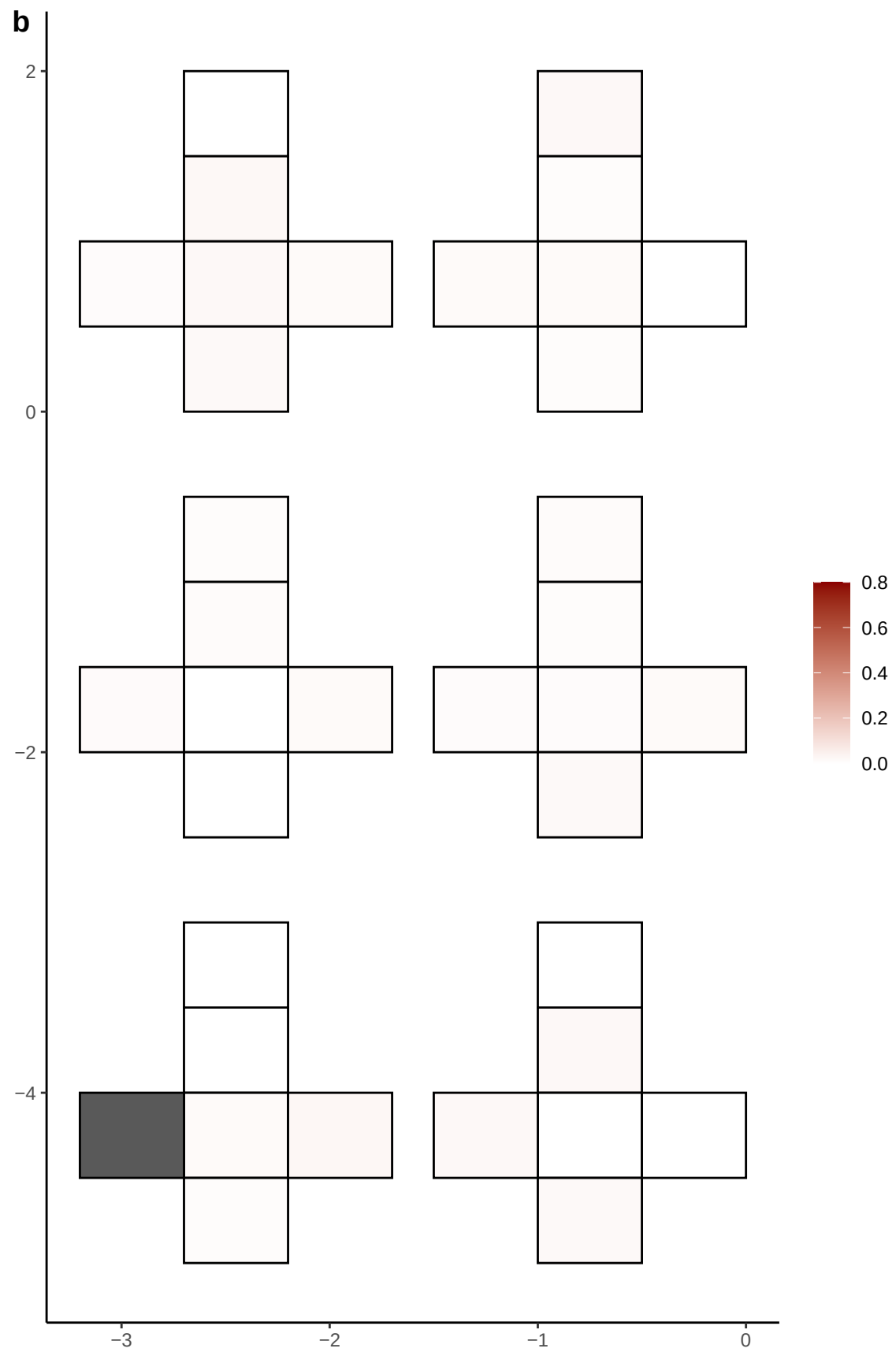

**a** ID01 4-33192479-C-A

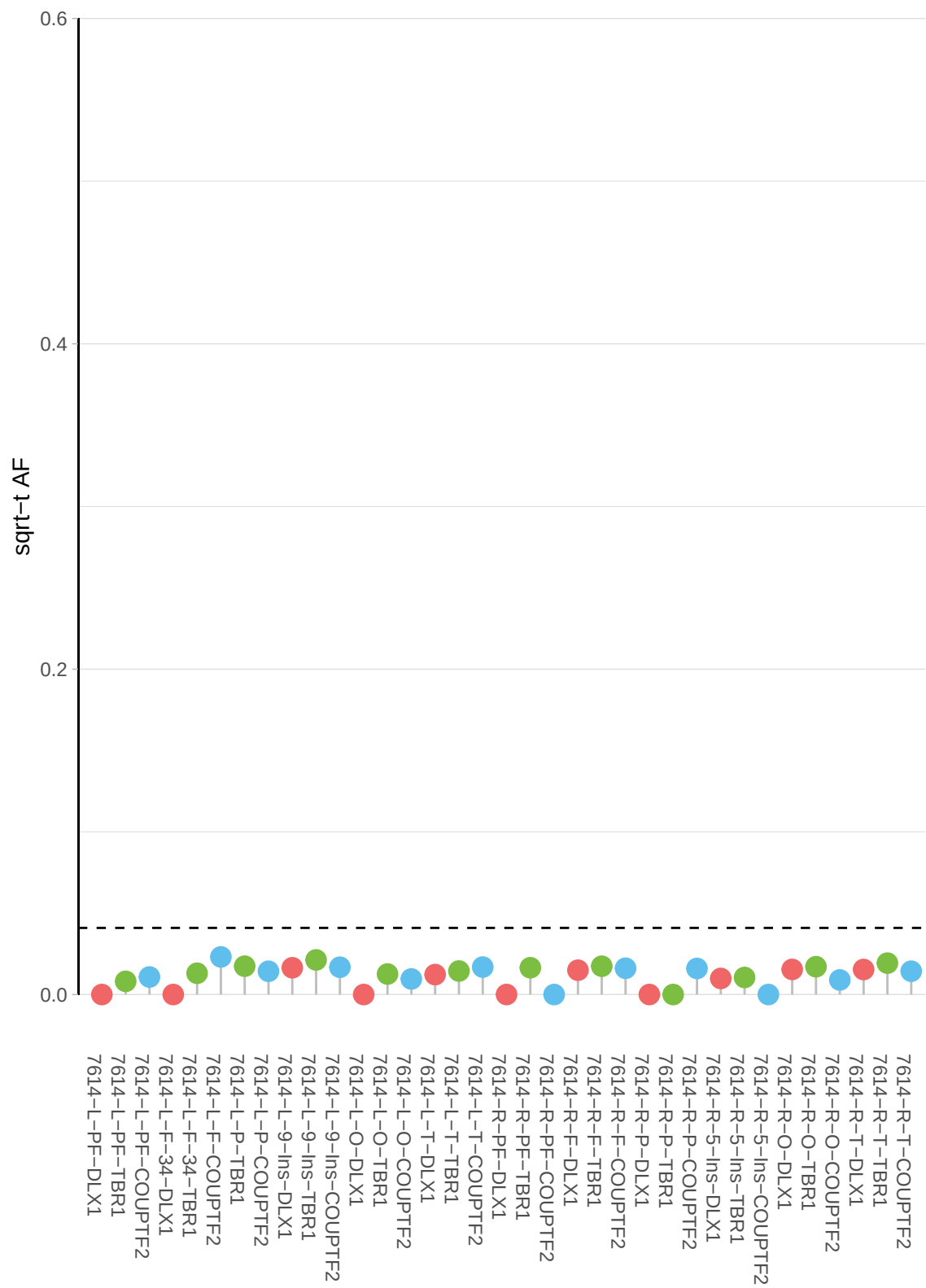

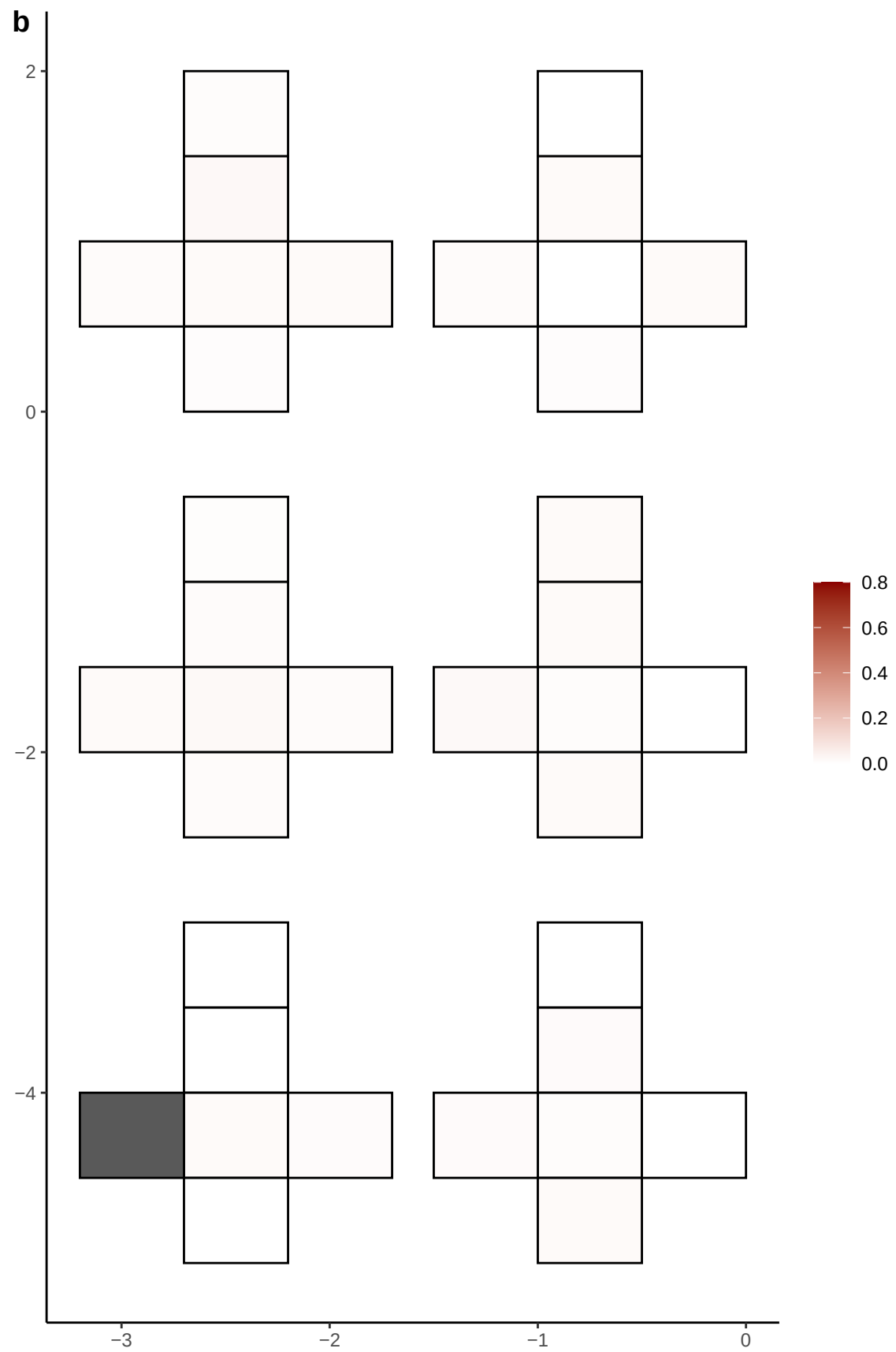

**a** ID01 4-55310609-G-A

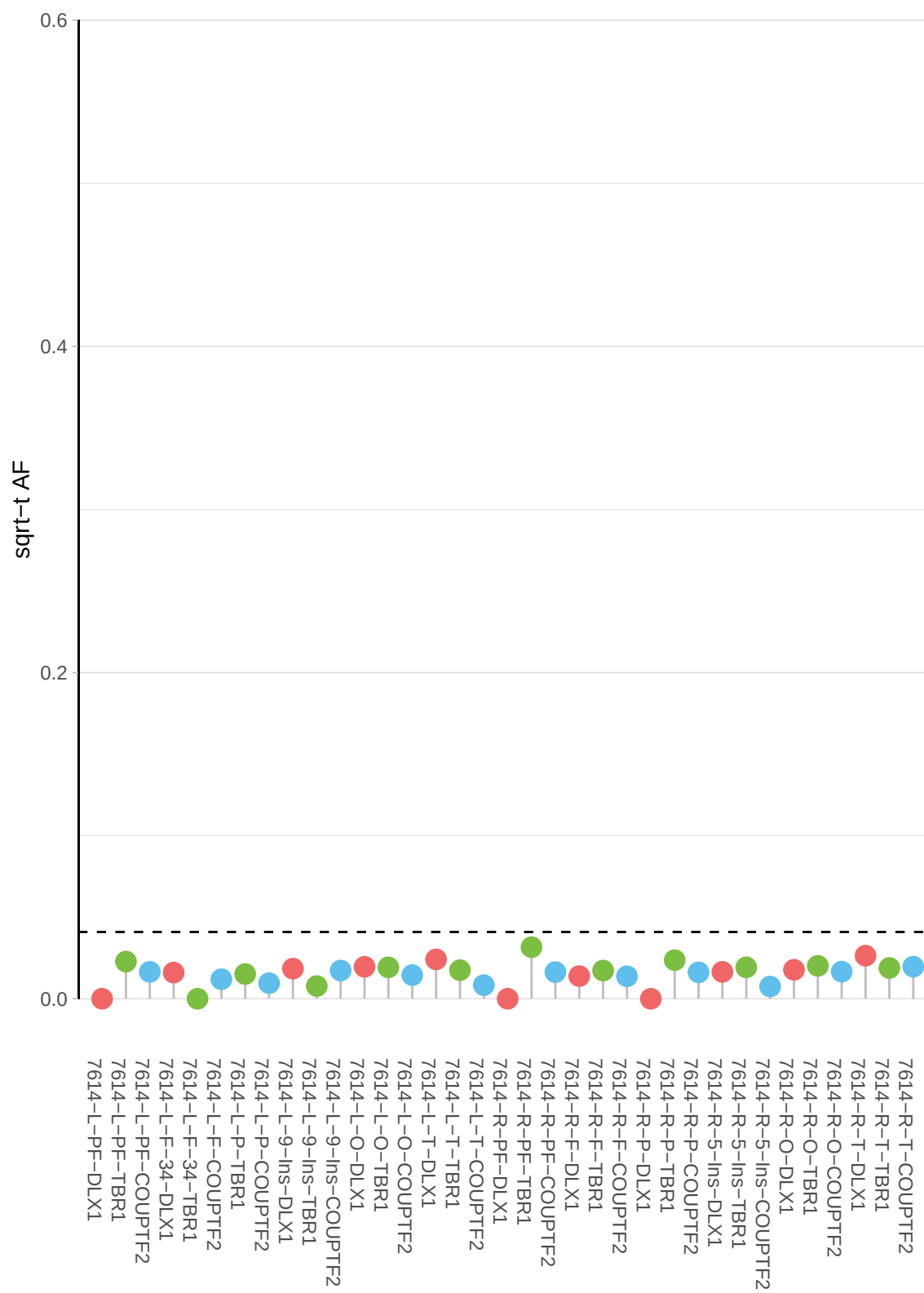

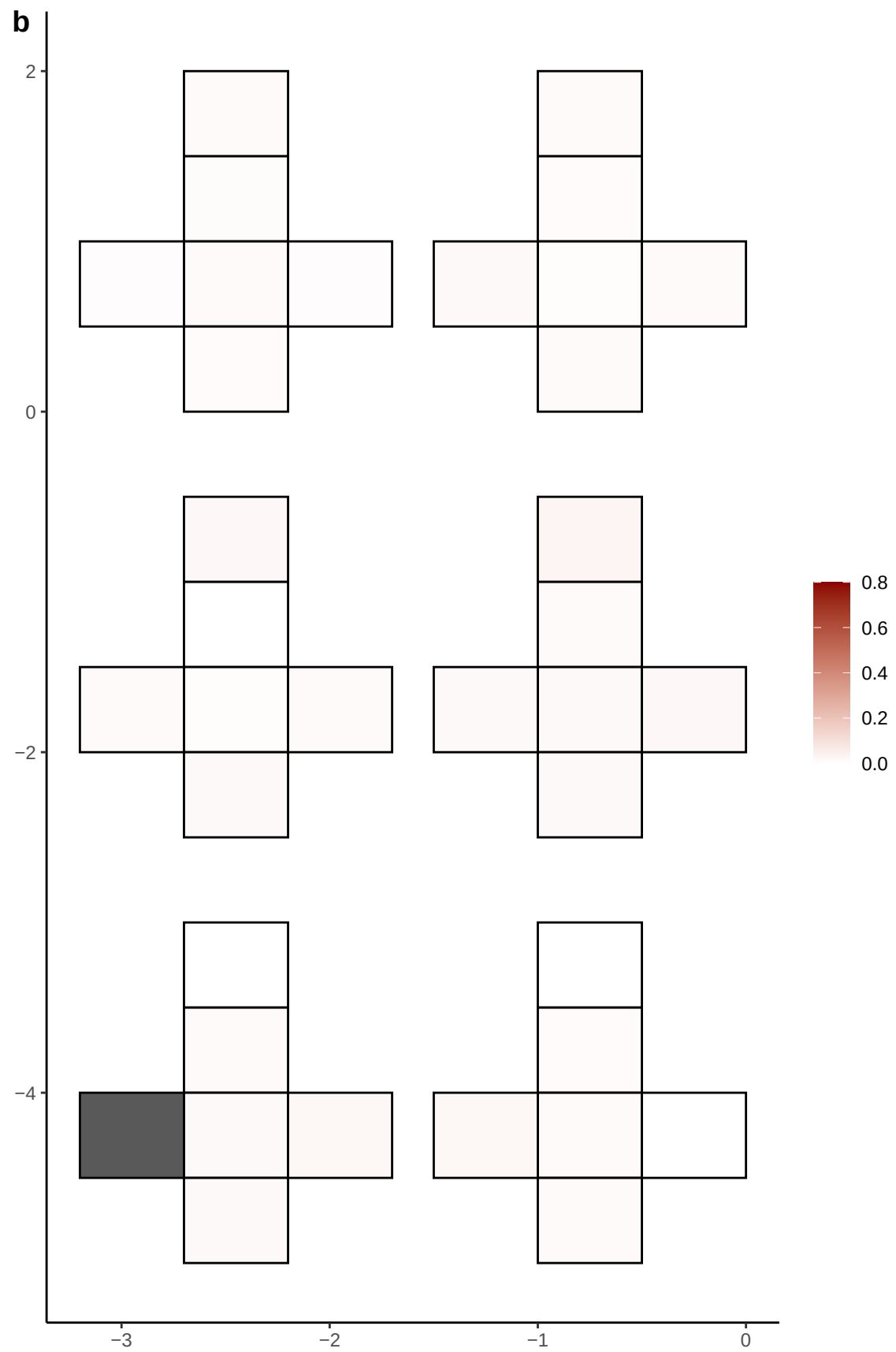

**a** ID01 4-55312148-T-C

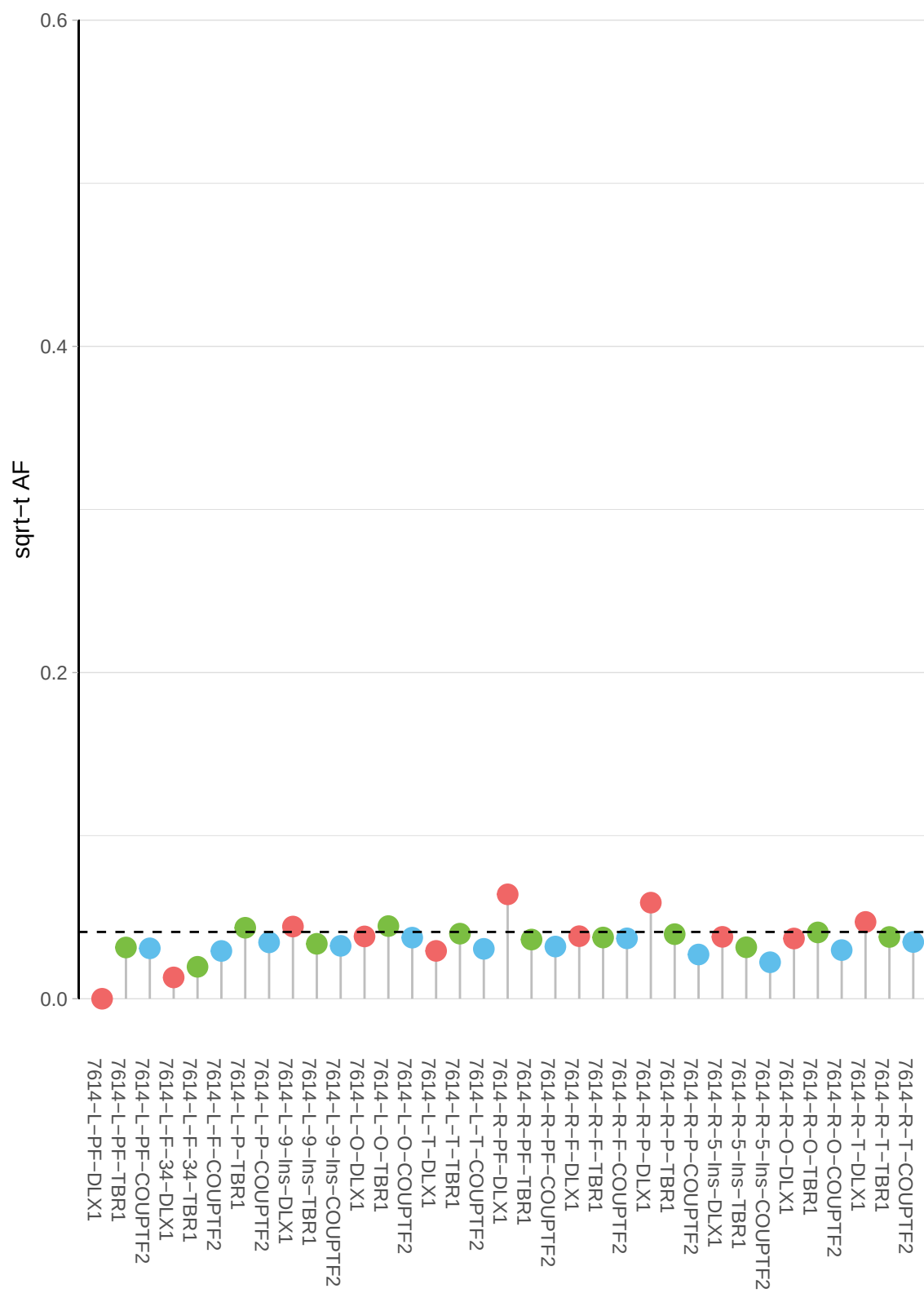

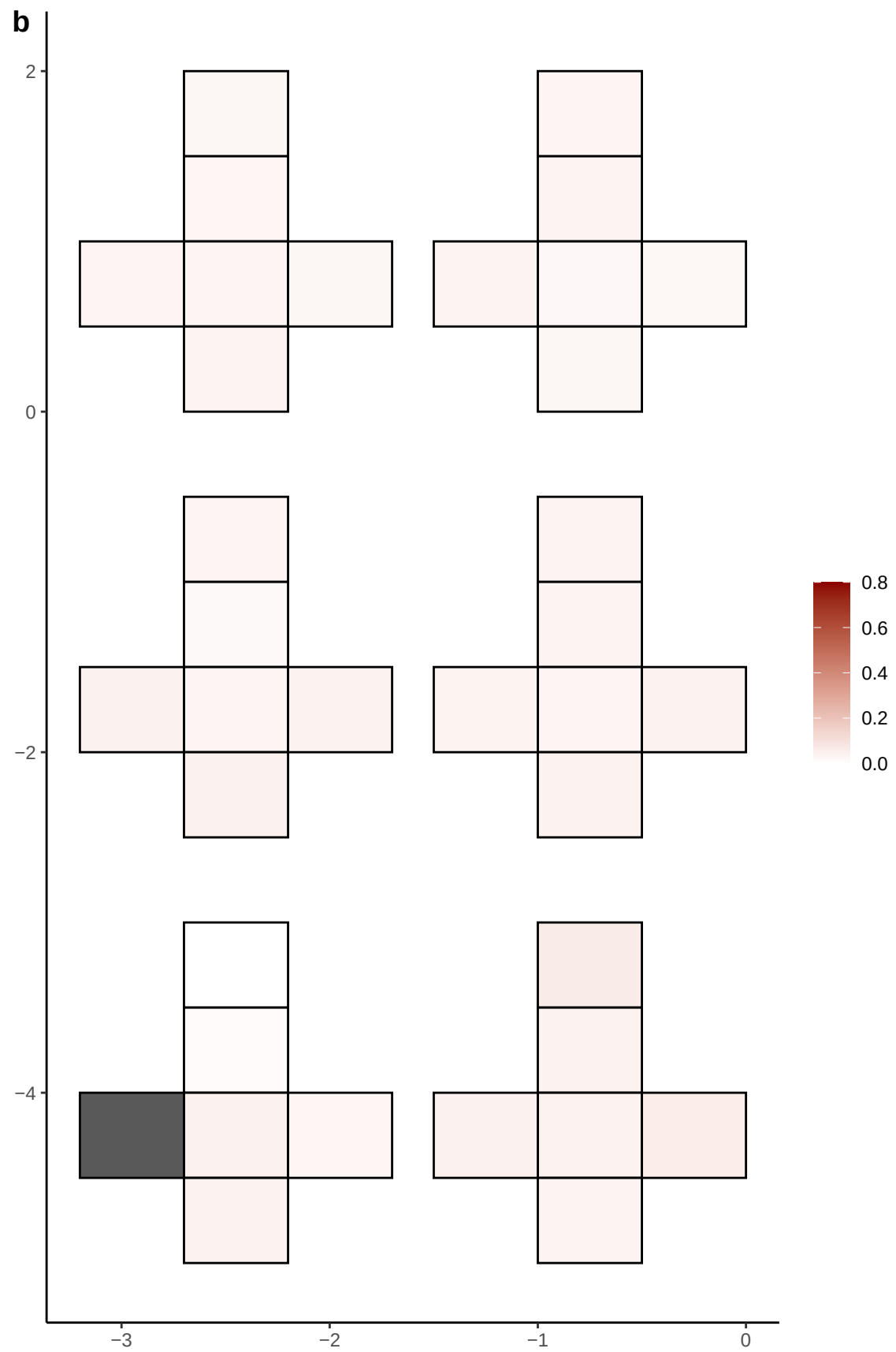

**a** ID01 4-66130457-C-T

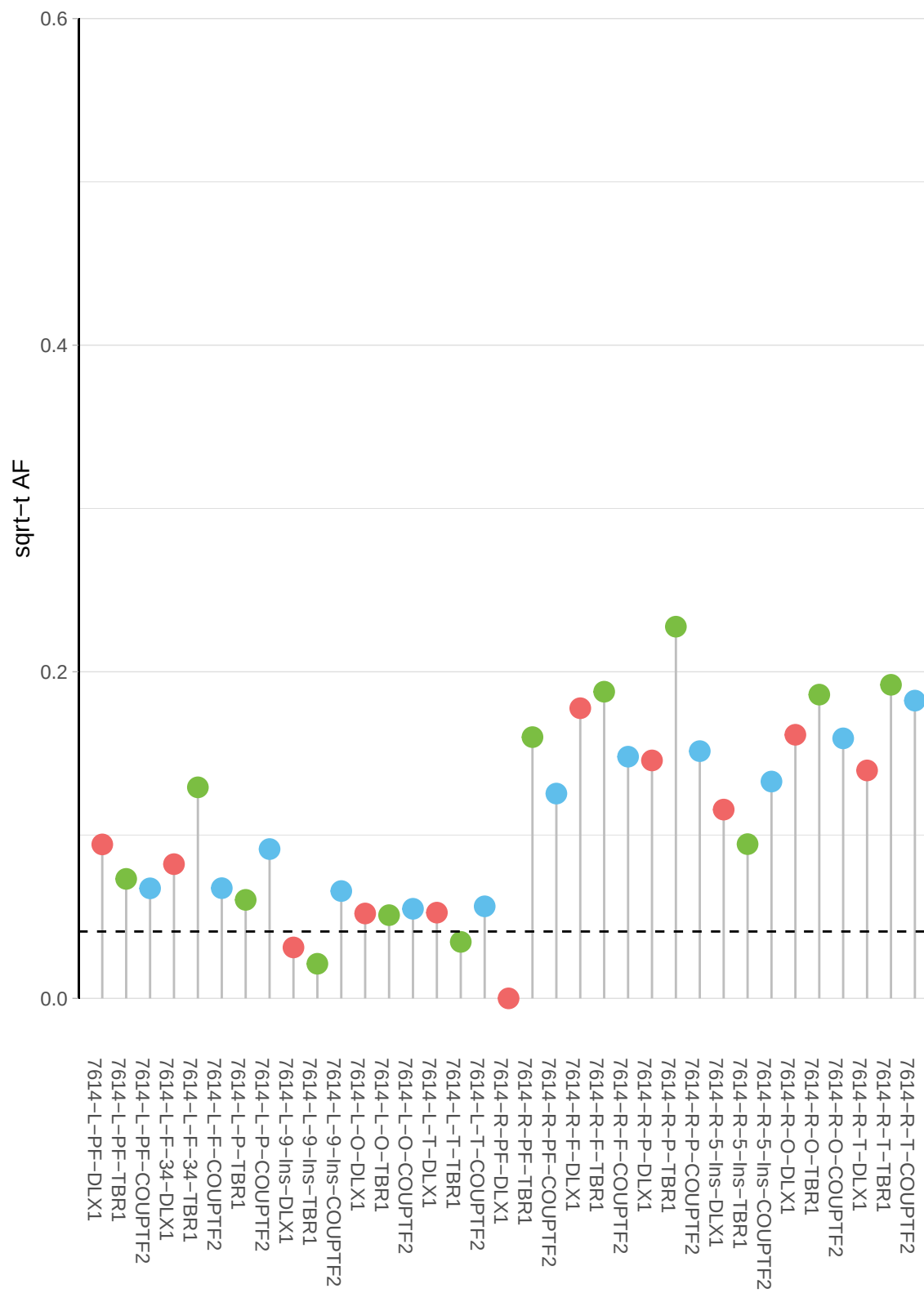

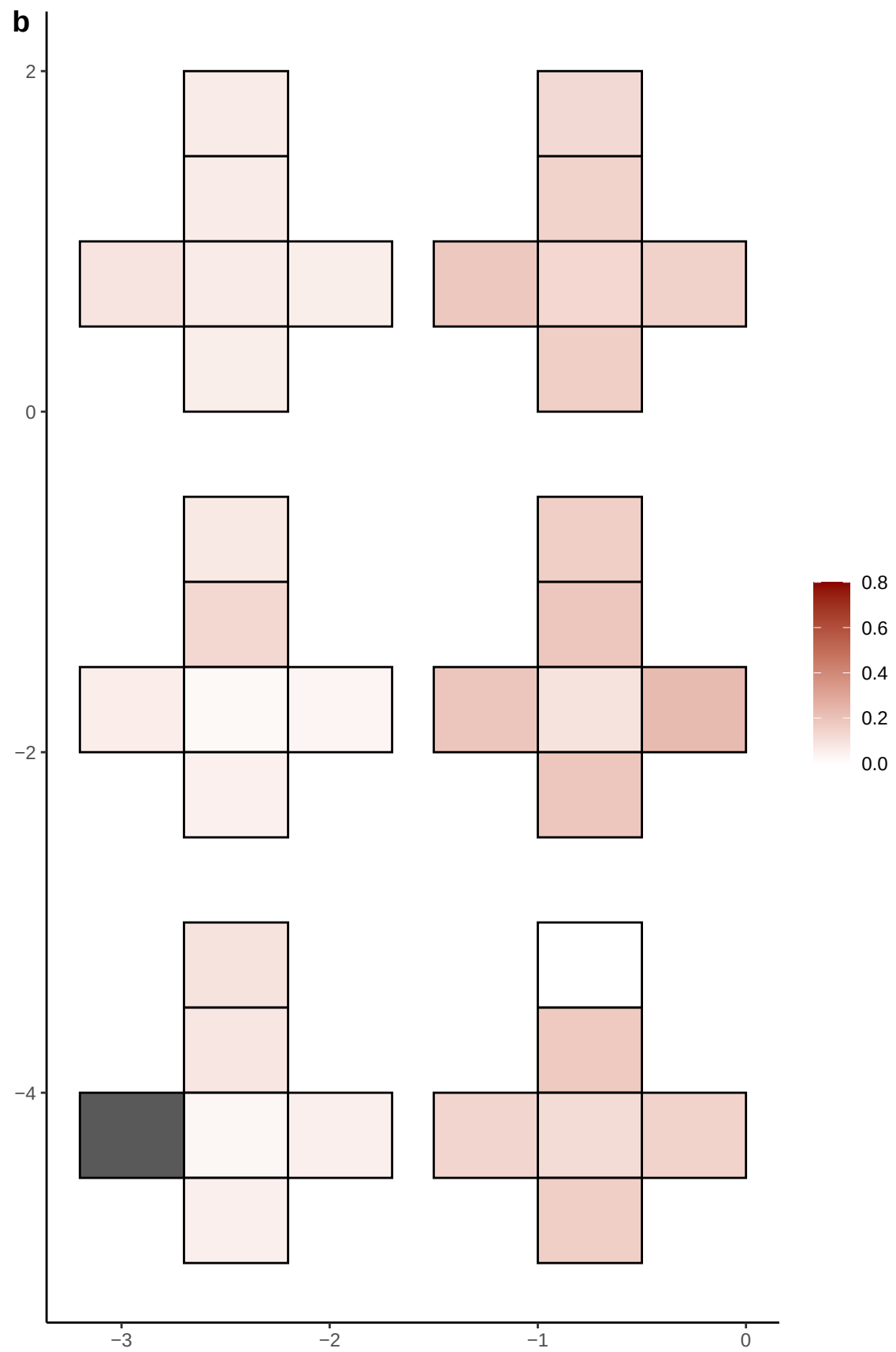

**a** ID01 4-67714666-G-A

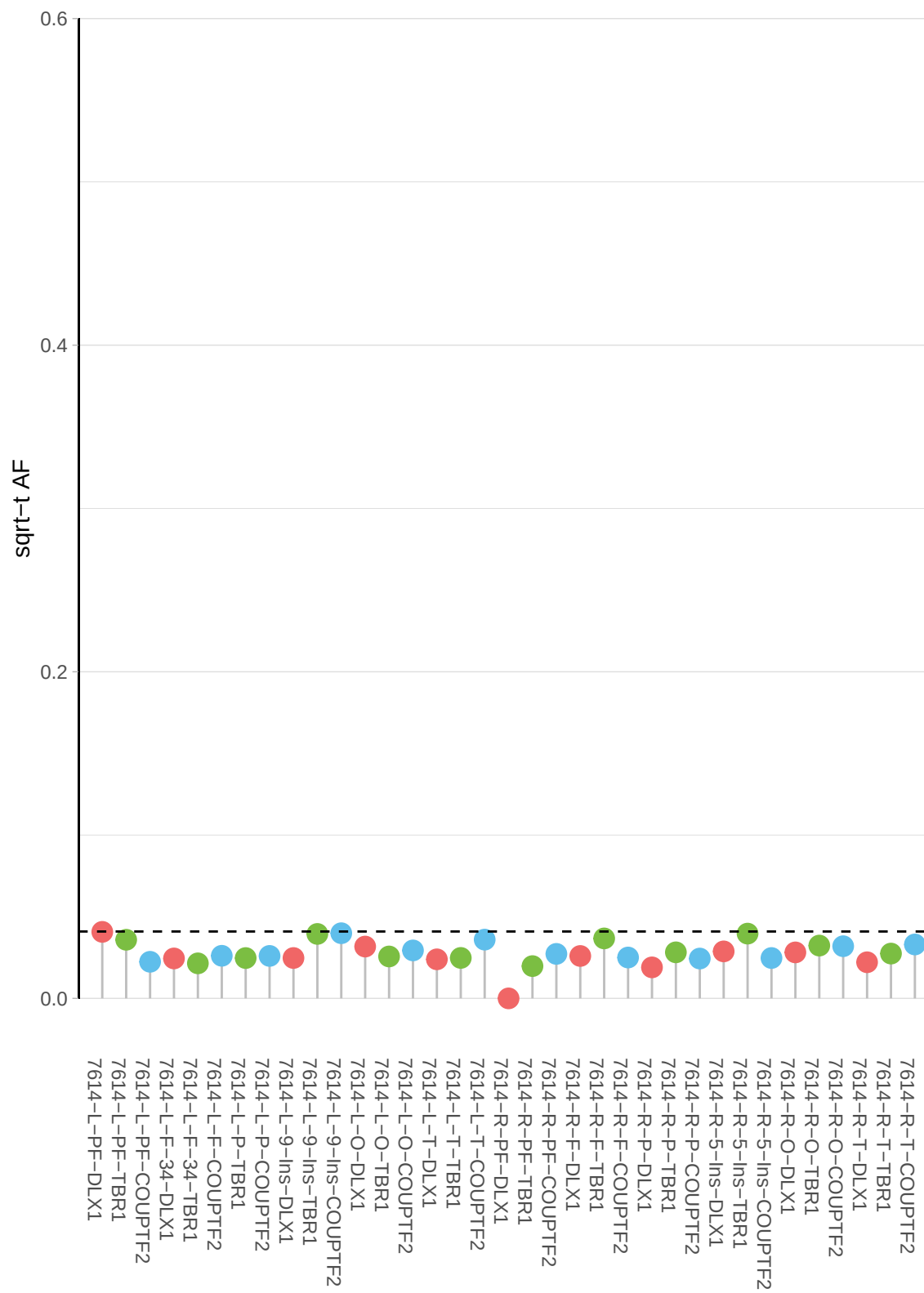

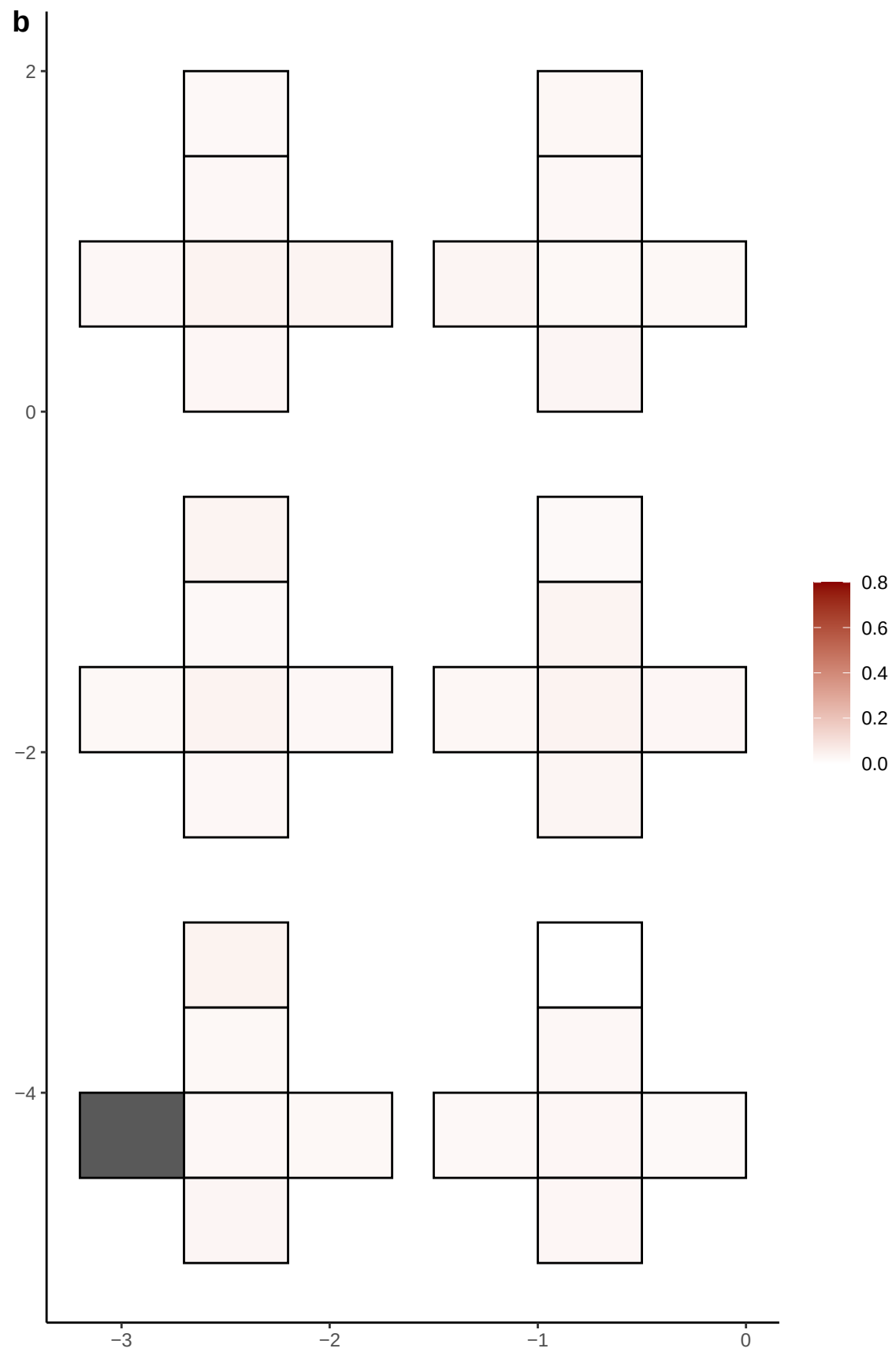

**a** ID01 4-71105167-A-T

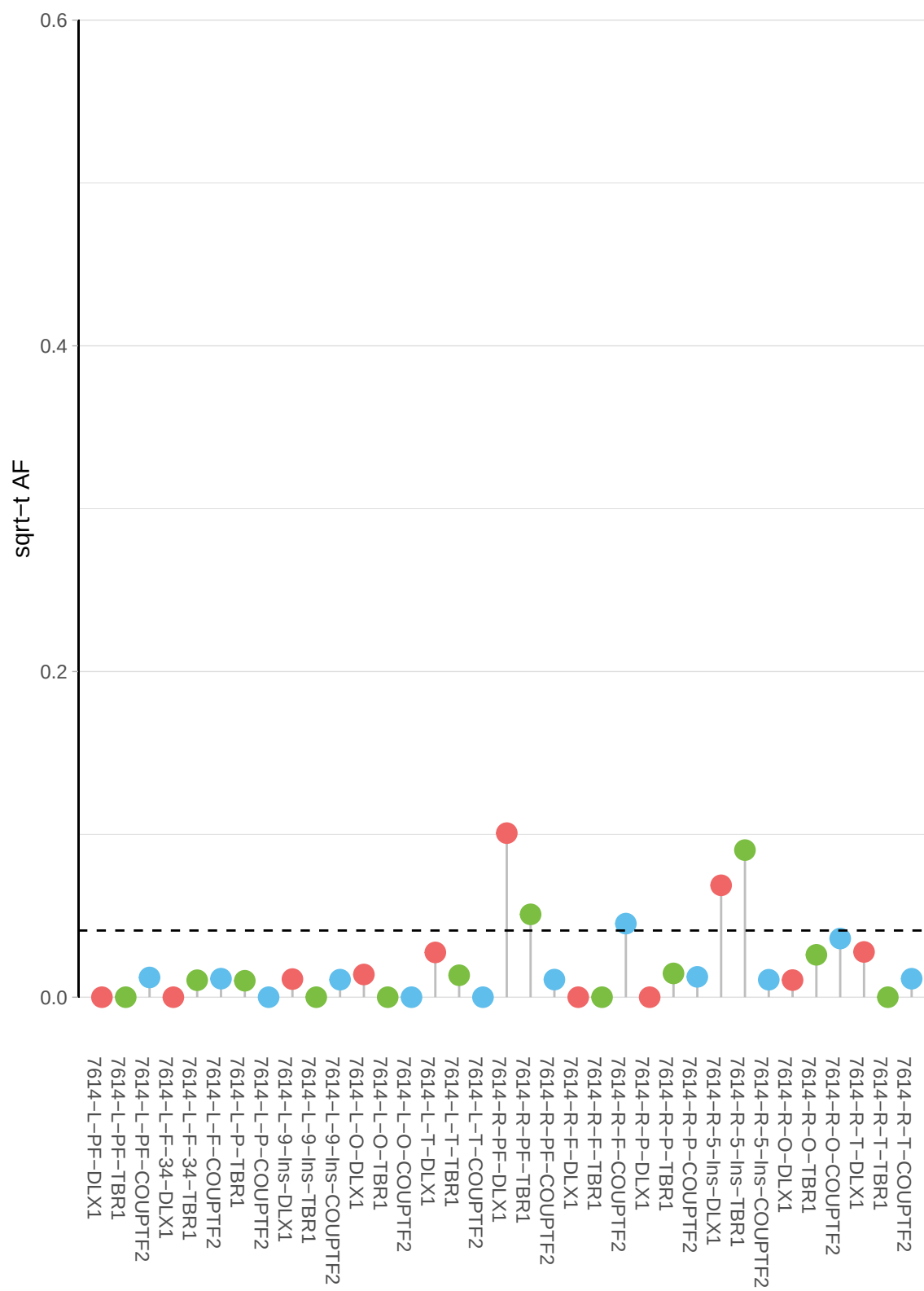

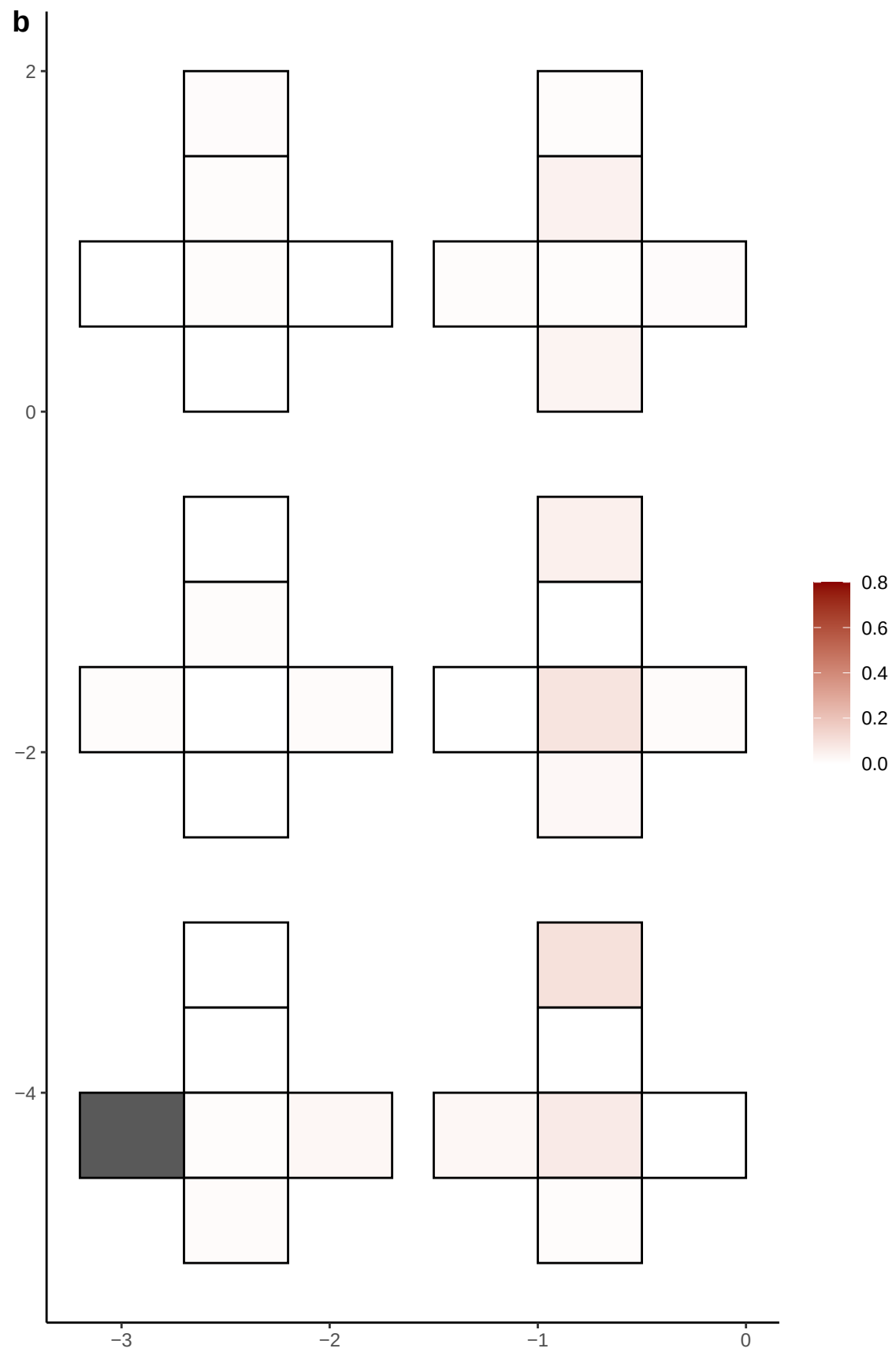

**a** ID01 4-74314197-C-G

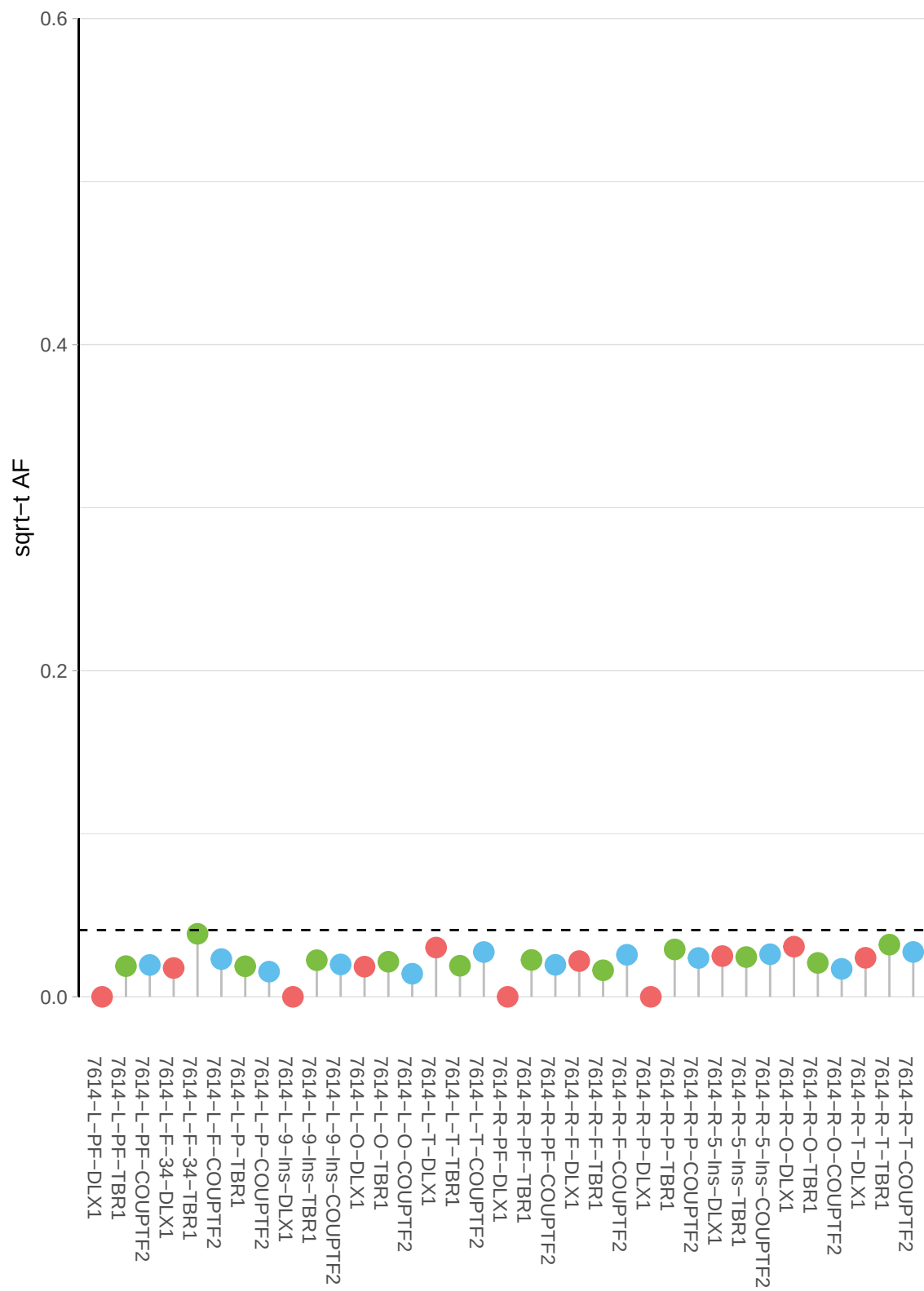

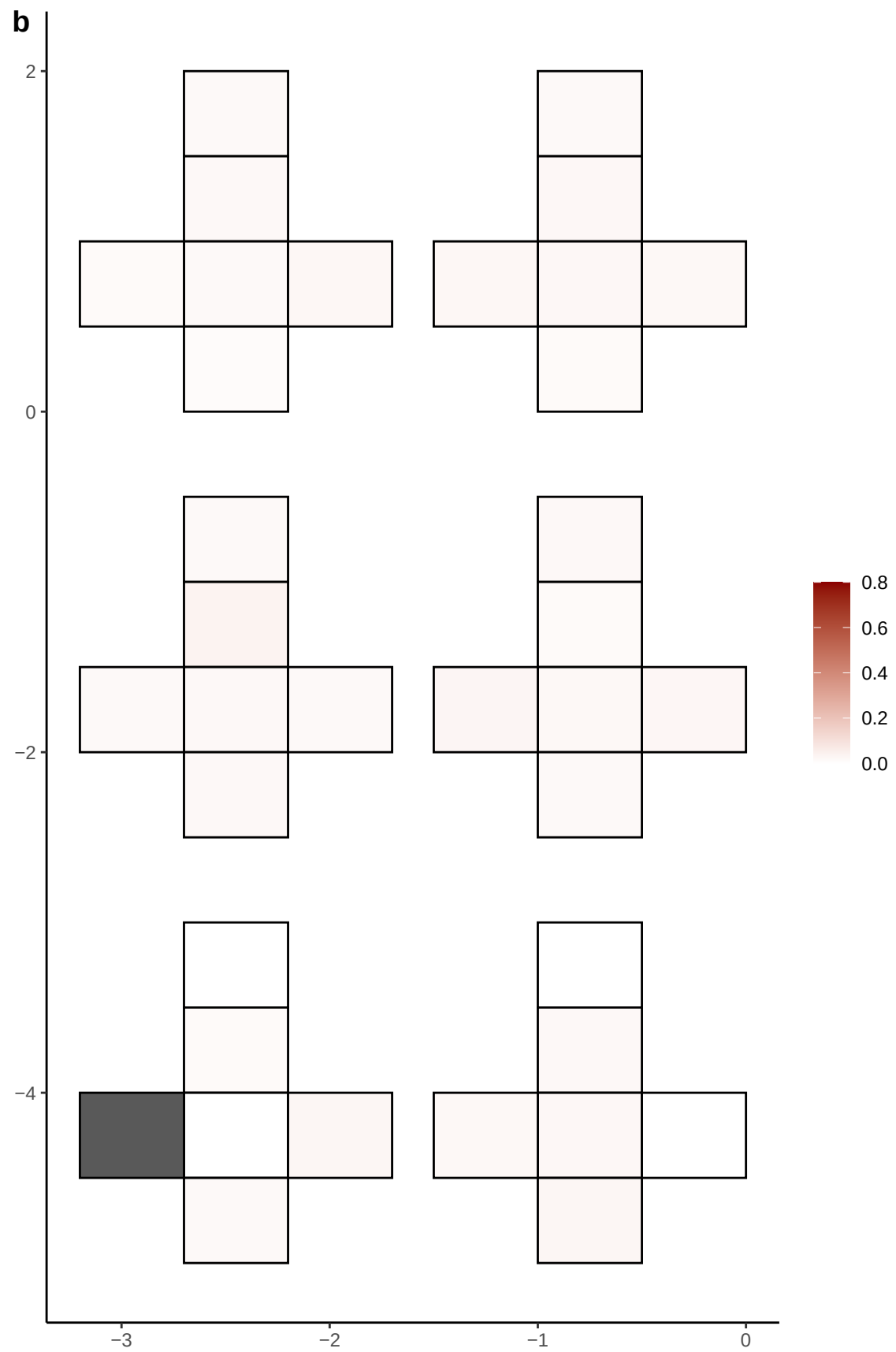

**a** ID01 4-83599250-T-C

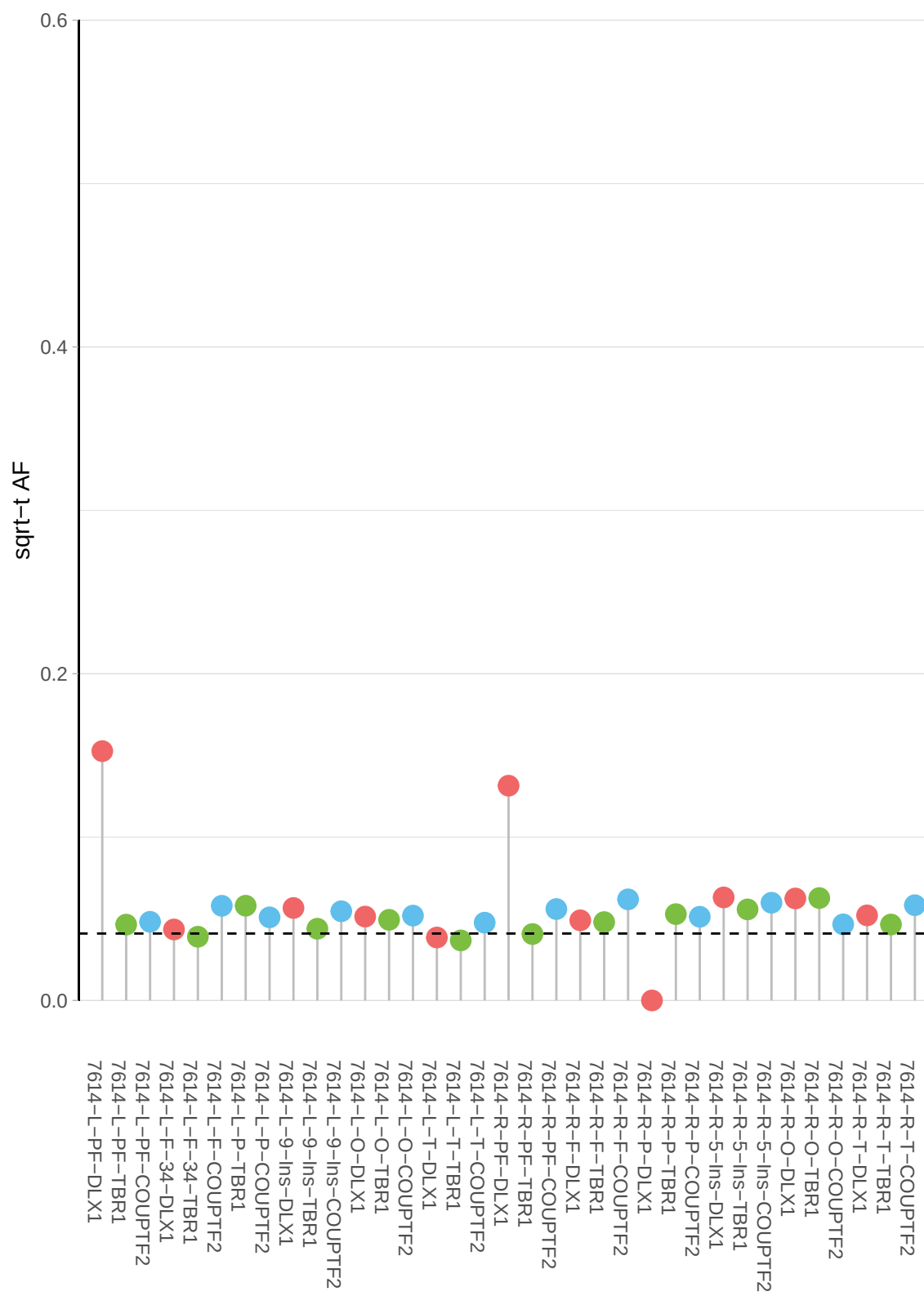

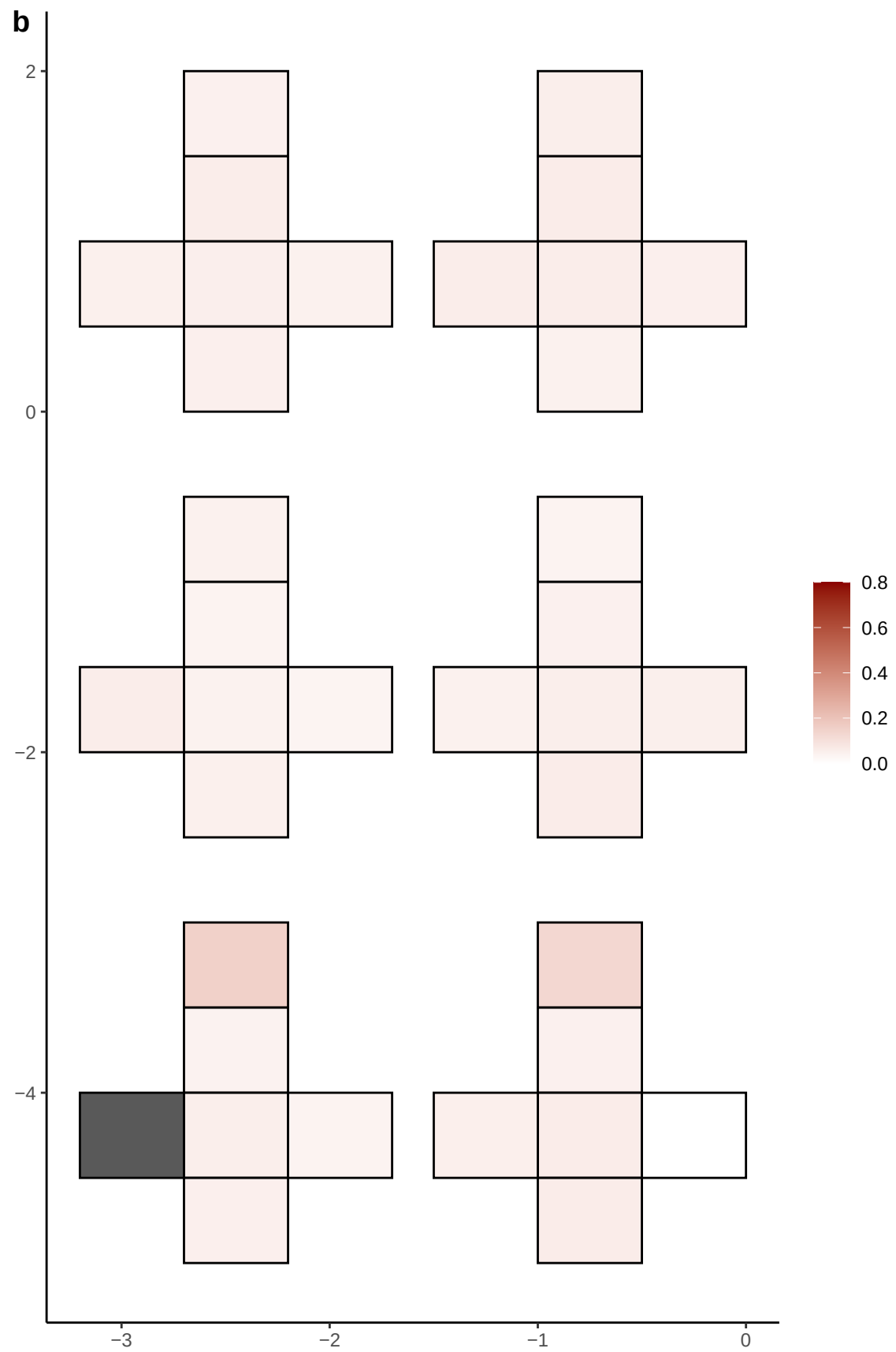

**a** ID01 4-93868224-A-C

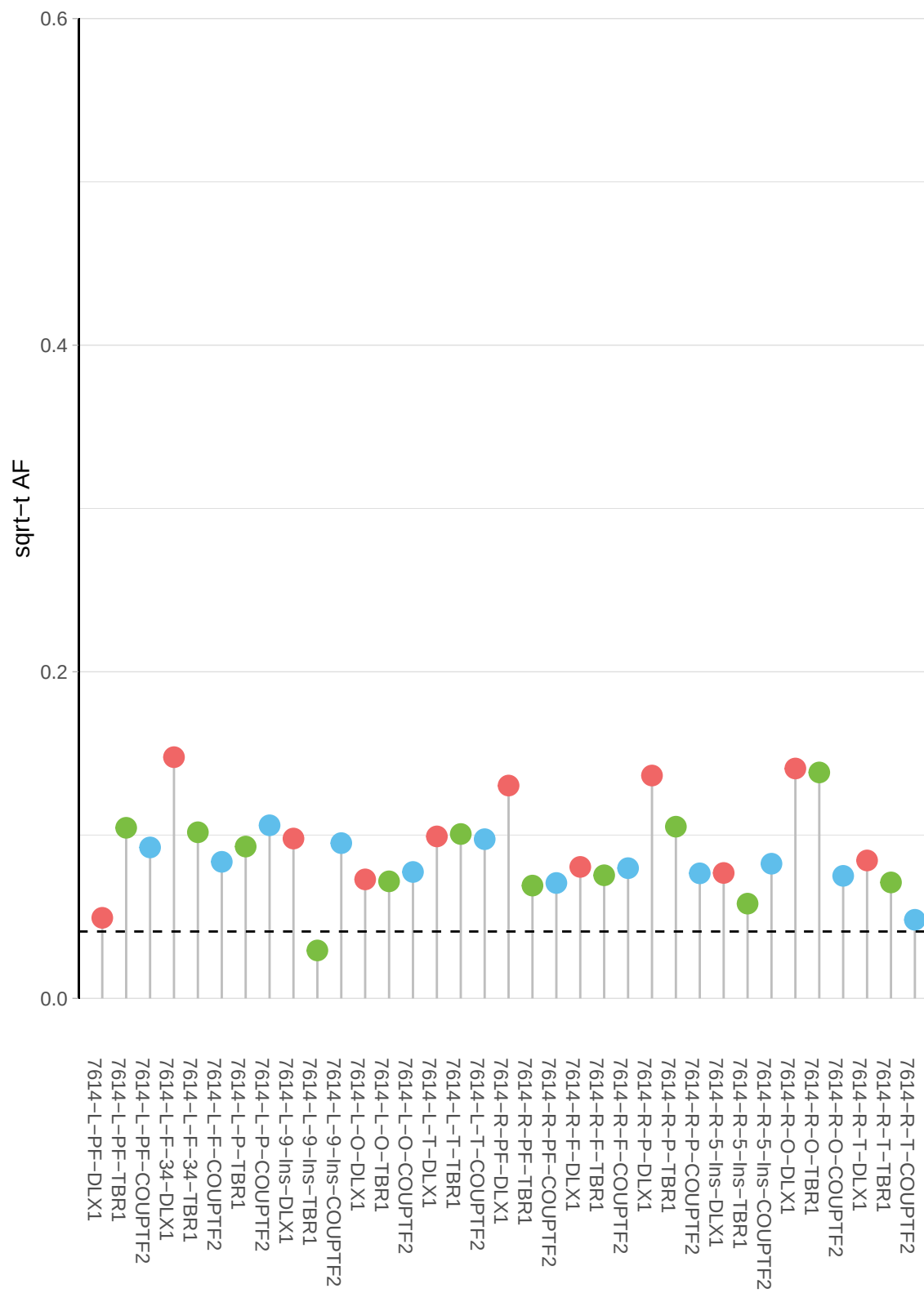

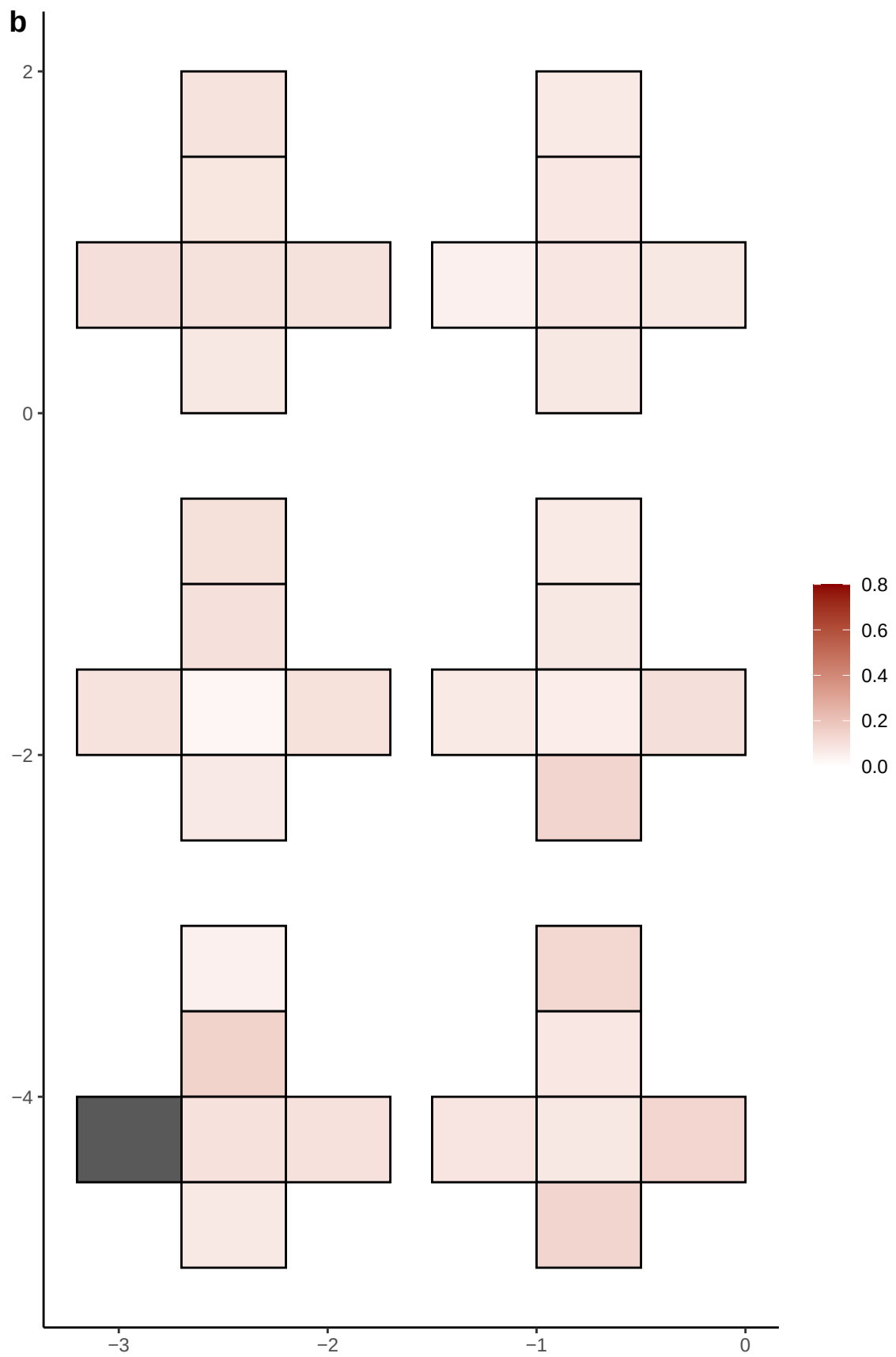

**a** ID01 5-109537368-T-C

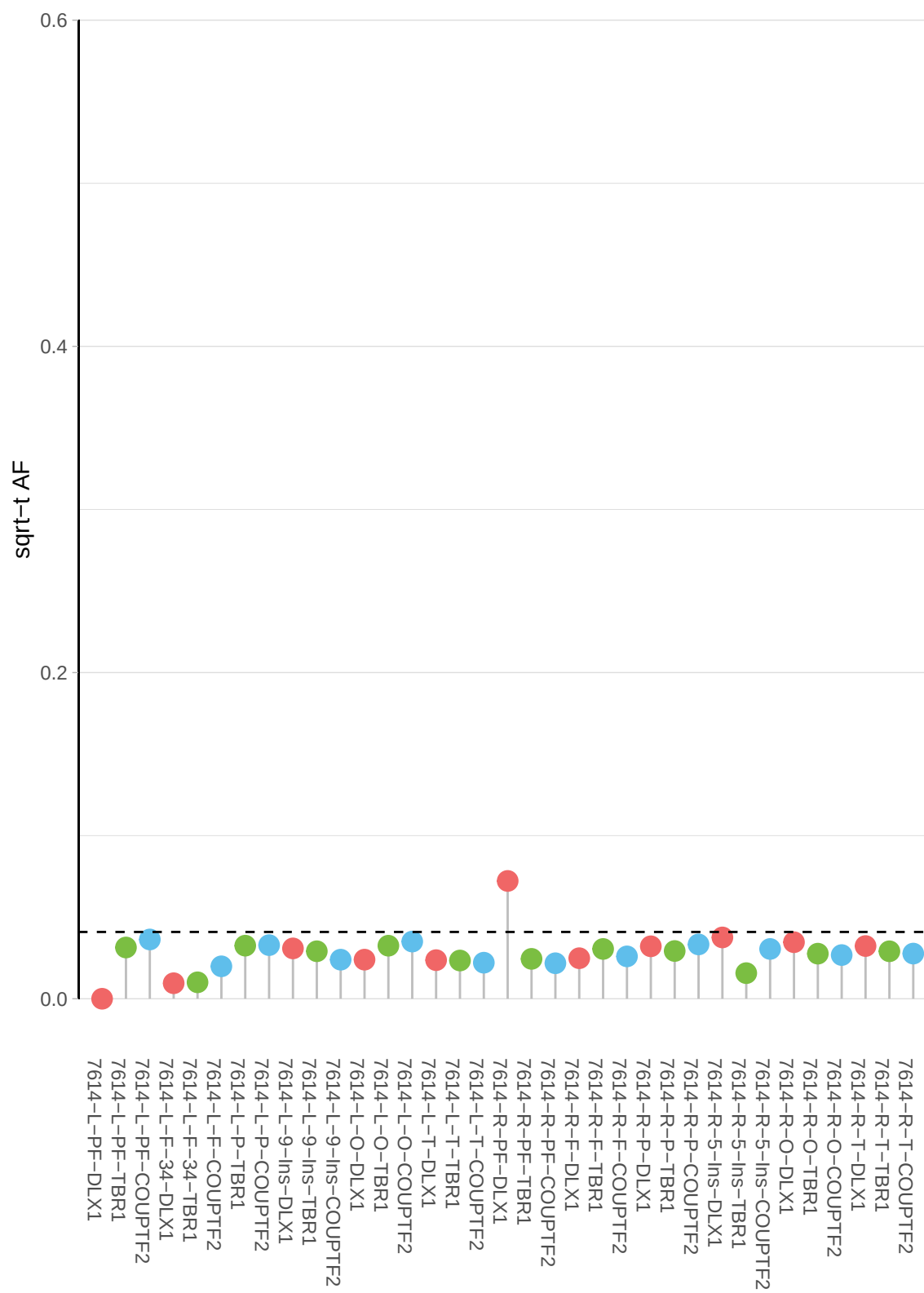

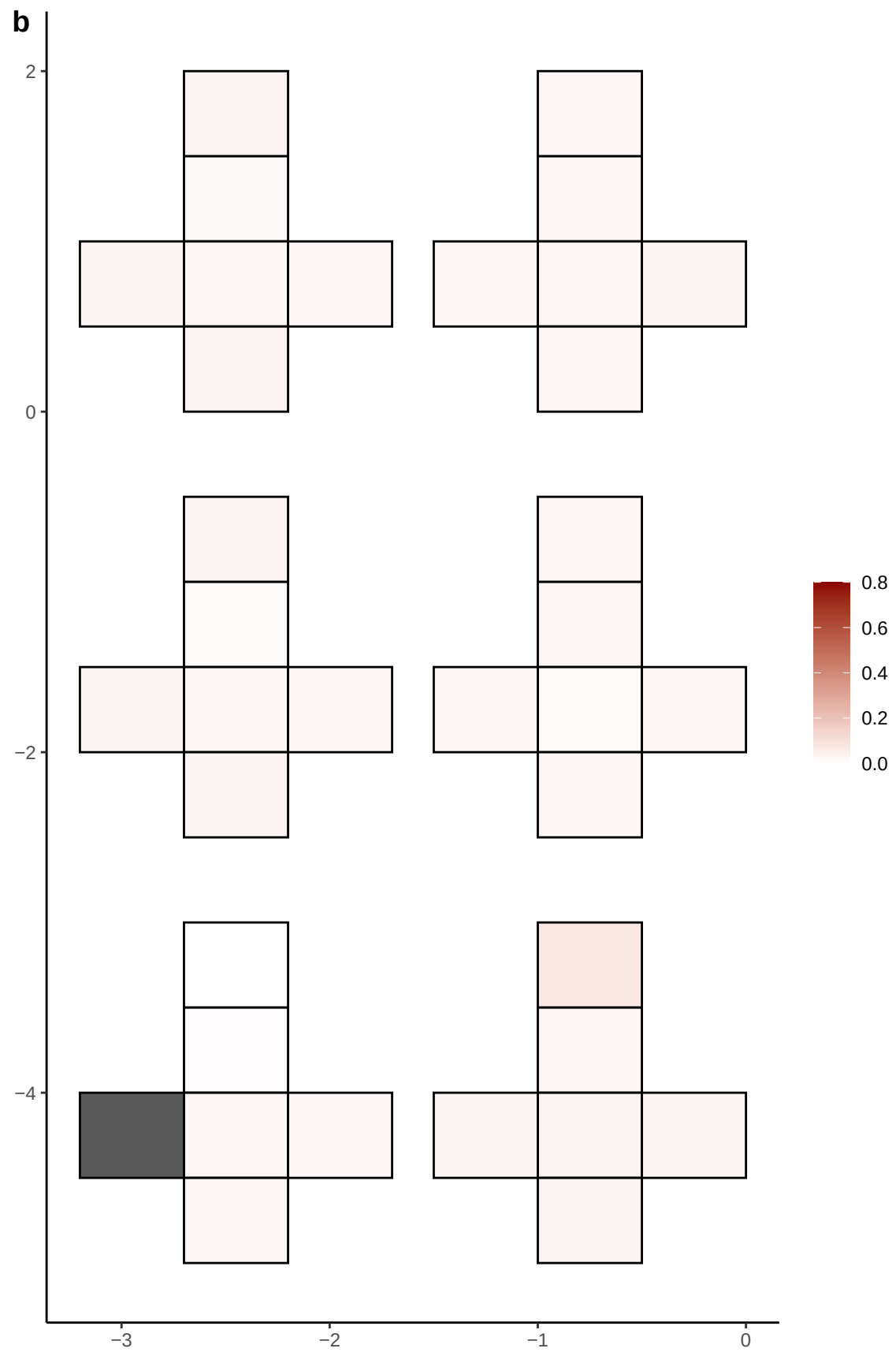

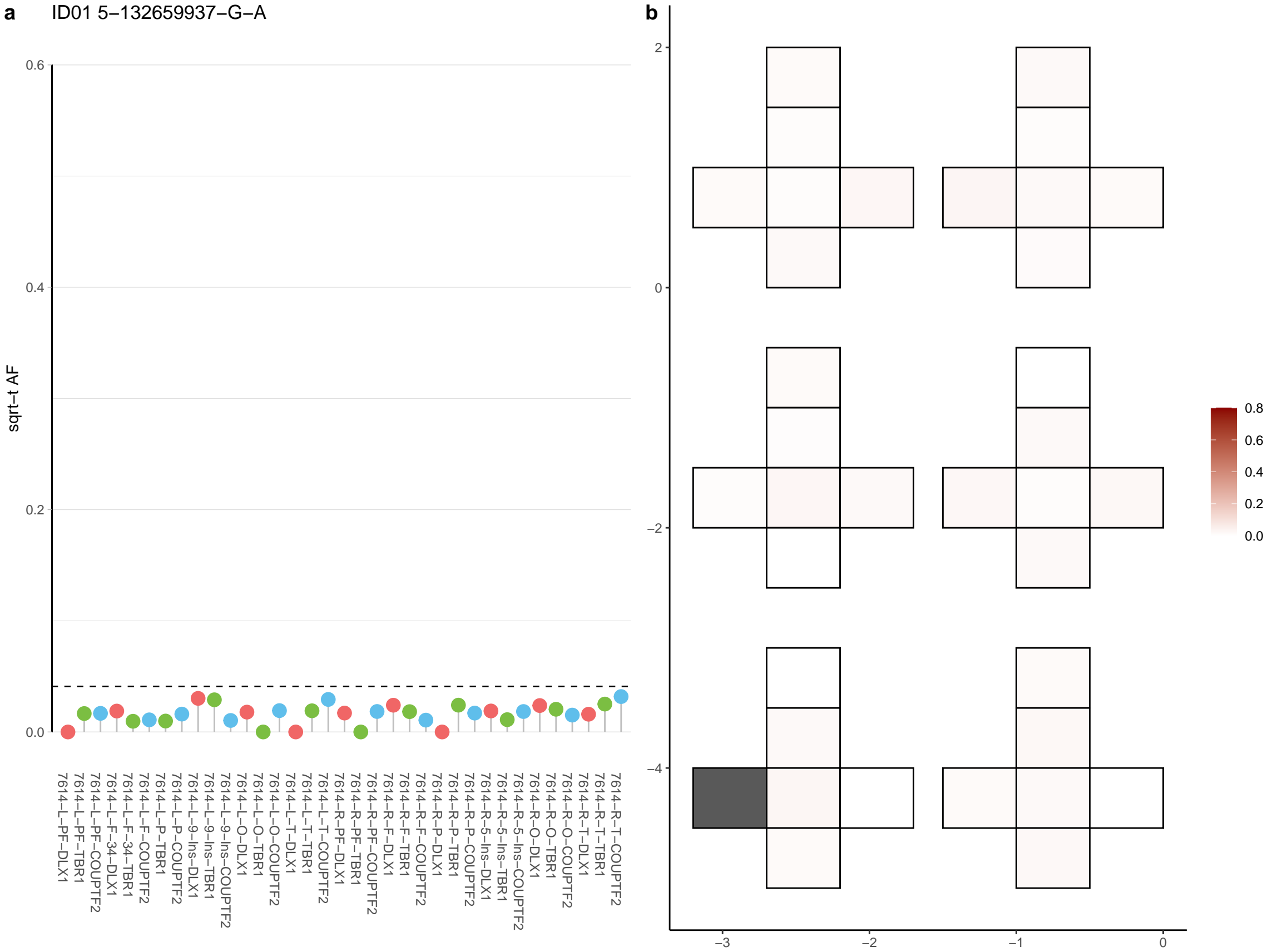

**a** ID01 5-146867769-A-G

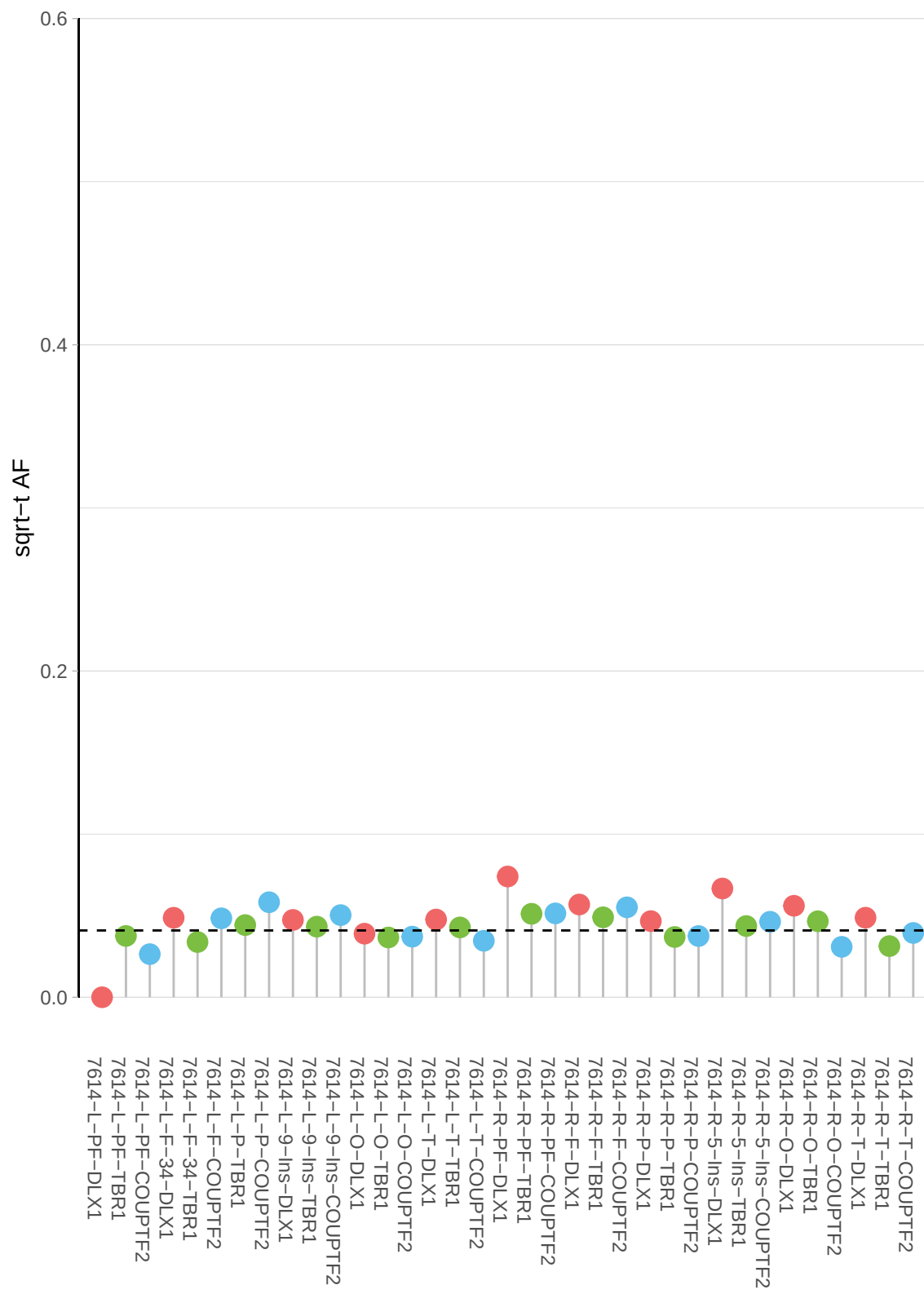

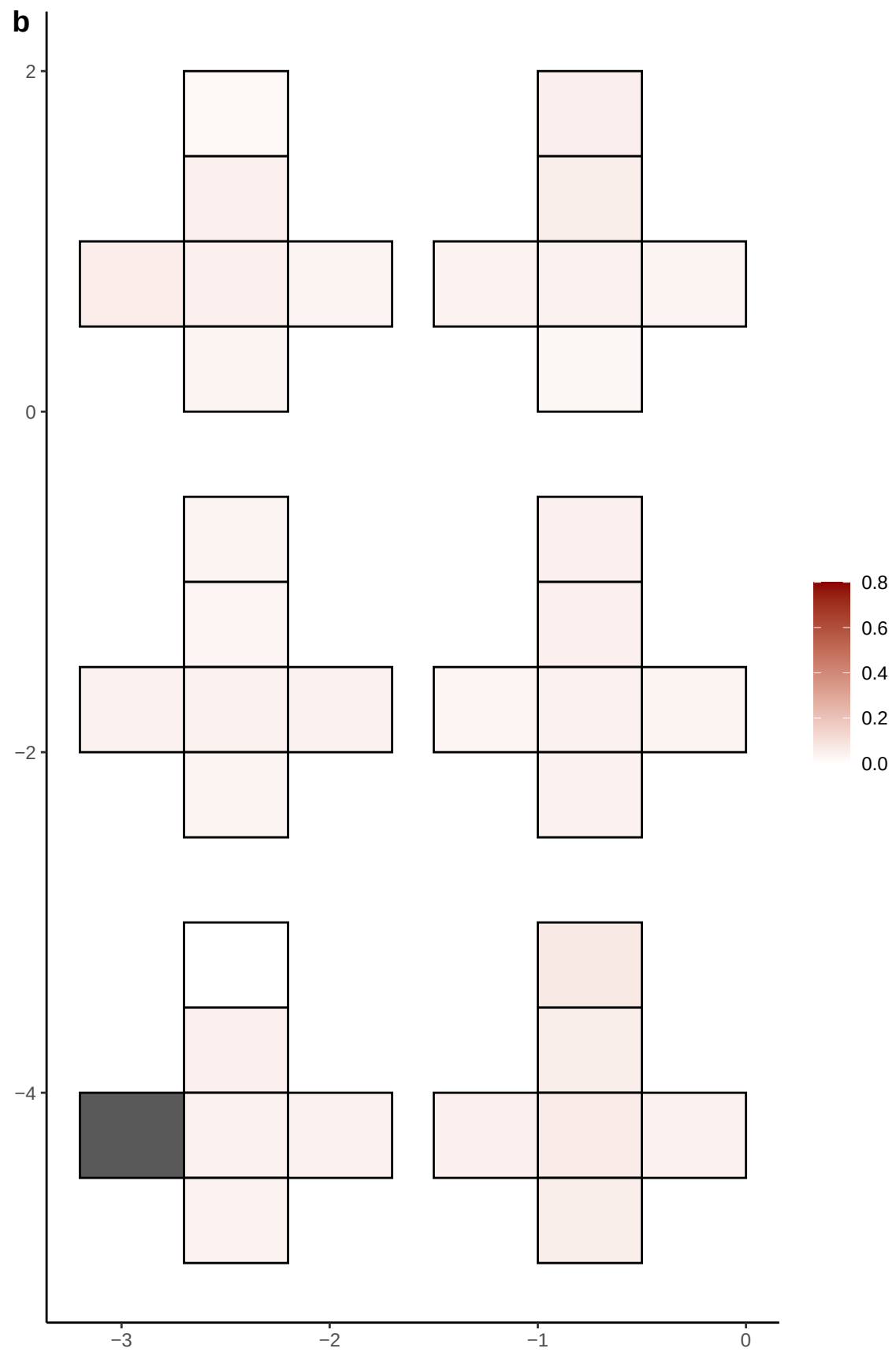

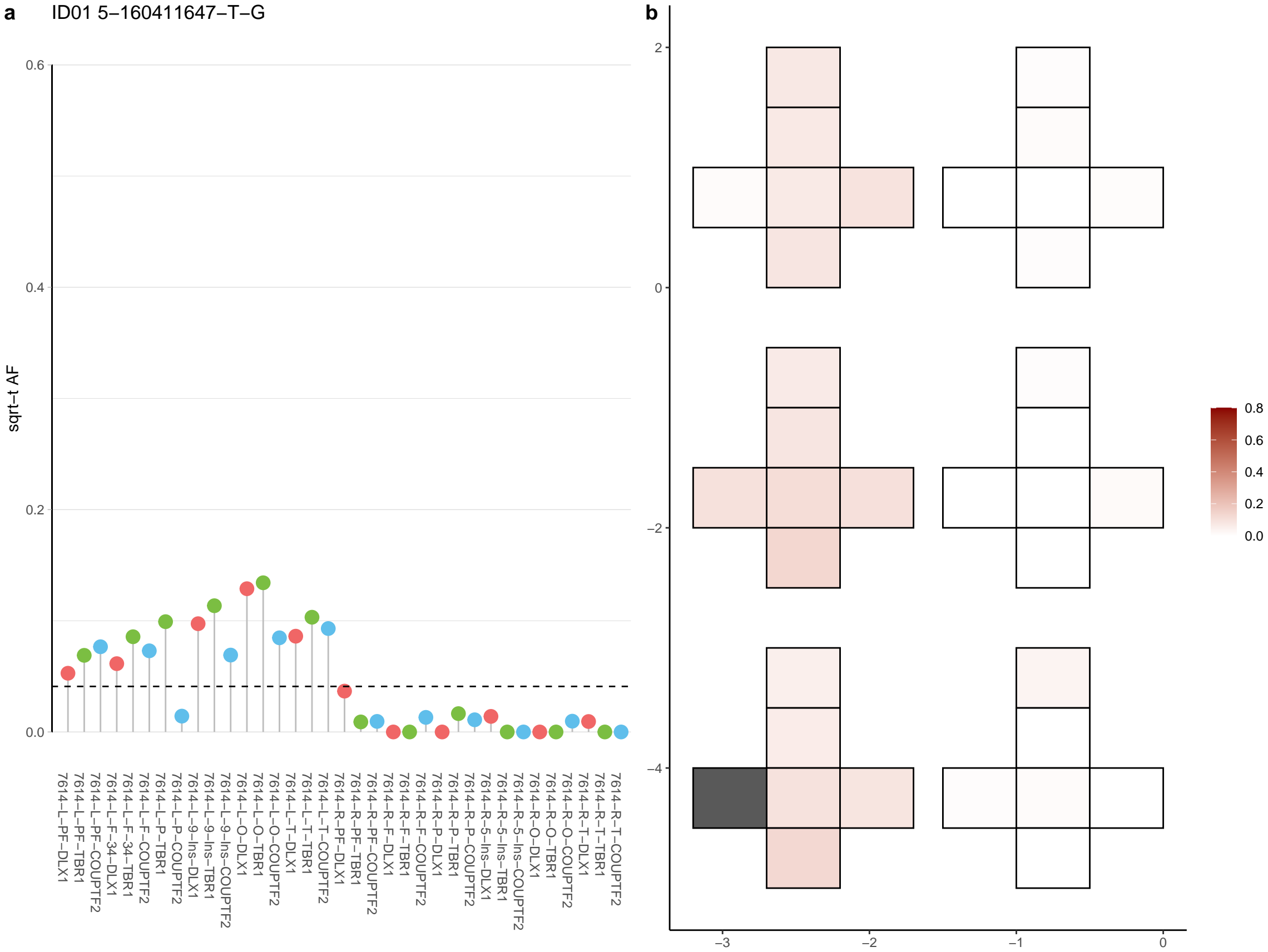

**a** ID01 5-17226551-A-G

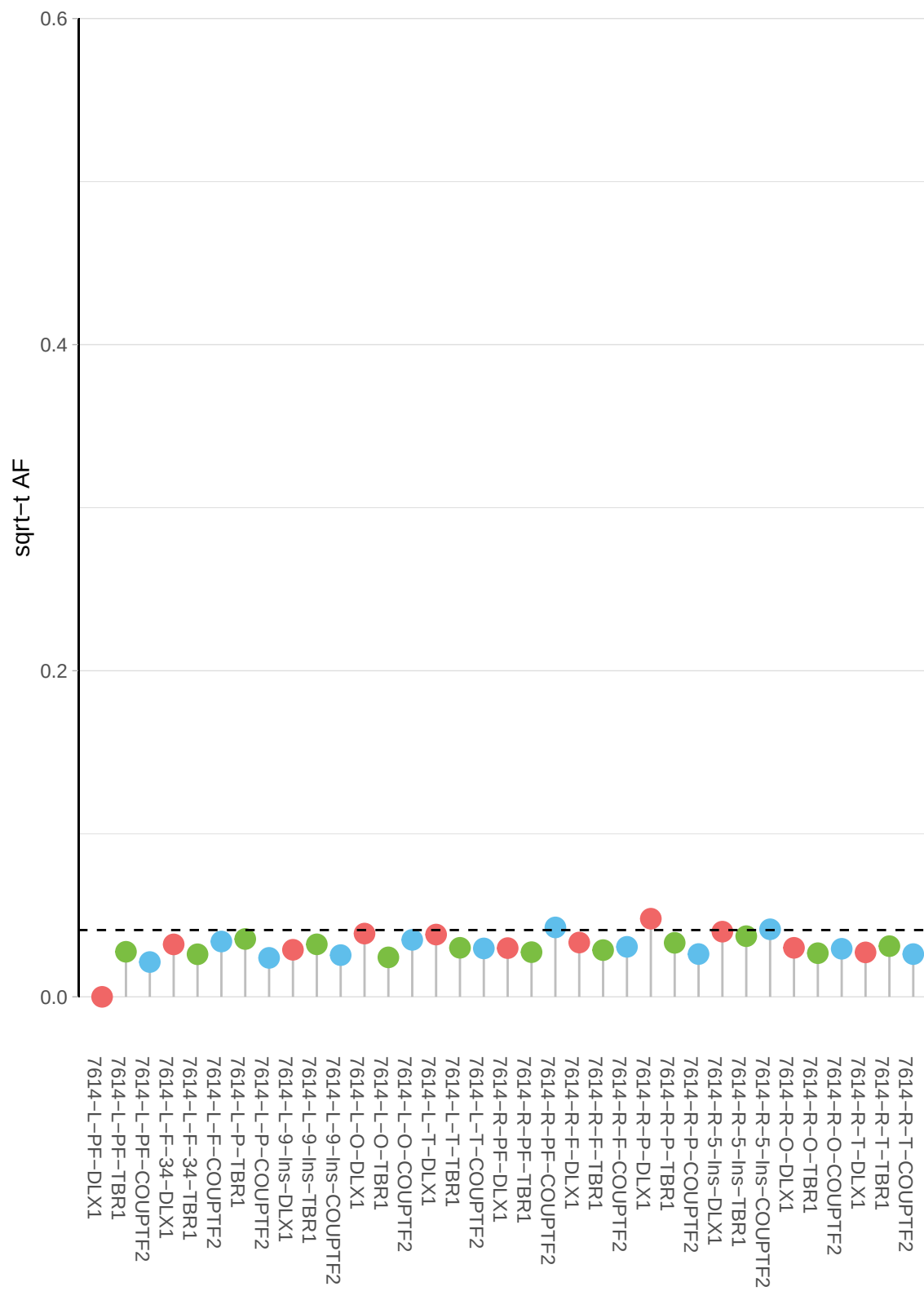

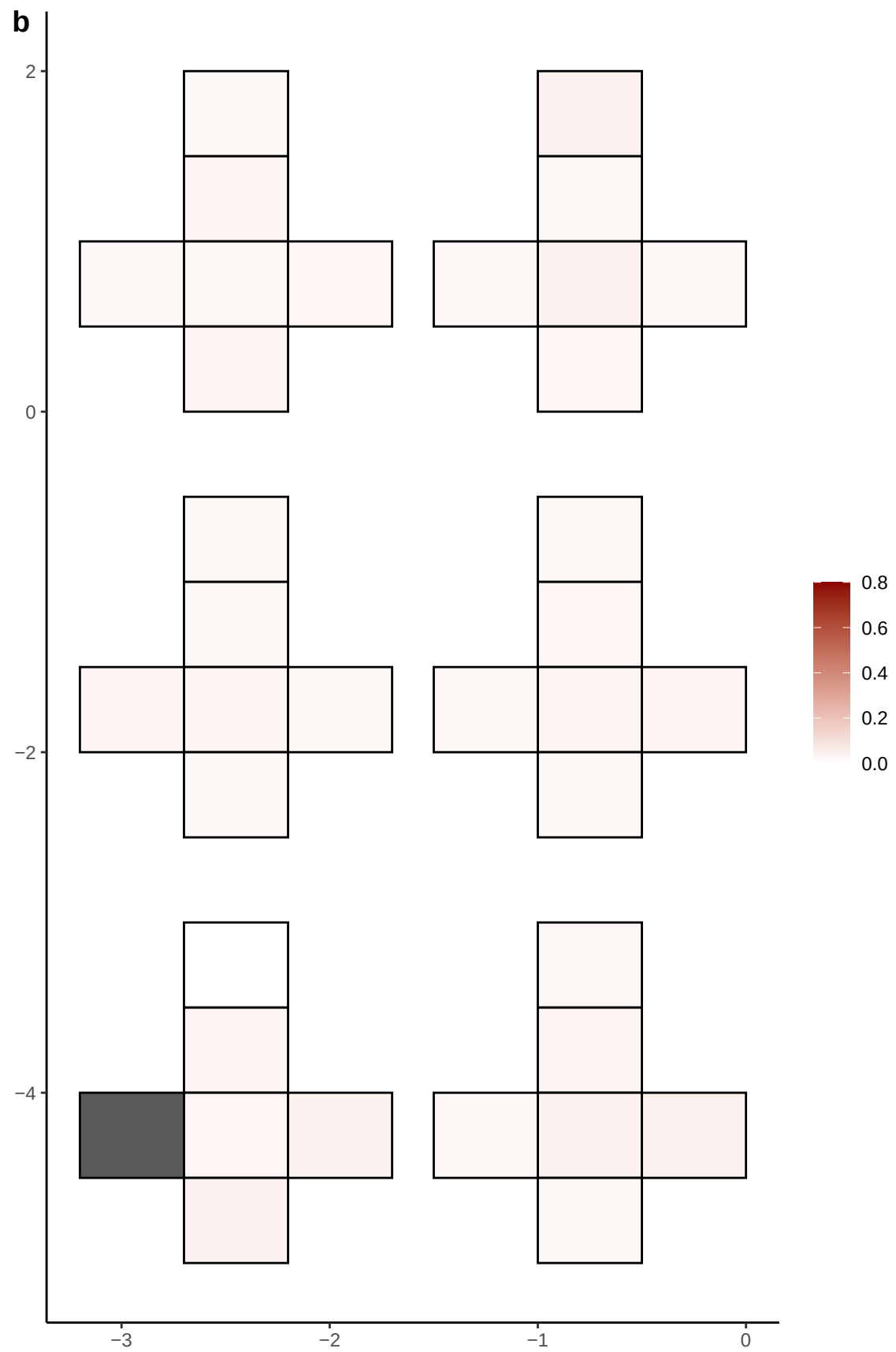

**a** ID01 5-172837236-C-T

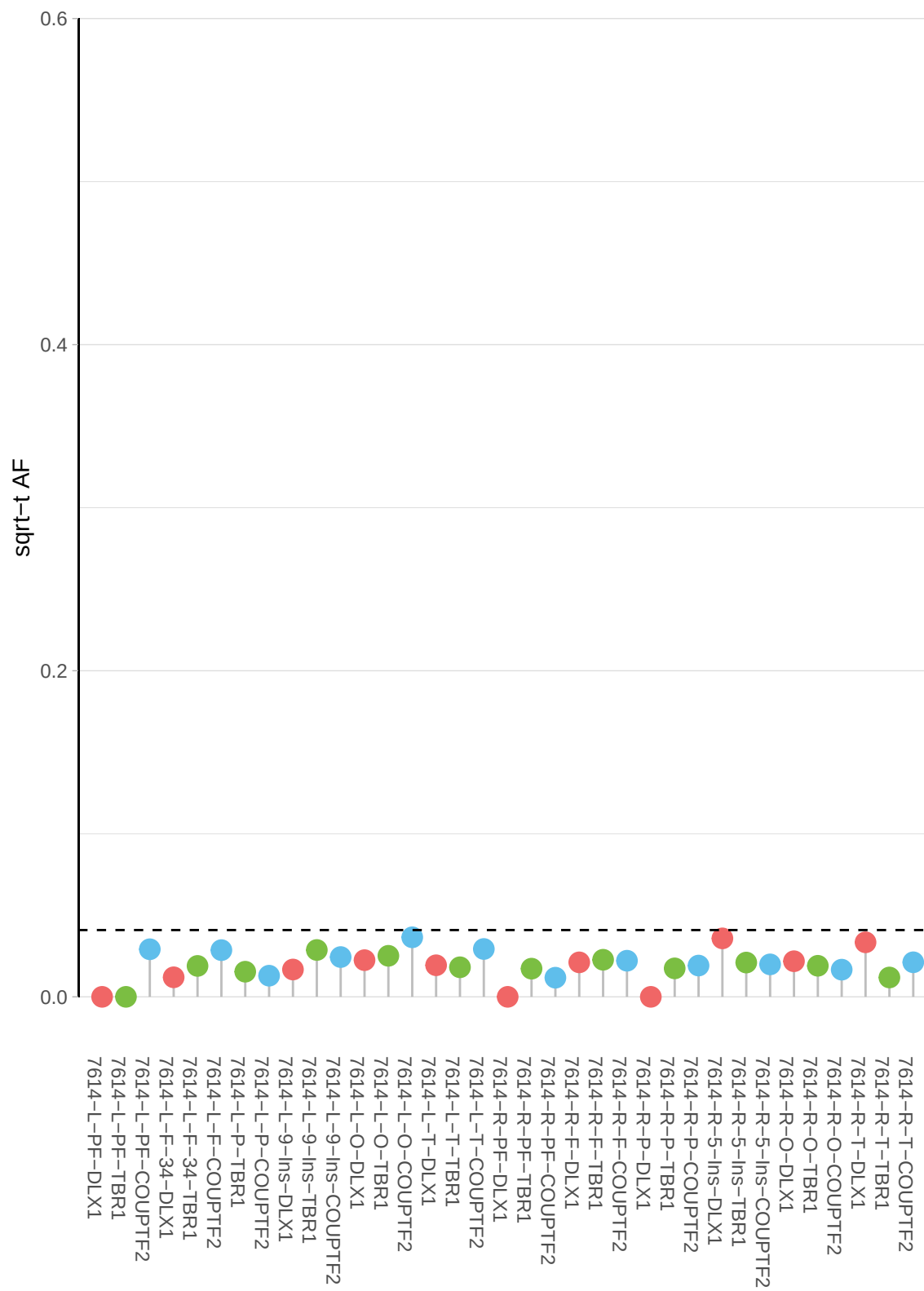

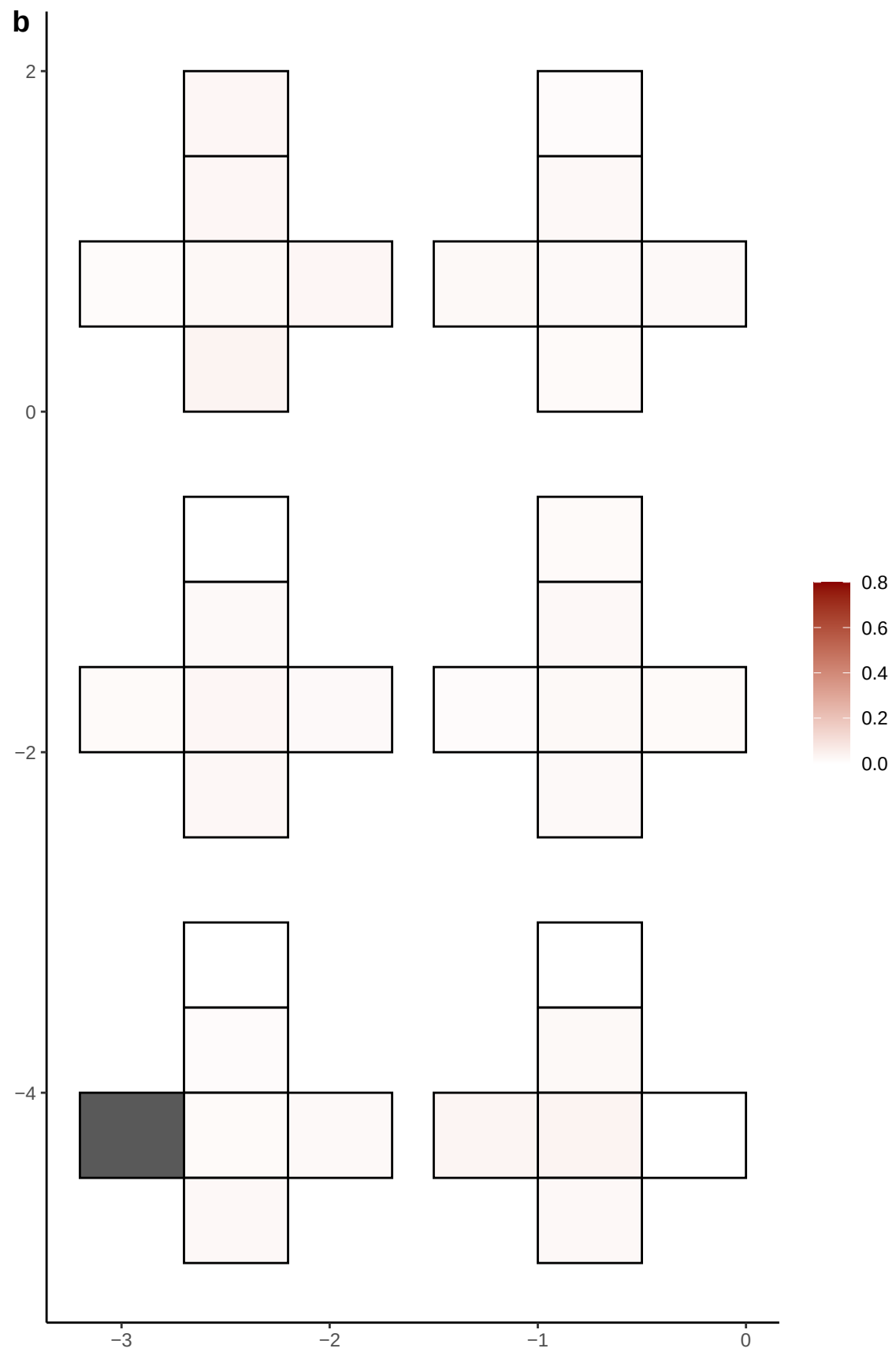

**a** ID01 5-179557321-C-A

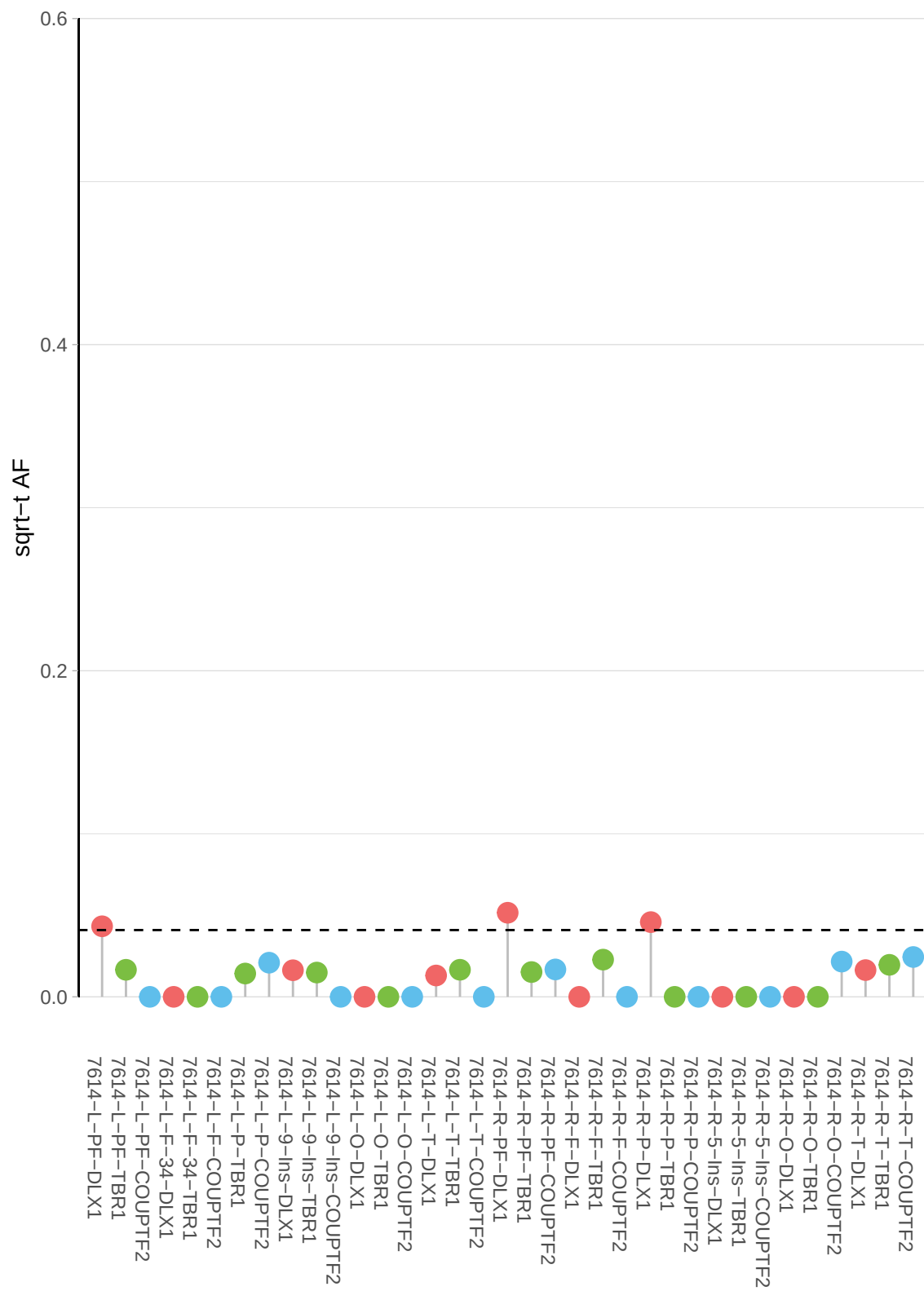

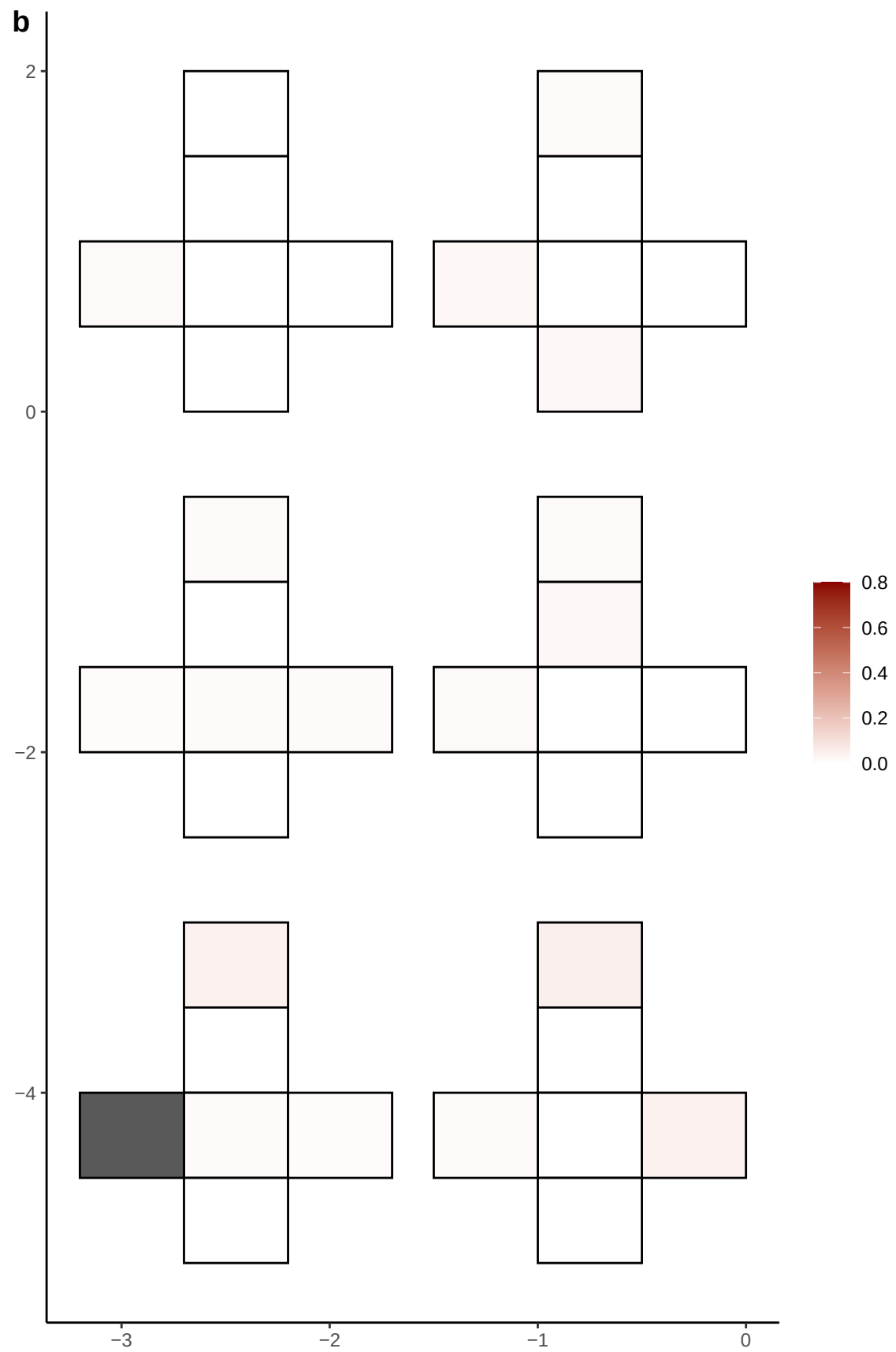

**a** ID01 5-44770209-G-A

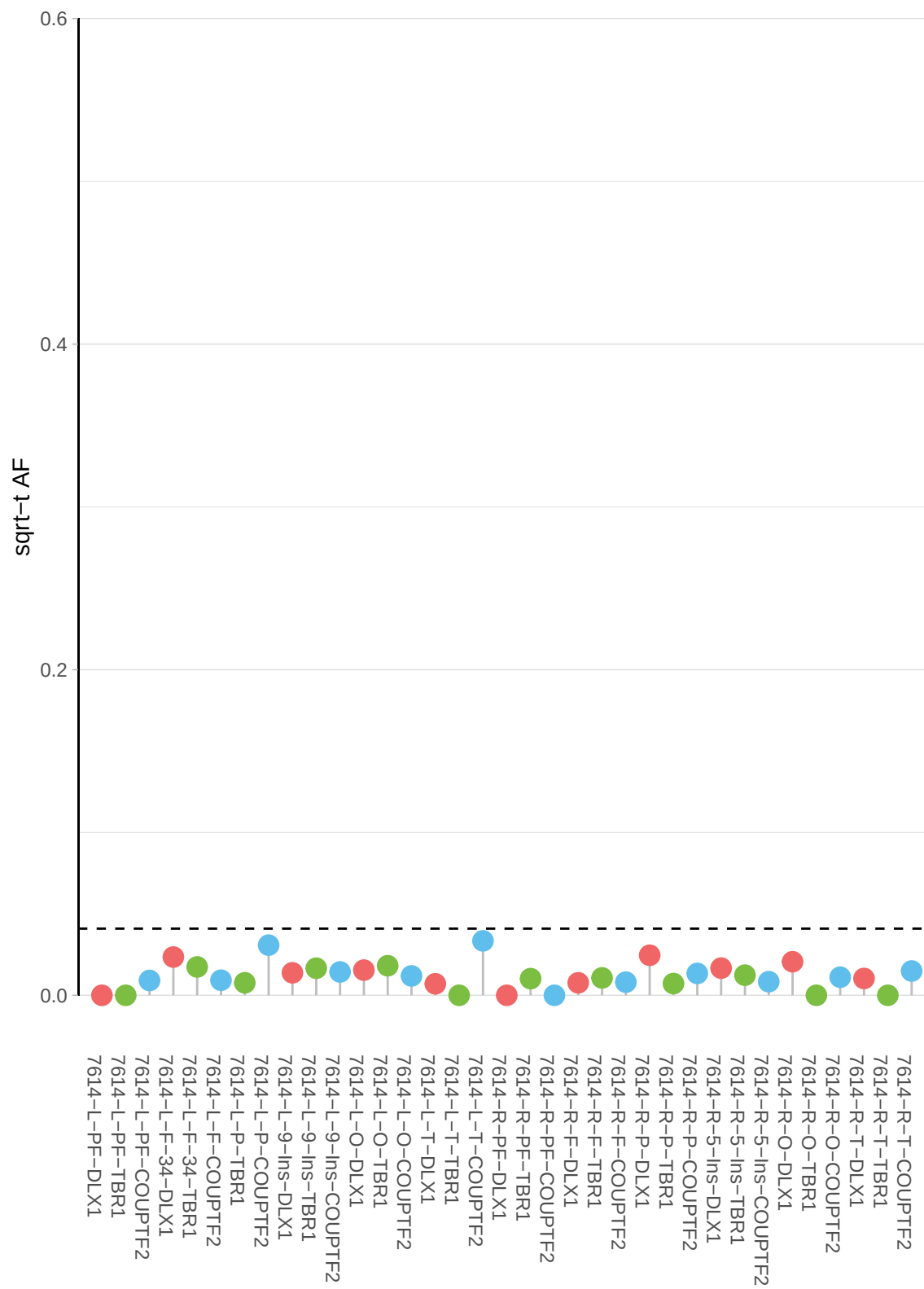

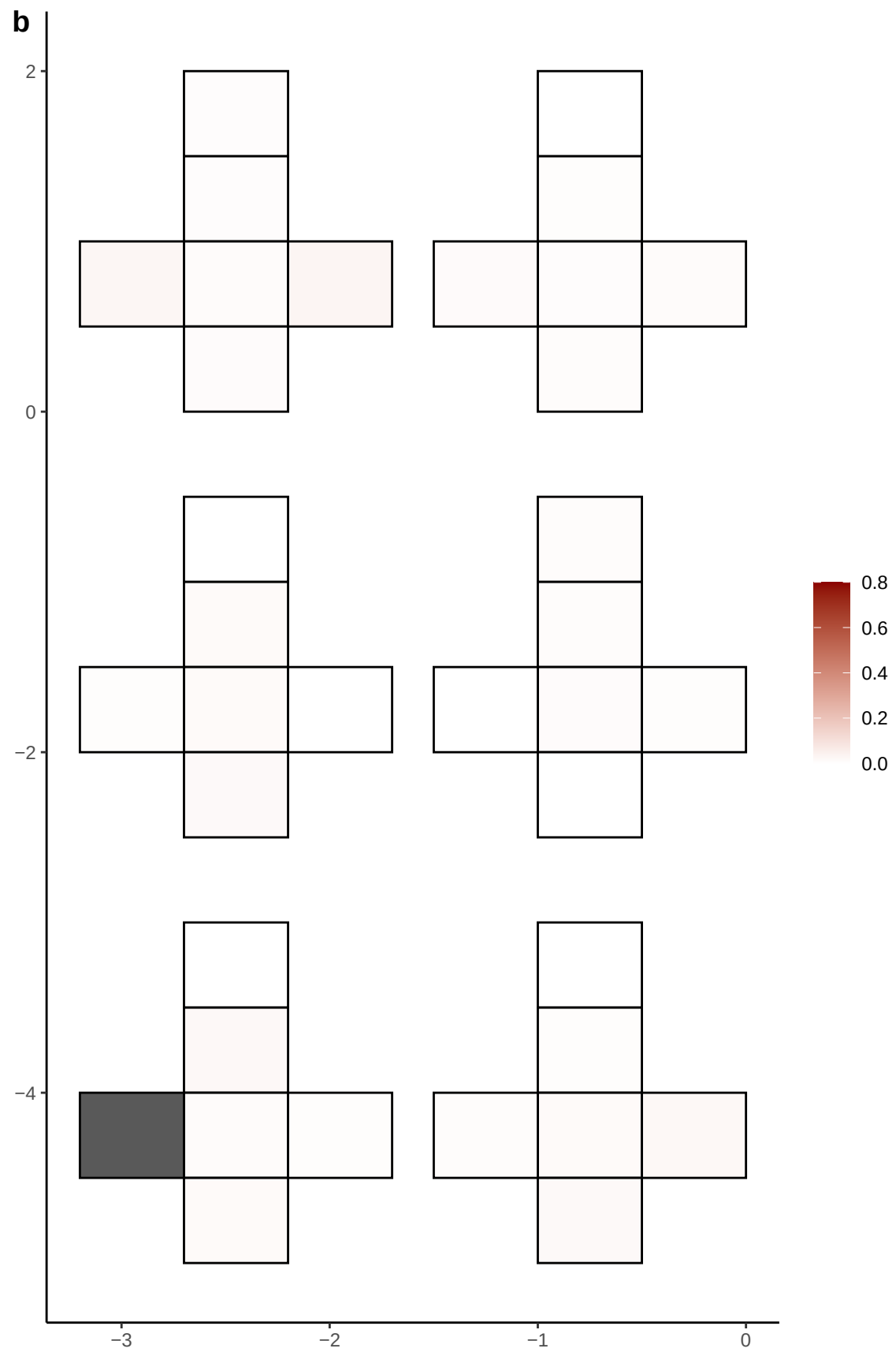

**a** ID01 5-51128677-C-A

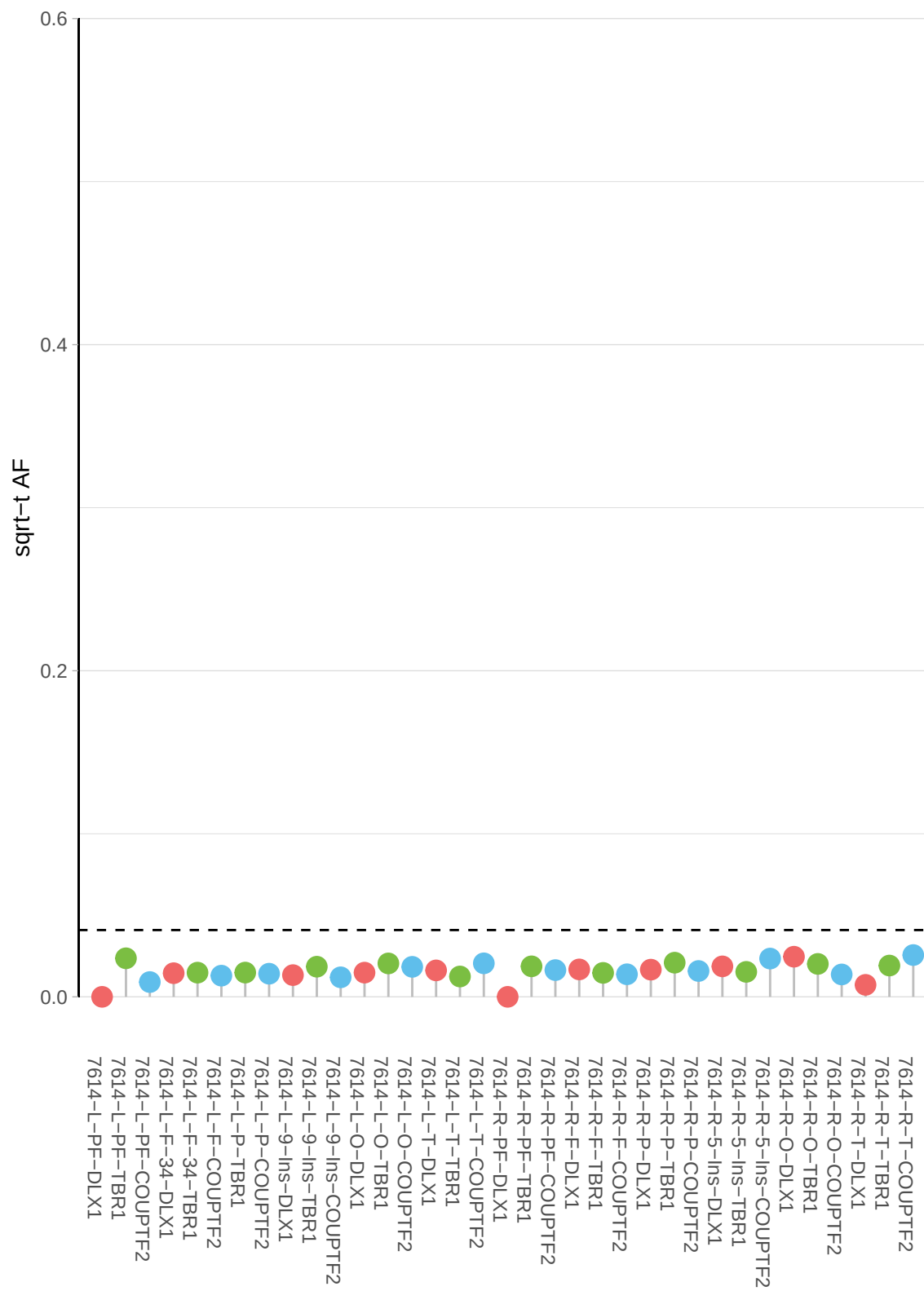

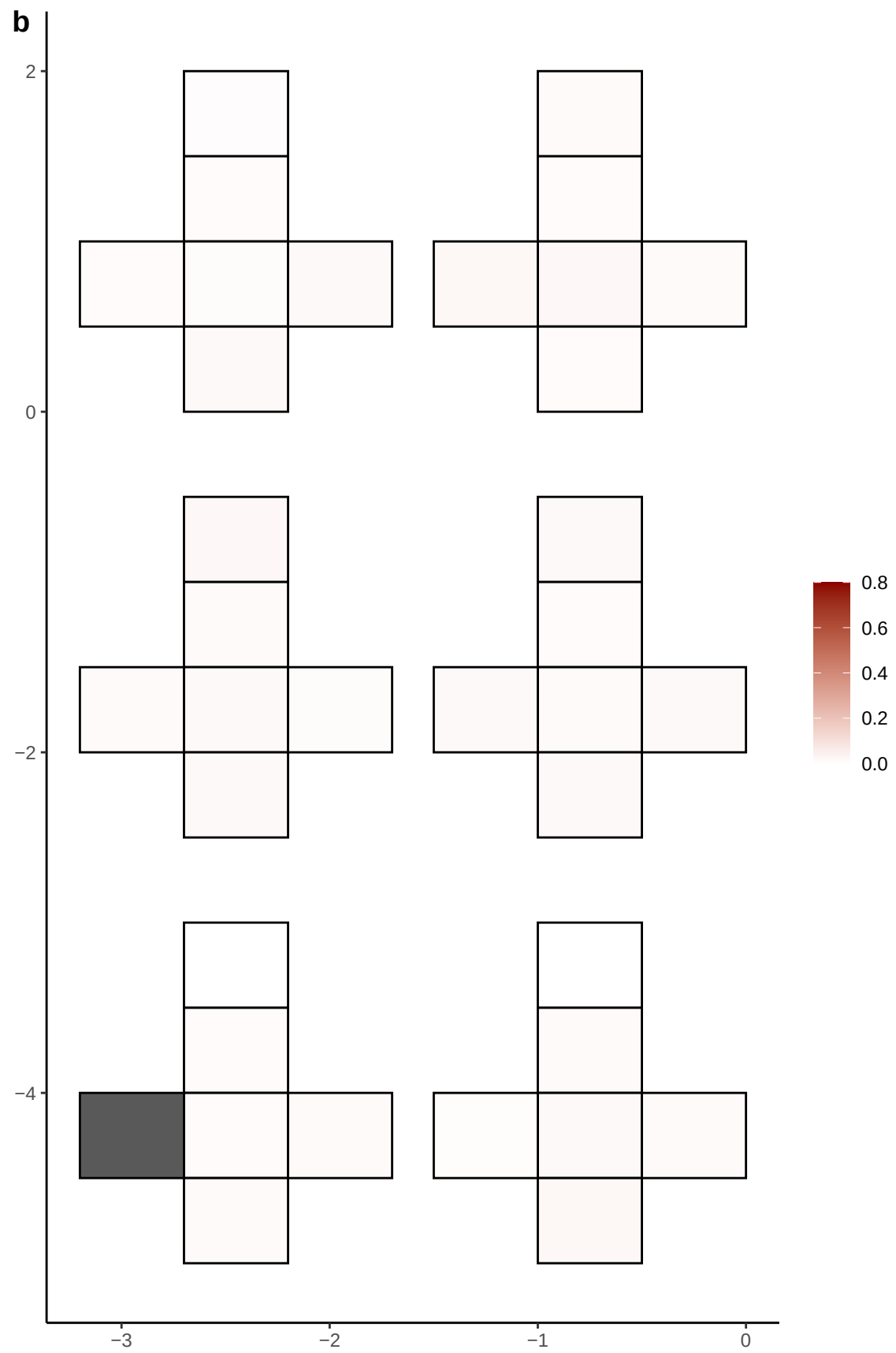

**a** ID01 5-65315424-G-A

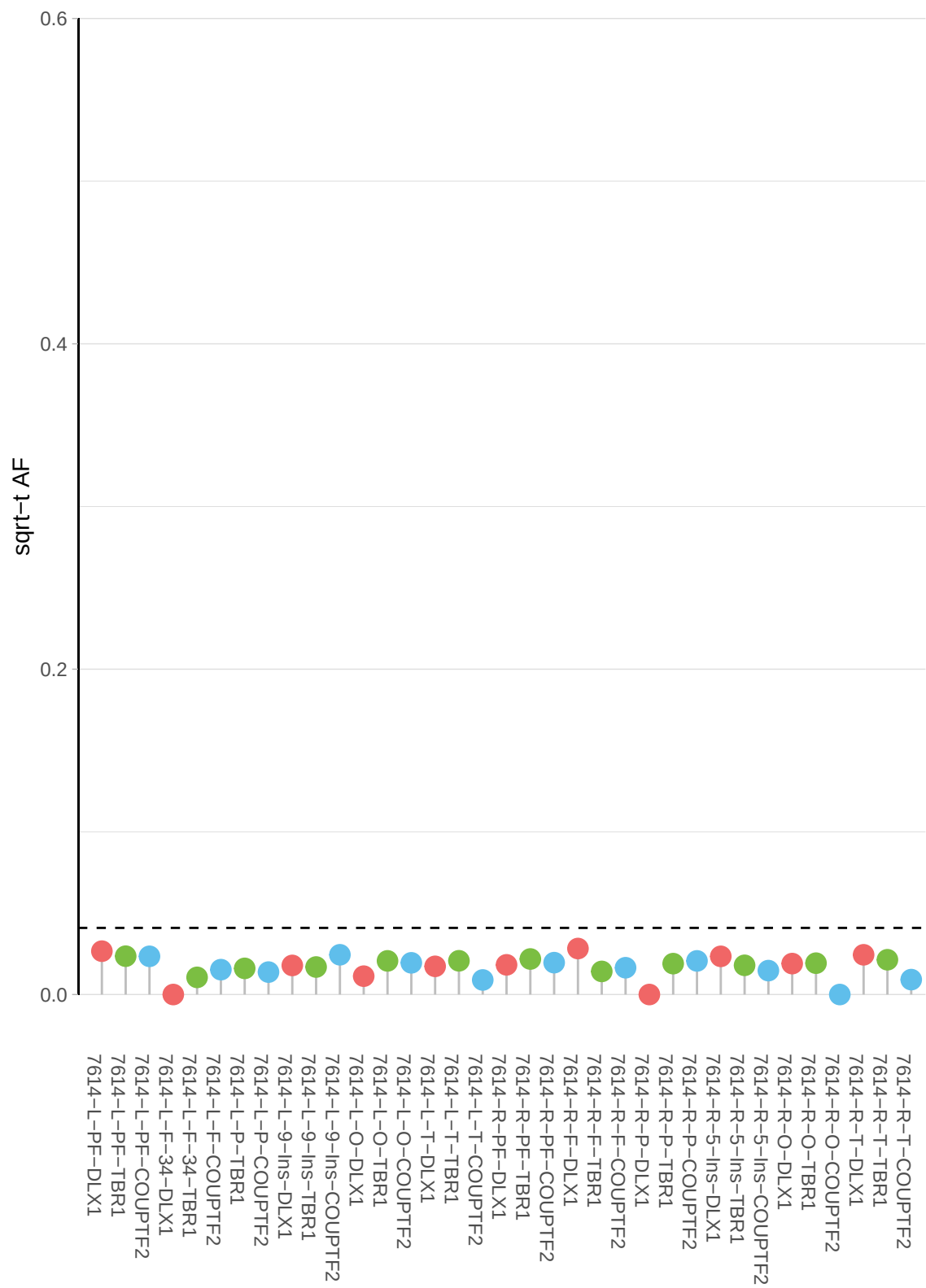

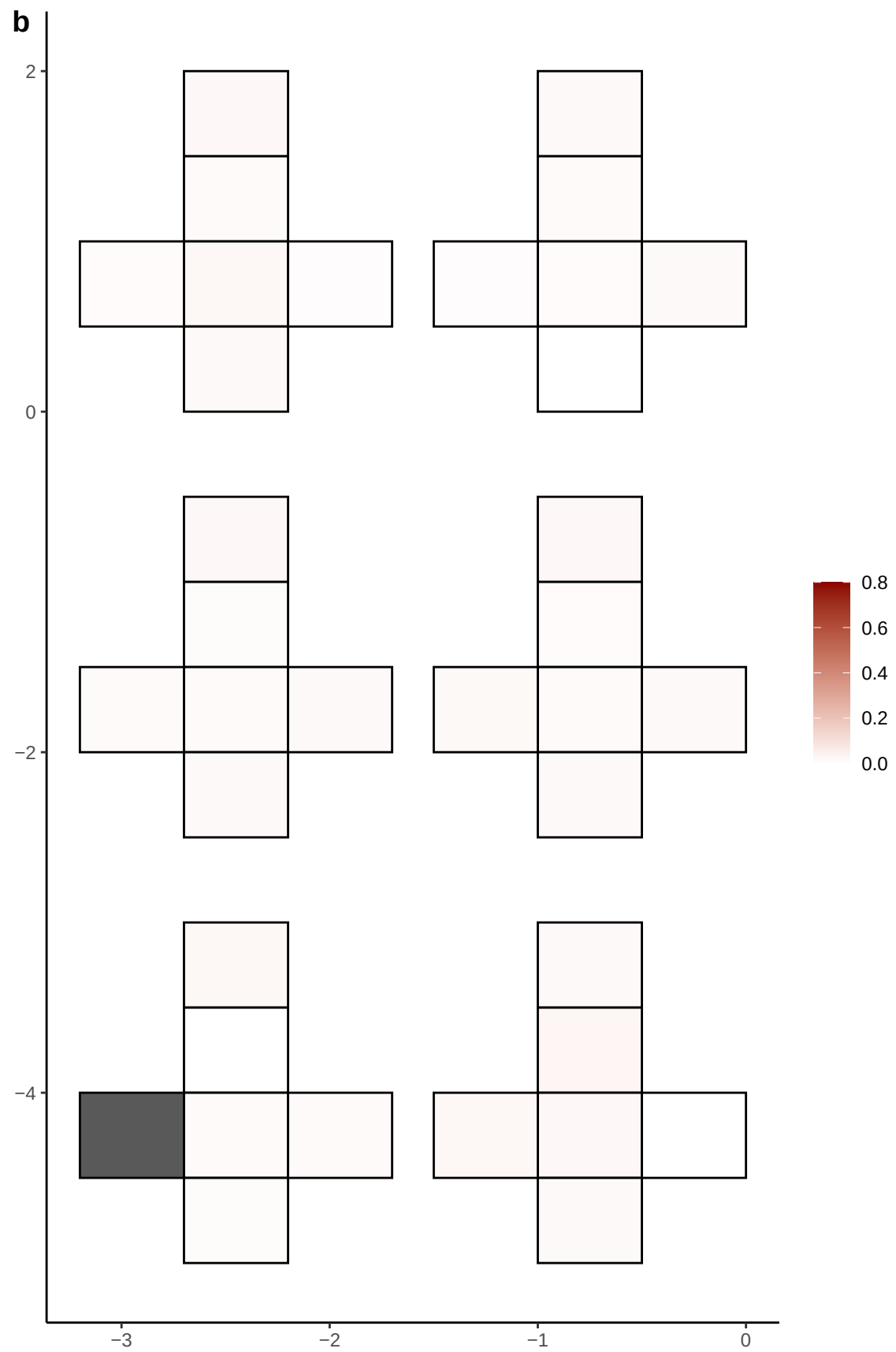

**a** ID01 5-72178944-ACG-A

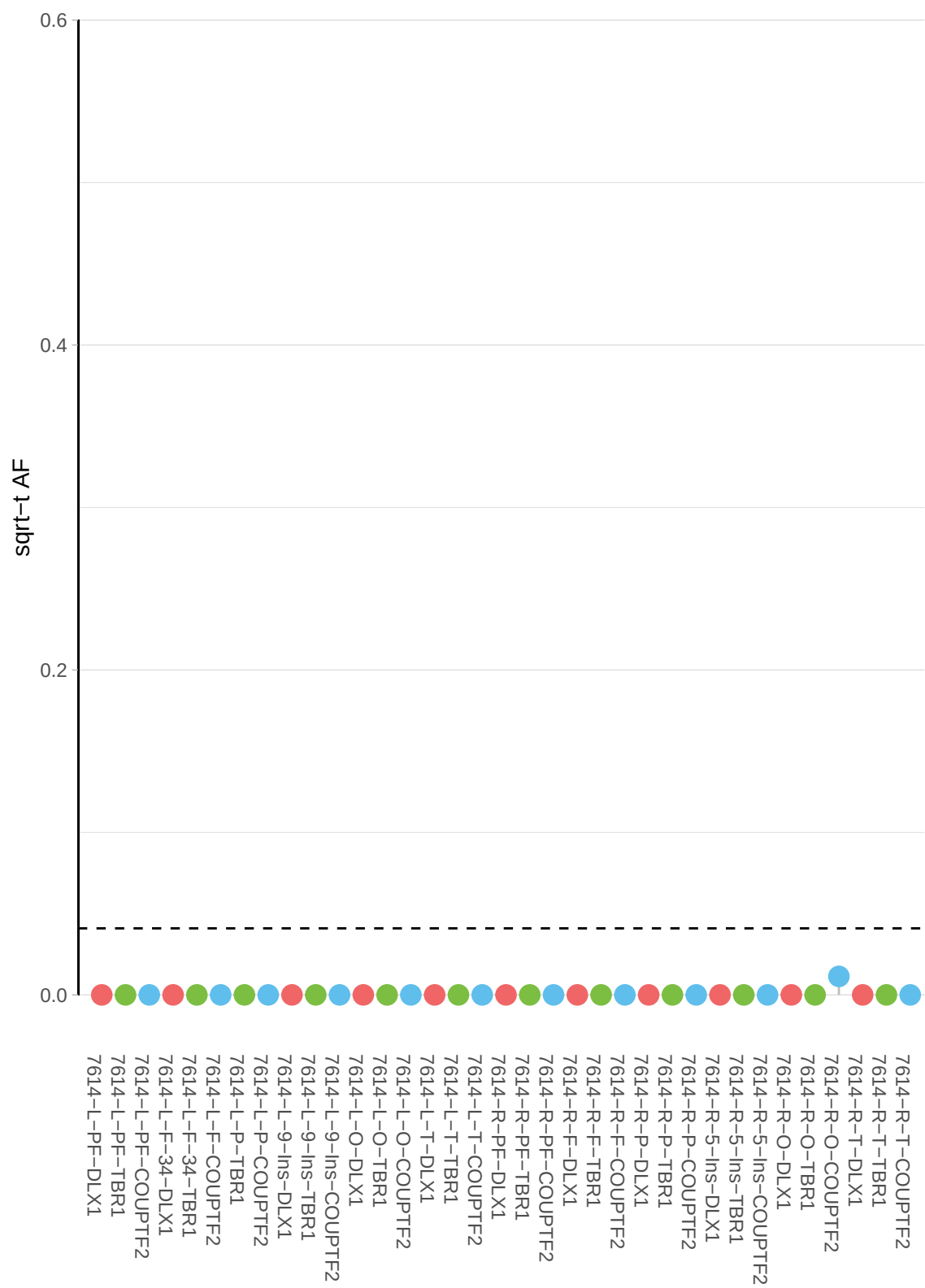

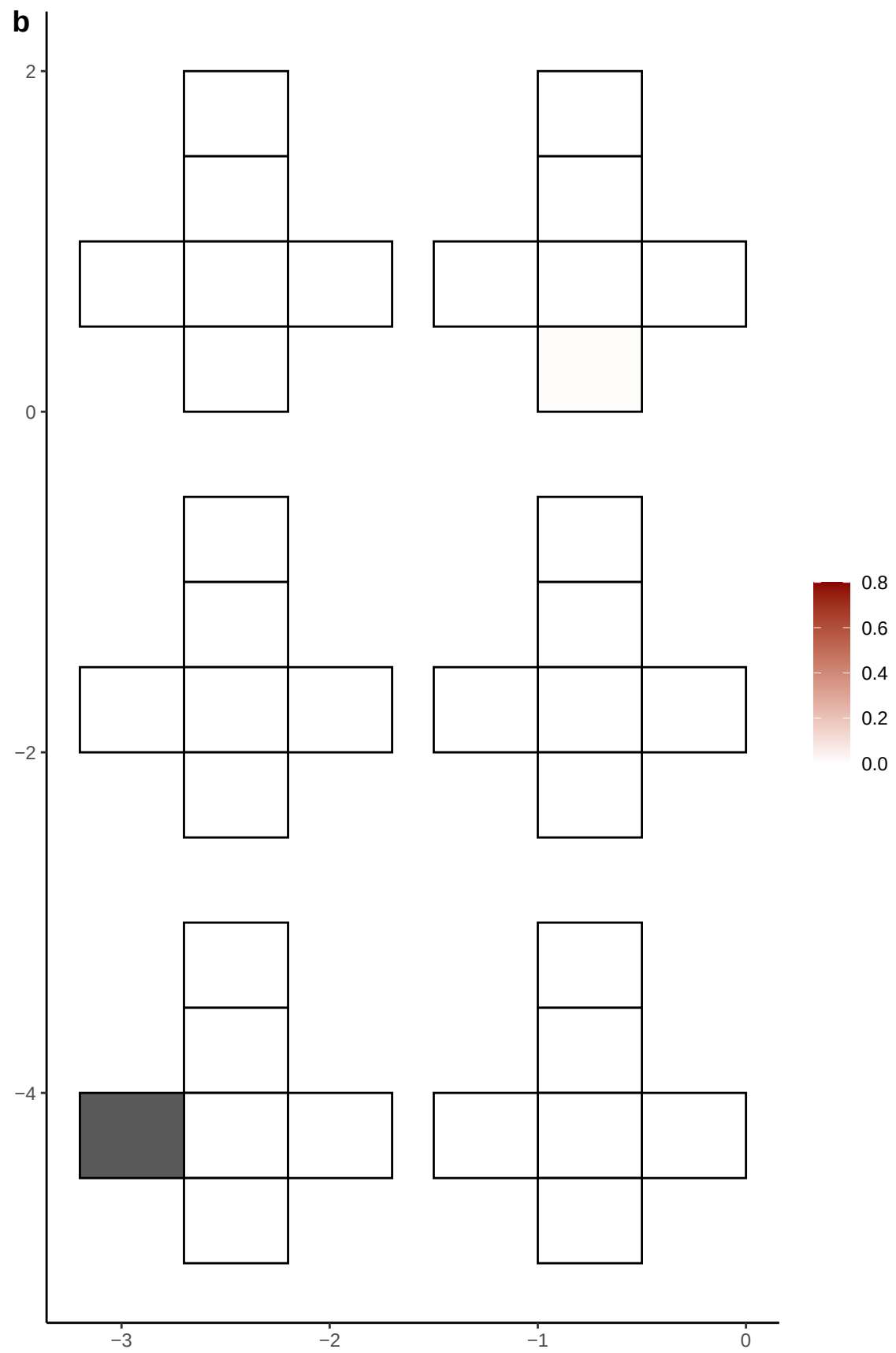

**a** ID01 5-72392587-A-G

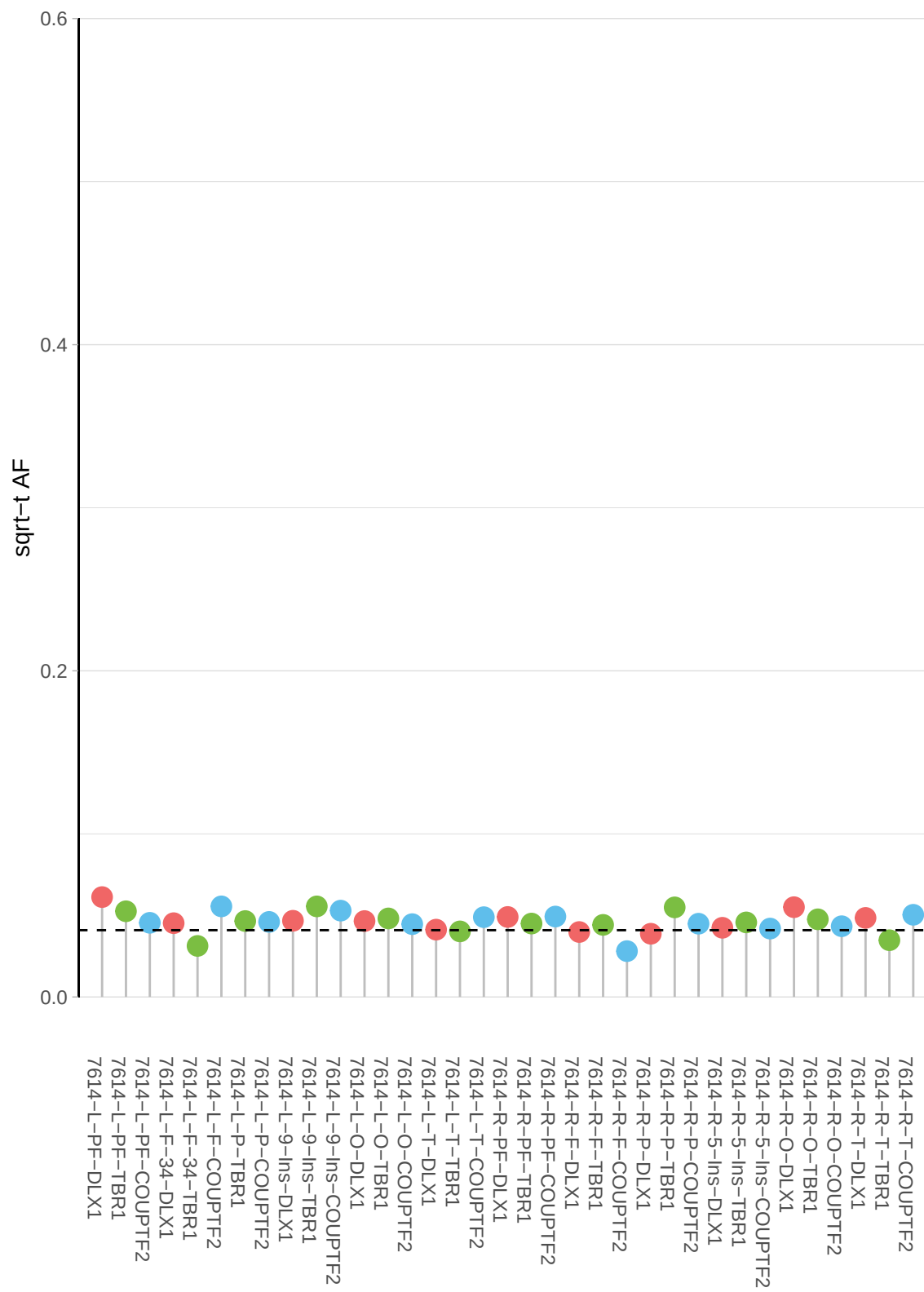

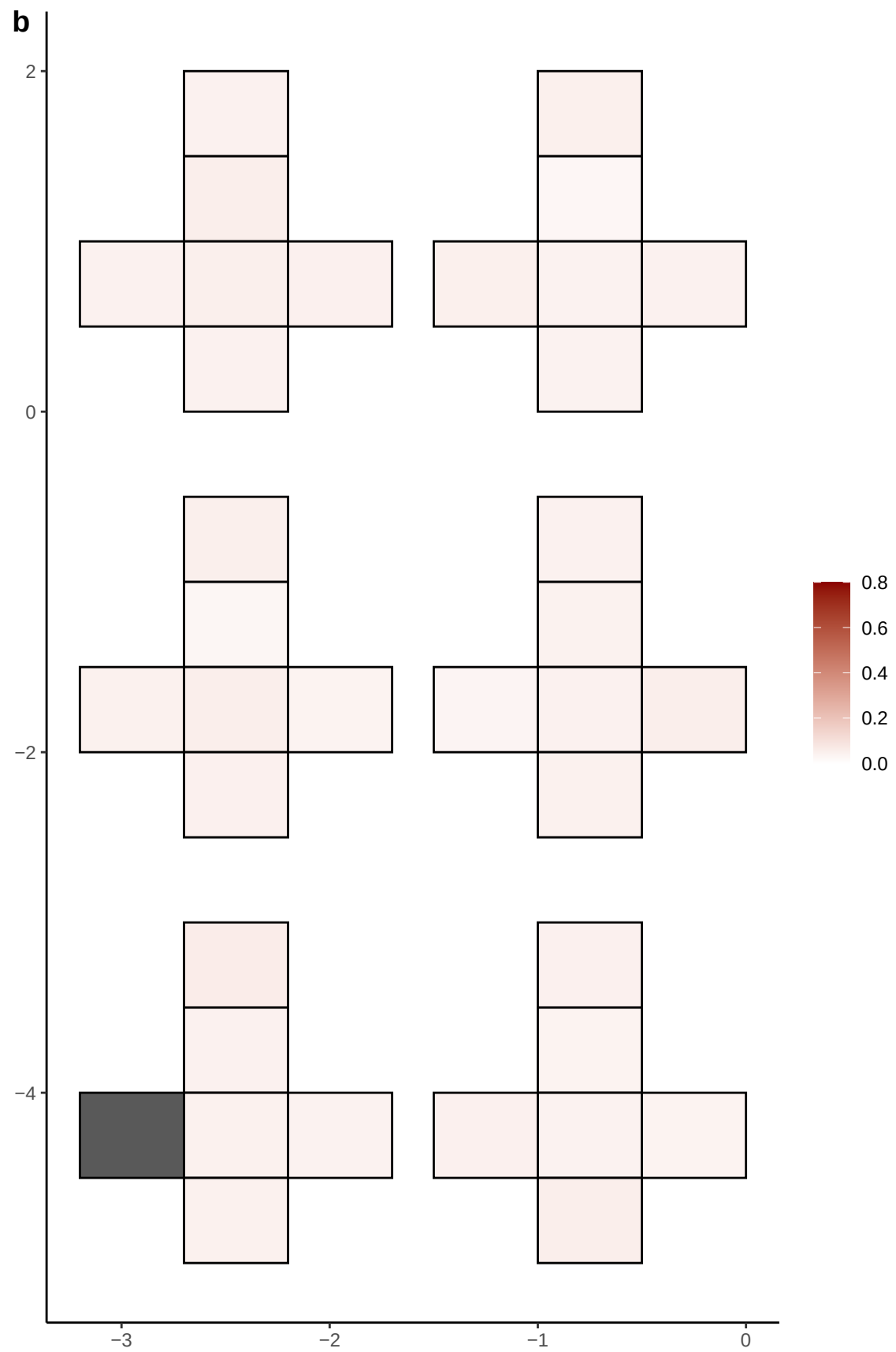

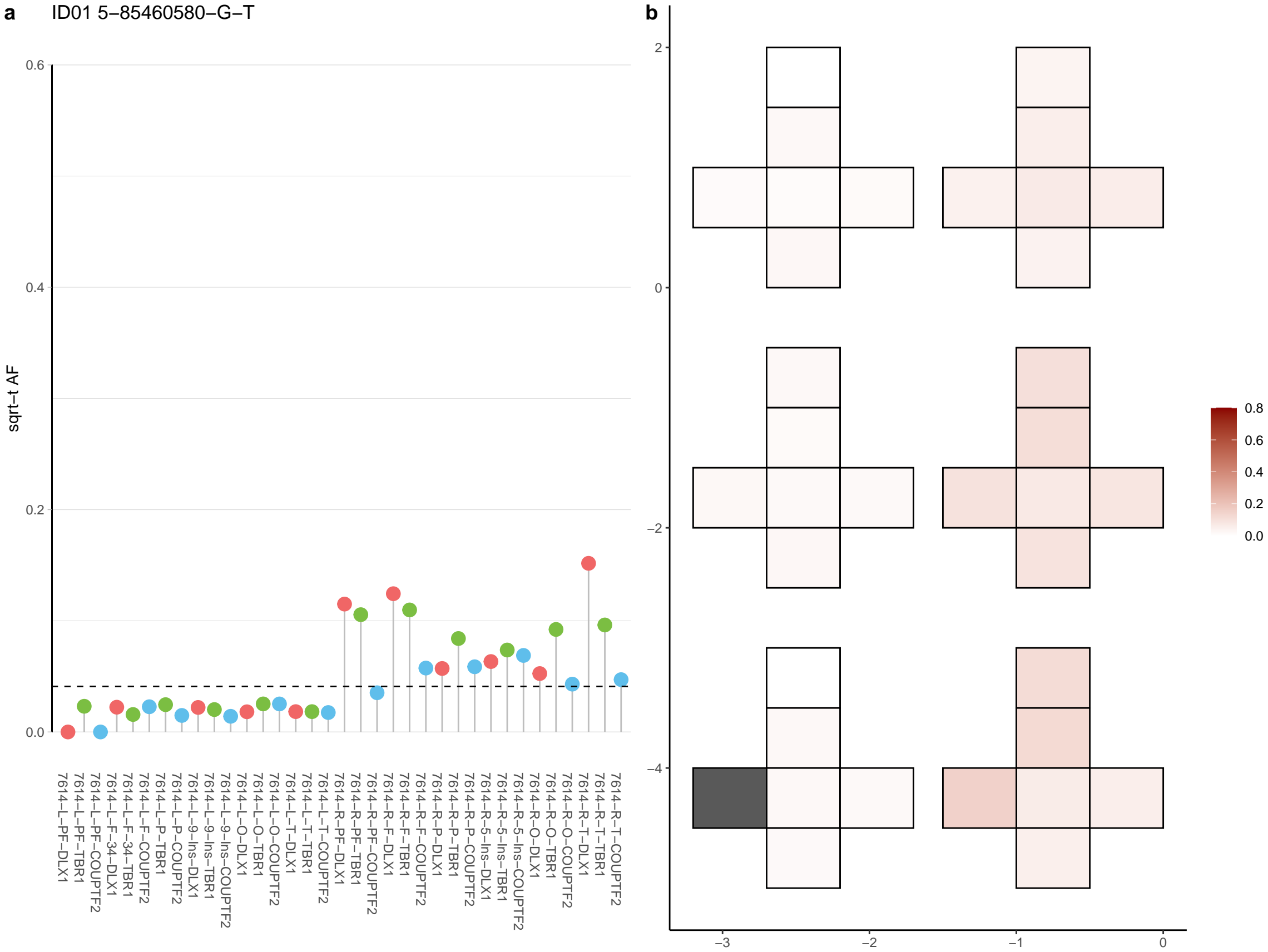

**a** ID01 6-101898717-G-A

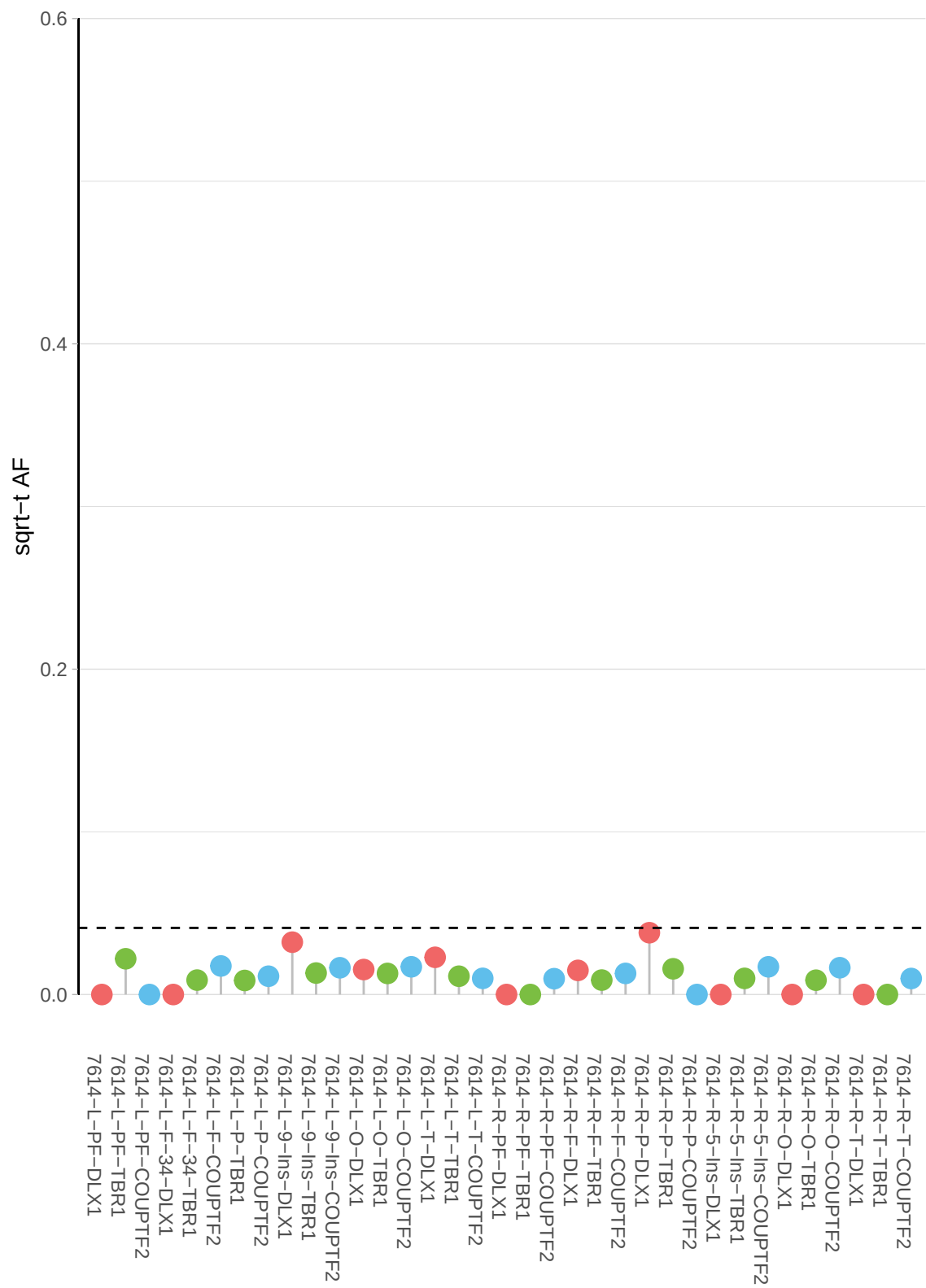

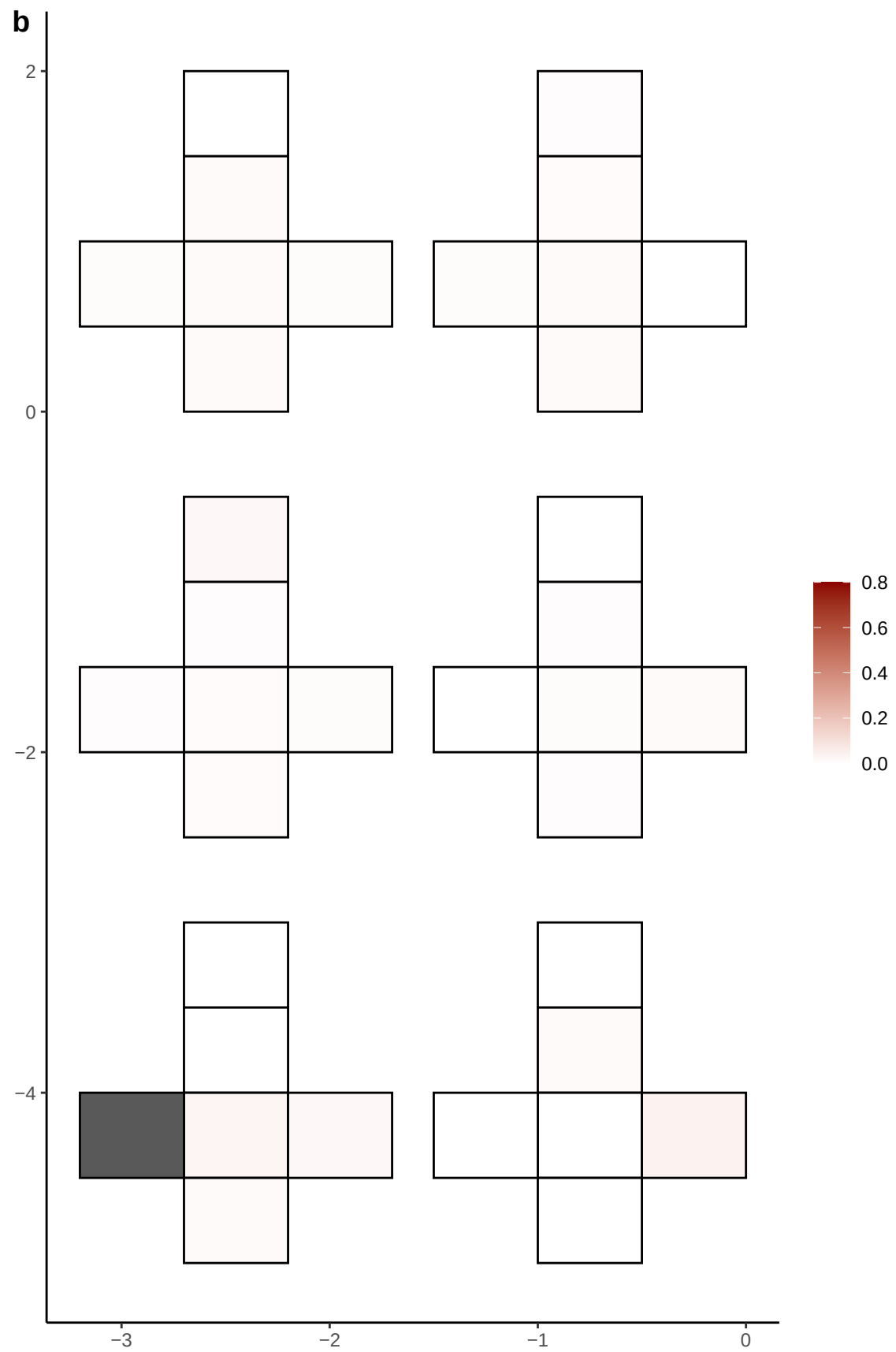

**a** ID01 6-110014503-C-T

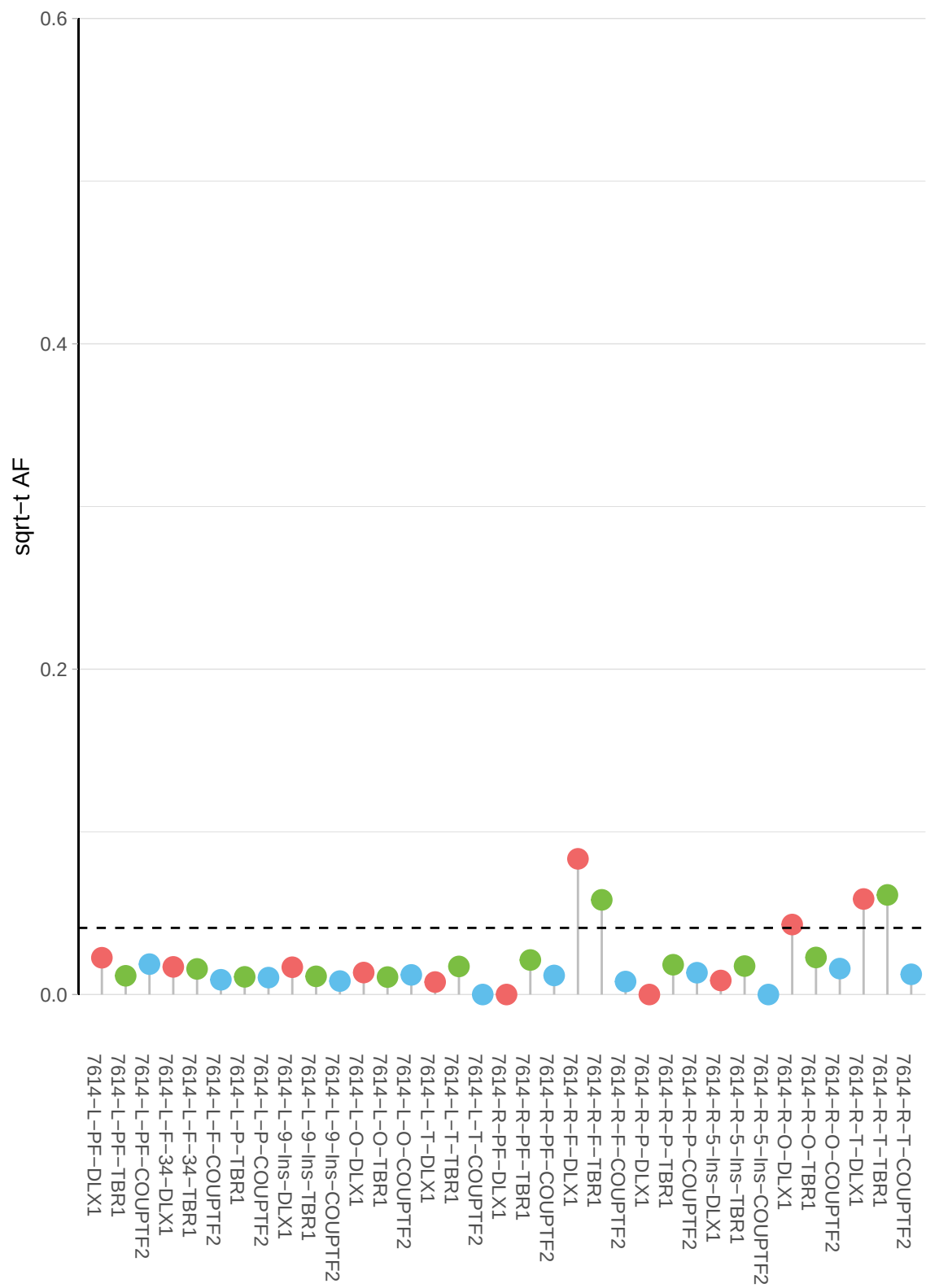

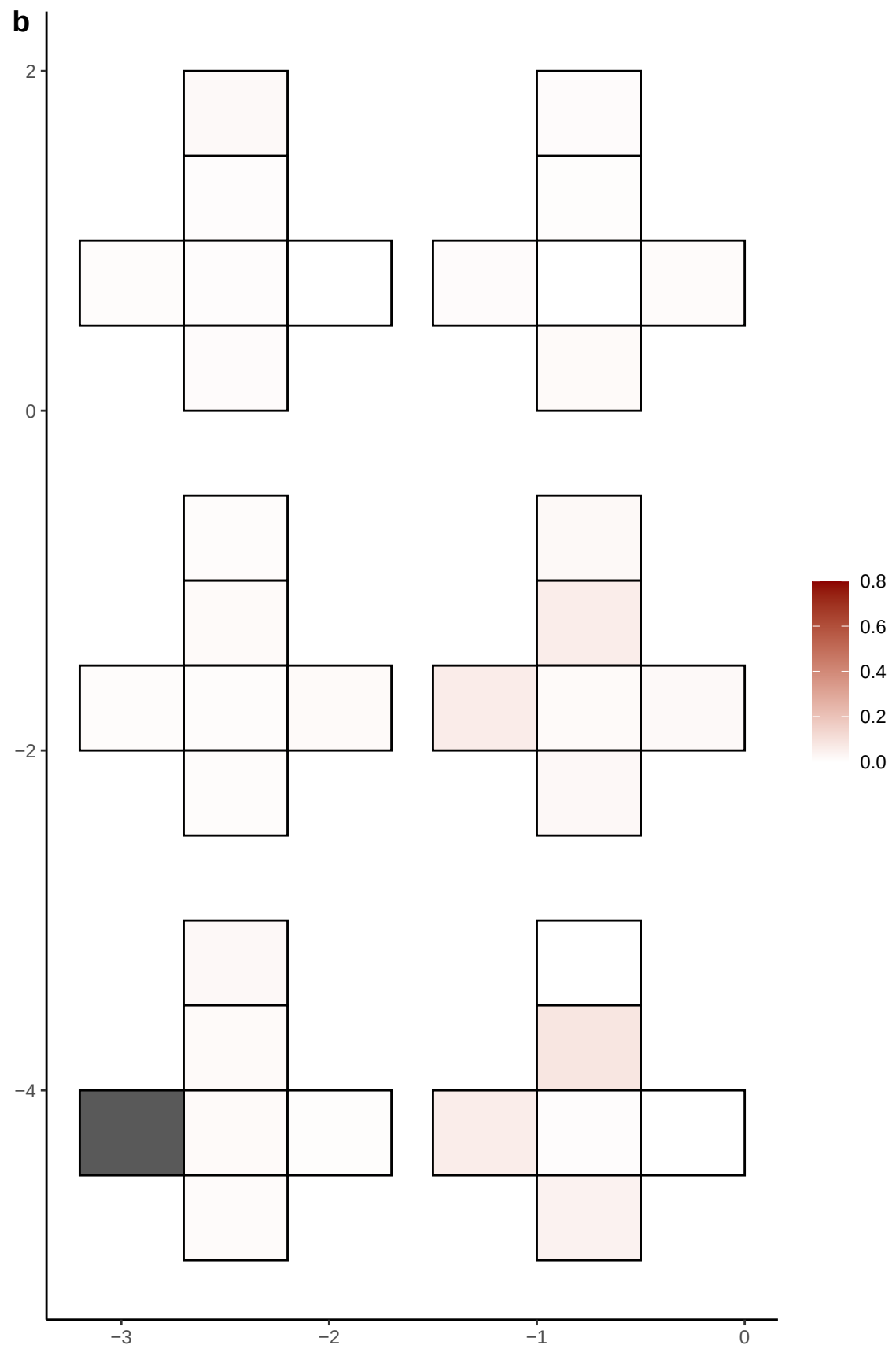

**a** ID01 6-114610552-GAACA-G

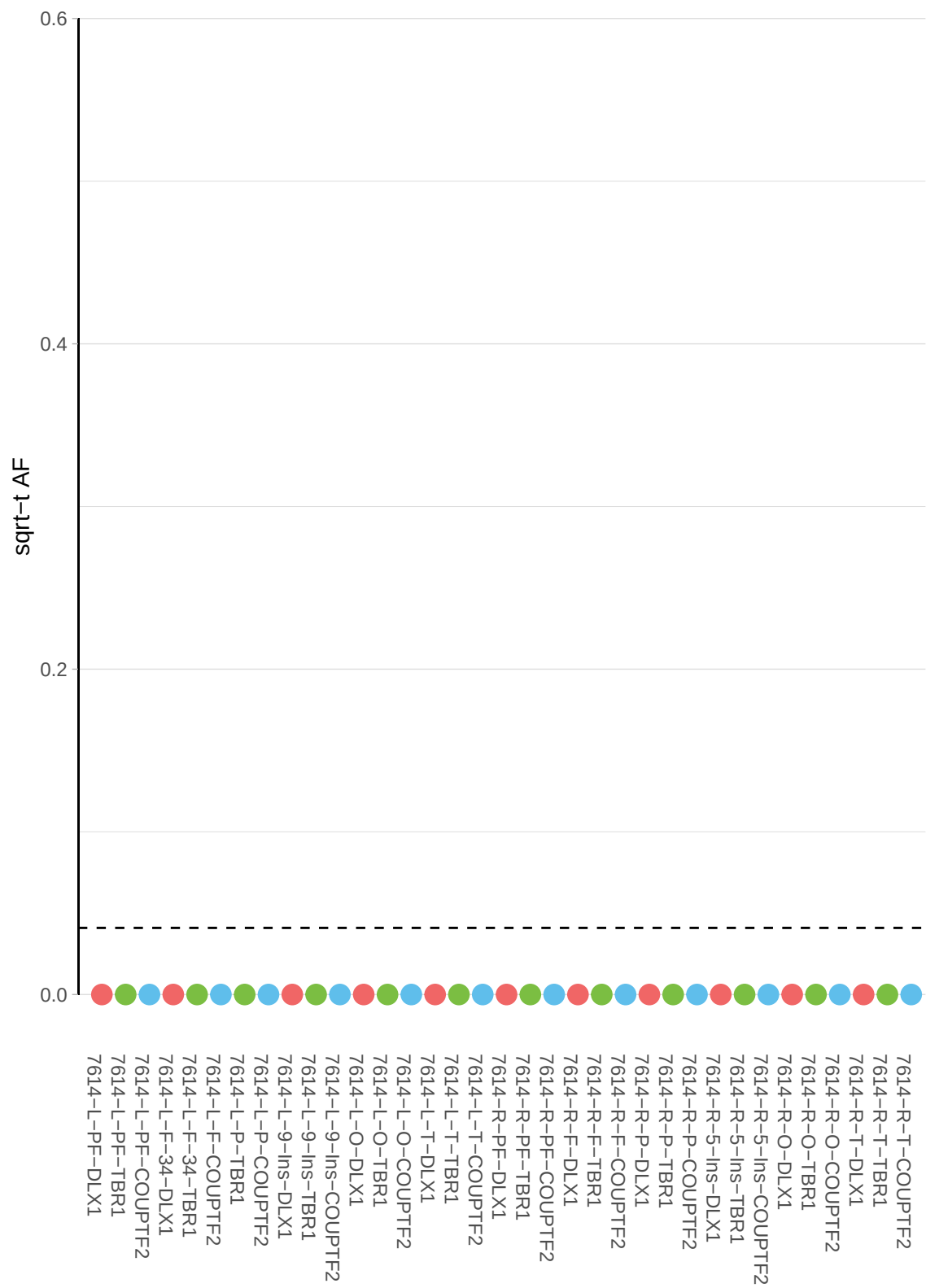

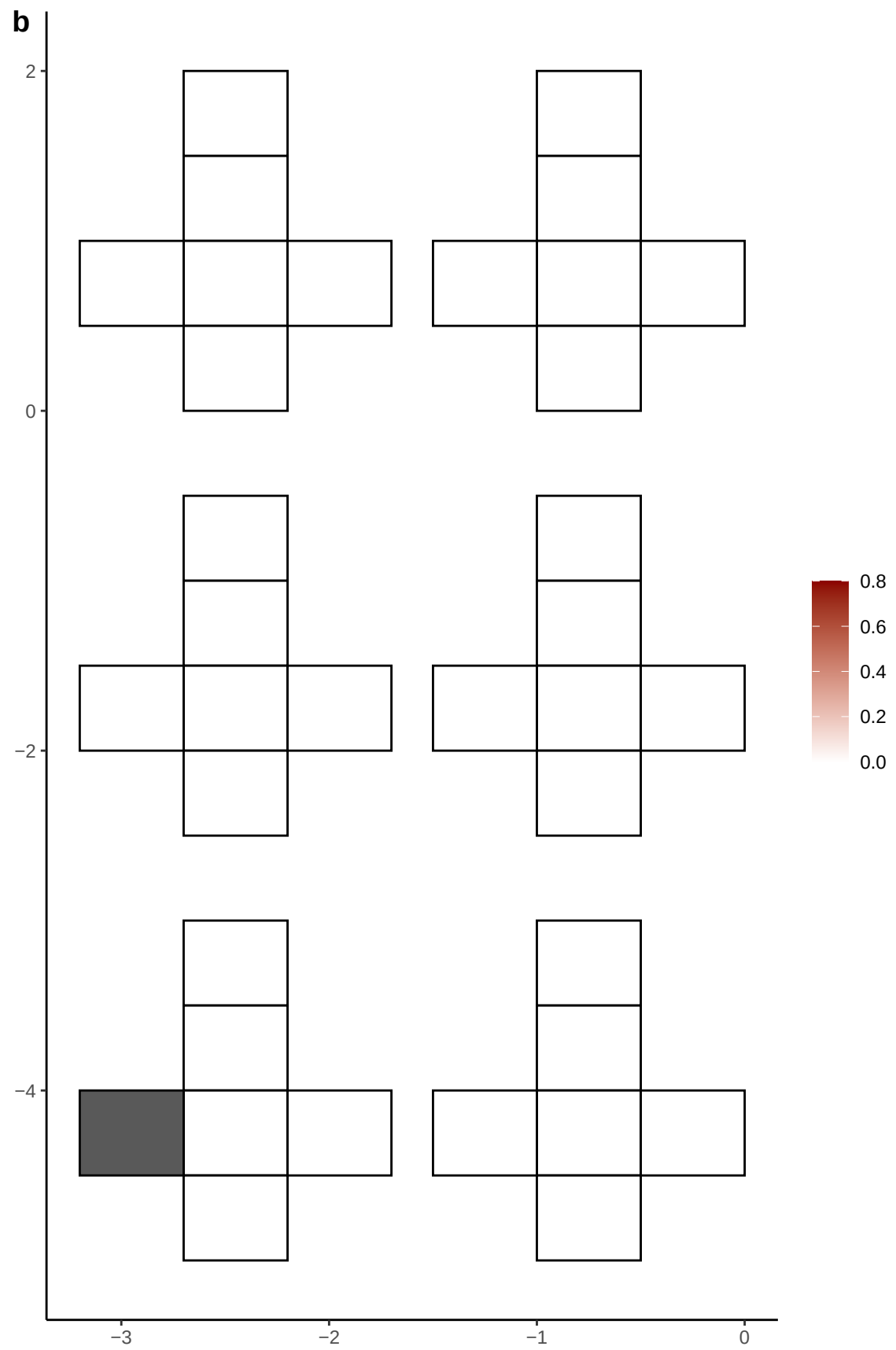

**a** ID01 6-121420086-C-T

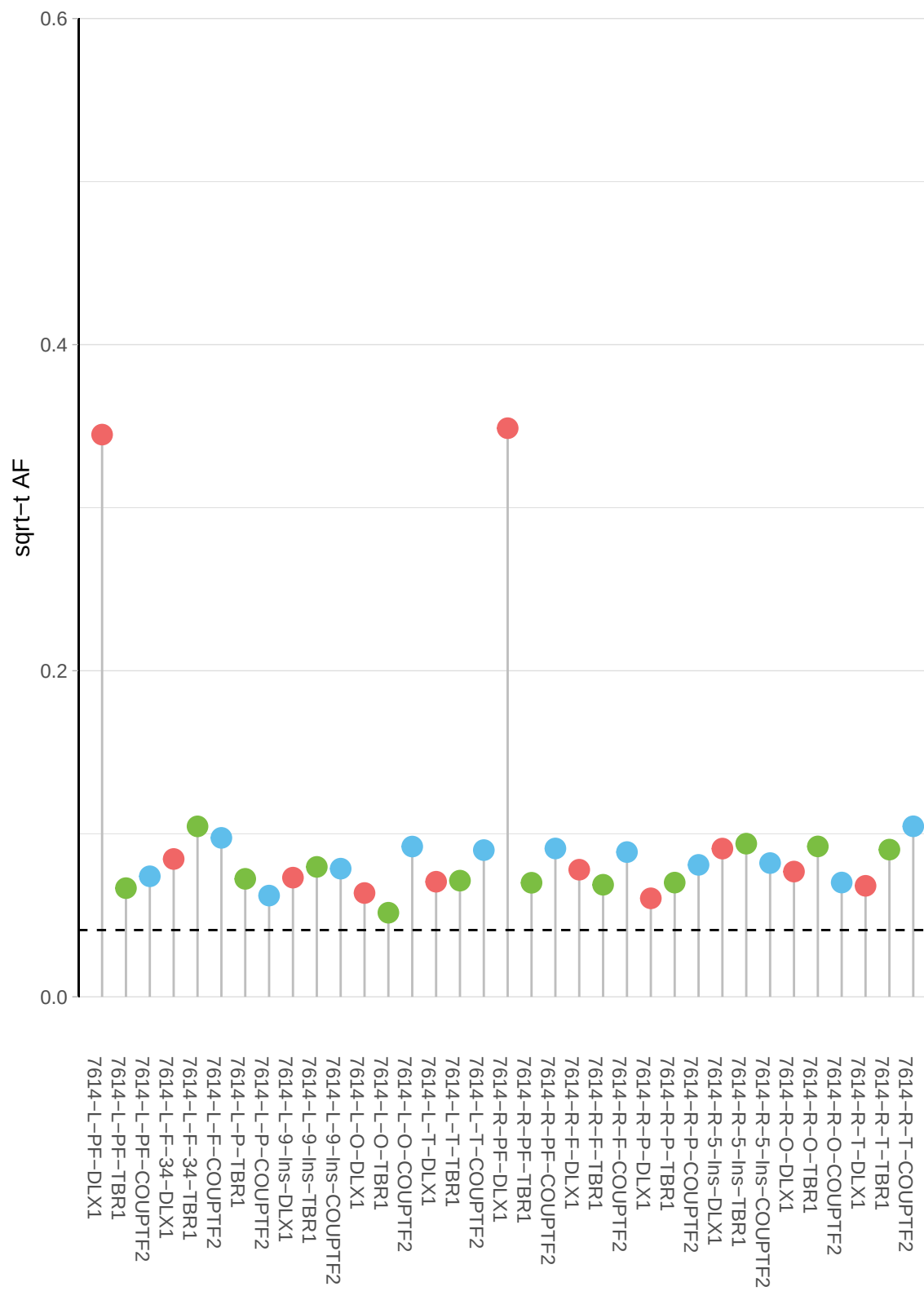

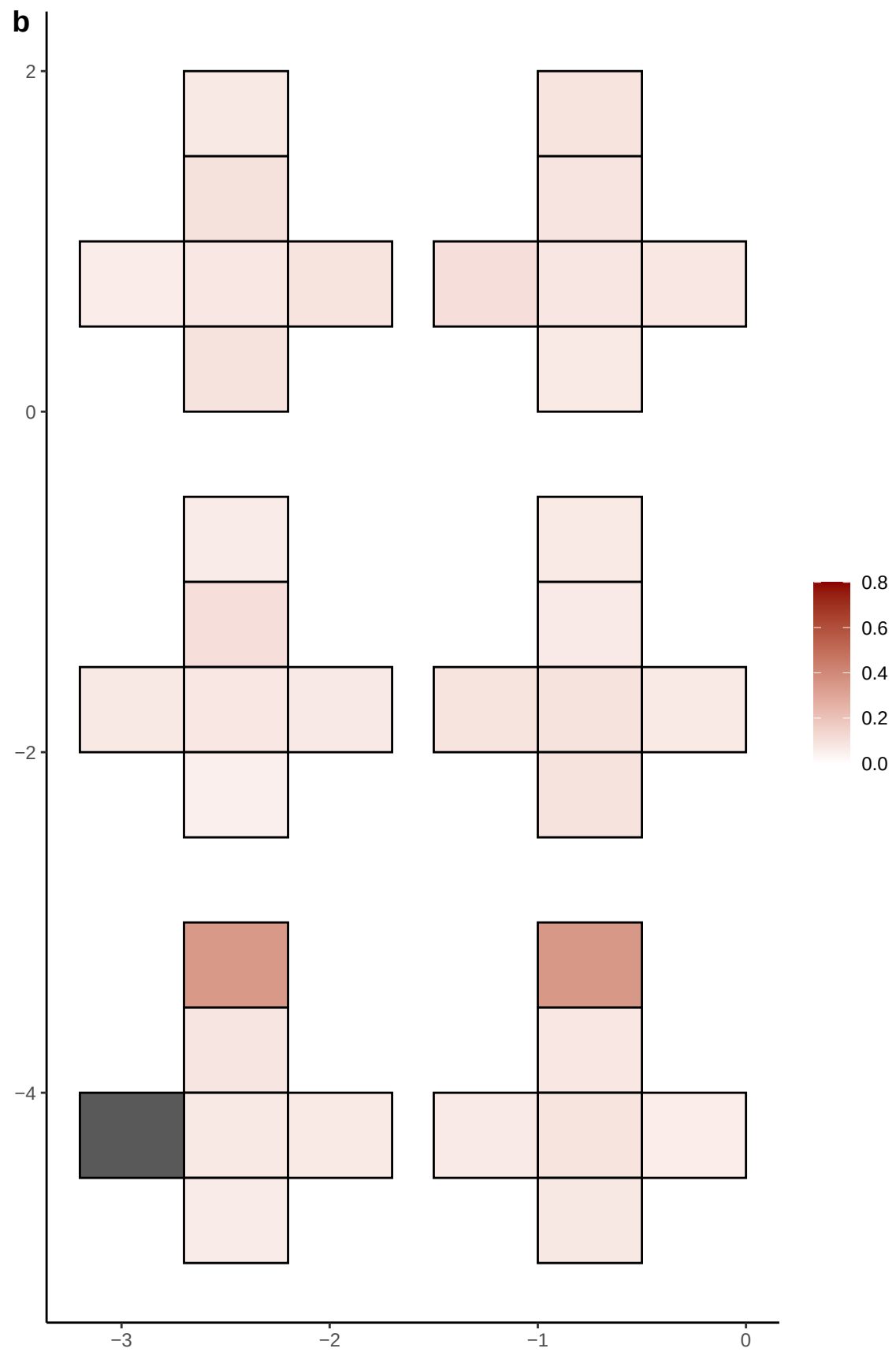

**a** ID01 6-122610884-C-T

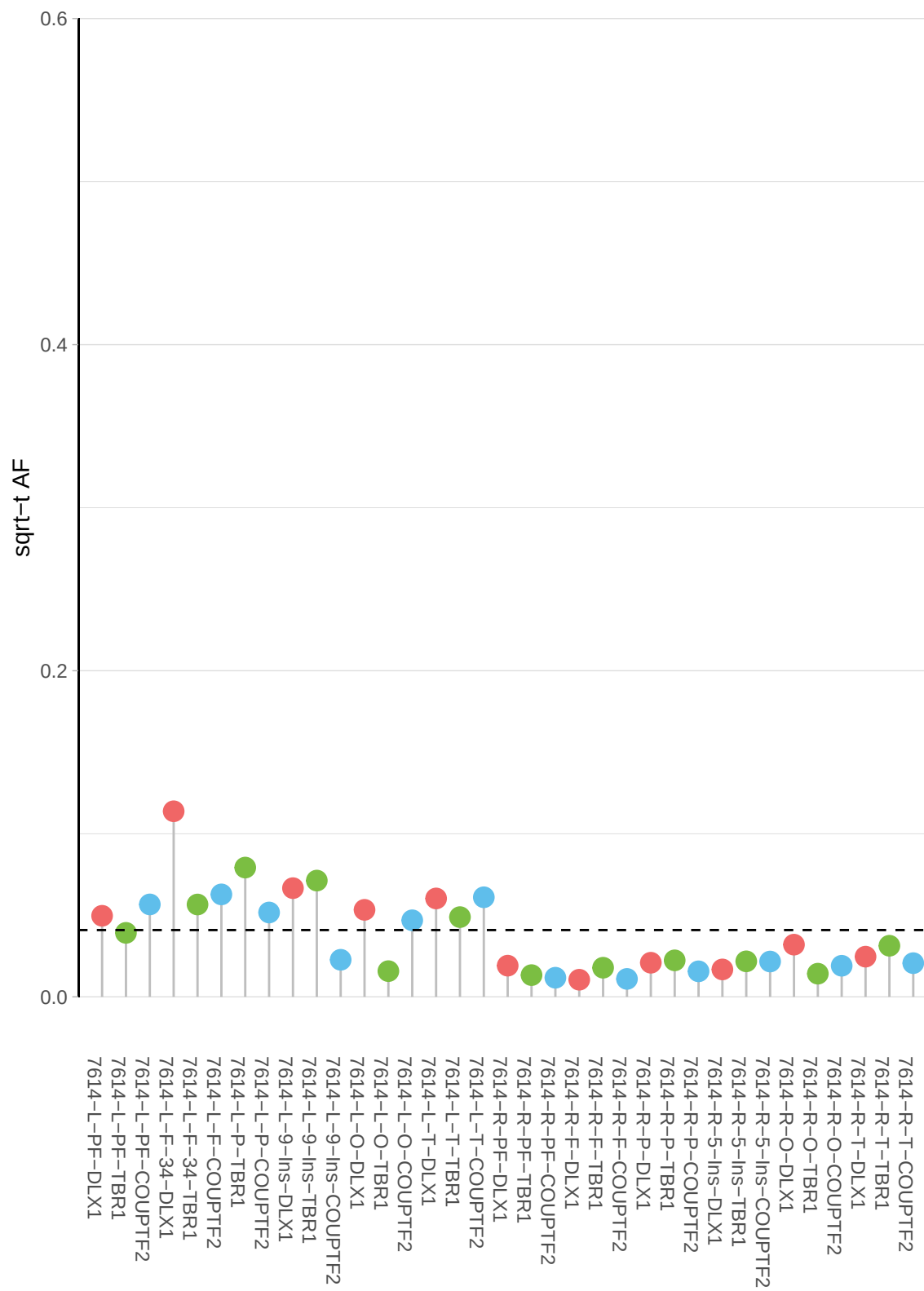

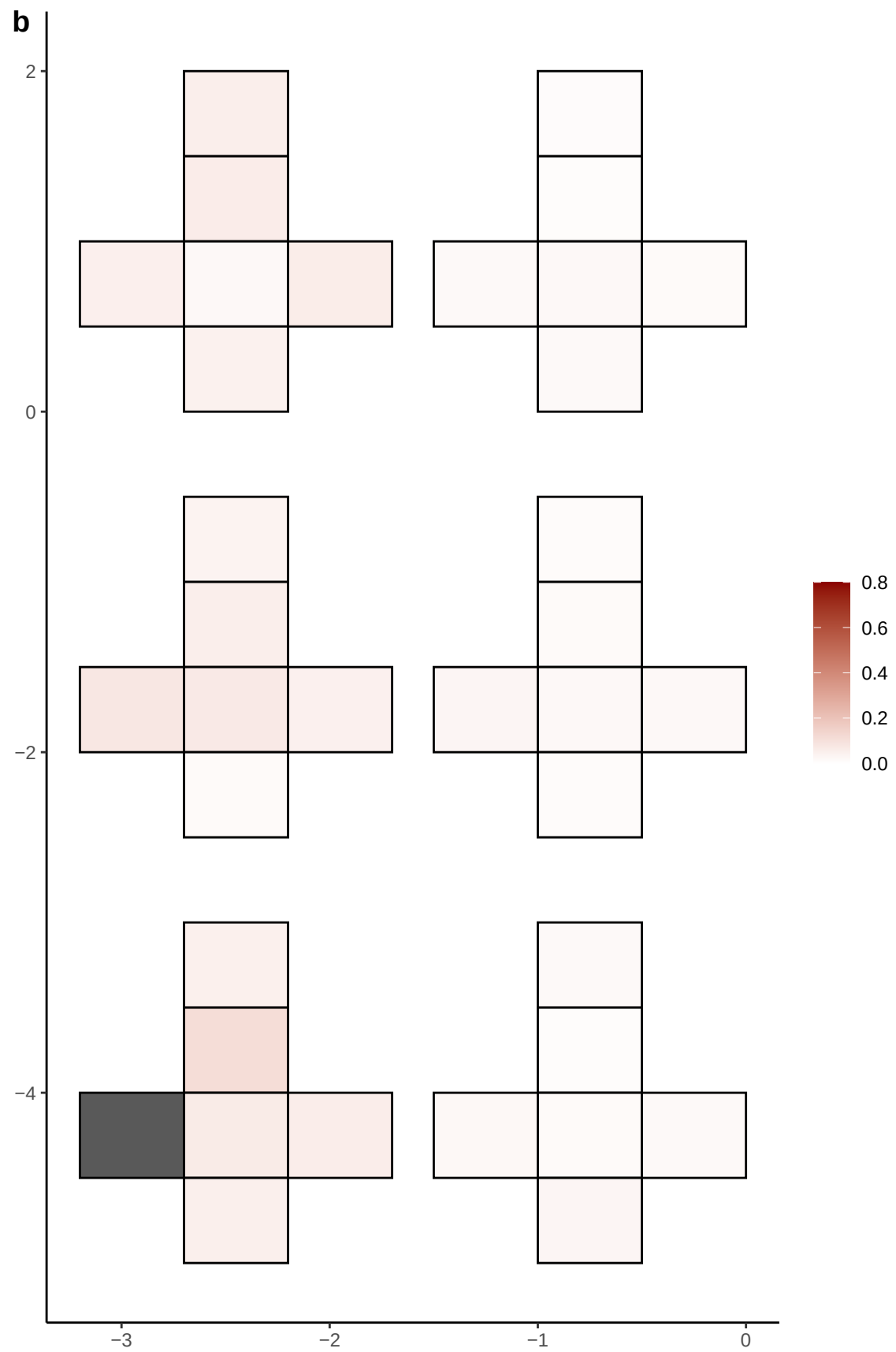

**a** ID01 6-125153224-A-C

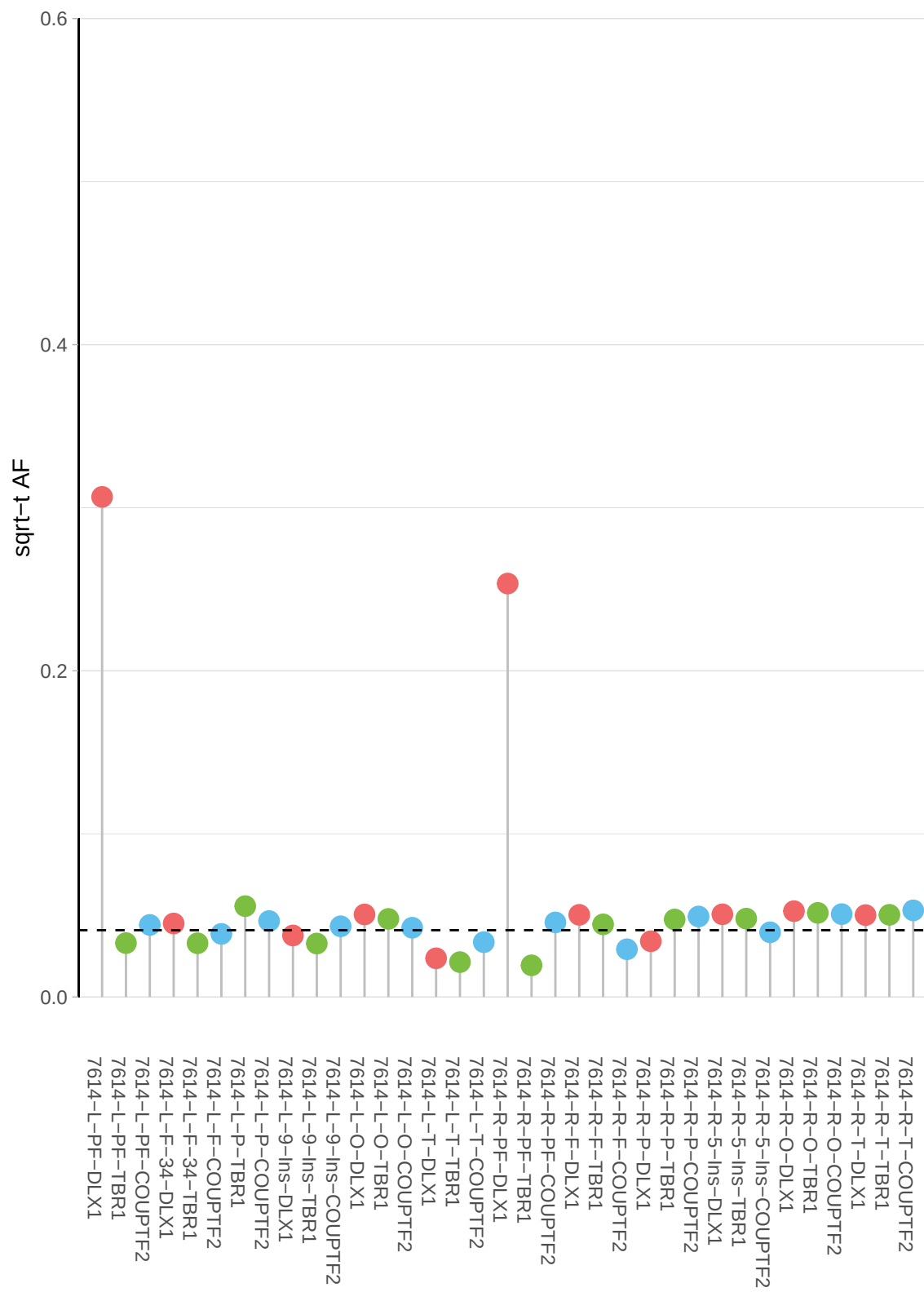

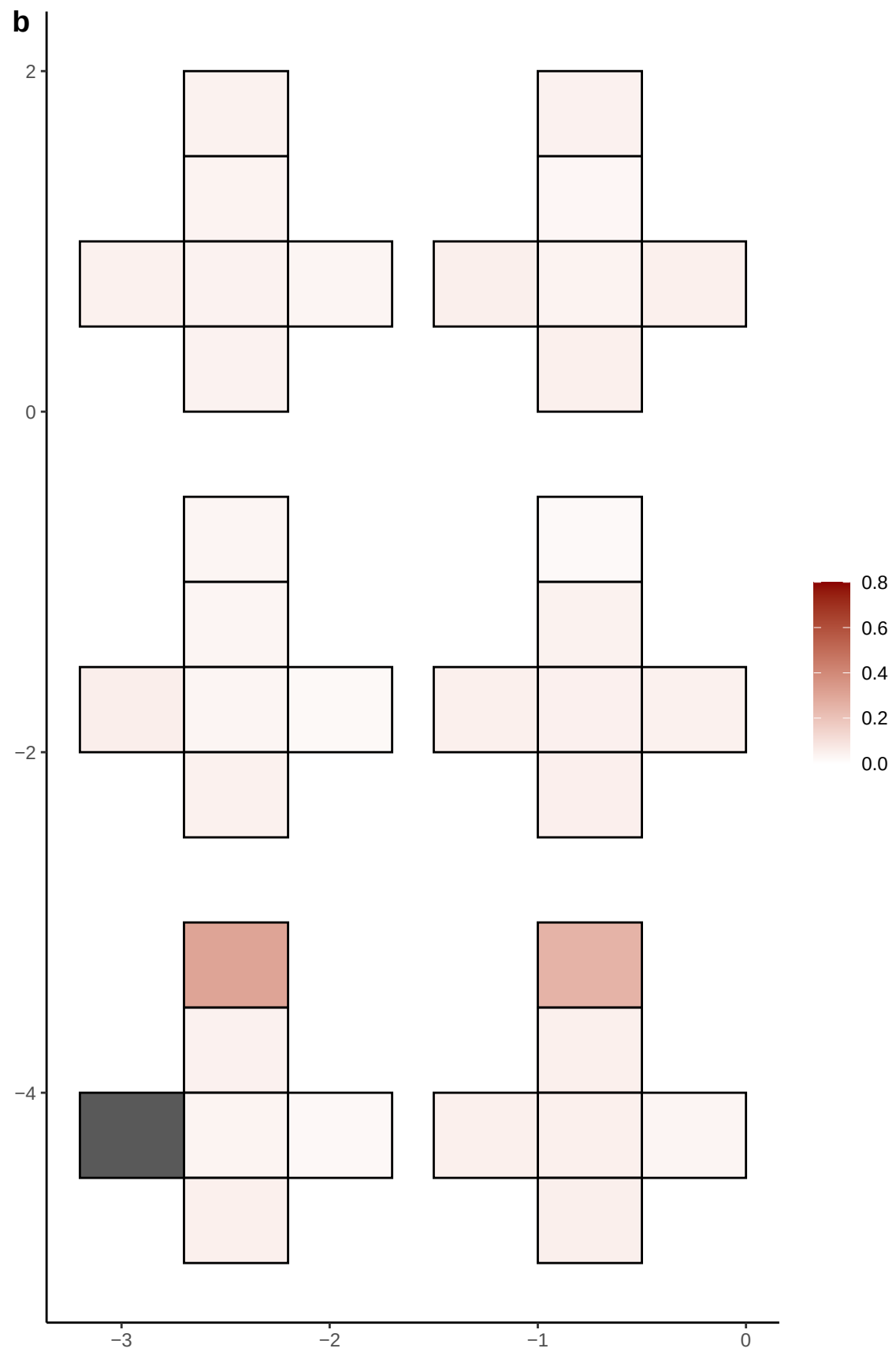

**a** ID01 6-125153225-G-T

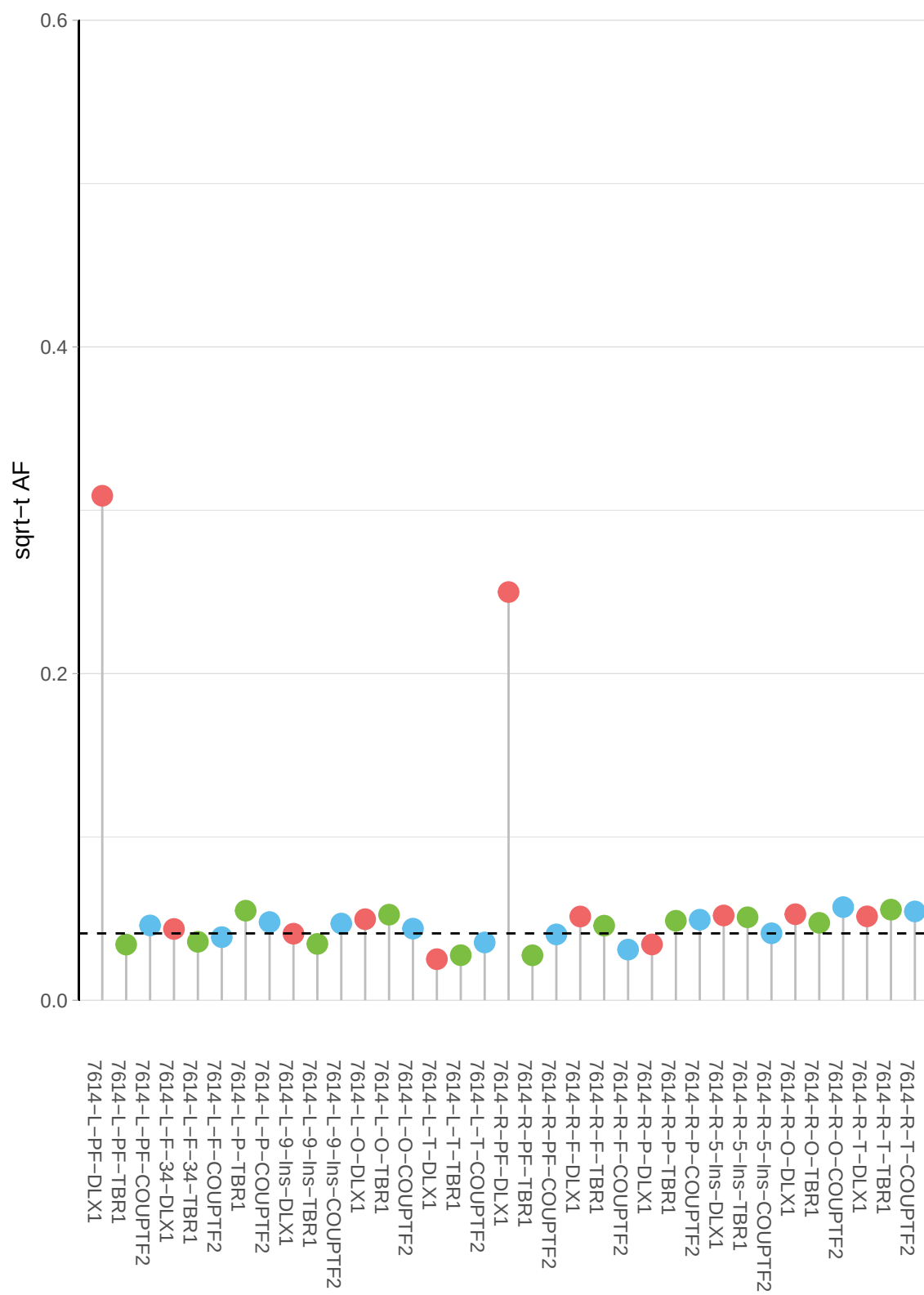

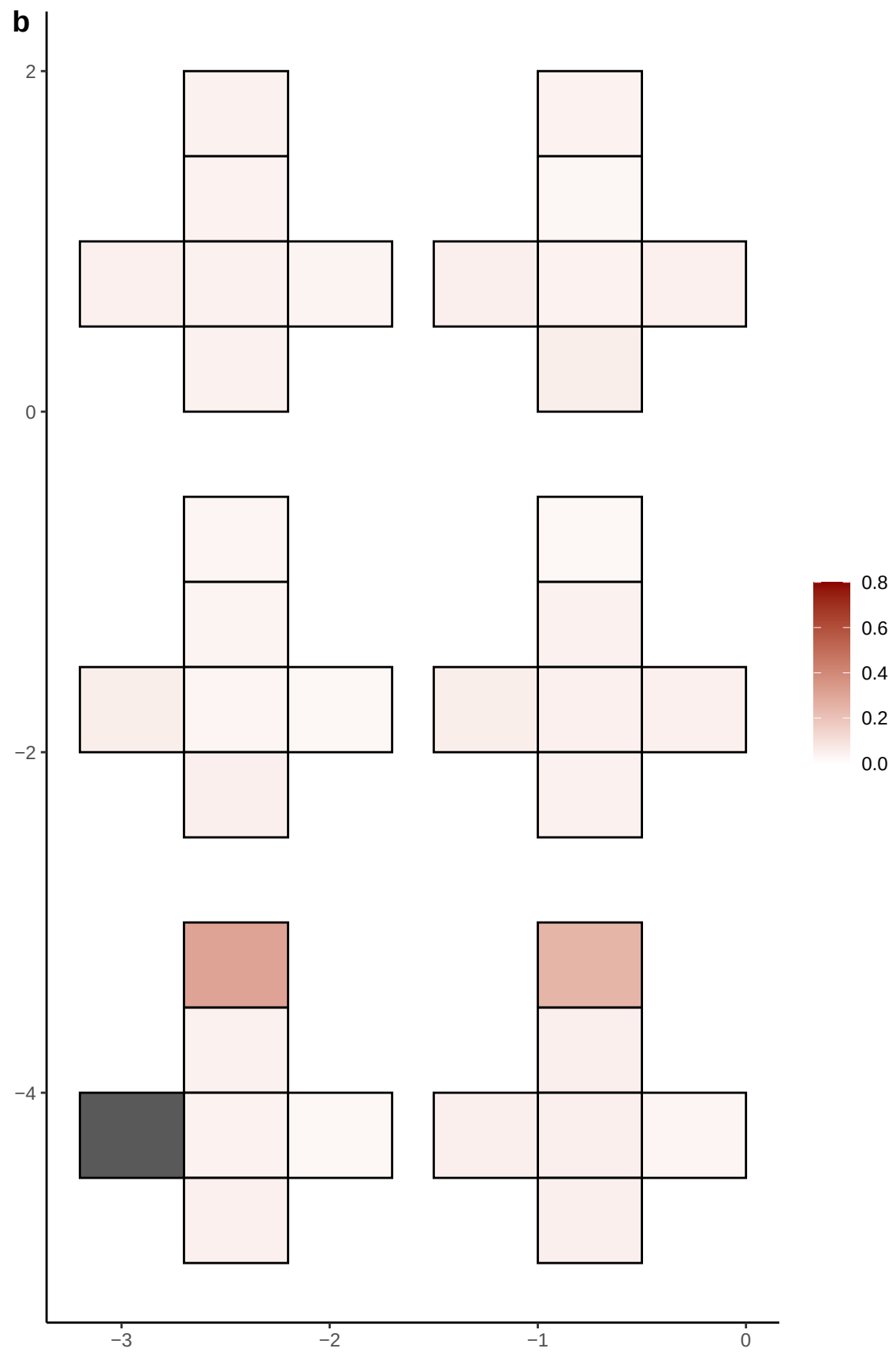

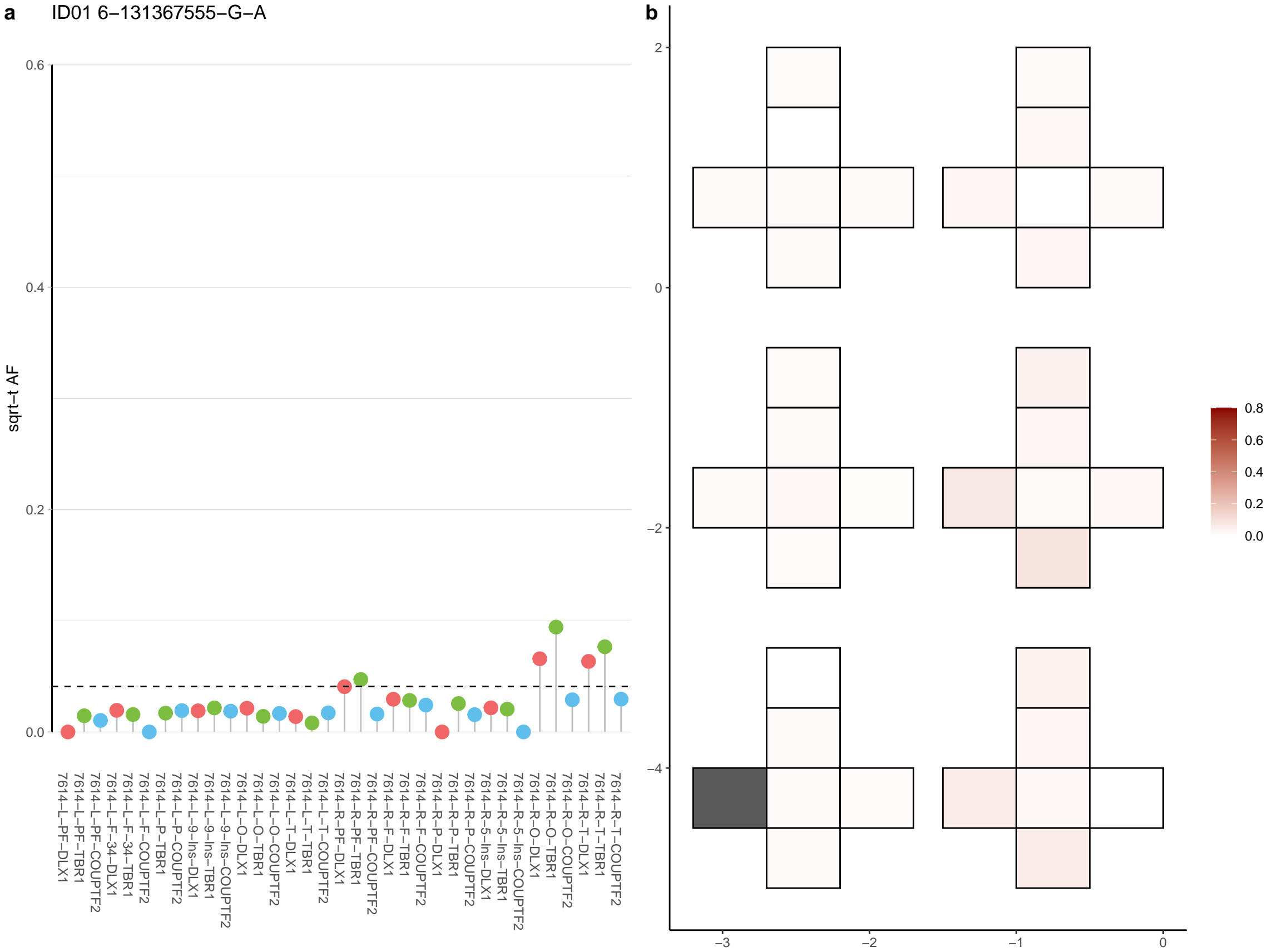

**a** ID01 6-154659392-C-T

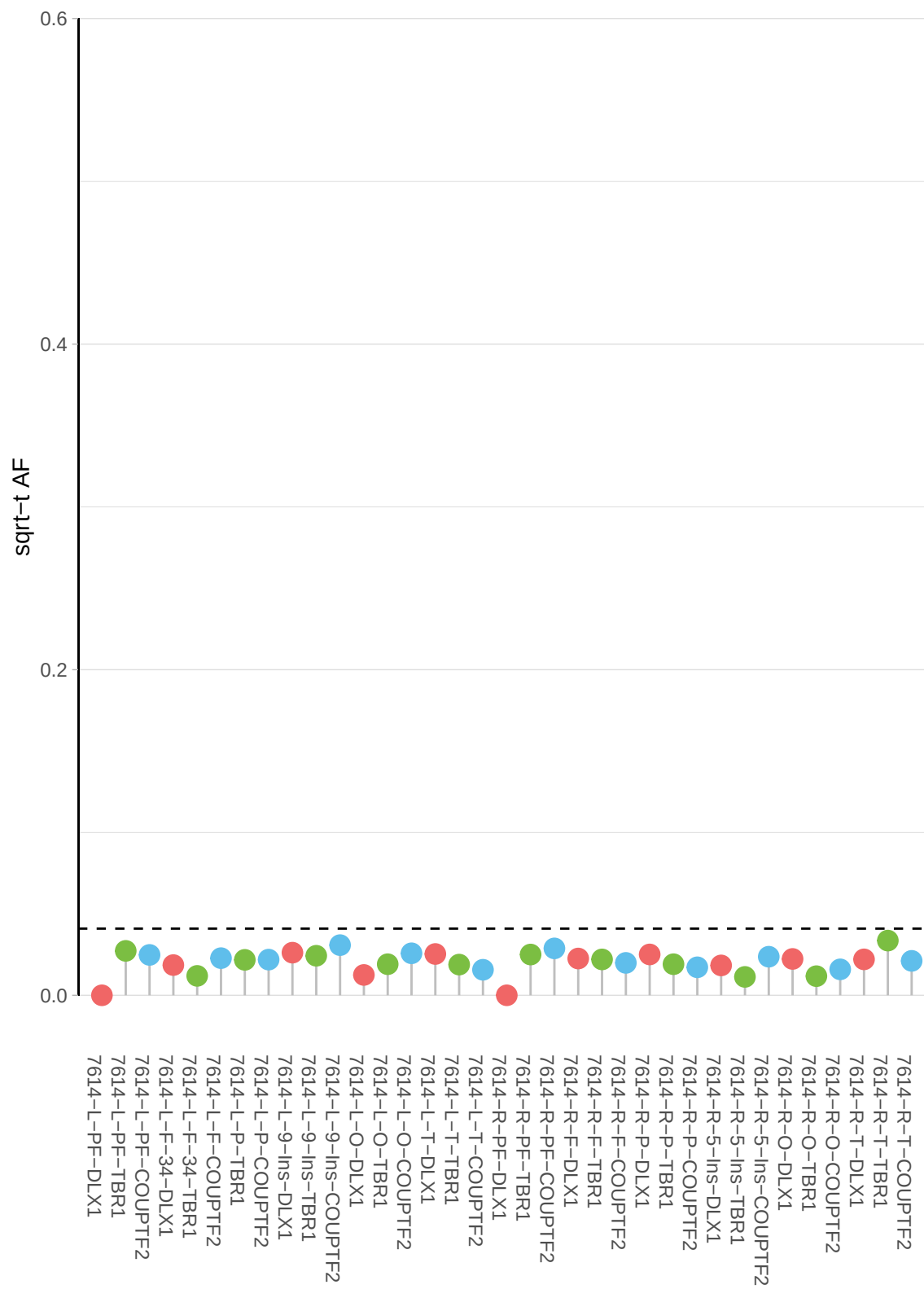

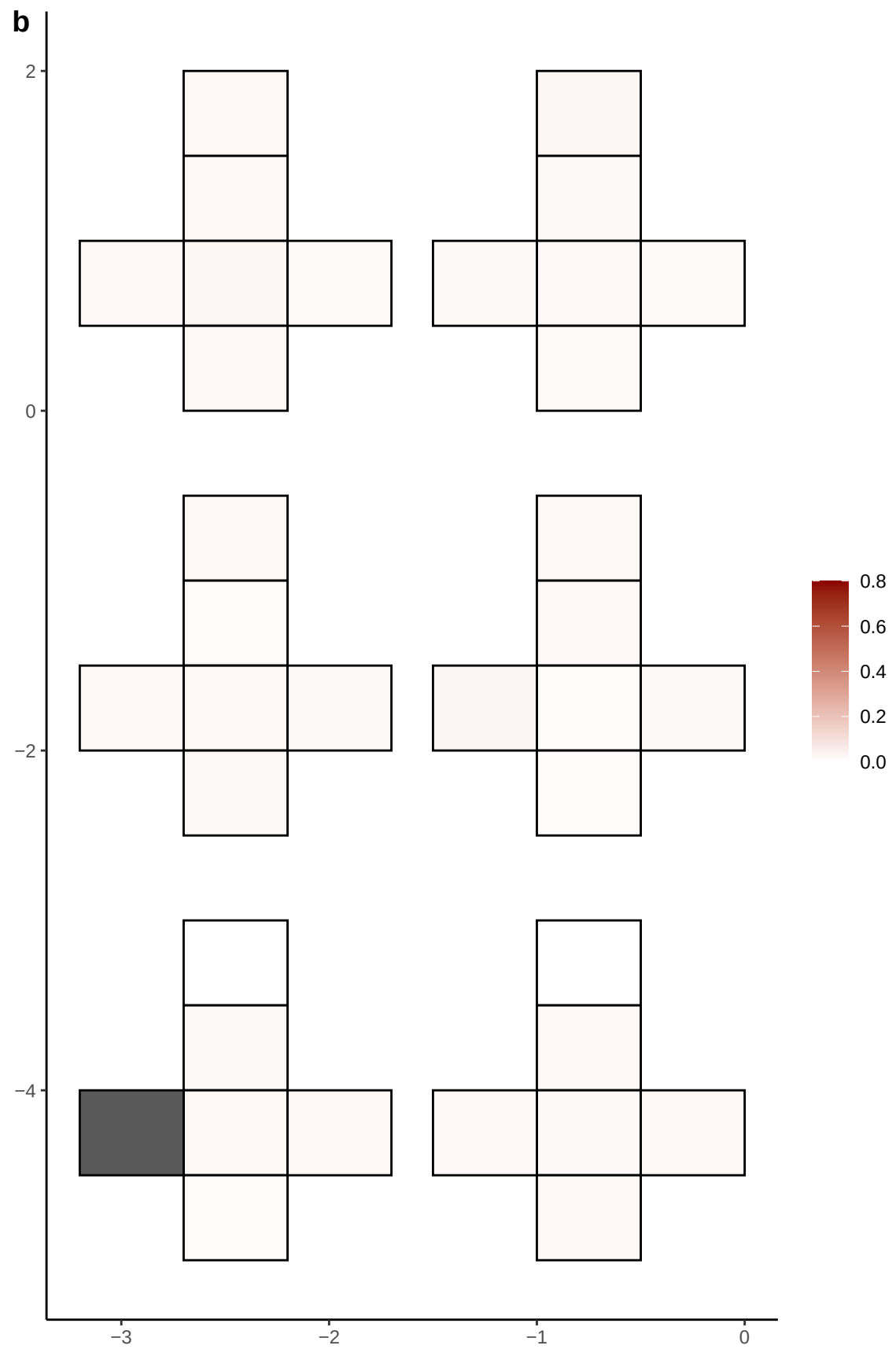

**a** ID01 6-156279892-A-G

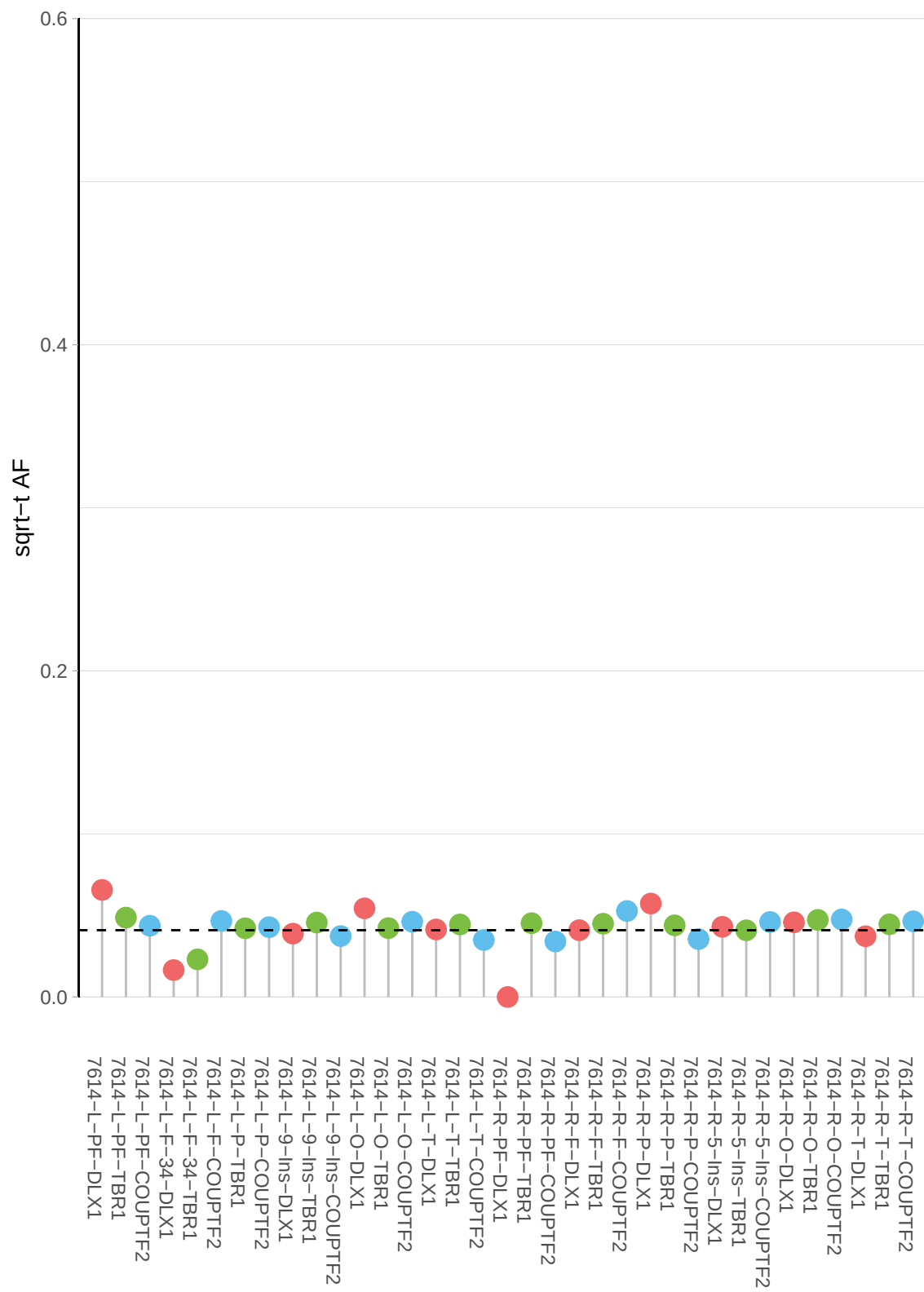

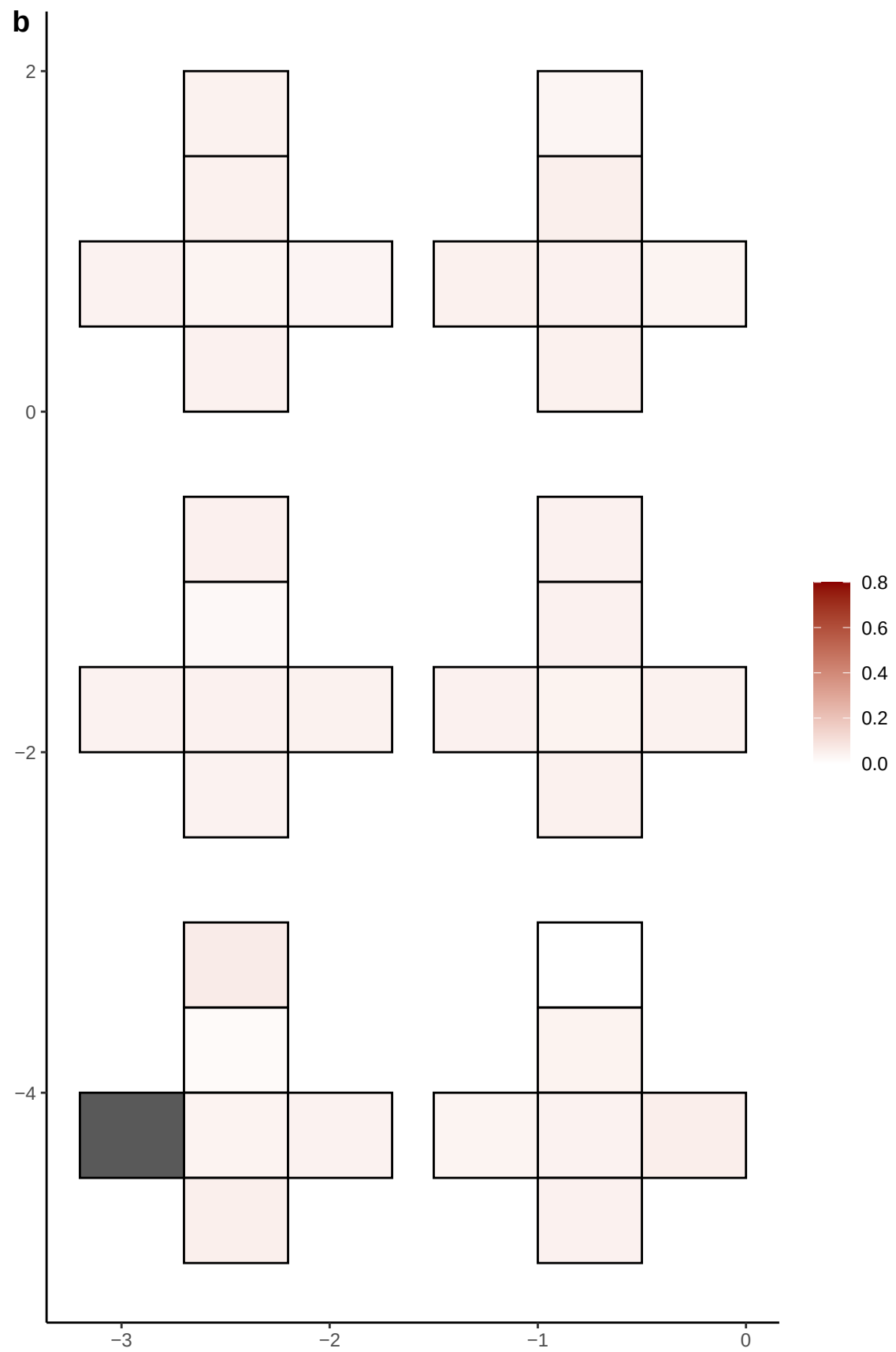

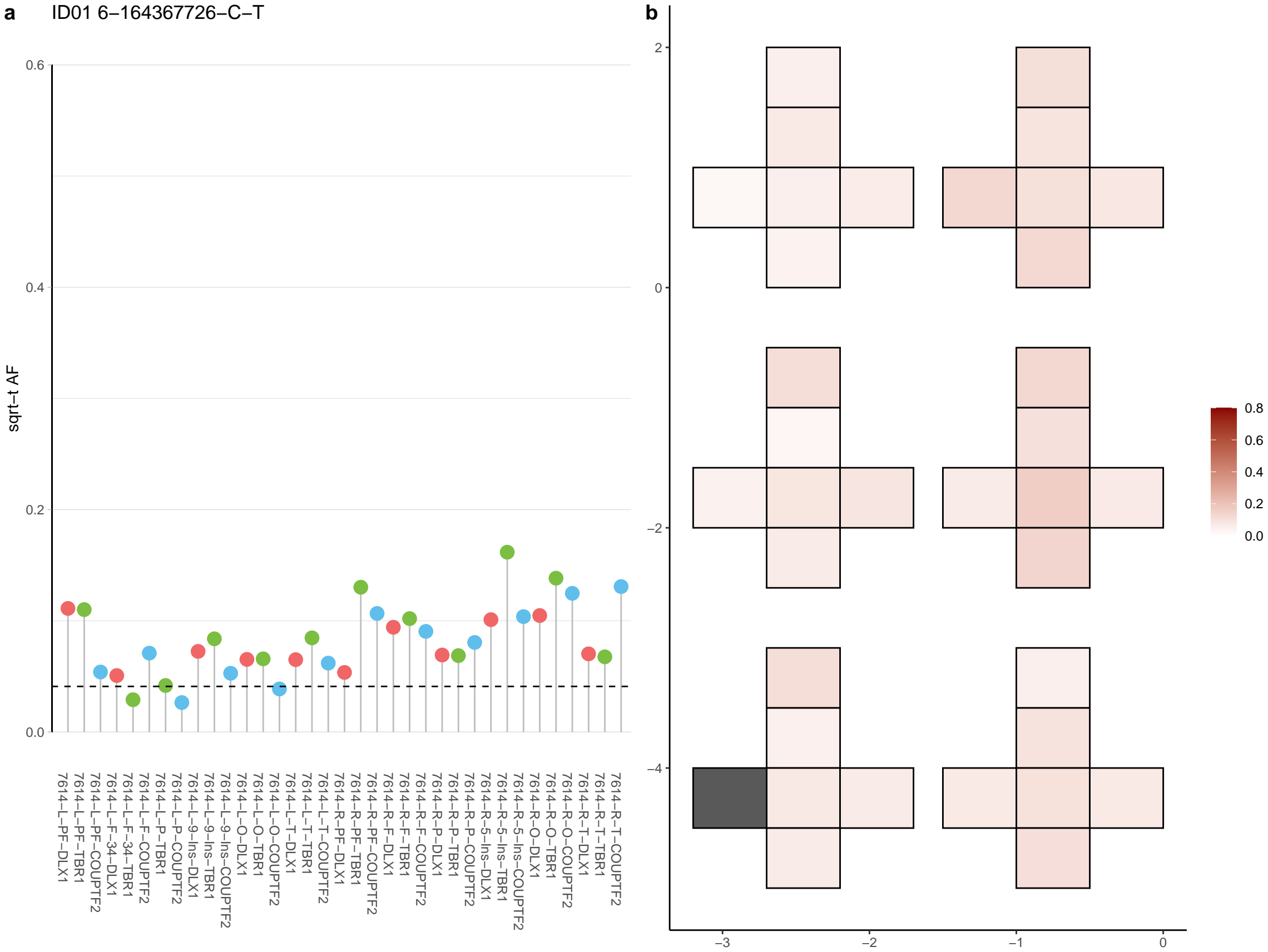

**a** ID01 6-21159973-A-C

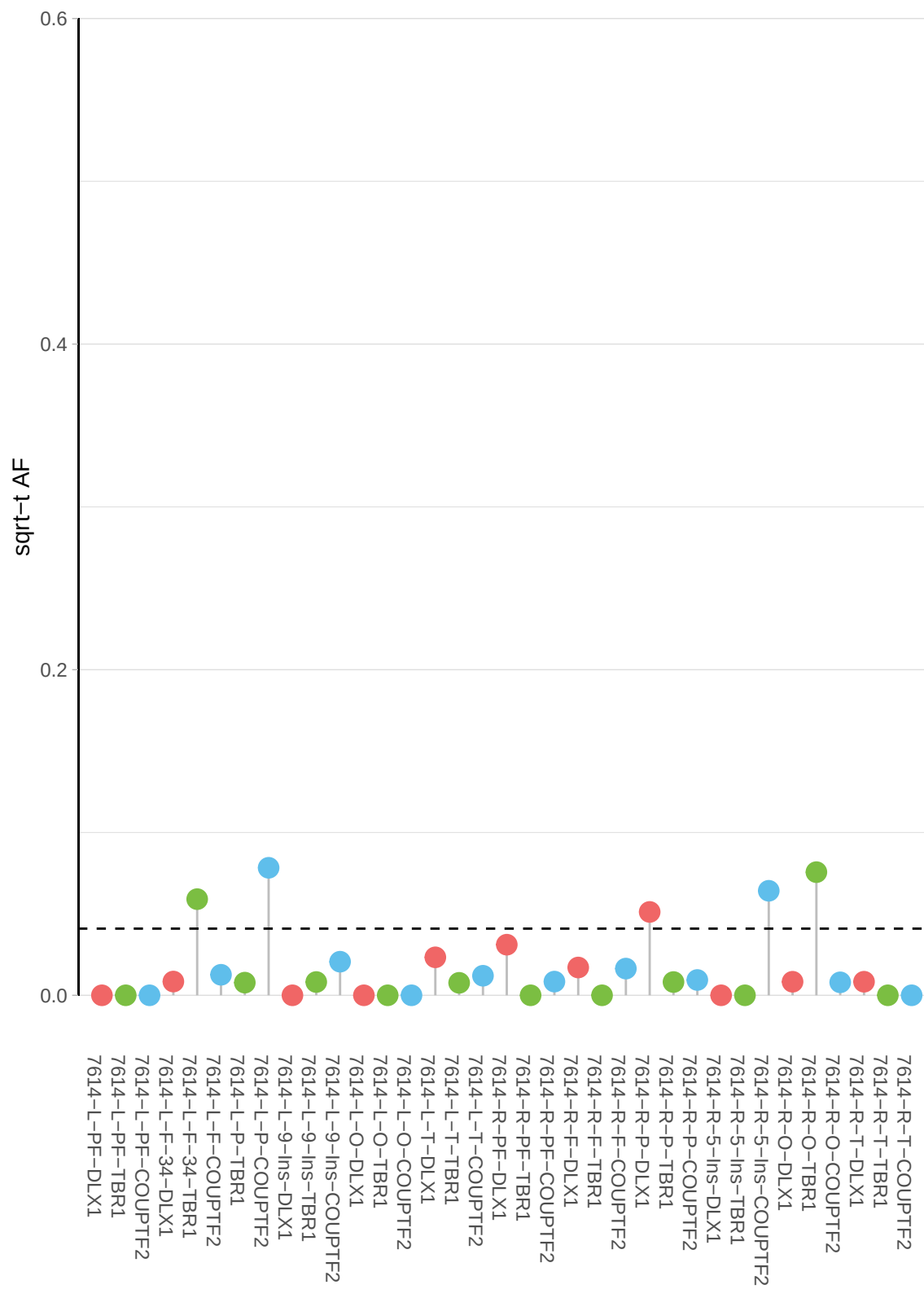

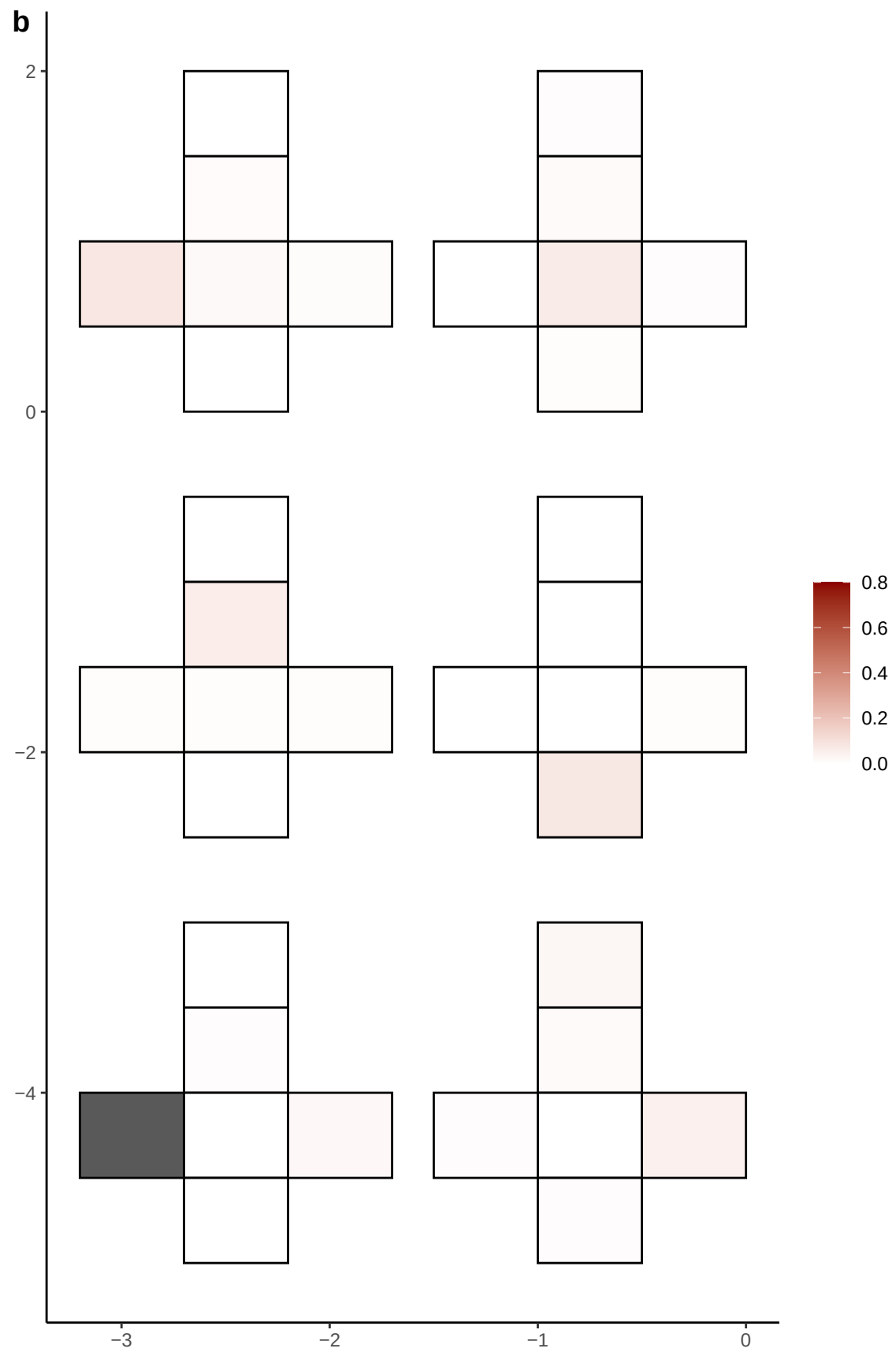

**a** ID01 6-23944712-G-T

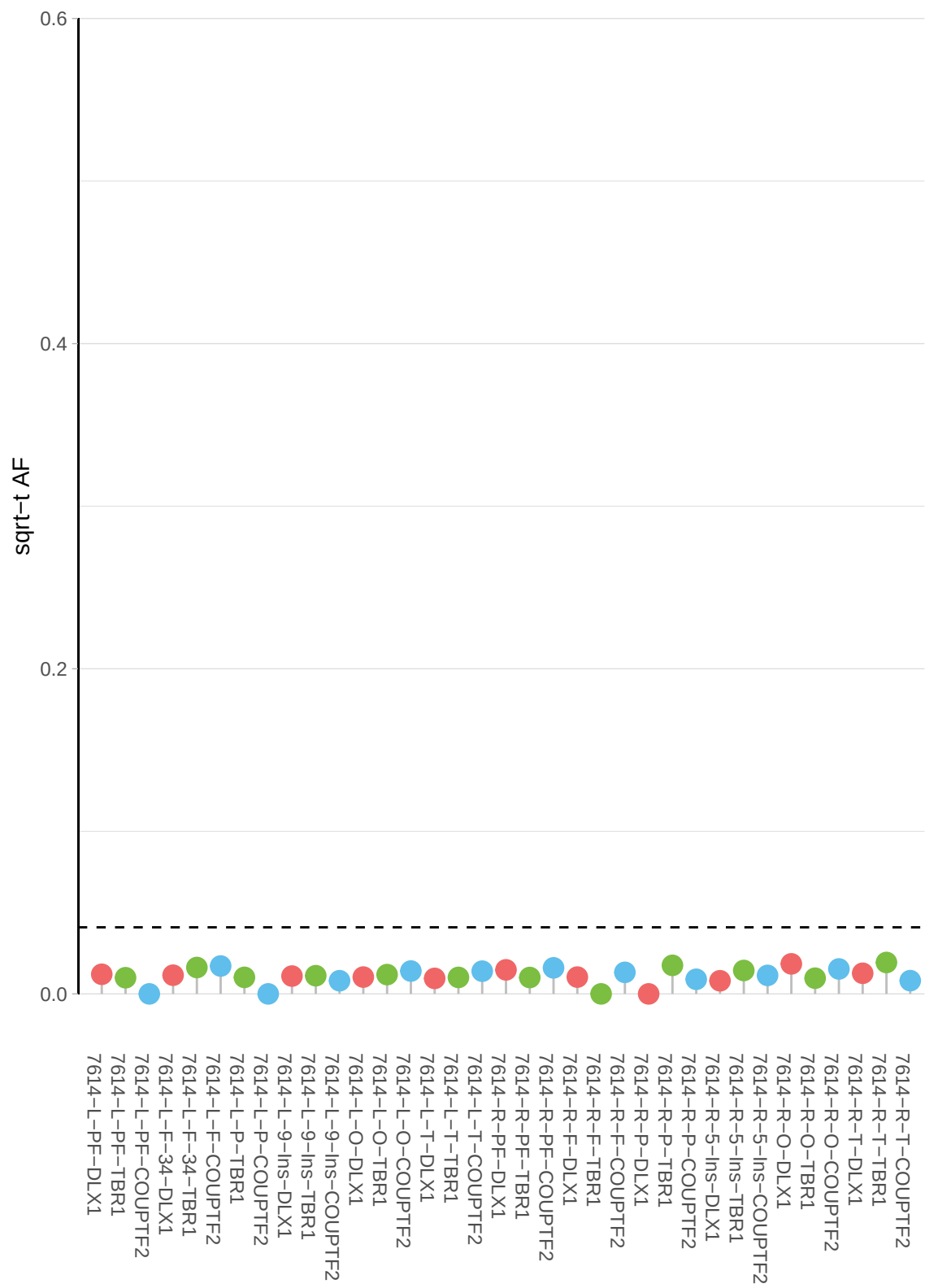

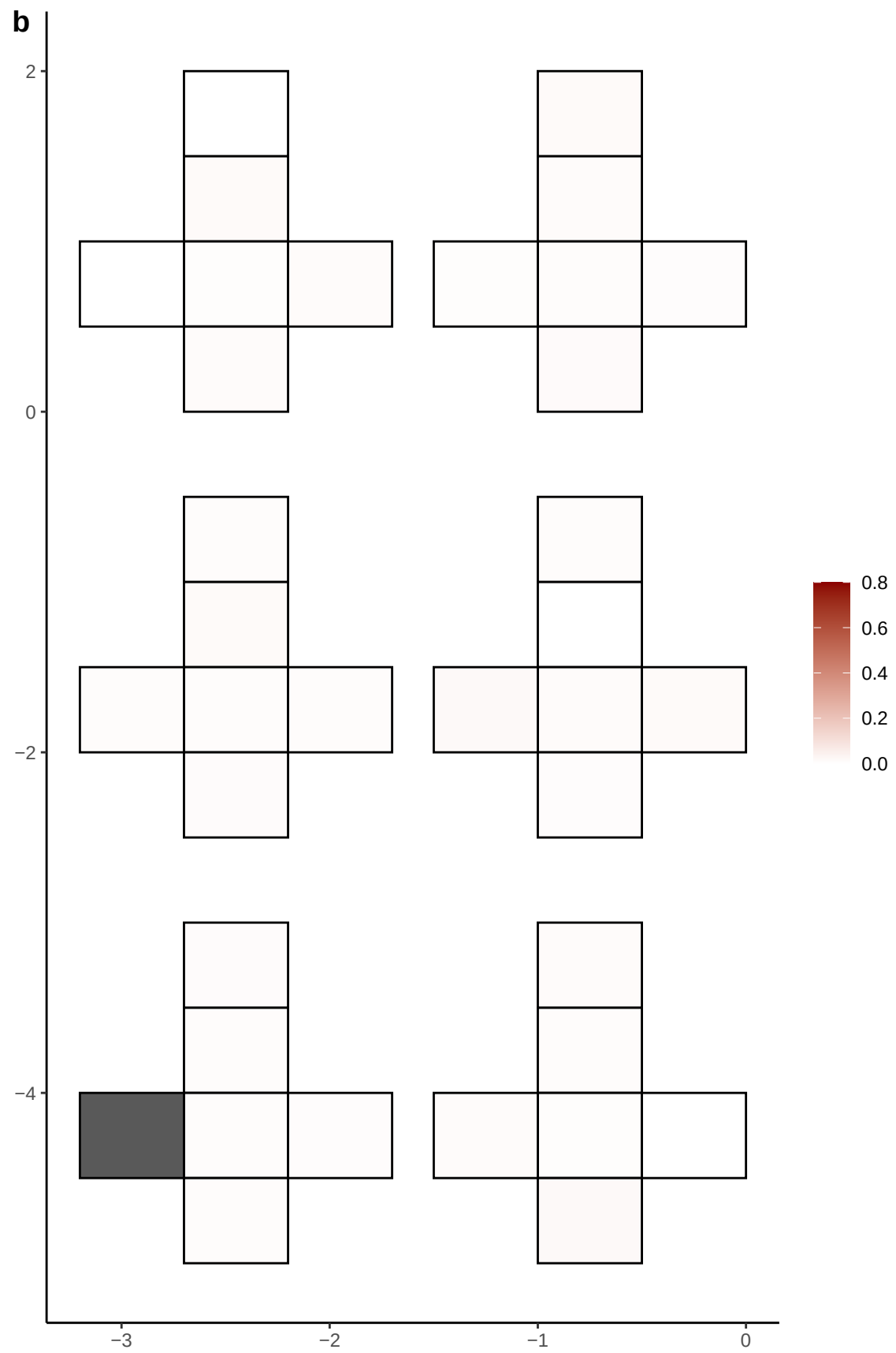

**a** ID01 6-27985938-C-T

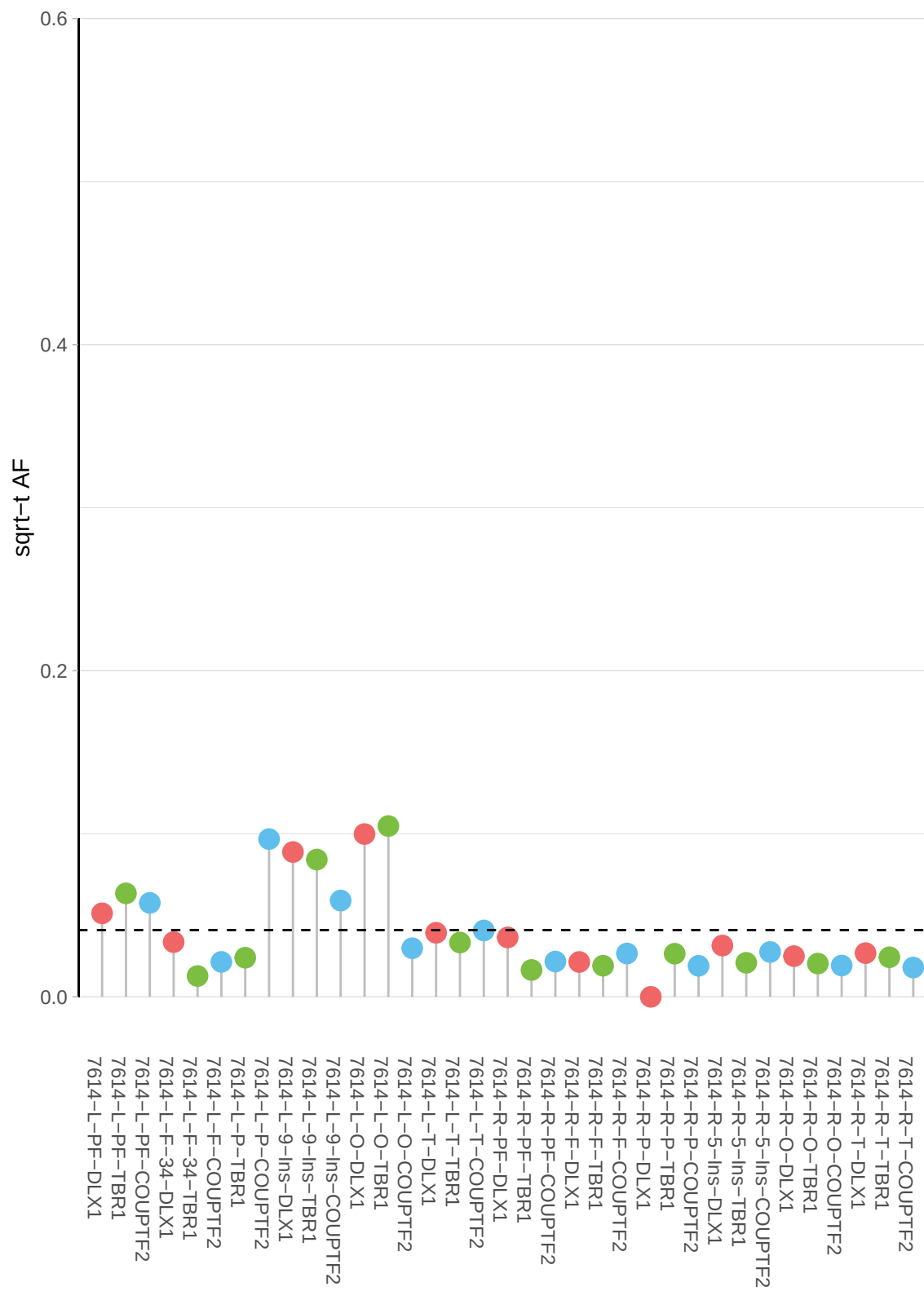

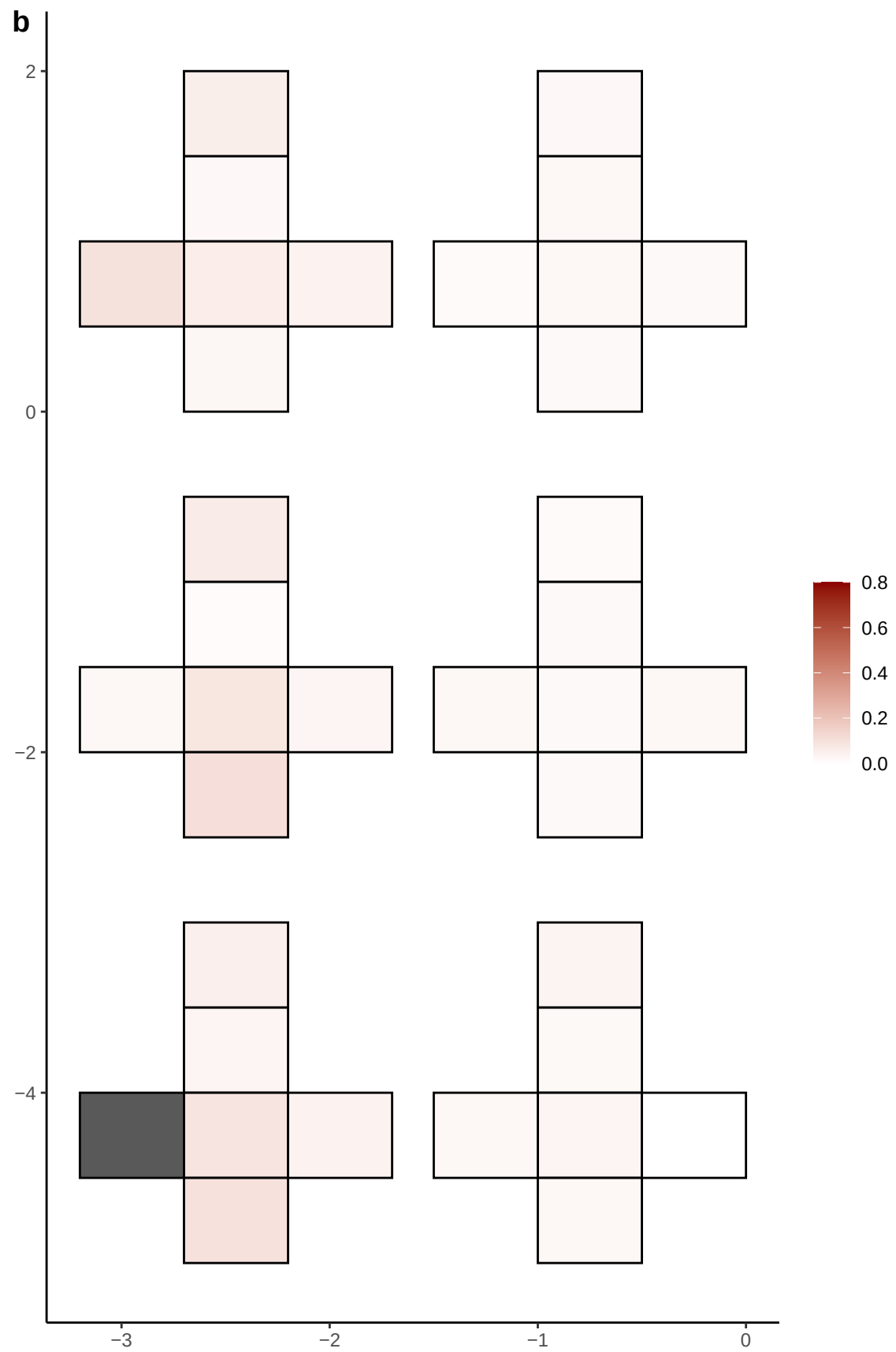

**a** ID01 6-40371400-C-T

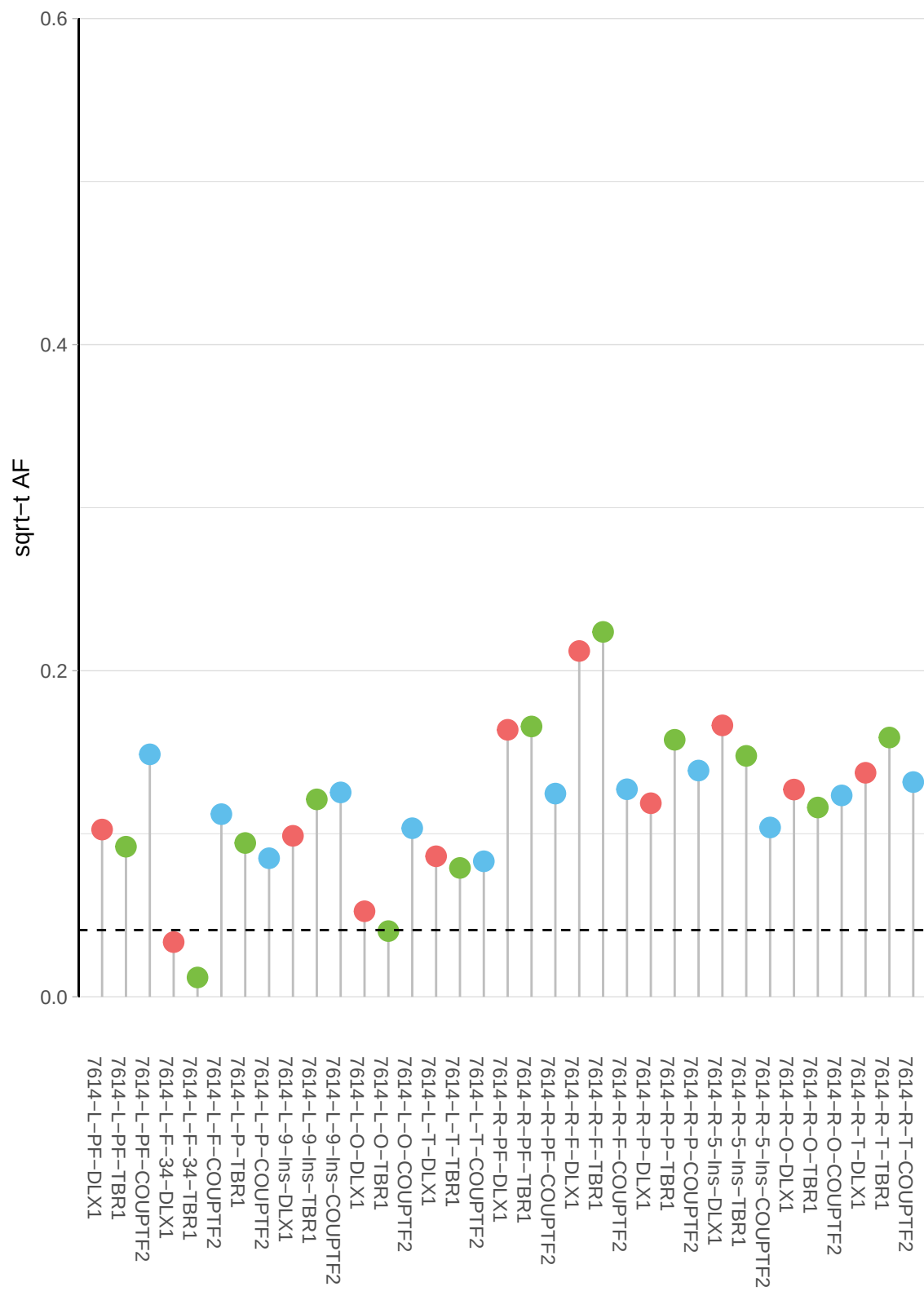

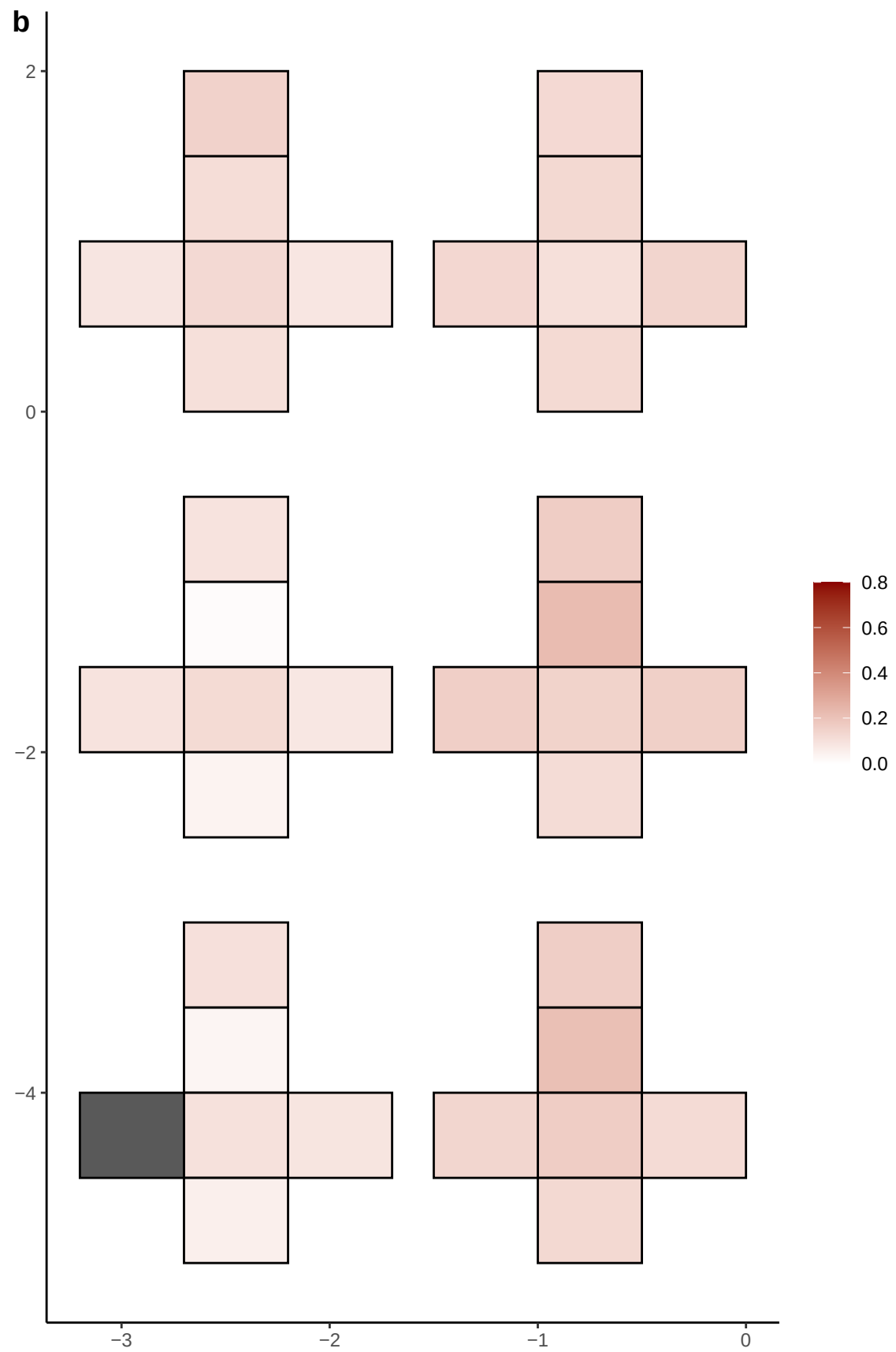

**a** ID01 6-45630553-G-A

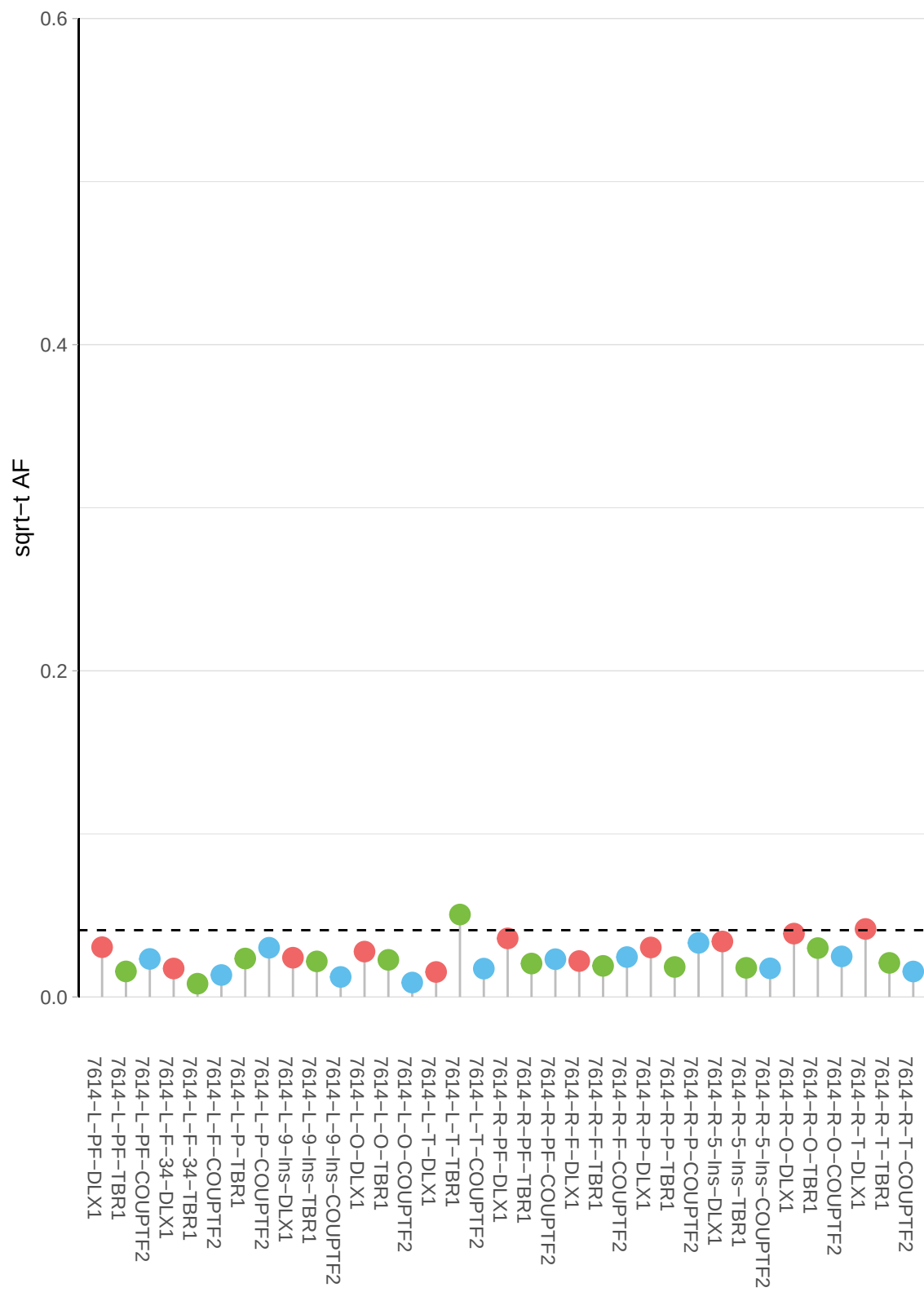

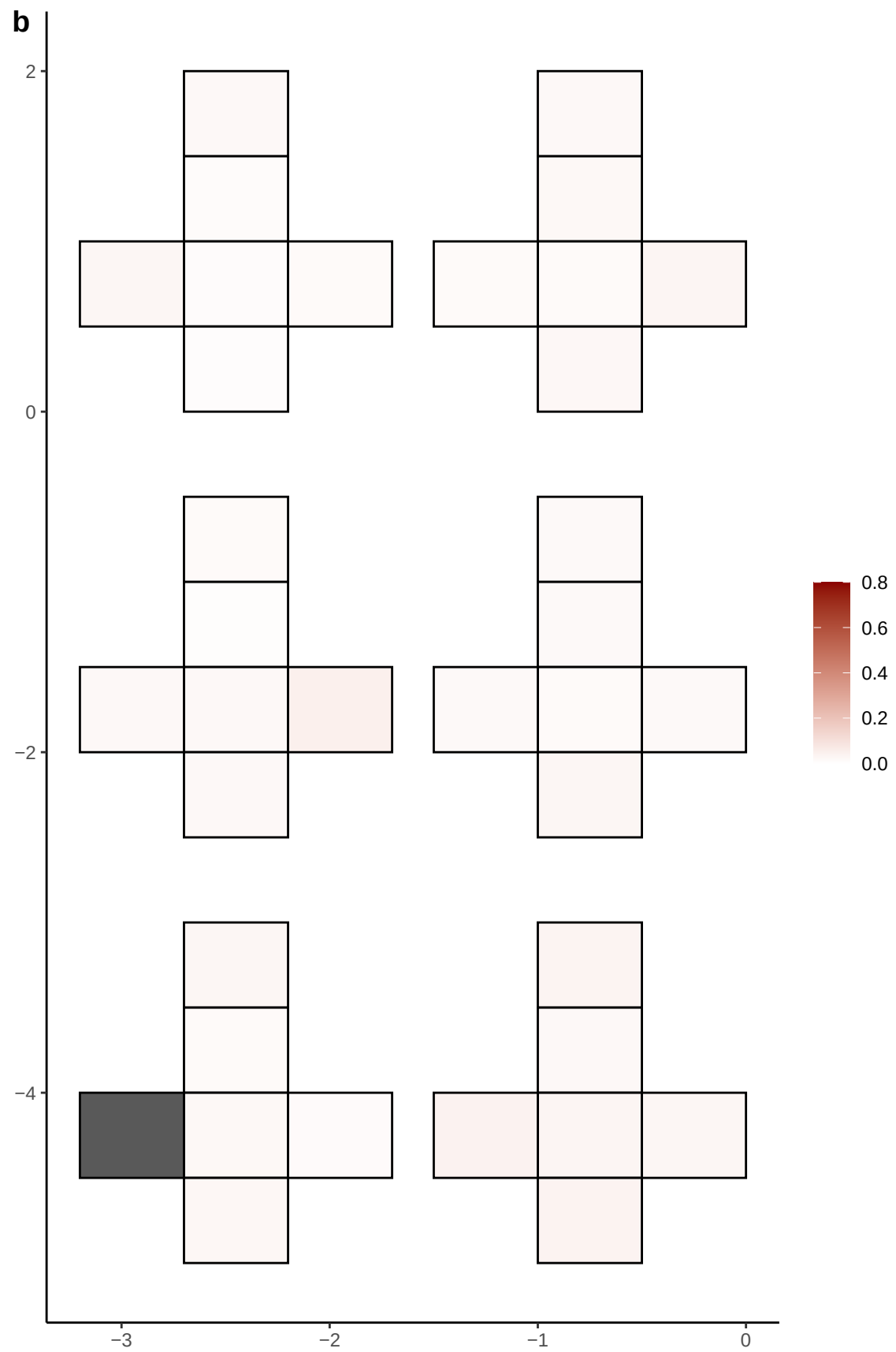

**a** ID01 6-55394736-C-T

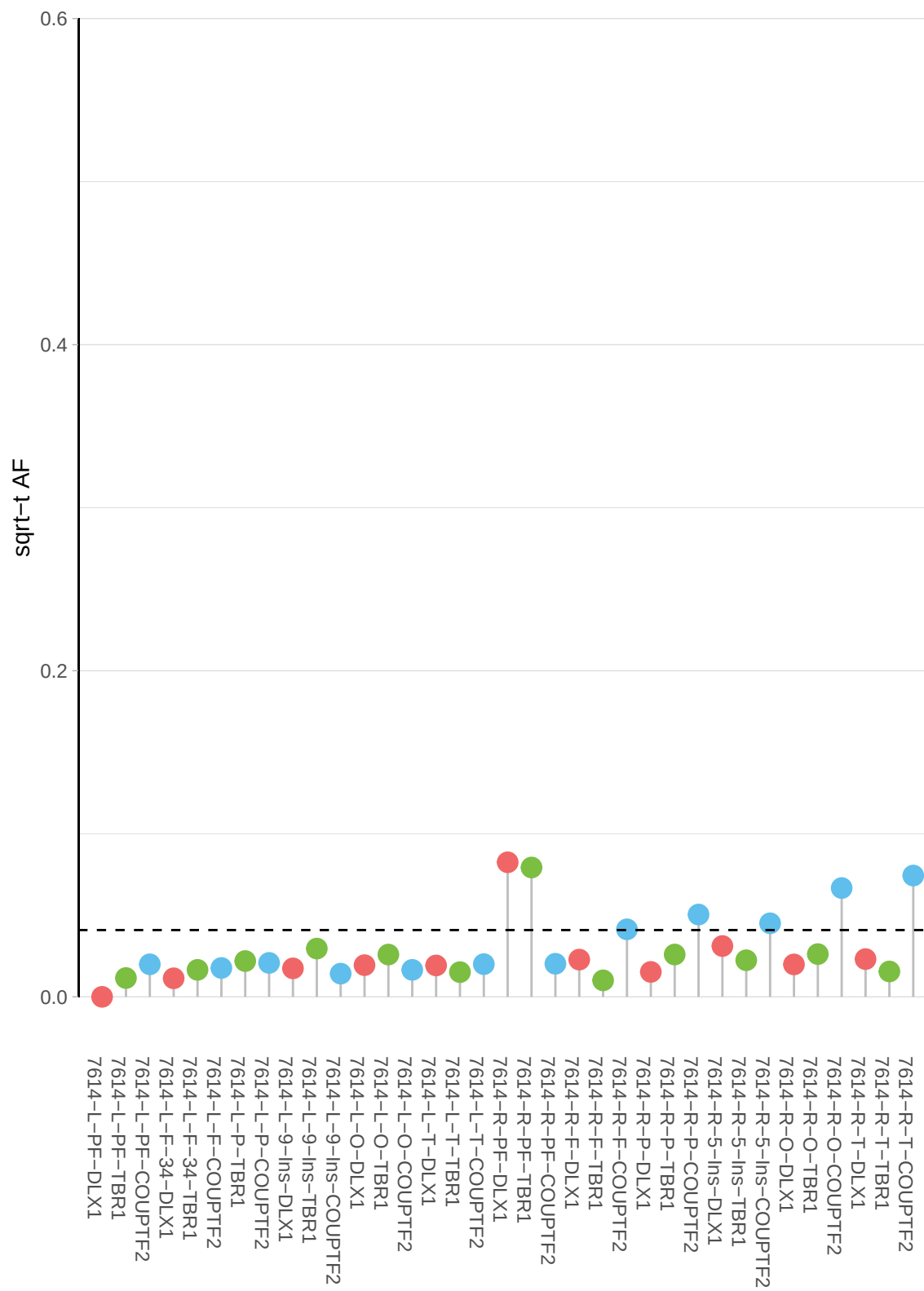

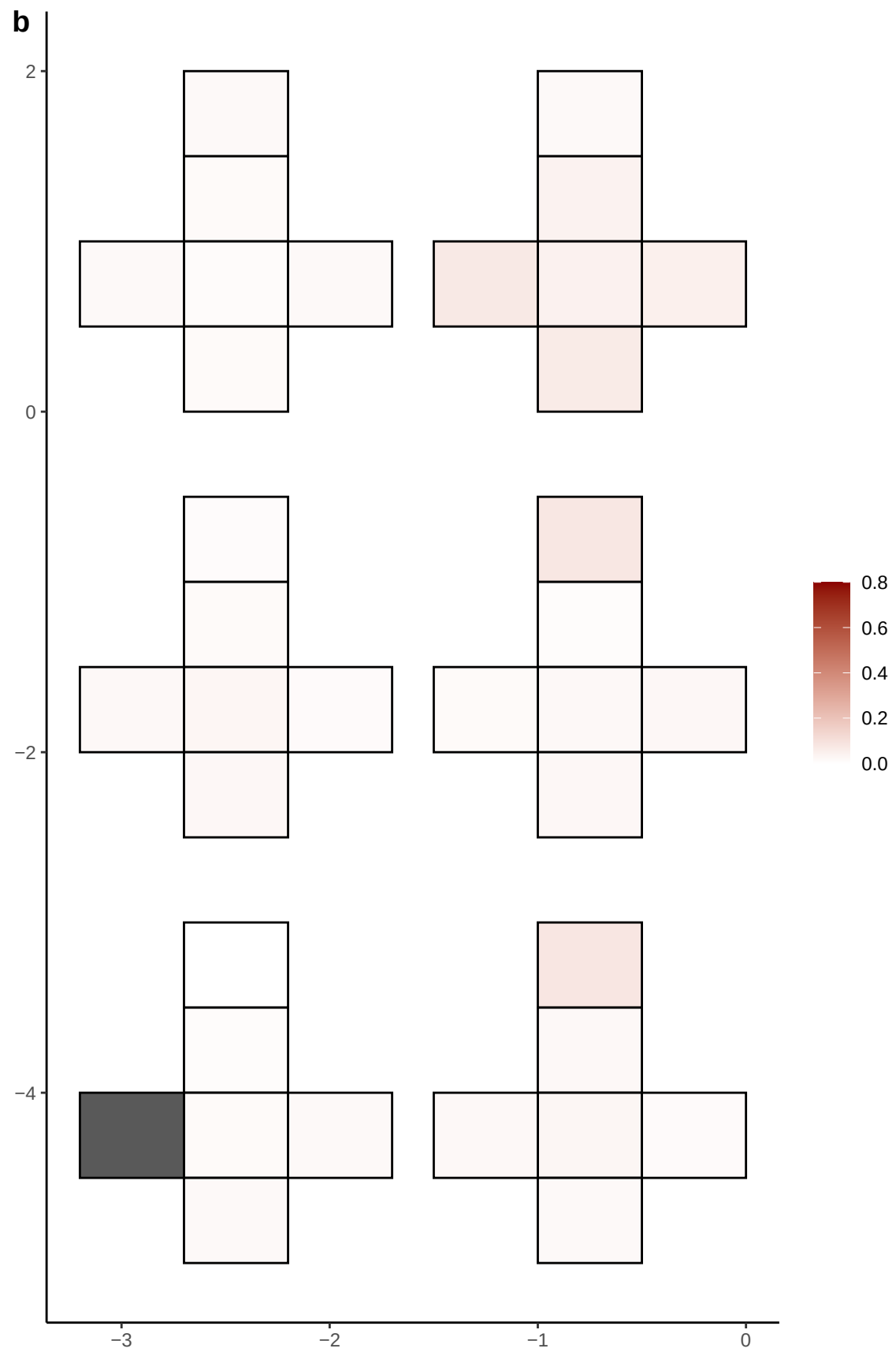

**a** ID01 6-57218601-A-G

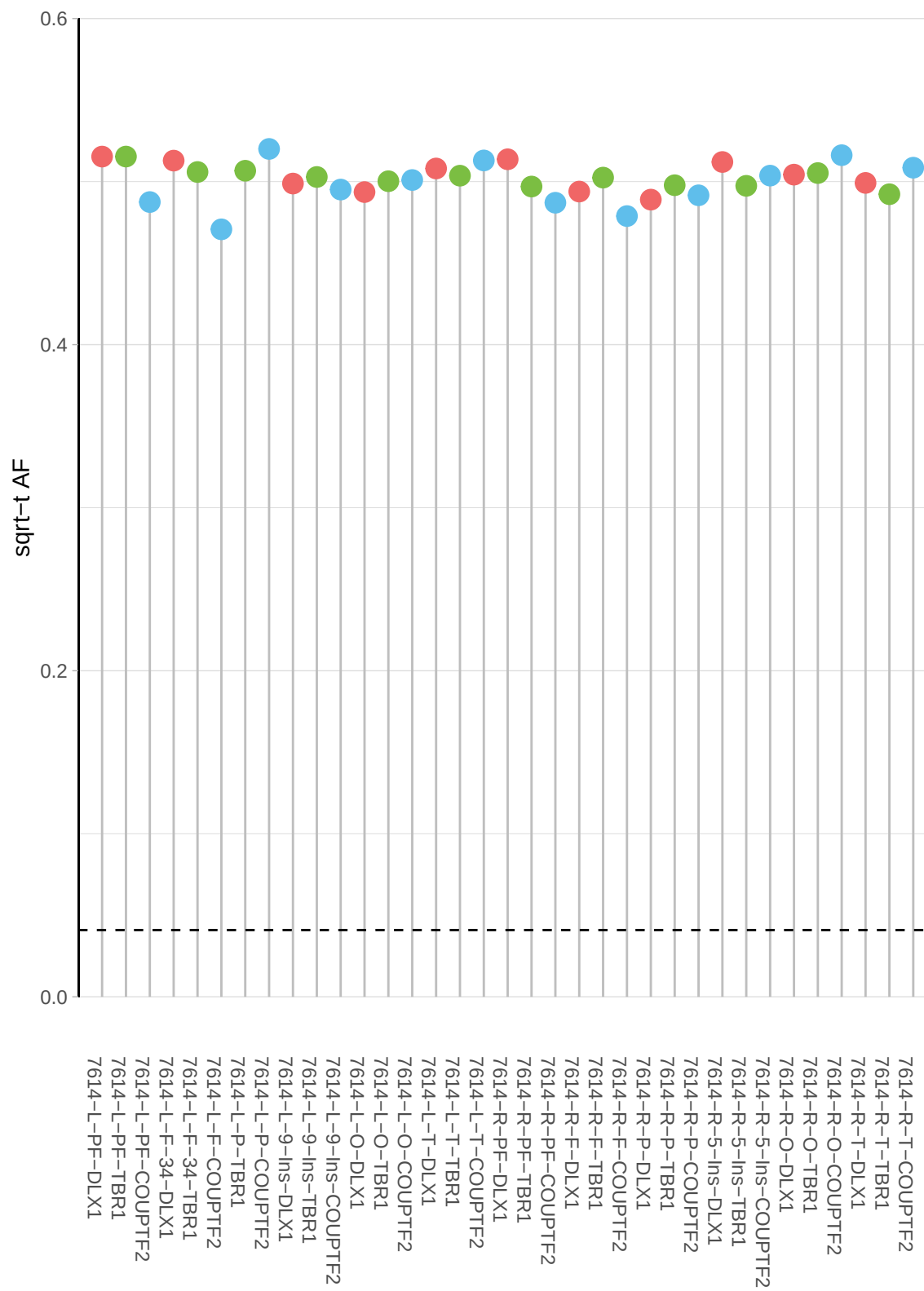

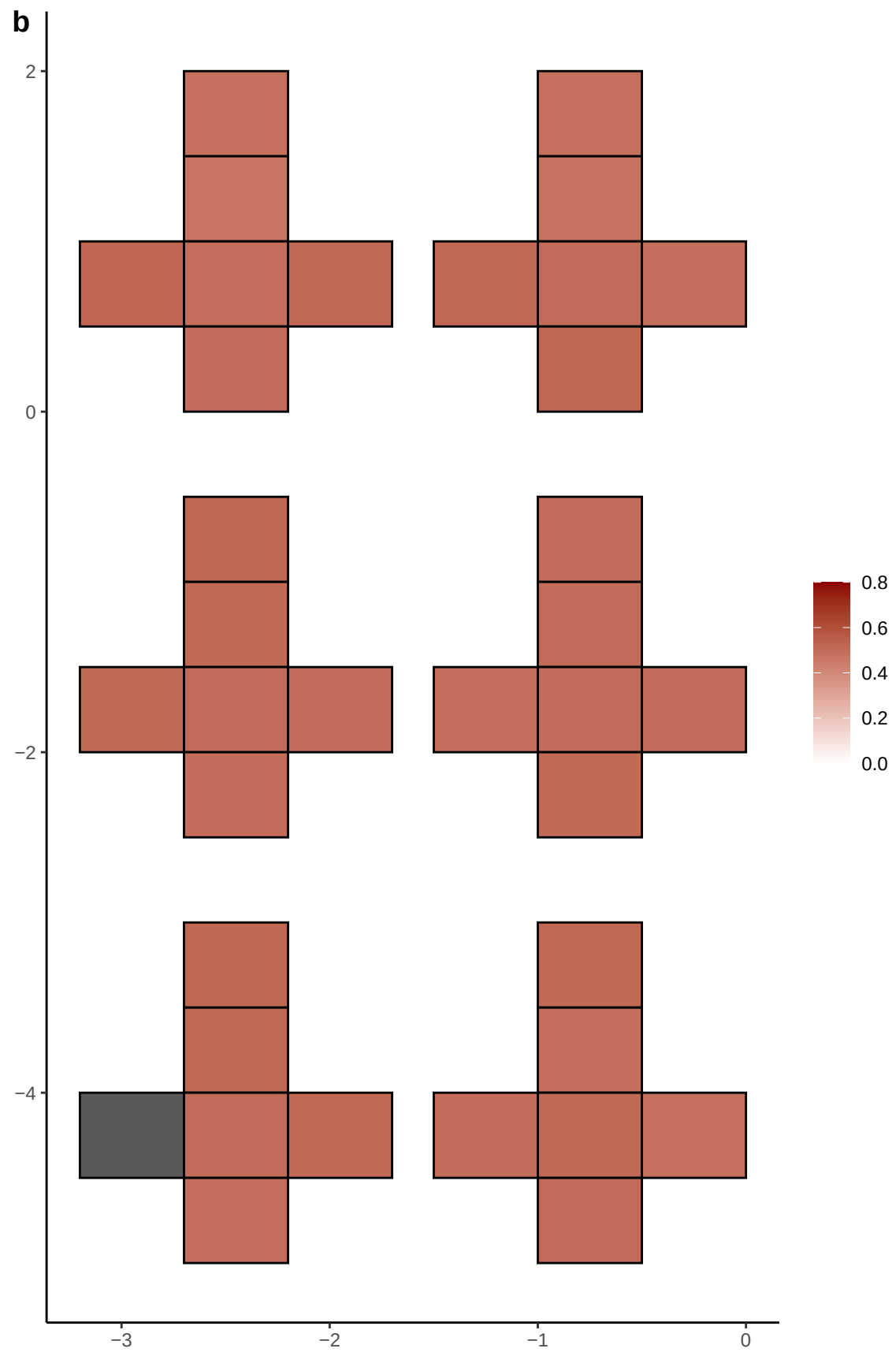

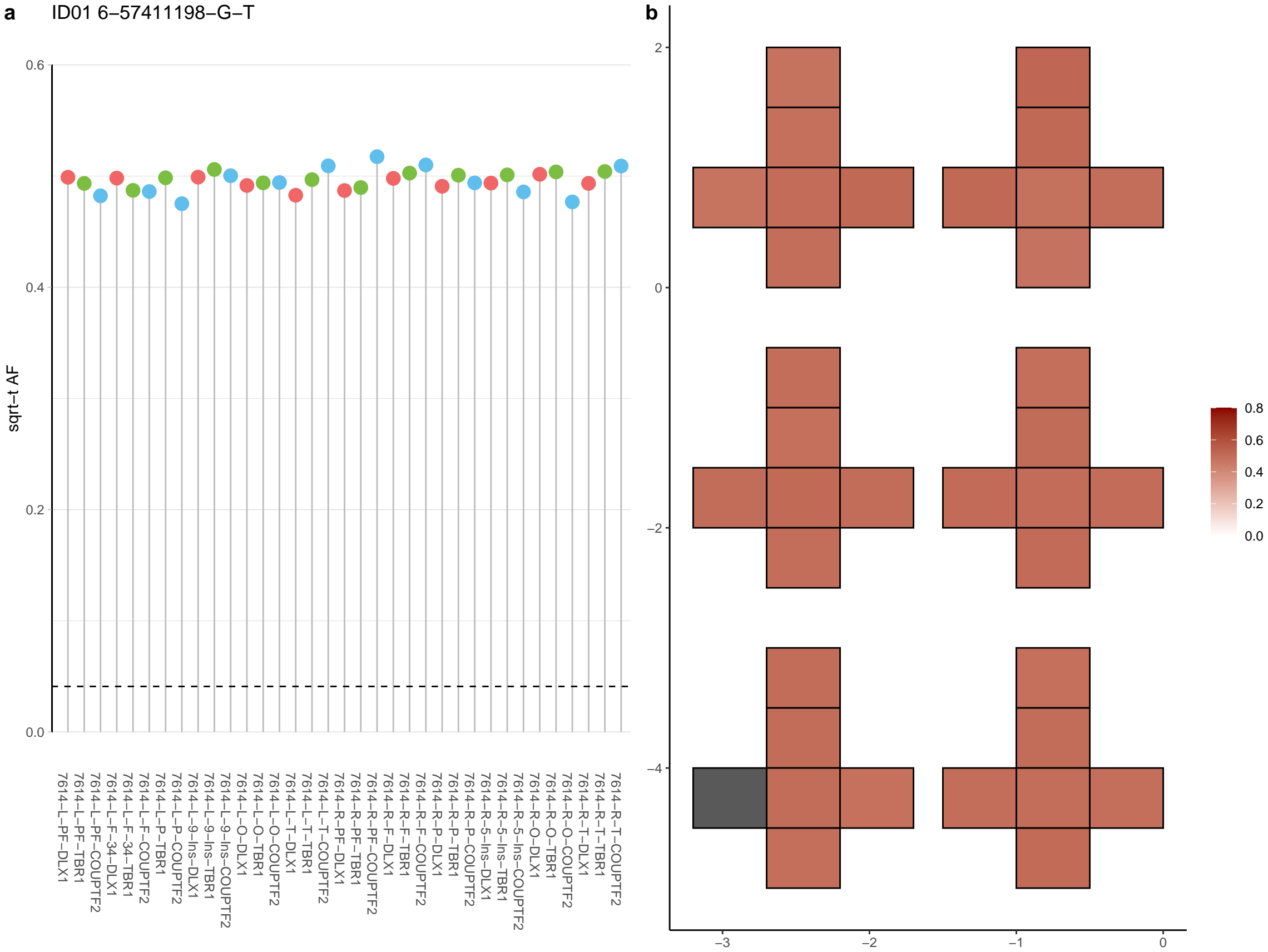

**a** ID01 6-7569605-G-A

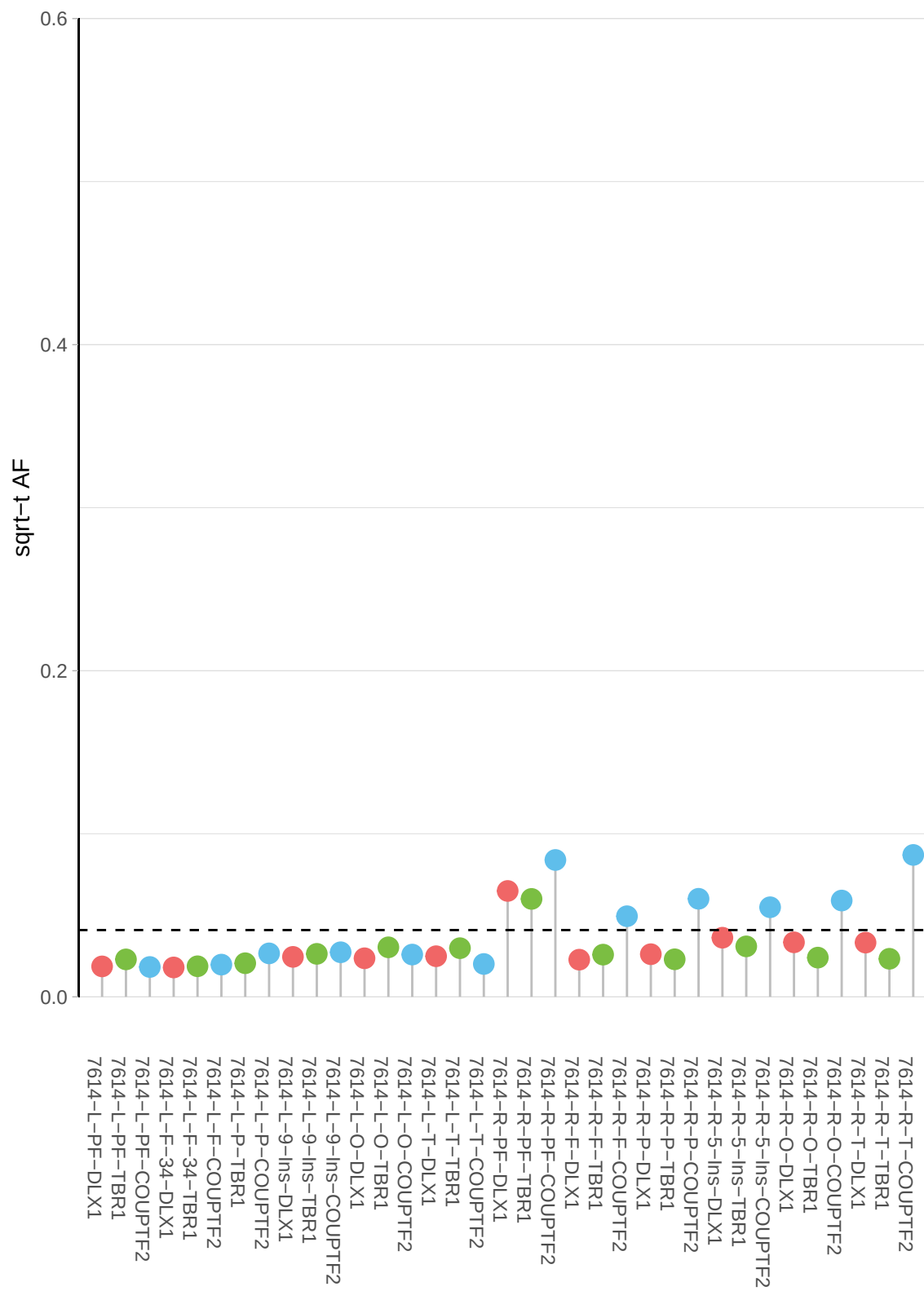

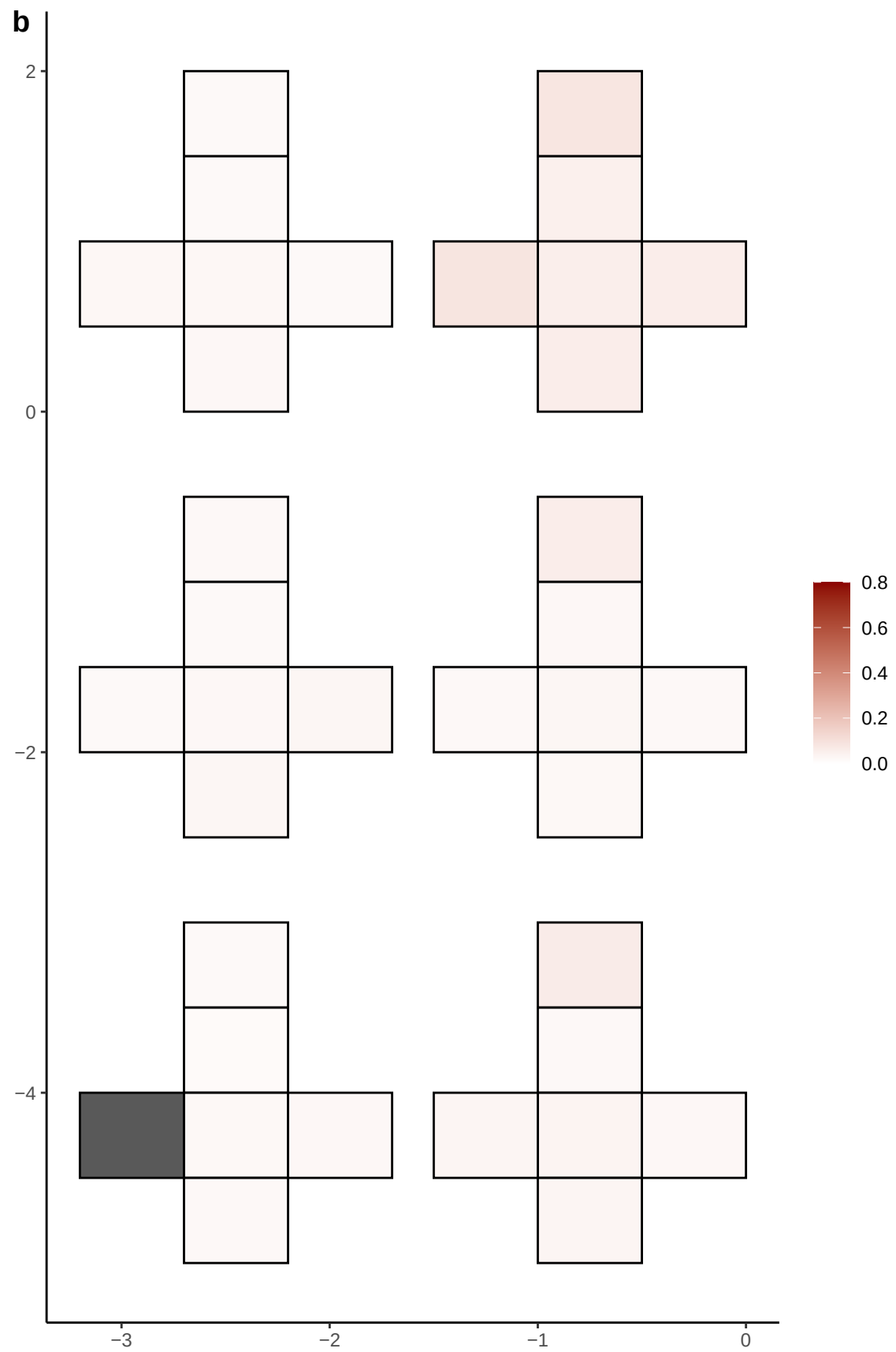

**a** ID01 6-80796617-T-G

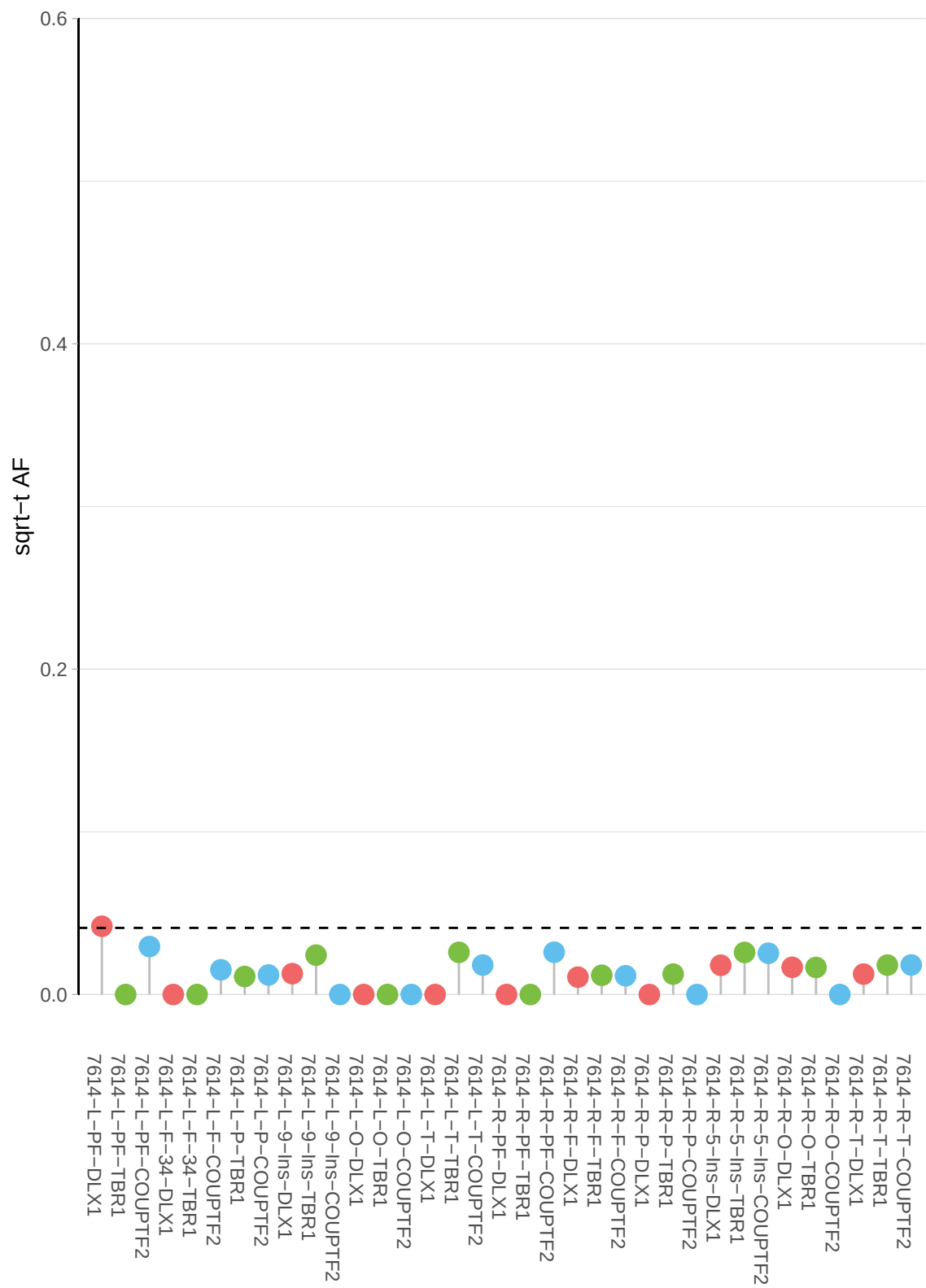

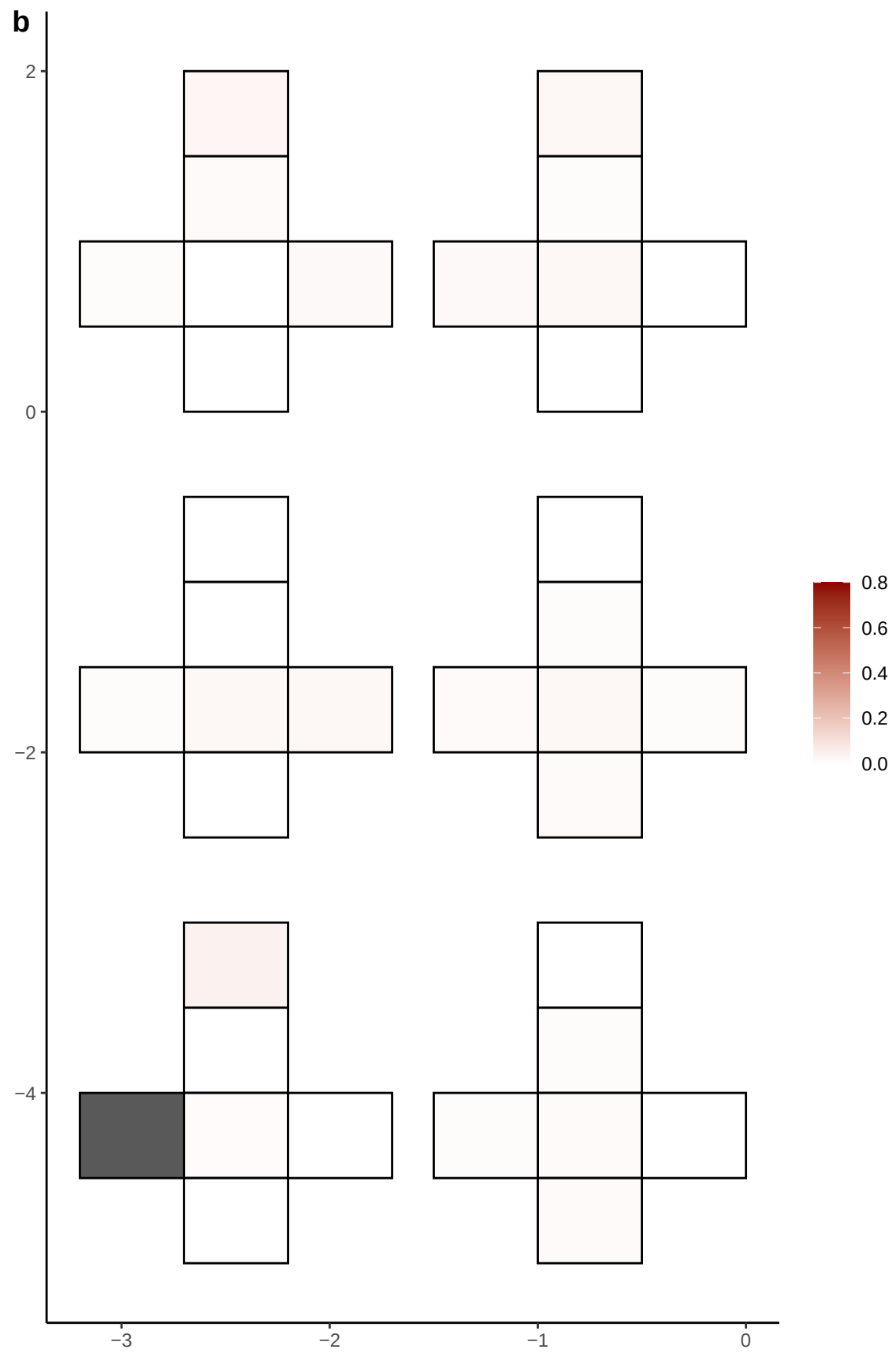

**a** ID01 7-121347585-T-G

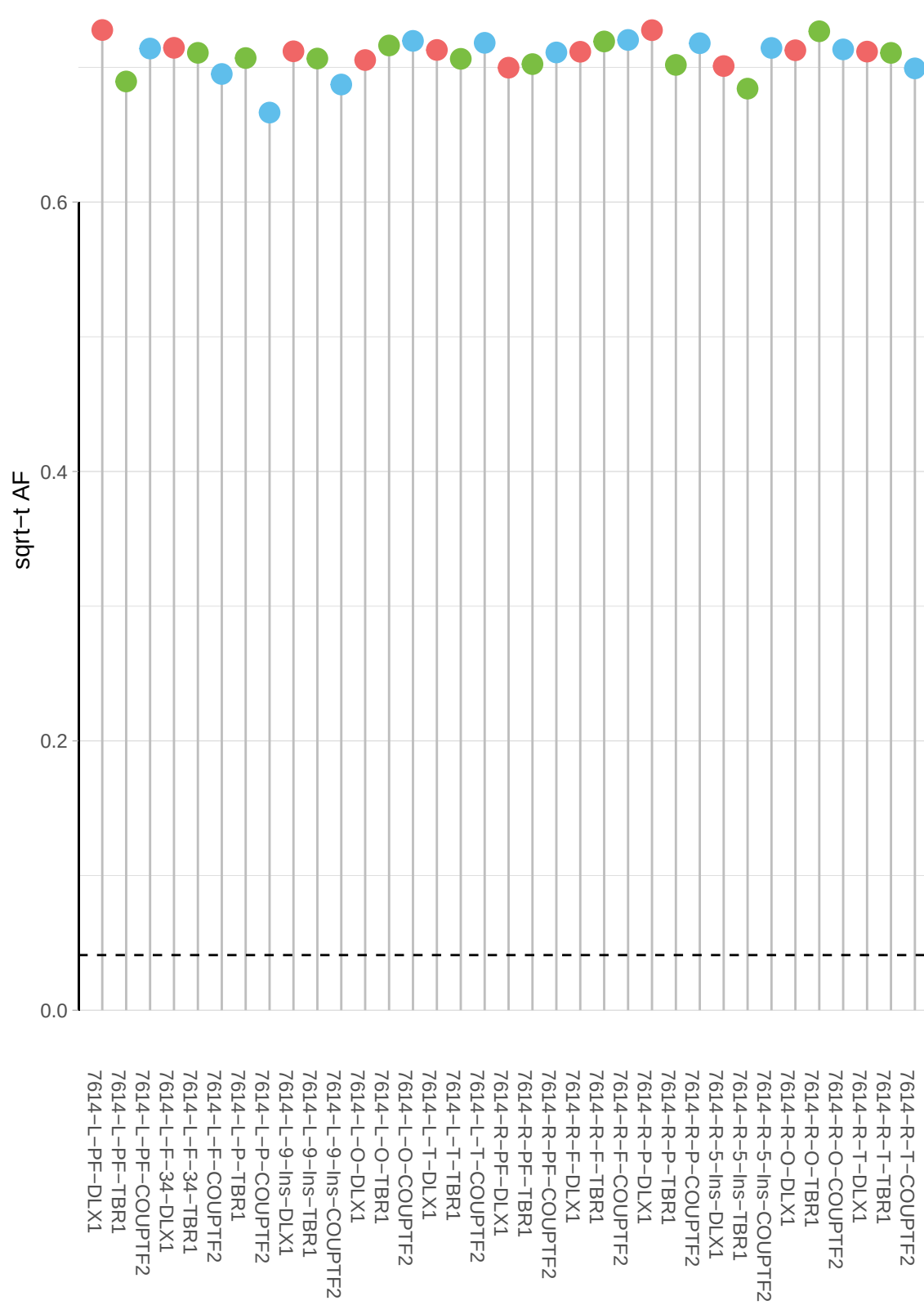

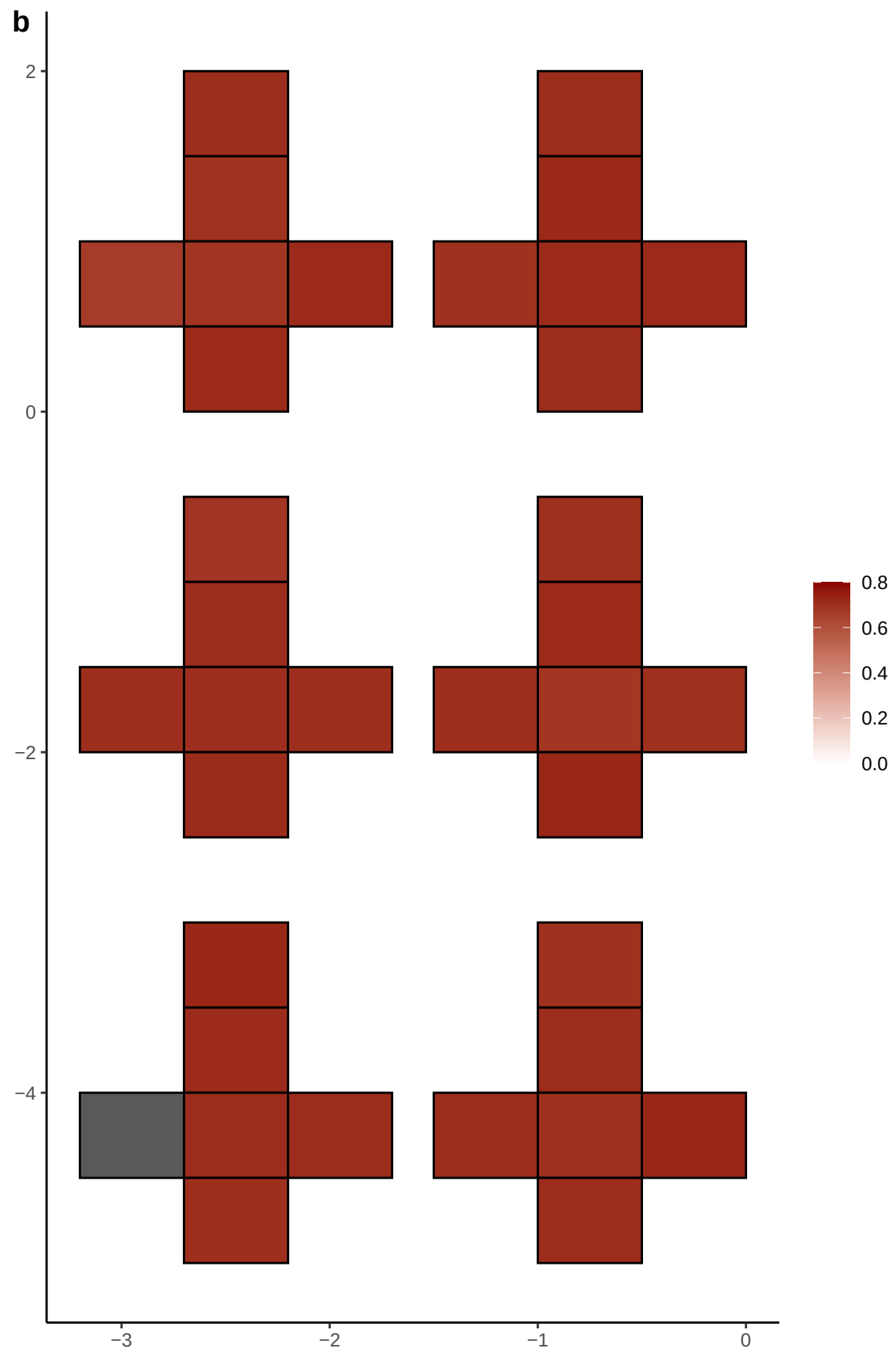

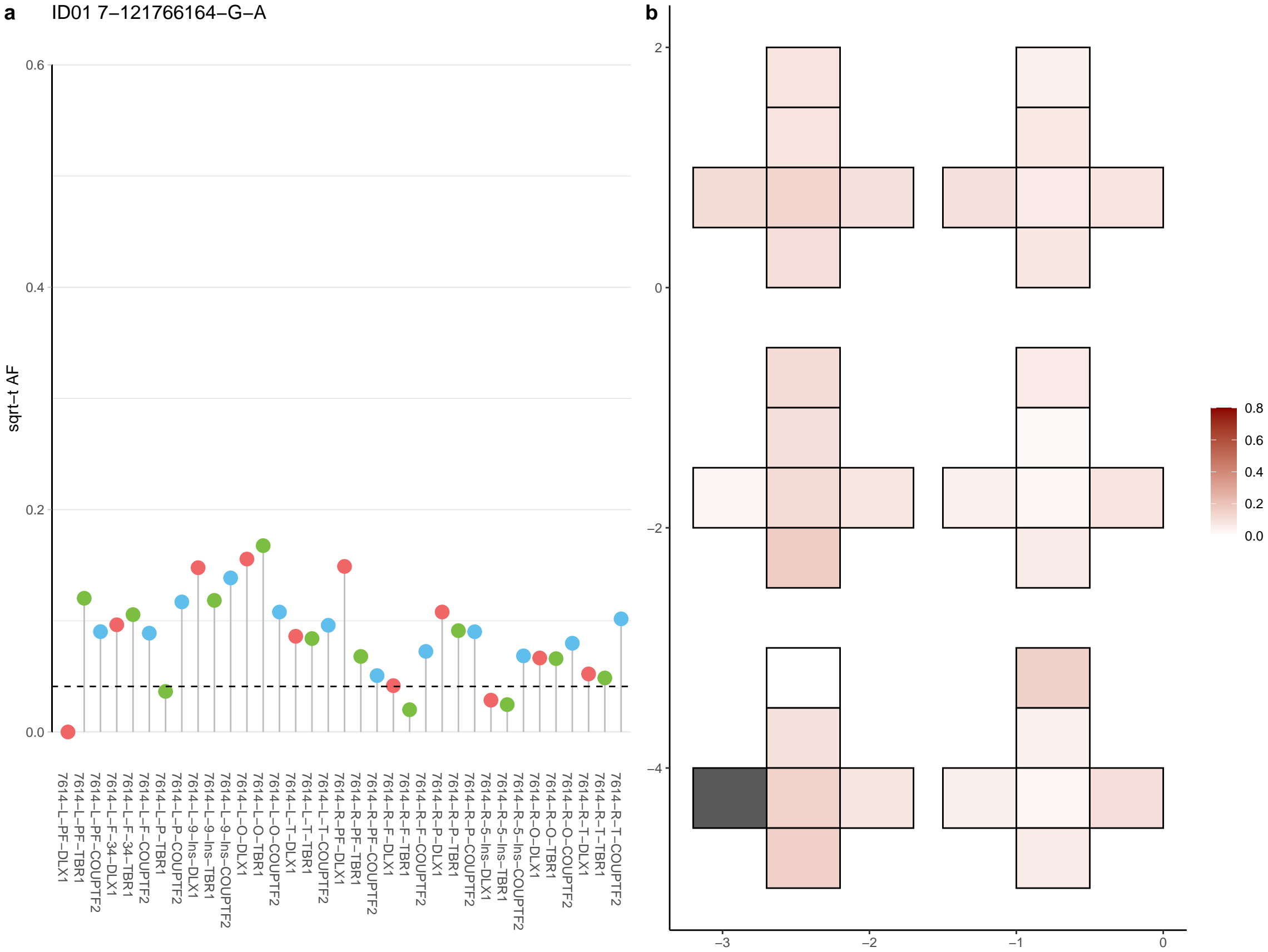

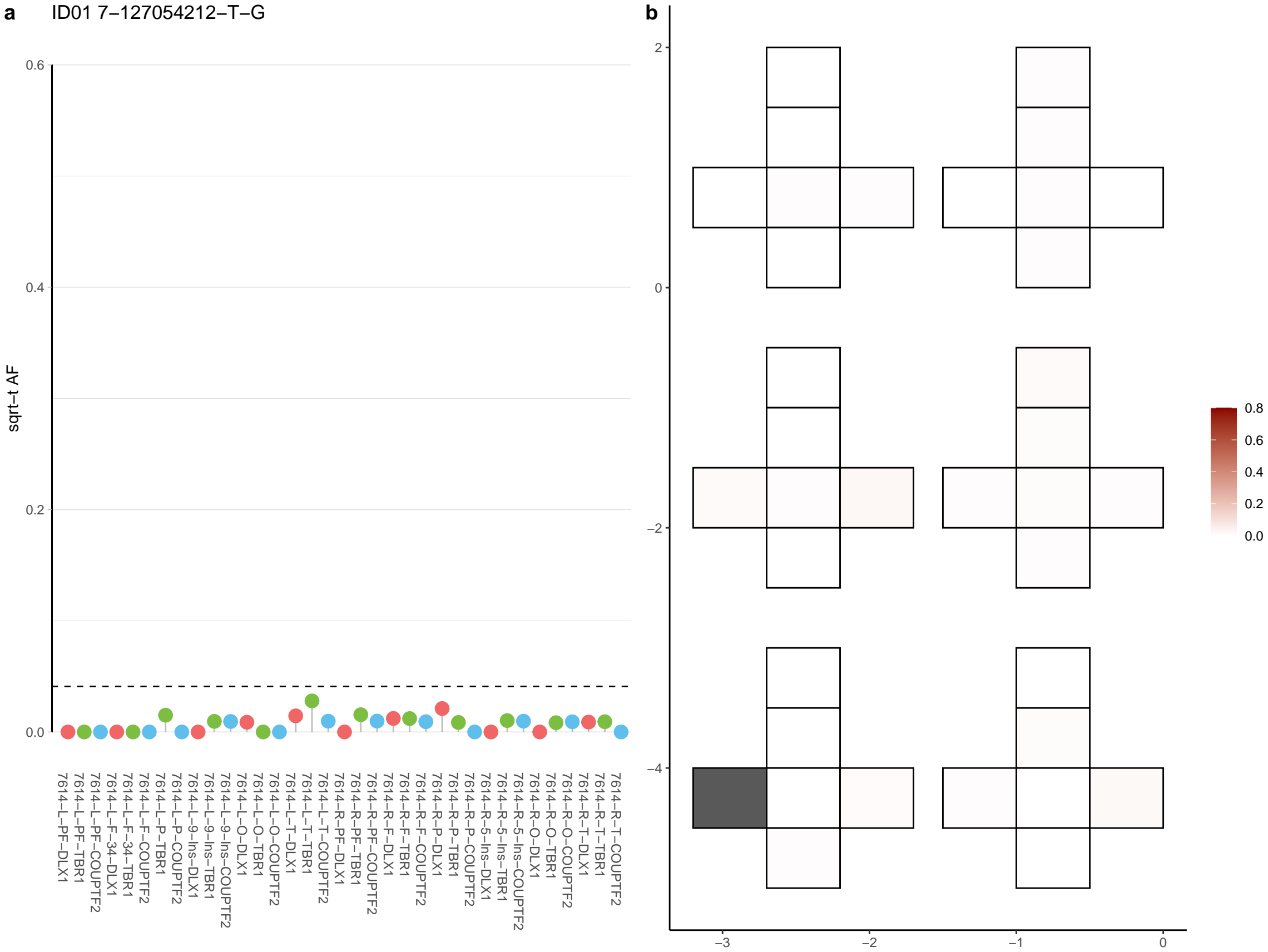

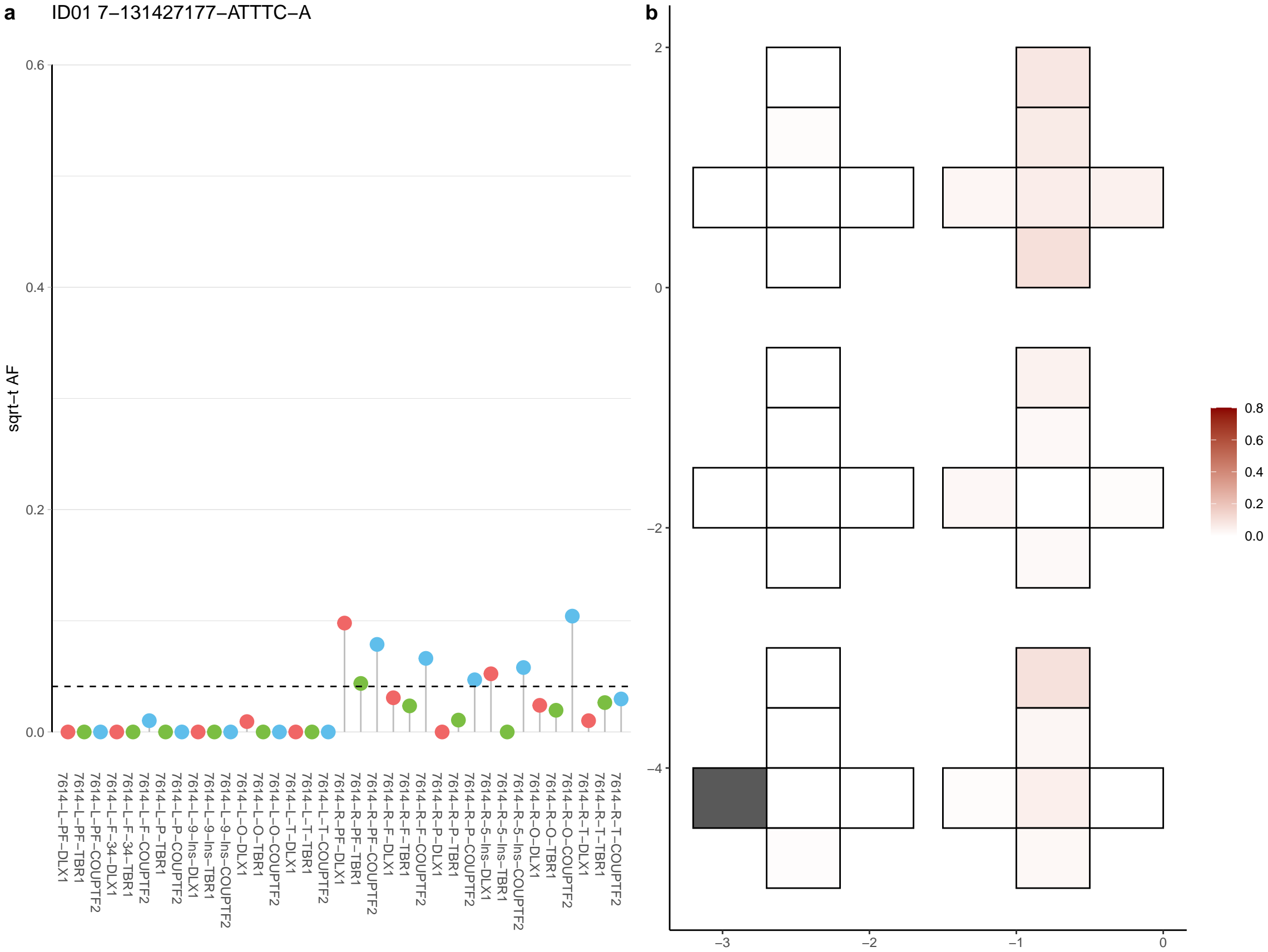

**a** ID01 7-141222076-G-A

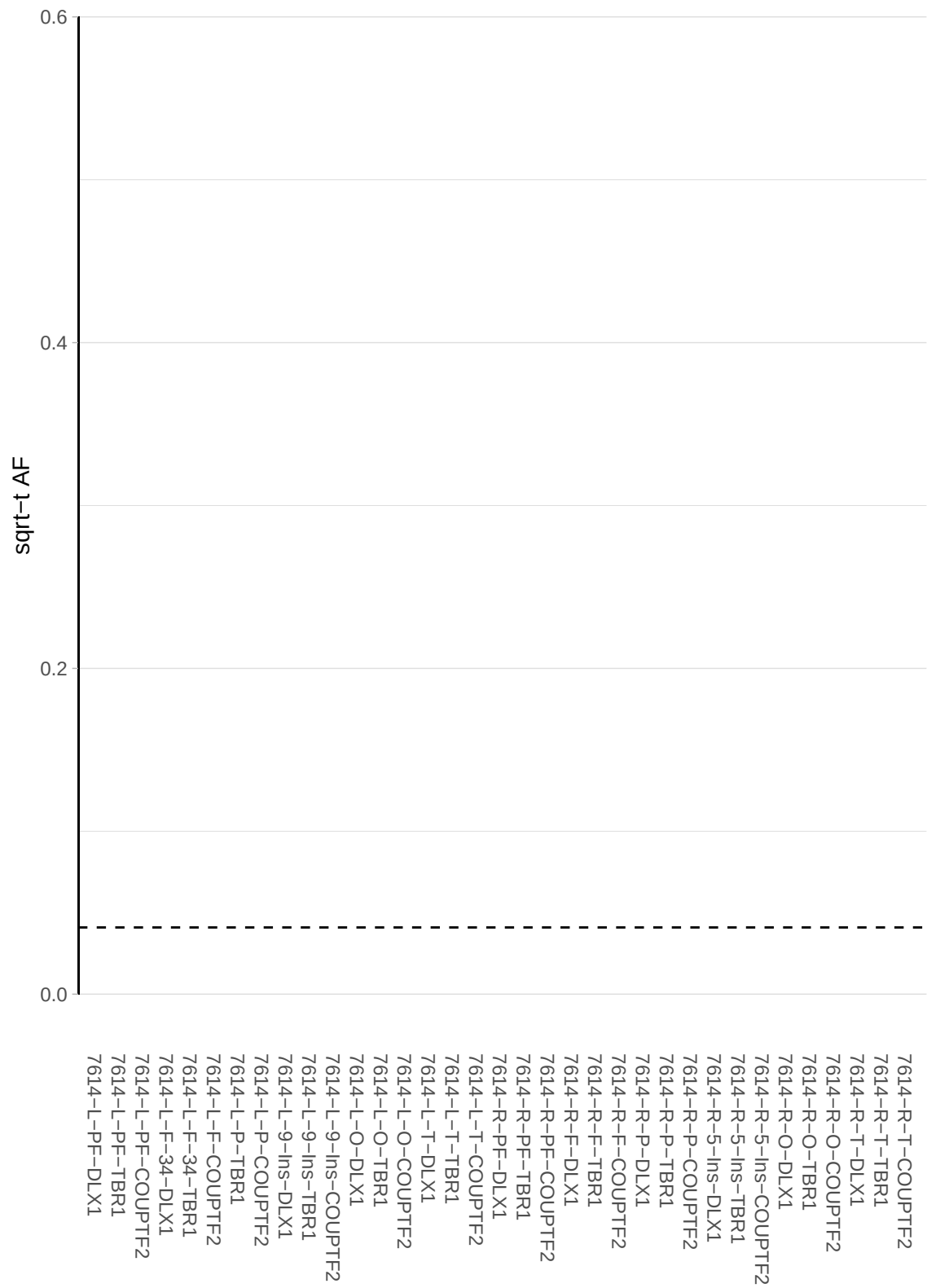

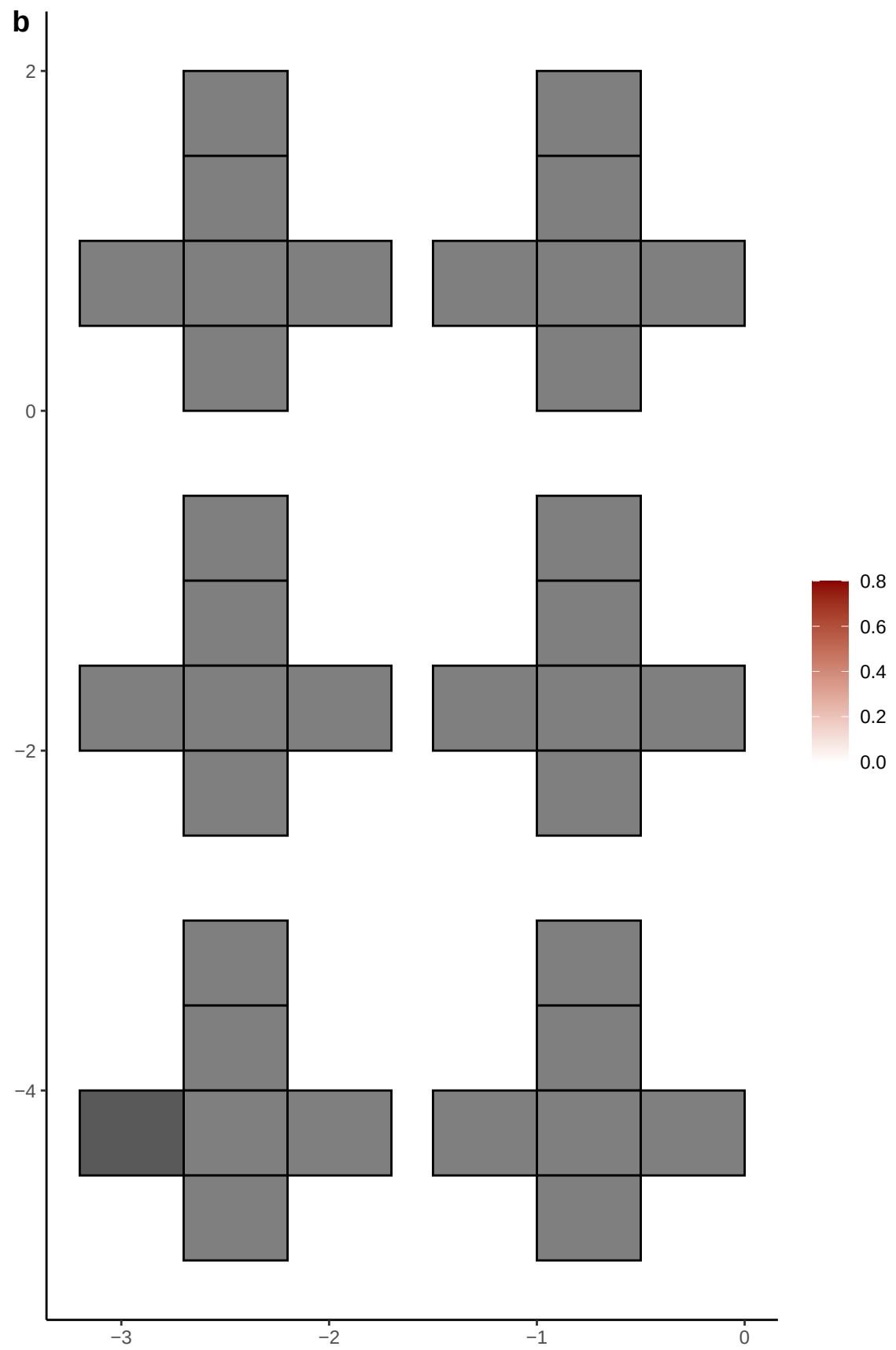

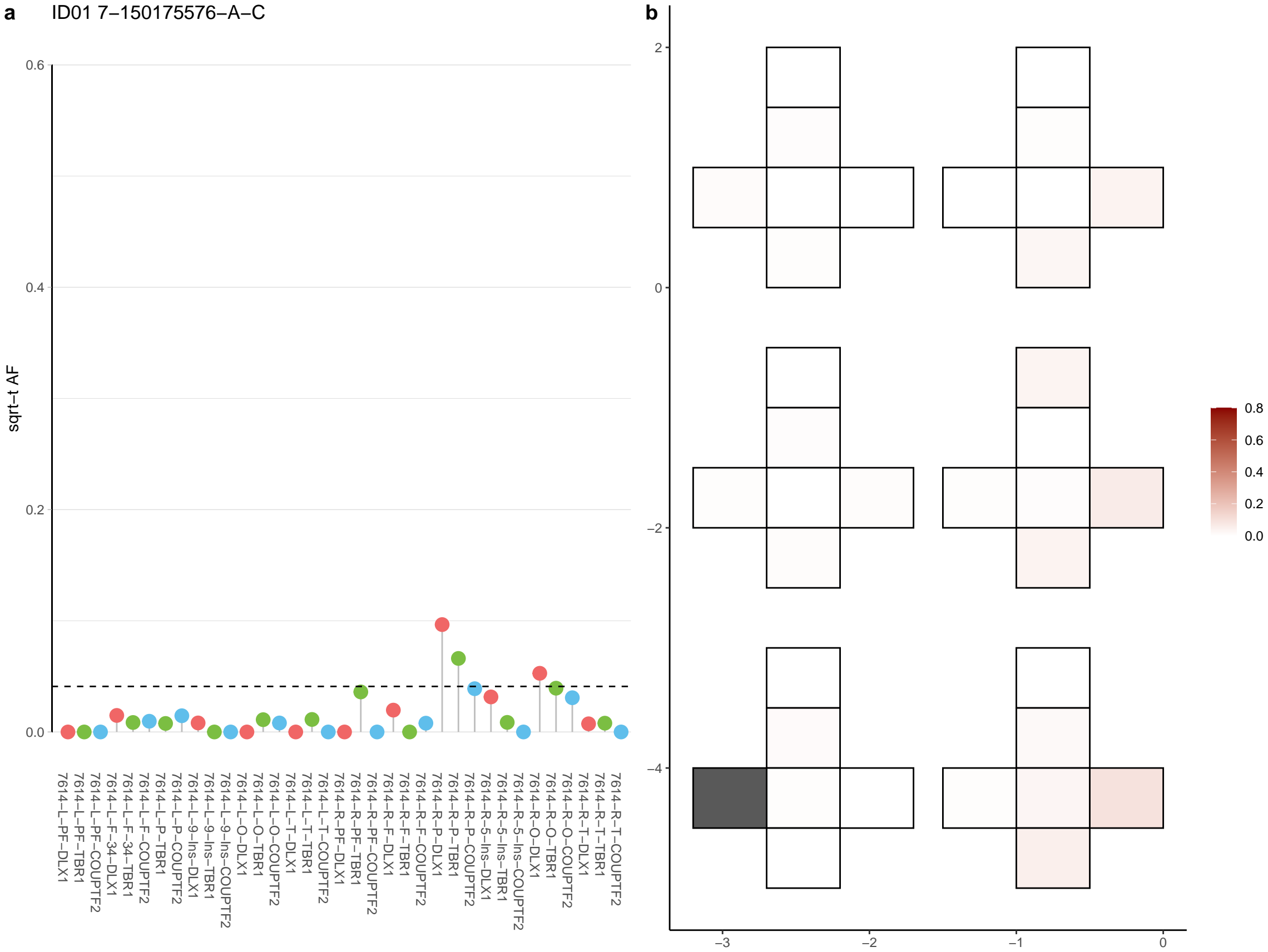

**a** ID01 7-31796412-T-G

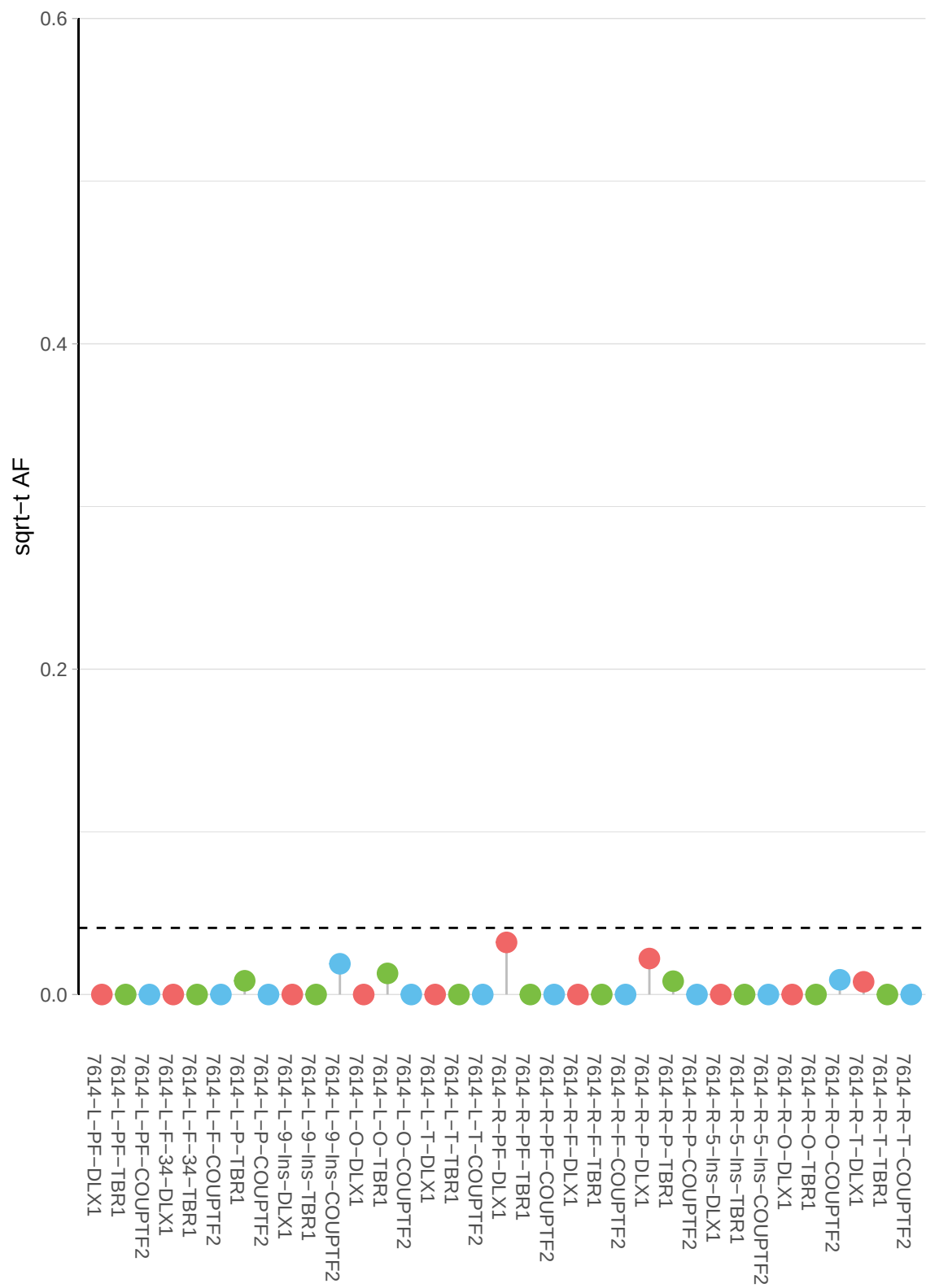

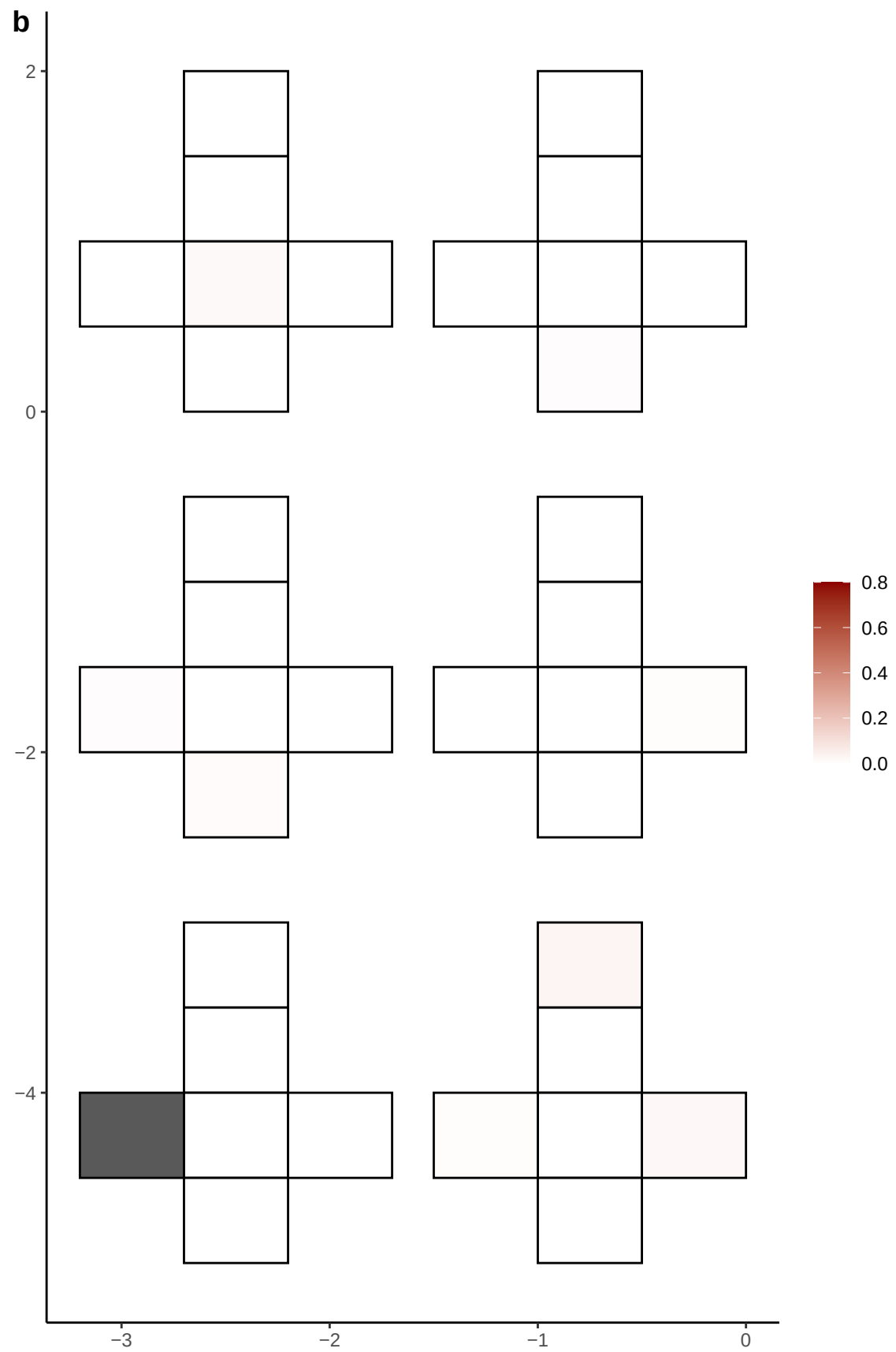

**a** ID01 7-33765368-A-G

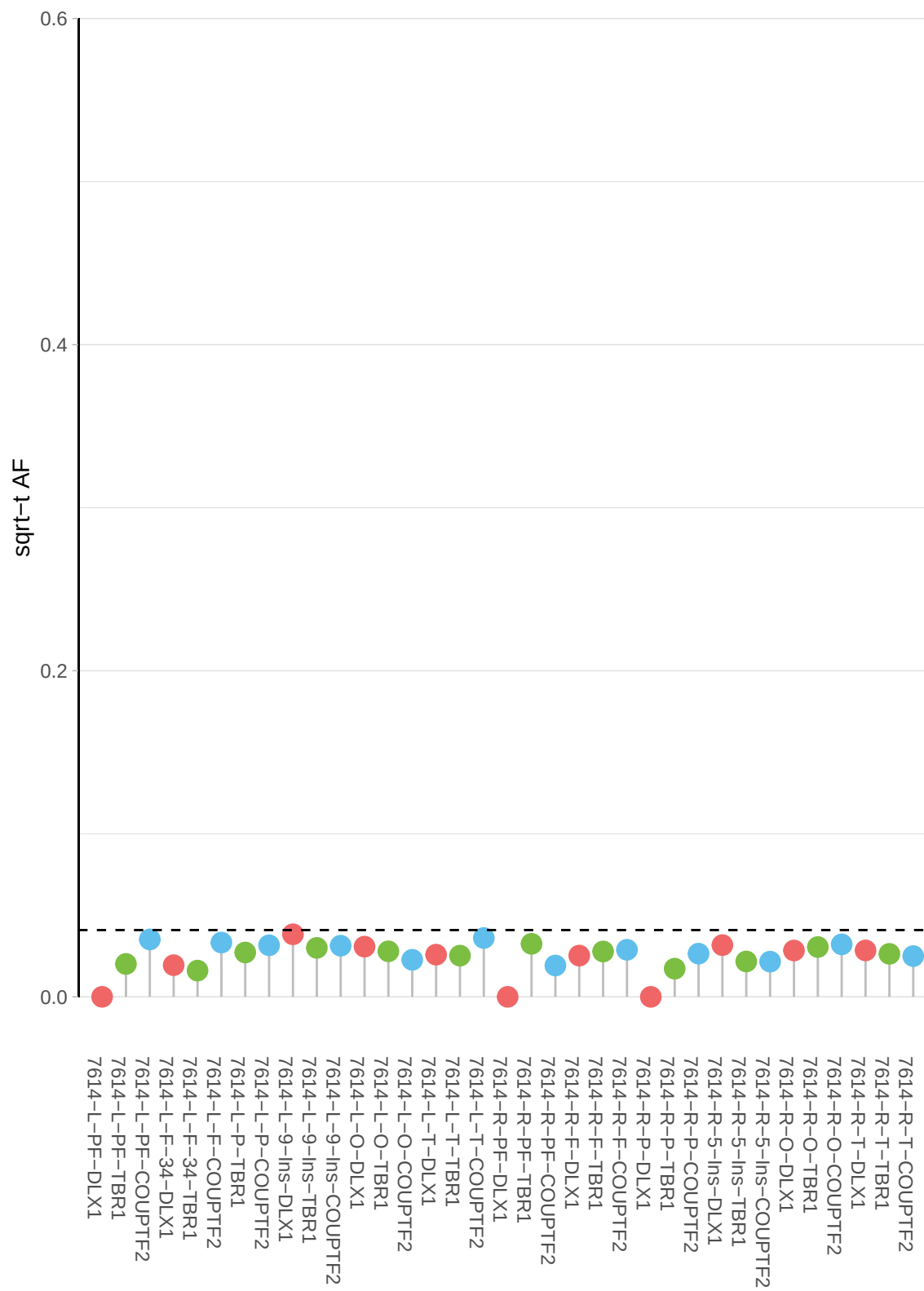

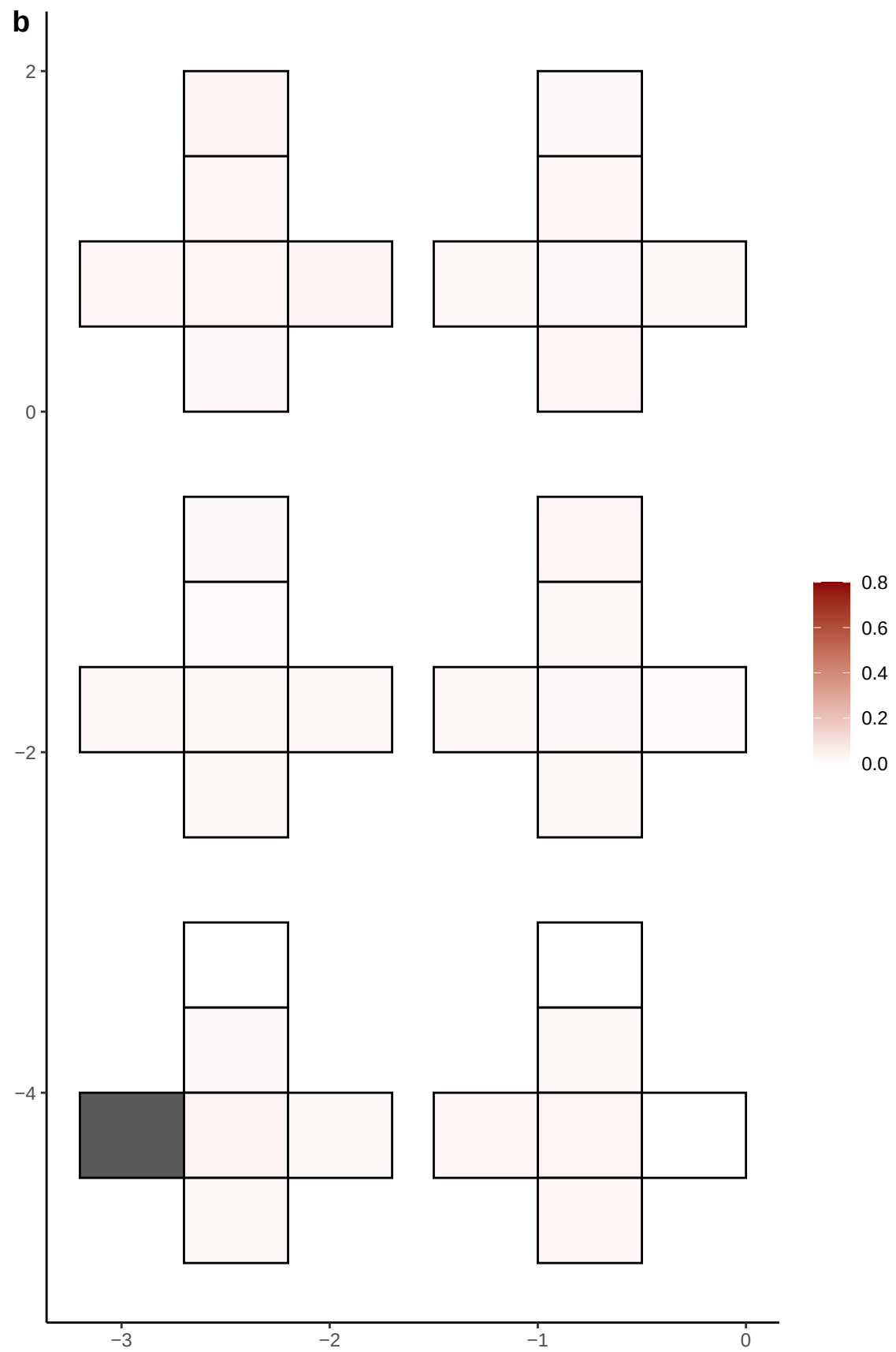

**a** ID01 7-37910766-C-T

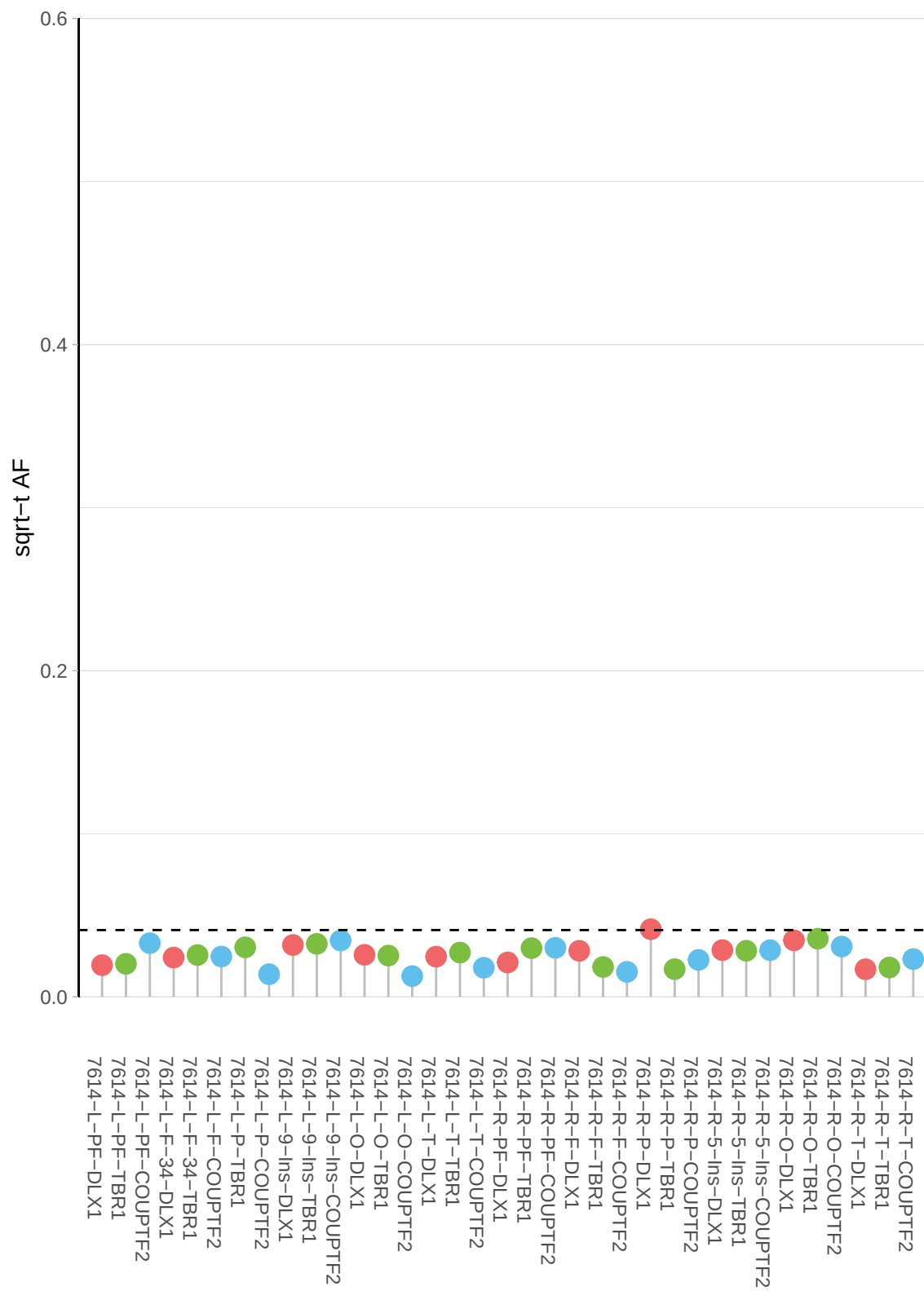

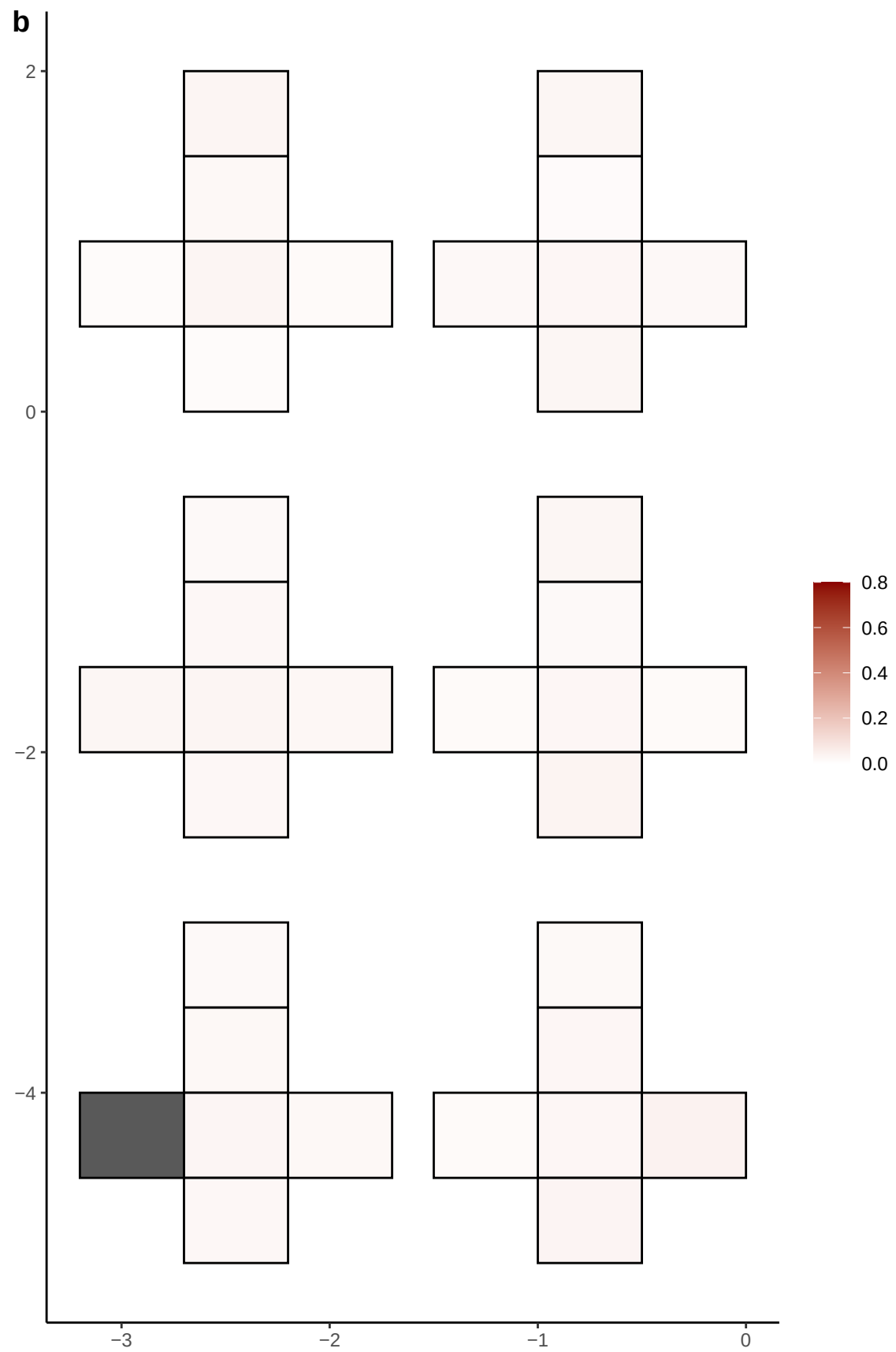

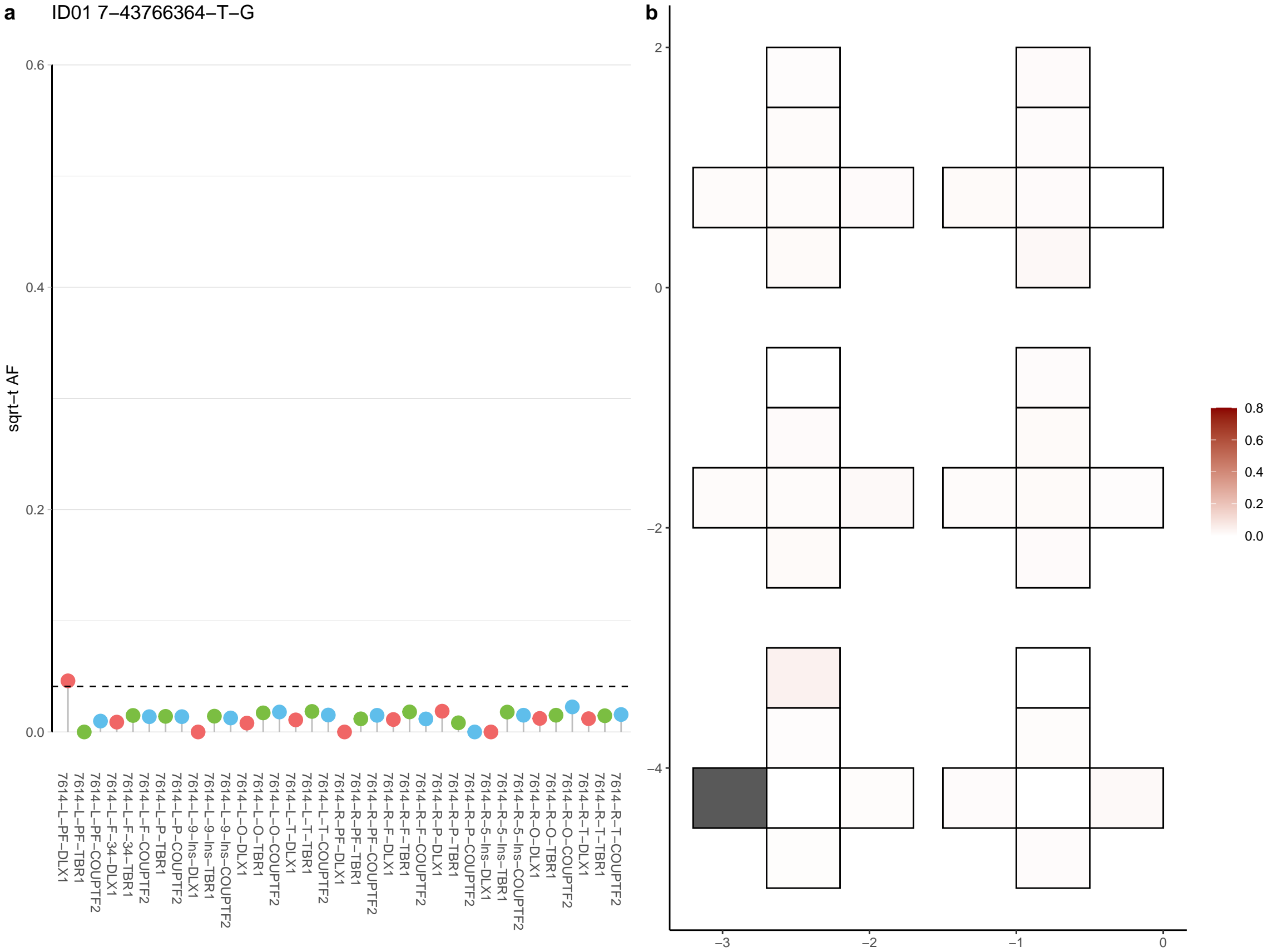

**a** ID01 7-67855805-C-T

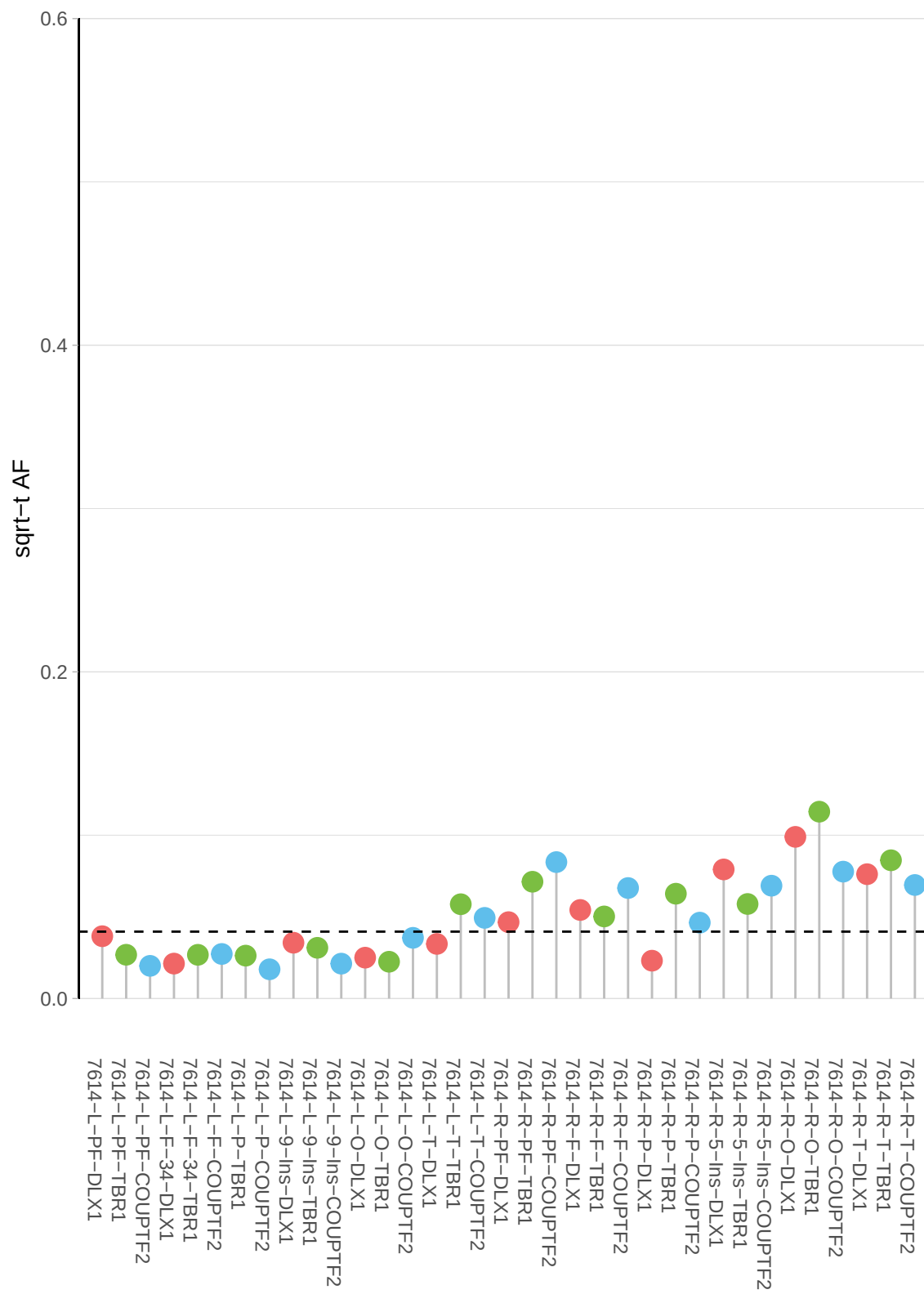

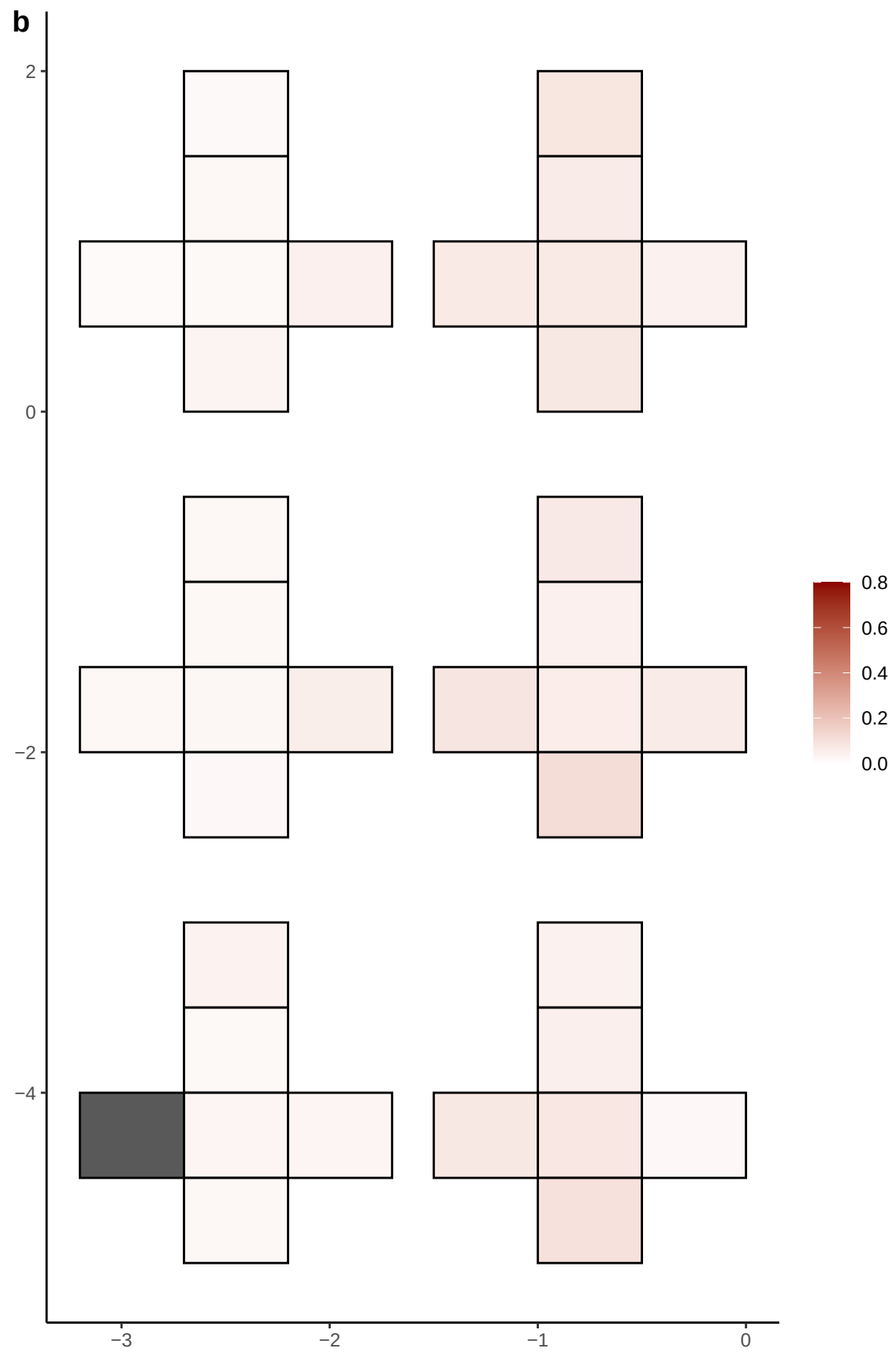

**a** ID01 7-80017095-C-T

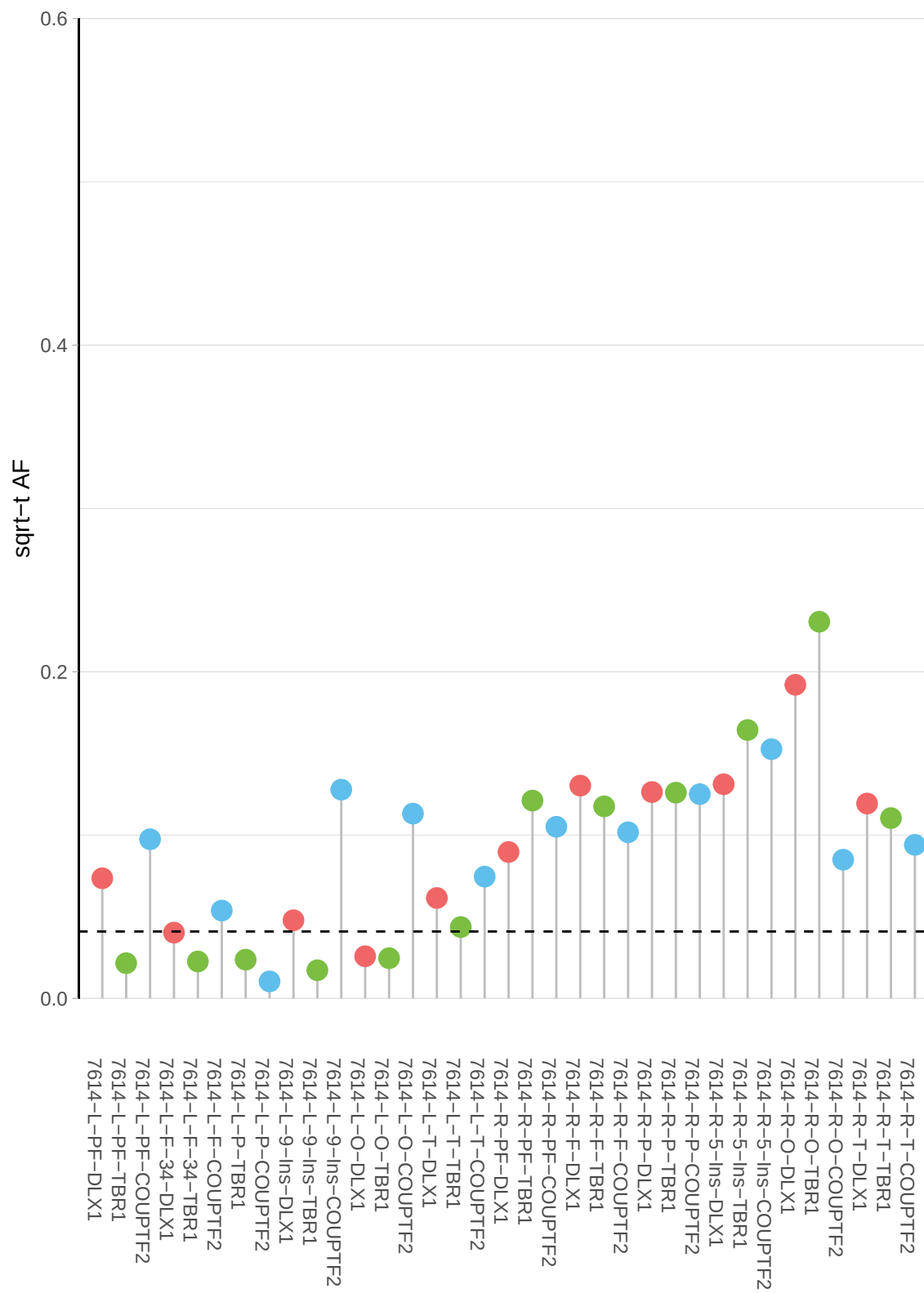

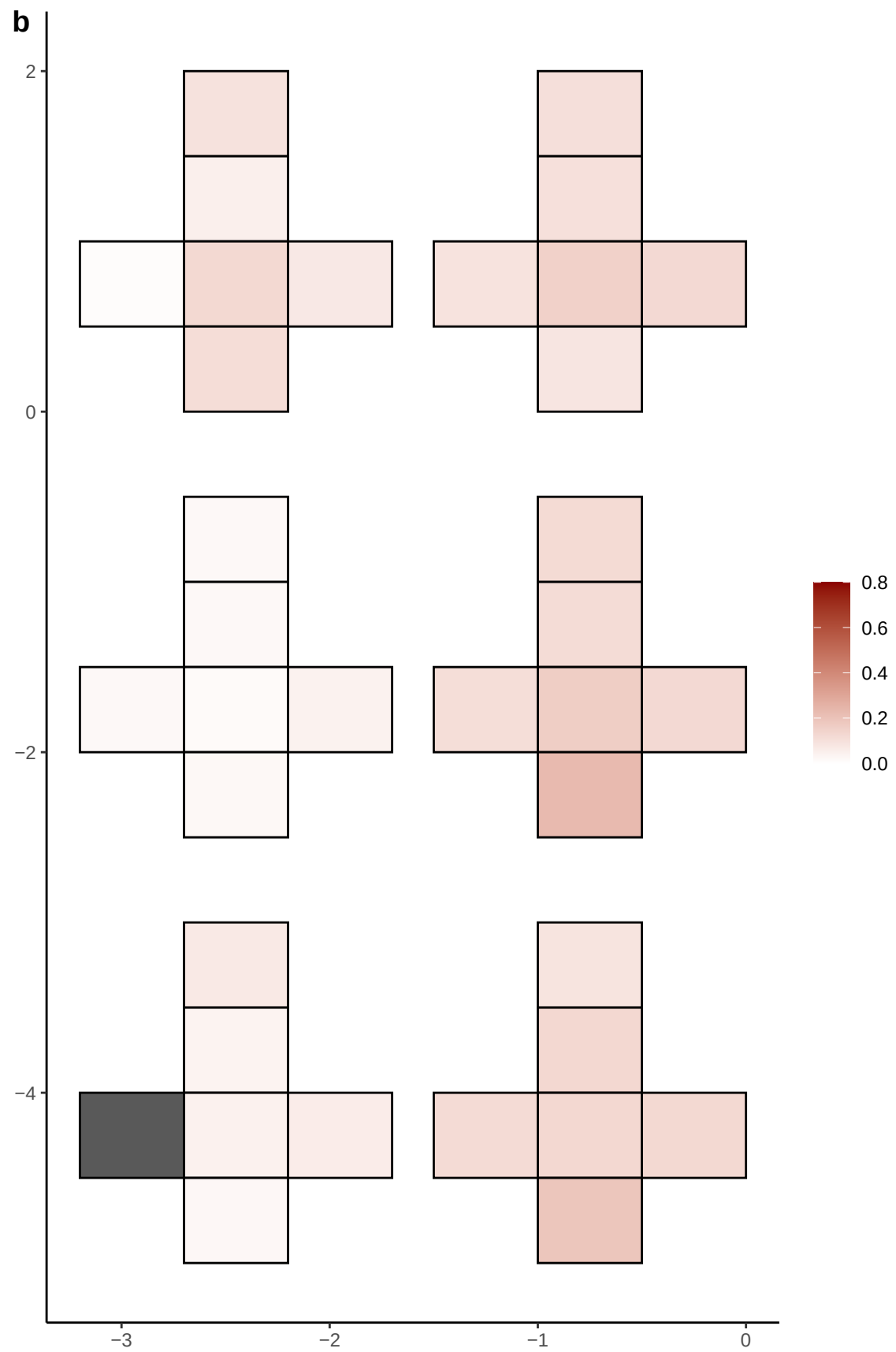

**a** ID01 7-87518026-C-T

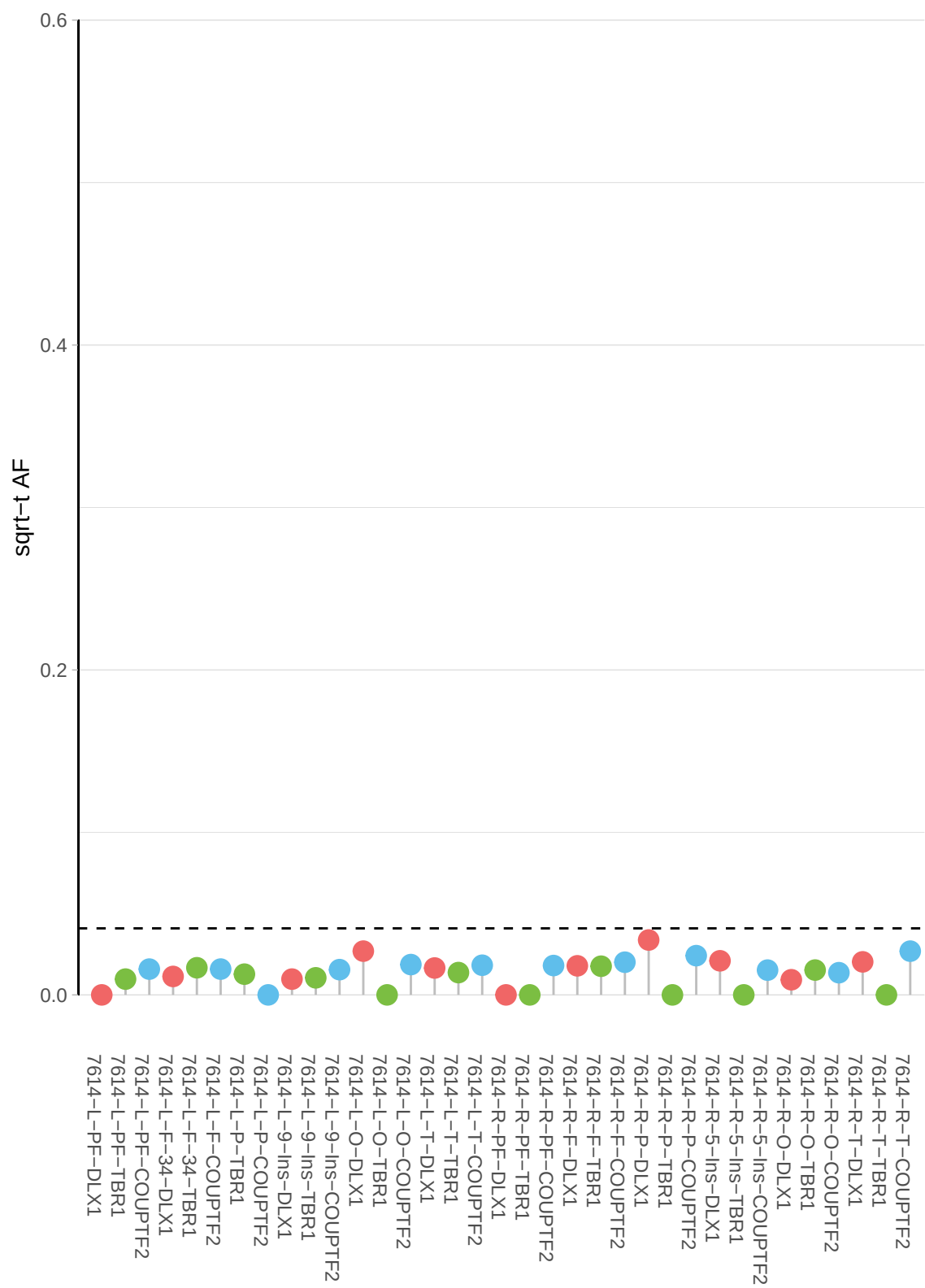

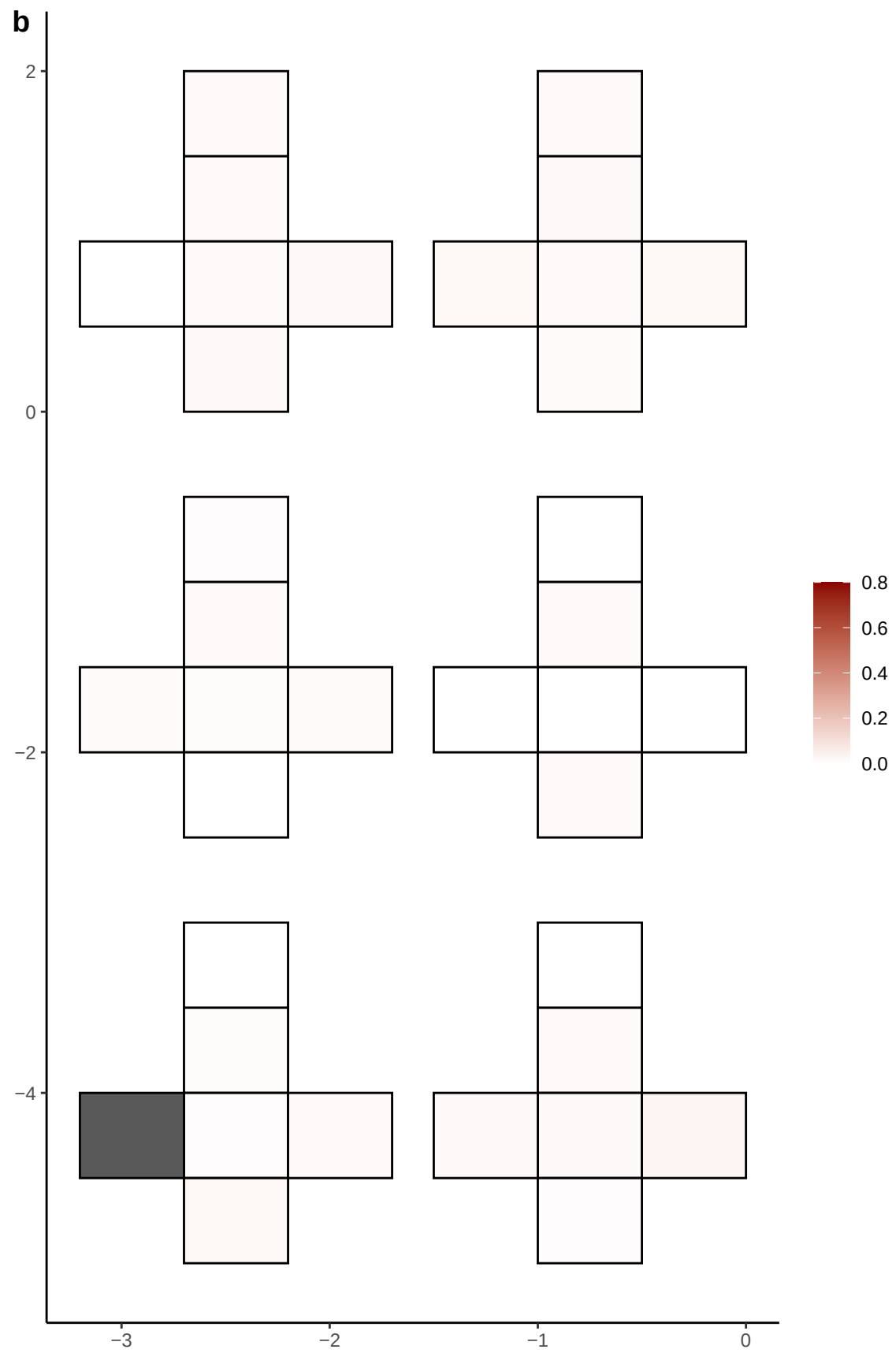

**a** ID01 8-100847587-A-AATGTGT

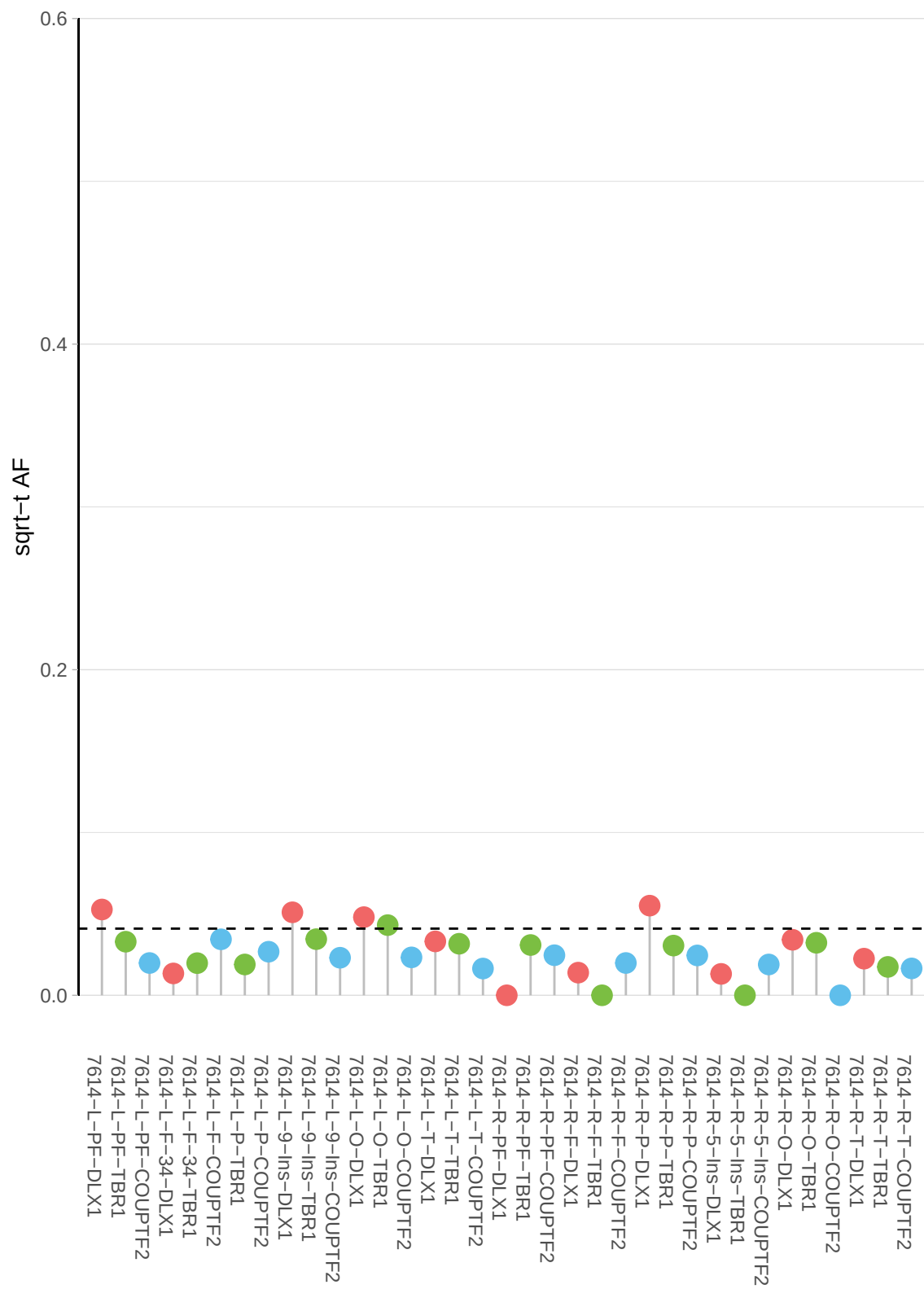

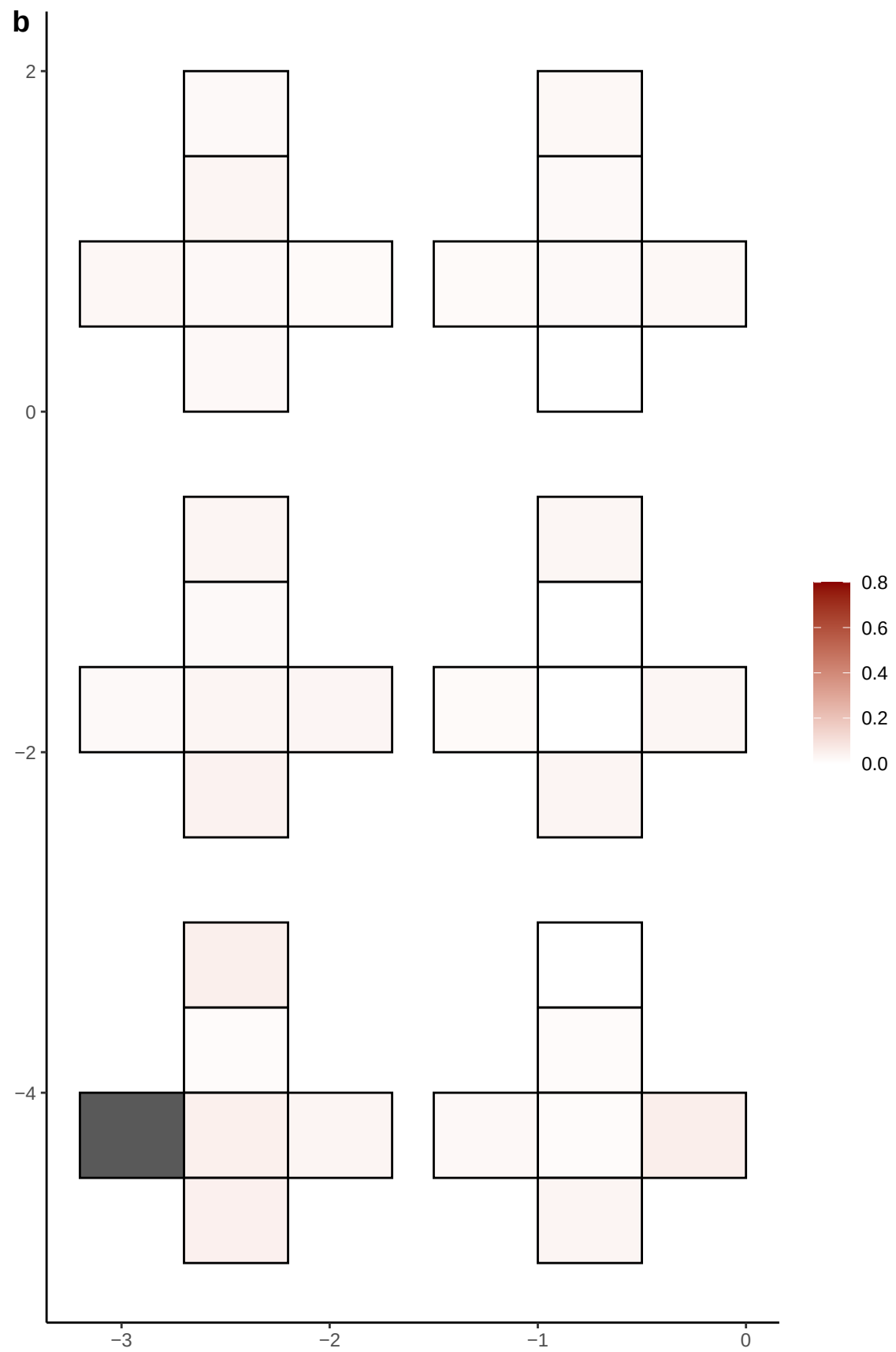

**a** ID01 8-116848693-T-A

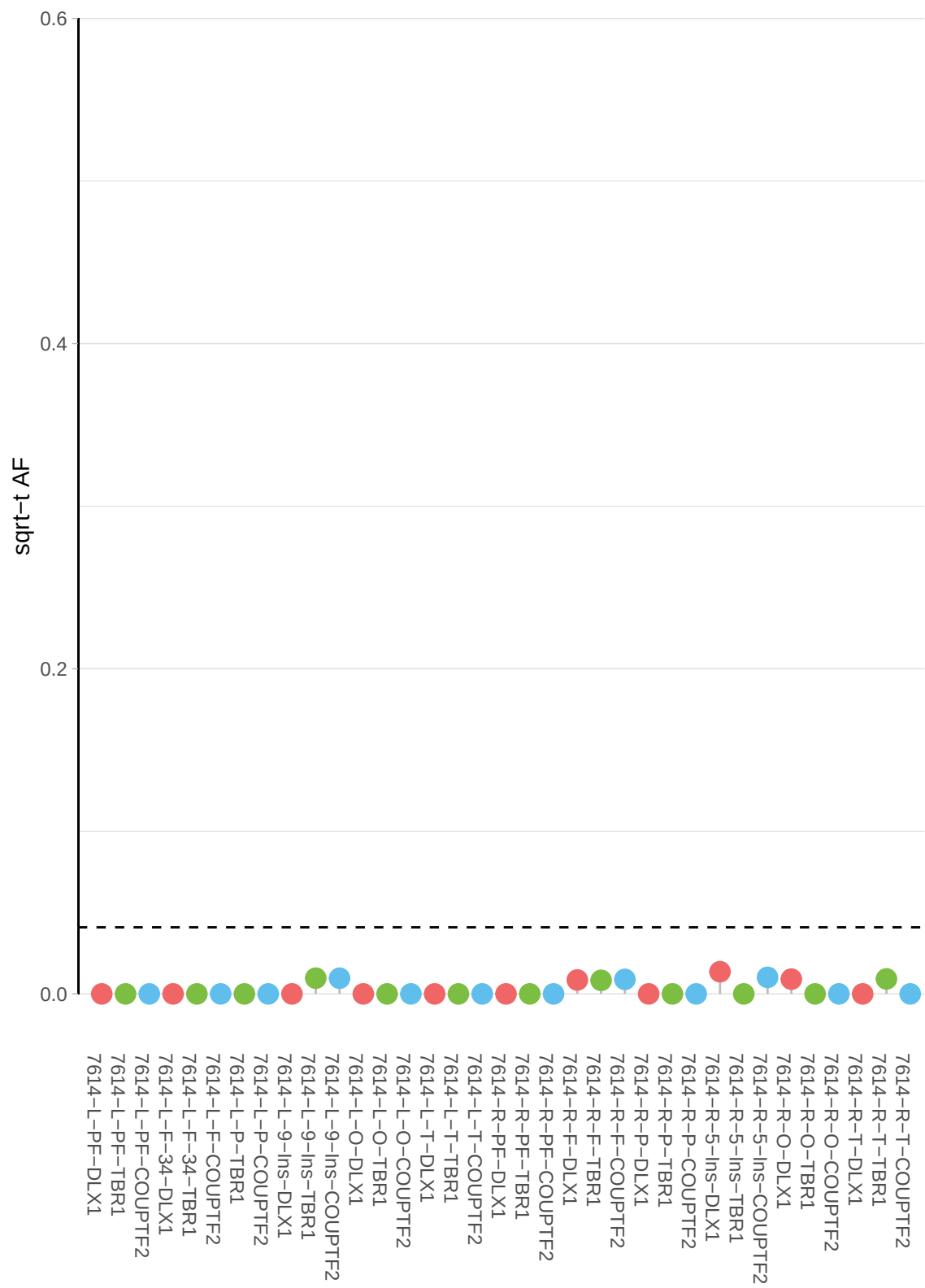

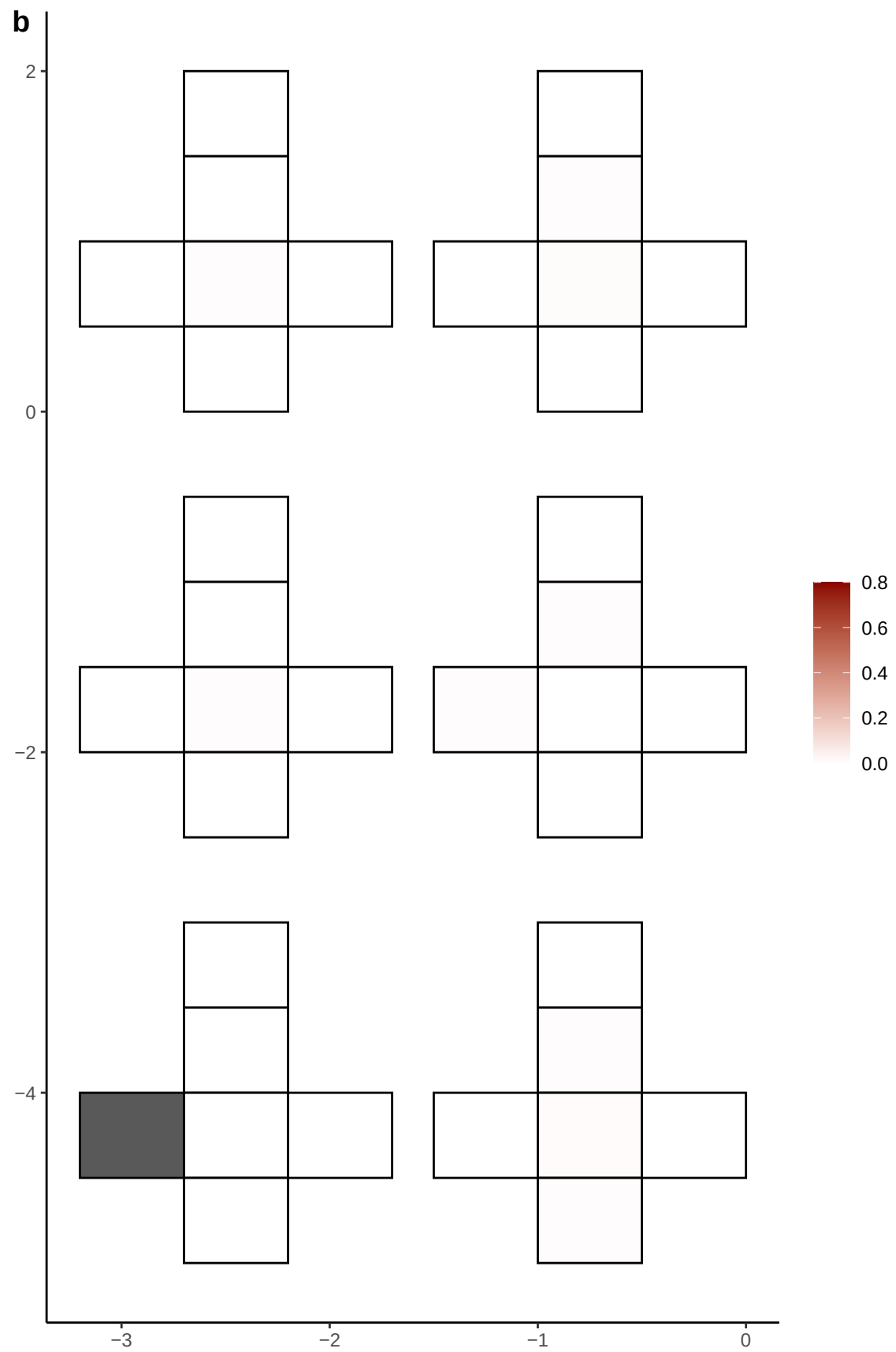

**a** ID01 8-121214447-G-A

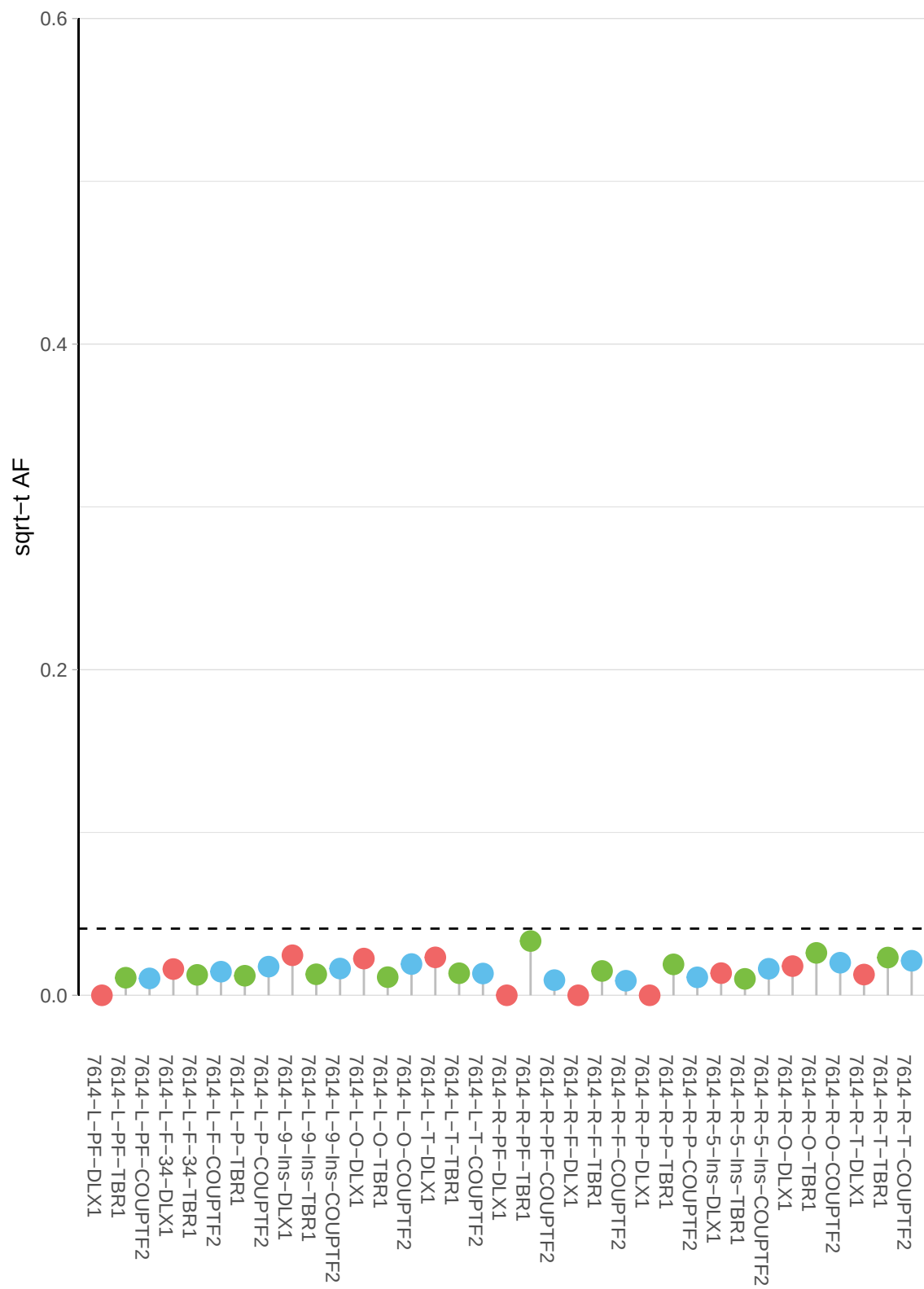

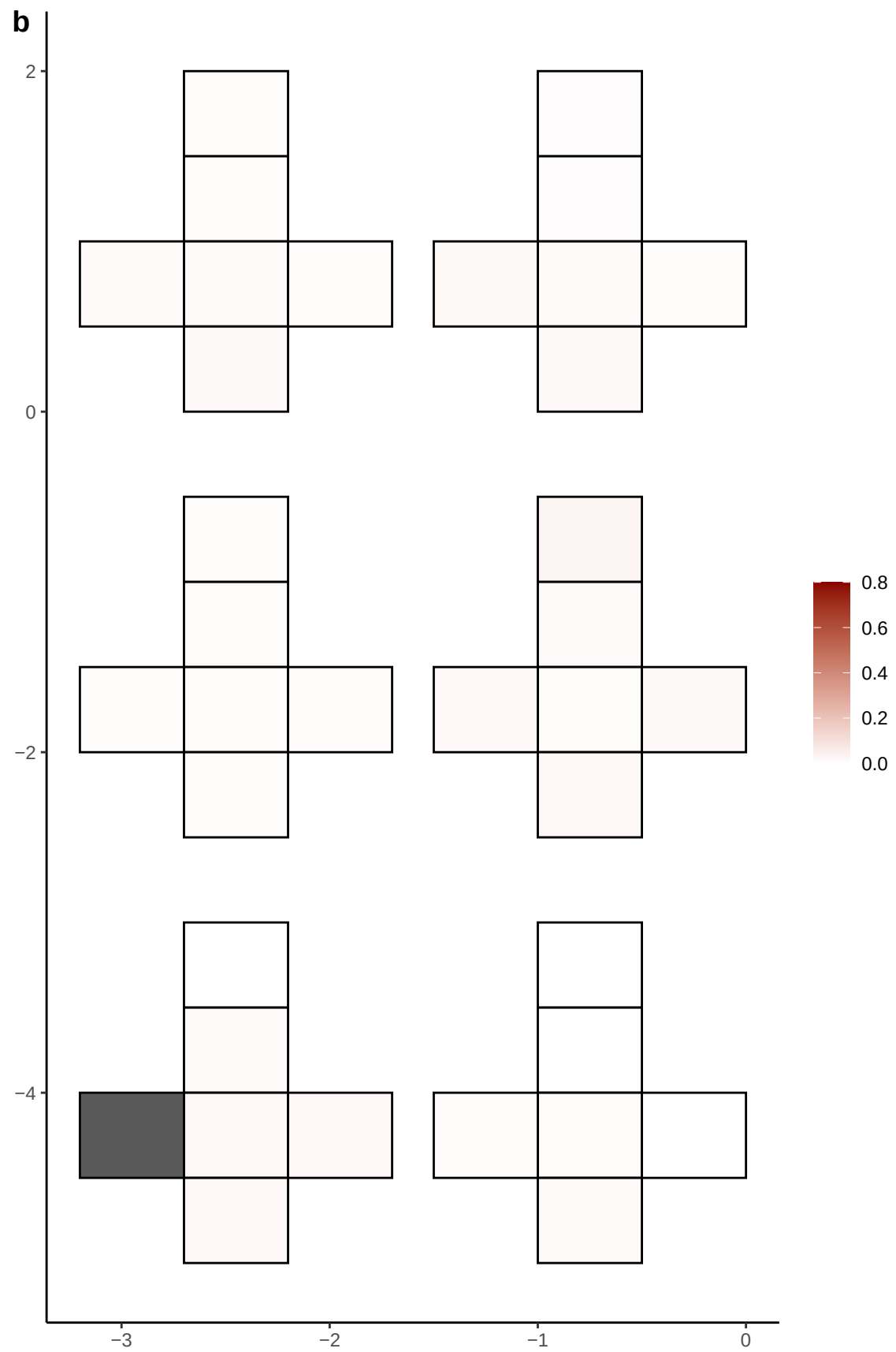

**a** ID01 8-122651370-C-A

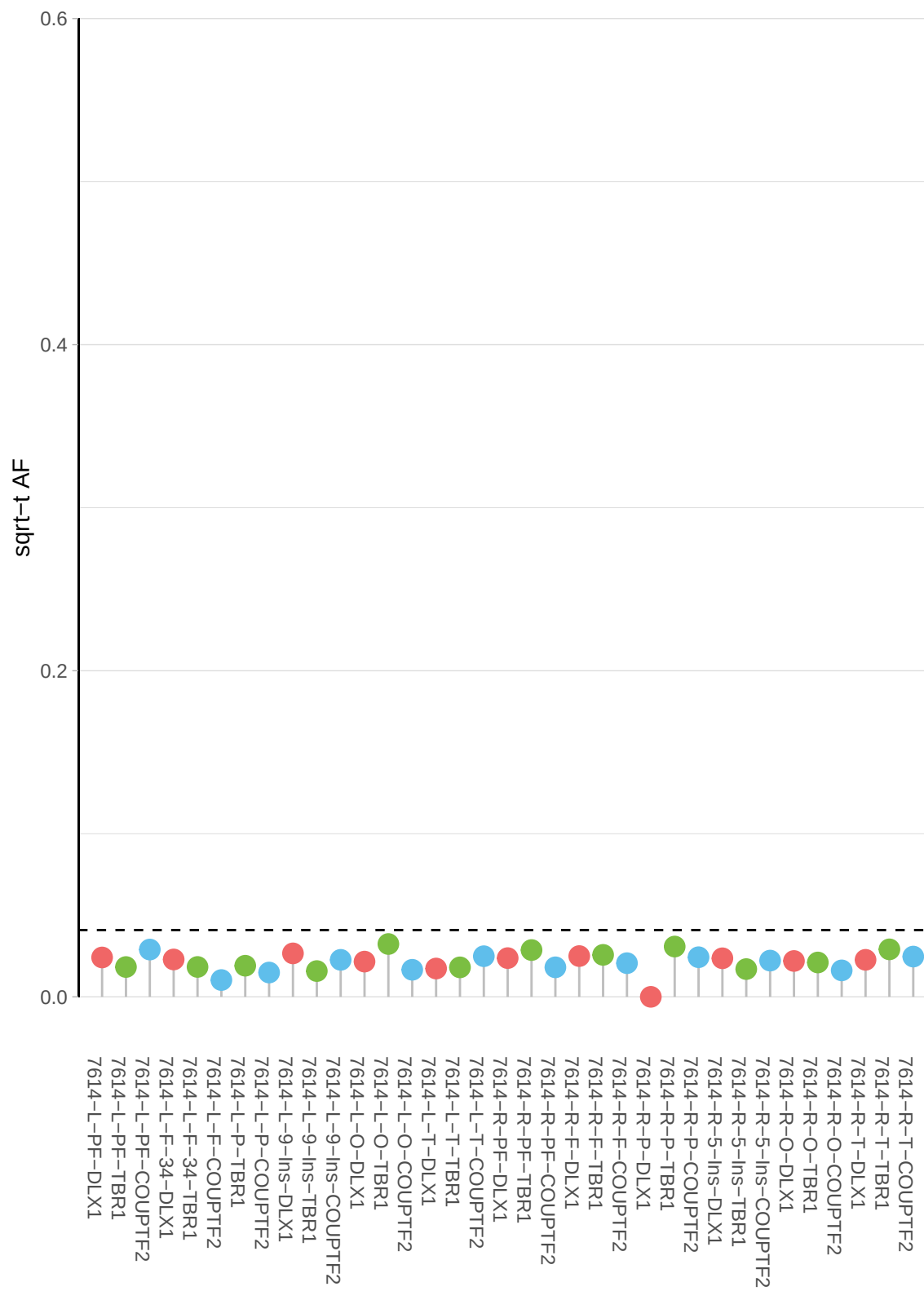

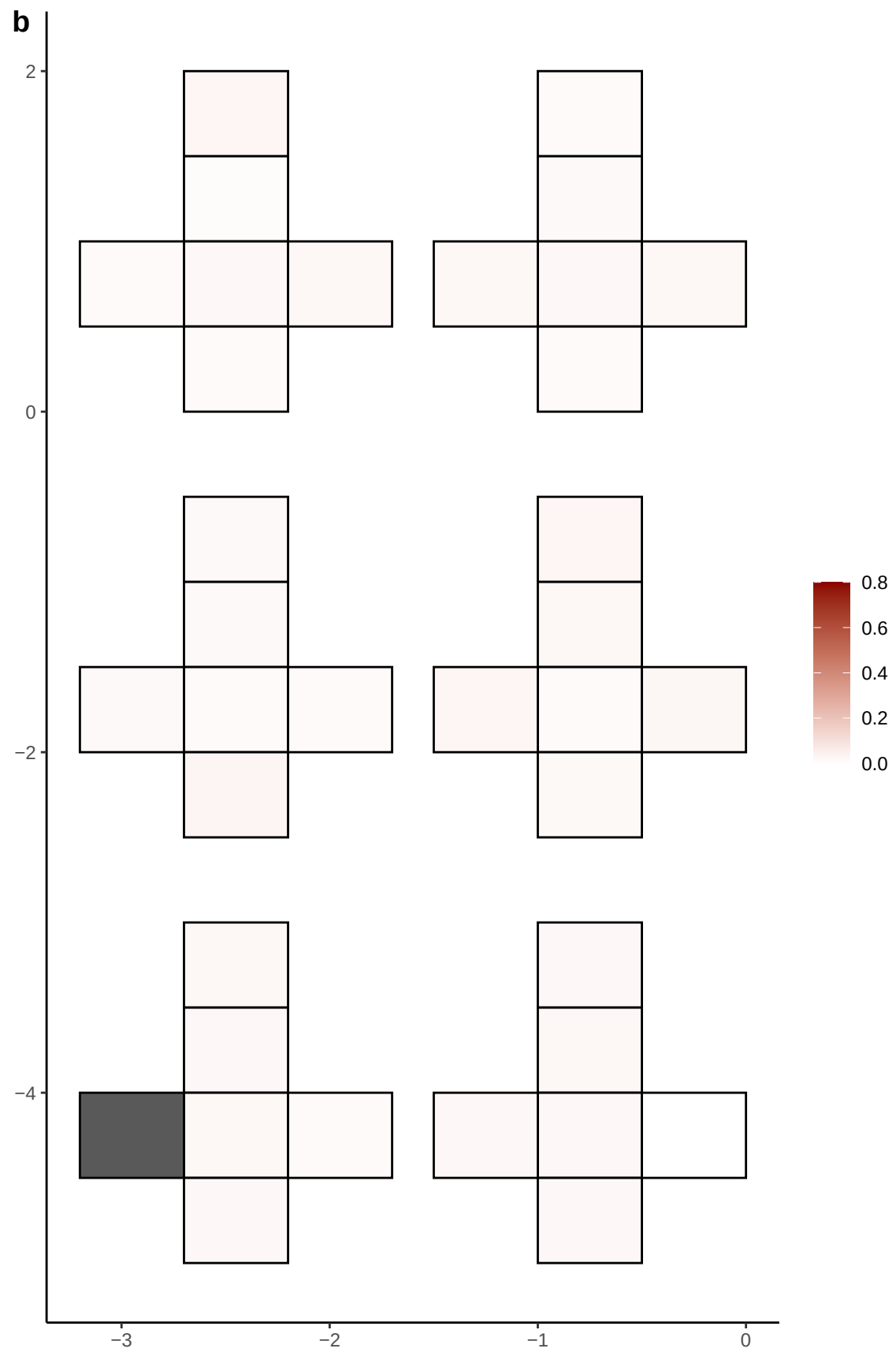

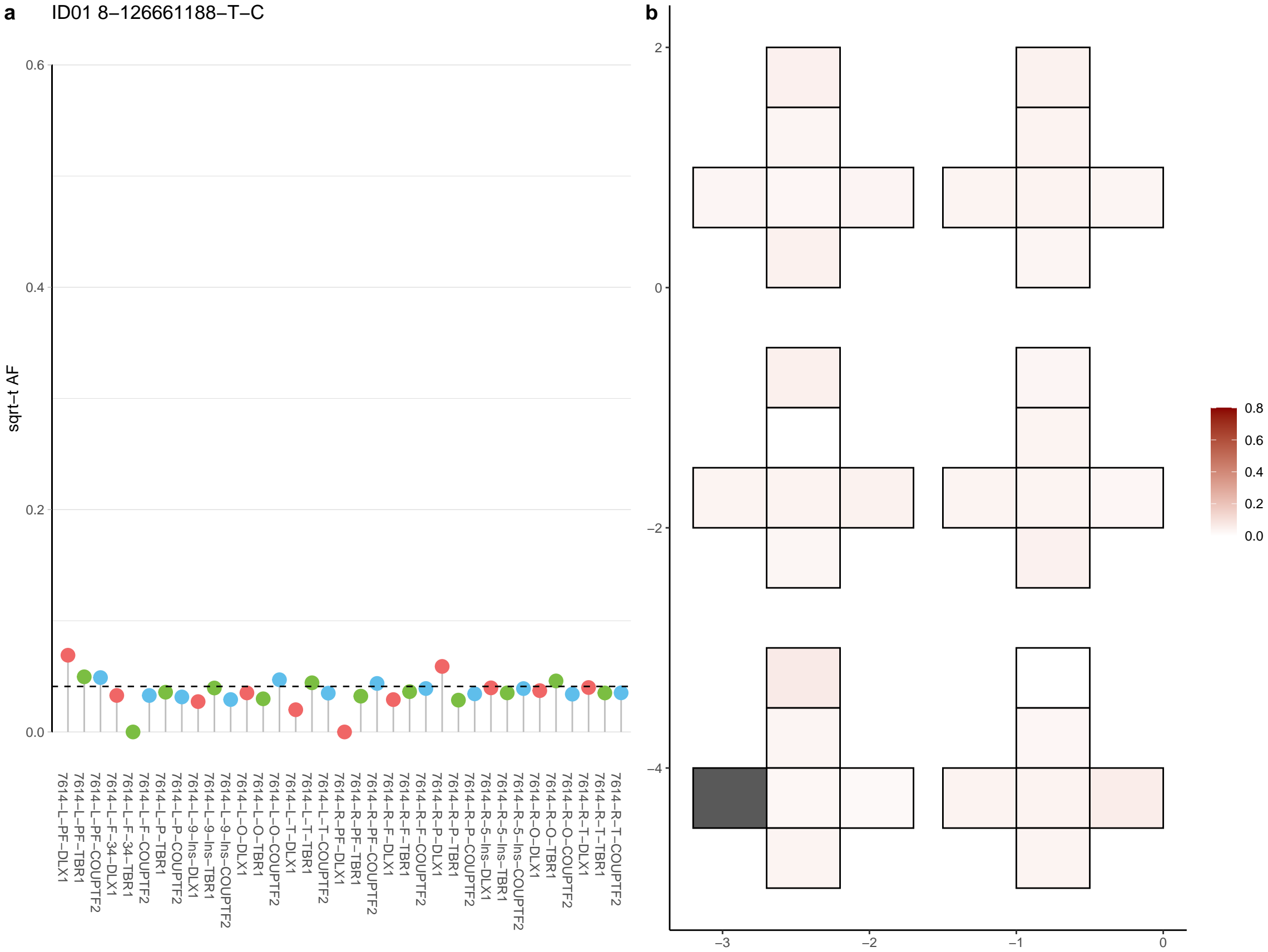

**a** ID01 8-138116363-A-G

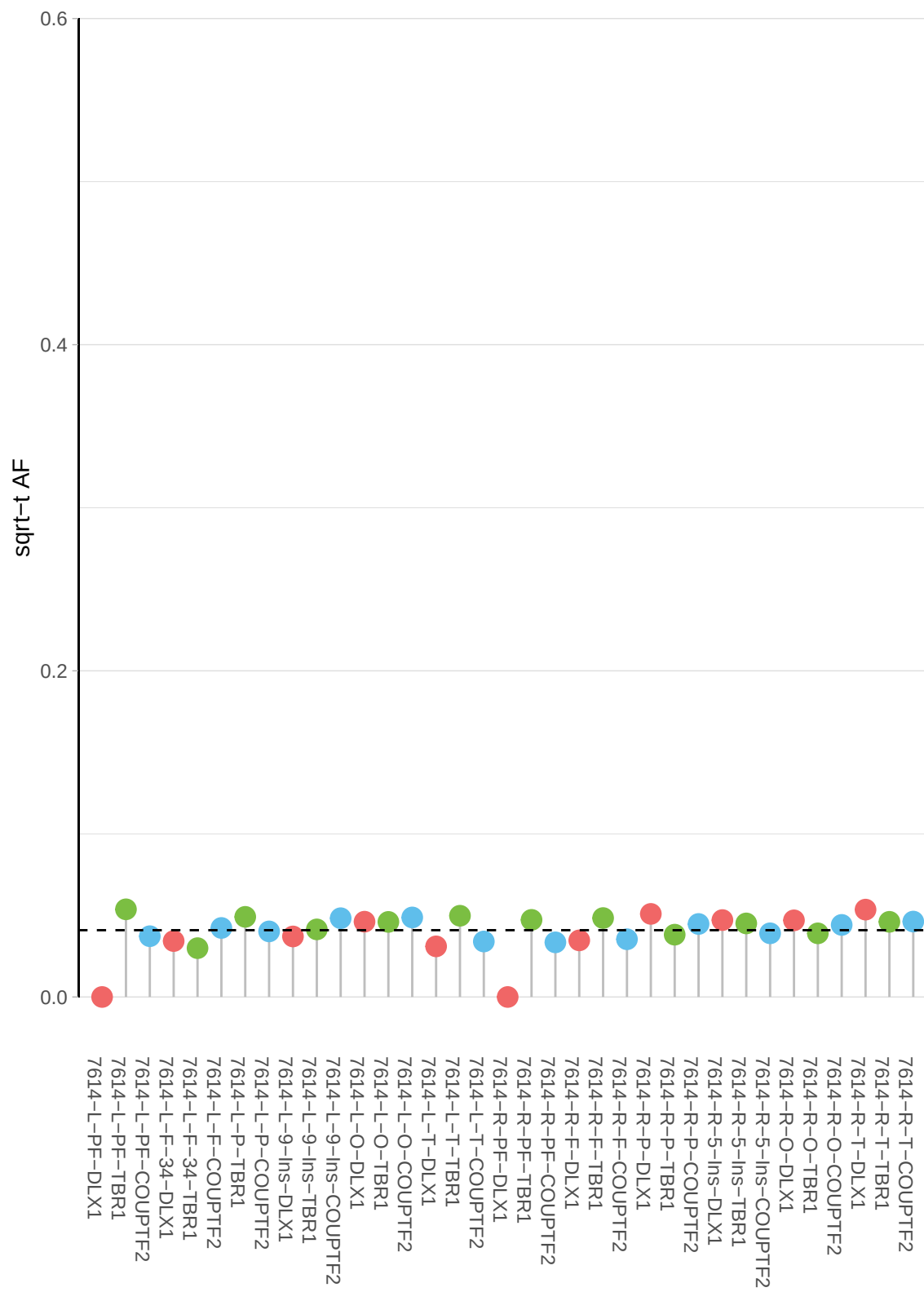

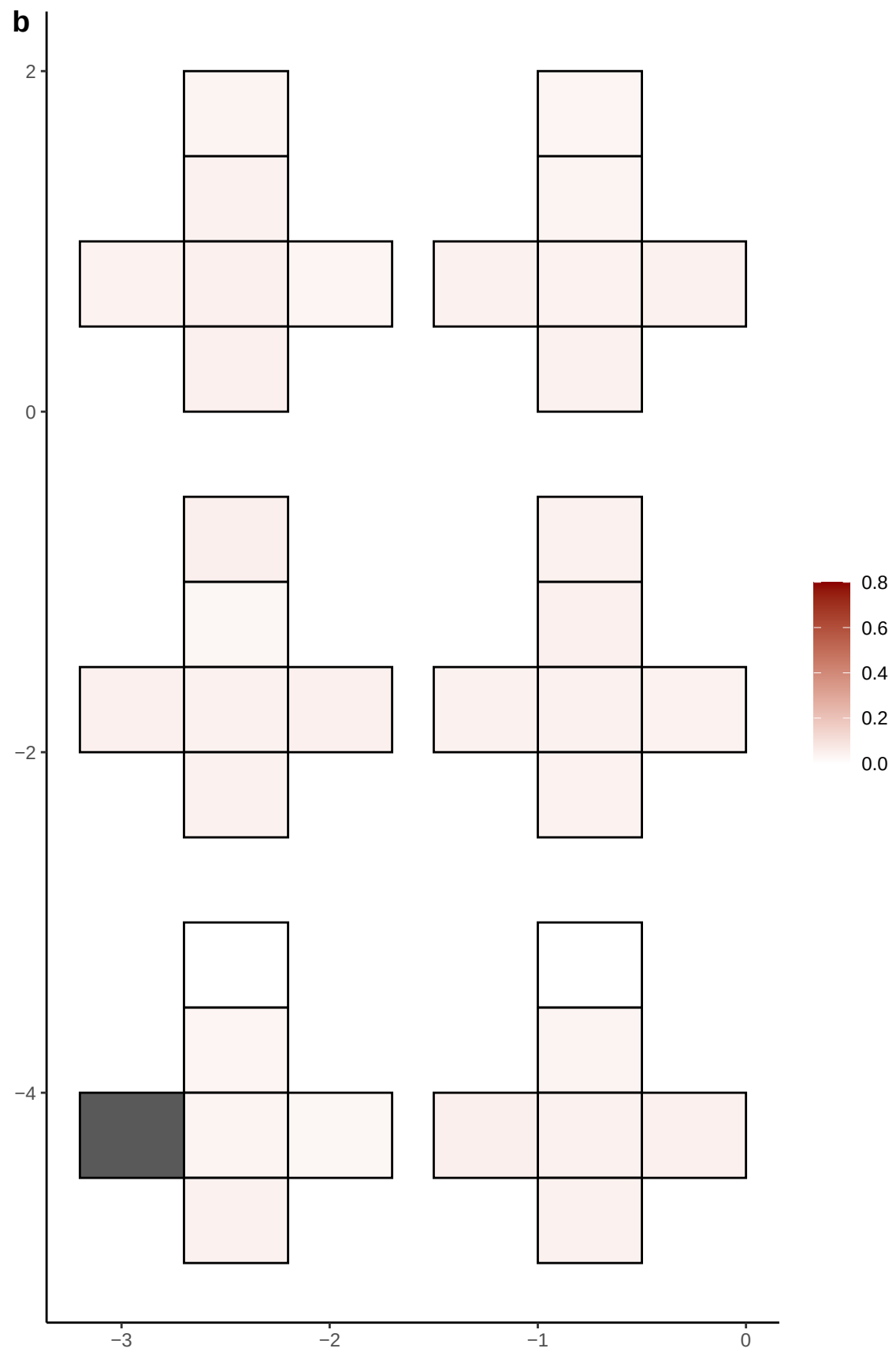

**a** ID01 8-13840424-T-C

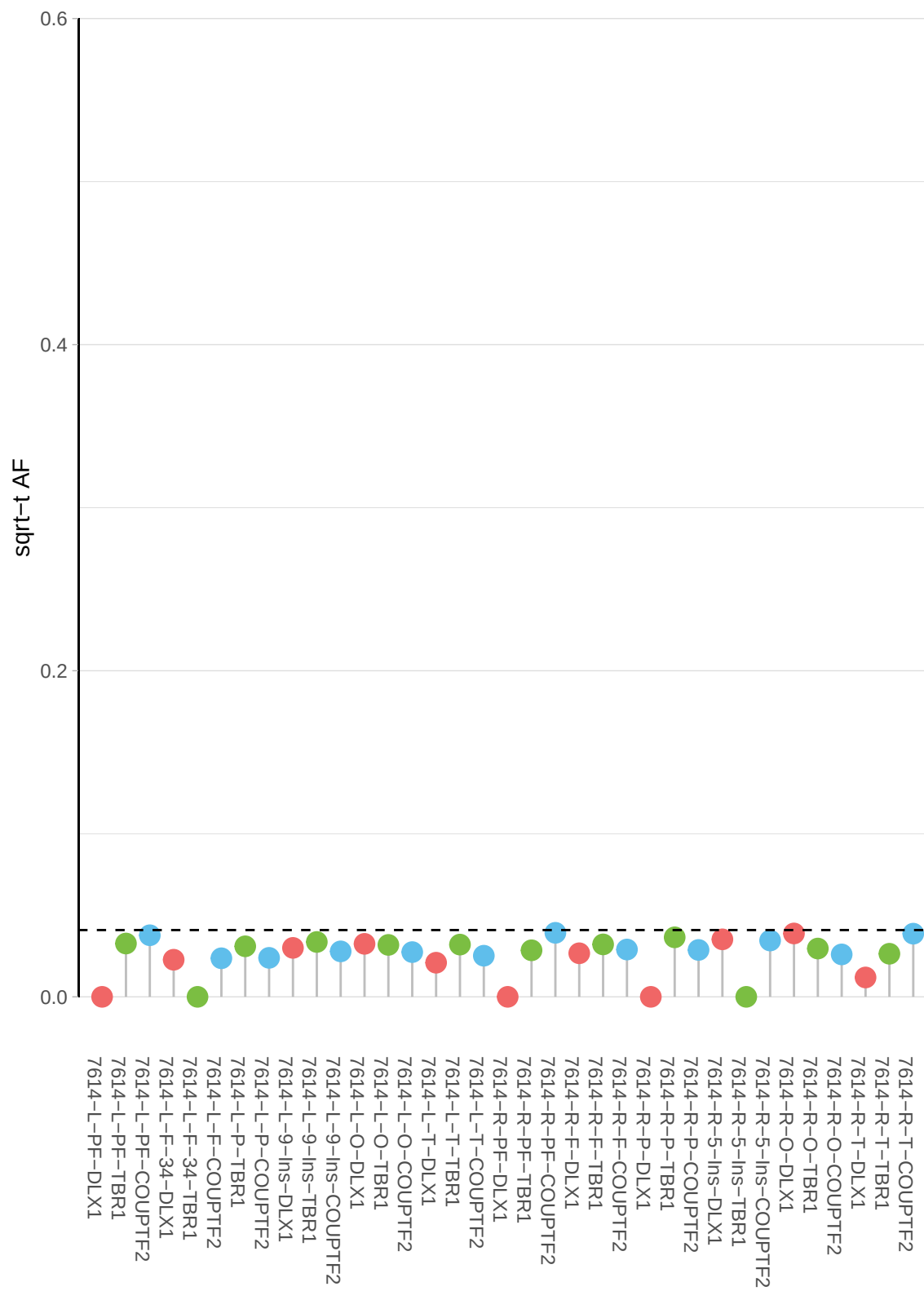

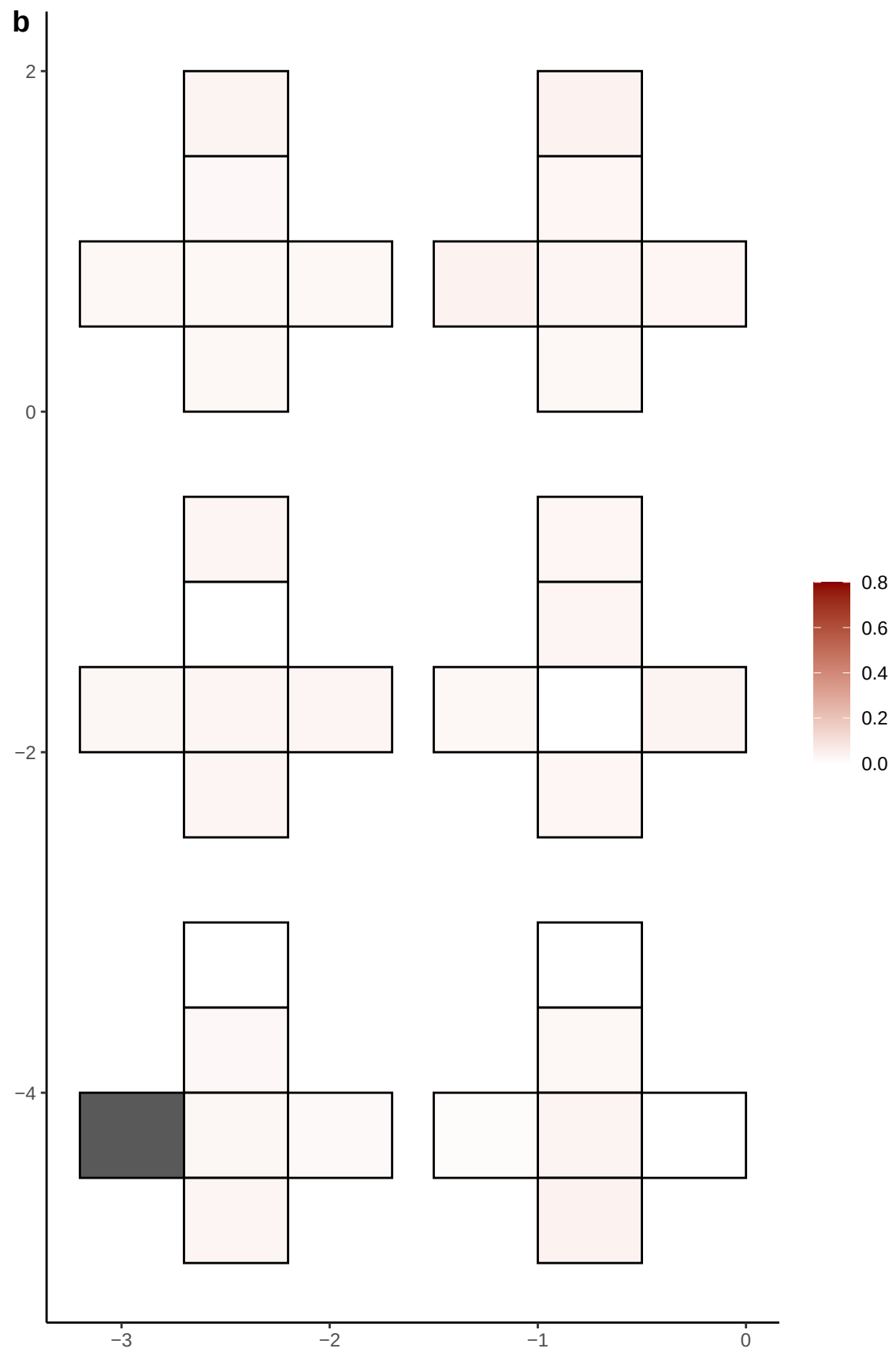

**a** ID01 8-14166252-C-T

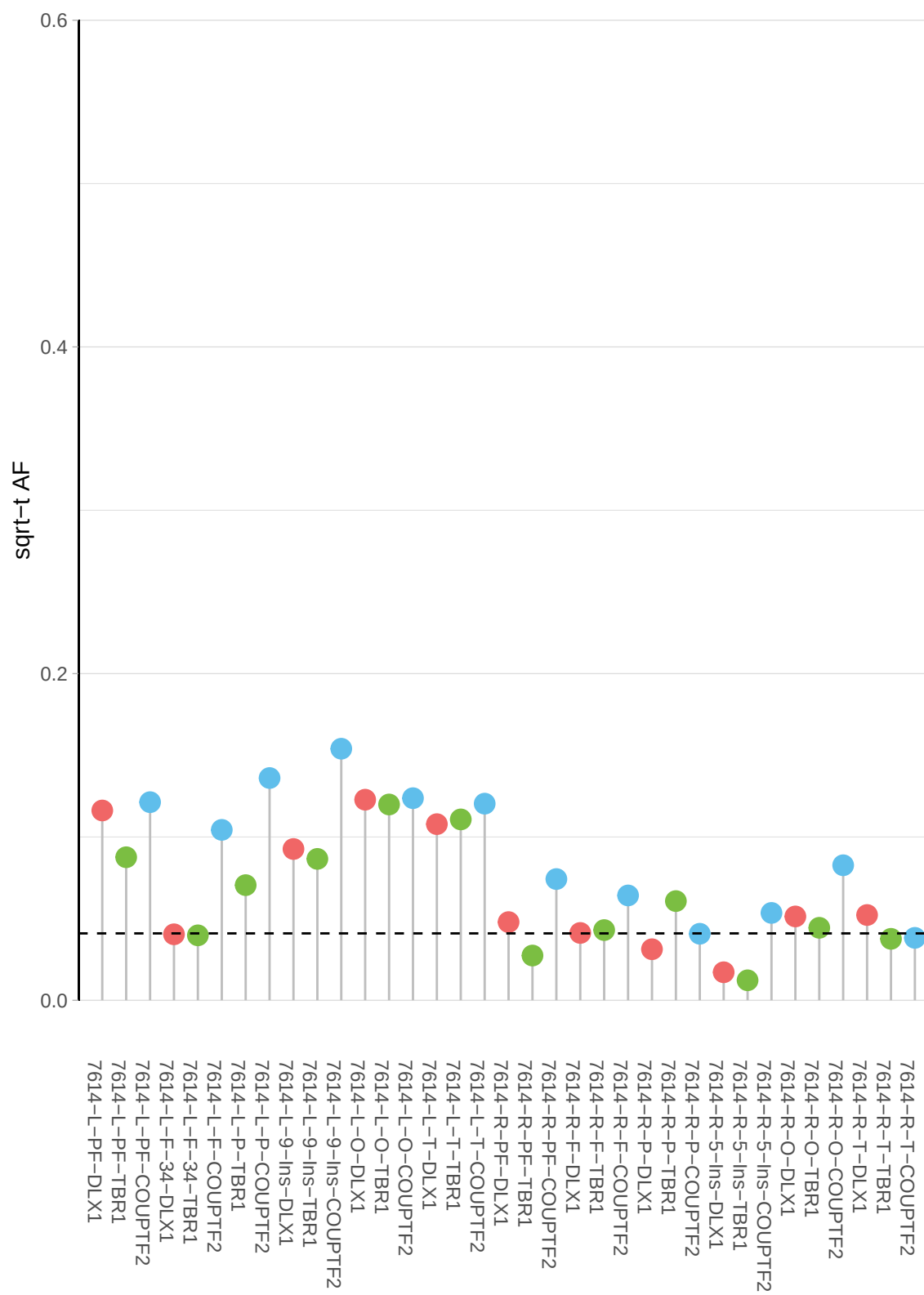

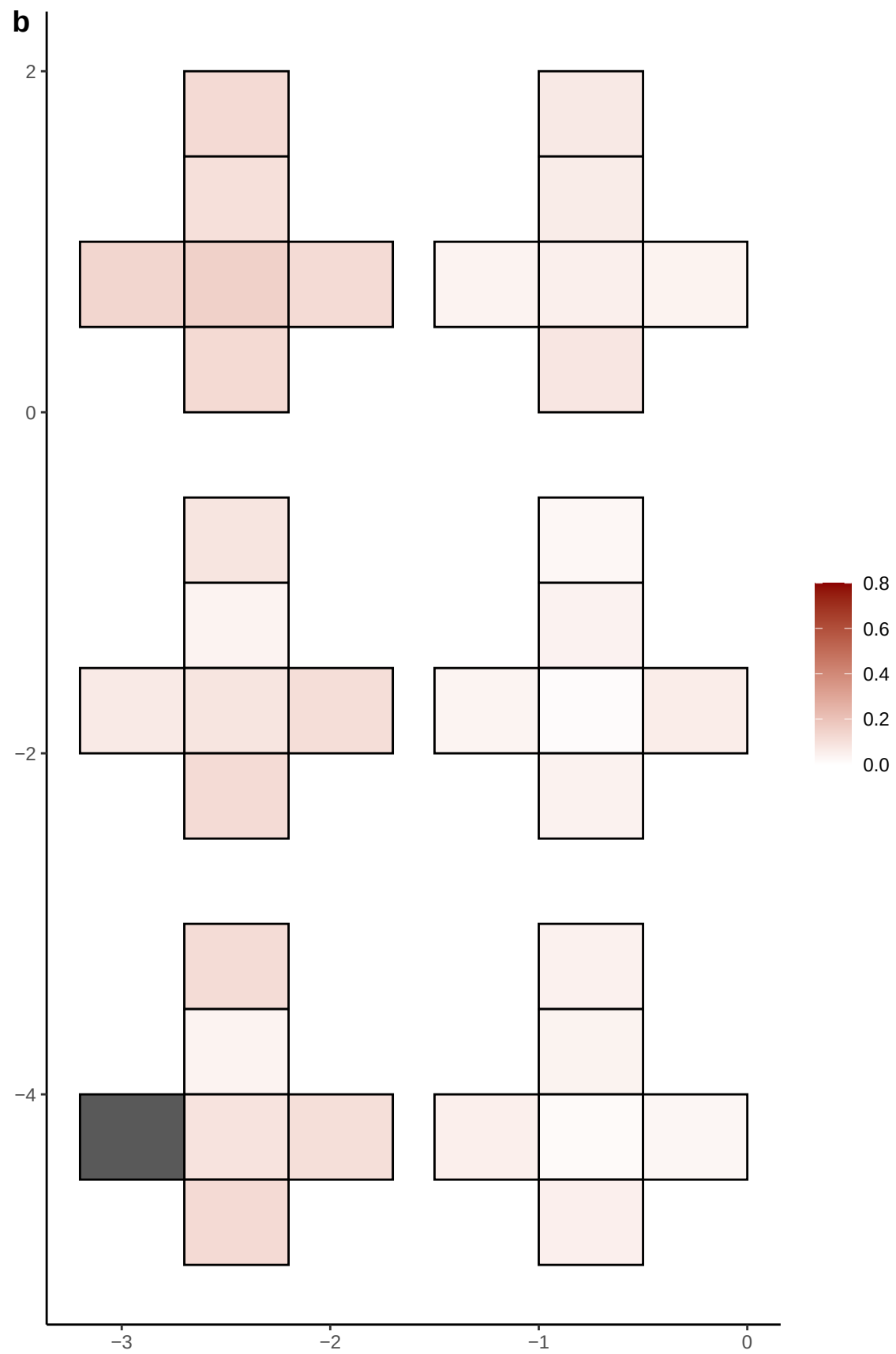

**a** ID01 8-14411299-CTT-C

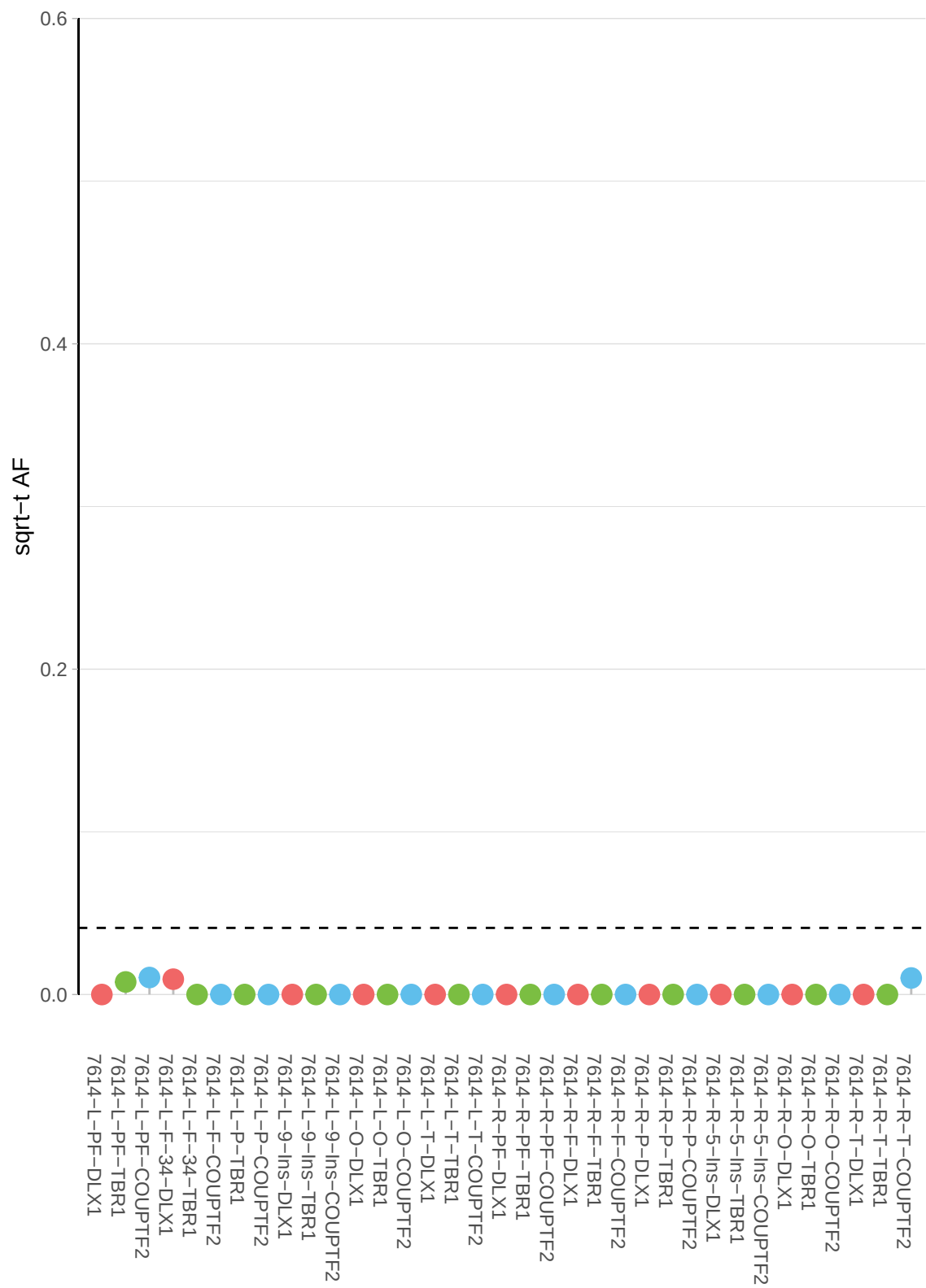

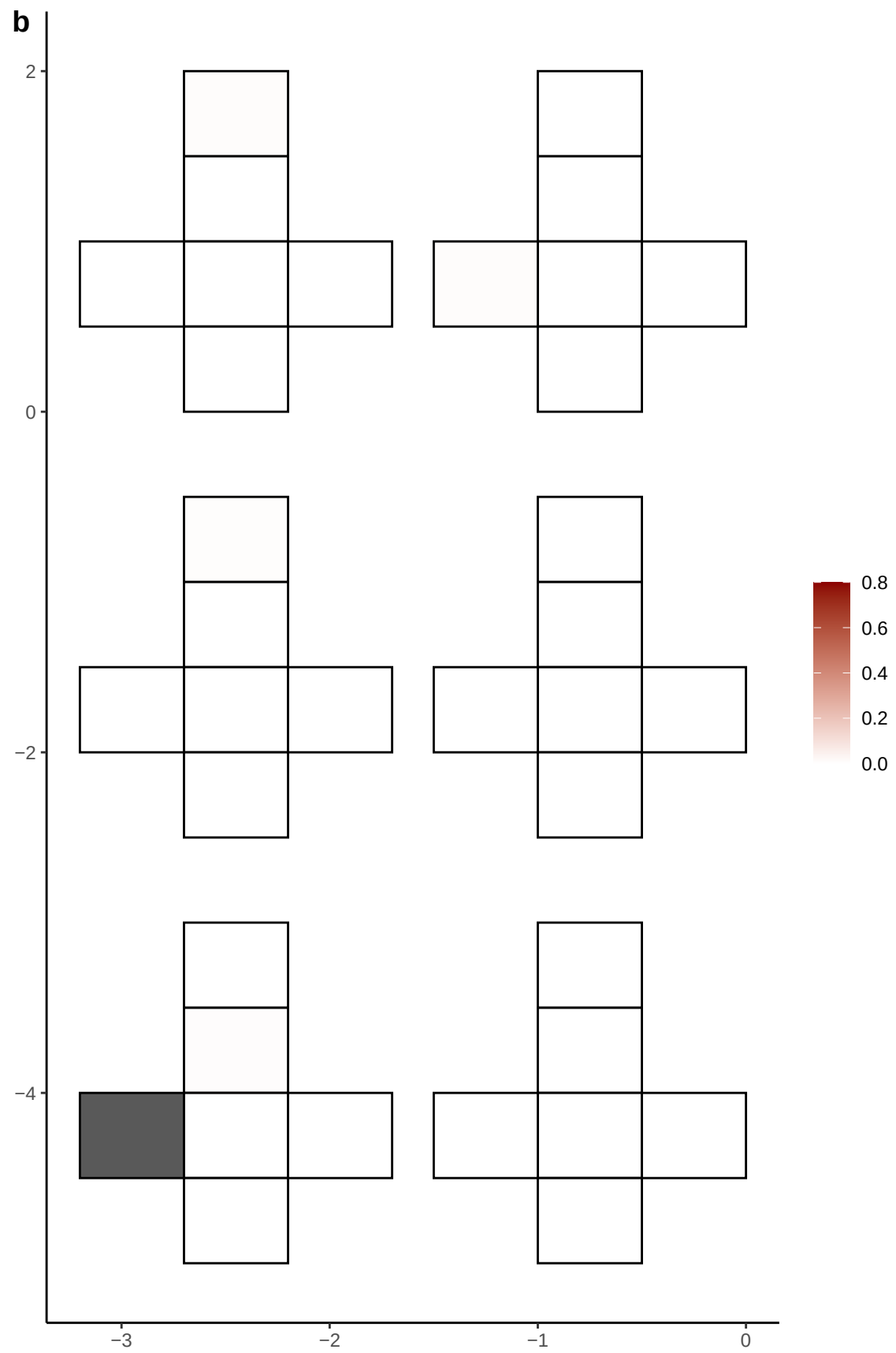

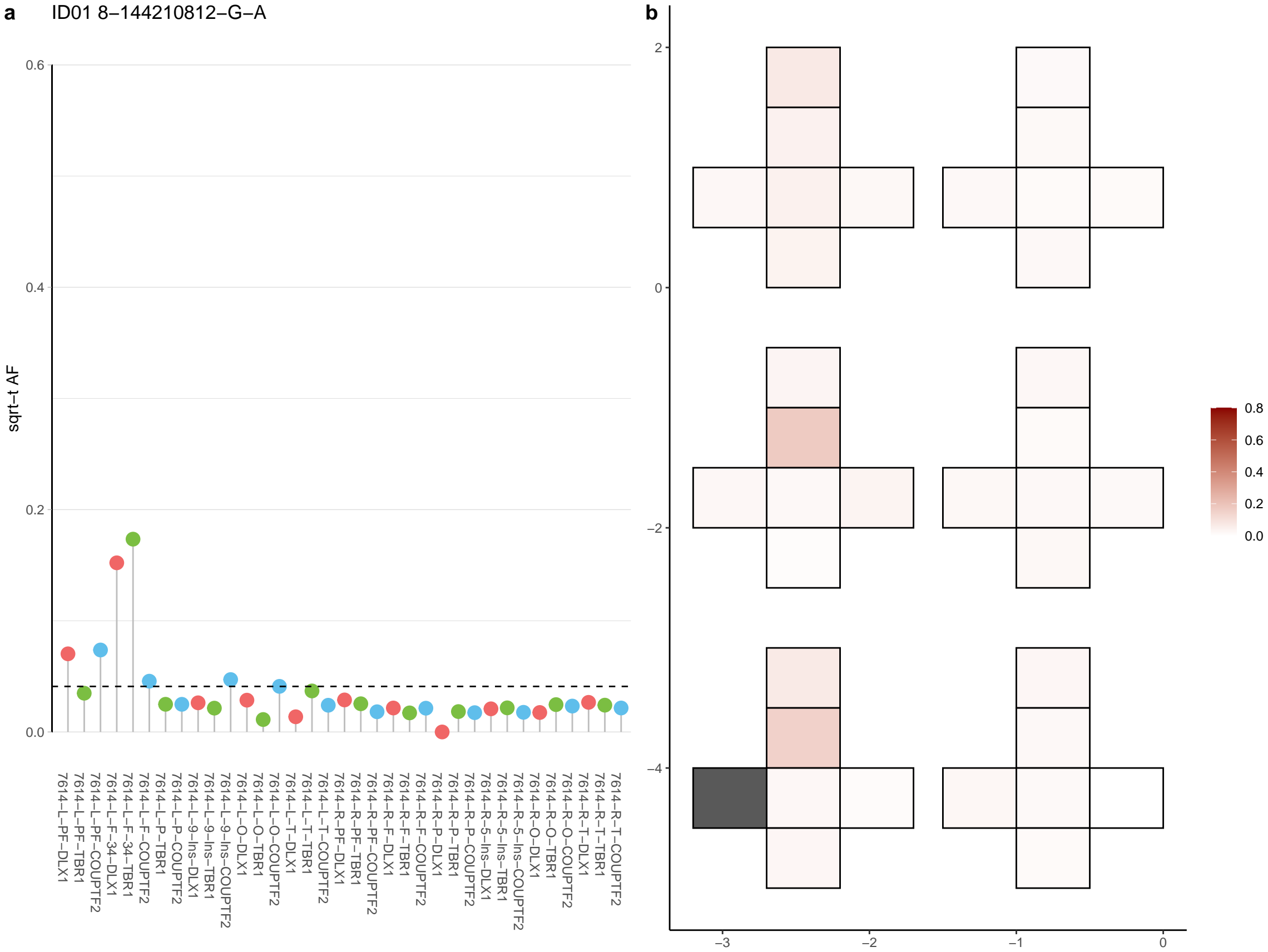

**a** ID01 8-2199684-G-A

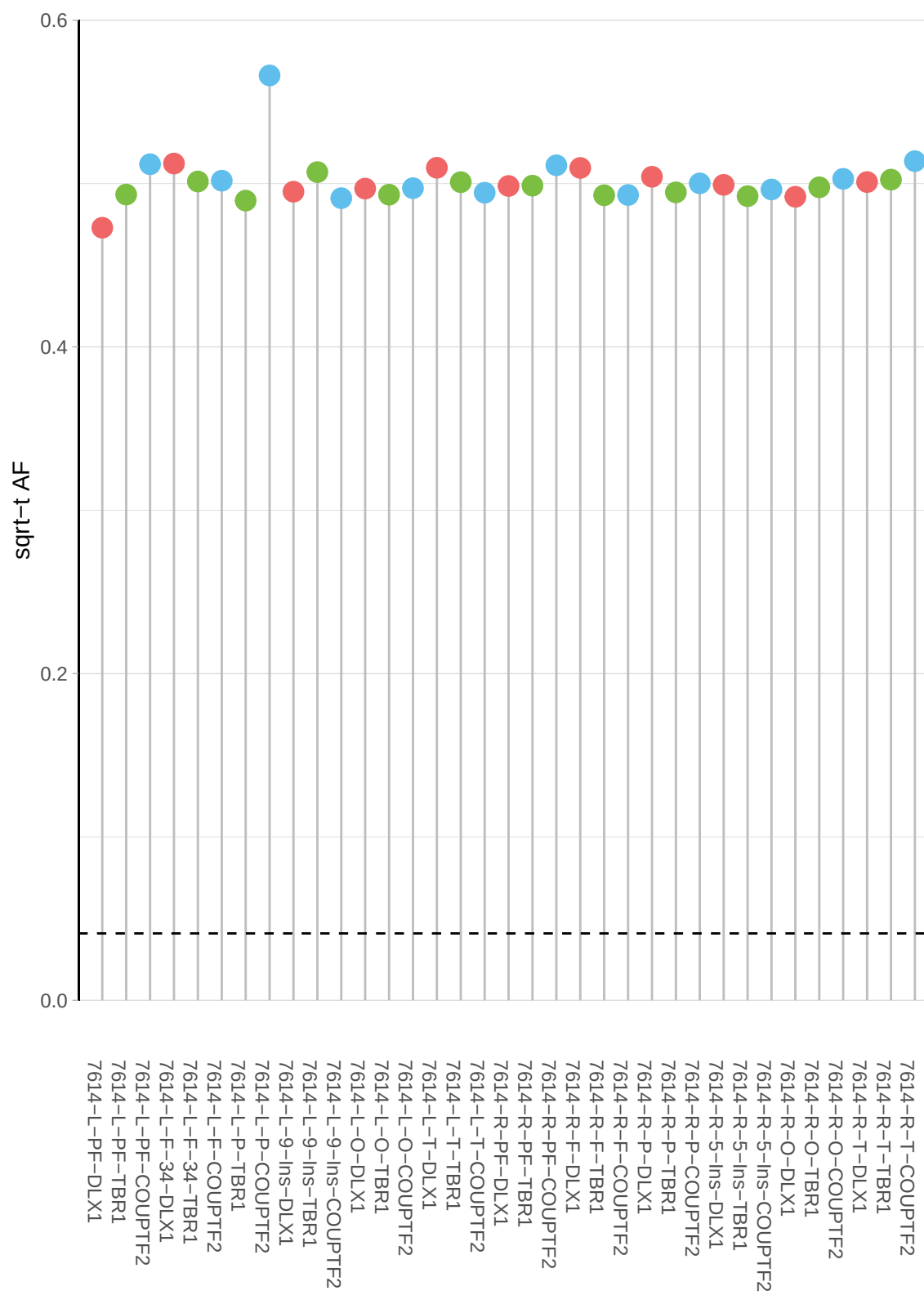

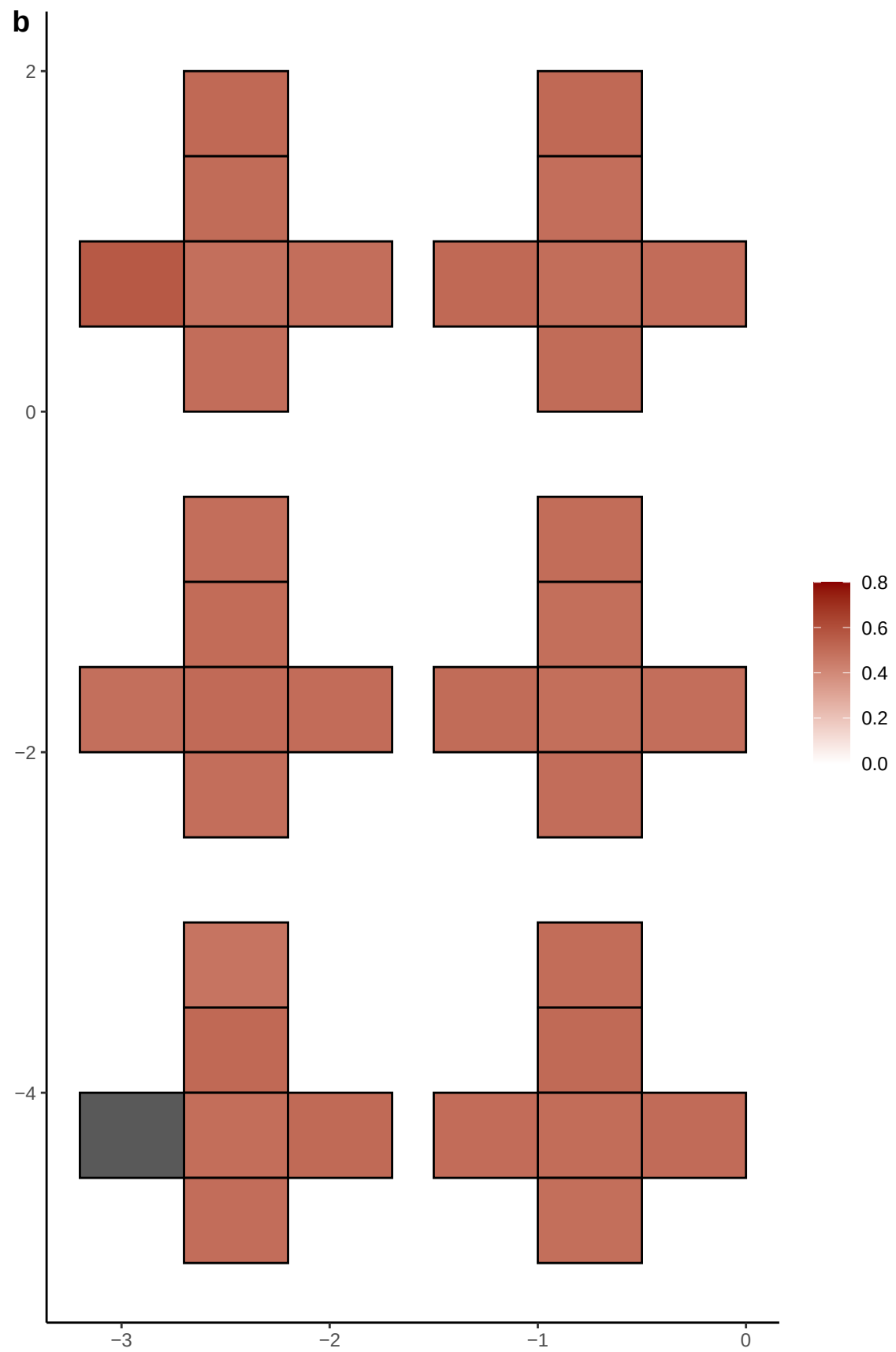

**a** ID01 8-221118-T-C

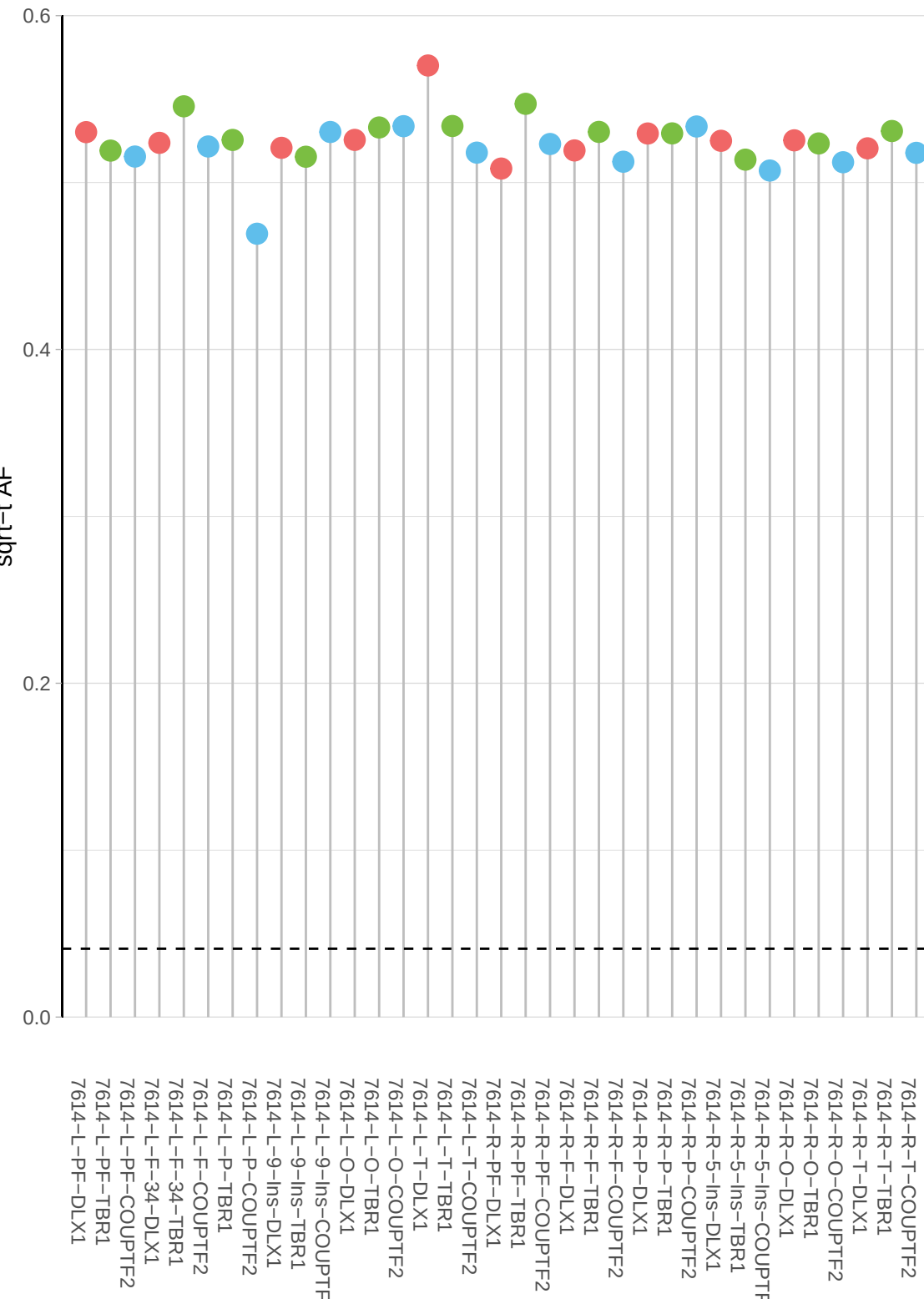

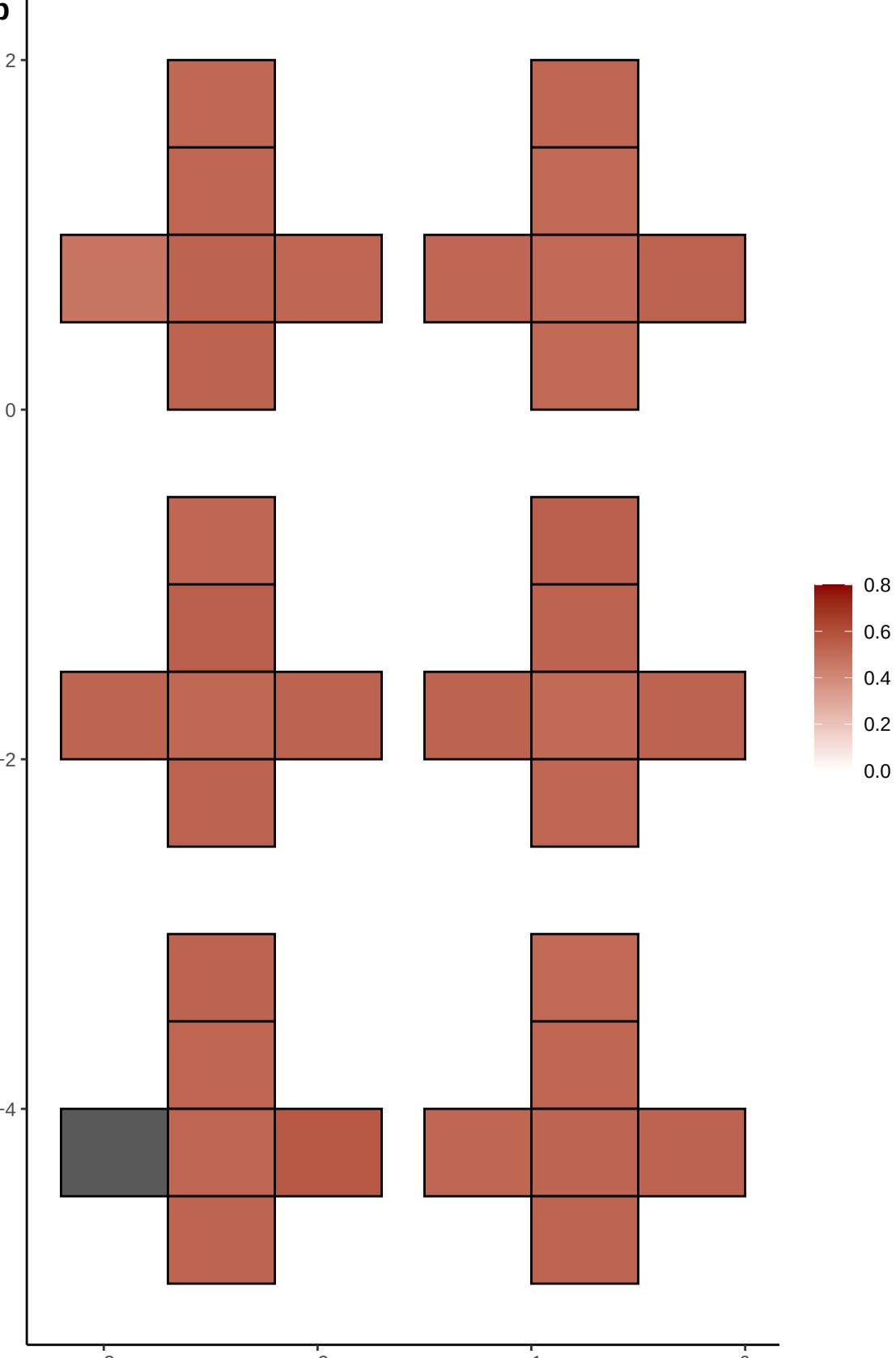

**a** ID01 8-2270450-T-C

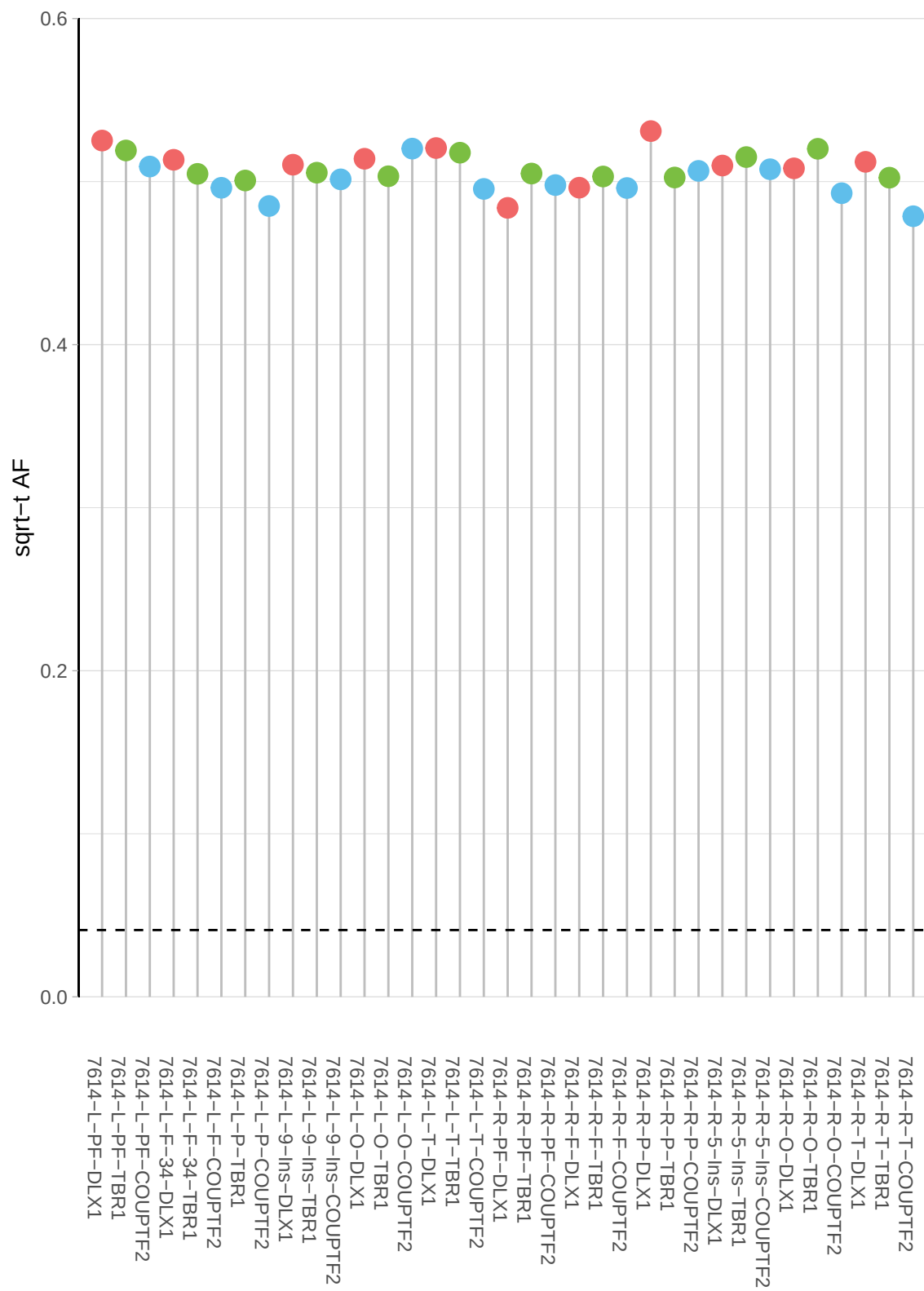

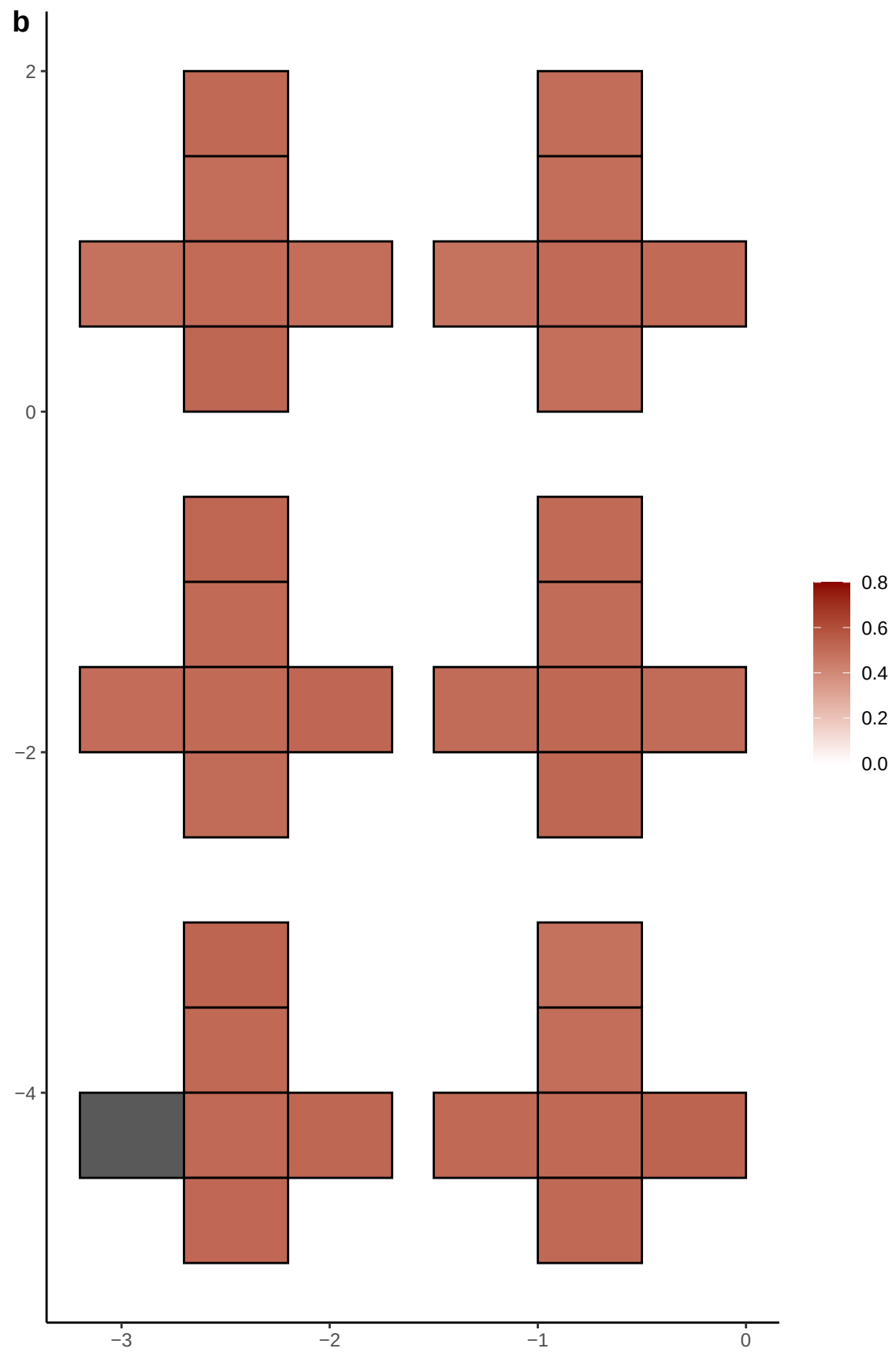

**a** ID01 8-25735478-CATA-C

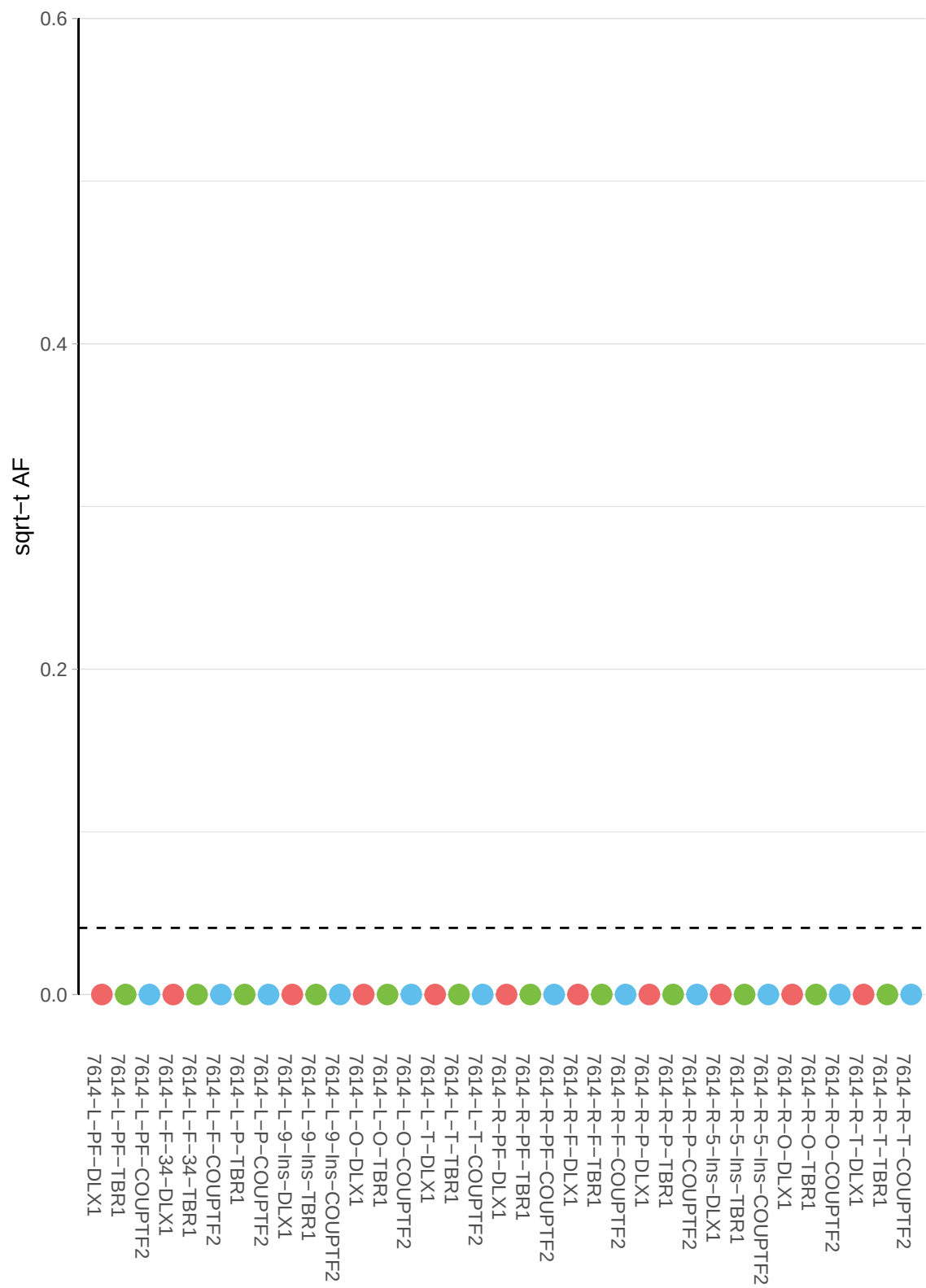

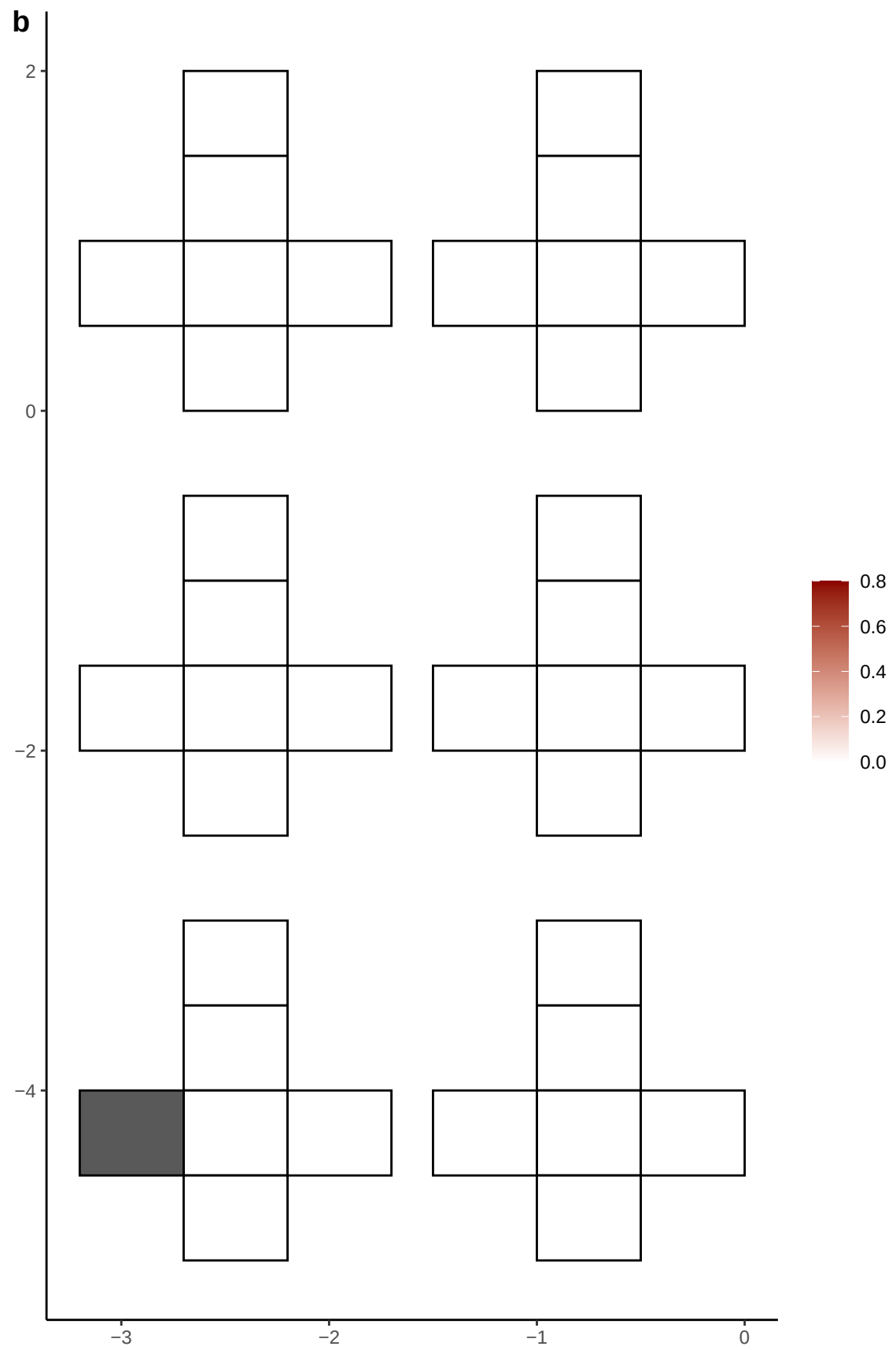

**a** ID01 8-27531157-G-A

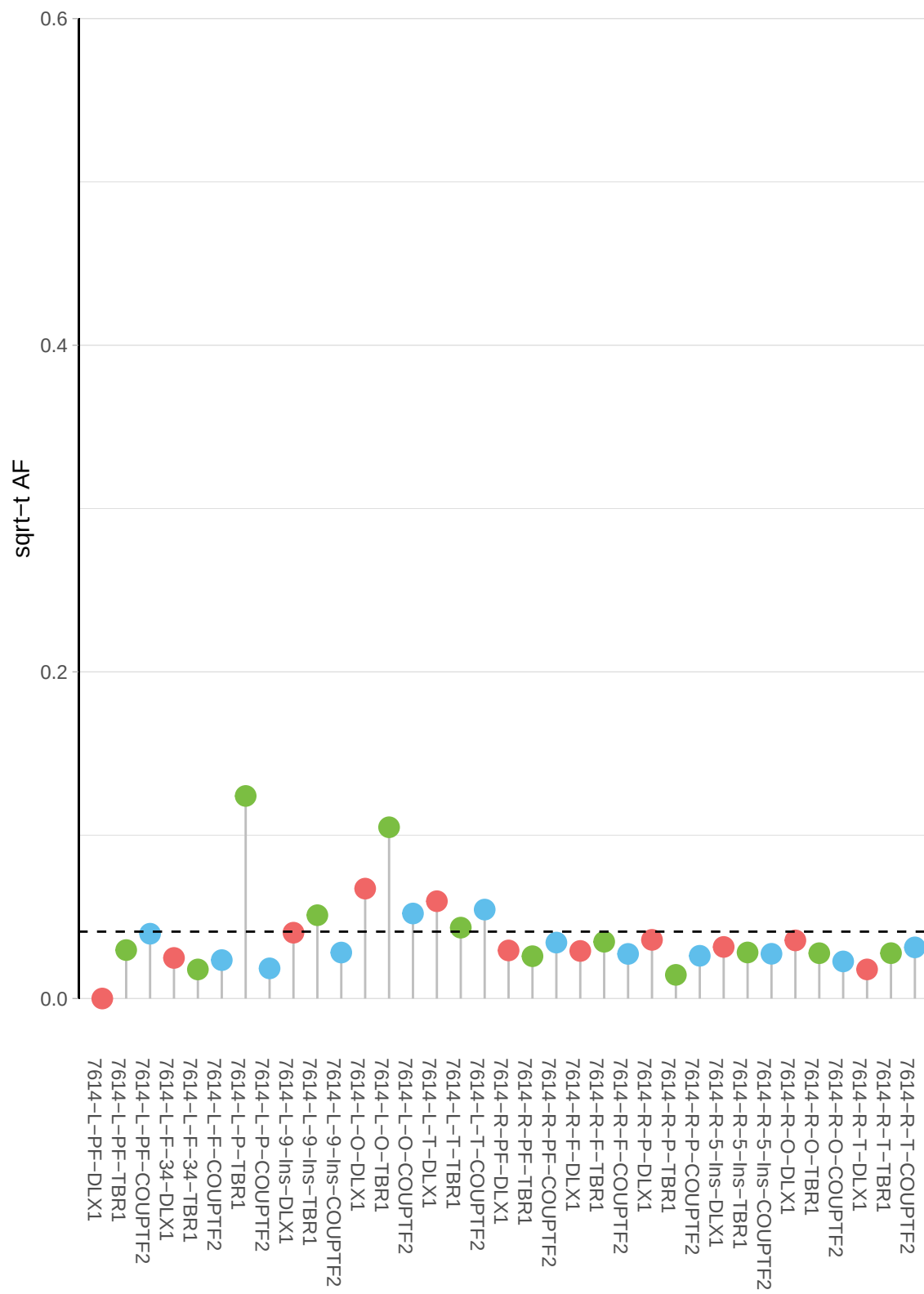

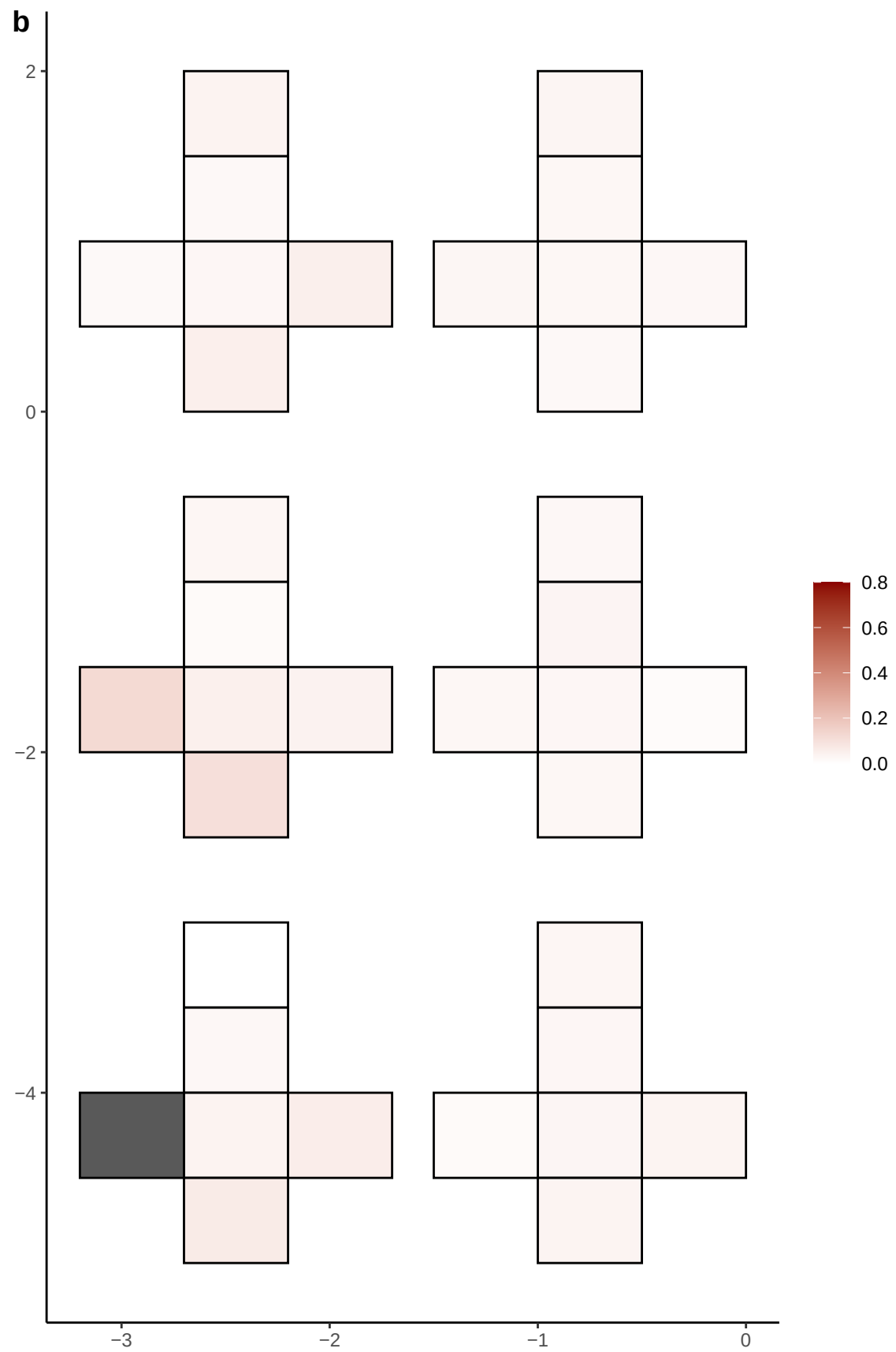

**a** ID01 8-28894088-T-G

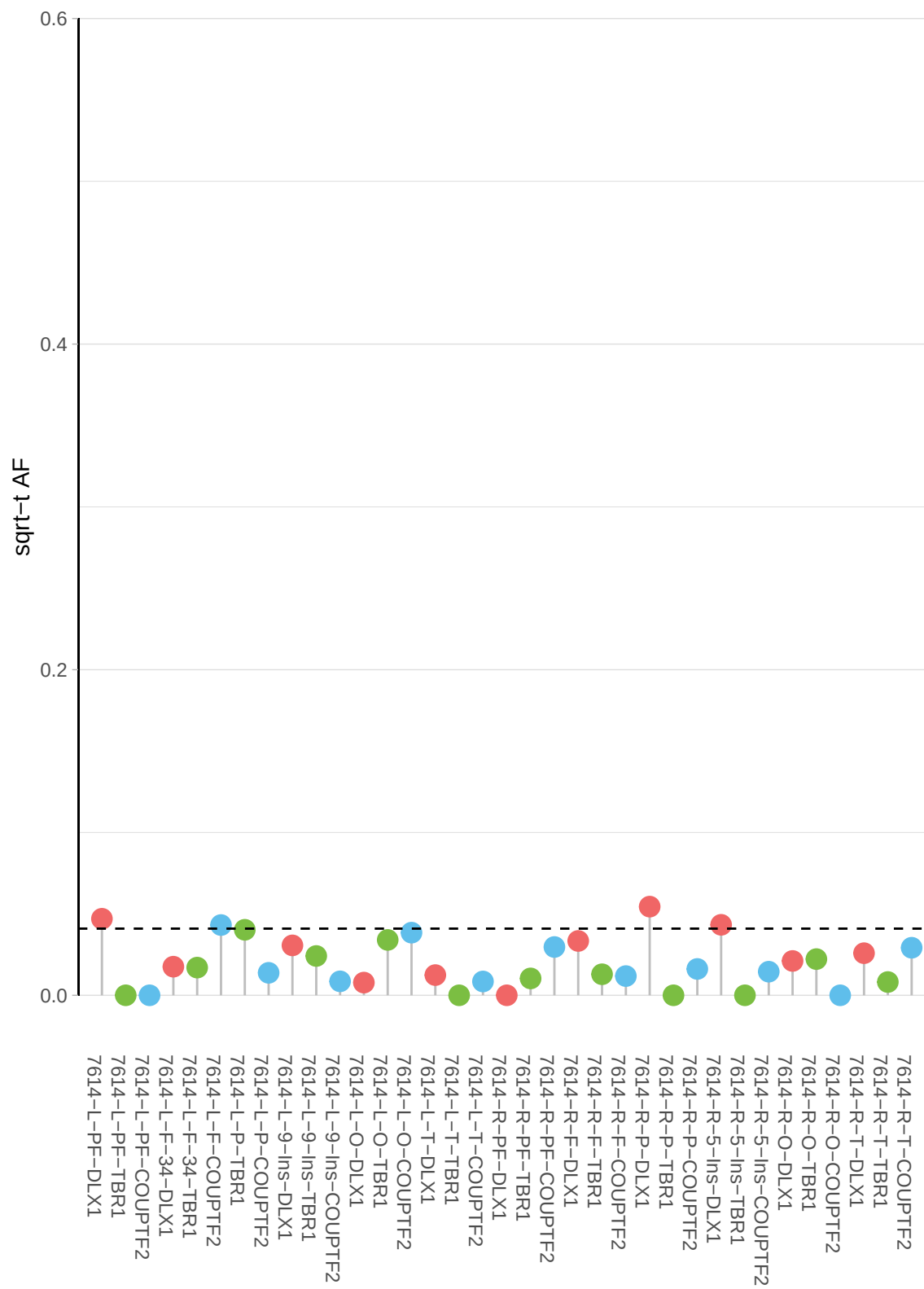

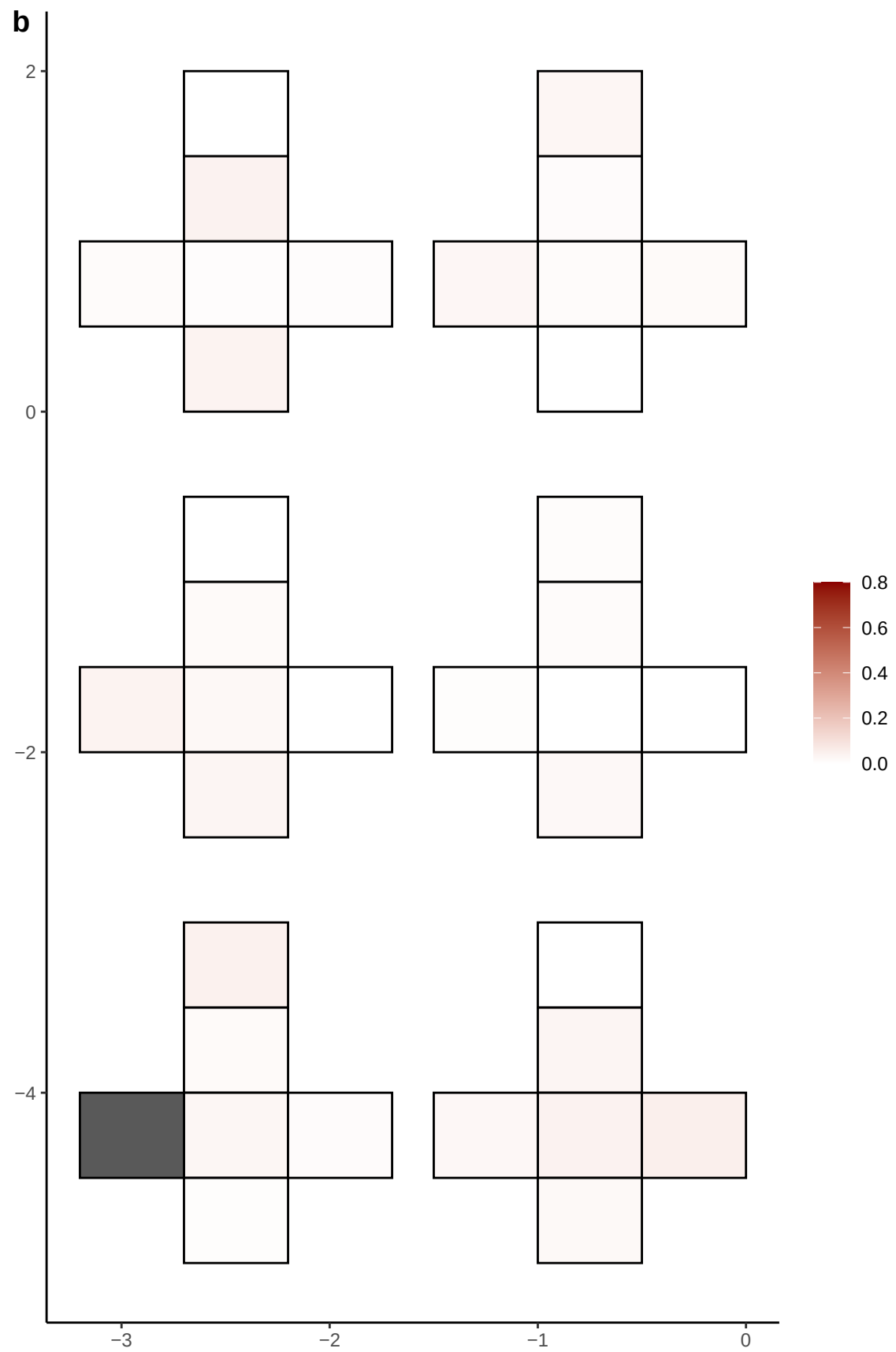

**a** ID01 8-63618020-A-C

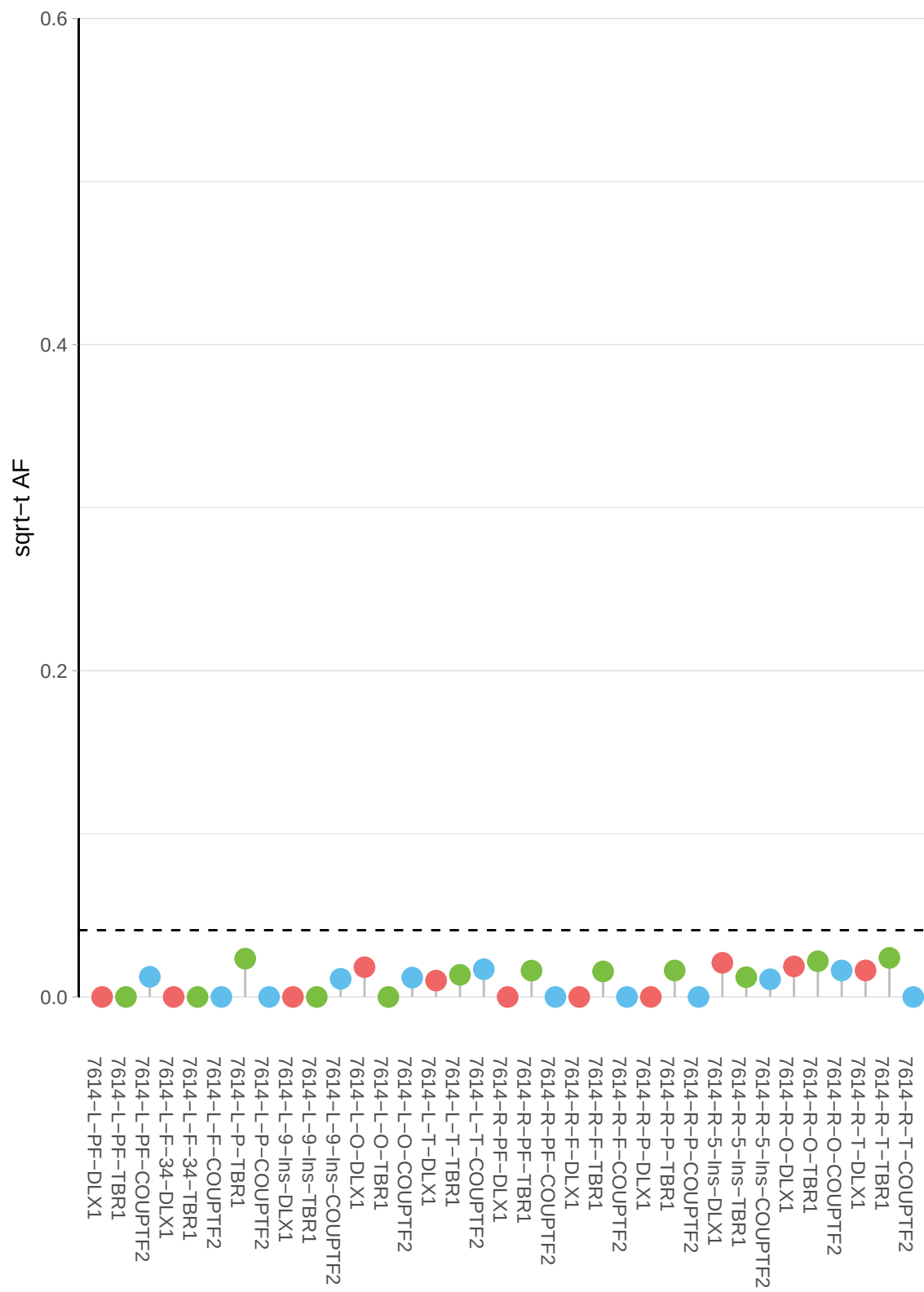

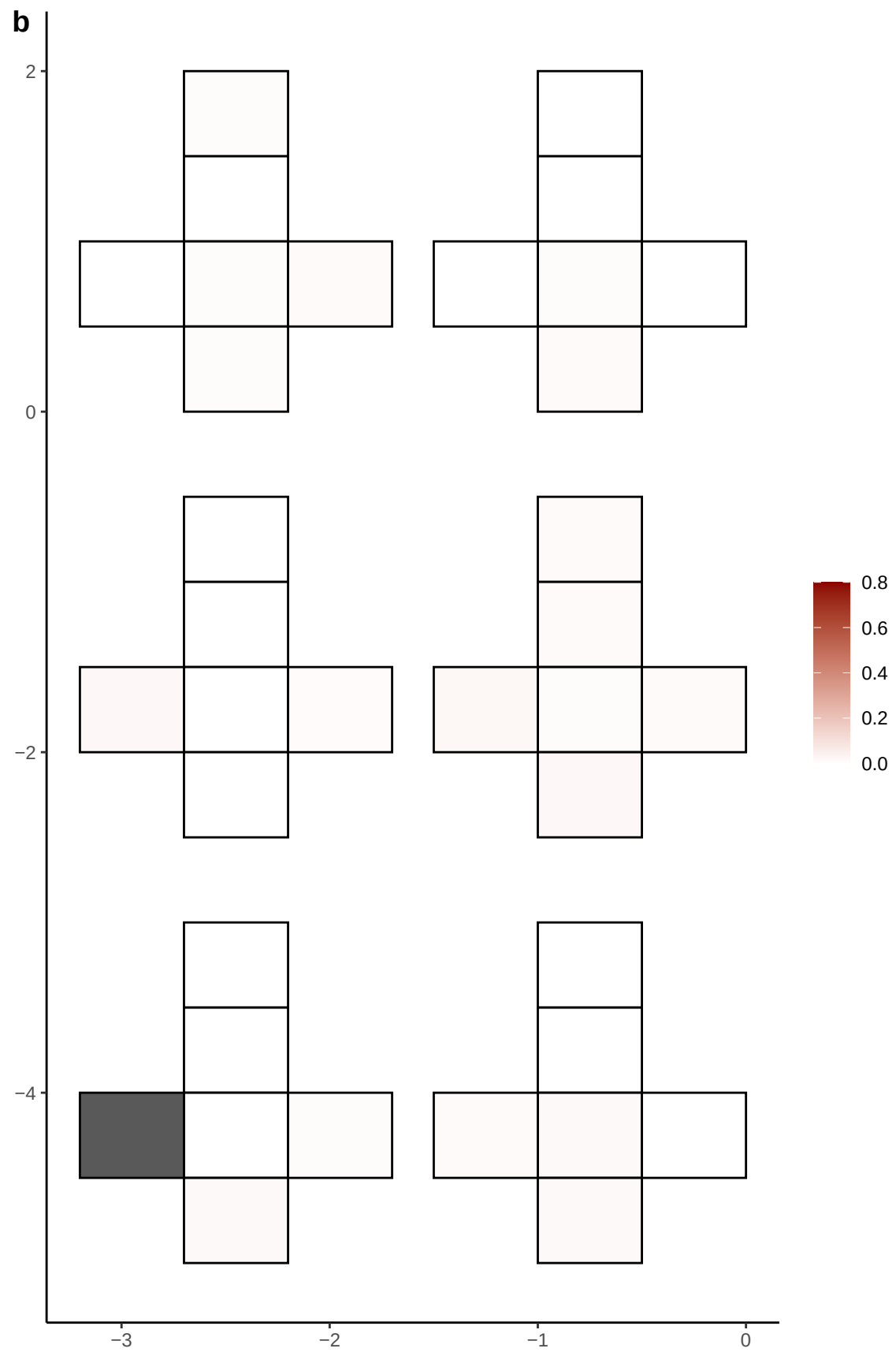

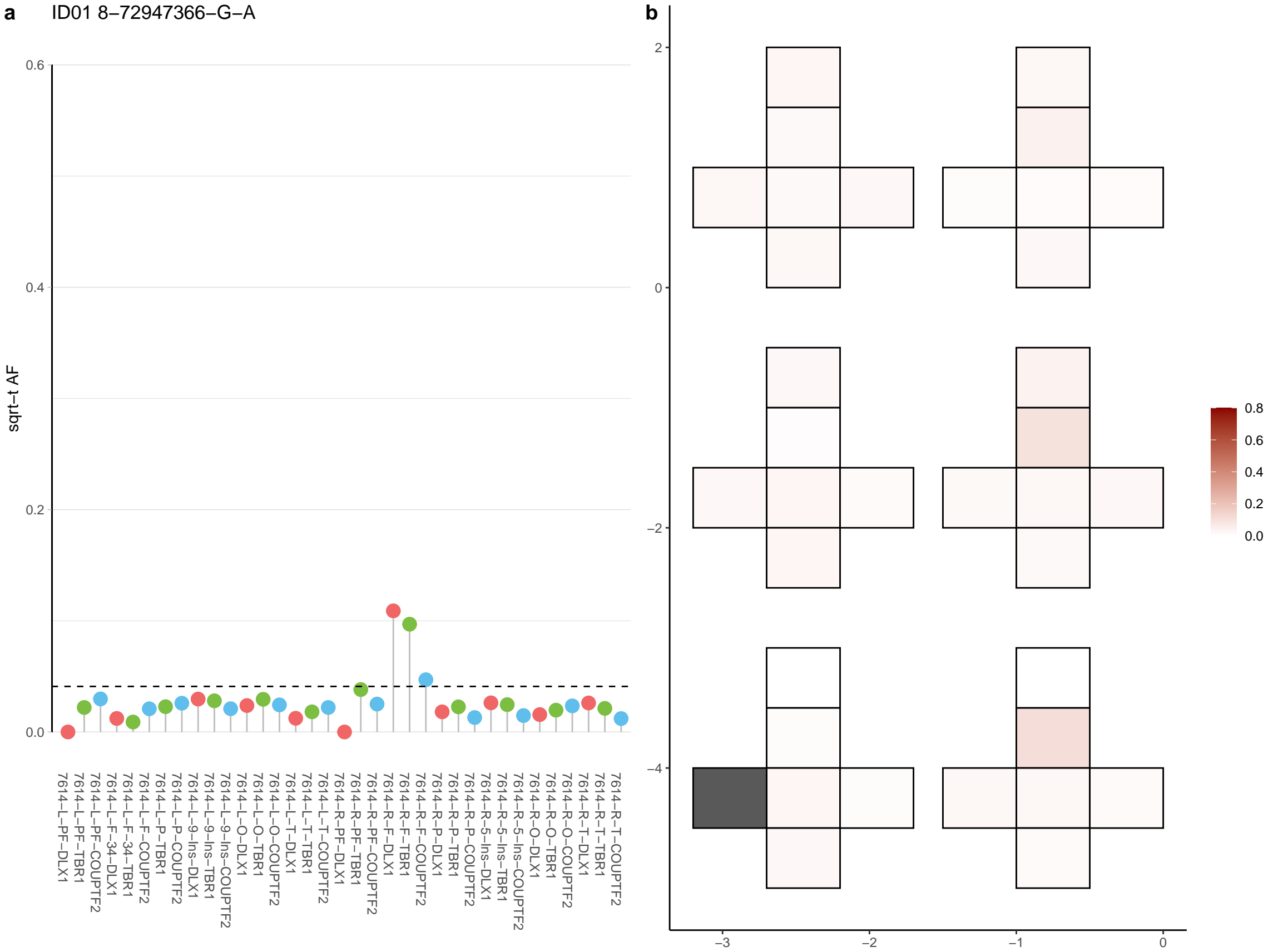

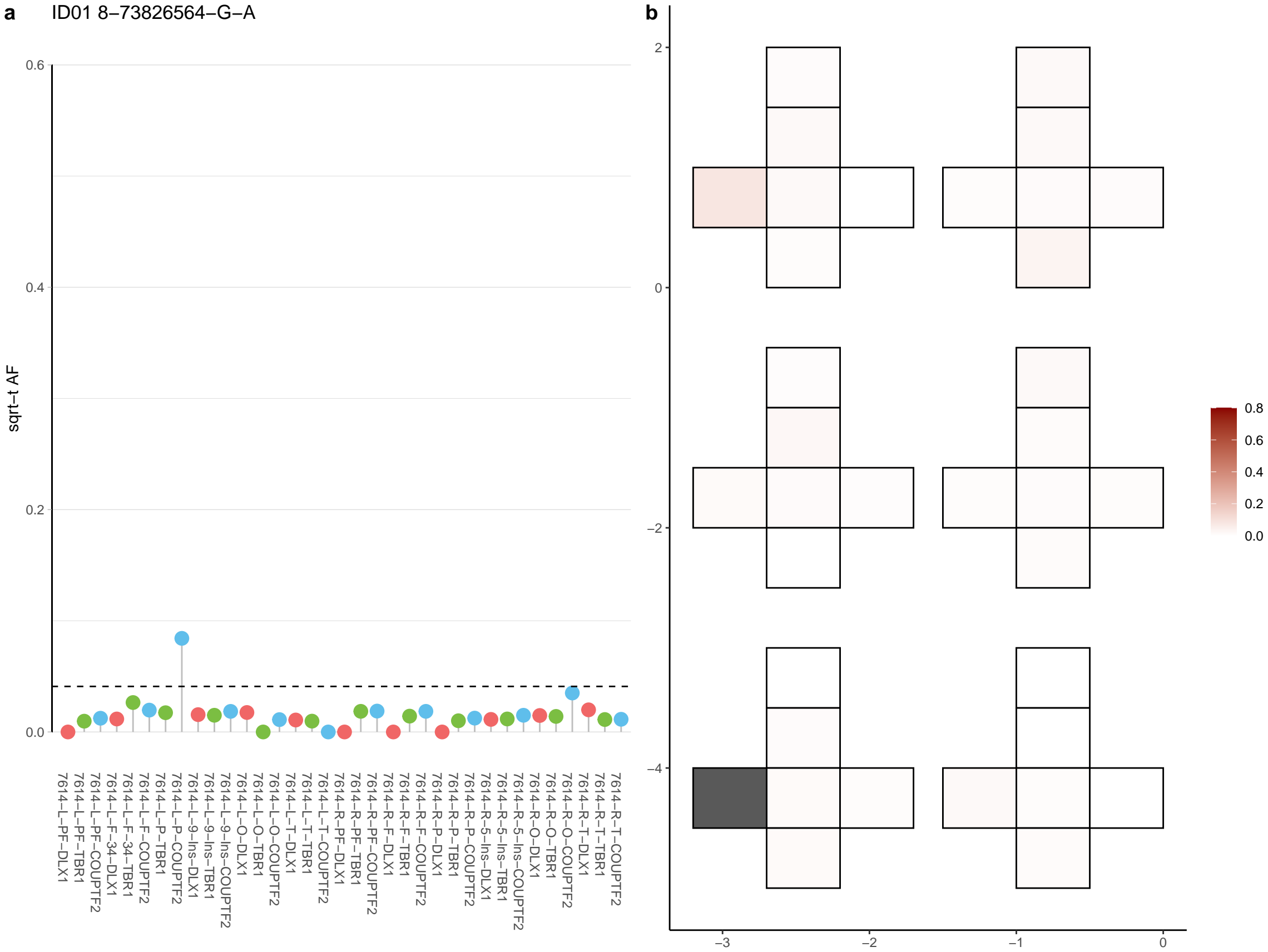

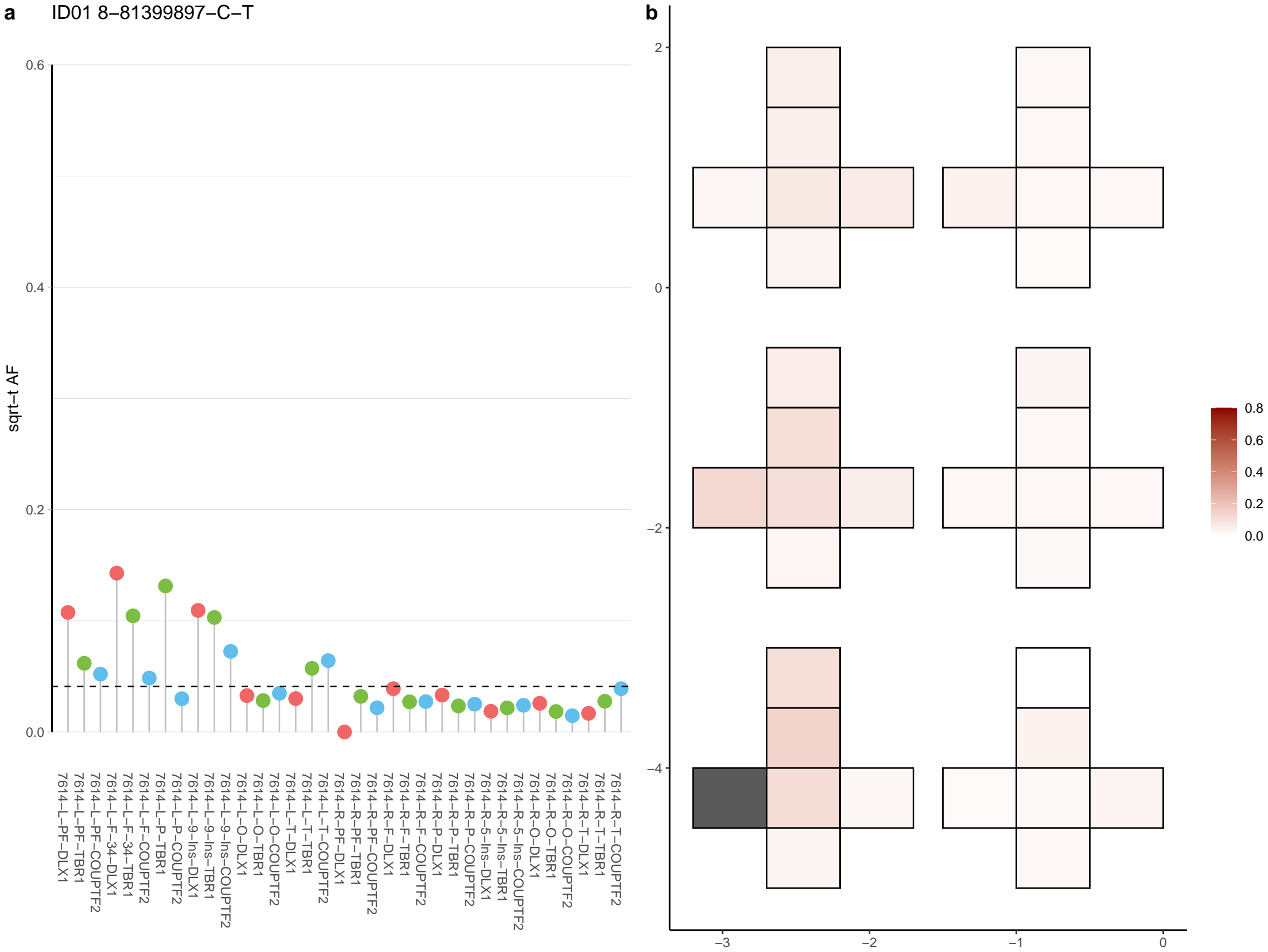

**a** ID01 9-133760767-C-T

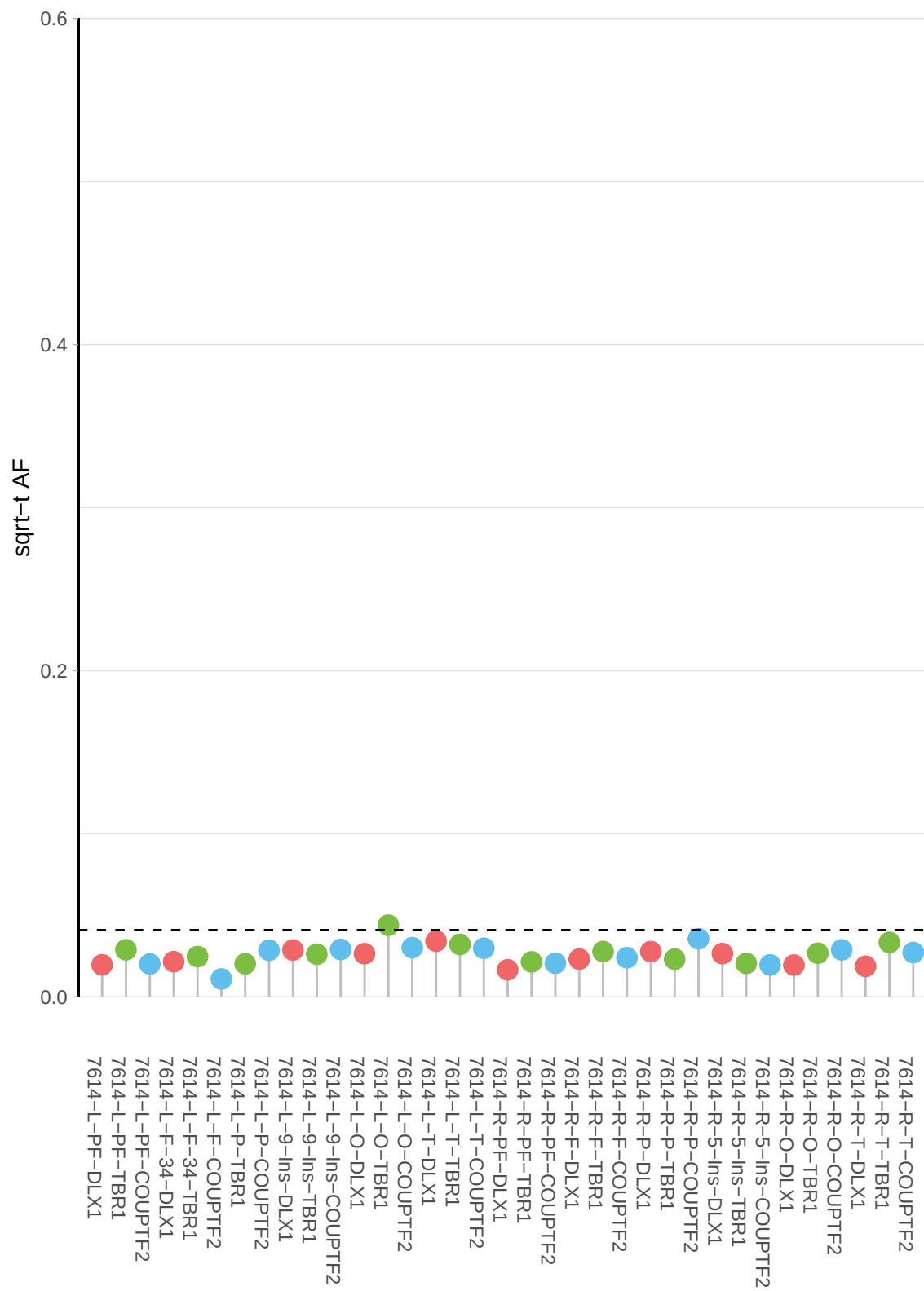

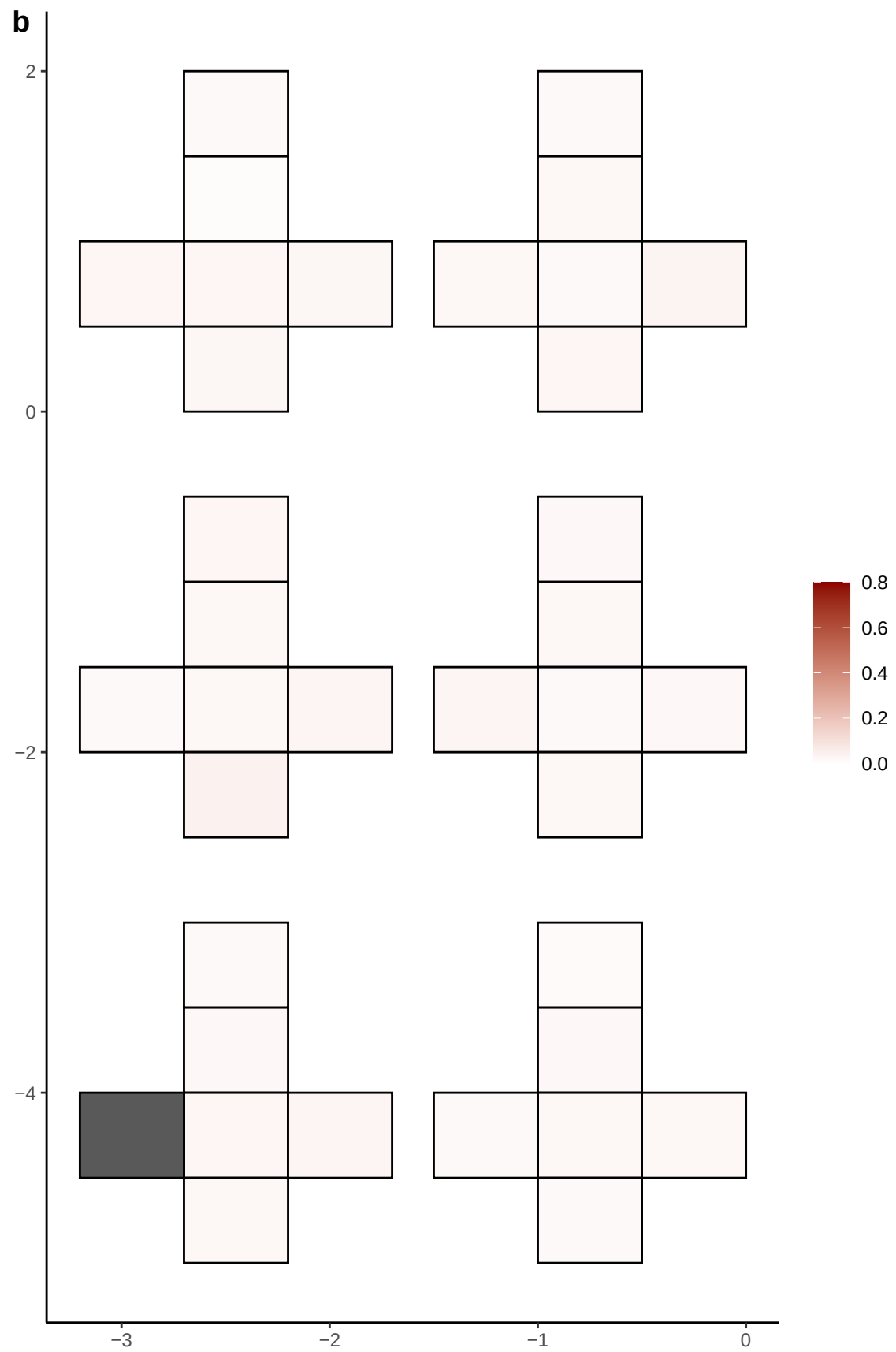

**a** ID01 9-134665838-G-A

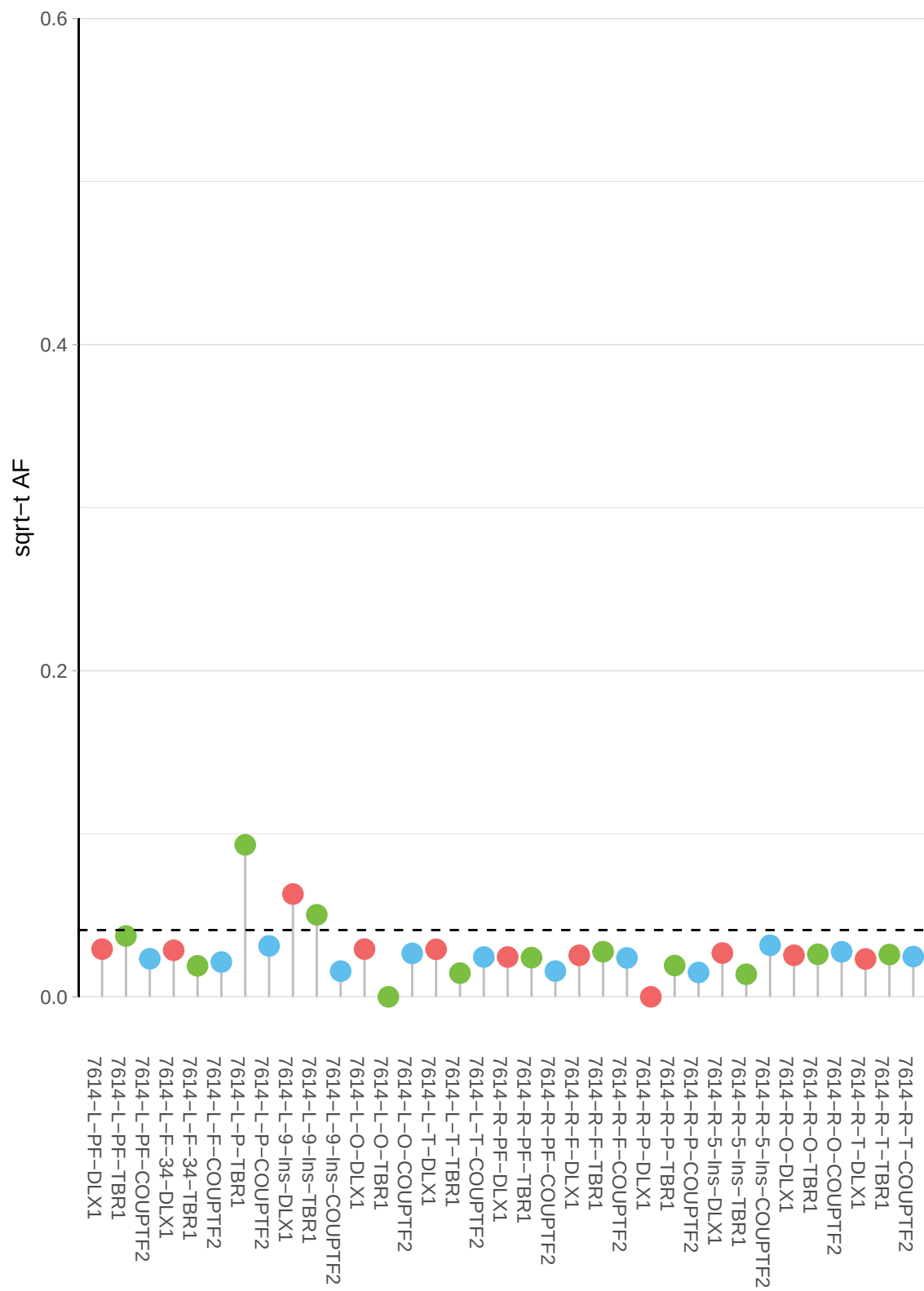

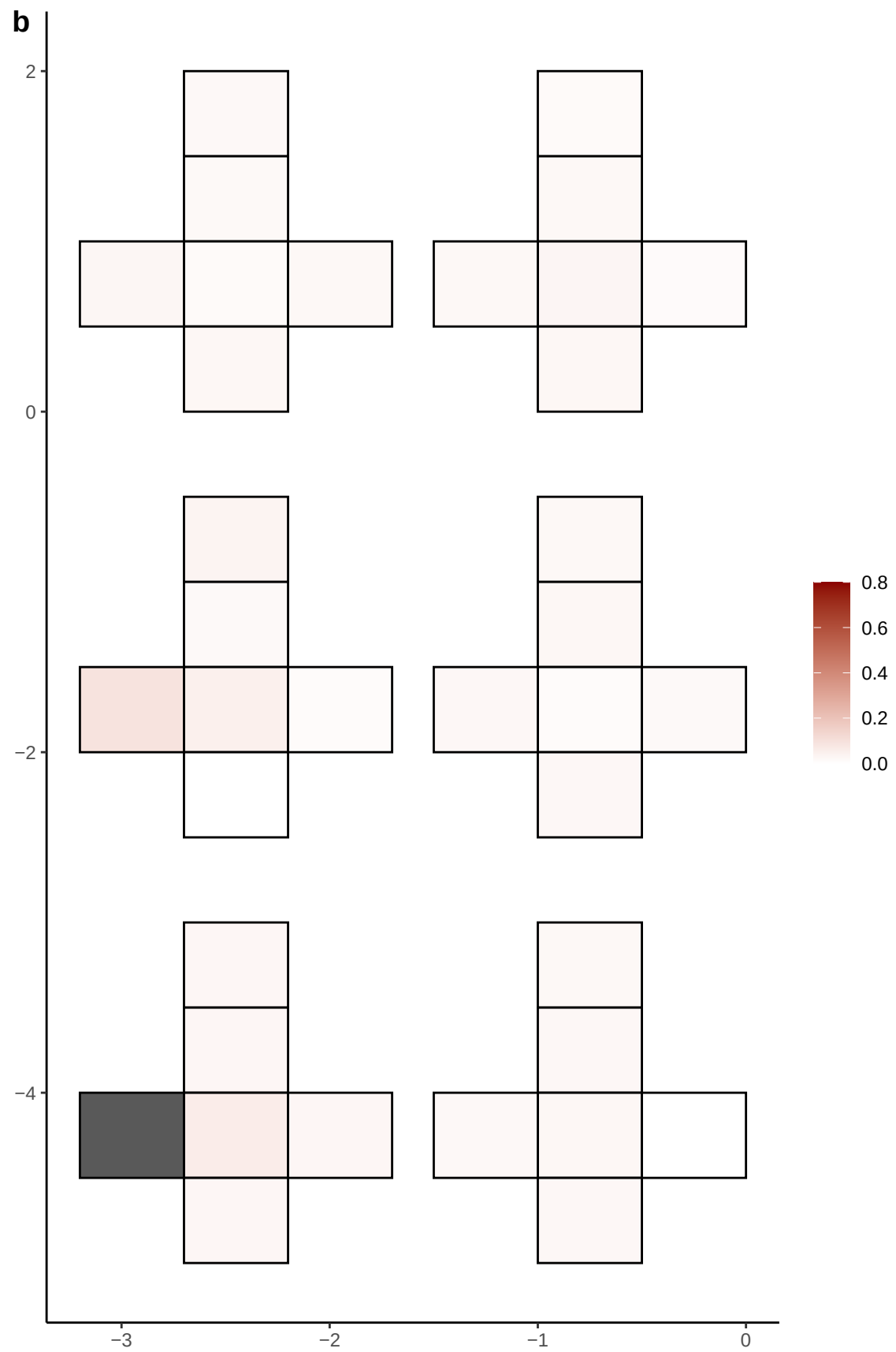

**a** ID01 9-134950488-G-T

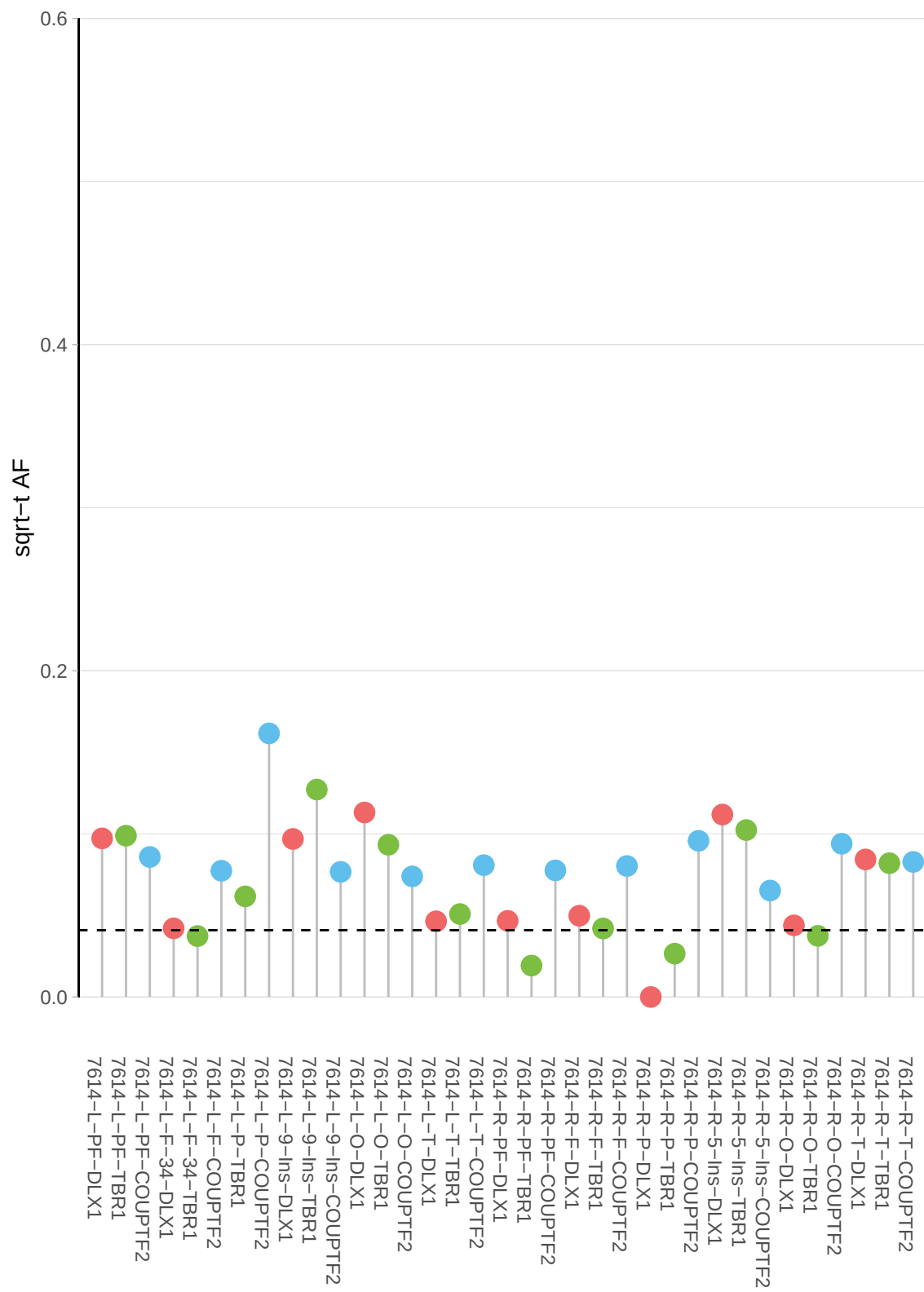

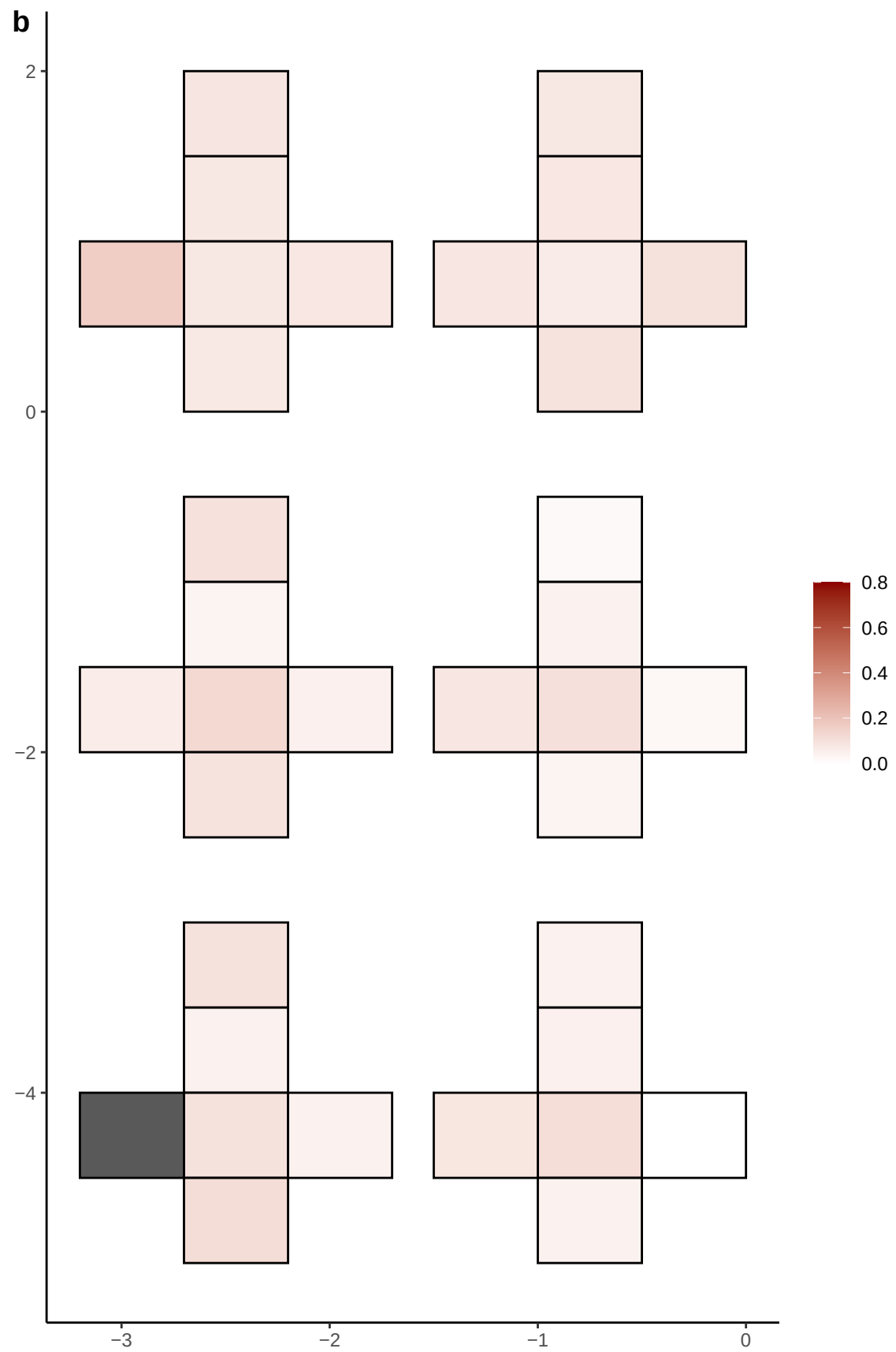

**a** ID01 9-1722340-A-G

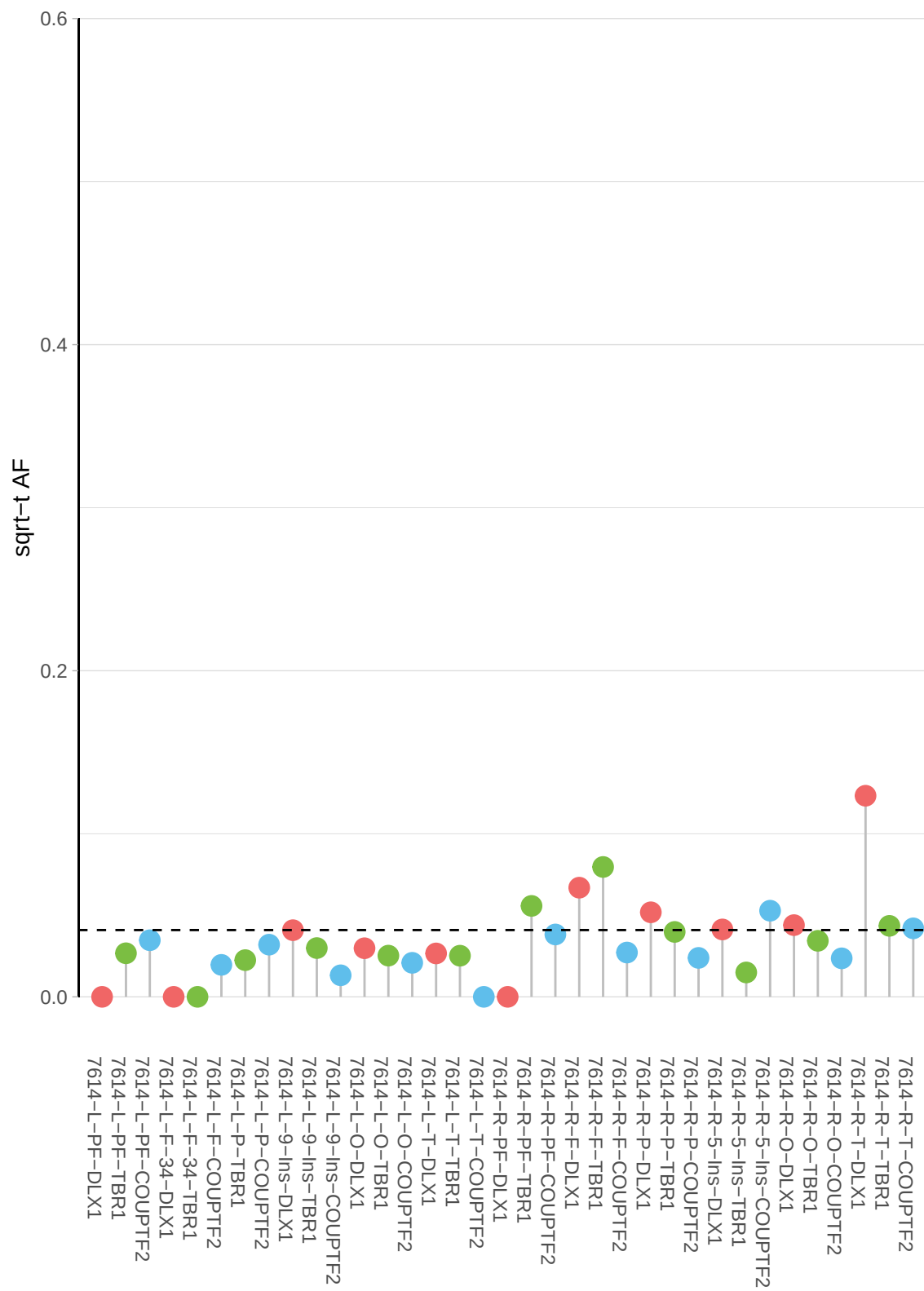

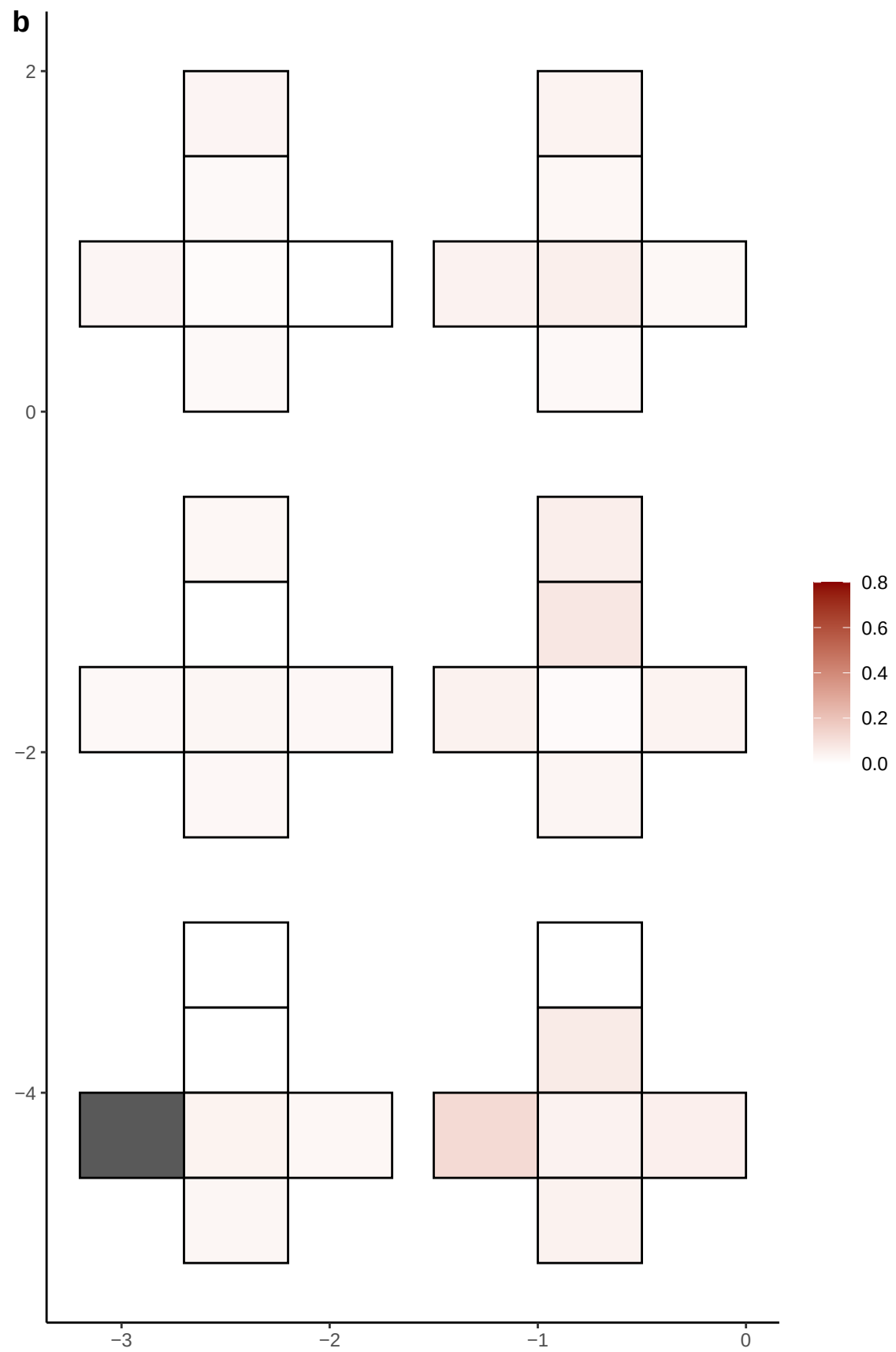

**a** ID01 9-38668272-C-T

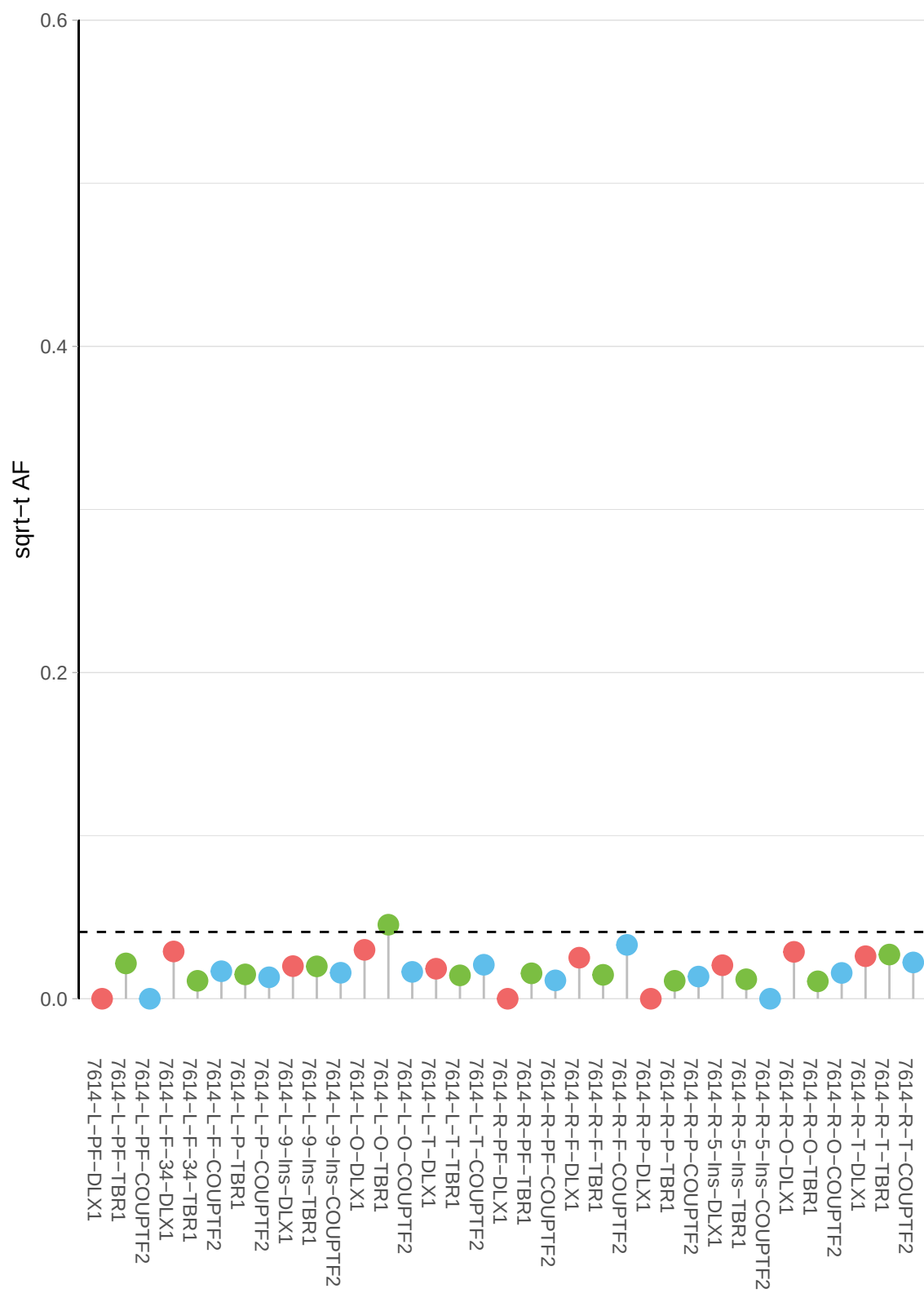

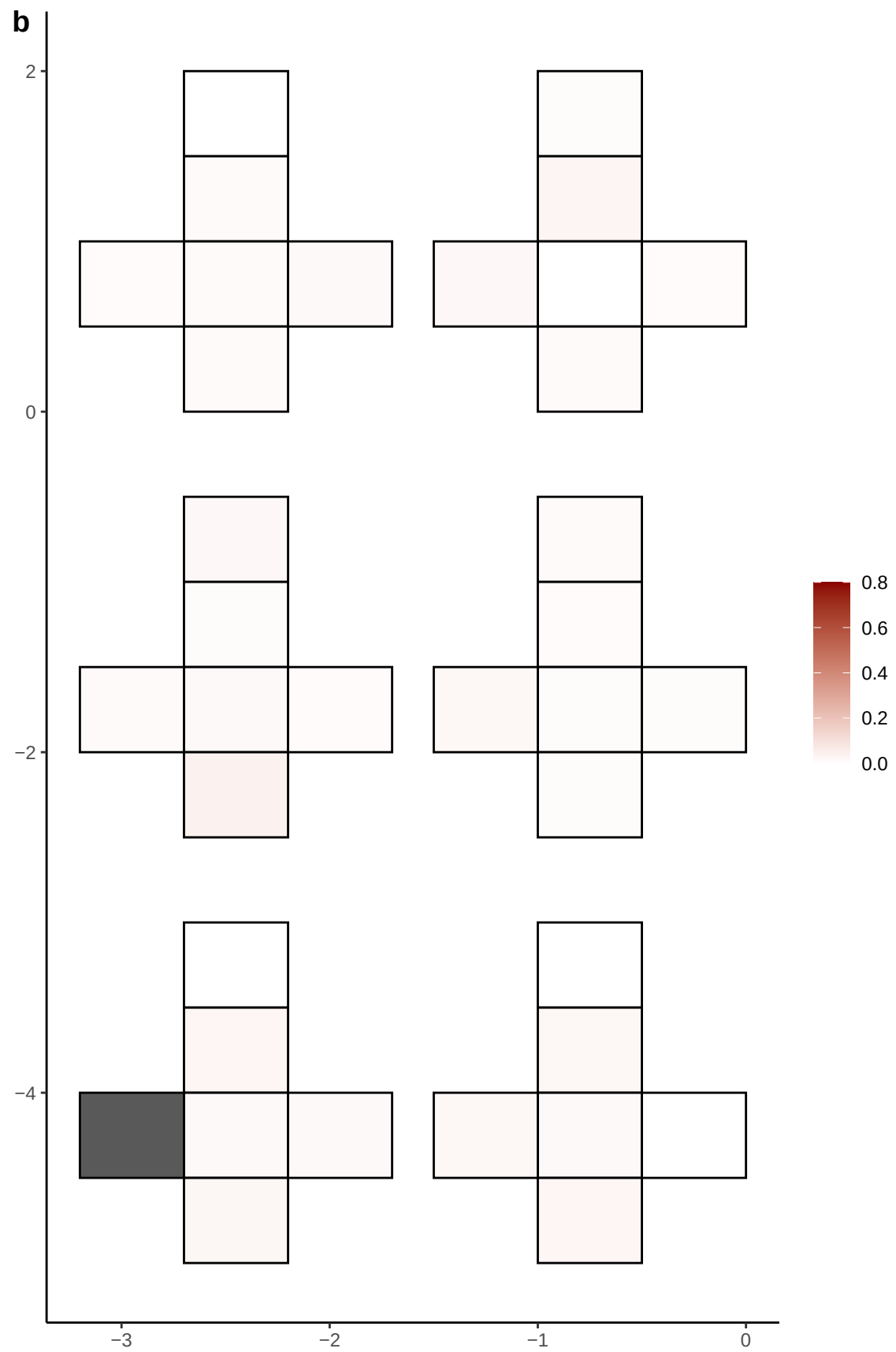

**a** ID01 9-76576921-A-G

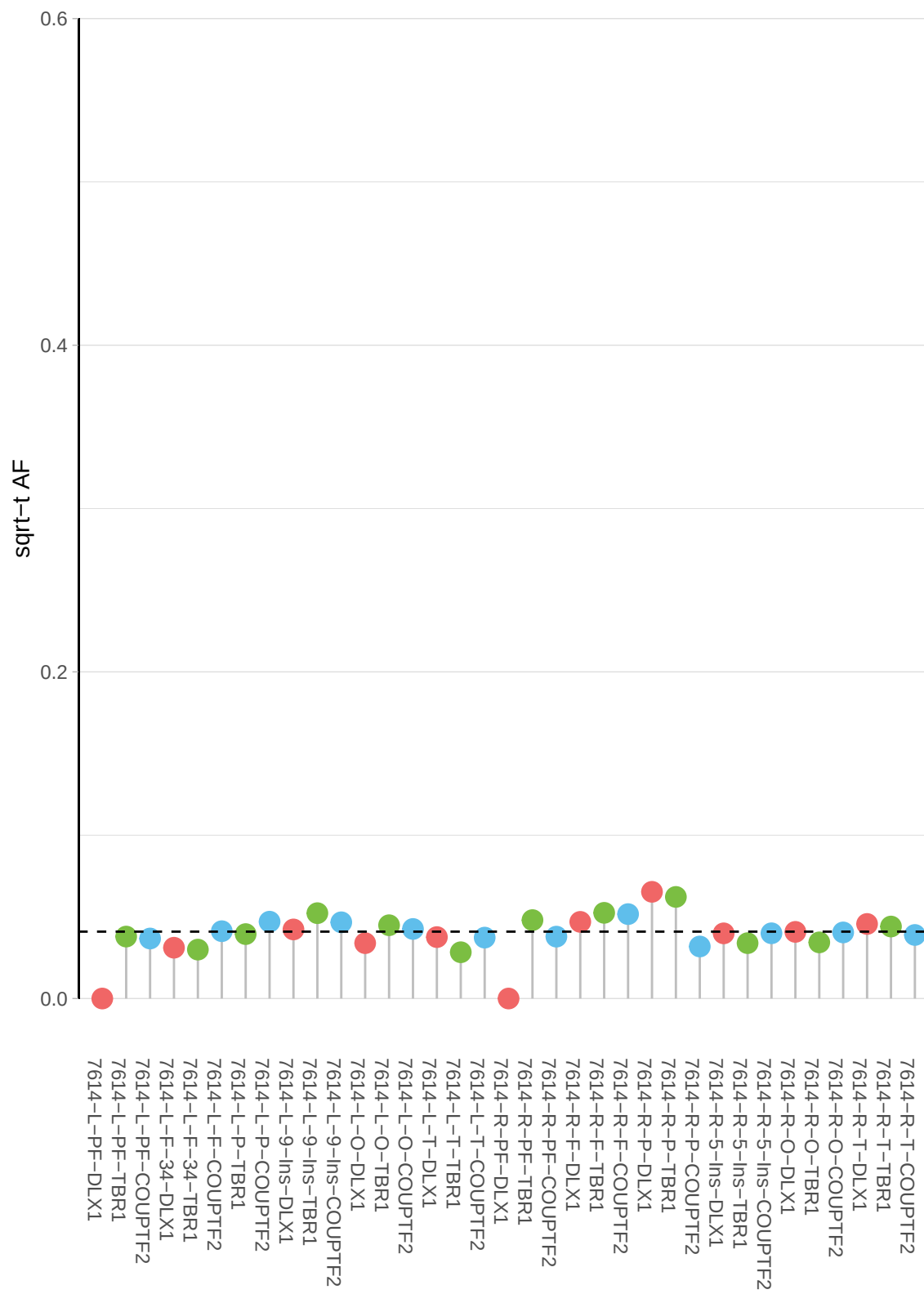

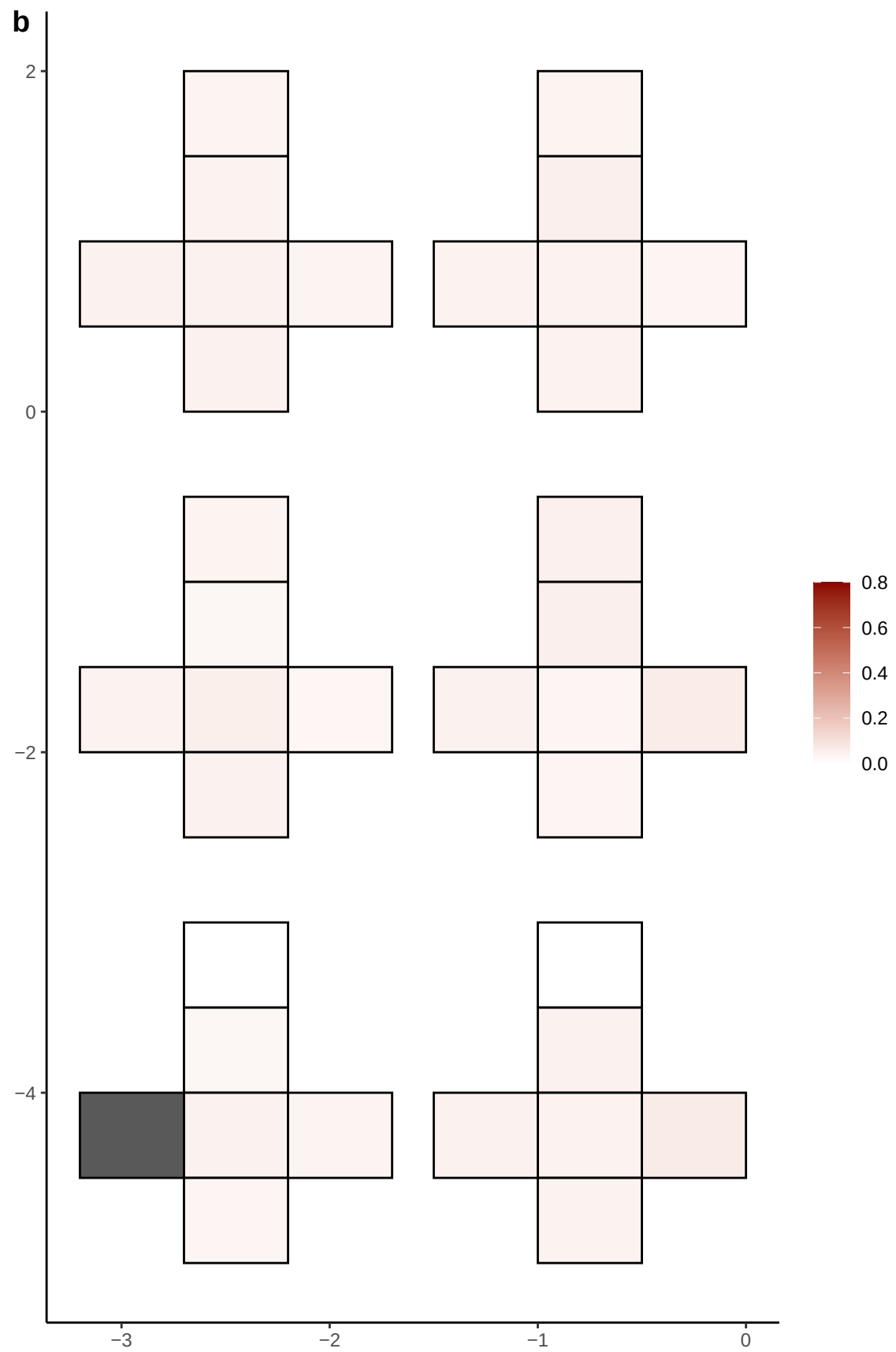

**a** ID01 9-78228121-C-G

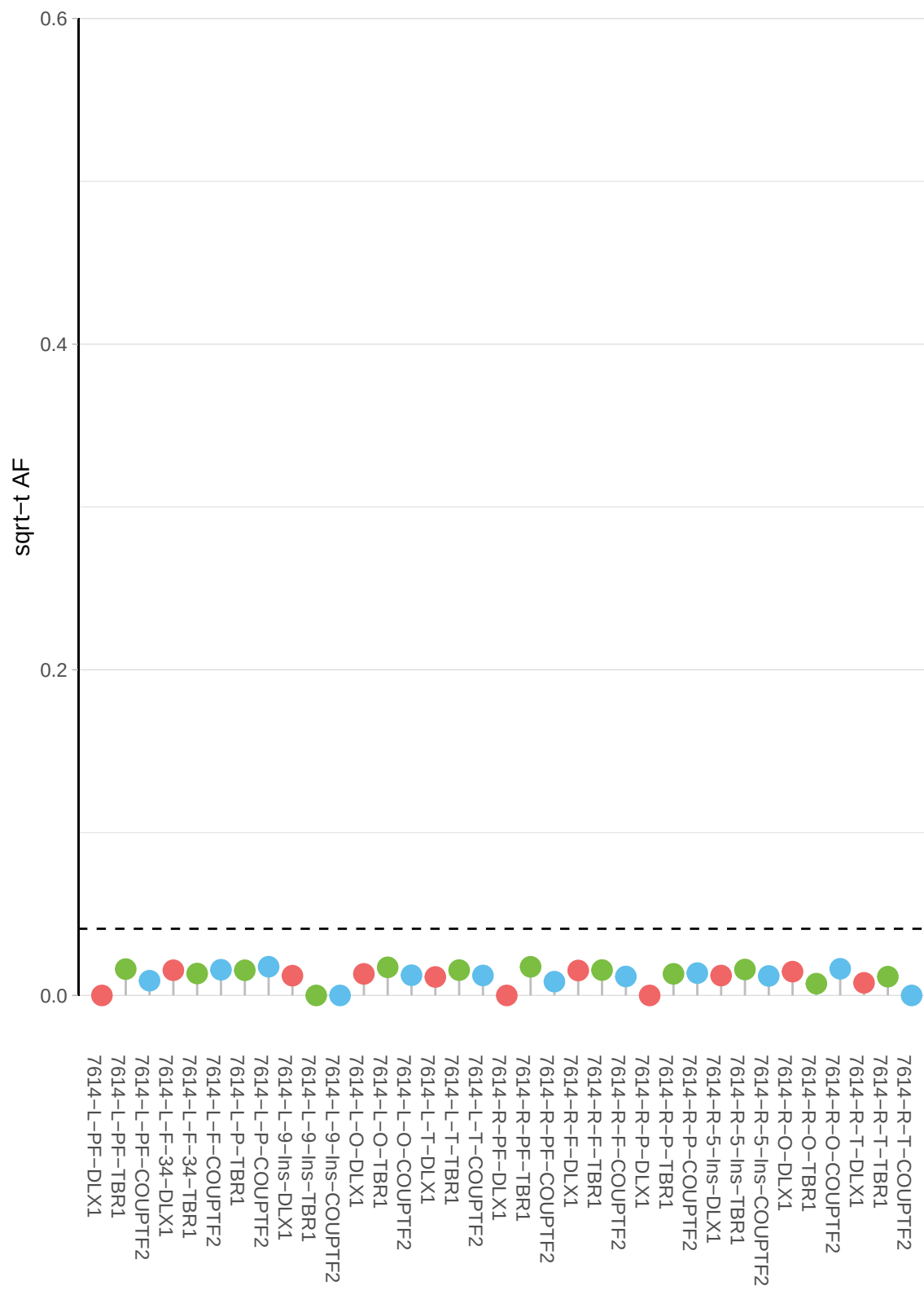

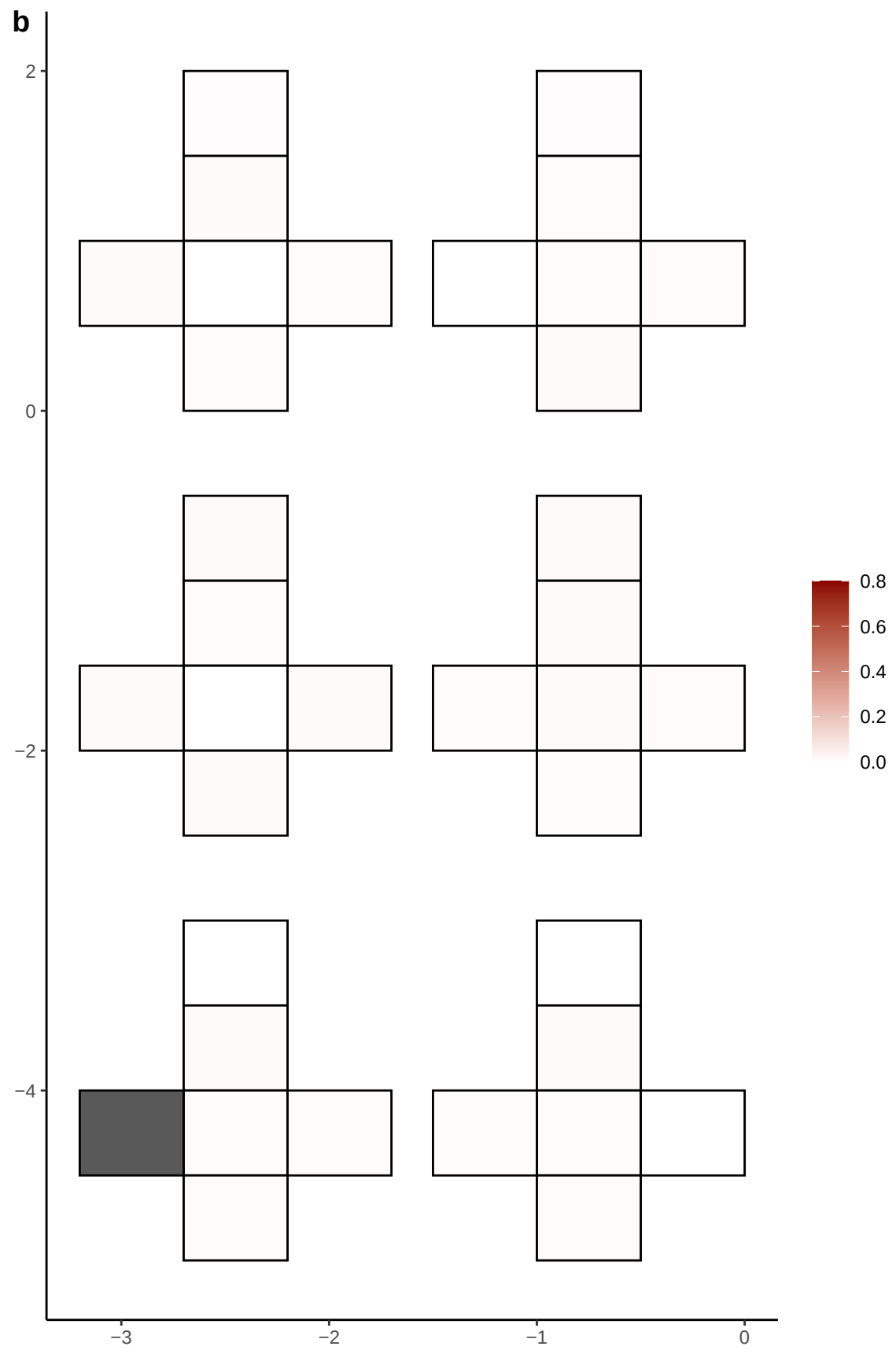

**a** ID01 9-7860755-C-T

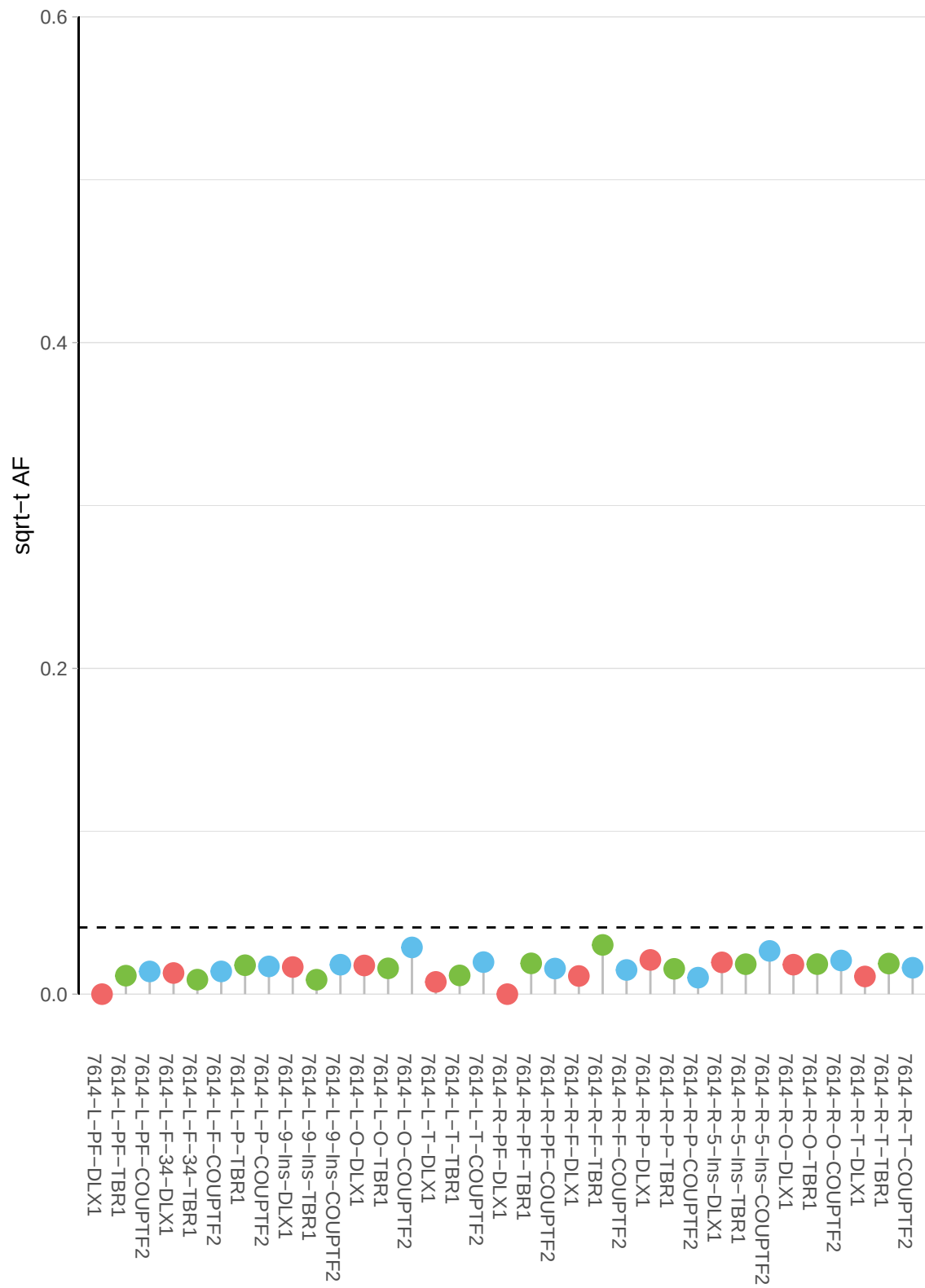

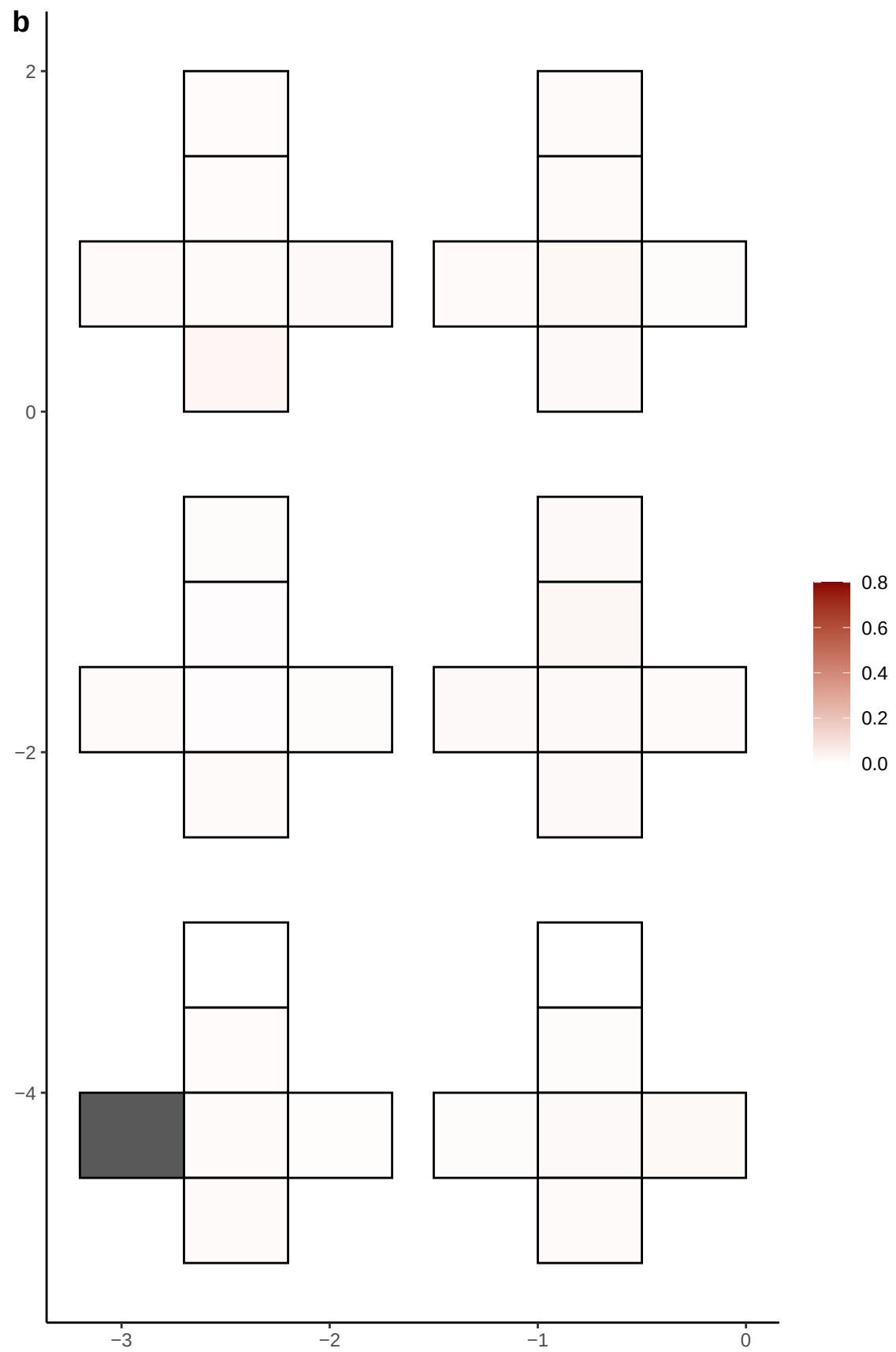

**a** ID01 9-78848802-T-G

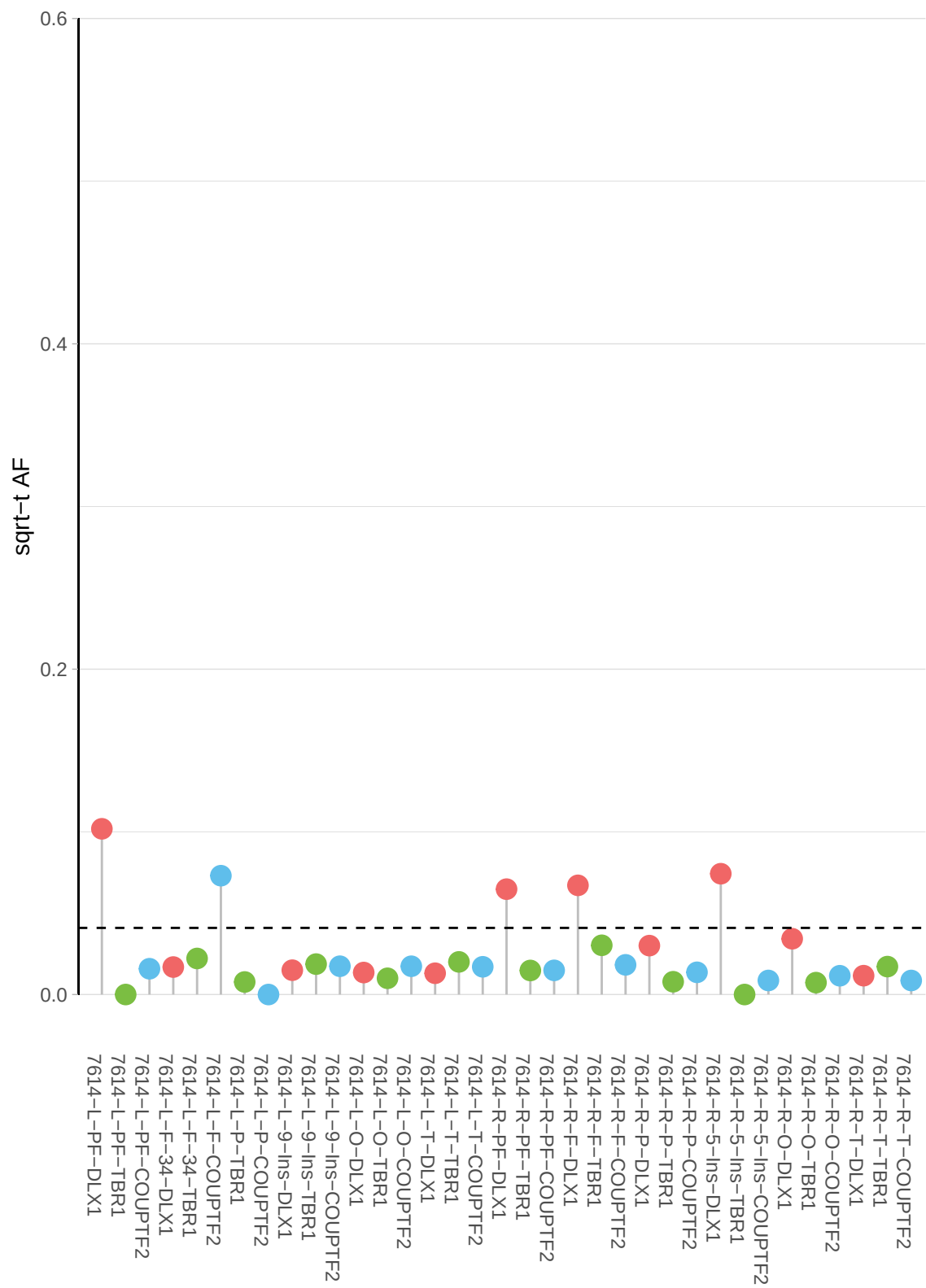

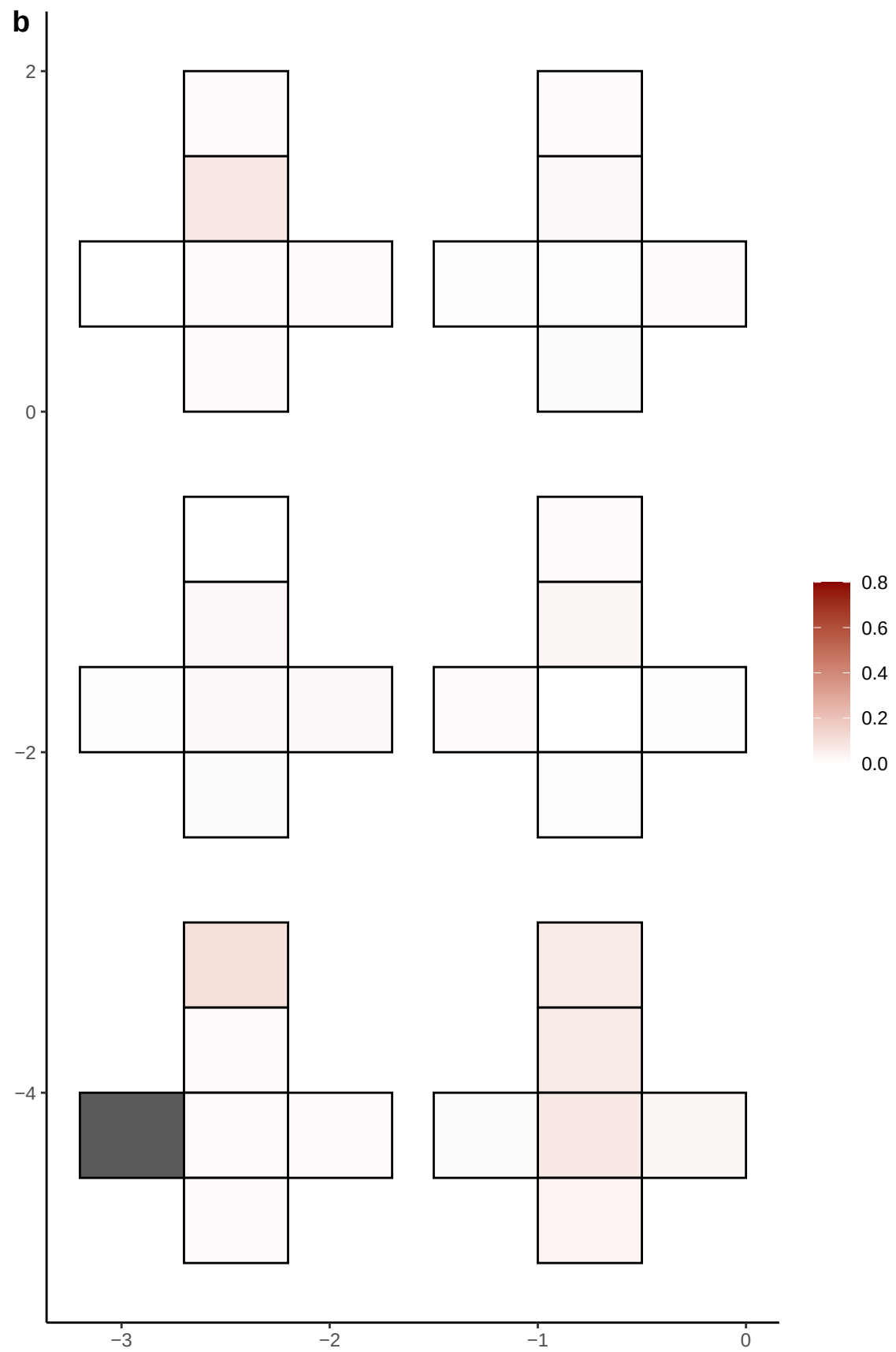

**a** ID01 9-88011667-A-C

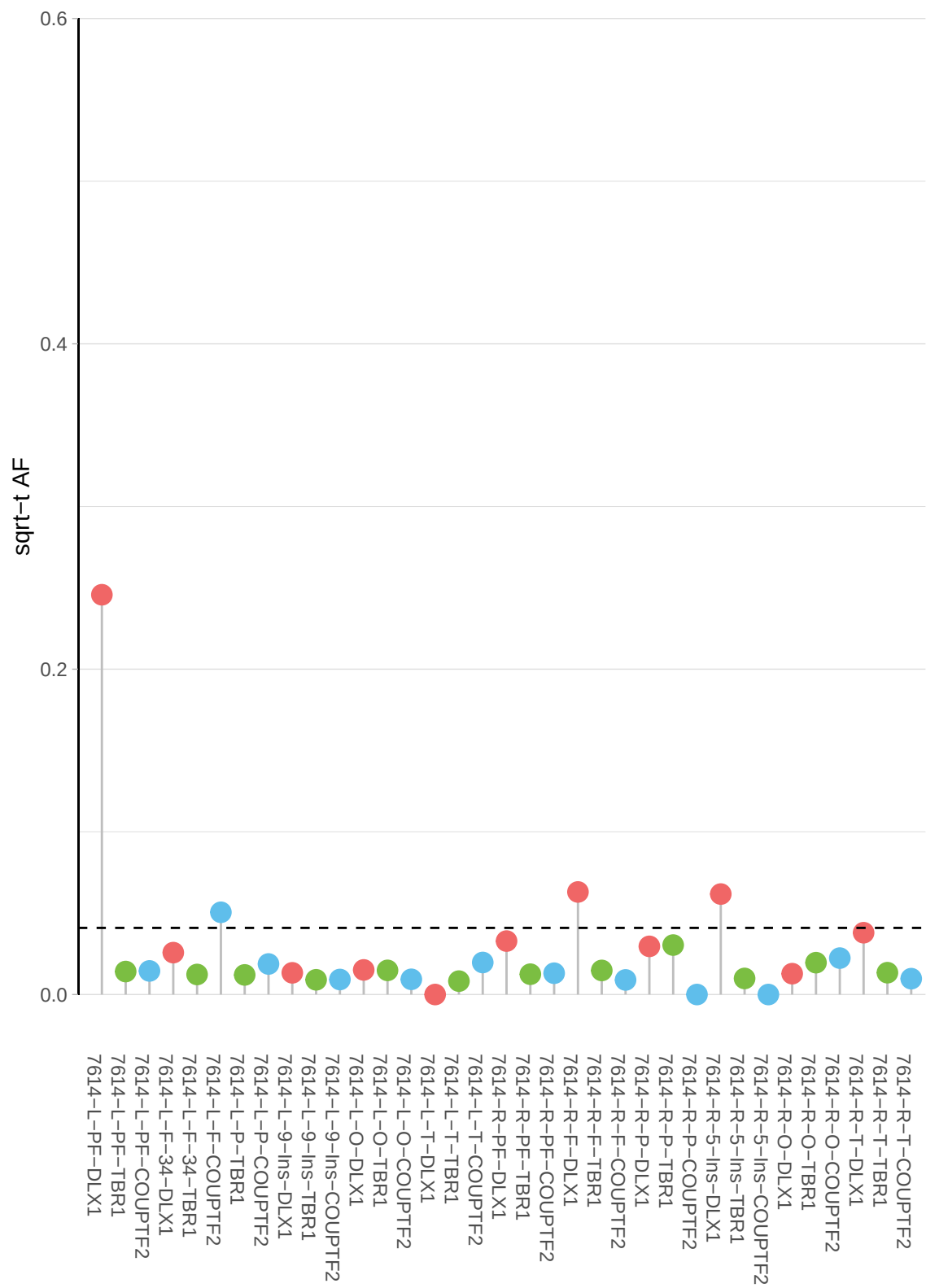

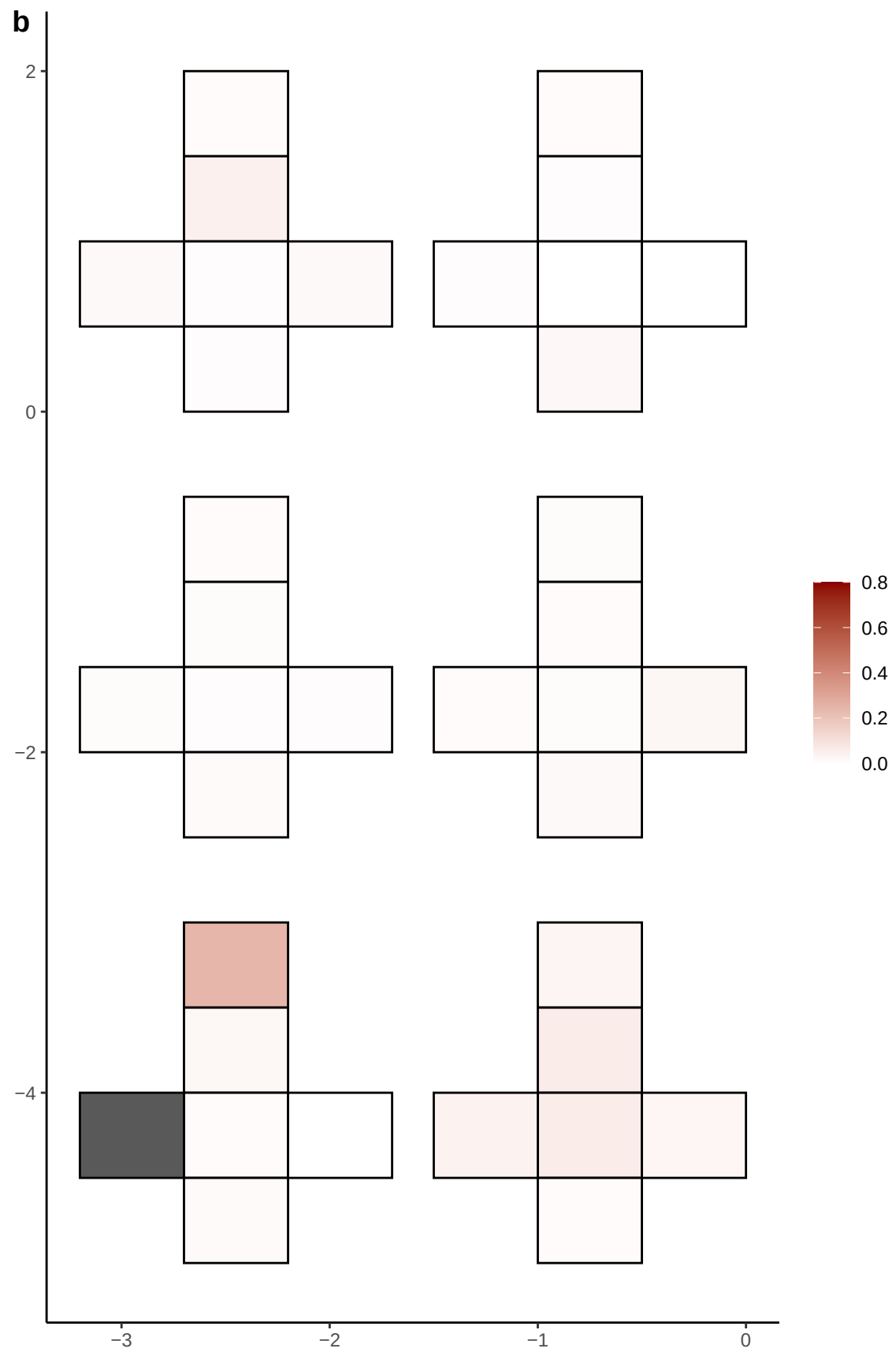

**a** ID01 9-9457115-T-A

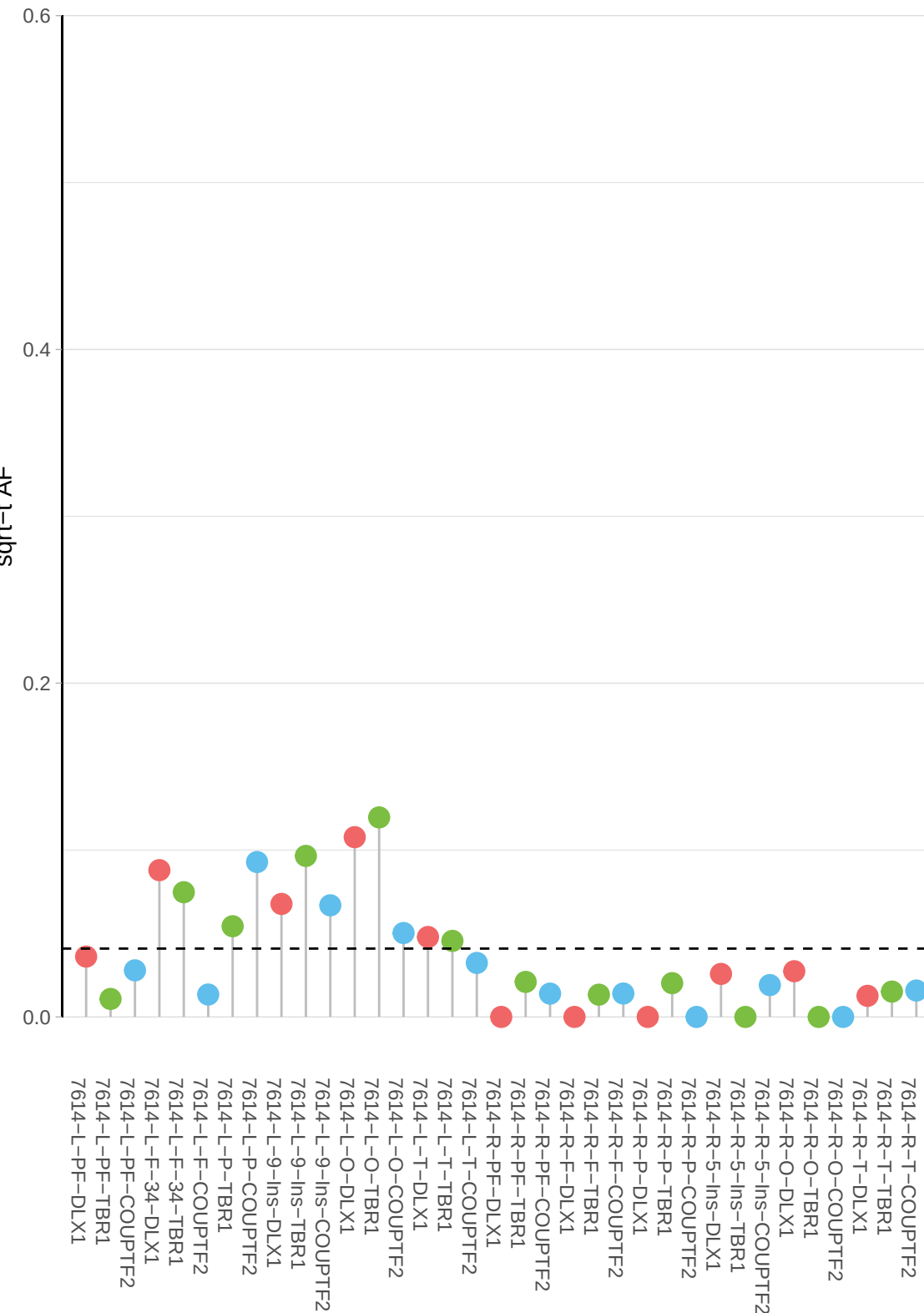

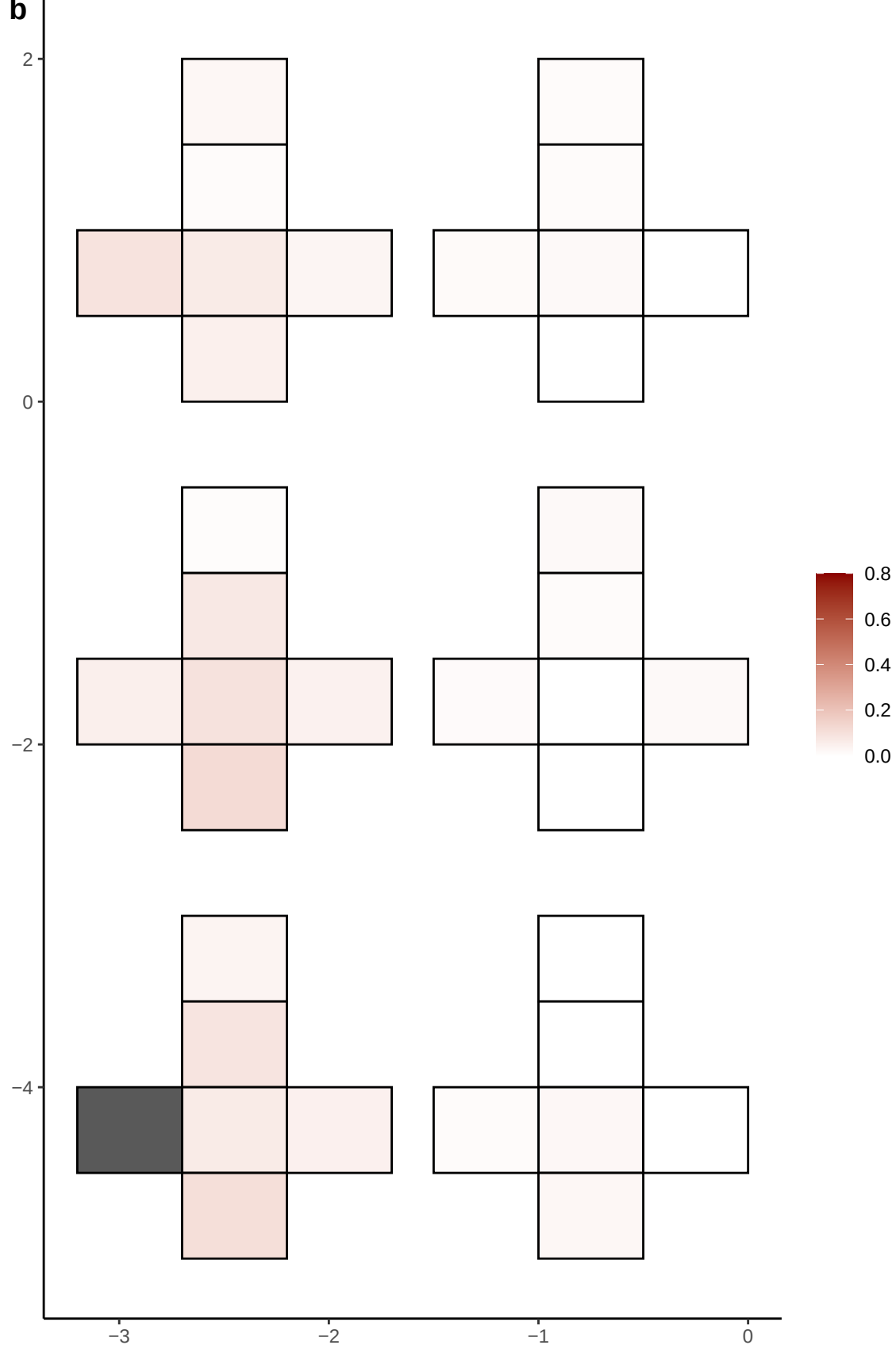

**a** ID01 X-10480747-T-A

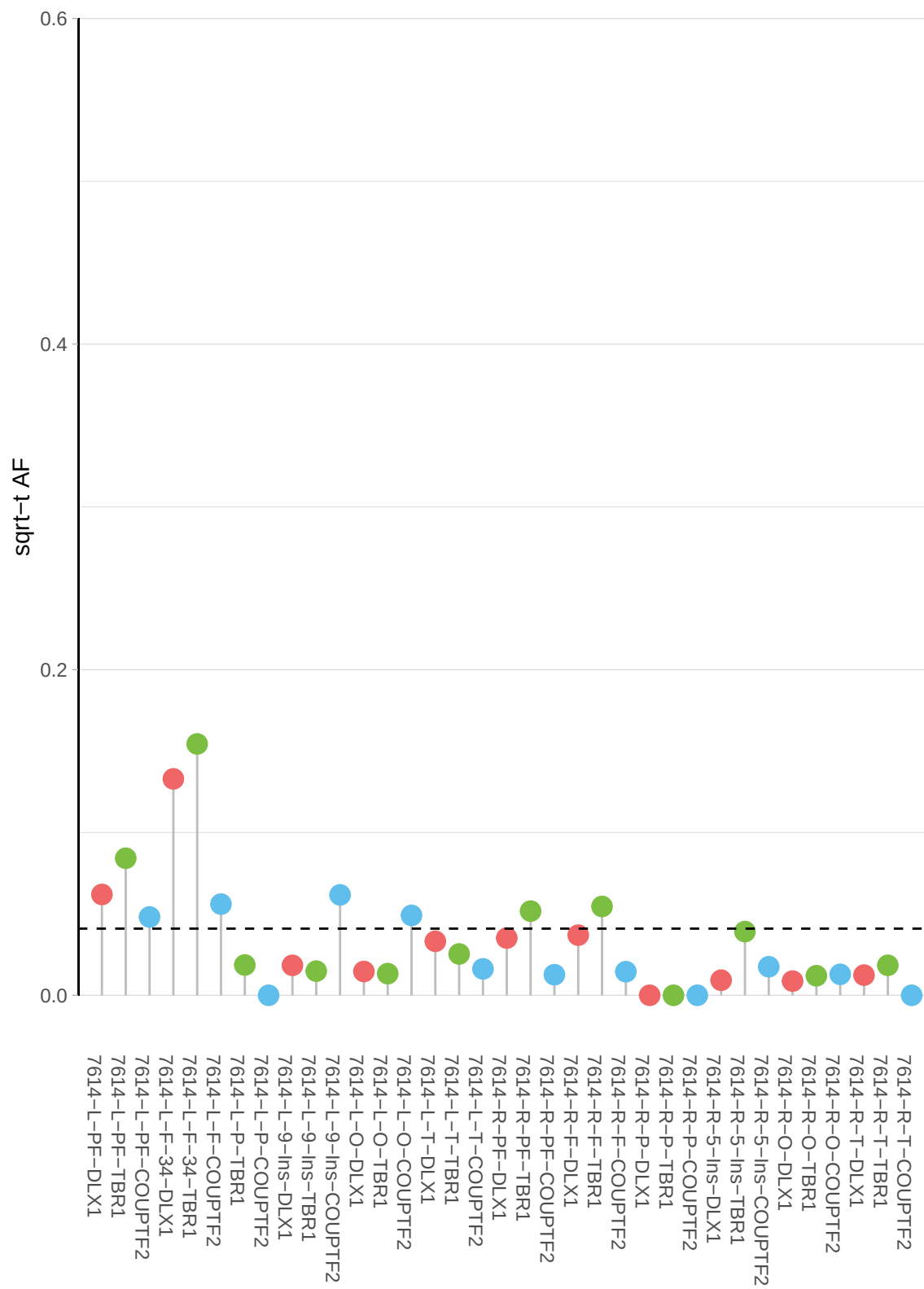

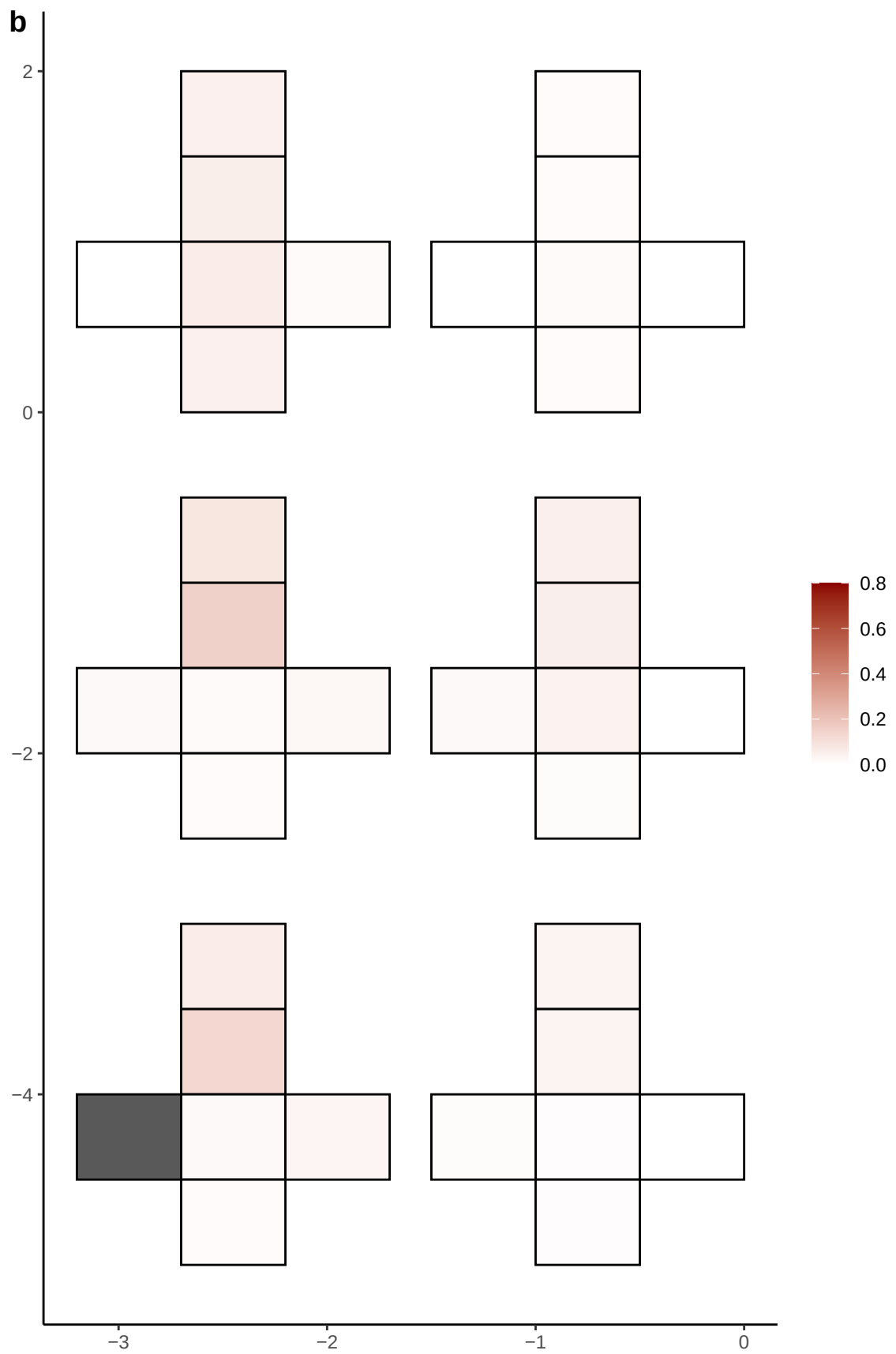

**a** ID01 X-117352547-A-T

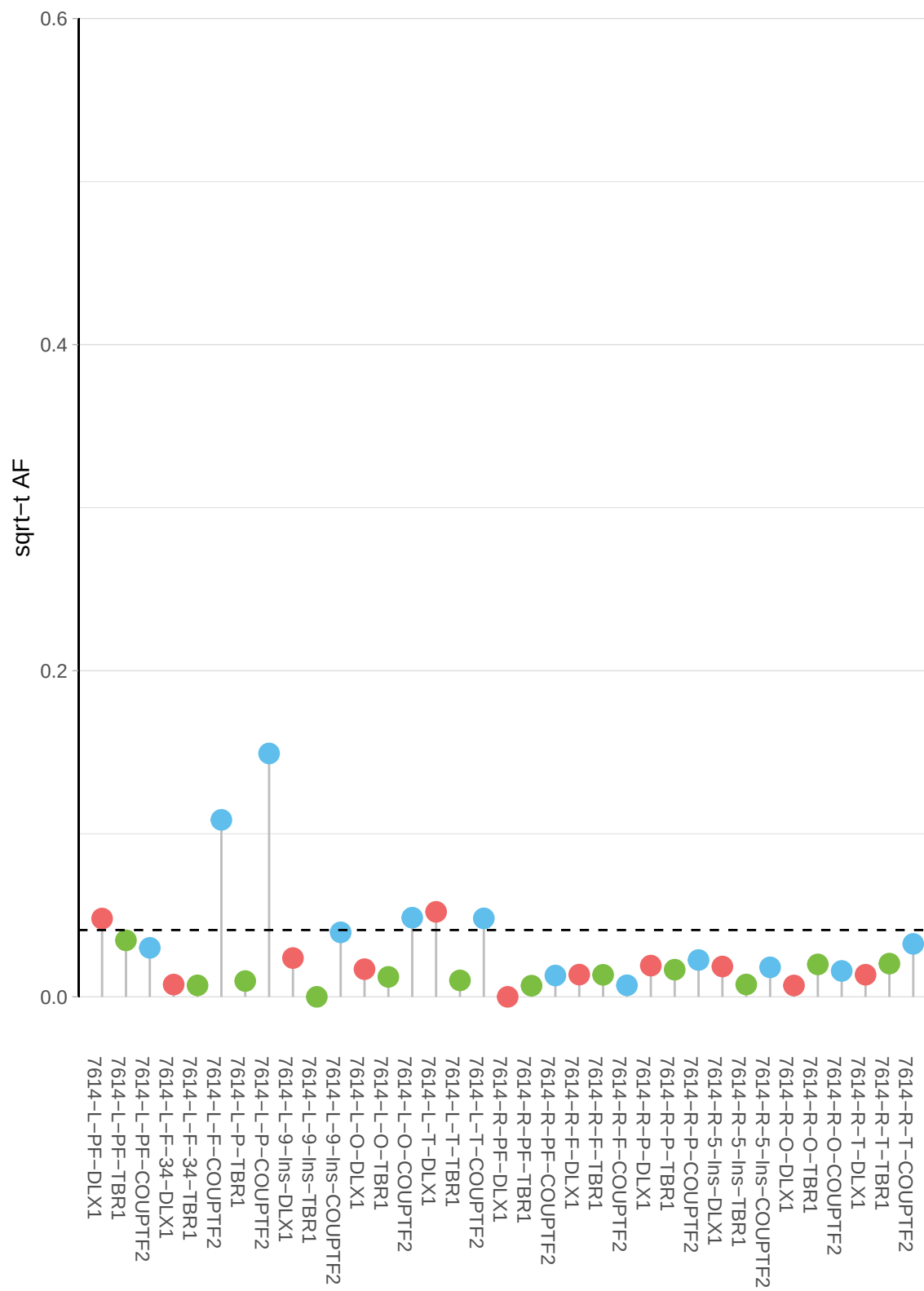

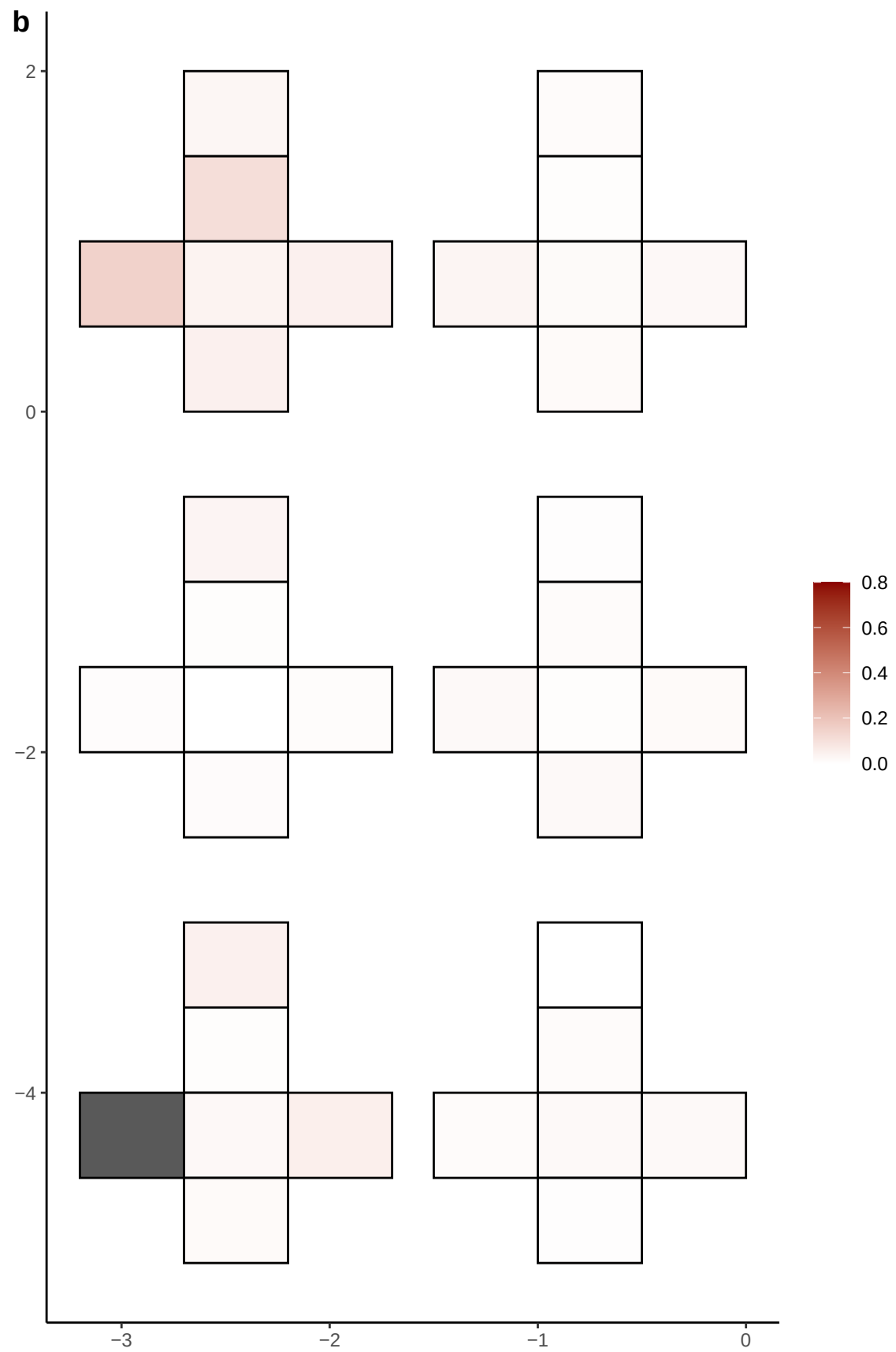

**a** ID01 X-124310752-C-T

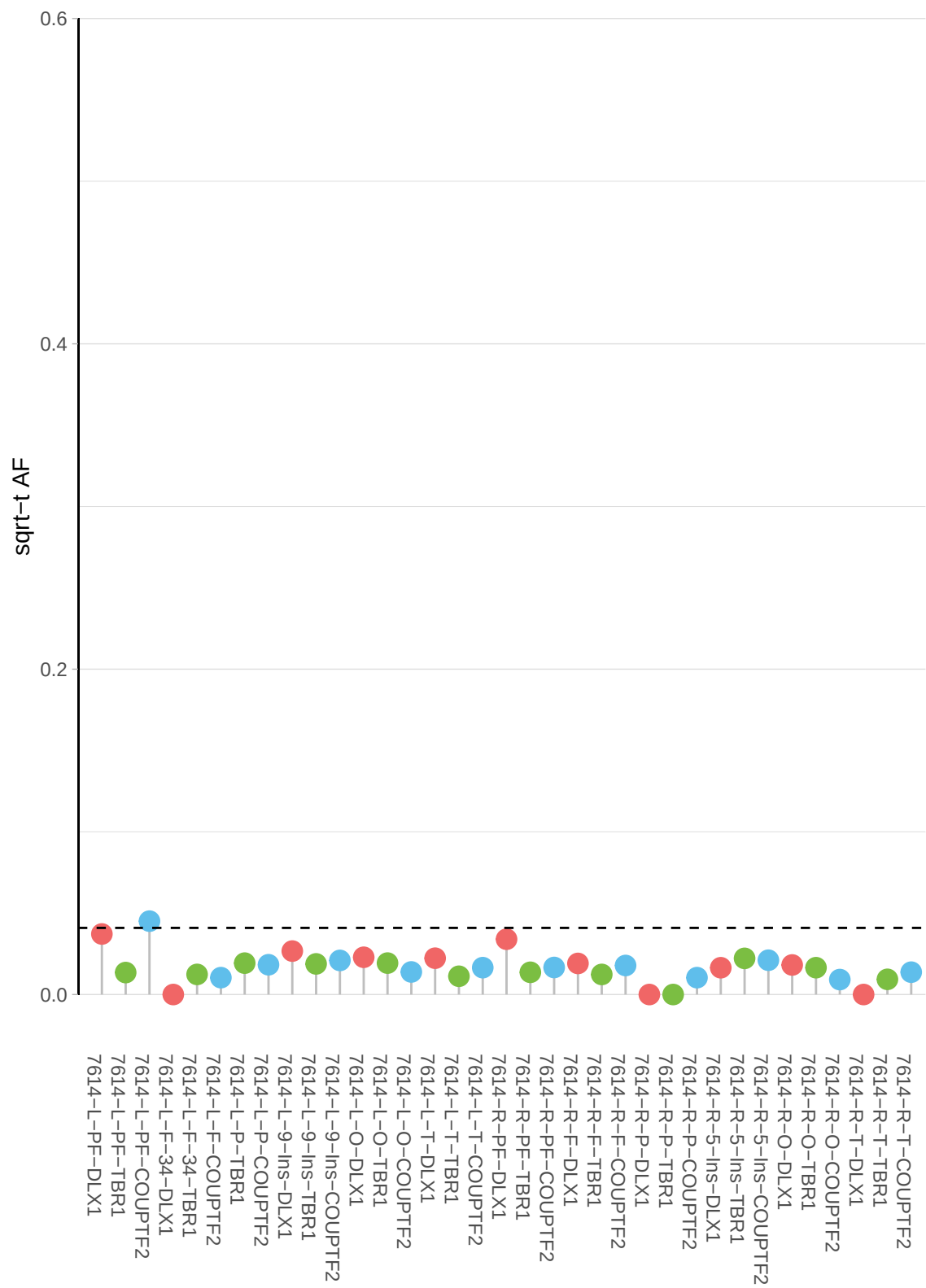

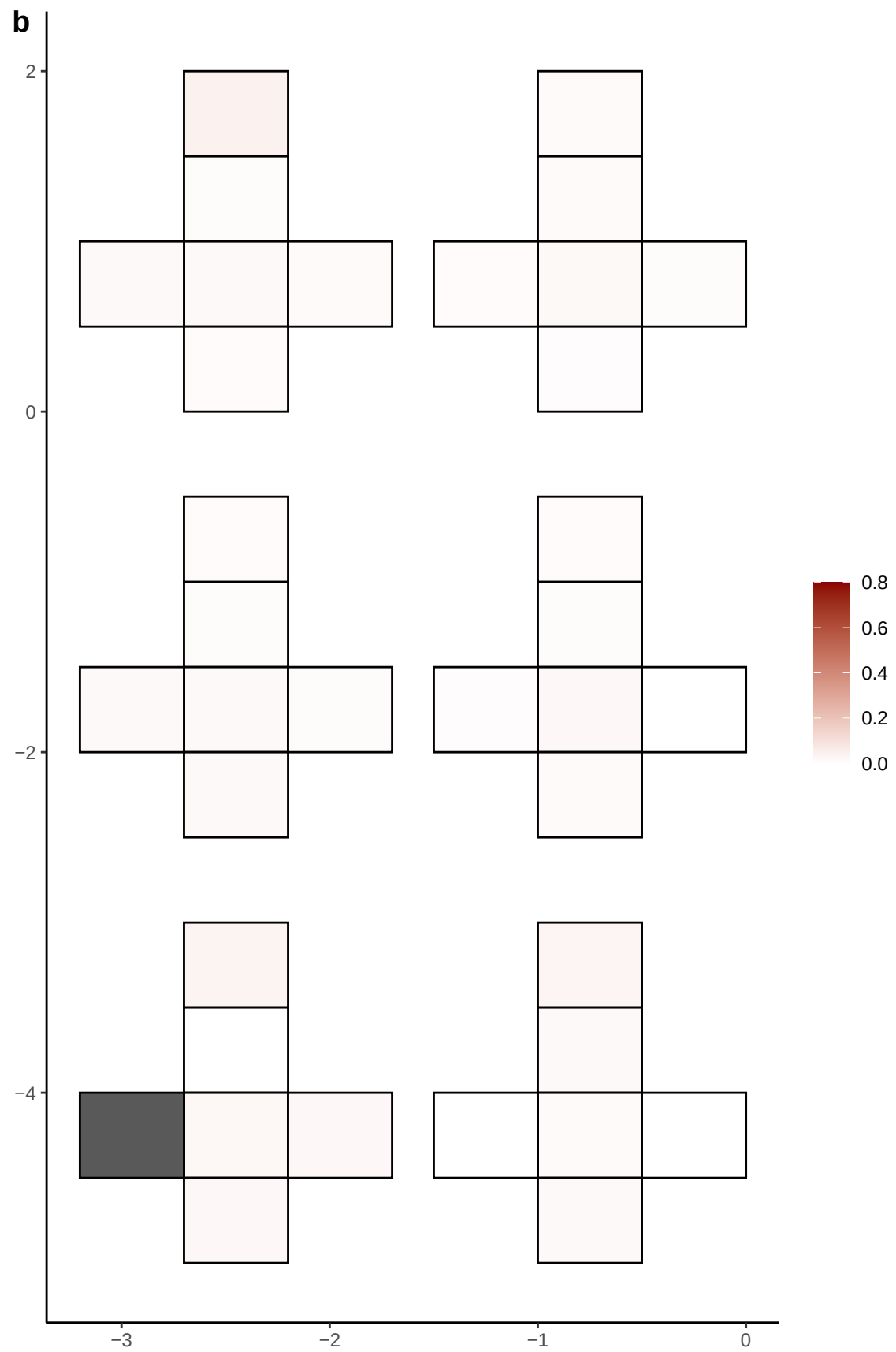

**a** ID01 X-130898293-A-G

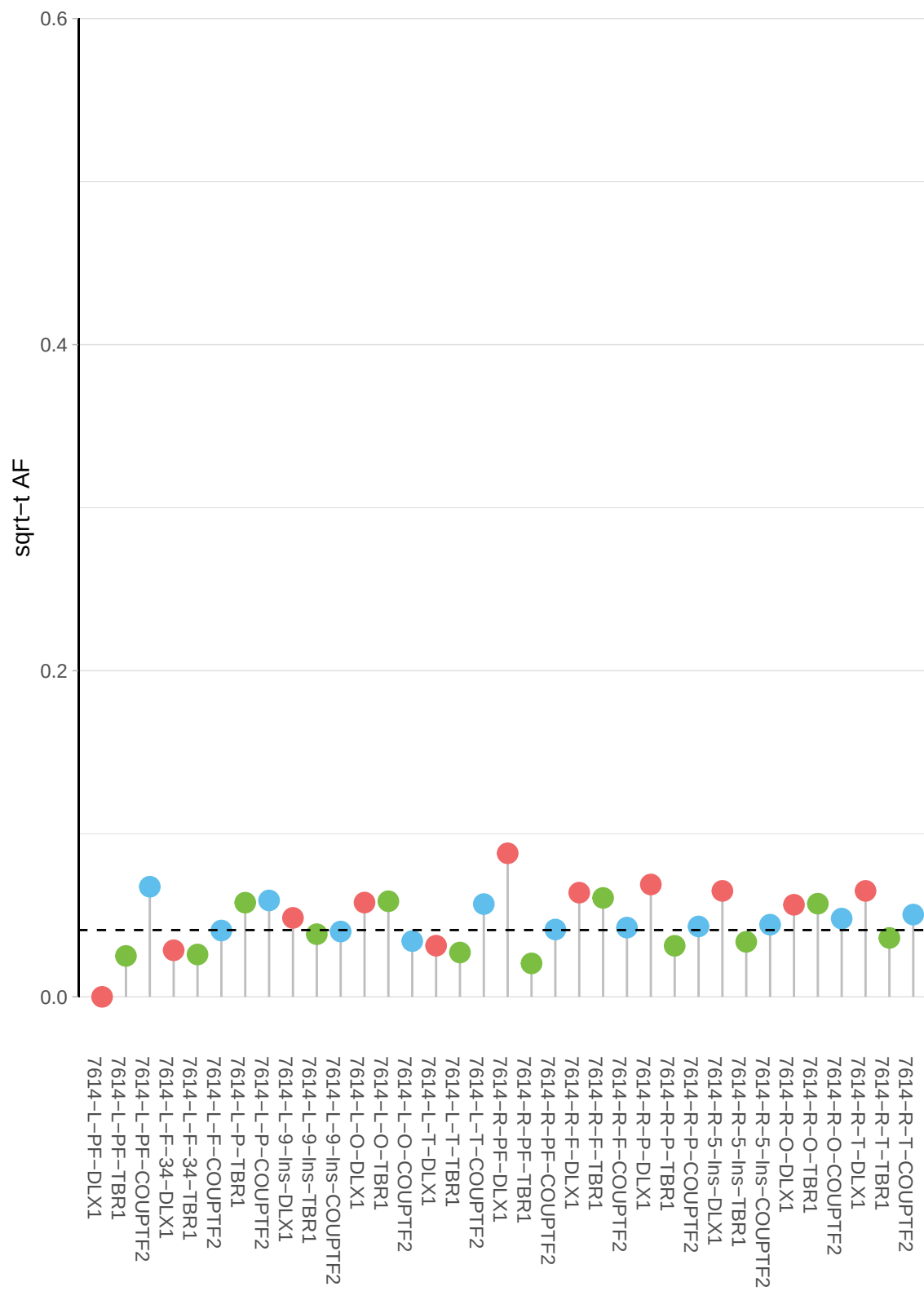

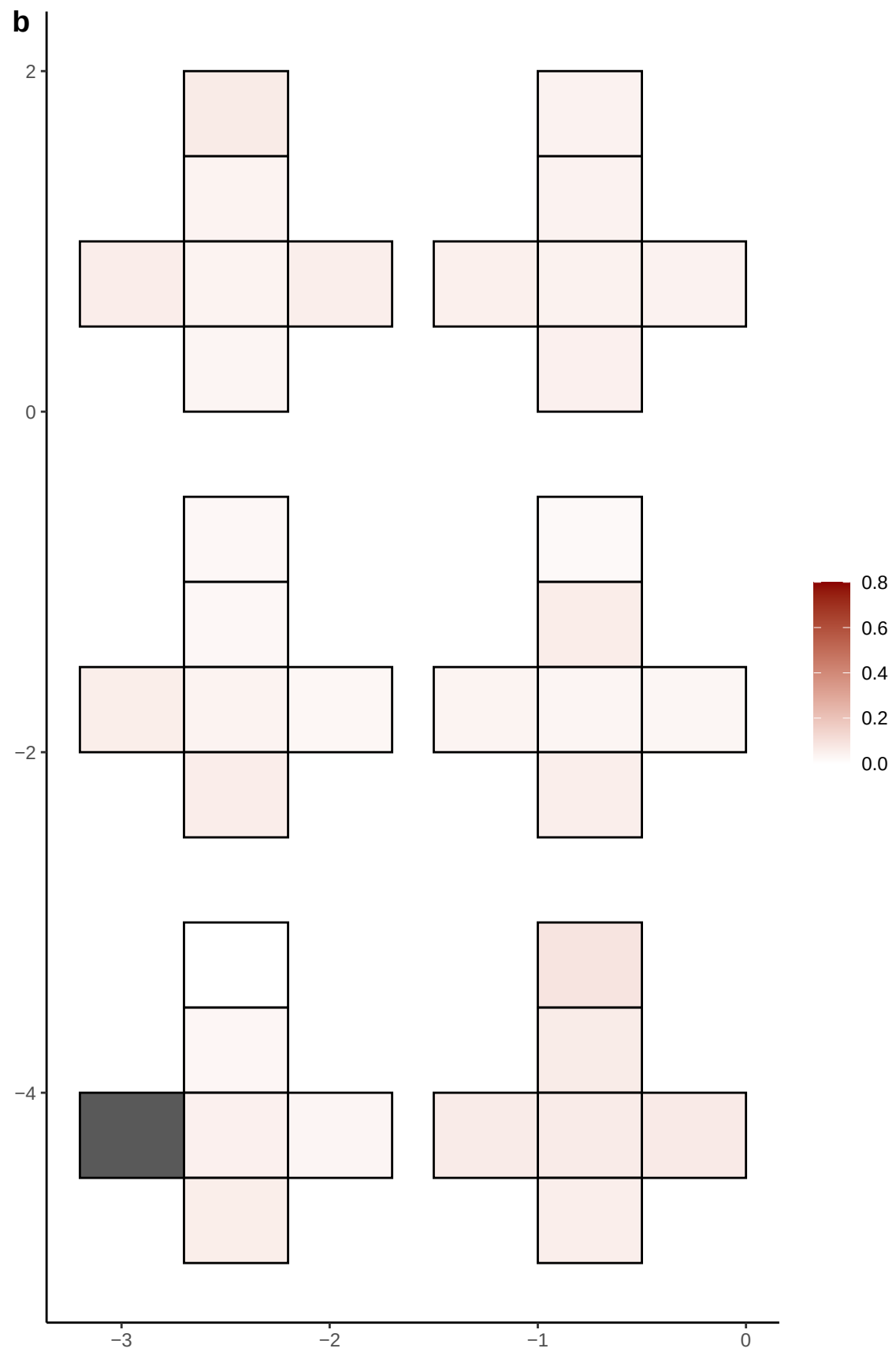

**a** ID01 X-130898294-C-T

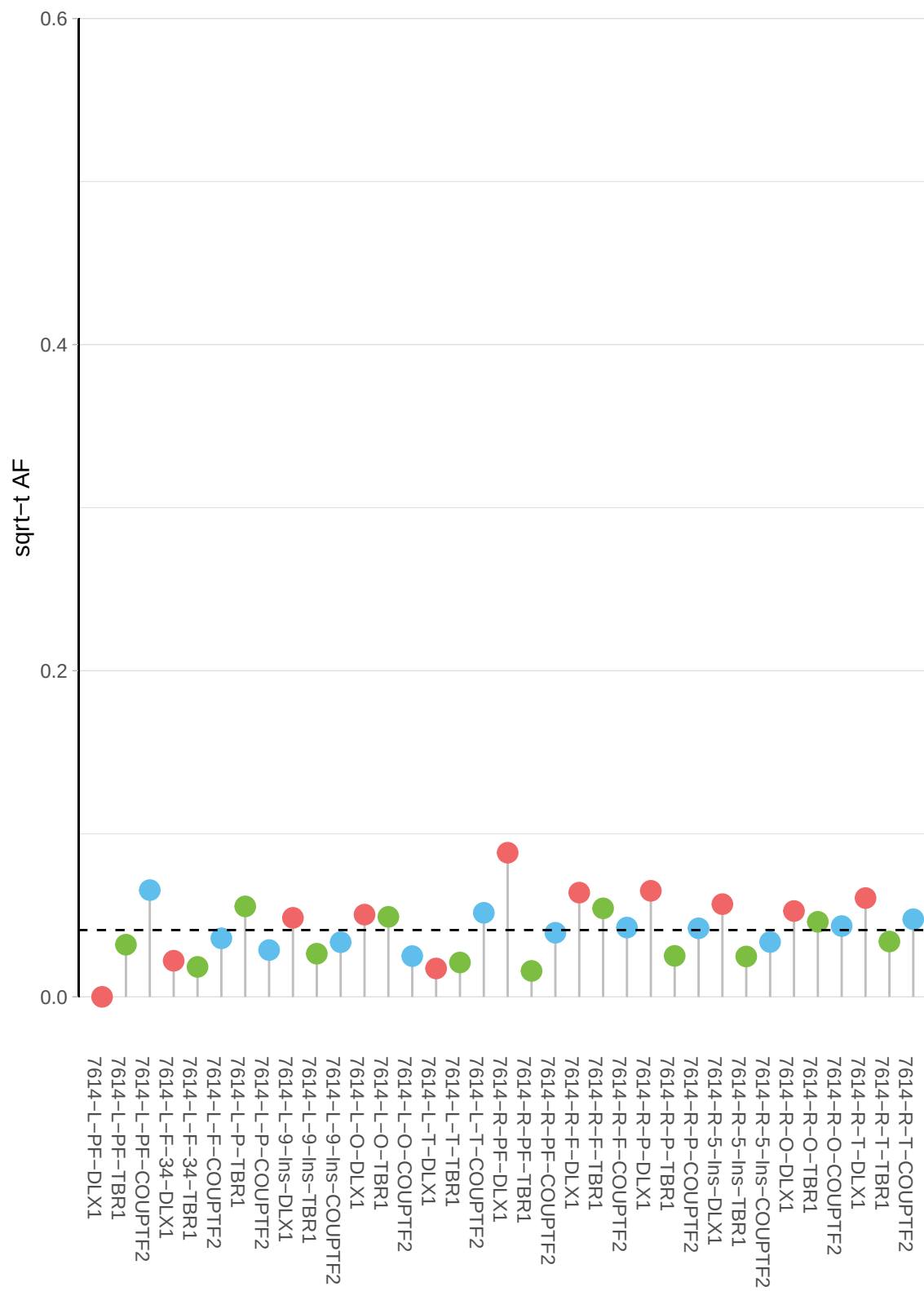

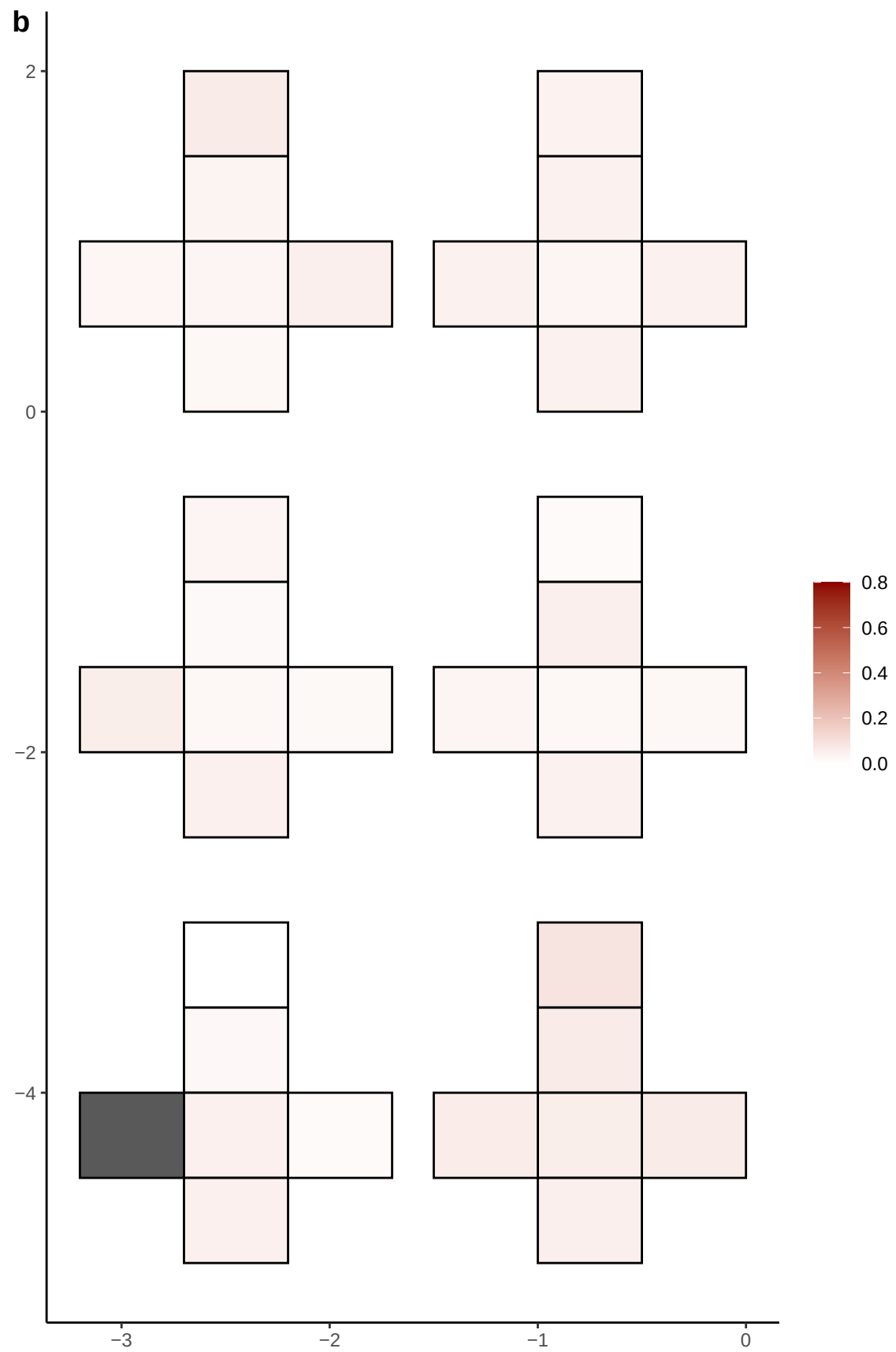

**a** ID01 X-138175923-C-T

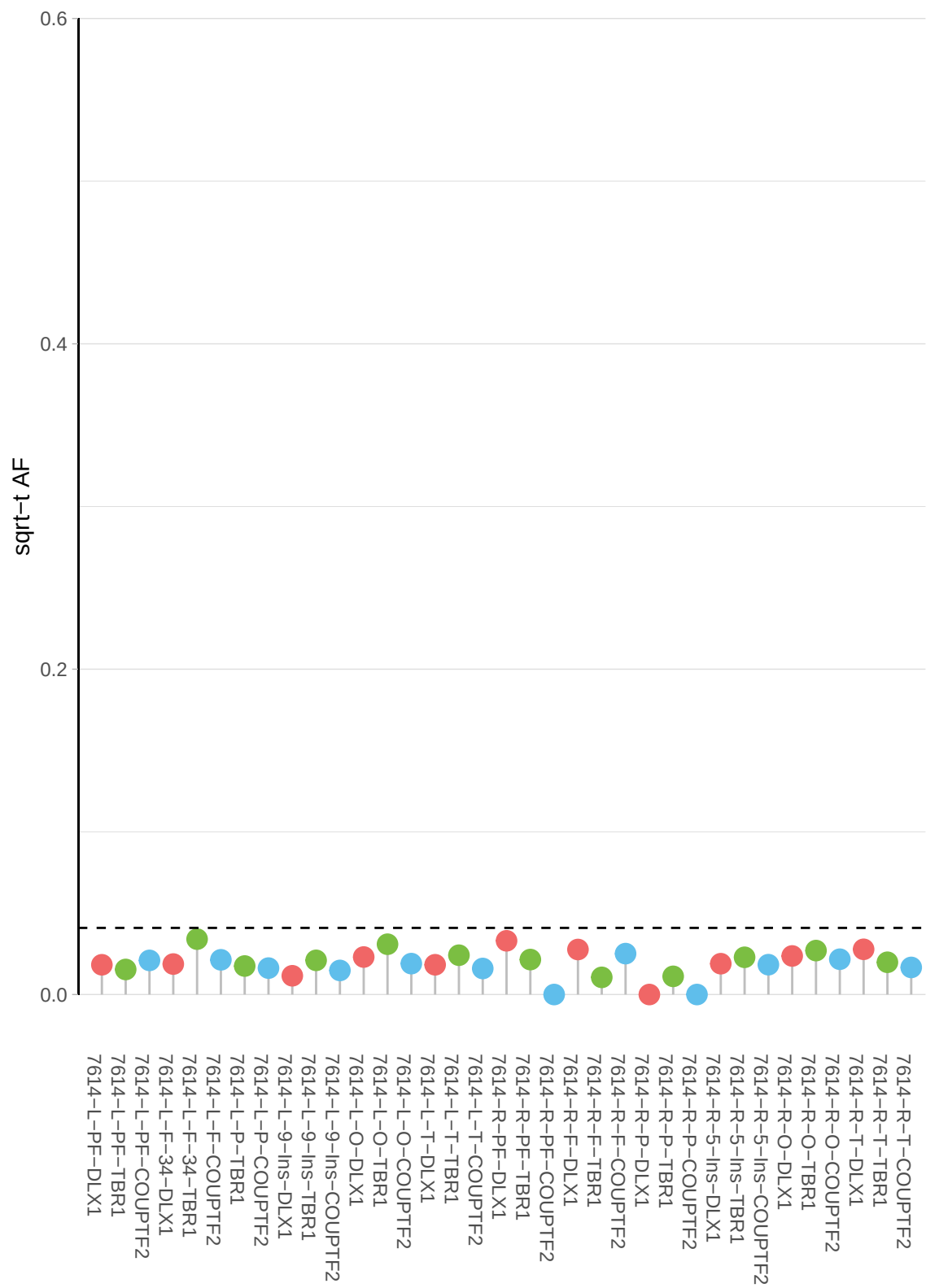

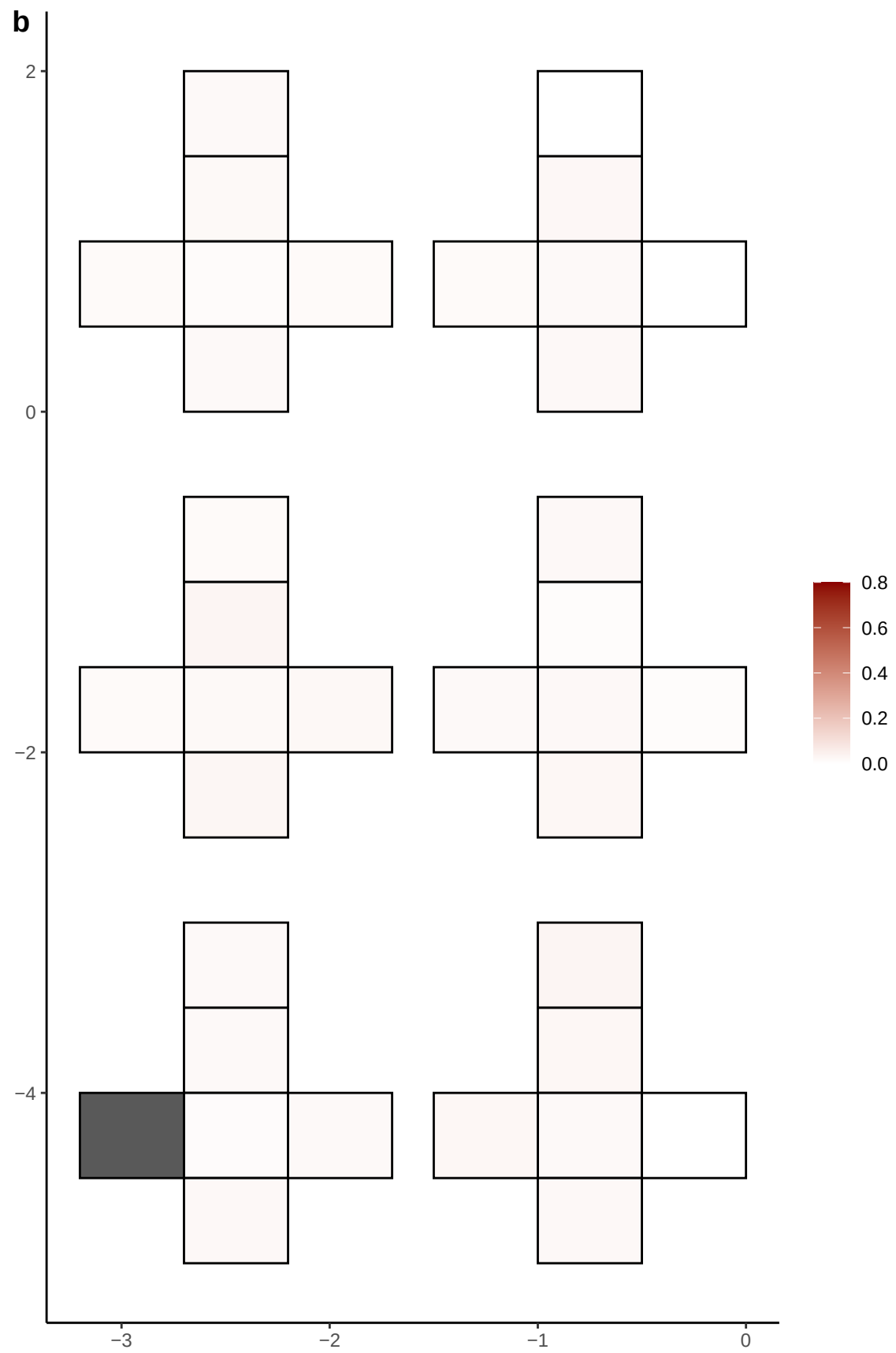

**a** ID01 X-139853112-G-A

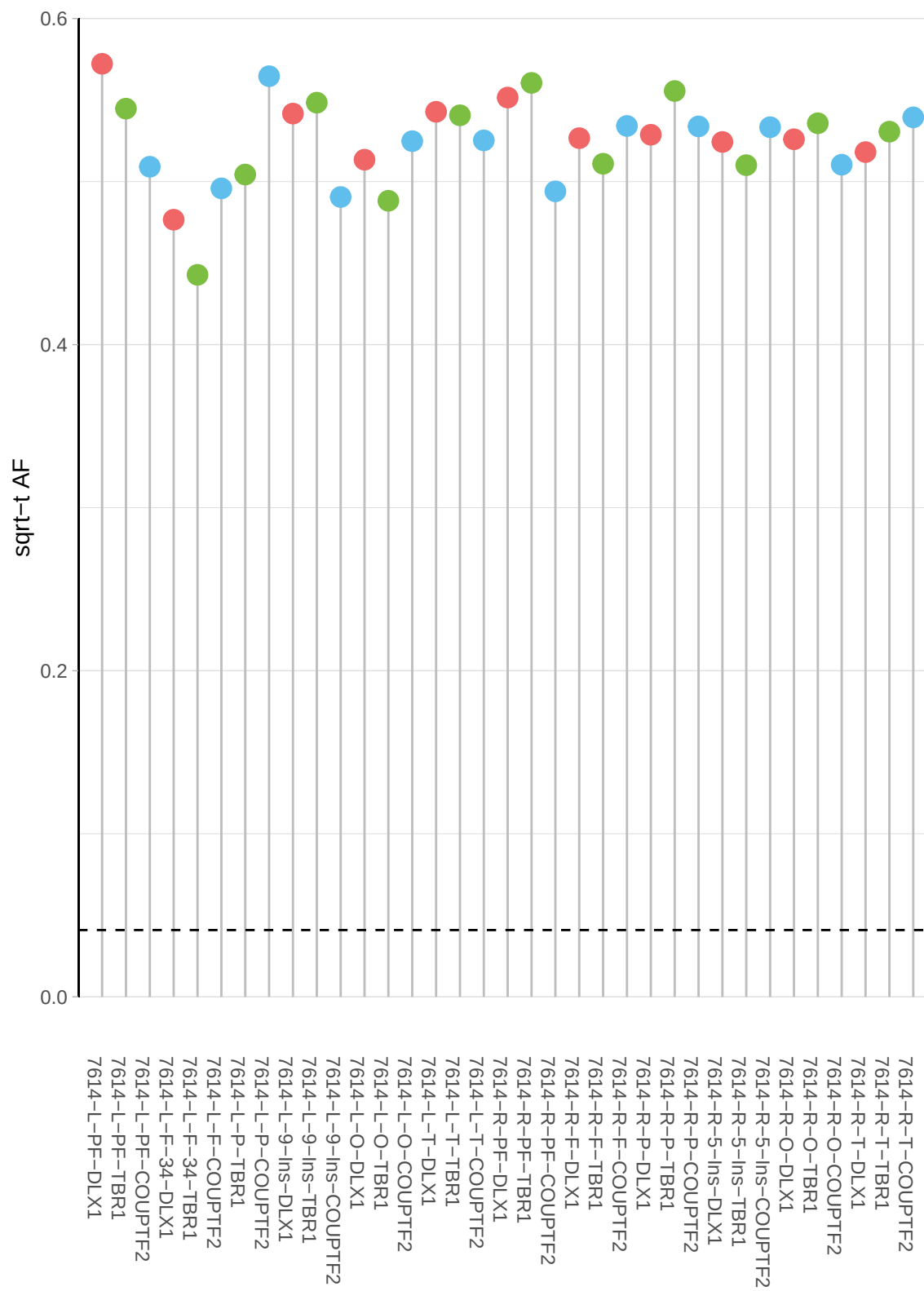

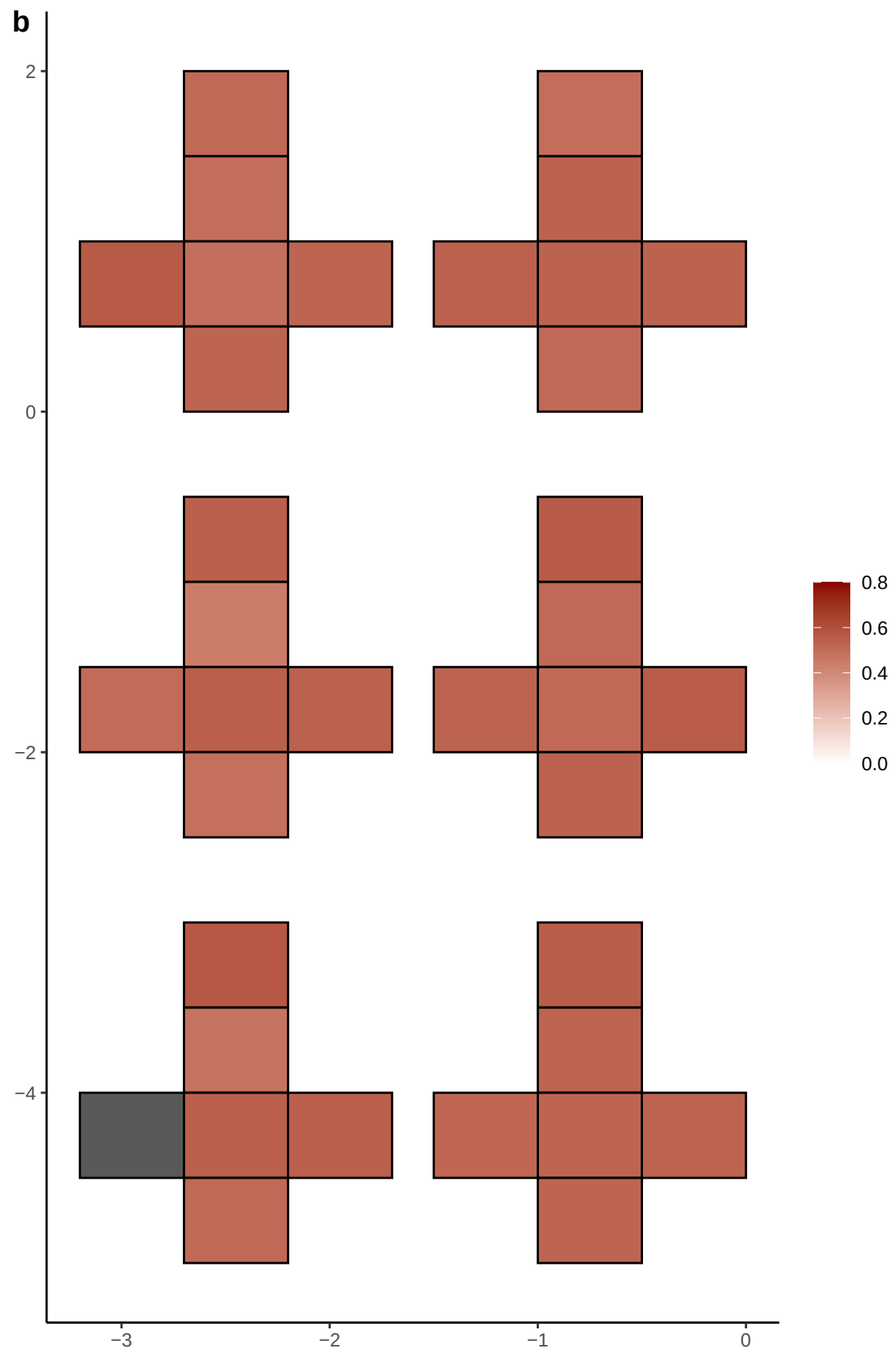

**a** ID01 X-141420247-C-G

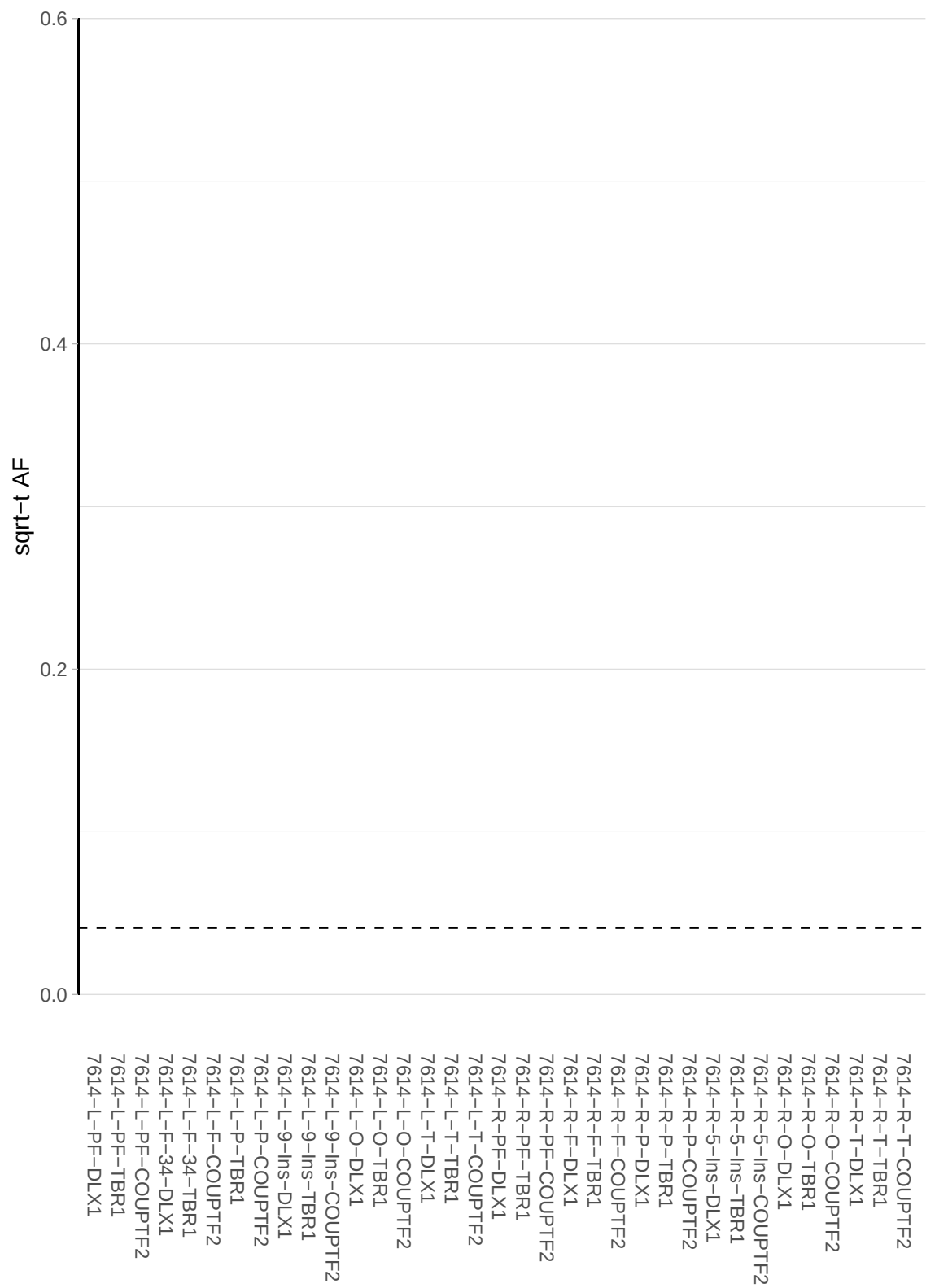

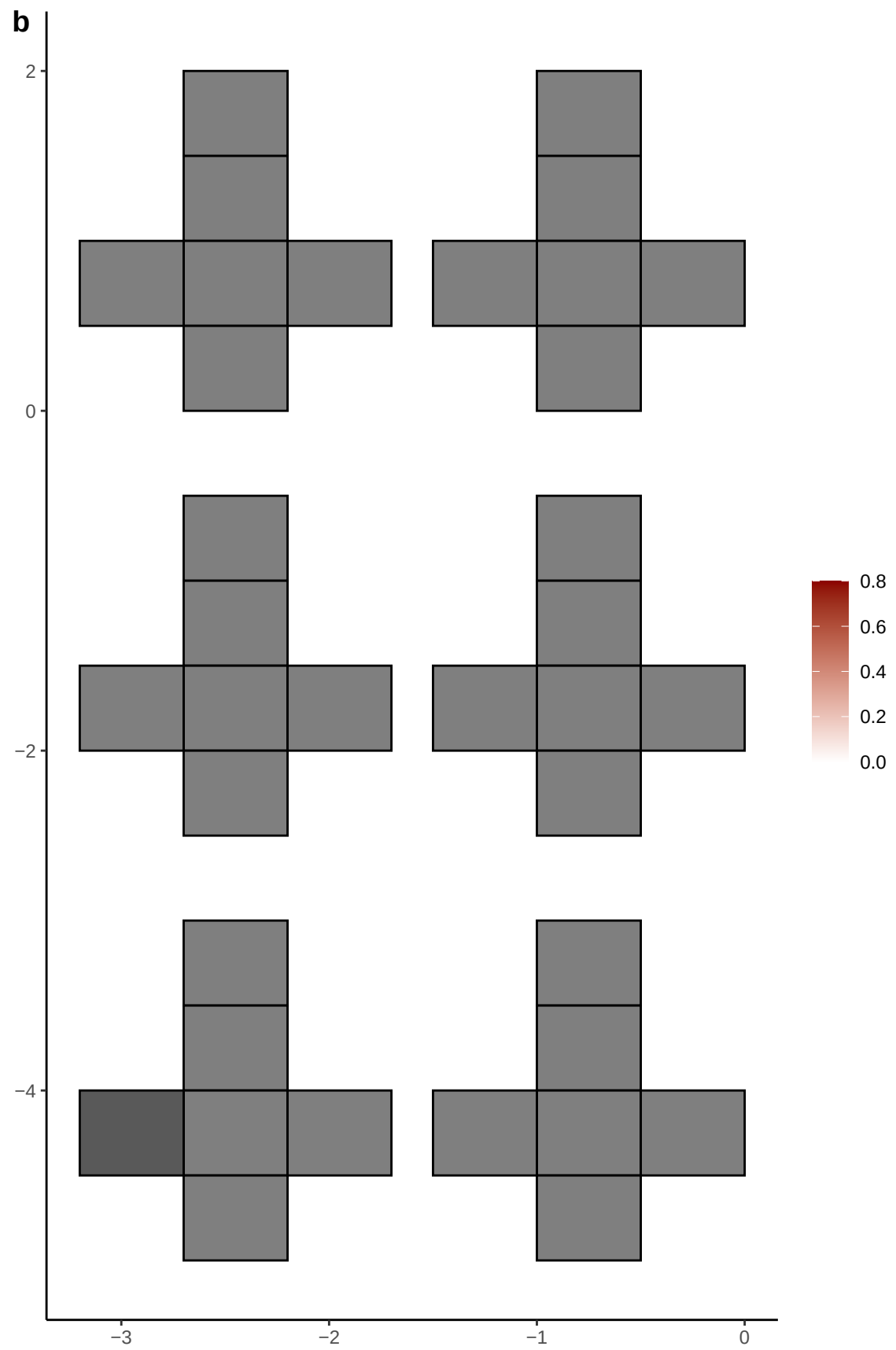

**a** ID01 X-141942973-G-A

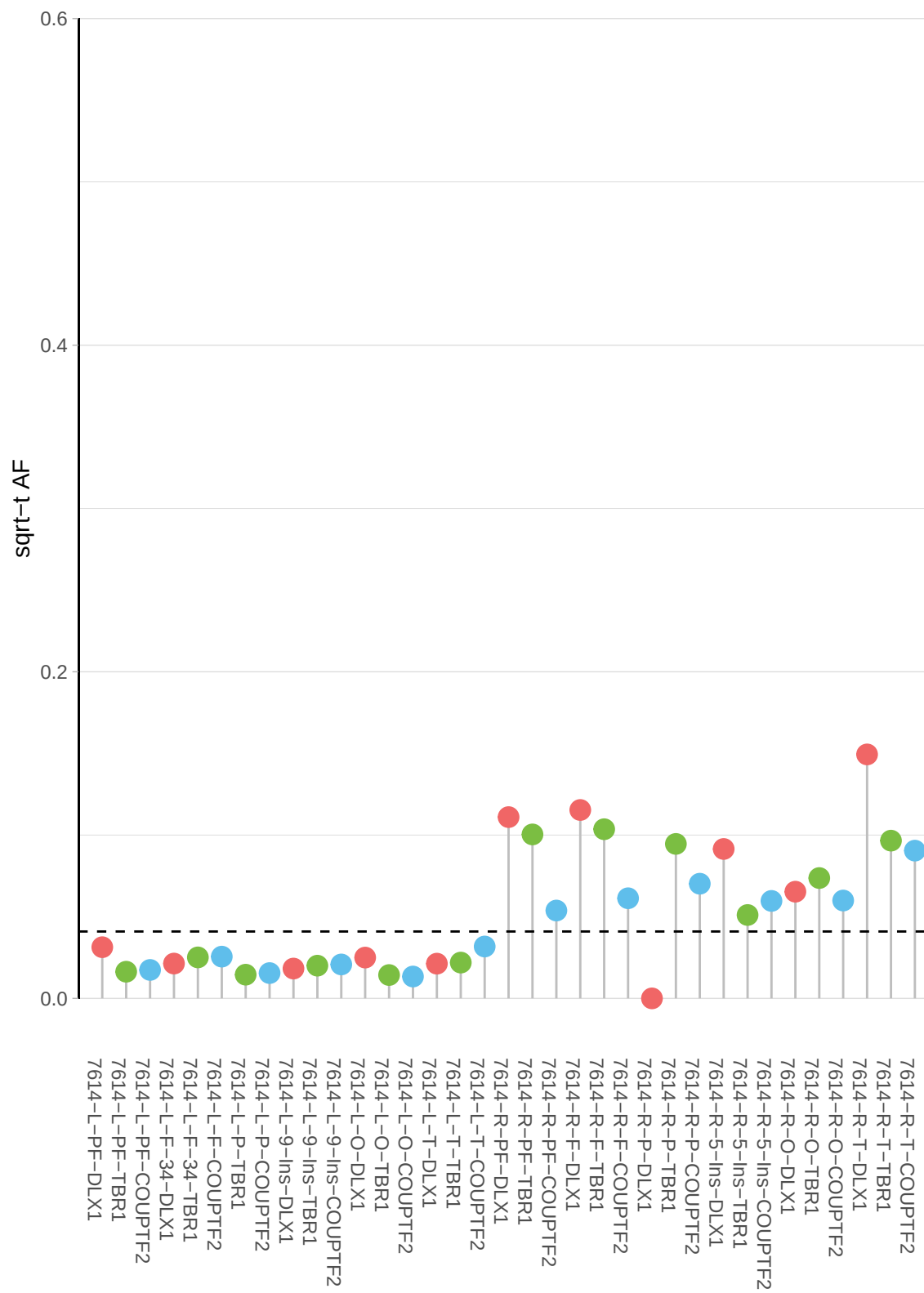

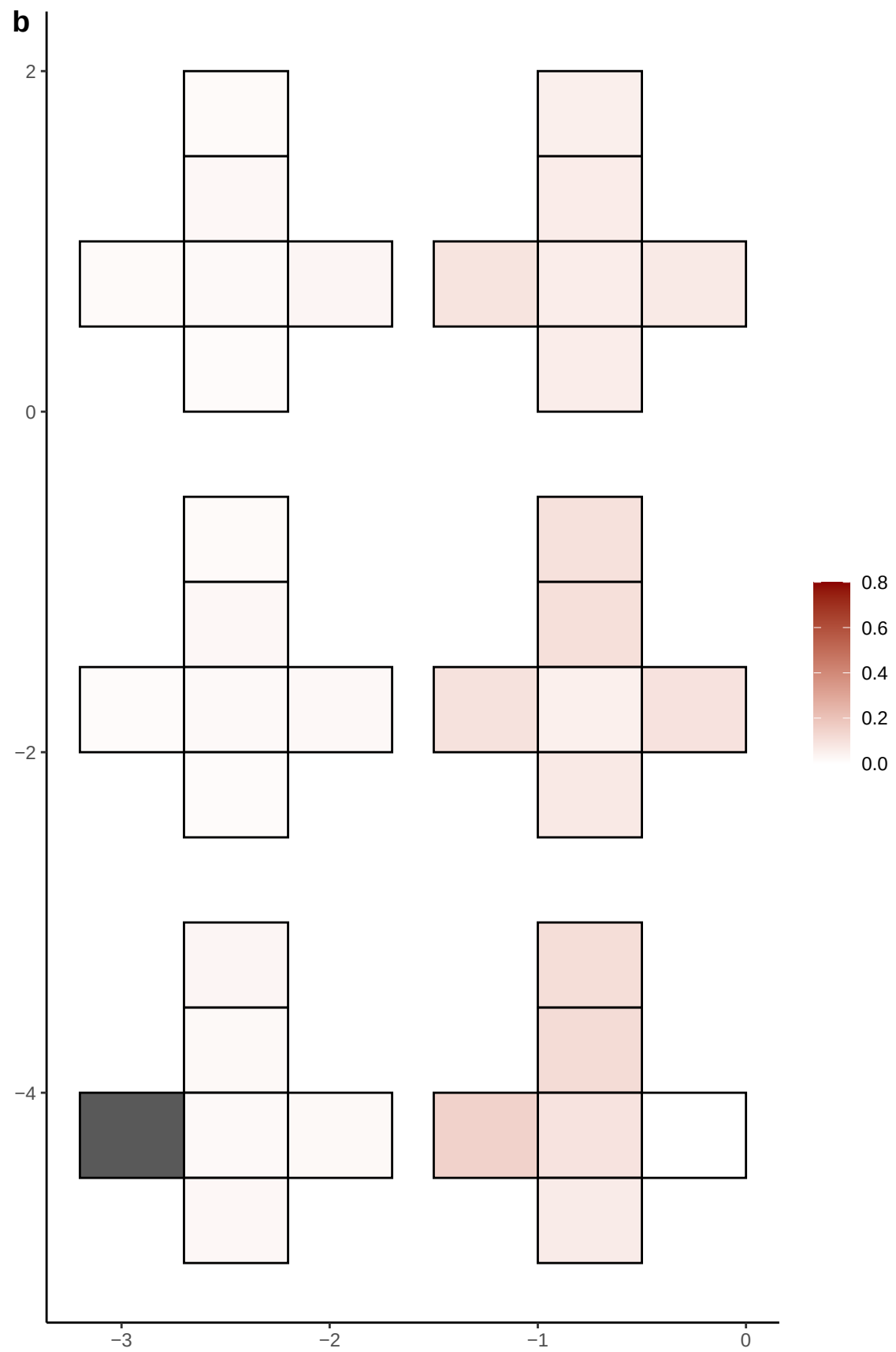

**a** ID01 X-51984755-C-T

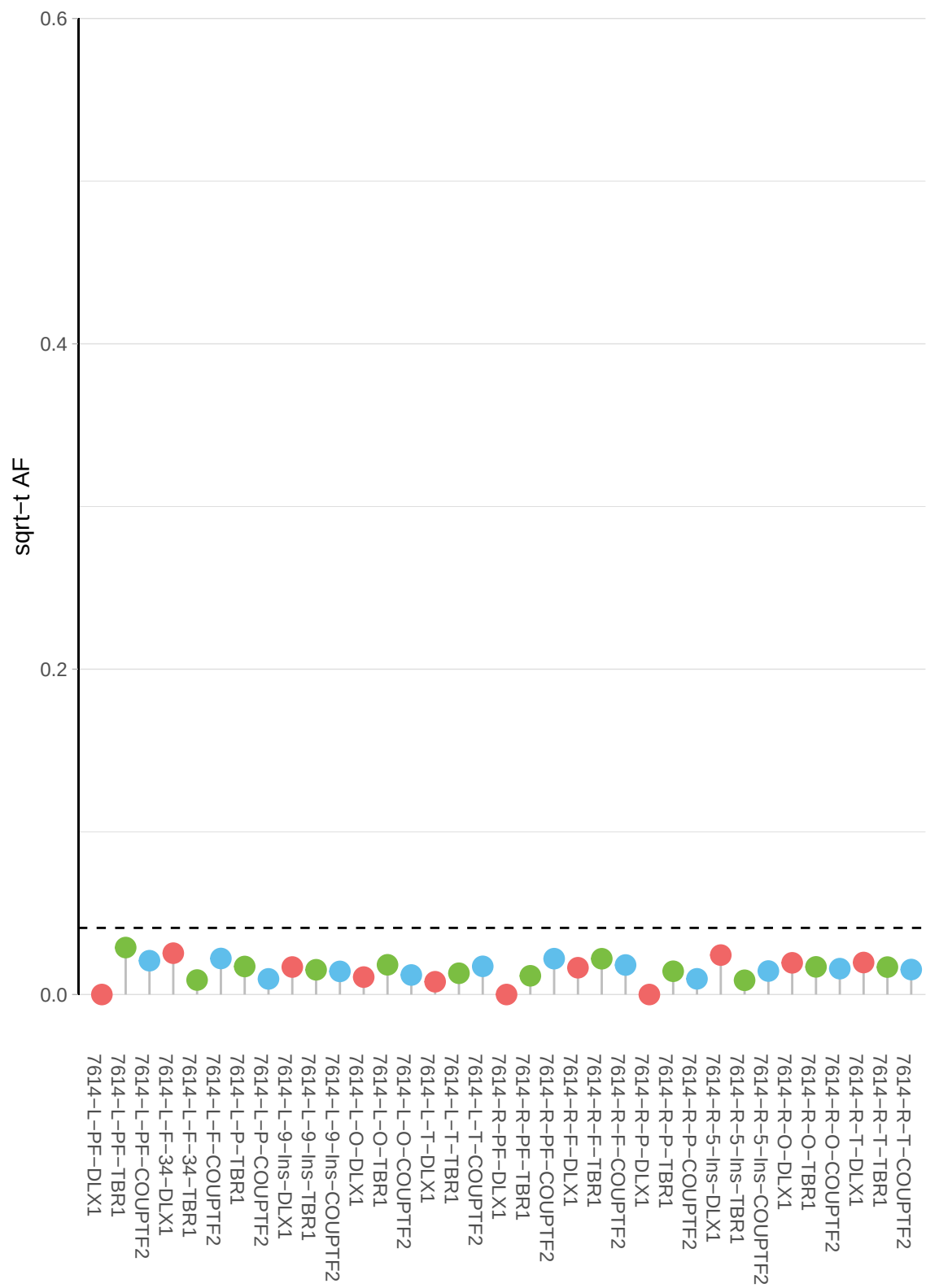

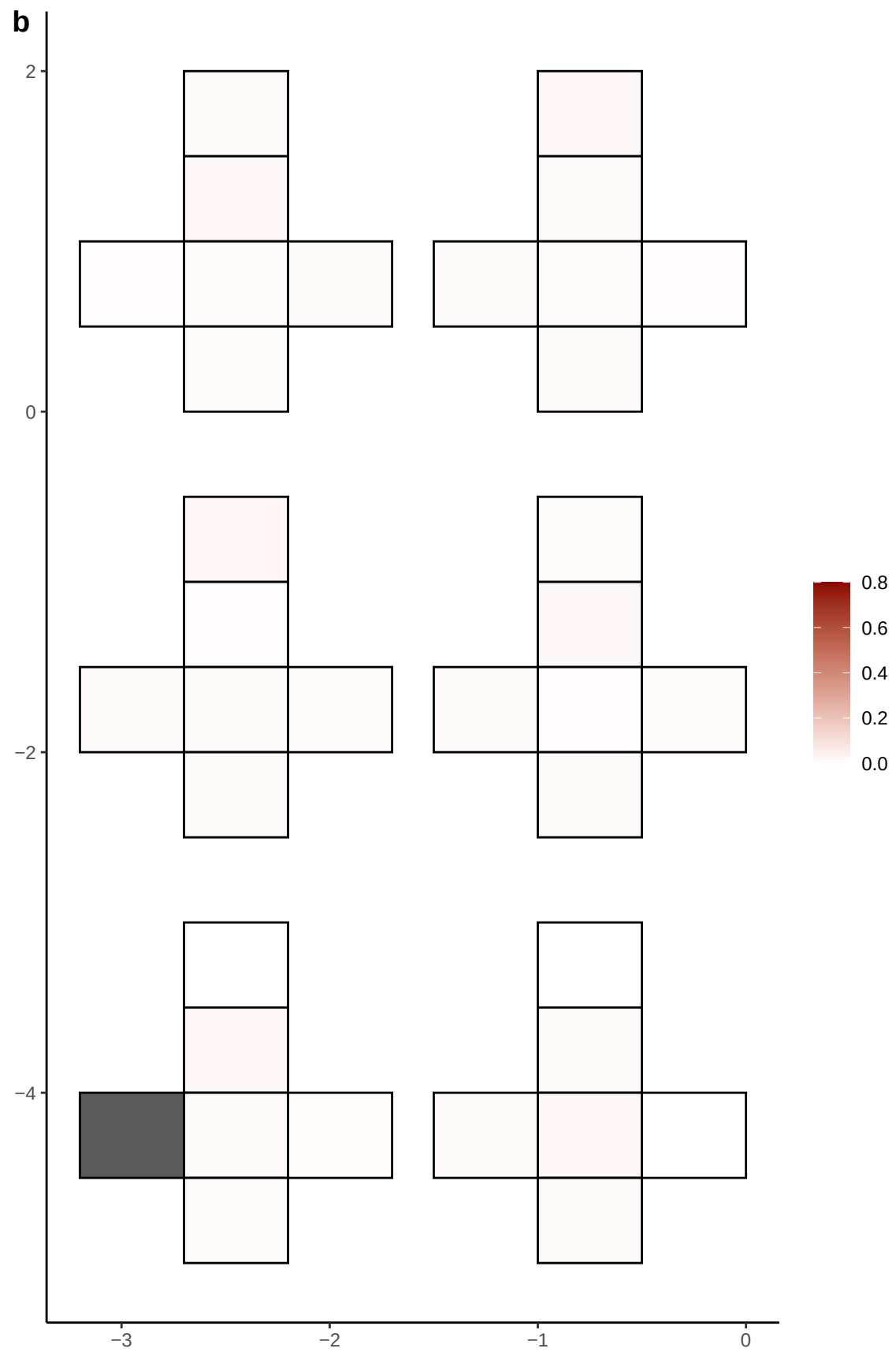

**a** ID01 X-54739596-C-G

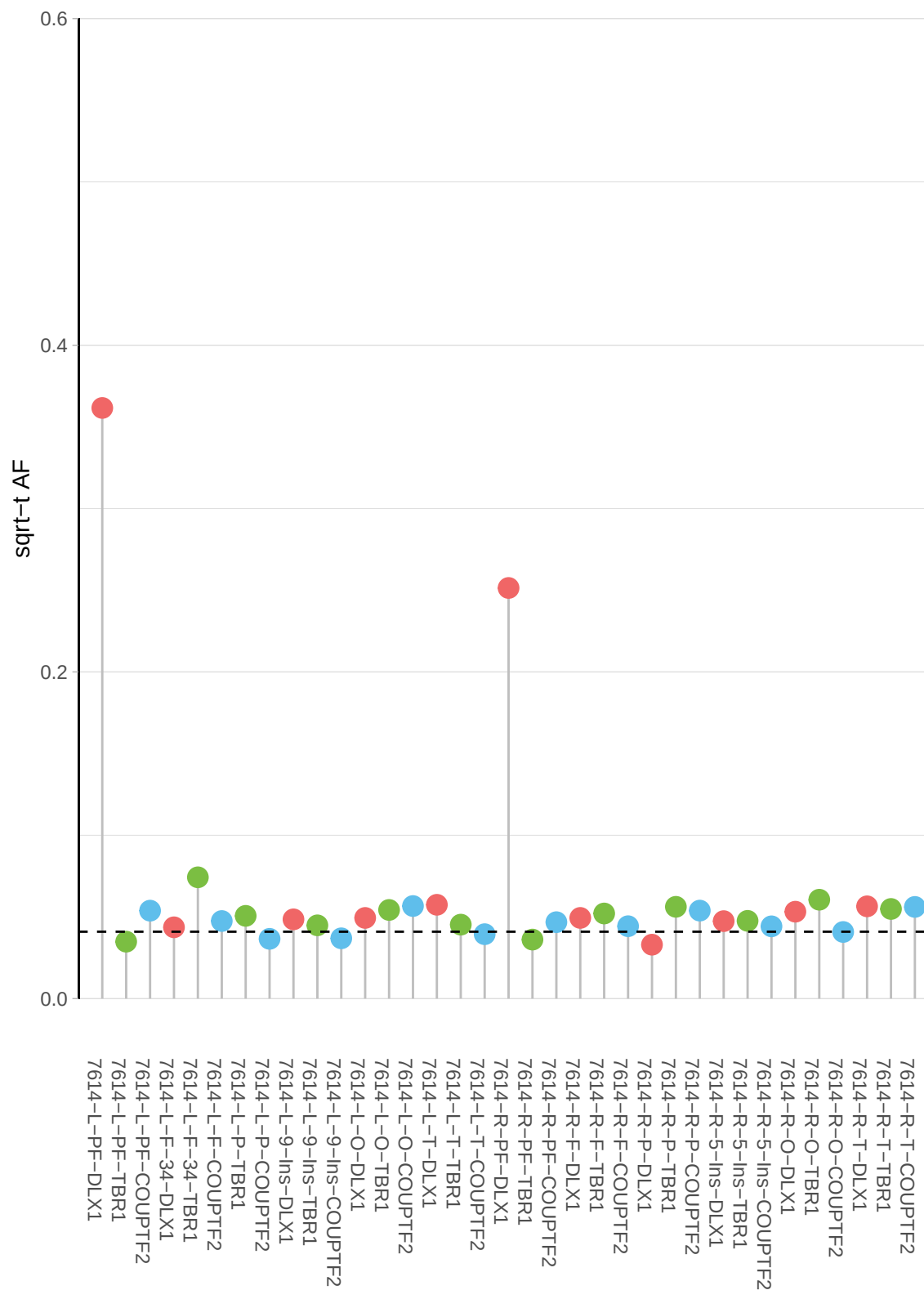

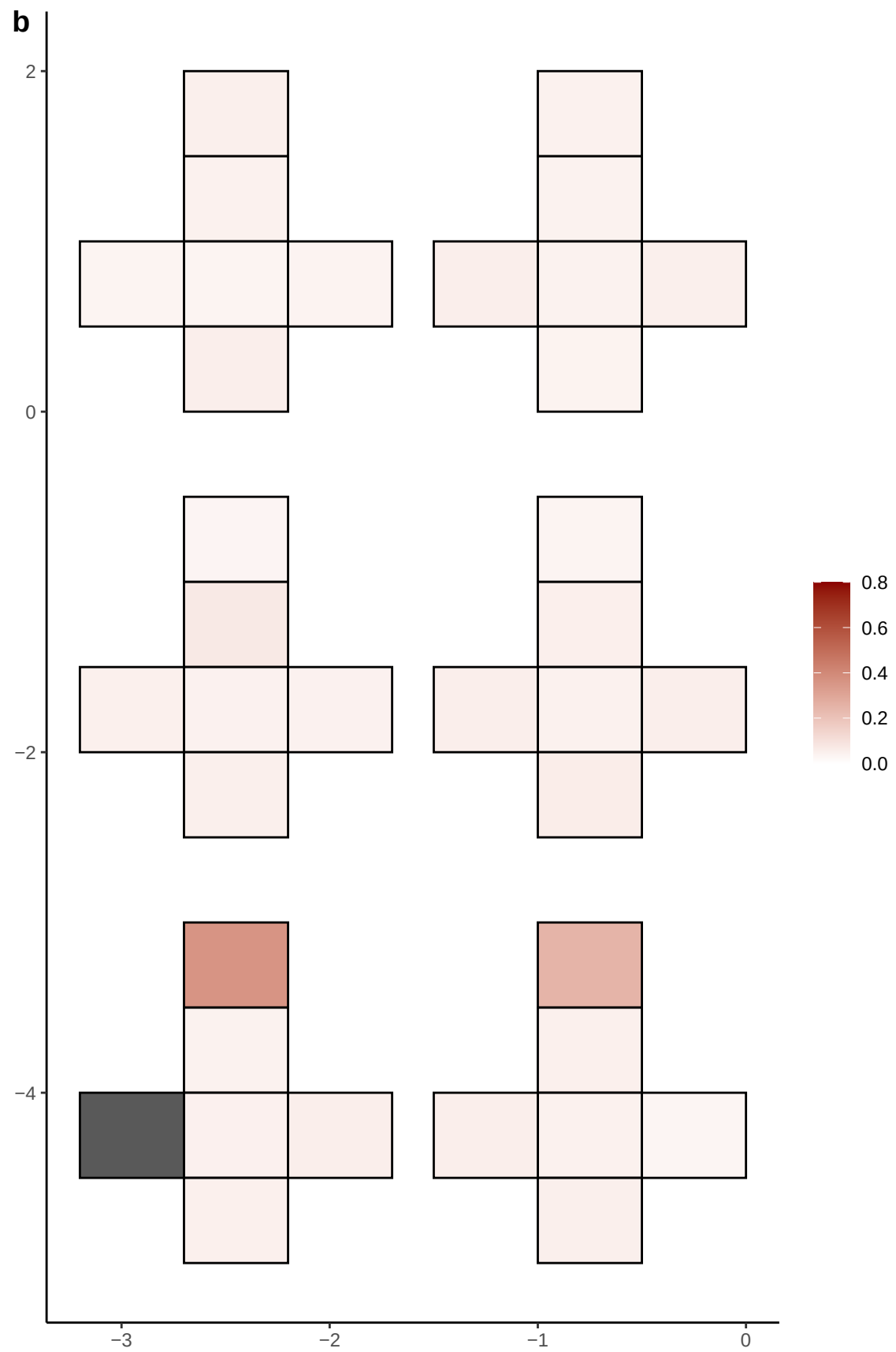

**a** ID01 X-62980946-T-C

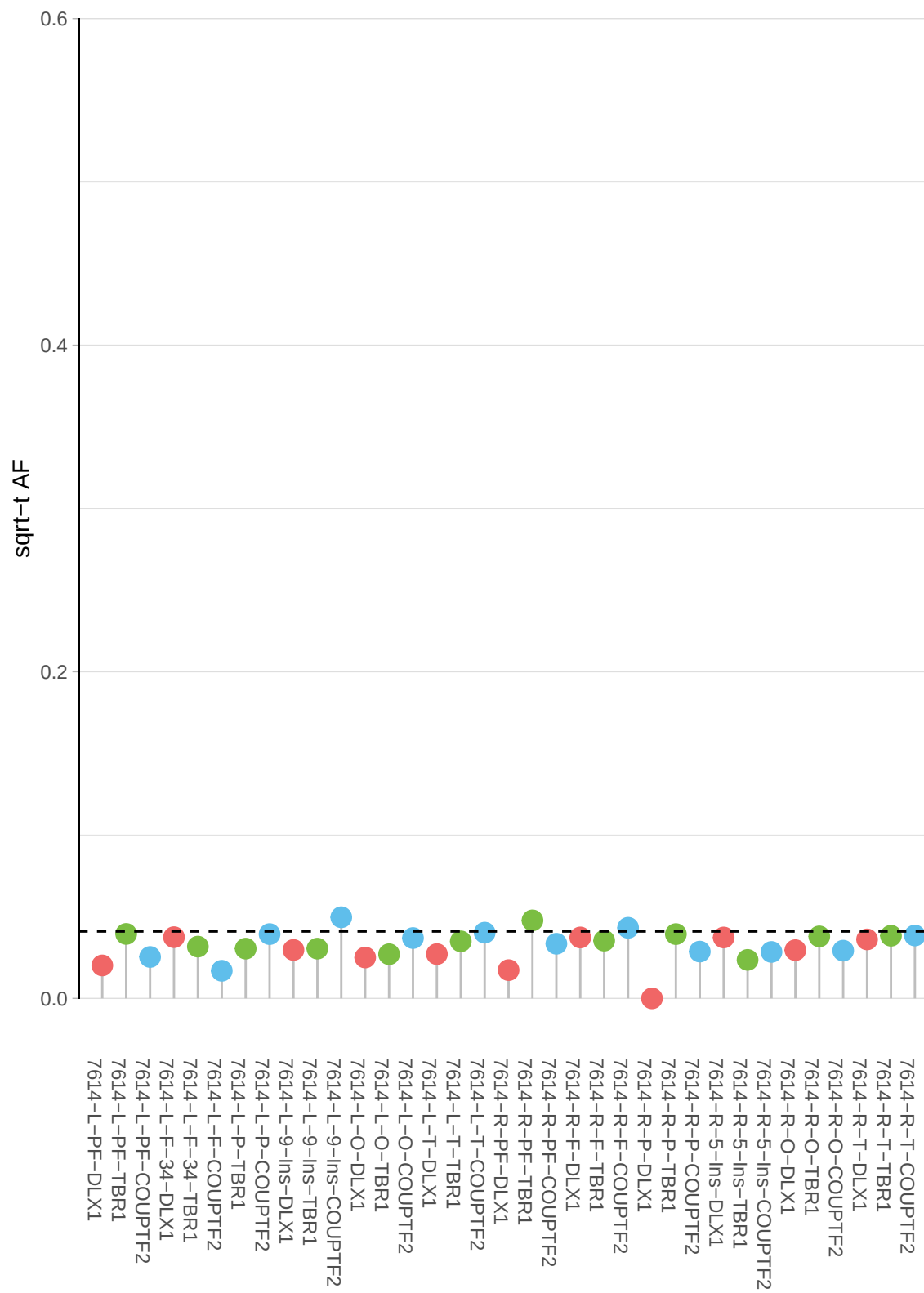

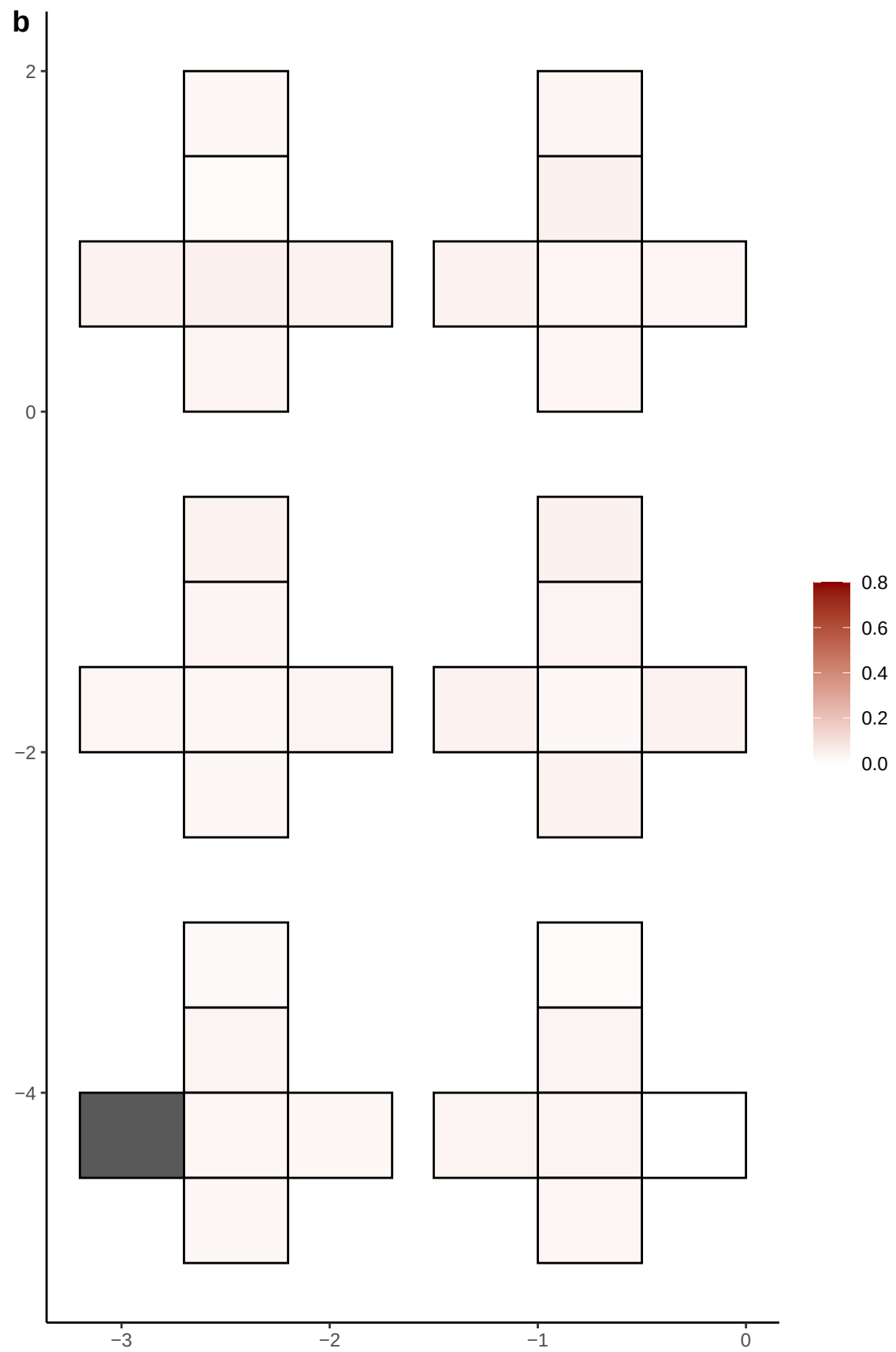

**a** ID01 X-77453233-C-T

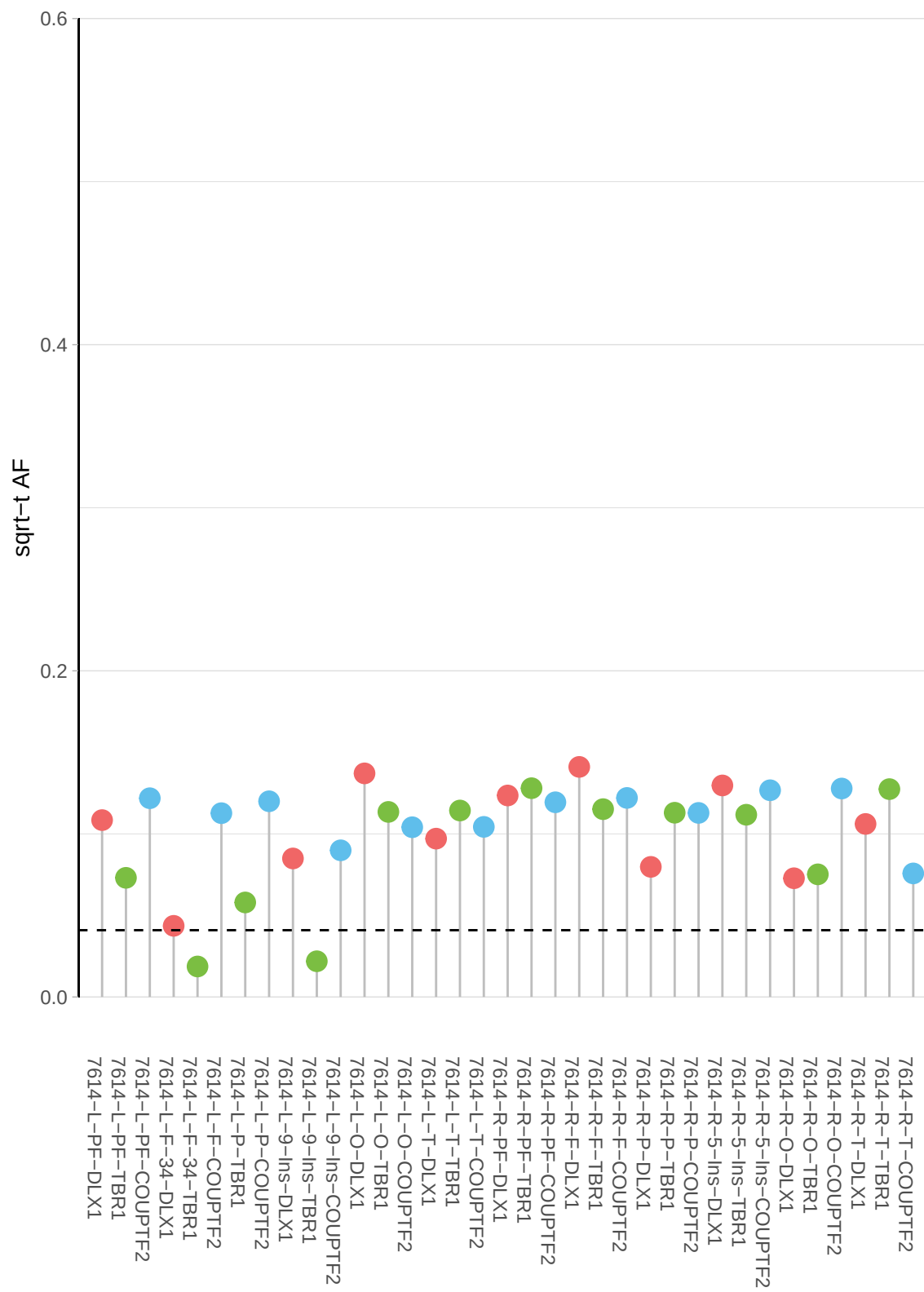

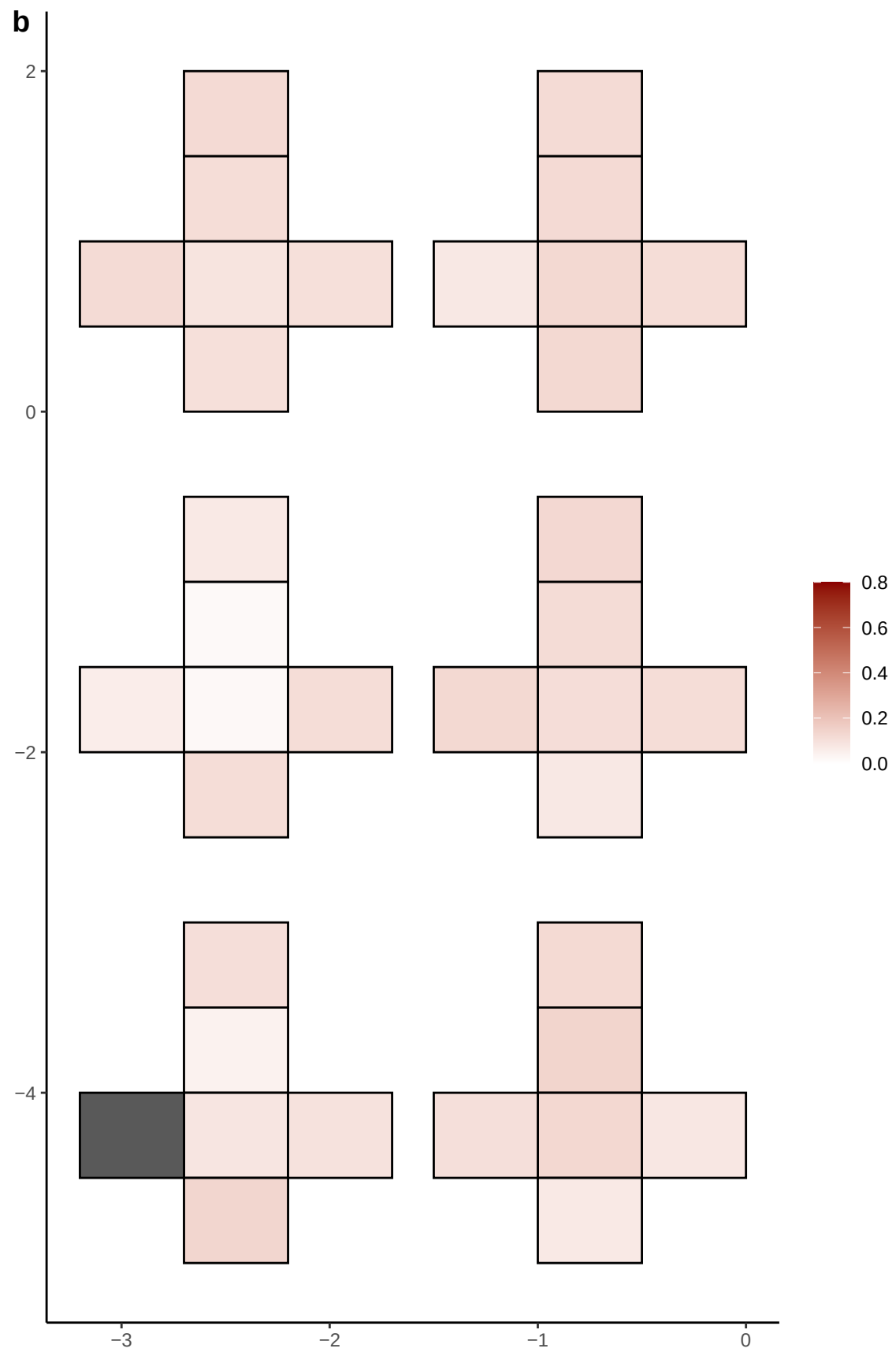

**a**

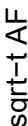

**b**

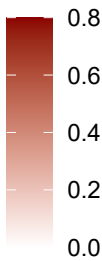

**a** ID05 1-102299198-G-T

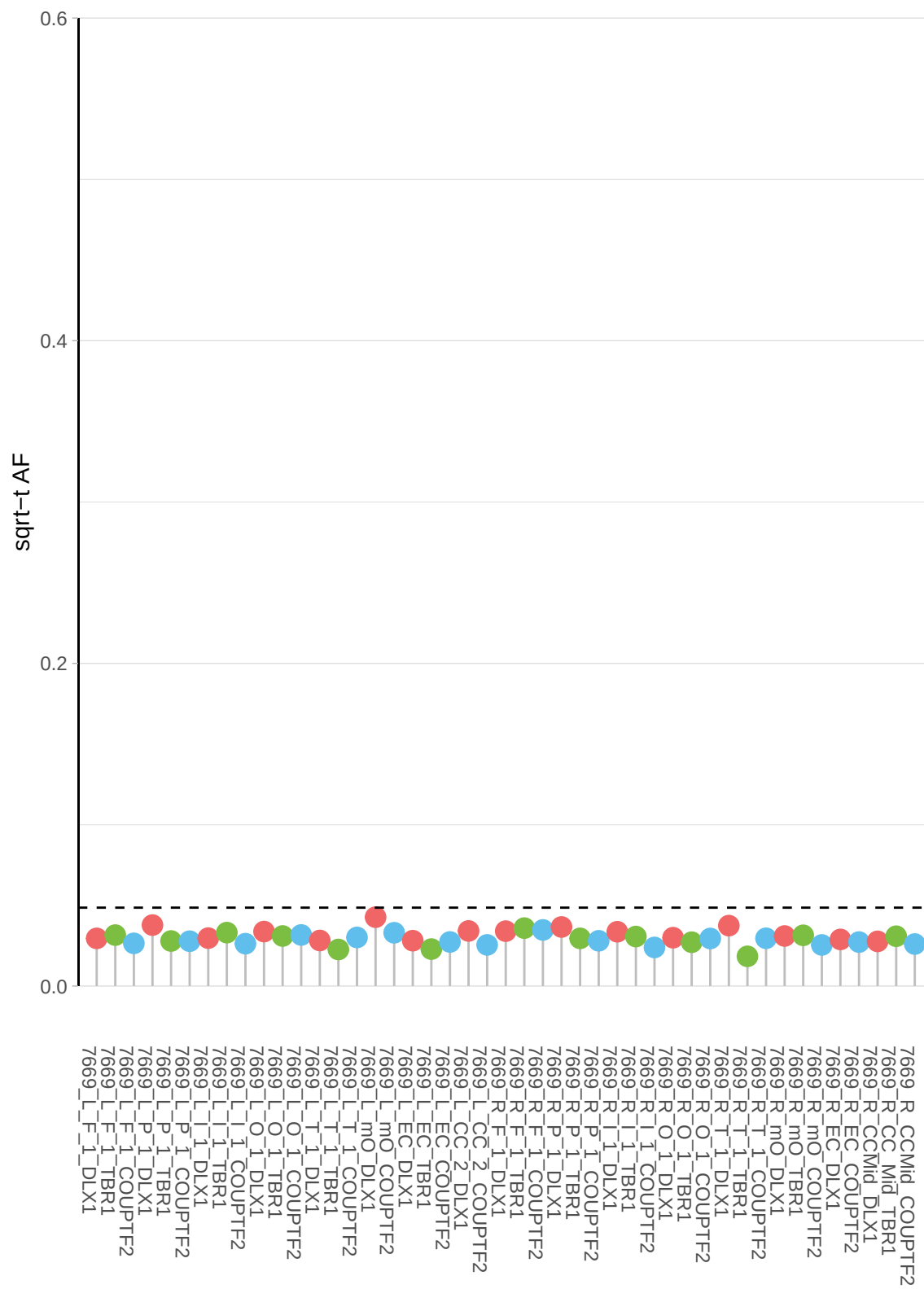

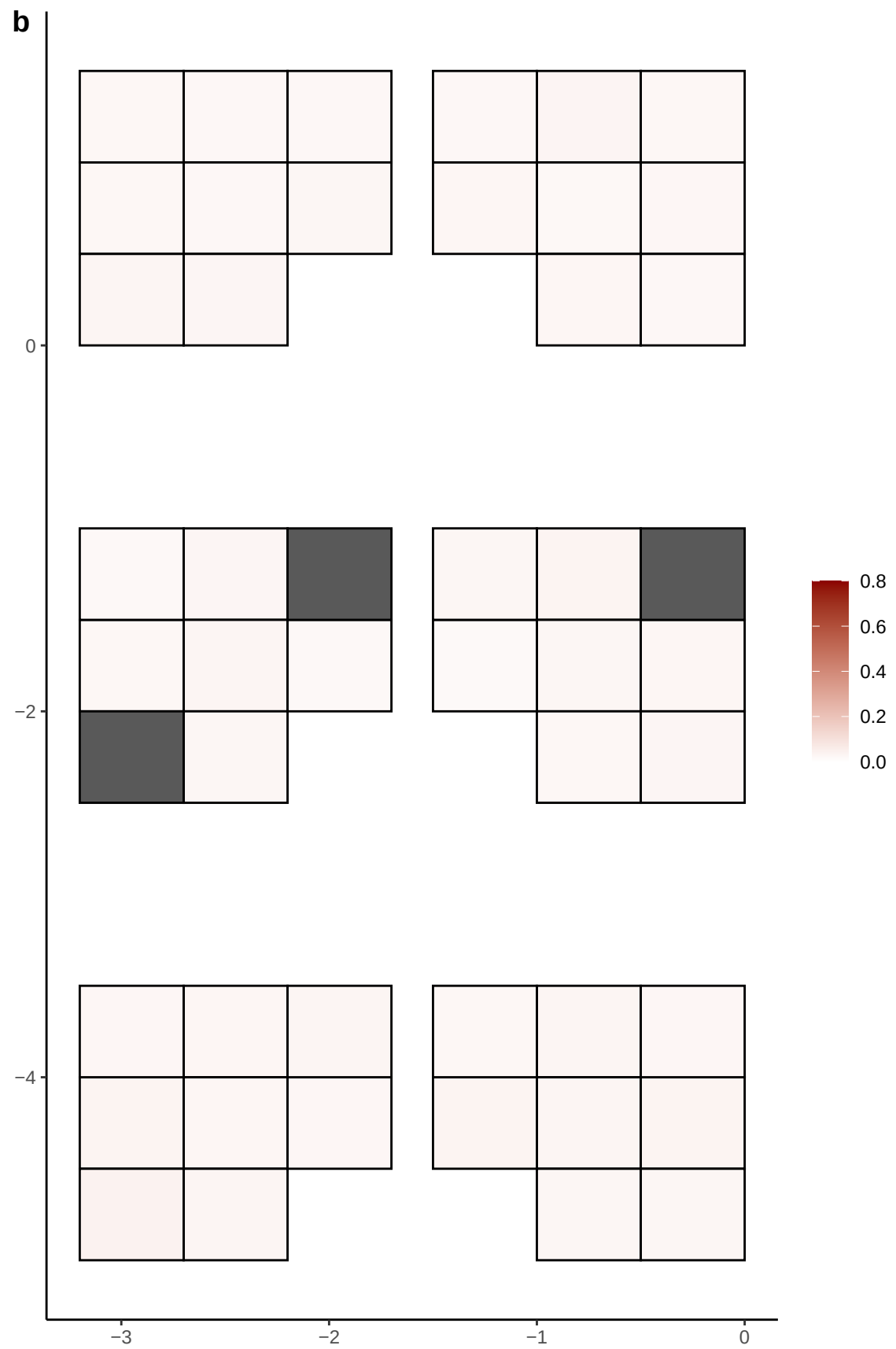

**a** ID05 1-103746984-A-T

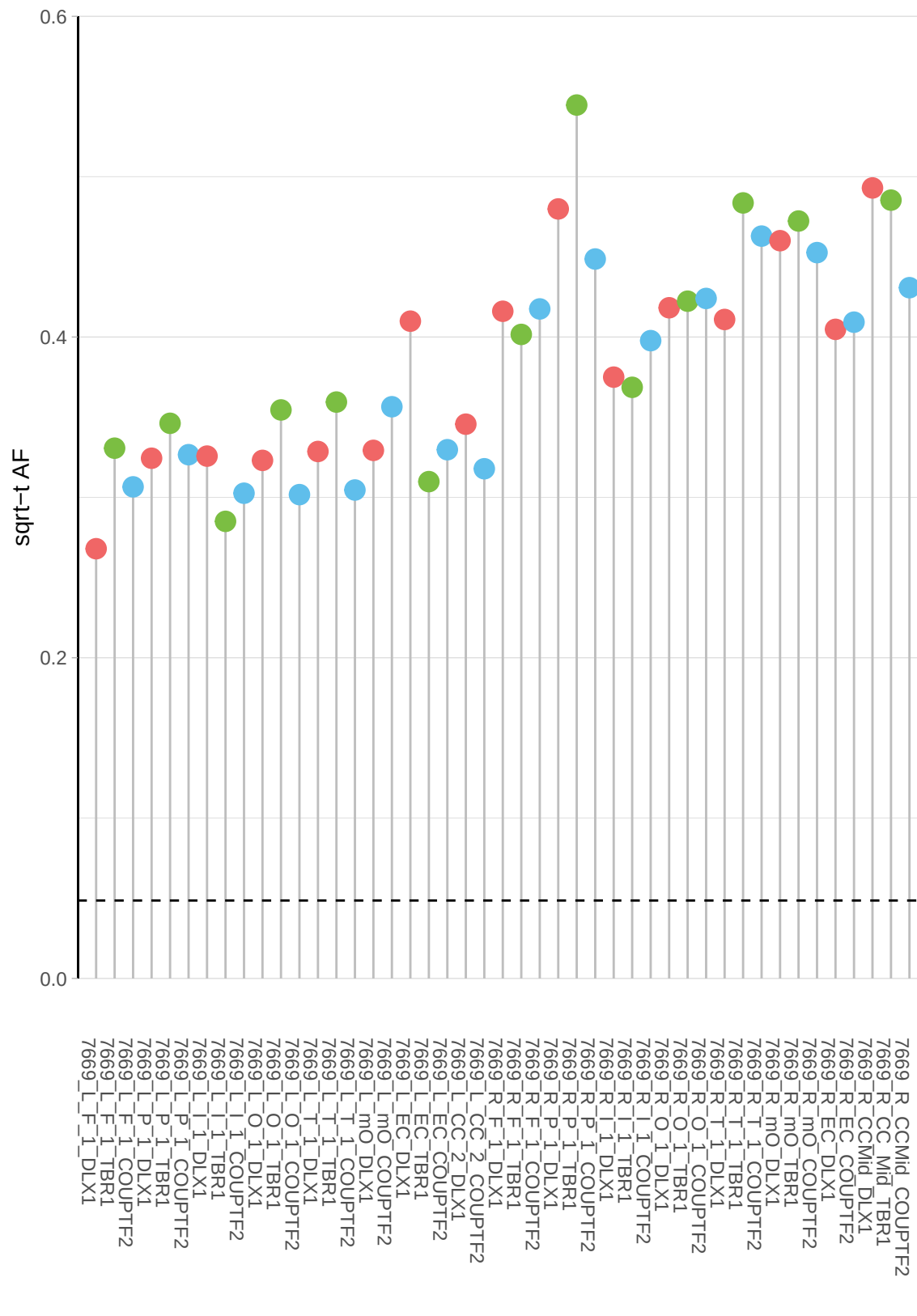

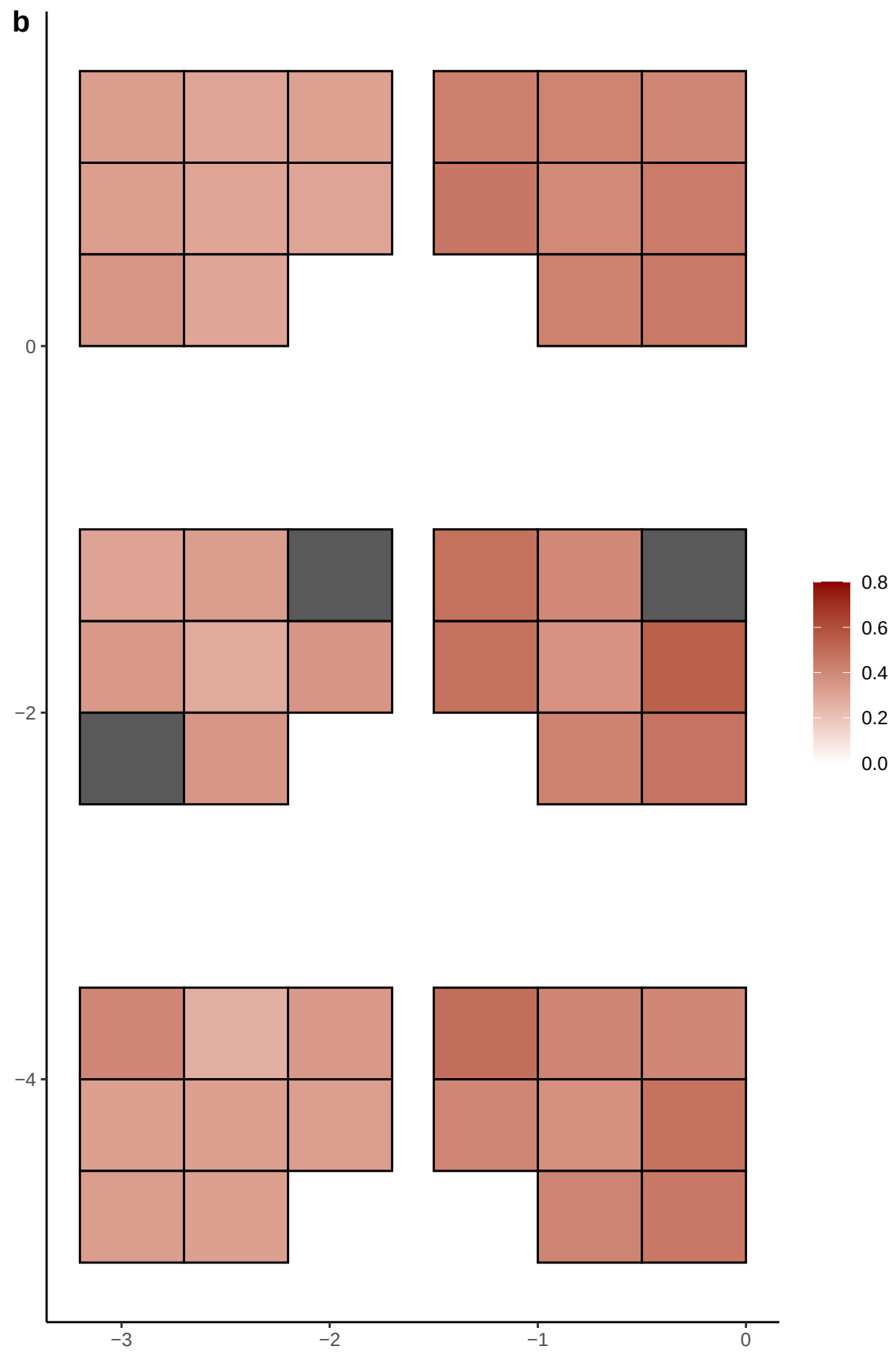

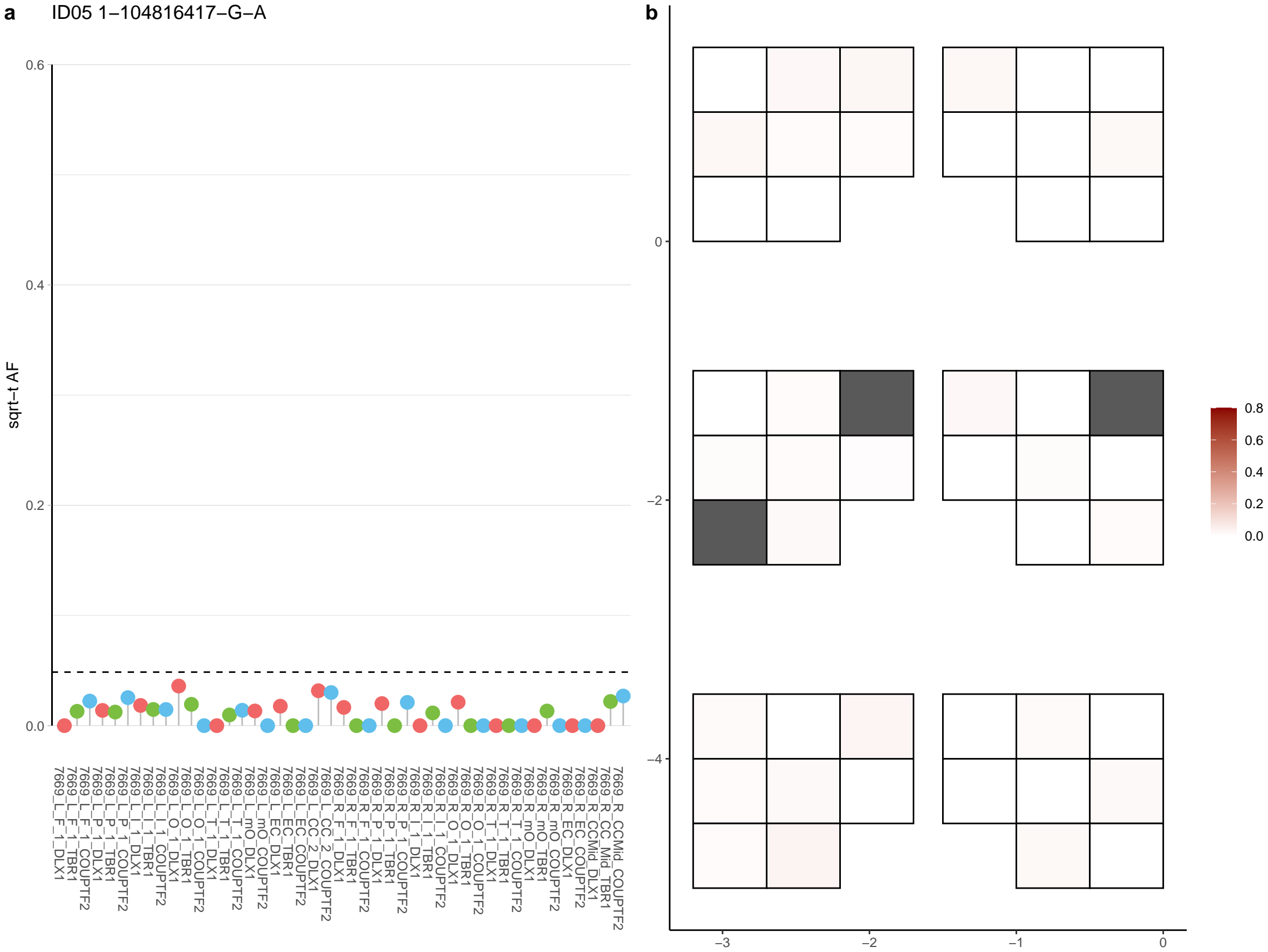

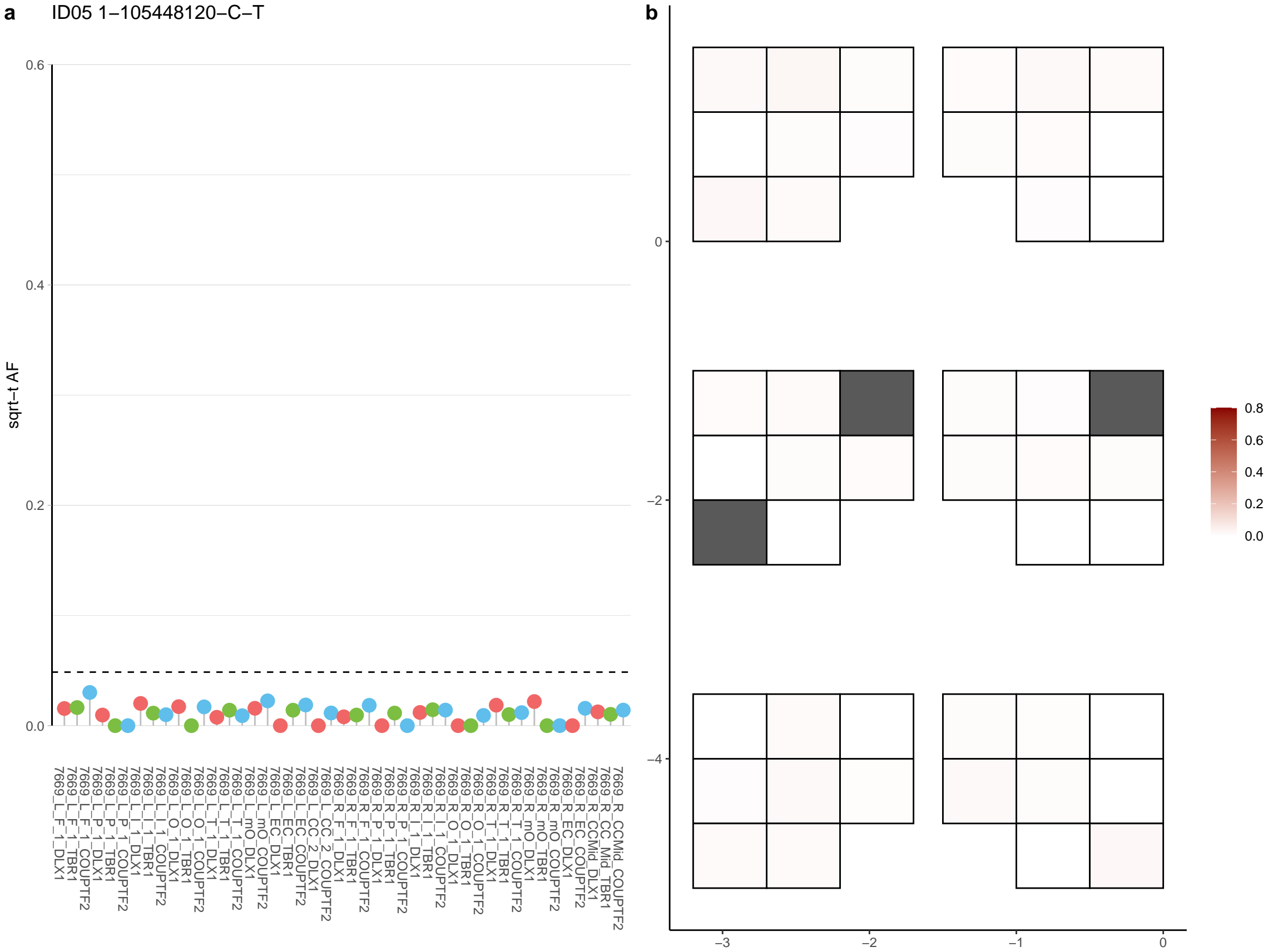

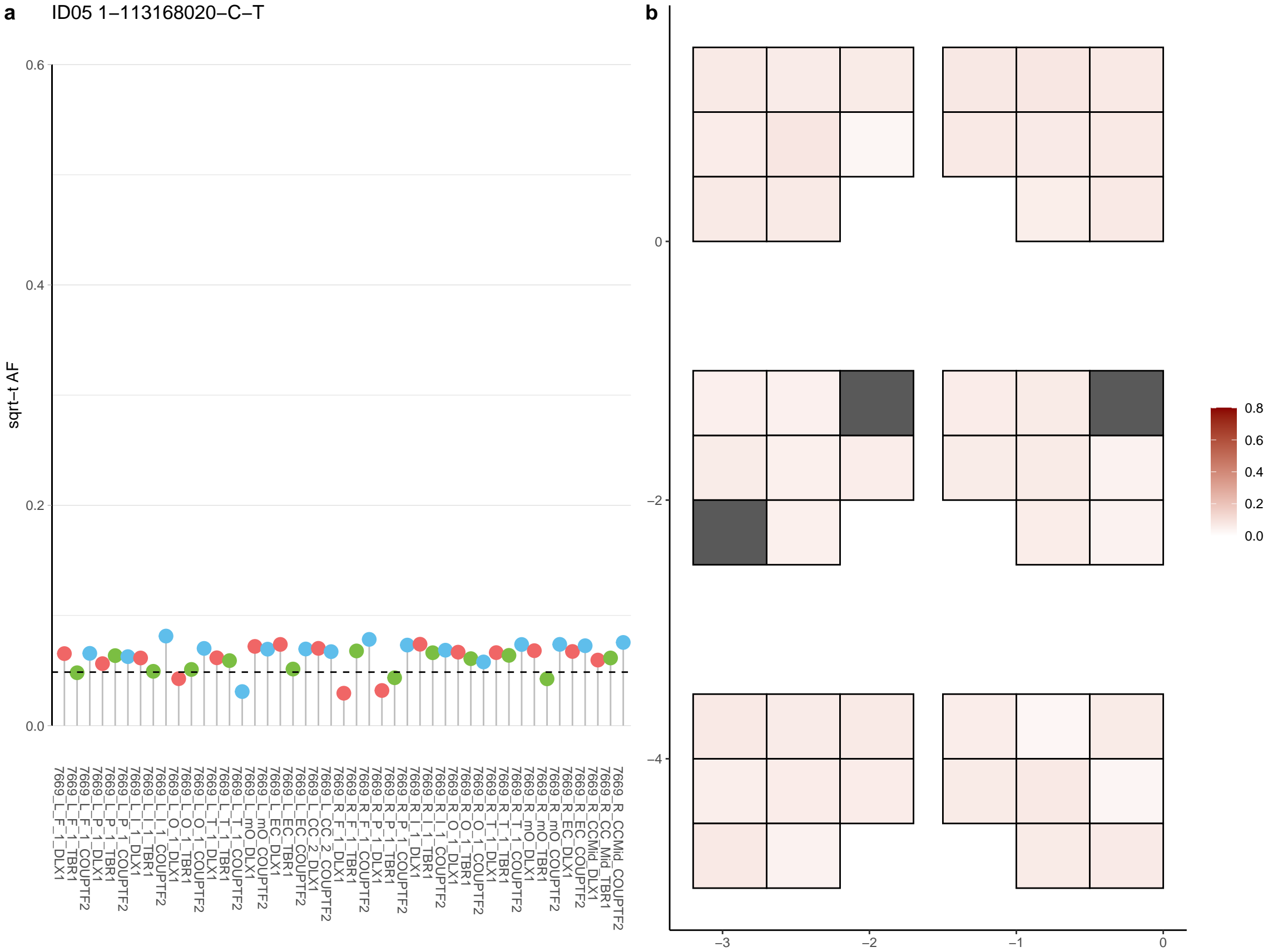

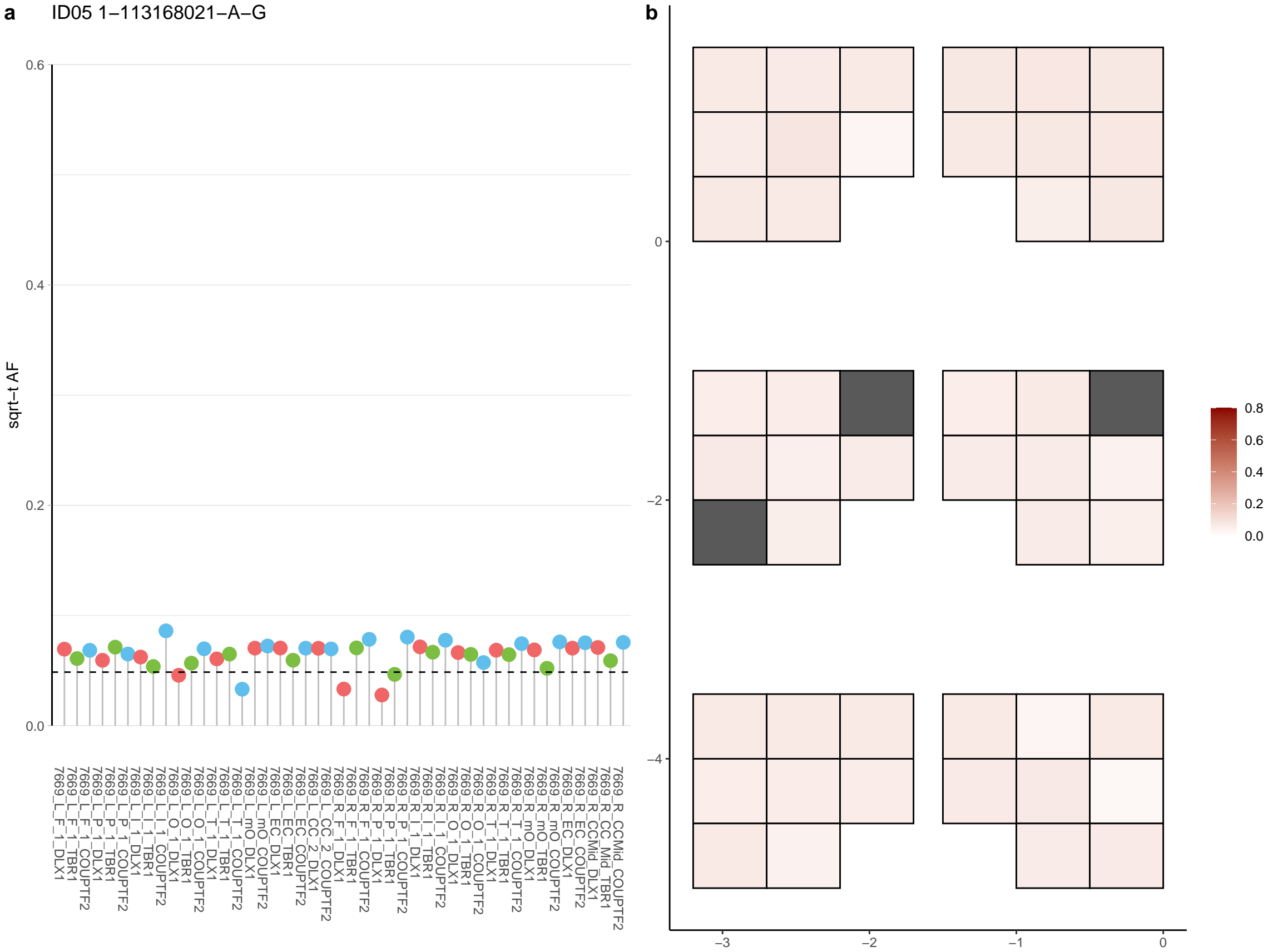

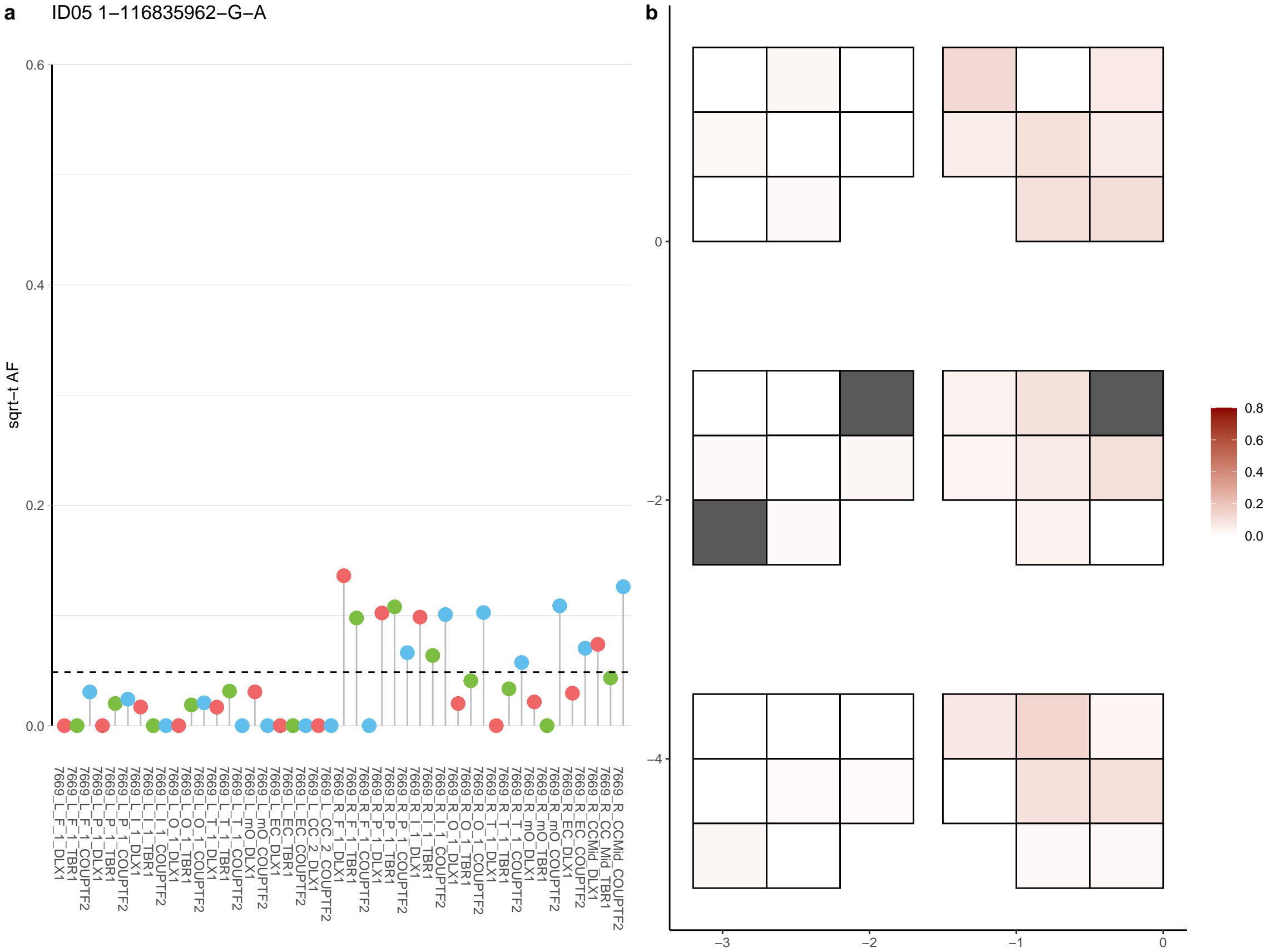

**a** ID05 1-117129159-C-T

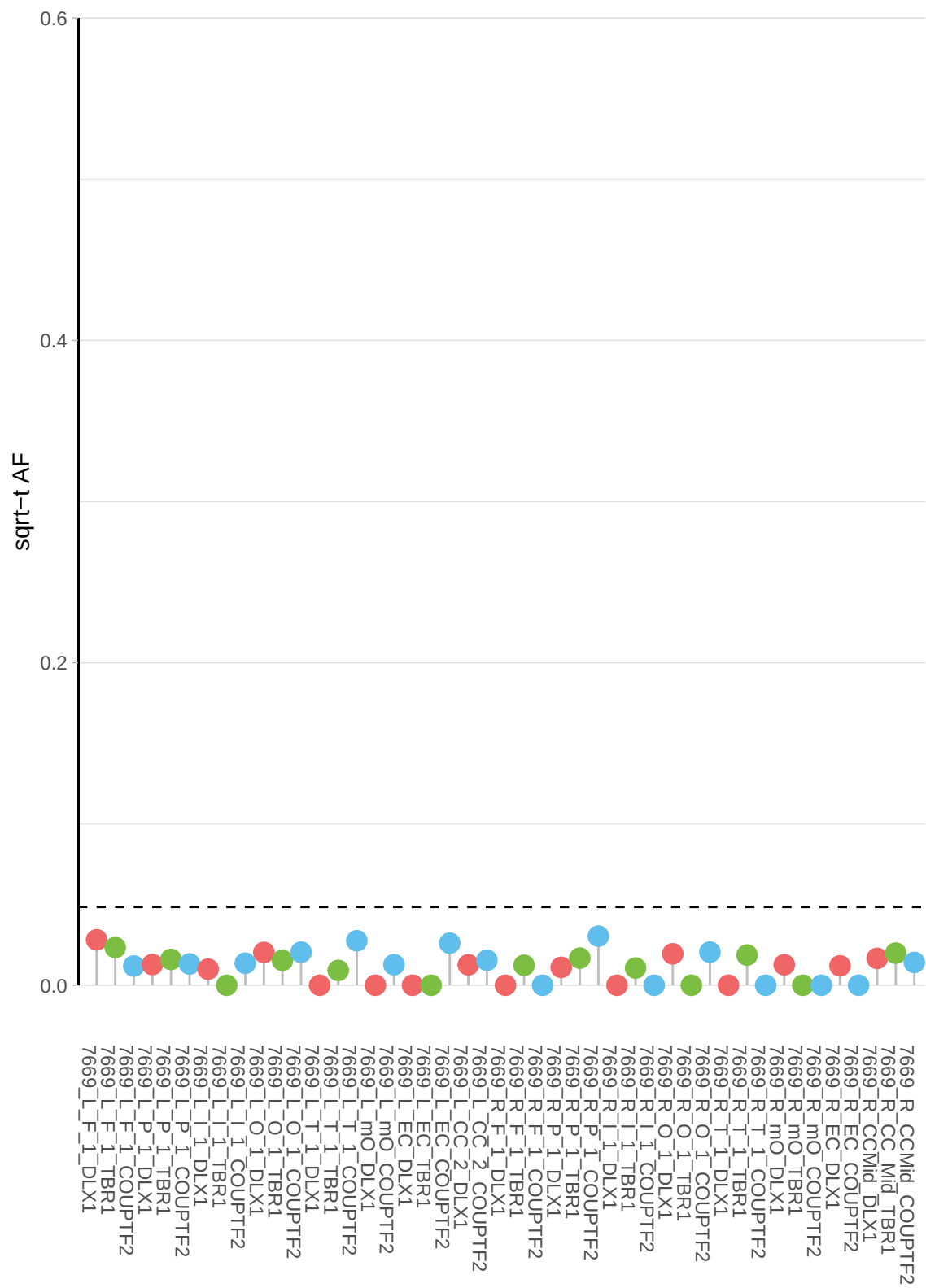

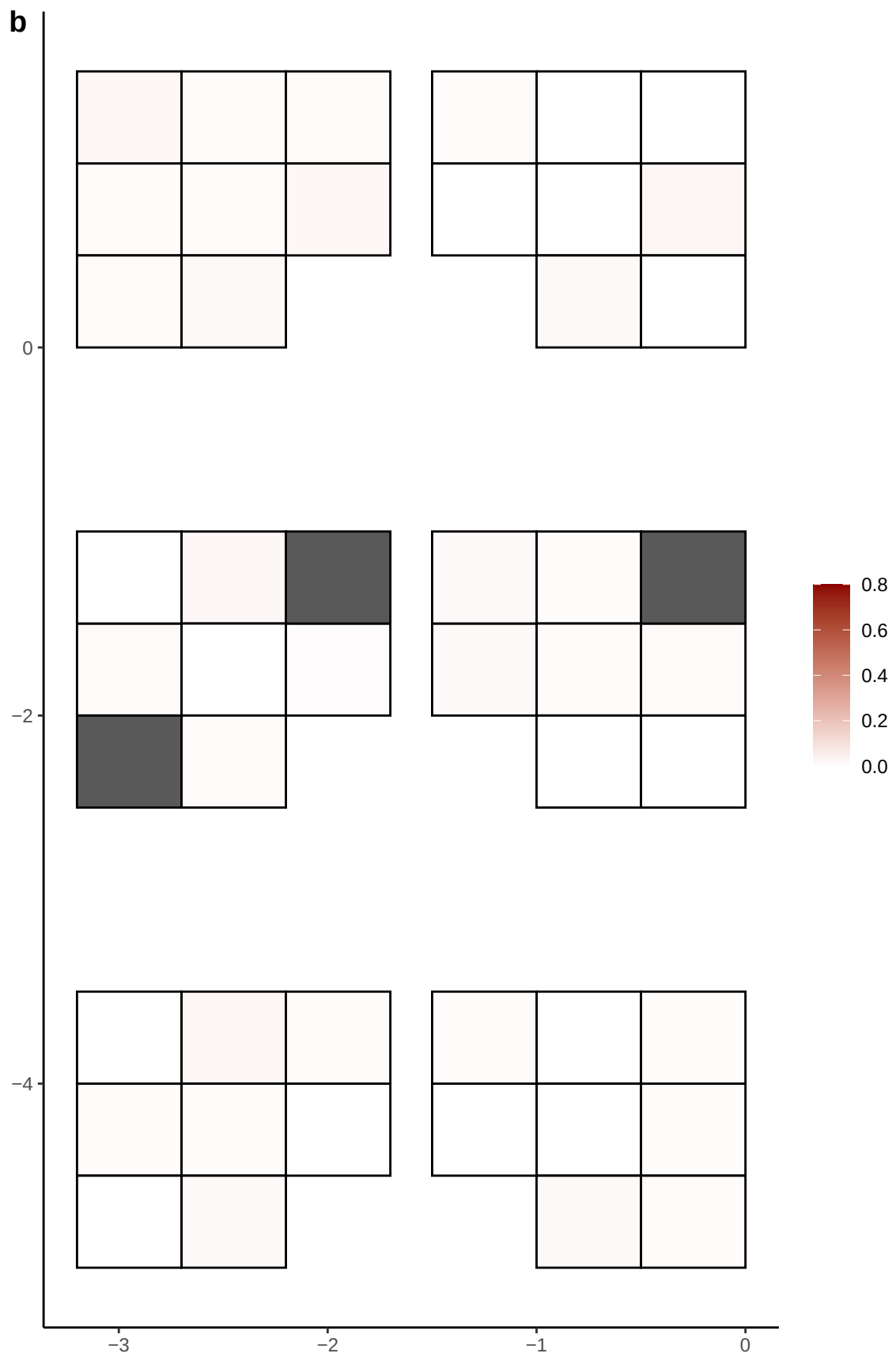

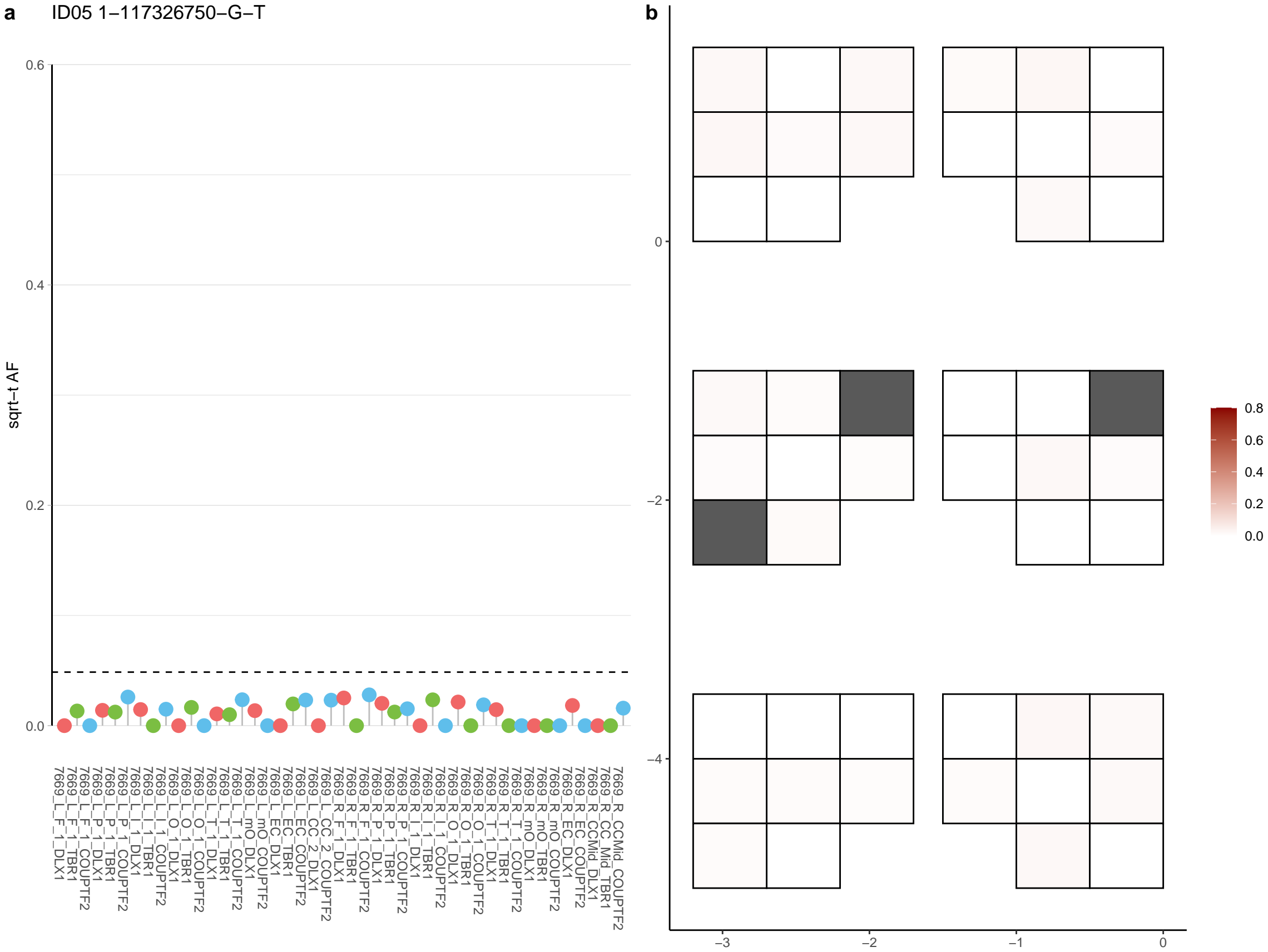

**a** ID05 1-1196724-A-T

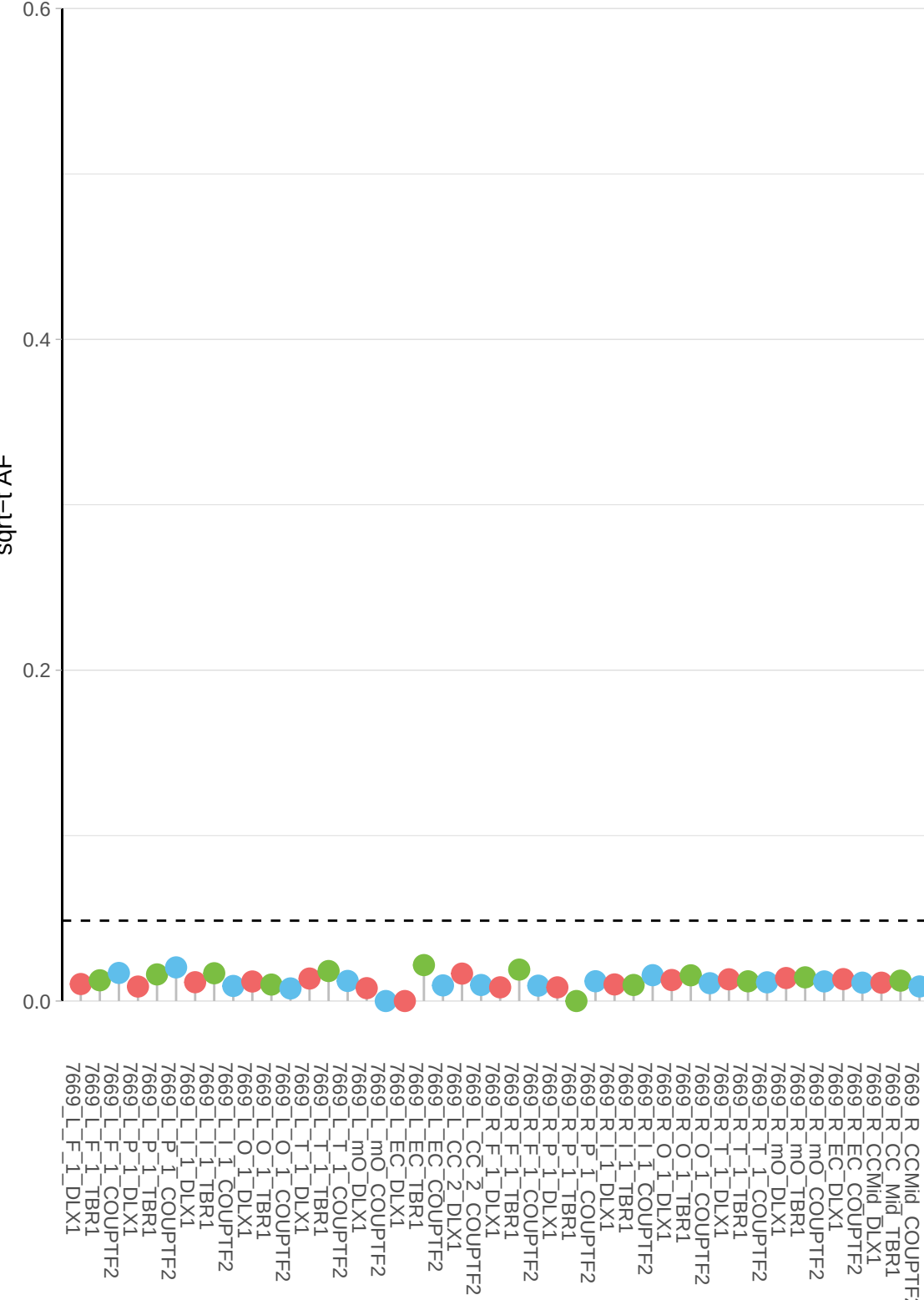

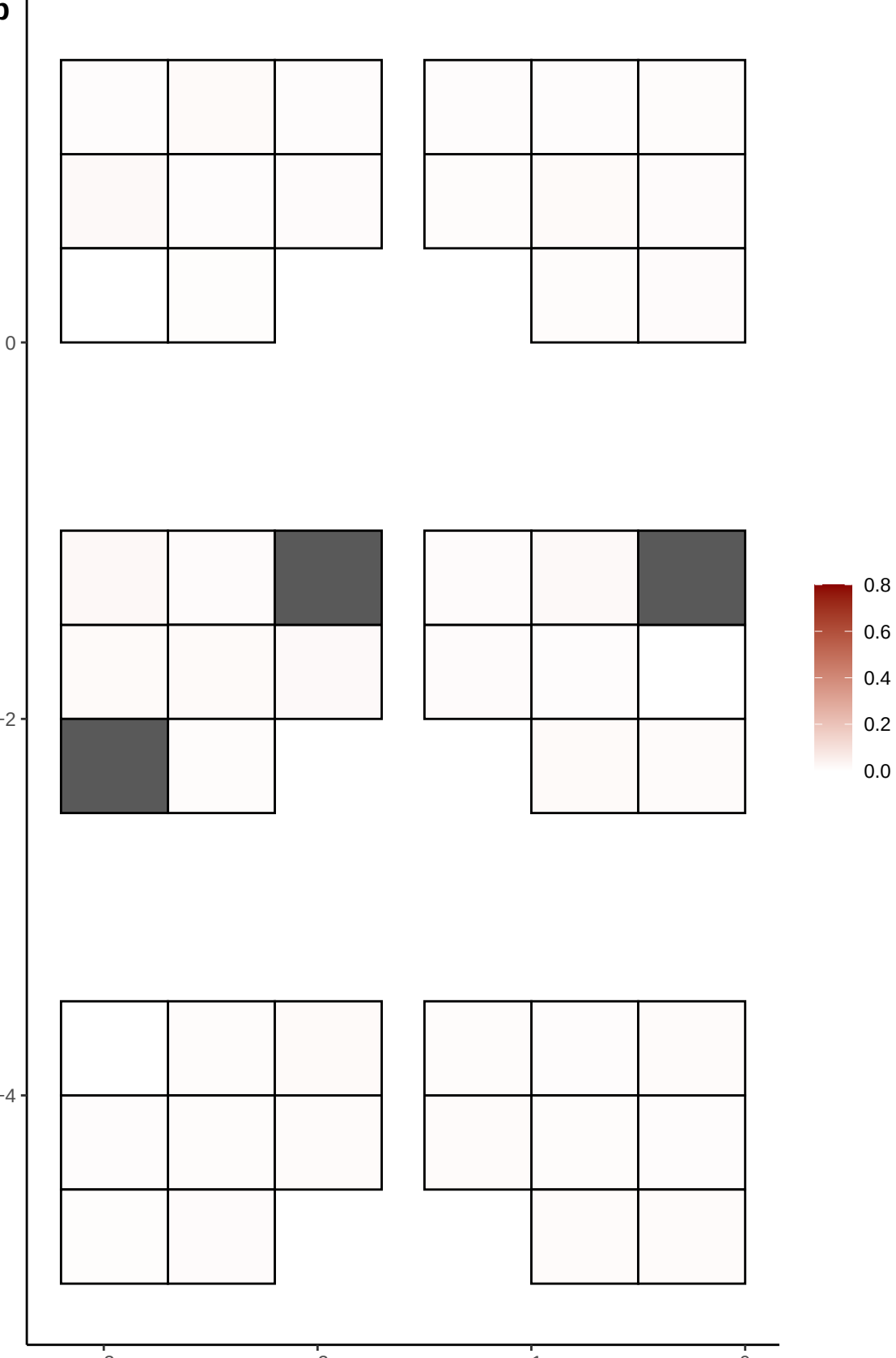

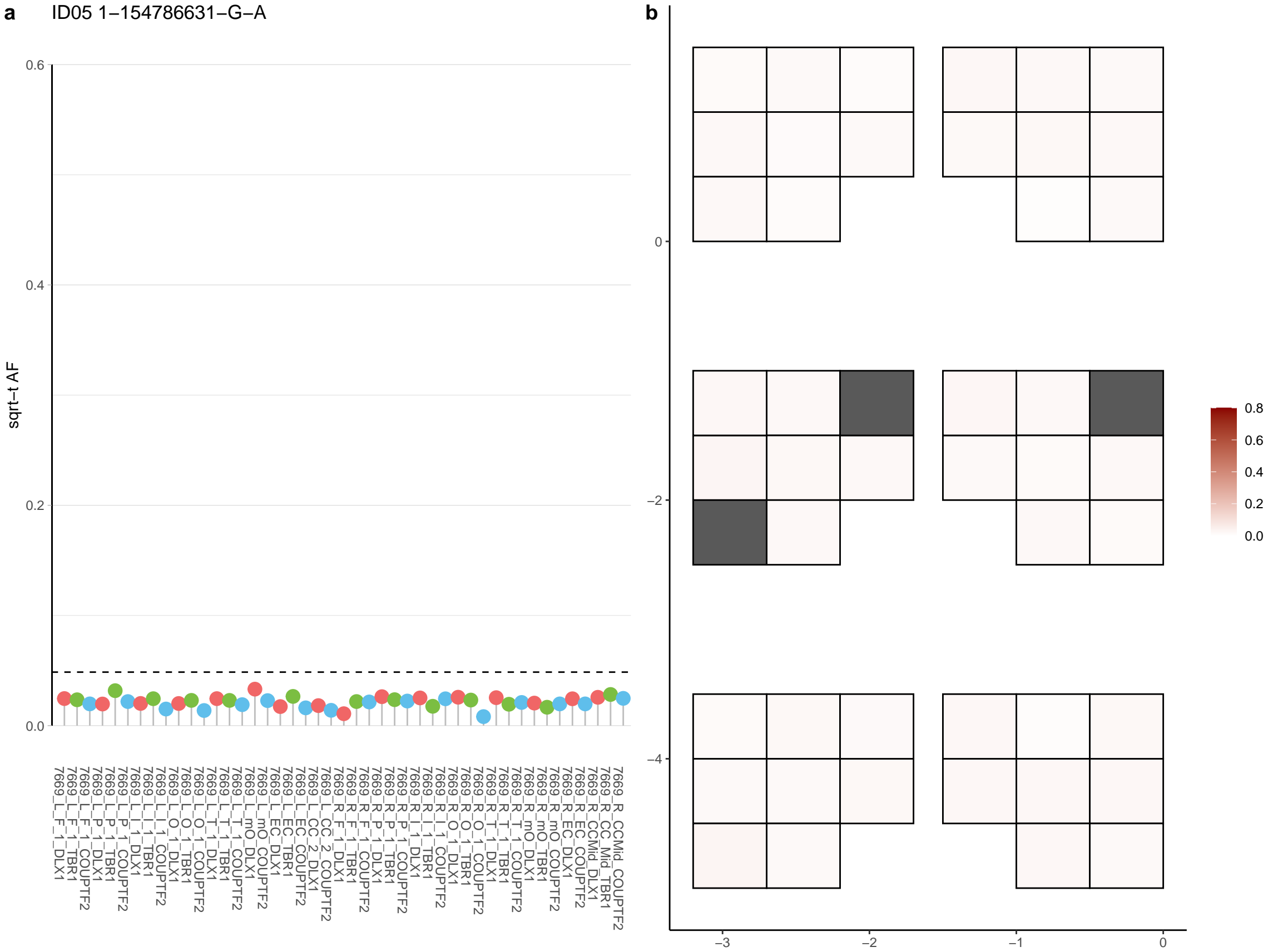

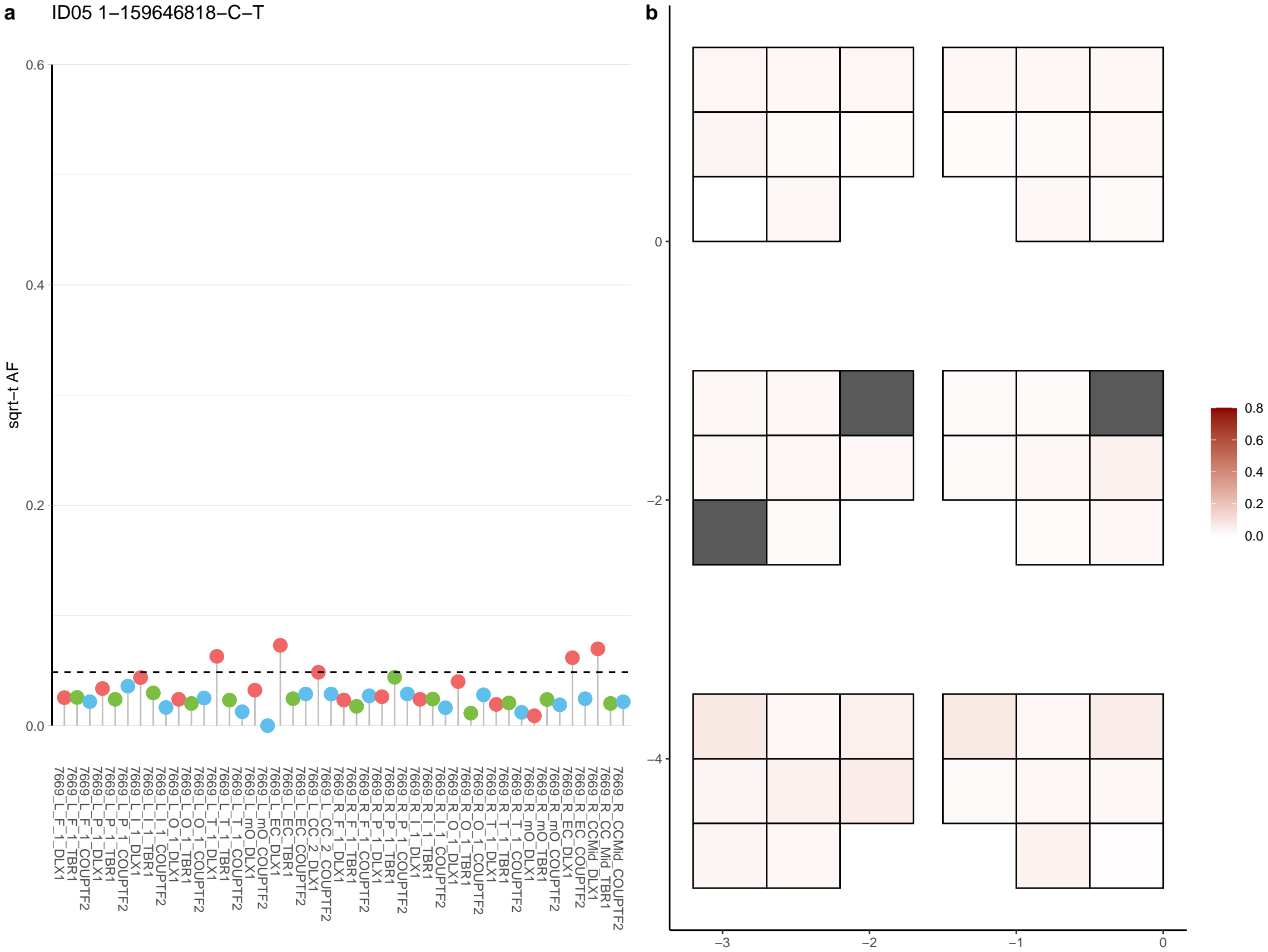

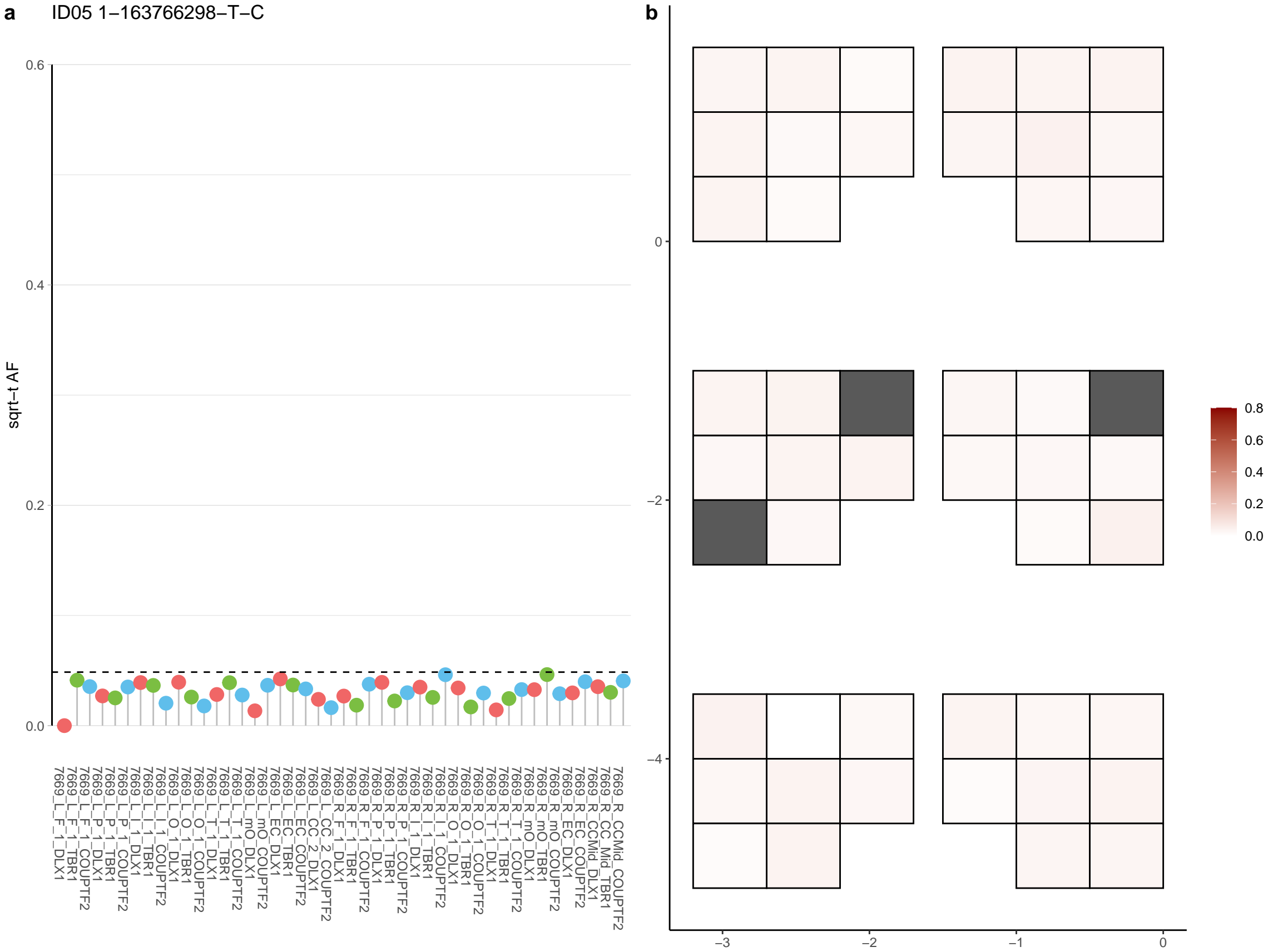

**a** ID05 1-163770532-T-A

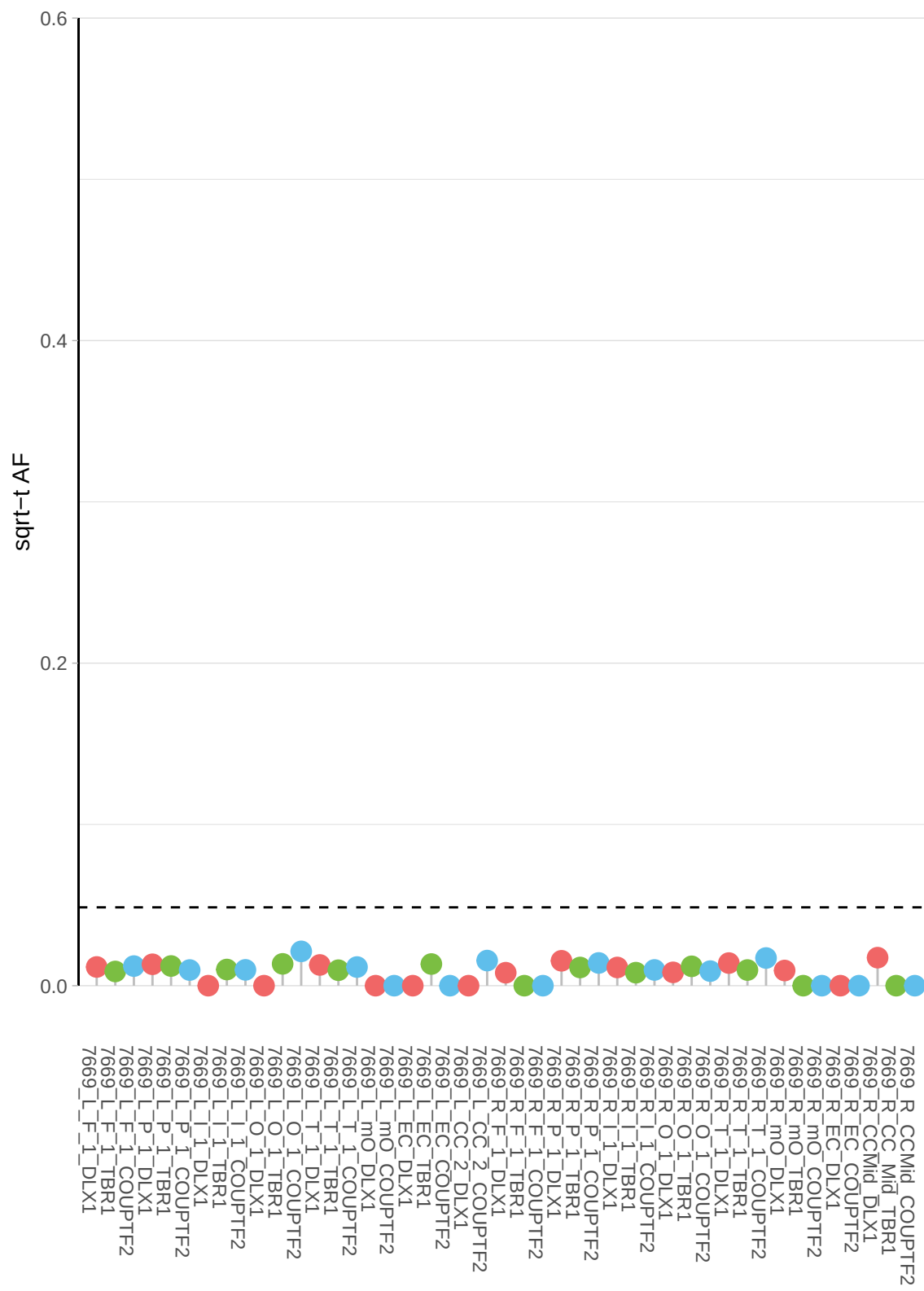

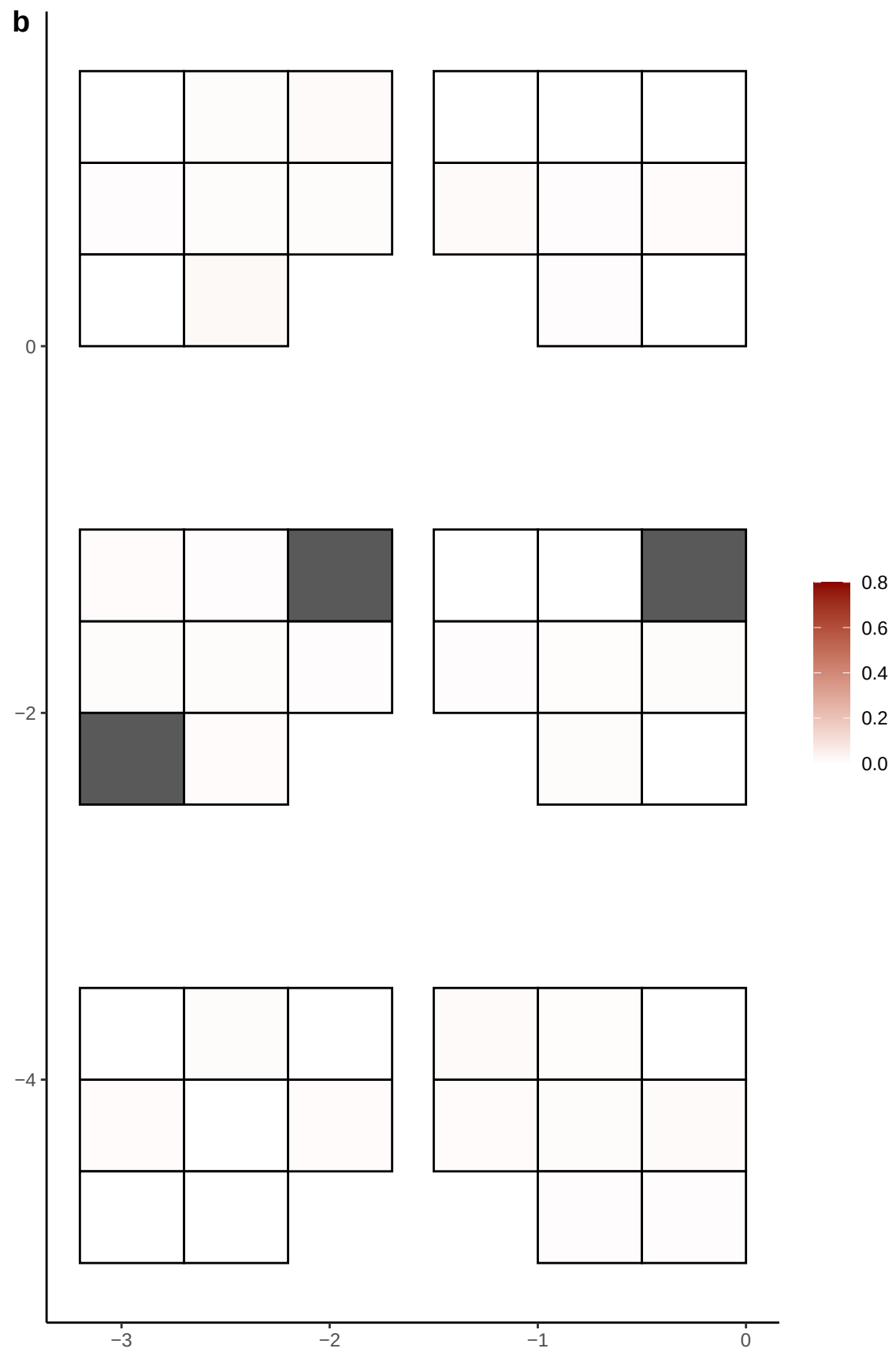

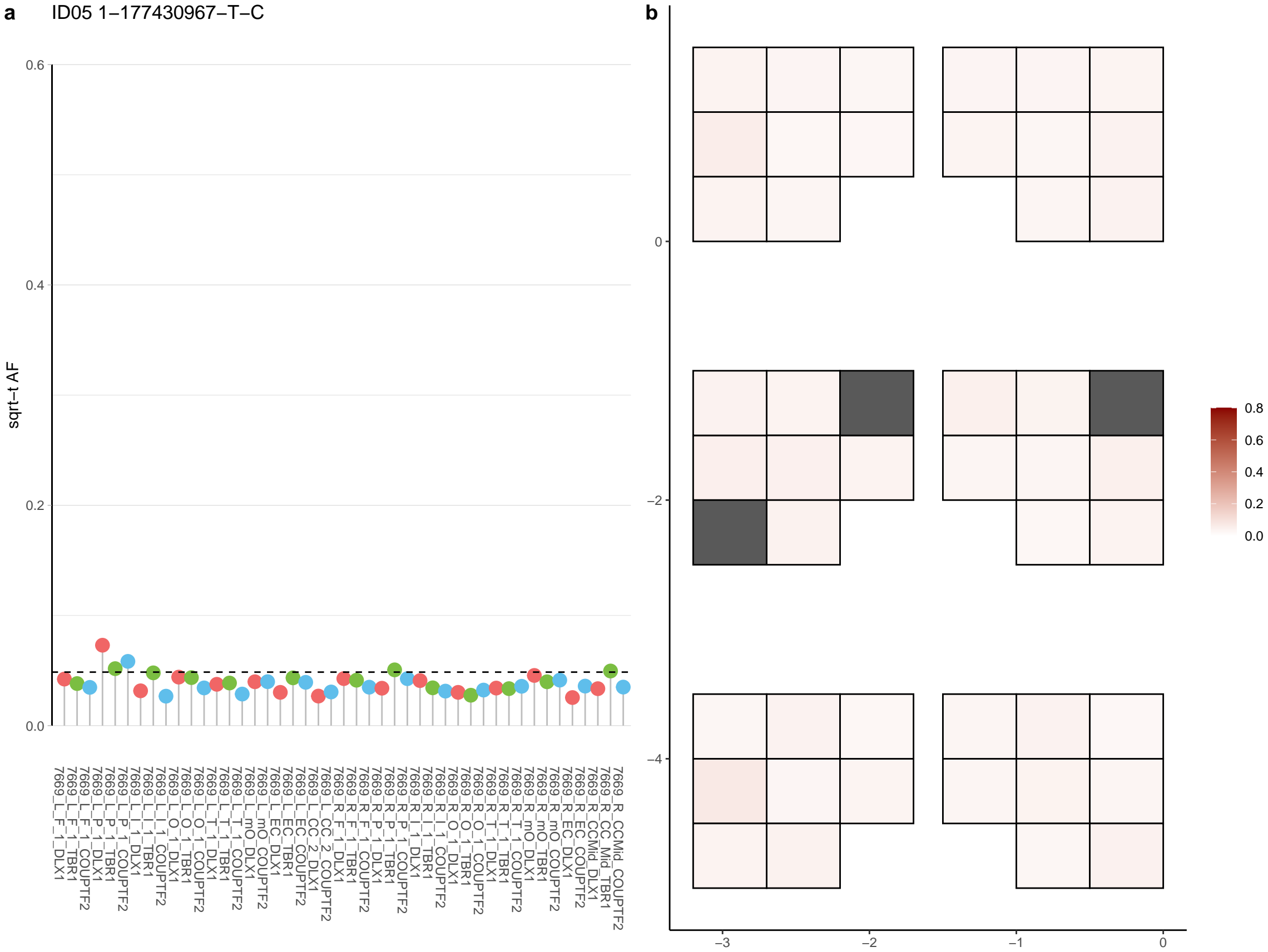

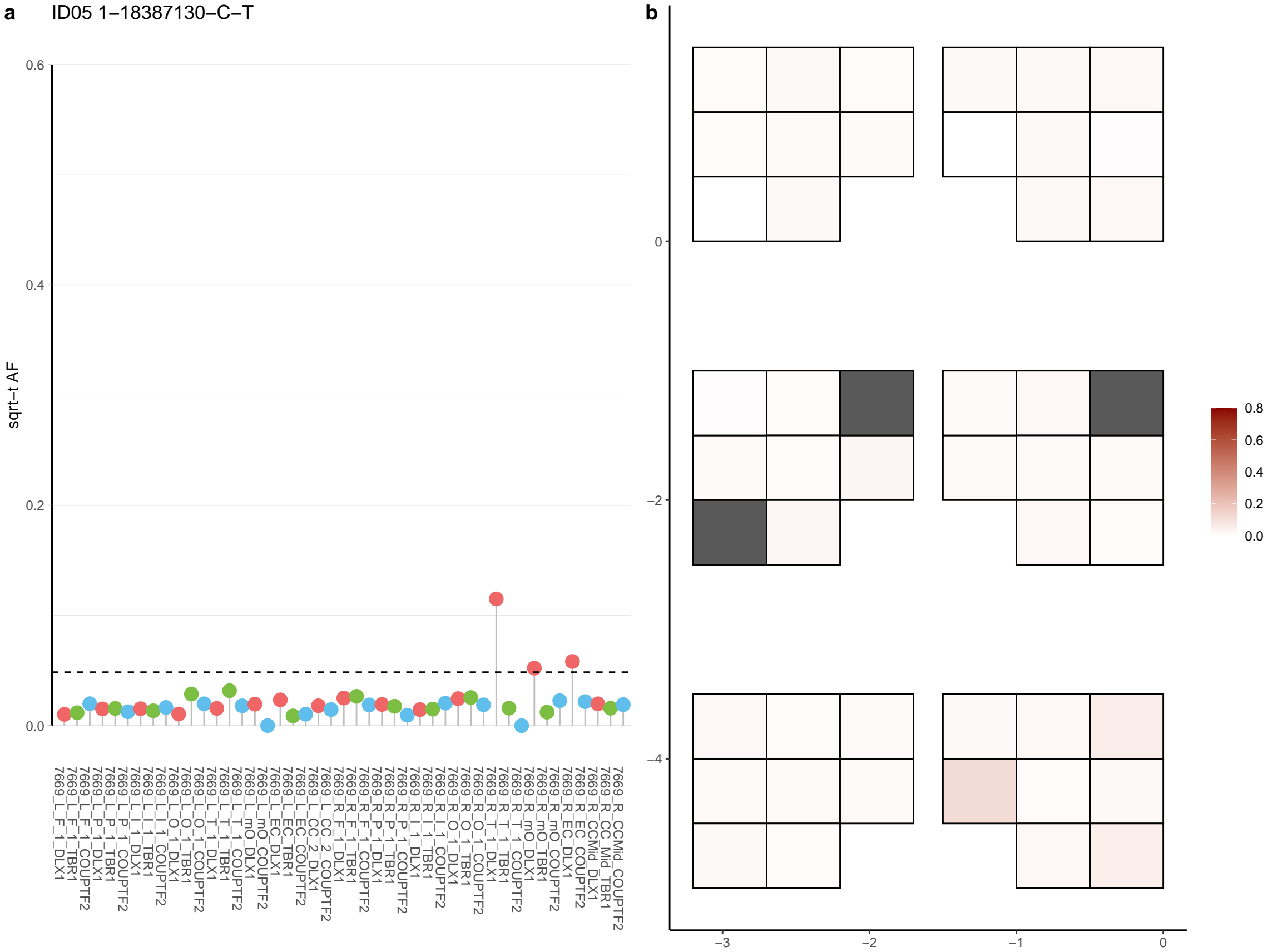

**a** ID05 1-18400310-C-T

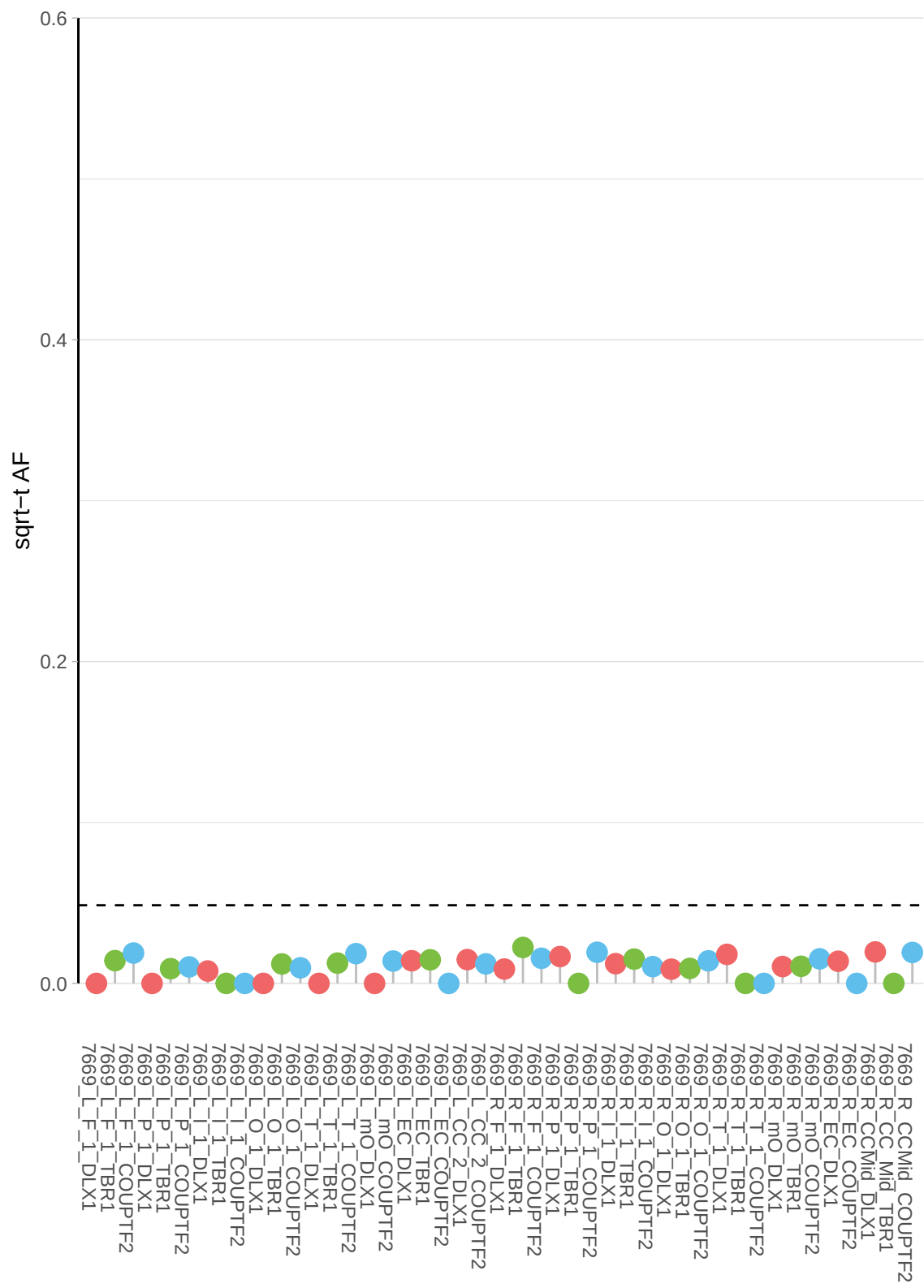

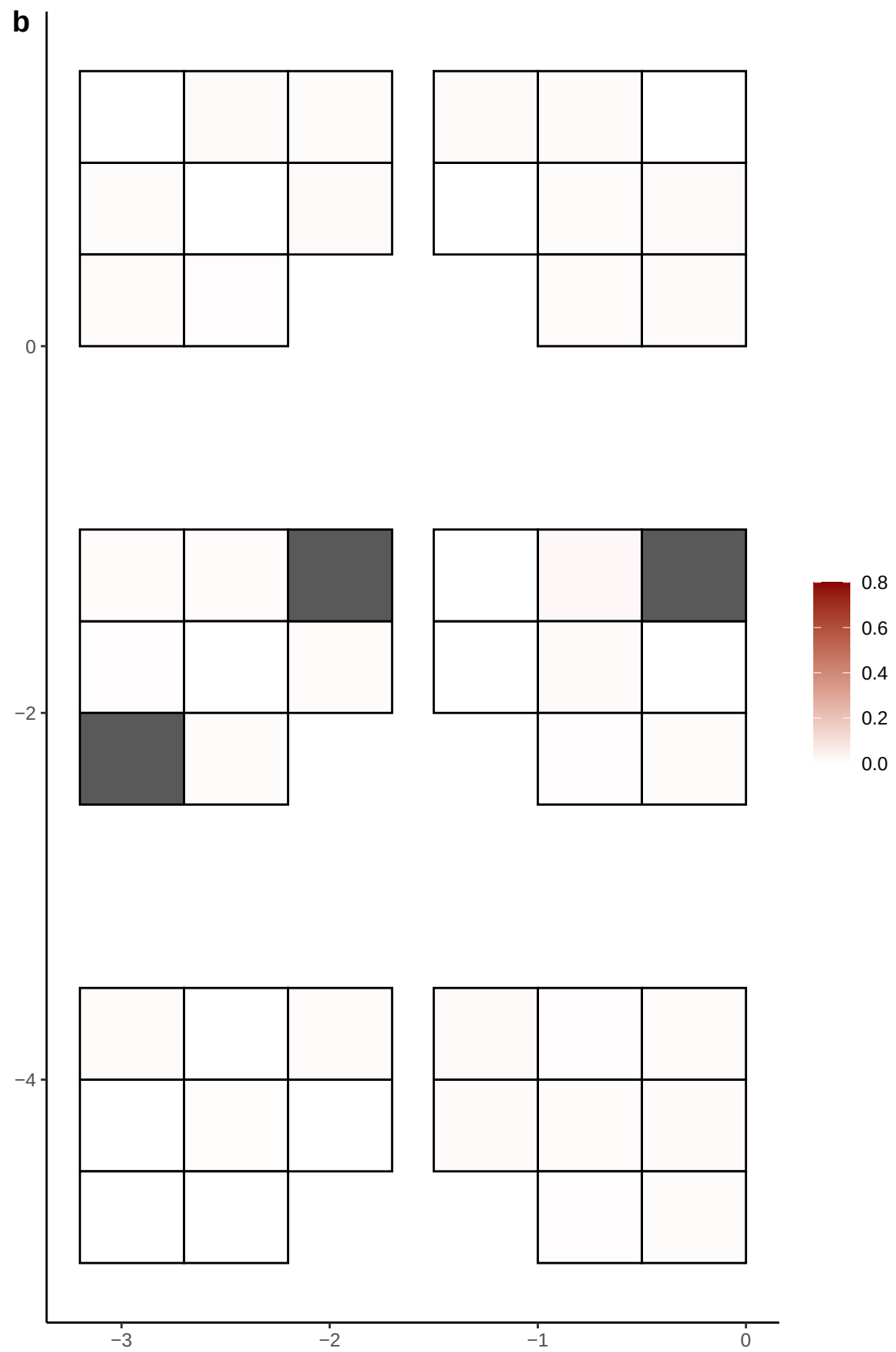

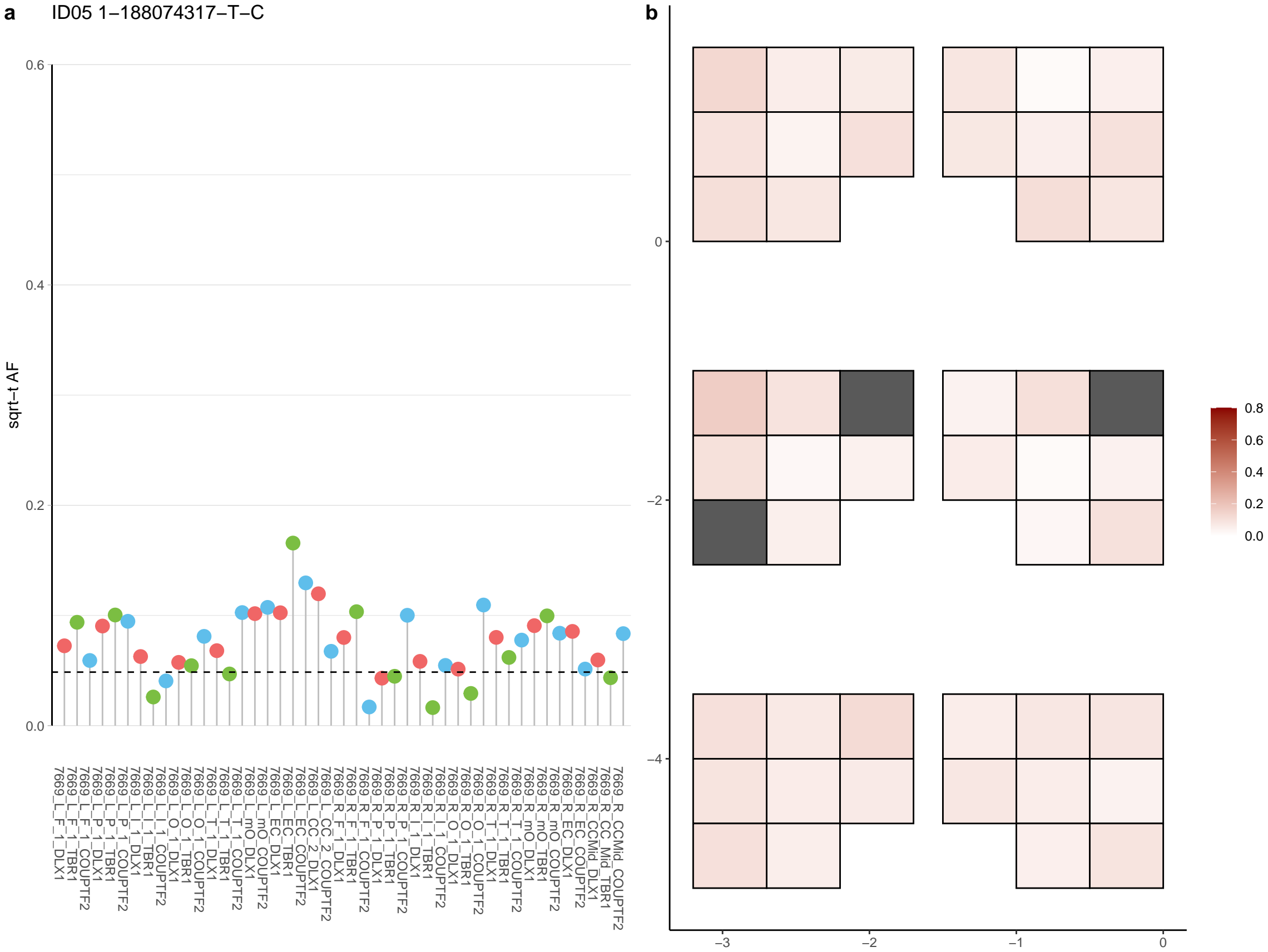

**a** ID05 1-191248571-G-A

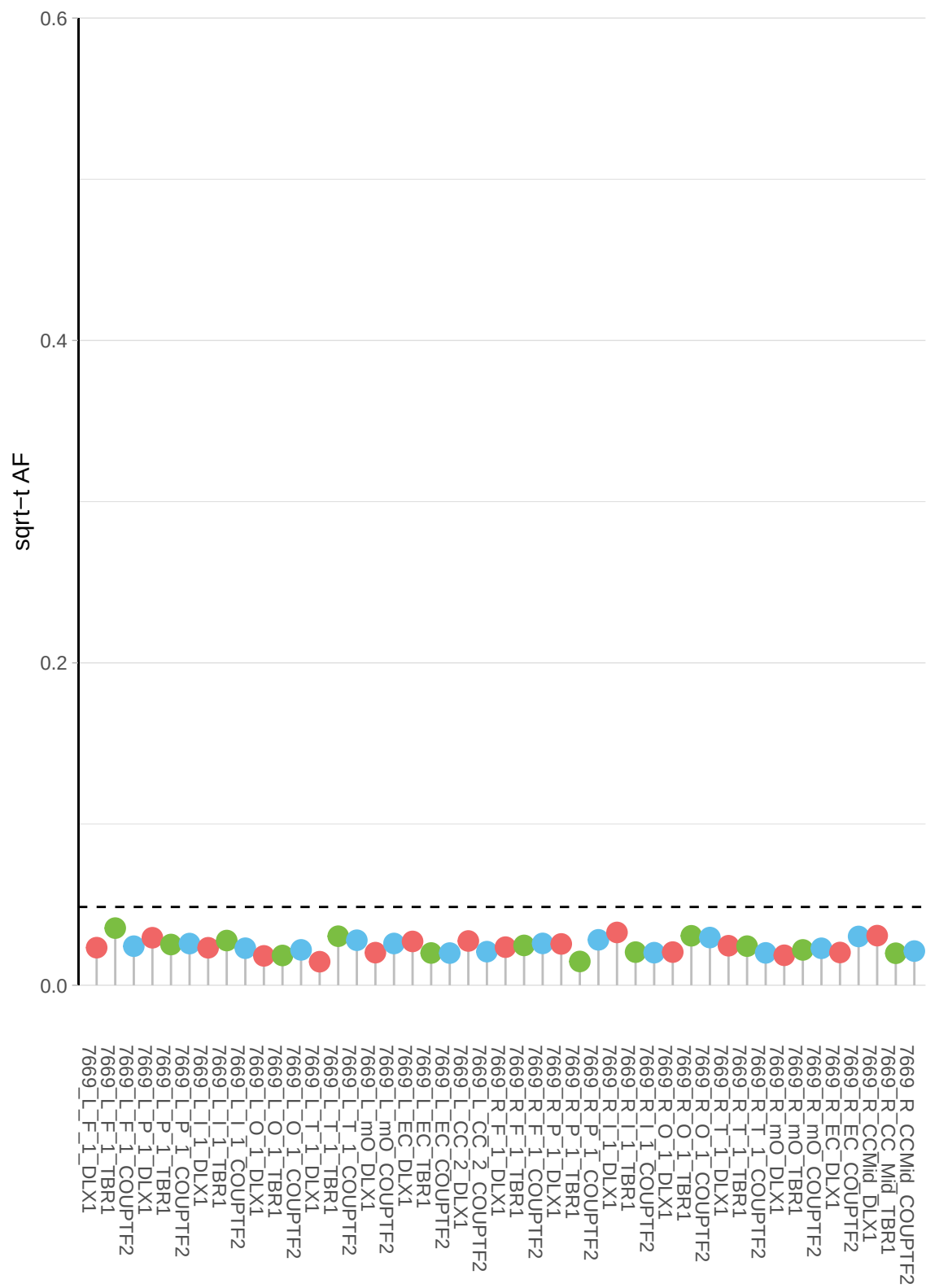

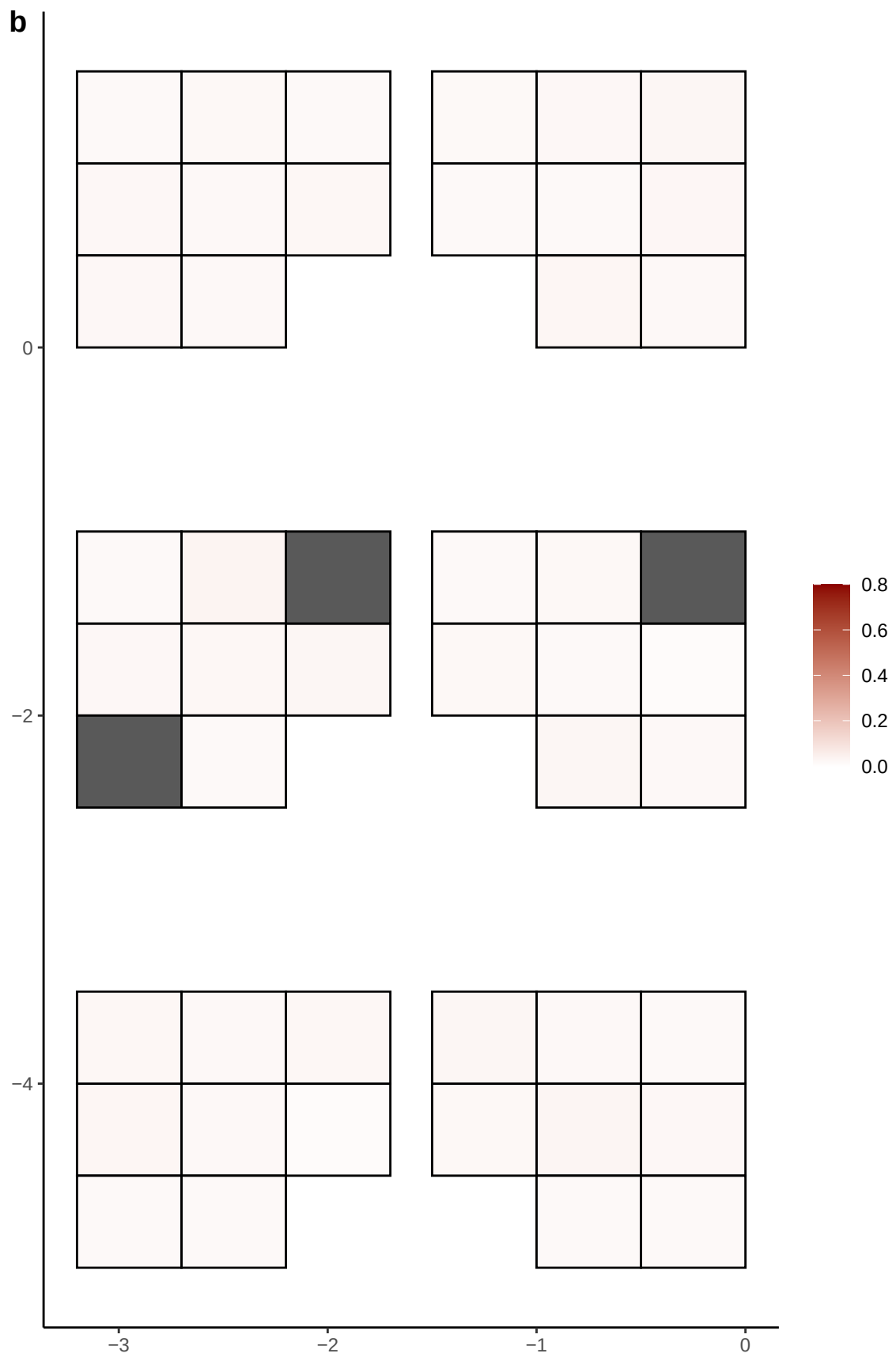

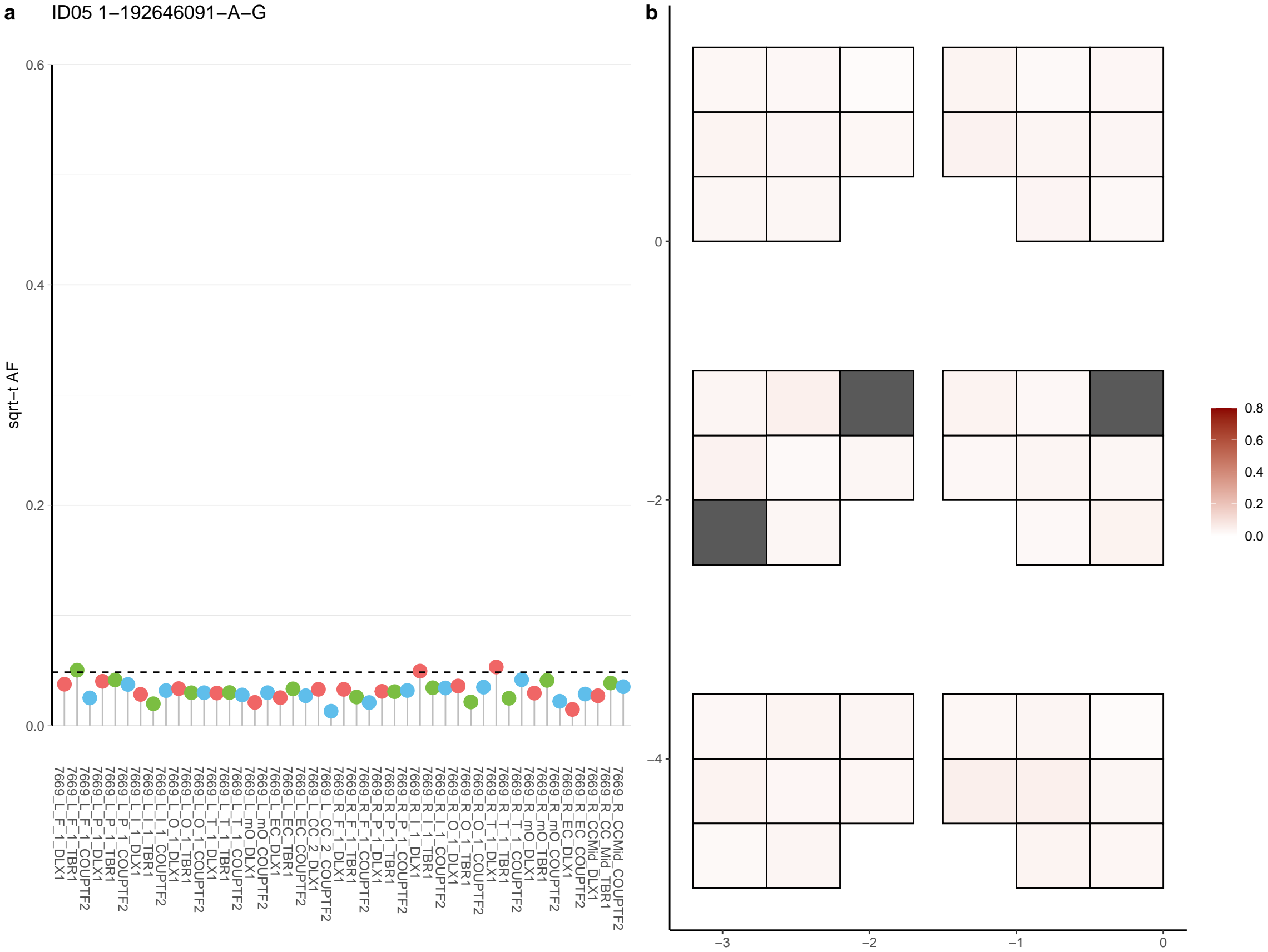

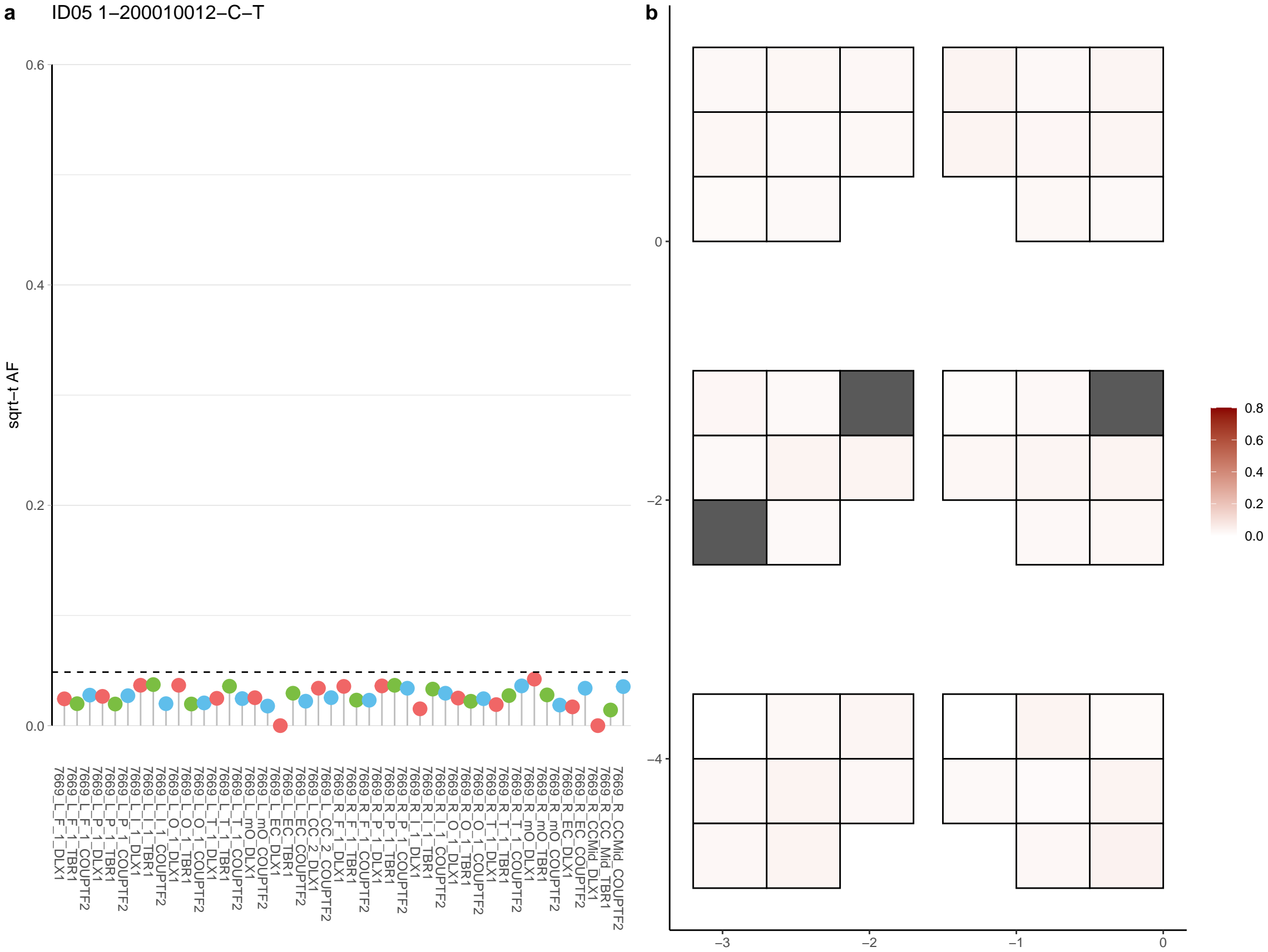

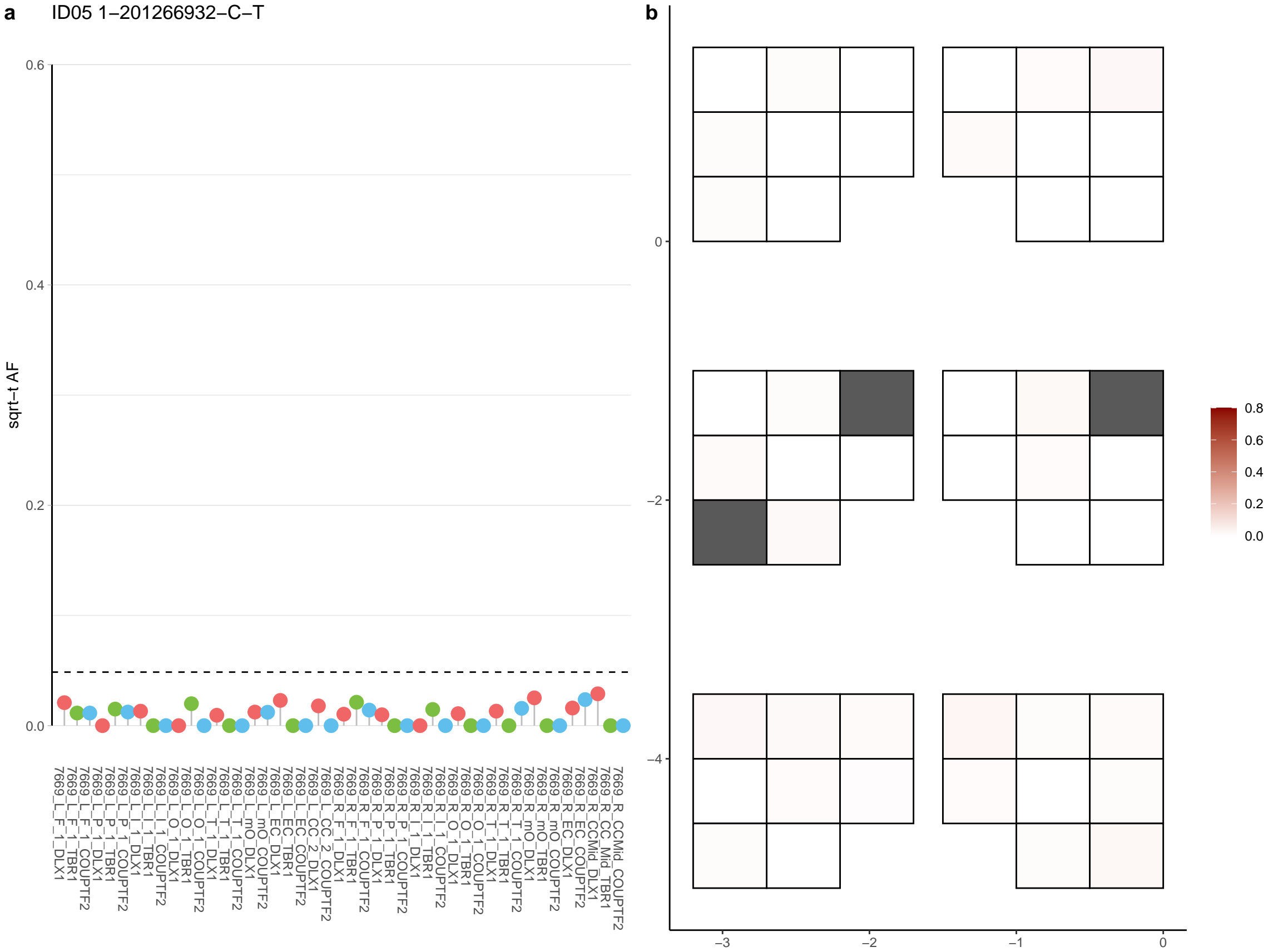

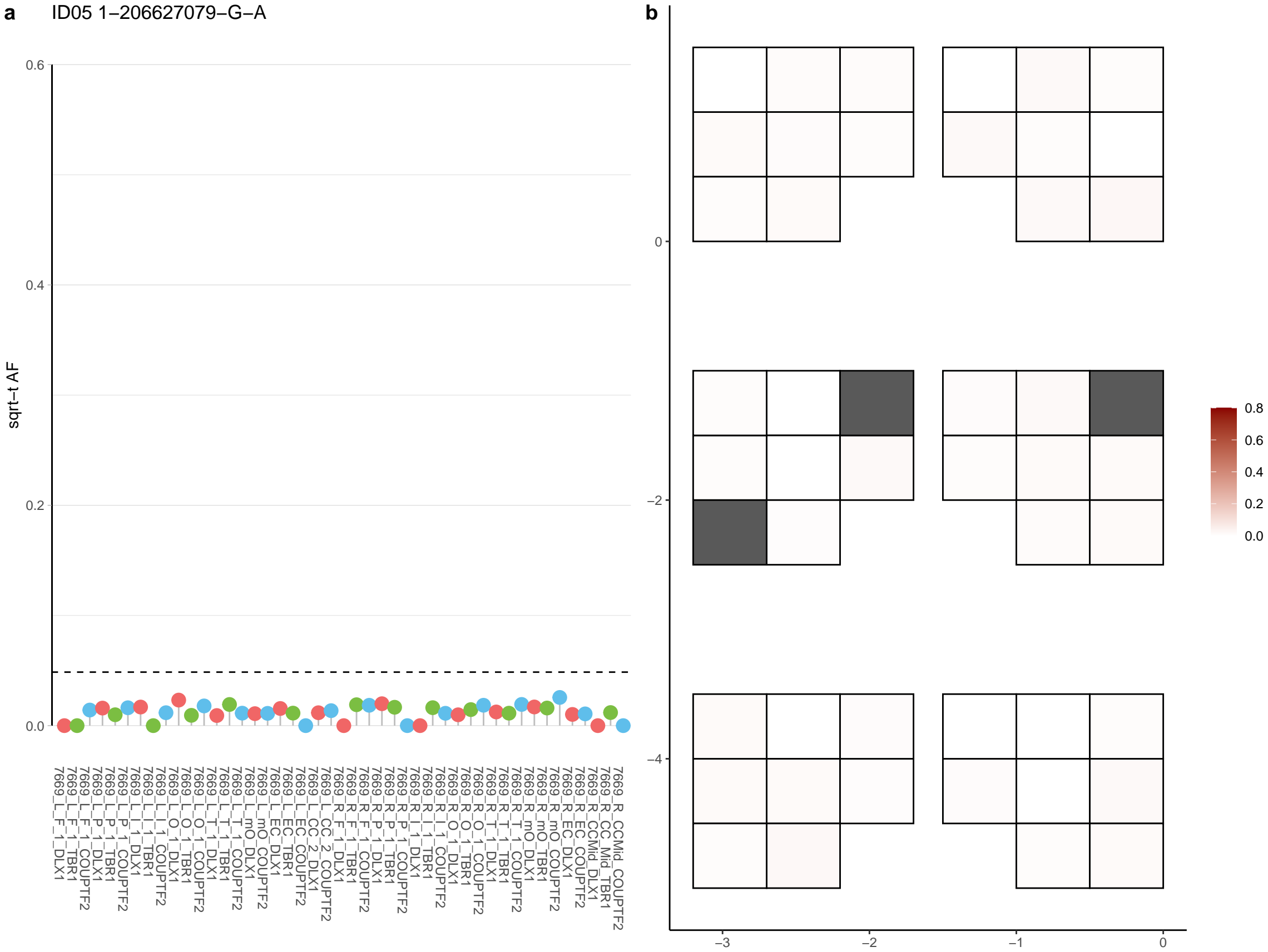

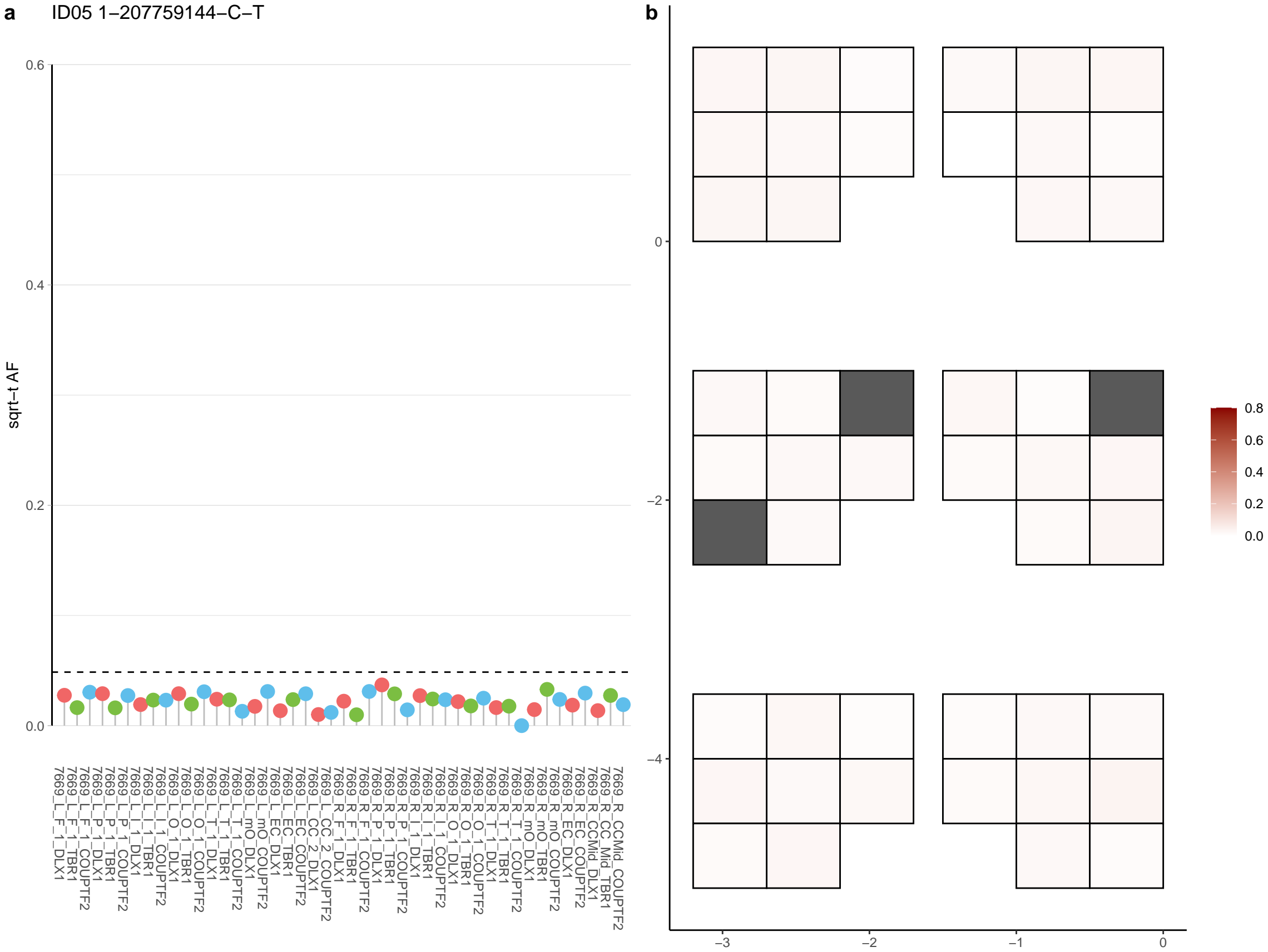

**a** ID05 1-210808944-G-A

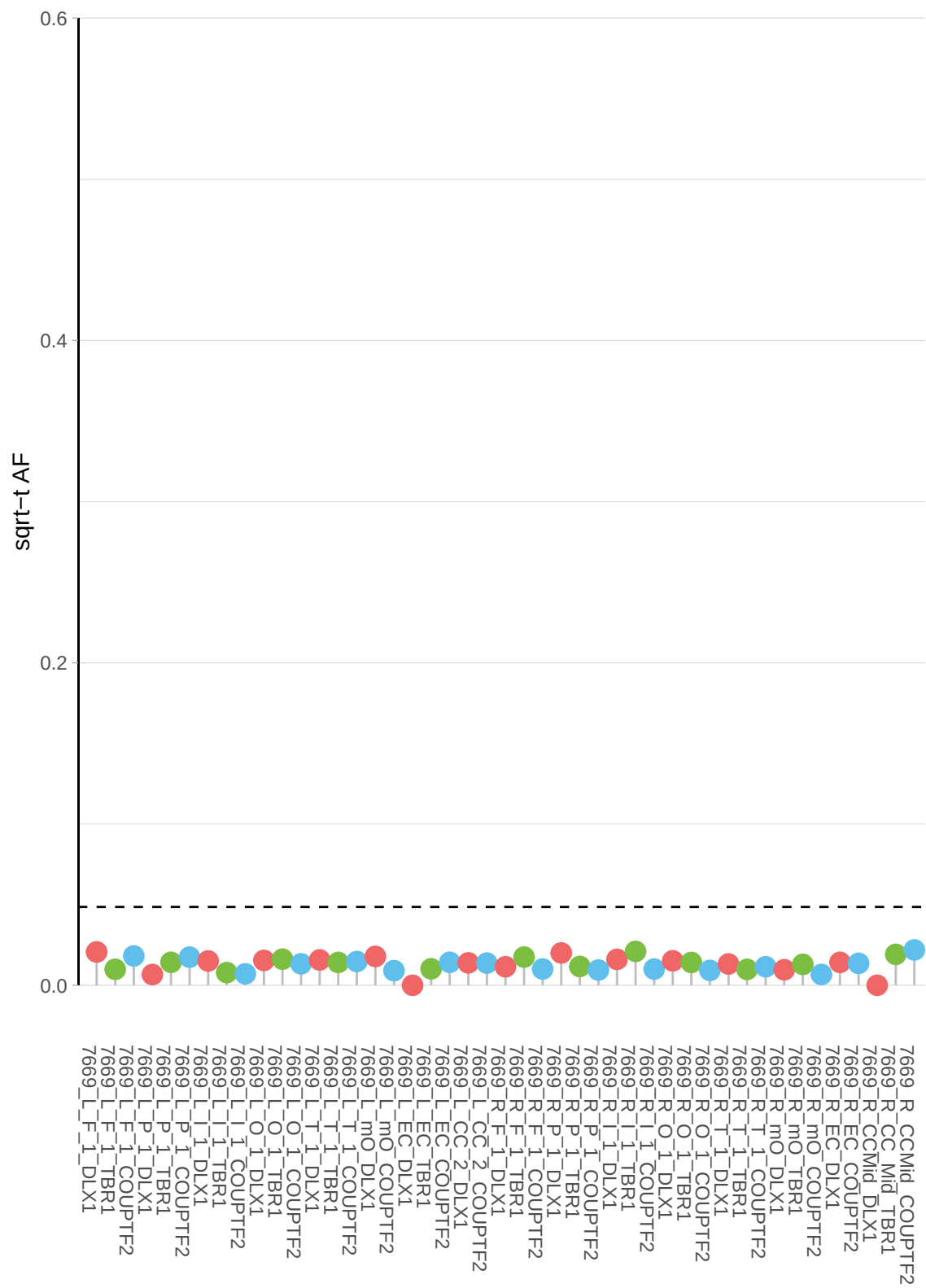

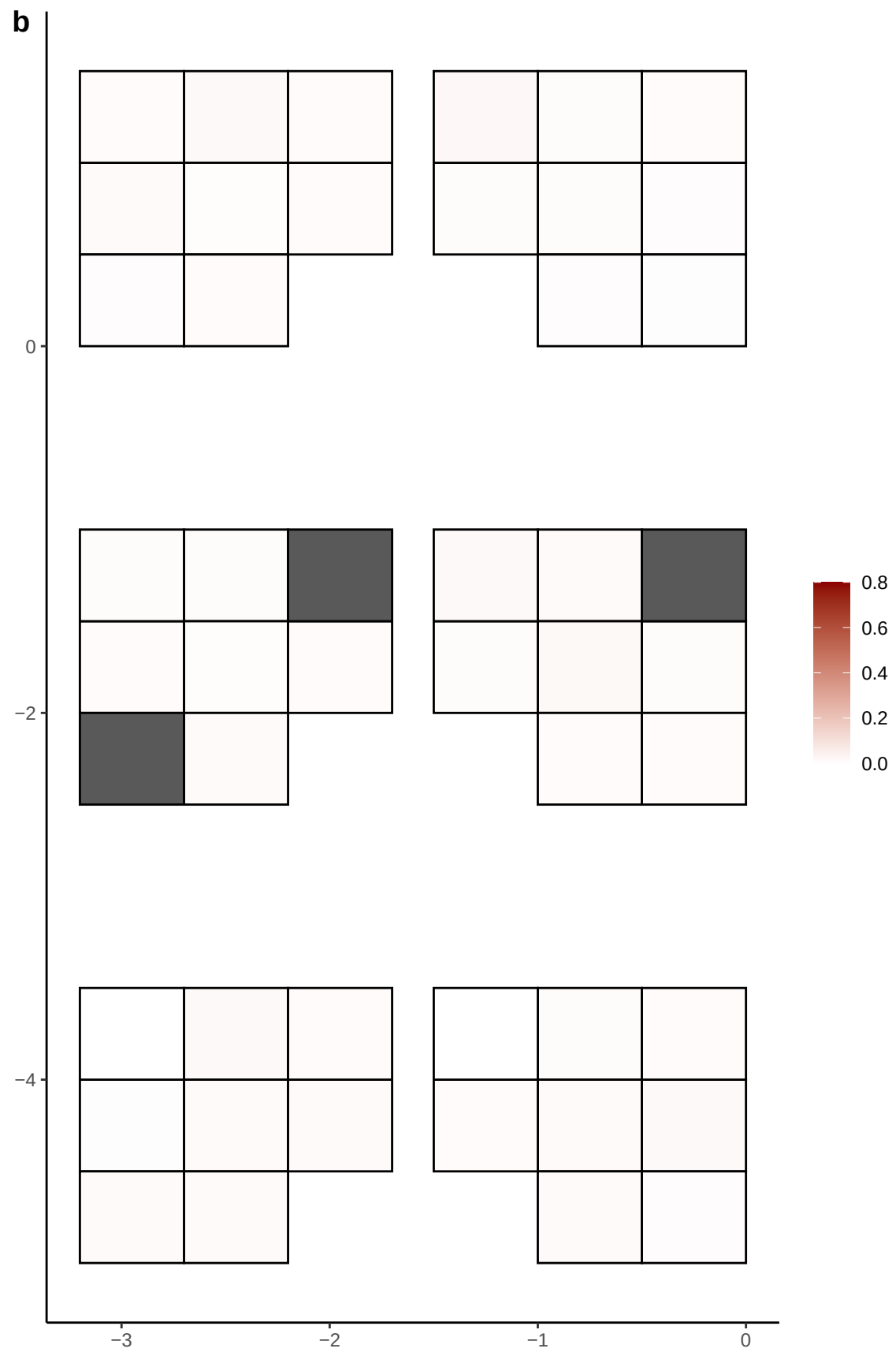

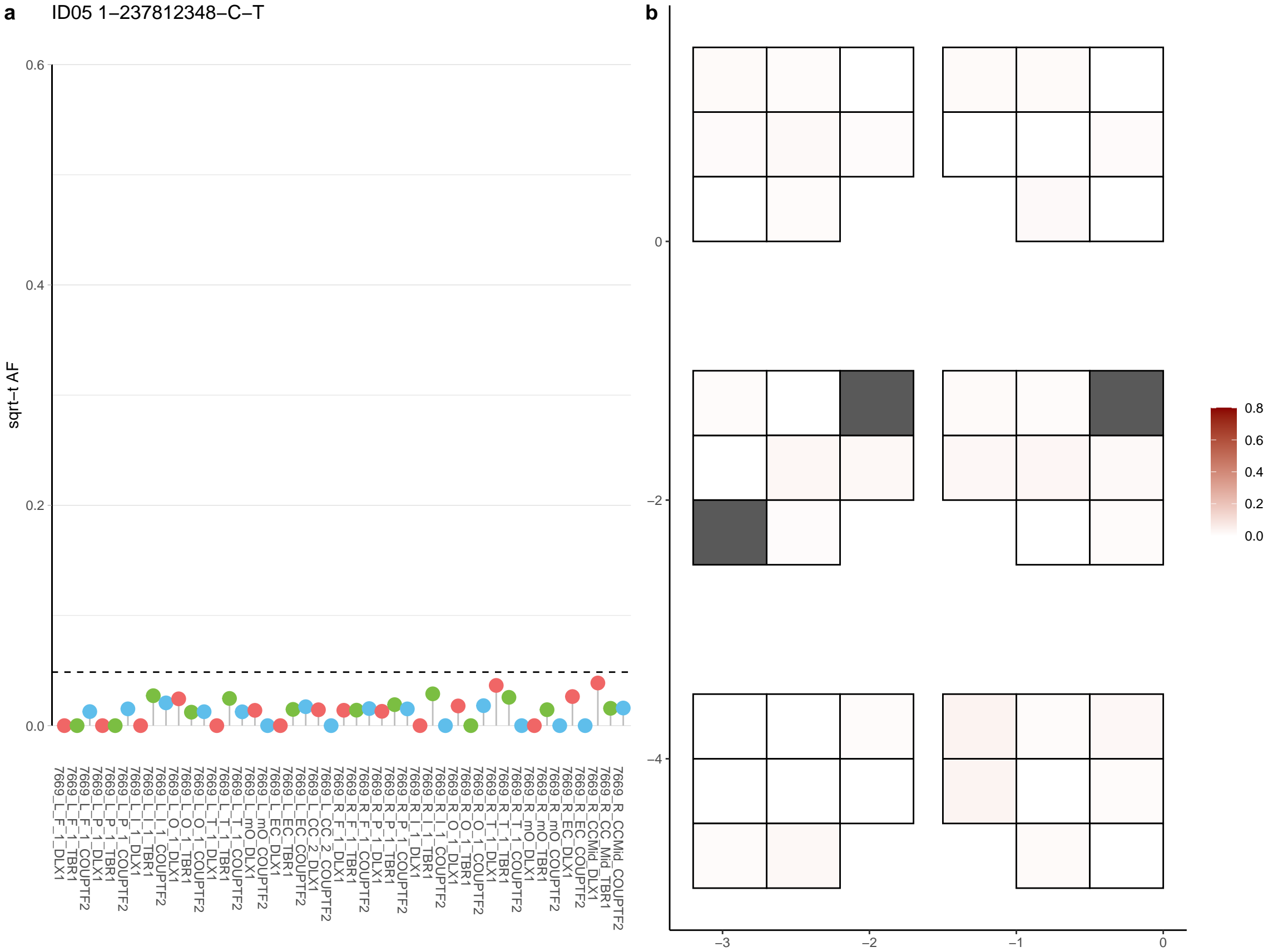

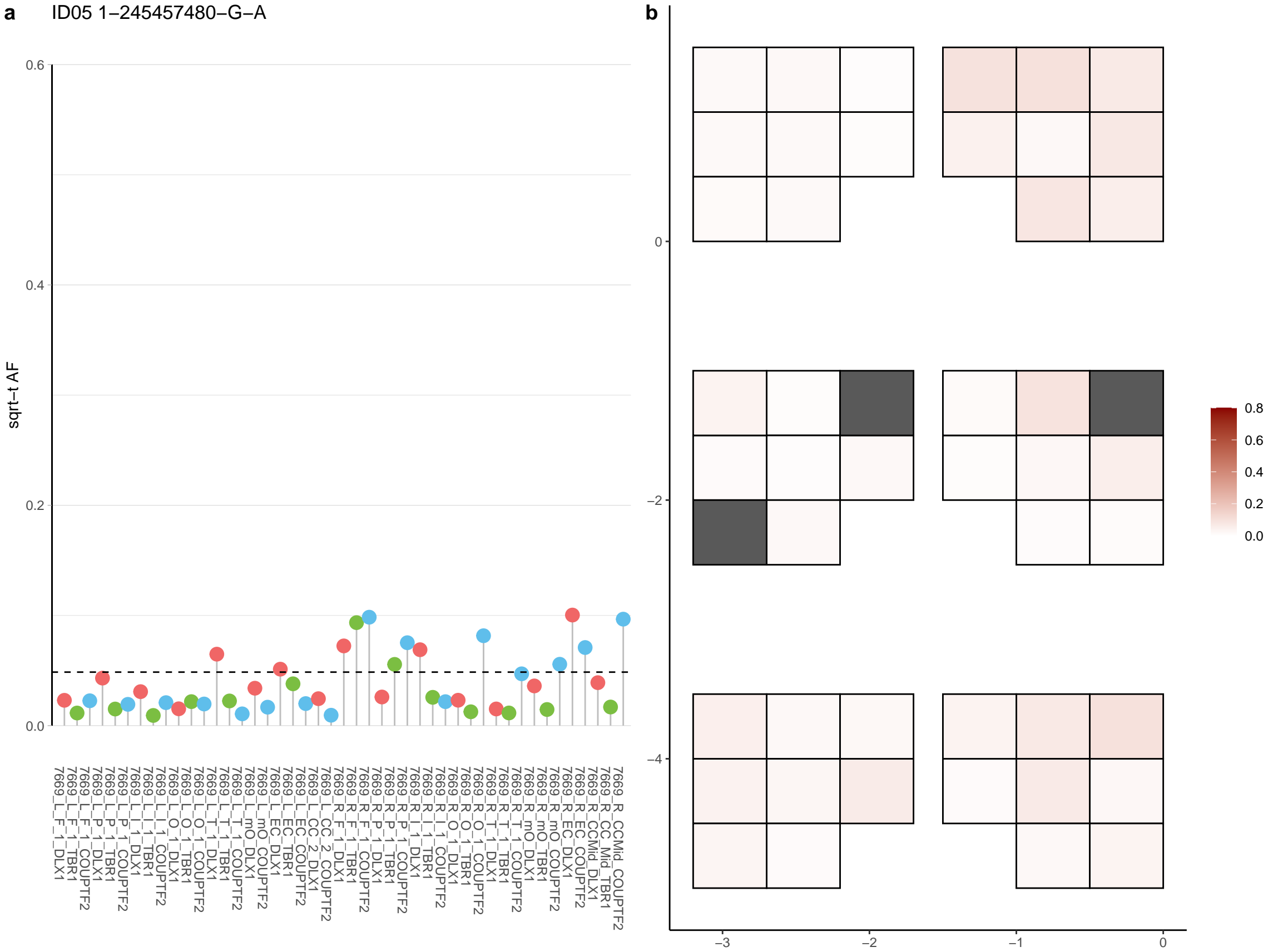

**a** ID05 1-248261416-C-T

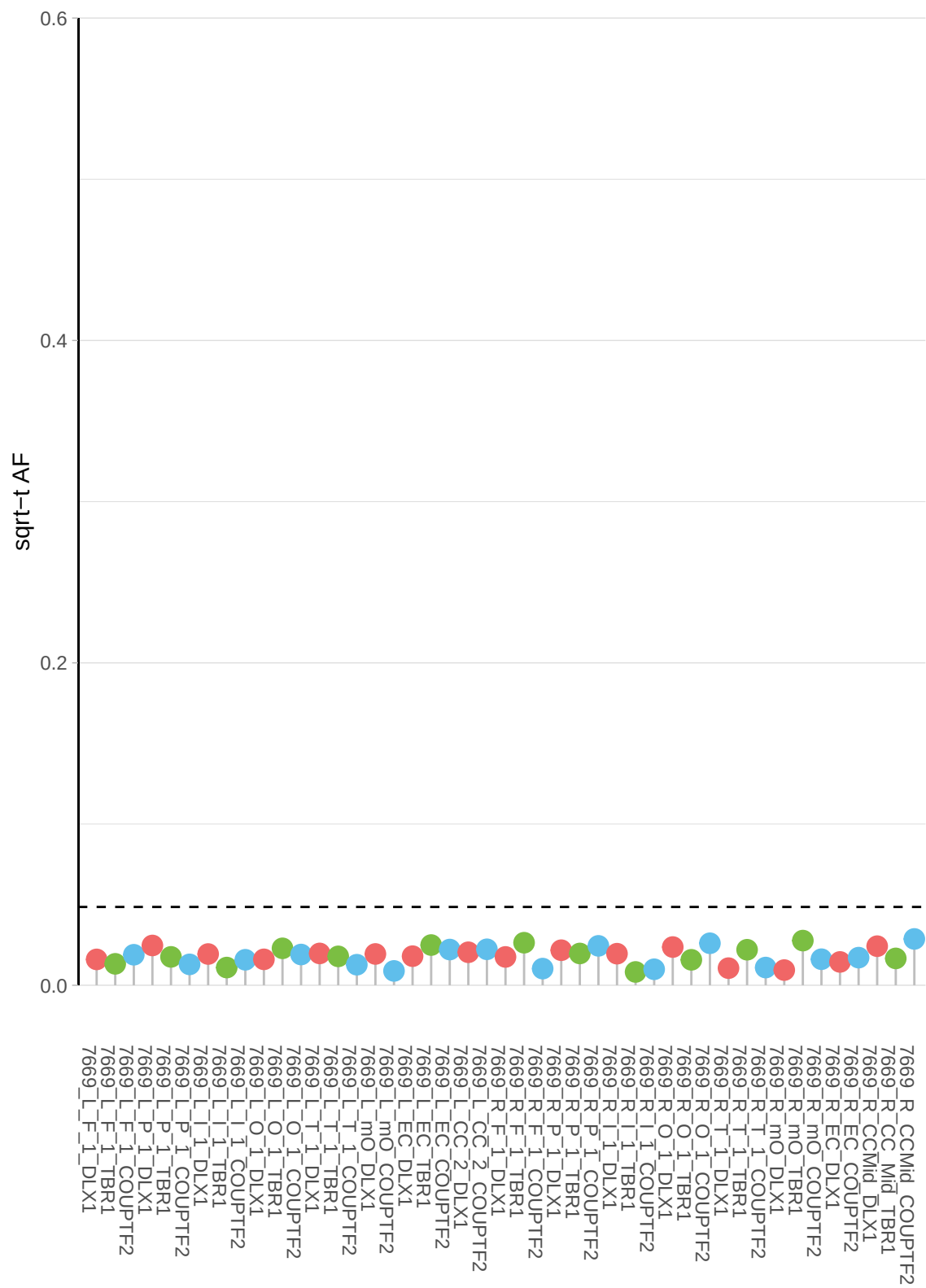

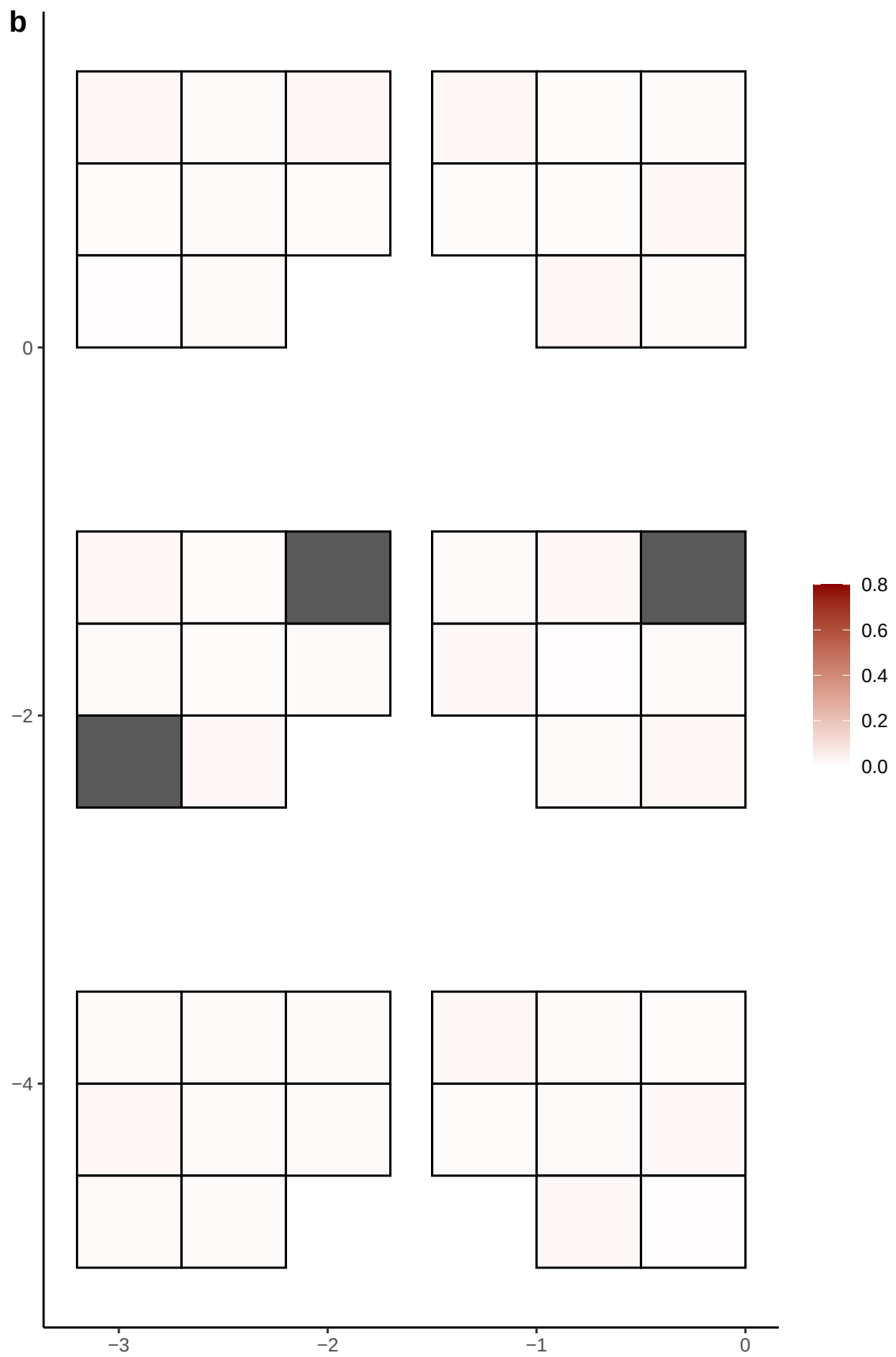

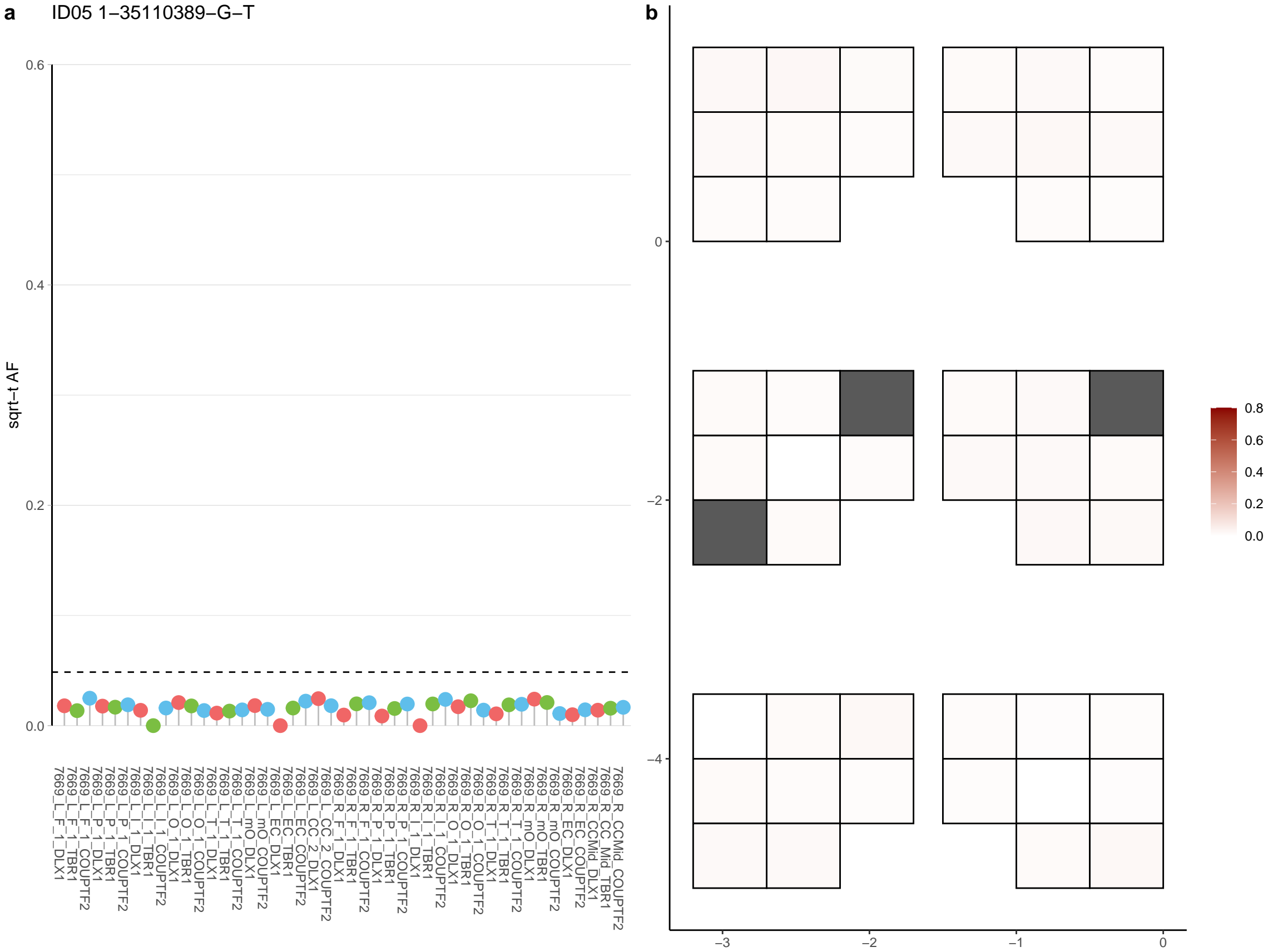

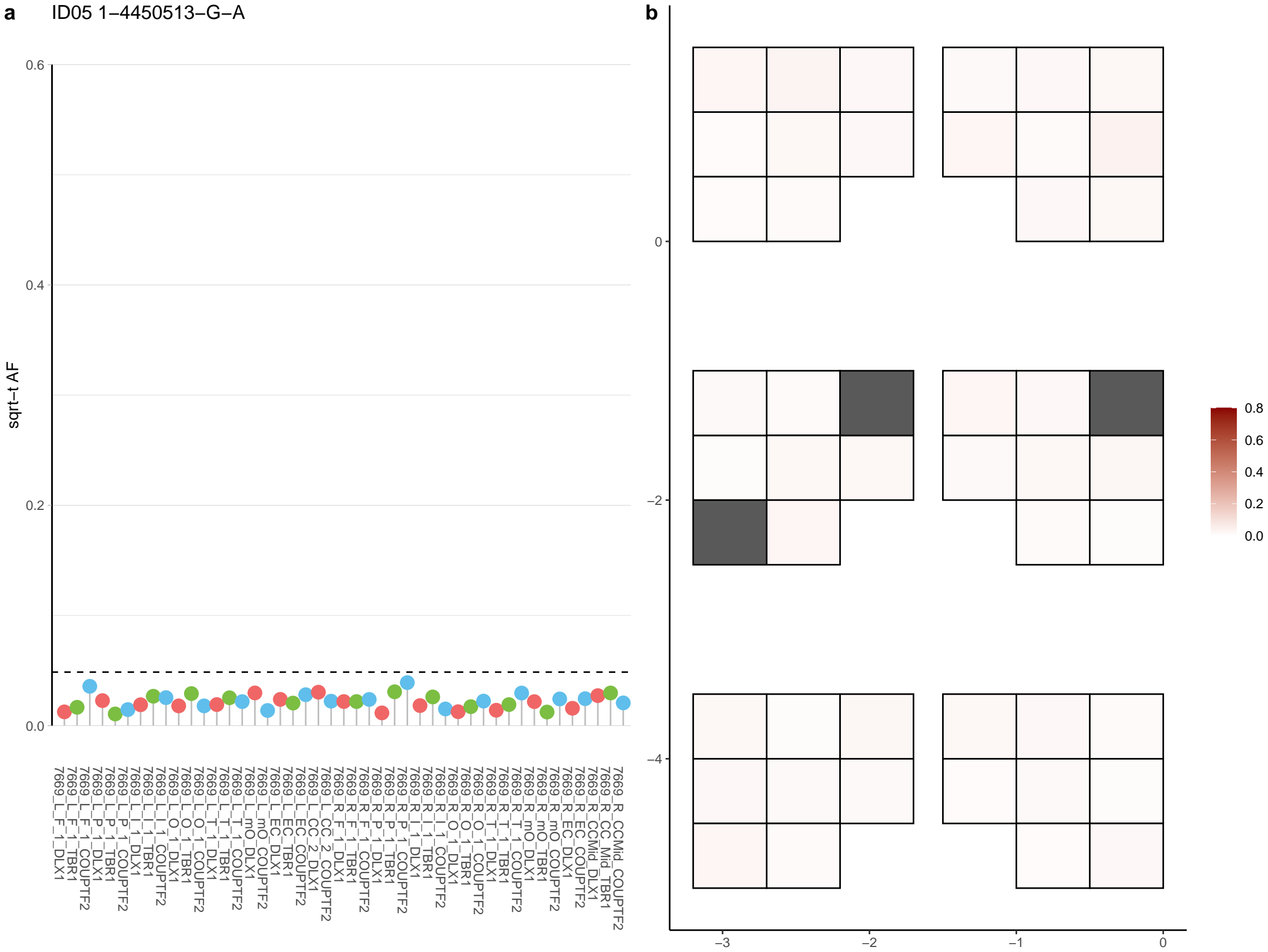

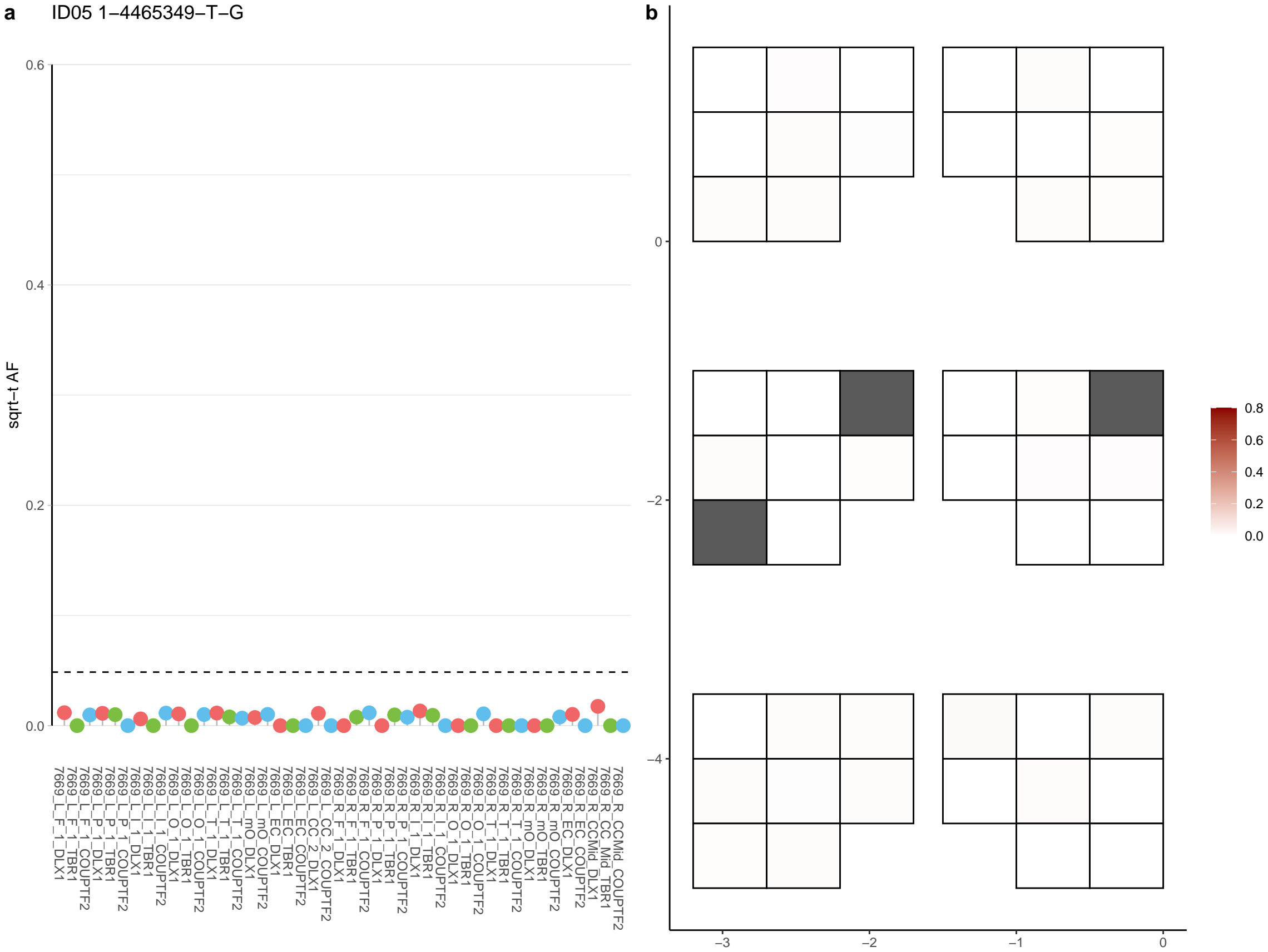

**a** ID05 1-50289297-G-A

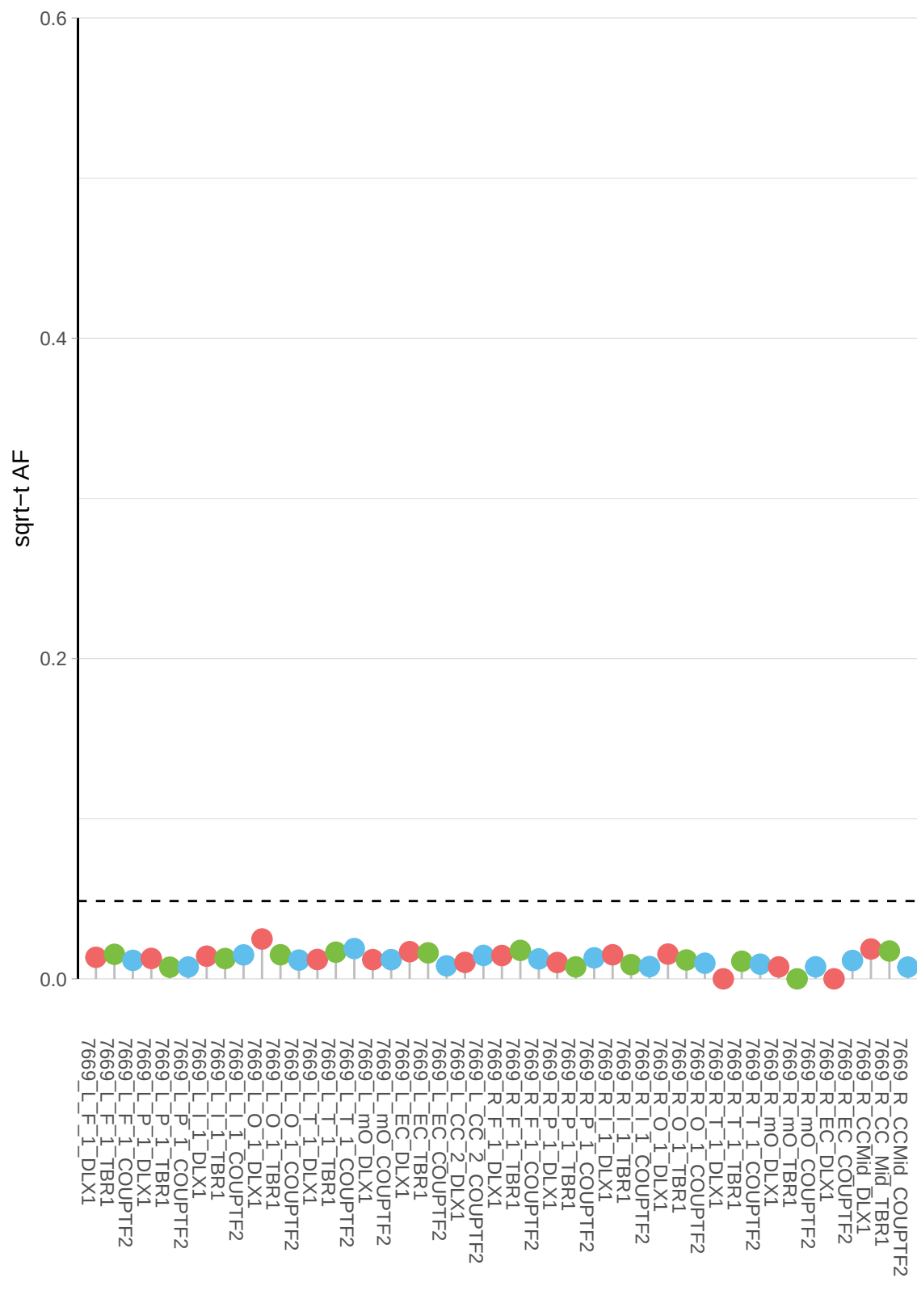

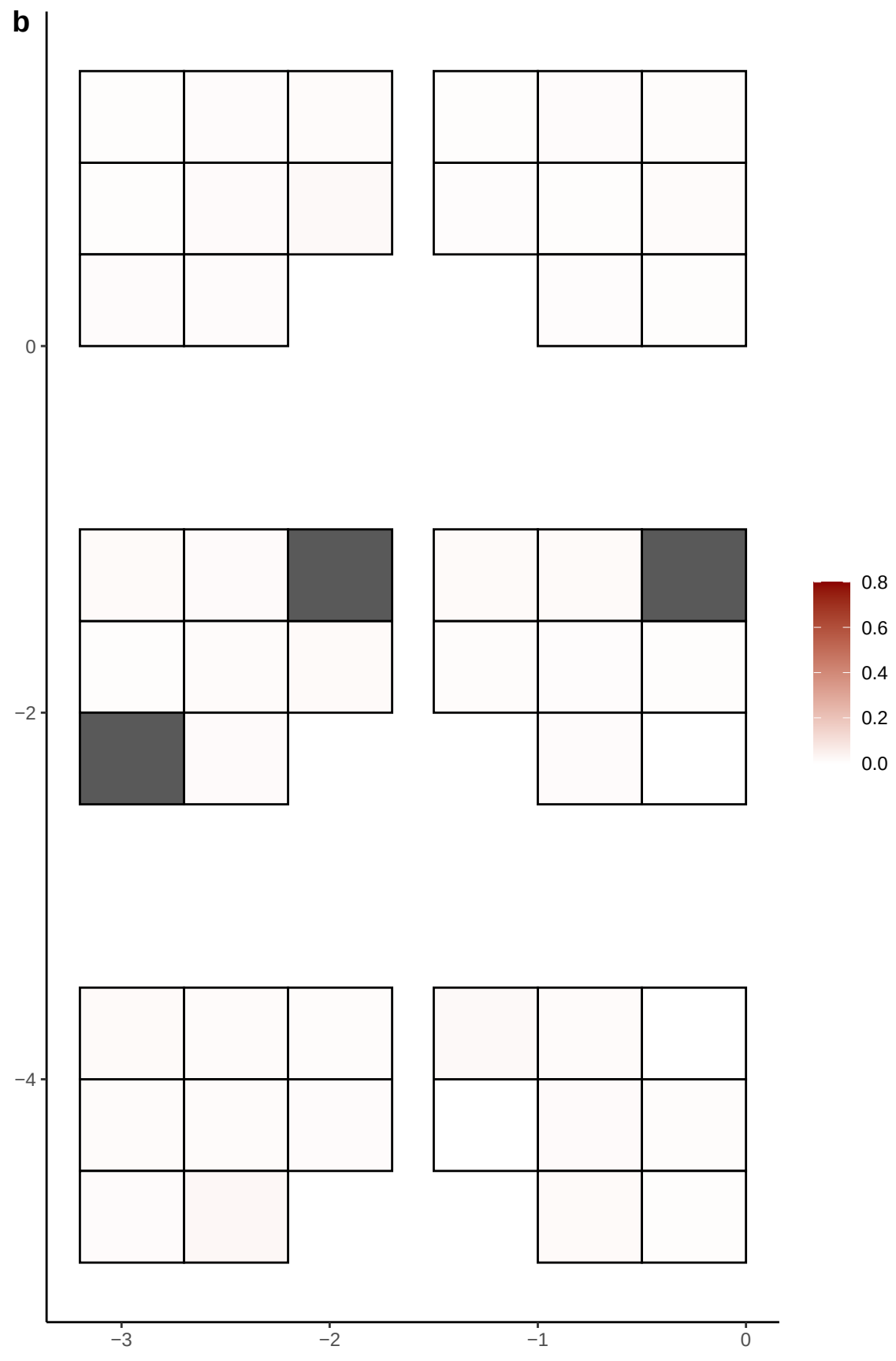

**a** ID05 1-53912650-G-A

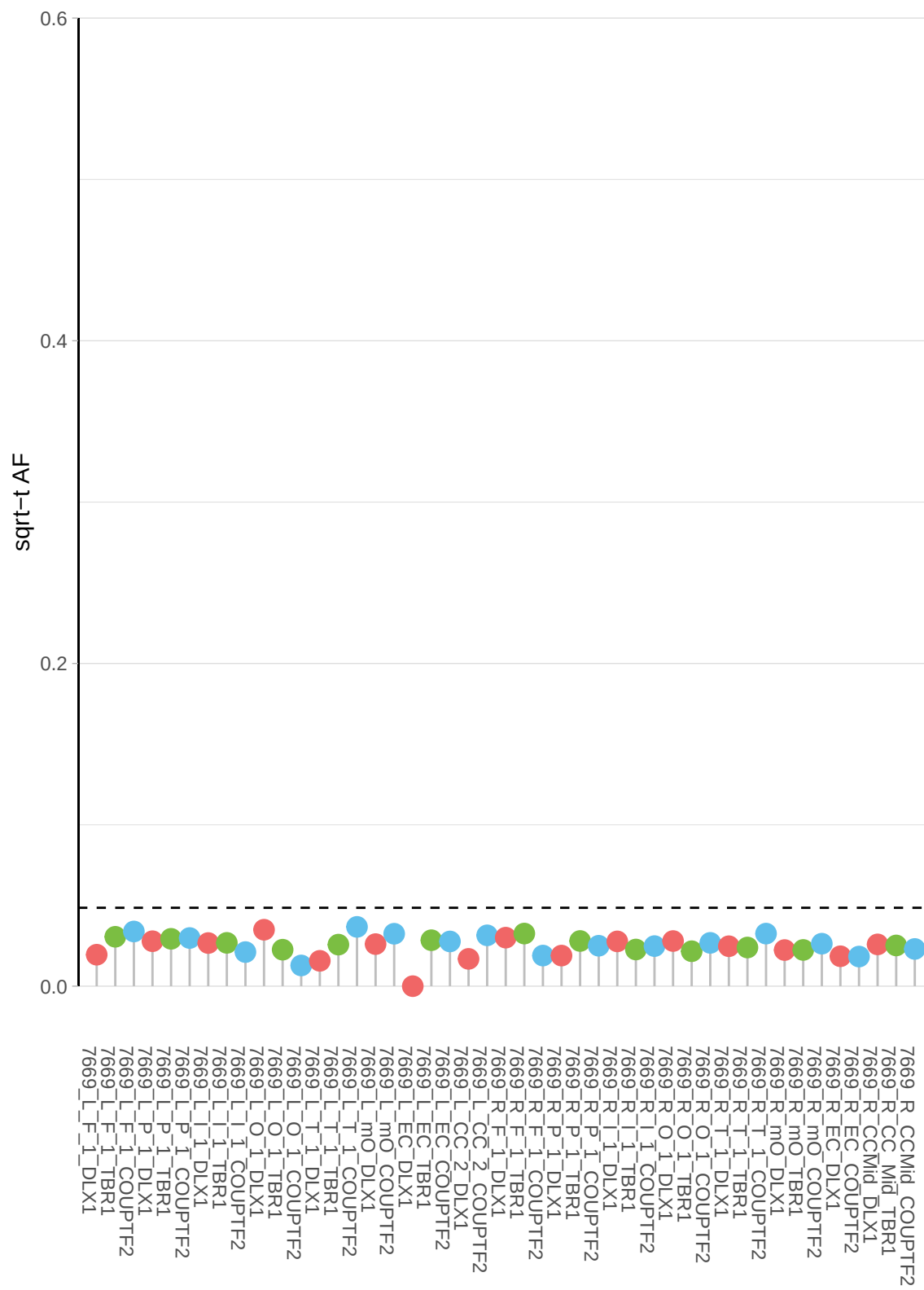

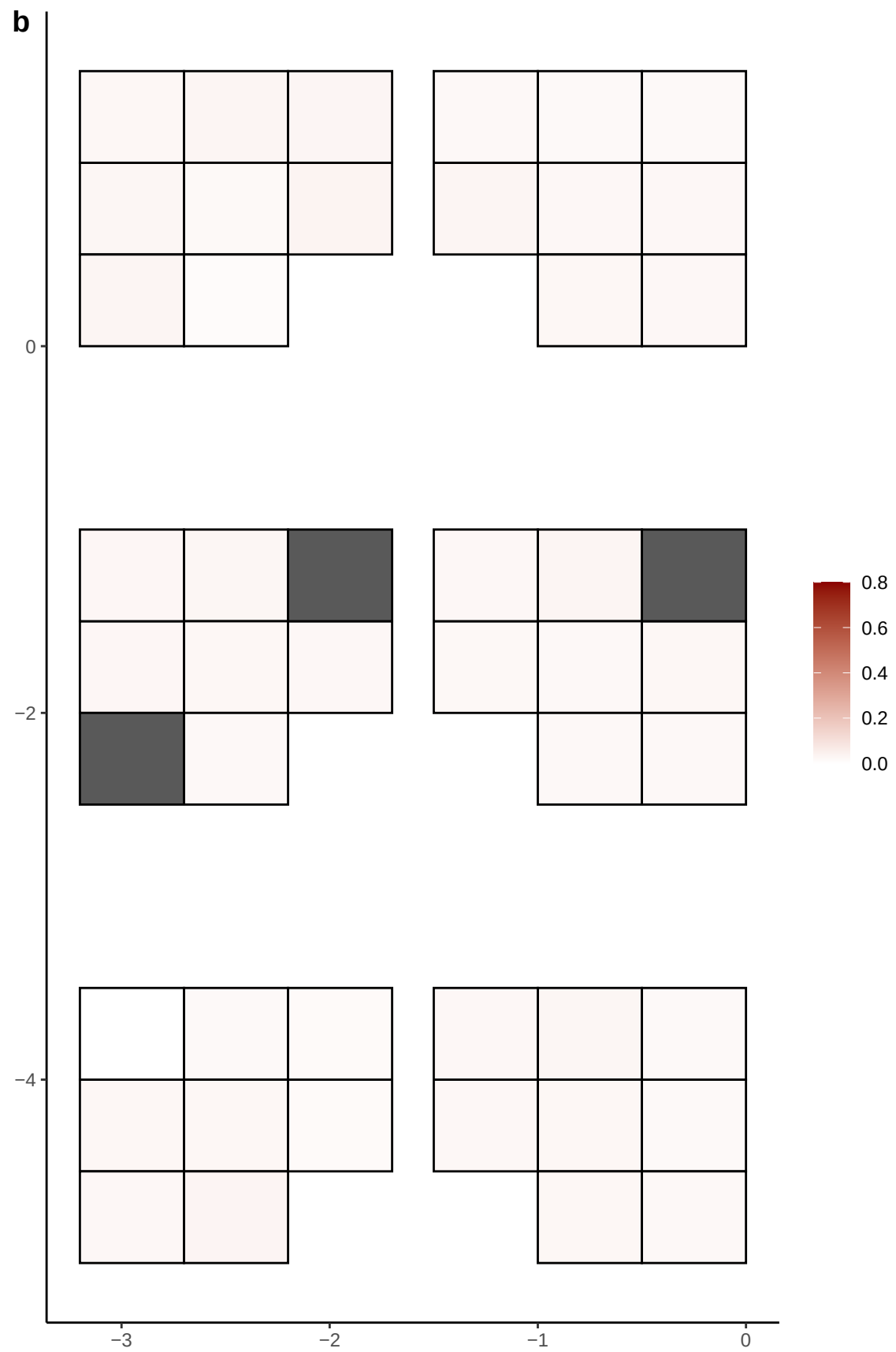

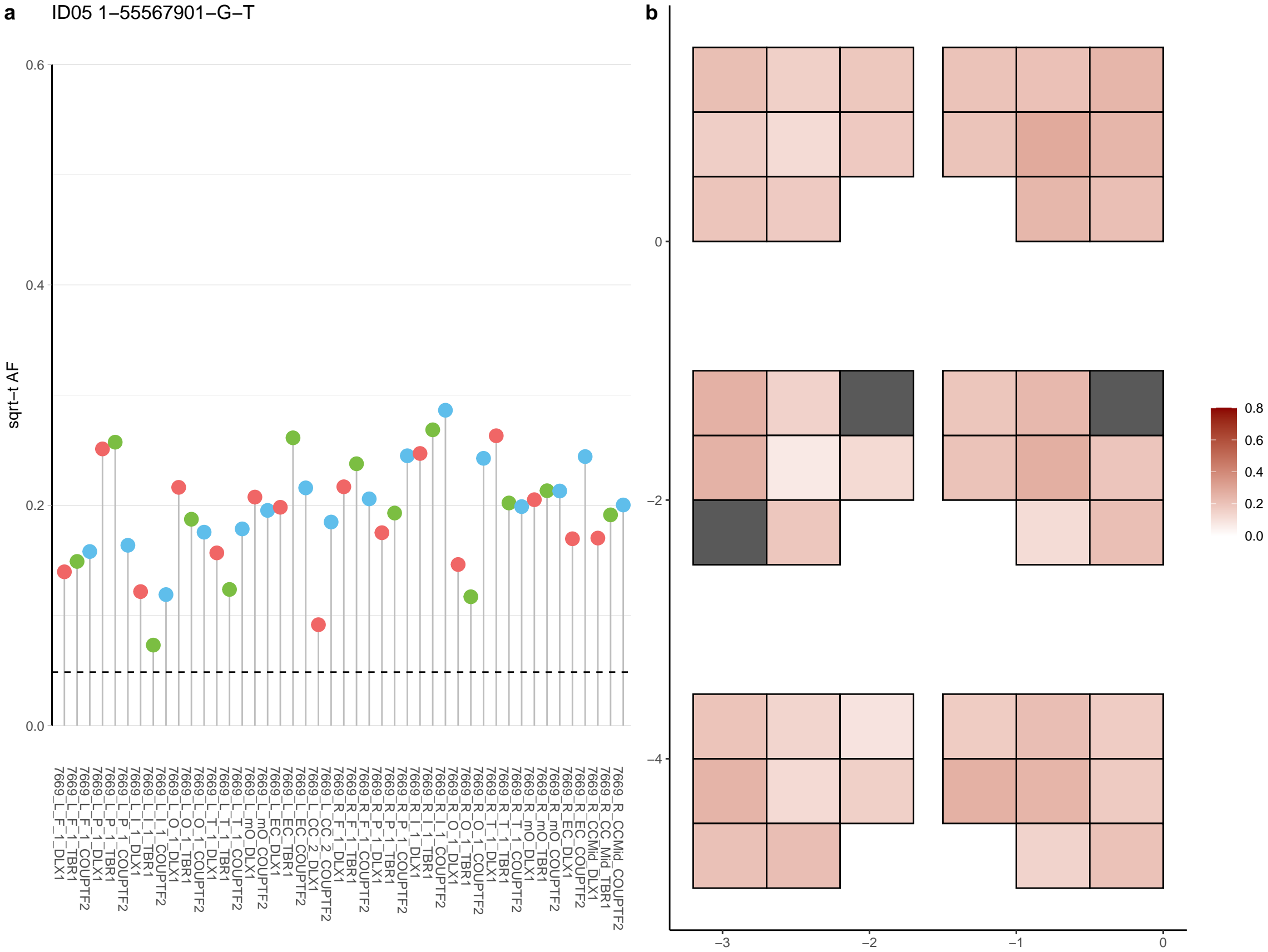

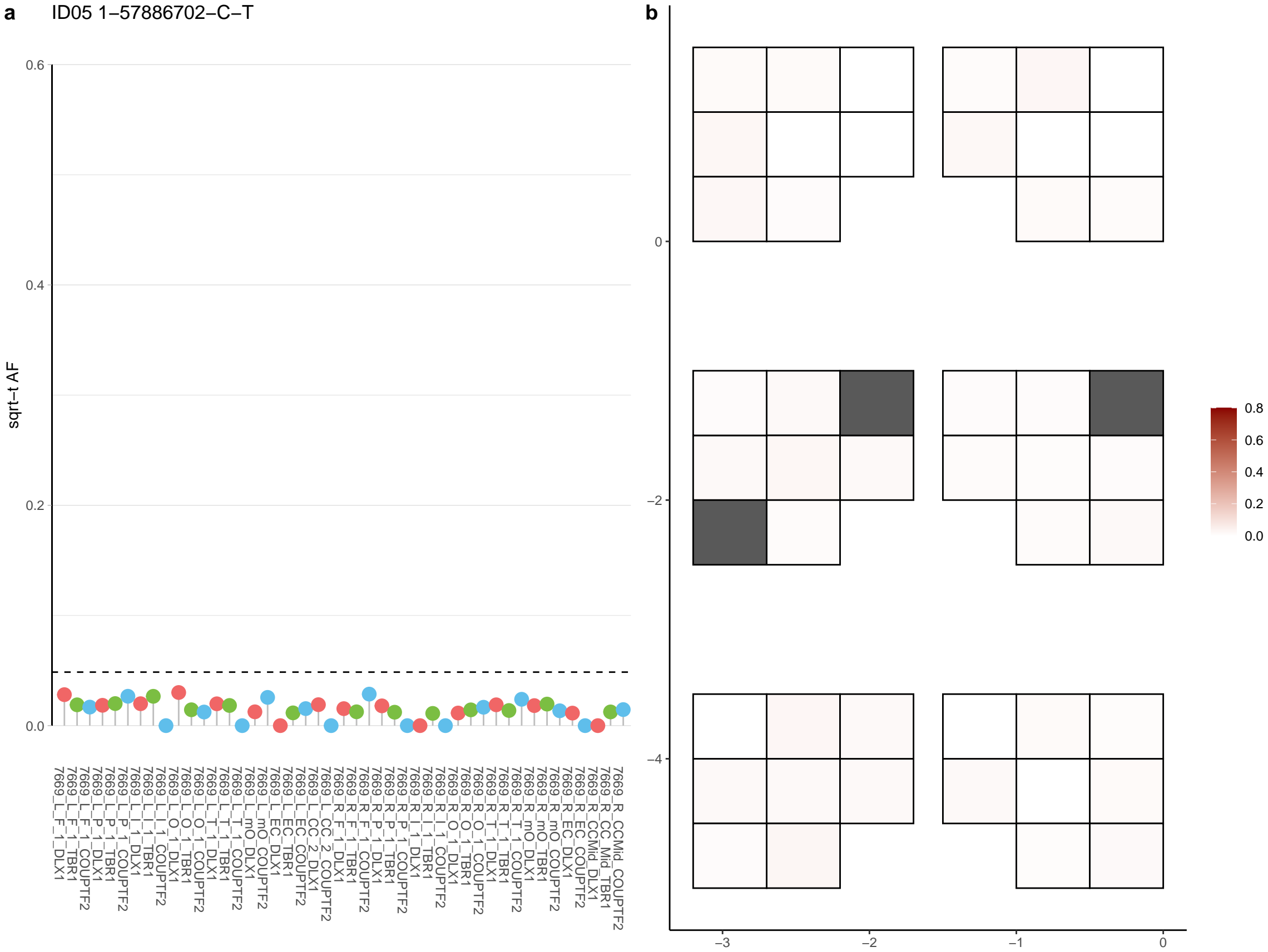

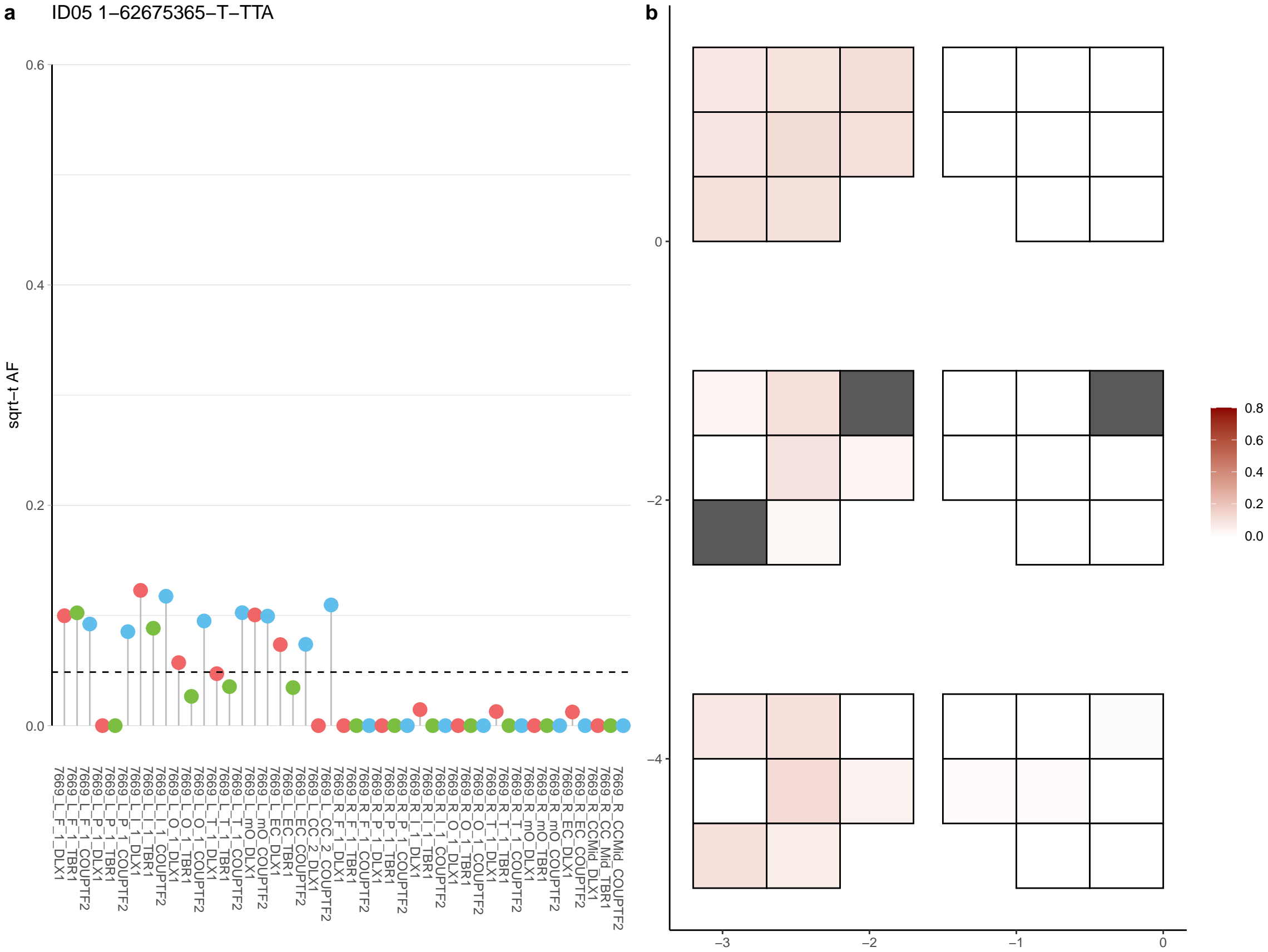

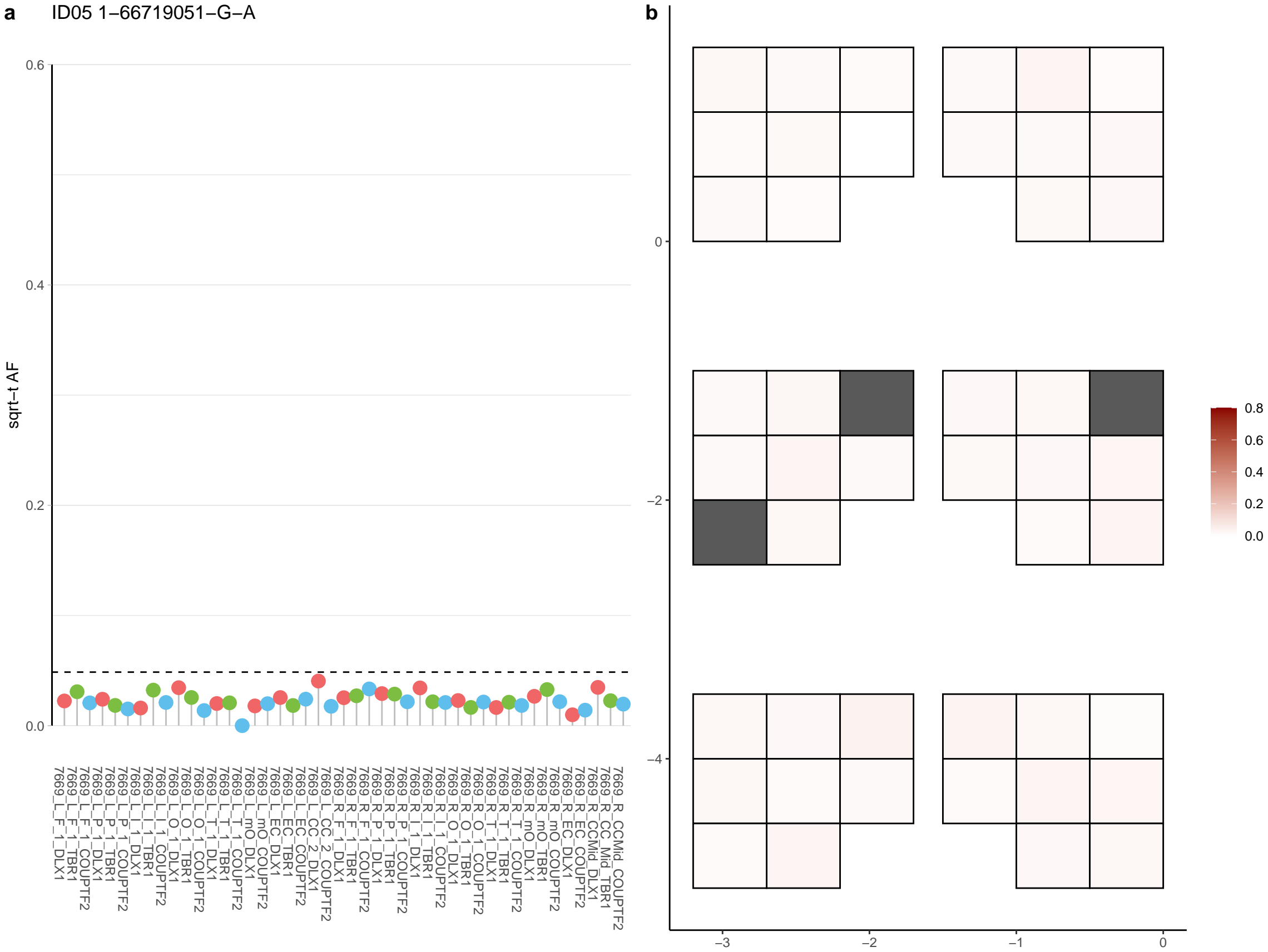

**a** ID05 1-67722227-C-A

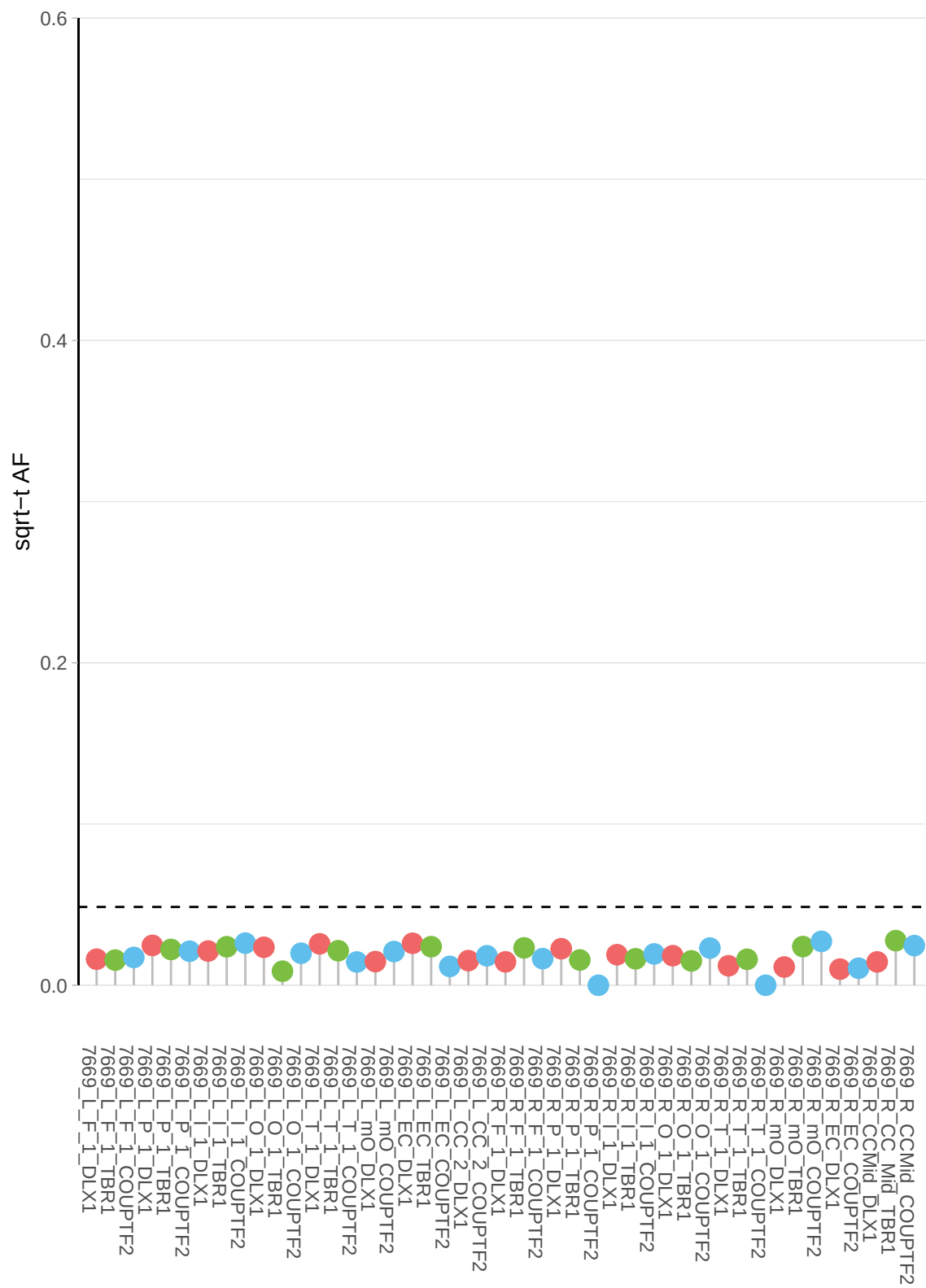

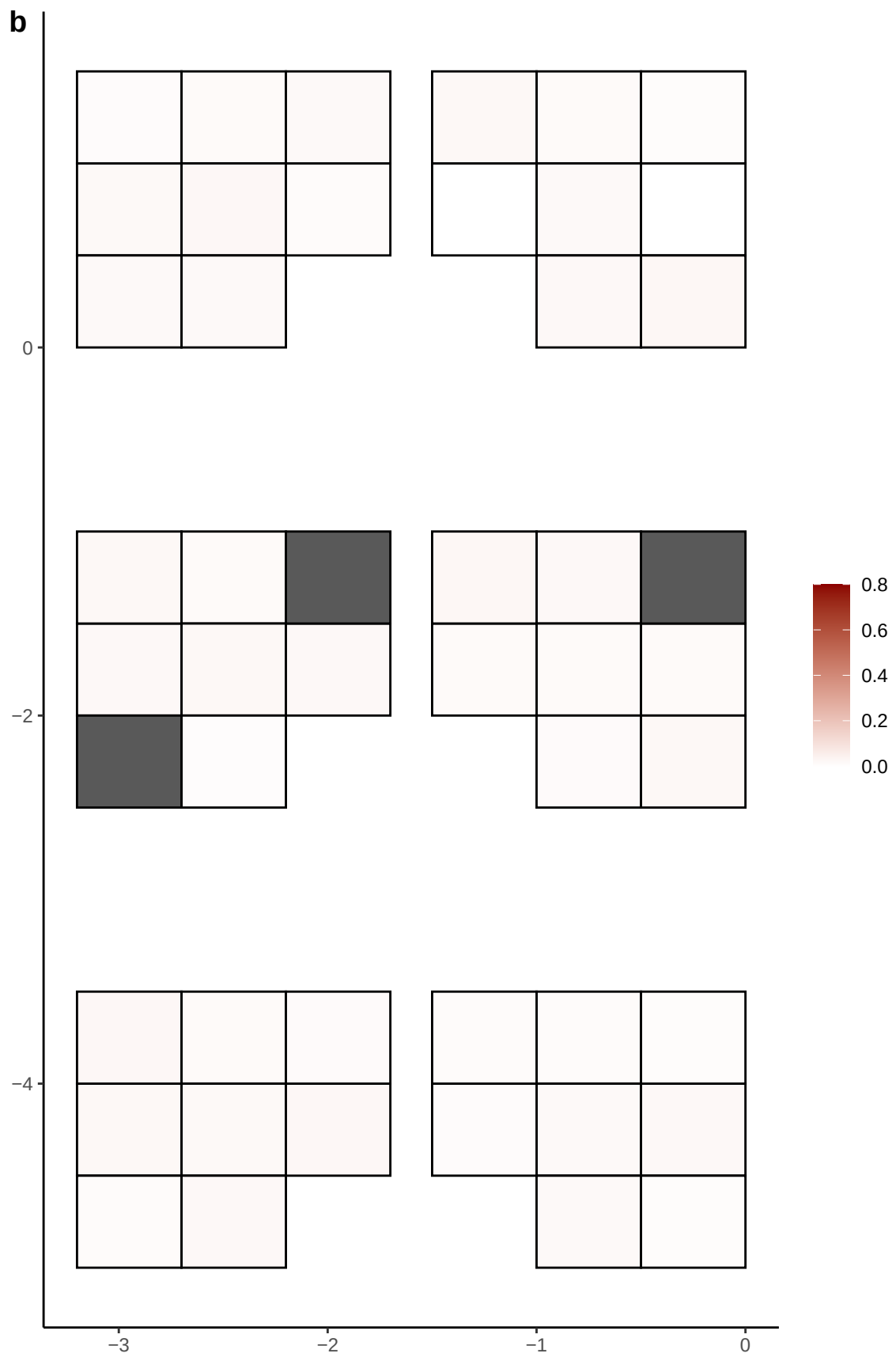

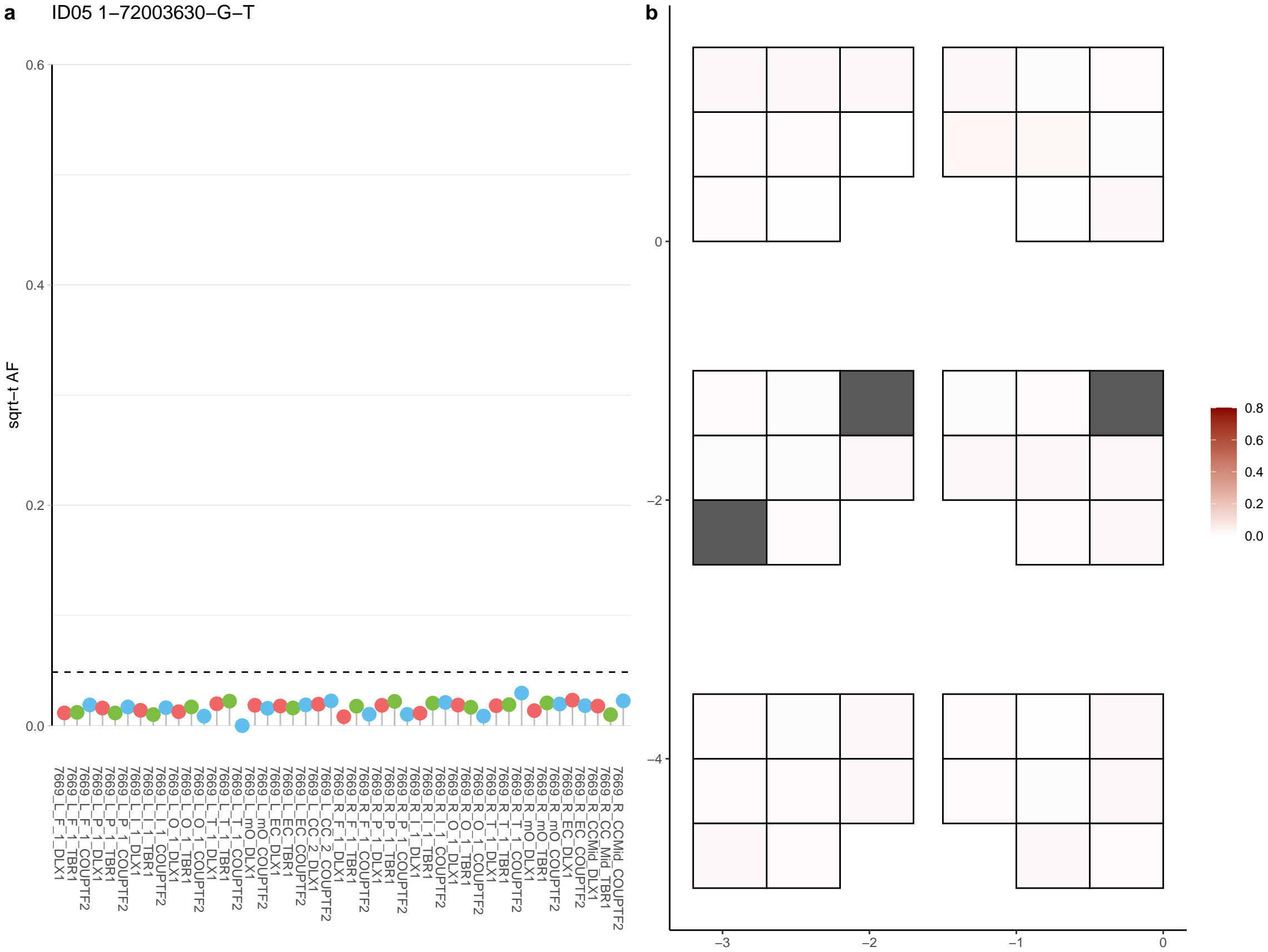

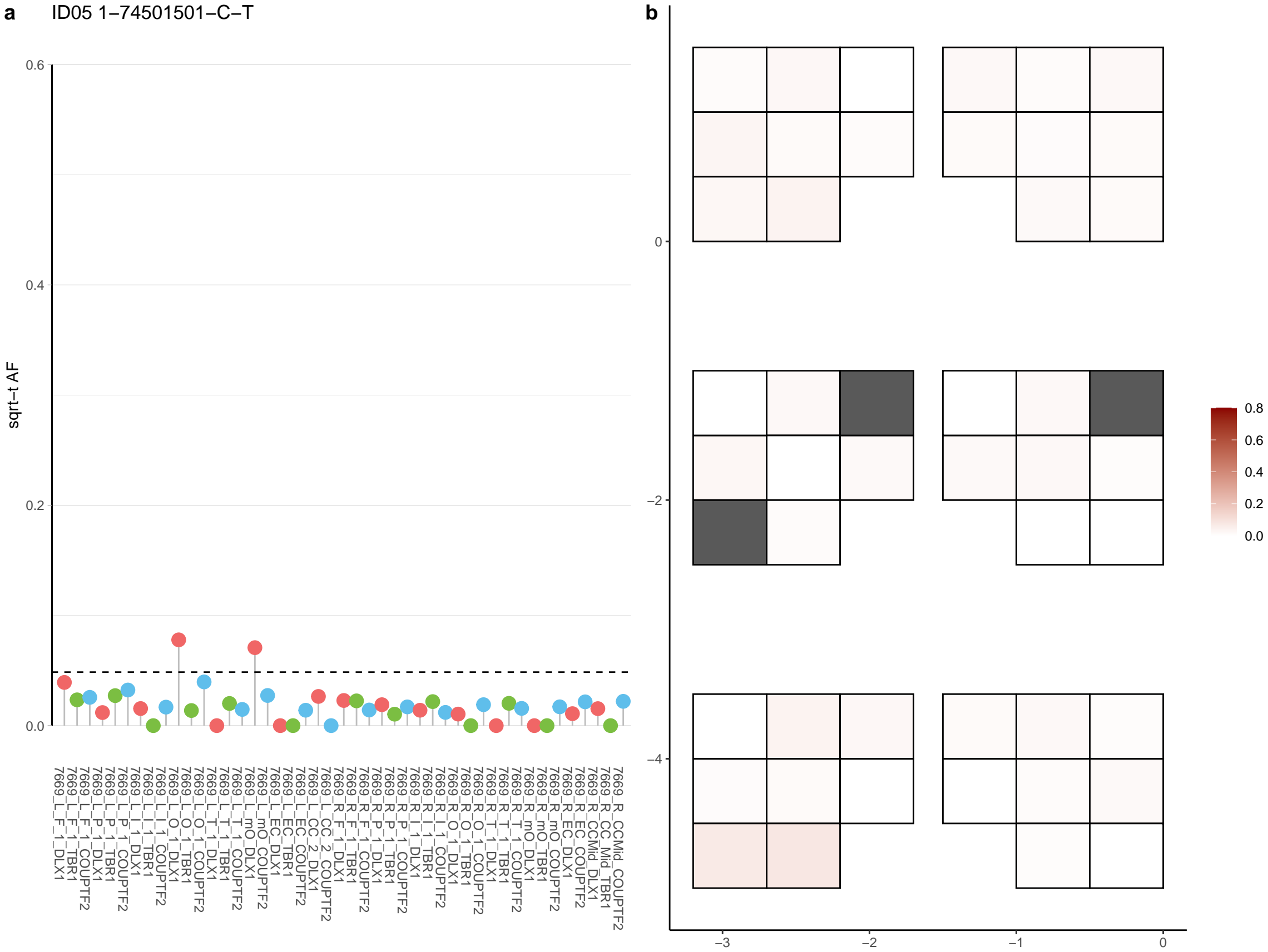

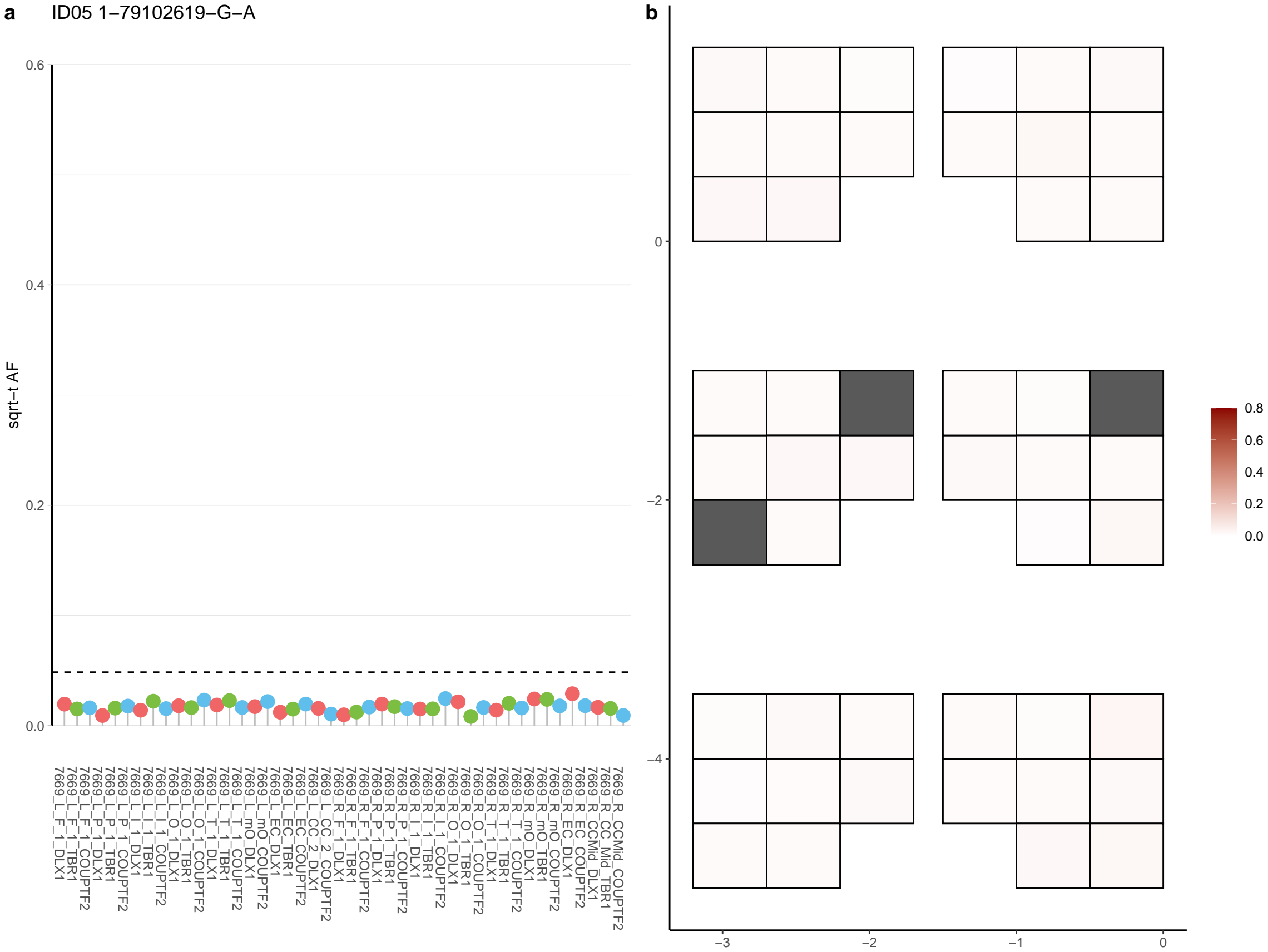

**a** ID05 1-83361707-GC-G

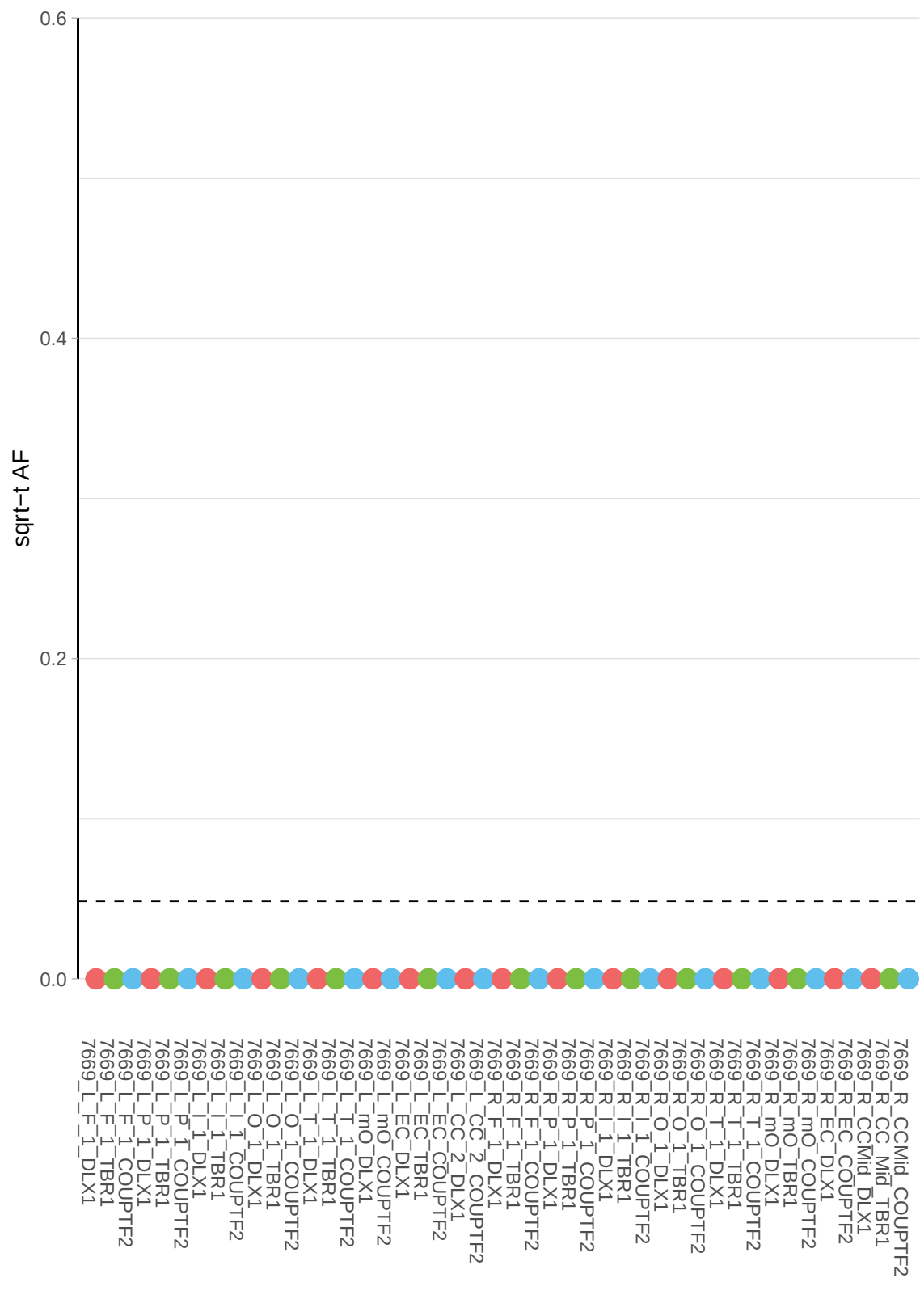

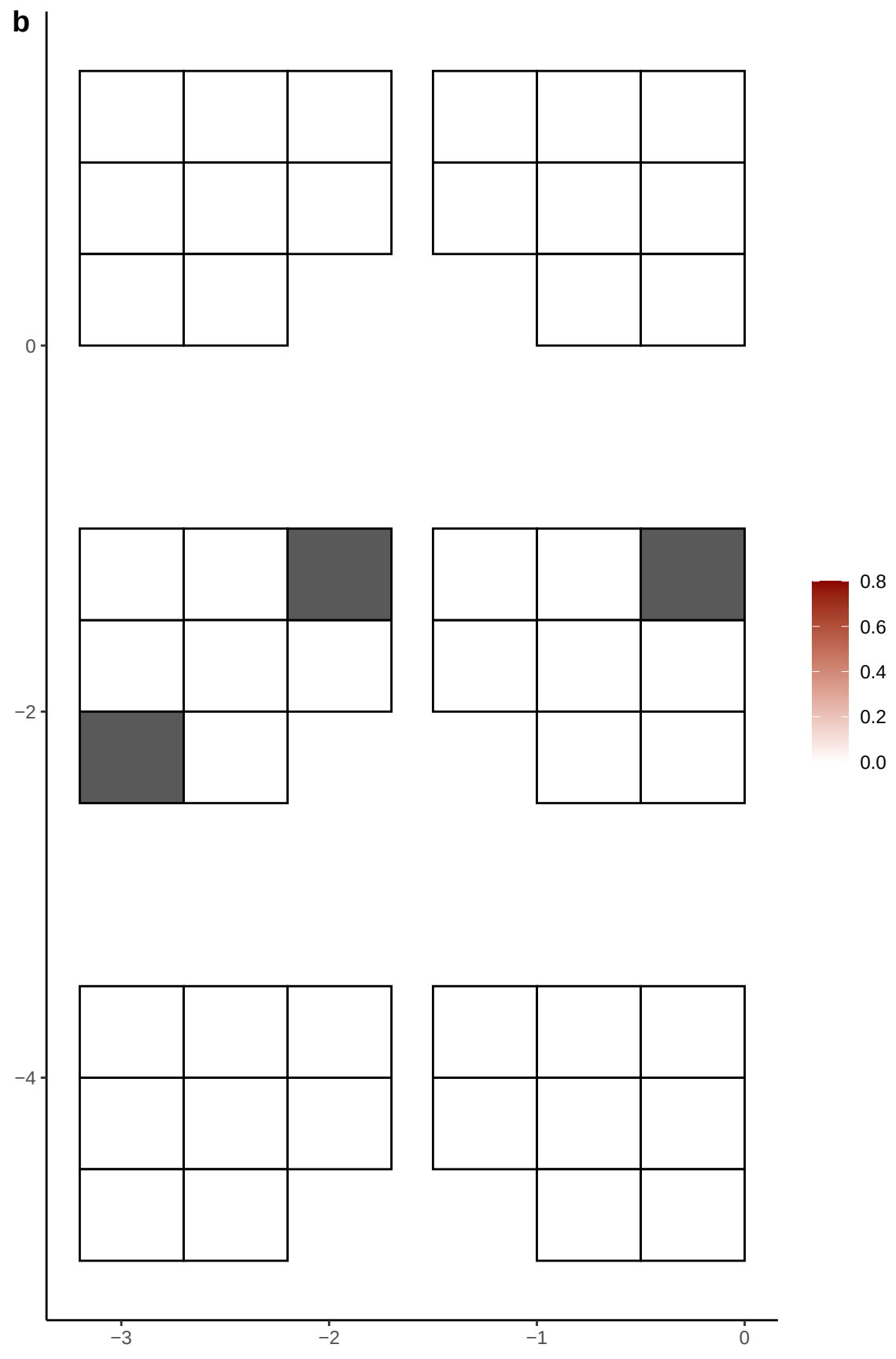

**a** ID05 1-9049289-T-C

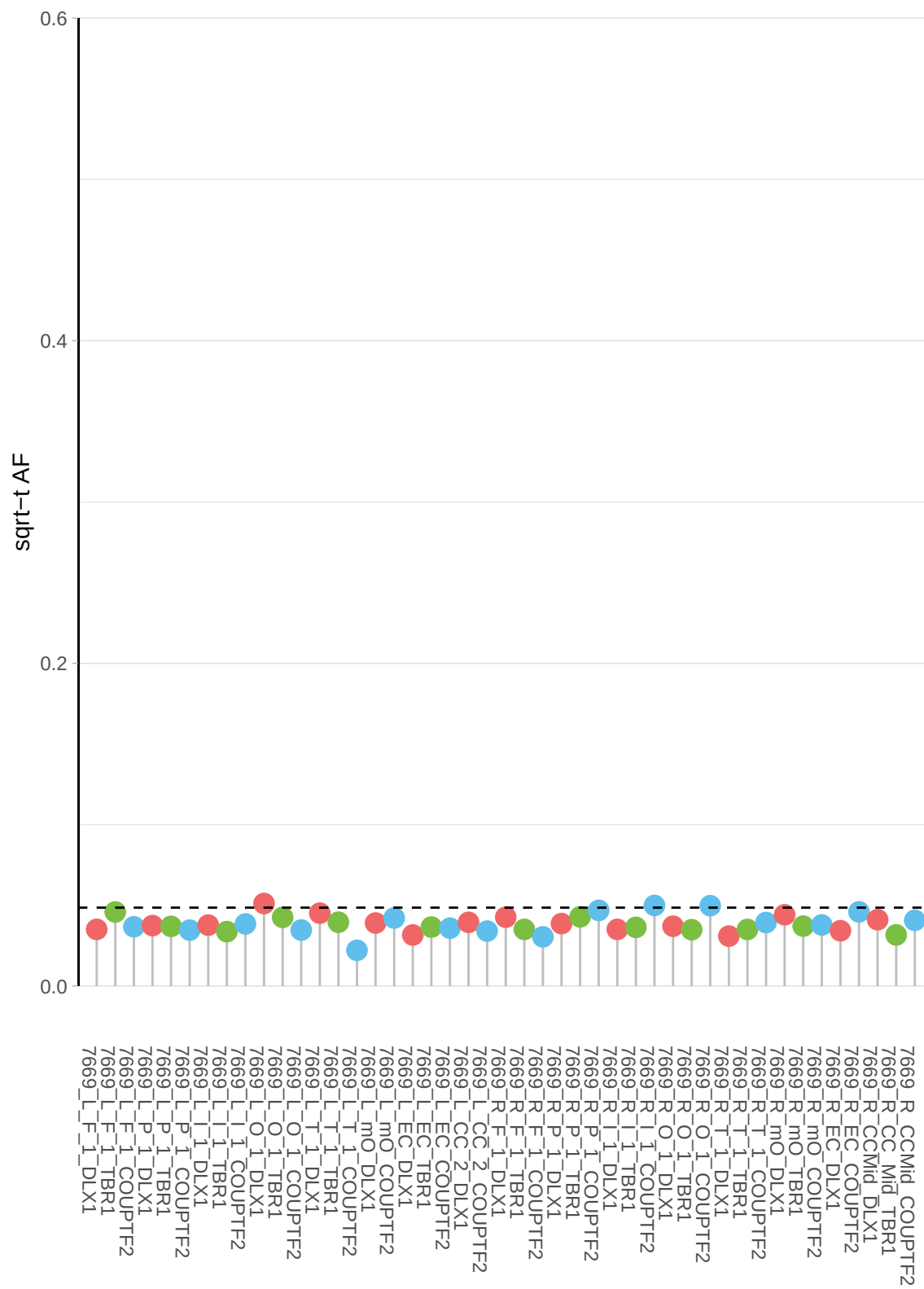

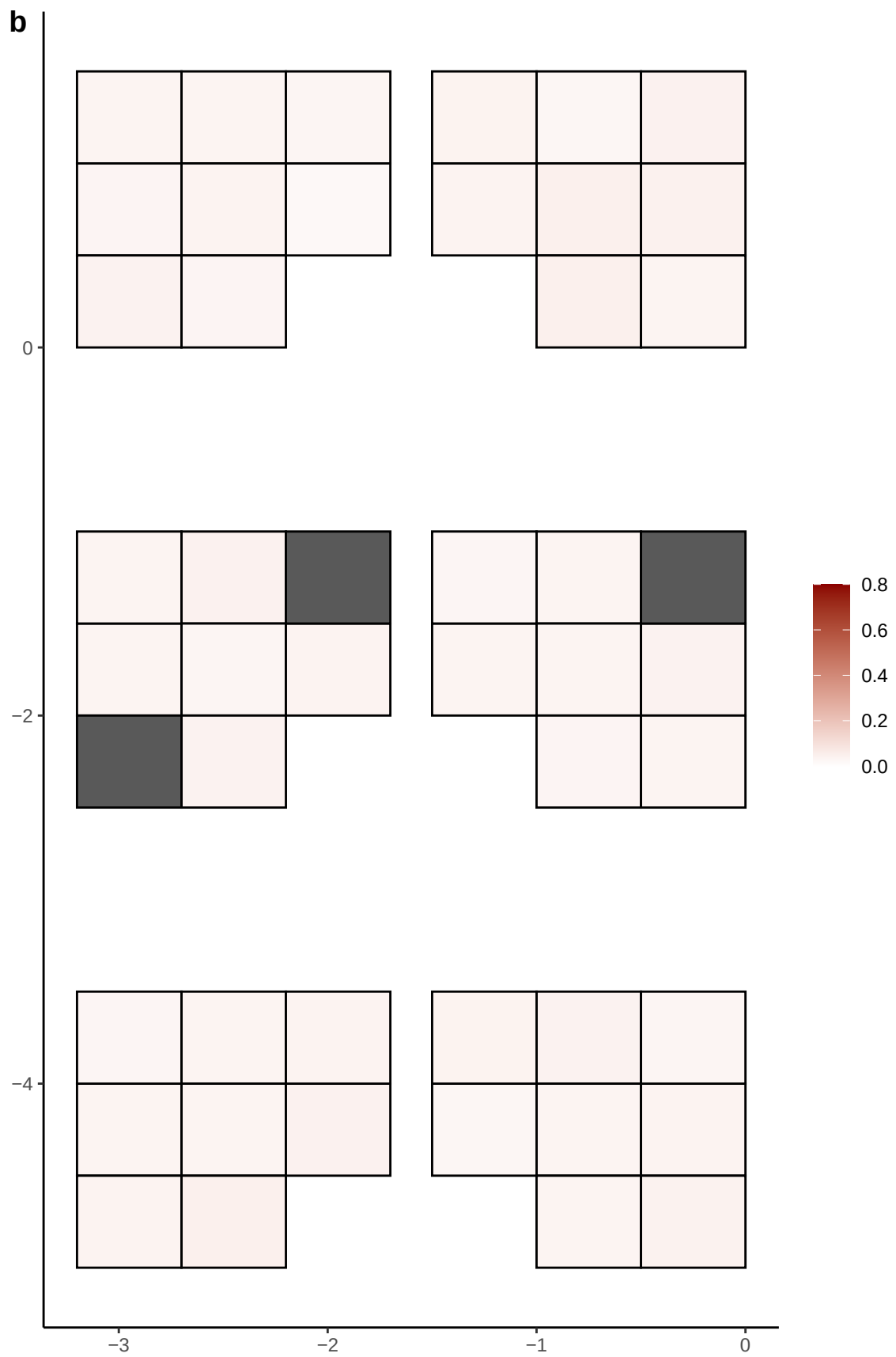

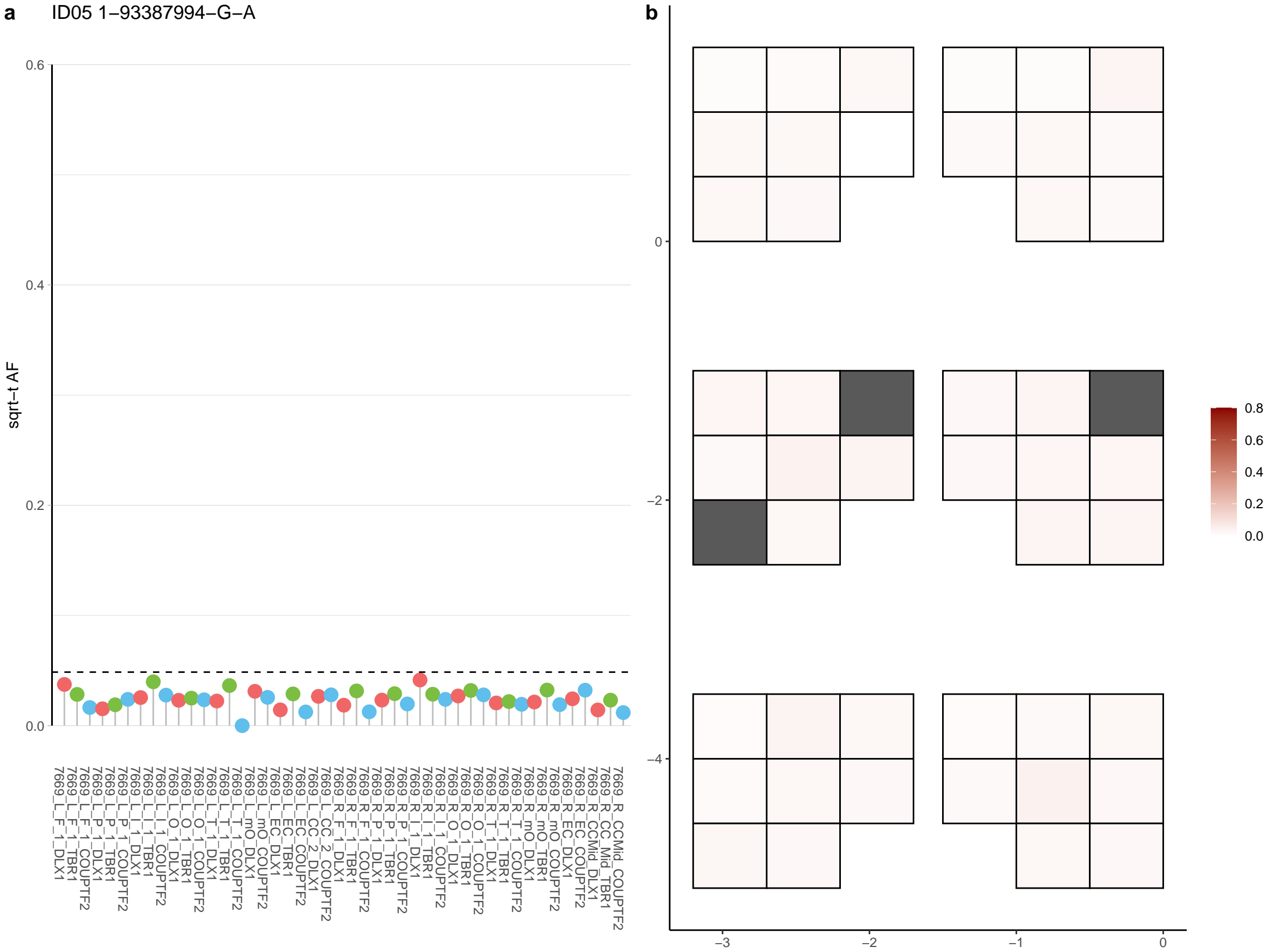

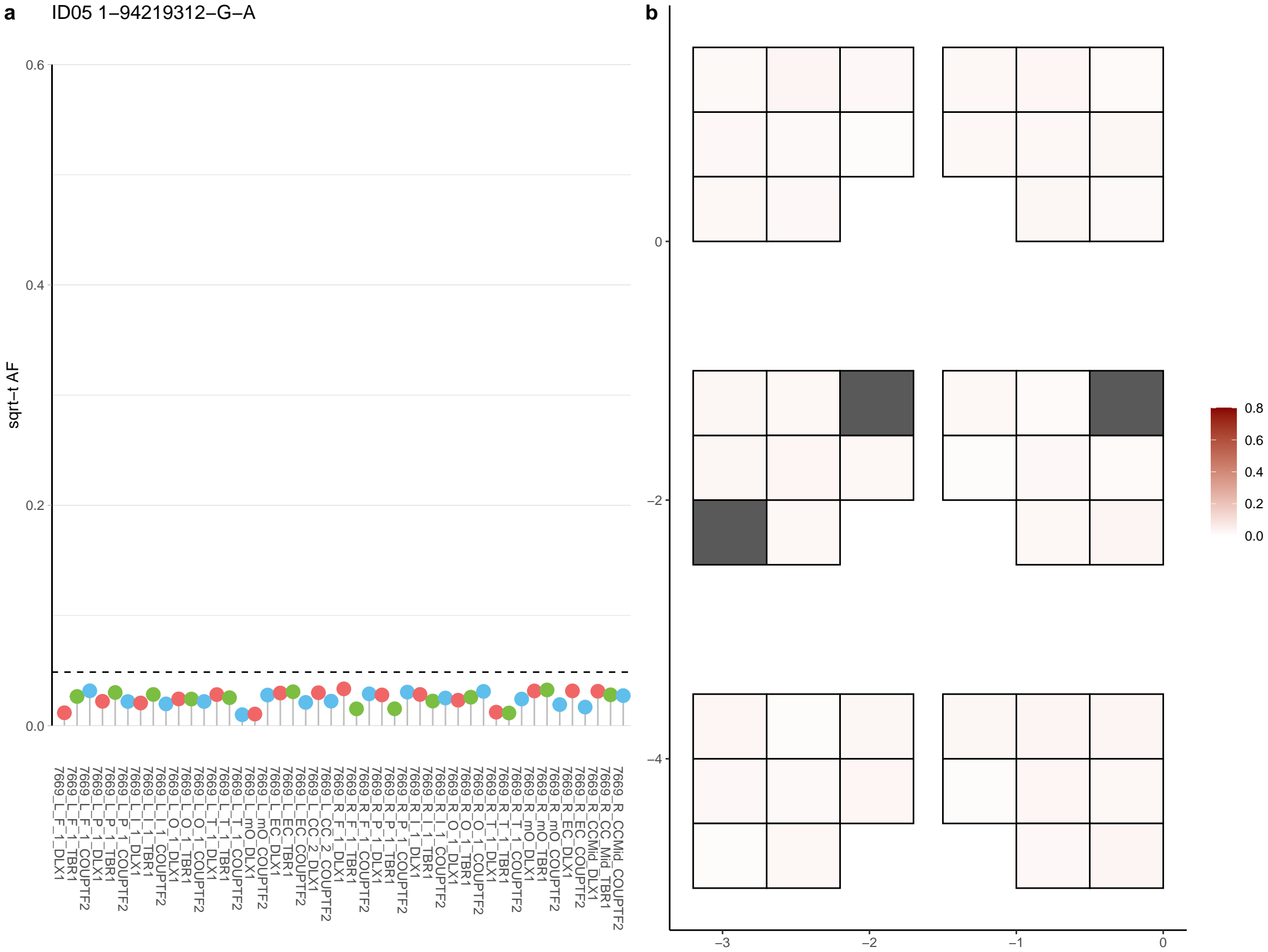

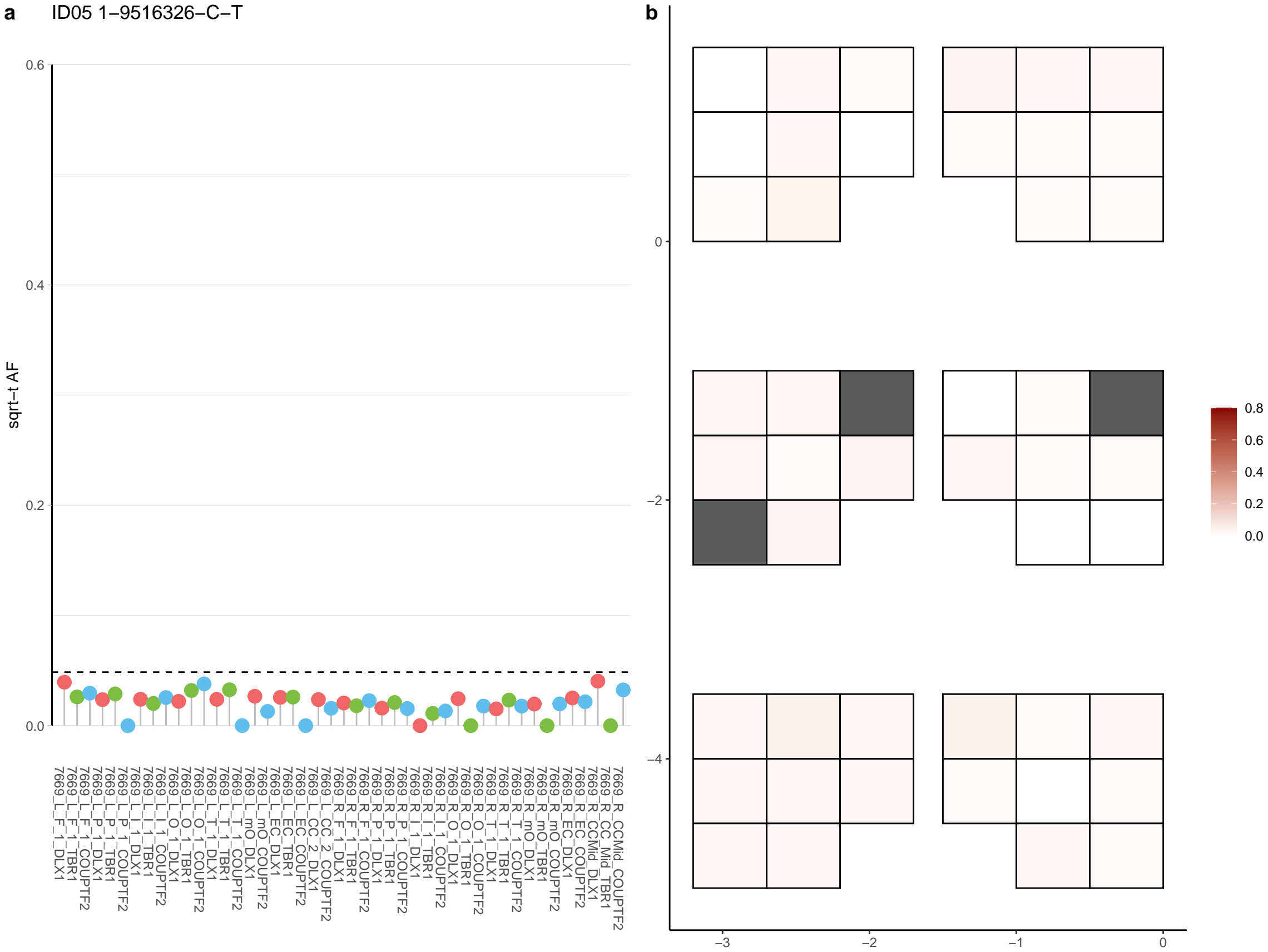

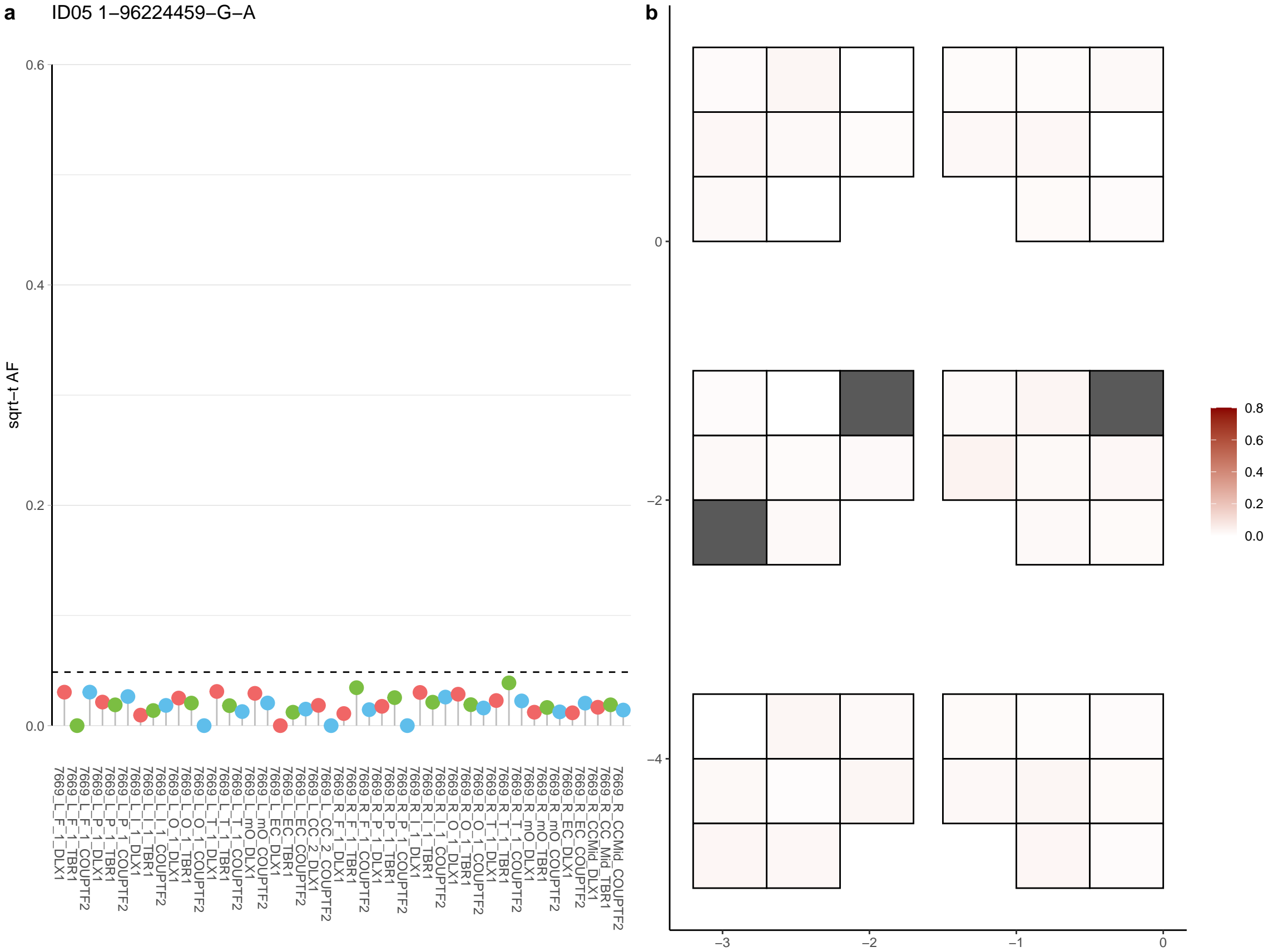

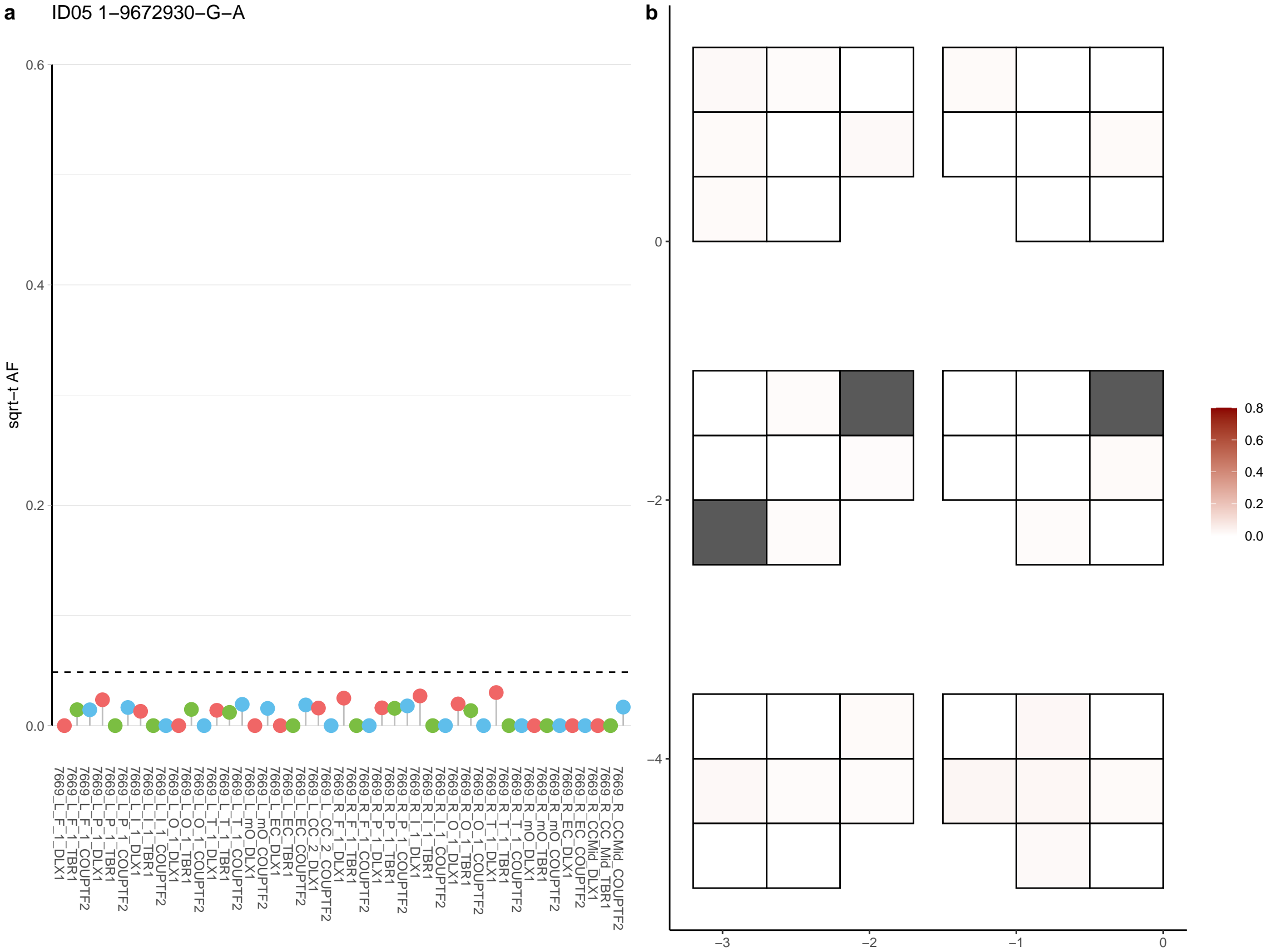

**a** ID05 1-98301896-T-C

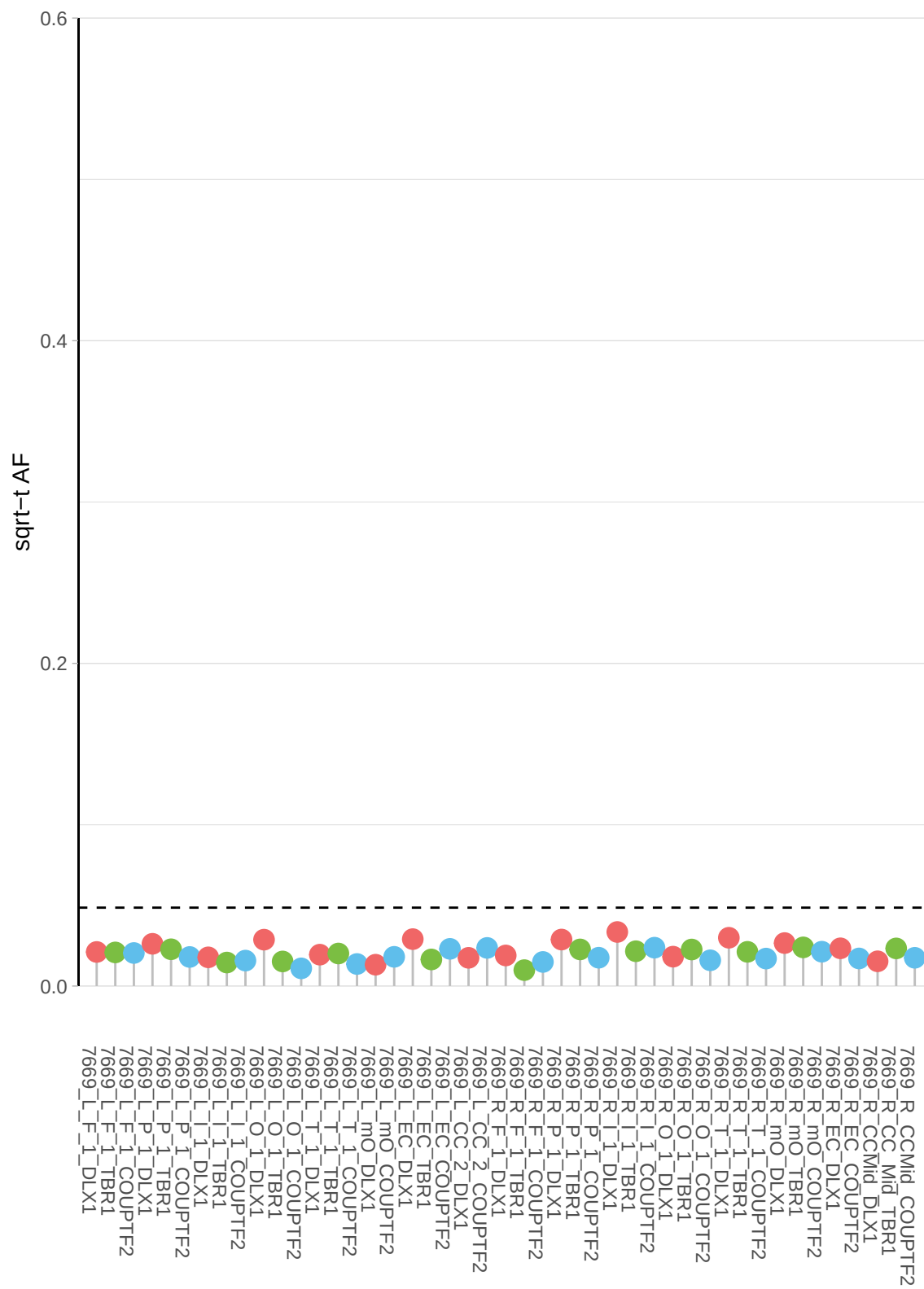

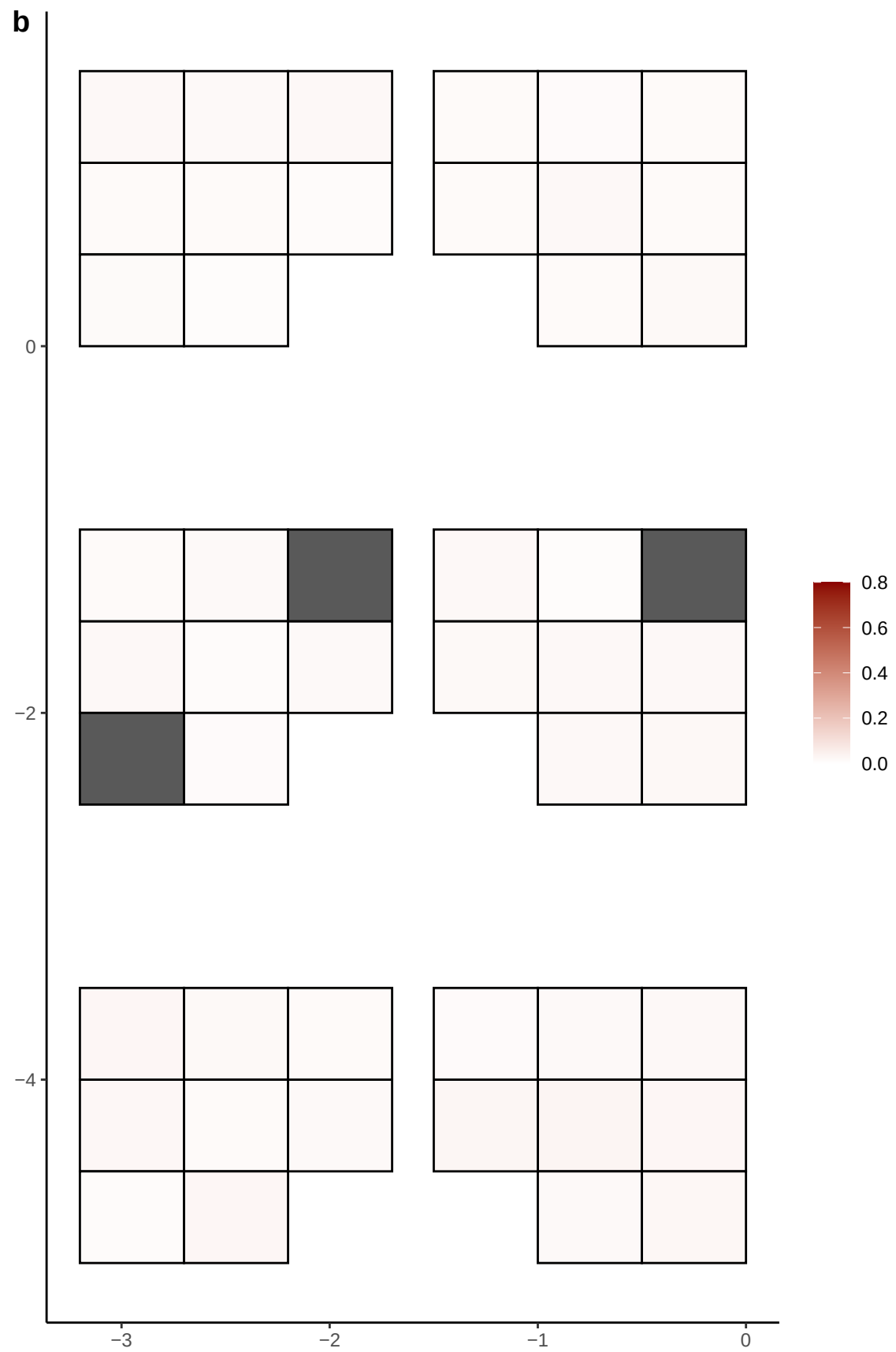

**a** ID05 10-105764174-A-G

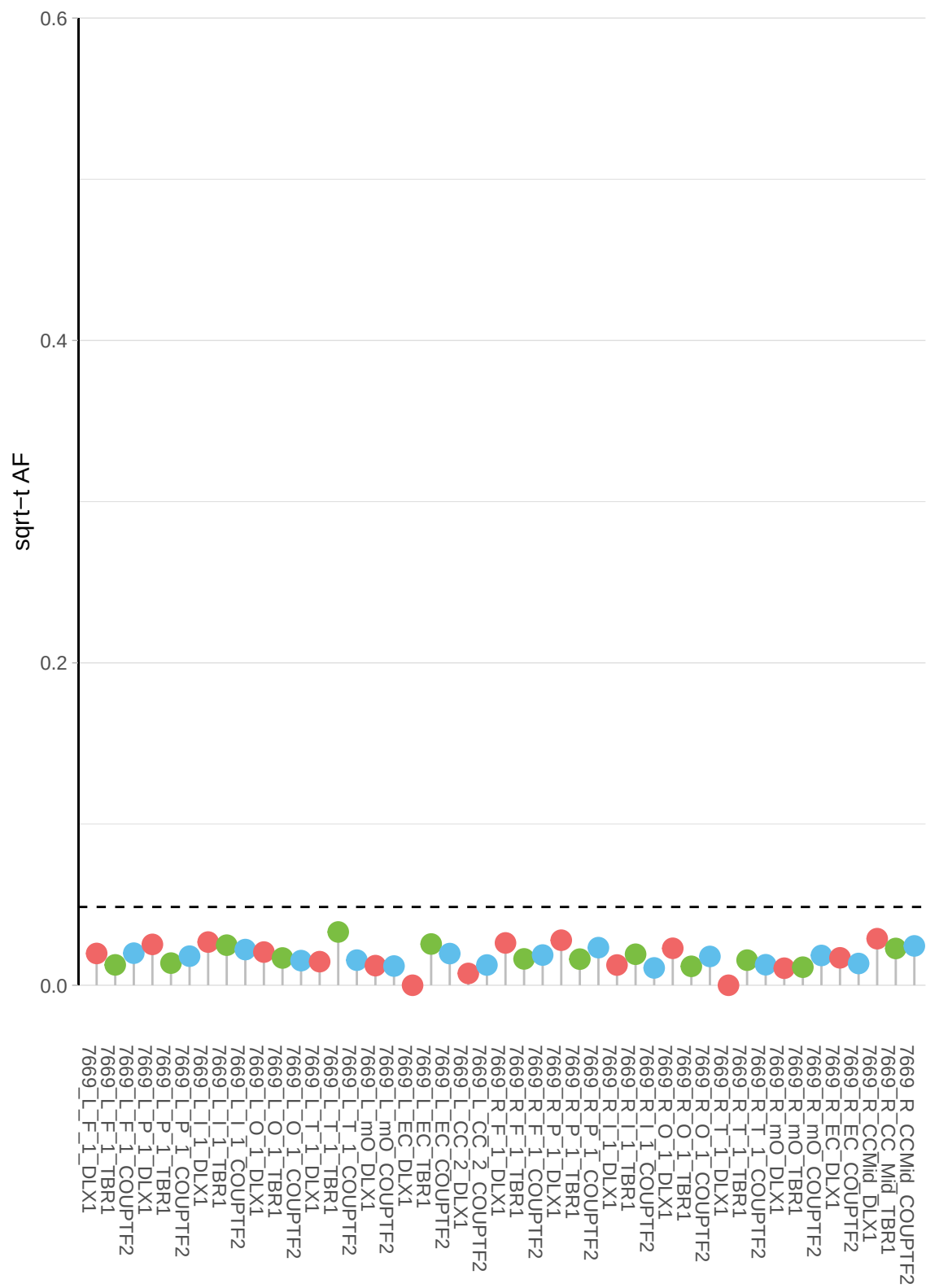

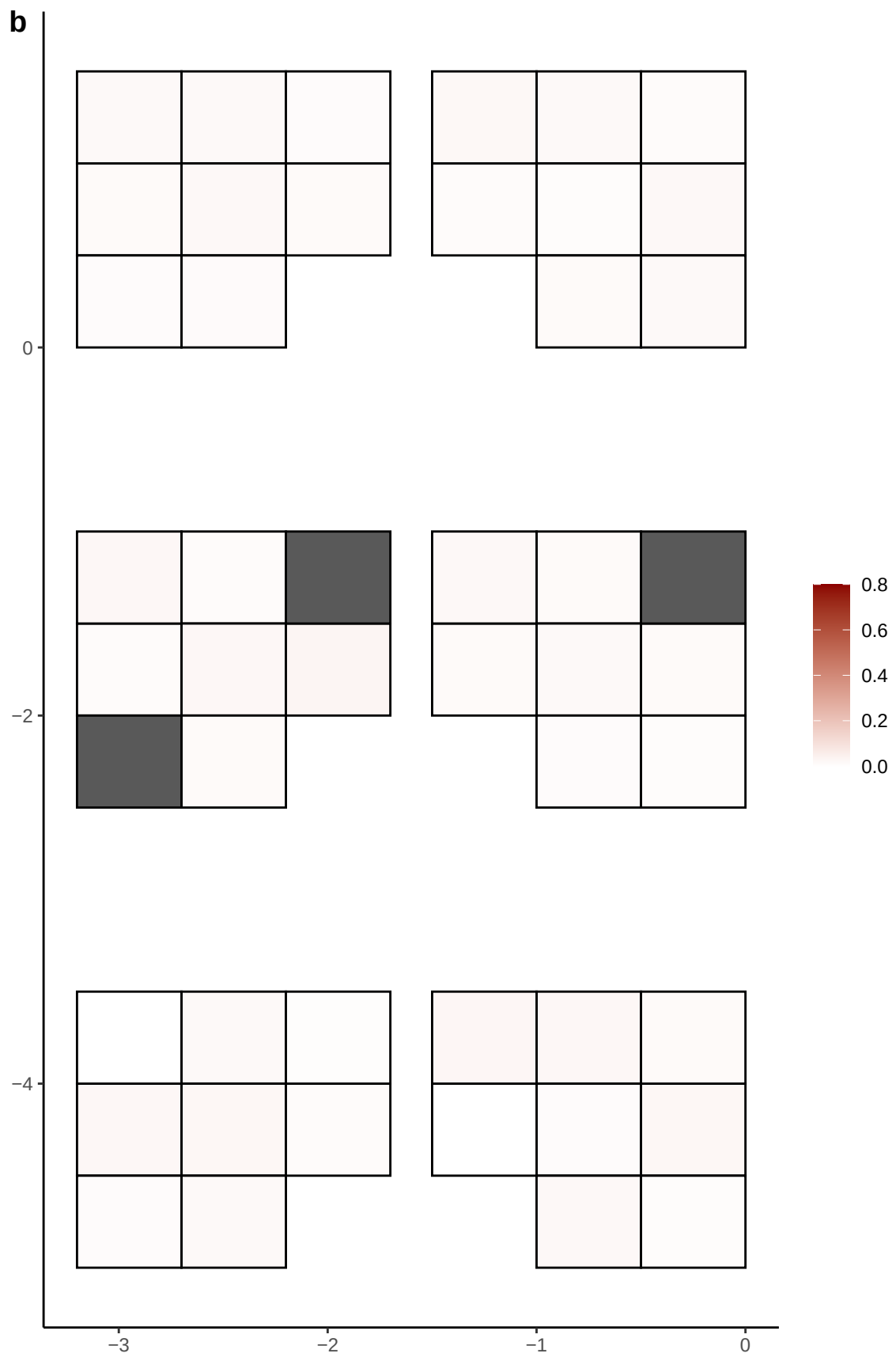

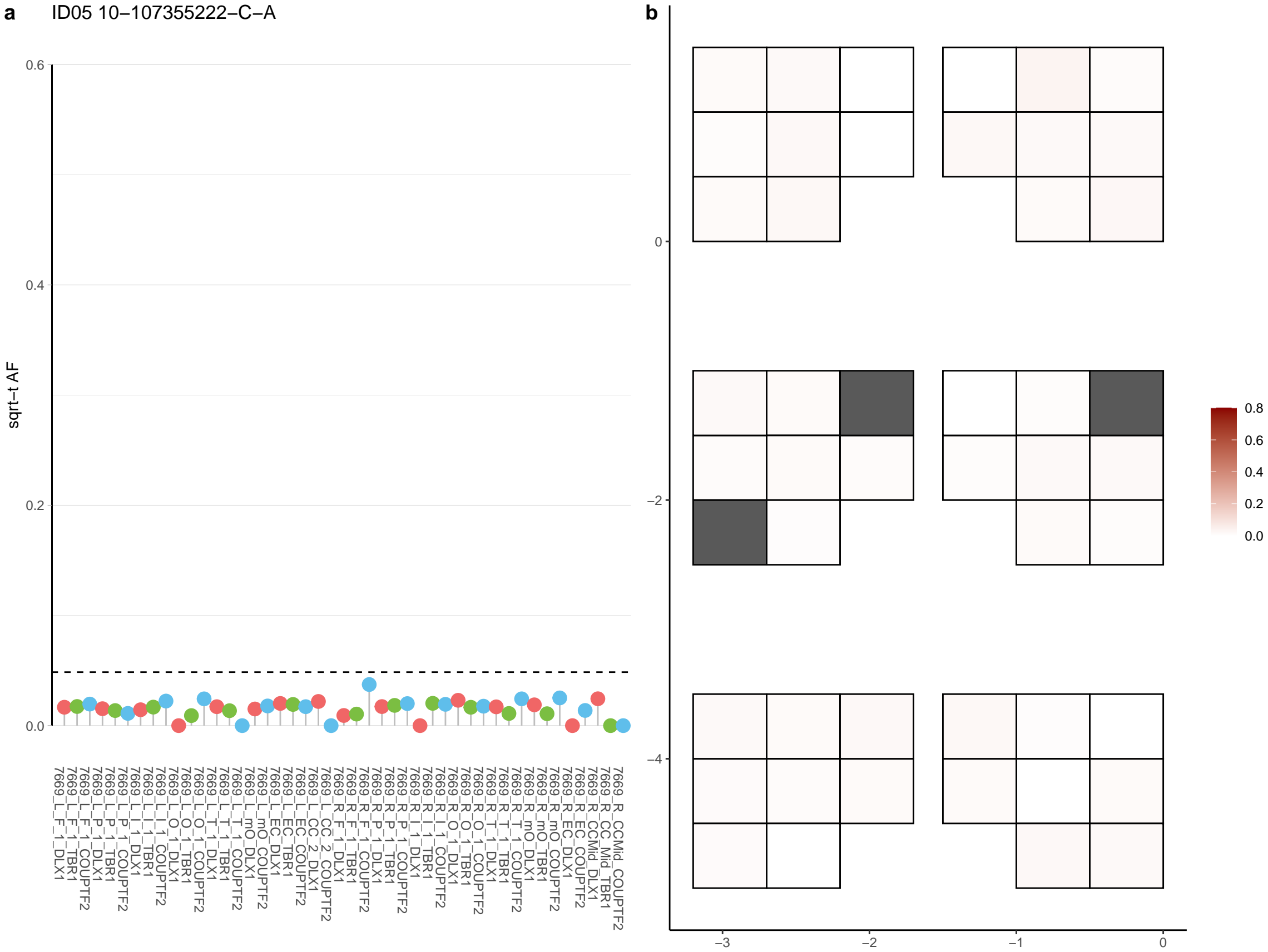

**a** ID05 10-108671103-C-T

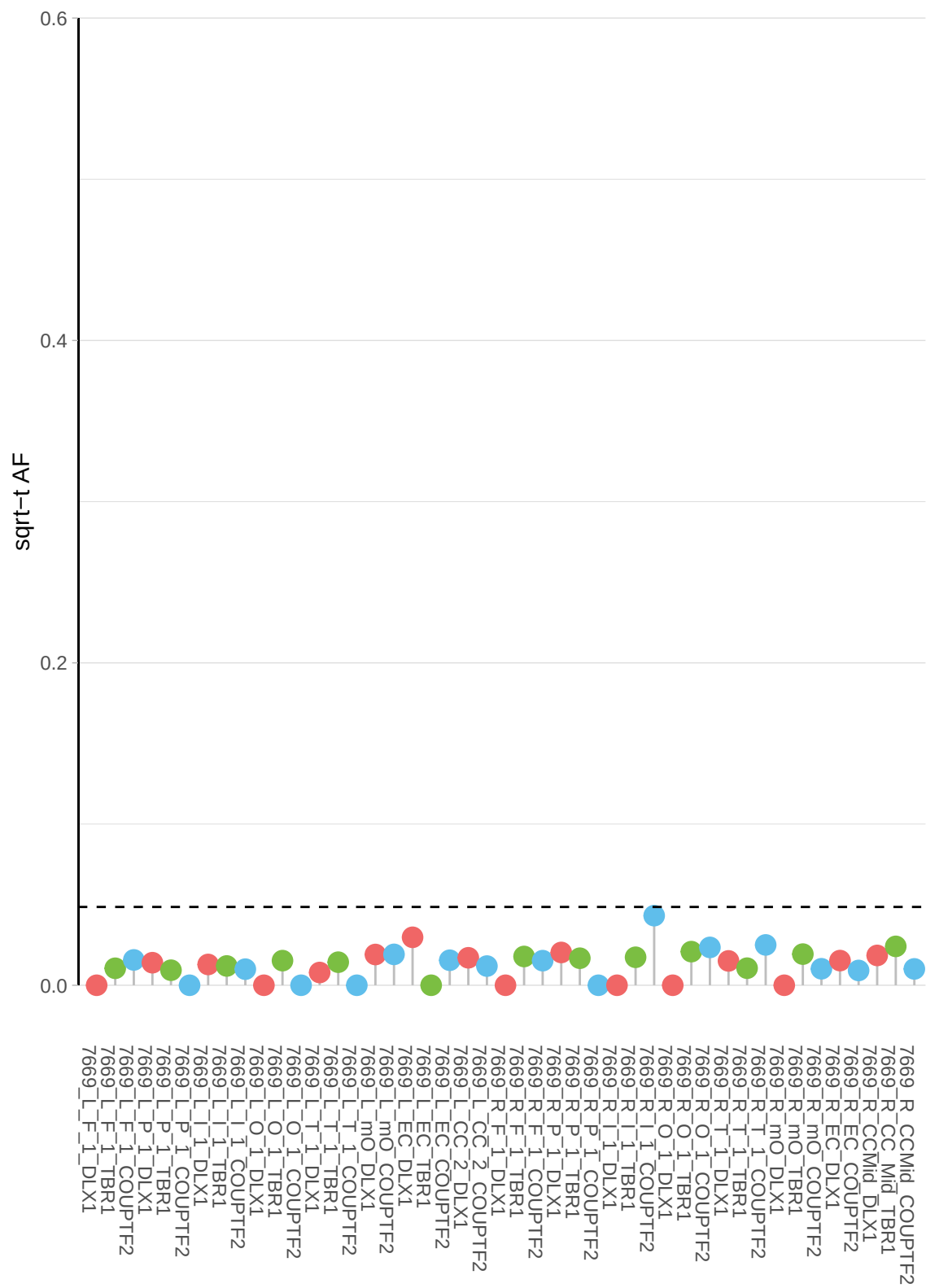

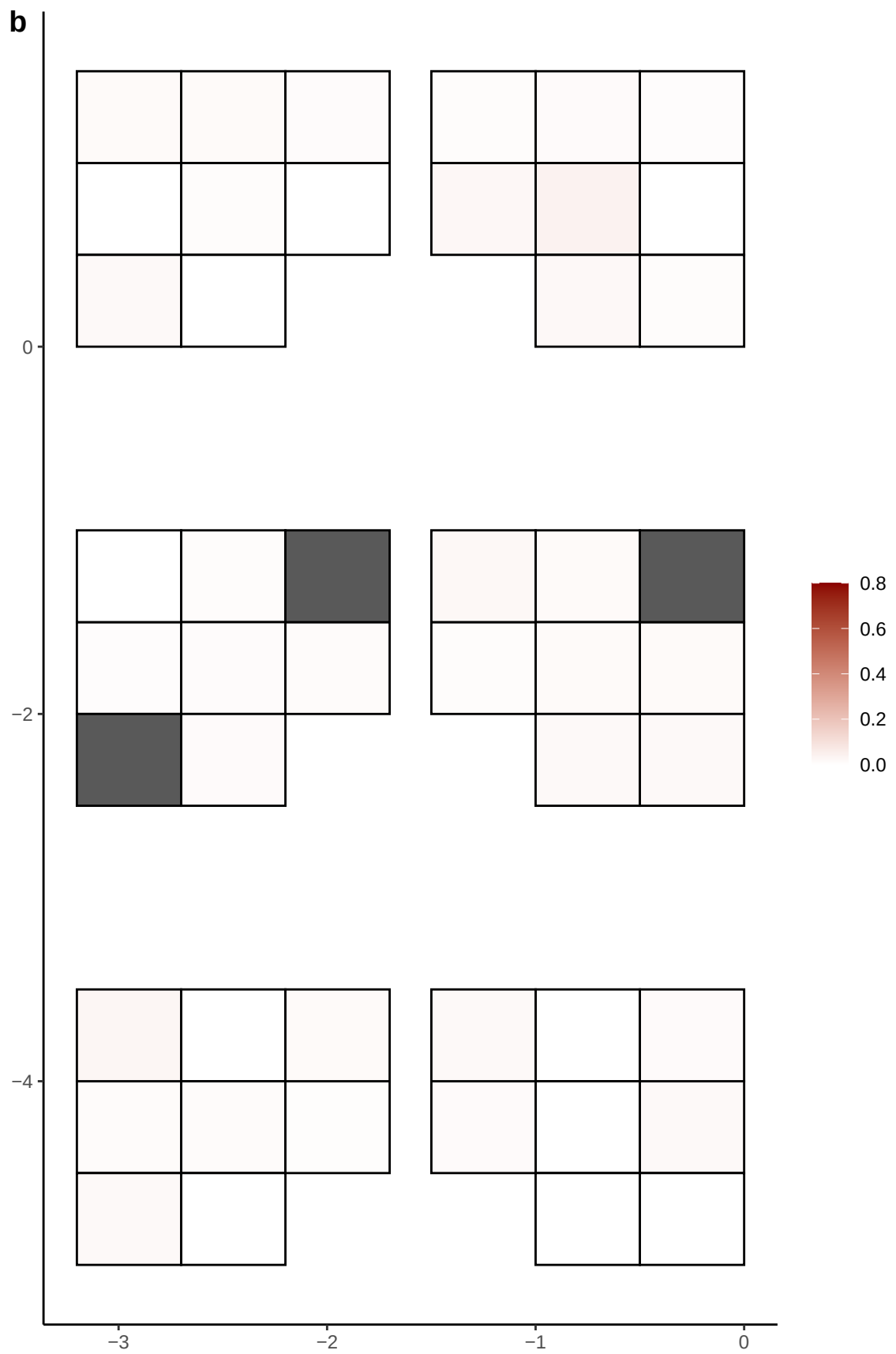

**a** ID05 10-109795130-C-G

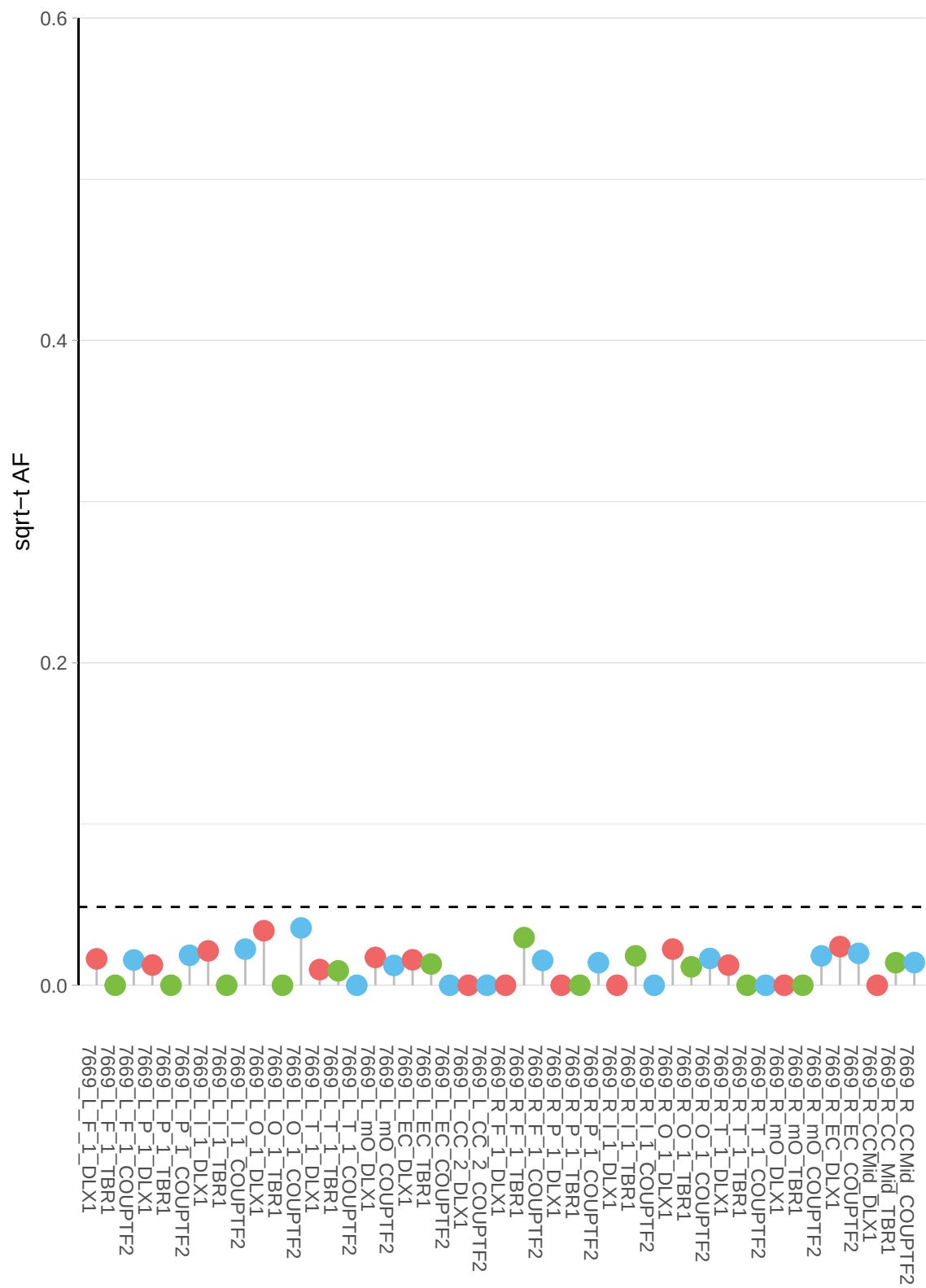

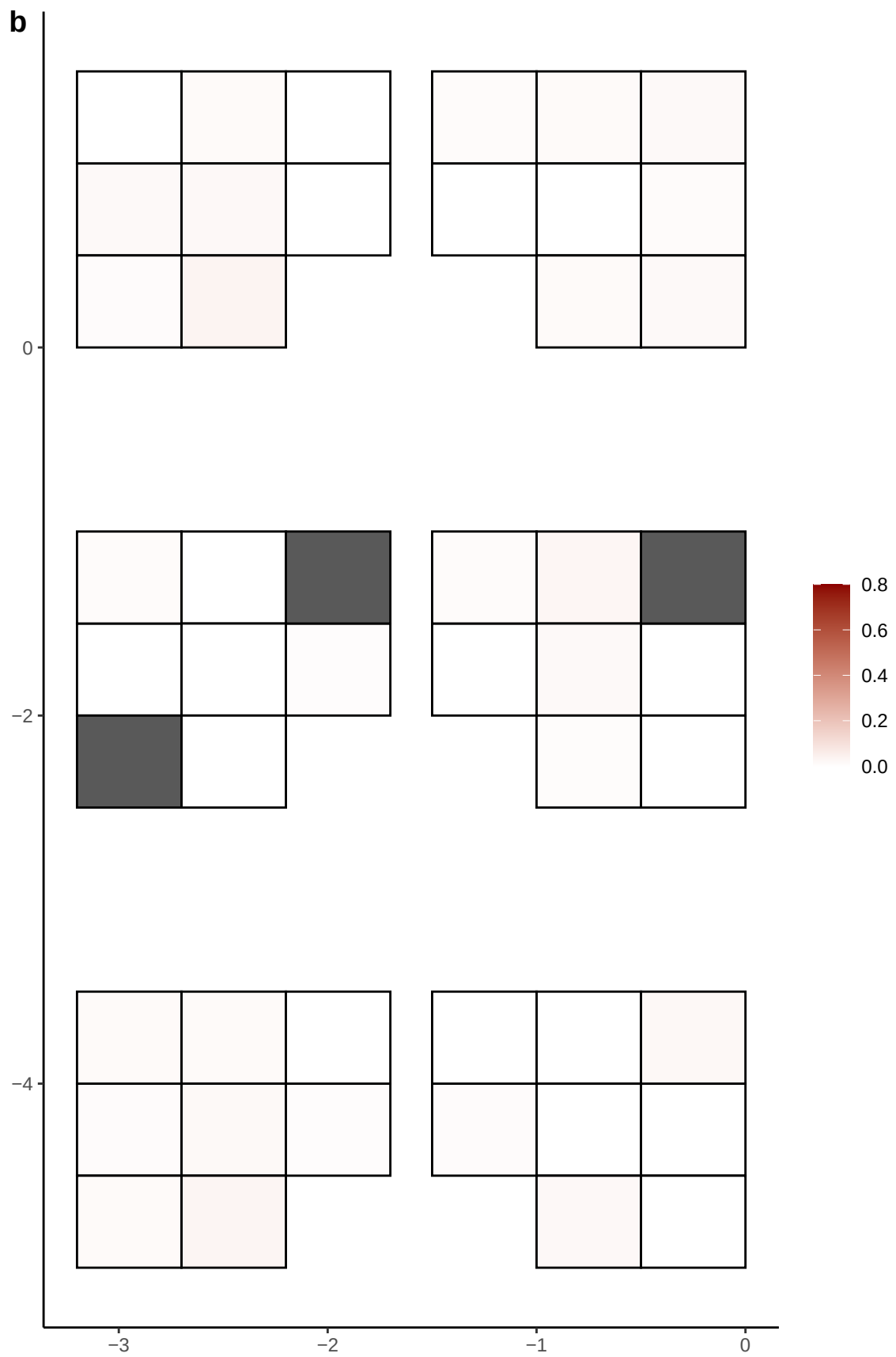

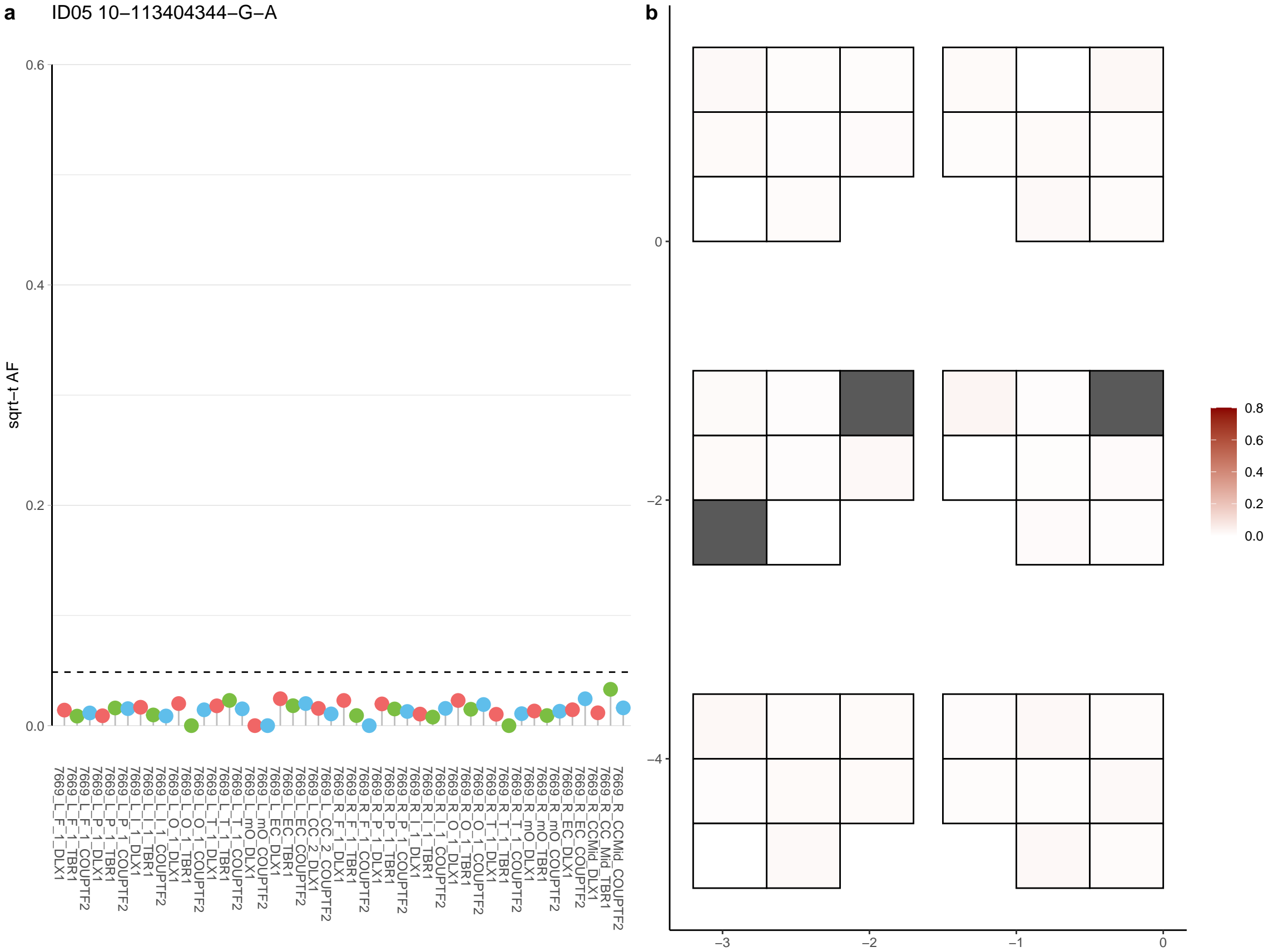

**a** ID05 10-116677604-C-T

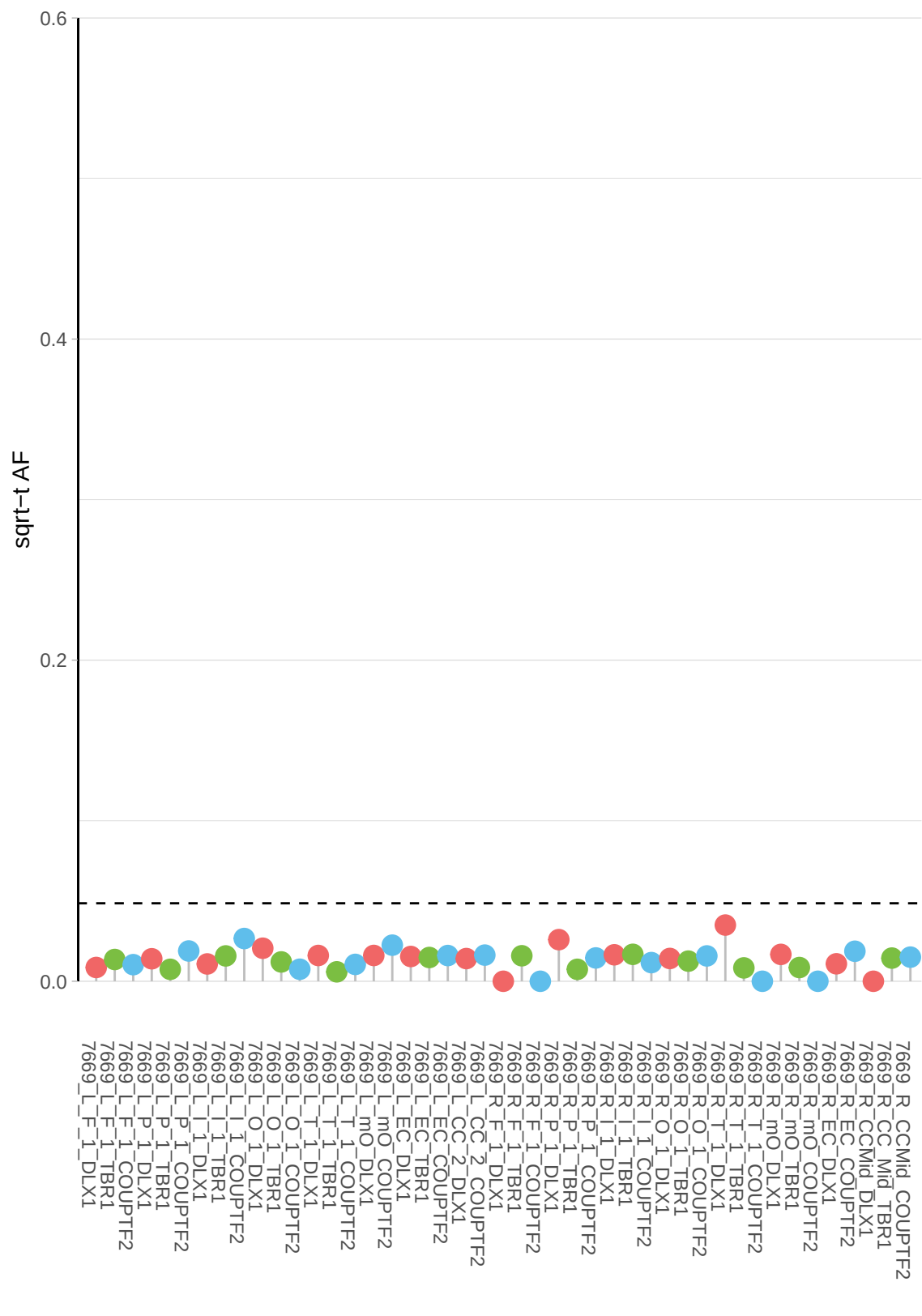

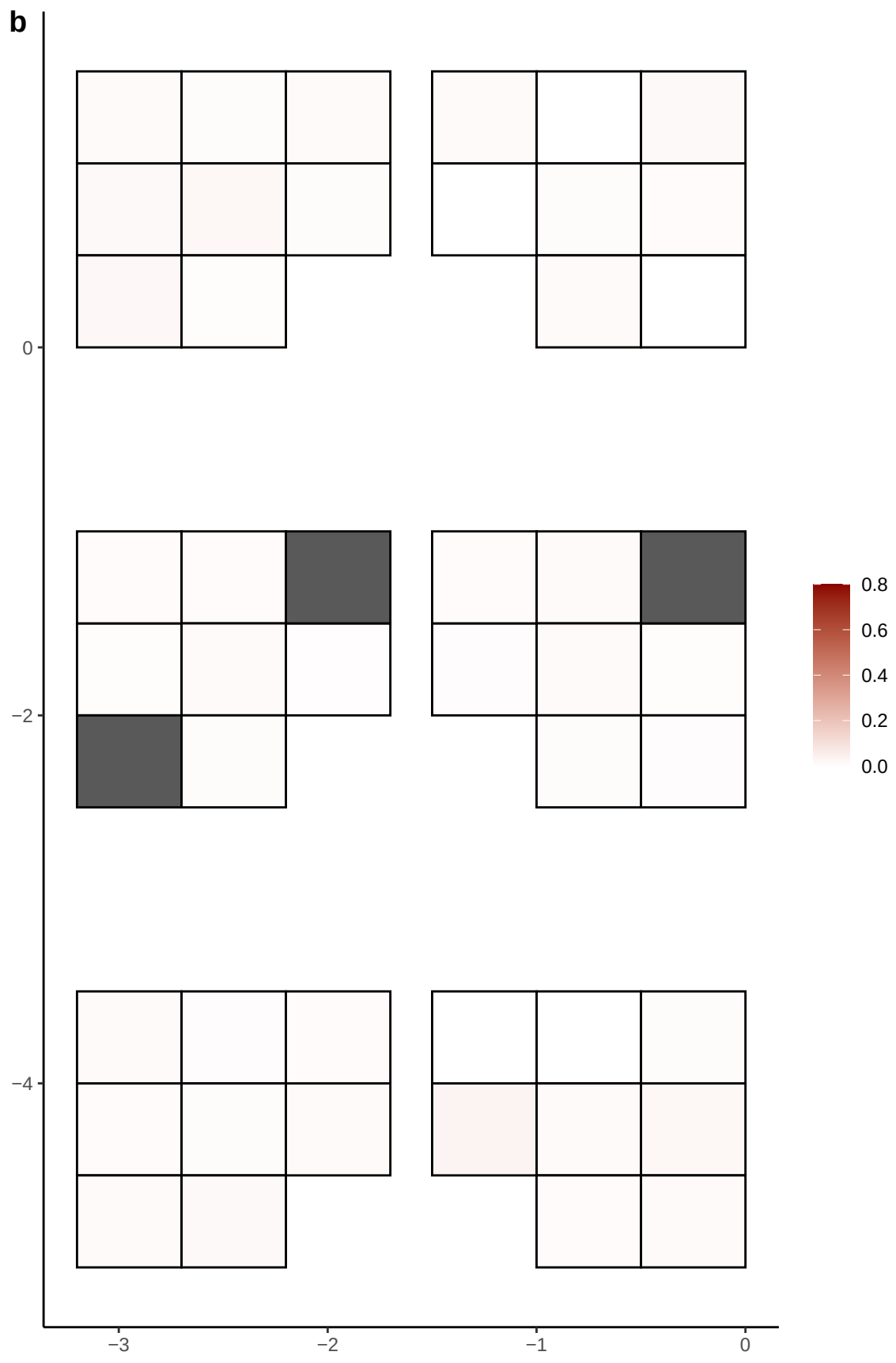

**a** ID05 10-117540461-G-A

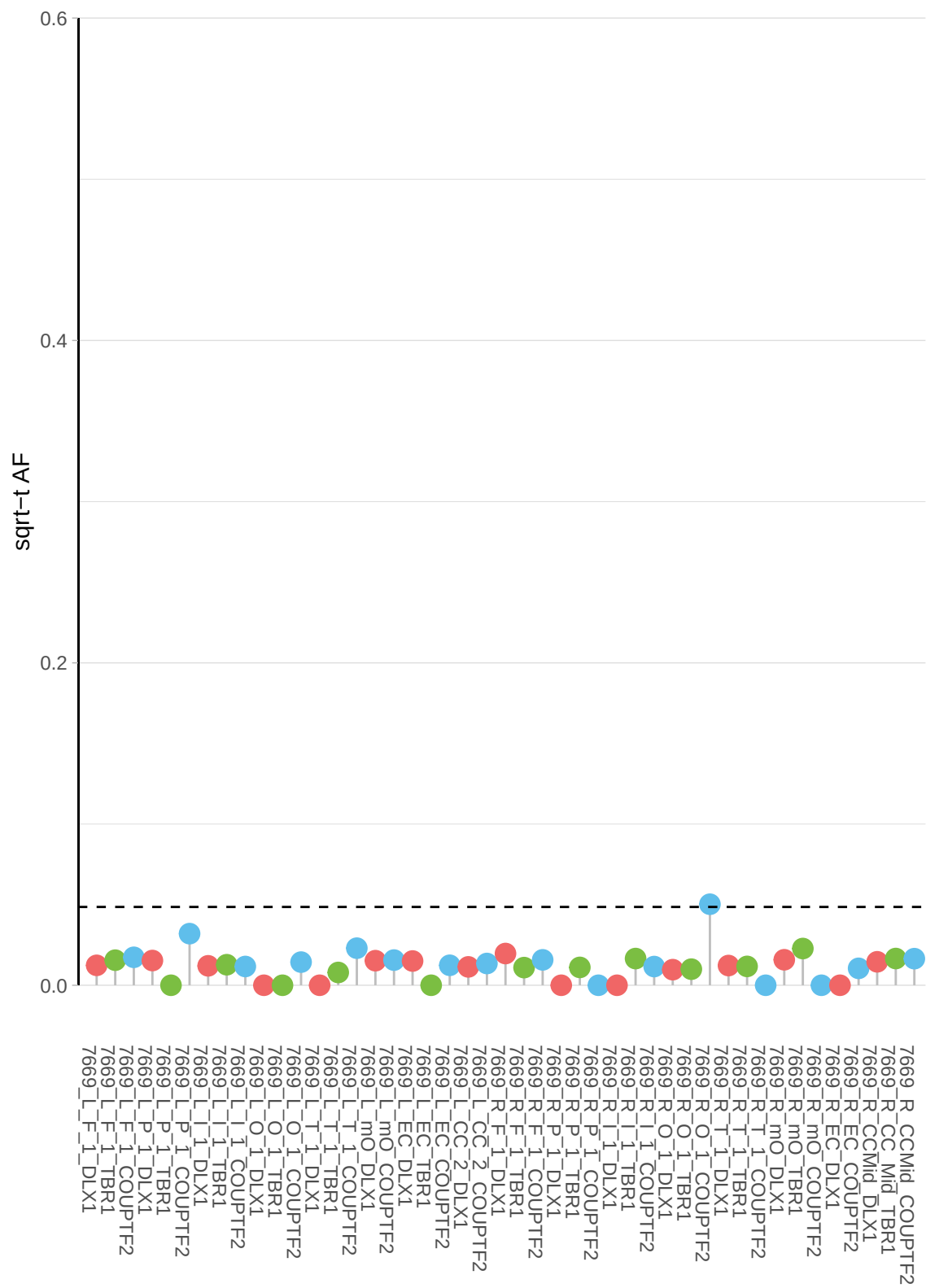

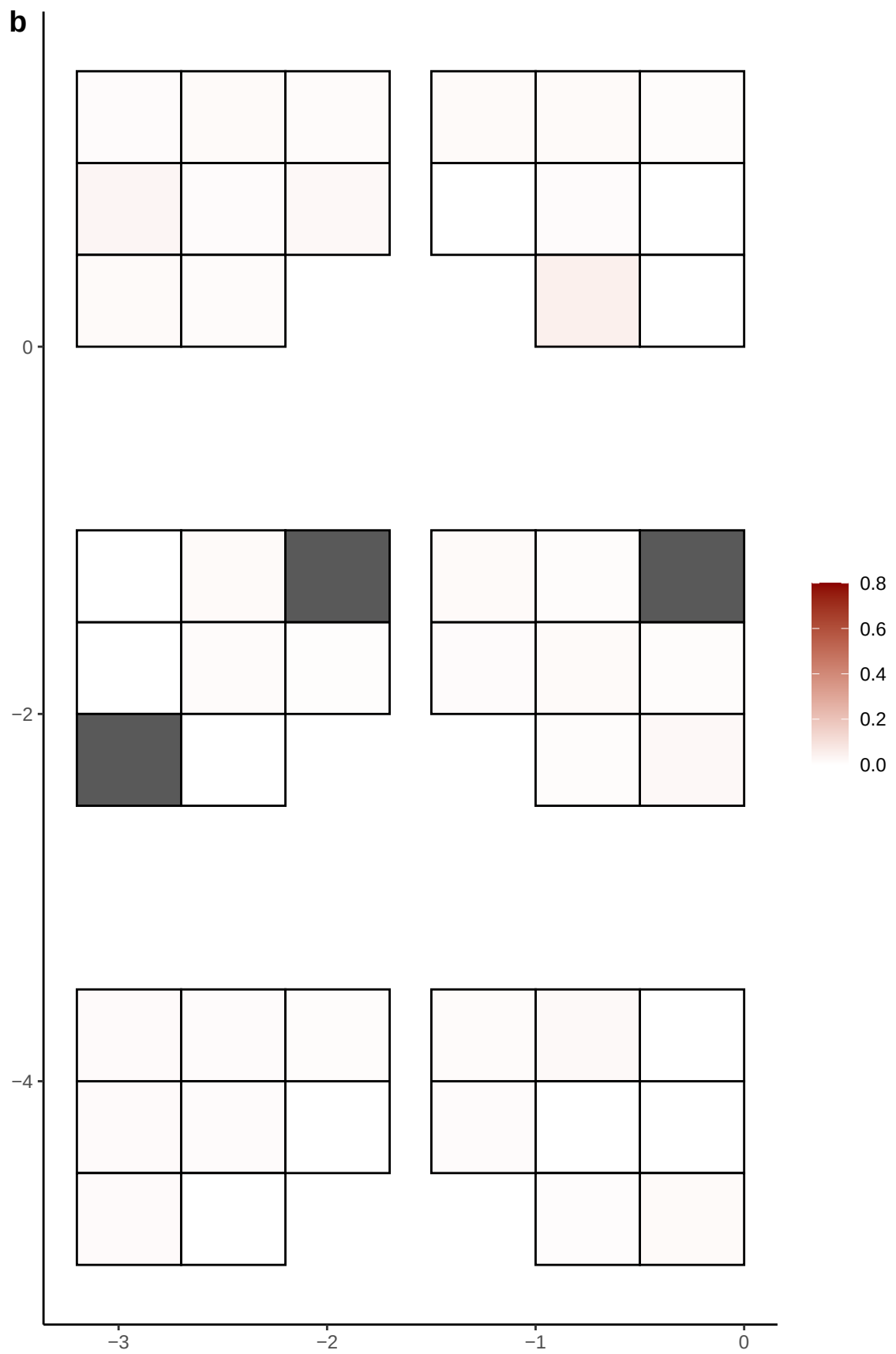

**a** ID05 10-121073761-C-T

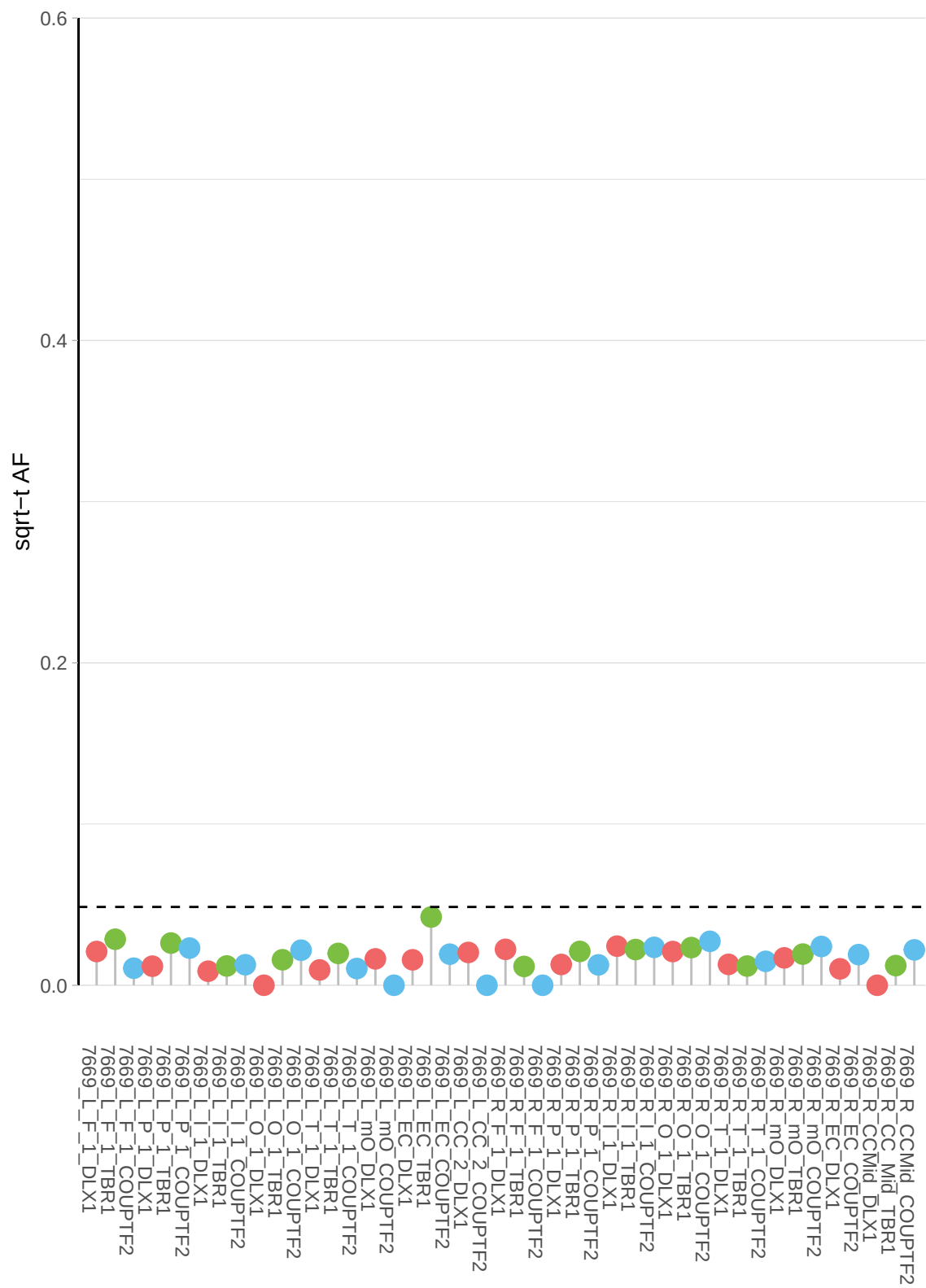

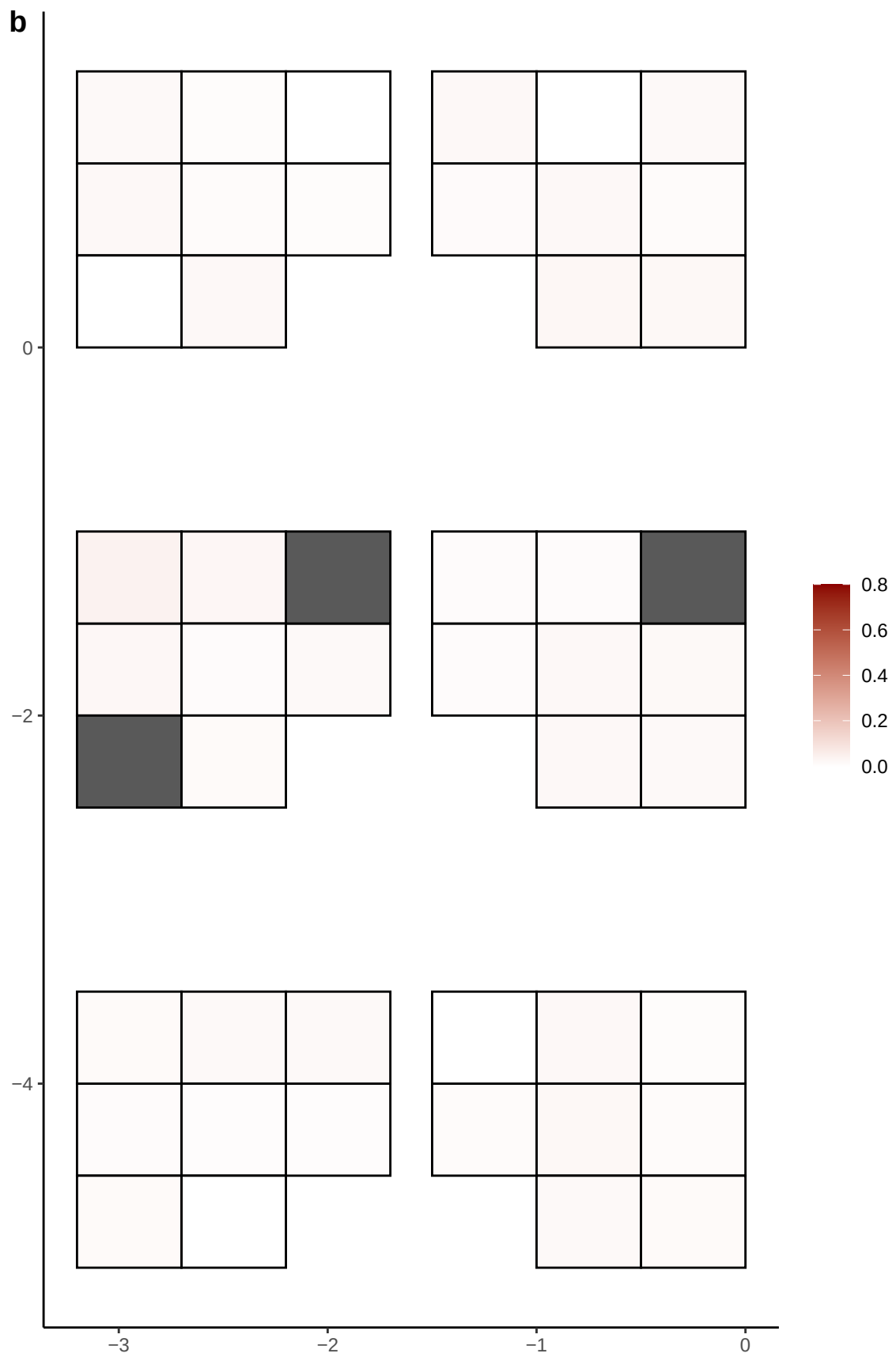

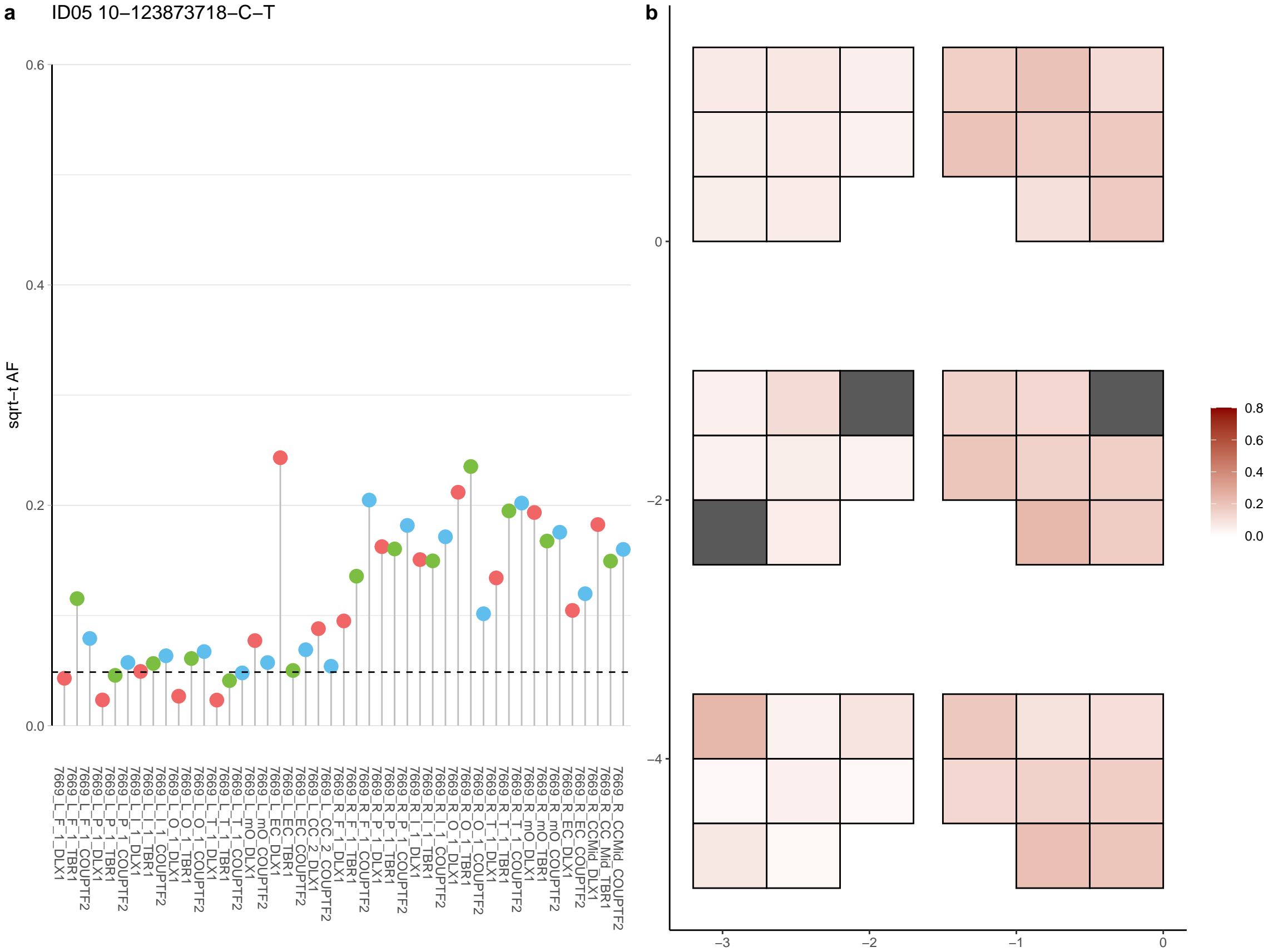

**a** ID05 10-125075590-G-A

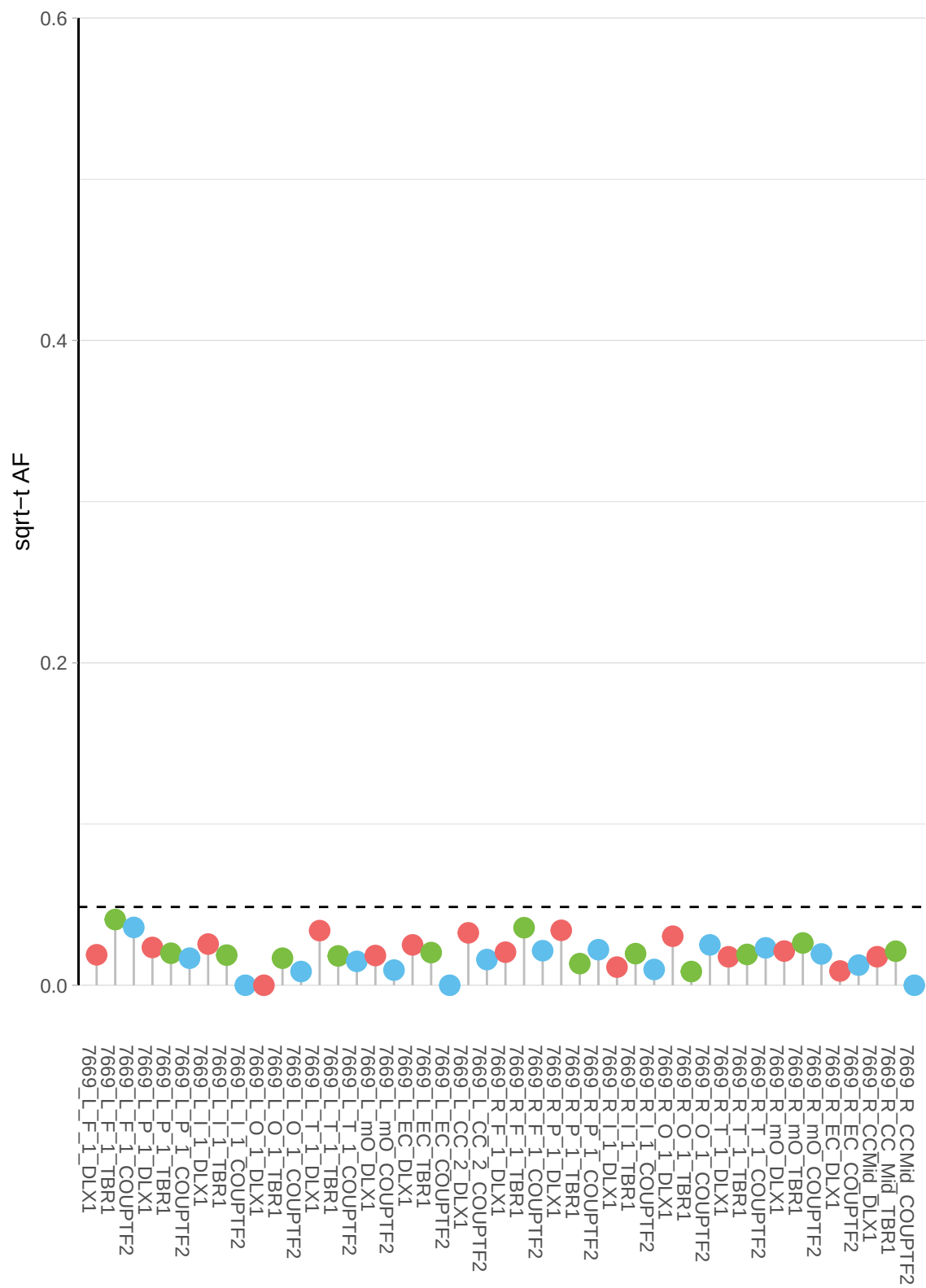

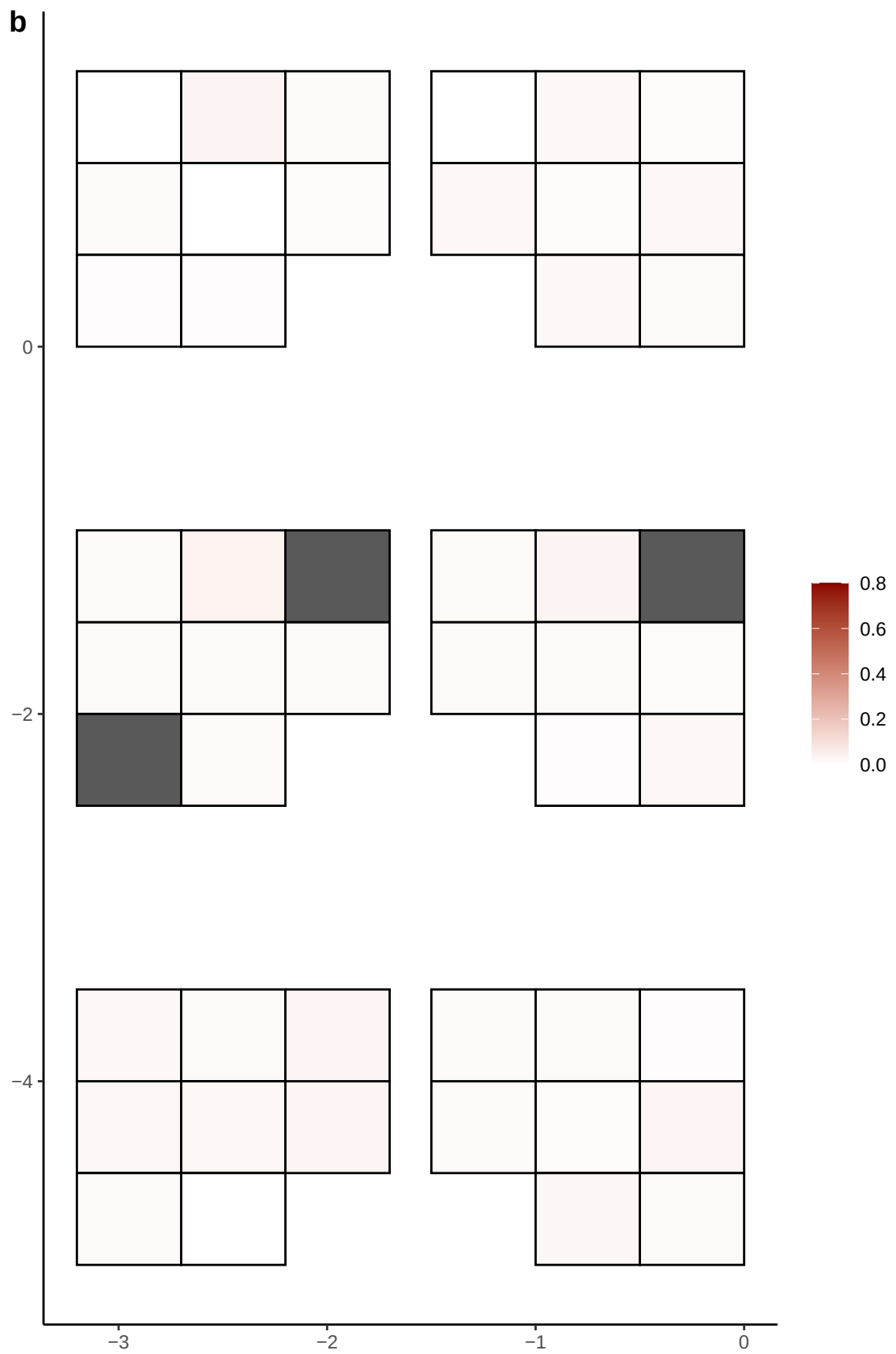

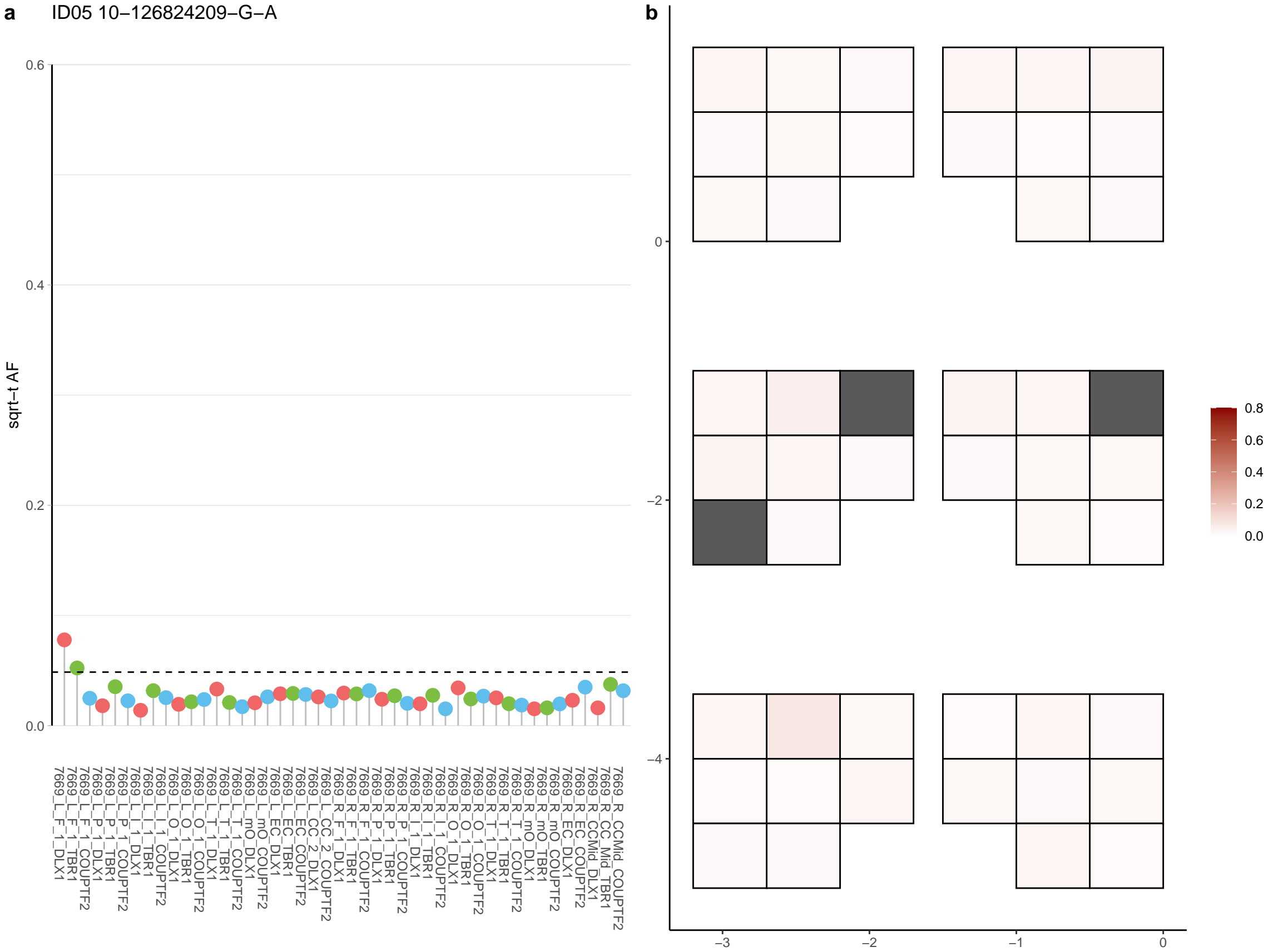

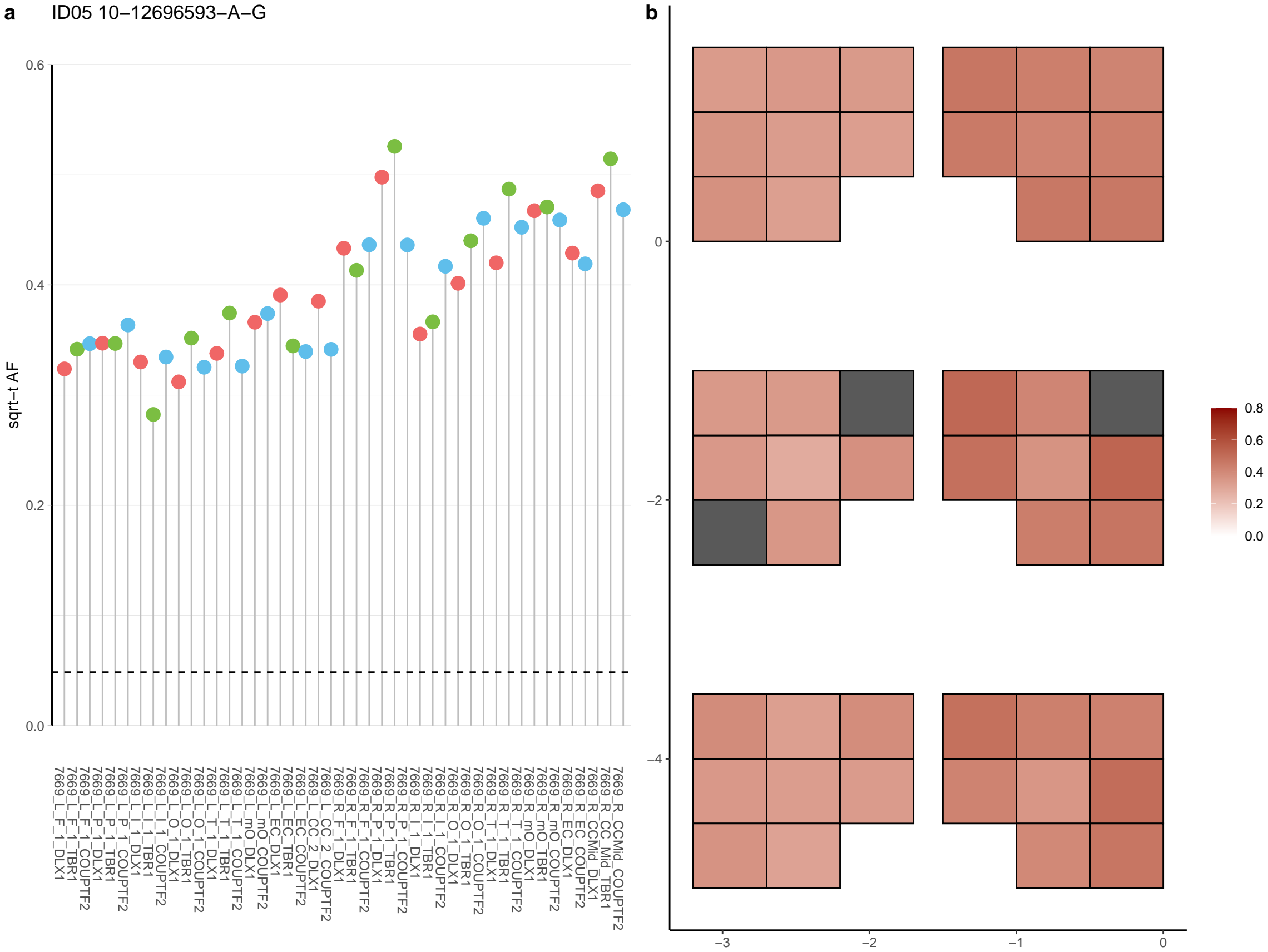

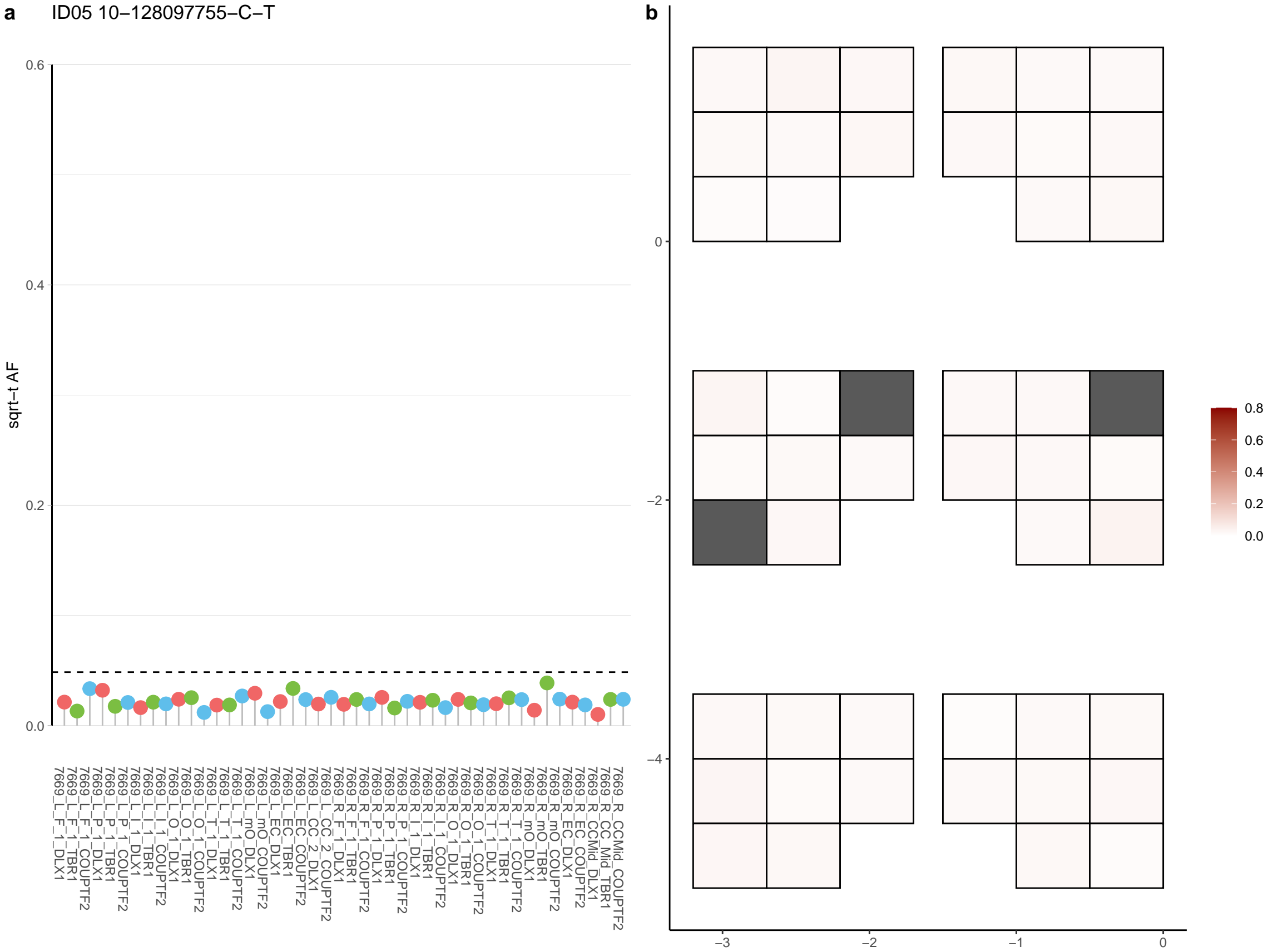

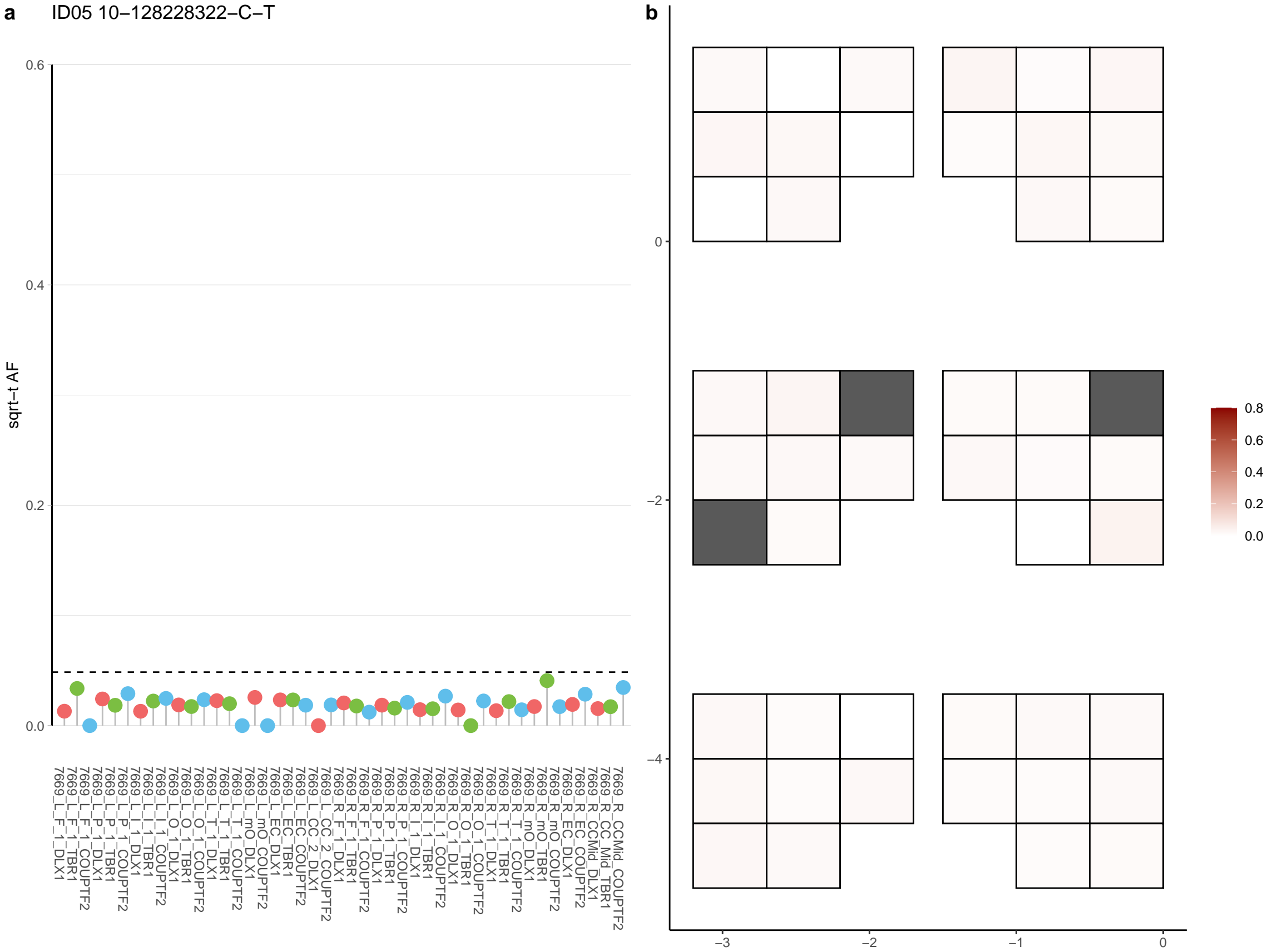

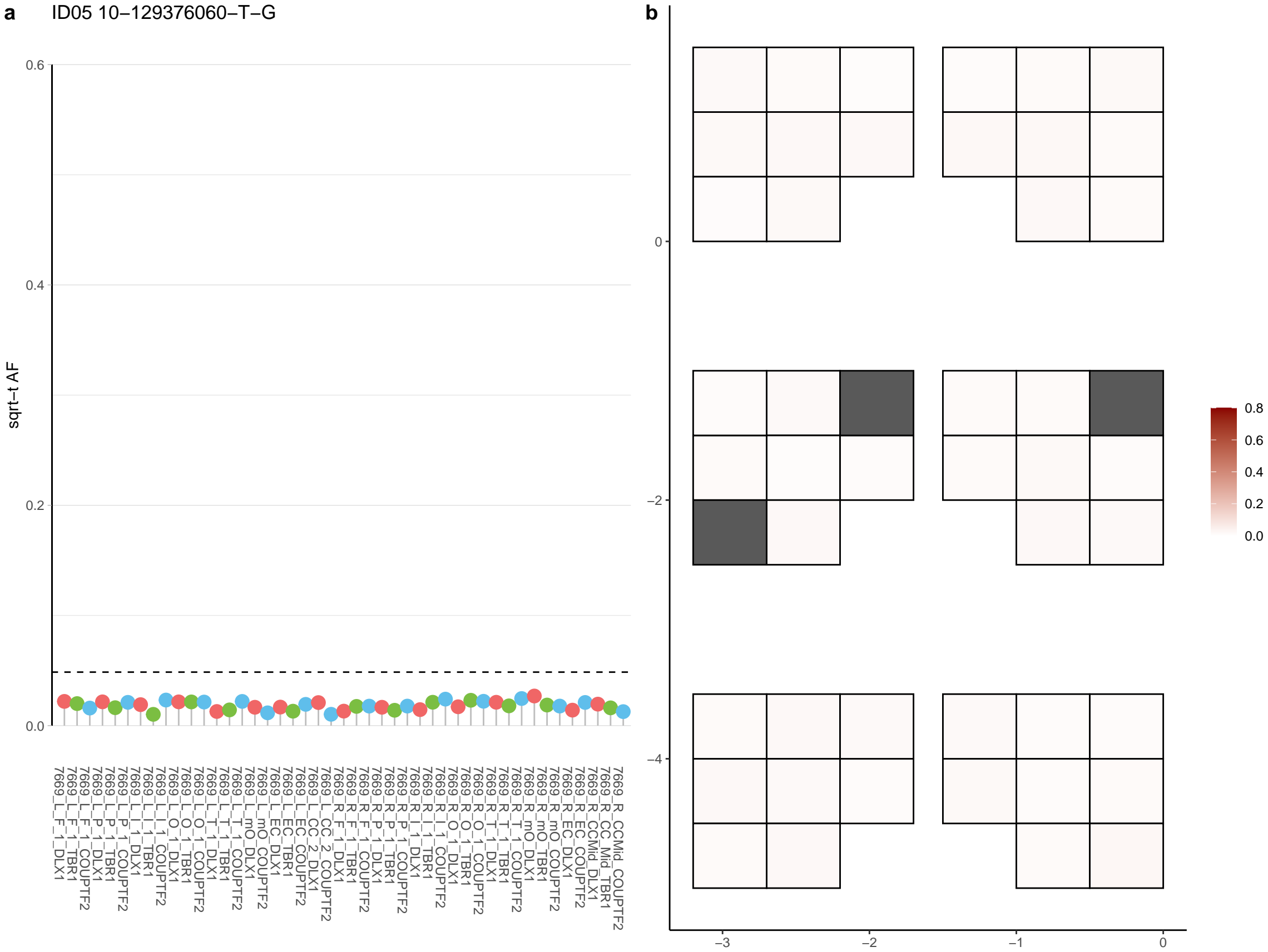

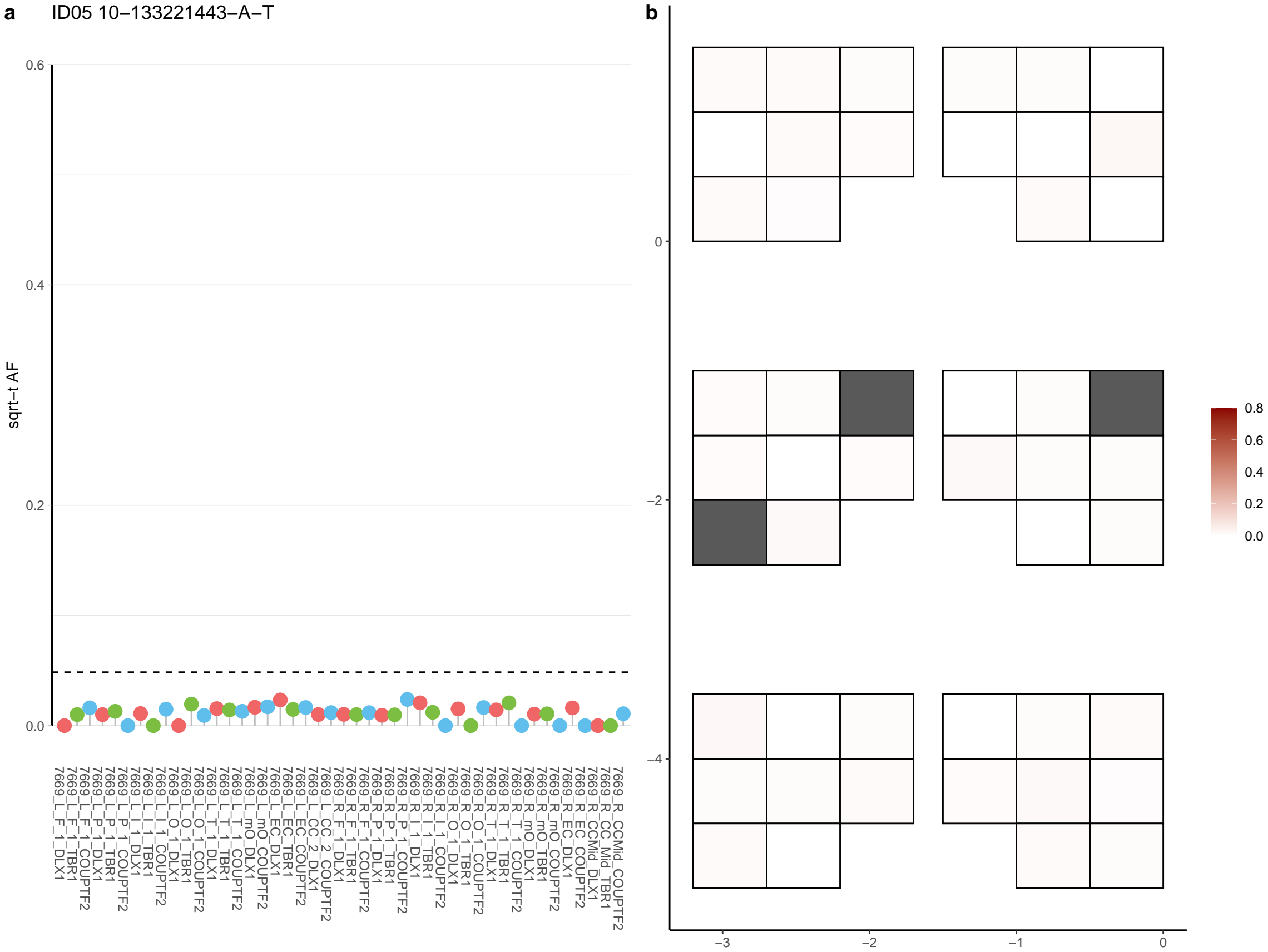

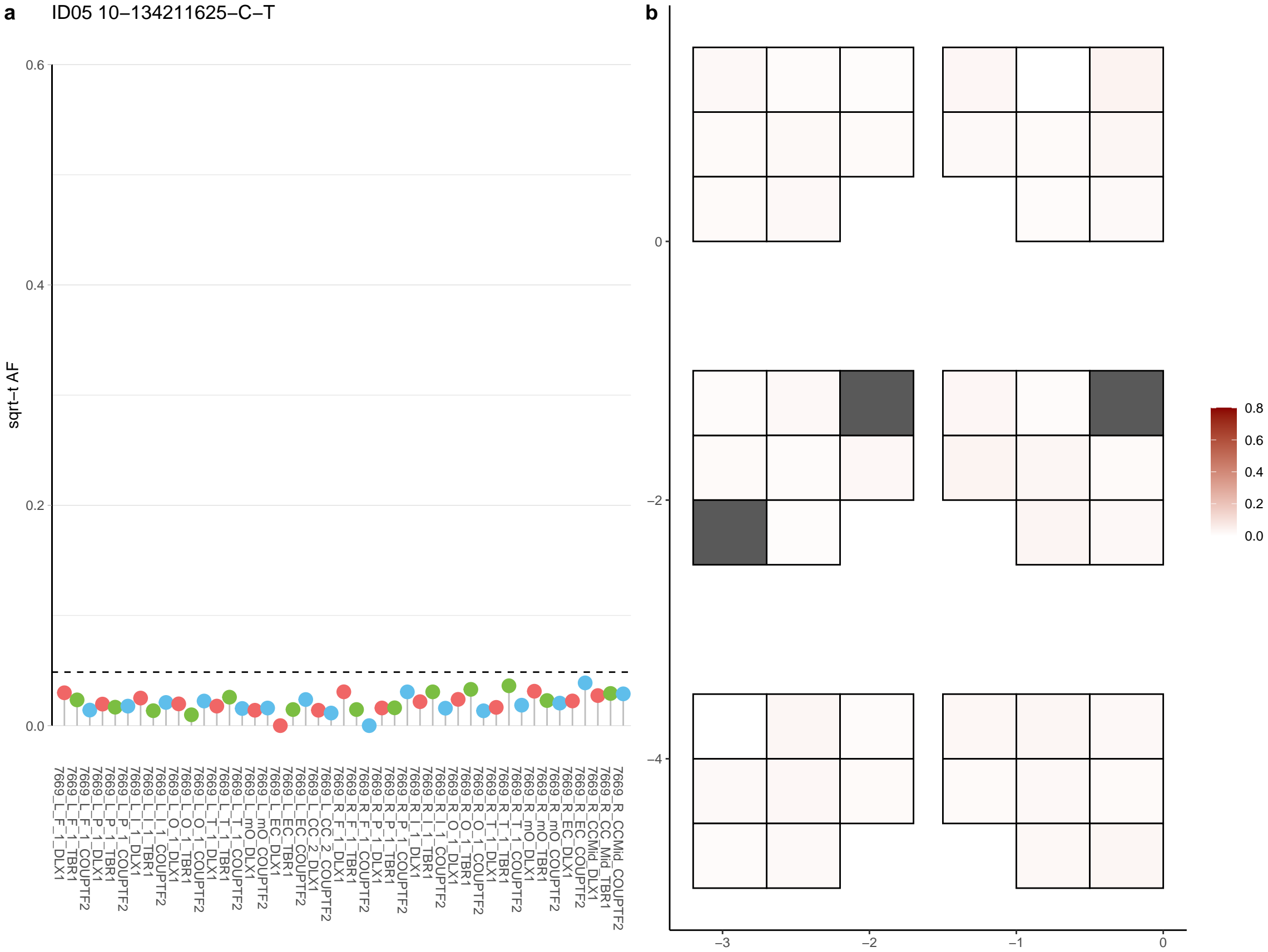

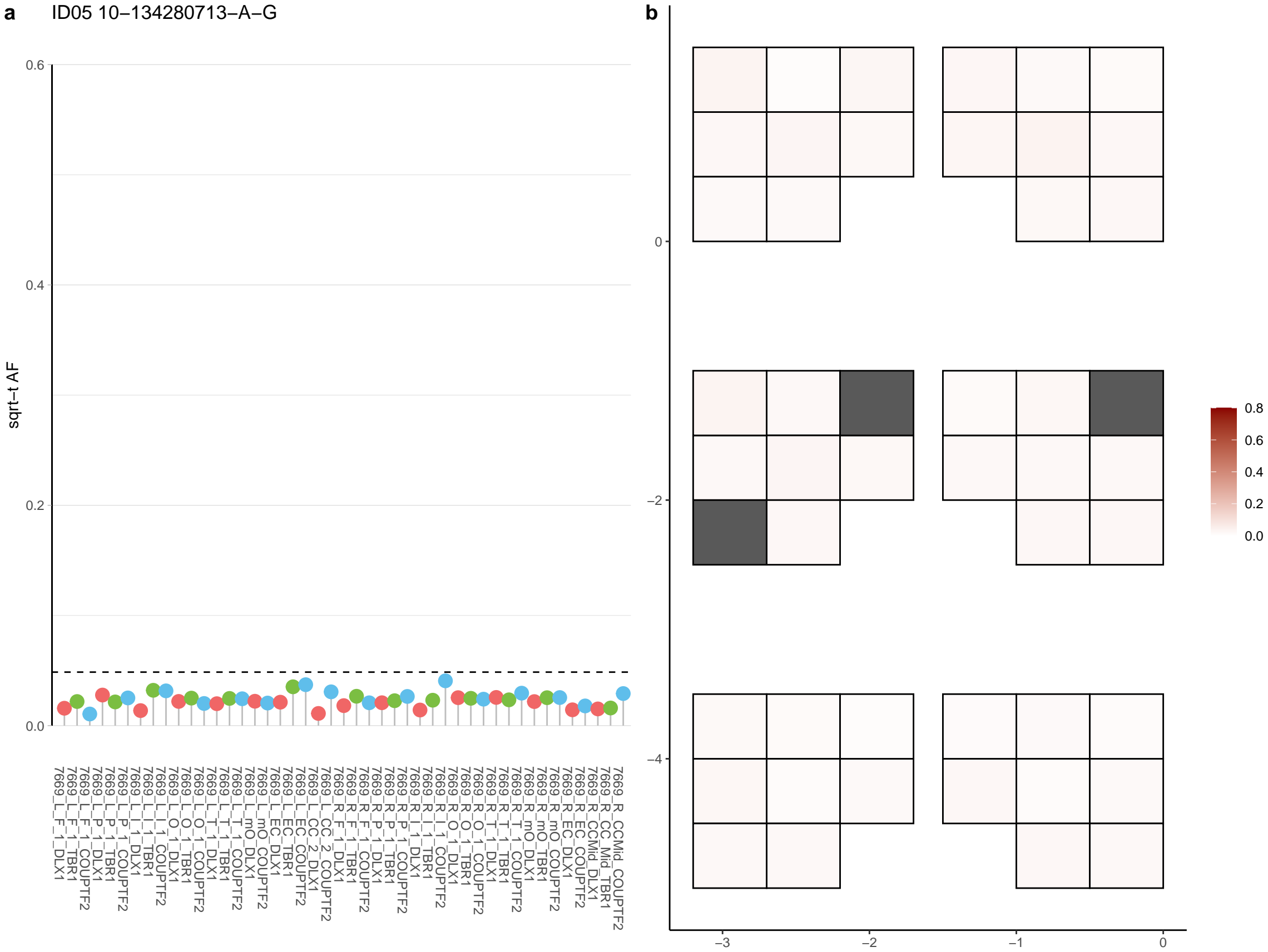

**a** ID05 10-13796054-C-T

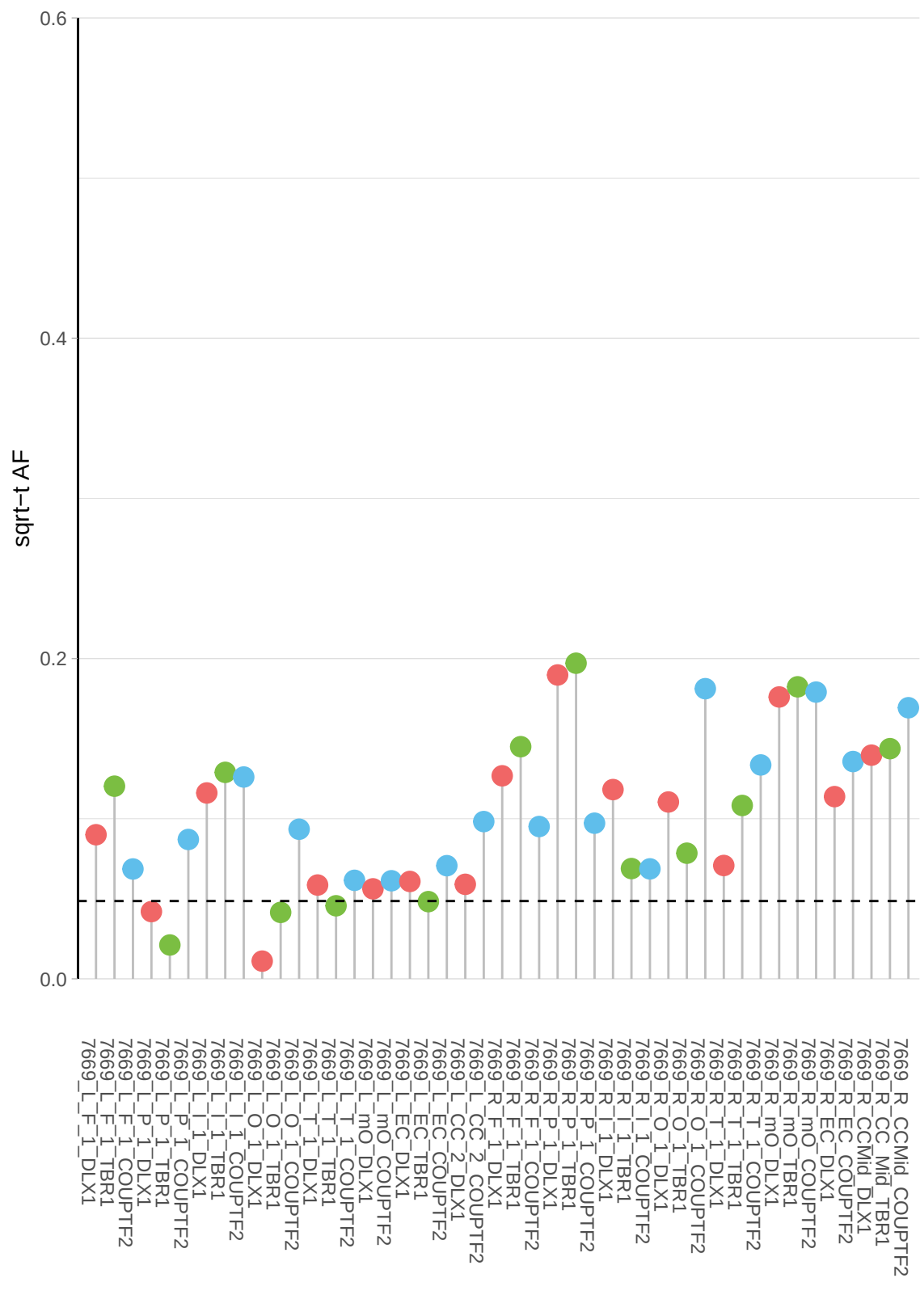

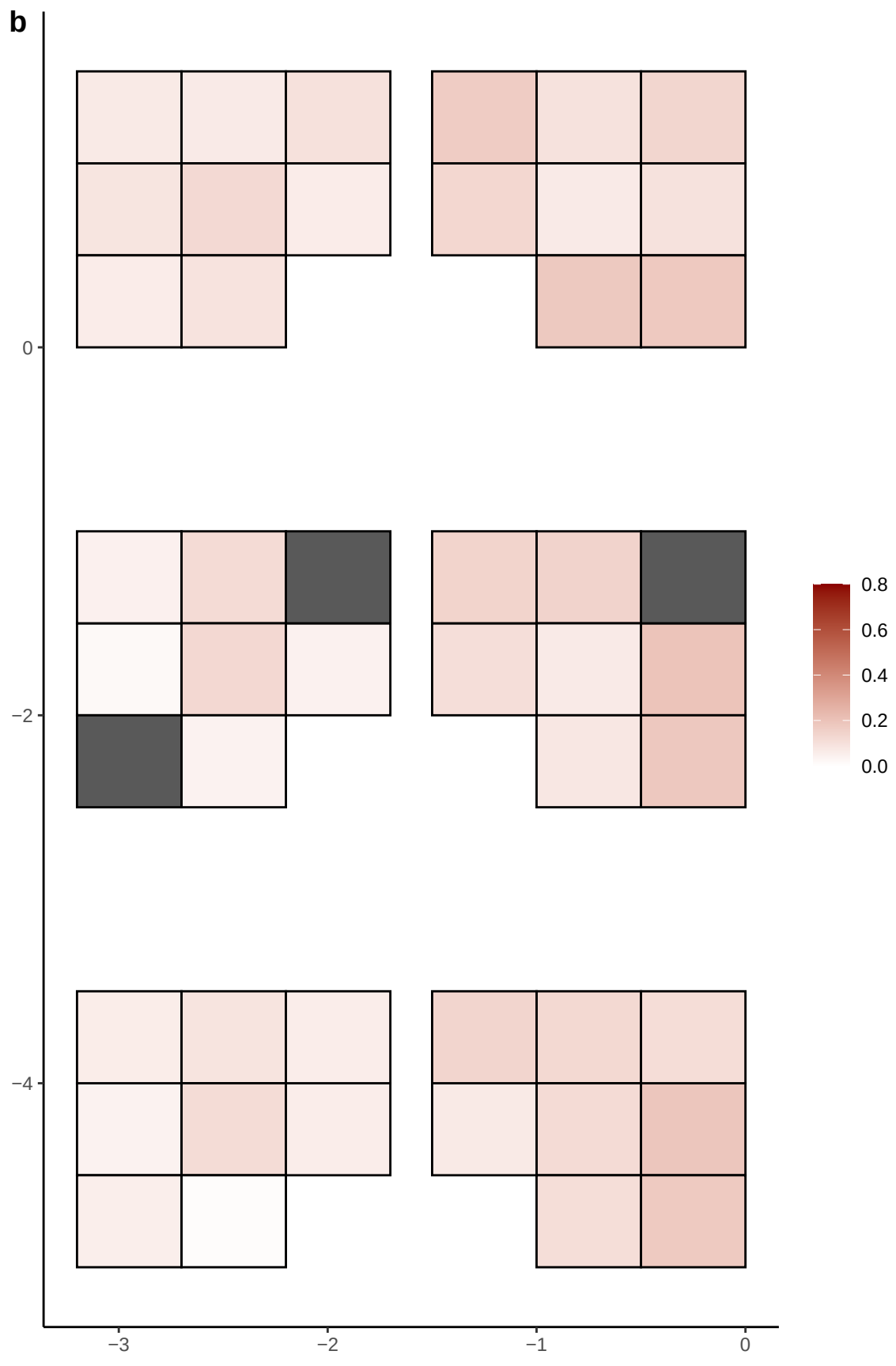

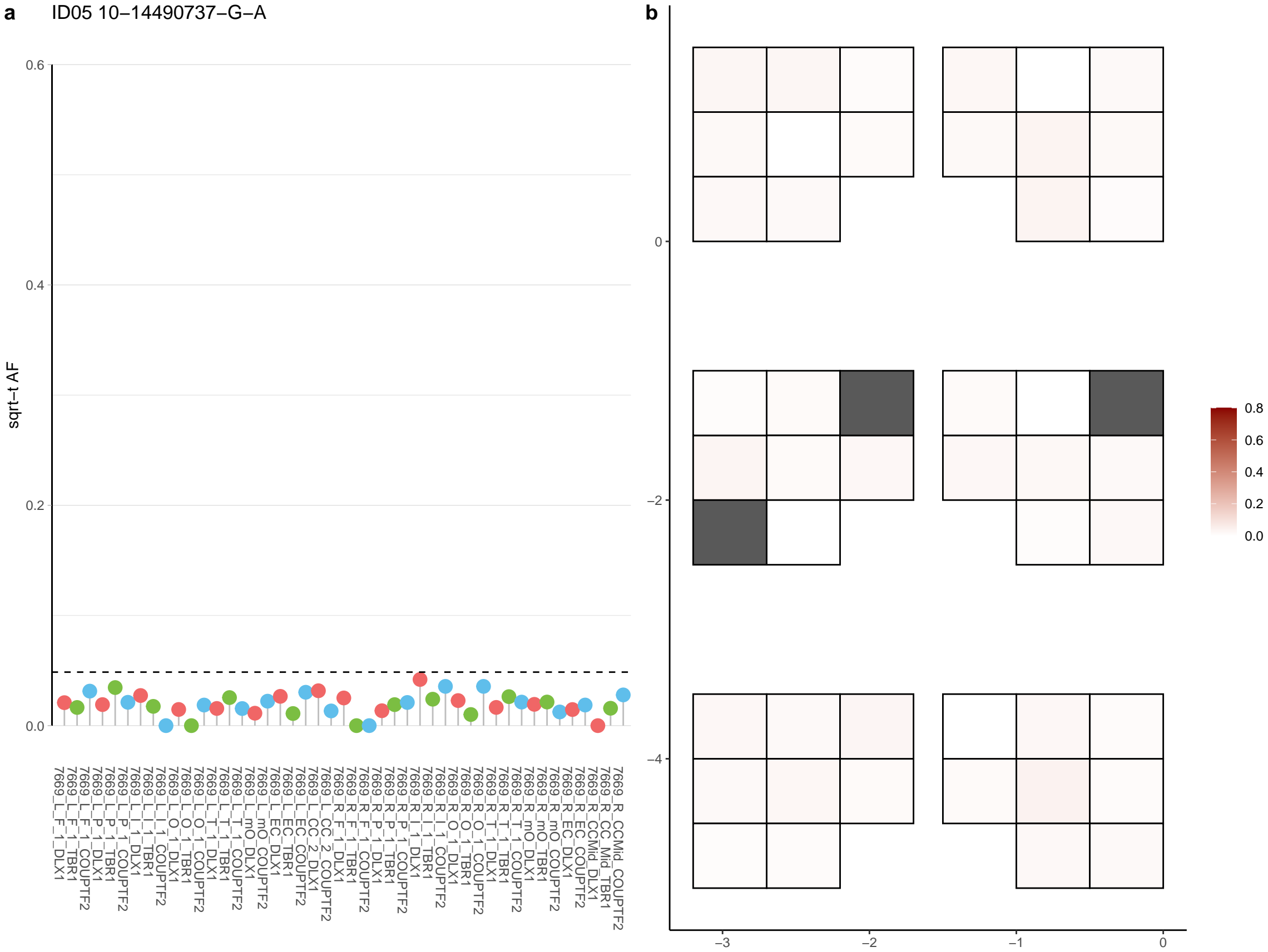

**a** ID05 10-18693563-A-G

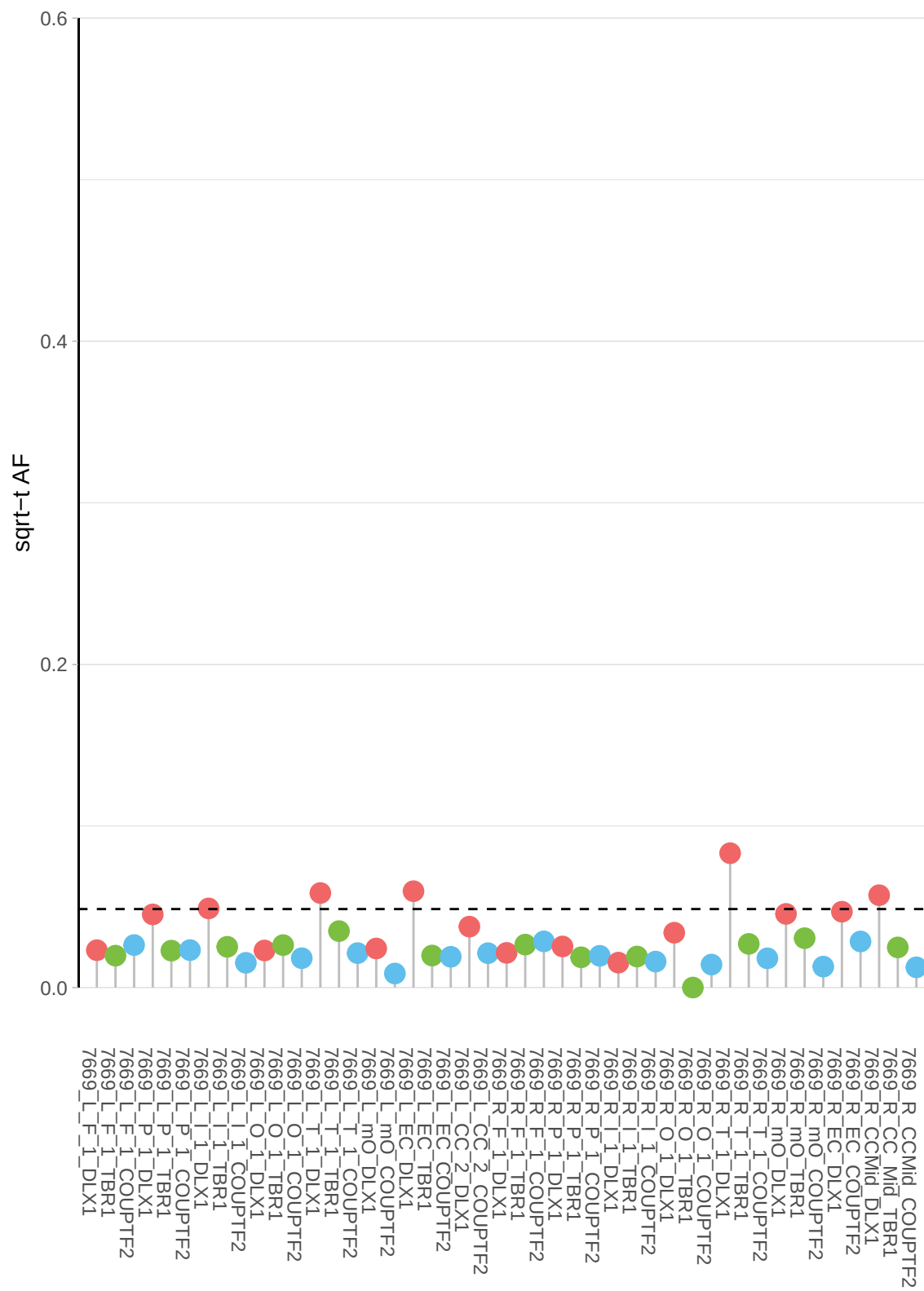

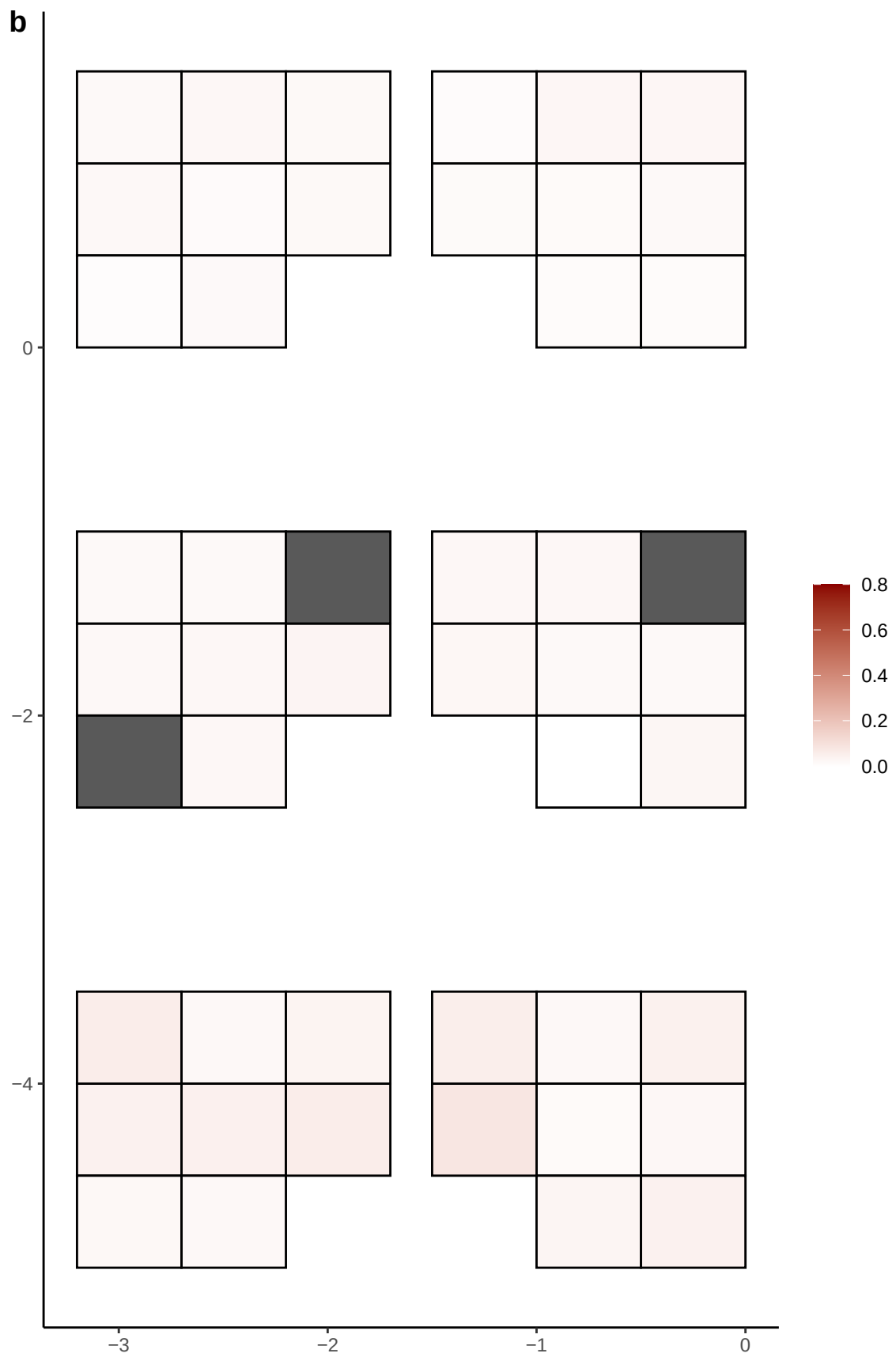

**a** ID05 10-18760480-C-T

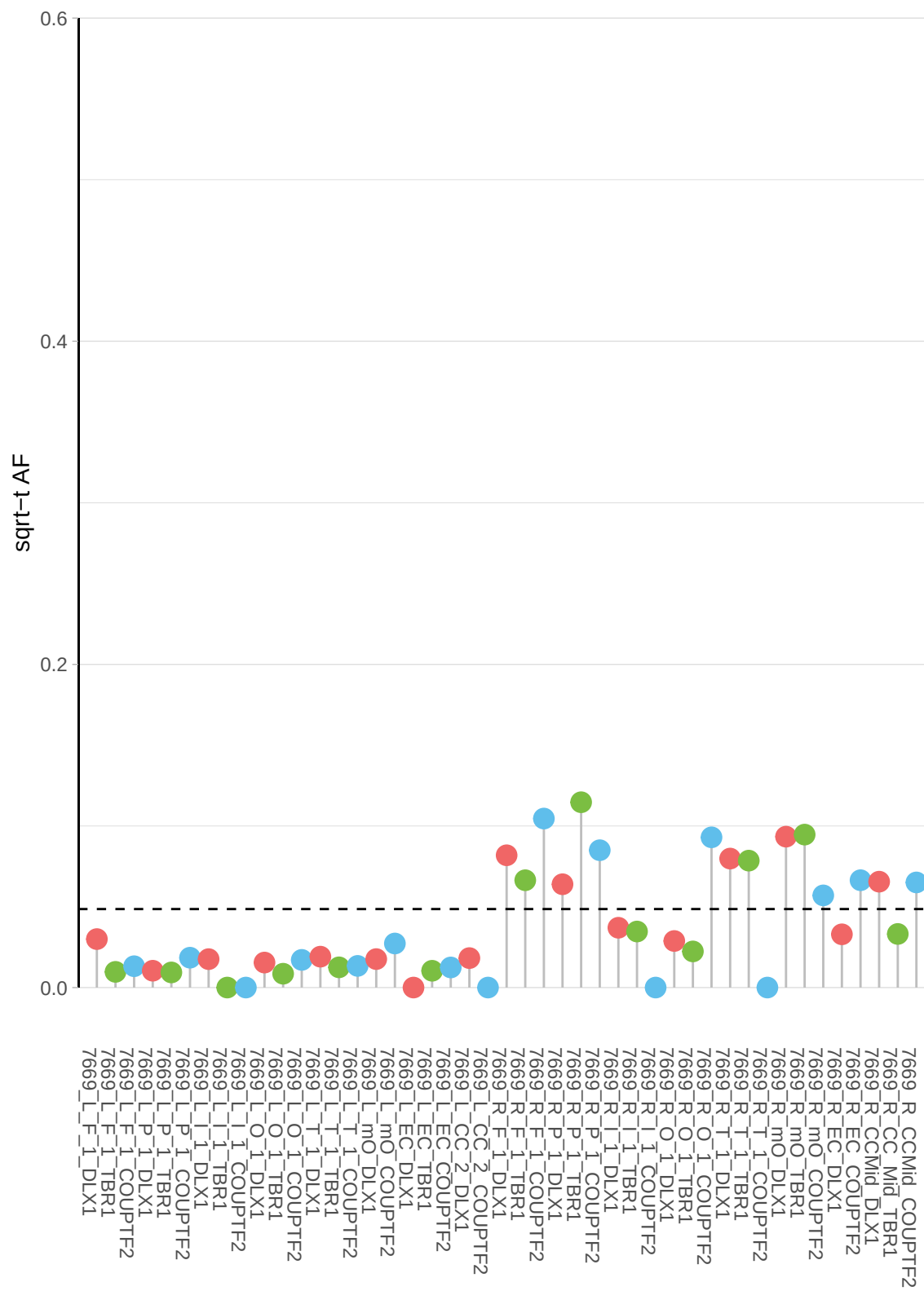

**b**

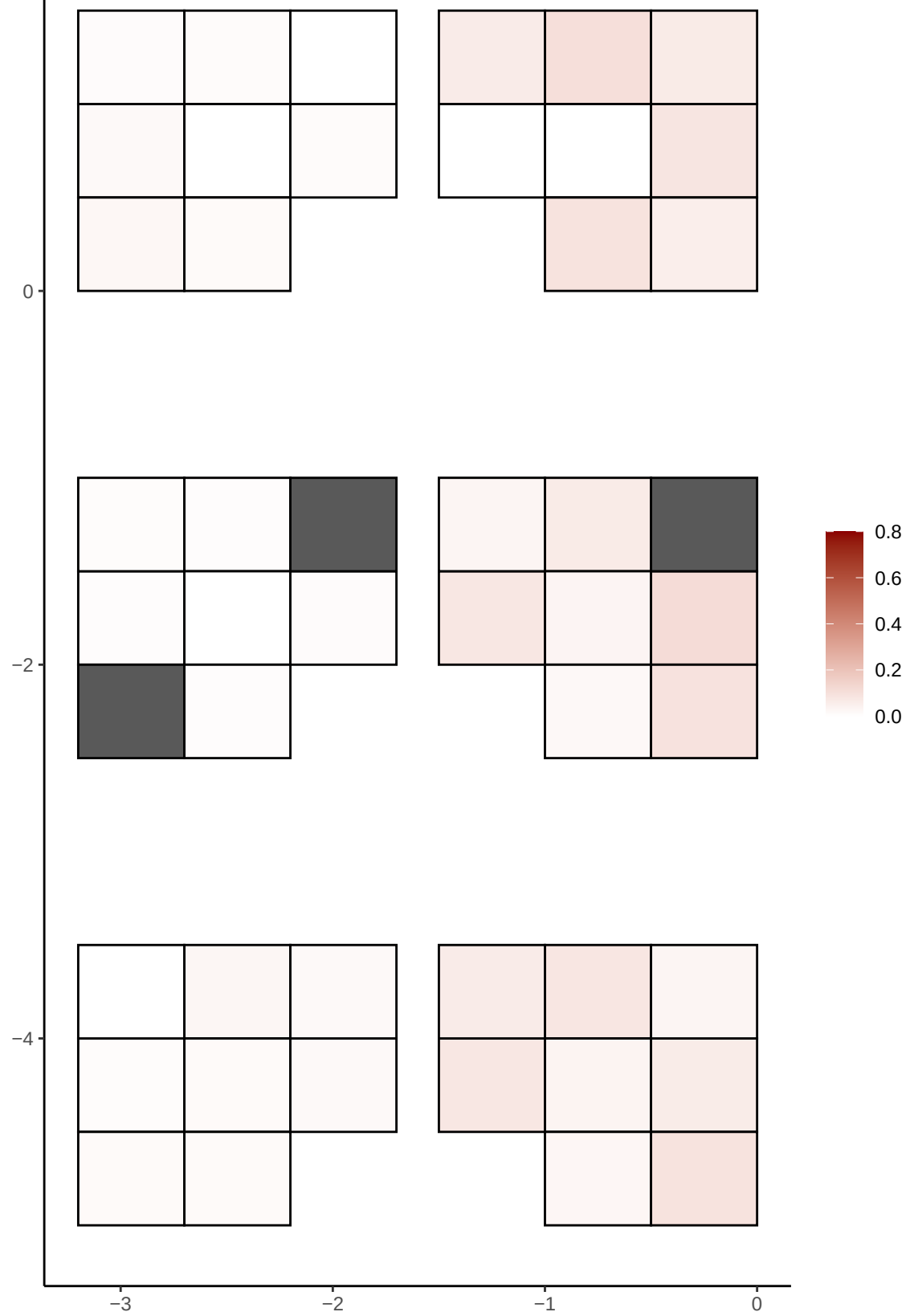

**a** ID05 10-2261697-A-C

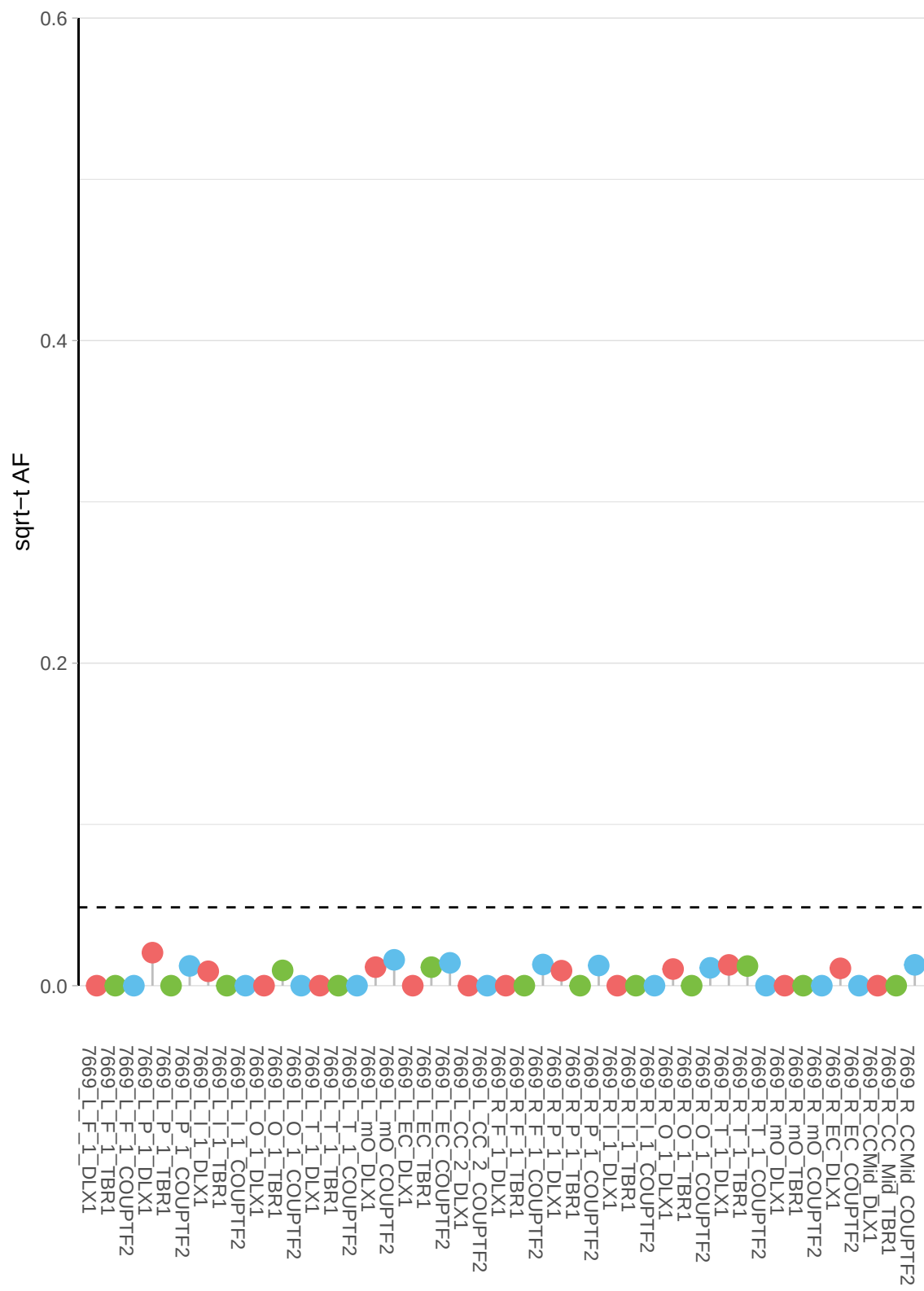

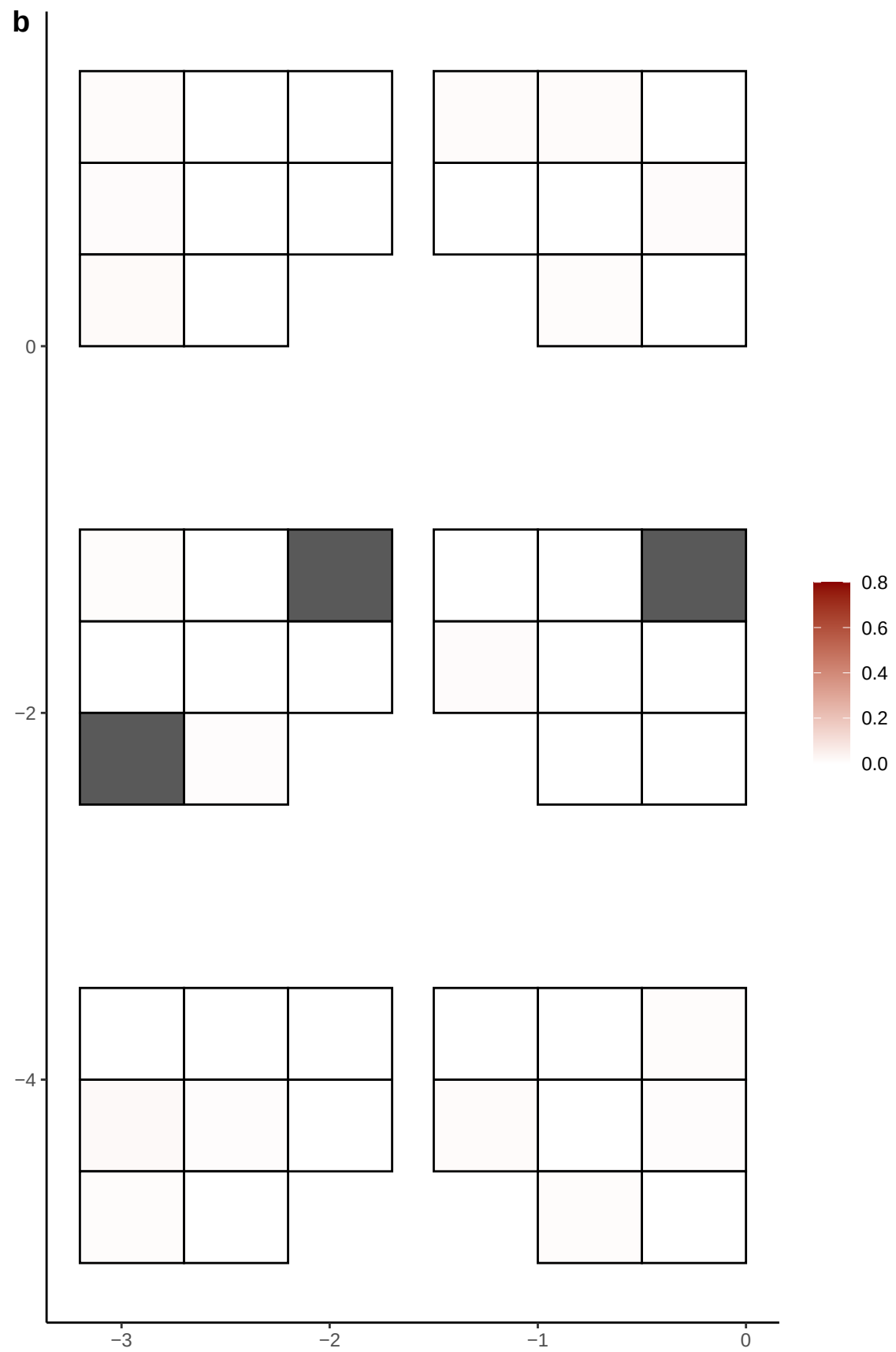

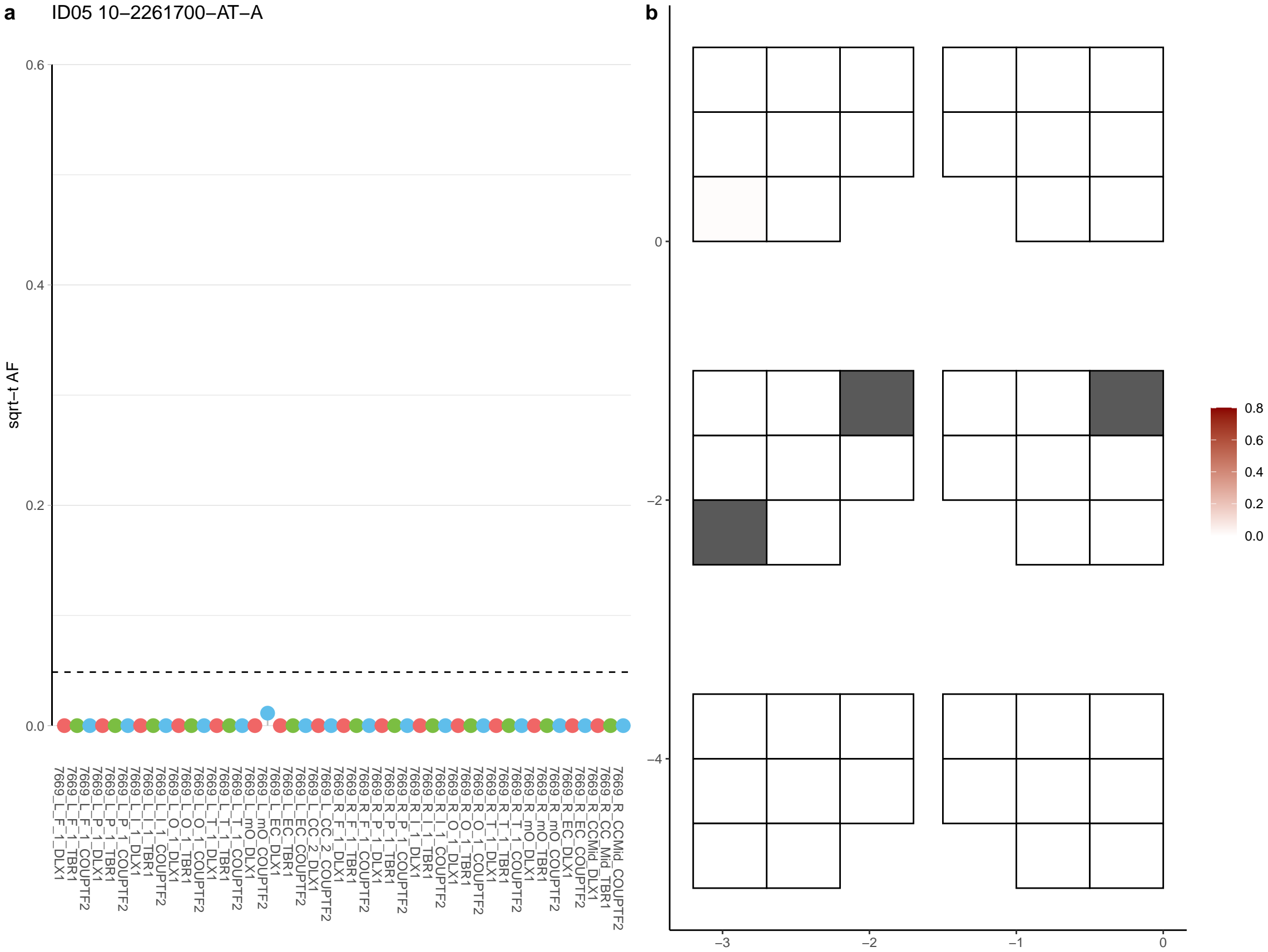

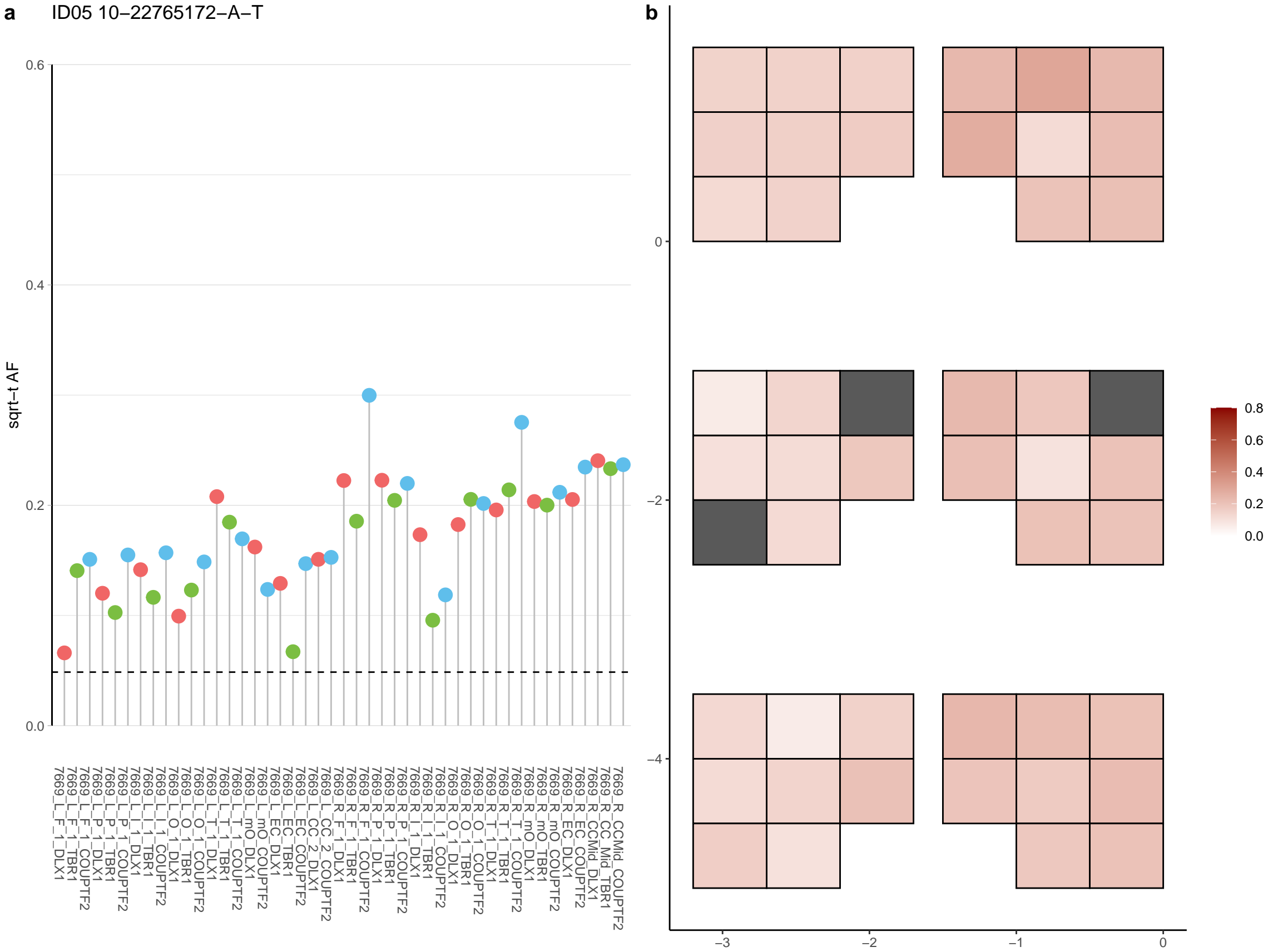

**a** ID05 10-23976083-C-A

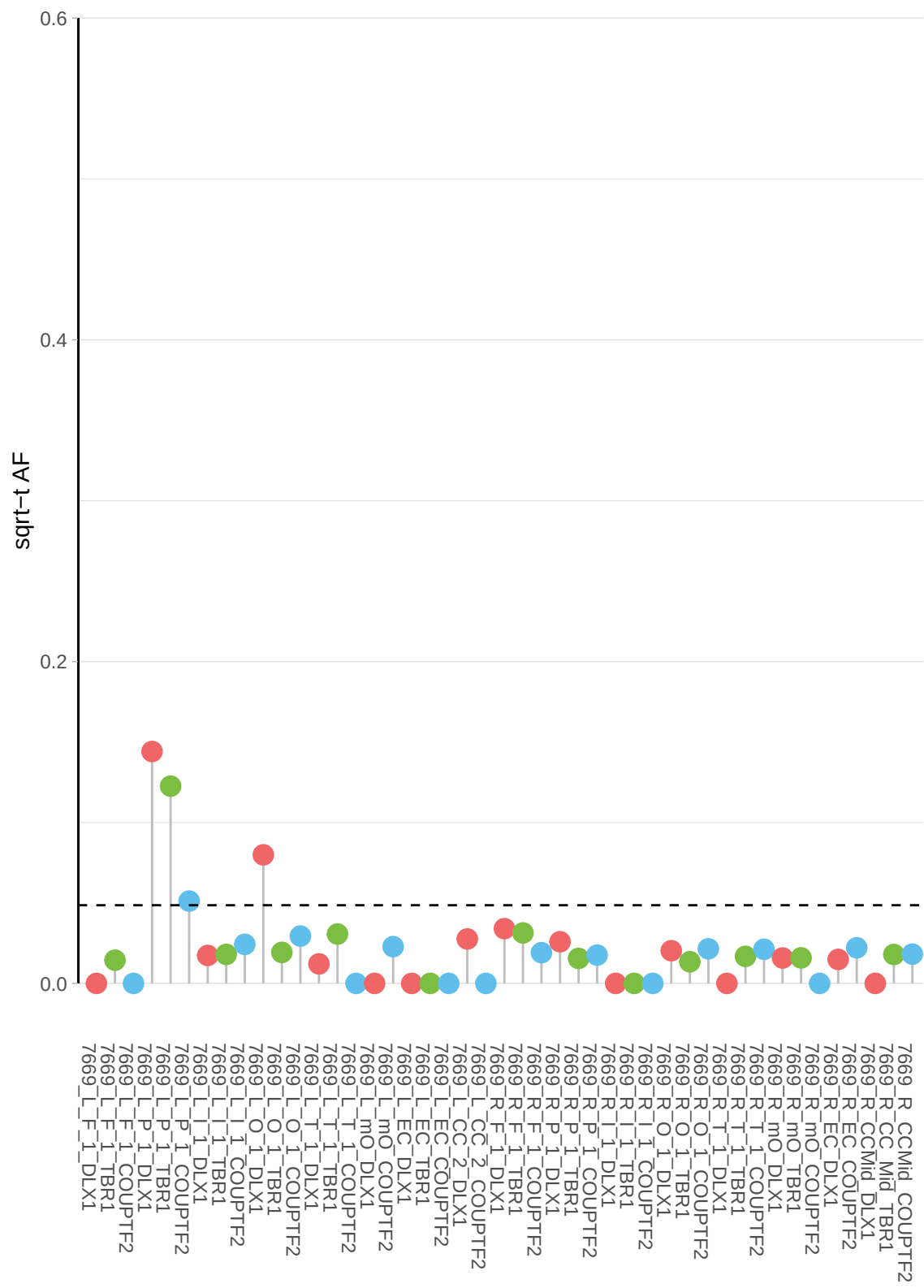

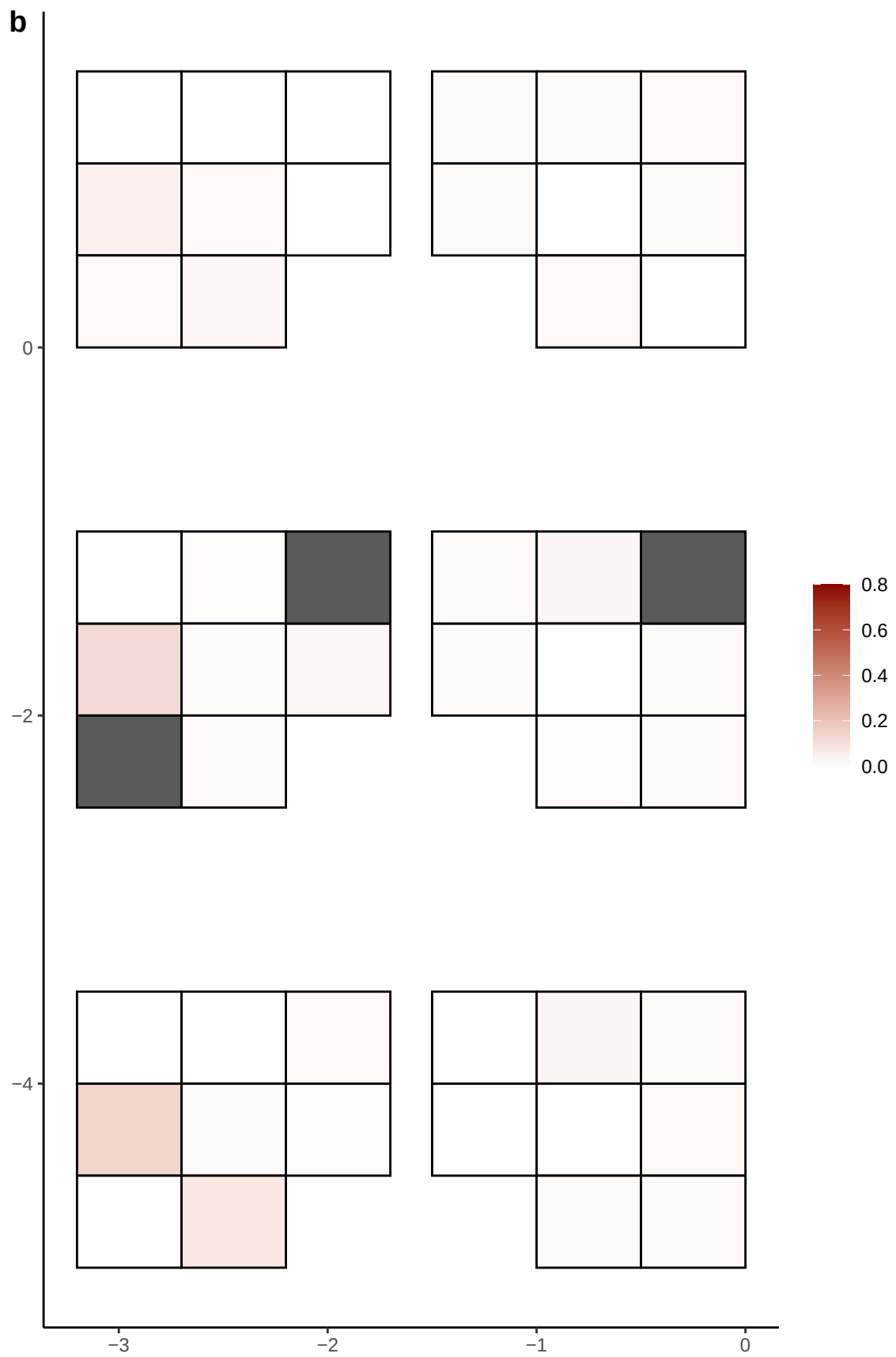

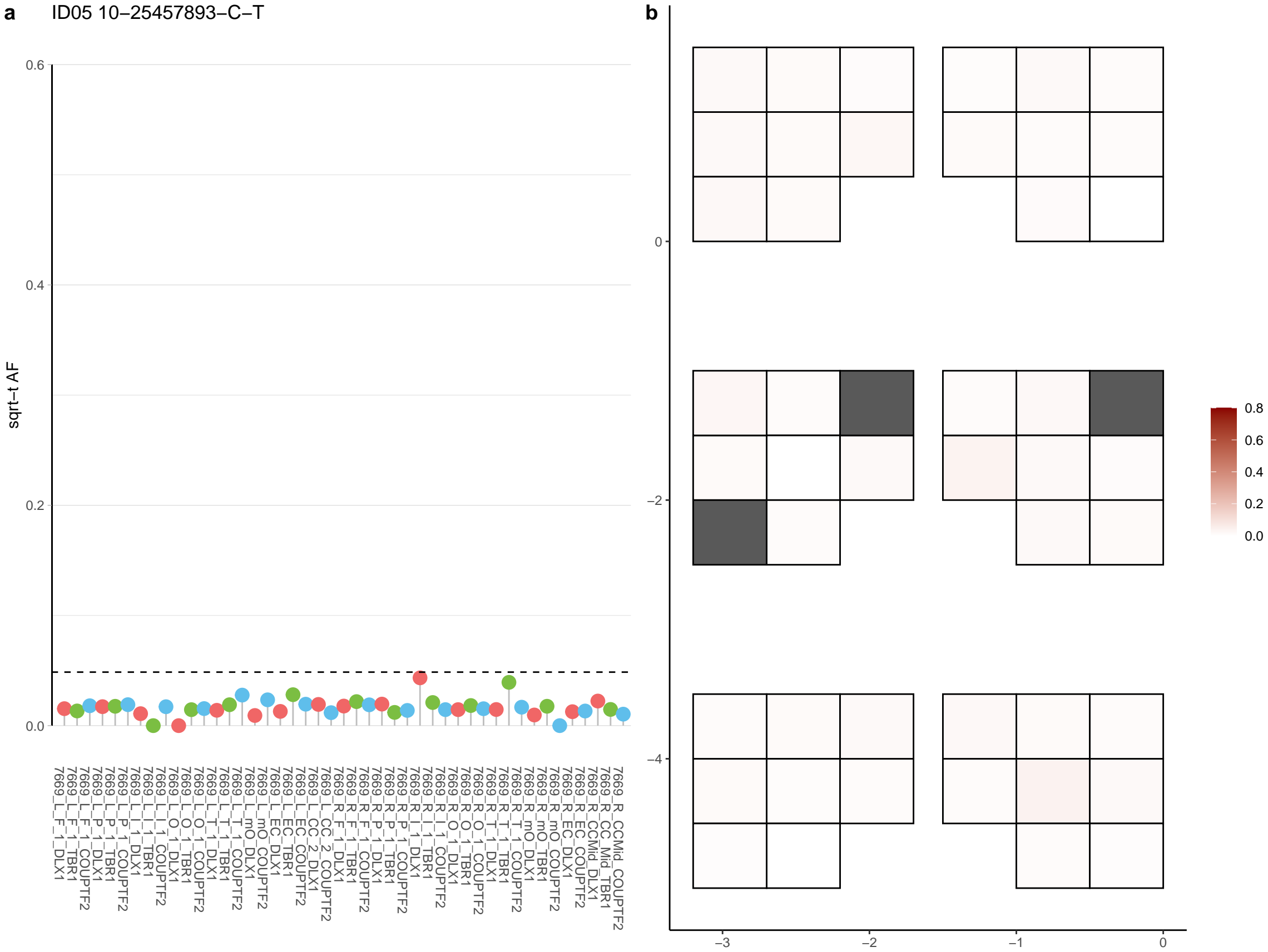

**a** ID05 10-25610159-G-A

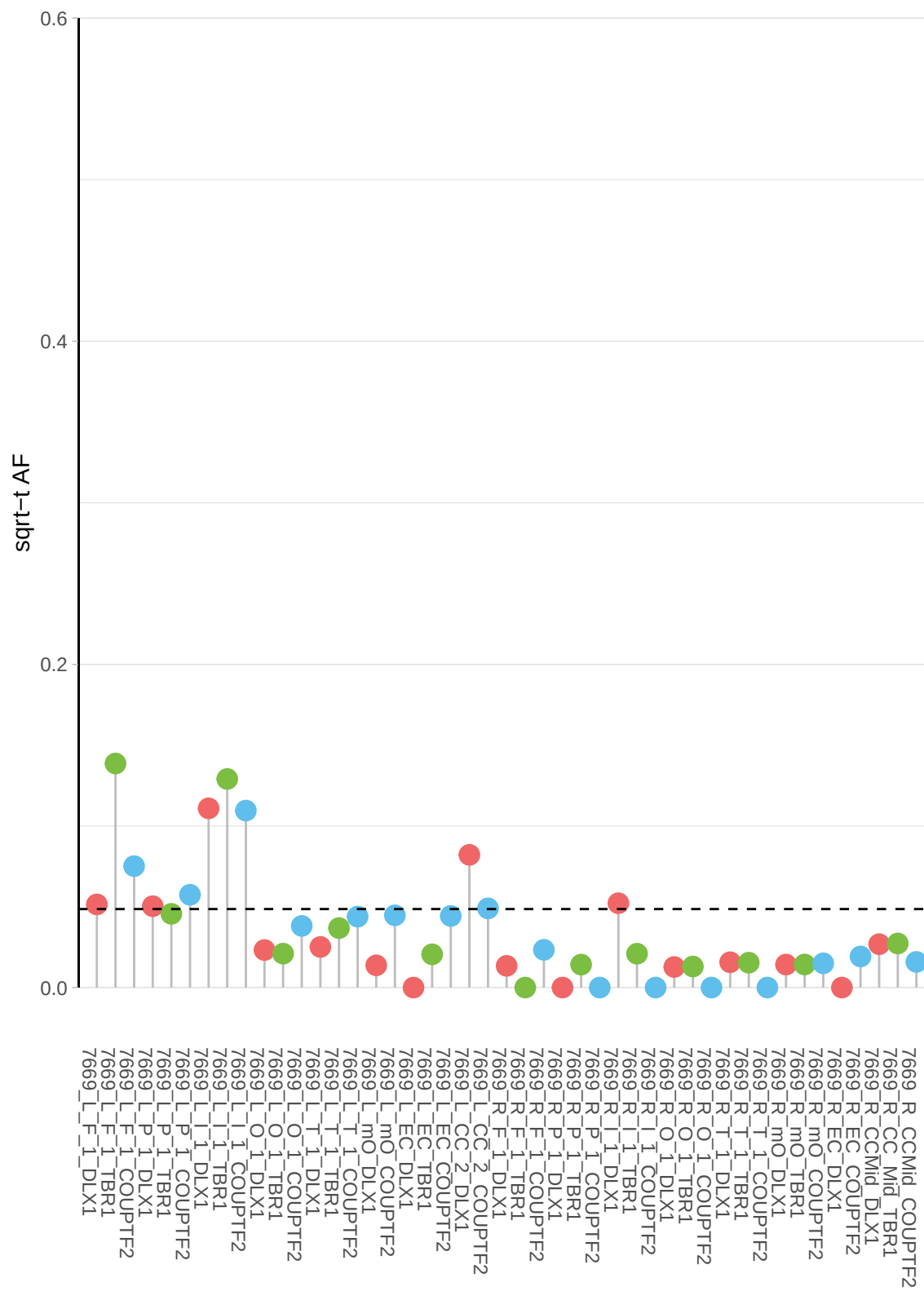

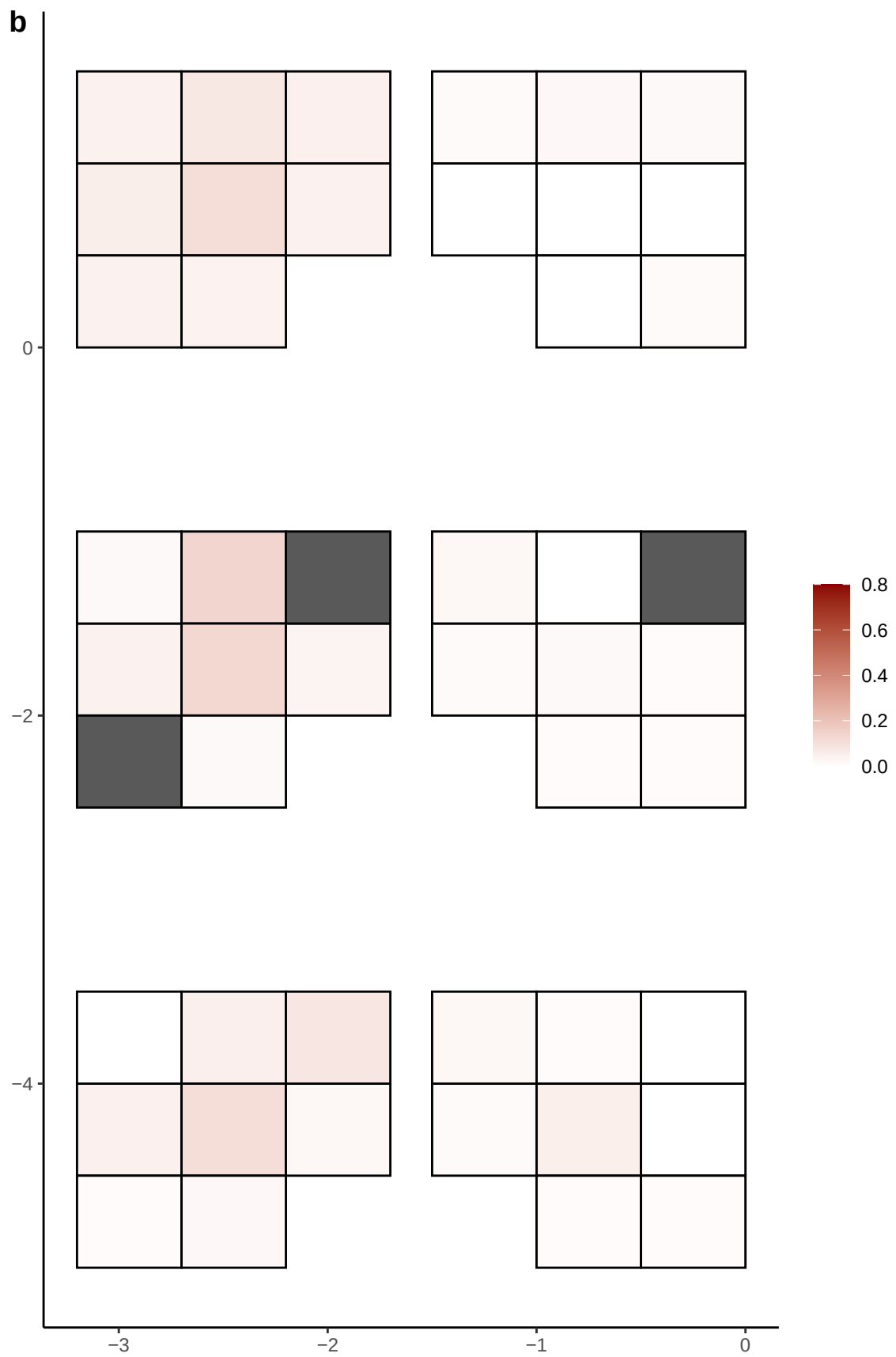

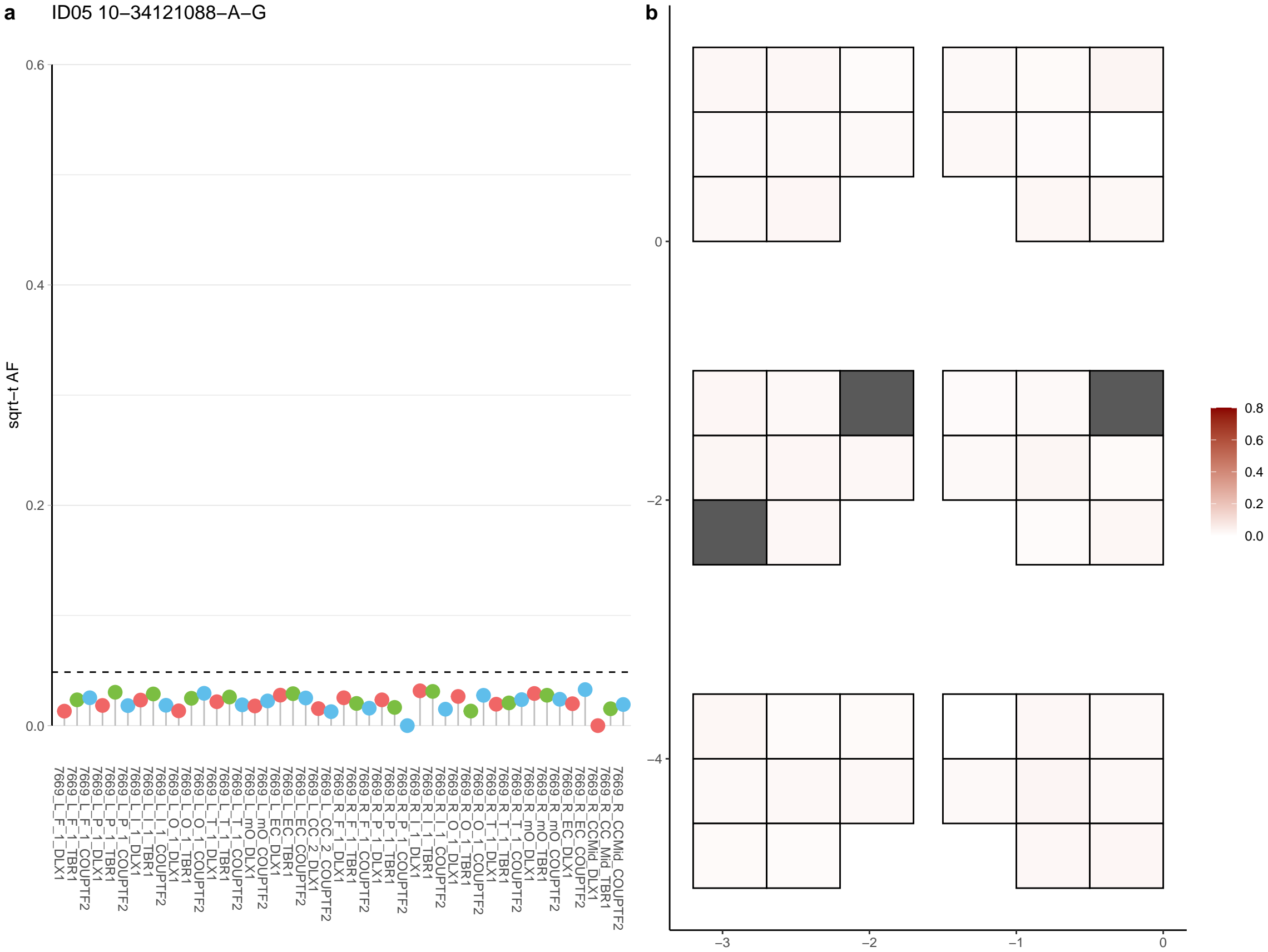

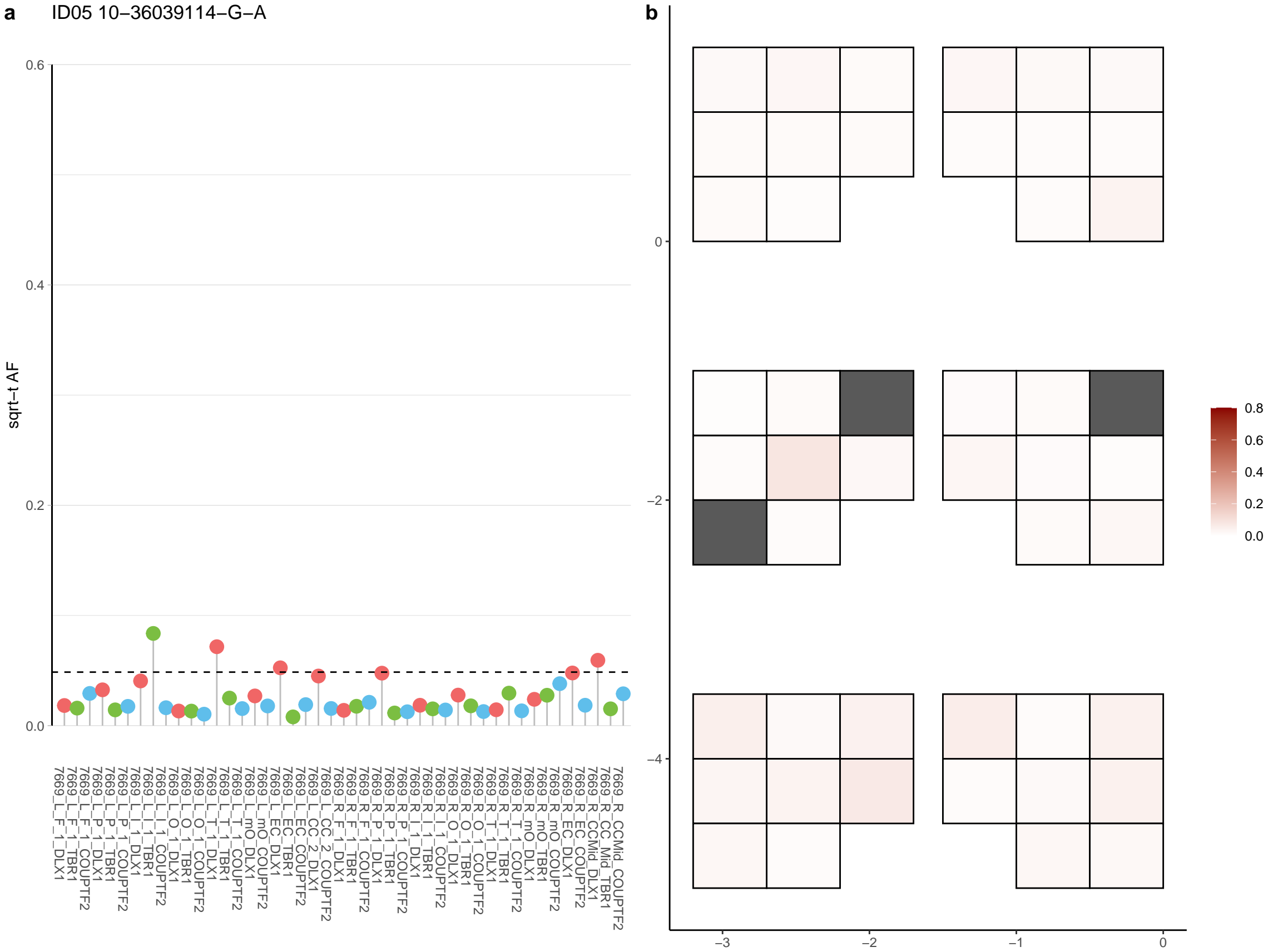

**a** ID05 10-58152659-C-T

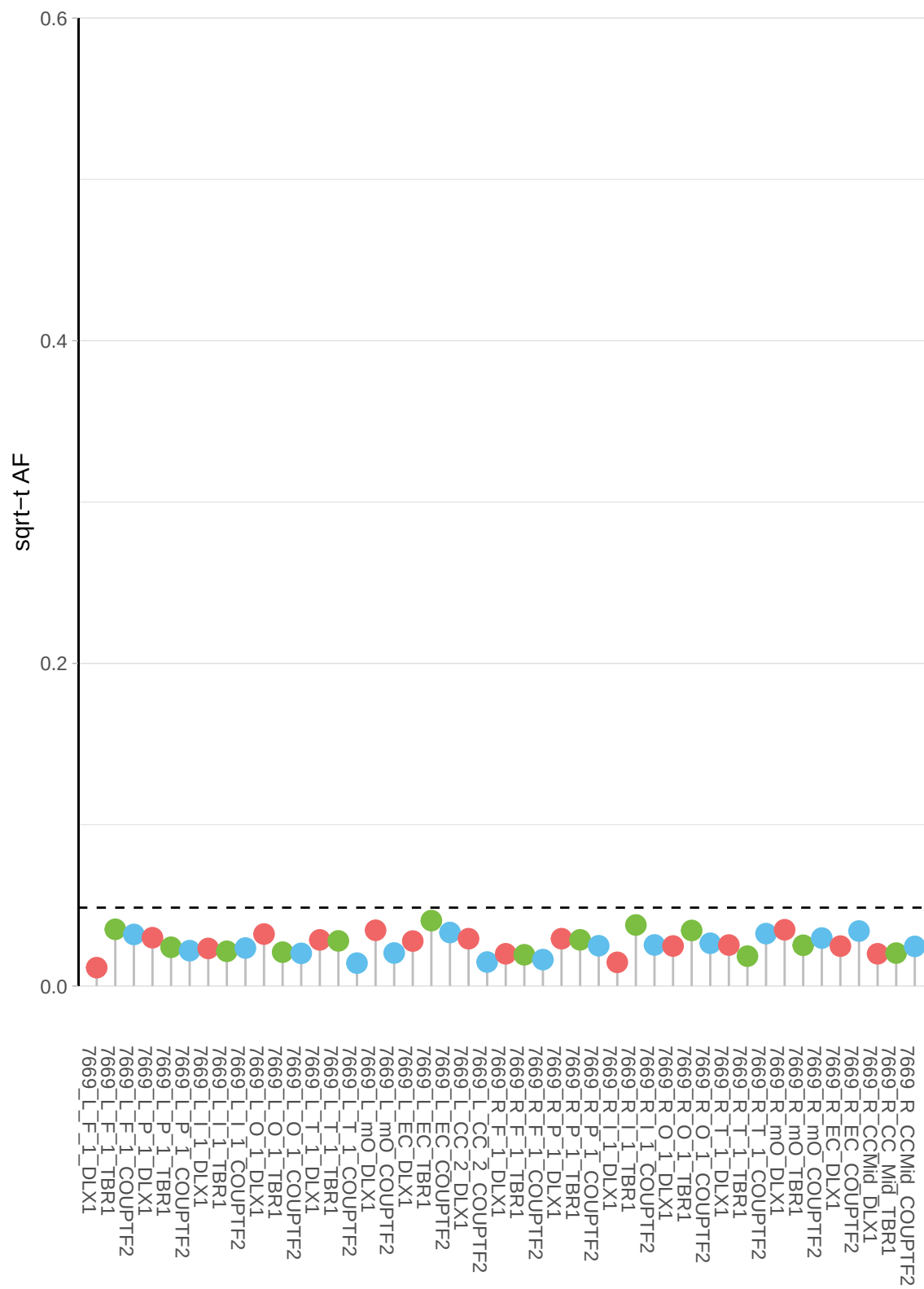

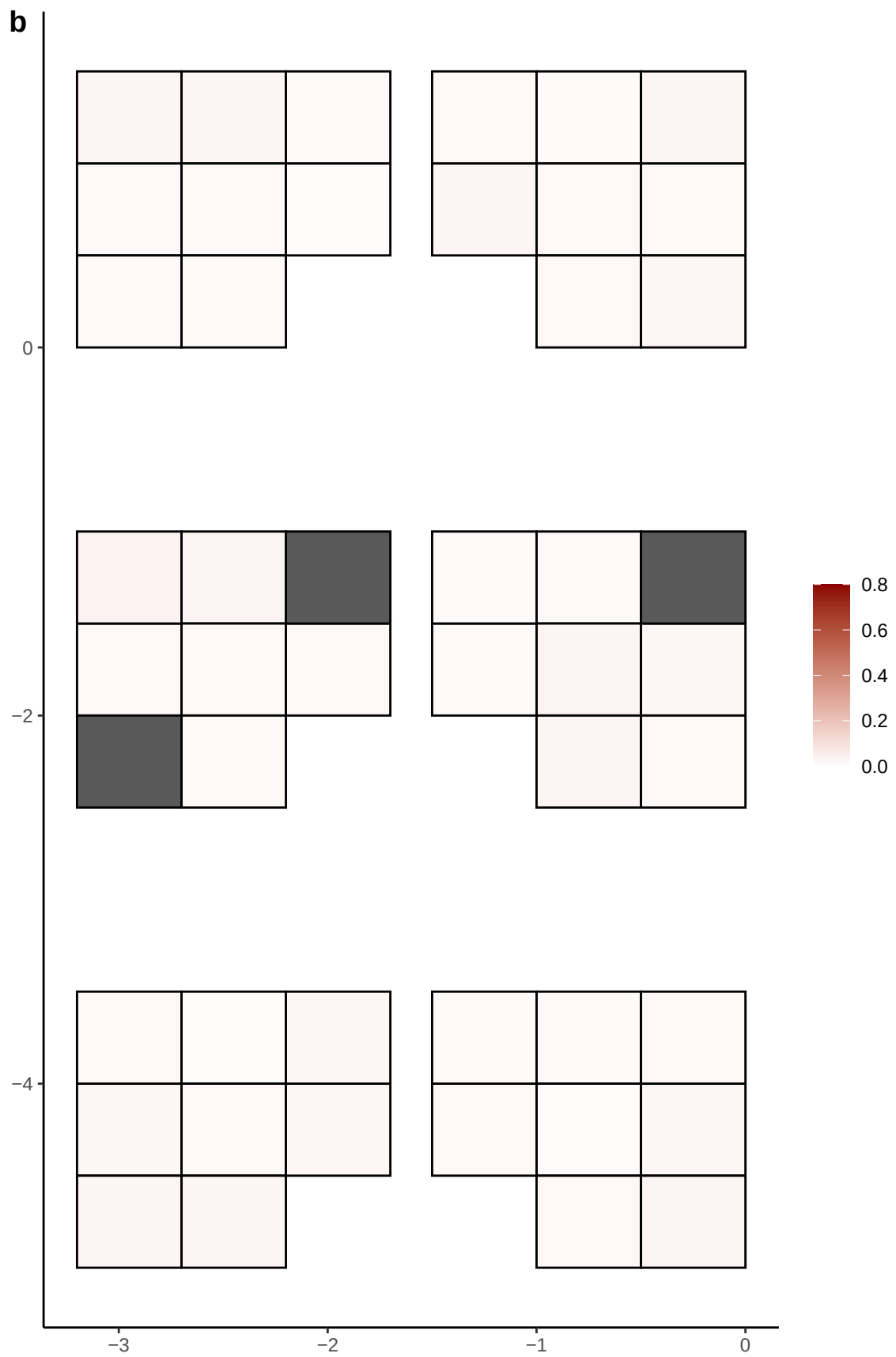

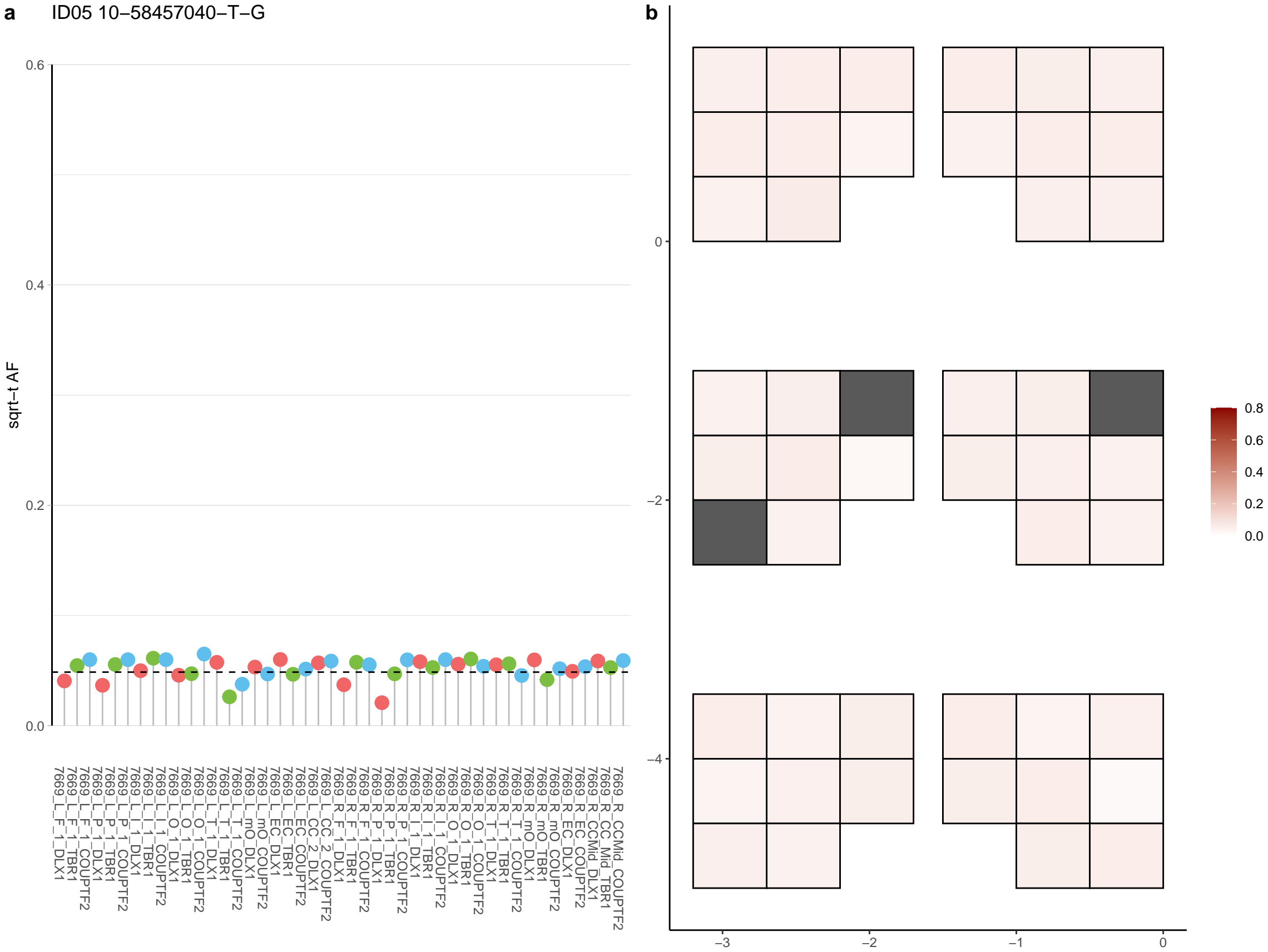

**a** ID05 10-60559957-C-T

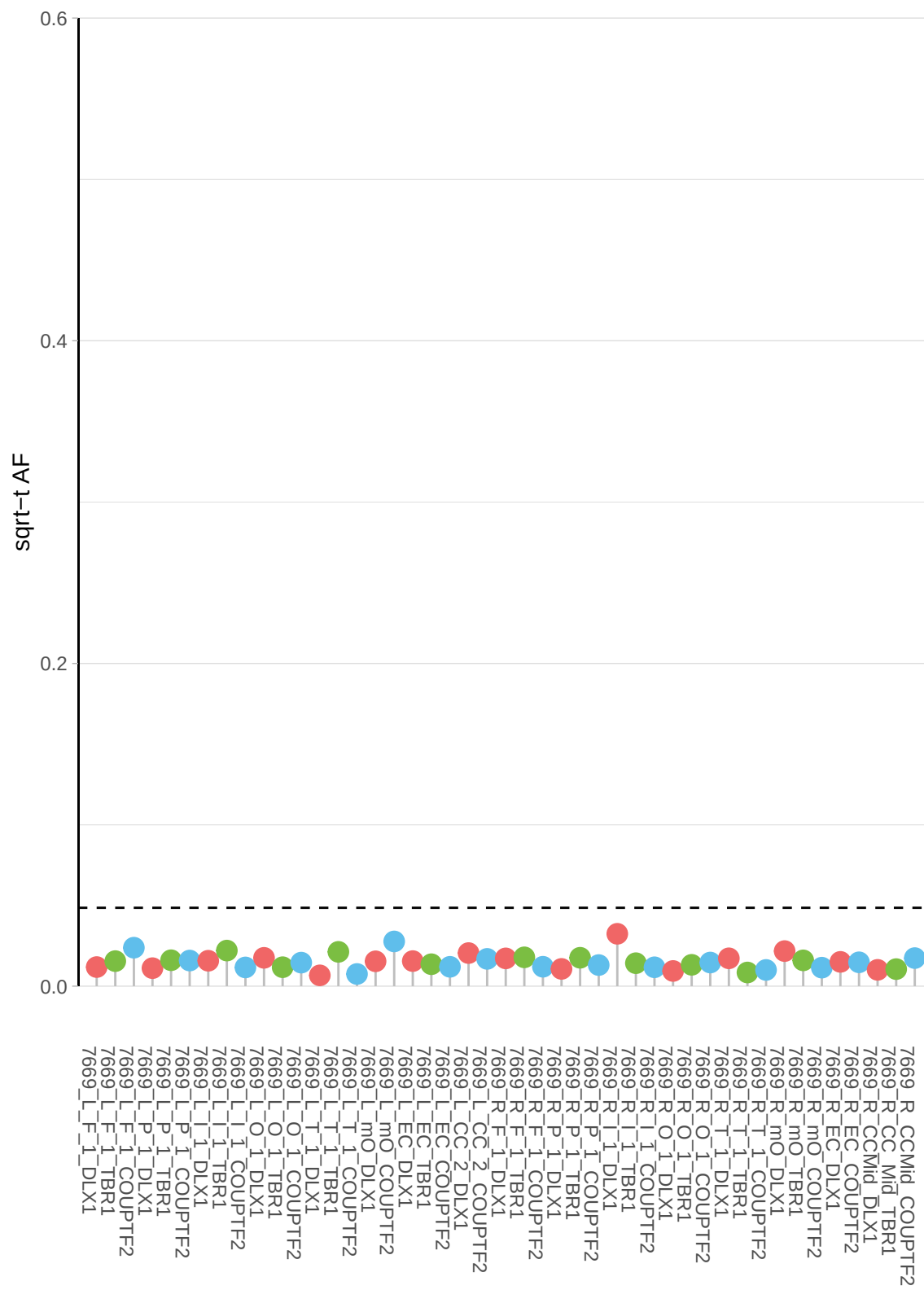

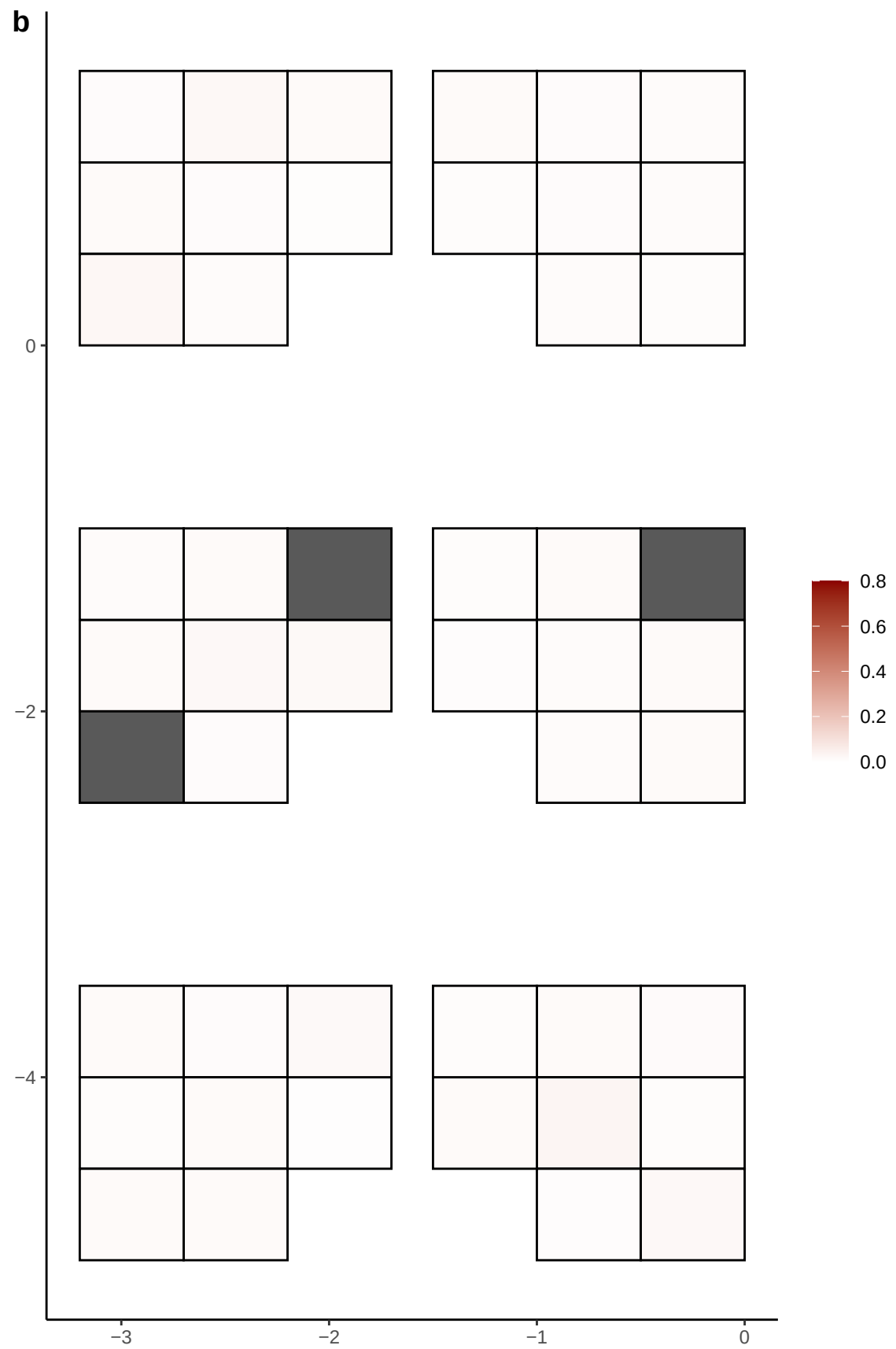

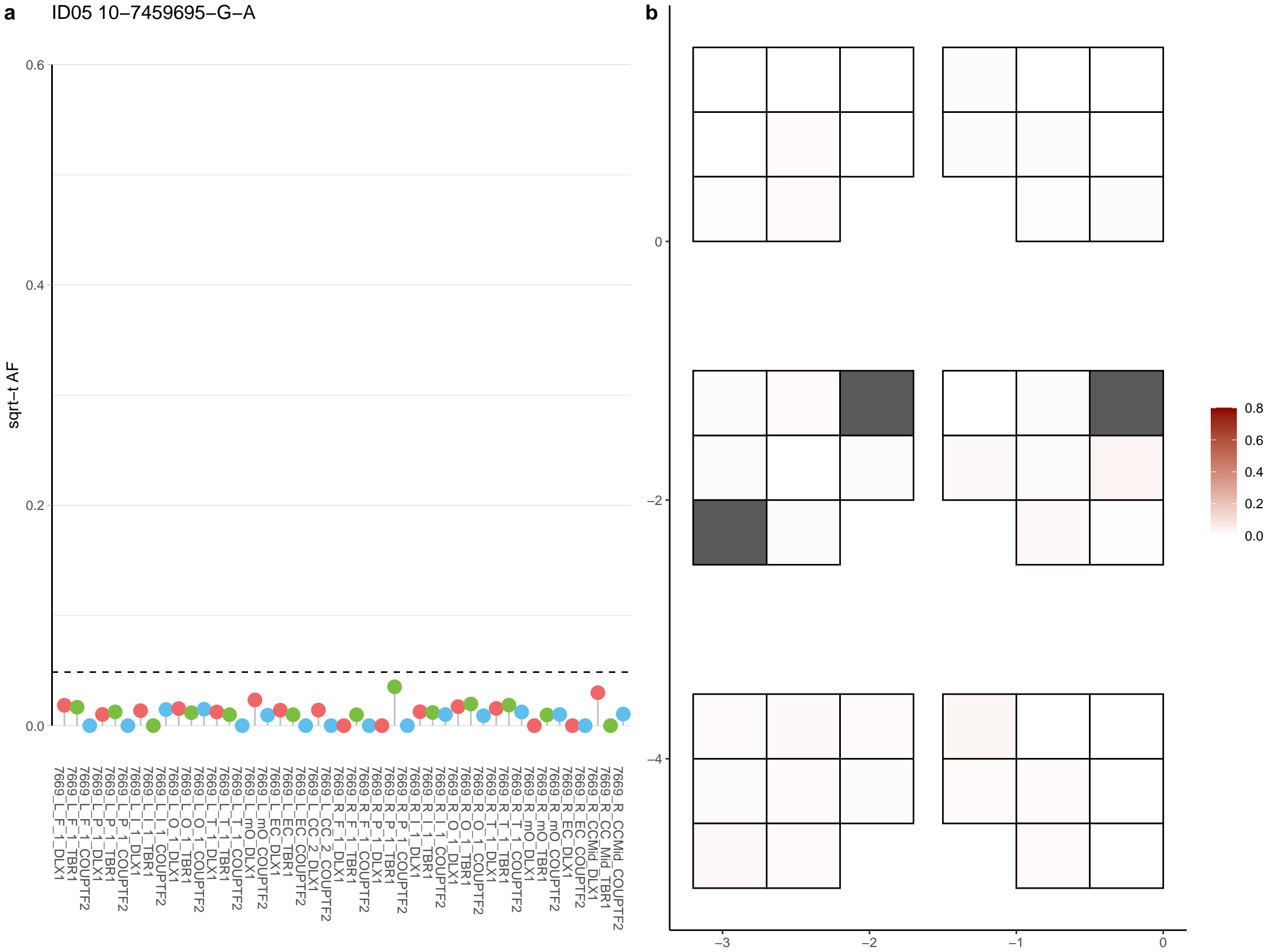

**a** ID05 10-79843135-G-A

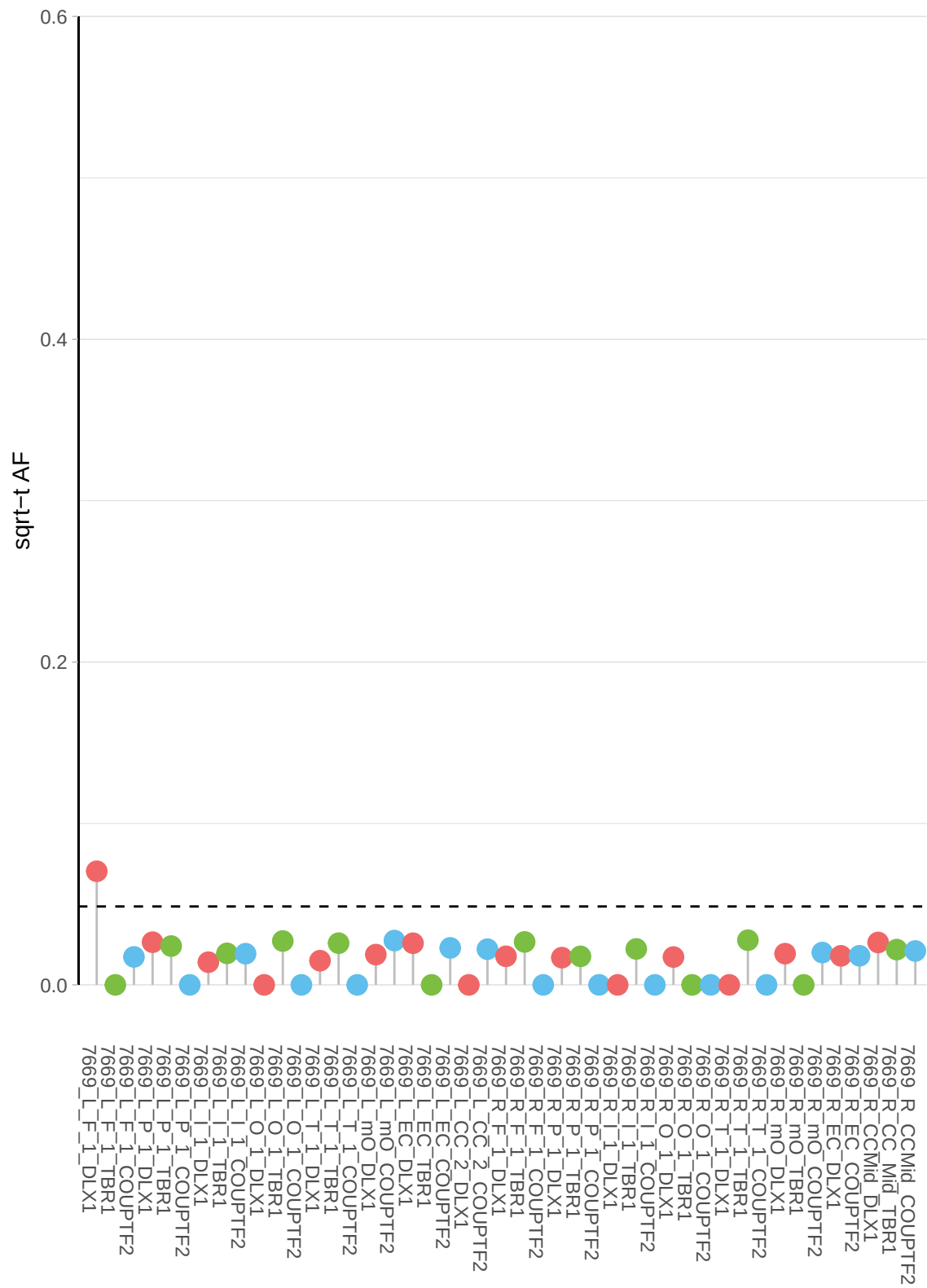

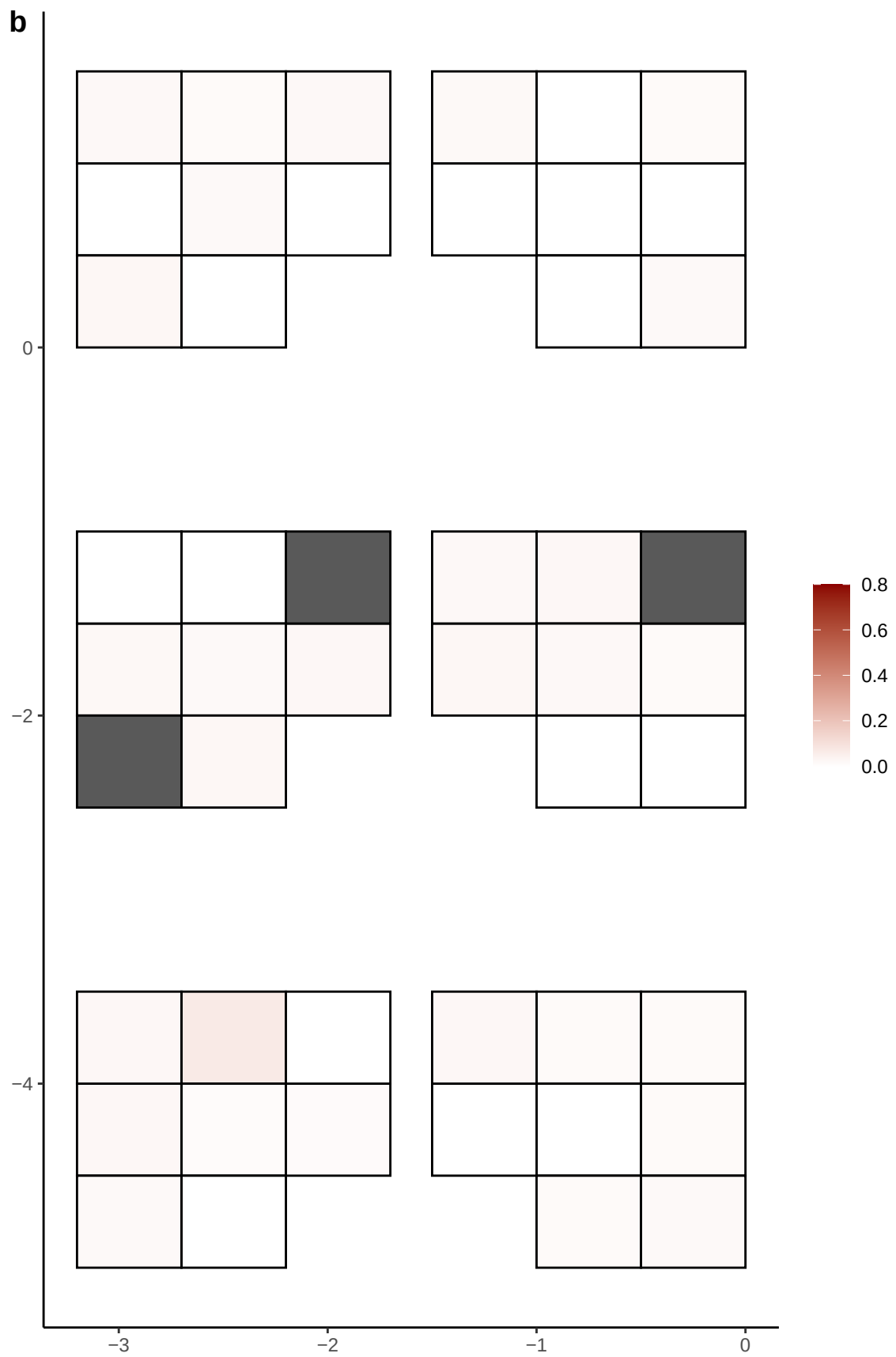

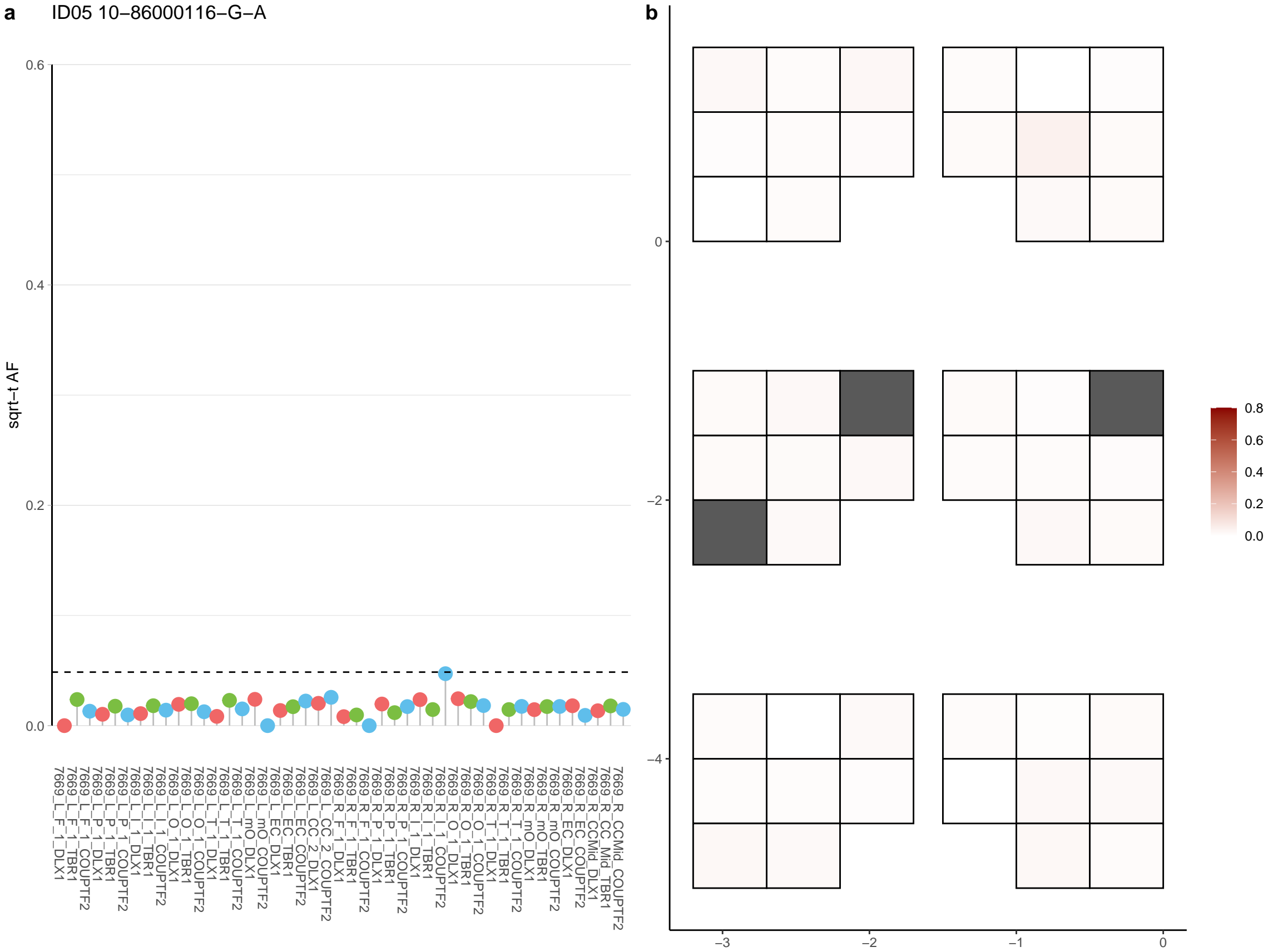

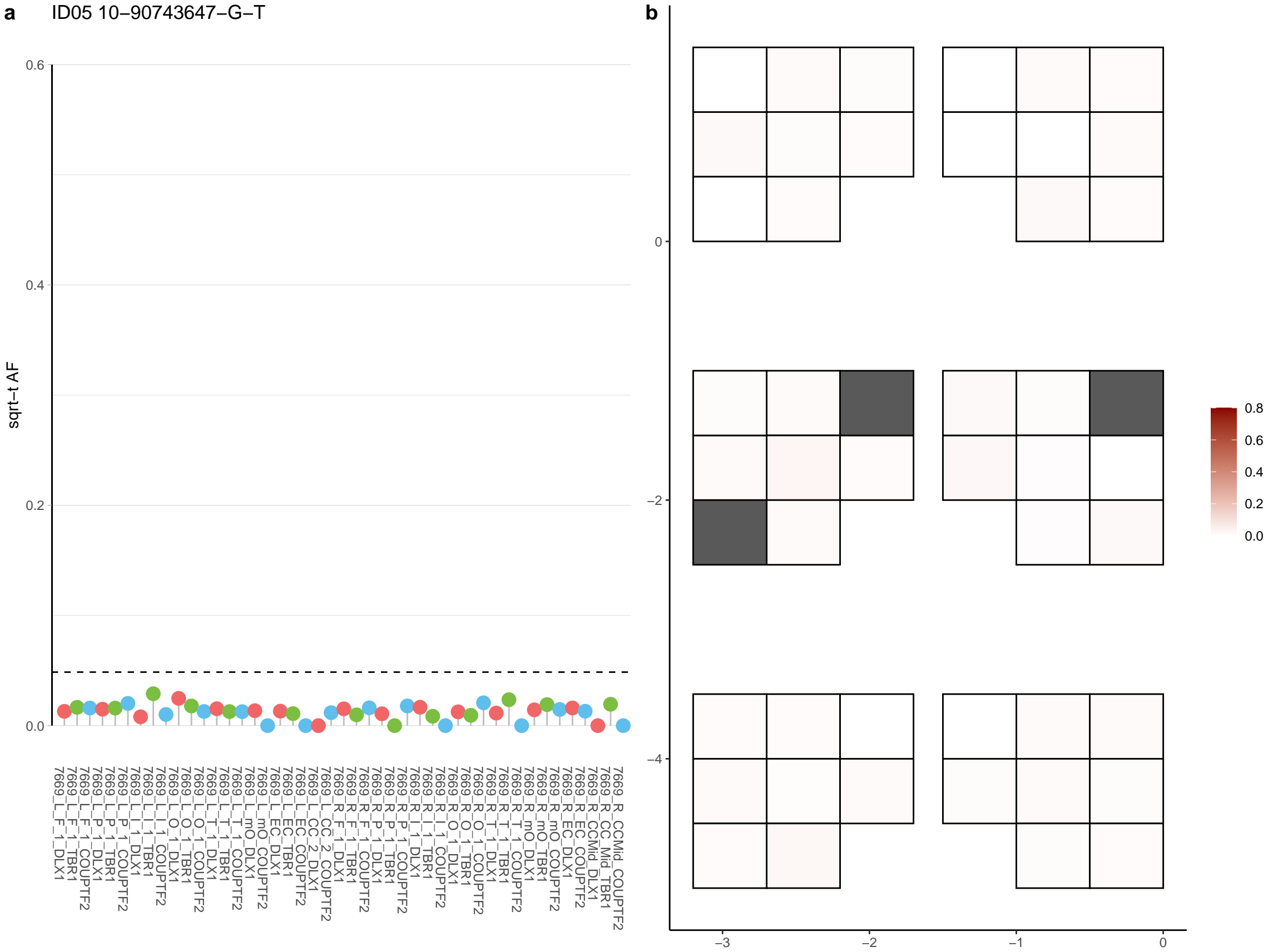

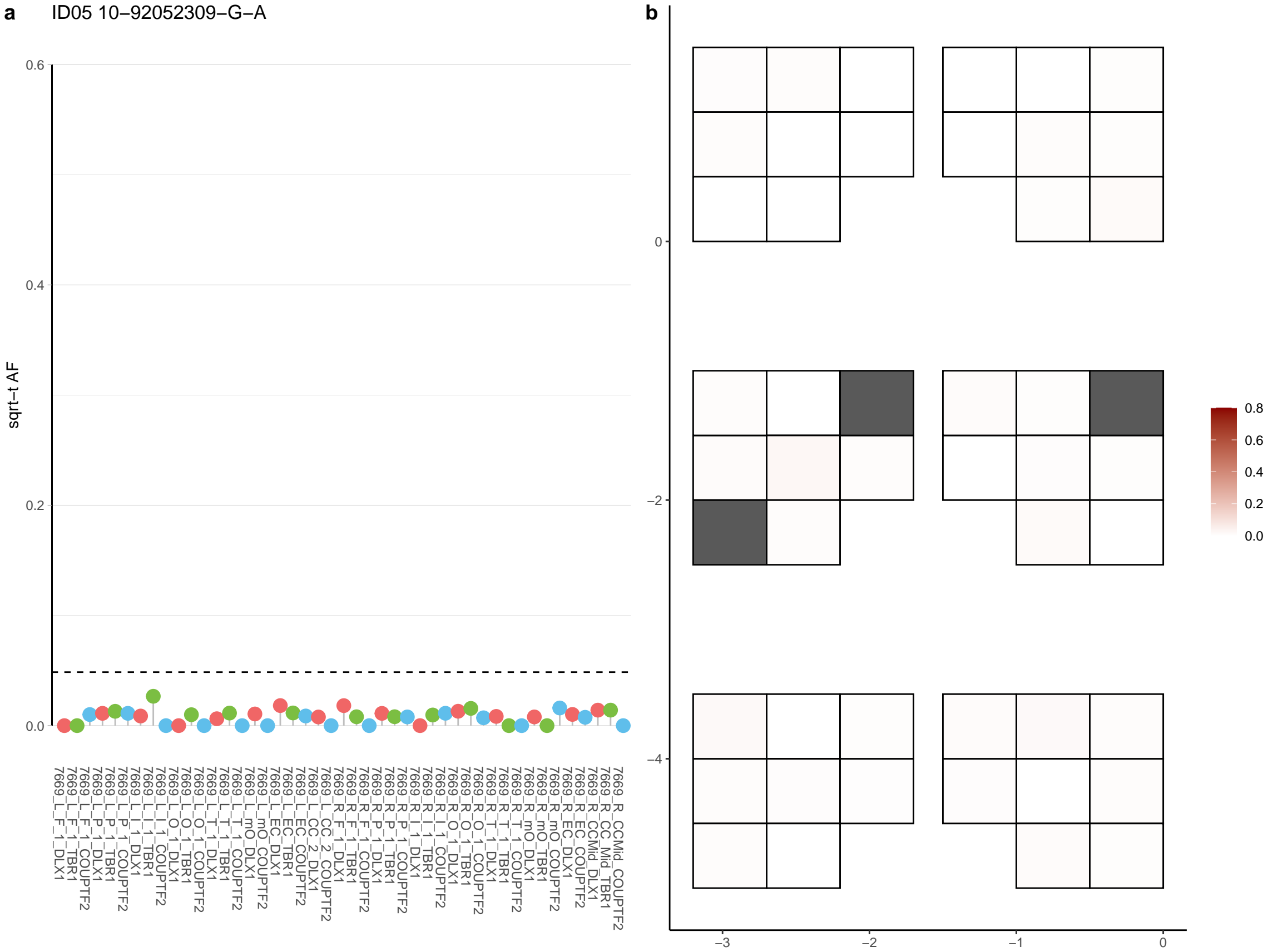

**a** ID05 10-98832334-C-T

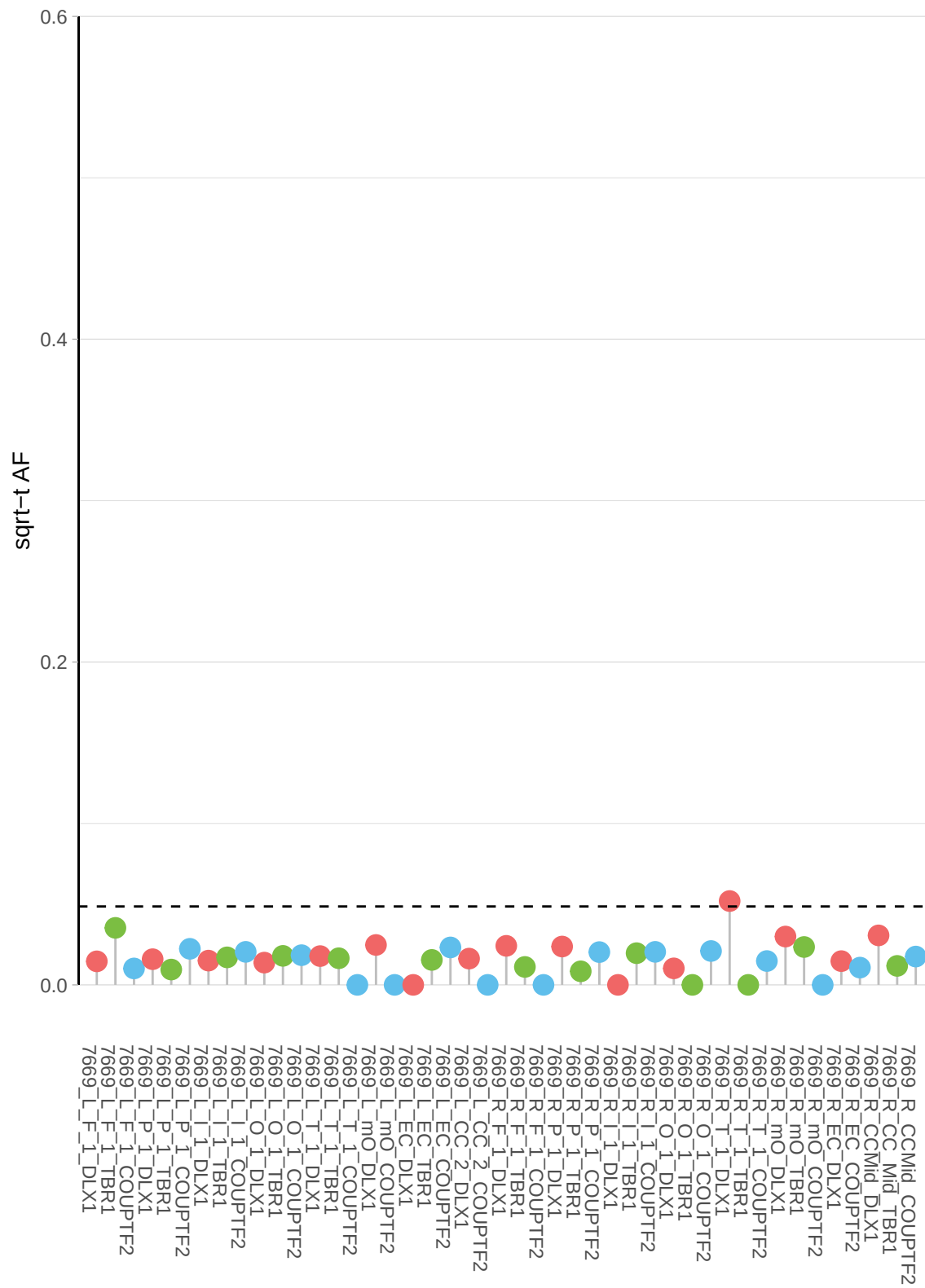

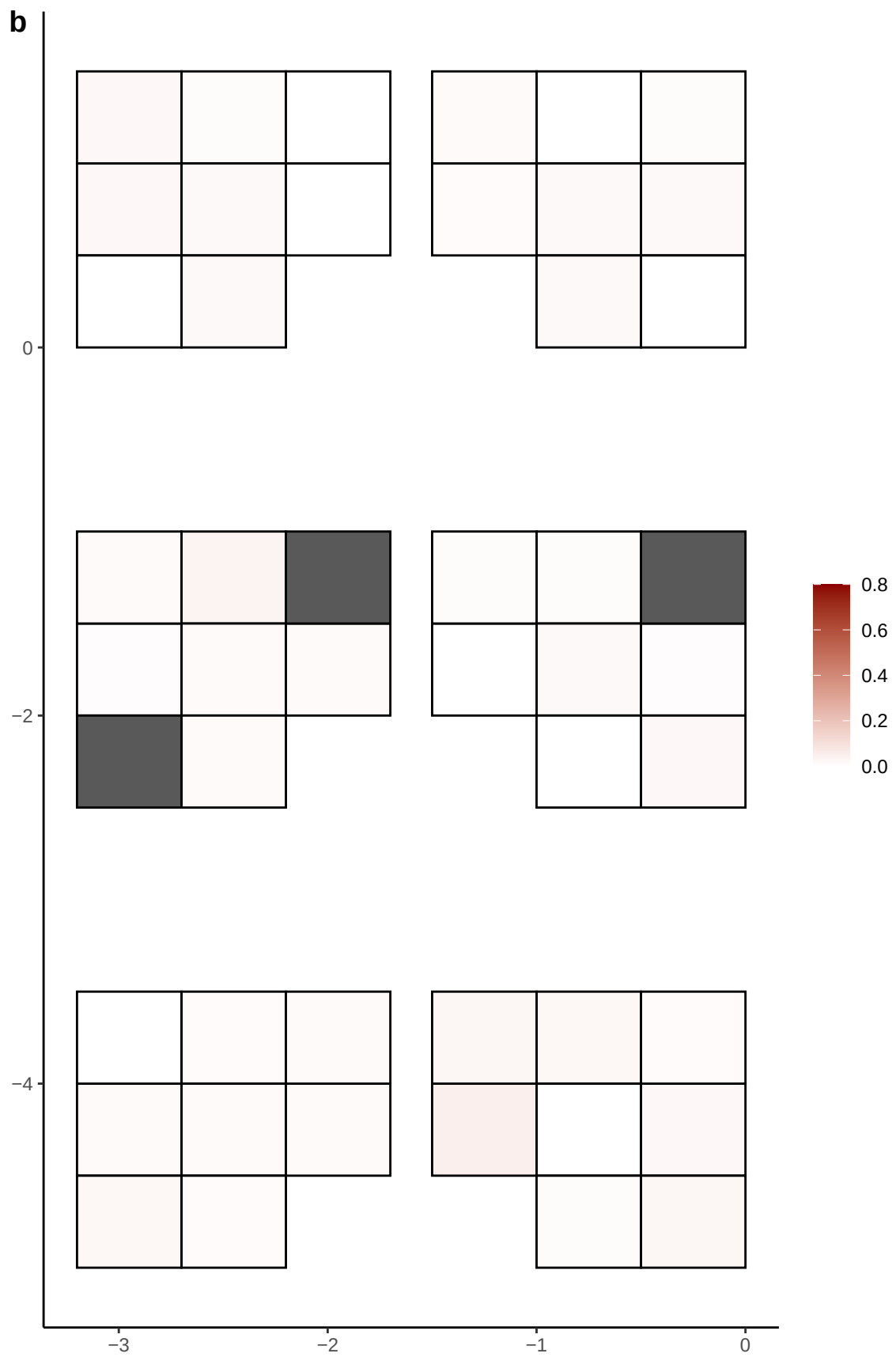

**a** ID05 10-98832335-C-A

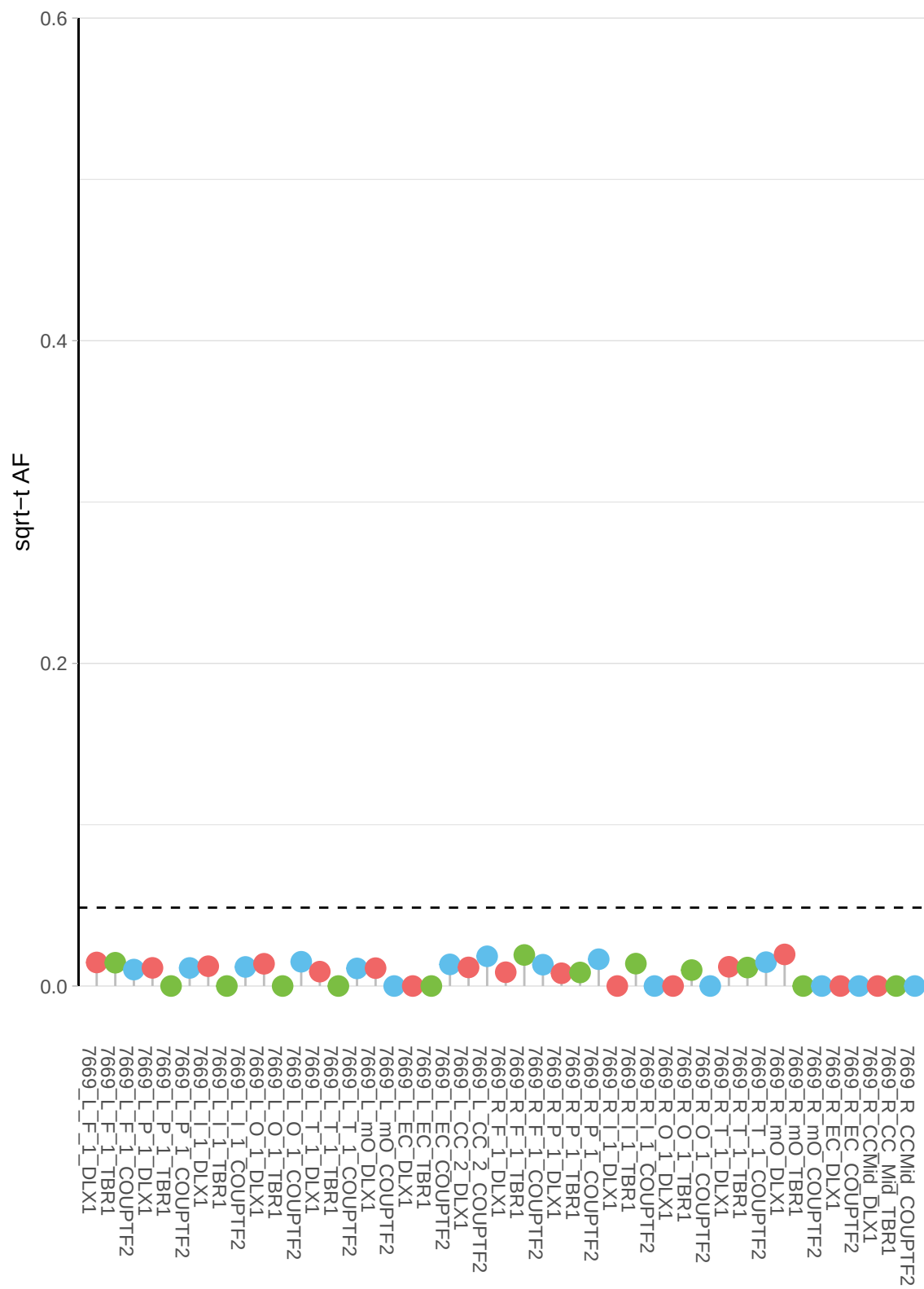

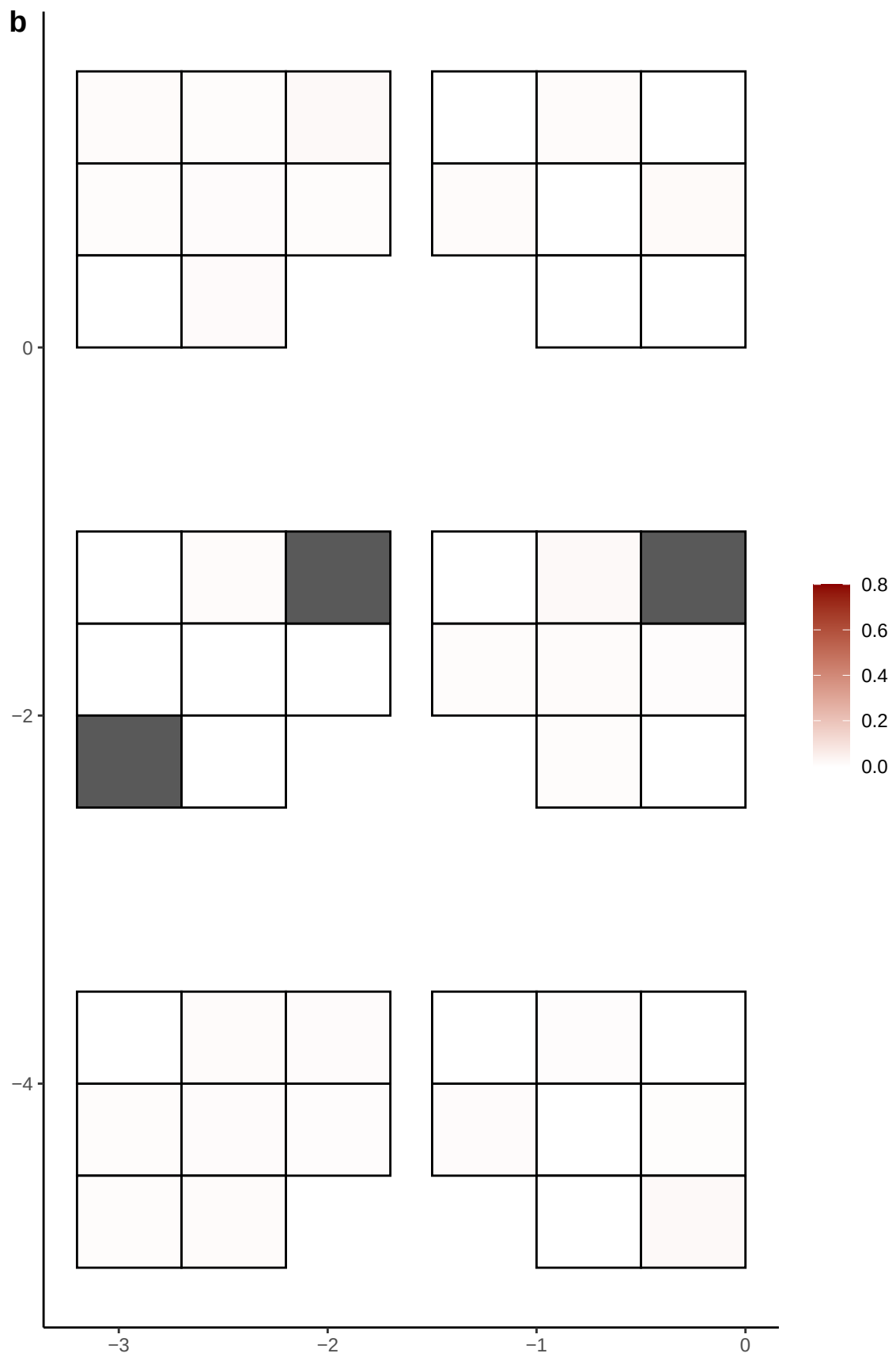

**a** ID05 11-105769654-T-C

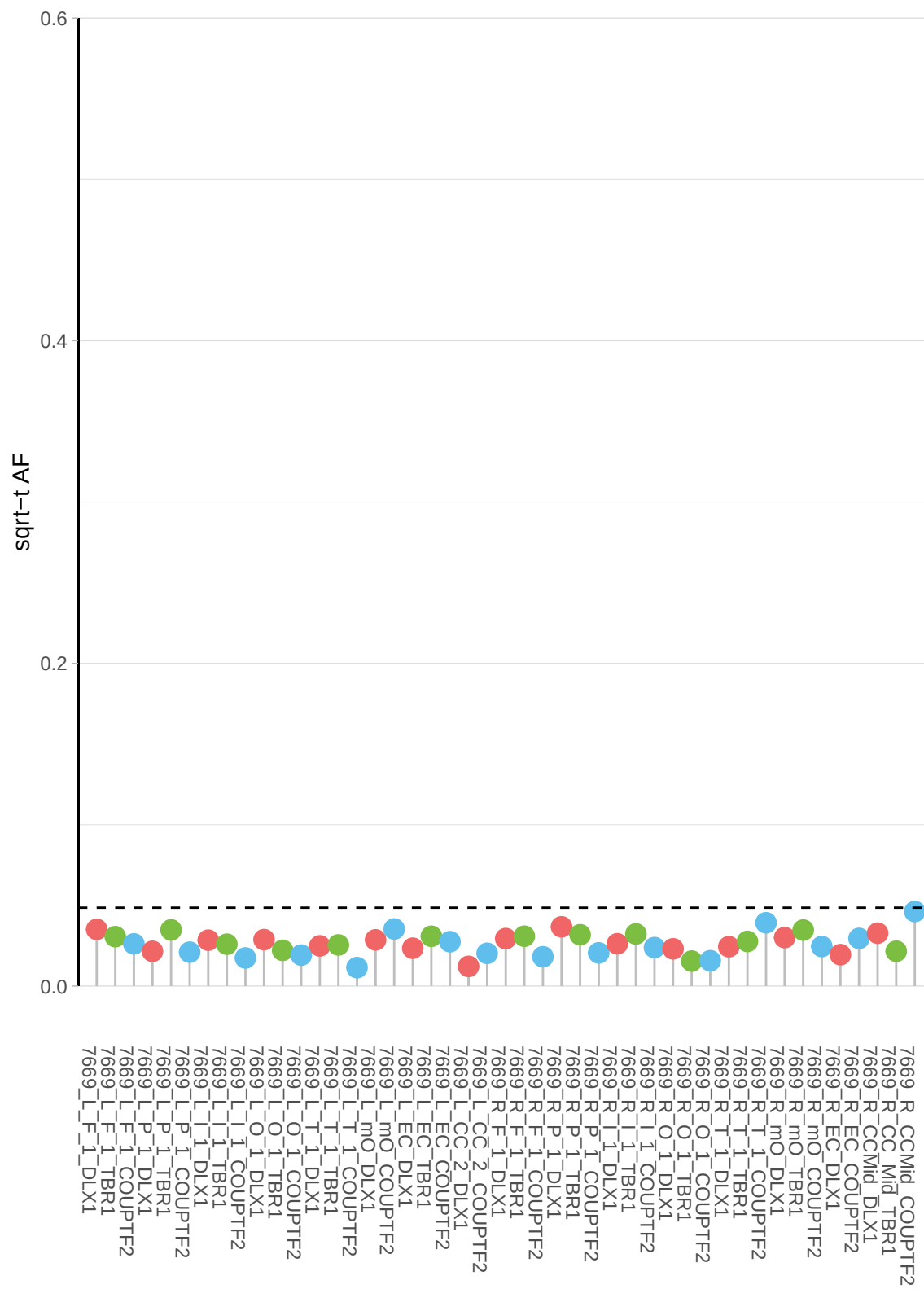

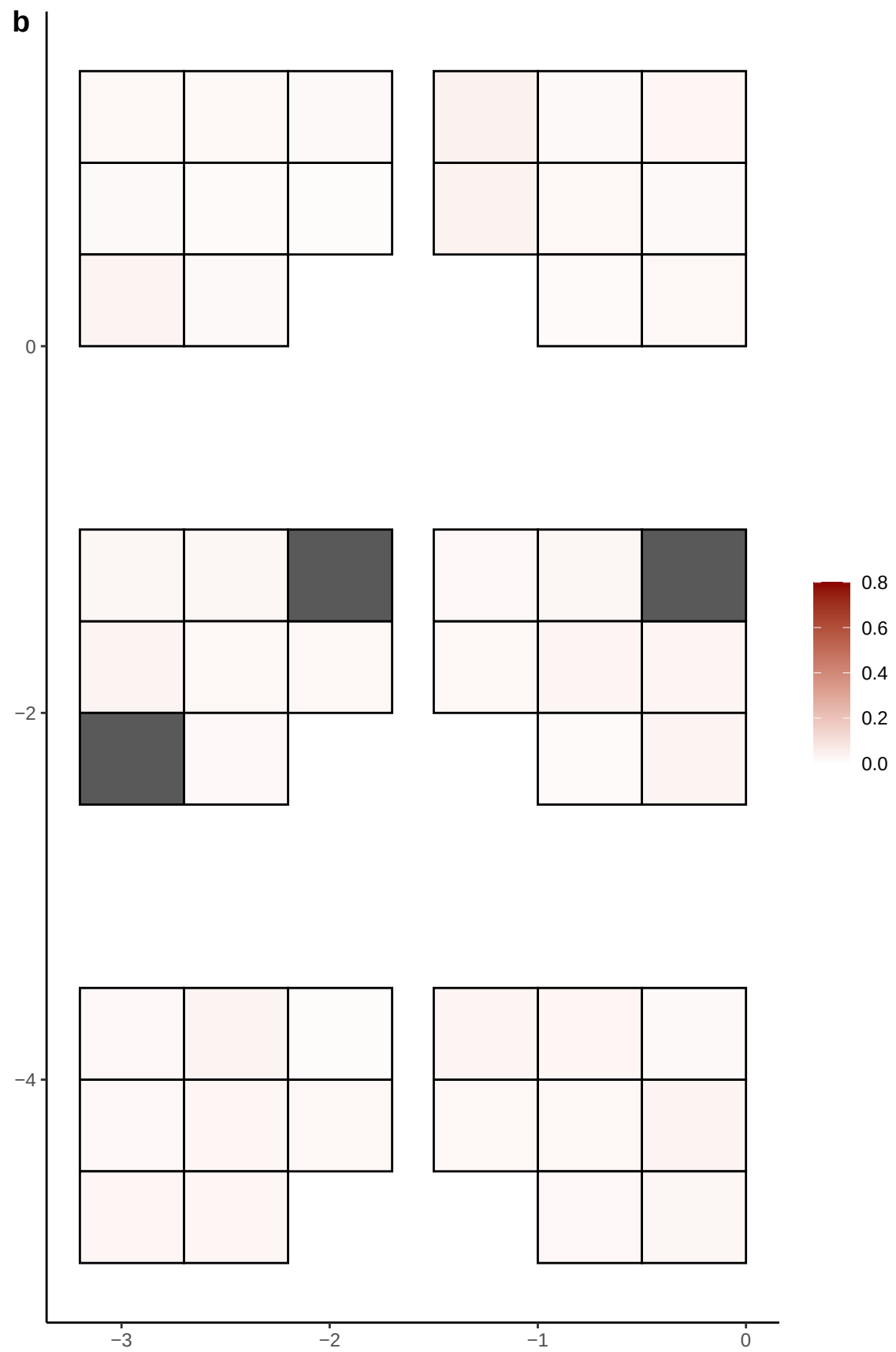

**a** ID05 11-109546001-G-C

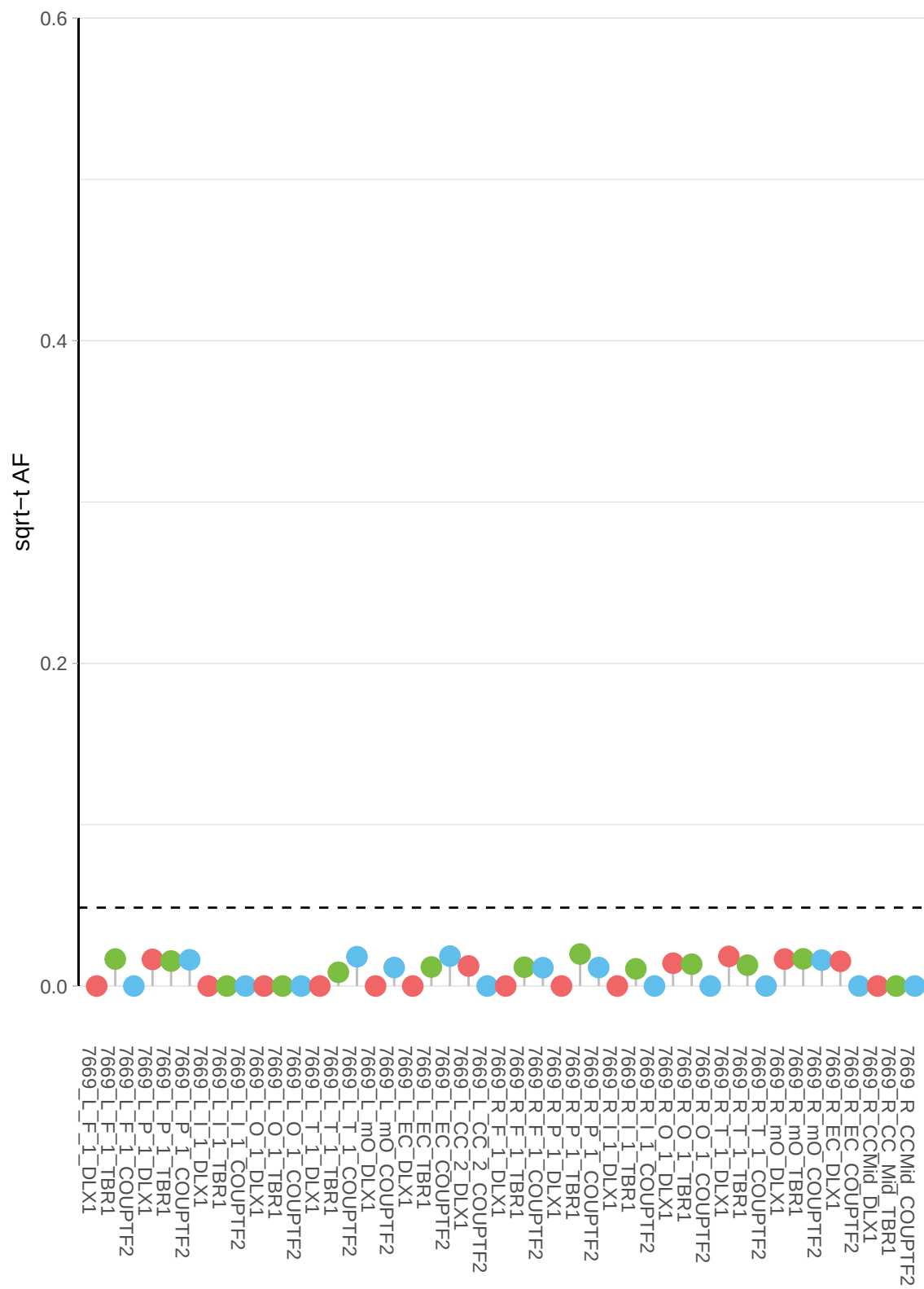

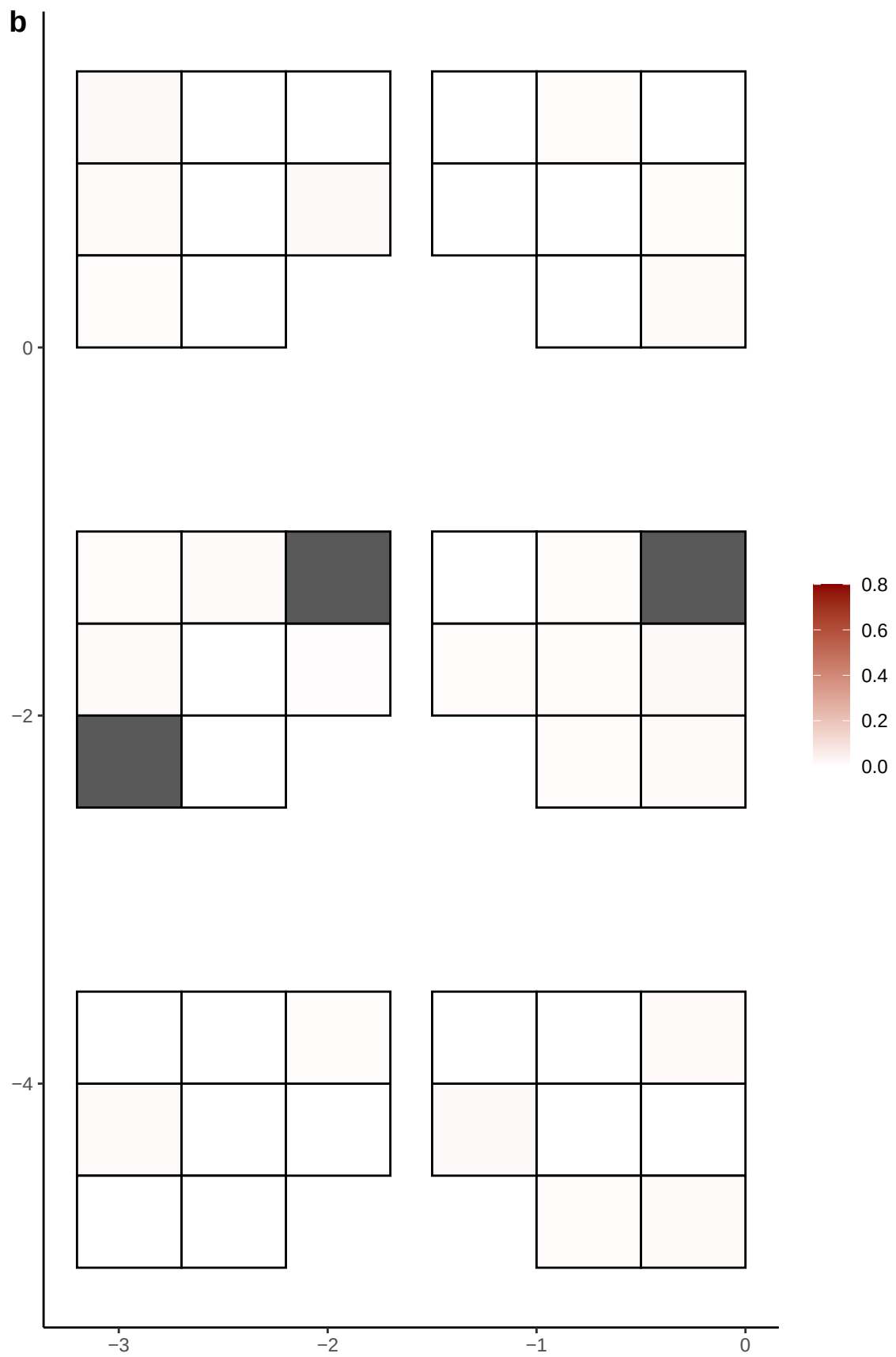

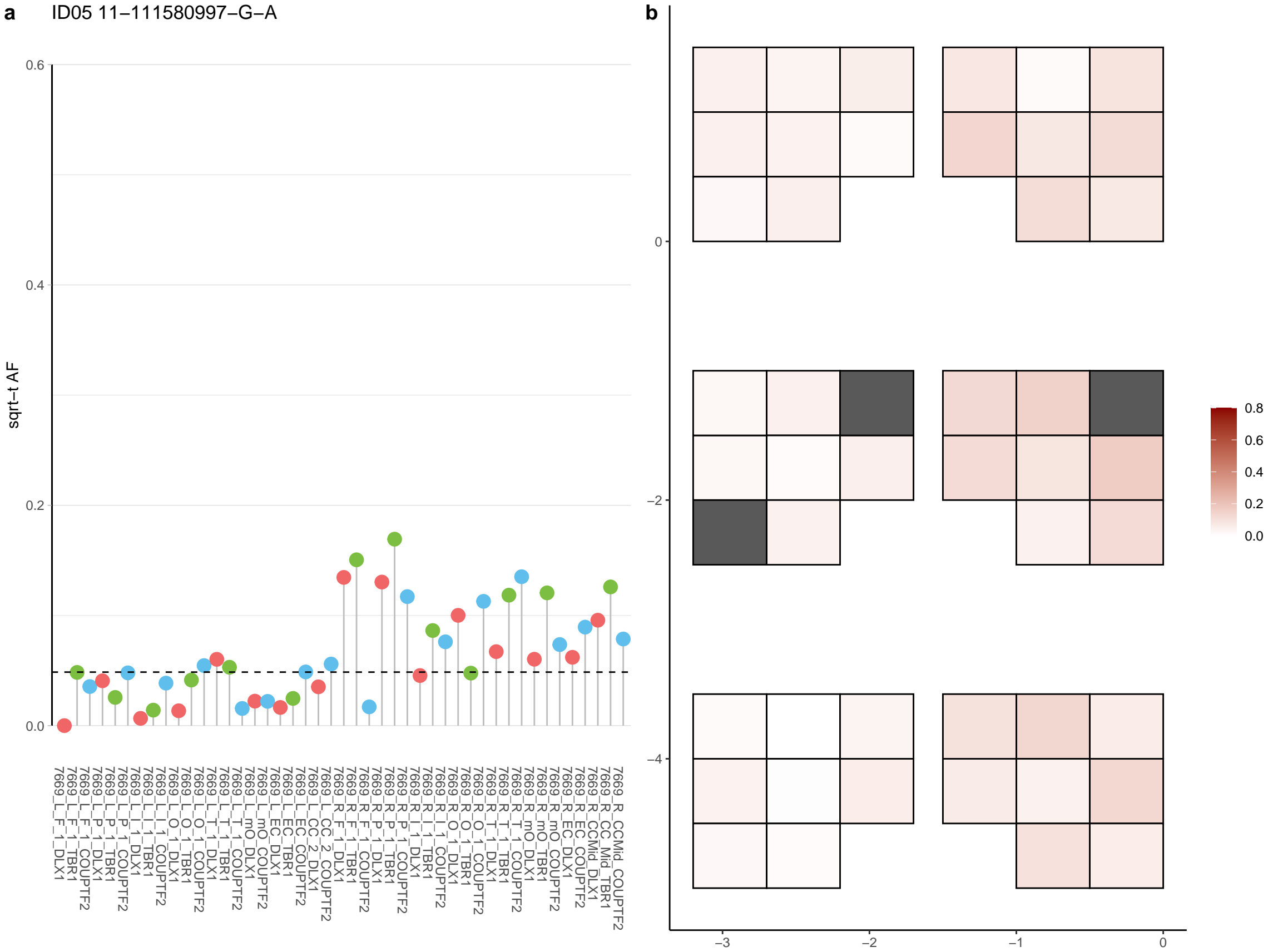

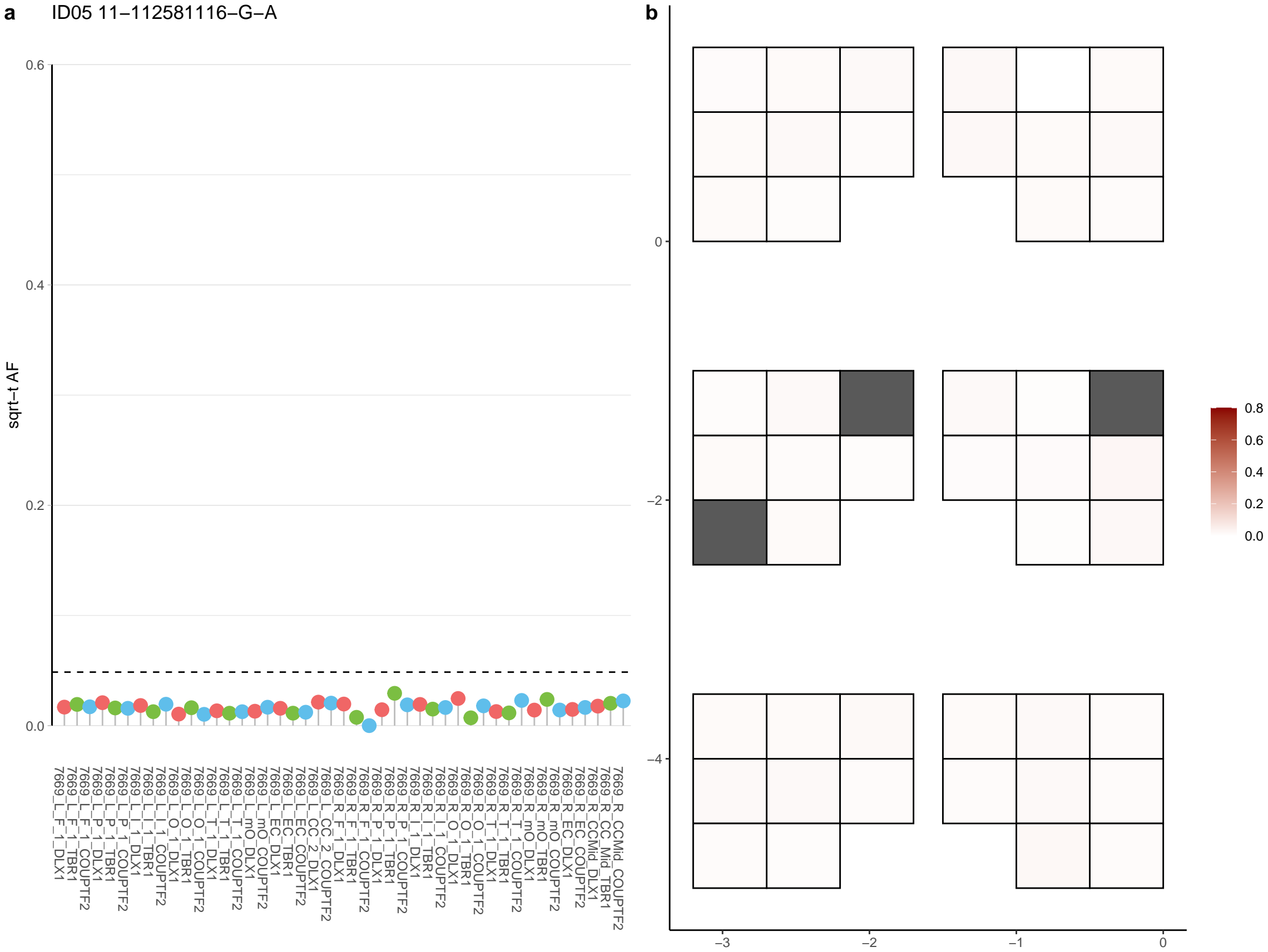

**a** ID05 11-112824453-C-T

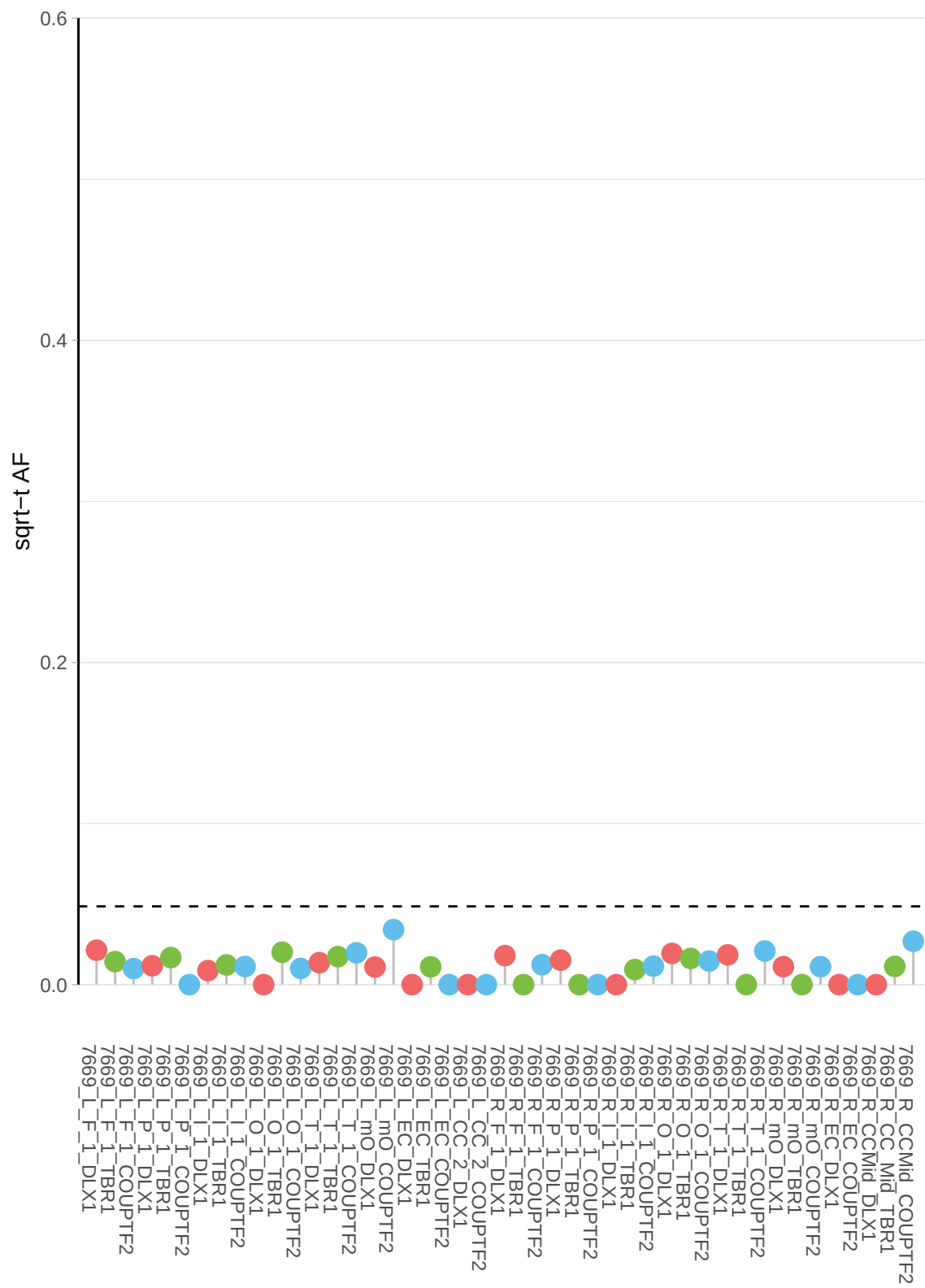

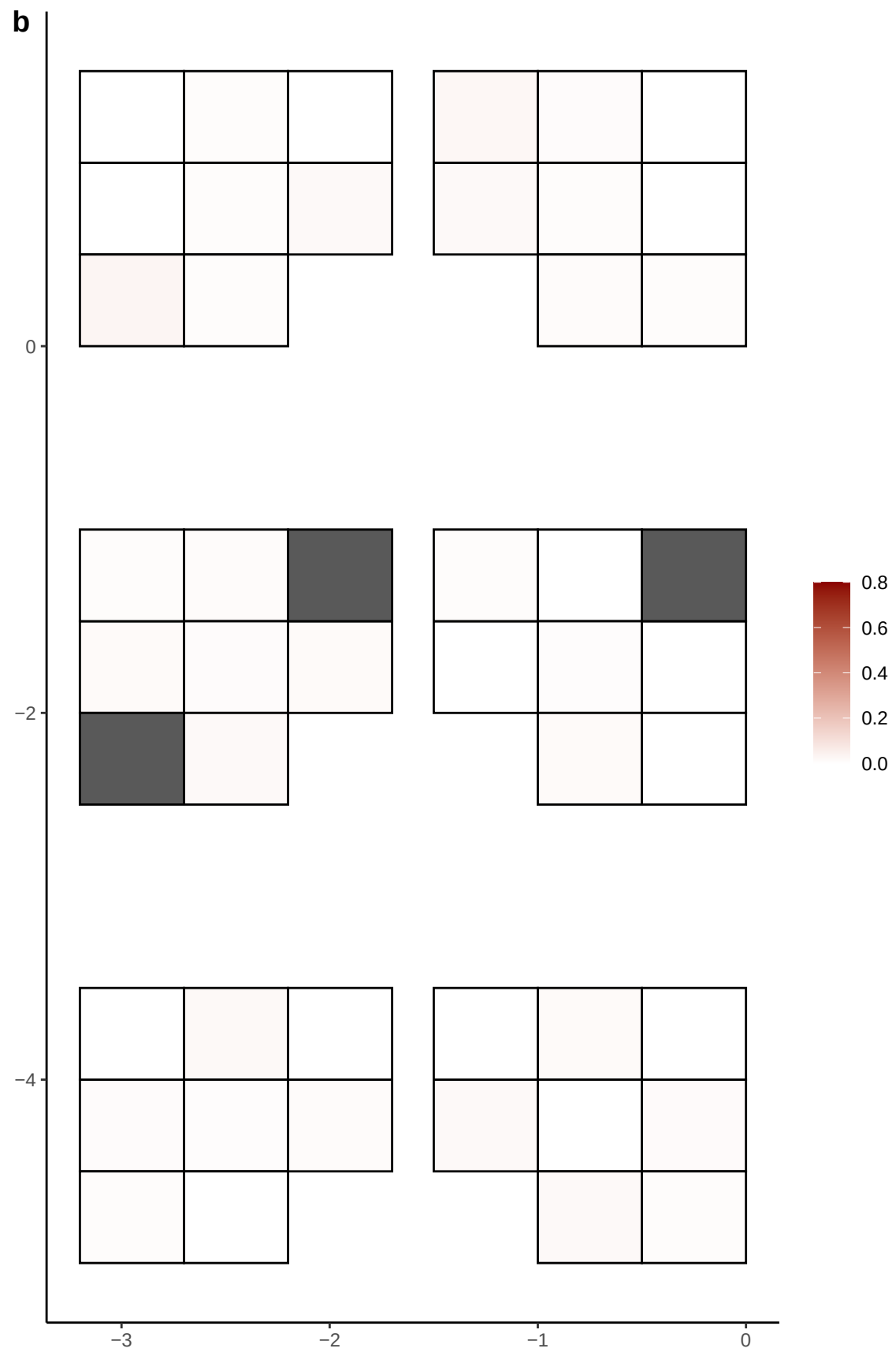

**a** ID05 11-11681099-G-C

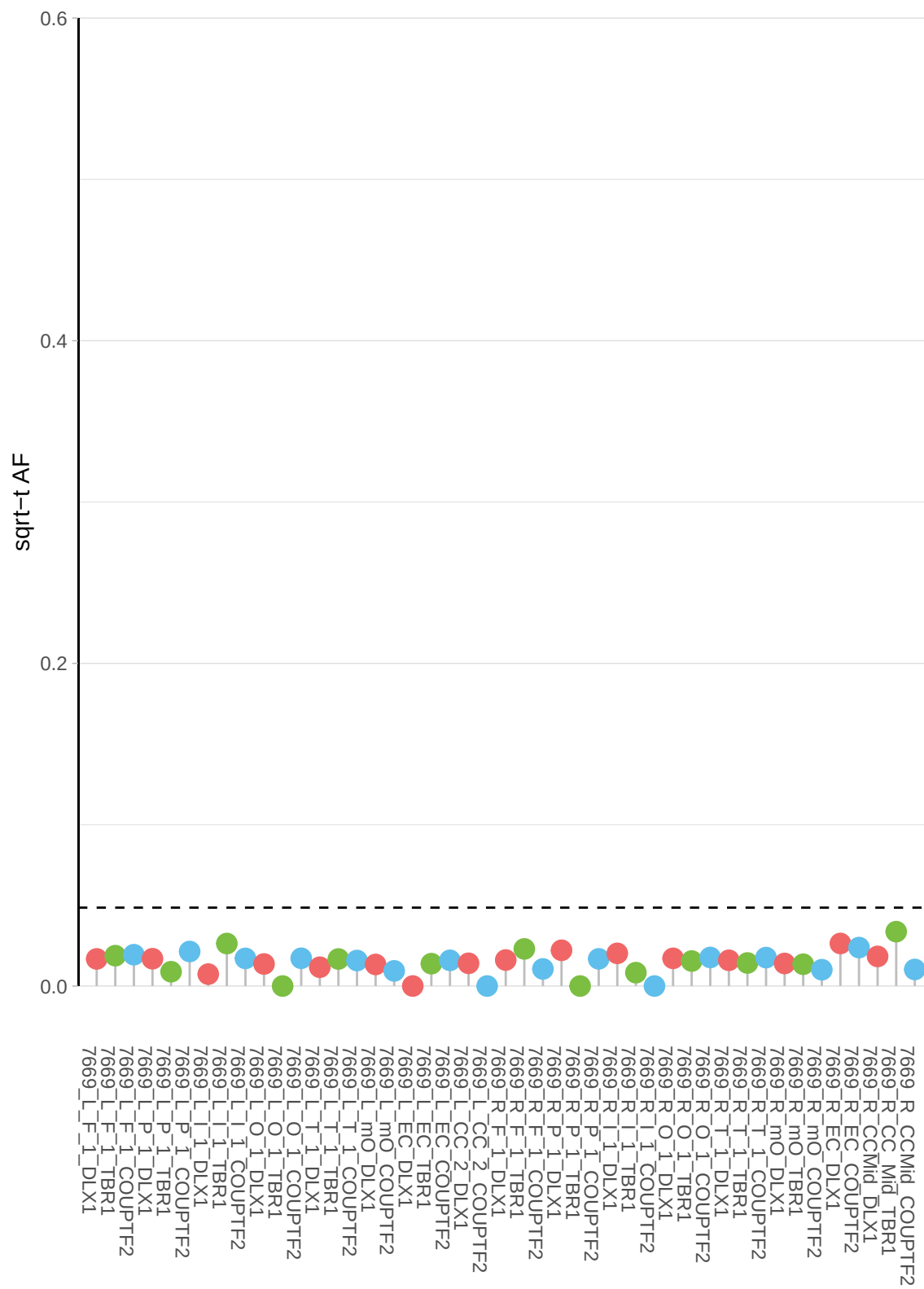

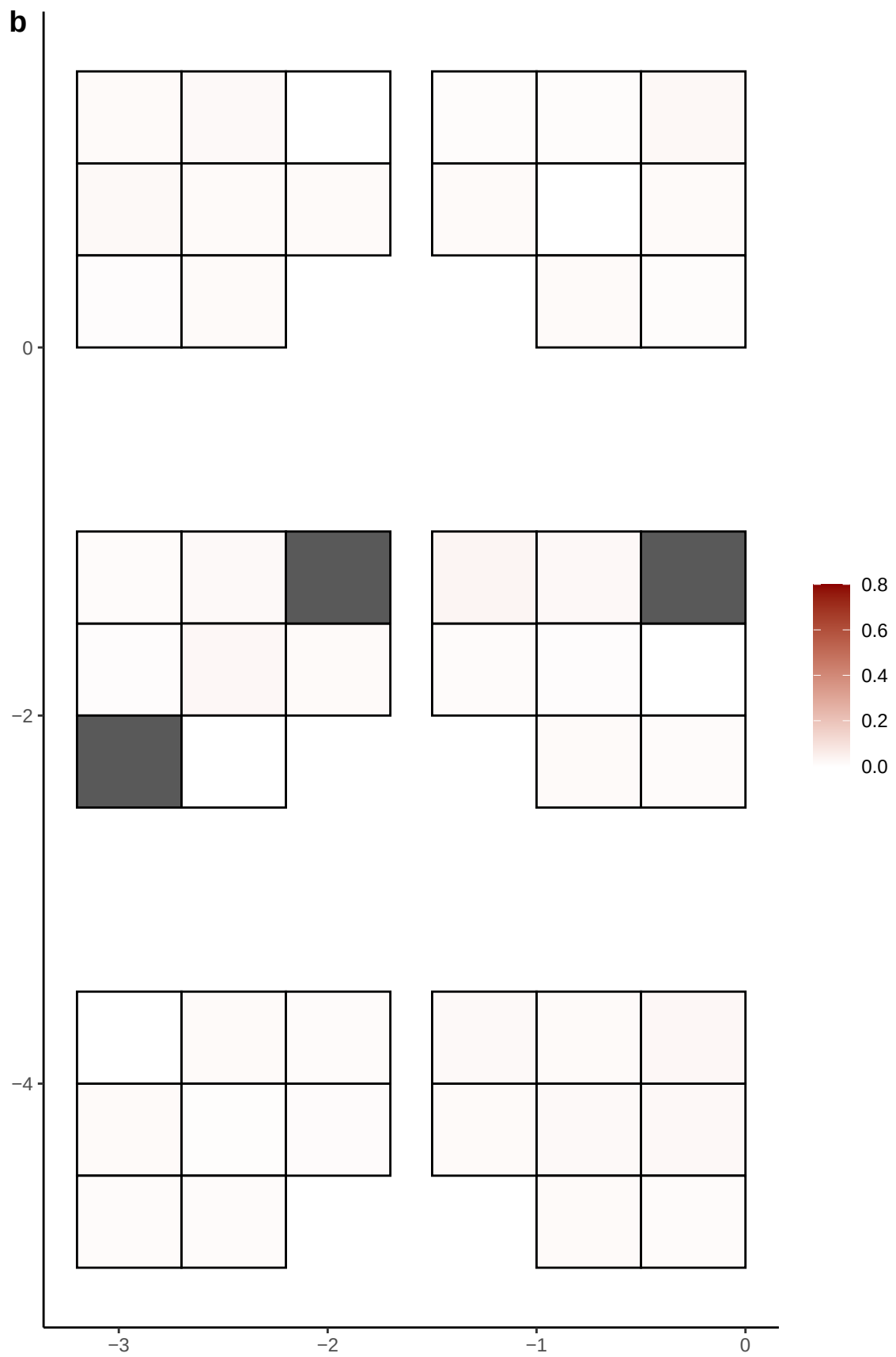

**a** ID05 11-123817821-G-A

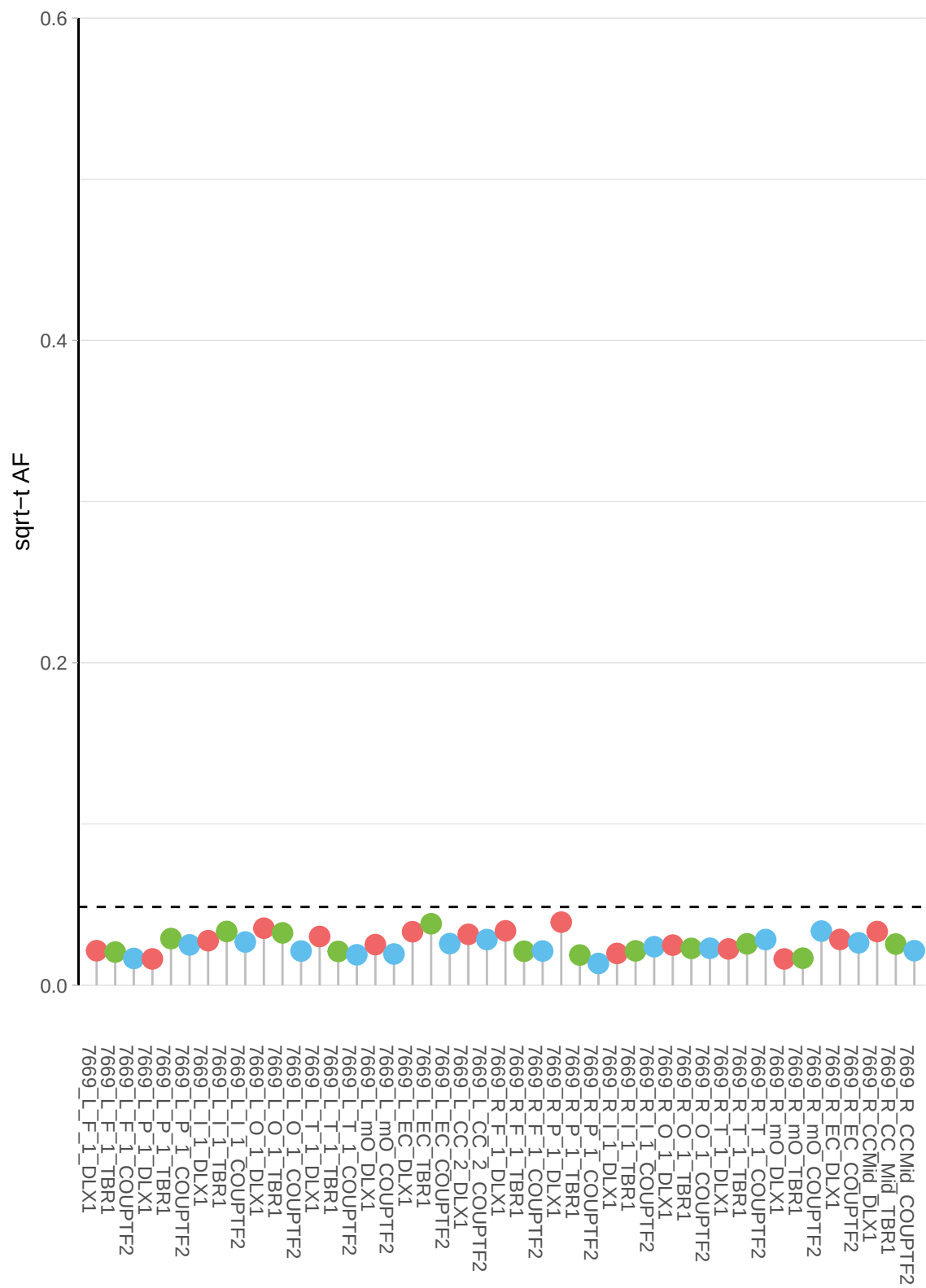

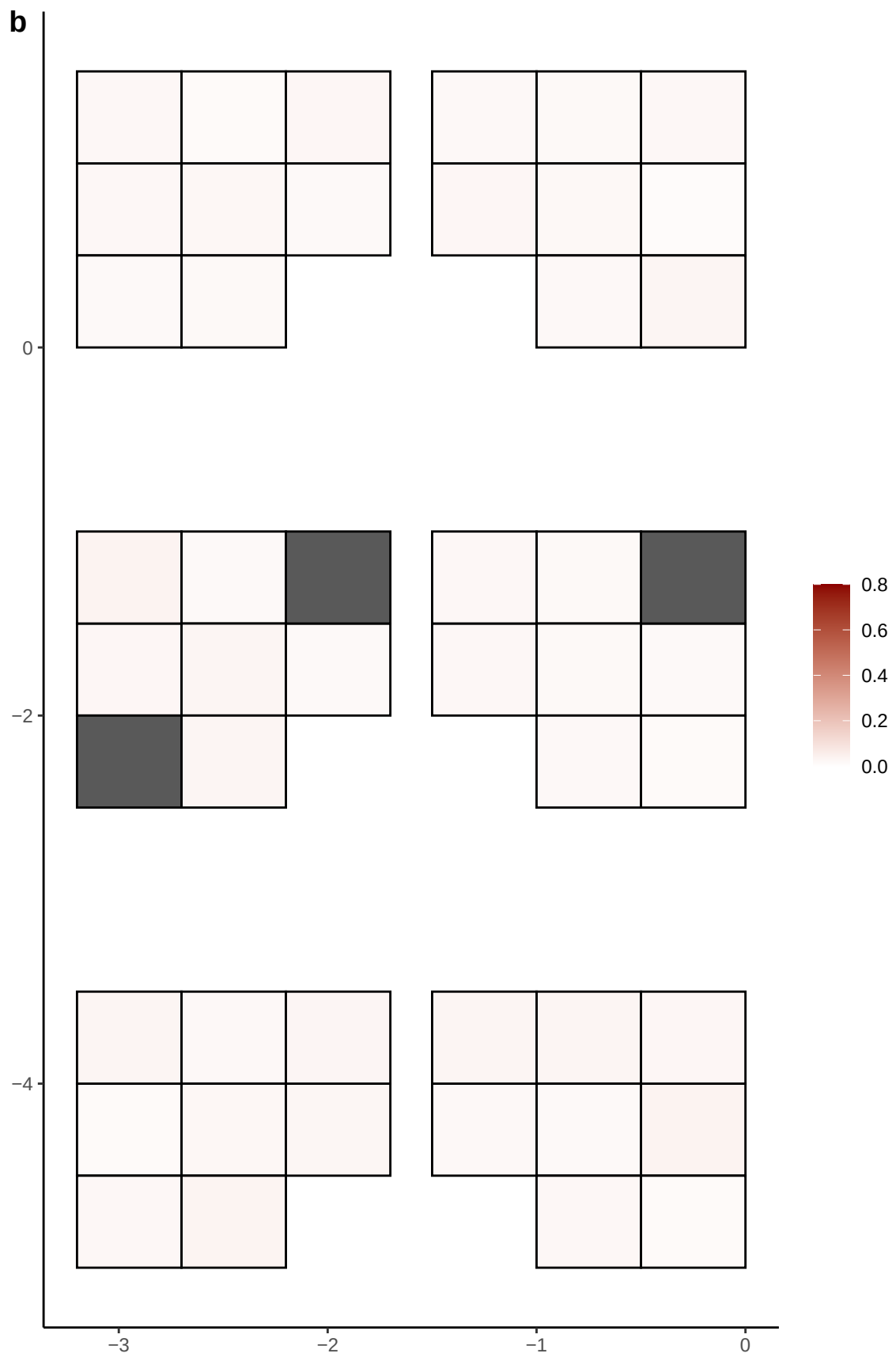

**a** ID05 11-125271170-G-C

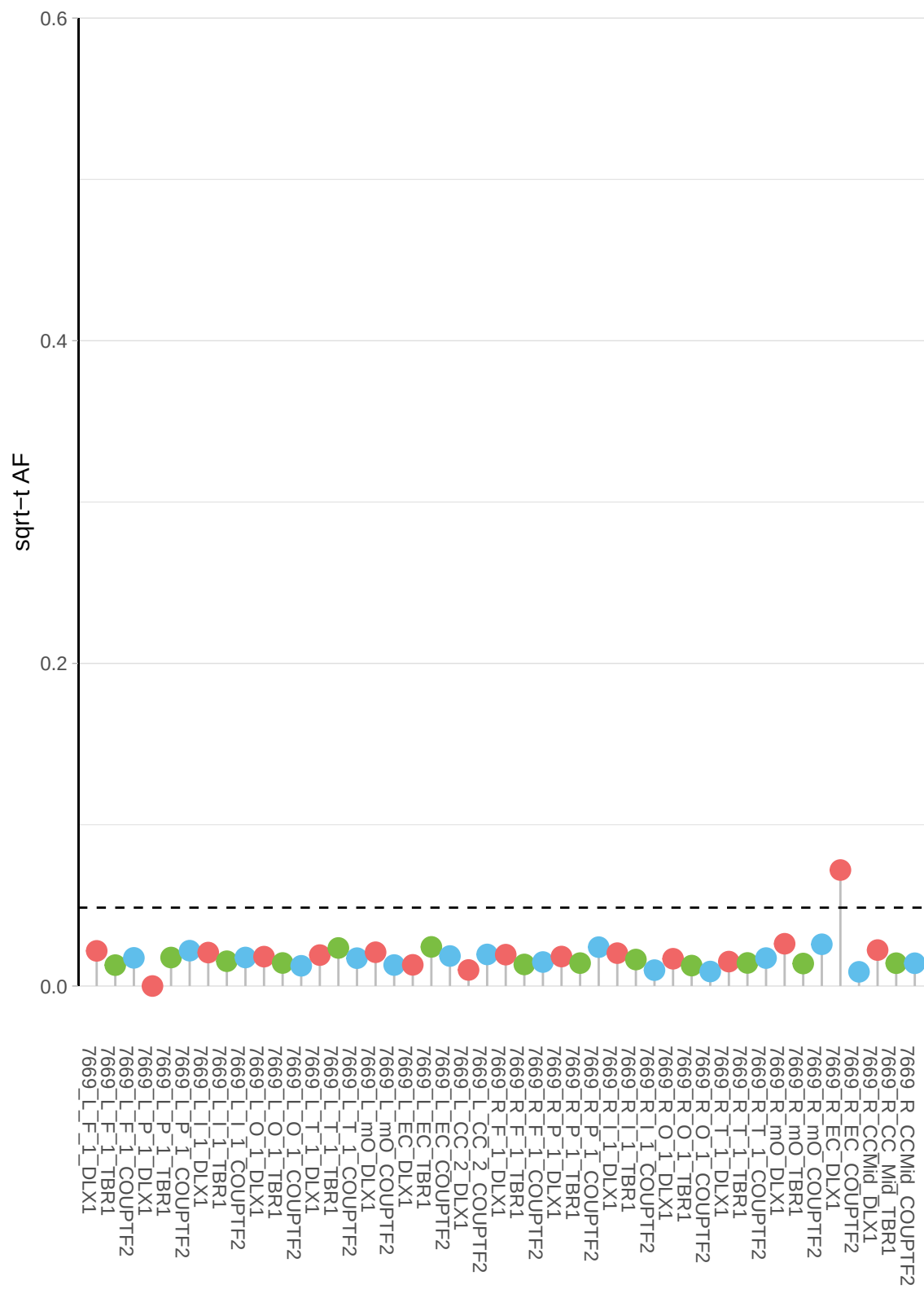

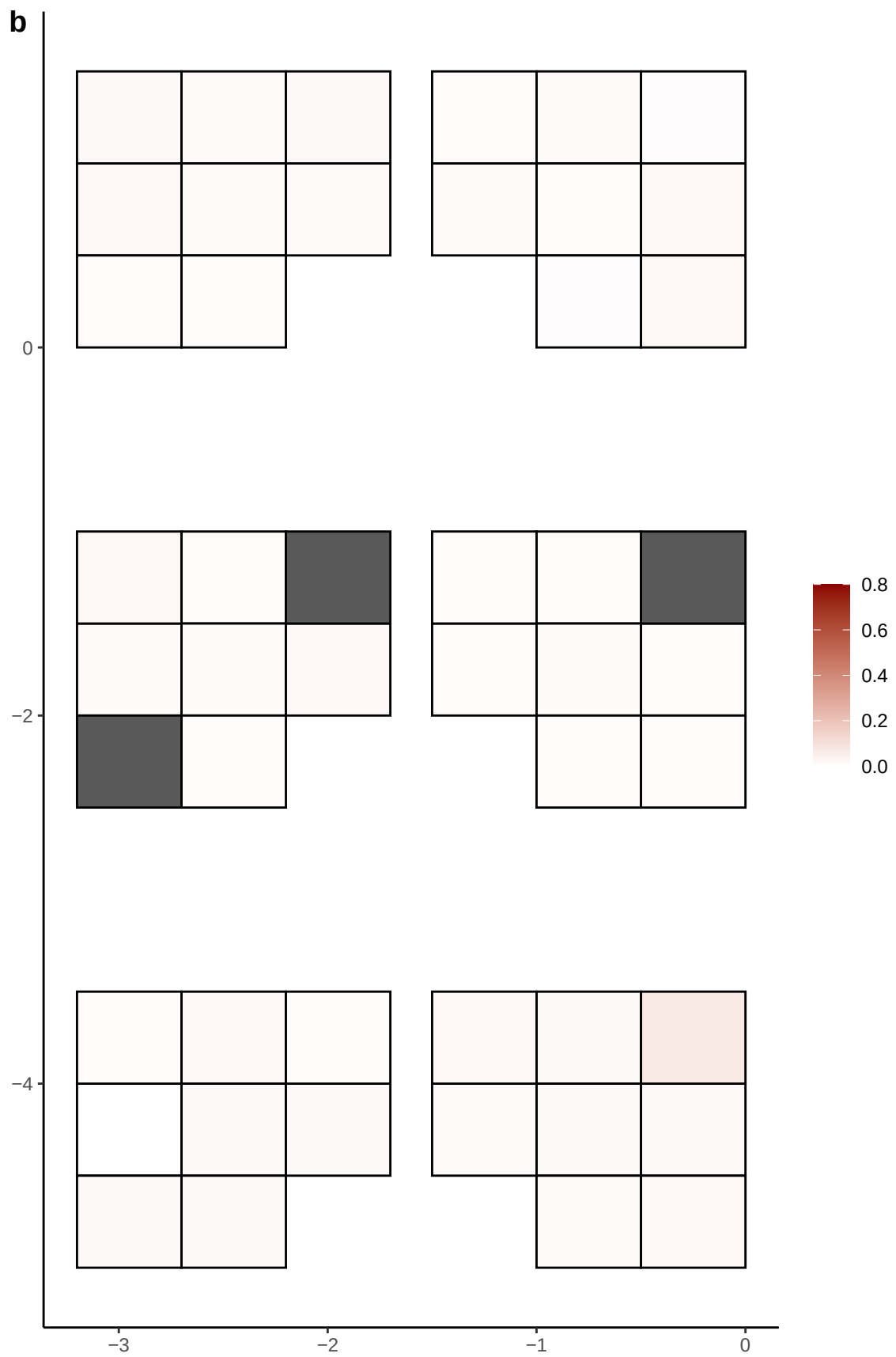

**a** ID05 11-127350815-C-T

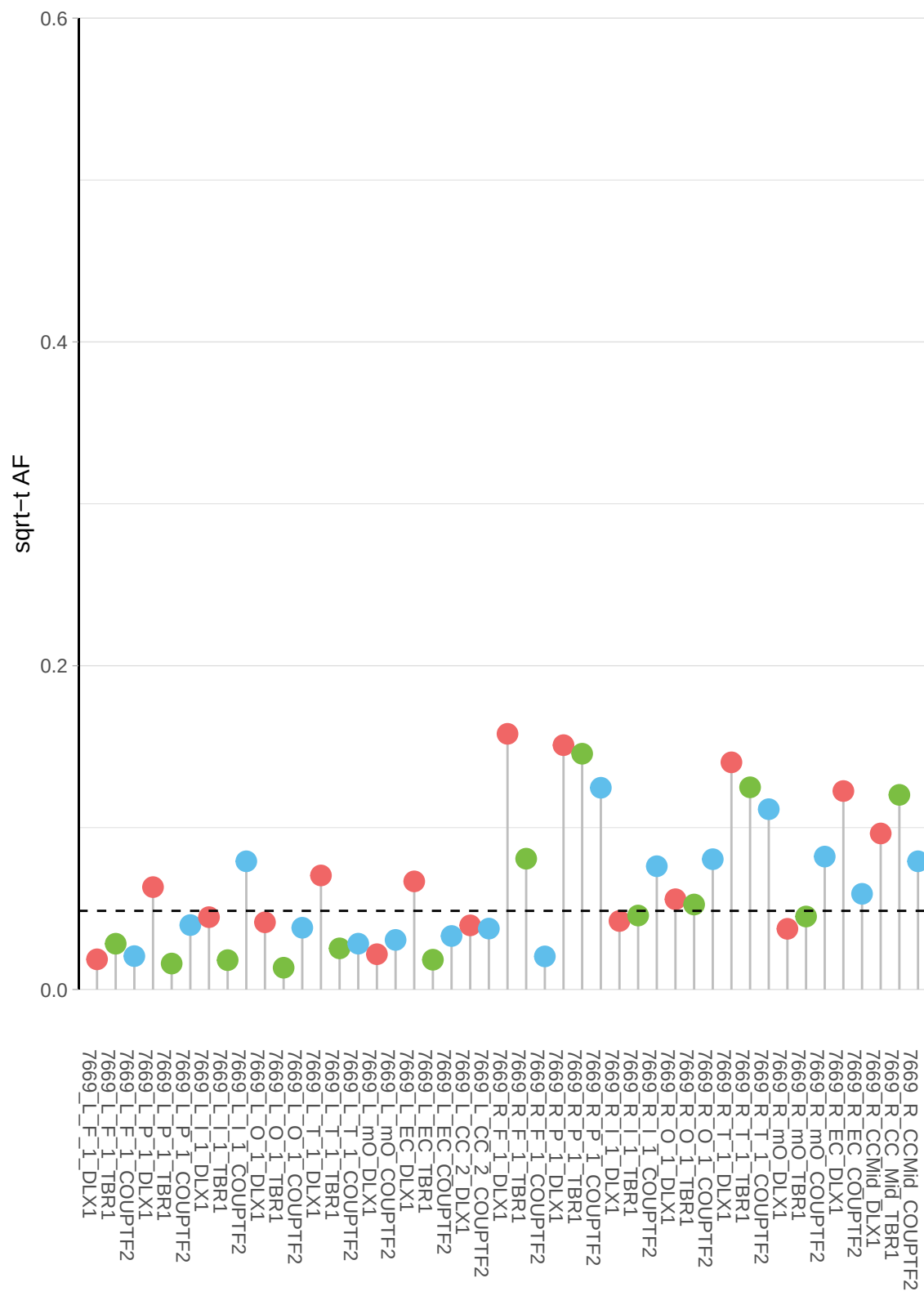

**b**

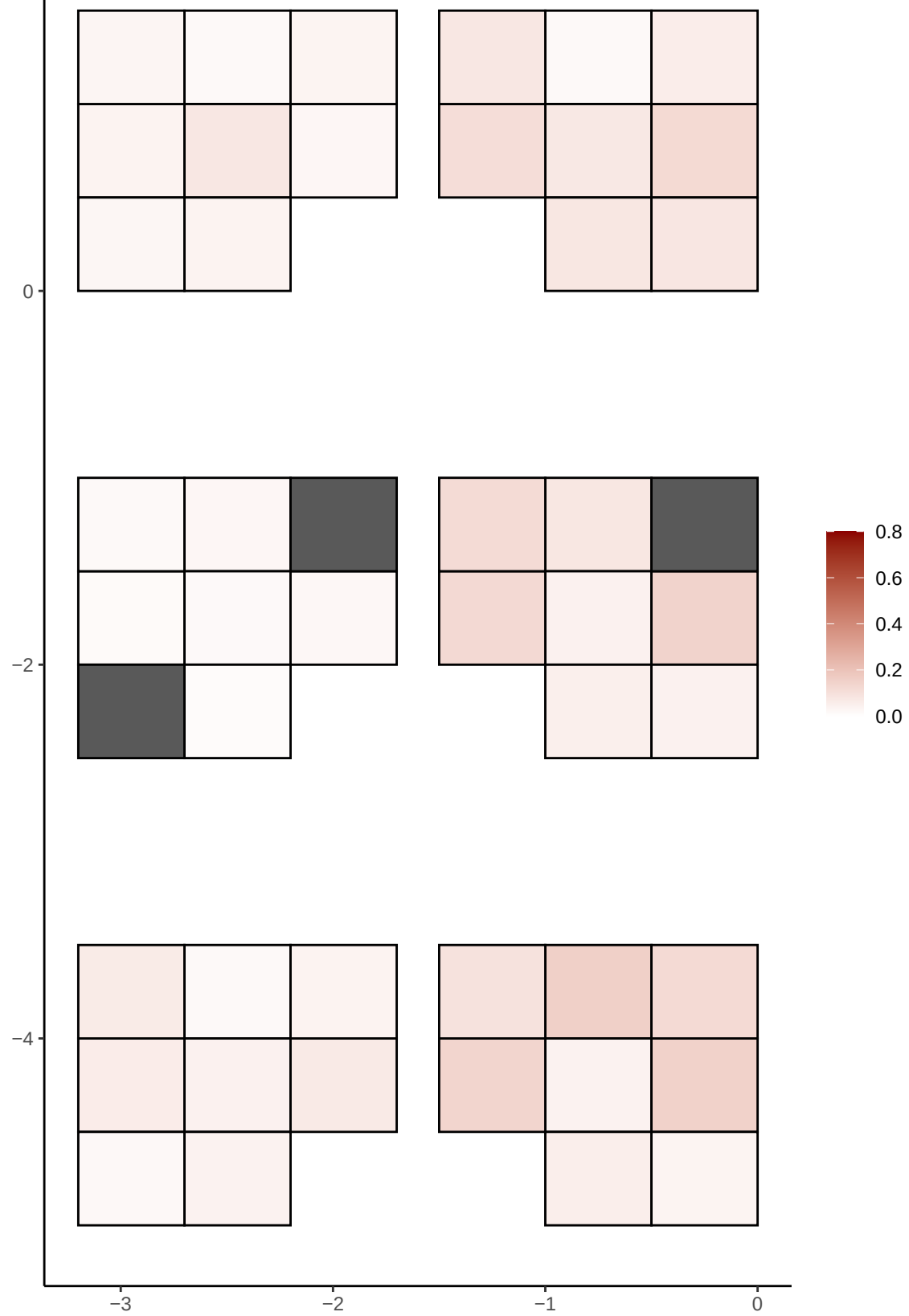

**a** ID05 11-133014880-G-C

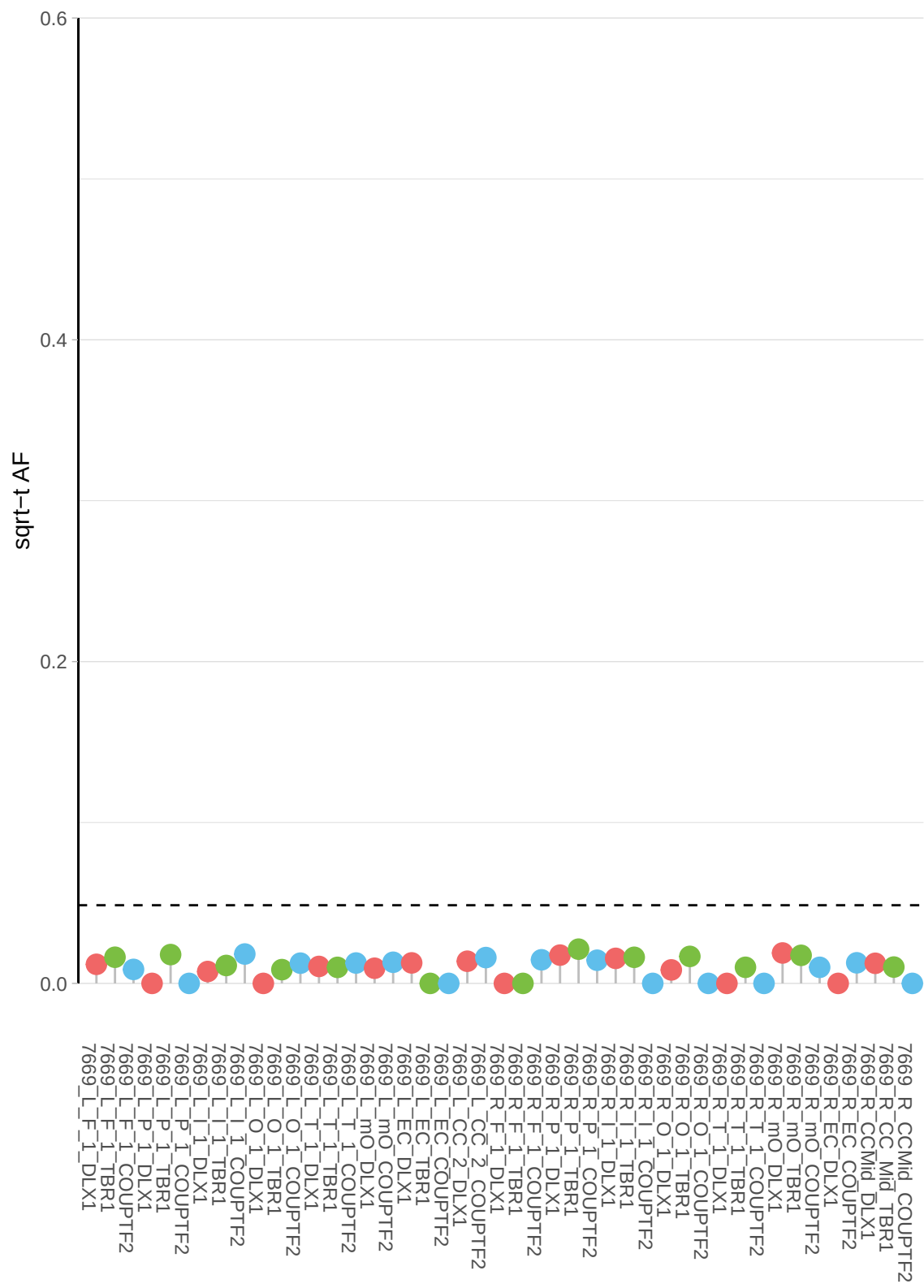

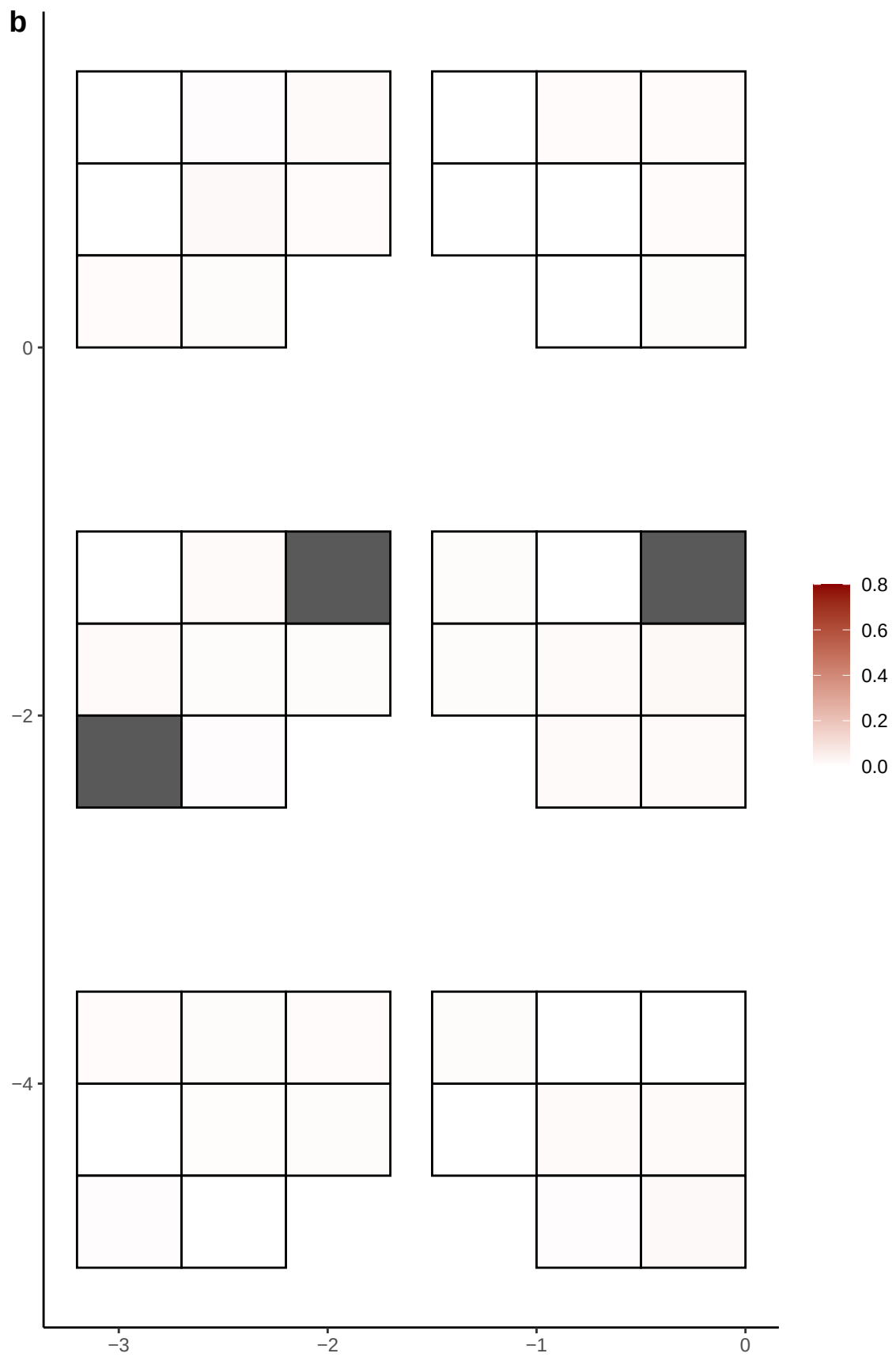

**a** ID05 11-21194438-G-A

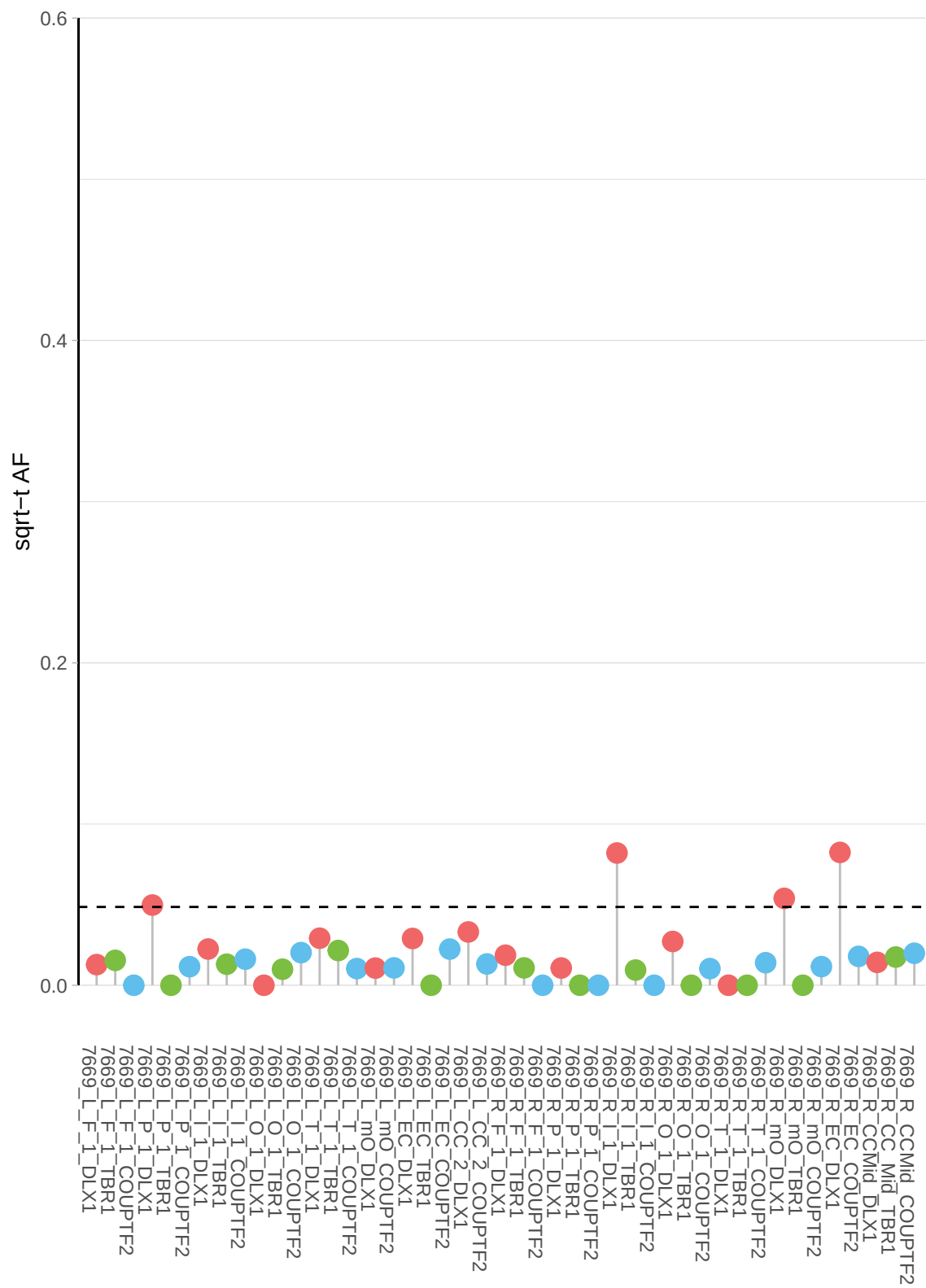

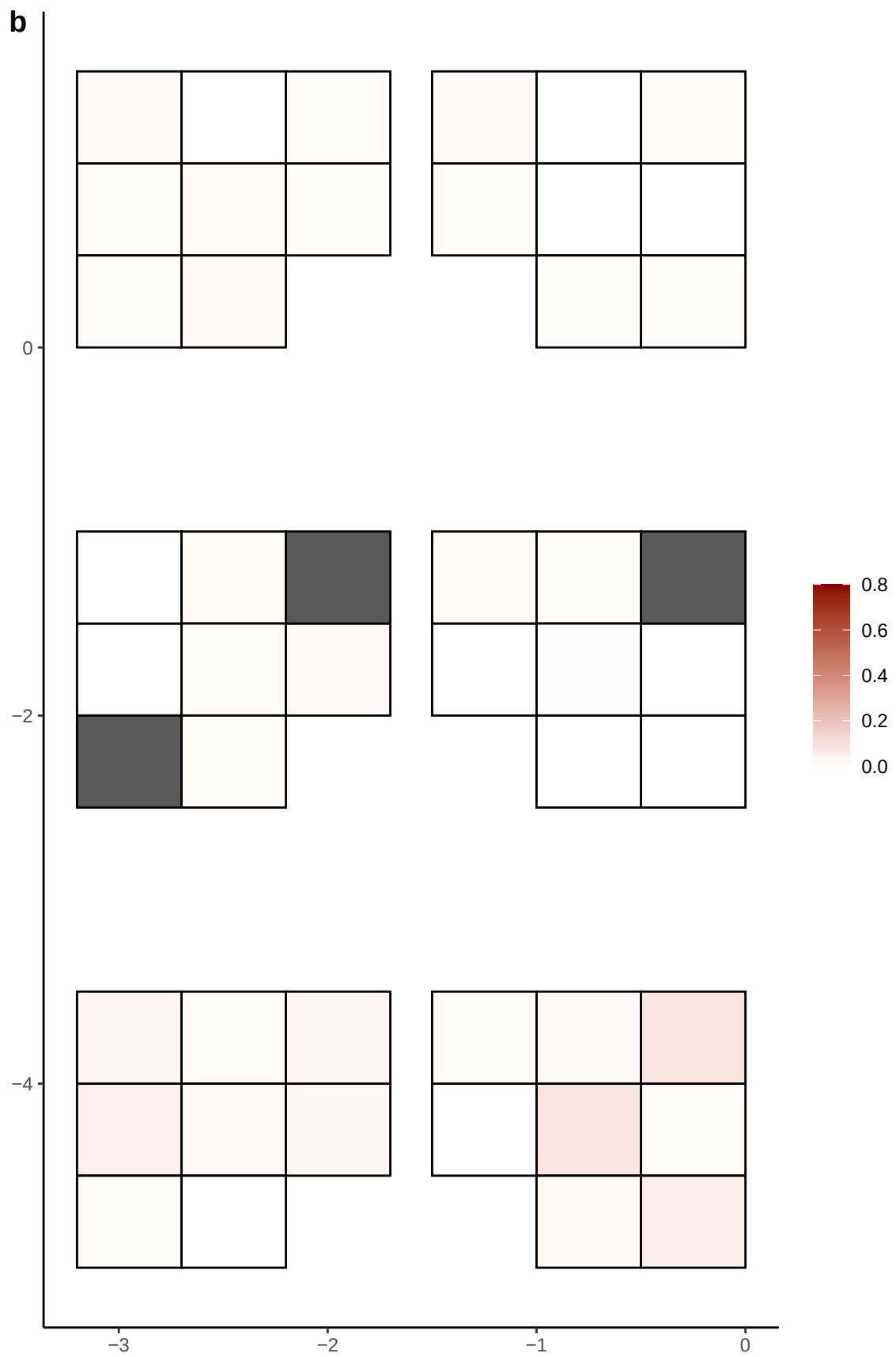

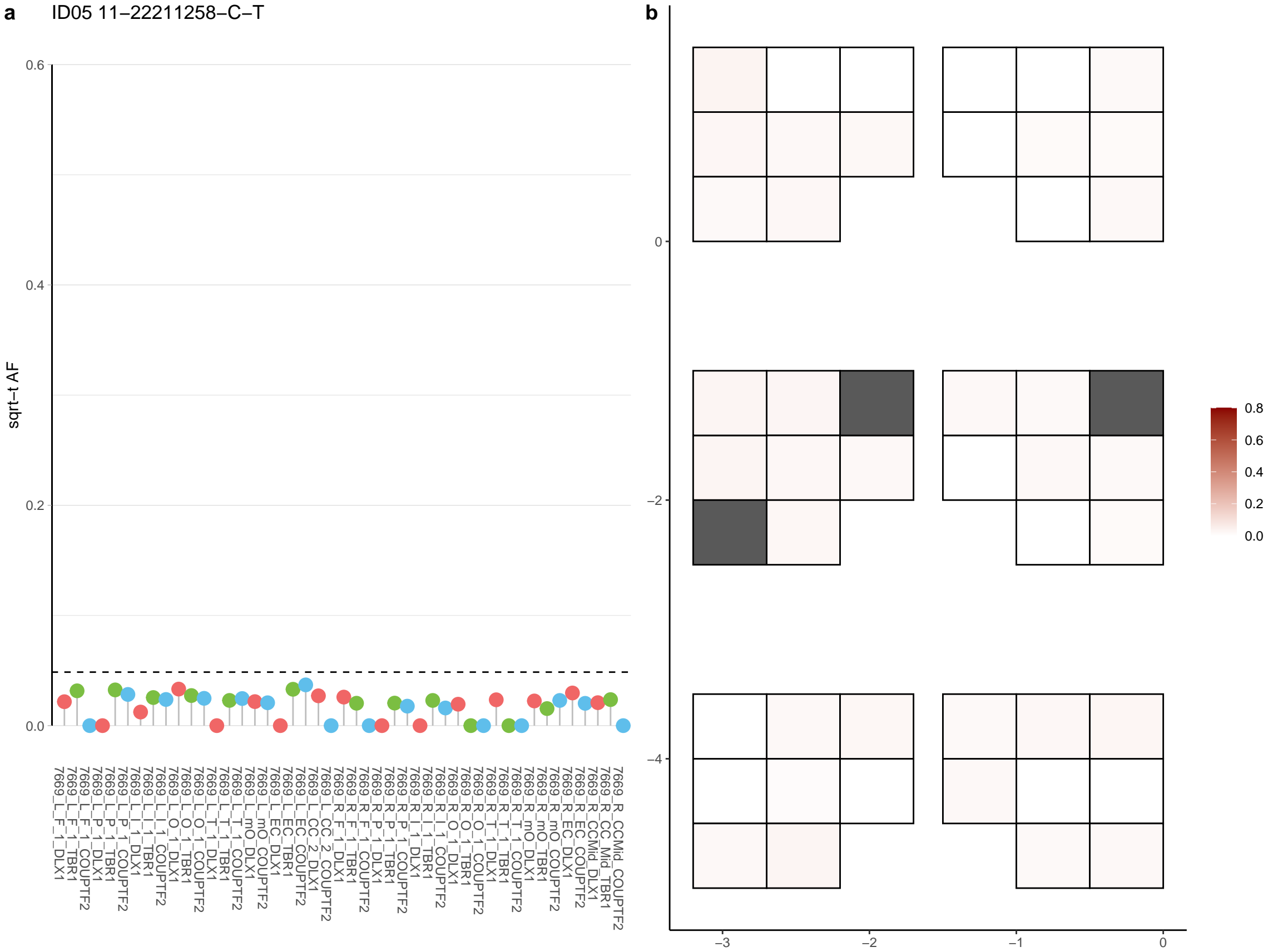

**a** ID05 11-25958939-C-T

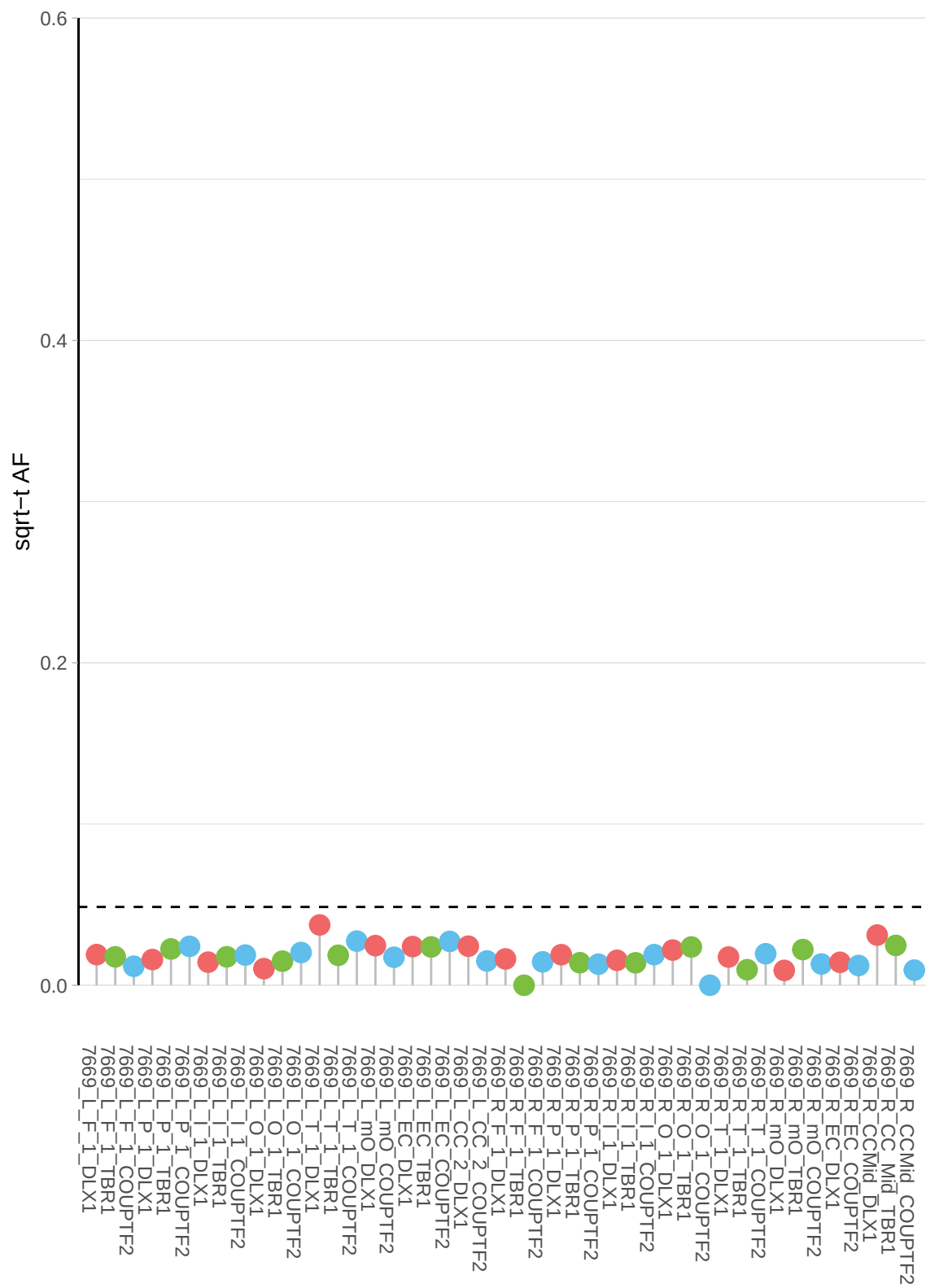

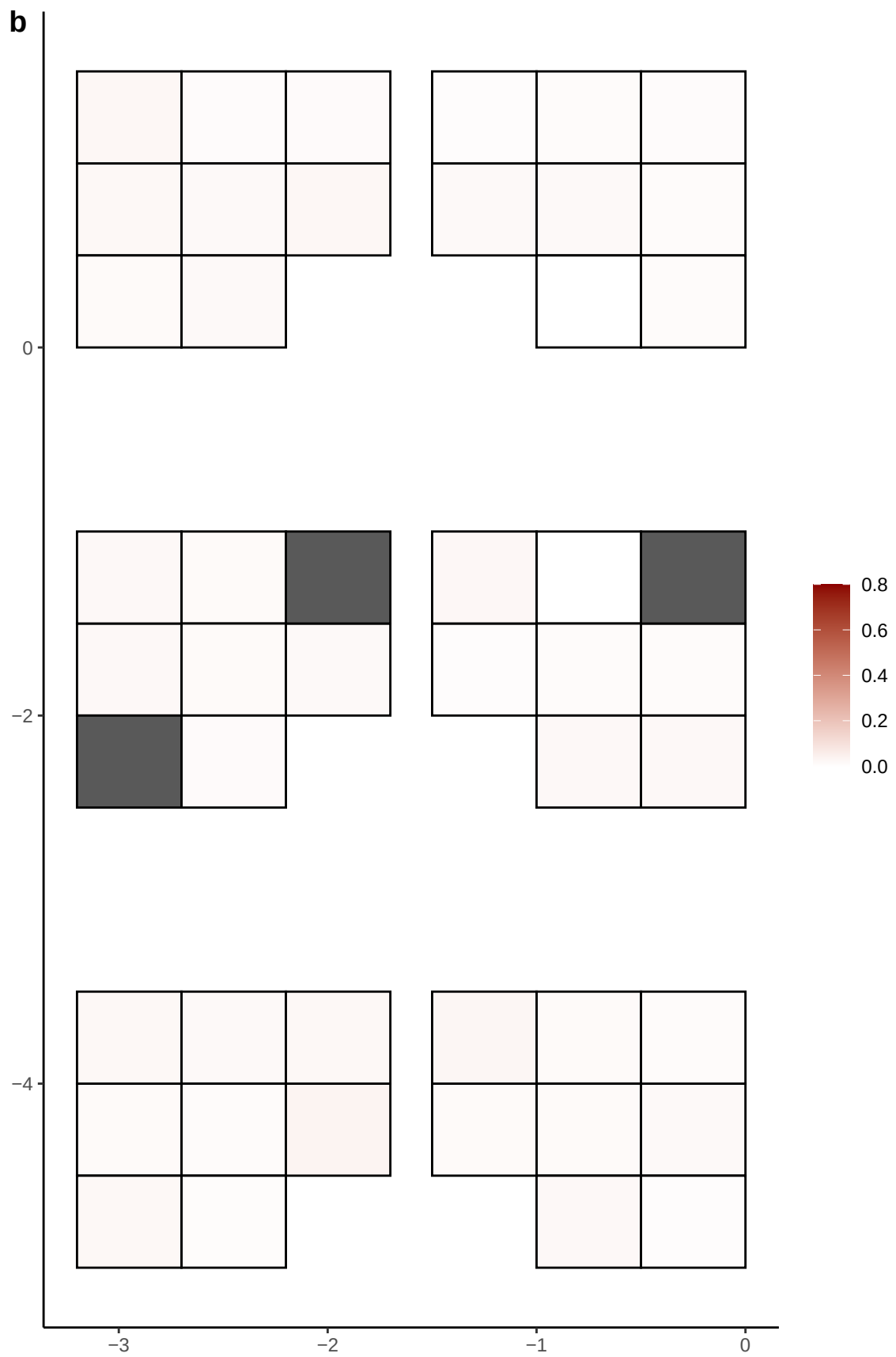

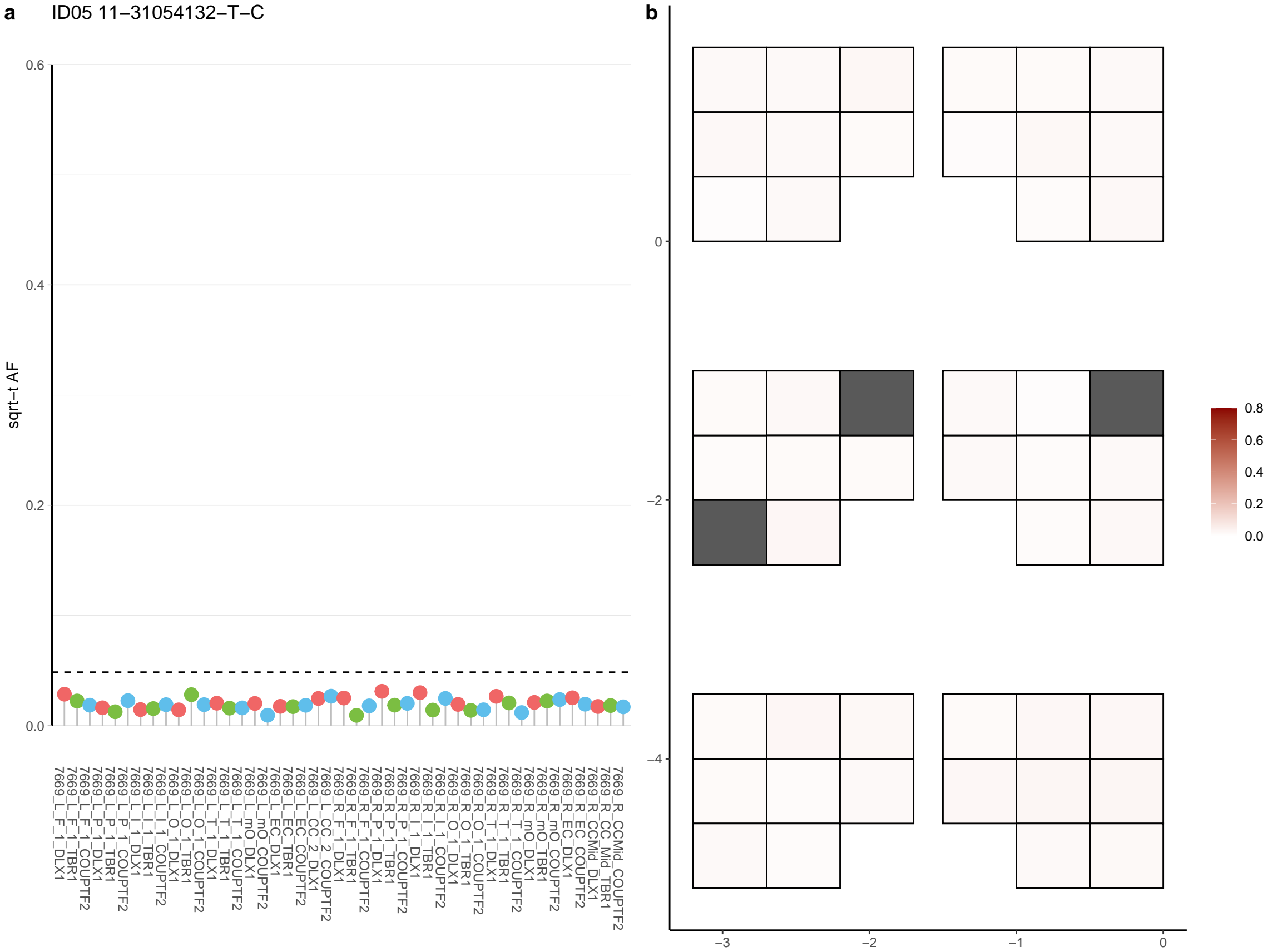

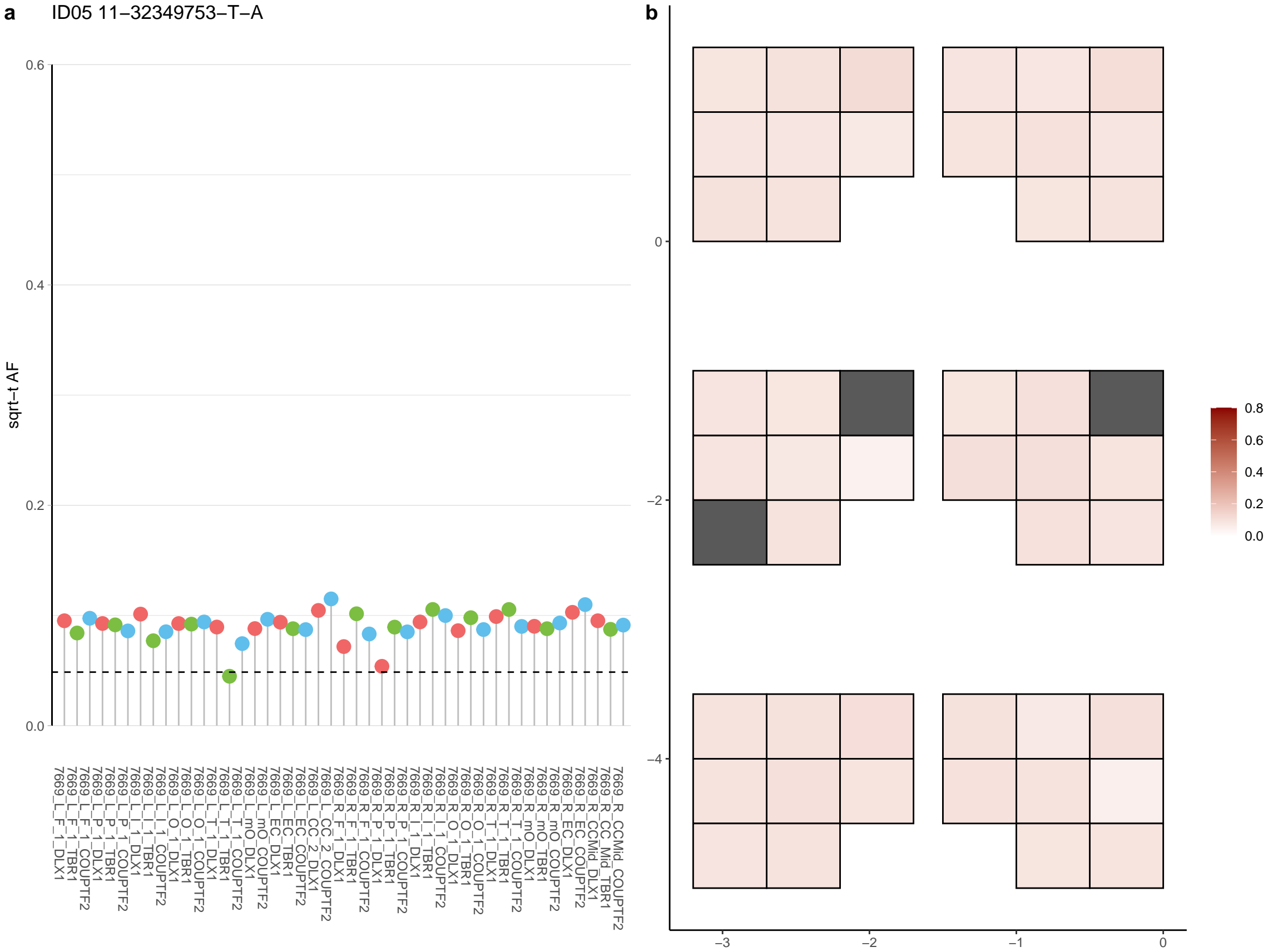

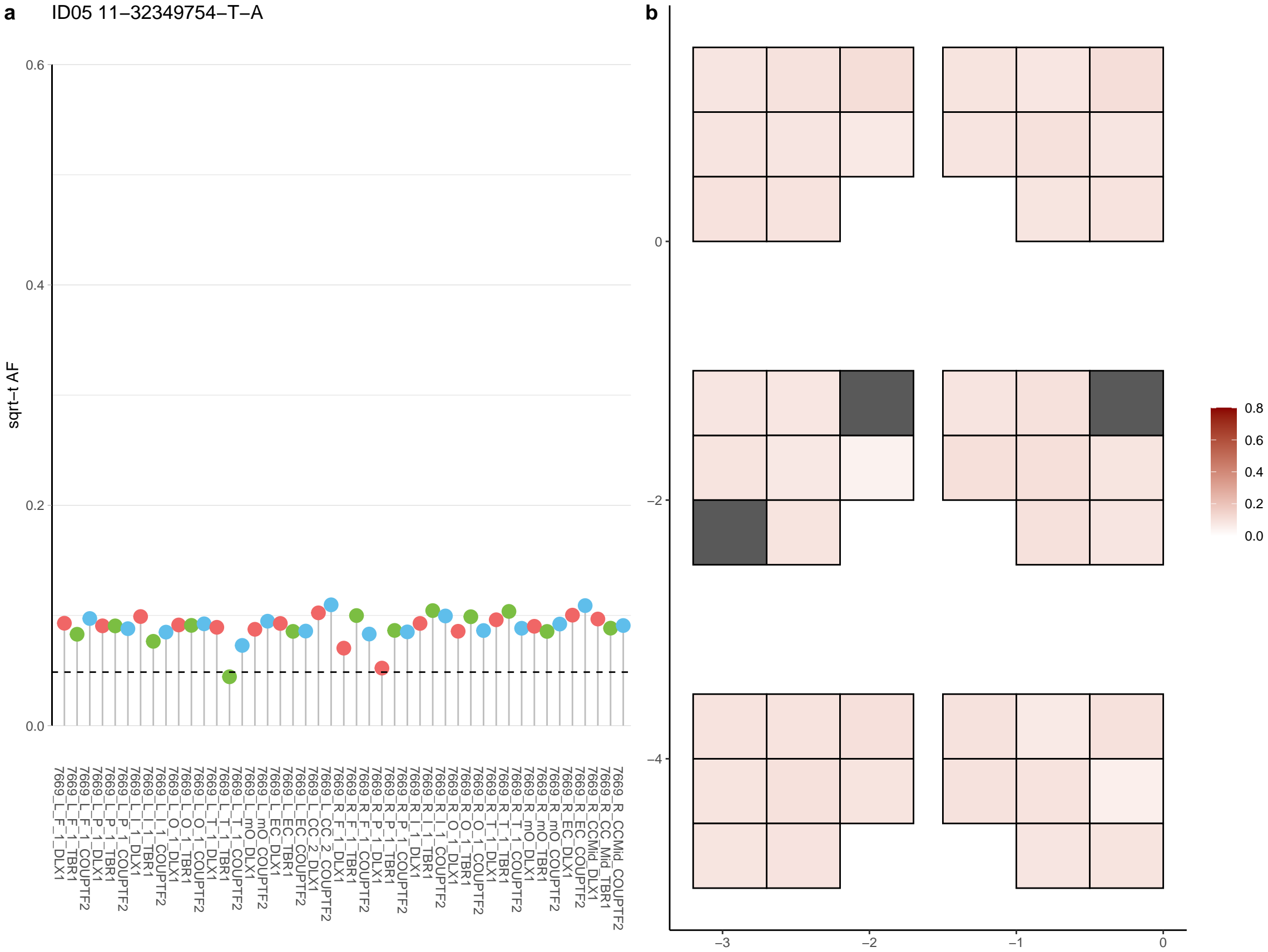

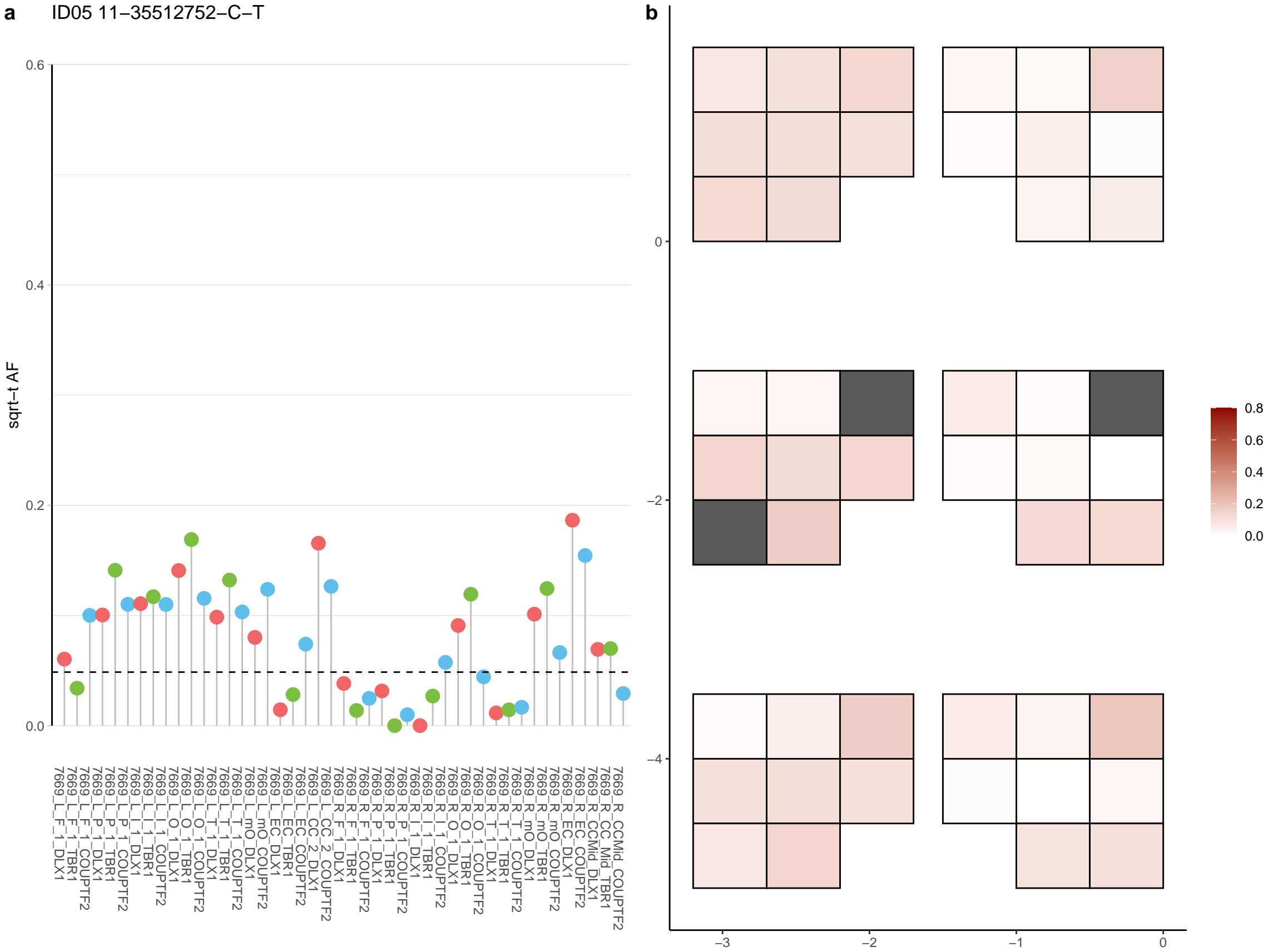

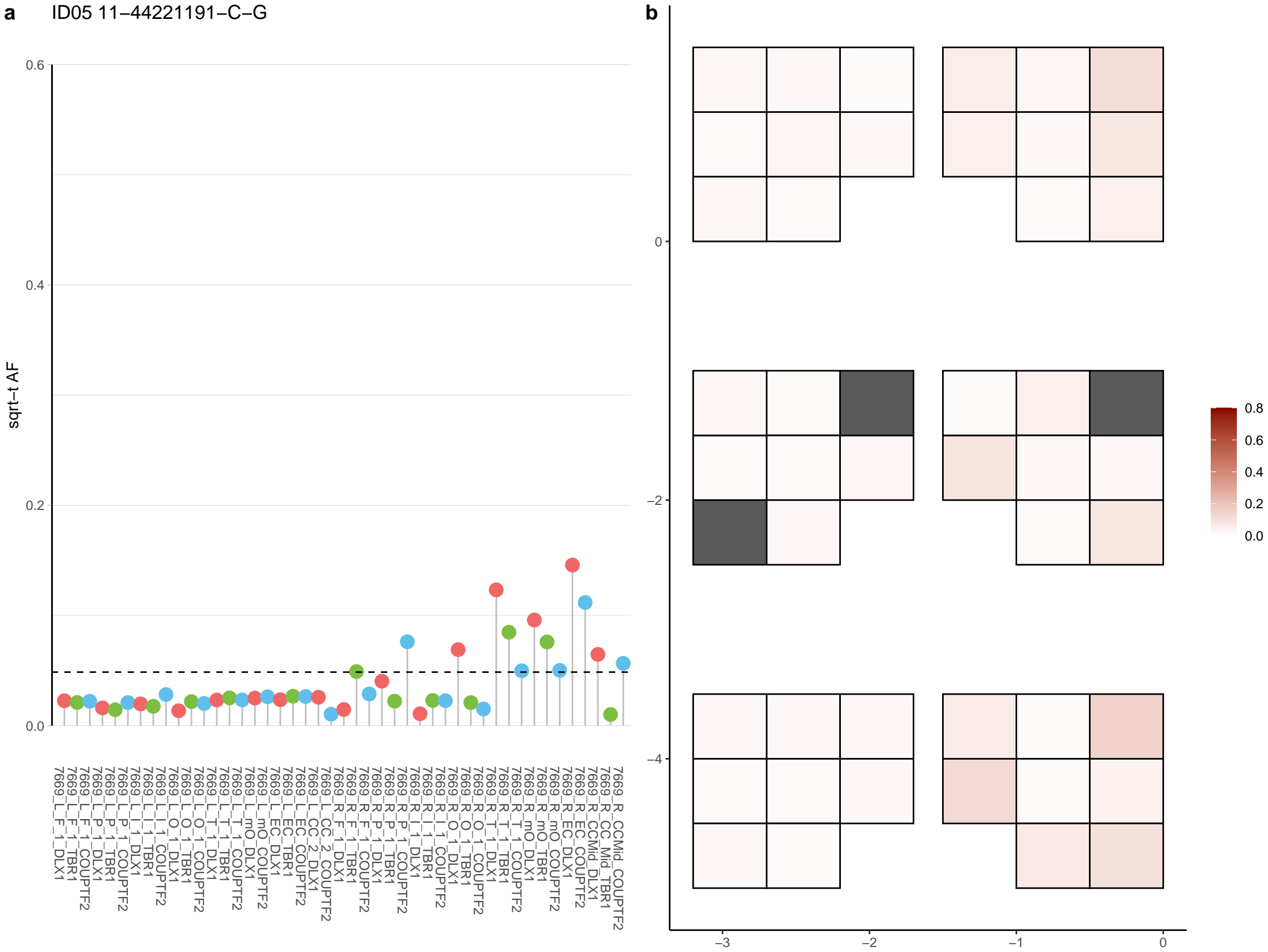

**a** ID05 11-45873419-G-A

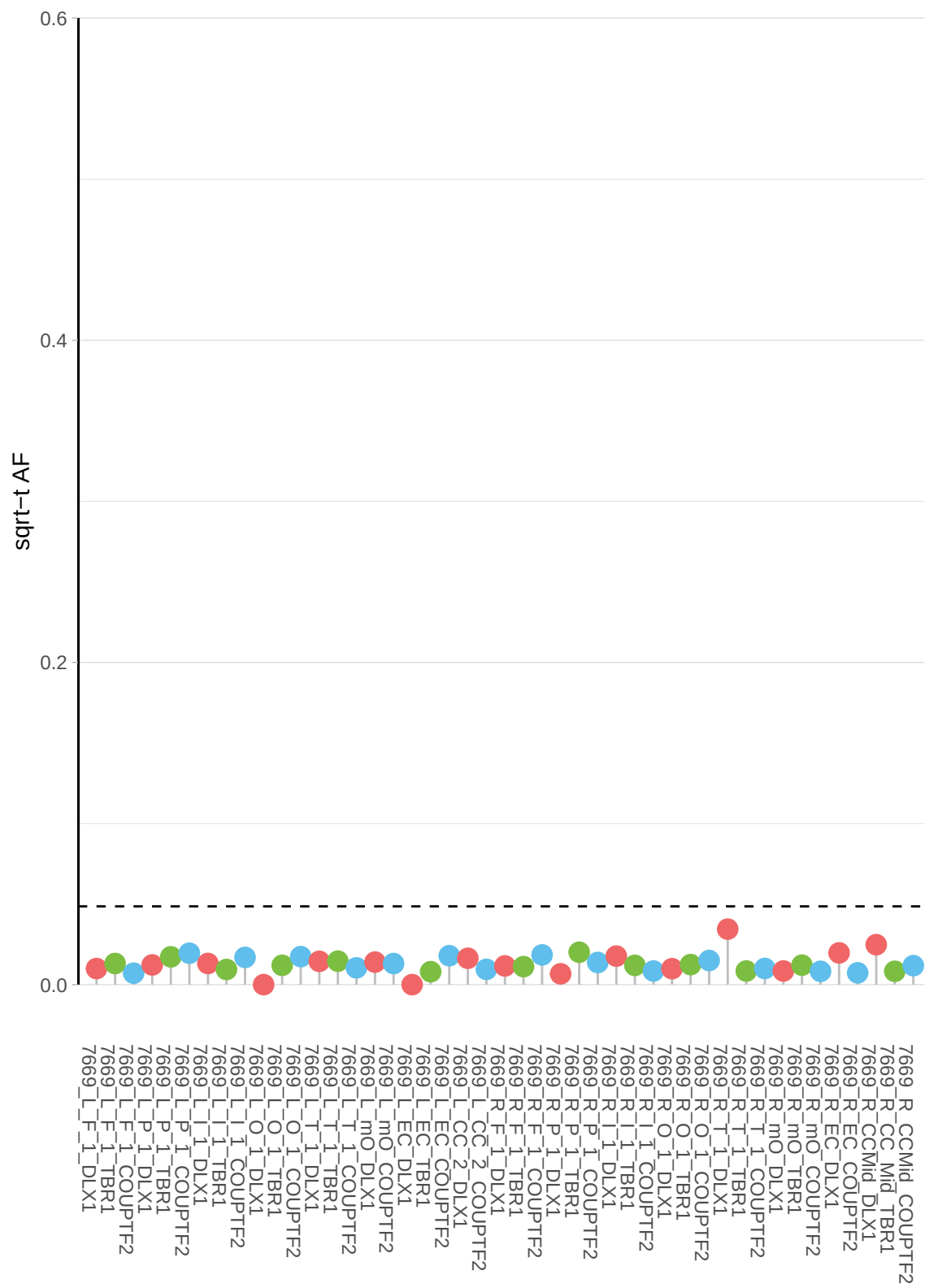

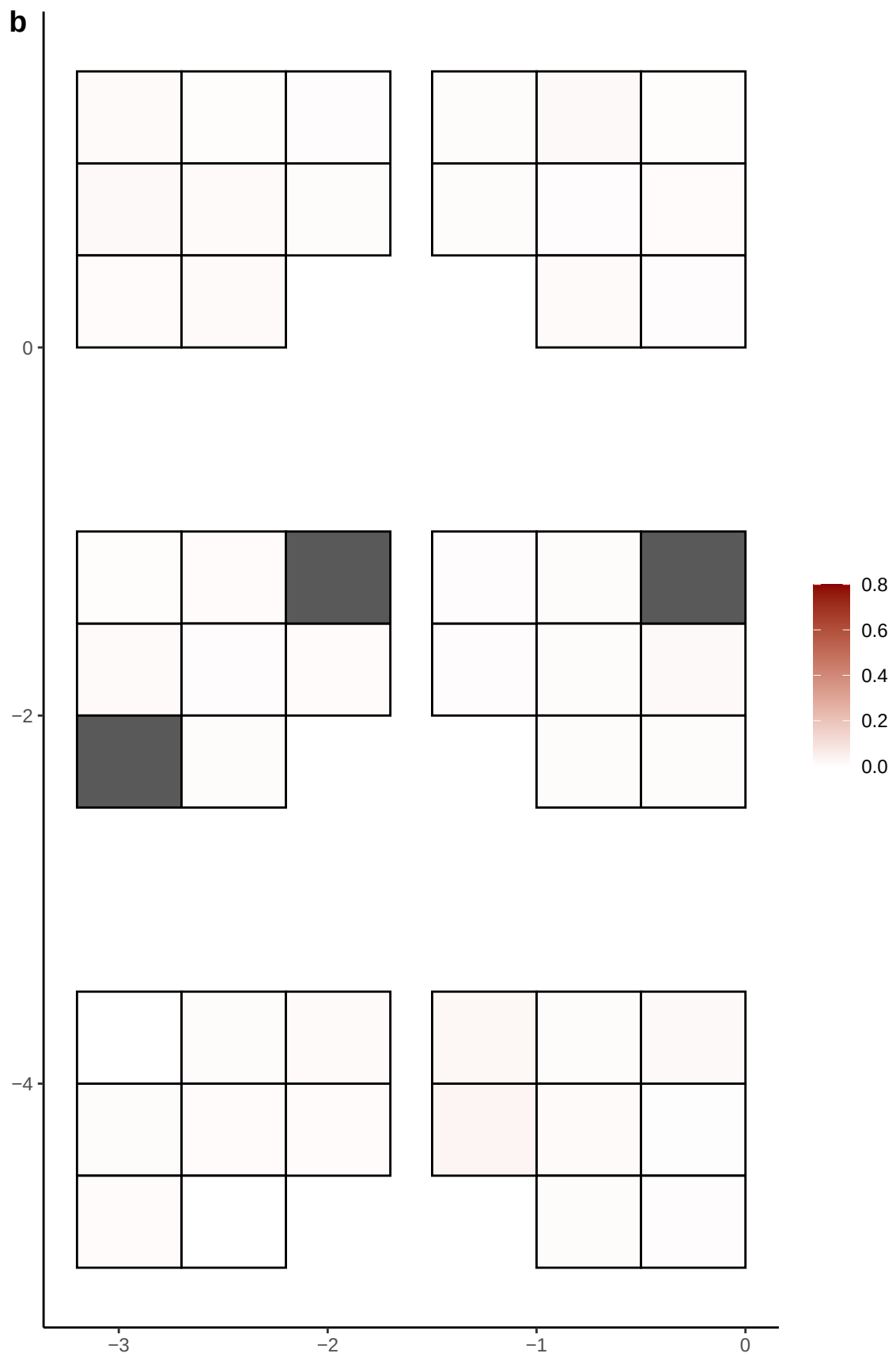

**a** ID05 11-55838876-G-T

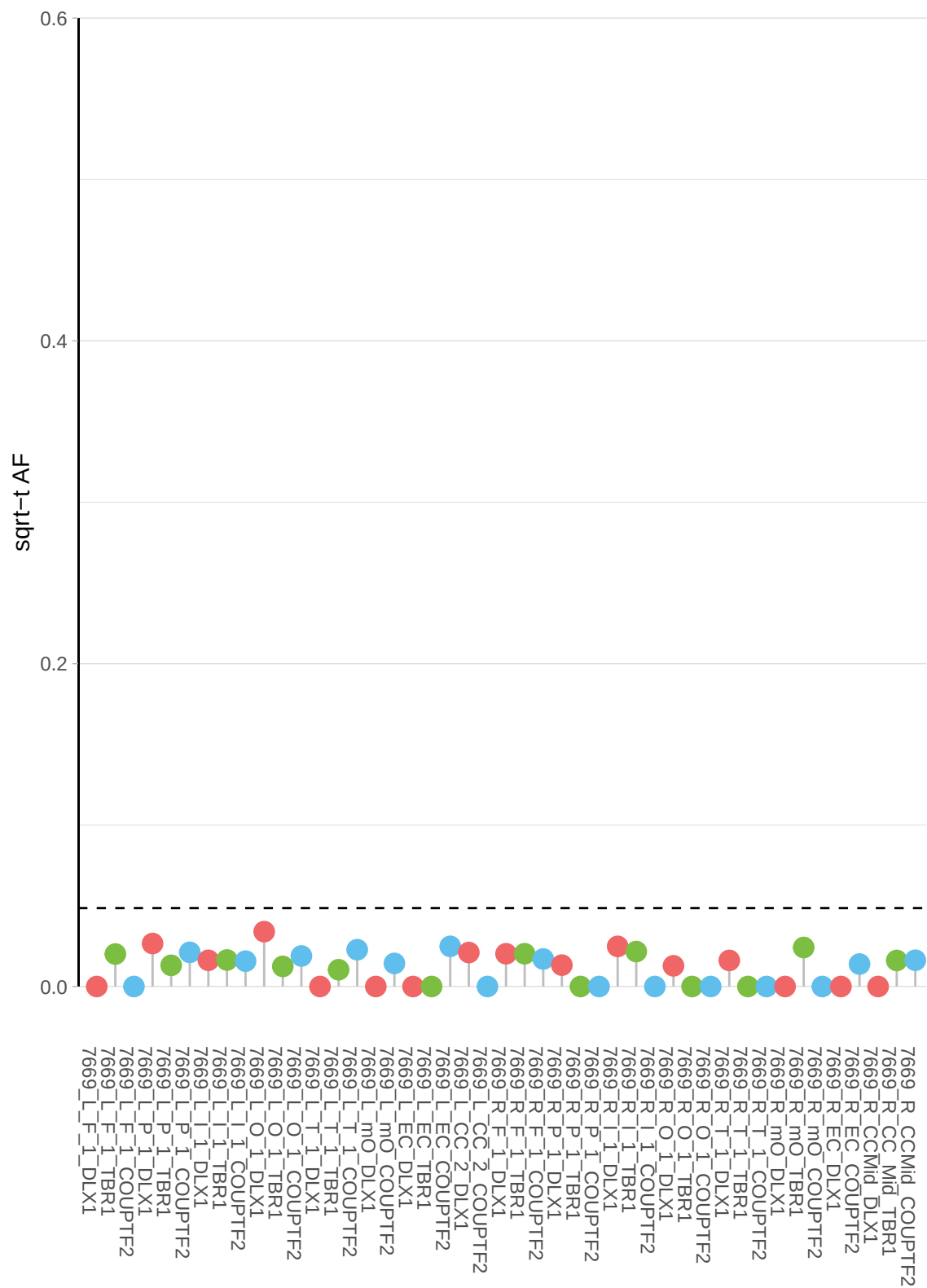

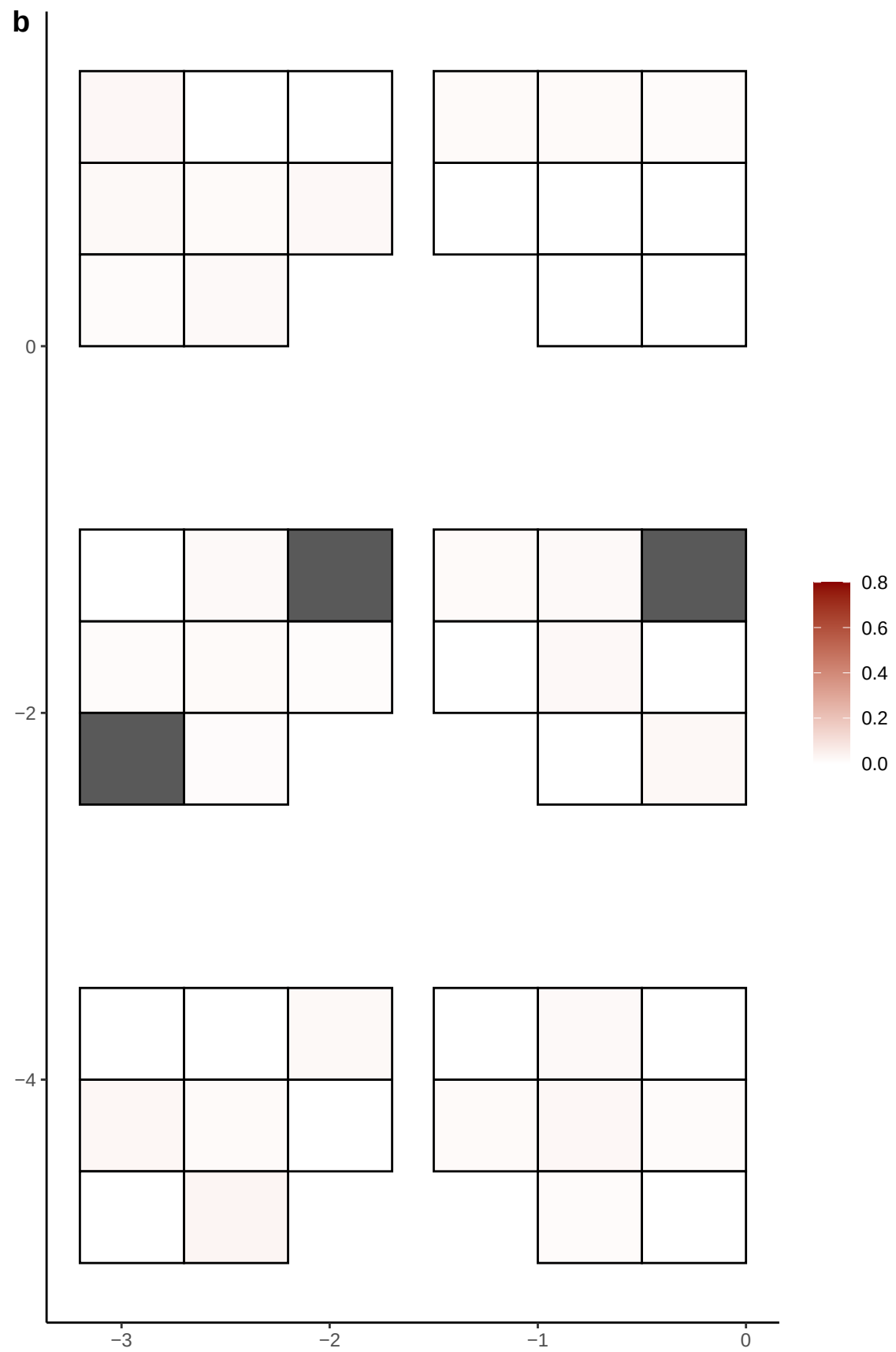

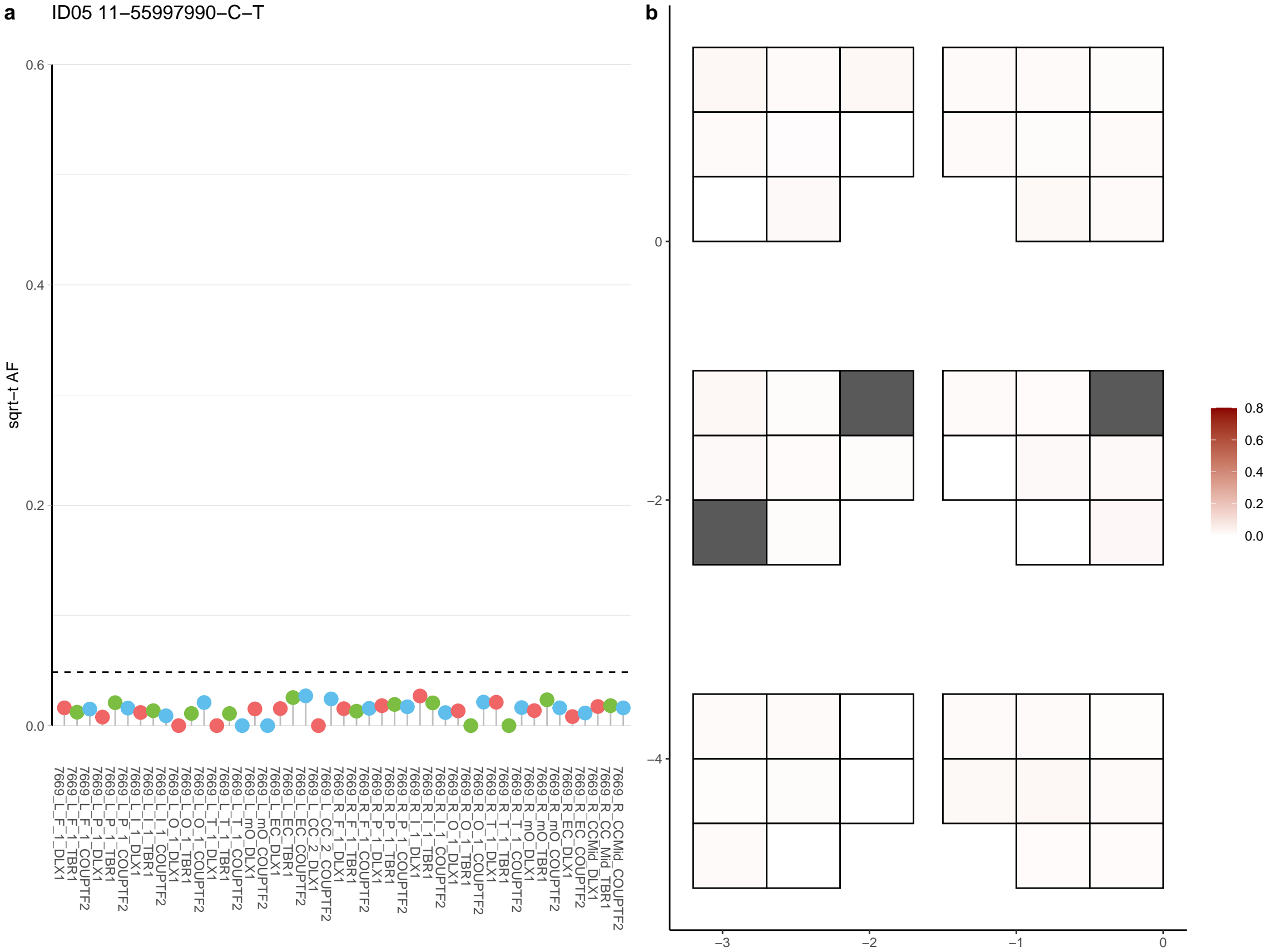

**a** ID05 11-56409116-C-T

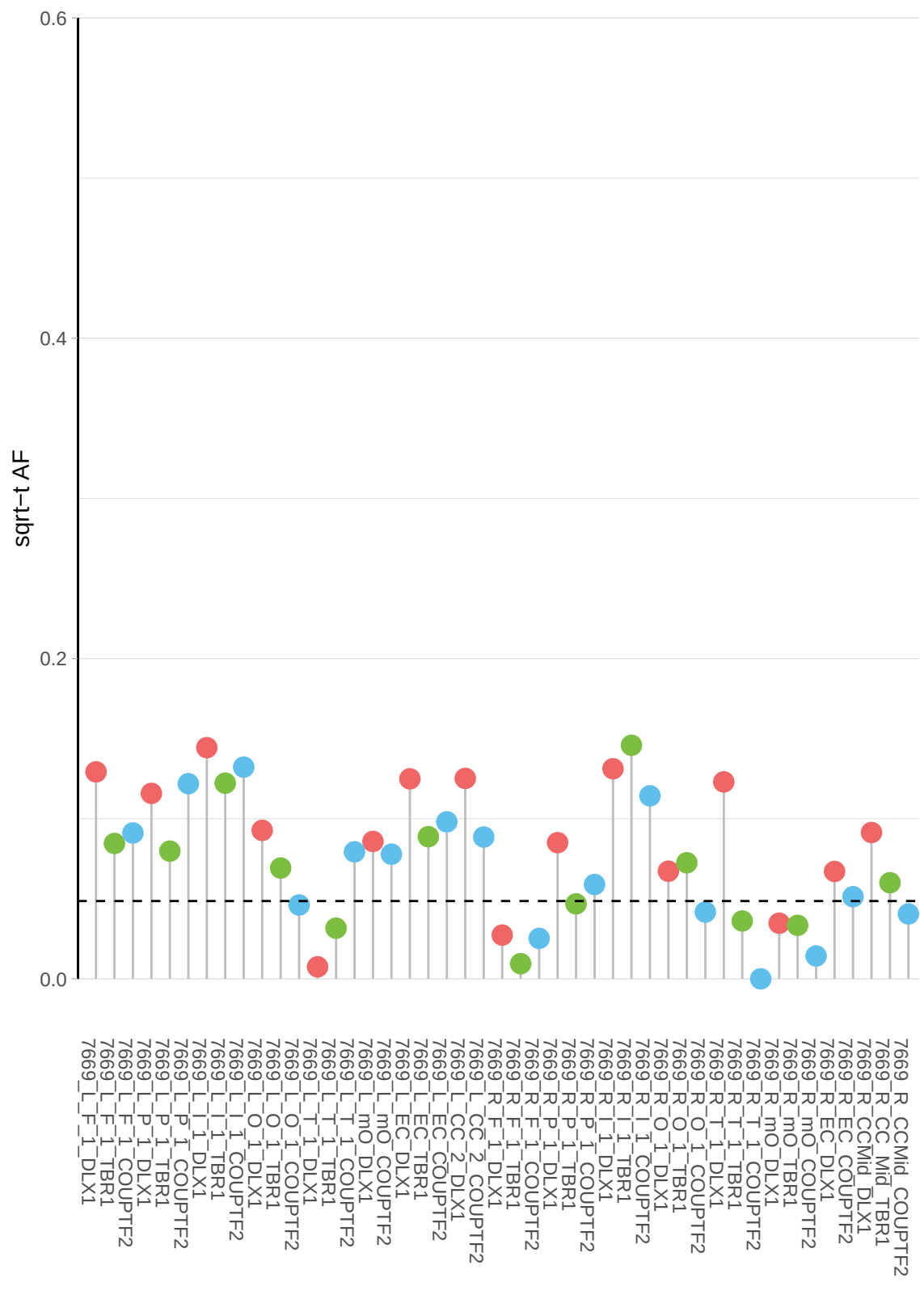

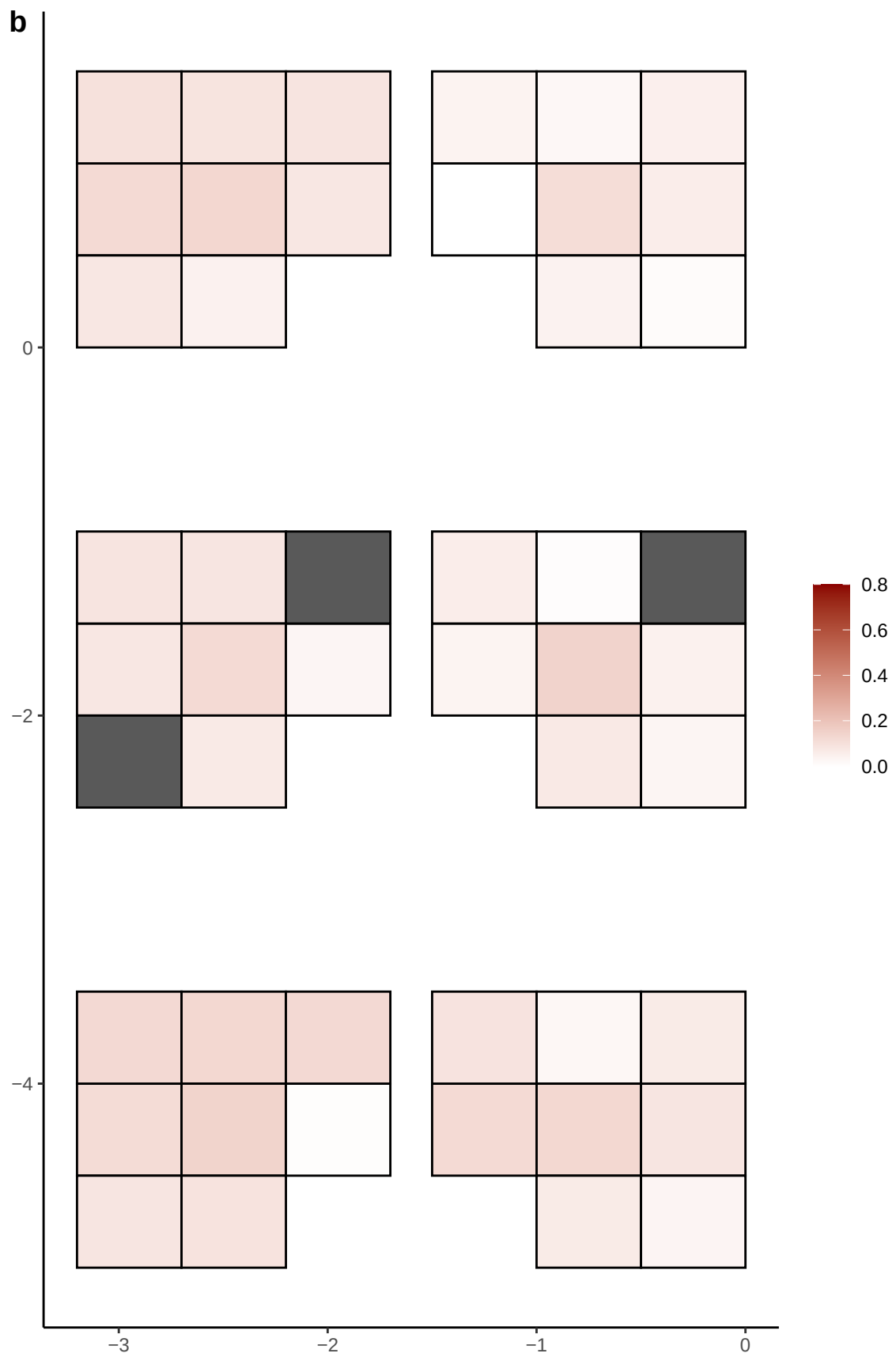

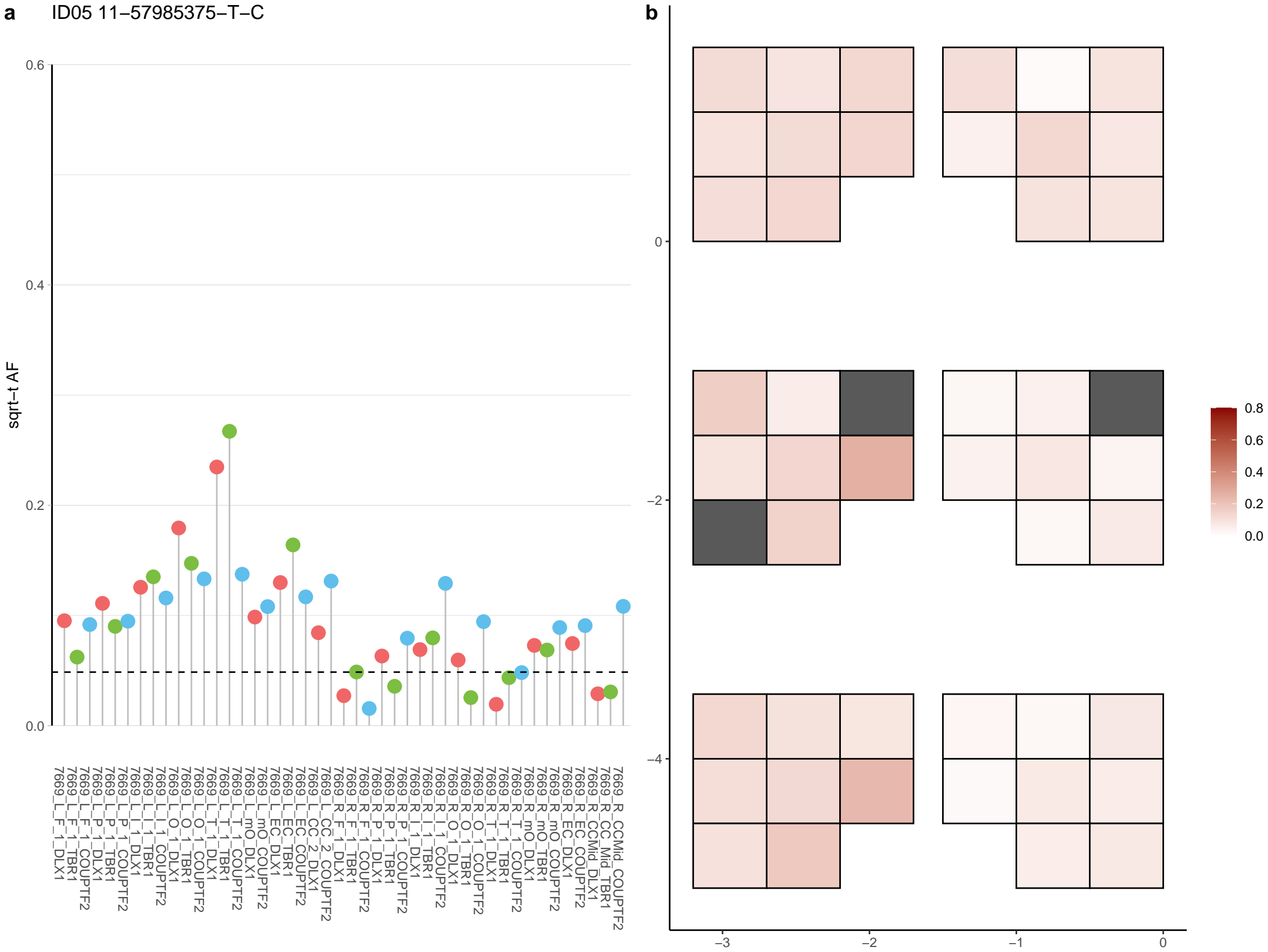

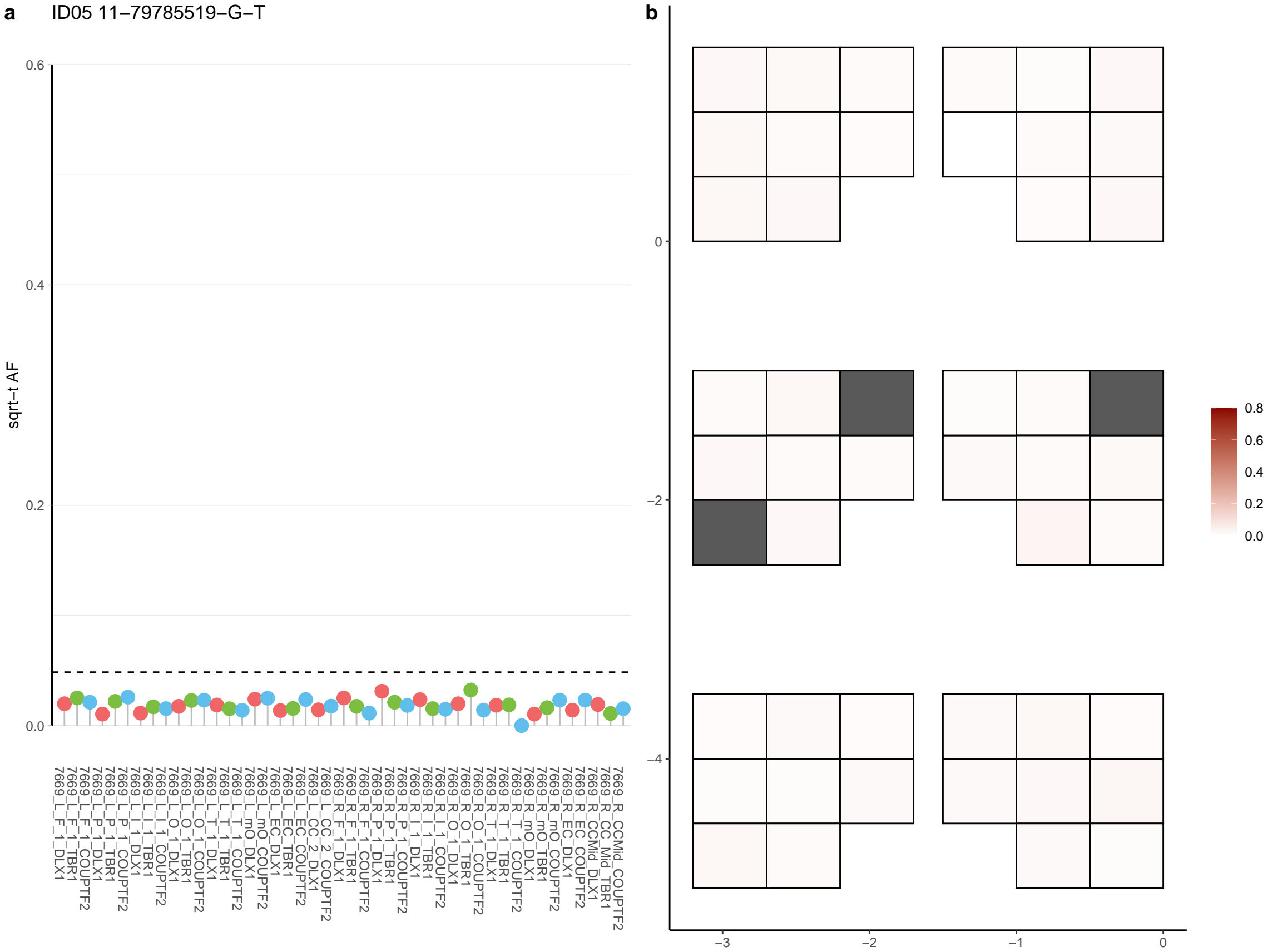

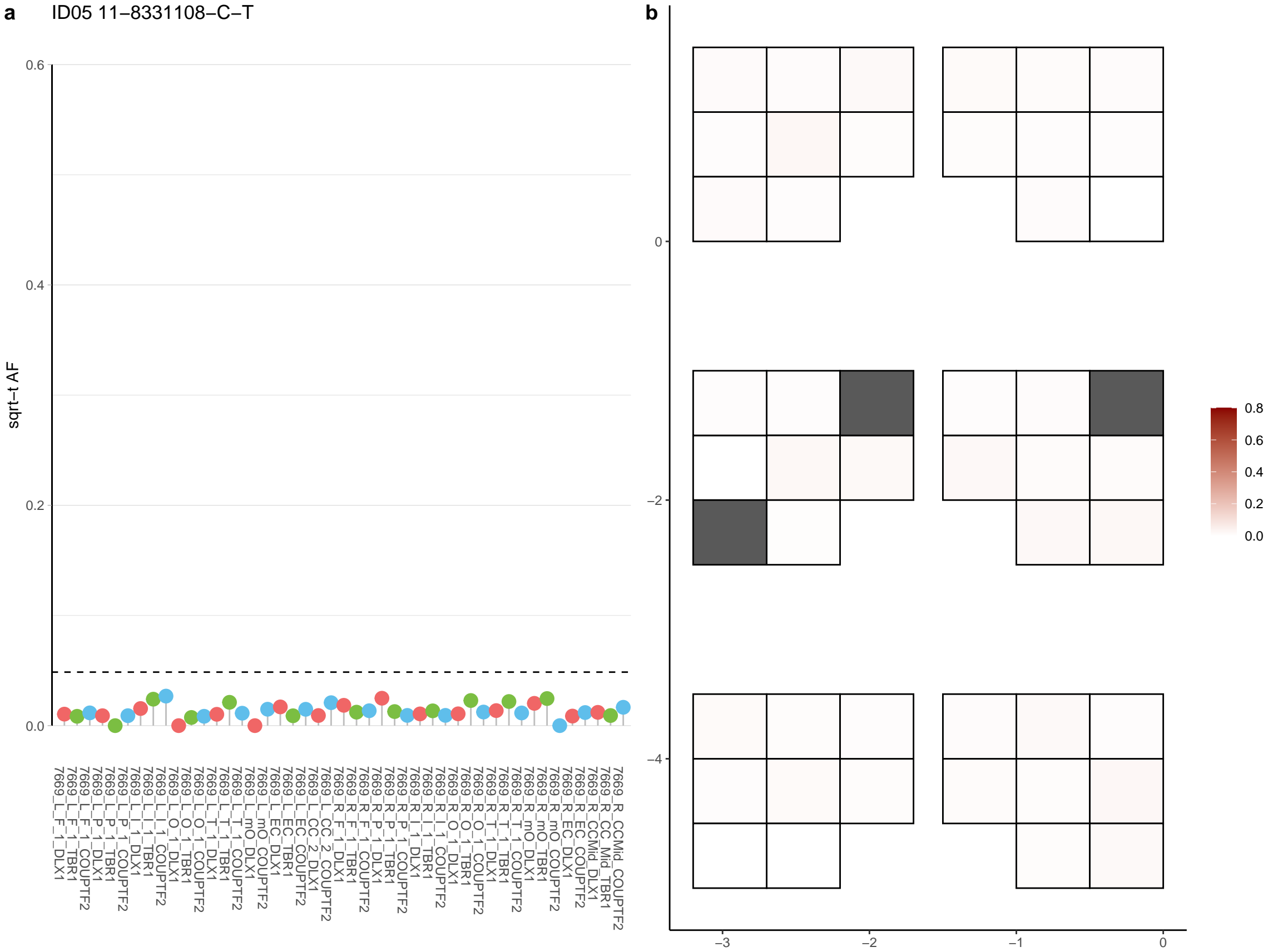

**a** ID05 11-91169265-C-T

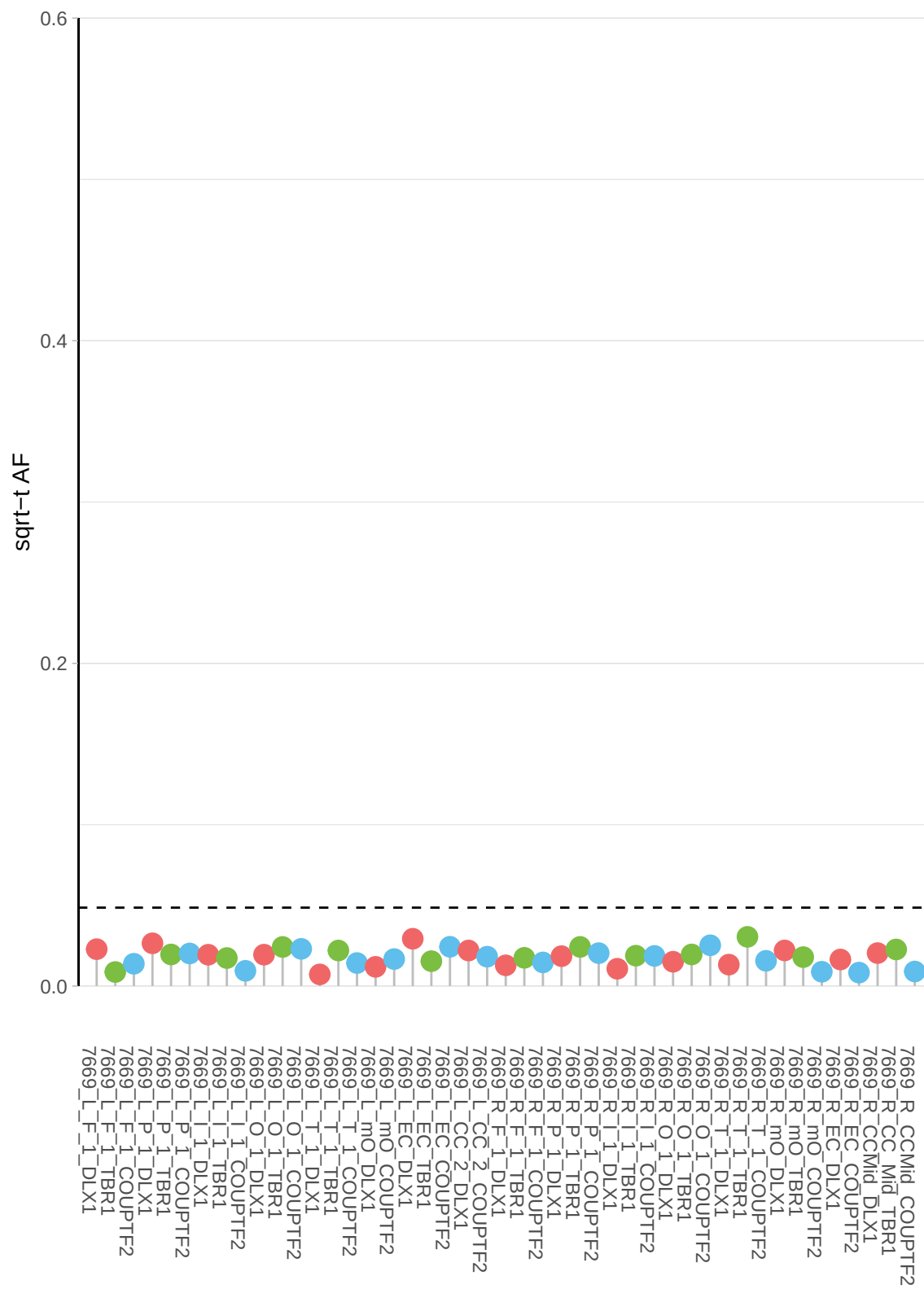

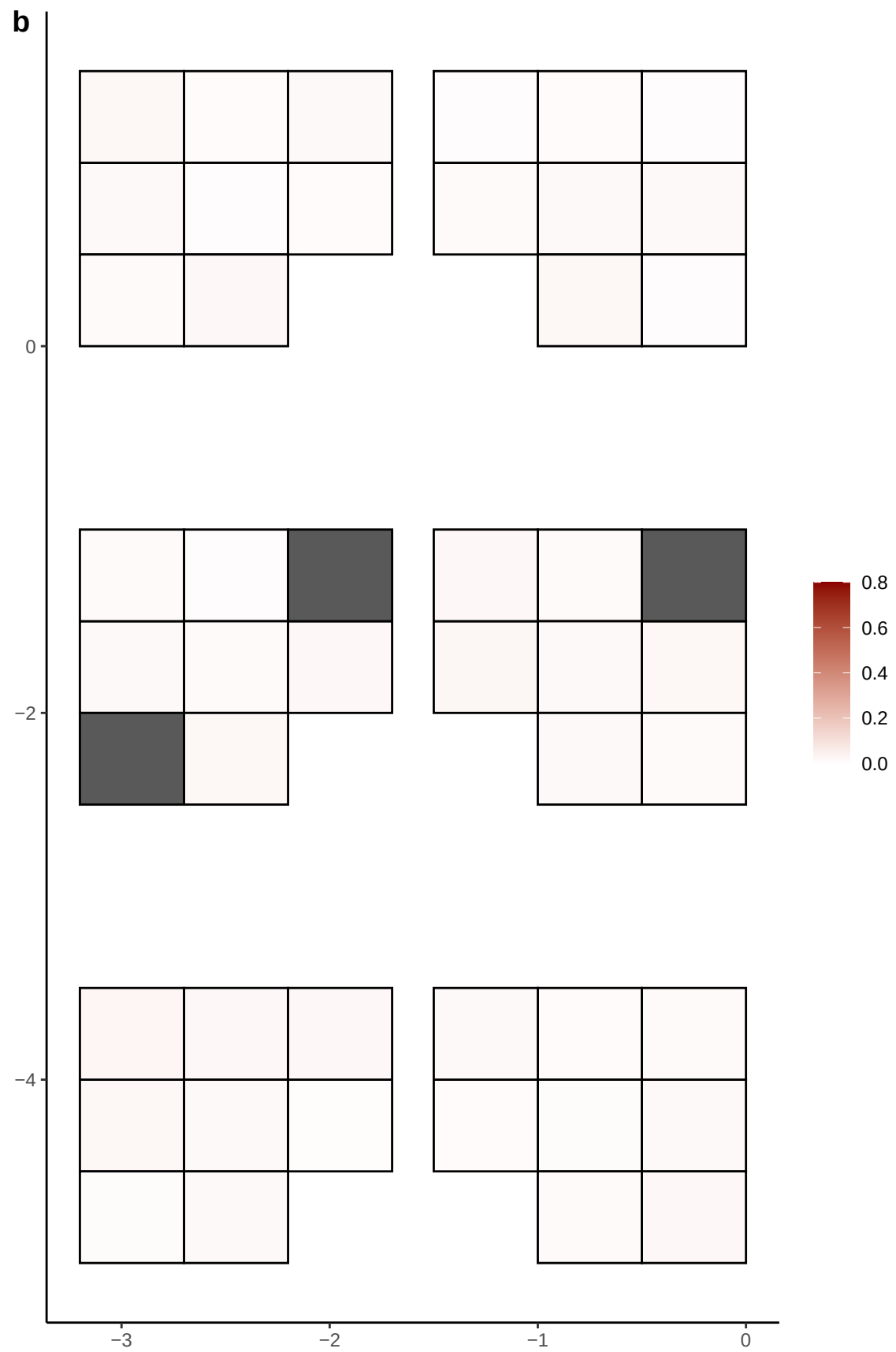

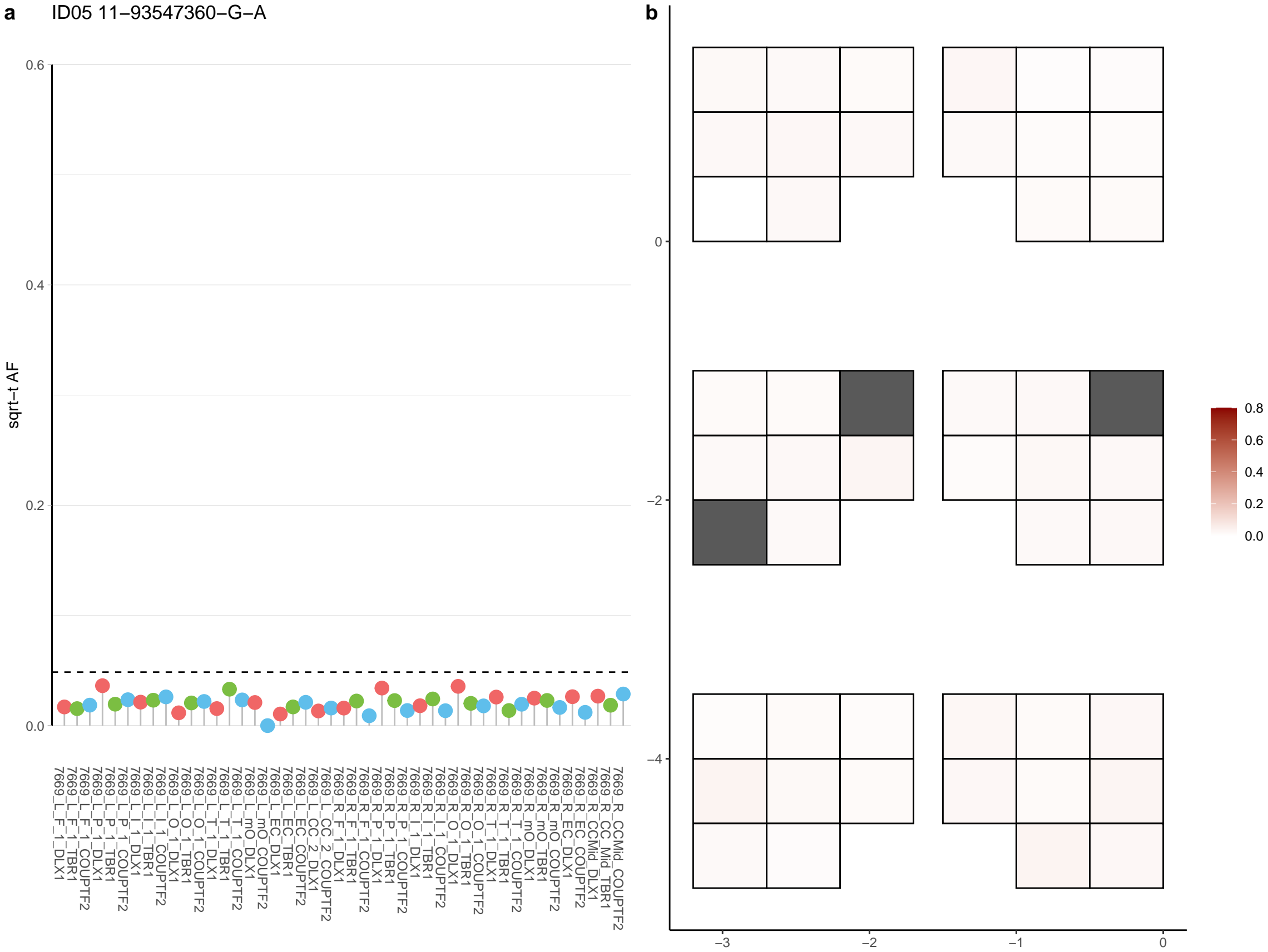

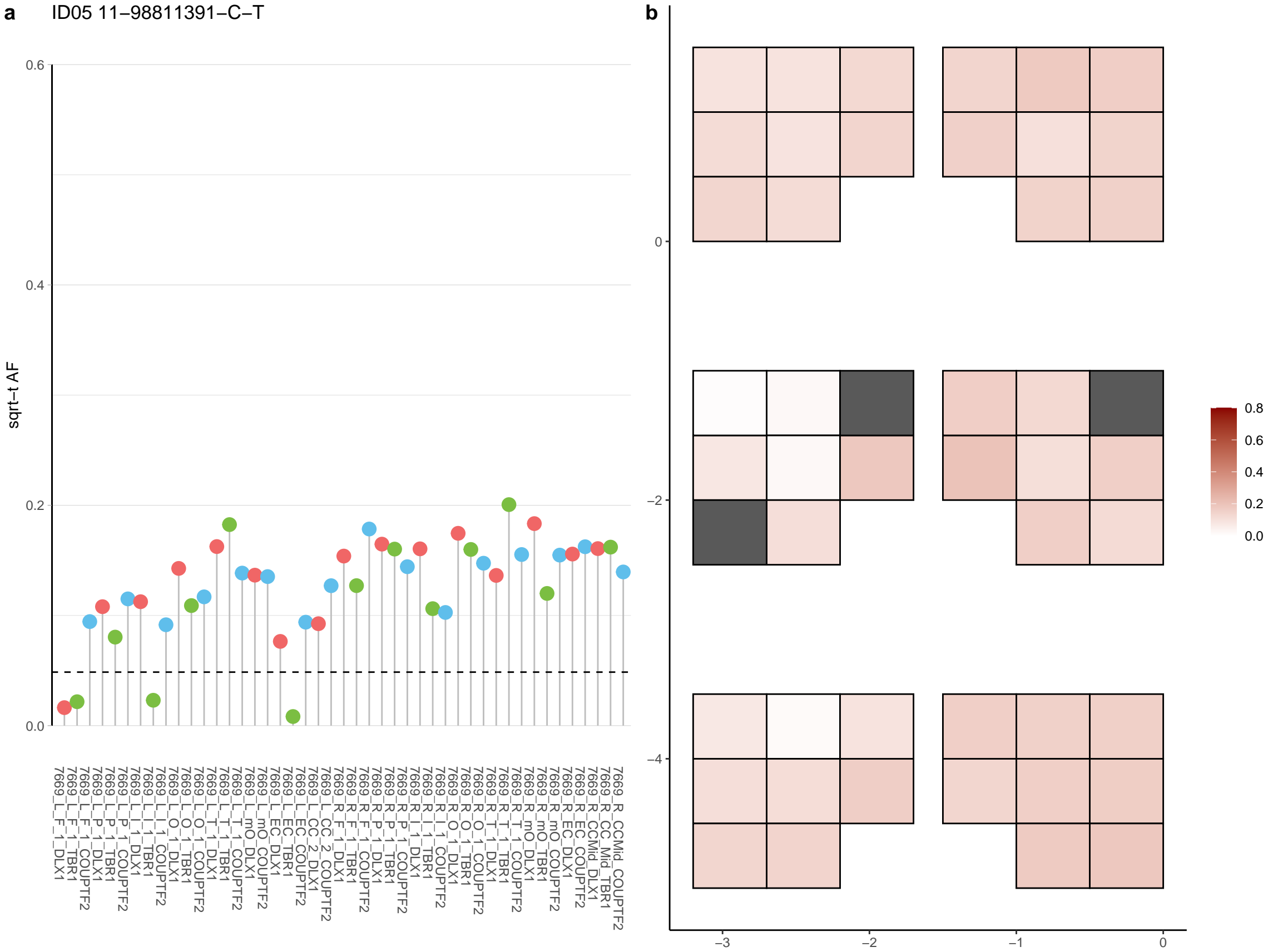

**a** ID05 11-99288625-T-A

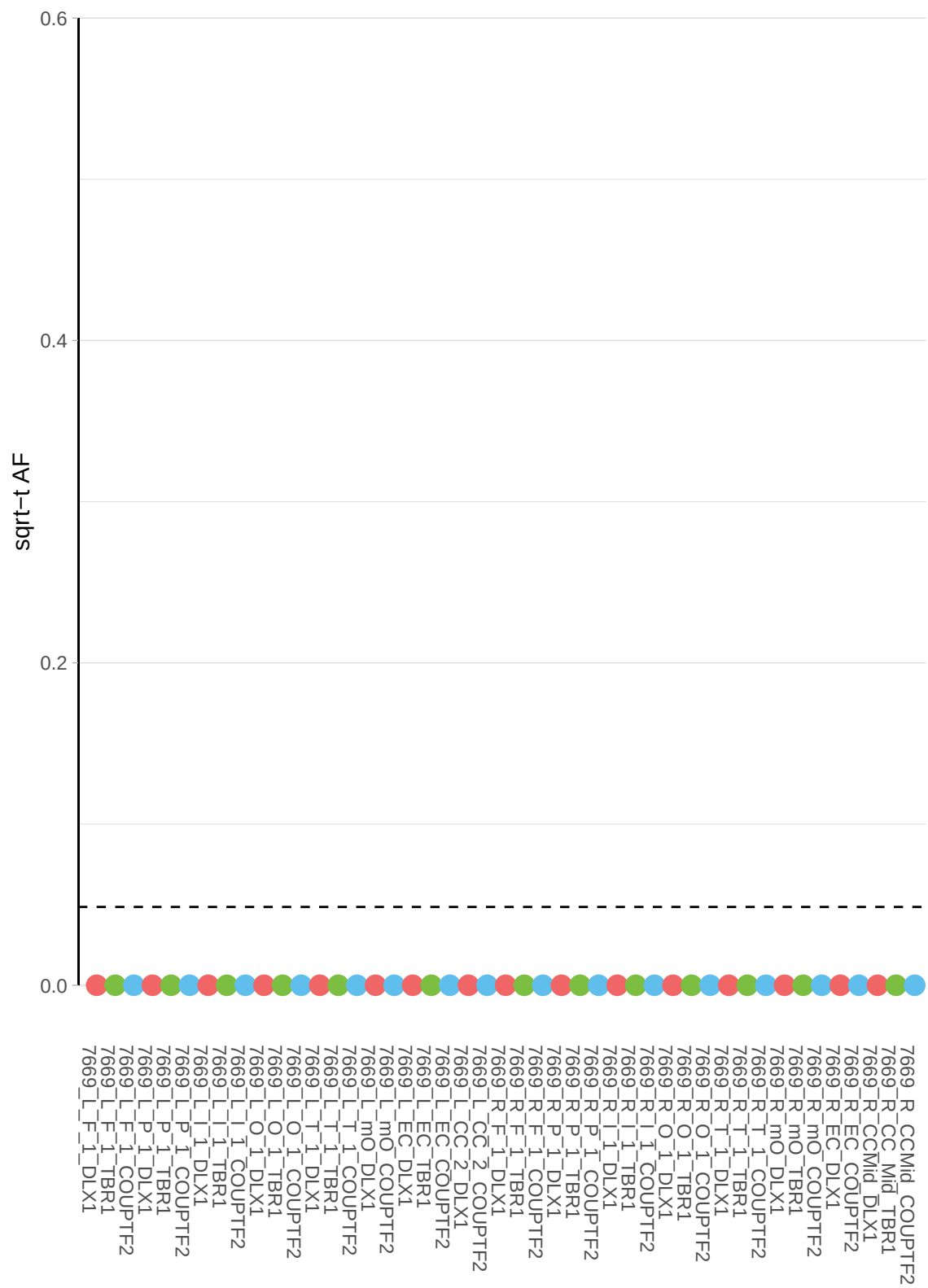

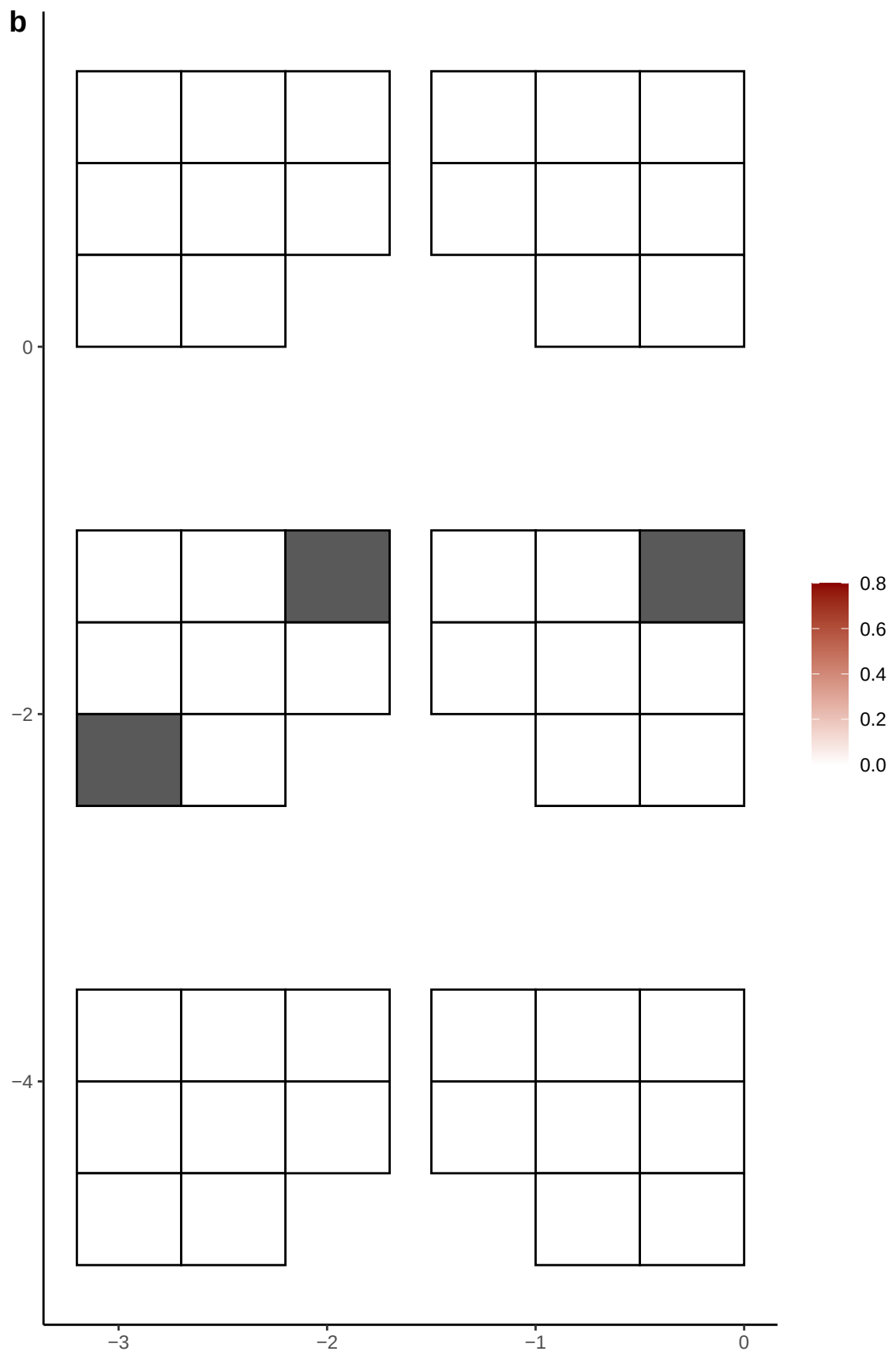

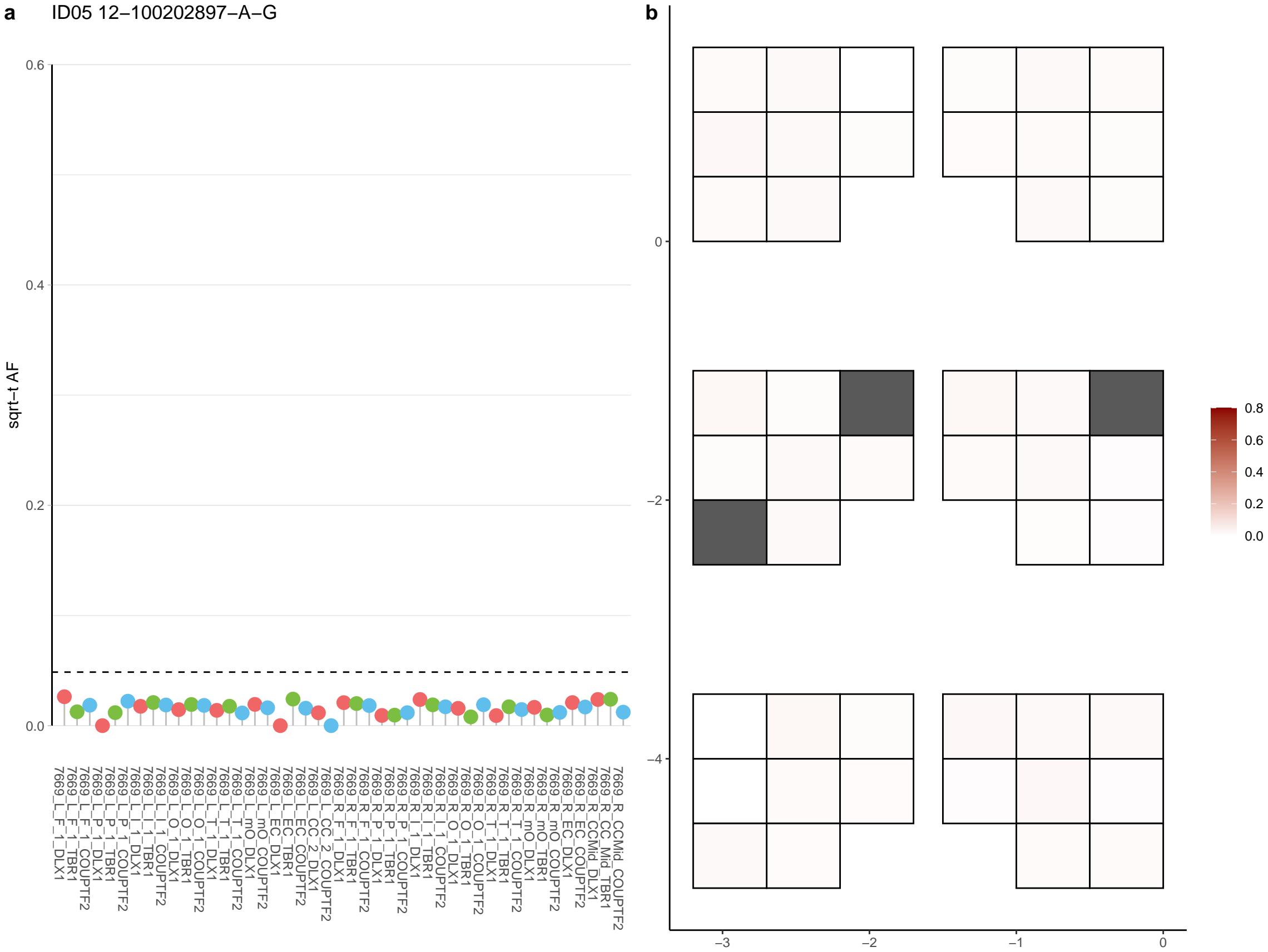

**a** ID05 12-110209050-G-A

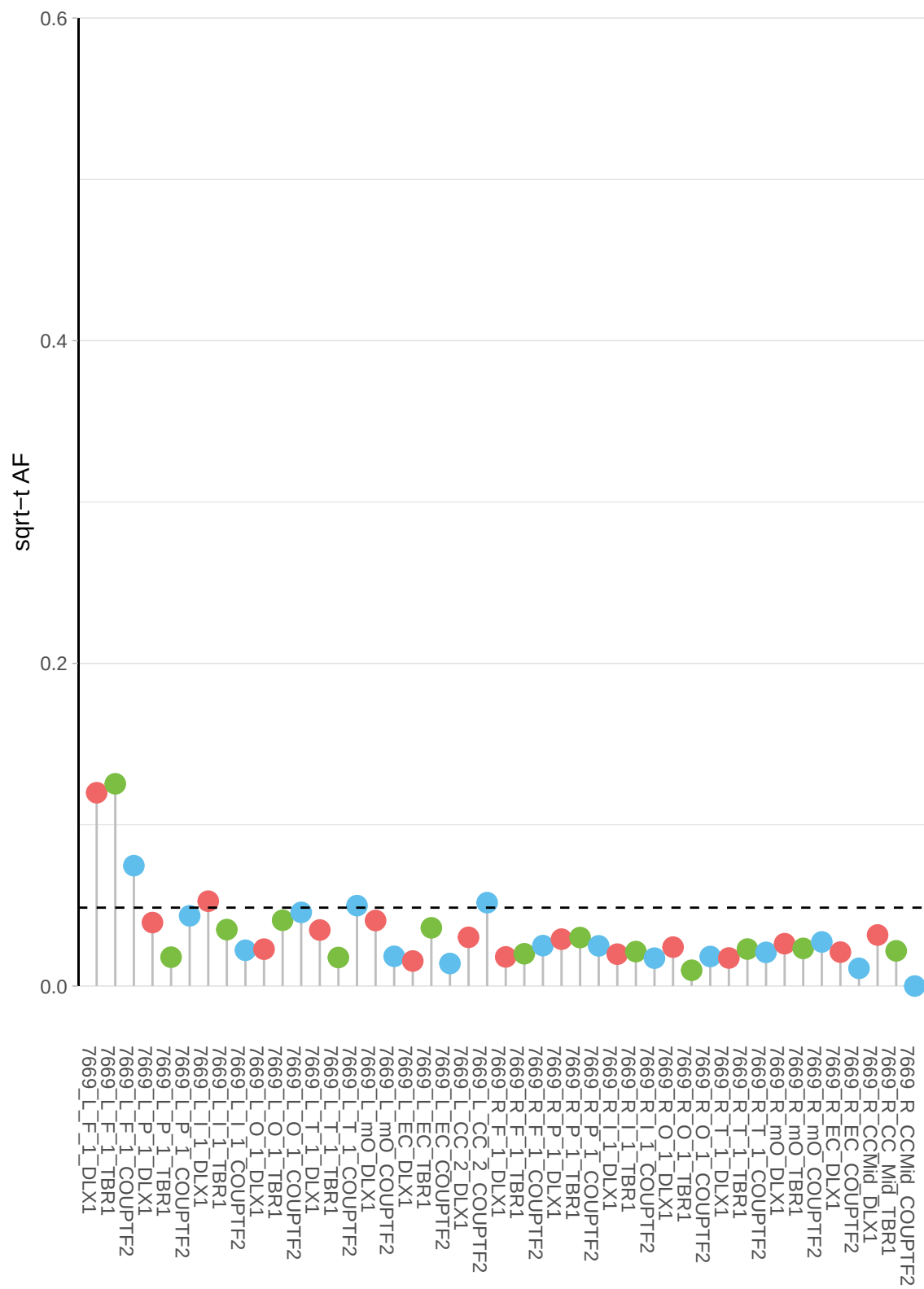

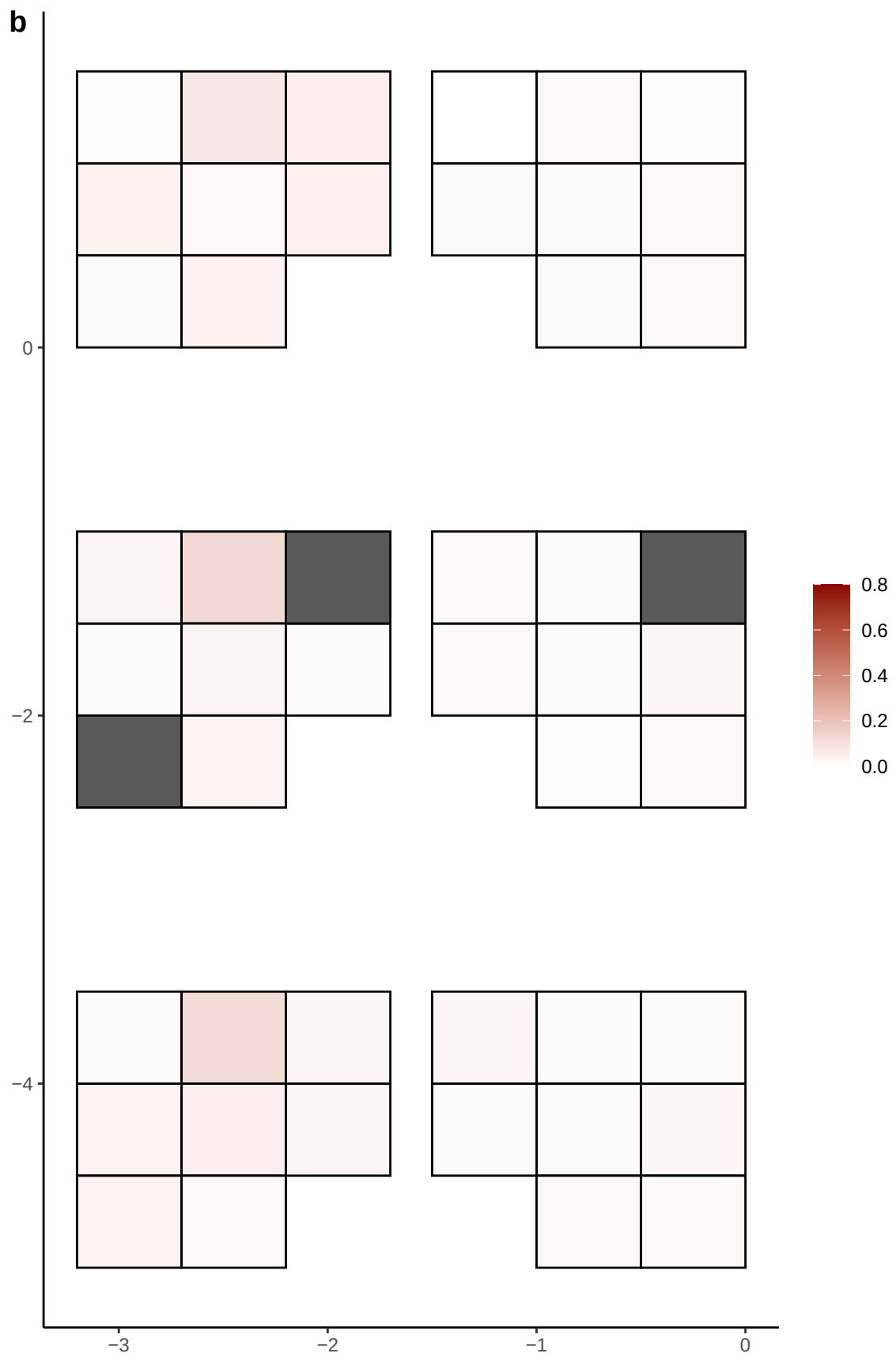

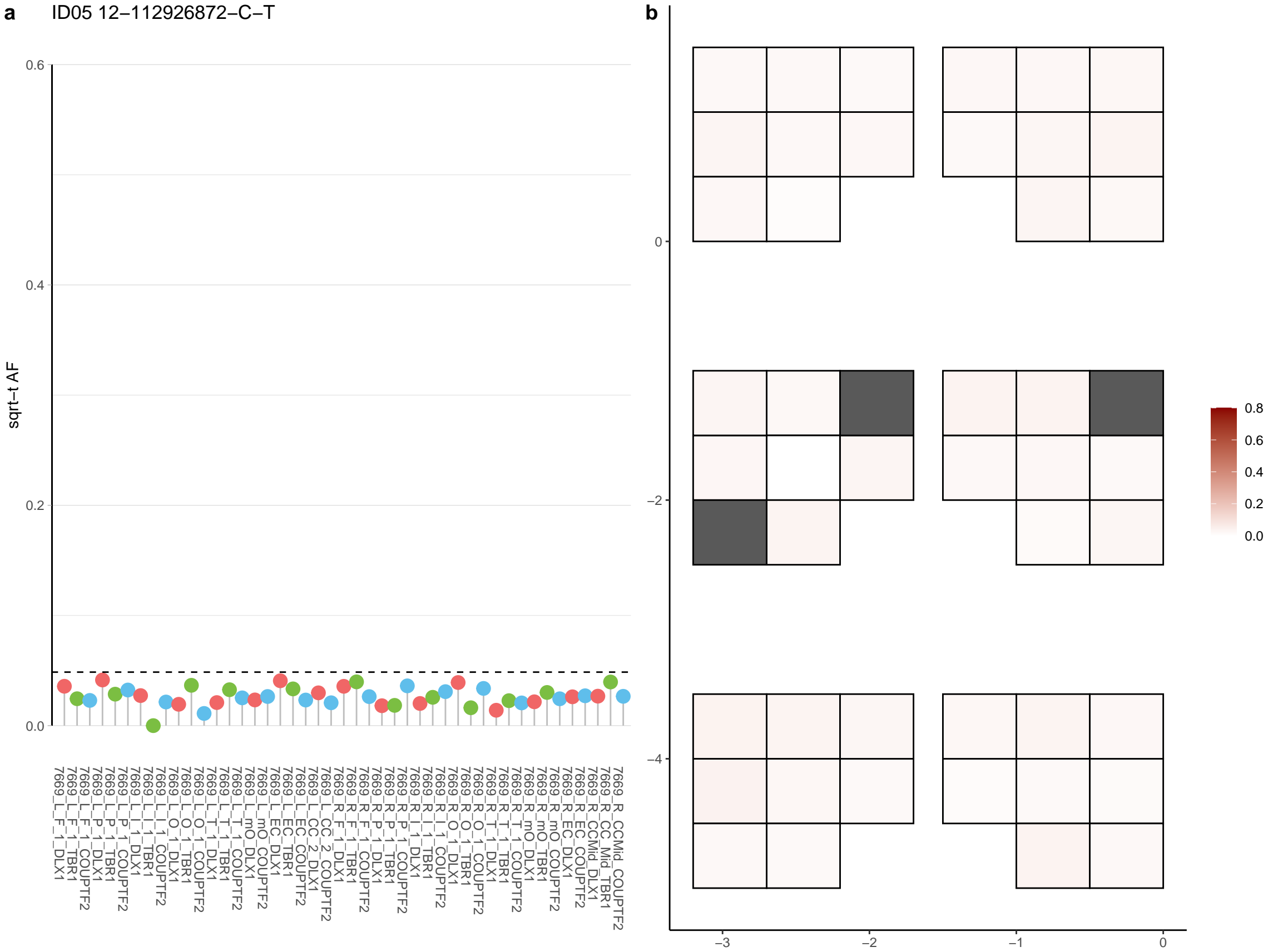

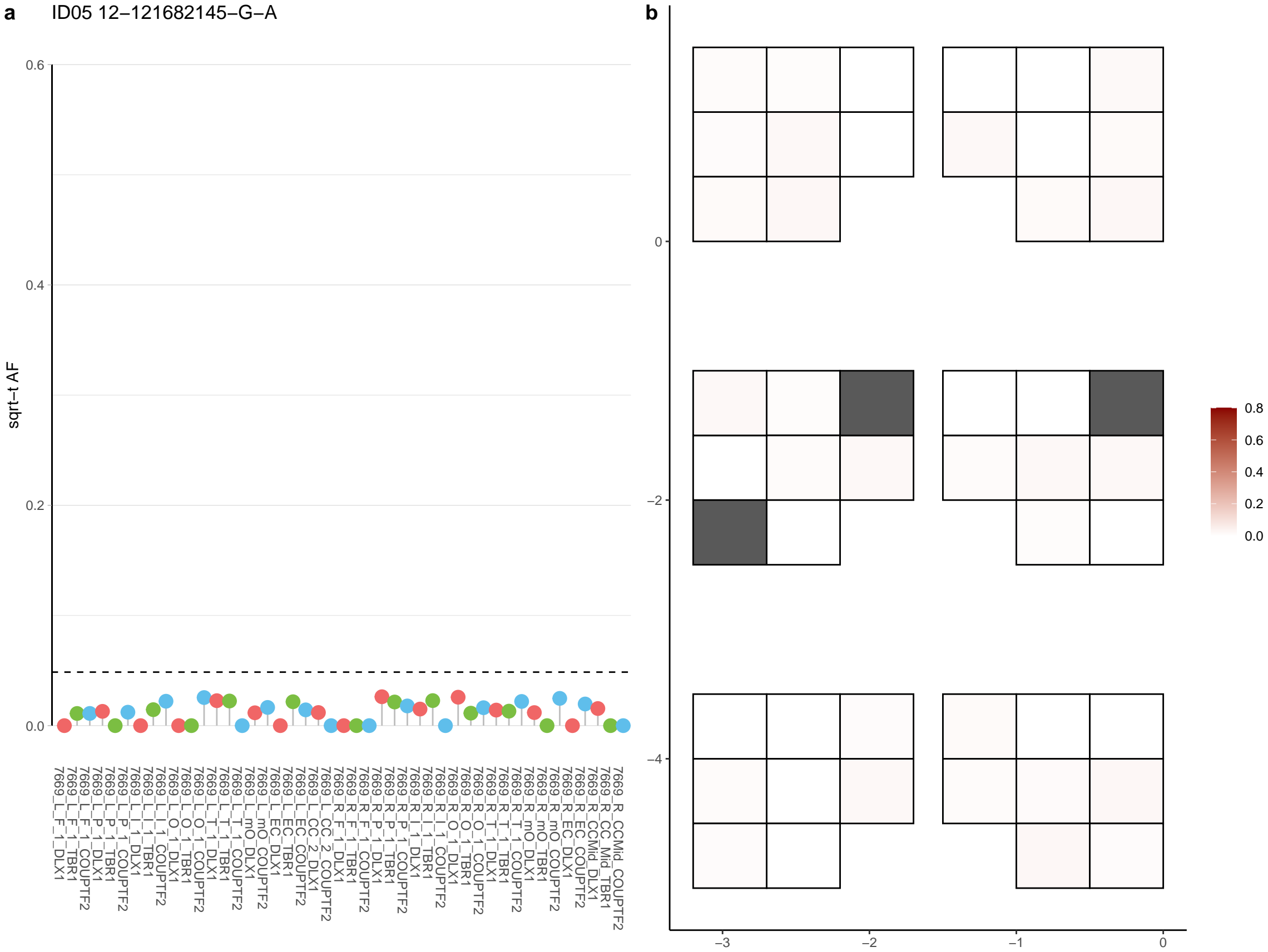

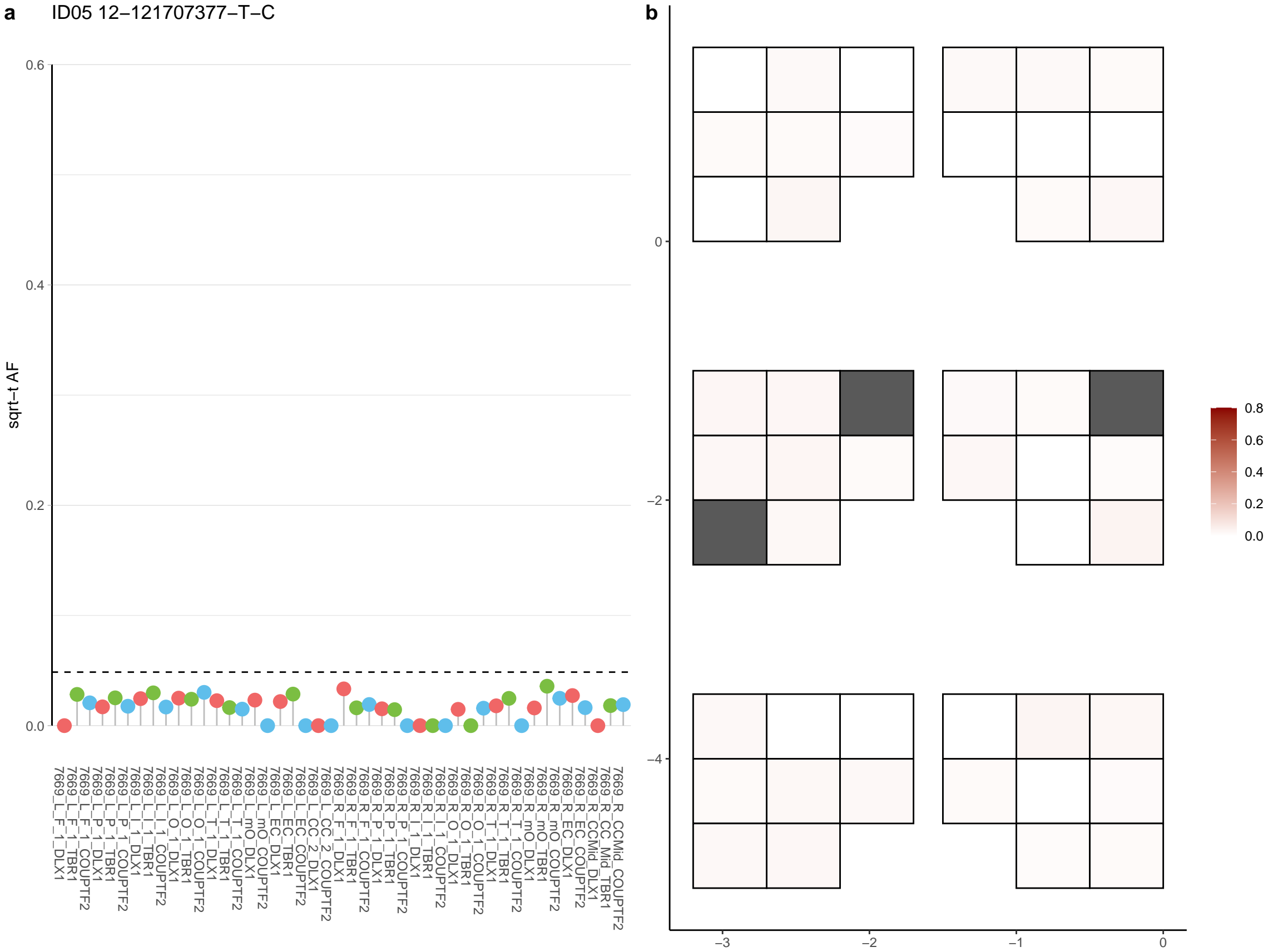

**a** ID05 12-127258051-G-T

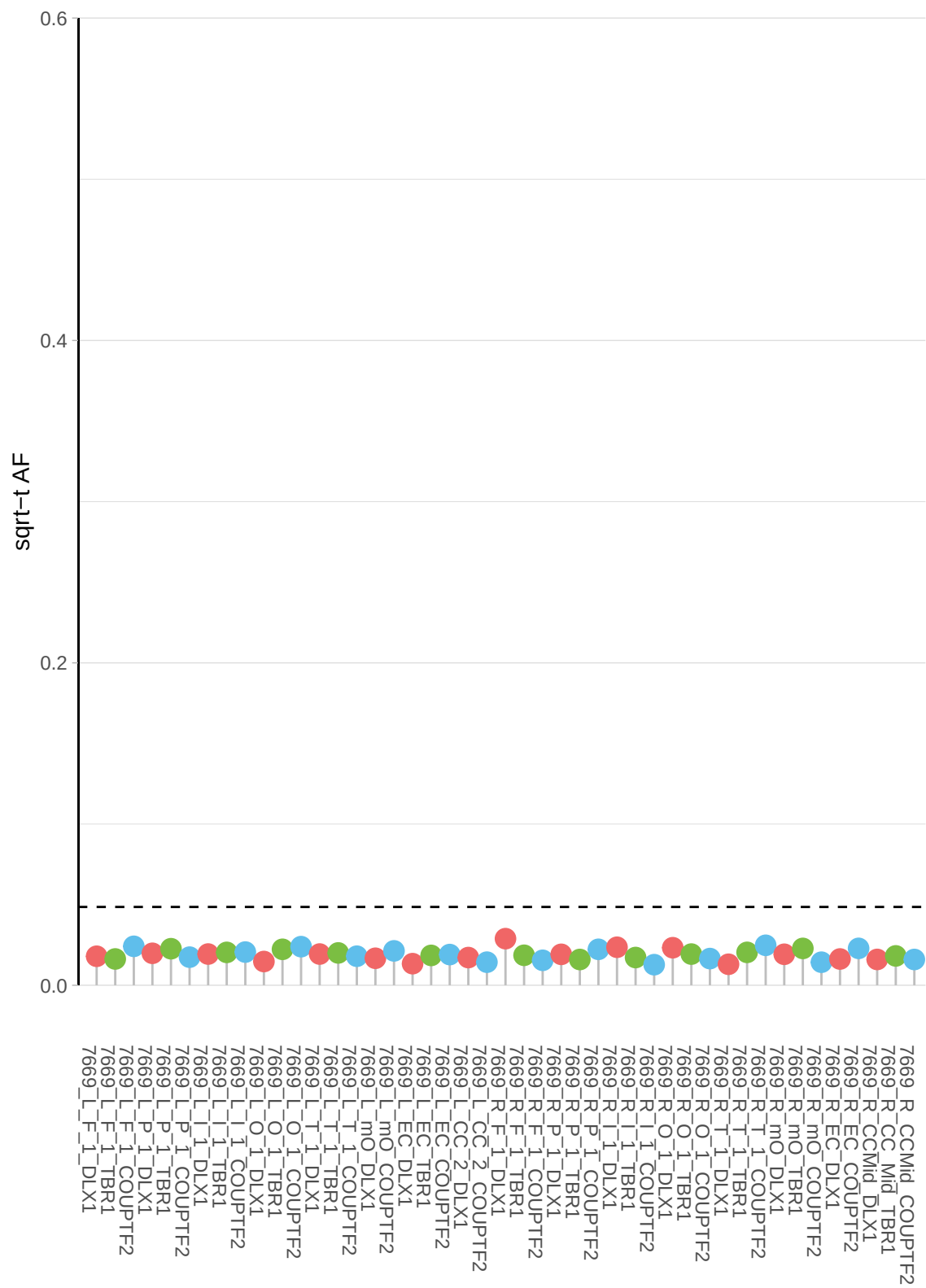

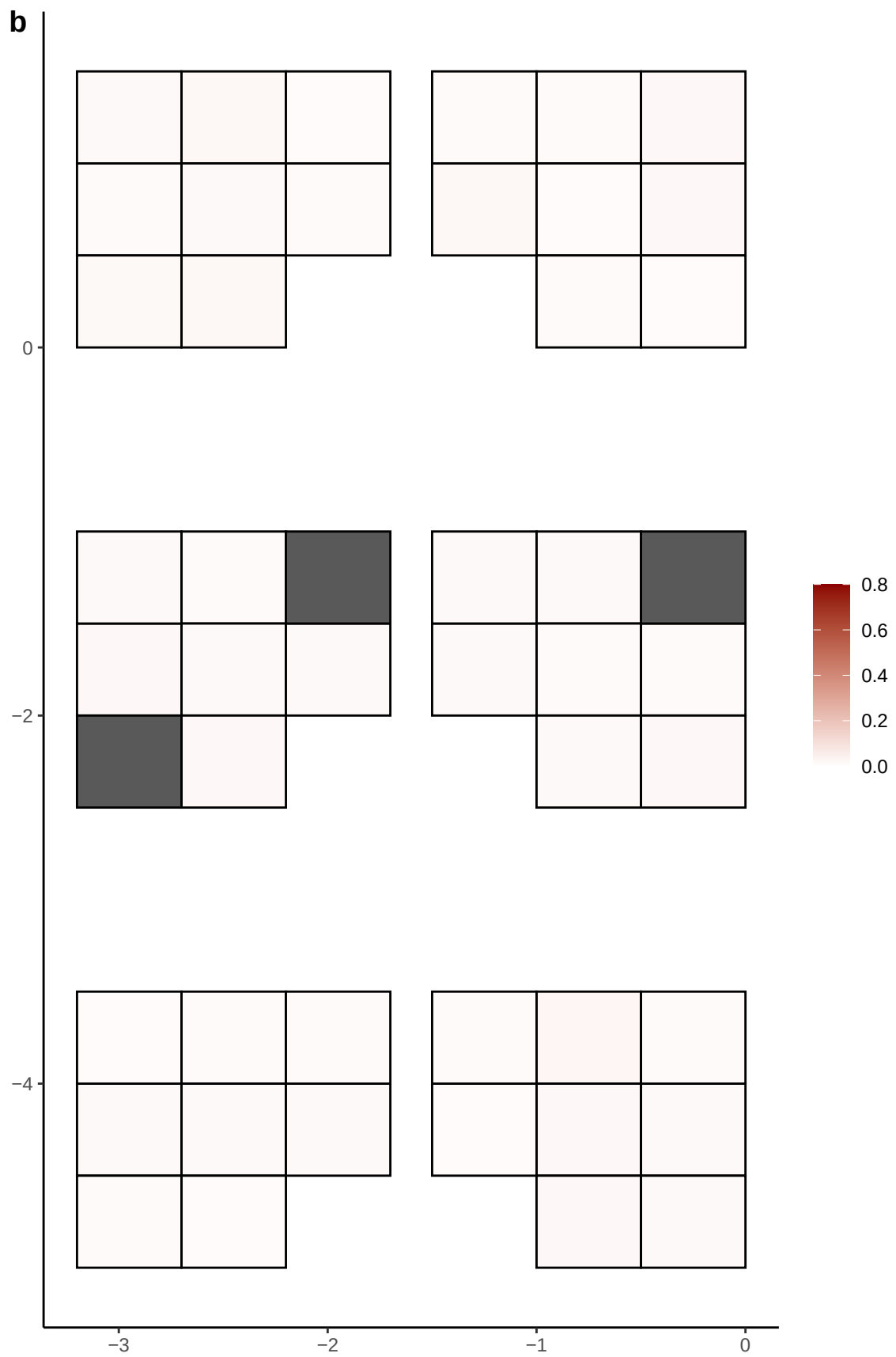

**a** ID05 12-128703875-C-T

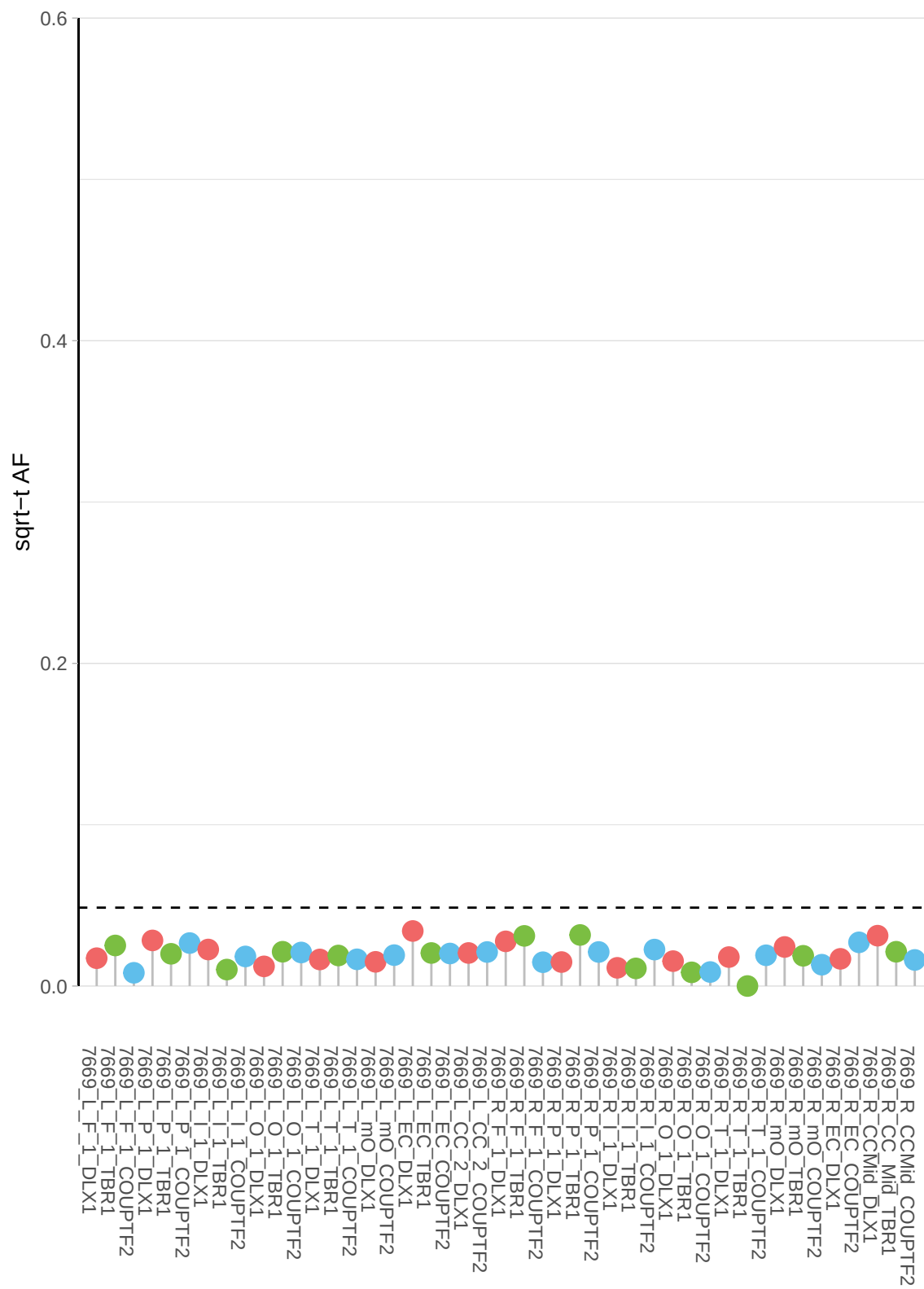

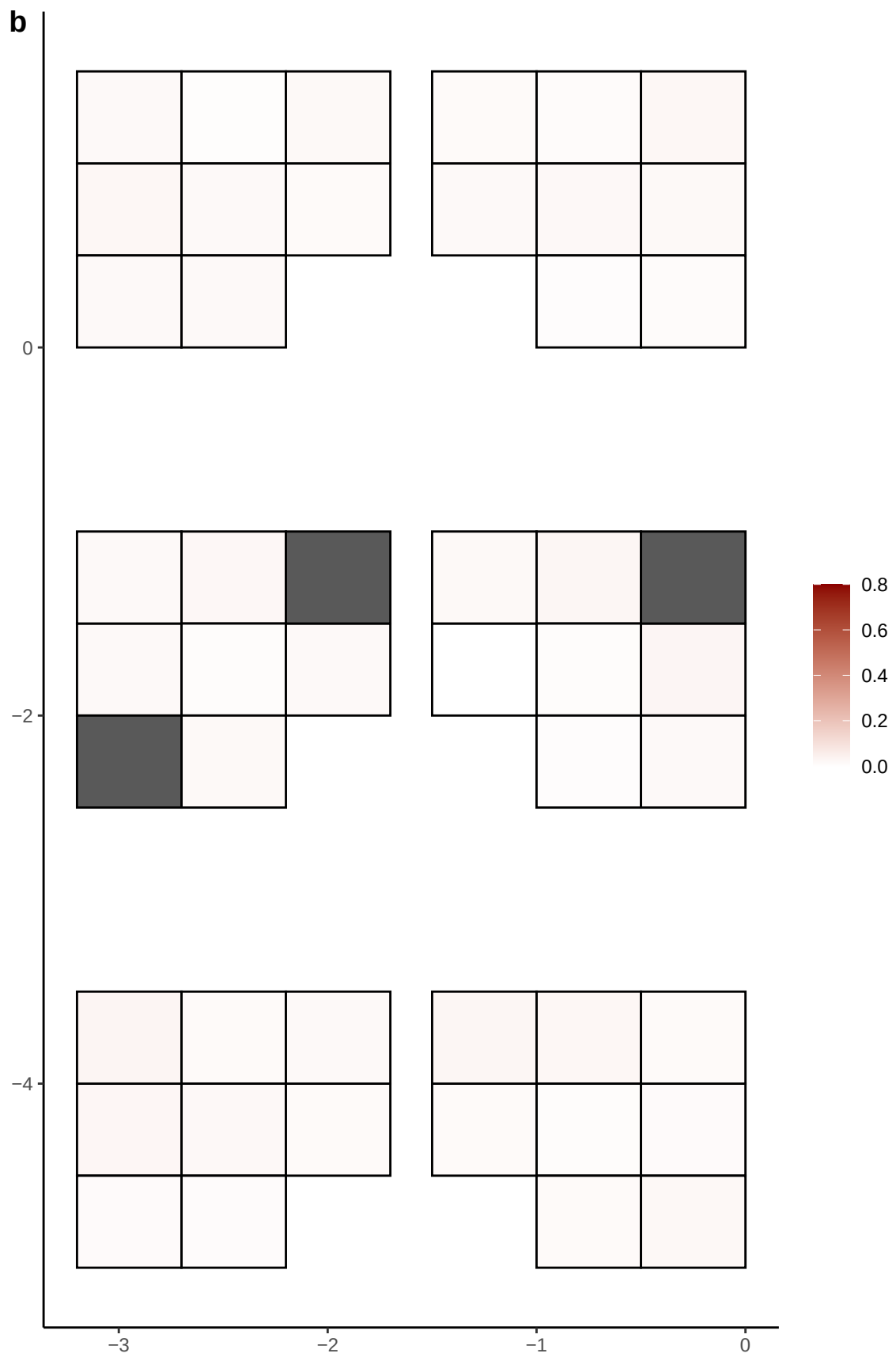

**a** ID05 12-130892713-C-T

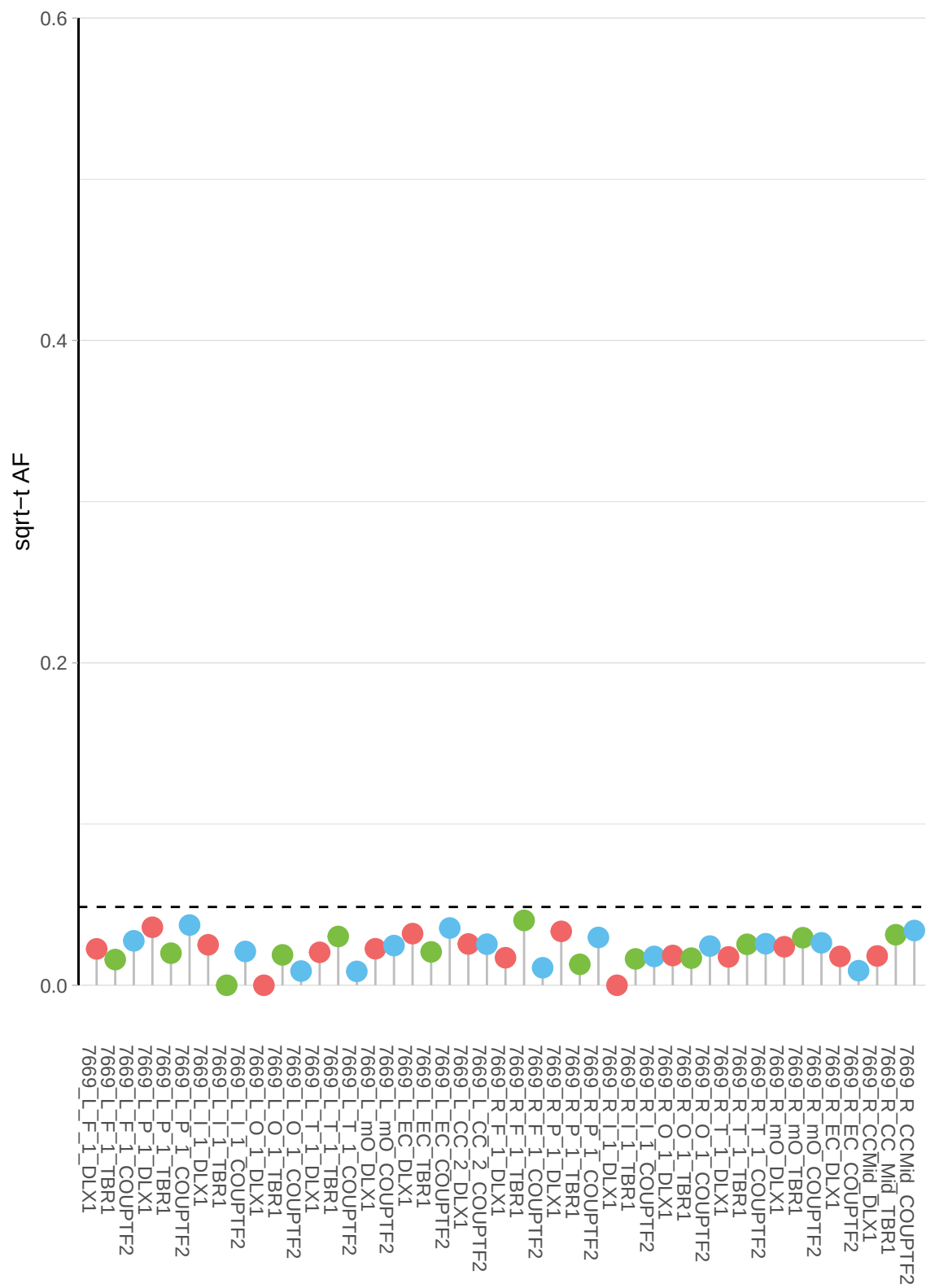

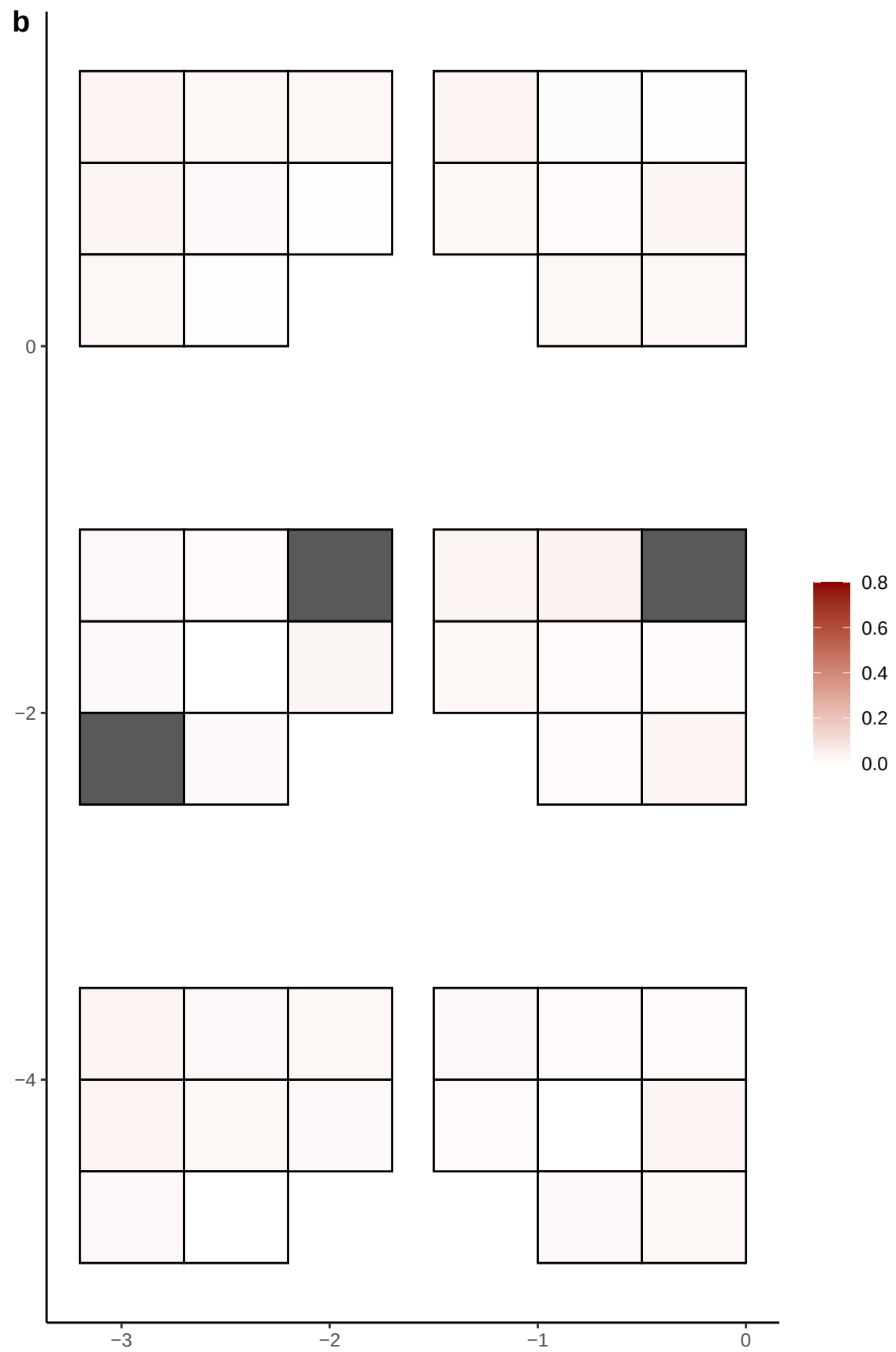

**a** ID05 12-131906235-G-A

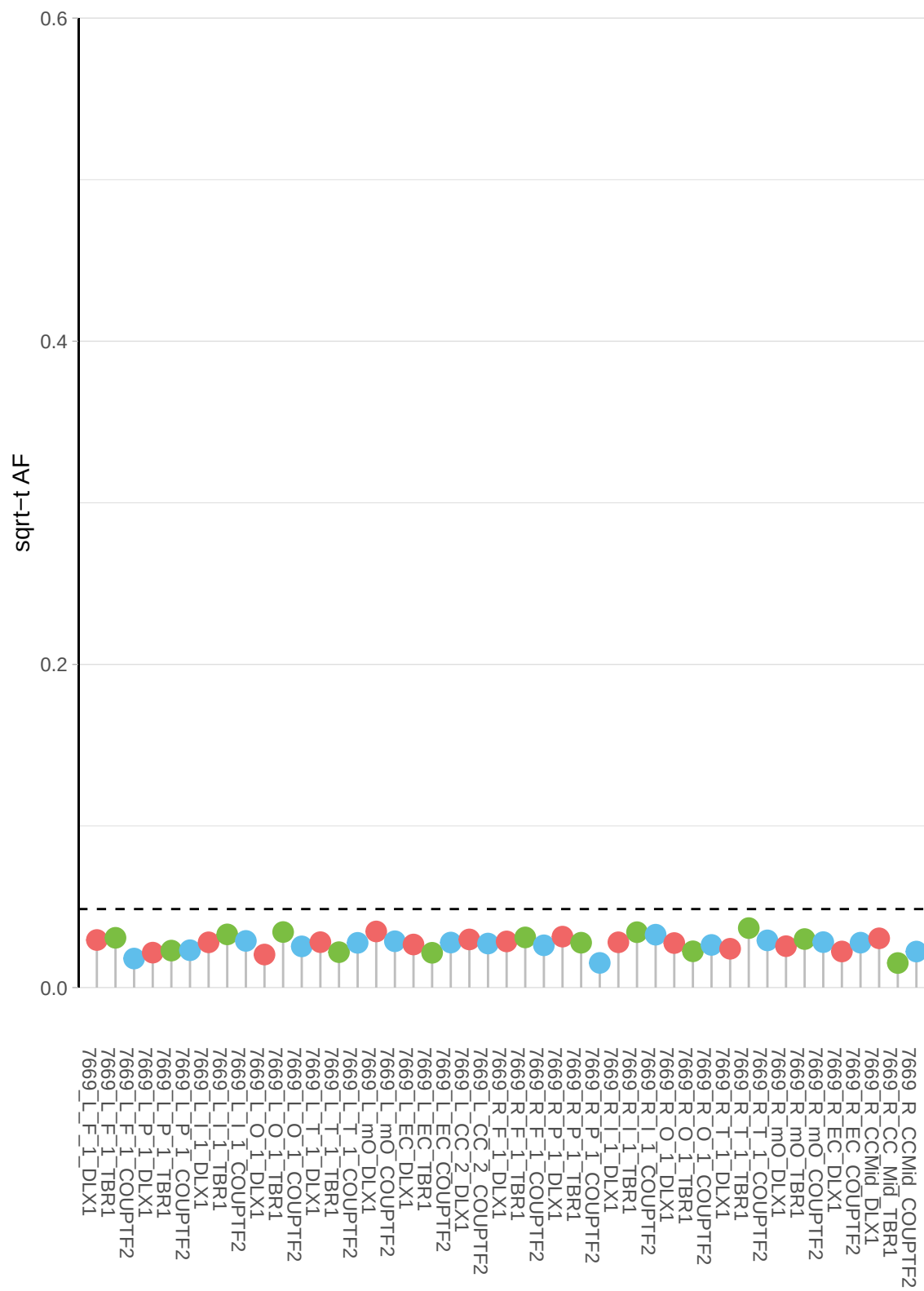

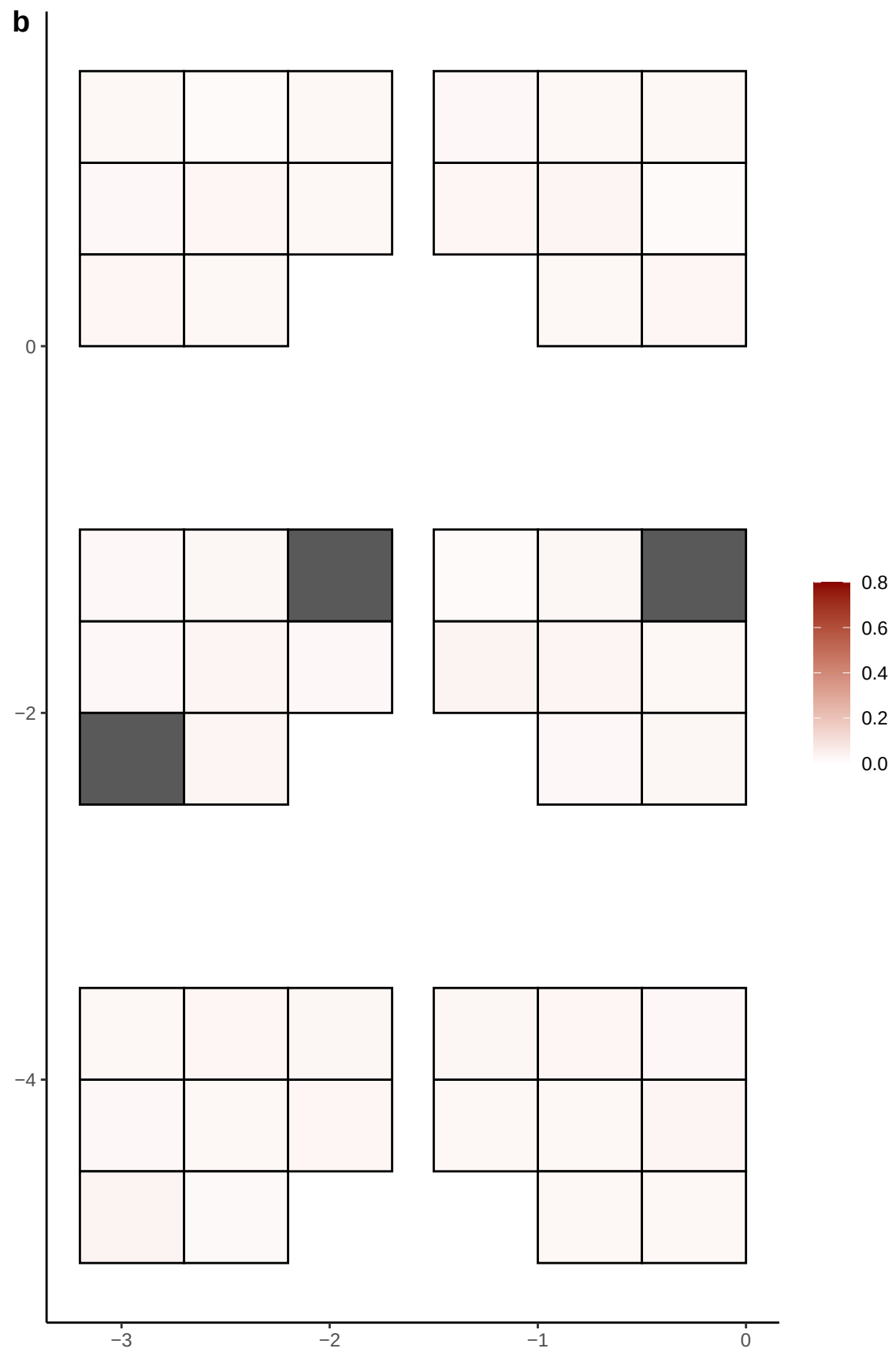

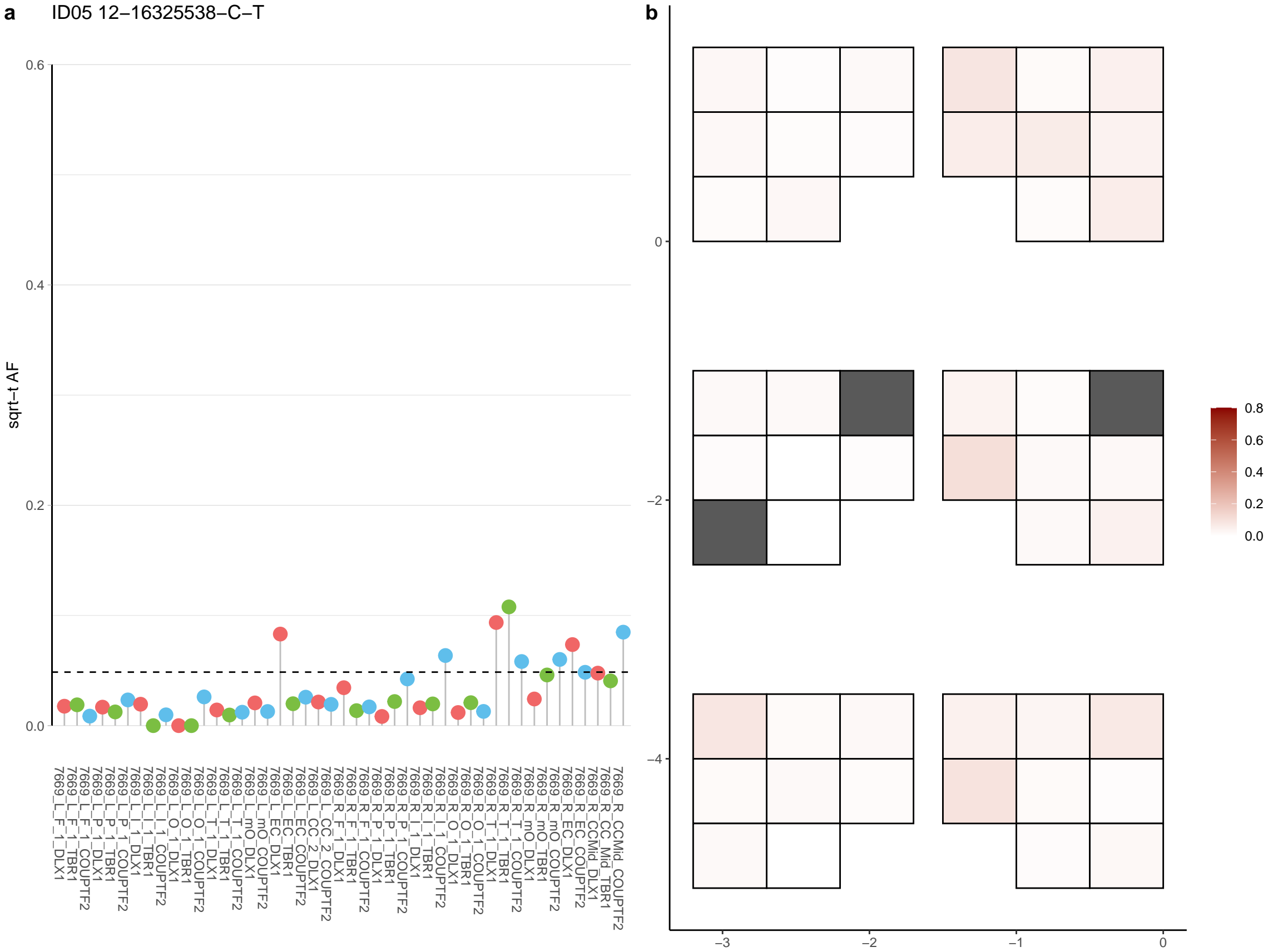

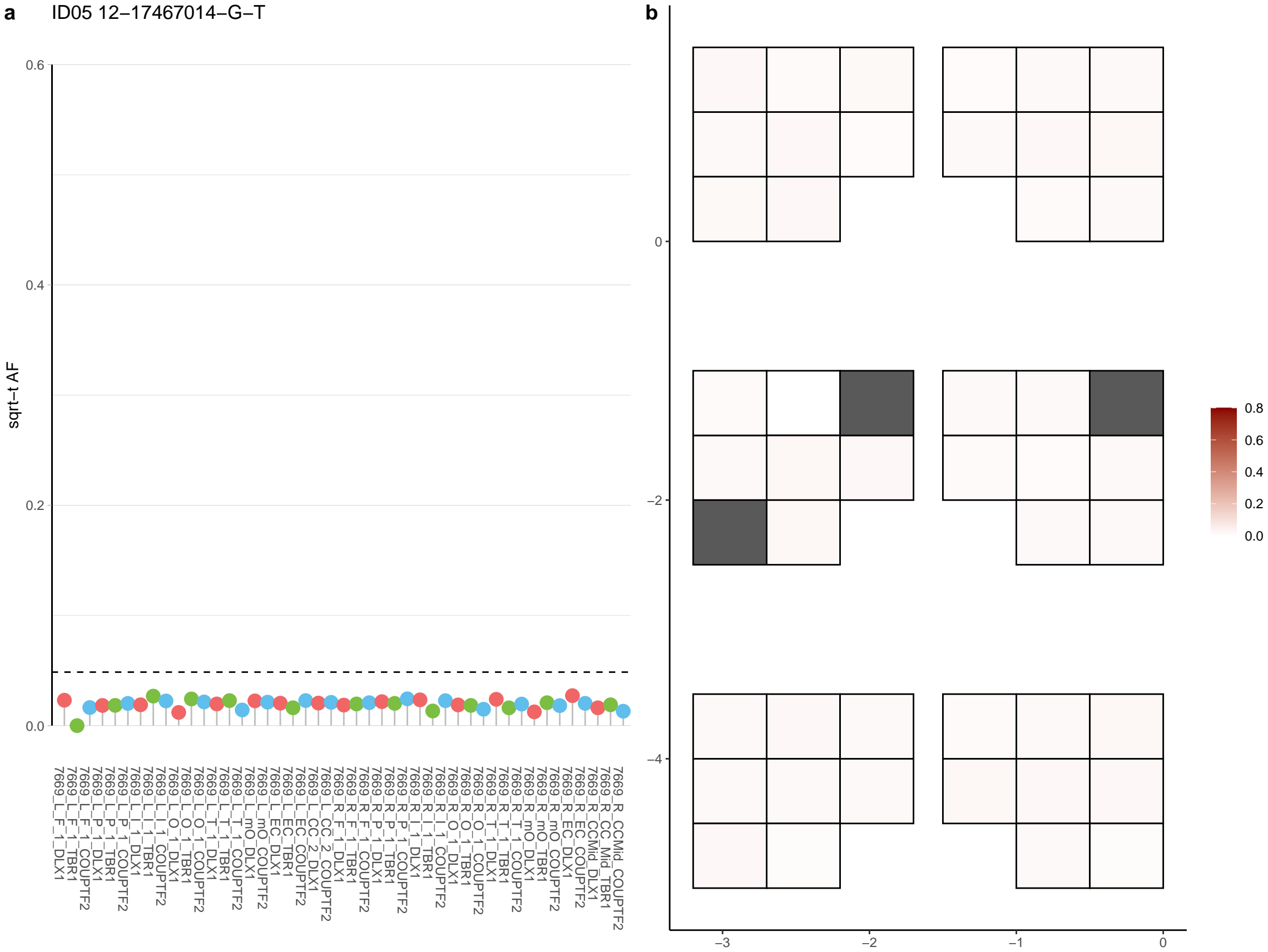

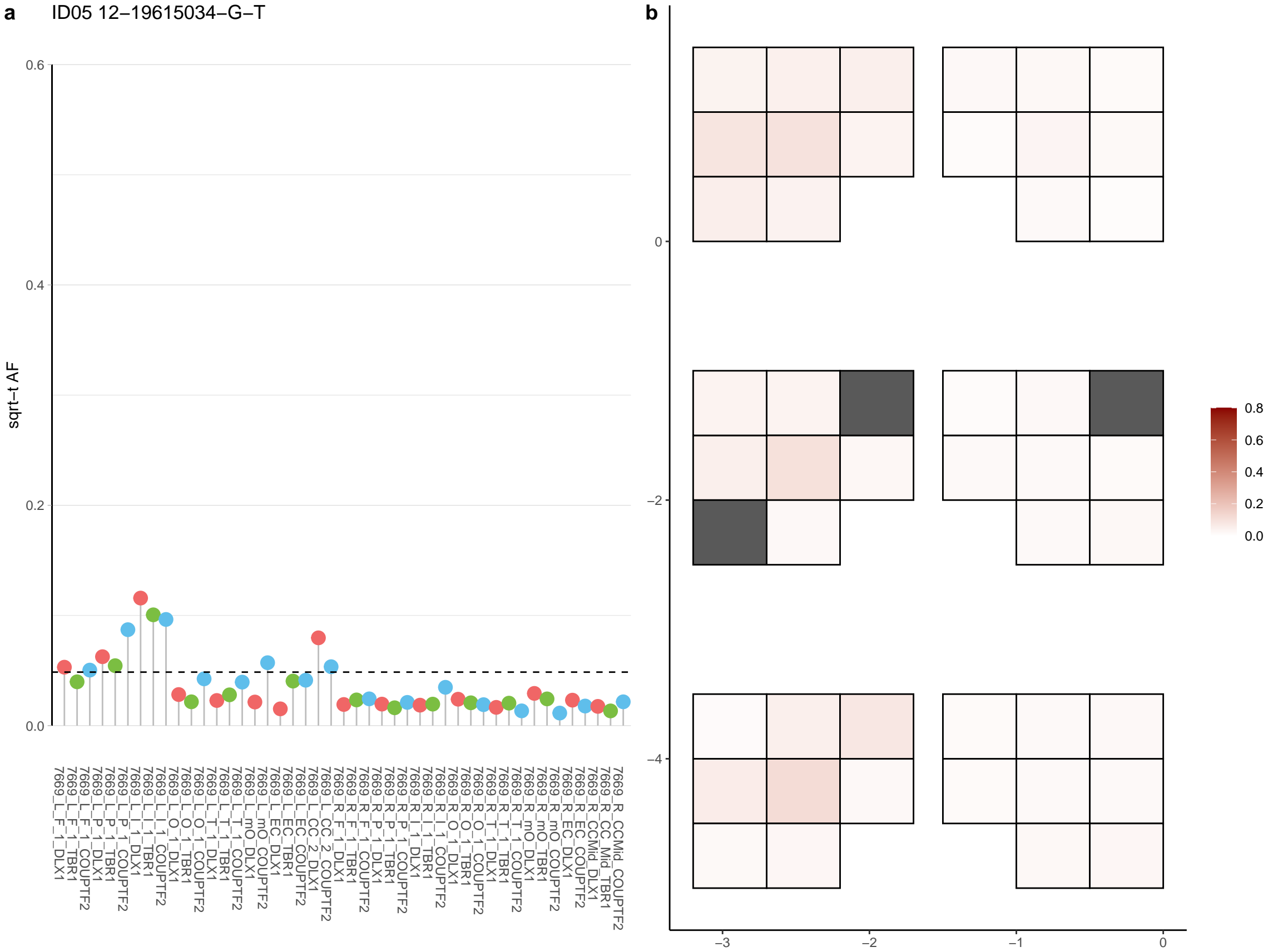

**a** ID05 12-20421175-G-A

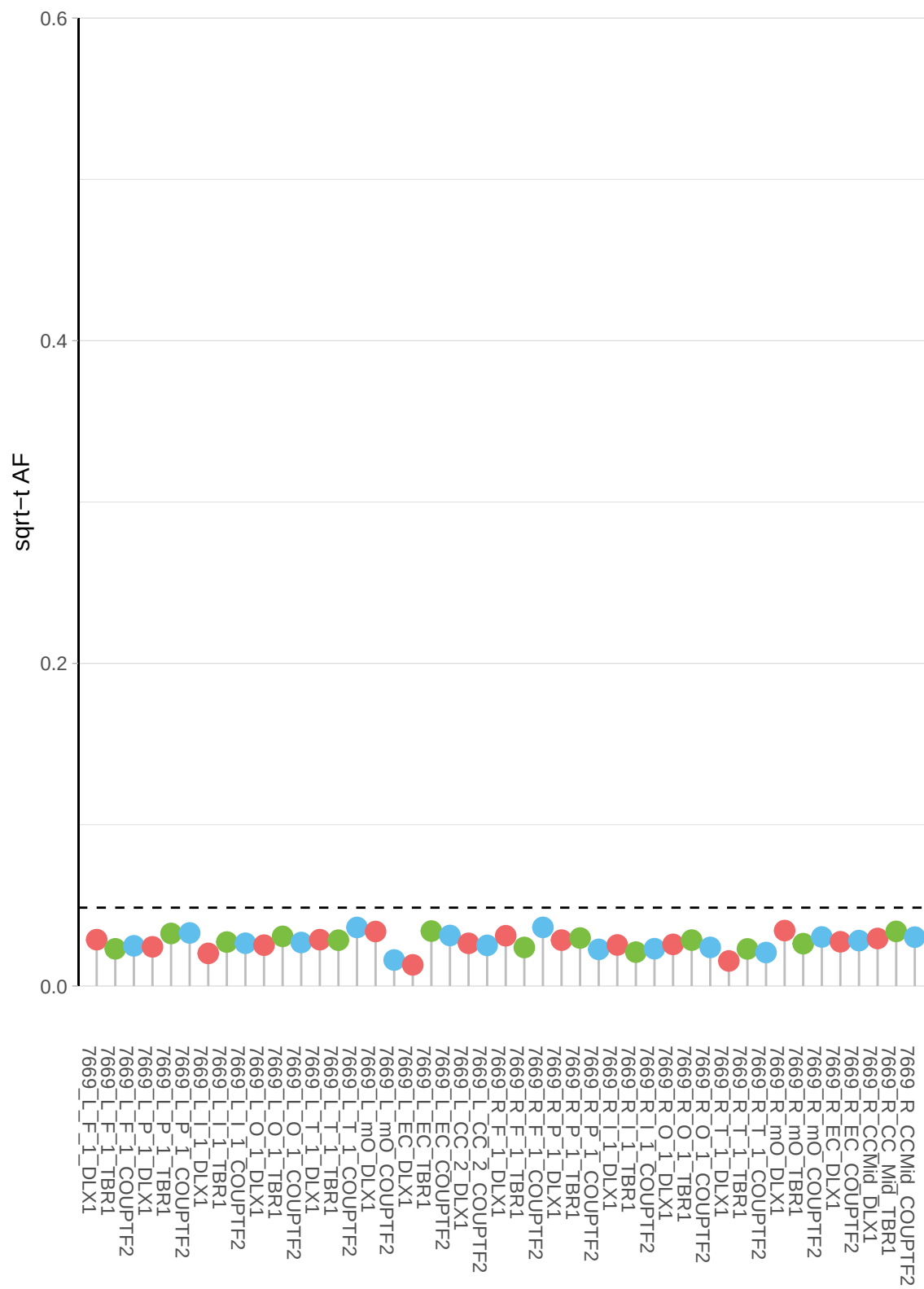

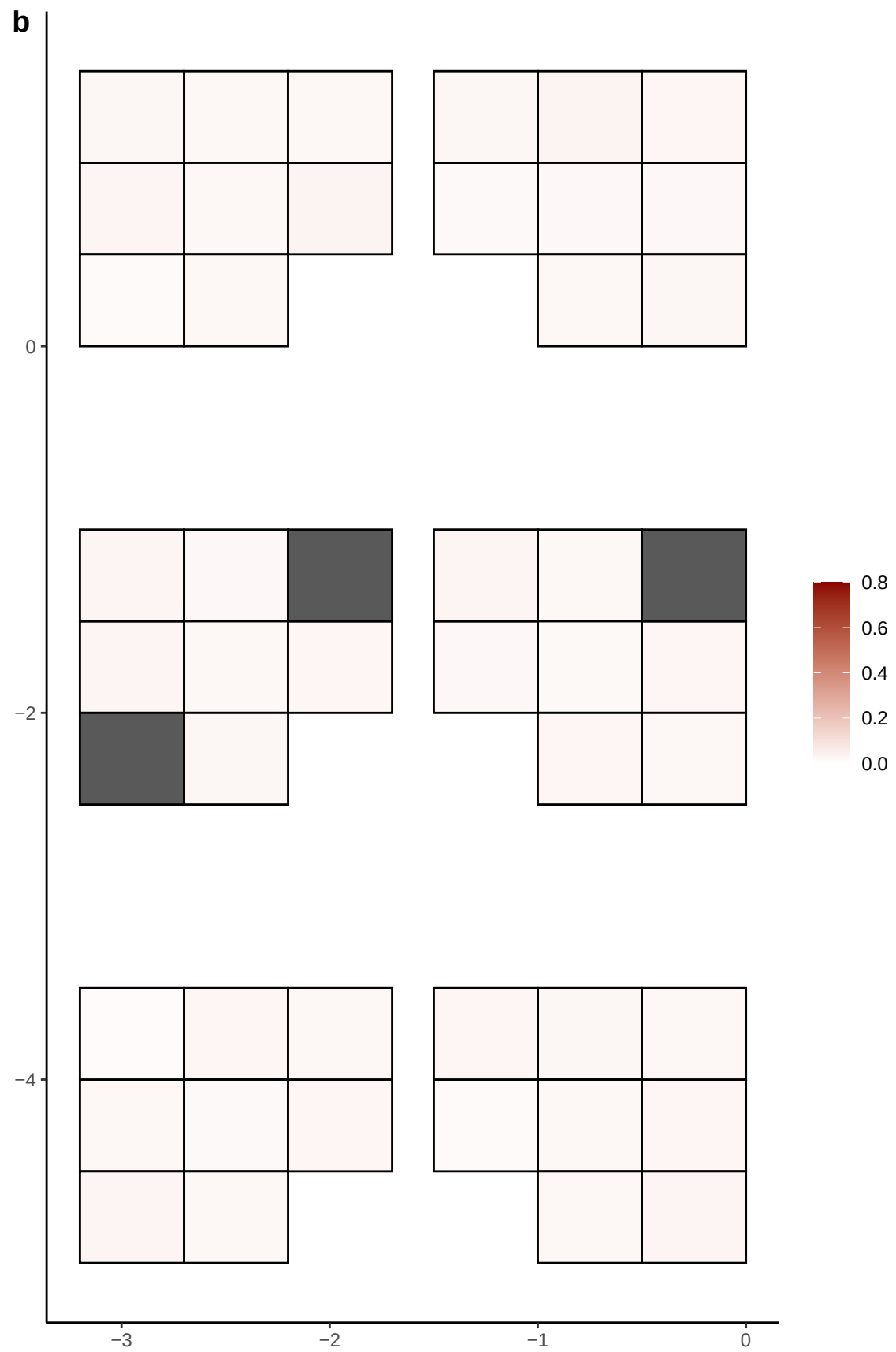

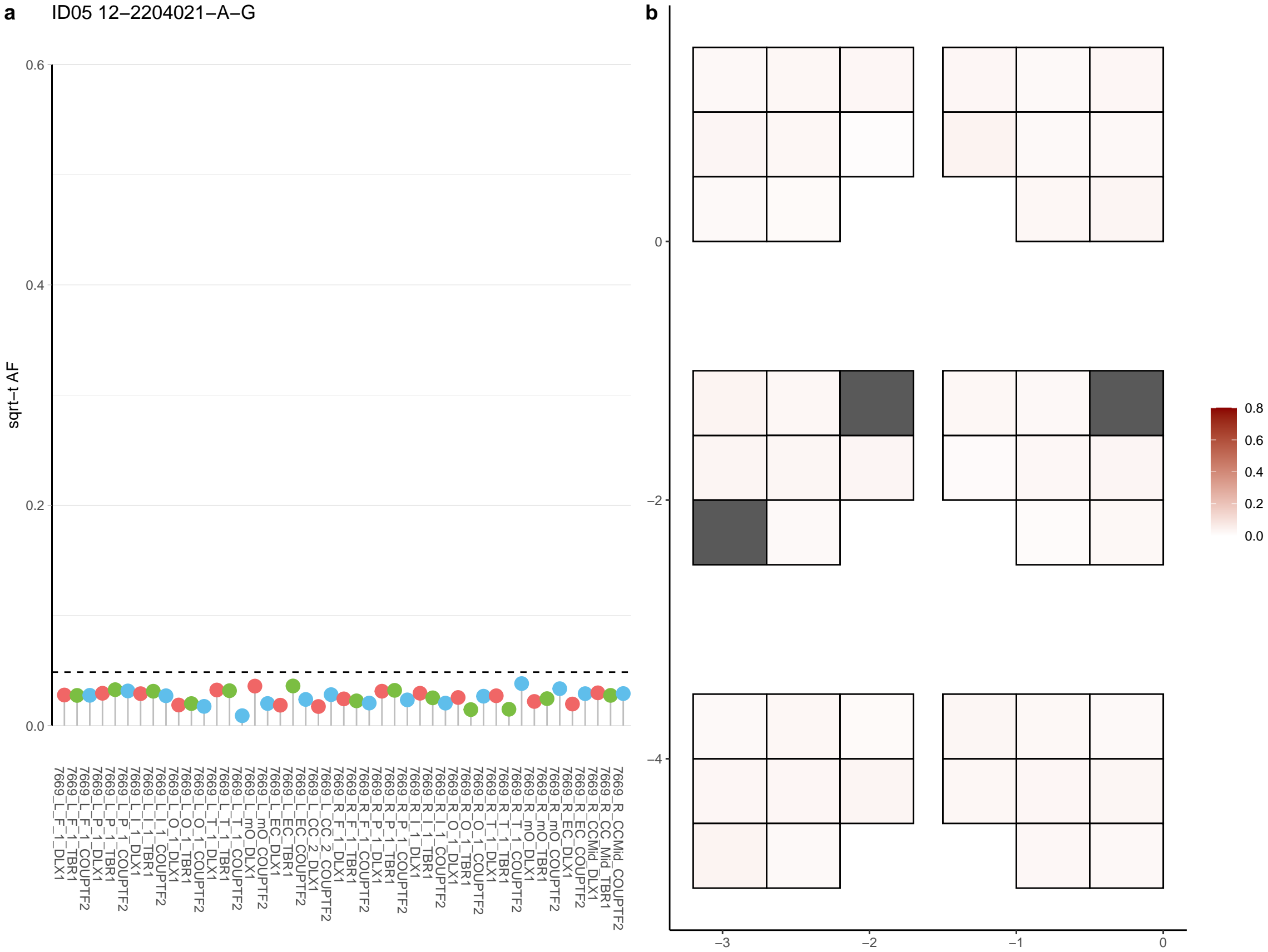

**a** ID05 12-24436884-C-T

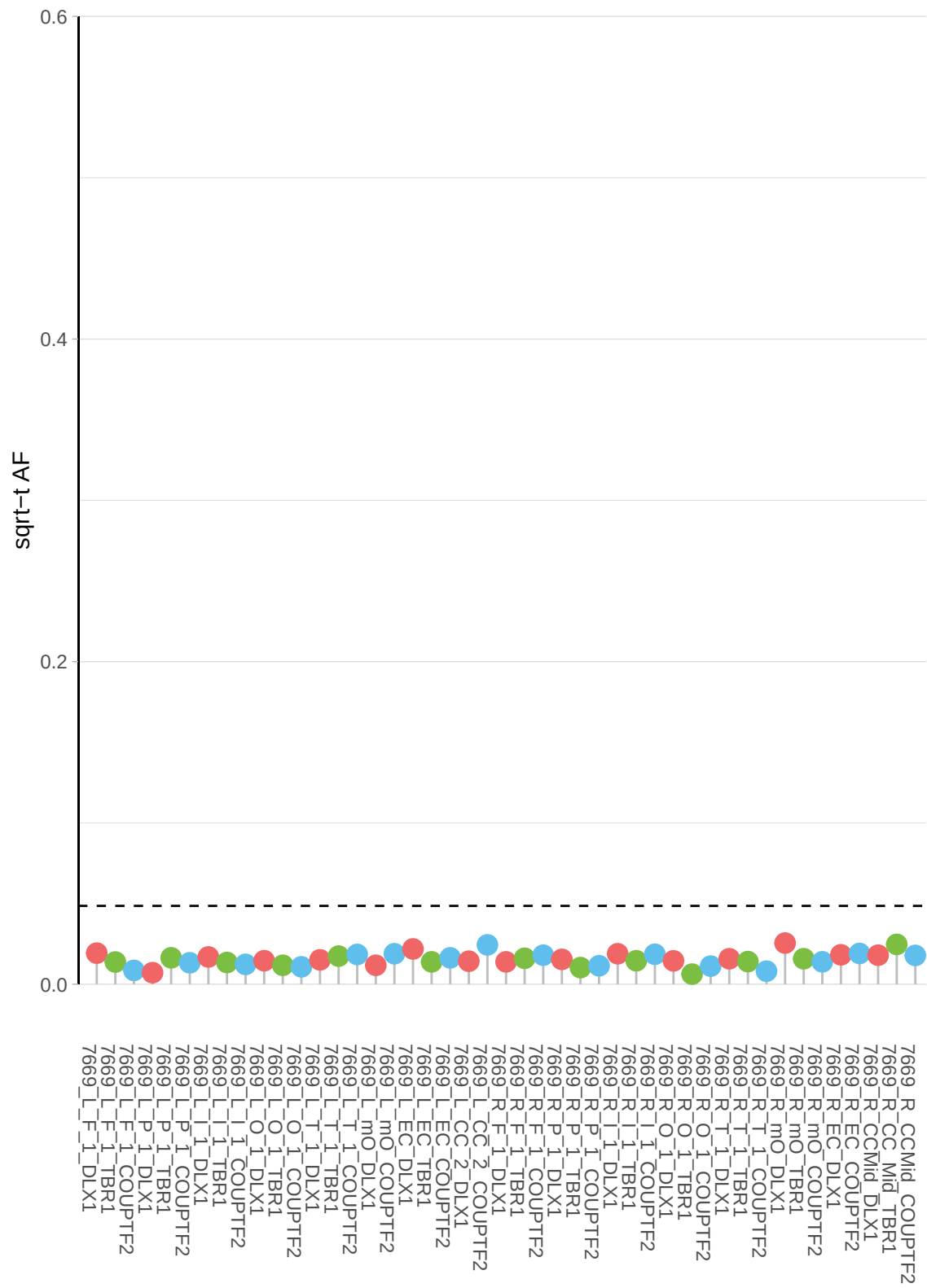

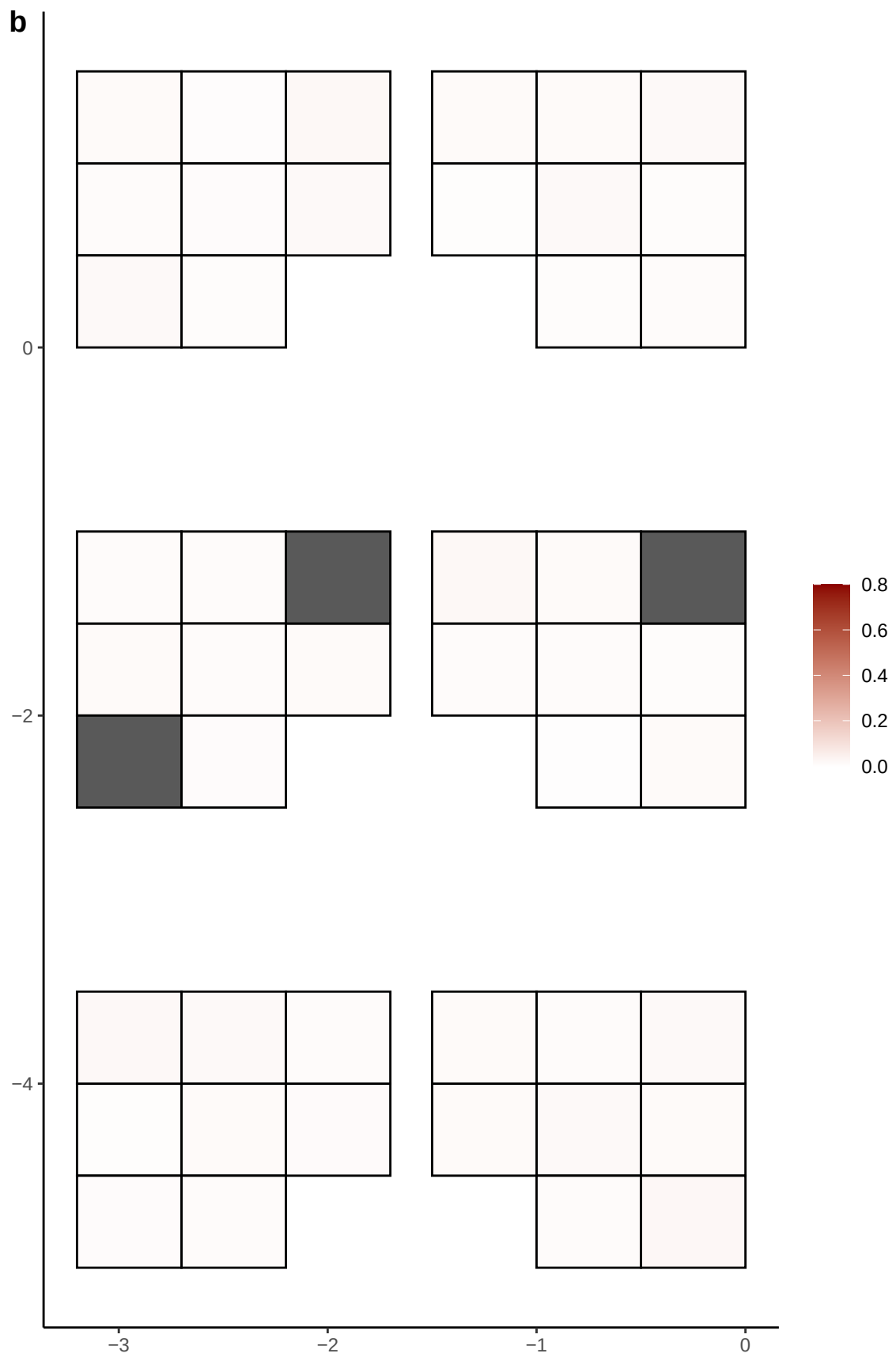

**a** ID05 12-55677268-C-A

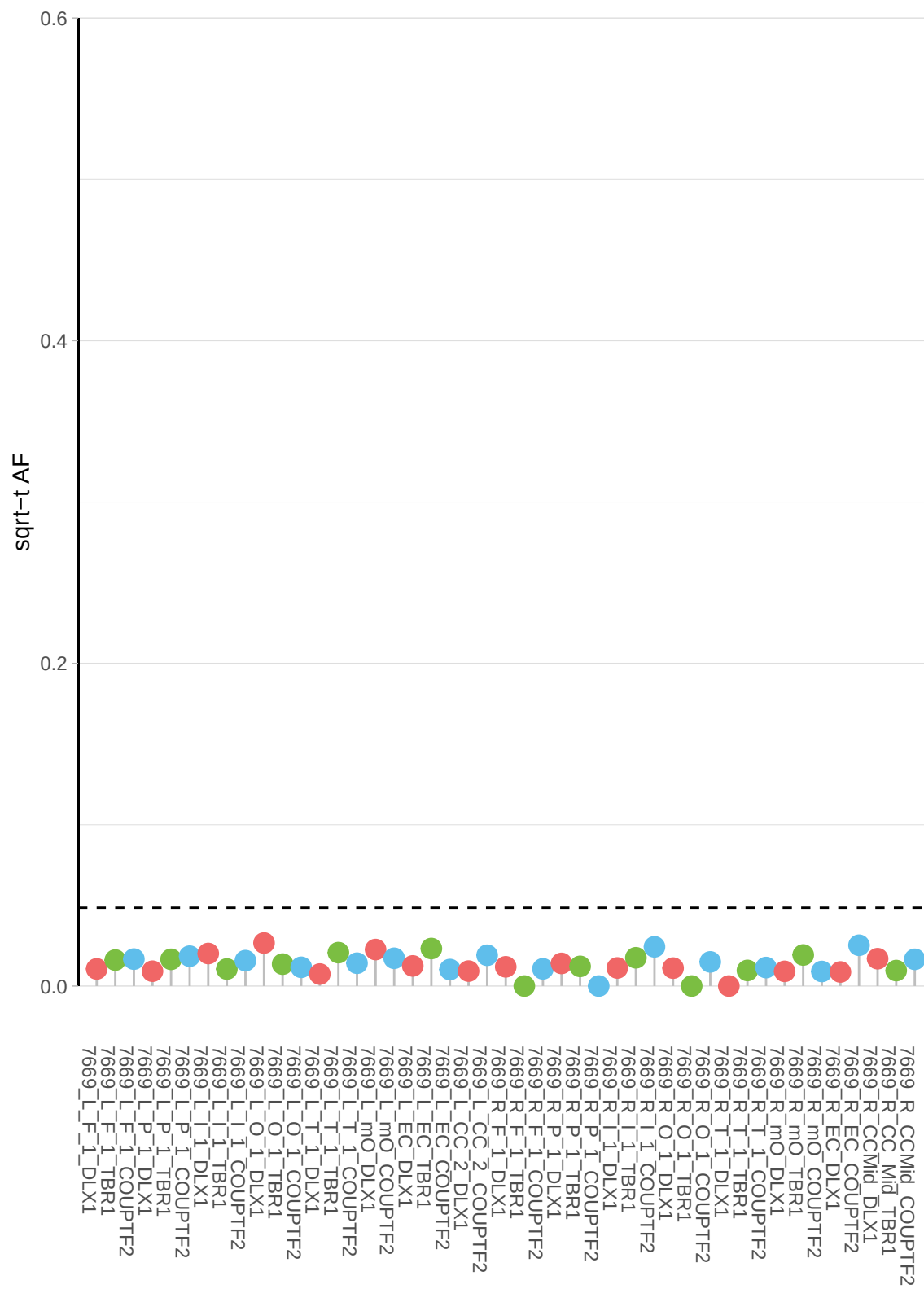

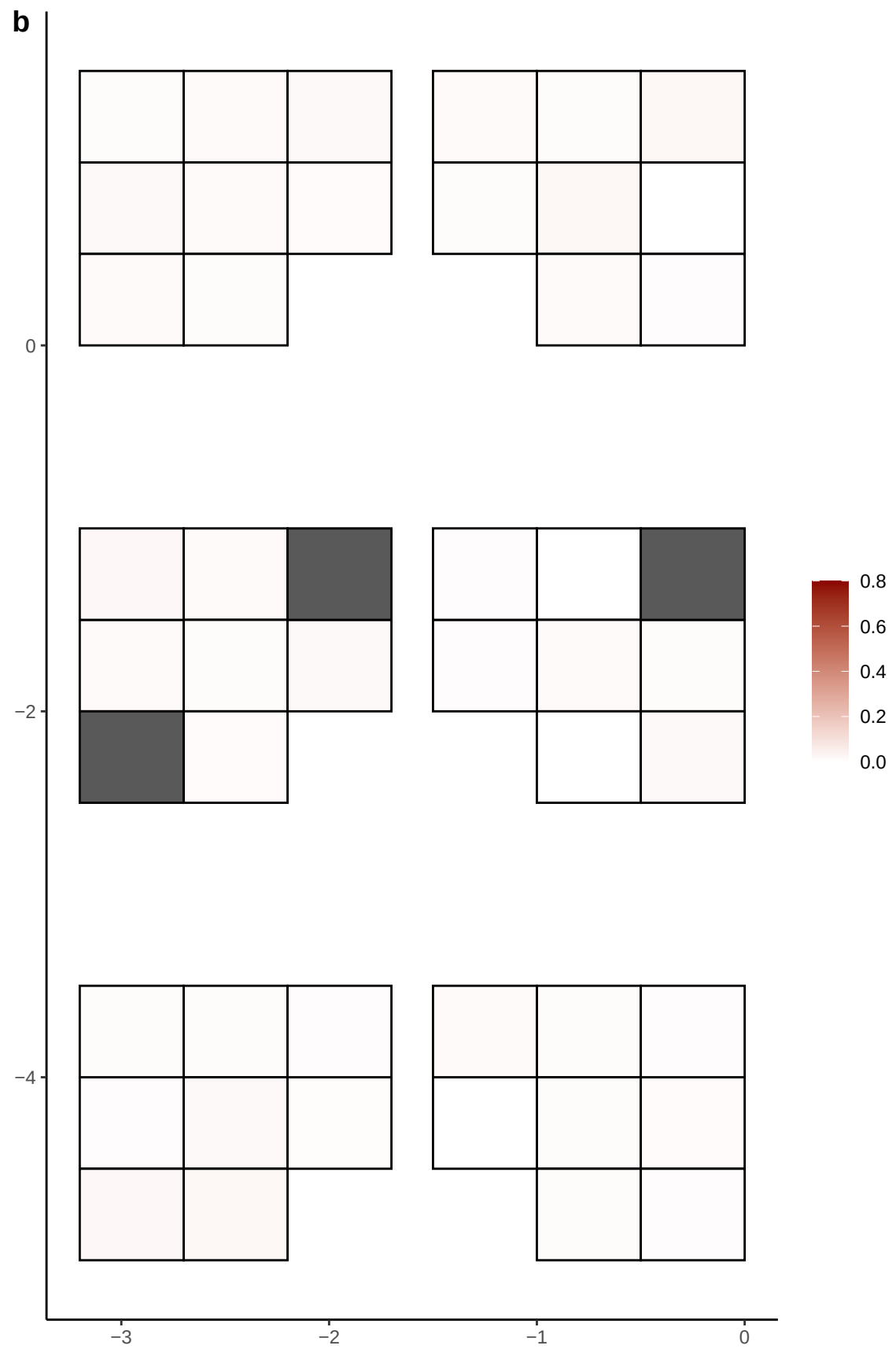

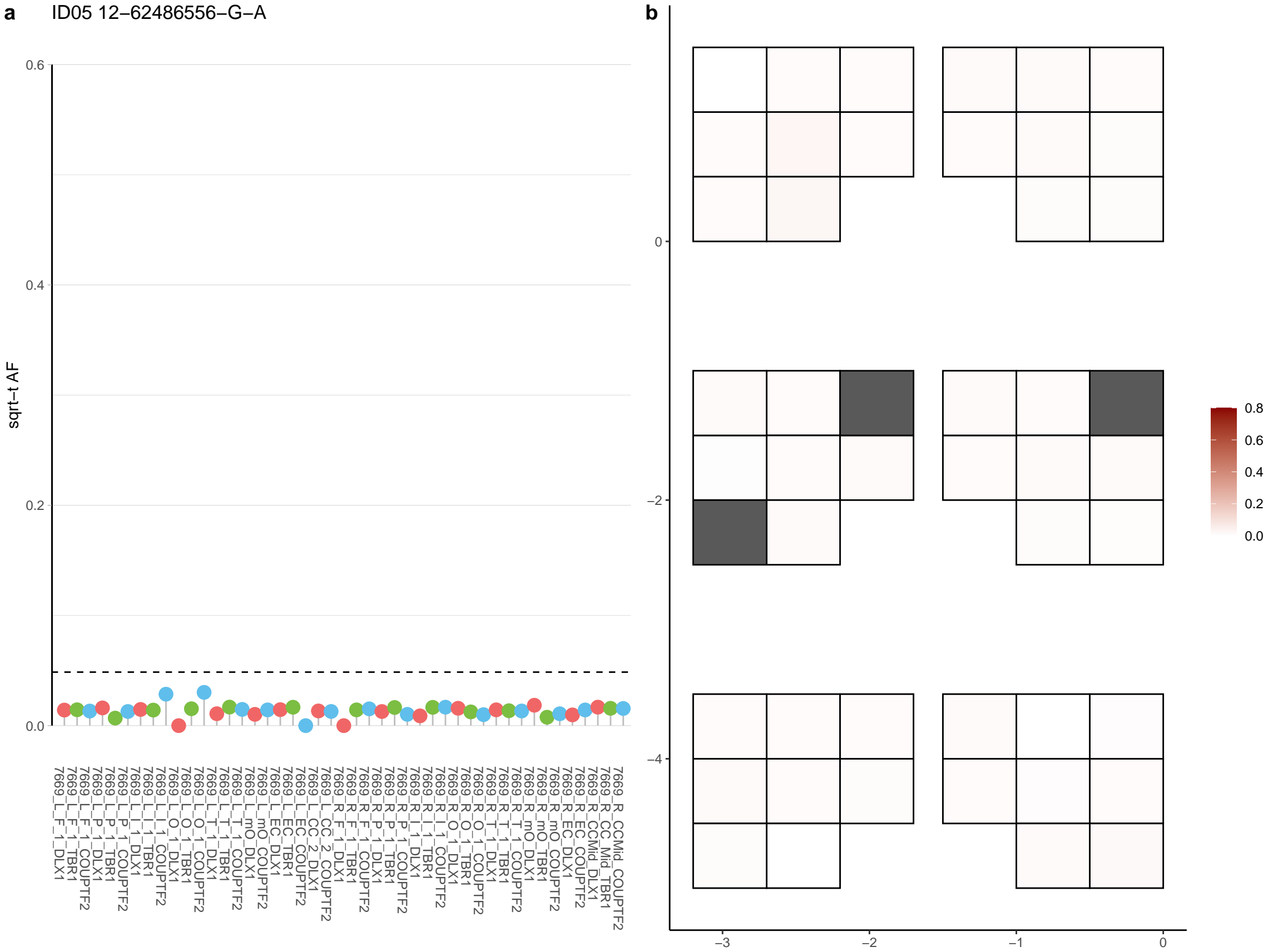

**a** ID05 12-73783719-C-T

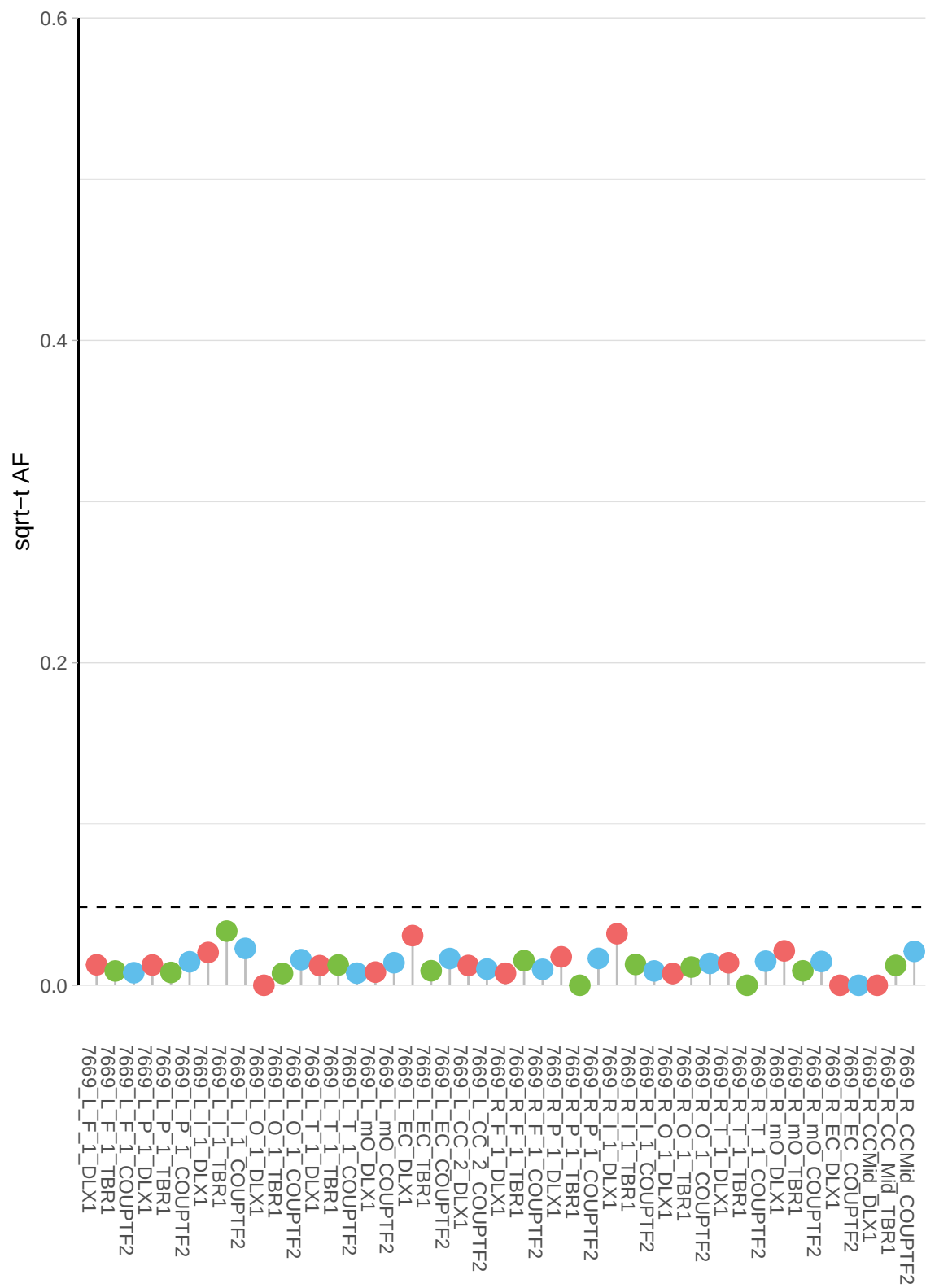

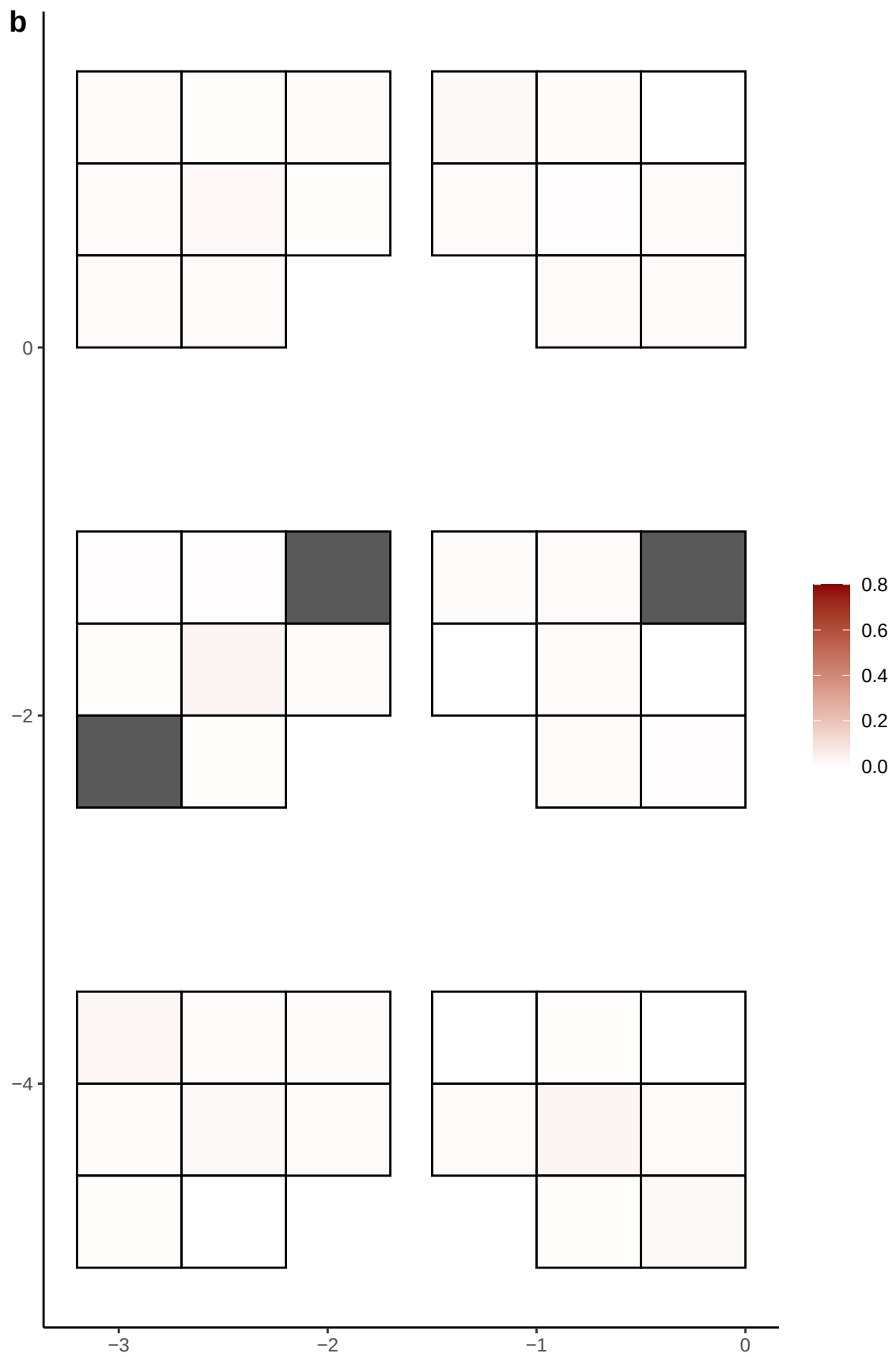

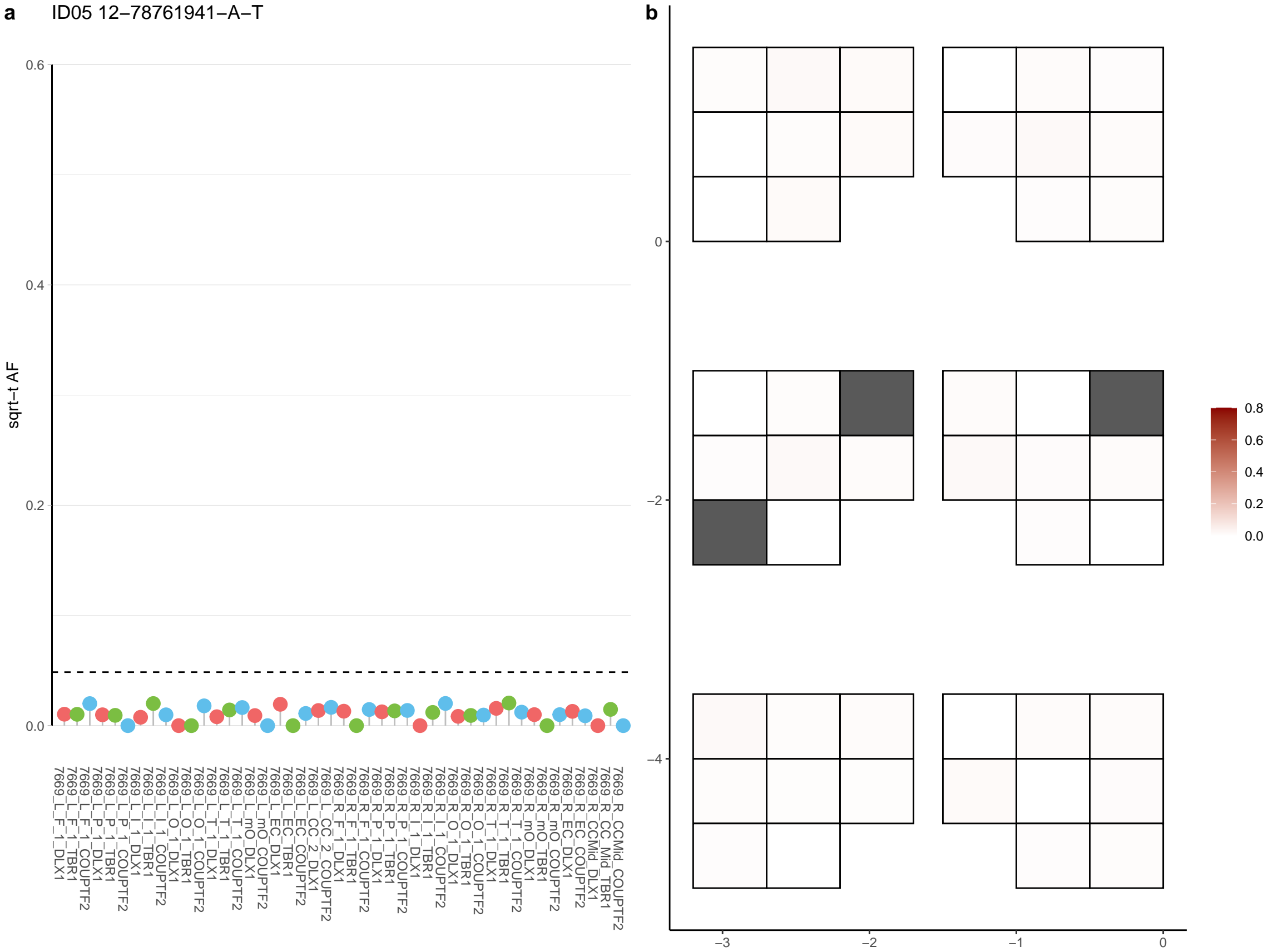

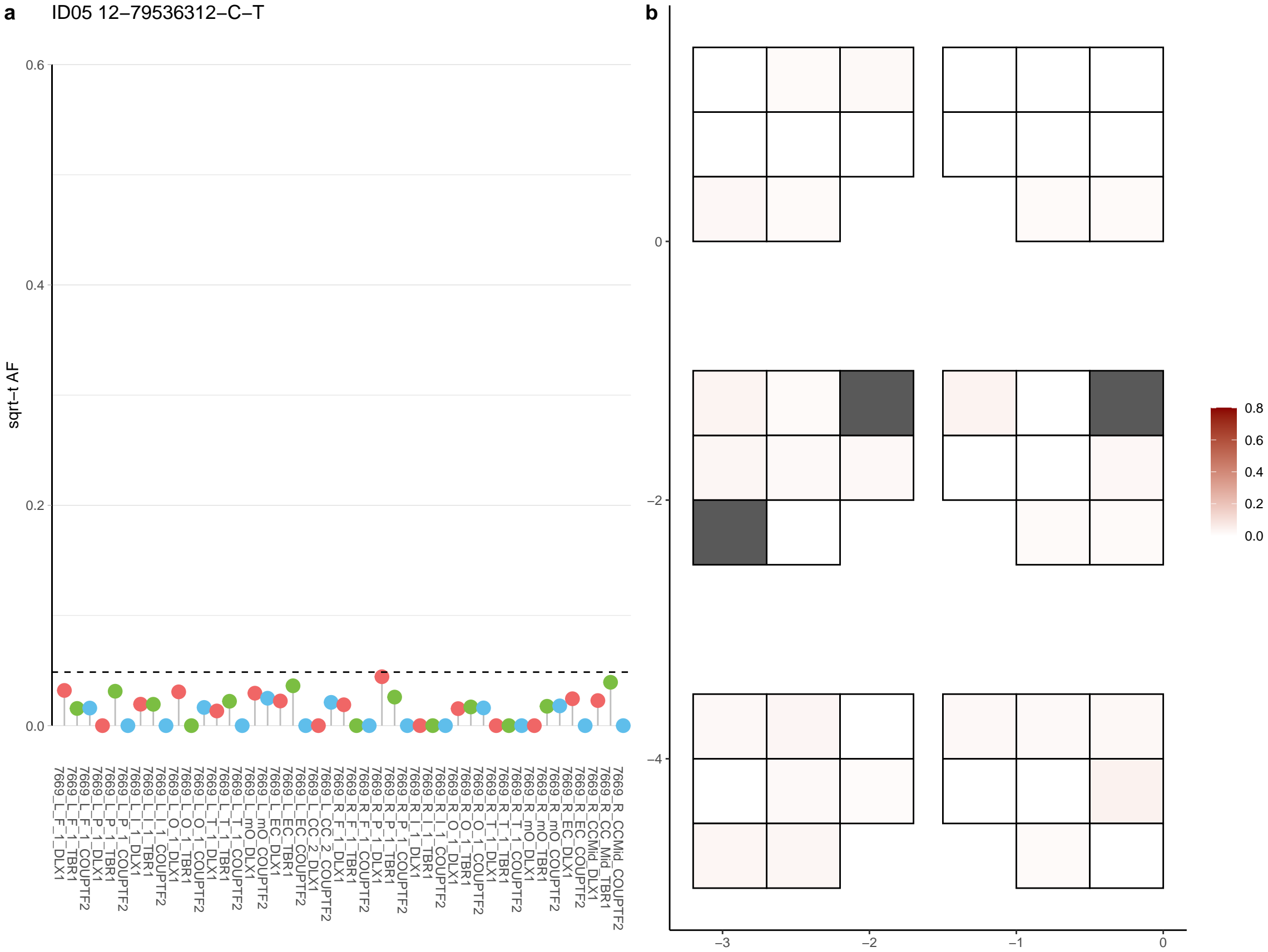

**a** ID05 12-85014452-T-A

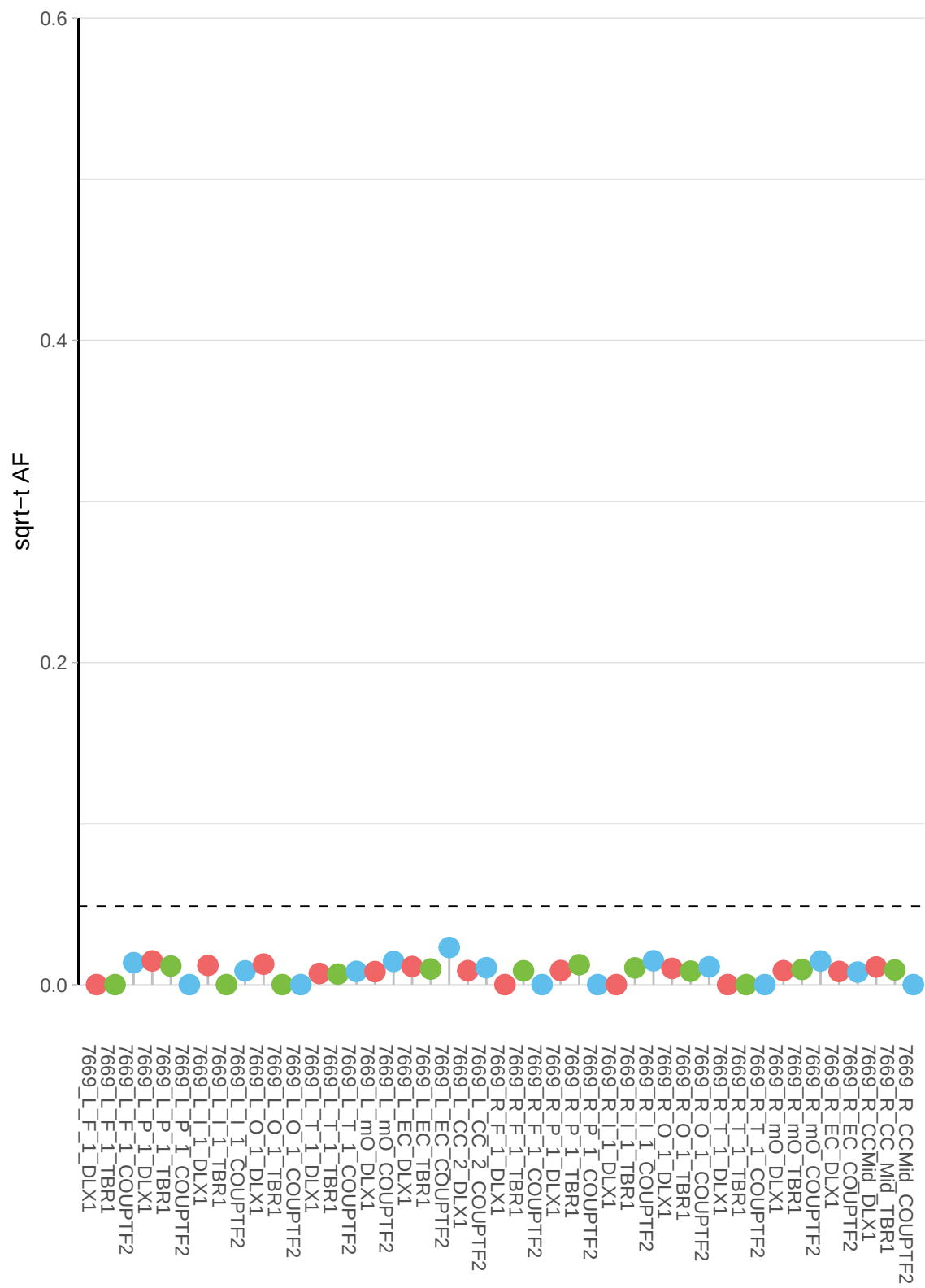

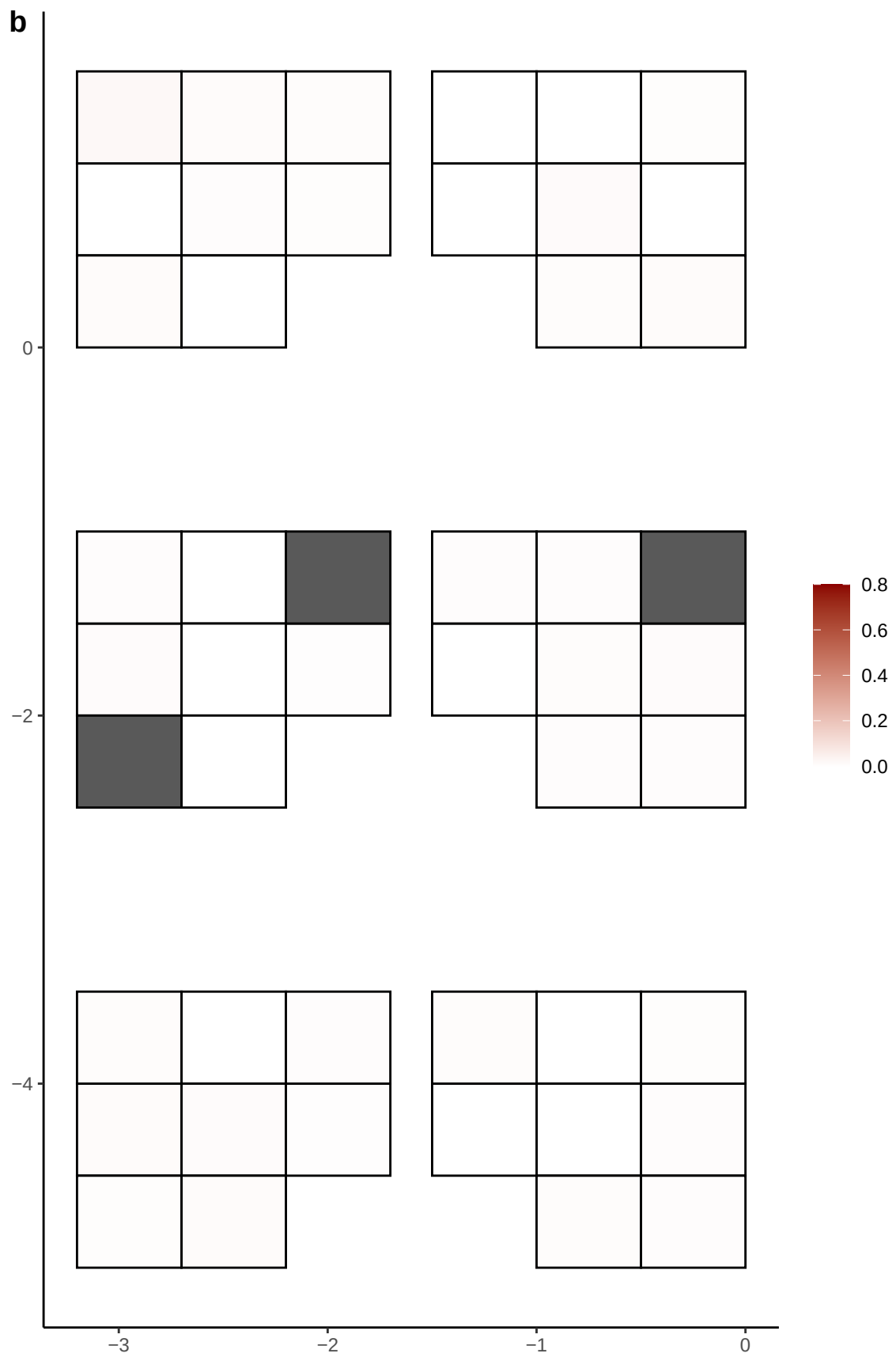

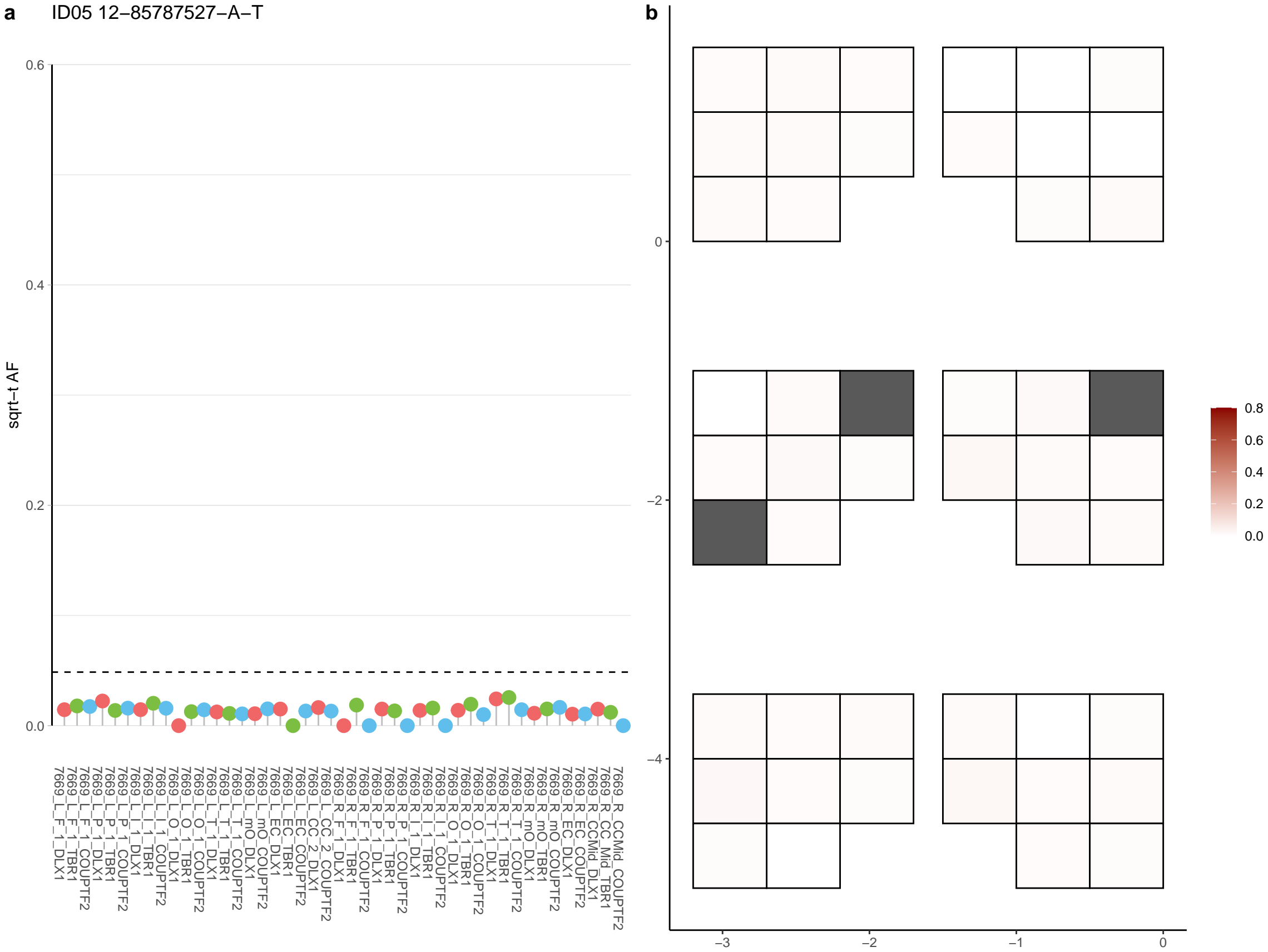

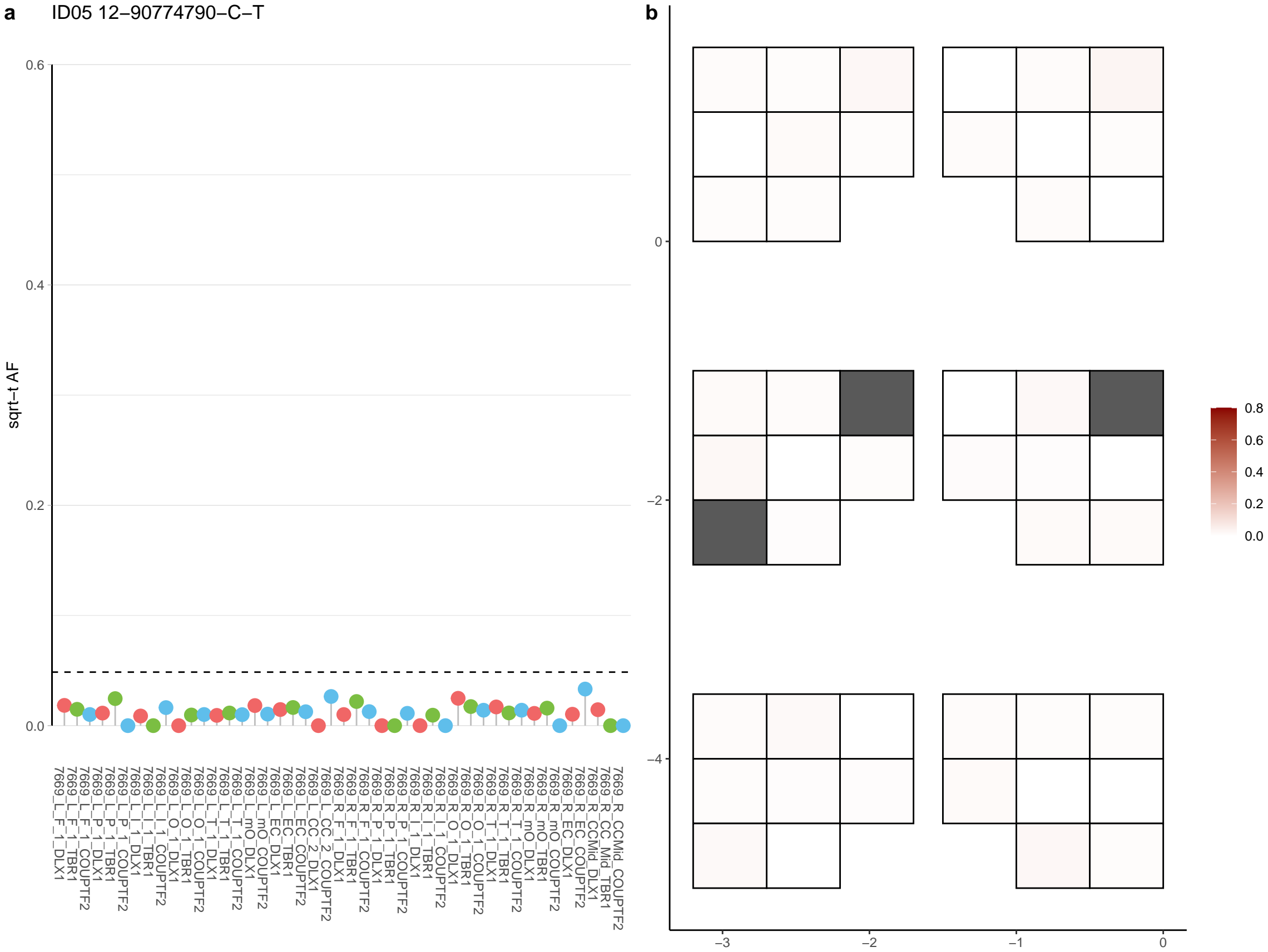

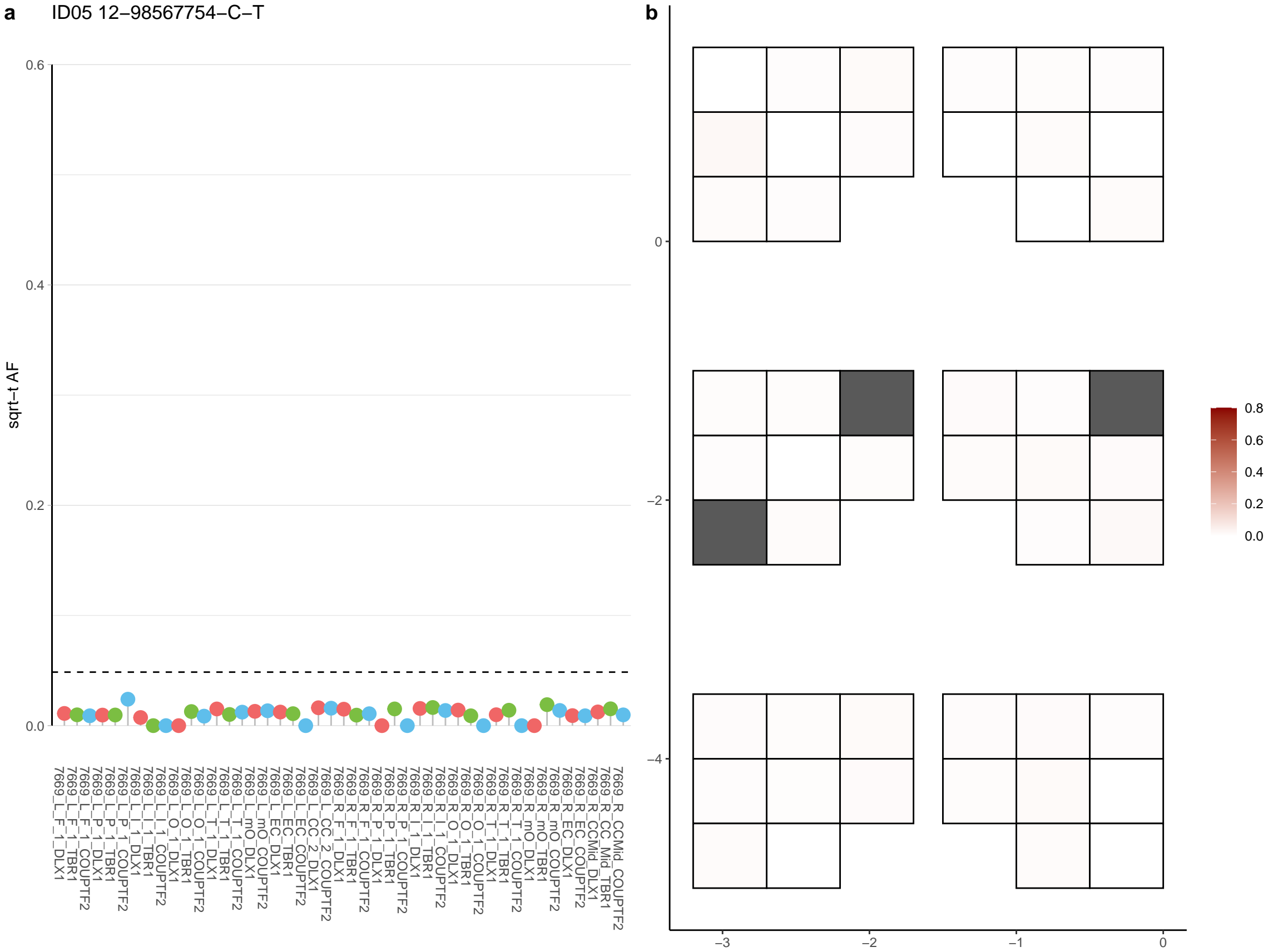

**a** ID05 12-98674630-C-T

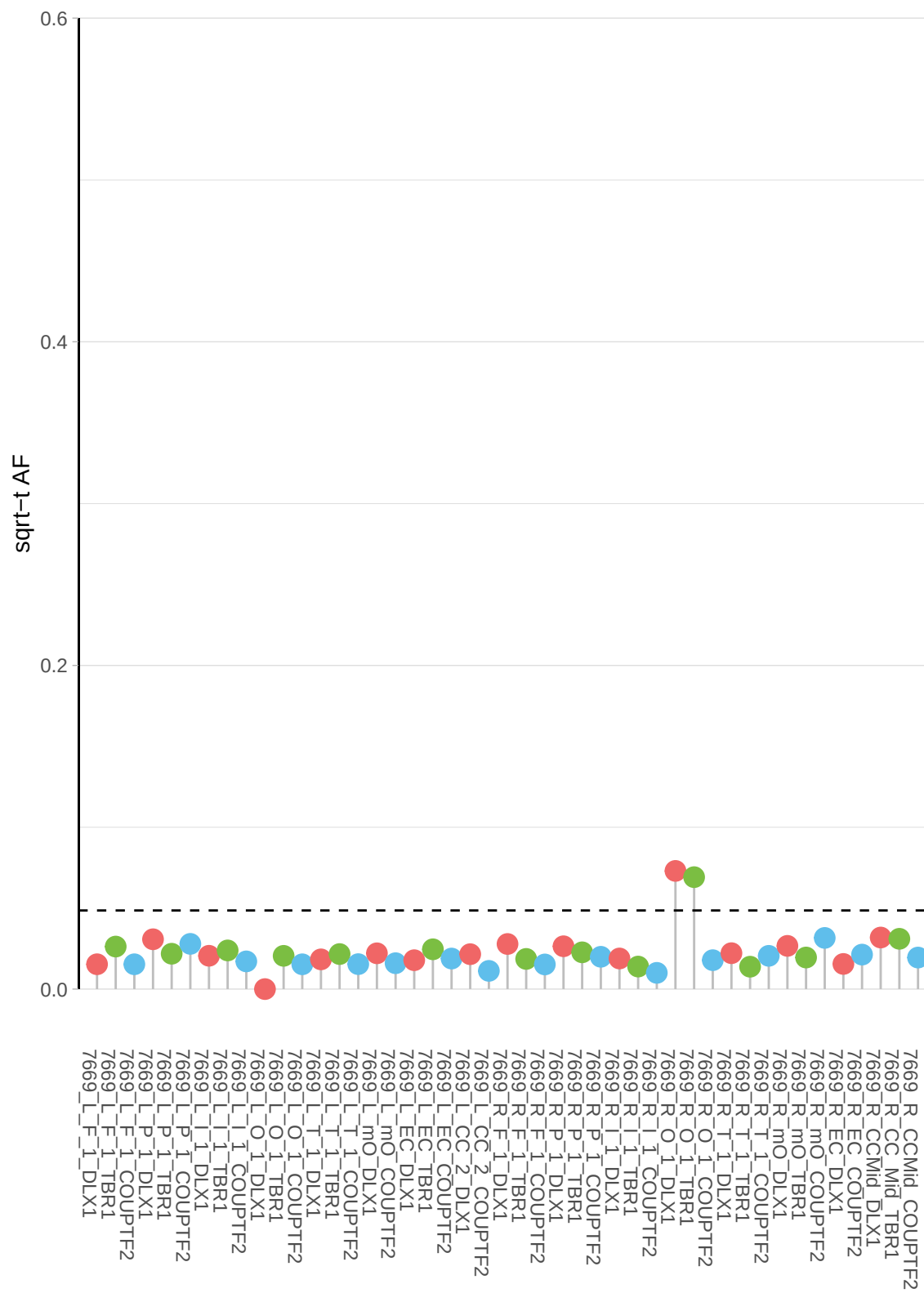

**b**

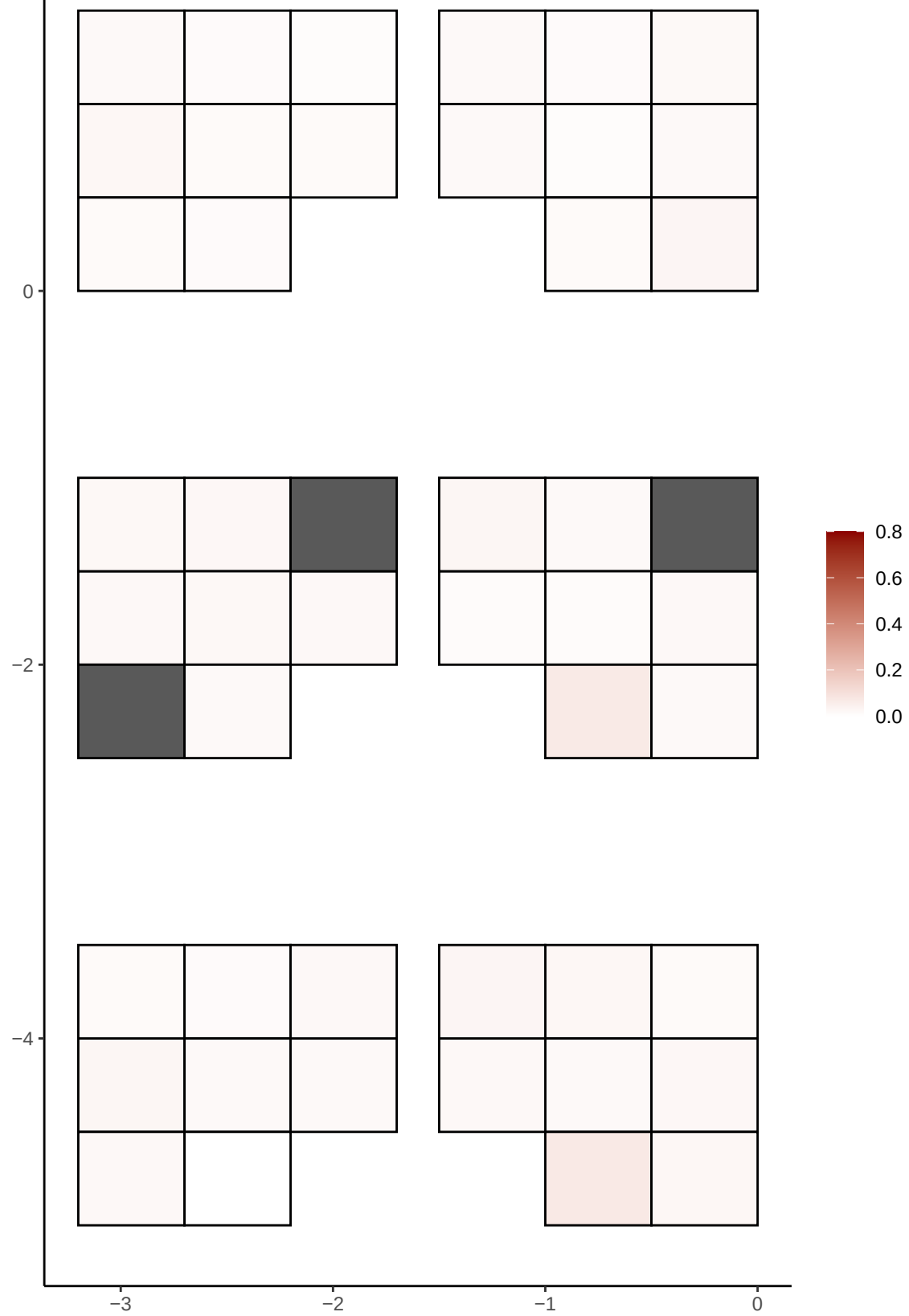

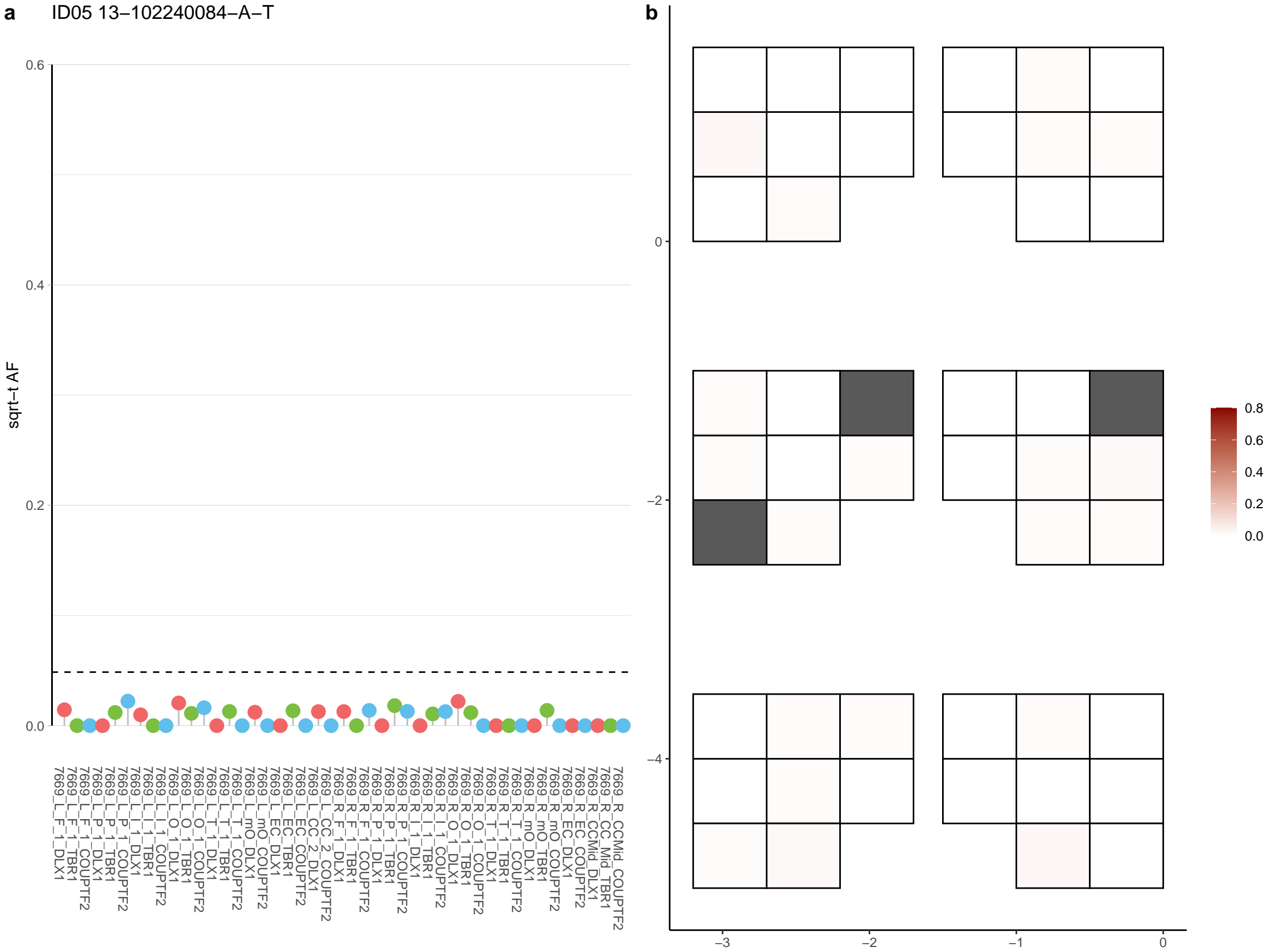

**a** ID05 13-106141548-C-T

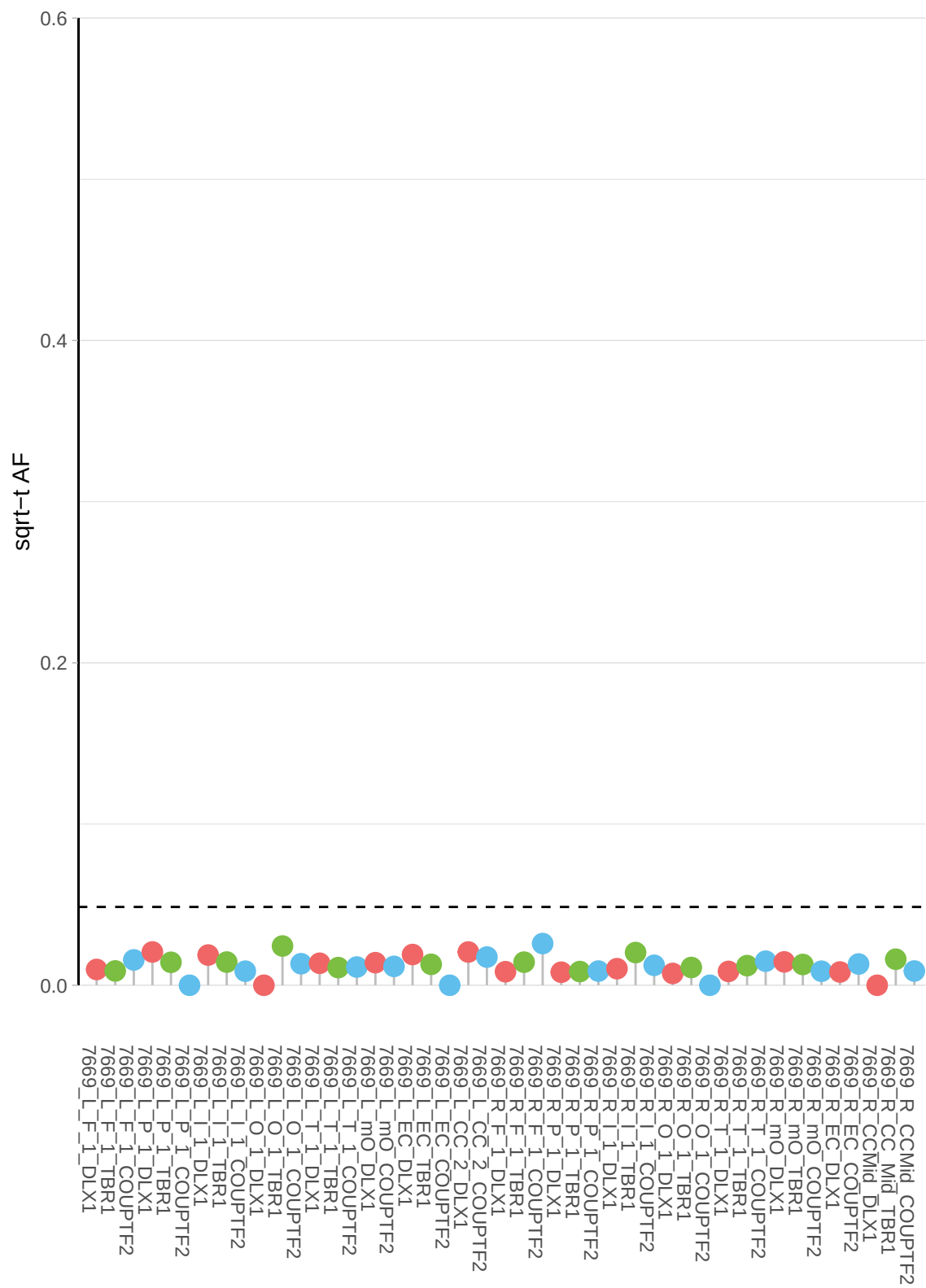

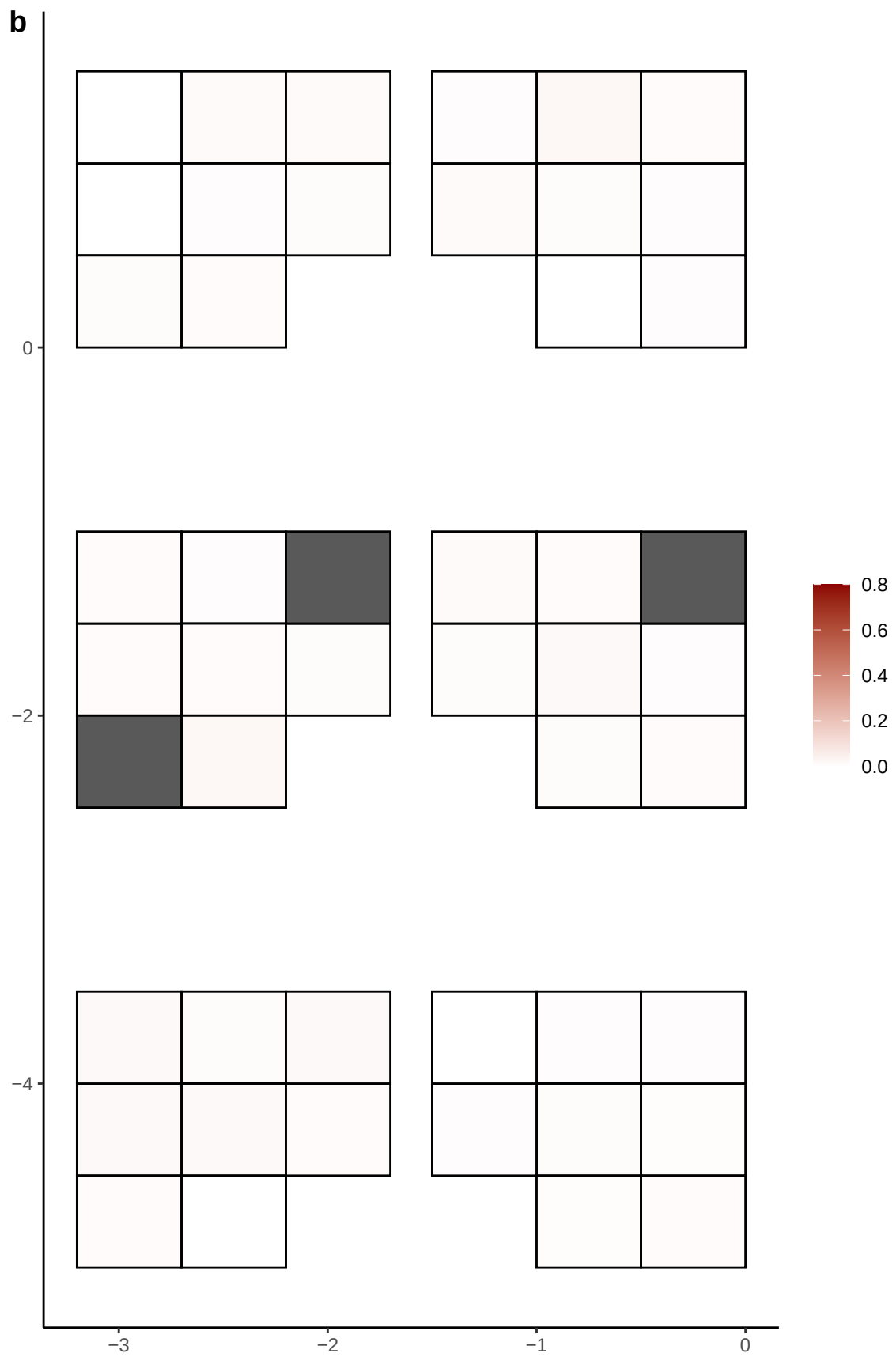

**a** ID05 13-109148397-G-A

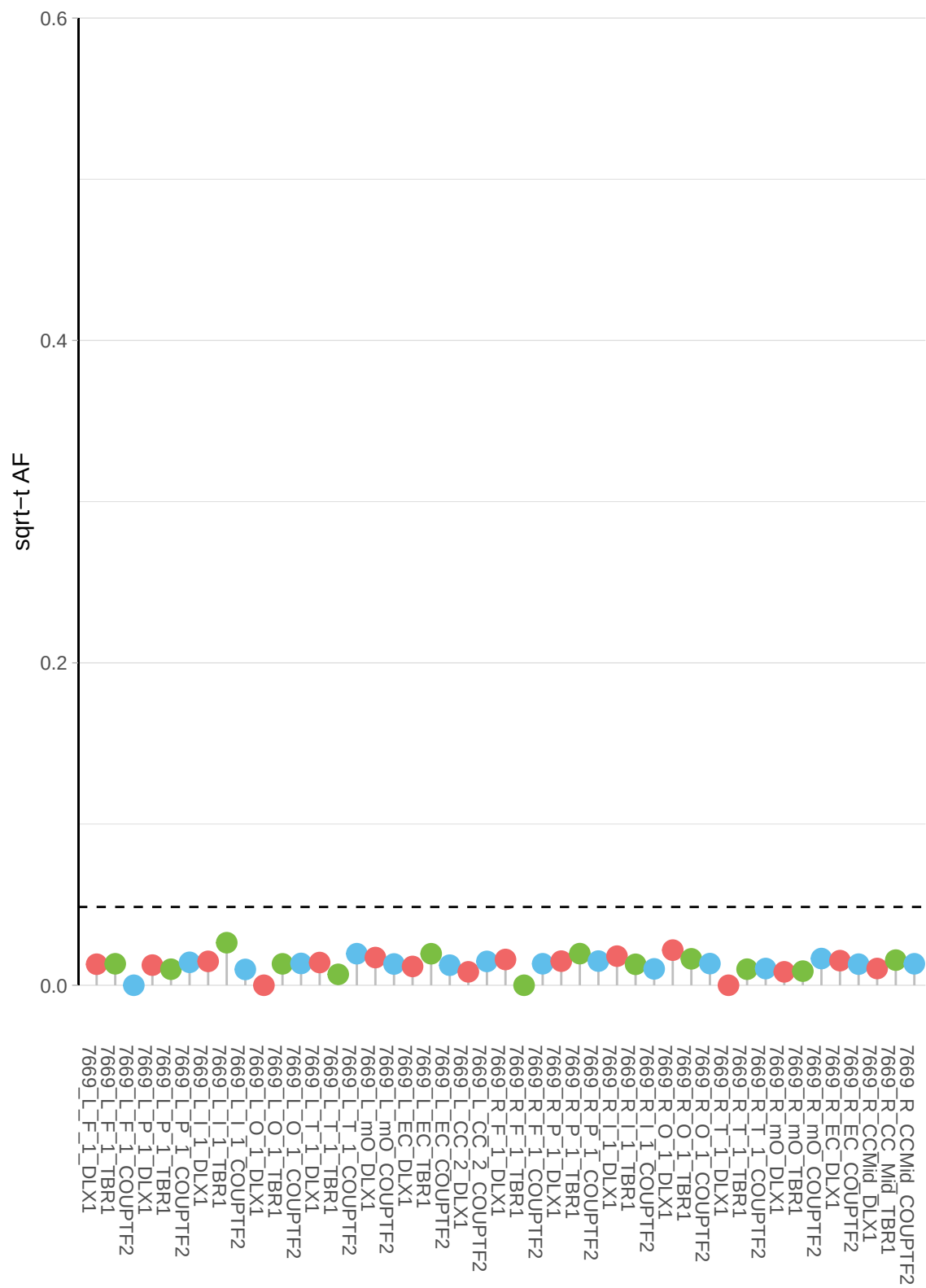

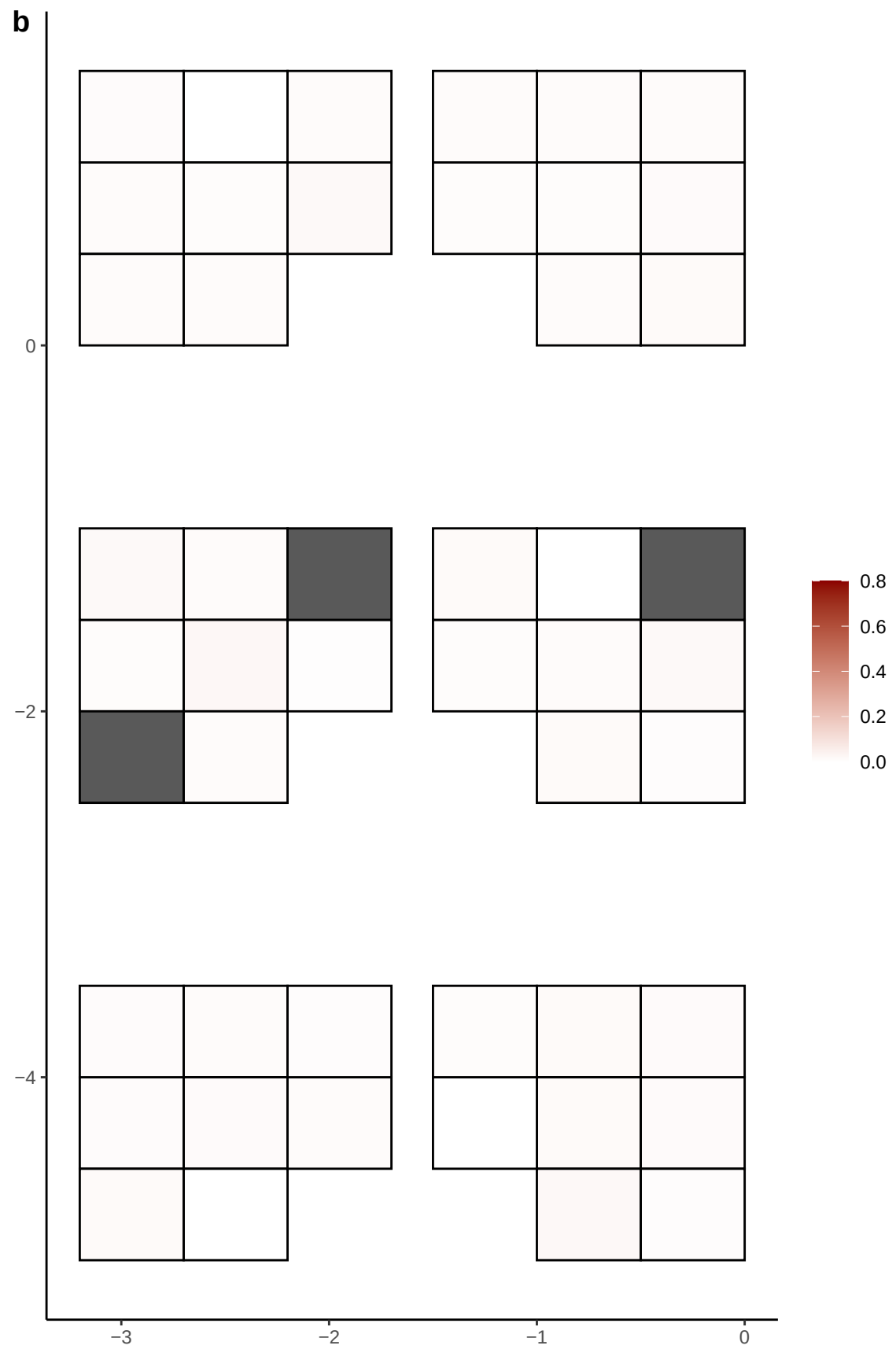

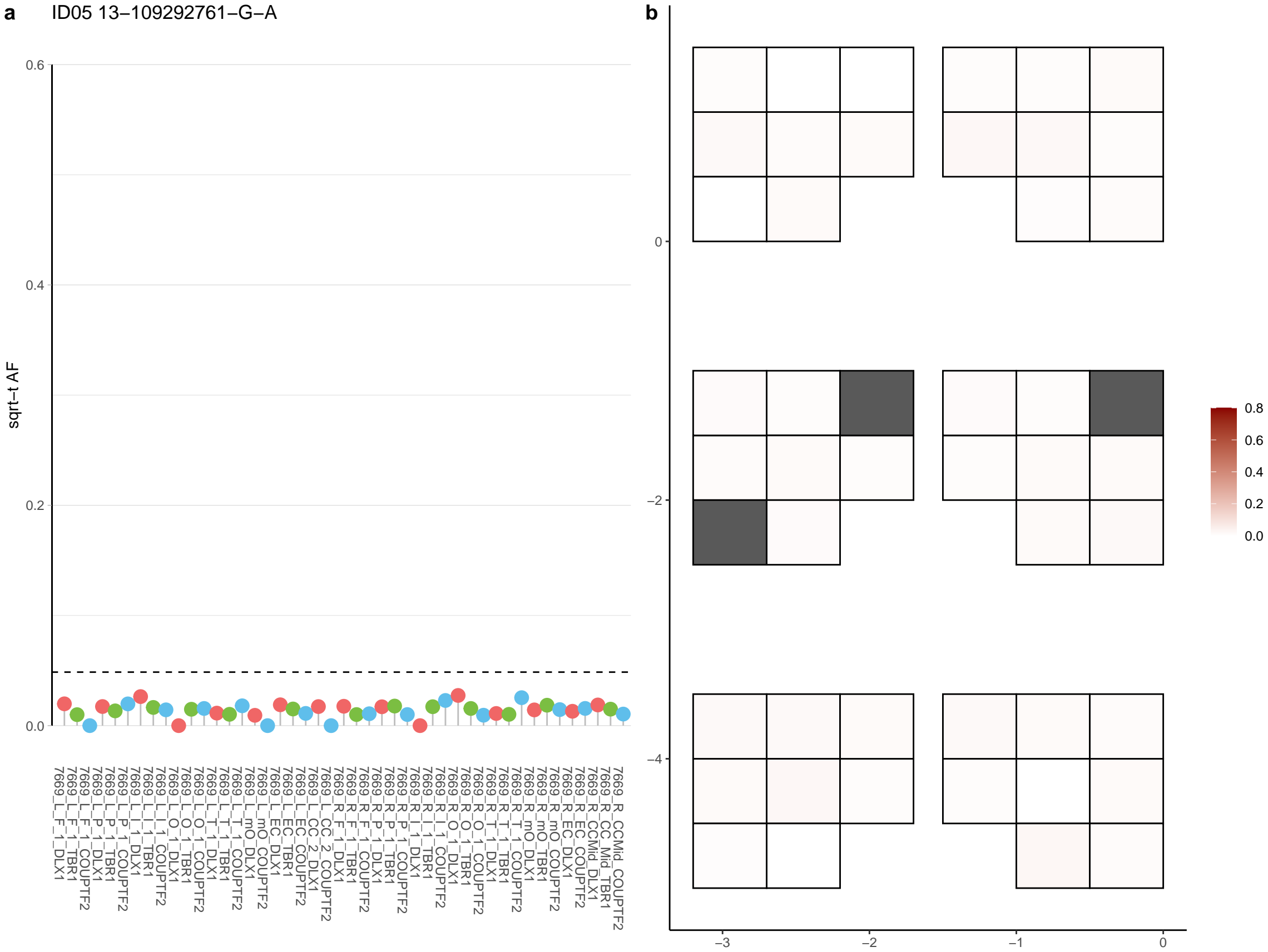

**a** ID05 13-111450669-C-T

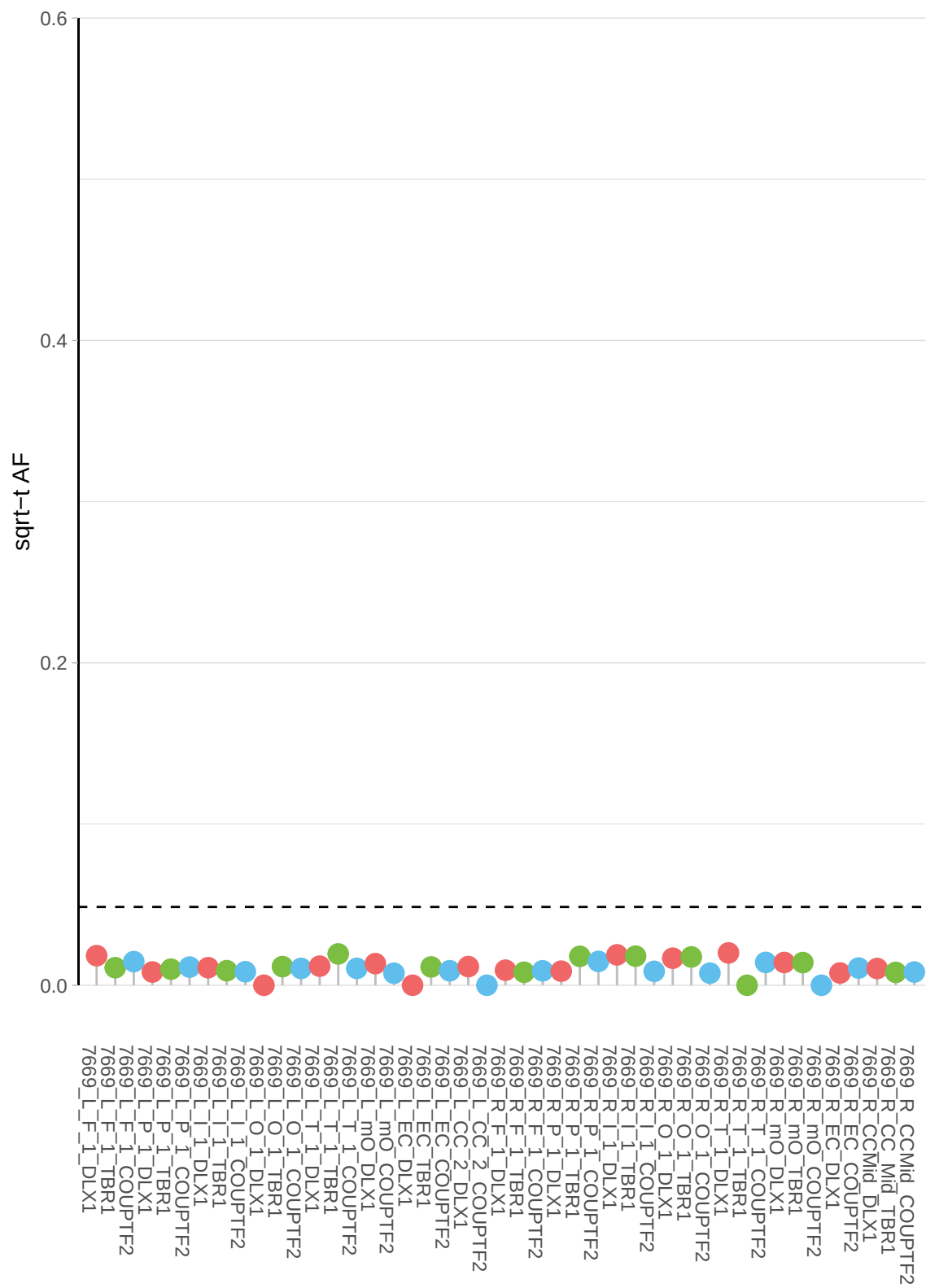

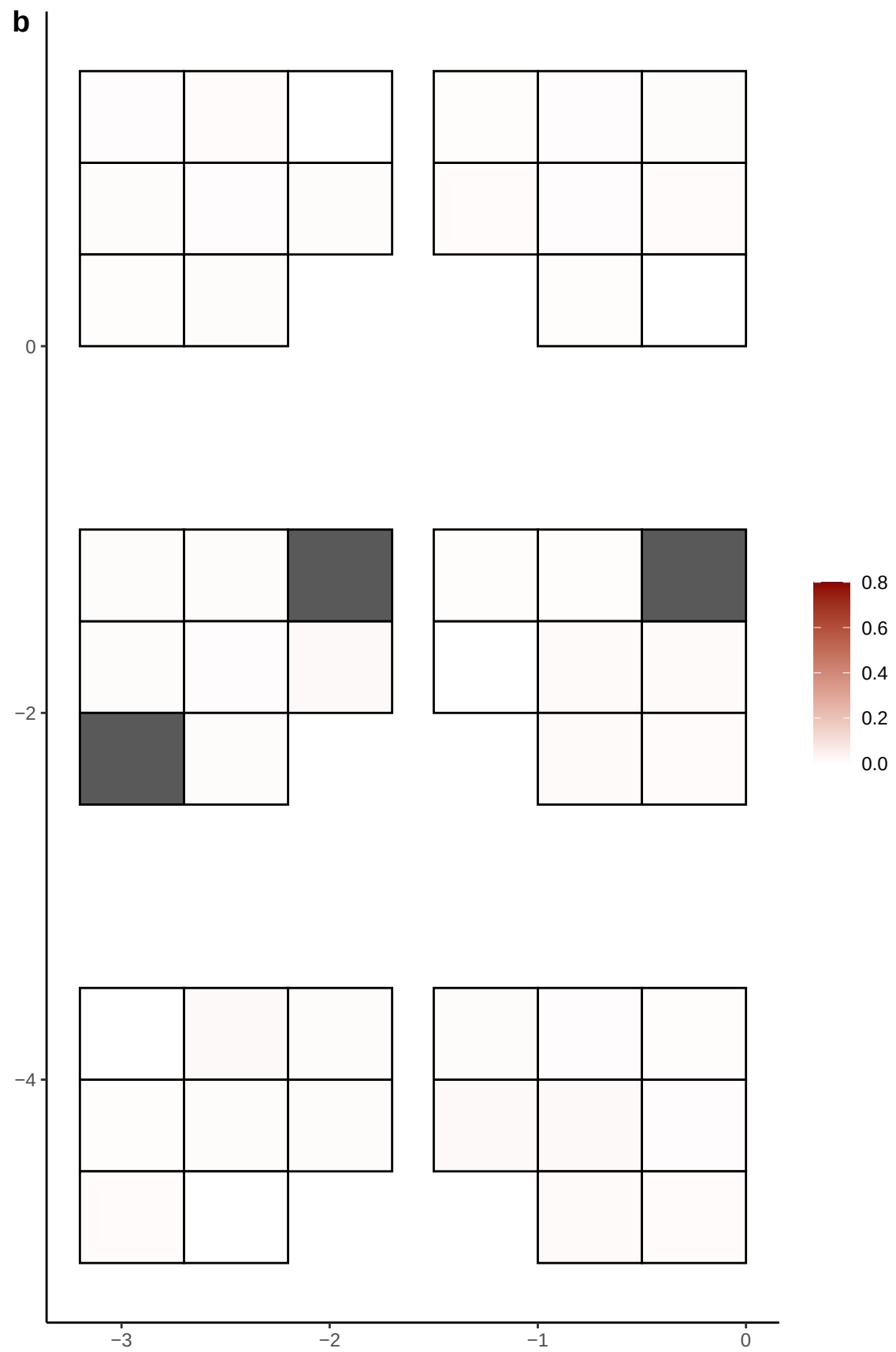

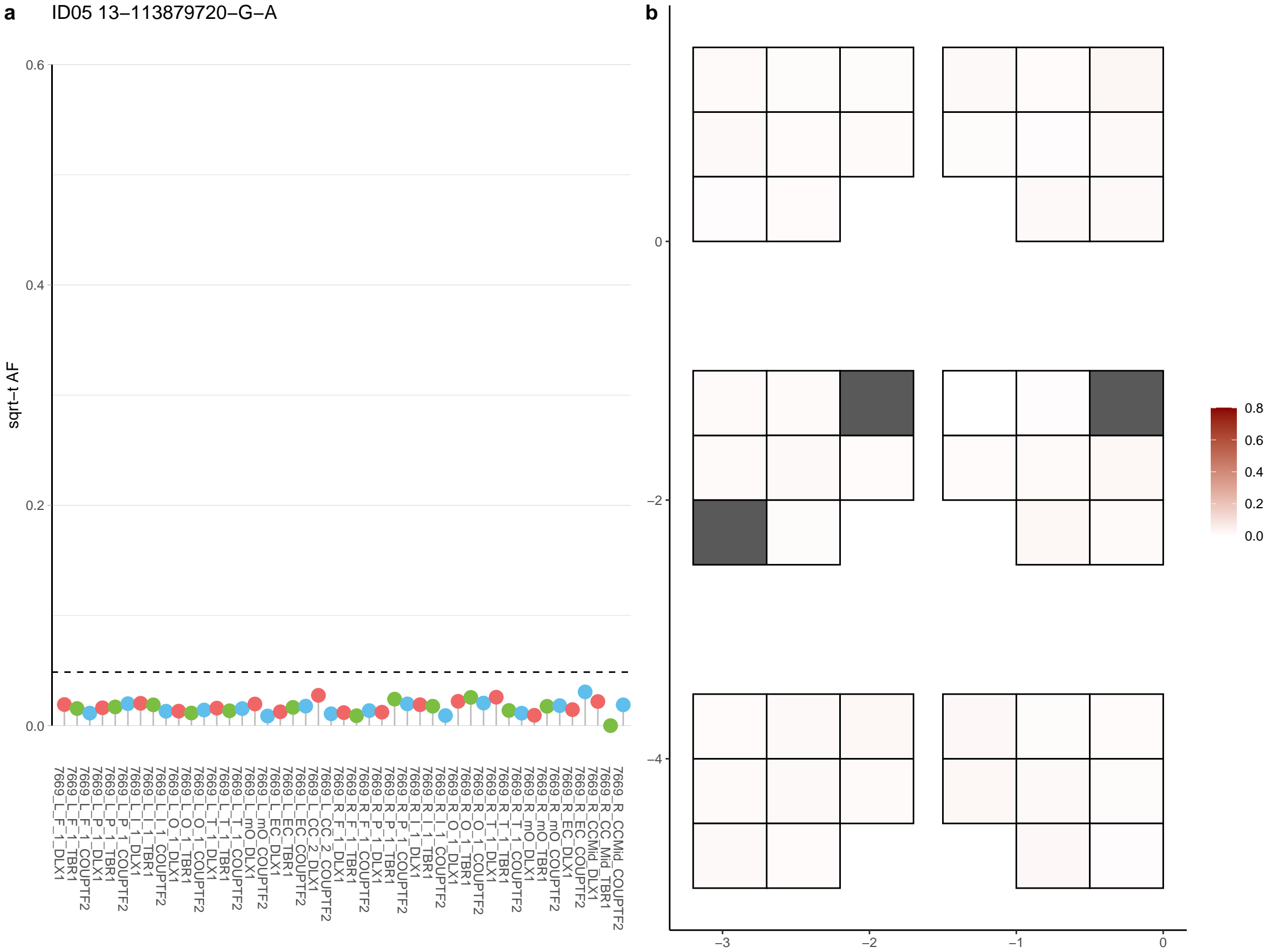

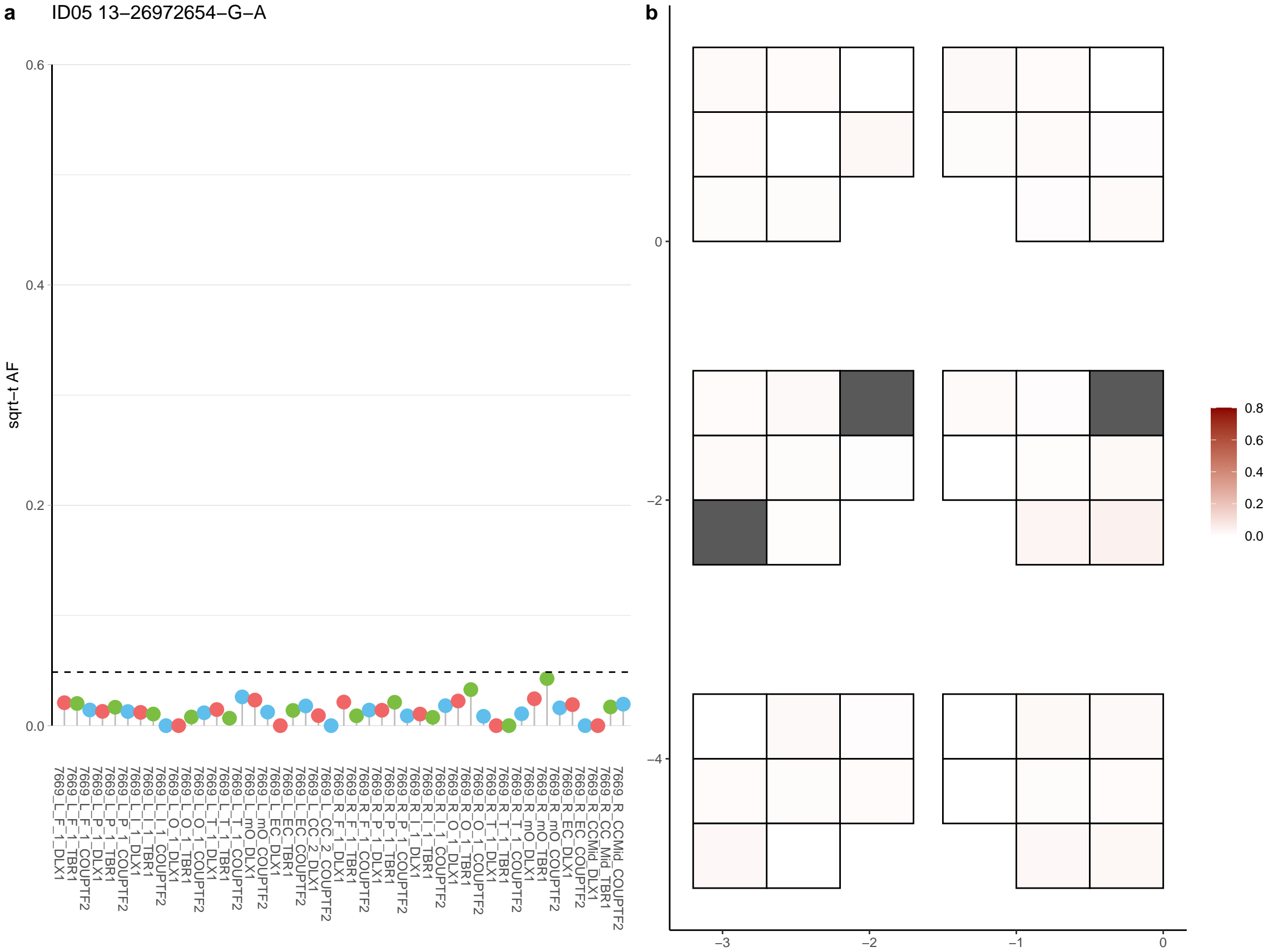

**a** ID05 13-26972655-G-A

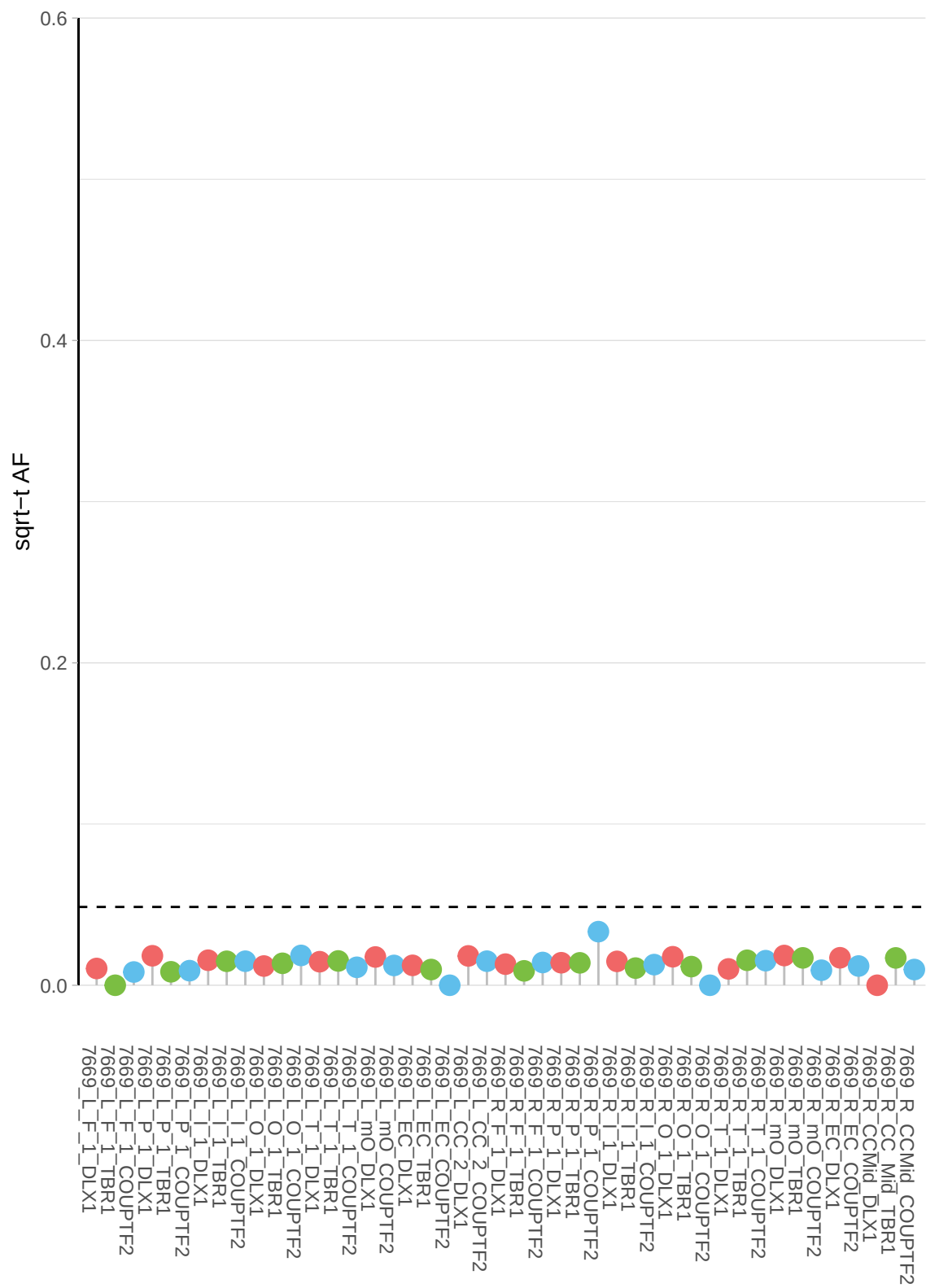

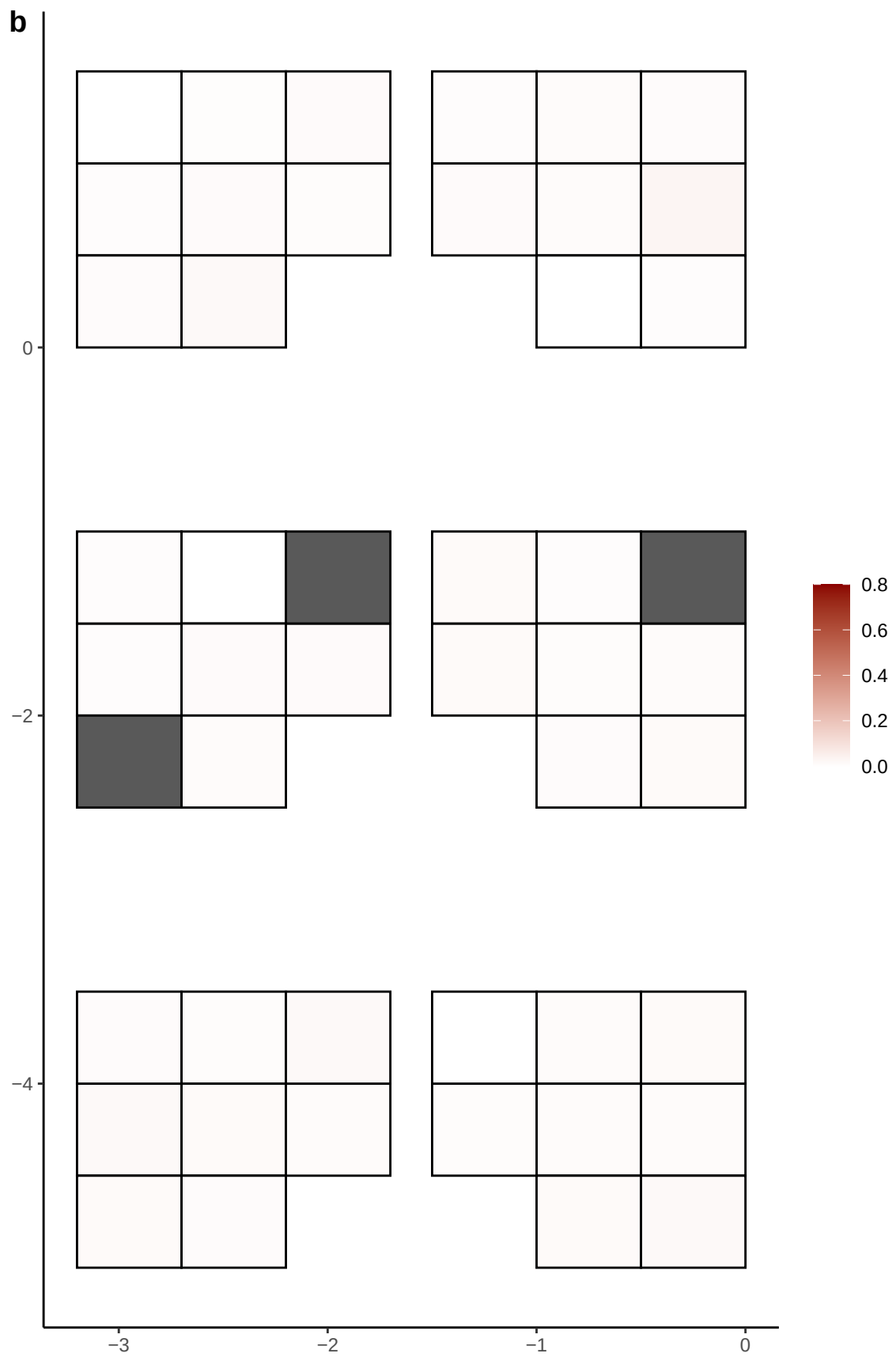

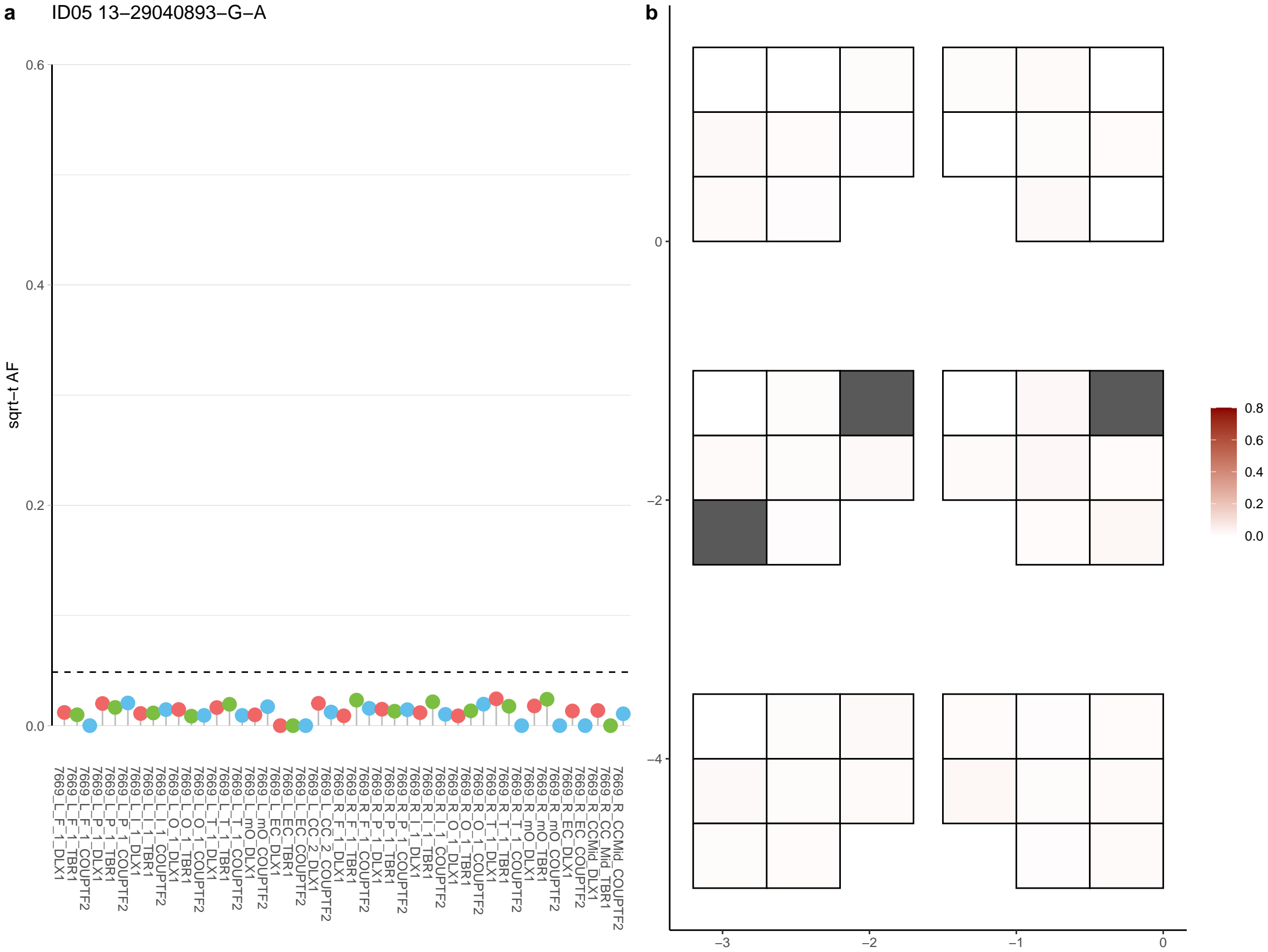

**a**

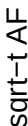

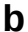

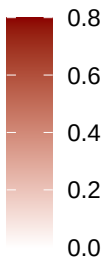

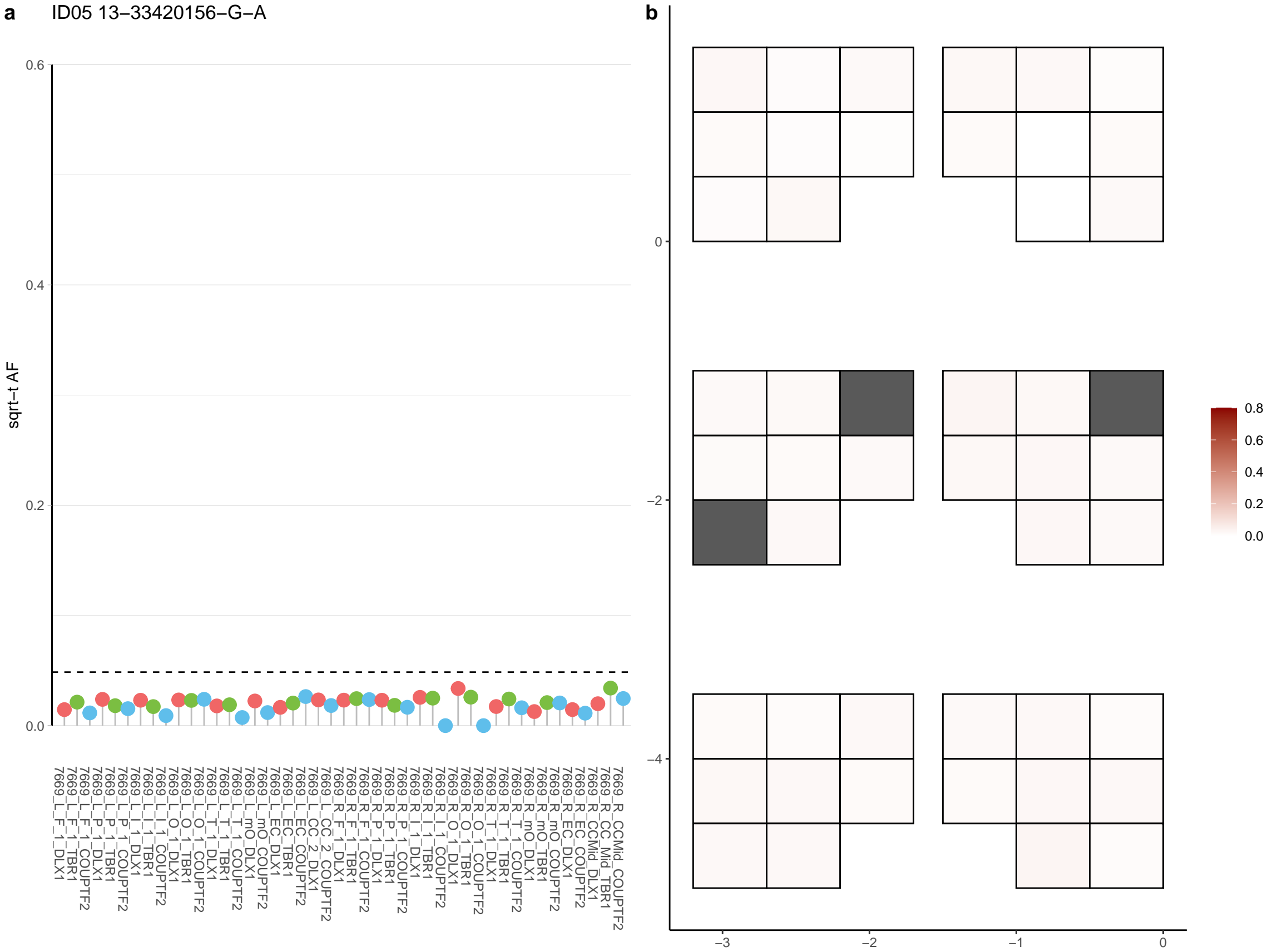

**a** ID05 13-41556737-G-C

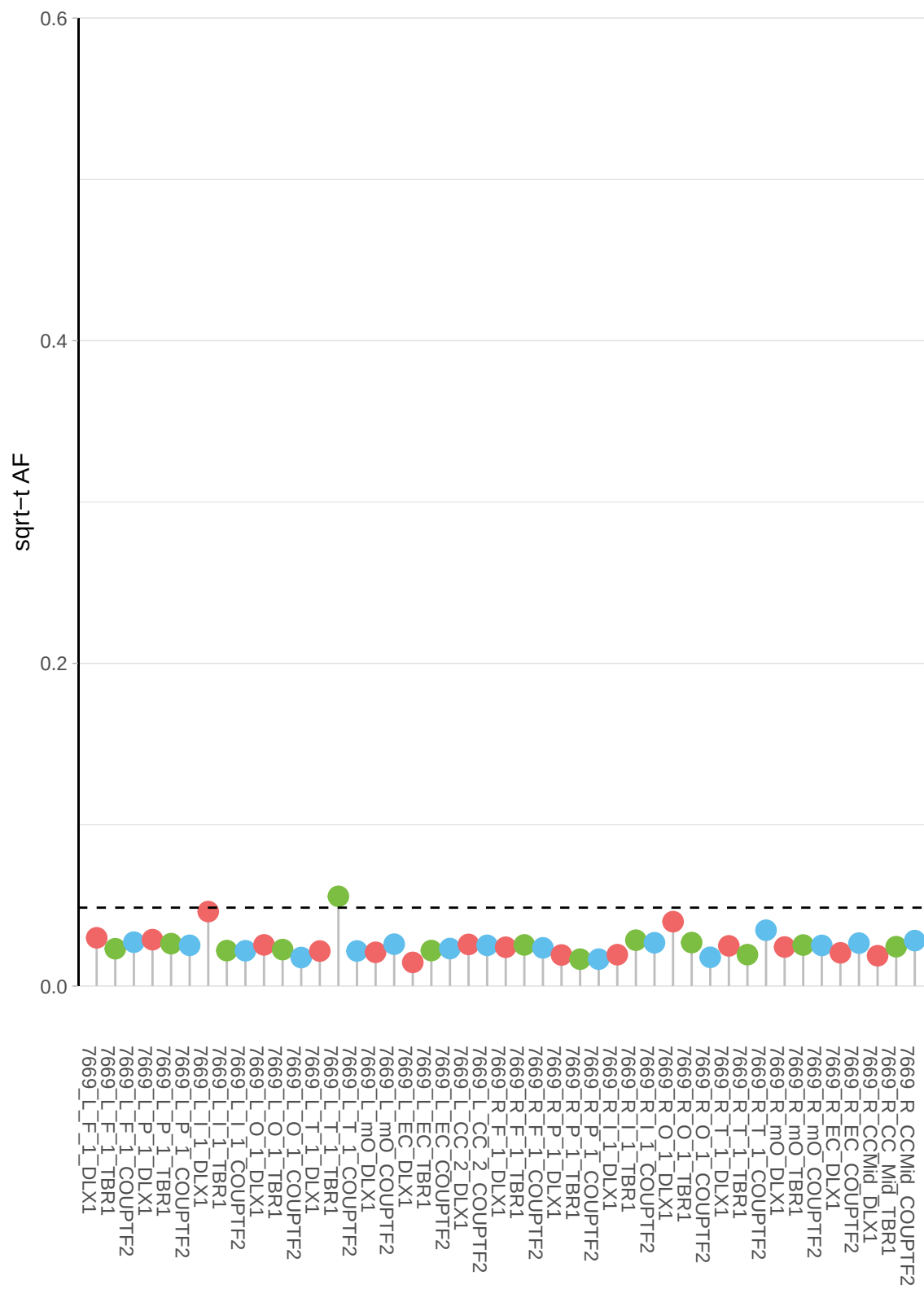

**b**

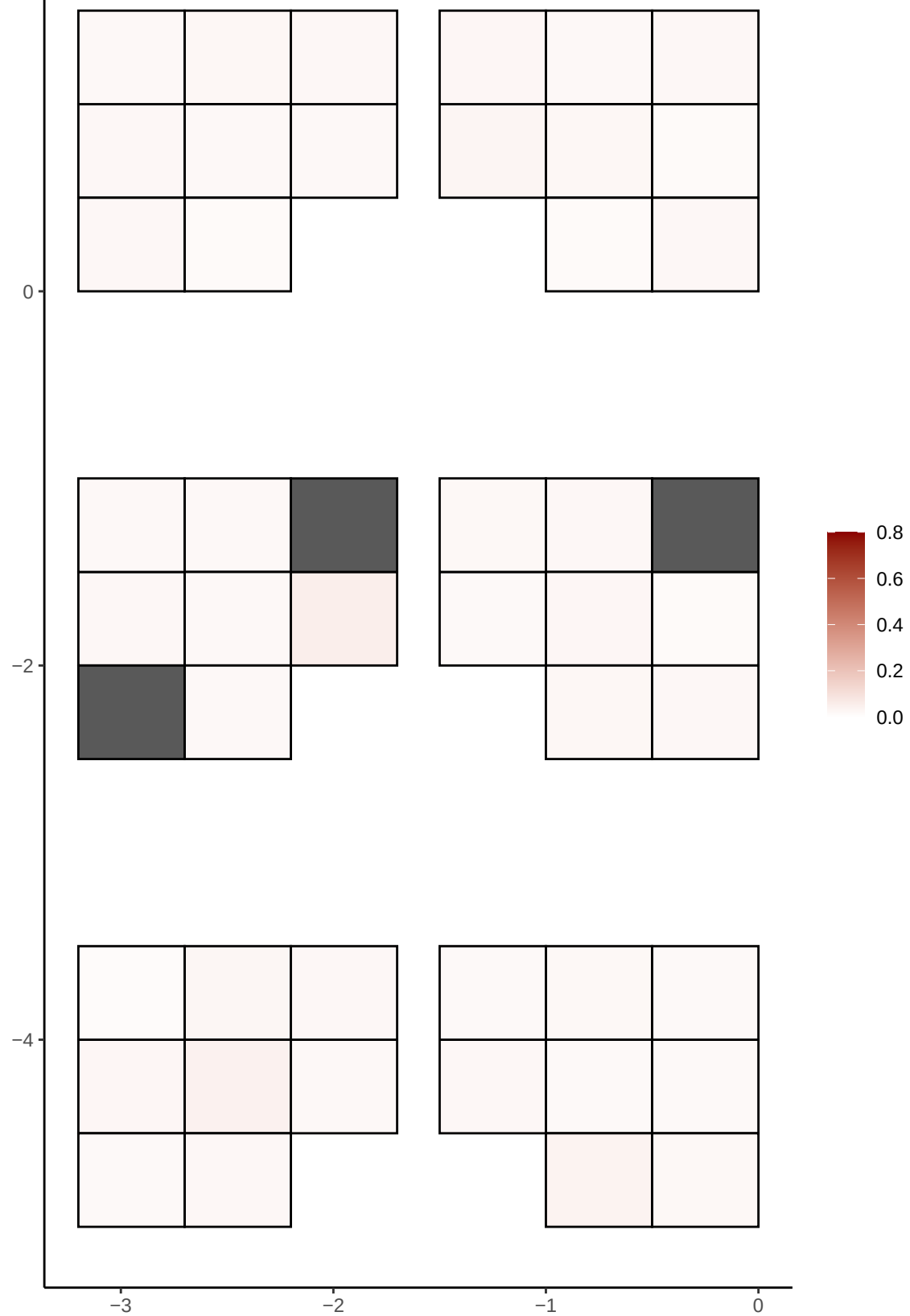

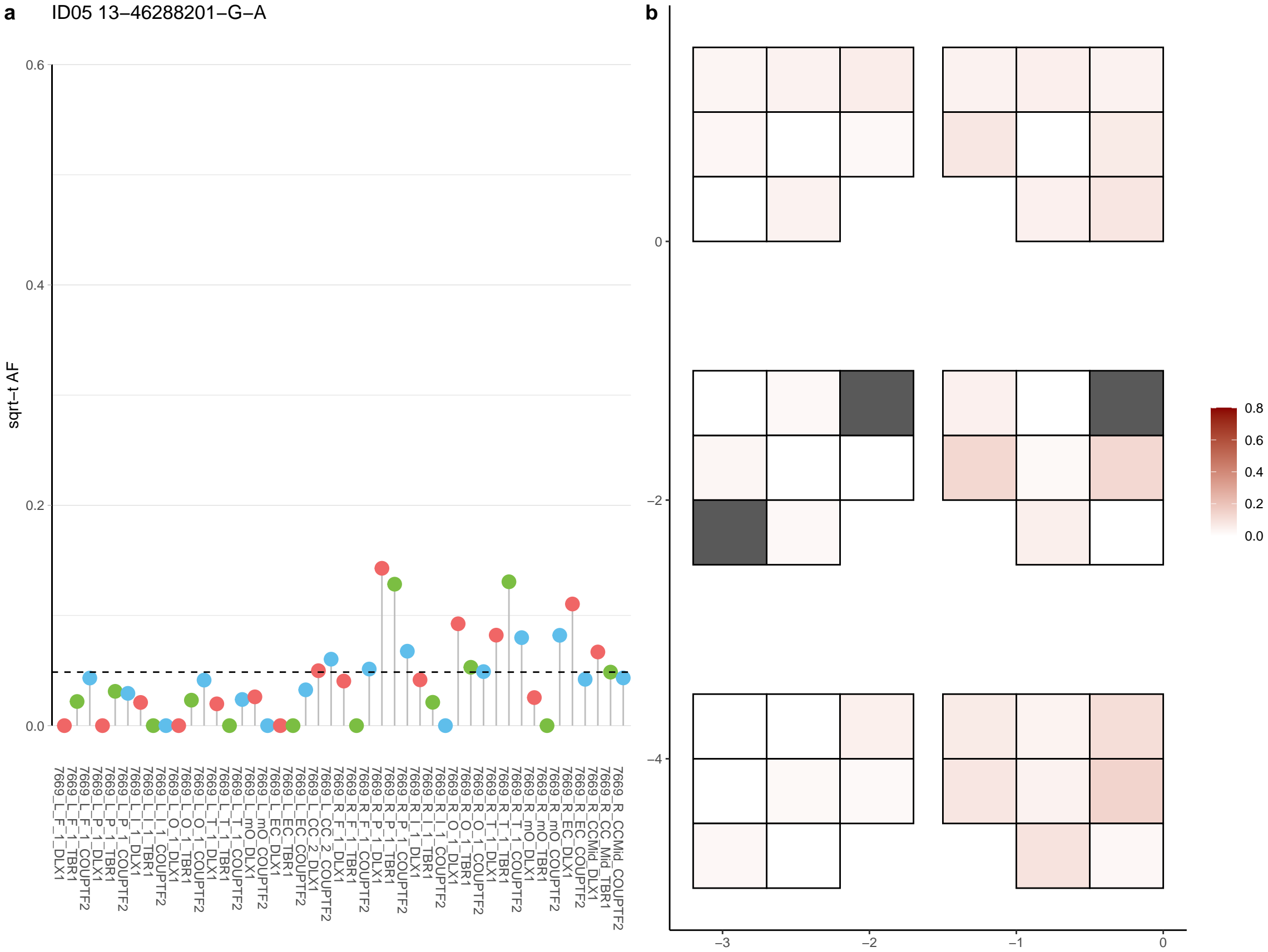

**a** ID05 13-55140512-C-T

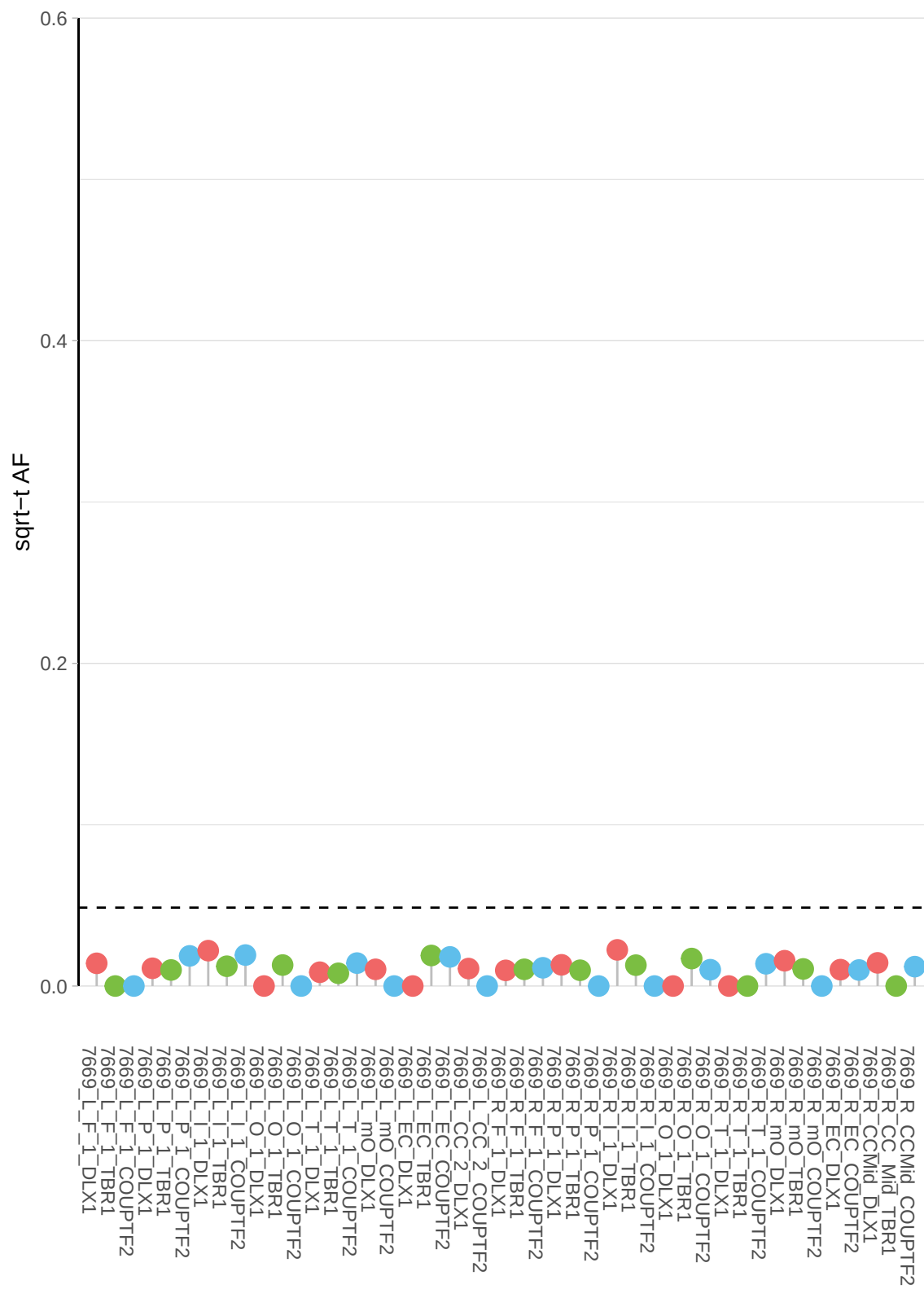

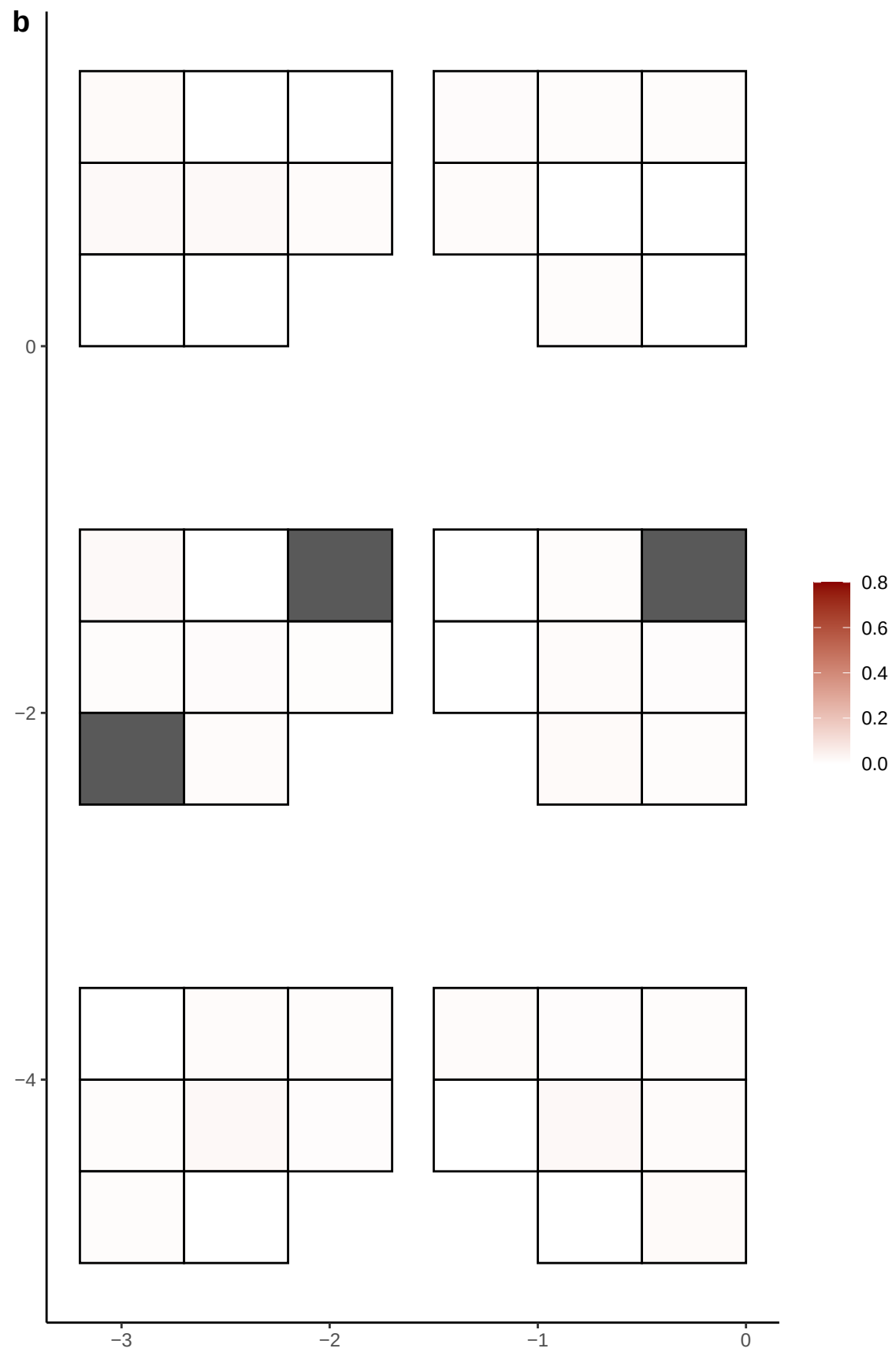

**a** ID05 13-56431067-T-A

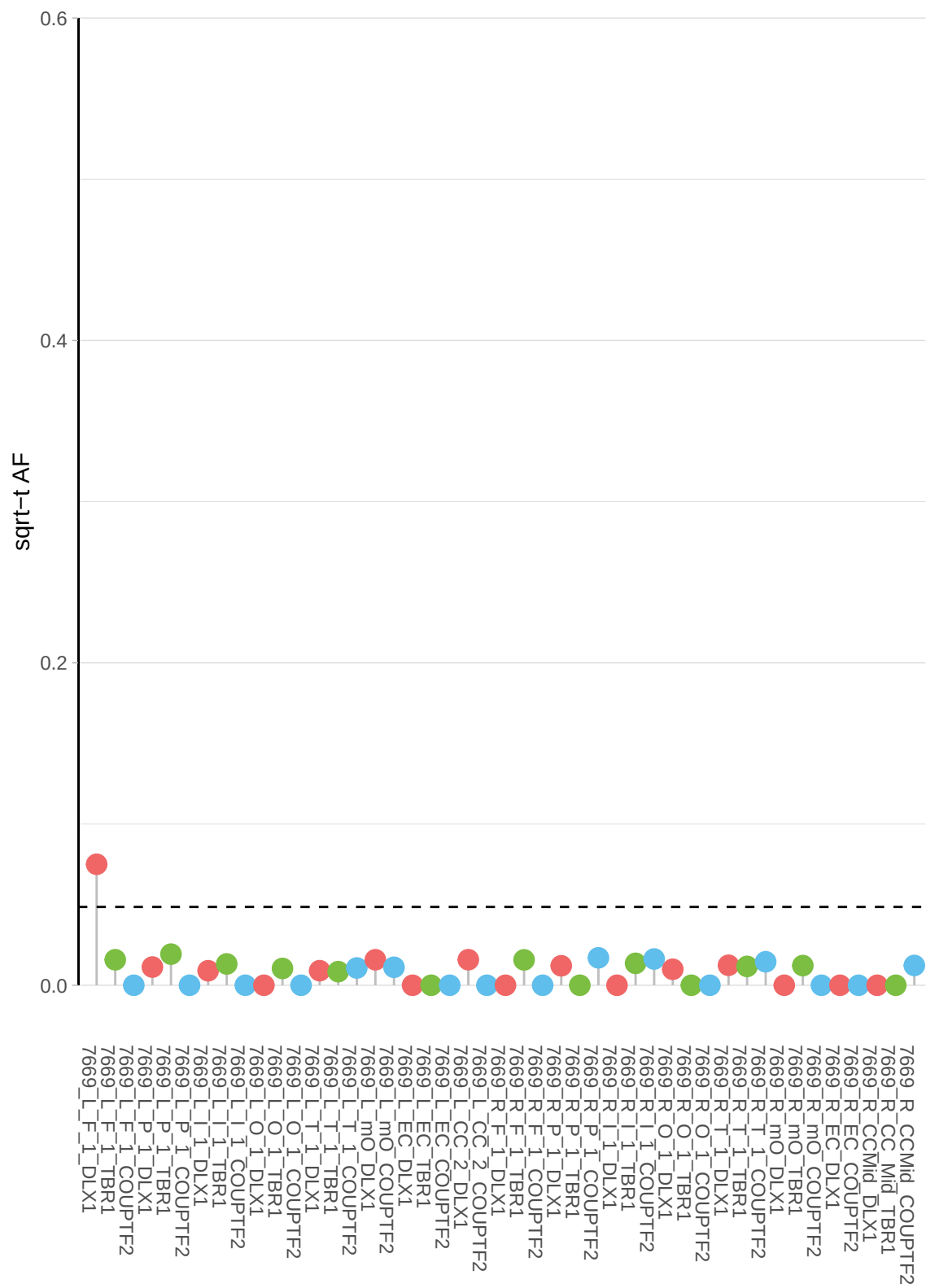

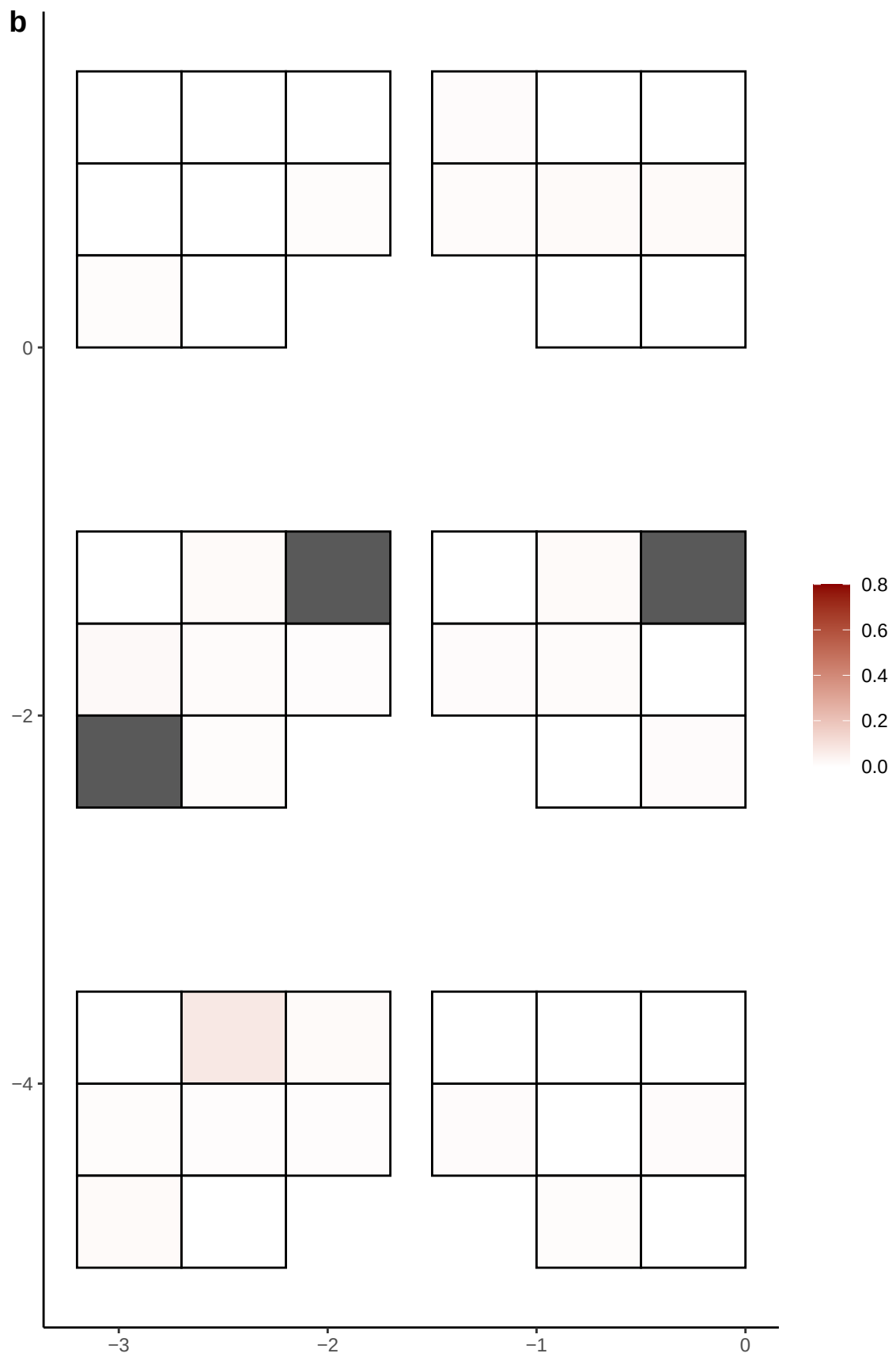

**a** ID05 13-59159499-G-A

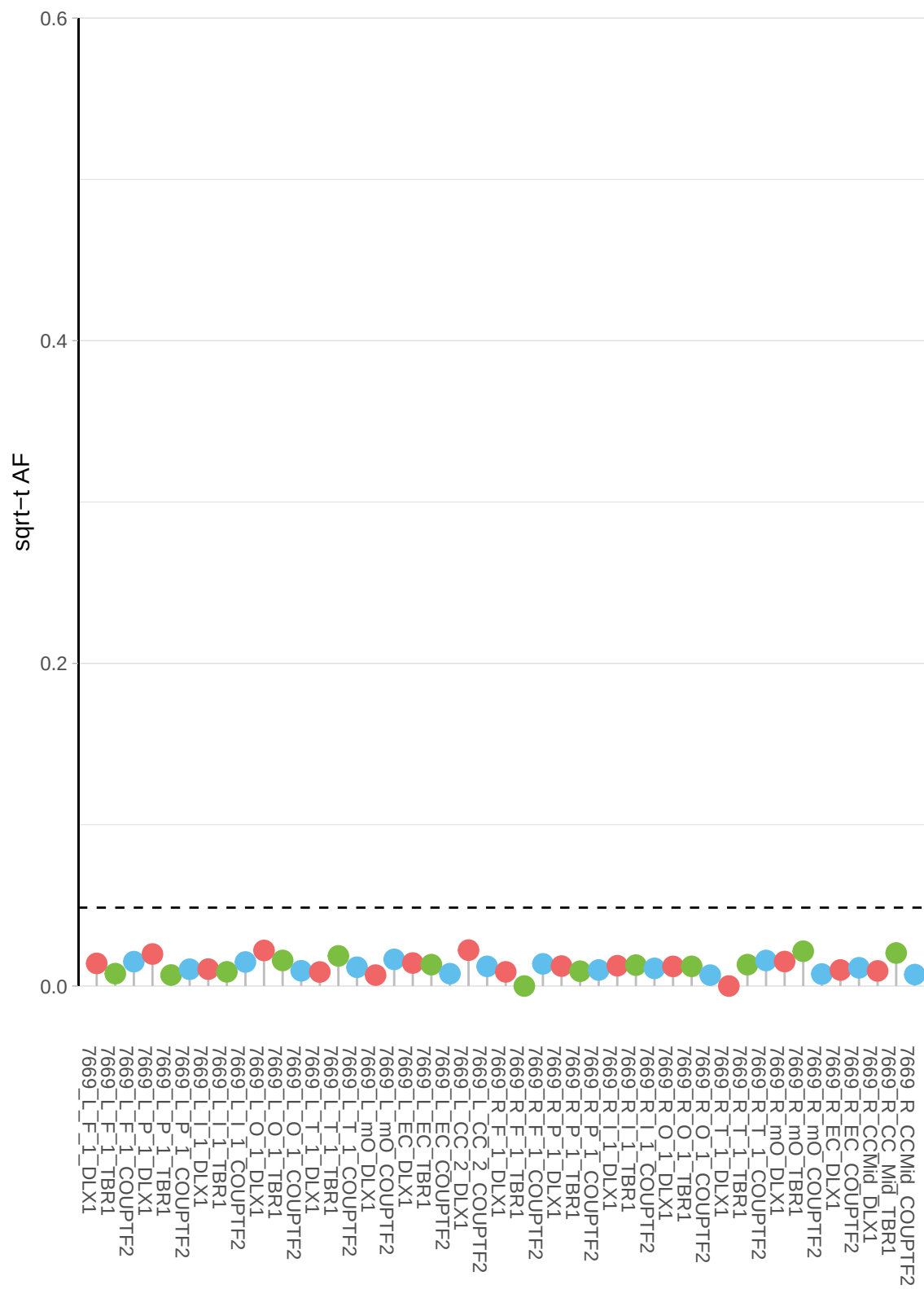

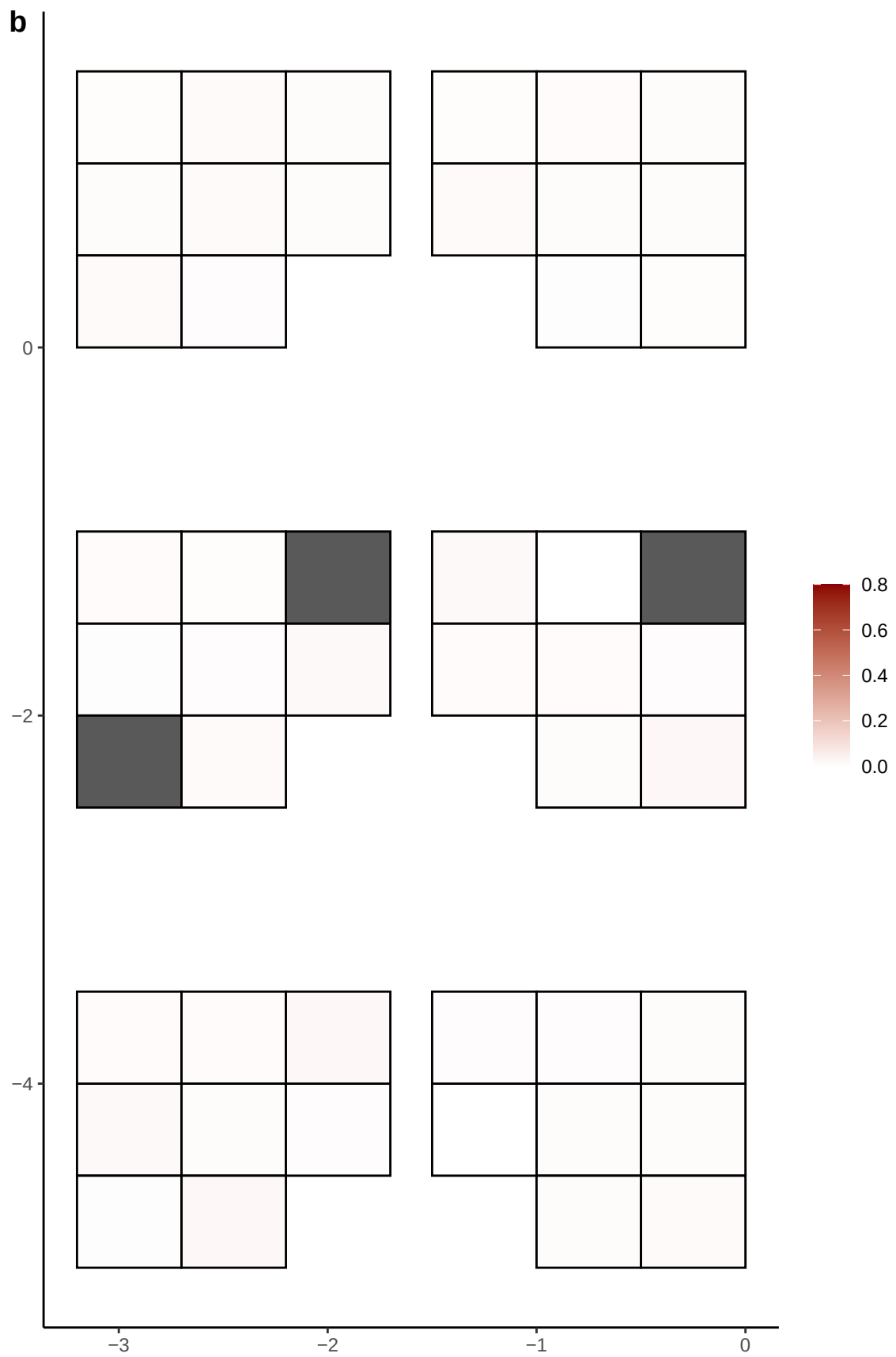

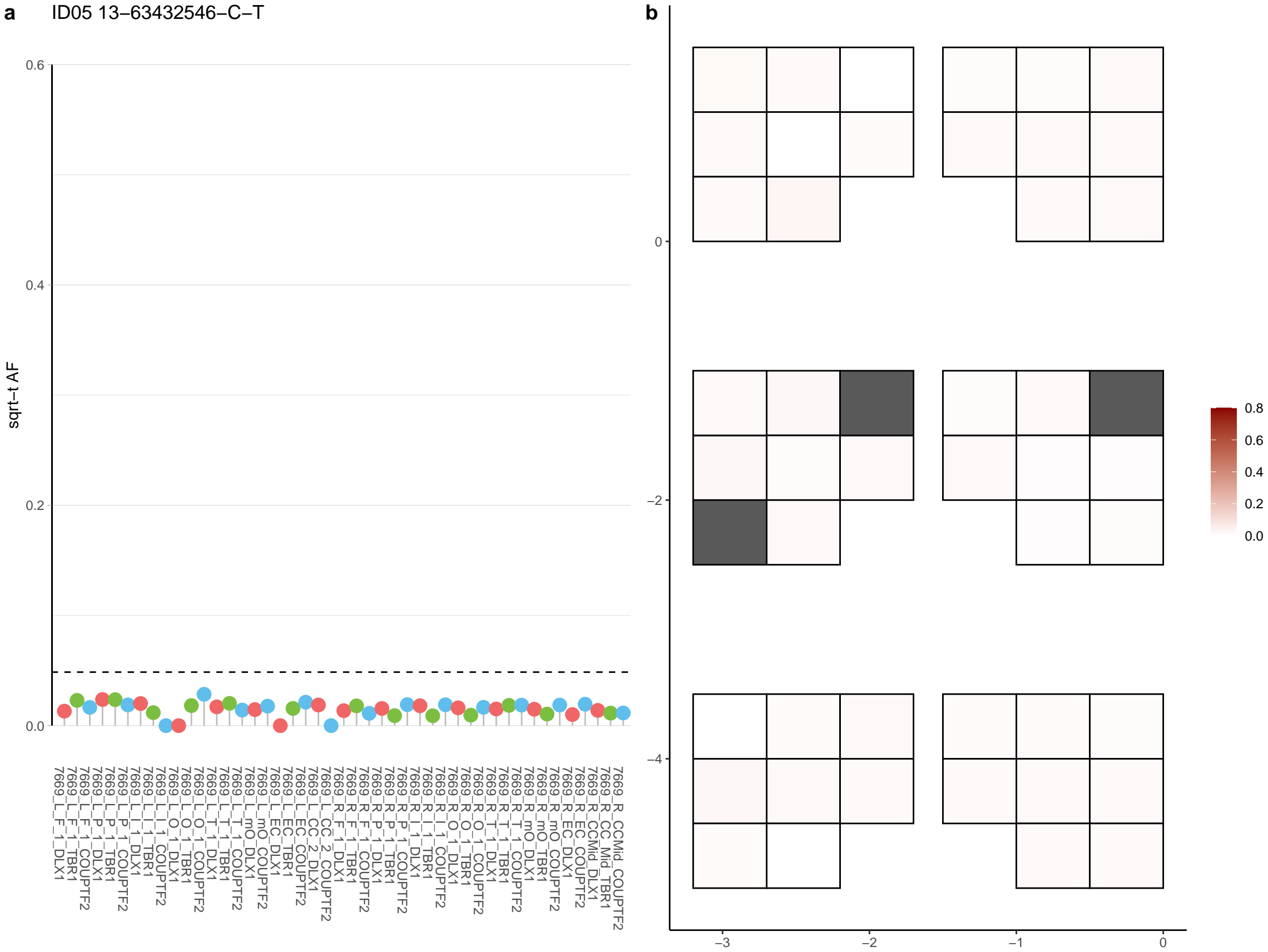

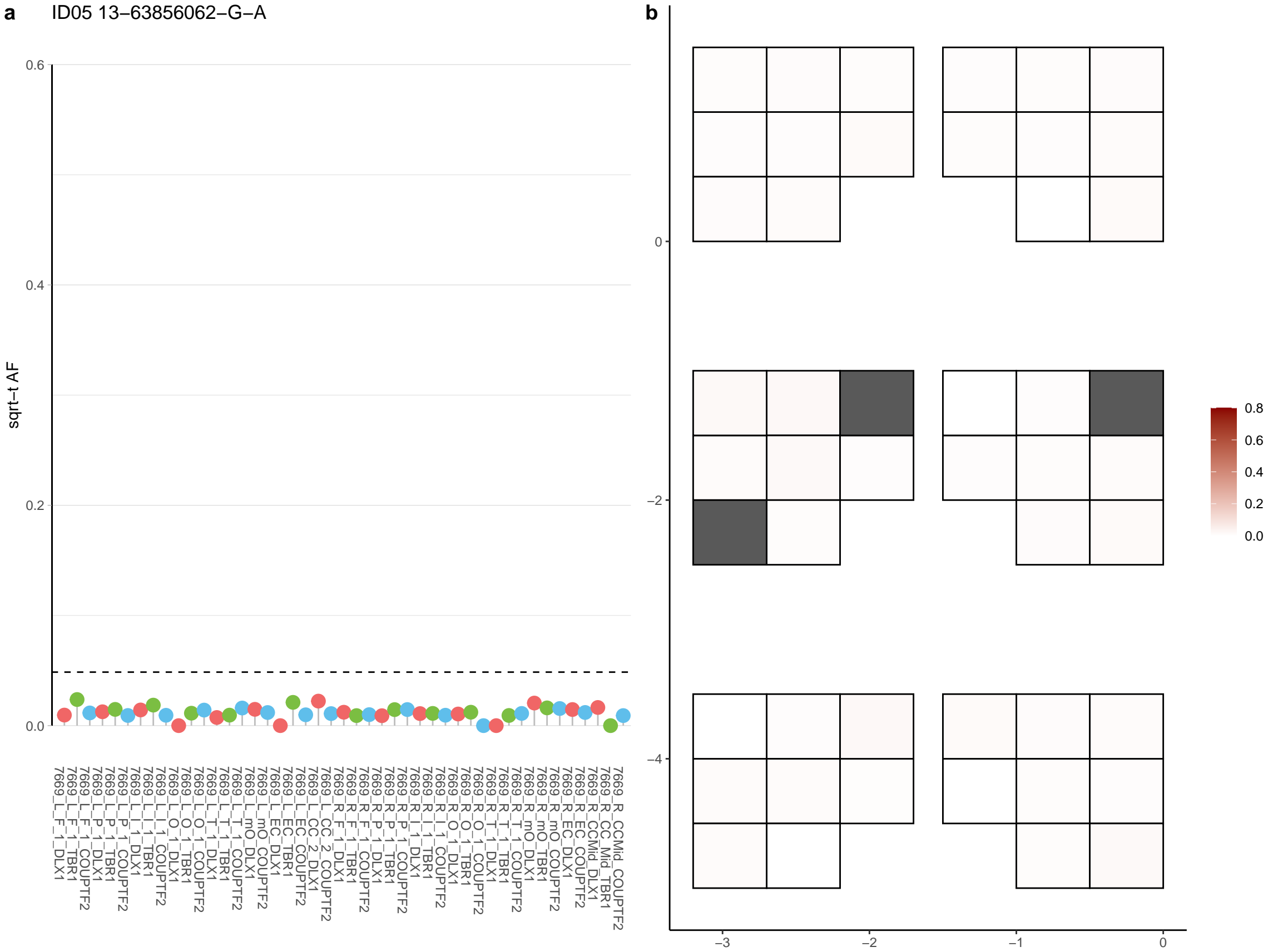

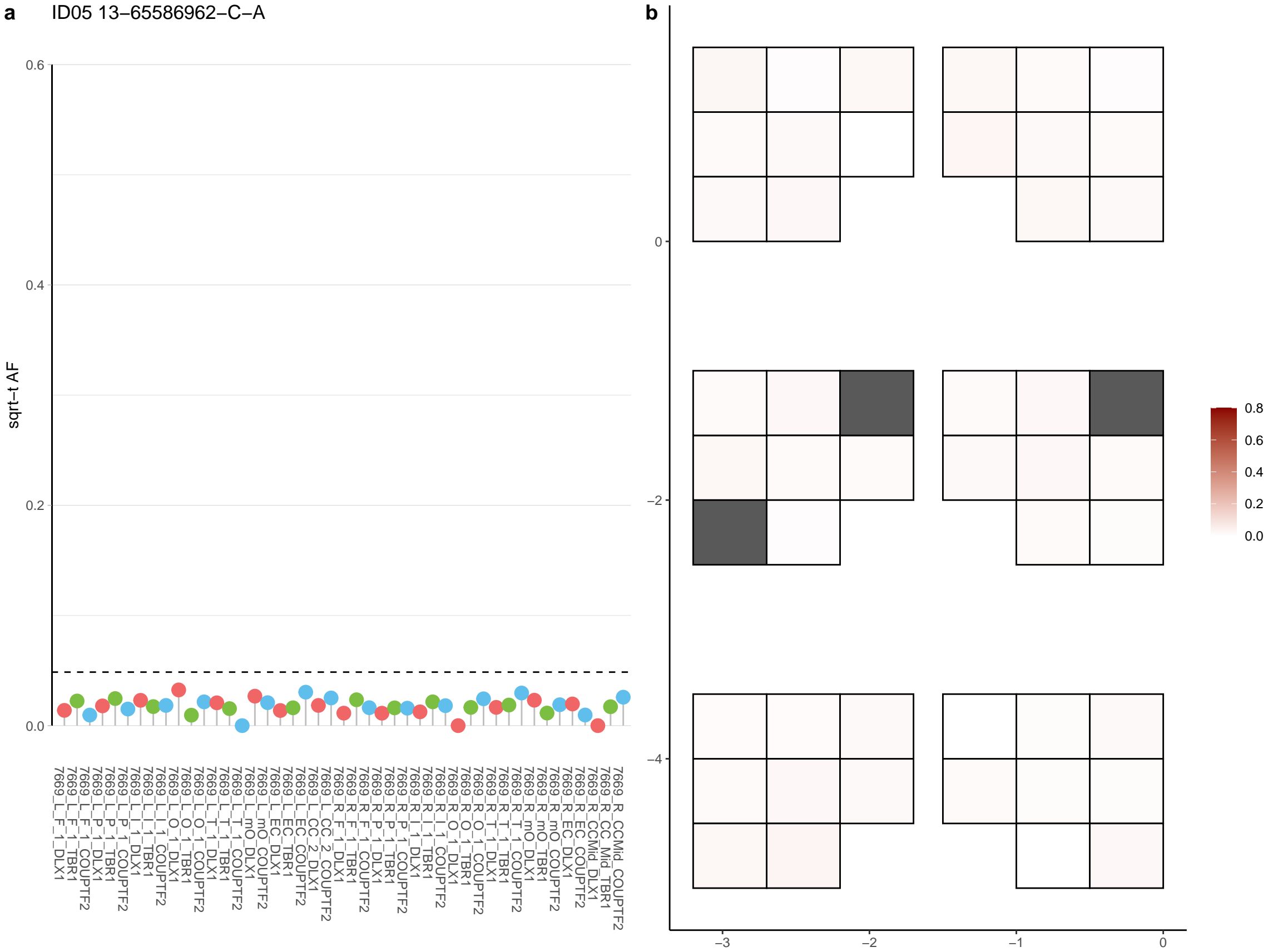

**a** ID05 13-67322807-C-A

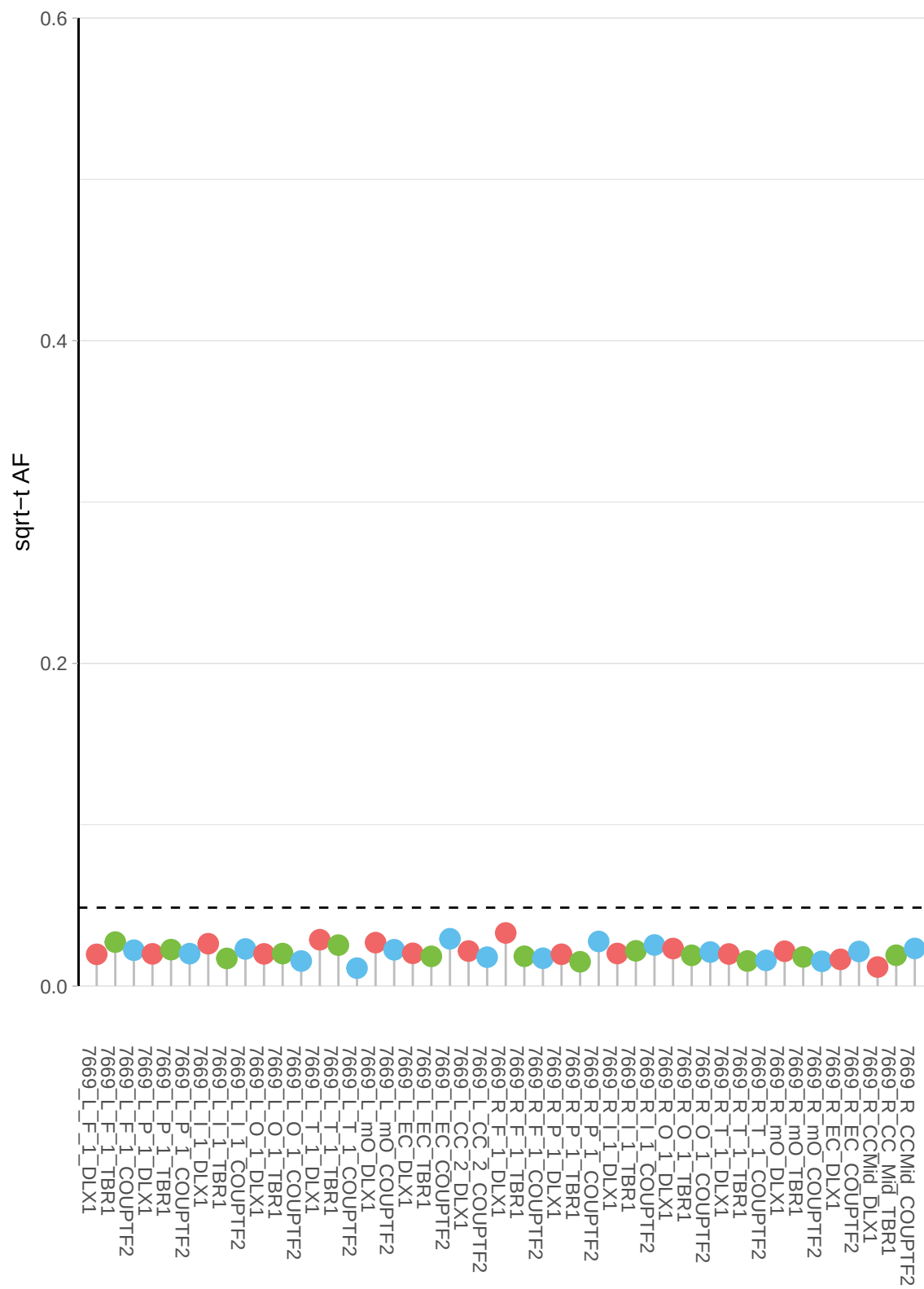

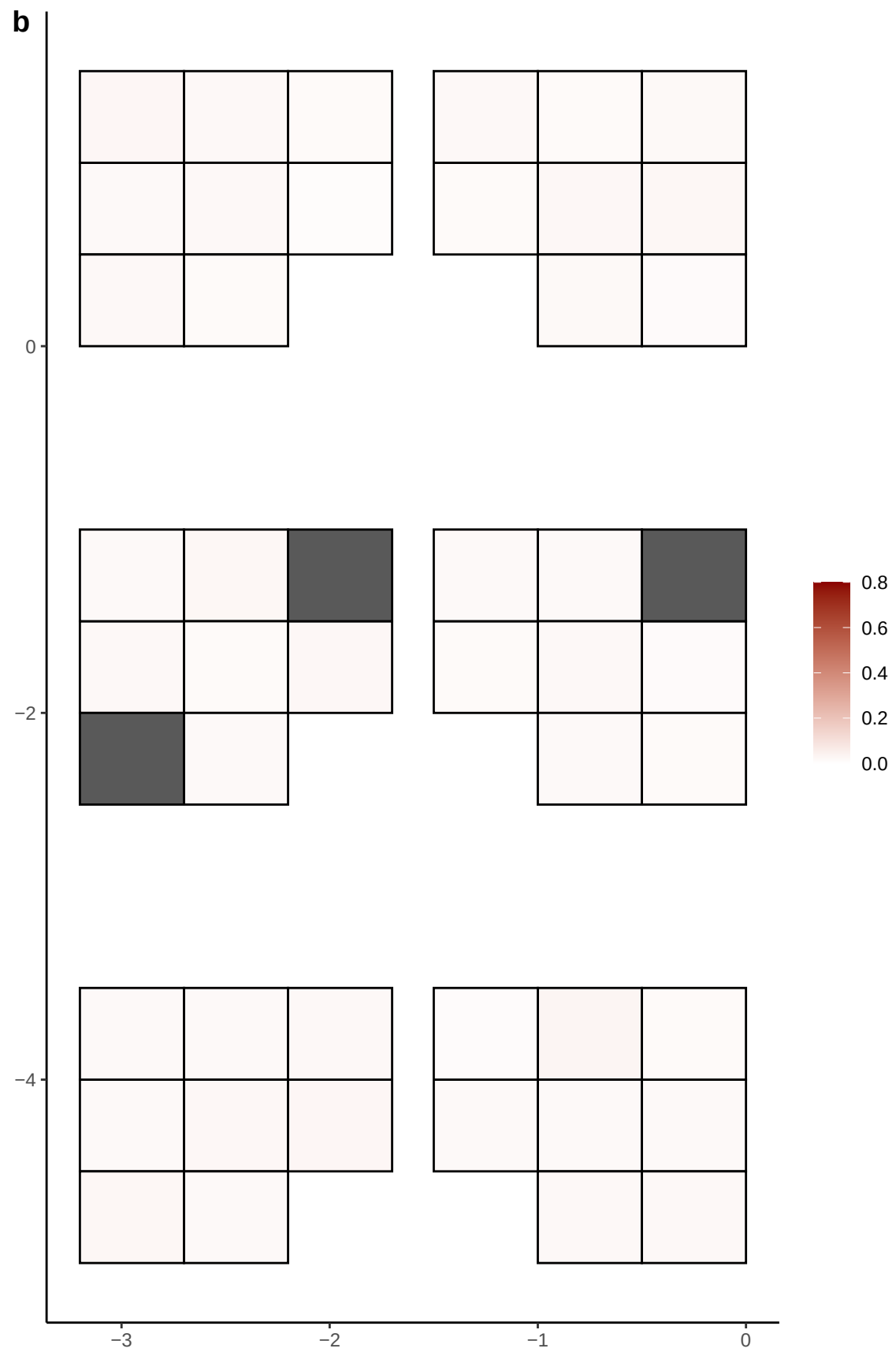

**a** ID05 13-67852564-G-A

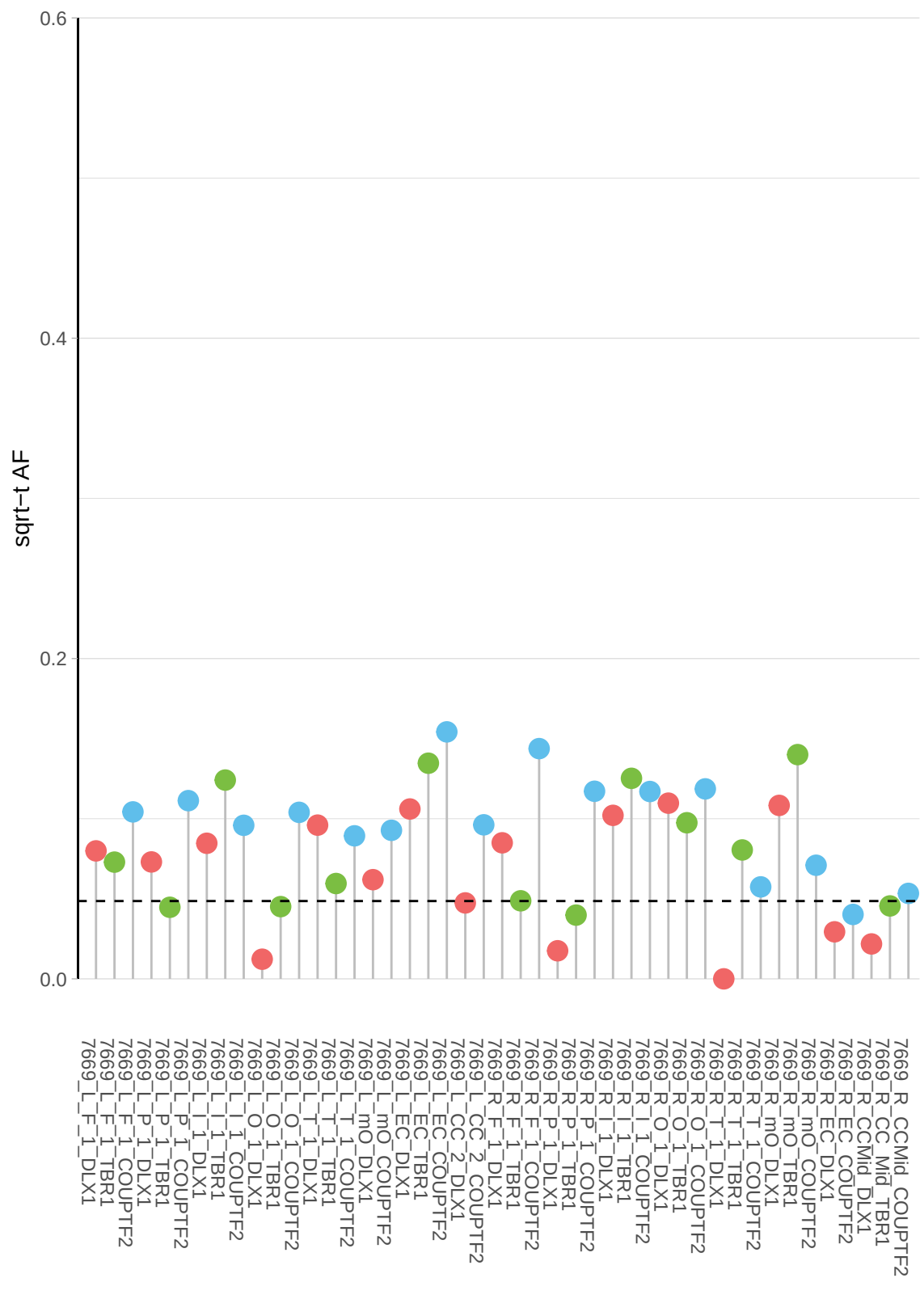

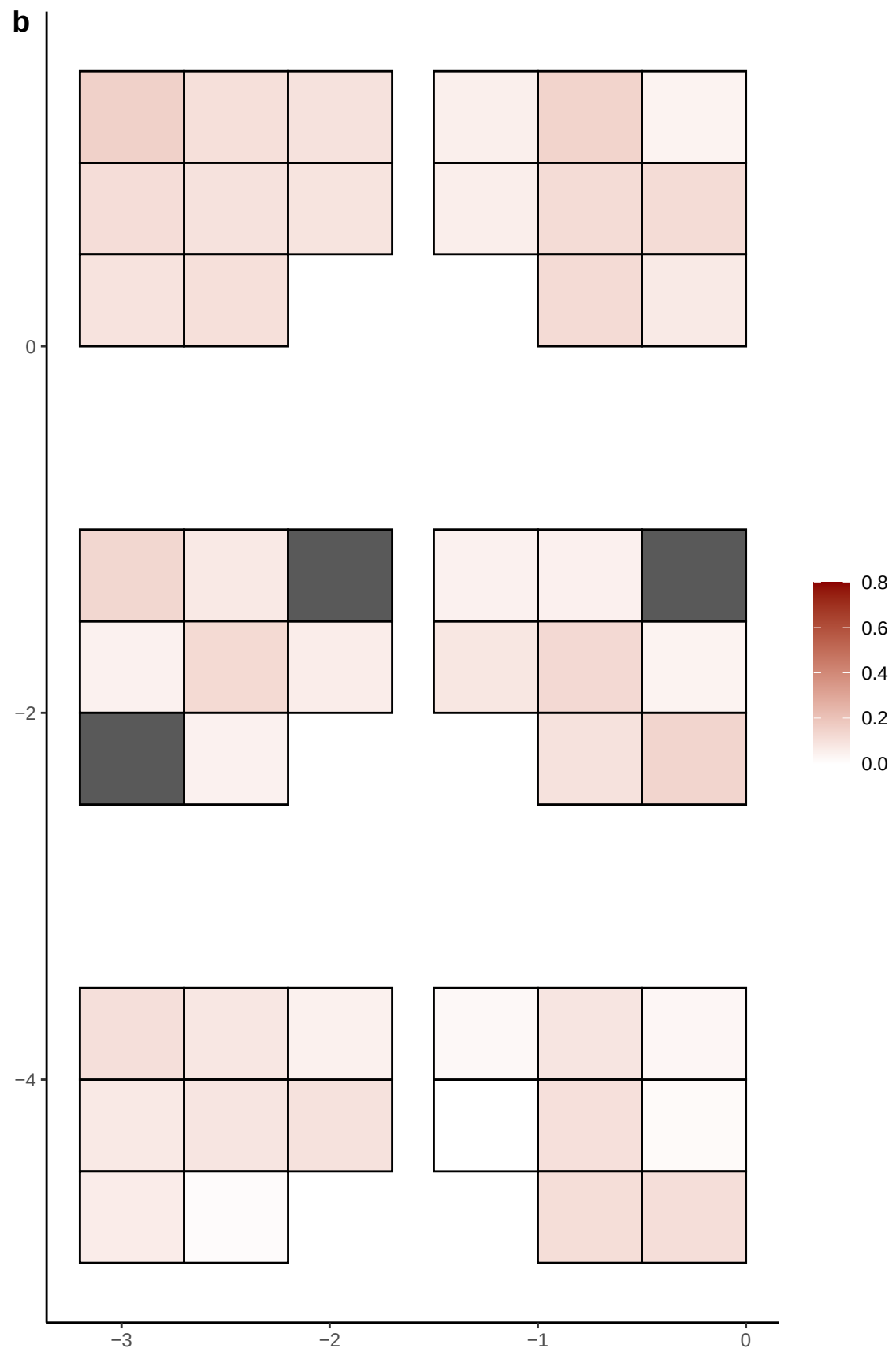

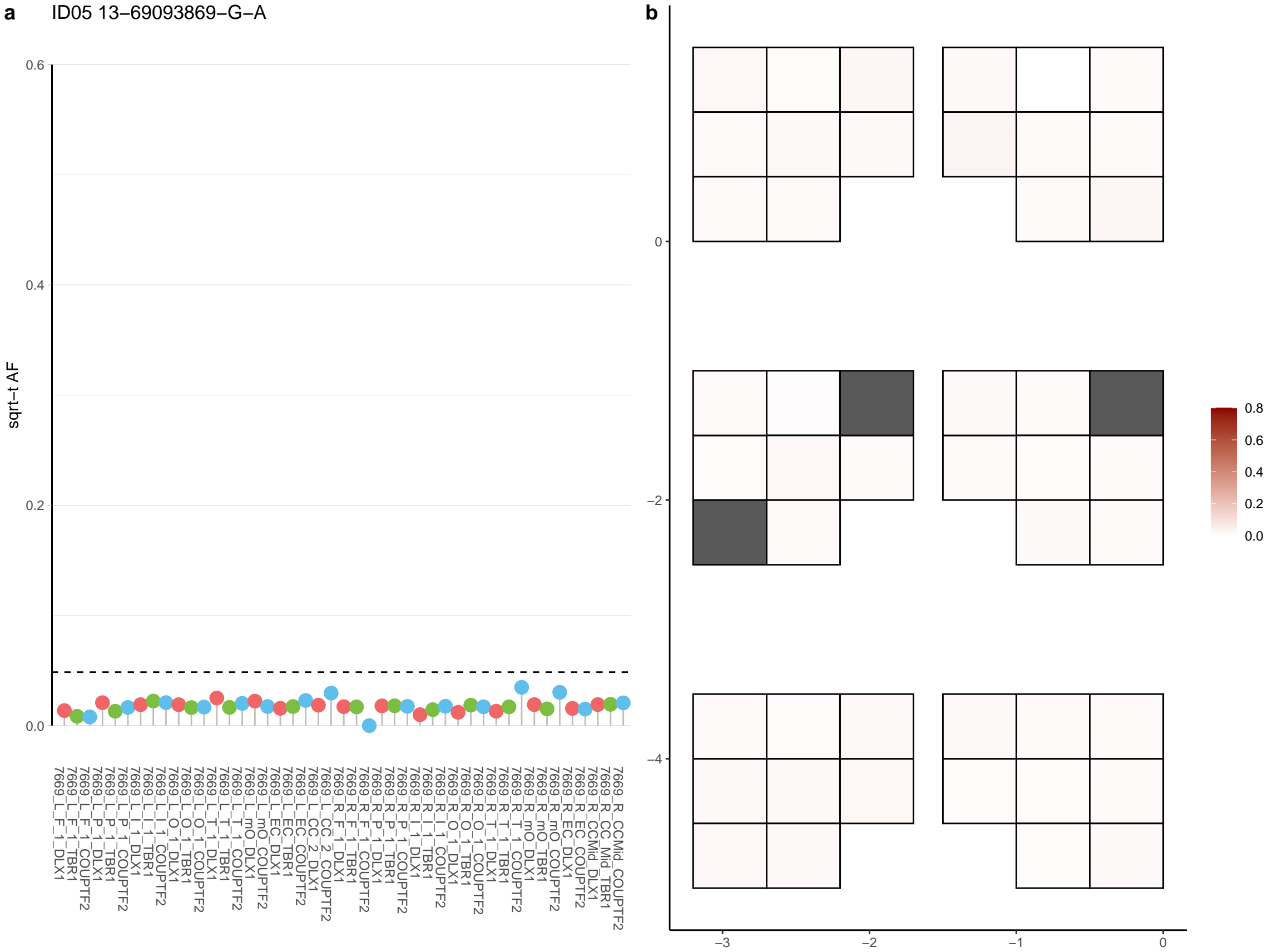

**a**

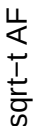

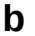

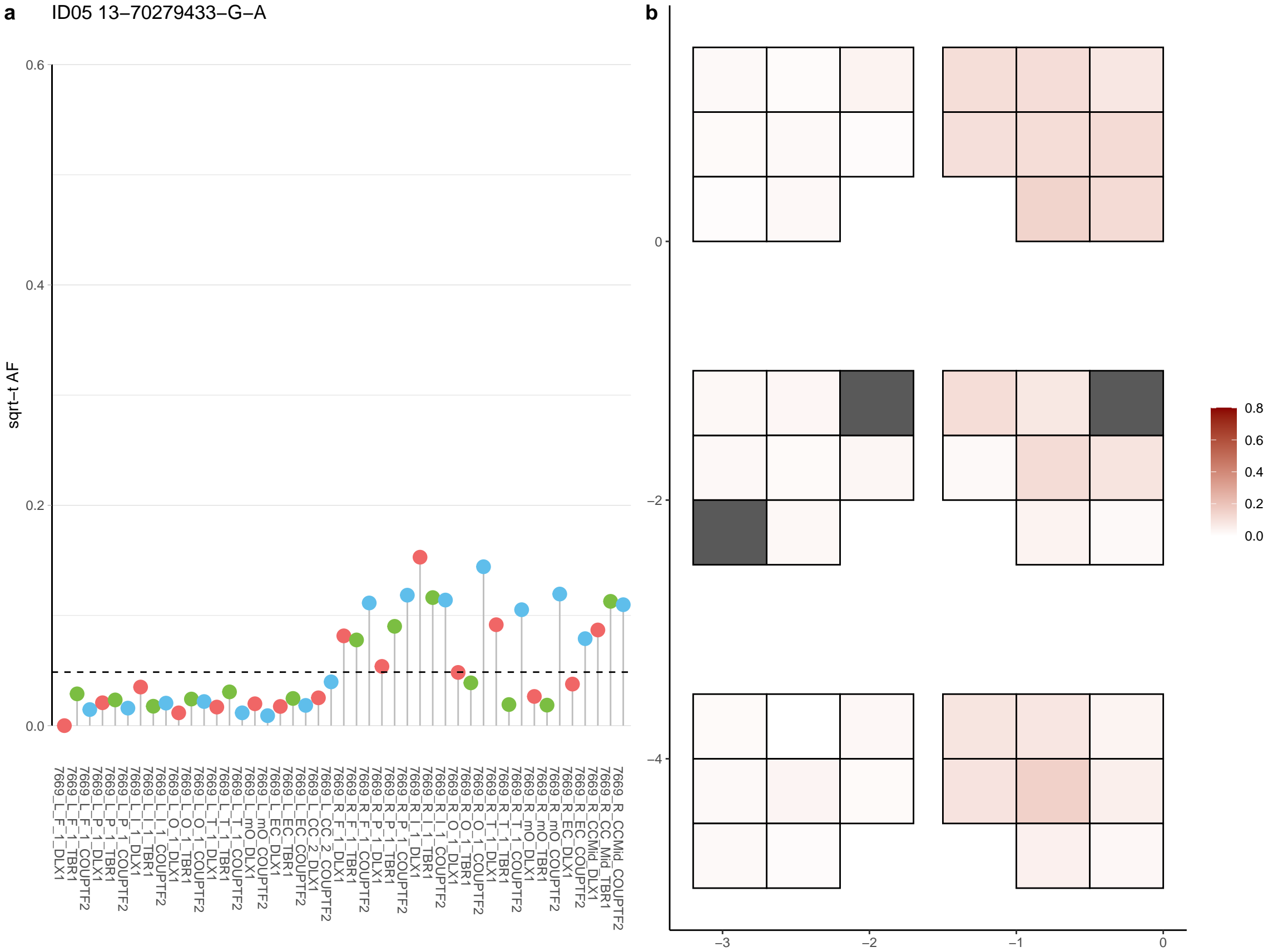

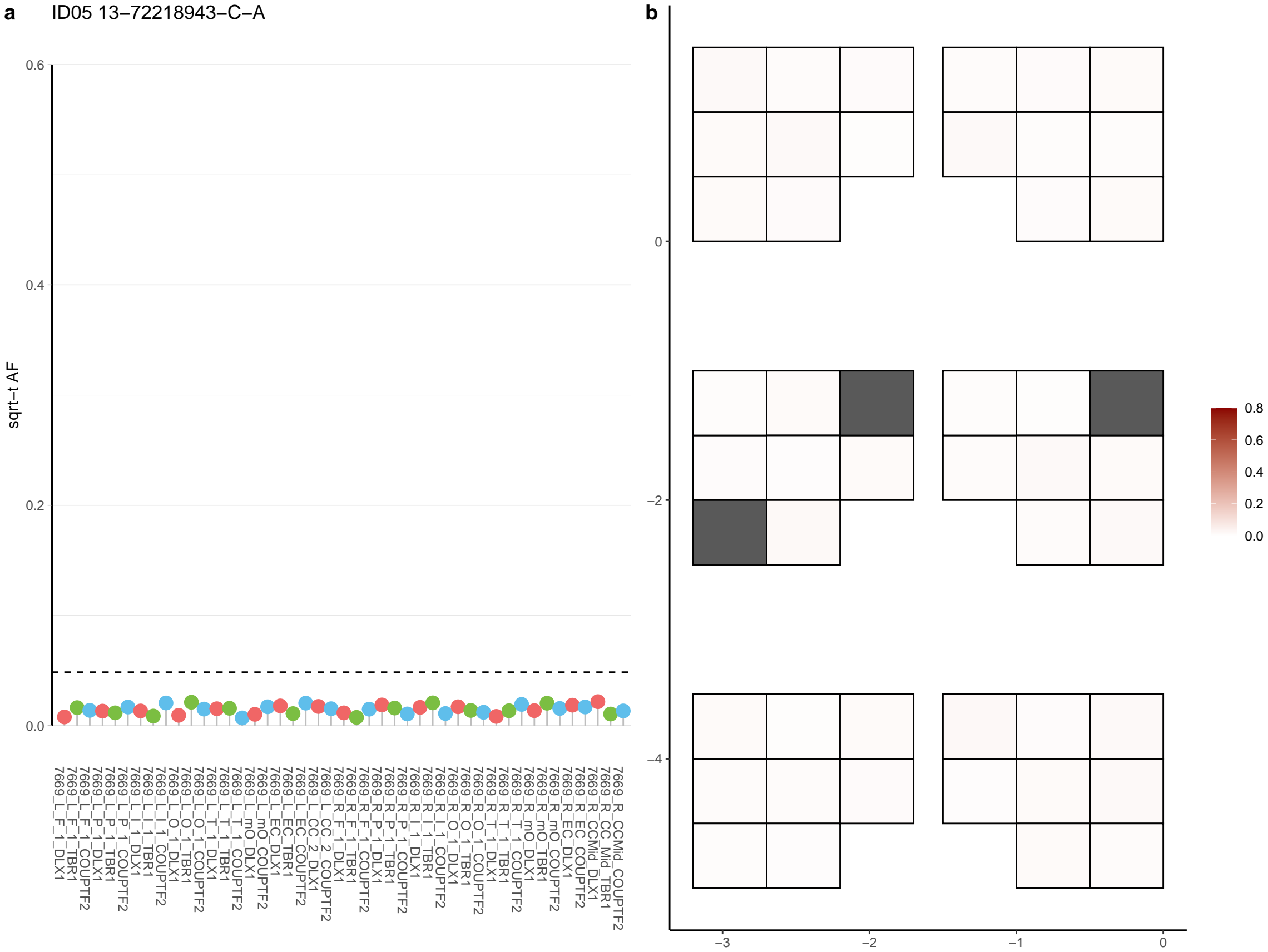

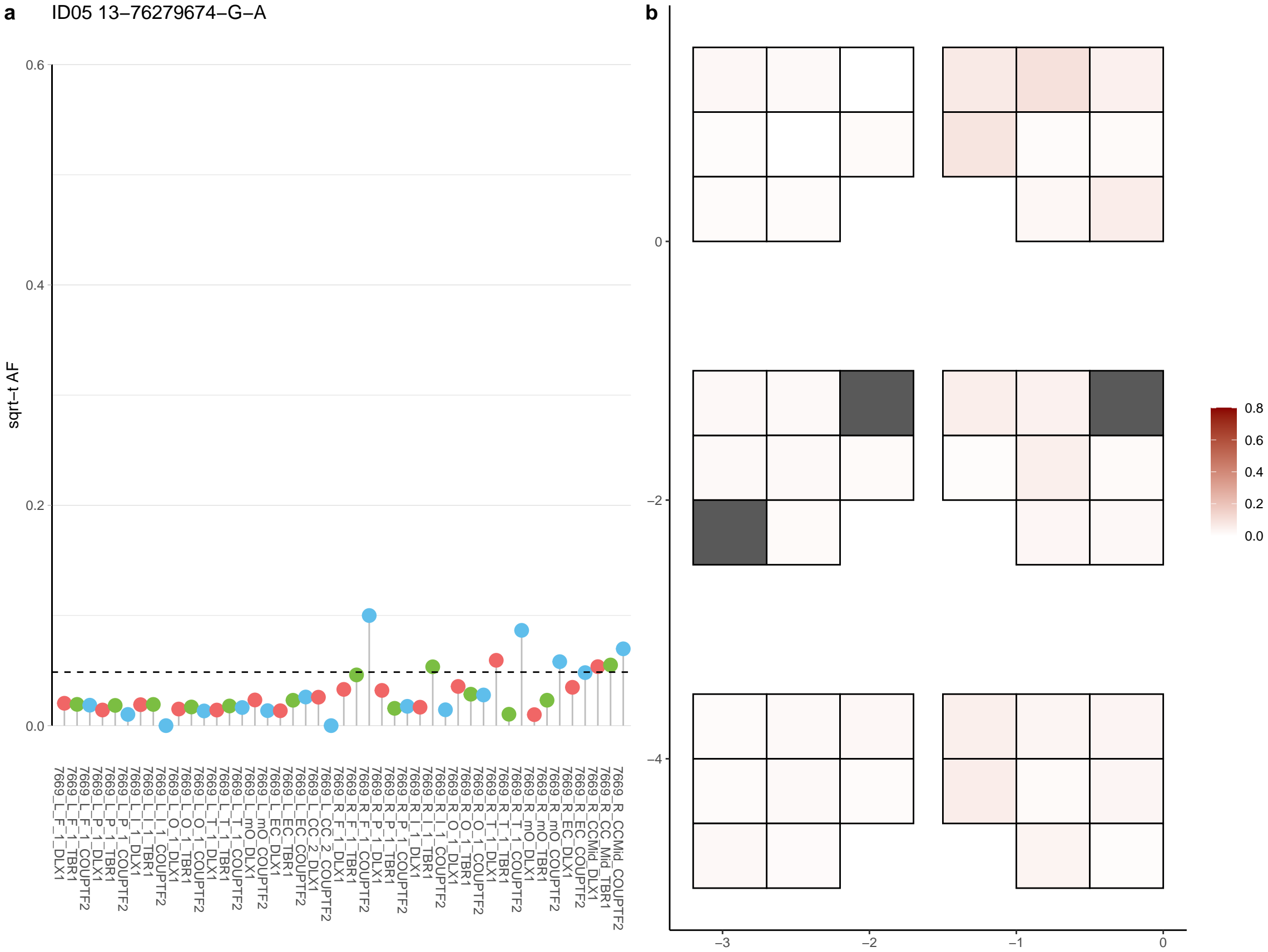

**a** ID05 13-76874198-C-T

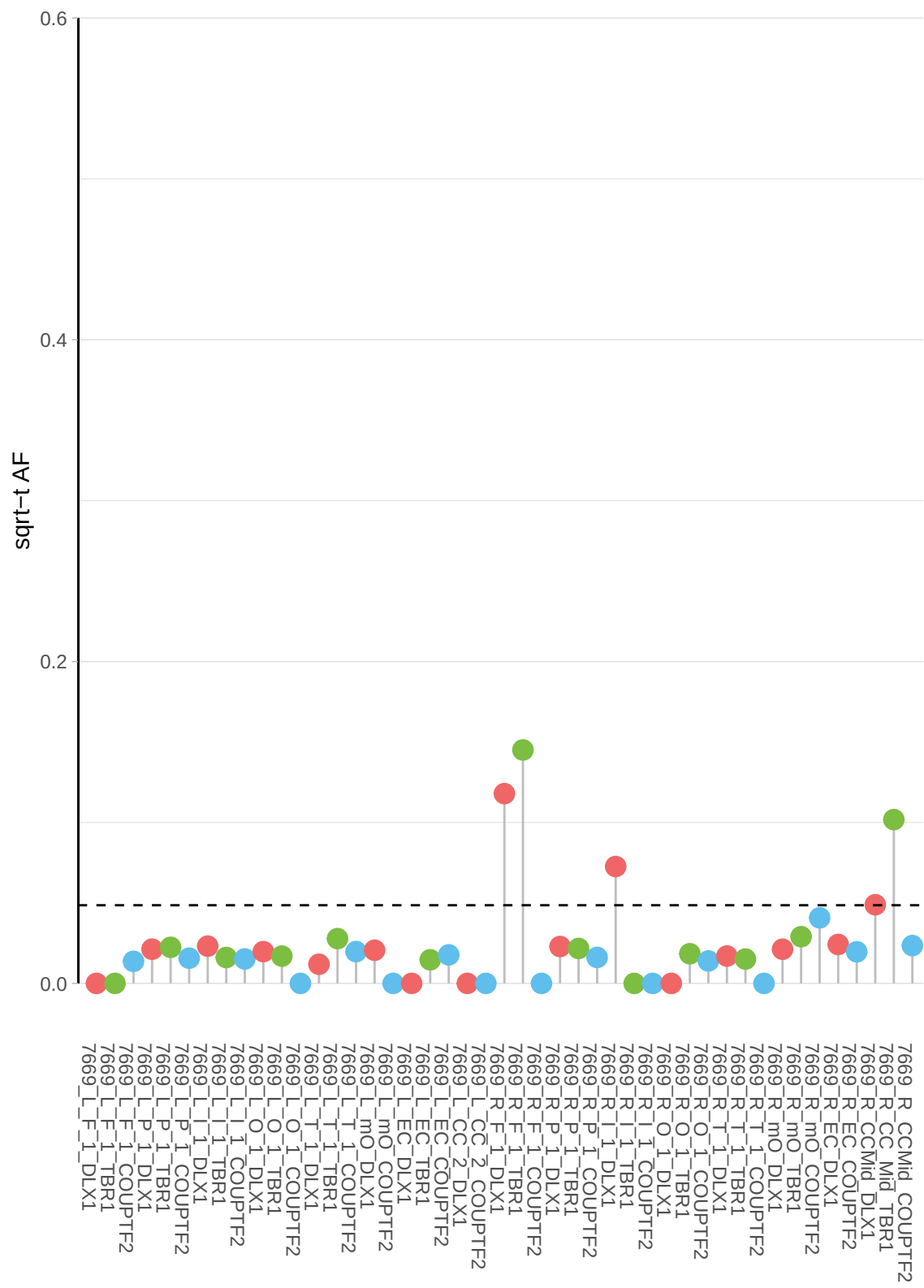

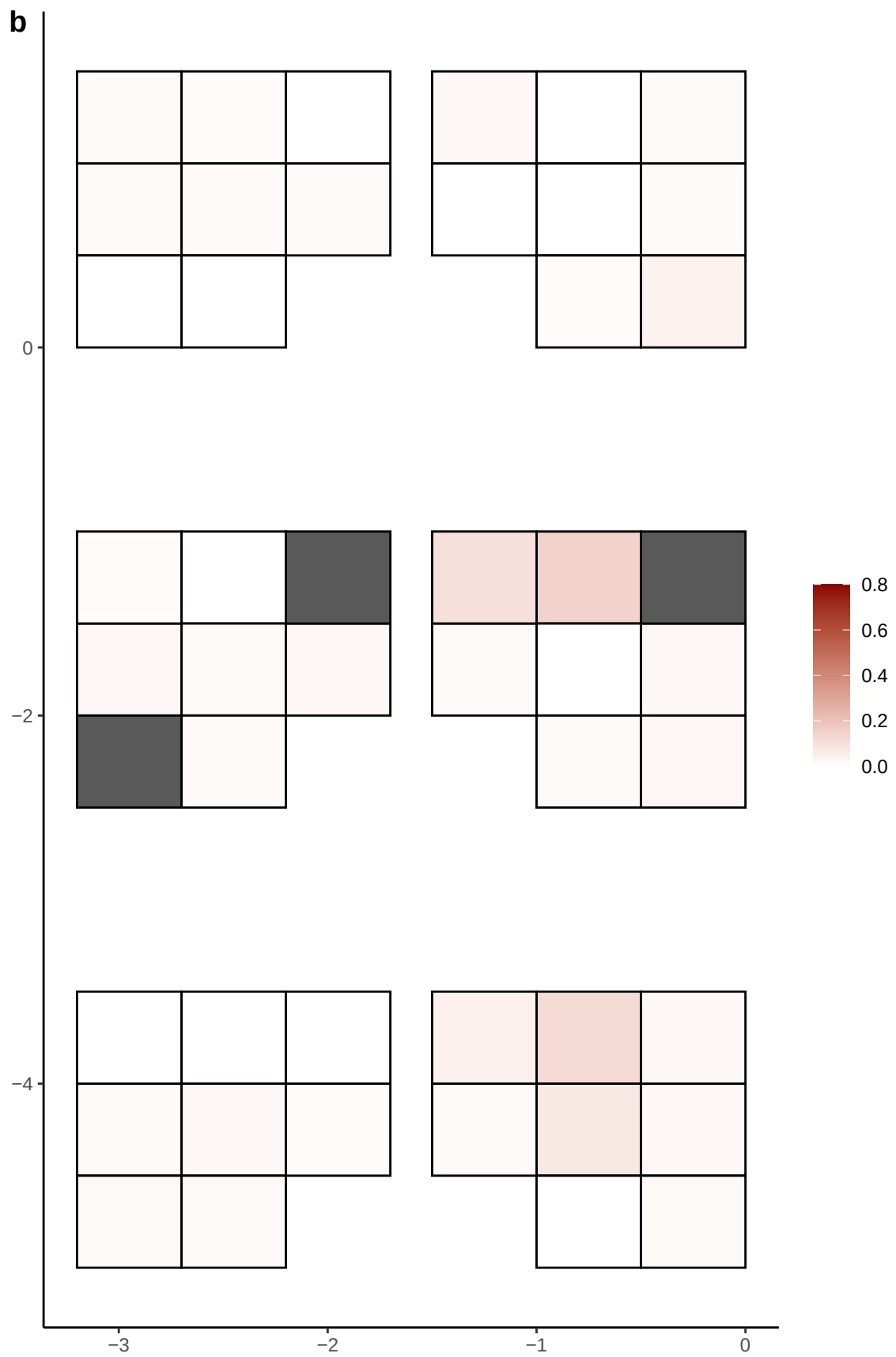

**a** ID05 13-77509041-G-T

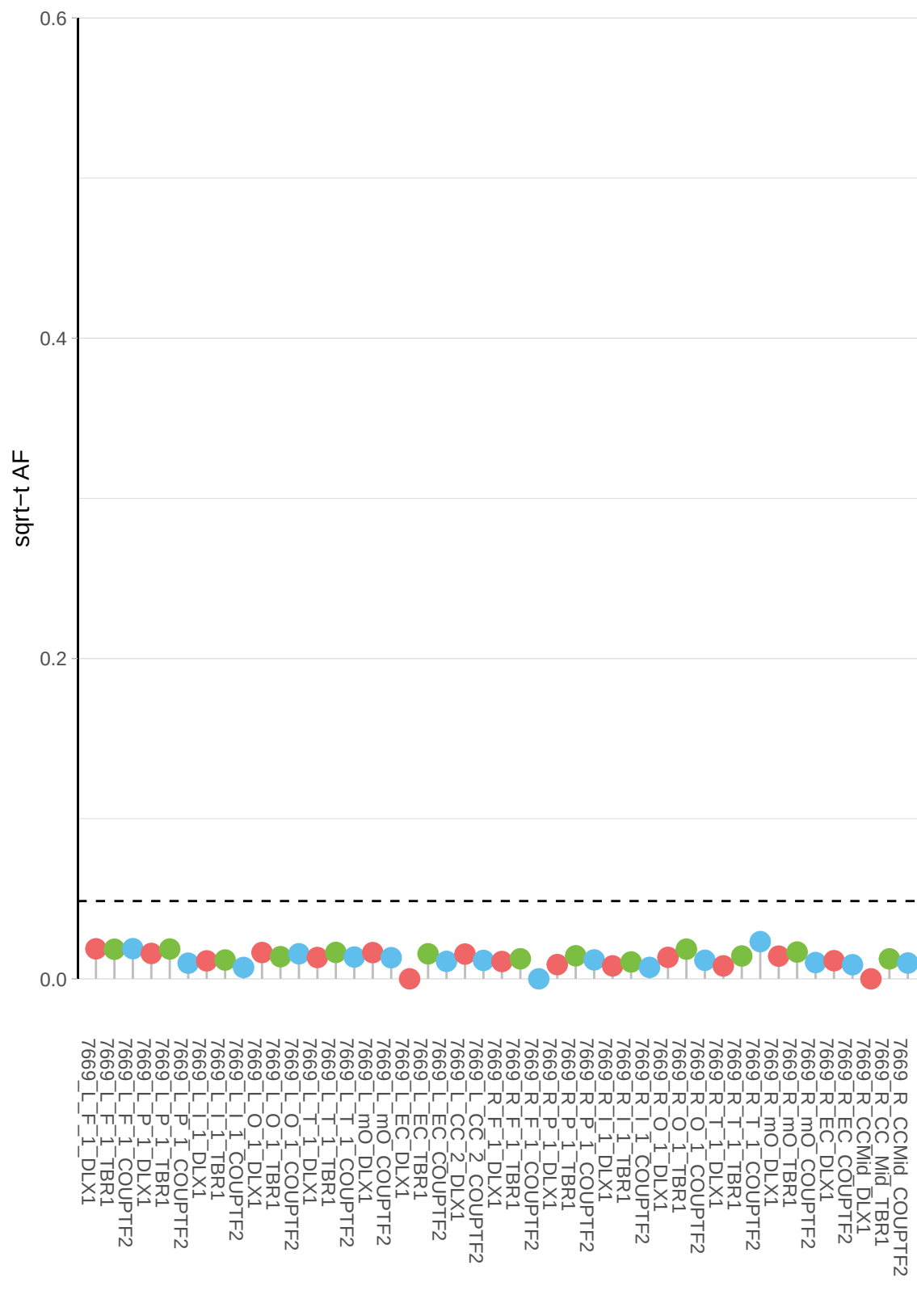

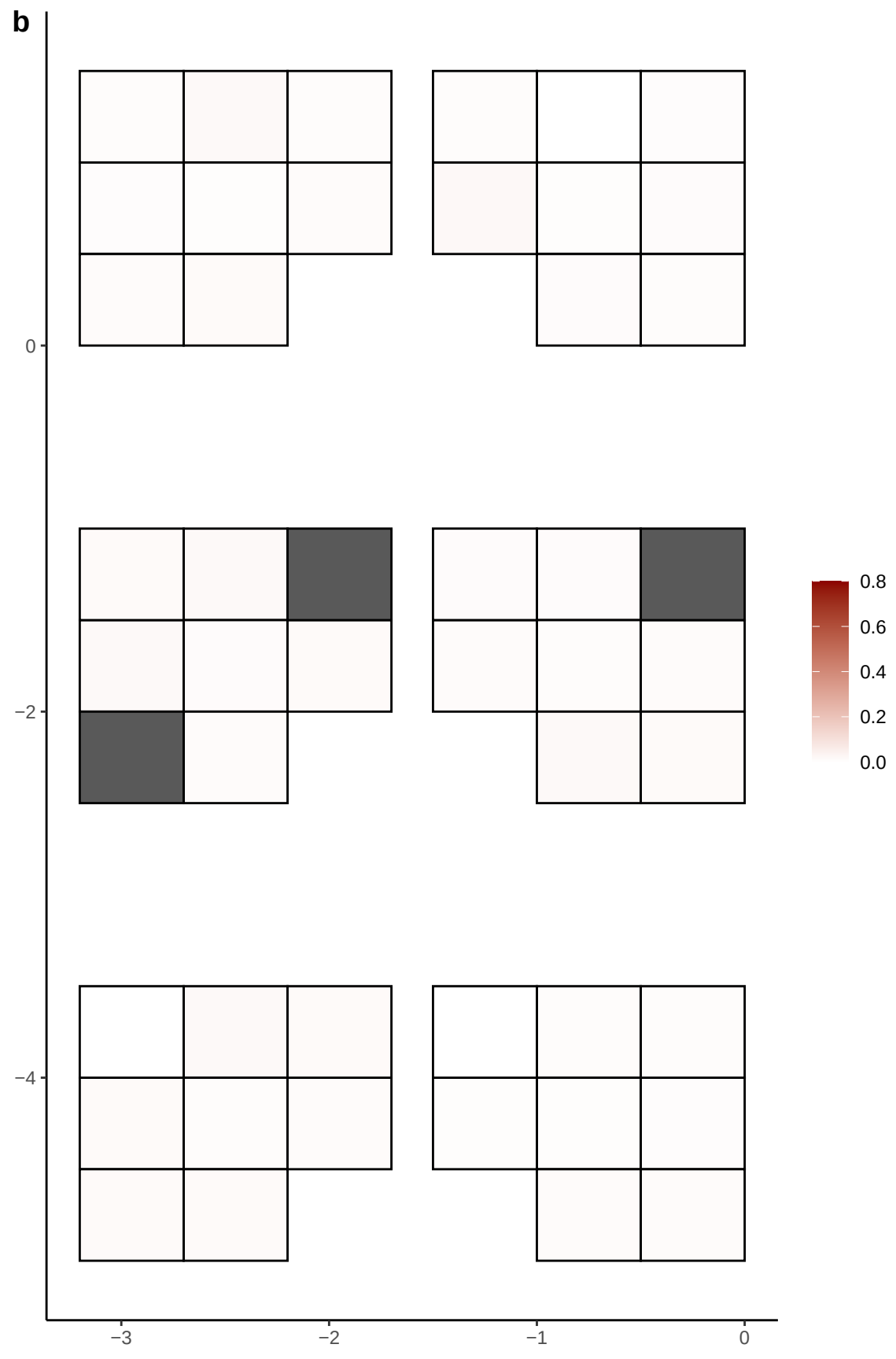

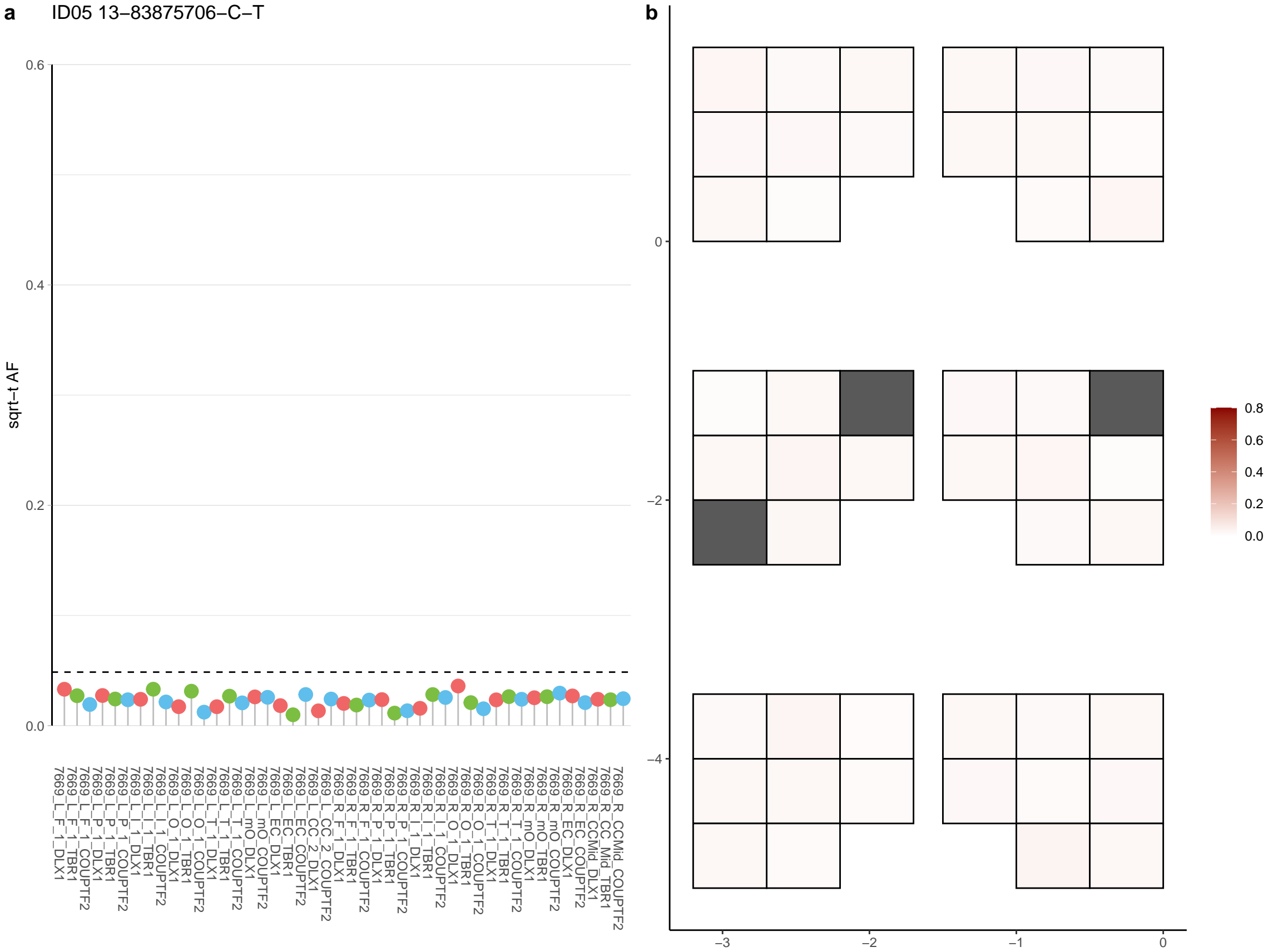

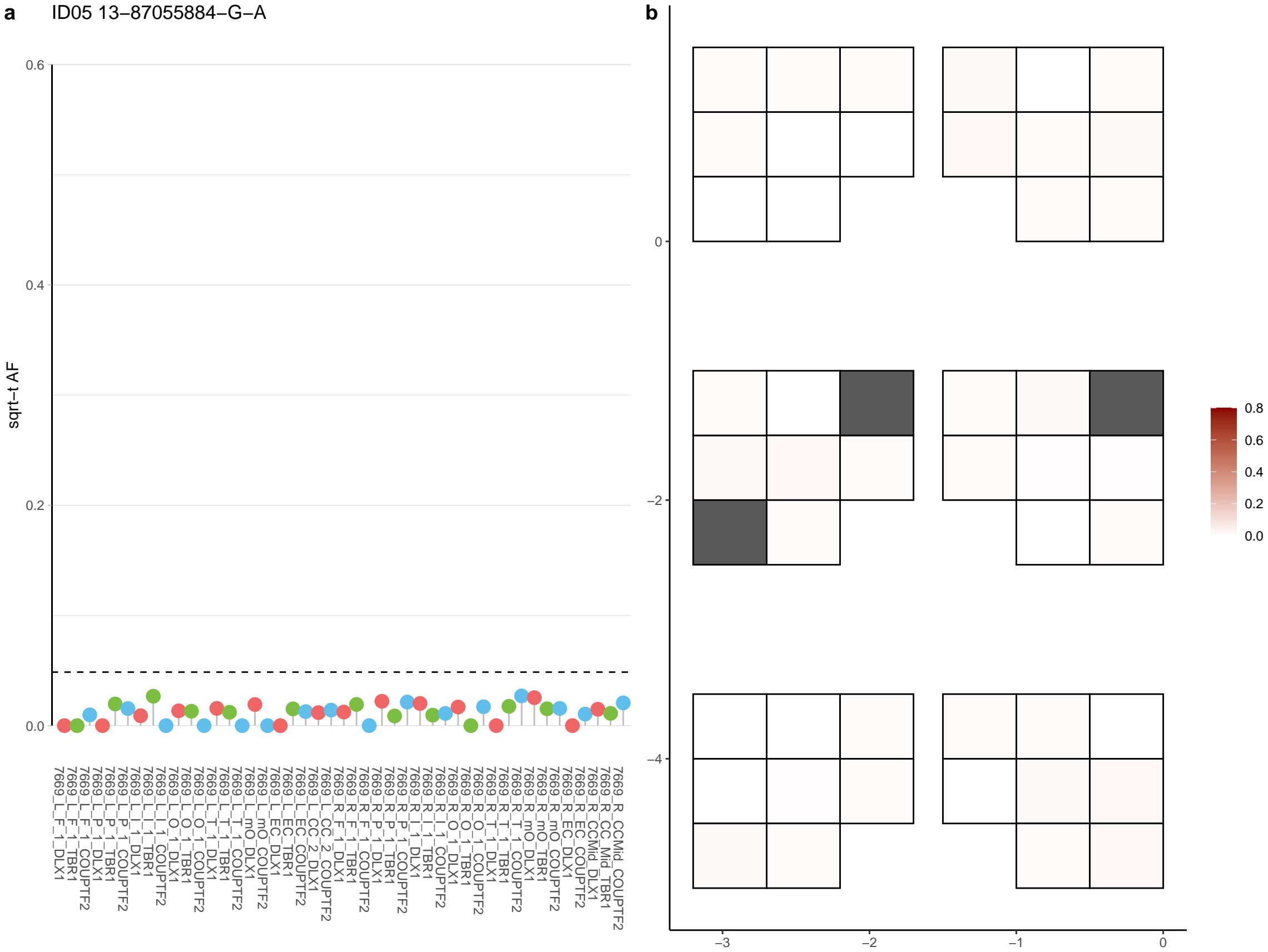

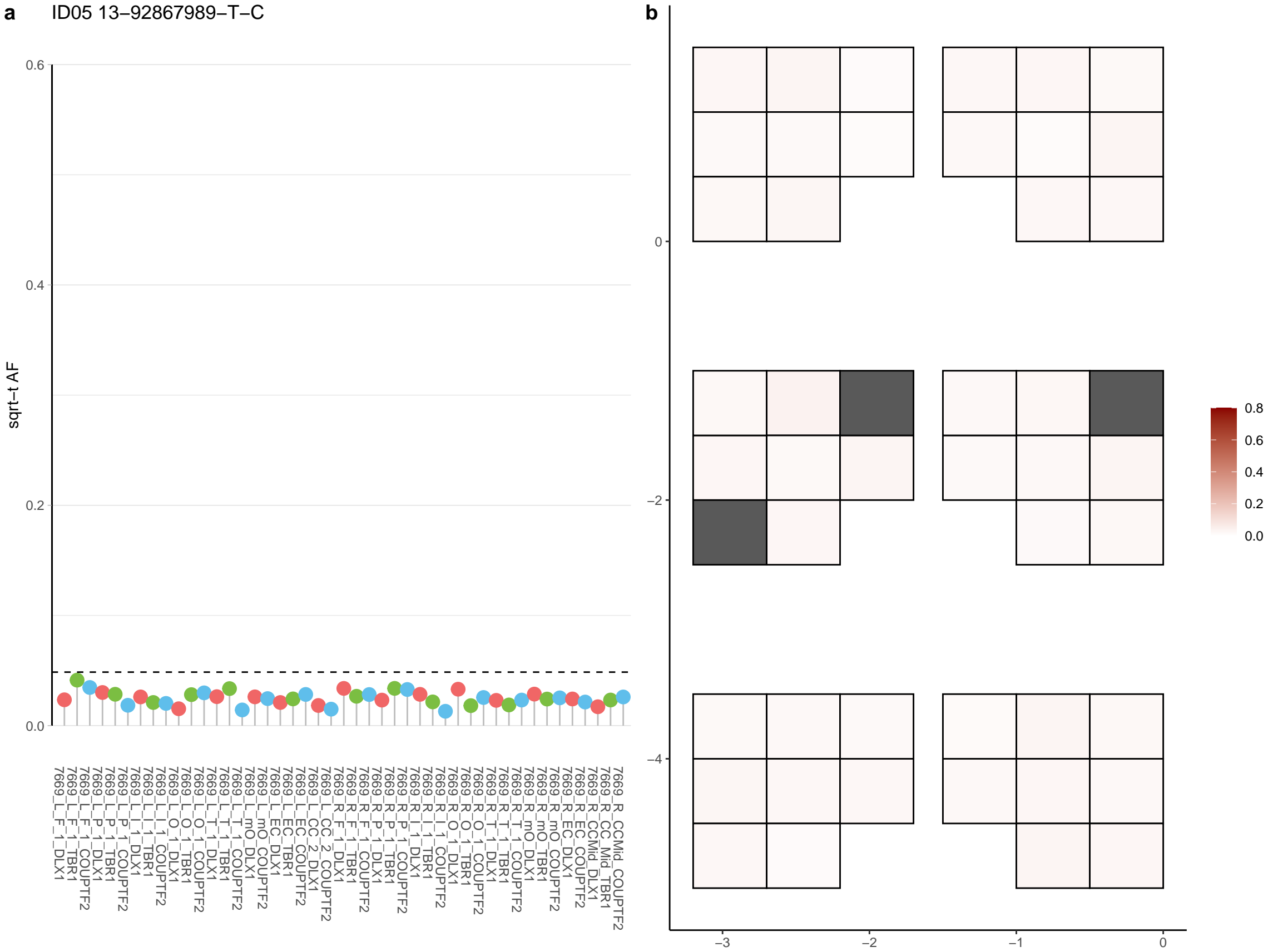

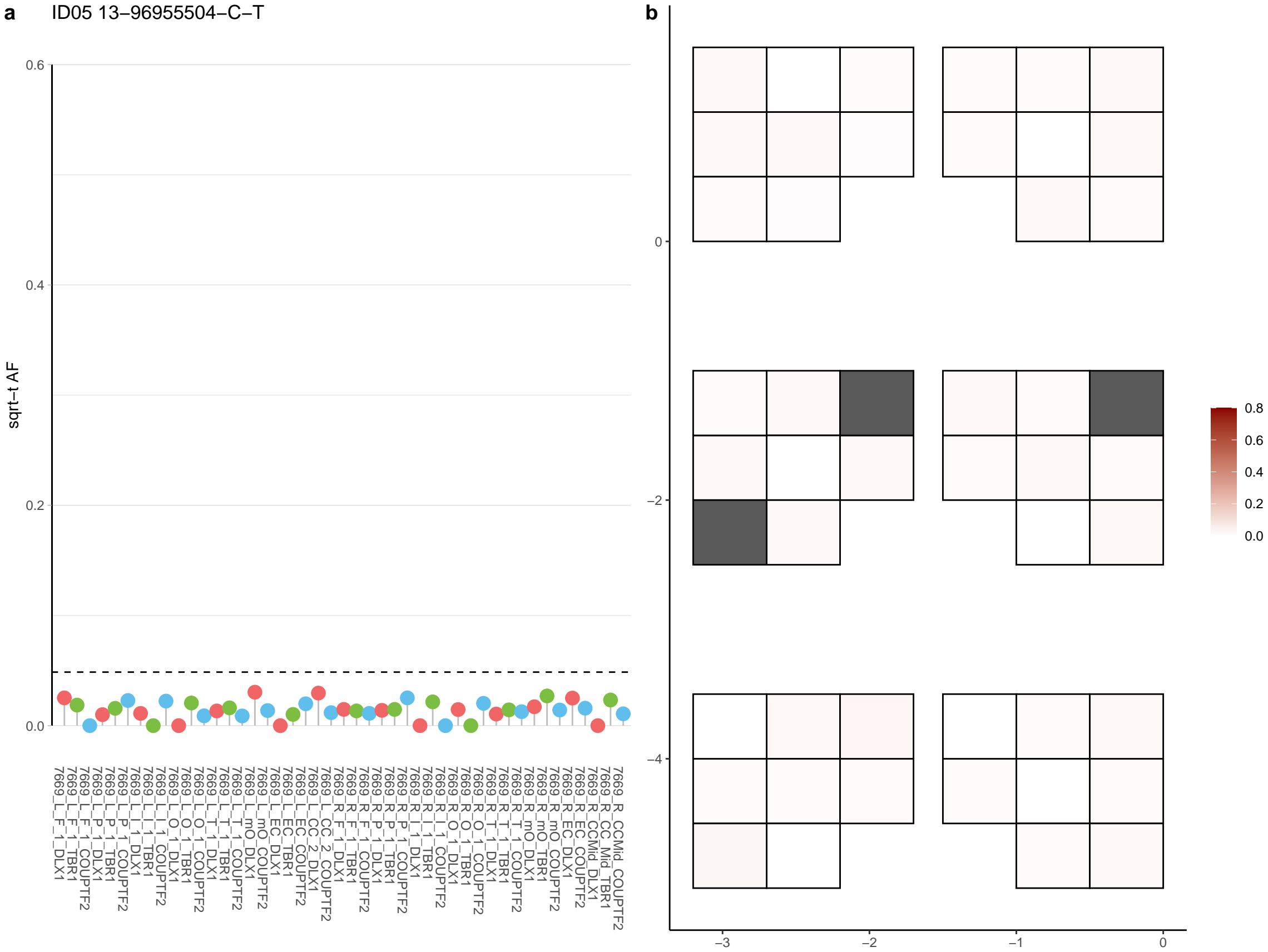

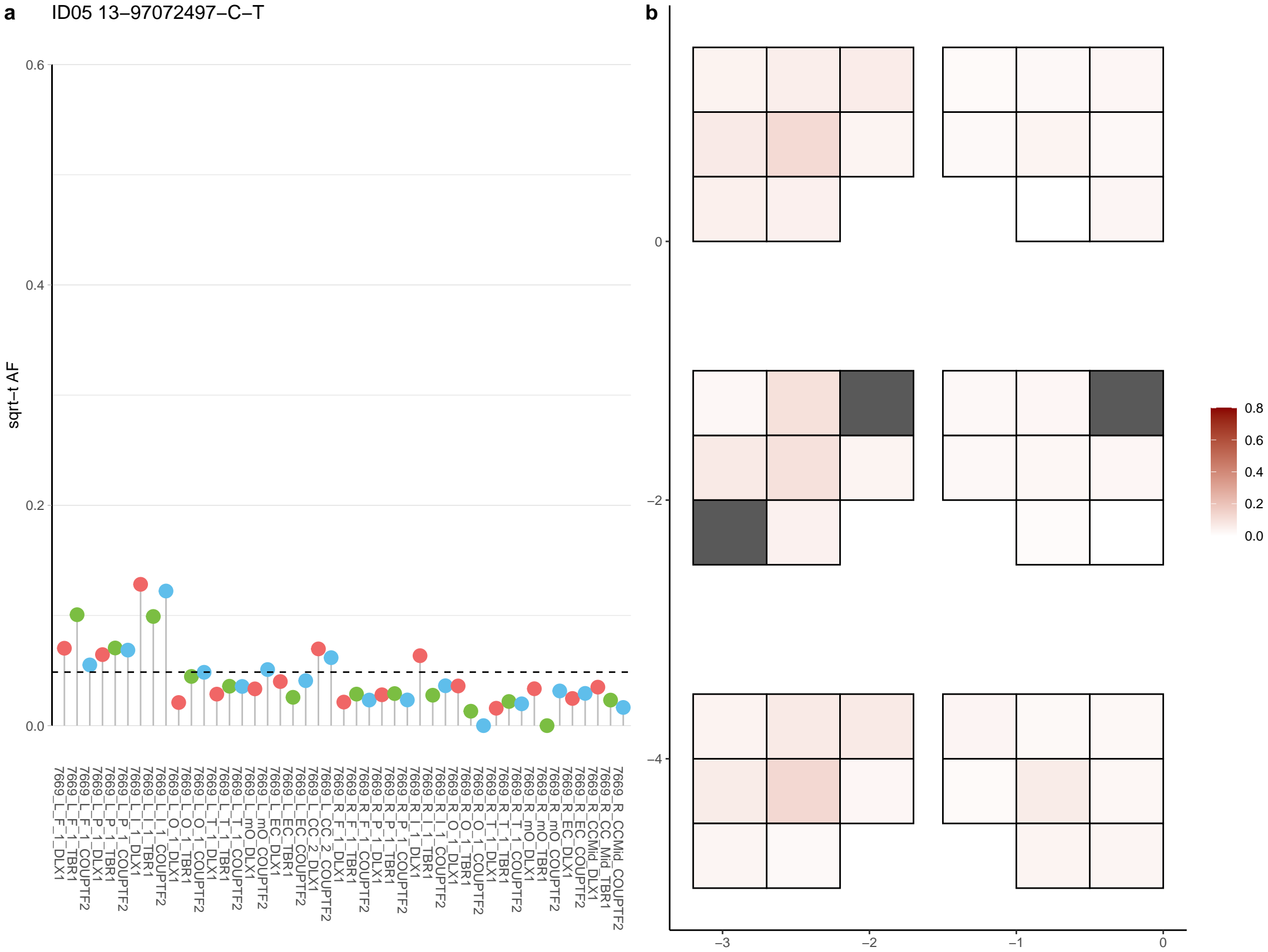

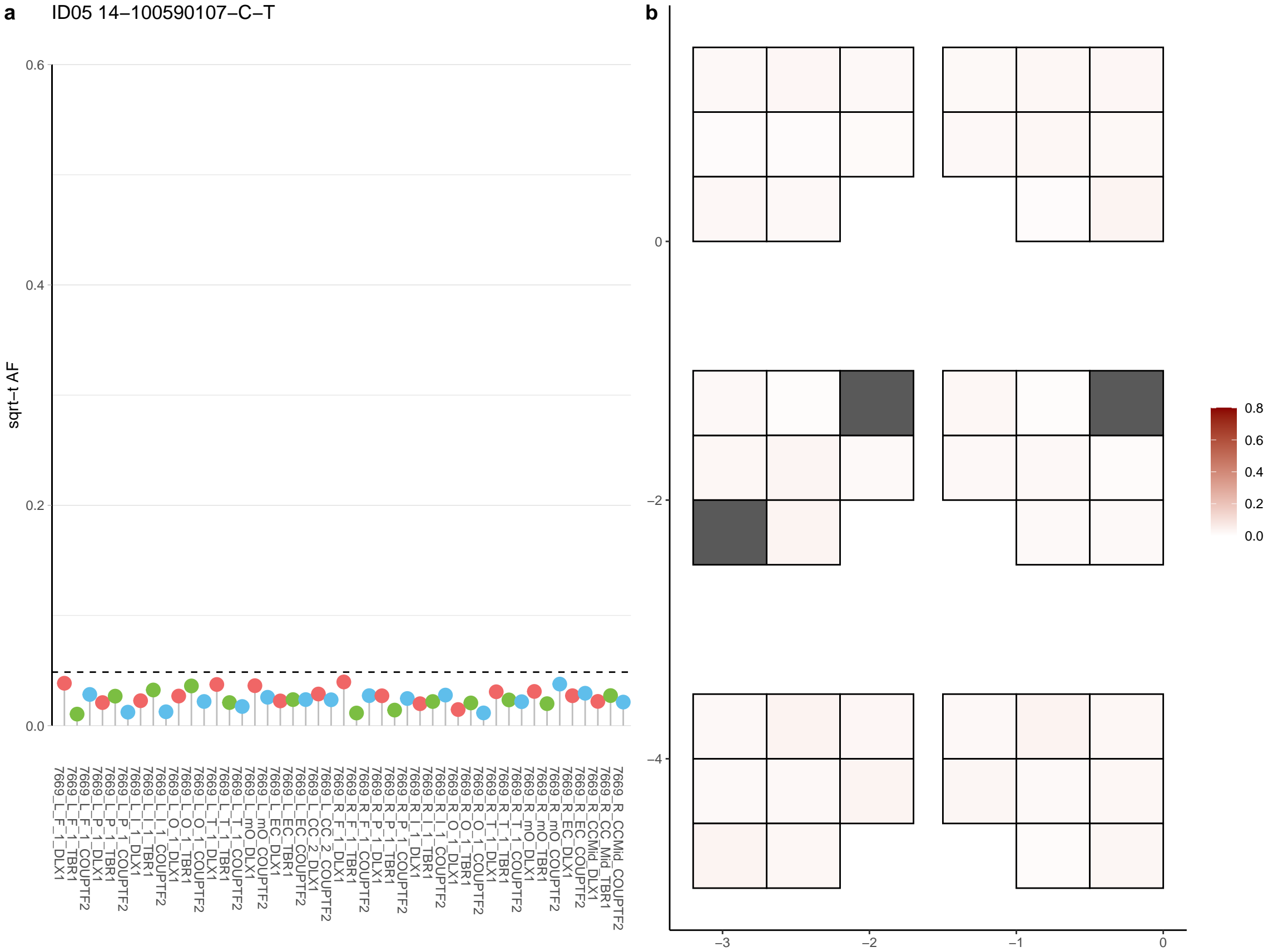

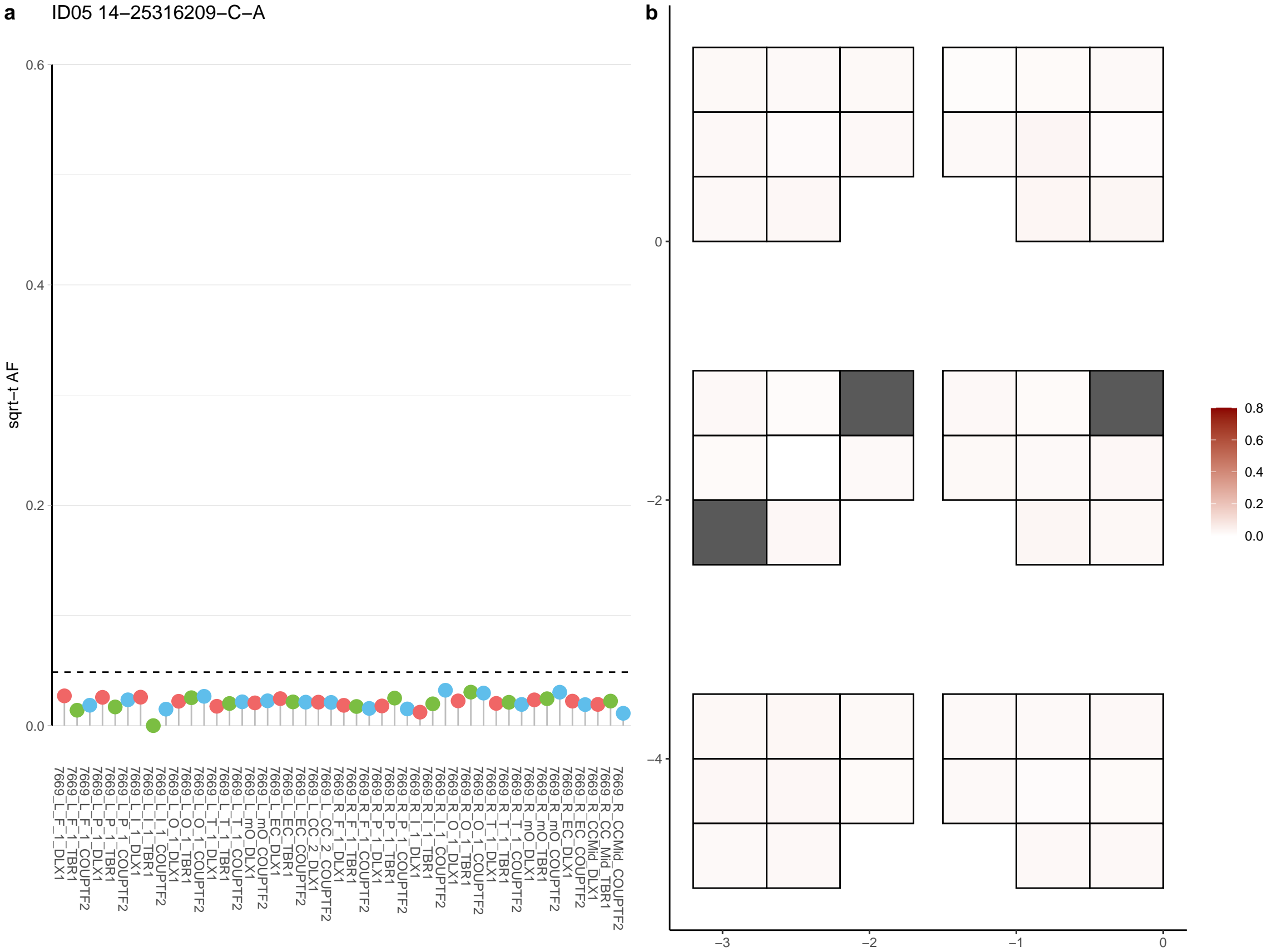

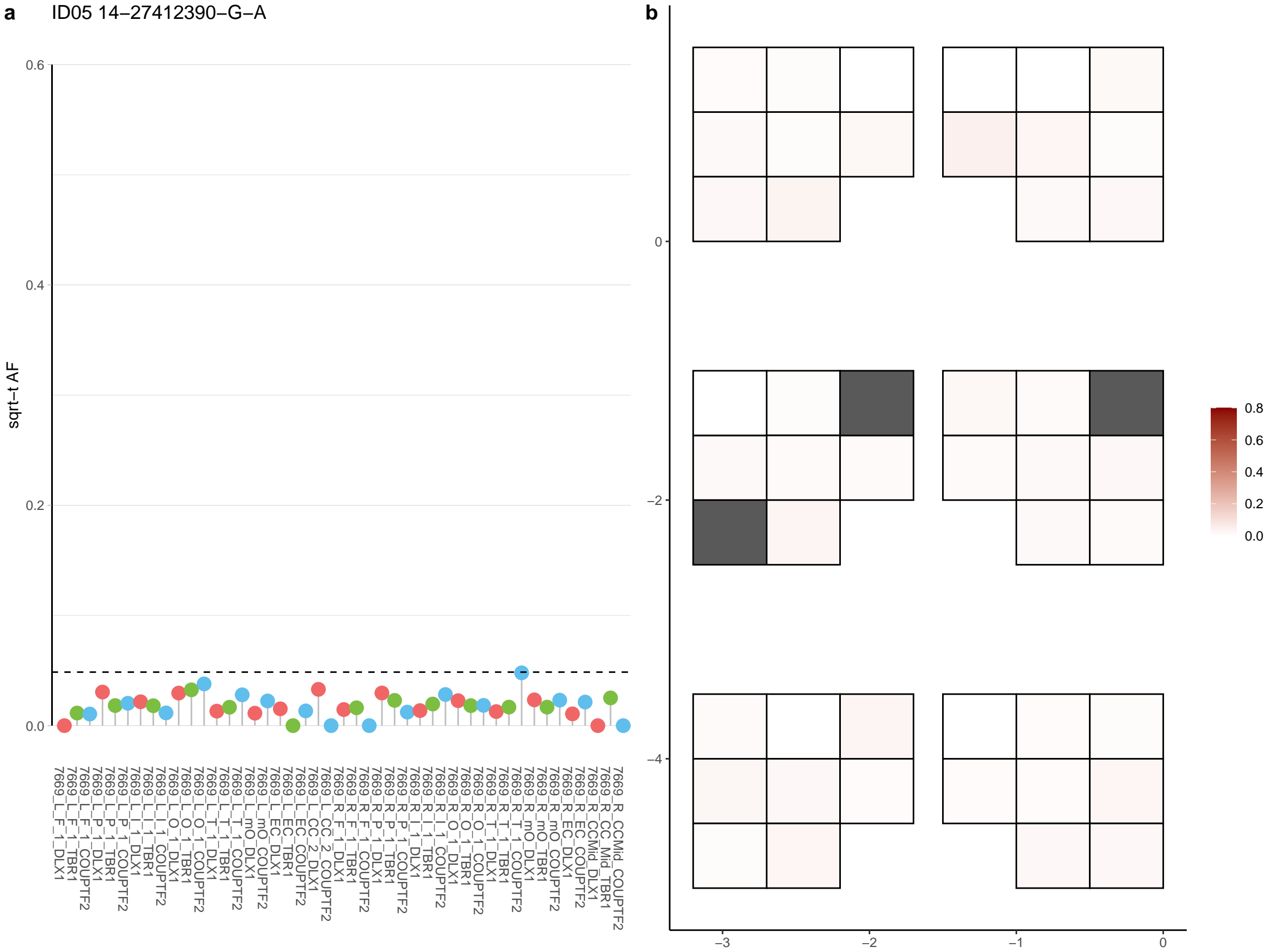

**a** ID05 14-27649824-C-T

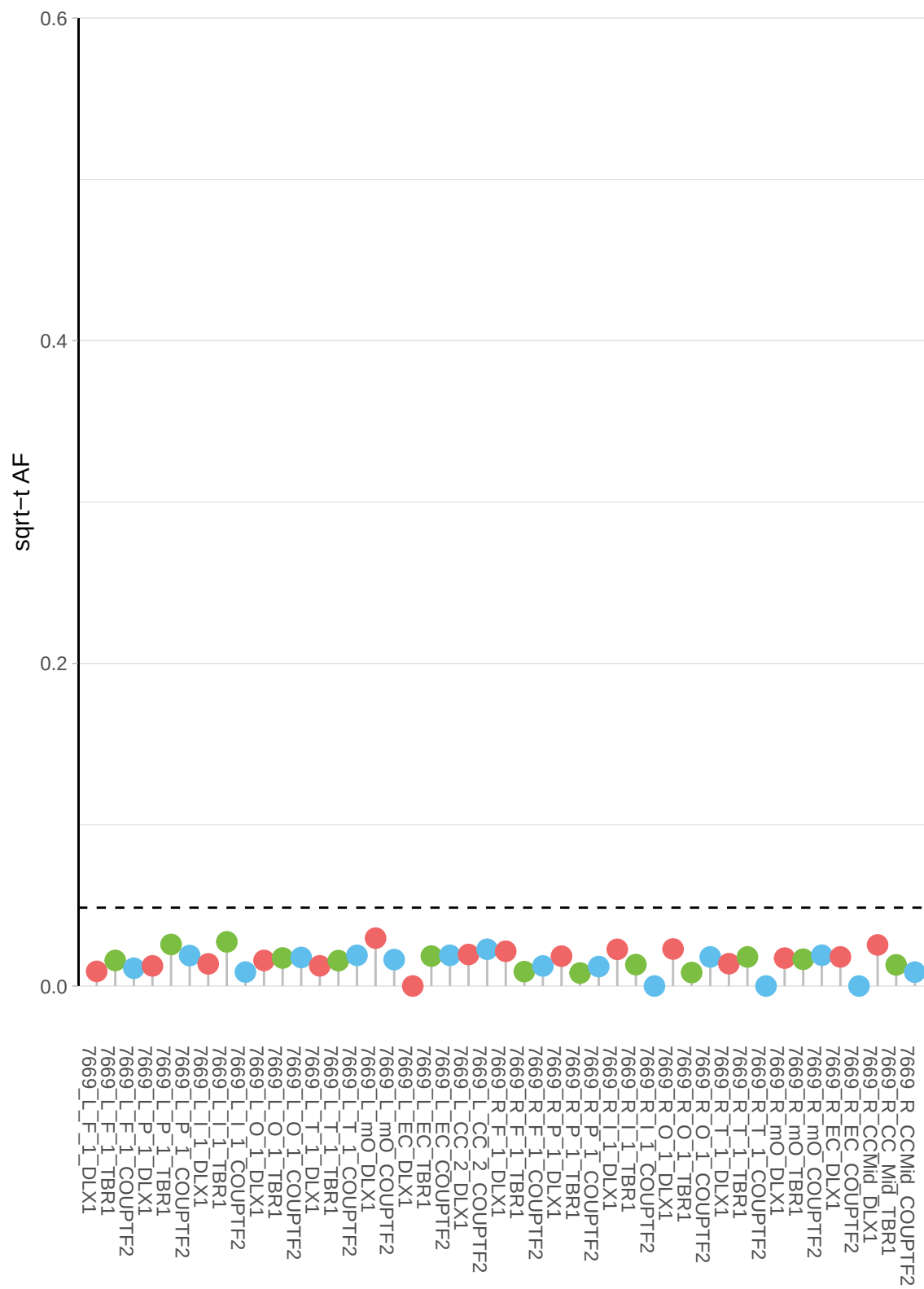

**b**

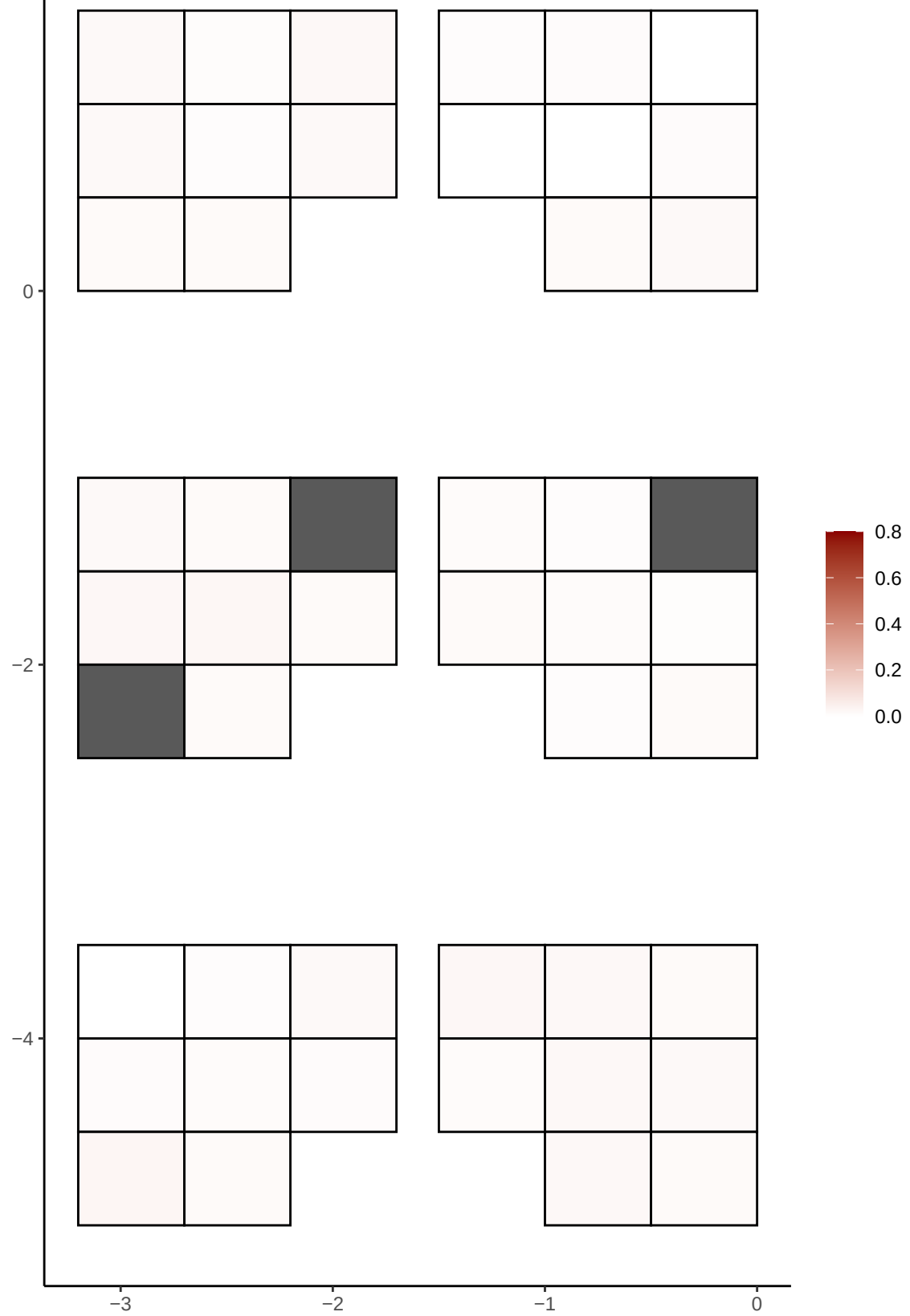

**a** ID05 14-36924791-G-C

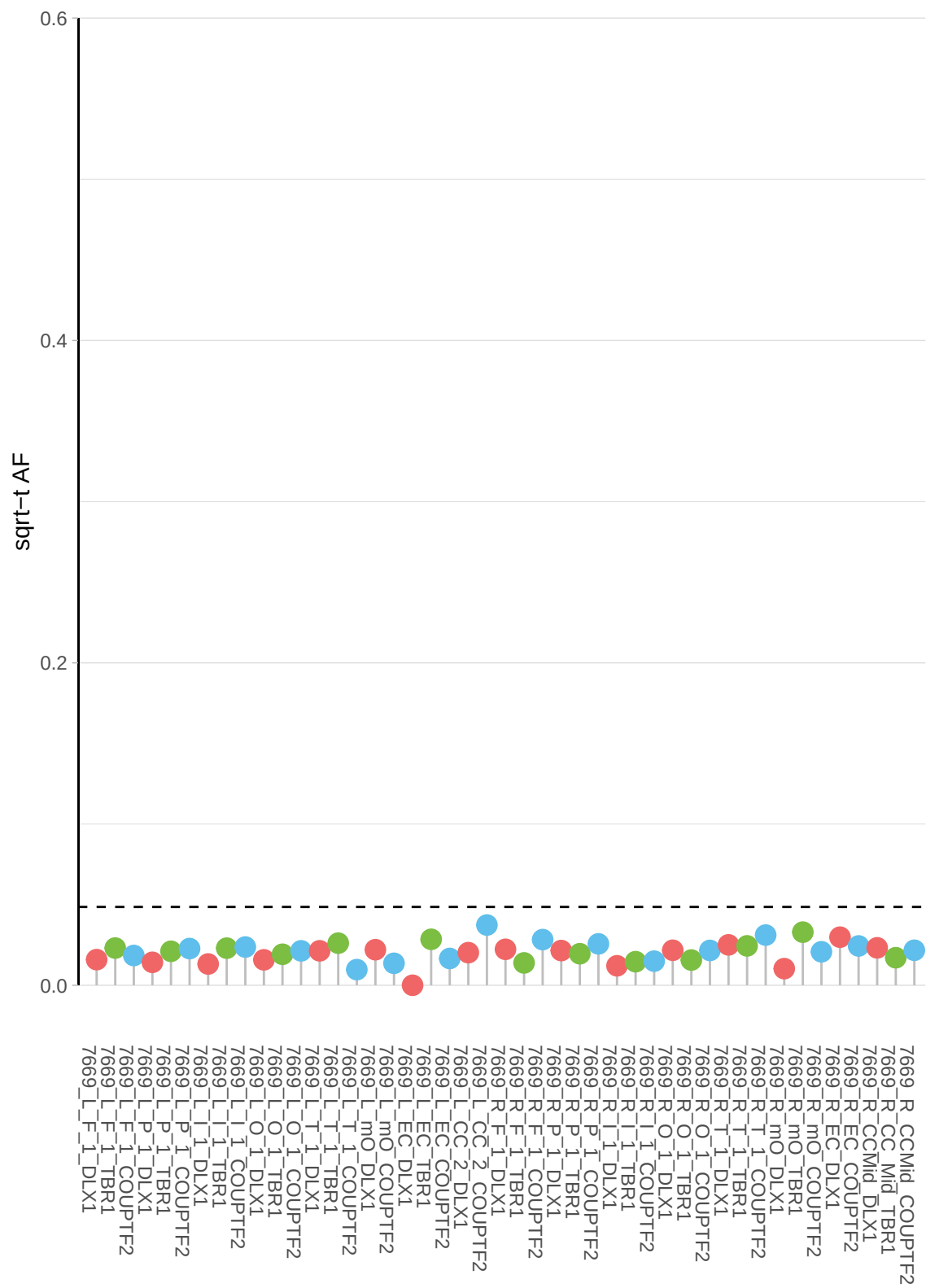

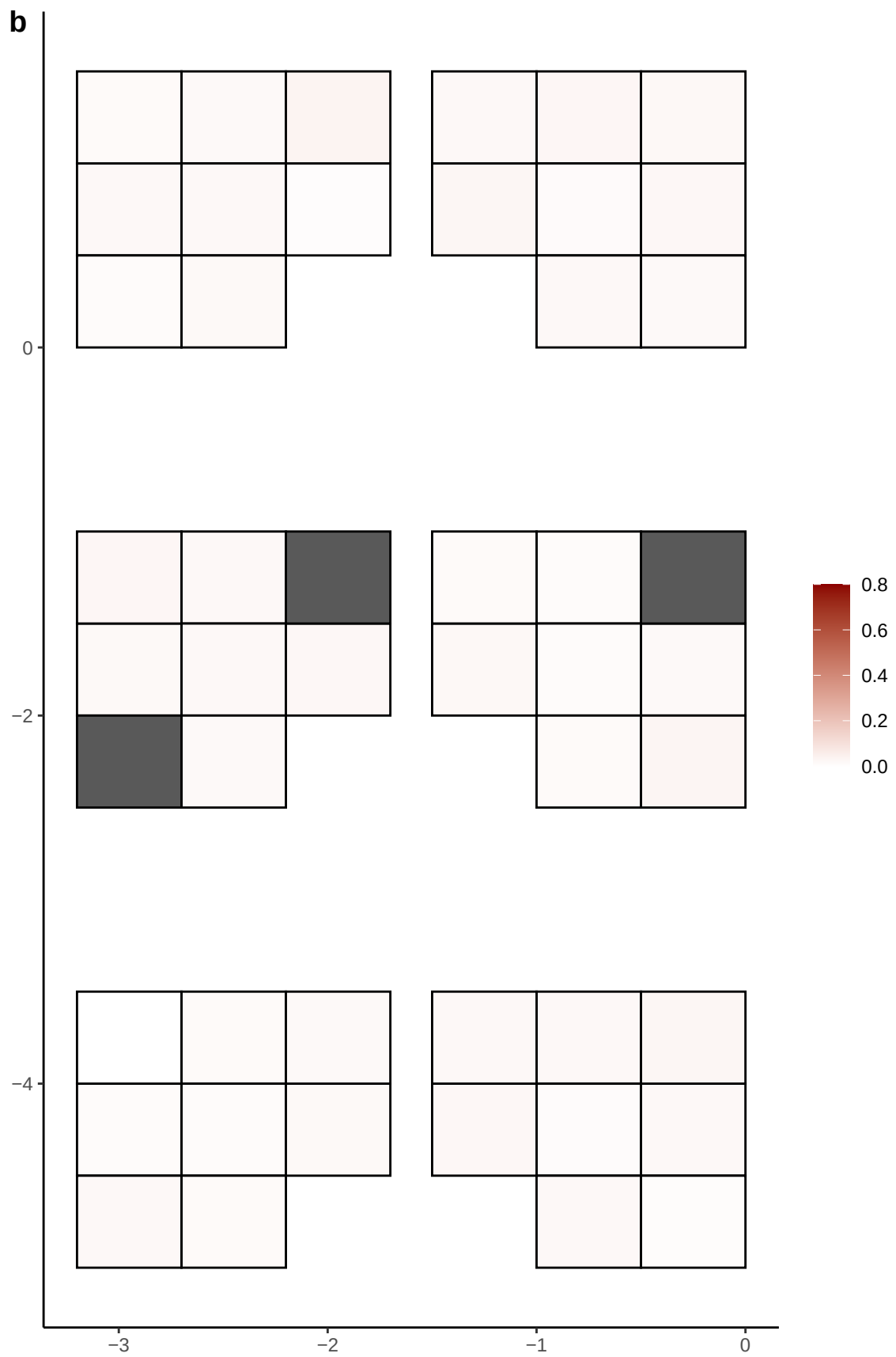

**a** ID05 14-41027959-C-T

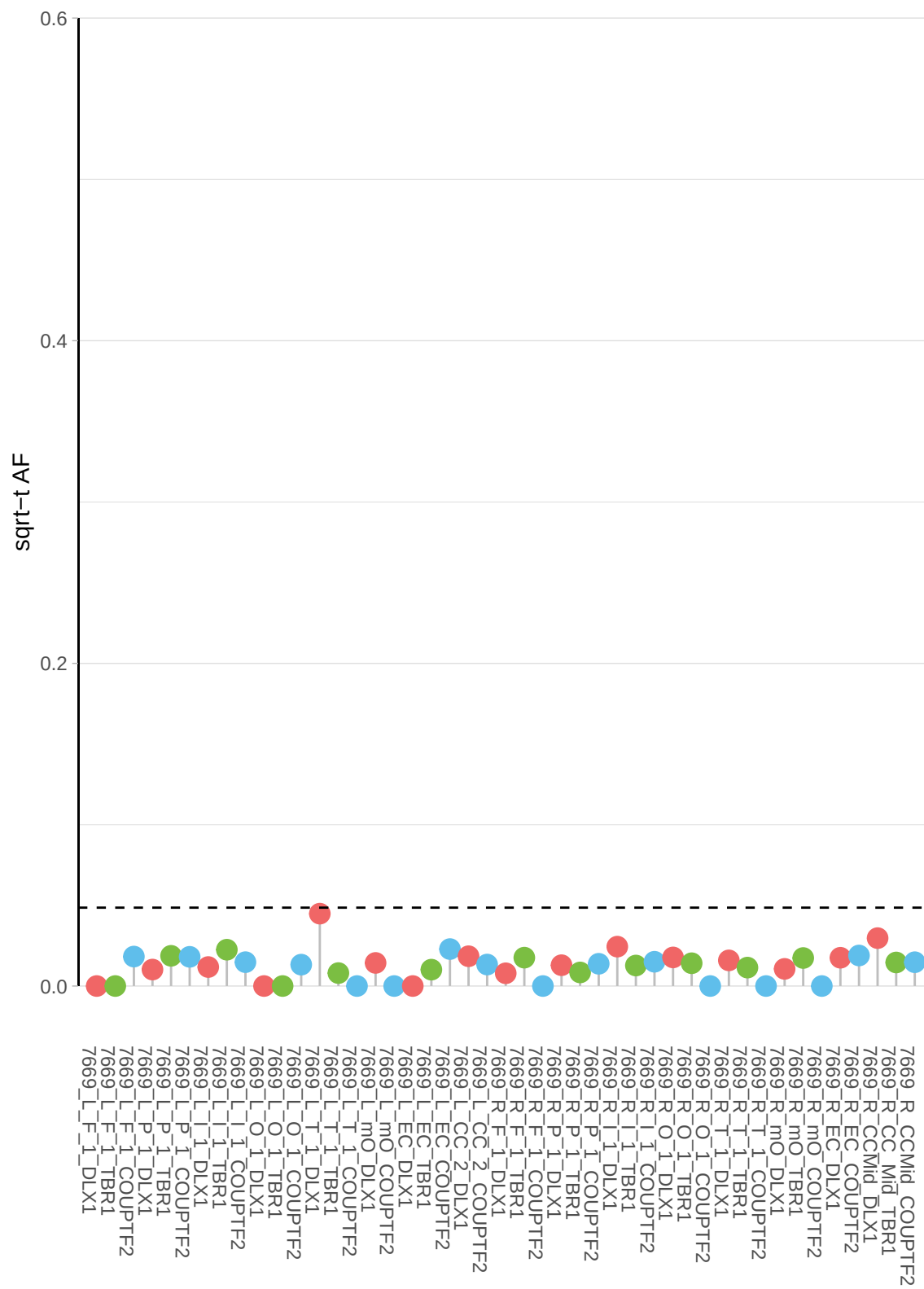

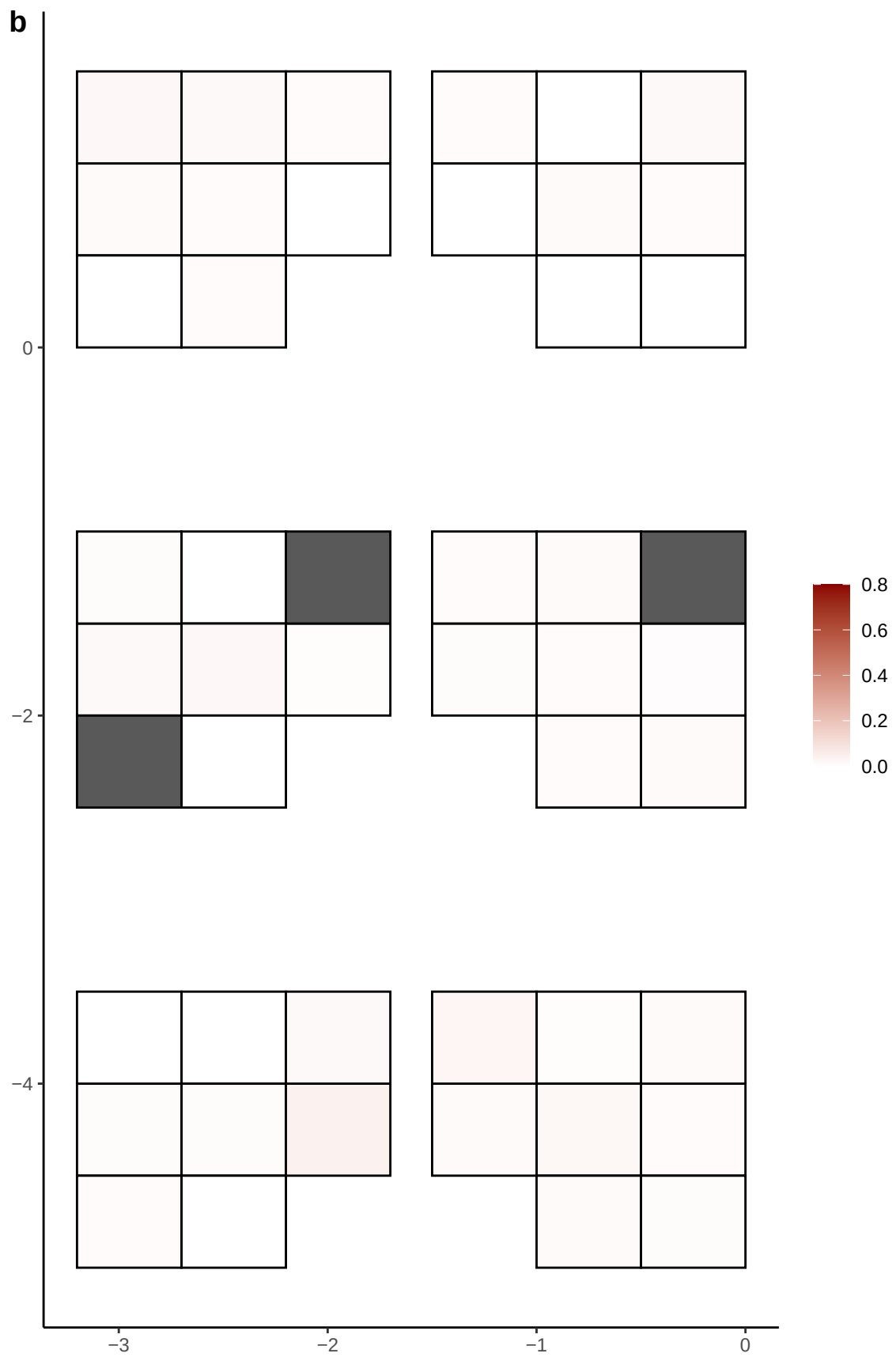

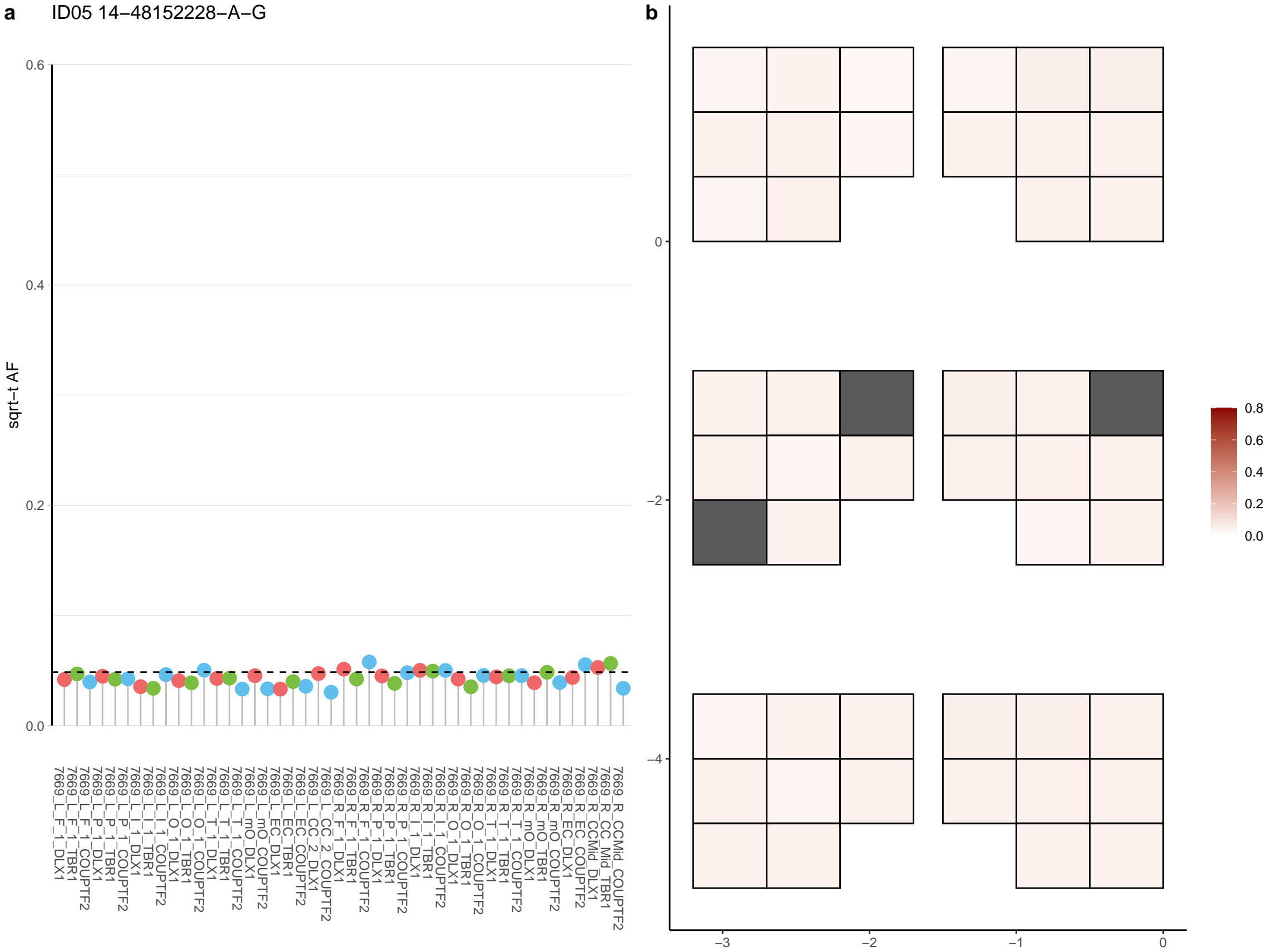

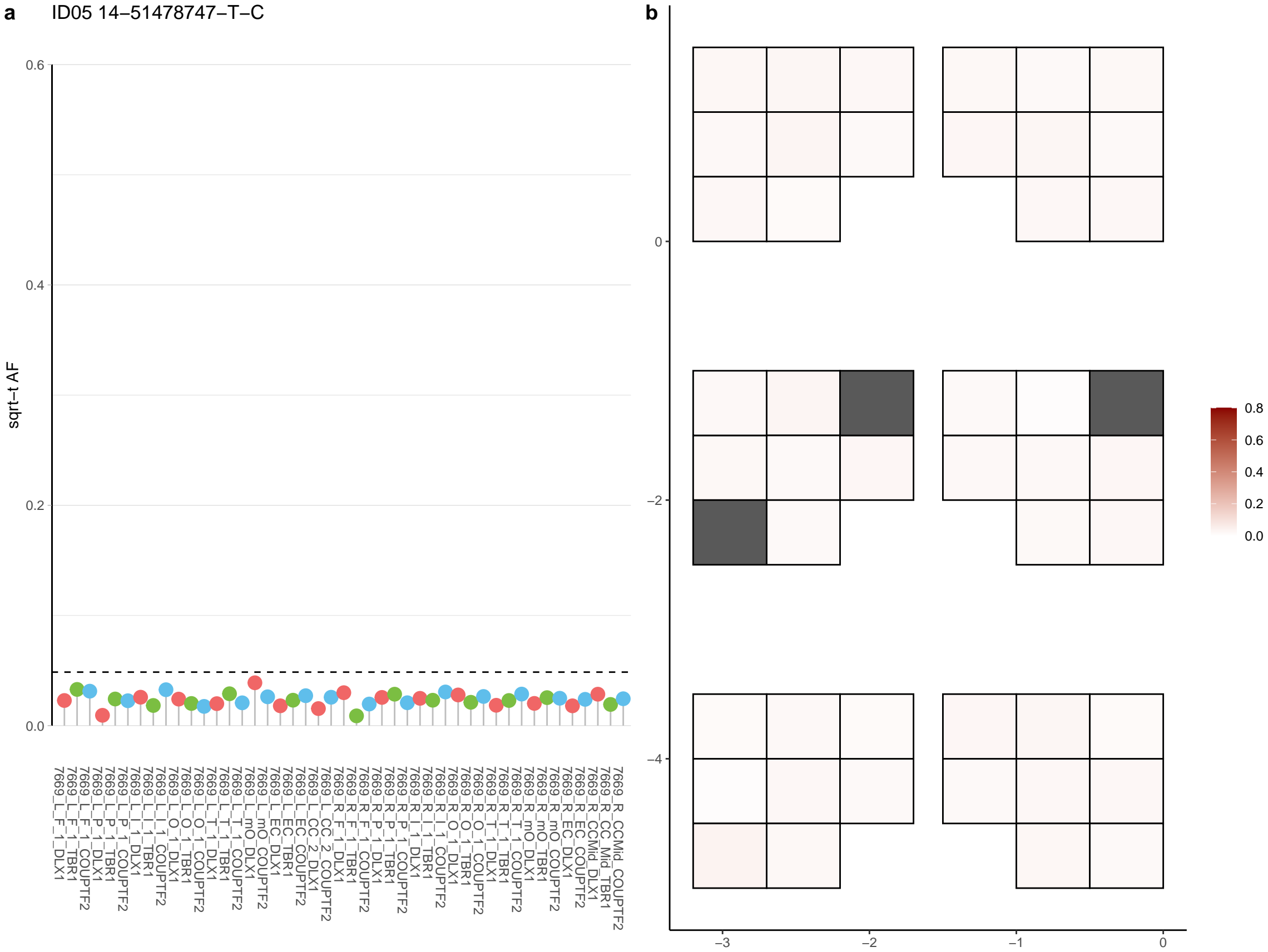

**a** ID05 14-55887203-A-G

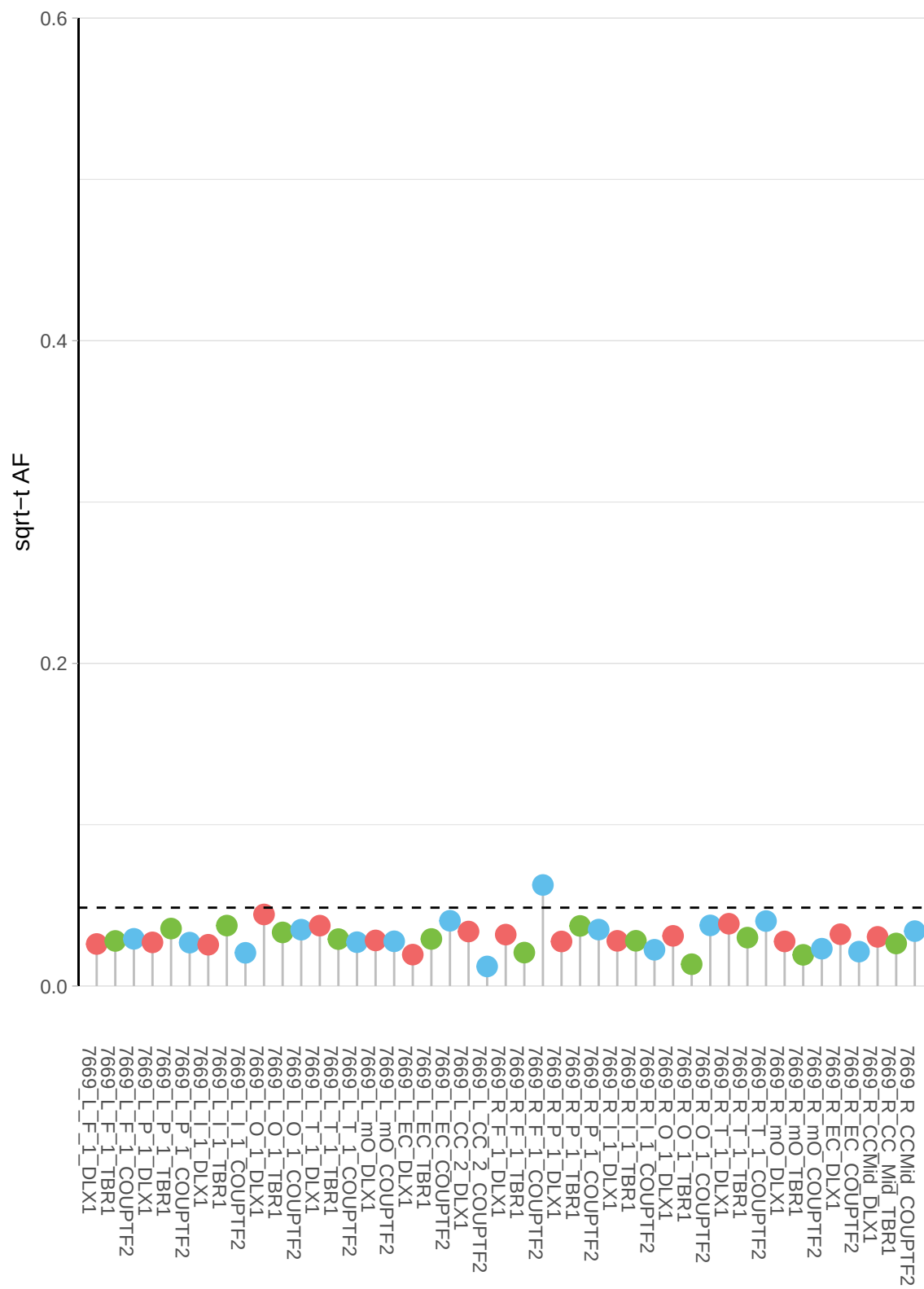

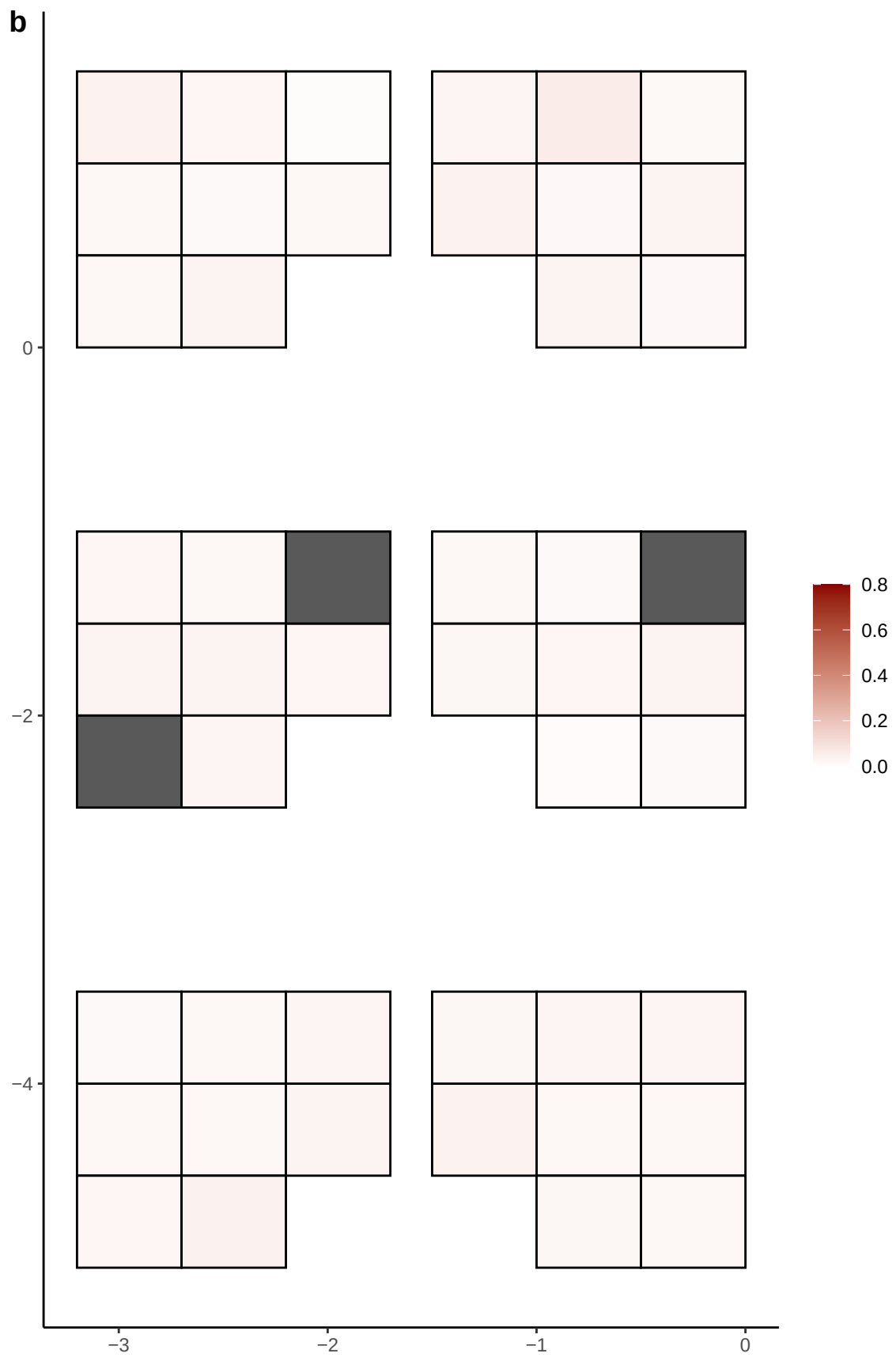

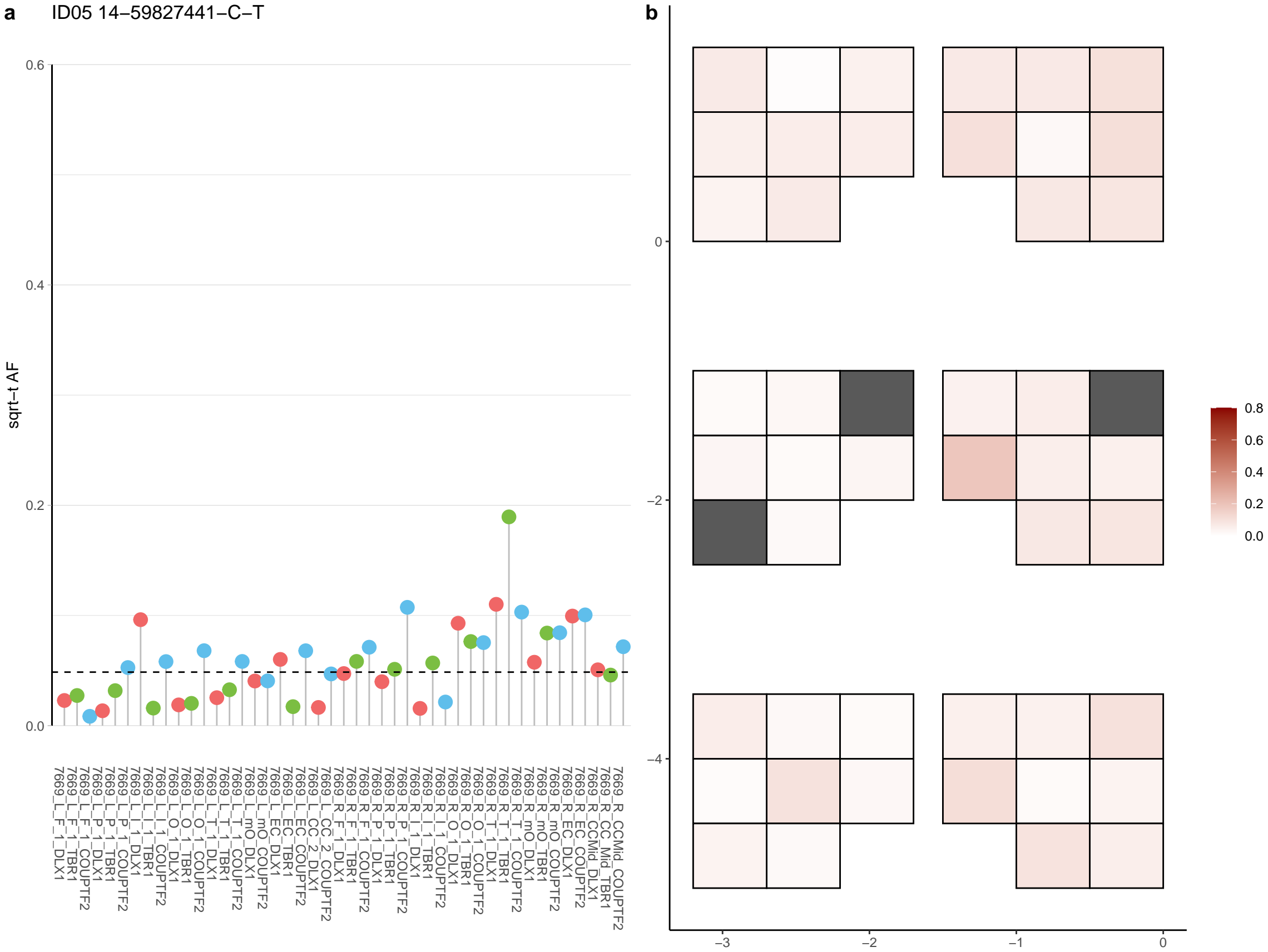

**a** ID05 14-65264866-C-T

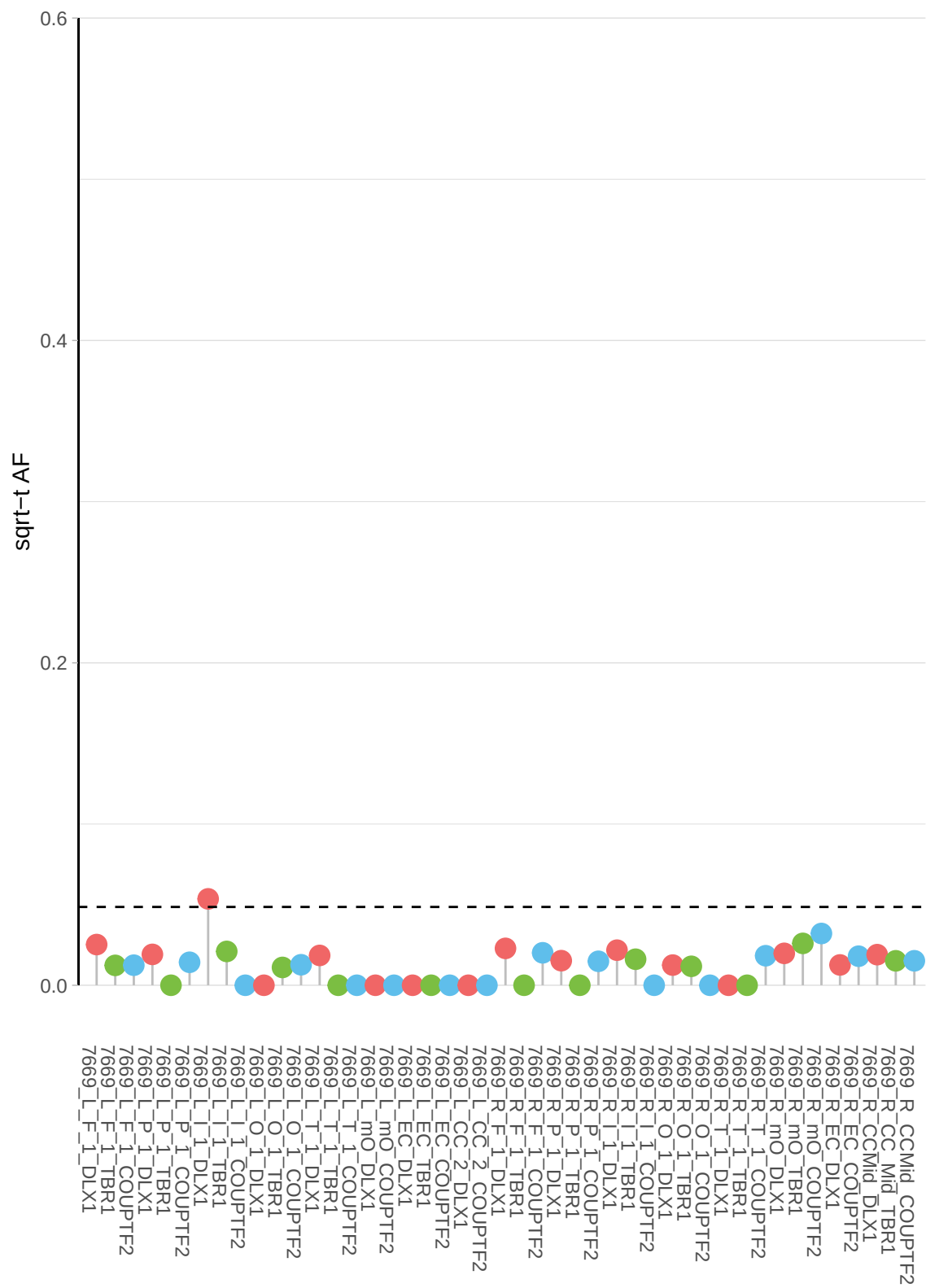

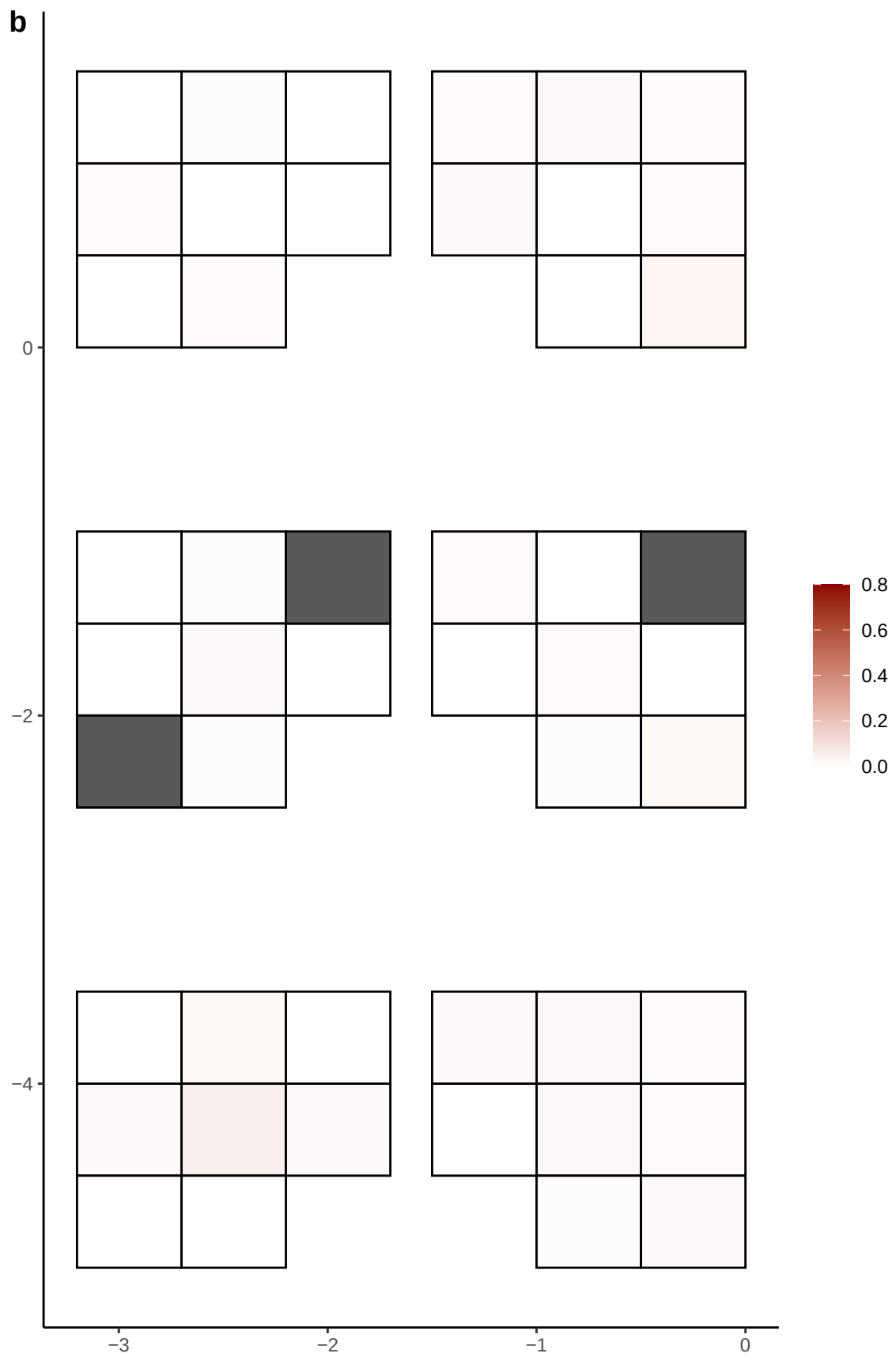

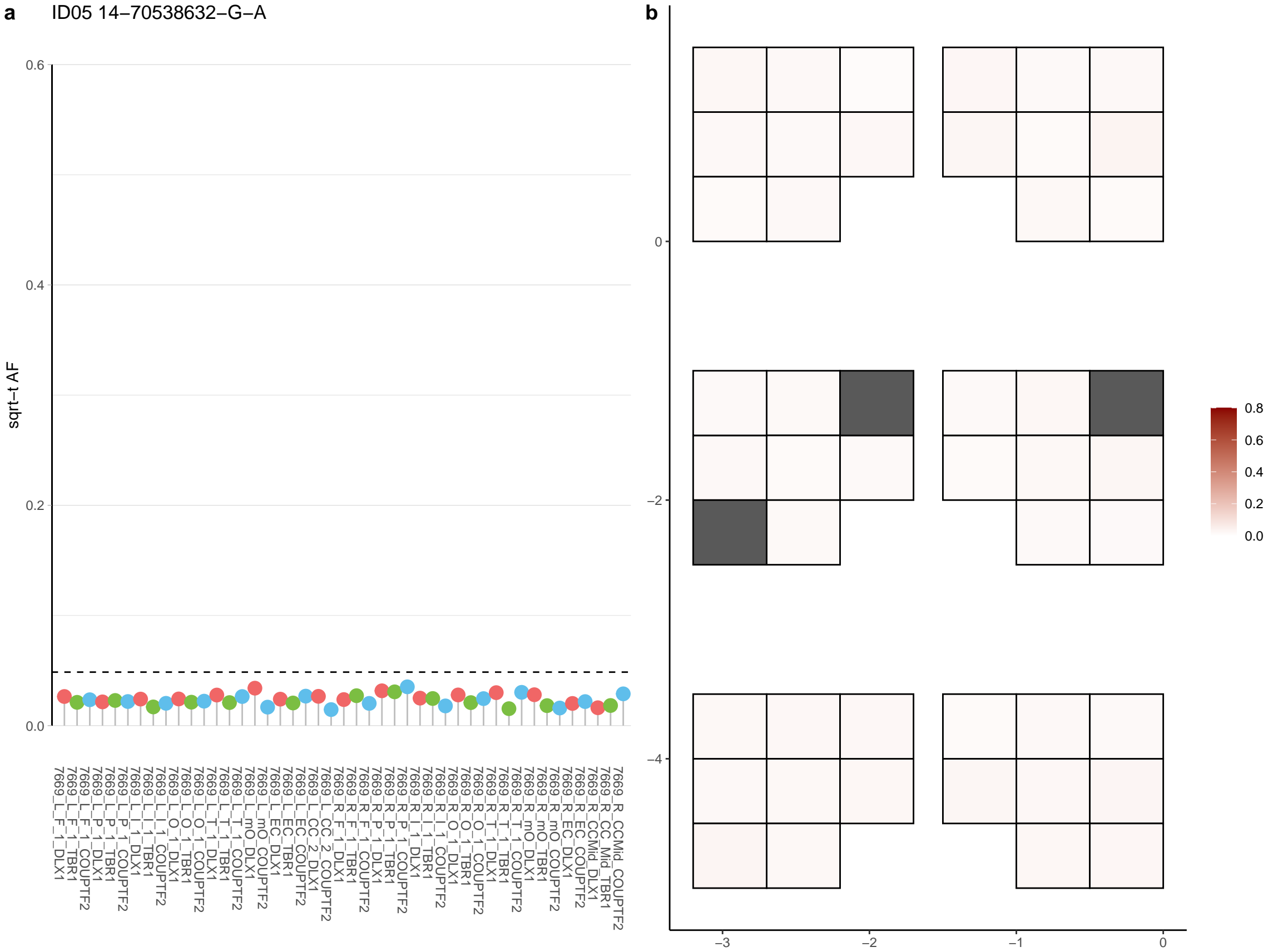

**a** ID05 14-77126705-G-A

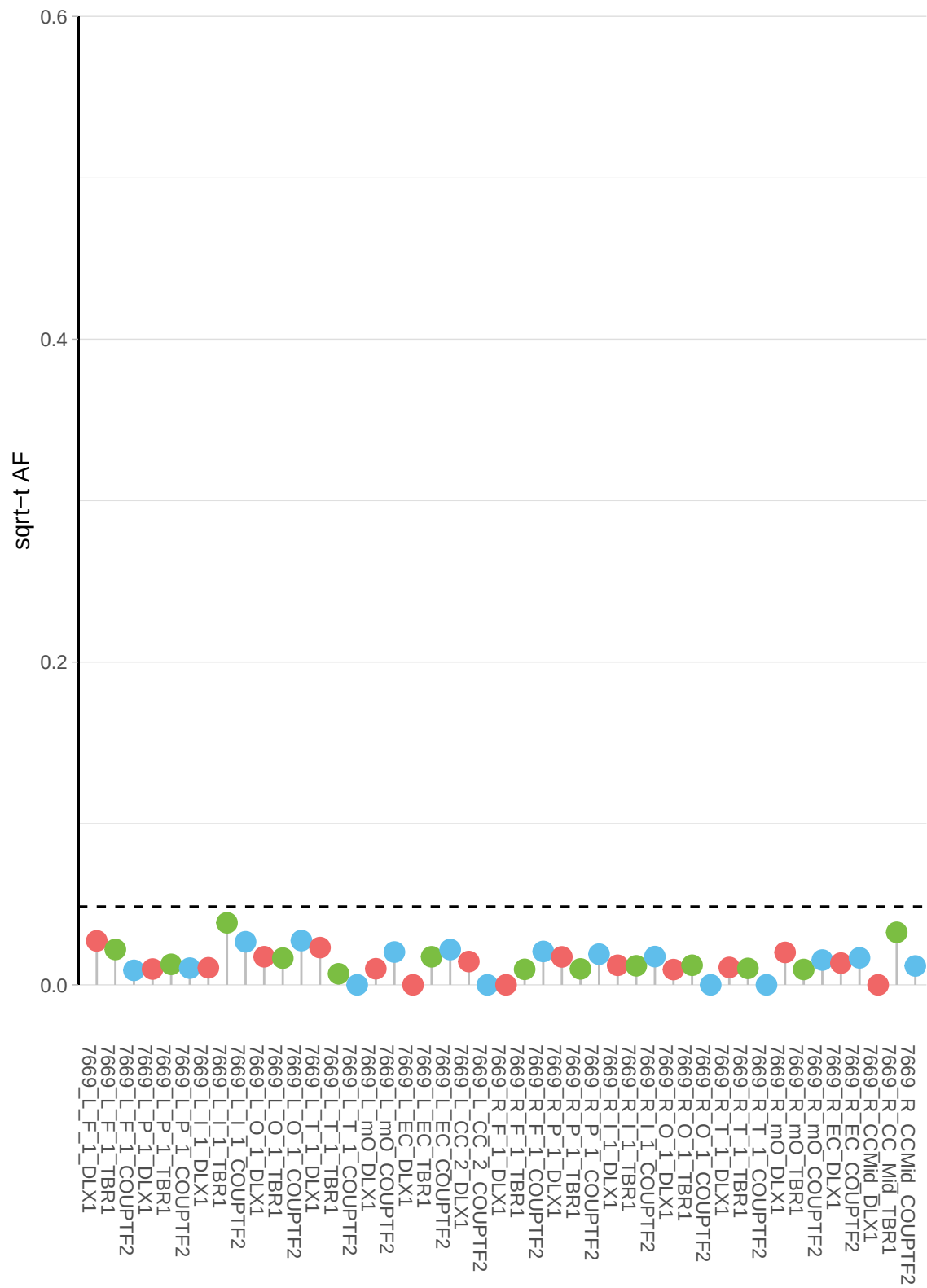

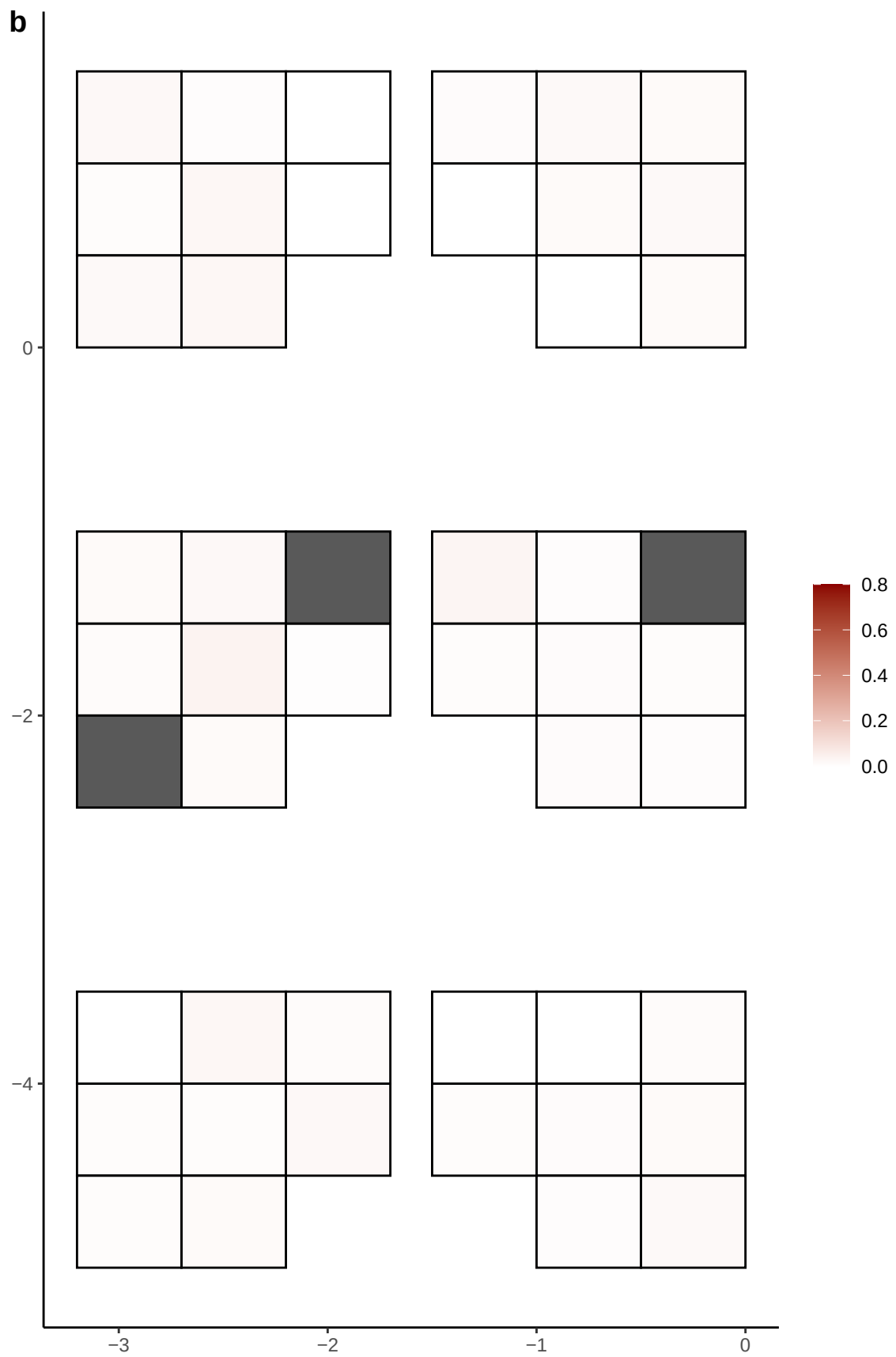

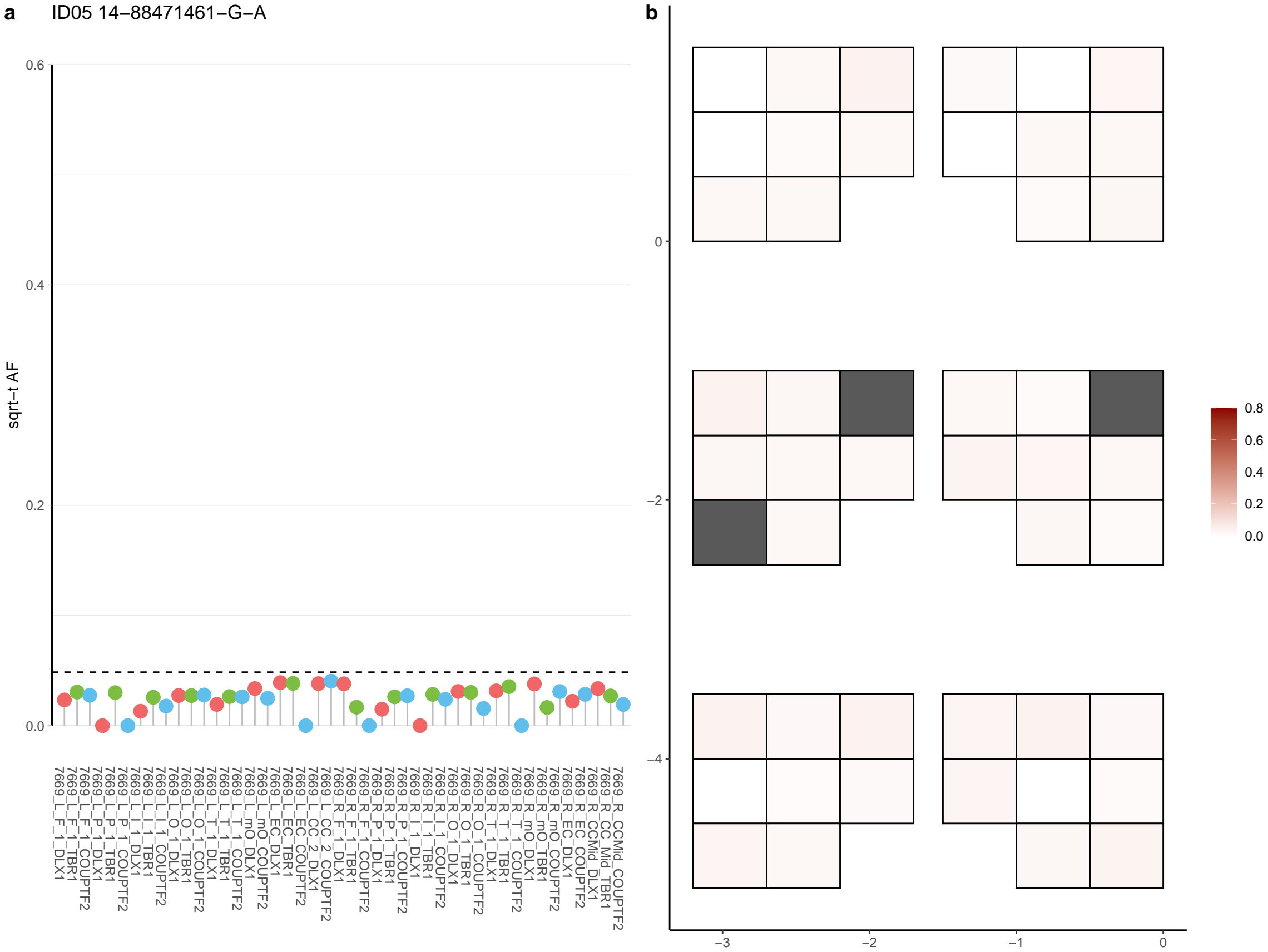

**a** ID05 14-89375012-G-T

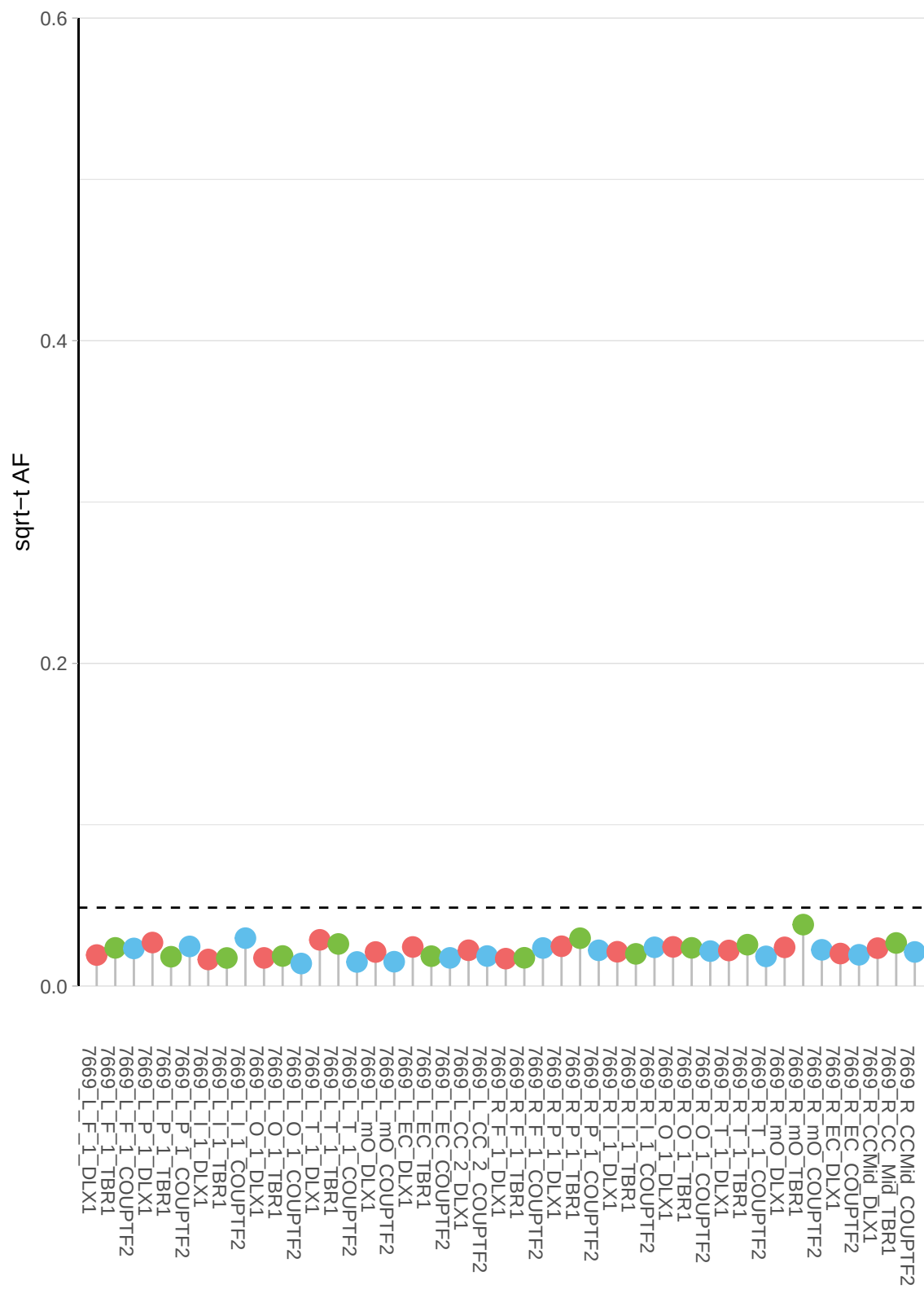

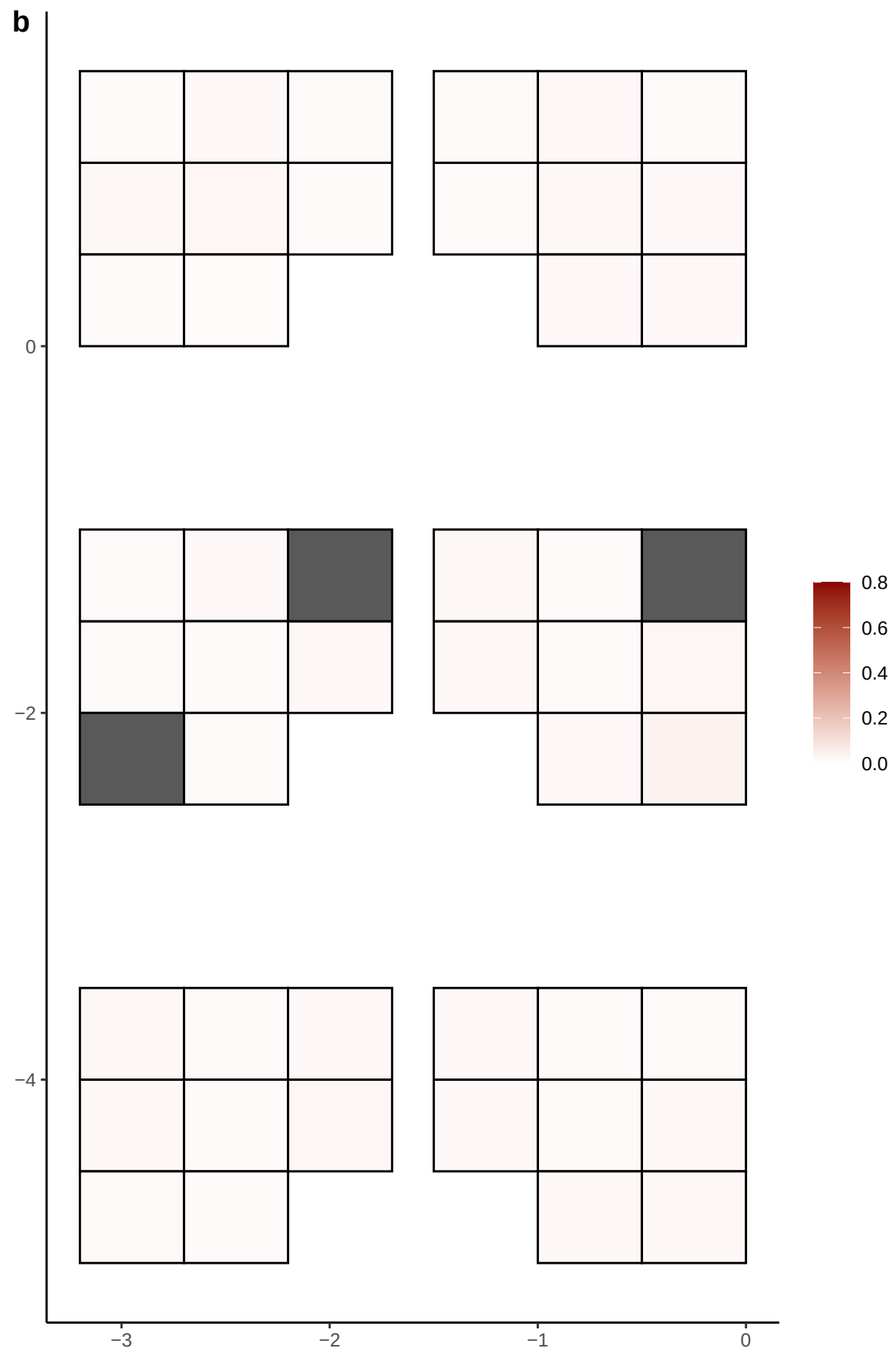

**a** ID05 14-93813737-C-T

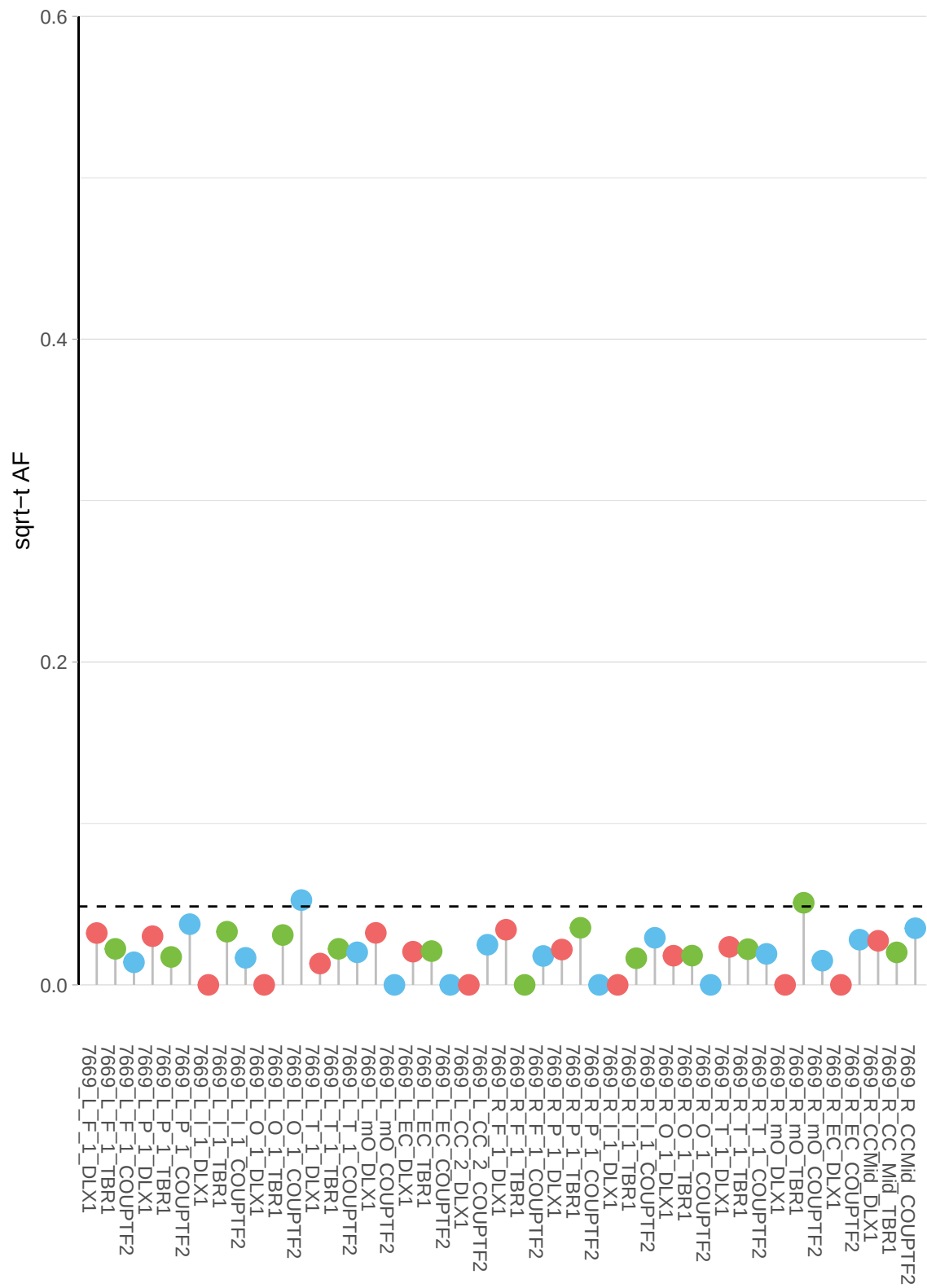

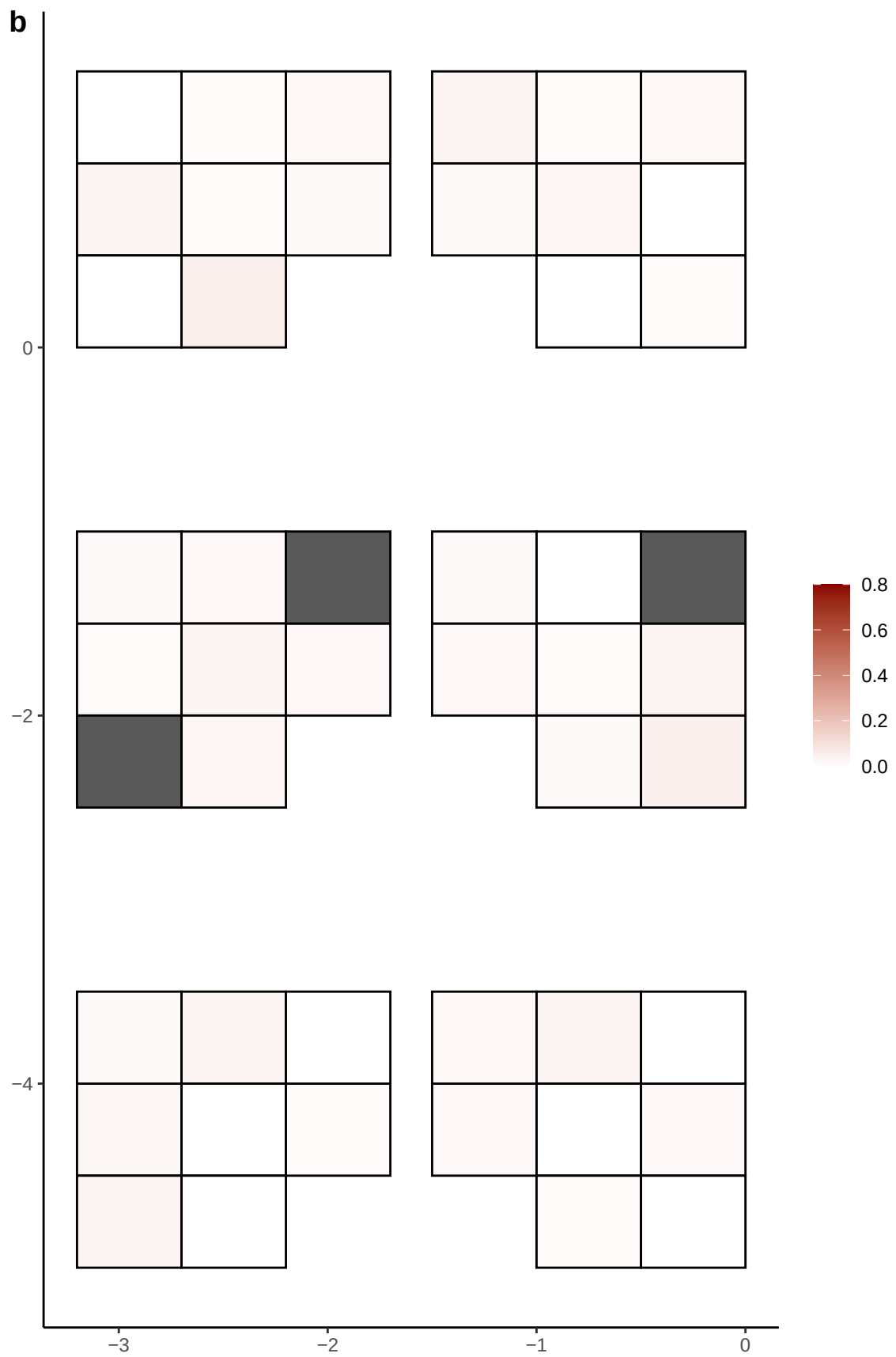

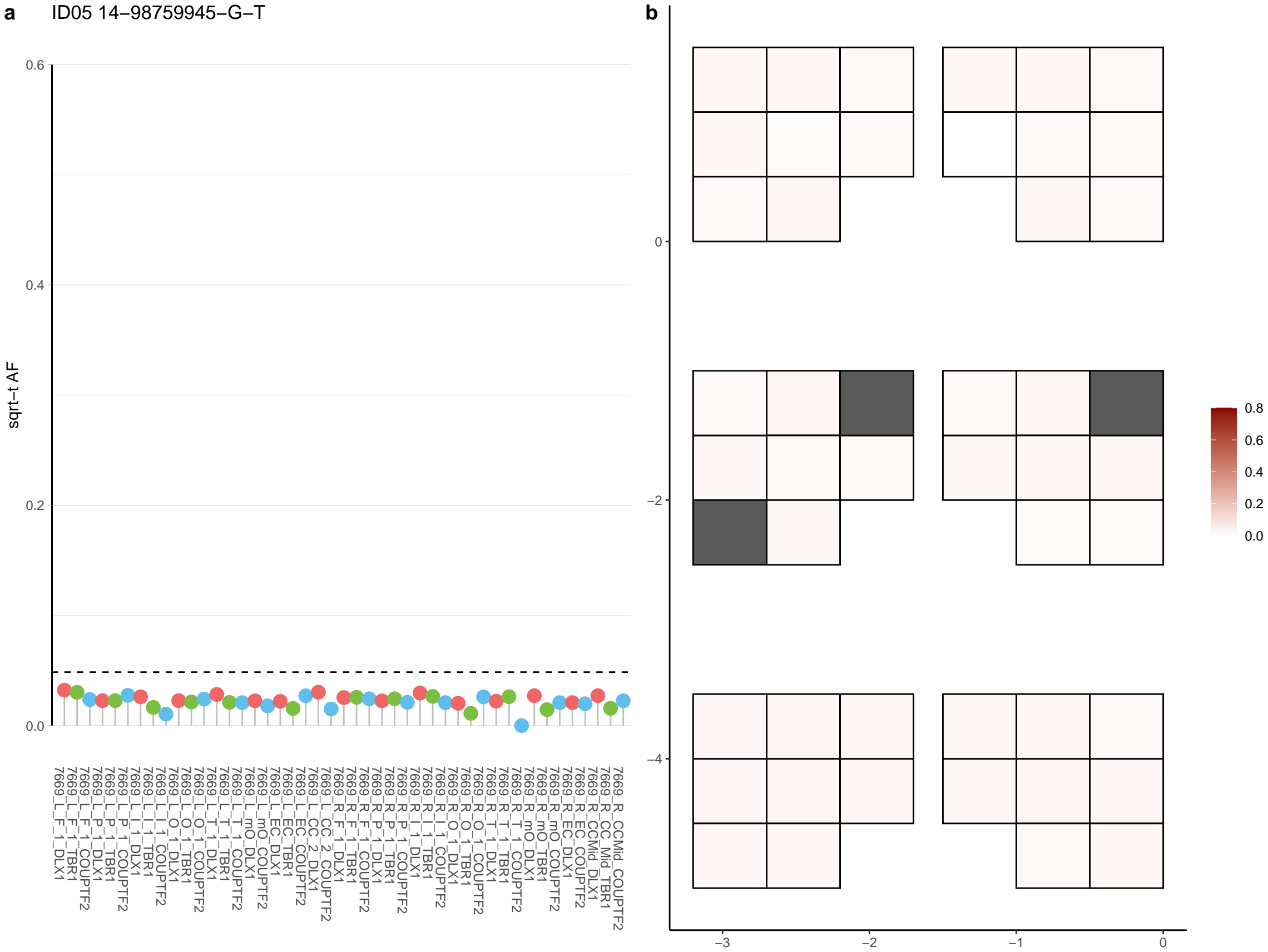

**a** ID05 15-26356007-C-T

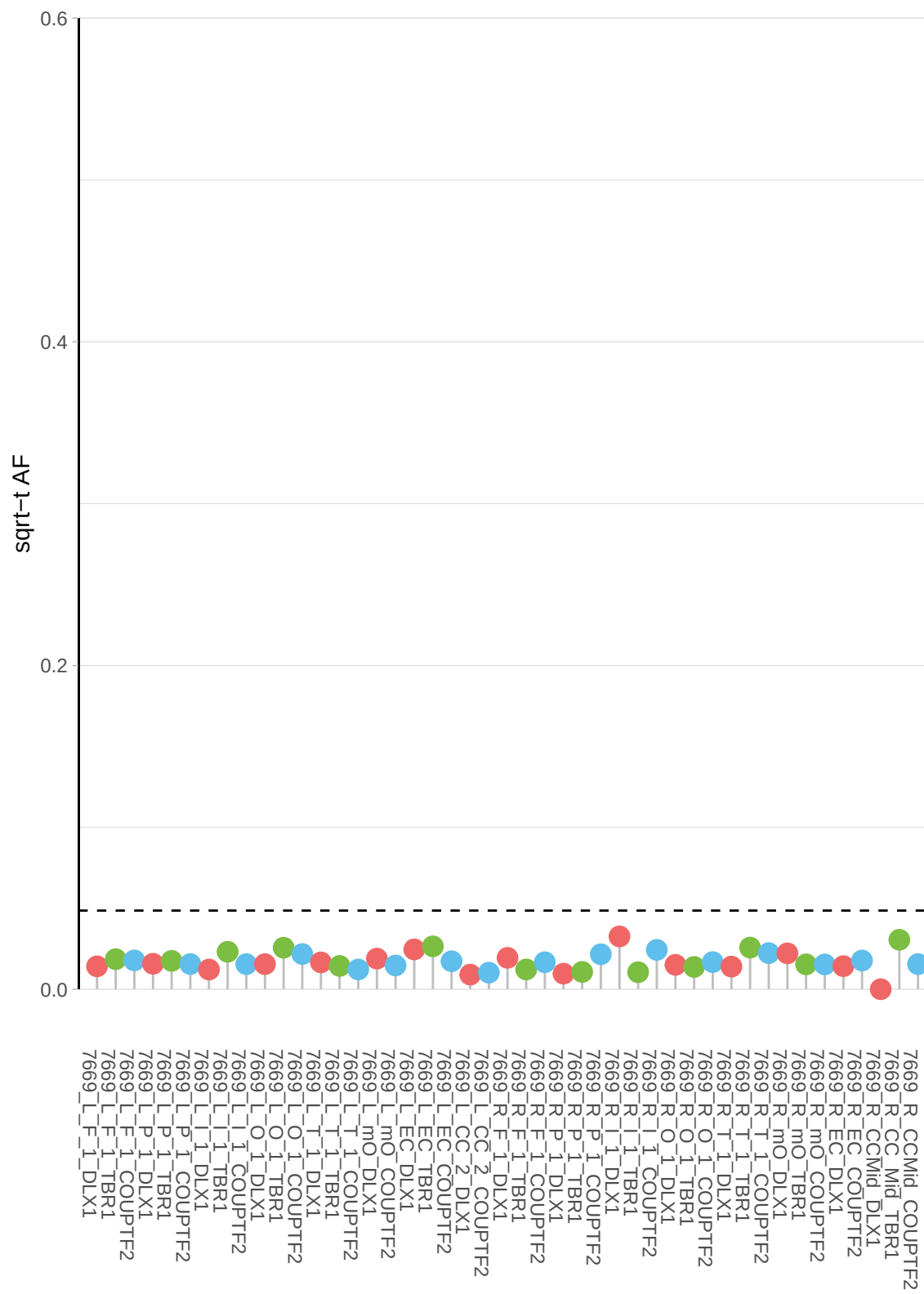

**b**

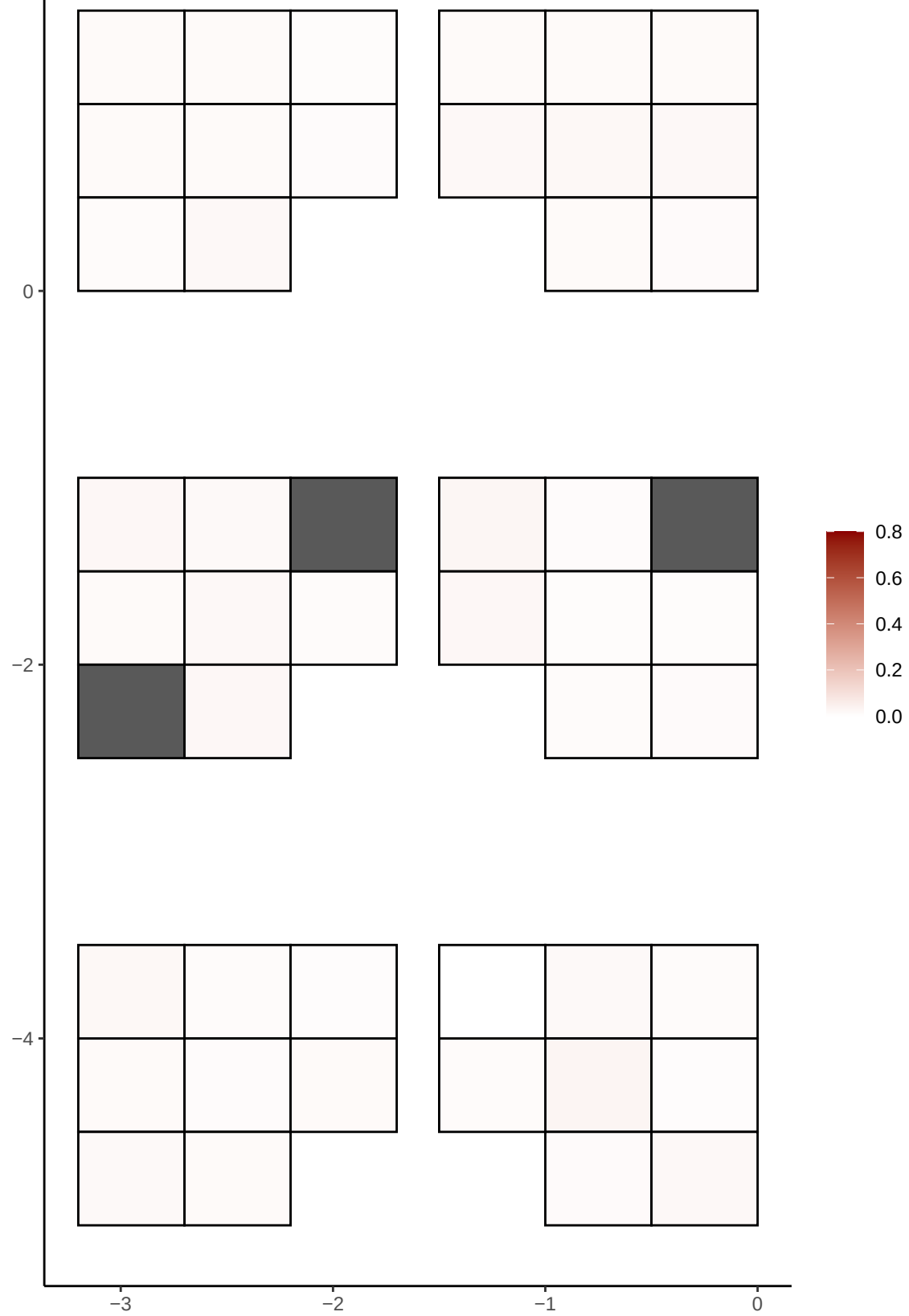

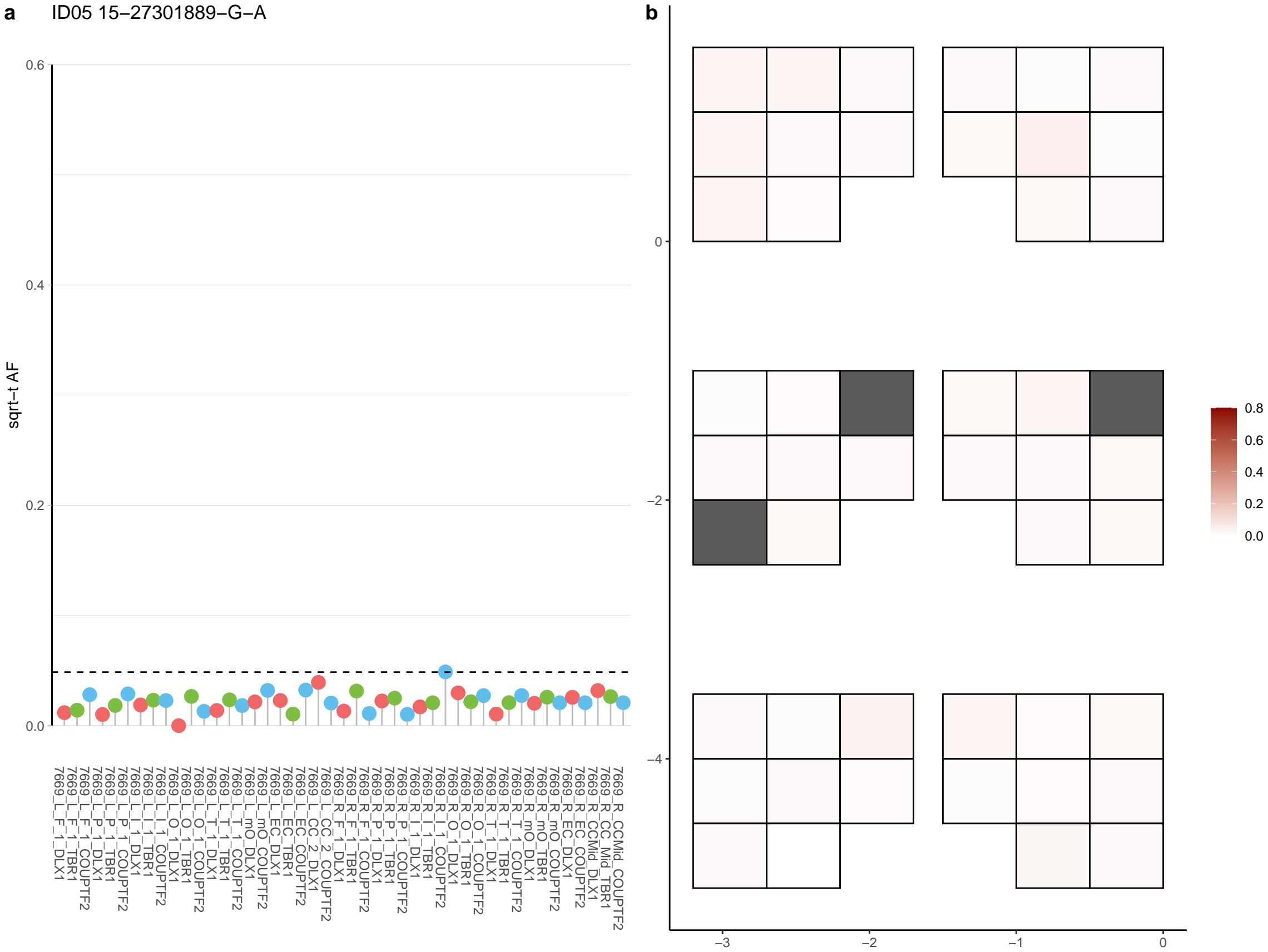

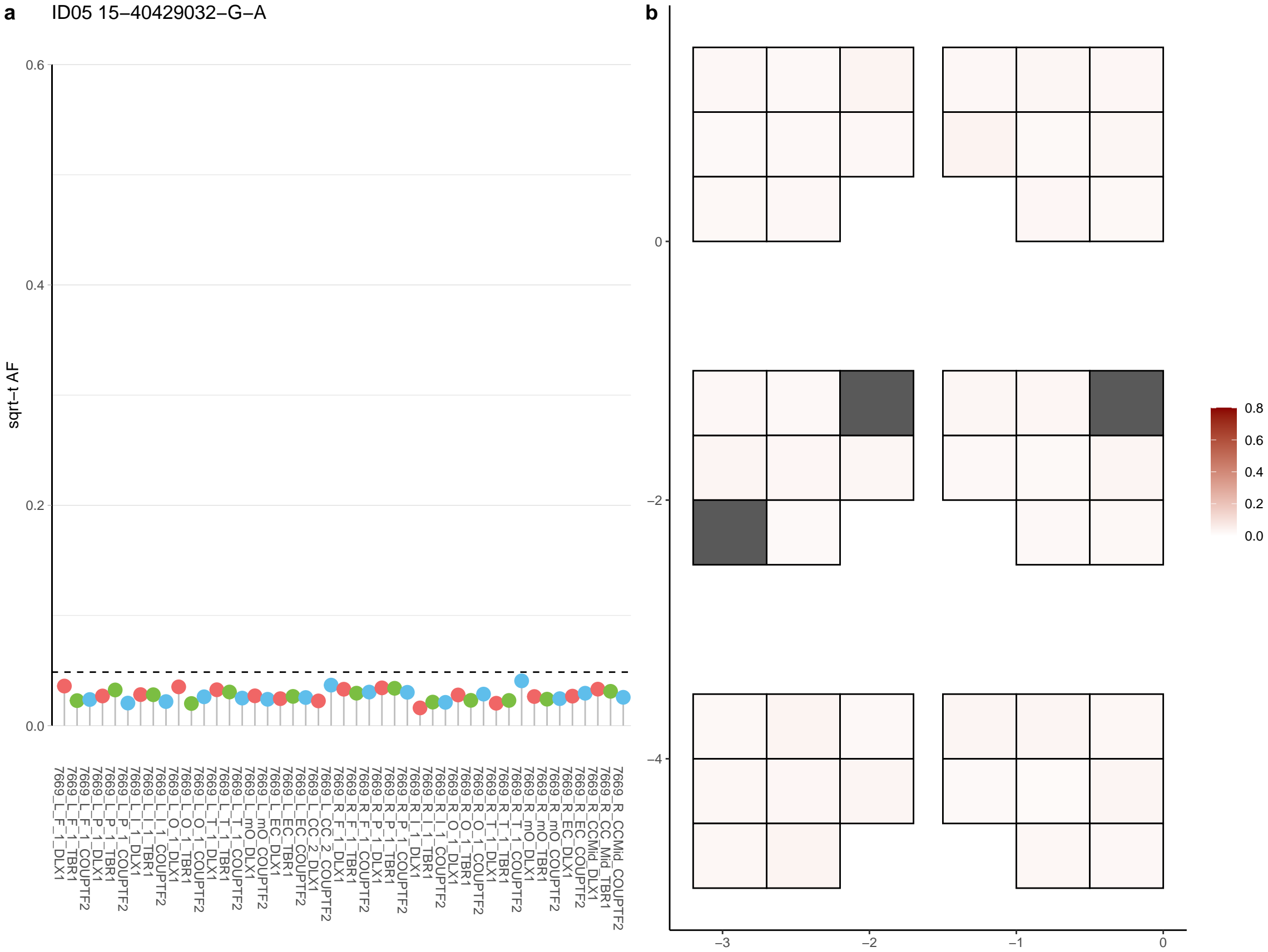

**a** ID05 15-46932376-G-A

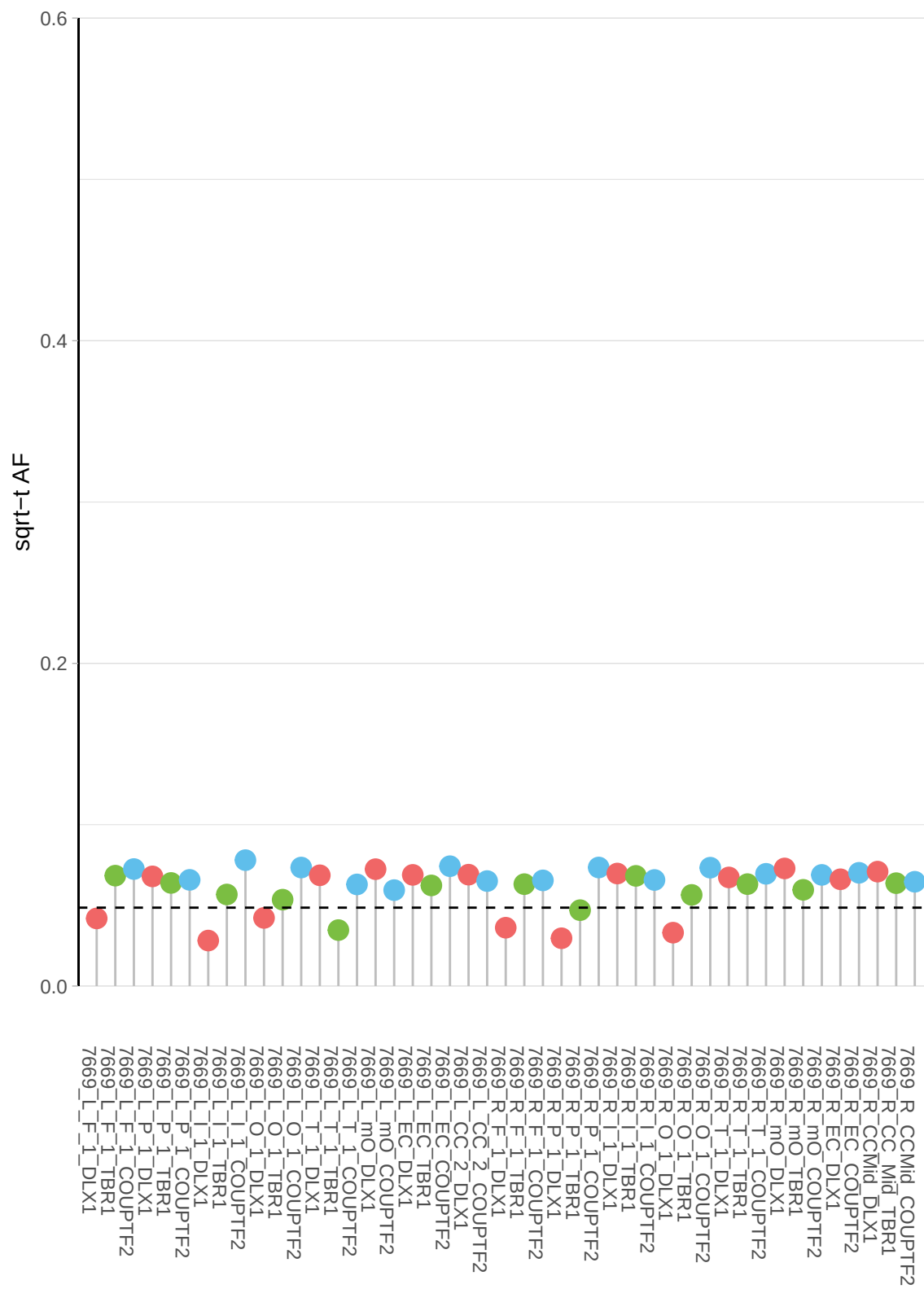

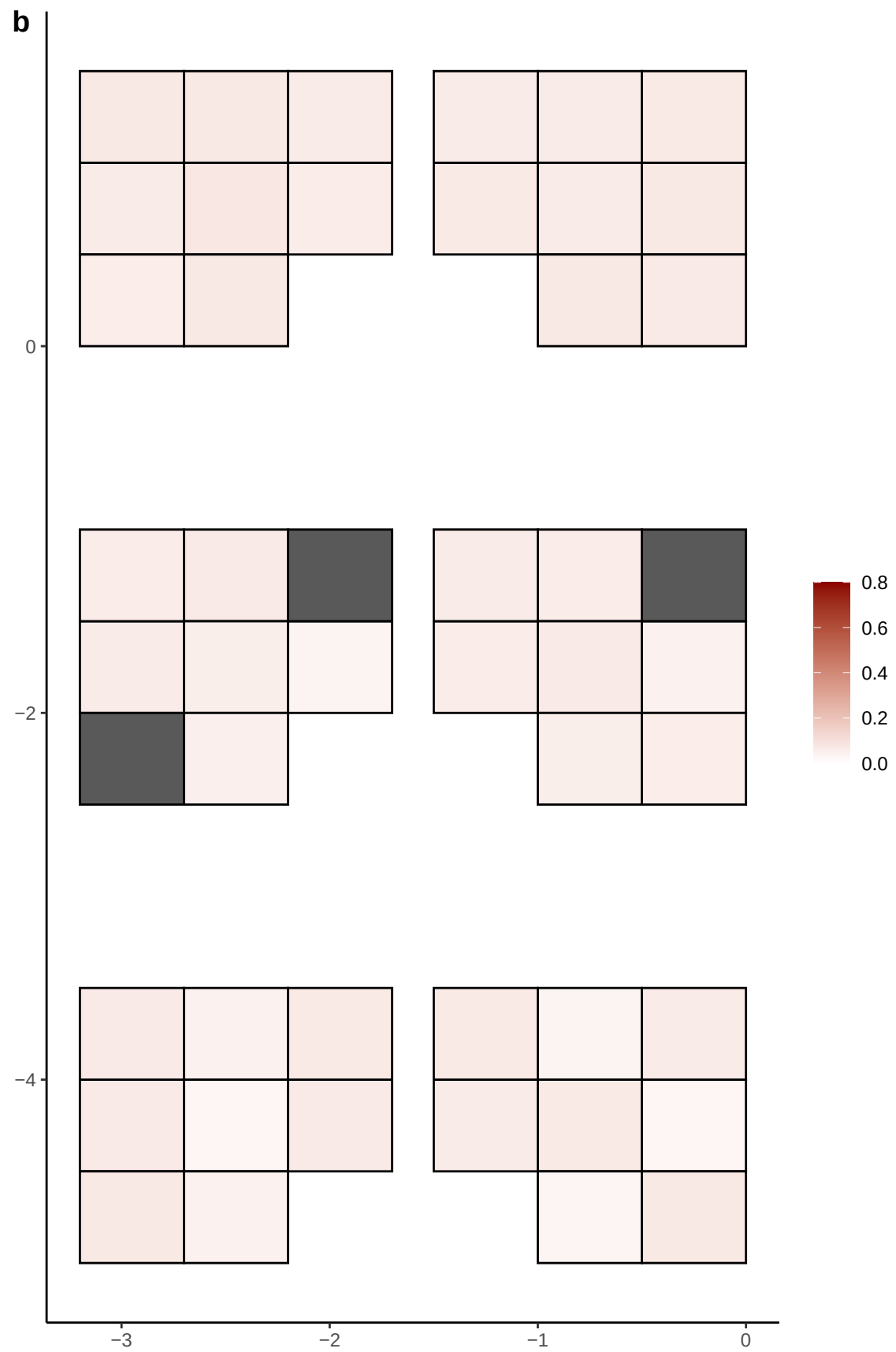

**a** ID05 15-46962188-T-A

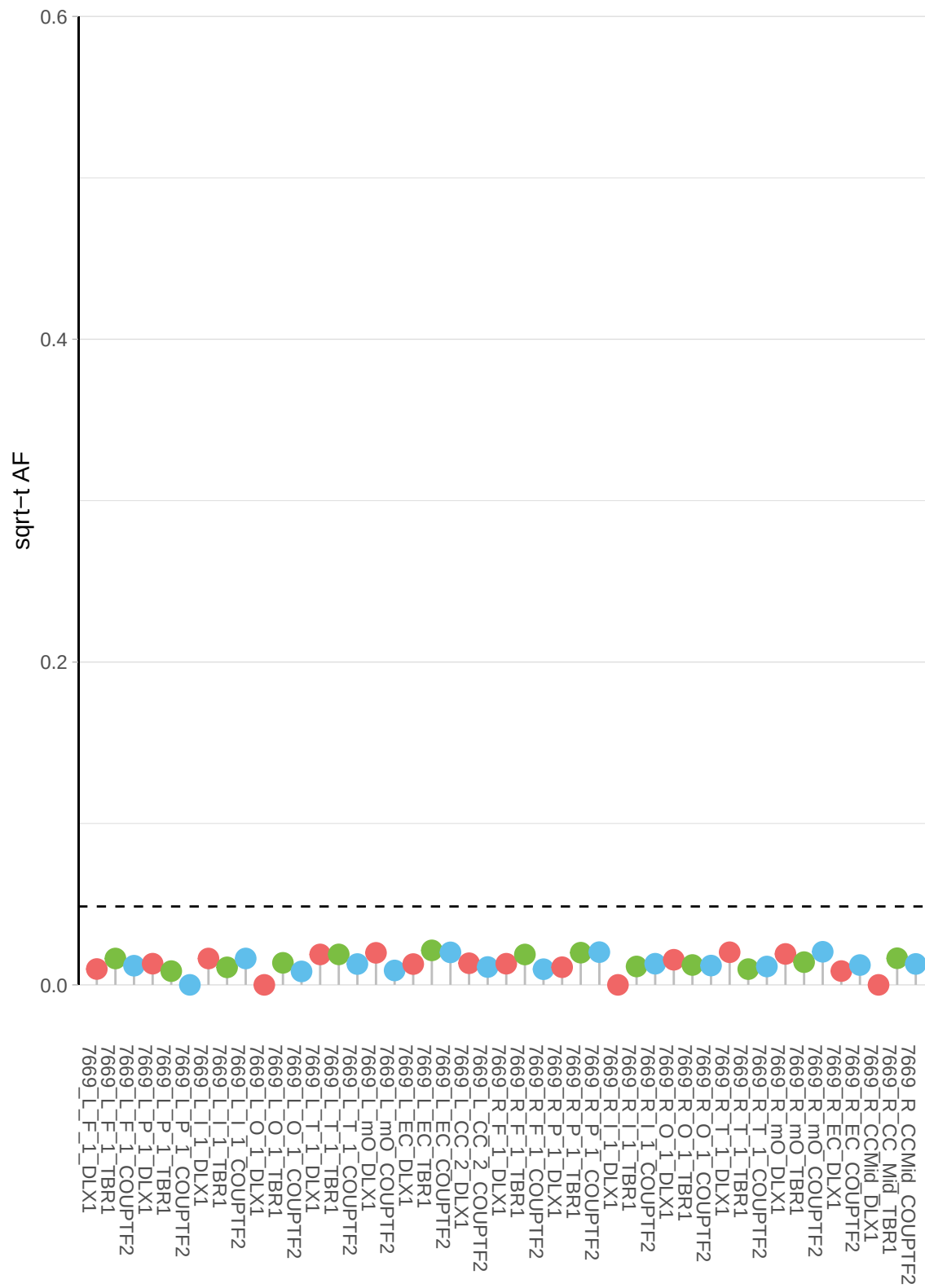

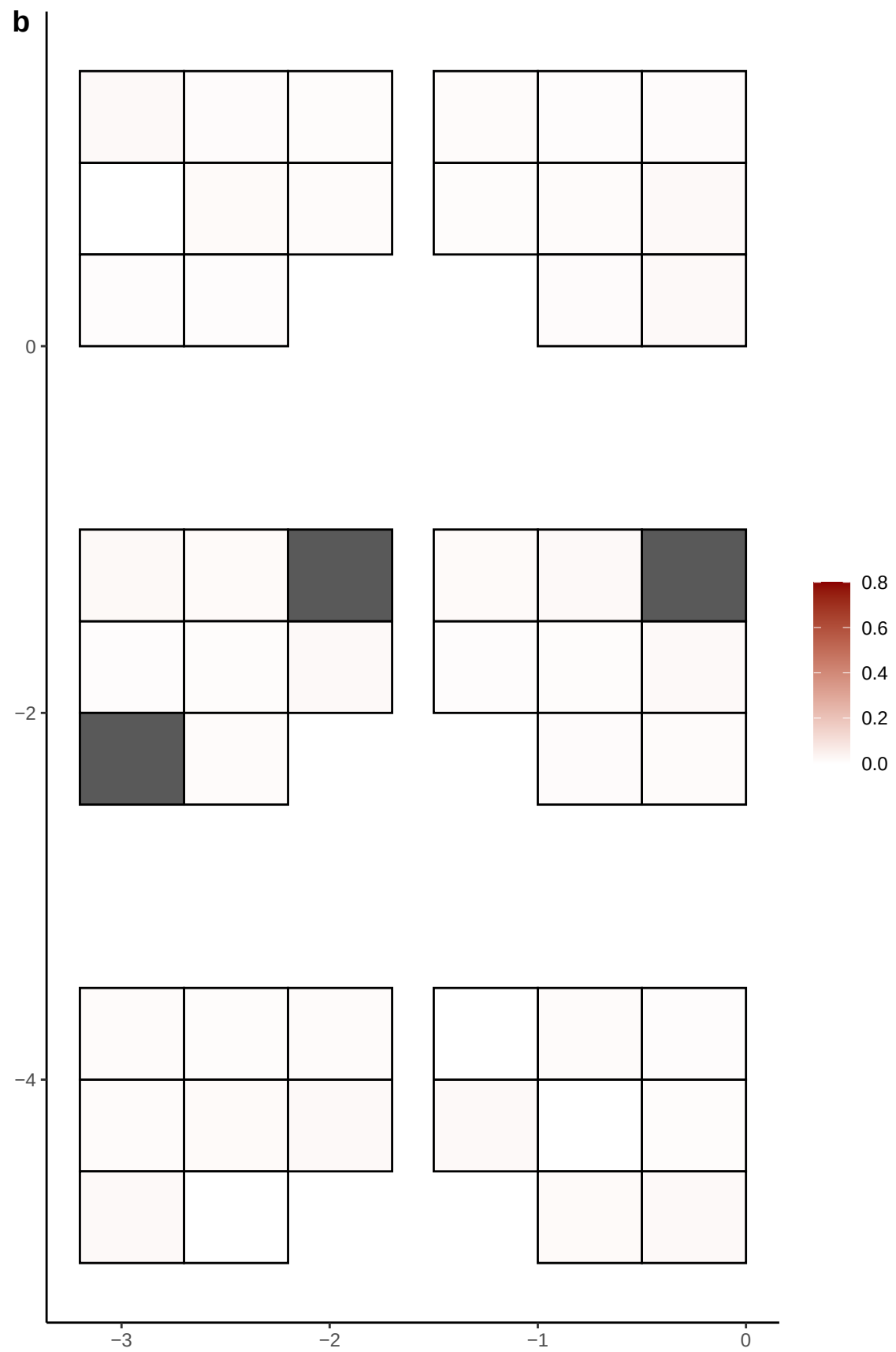

**a** ID05 15-51489592-C-T

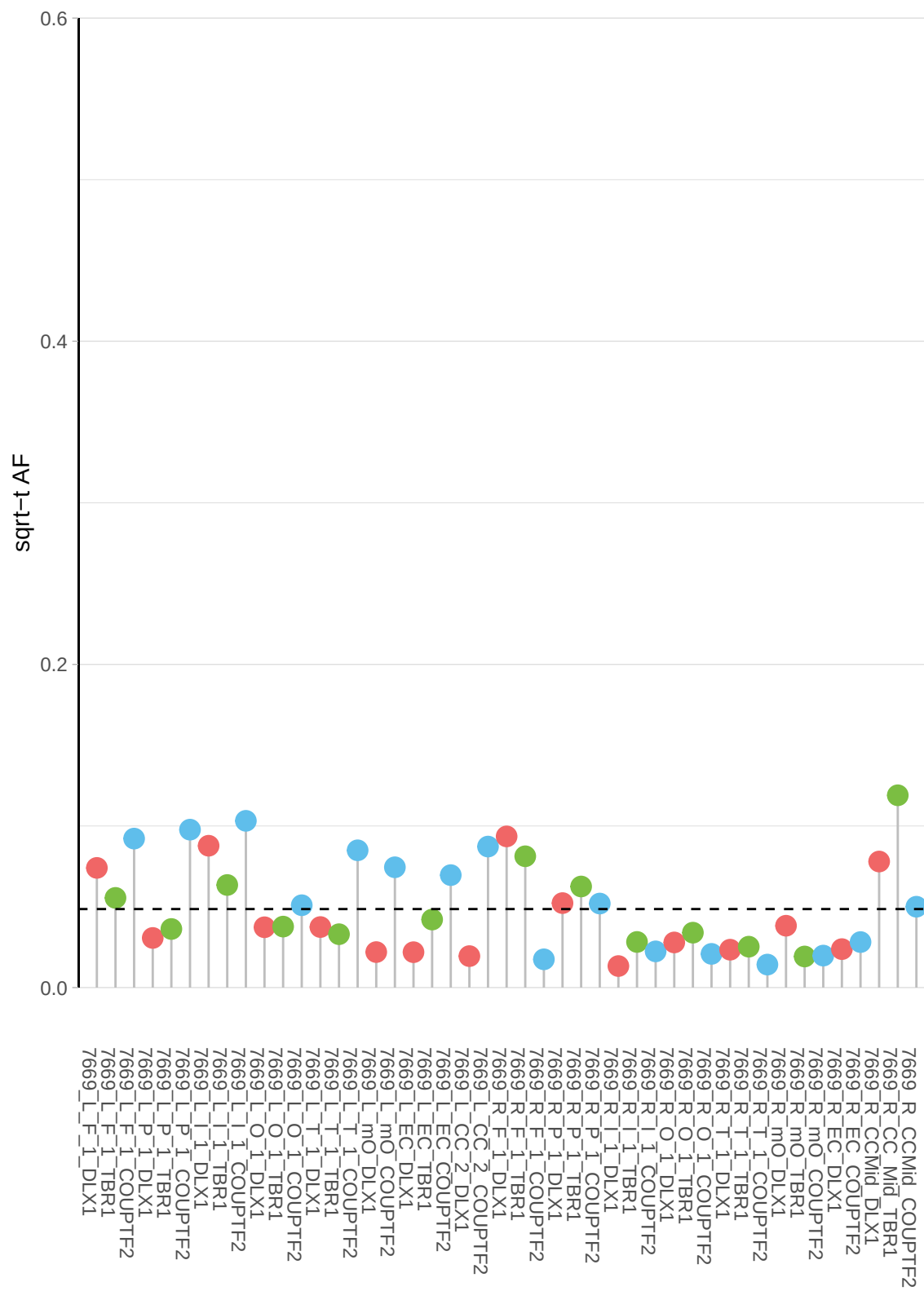

**b**

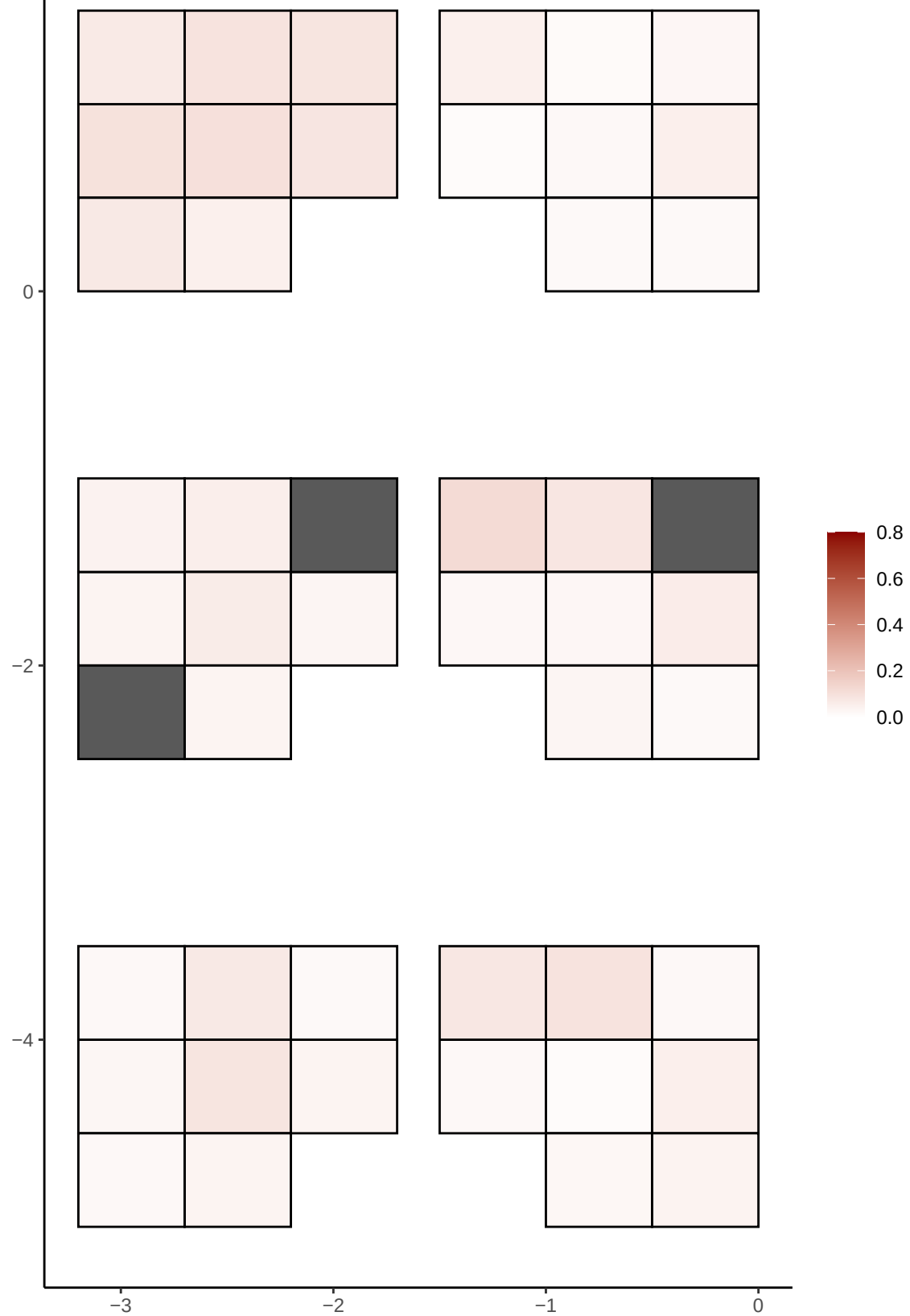

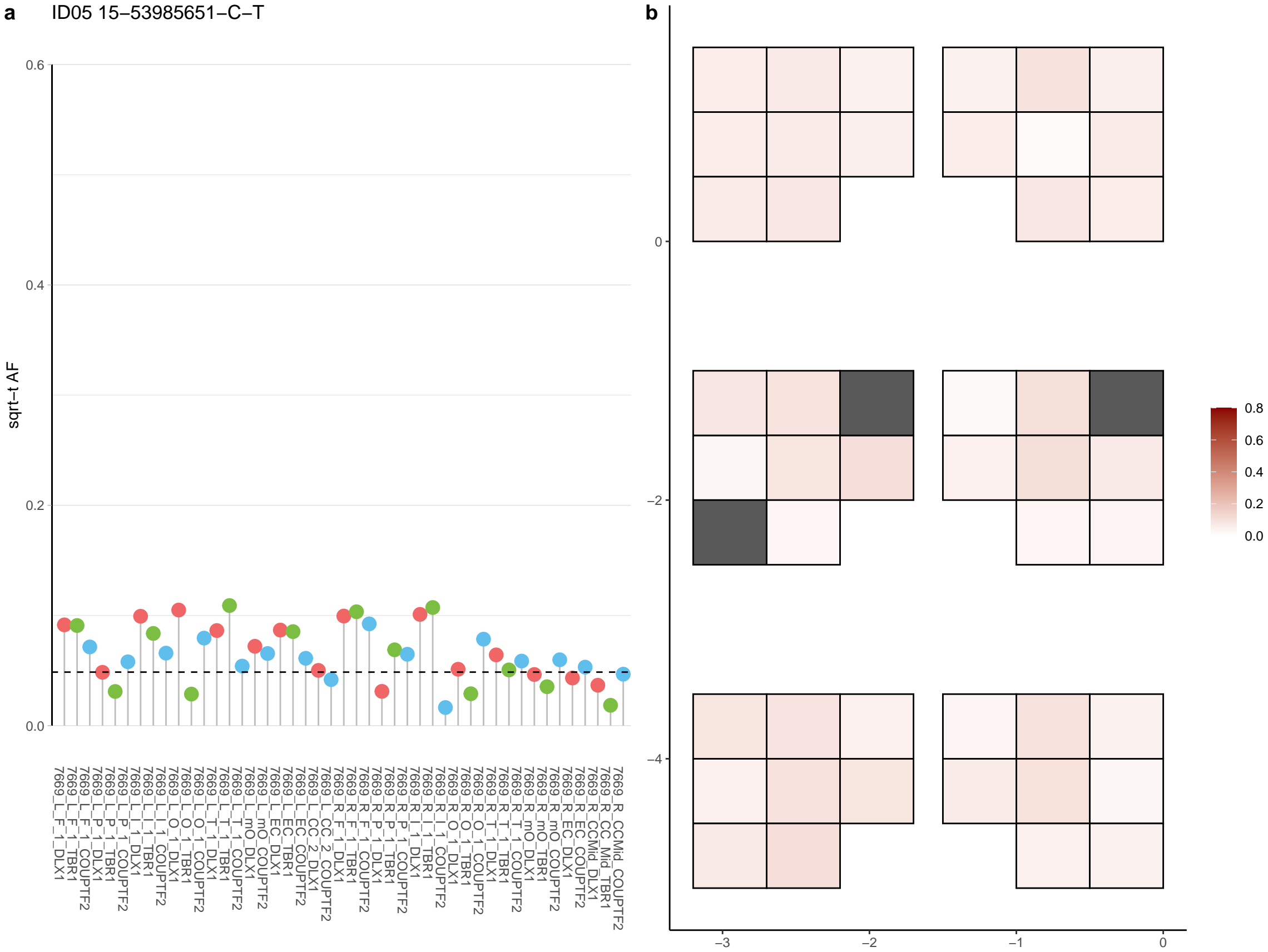

**a** ID05 15-60164292-C-T

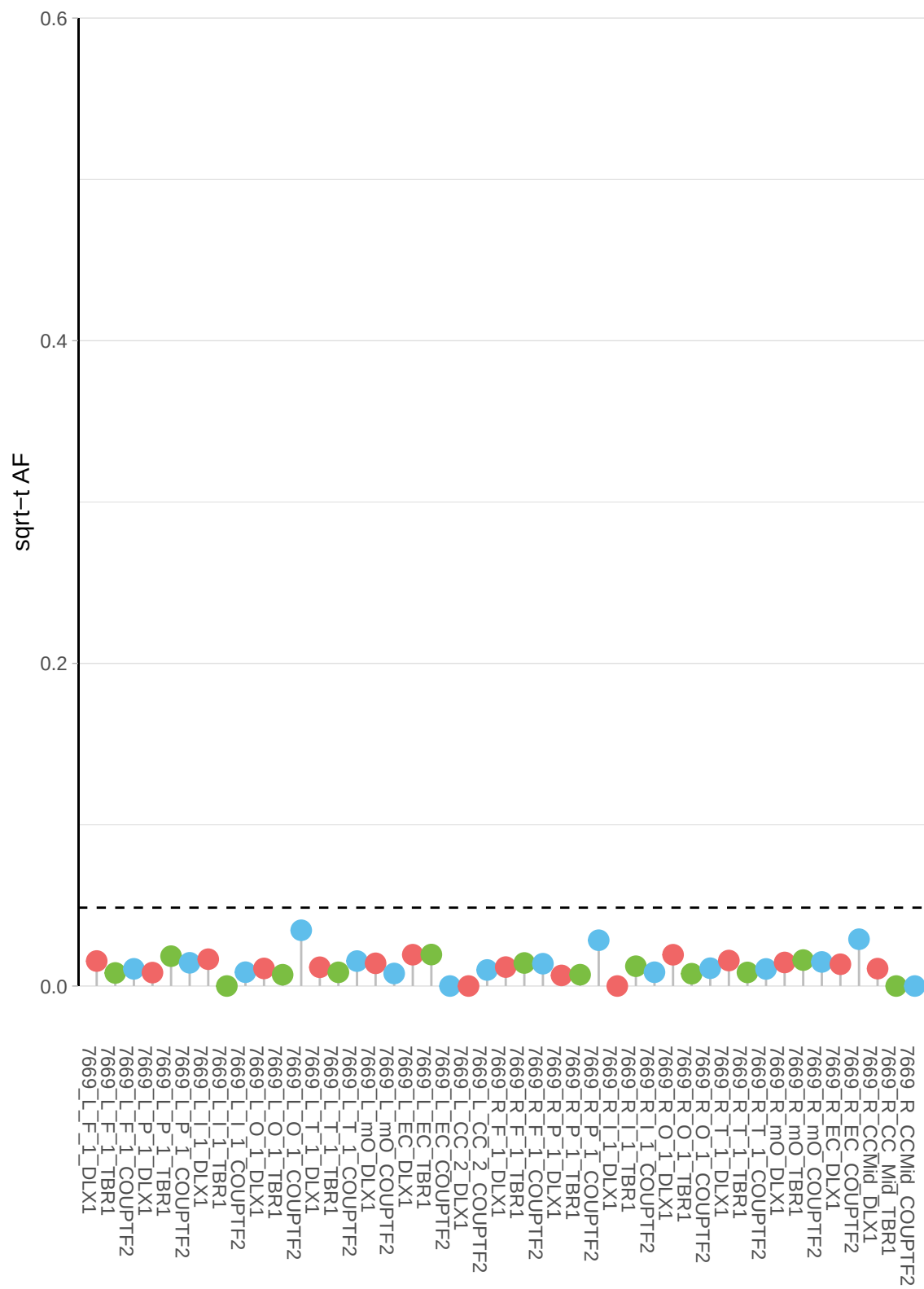

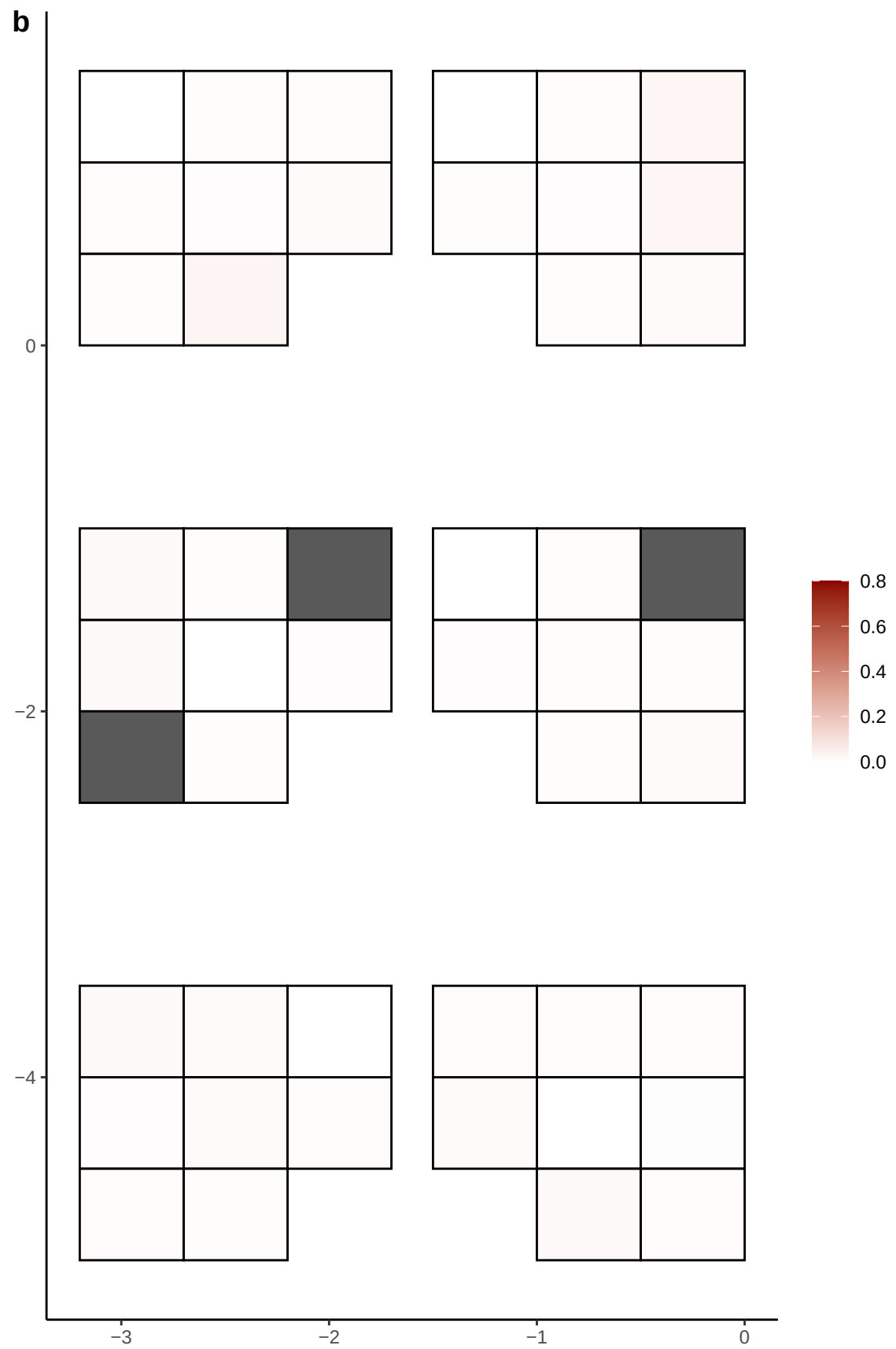

**a** ID05 15-68132566-T-C

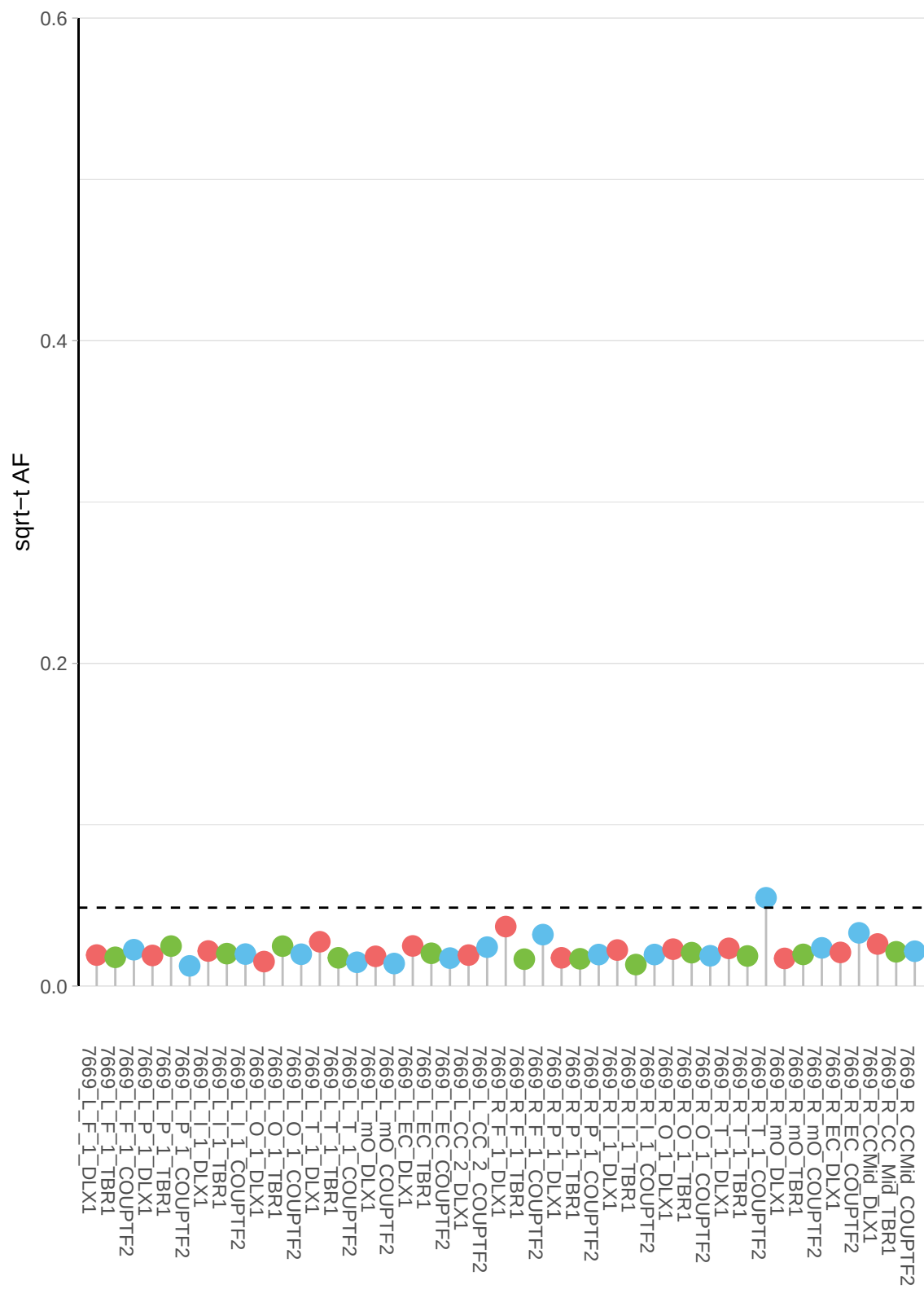

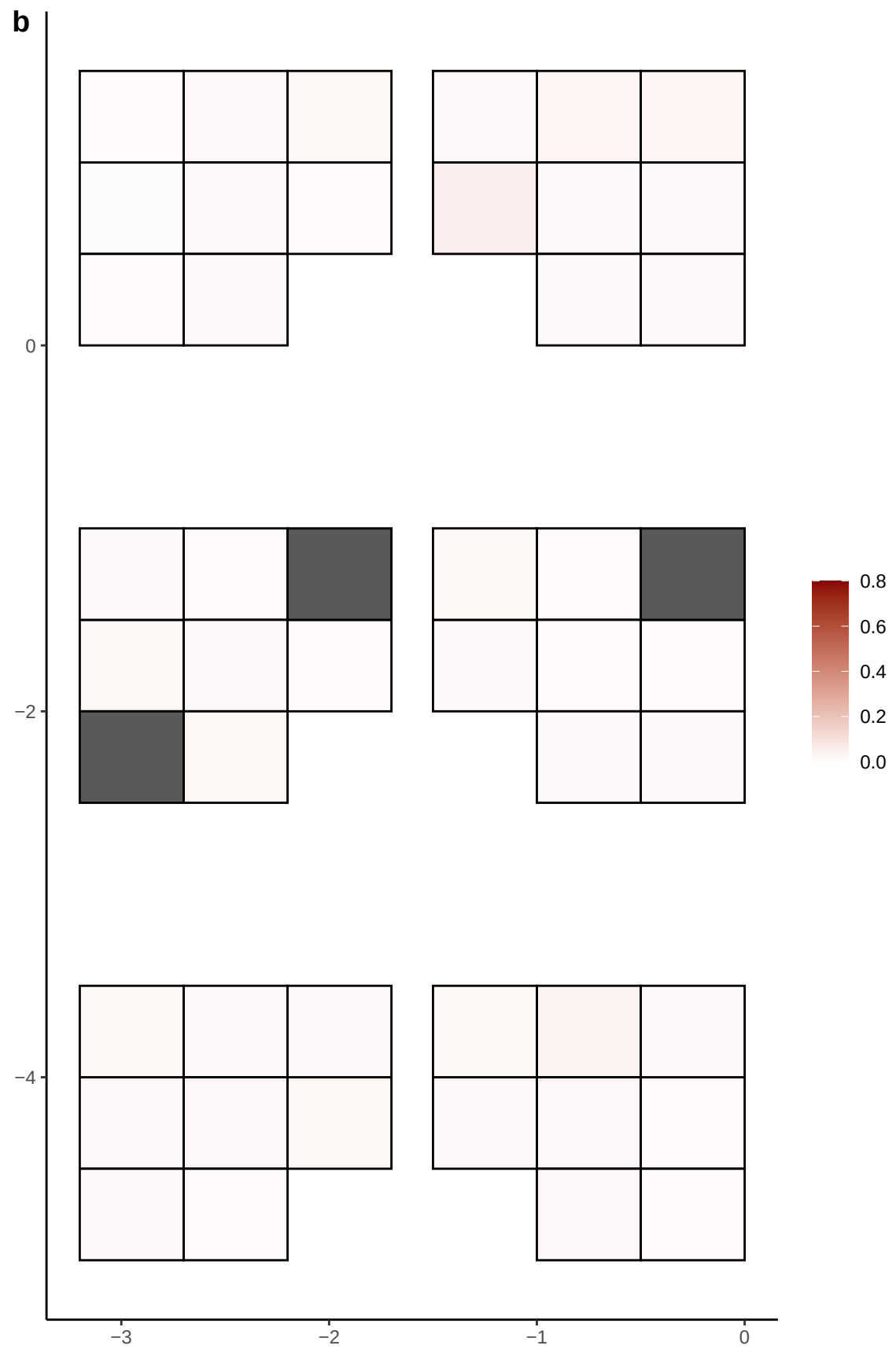

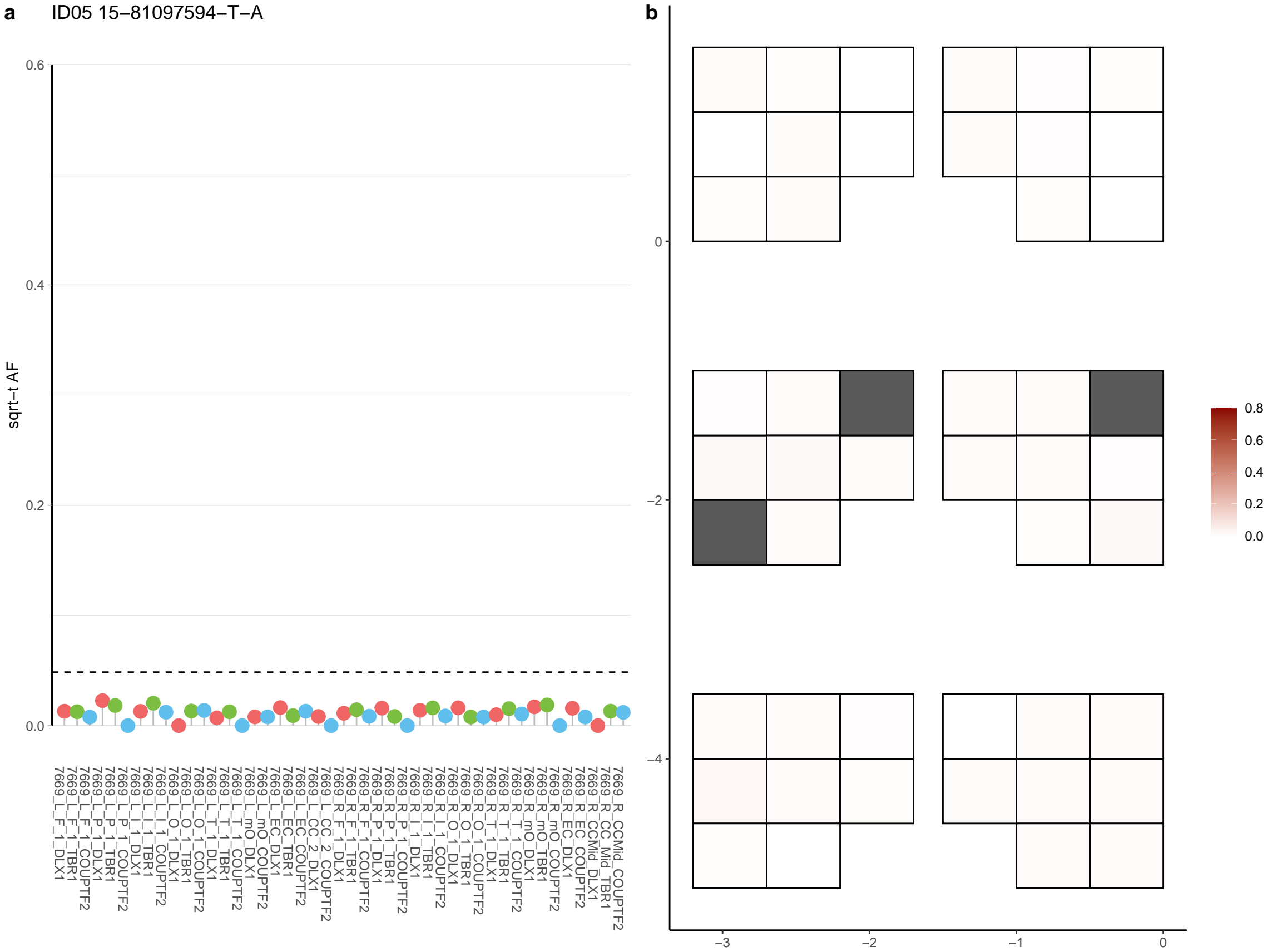

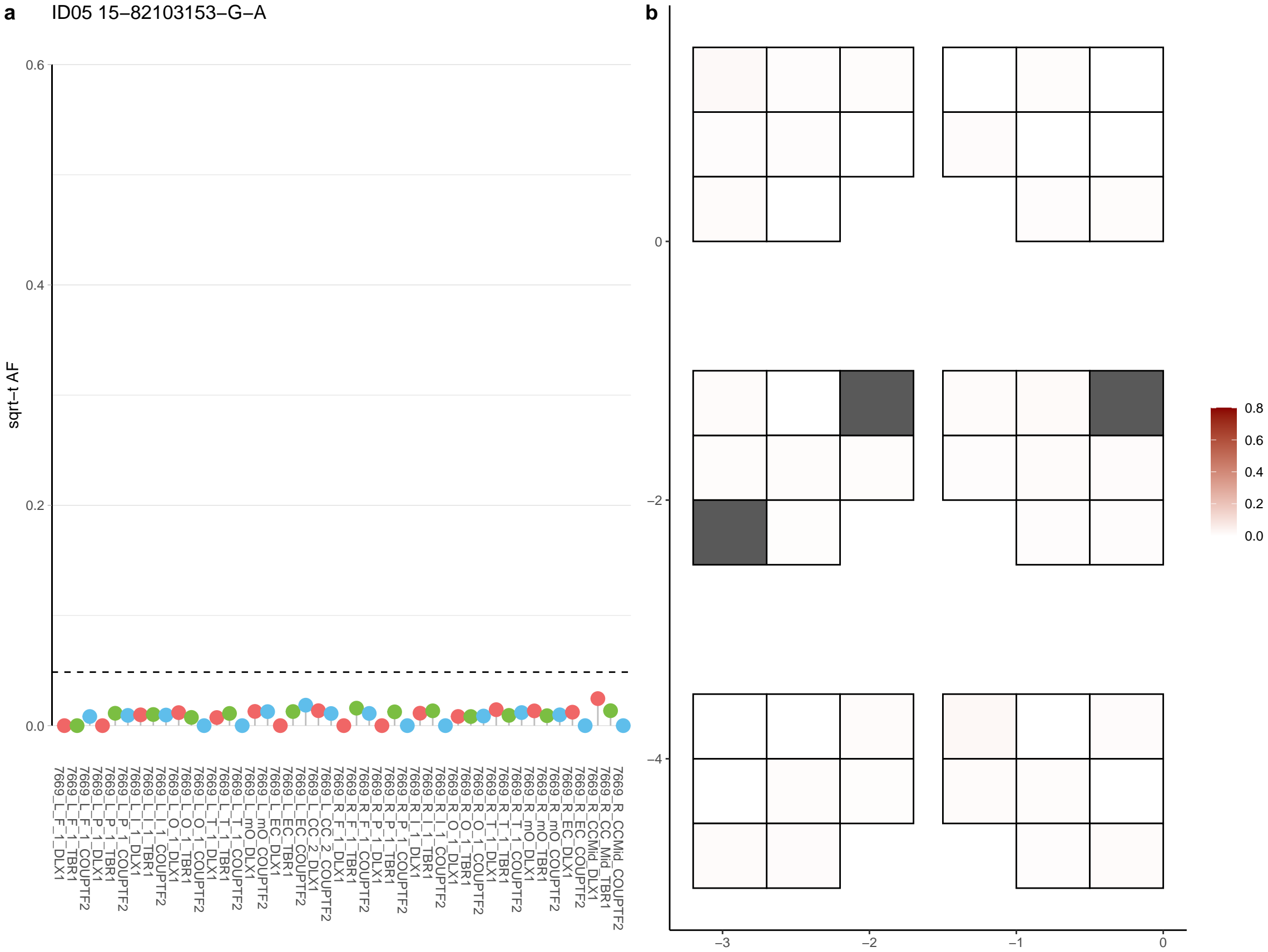

**a** ID05 15-89449716-CT-C

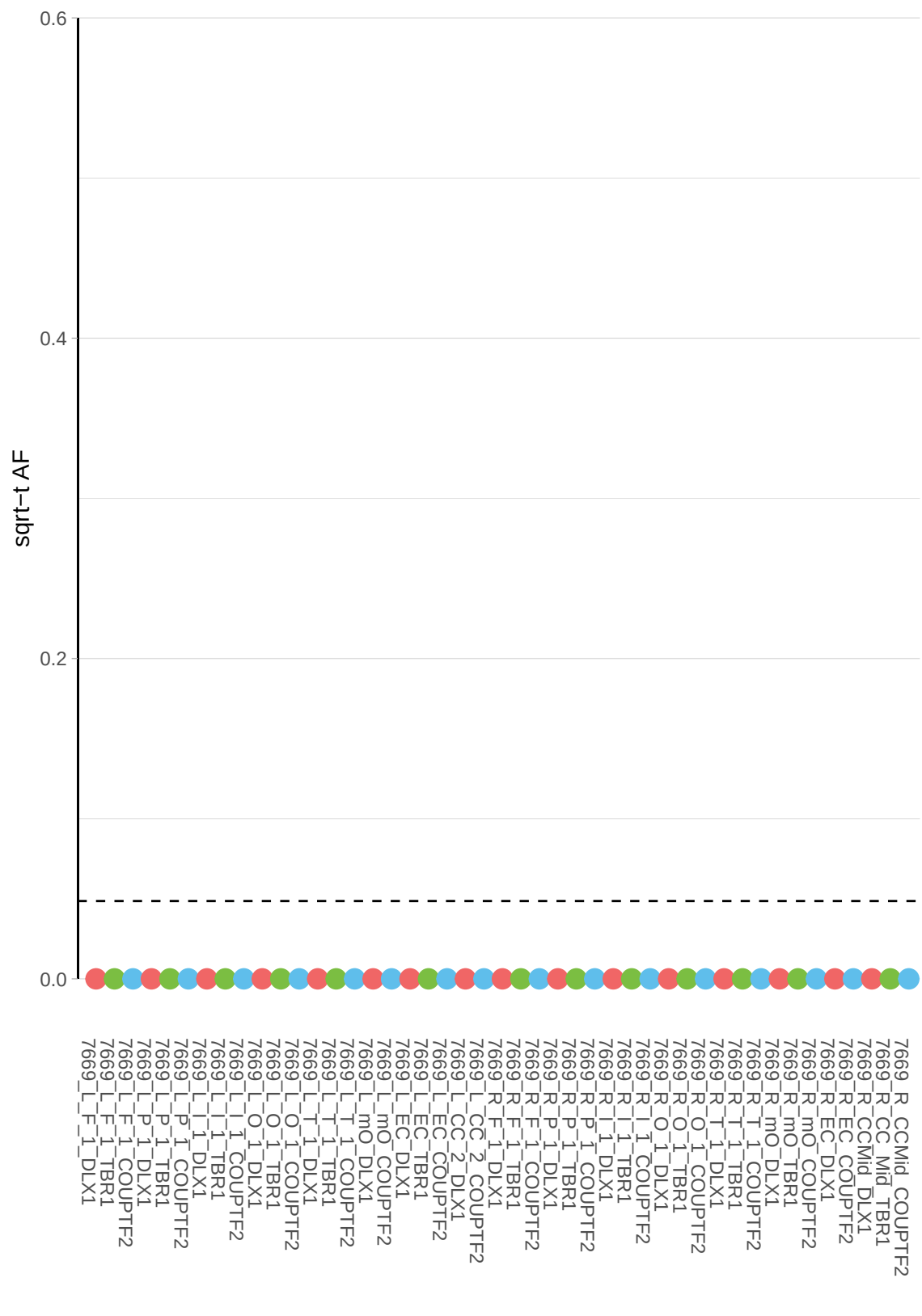

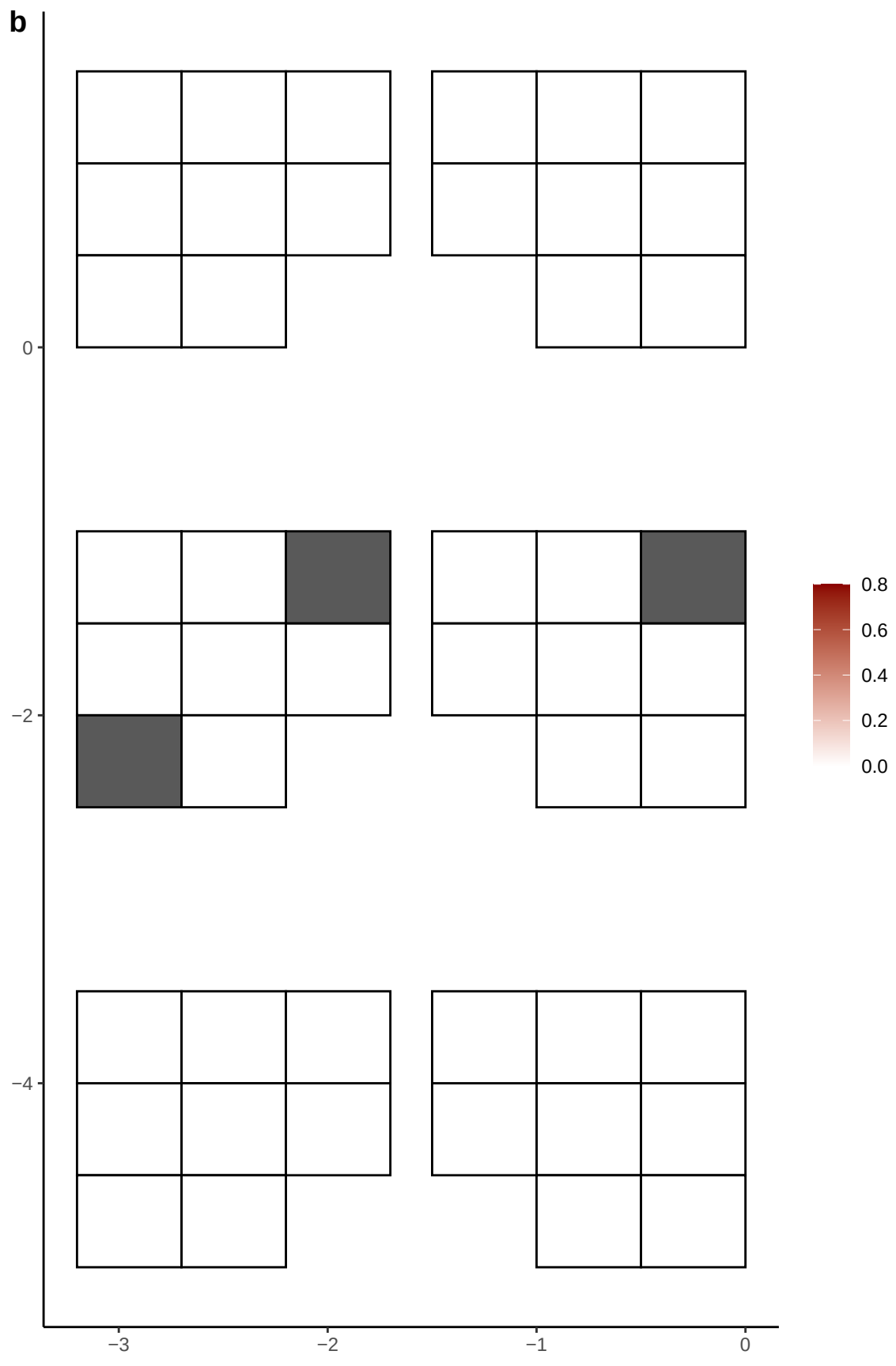

**a** ID05 15-92865289-G-A

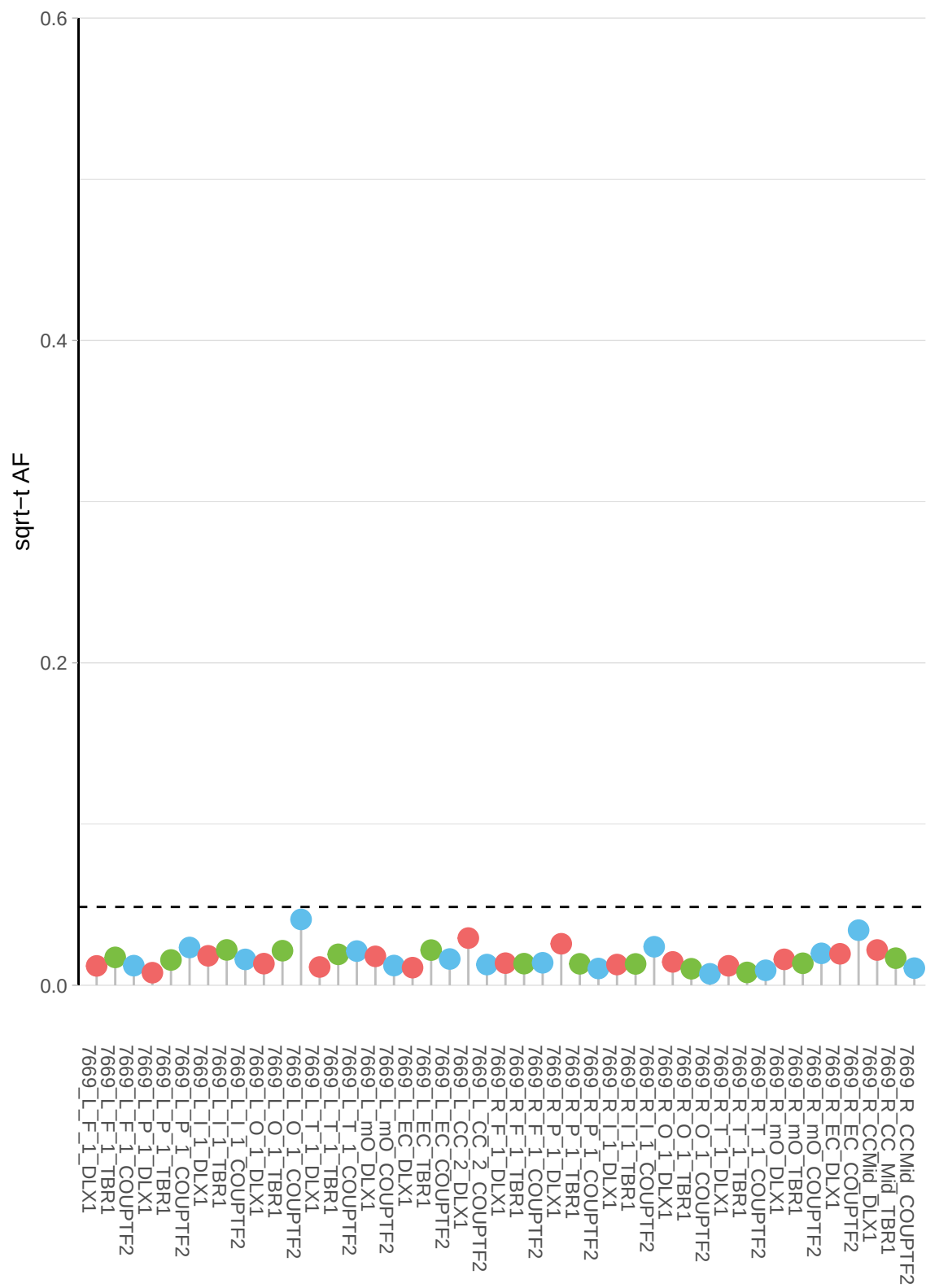

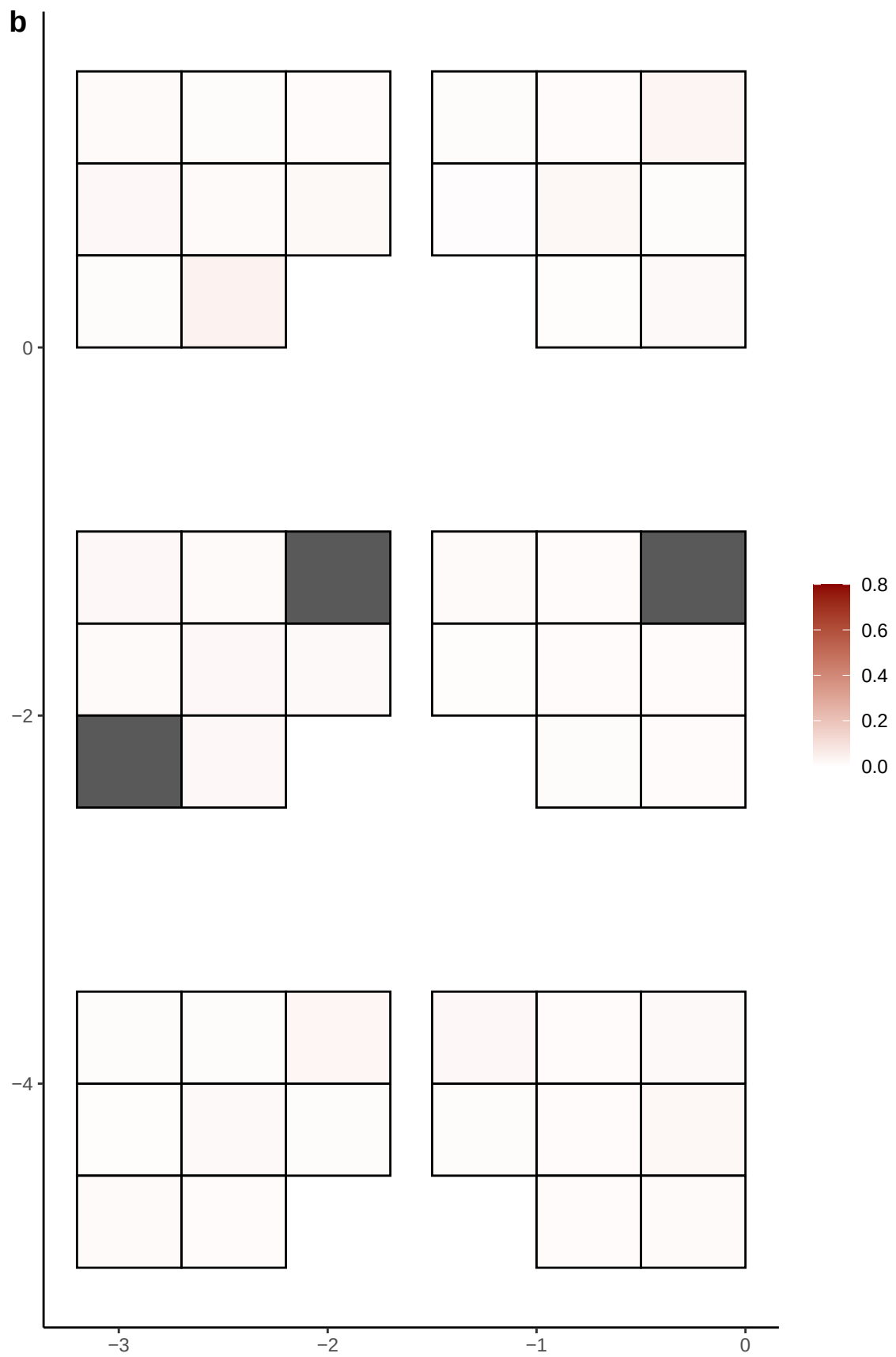

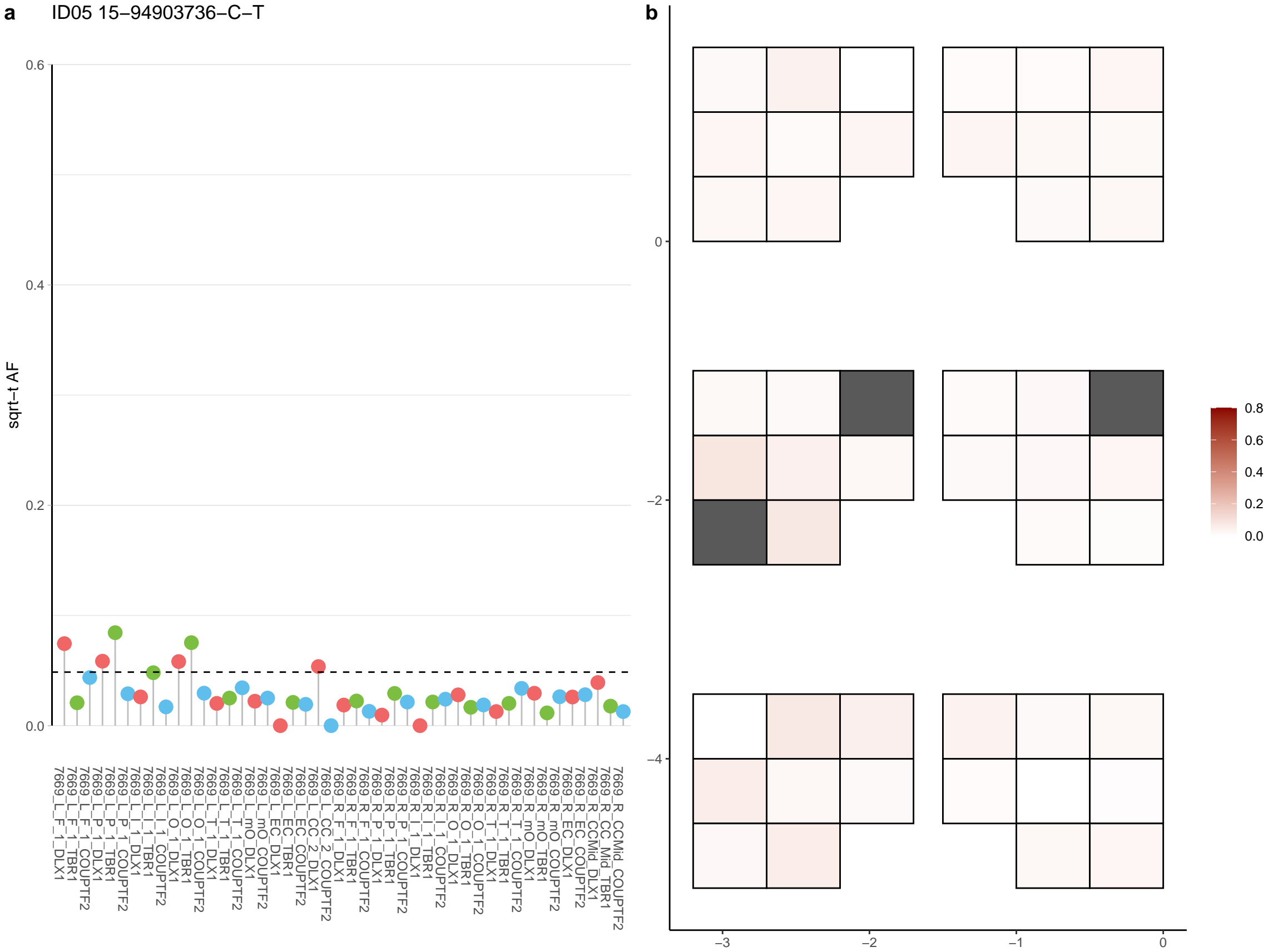

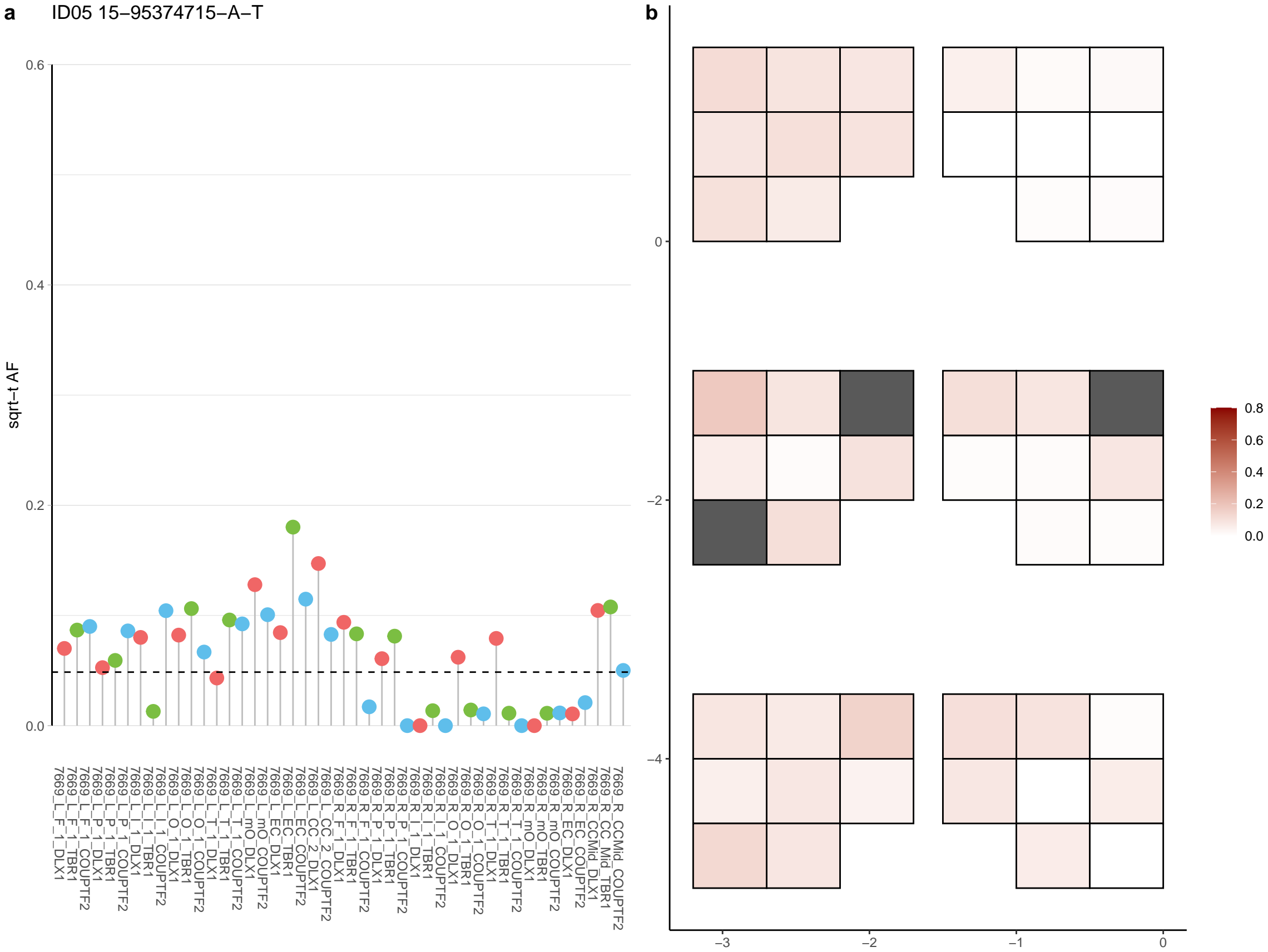

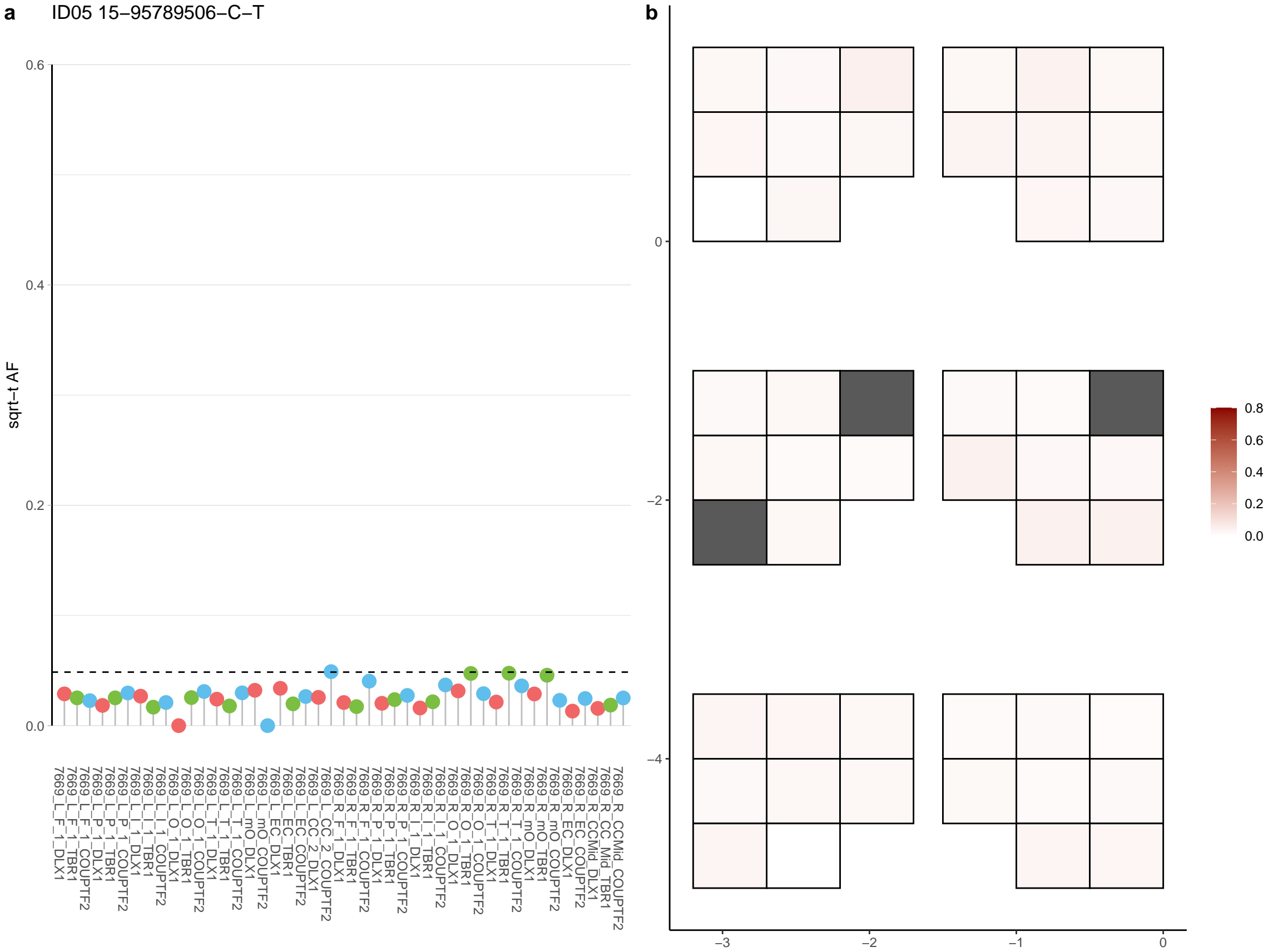

**a** ID05 15-95947987-C-G

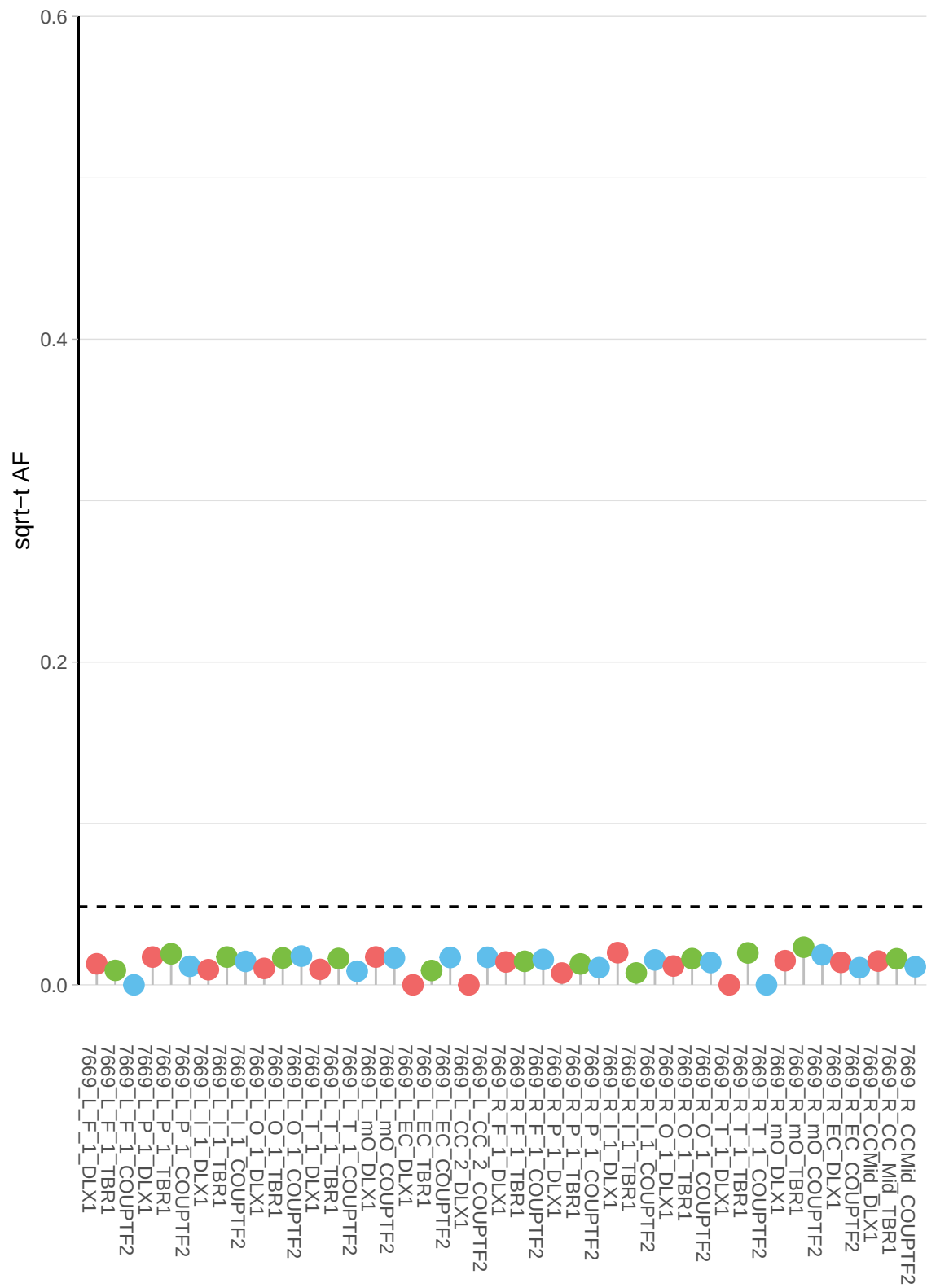

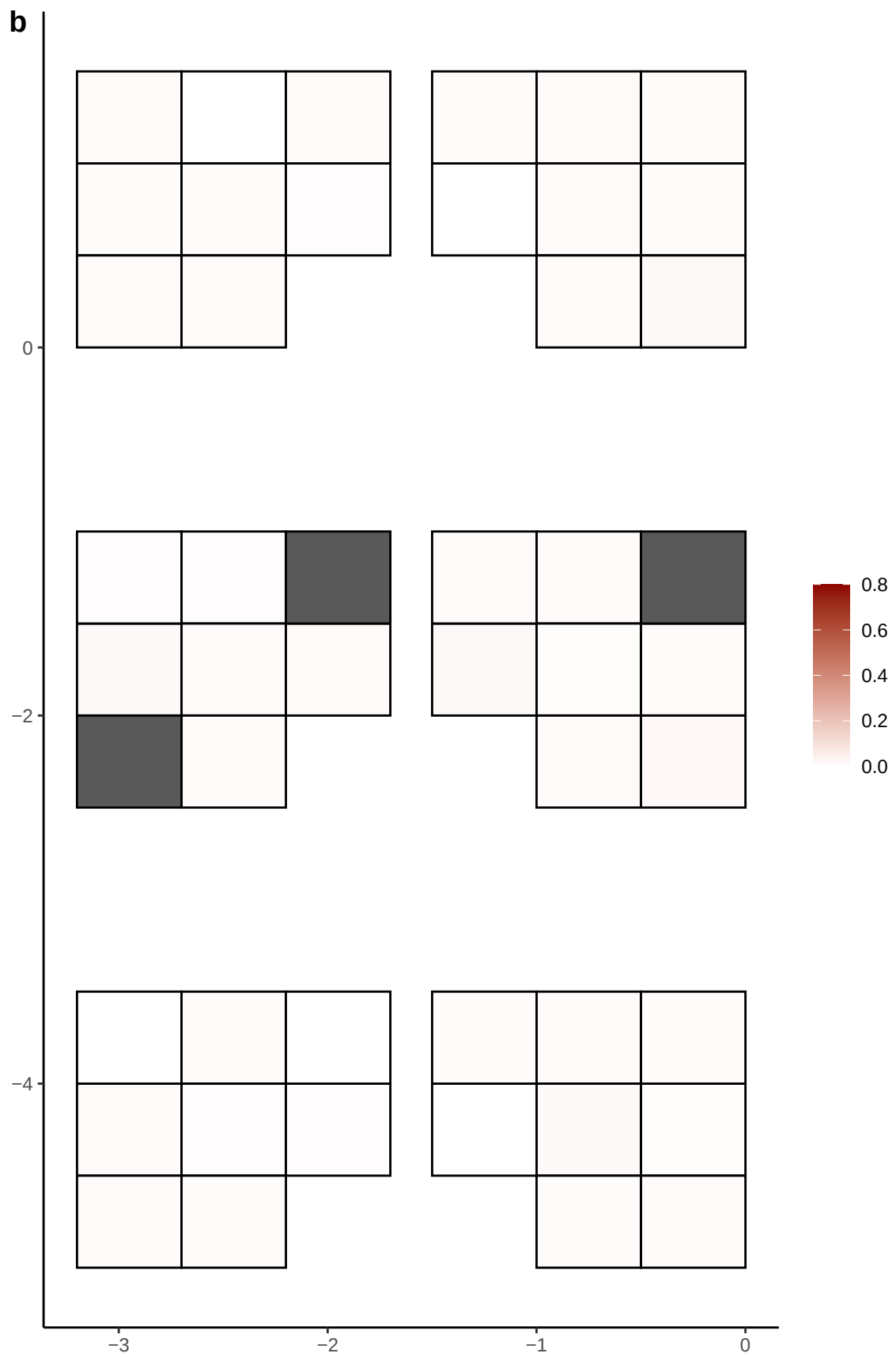

**a**

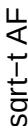

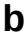

**a** ID05 16-24164513-T-C

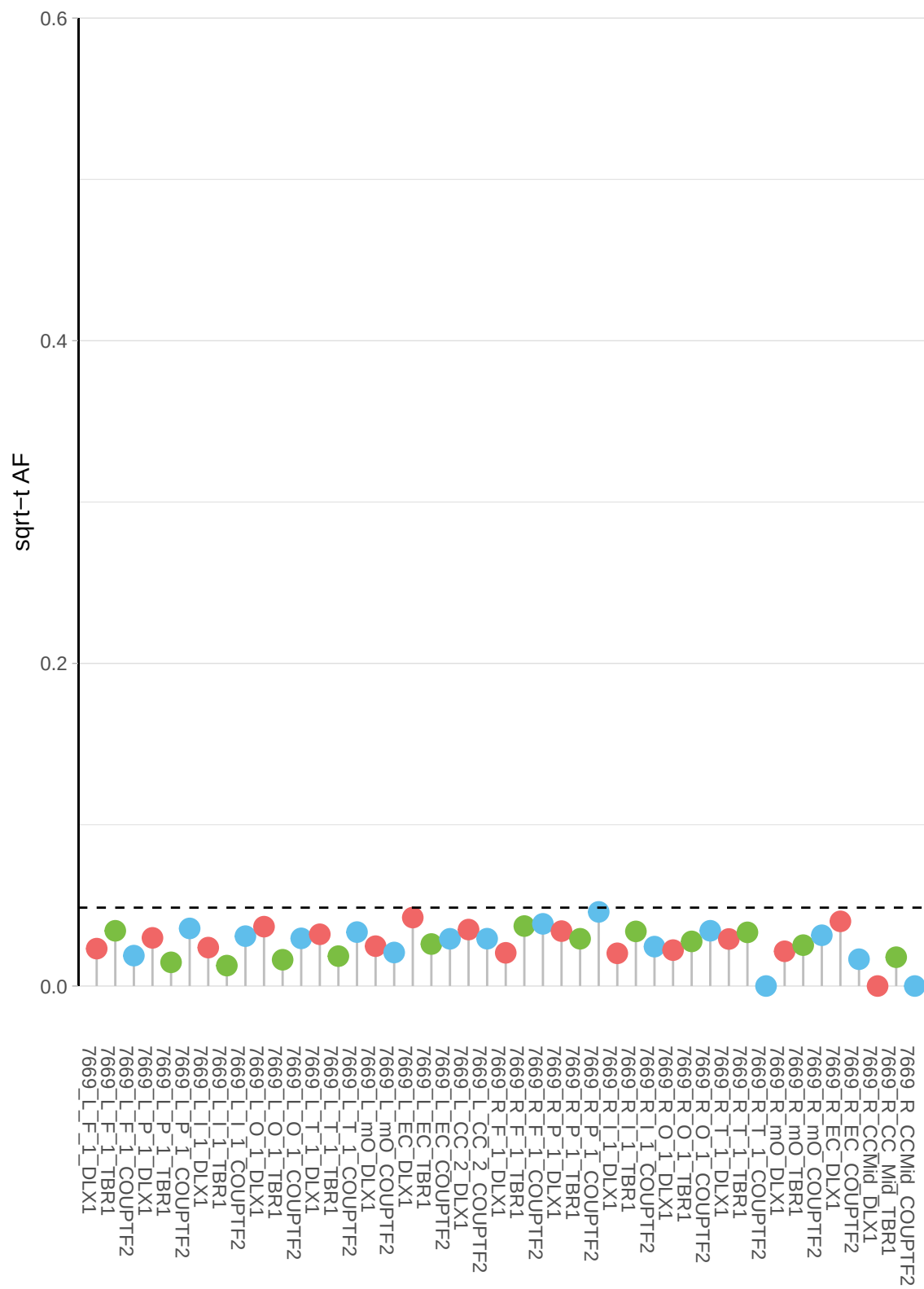

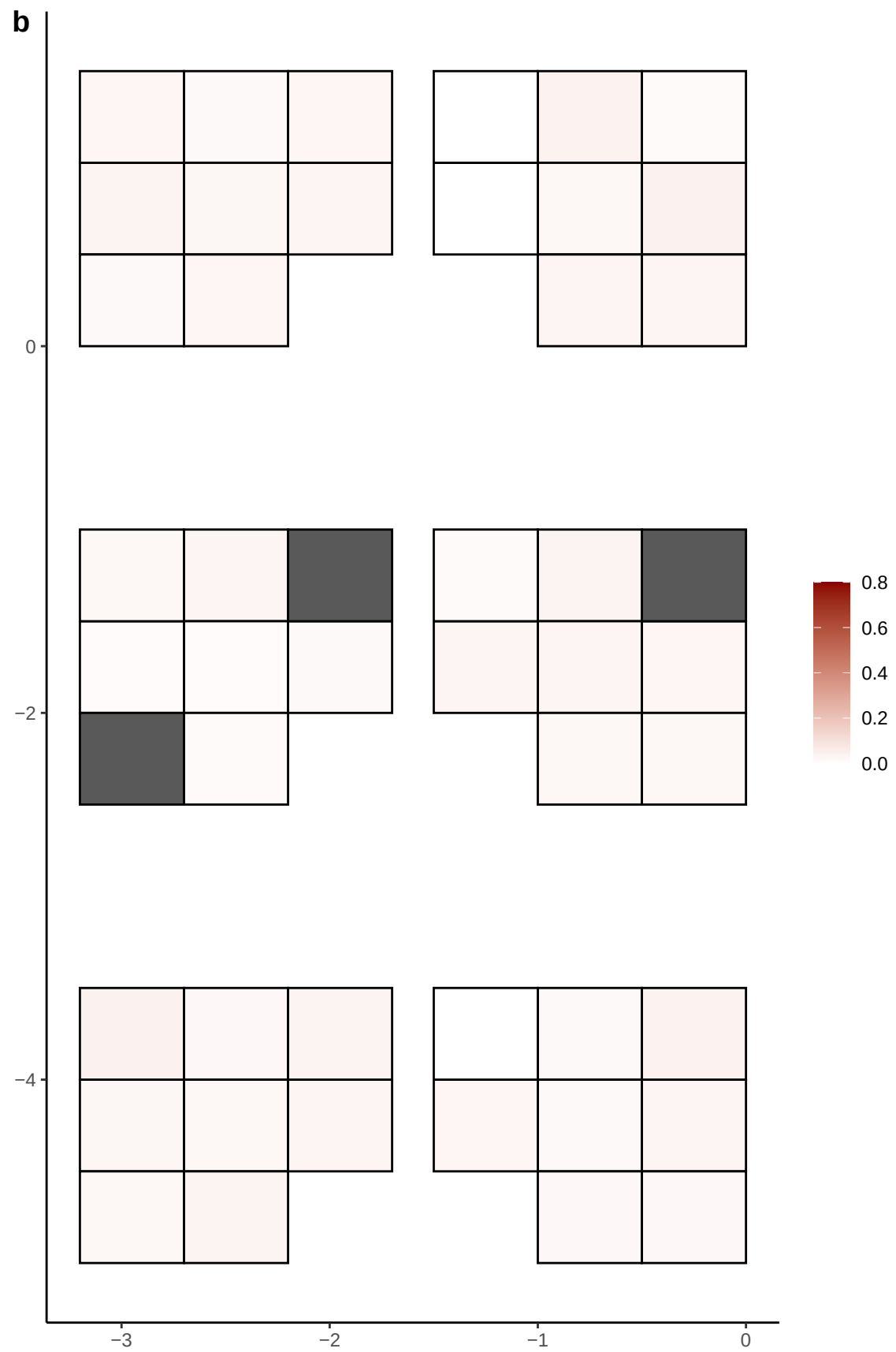

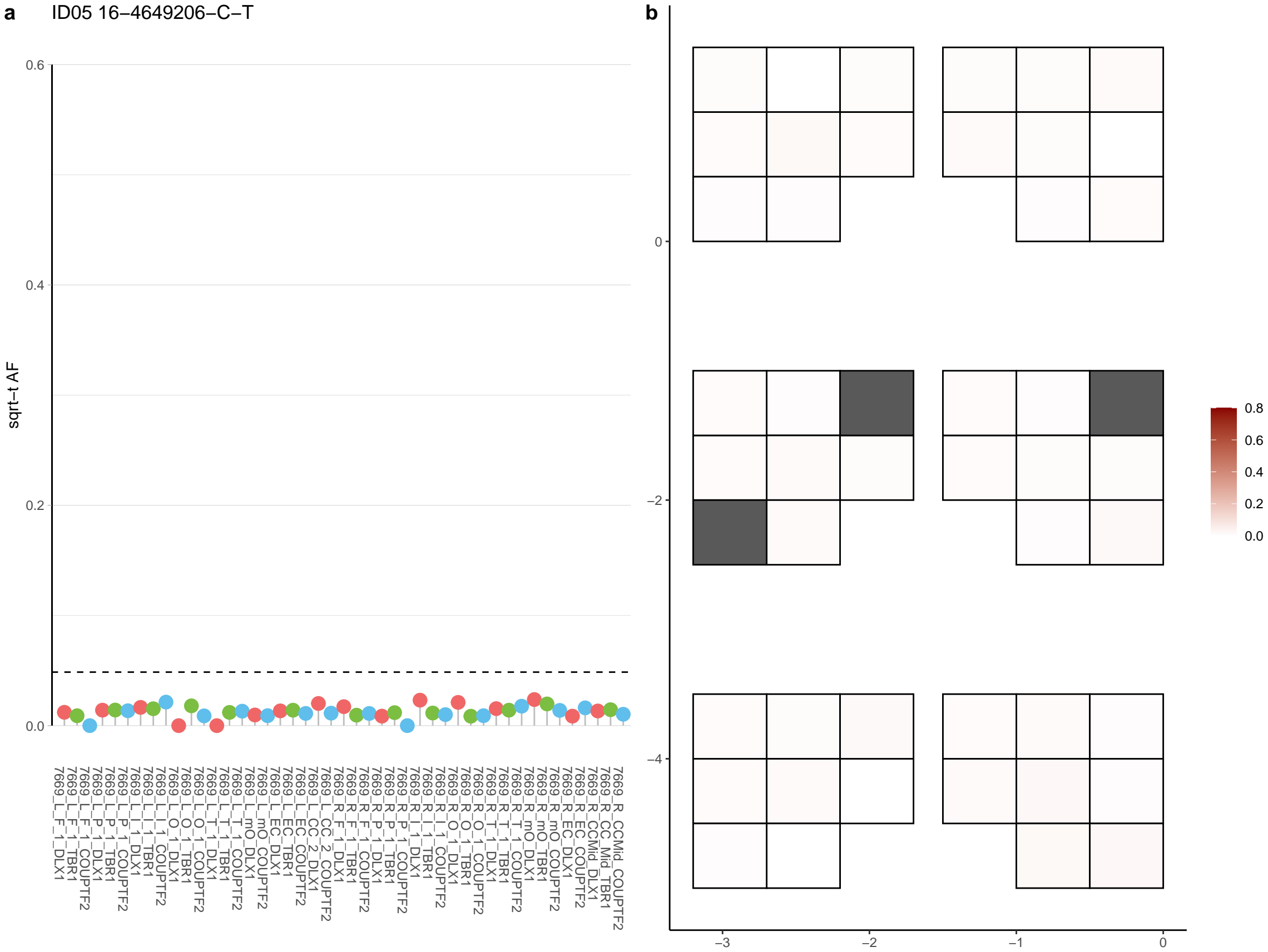

**a** ID05 16-47975657-G-A

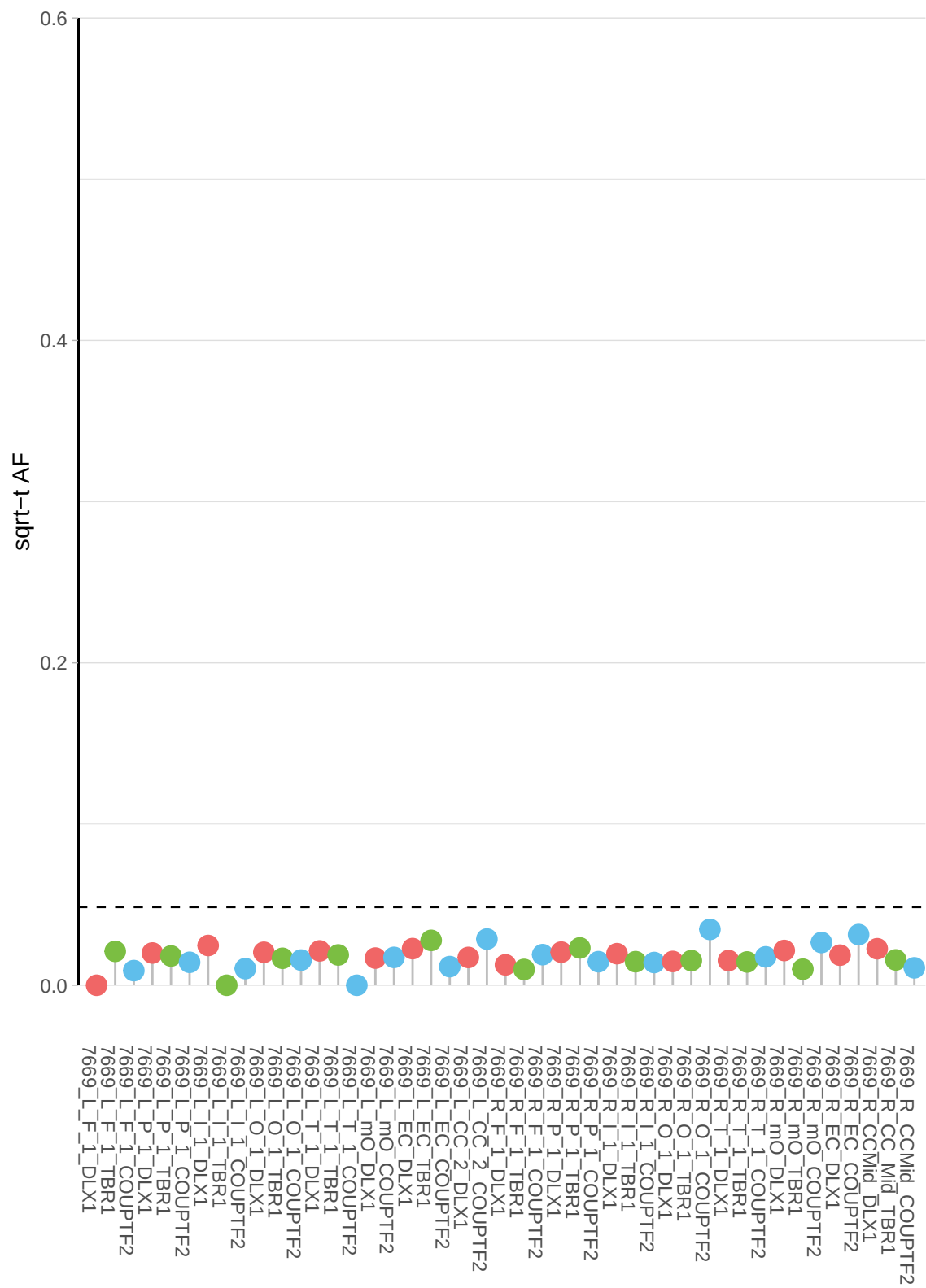

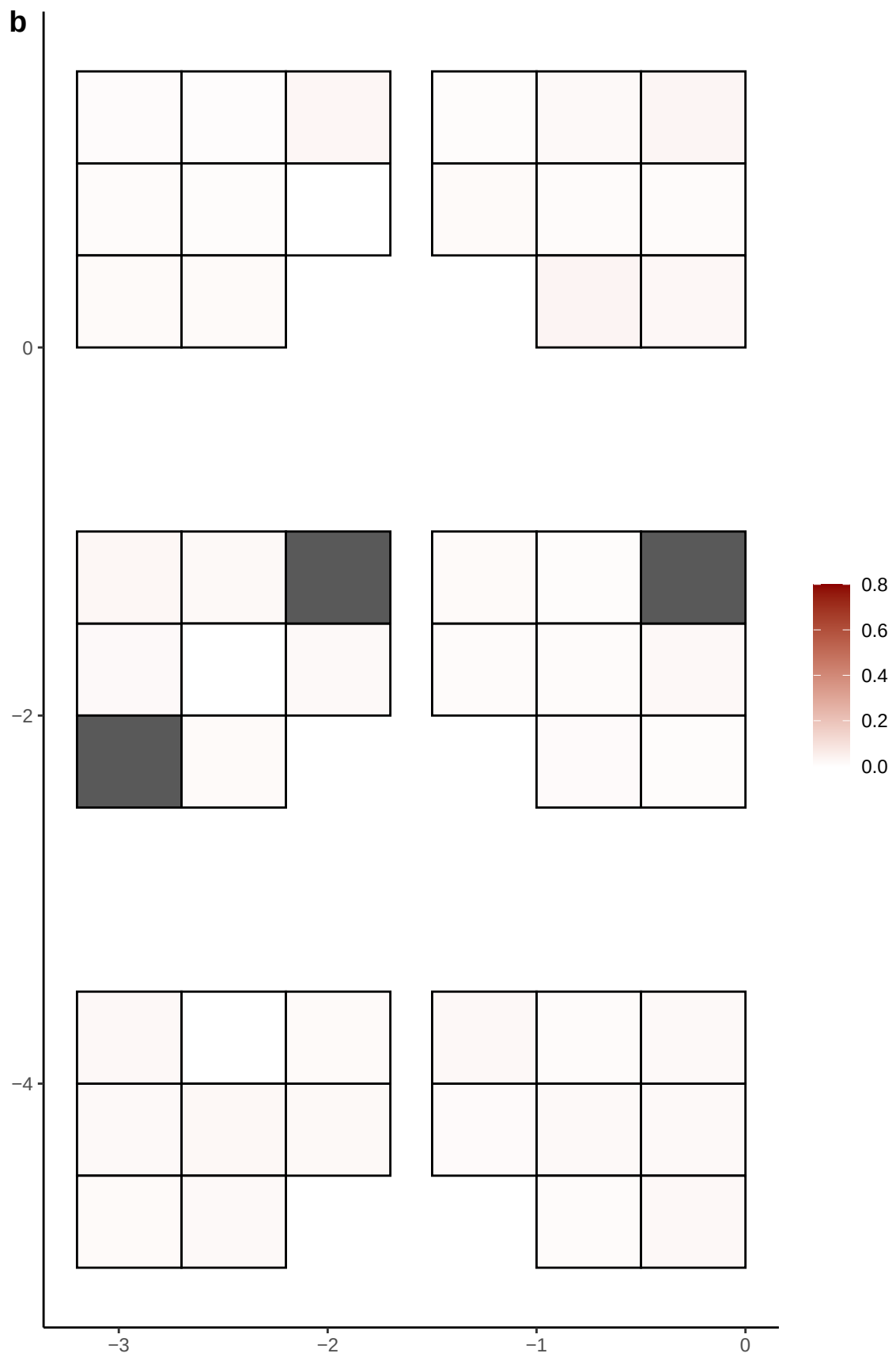

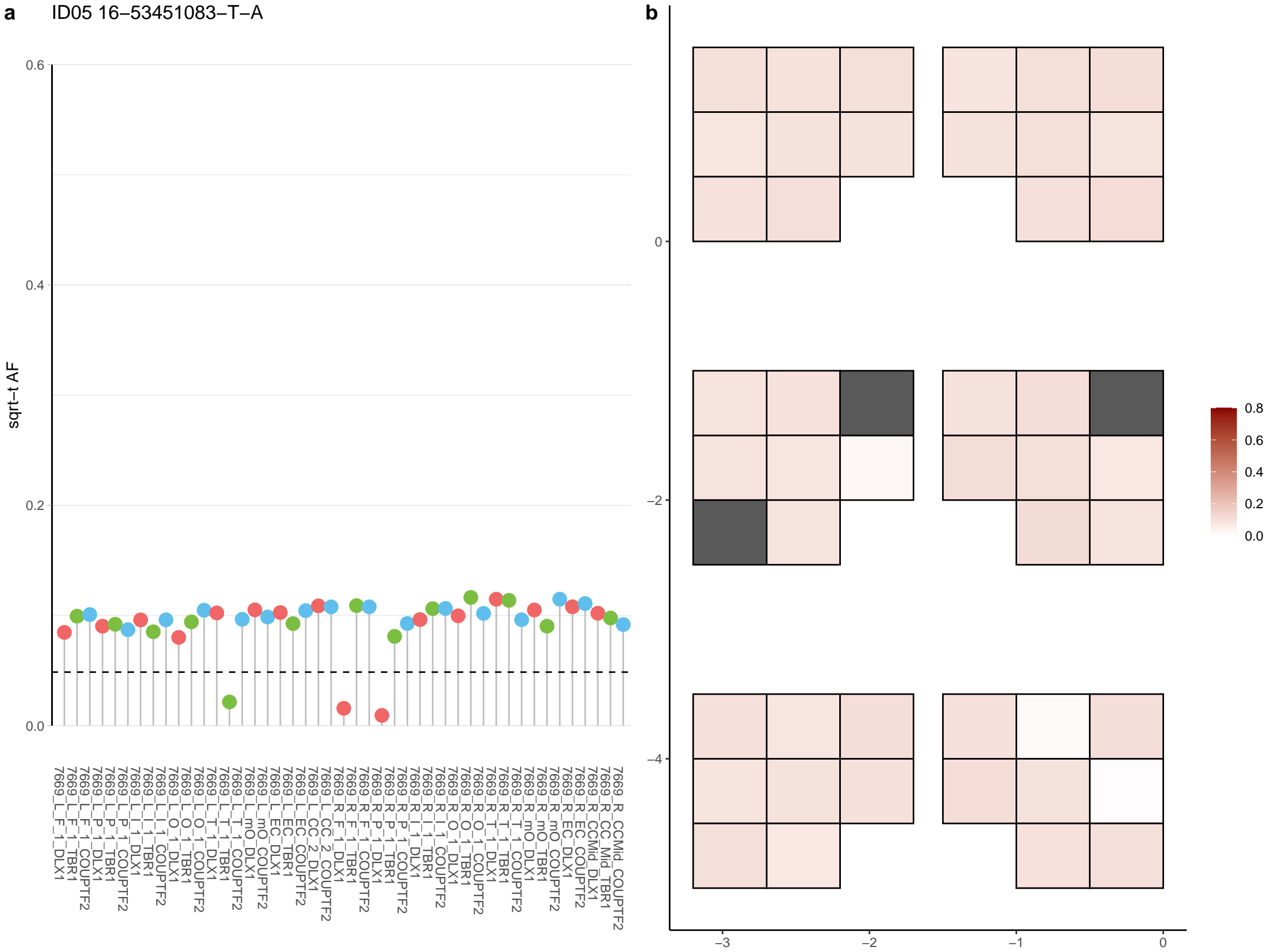

**a** ID05 16-53451084-T-A

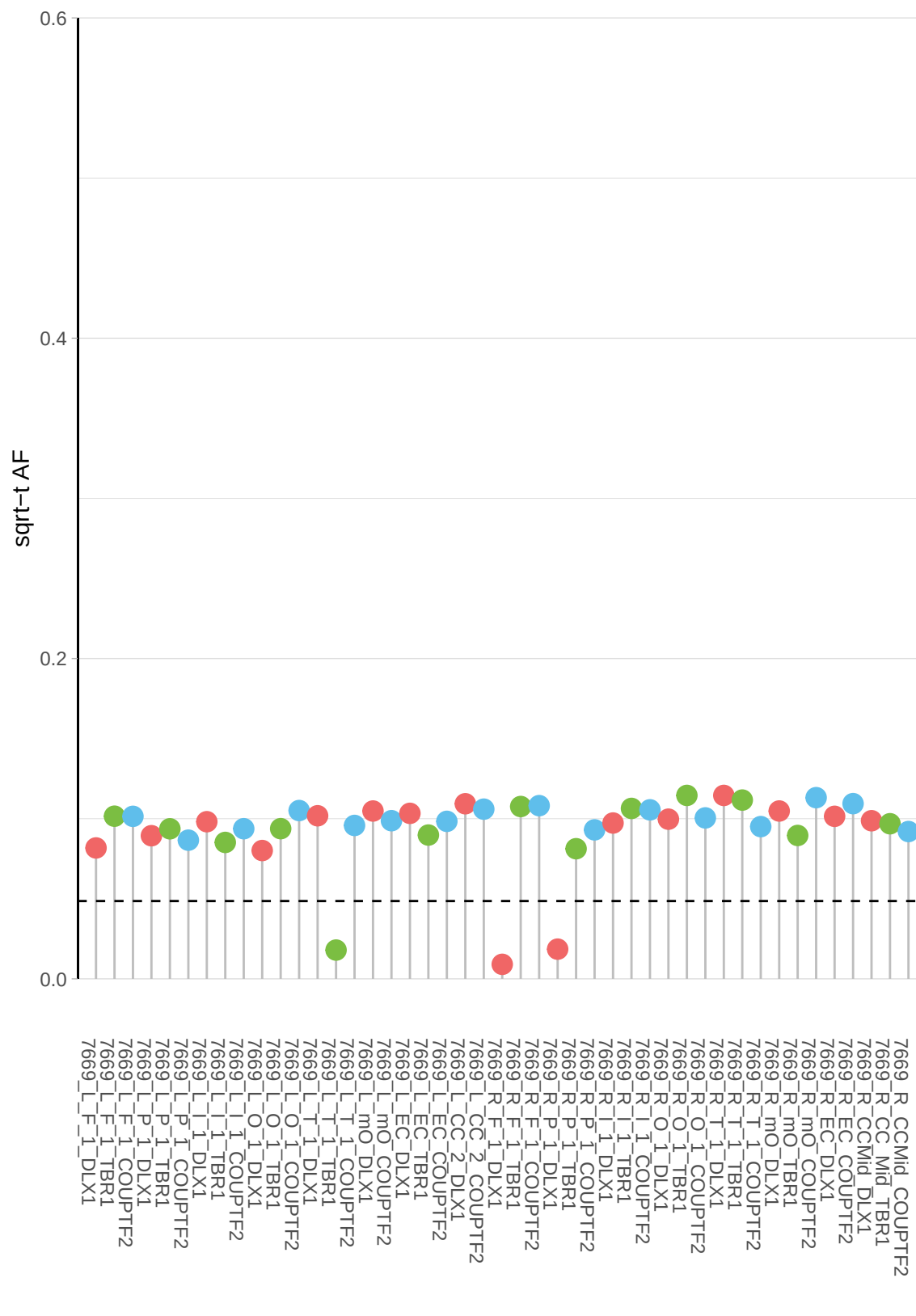

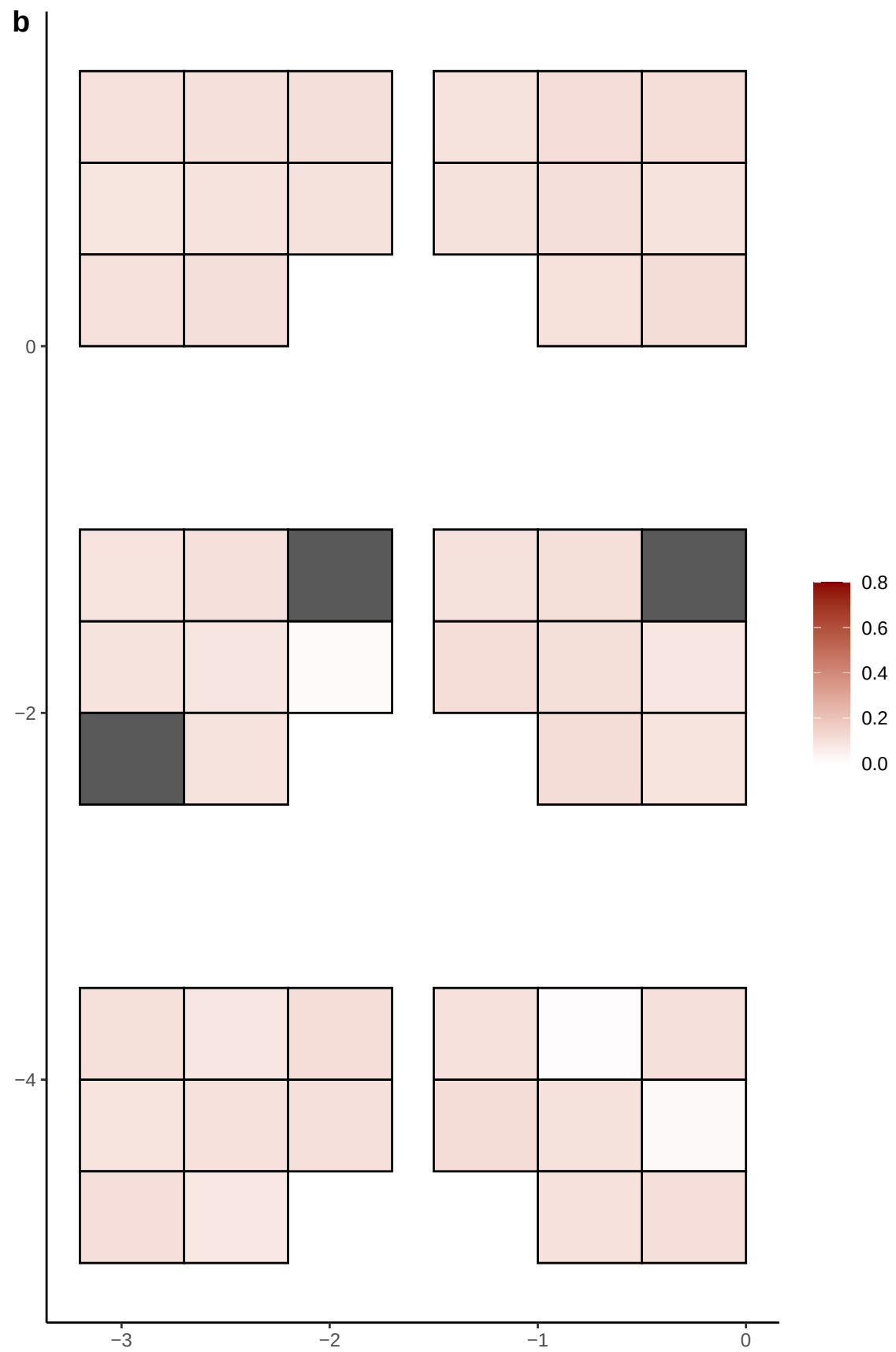

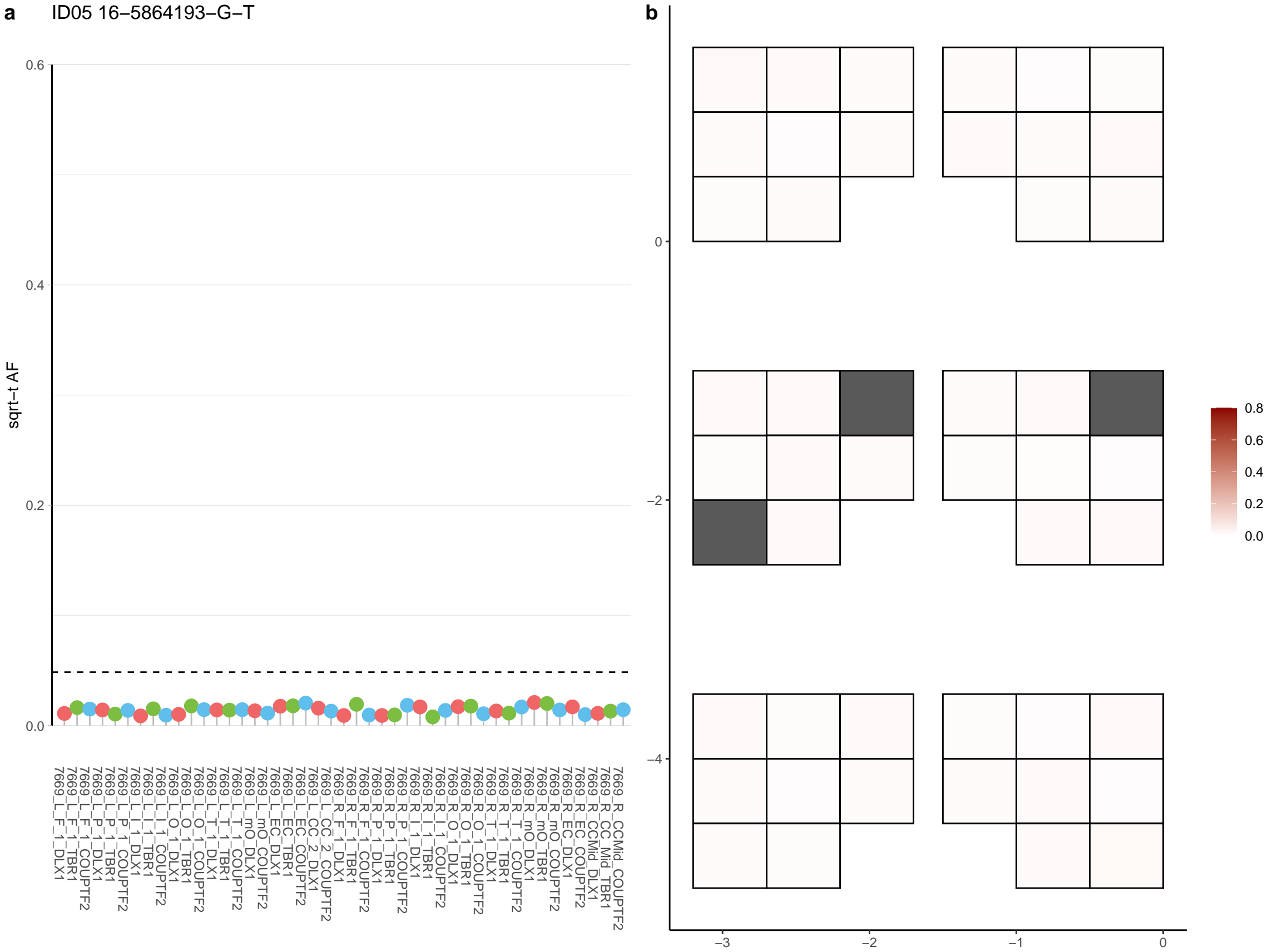

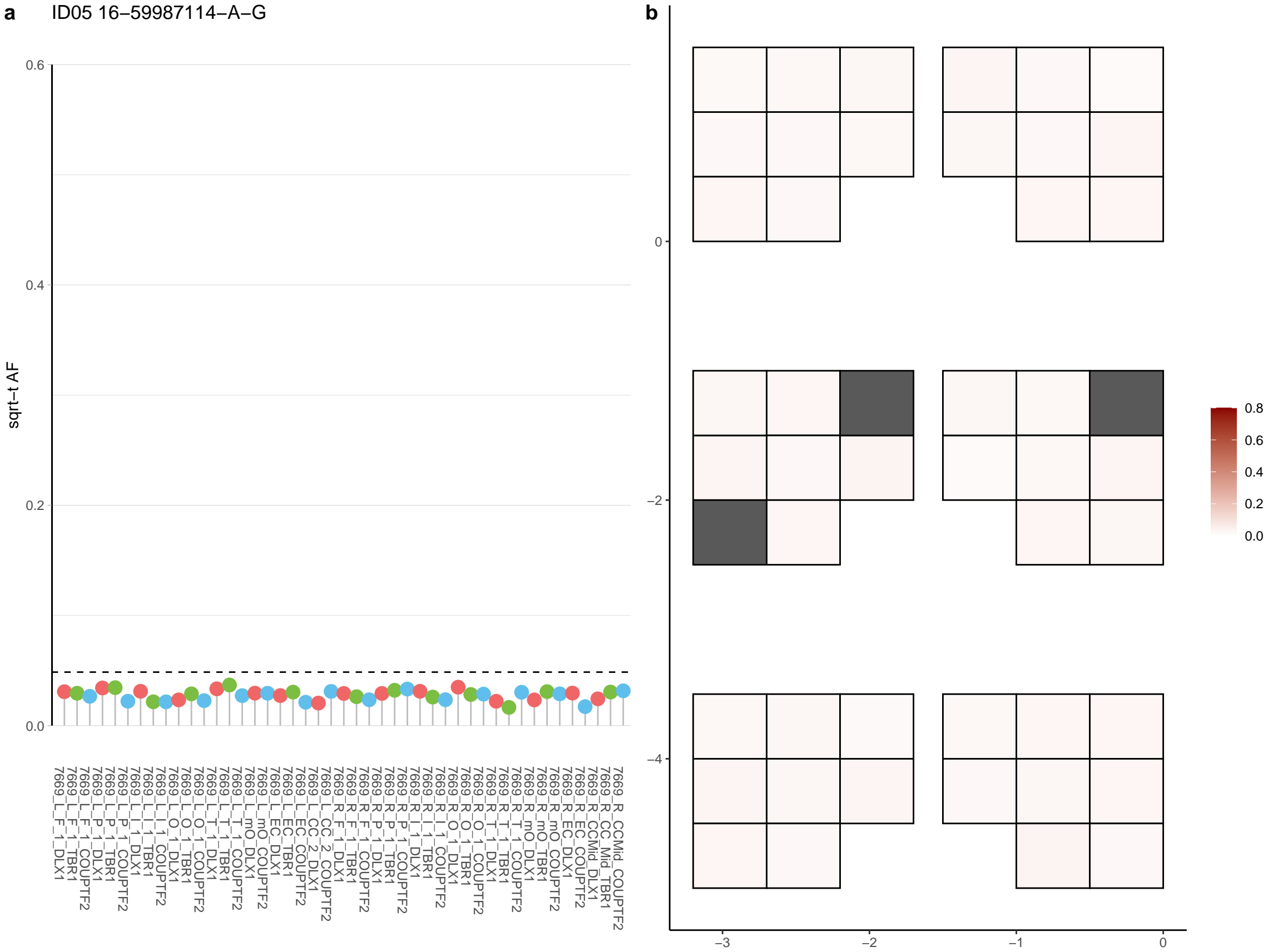

**a** ID05 16-60721807-G-T

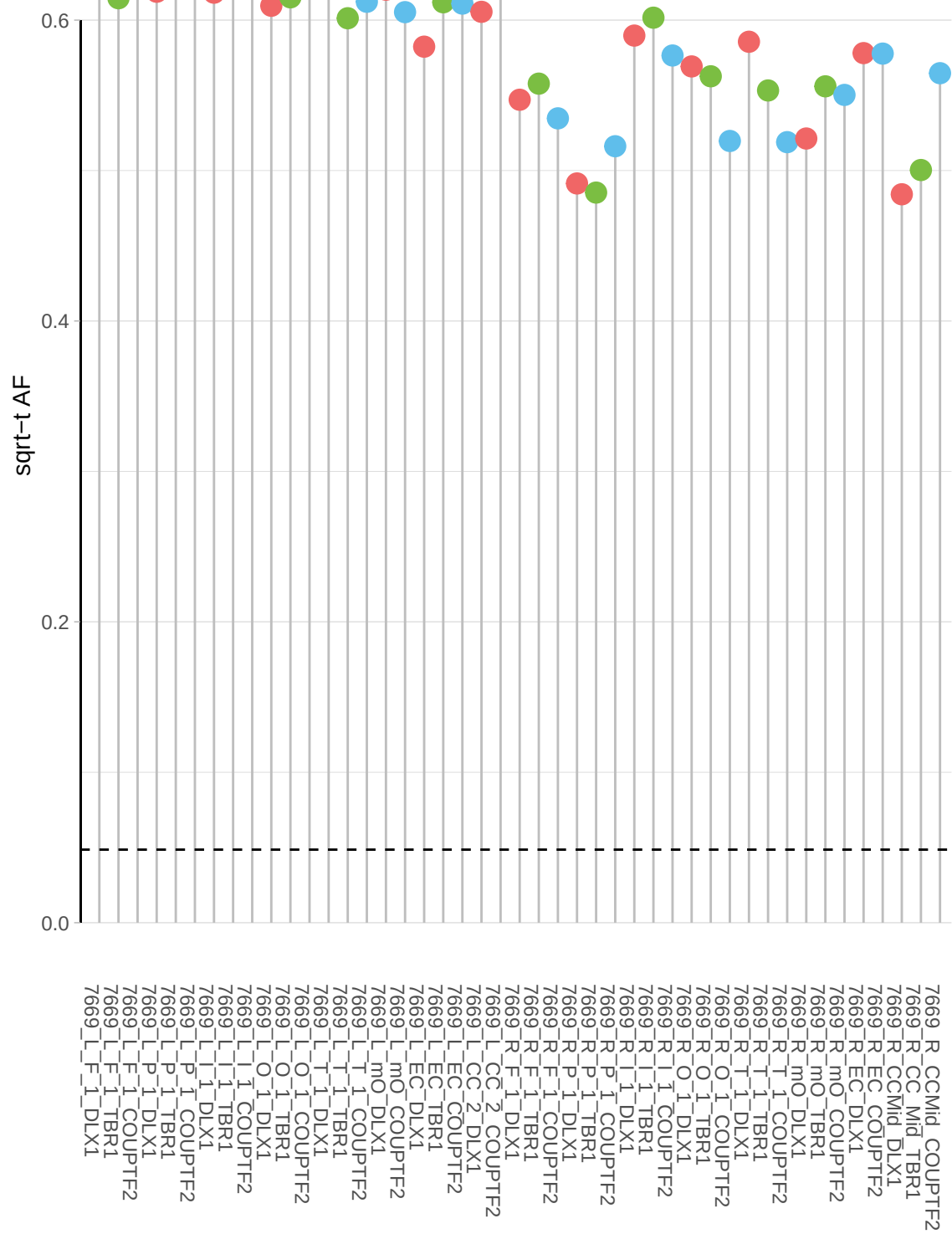

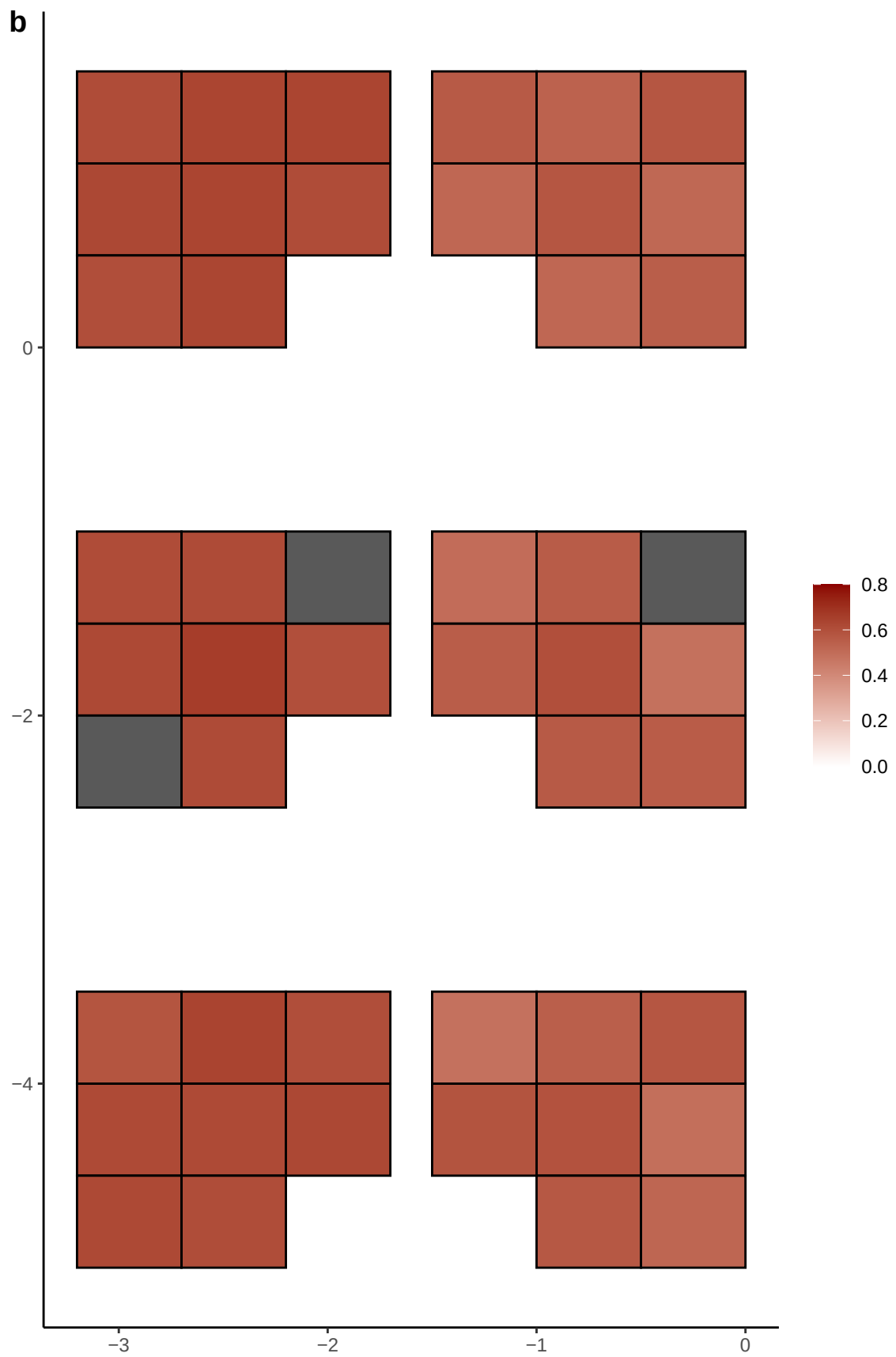

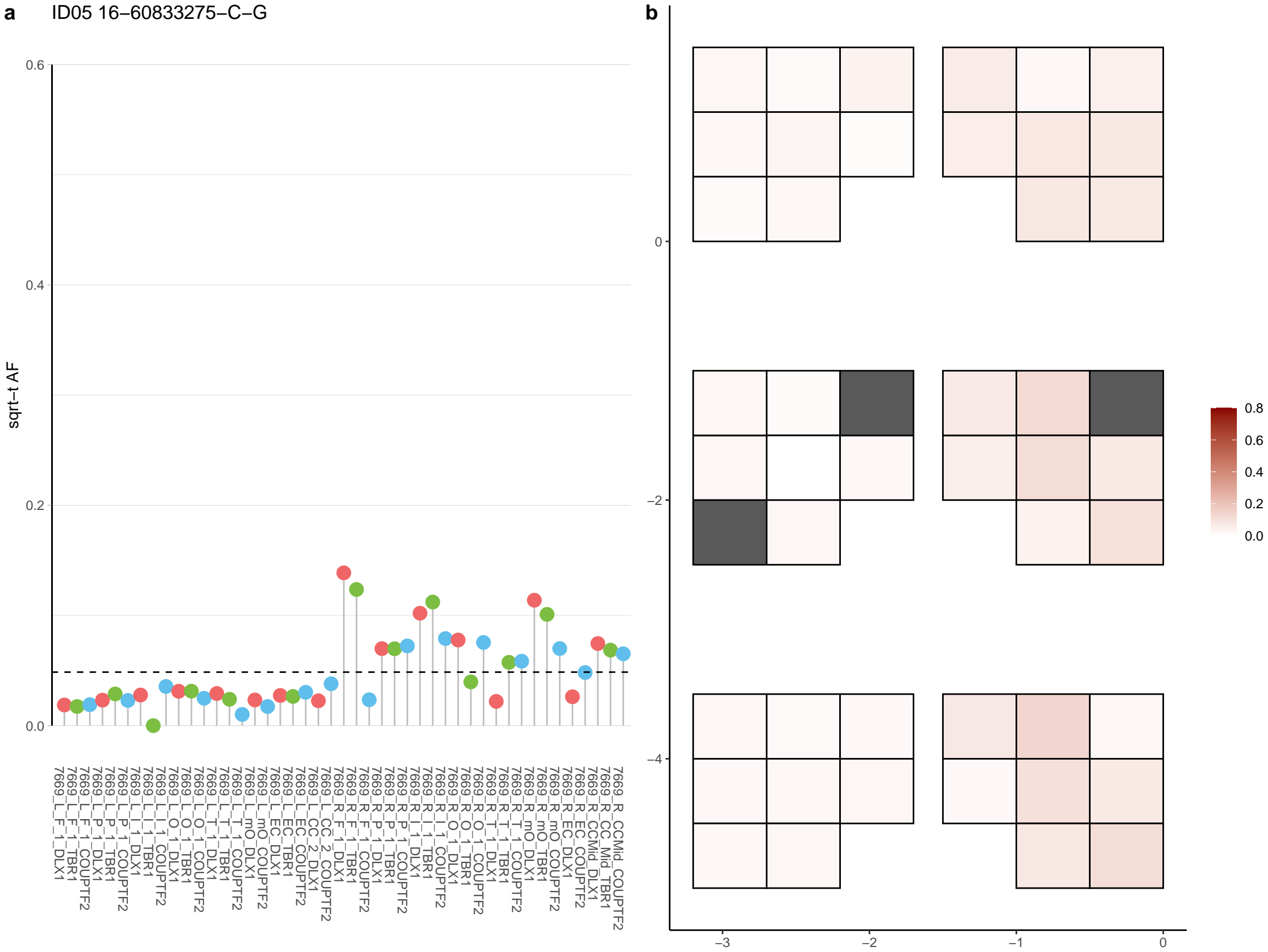

**a** ID05 16-63481903-A-T

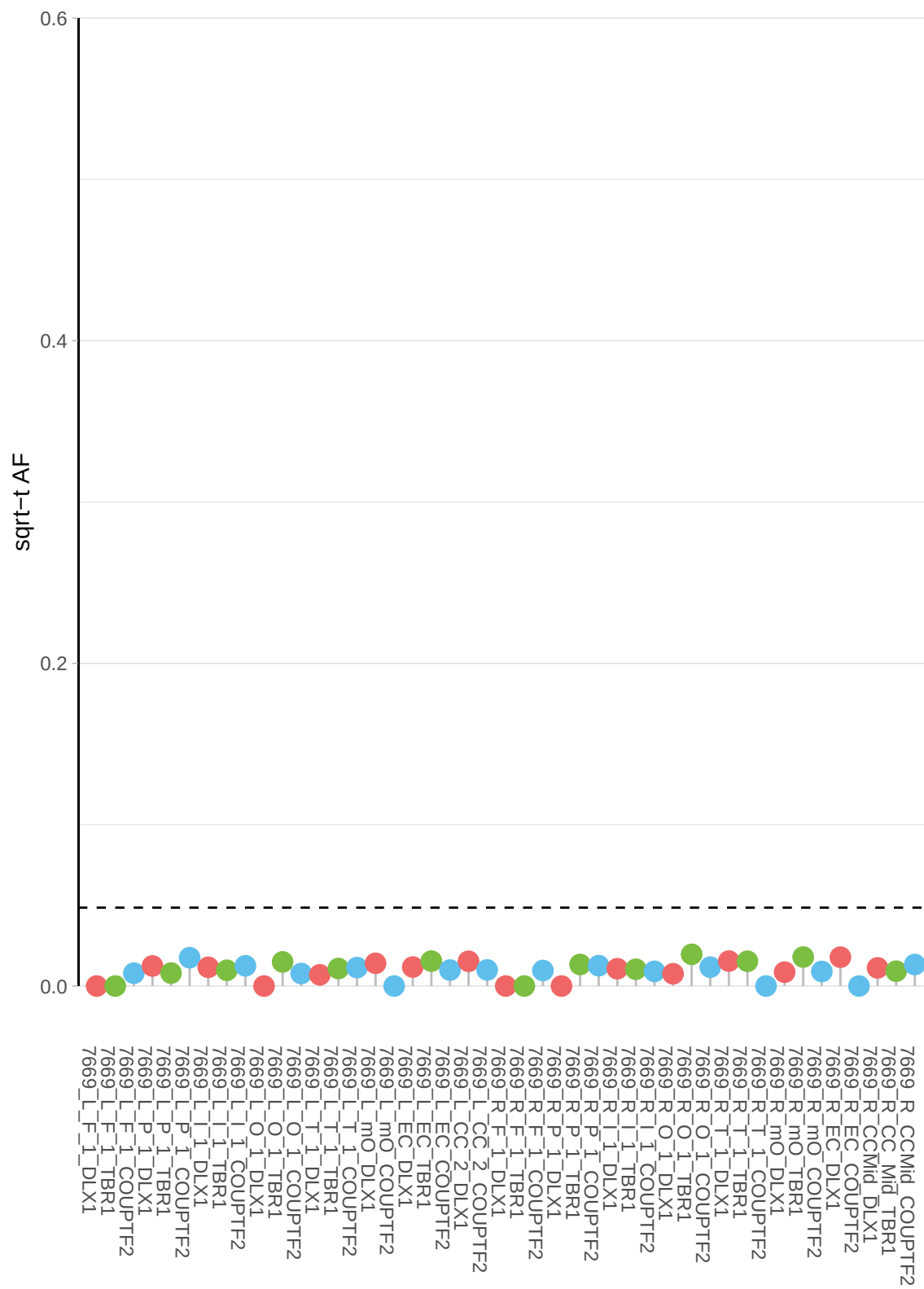

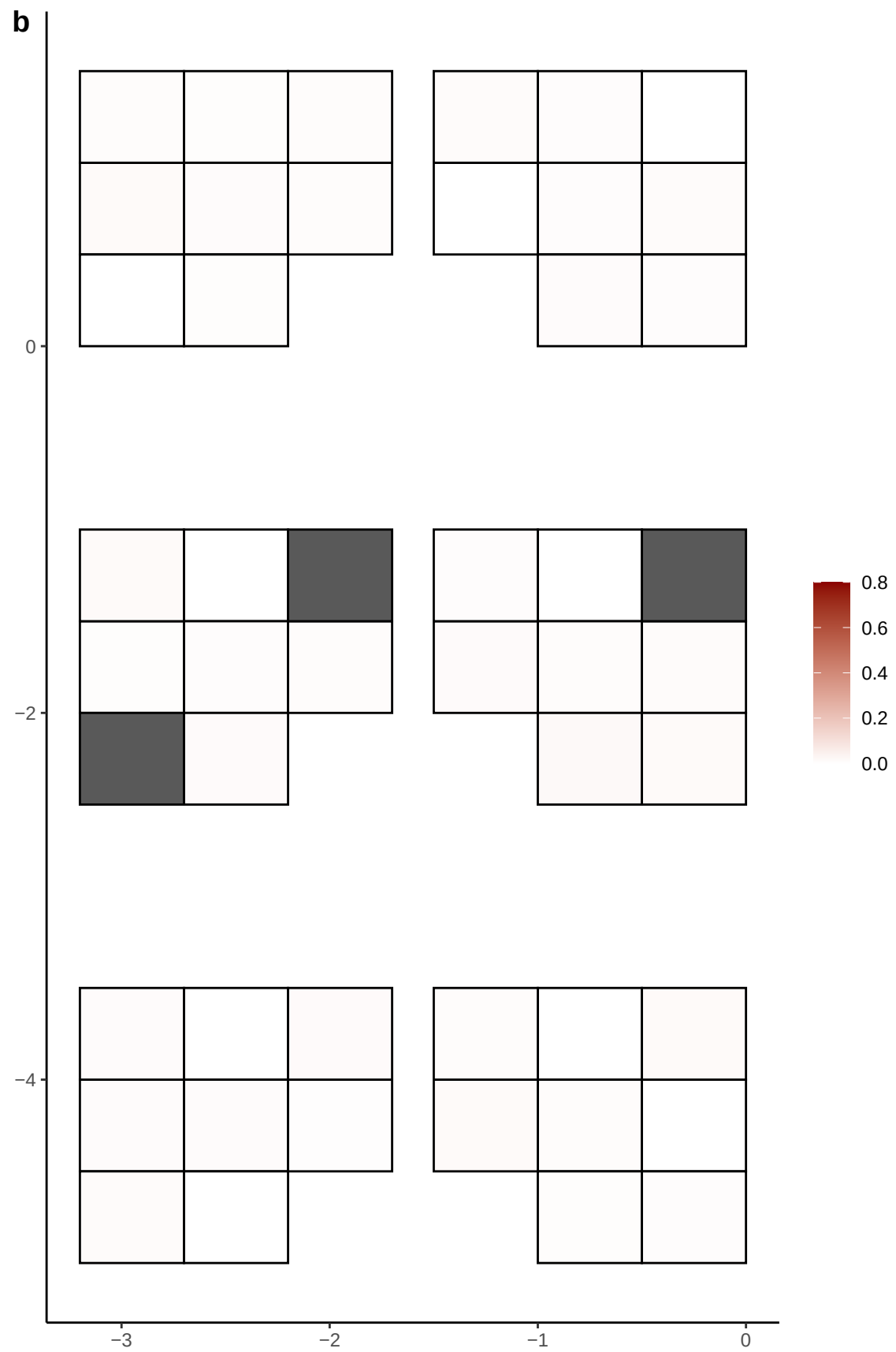

**a** ID05 16-64667272-C-T

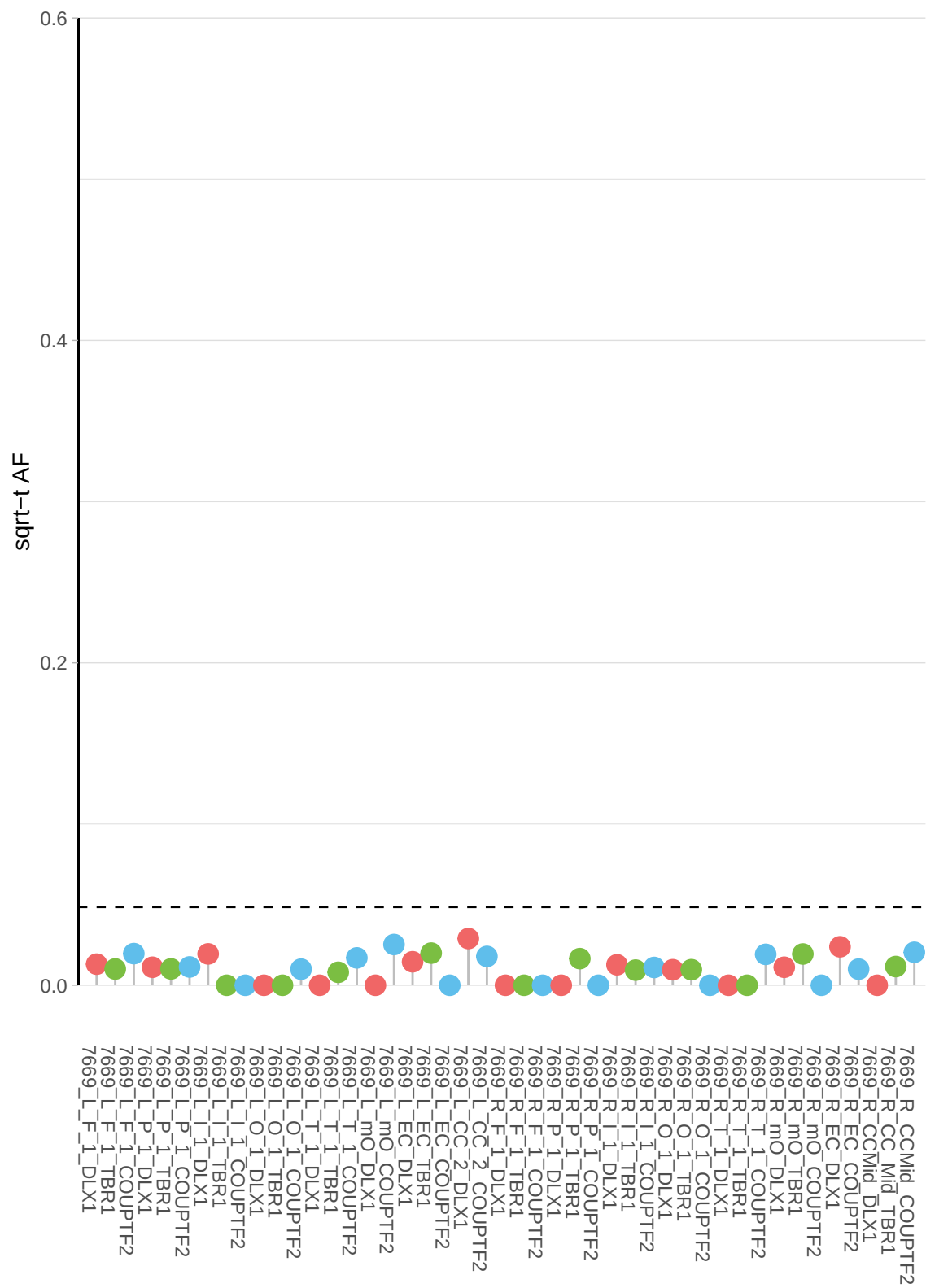

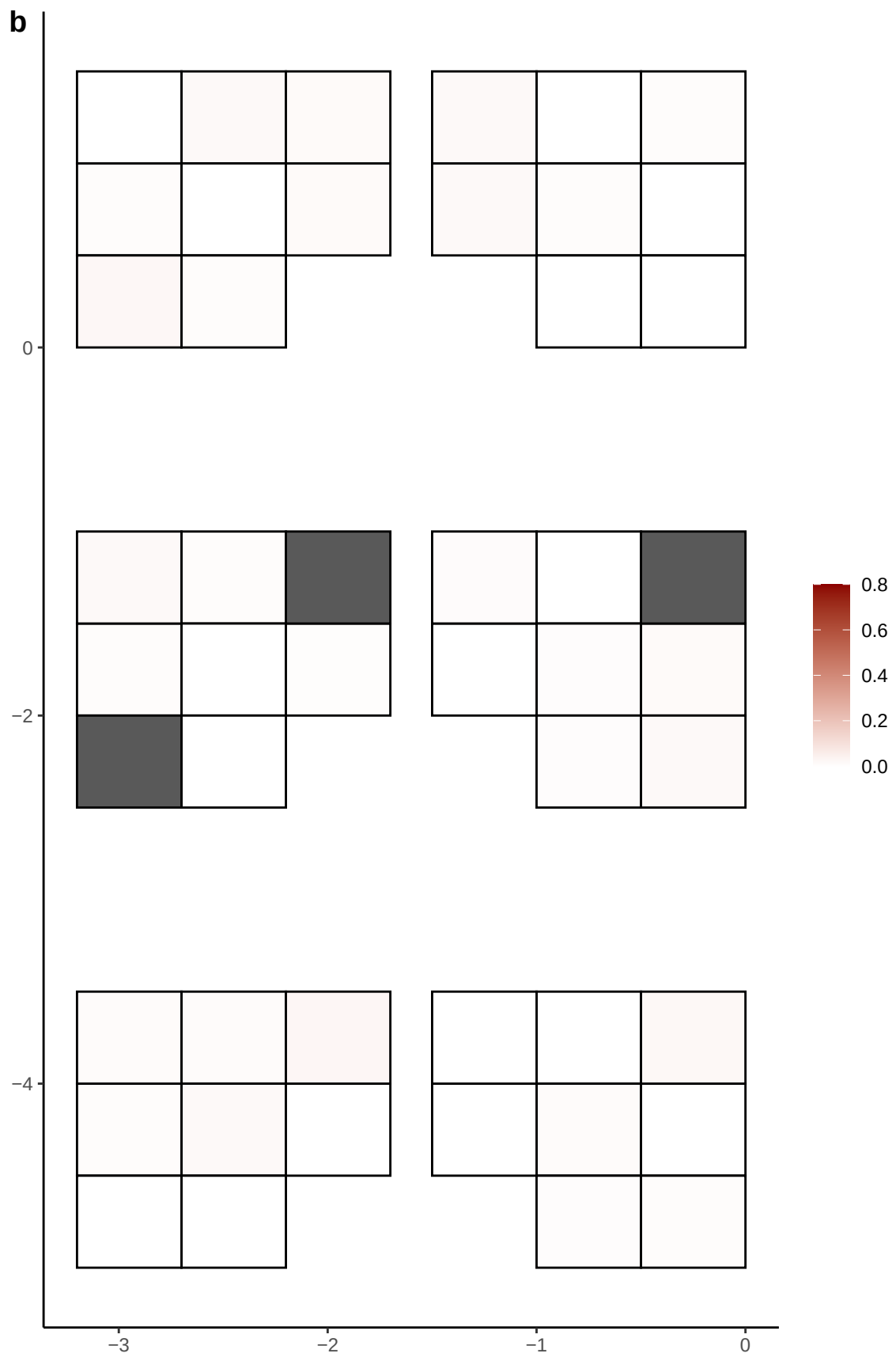

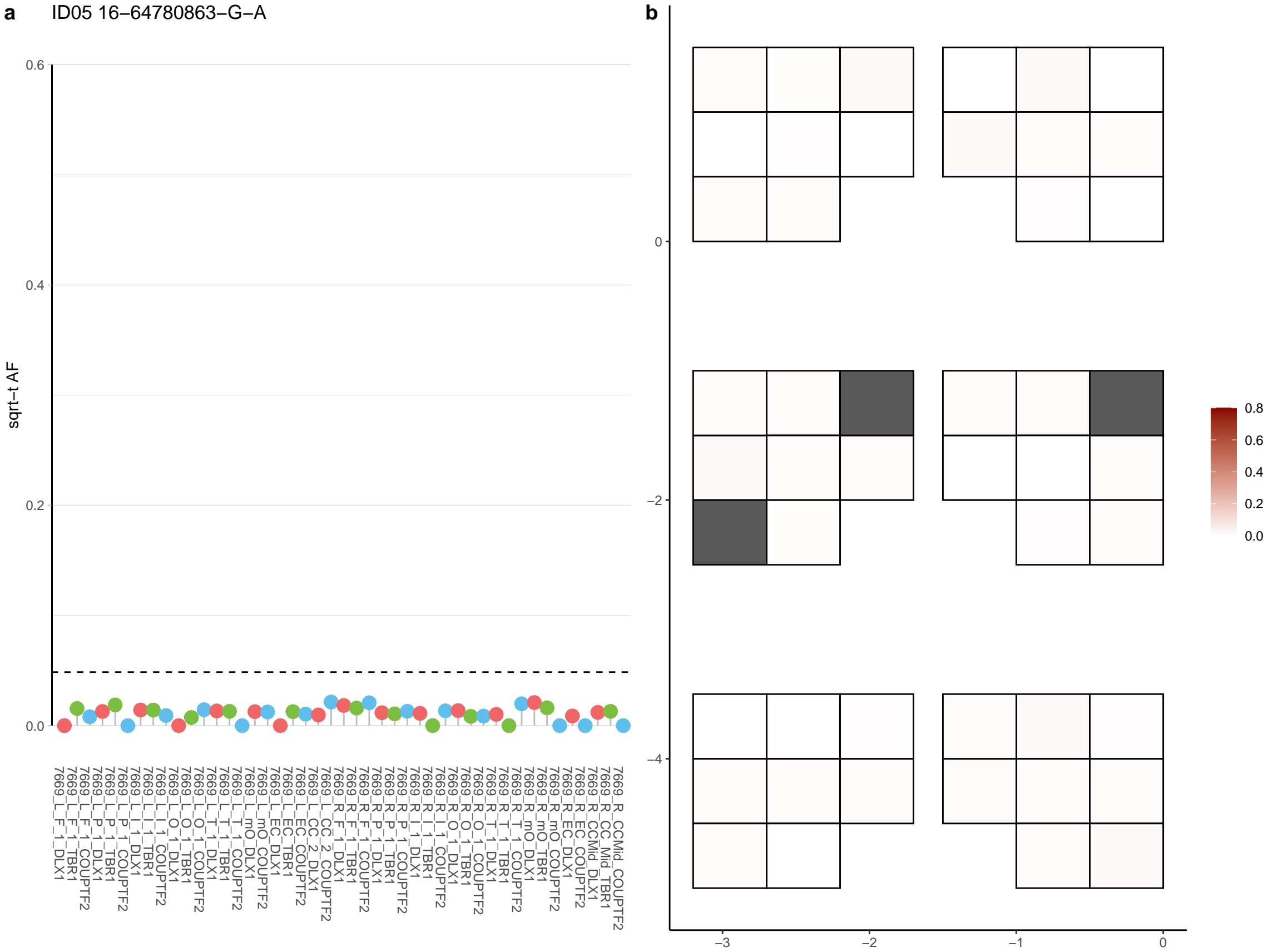

**a** ID05 16-71798192-C-A

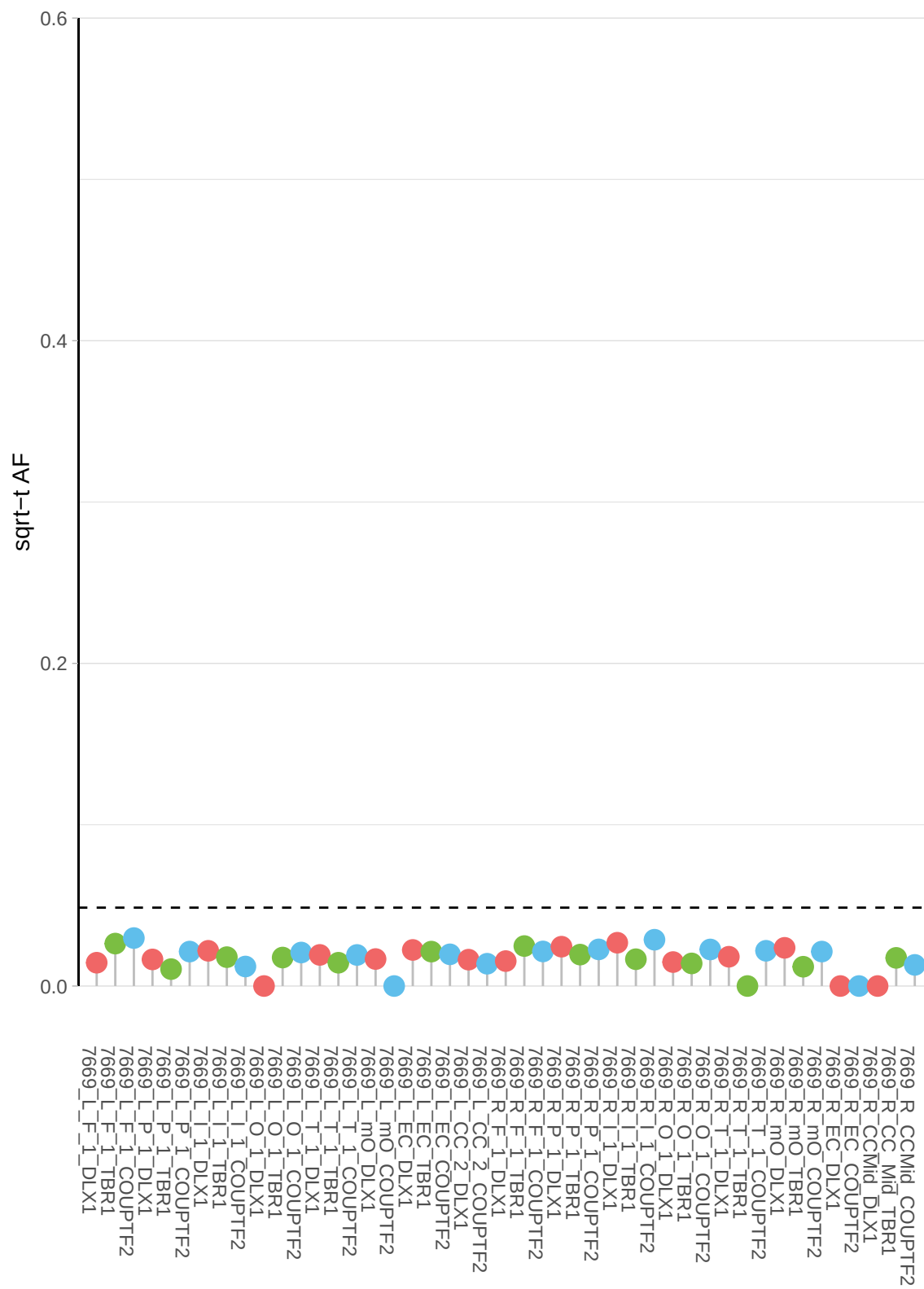

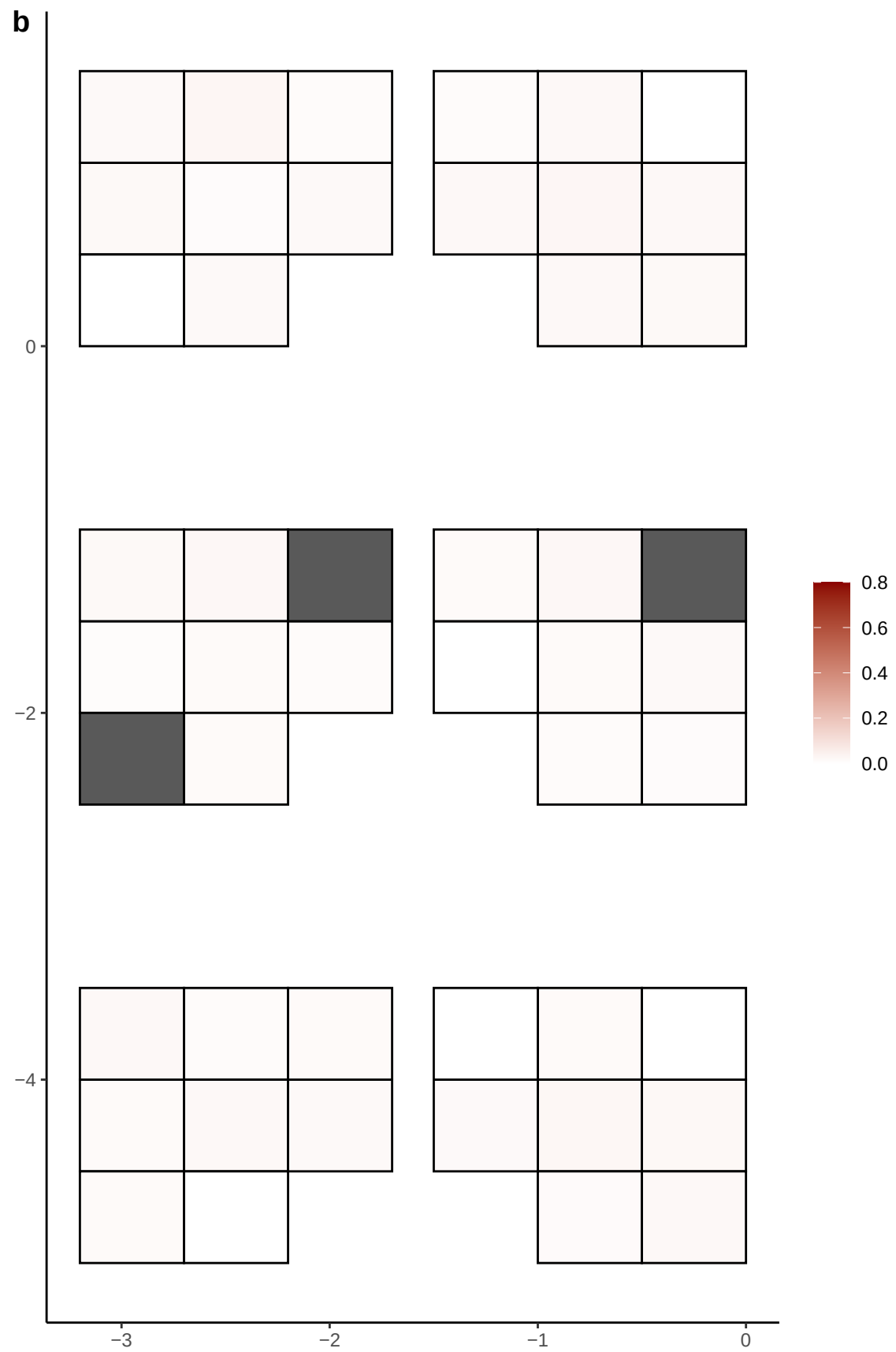

**a** ID05 16-73492923-G-T

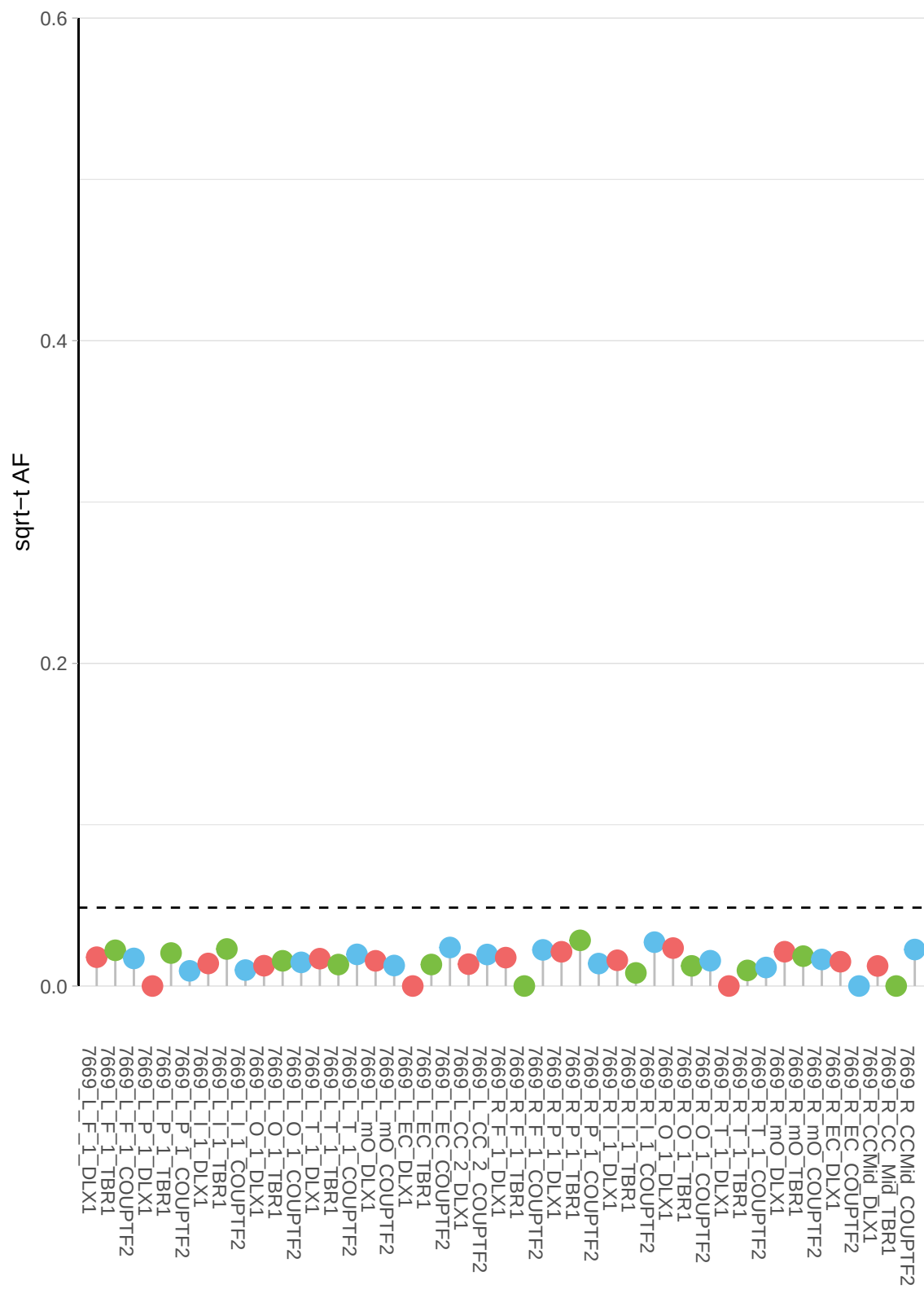

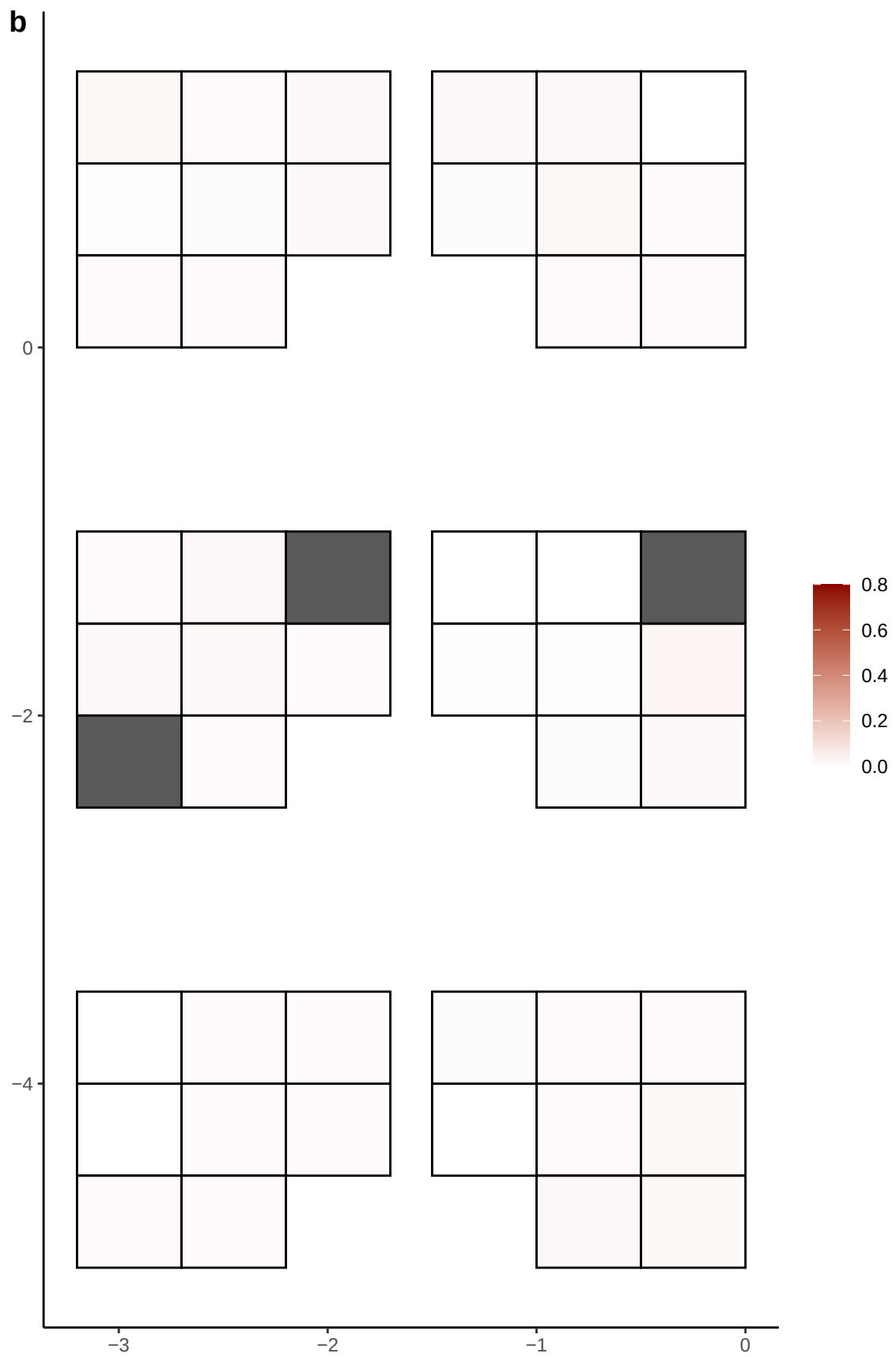

**a** ID05 16-77437520-G-T

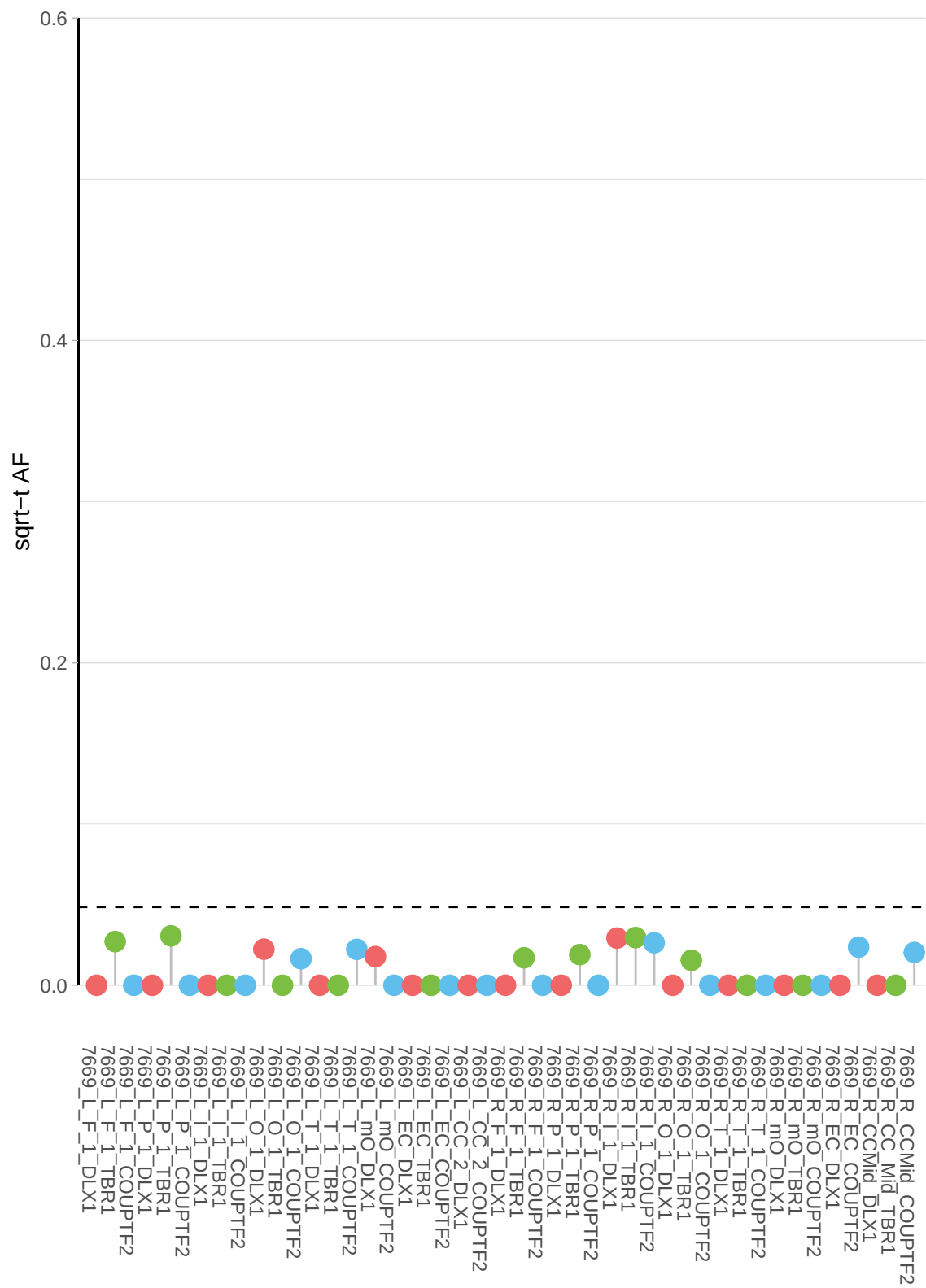

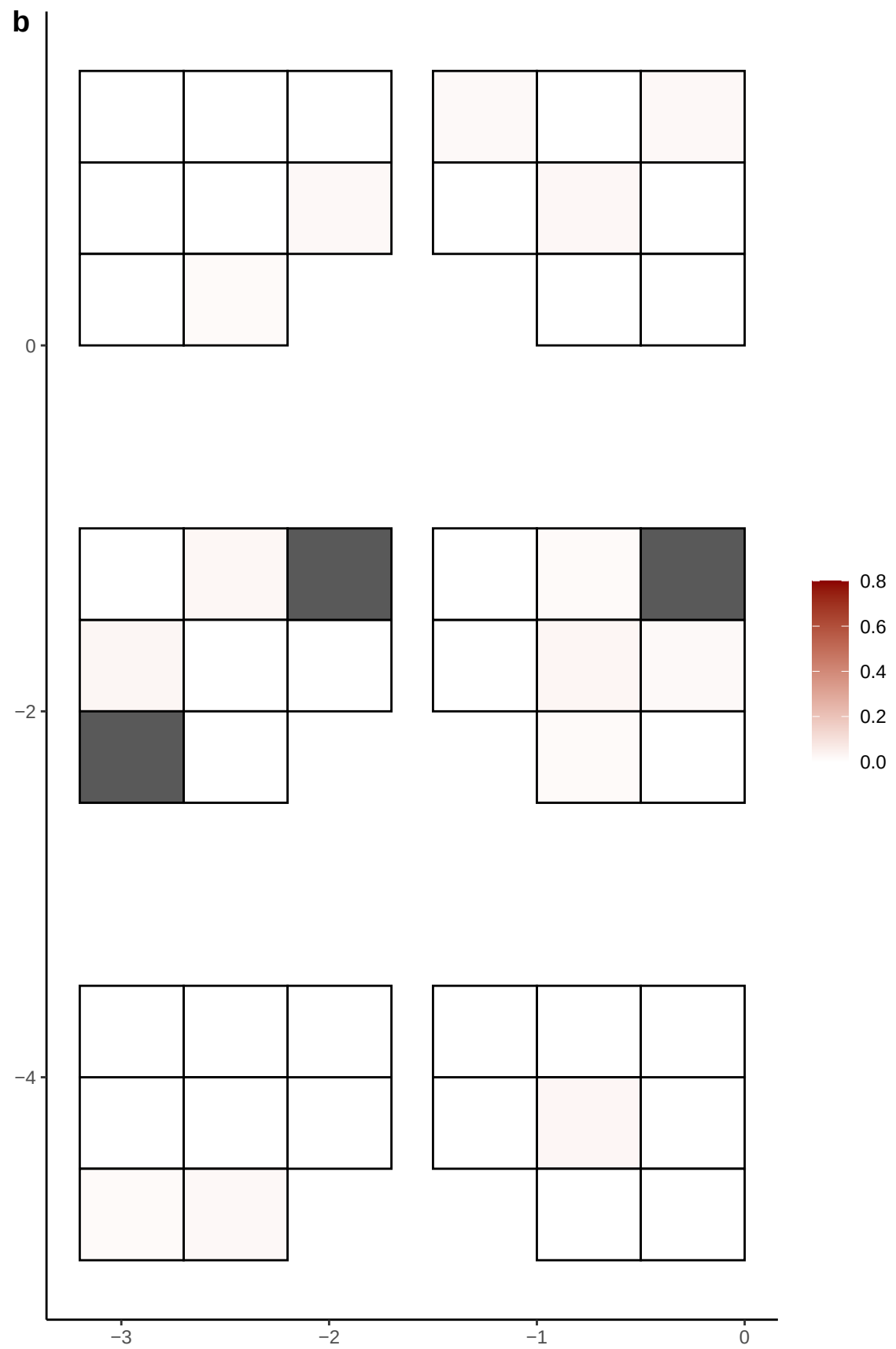

**a** ID05 16-83153418-C-T

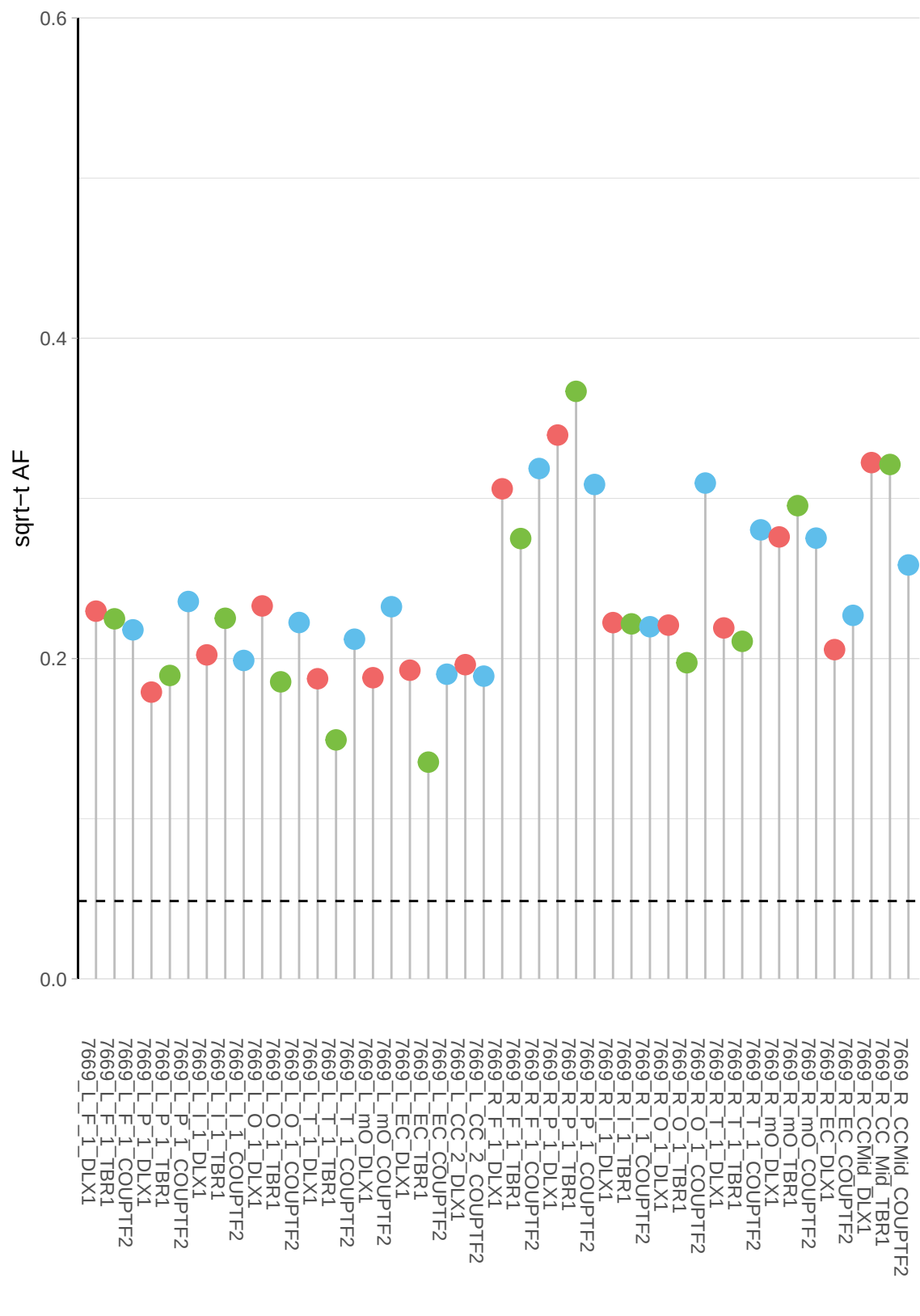

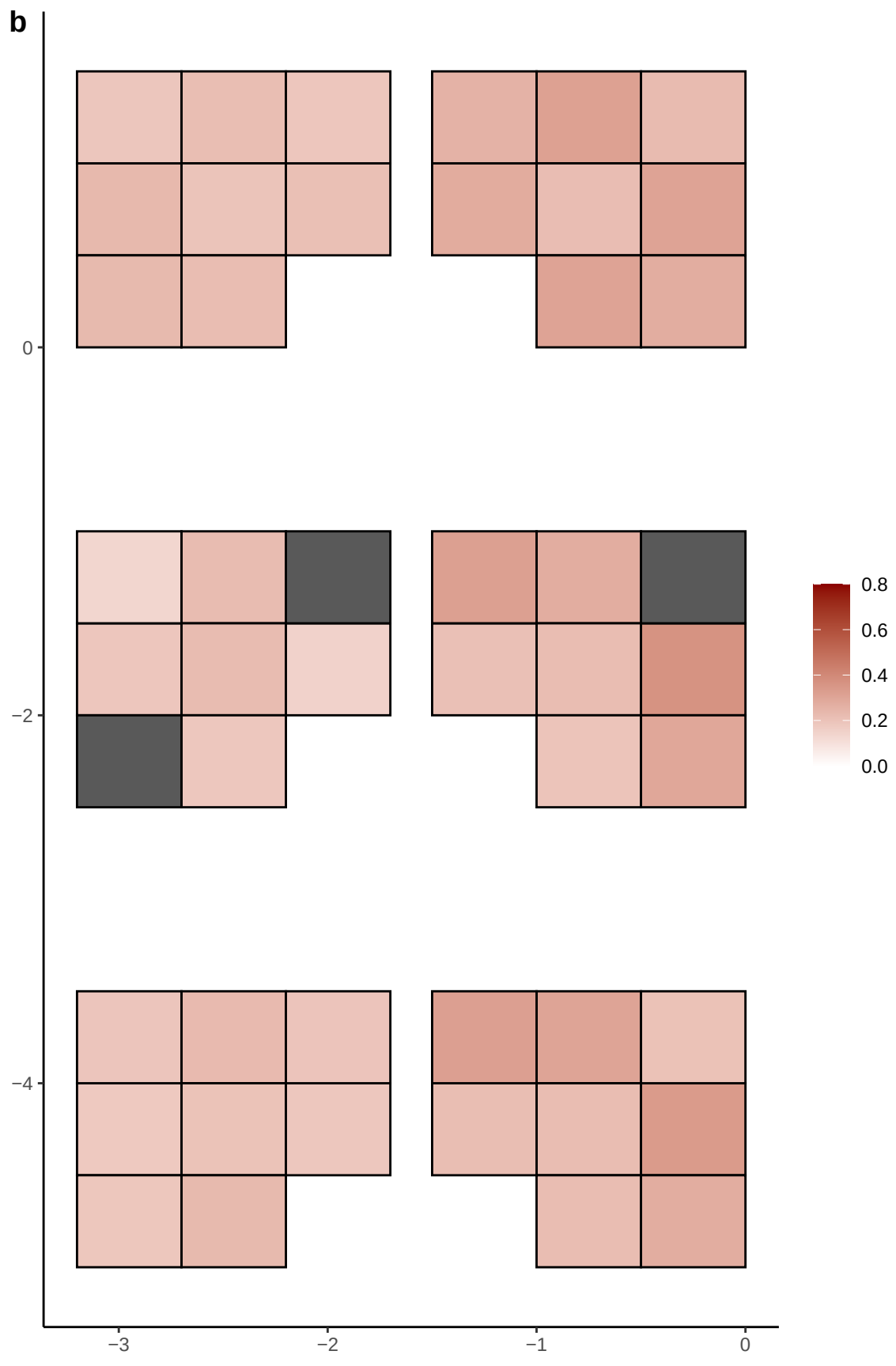

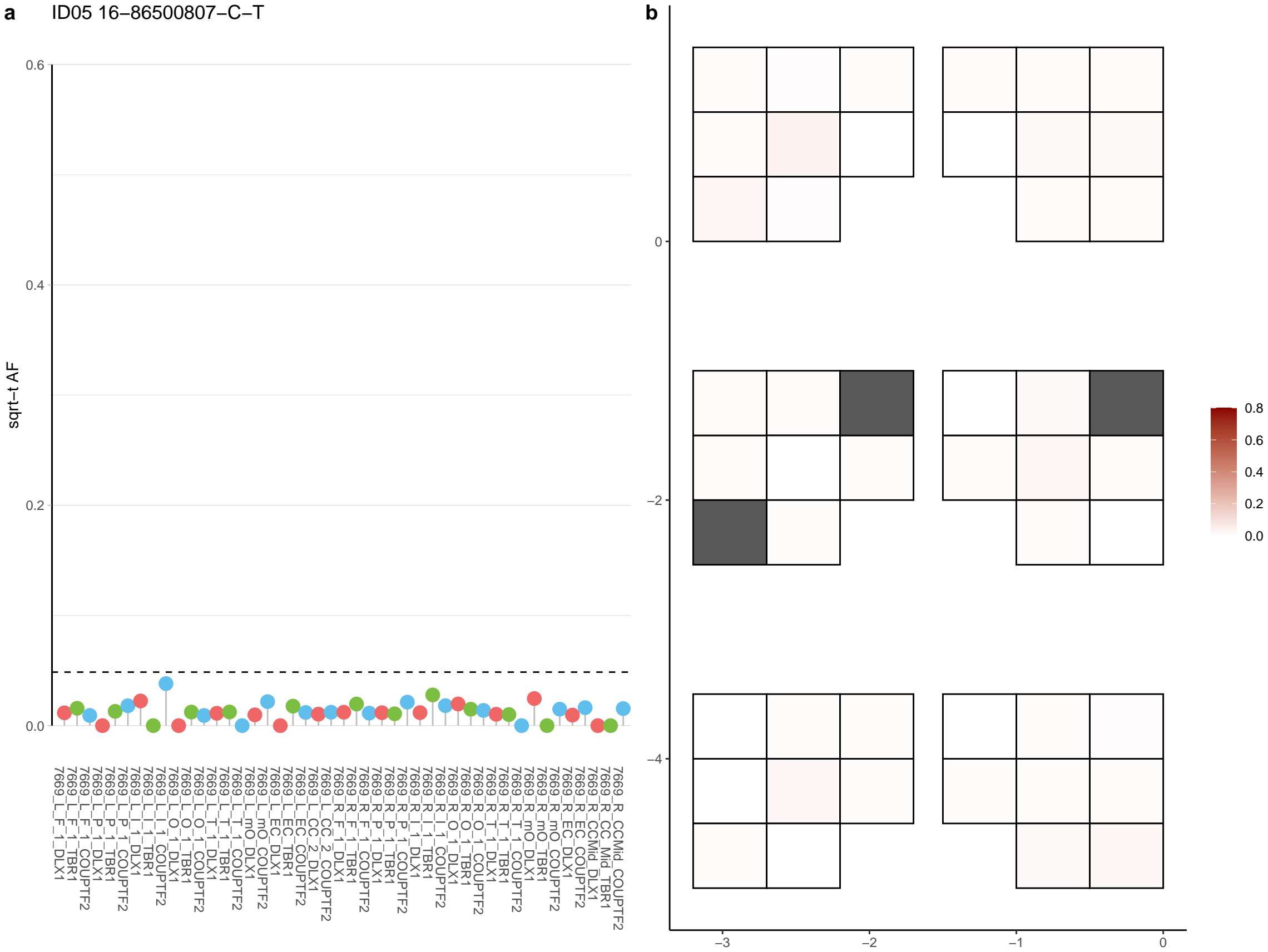

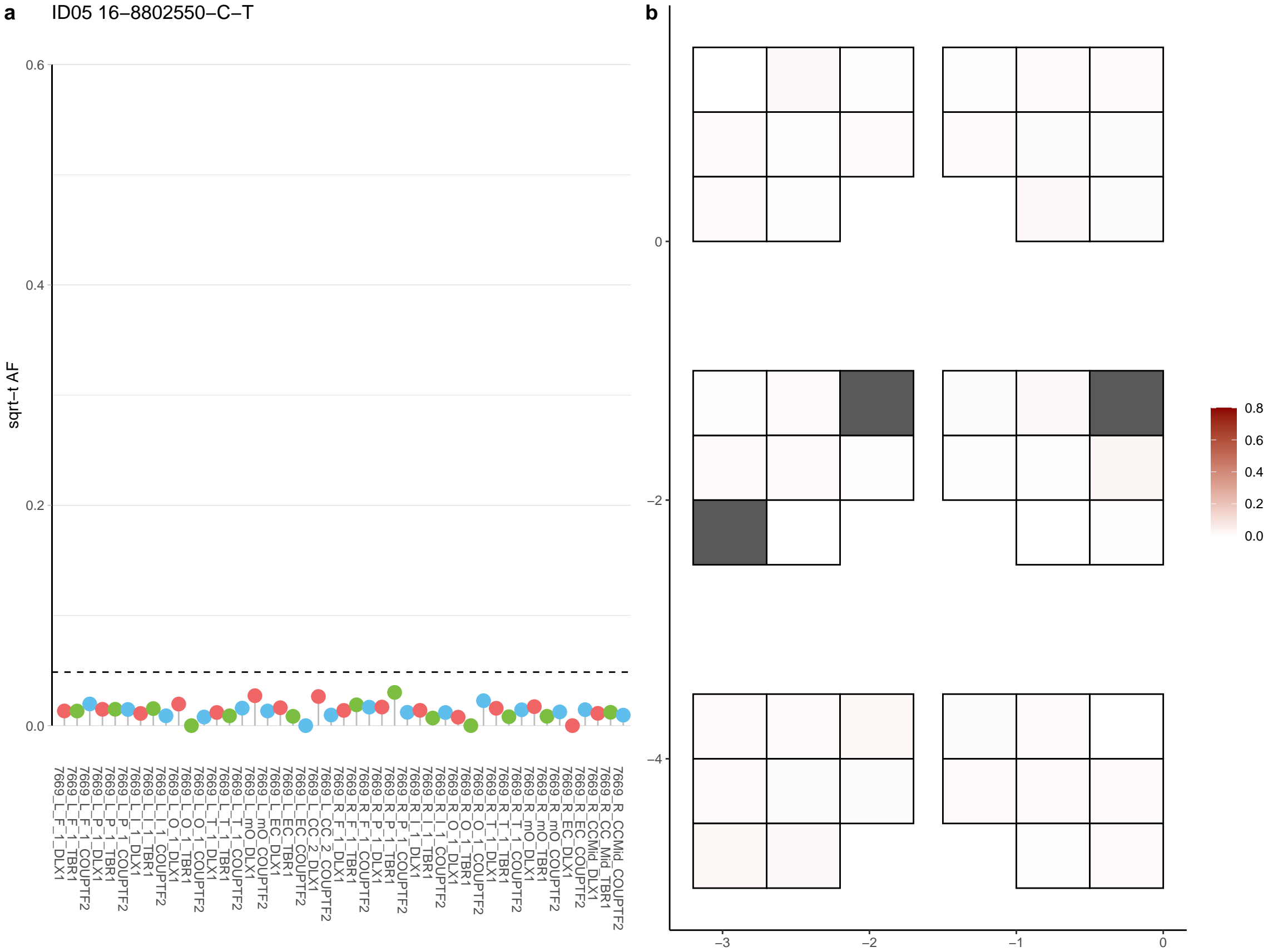

**a** ID05 16-925921-C-T

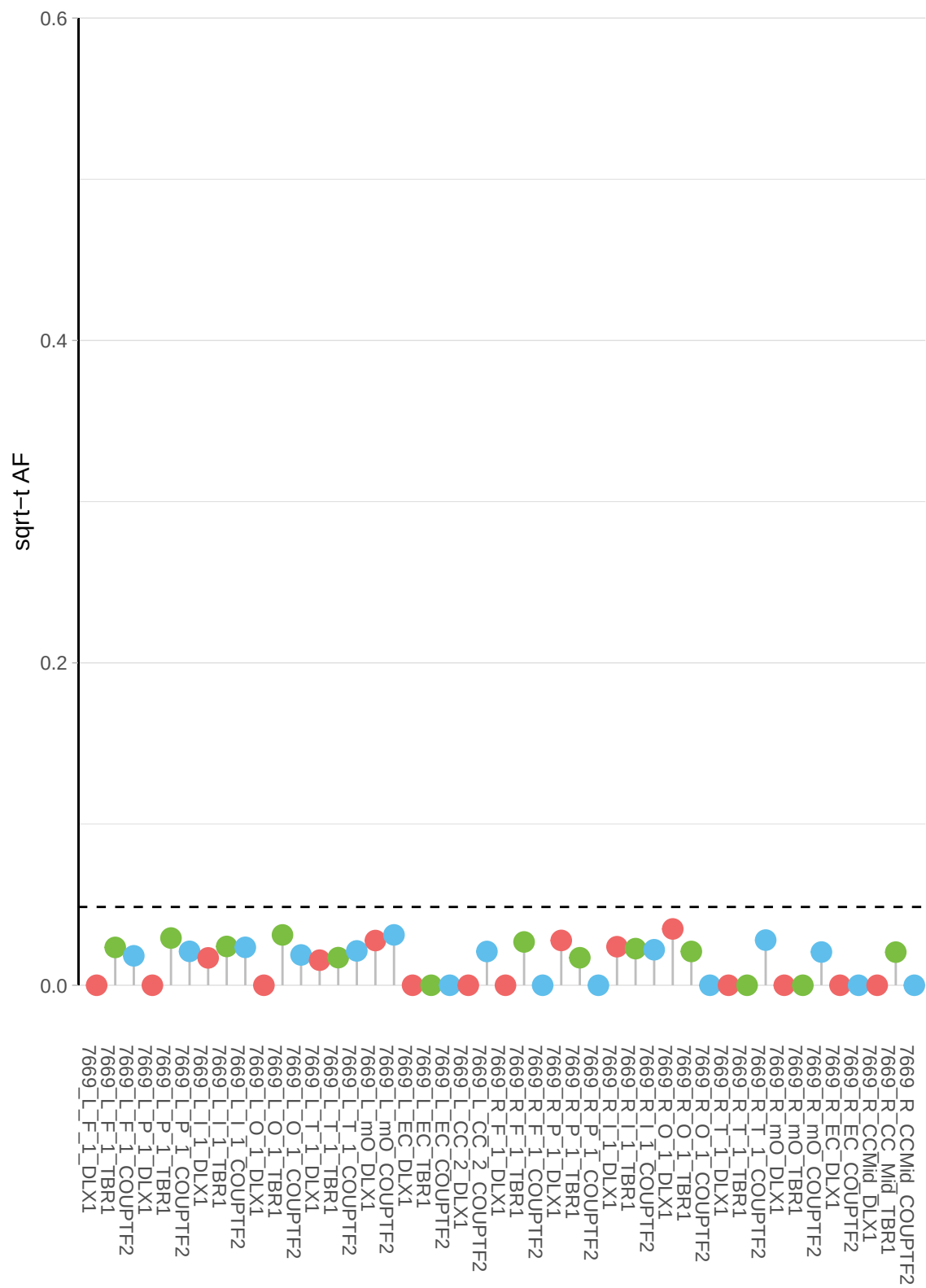

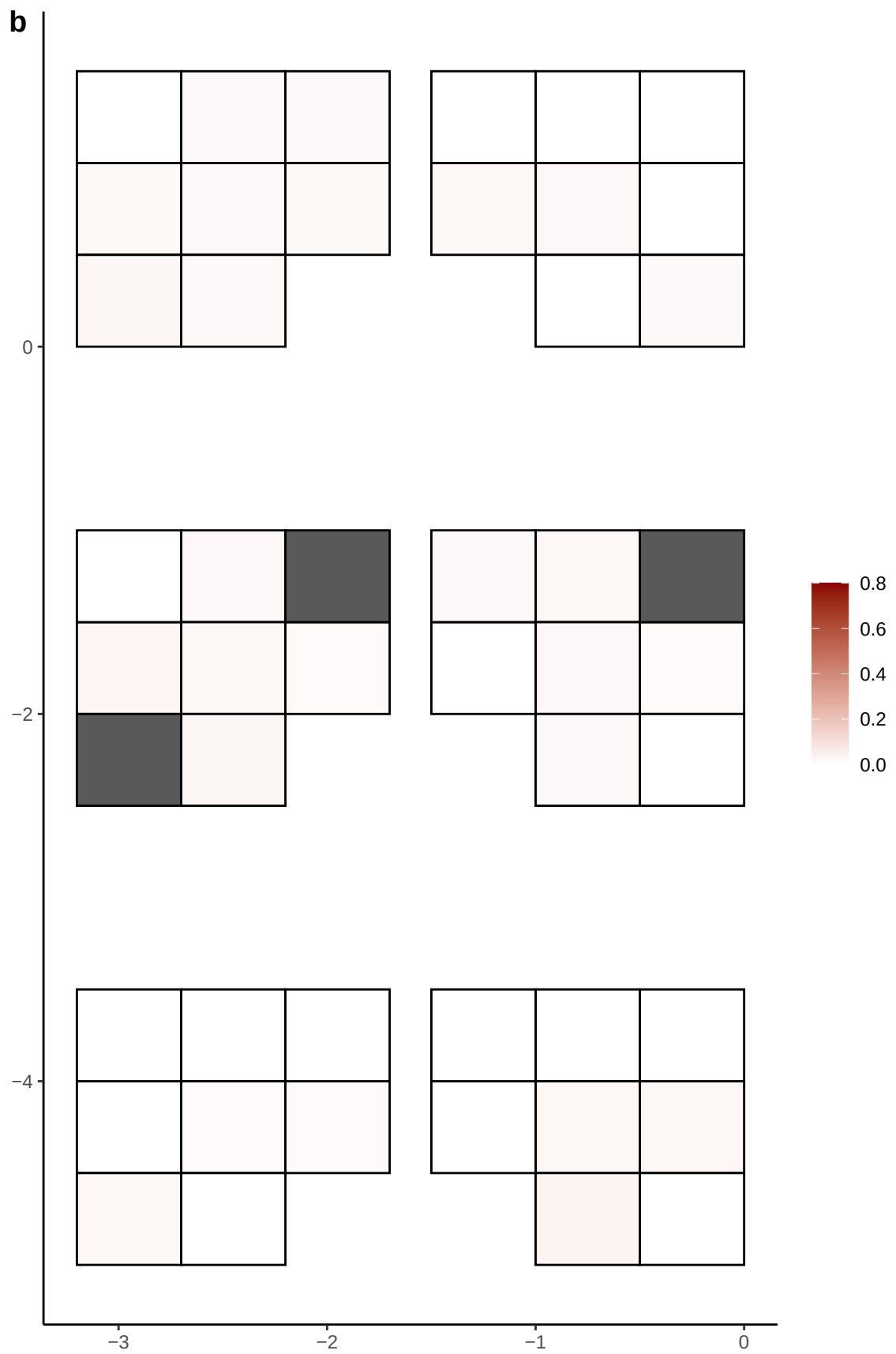

**a** ID05 16-9382504-C-T

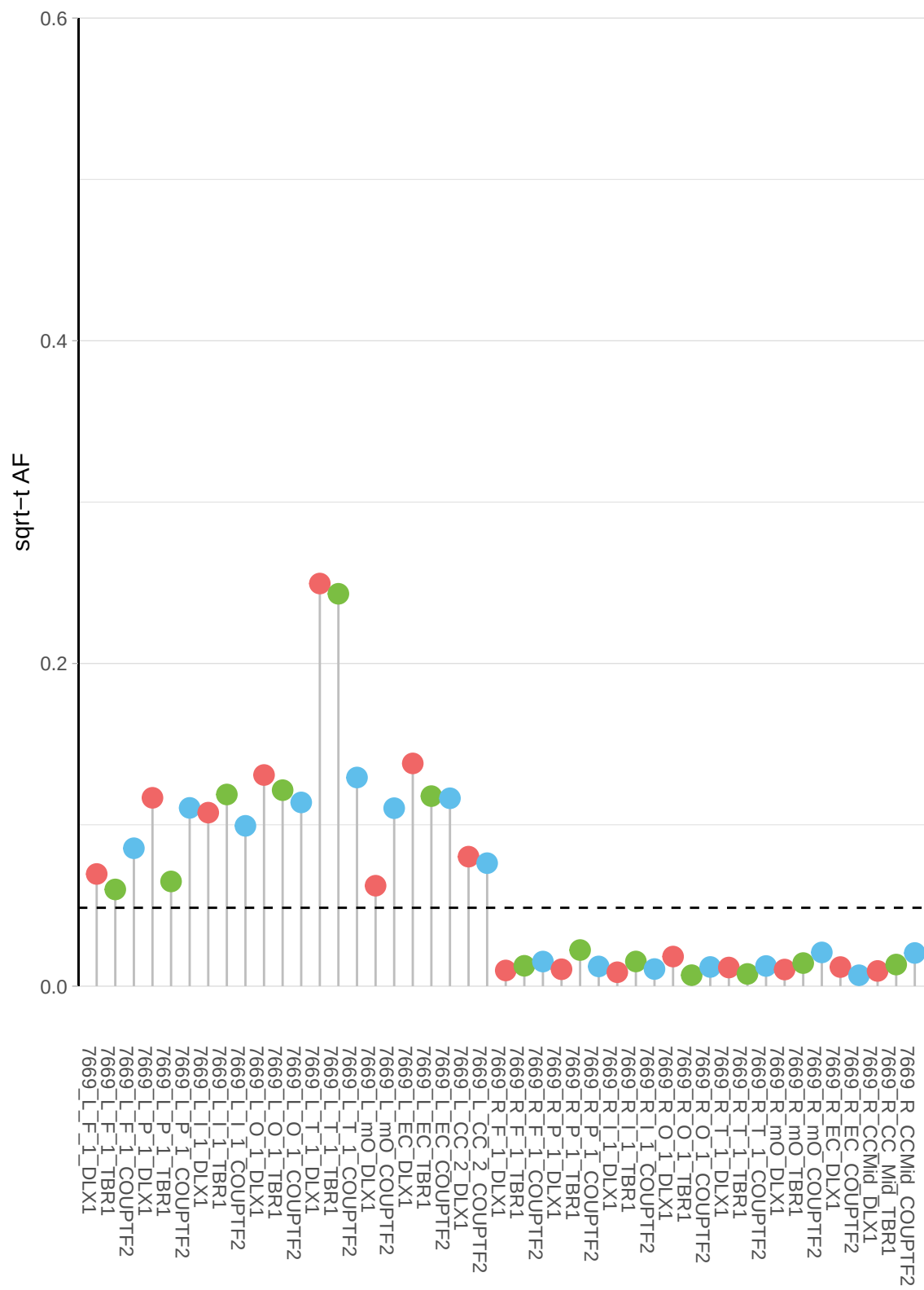

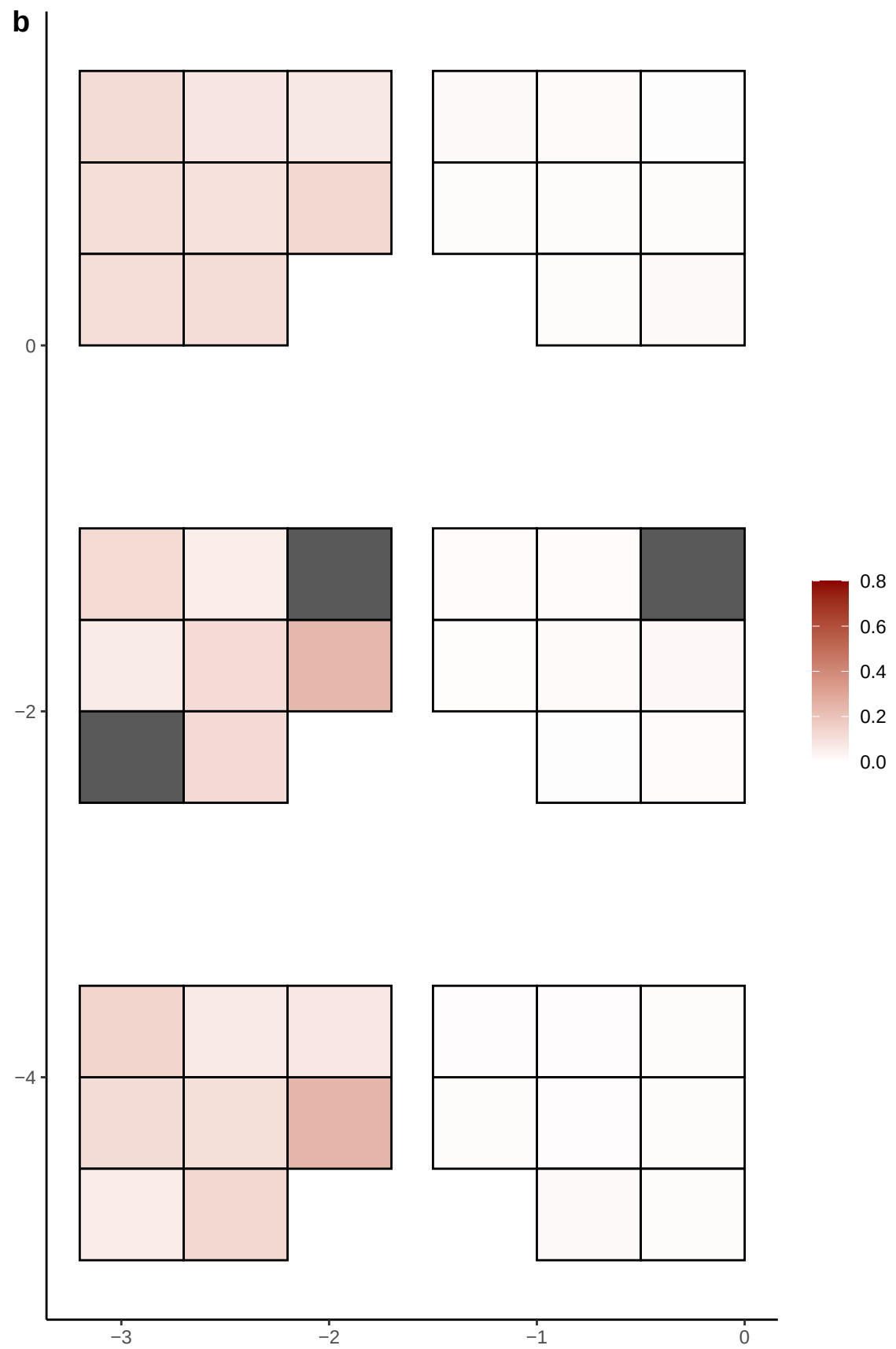

**a** ID05 17-10283611-C-T

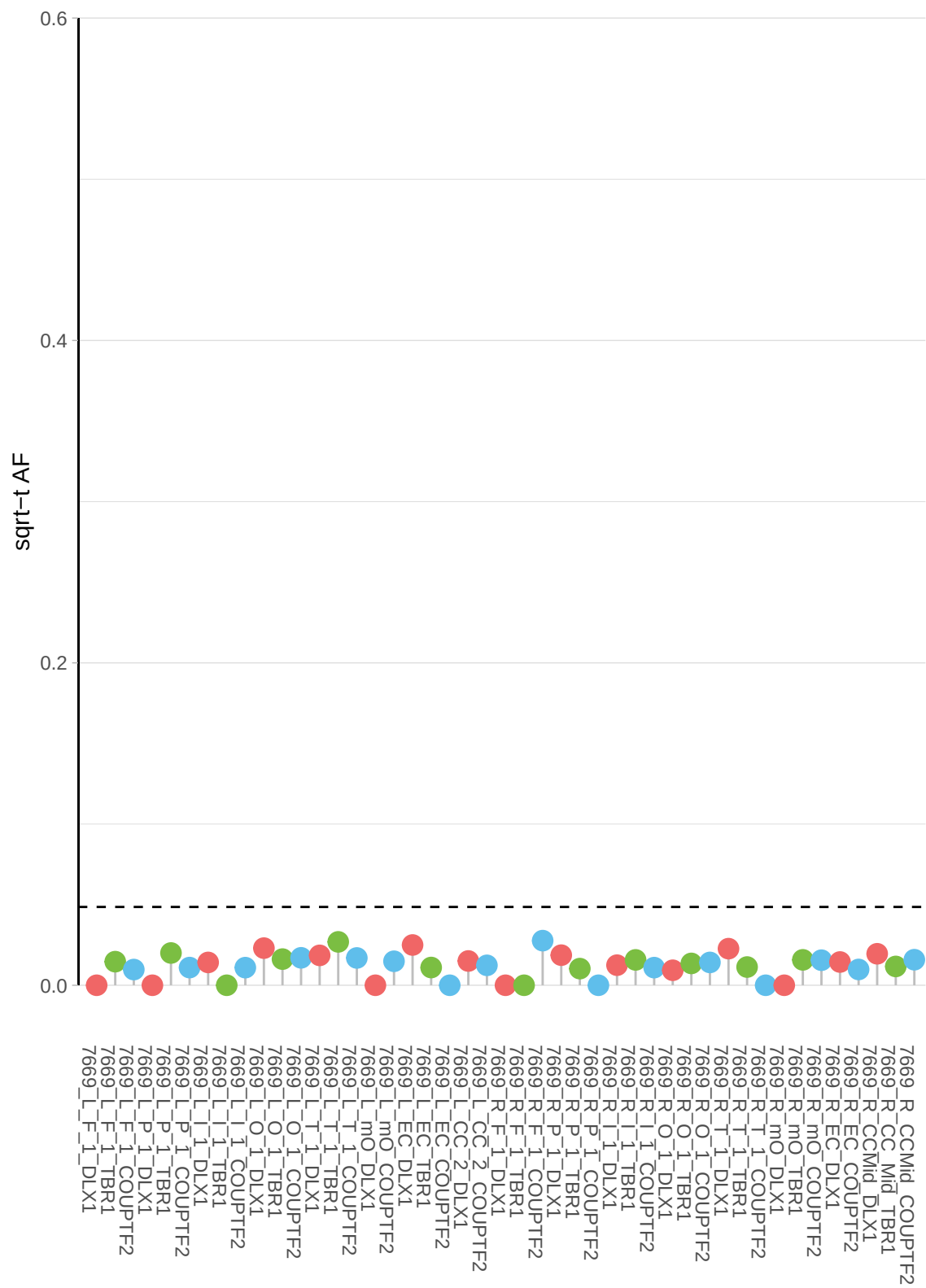

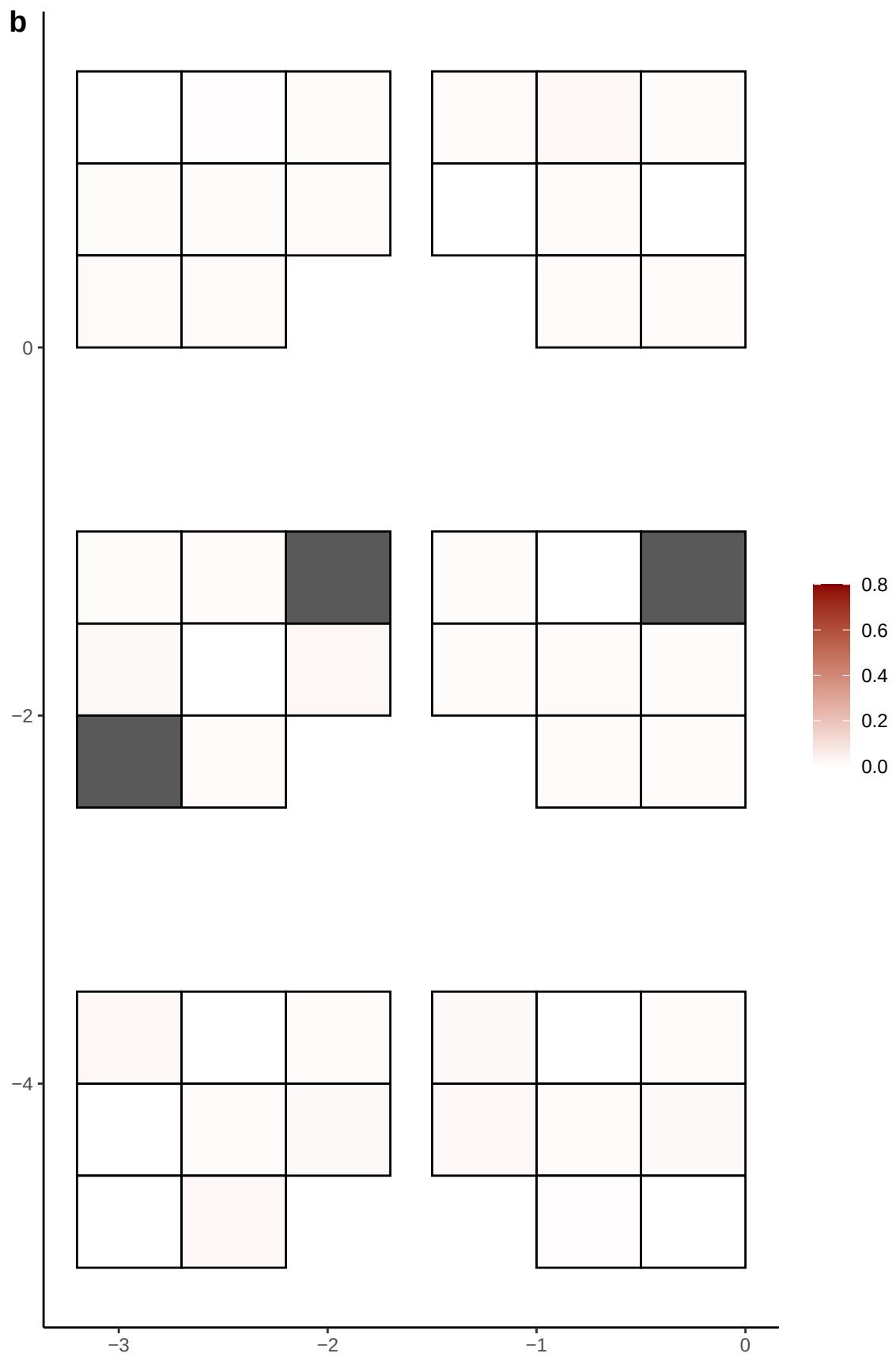

**a** ID05 17-12534415-G-T

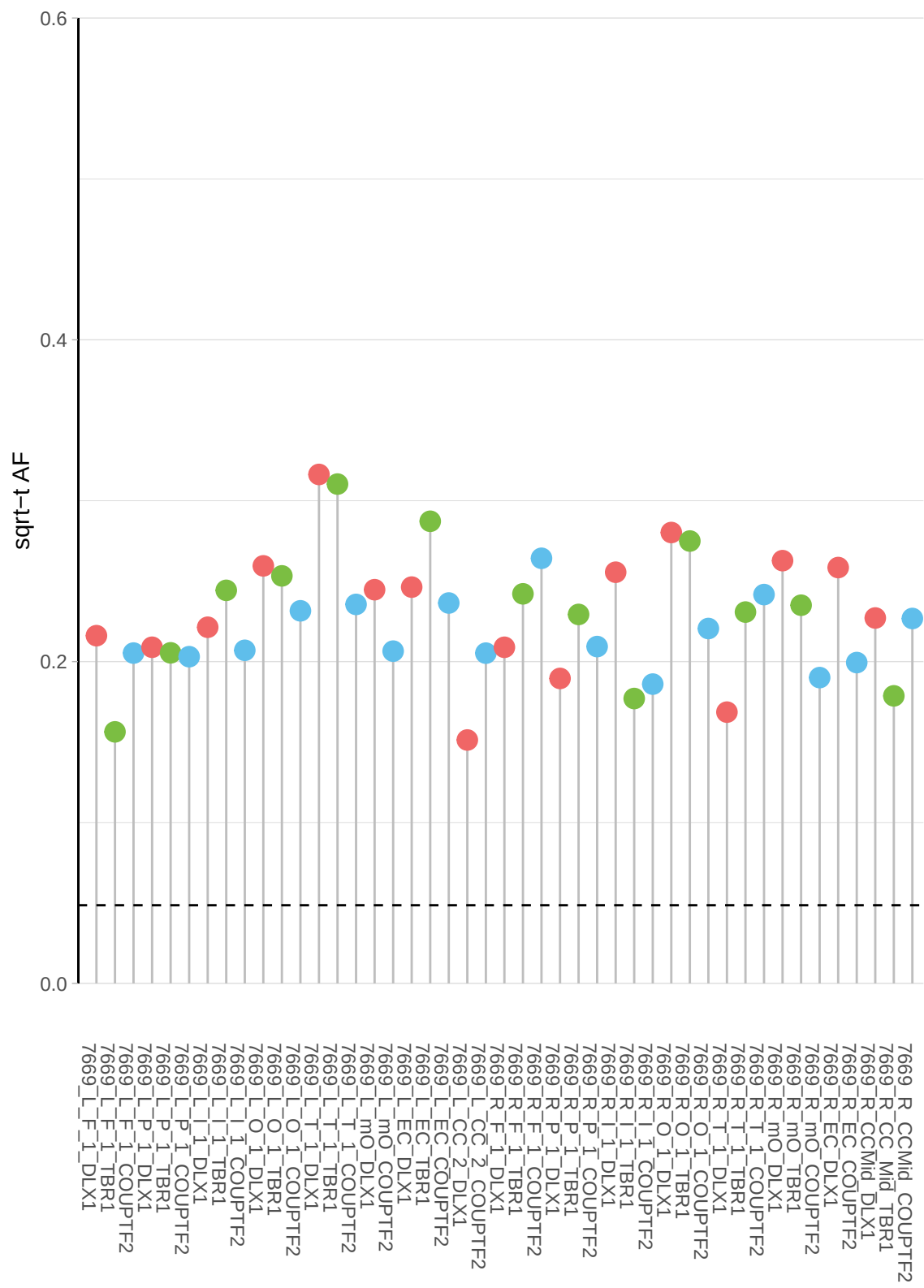

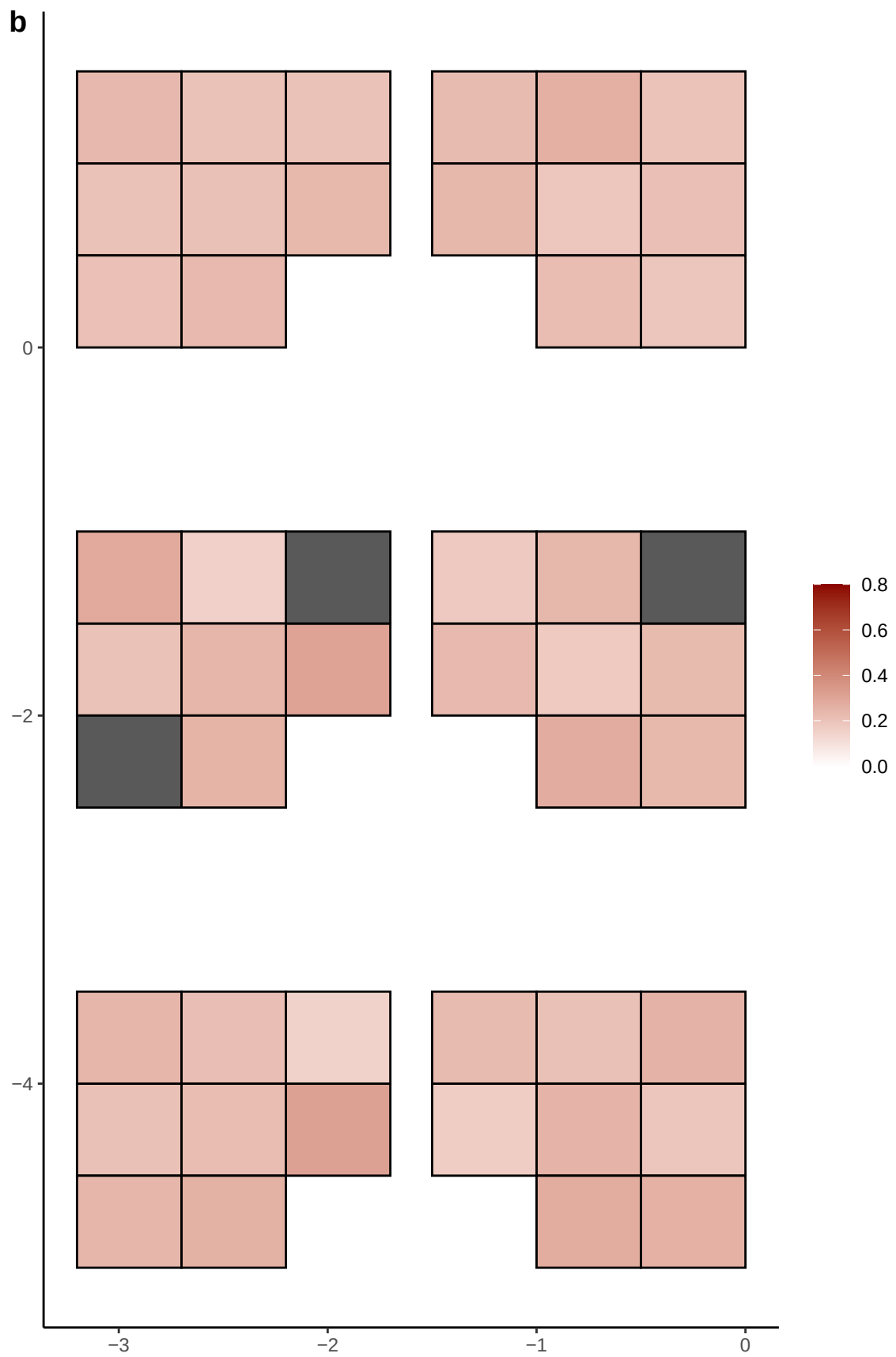

**a** ID05 17-1478793-G-A

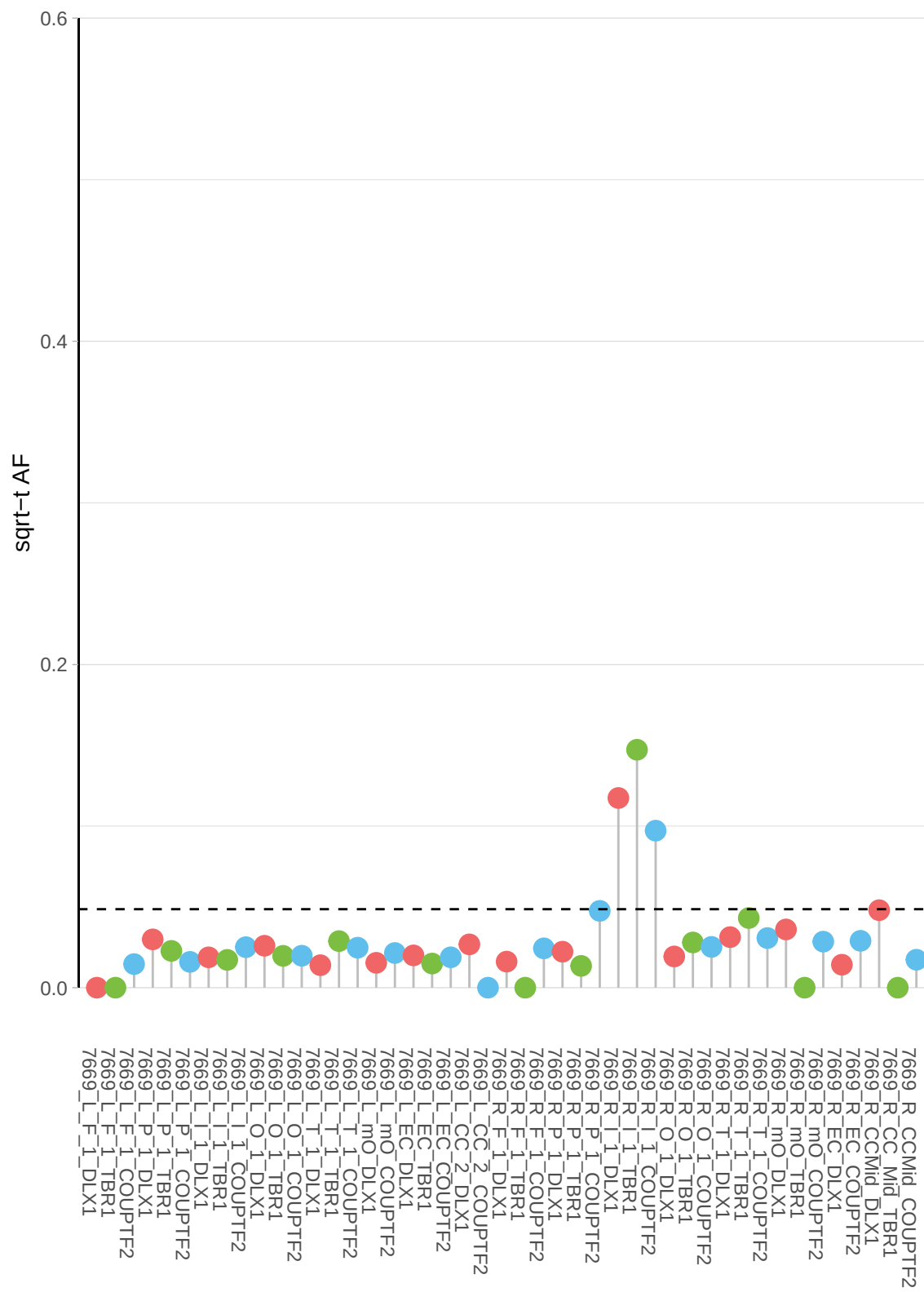

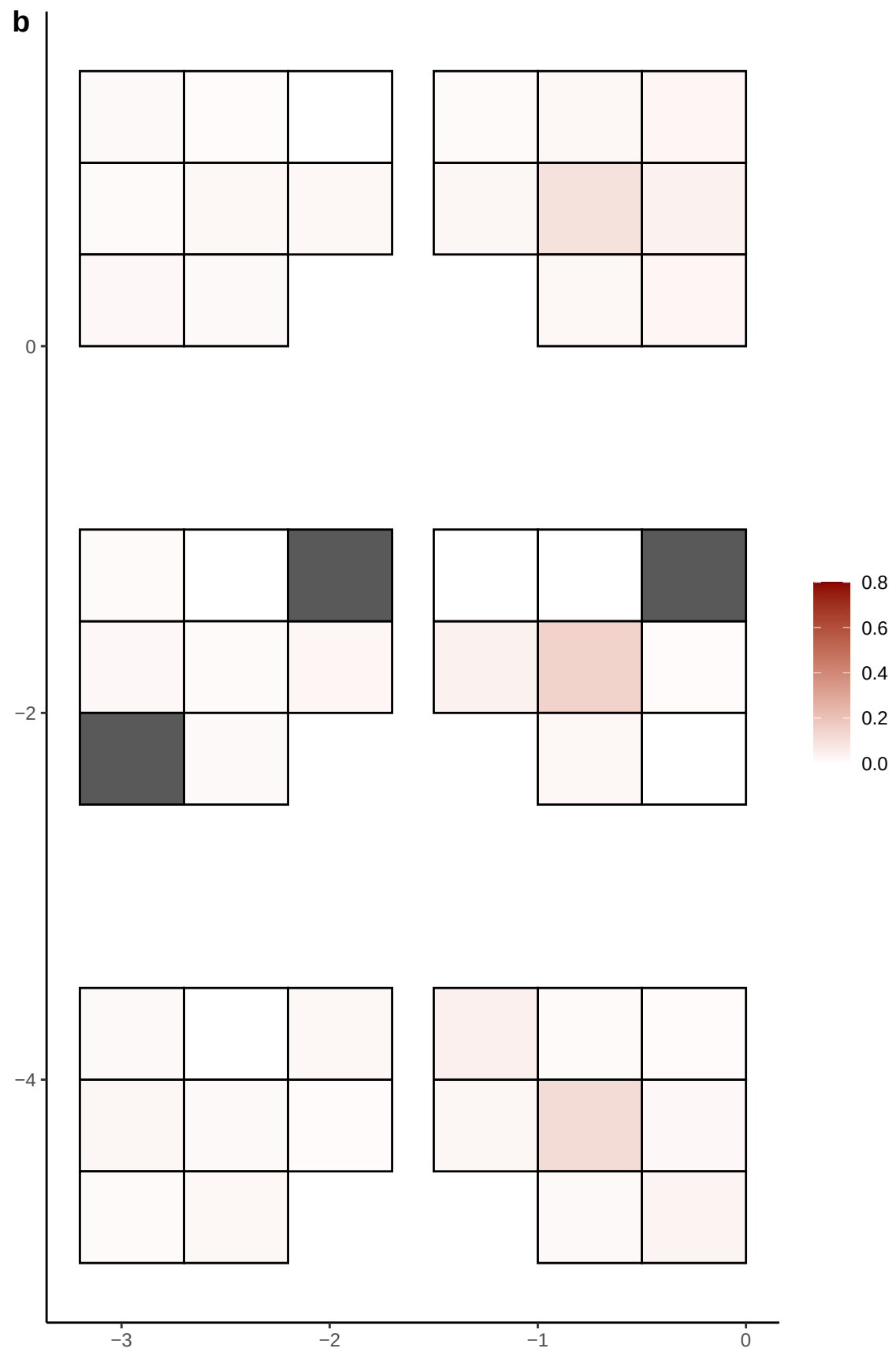

**a** ID05 17-1620800-G-T

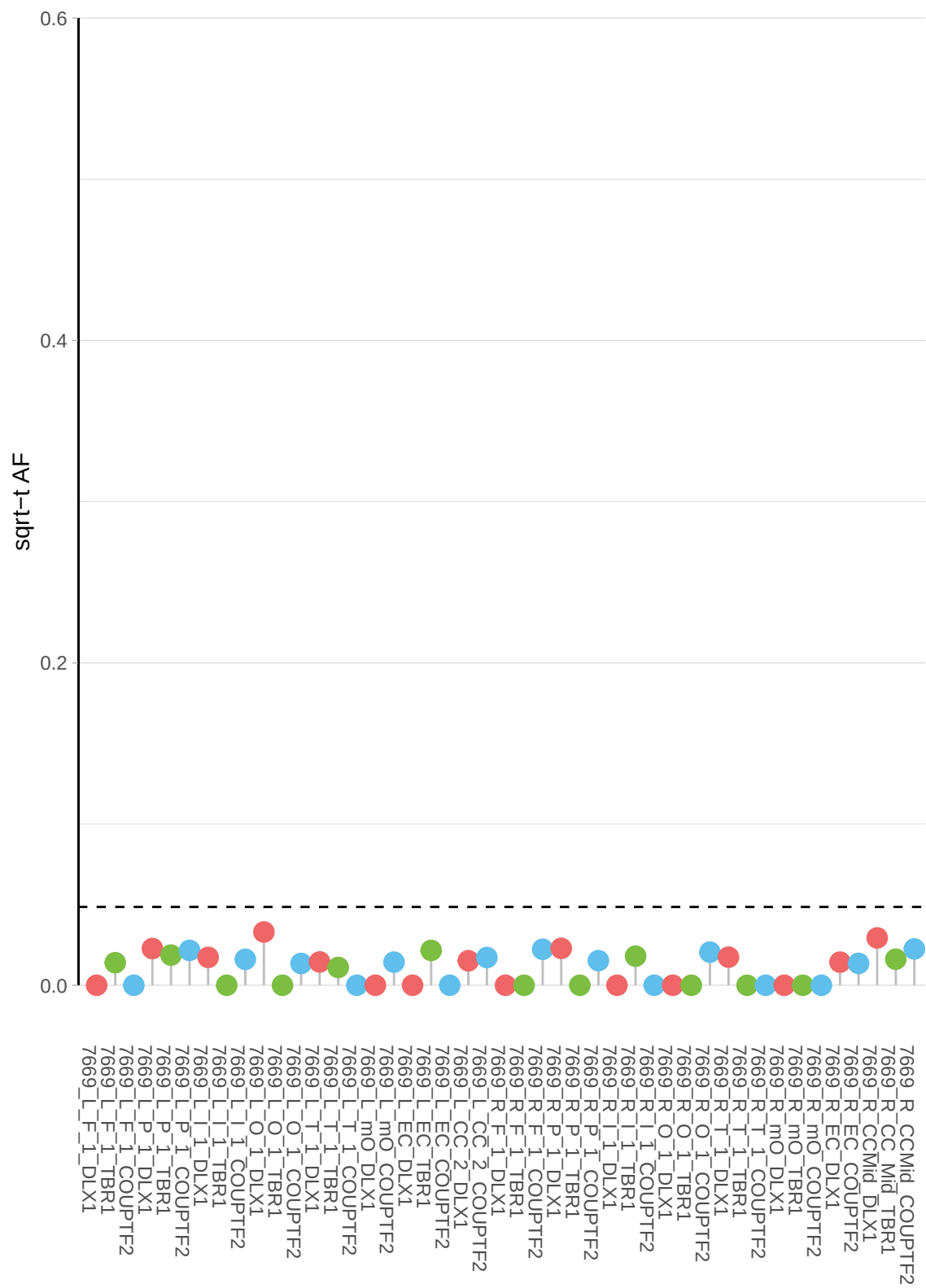

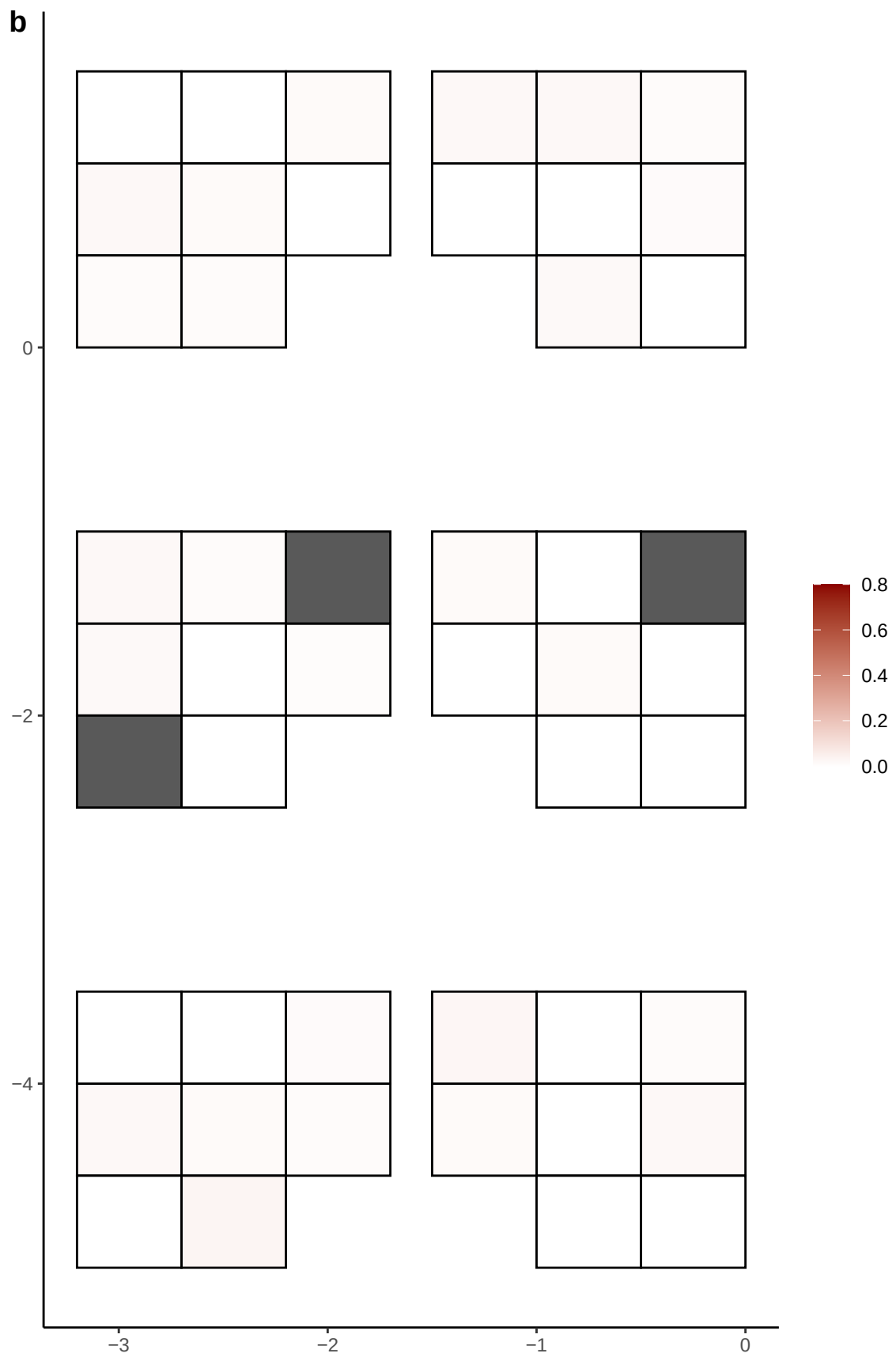

**a** ID05 17-17411221-G-A

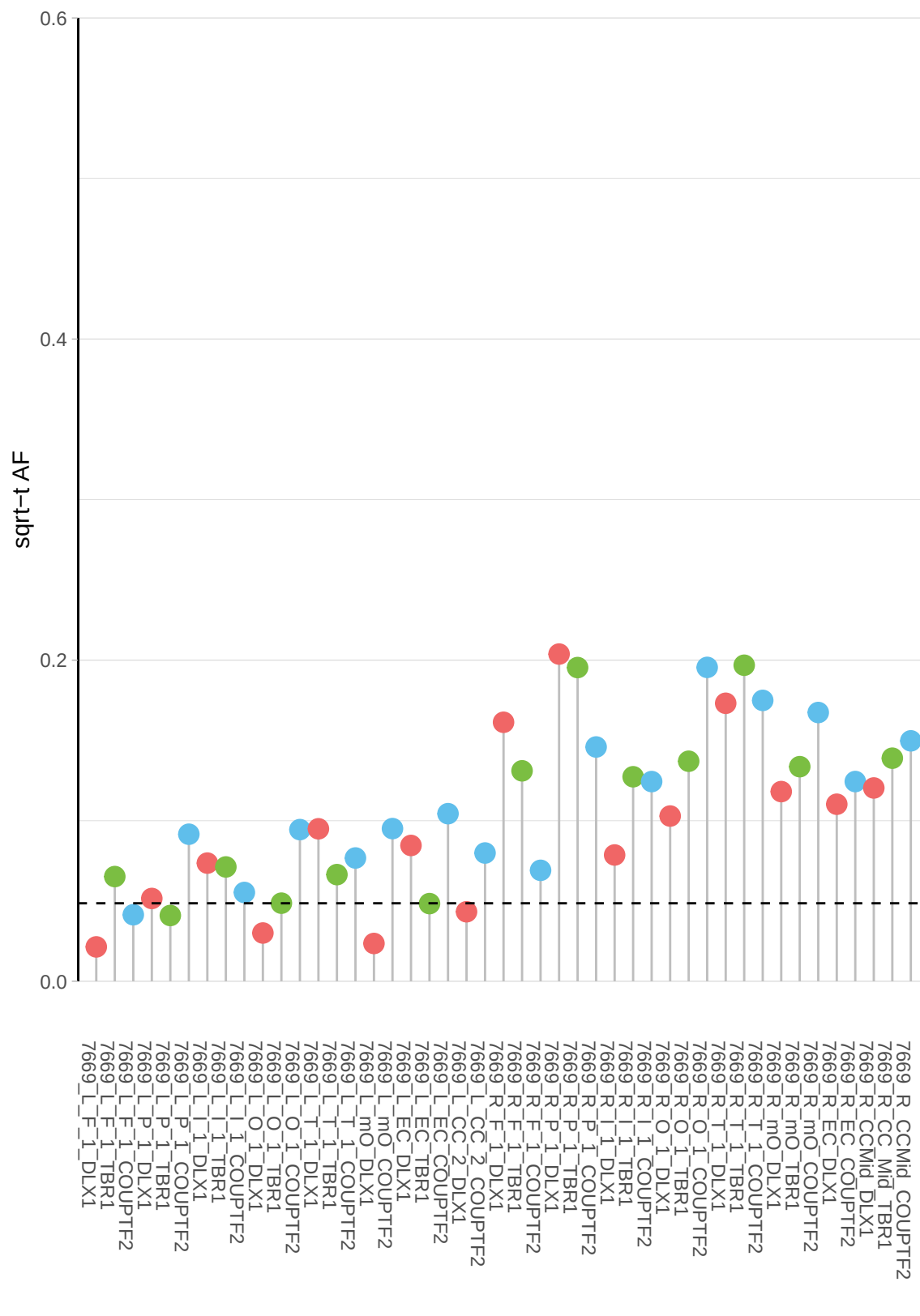

**b**

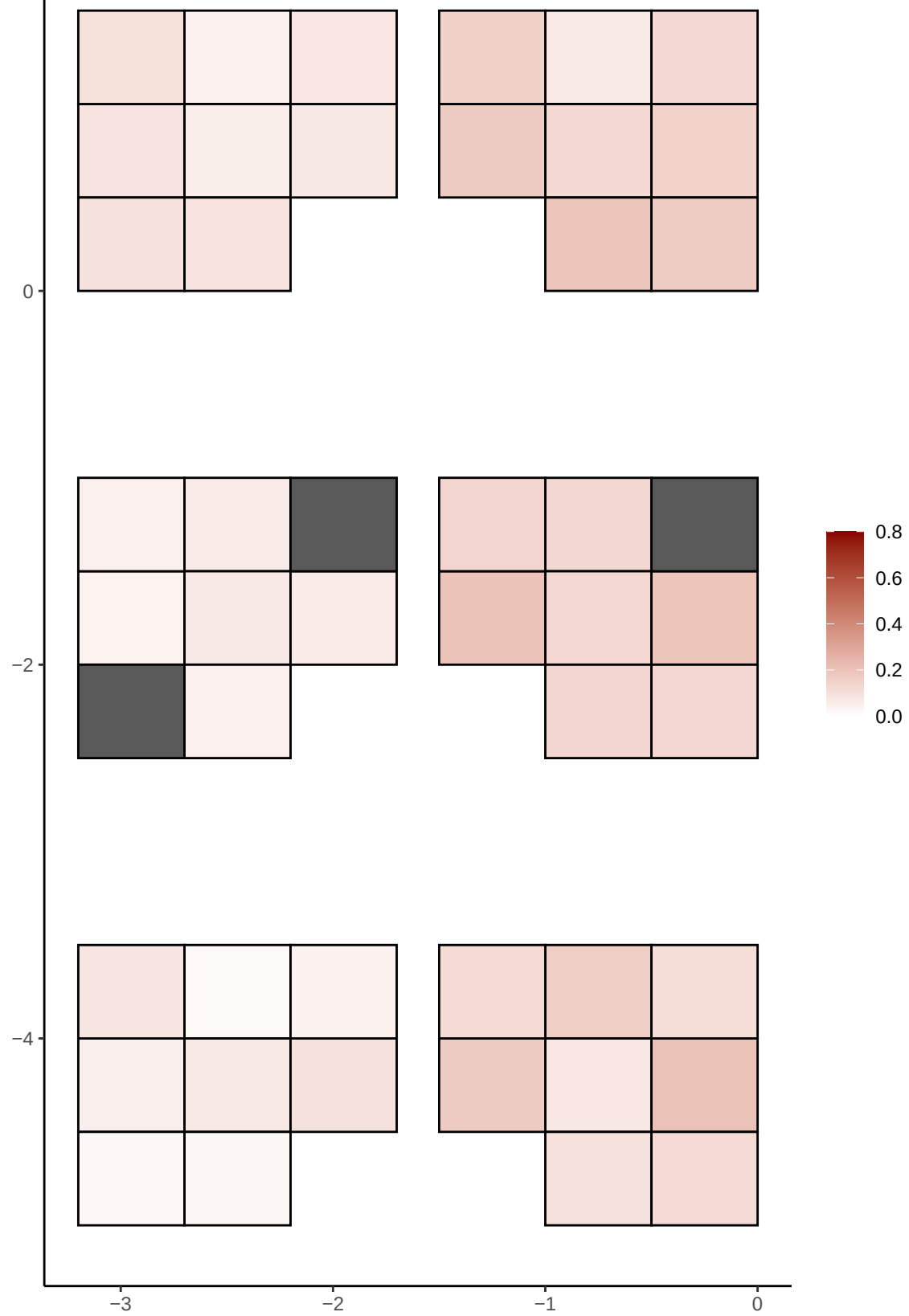

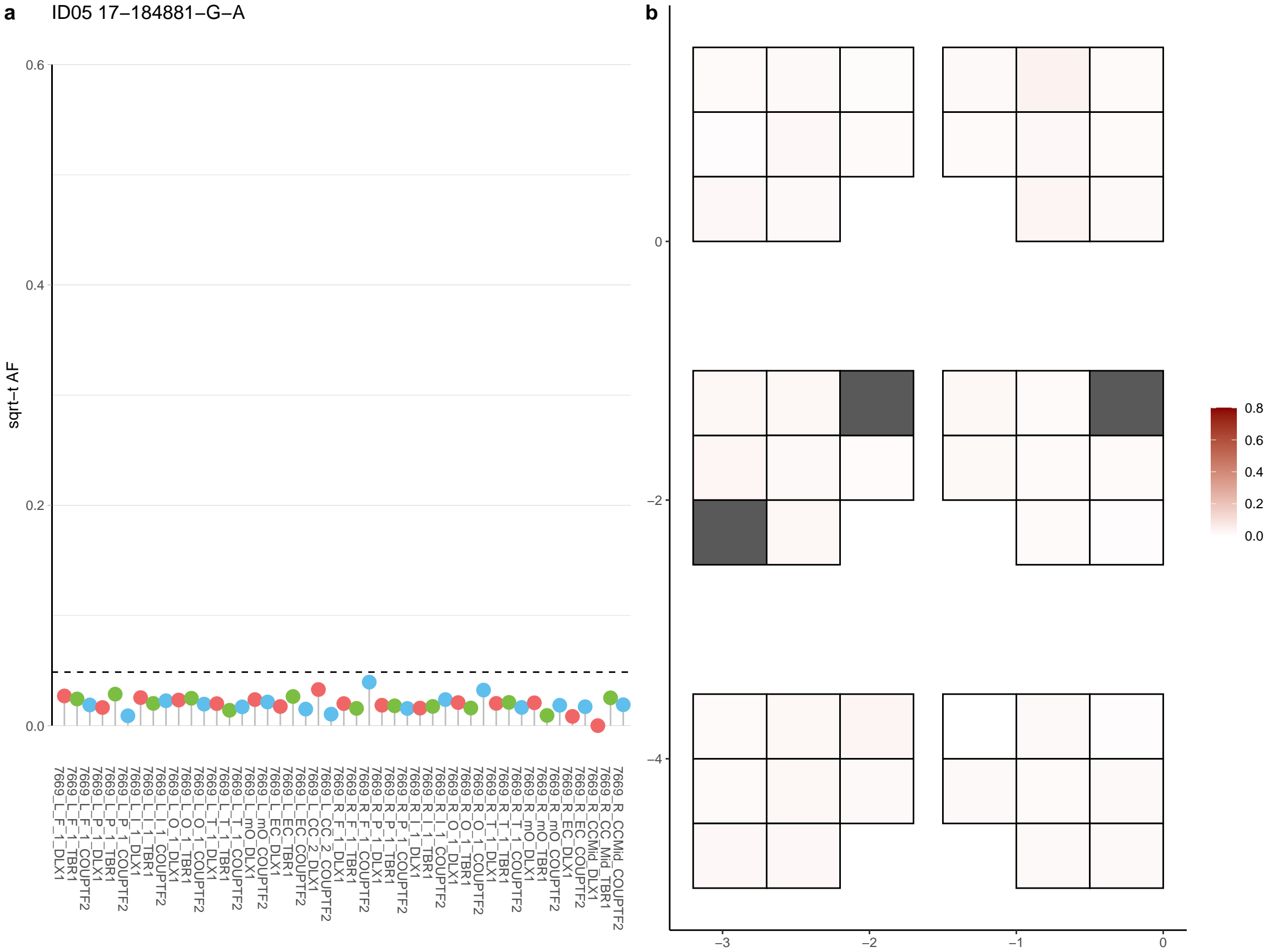

**a** ID05 17-20005504-G-A

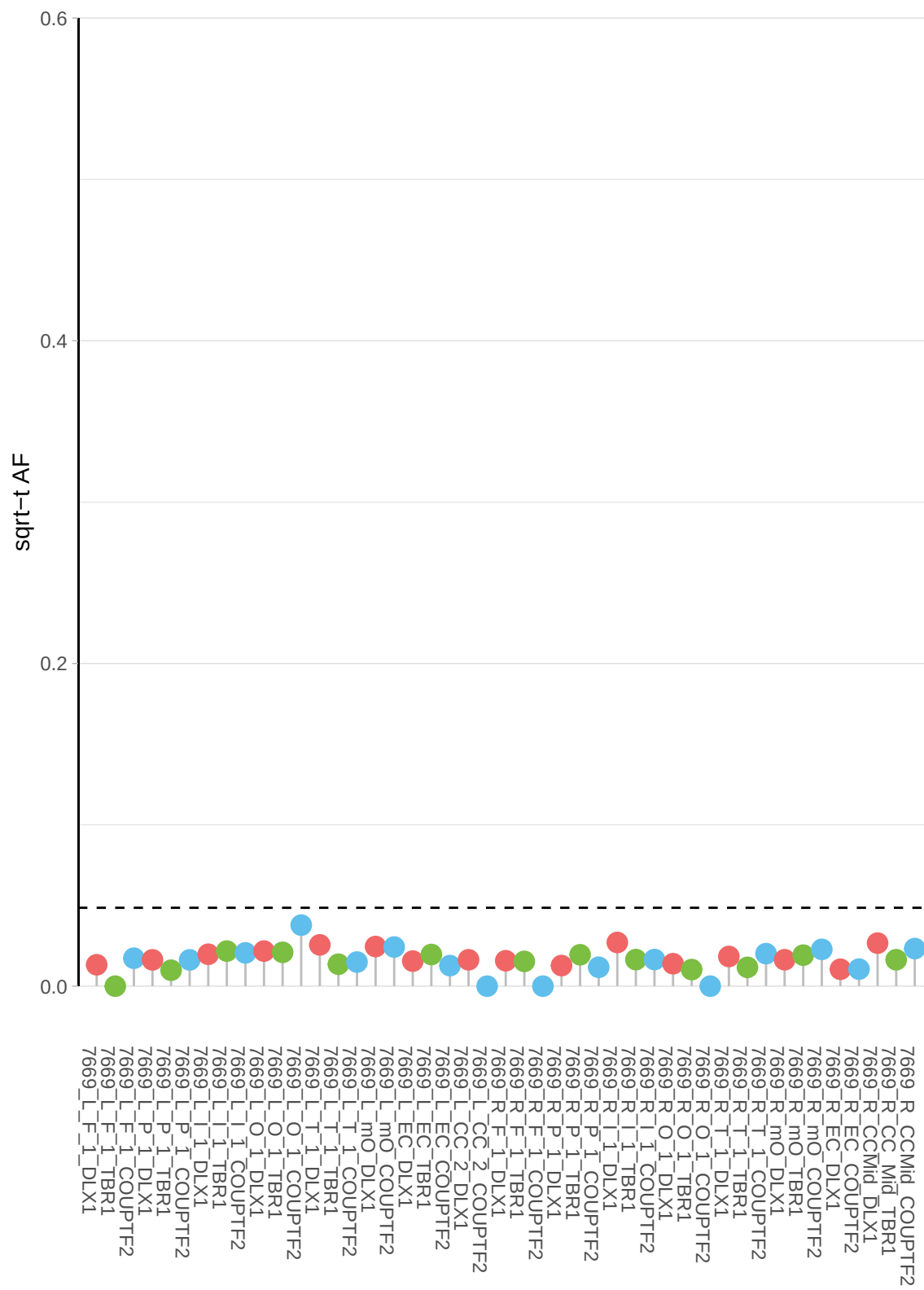

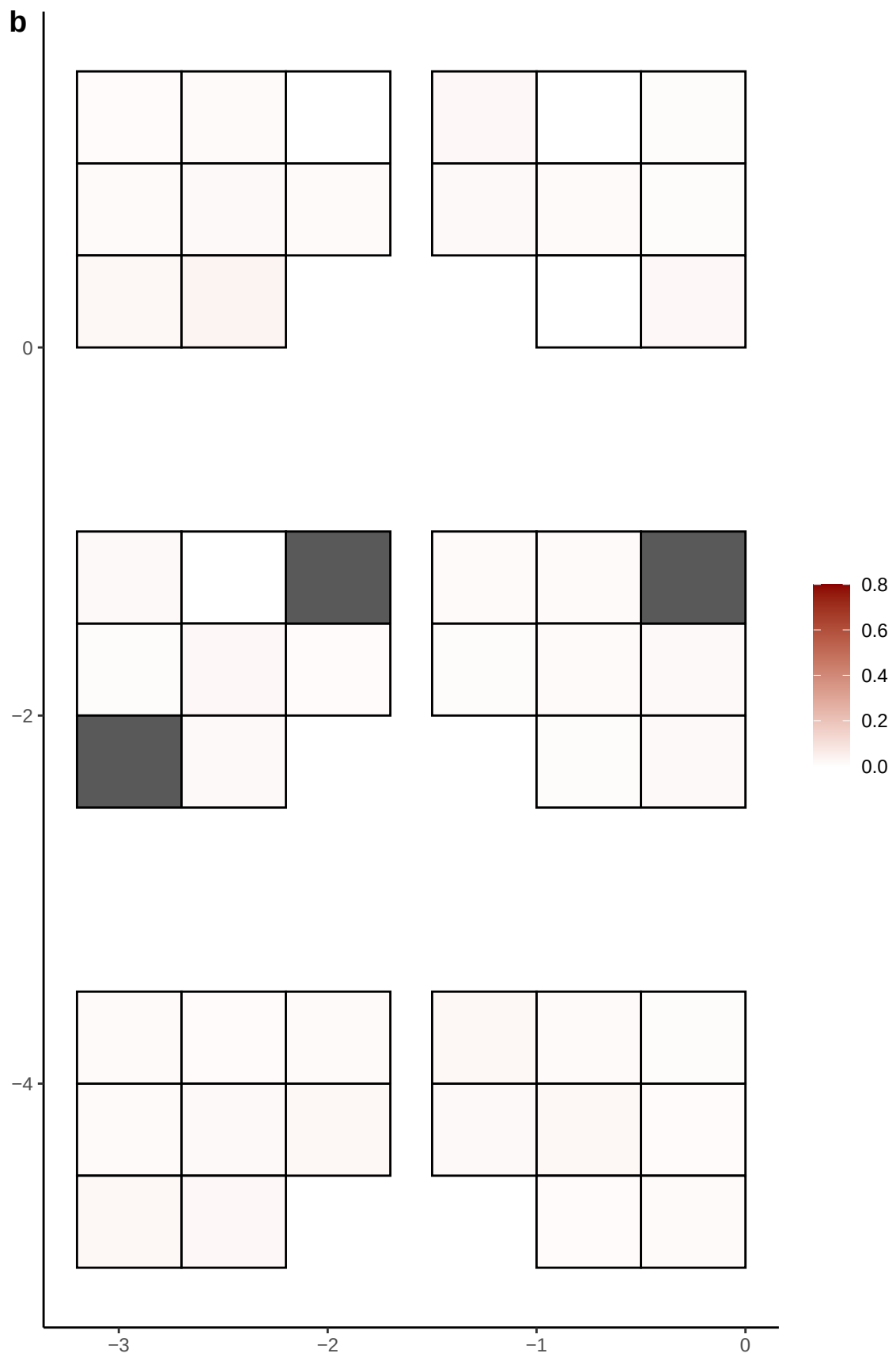

**a** ID05 17-20005505-G-A

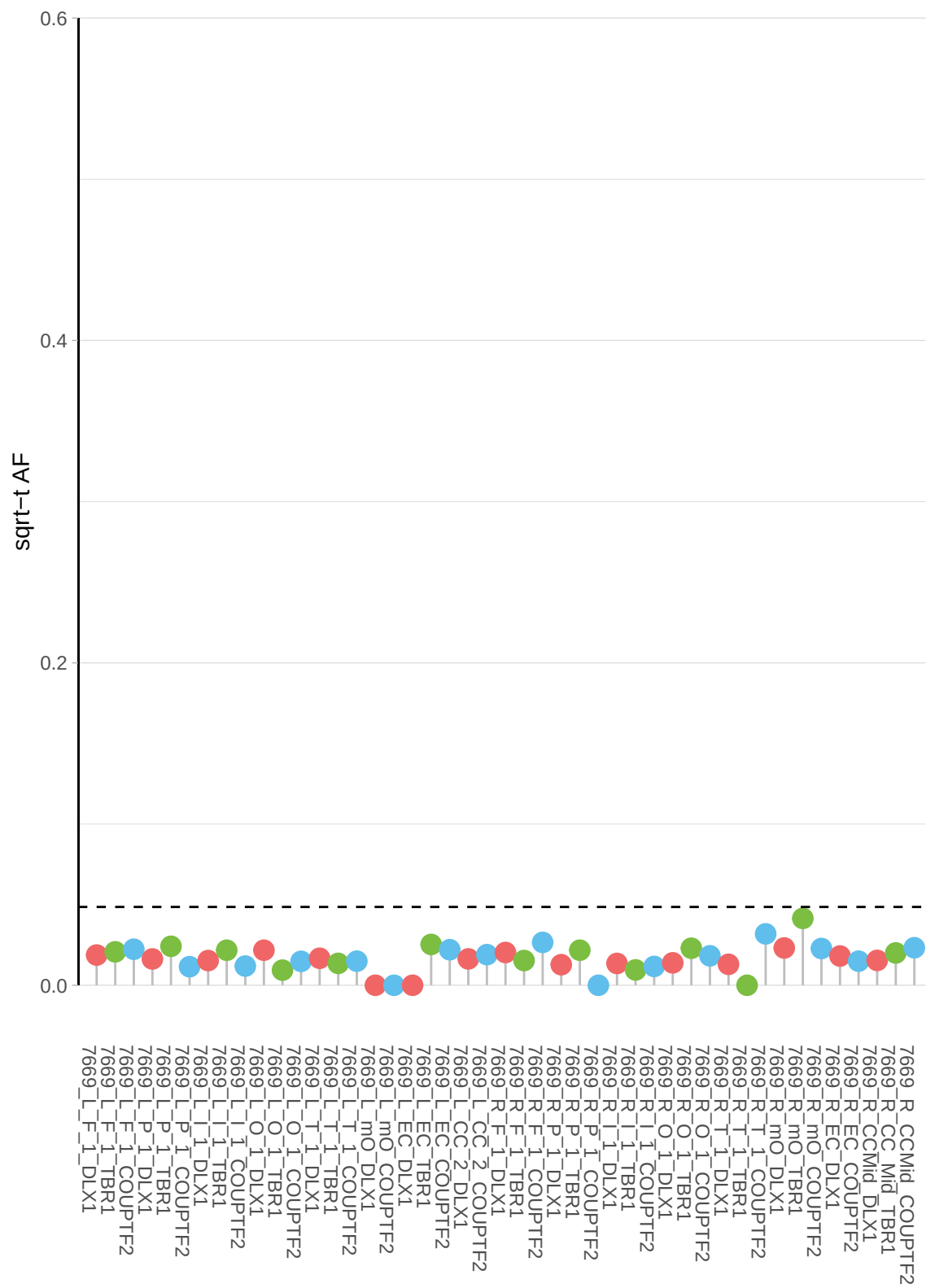

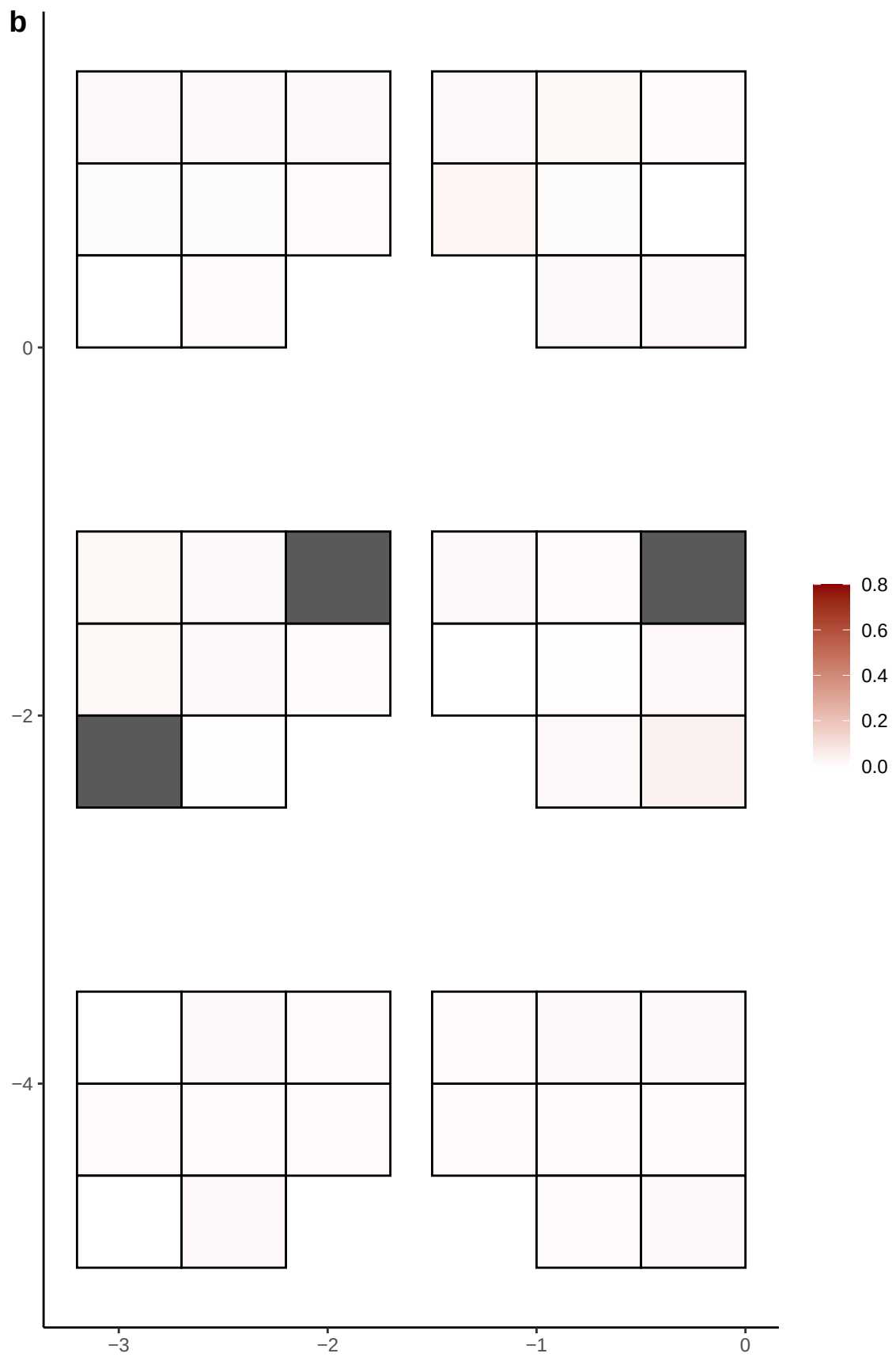

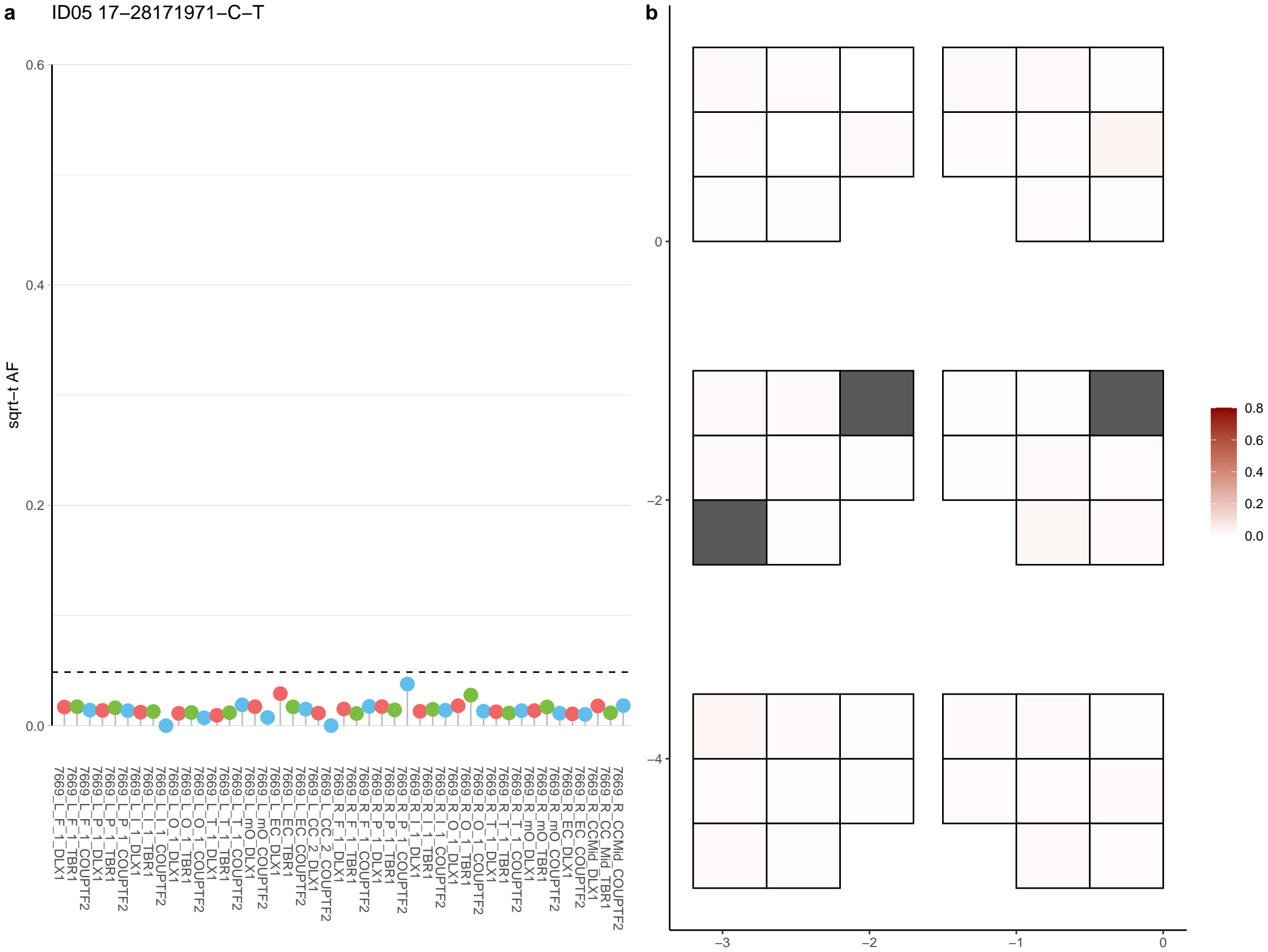

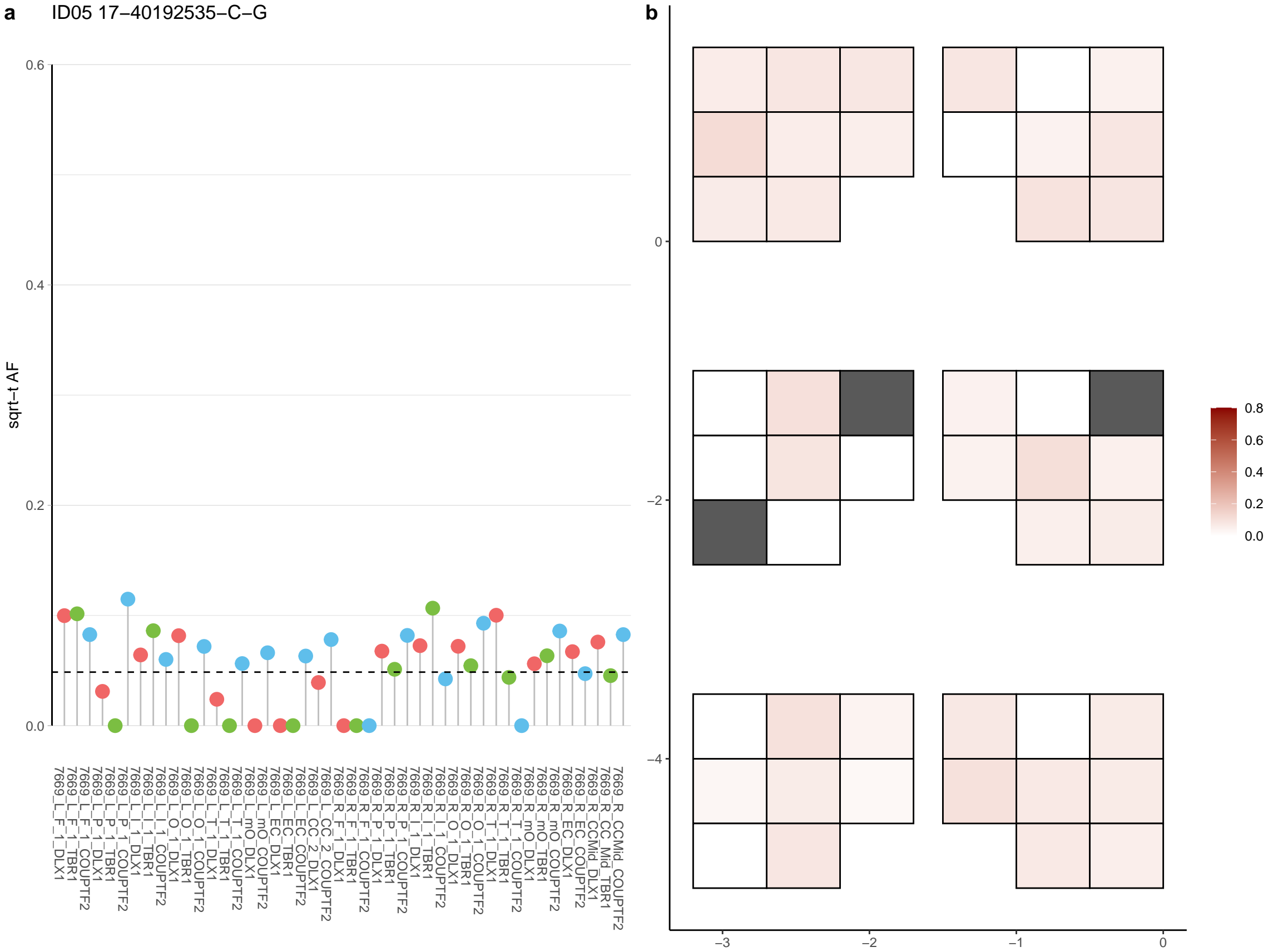

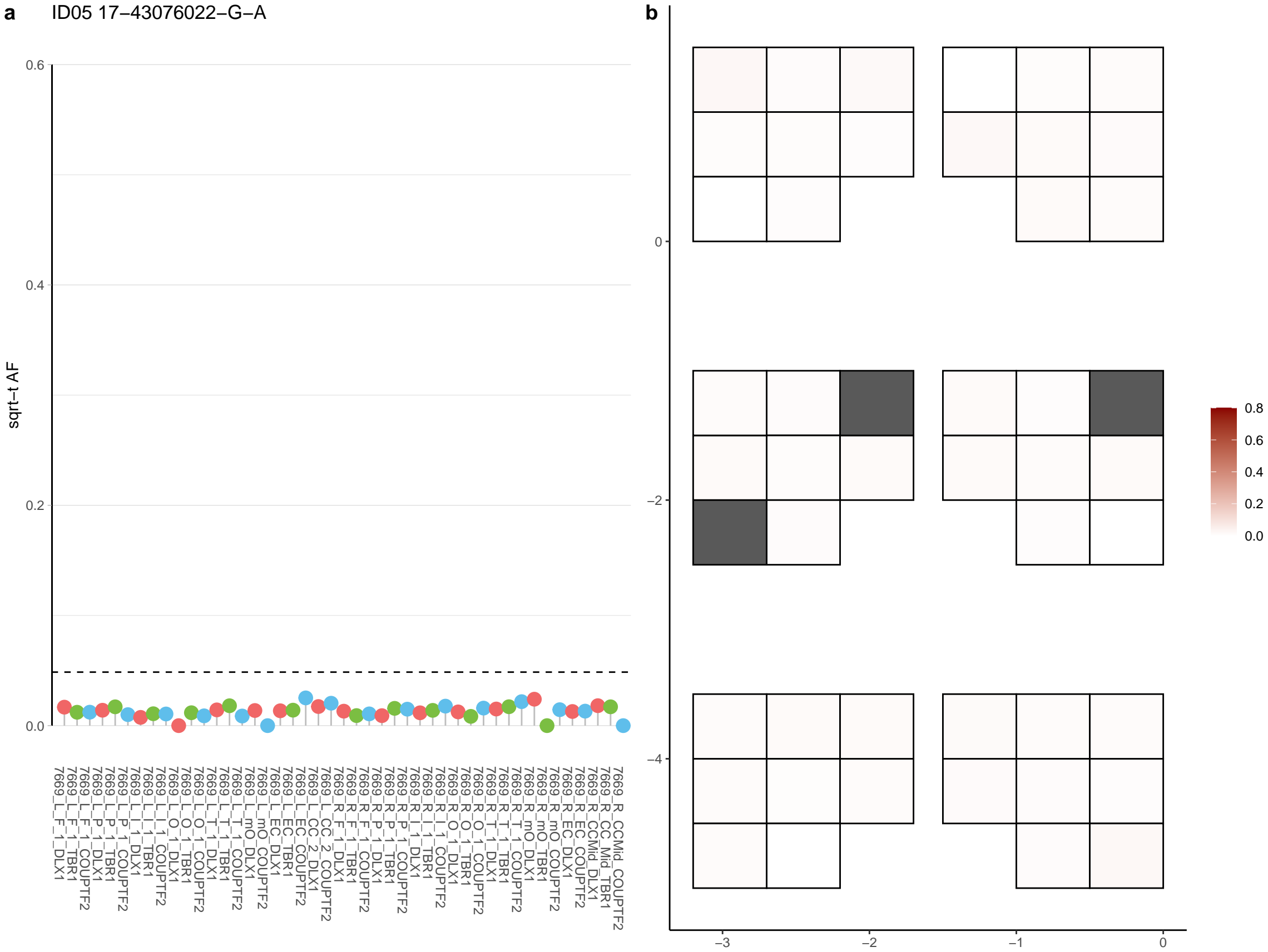

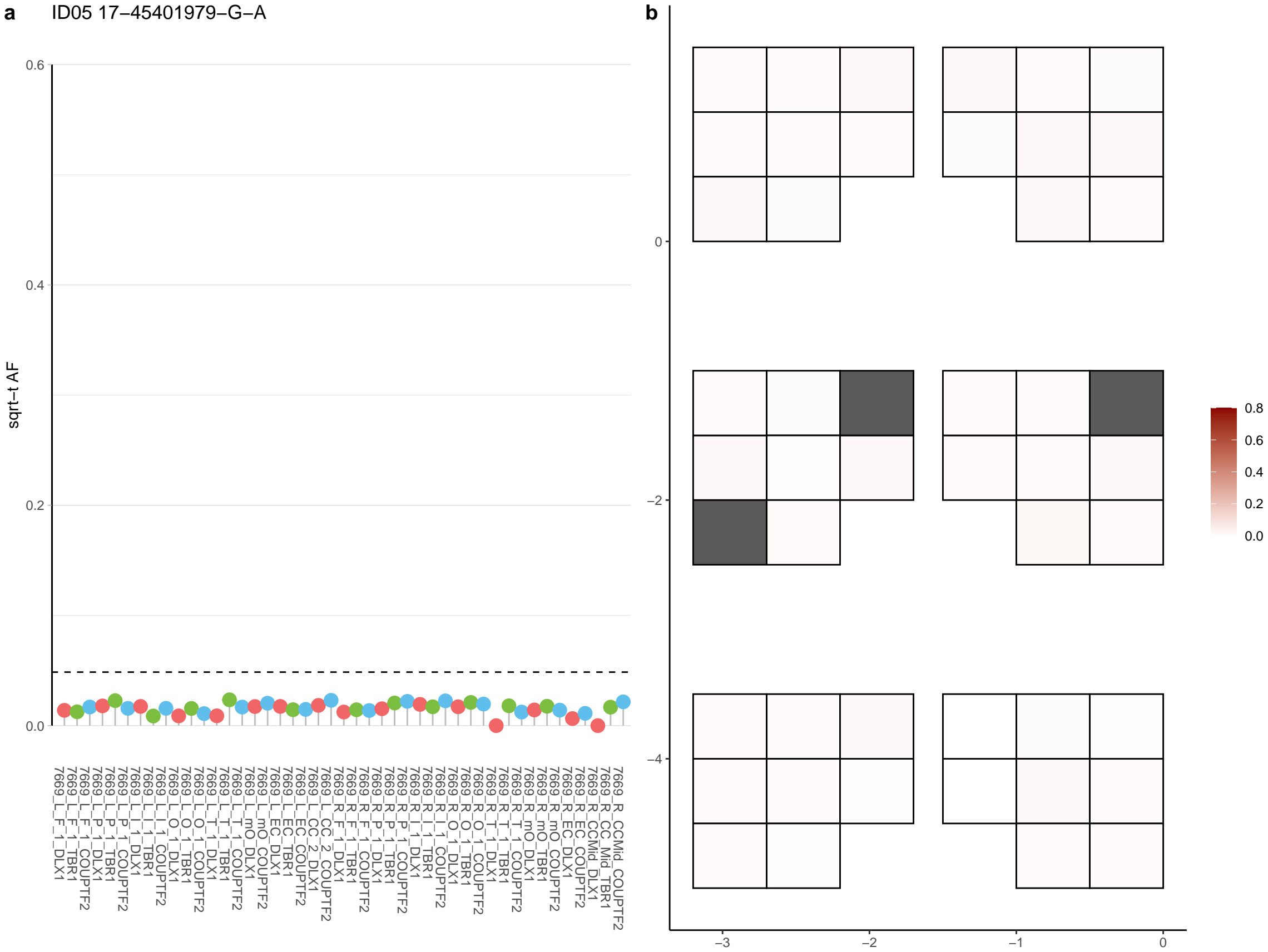

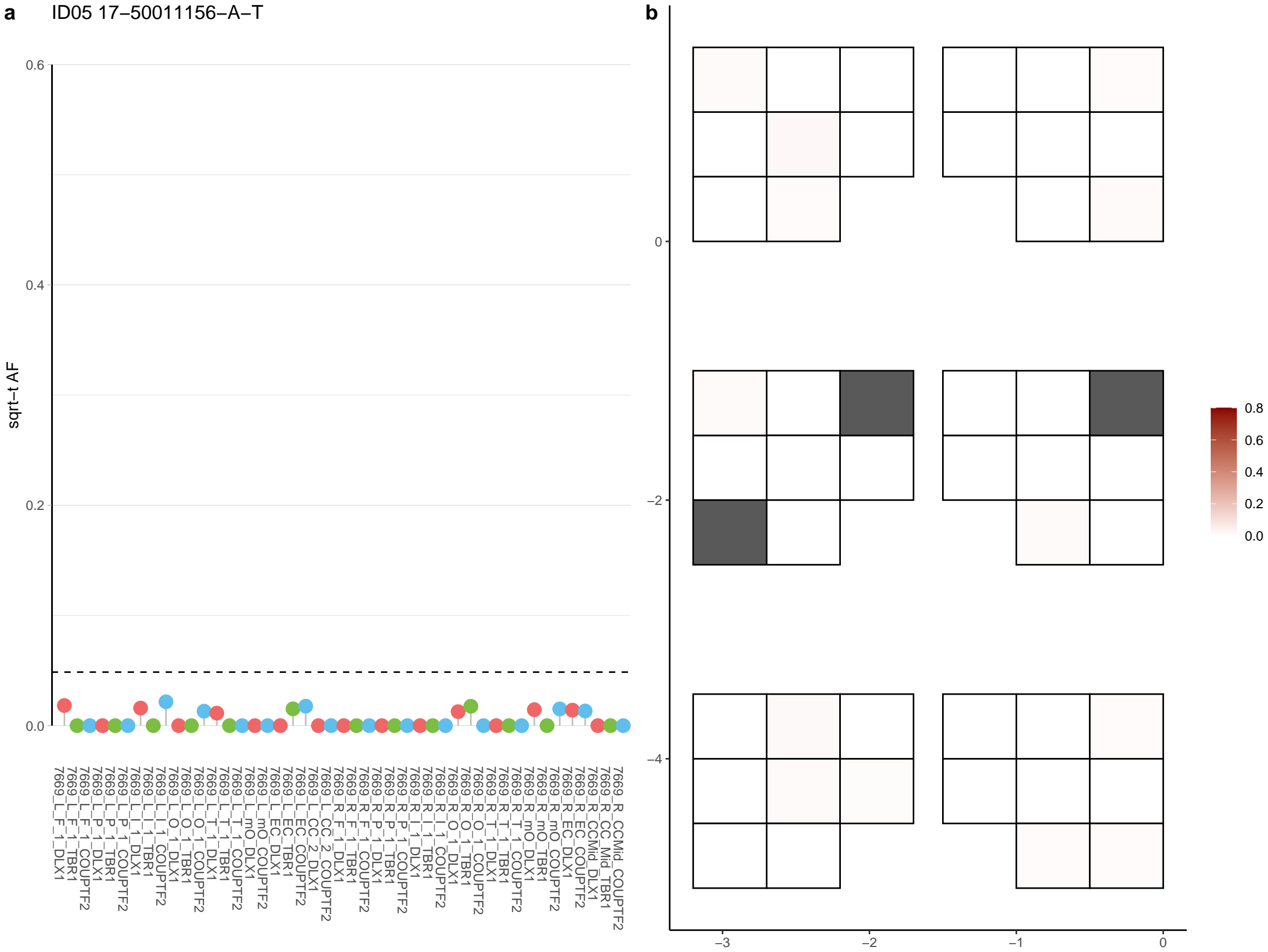

**a** ID05 17-59031927-C-T

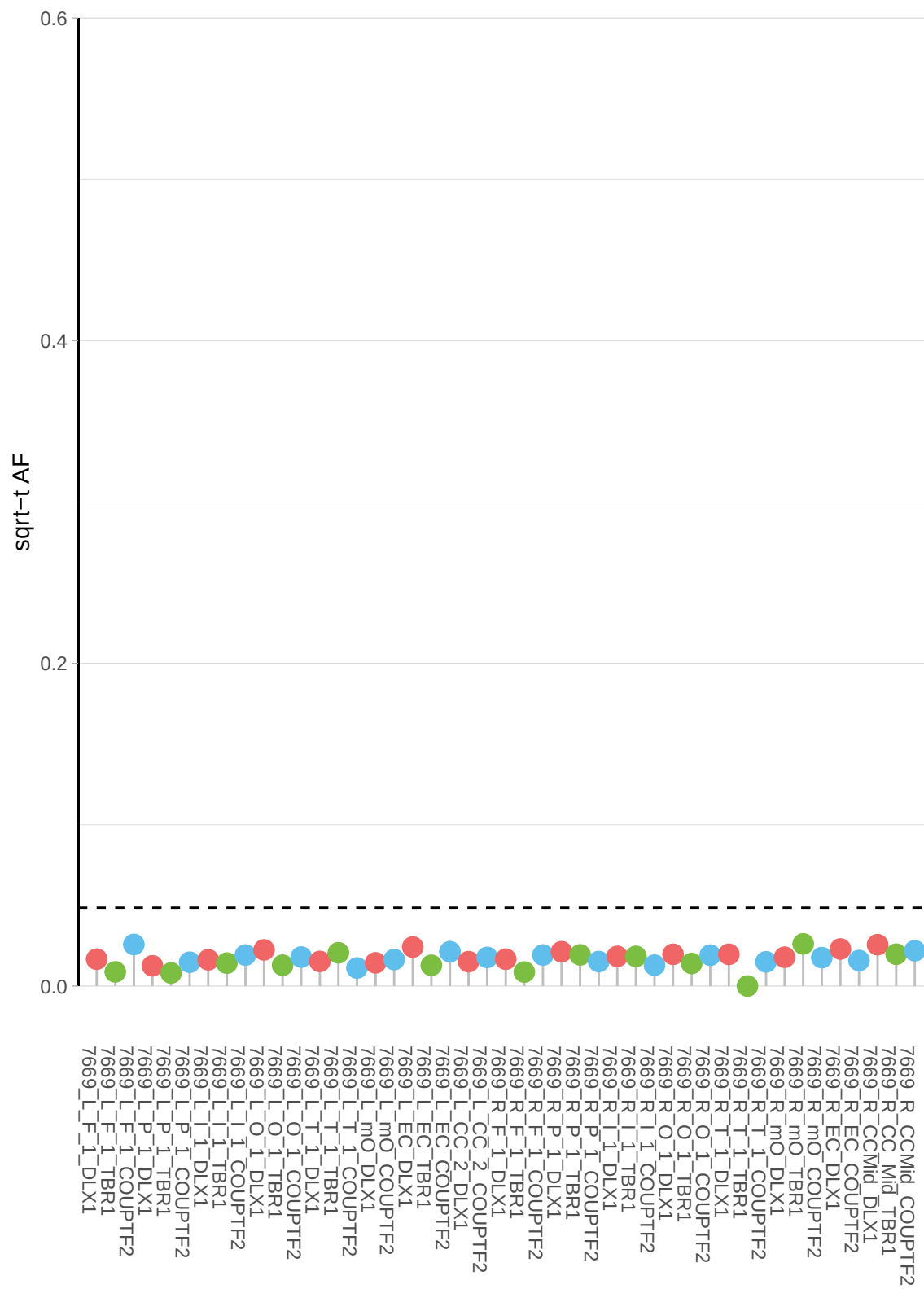

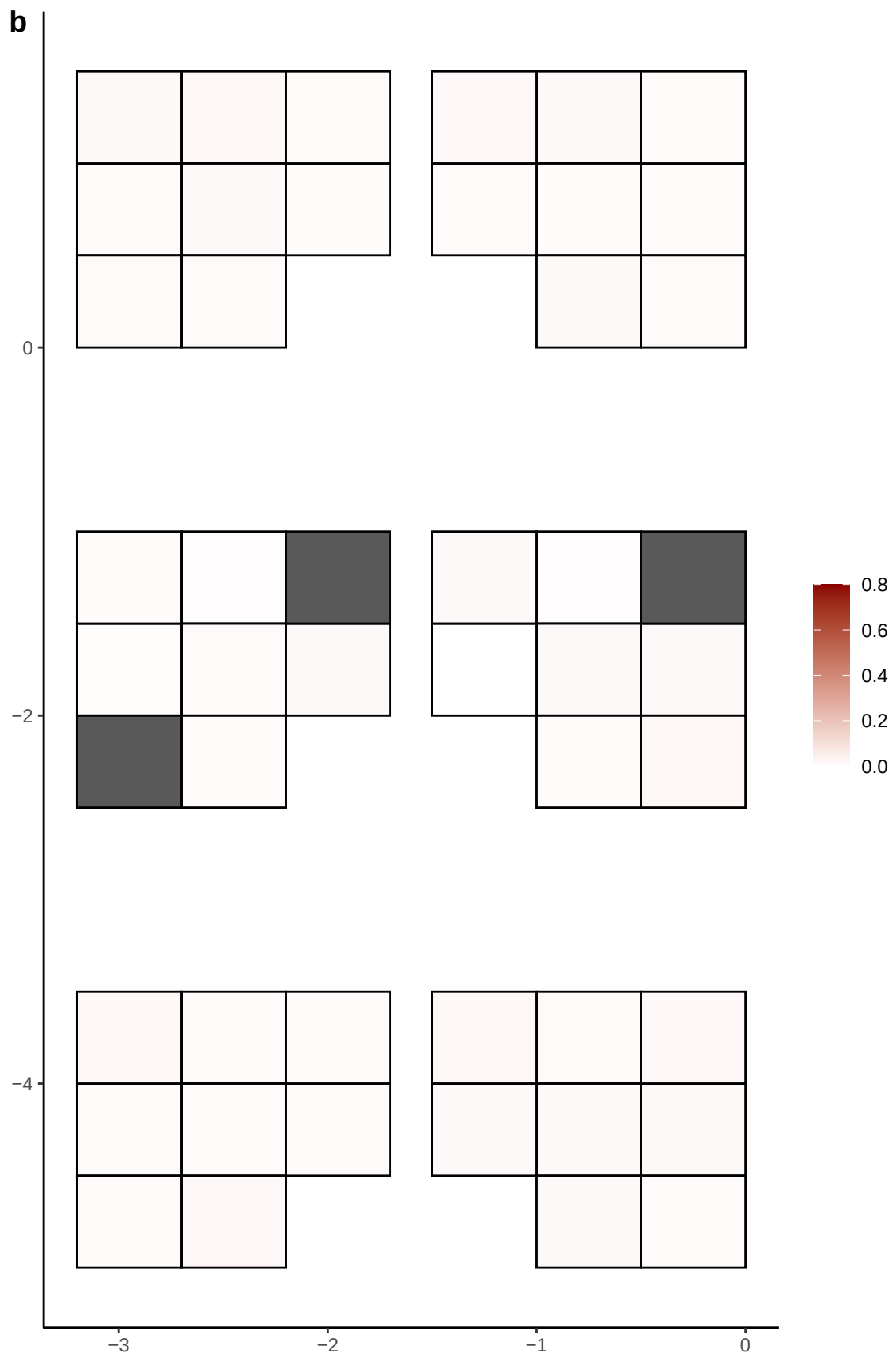

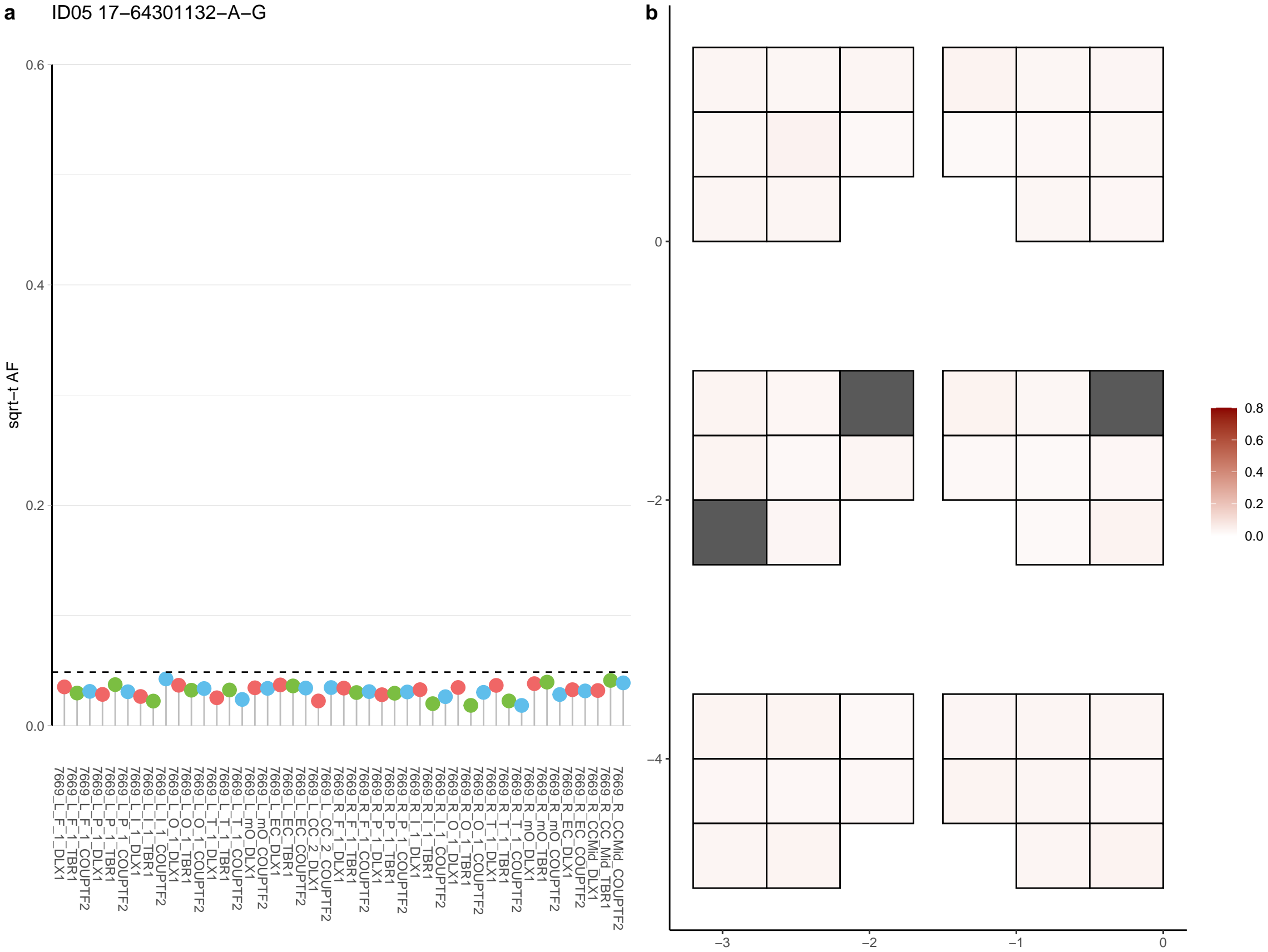

**a** ID05 17-66742030-T-G

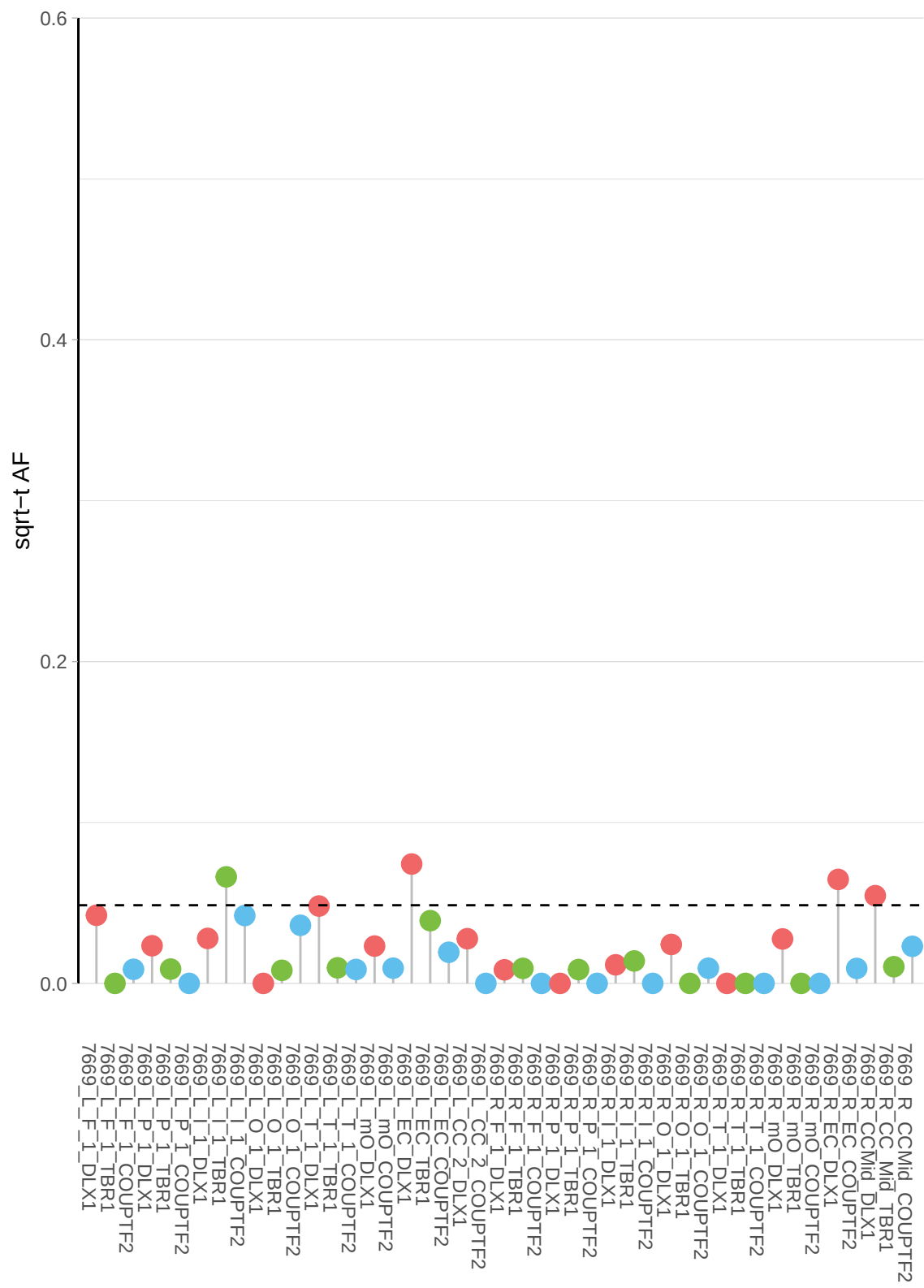

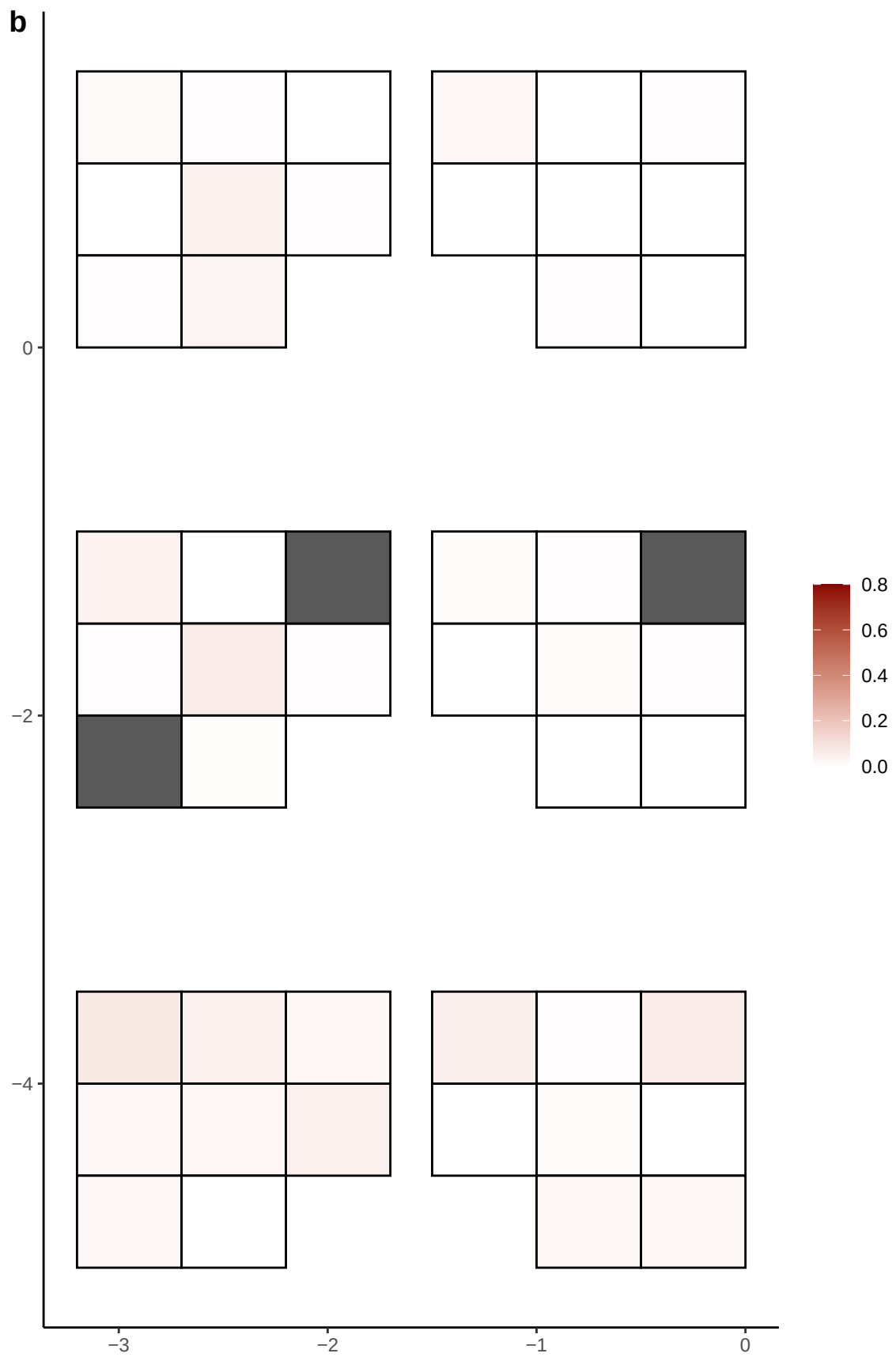

**a** ID05 17-73327453-G-A

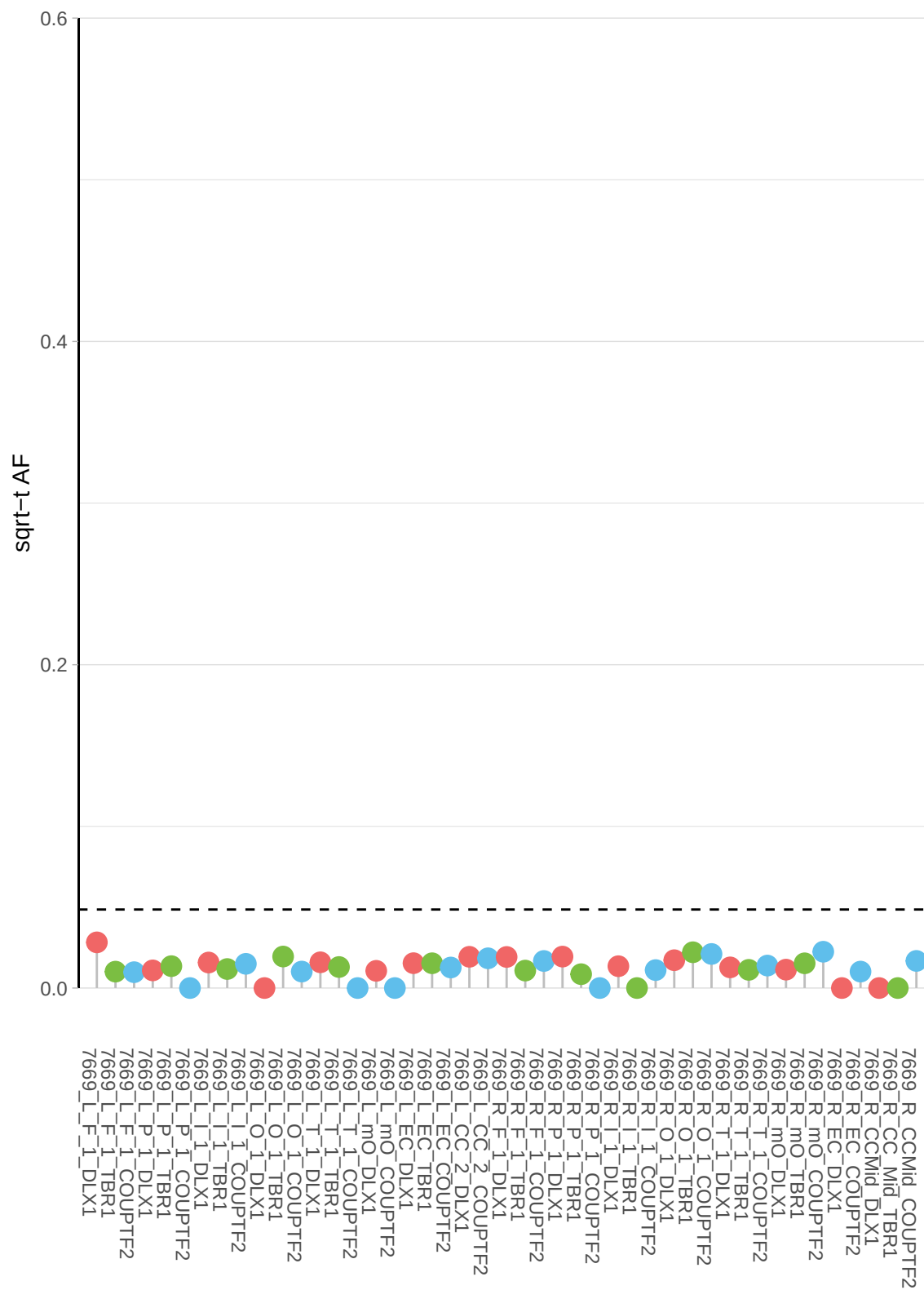

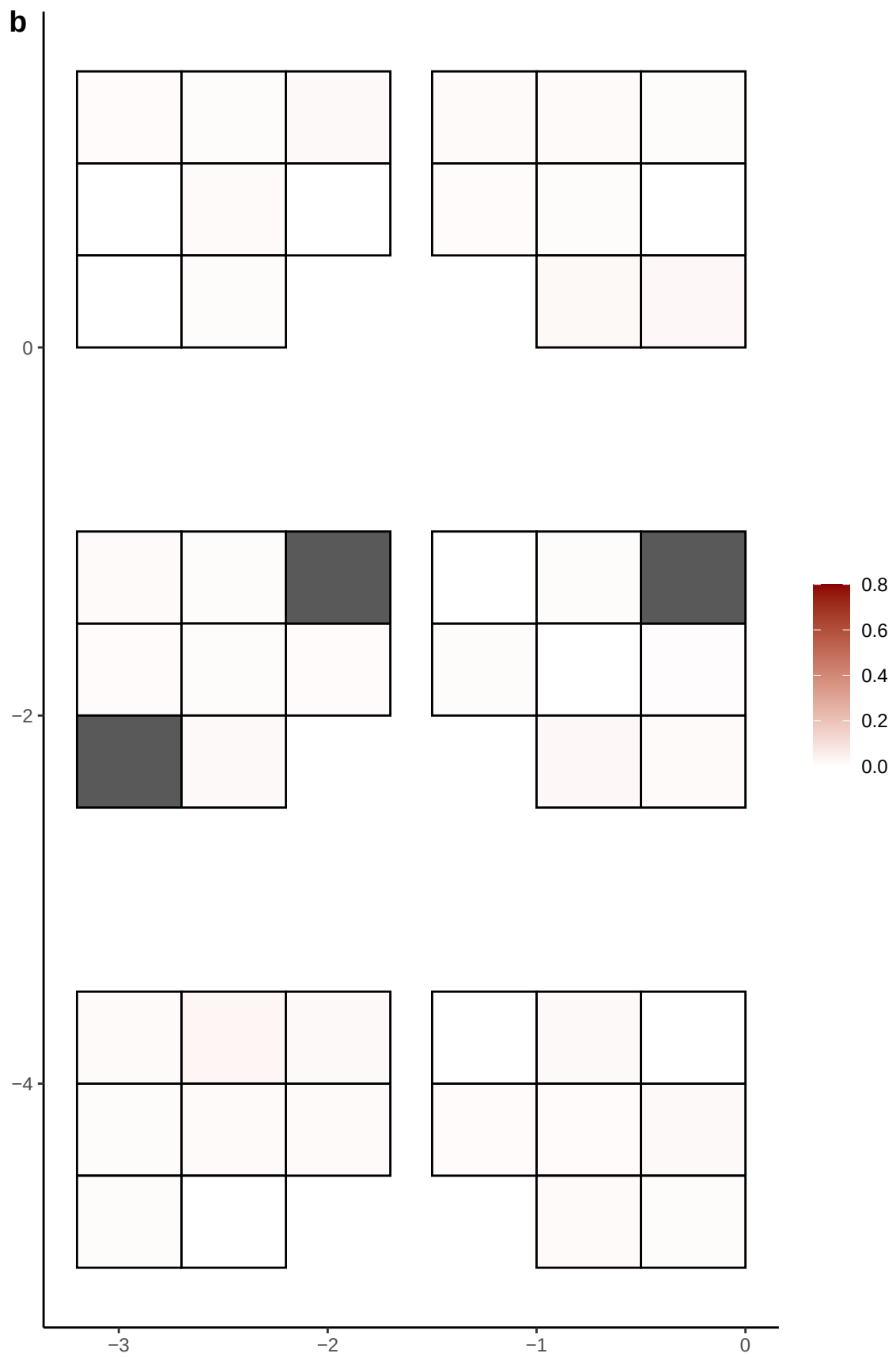

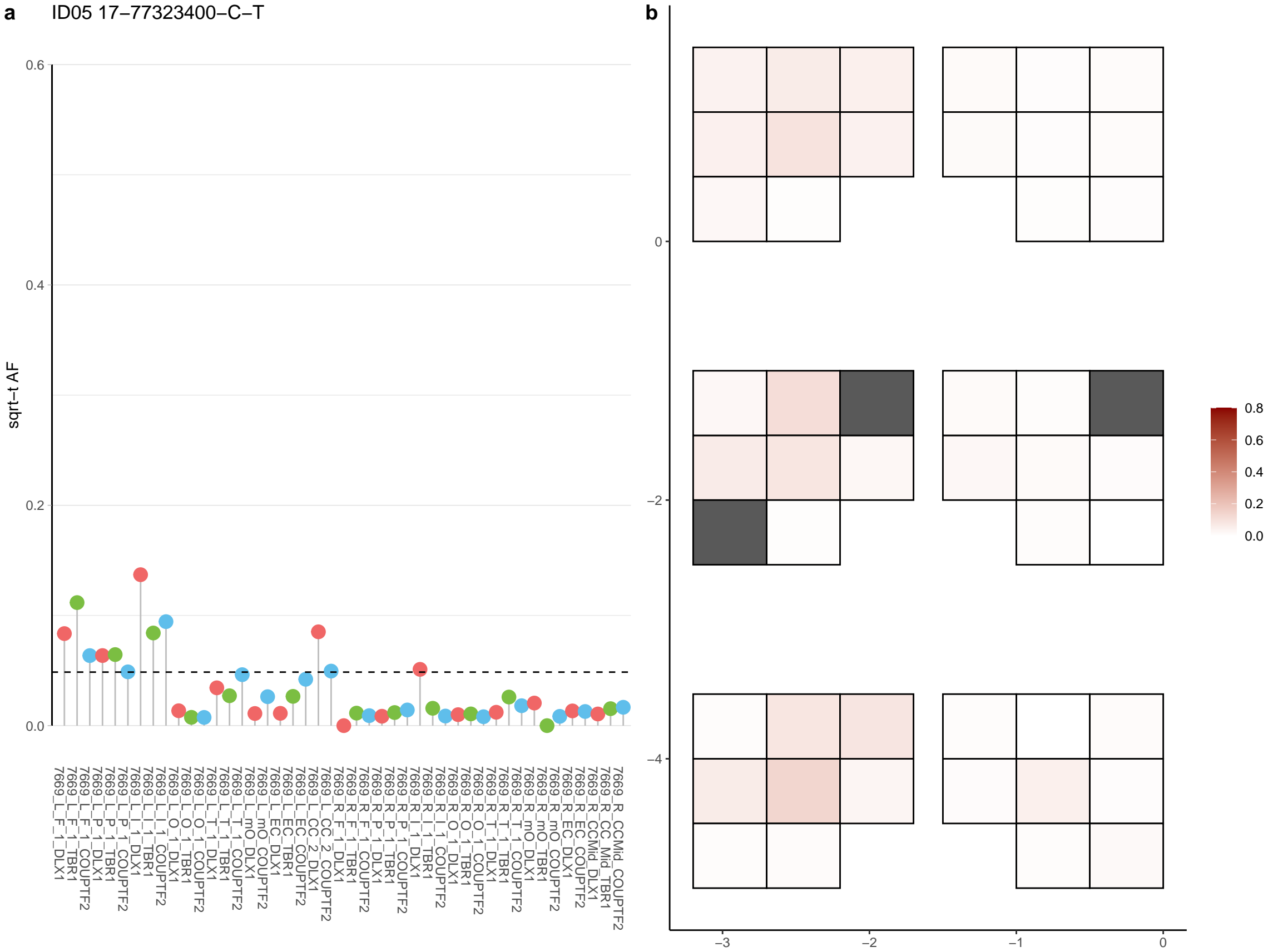

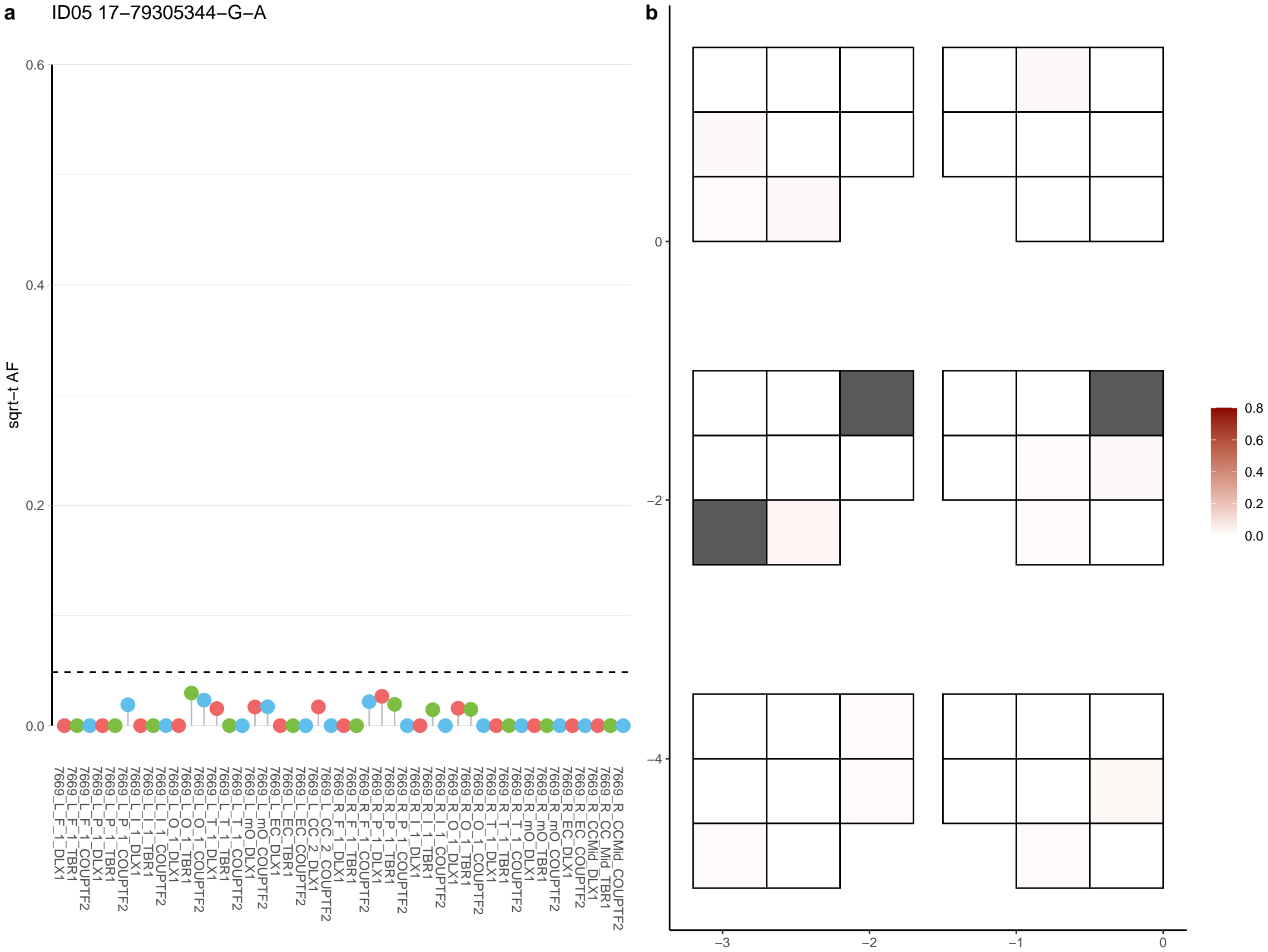

**a** ID05 17-80207448-G-A

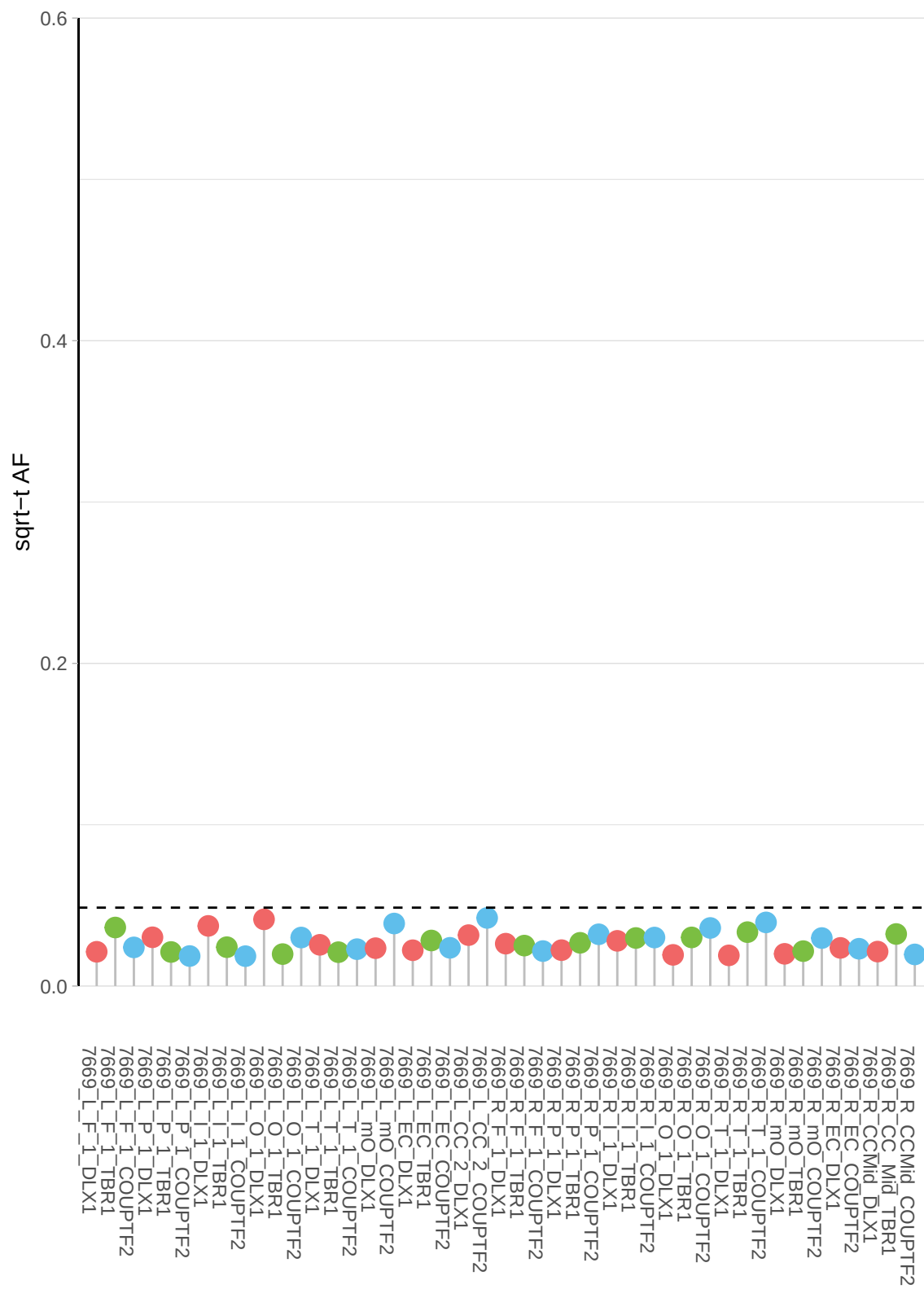

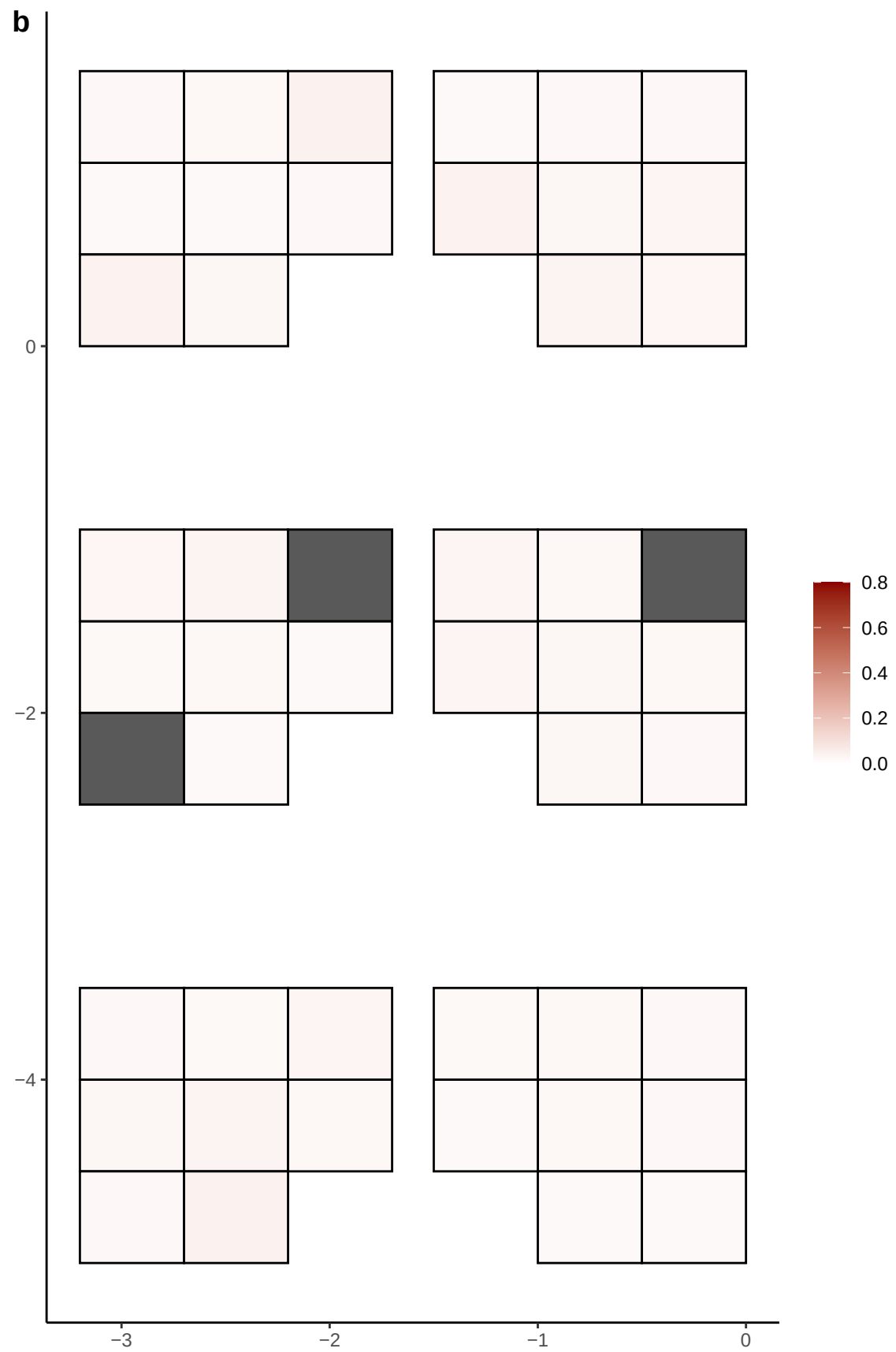

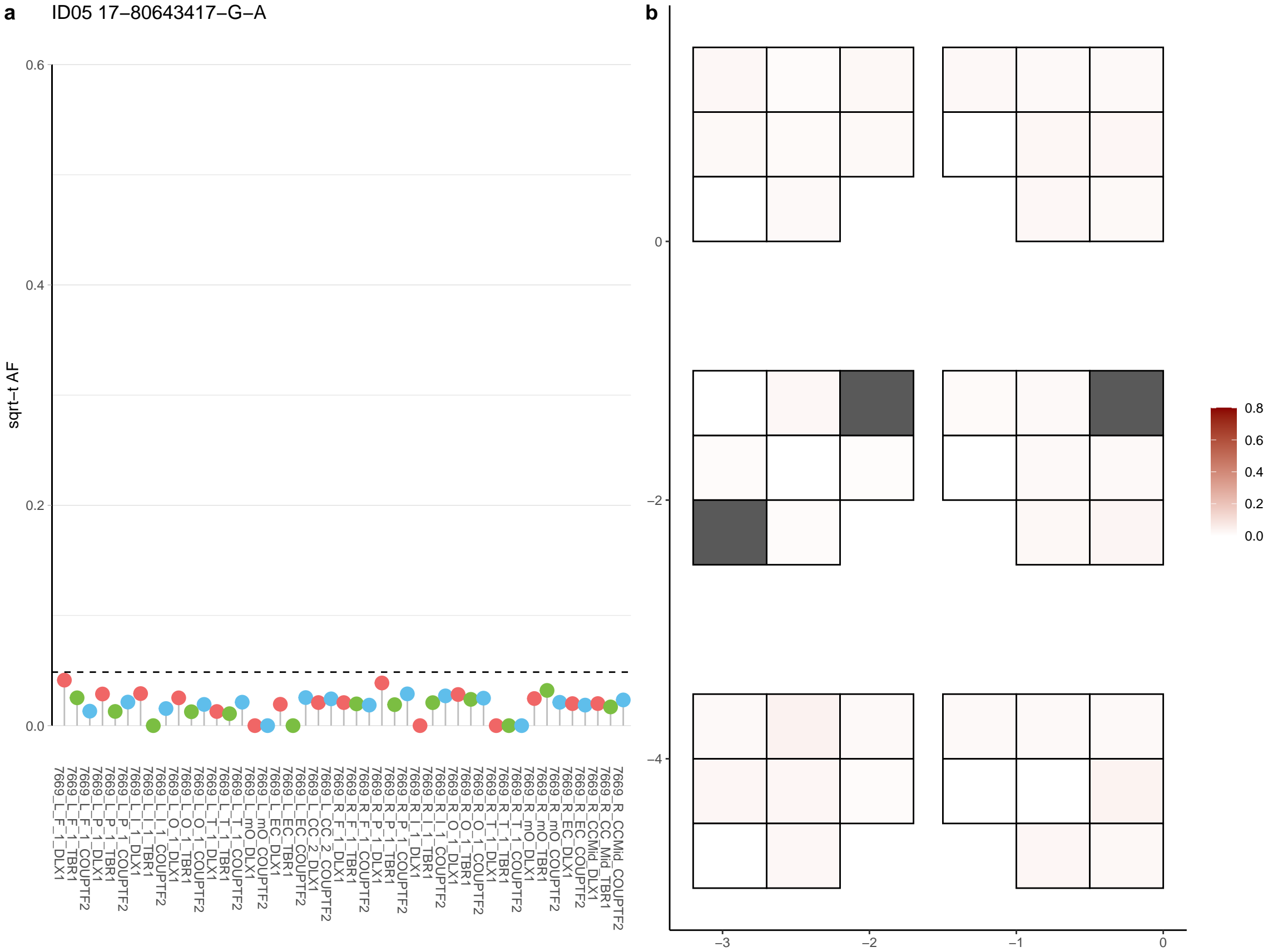

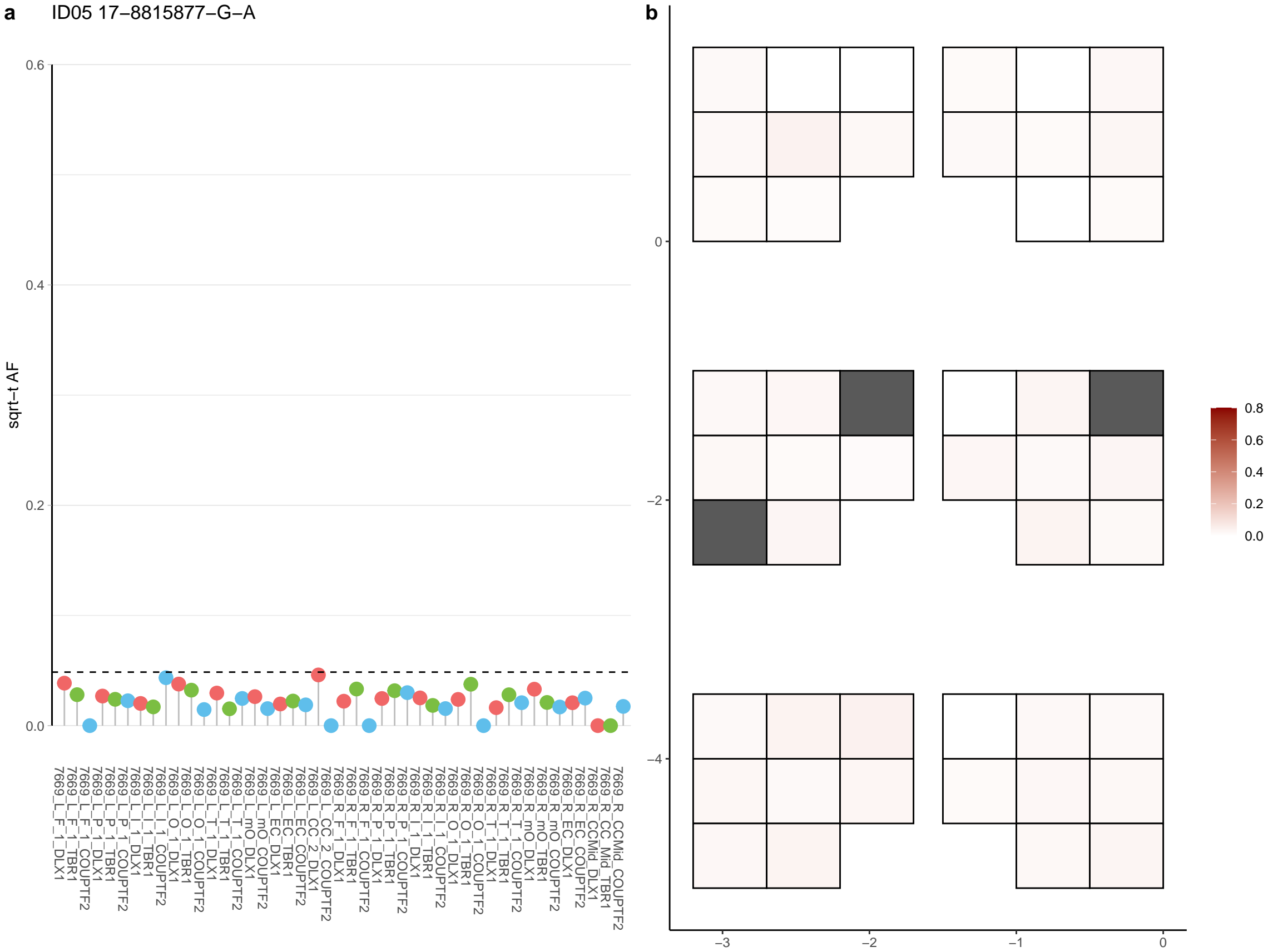

**a** ID05 17-9084127-C-T

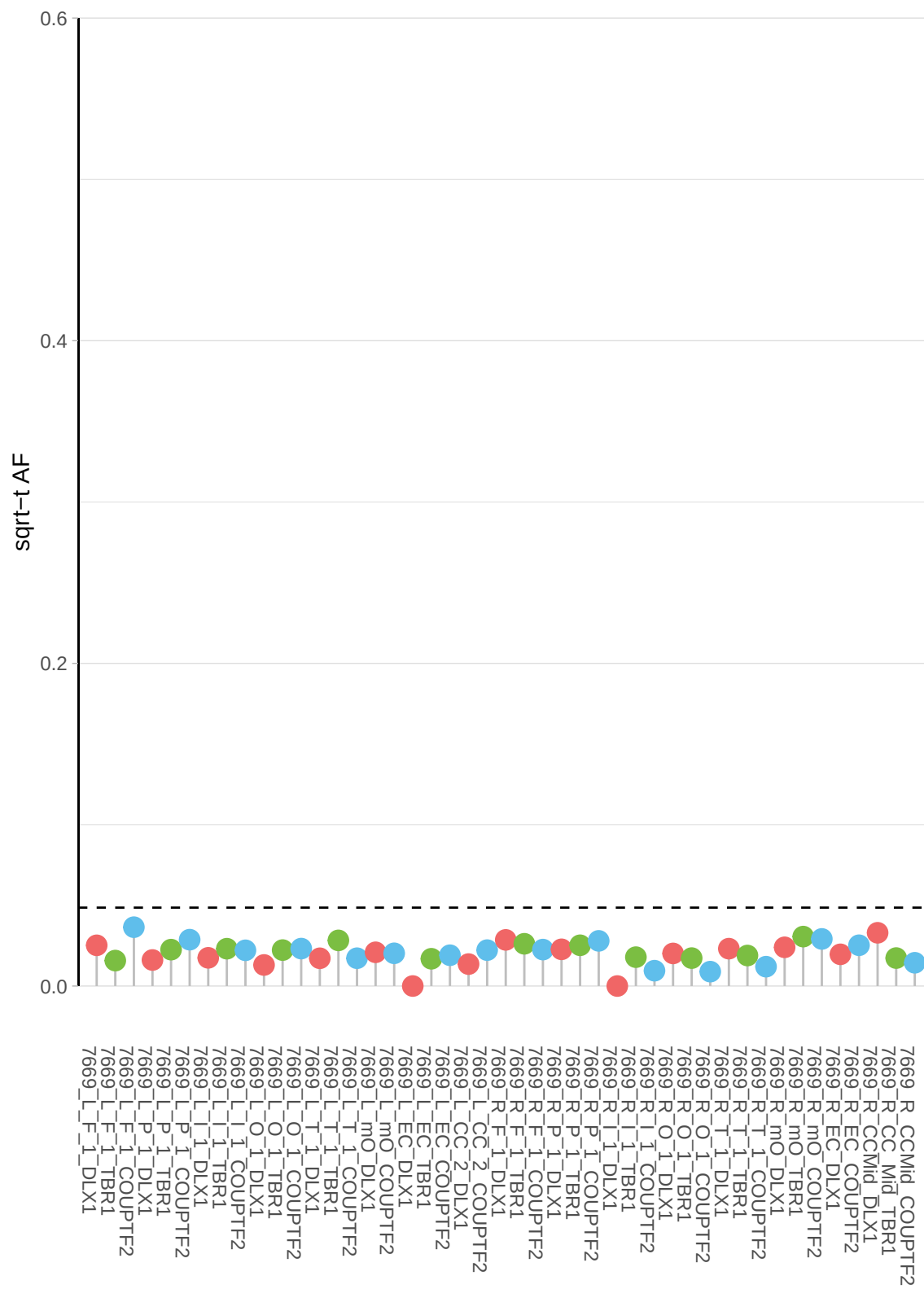

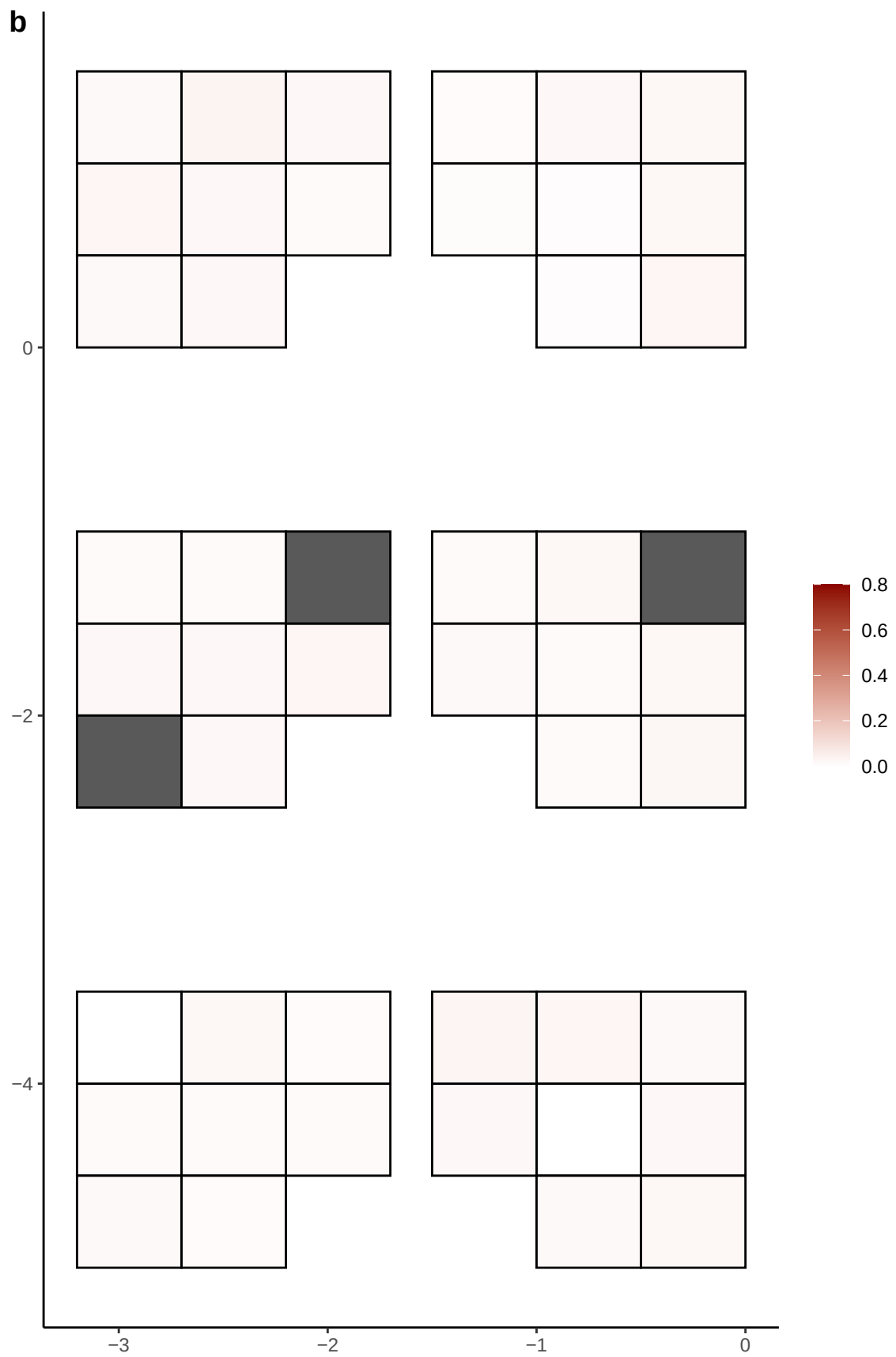

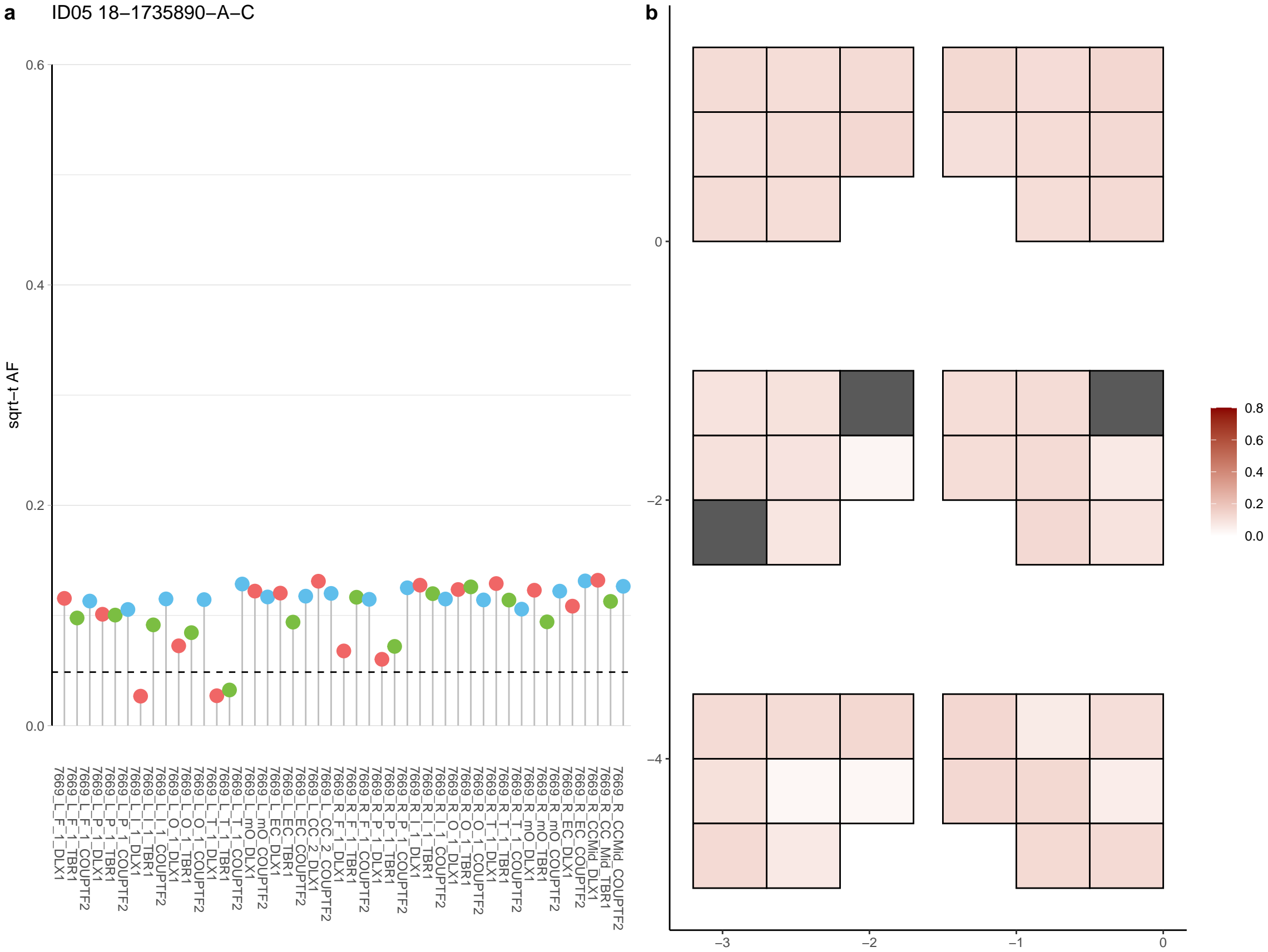

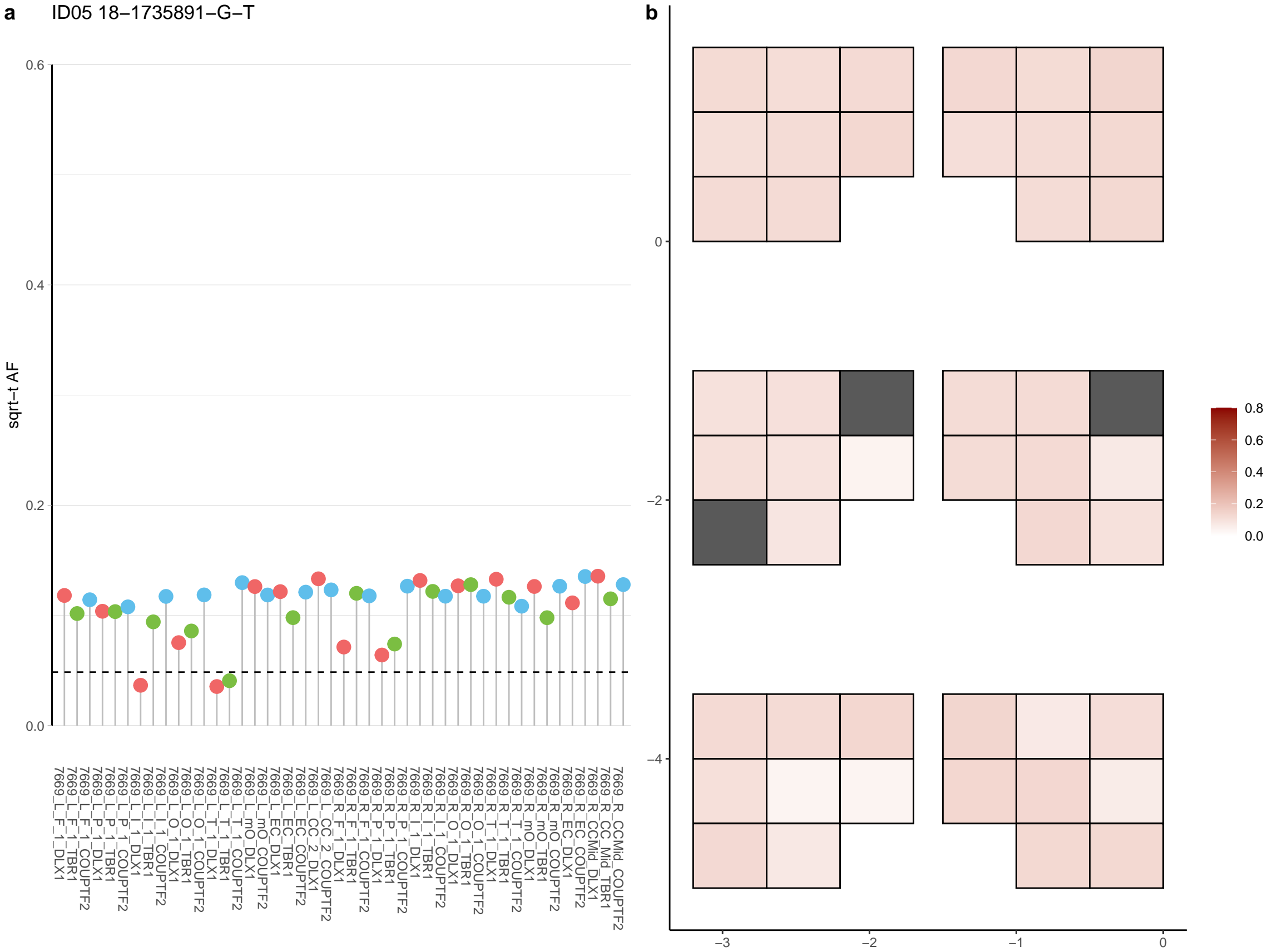

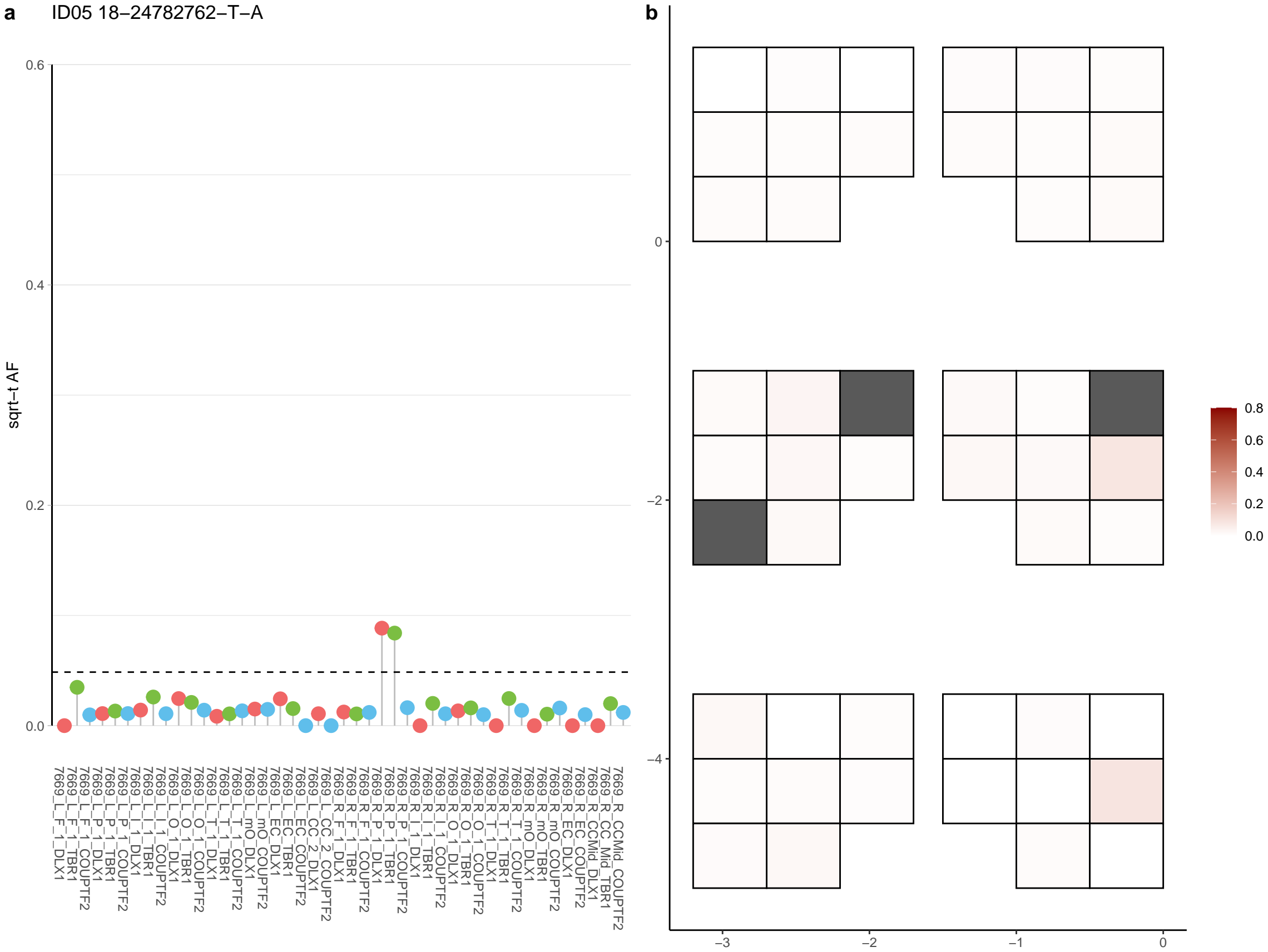

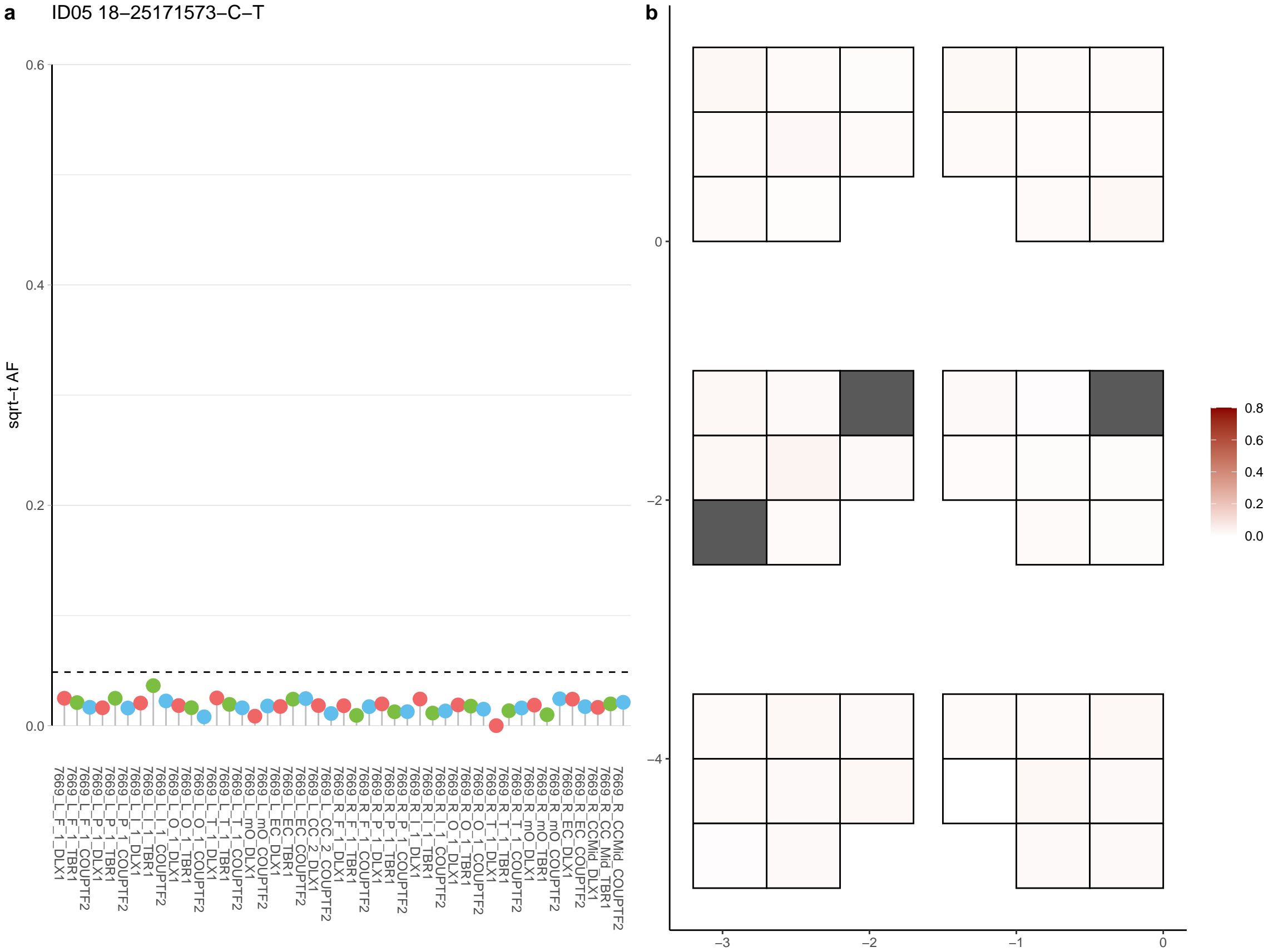

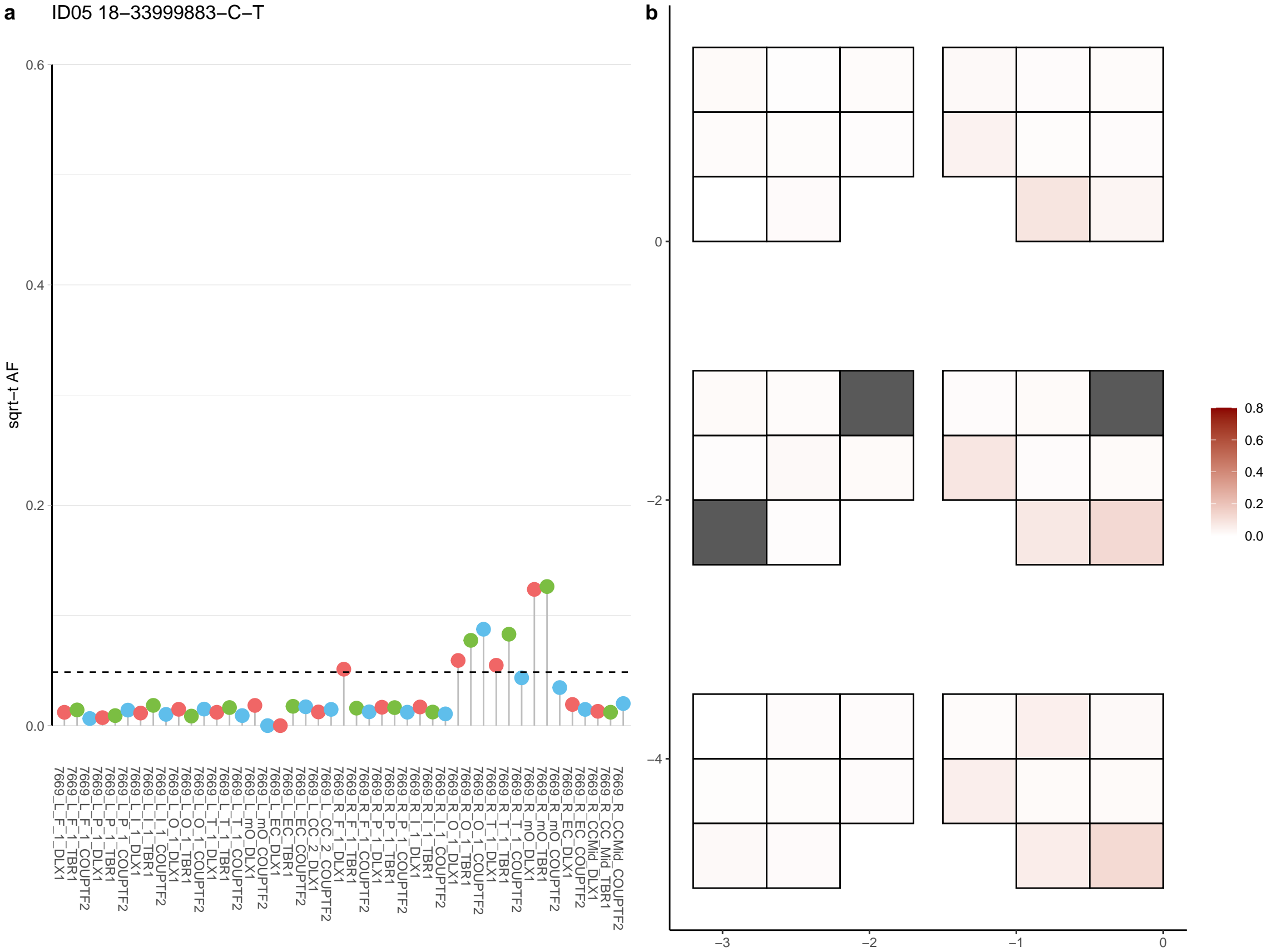

**a** ID05 18-36090085-A-C

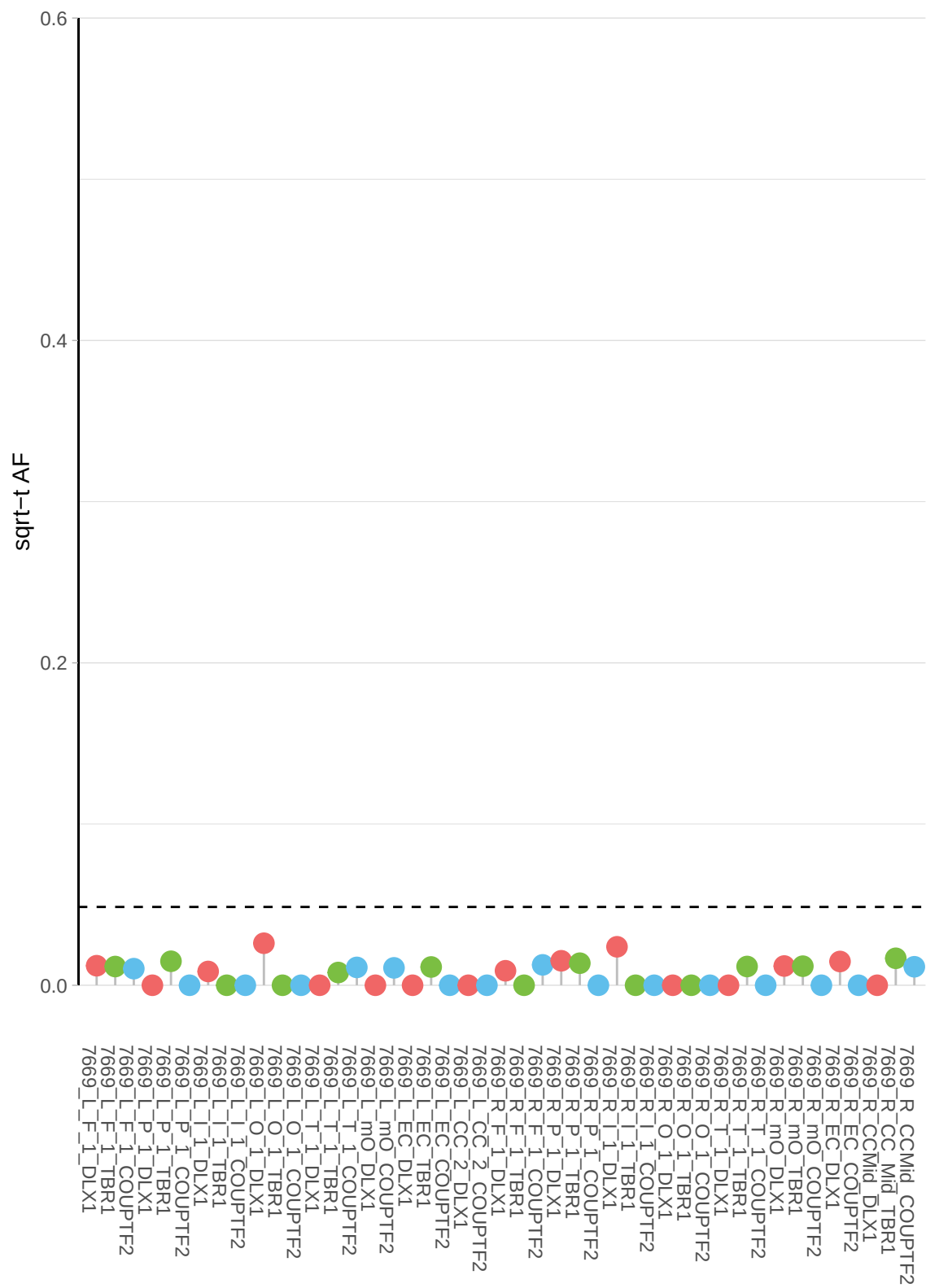

**b**

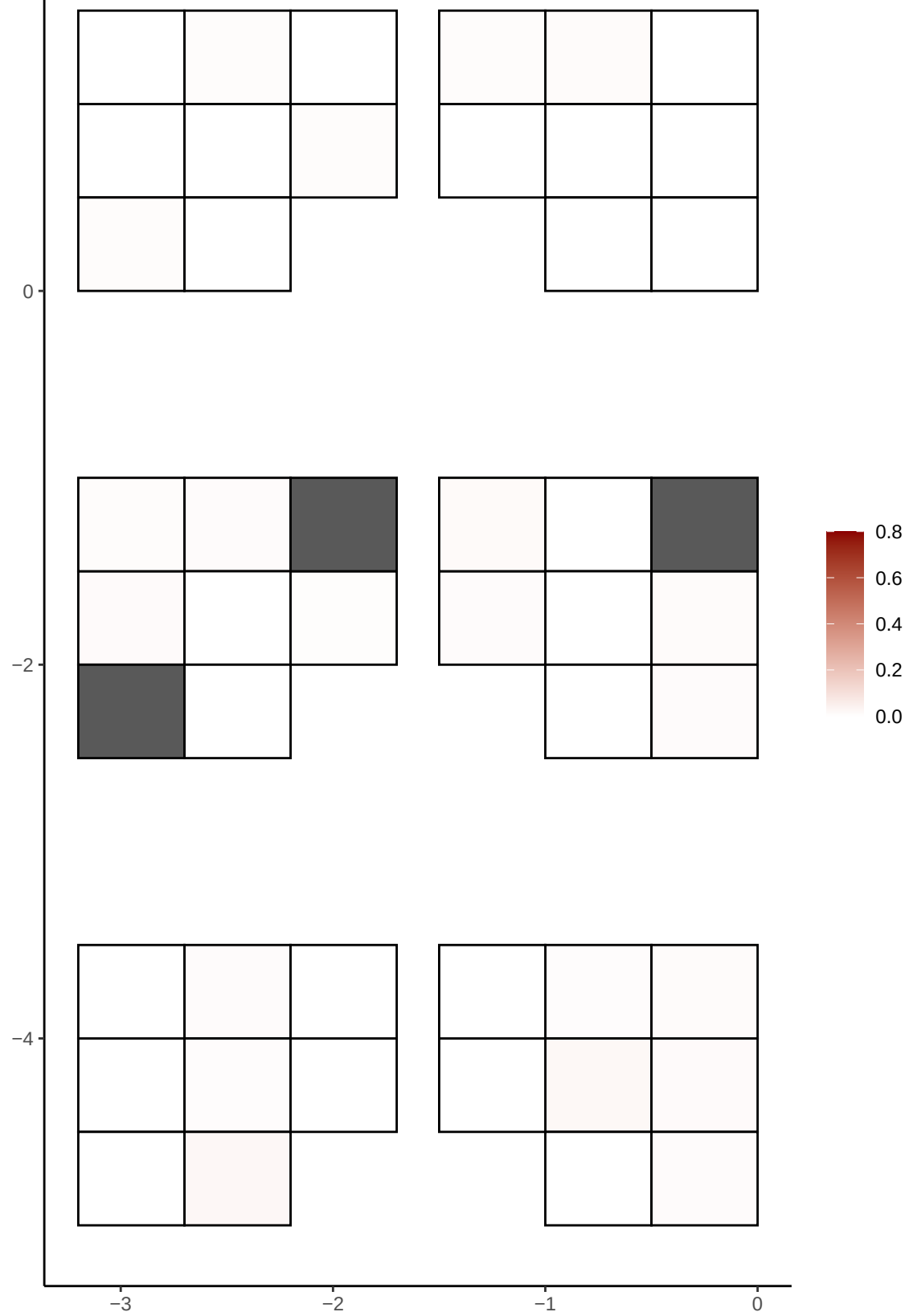

**a** ID05 18-36334609-T-C

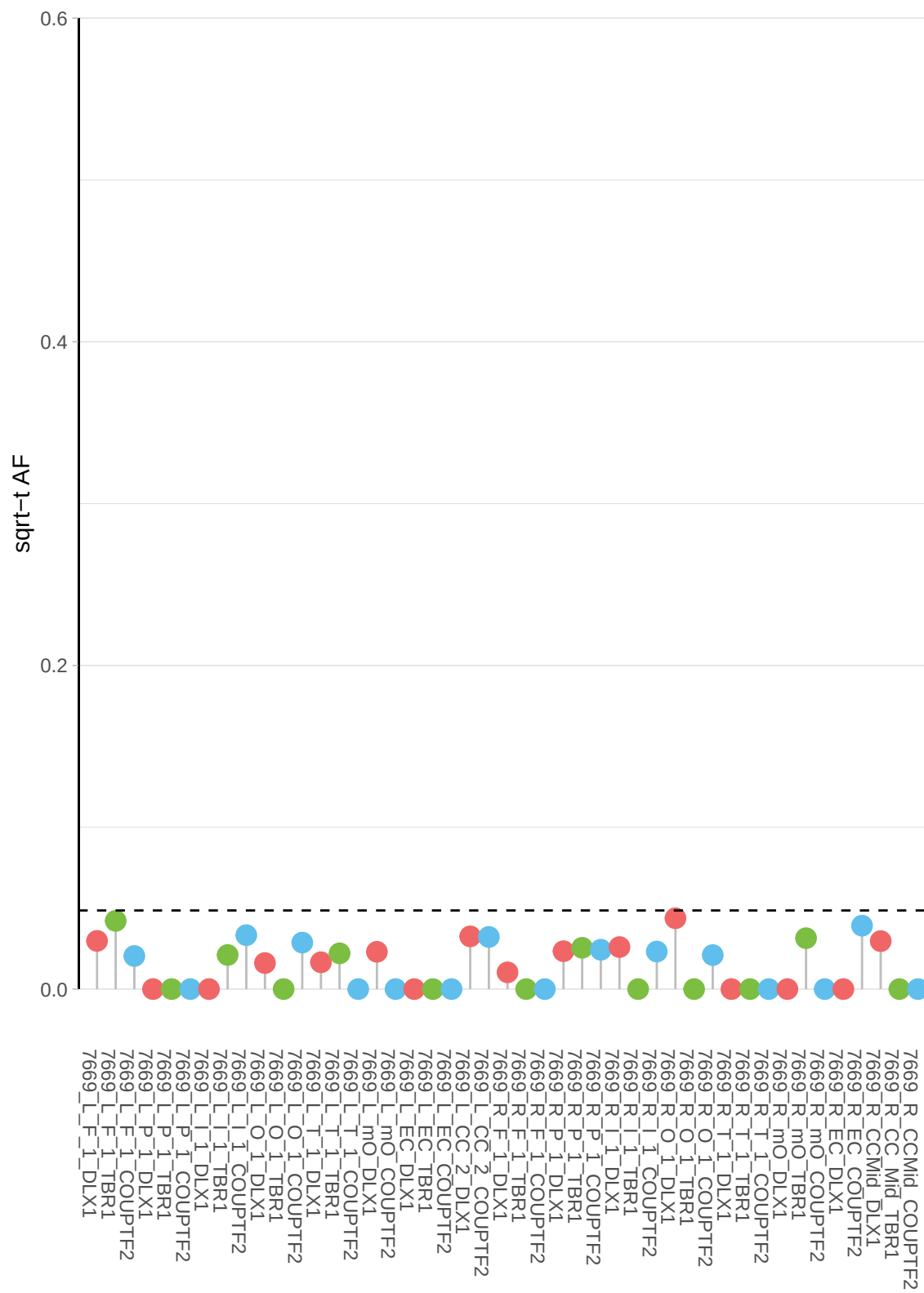

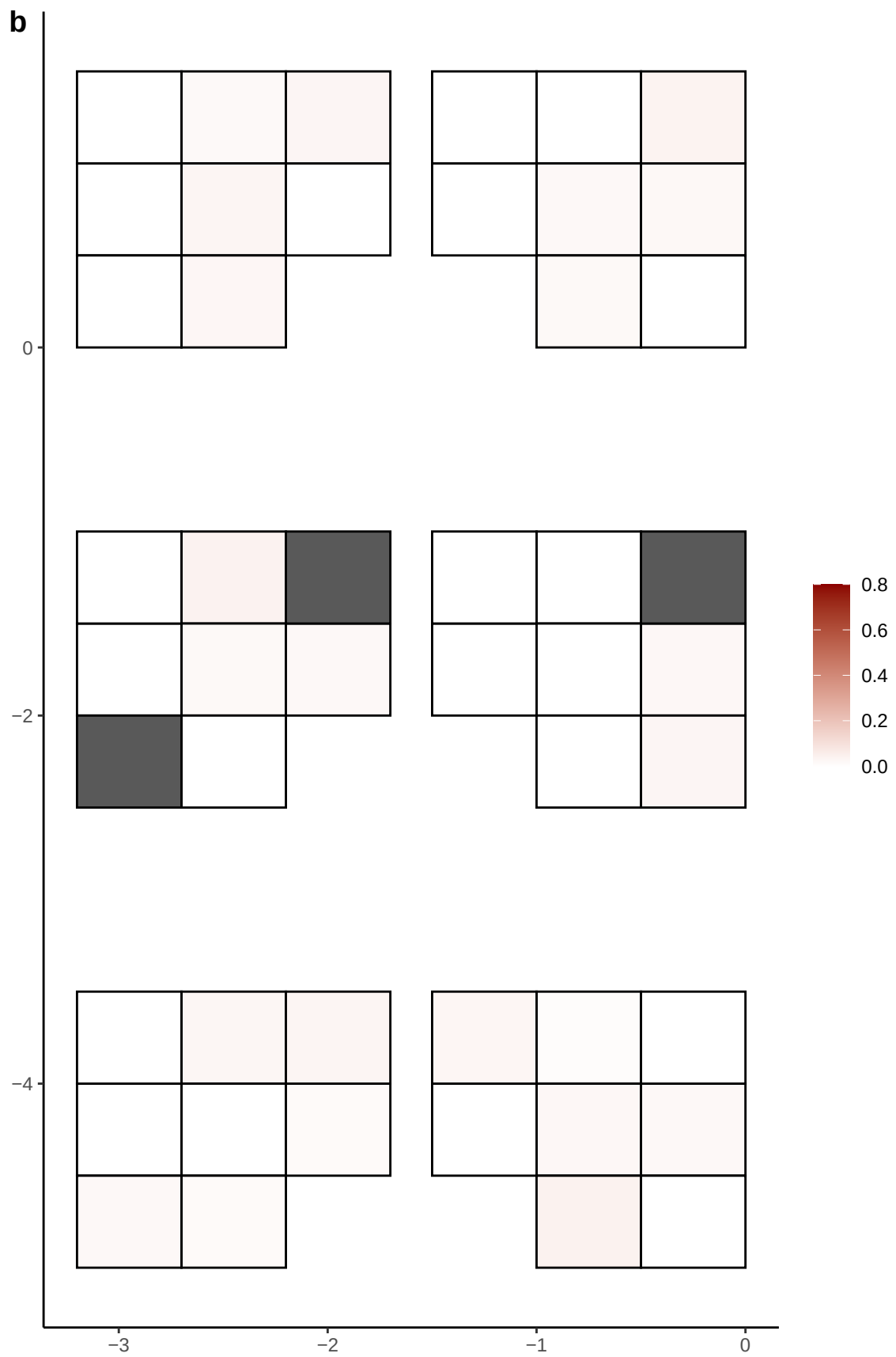

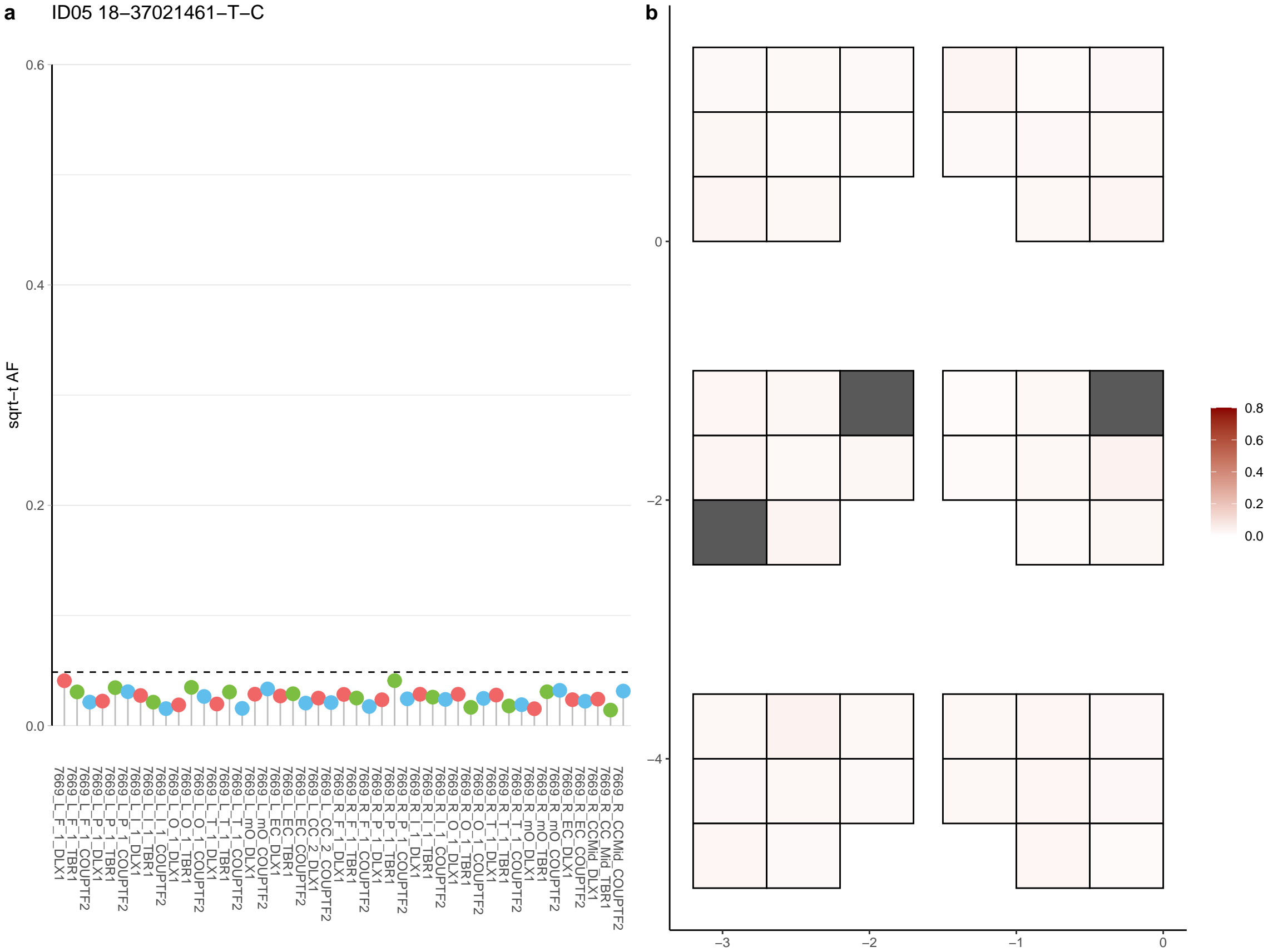

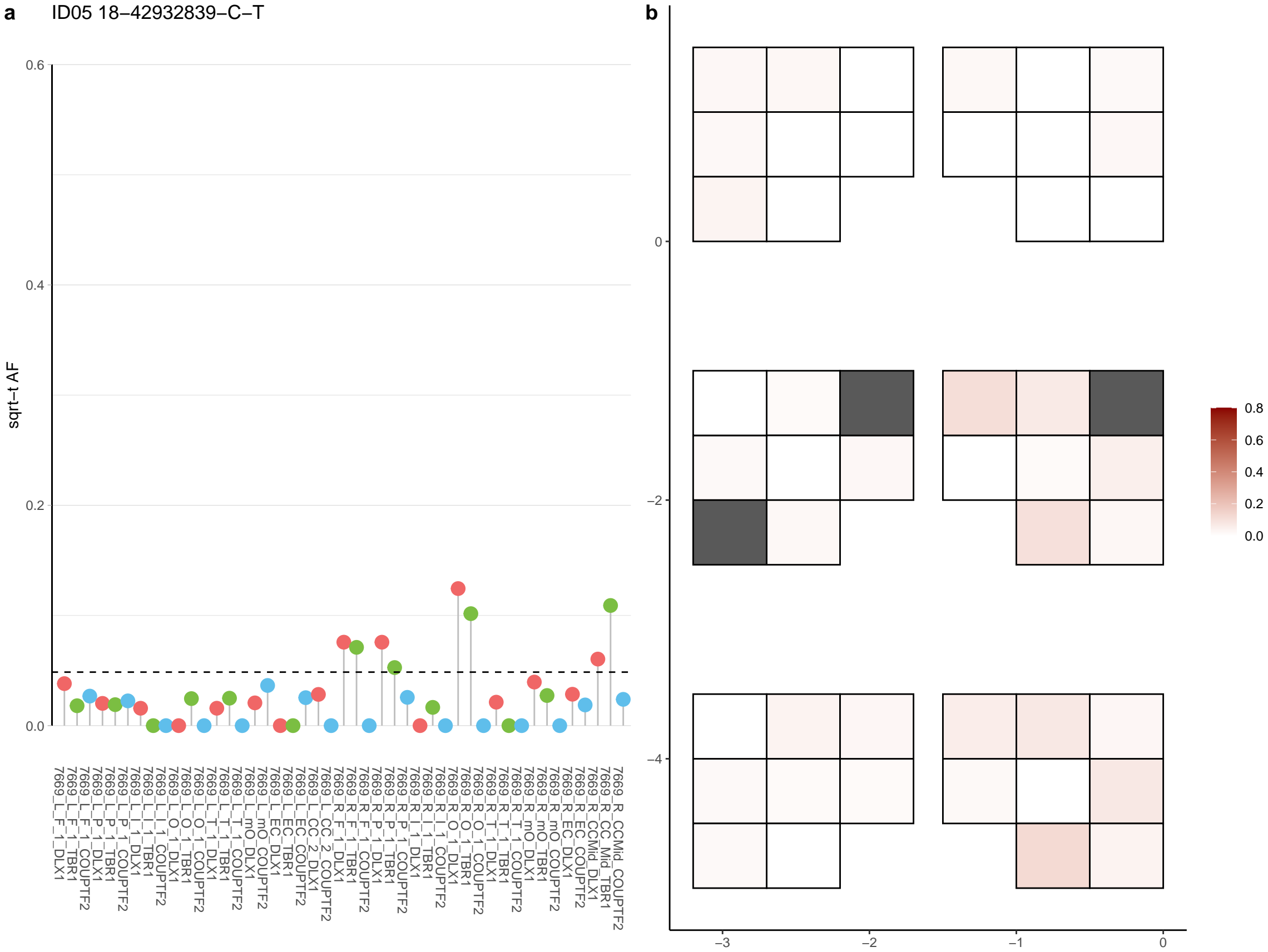

**a** ID05 18-49194474-G-A

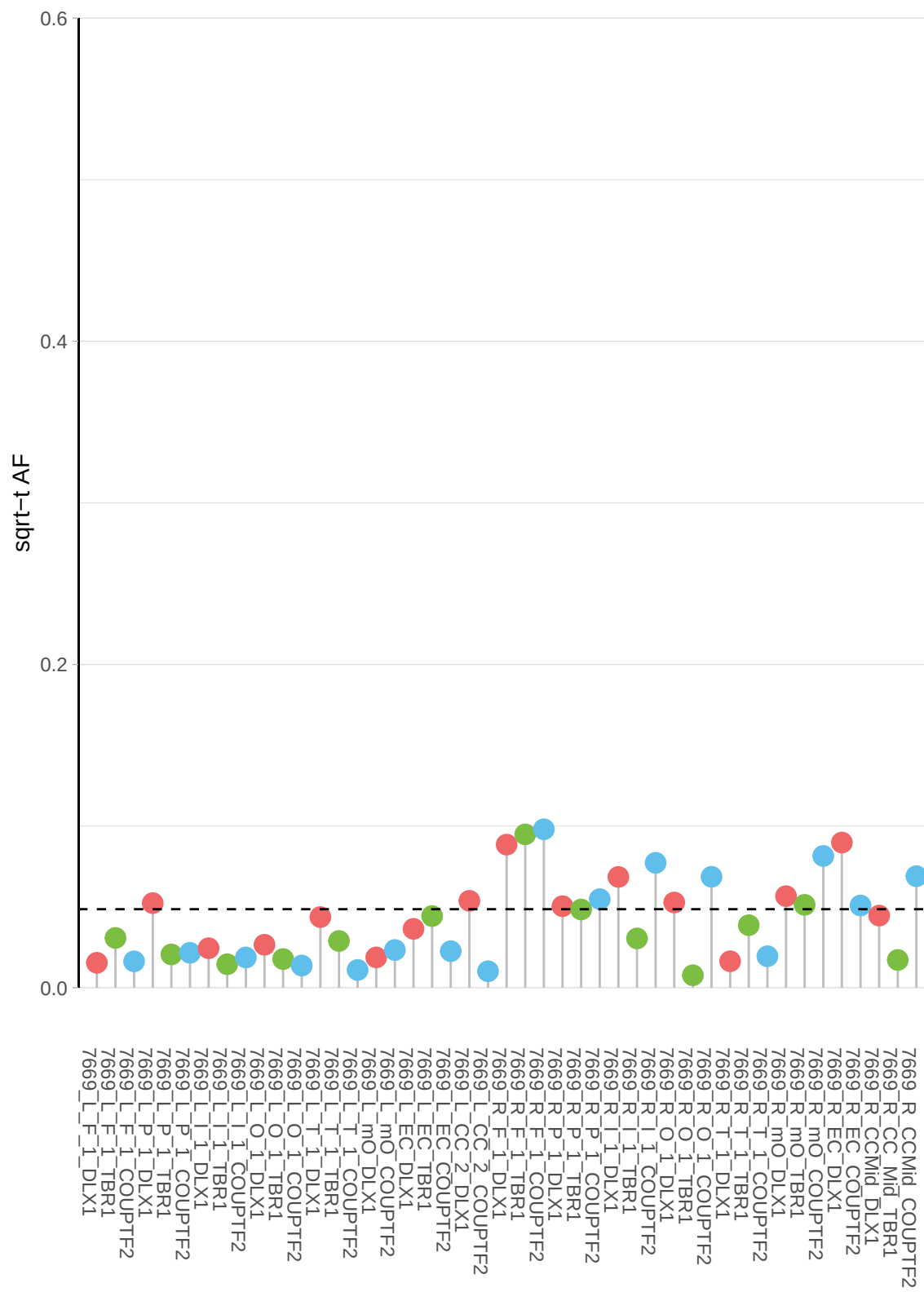

**b**

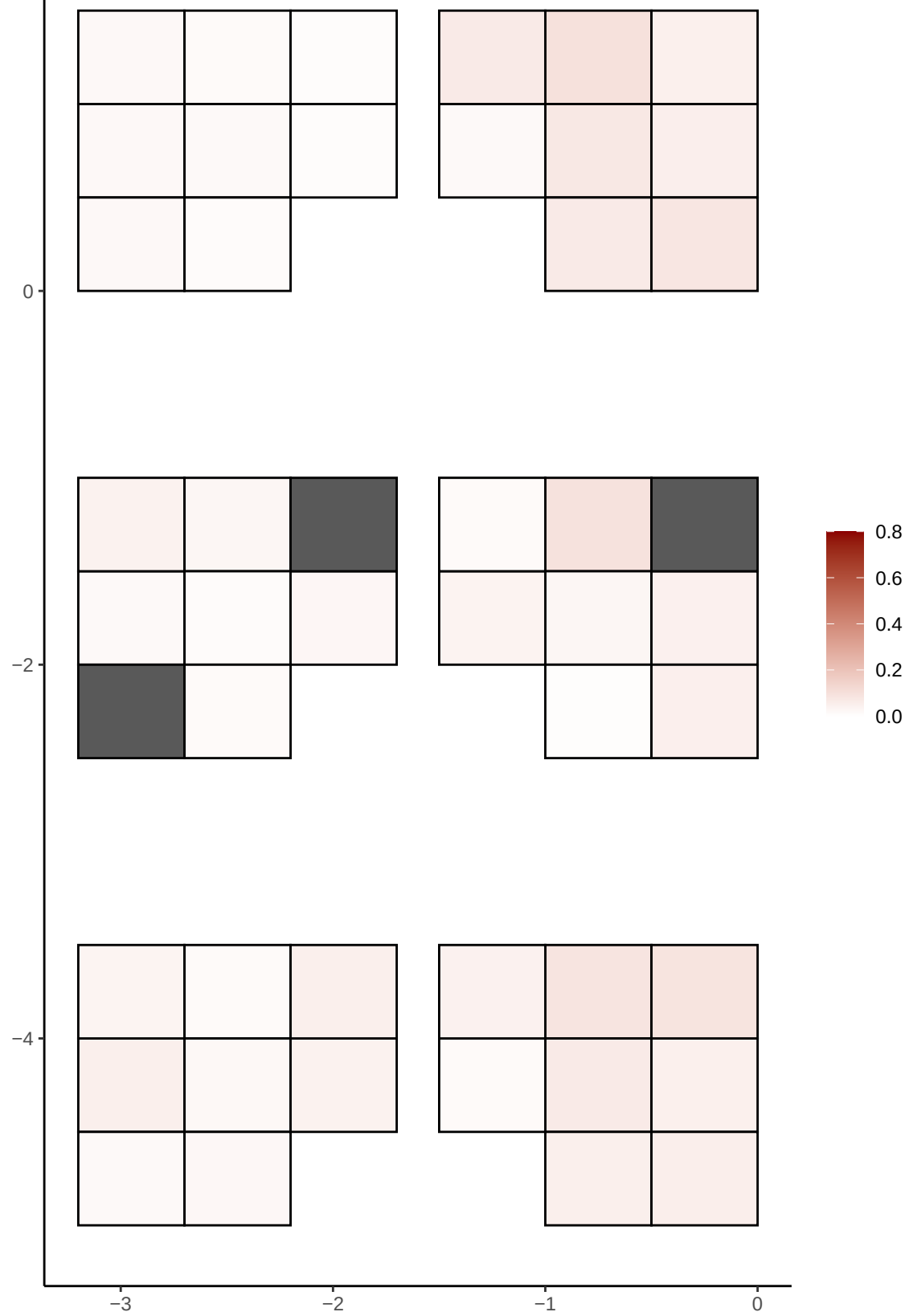

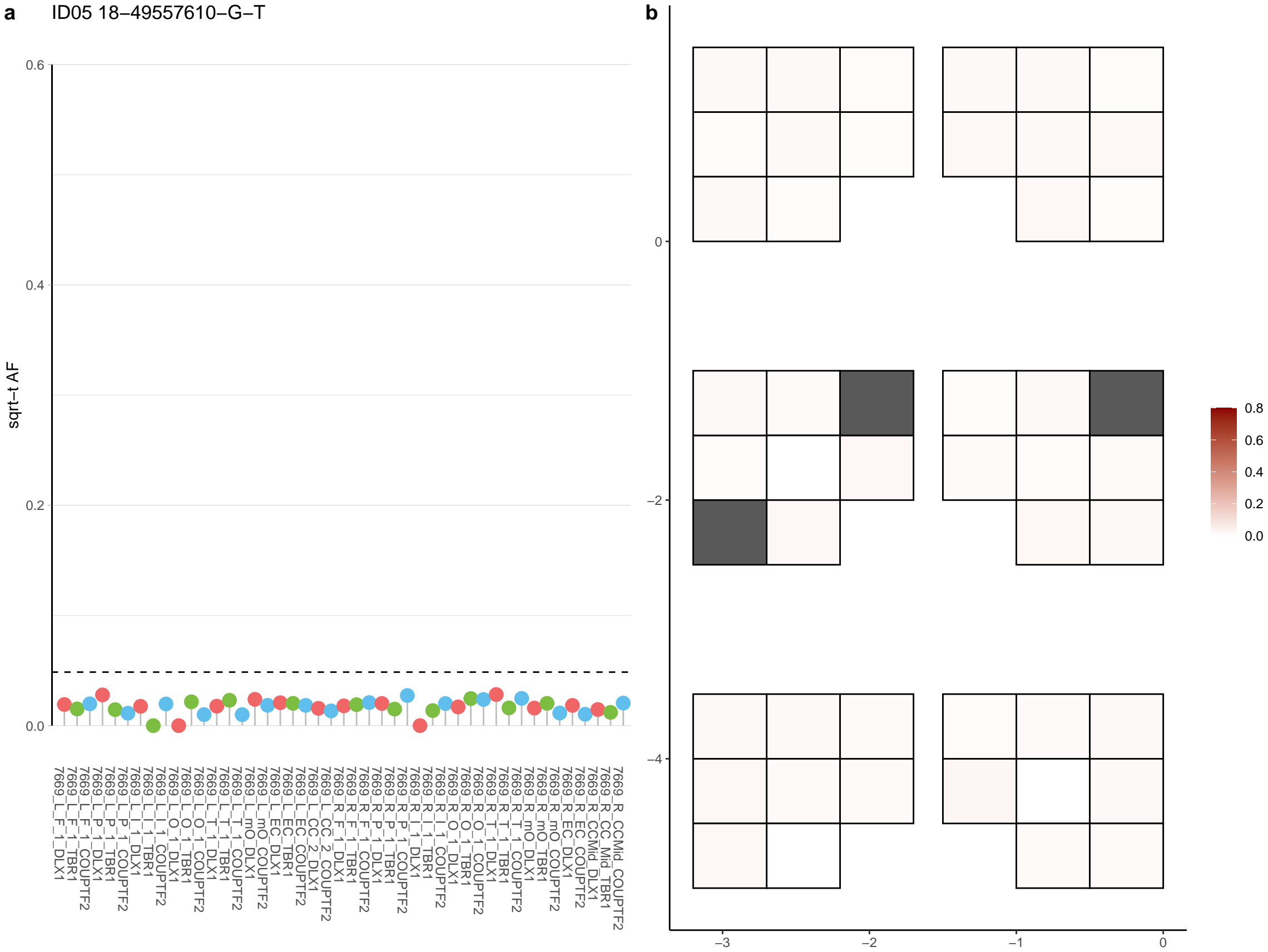

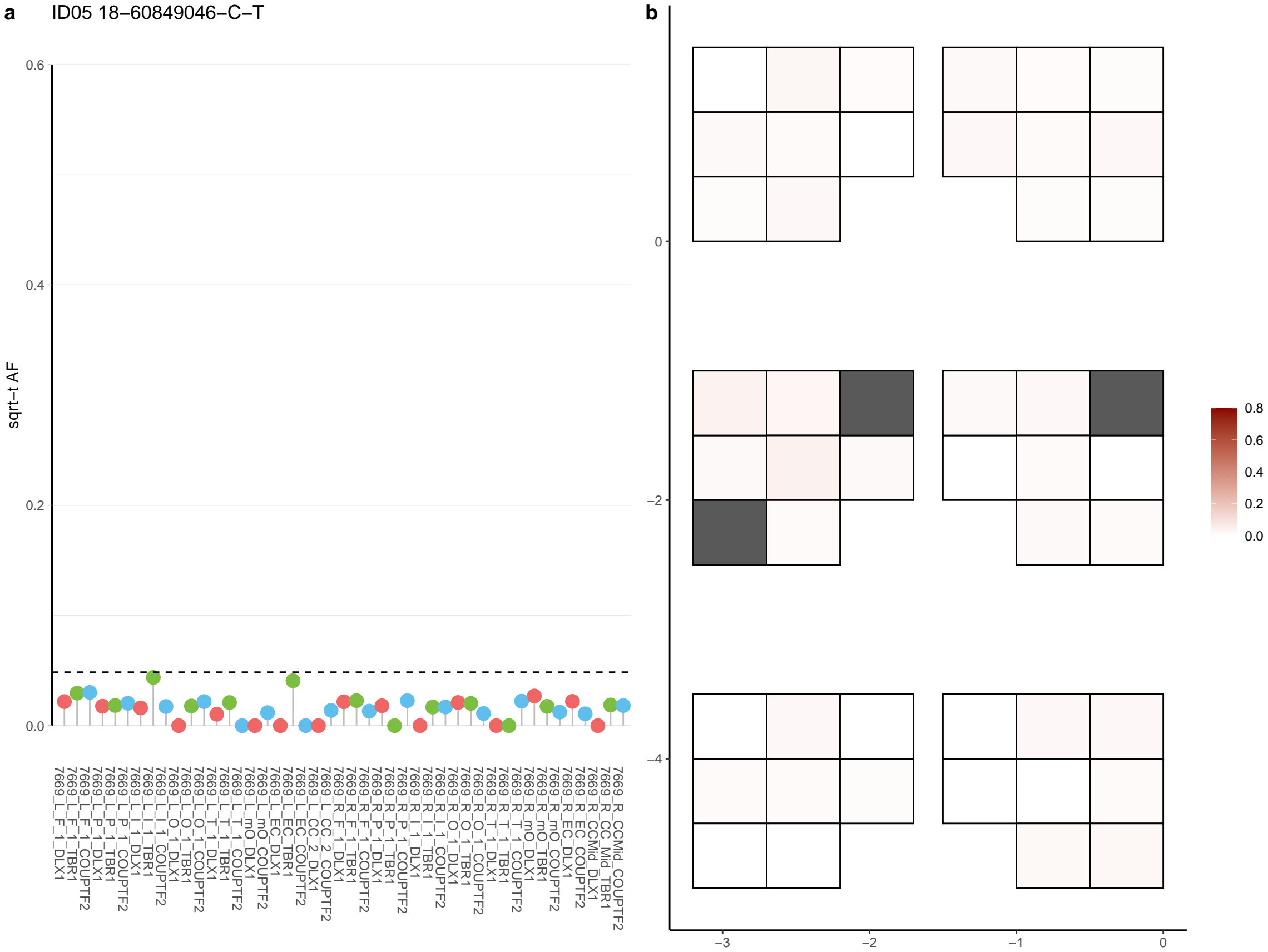

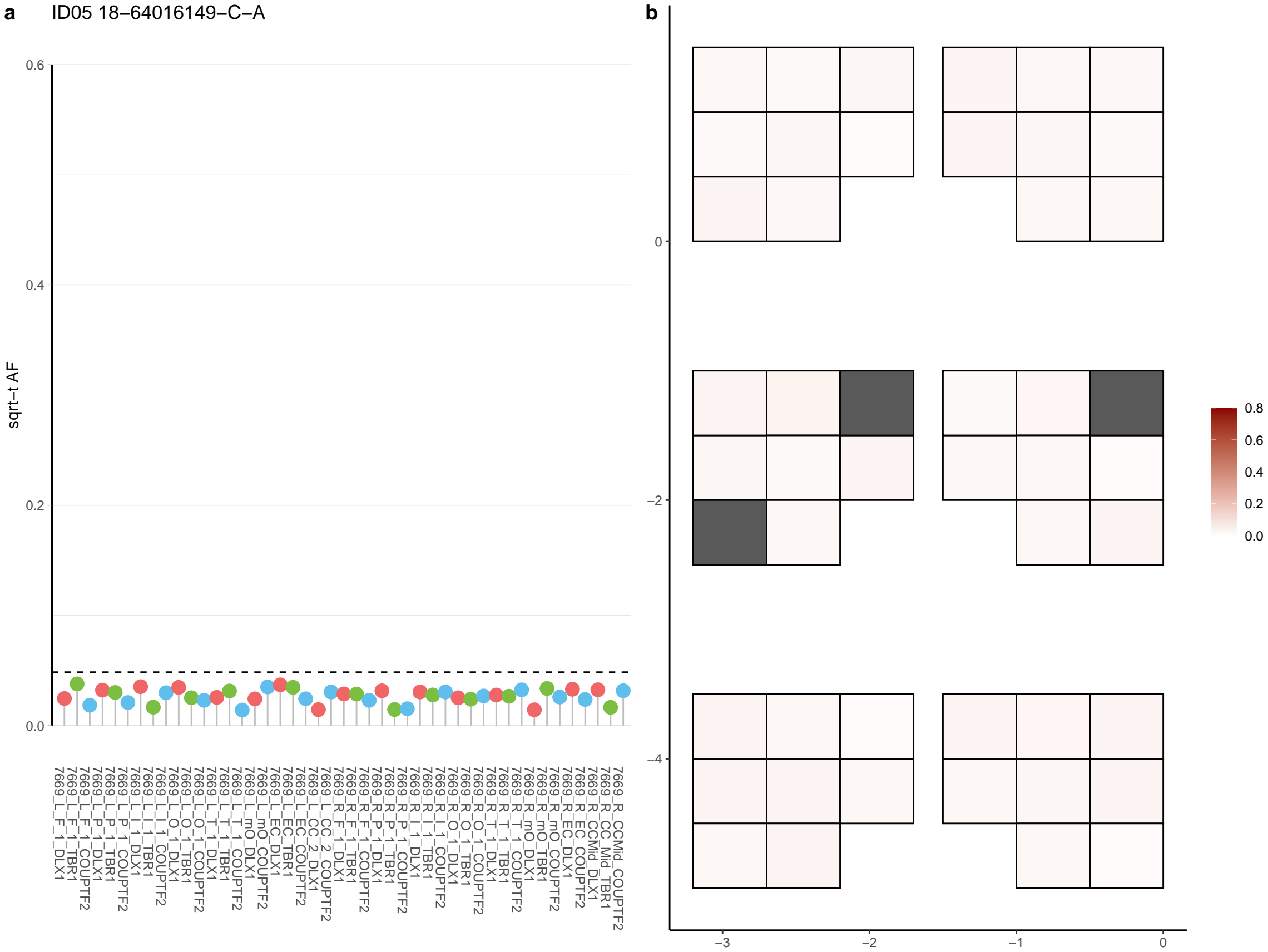

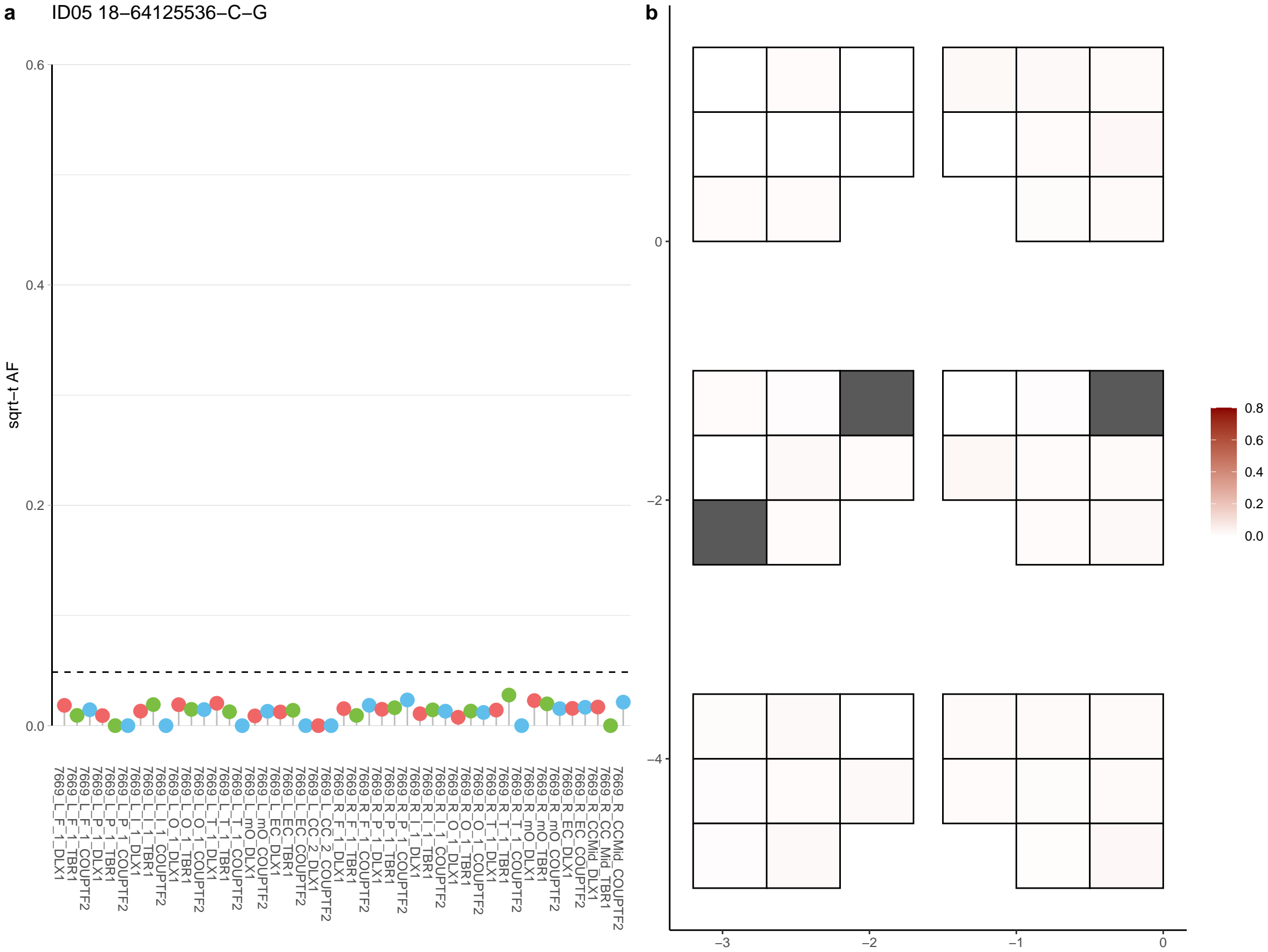

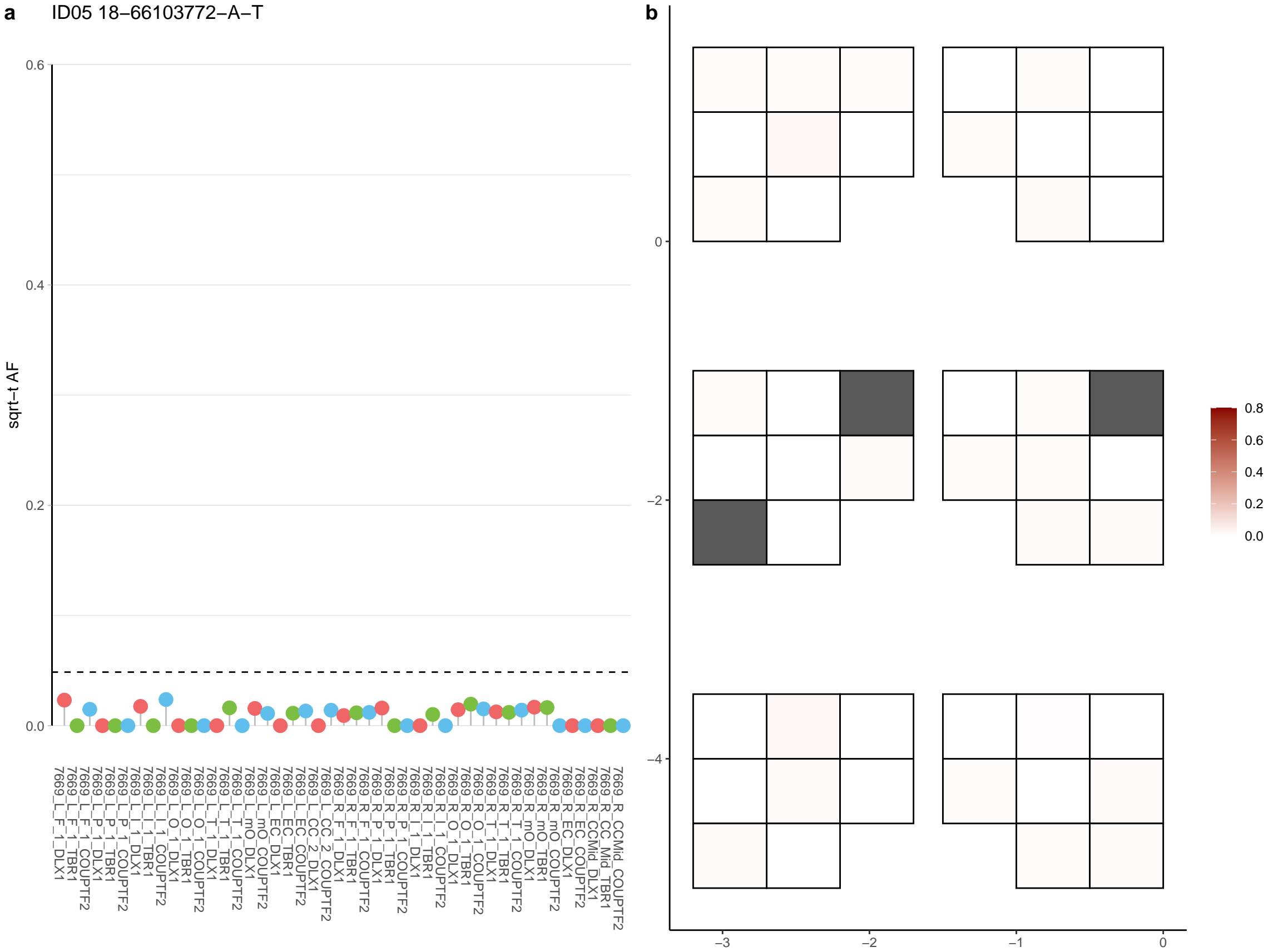

**a** ID05 18-68768921-C-T

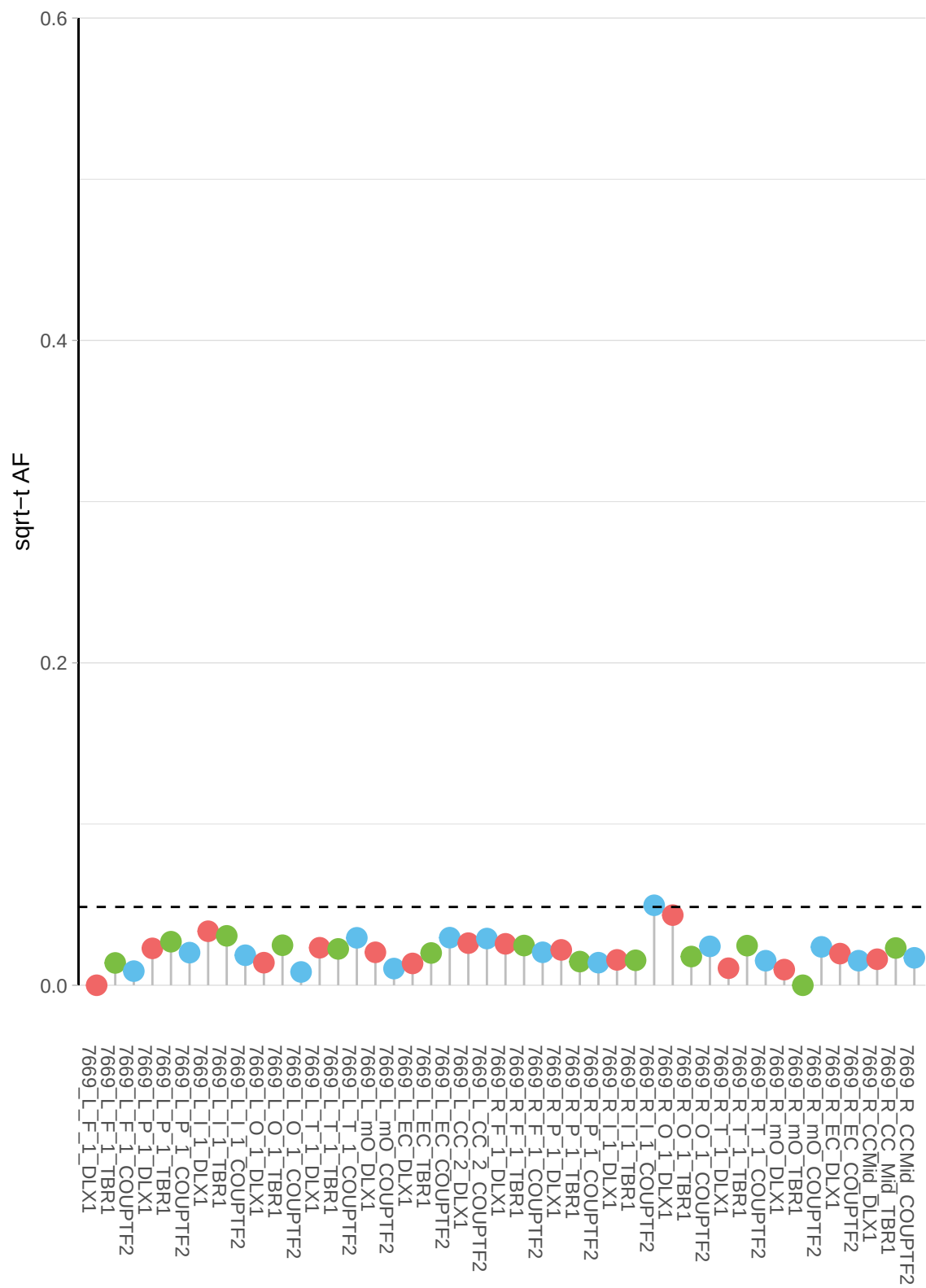

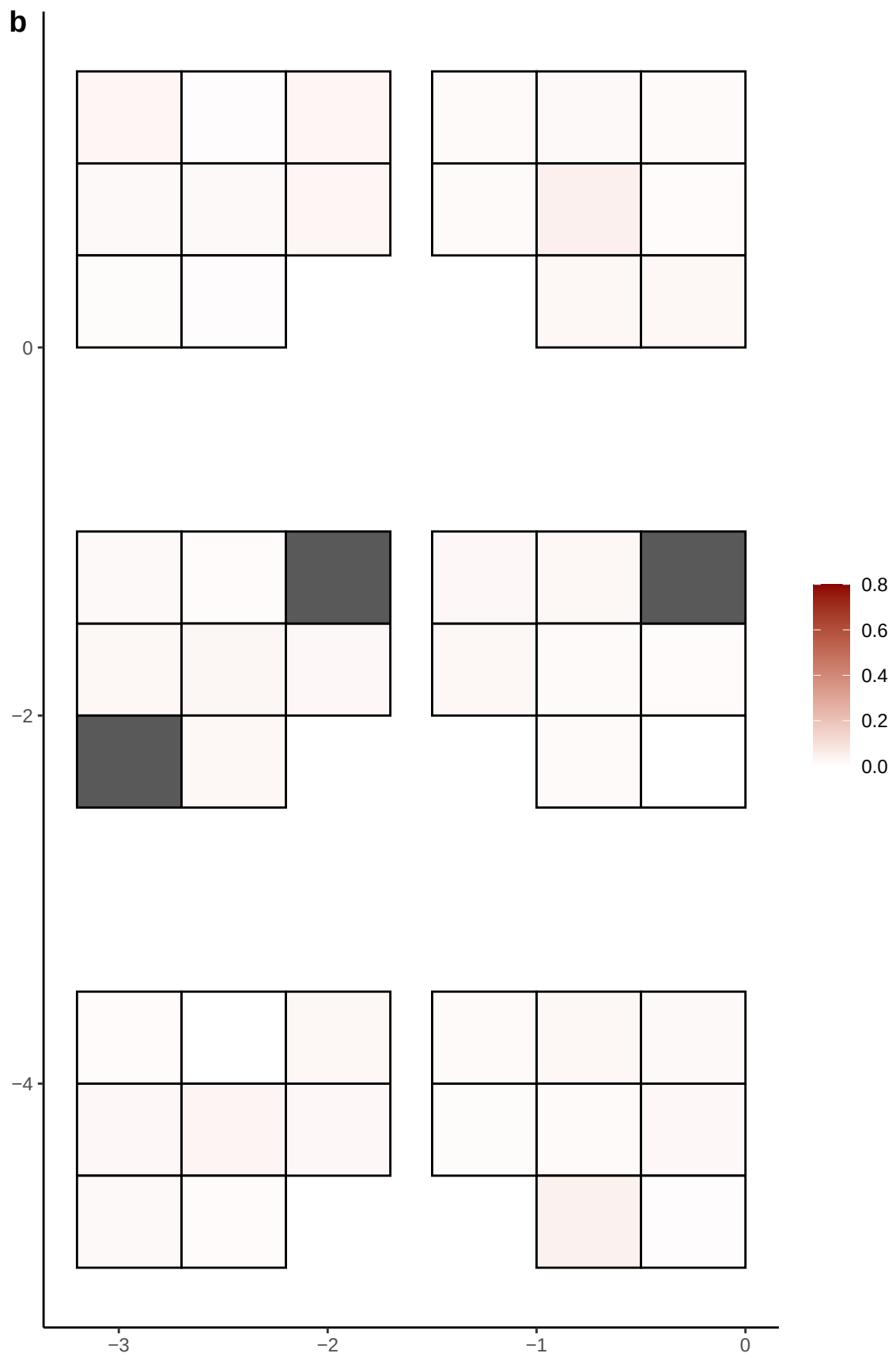

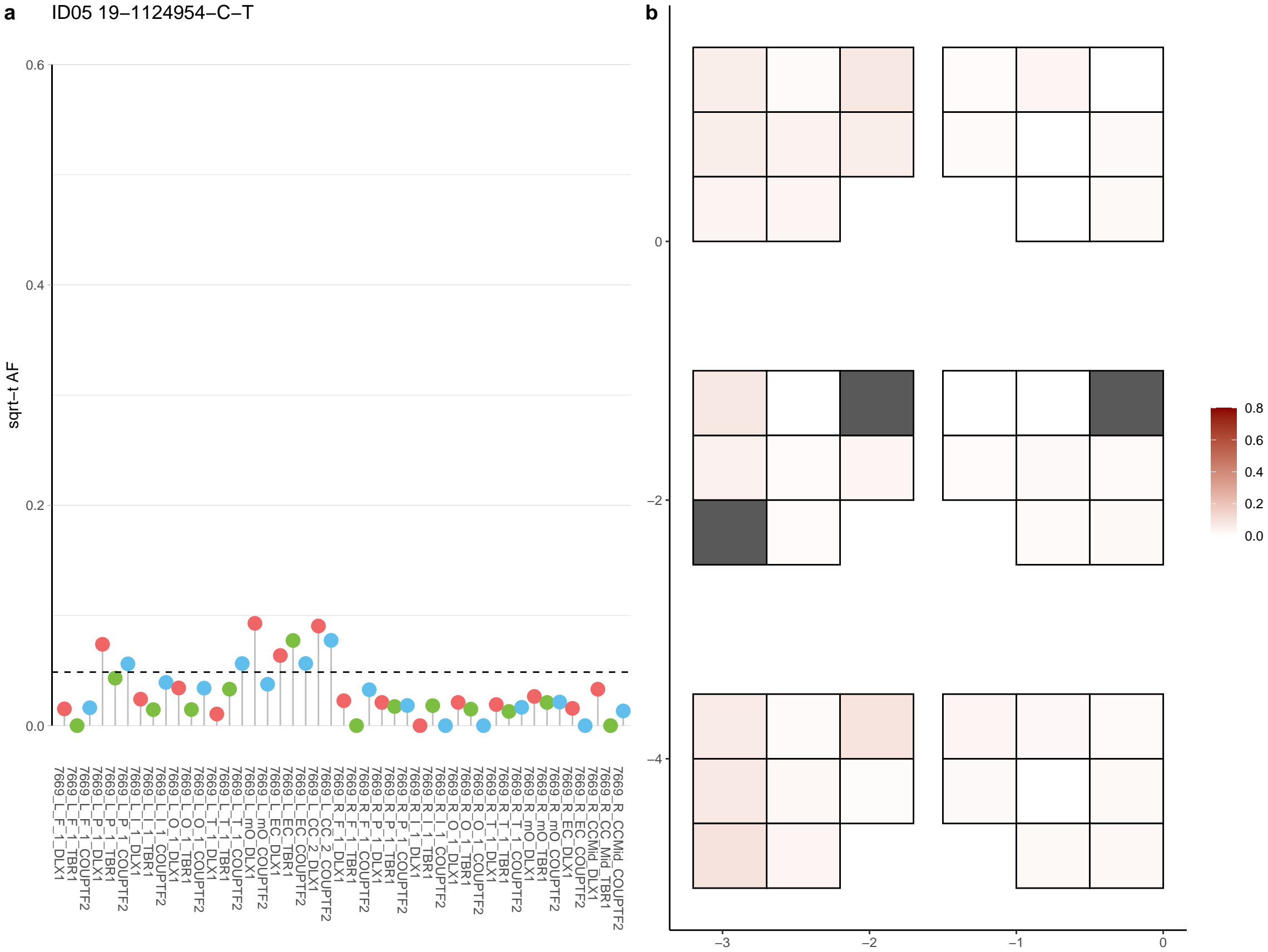

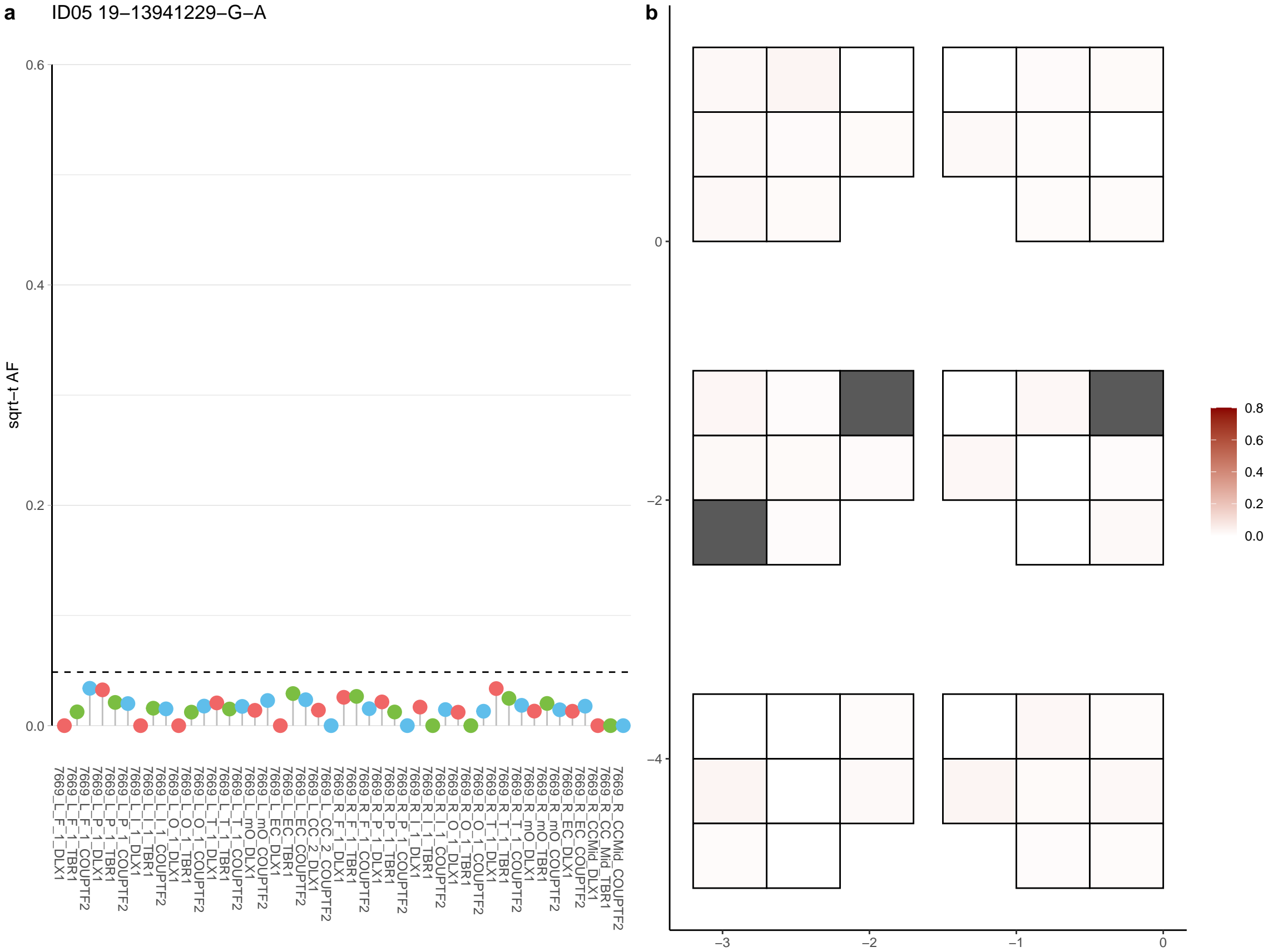

**a** ID05 19-15010579-G-A

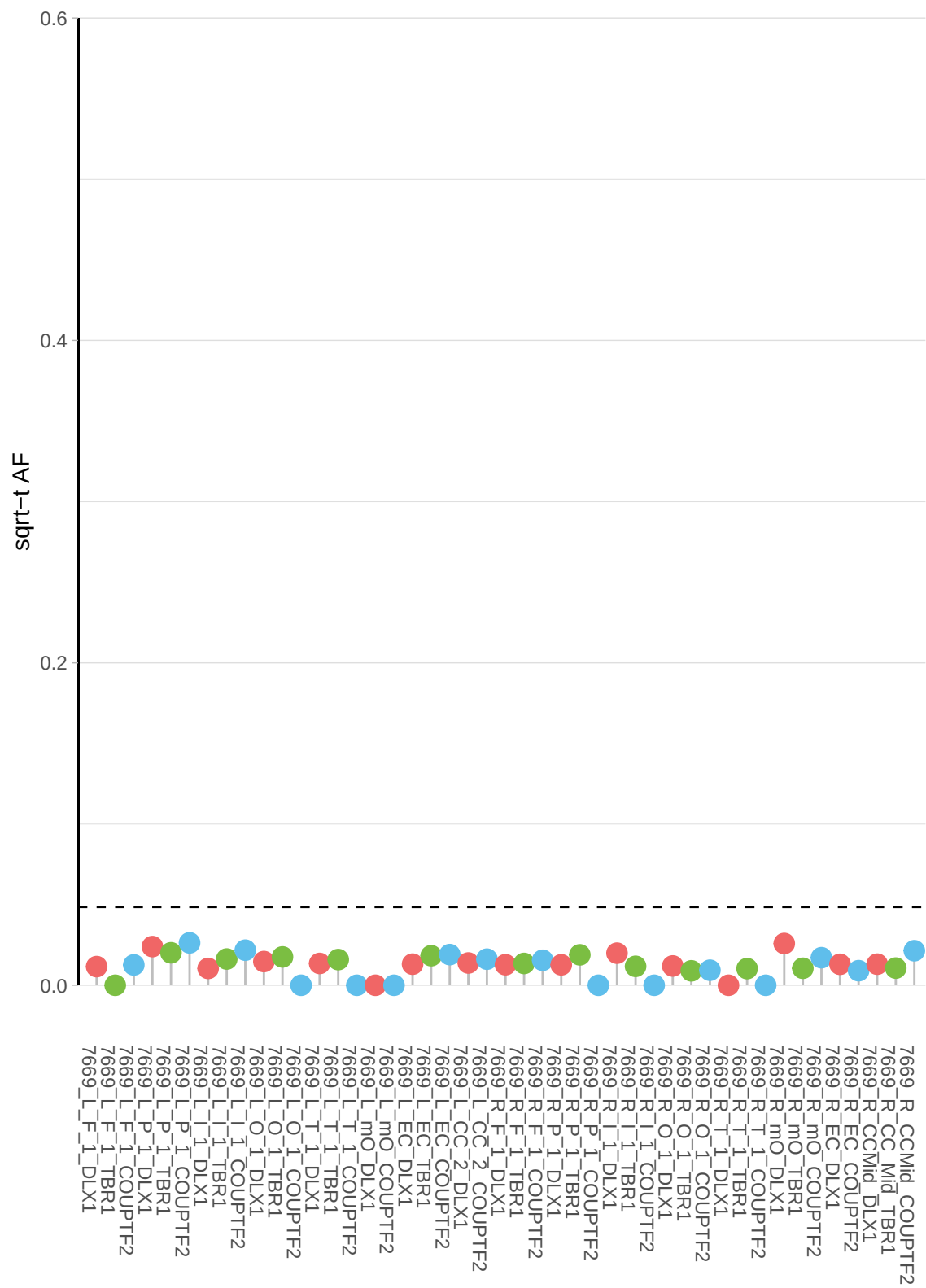

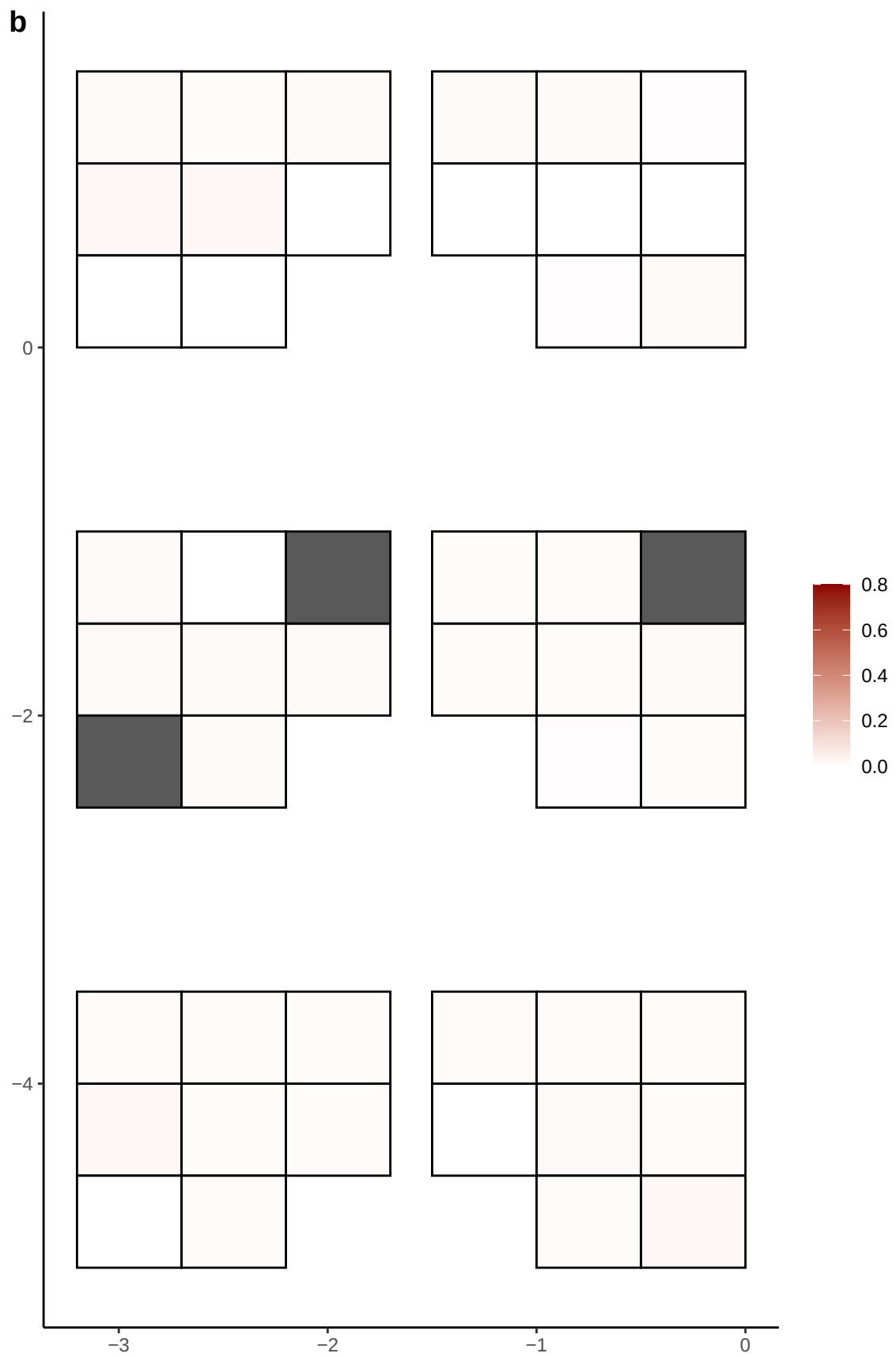

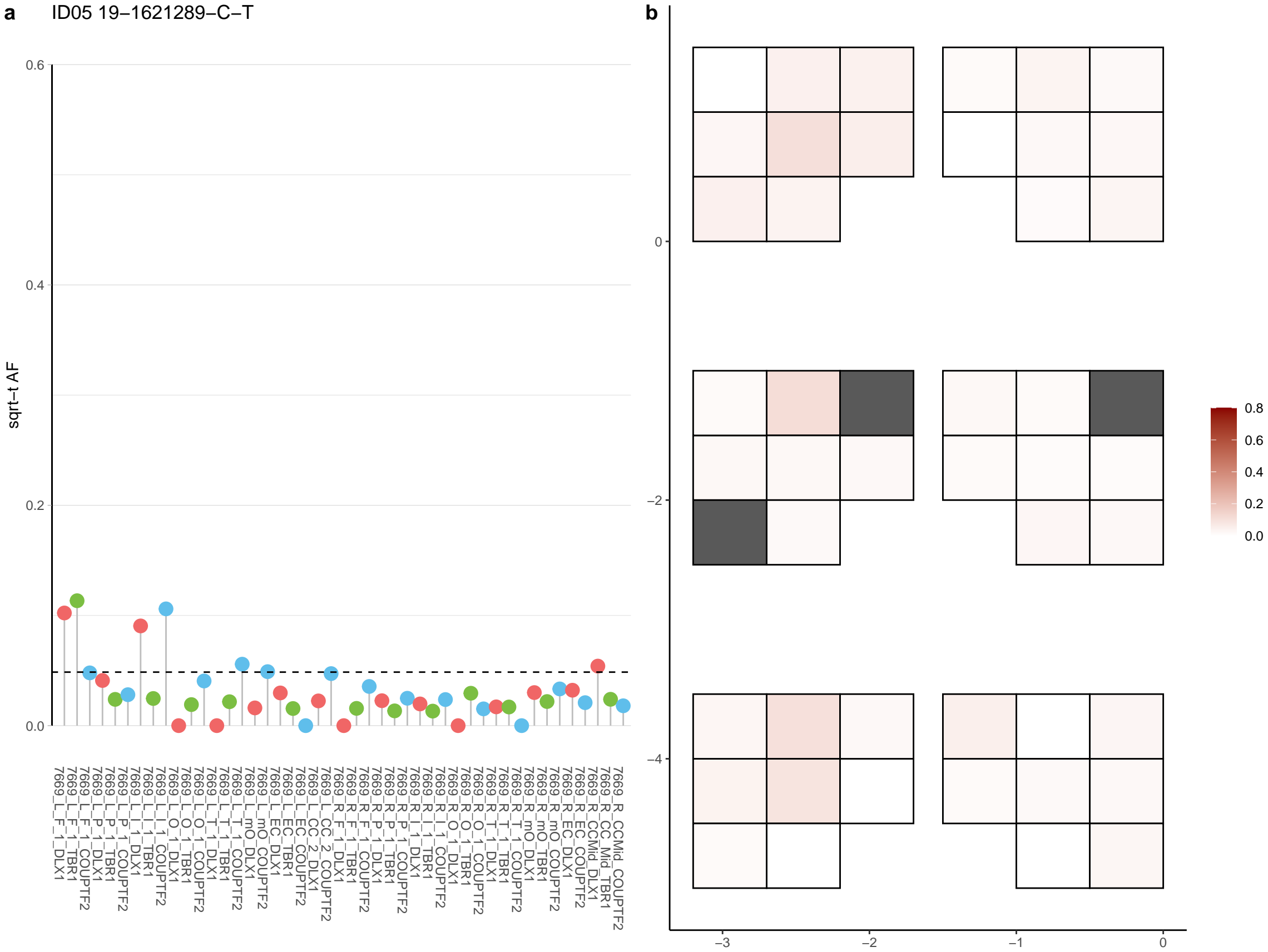

**a** ID05 19-30808081-C-T

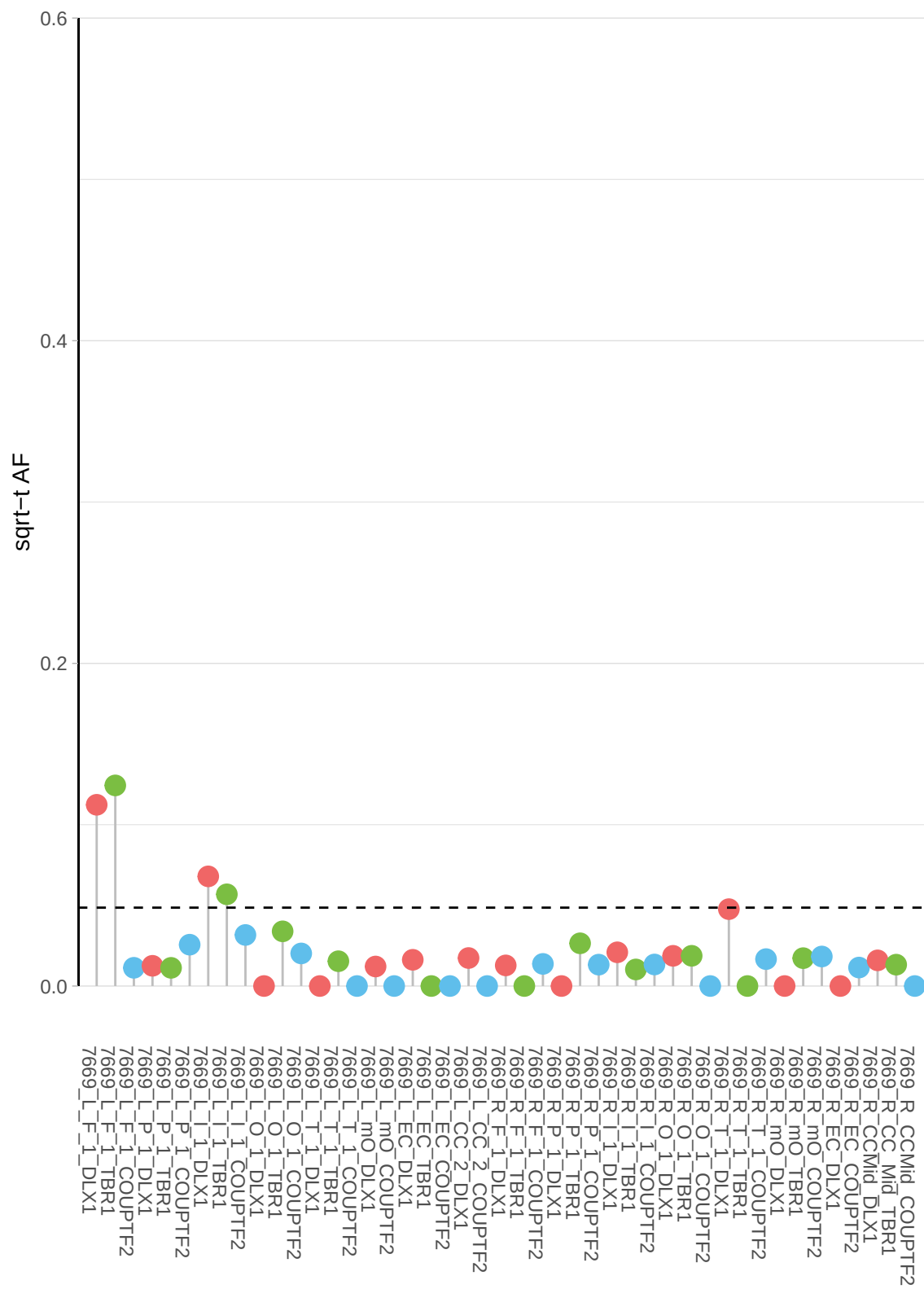

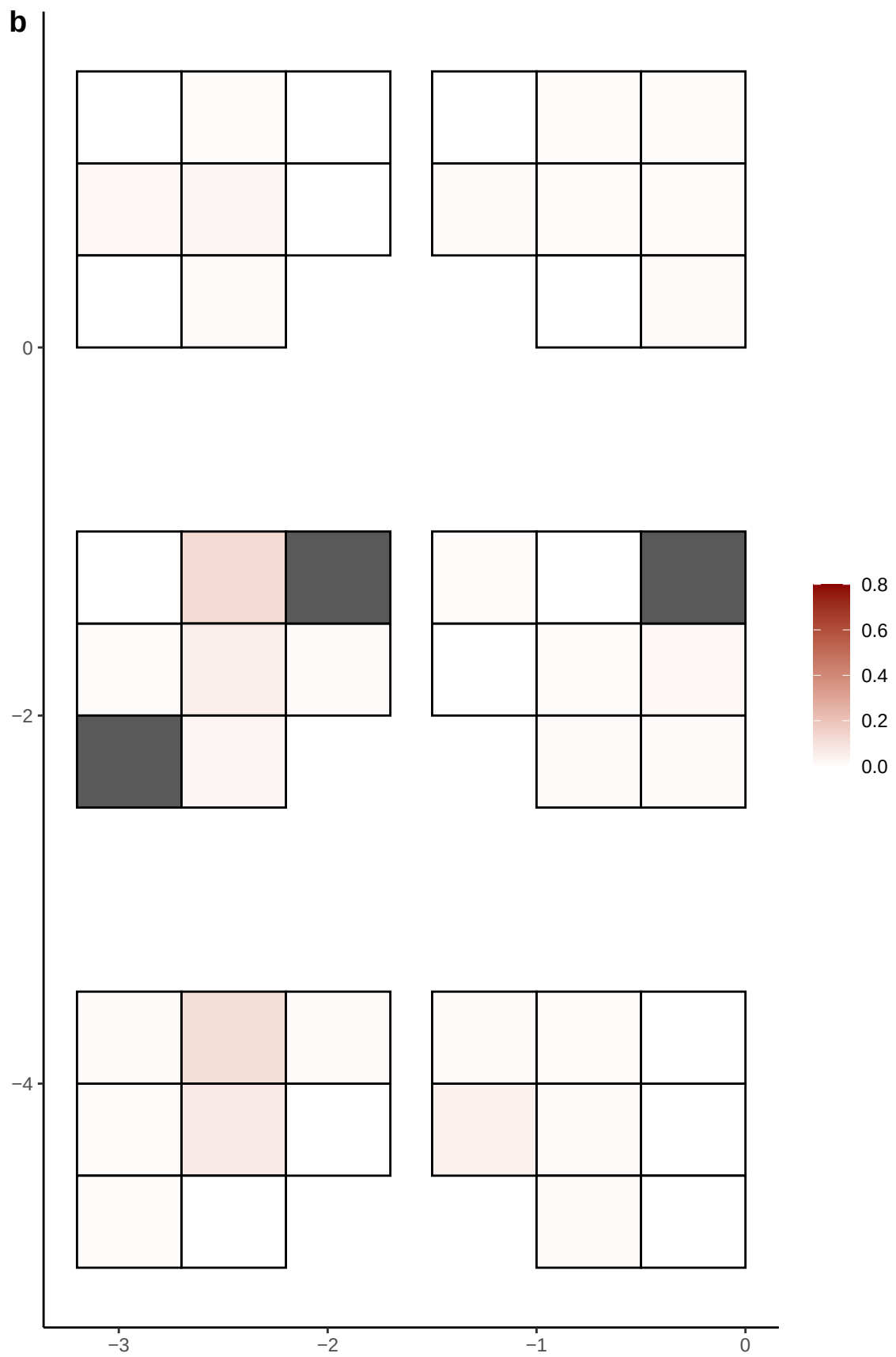

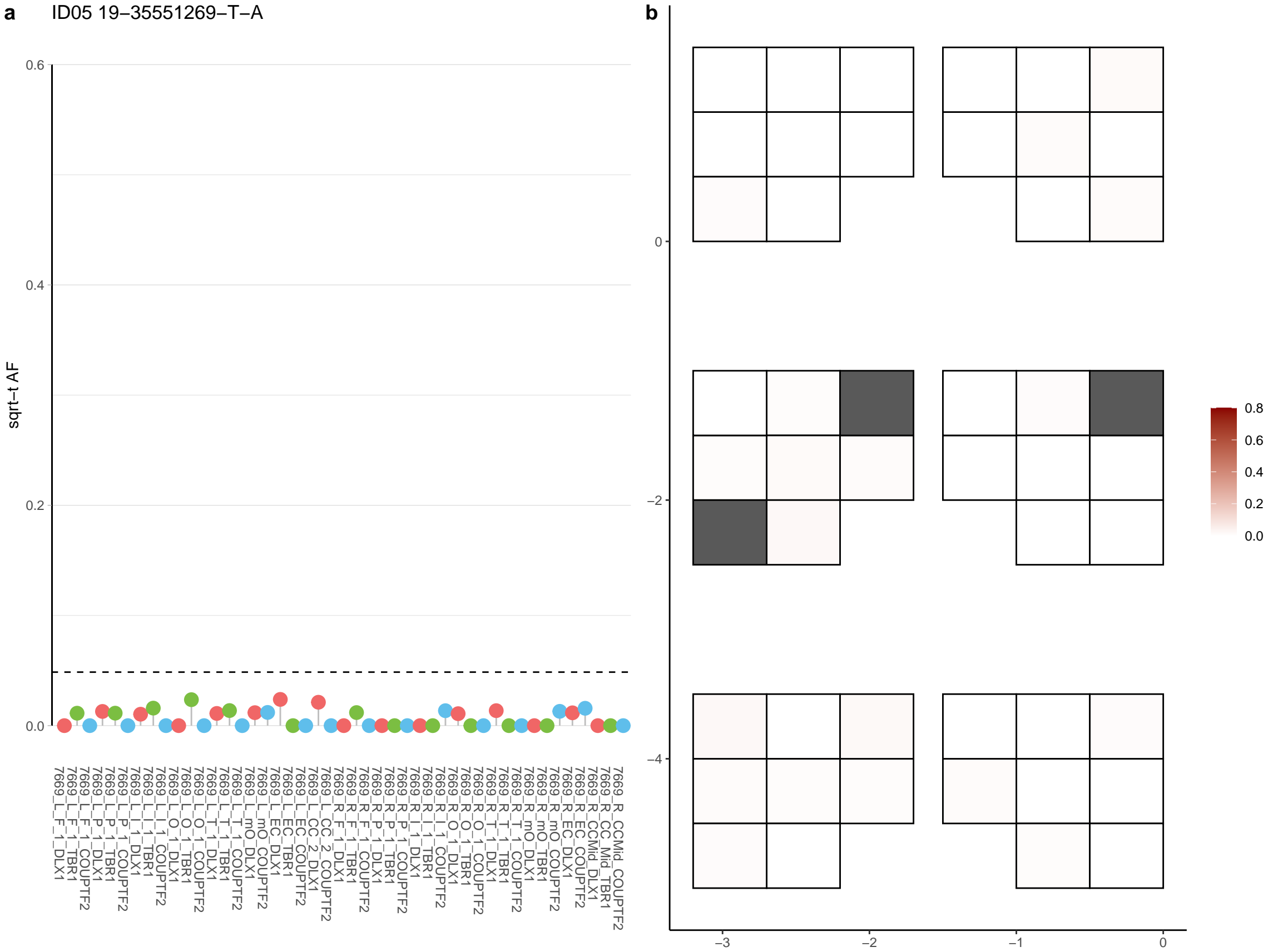

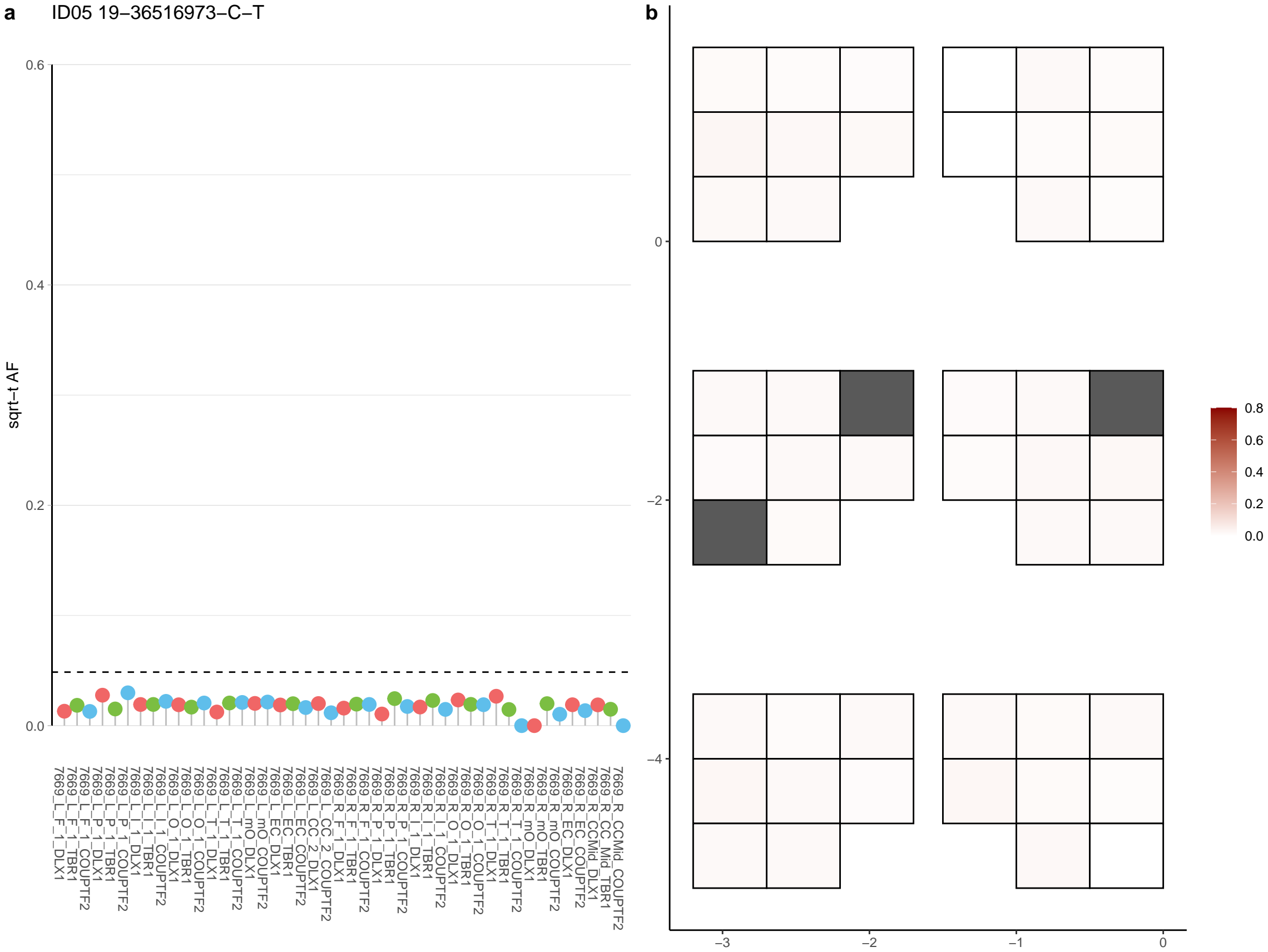

**a** ID05 19-46596282-C-T

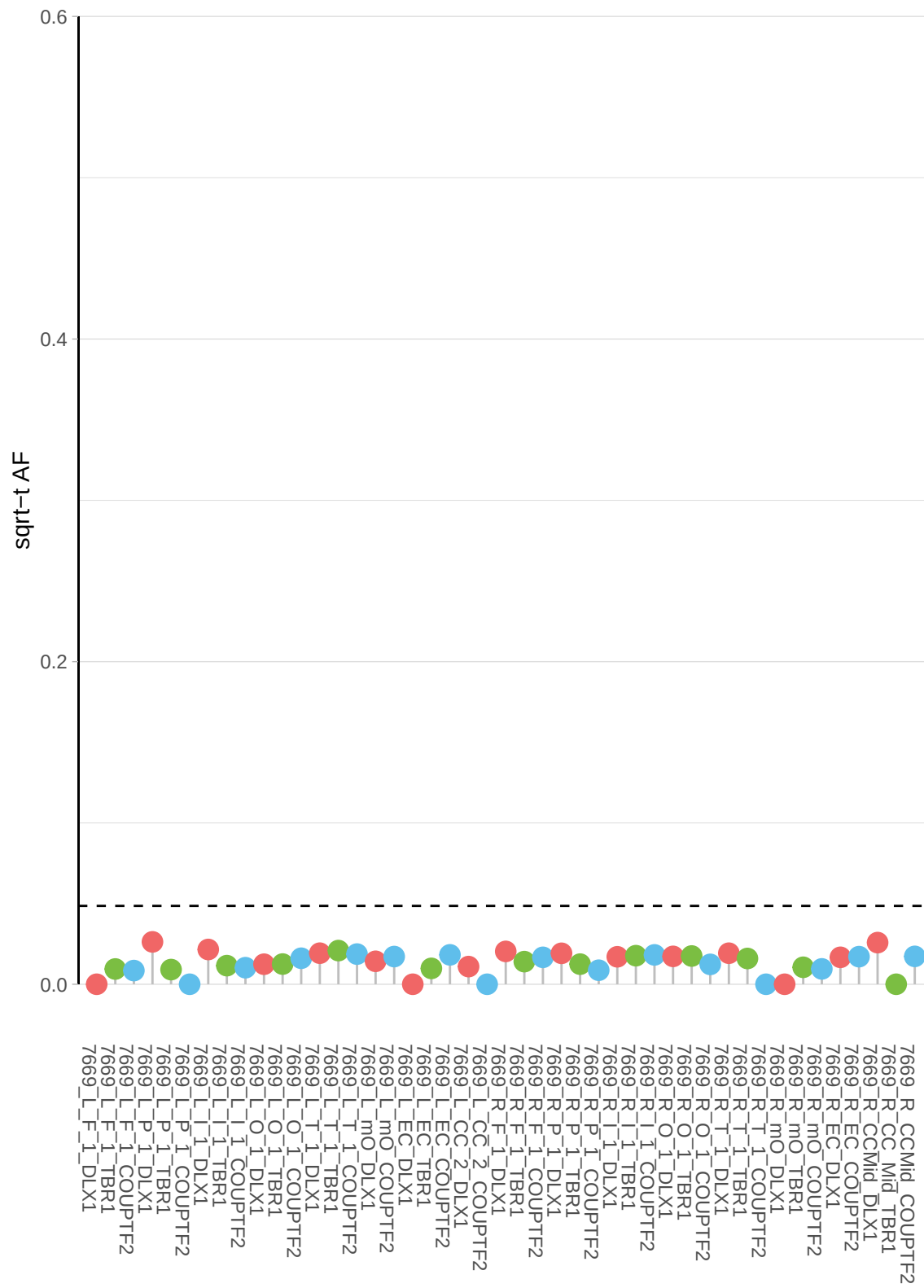

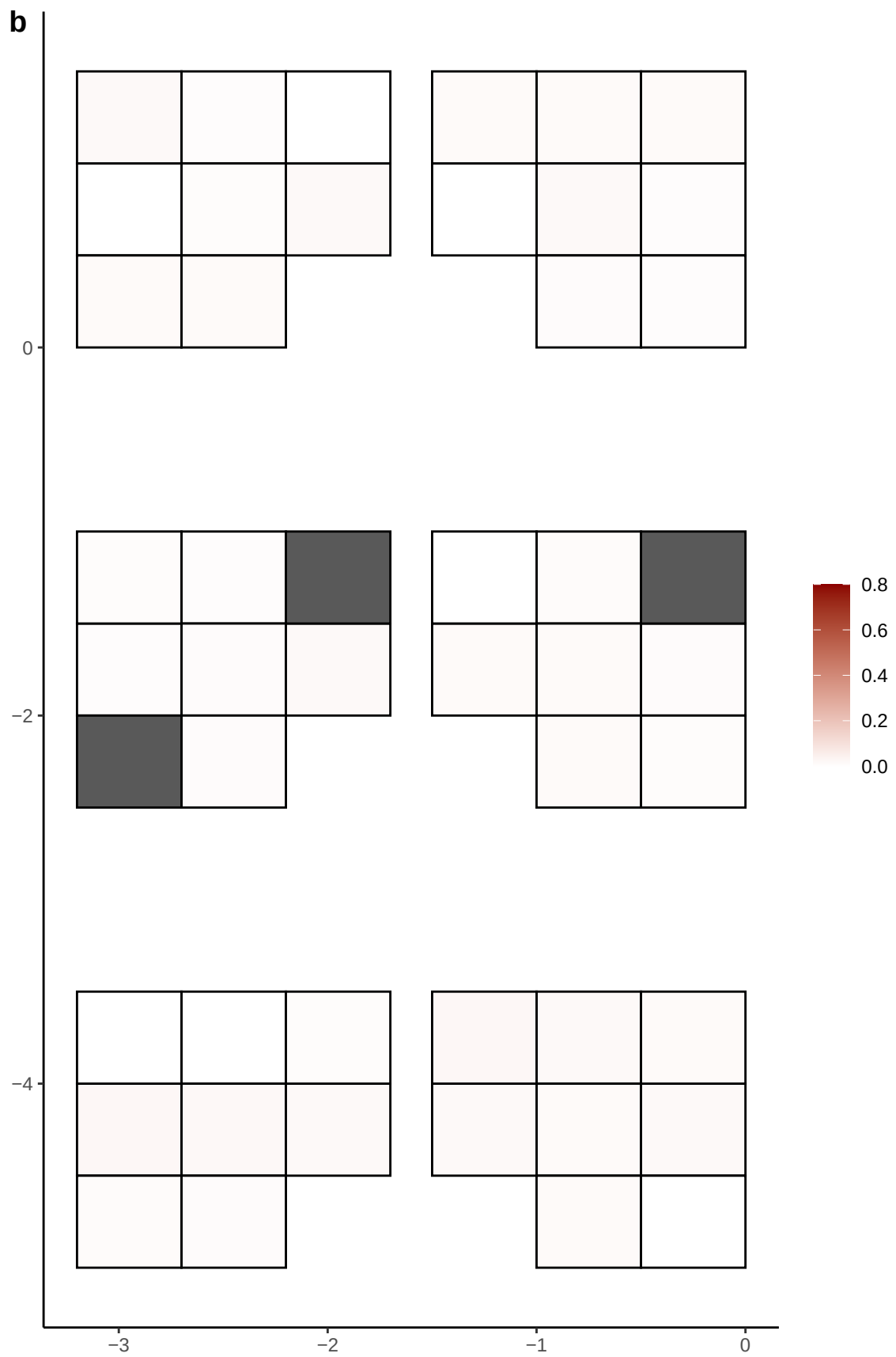

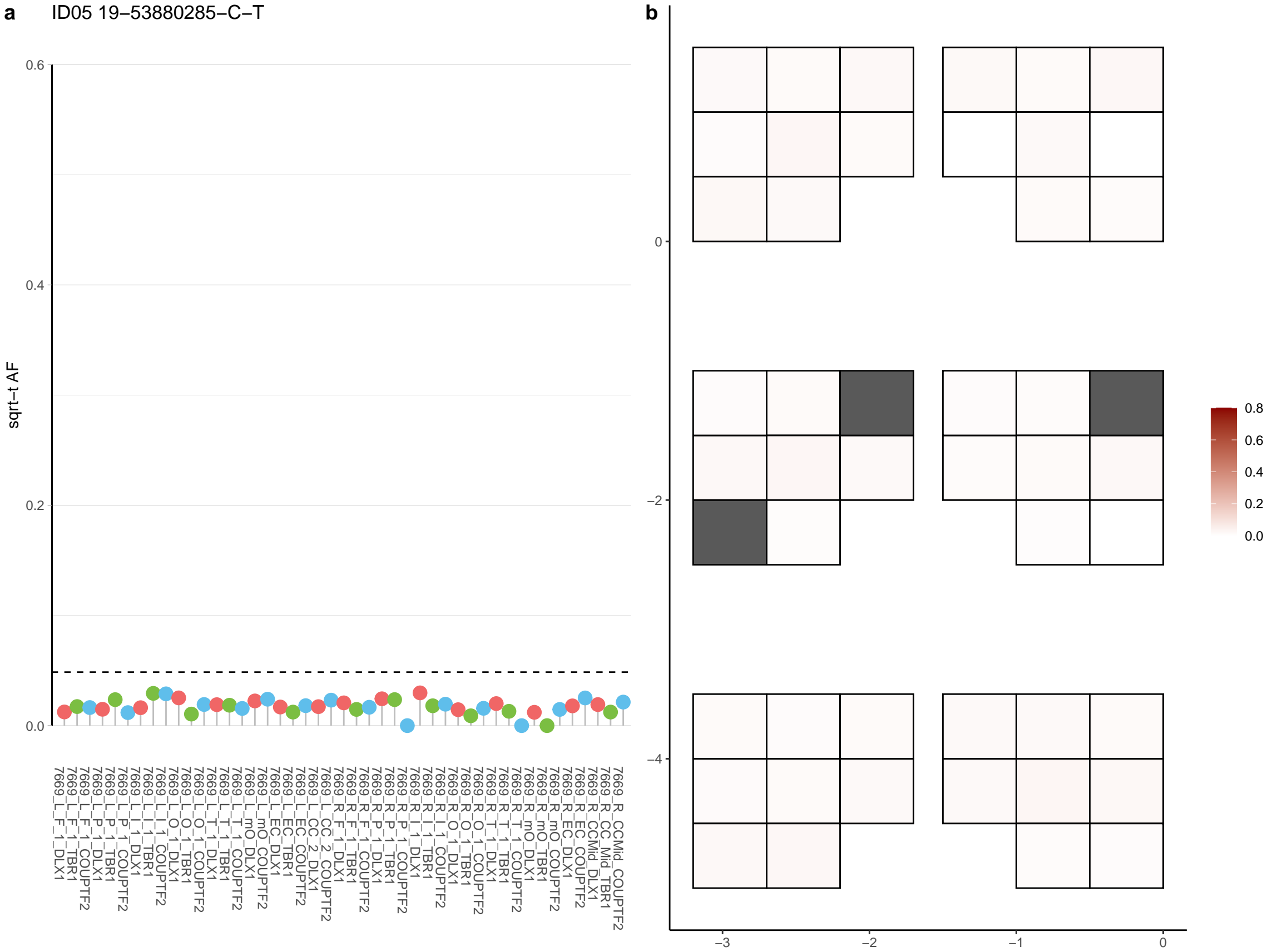

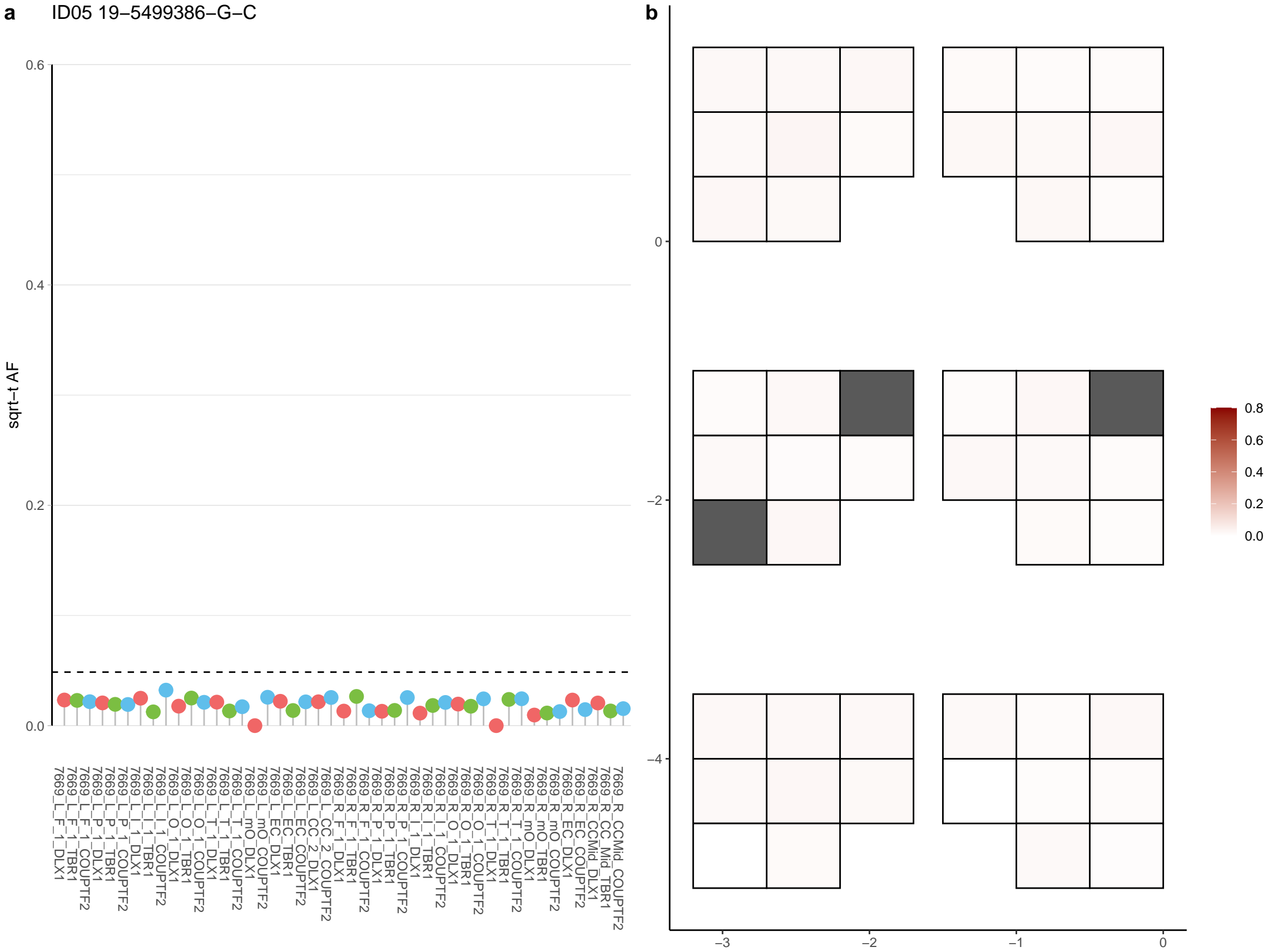

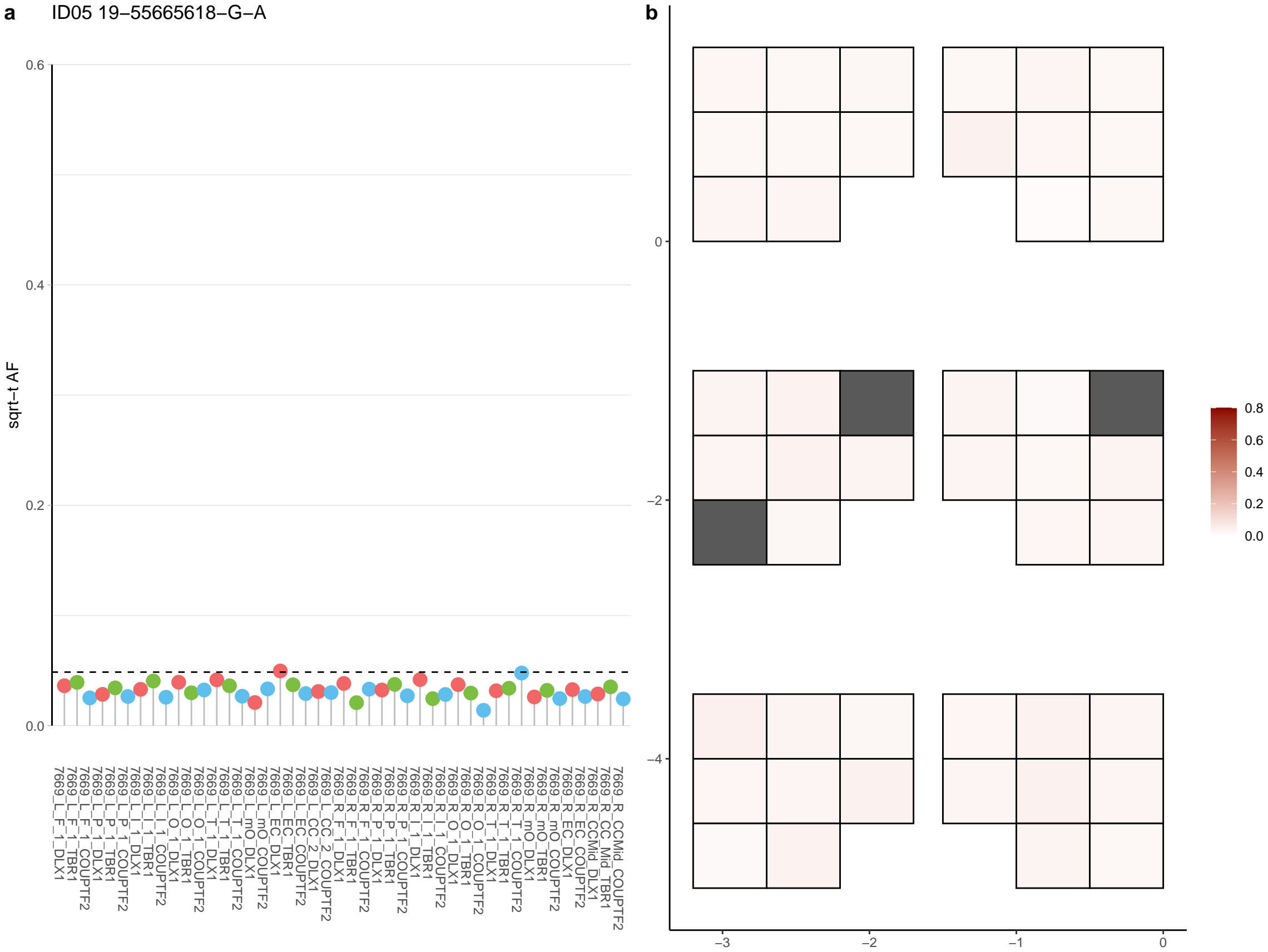

**a** ID05 19-57342142-G-A

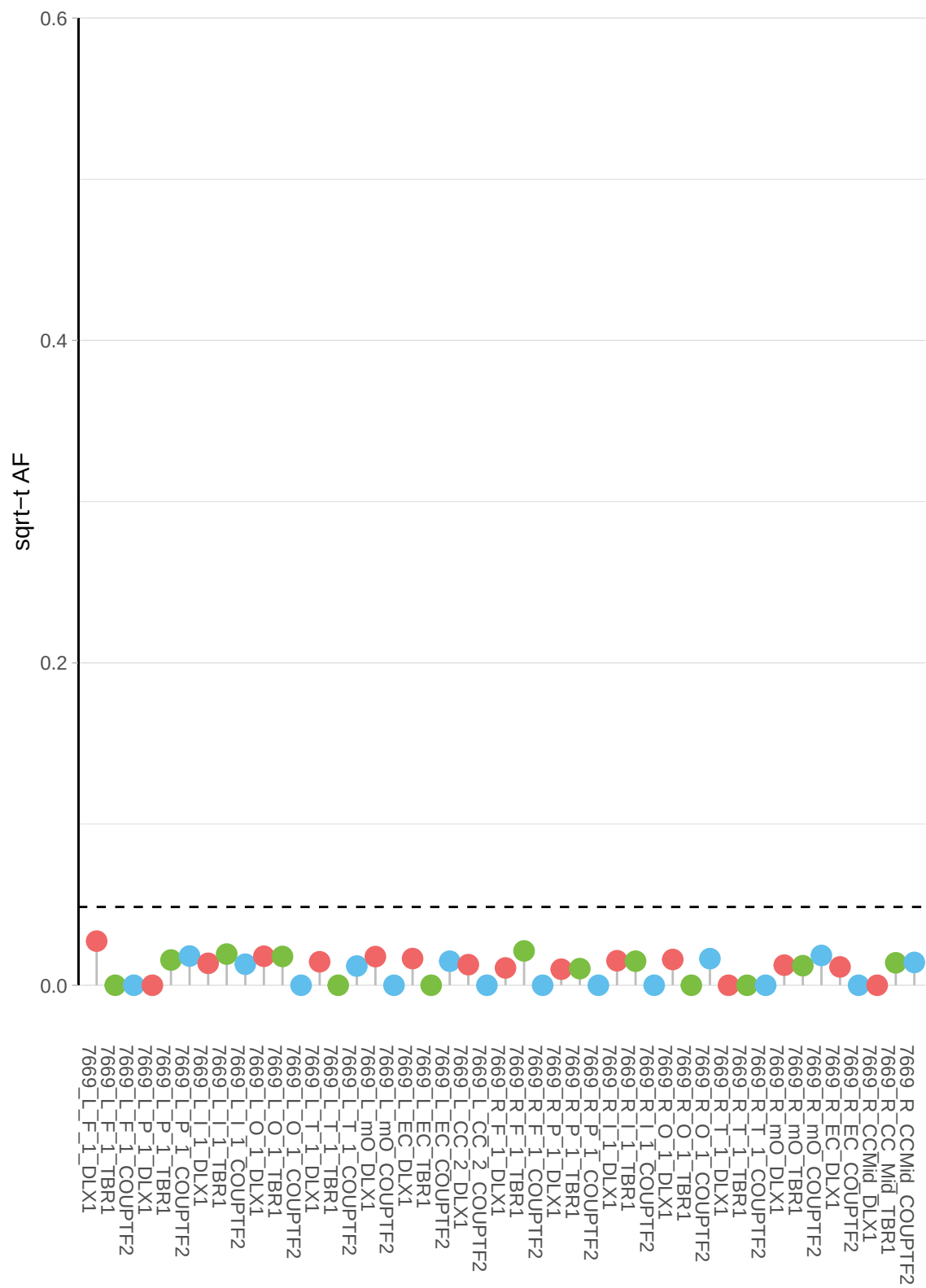

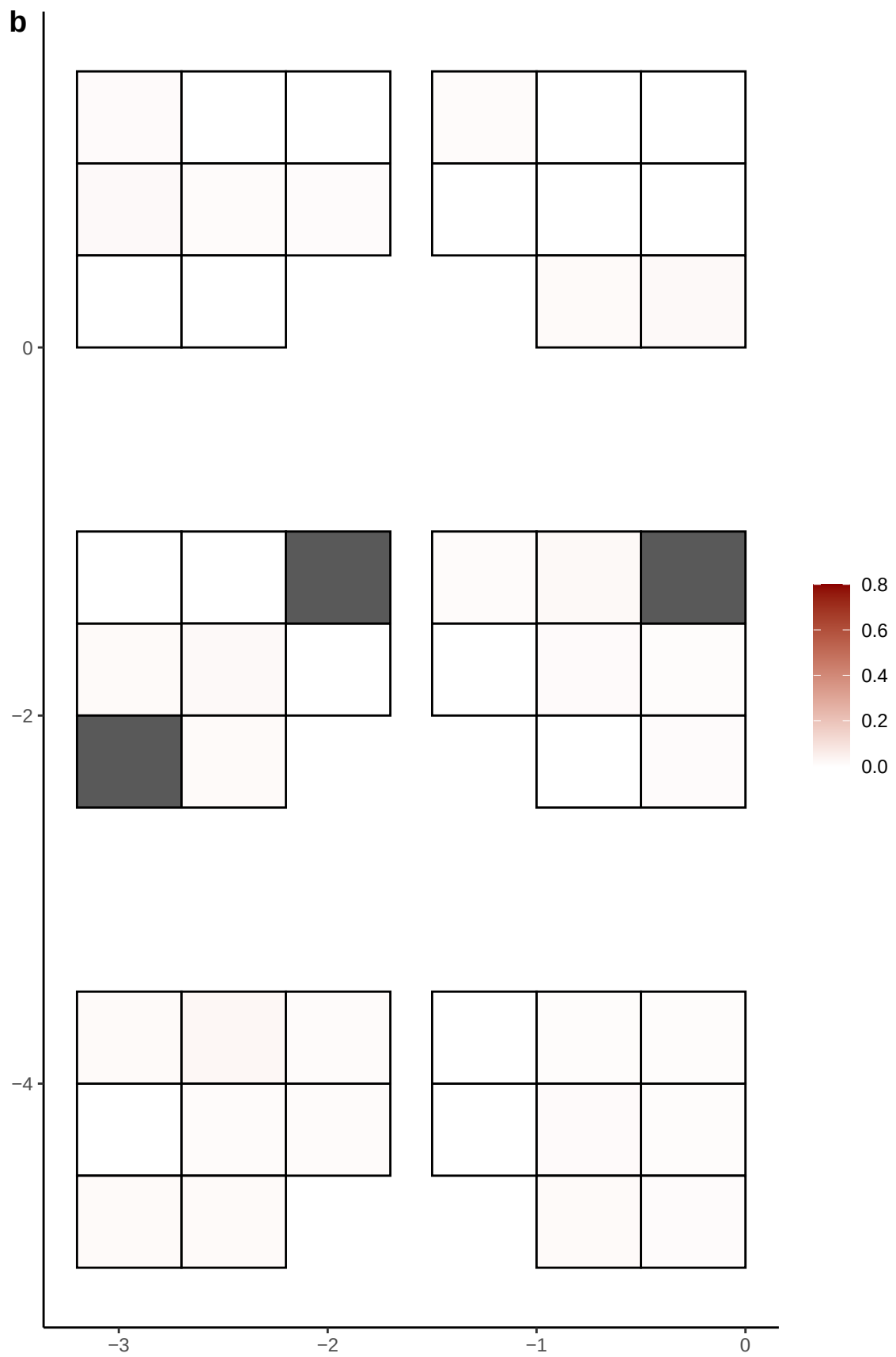

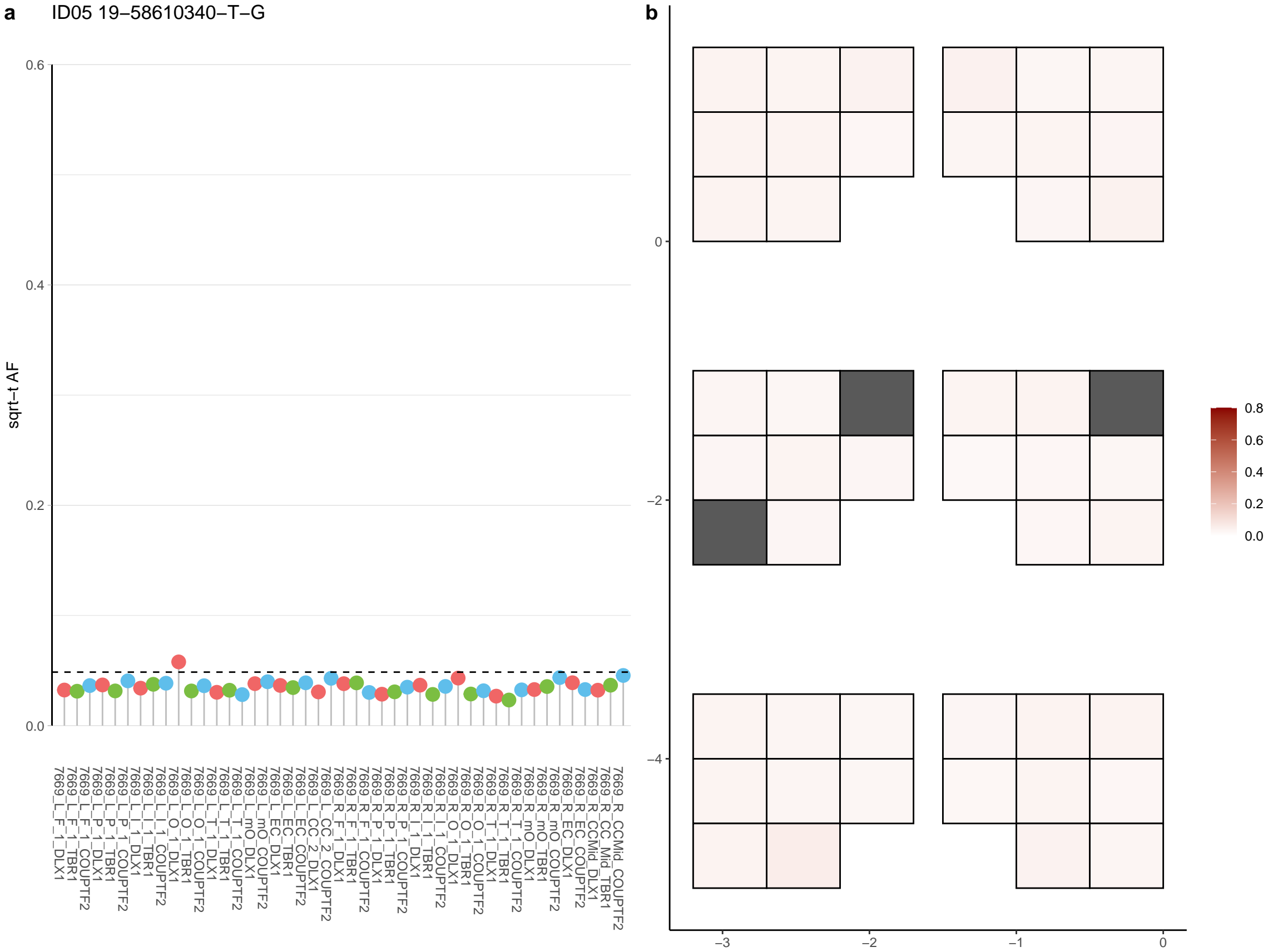

**a** ID05 19-5879037-C-T

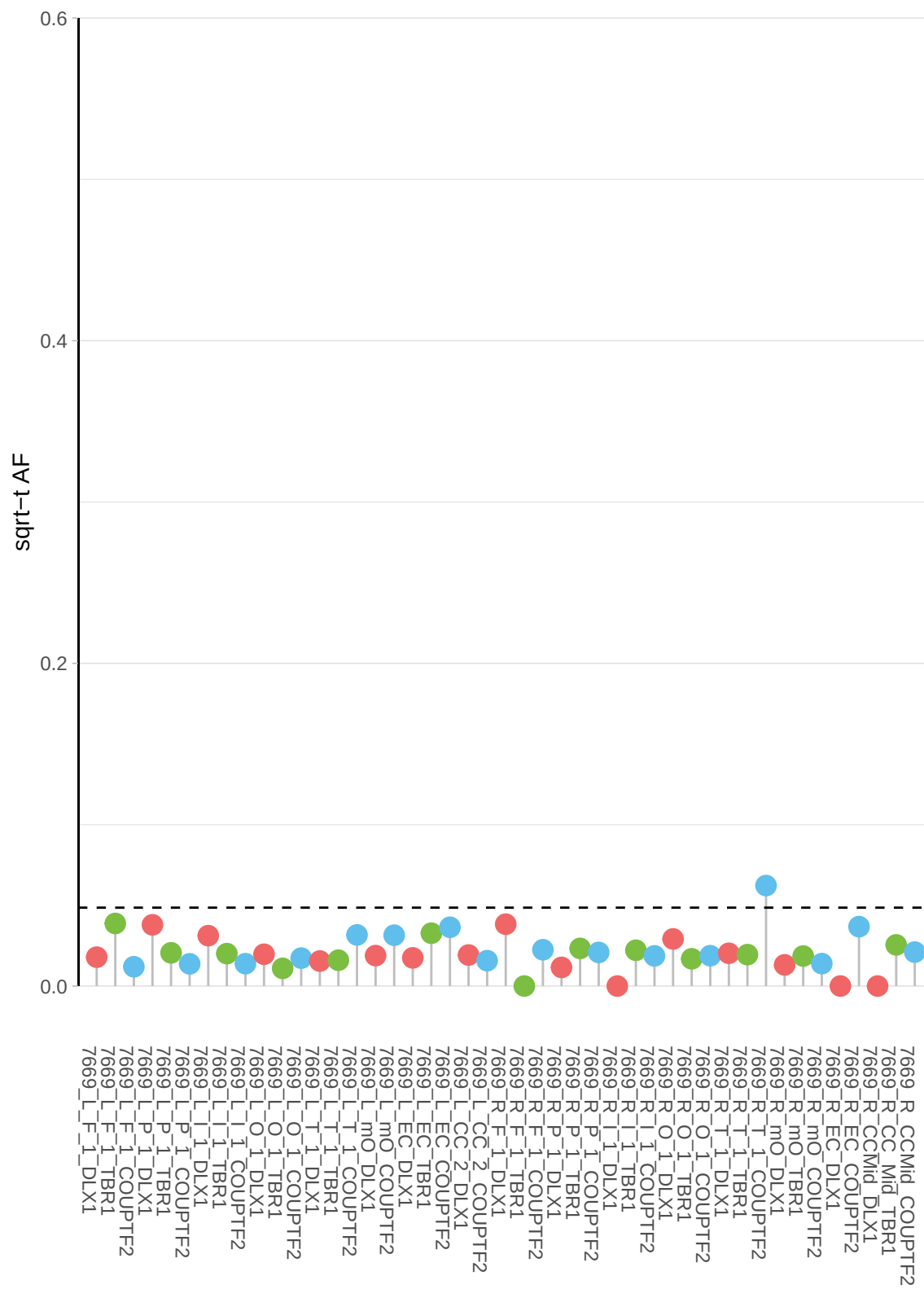

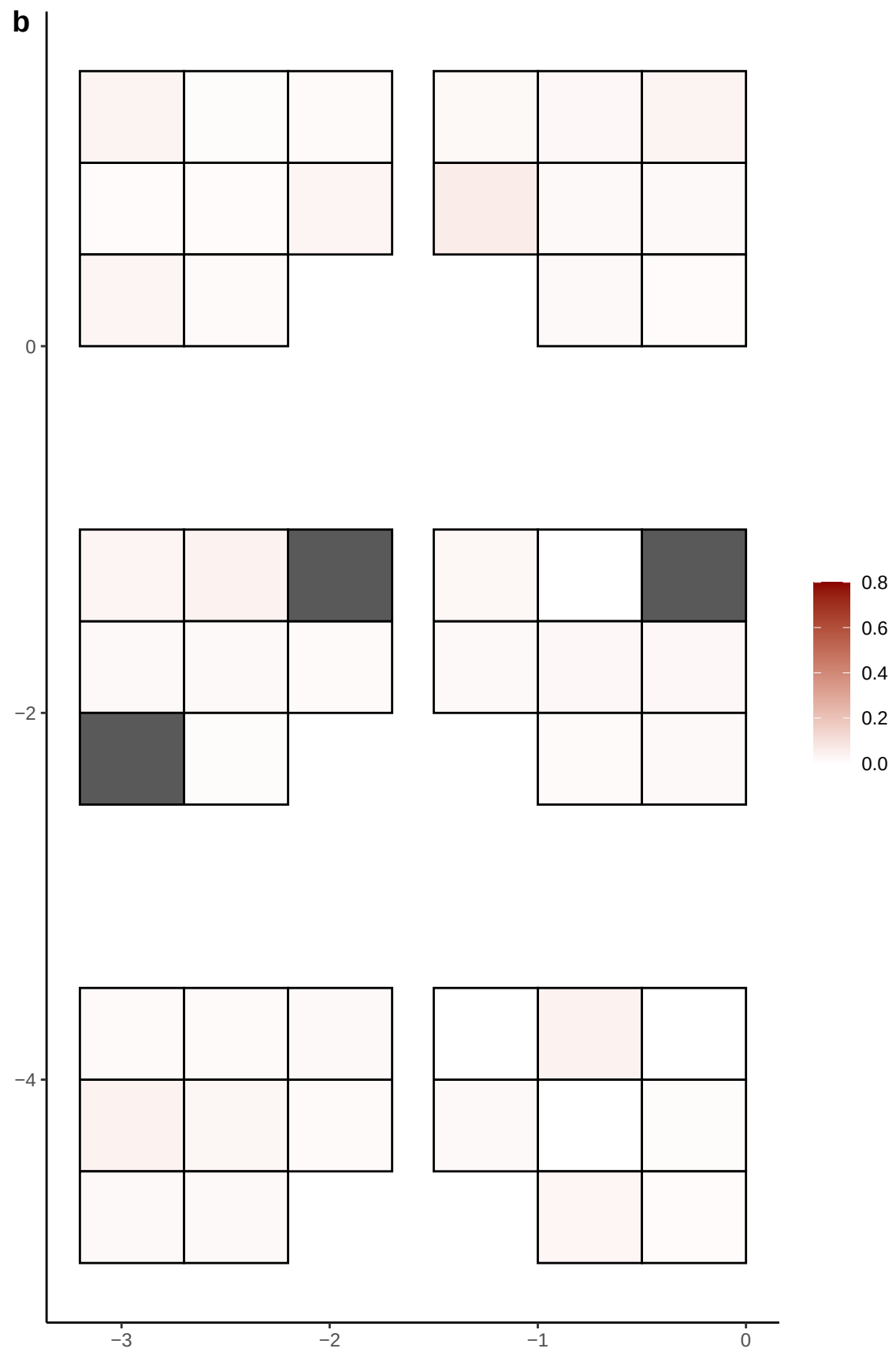

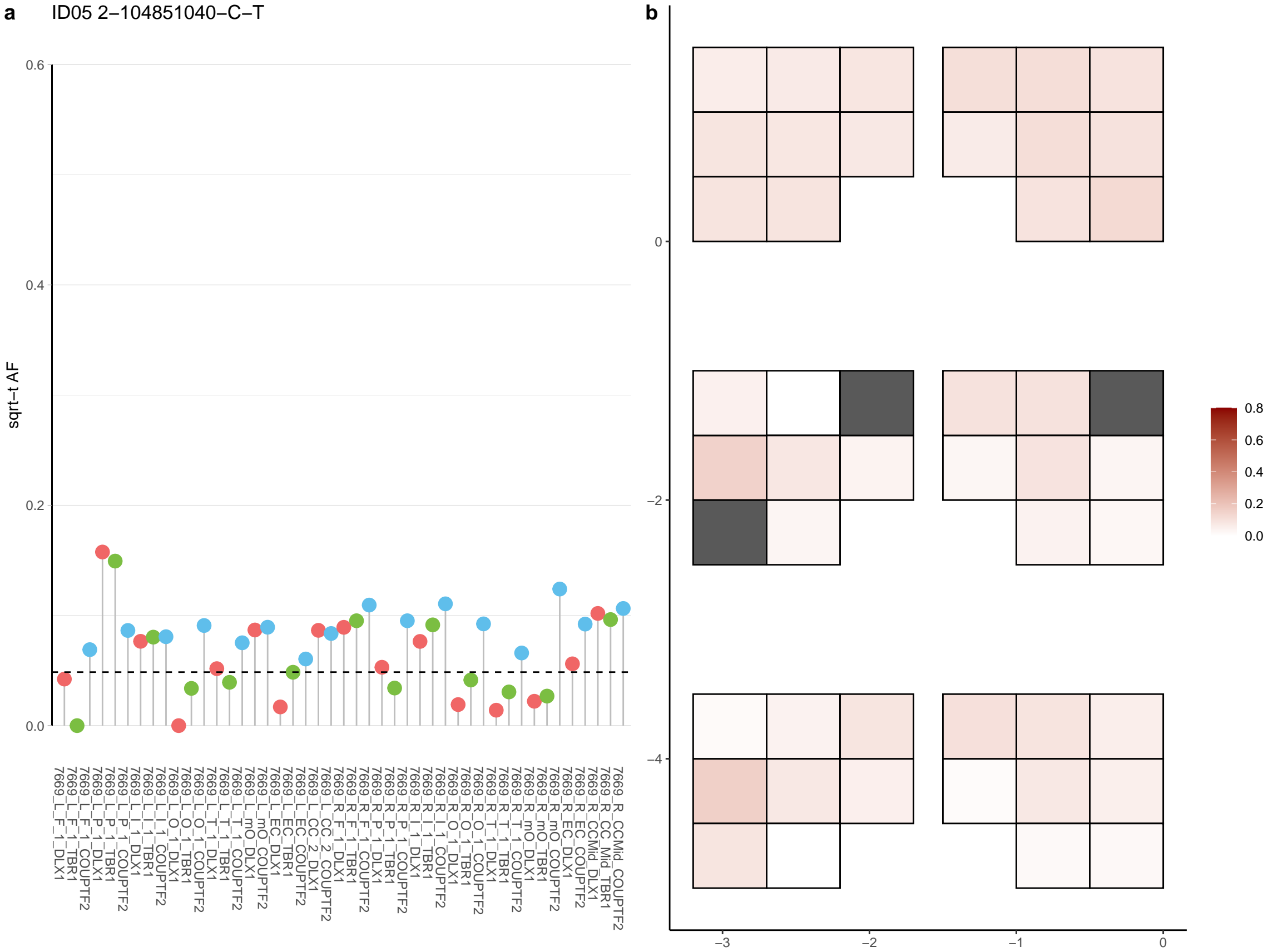

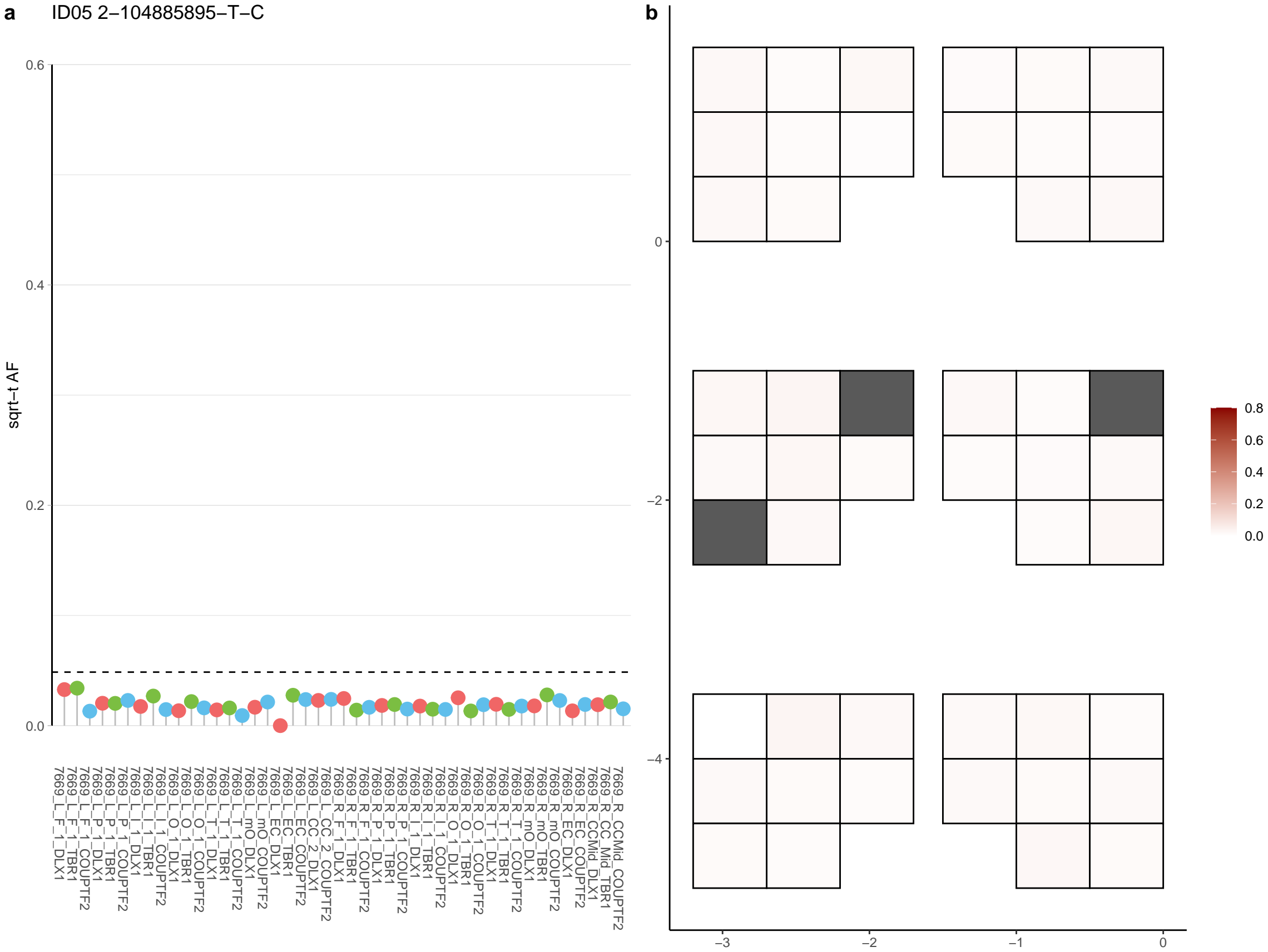

**a** ID05 2-105083647-G-A

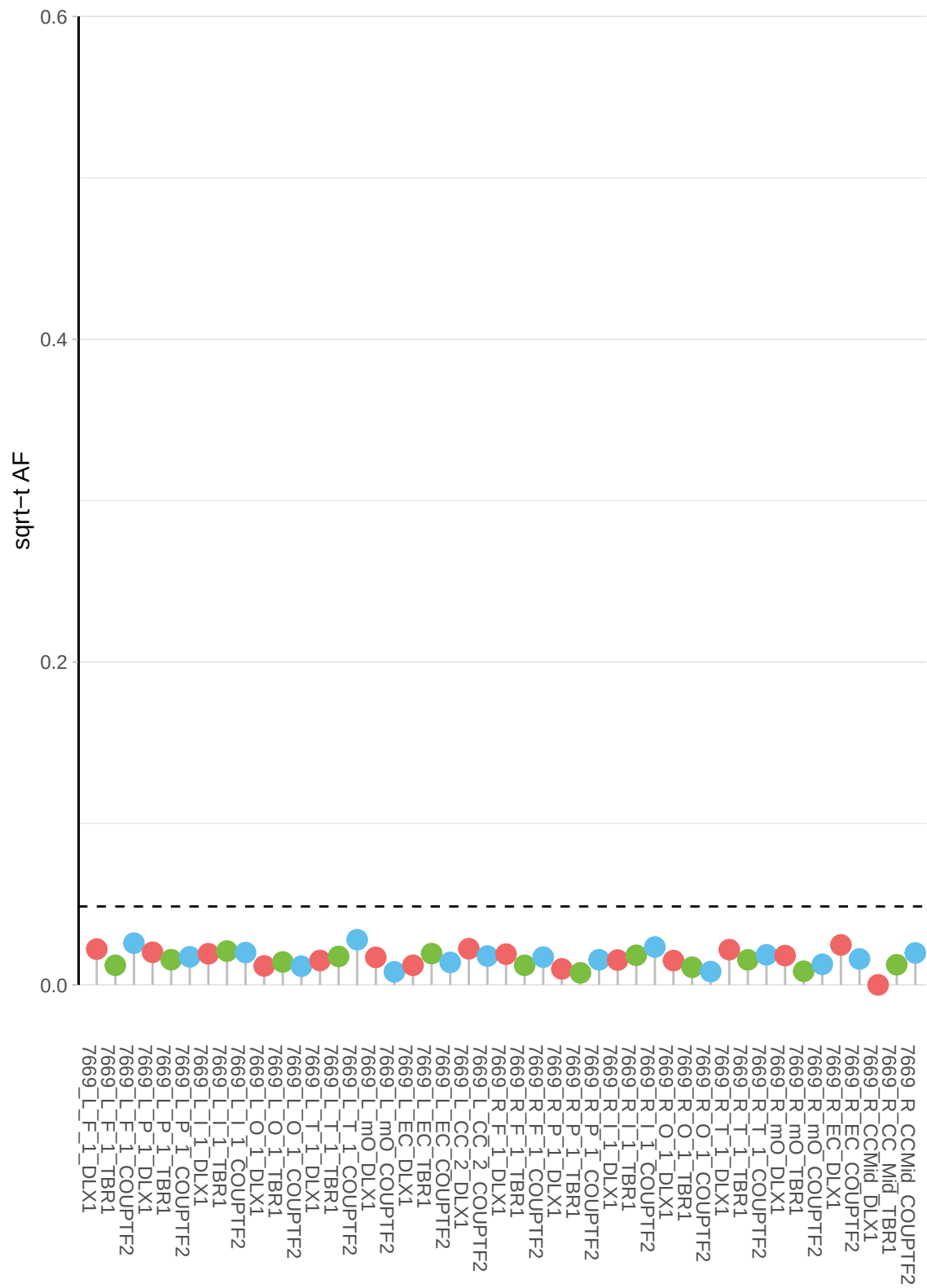

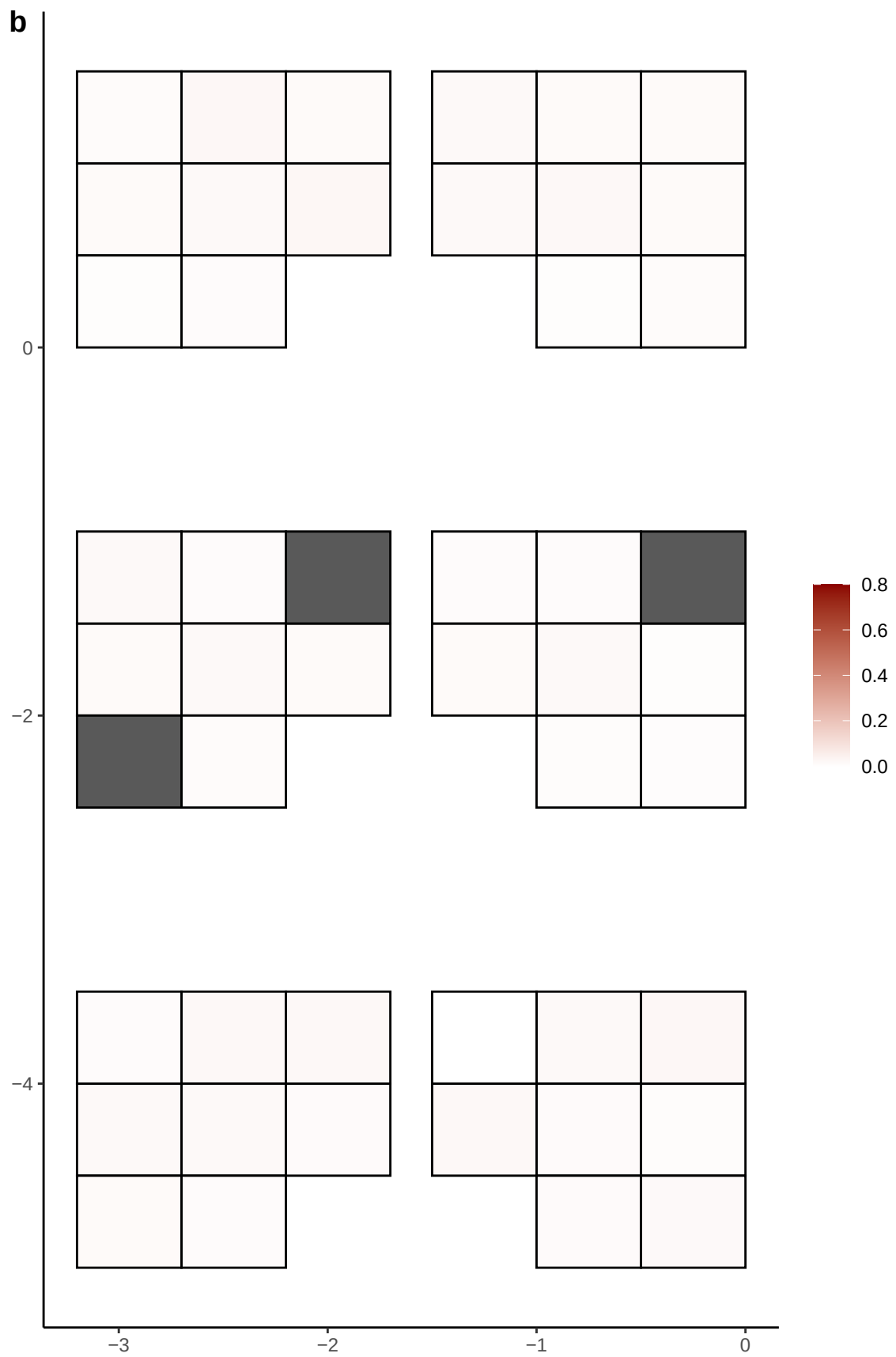

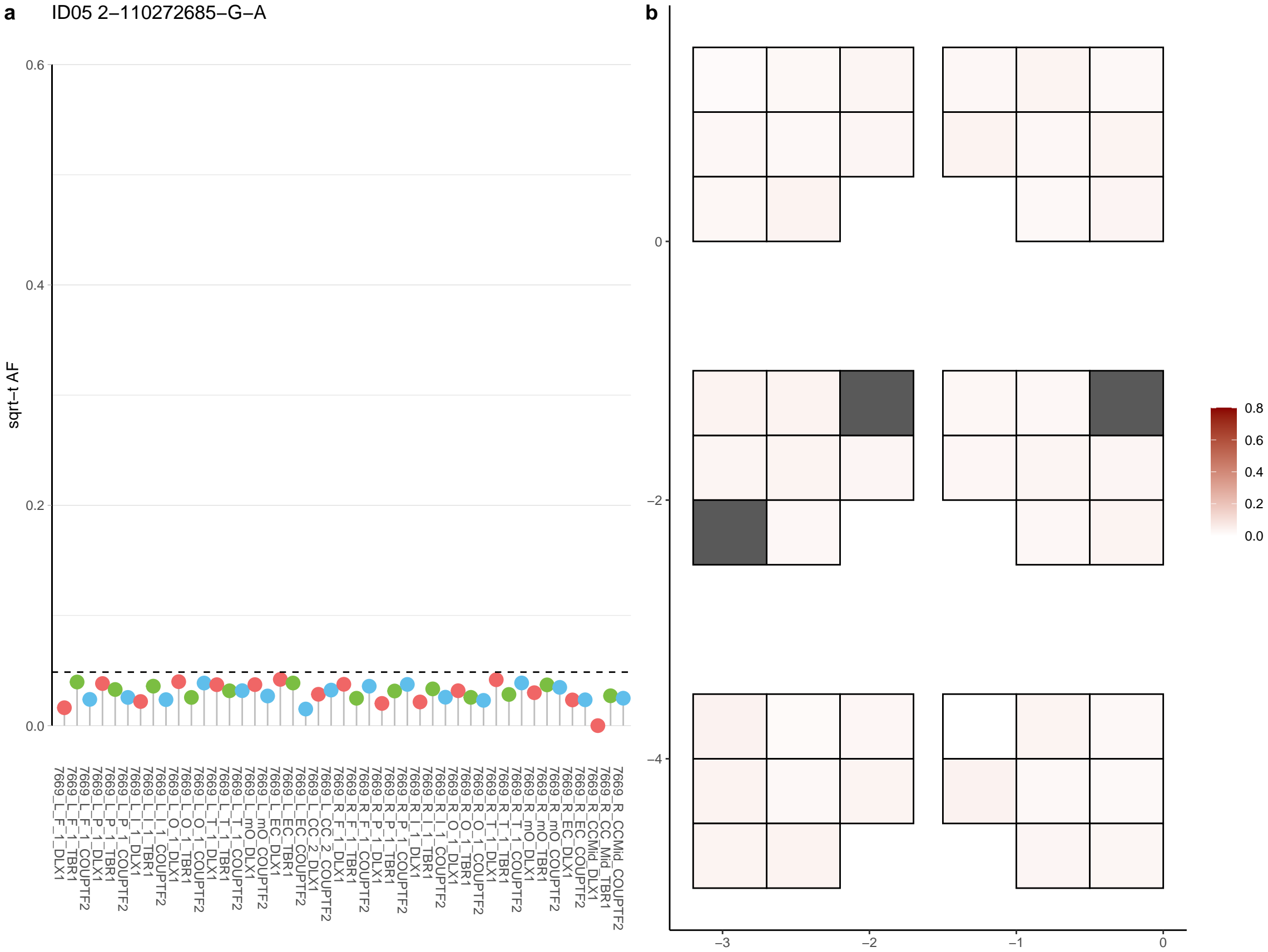

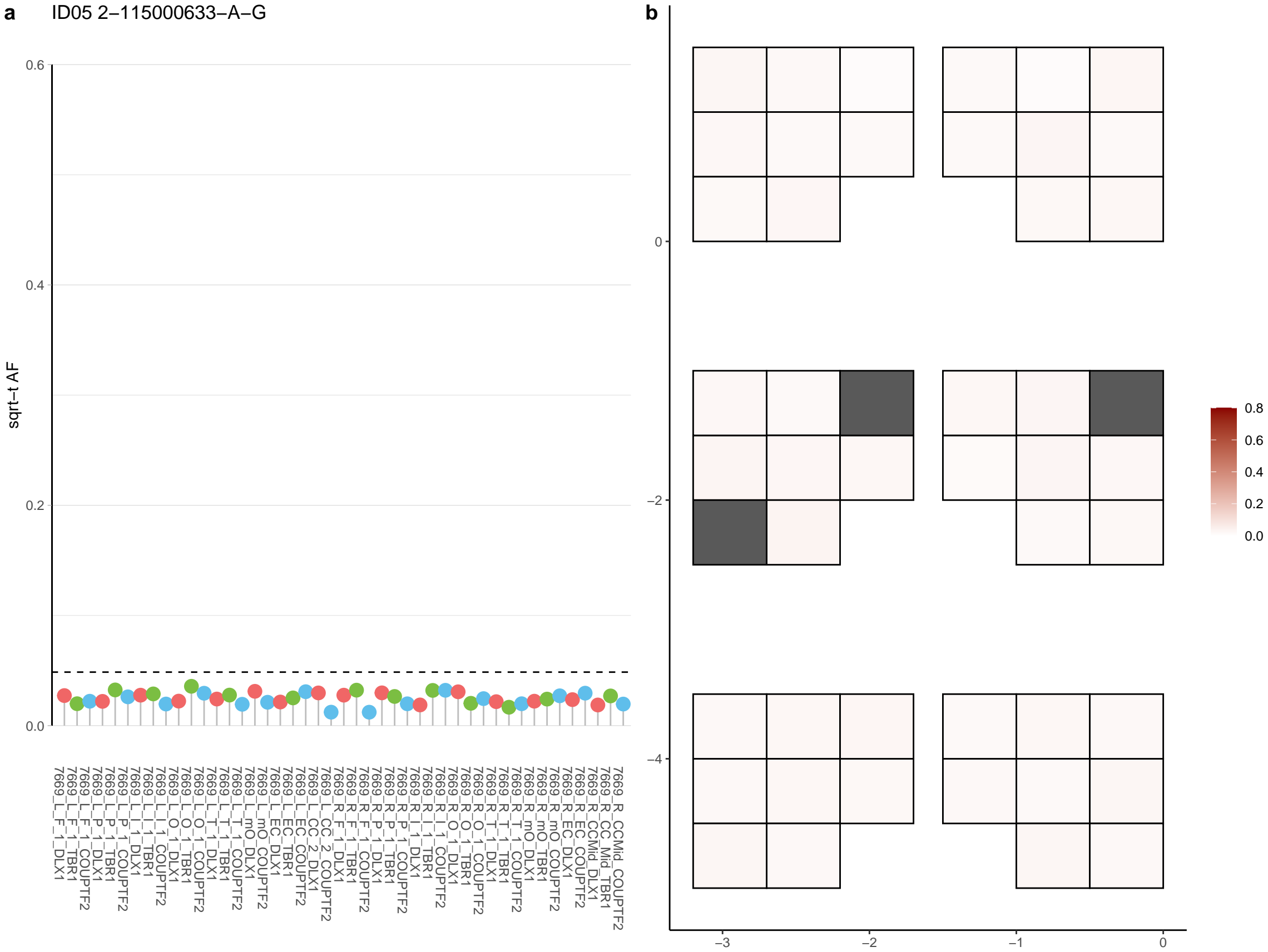

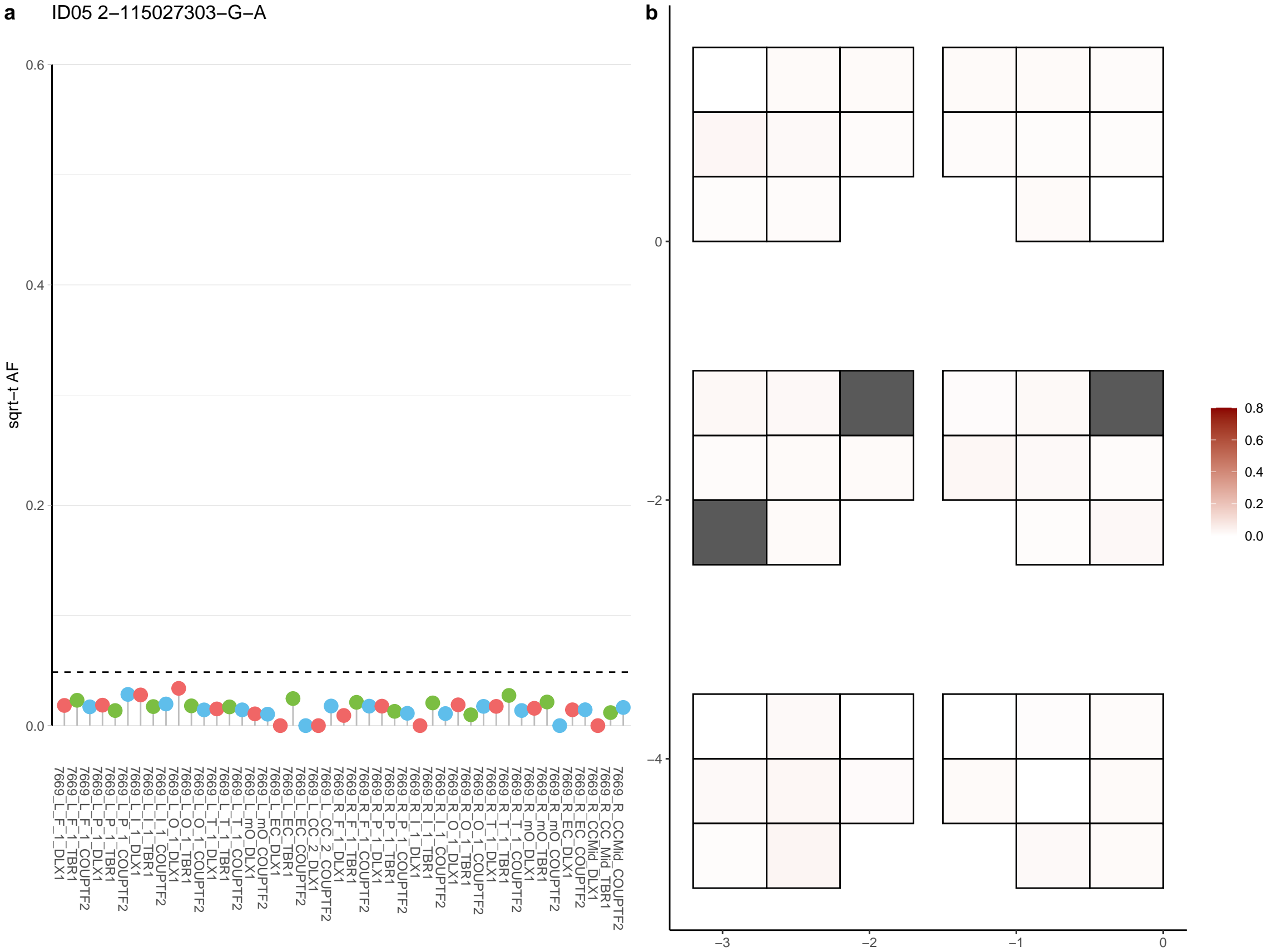

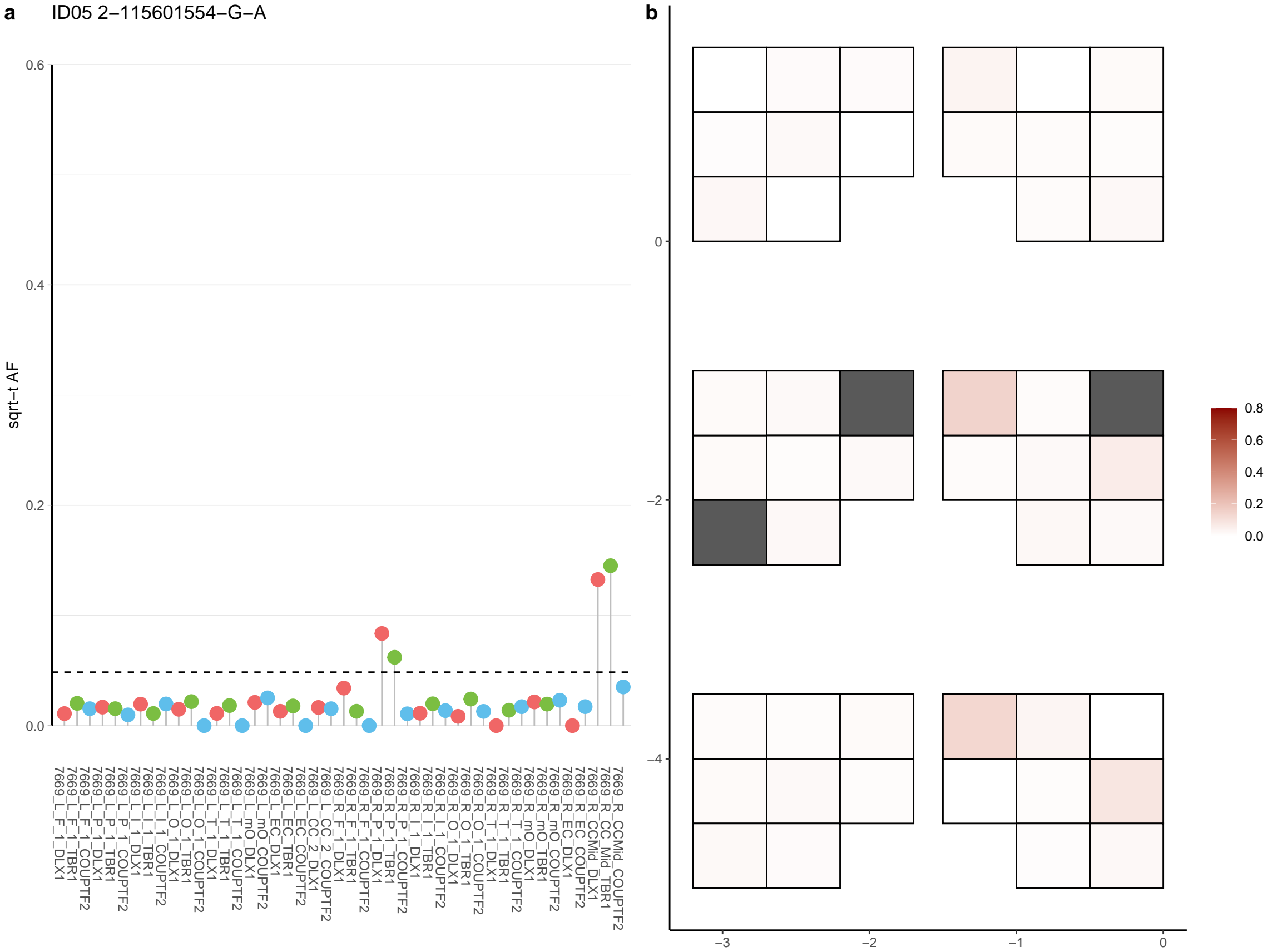

**a** ID05 2-115708162-C-T

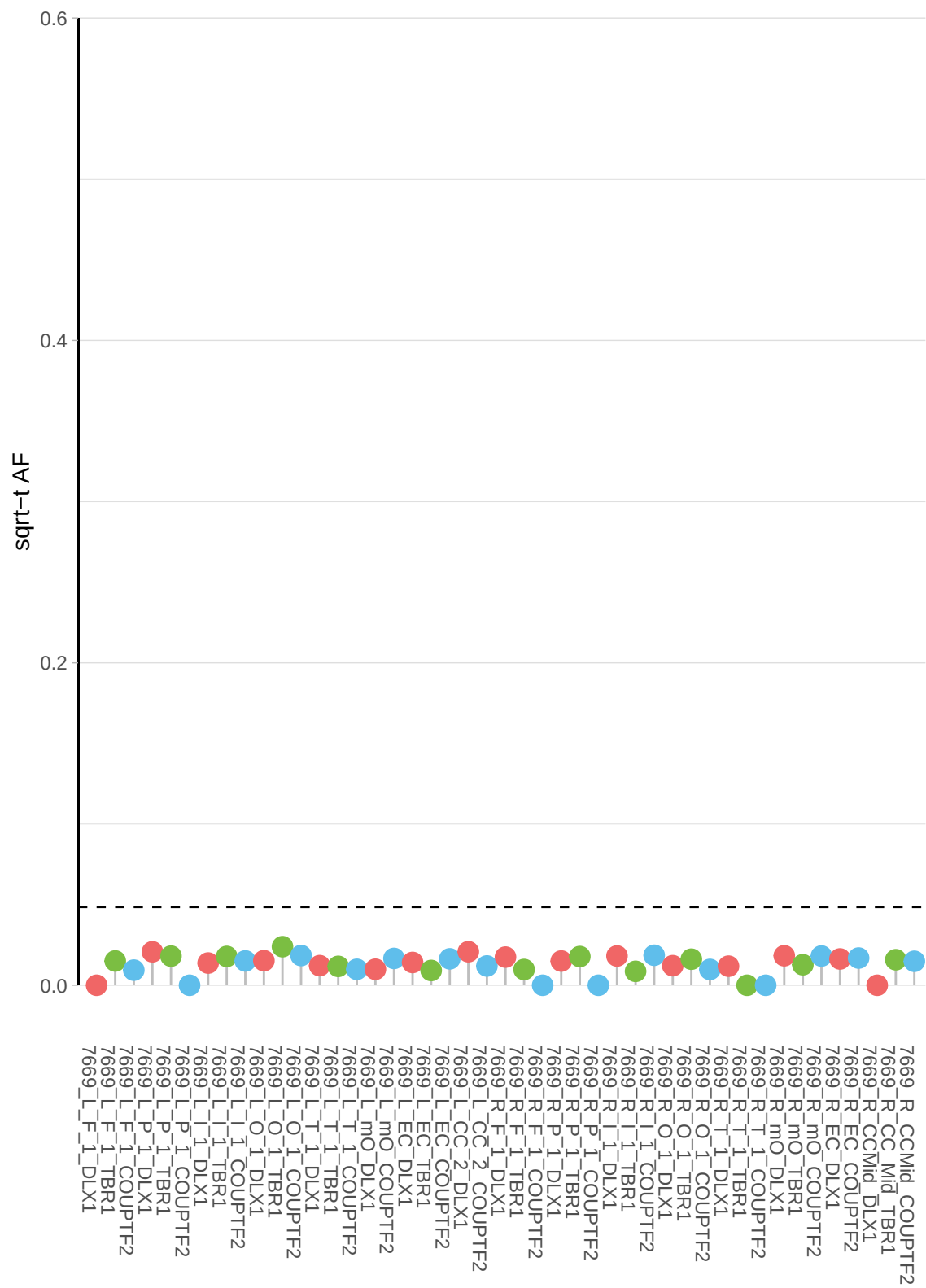

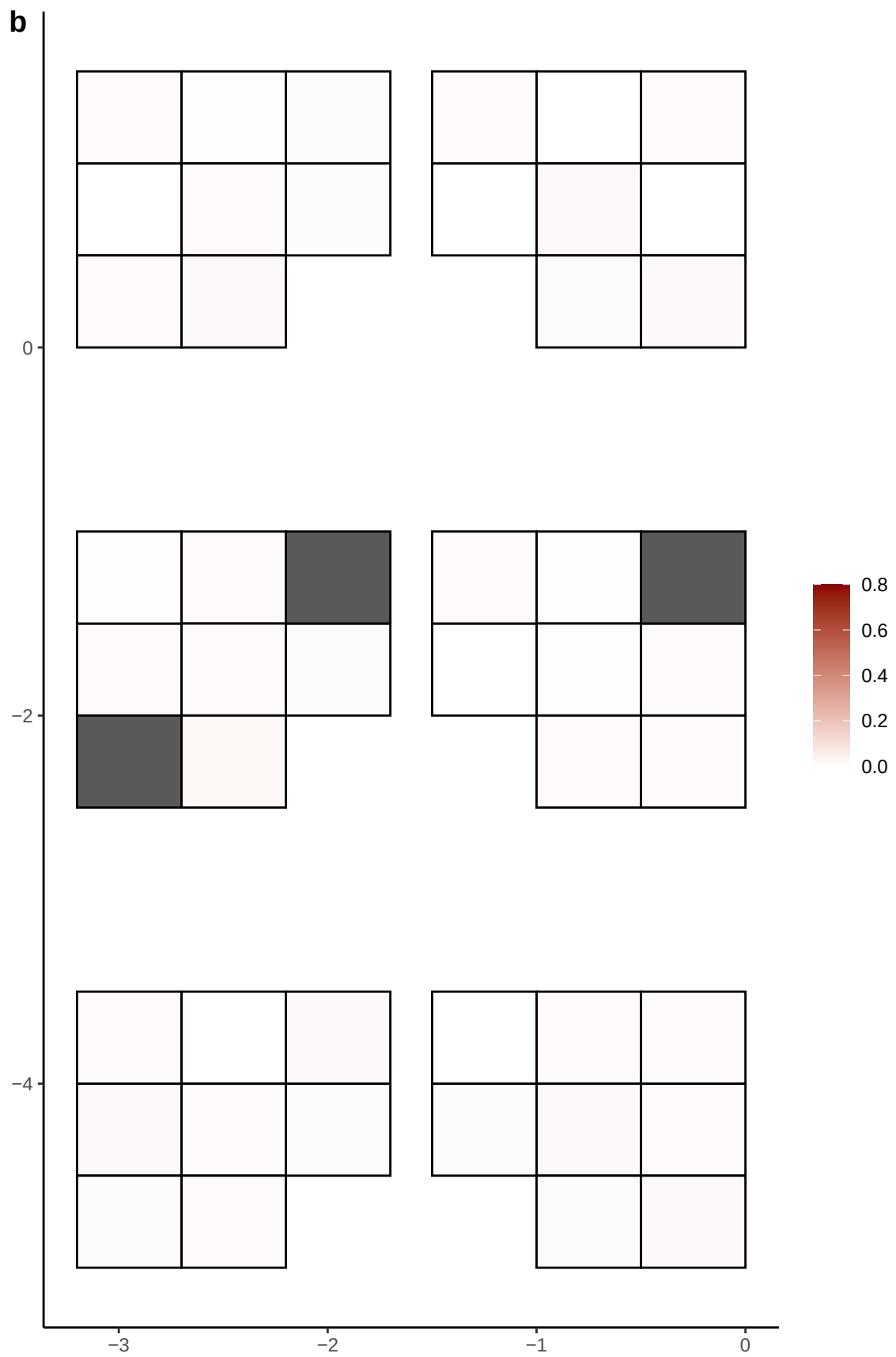

**a** ID05 2-116395374-G-T

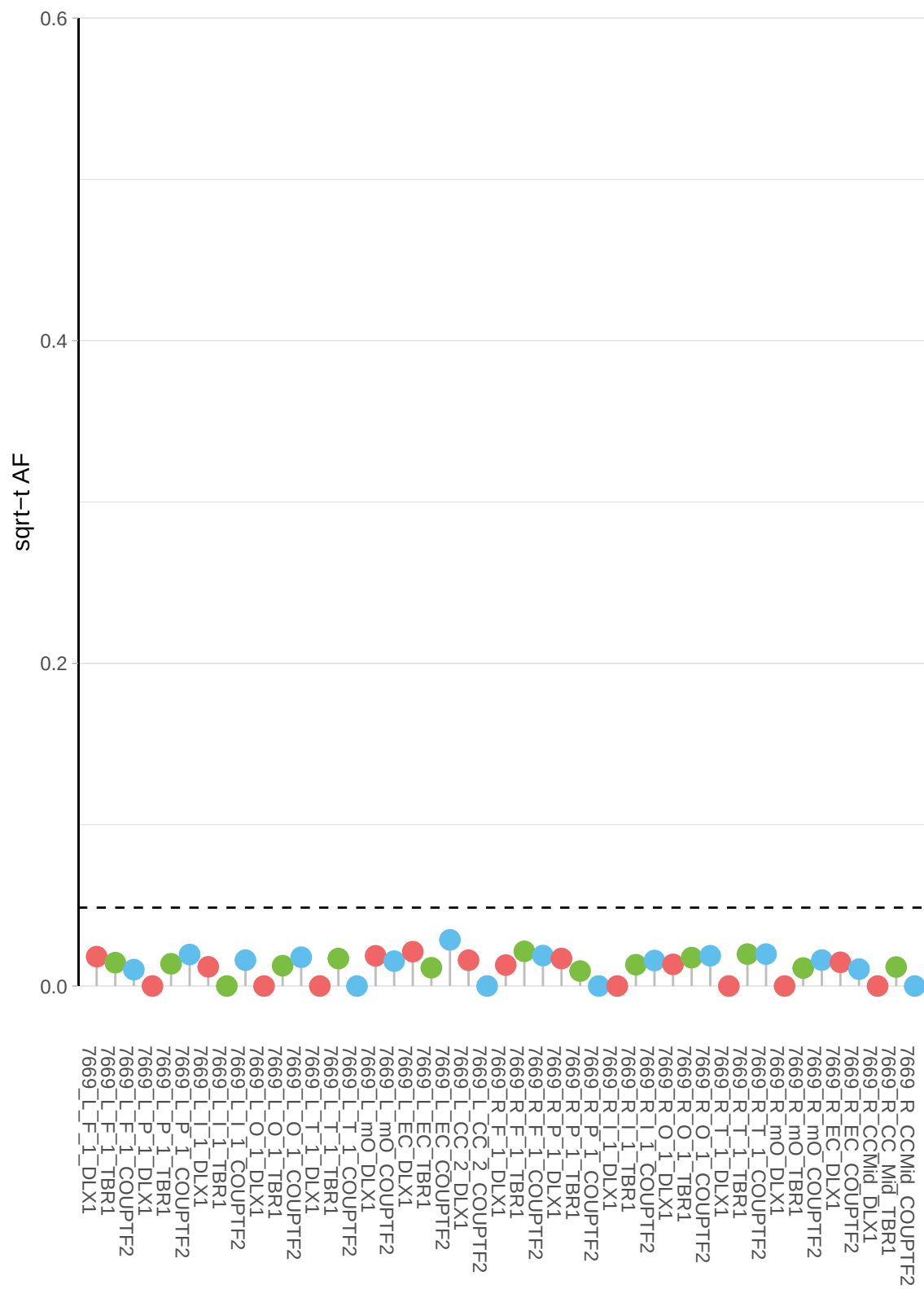

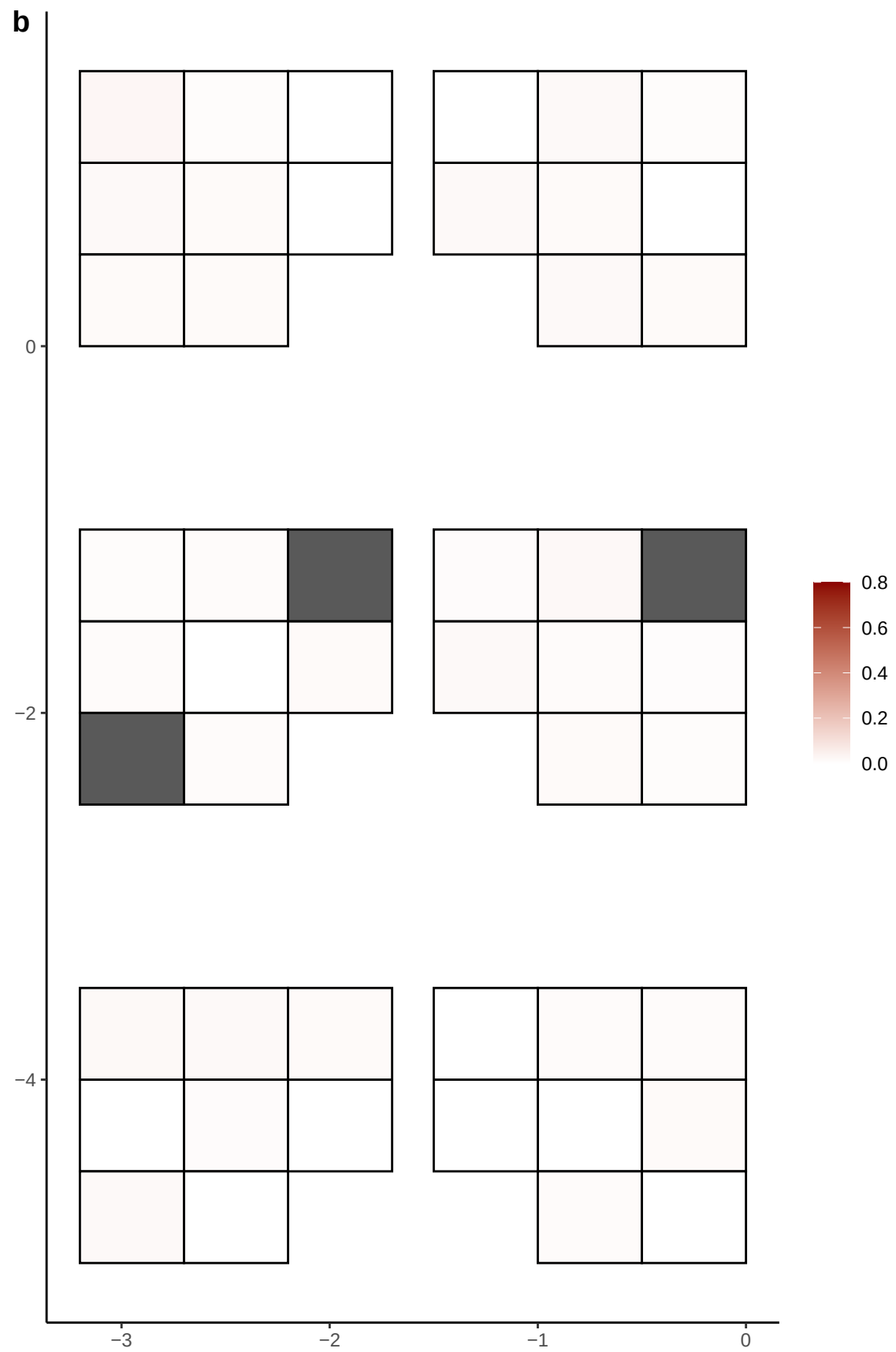

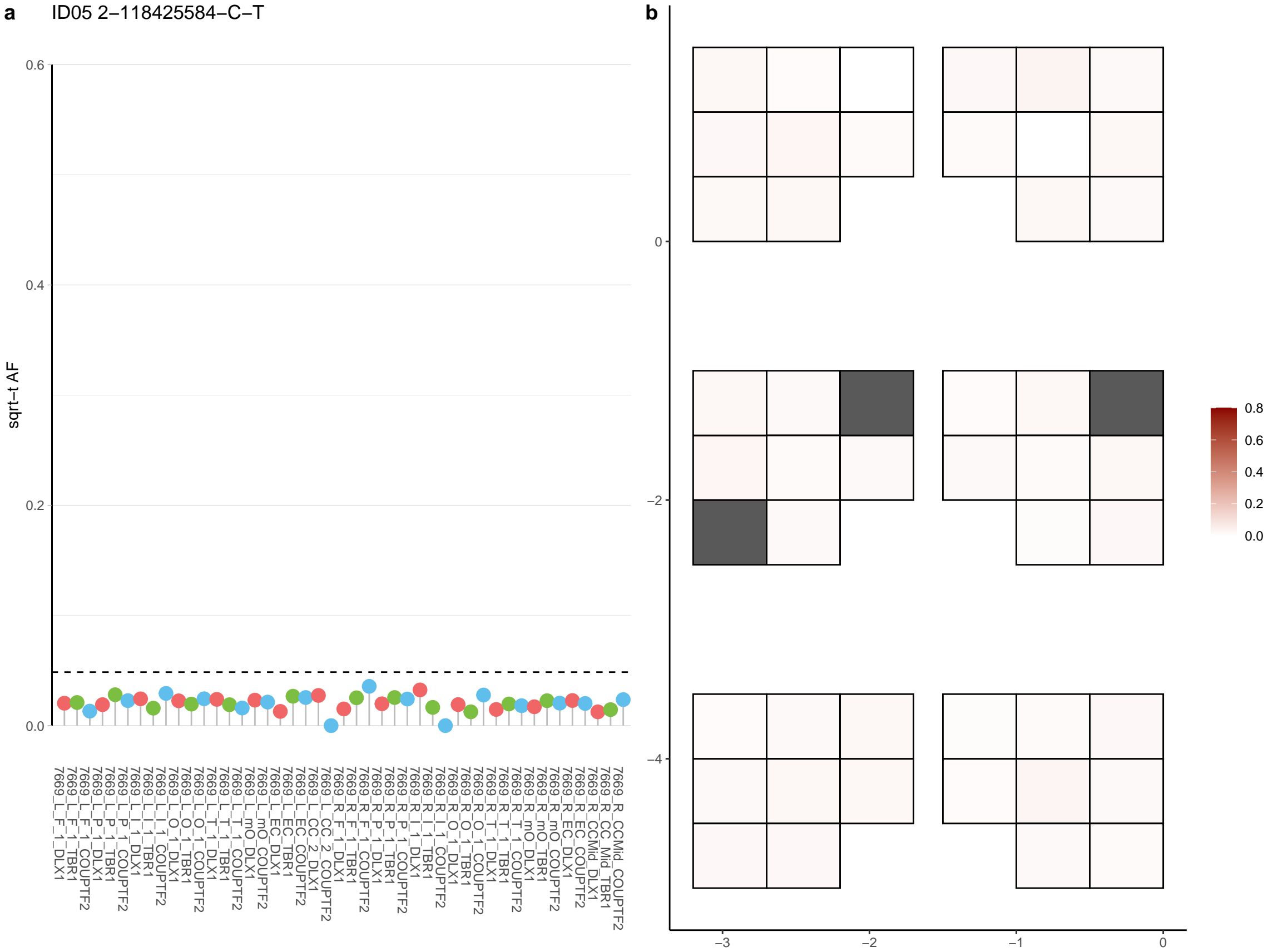

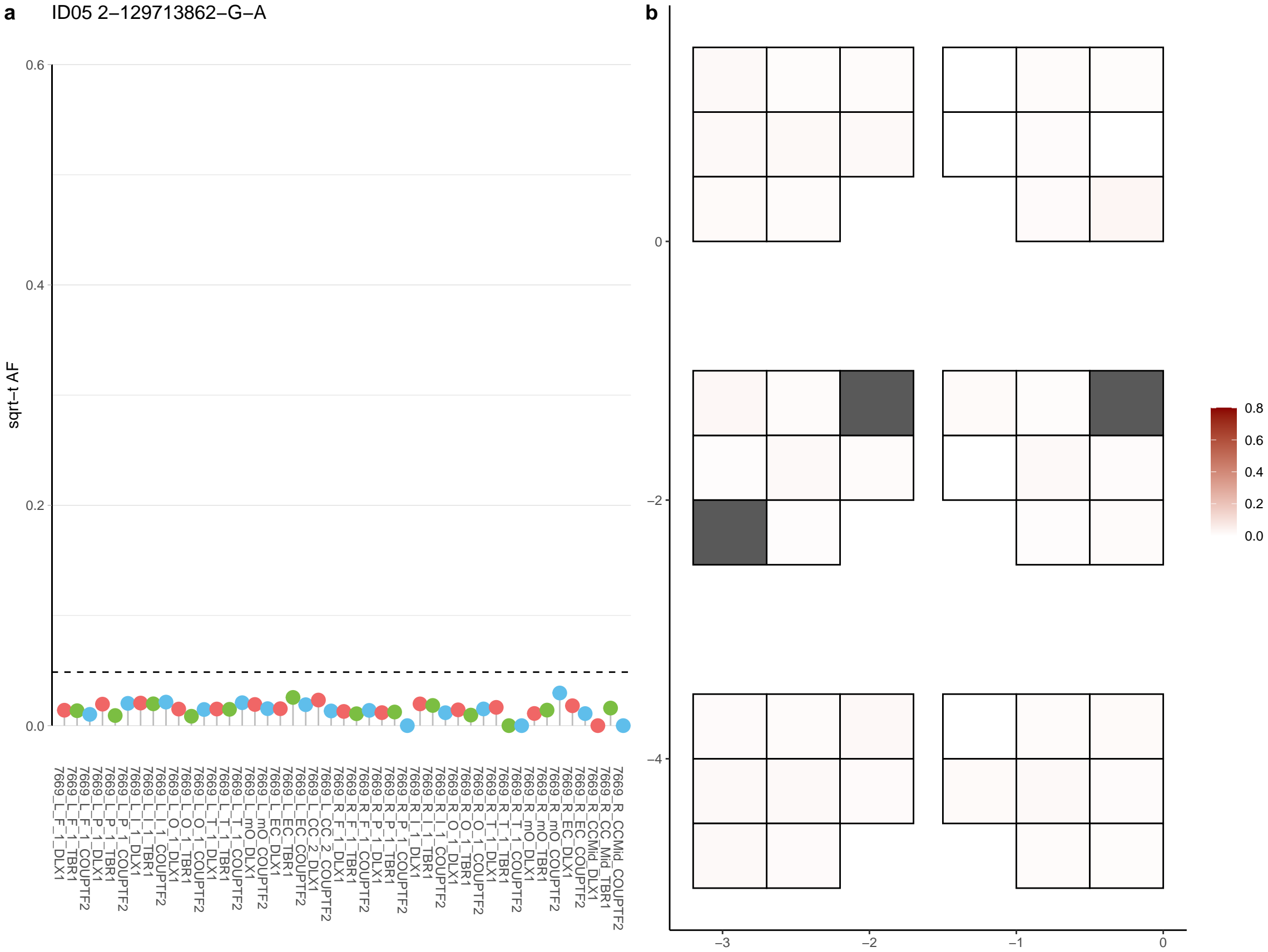

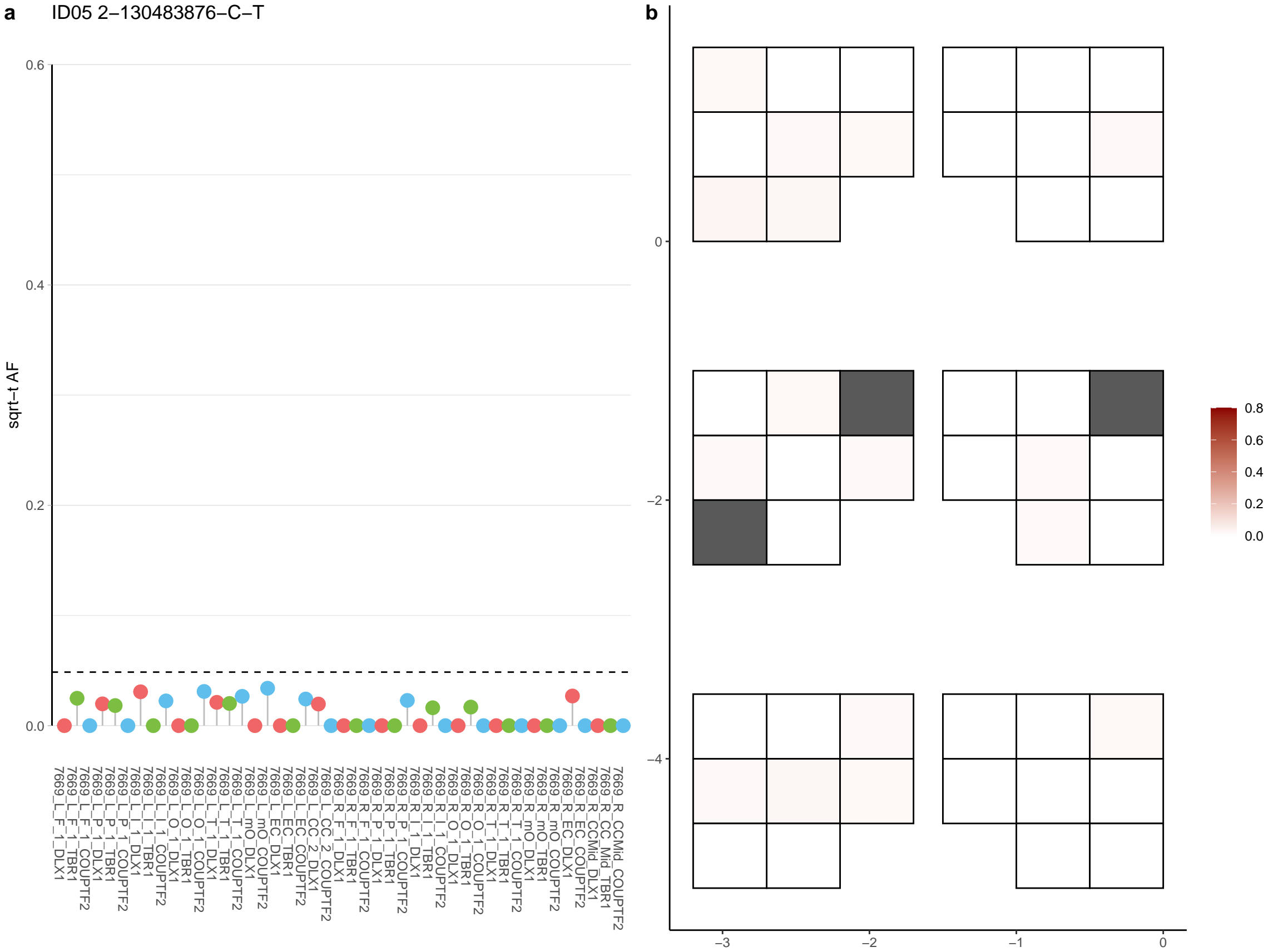

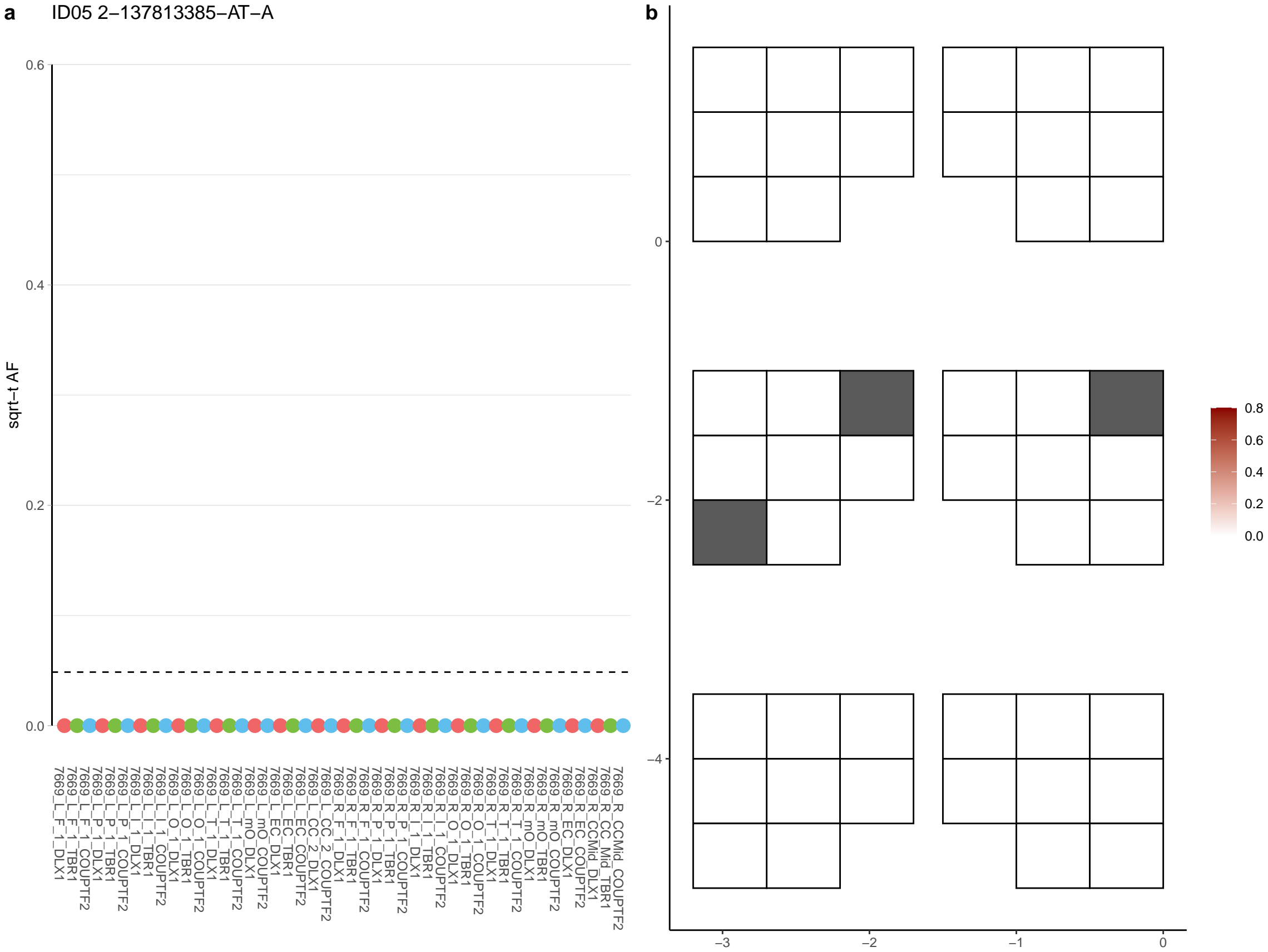

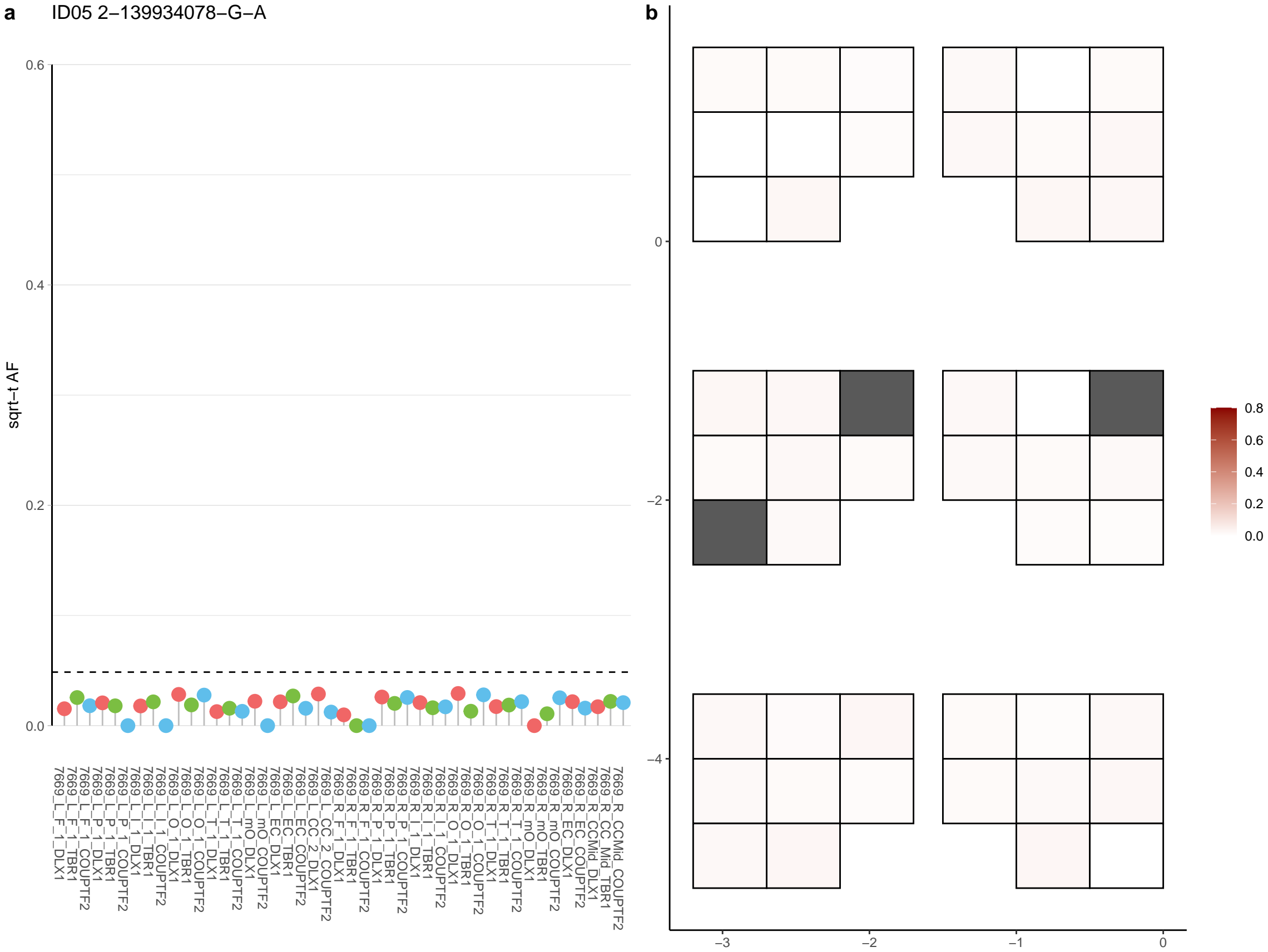

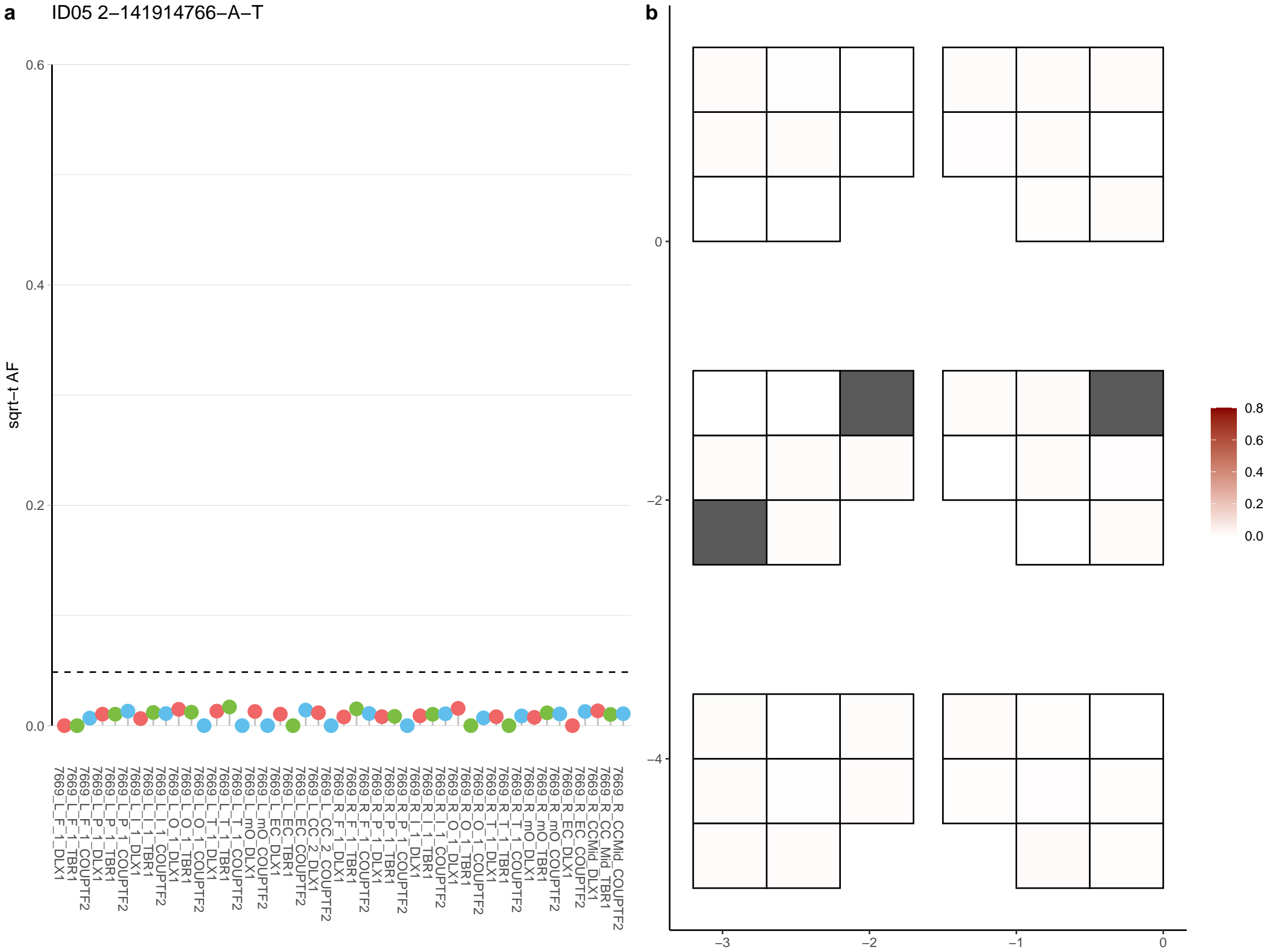

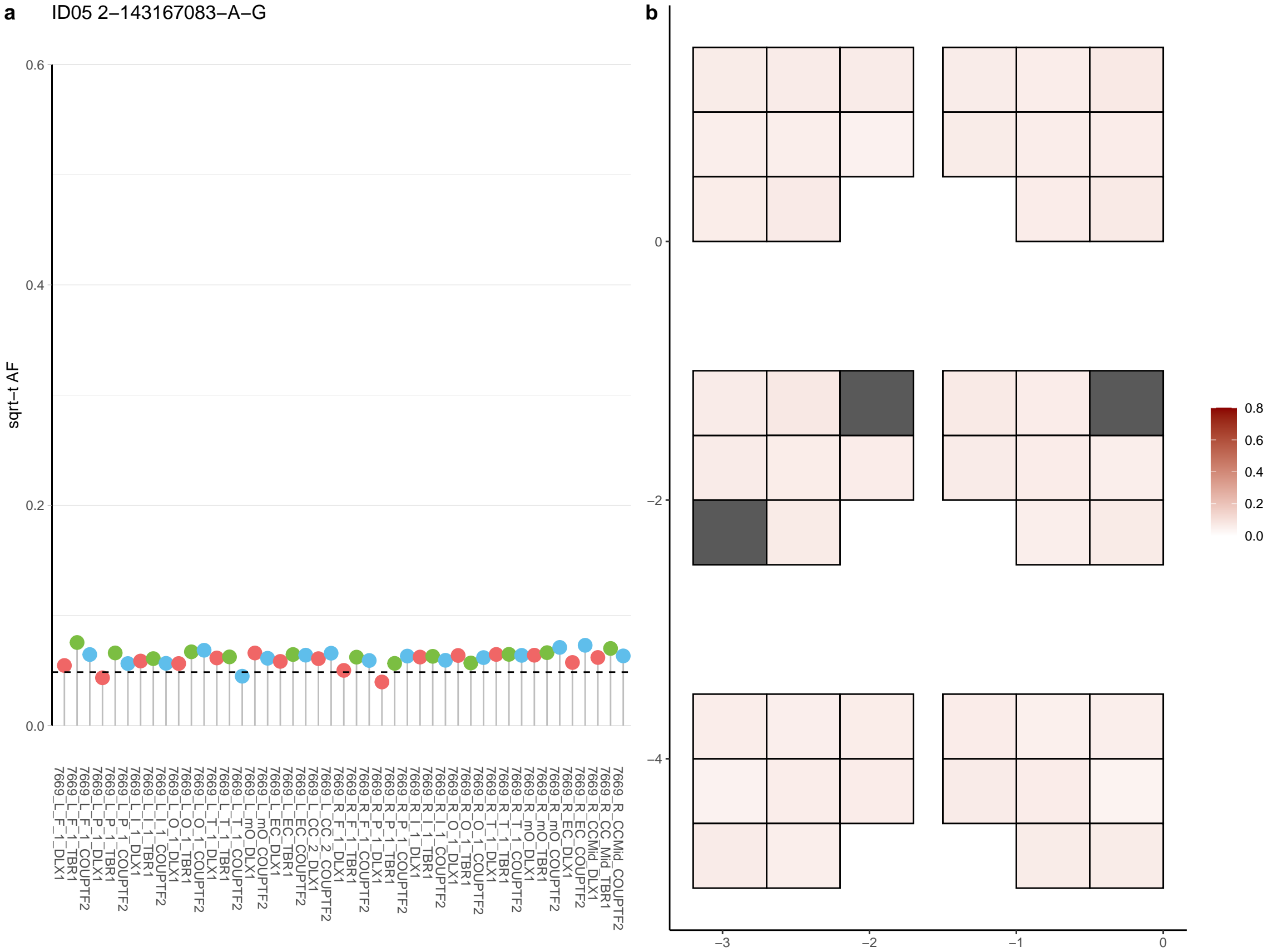

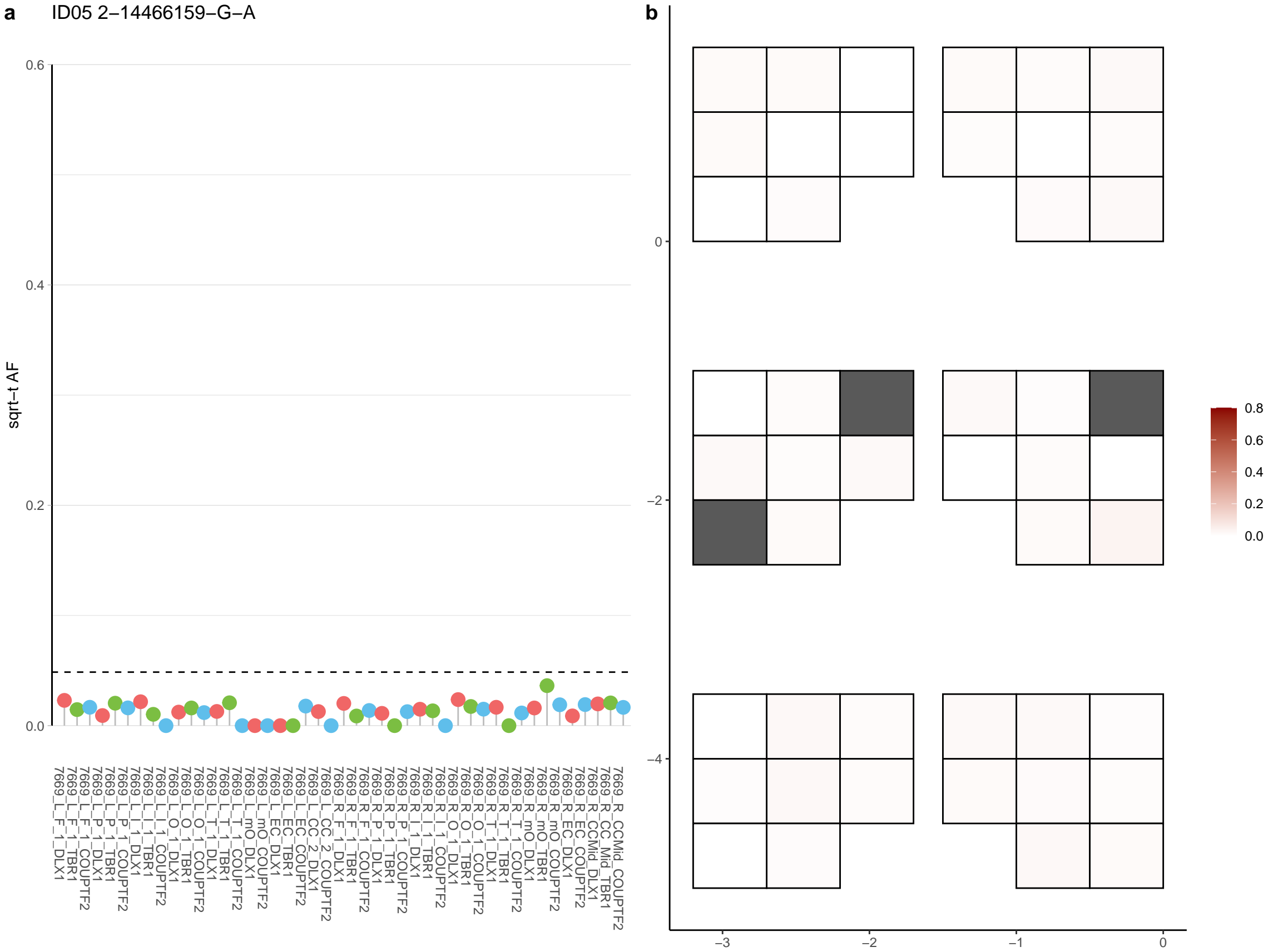

**a** ID05 2-159760148-C-T

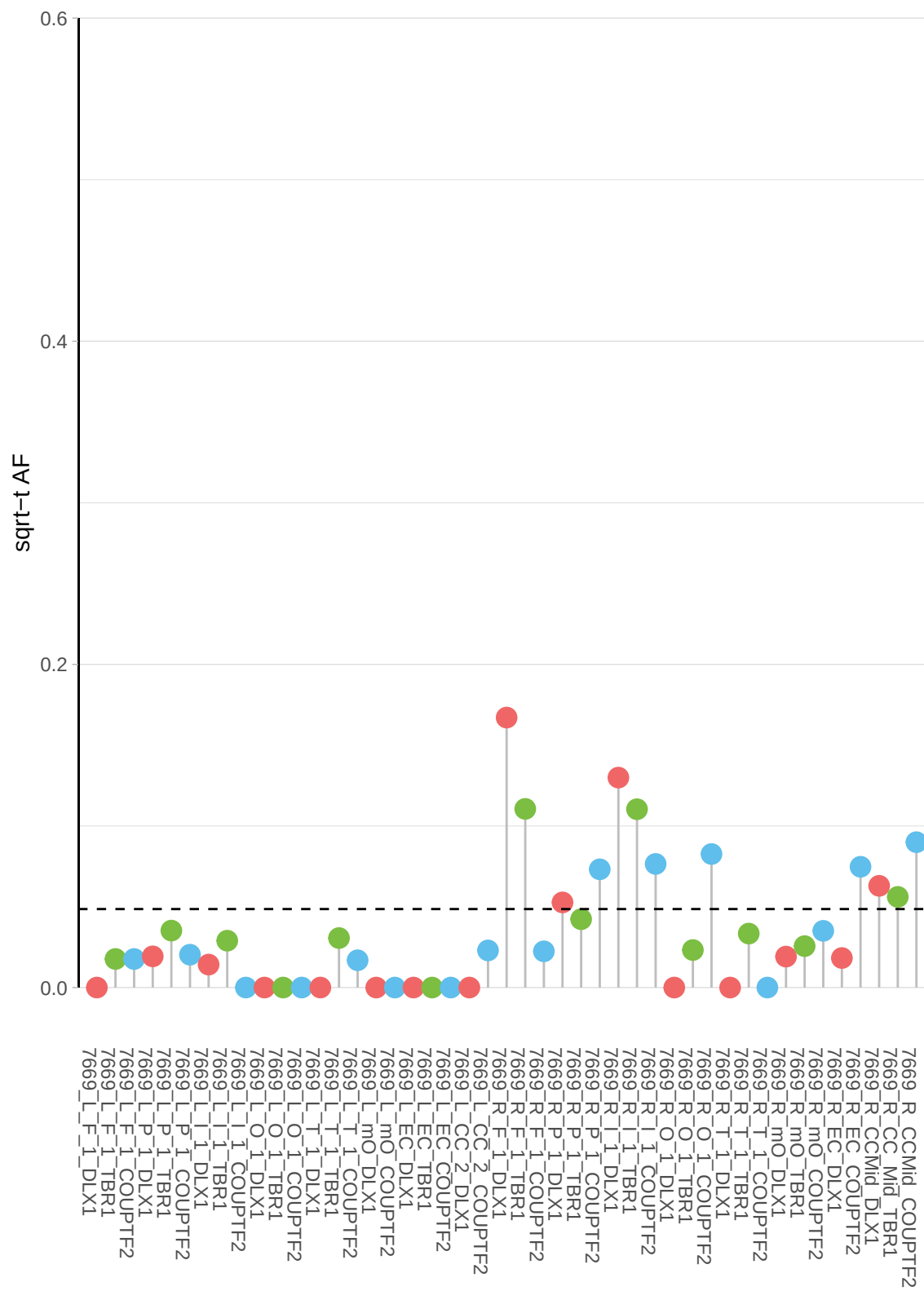

**b**

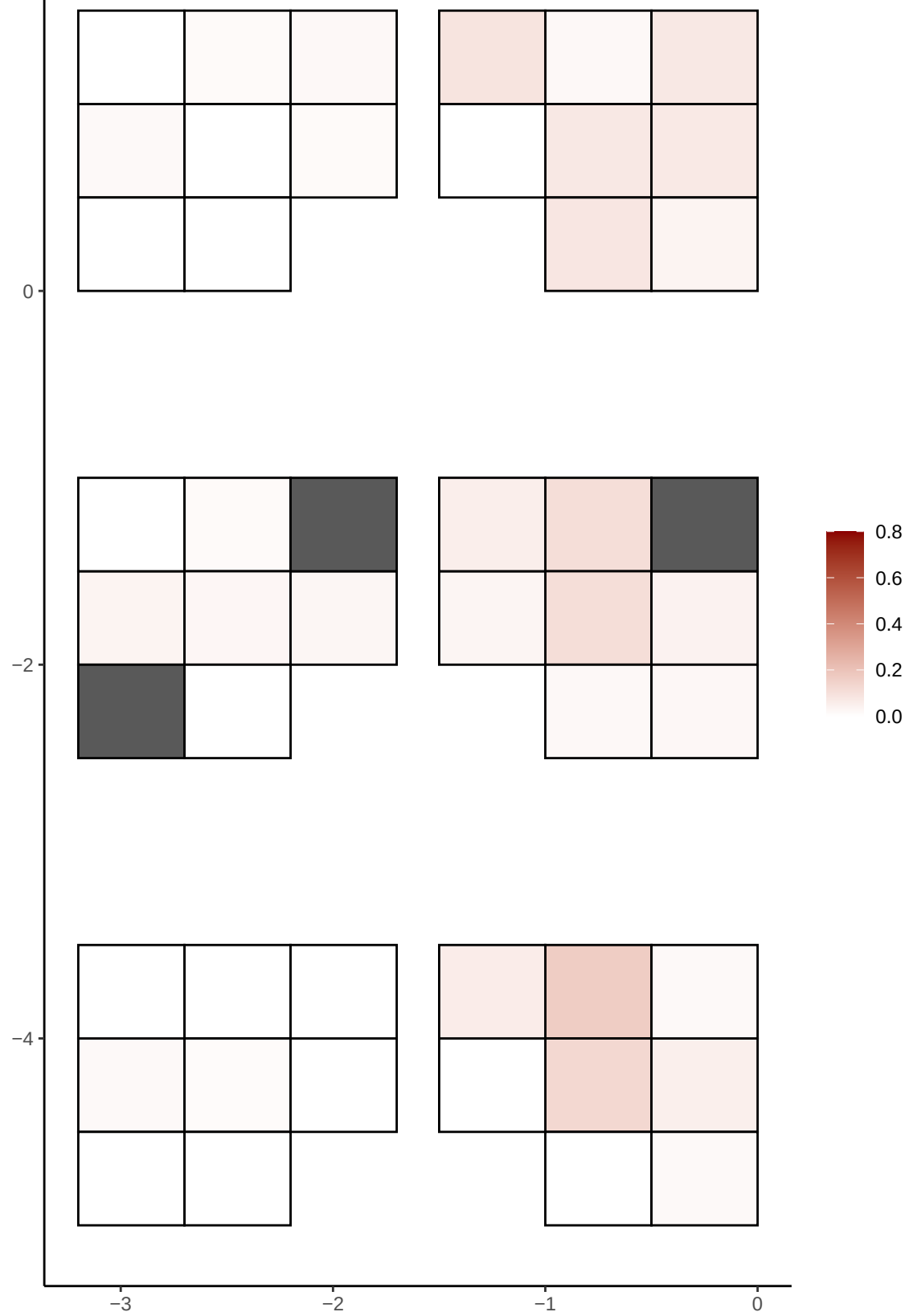

**a** ID05 2-163073318-G-A

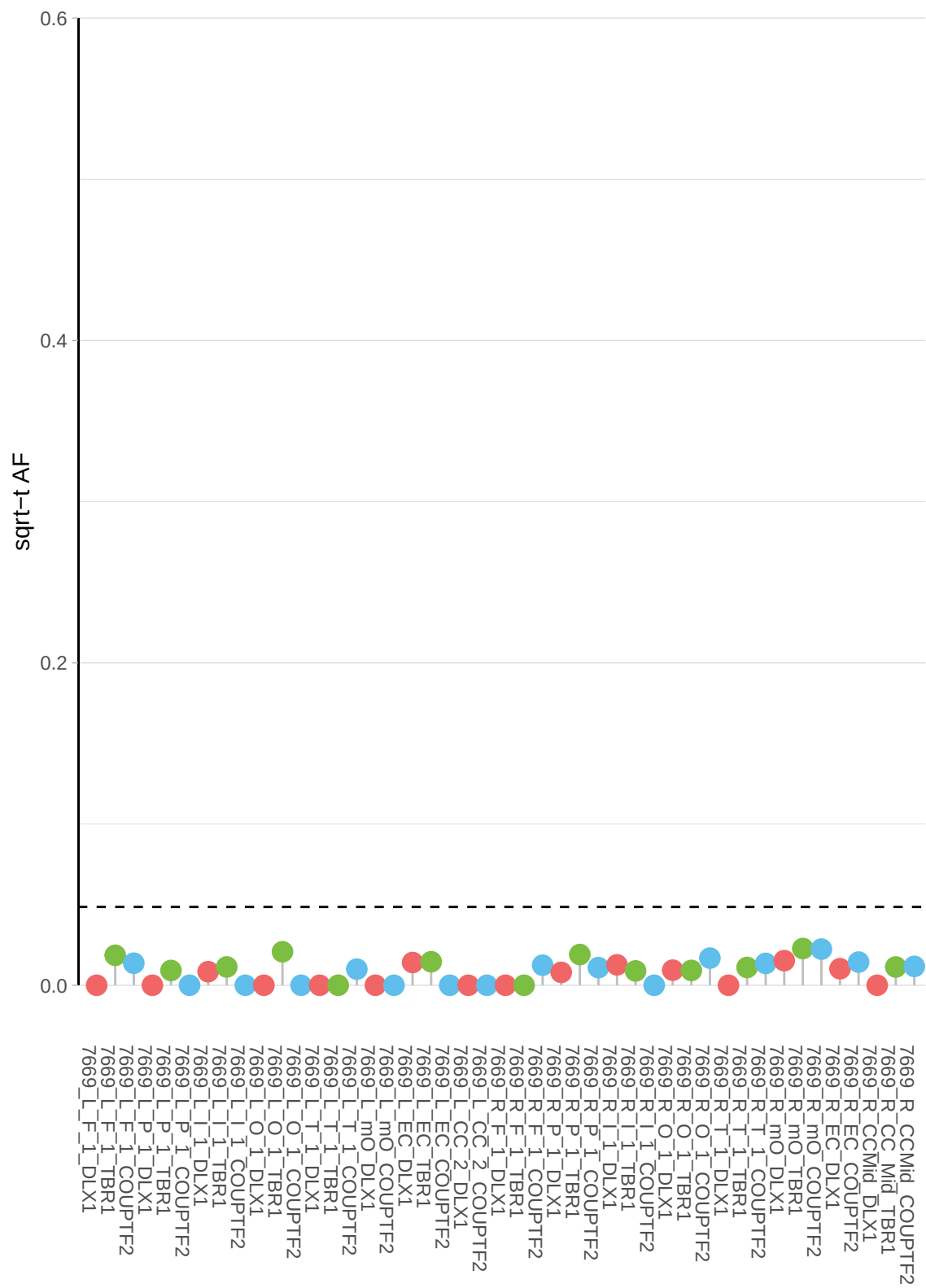

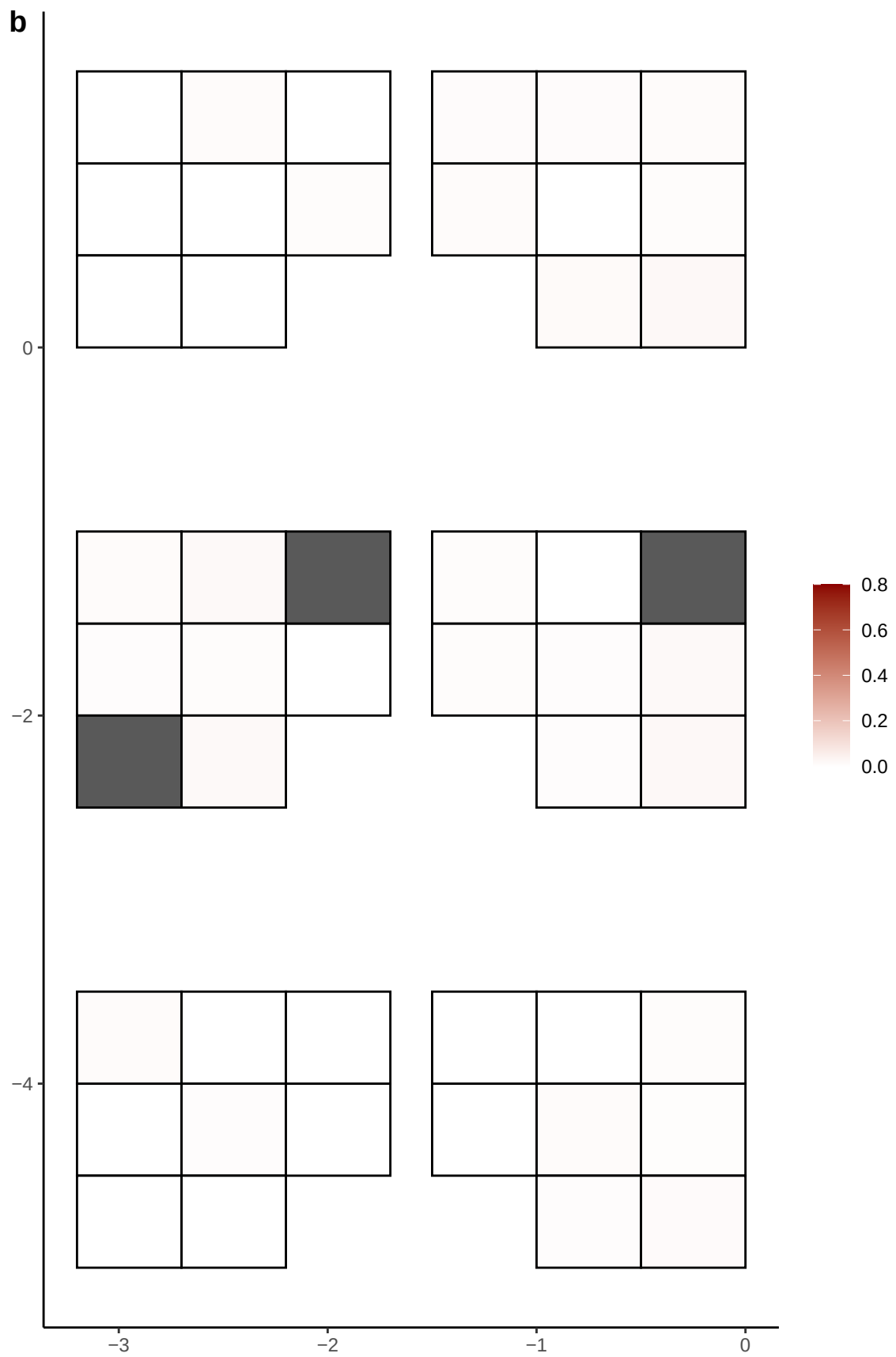

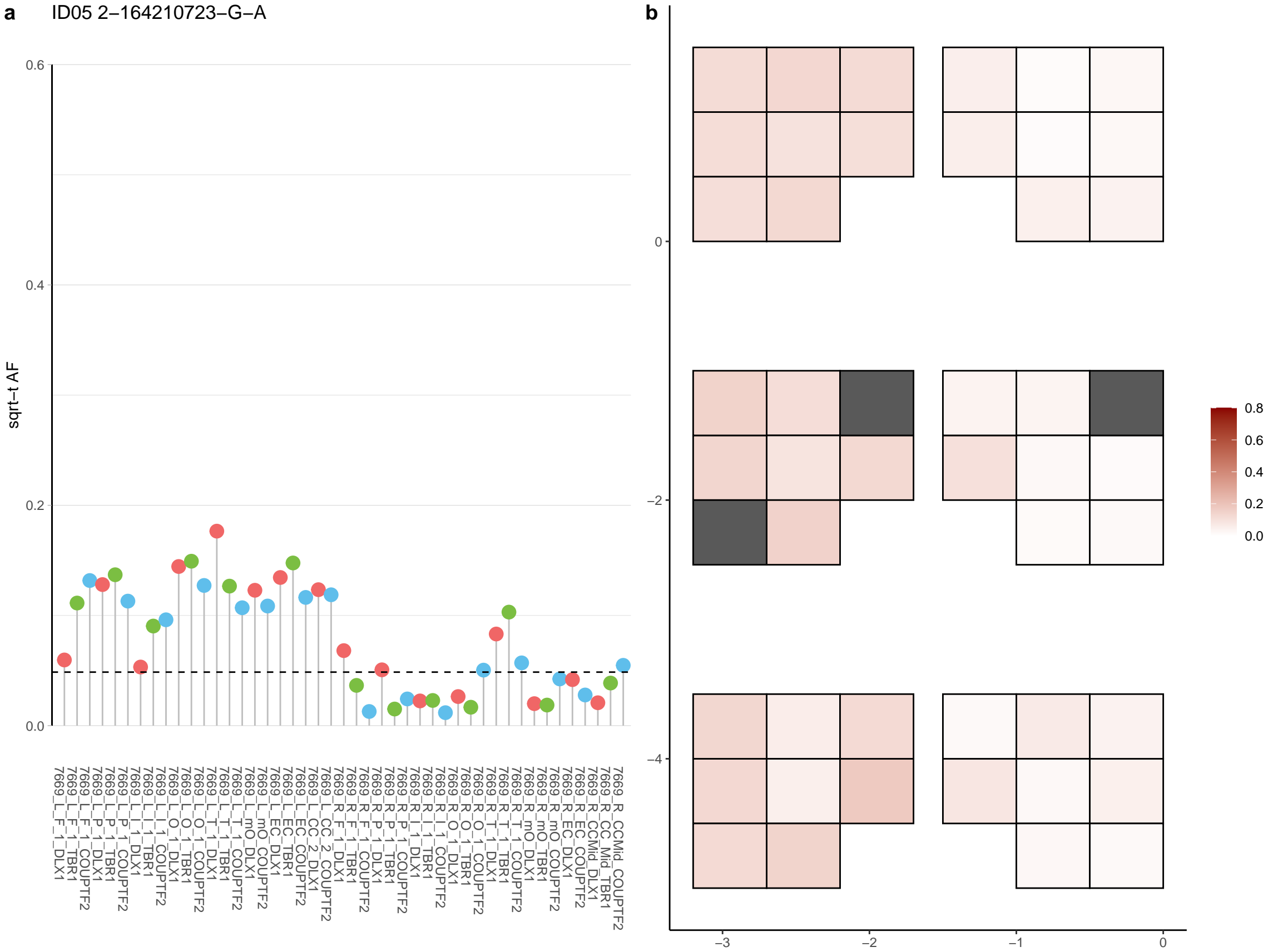

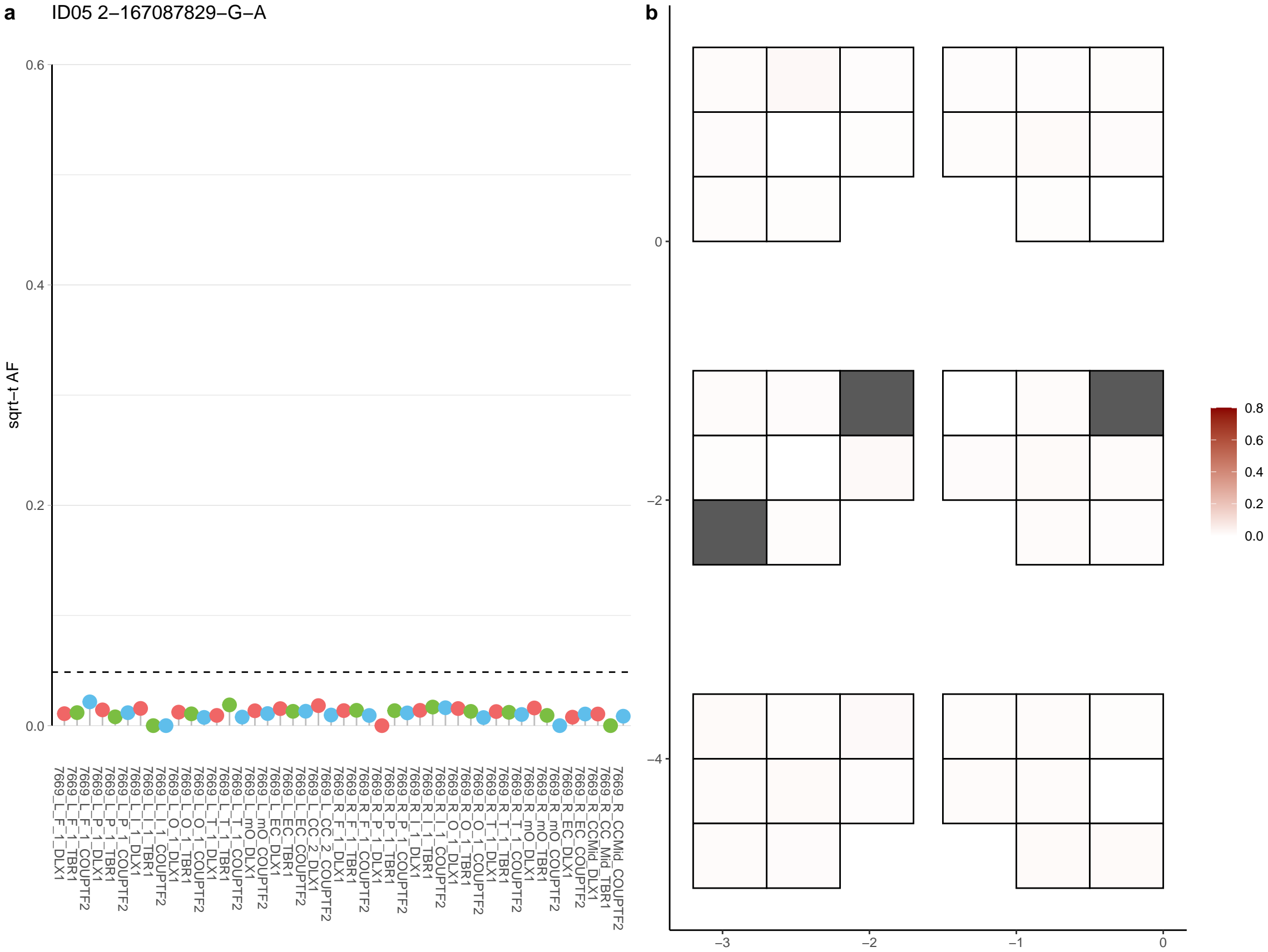

**a** ID05 2-167342653-C-T

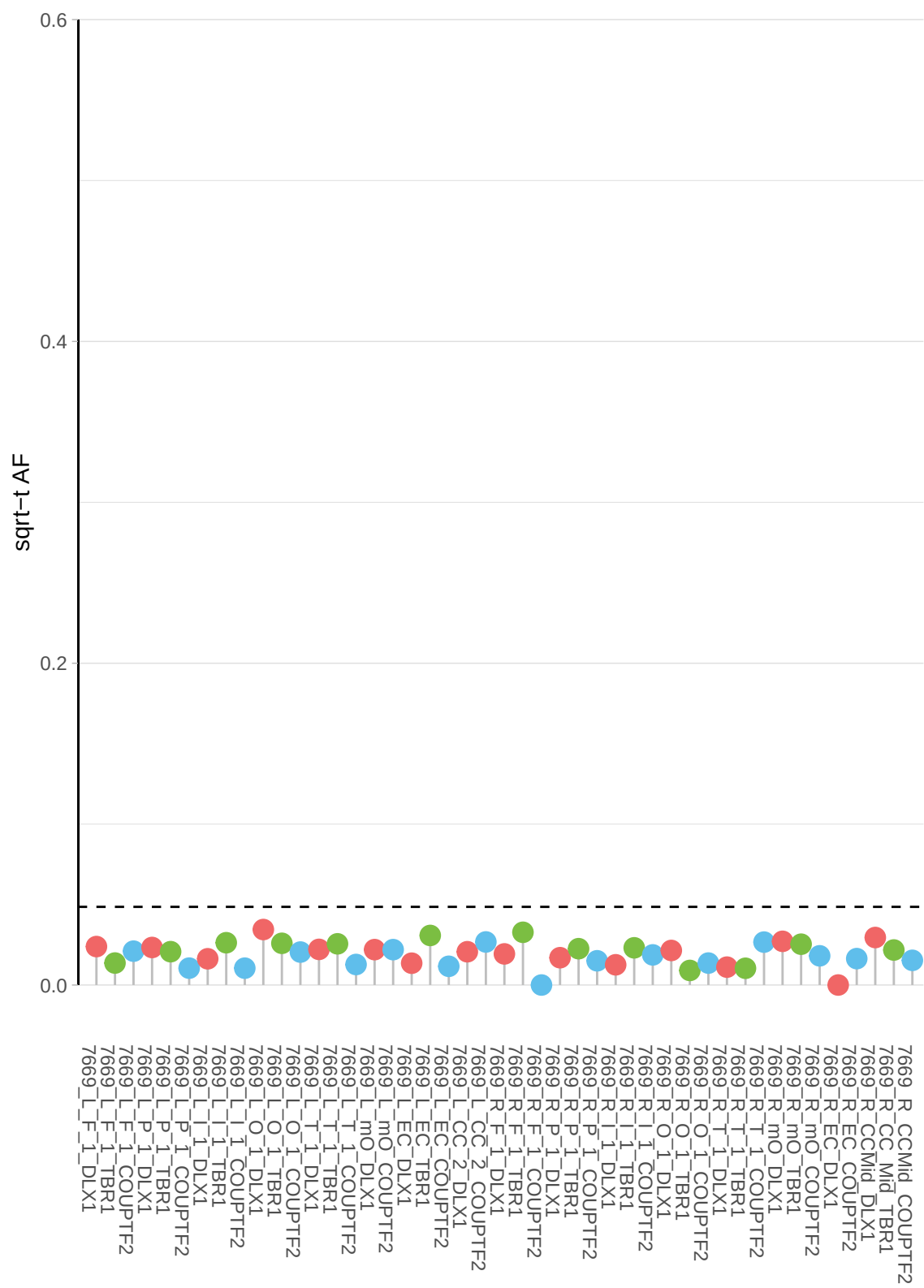

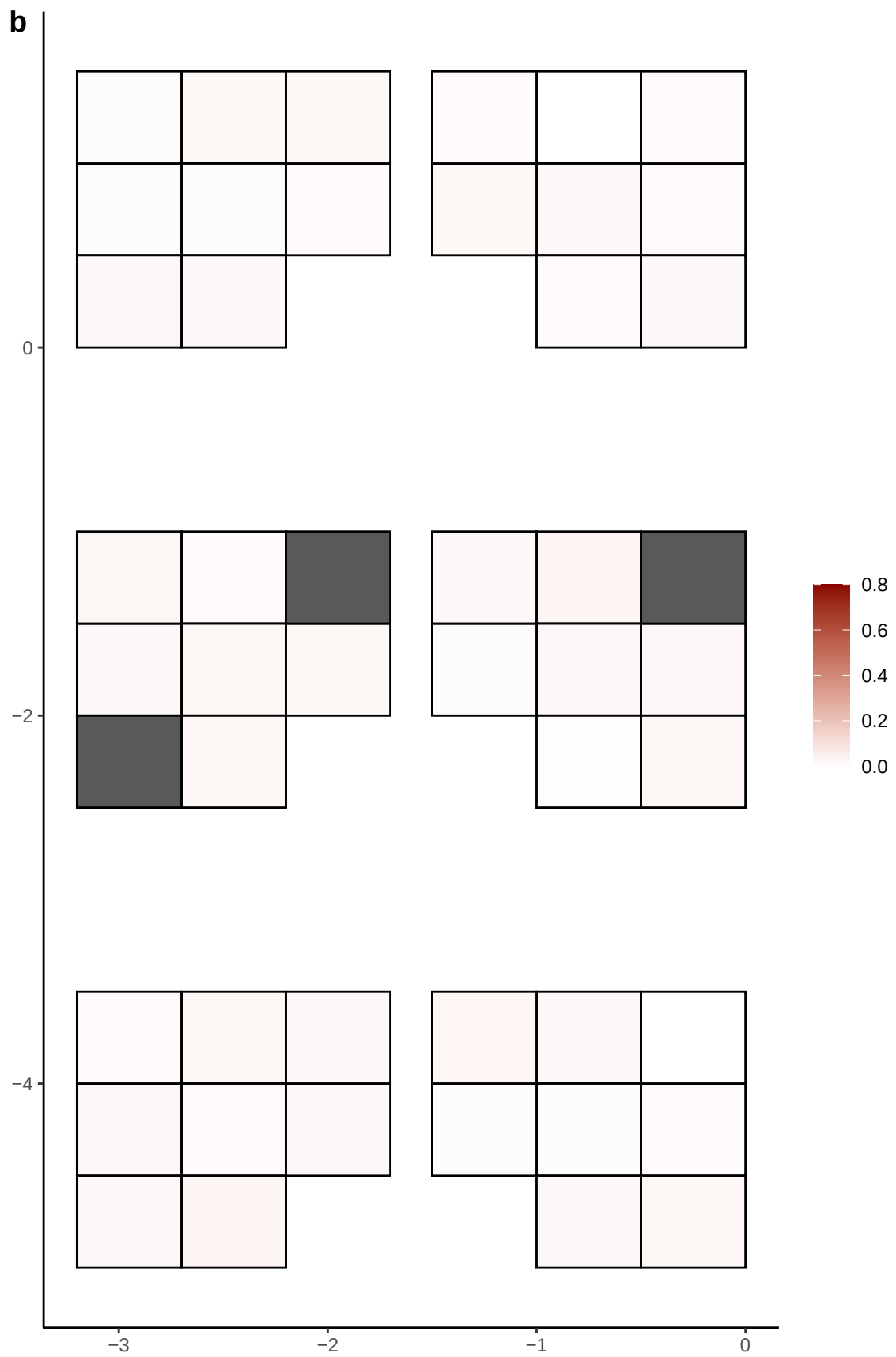

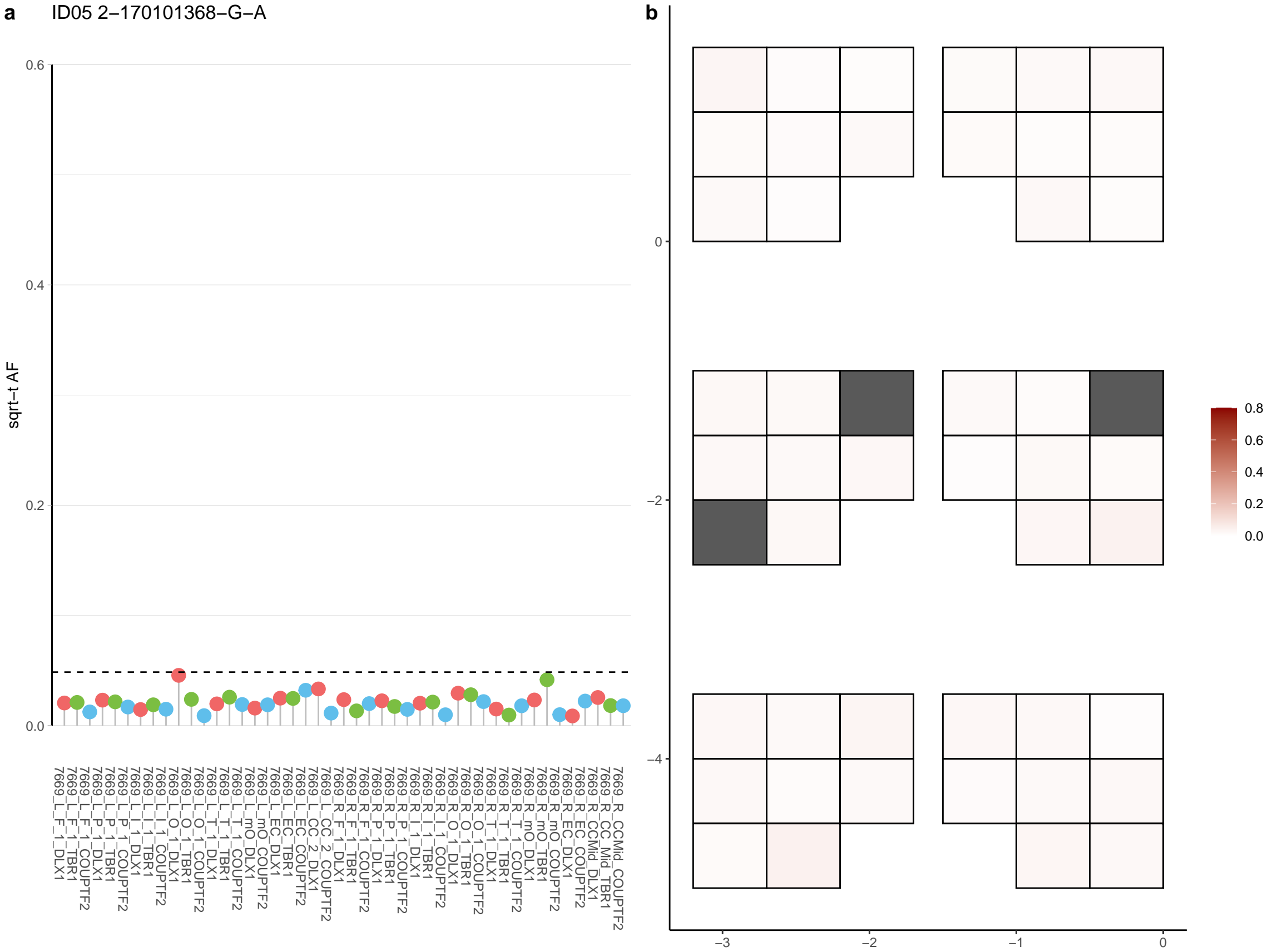

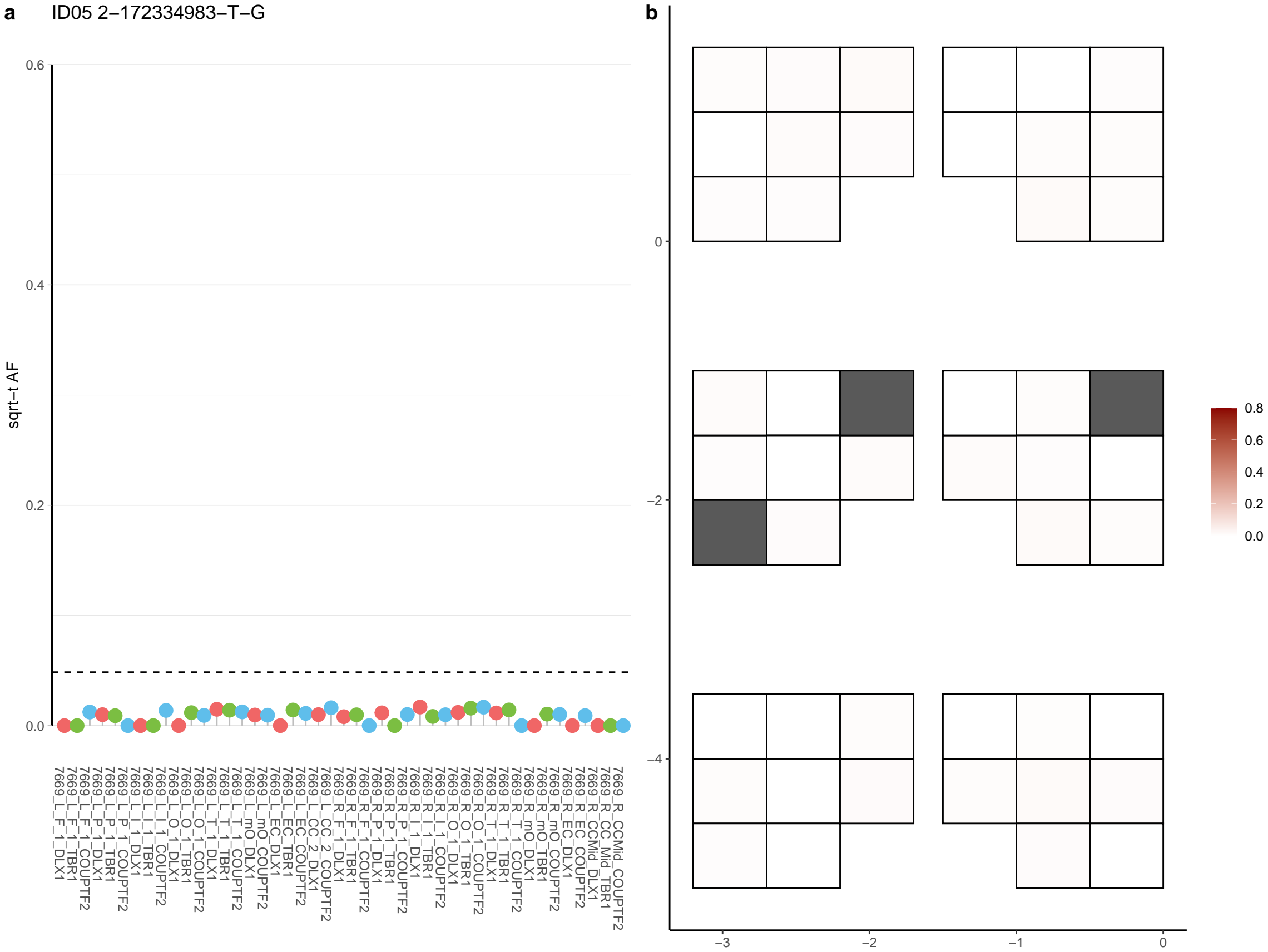

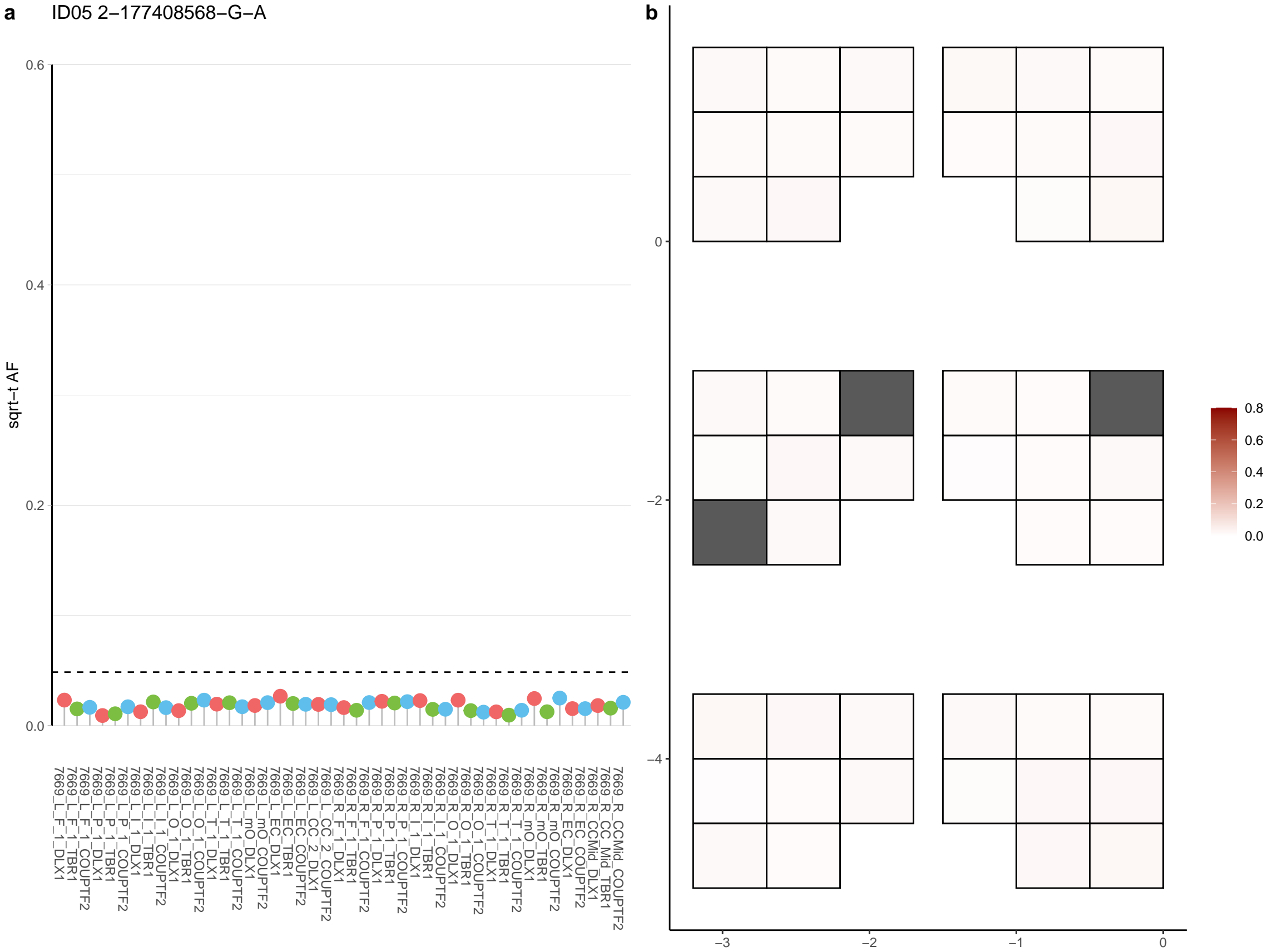

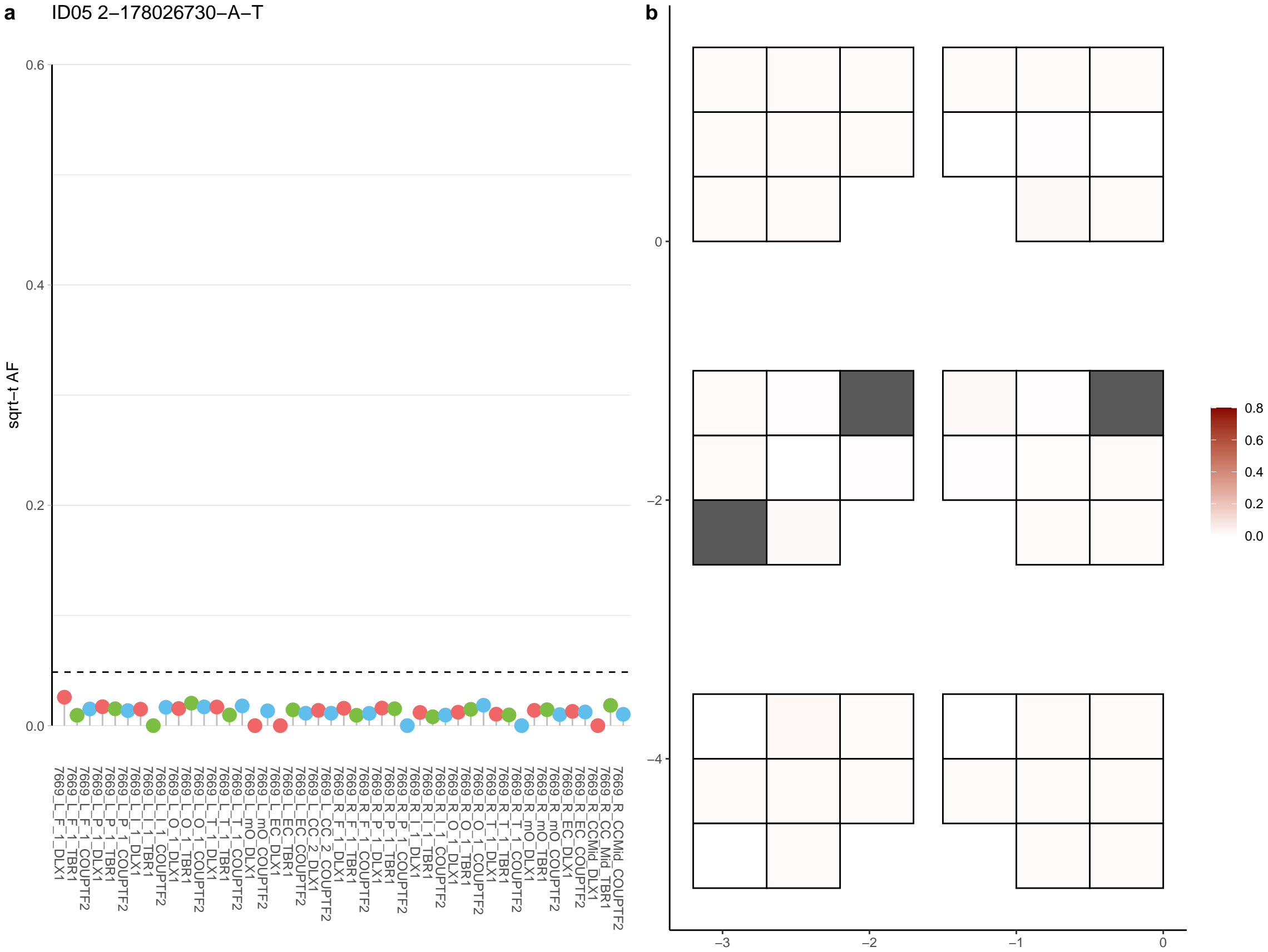

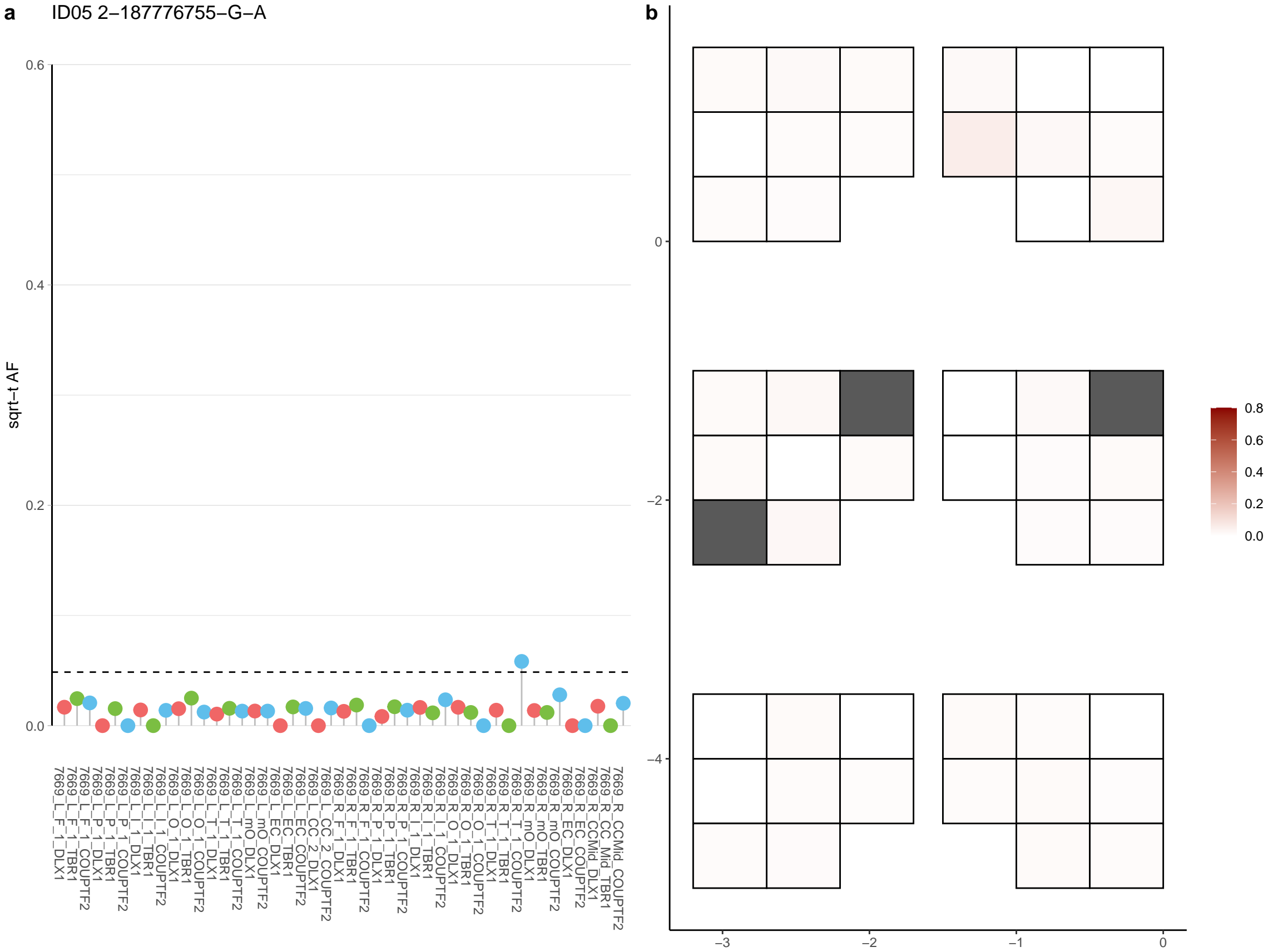

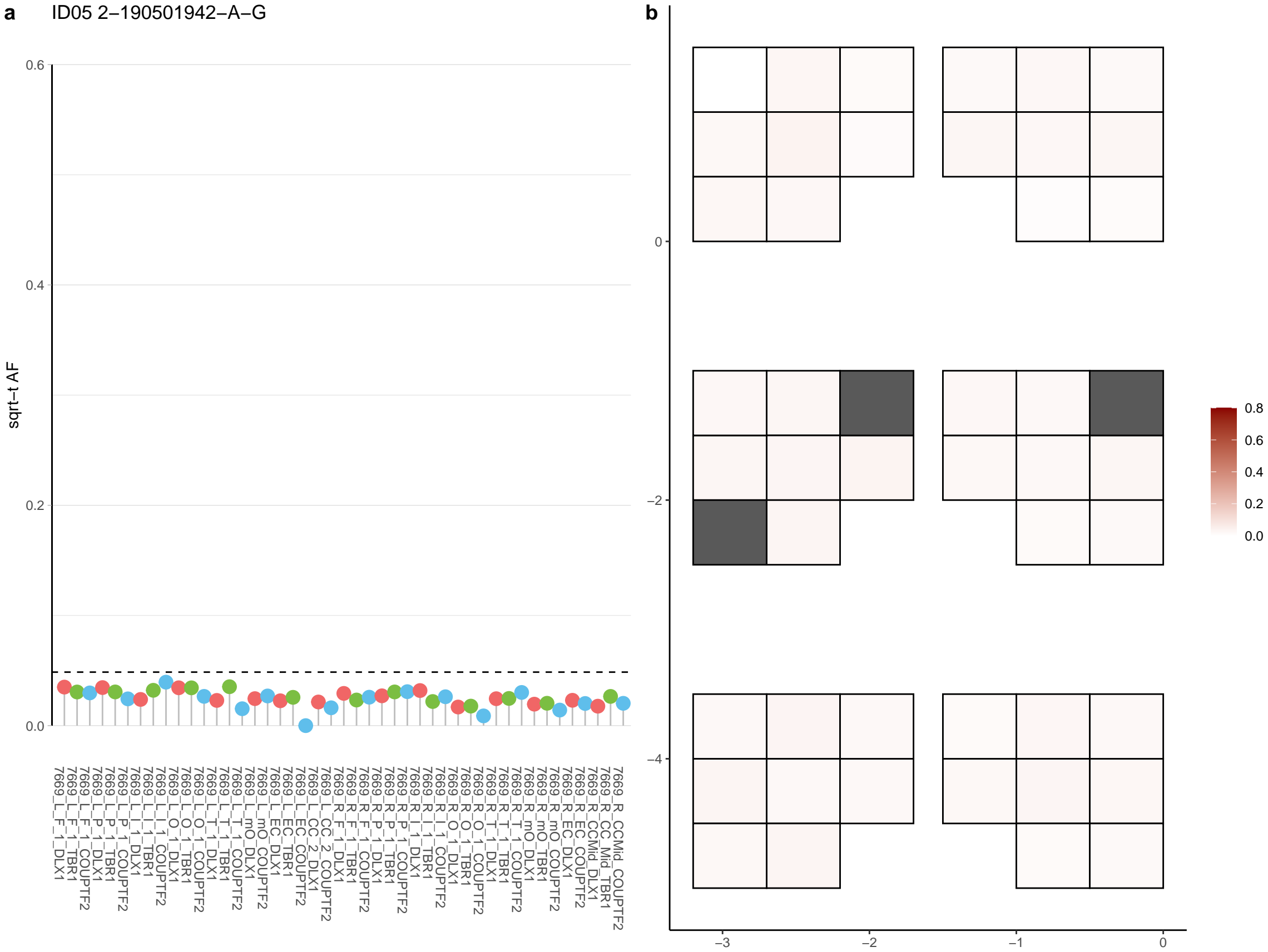

**a** ID05 2-193626603-C-A

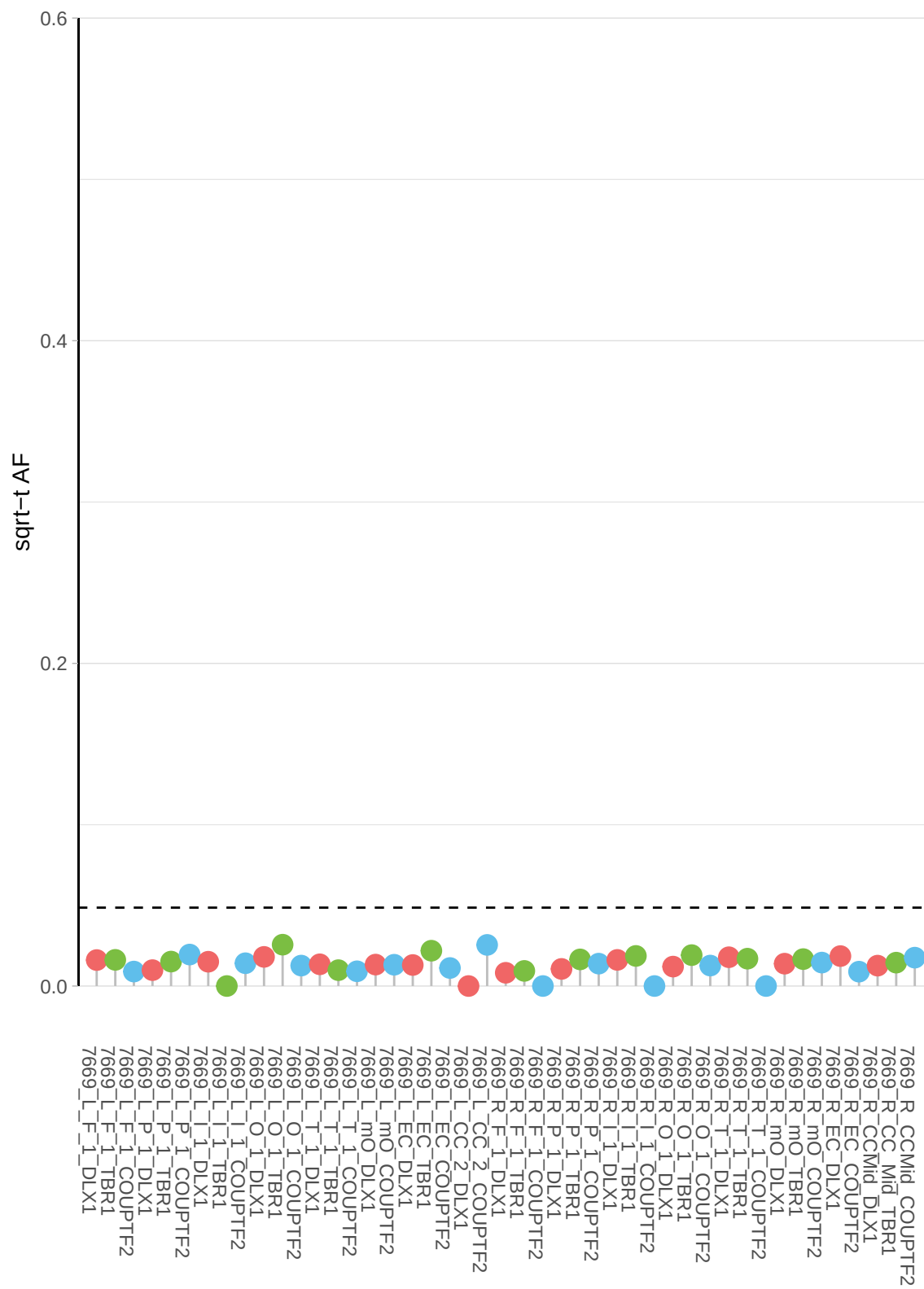

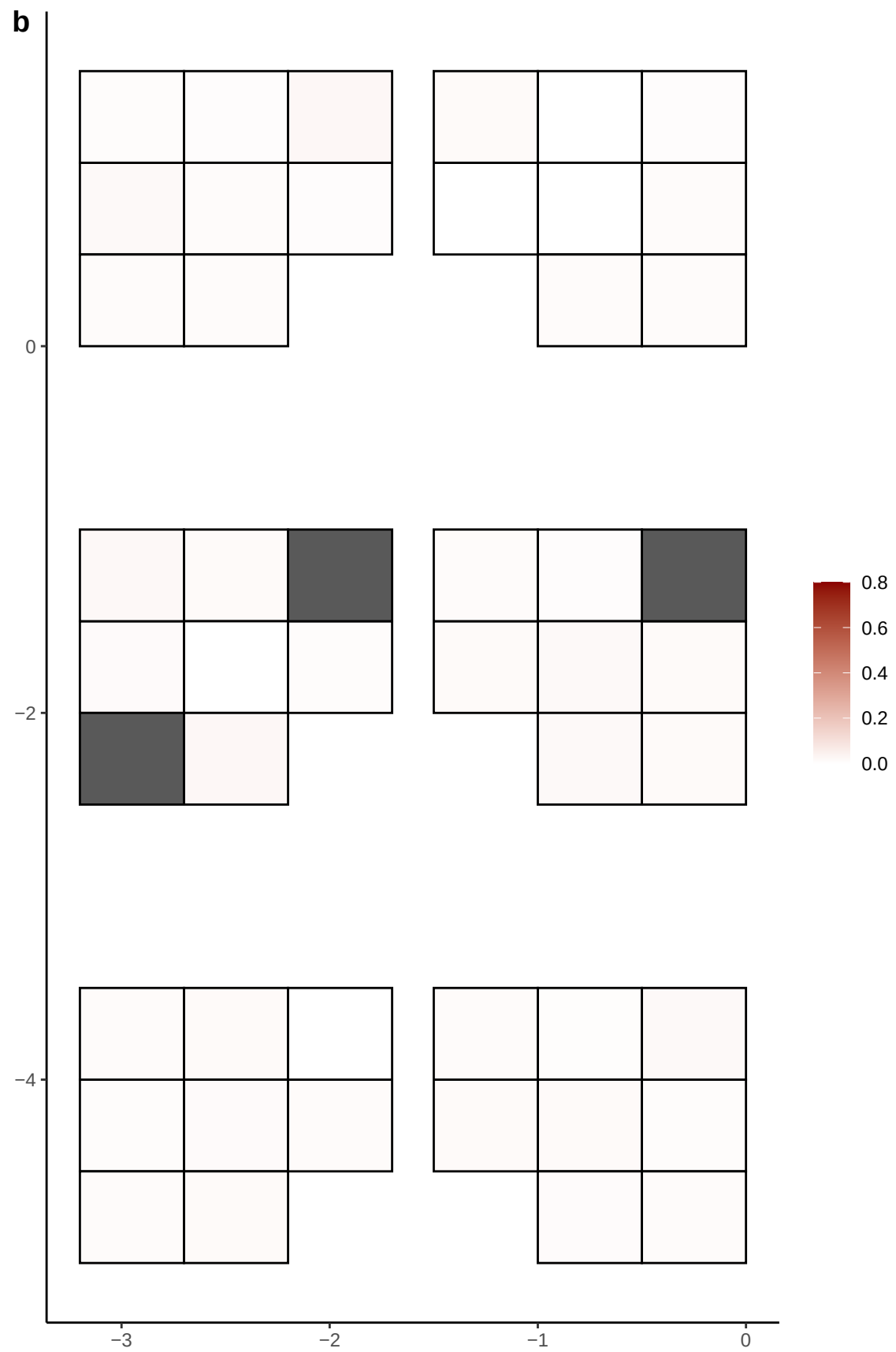

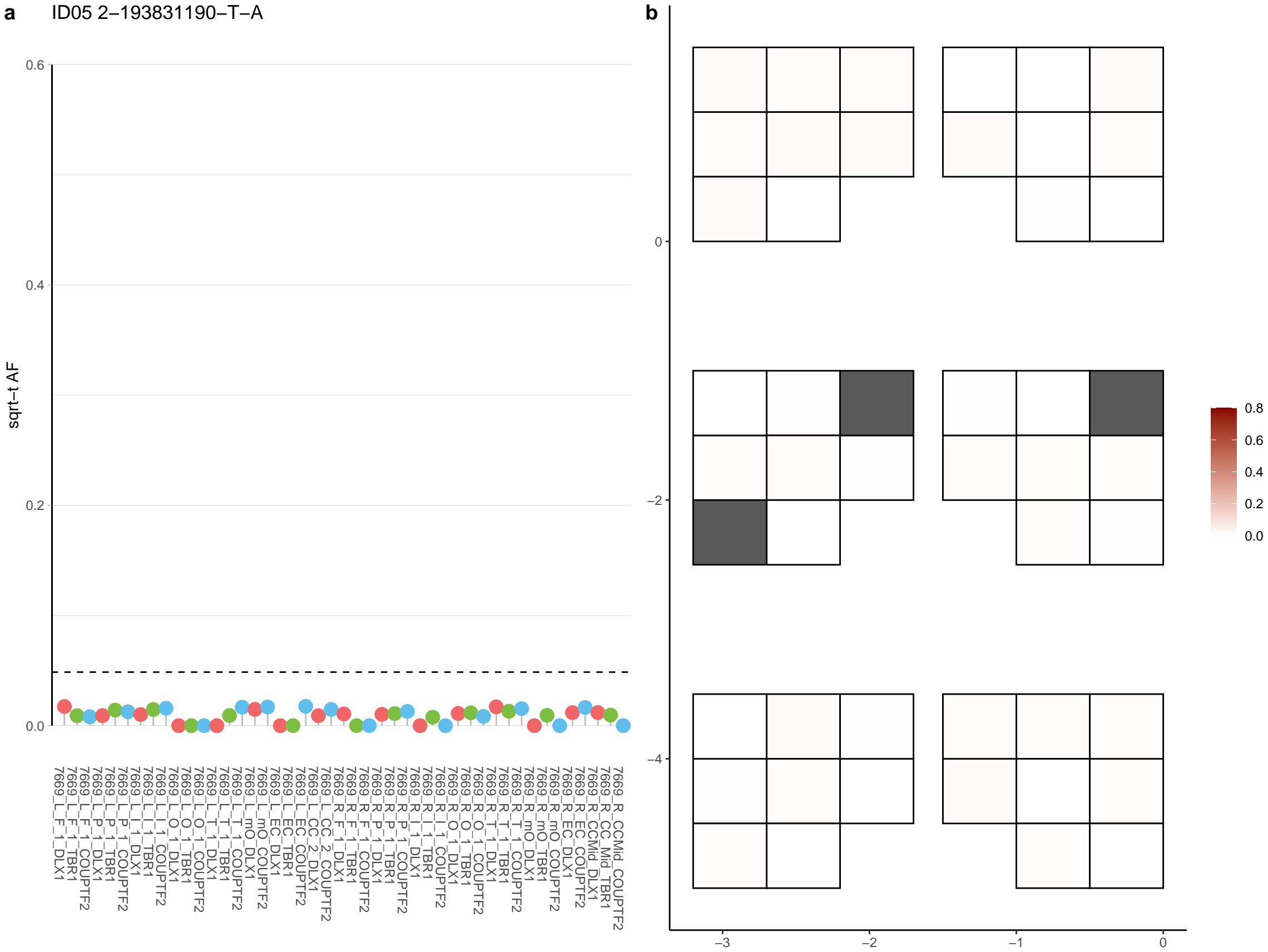

**a** ID05 2-198773820-A-G

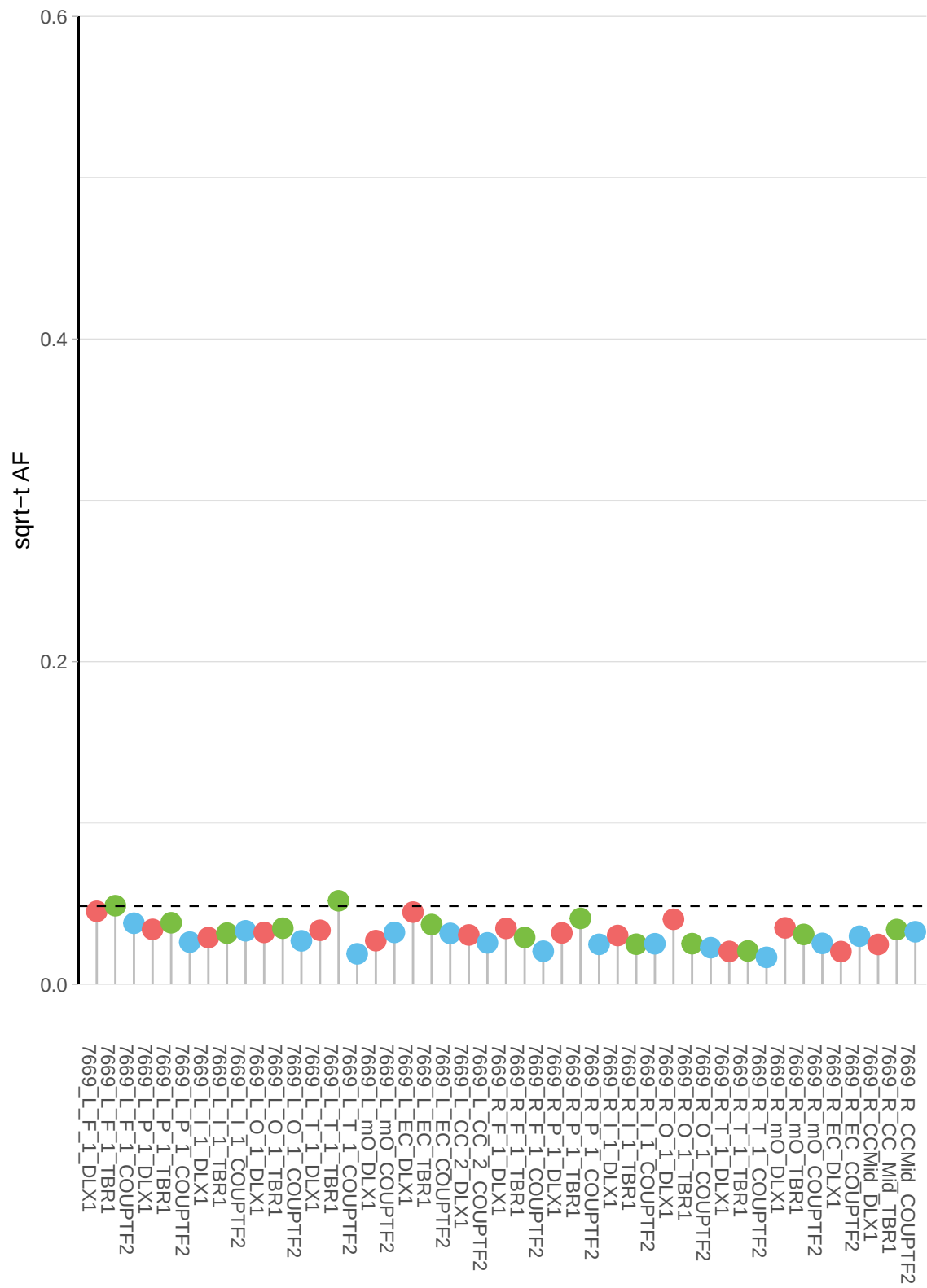

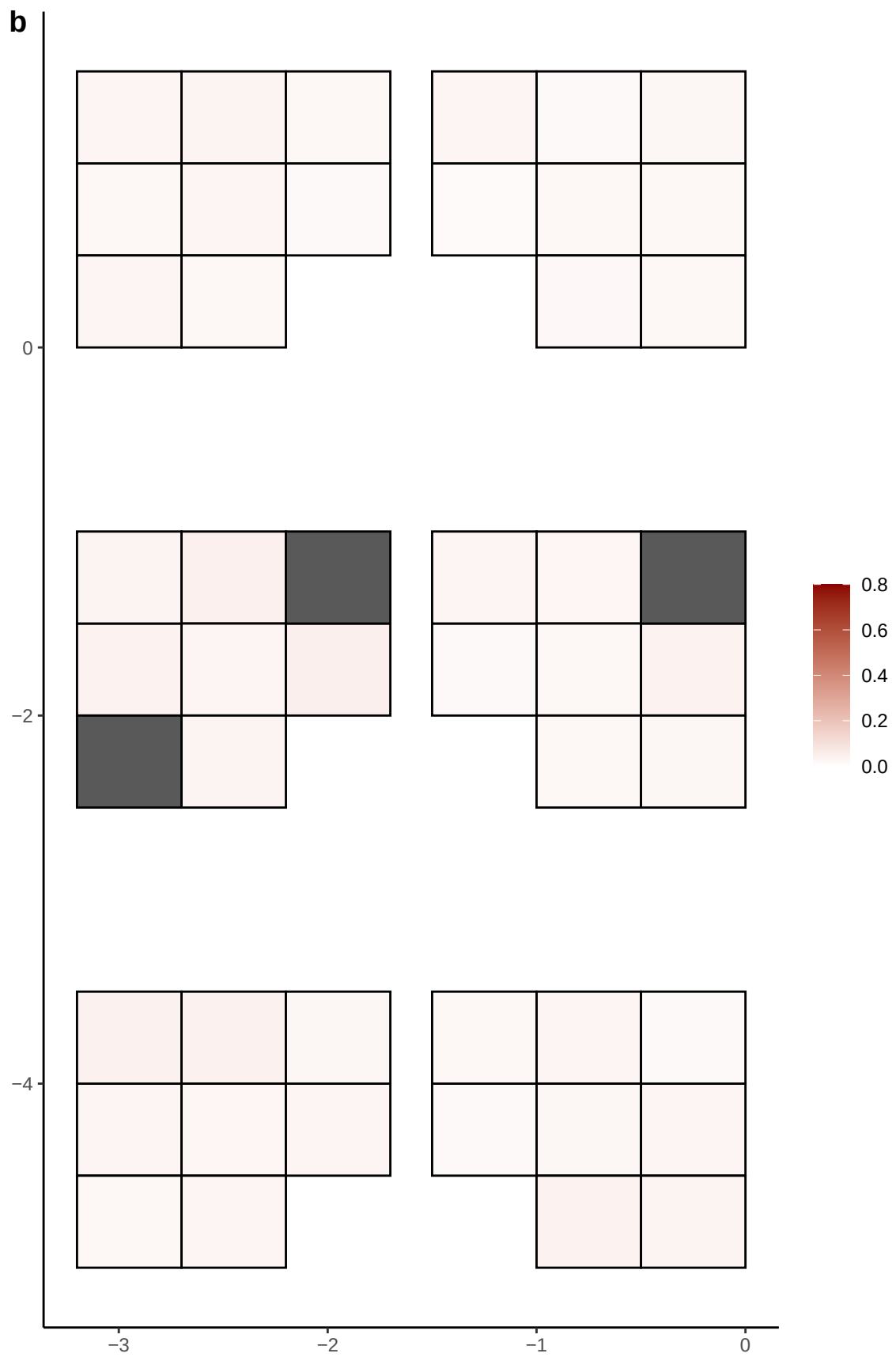

**a**

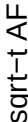

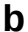

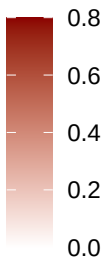

**a** ID05 2-204722237-C-T

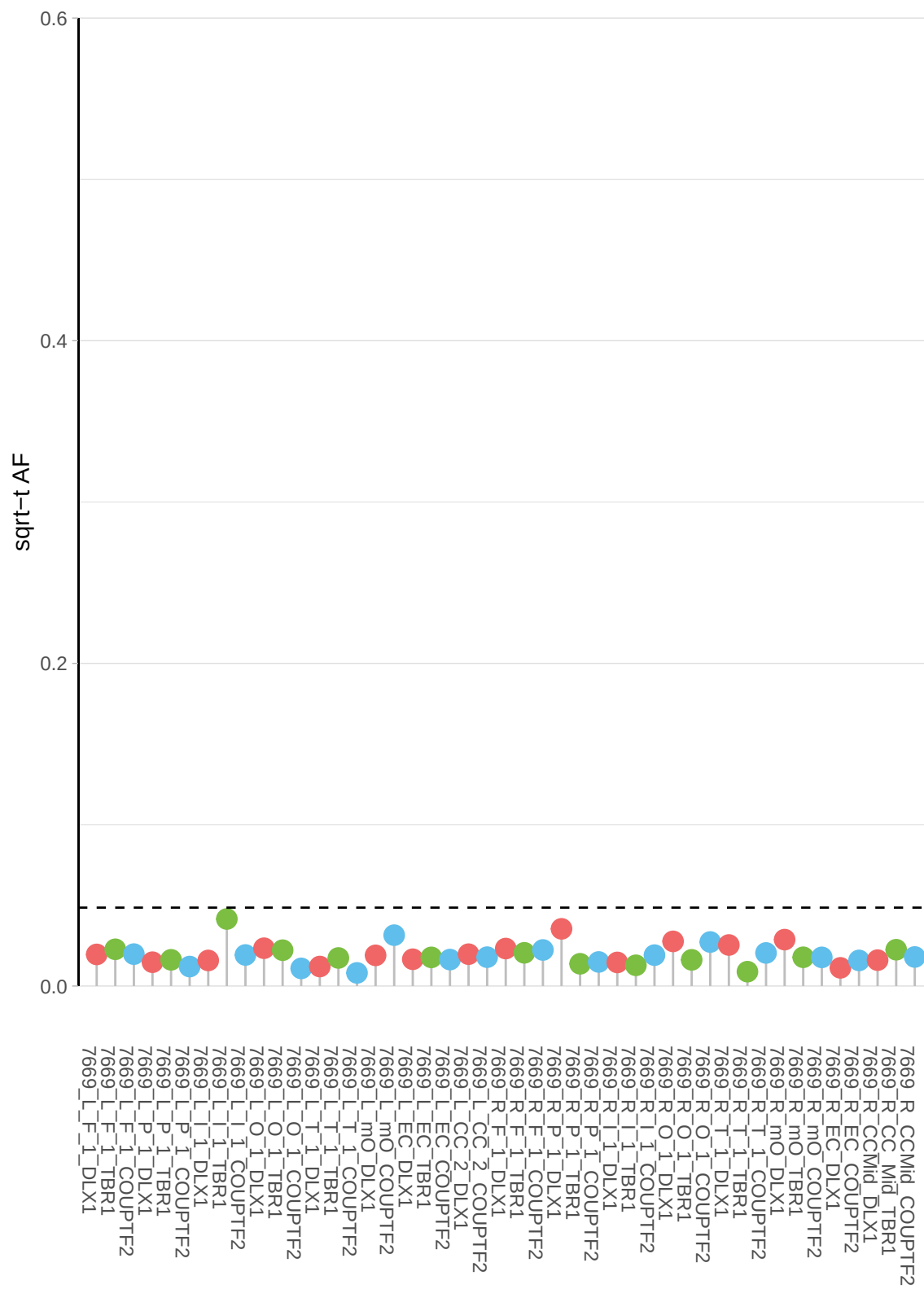

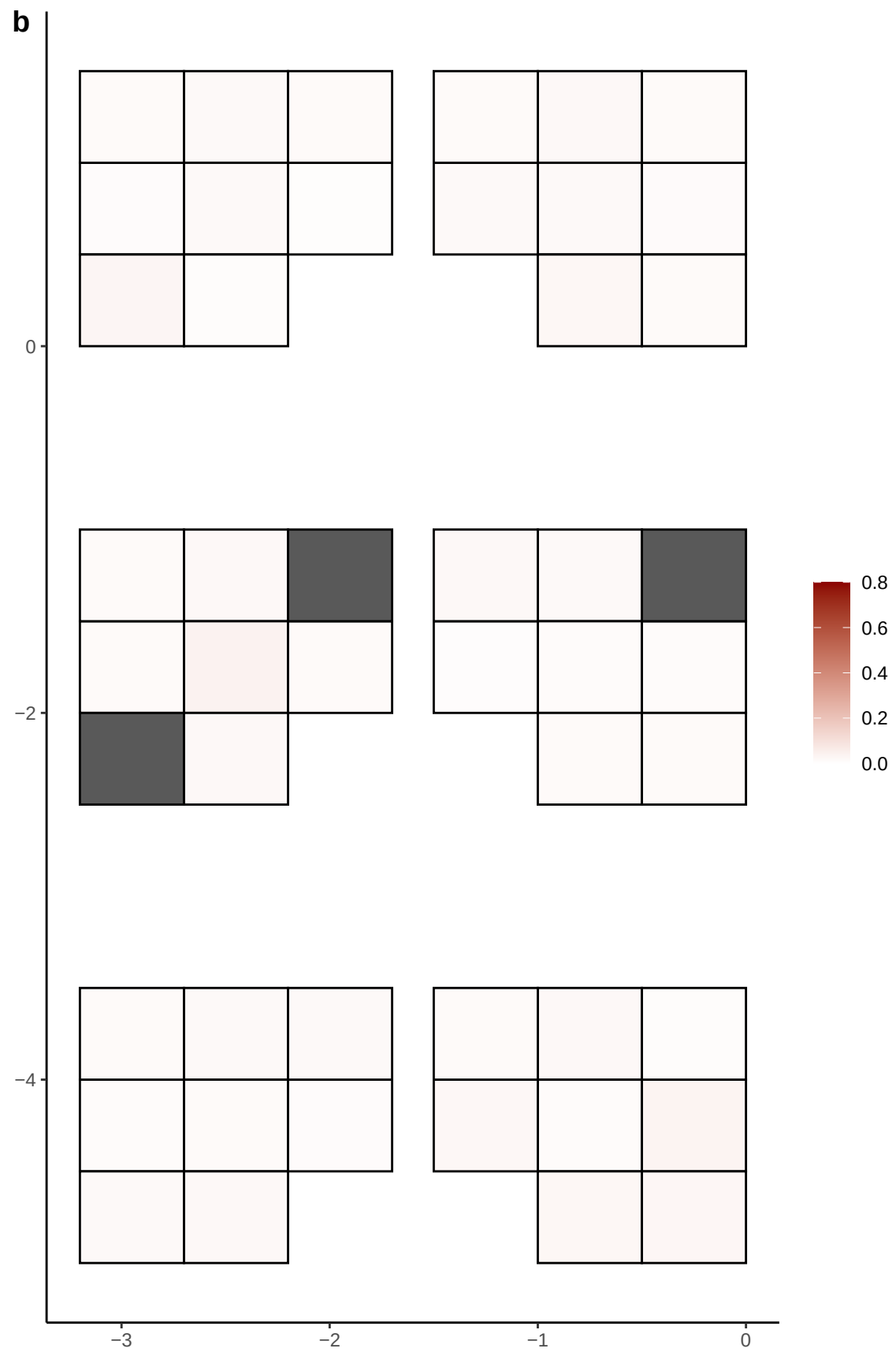

**a** ID05 2-209569113-C-T

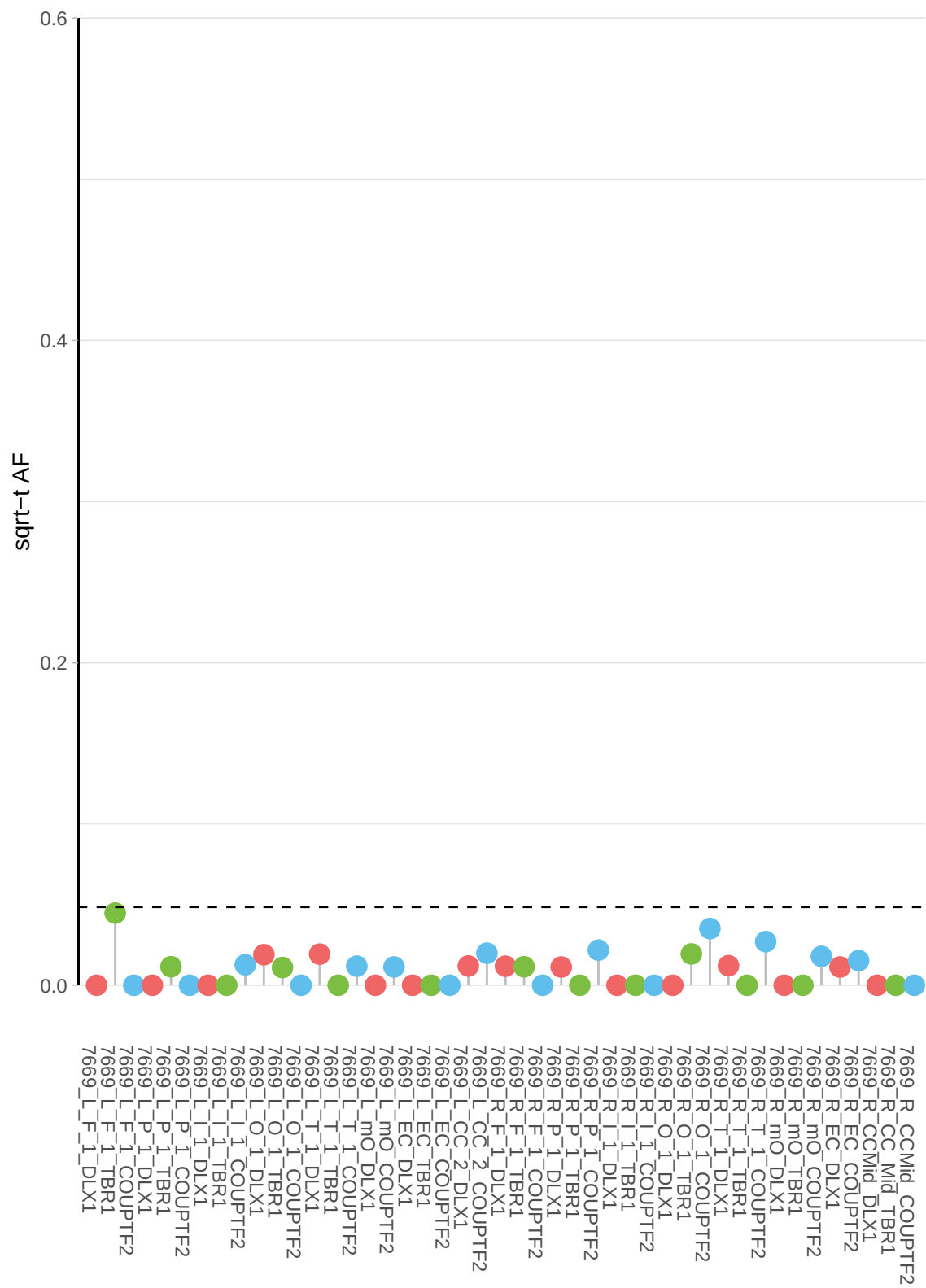

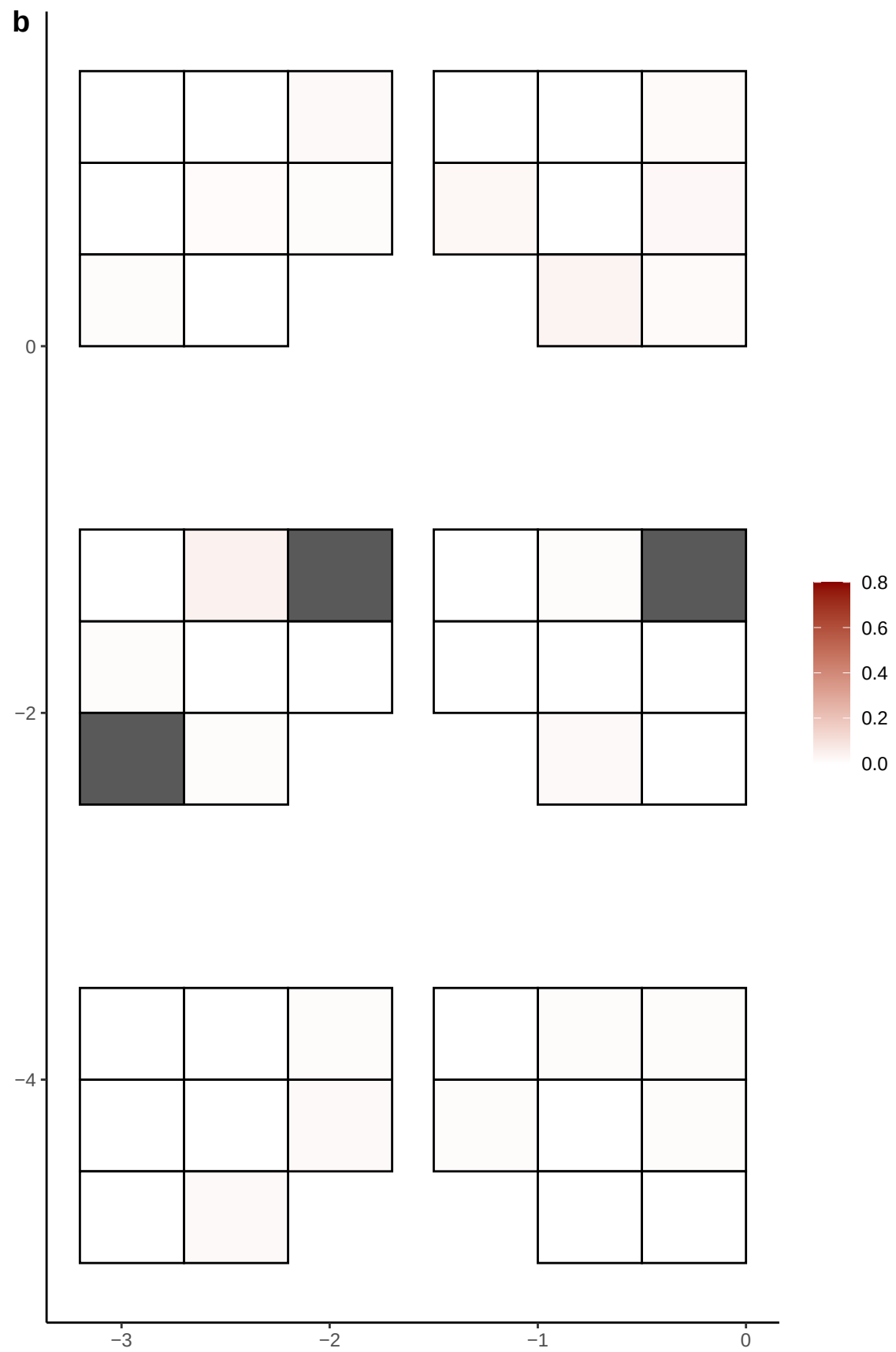

**a** ID05 2-213063972-C-T

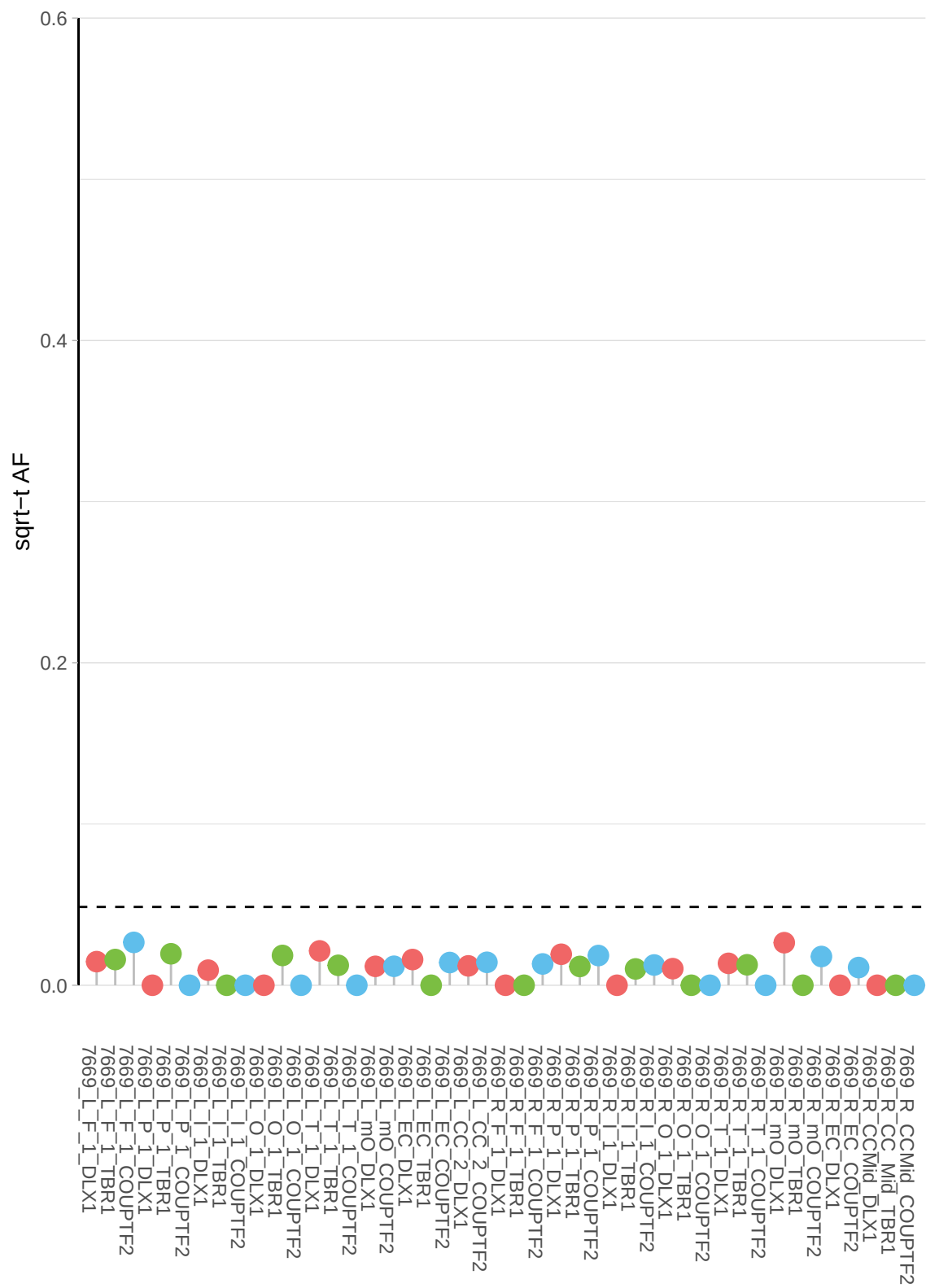

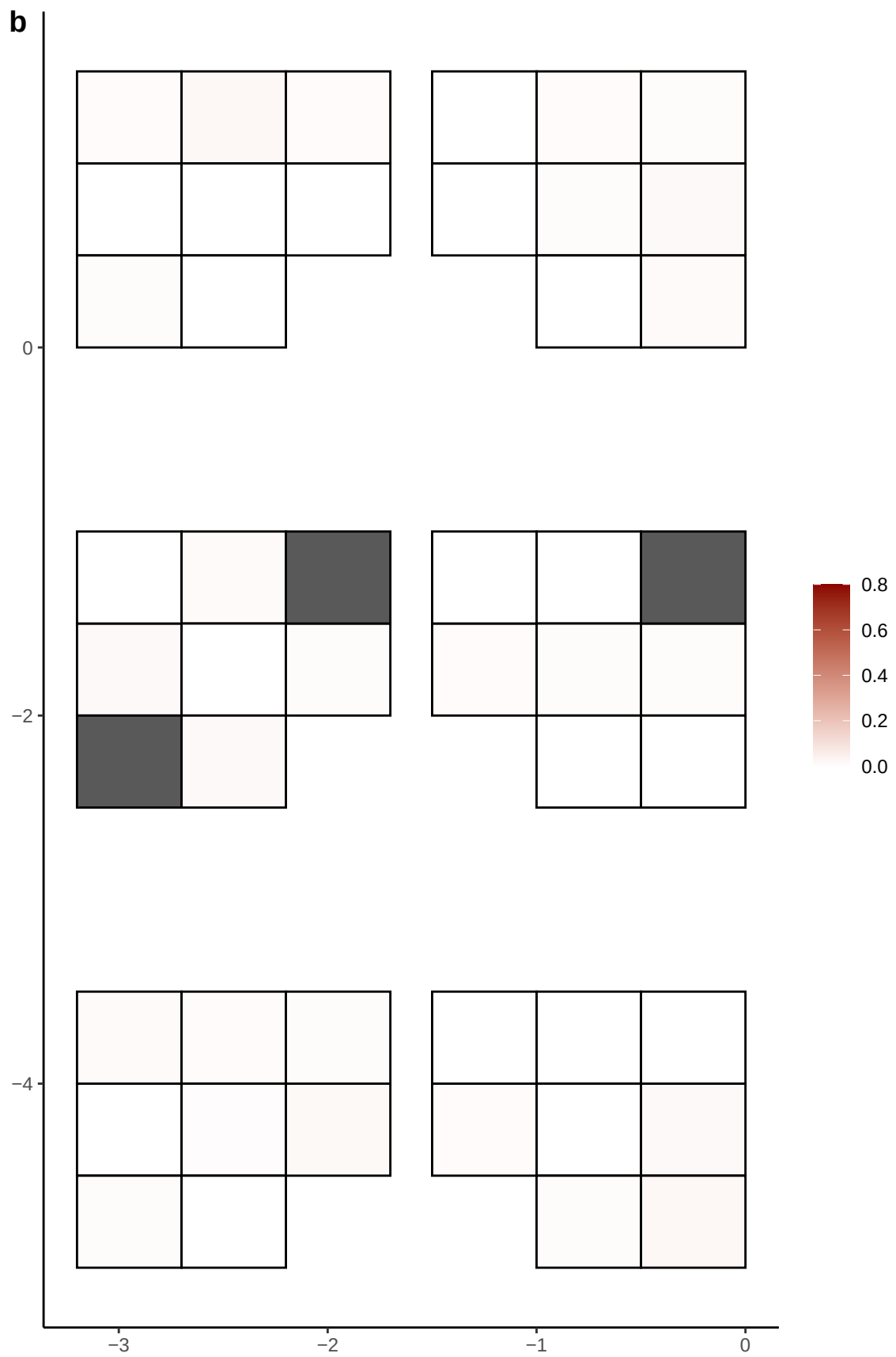

**a** ID05 2-22209916-G-A

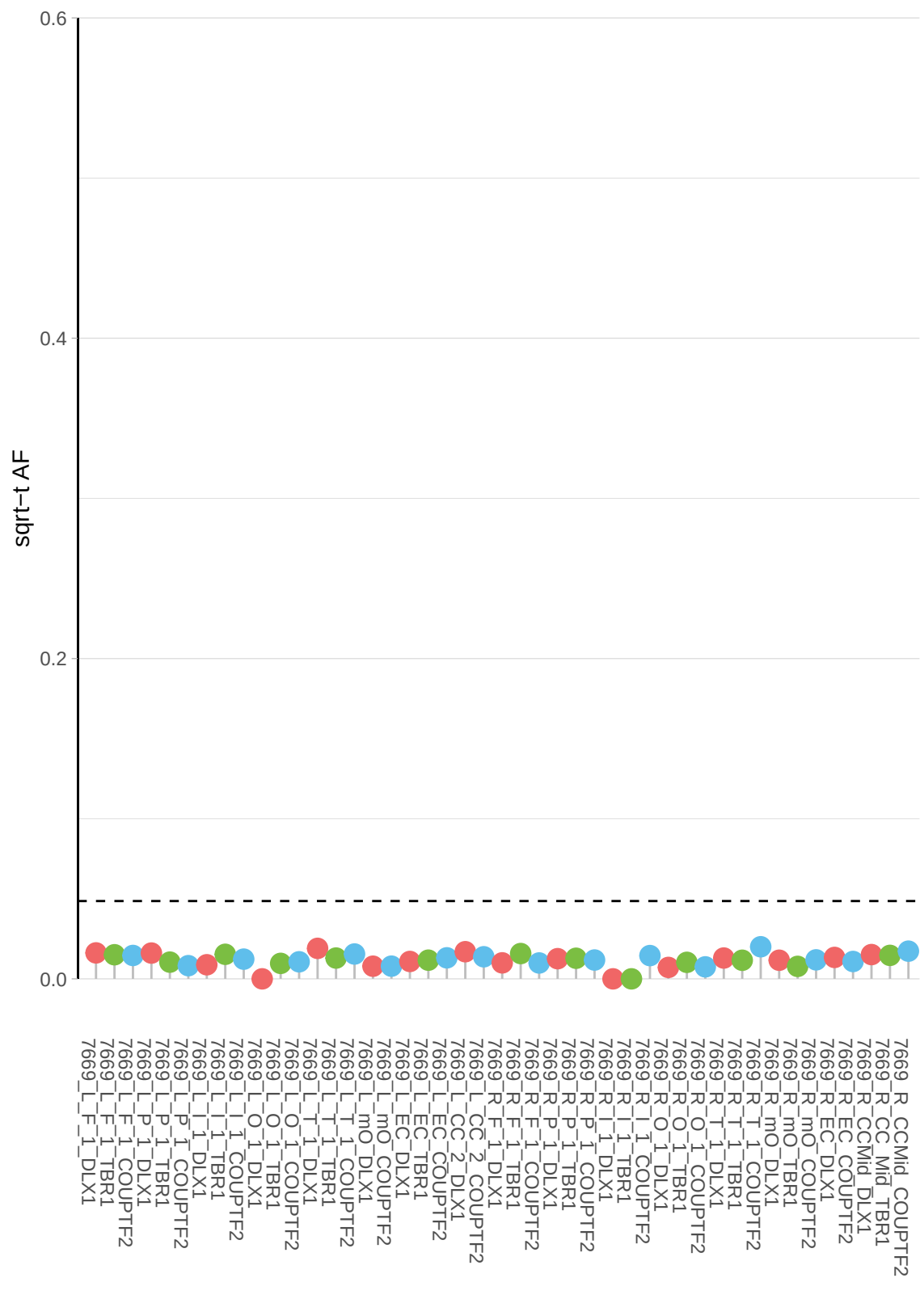

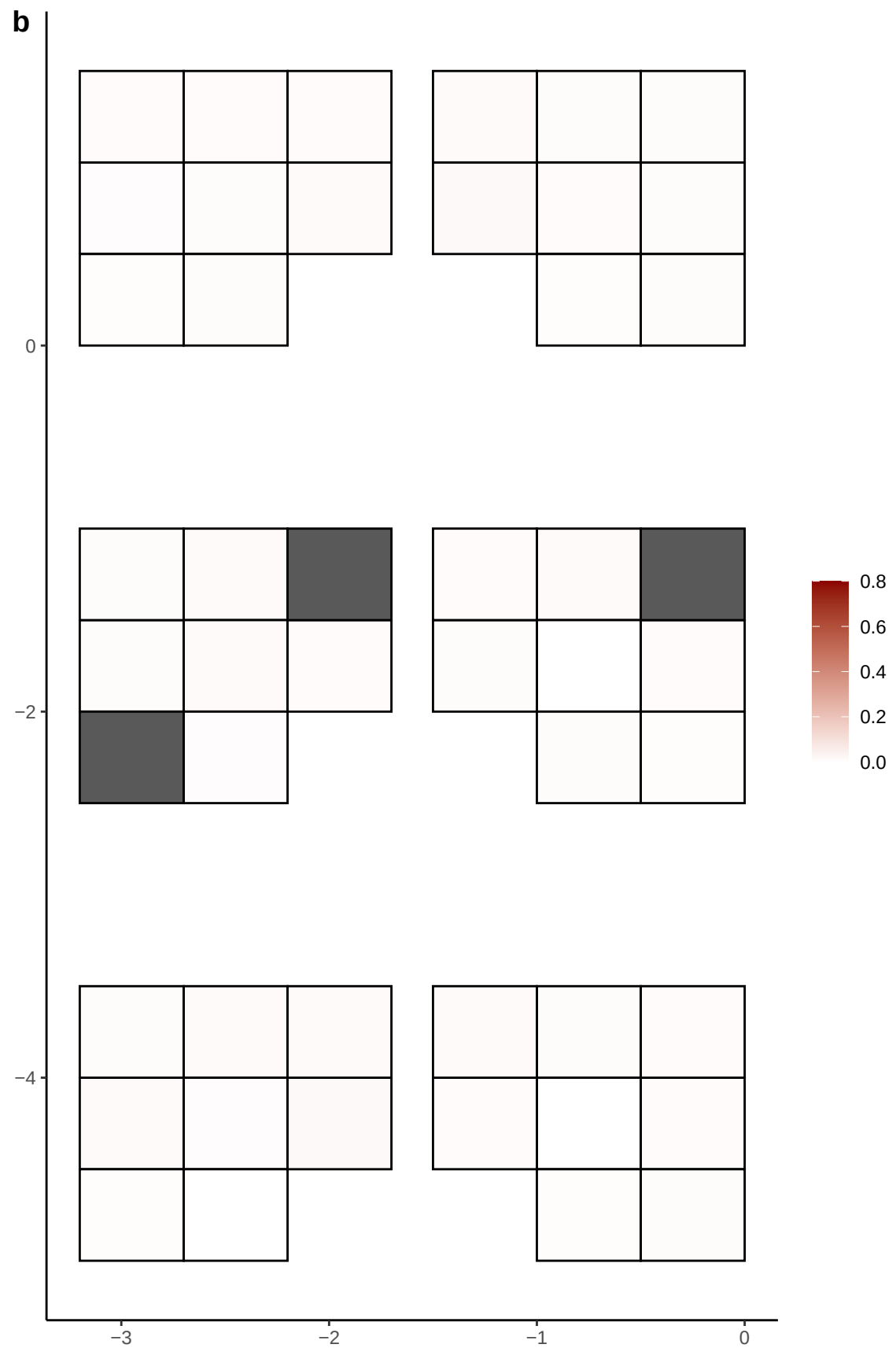

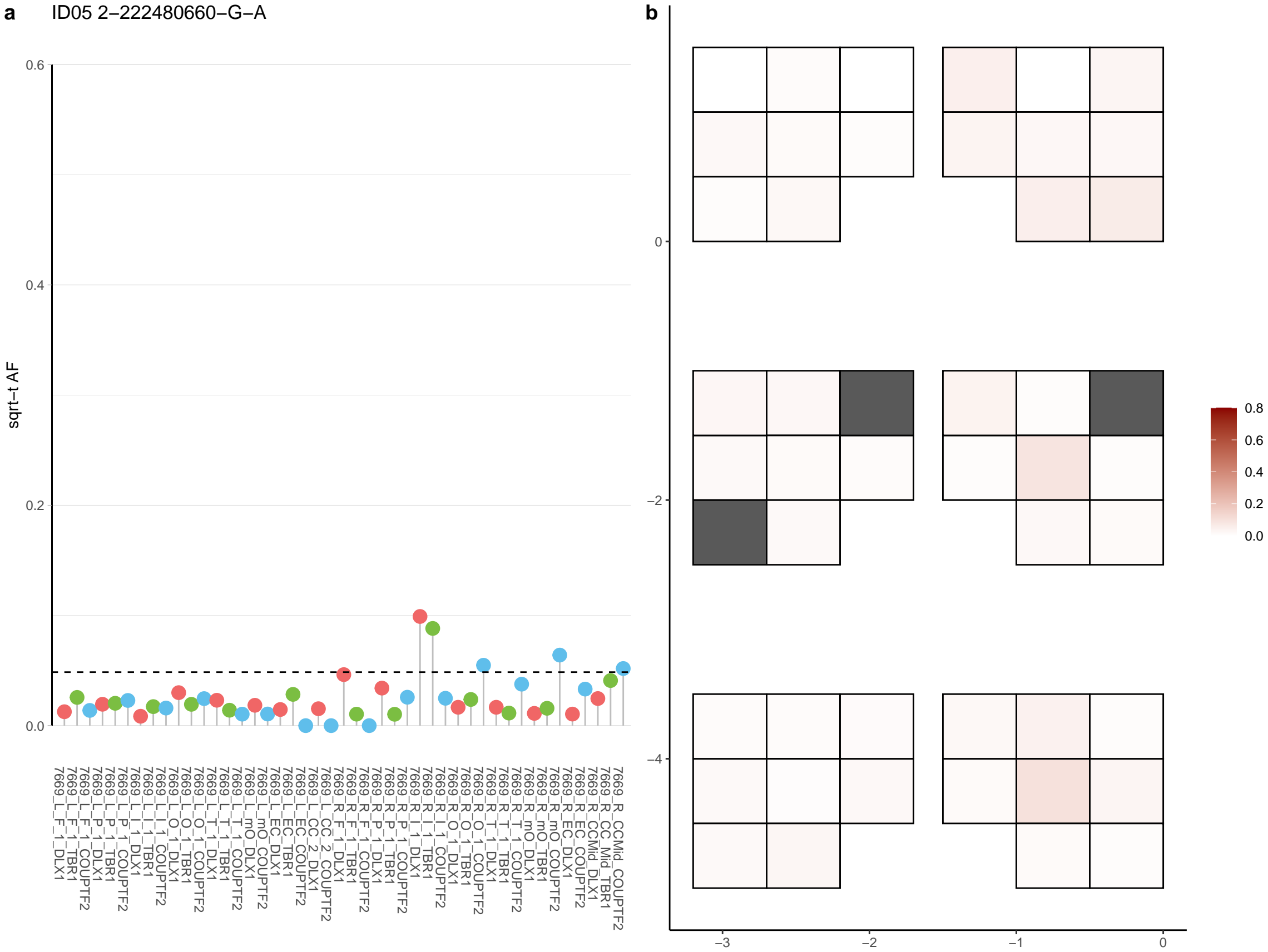

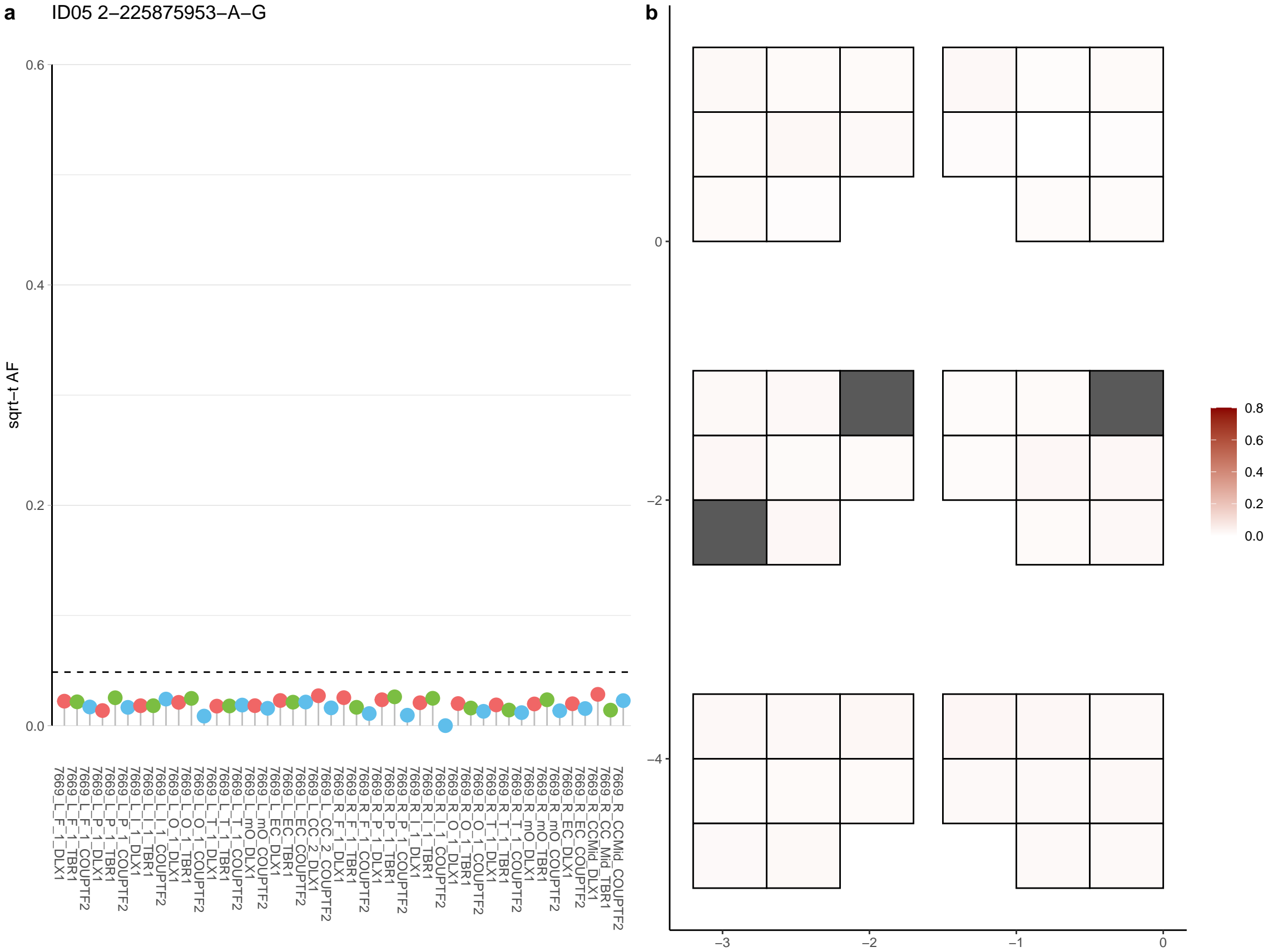

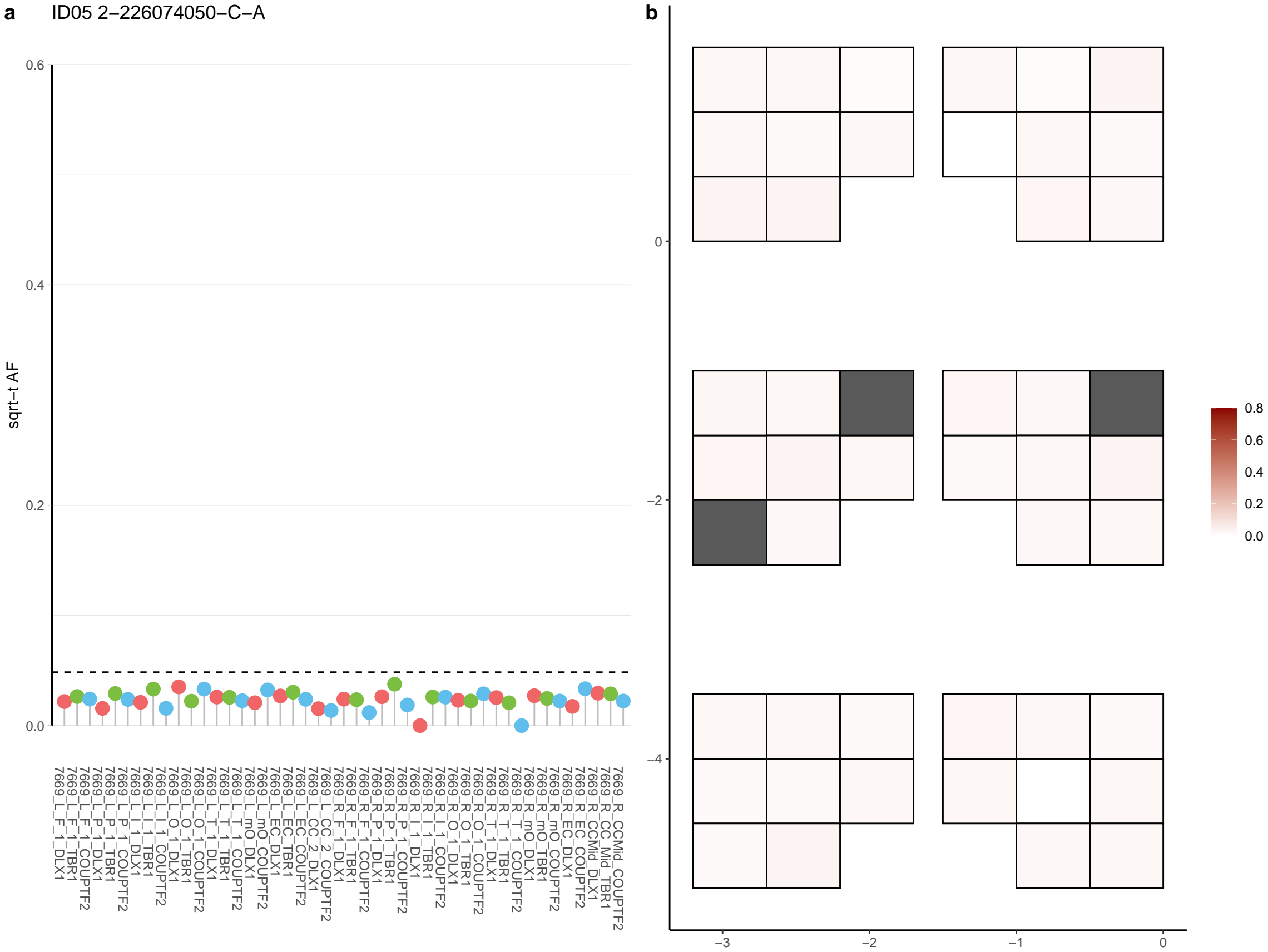

**a** ID05 2-226299479-C-T

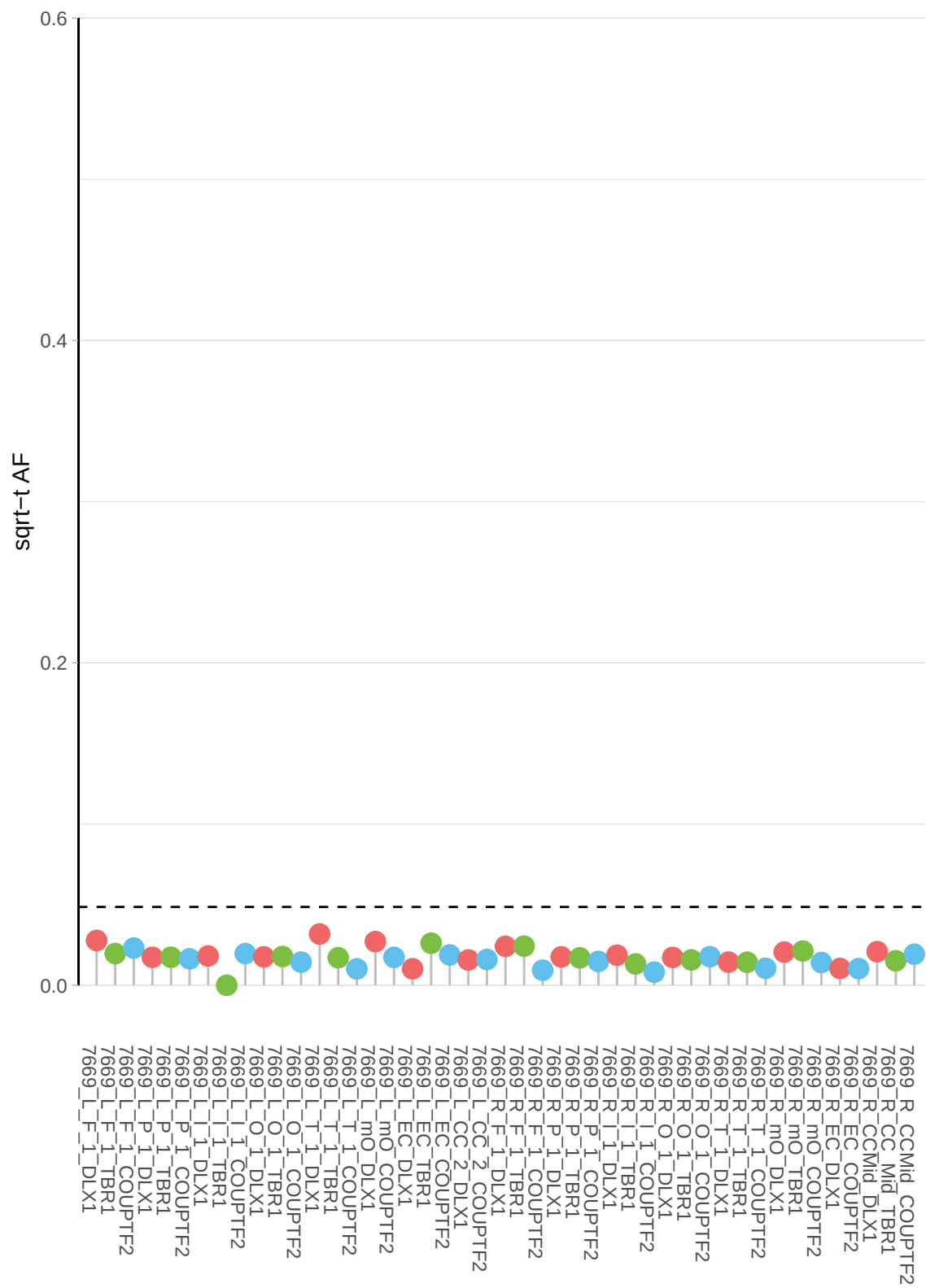

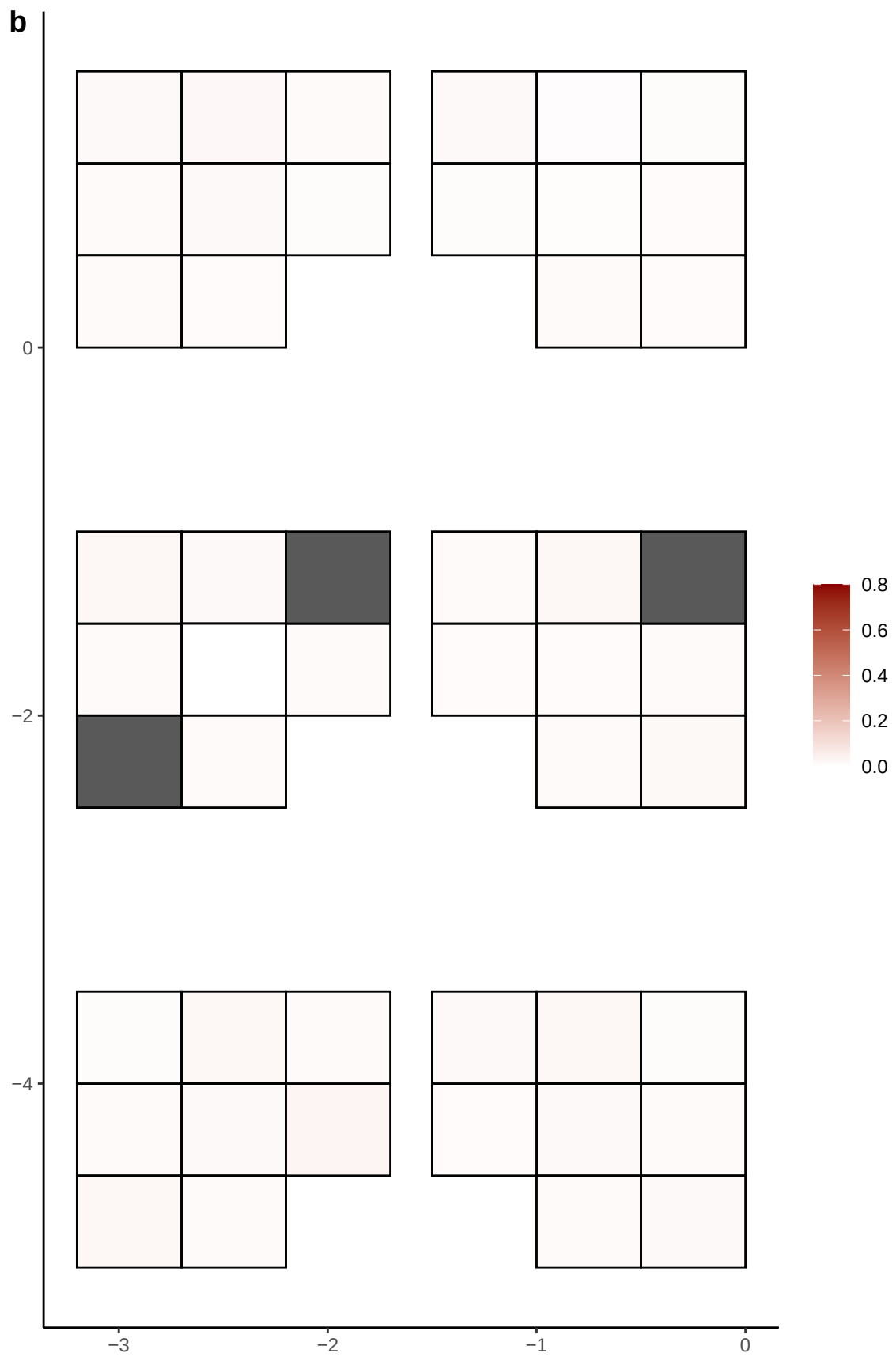

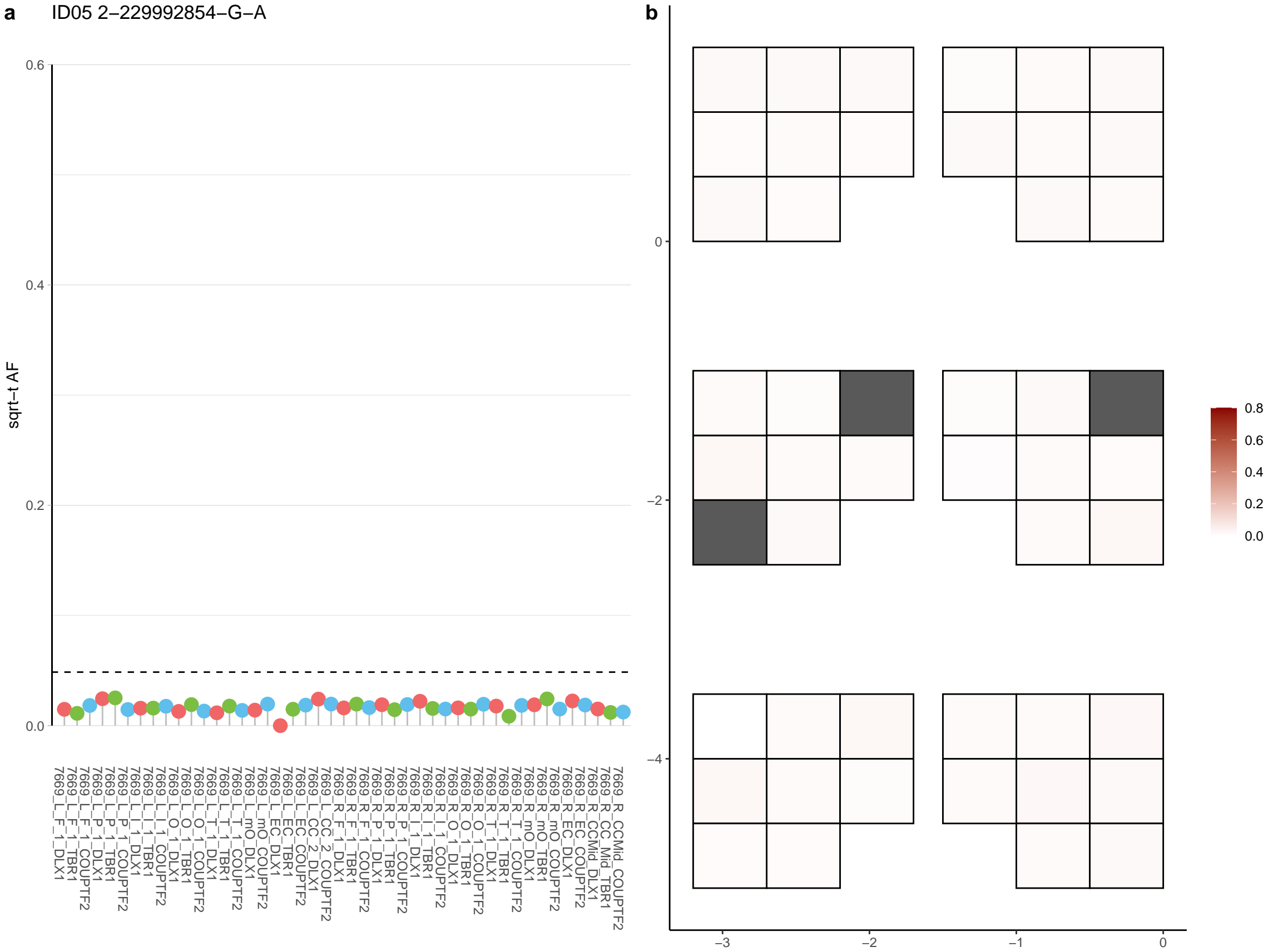

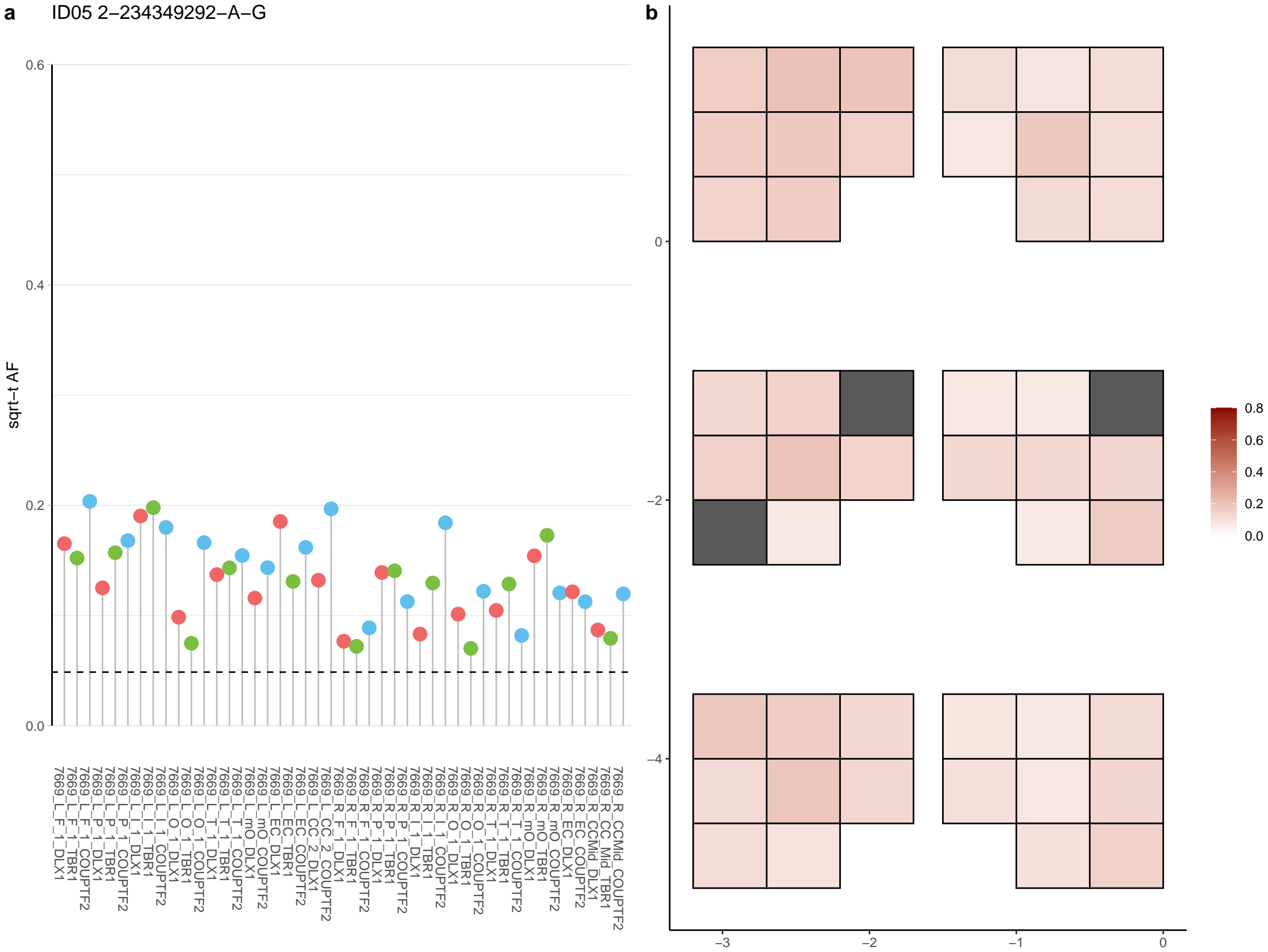

**a** ID05 2-240582791-G-A

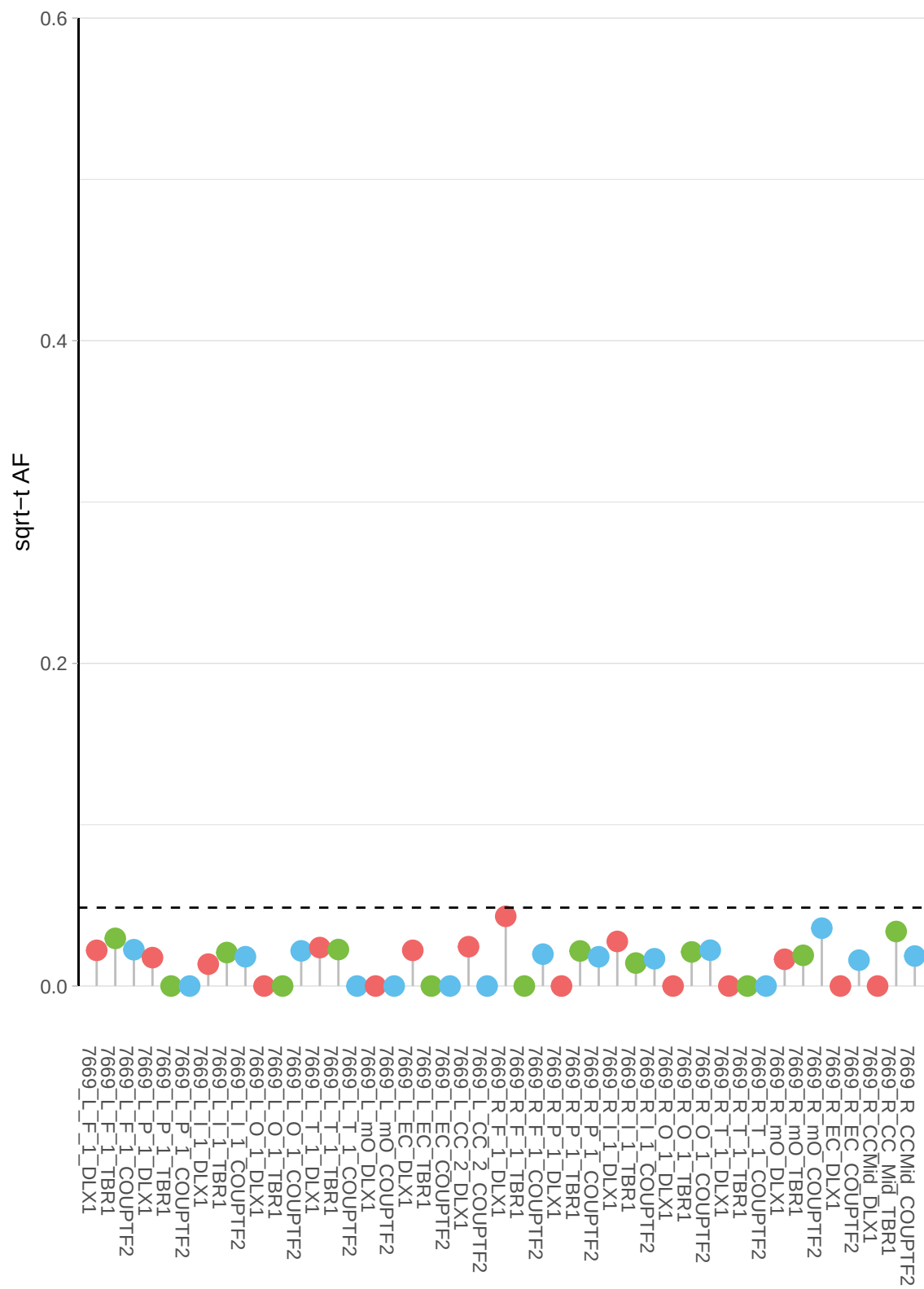

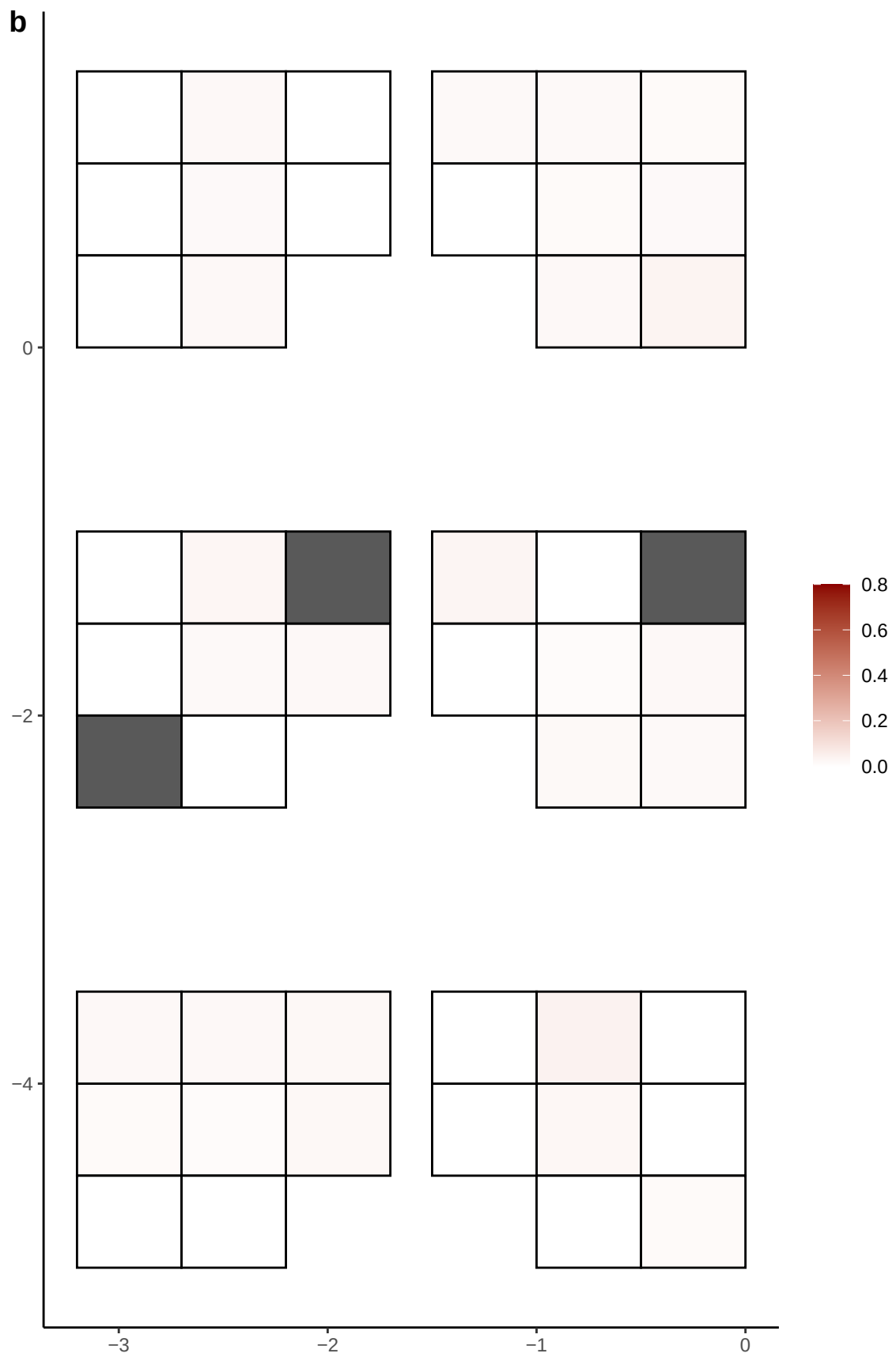

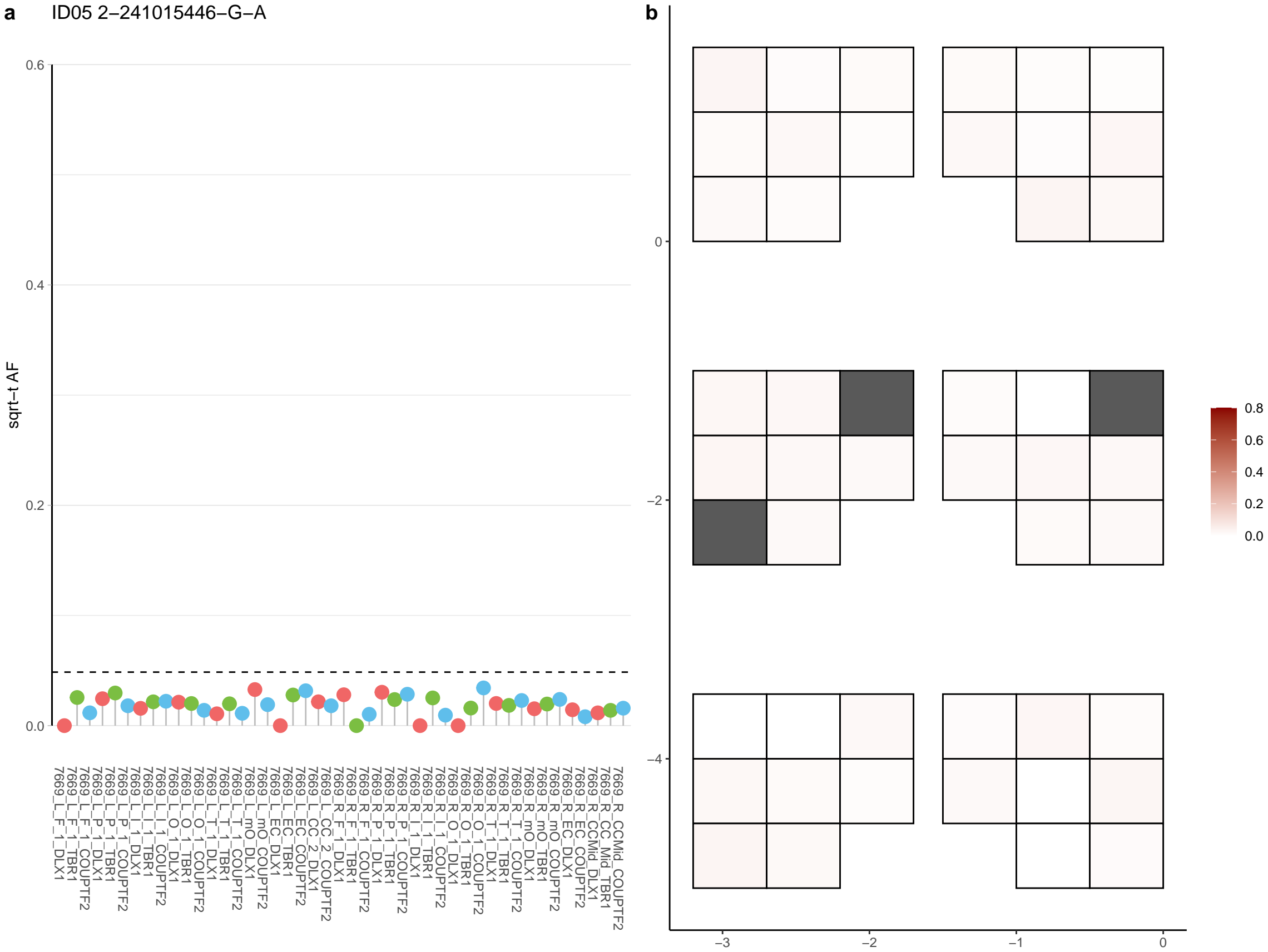

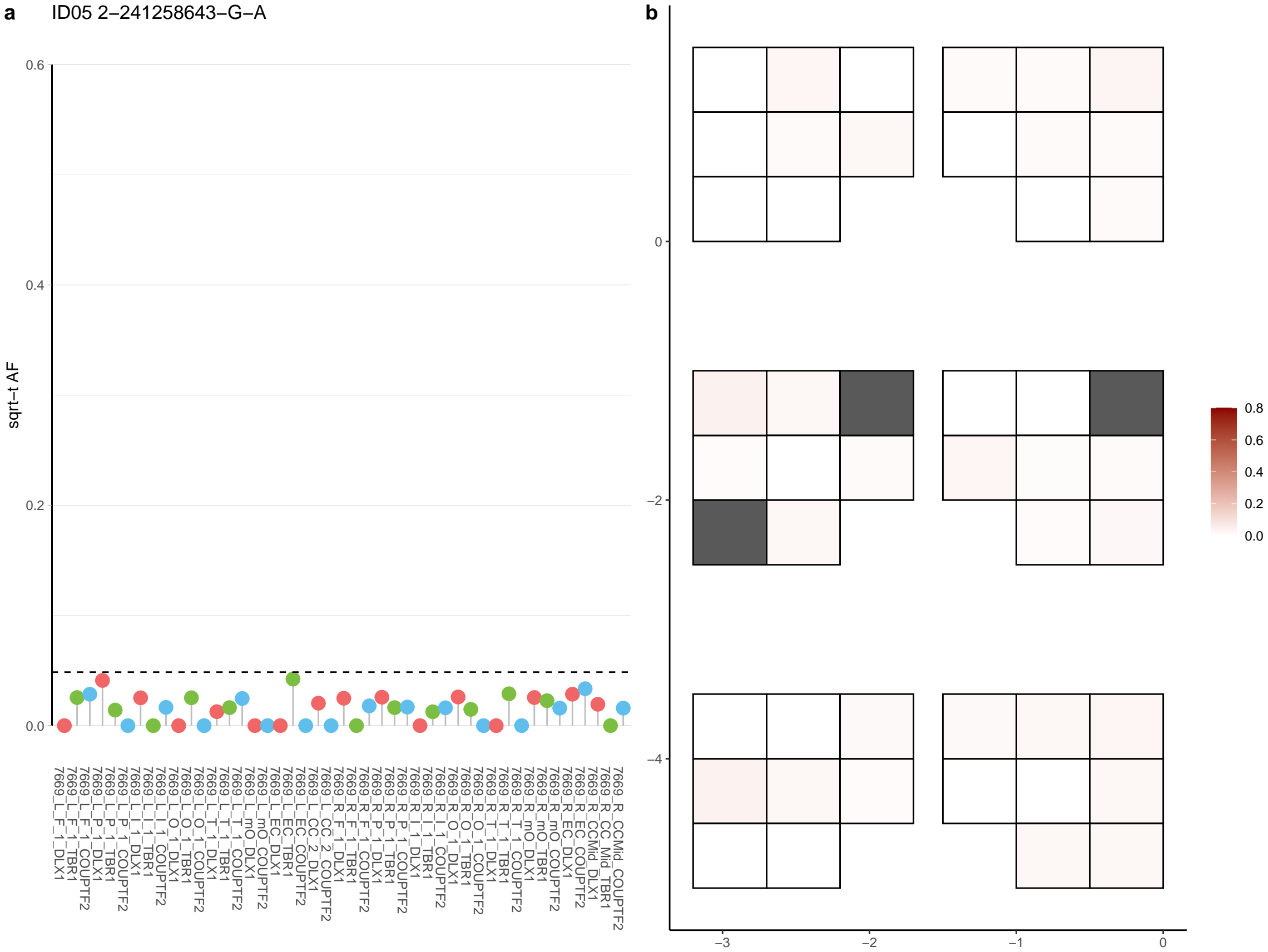

**a** ID05 2-27102490-G-A

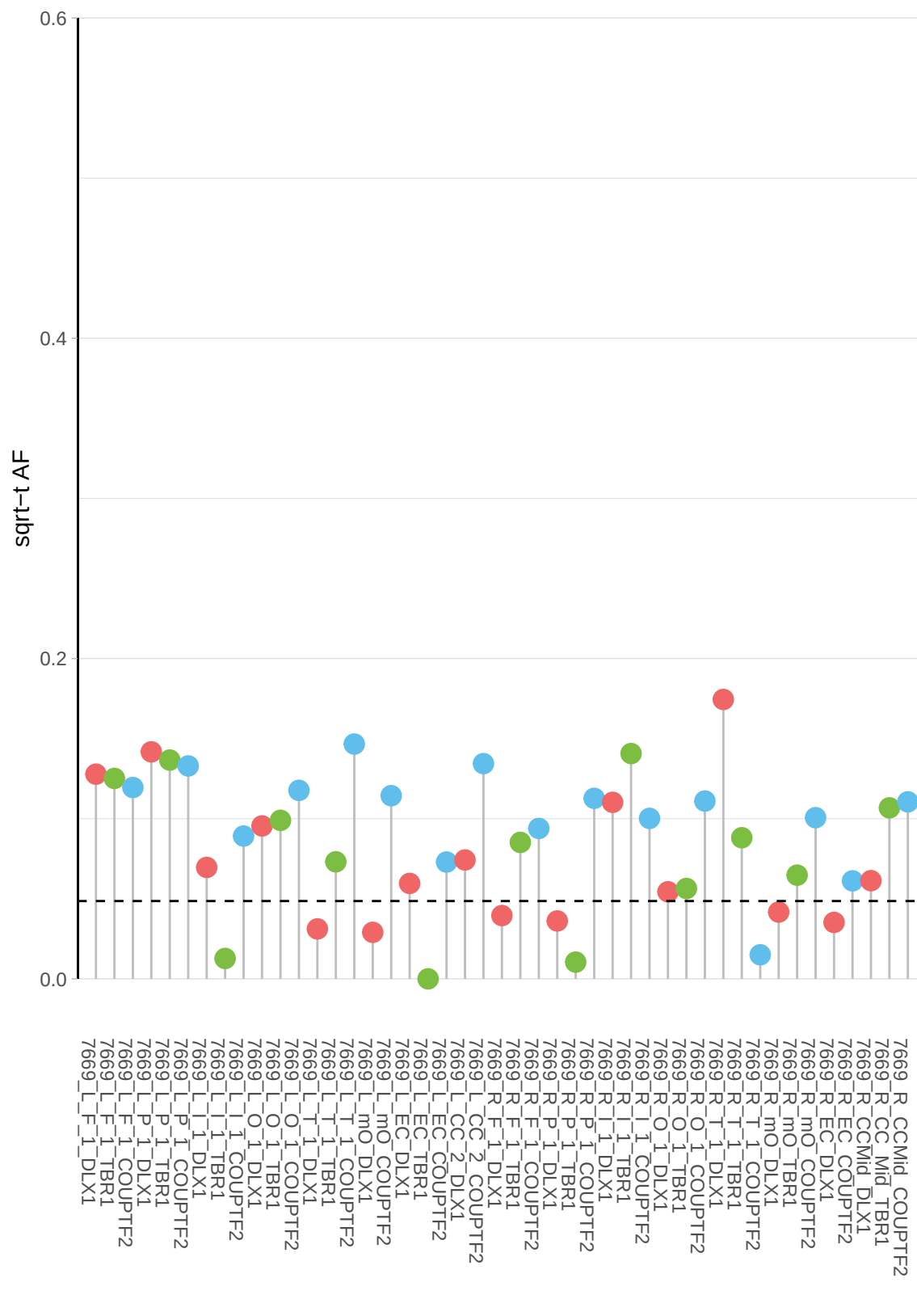

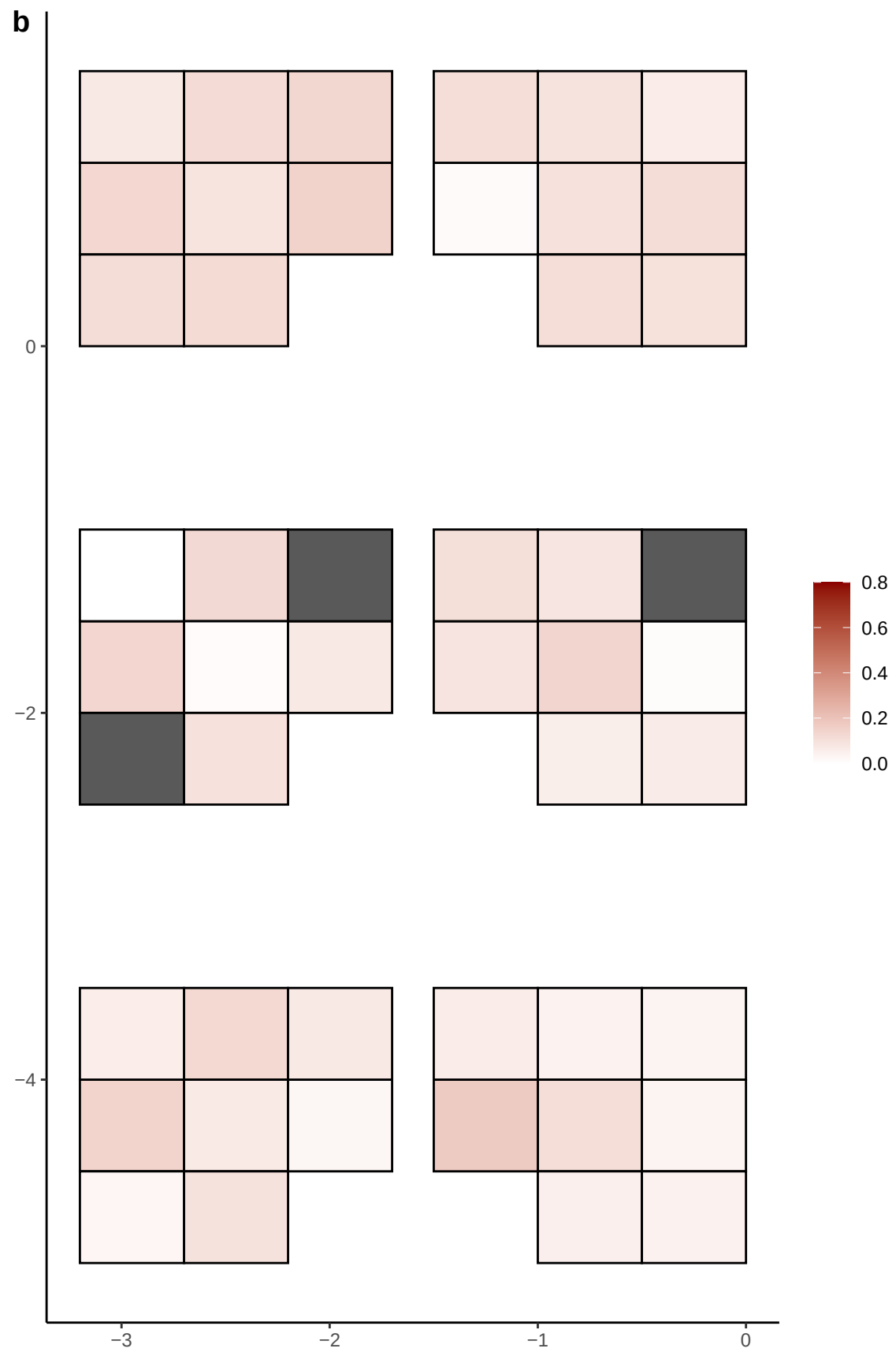

**a** ID05 2-3282940-C-T

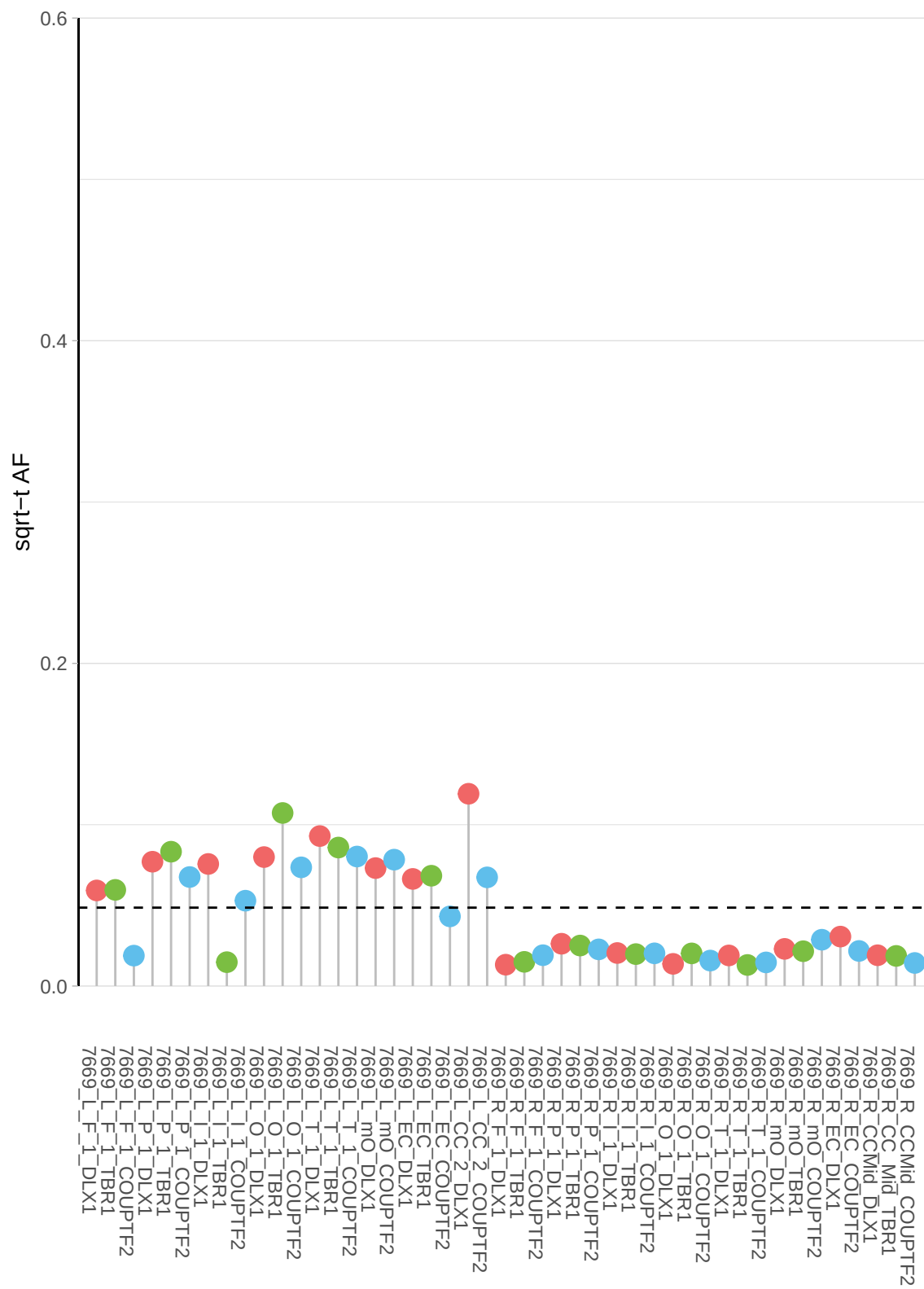

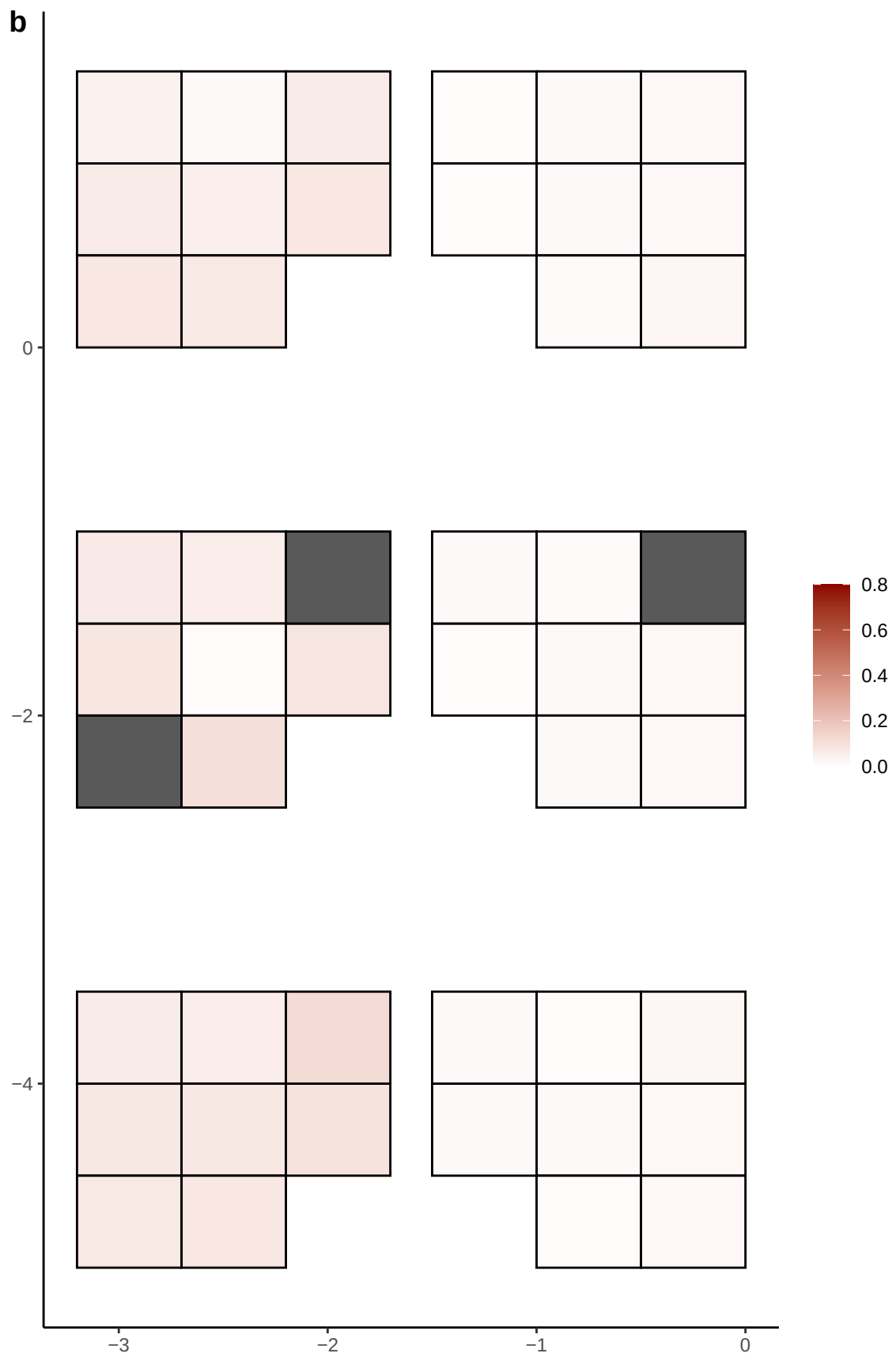

**a** ID05 2-34945794-T-C

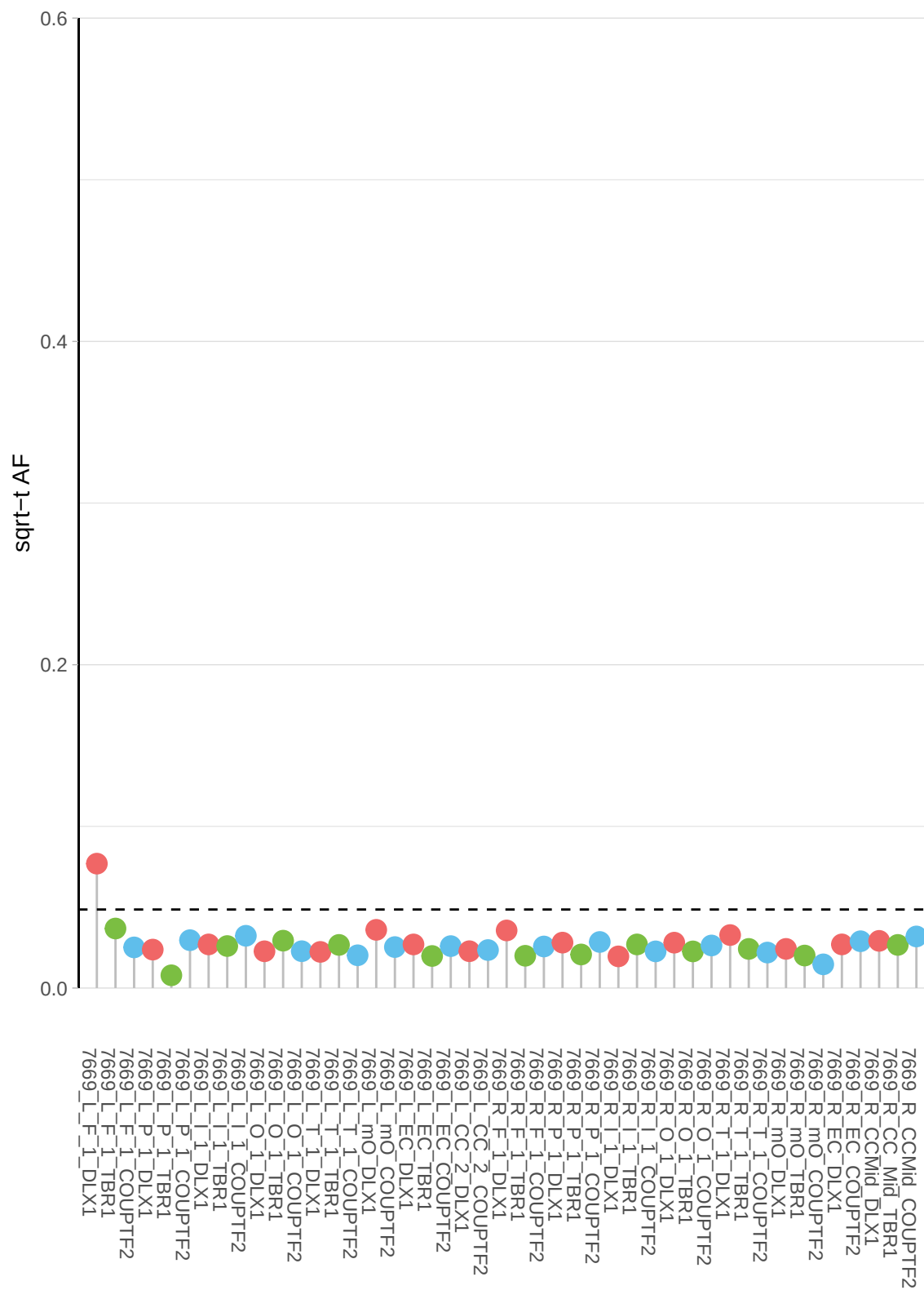

**b**

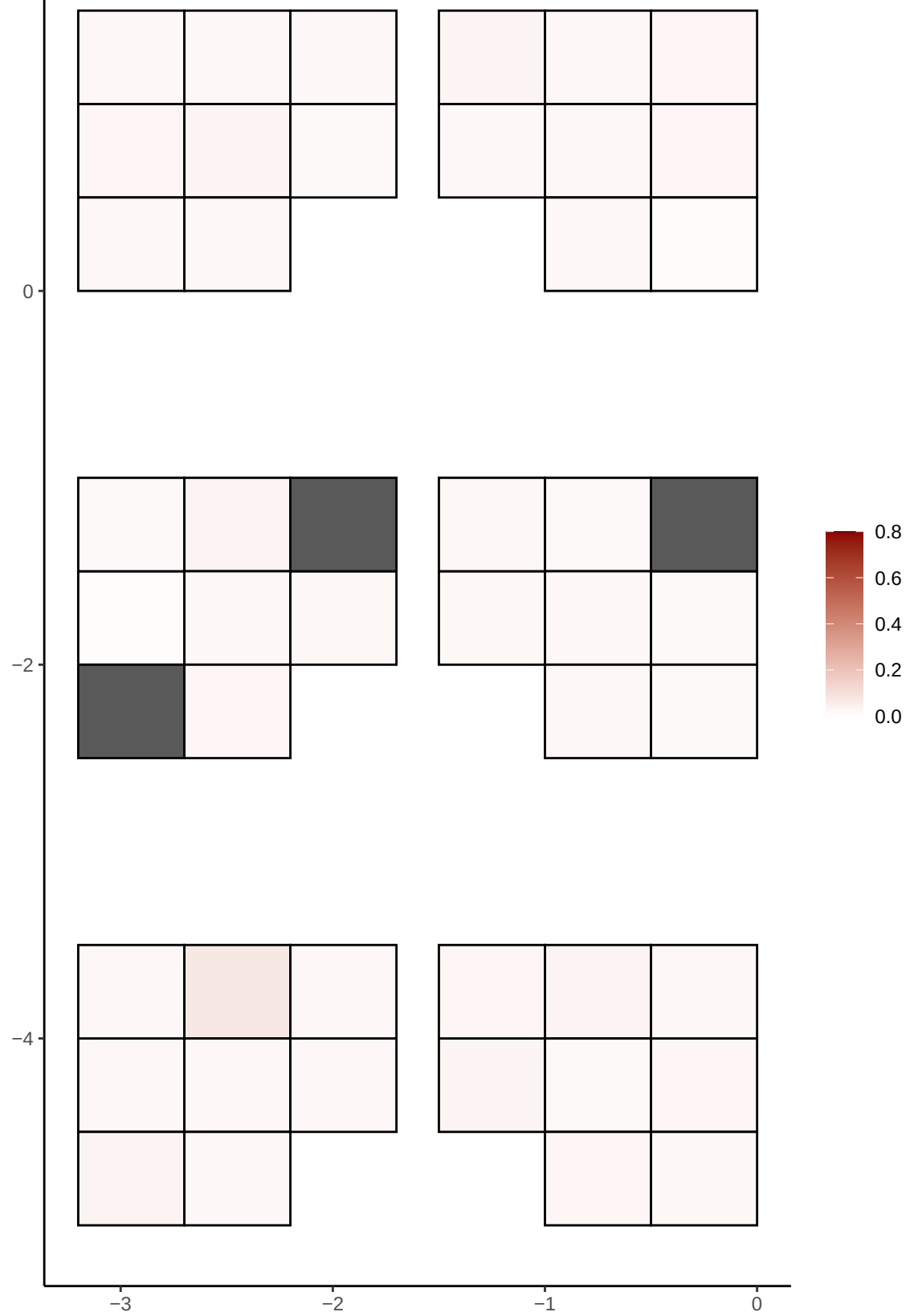

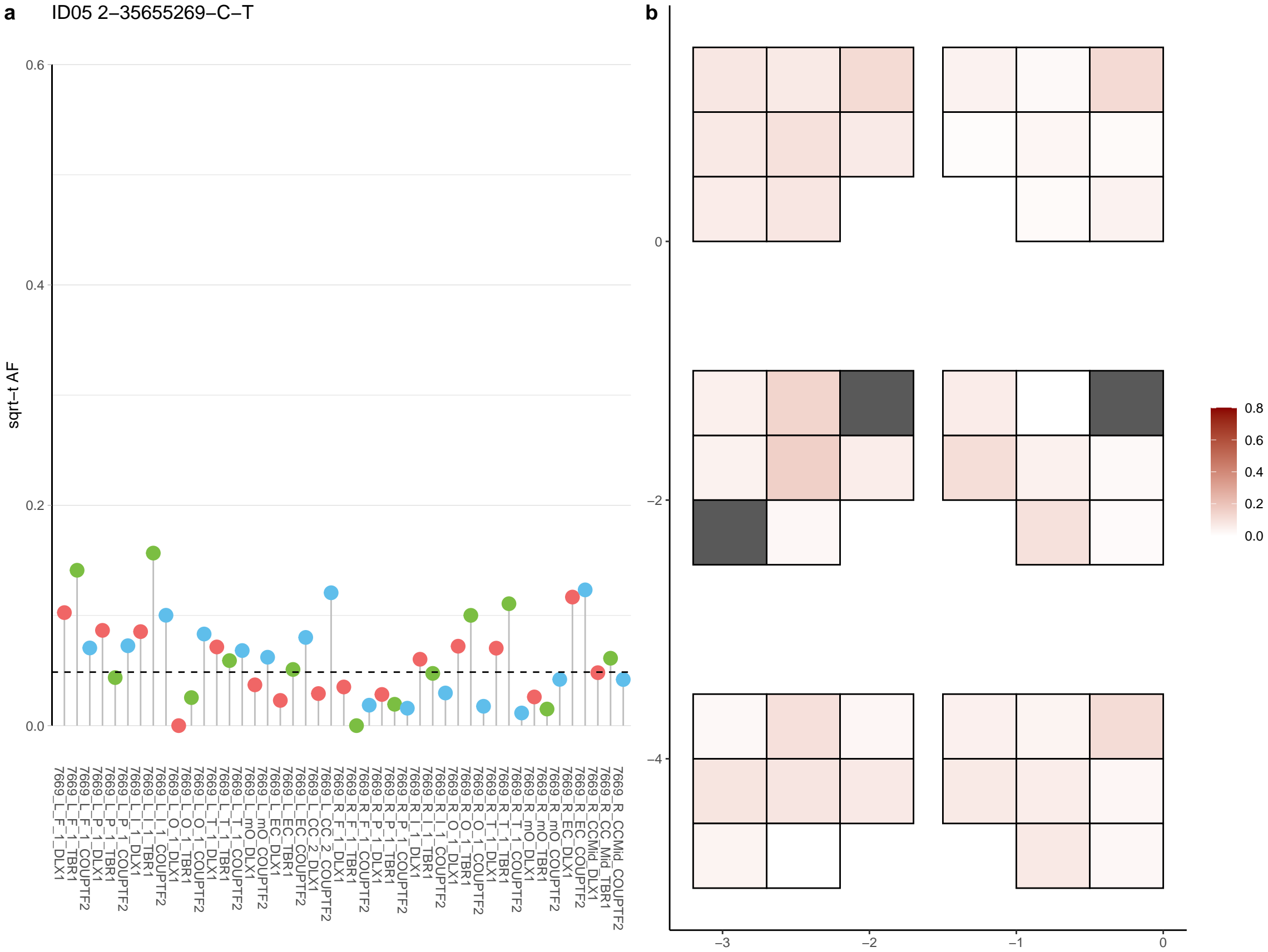

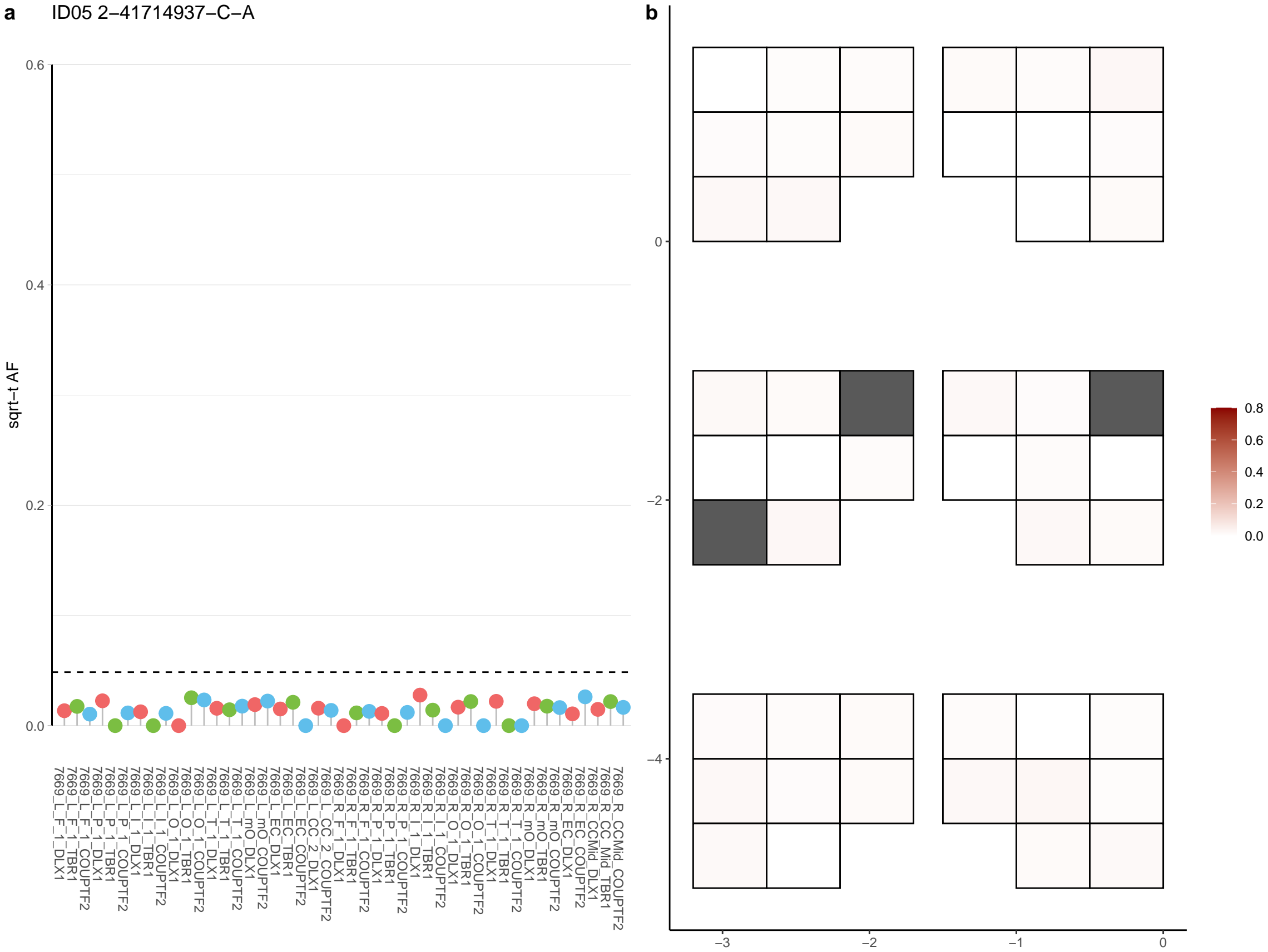

a ID05 2-424135-C-T

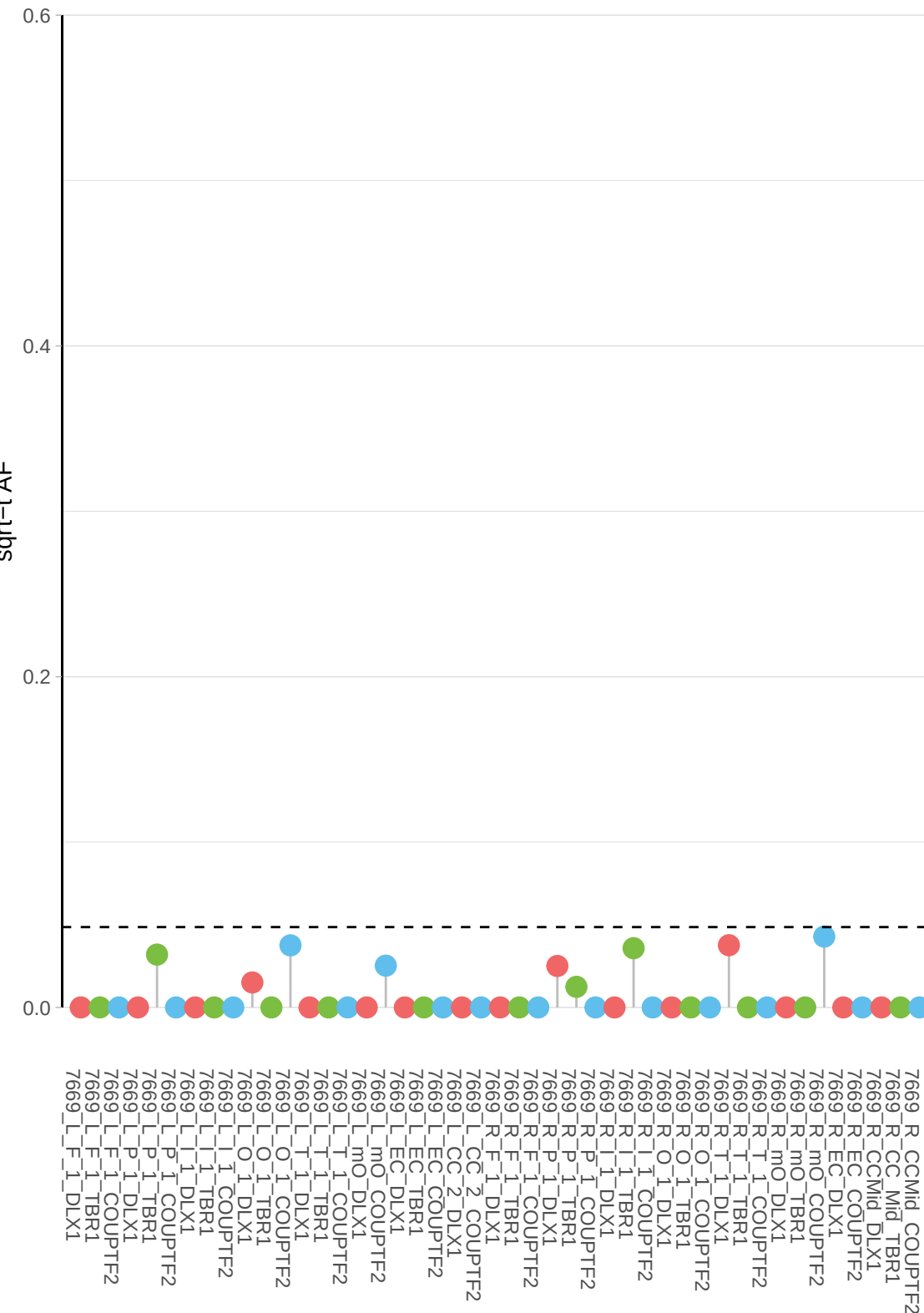

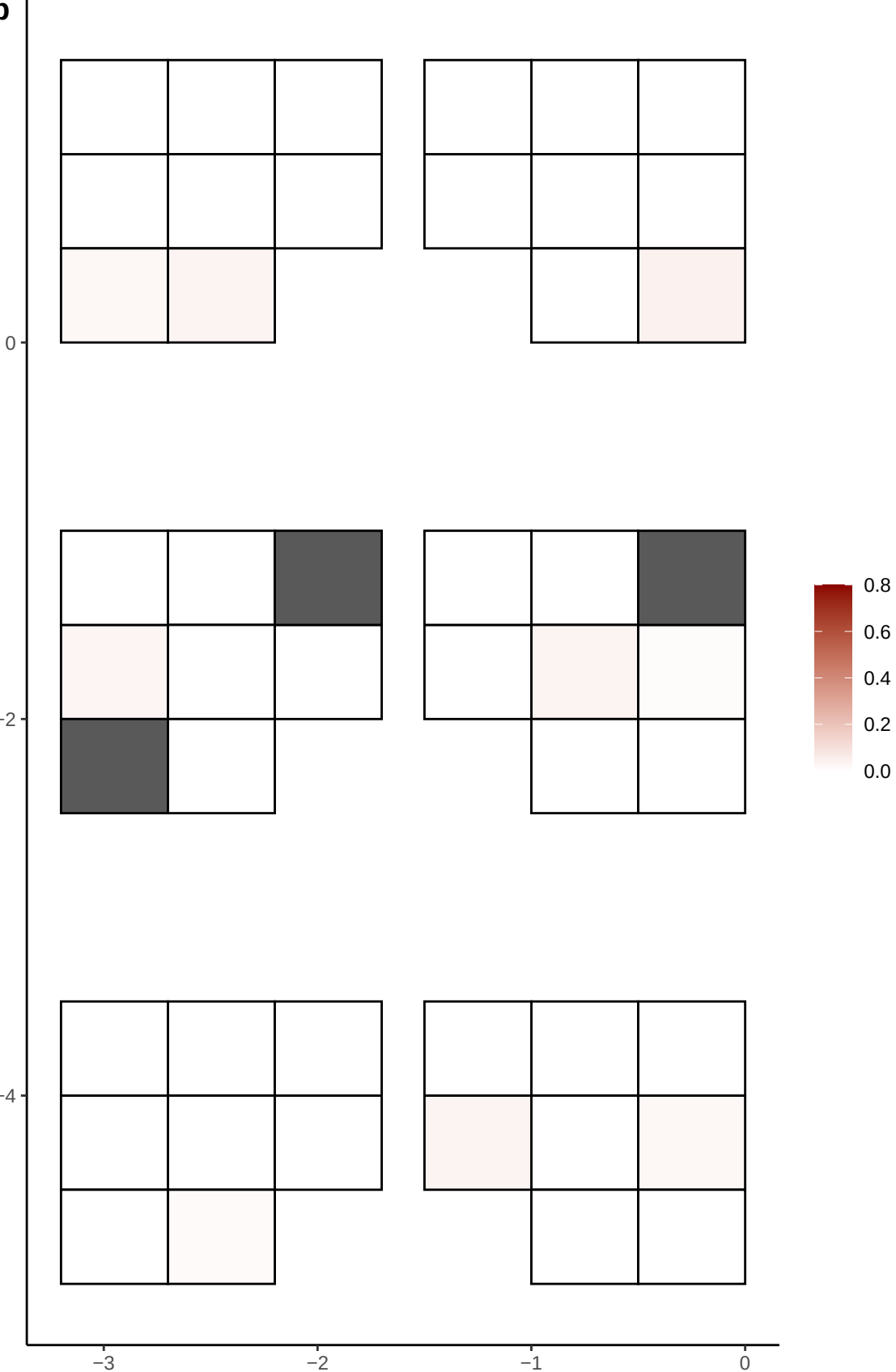

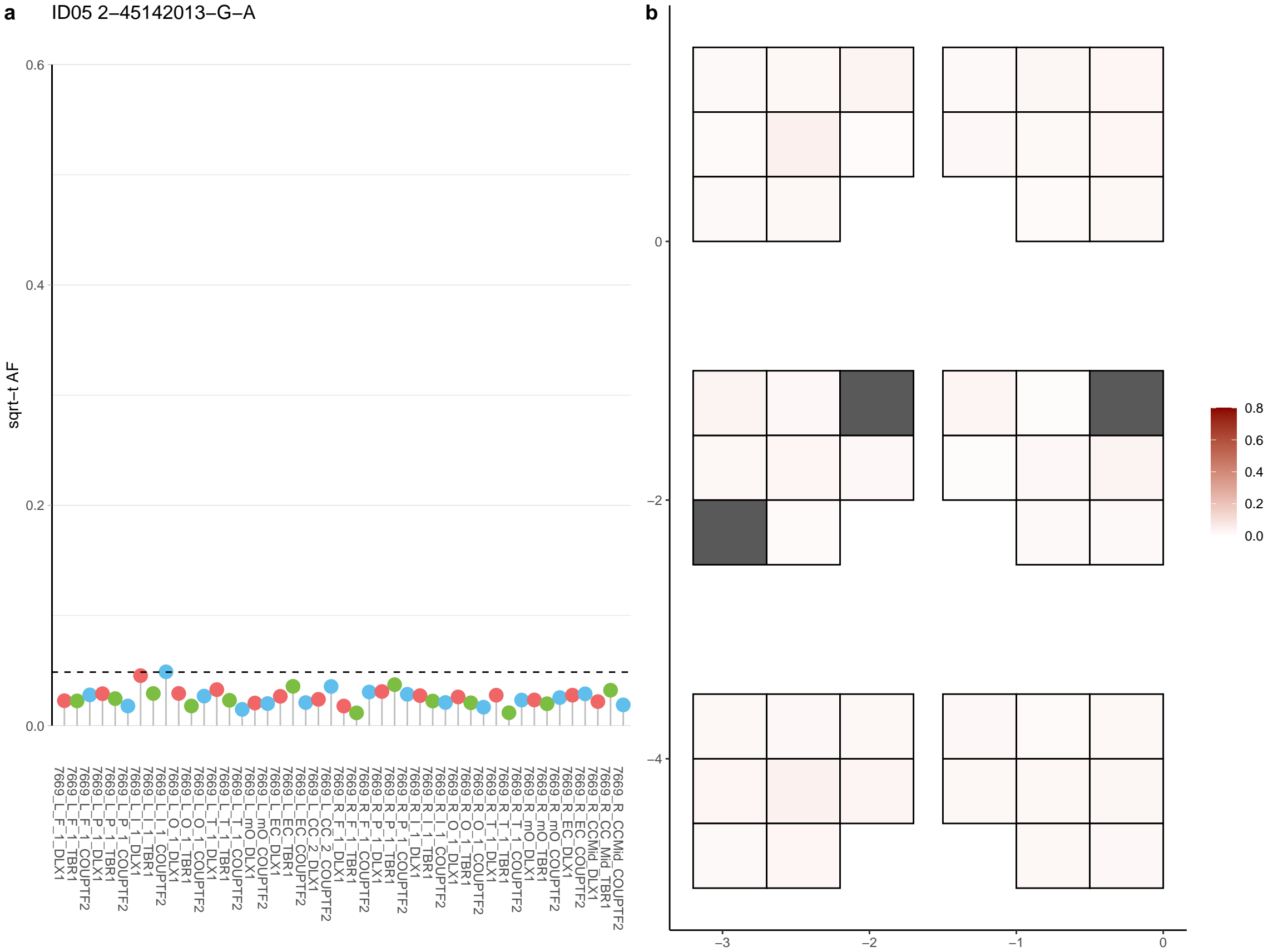

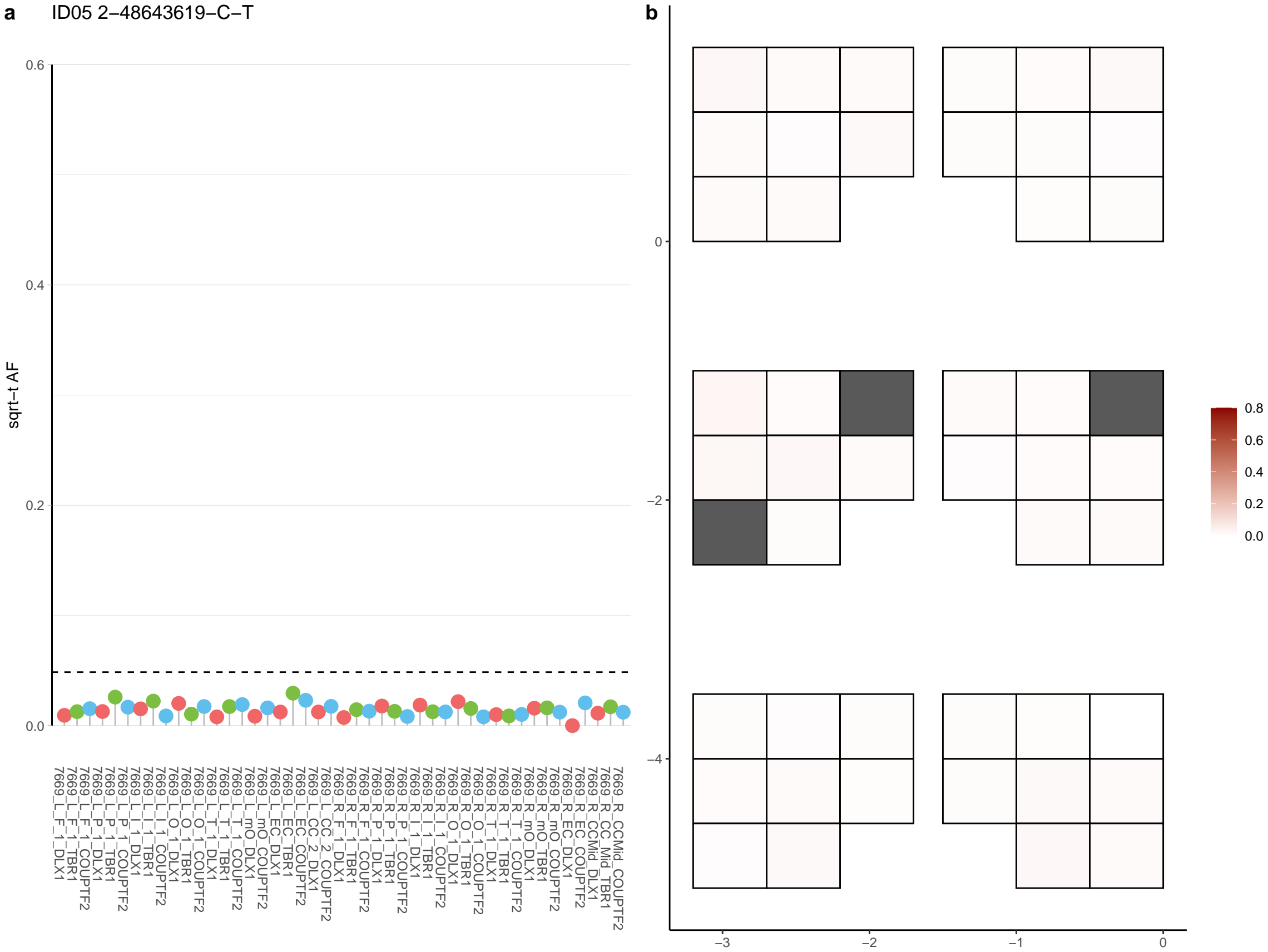

**a** ID05 2-52452769-G-A

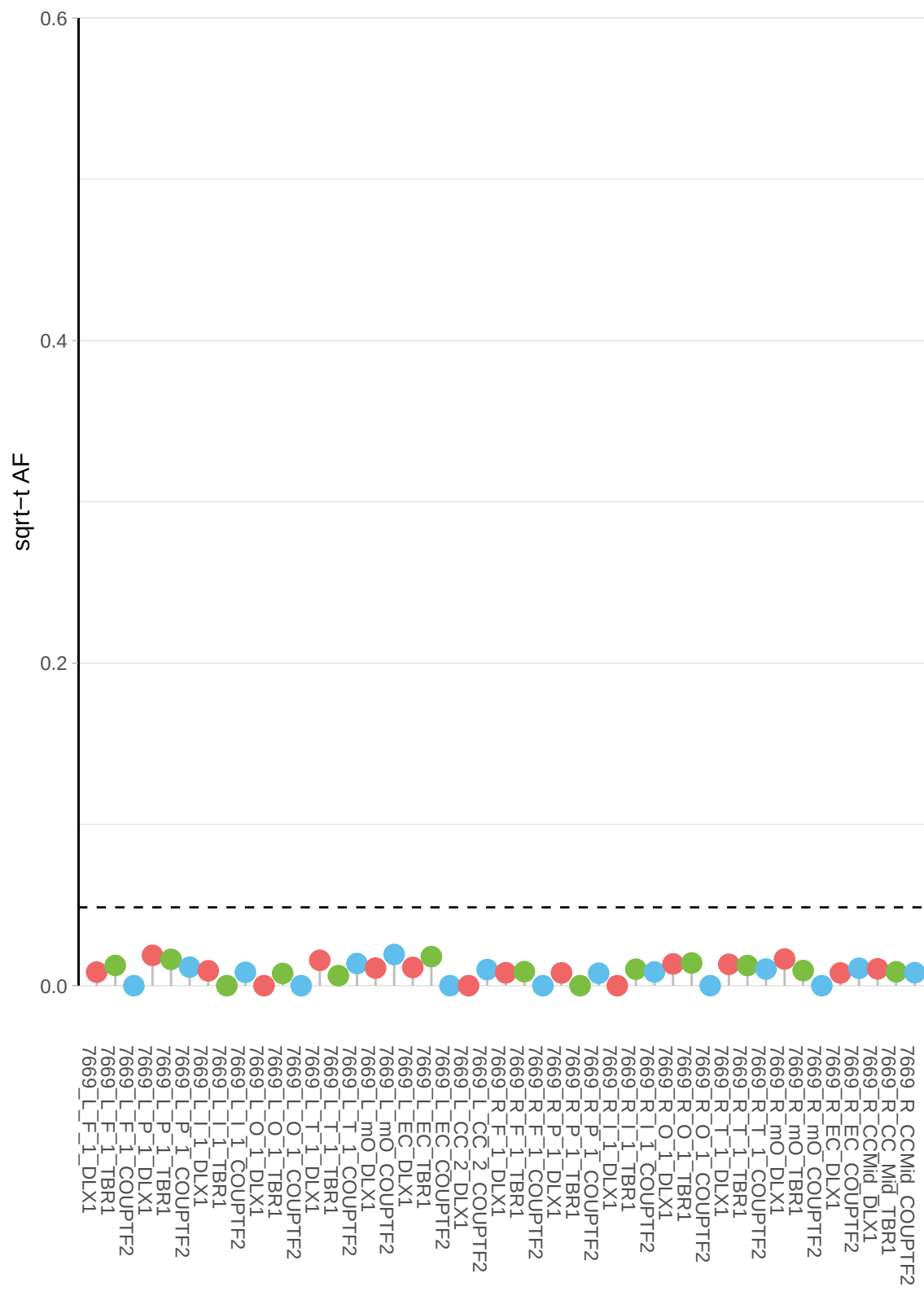

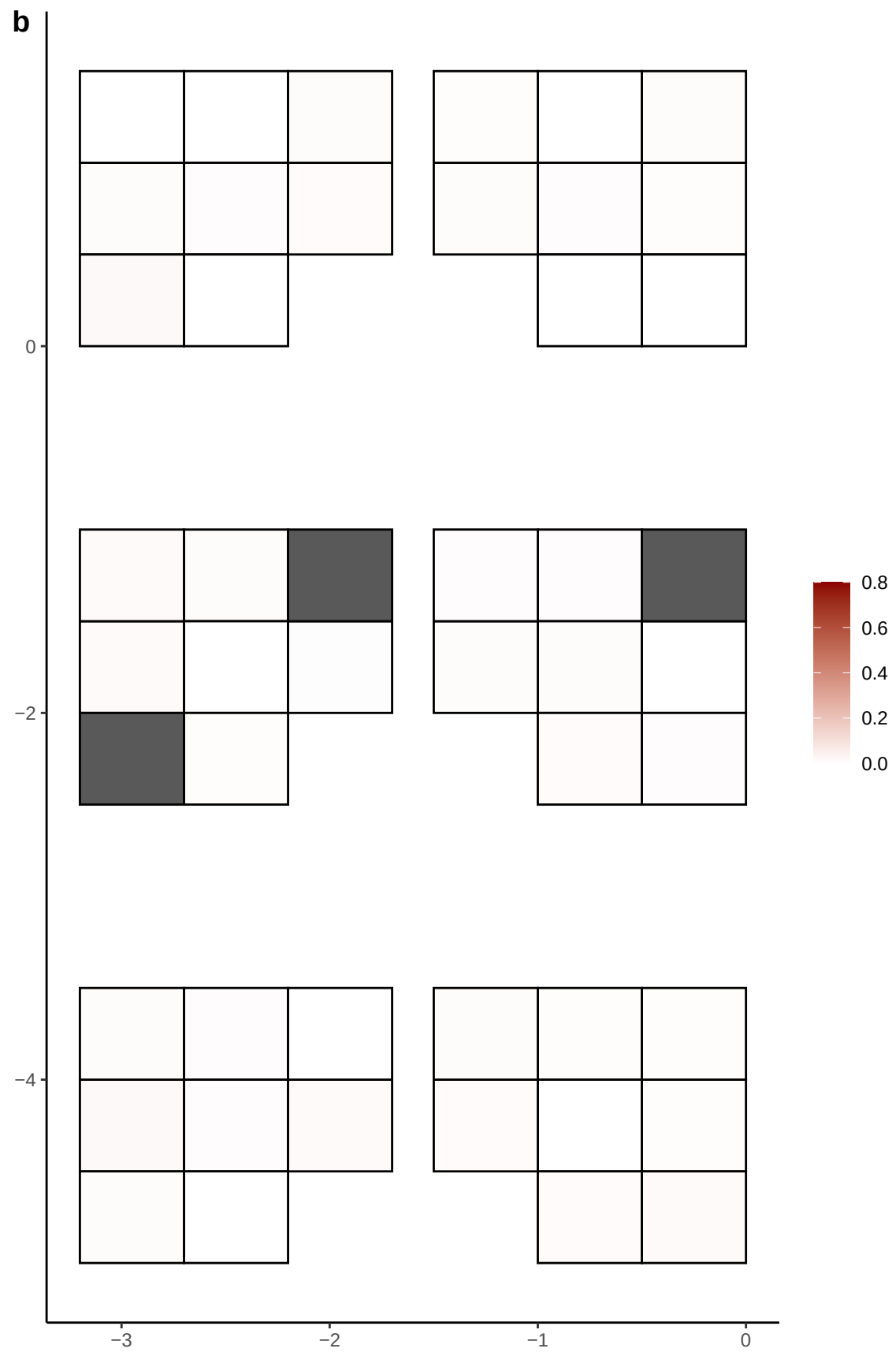

**a** ID05 2-53839419-G-A

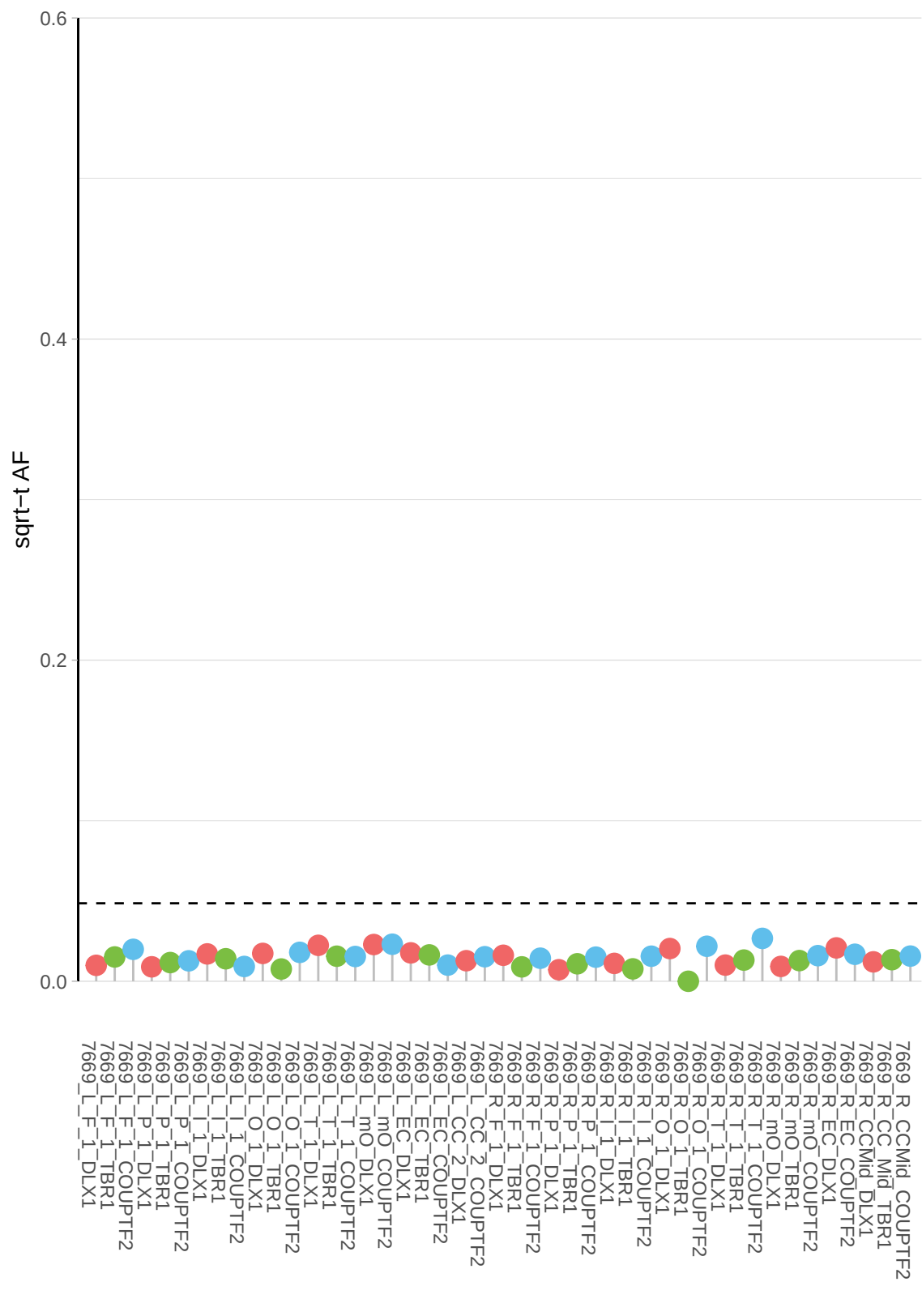

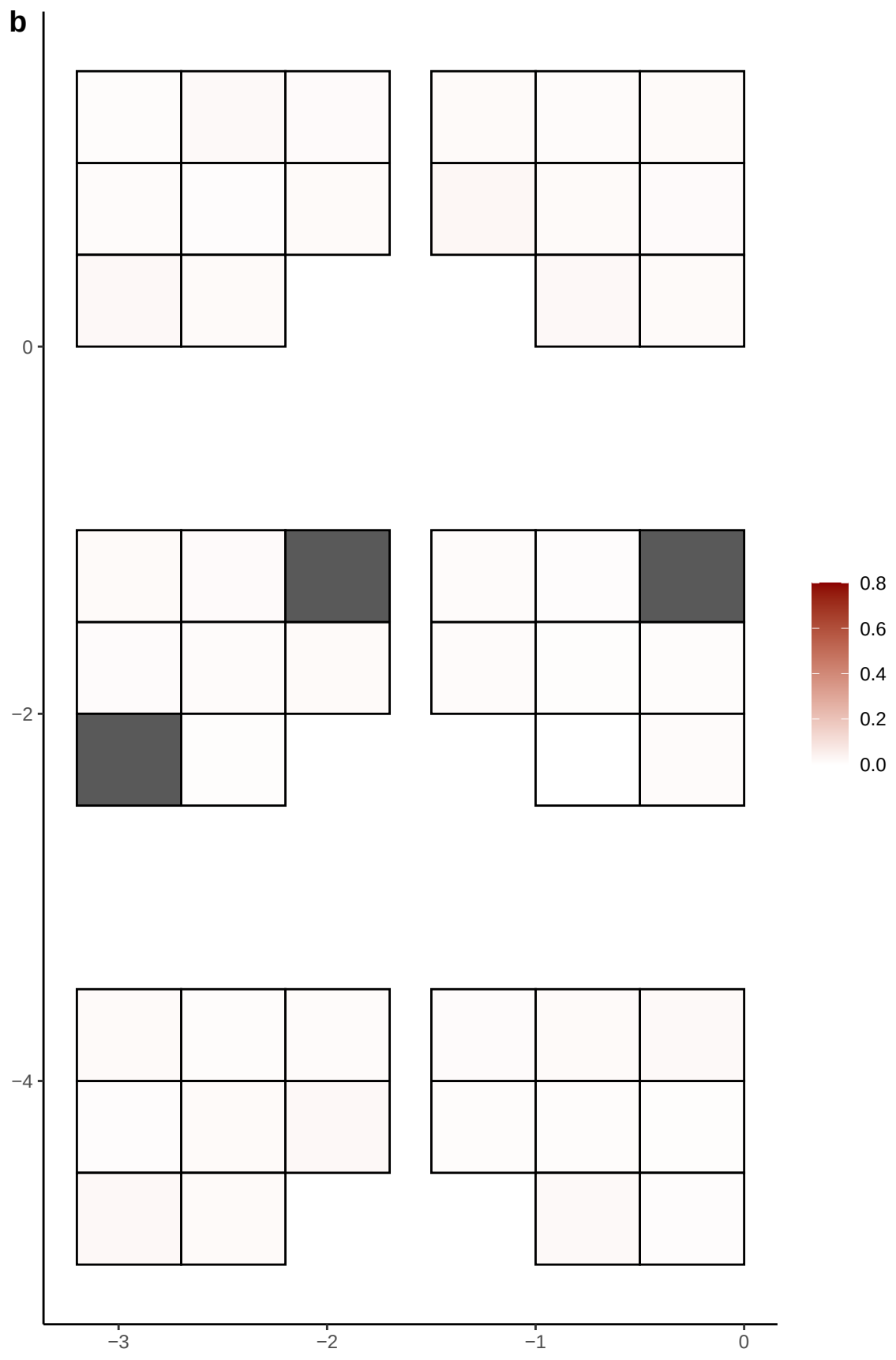

**a** ID05 2-57021433-T-C

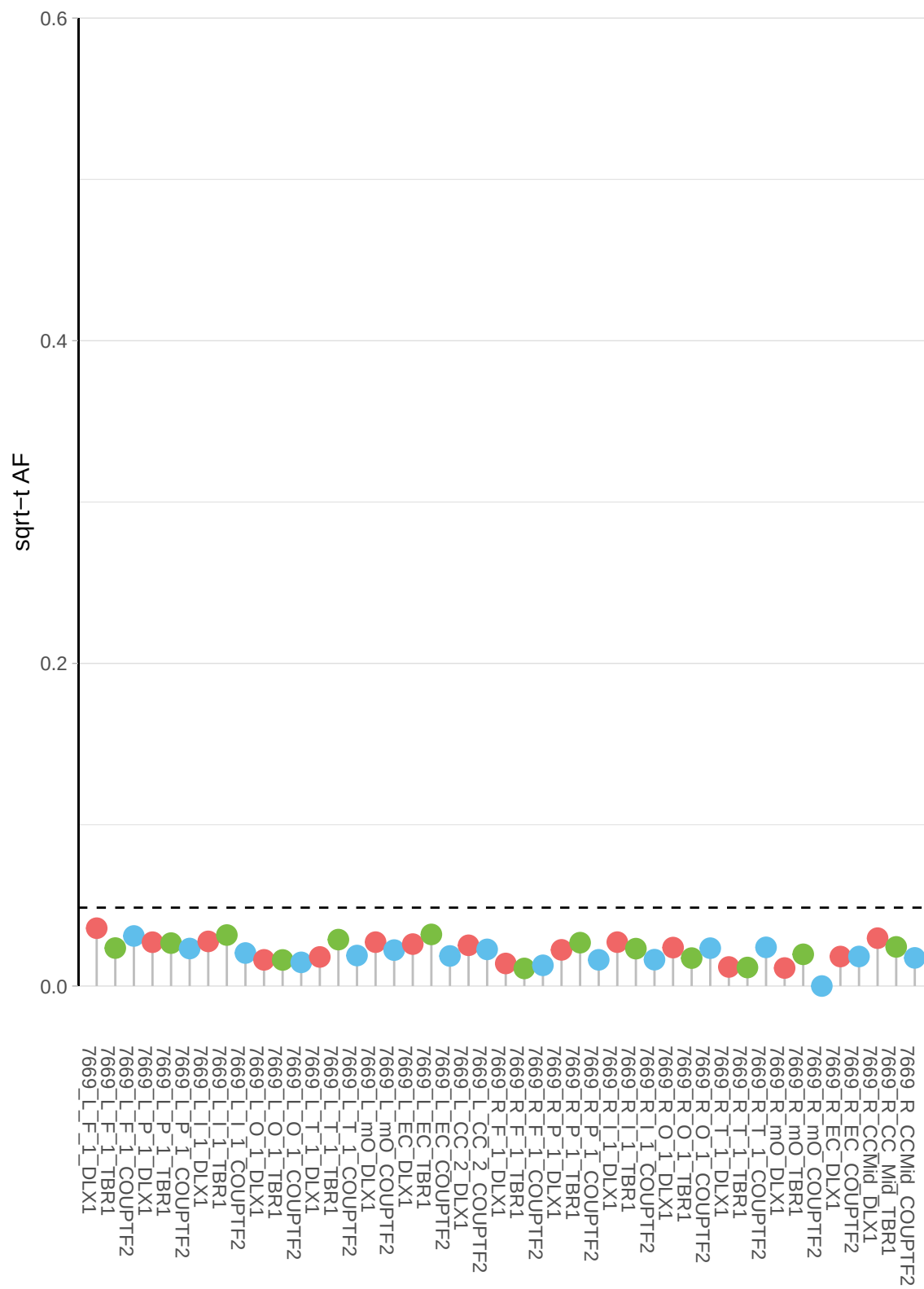

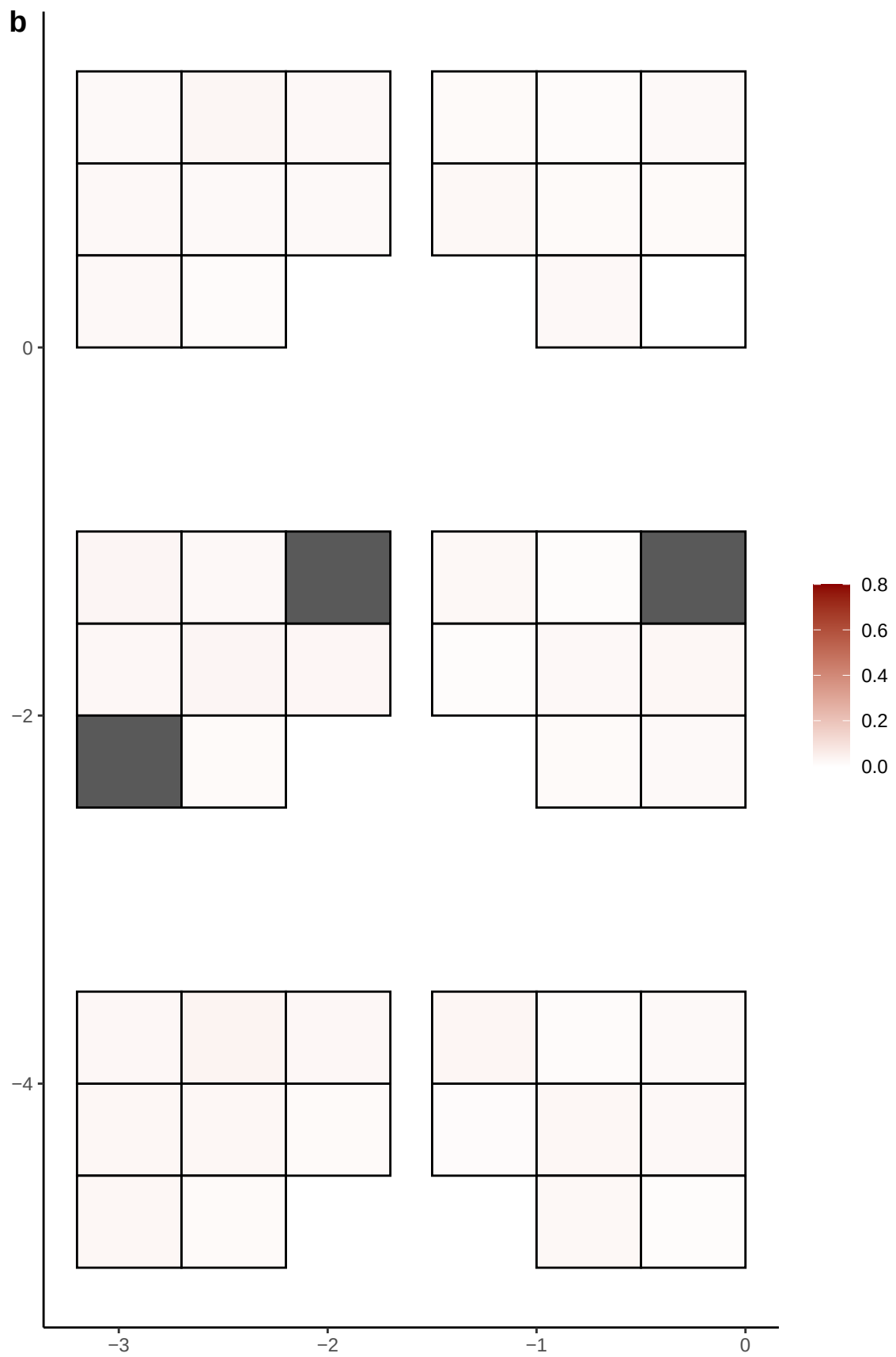

**a** ID05 2-64648189-C-T

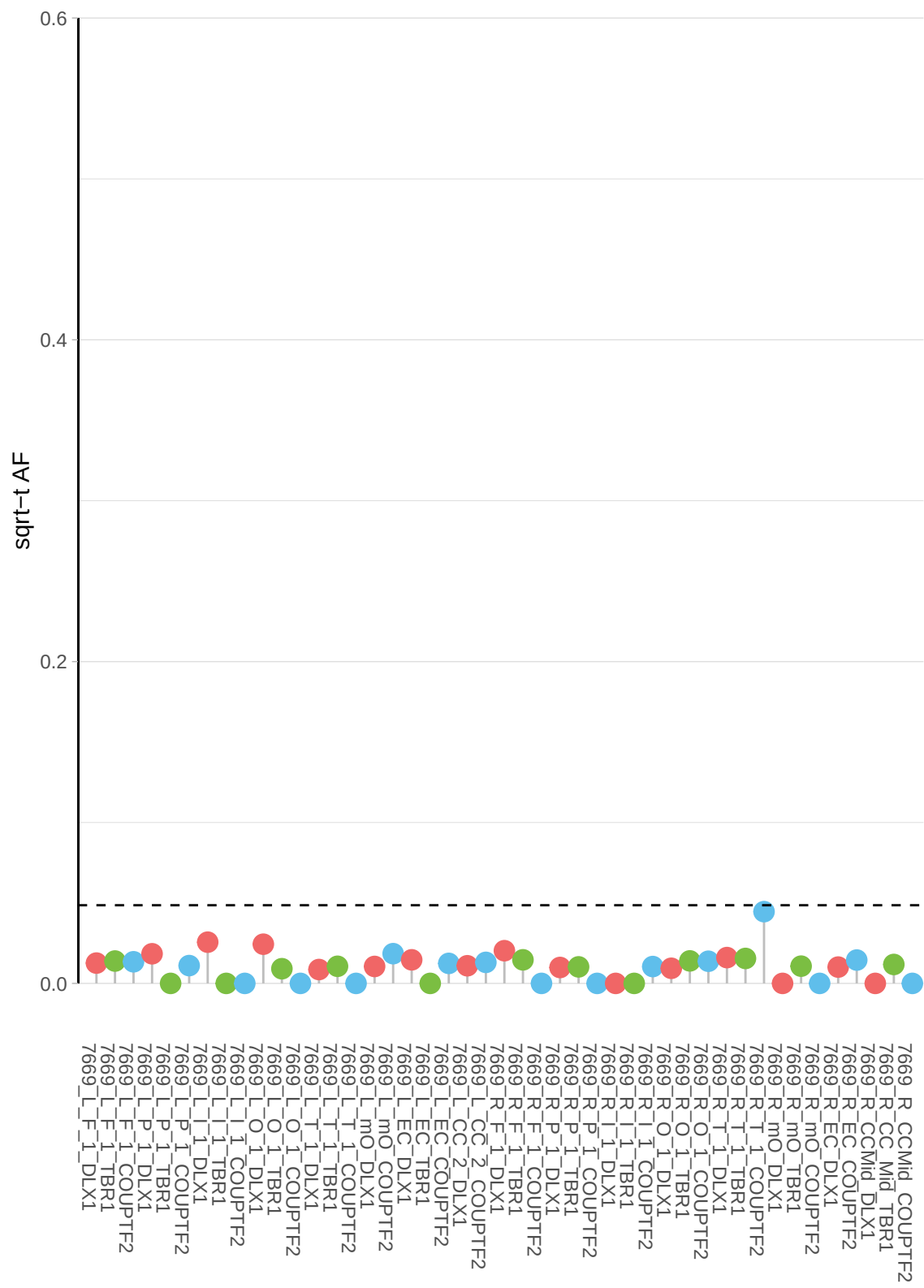

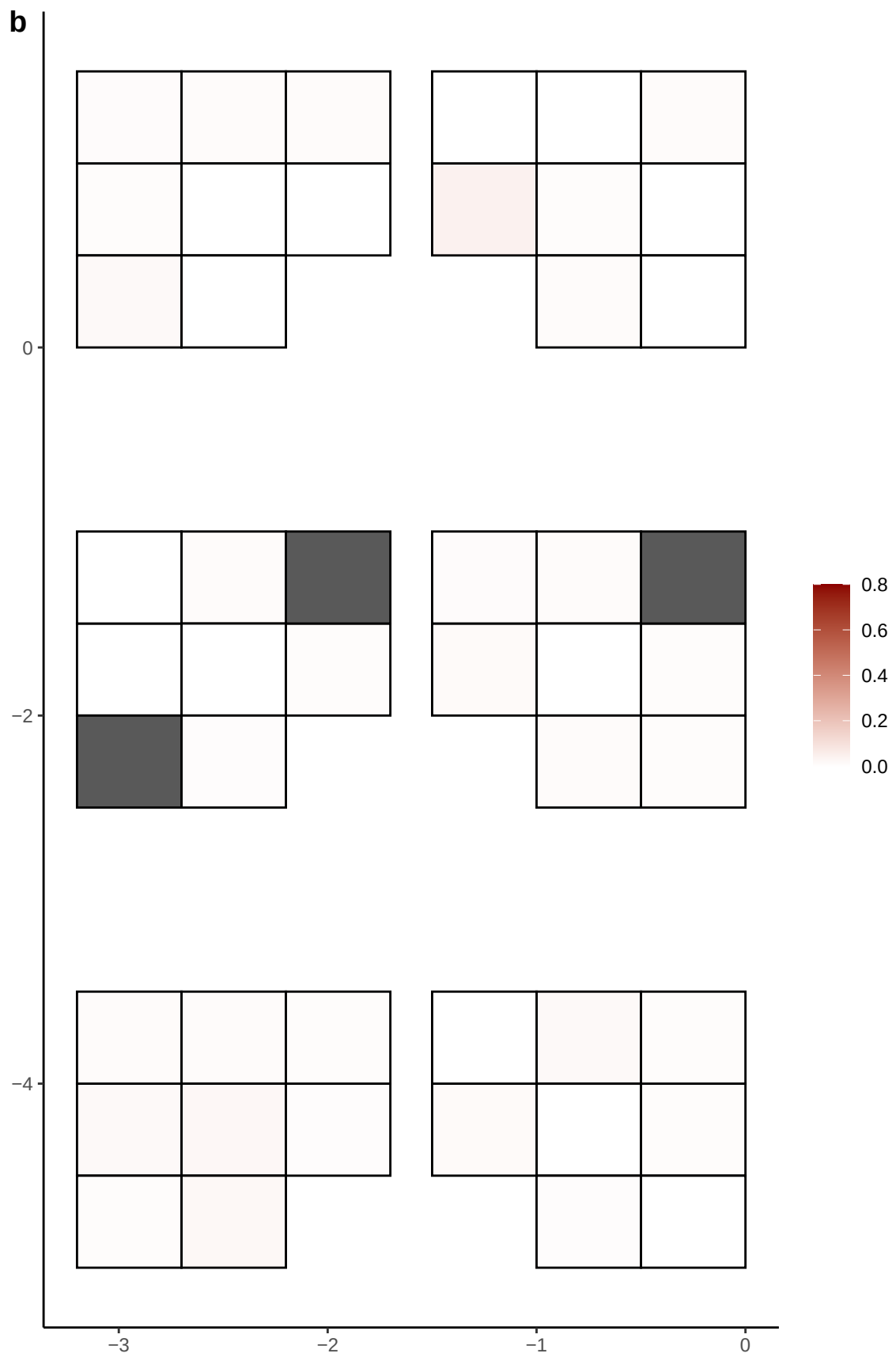

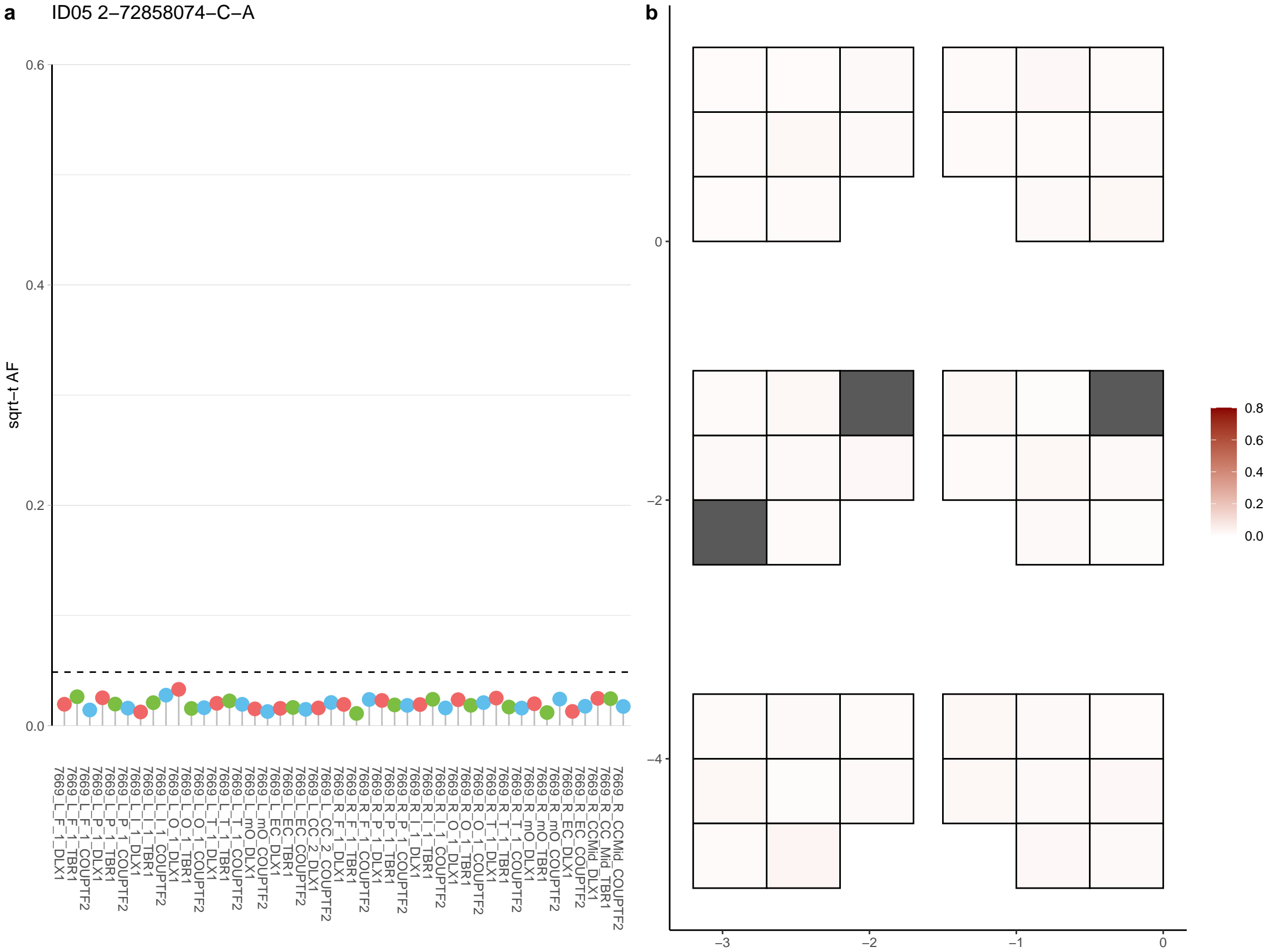

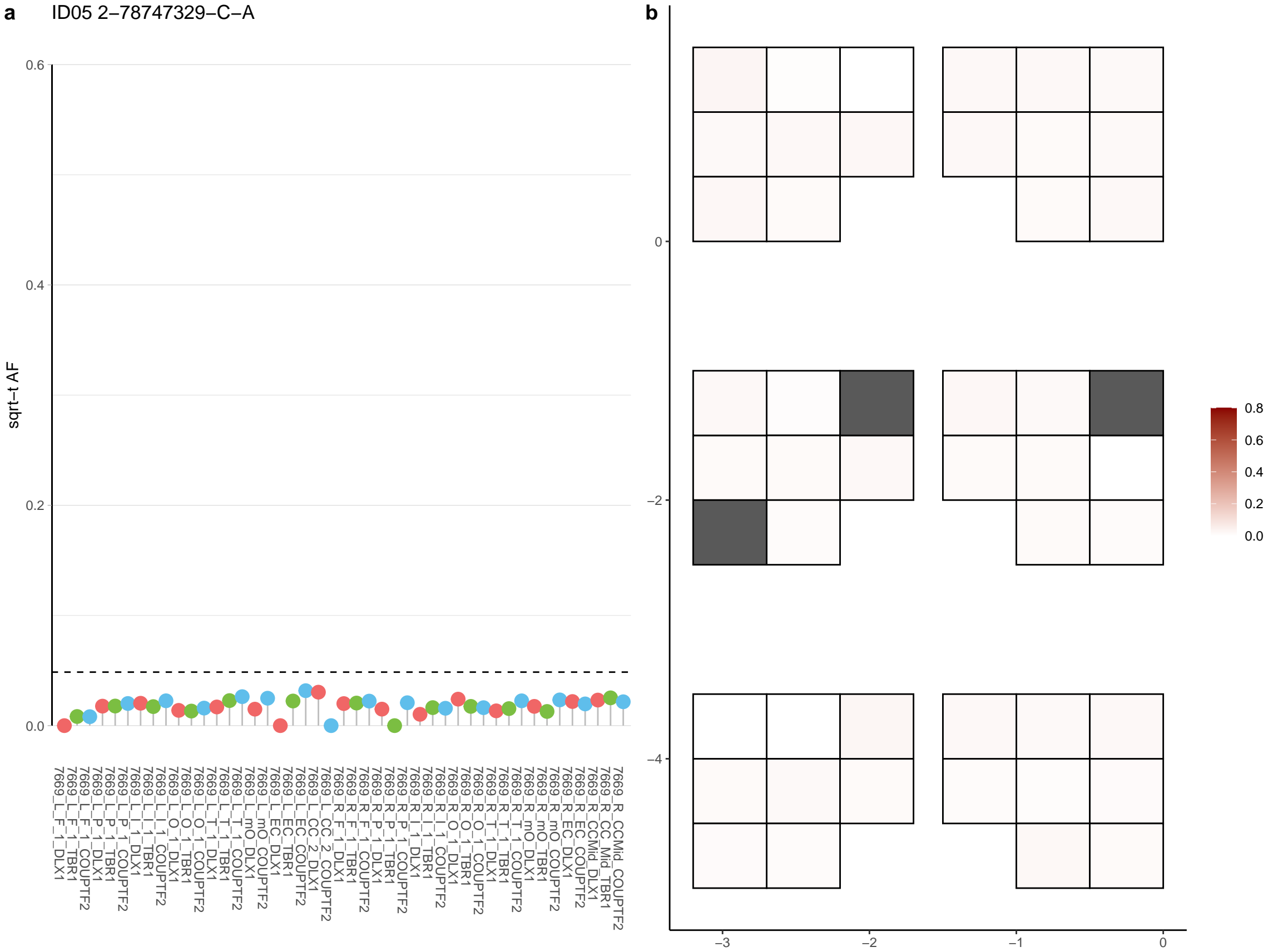

**a** ID05 2-82189643-C-A

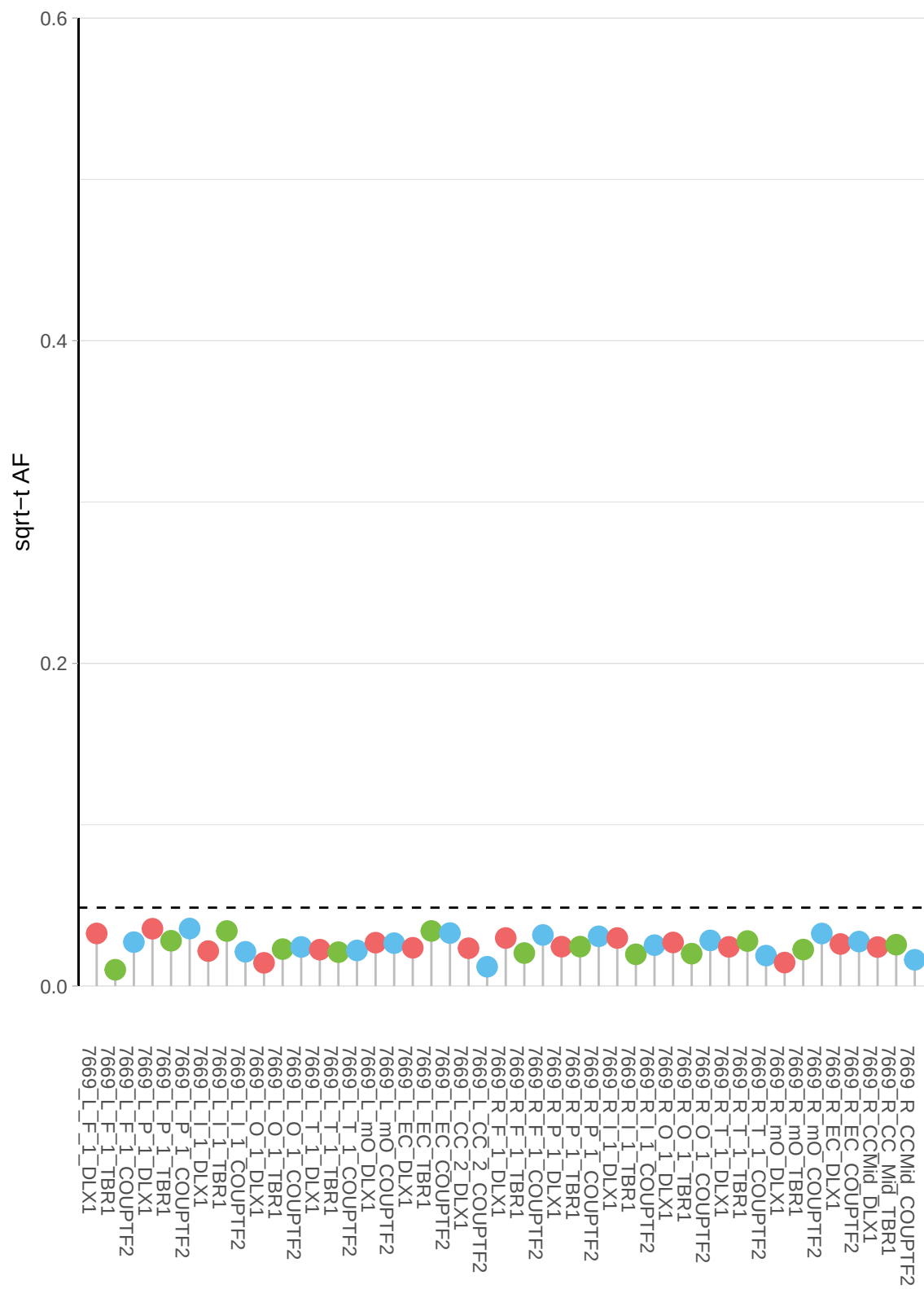

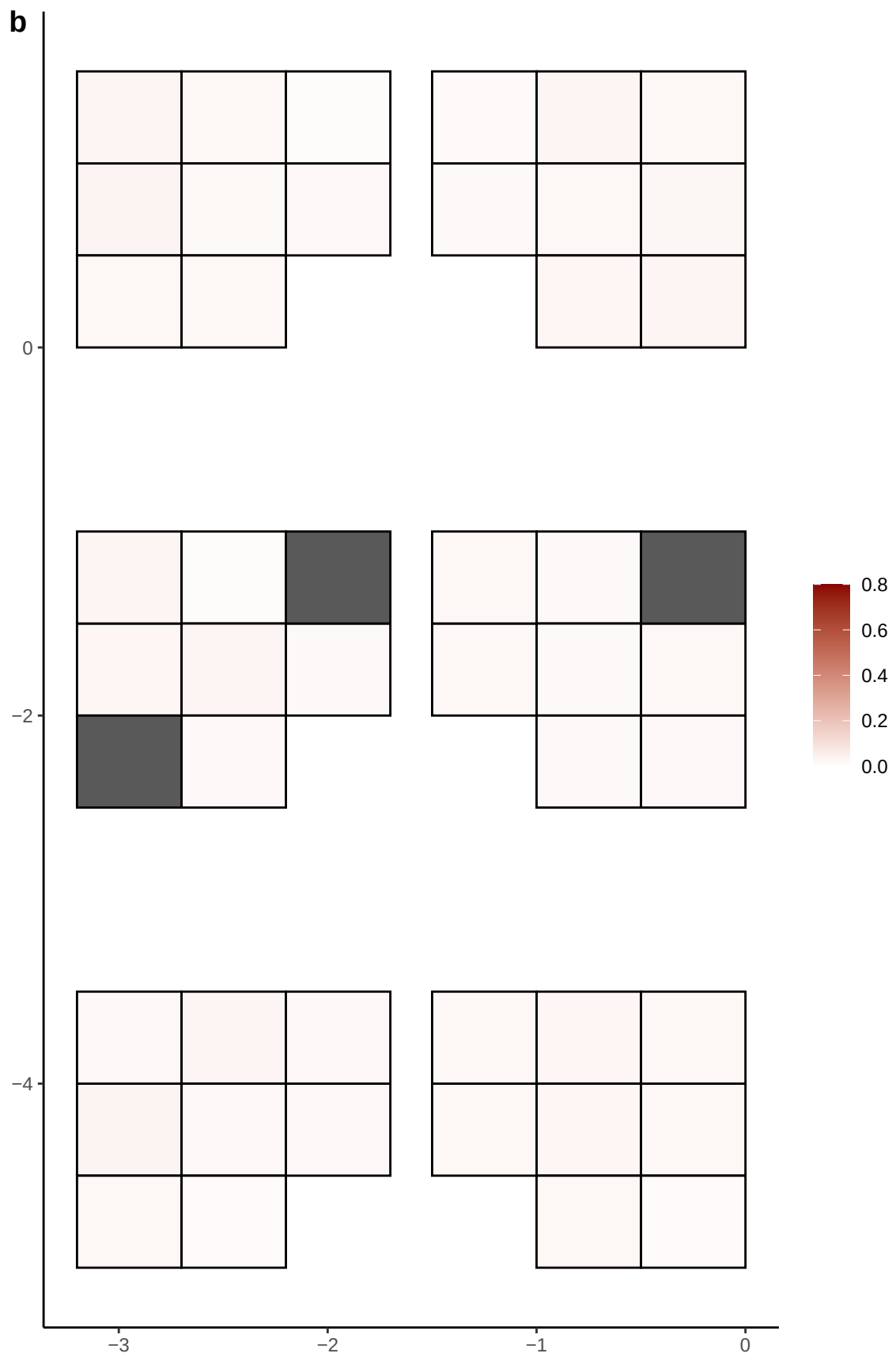

**a** ID05 2-86260205-G-A

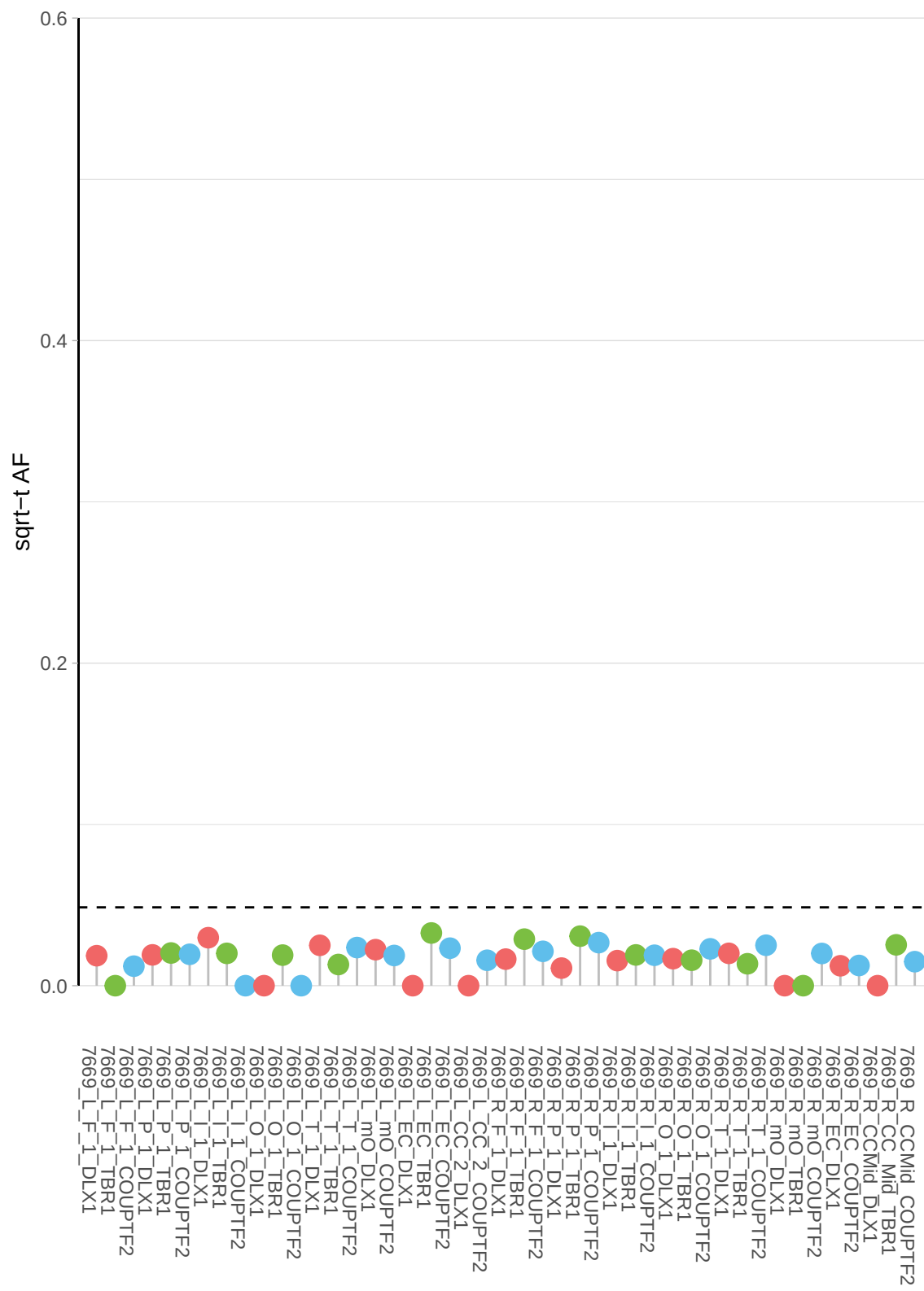

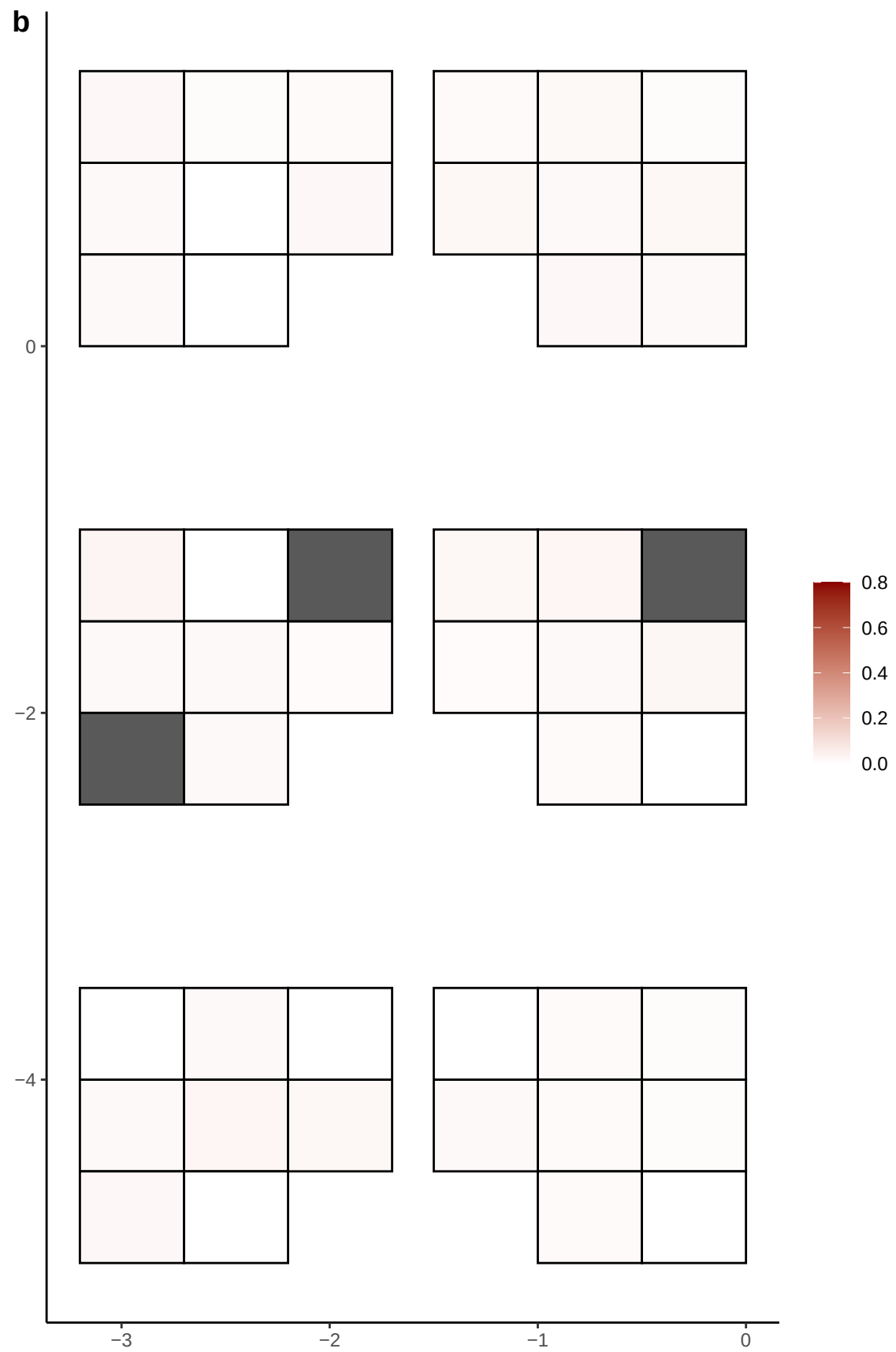

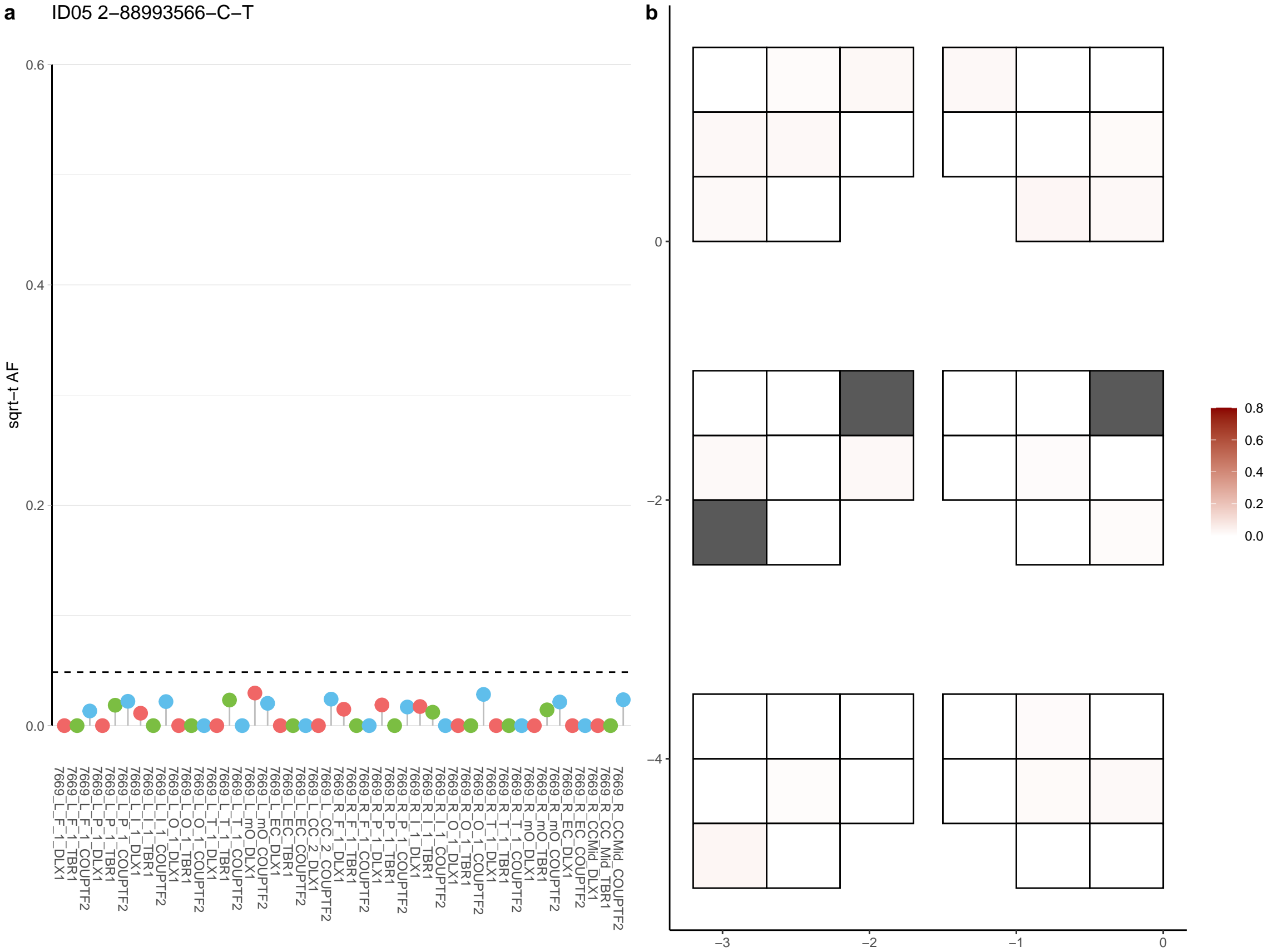

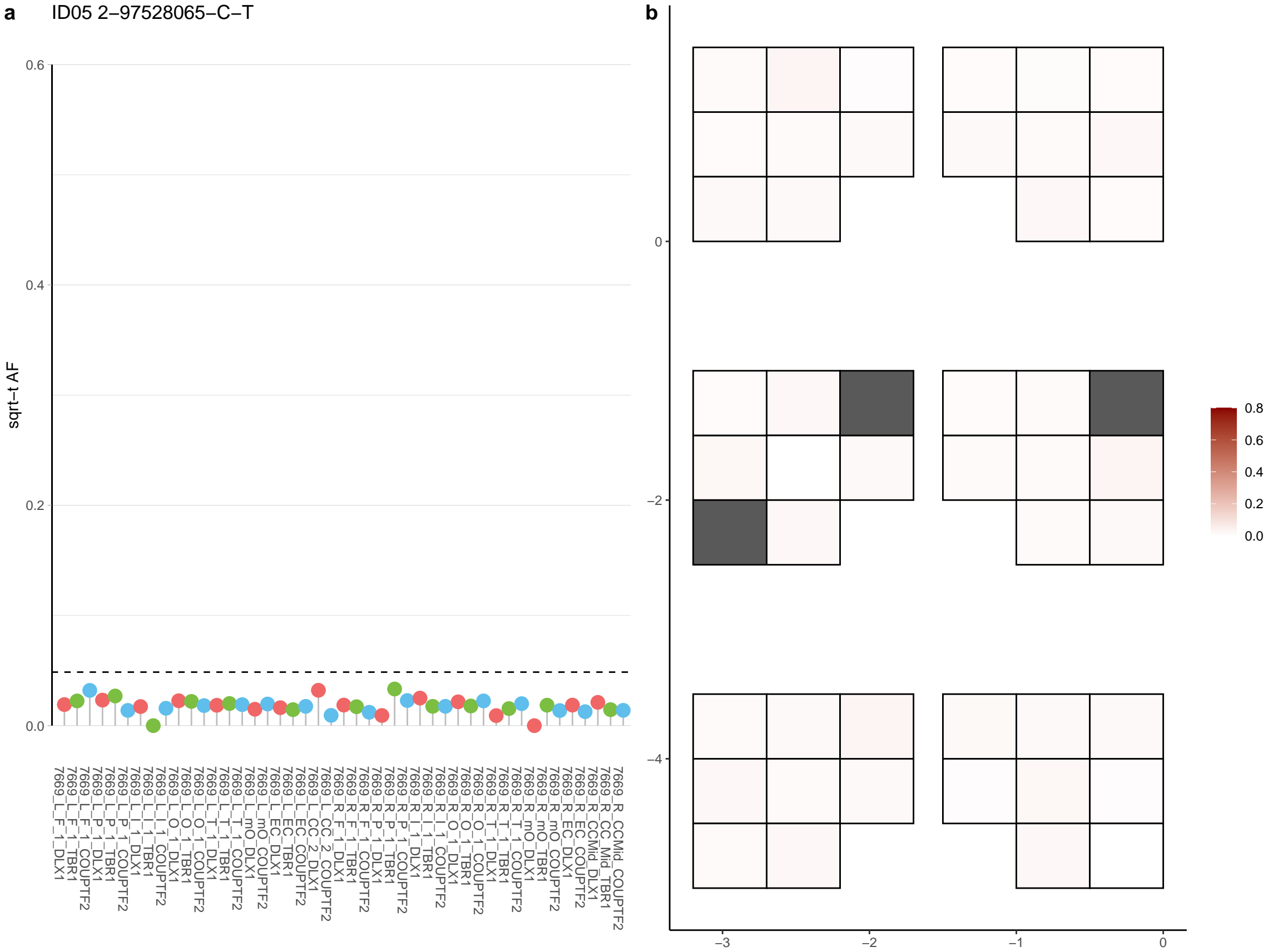

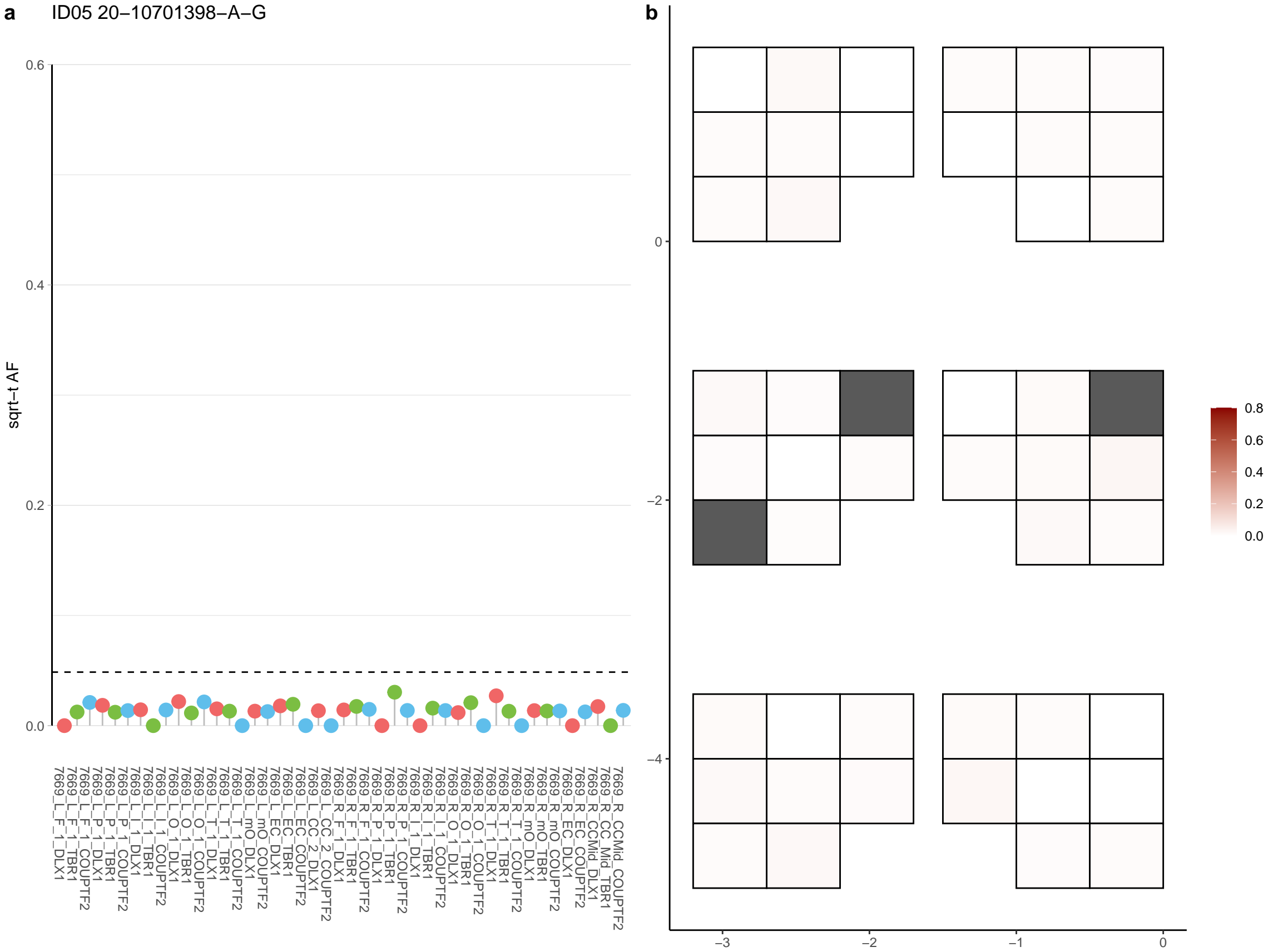

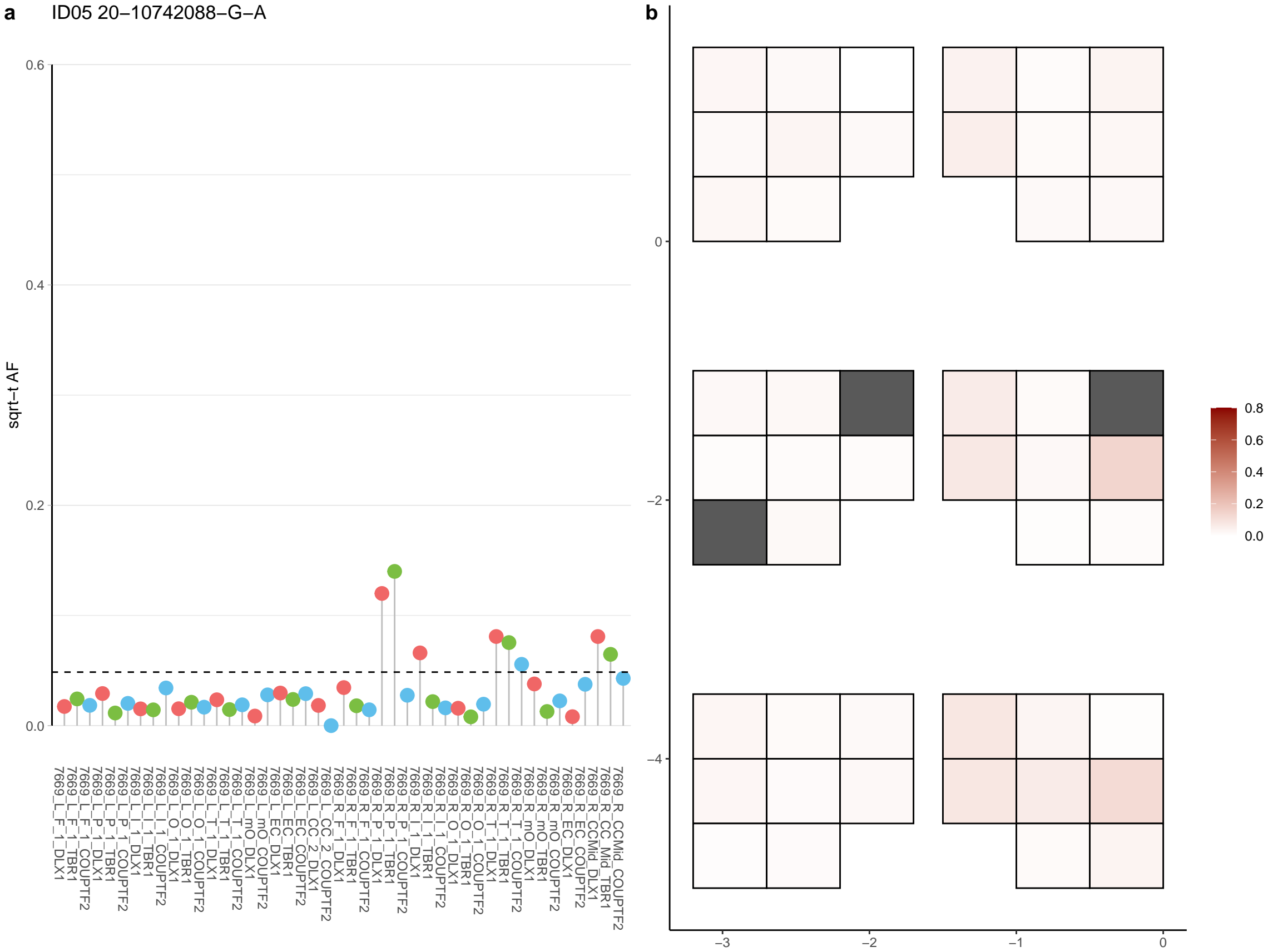

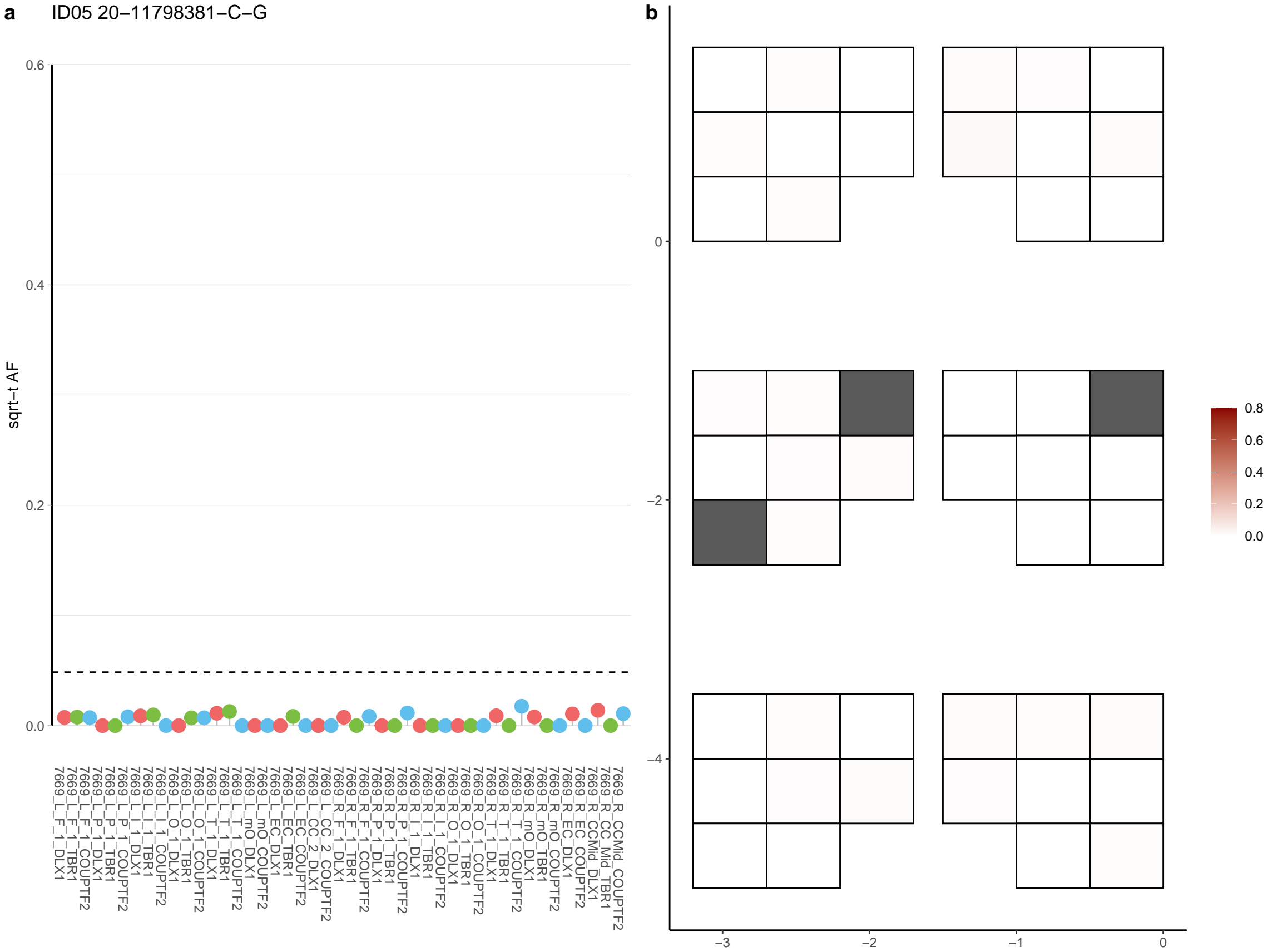

**a** ID05 20-12514966-T-G

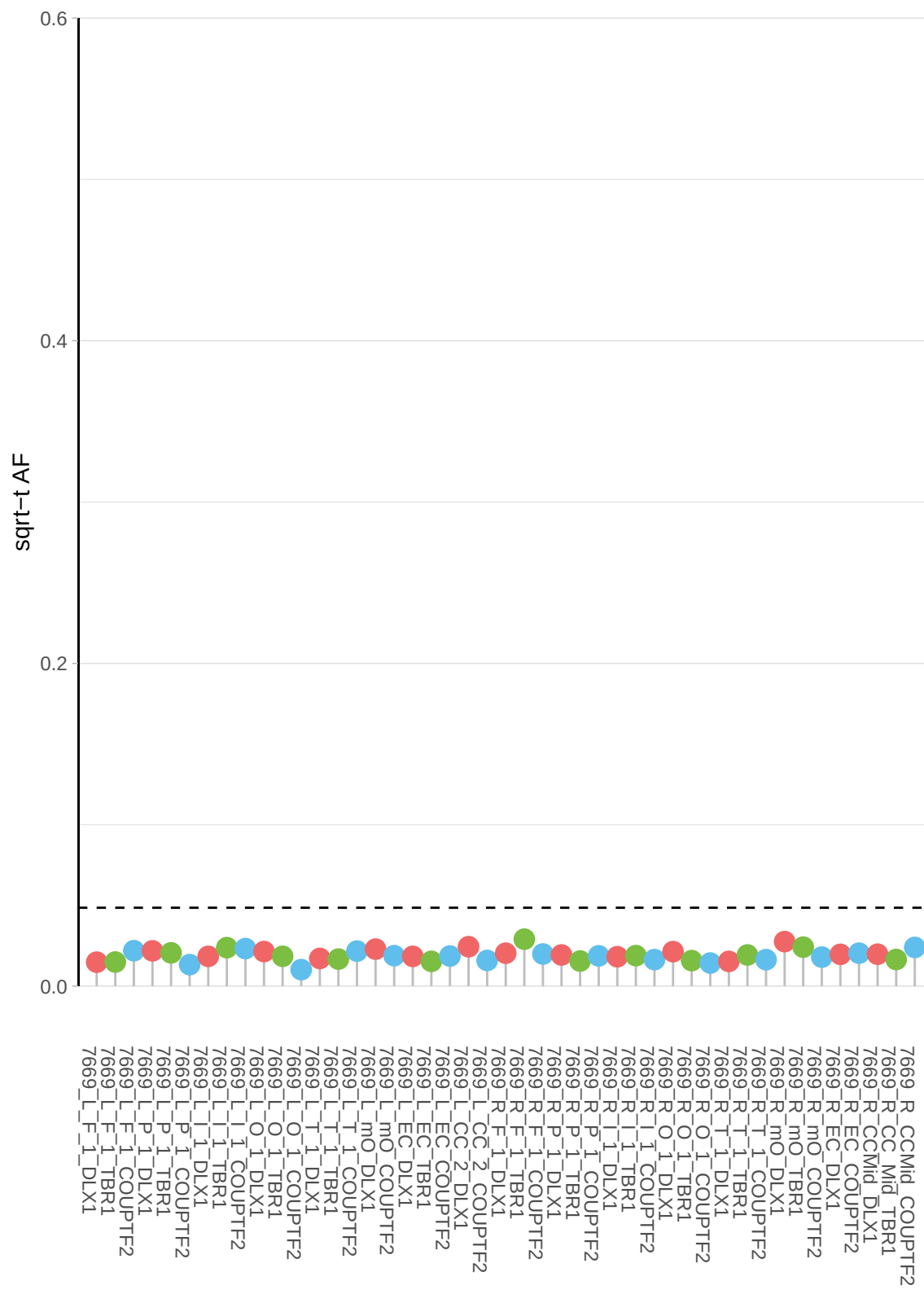

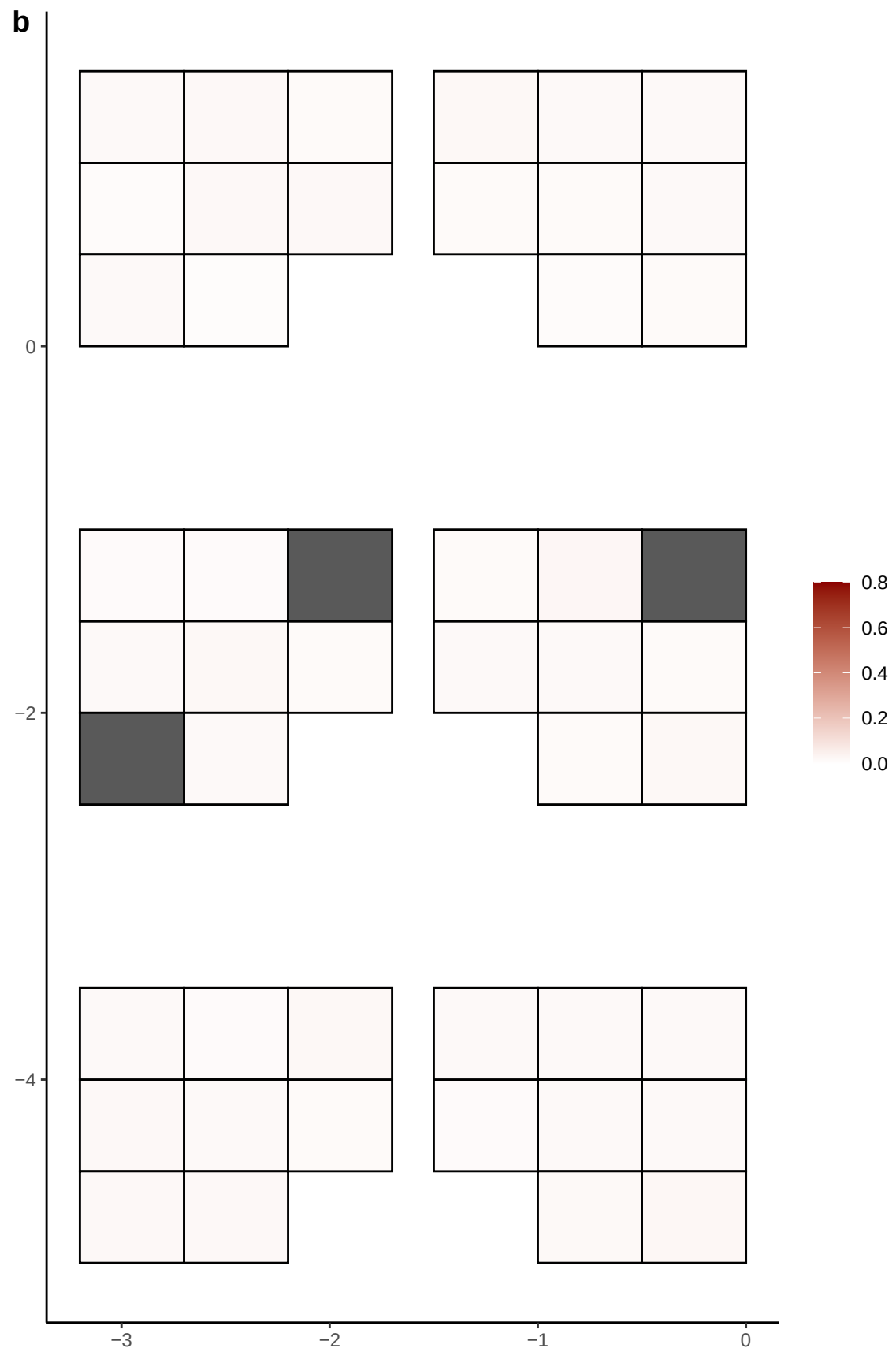

**a** ID05 20-15344074-C-T

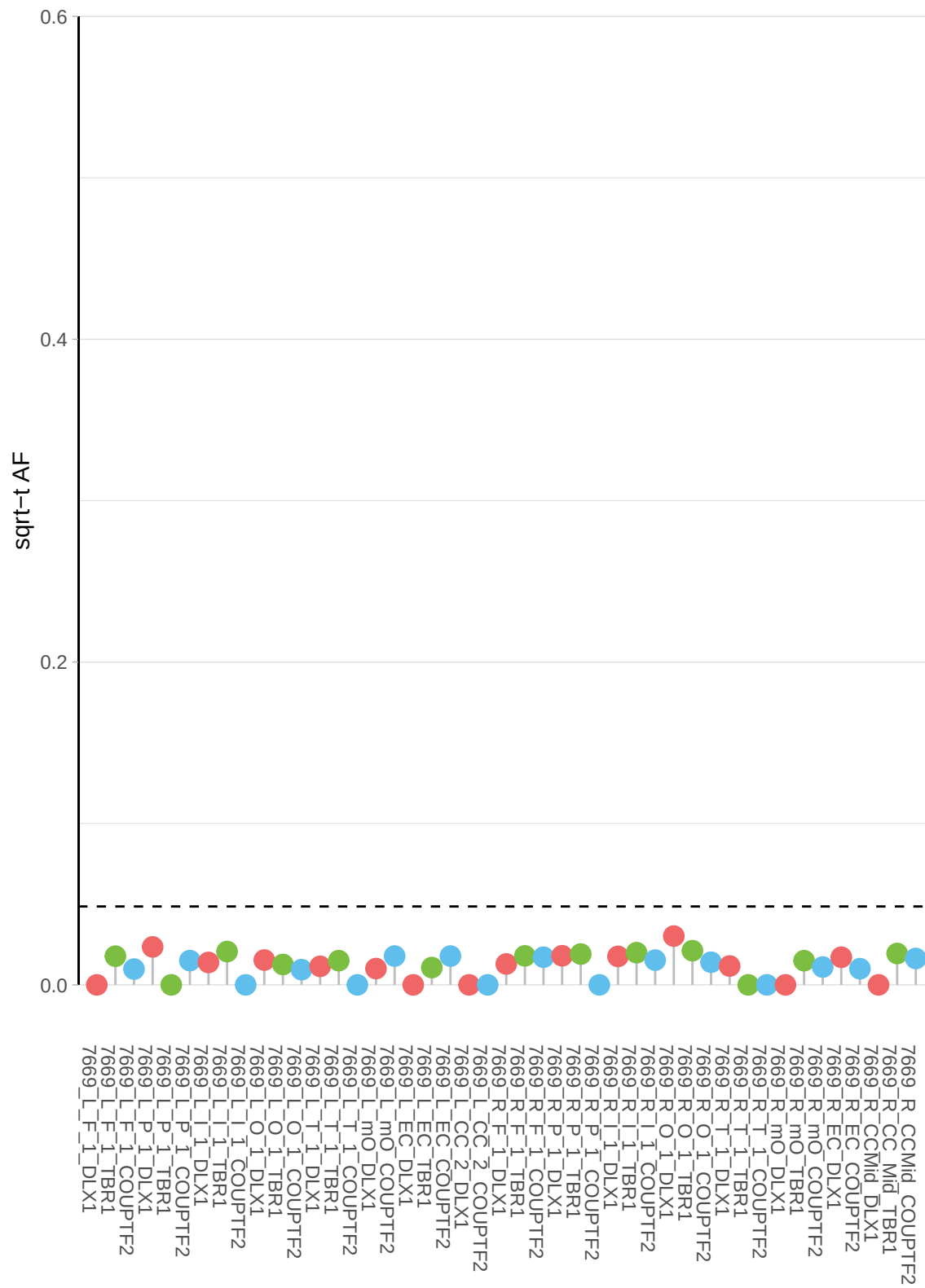

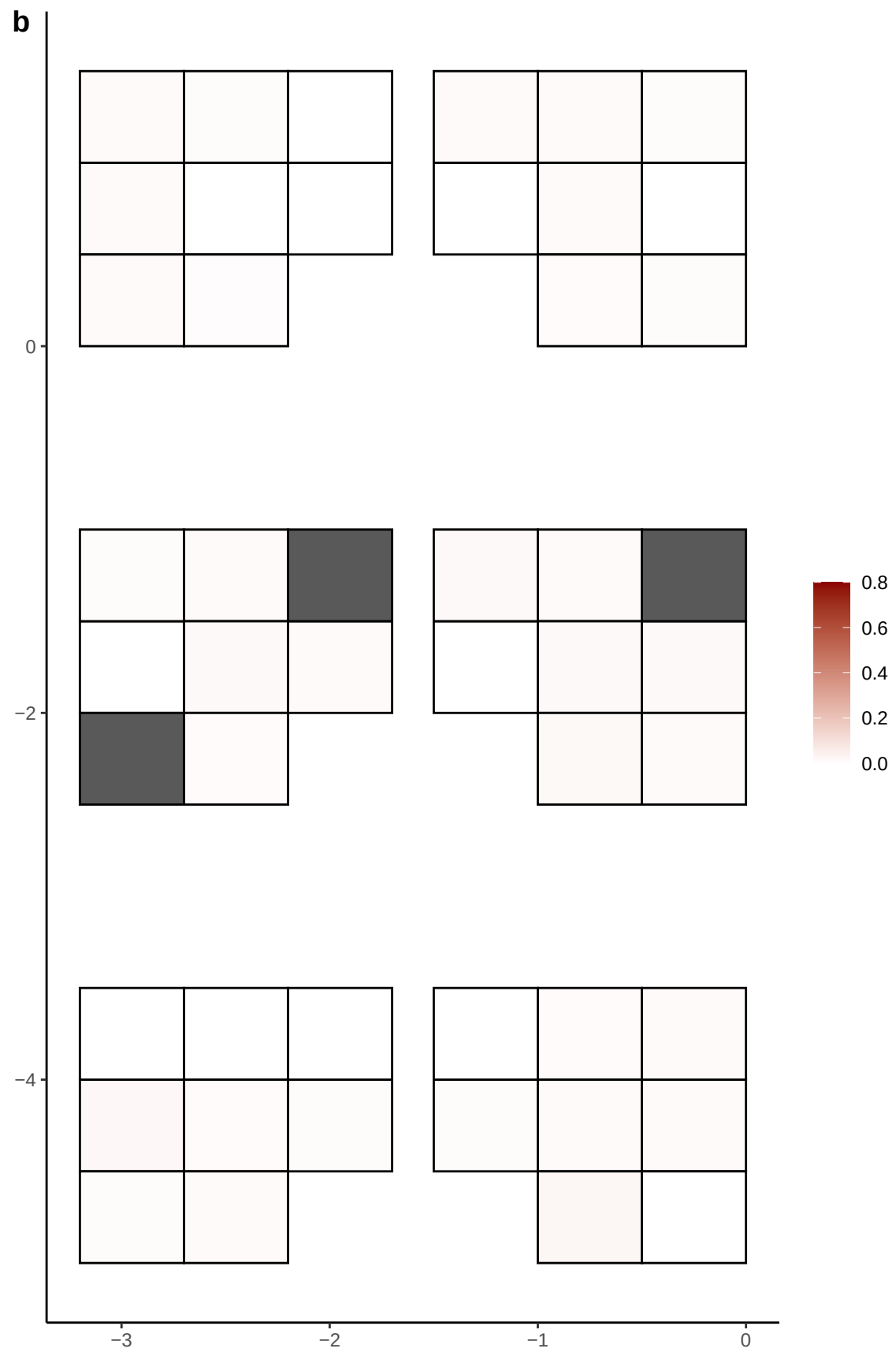

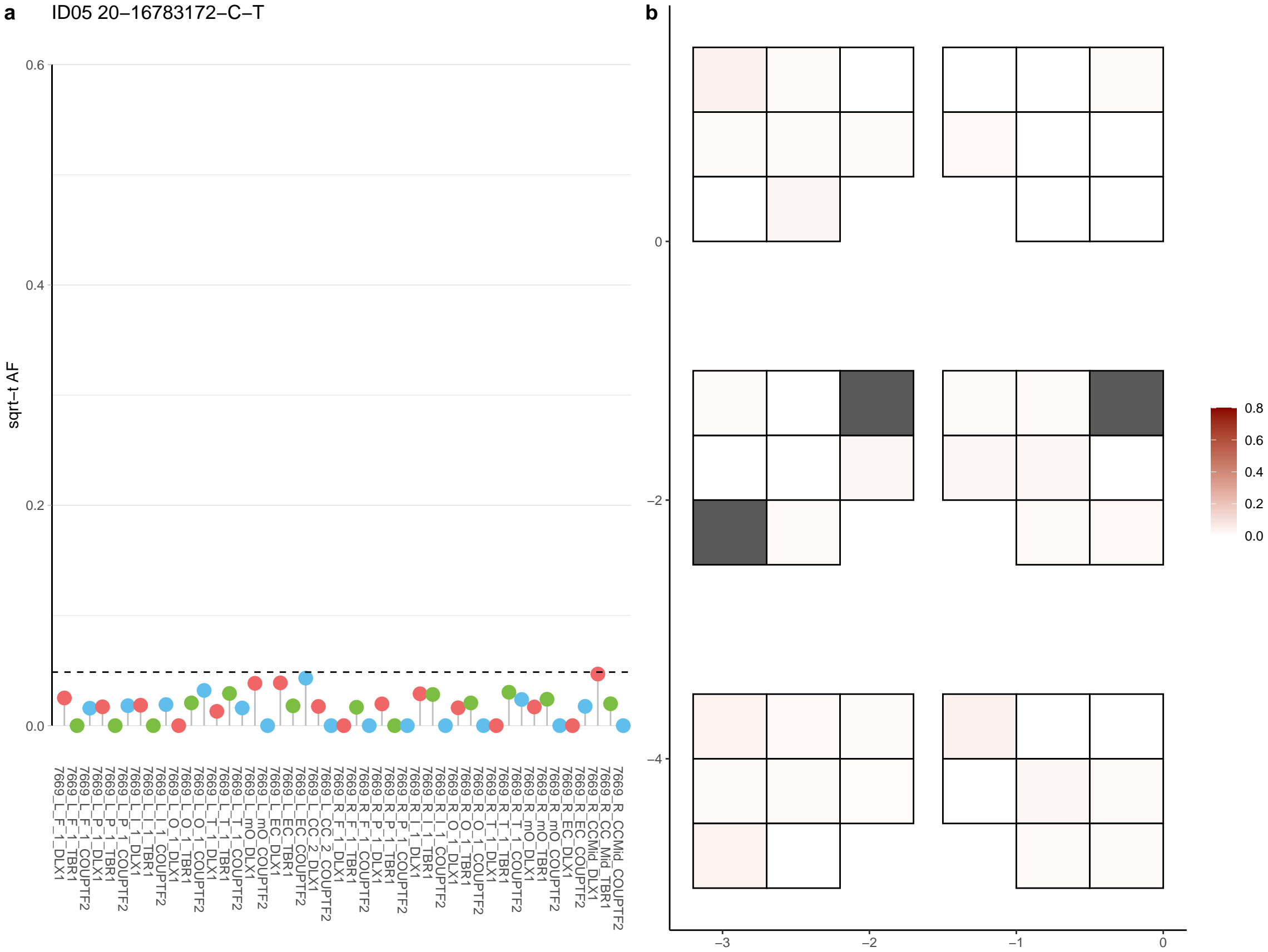

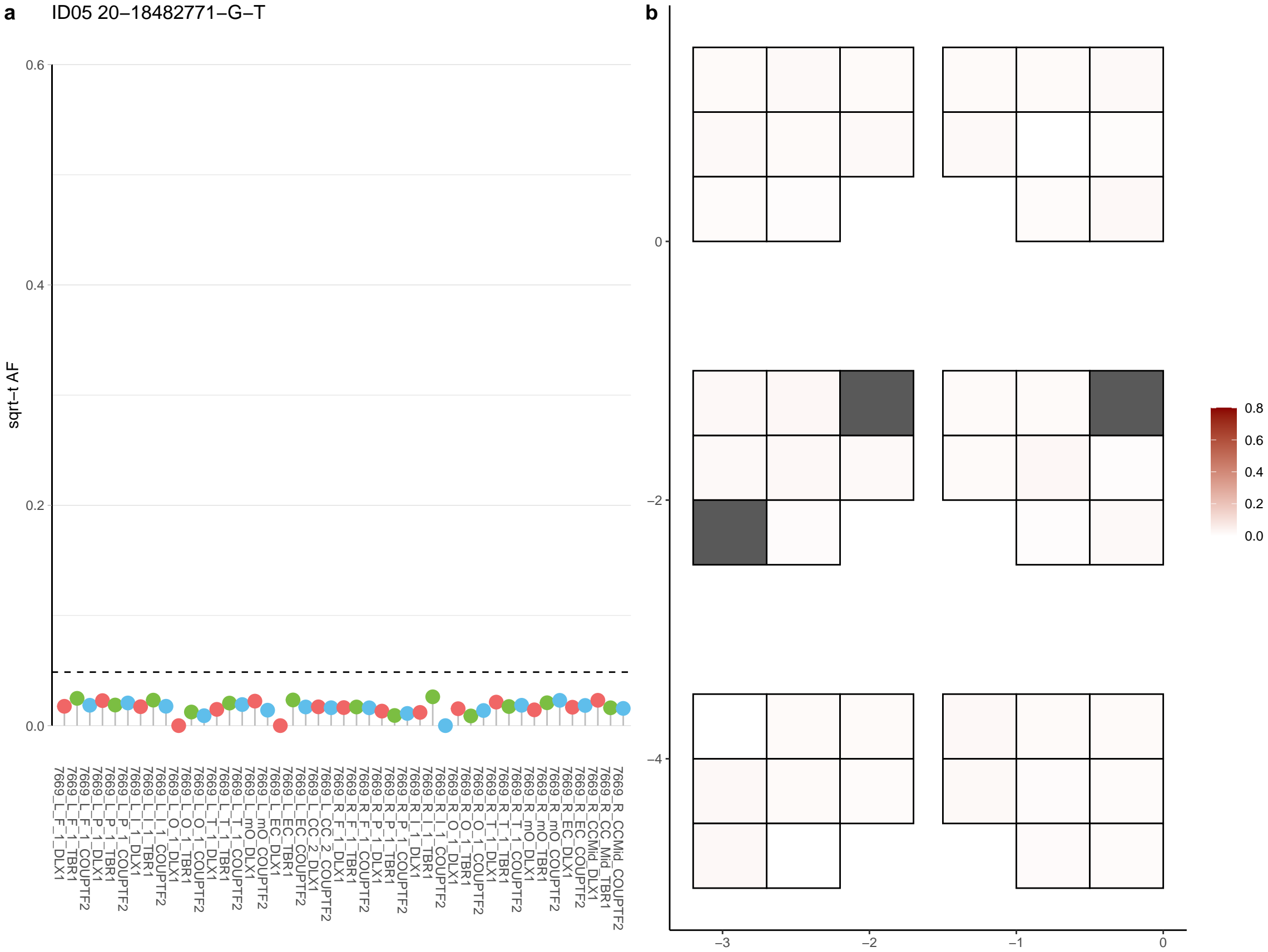

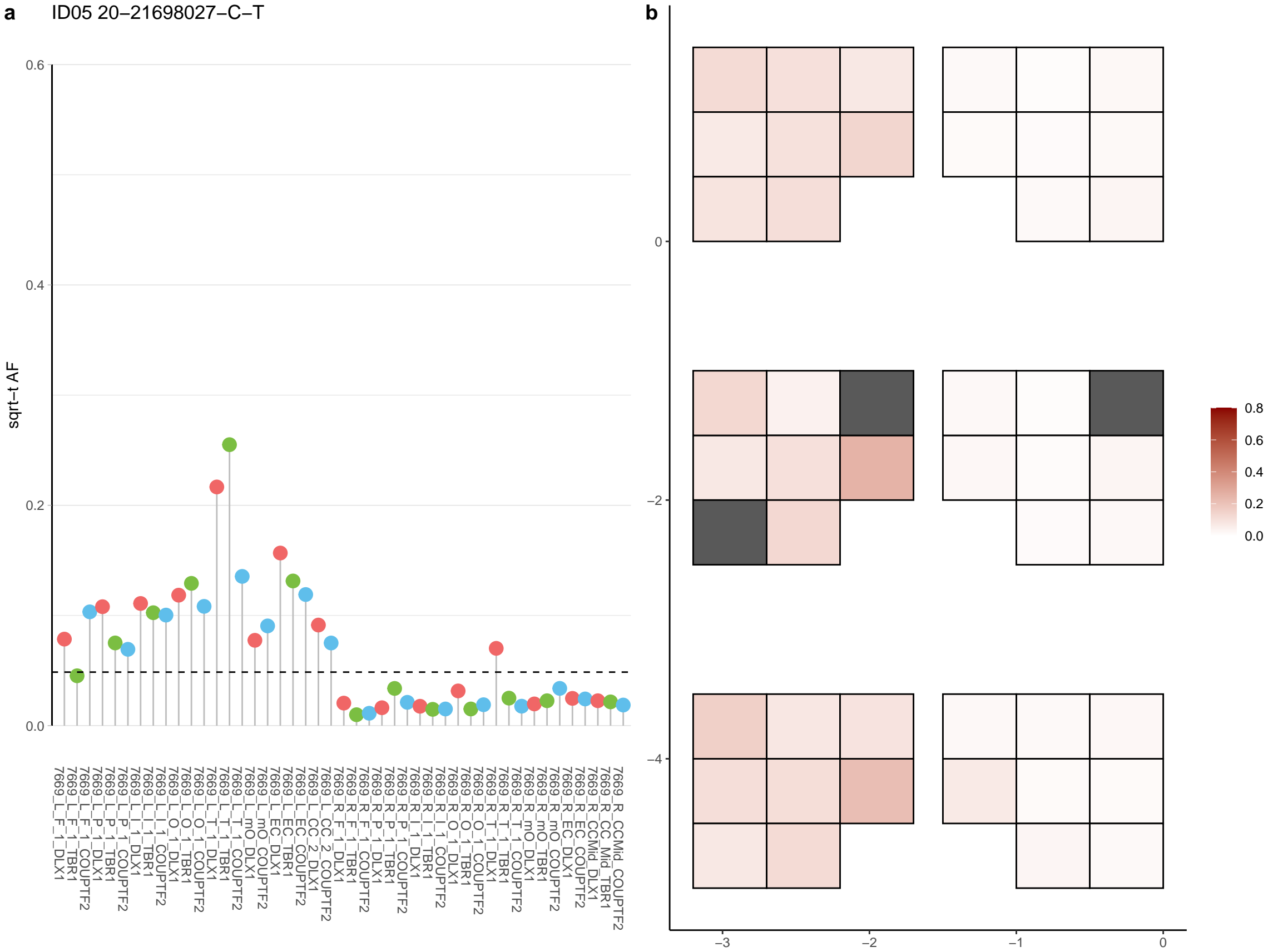

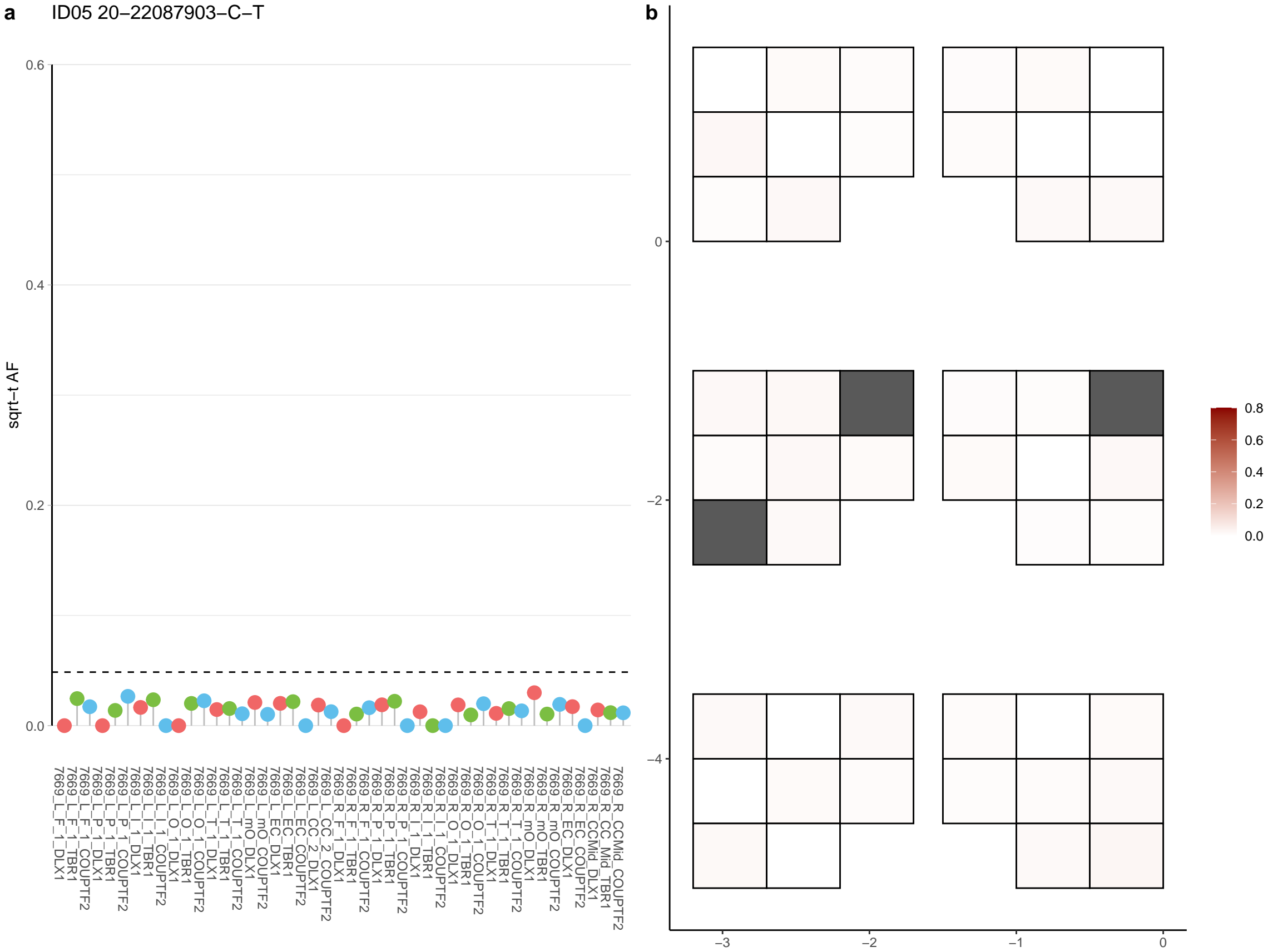

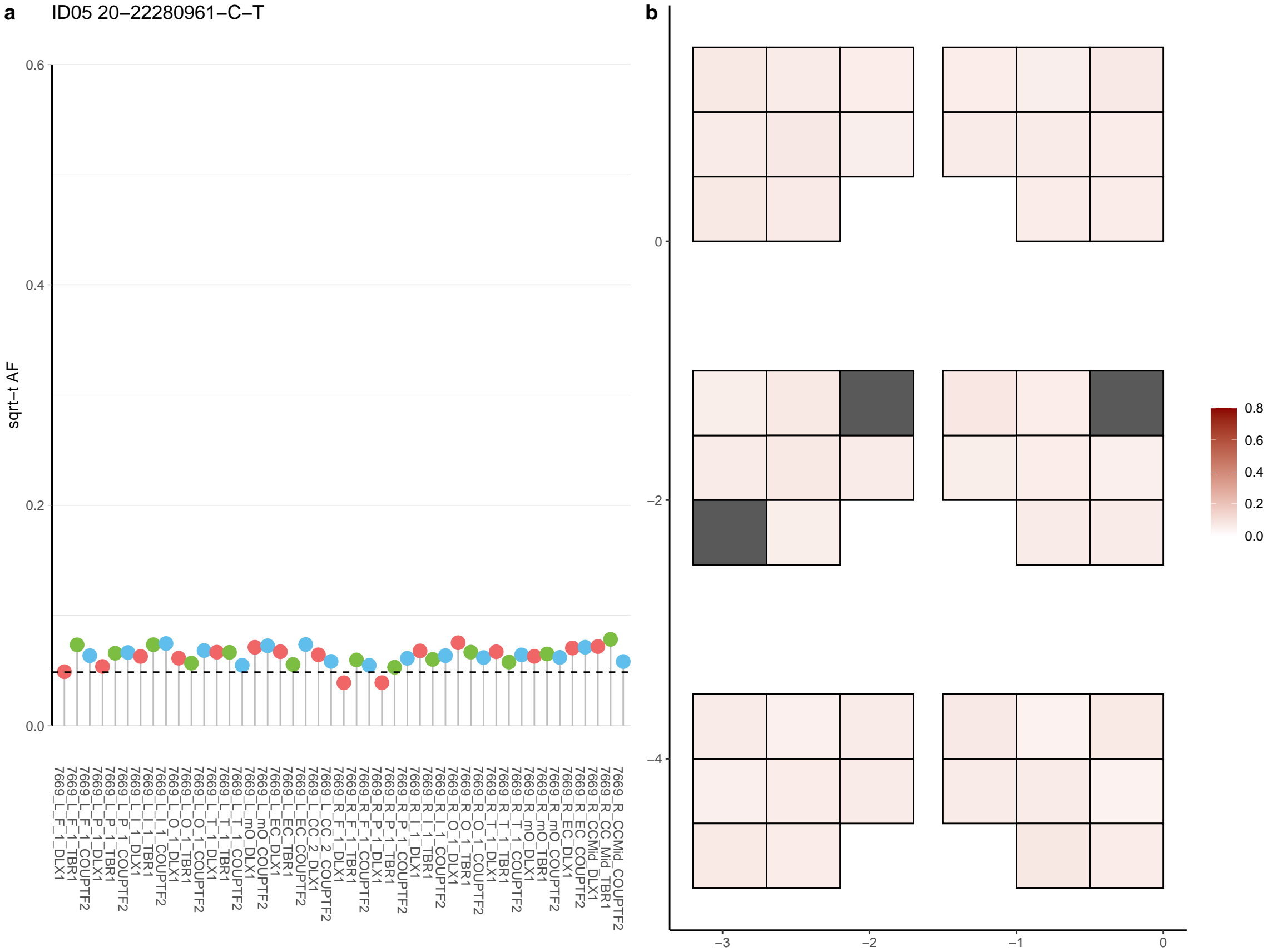

**a** ID05 20-22383923-C-A

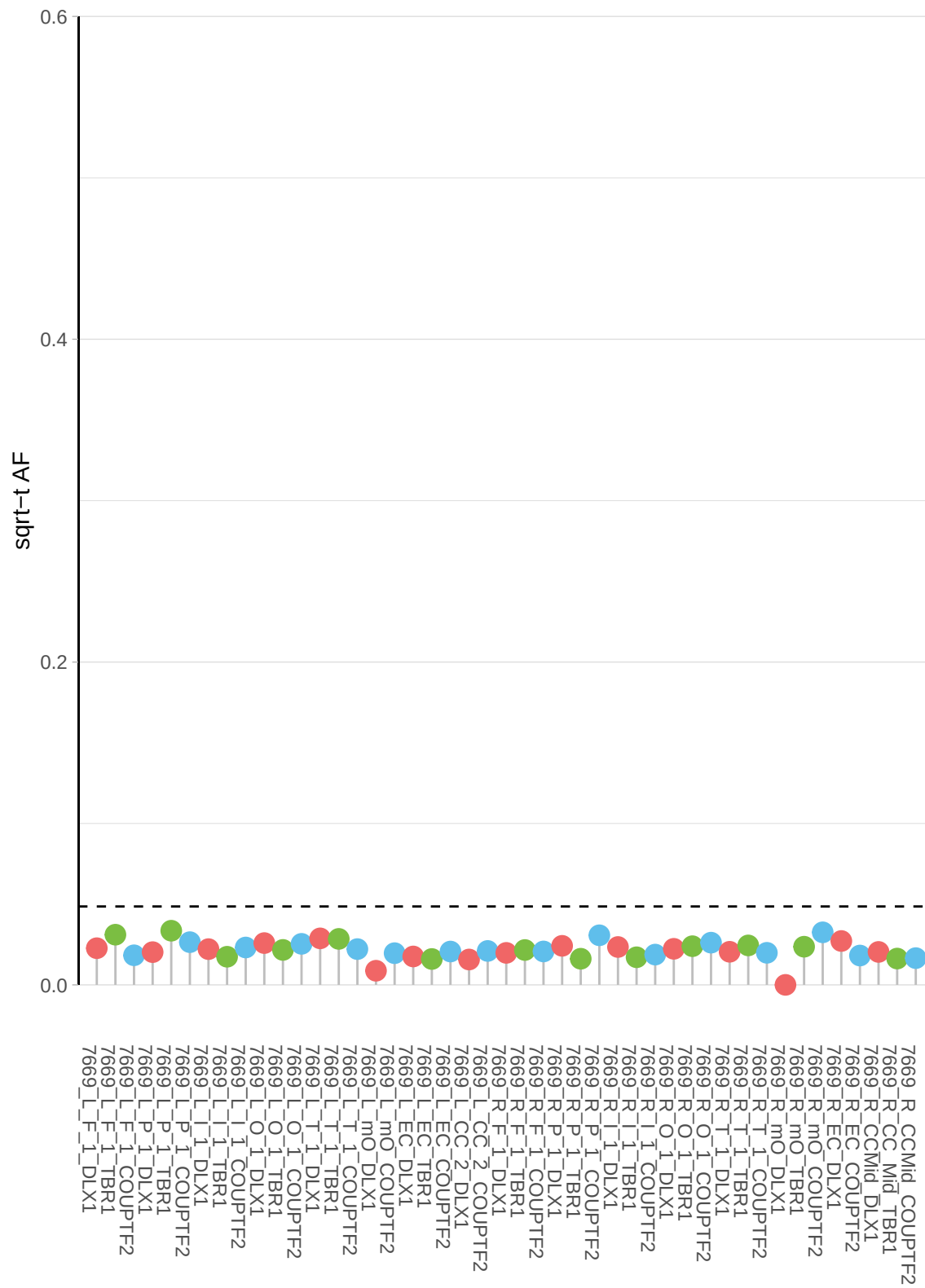

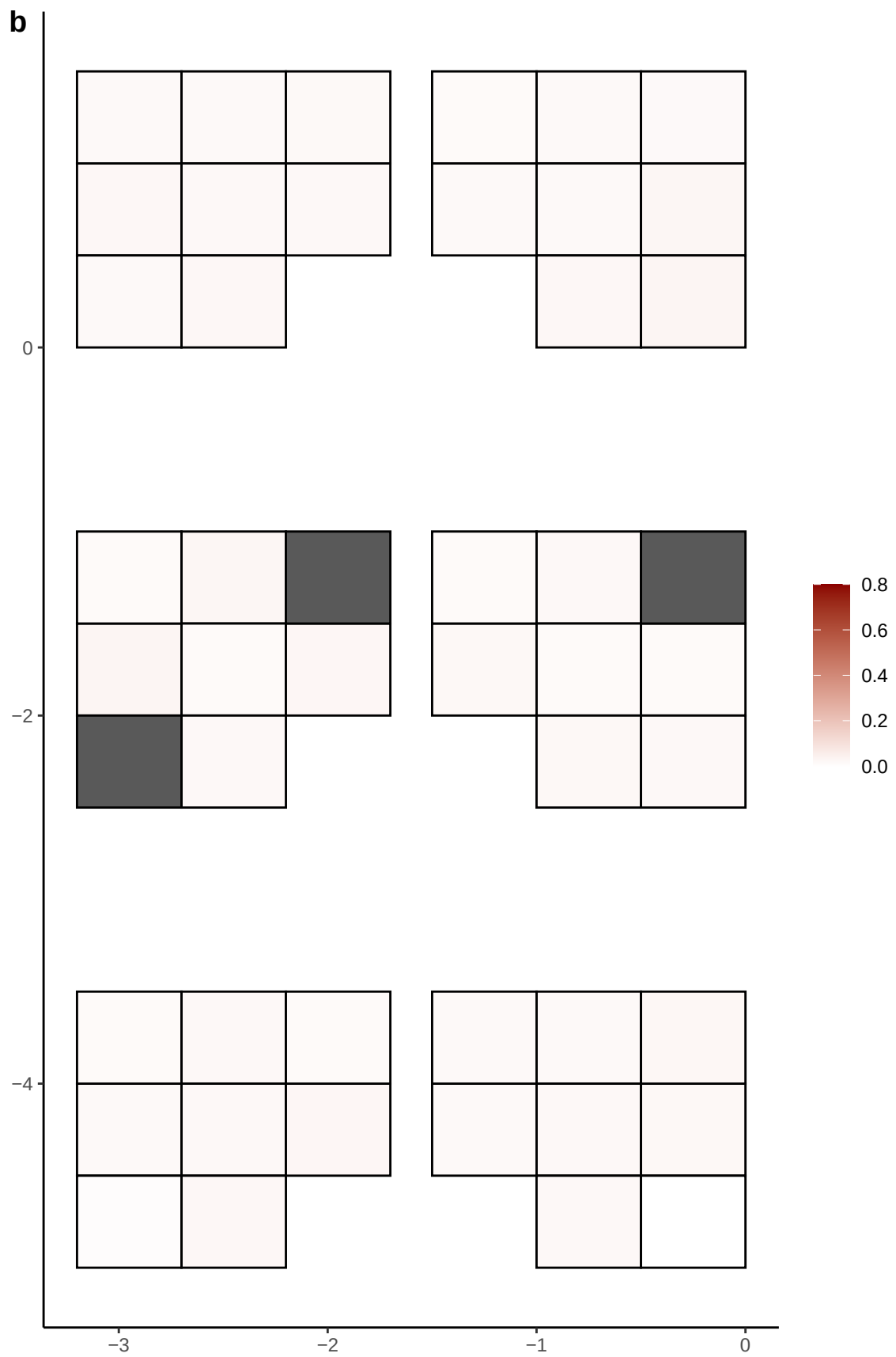

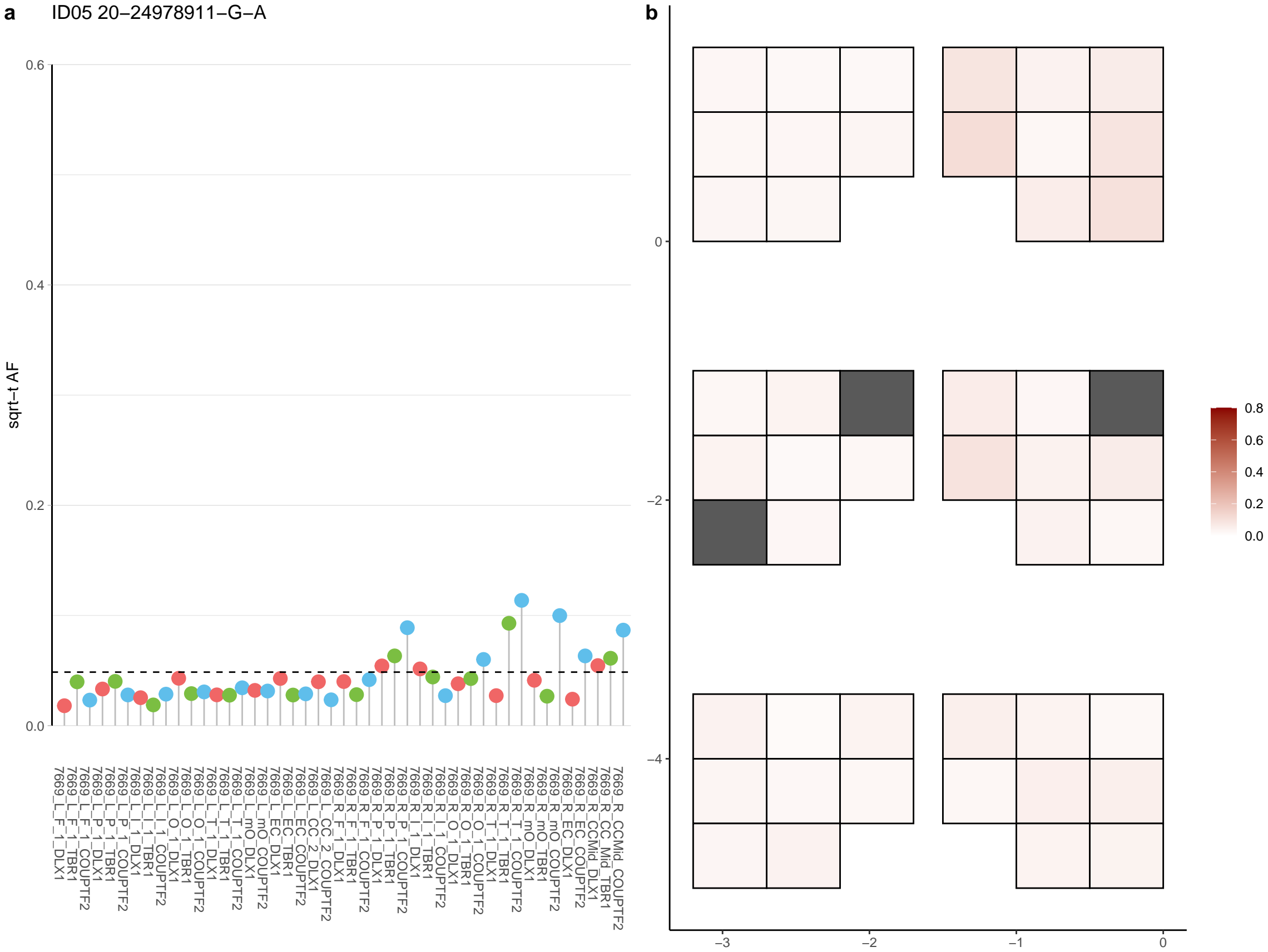

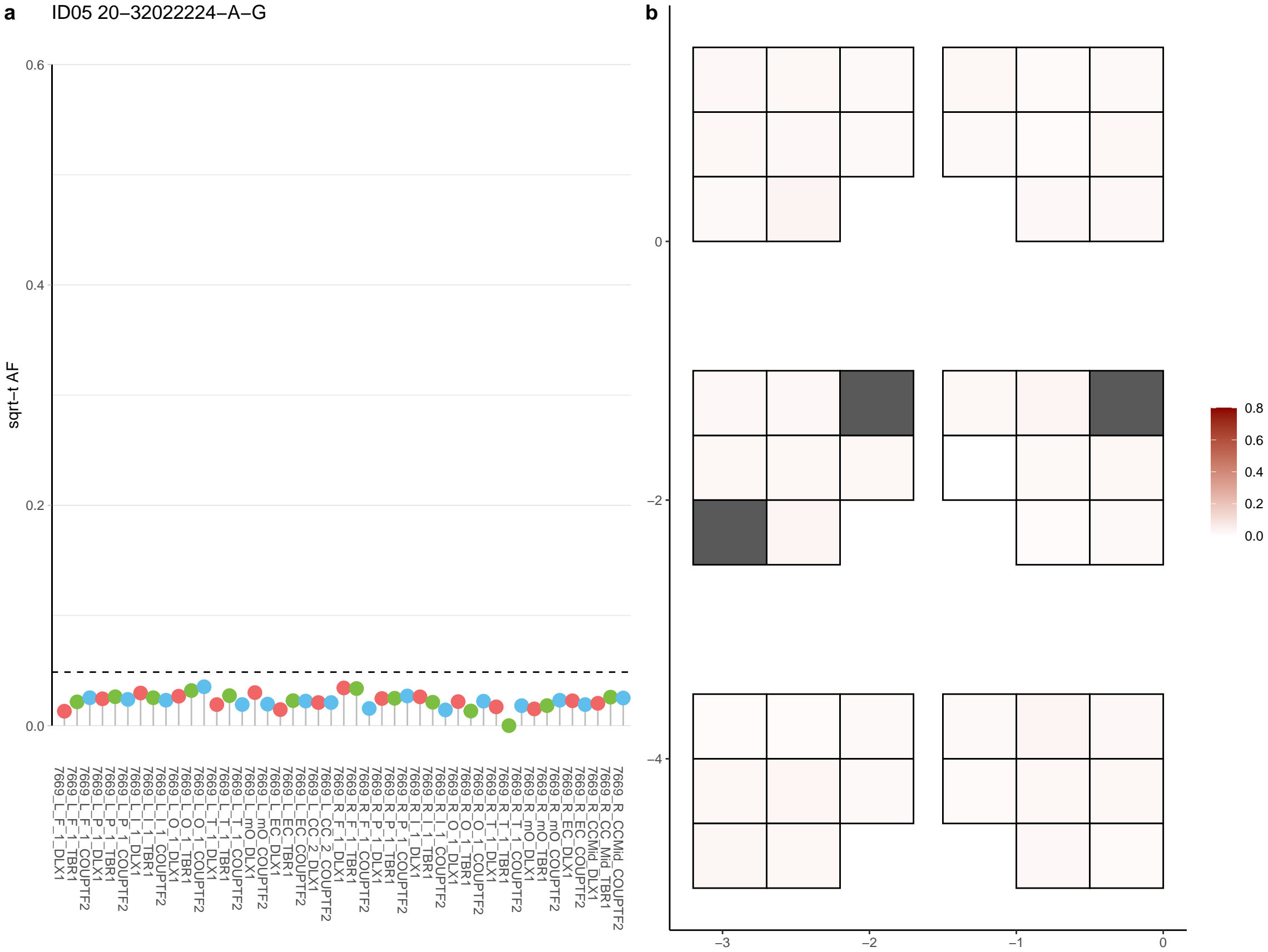

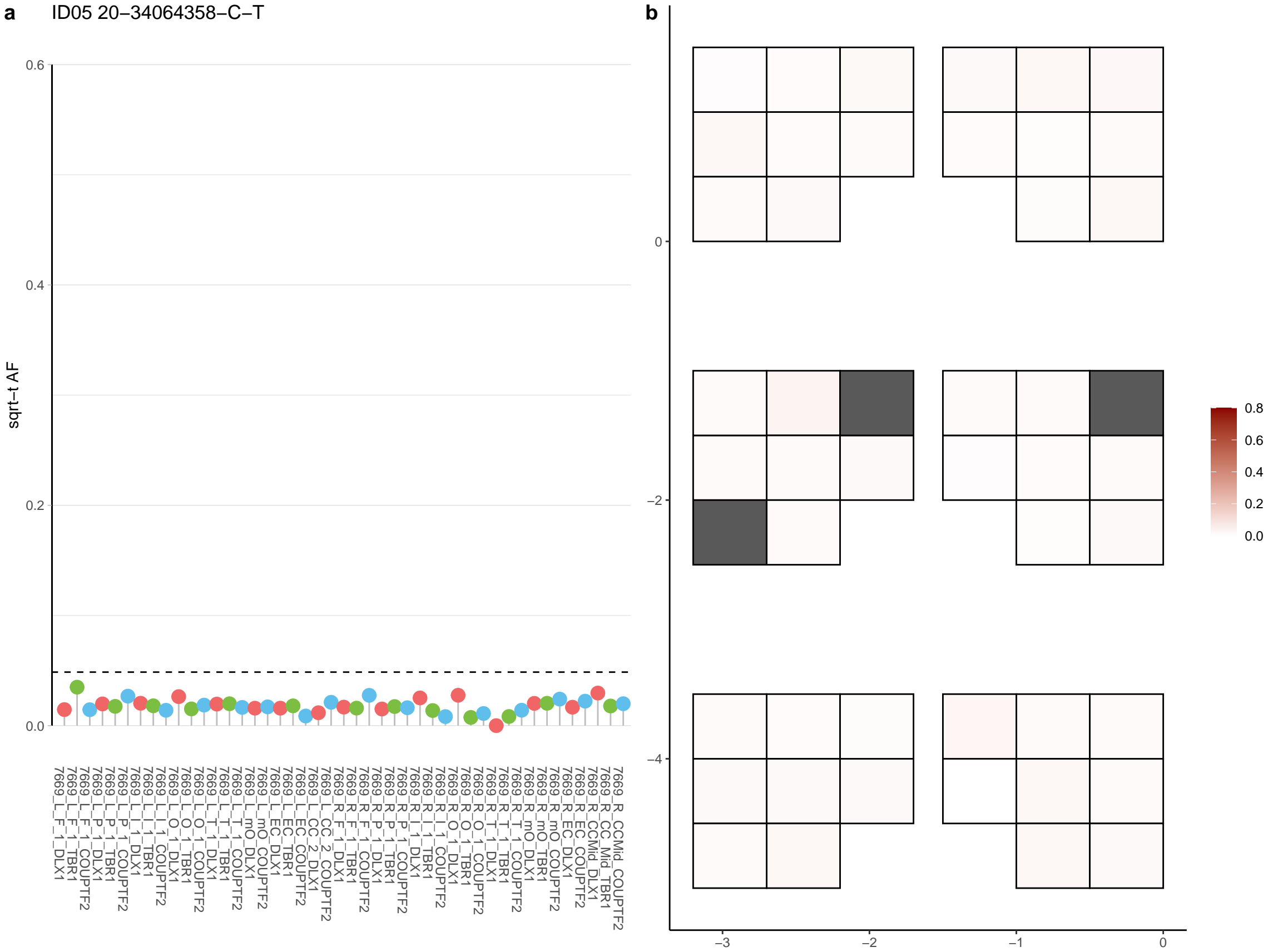

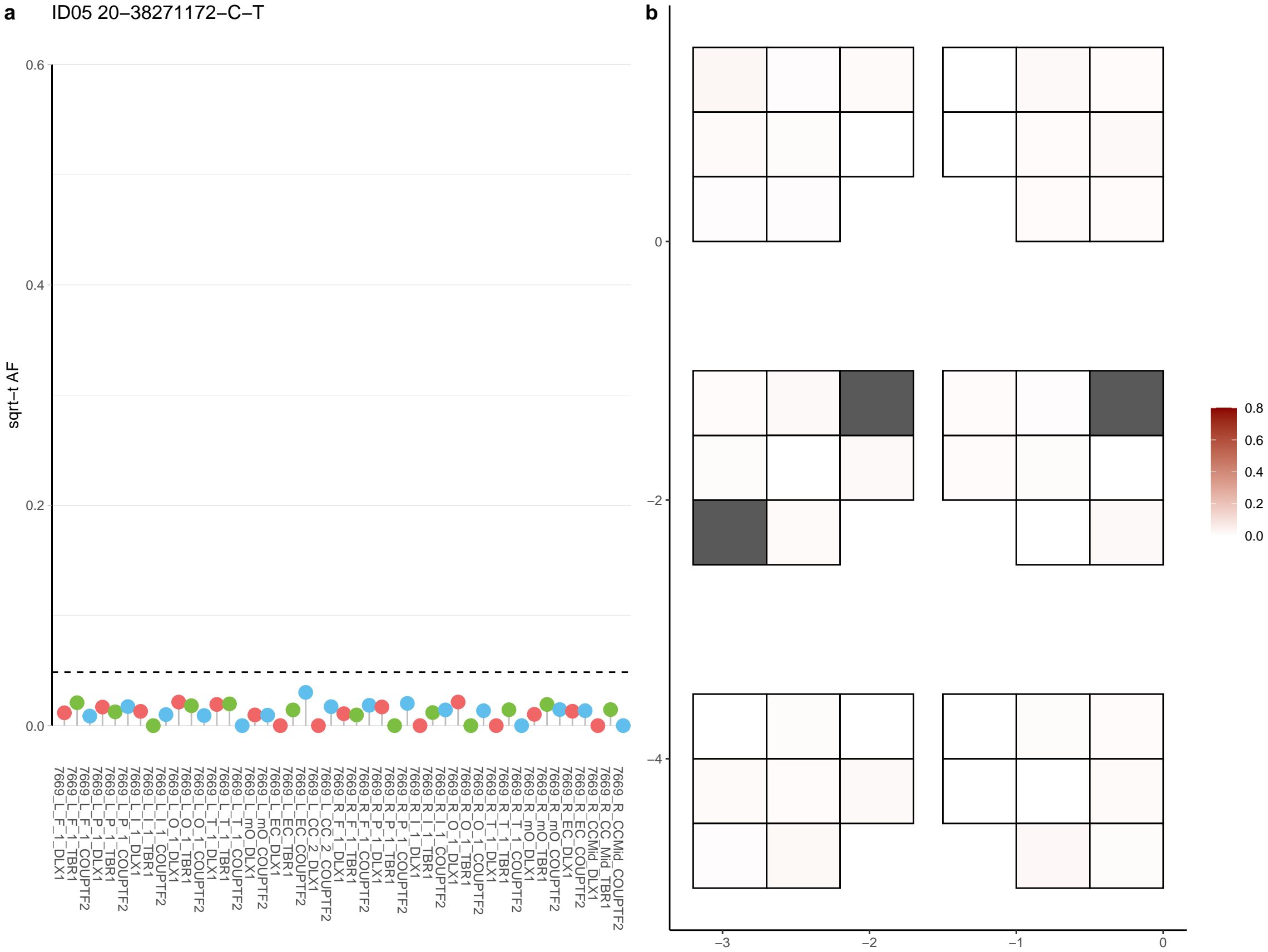

**a** ID05 20-39222576-G-T

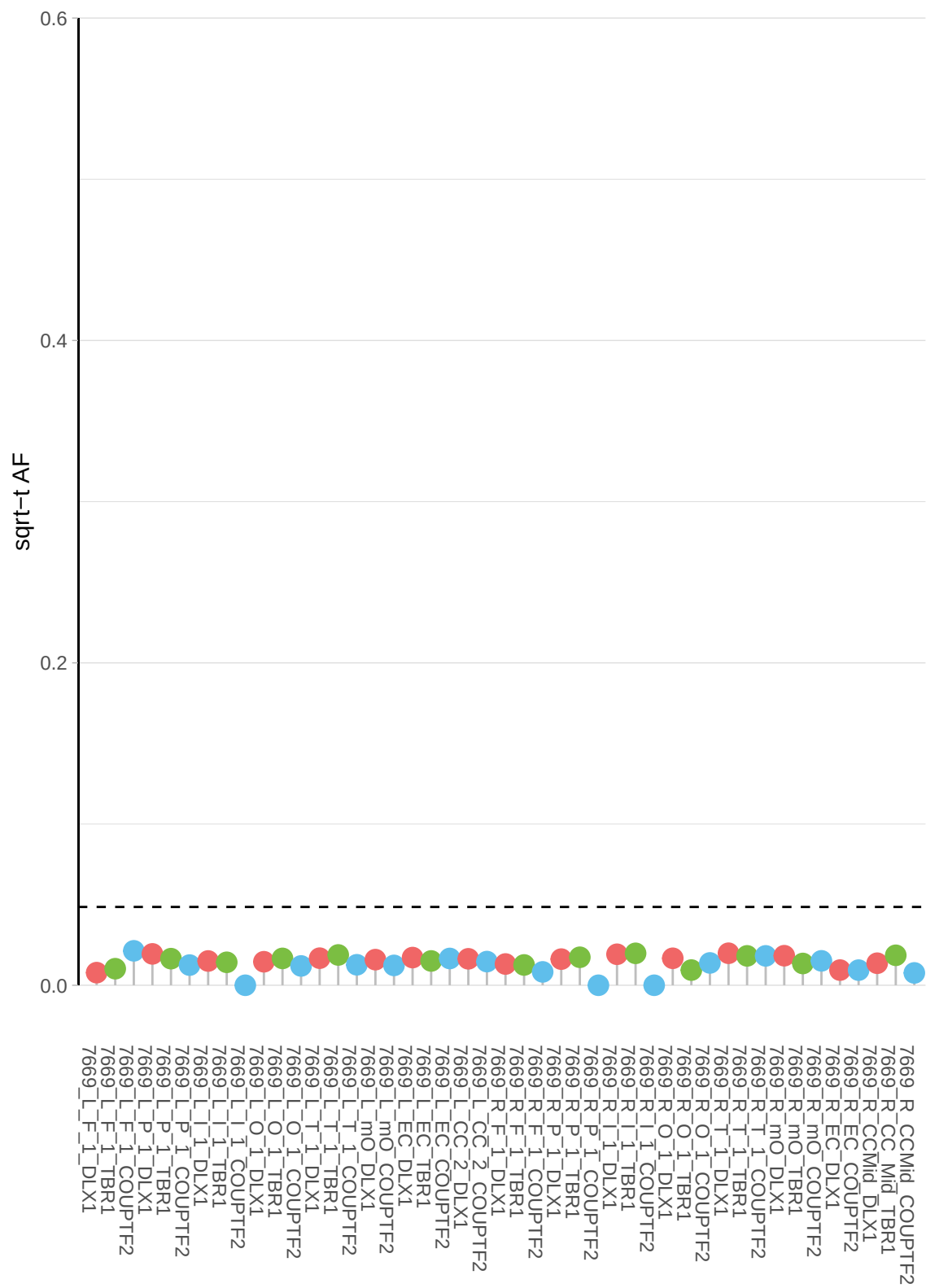

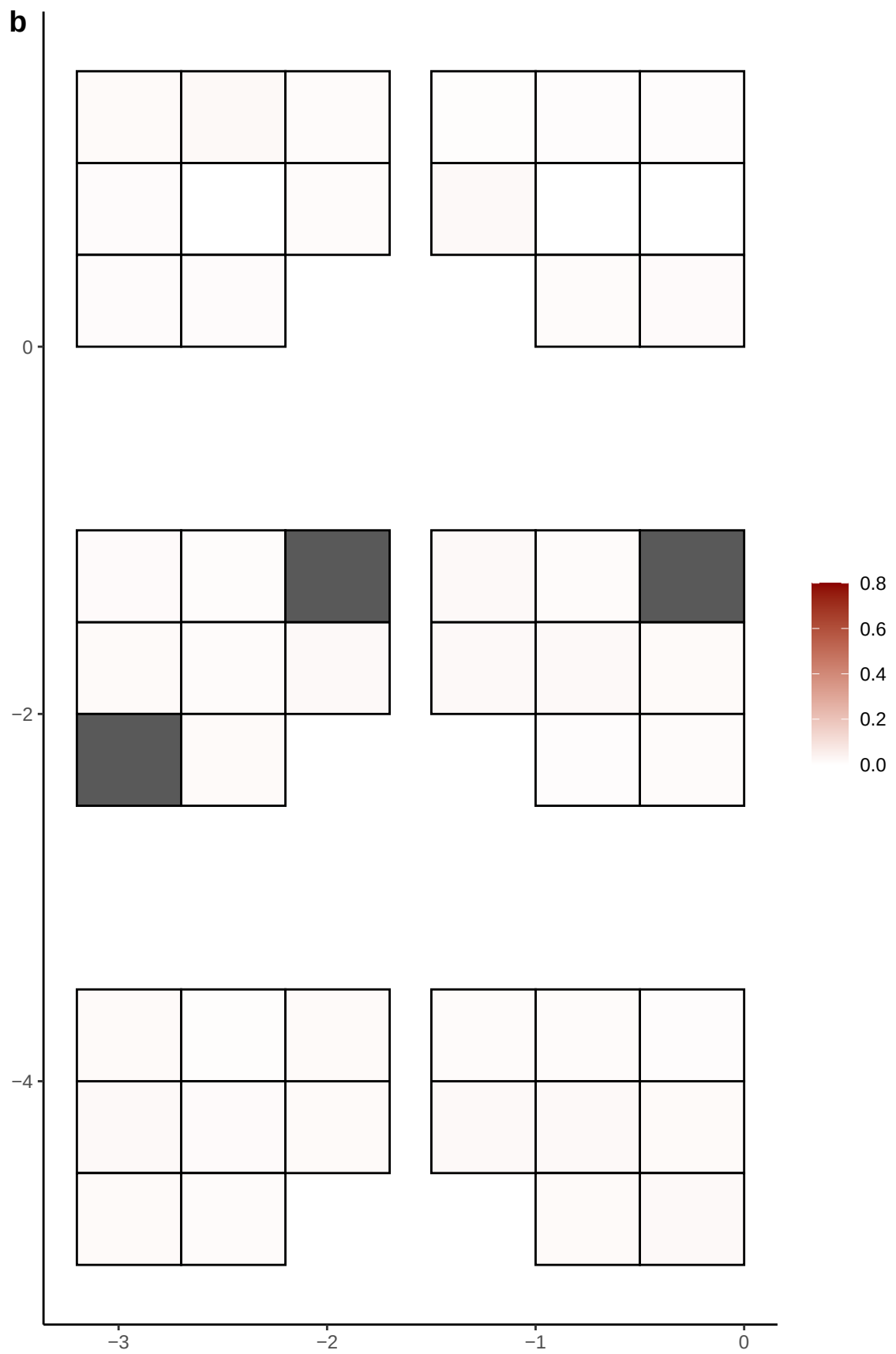

**a** ID05 20-39714075-G-A

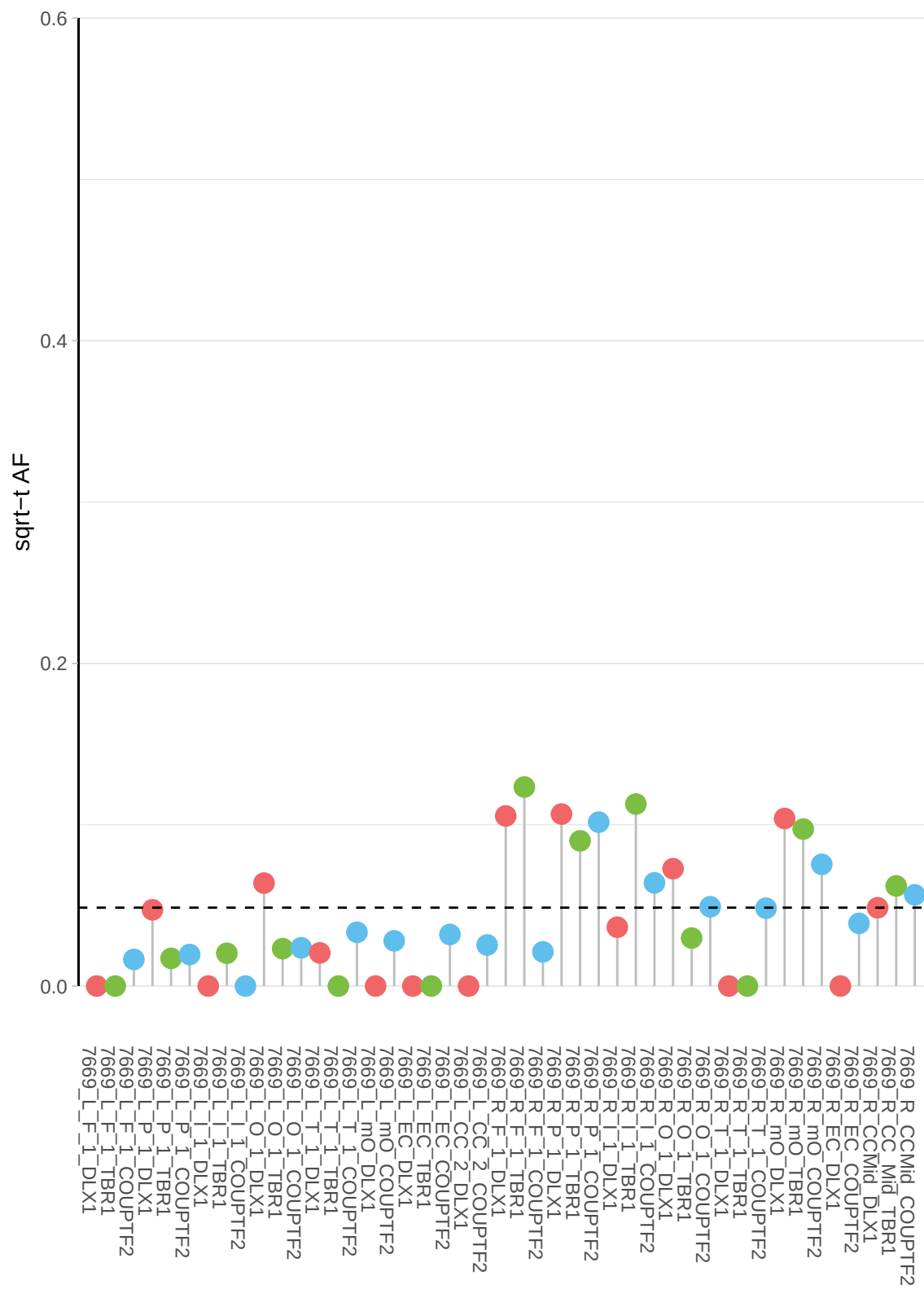

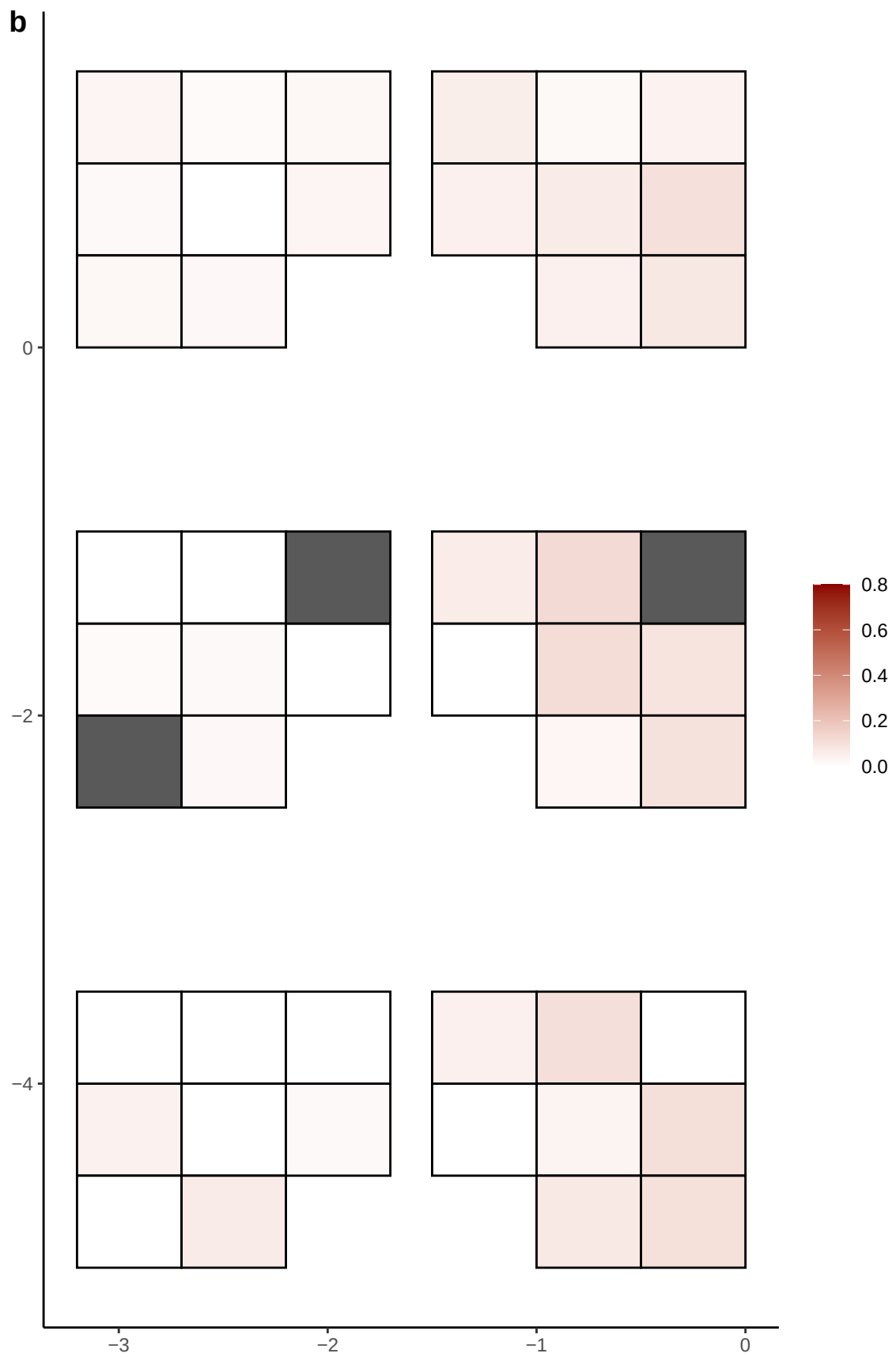

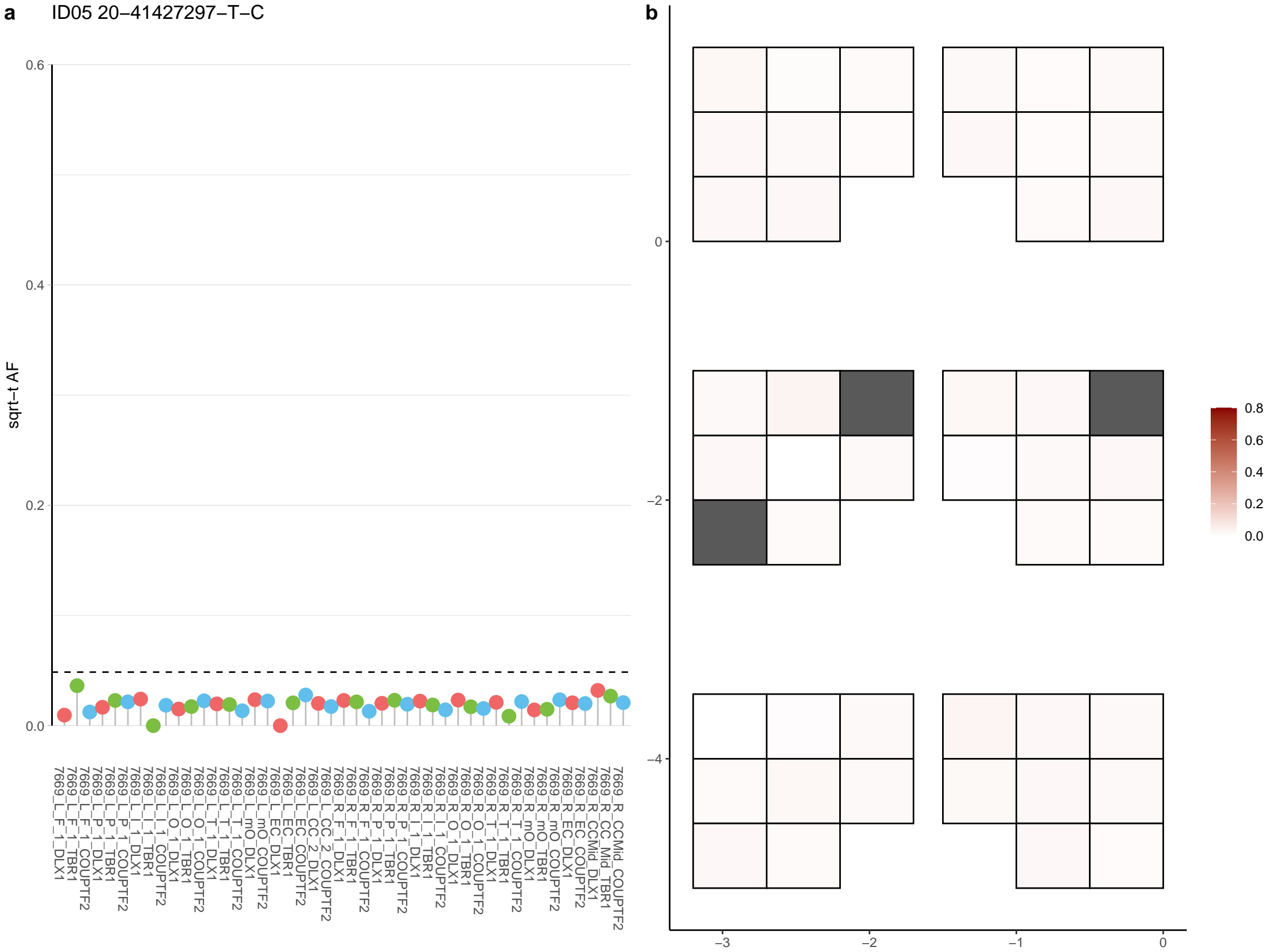

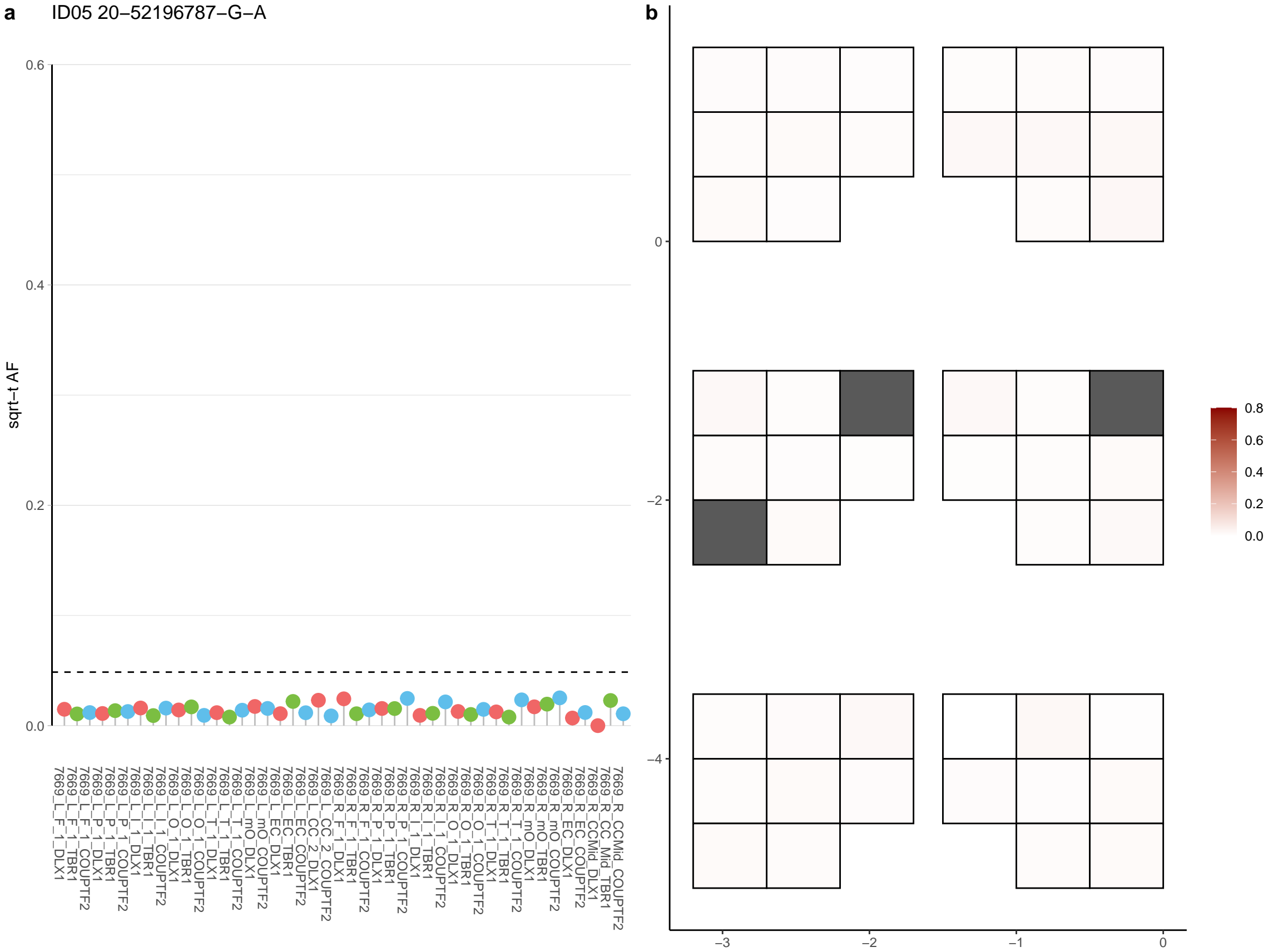

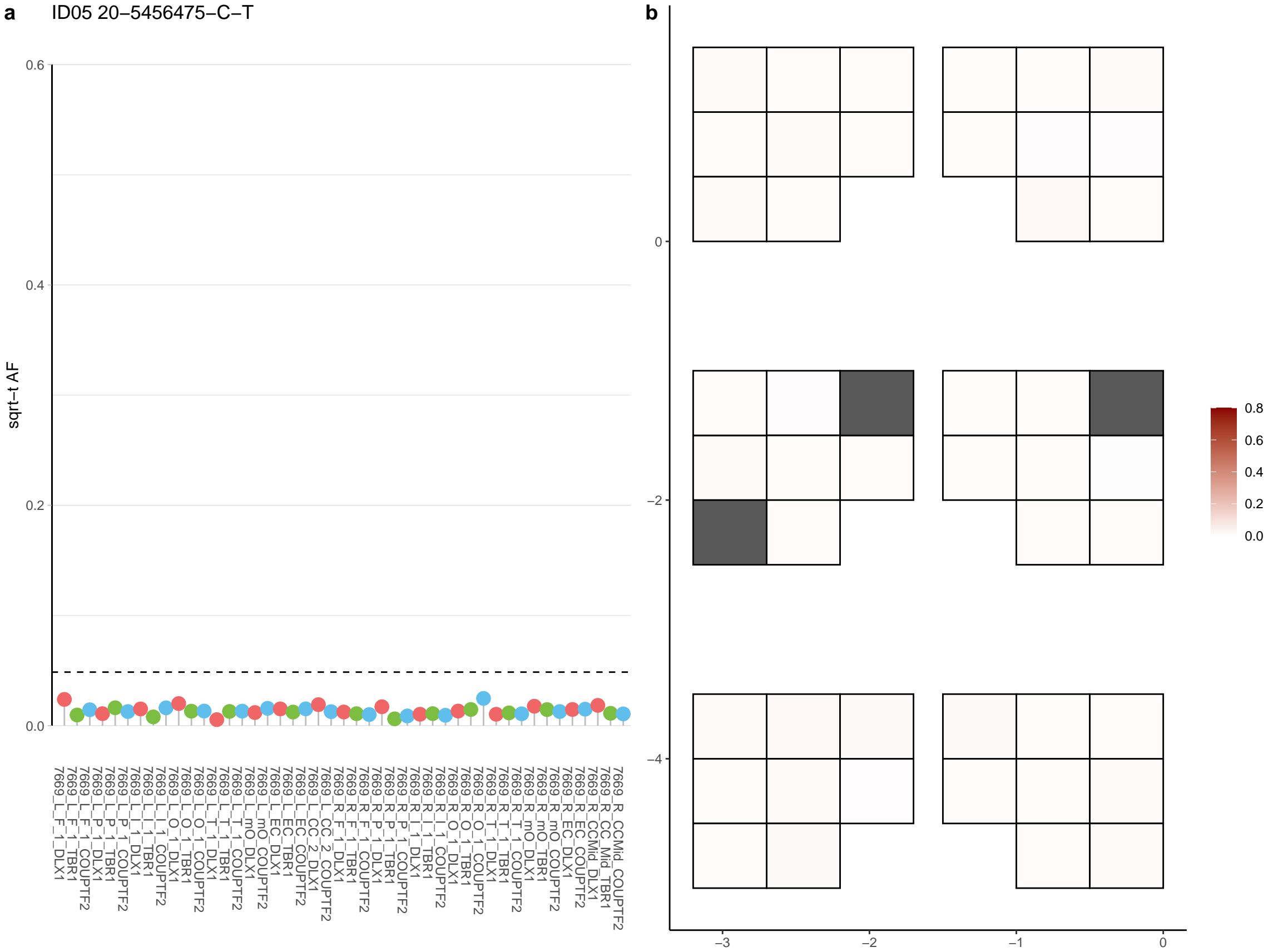

**a** ID05 20-56452953-A-G

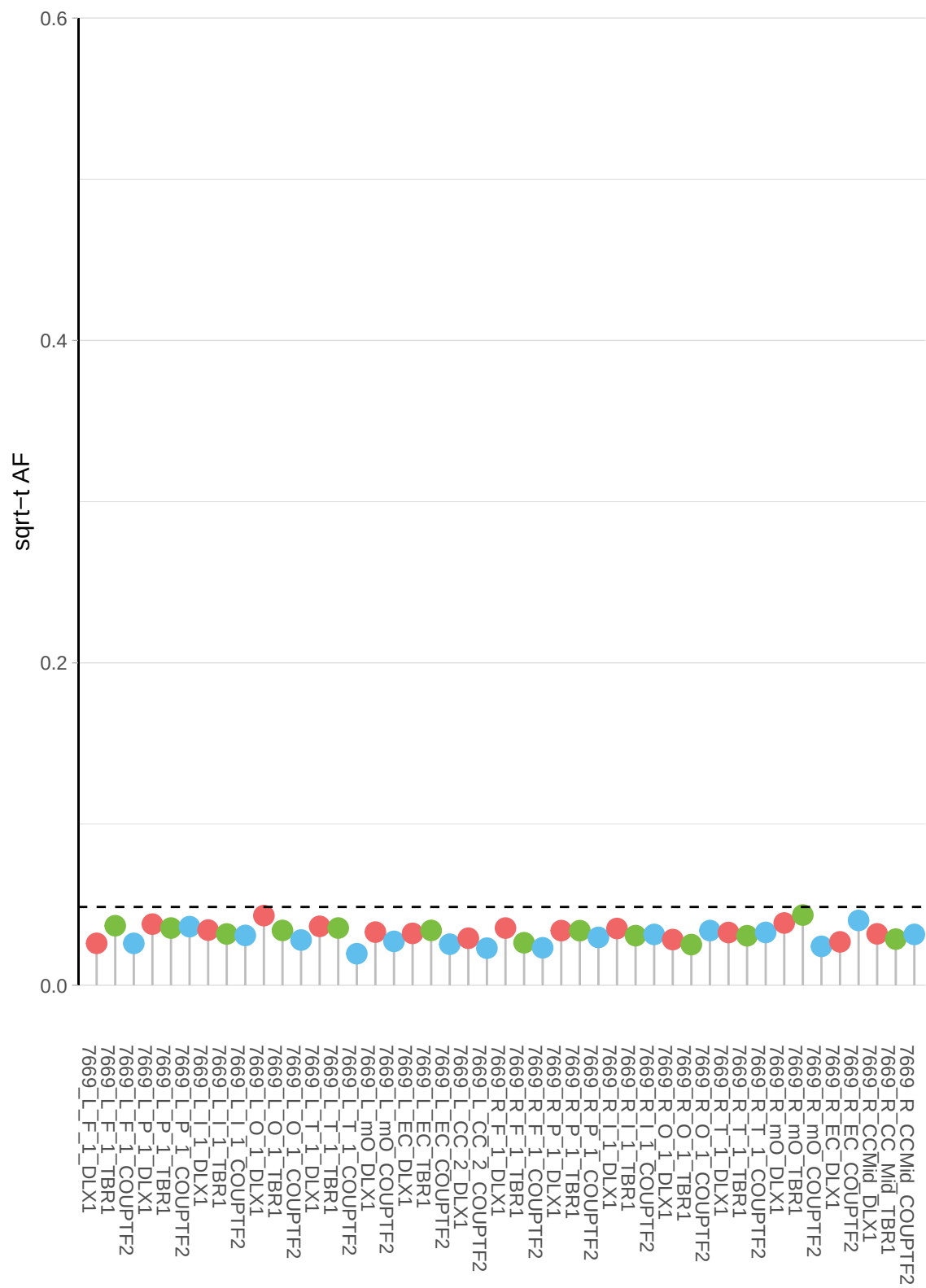

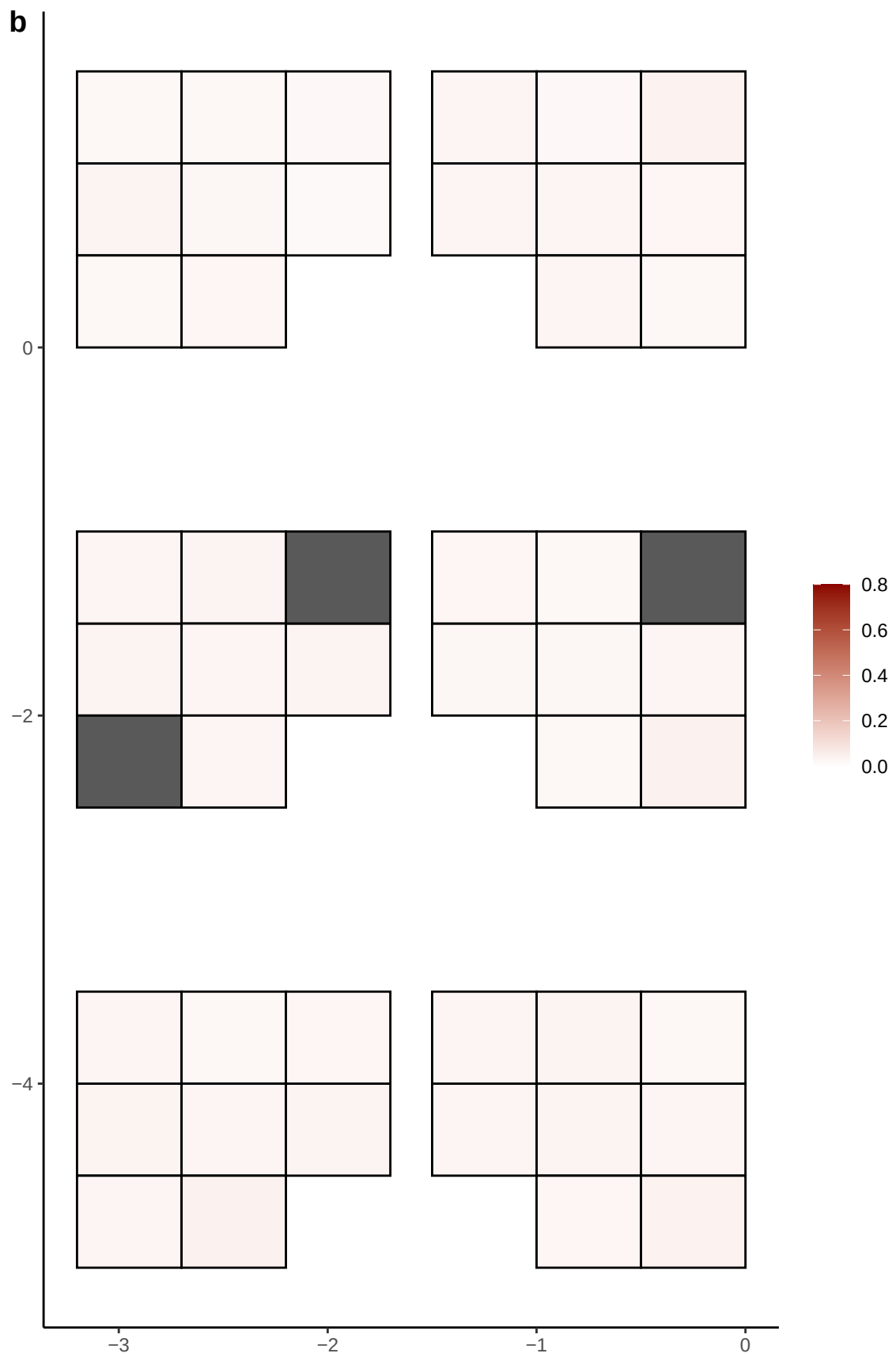

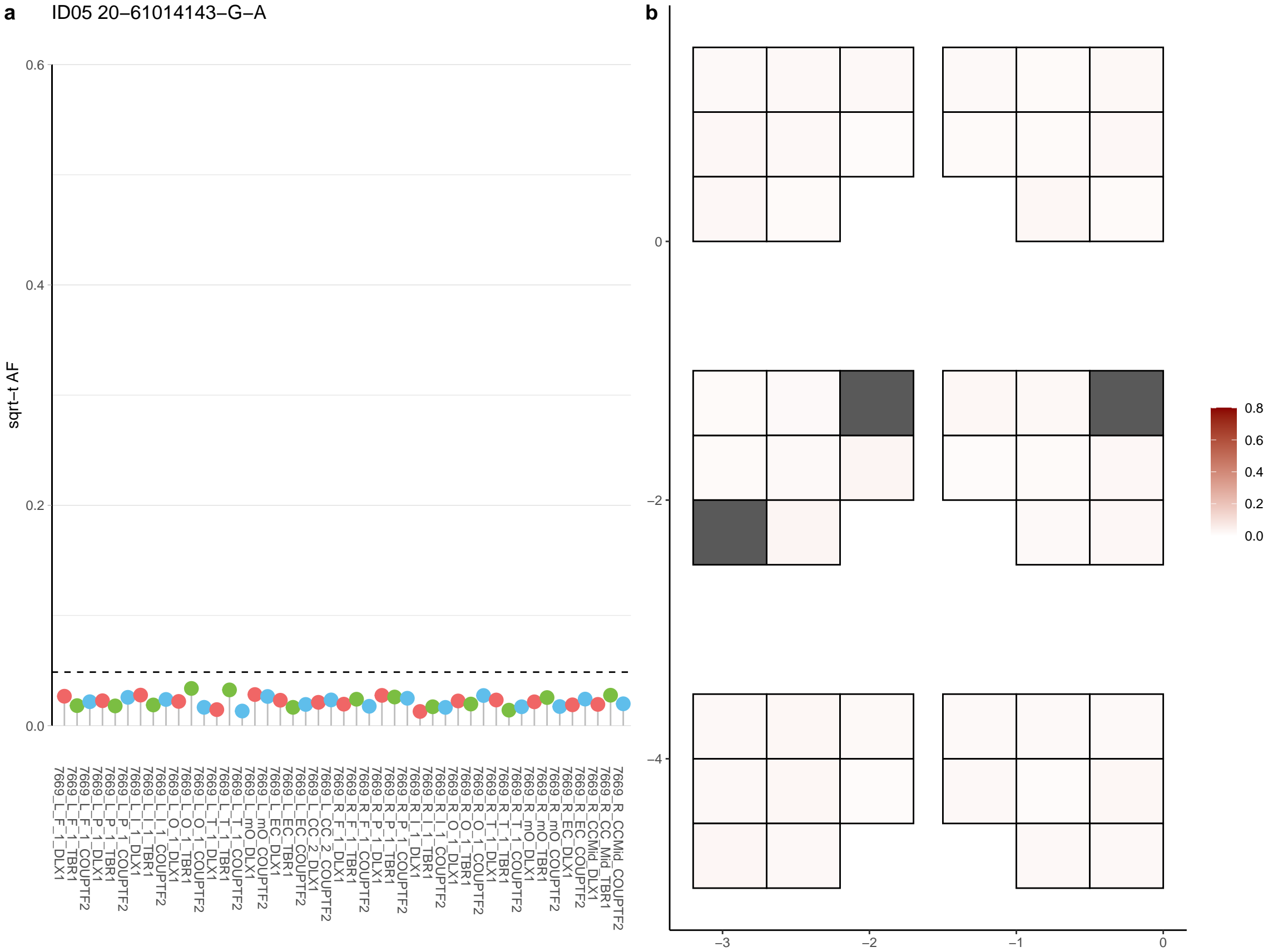

**a** ID05 20-9698158-A-G

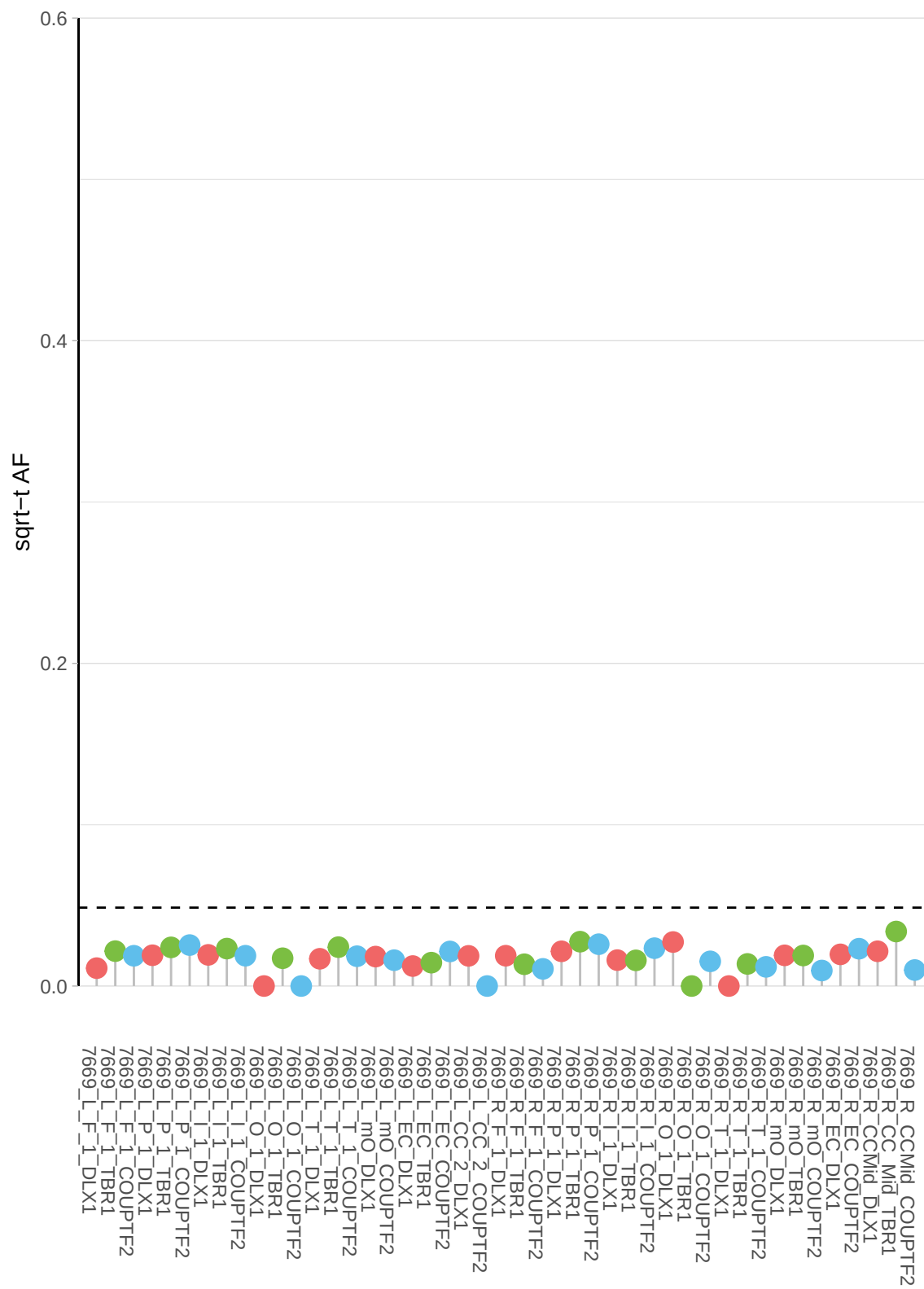

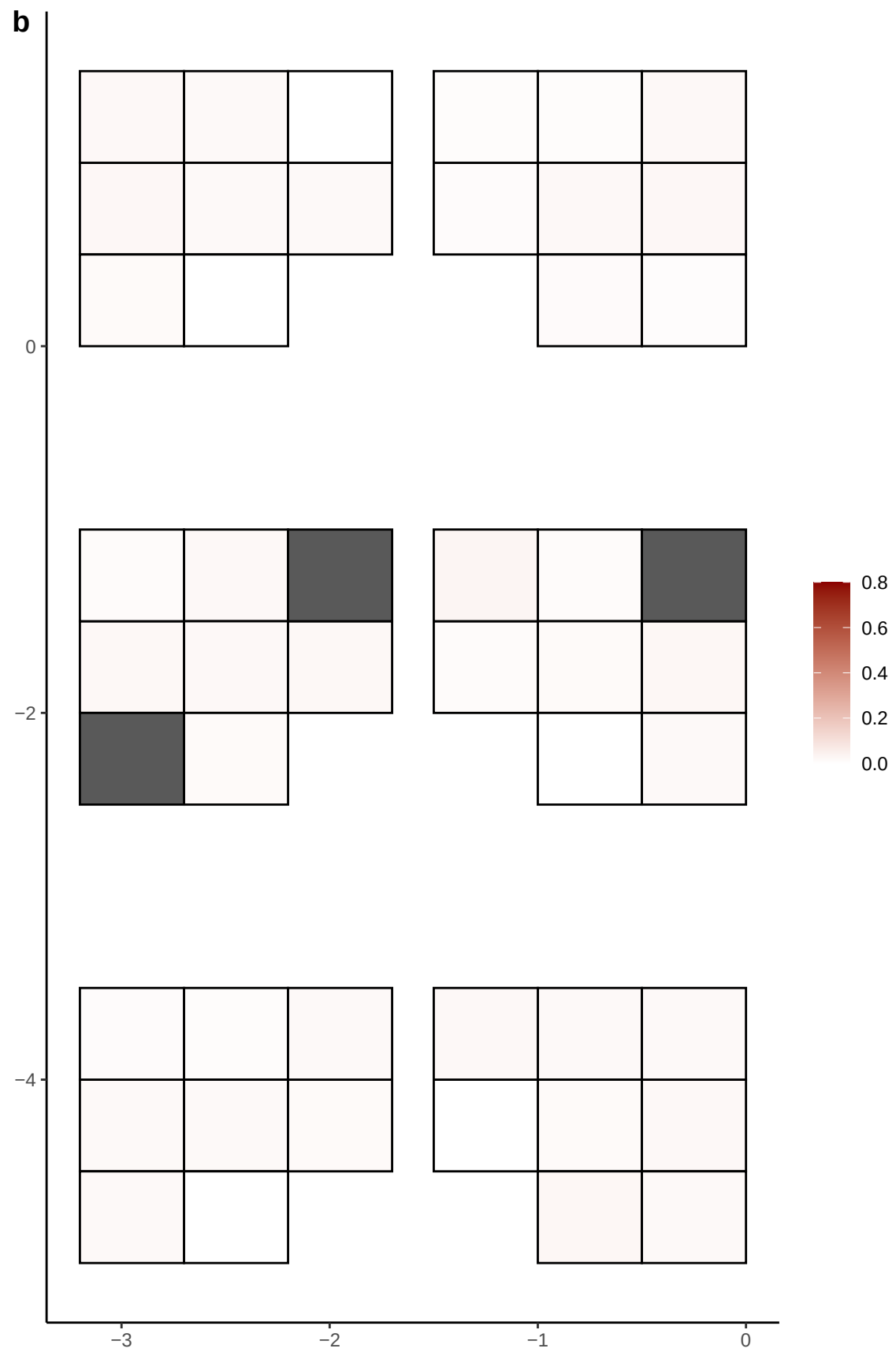

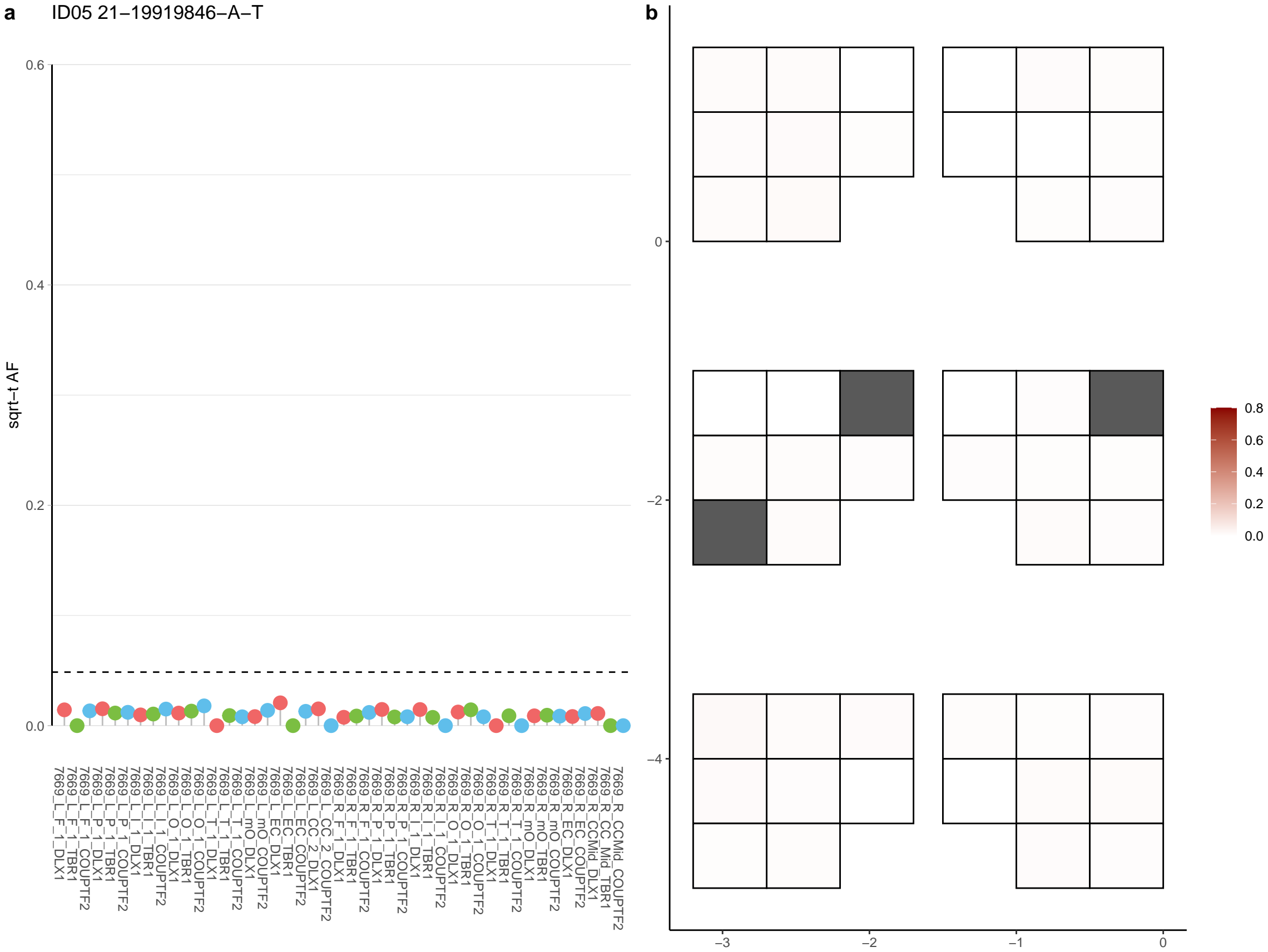

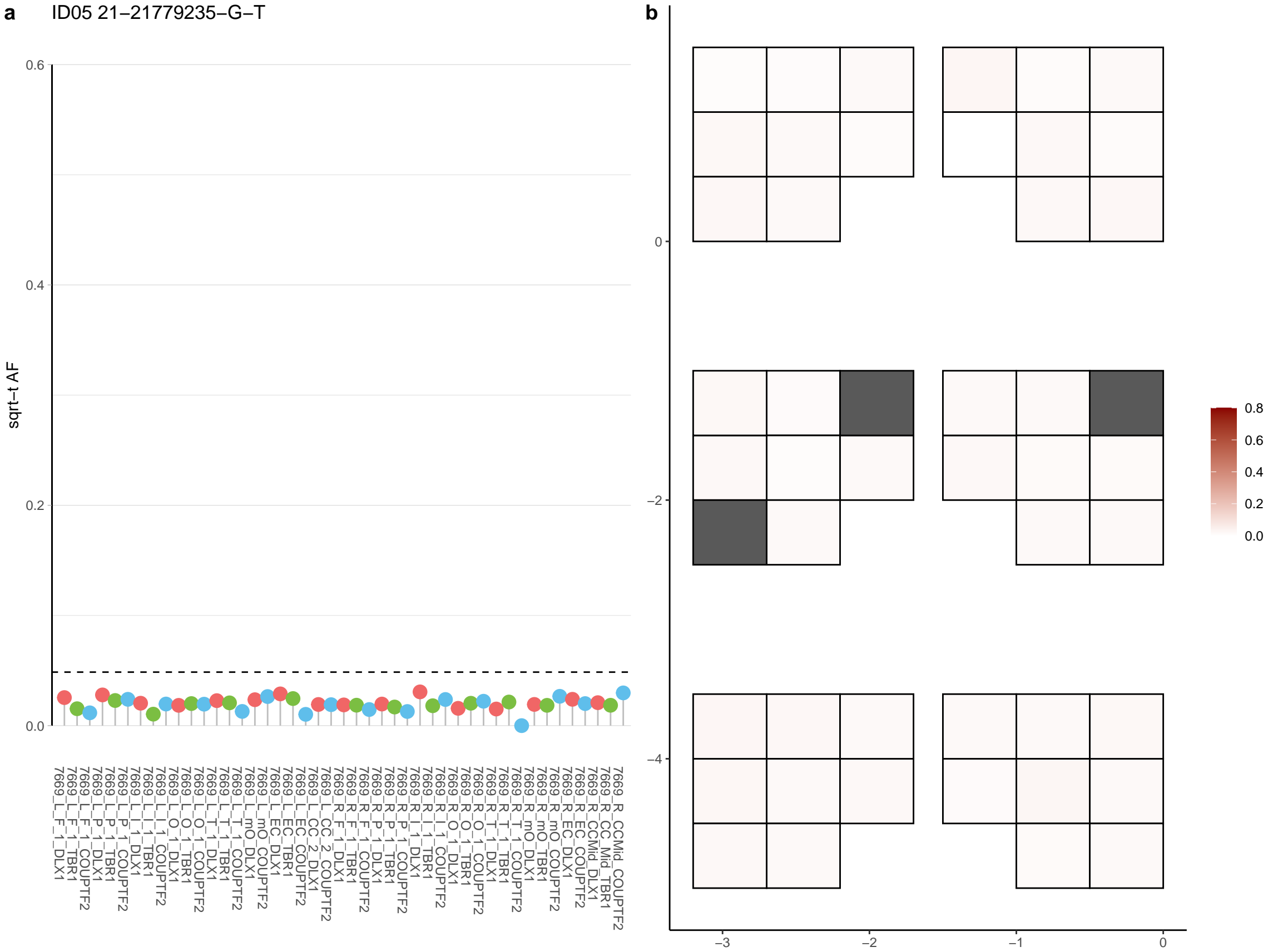

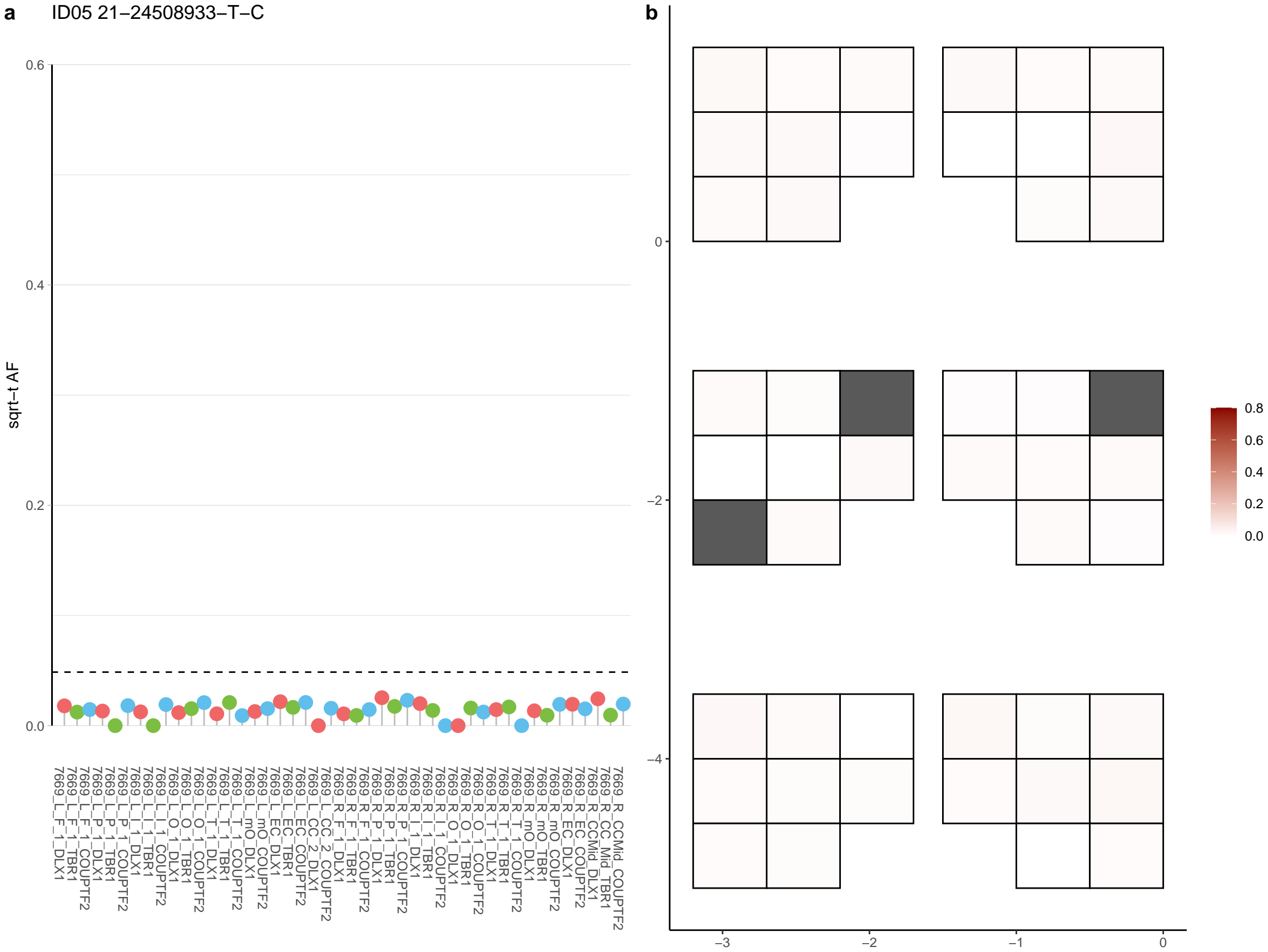

**a** ID05 21-28769343-G-A

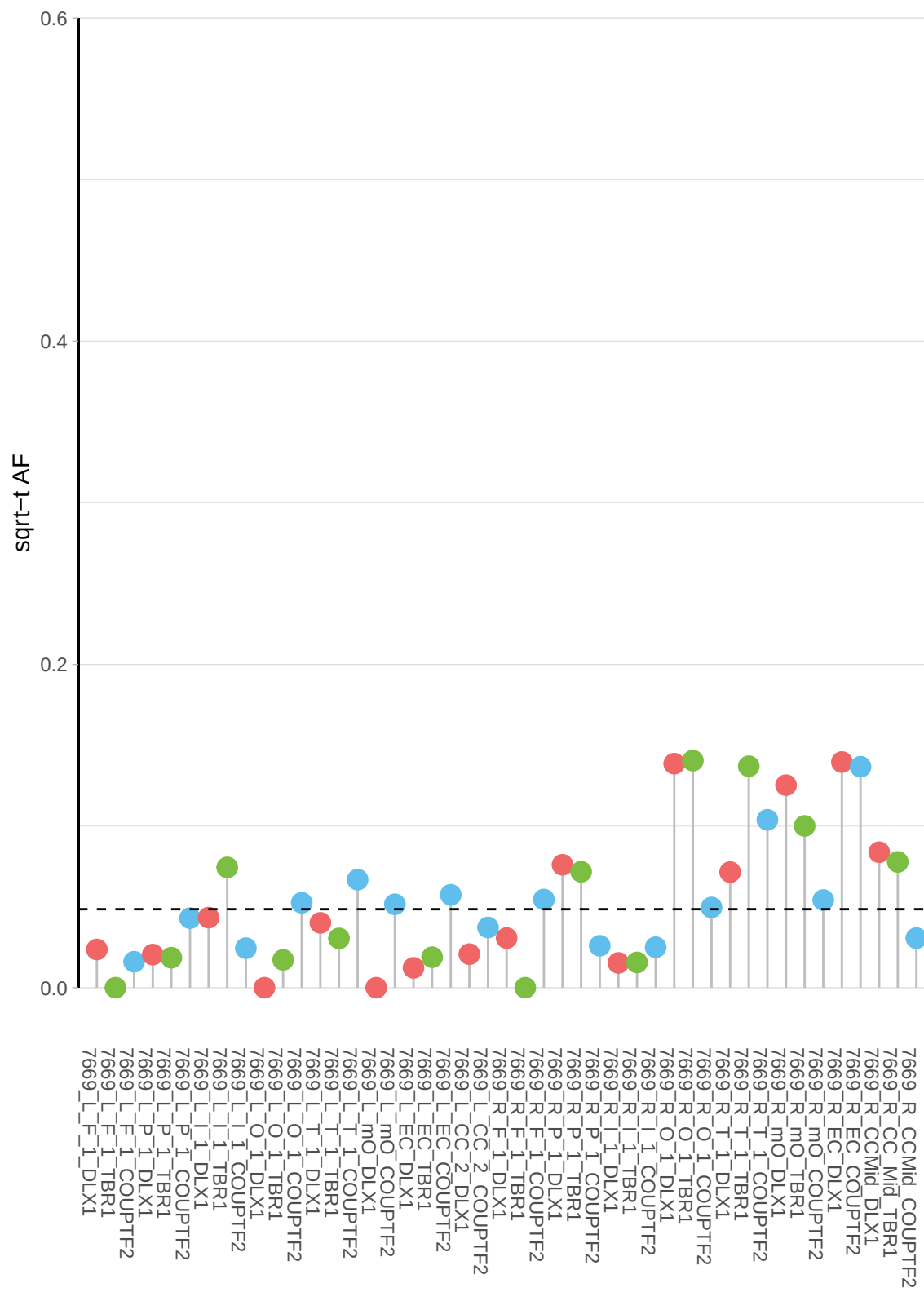

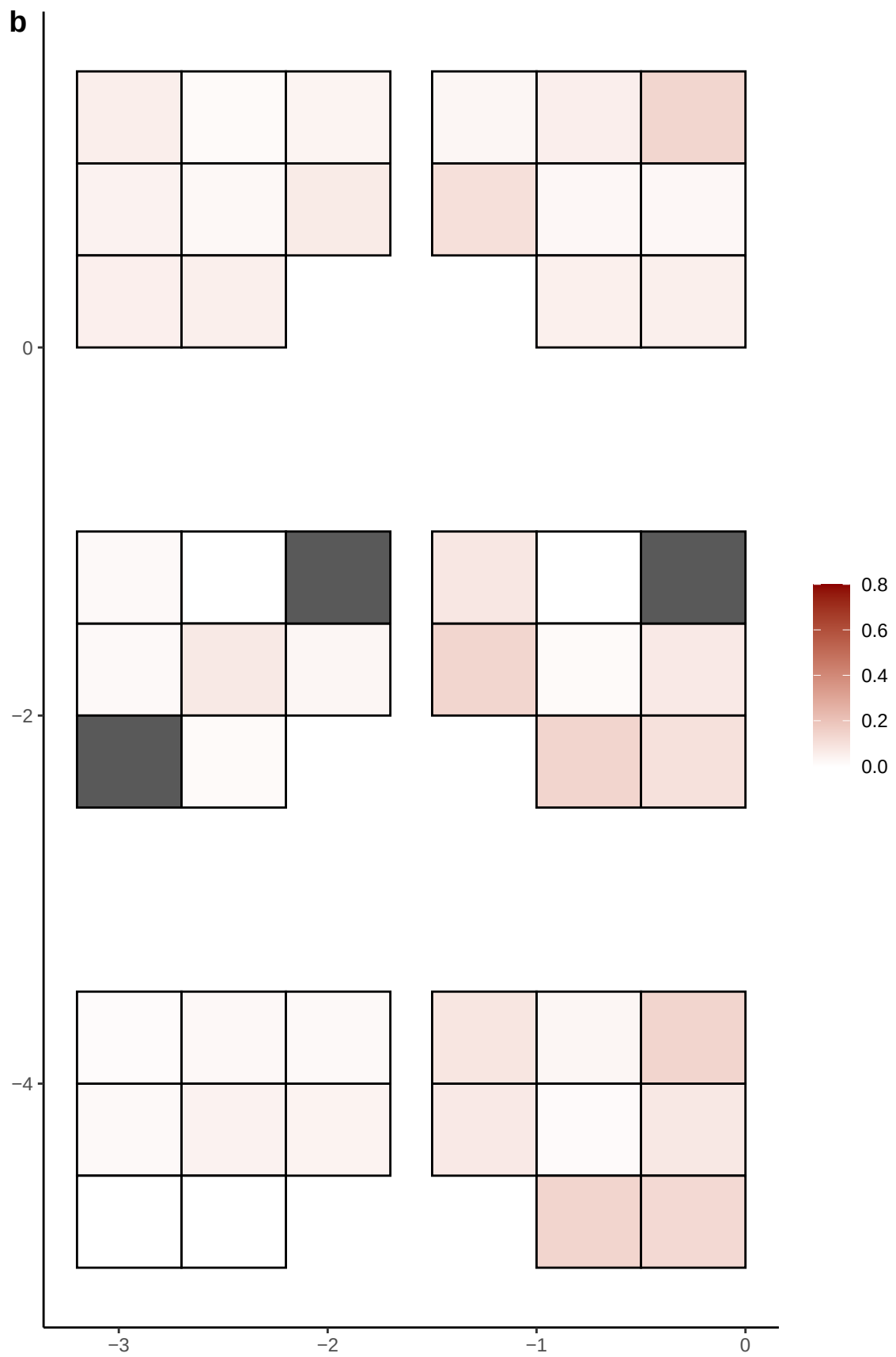

**a** ID05 21-32316236-C-T

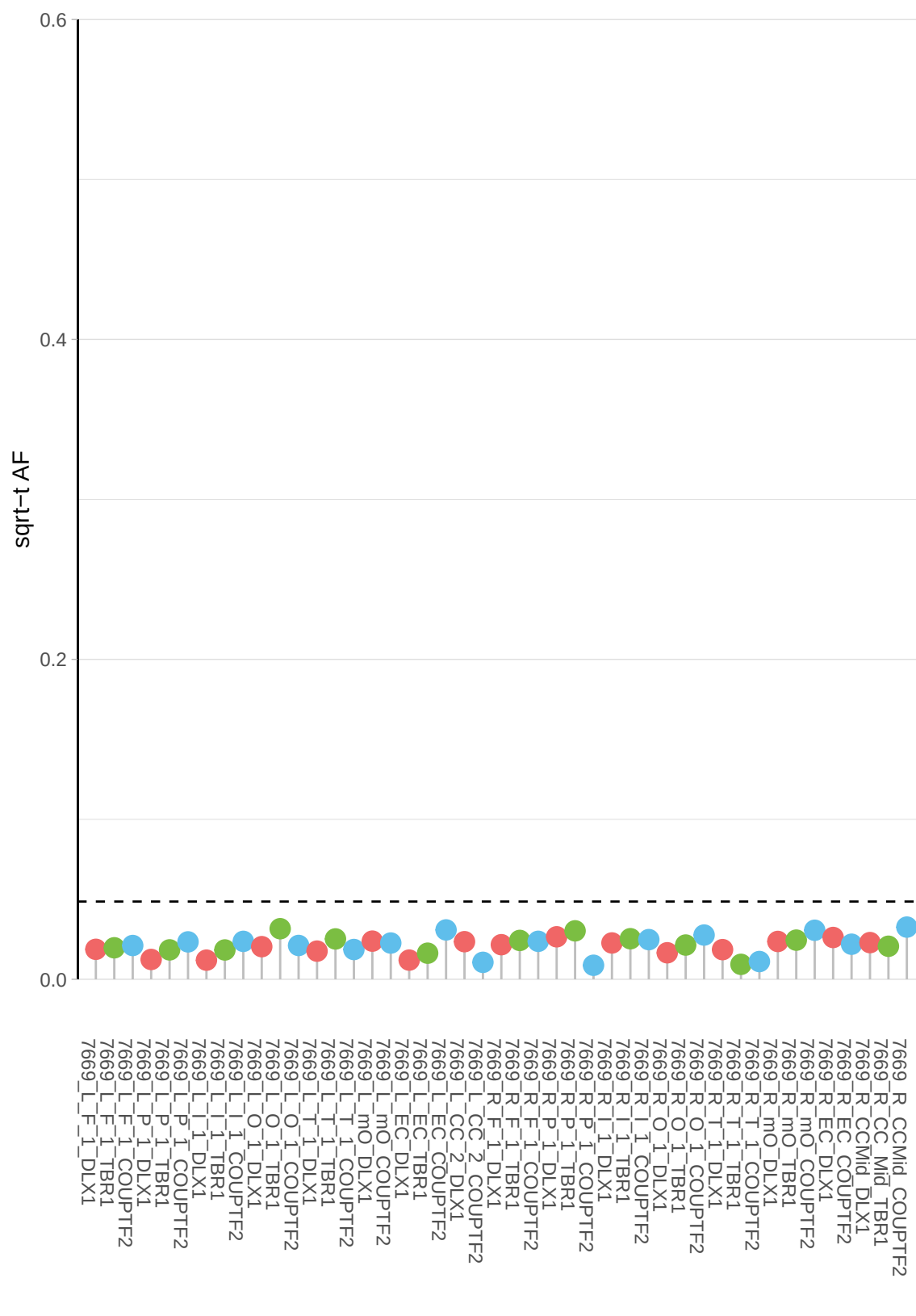

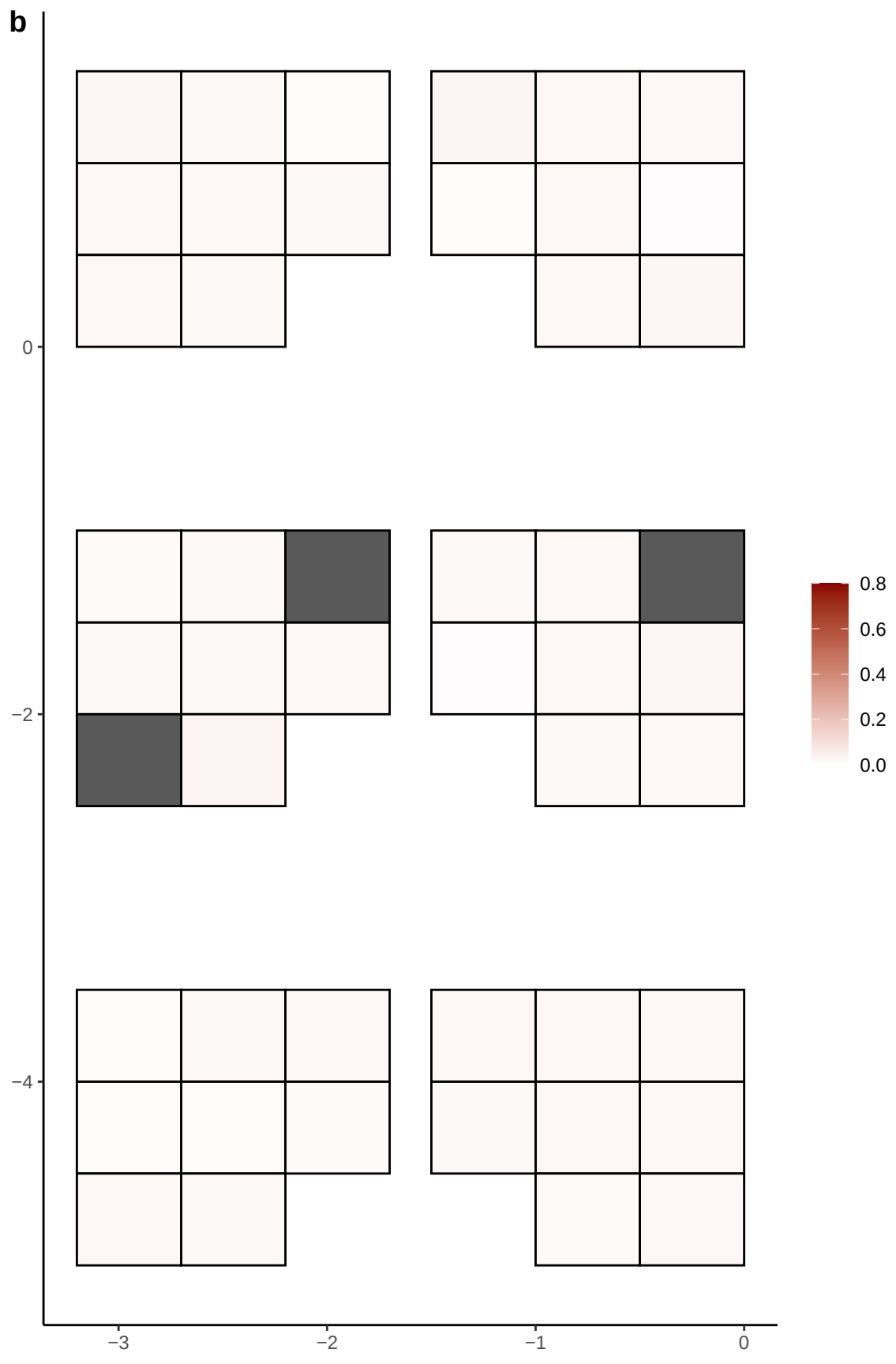

**a** ID05 21-32682912-C-T

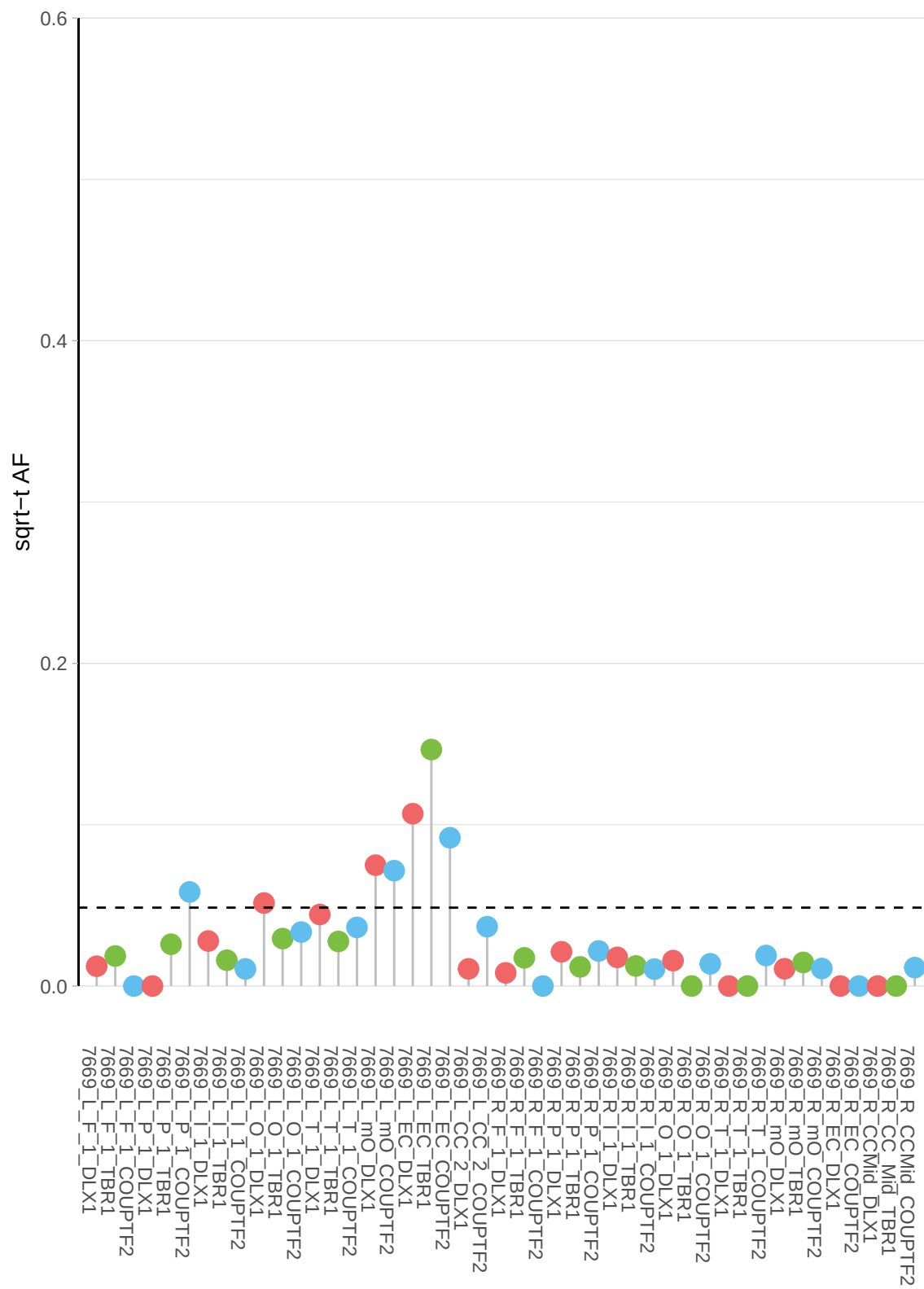

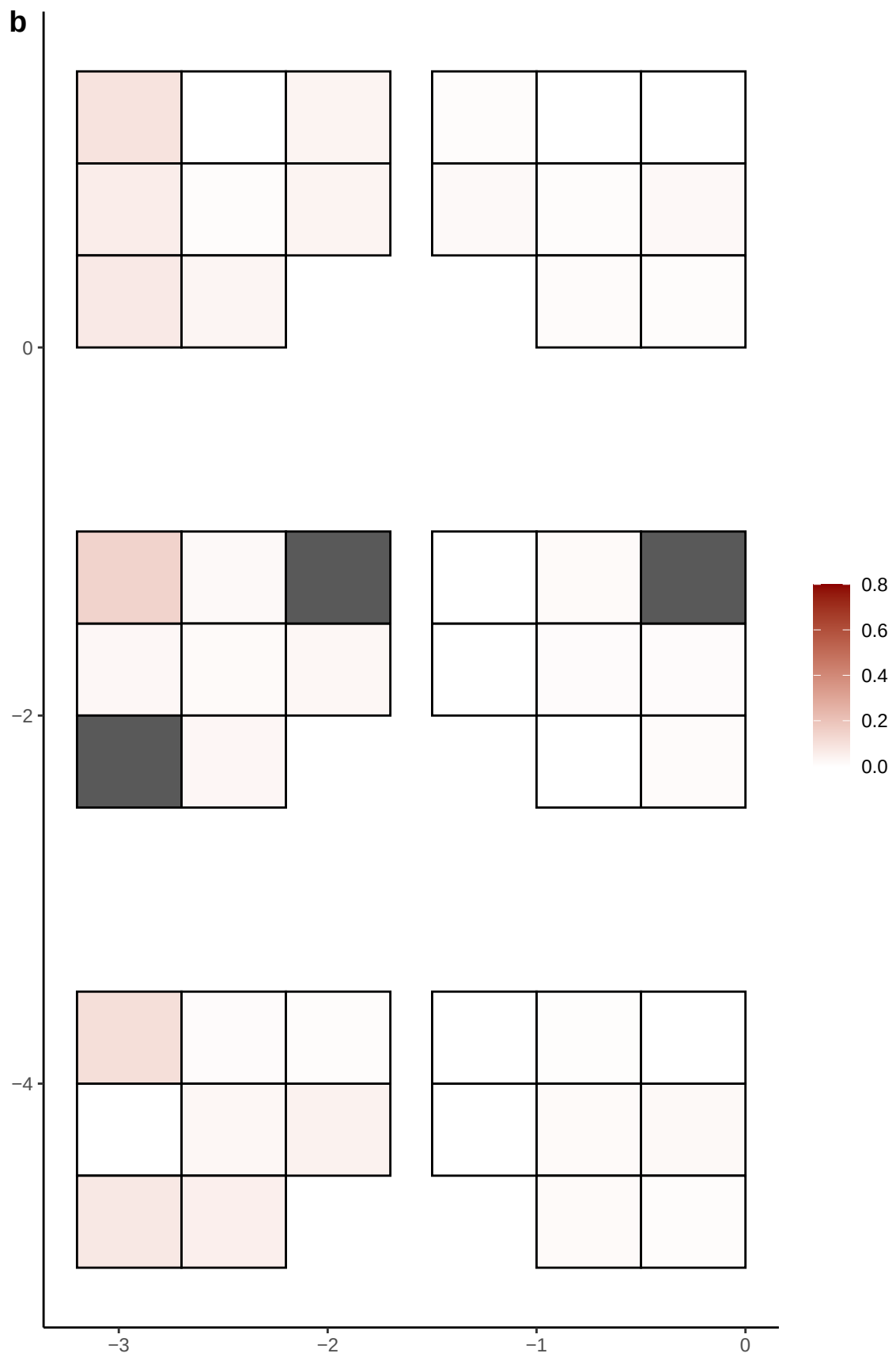

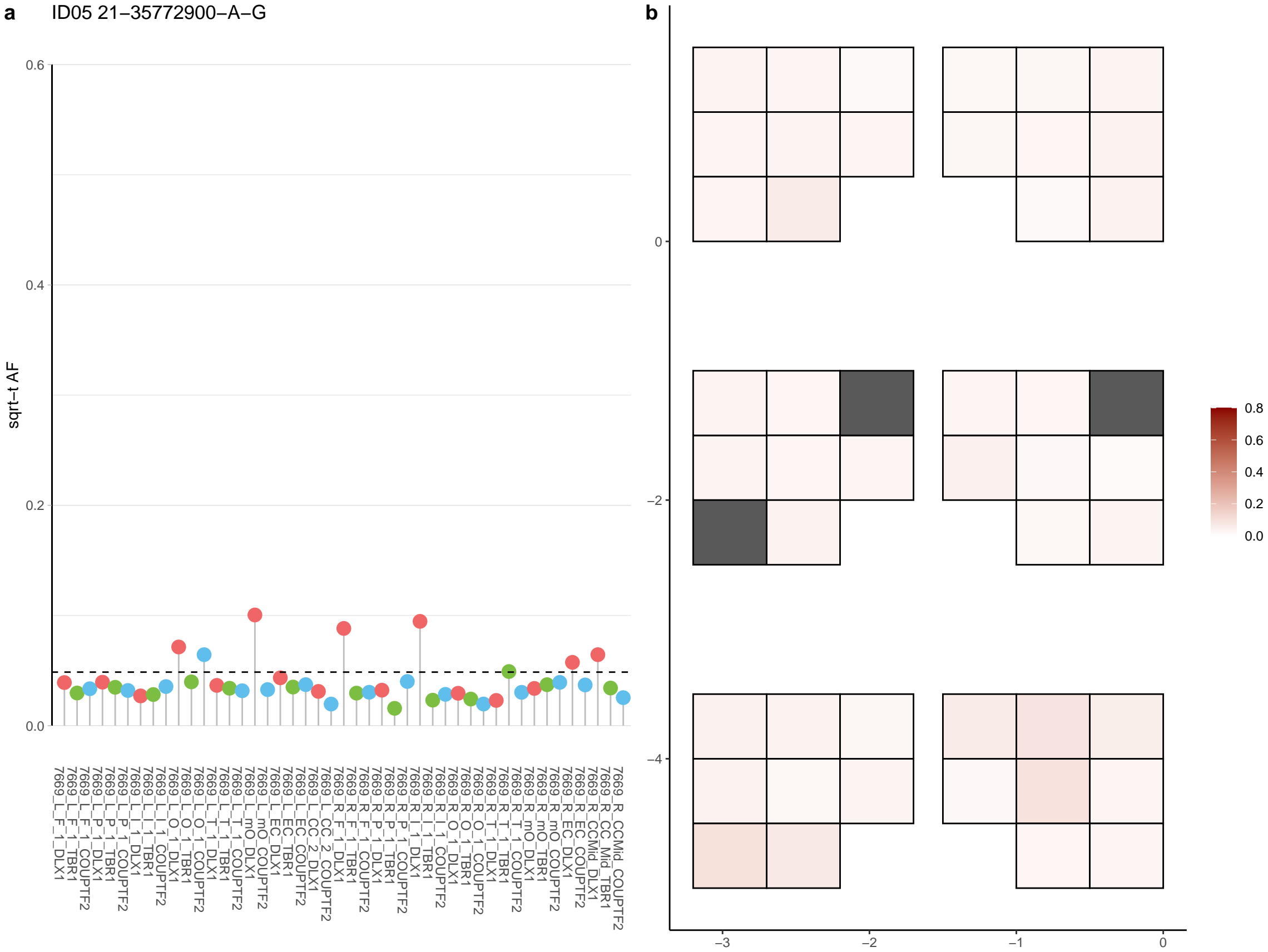

**a** ID05 21-41023573-C-T

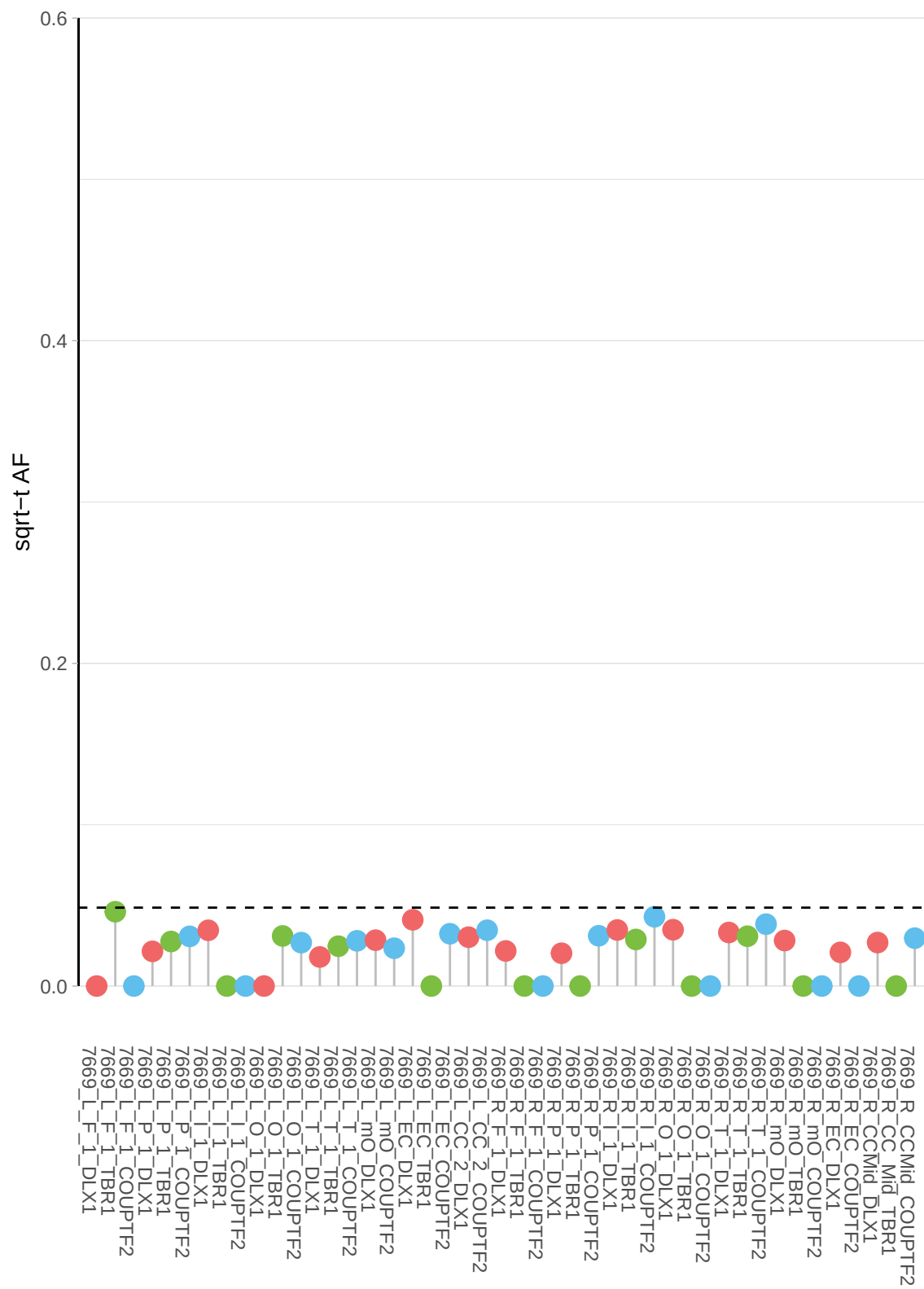

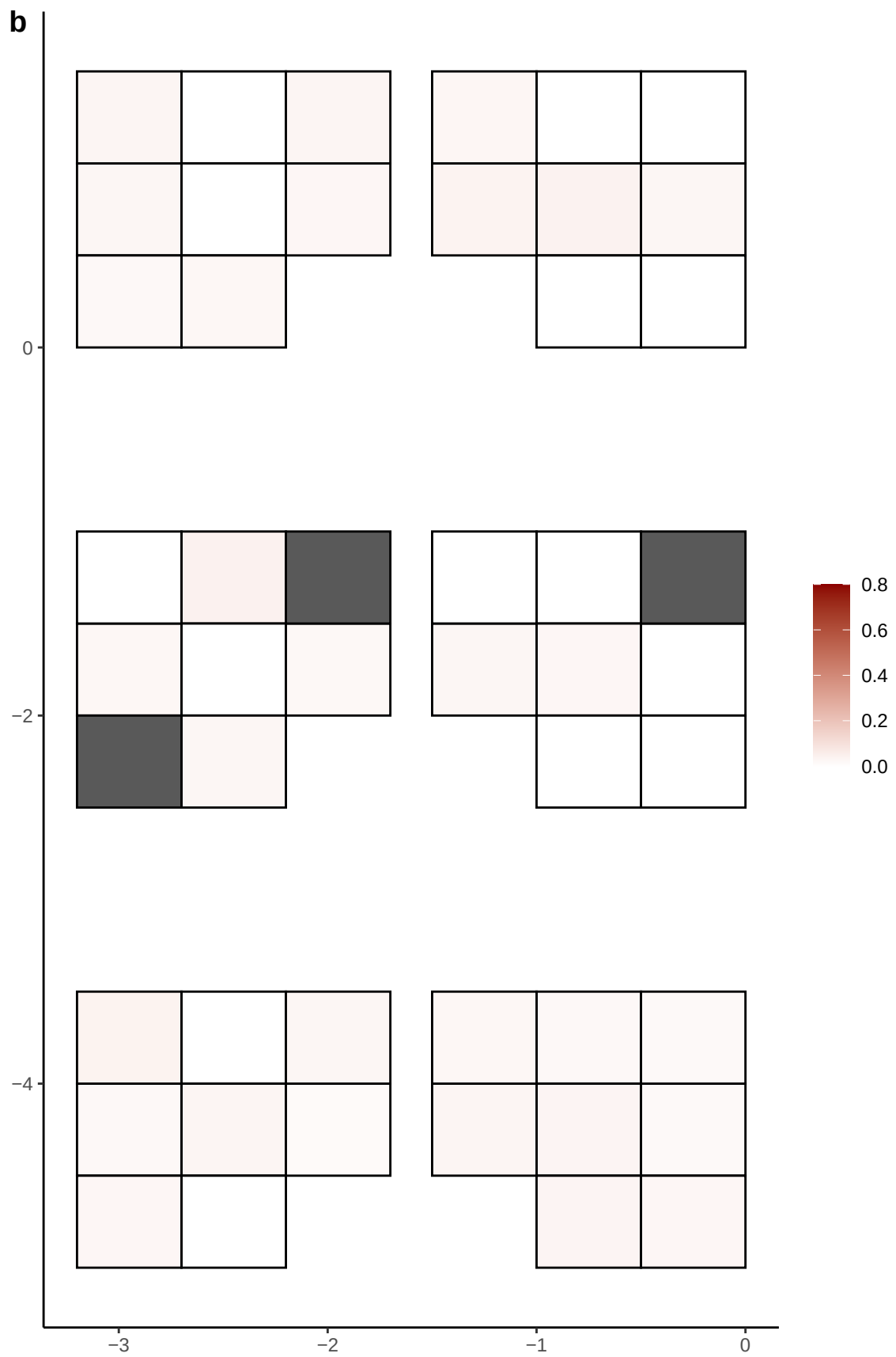

**a** ID05 21-41236582-C-T

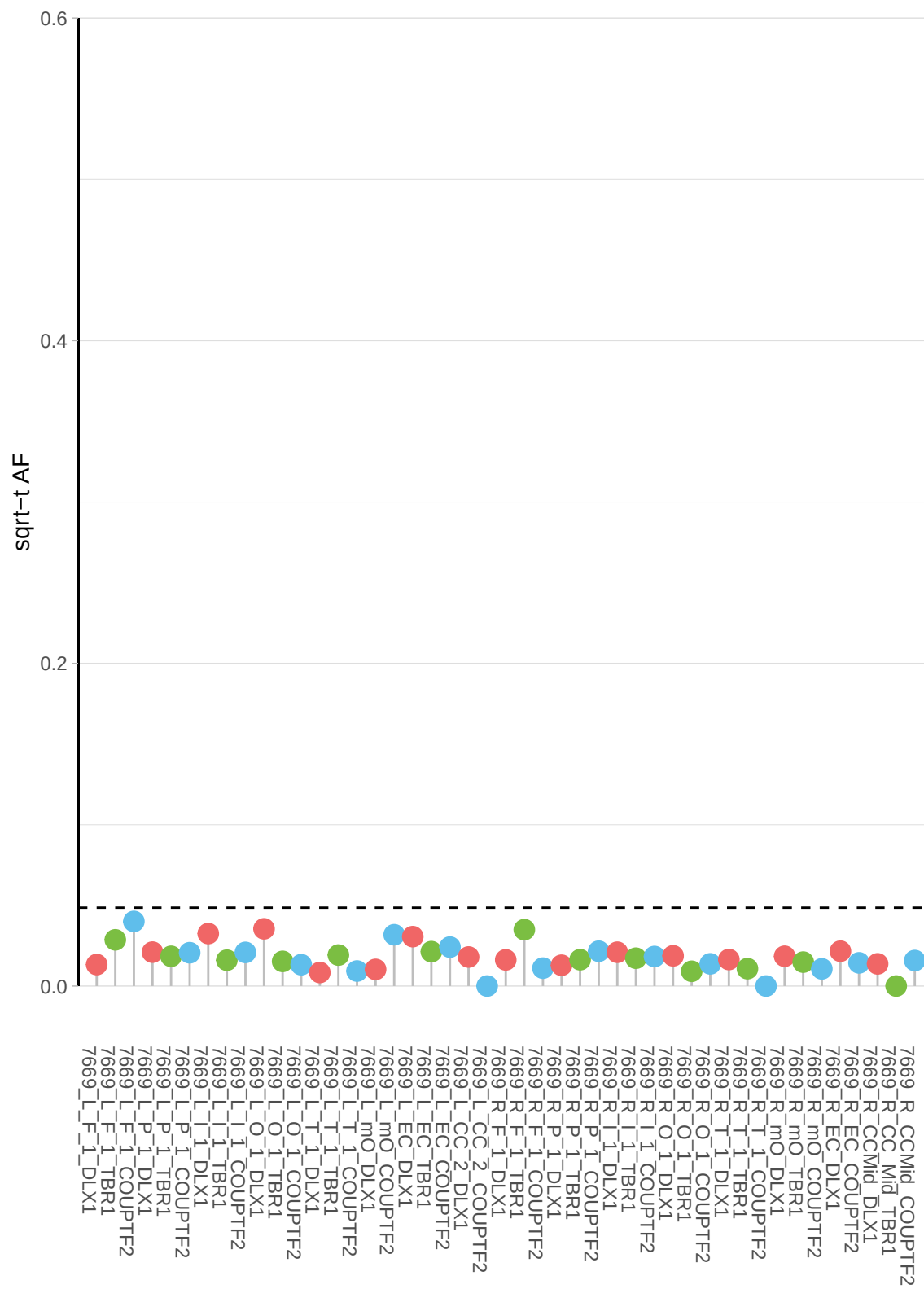

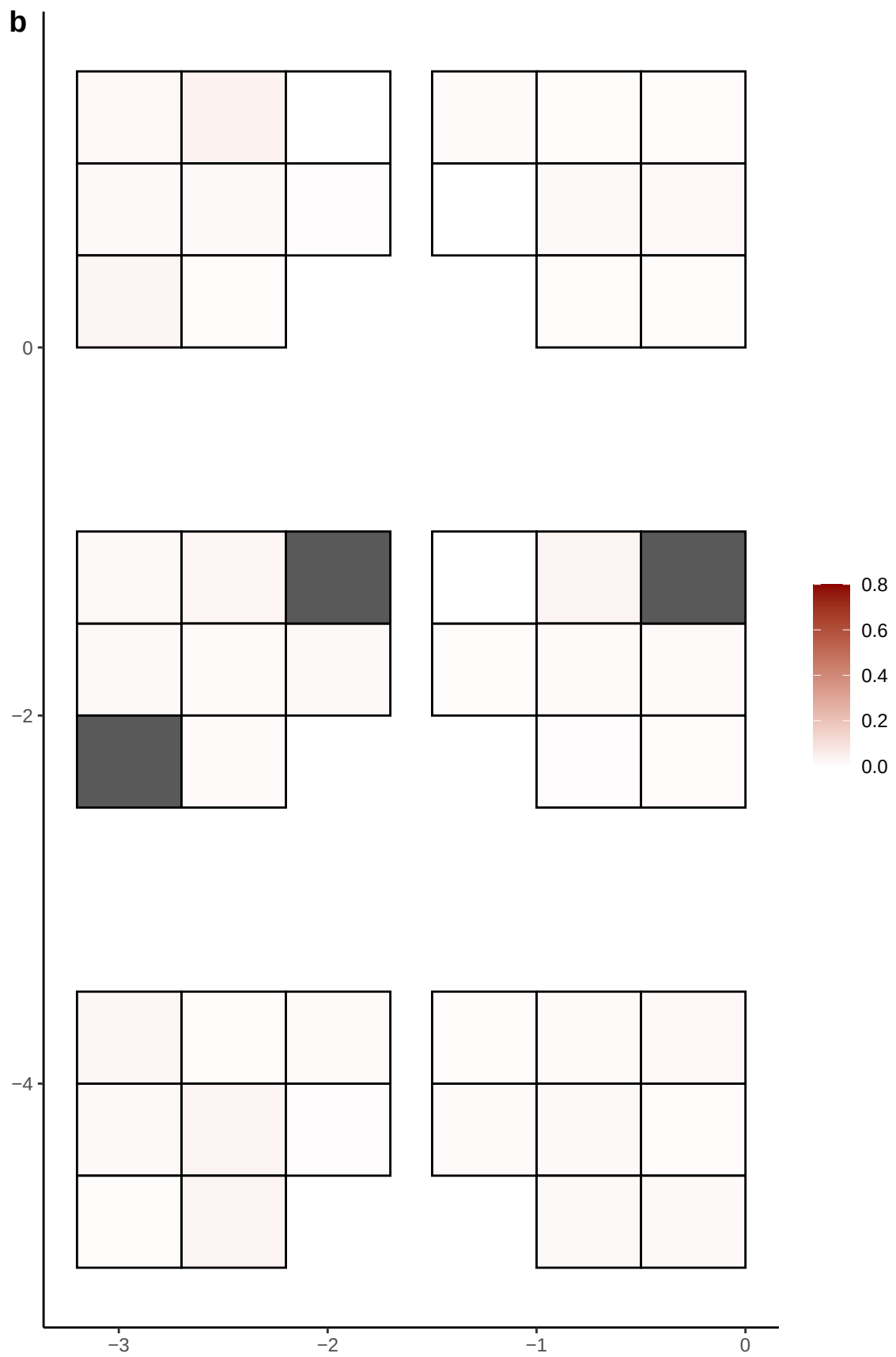

**a** ID05 21-41370607-T-C

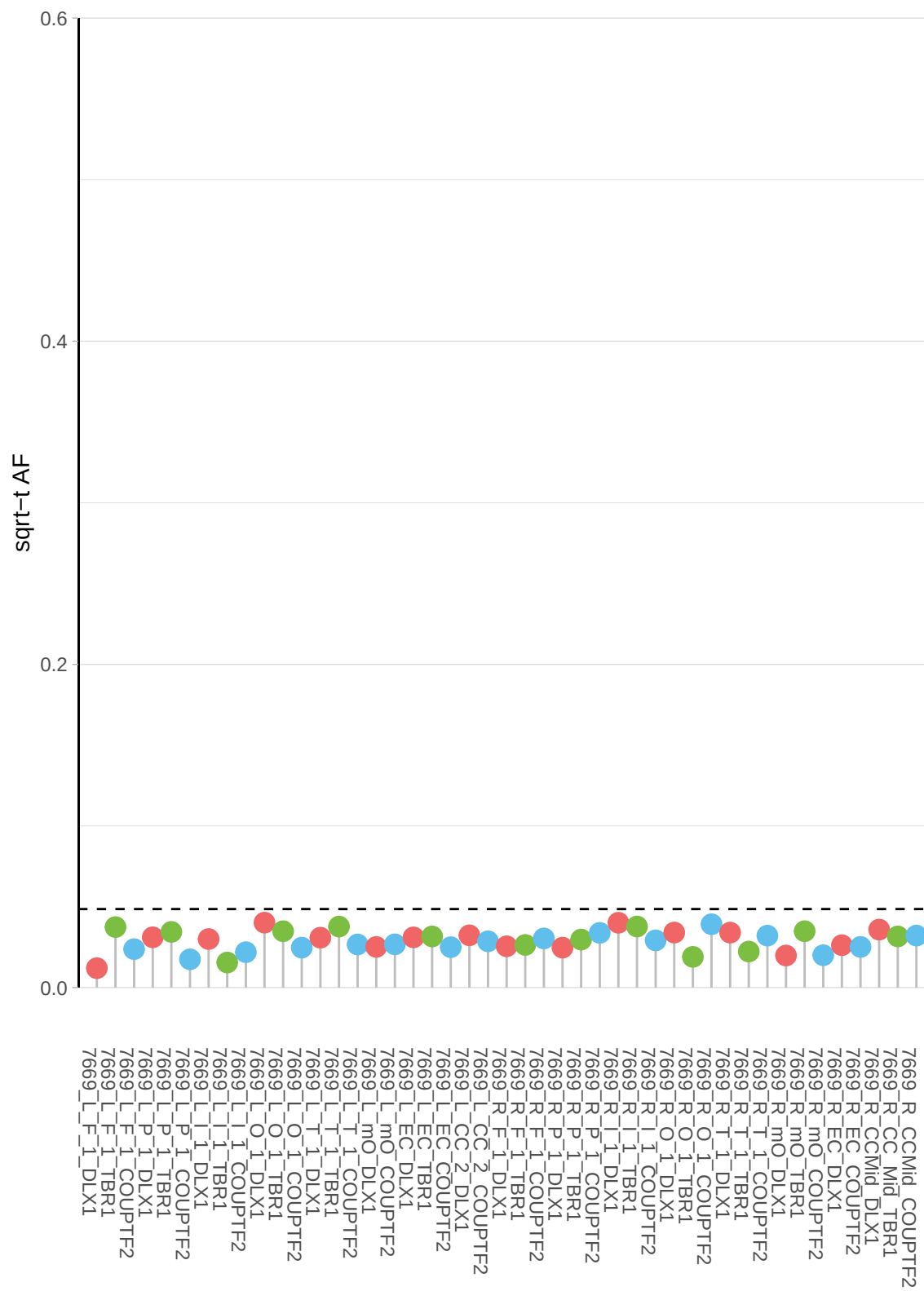

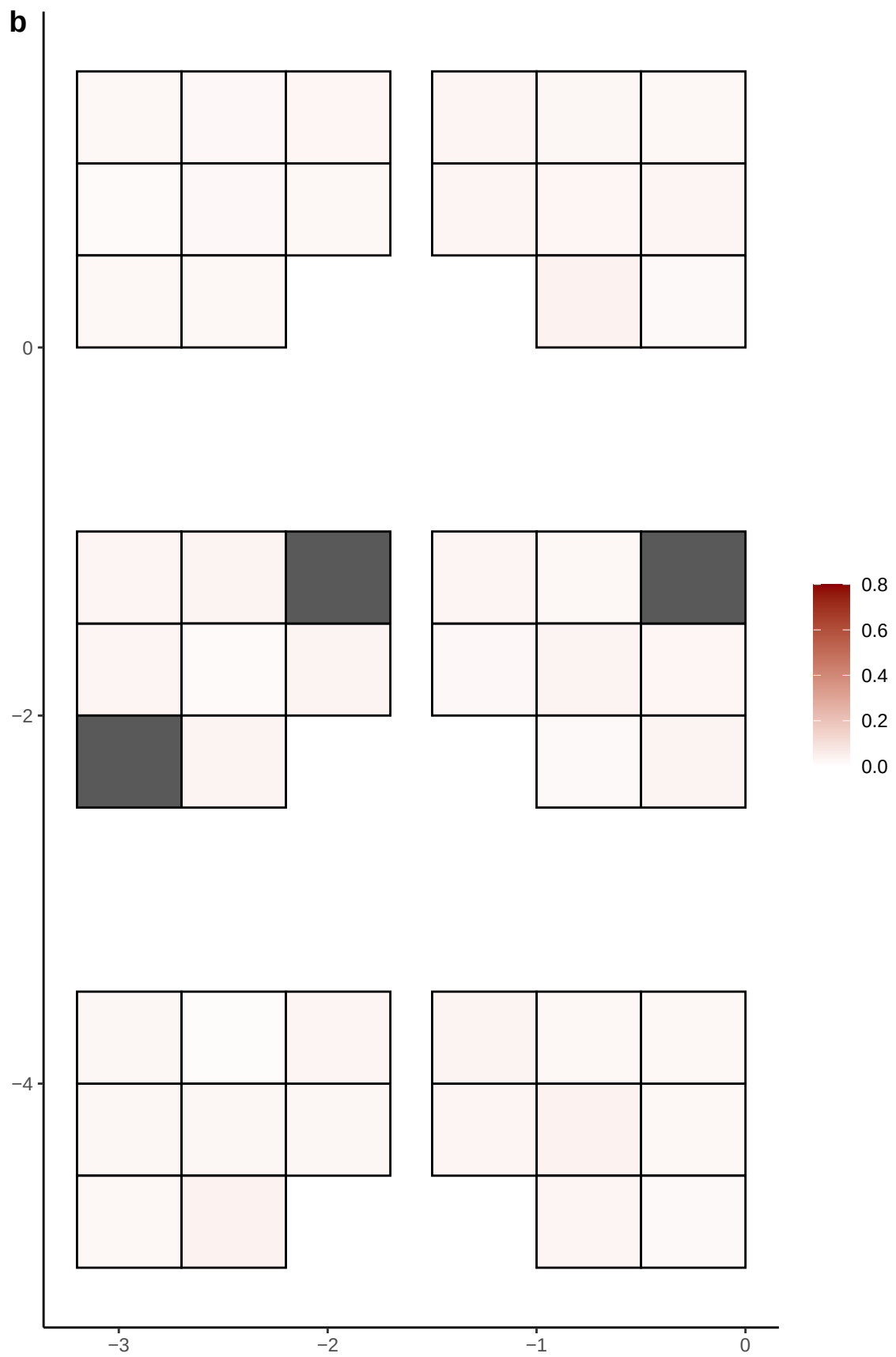

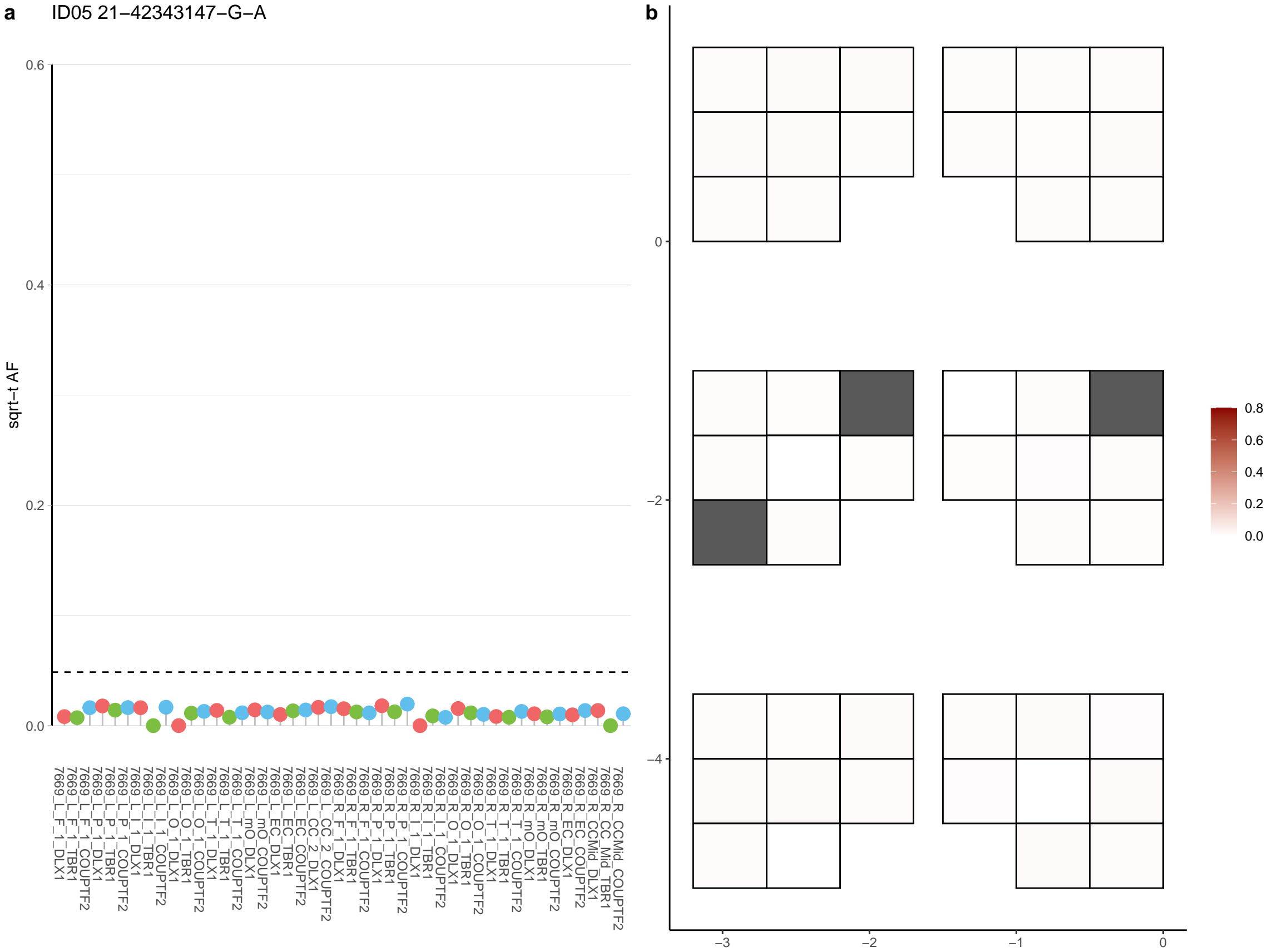

**a** ID05 21-42676569-G-A

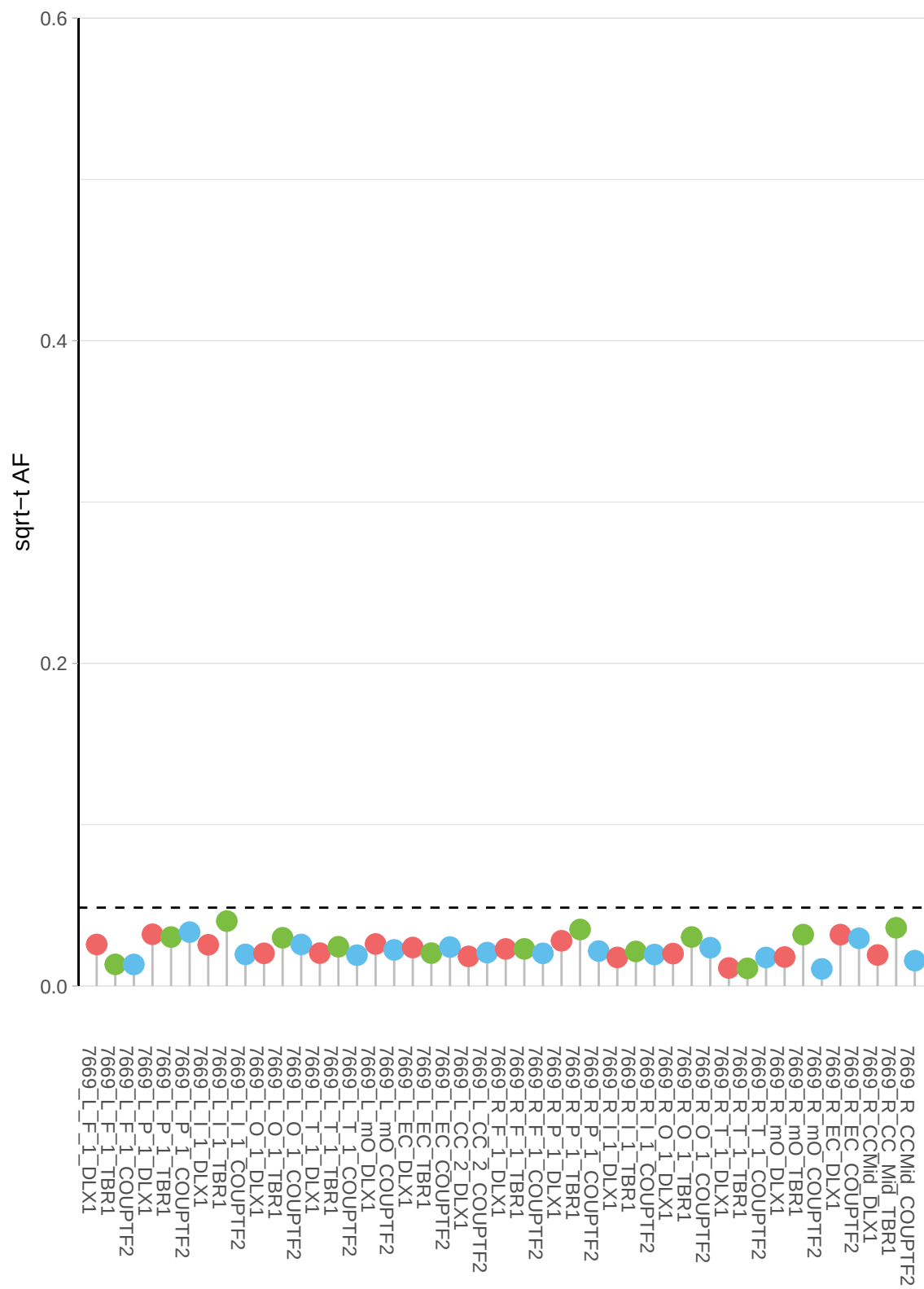

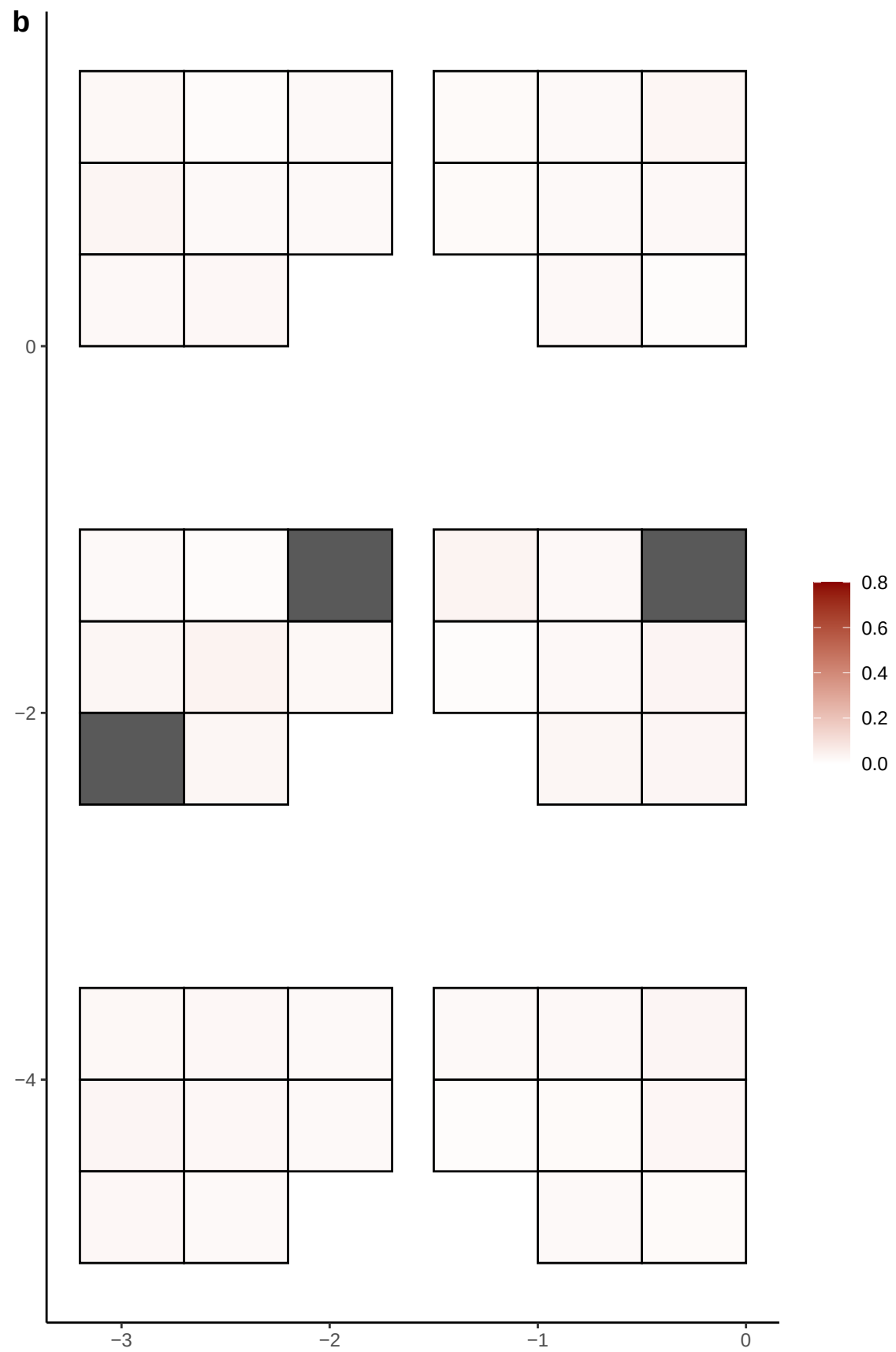

**a** ID05 21-43768376-C-T

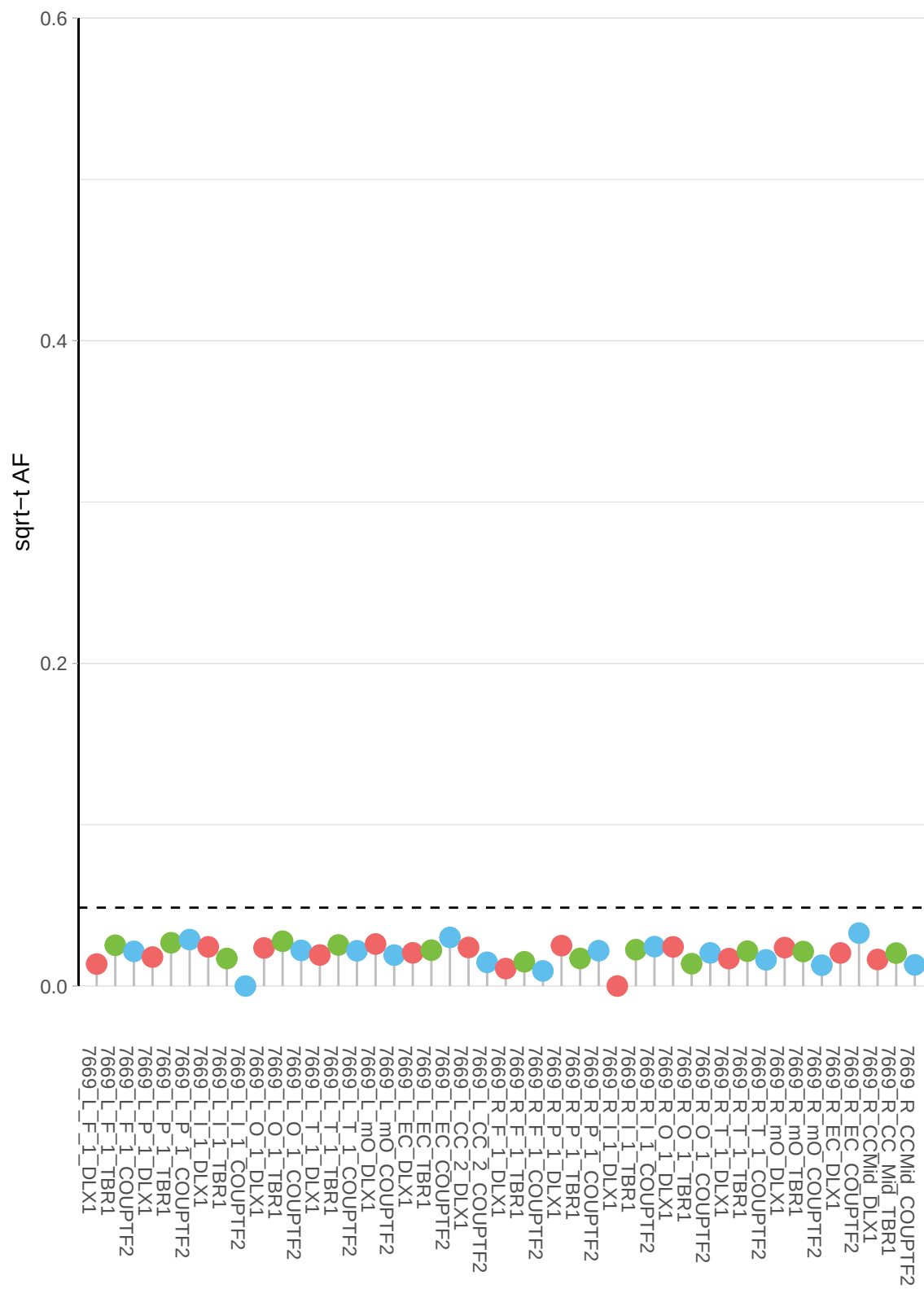

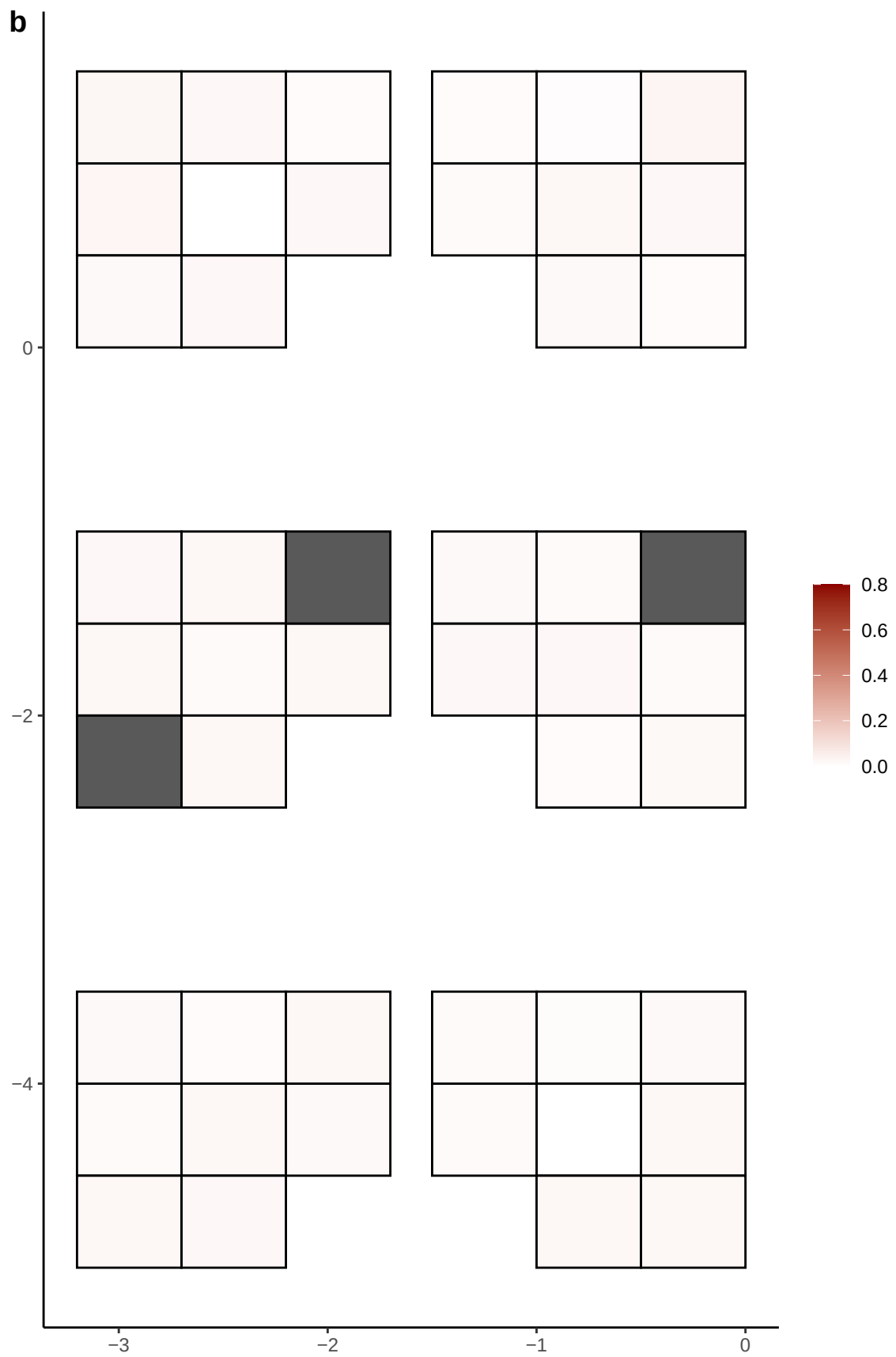

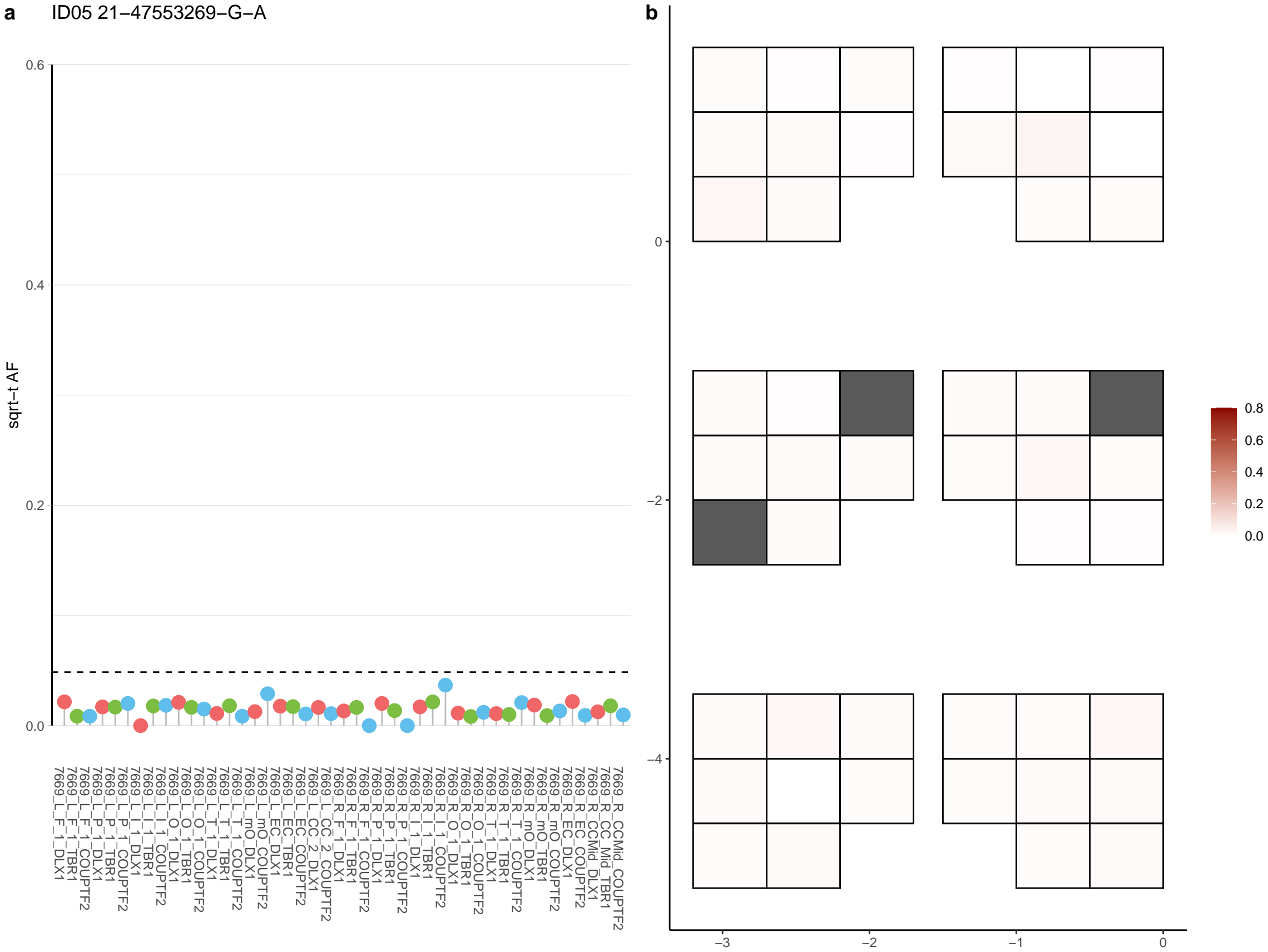

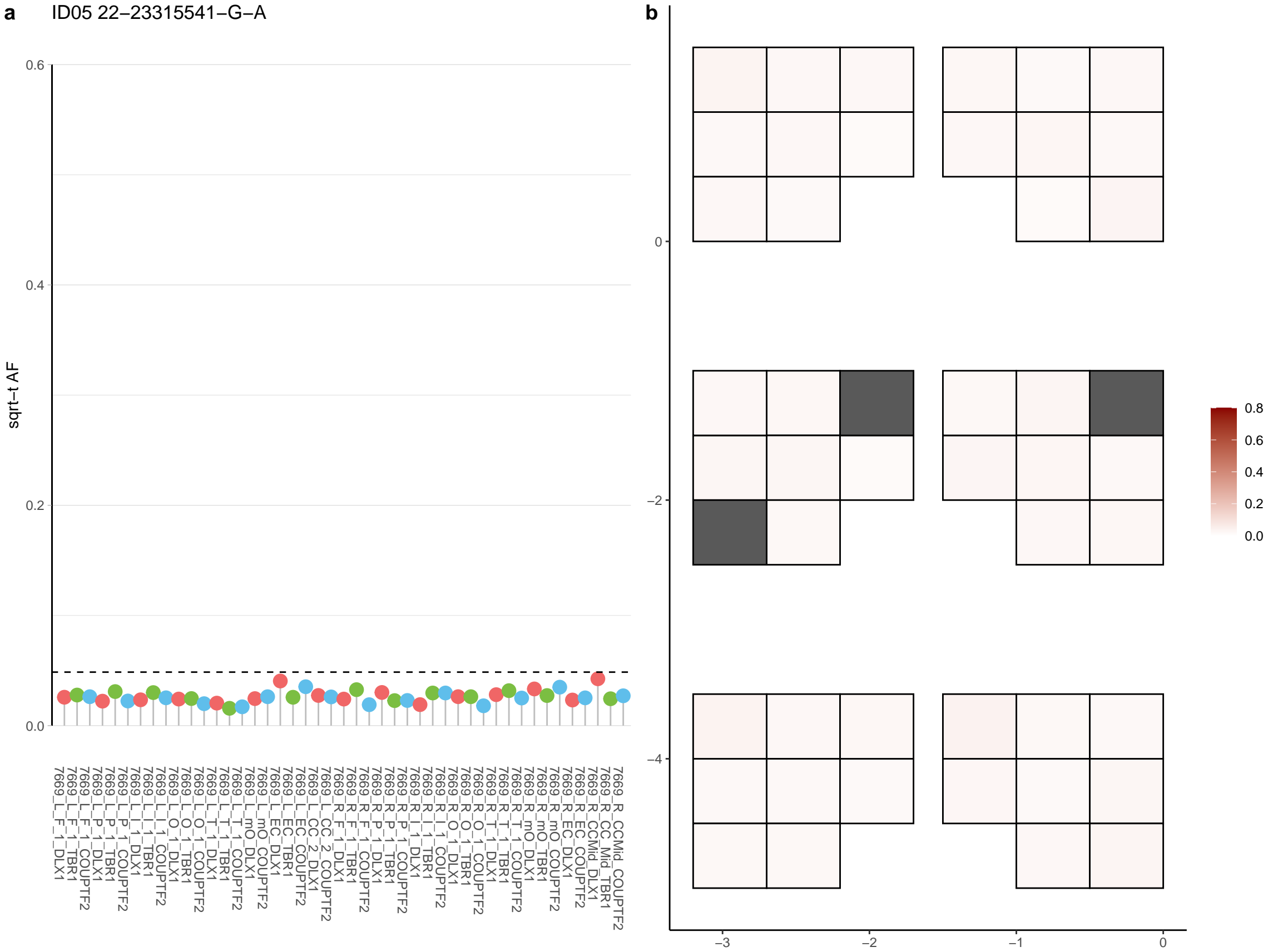

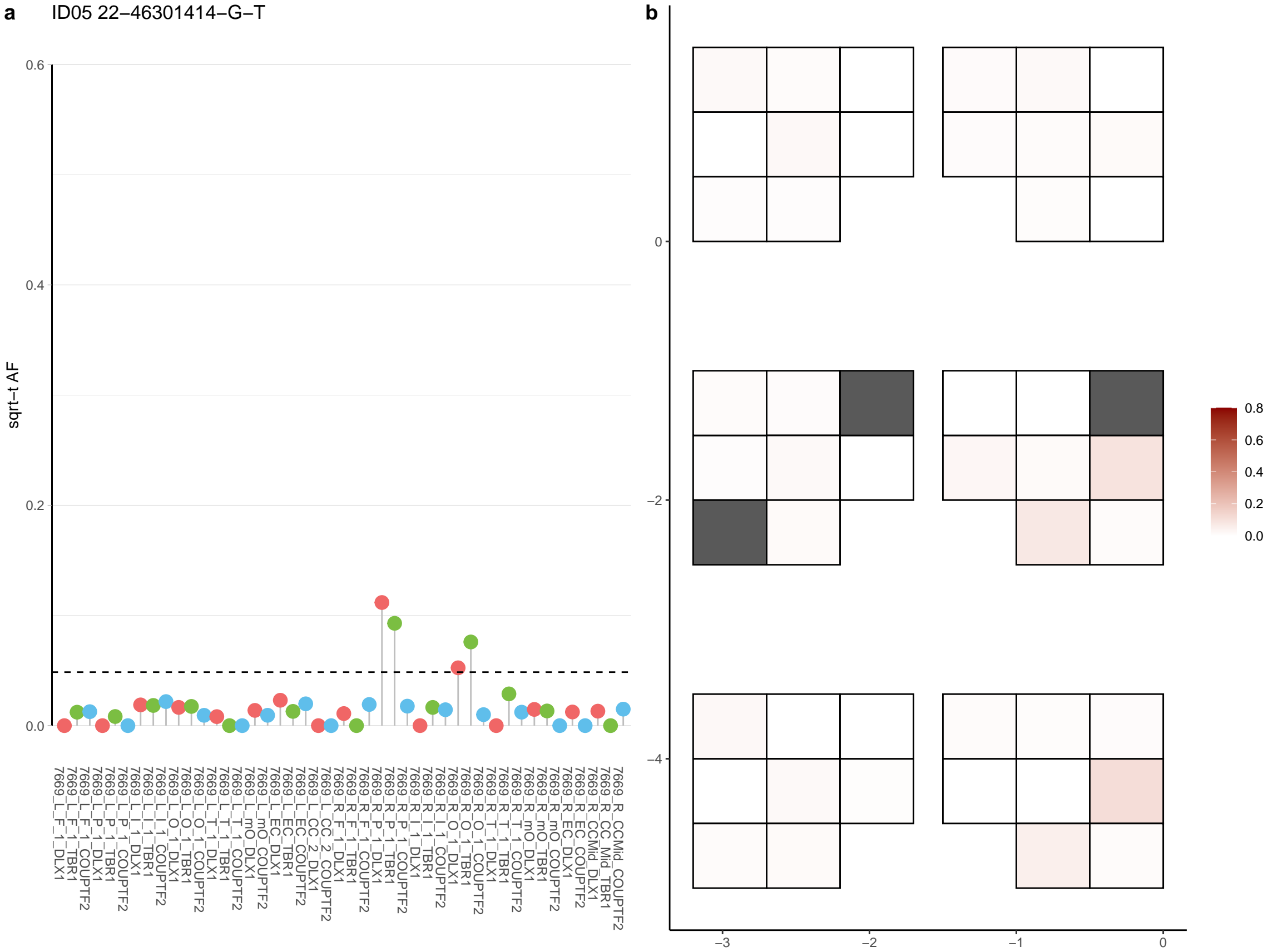

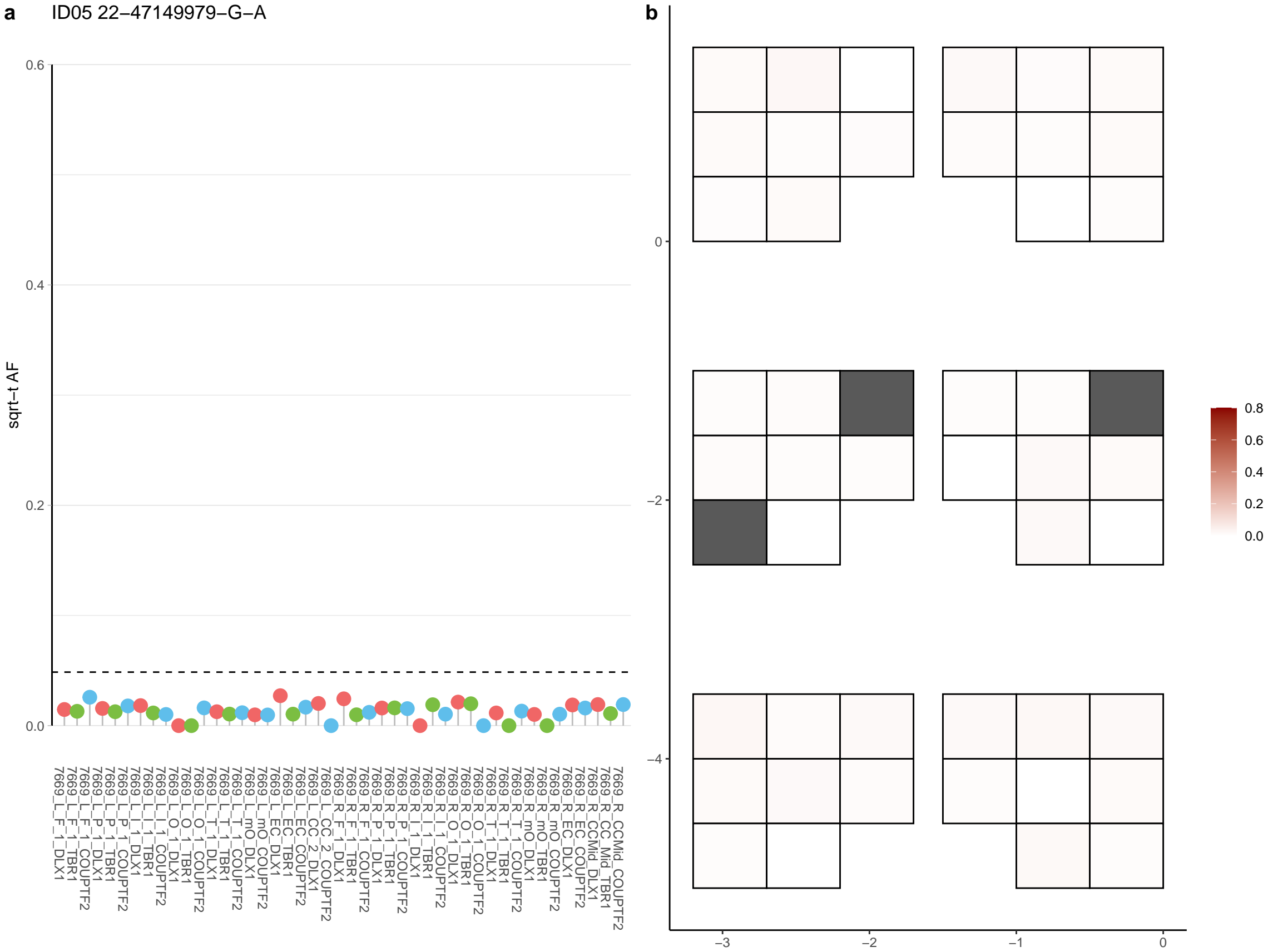

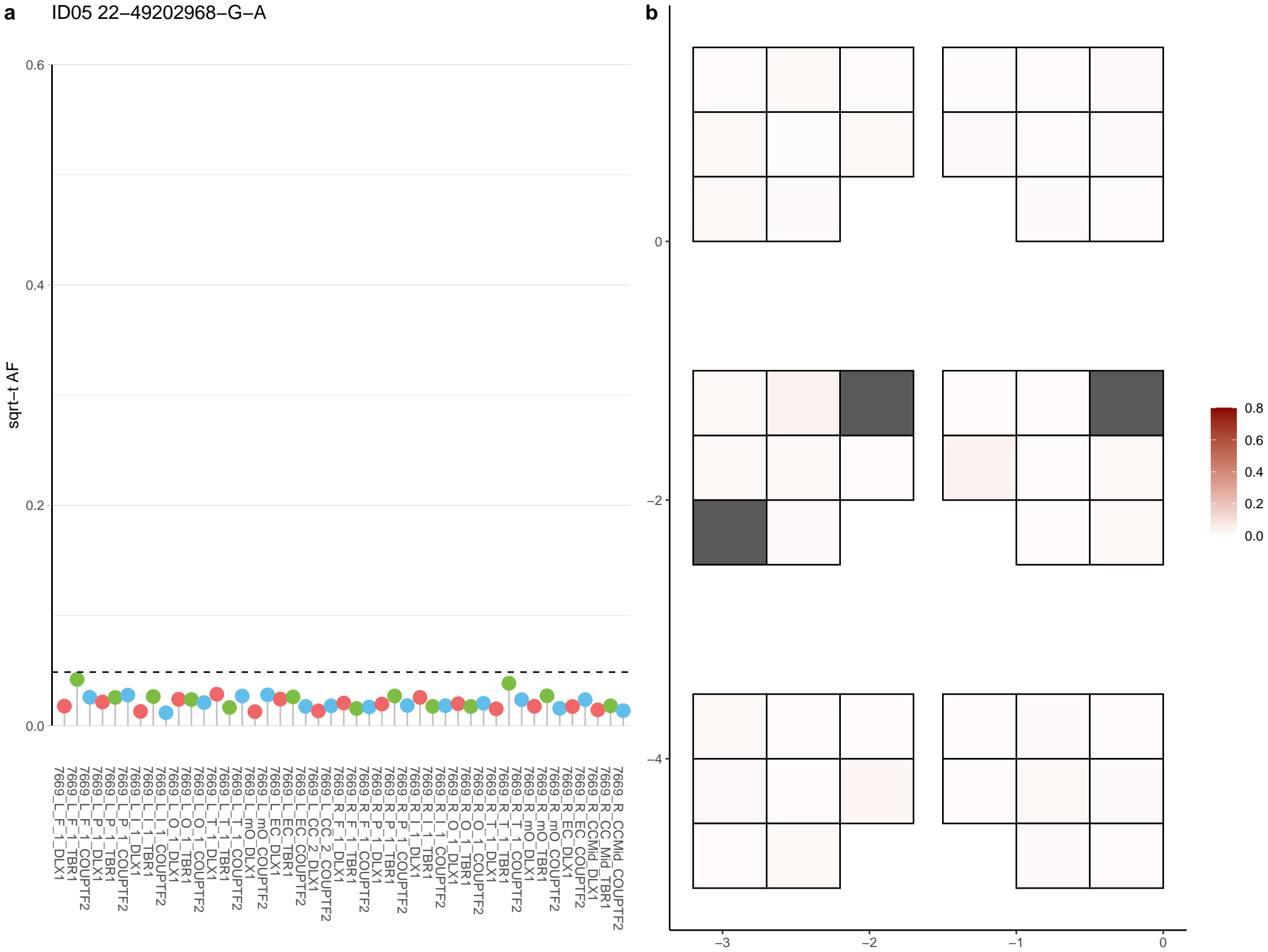

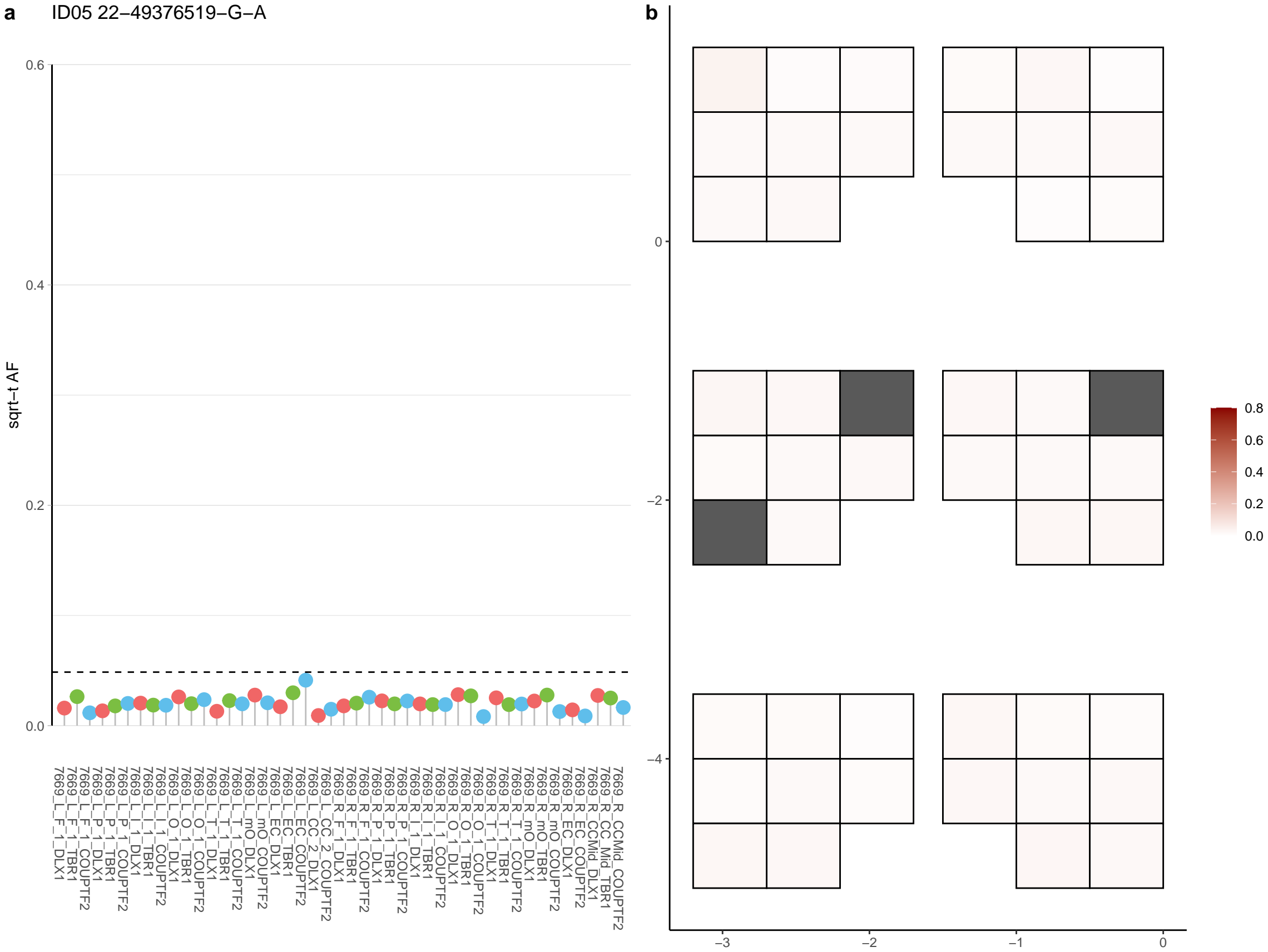

**a** ID05 22-49717540-G-A

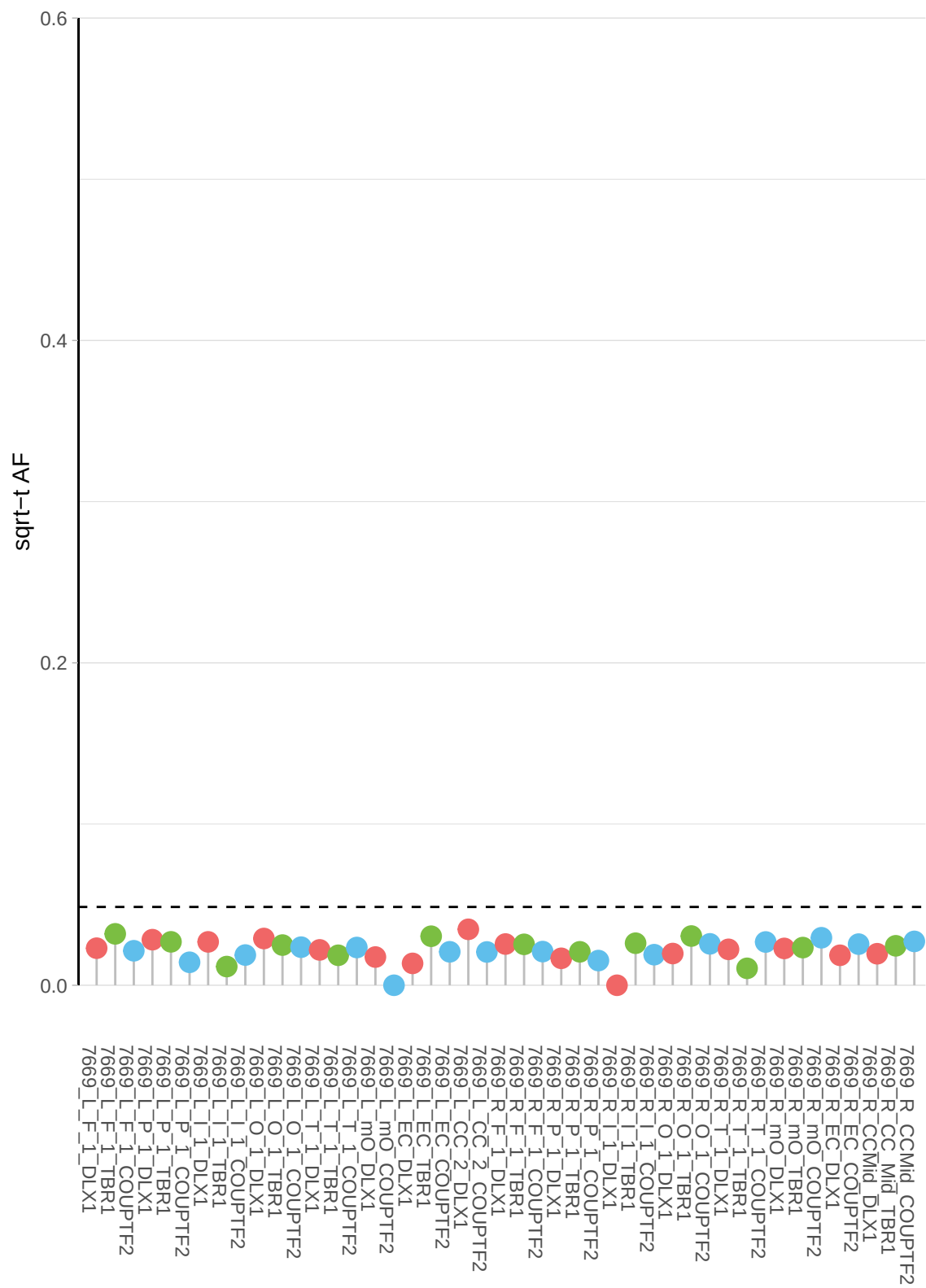

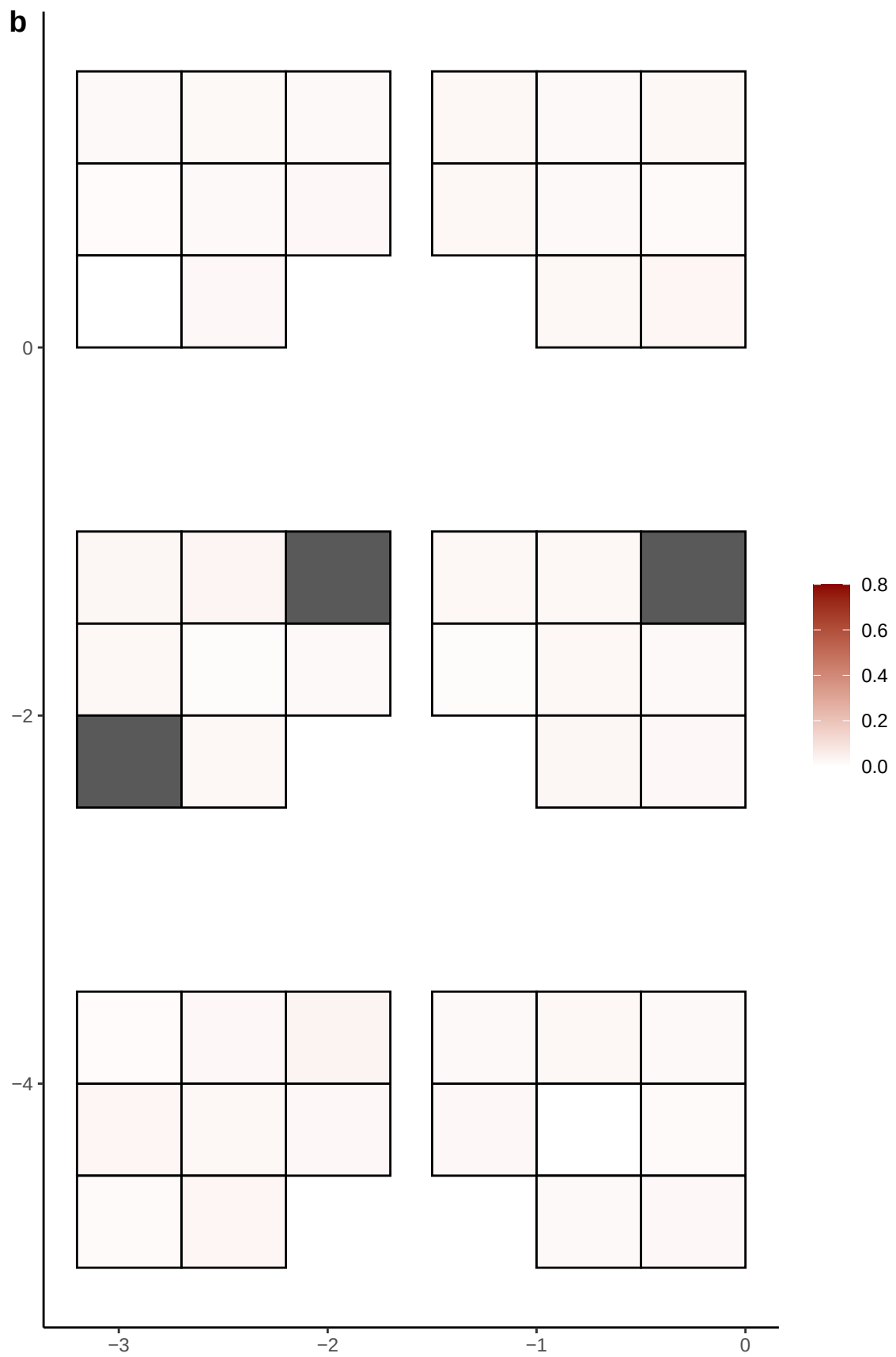

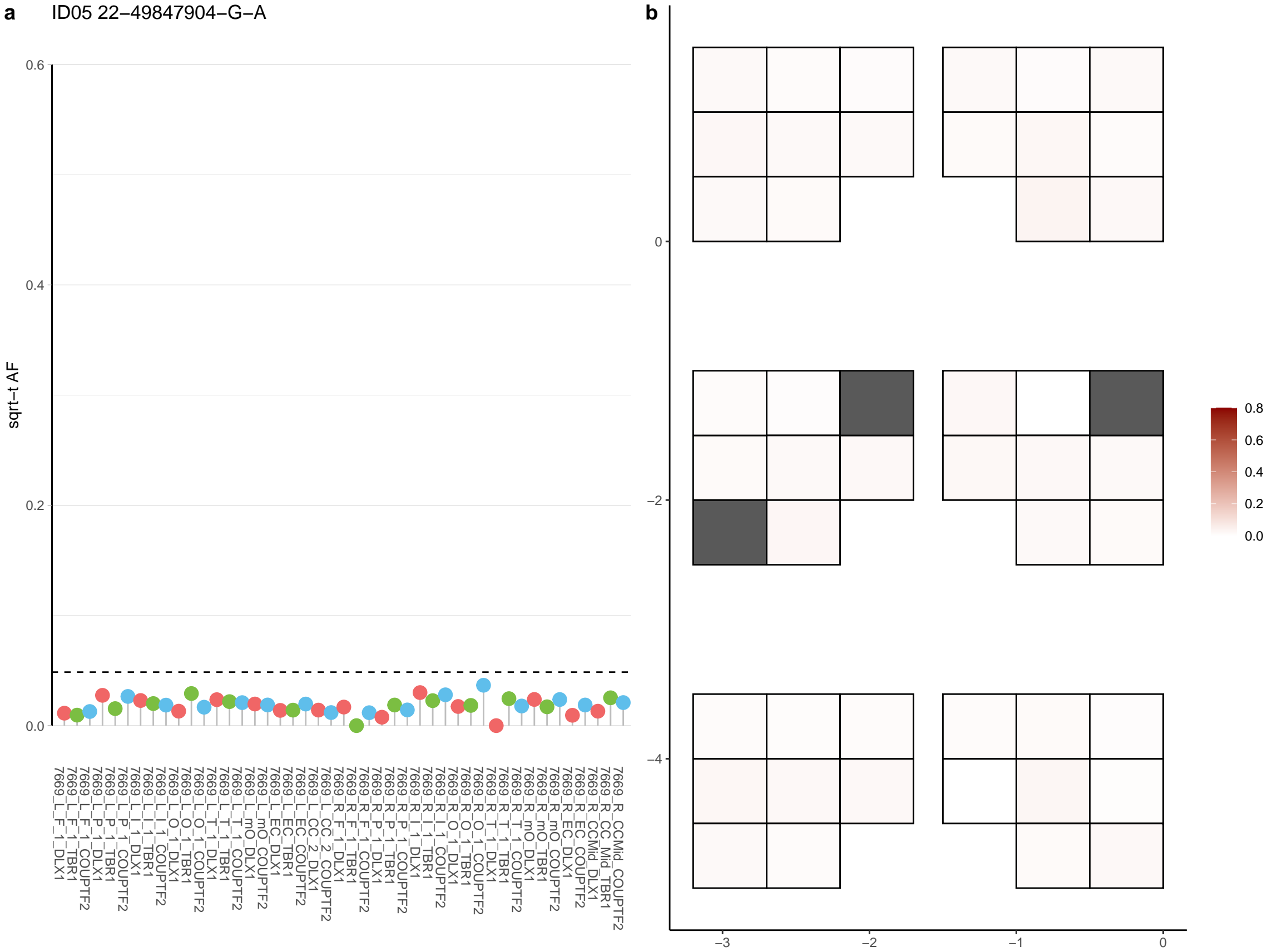

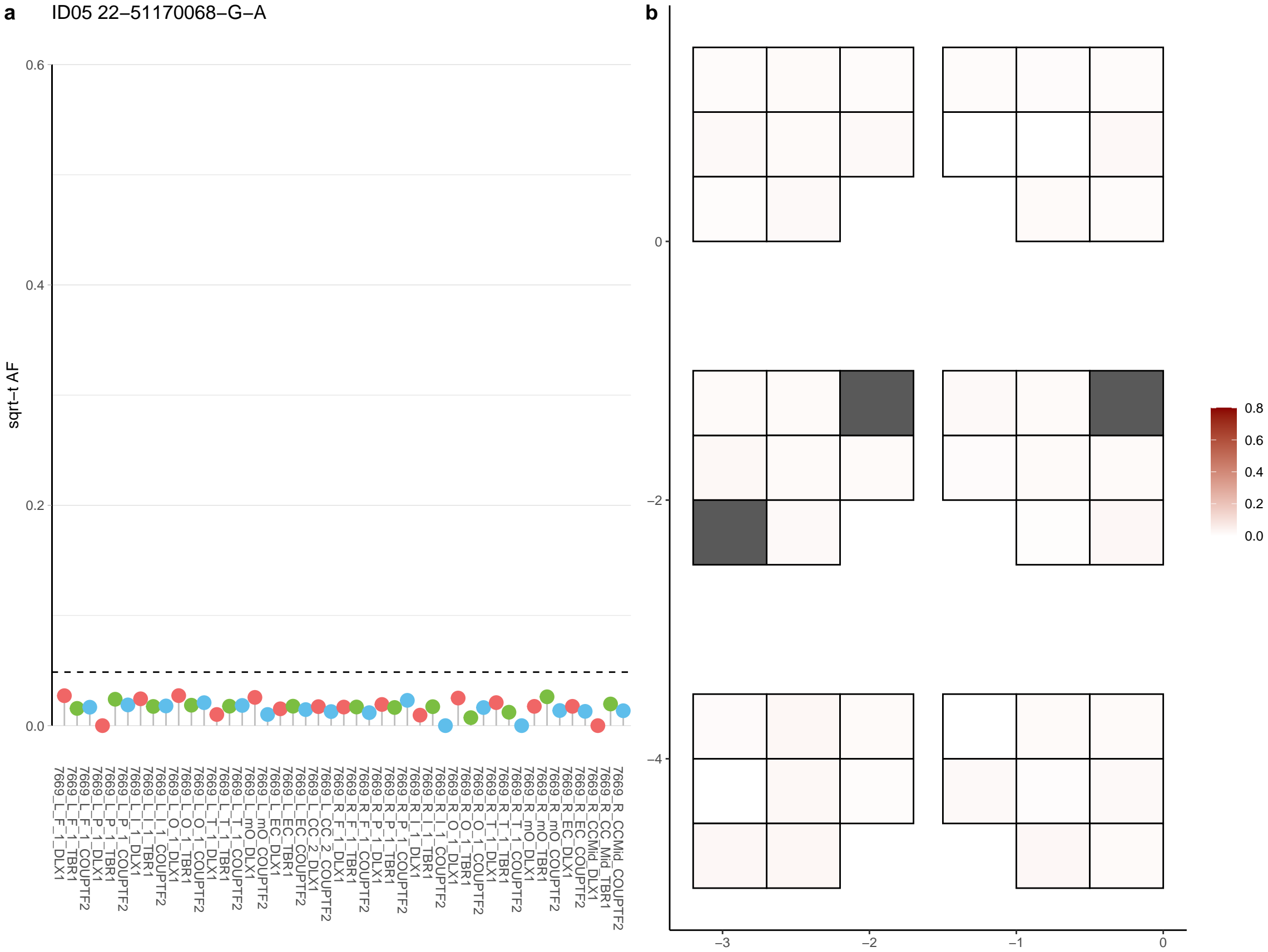

**a** ID05 3-101156372-T-C

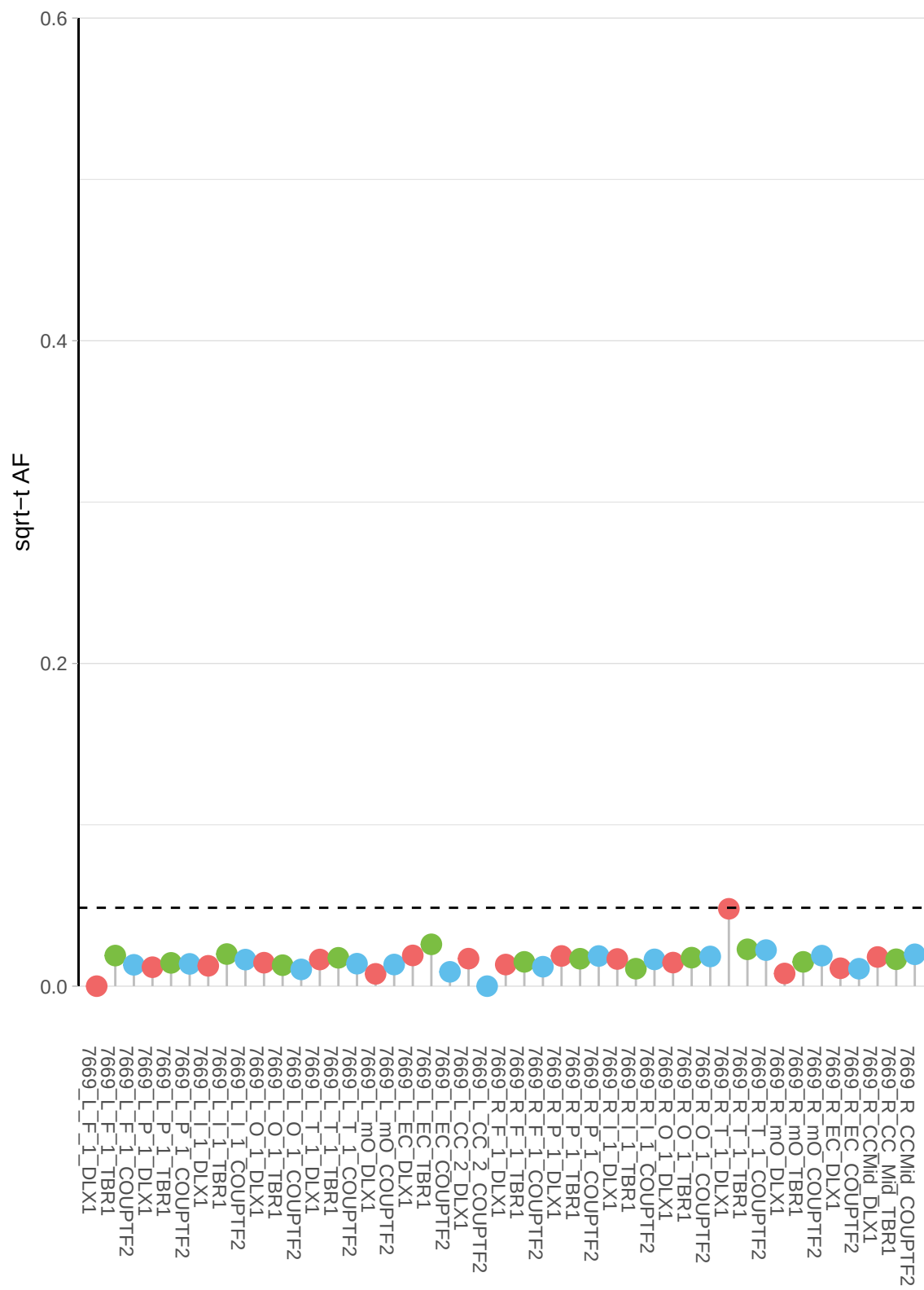

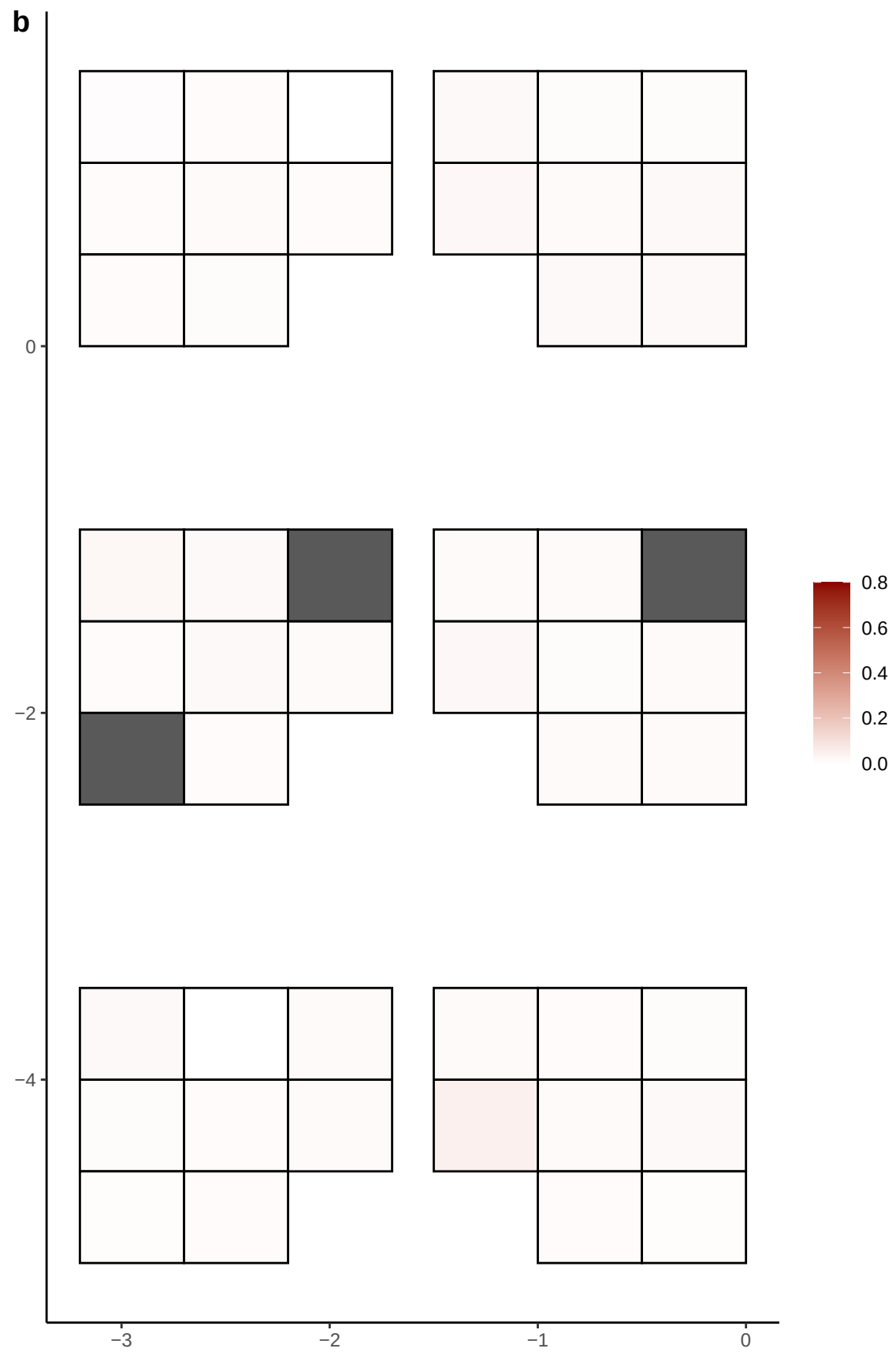

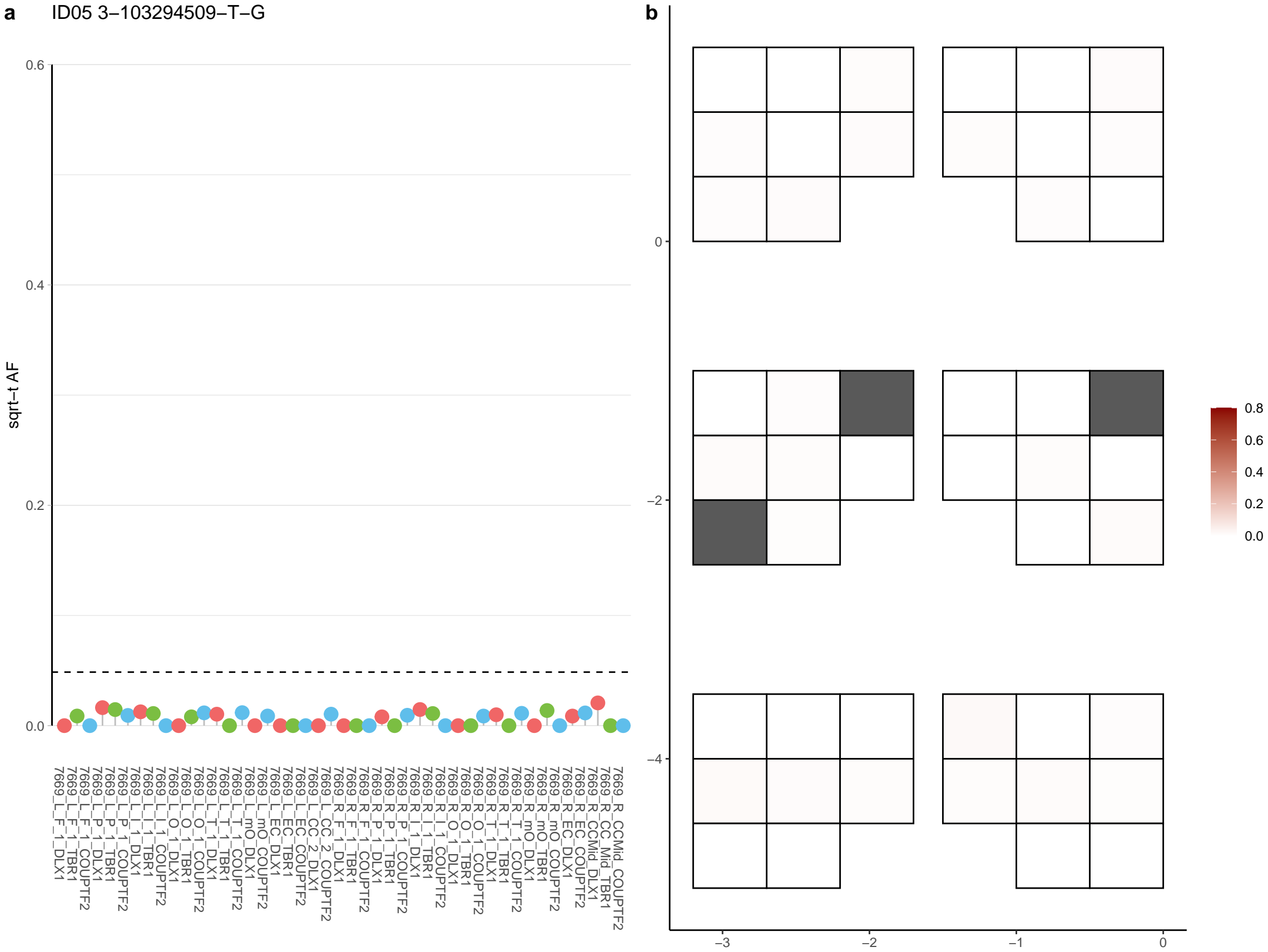

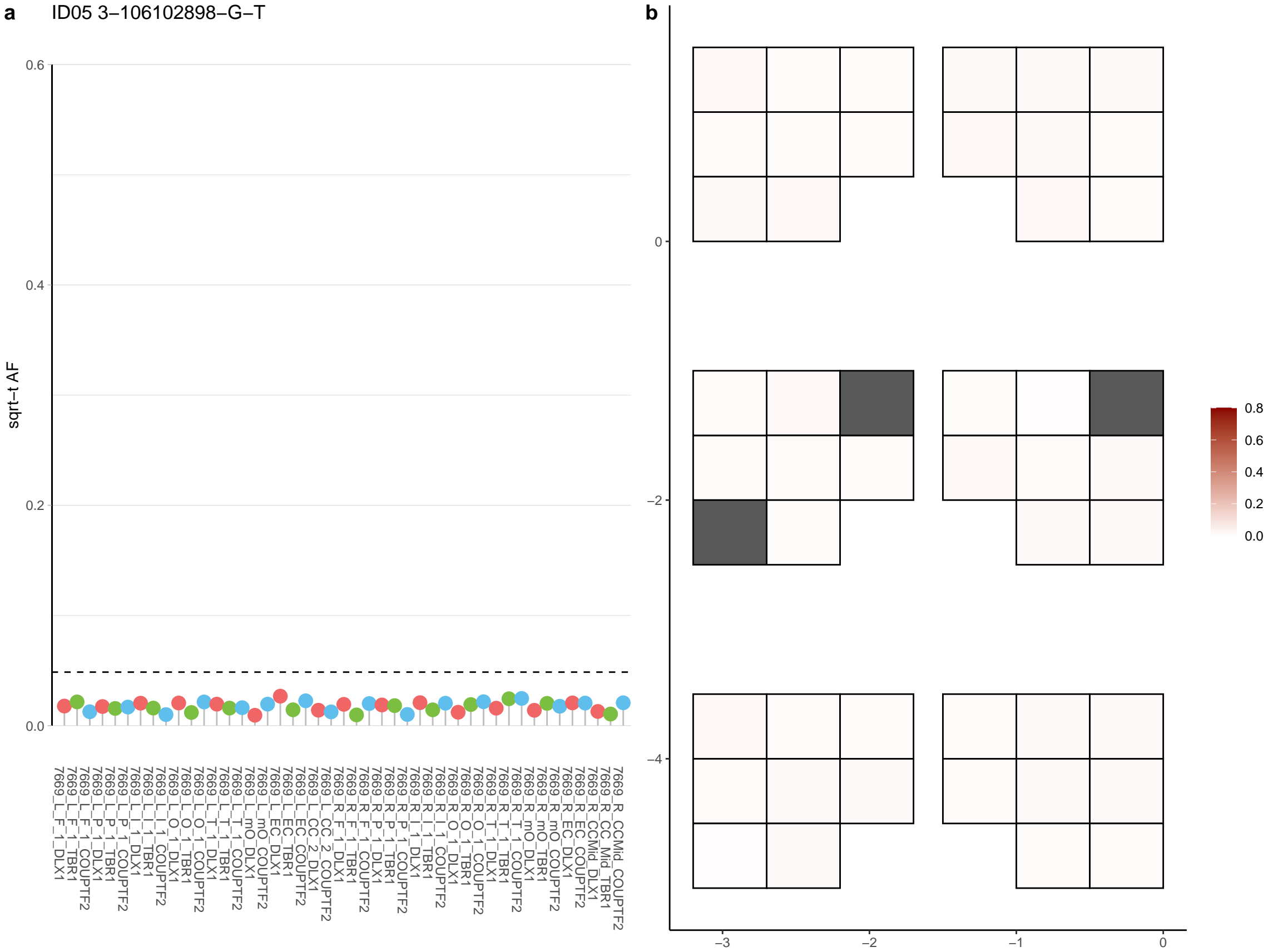

**a** ID05 3-107821326-C-T

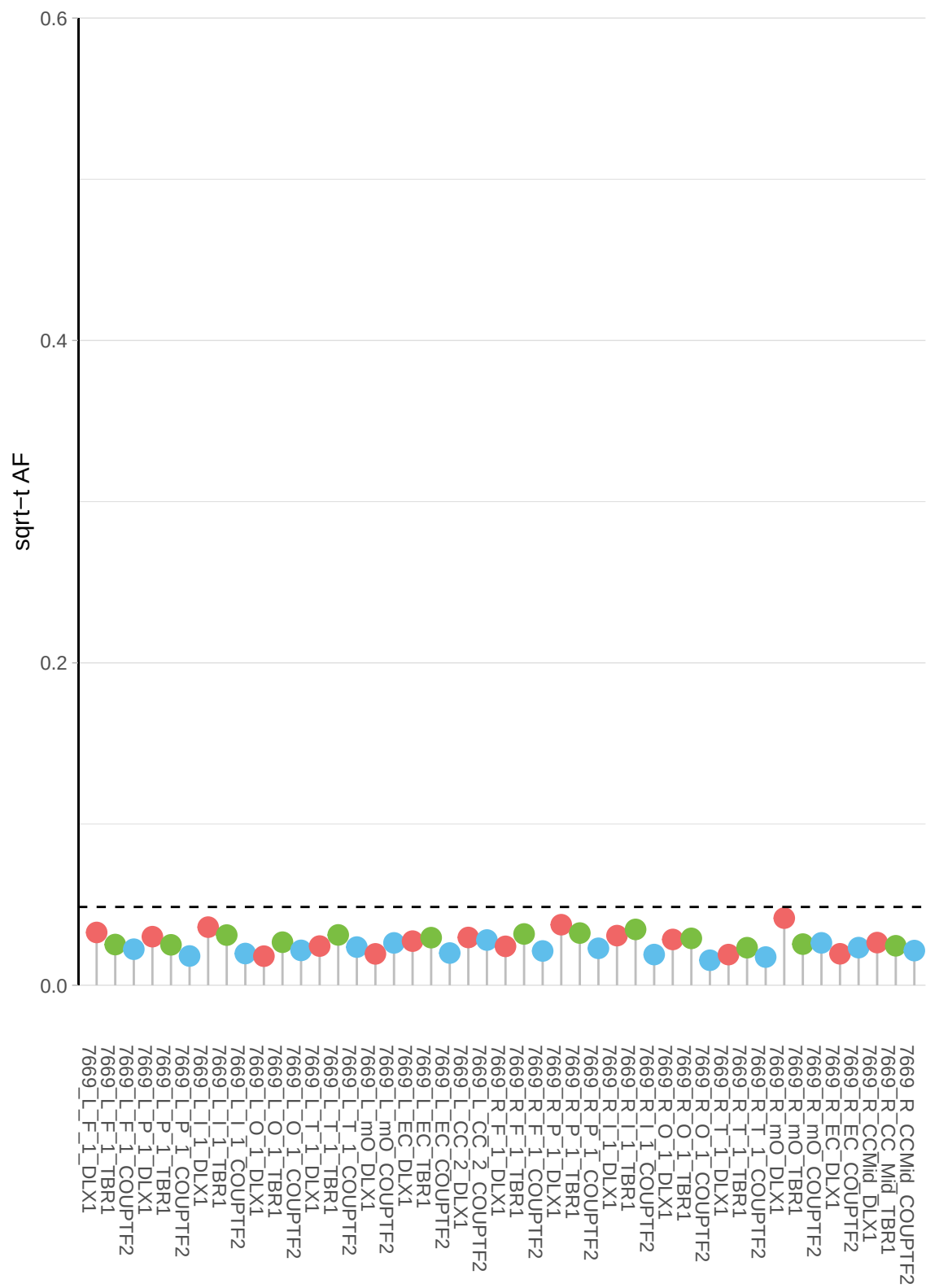

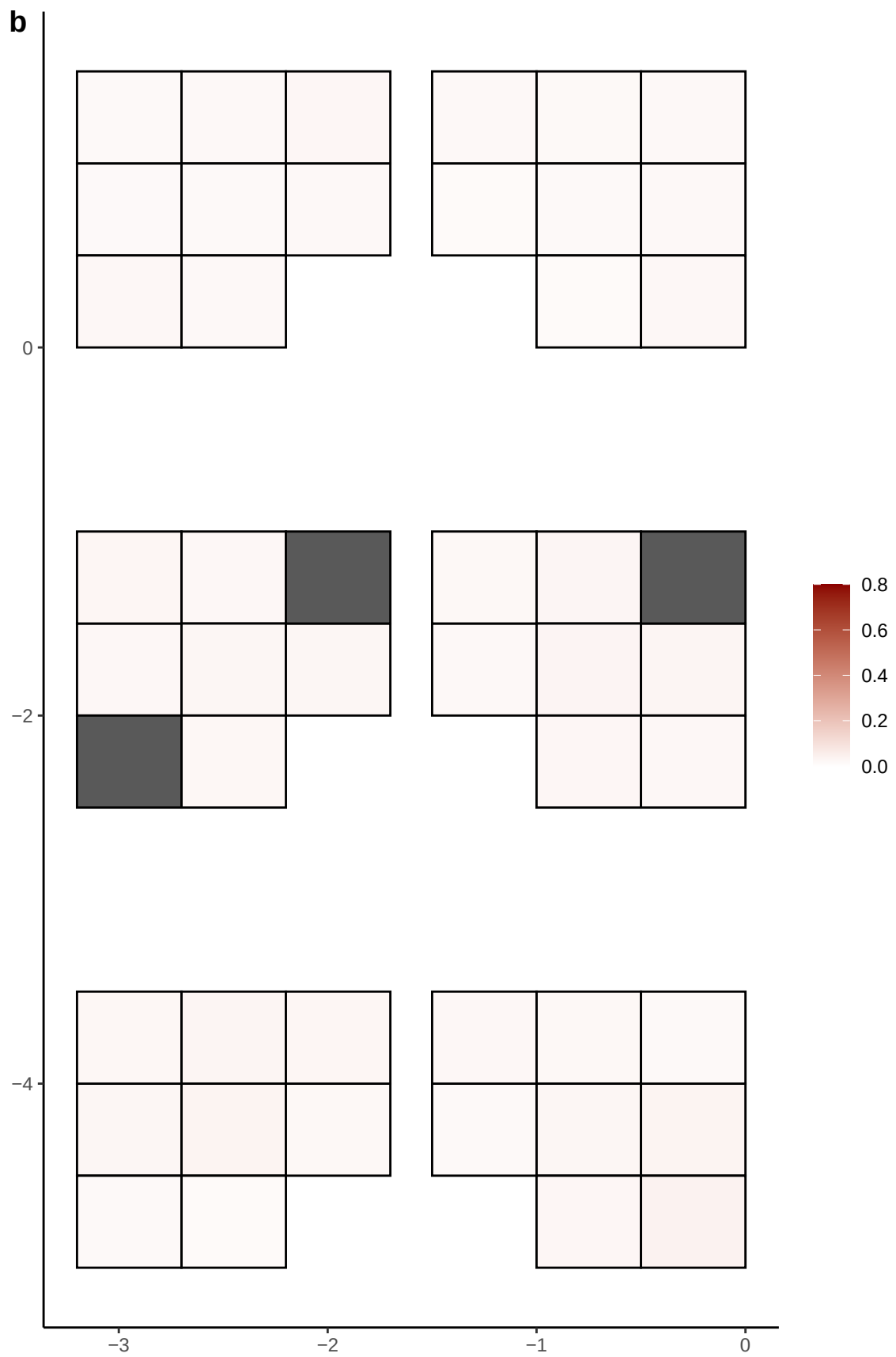

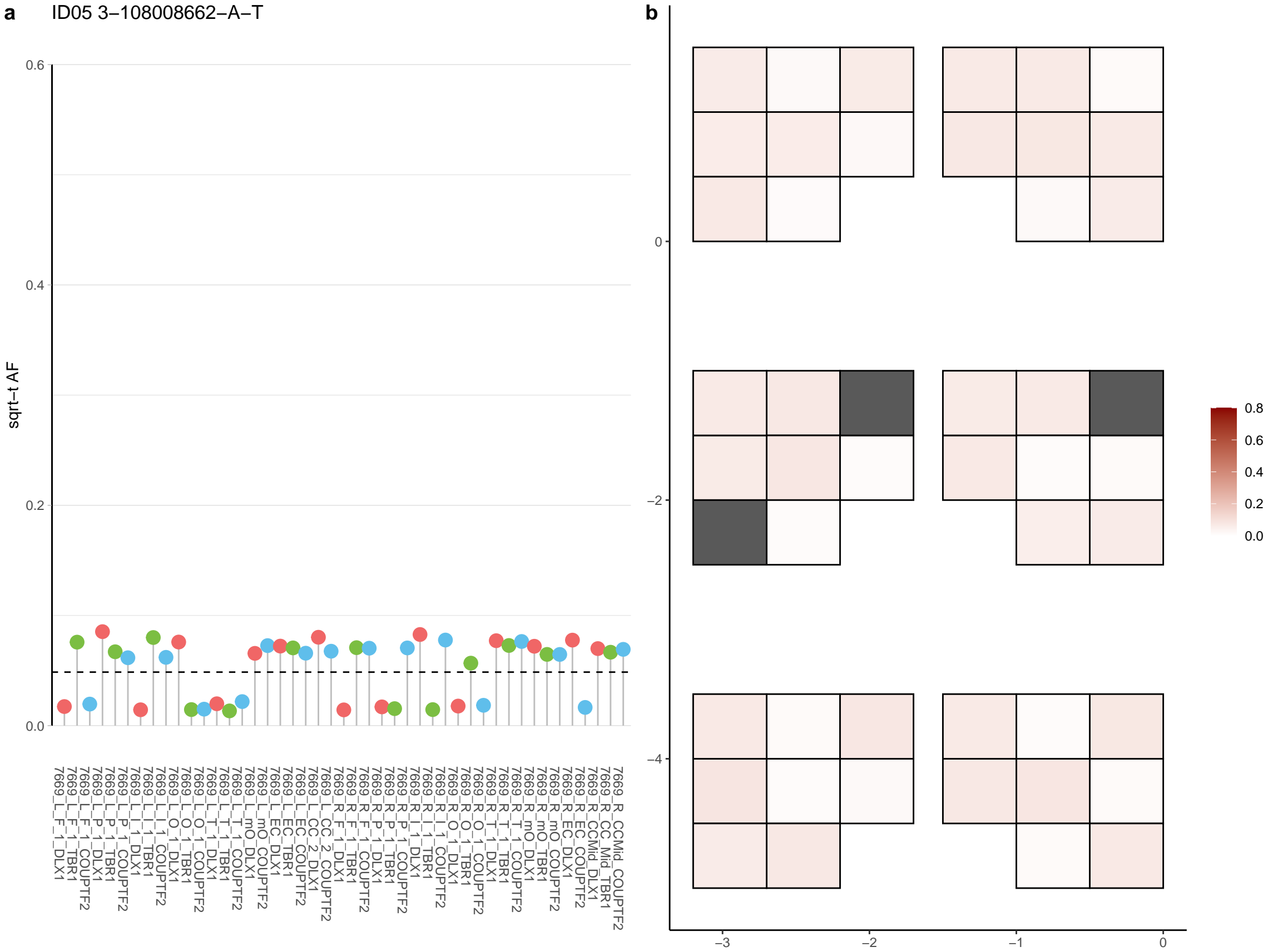

**a** ID05 3-109900234-C-T

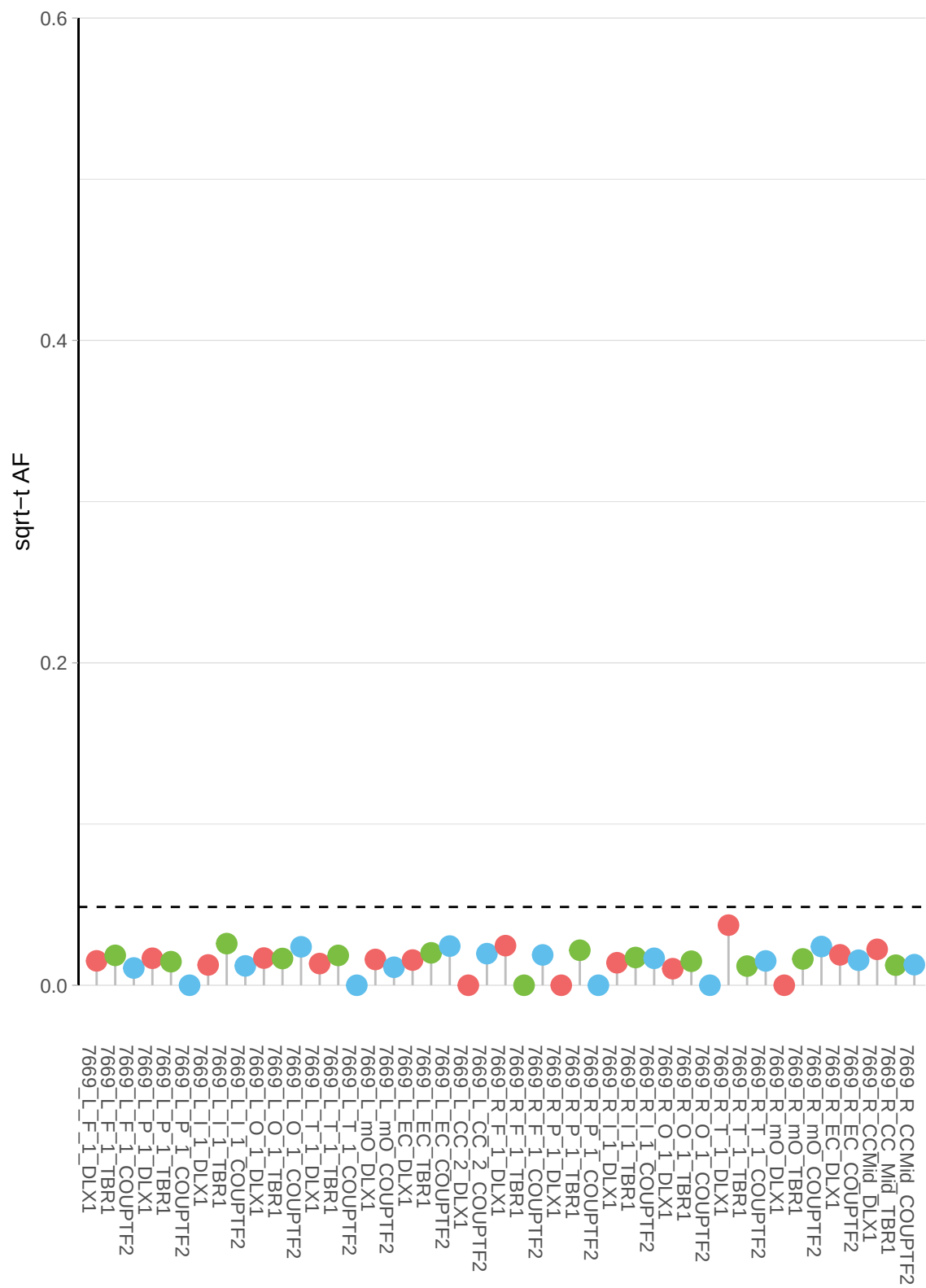

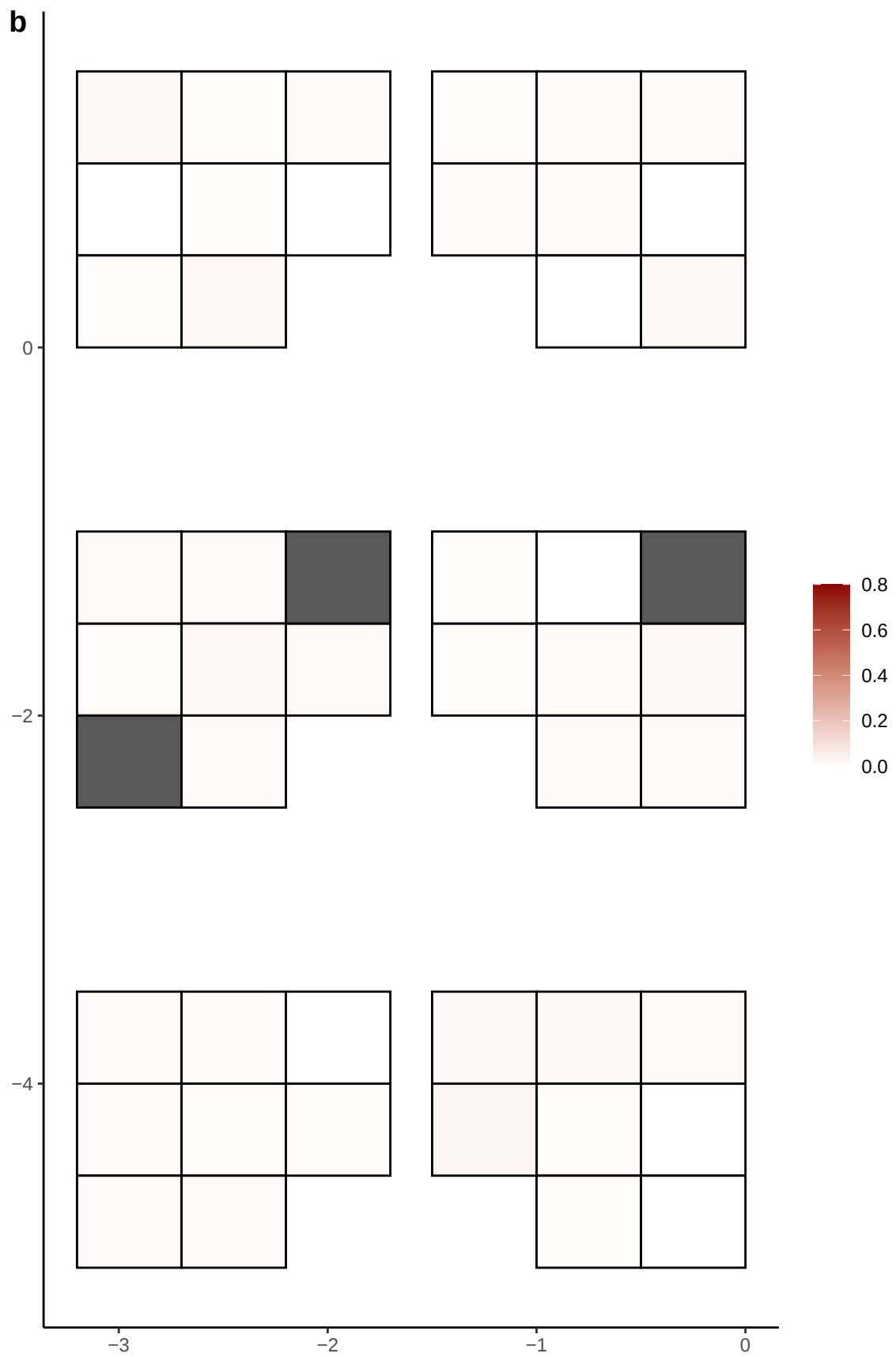

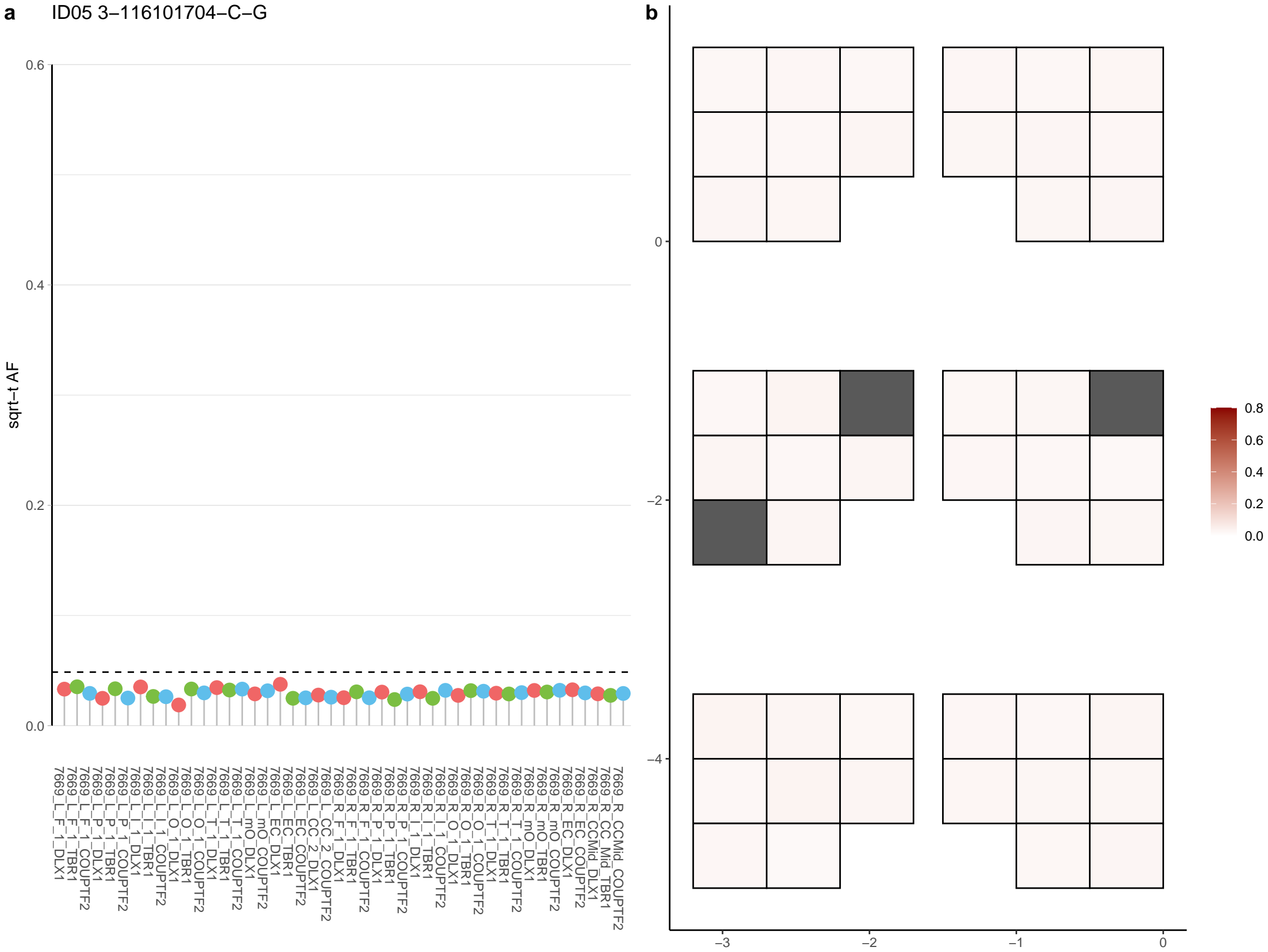

**a** ID05 3-116224135-C-T

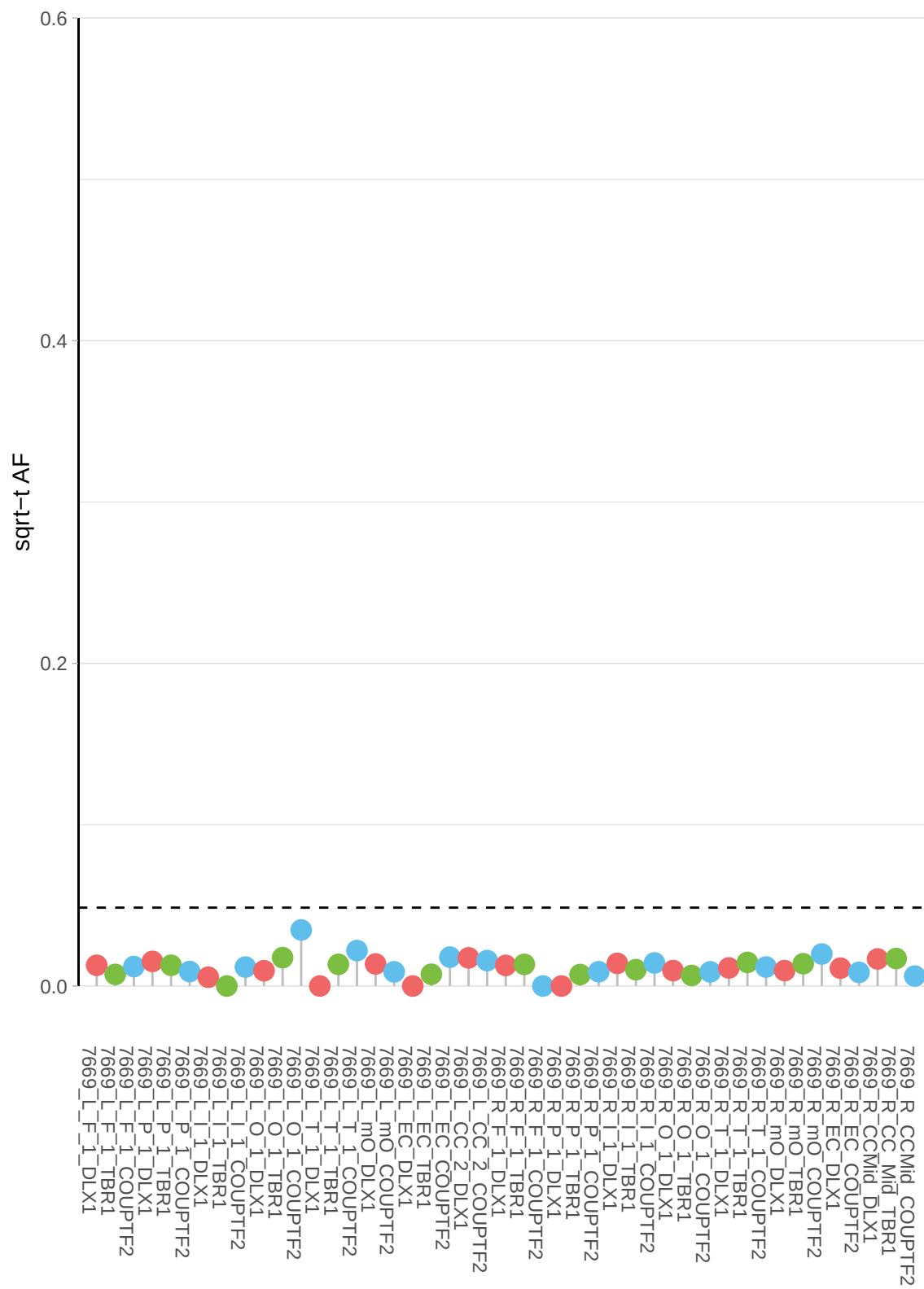

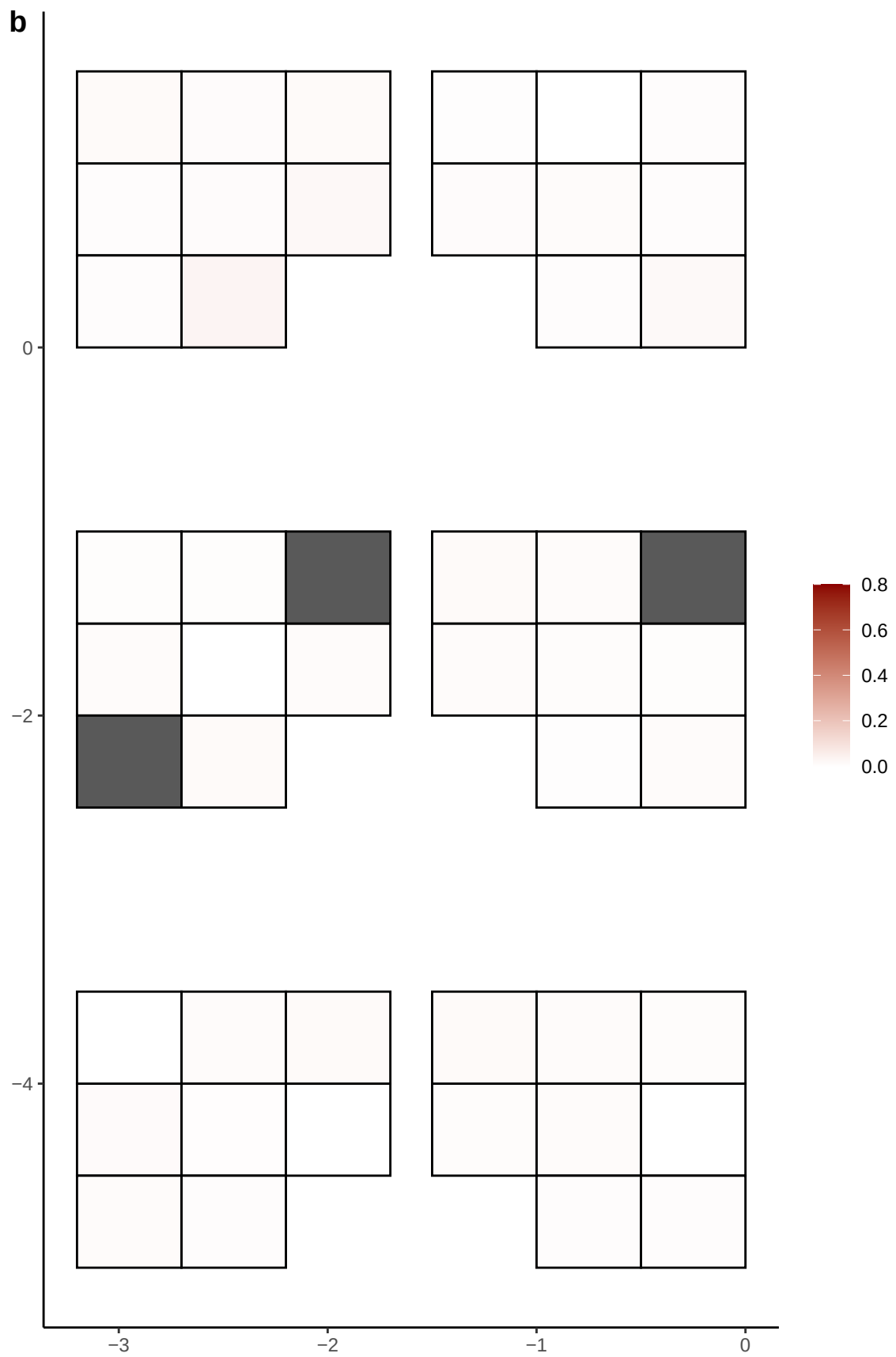

**a**

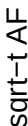

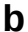

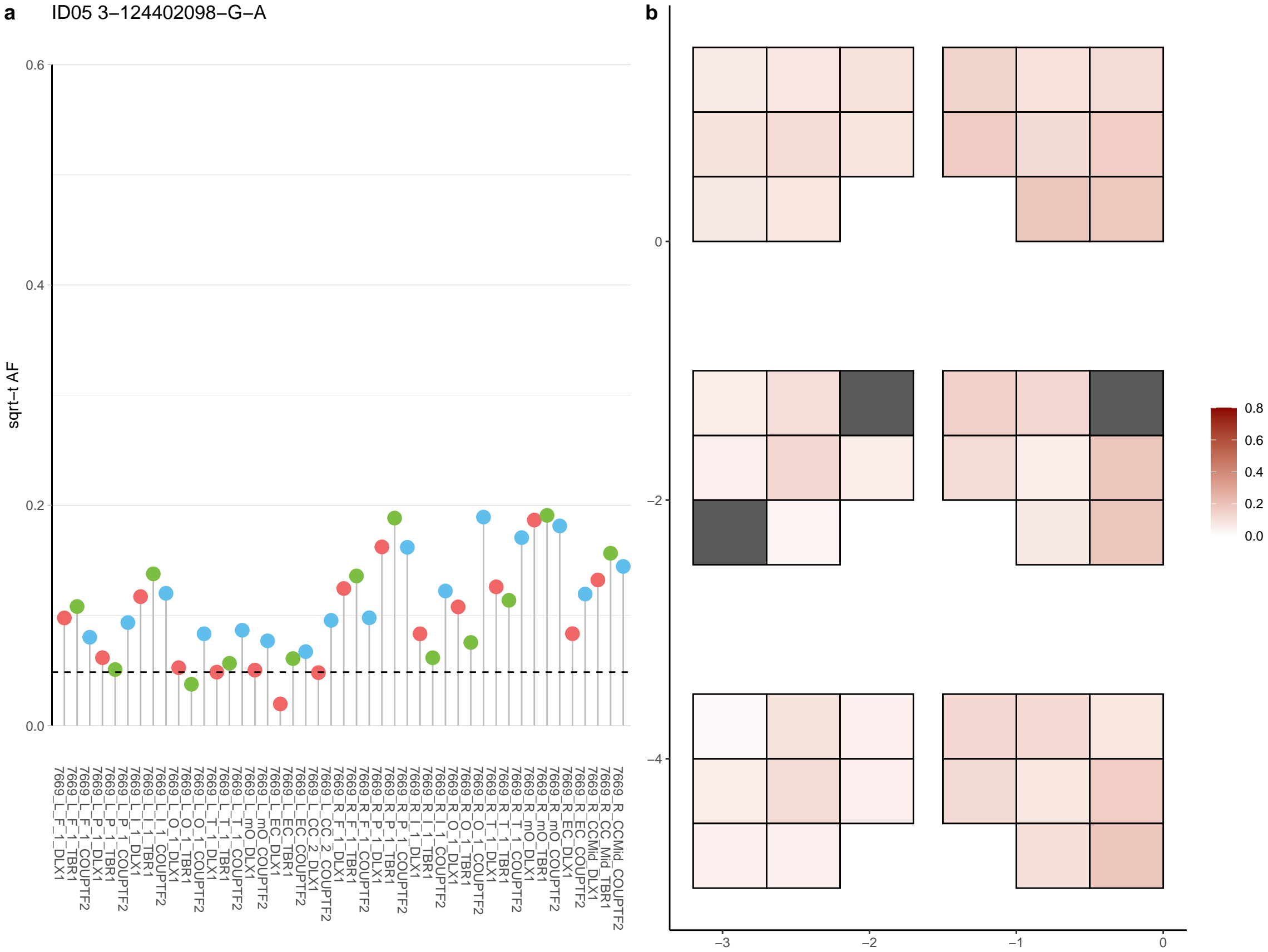

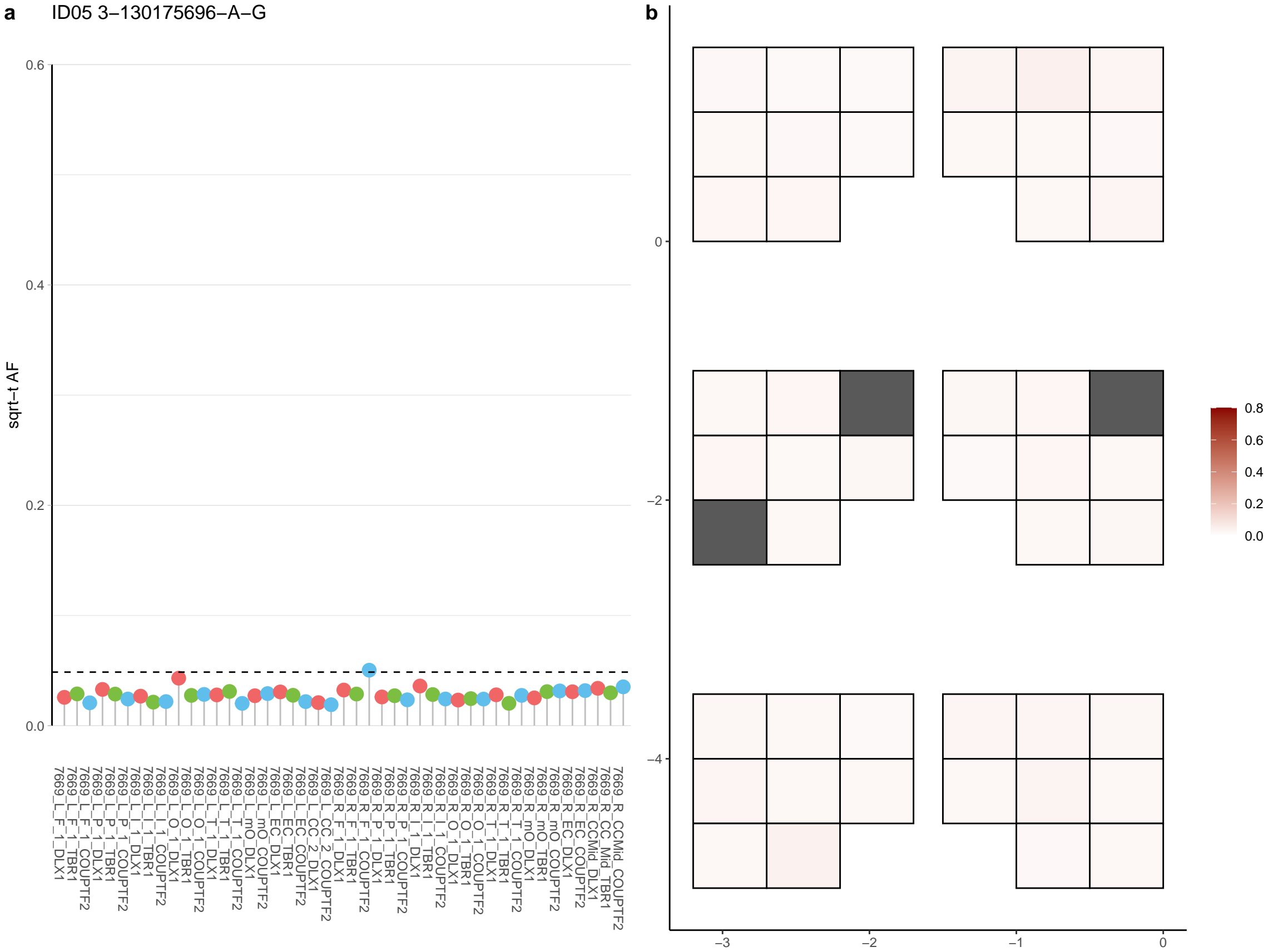

**a** ID05 3-137569791-G-A

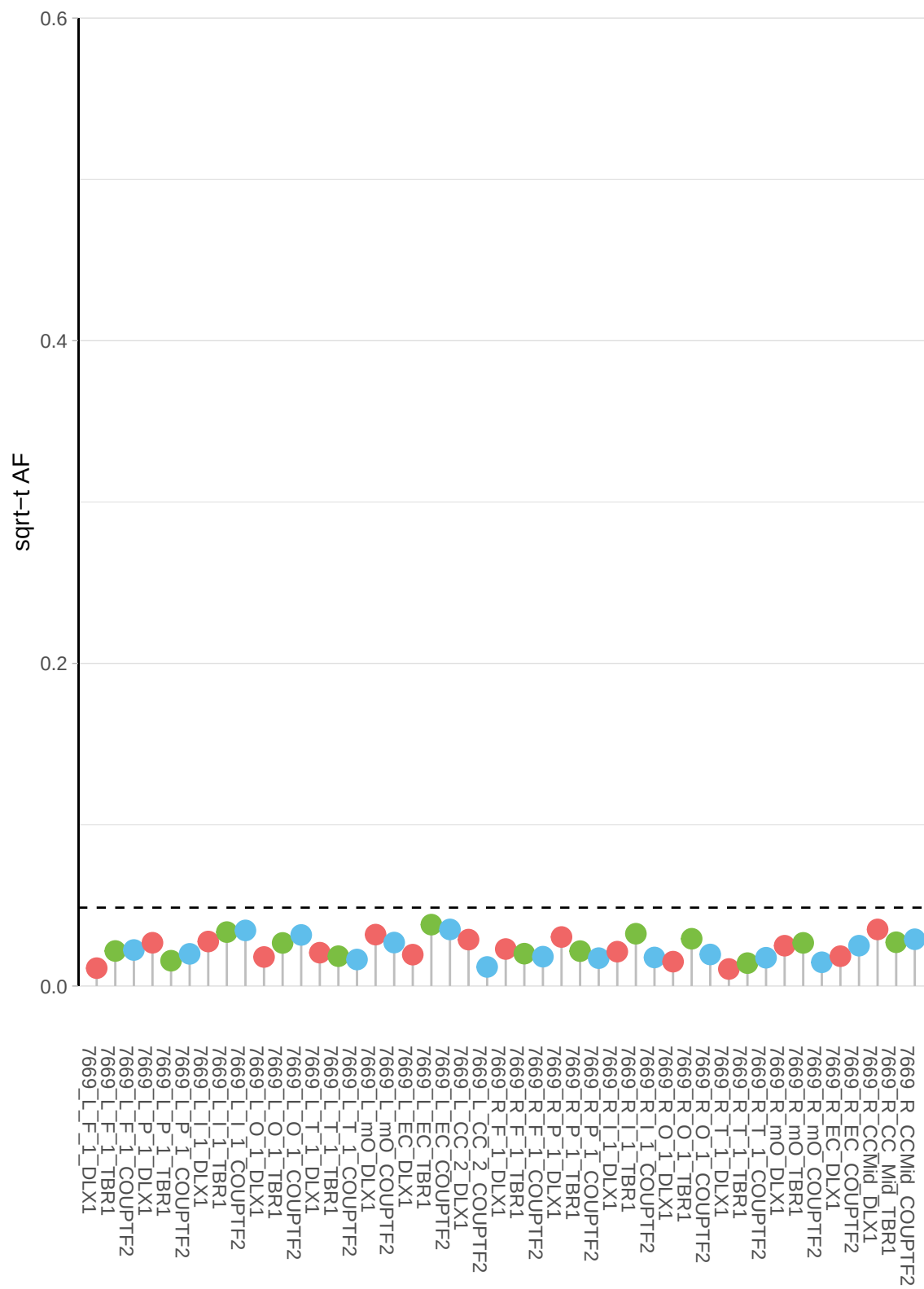

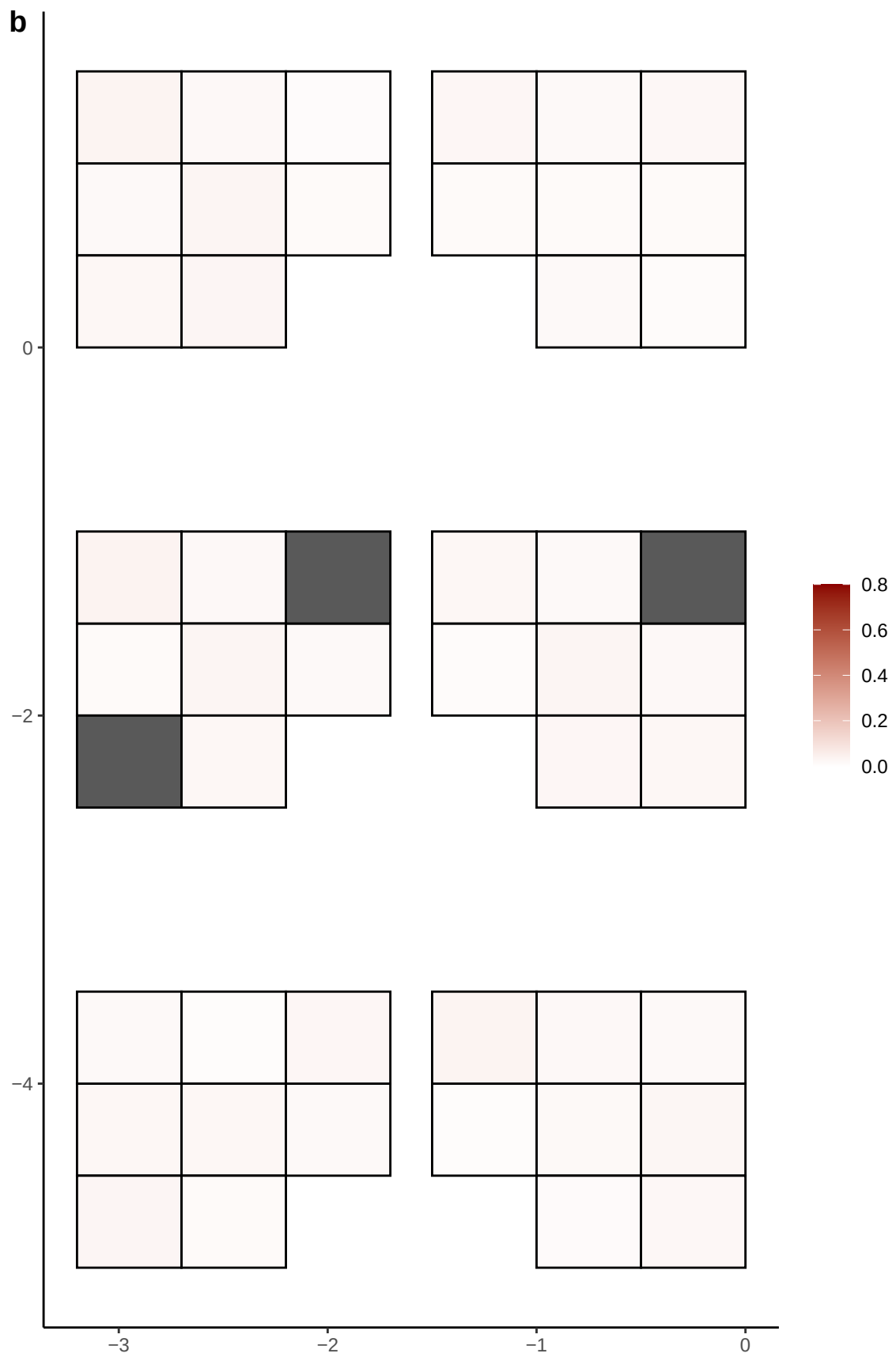

**a** ID05 3-145391171-A-G

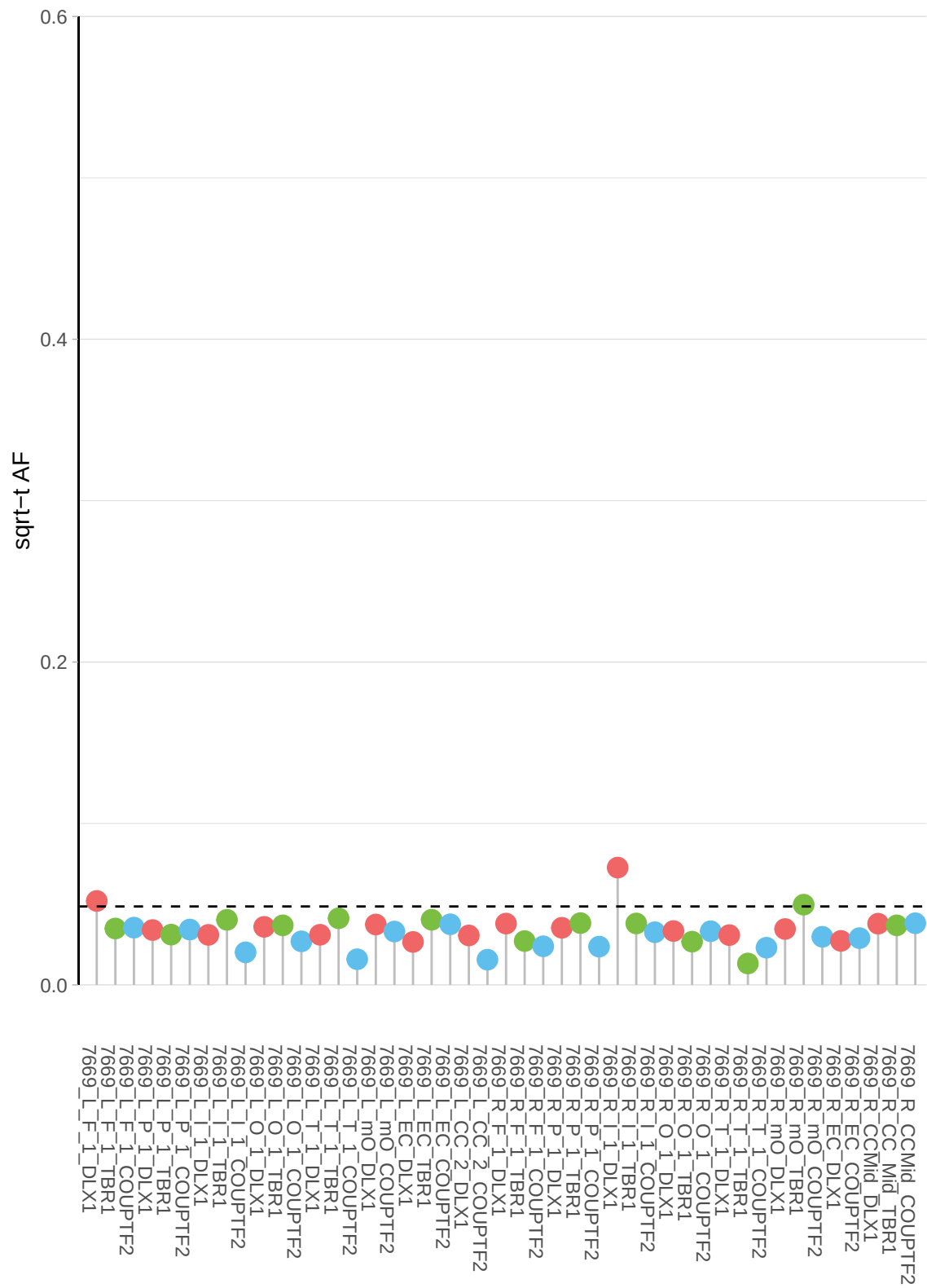

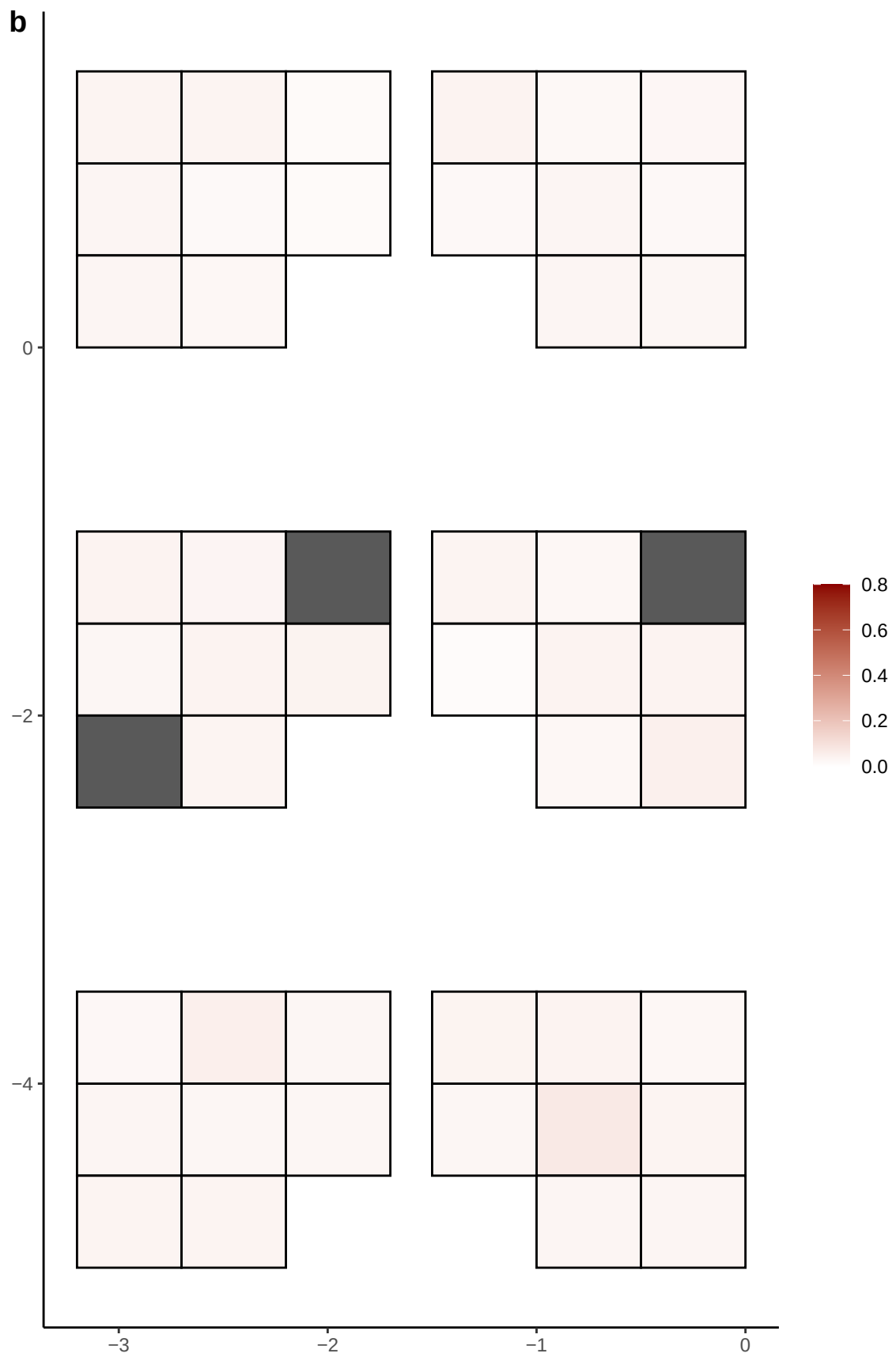

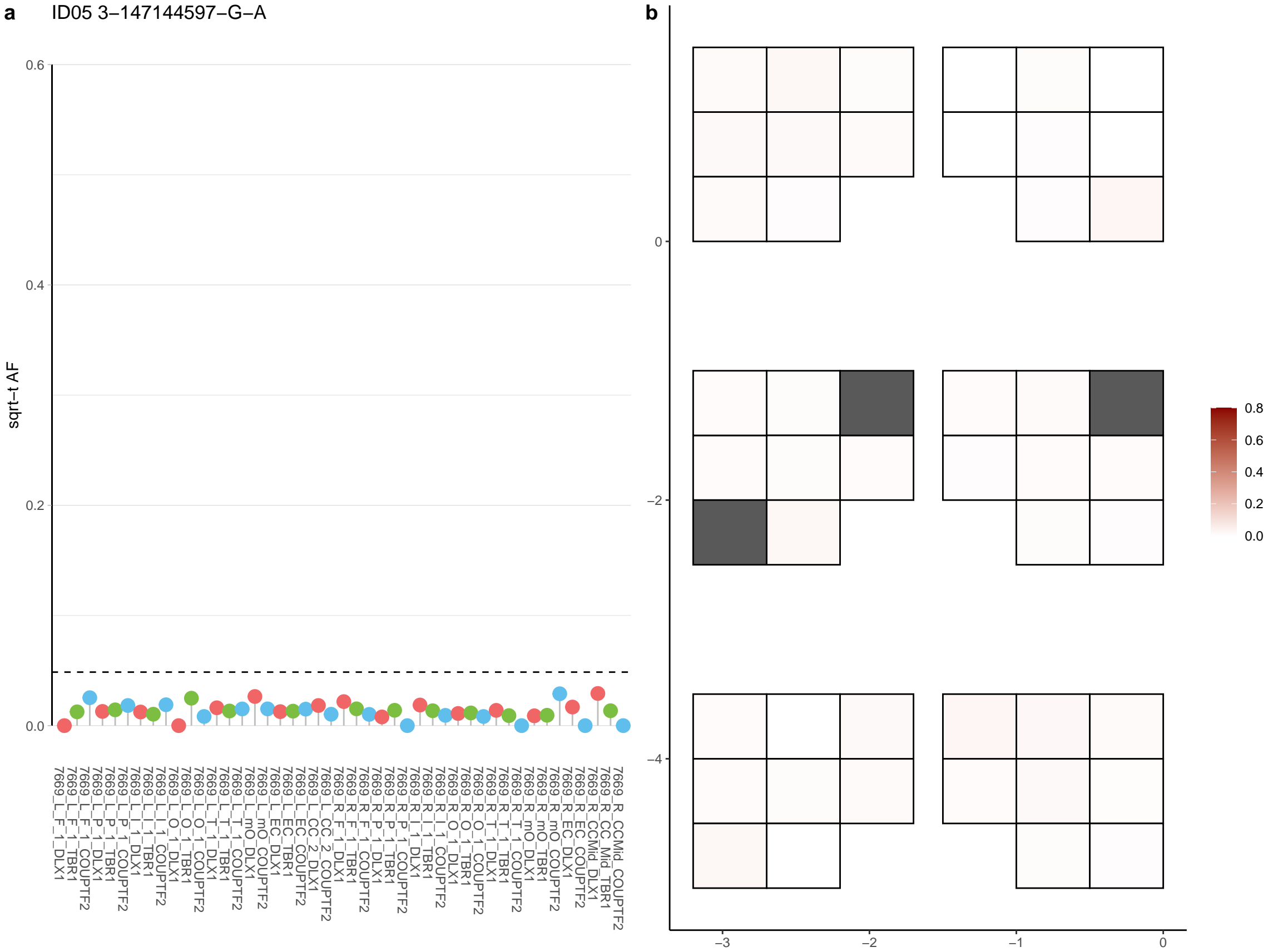

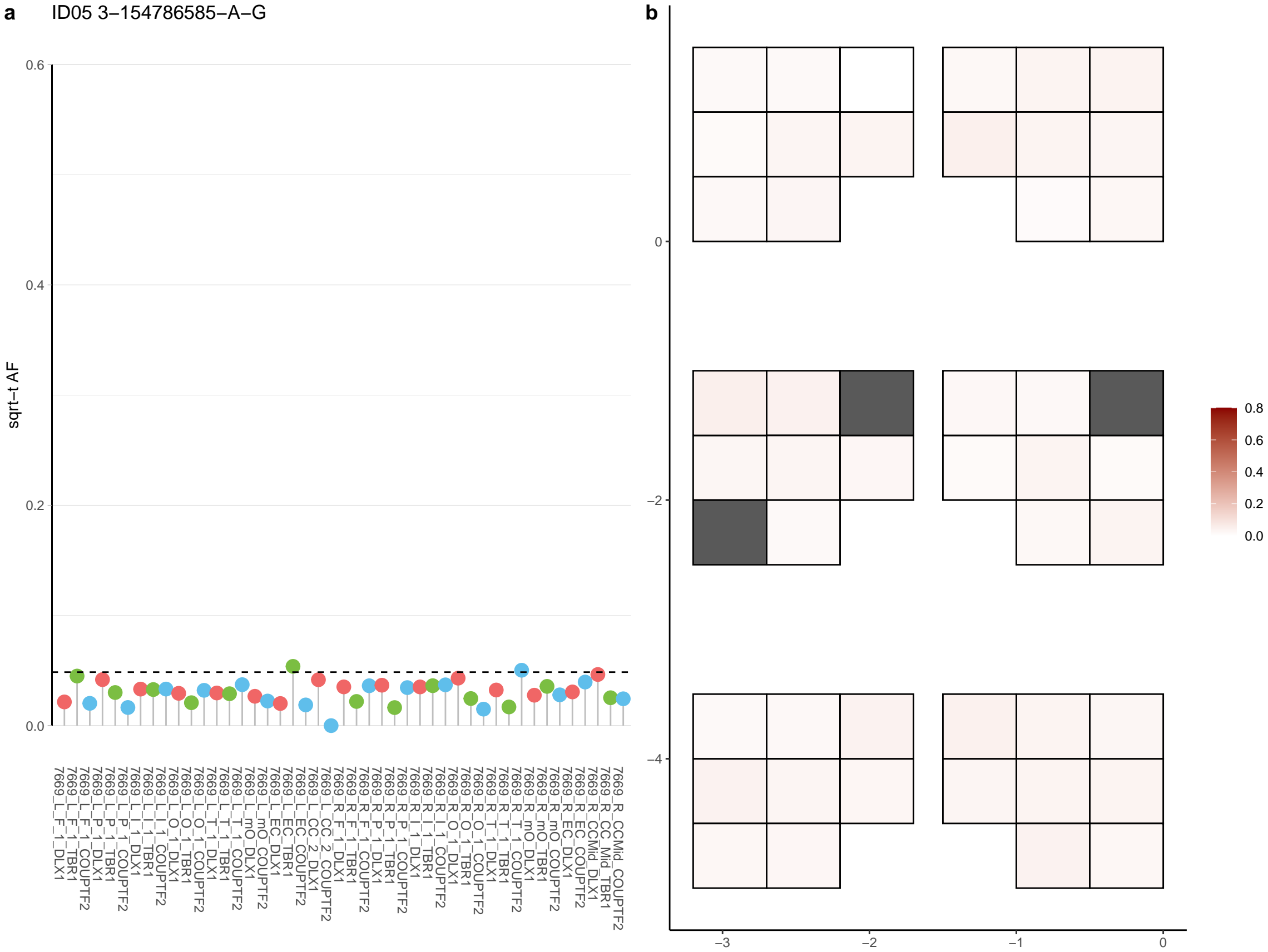

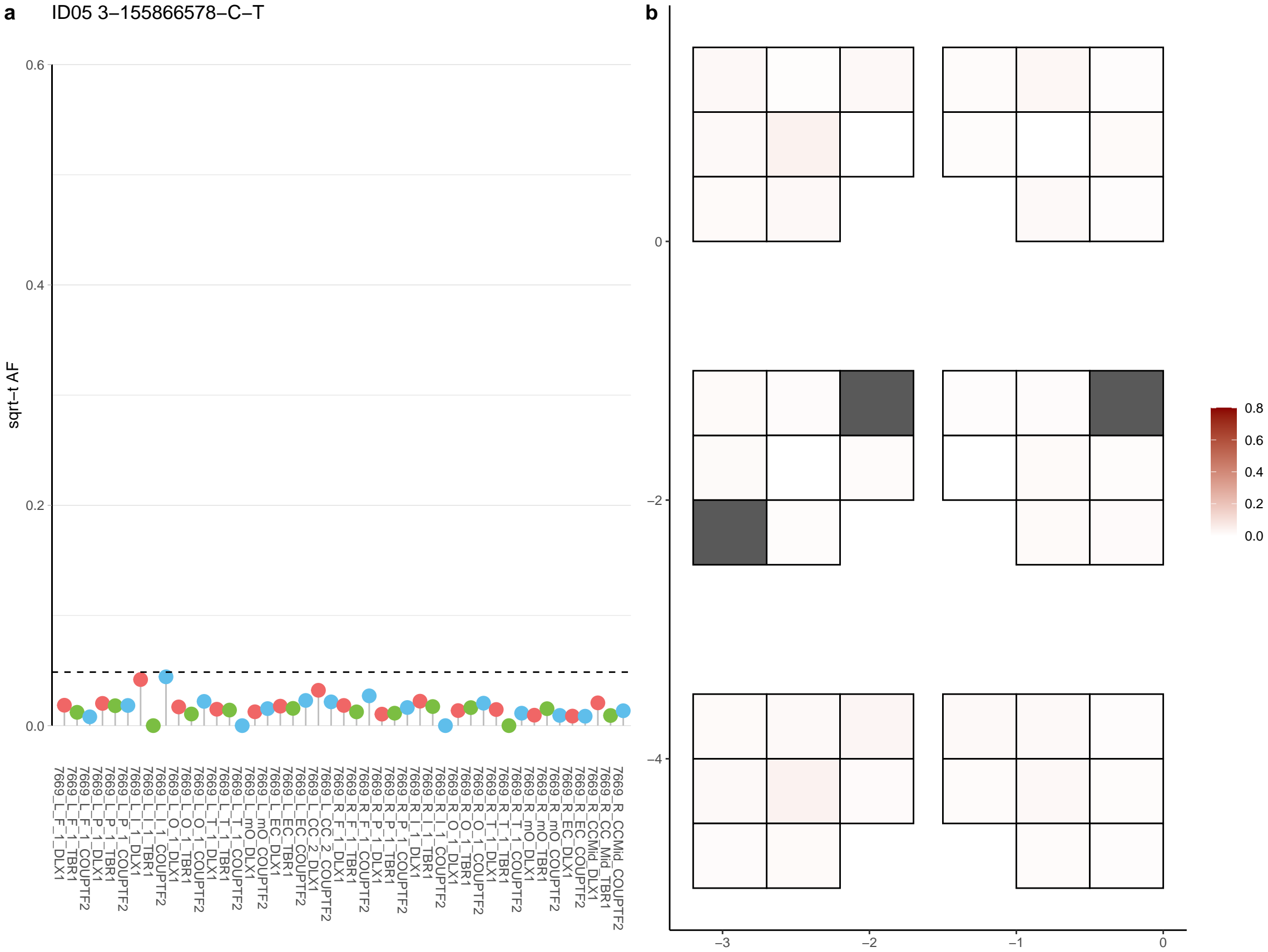

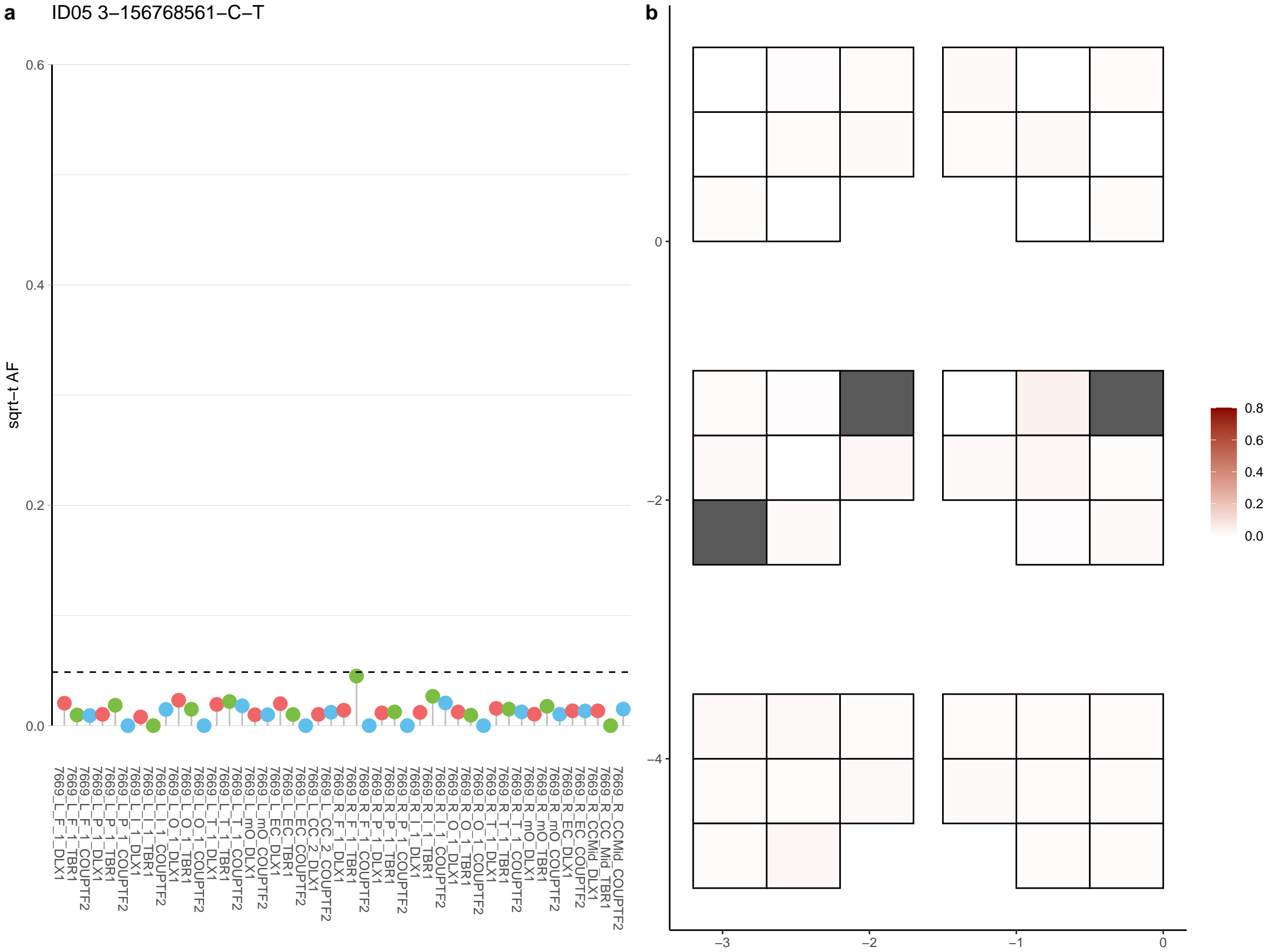

**a** ID05 3-158256477-C-T

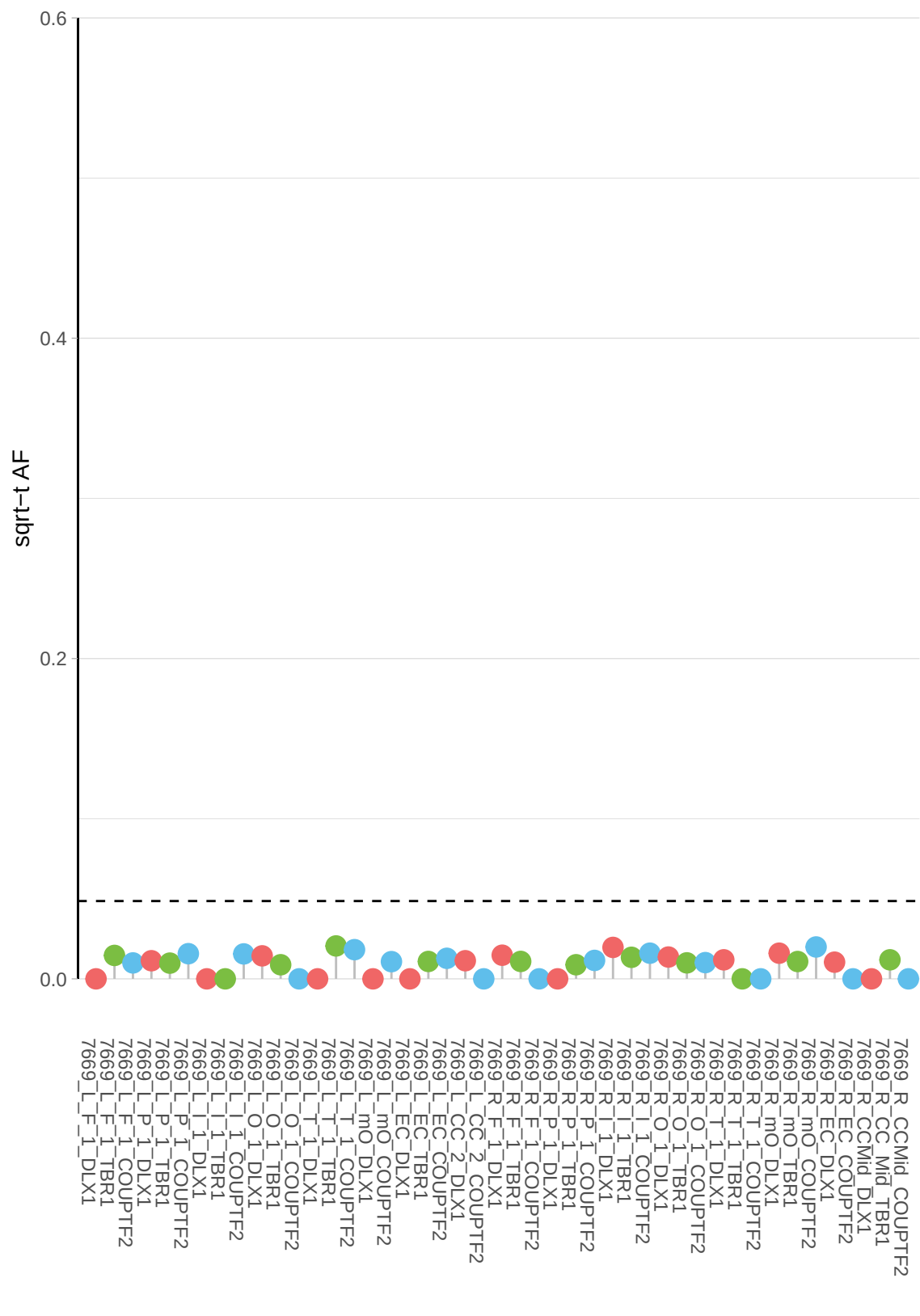

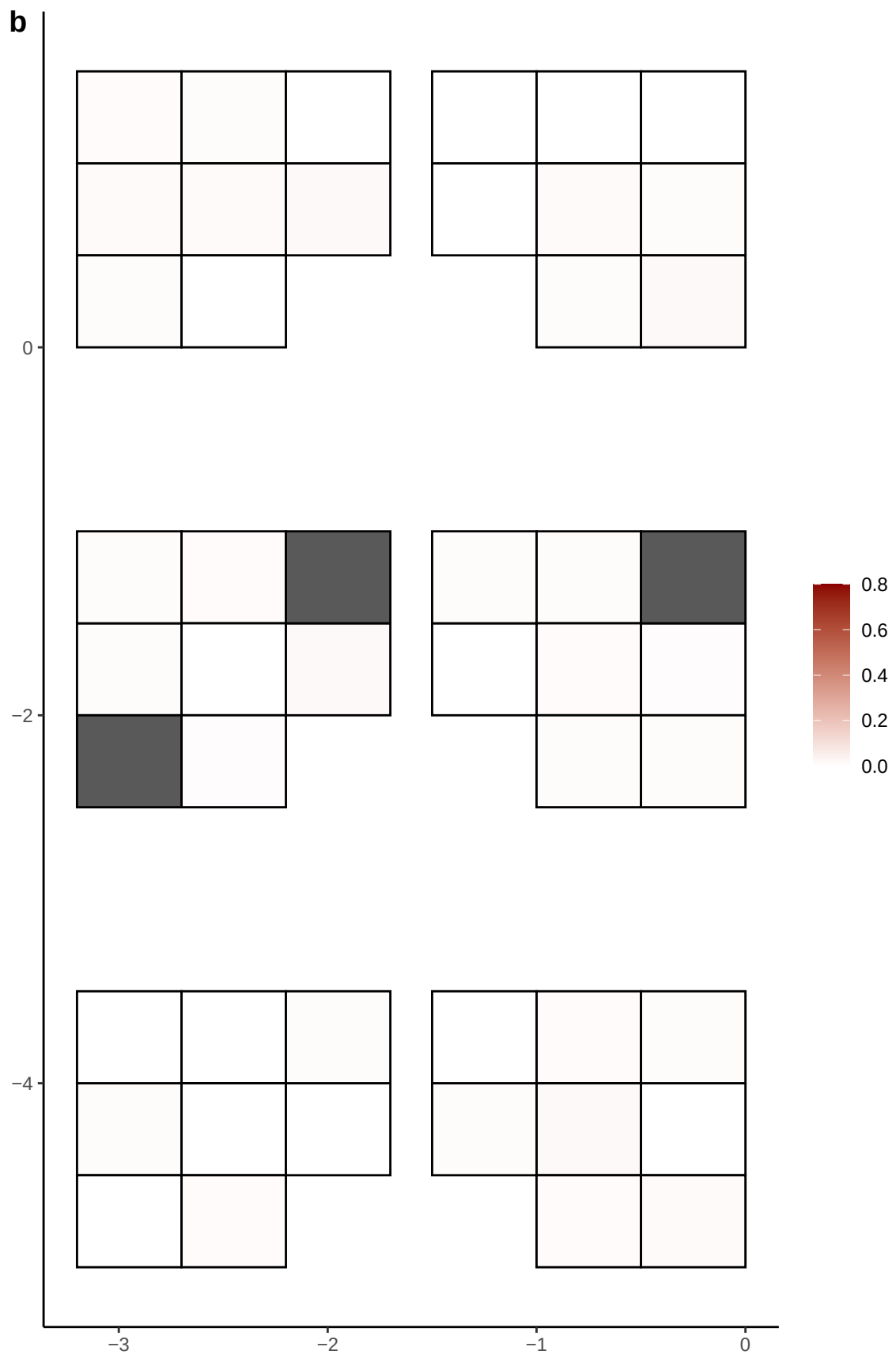

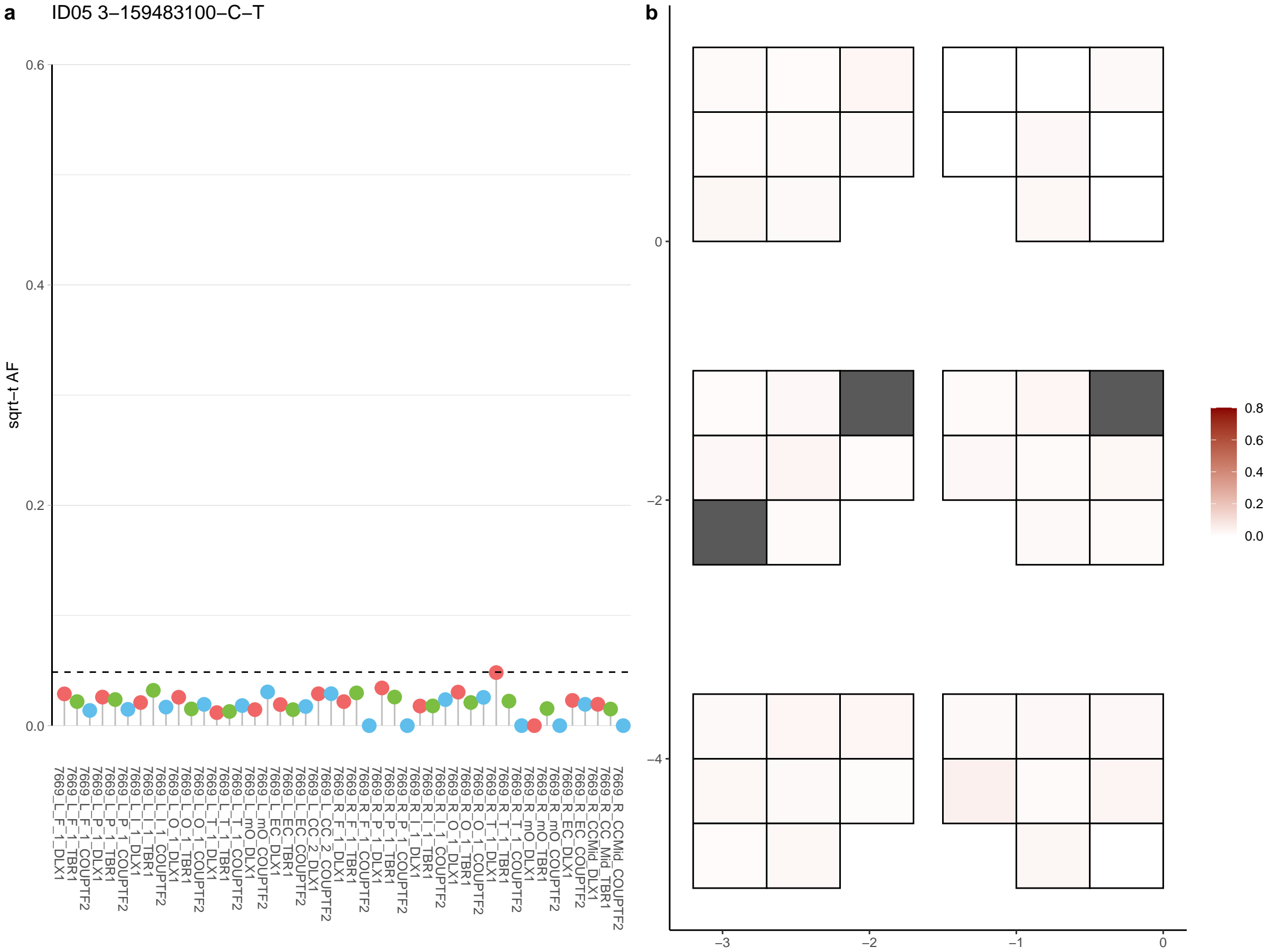

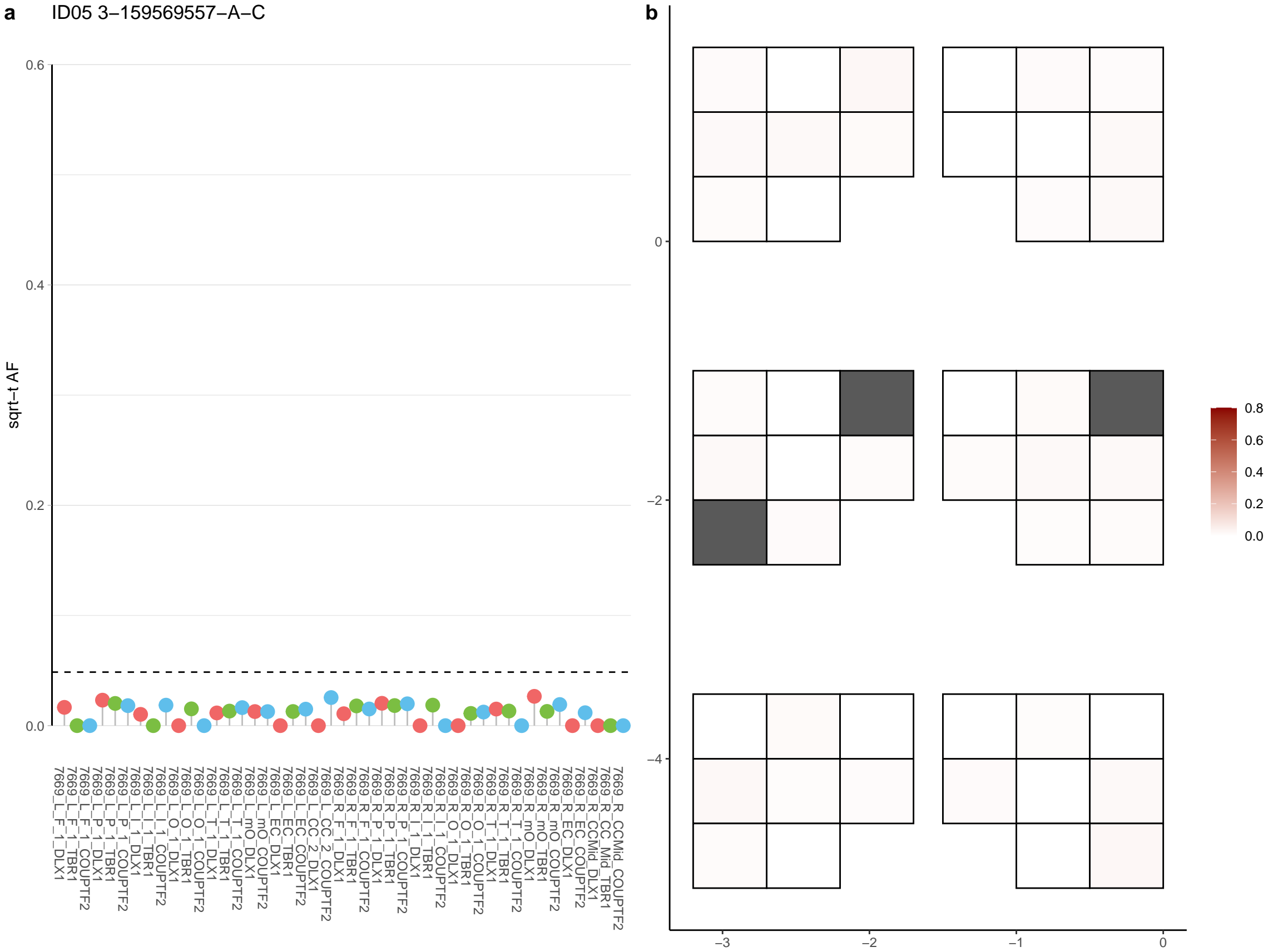

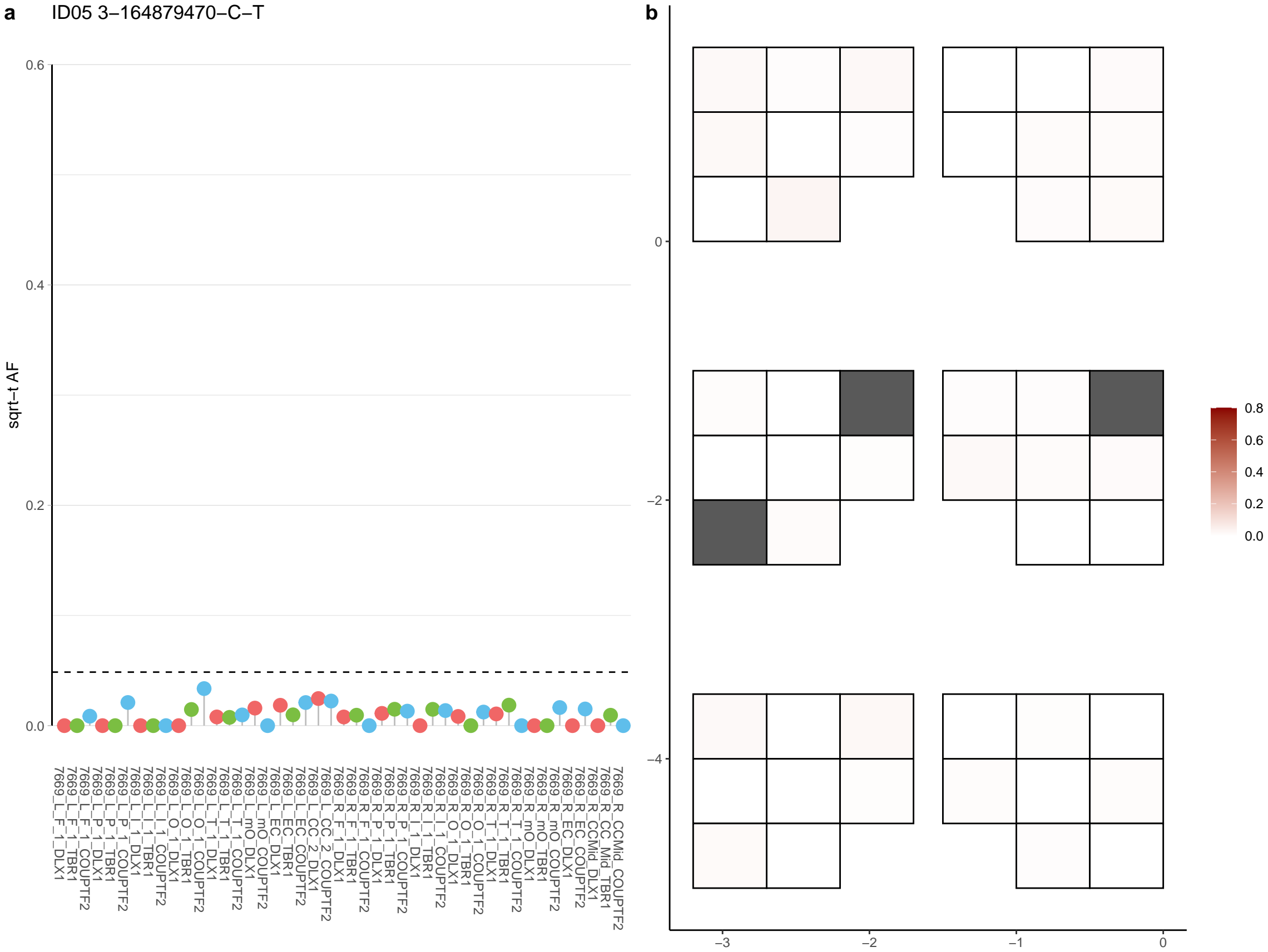

**a** ID05 3-16511350-A-C

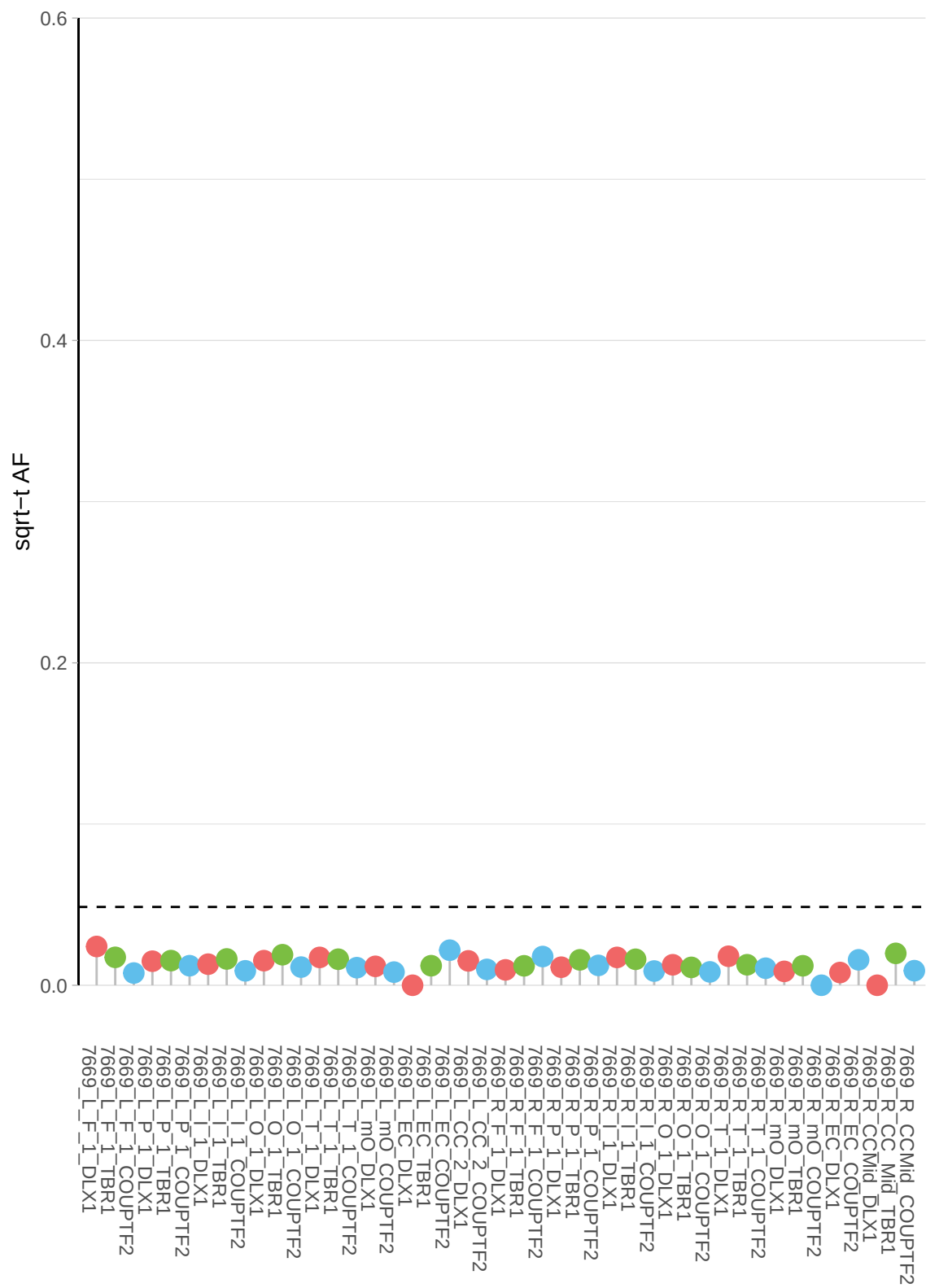

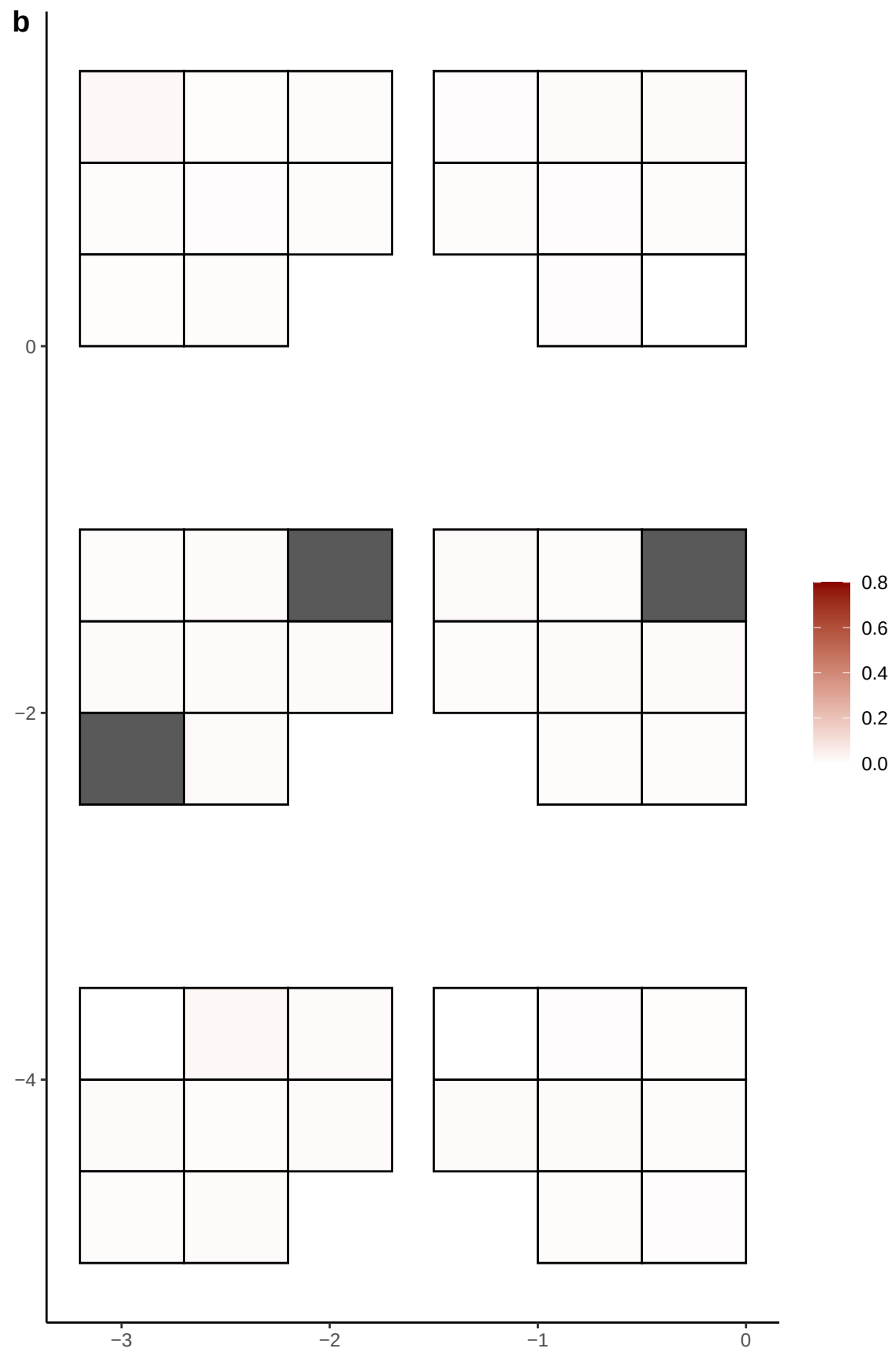

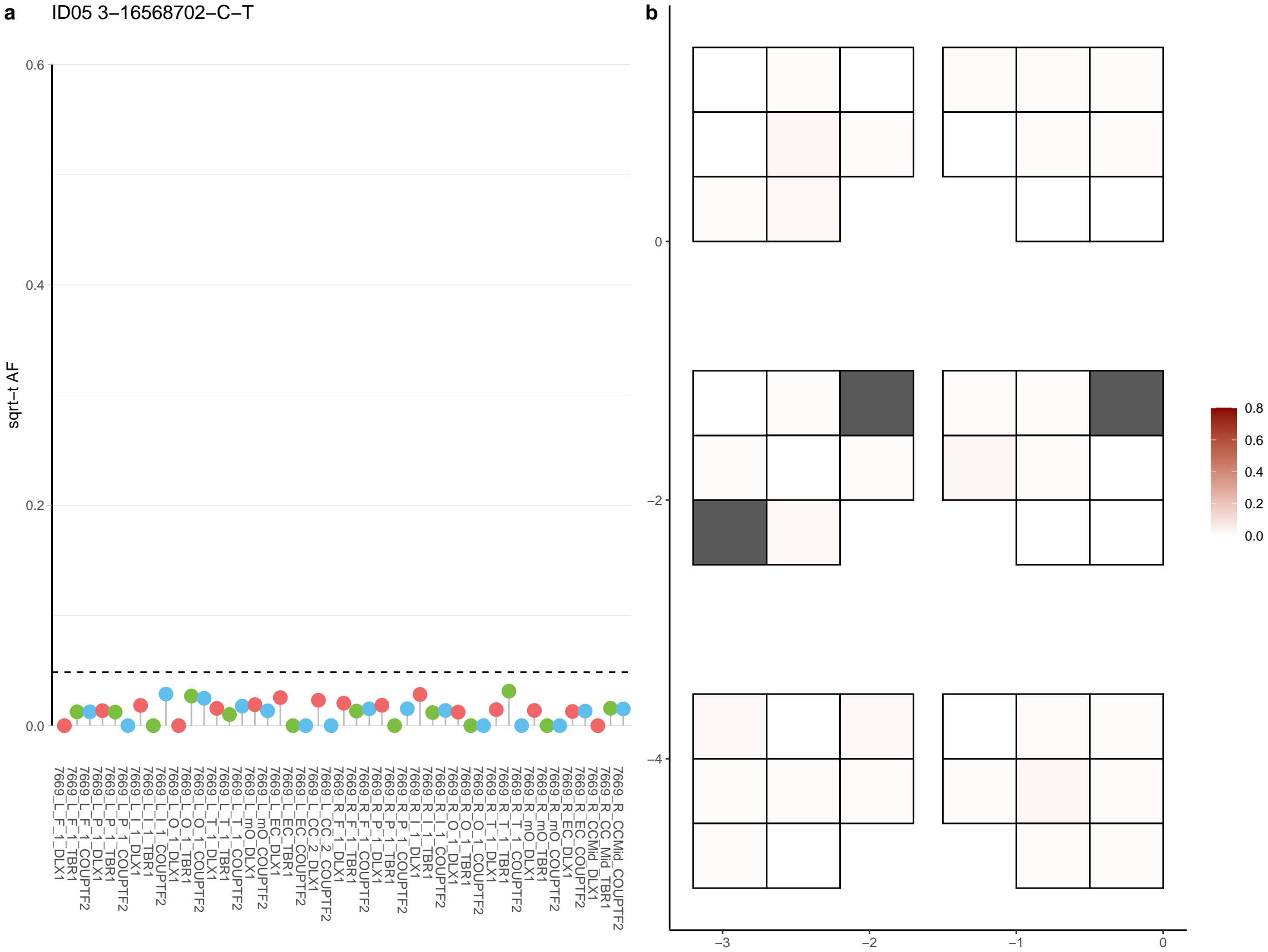

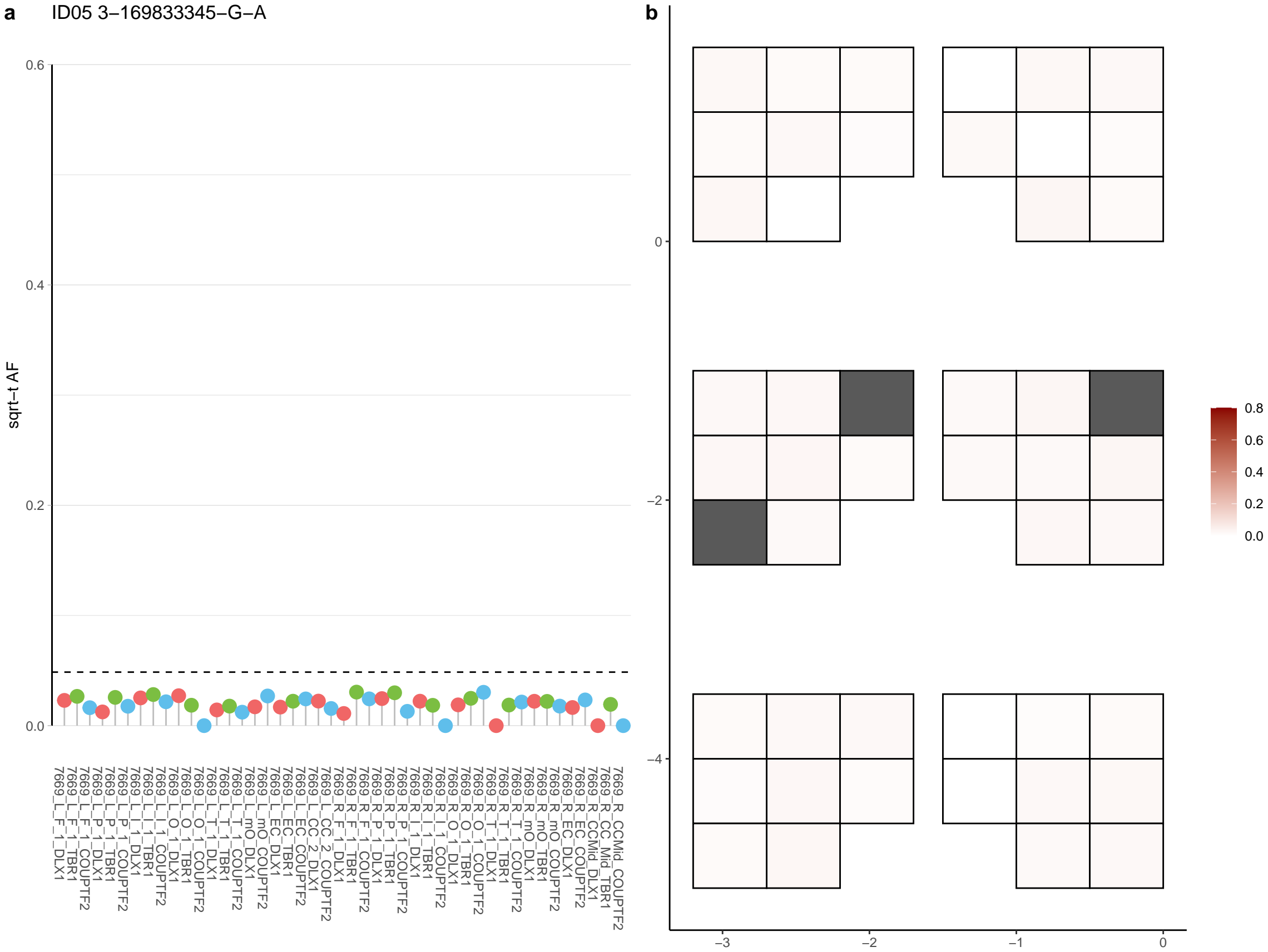

**a** ID05 3-172635725-G-A

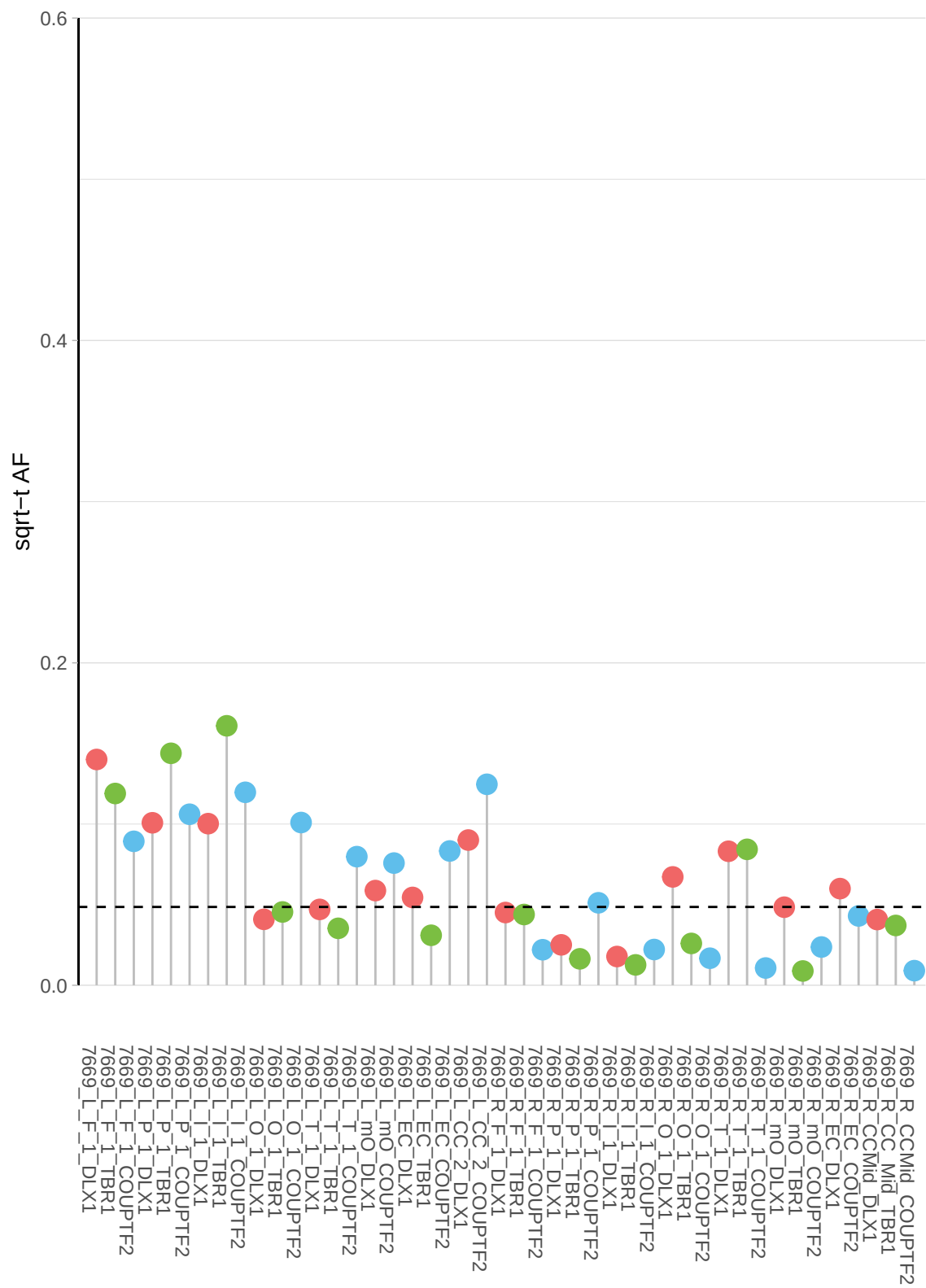

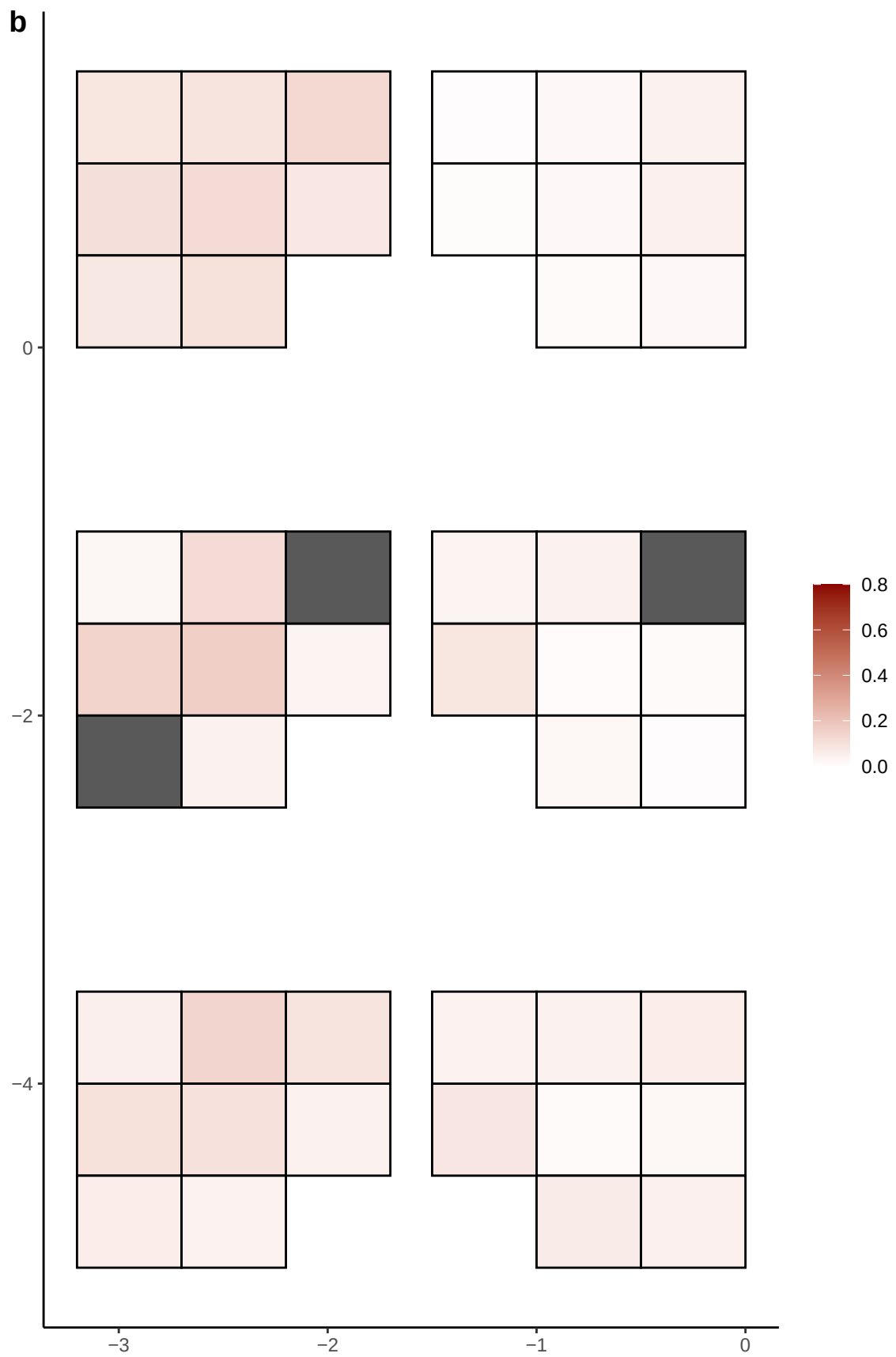

**a** ID05 3-173423567-C-A

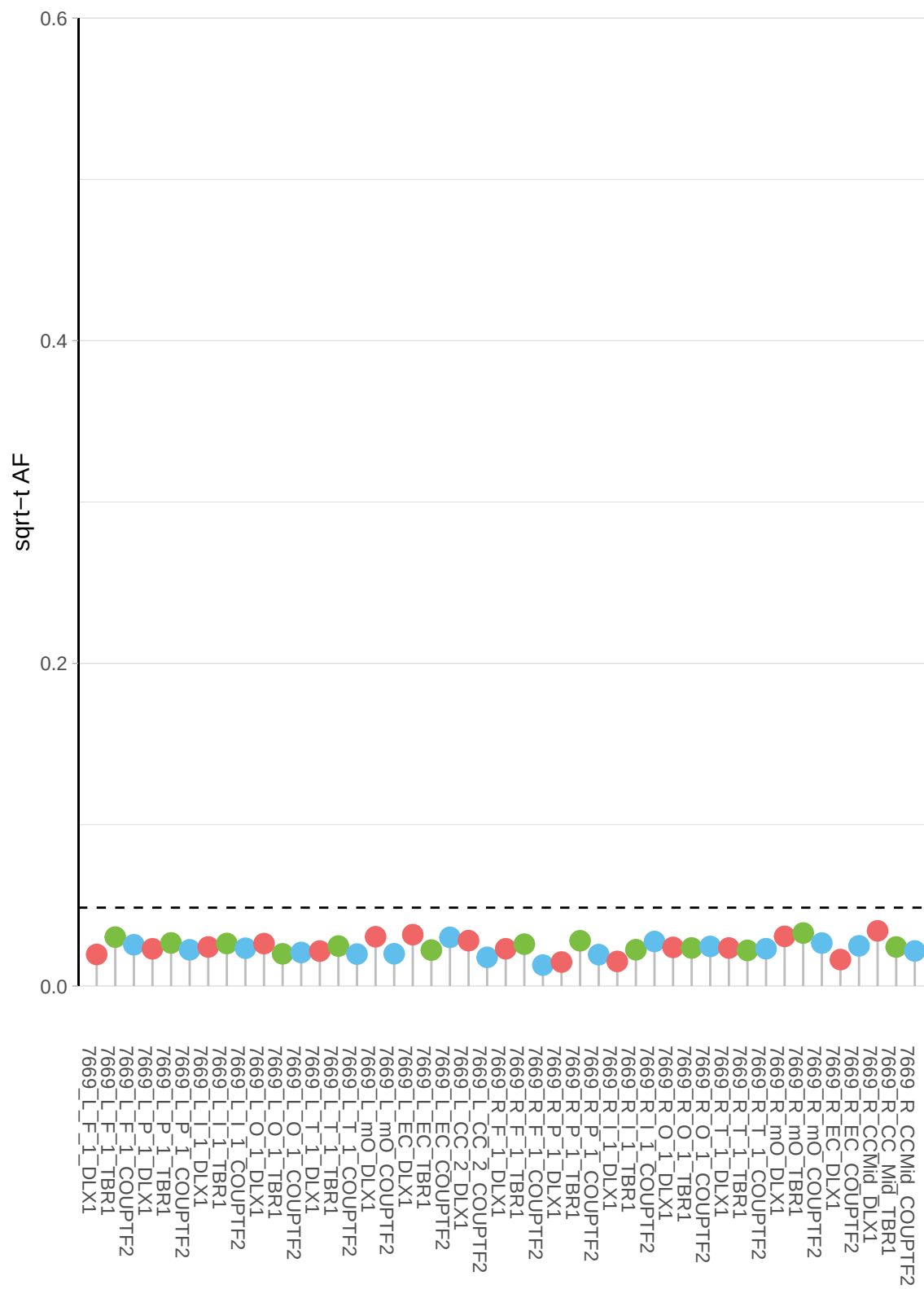

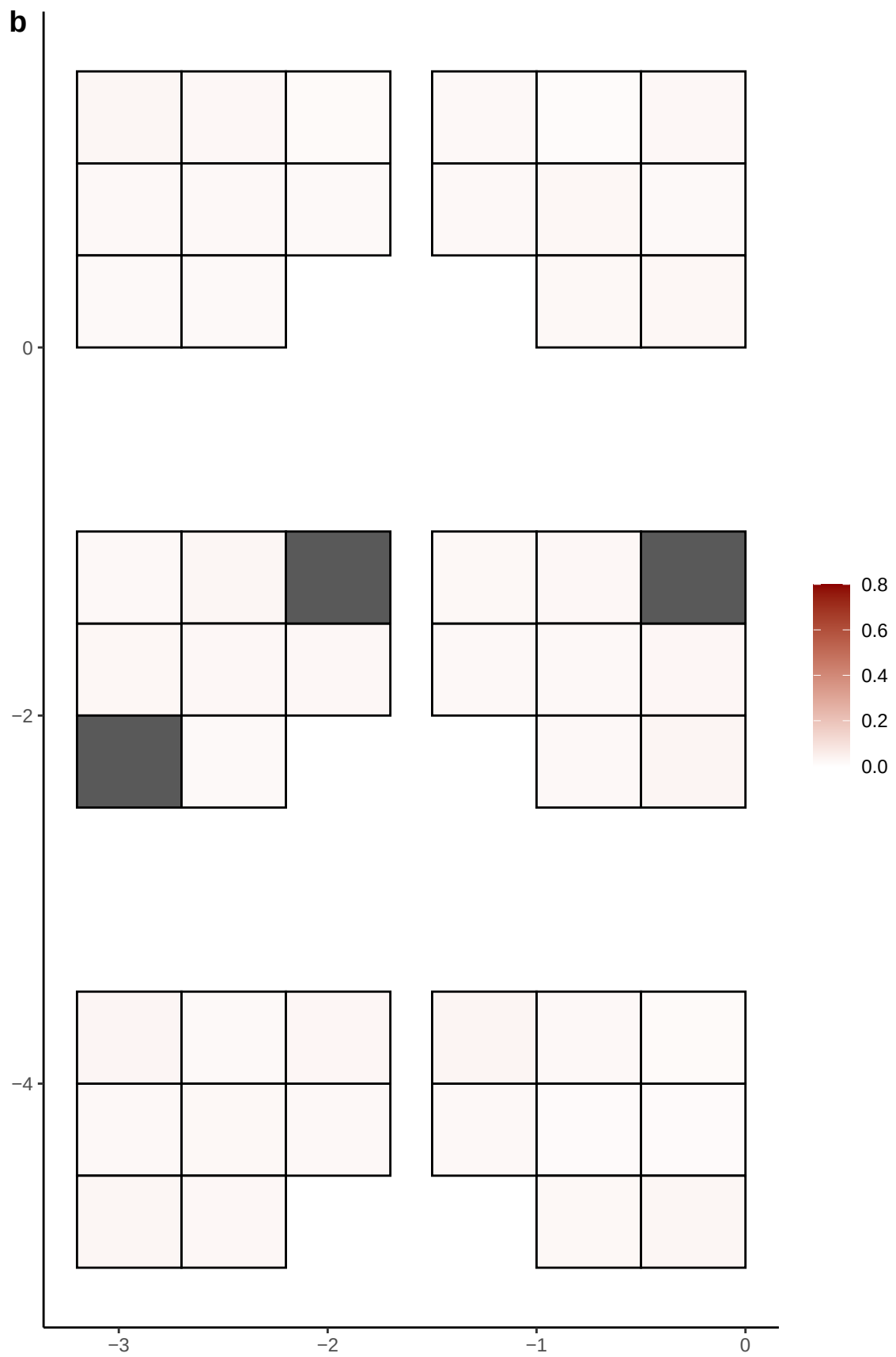

**a** ID05 3-181259475-C-T

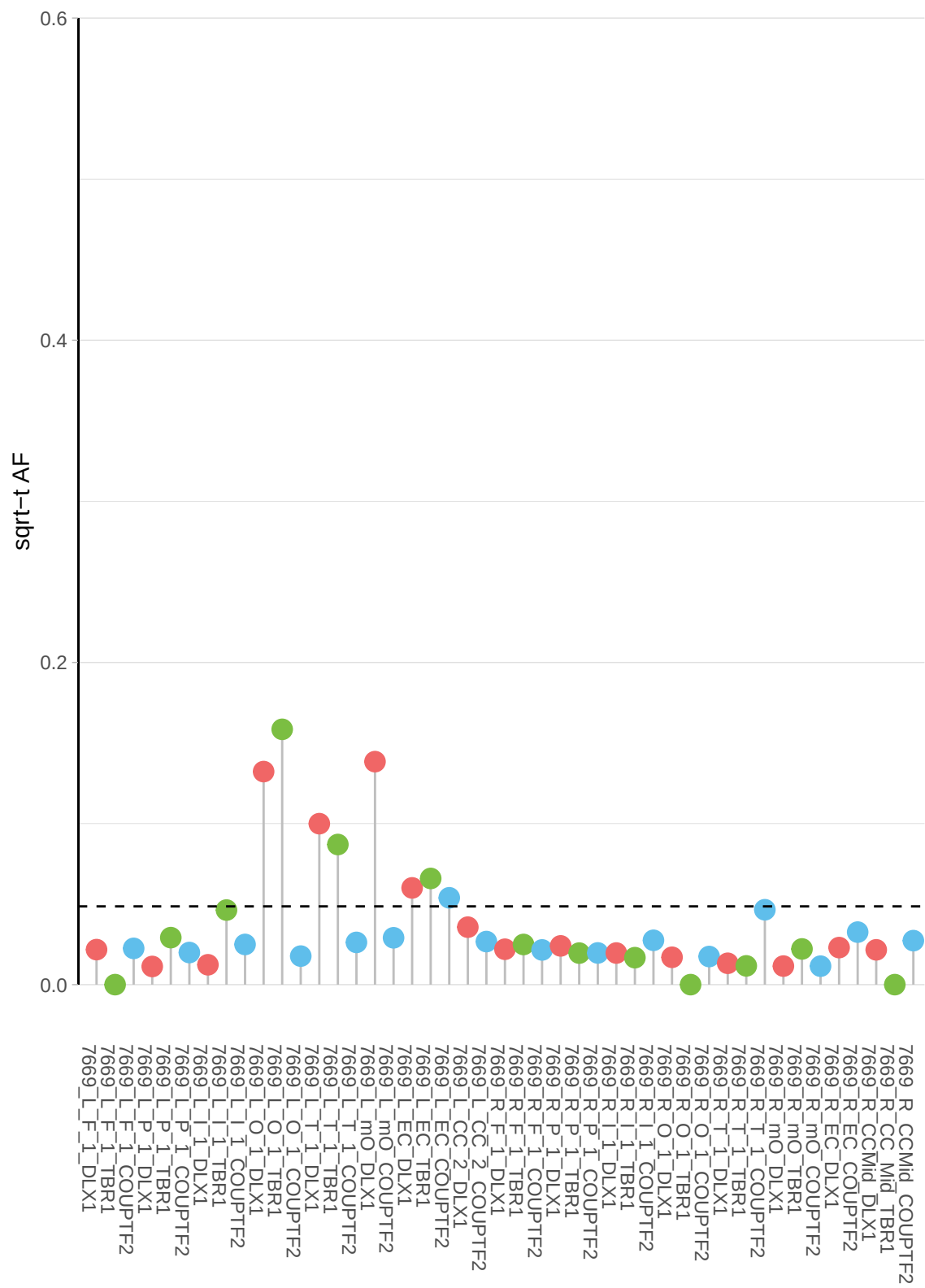

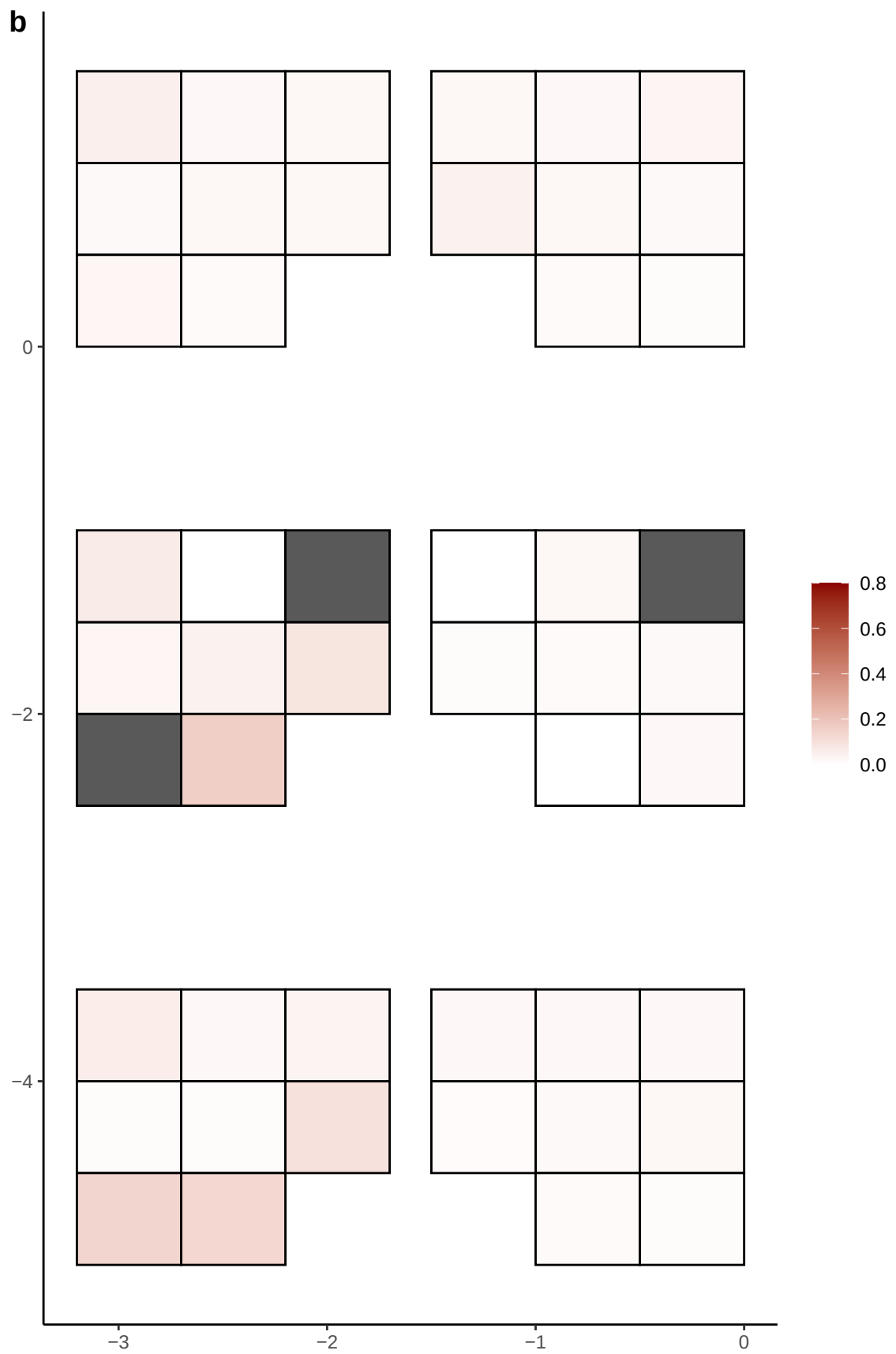

**a** ID05 3-185077743-C-T

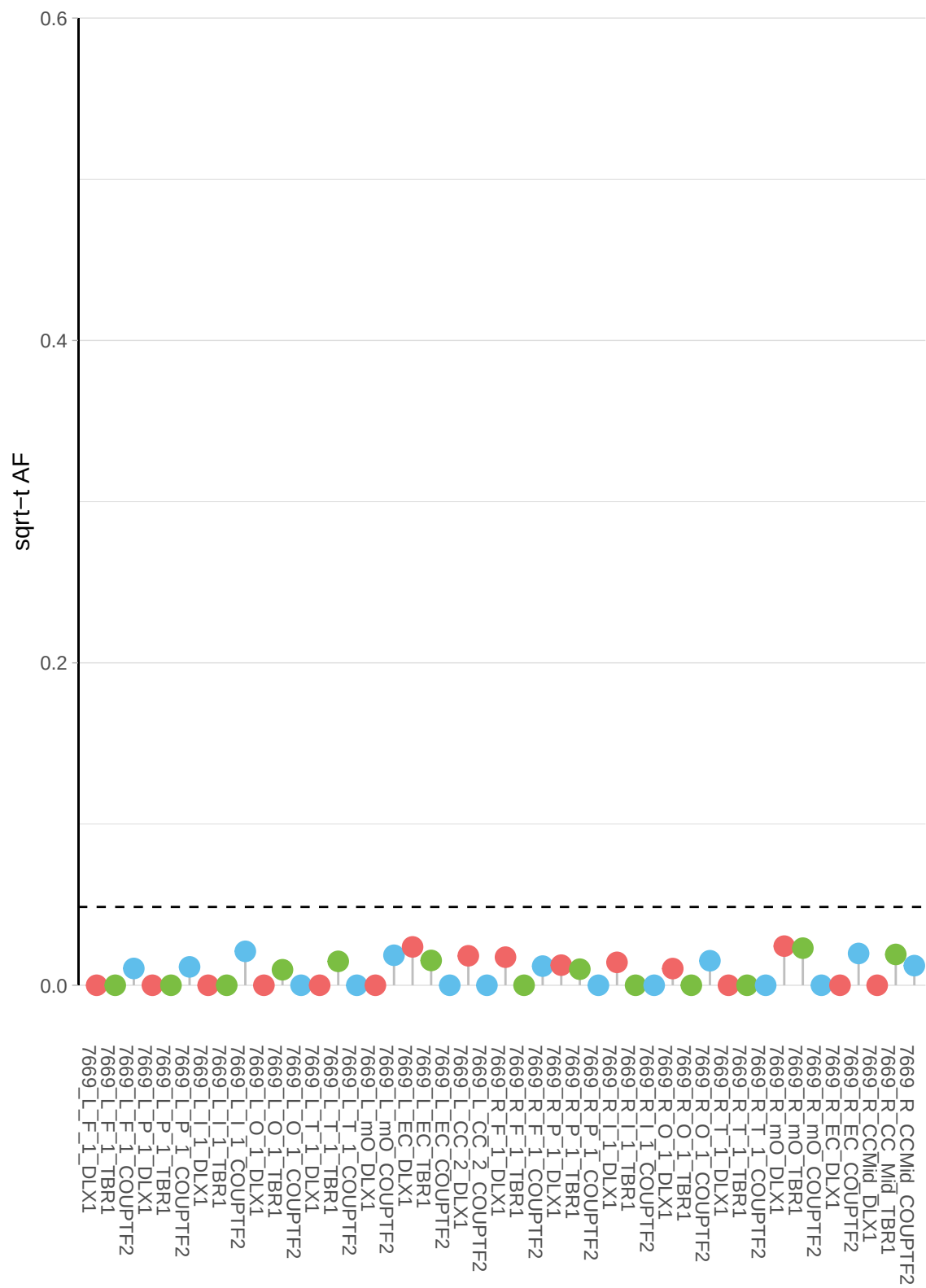

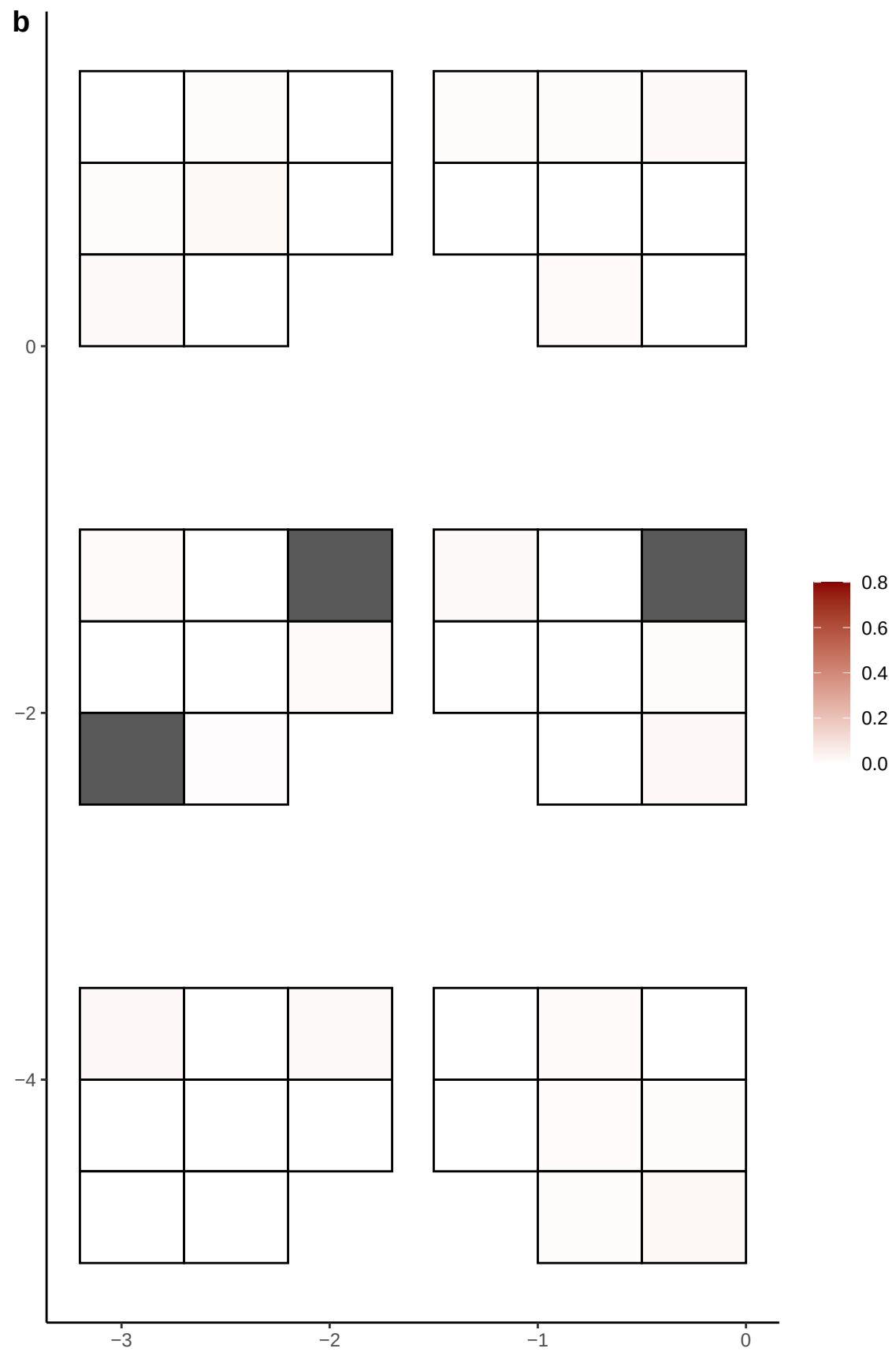

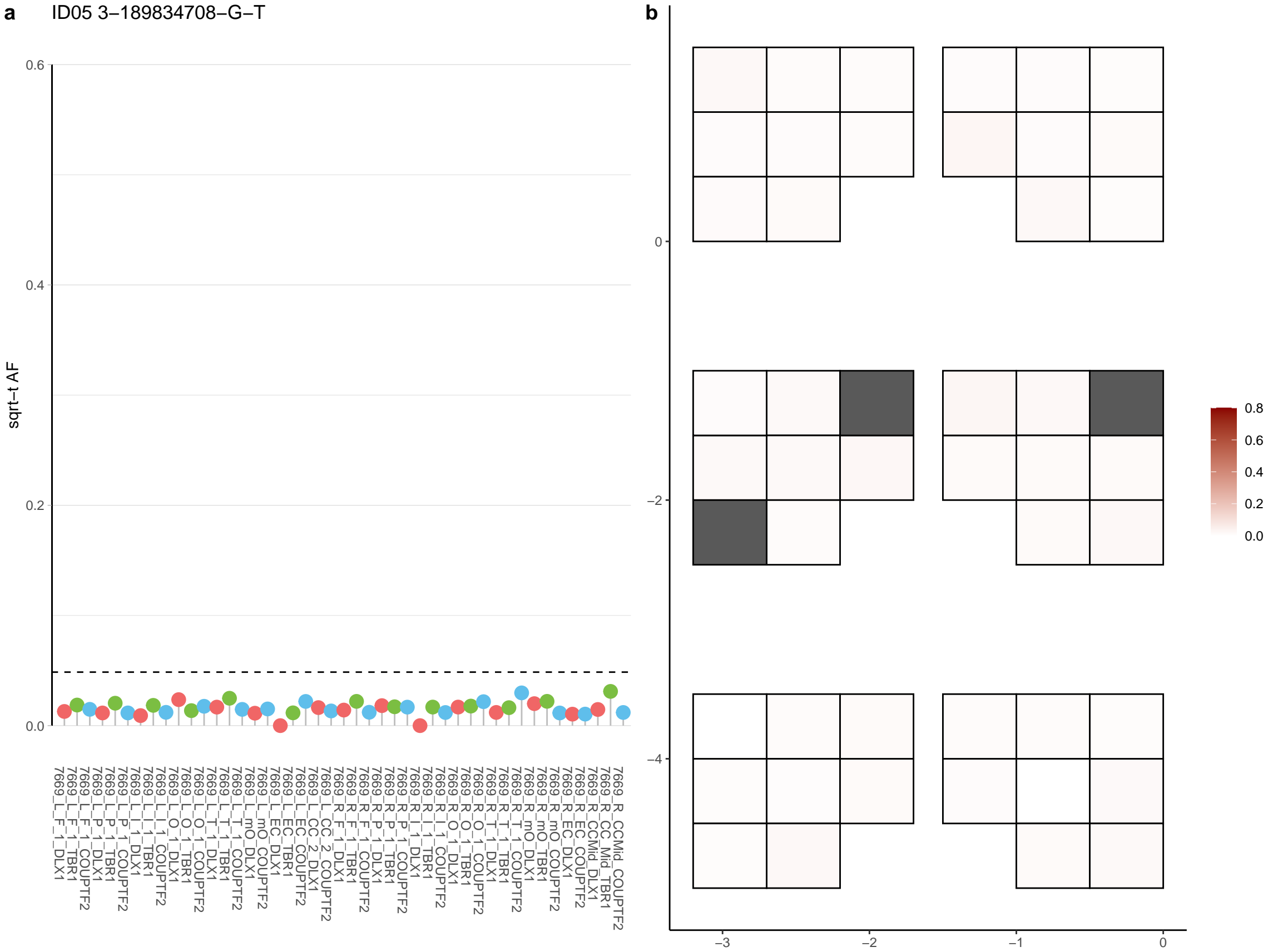

**a** ID05 3-190125961-G-A

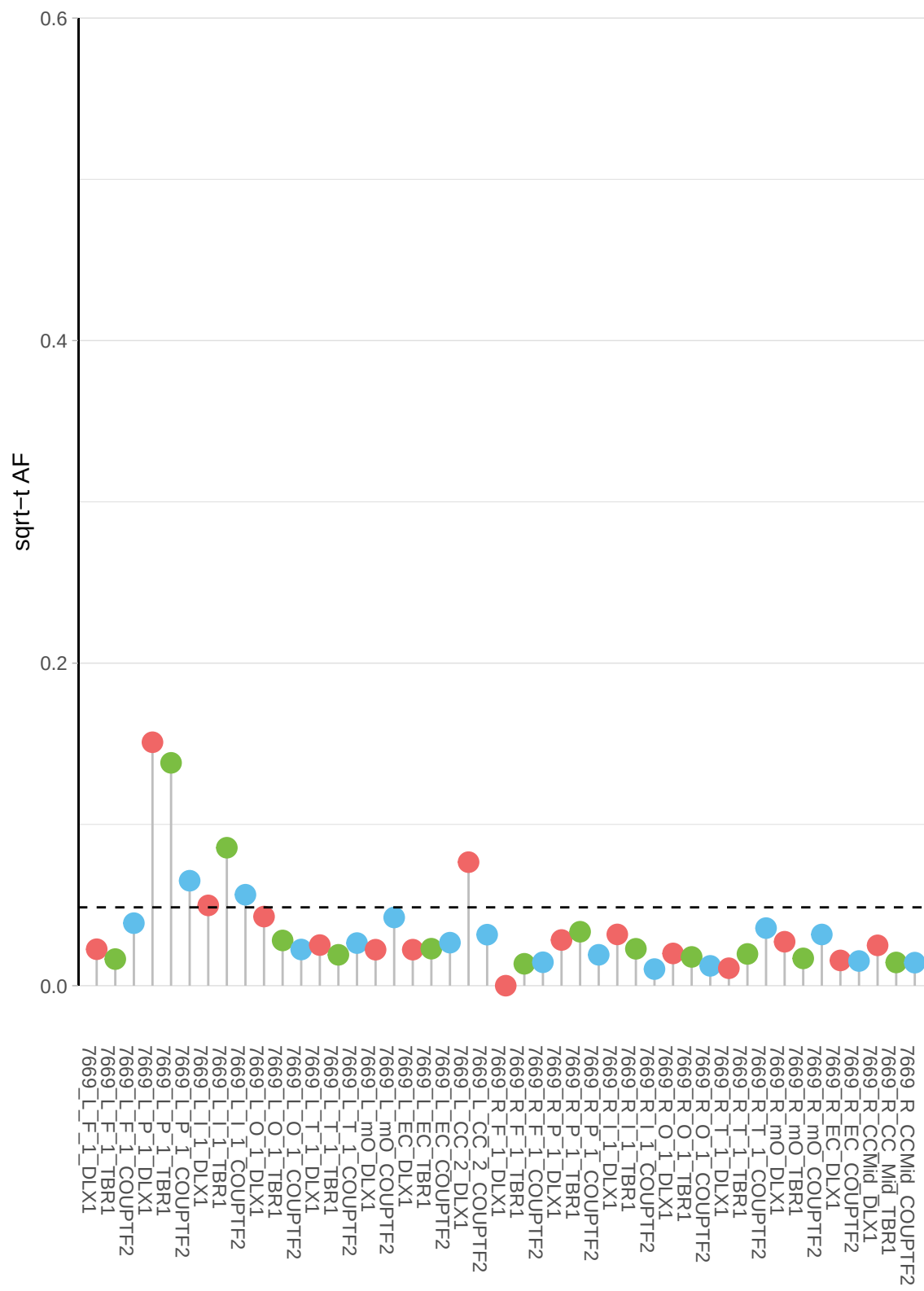

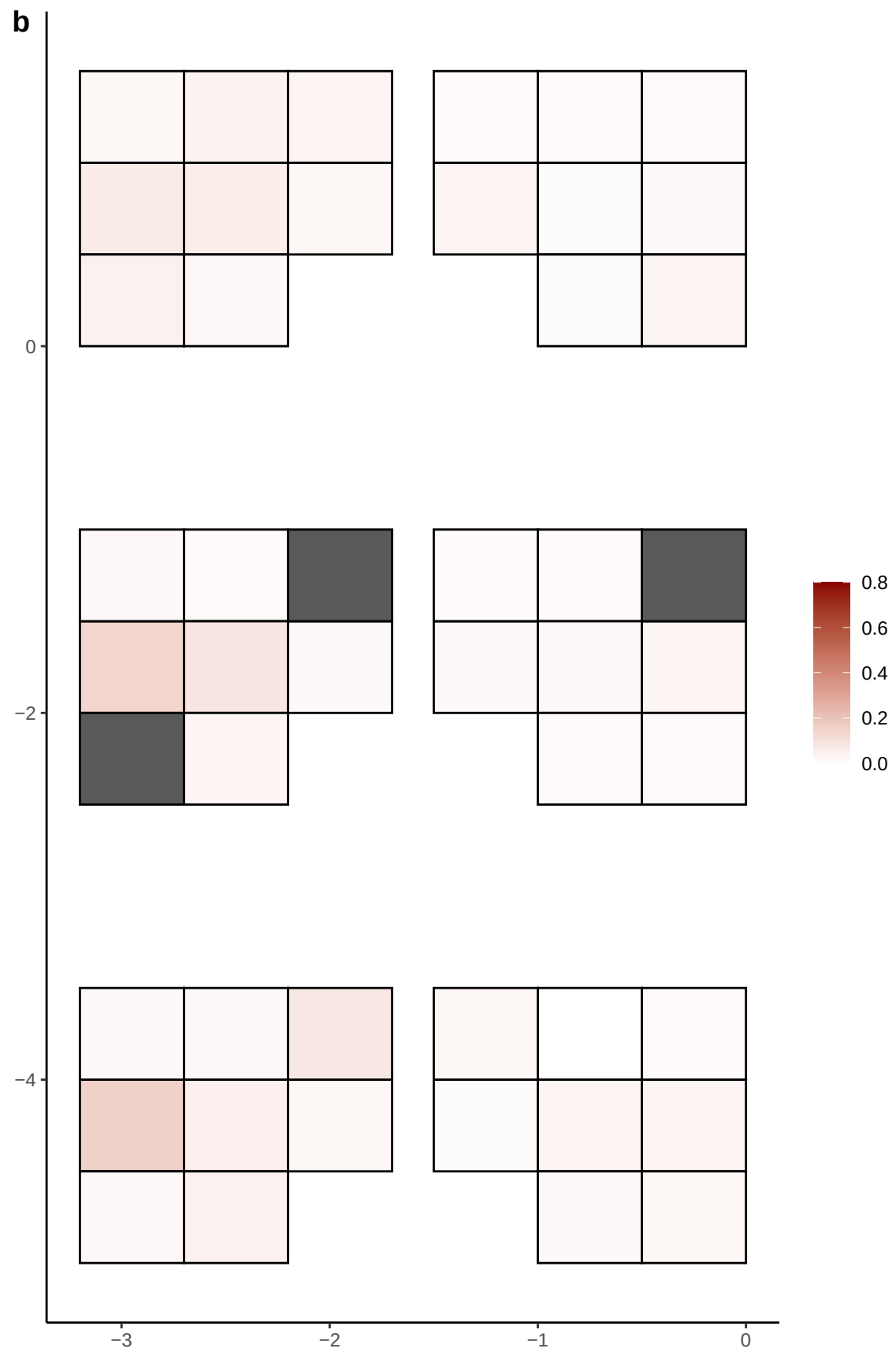

**a** ID05 3-193720234-C-T

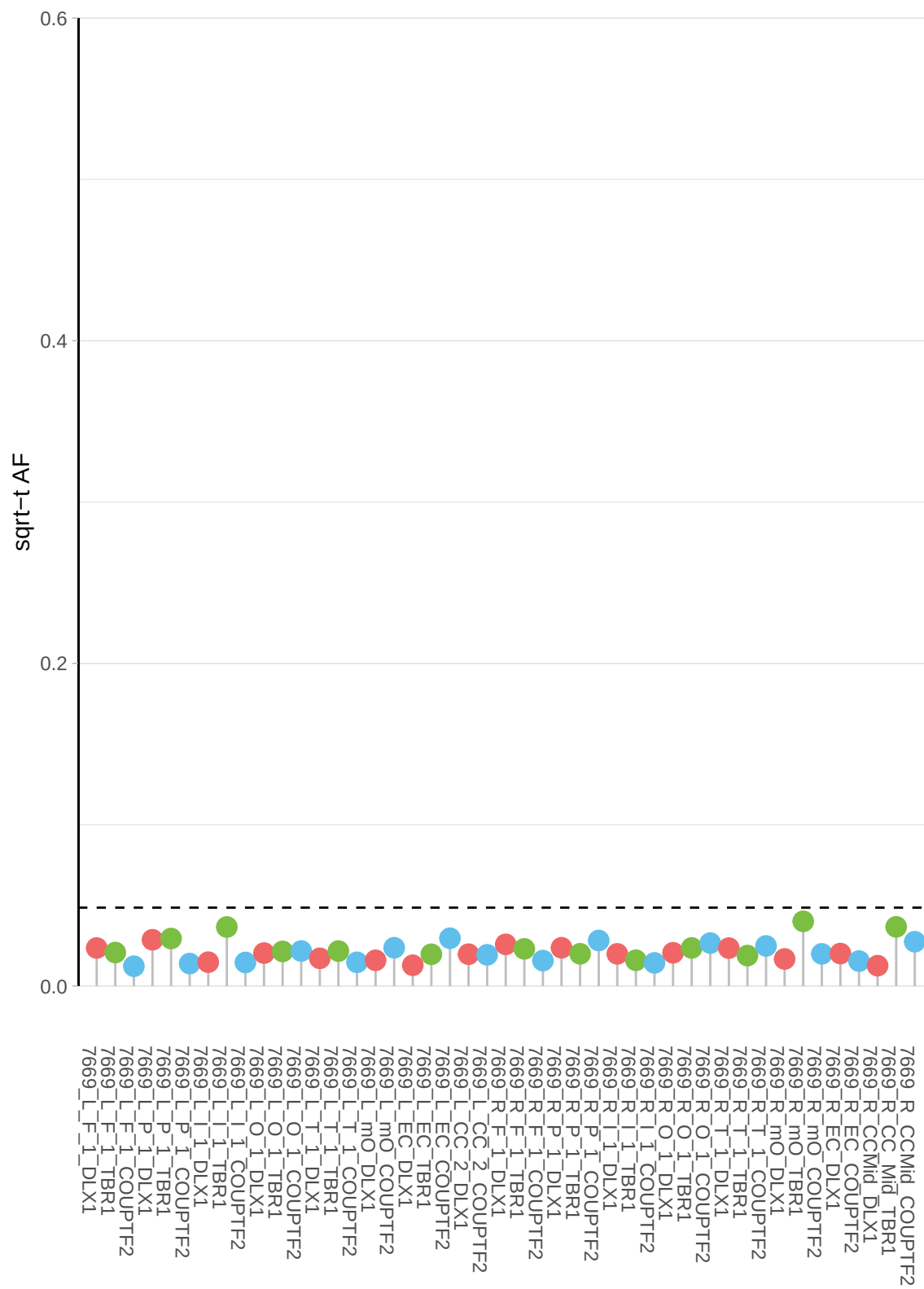

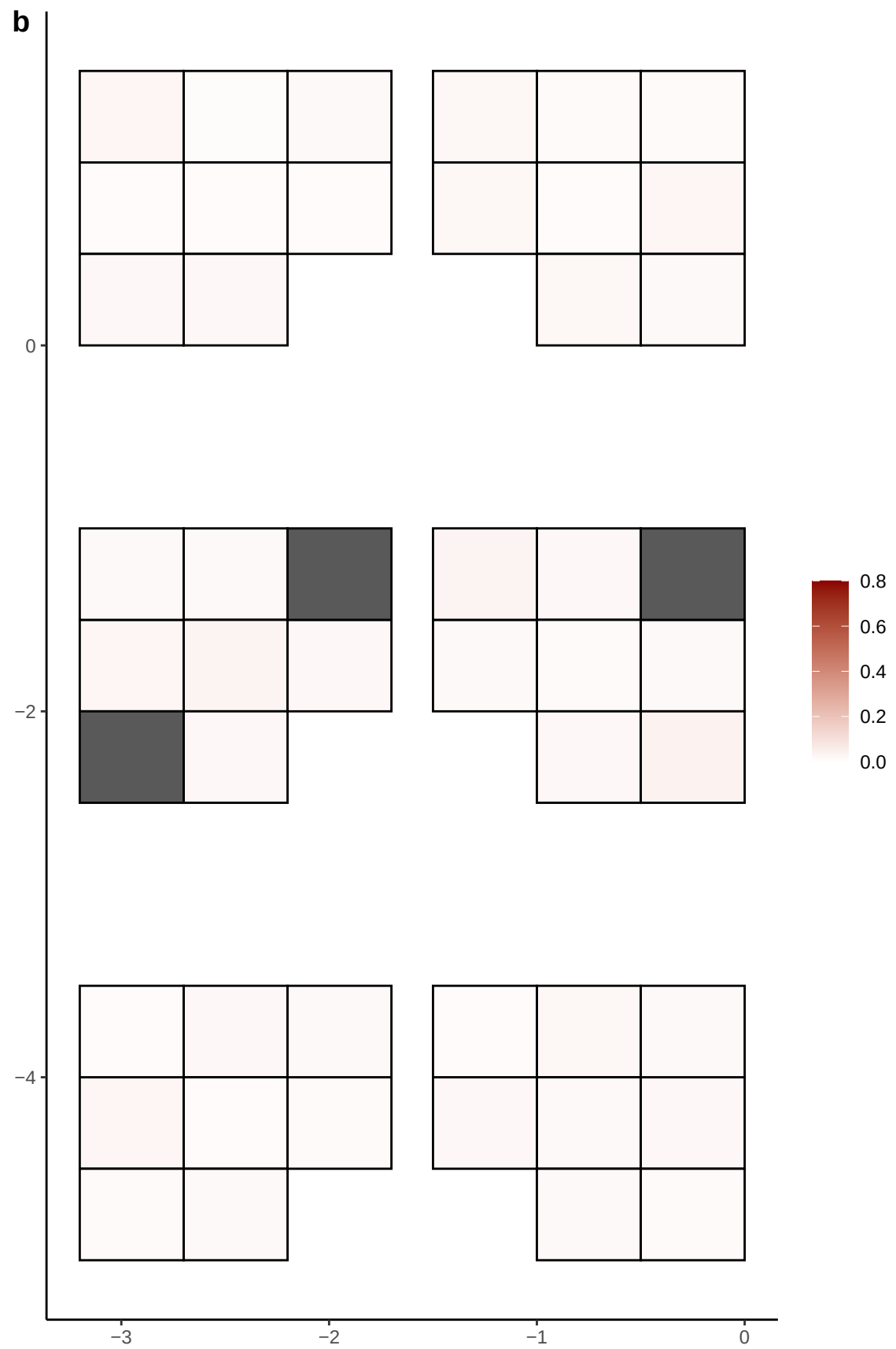

**a** ID05 3-195270223-T-G

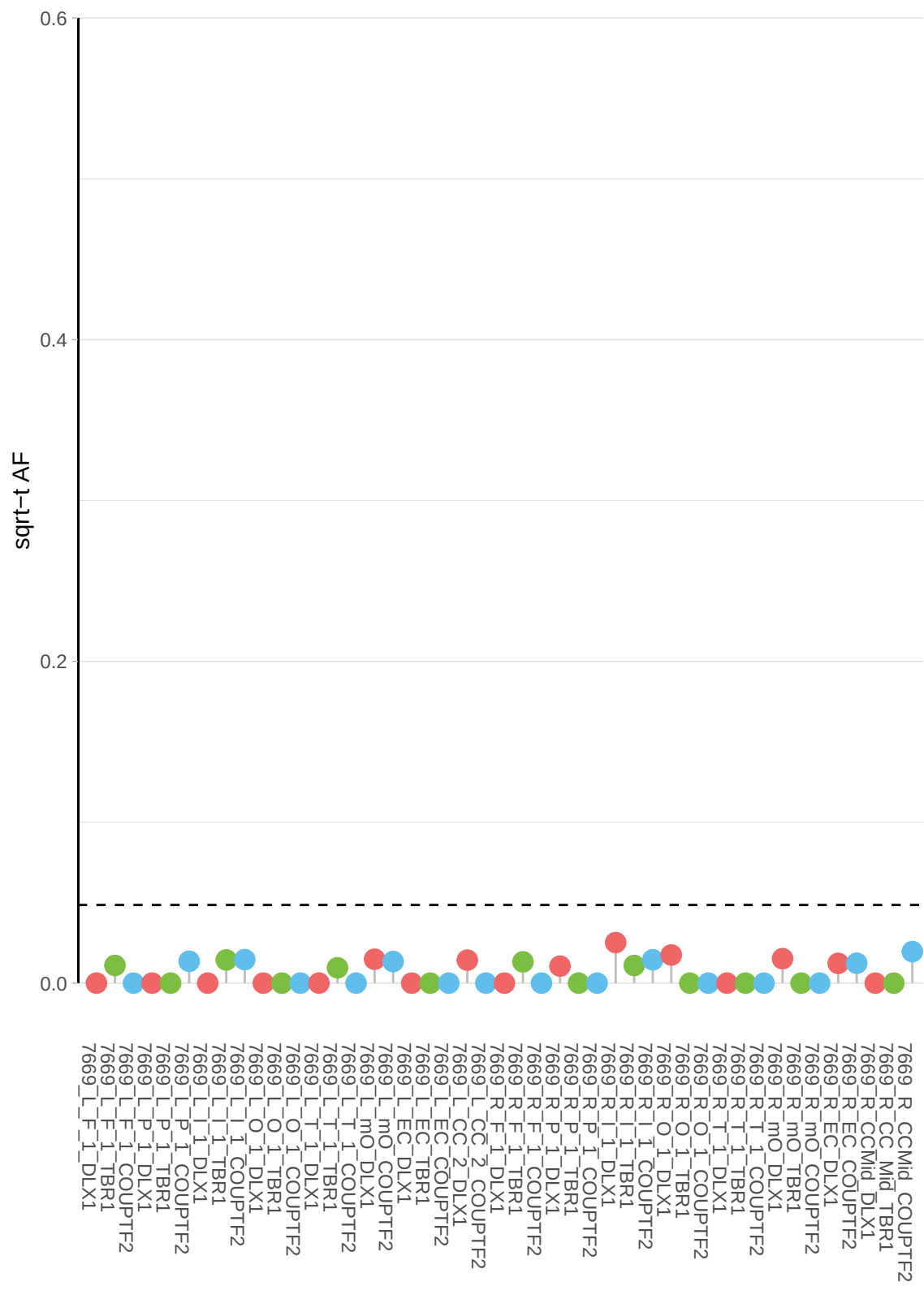

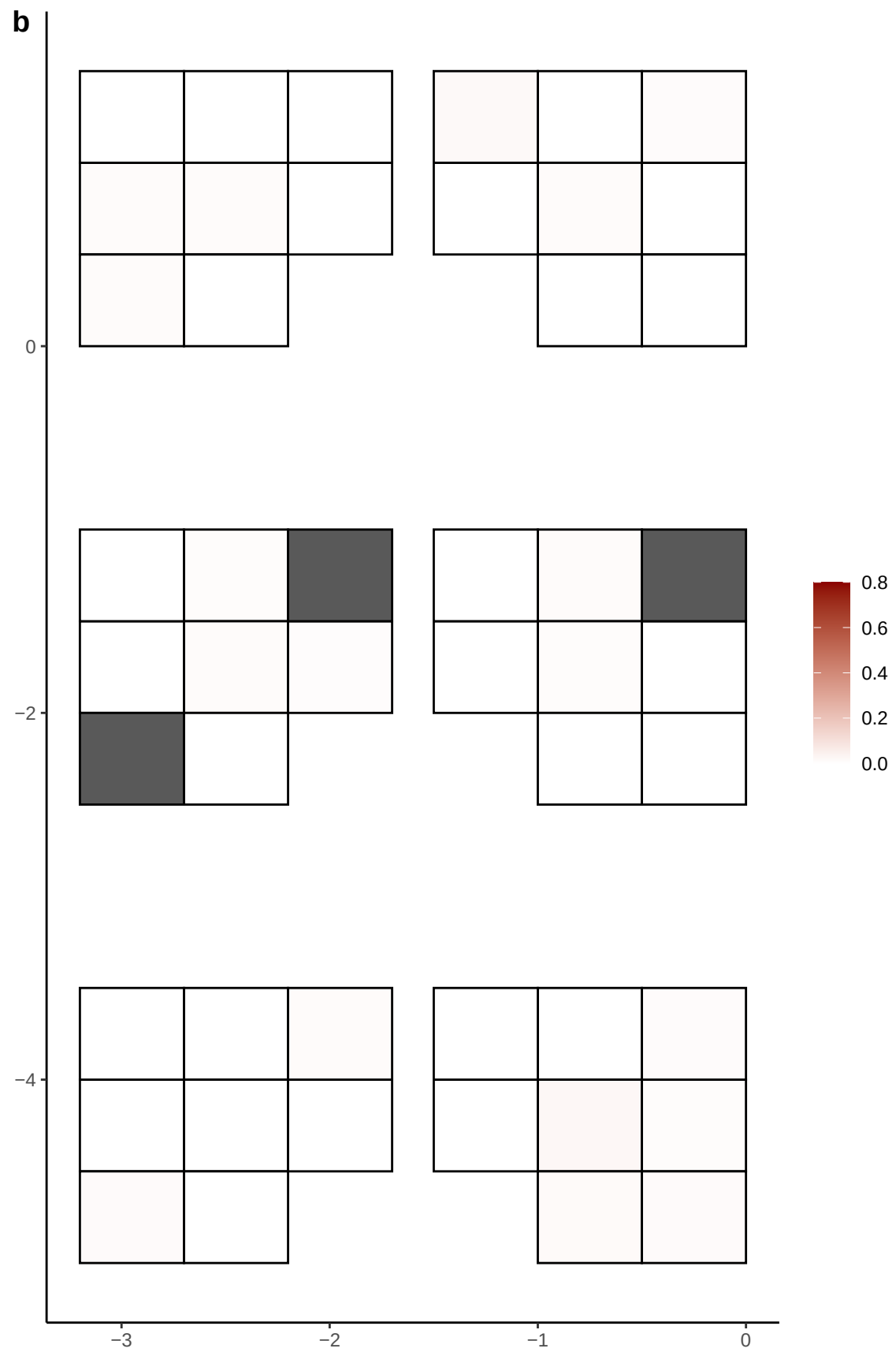

**a** ID05 3-20053569-C-T

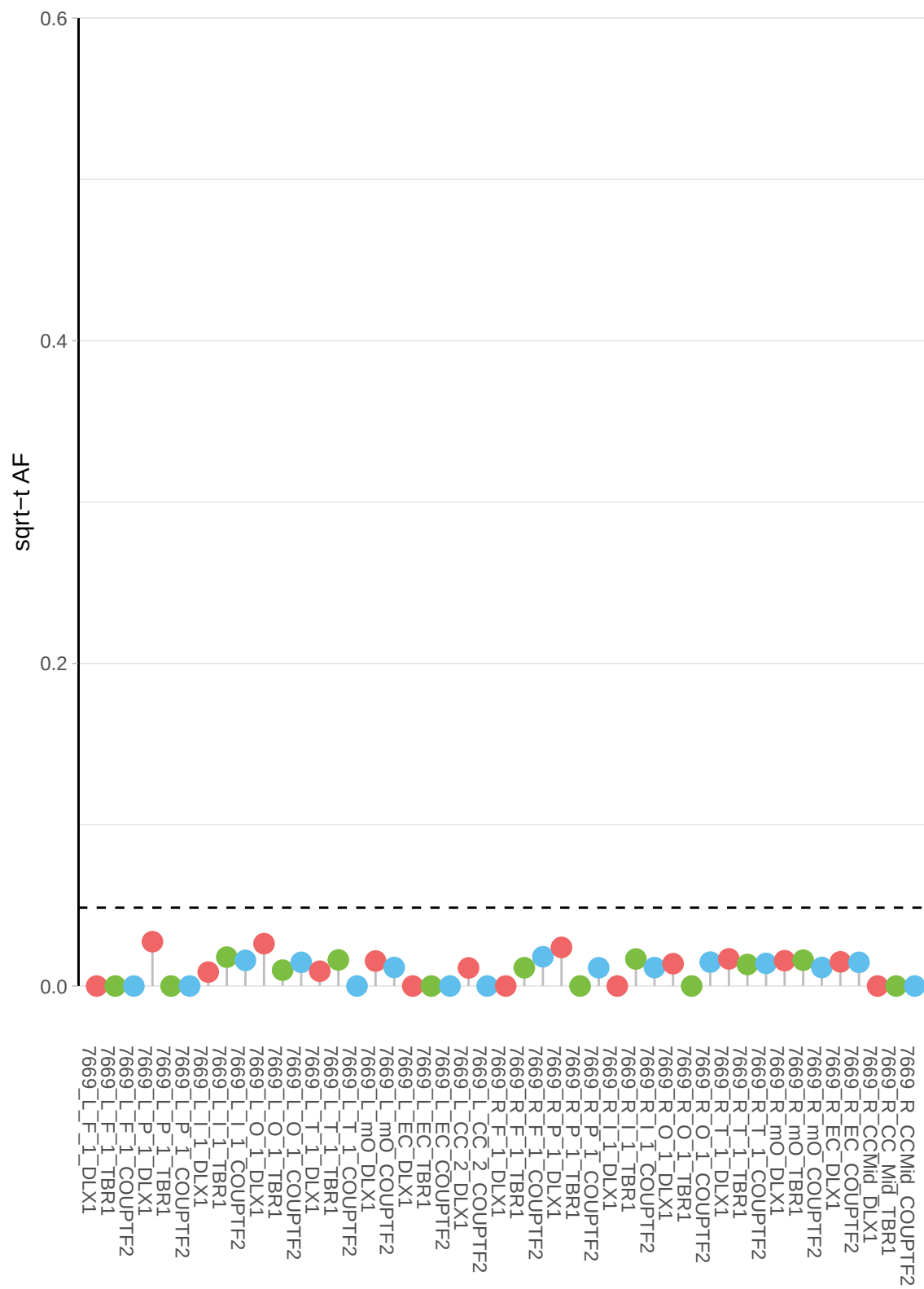

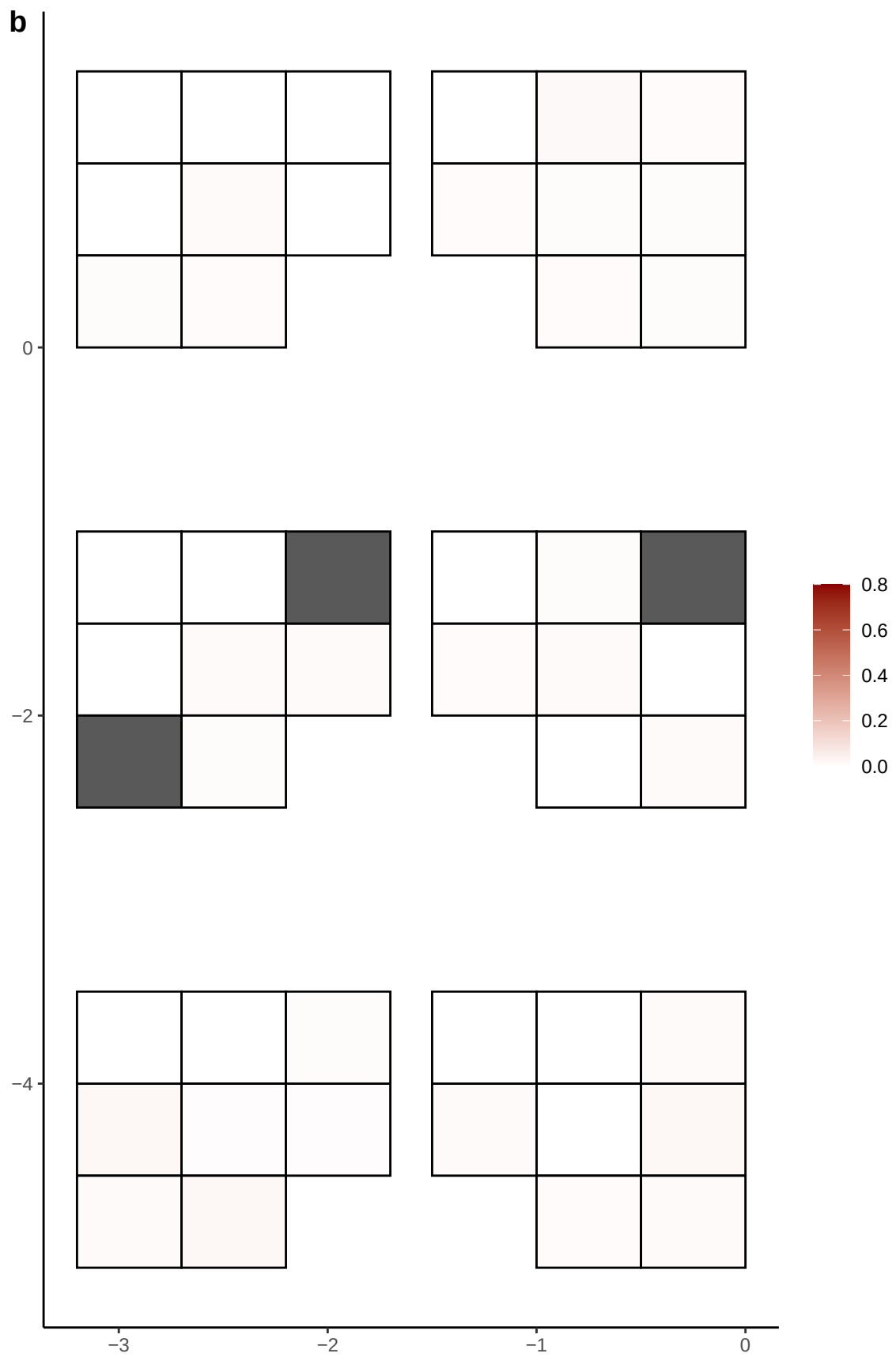

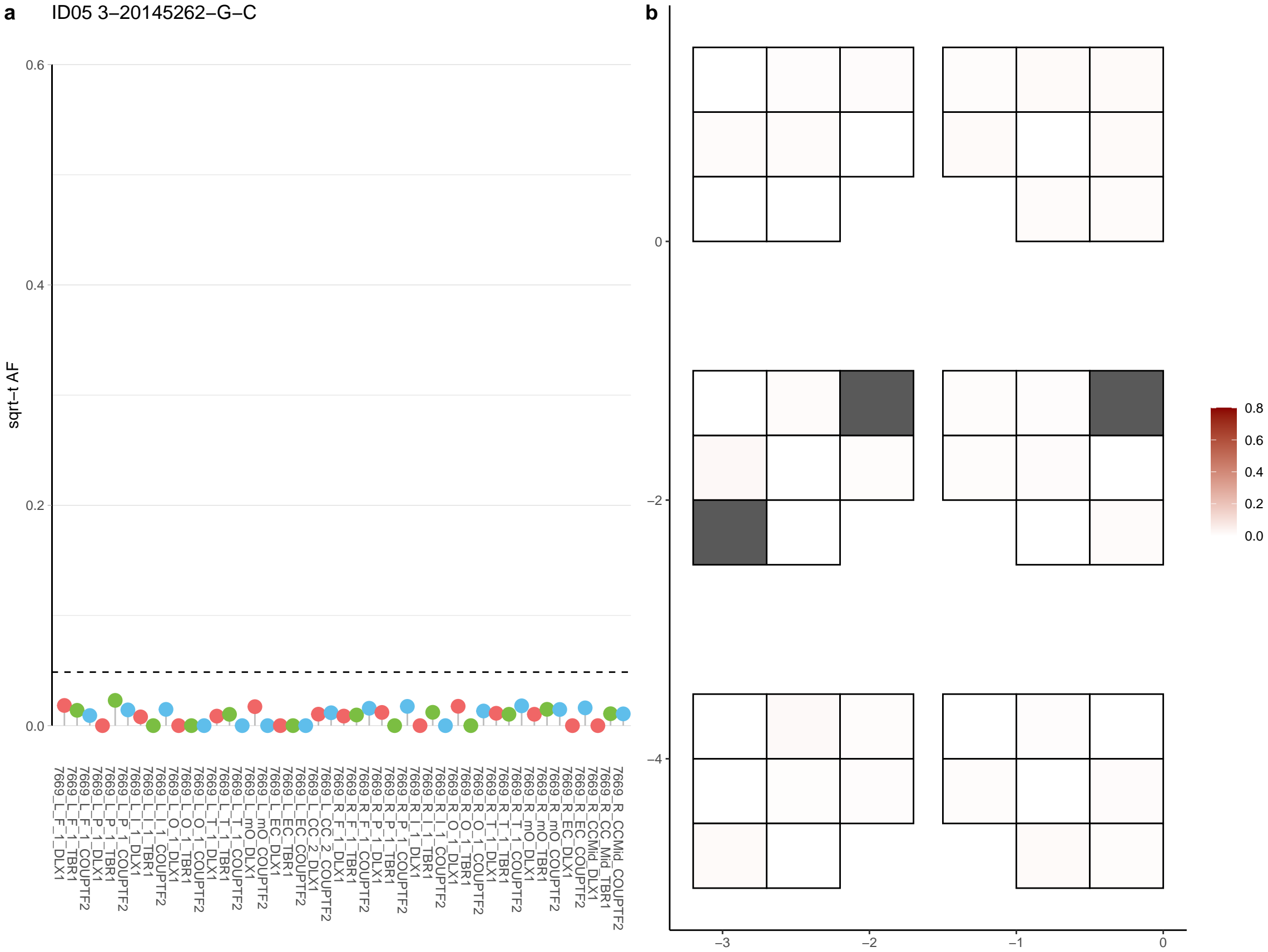

**a** ID05 3-20222654-C-T

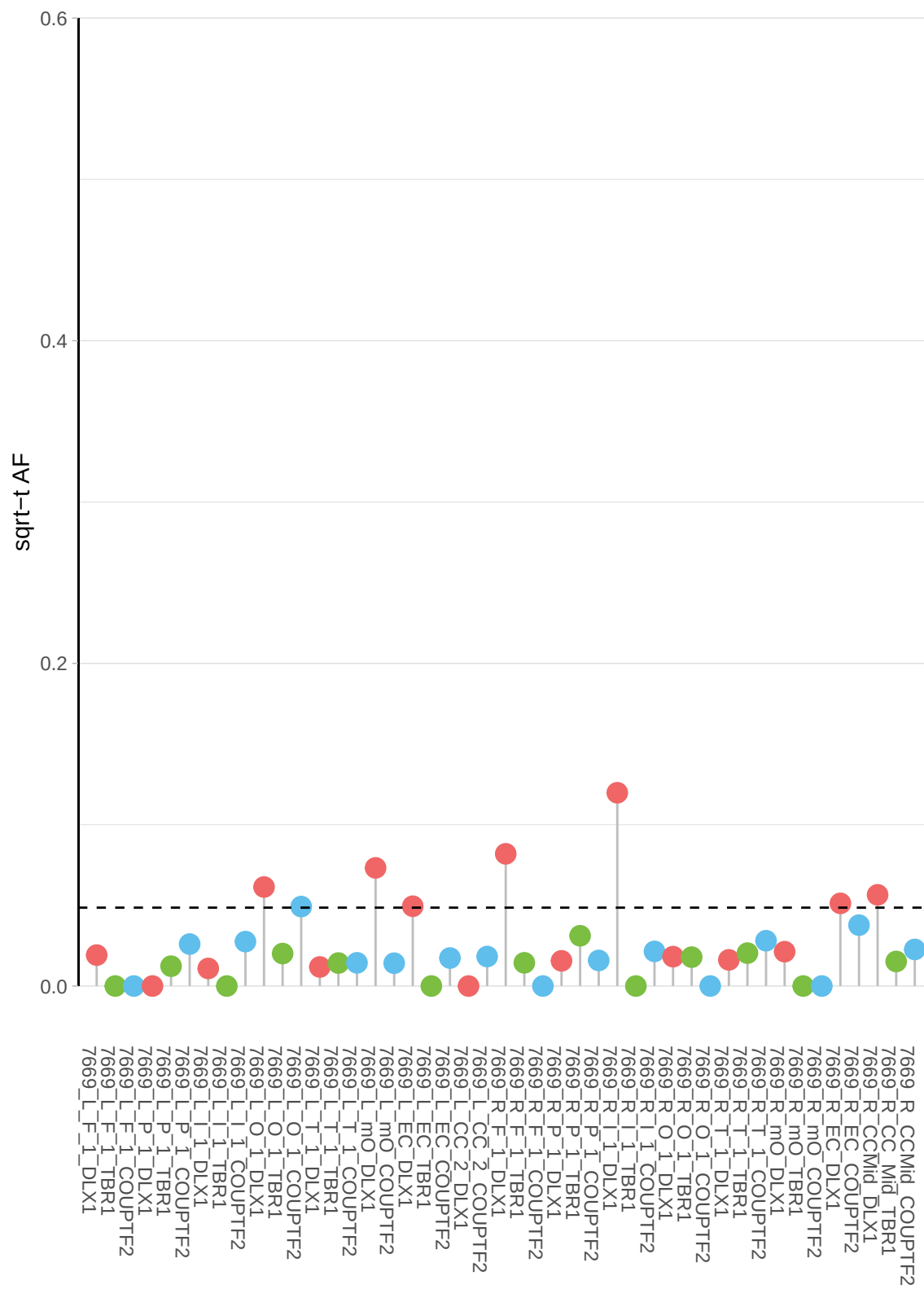

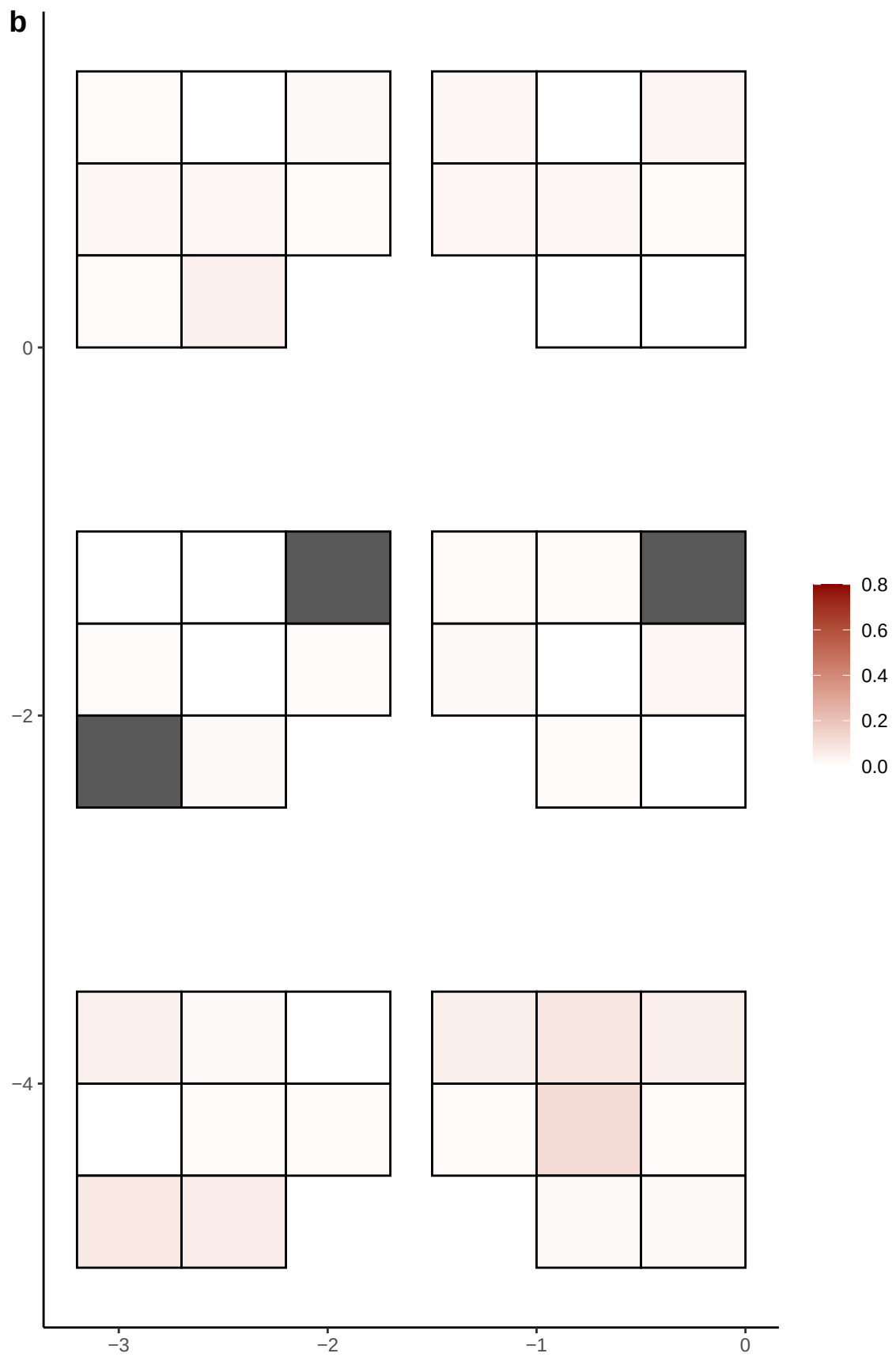

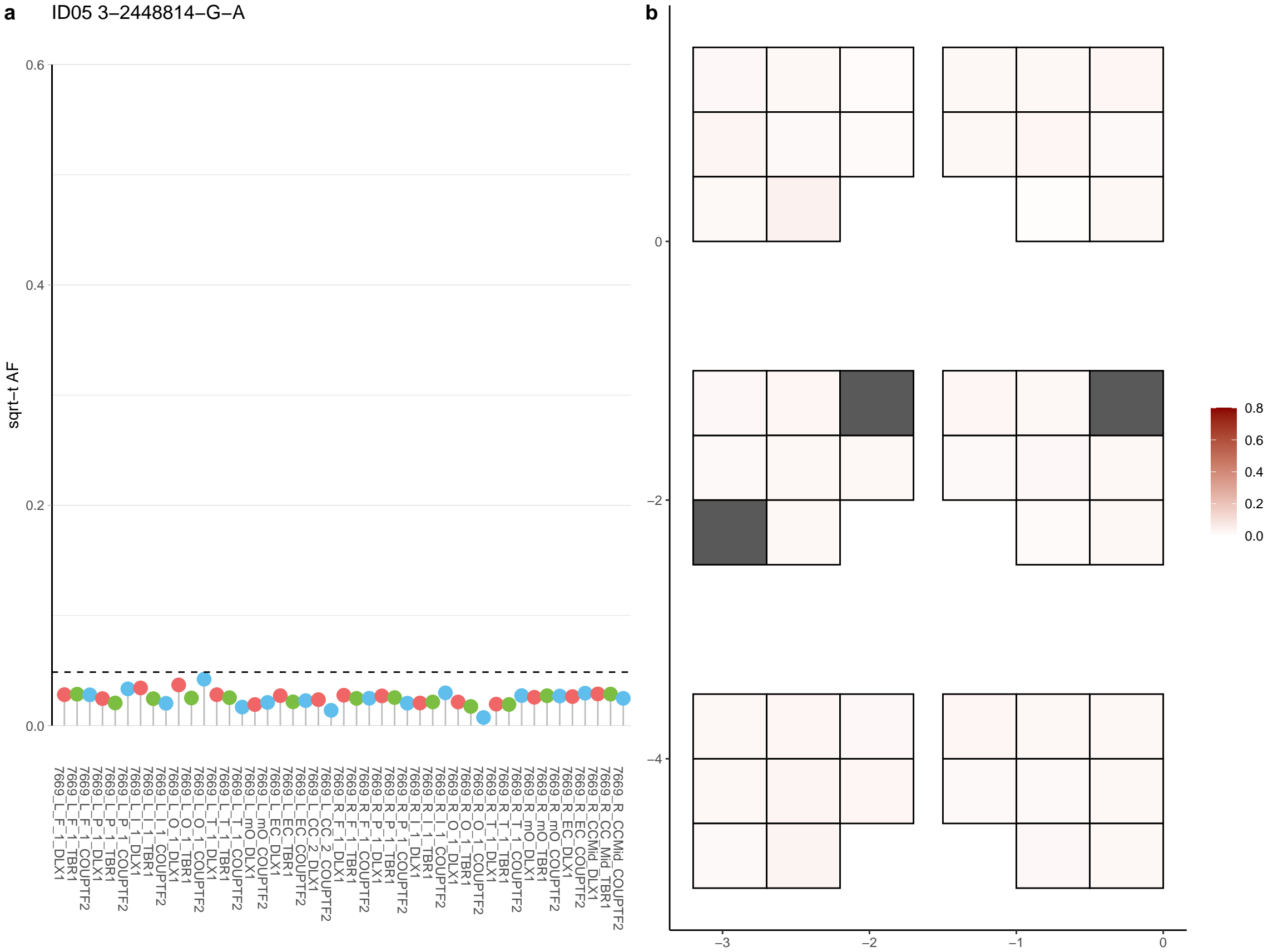

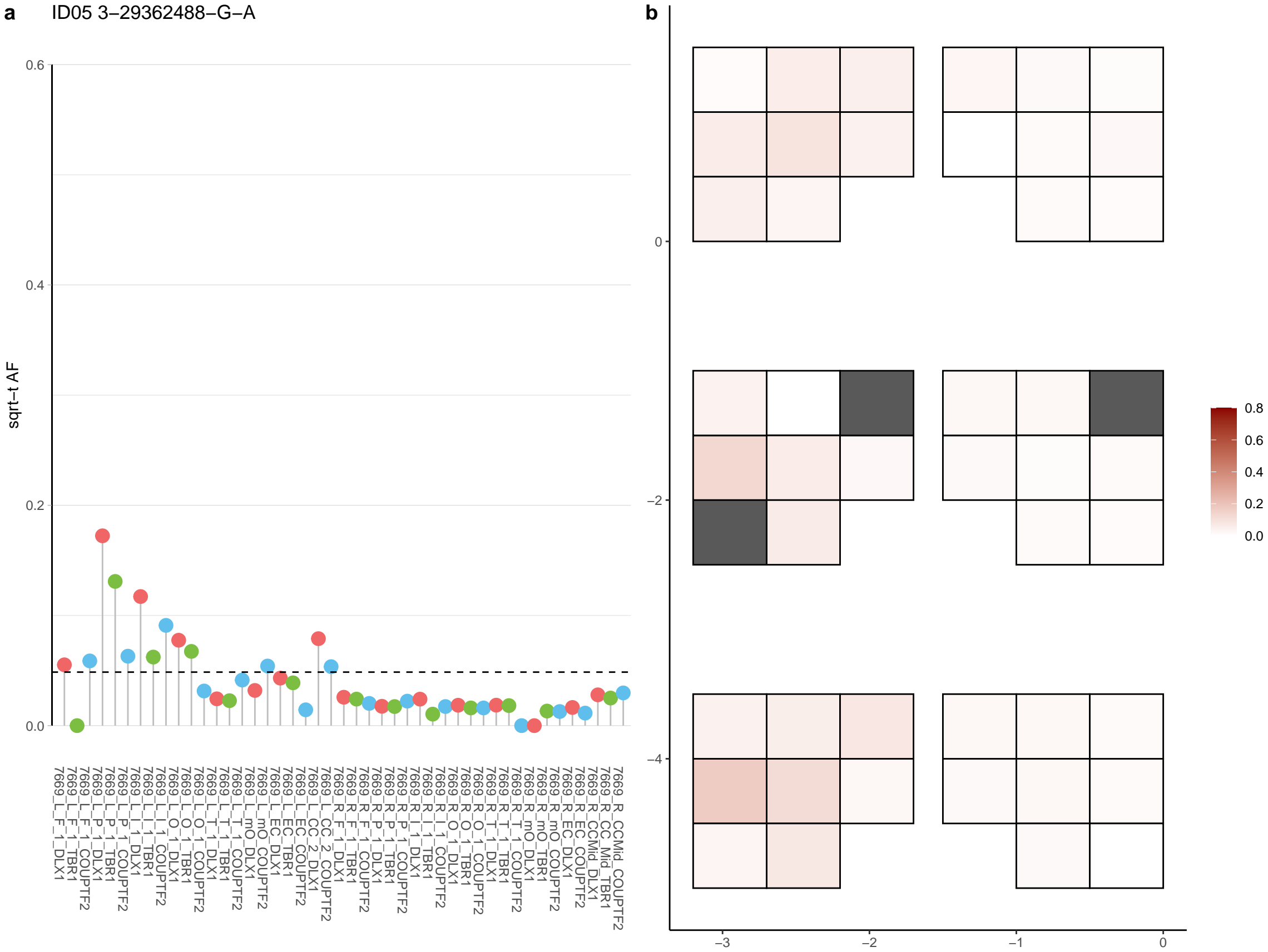

**a** ID05 3-29763350-T-C

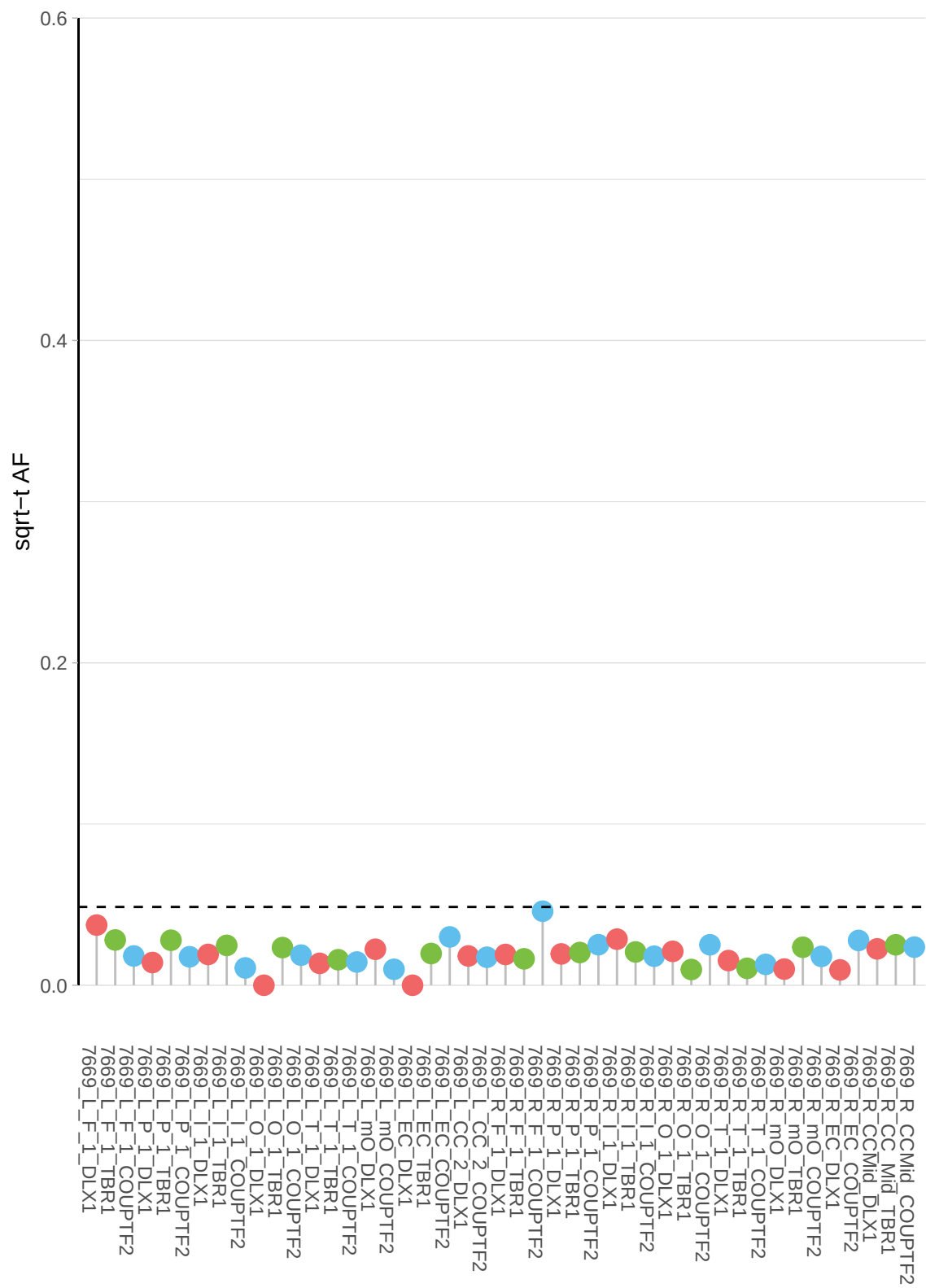

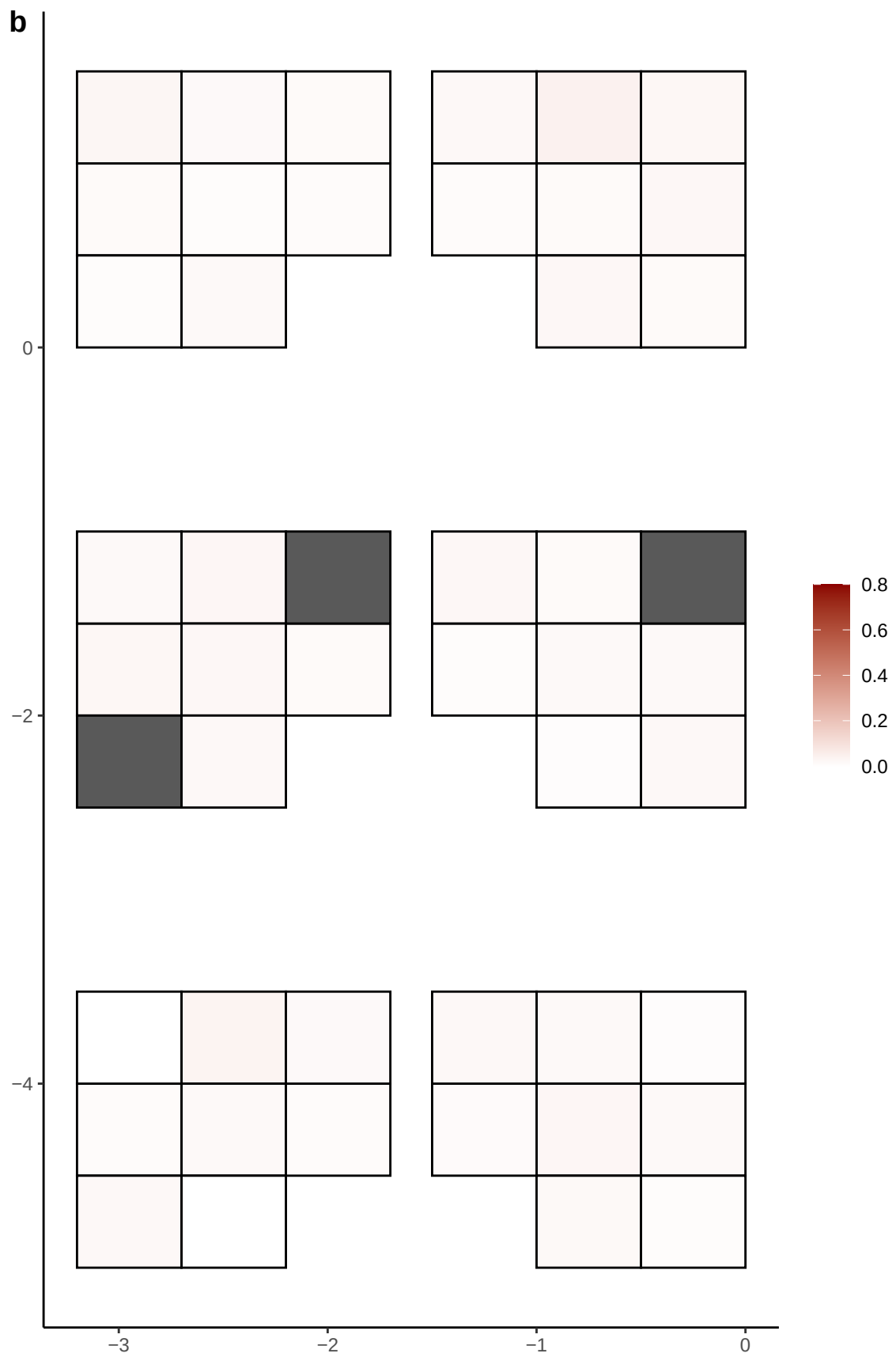

**a** ID05 3-32863852-G-A

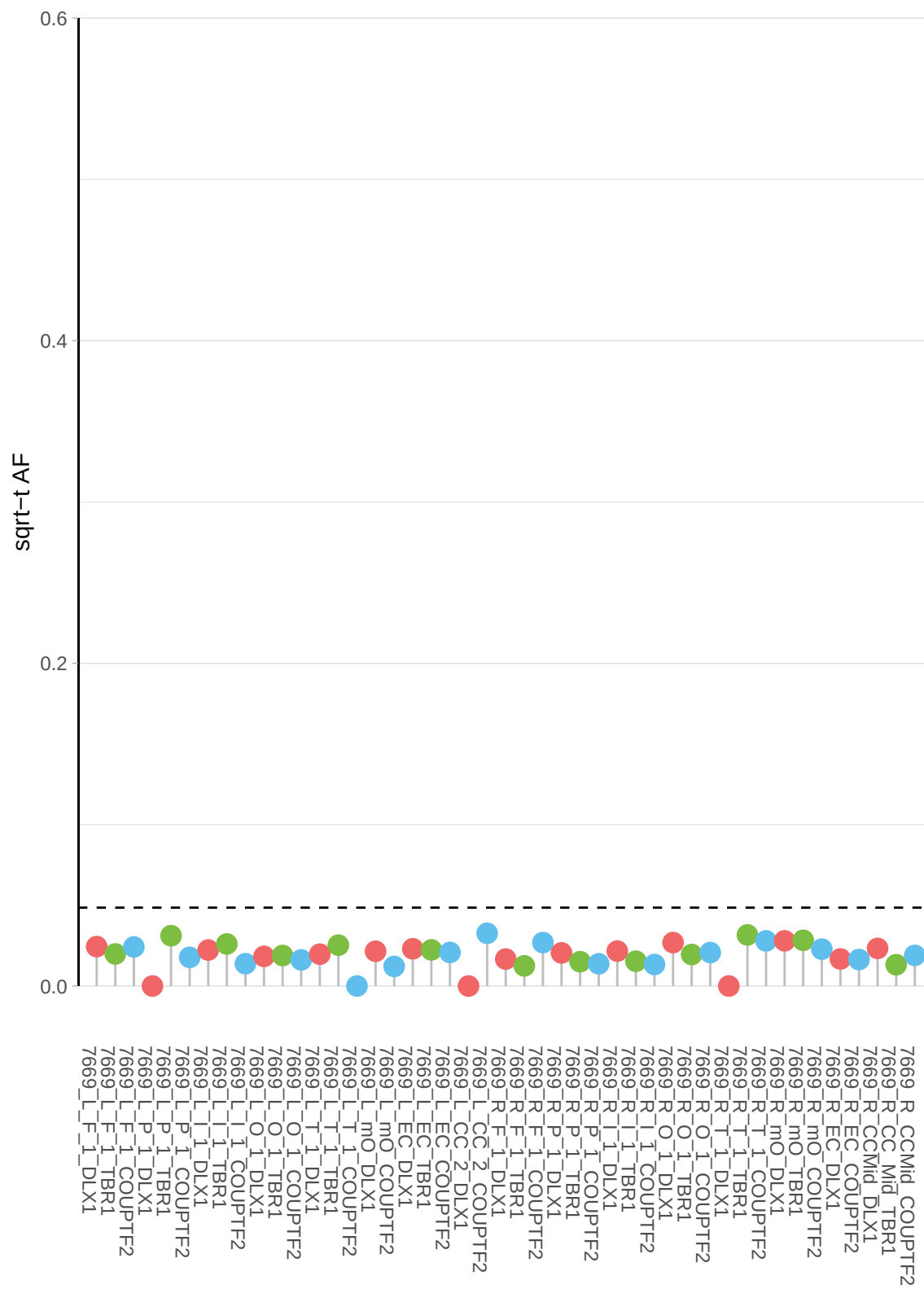

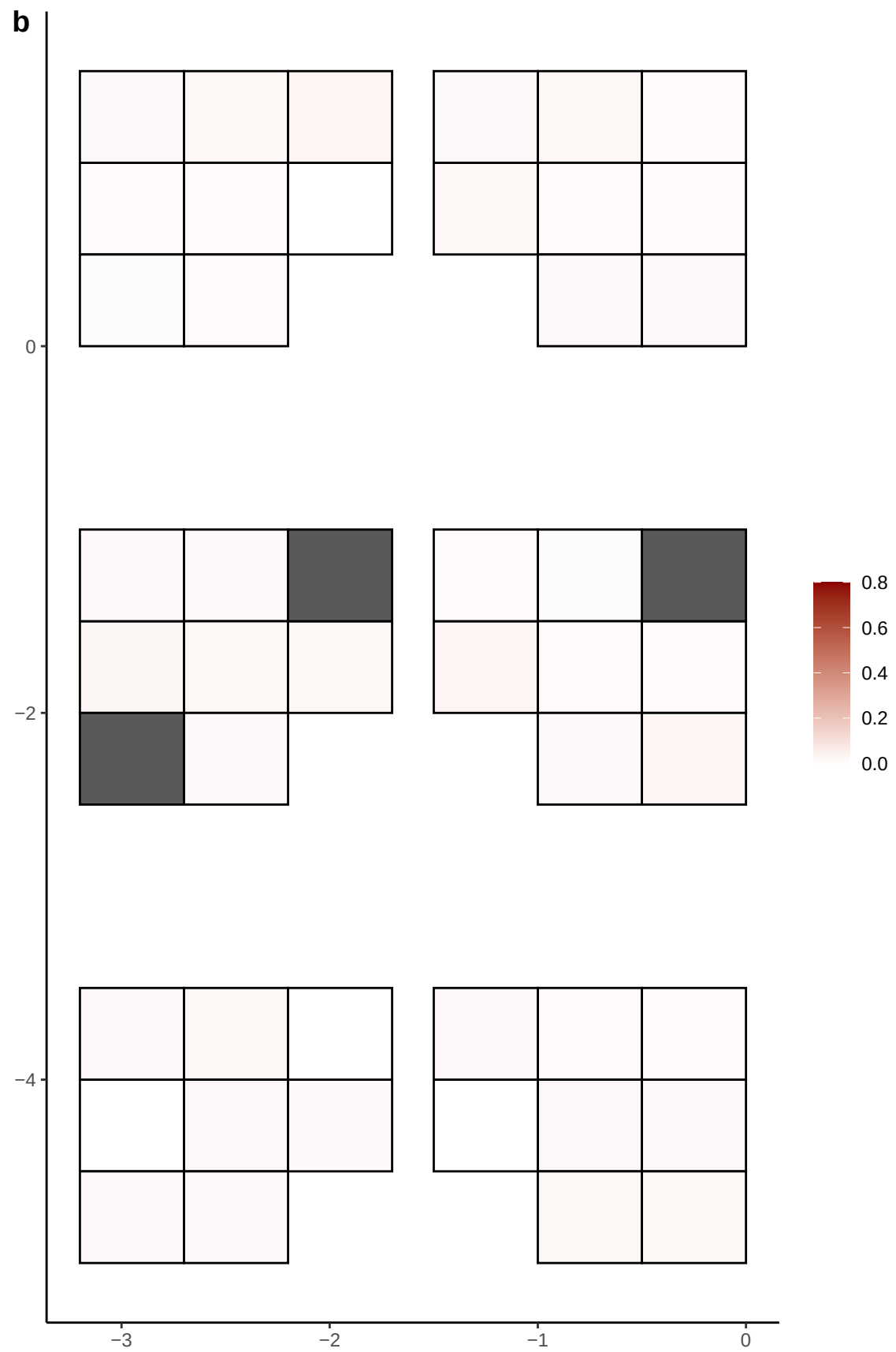

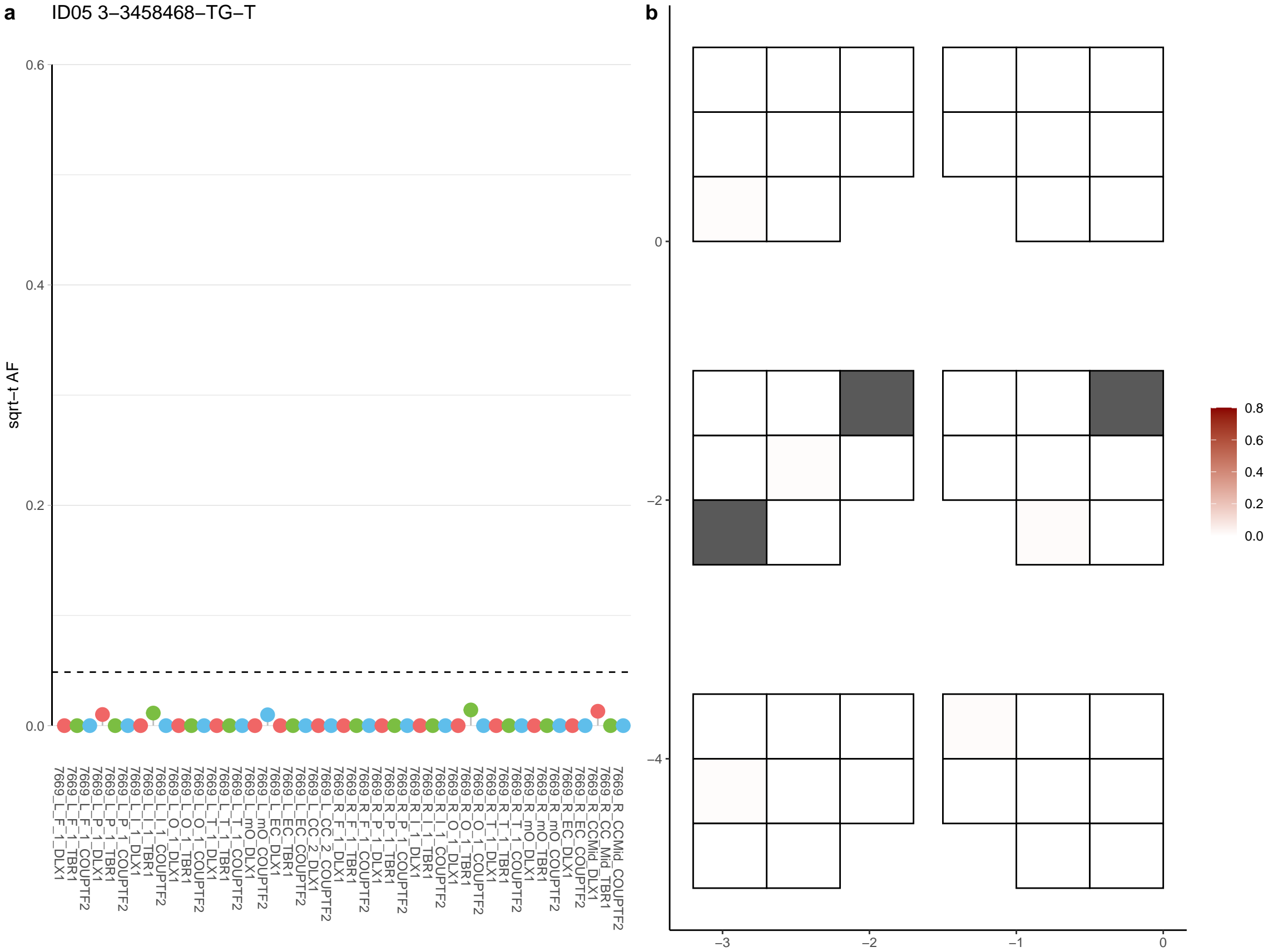

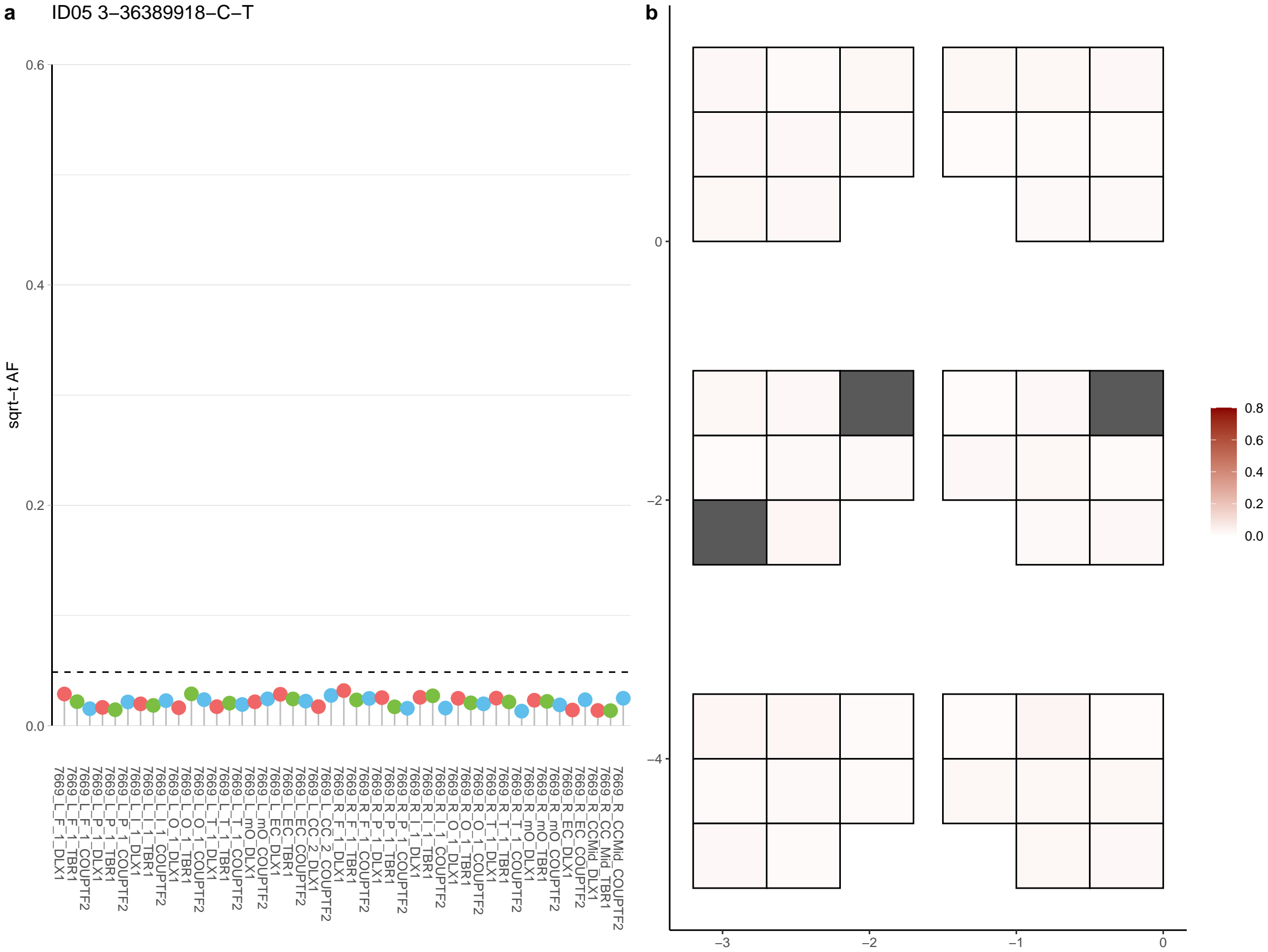

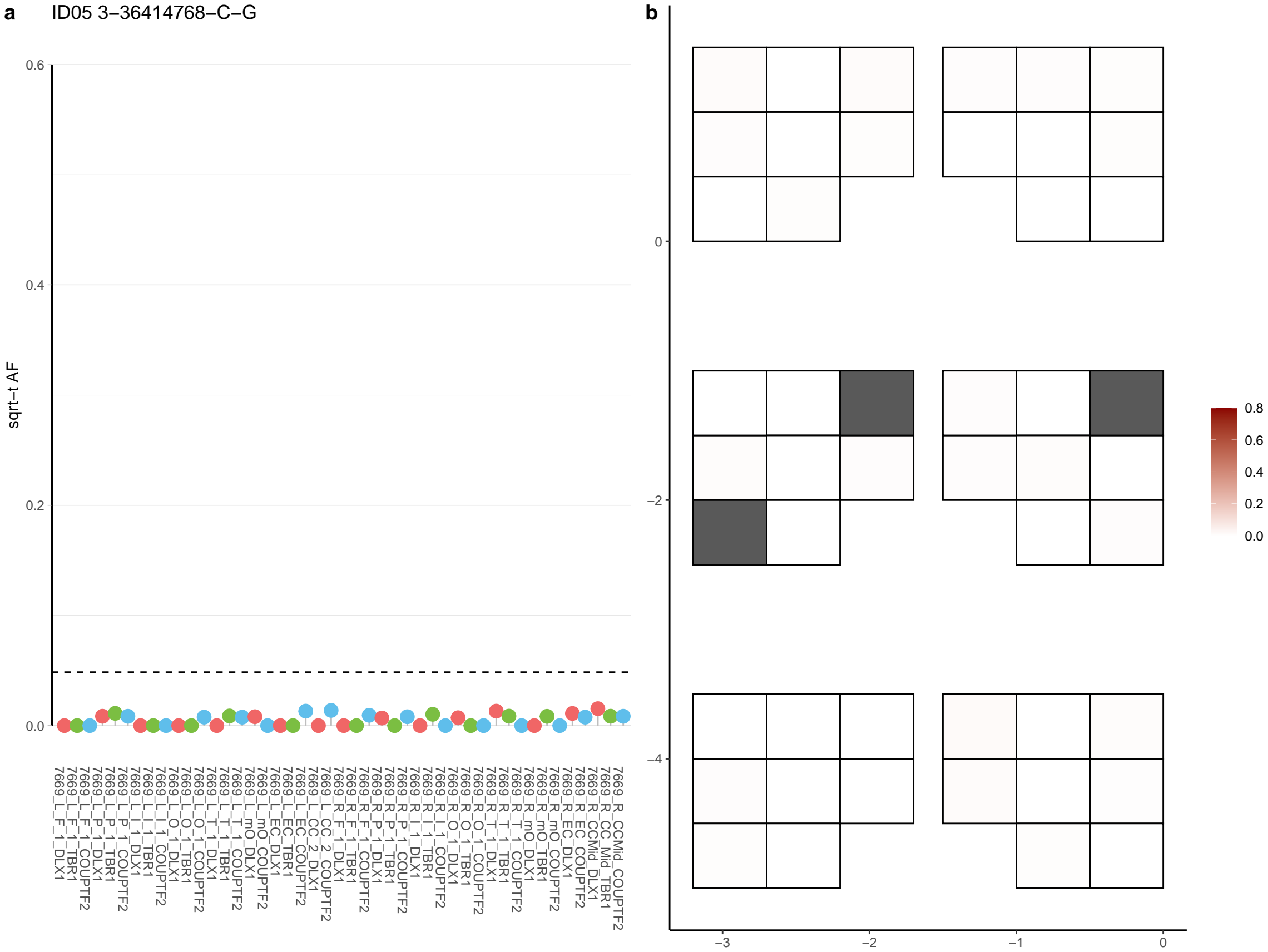

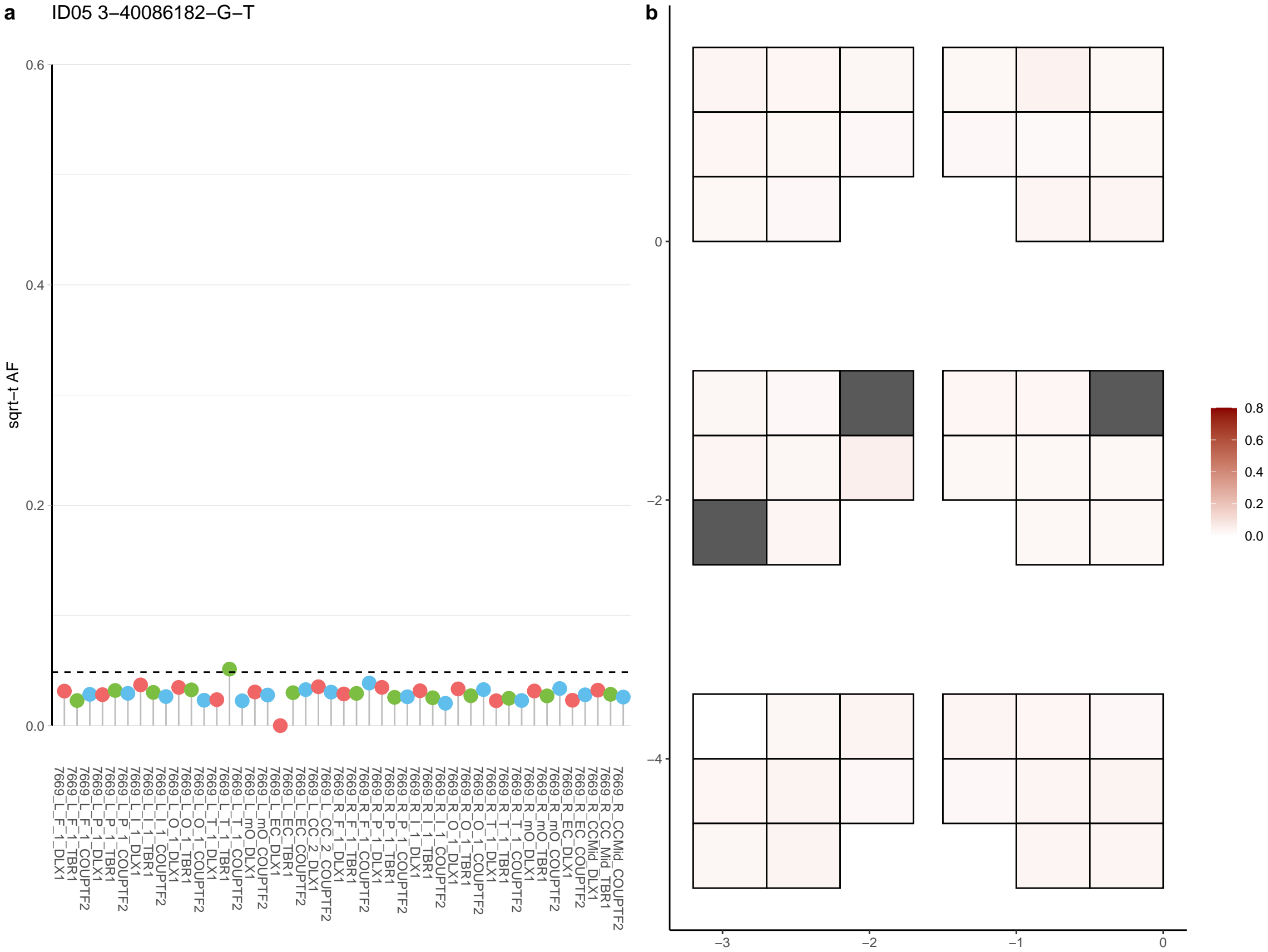

**a** ID05 3-44078495-G-A

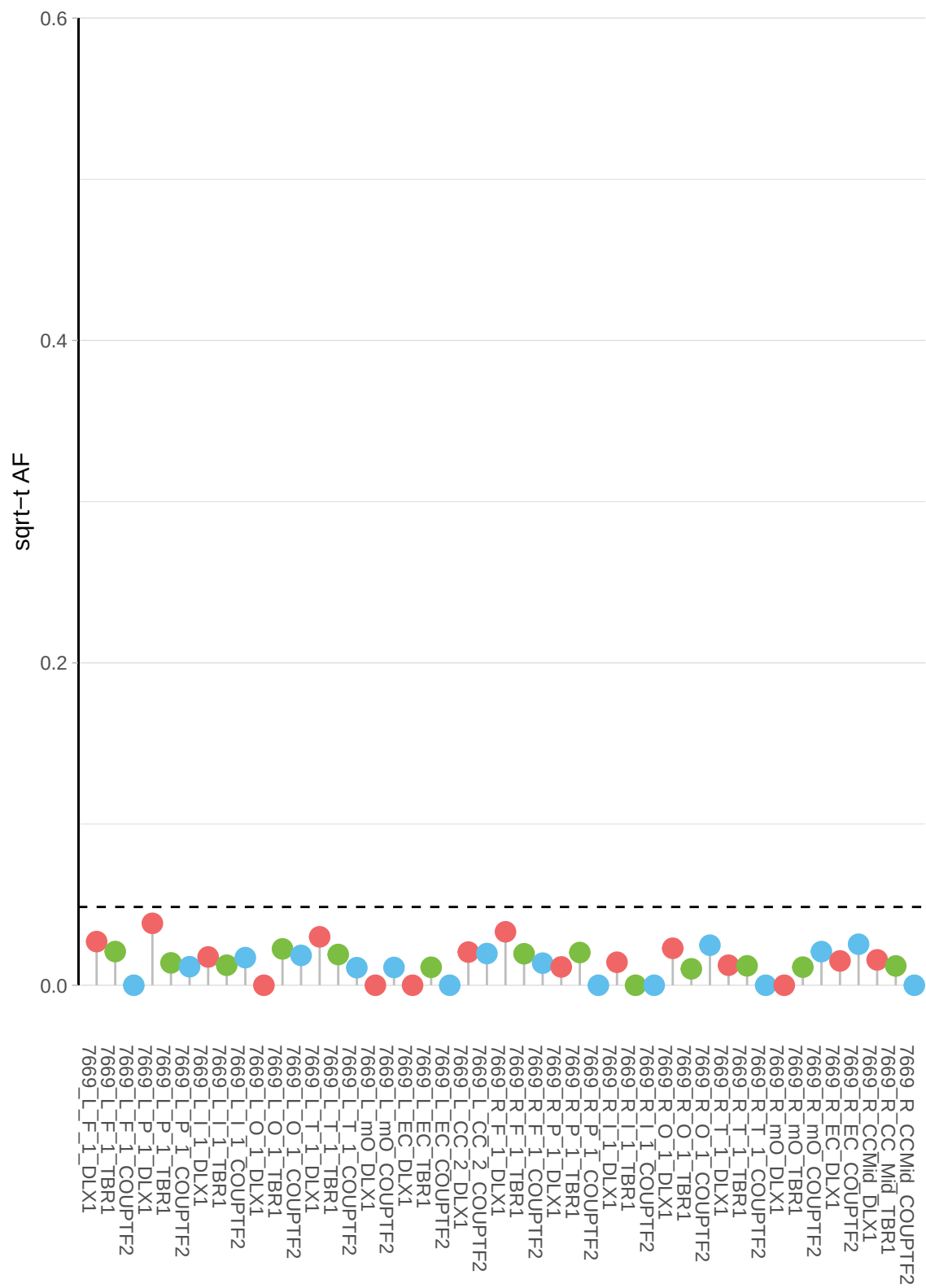

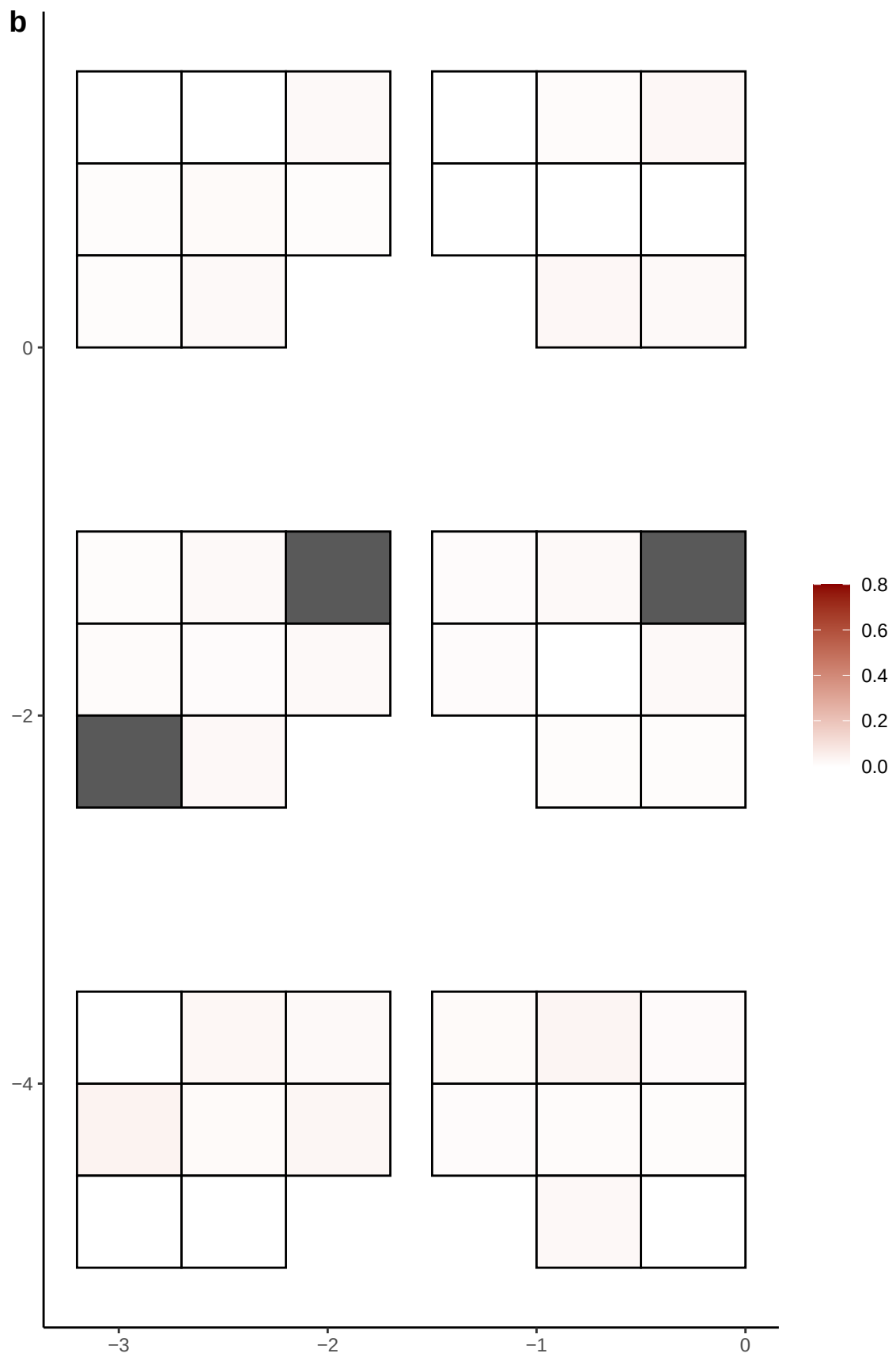

**a** ID05 3-492626-A-T

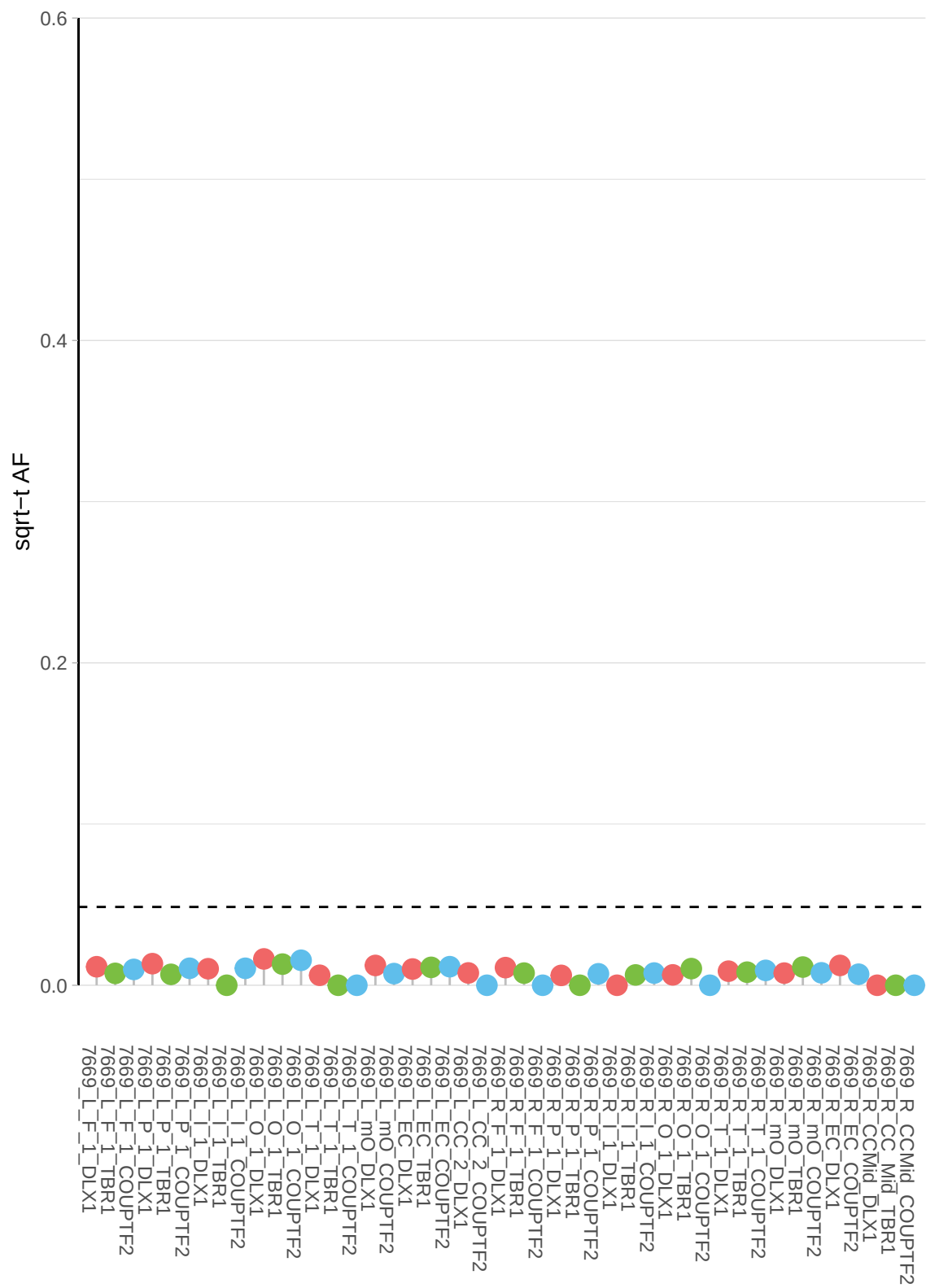

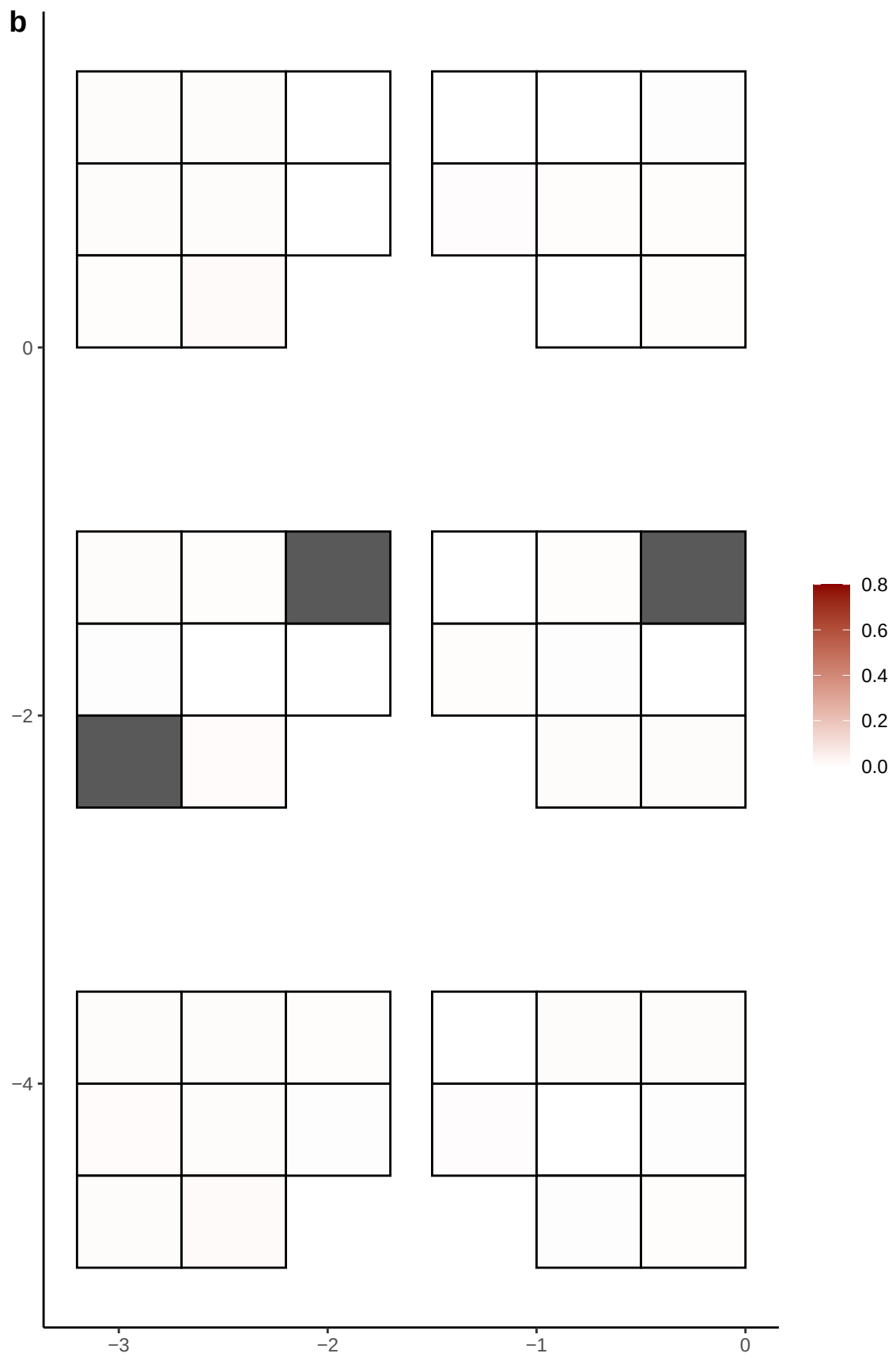

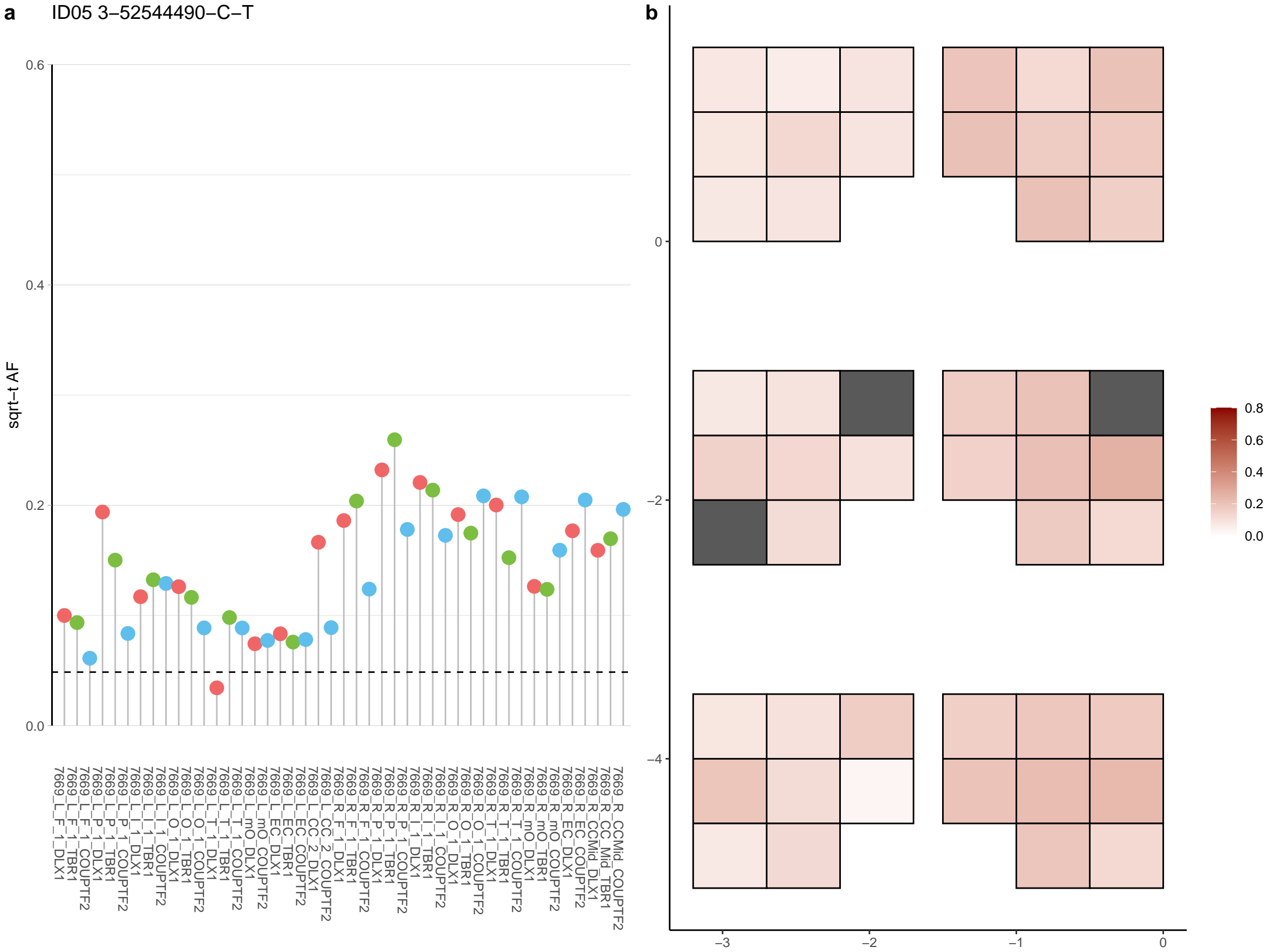

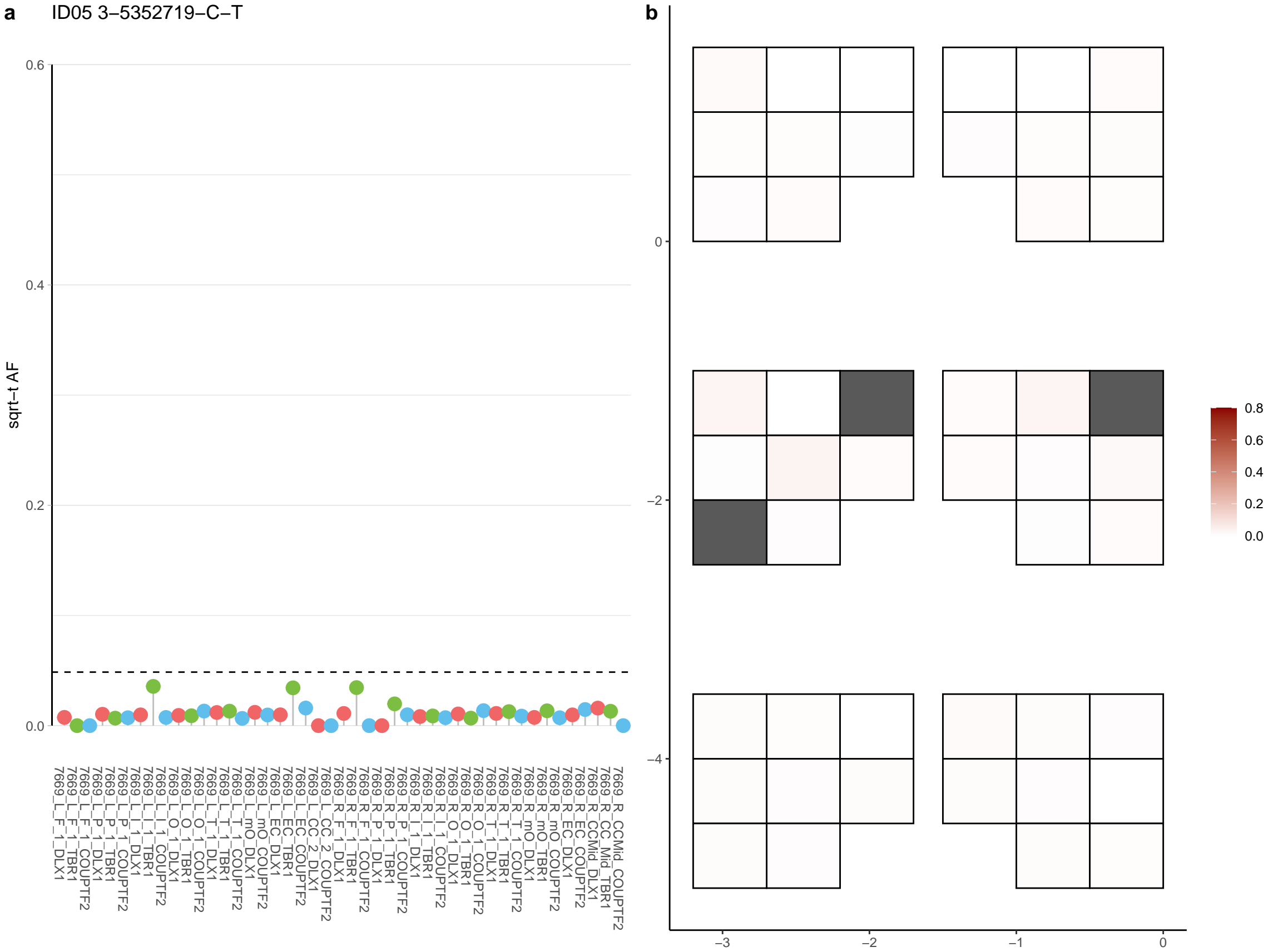

**a** ID05 3-54233044-C-T

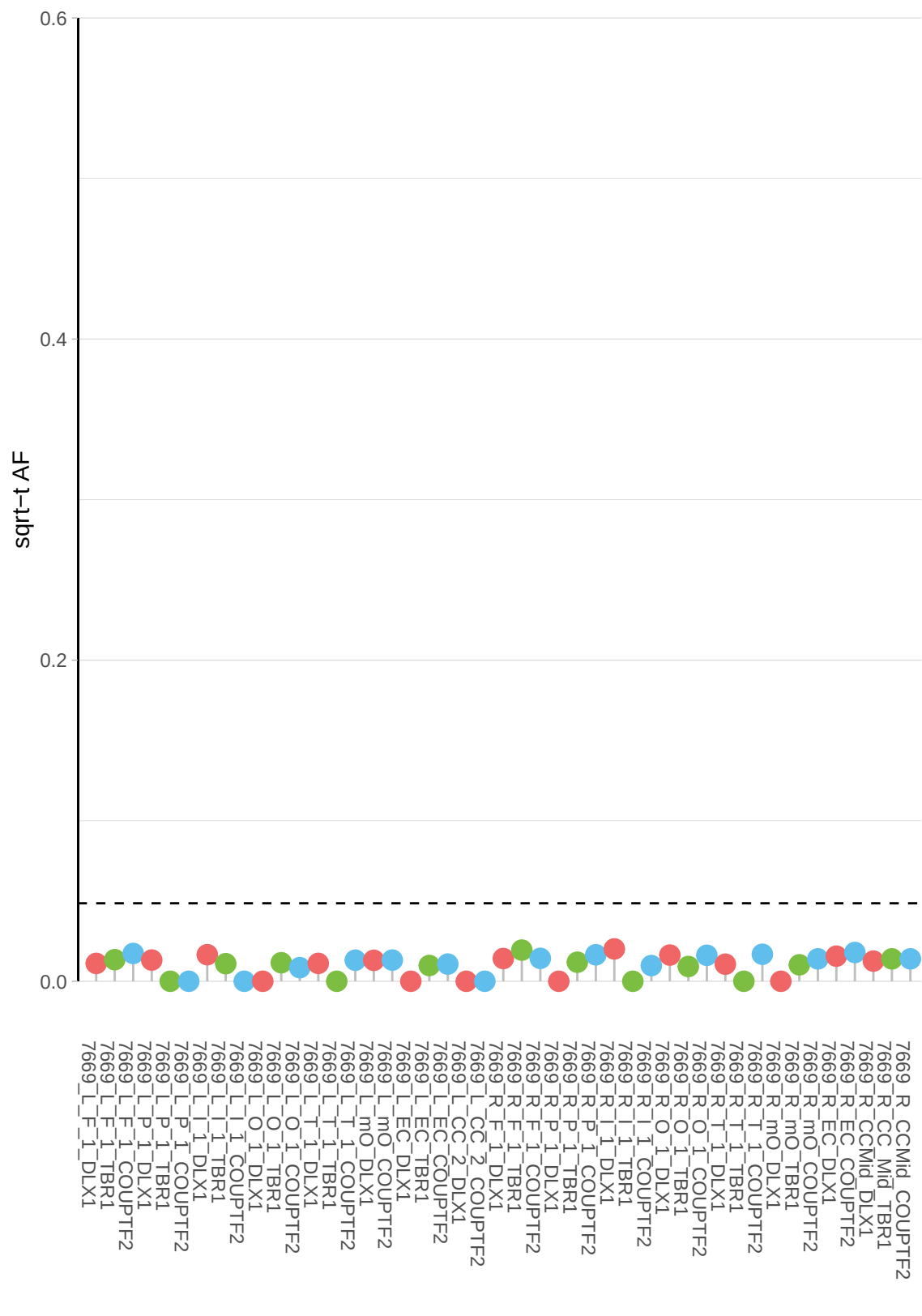

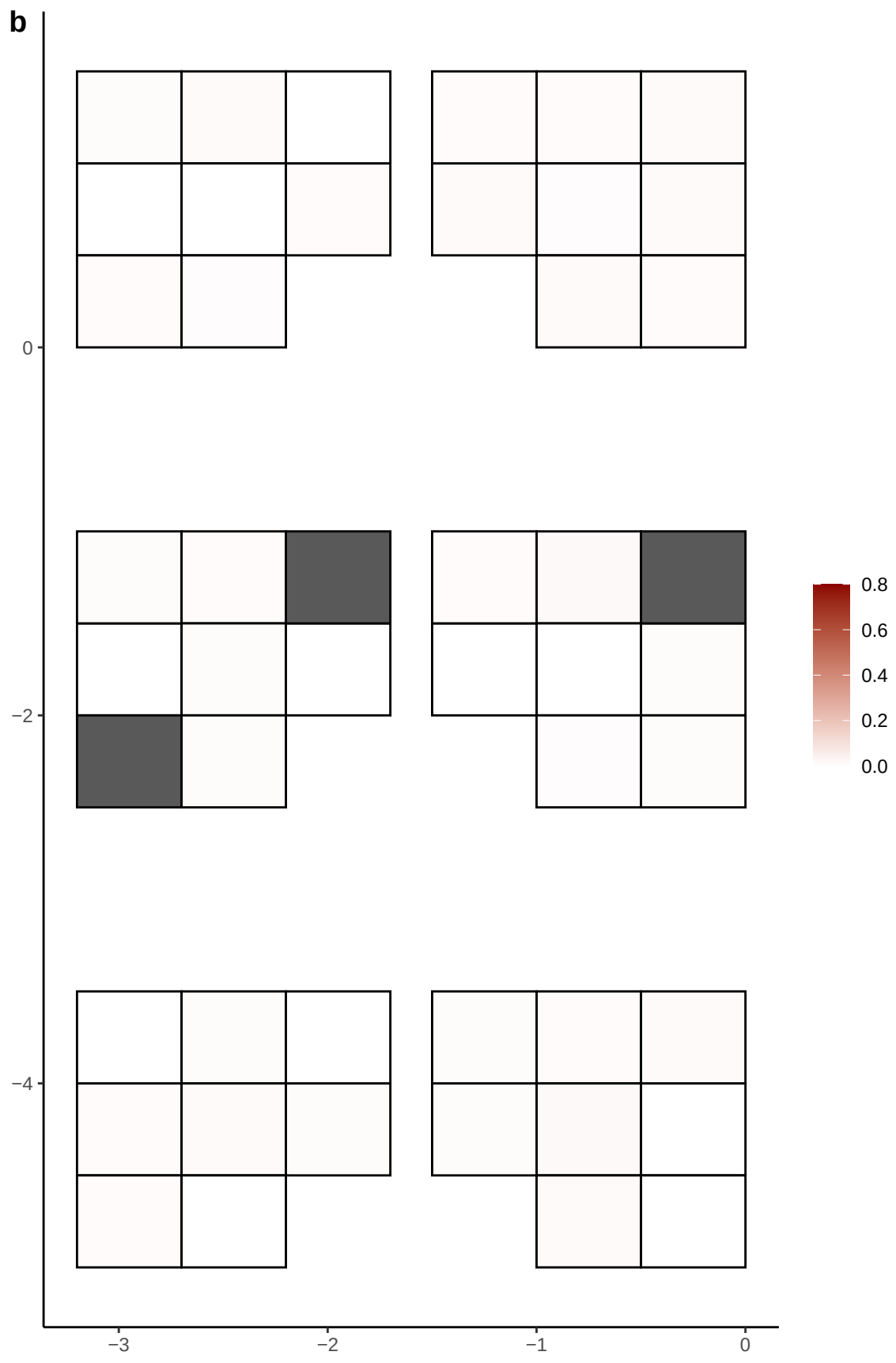

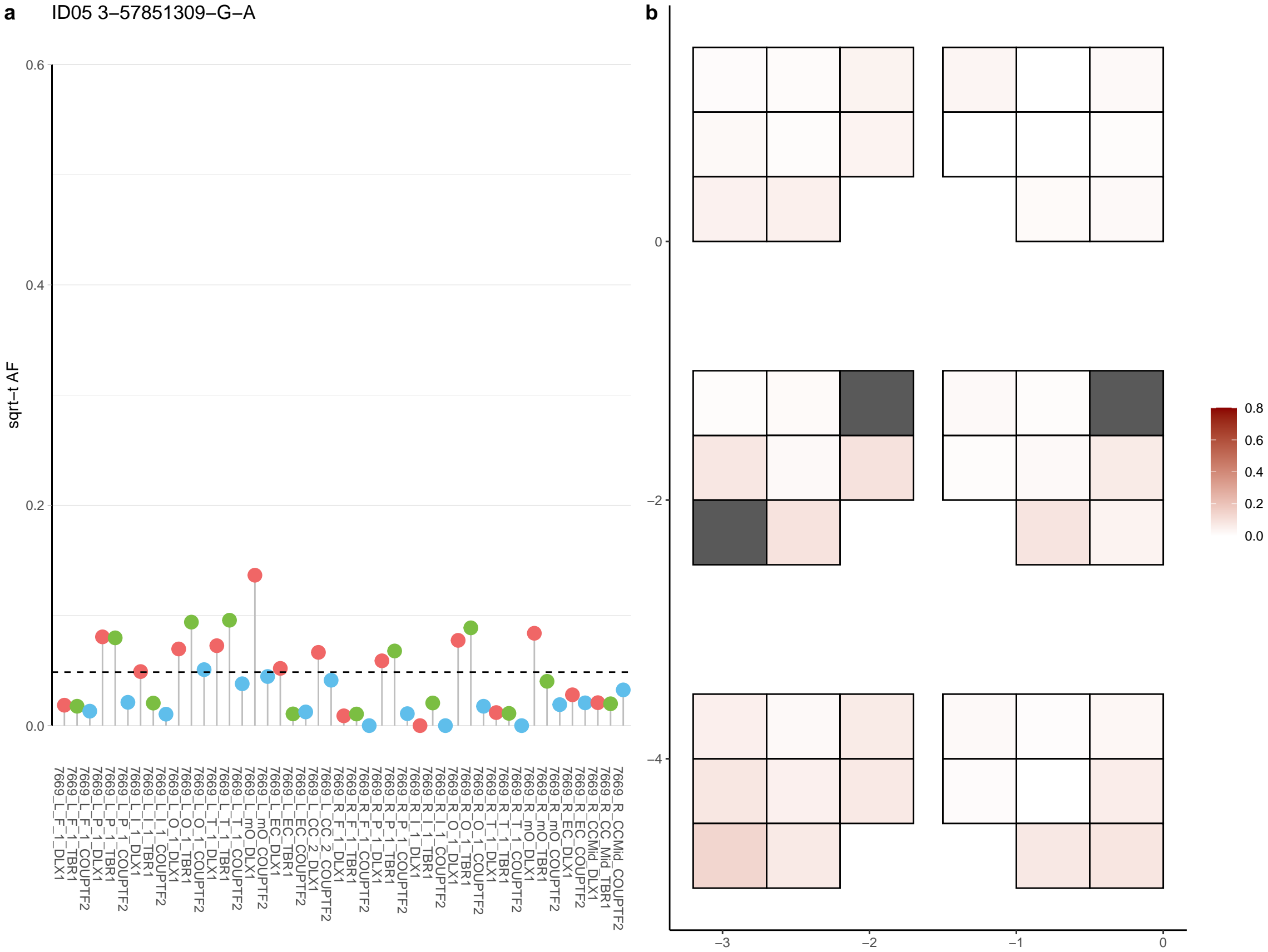

**a** ID05 3-58915984-C-A

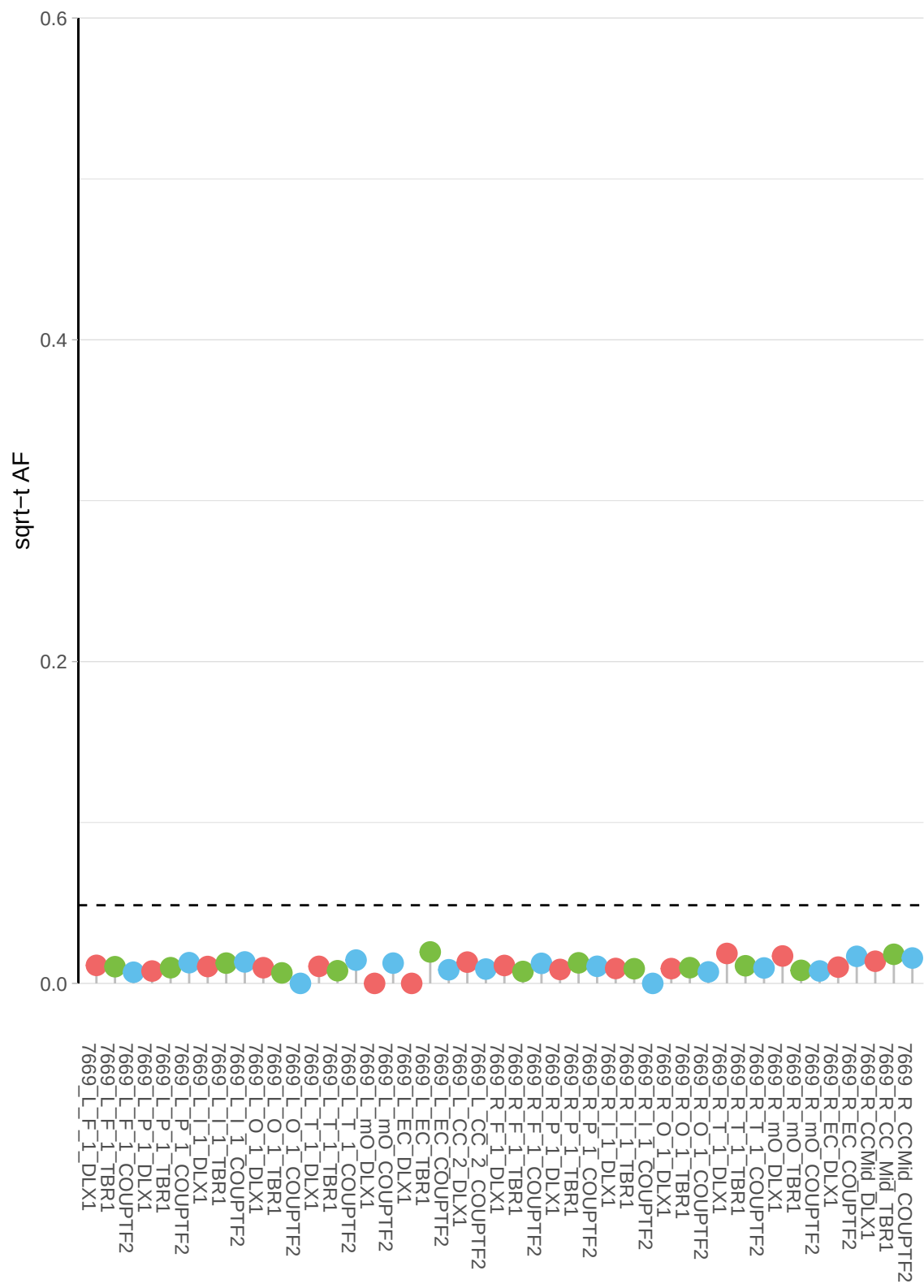

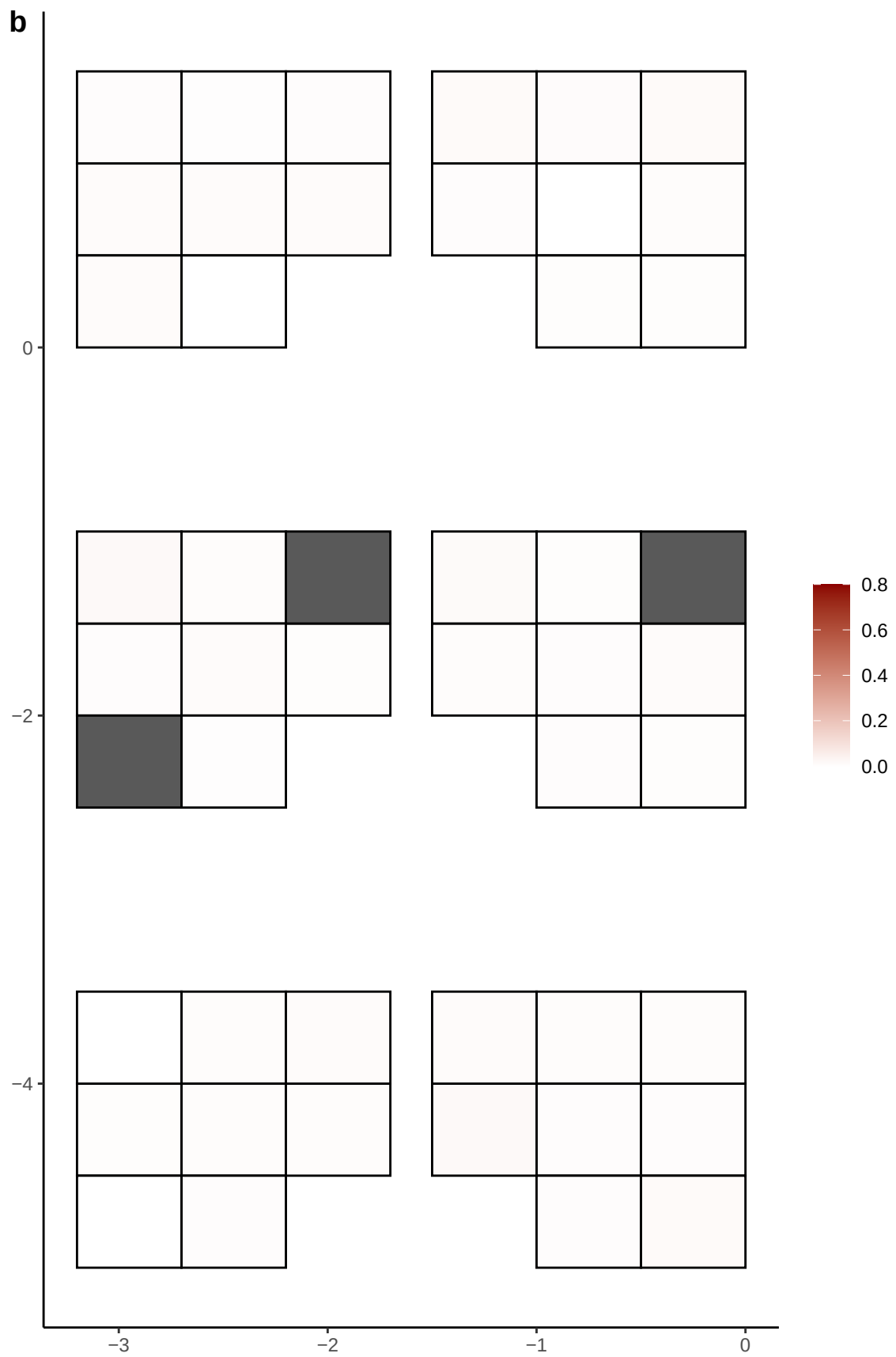

**a** ID05 3-61853127-C-T

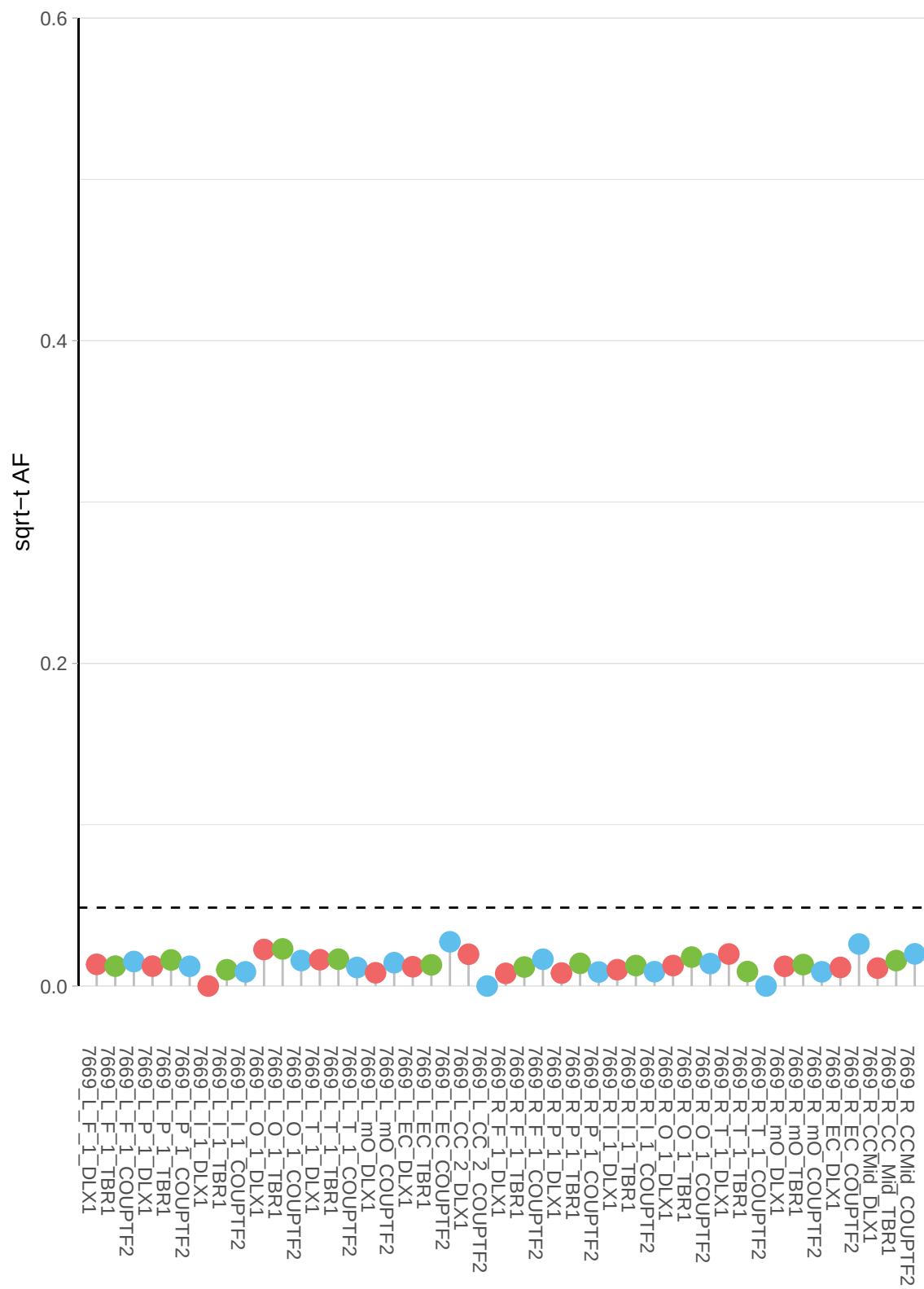

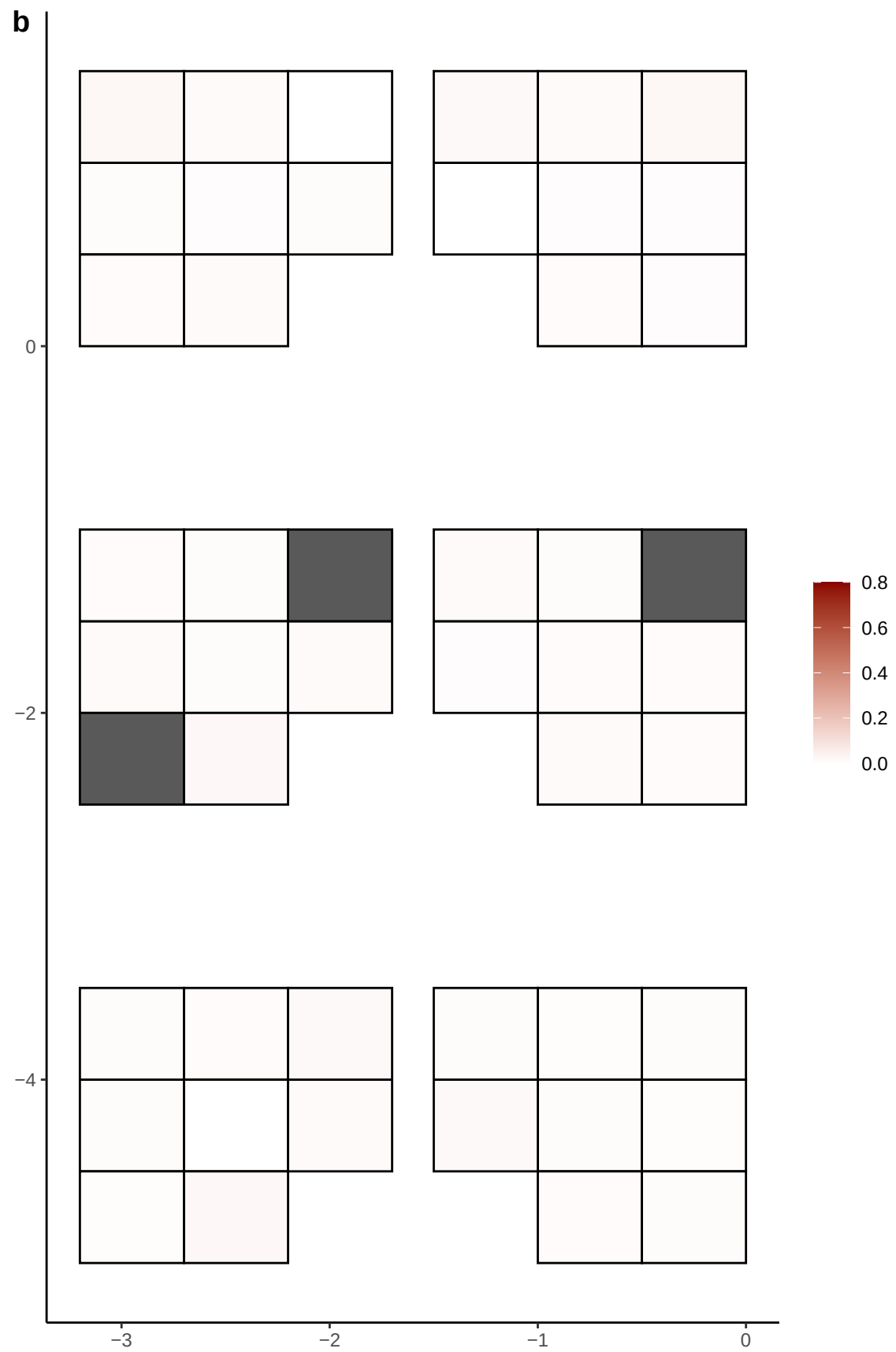

**a** ID05 3-69001163-C-T

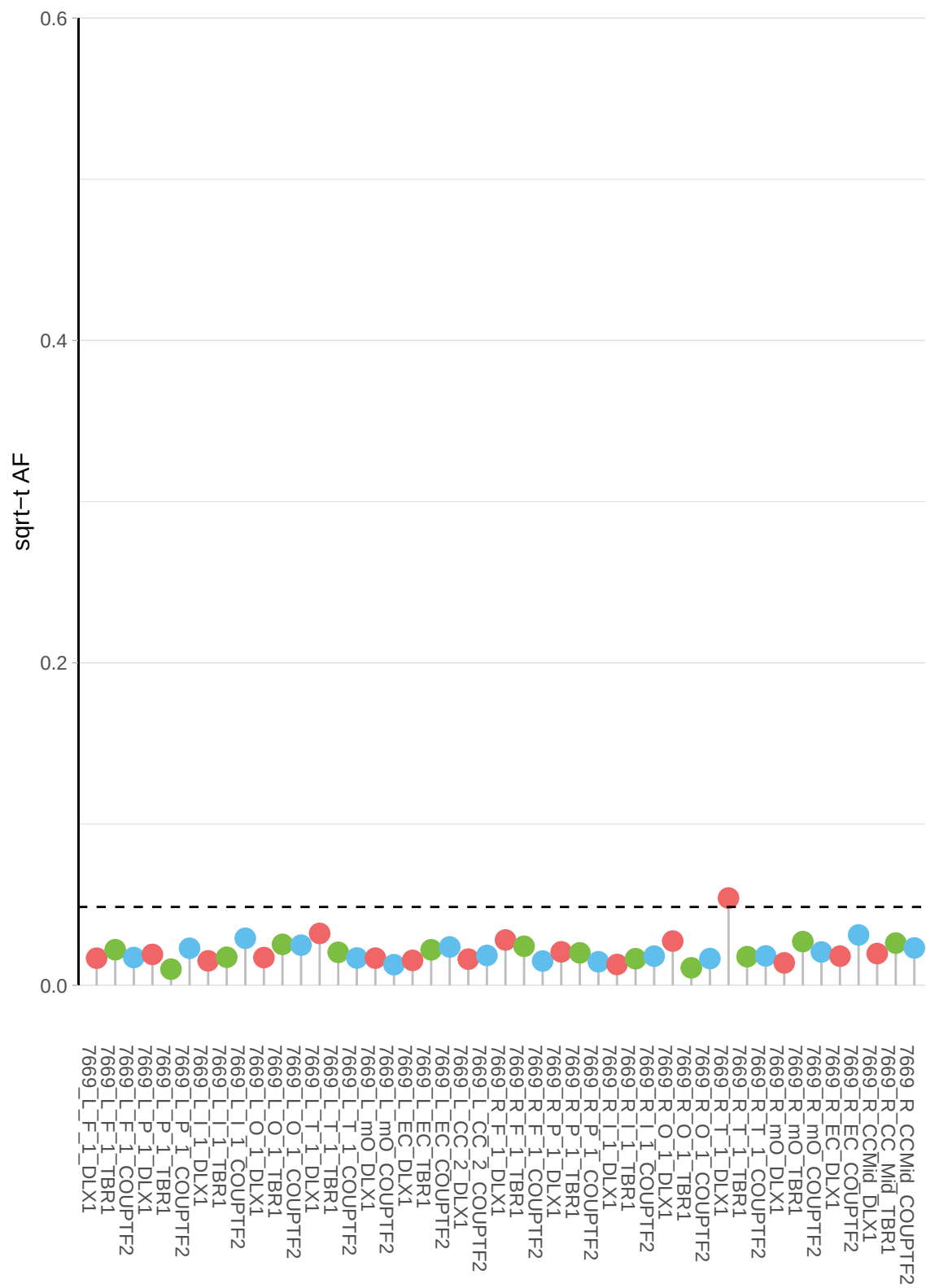

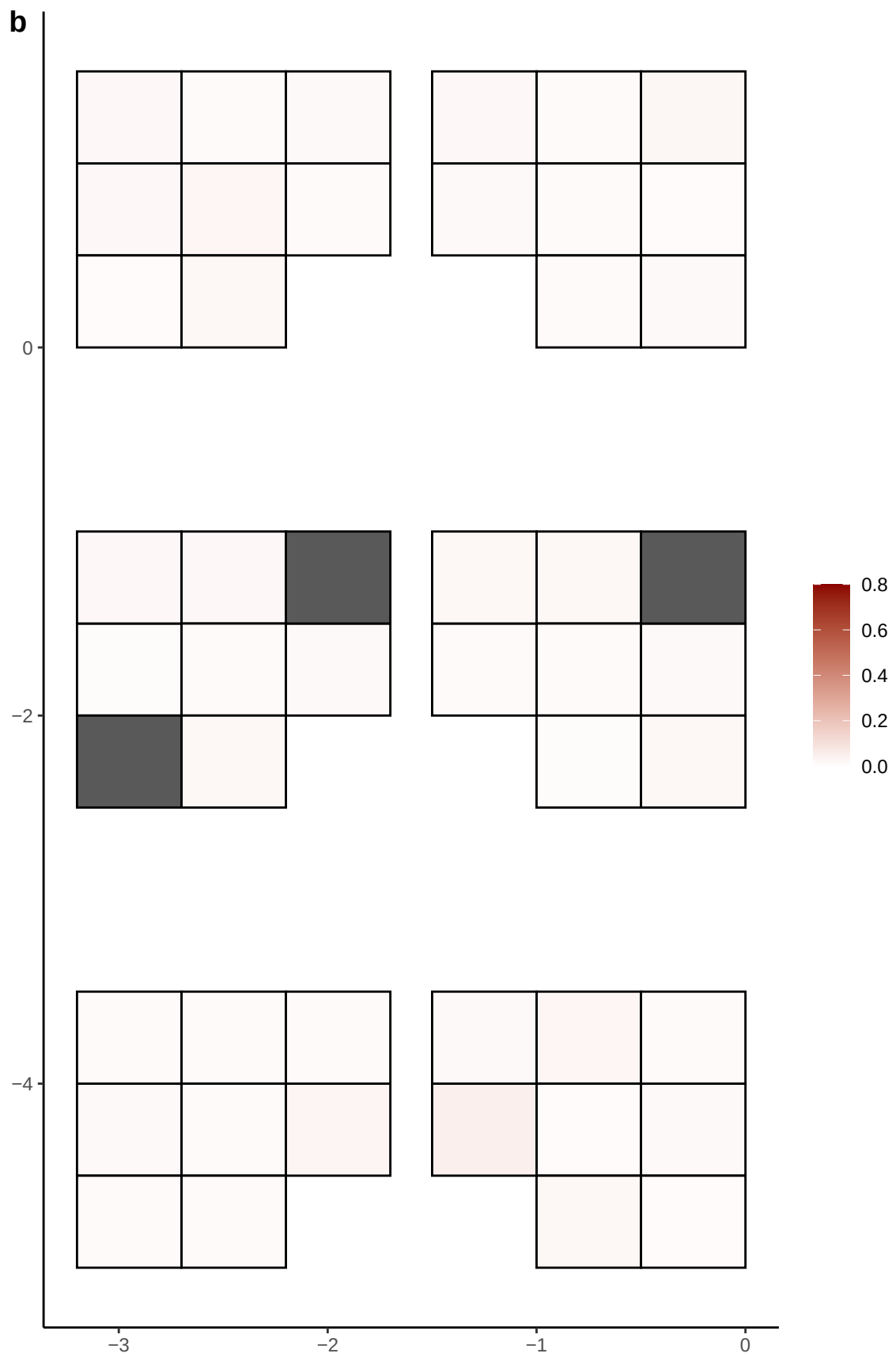

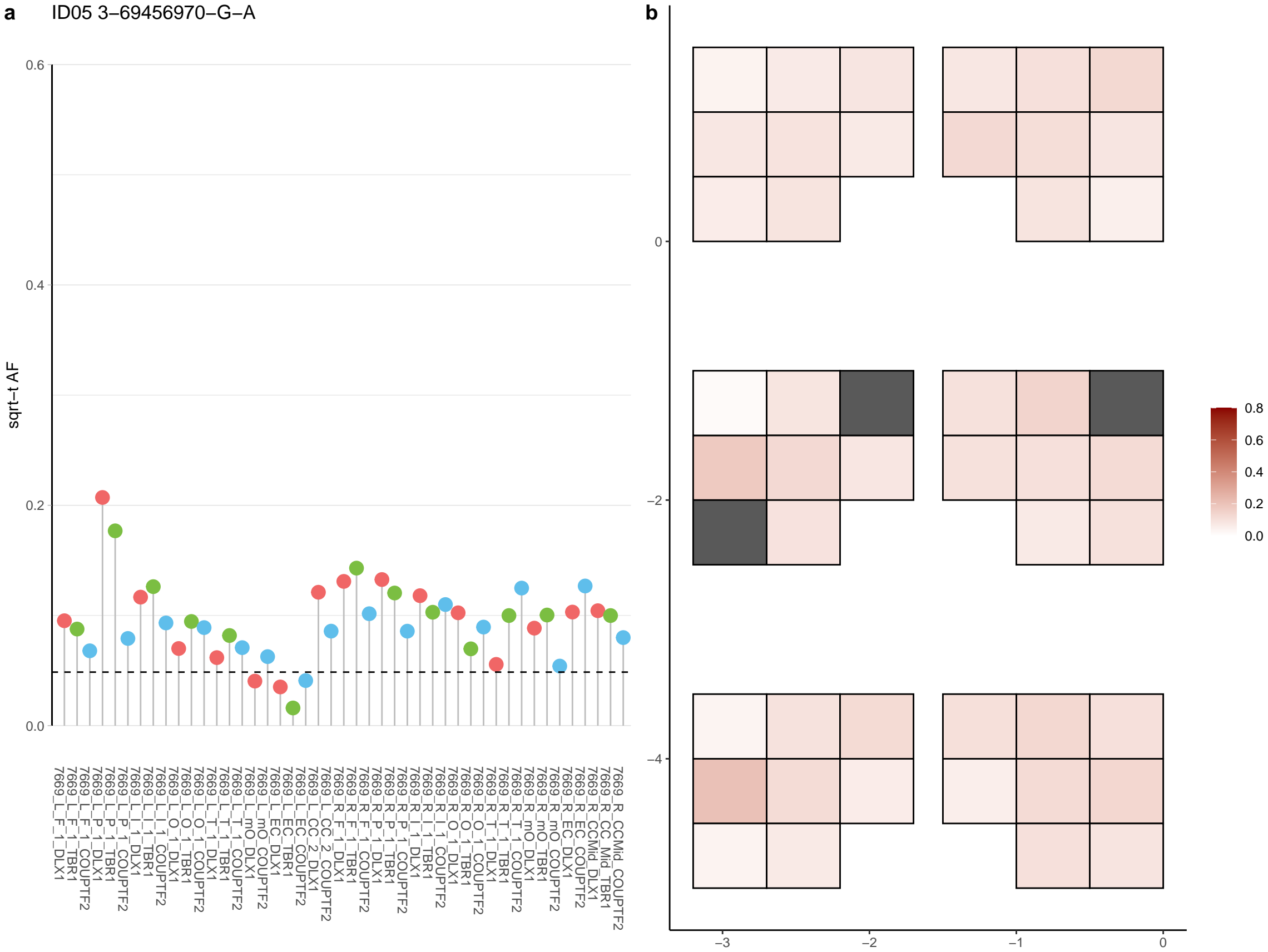

**a** ID05 3-78738978-T-C

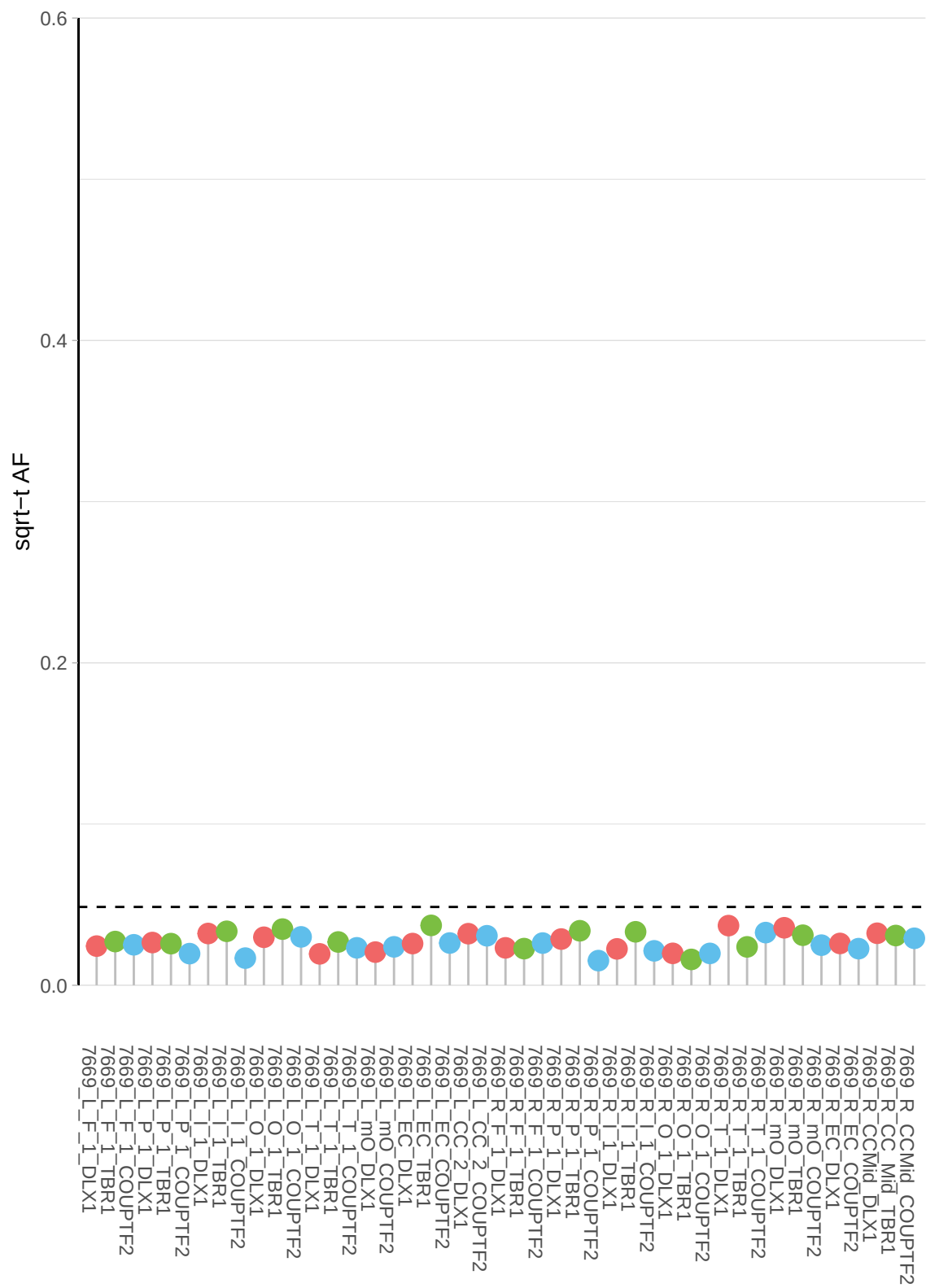

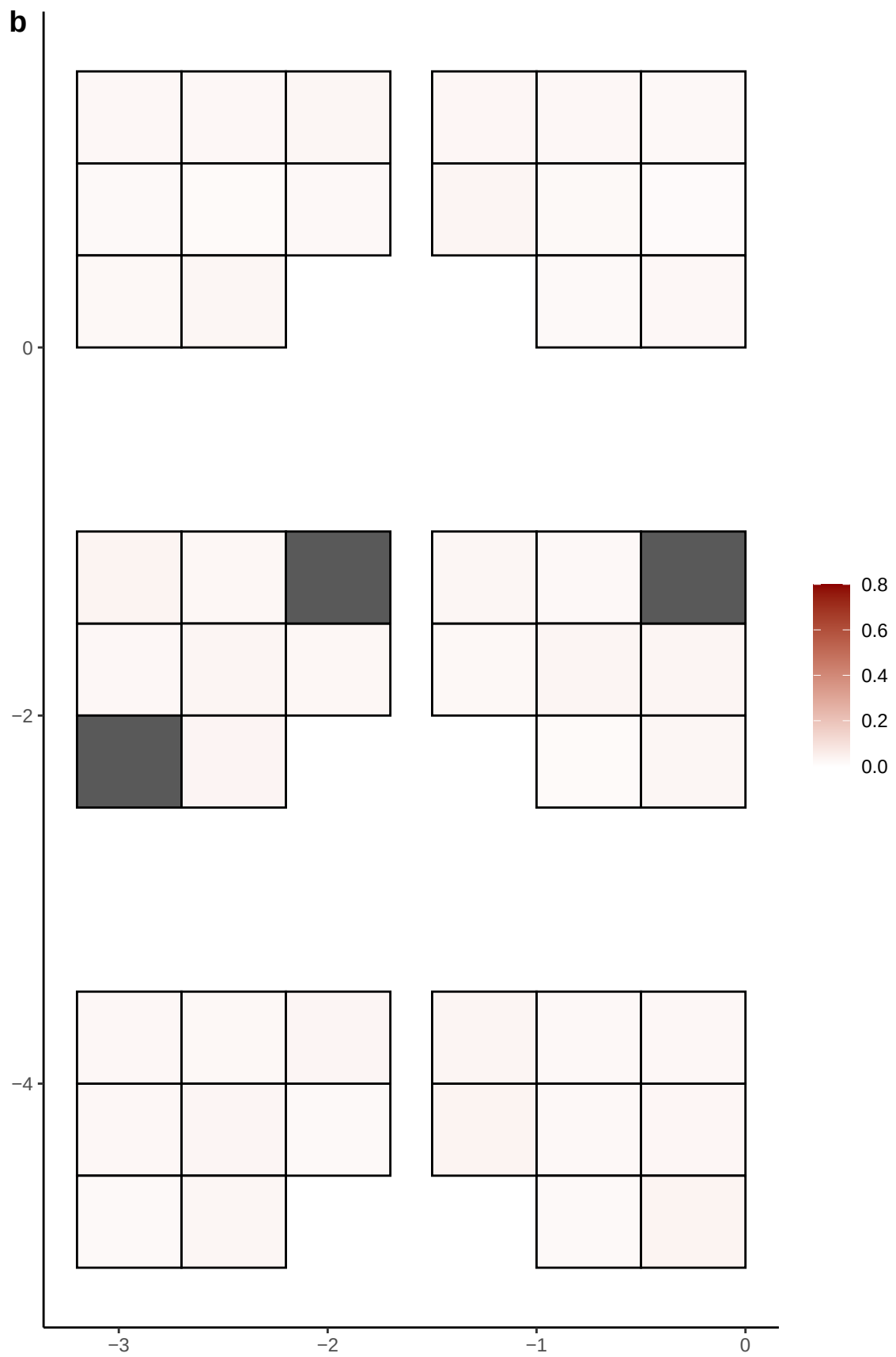

**a** ID05 3-79357944-G-C

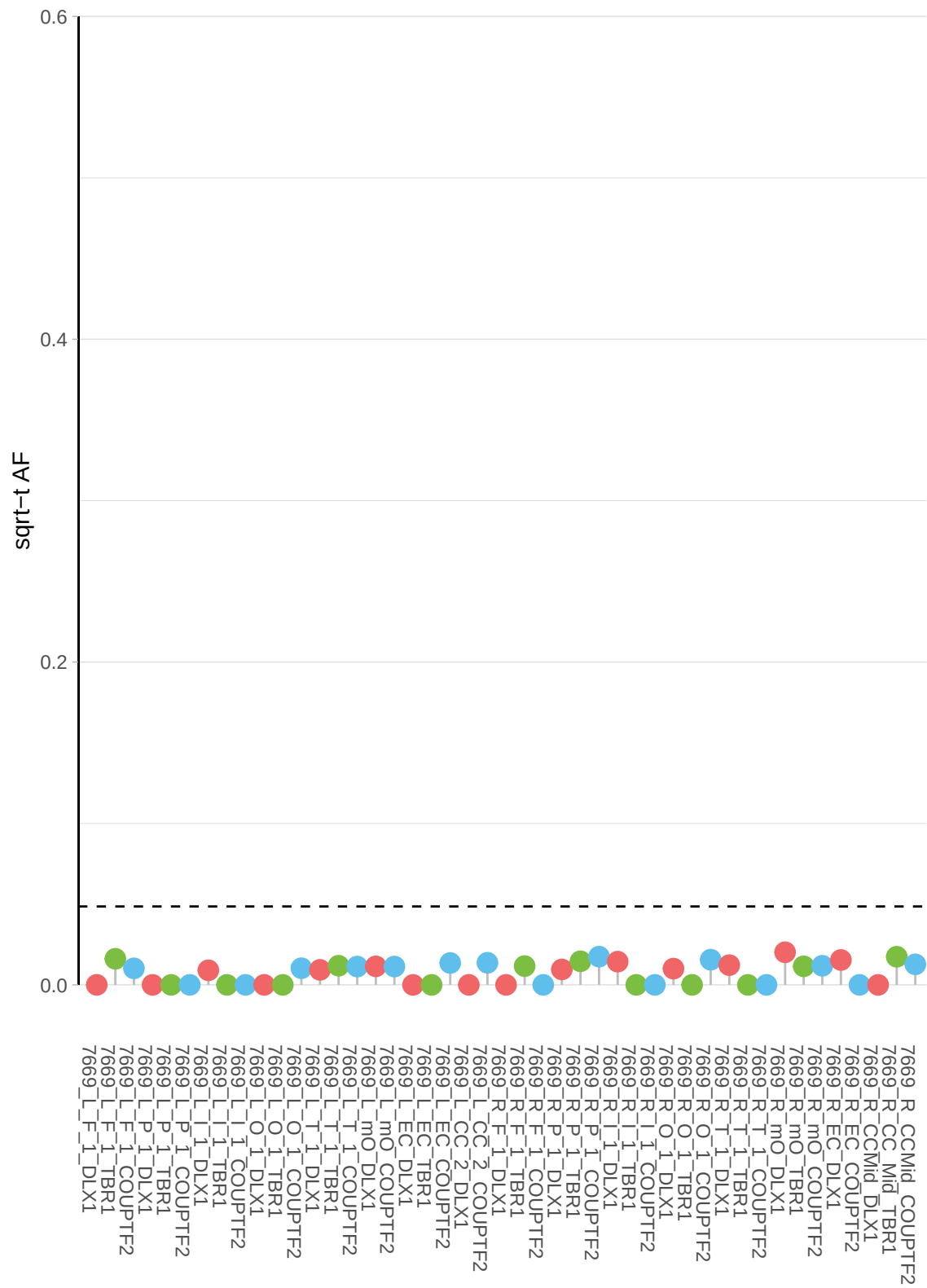

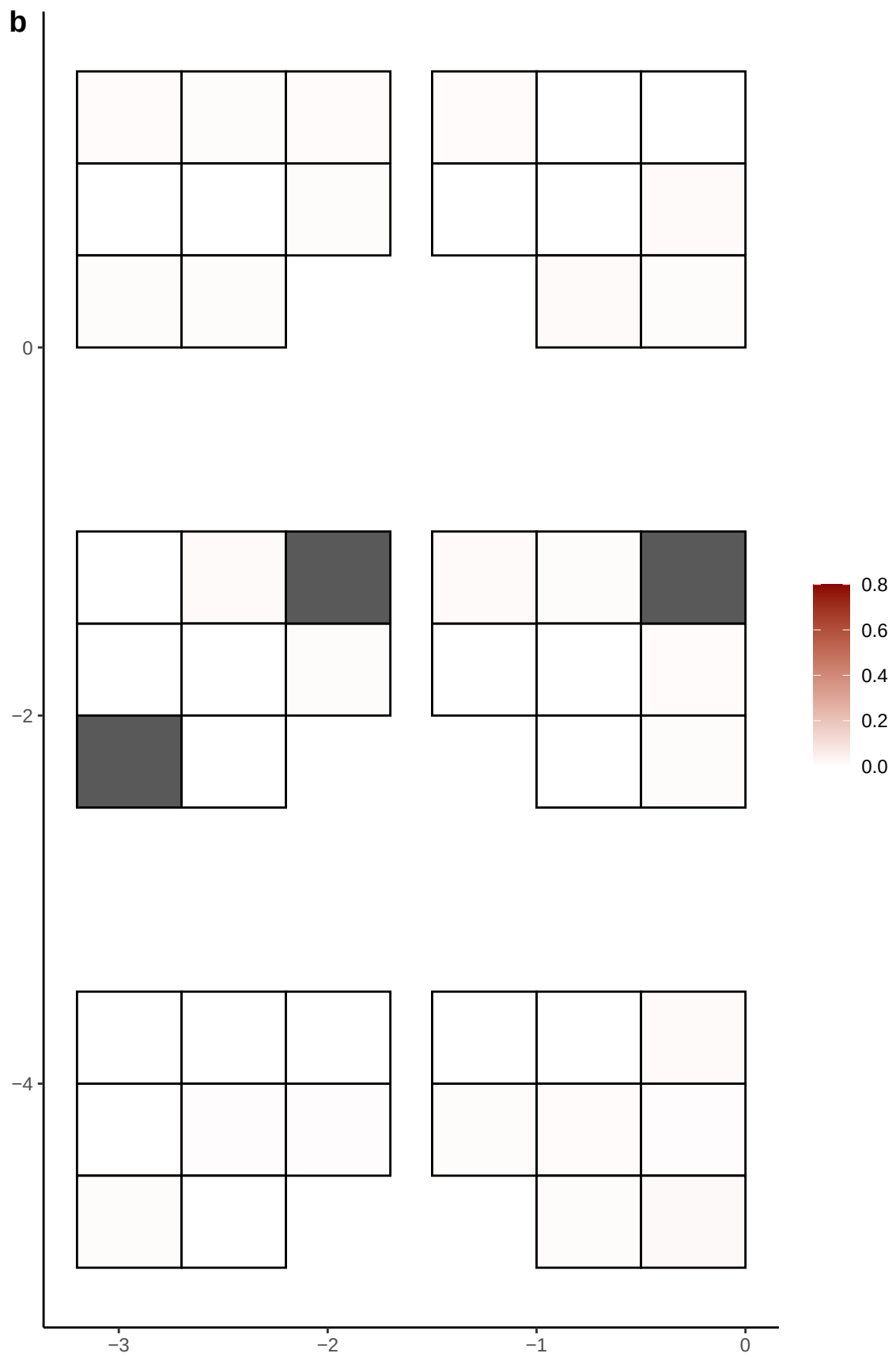

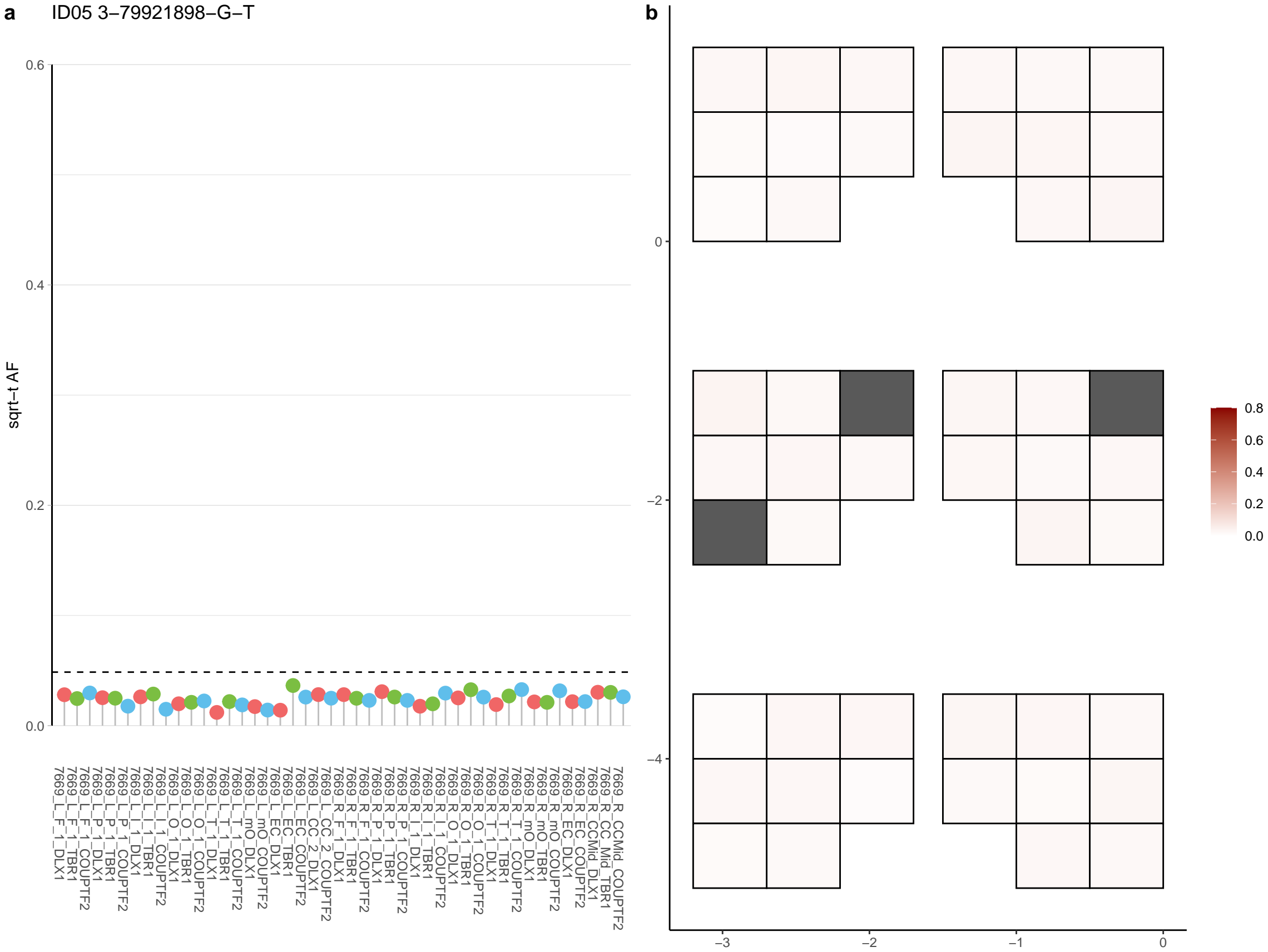

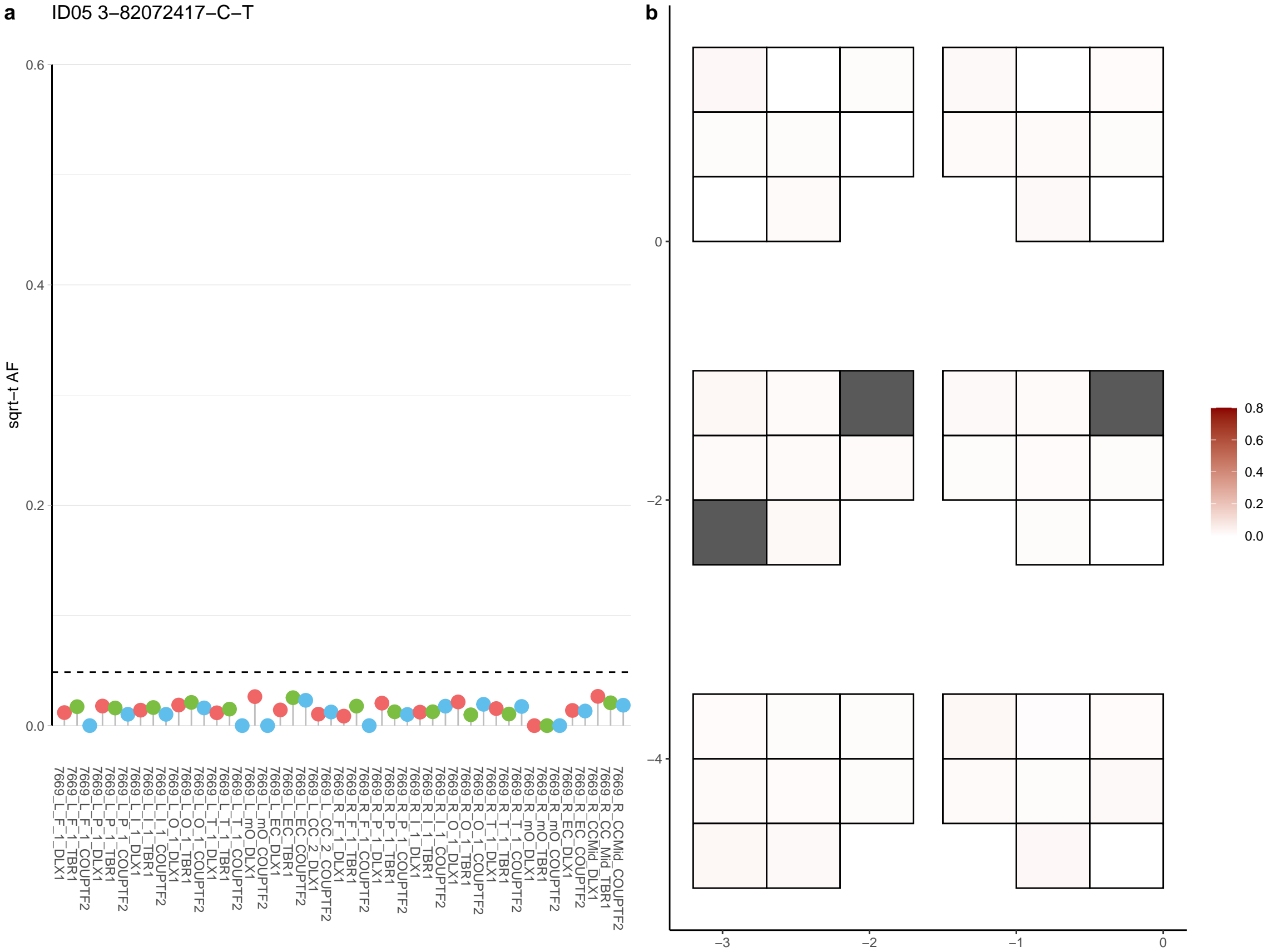

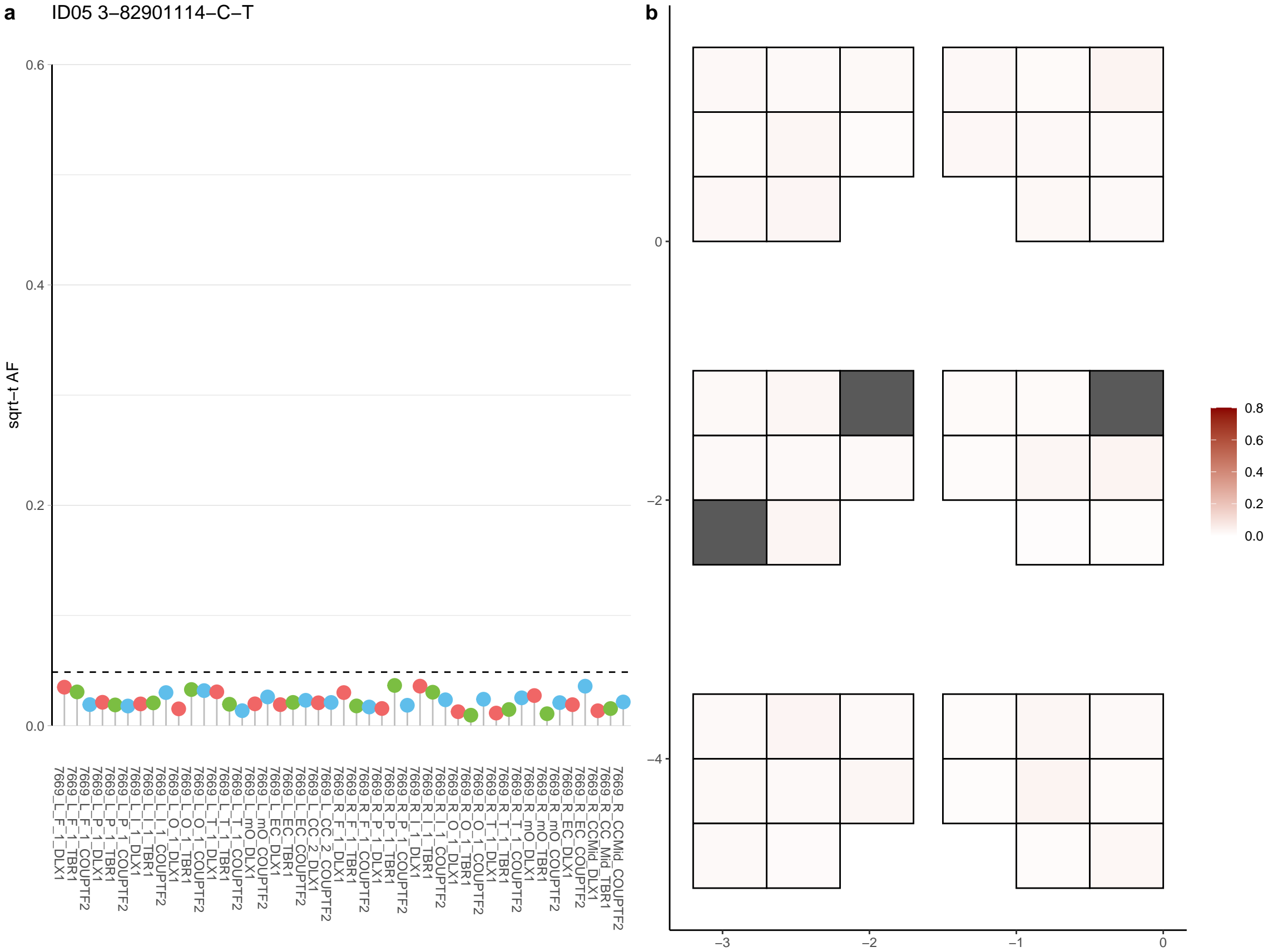

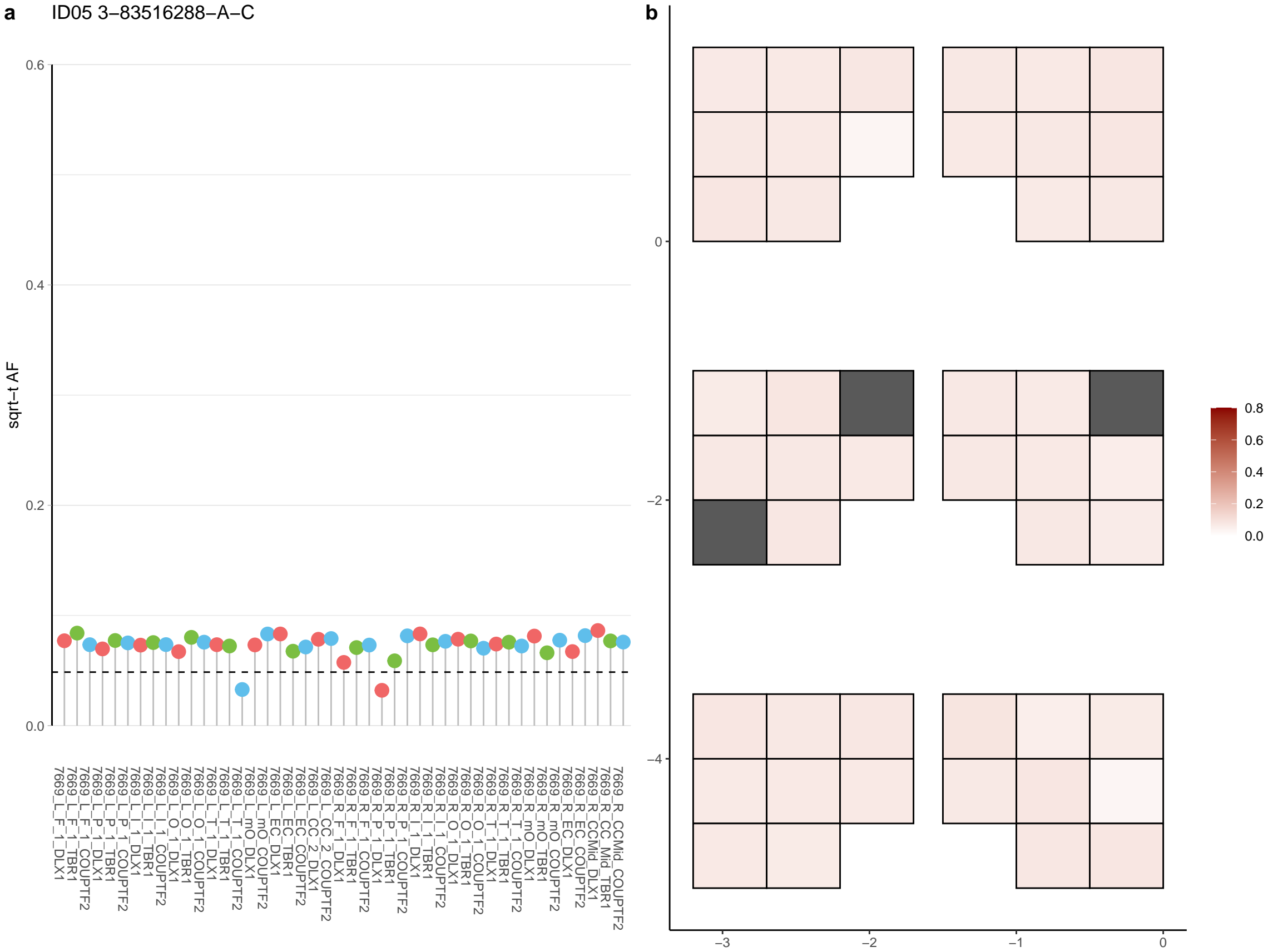

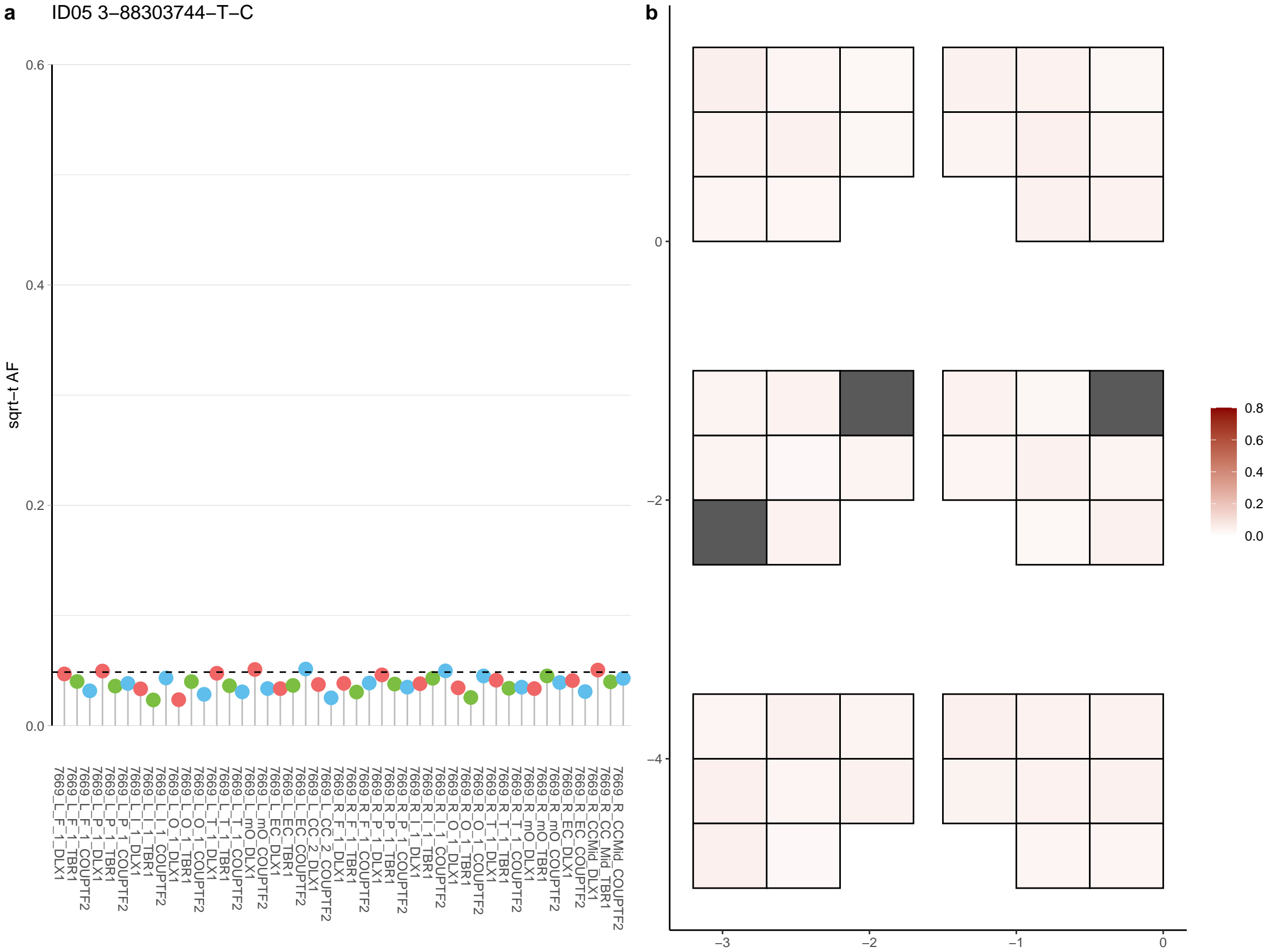

**a** ID05 3-9195857-C-T

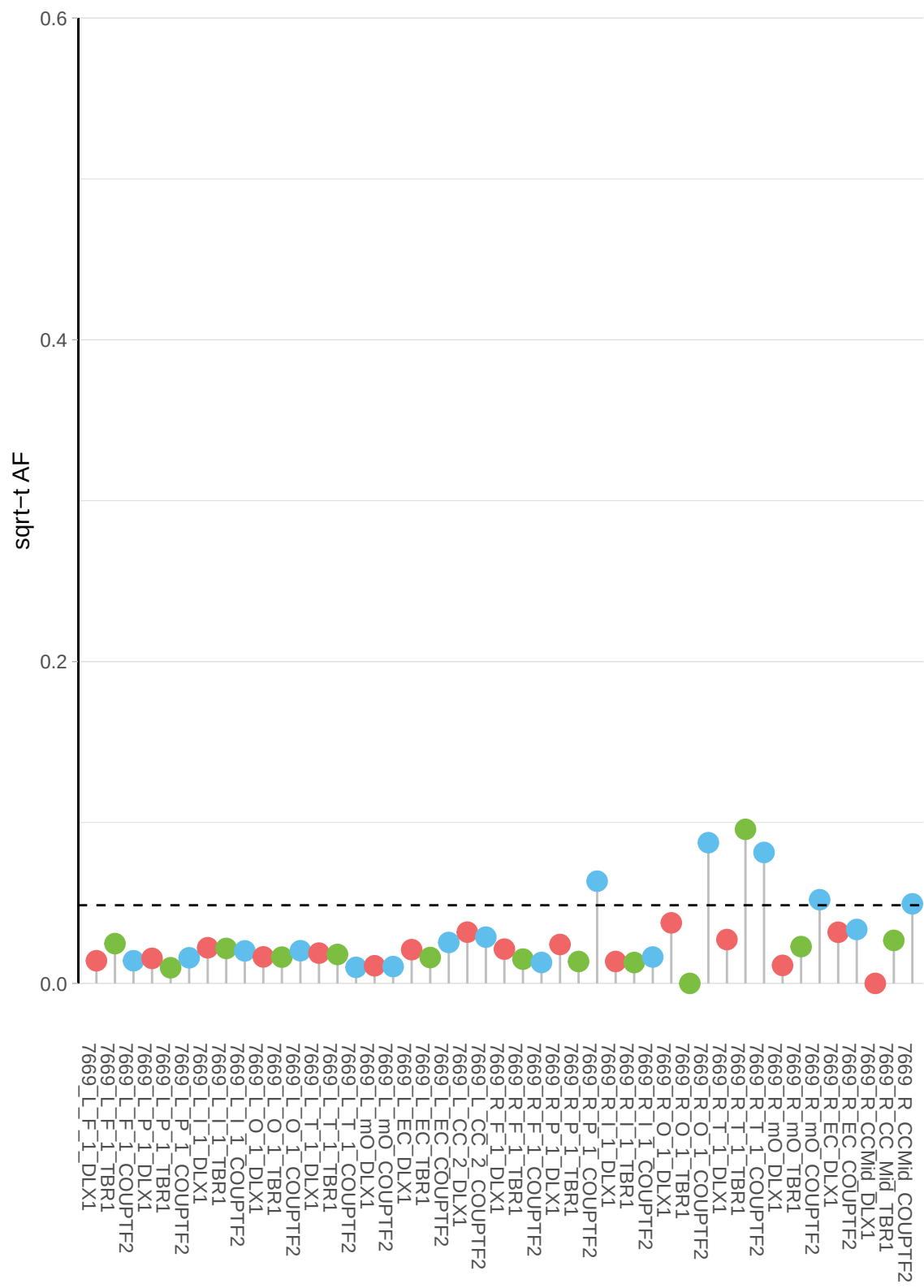

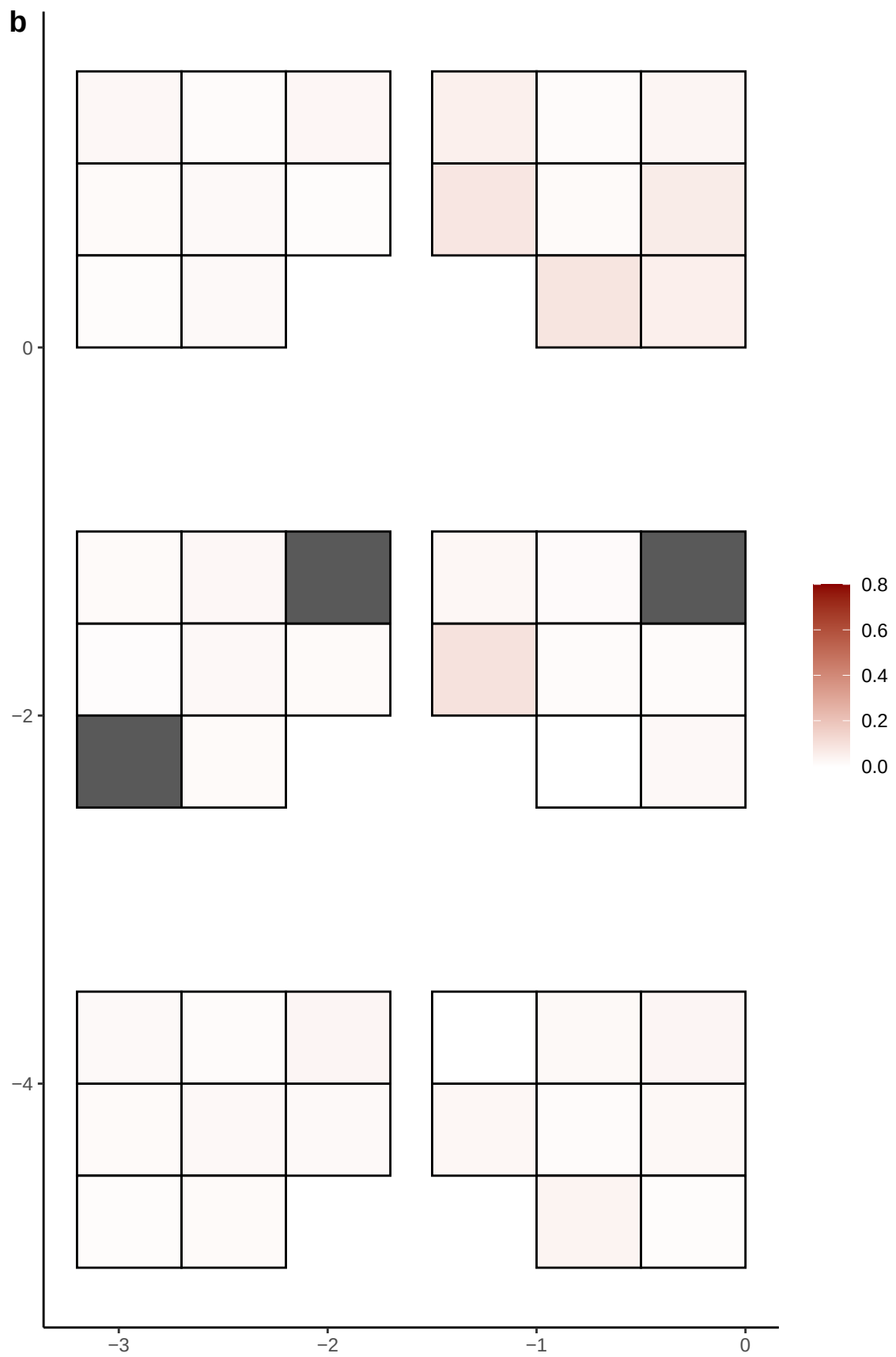

**a** ID05 3-9291135-G-A

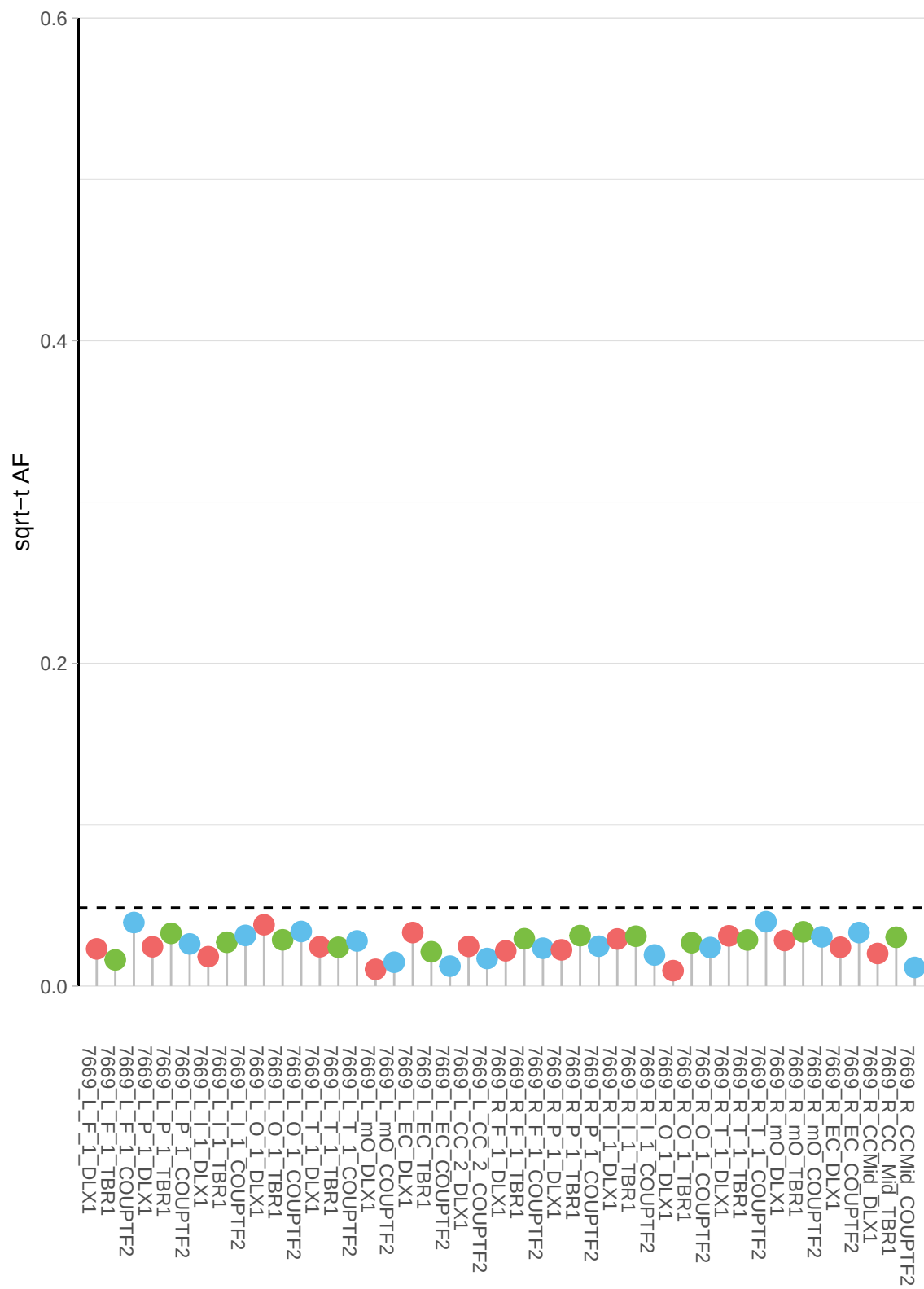

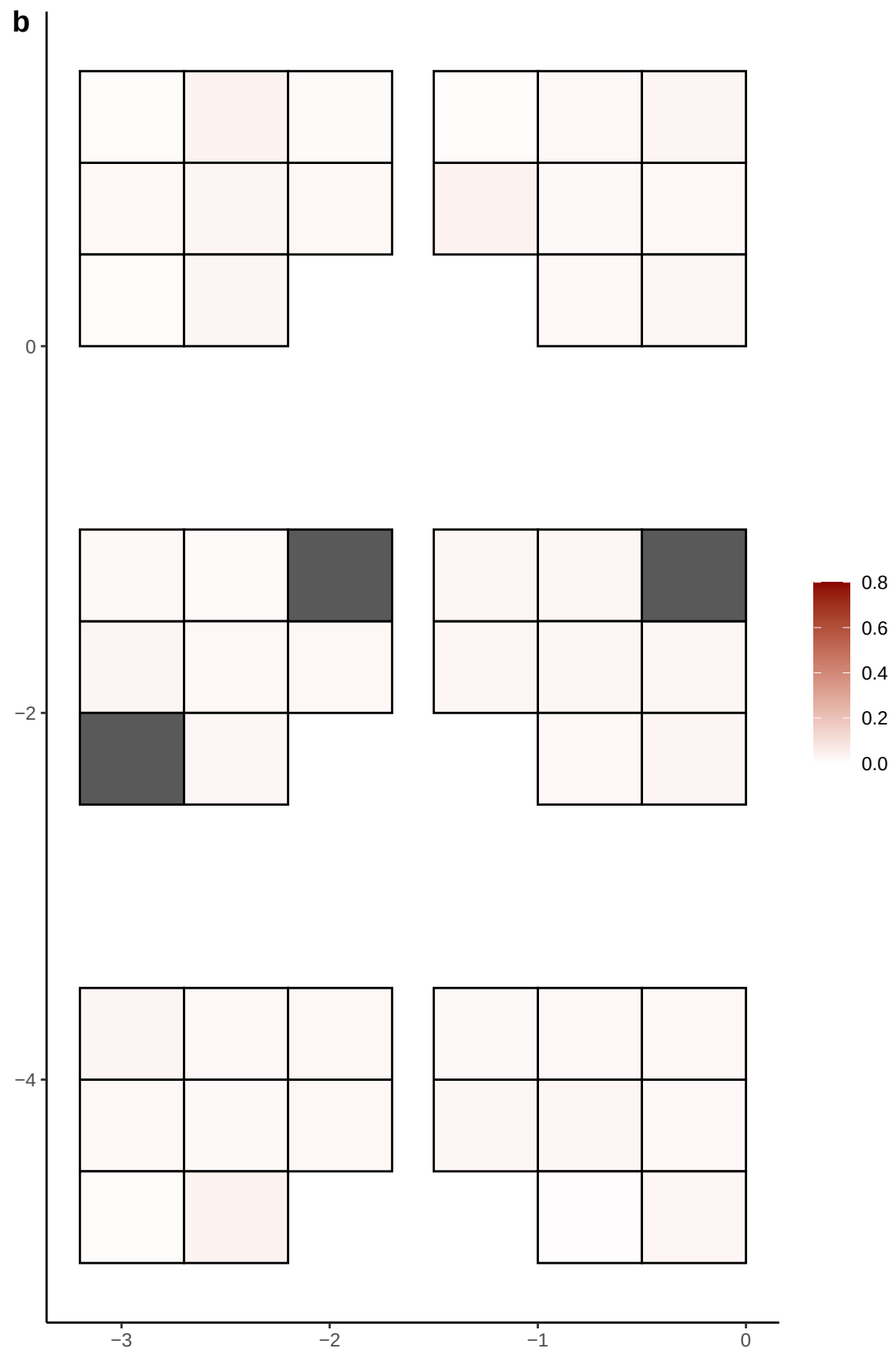

**a** ID05 3-9859360-C-T

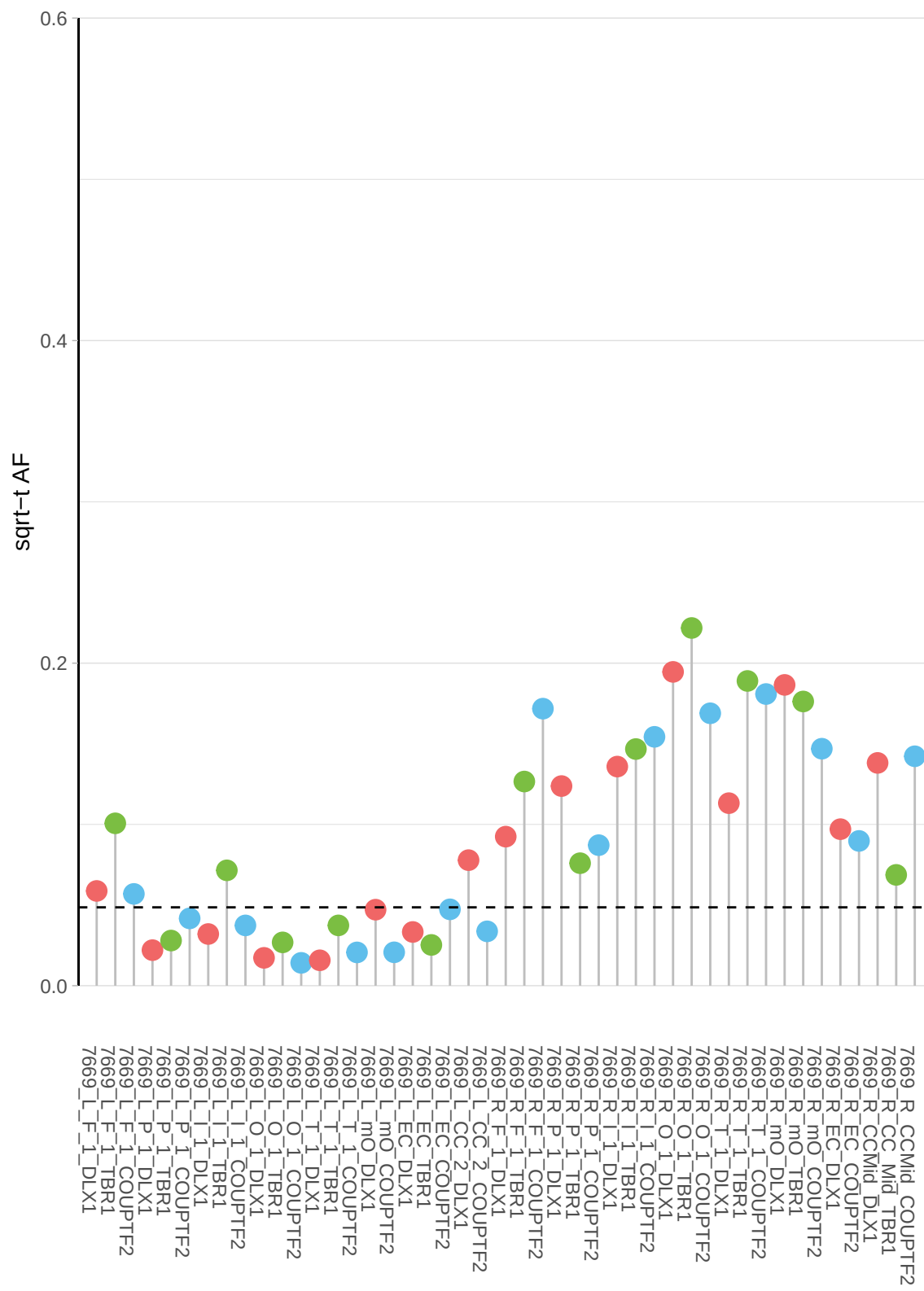

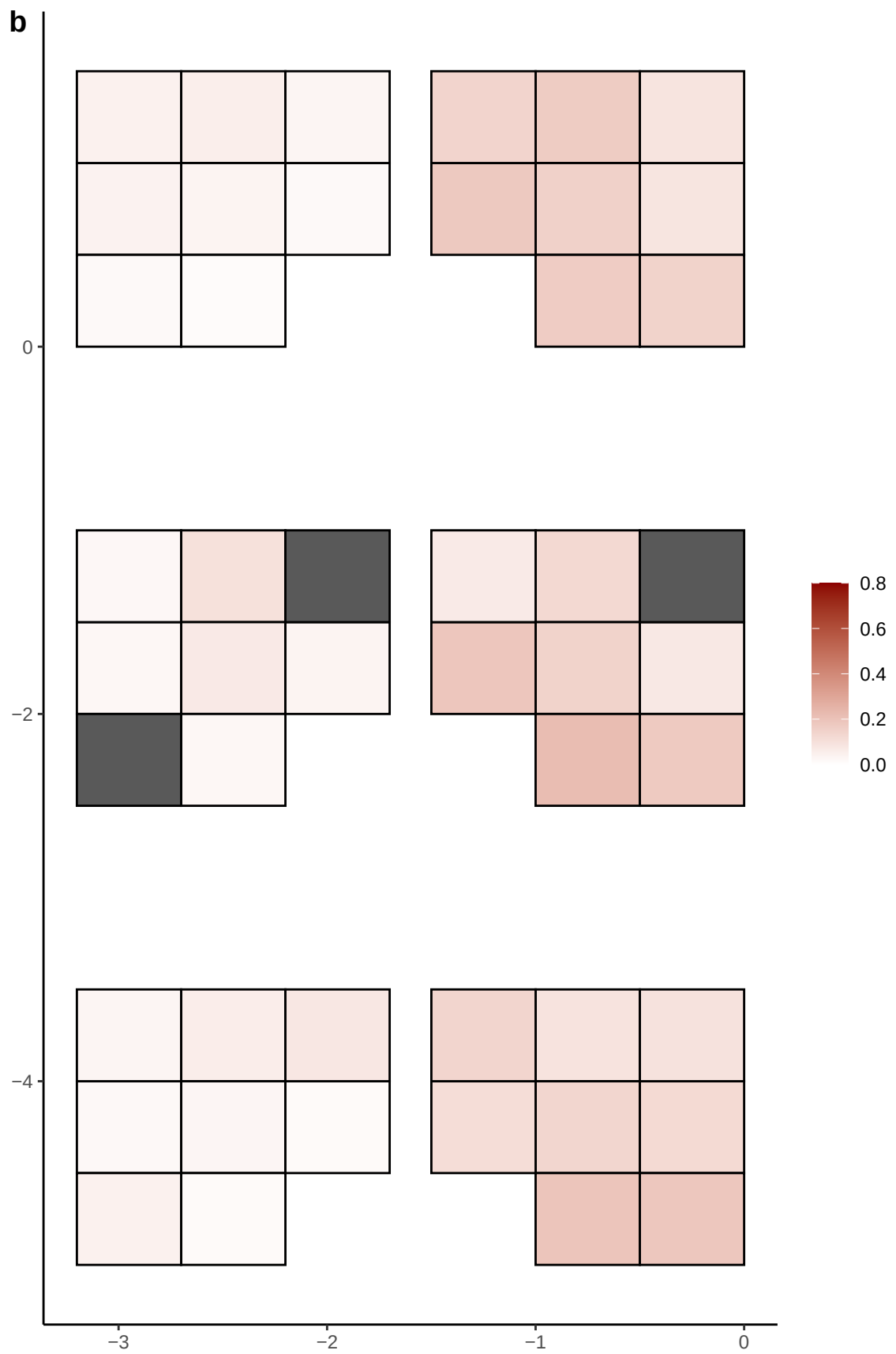

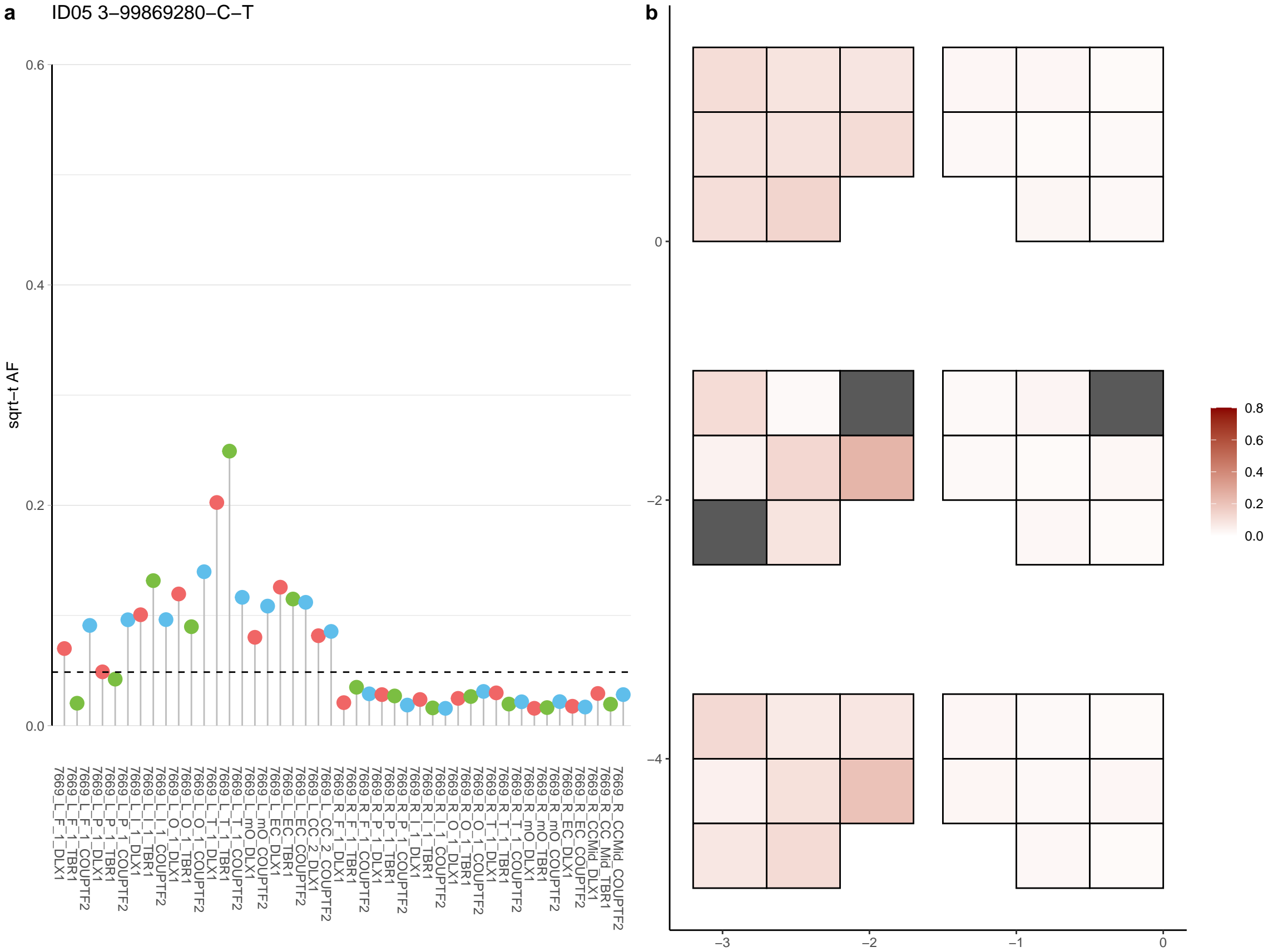

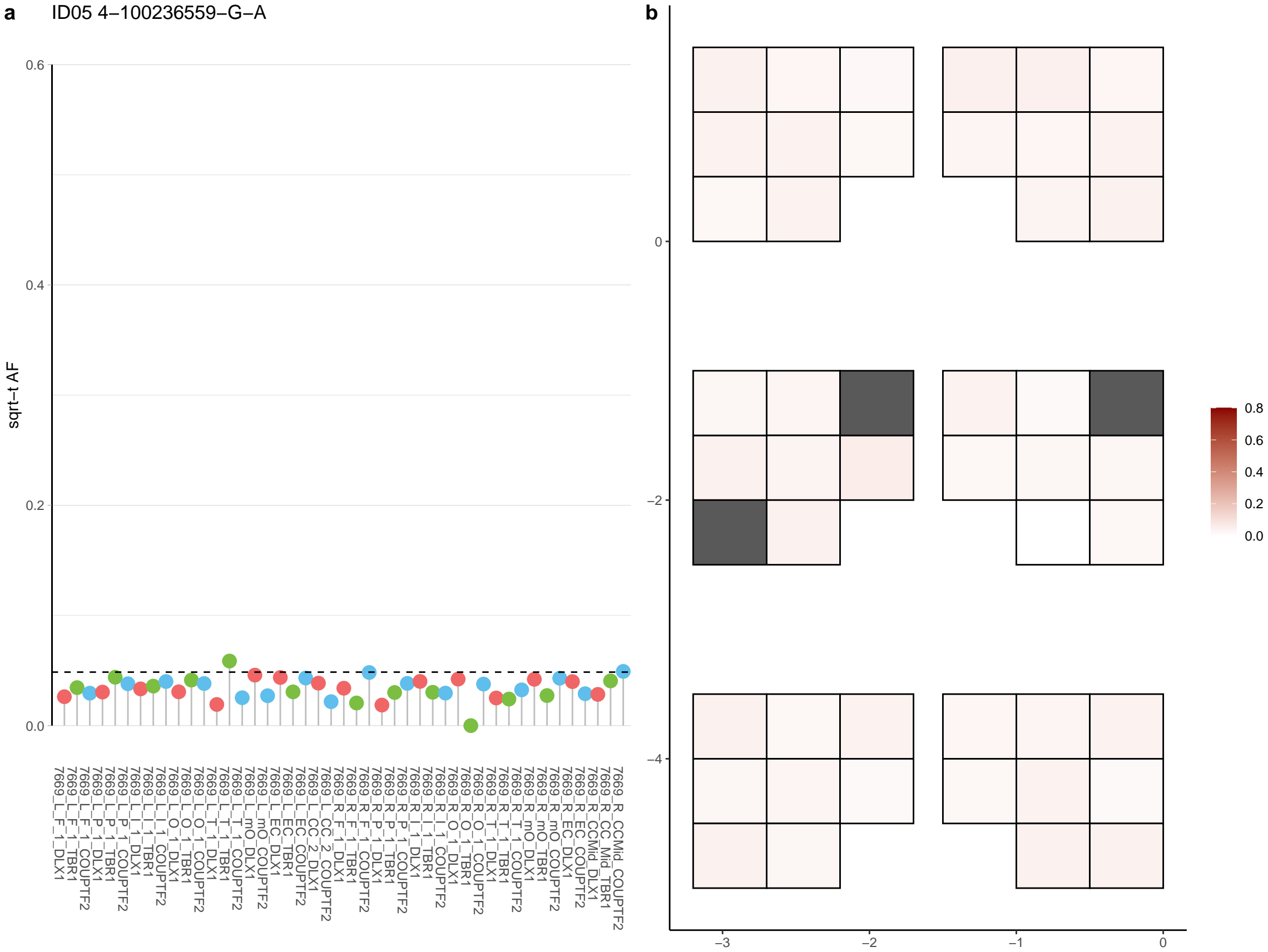

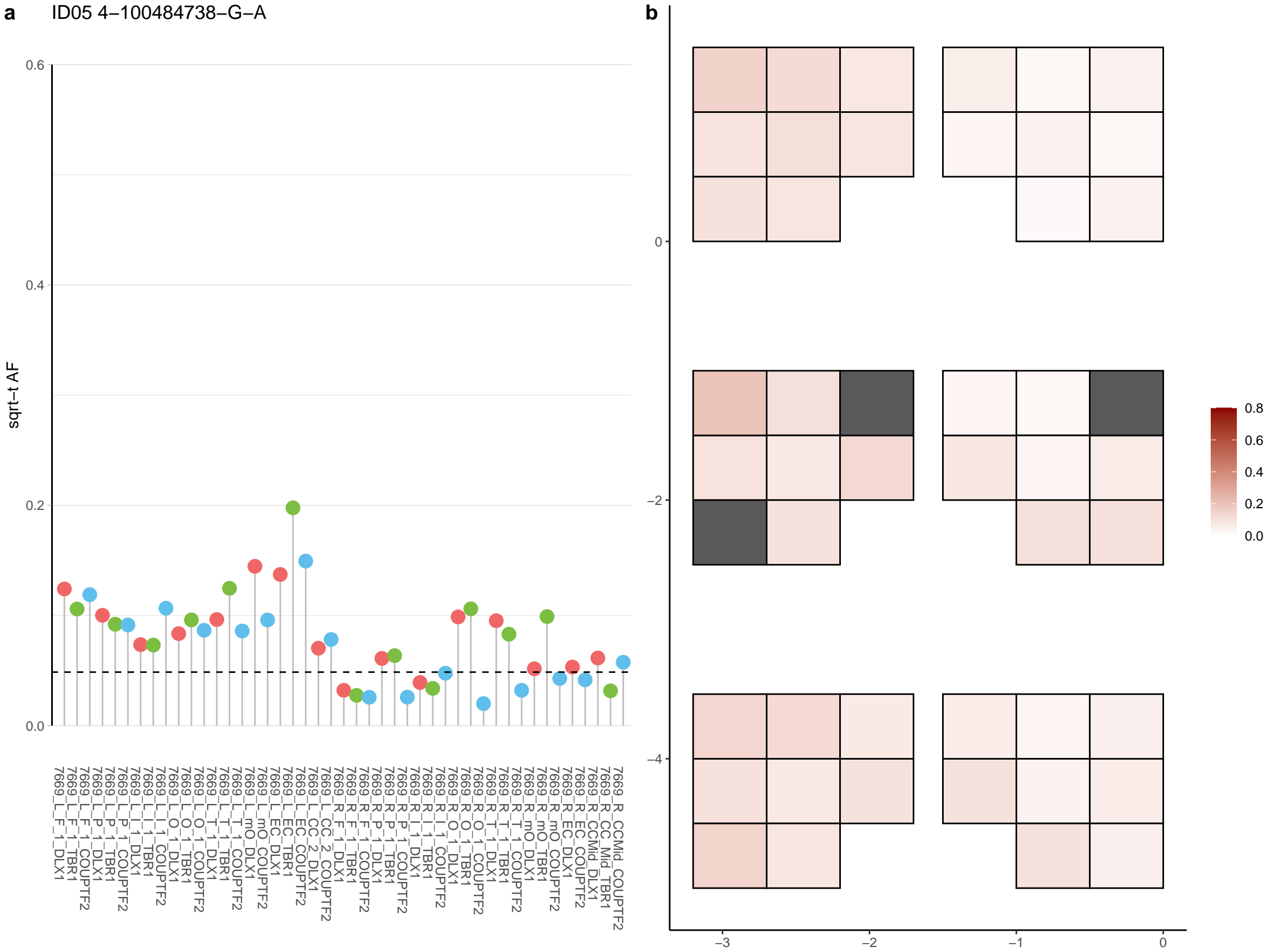

**a**

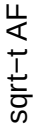

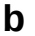

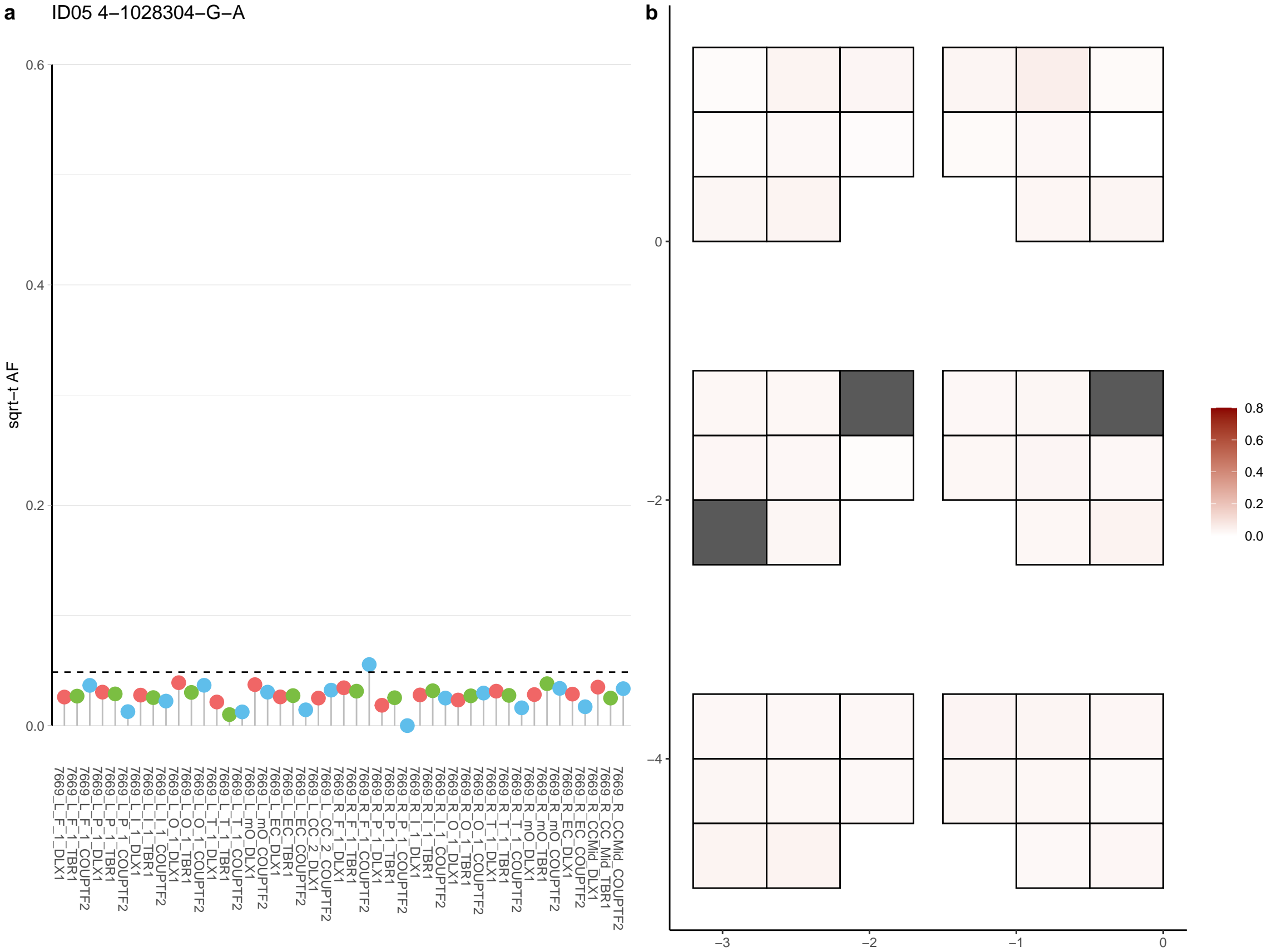

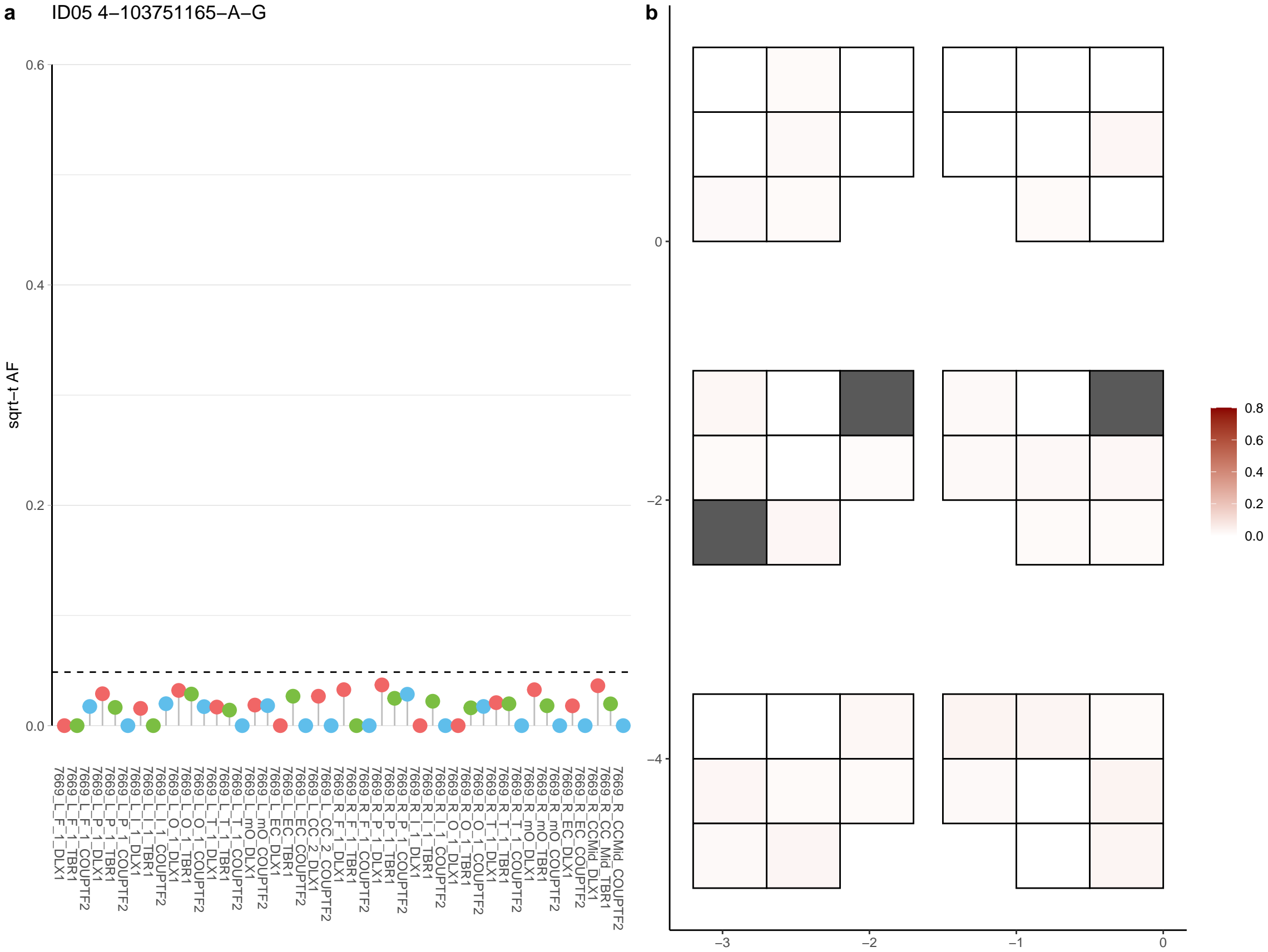

**a** ID05 4-104214112-C-T

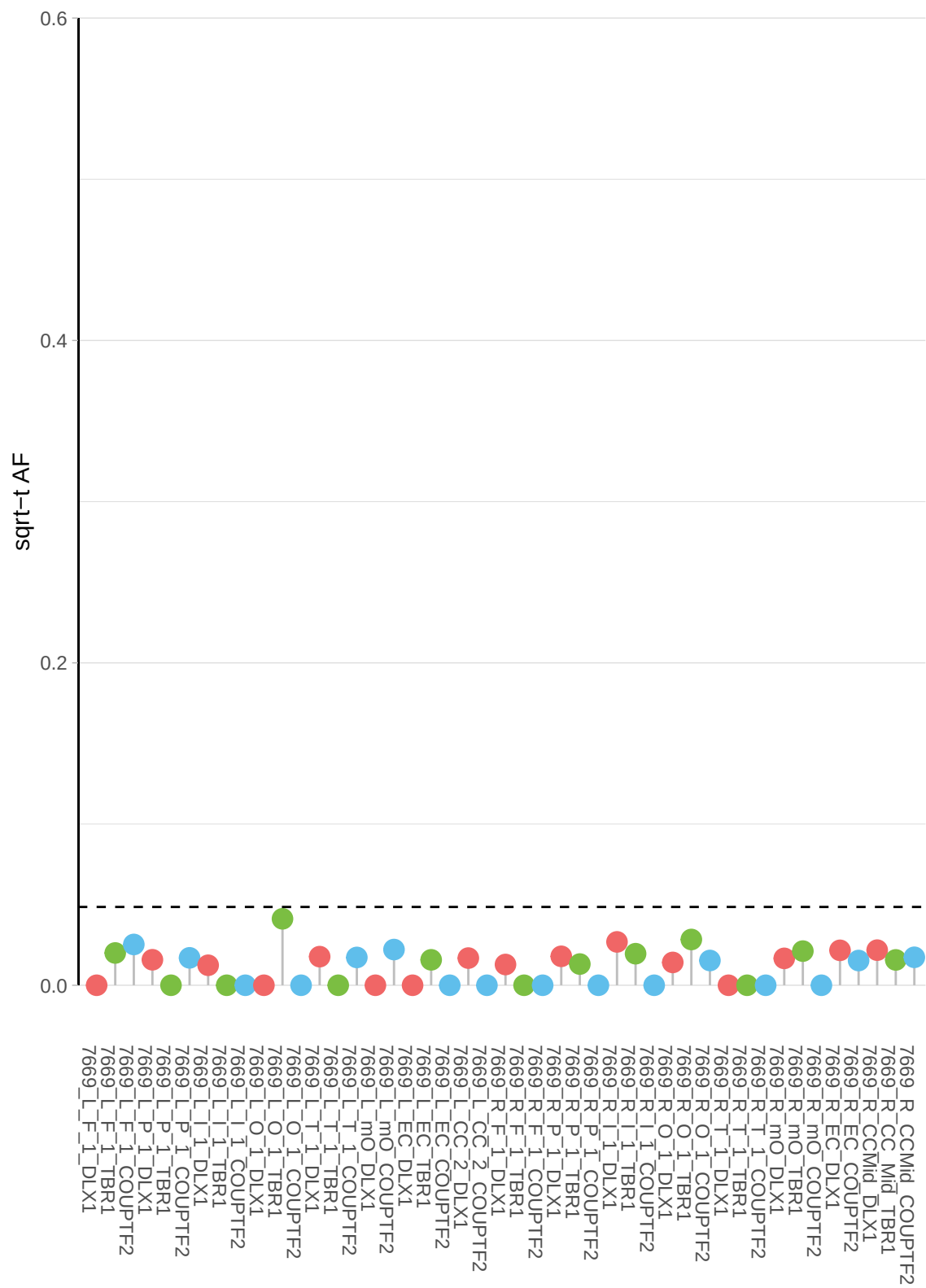

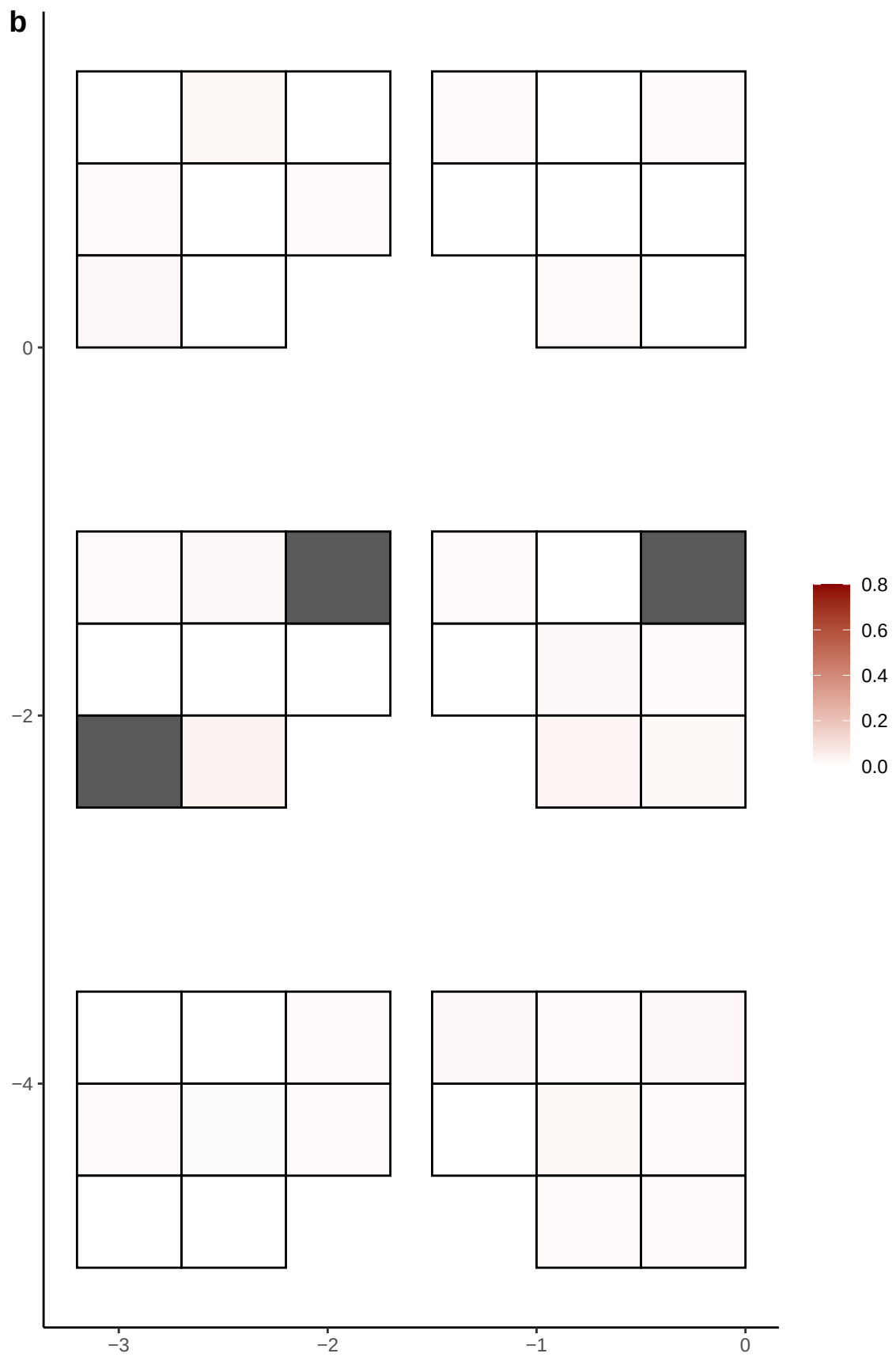

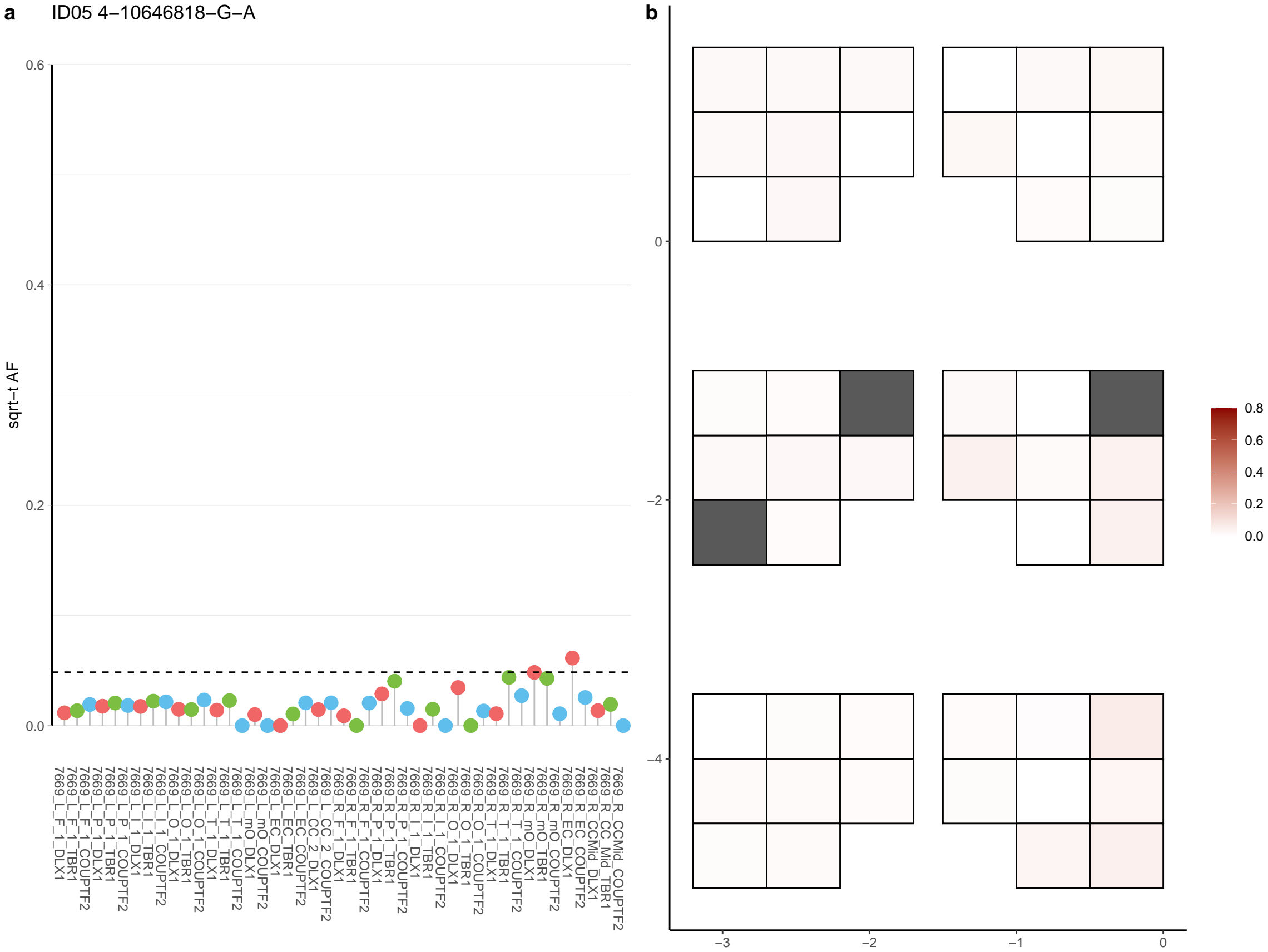

**a** ID05 4-109178107-T-A

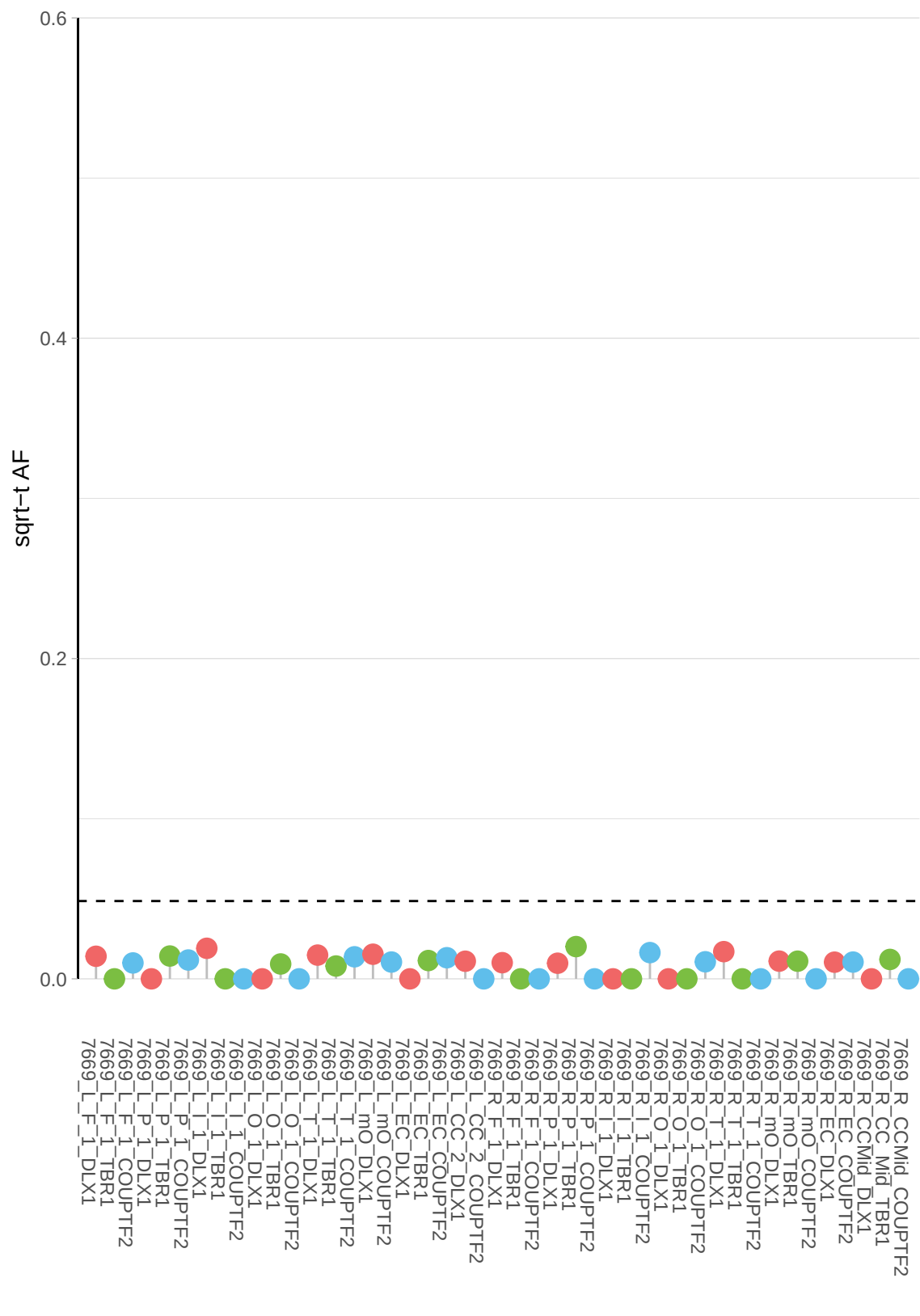

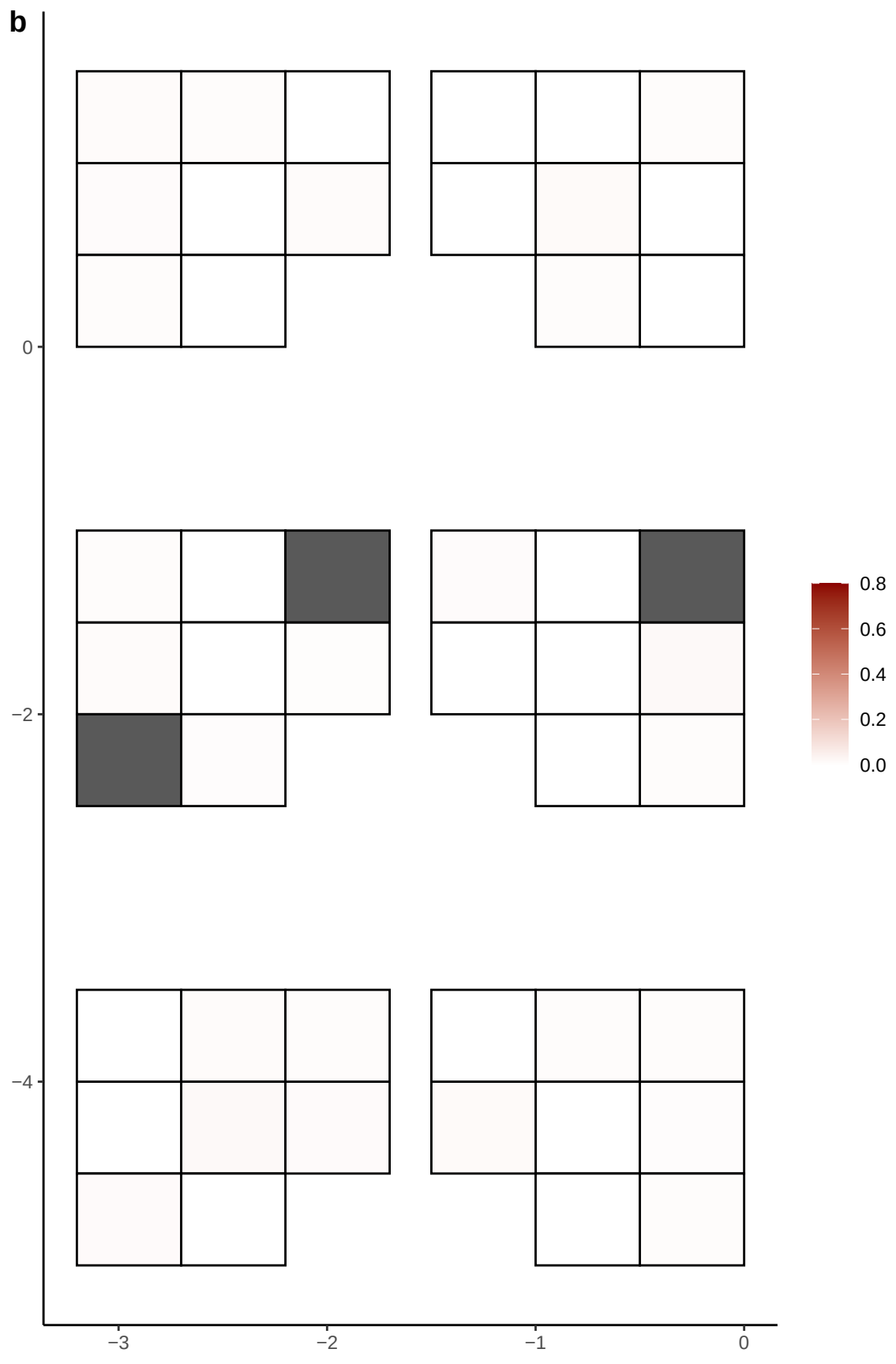

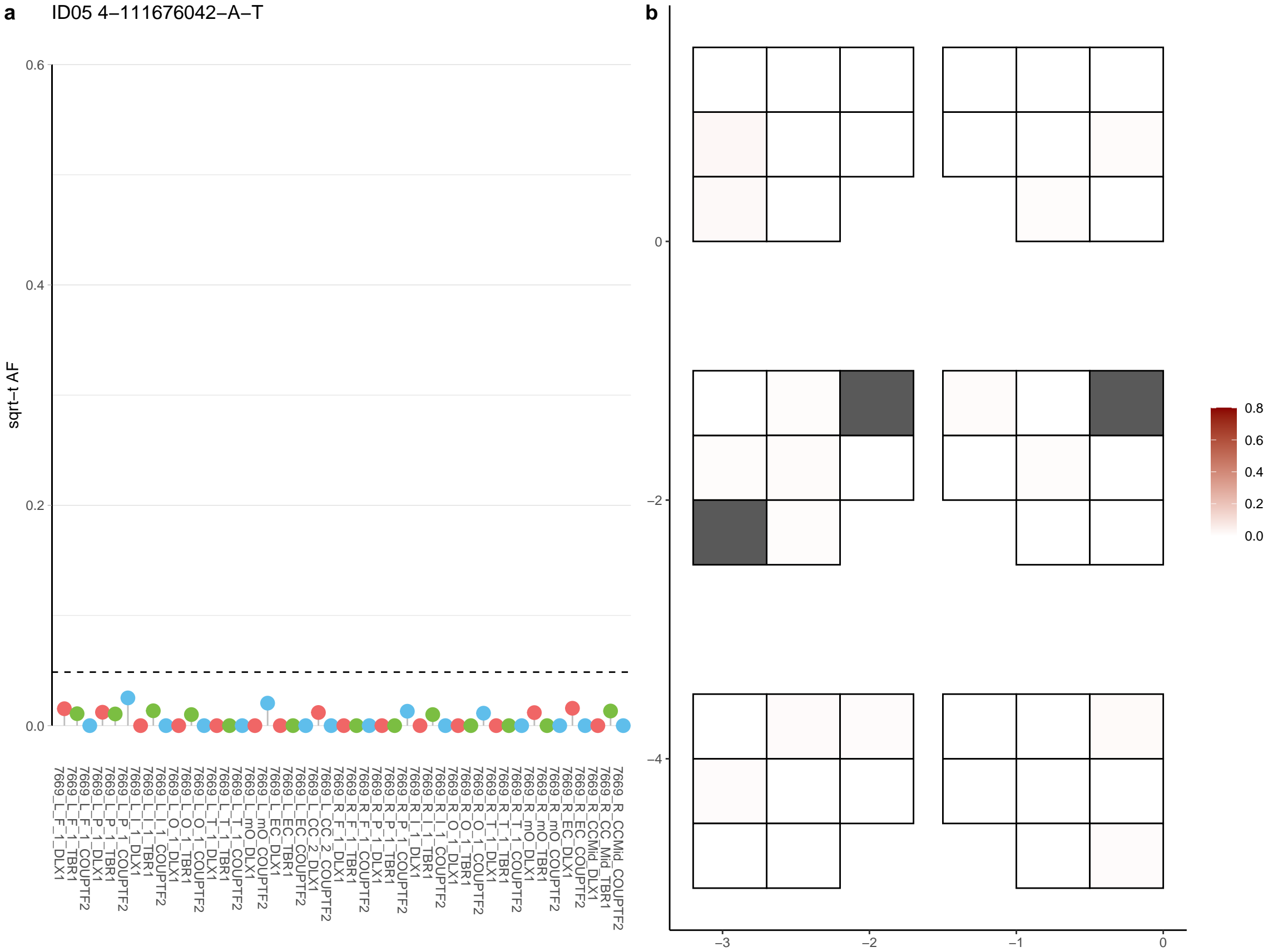

**a** ID05 4-111915329-C-T

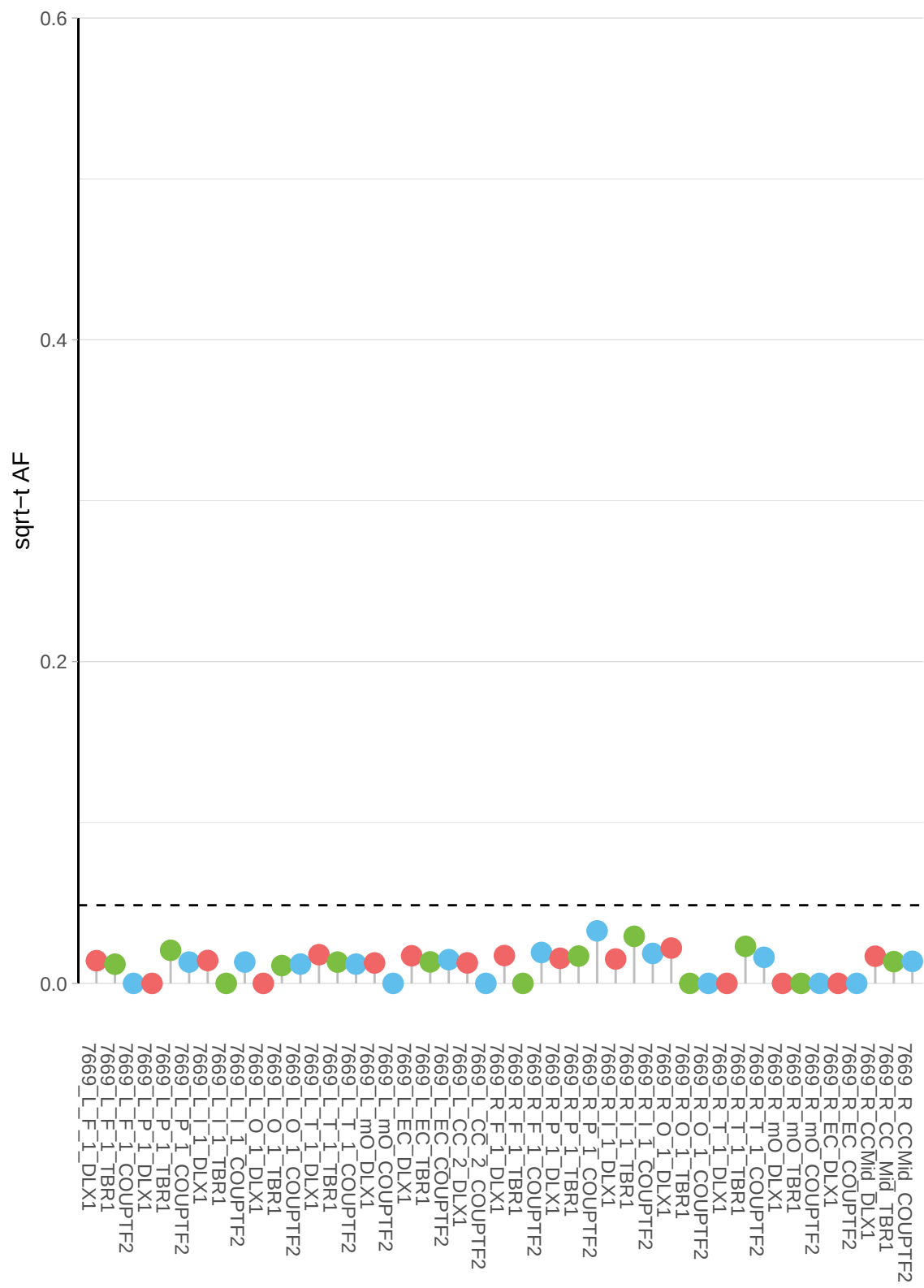

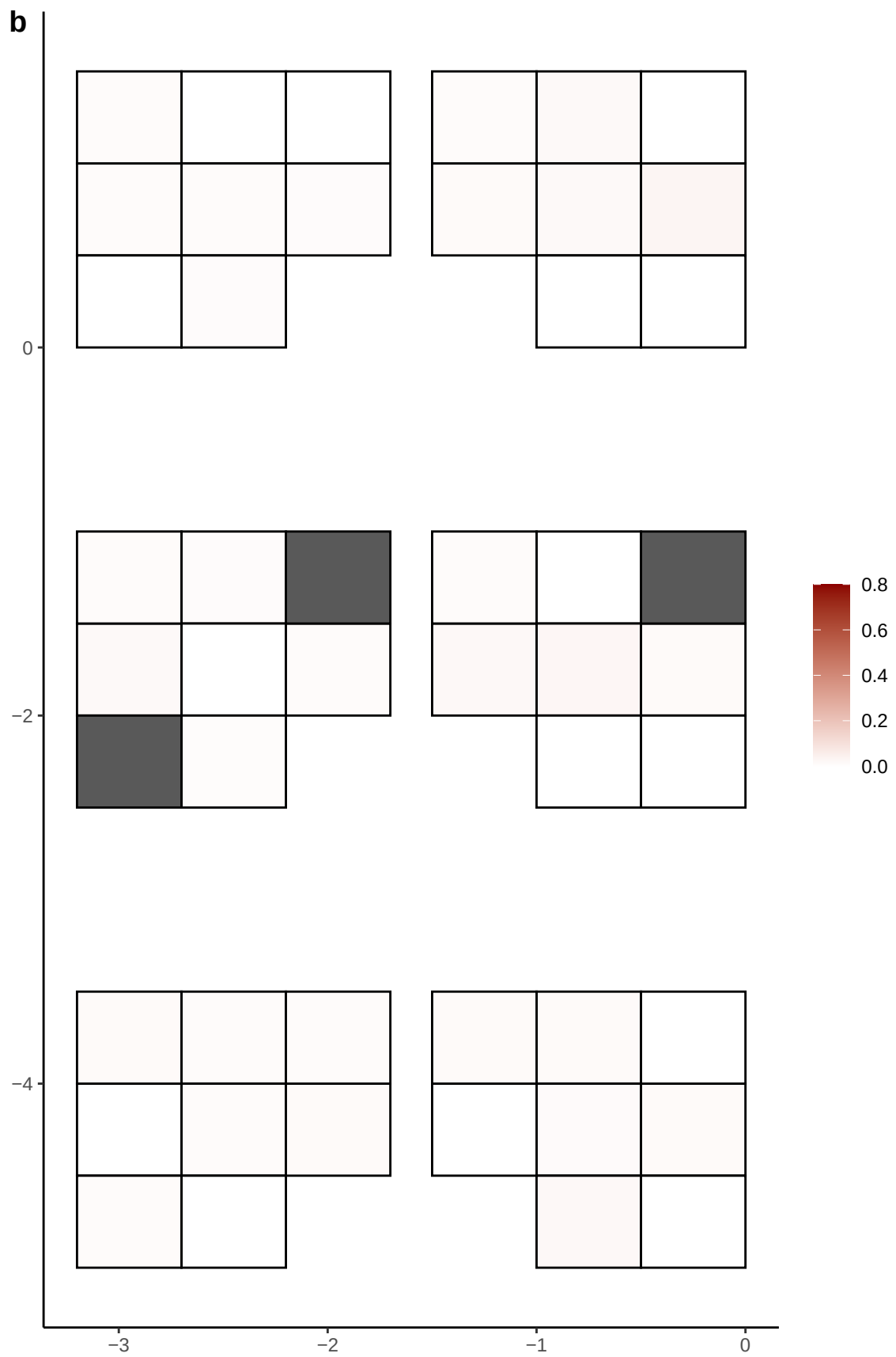

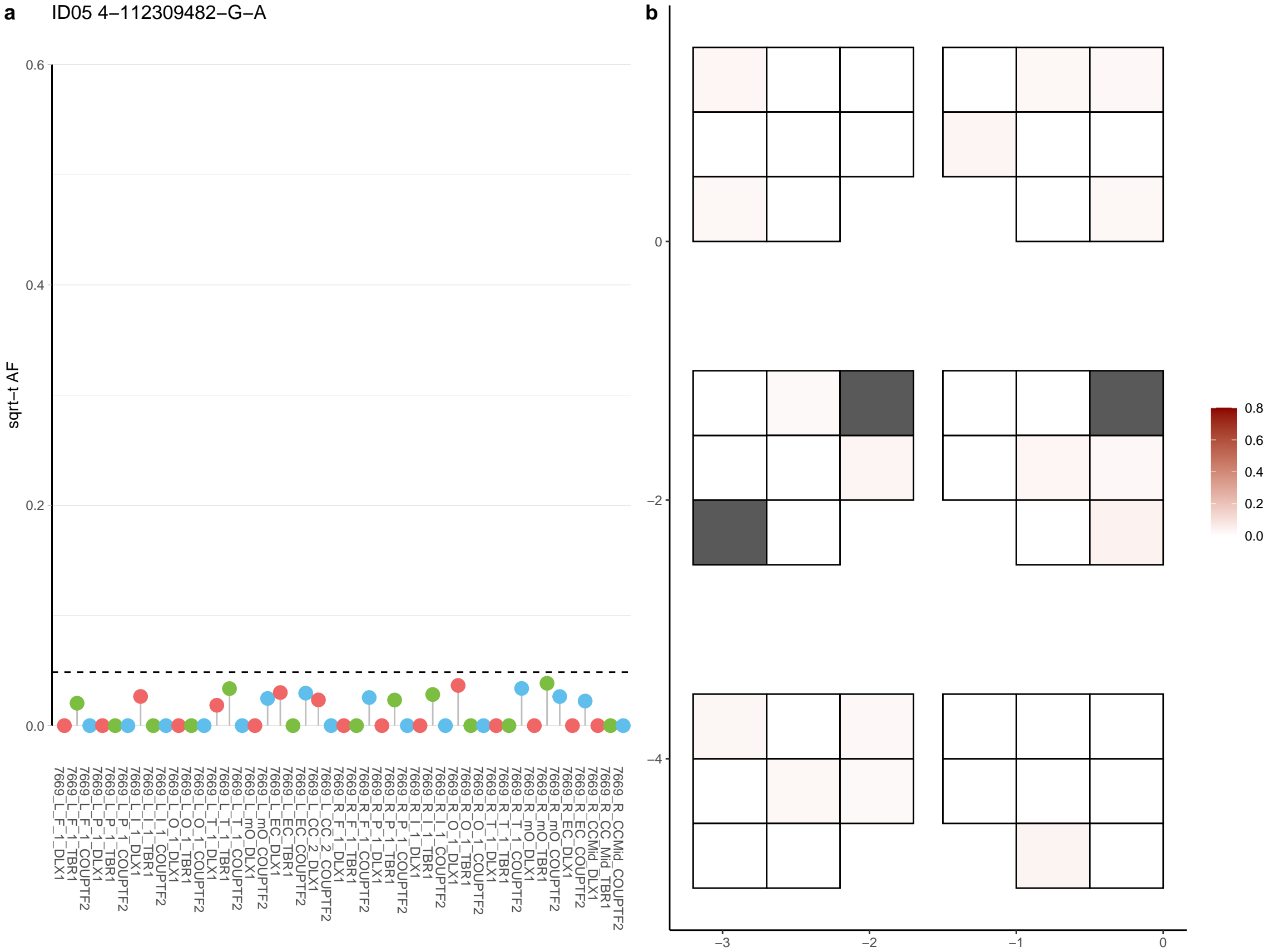

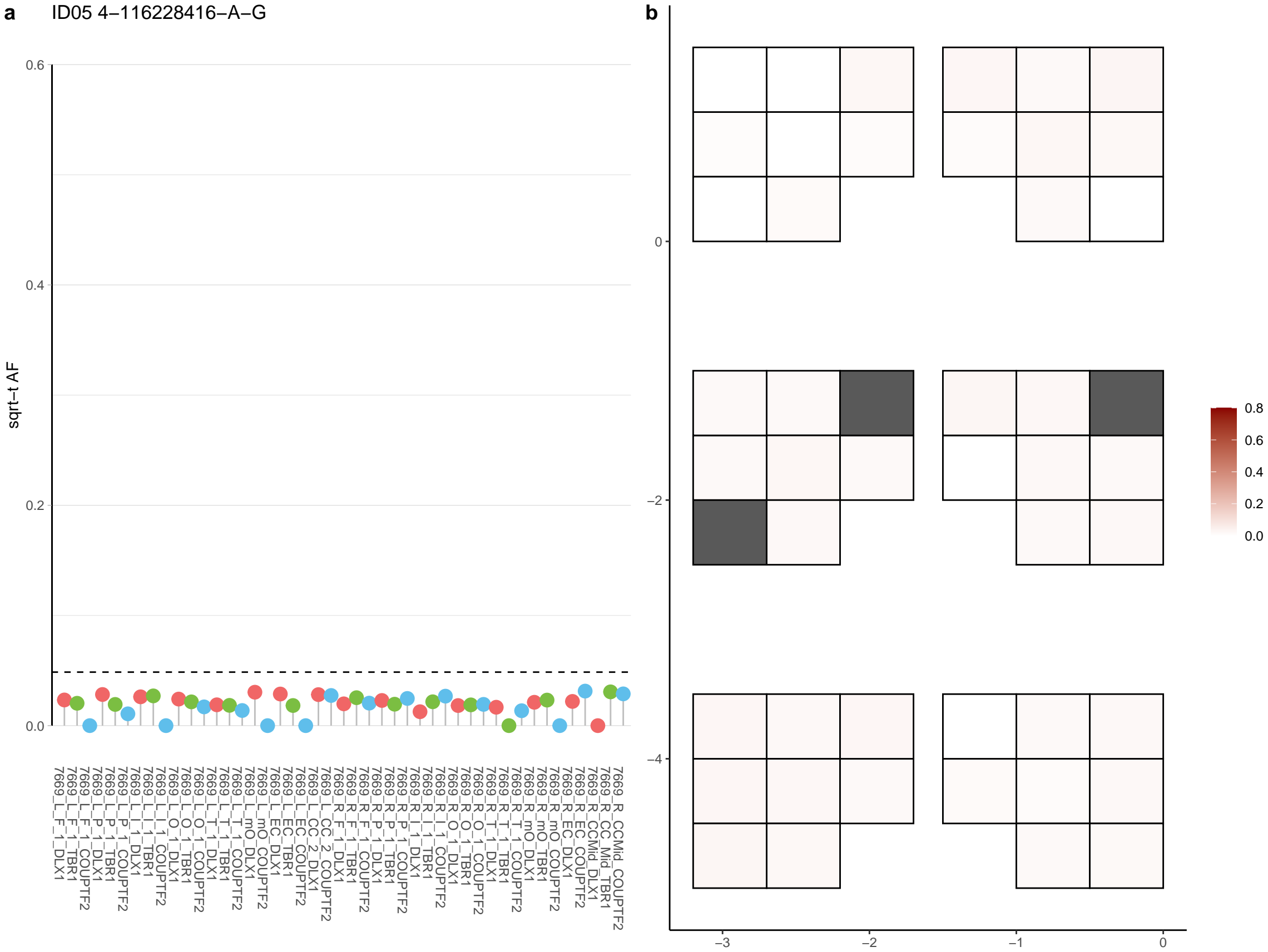

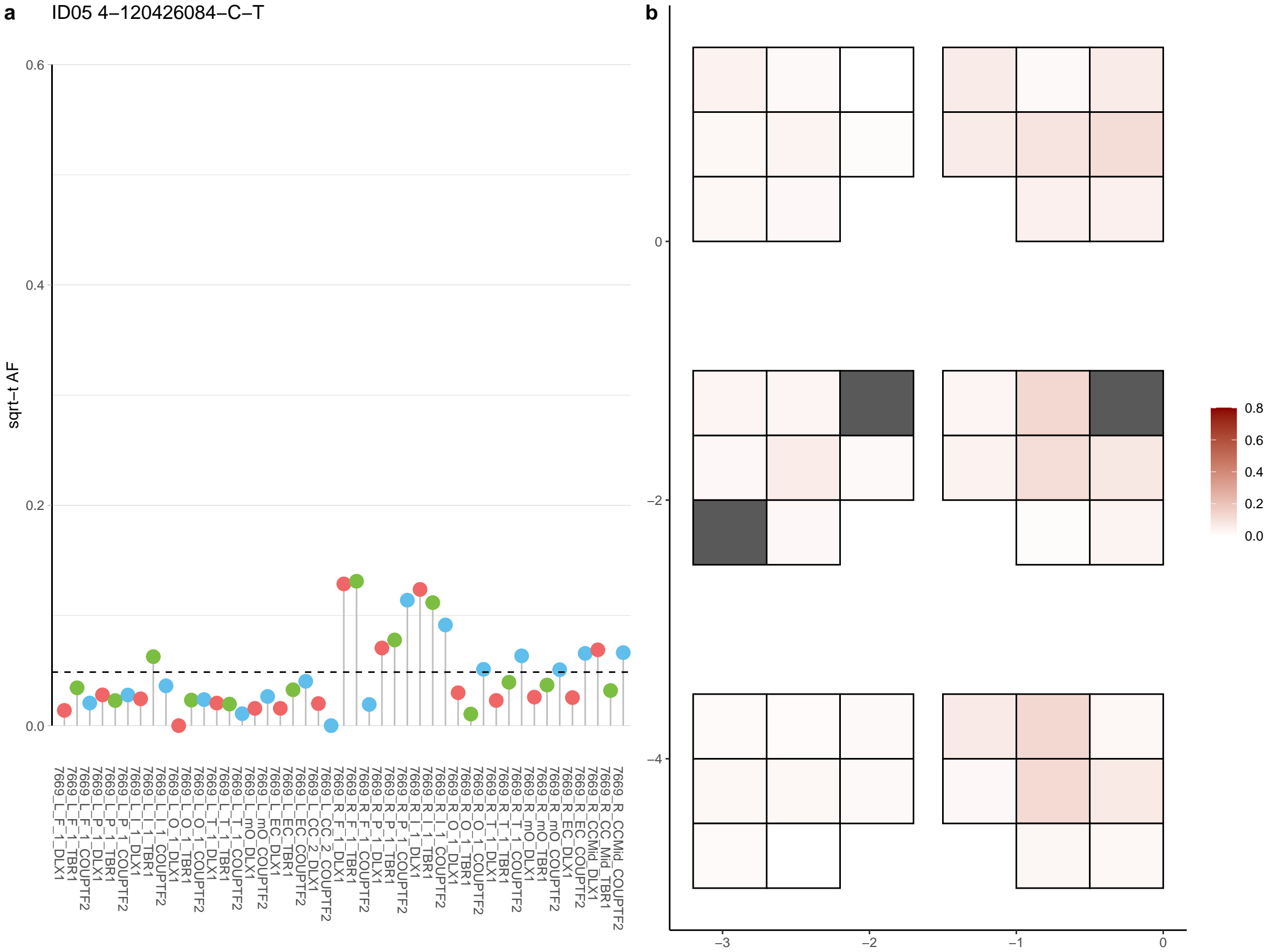

**a** ID05 4-121359392-T-G

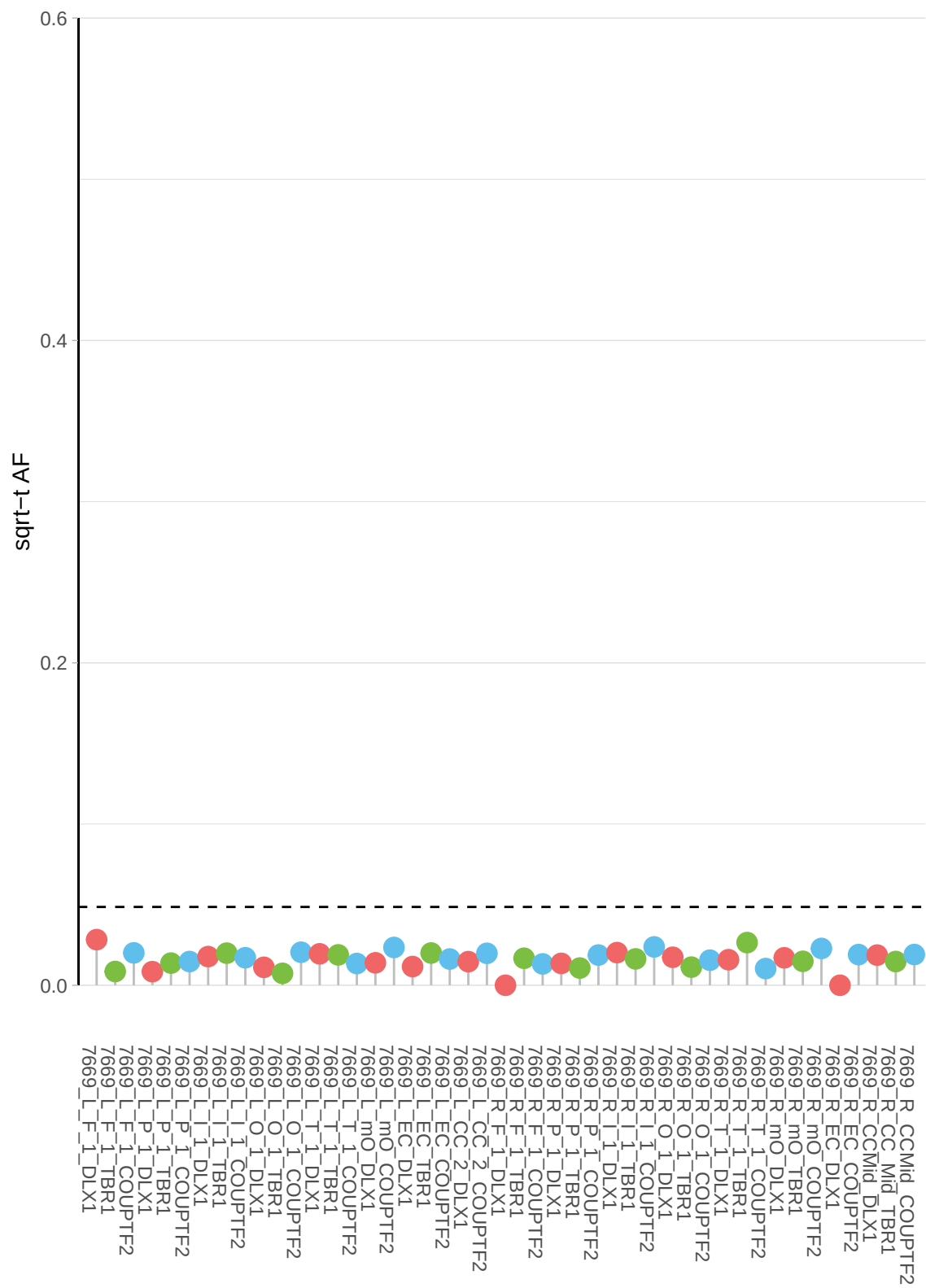

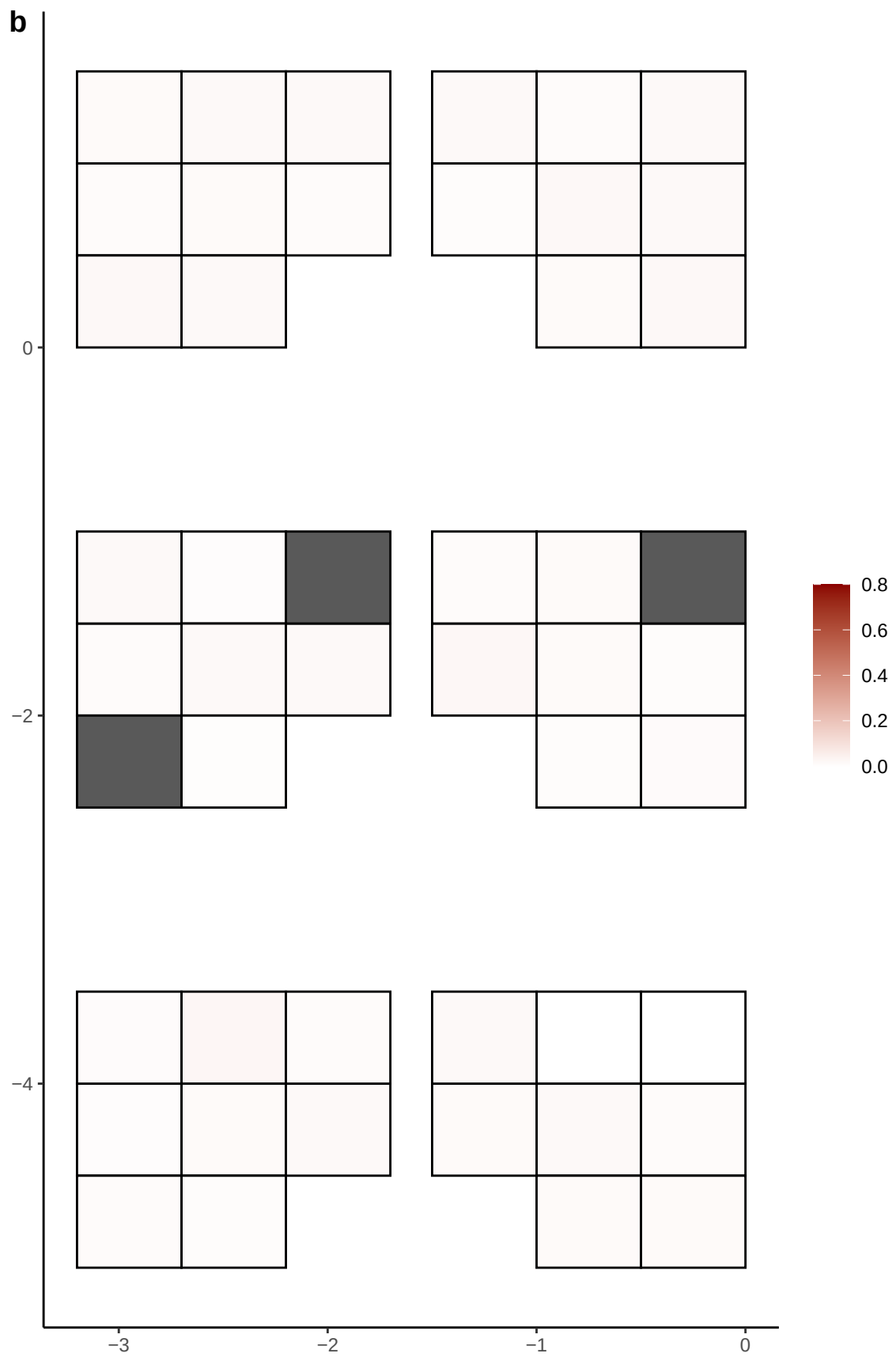

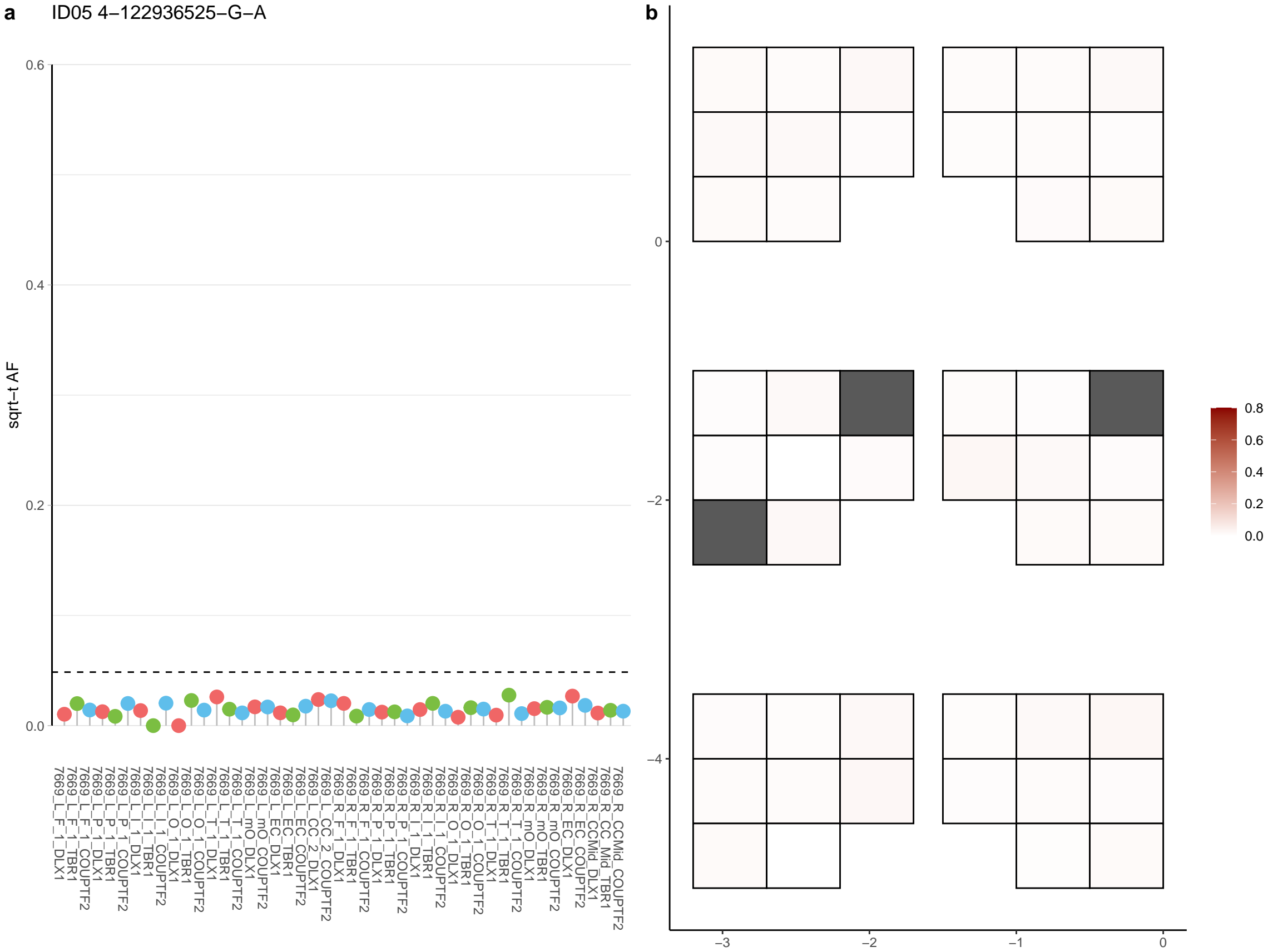

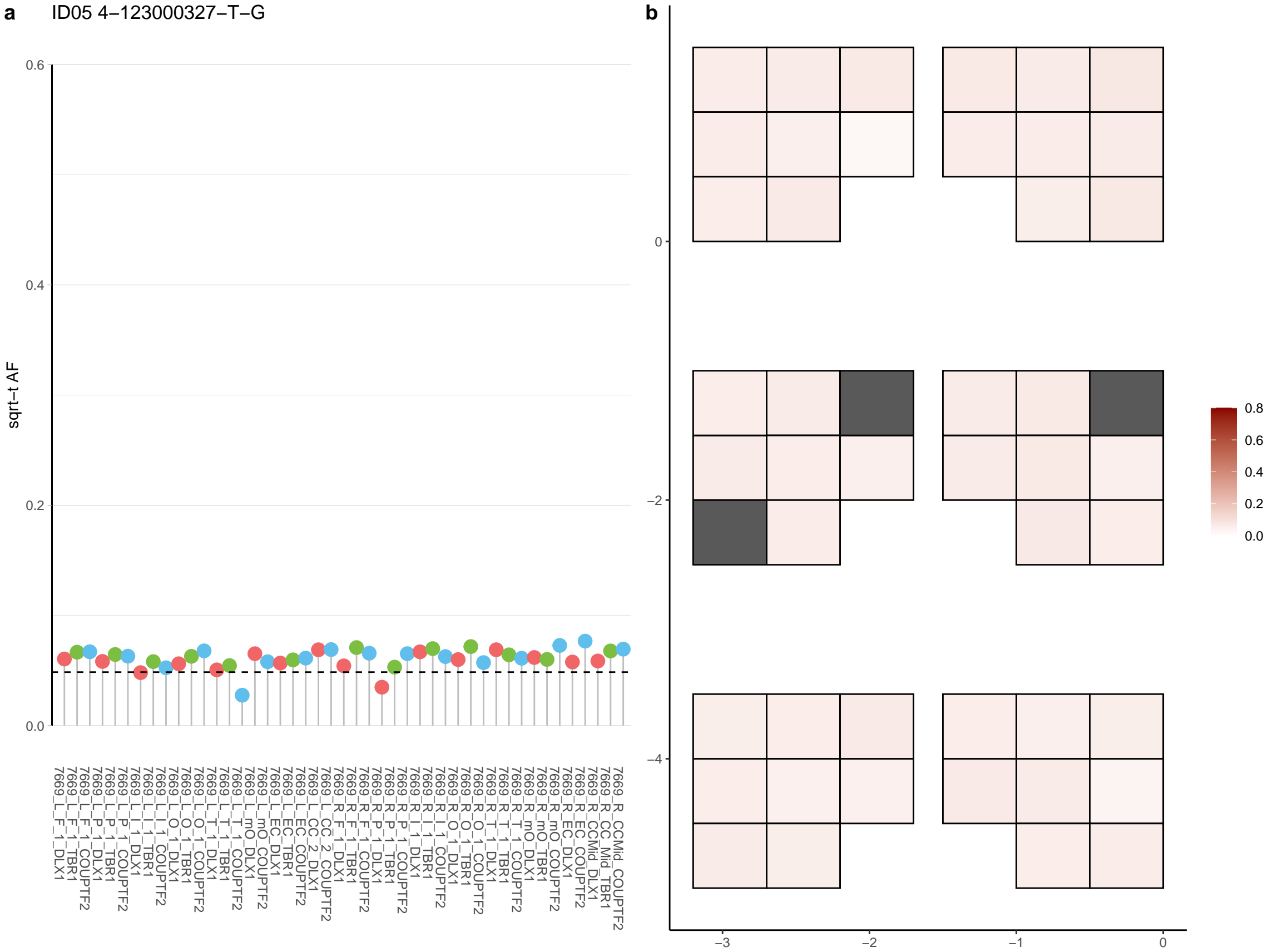

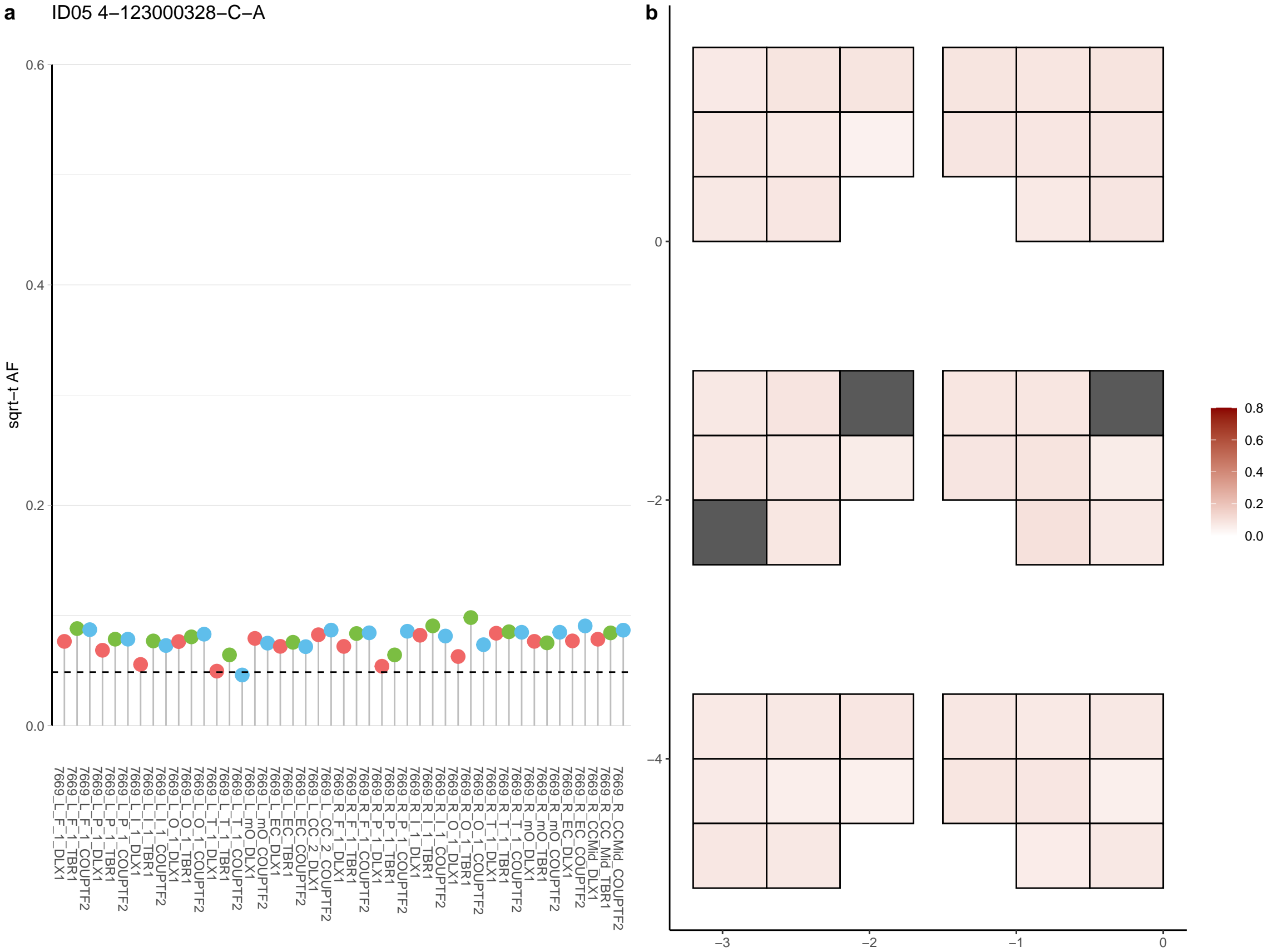

**a** ID05 4-125806400-T-C

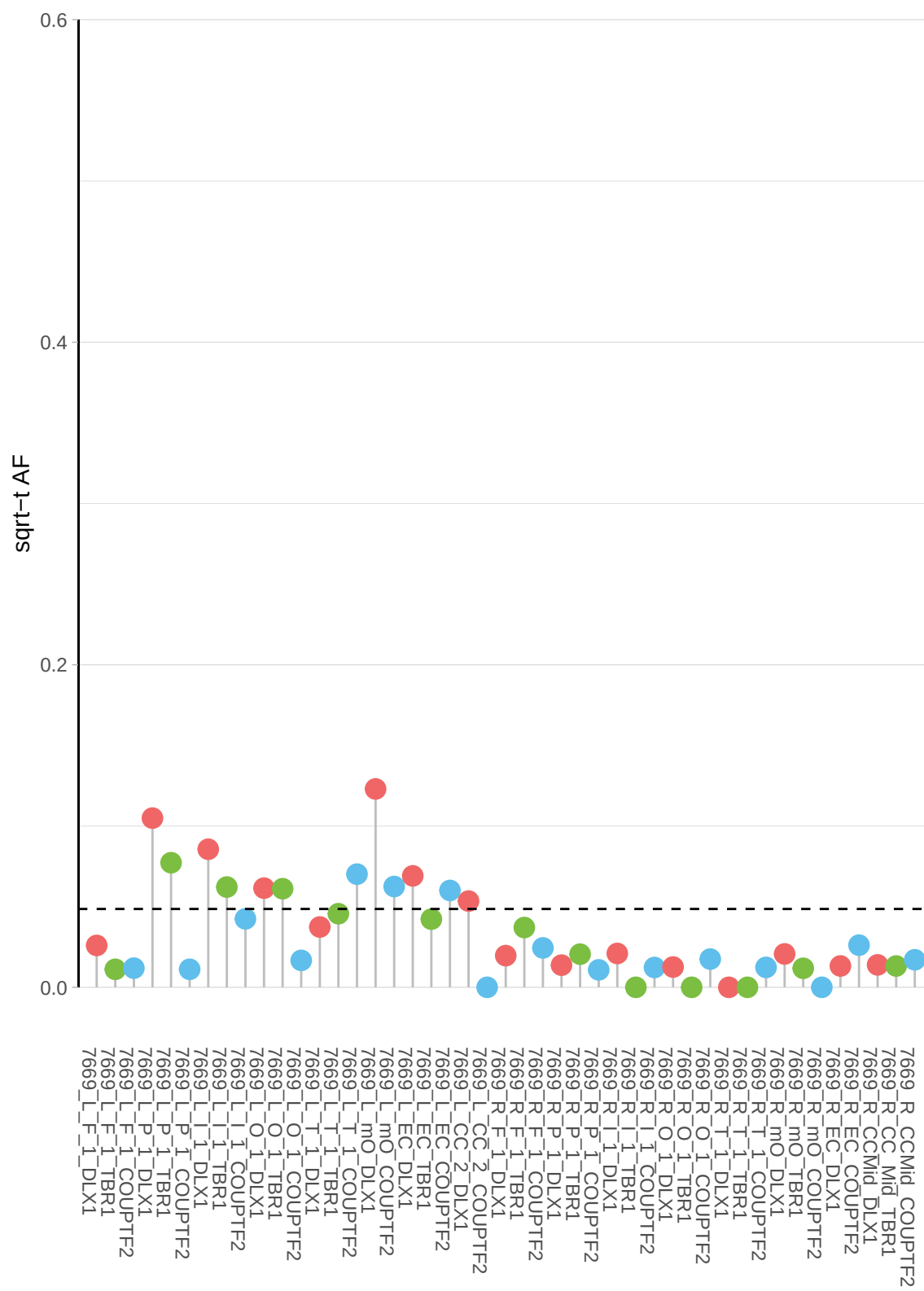

**b**

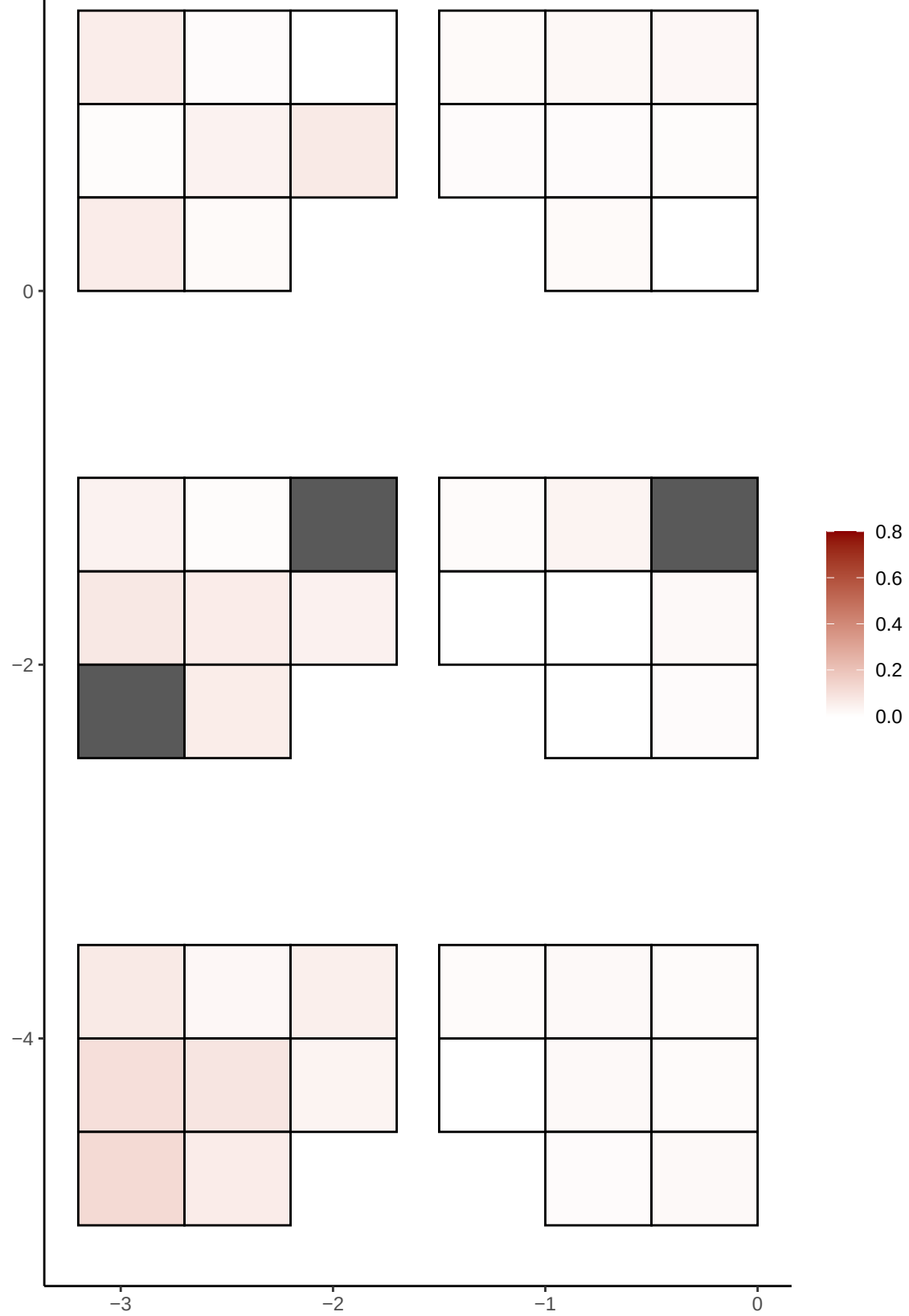

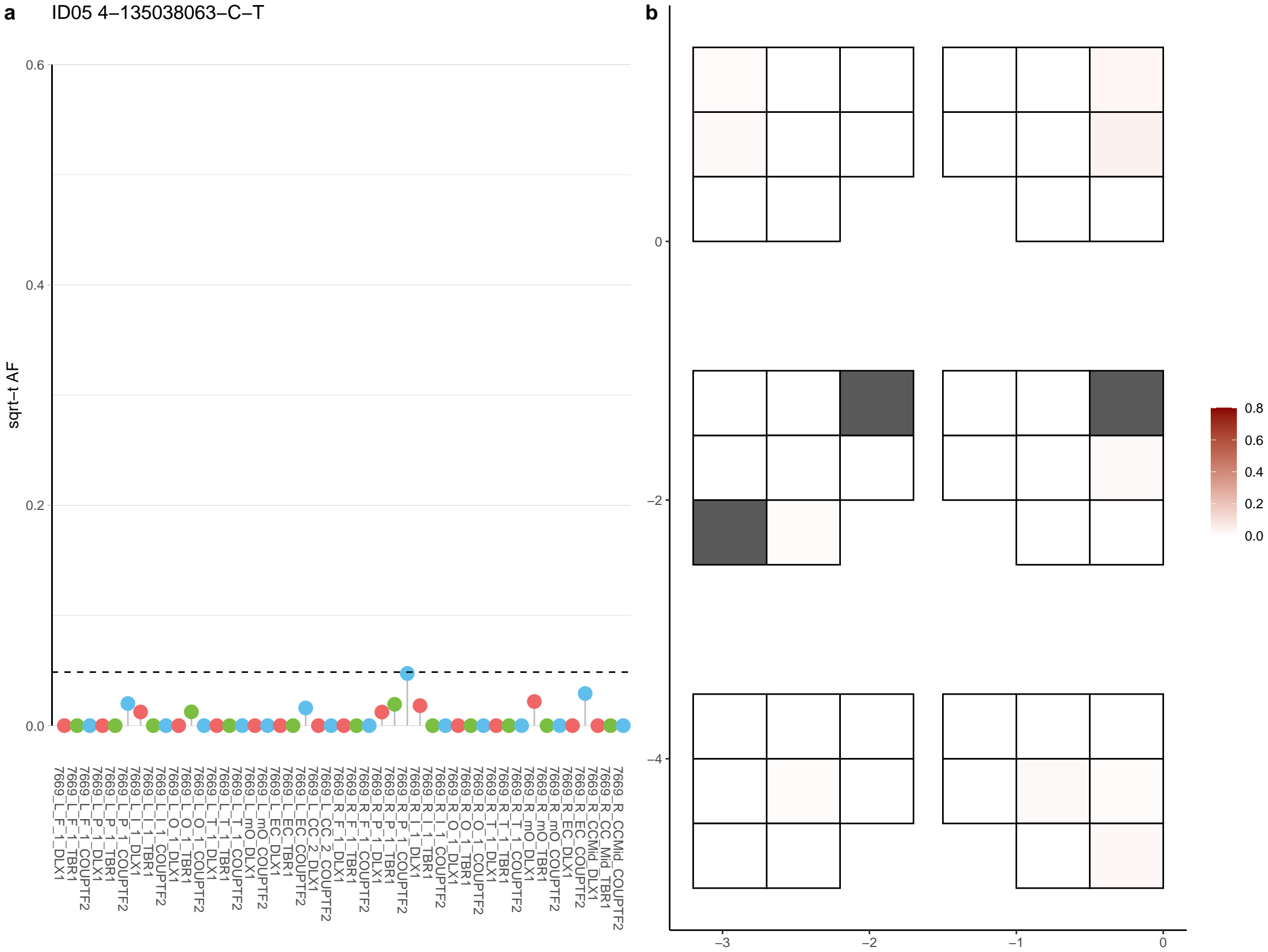

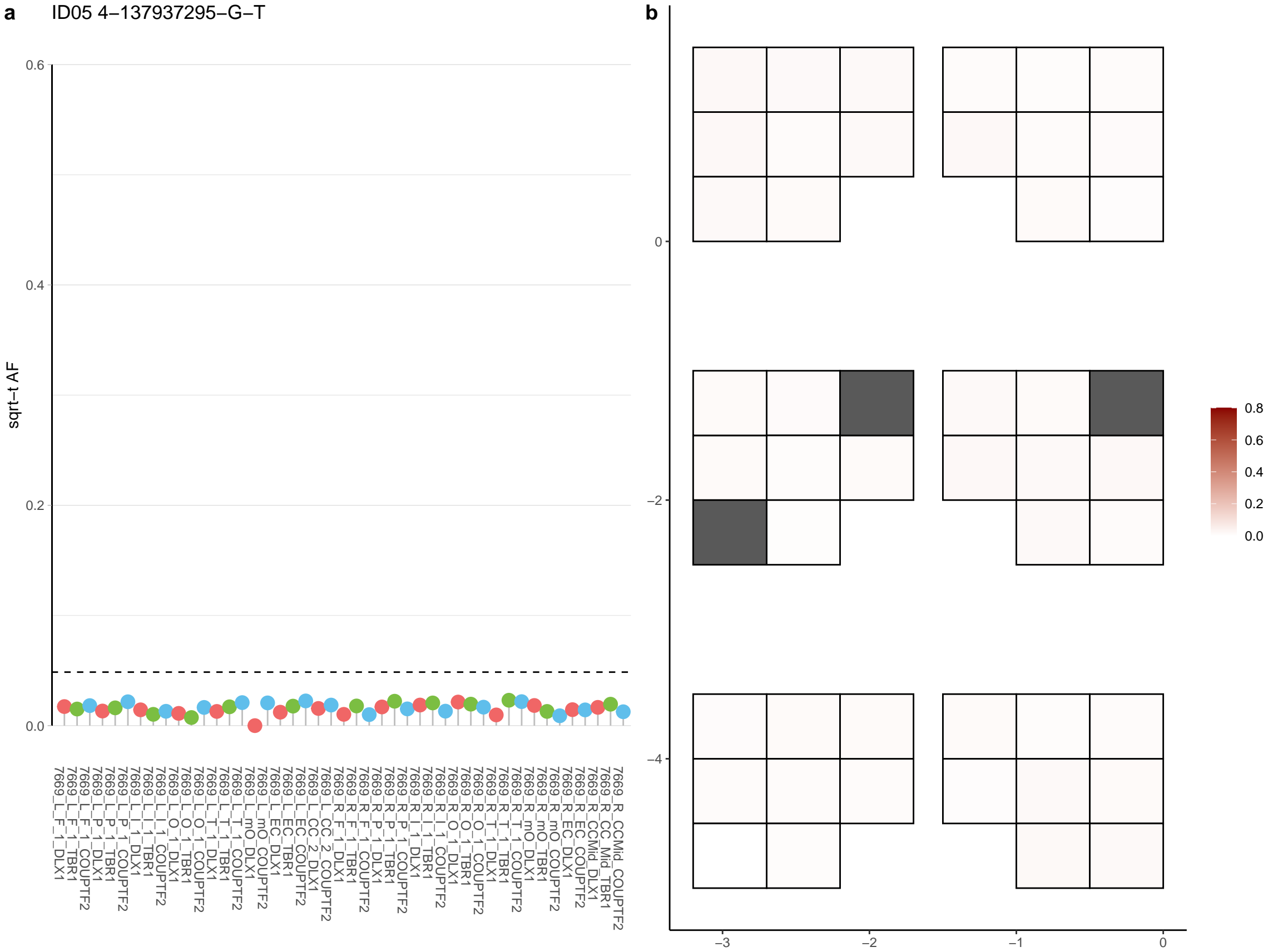

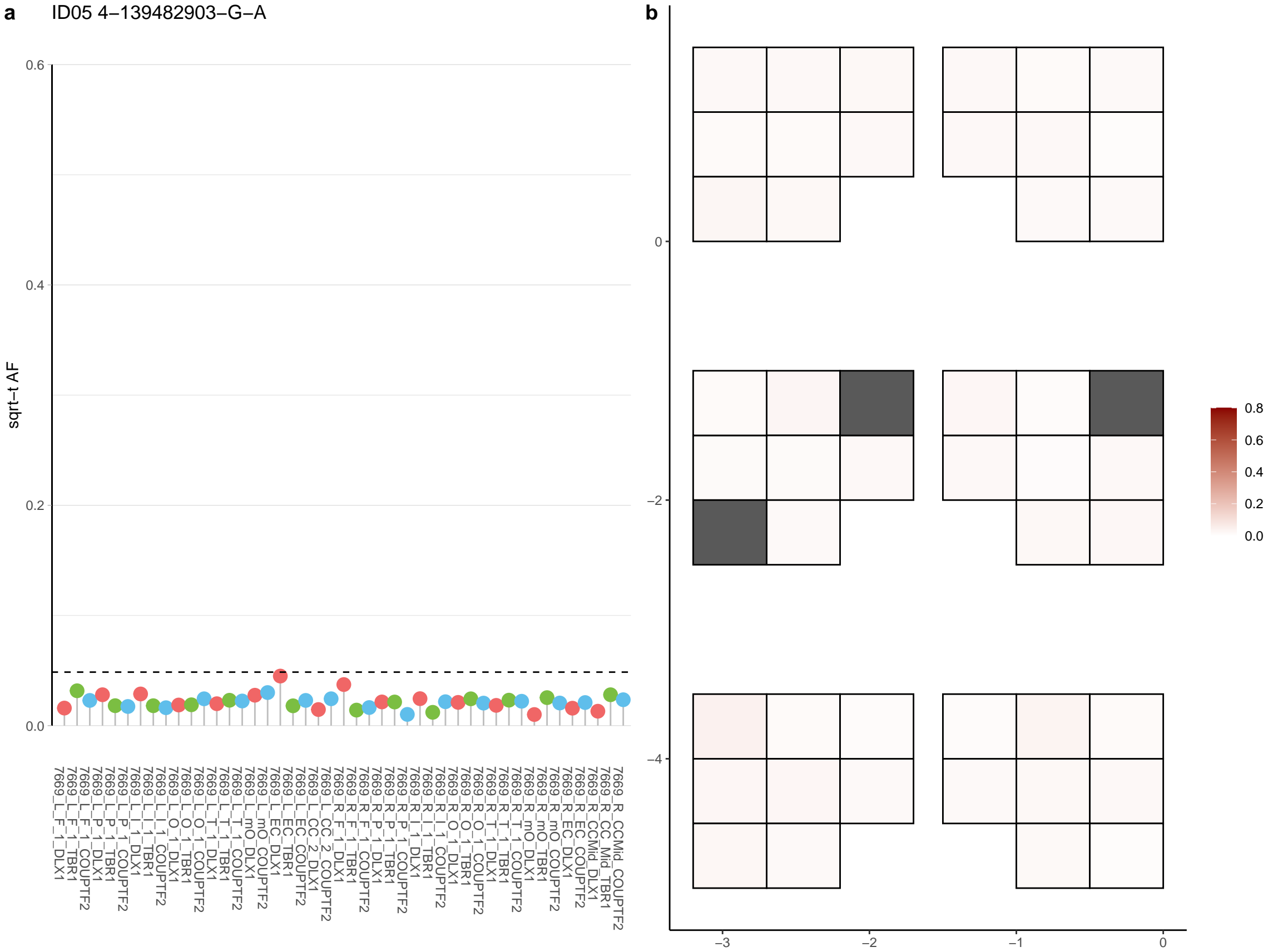

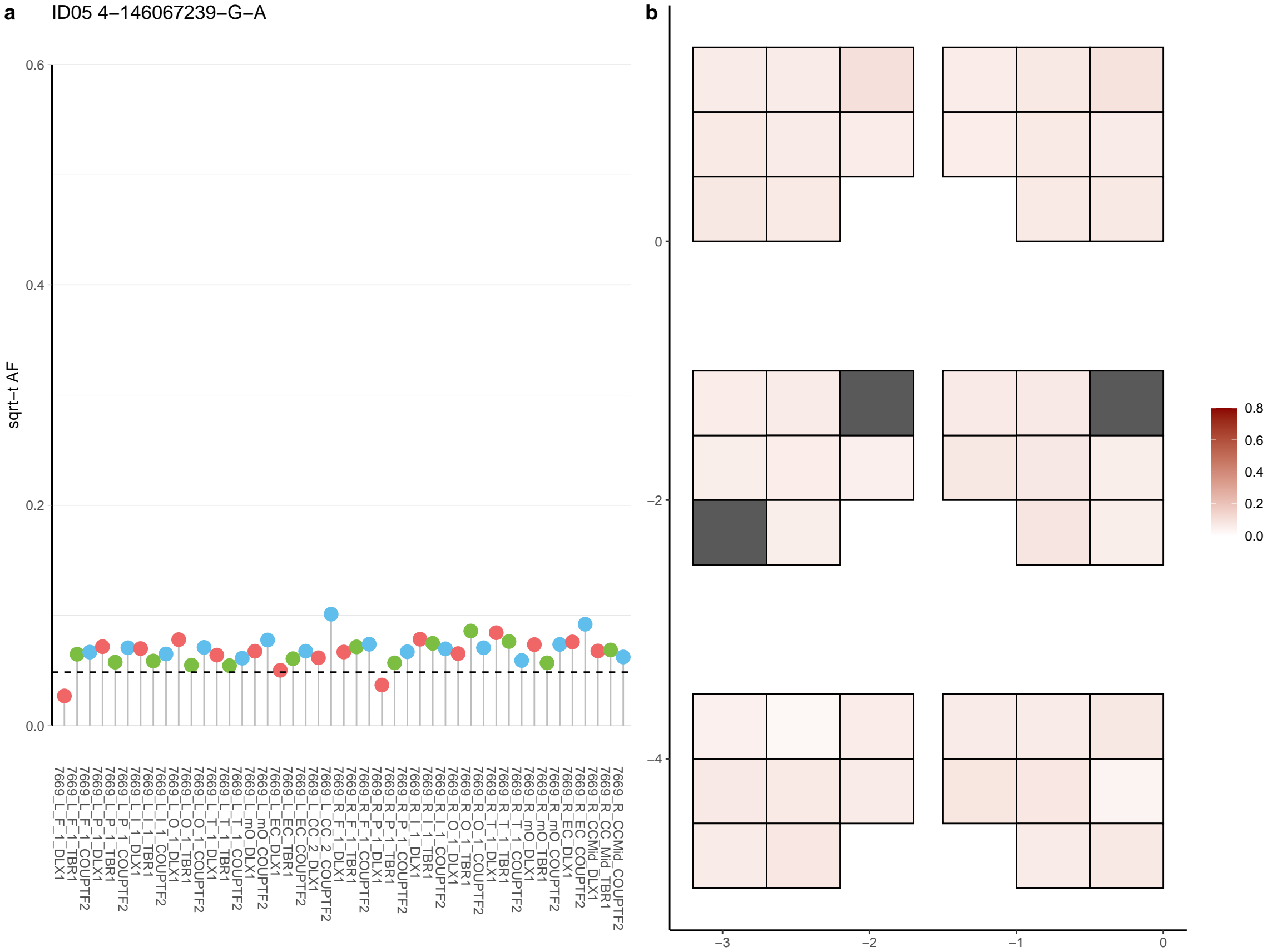

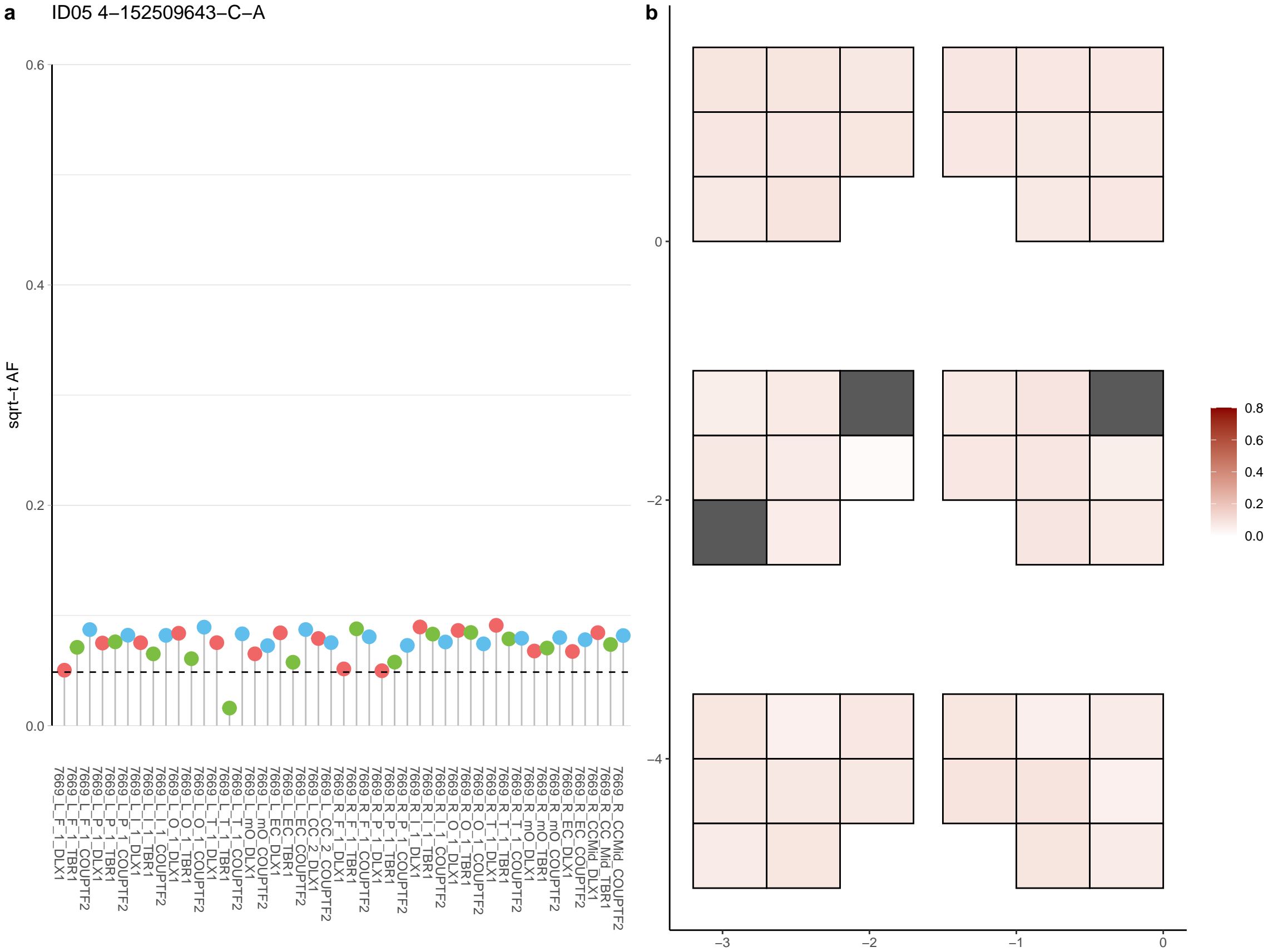

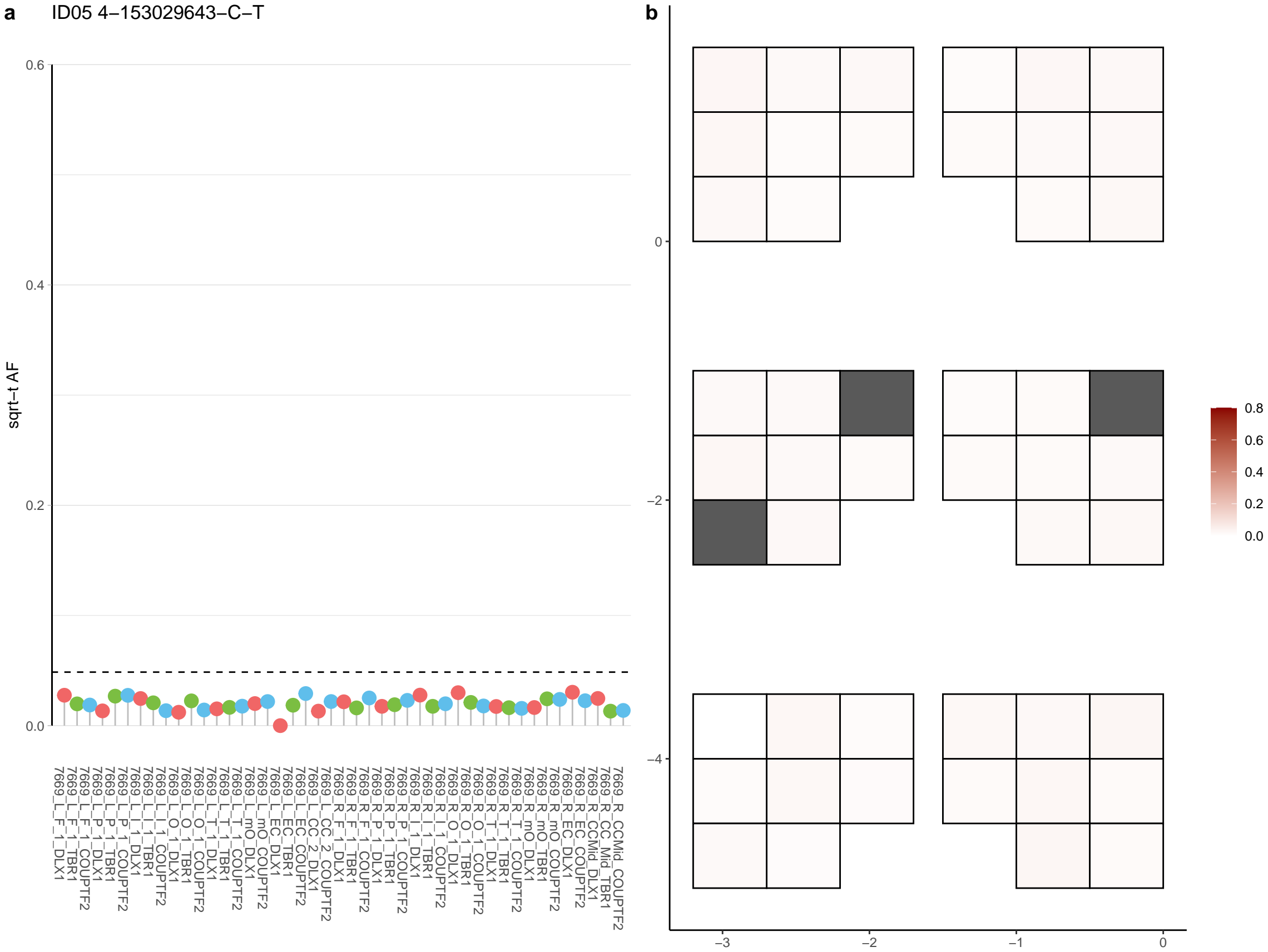

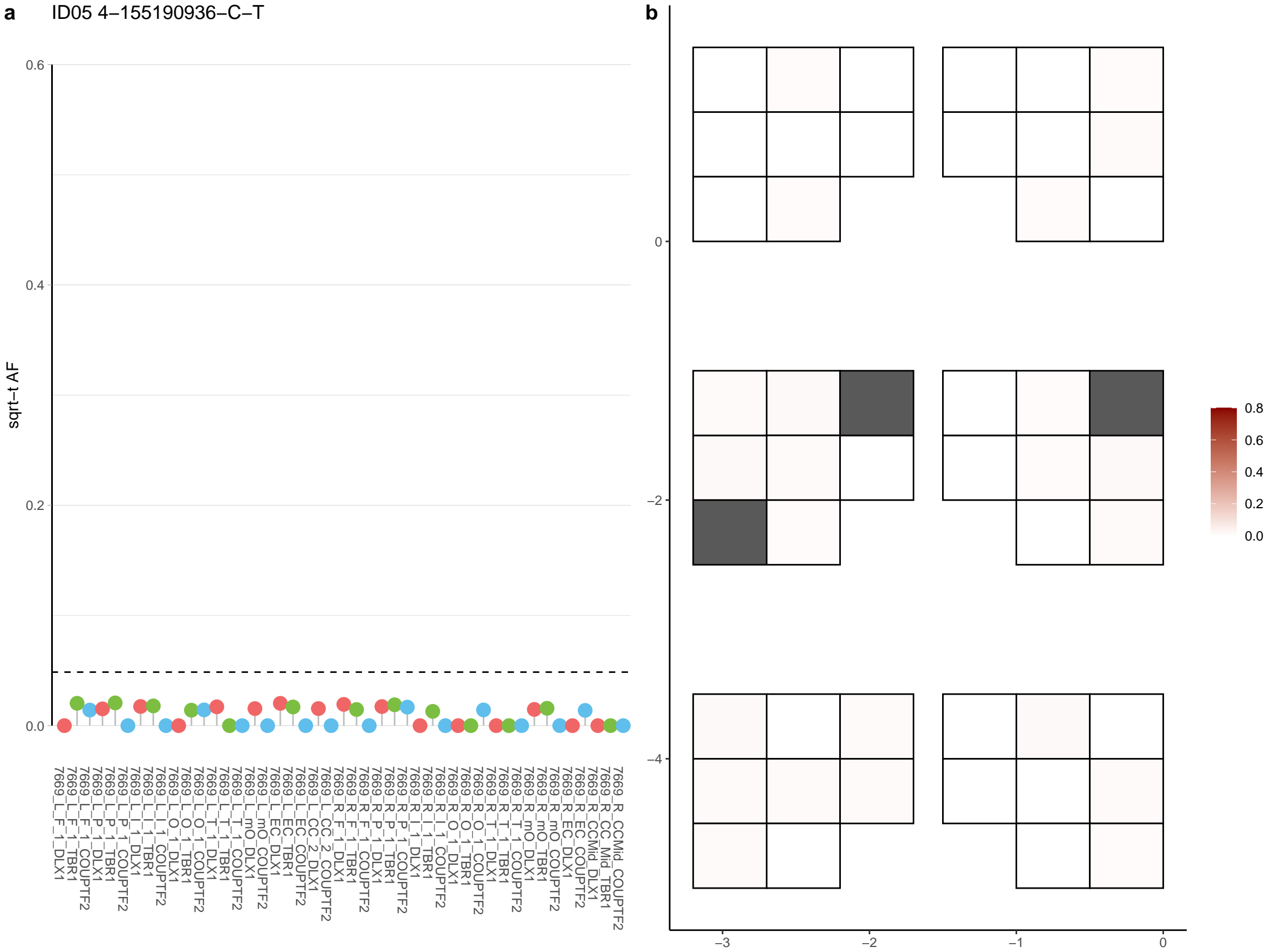

**a** ID05 4-155889870-C-T

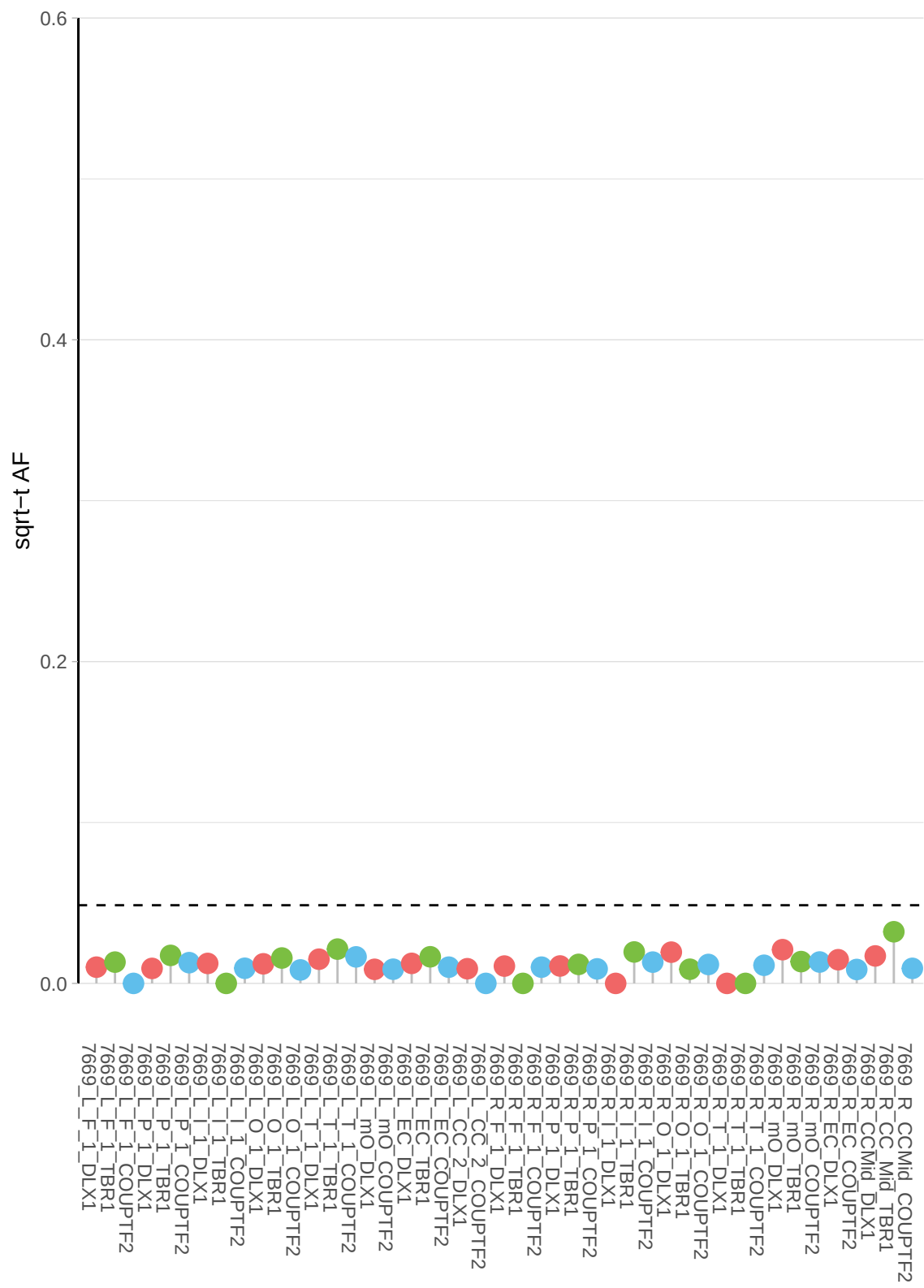

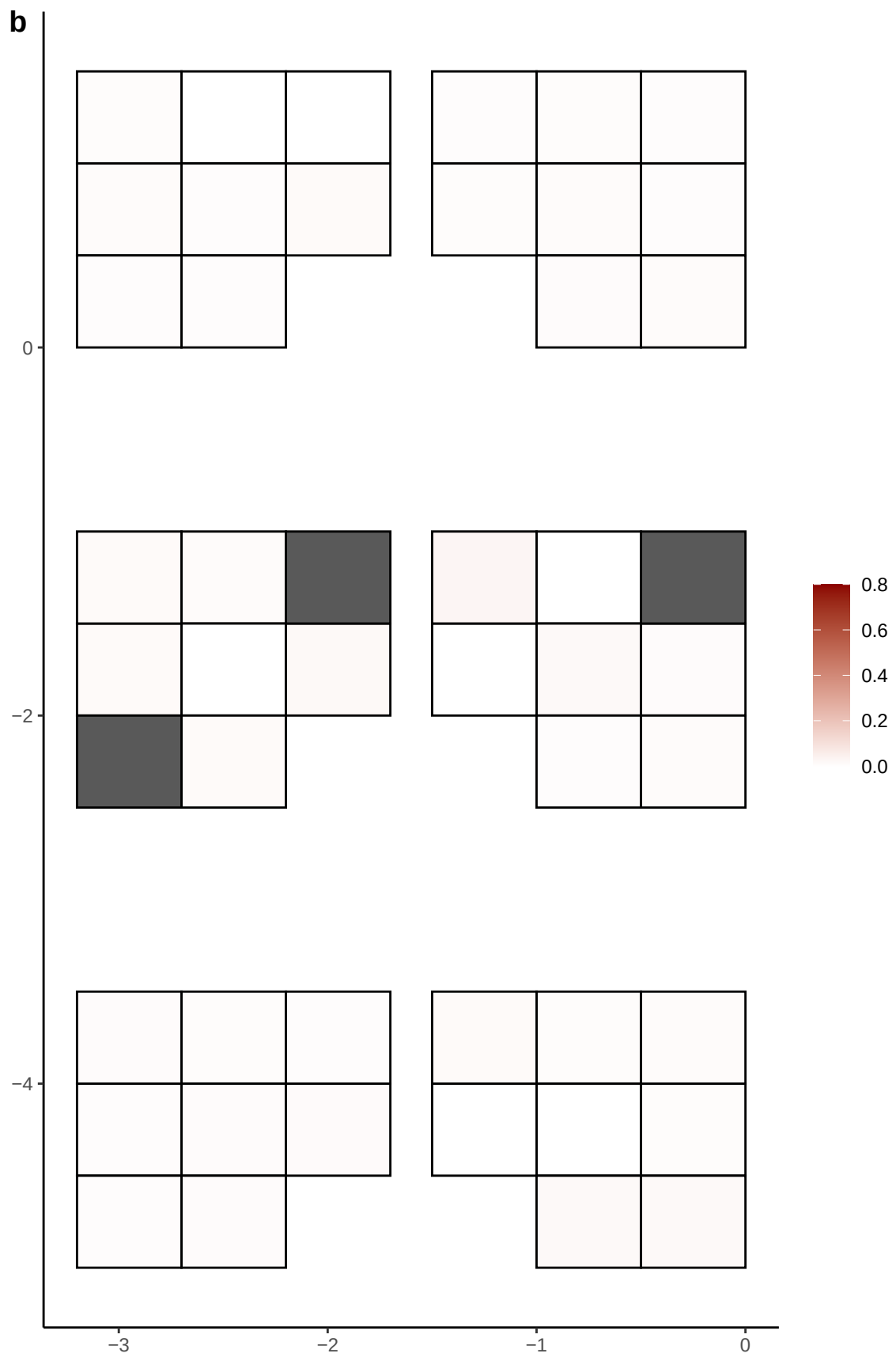

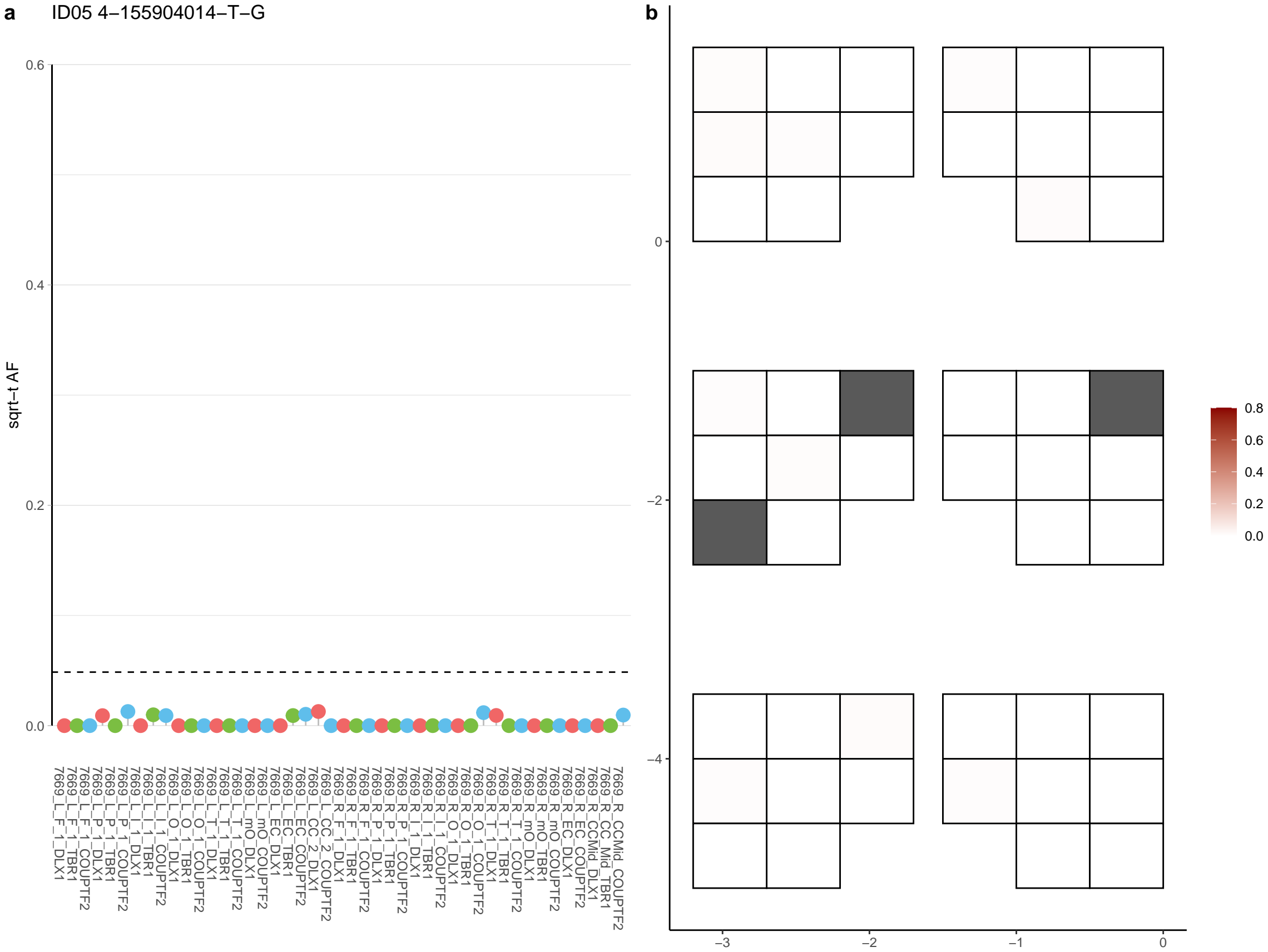

**a** ID05 4-165952565-A-G

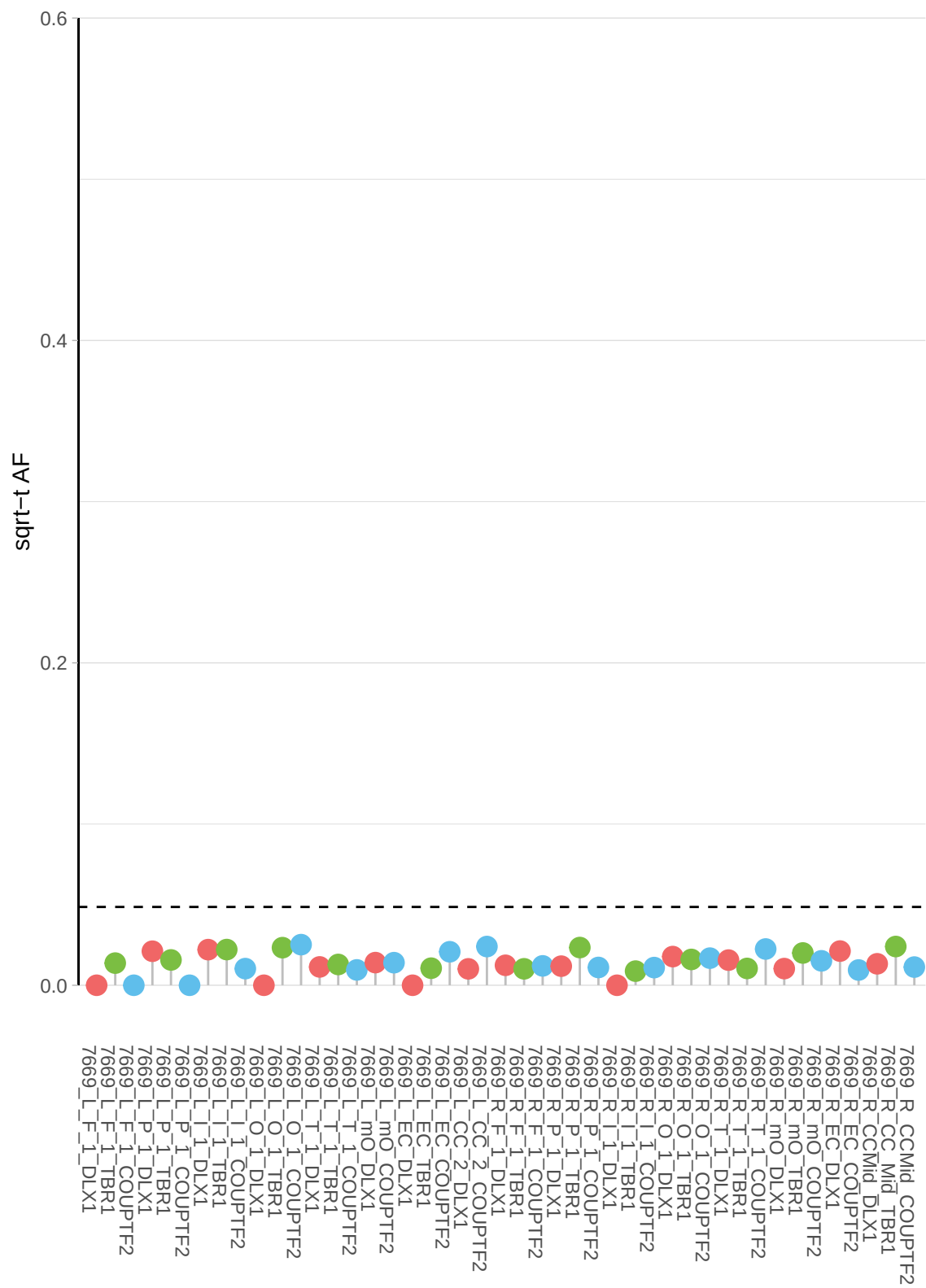

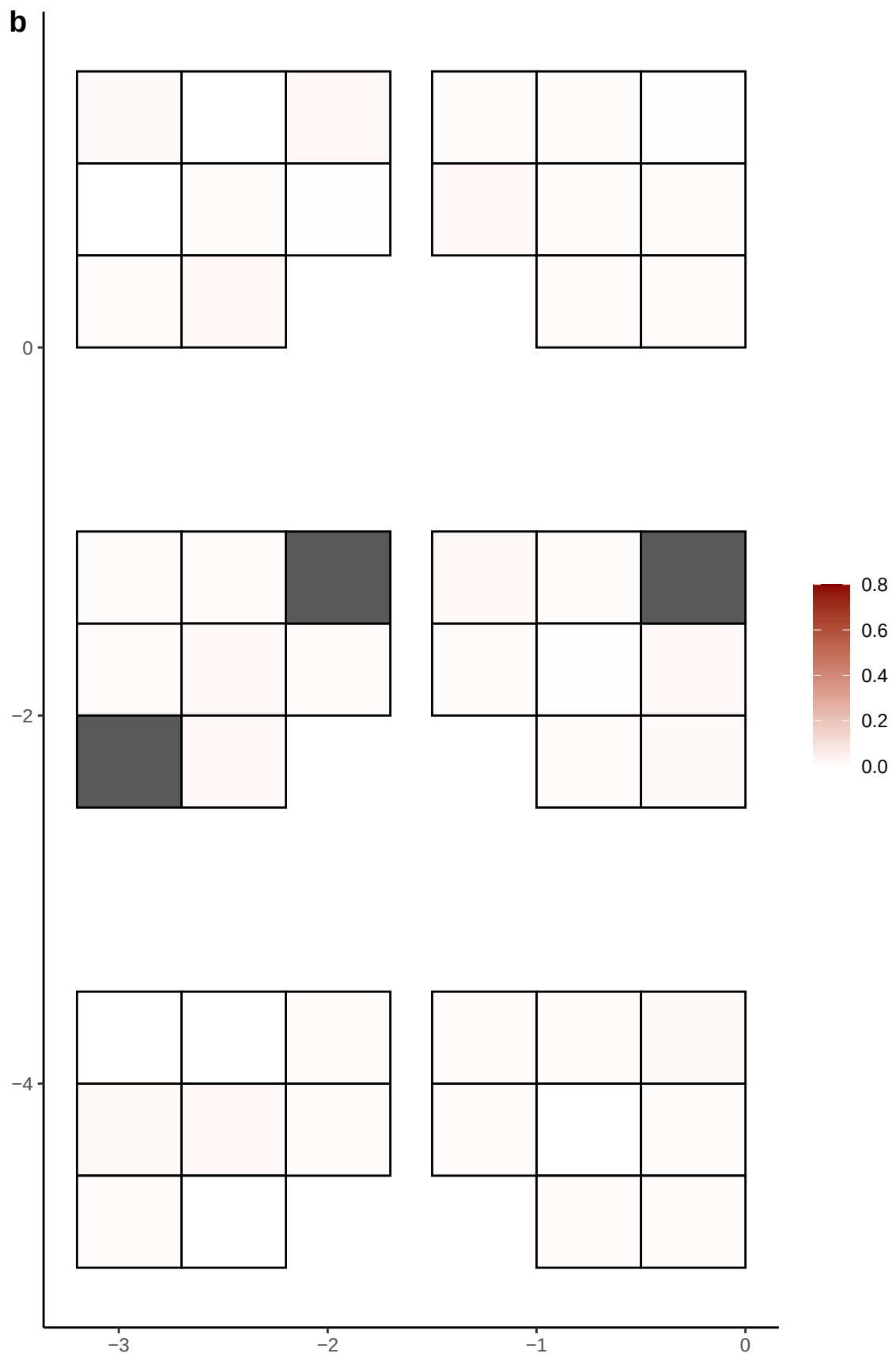

**a** ID05 4-168050168-C-A

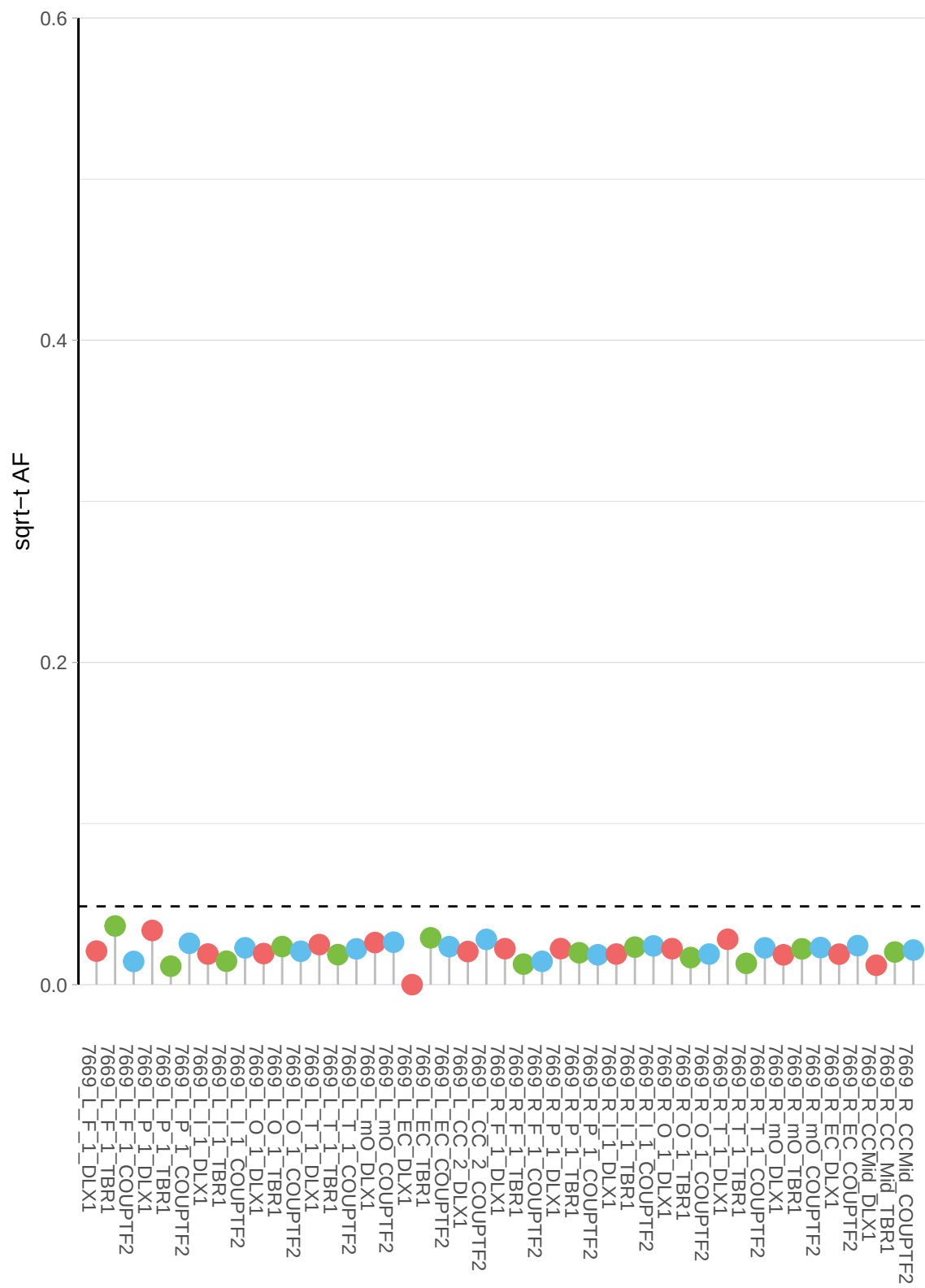

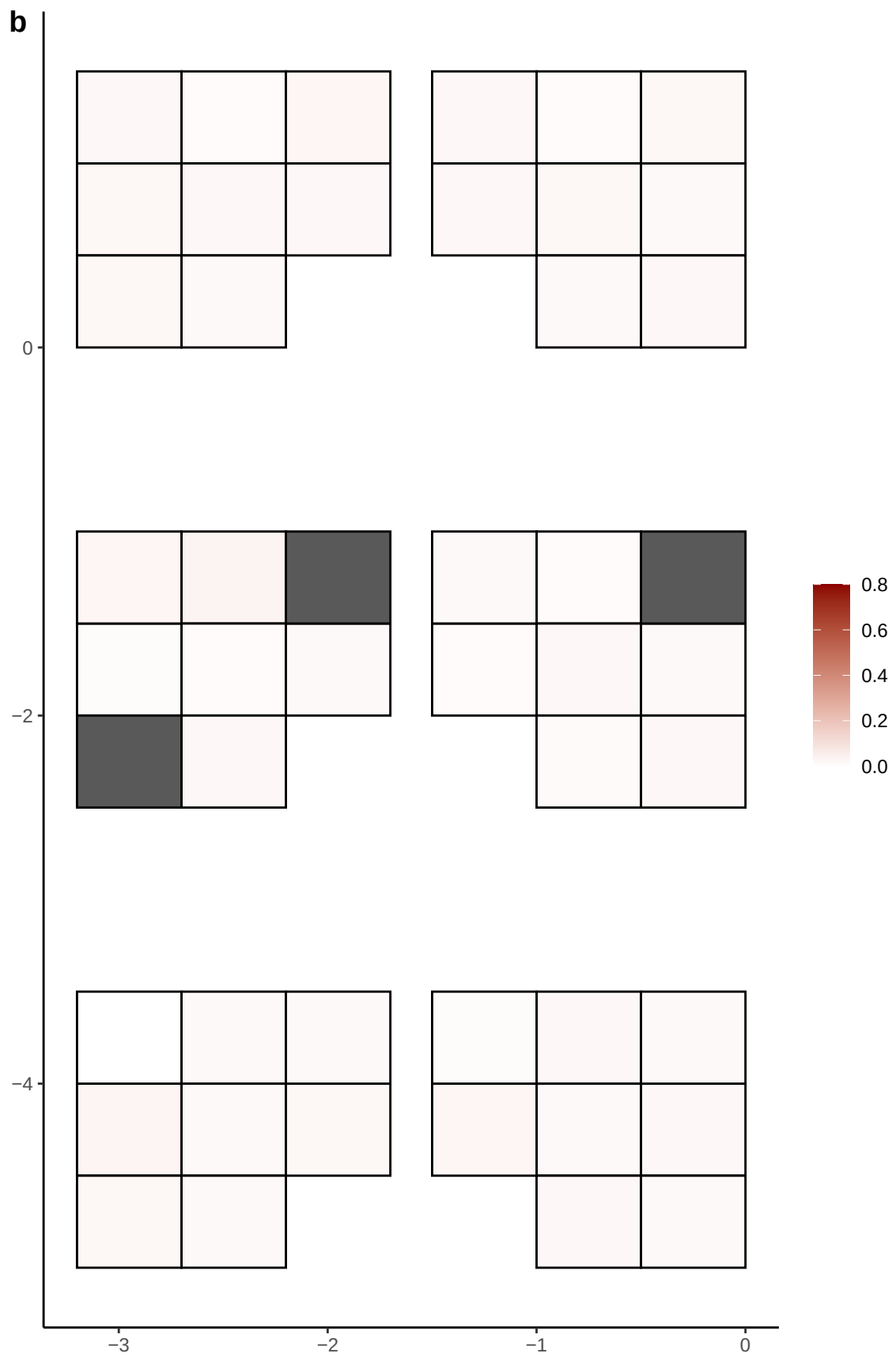

**a** ID05 4-183244212-C-T

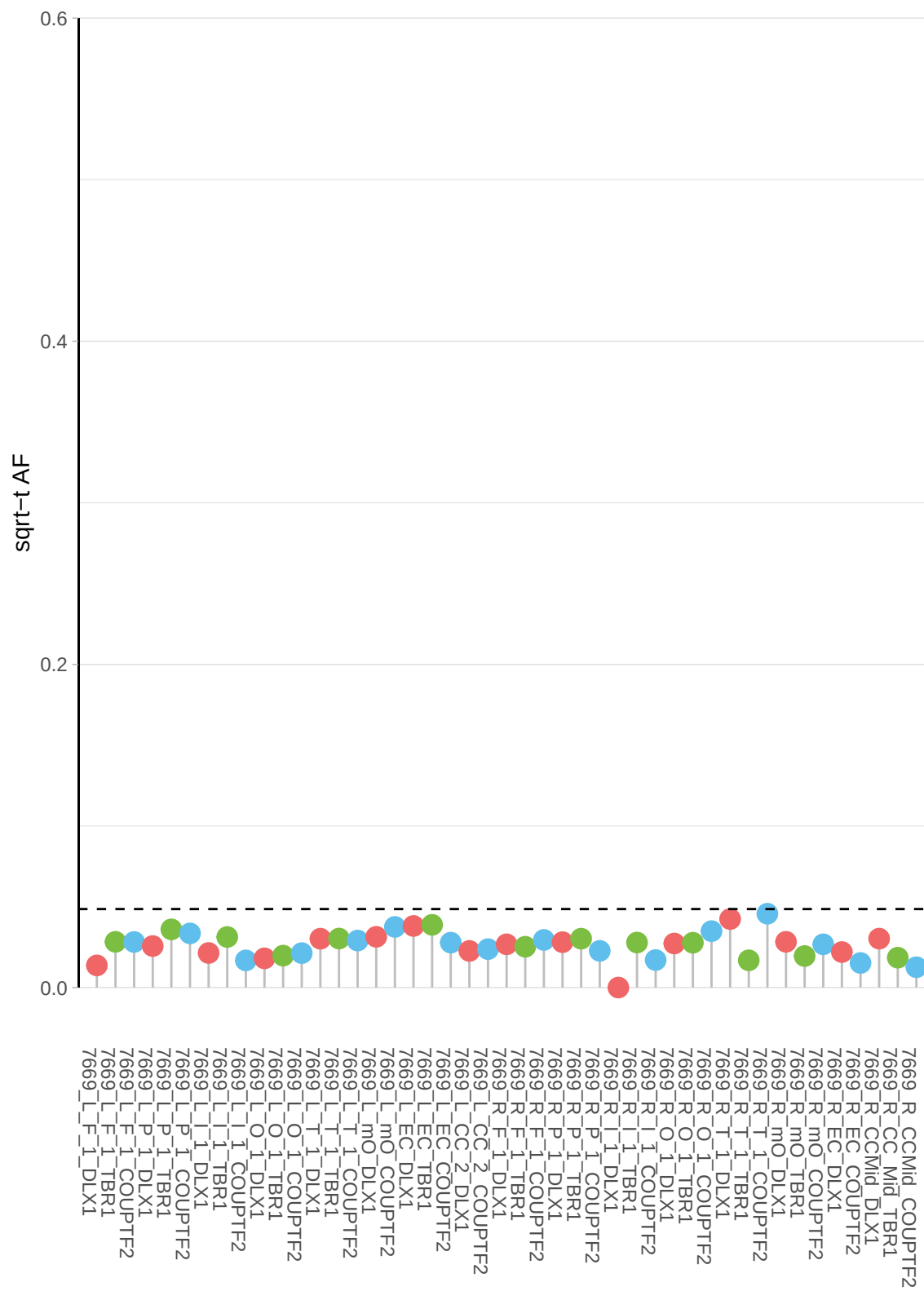

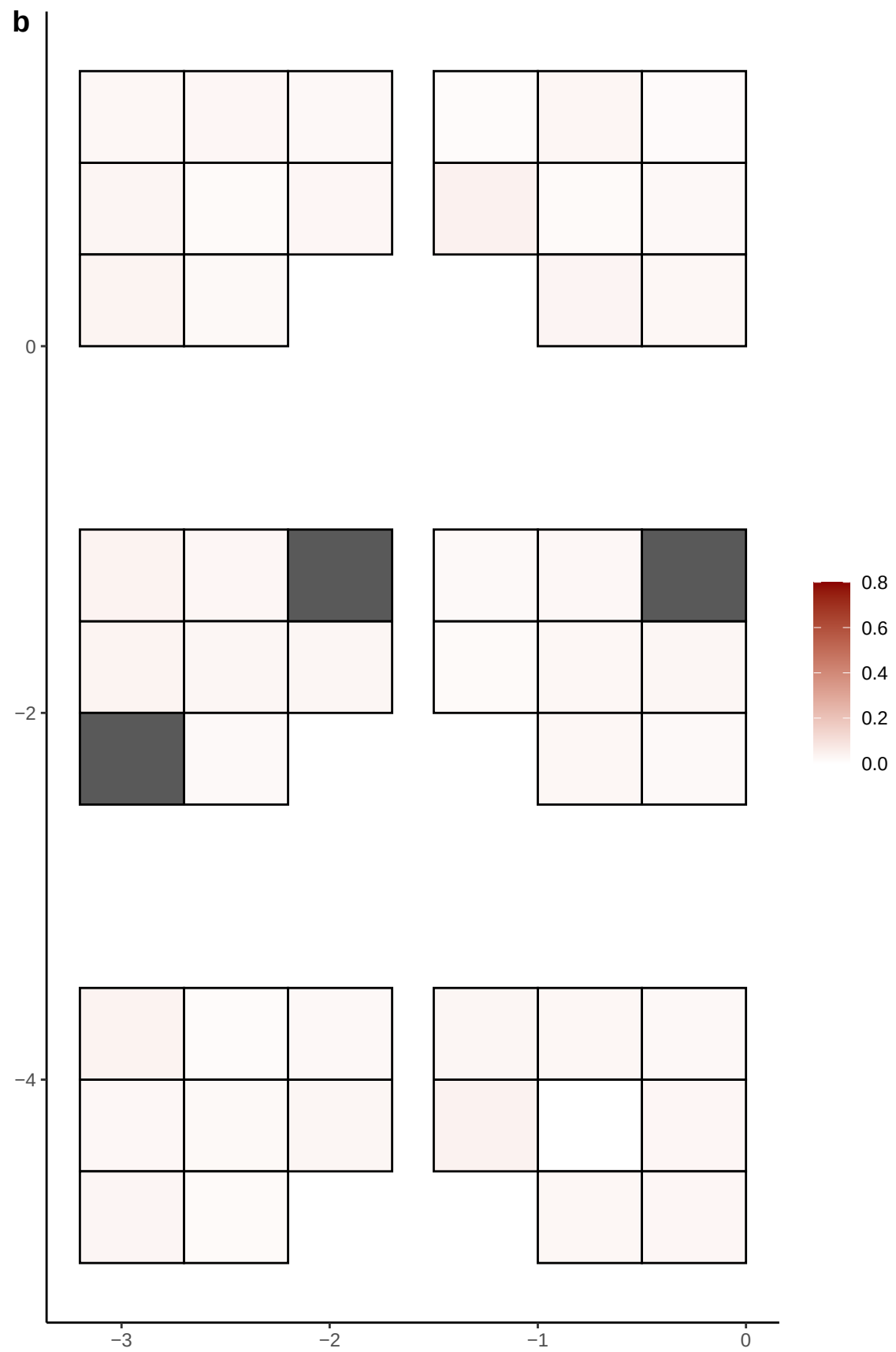

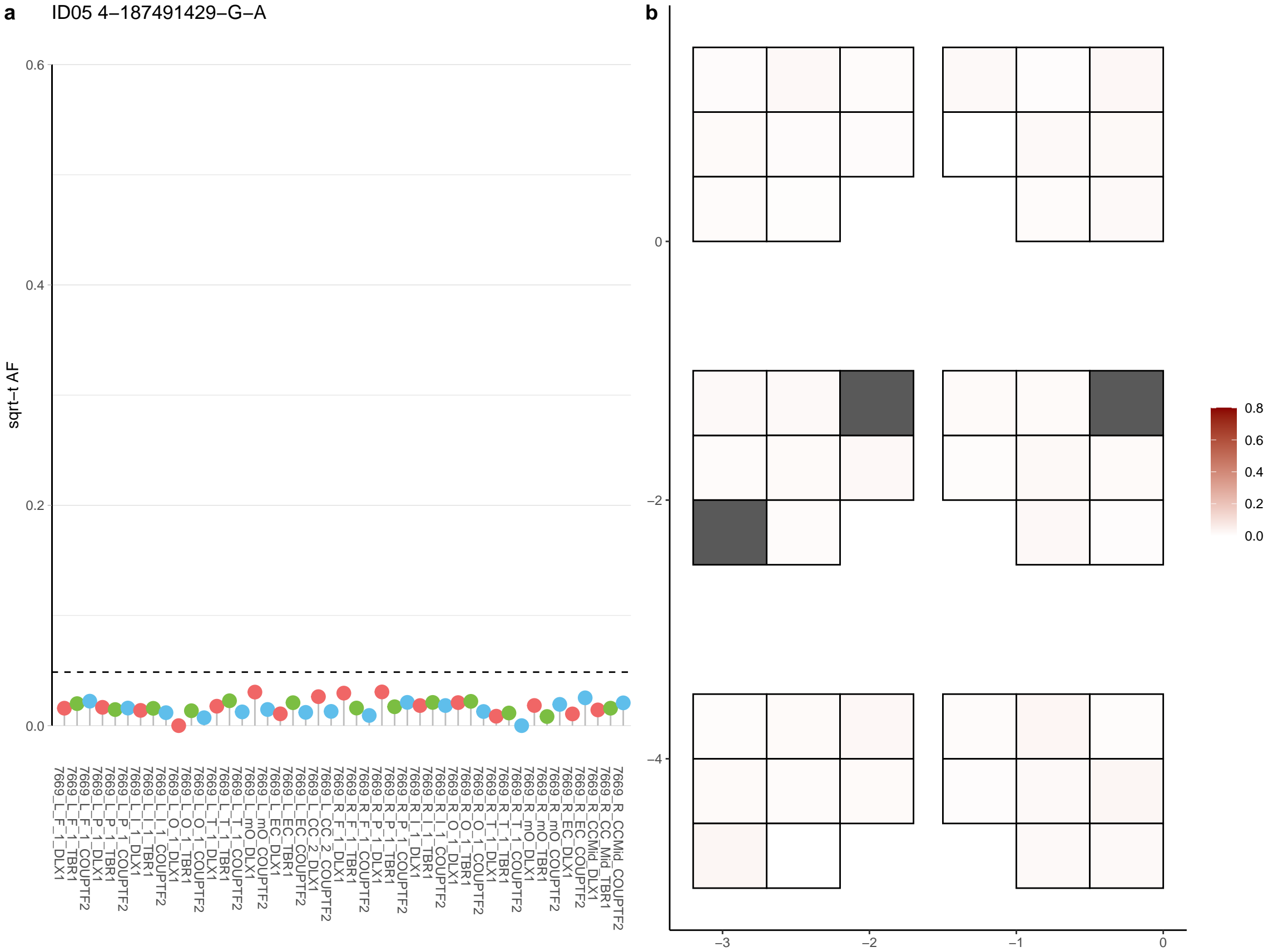

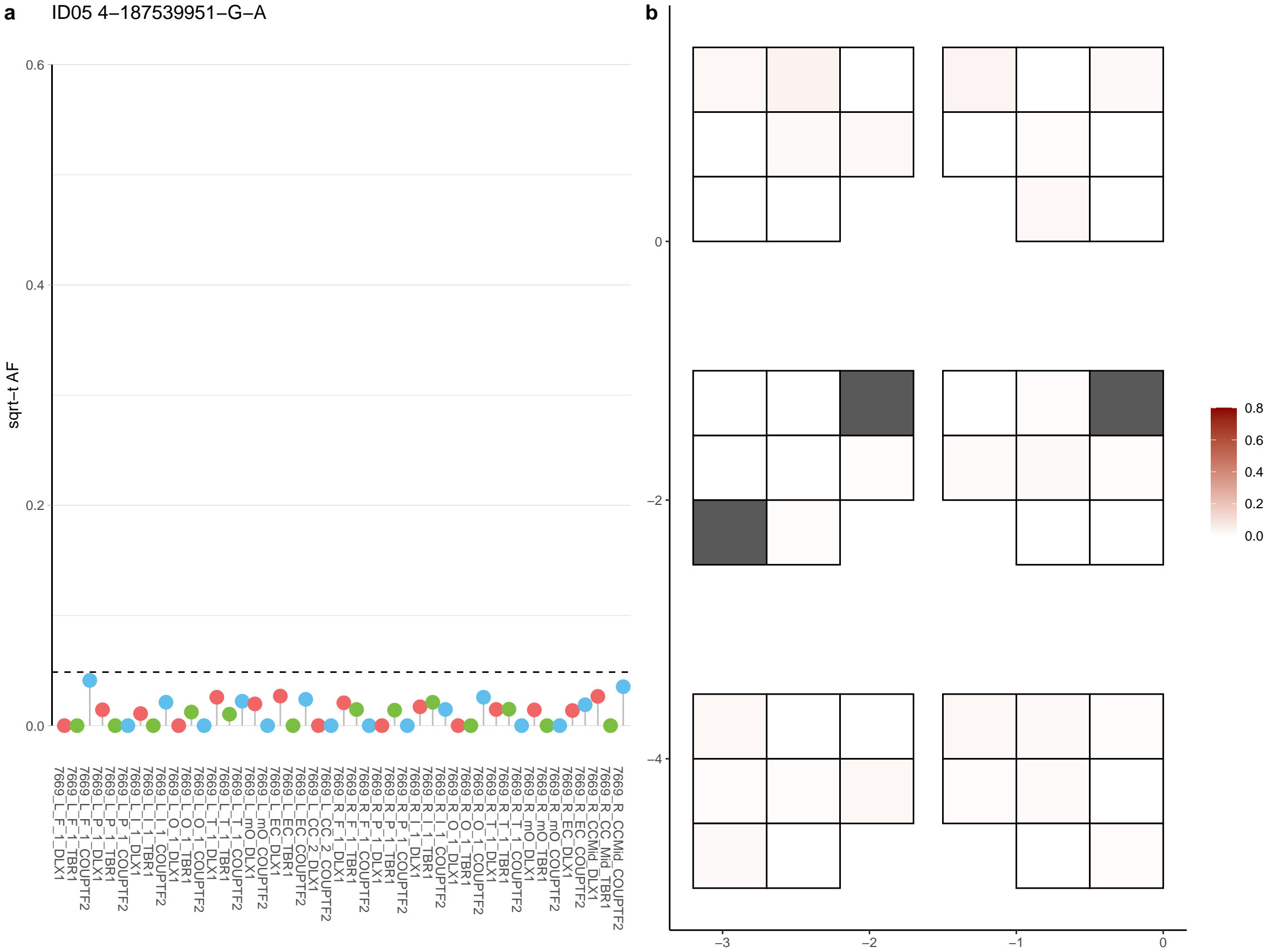

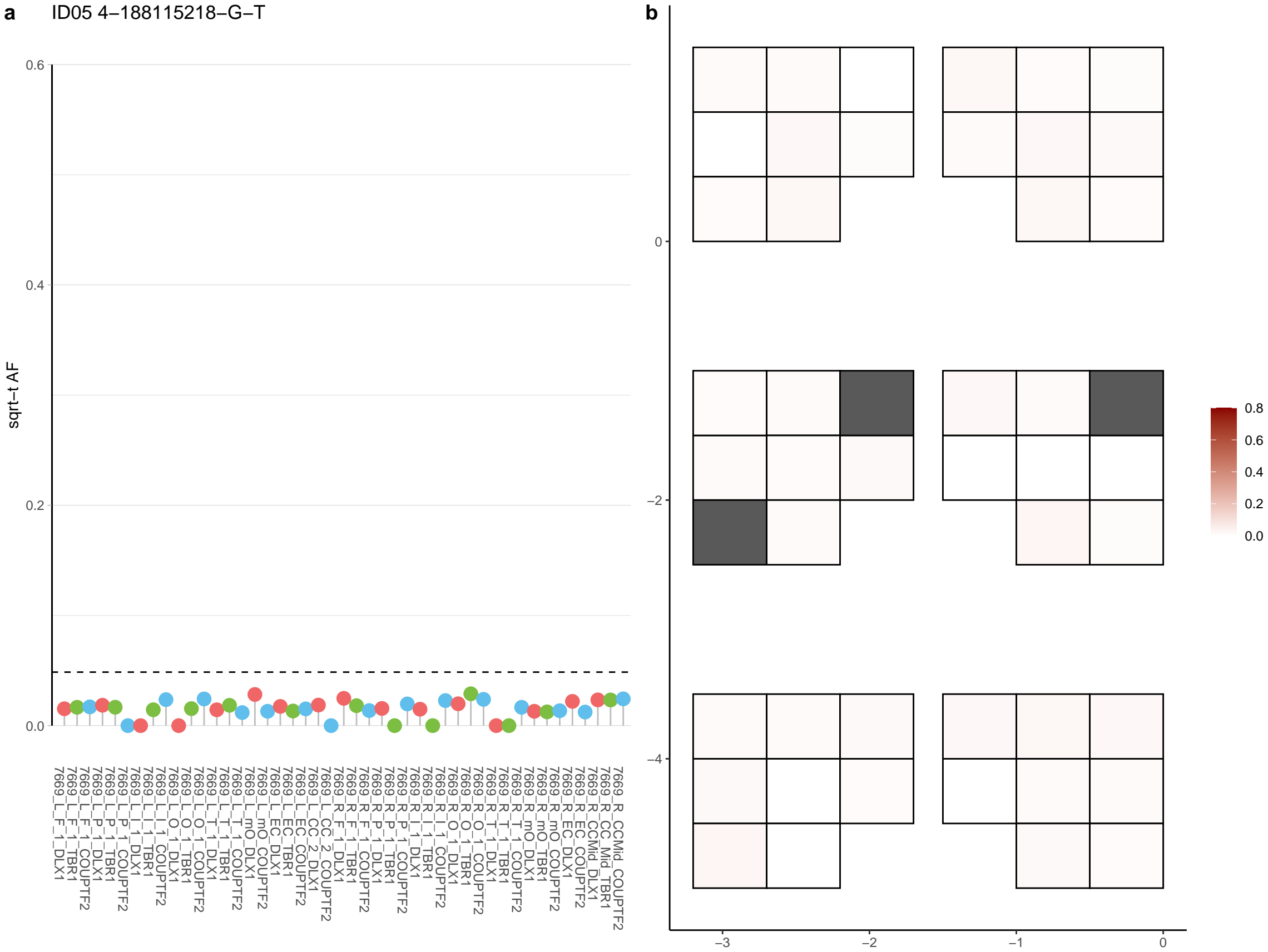

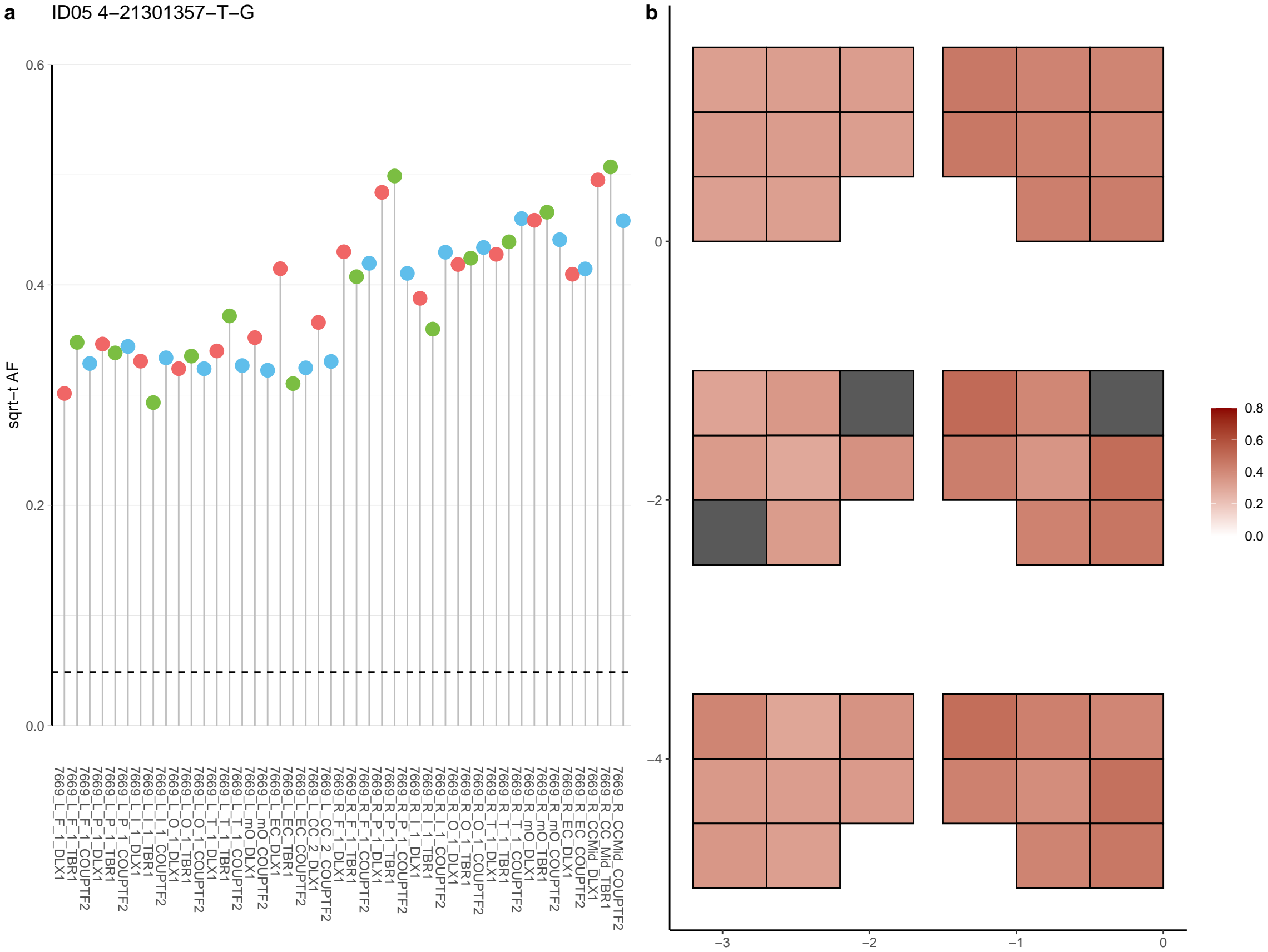

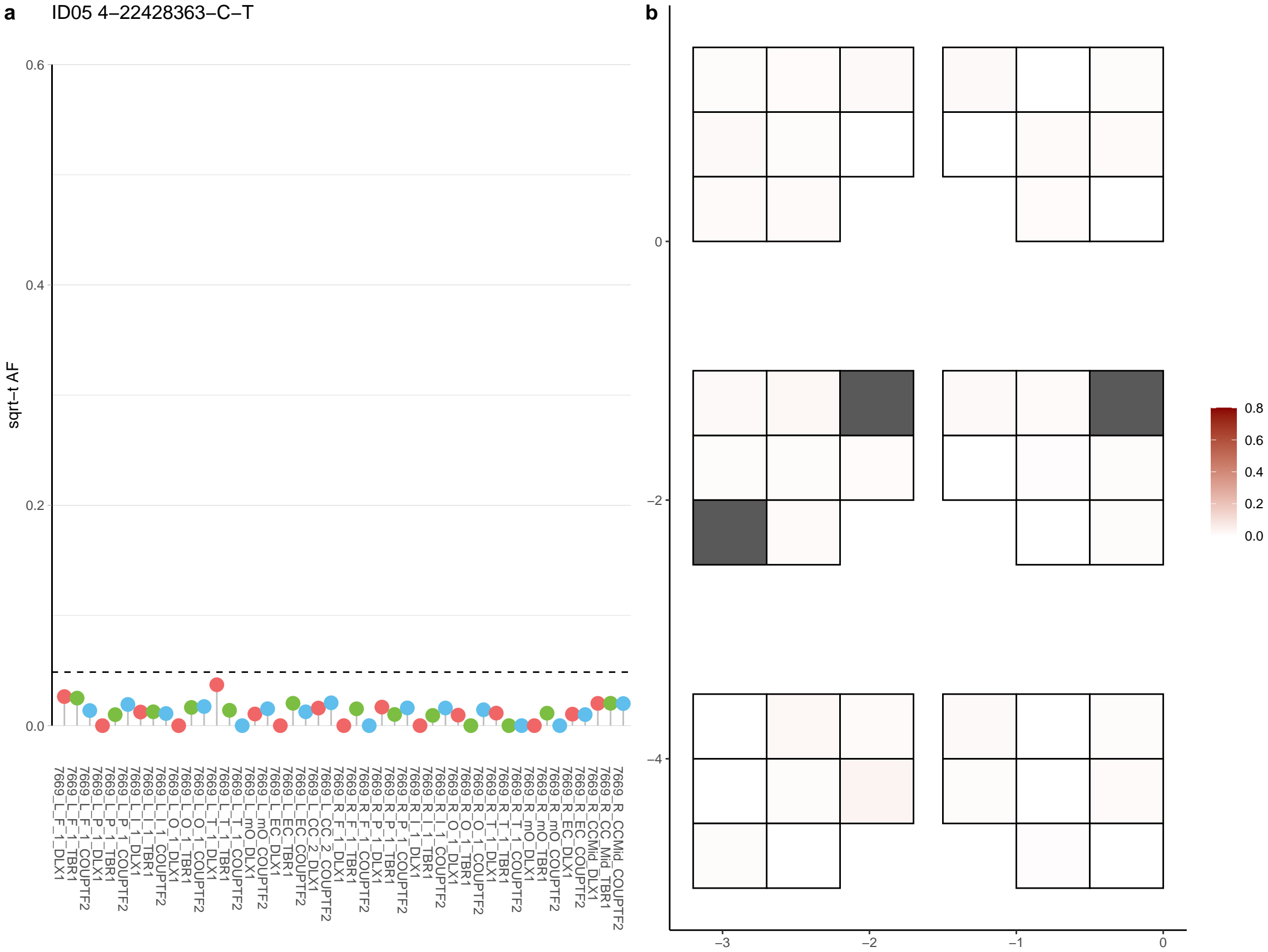

**a** ID05 4-23552226-G-A

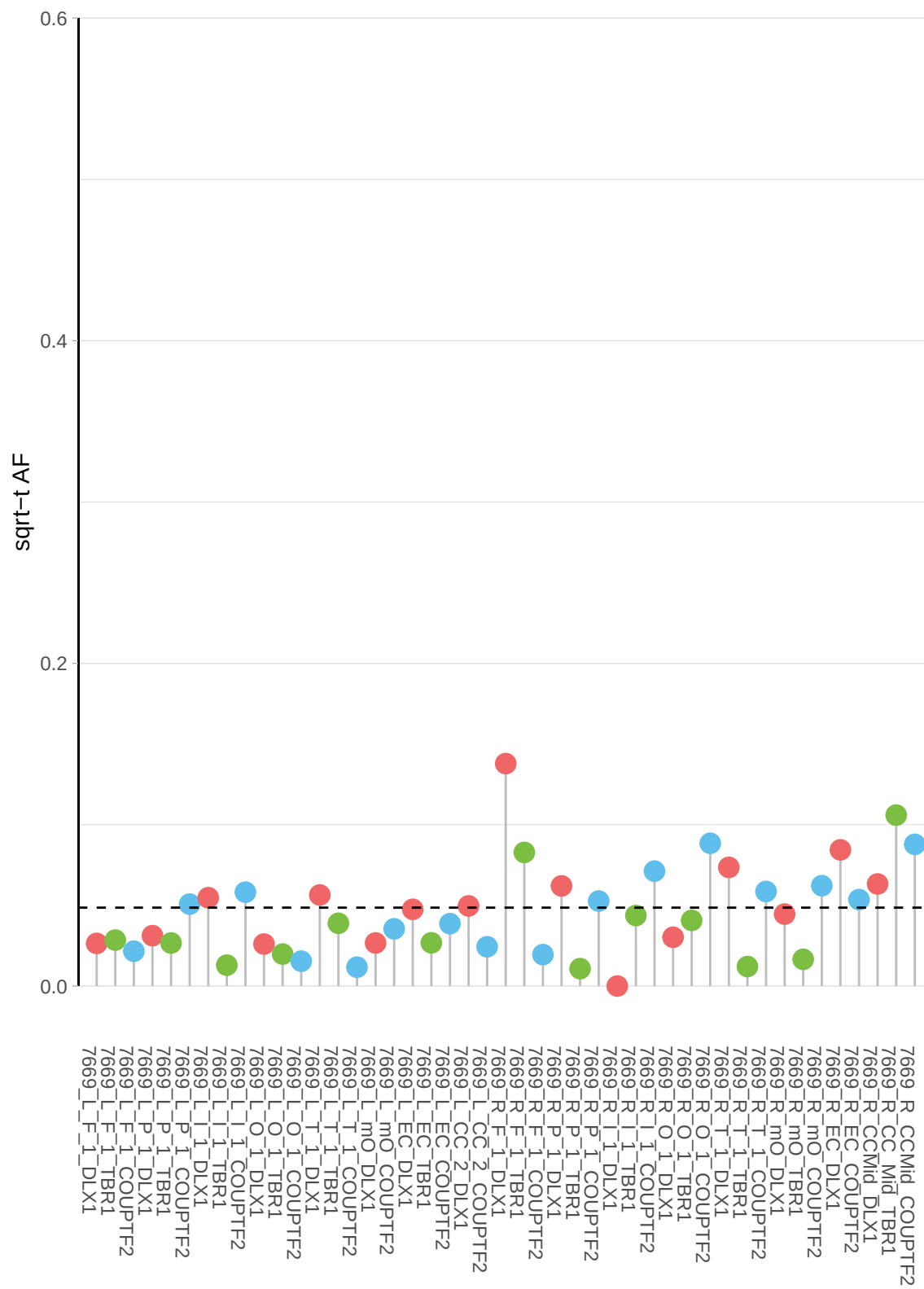

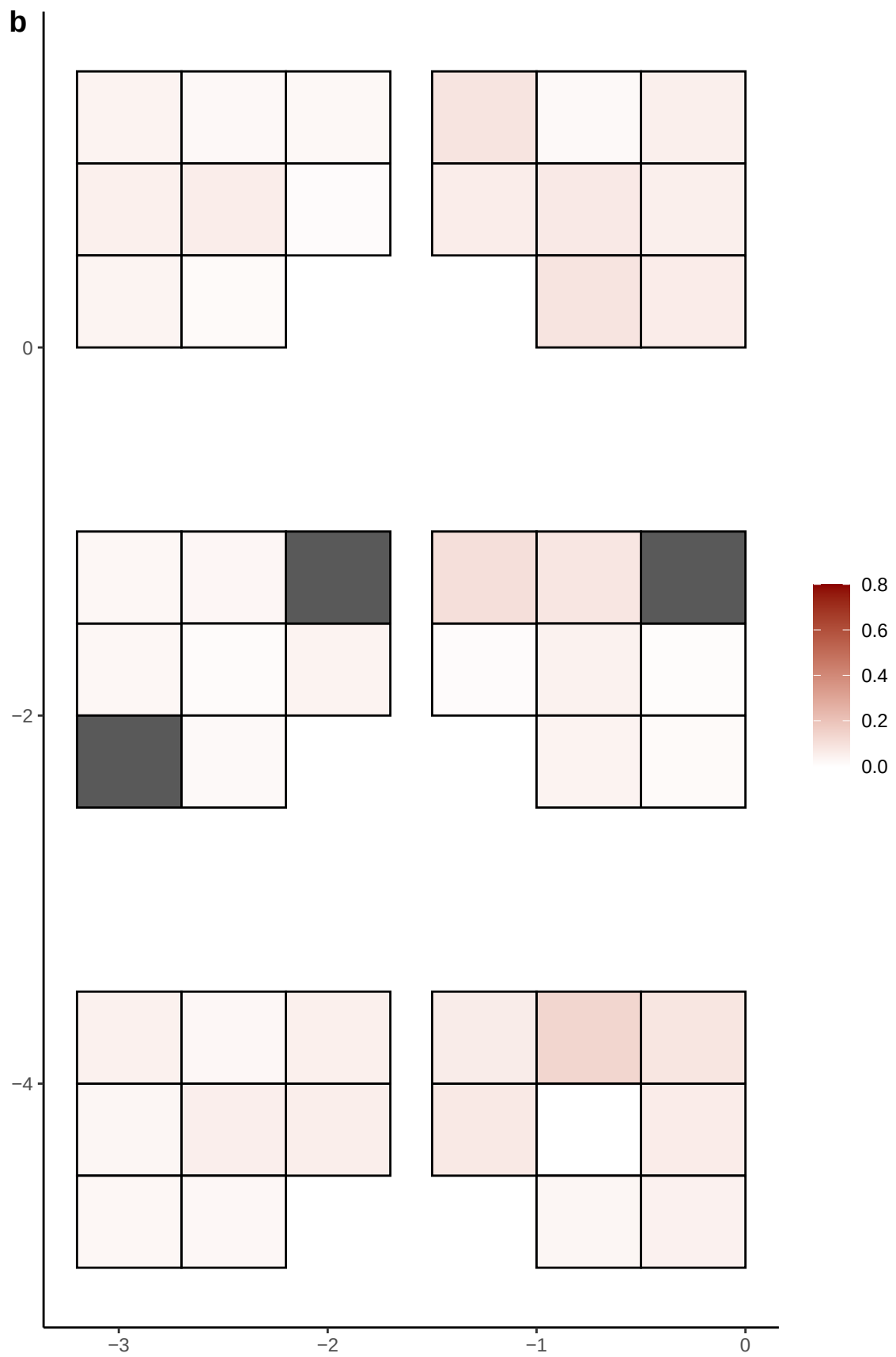

**a** ID05 4-25107945-C-T

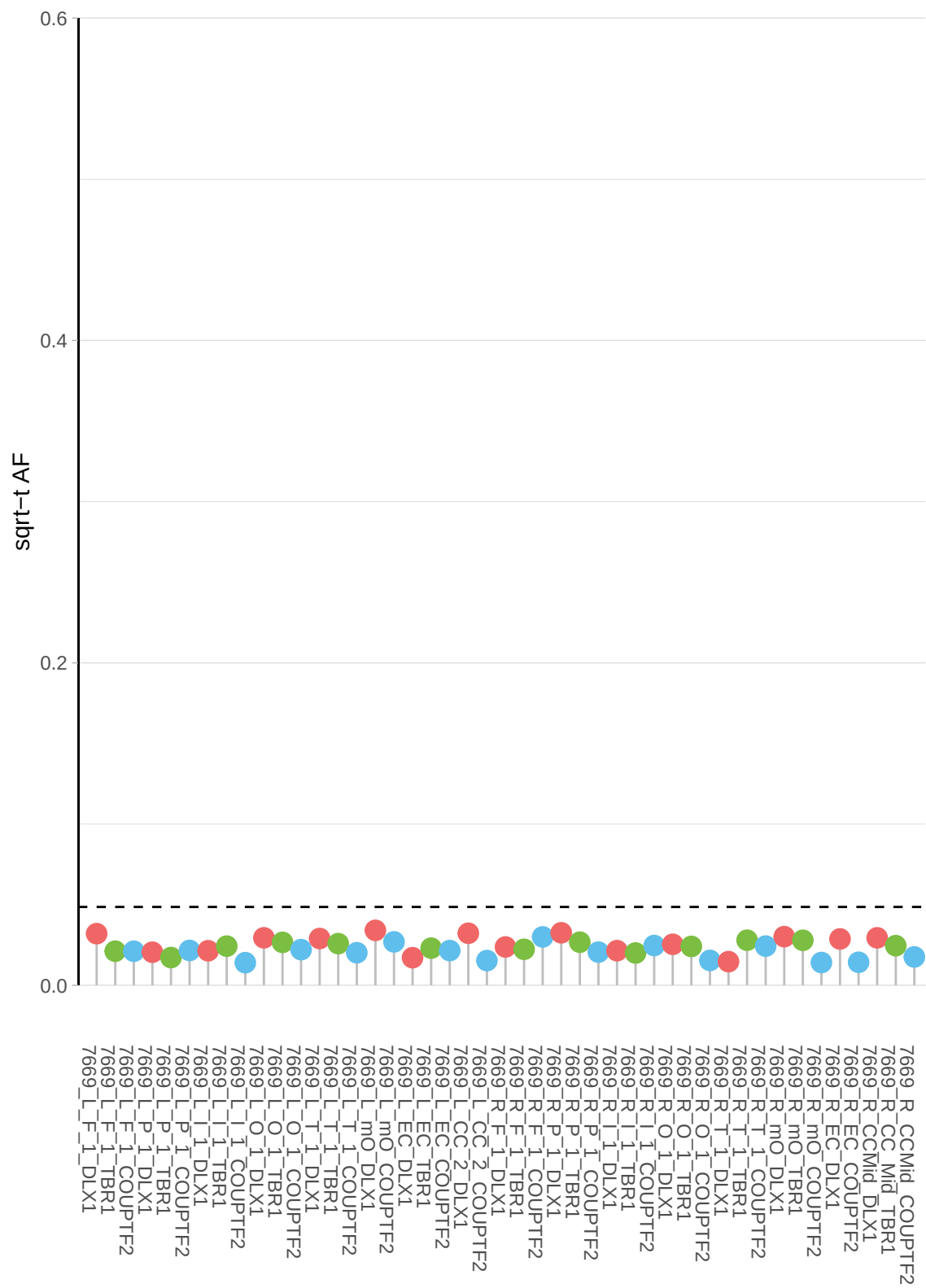

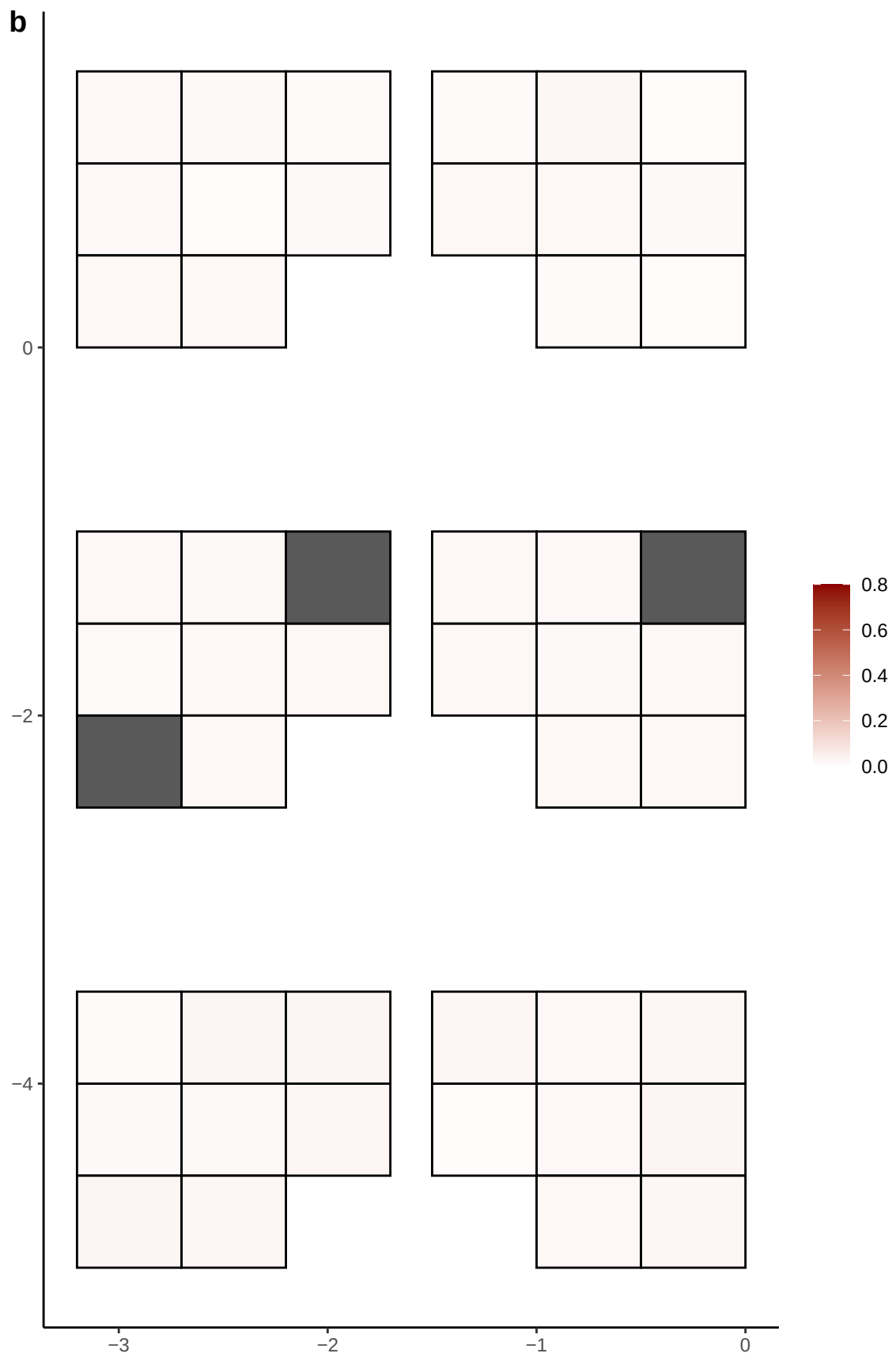

**a** ID05 4-26808694-C-T

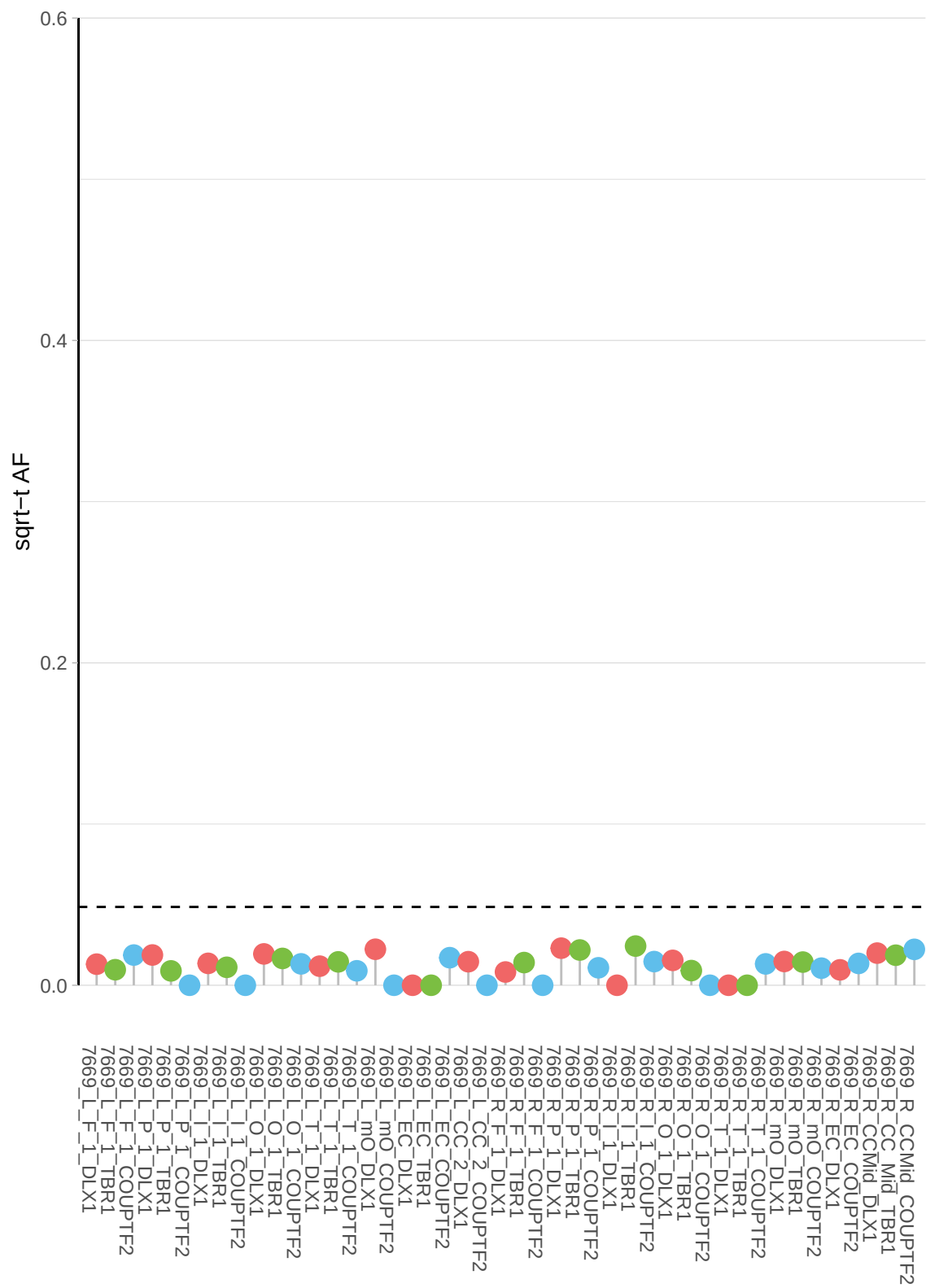

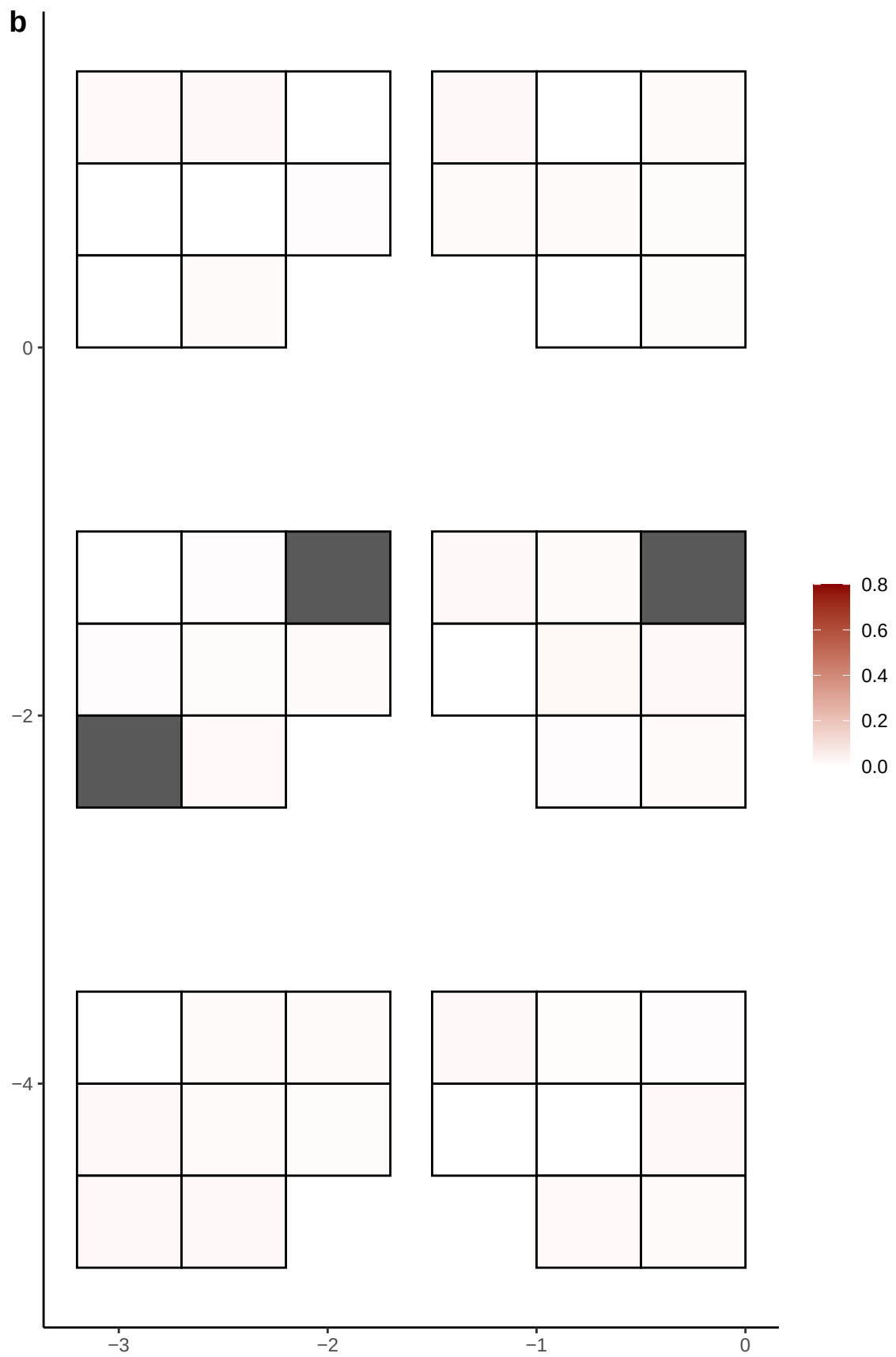

**a** ID05 4-30123079-T-G

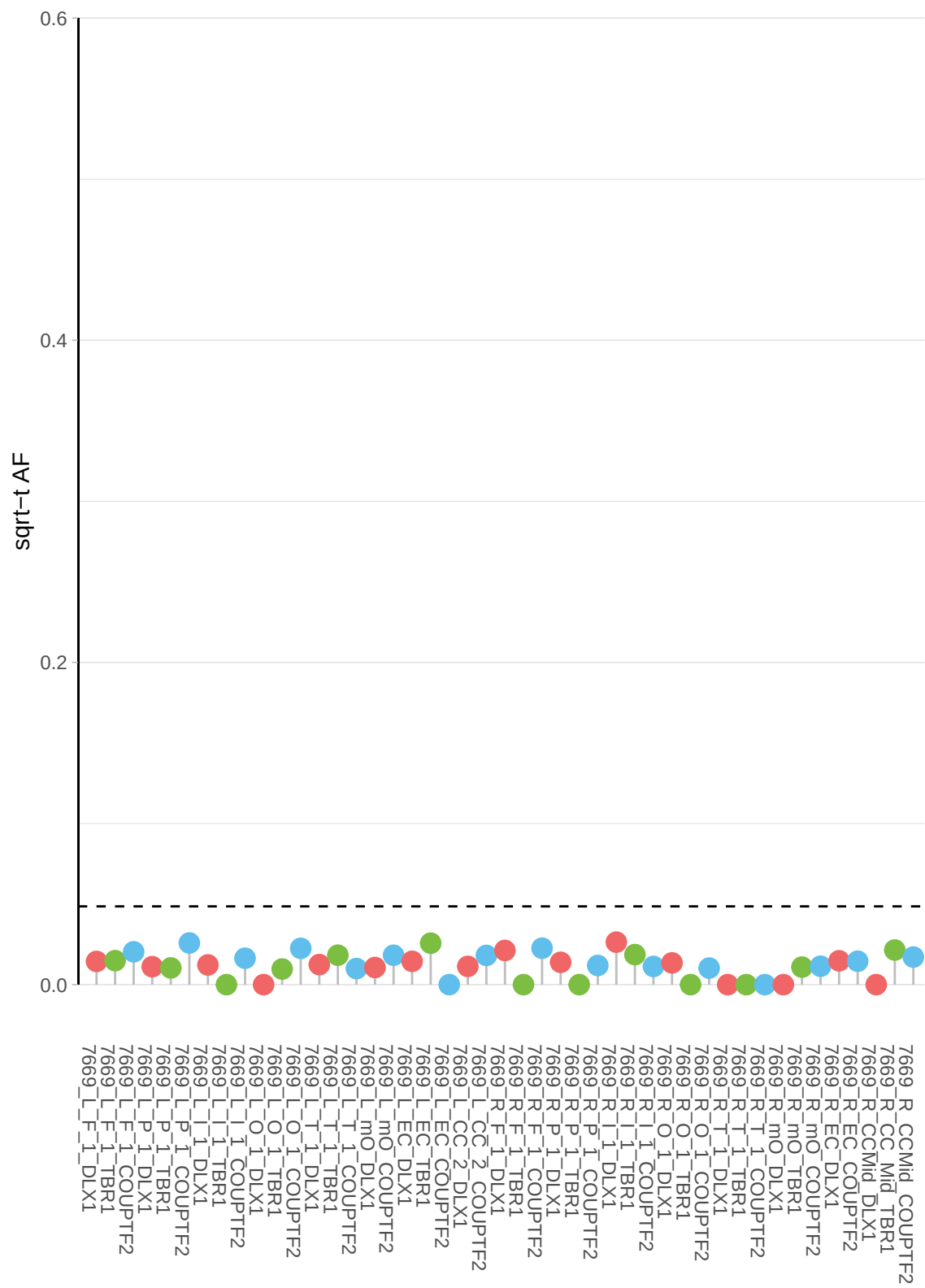

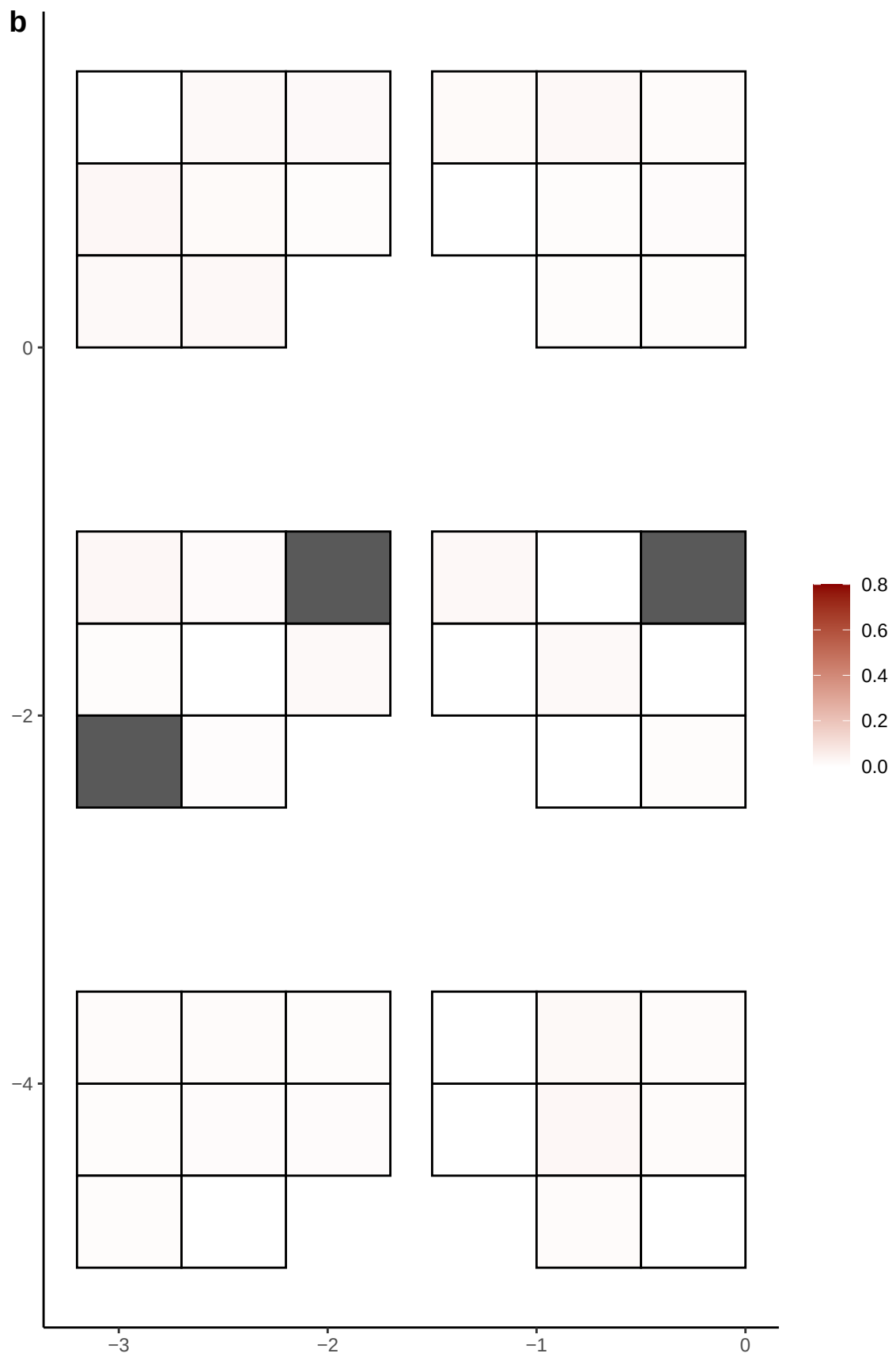

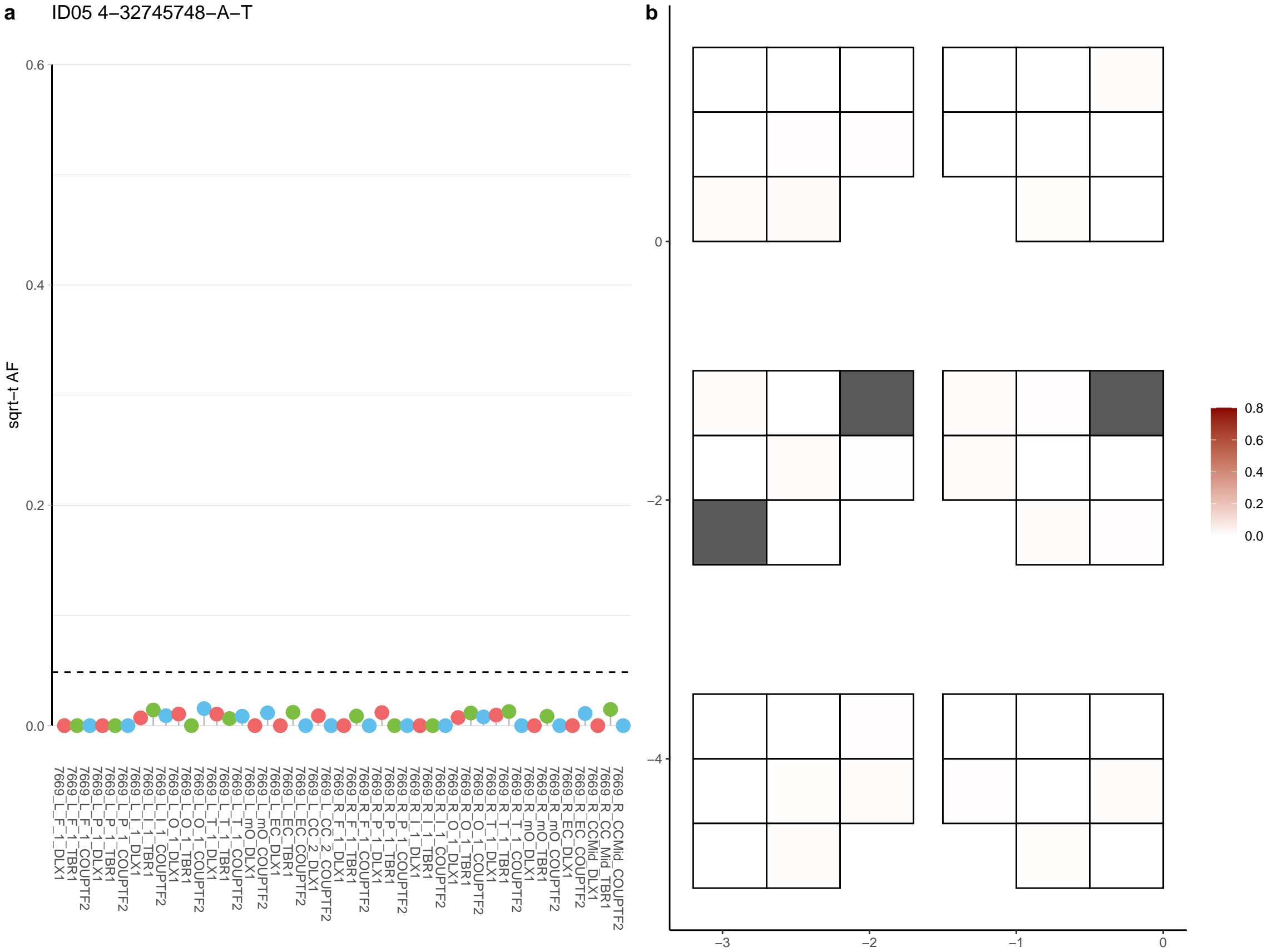

**a** ID05 4-34386270-C-T

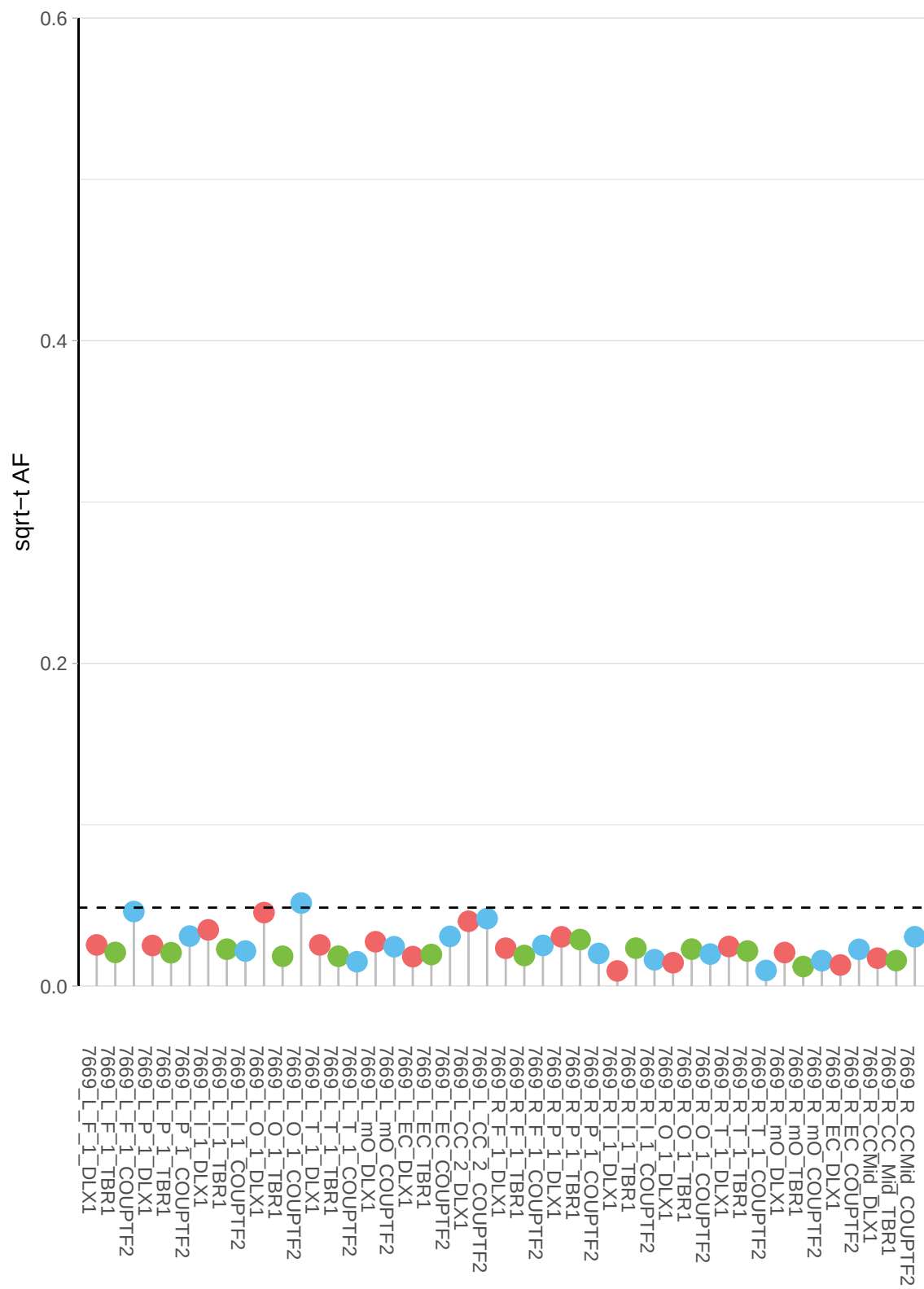

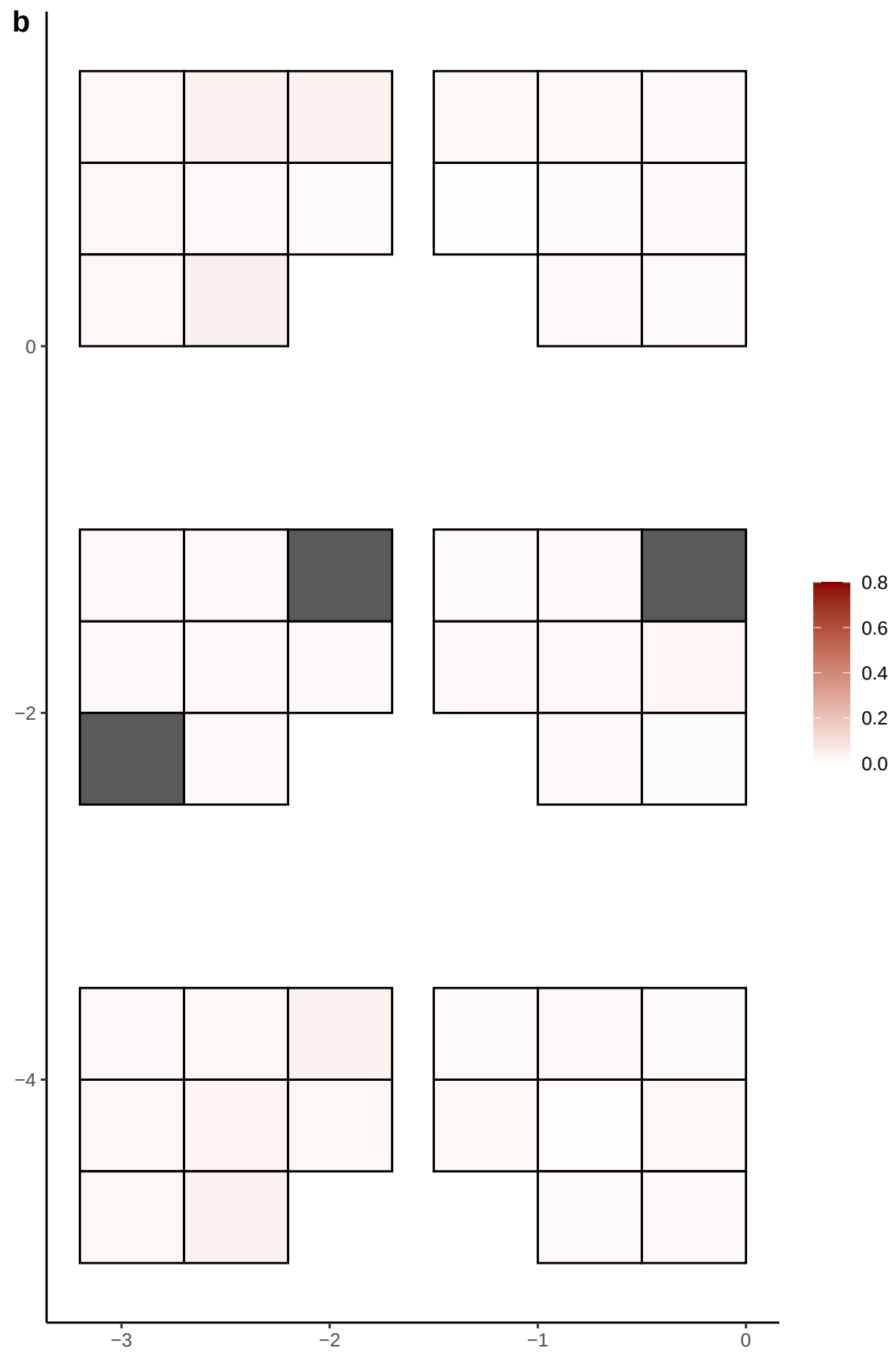

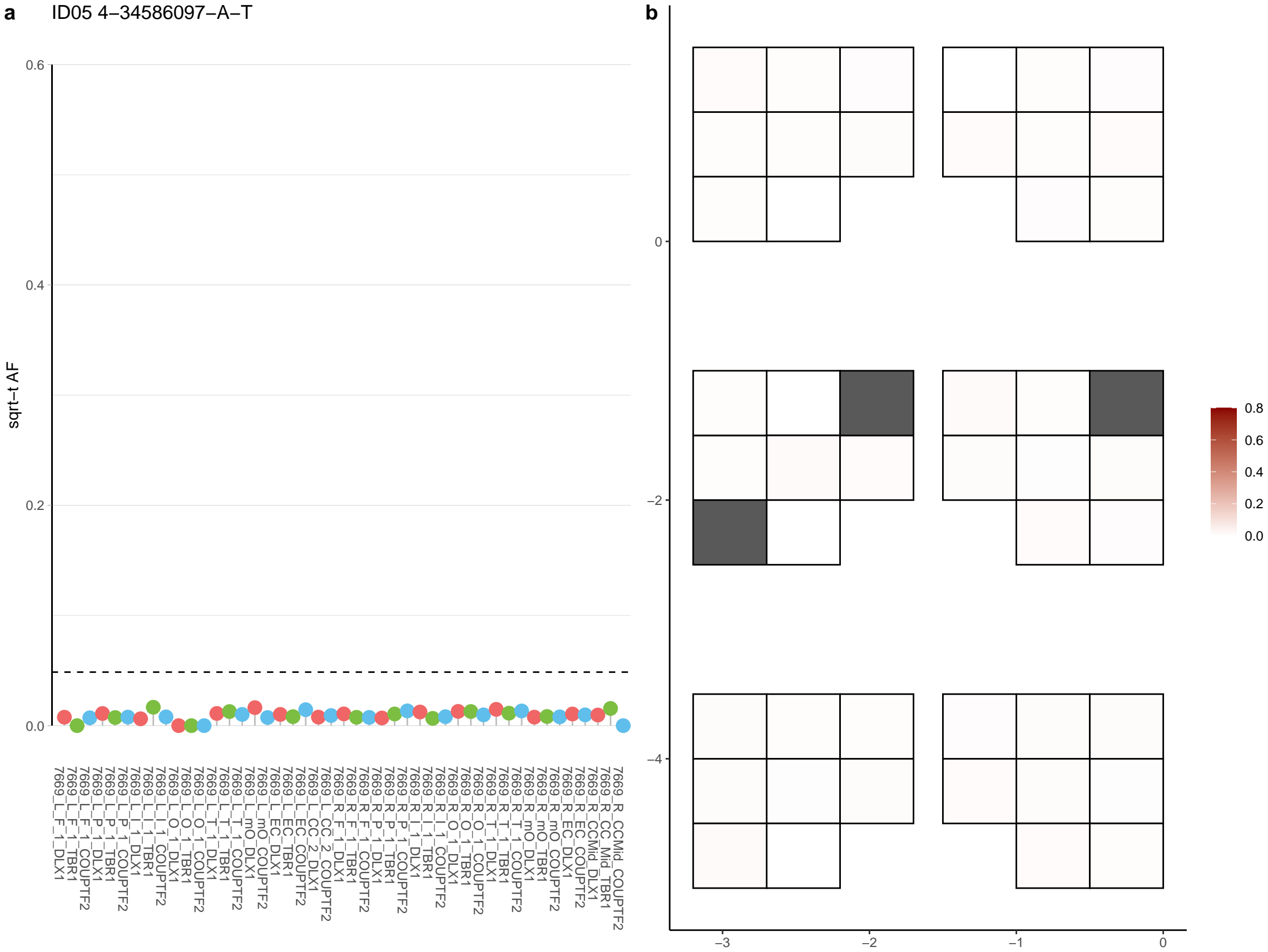

**a** ID05 4-36163098-C-T

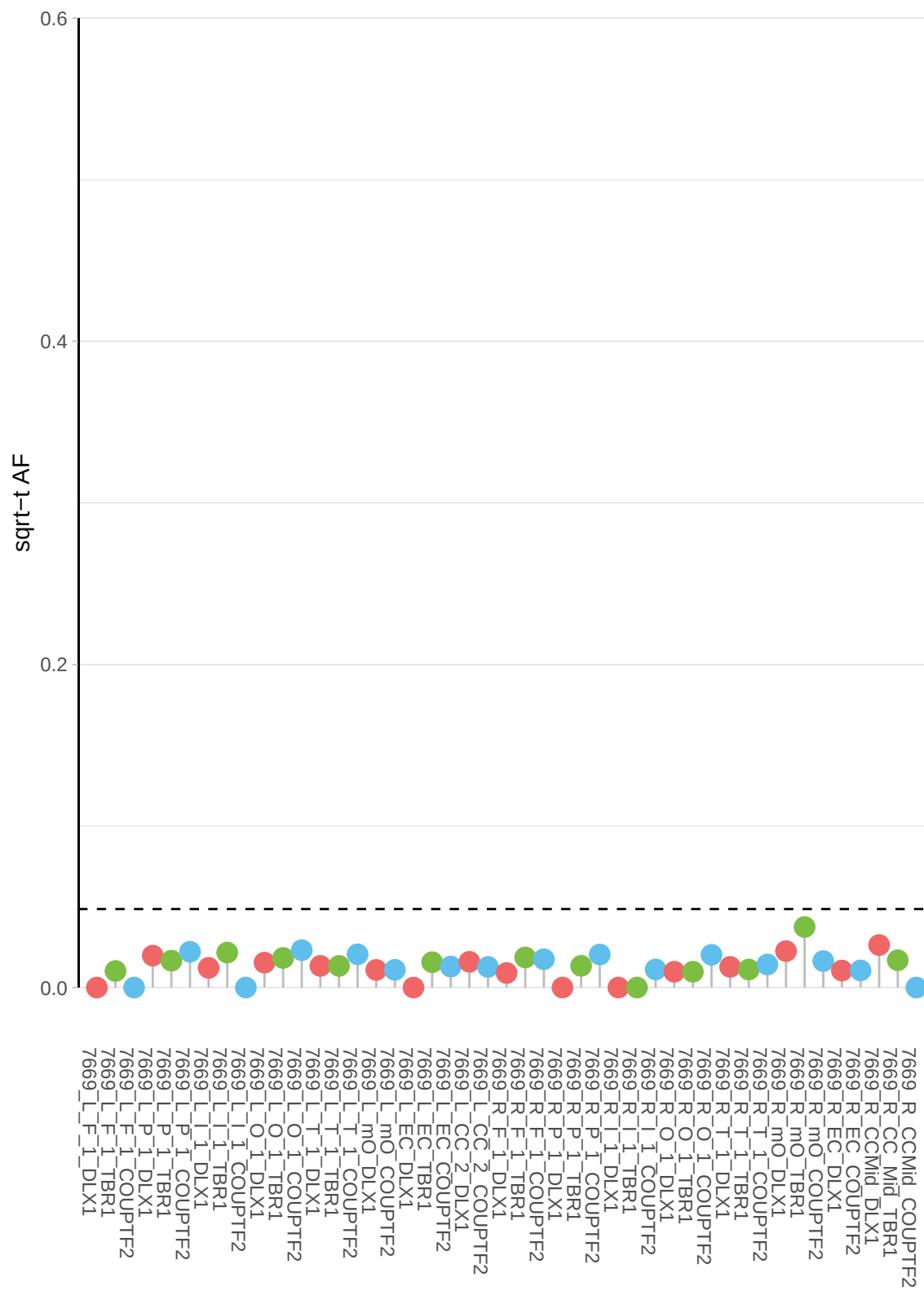

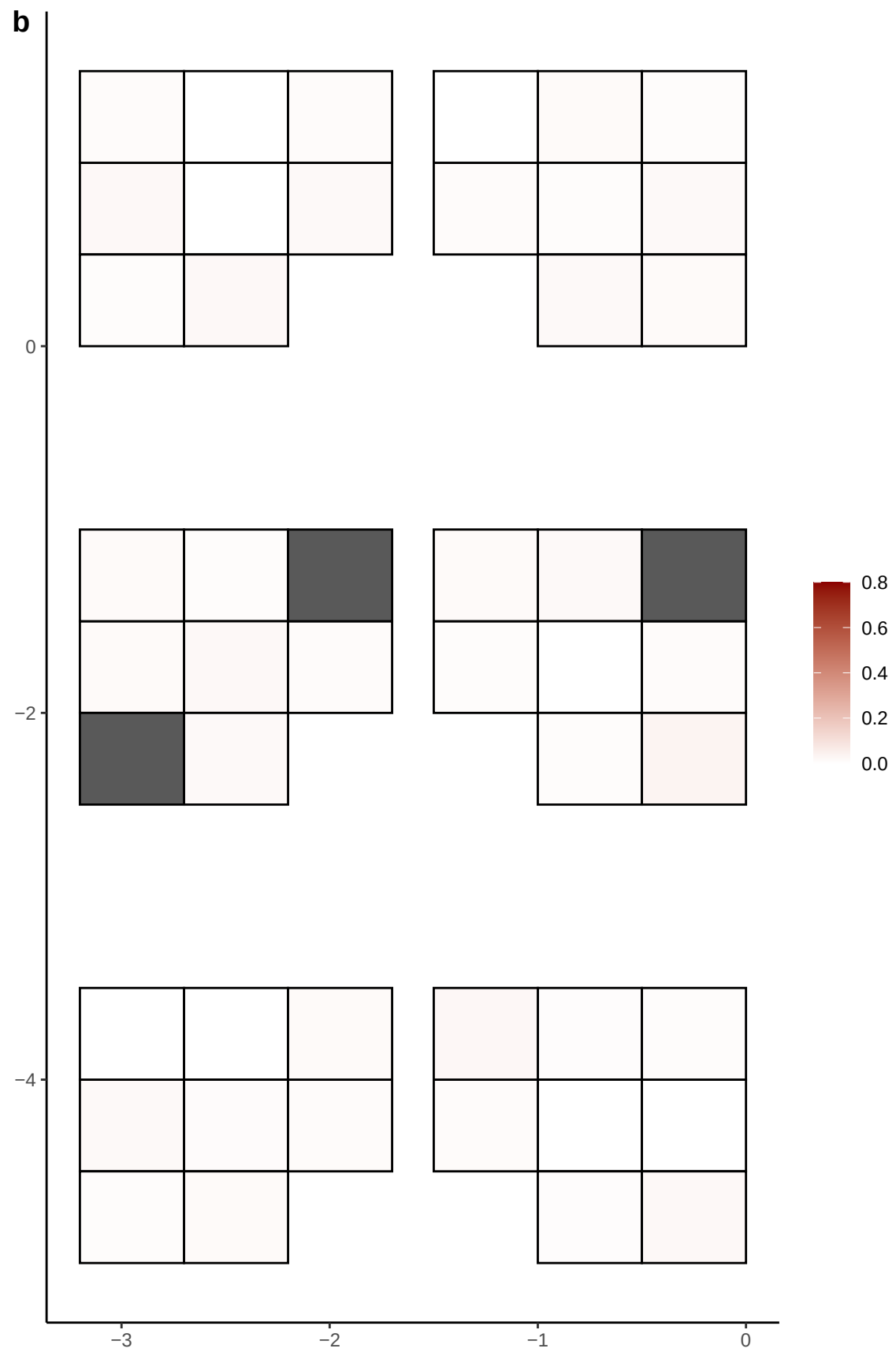

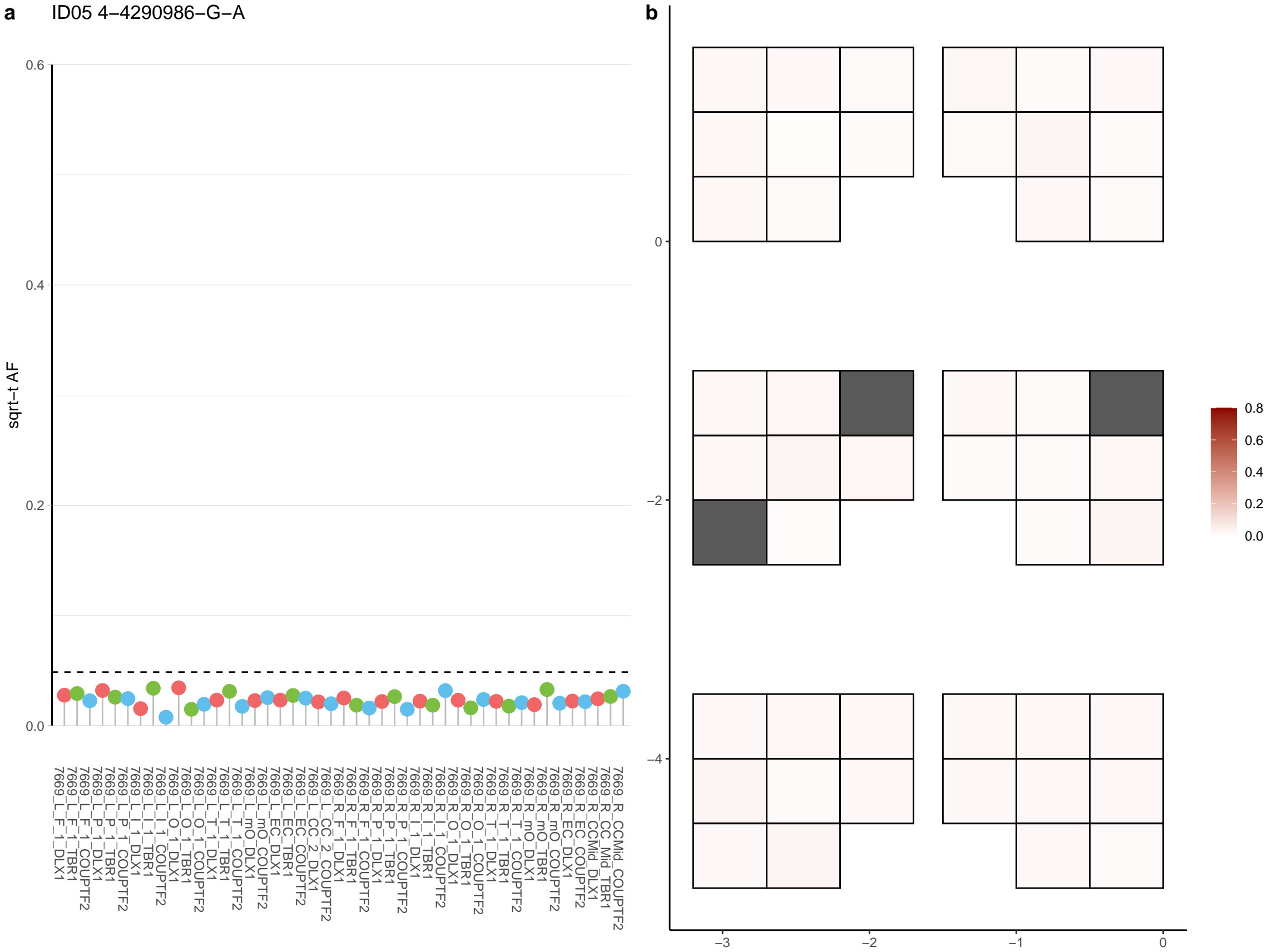

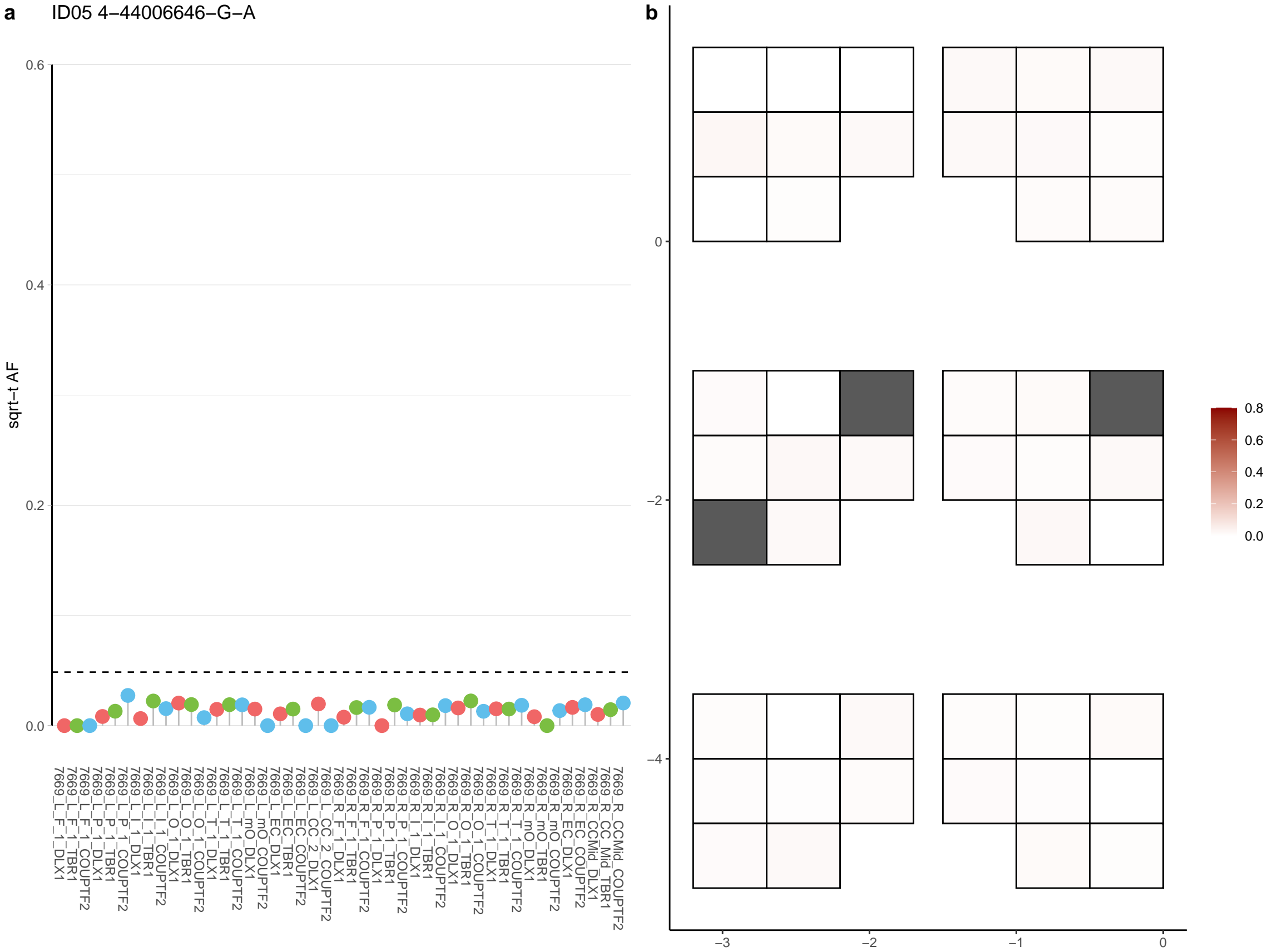

**a** ID05 4-5158885-G-A

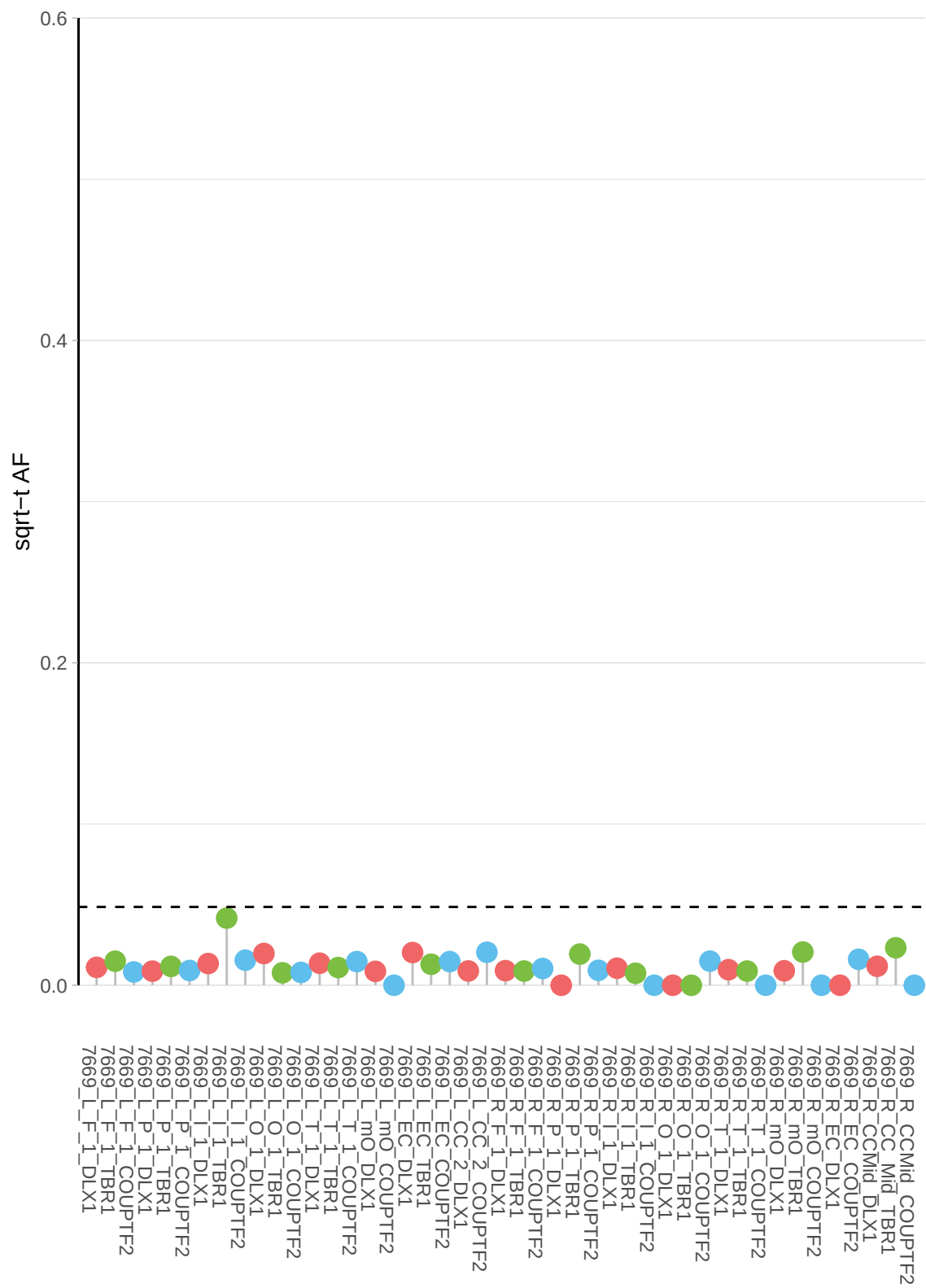

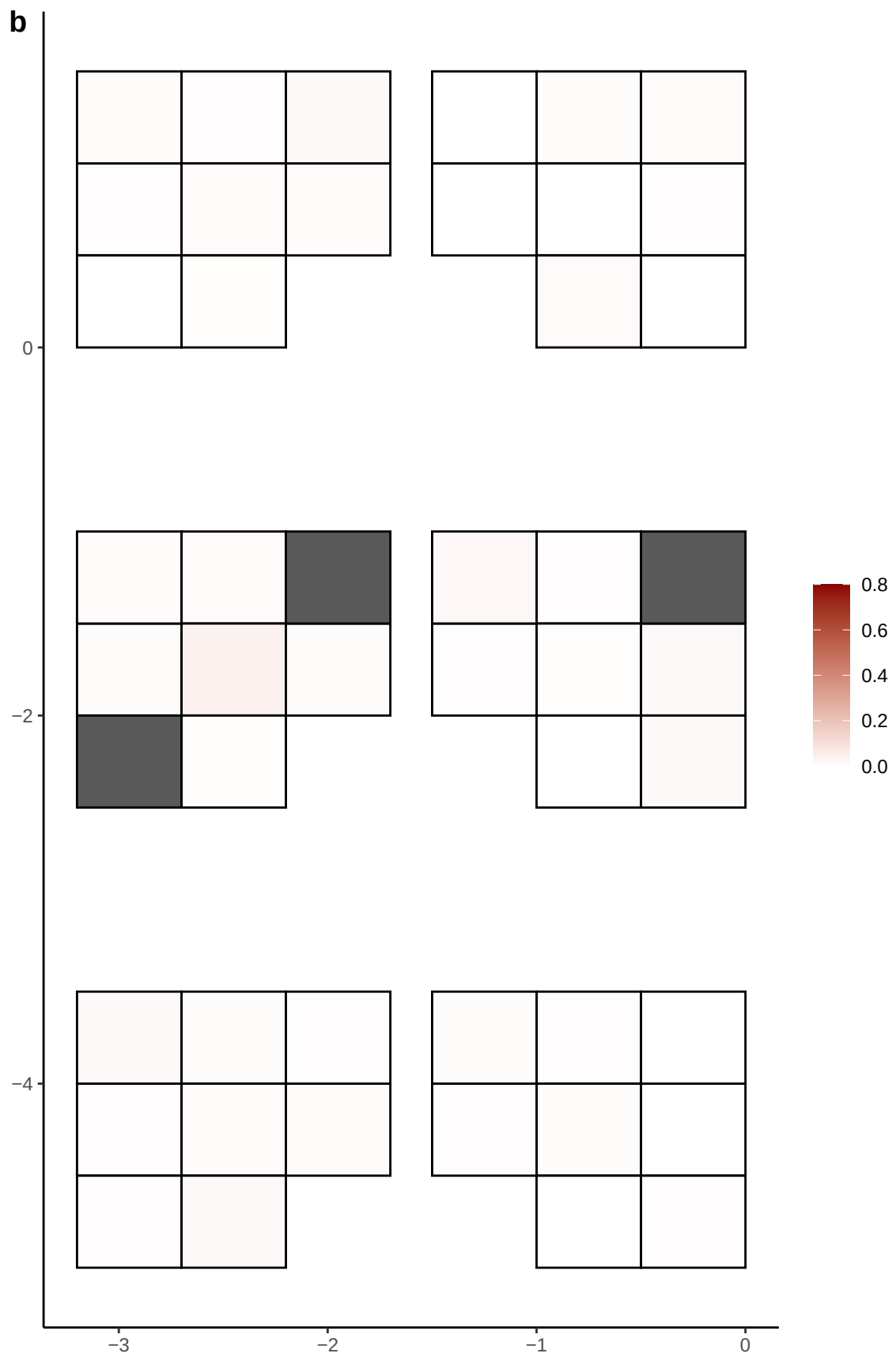

**a** ID05 4-52773-C-T

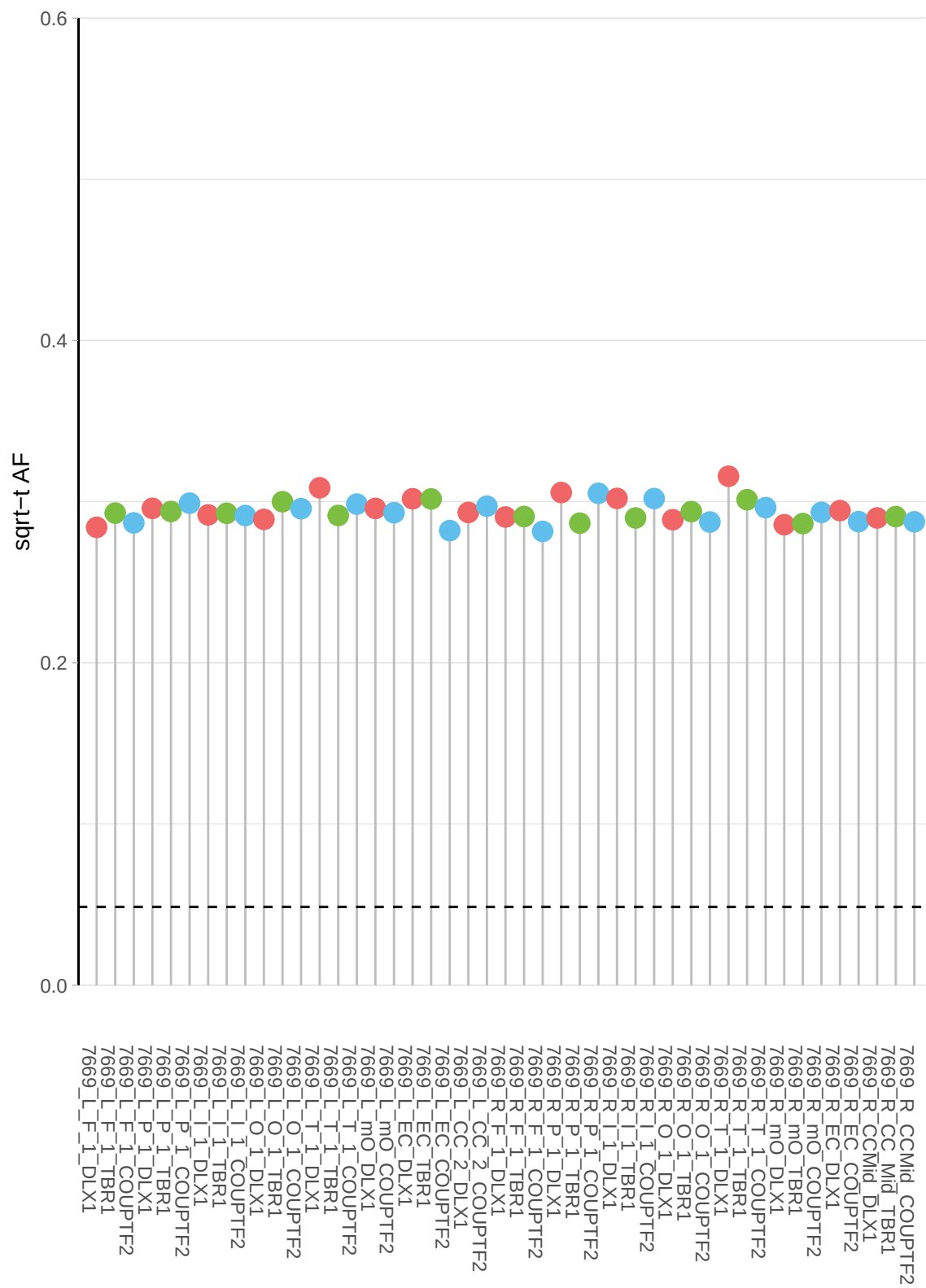

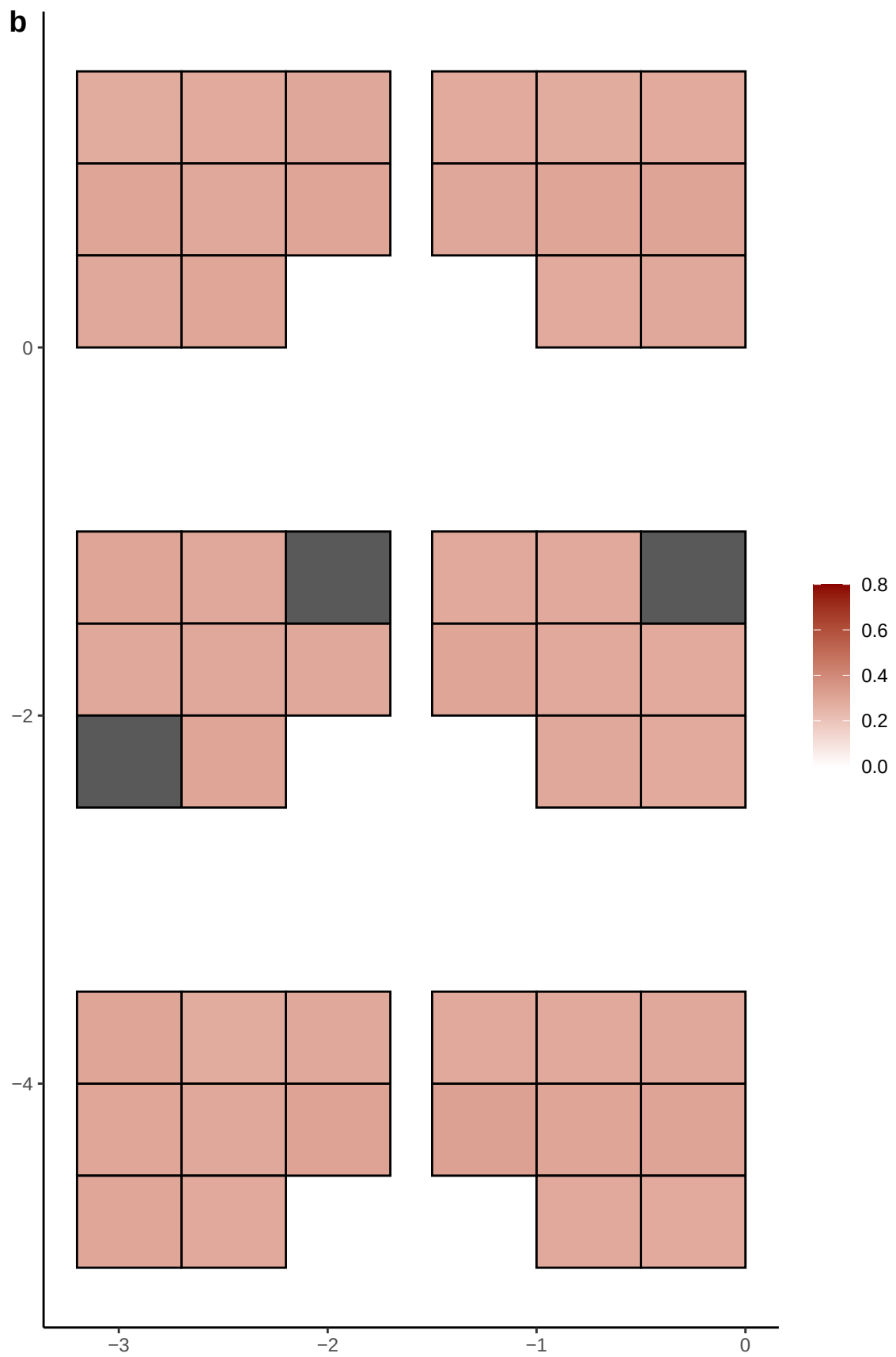

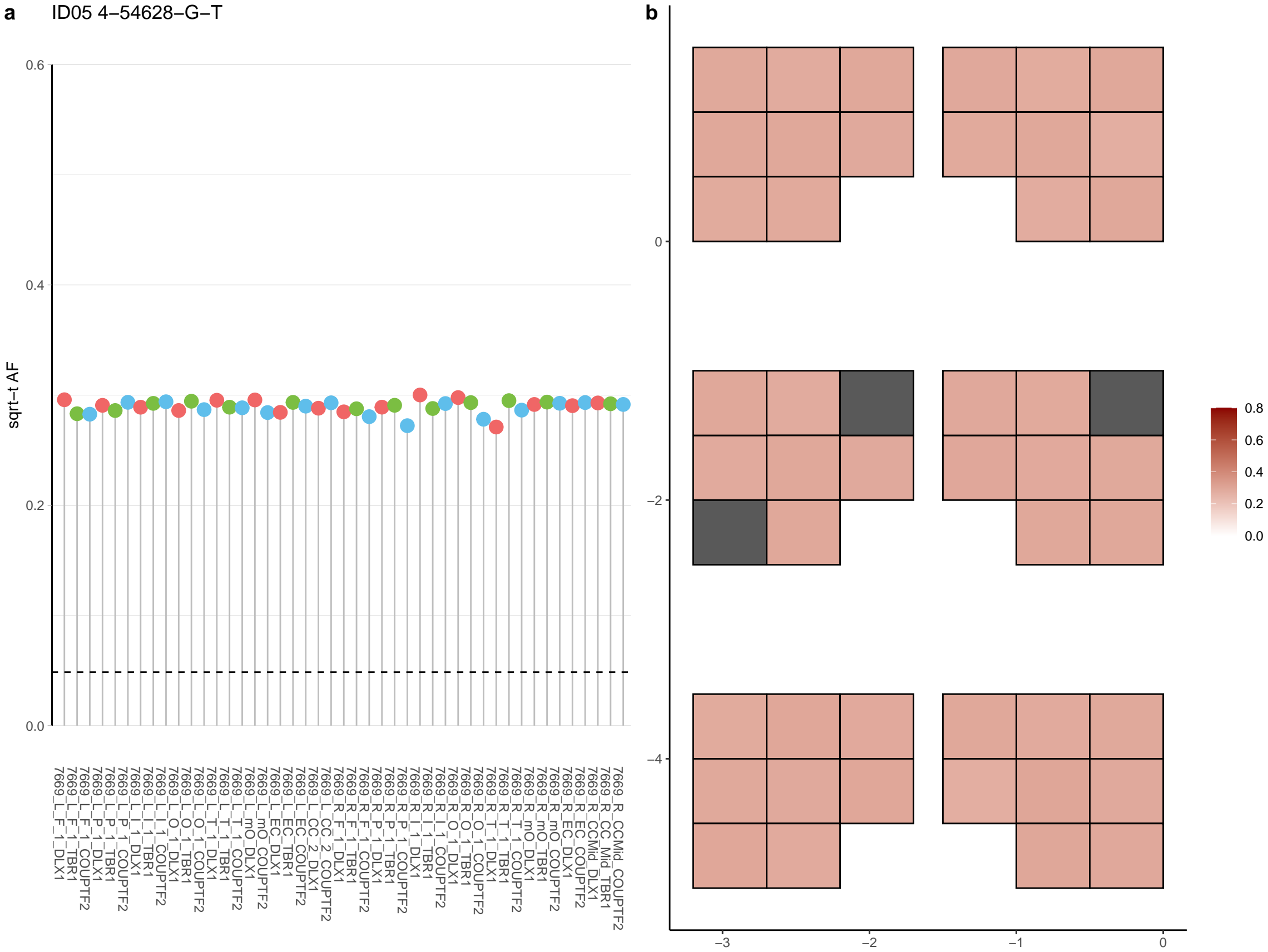

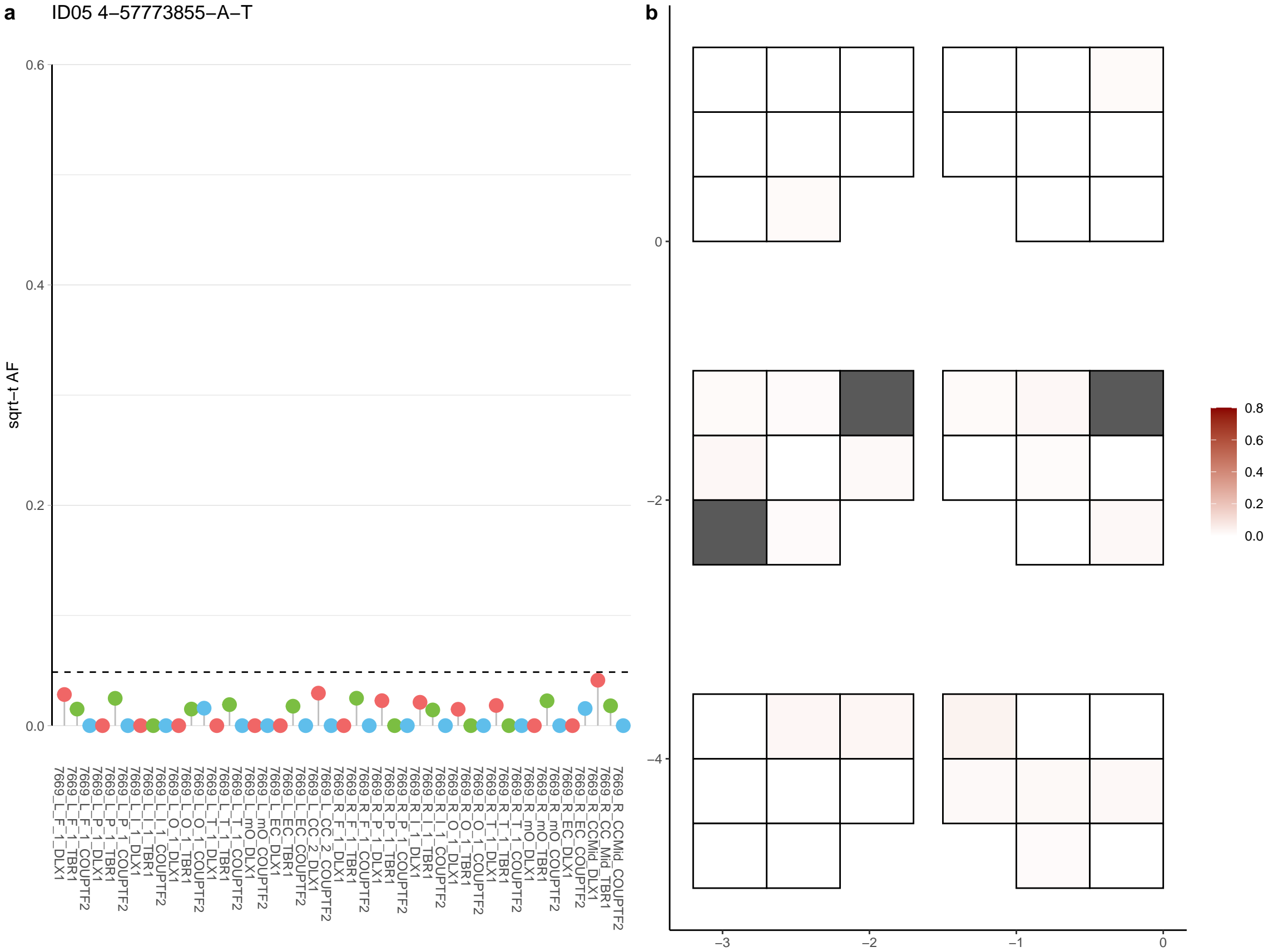

**a** ID05 4-58638686-G-A

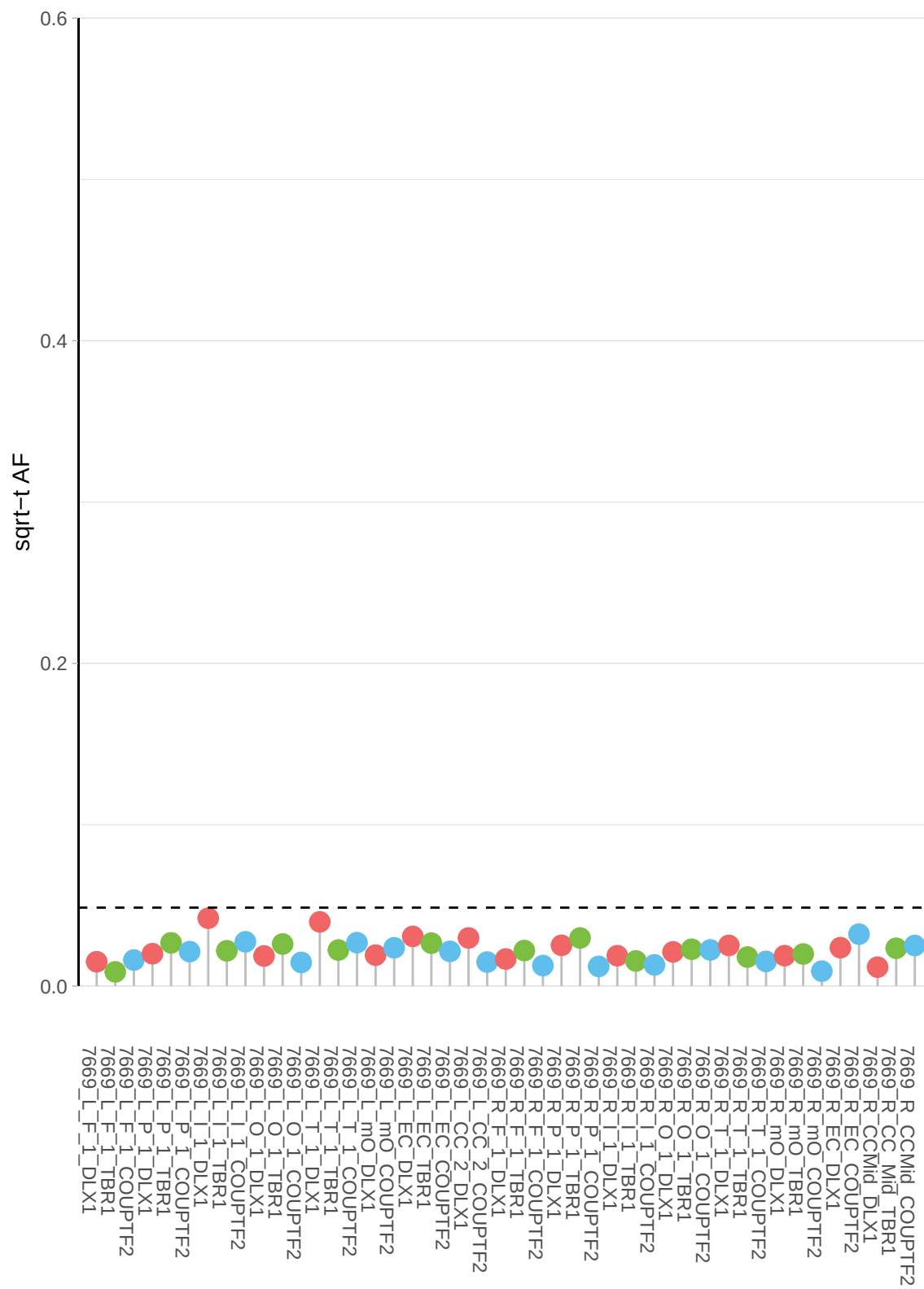

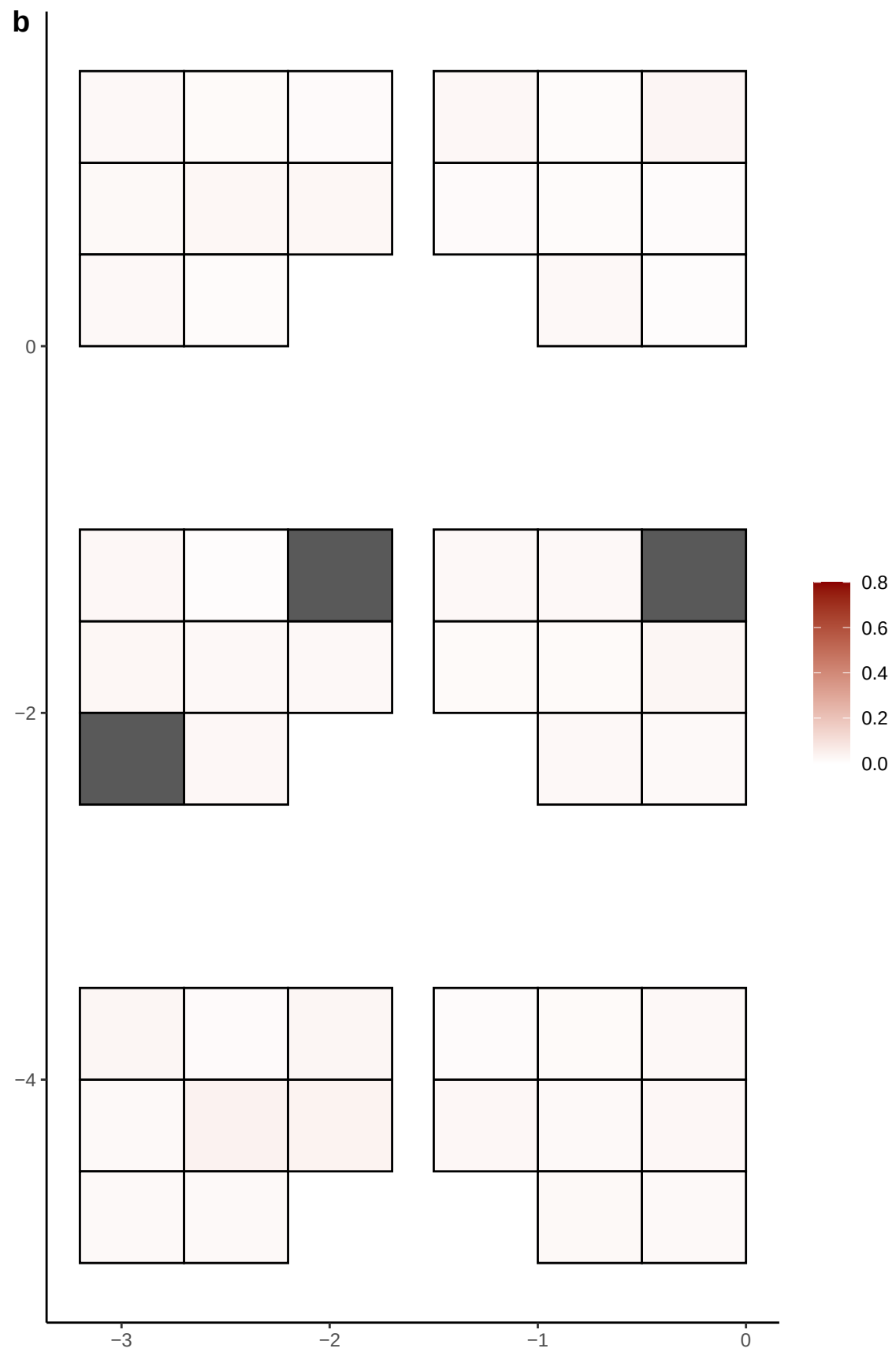

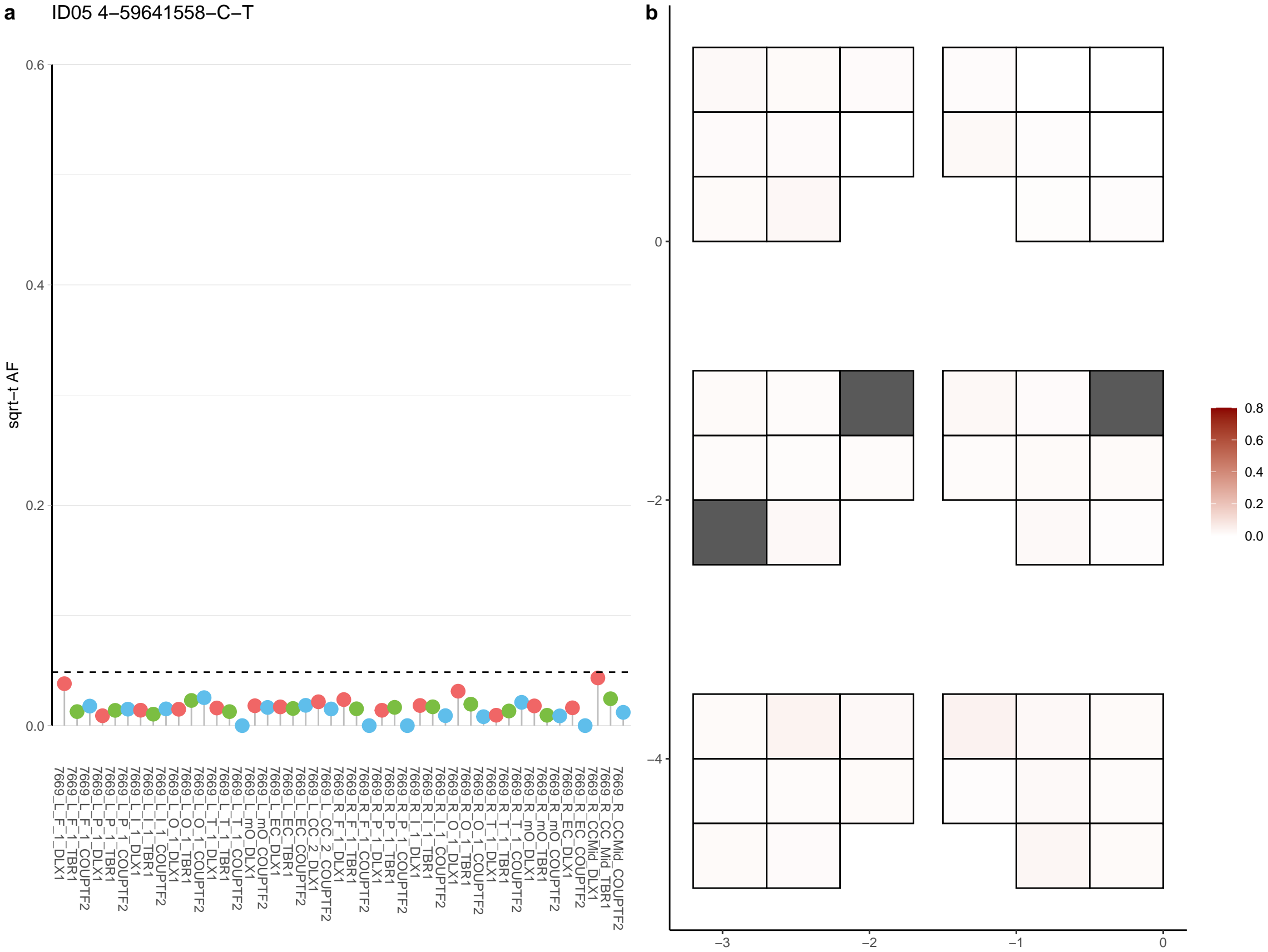

**a** ID05 4-60968373-G-A

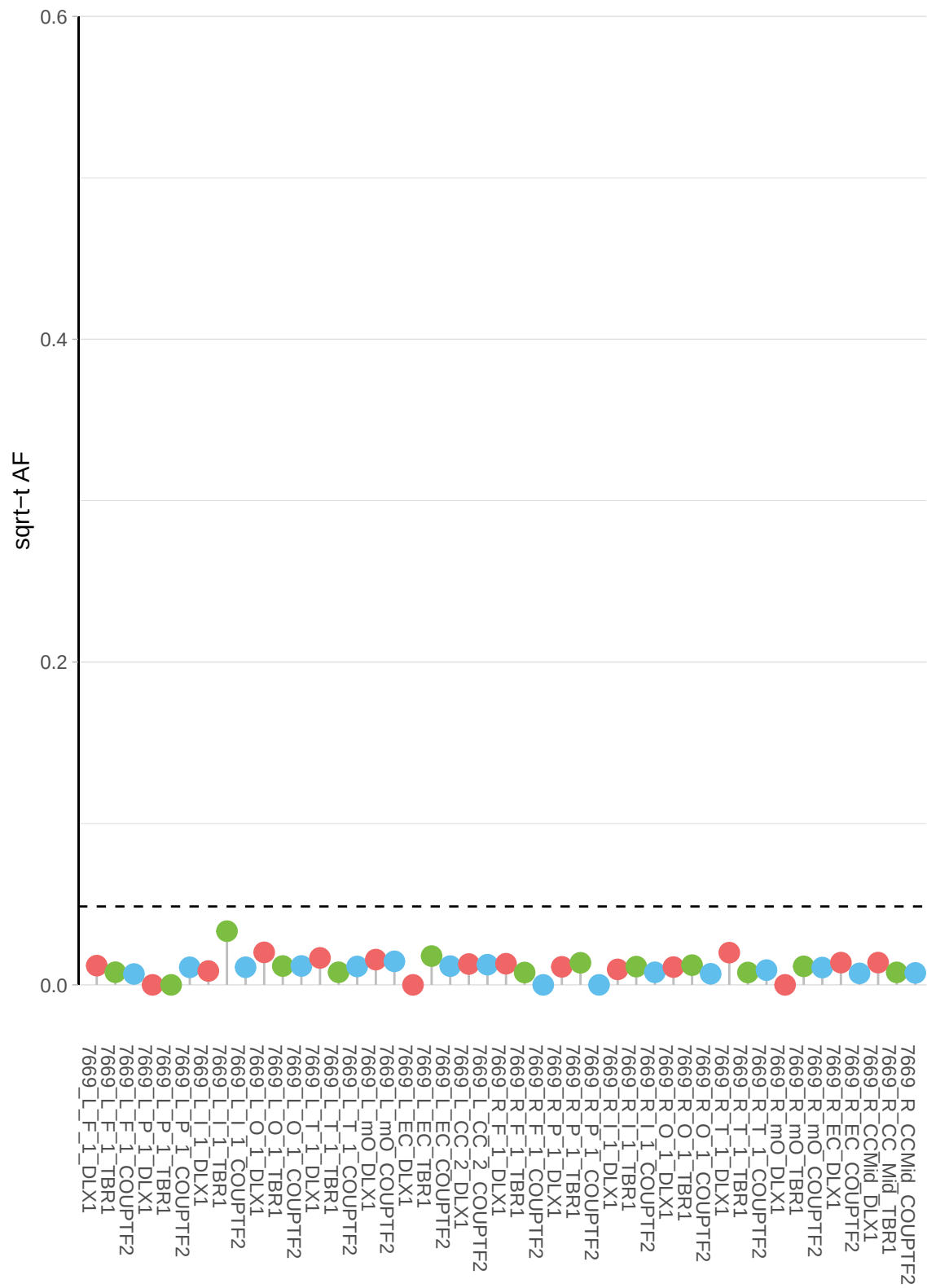

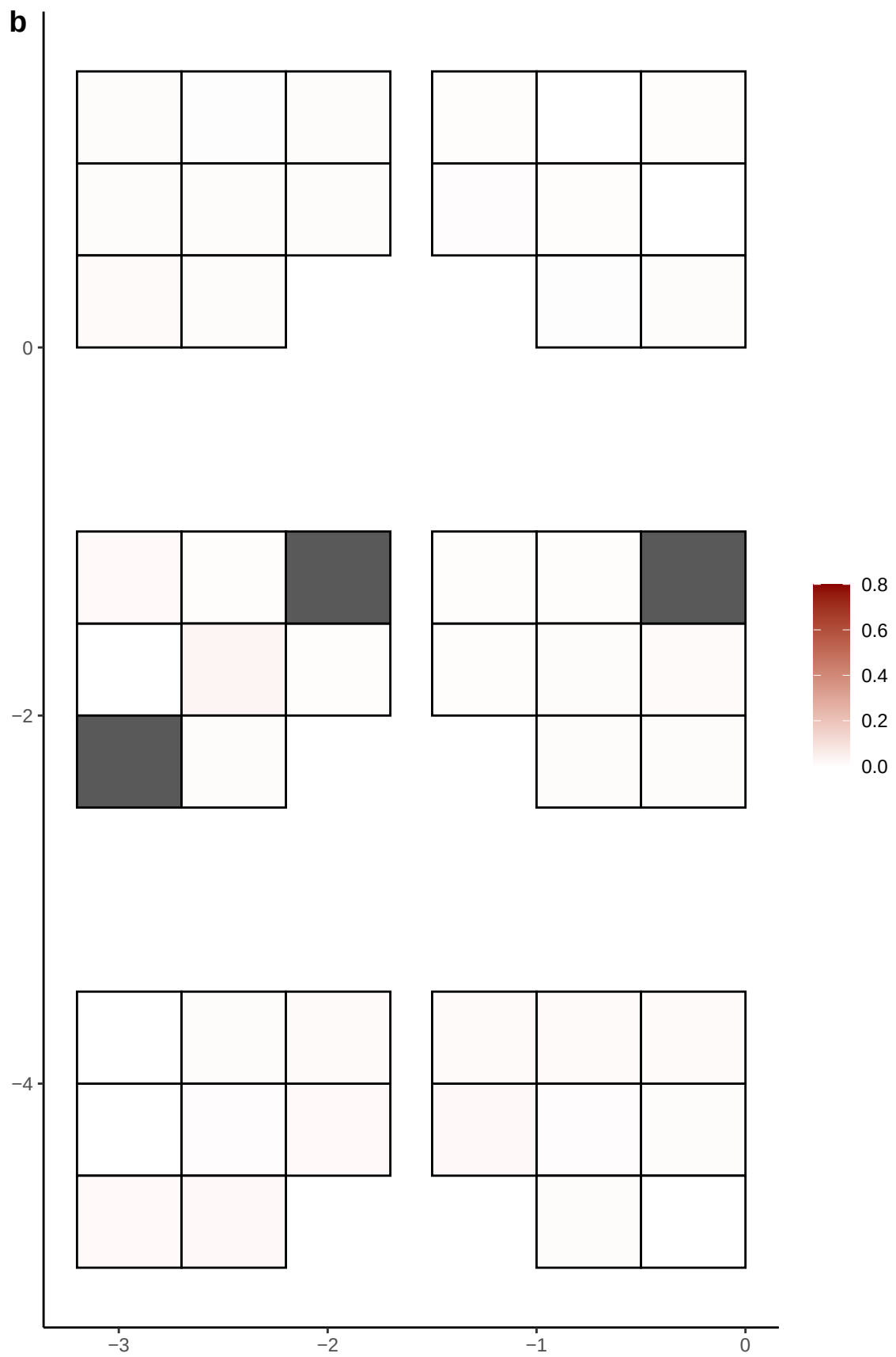

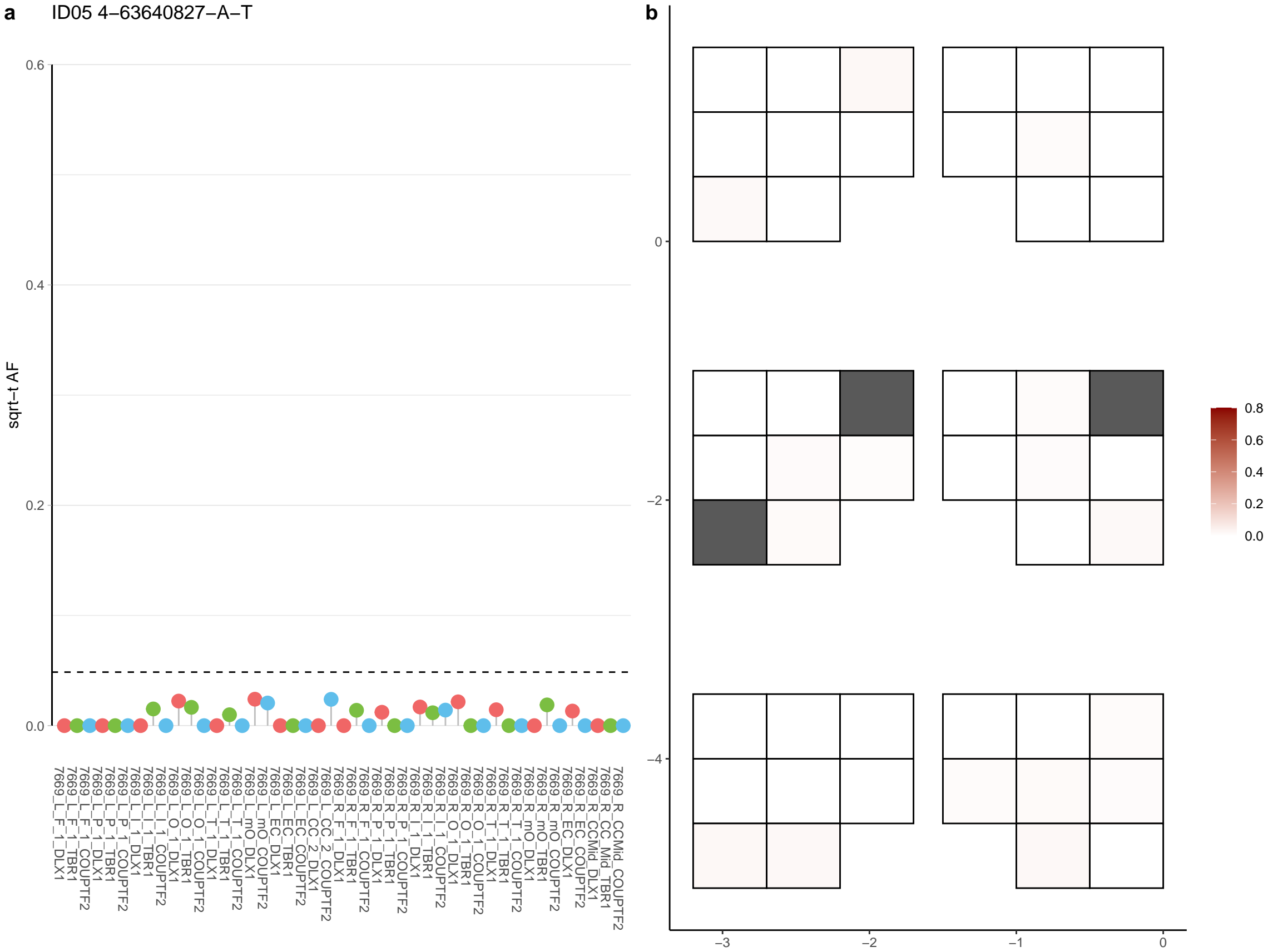

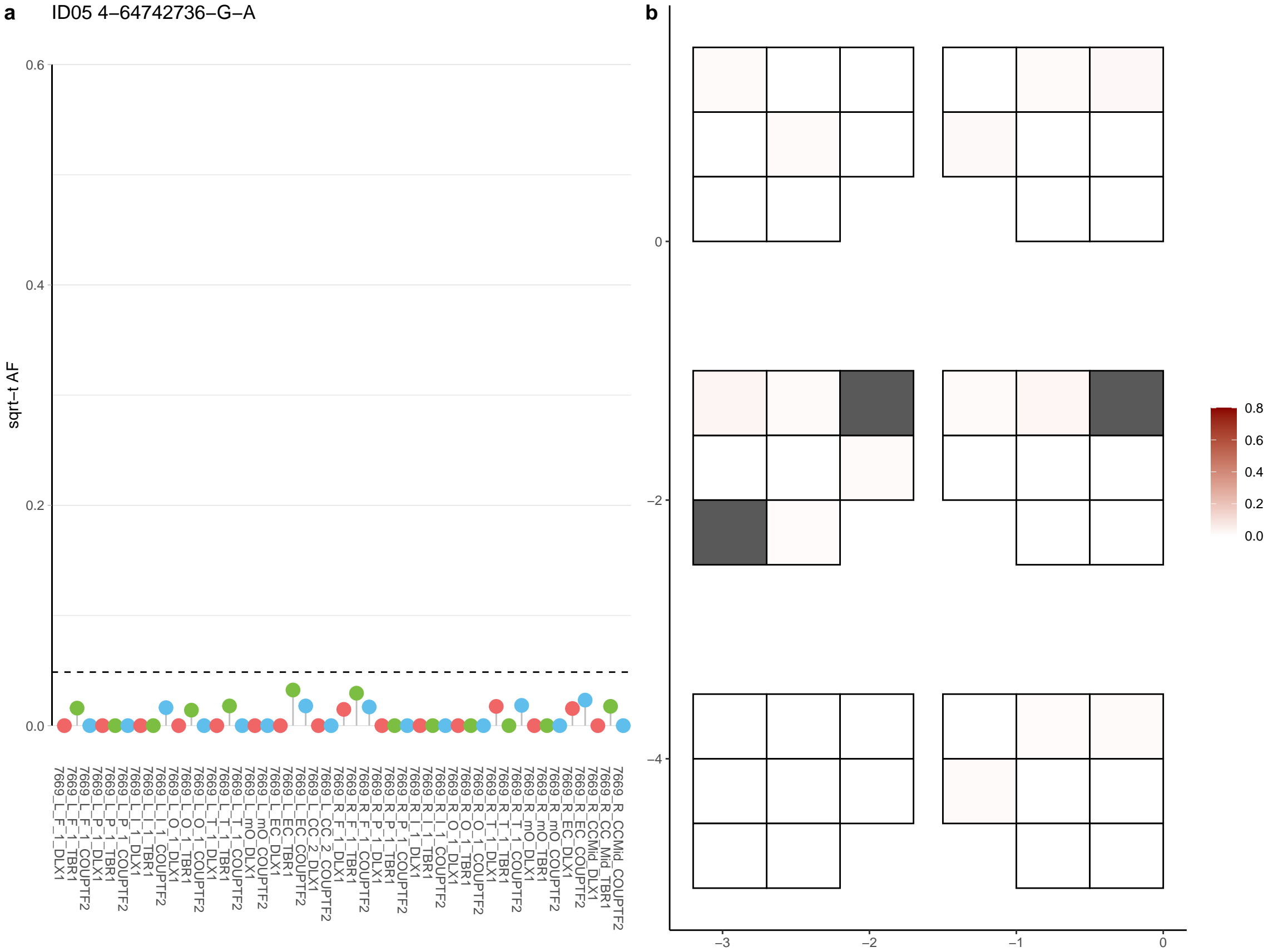

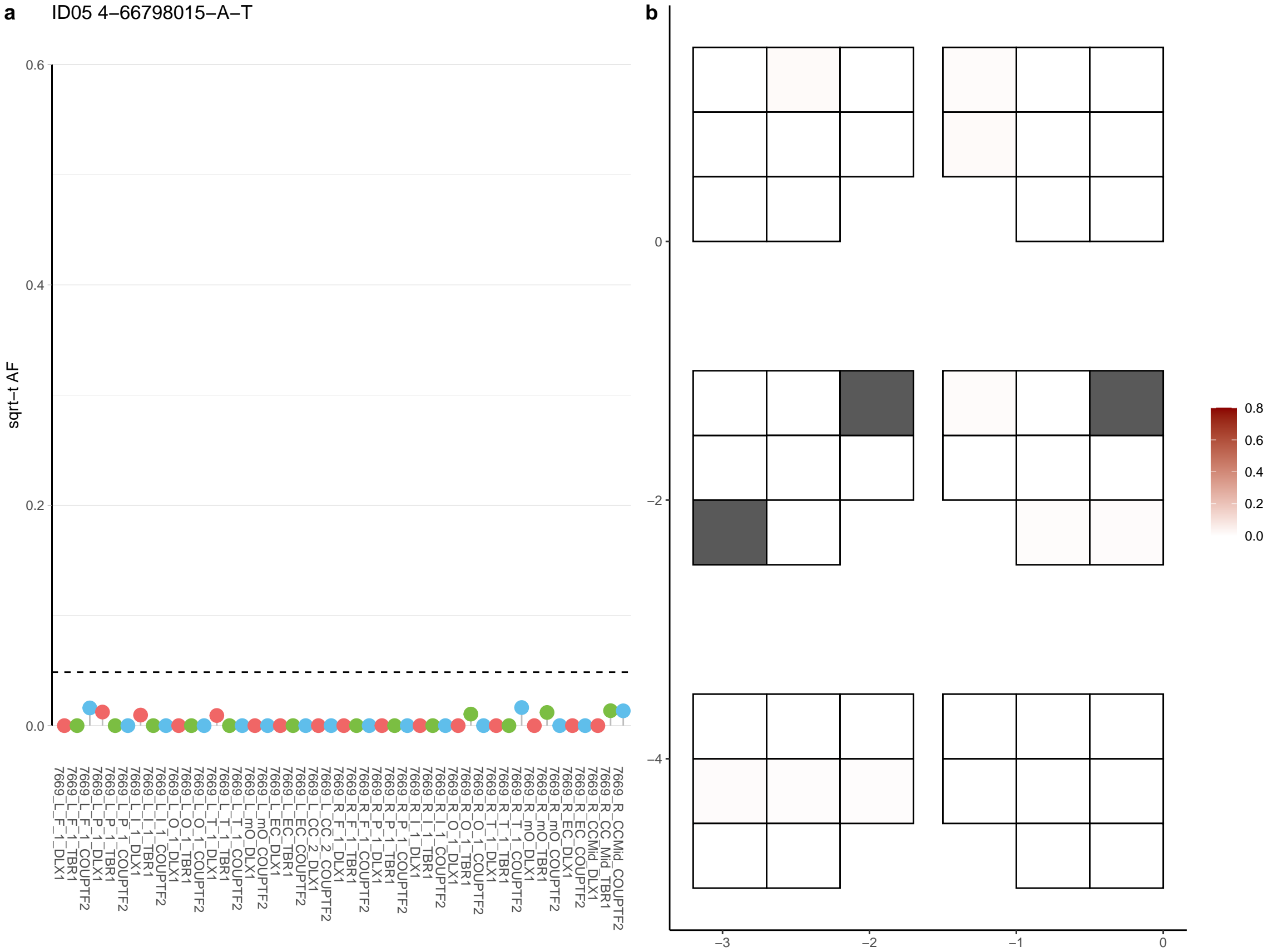

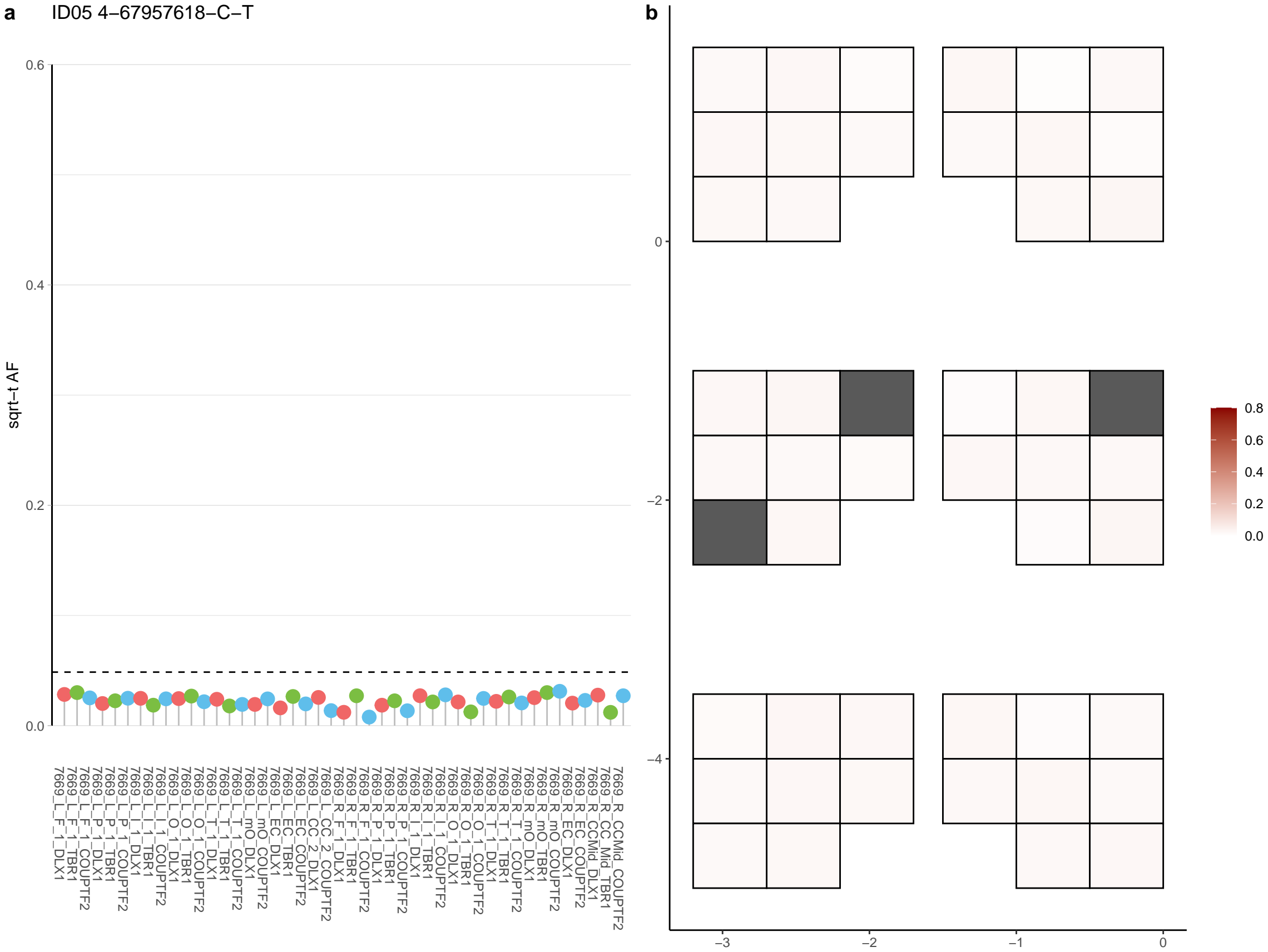

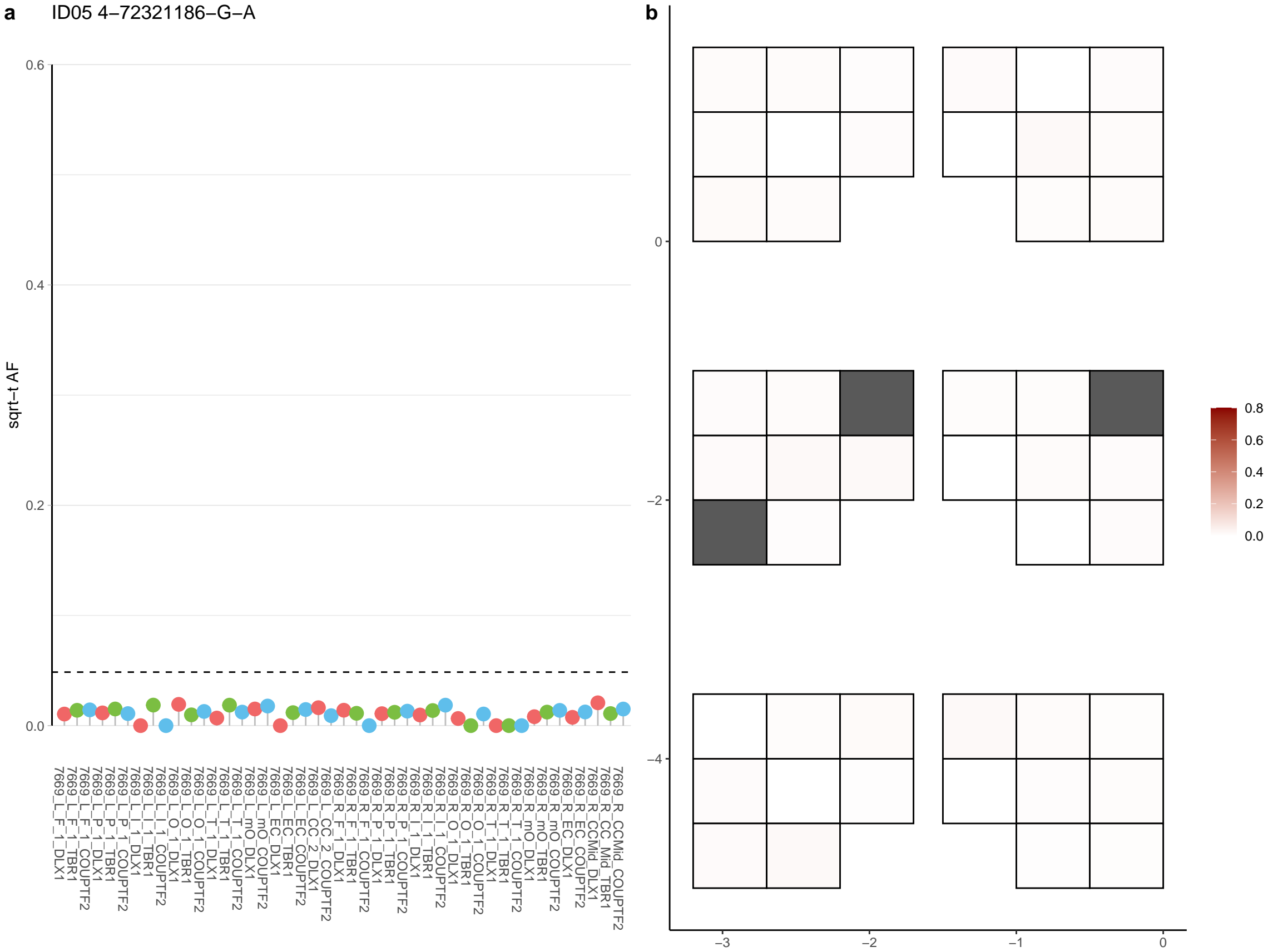

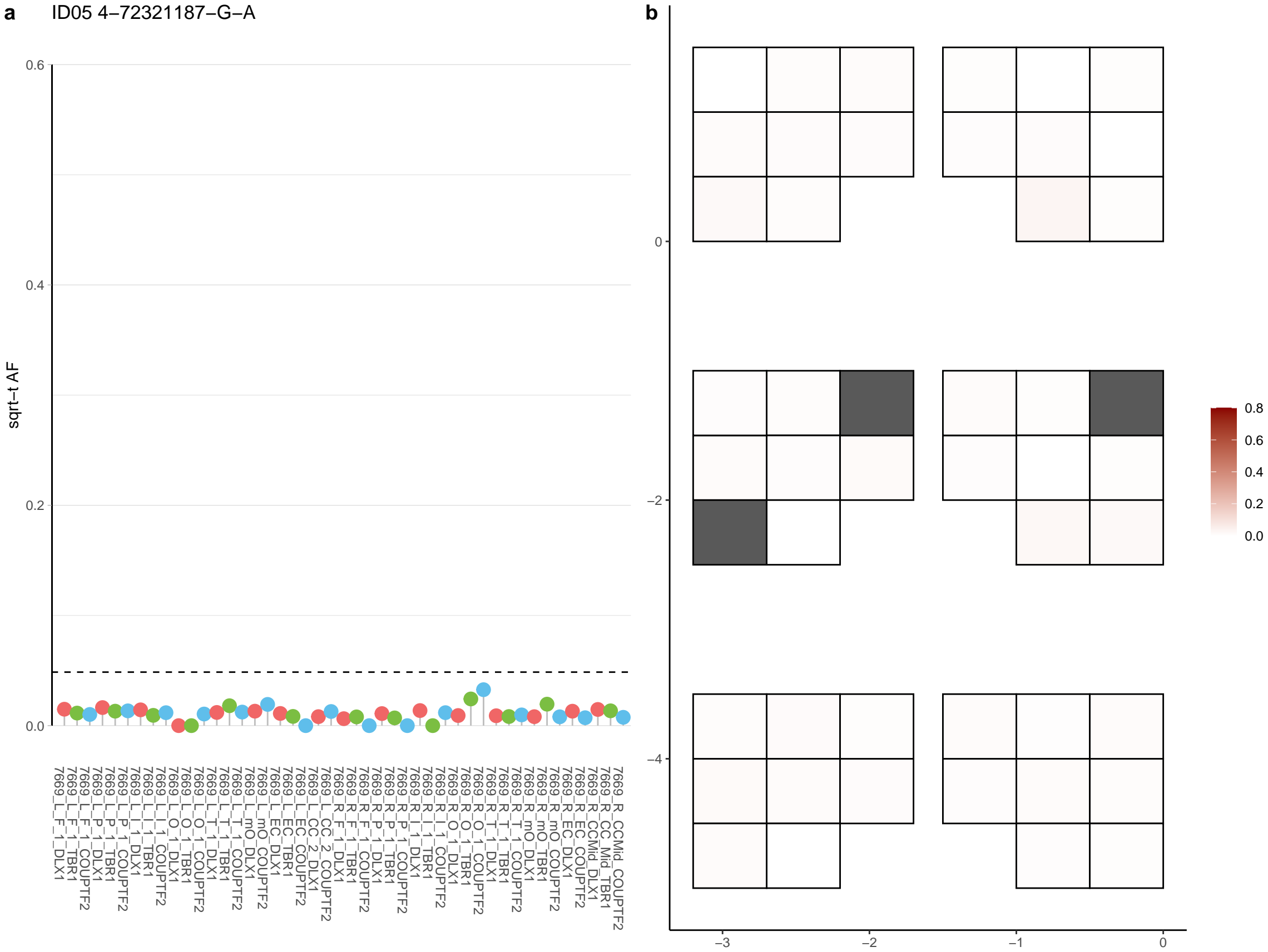

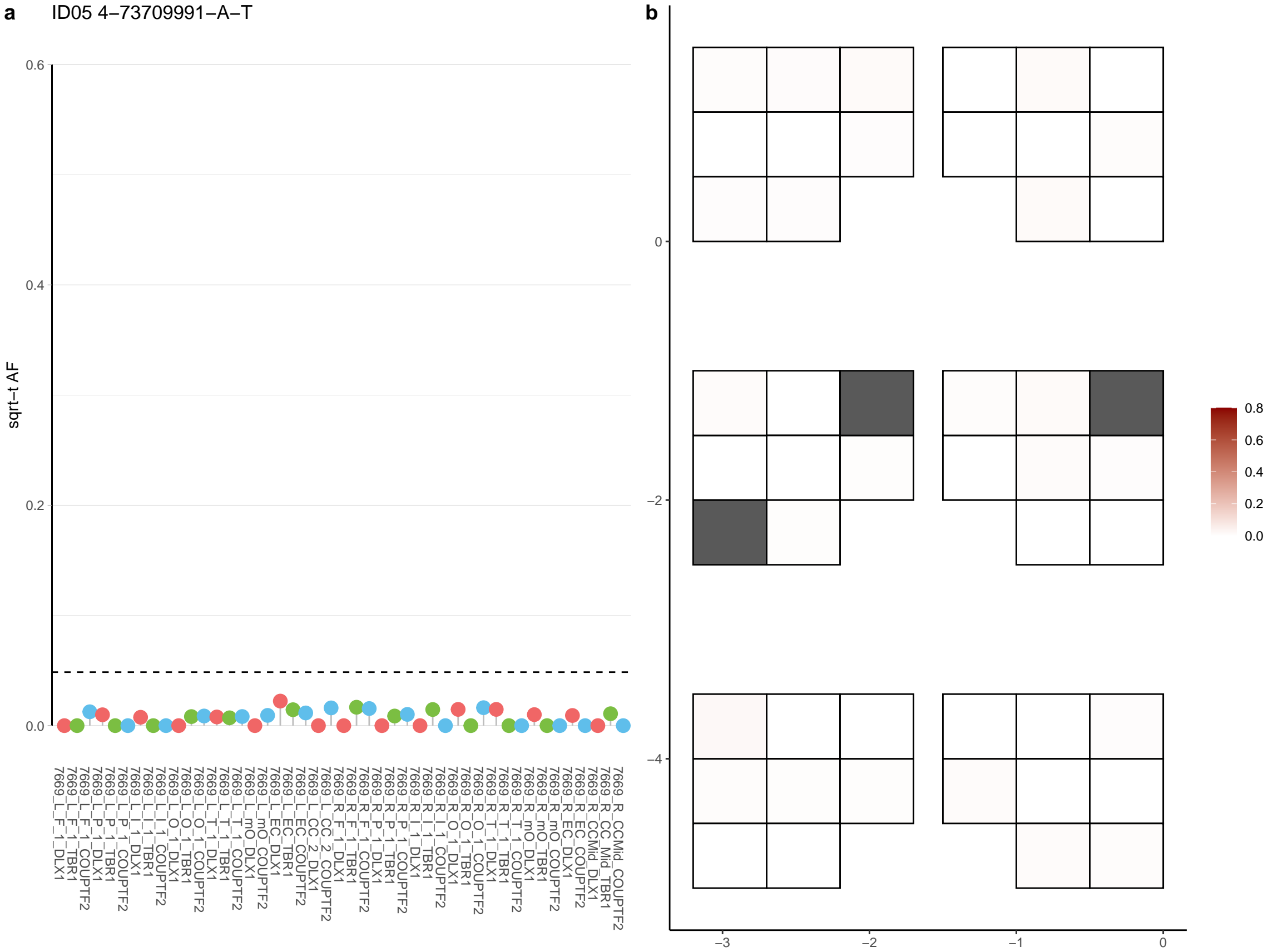

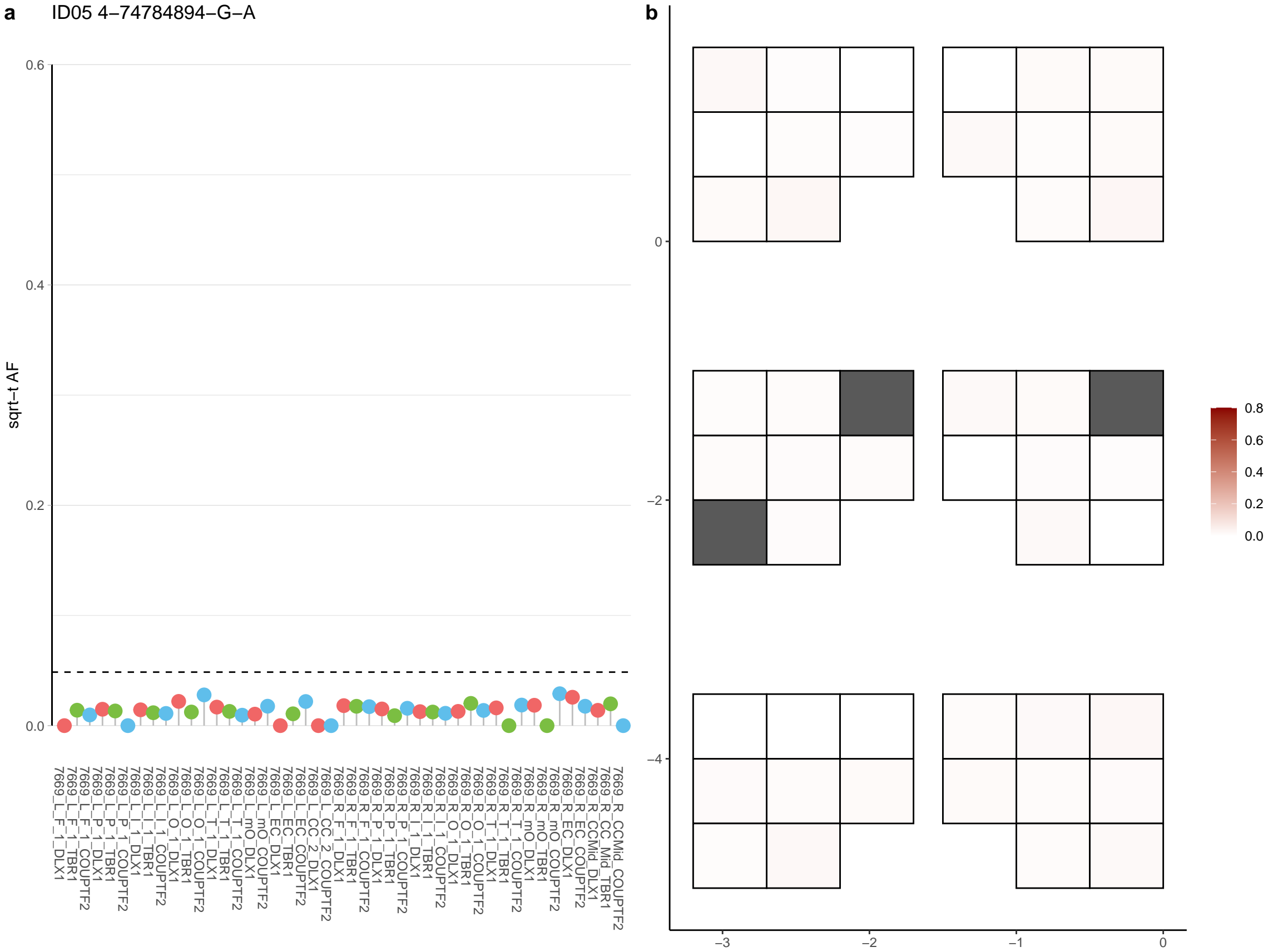

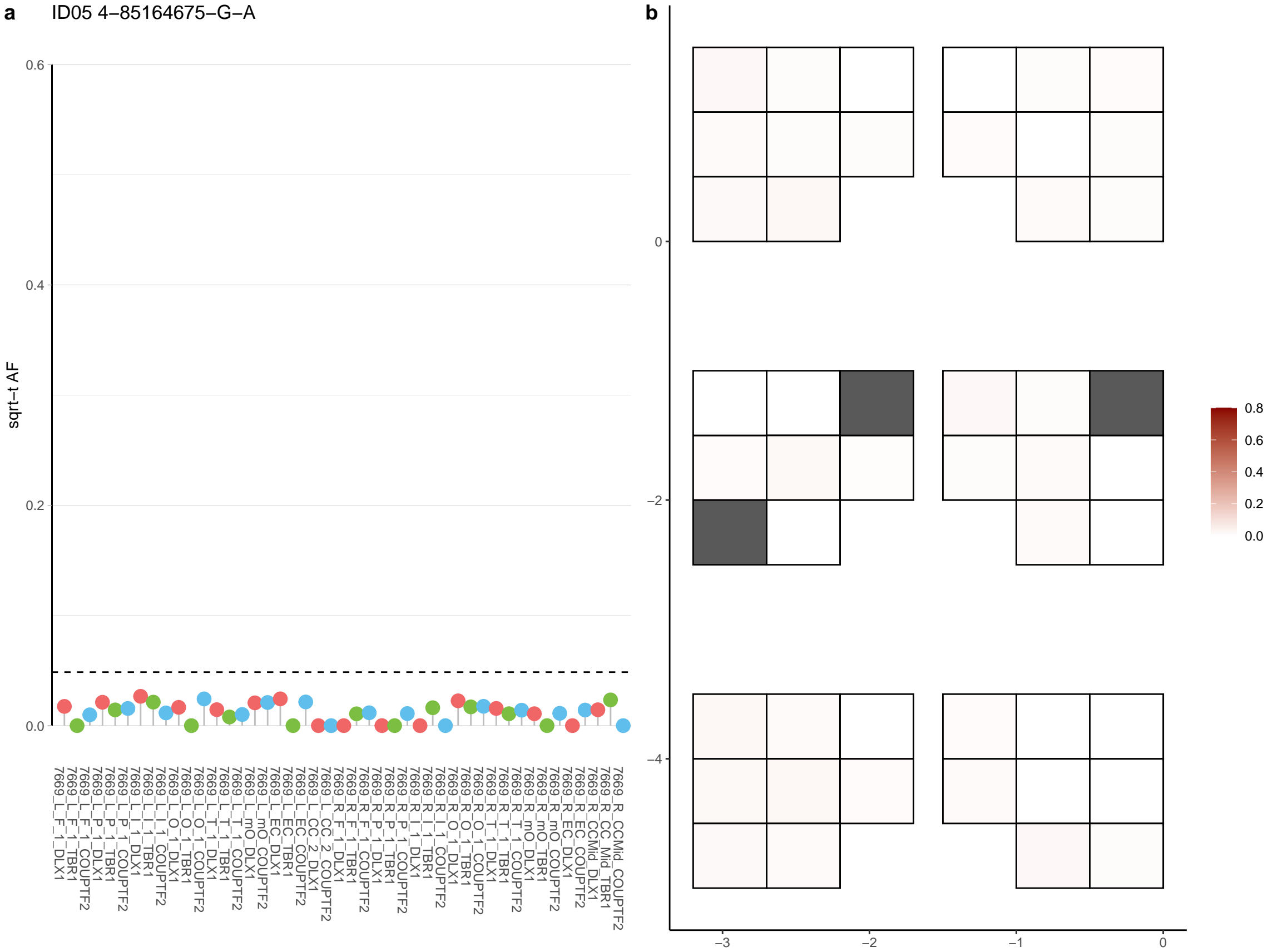

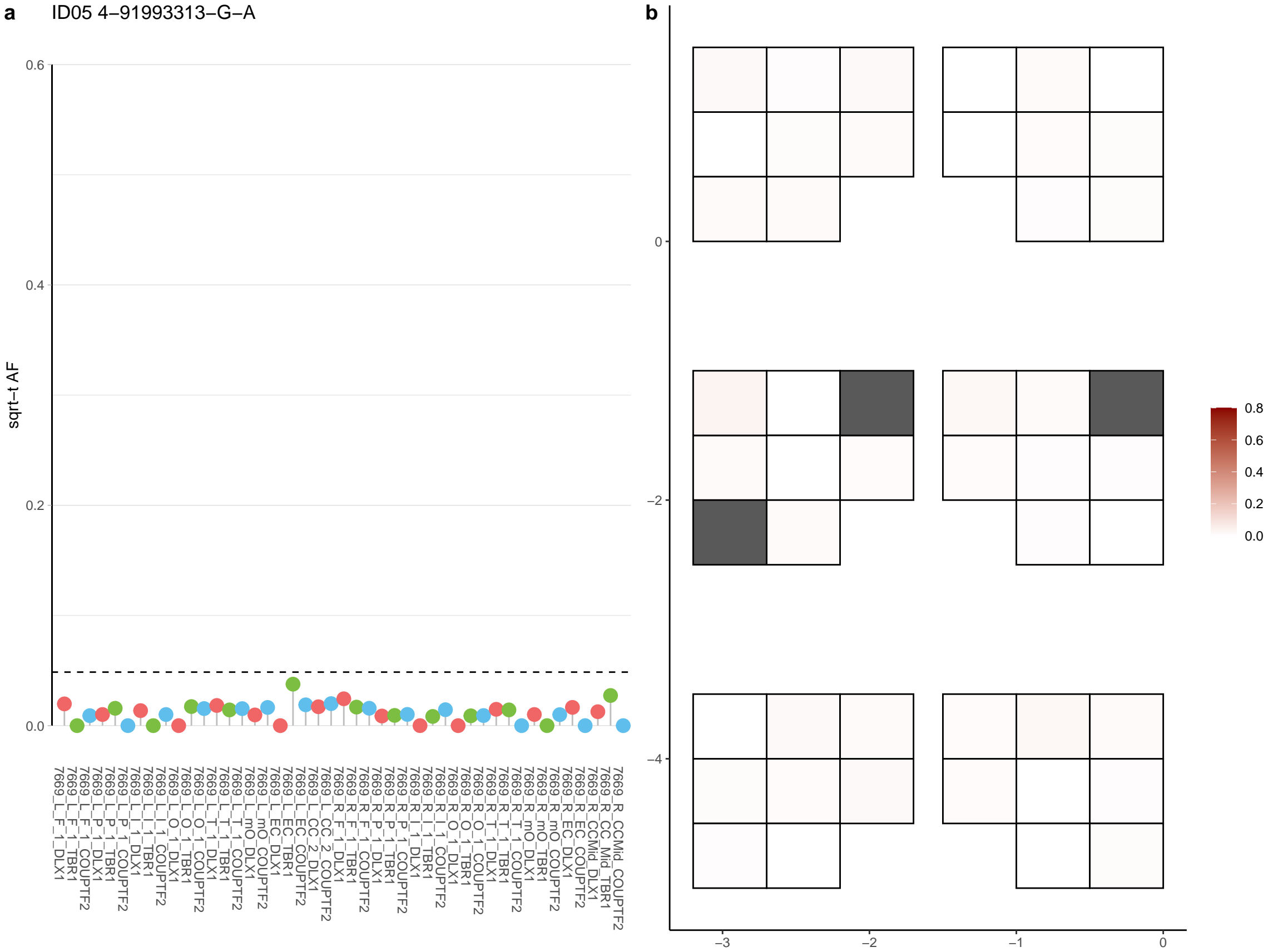

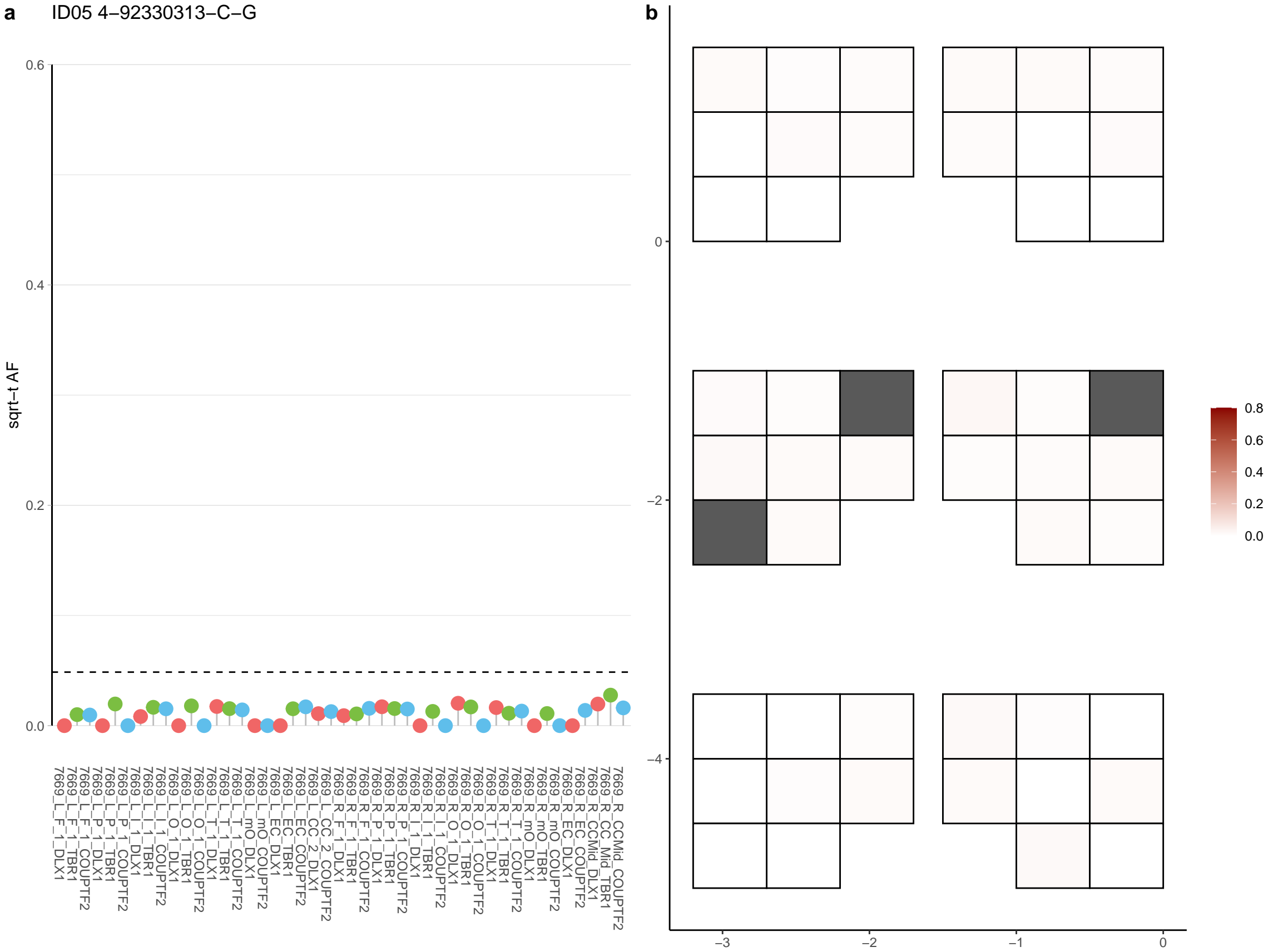

**a** ID05 4-92660887-G-A

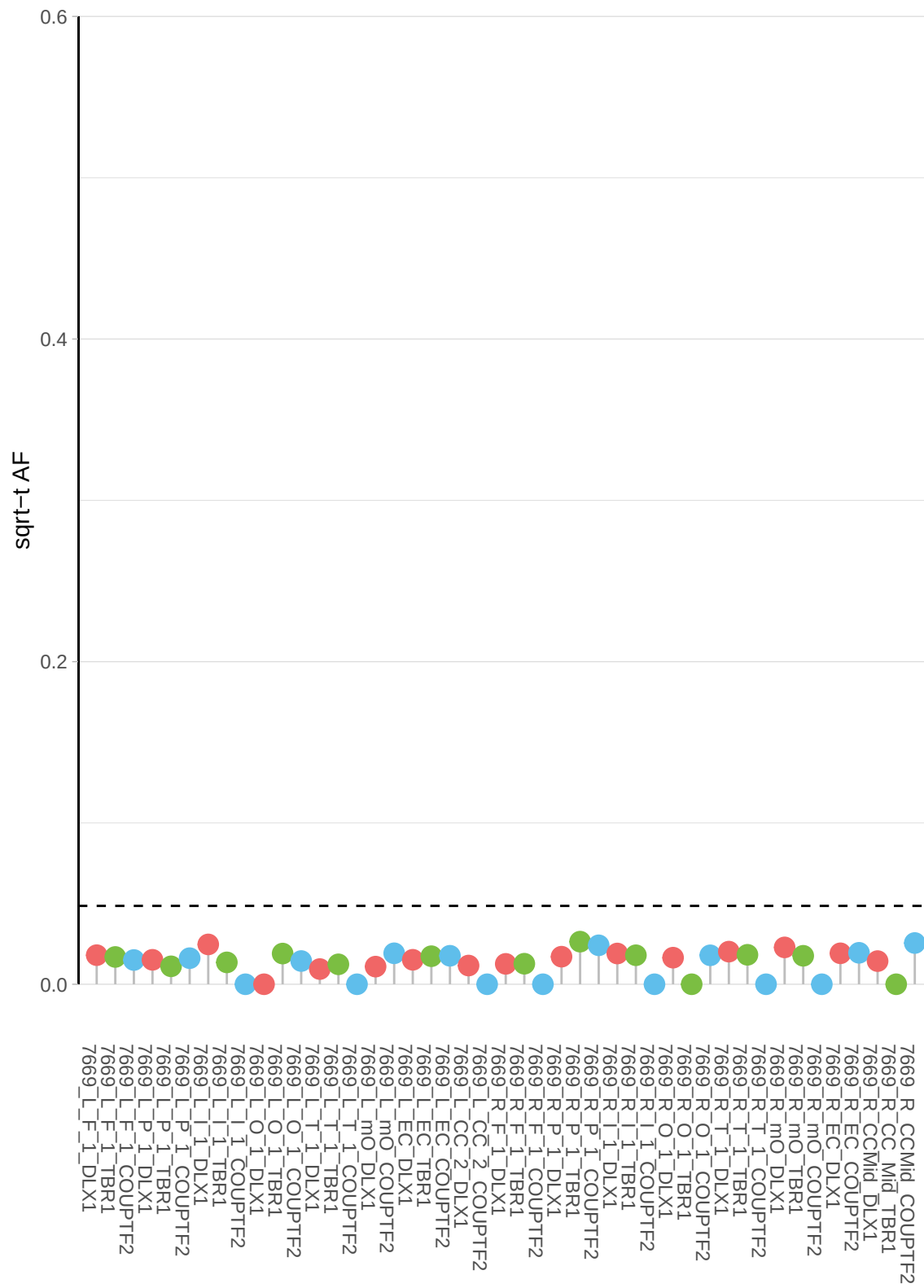

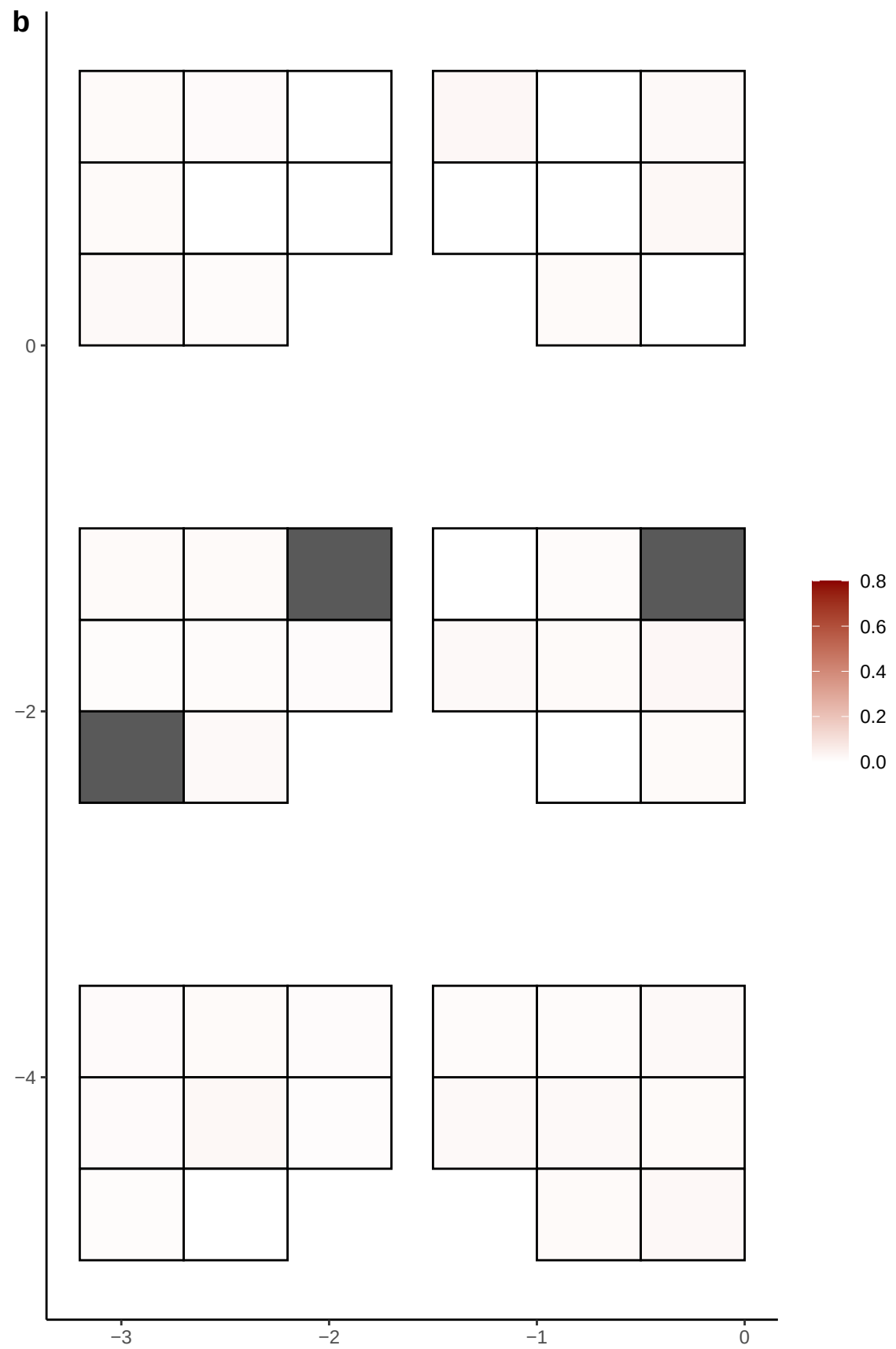

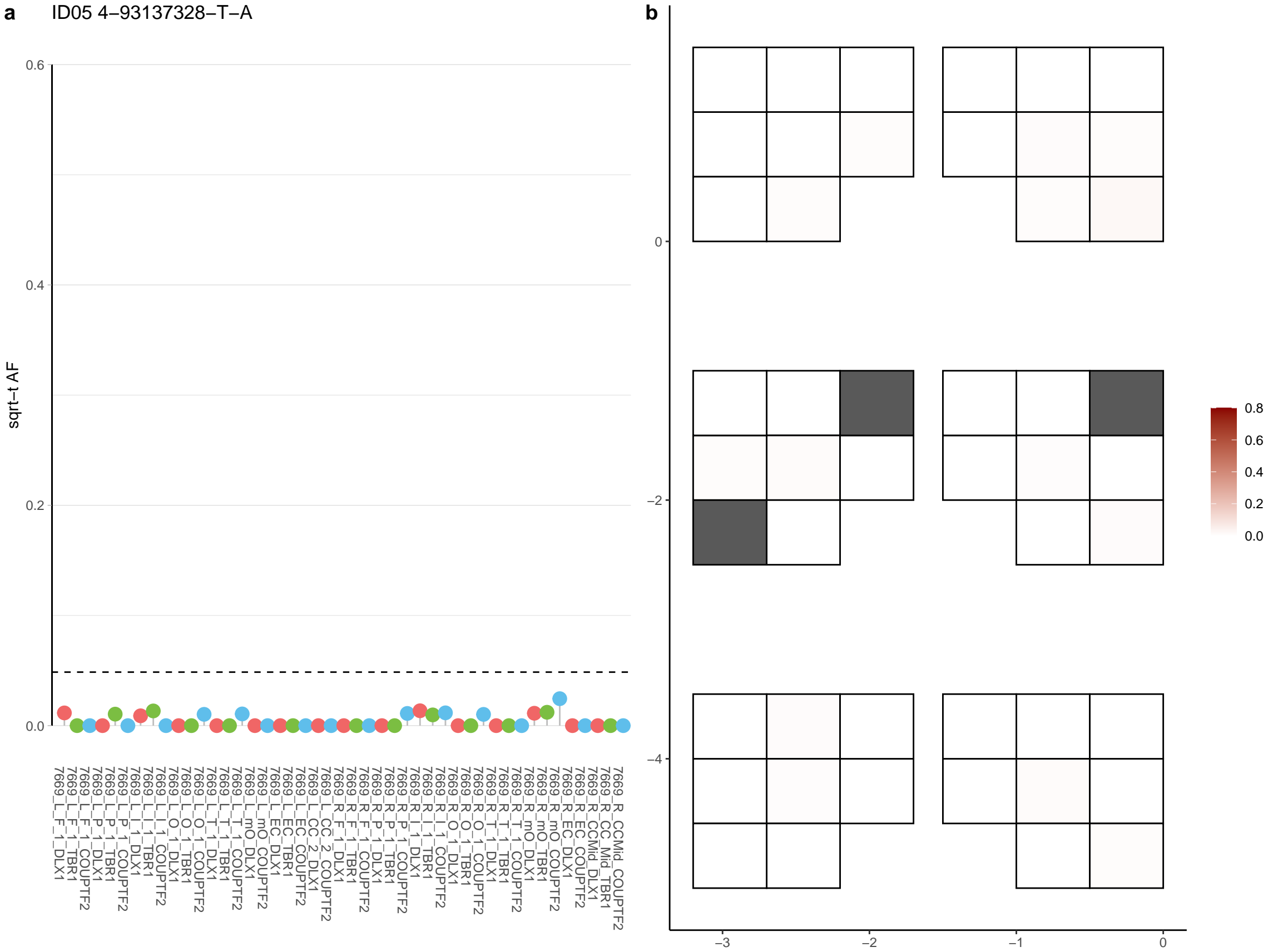

**a** ID05 4-94277150-C-T

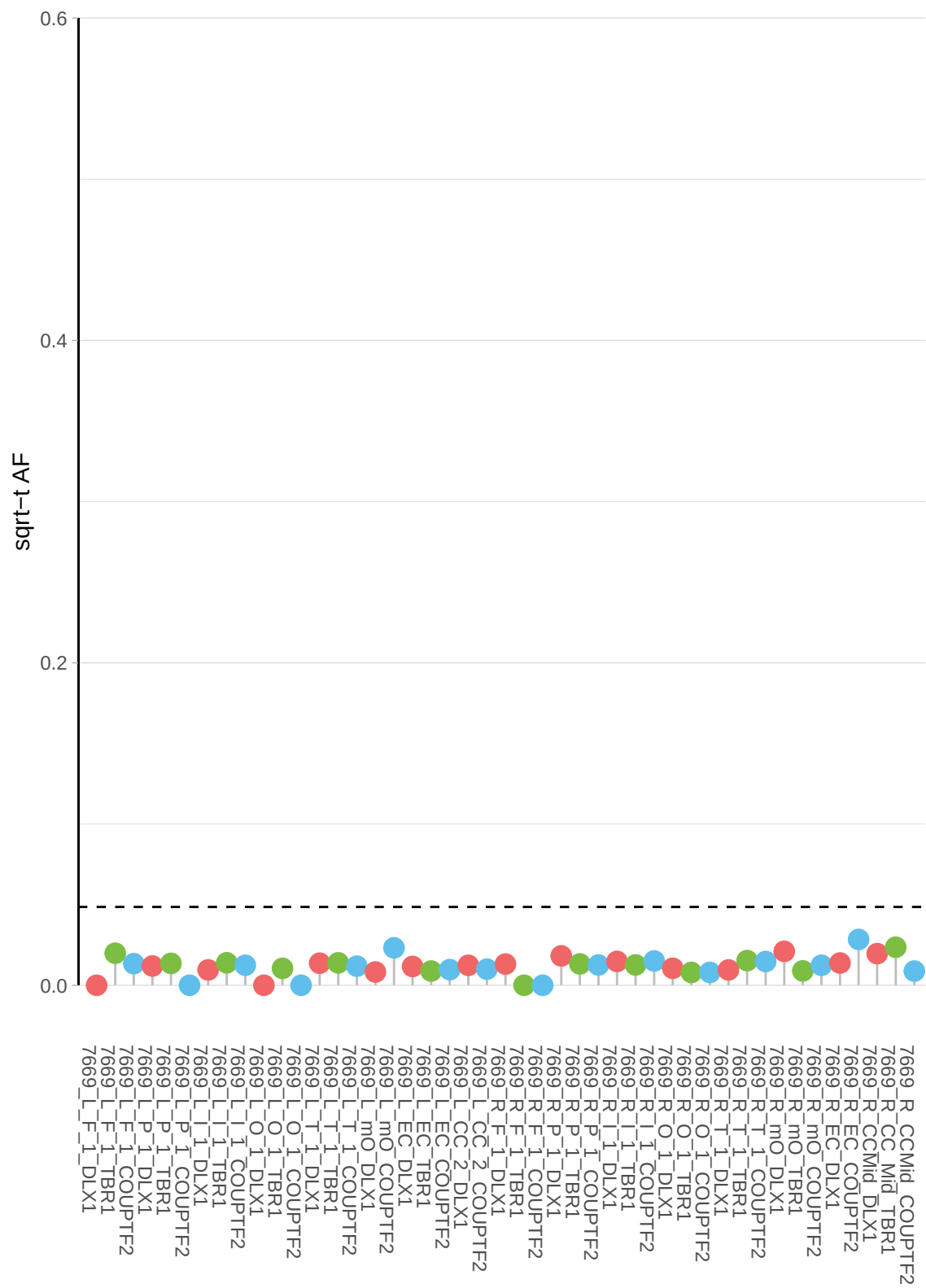

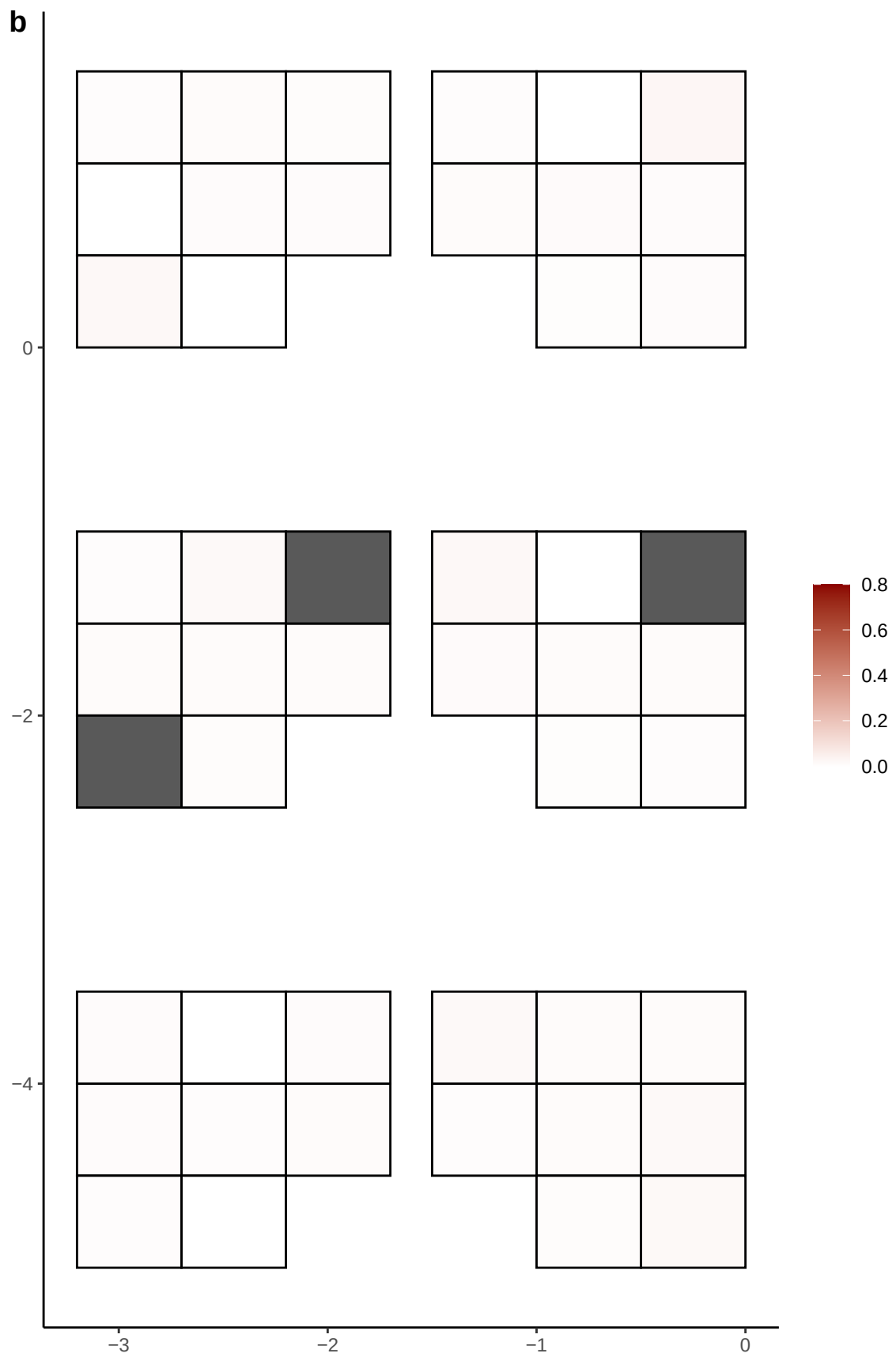

**a** ID05 4-96601800-C-T

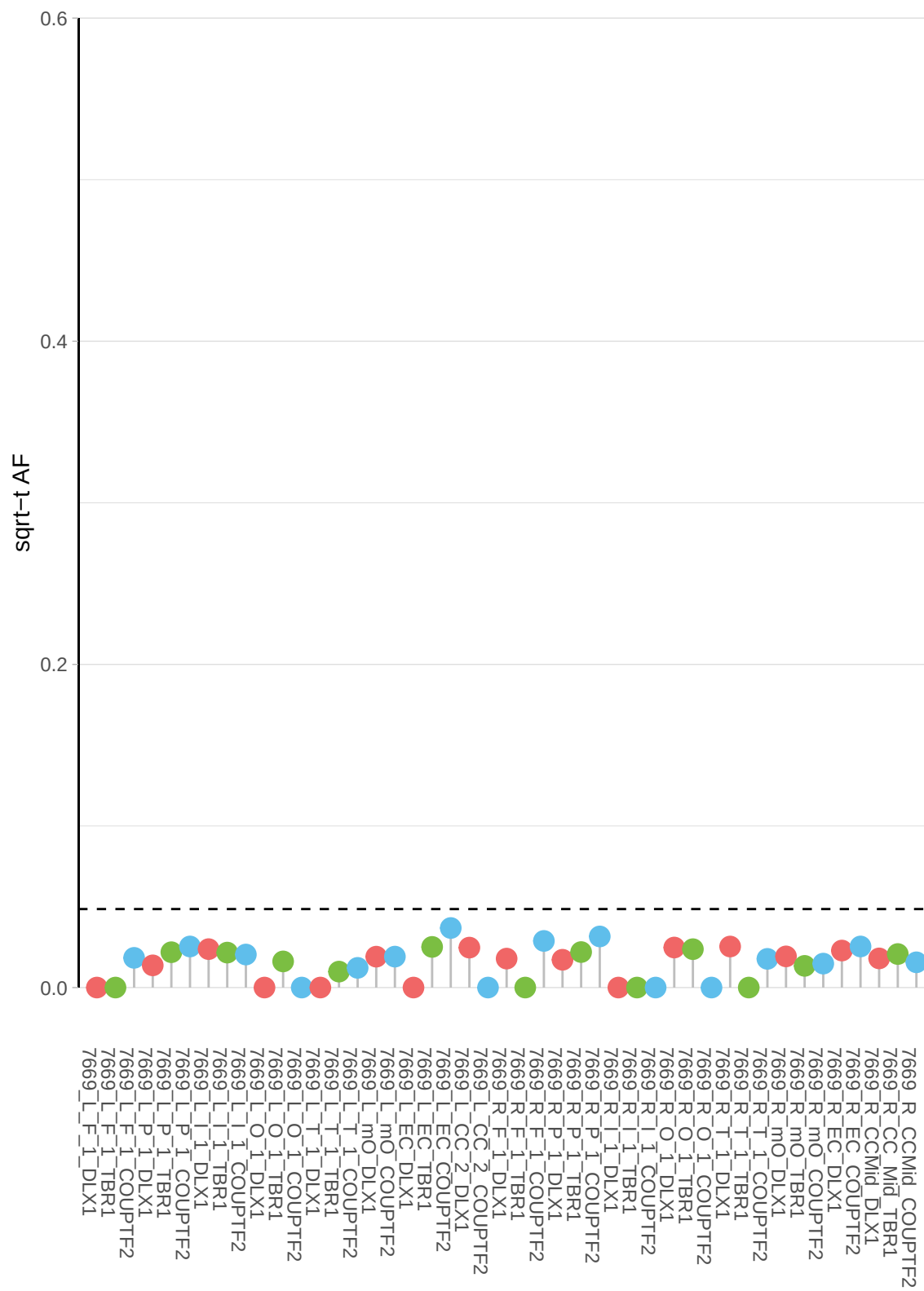

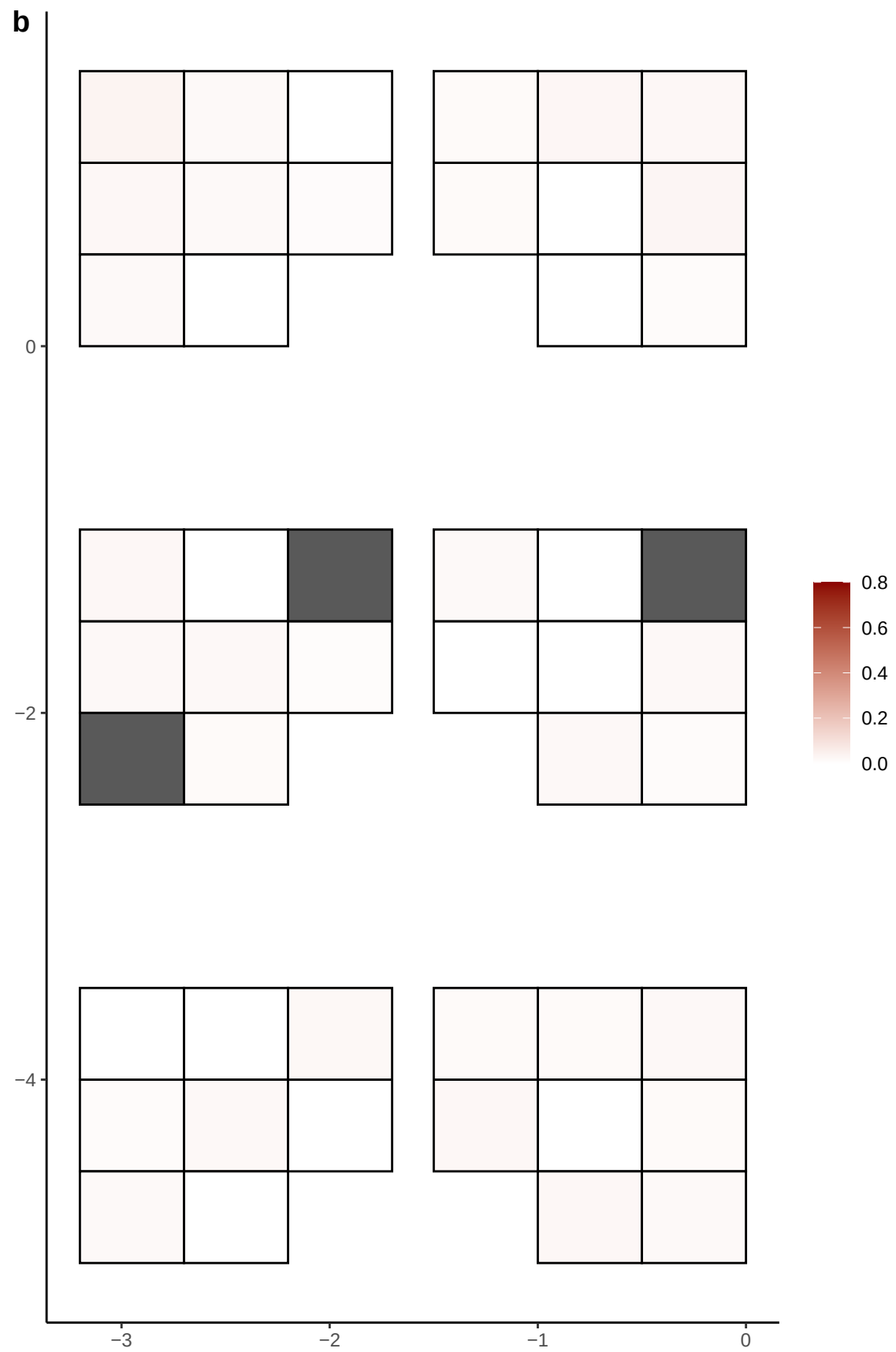

**a** ID05 5-10808399-G-A

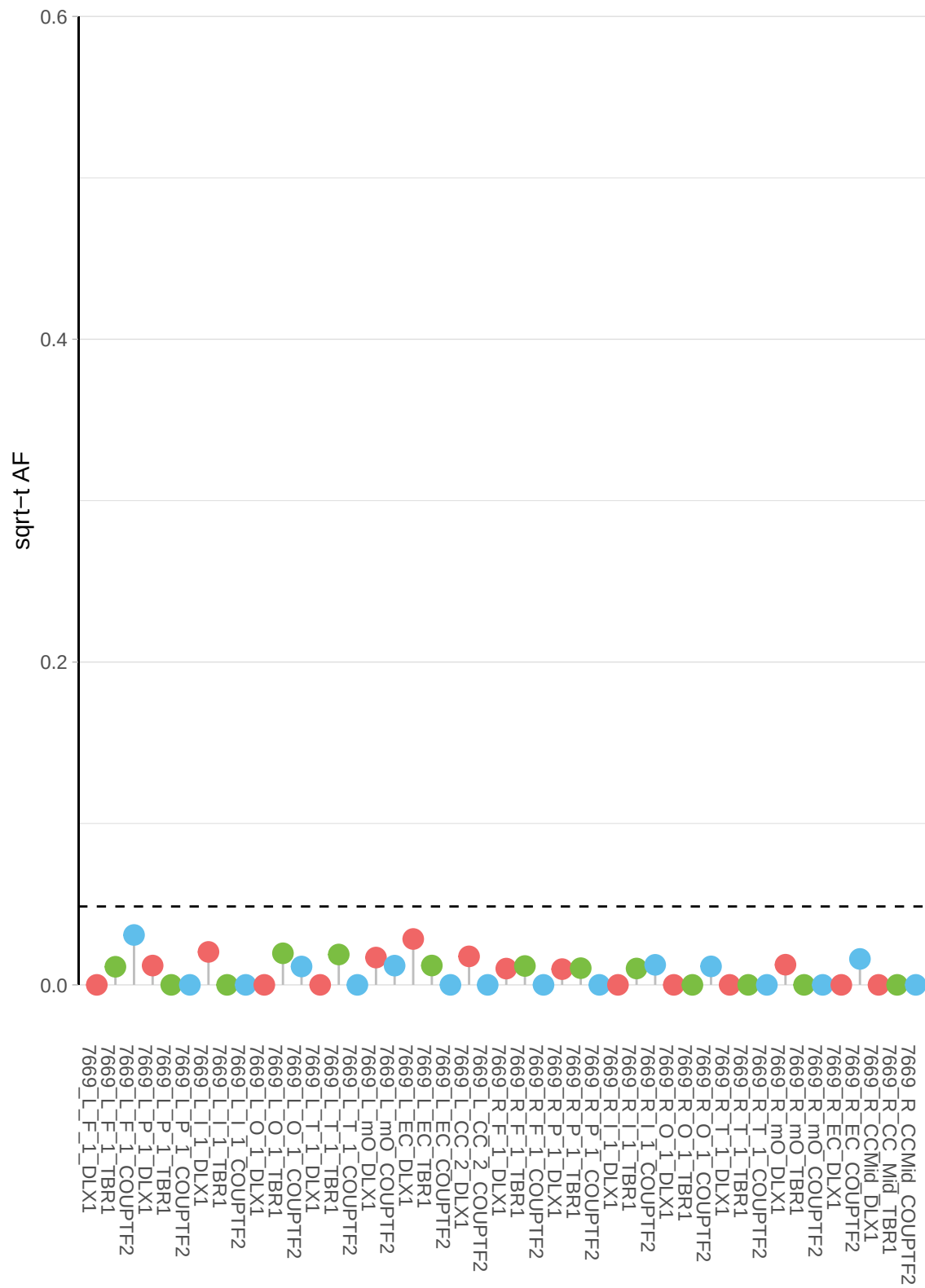

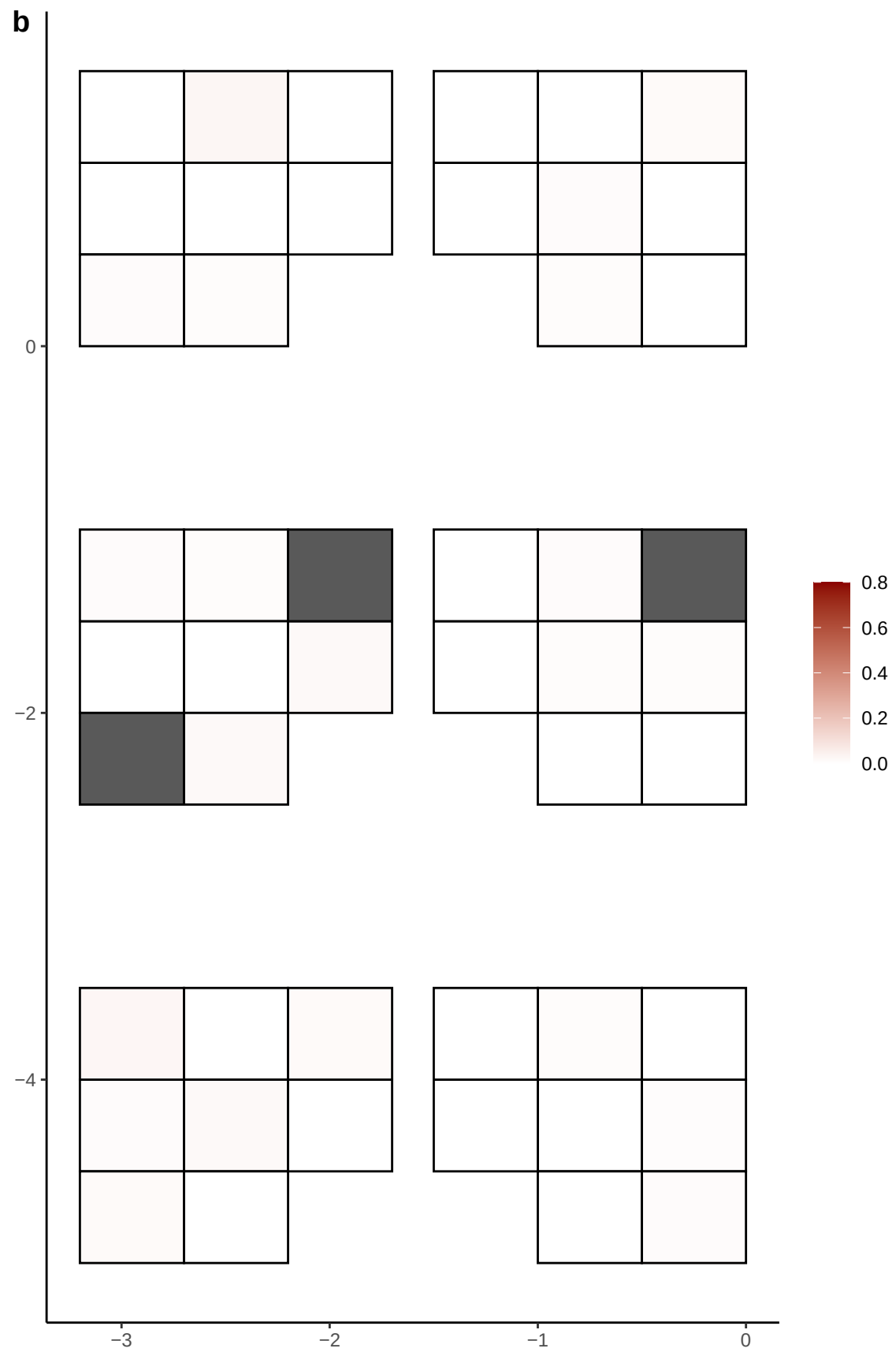

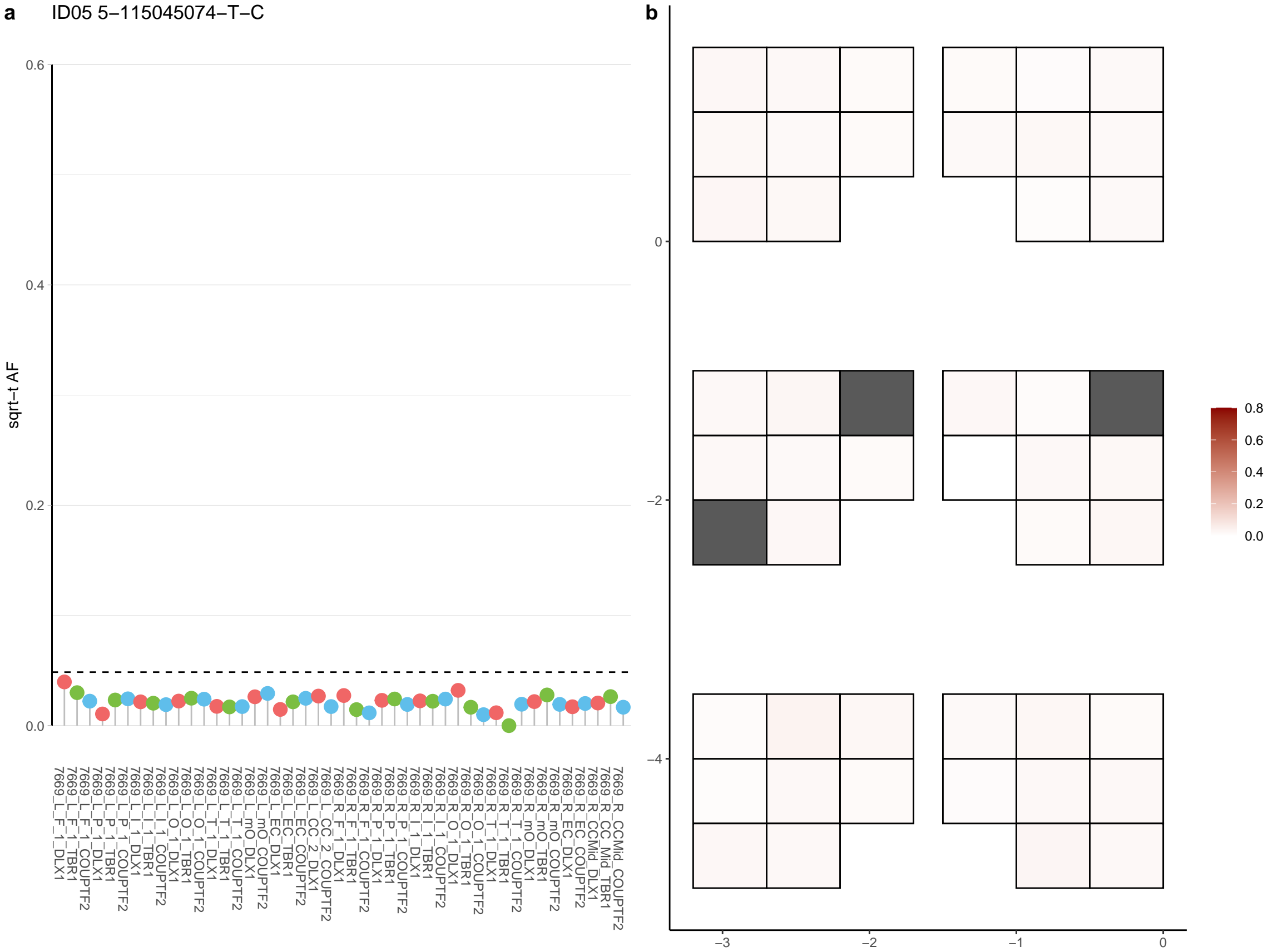

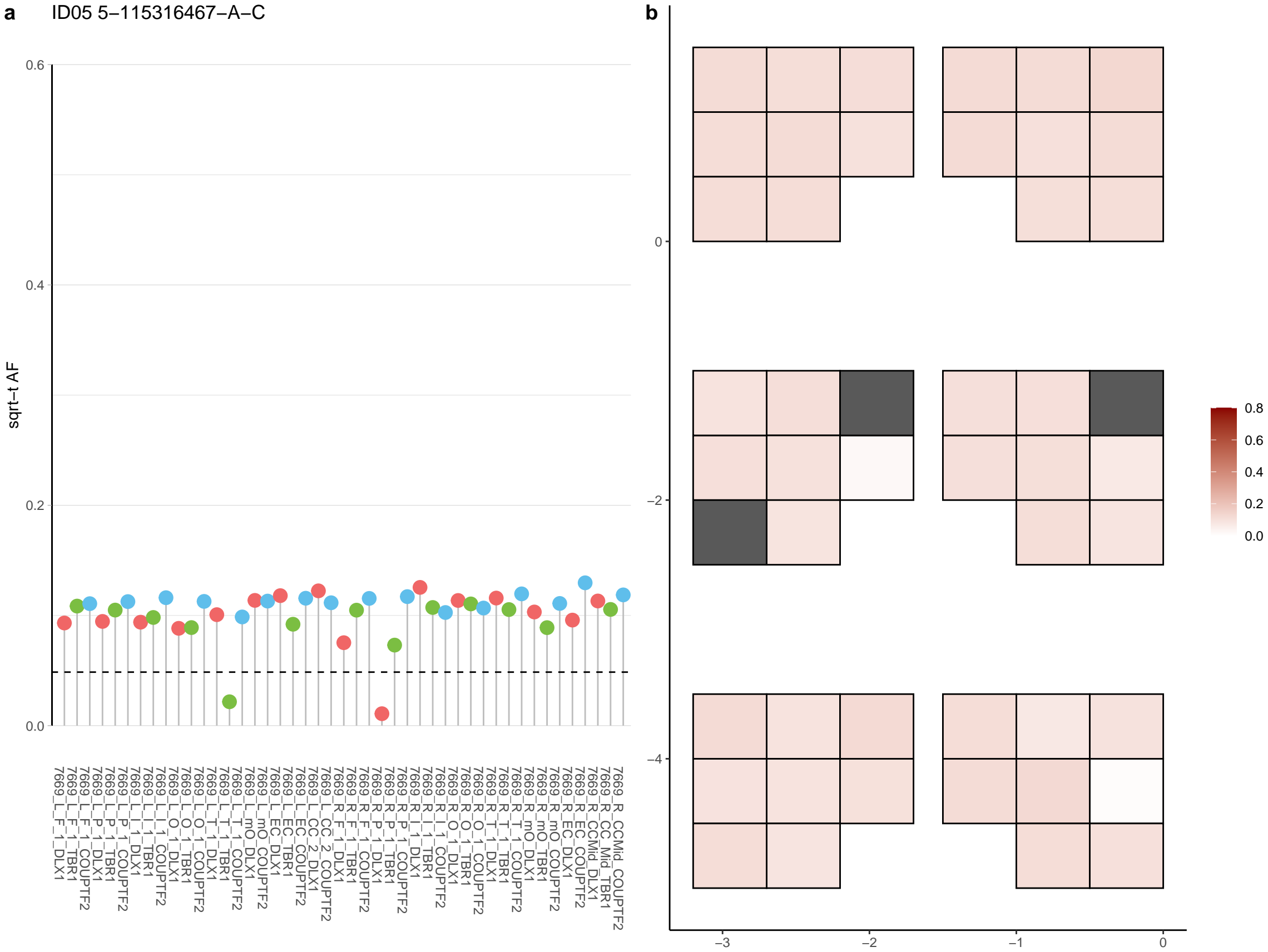

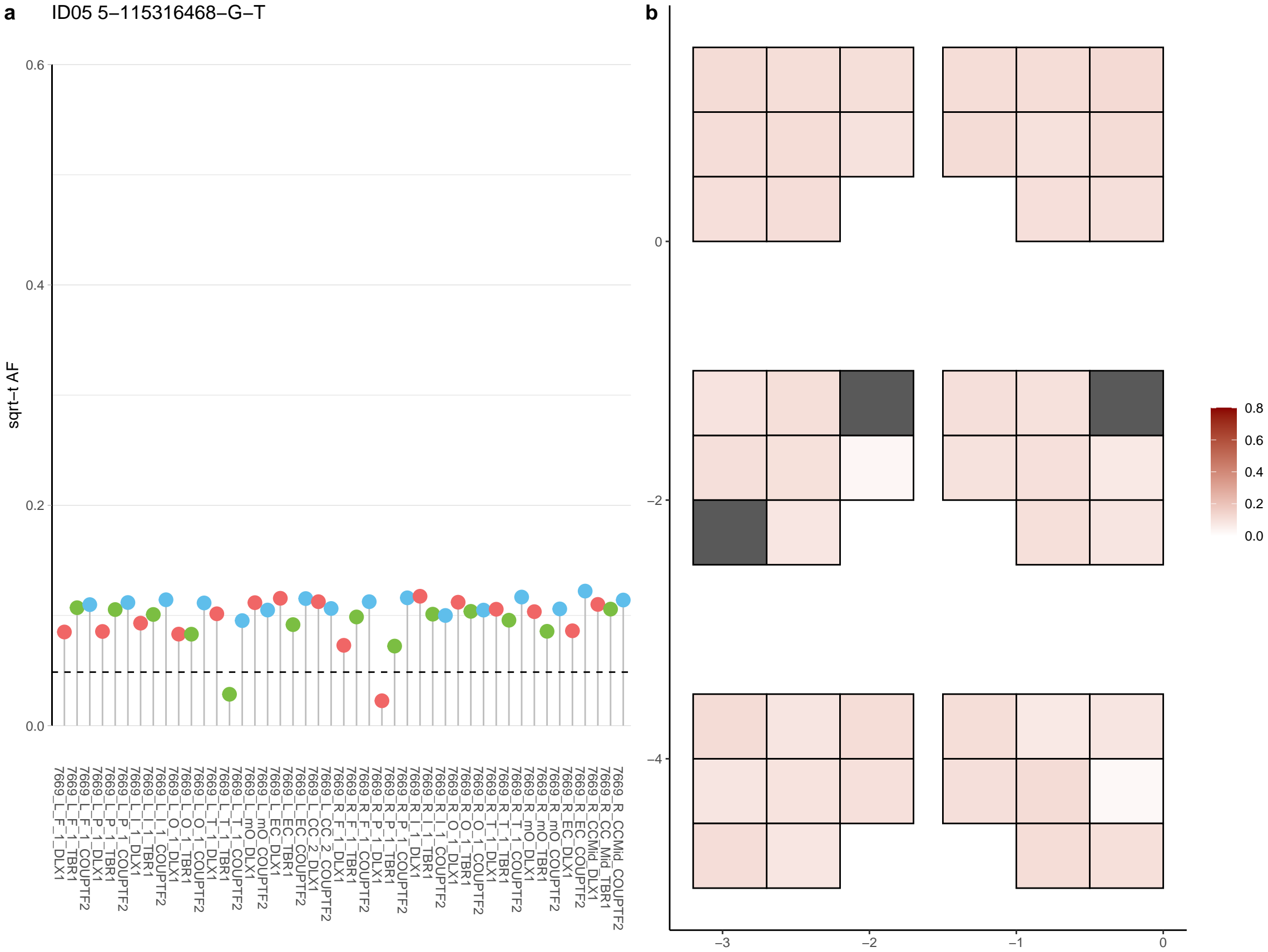

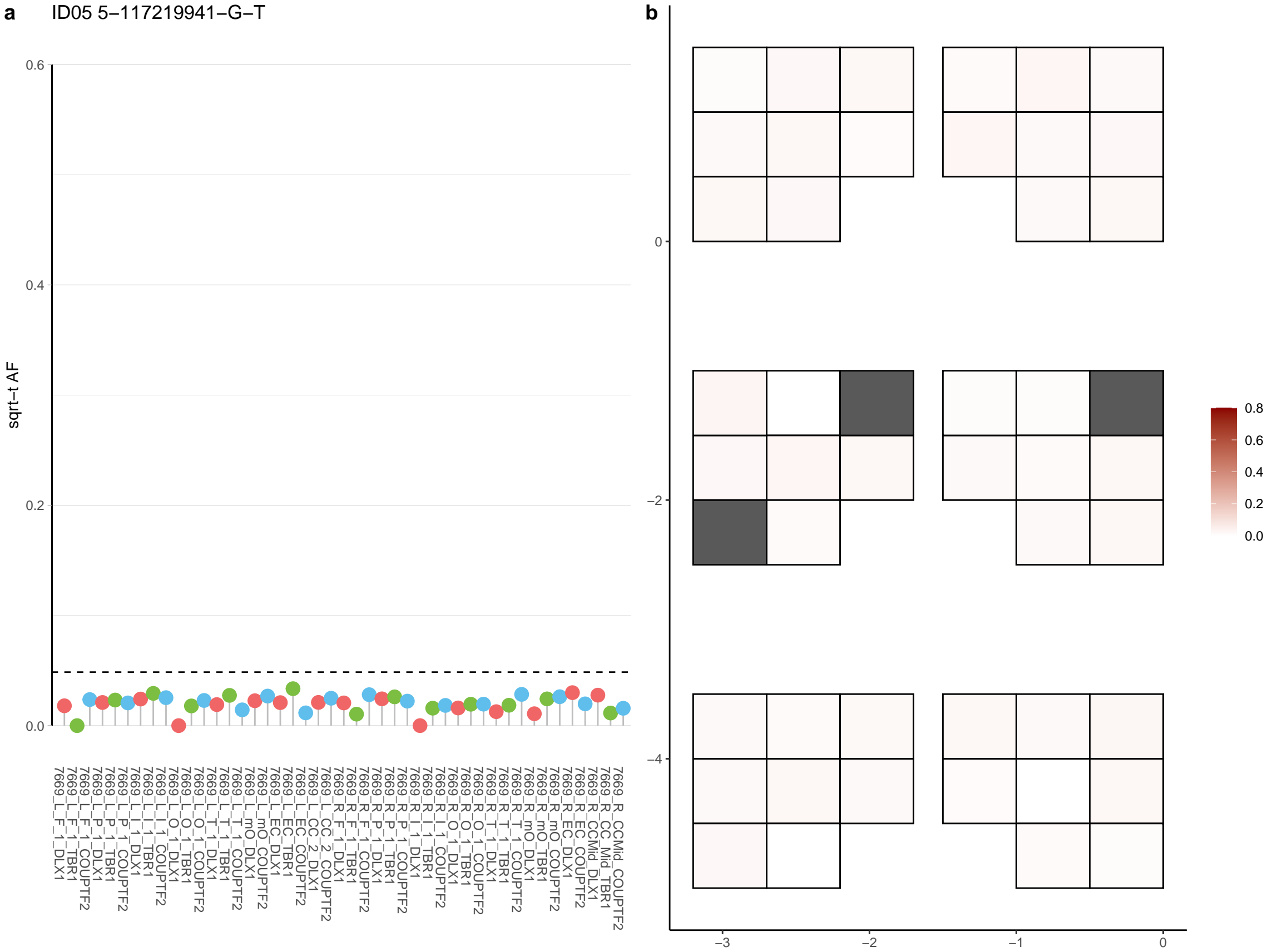

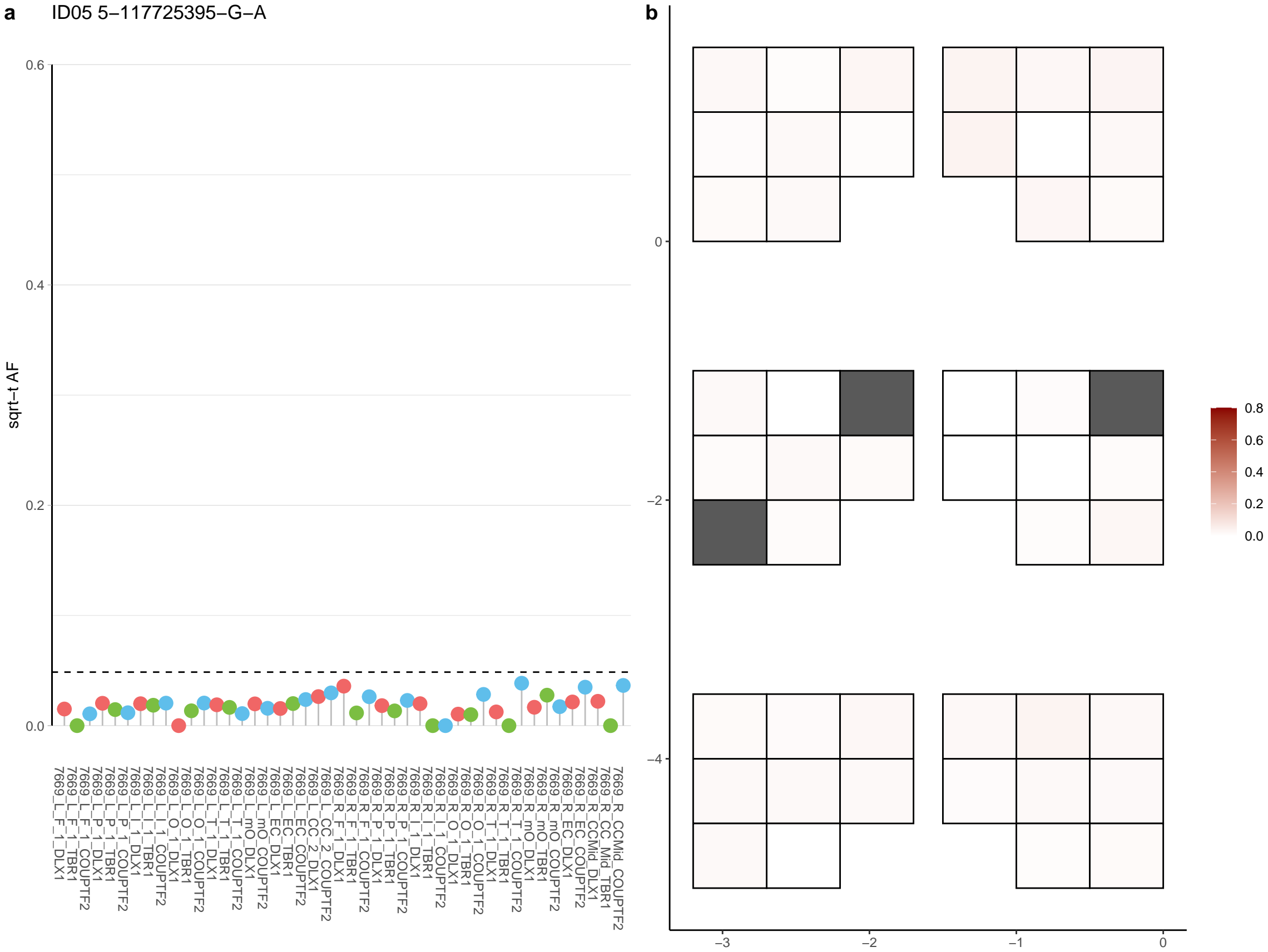

**a** ID05 5-121446225-C-T

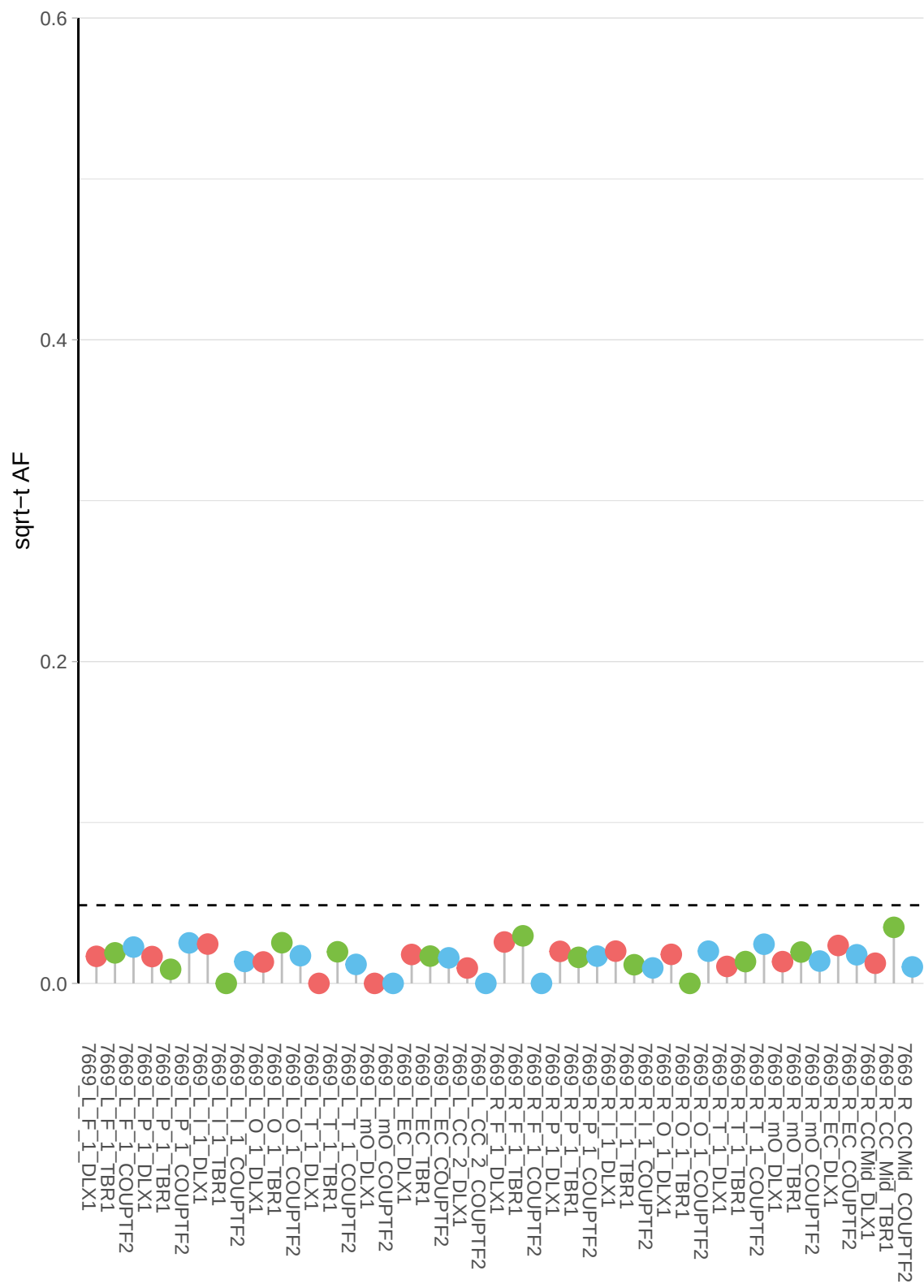

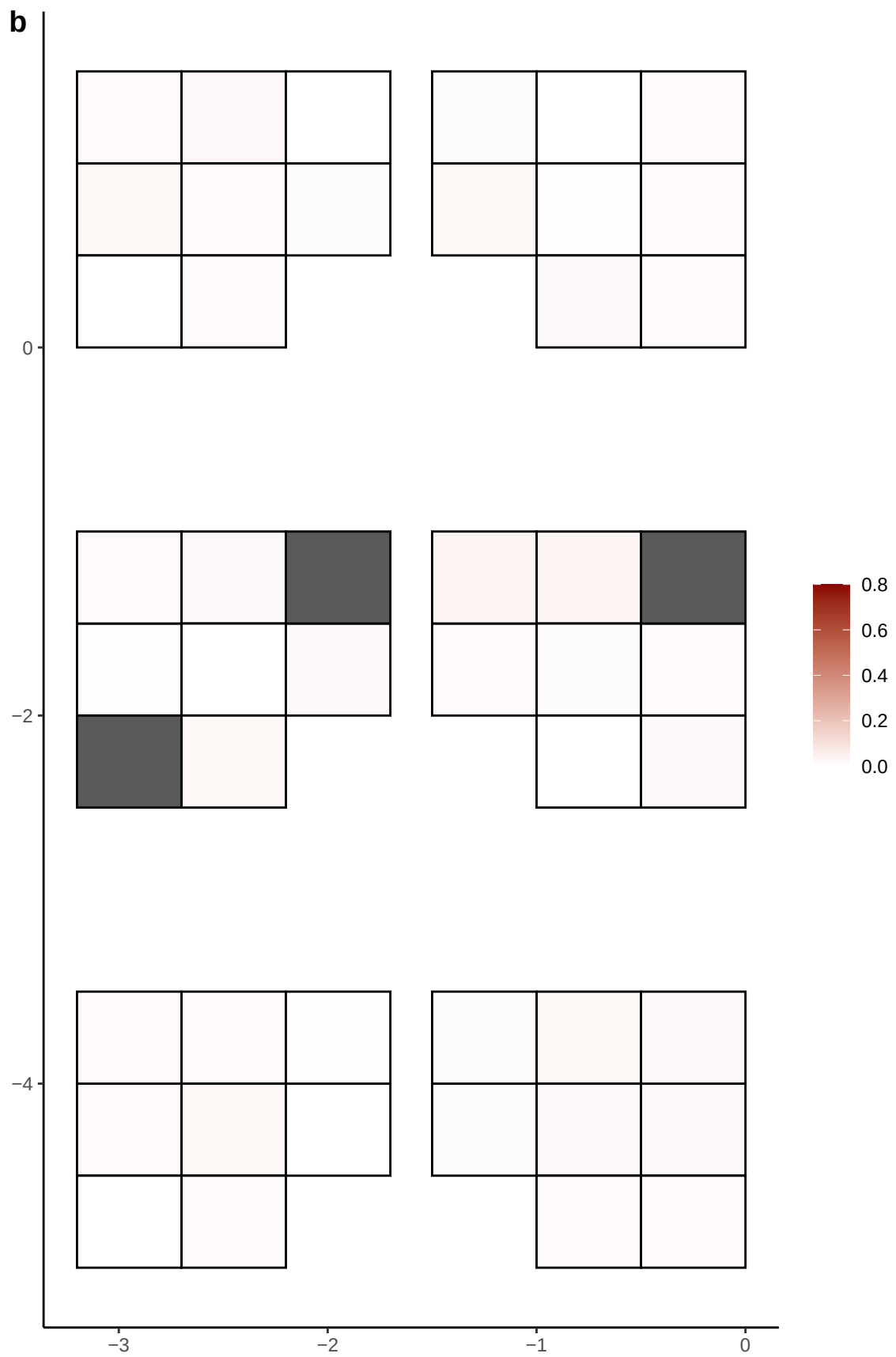

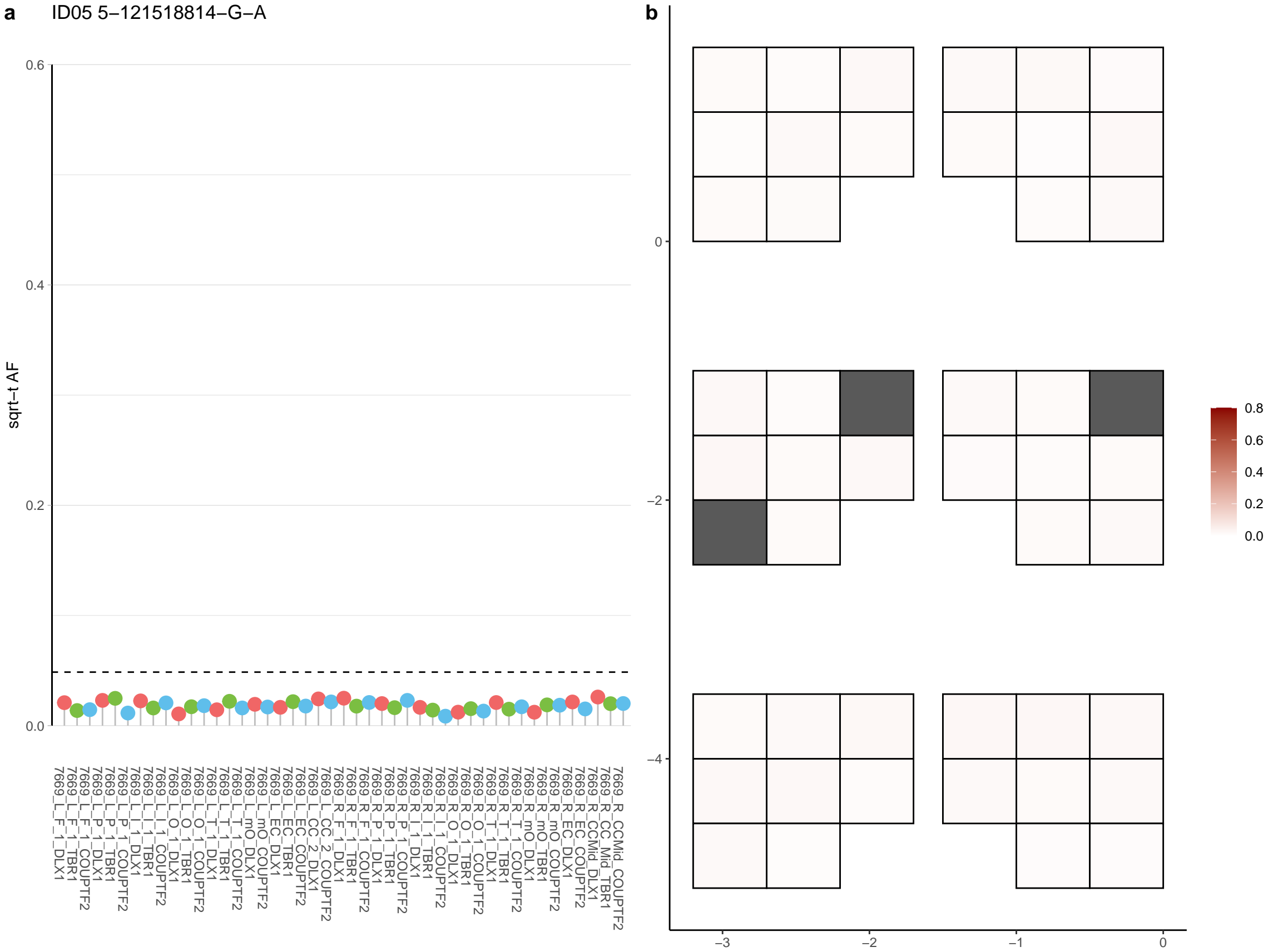

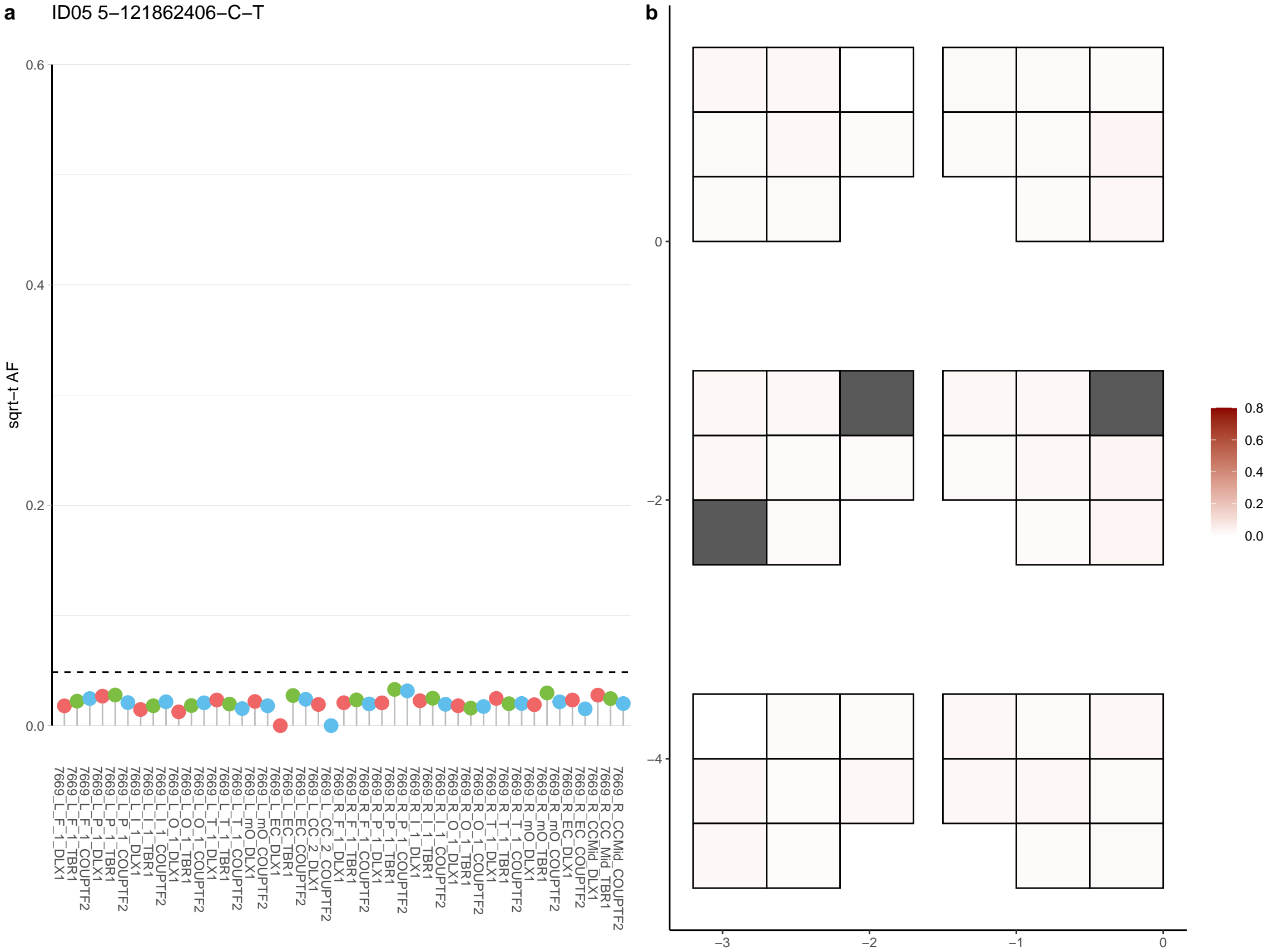

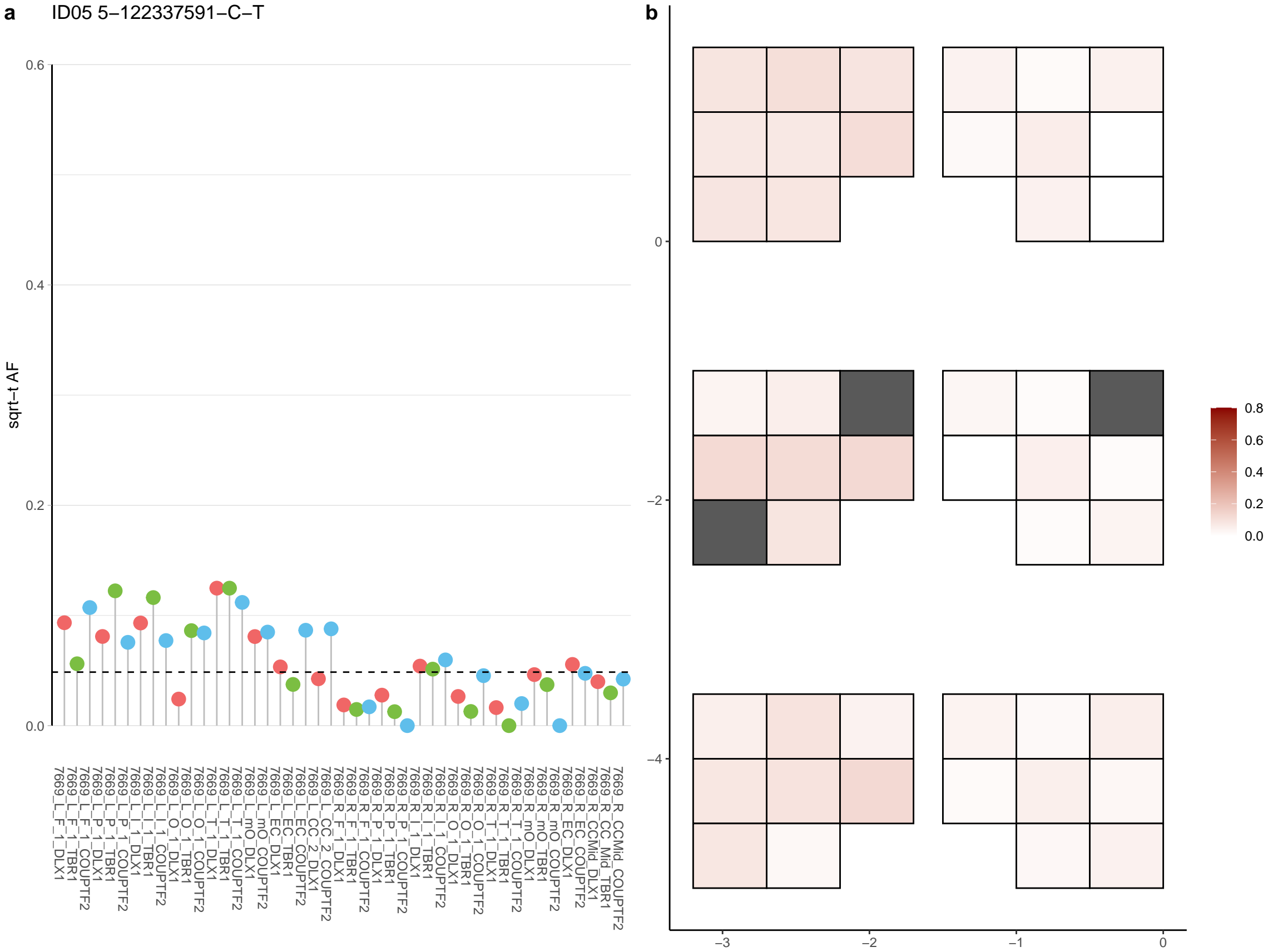

**a** ID05 5-13021088-C-T

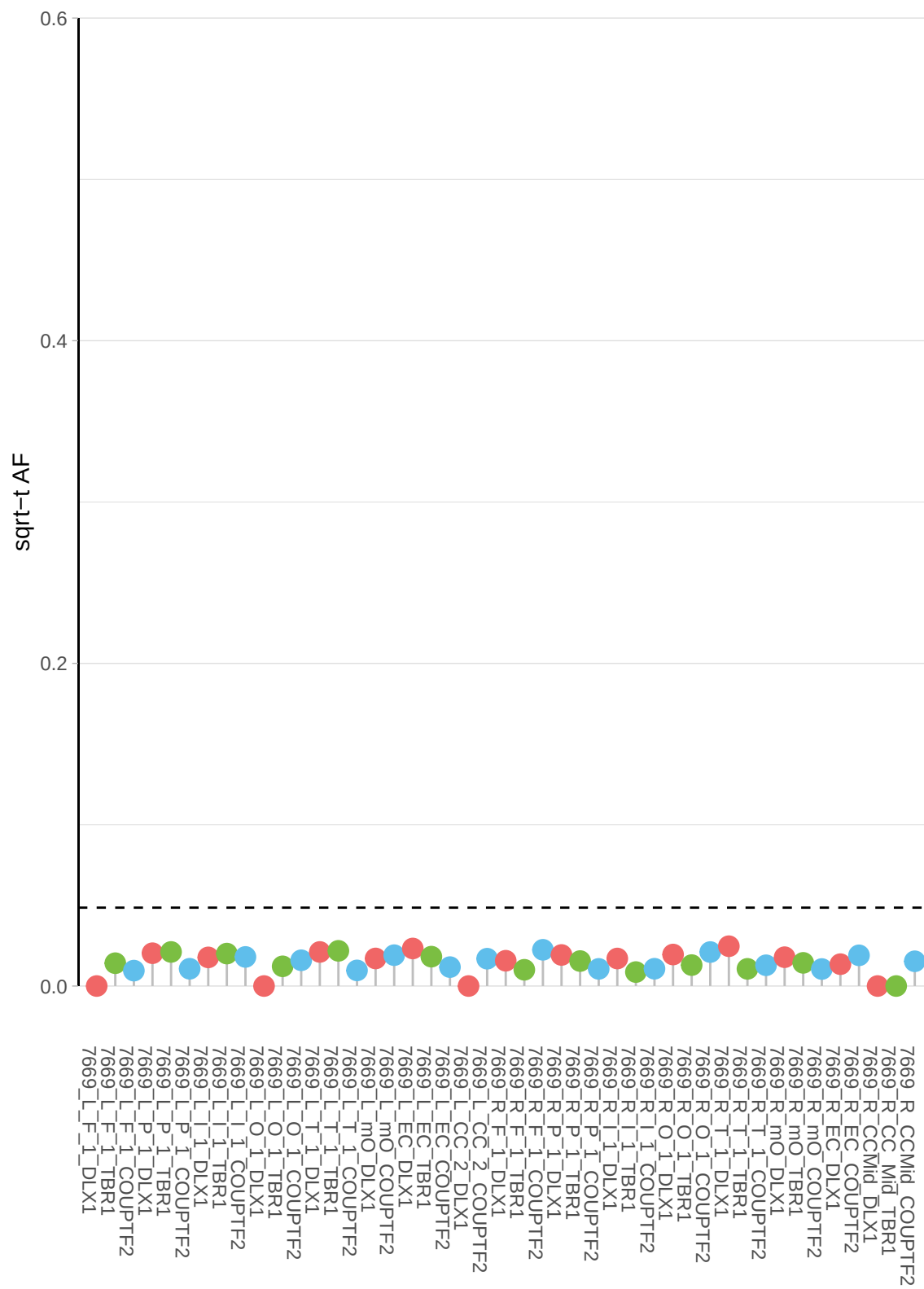

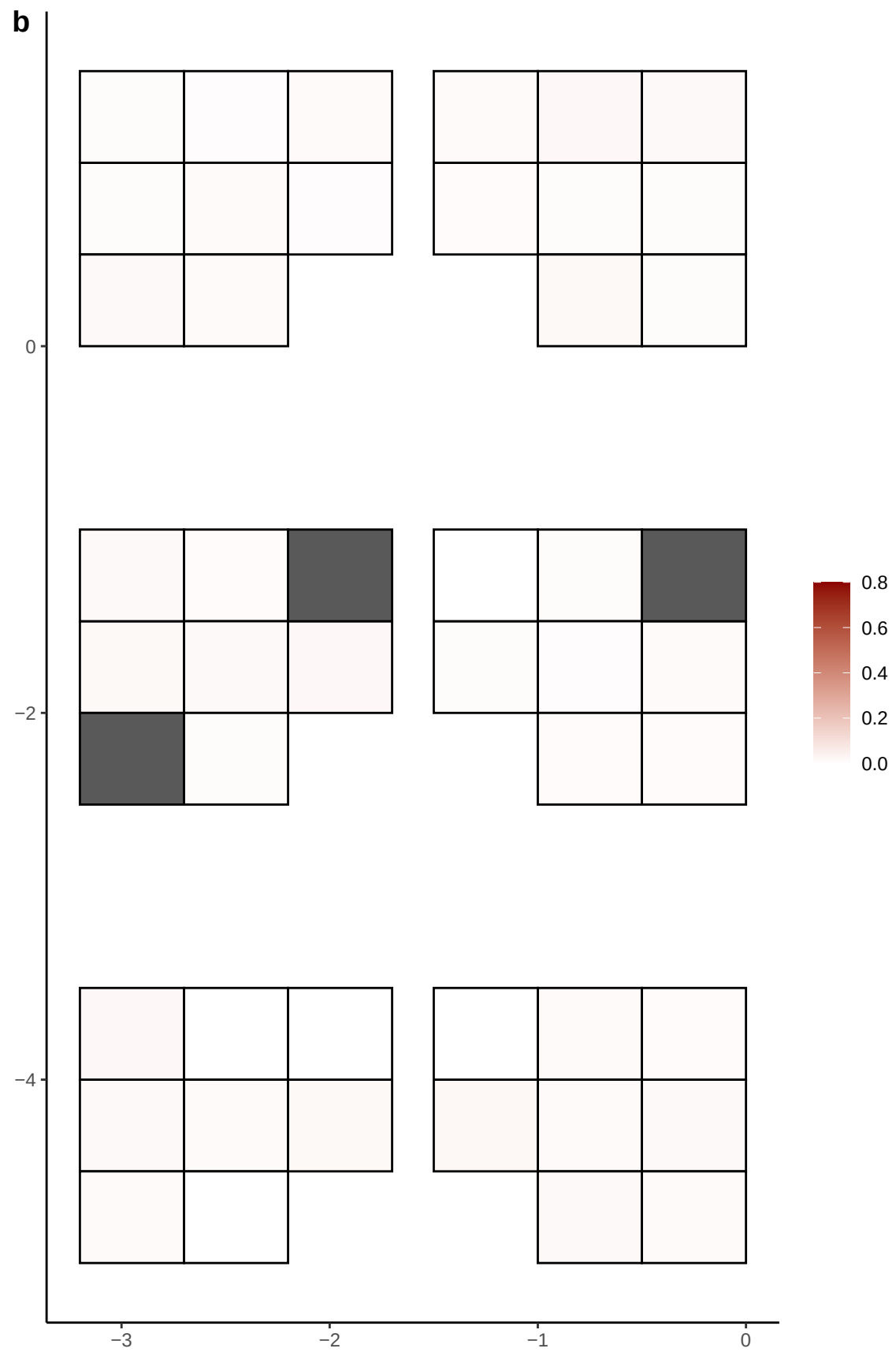

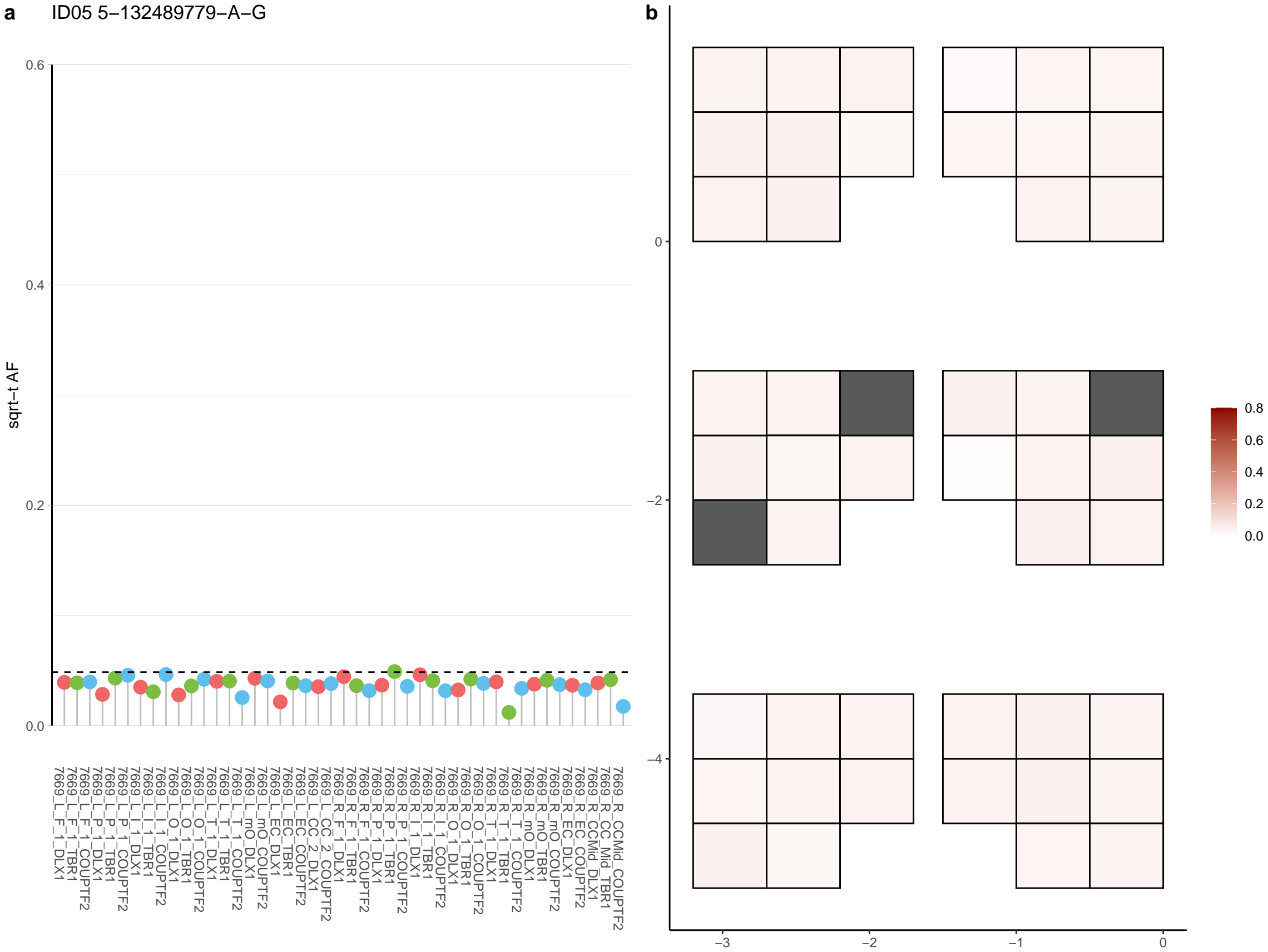

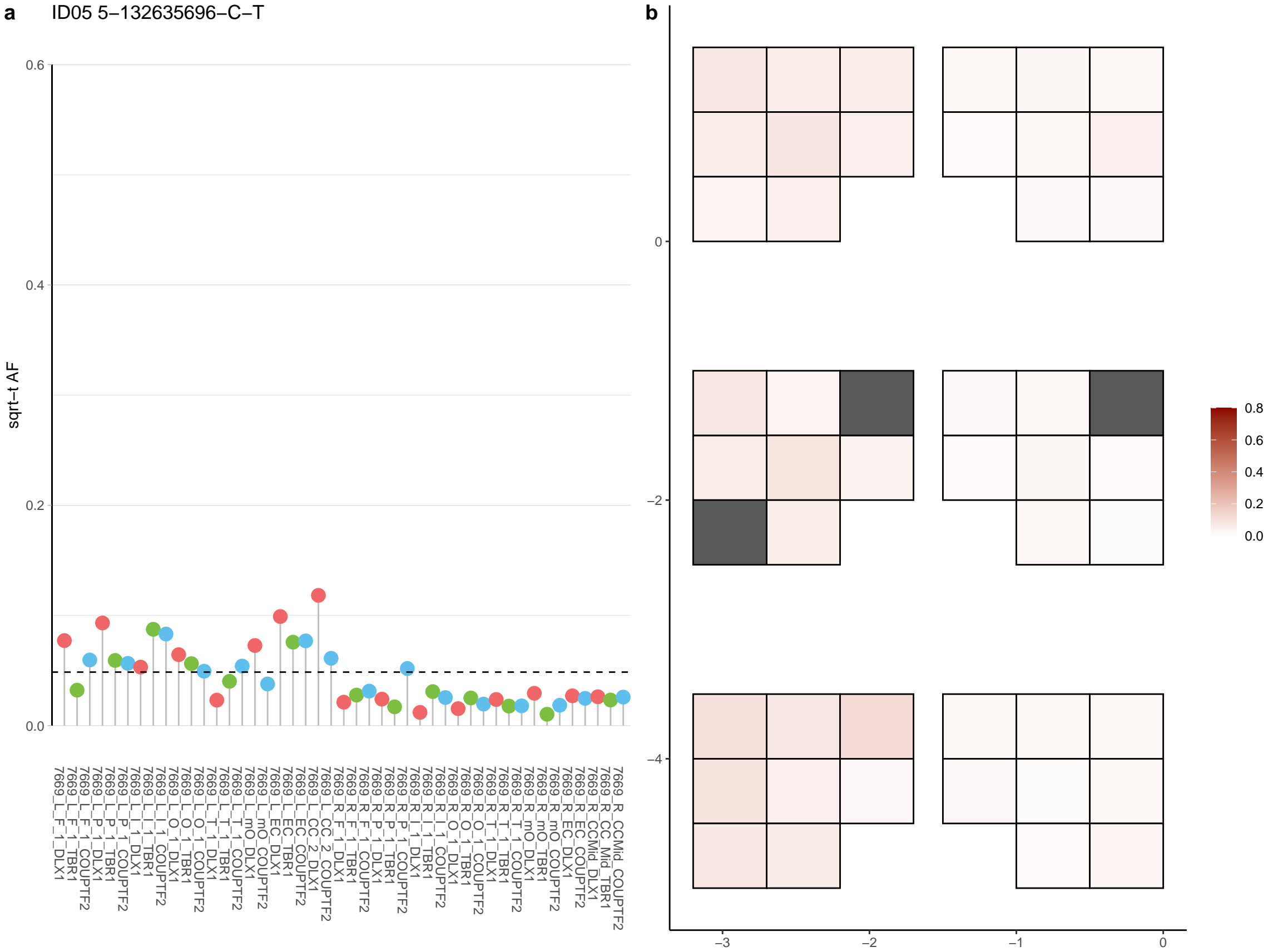

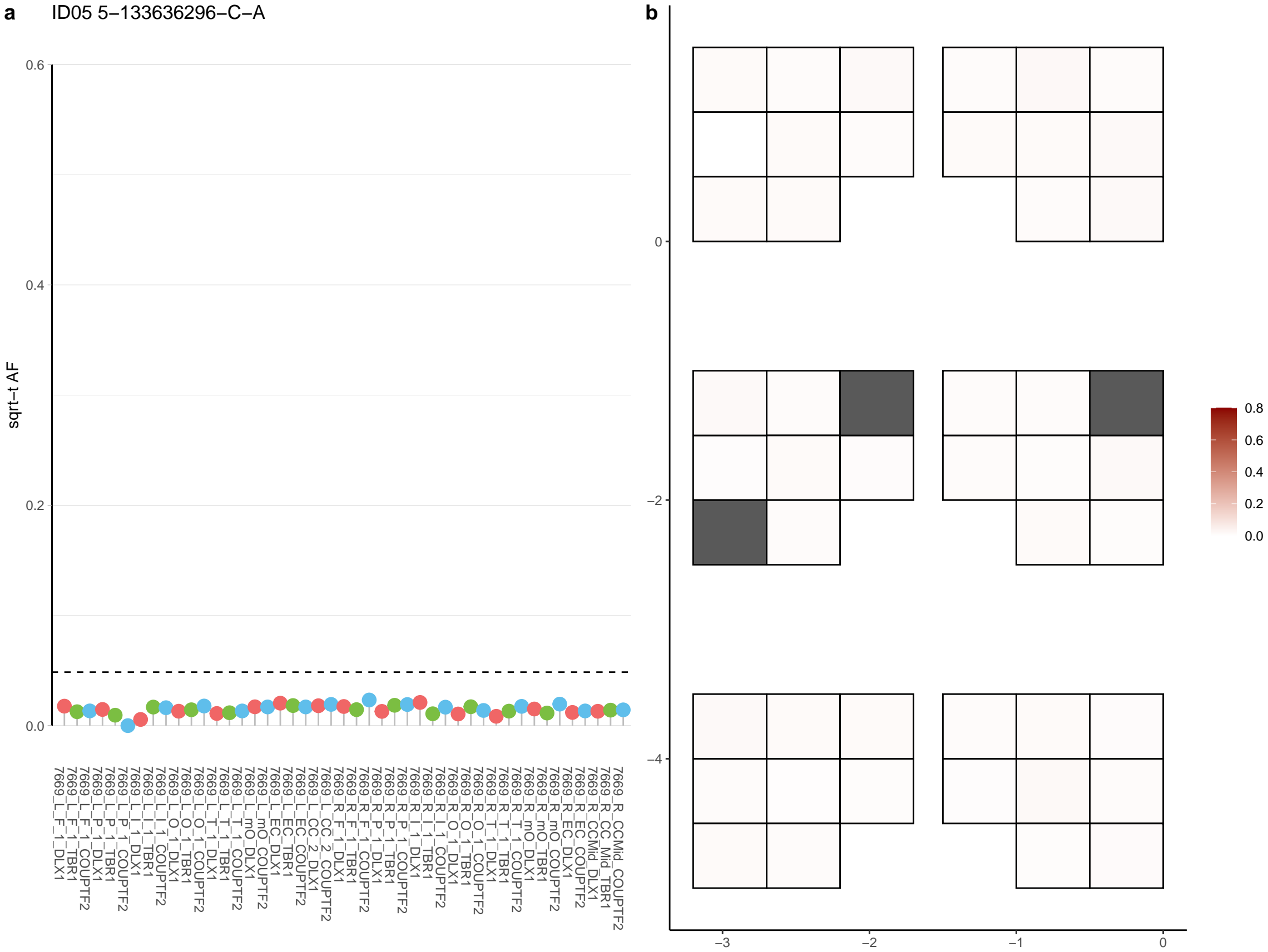

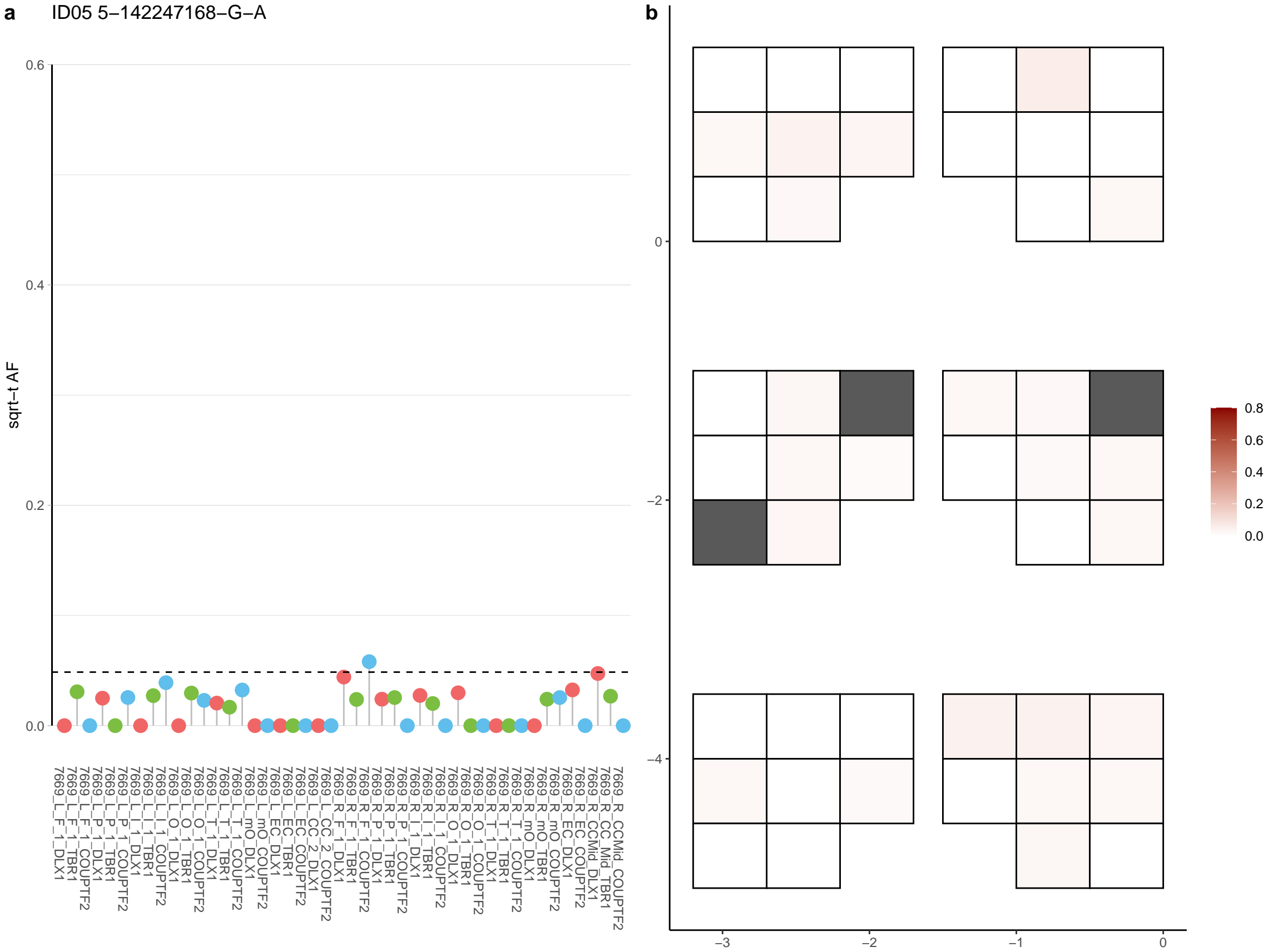

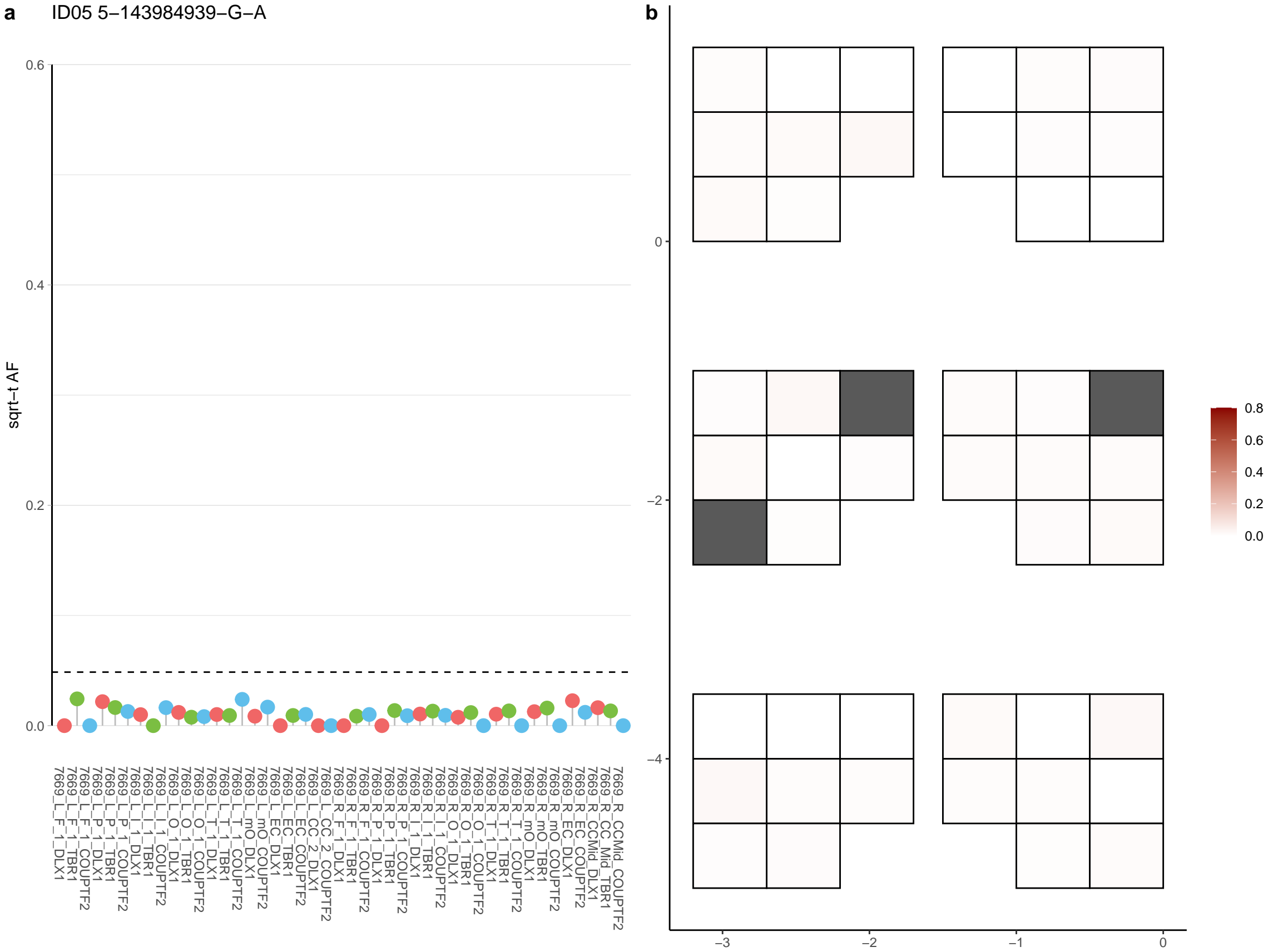

**a** ID05 5-155350078-C-T

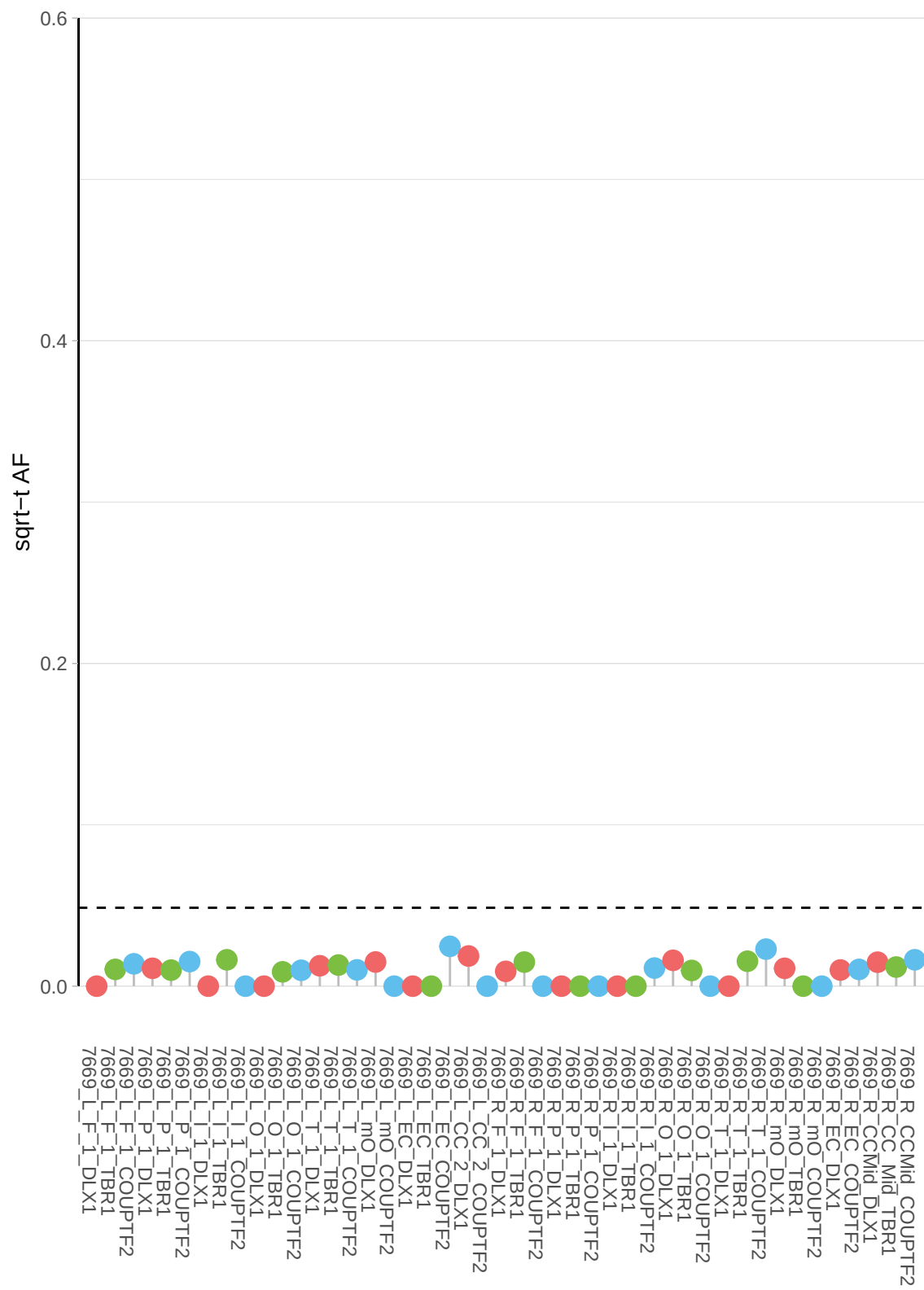

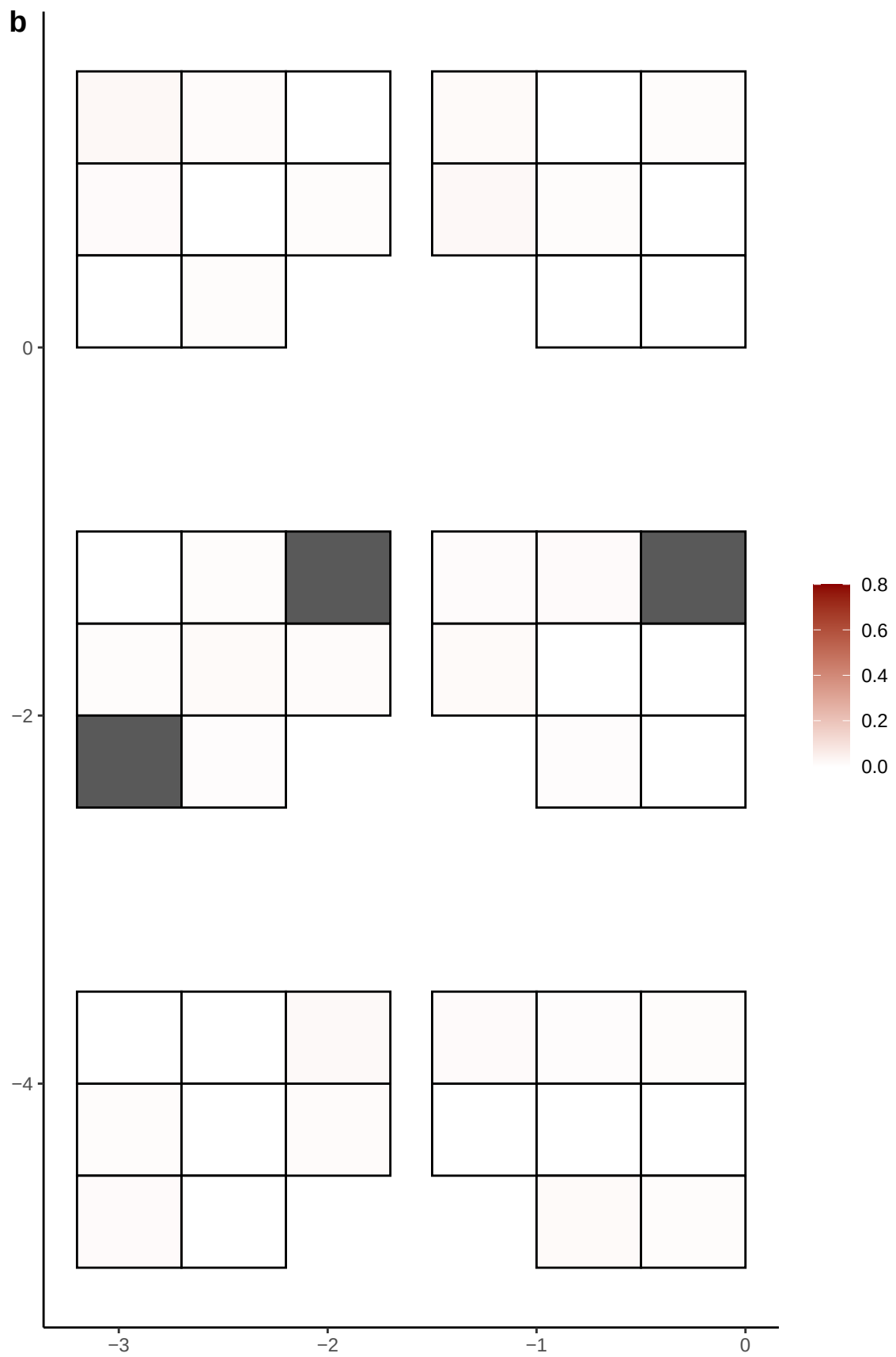

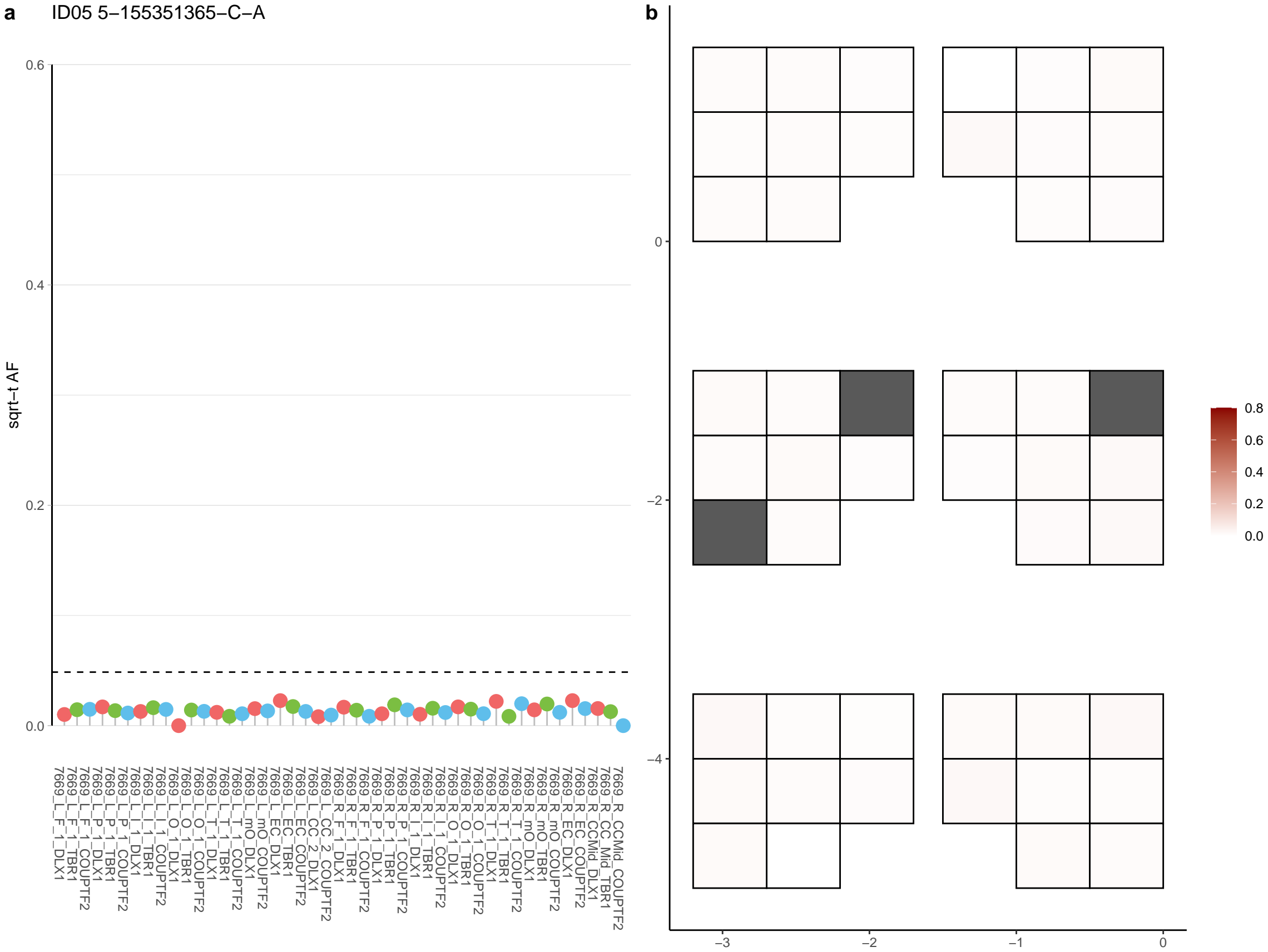

**a** ID05 5-158180215-G-A

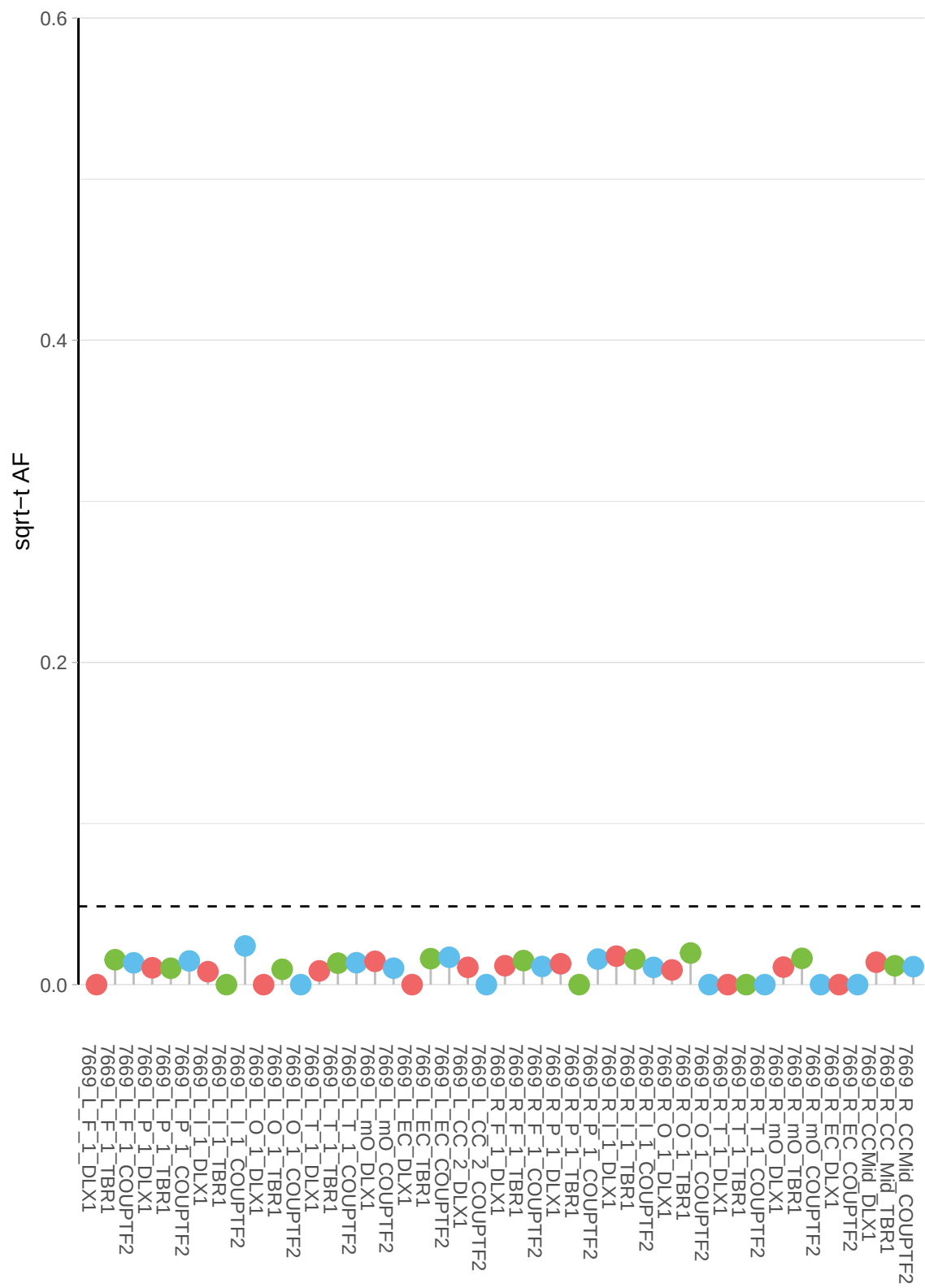

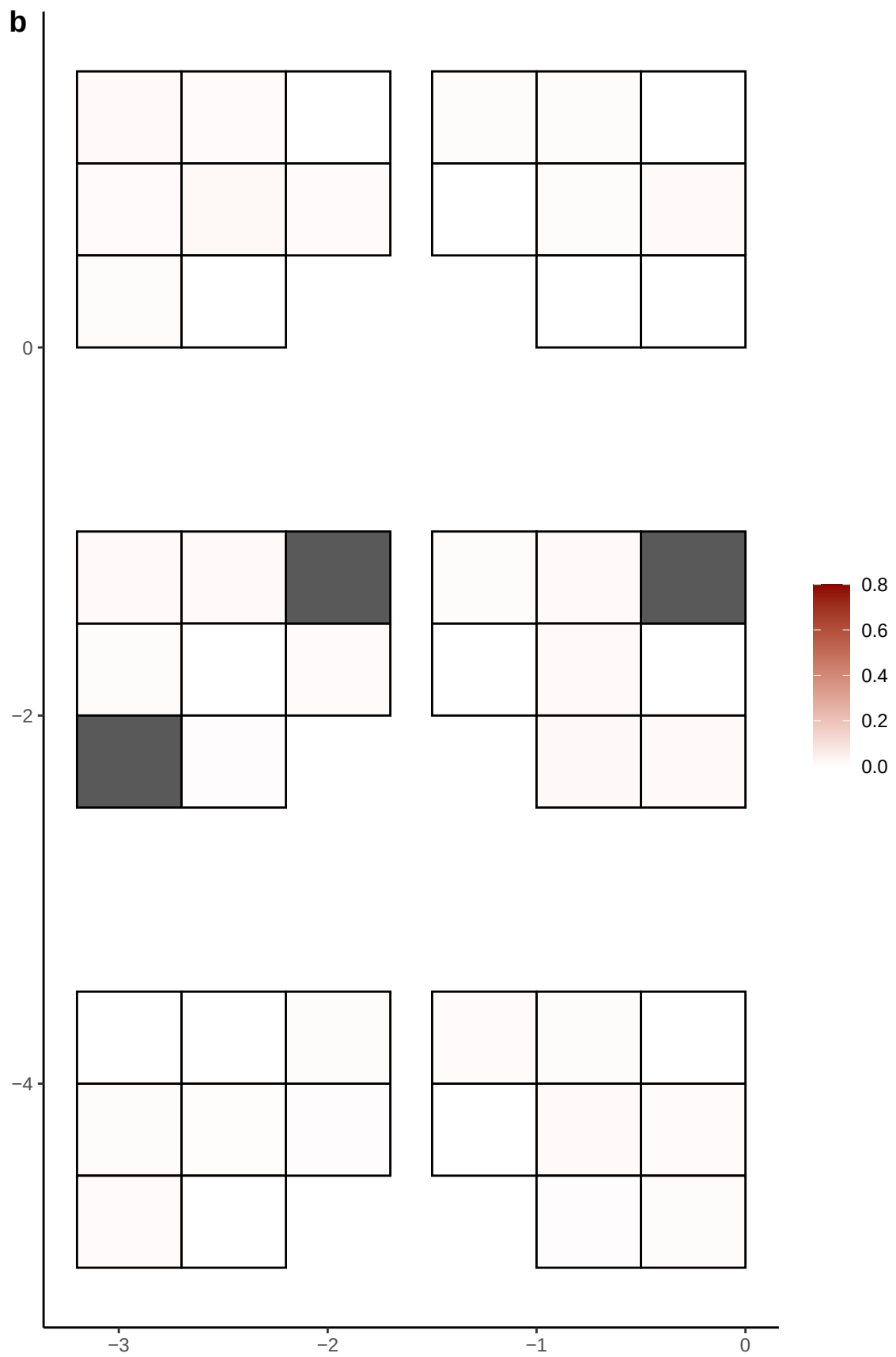

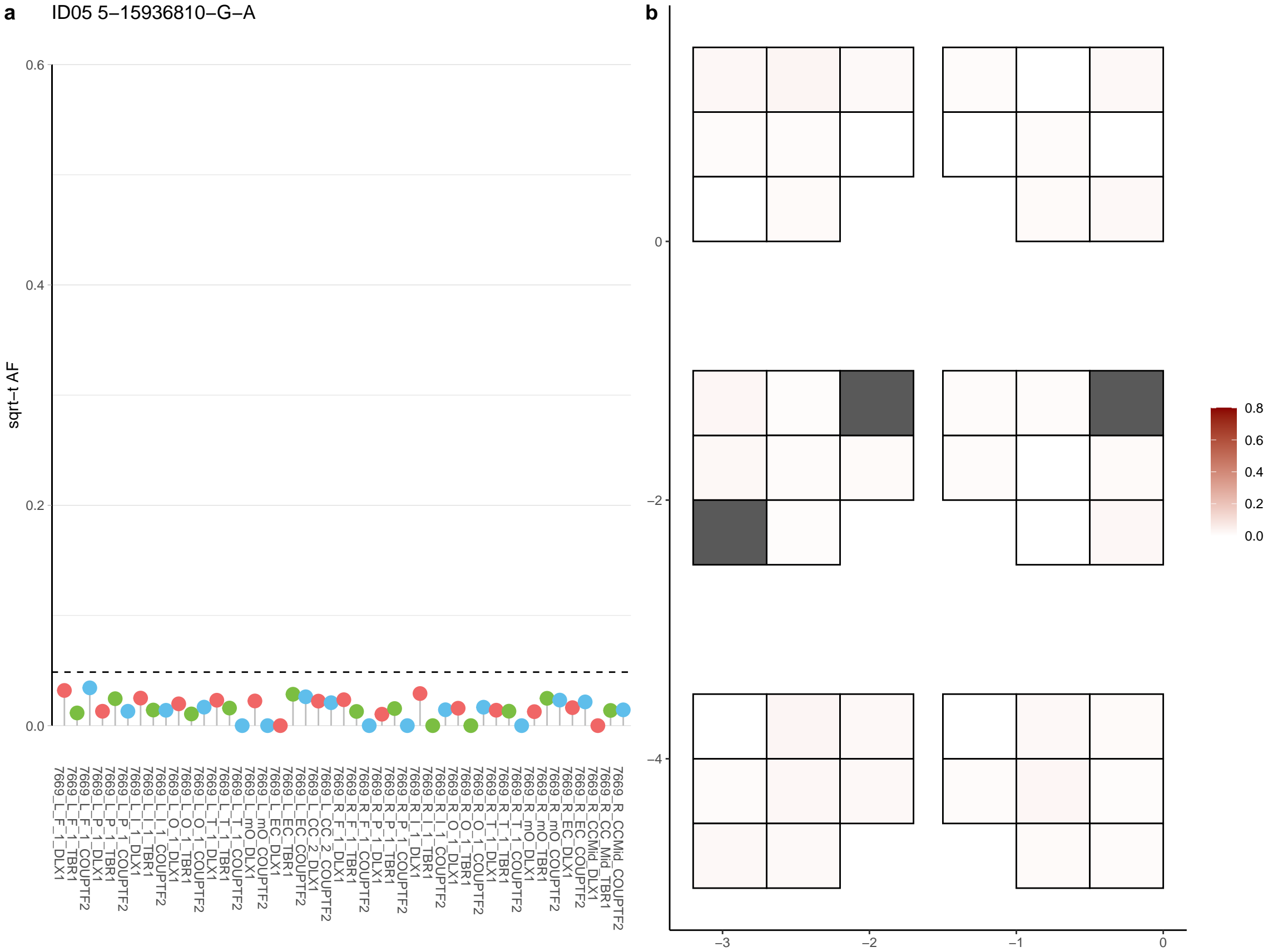

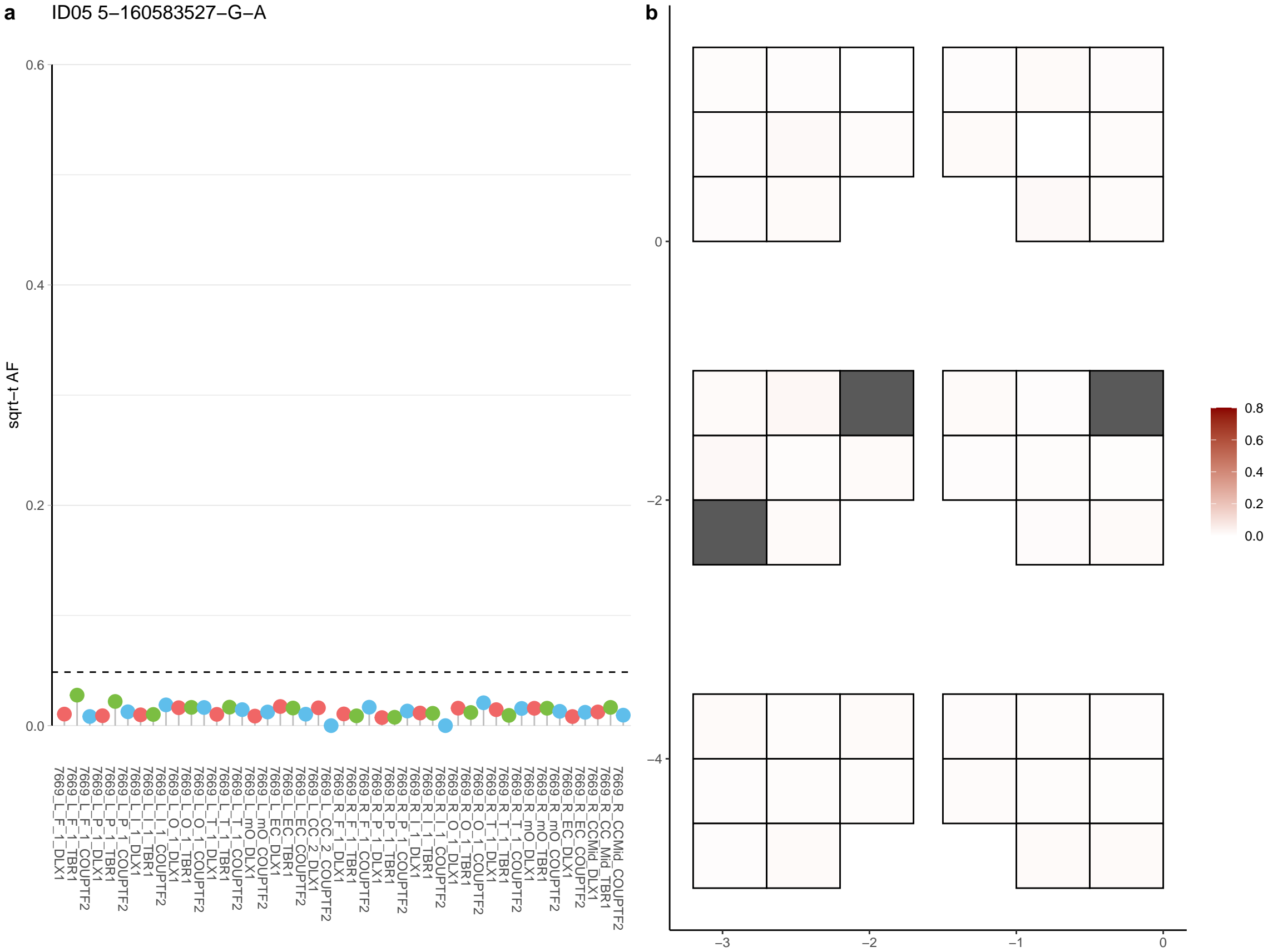

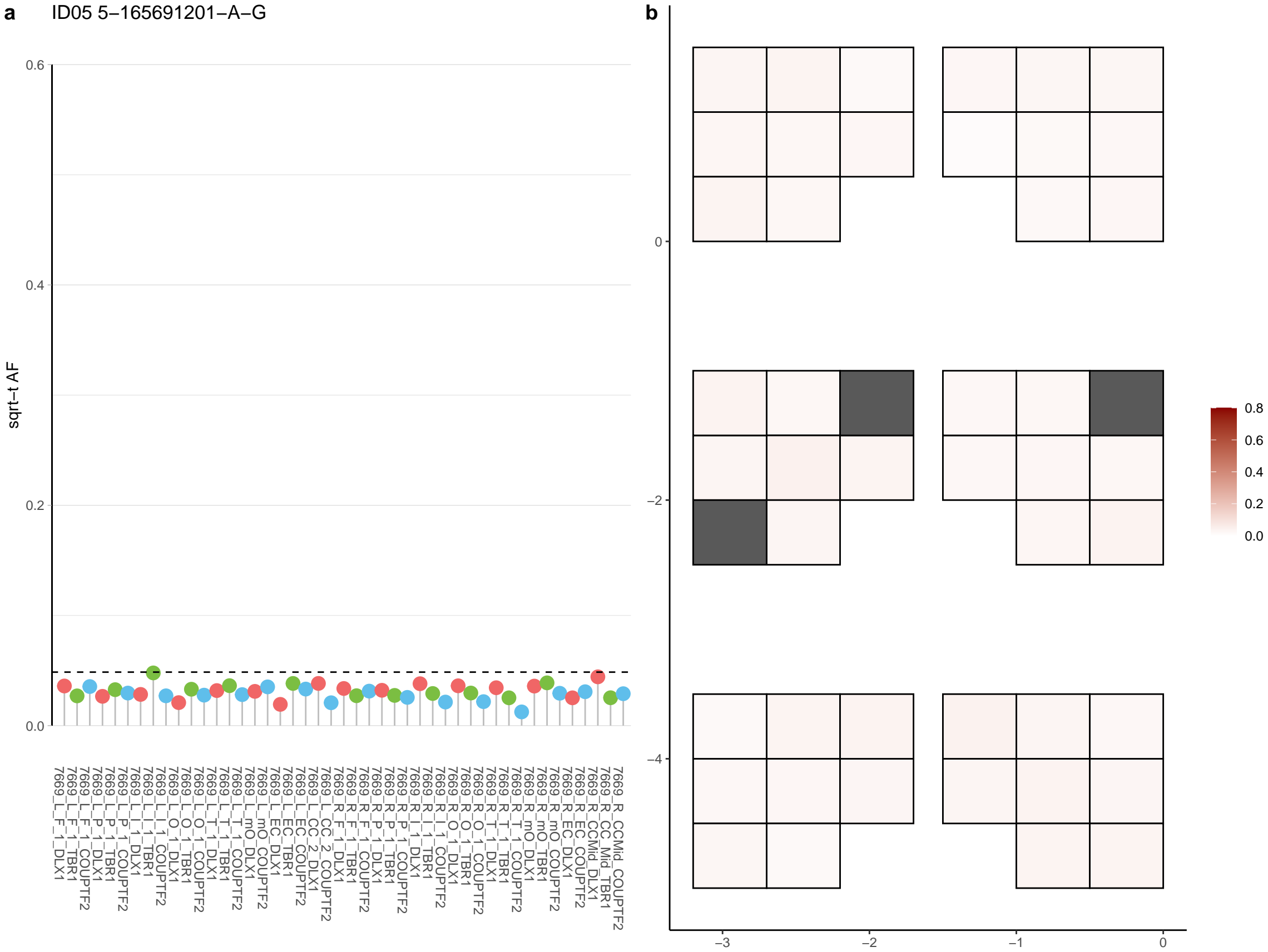

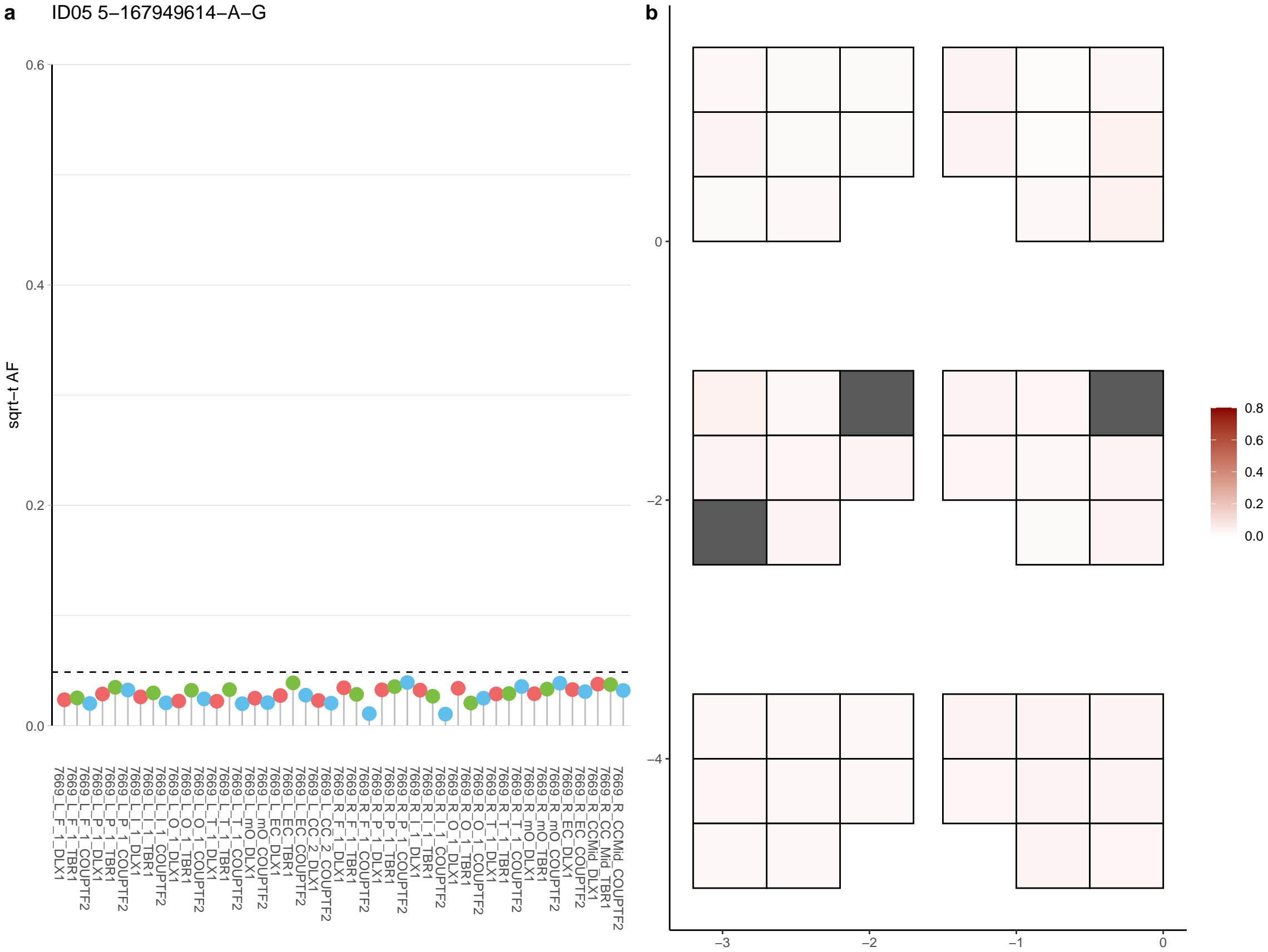

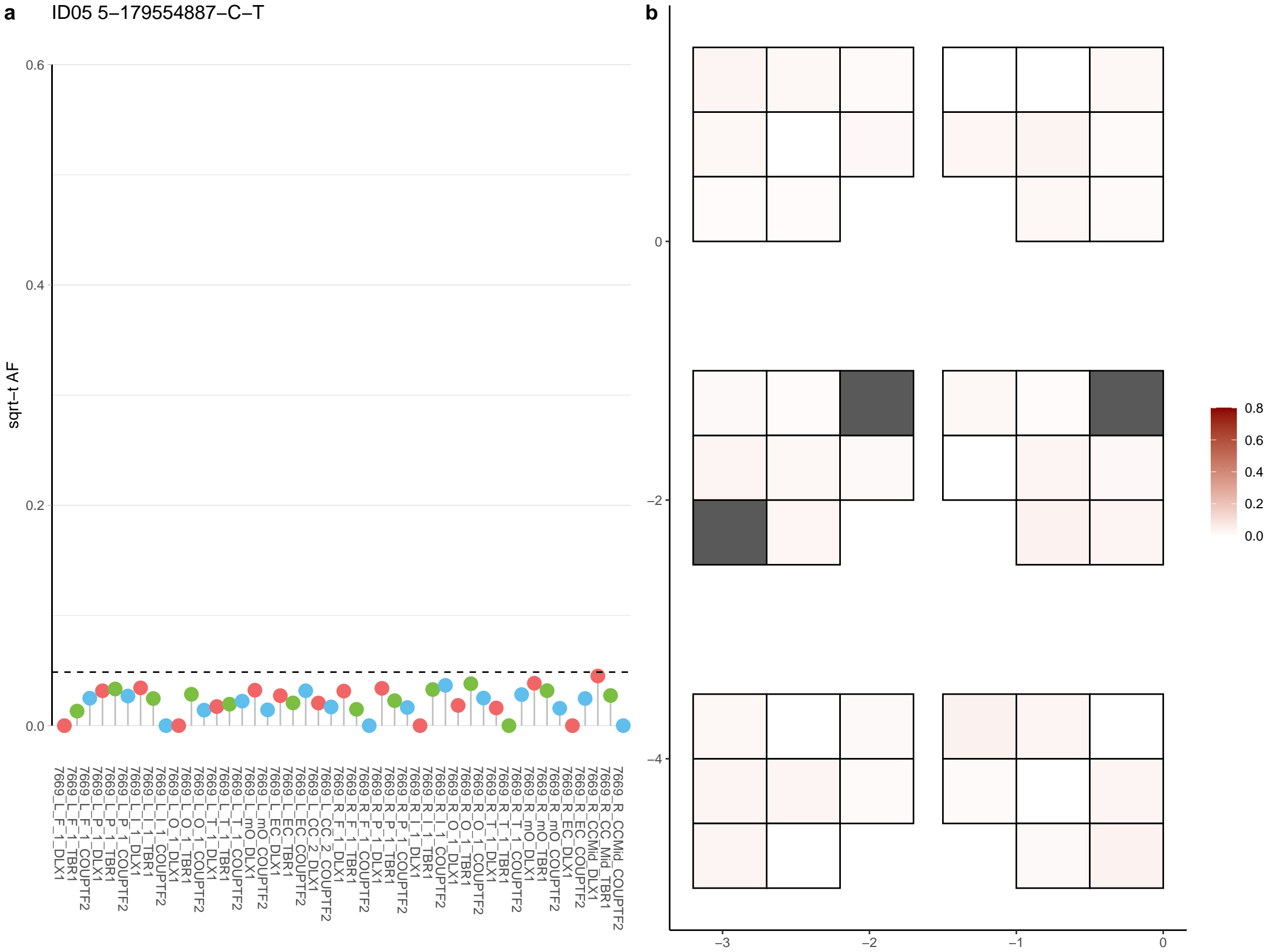

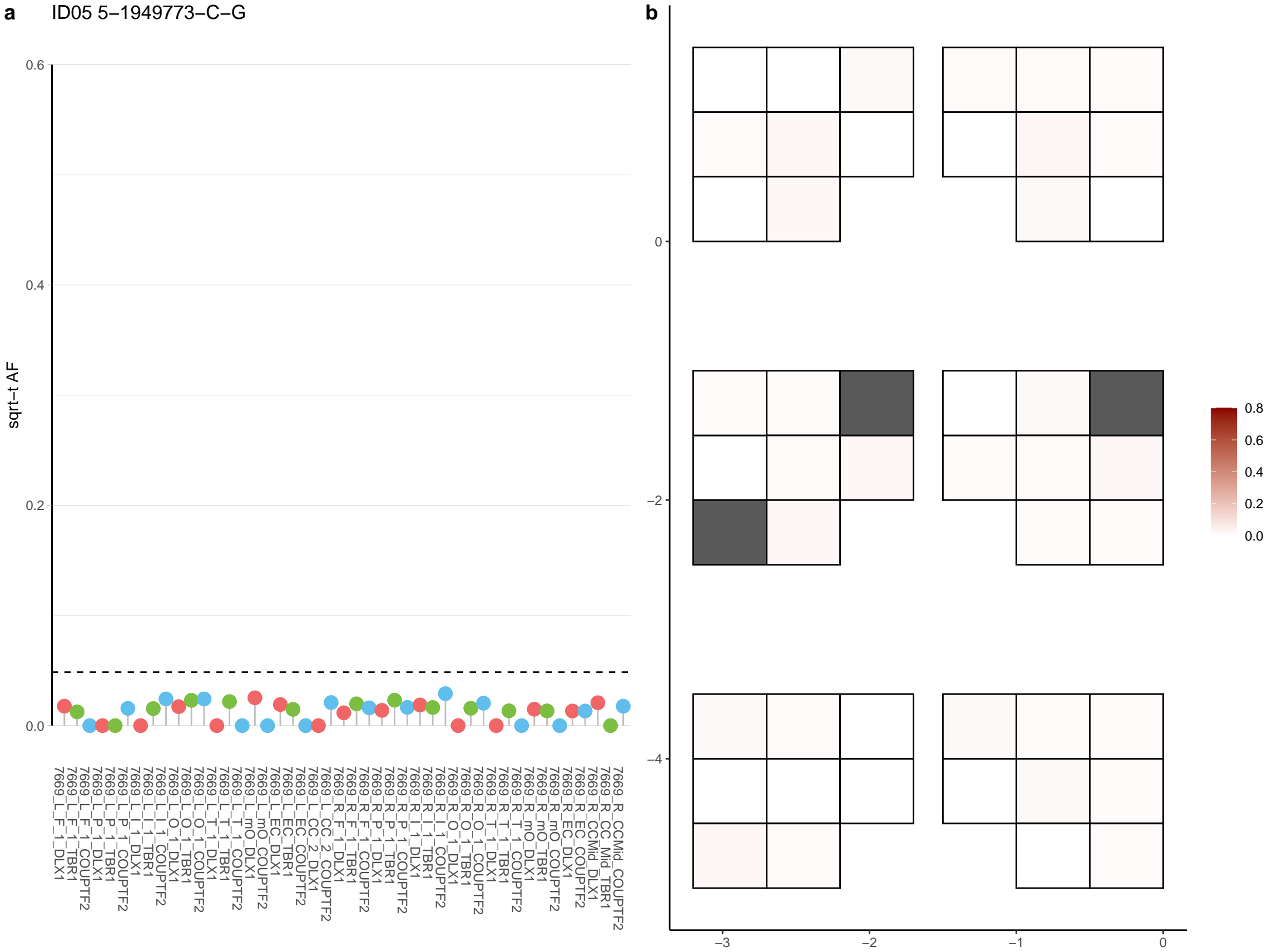

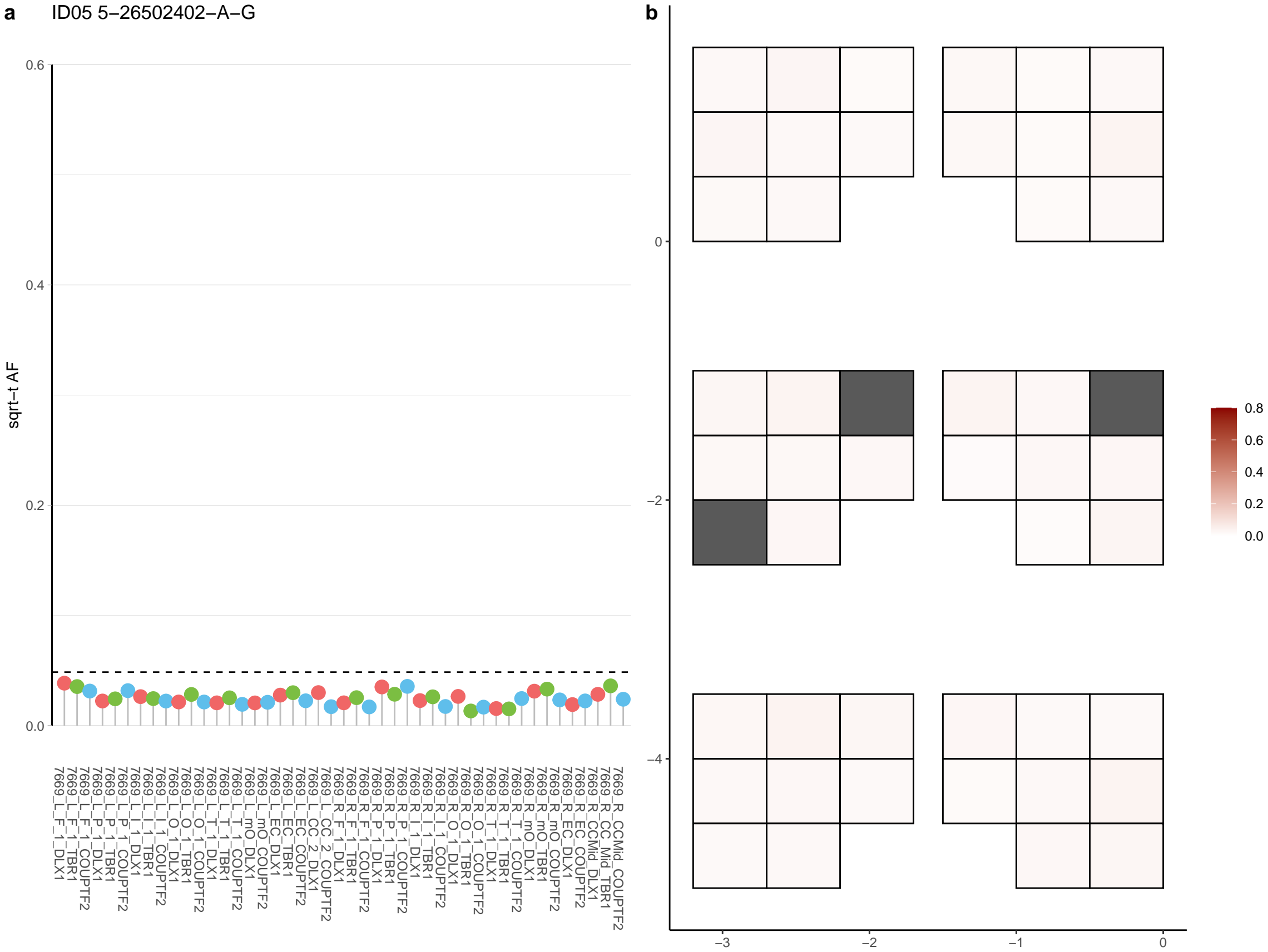

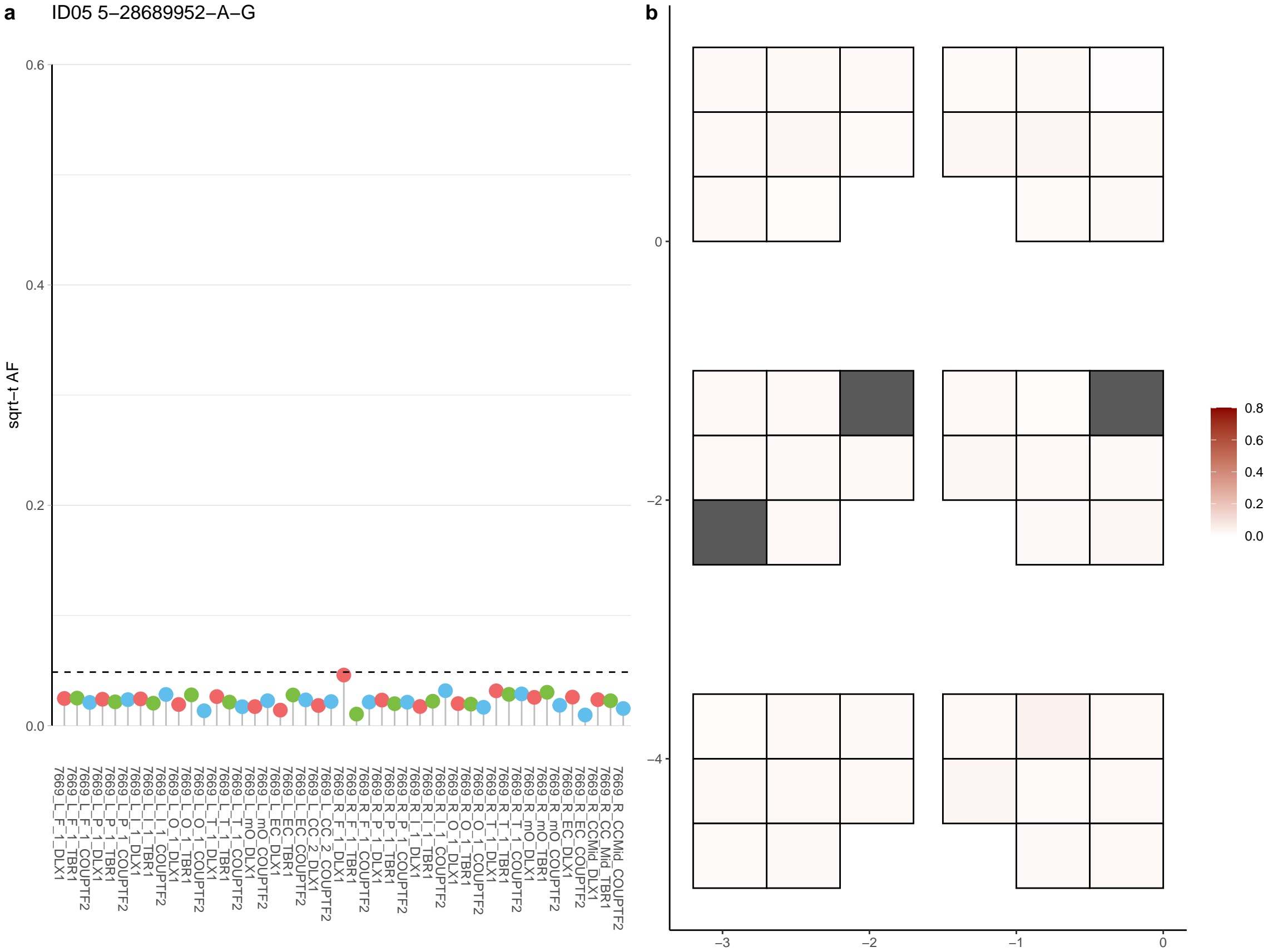

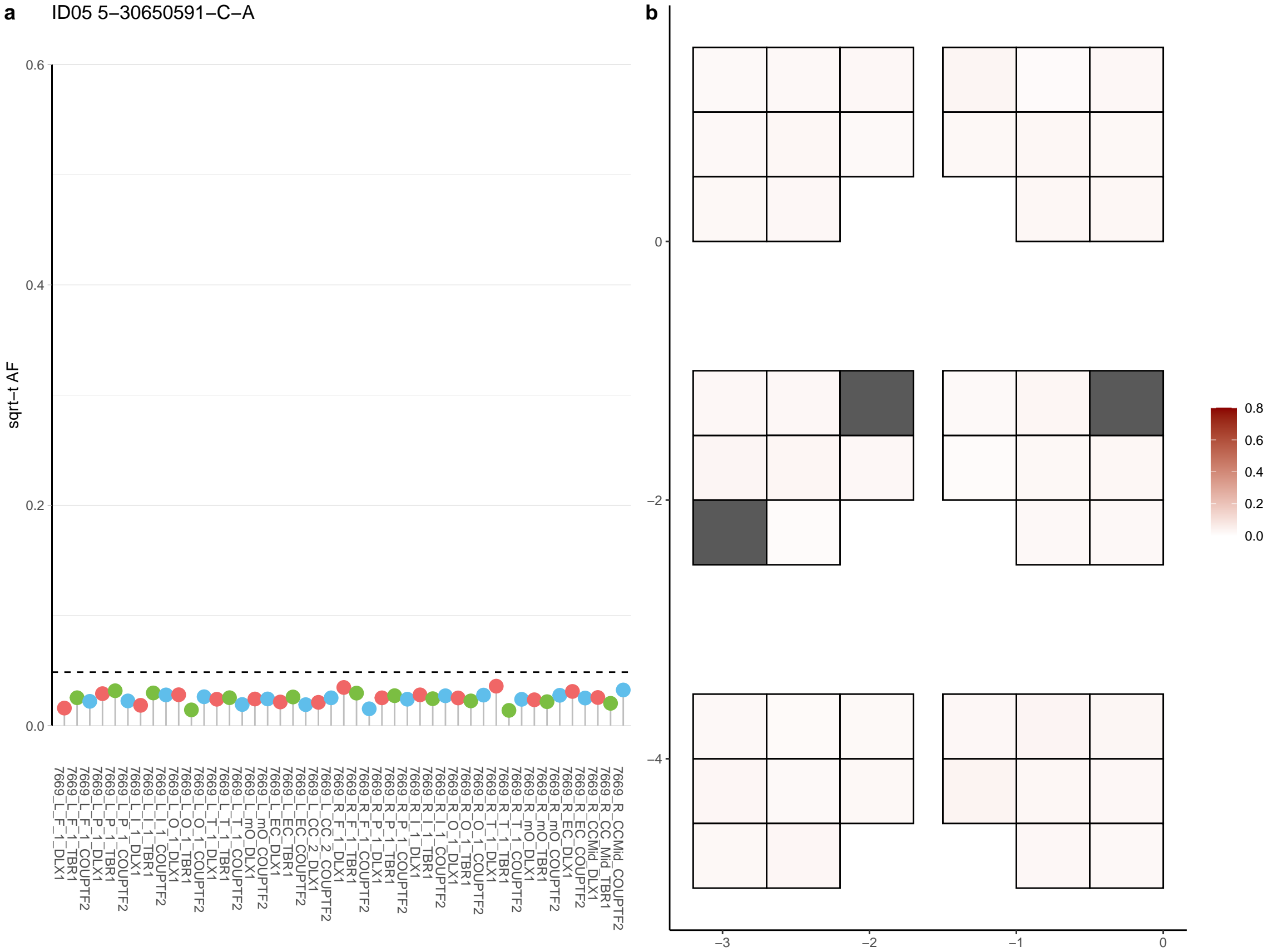

**a** ID05 5-30923233-C-T

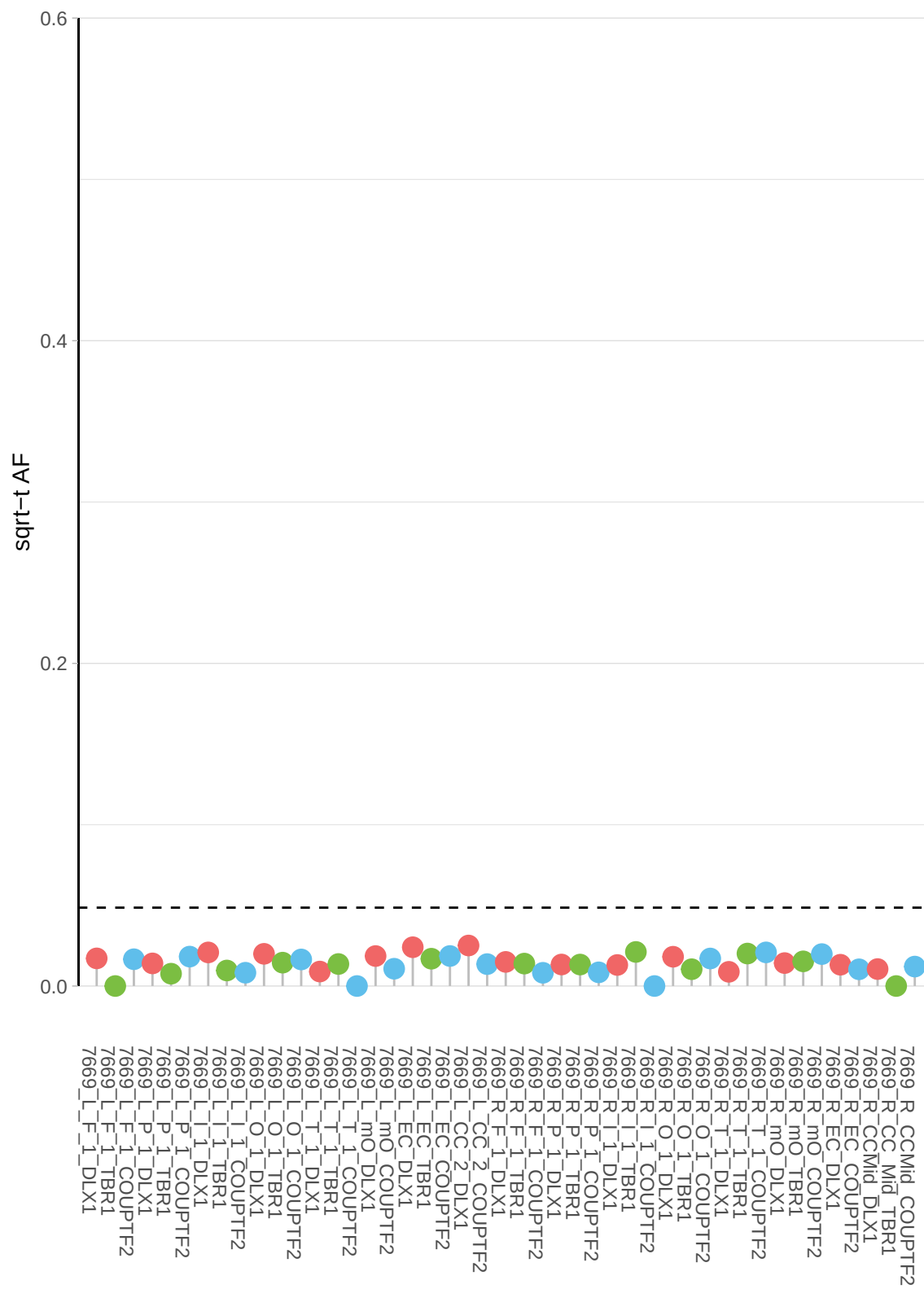

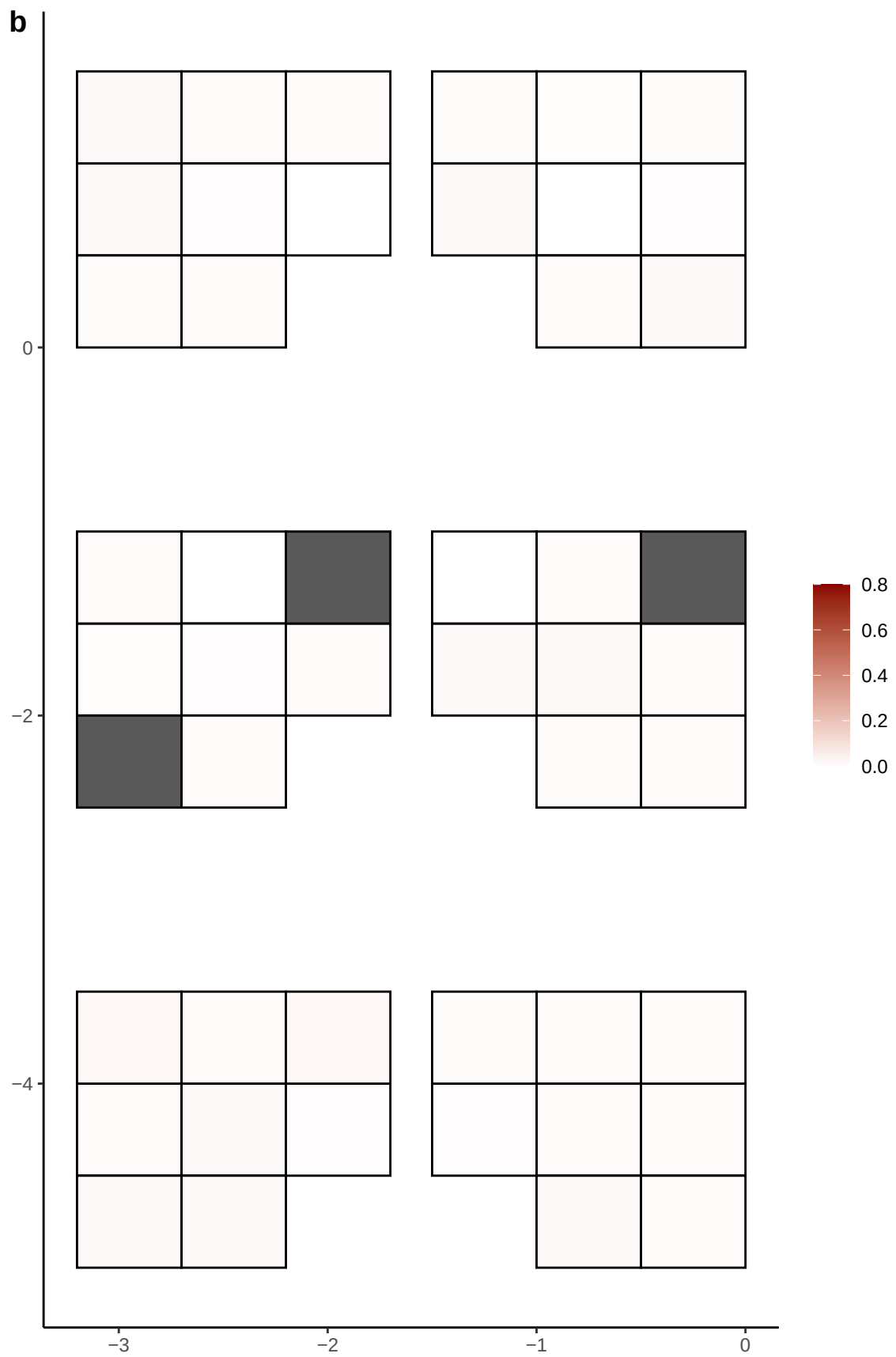

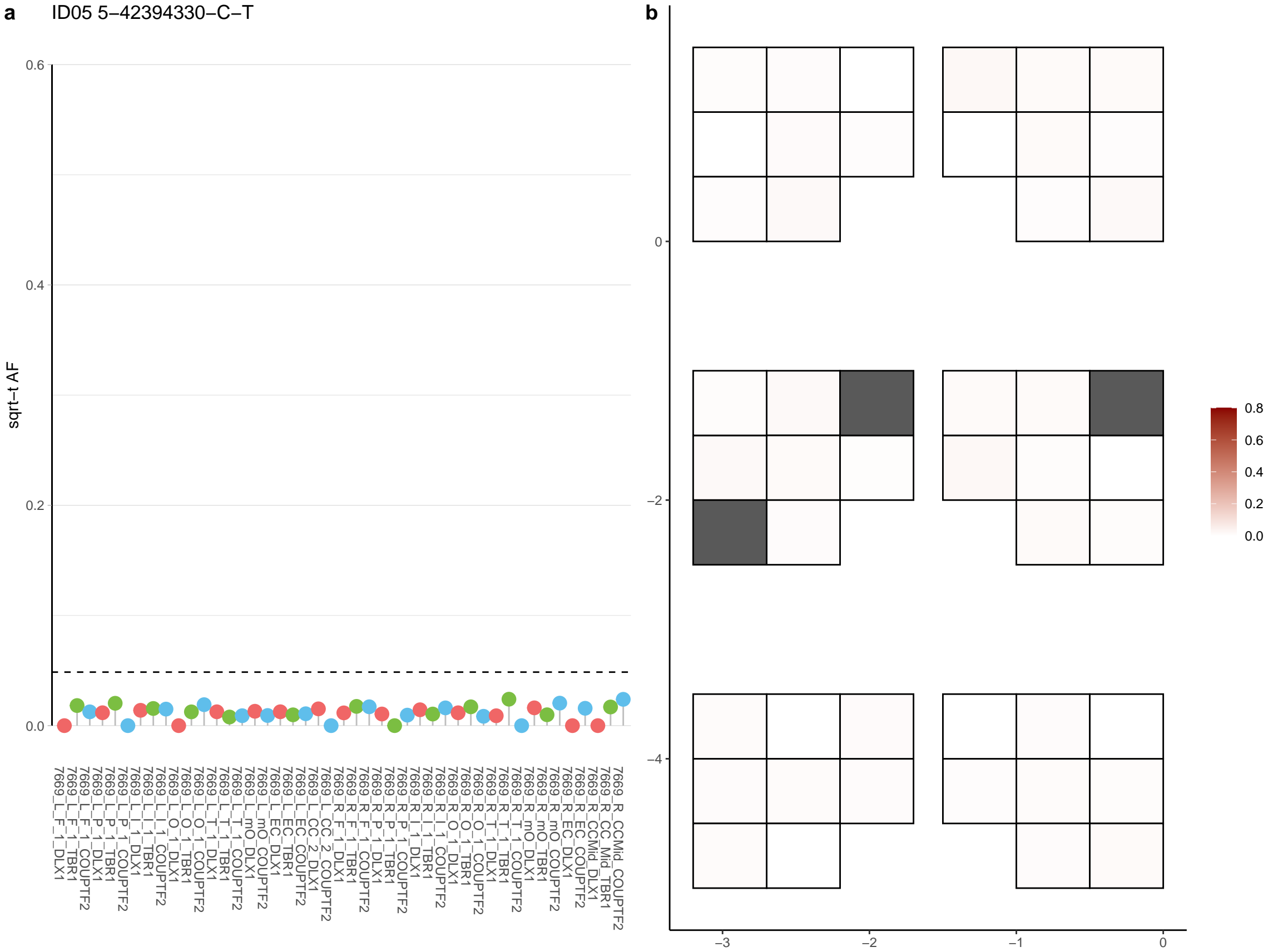

**a** ID05 5-42474677-G-T

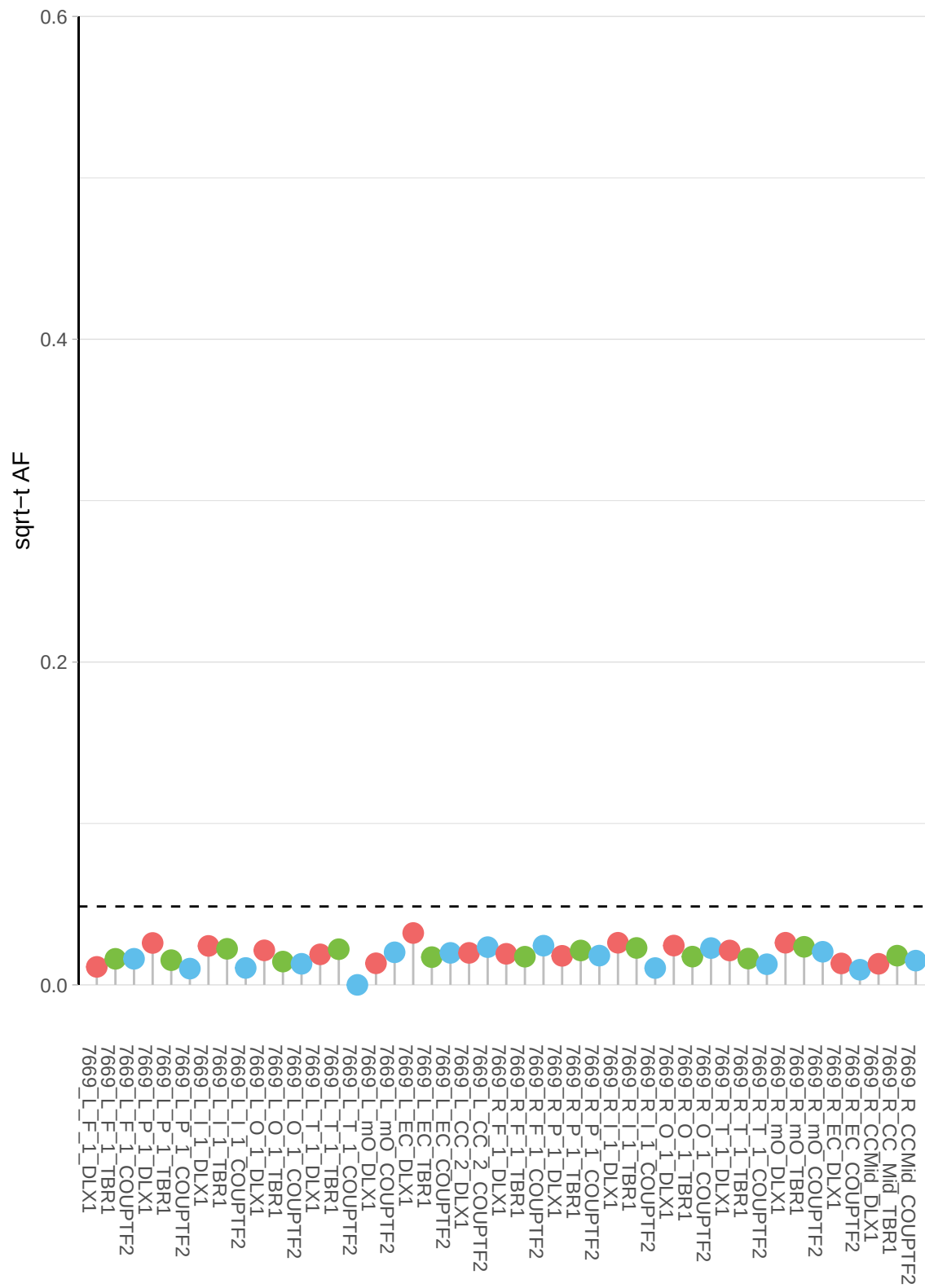

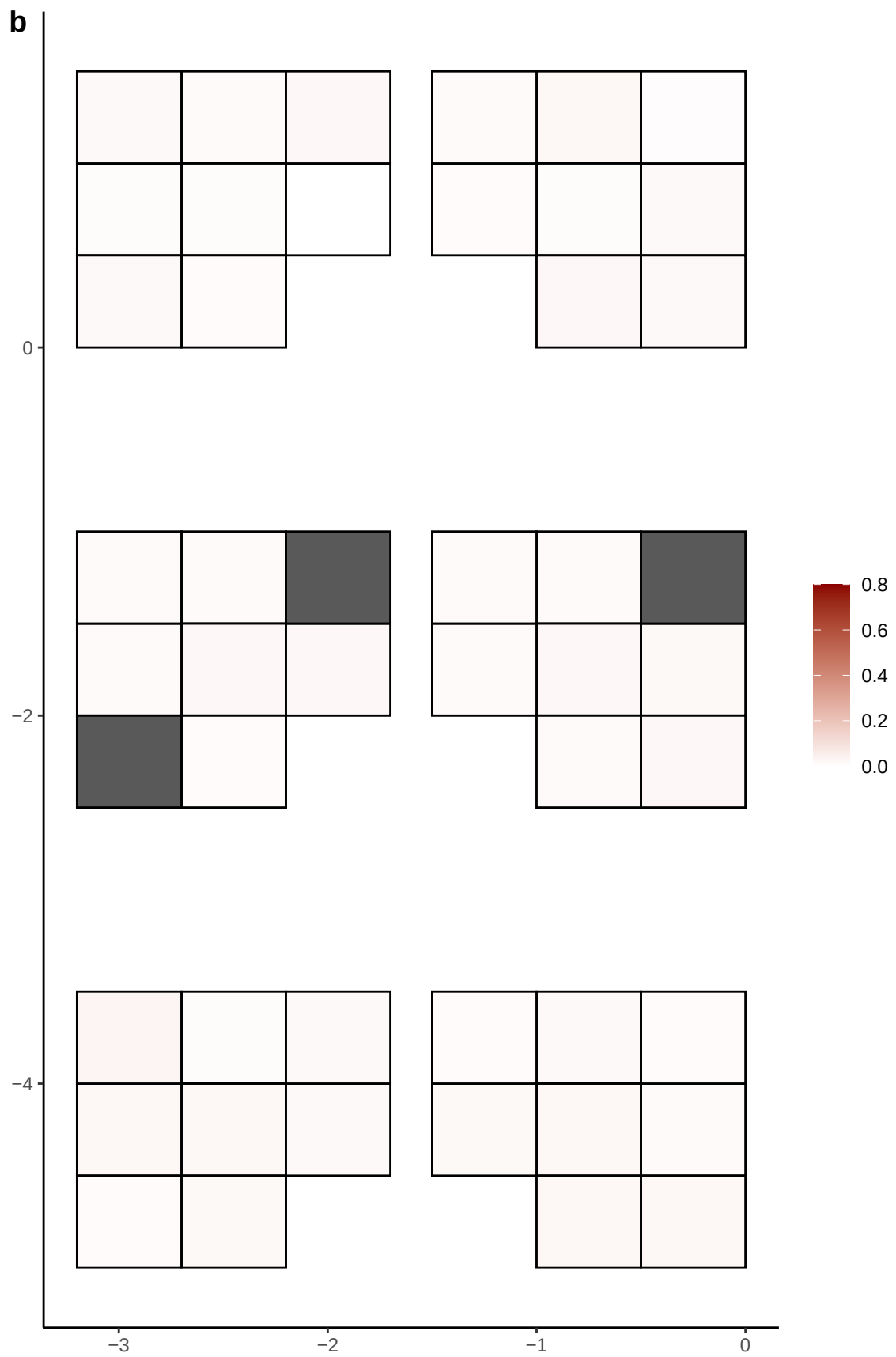

**a** ID05 5-50102837-T-C

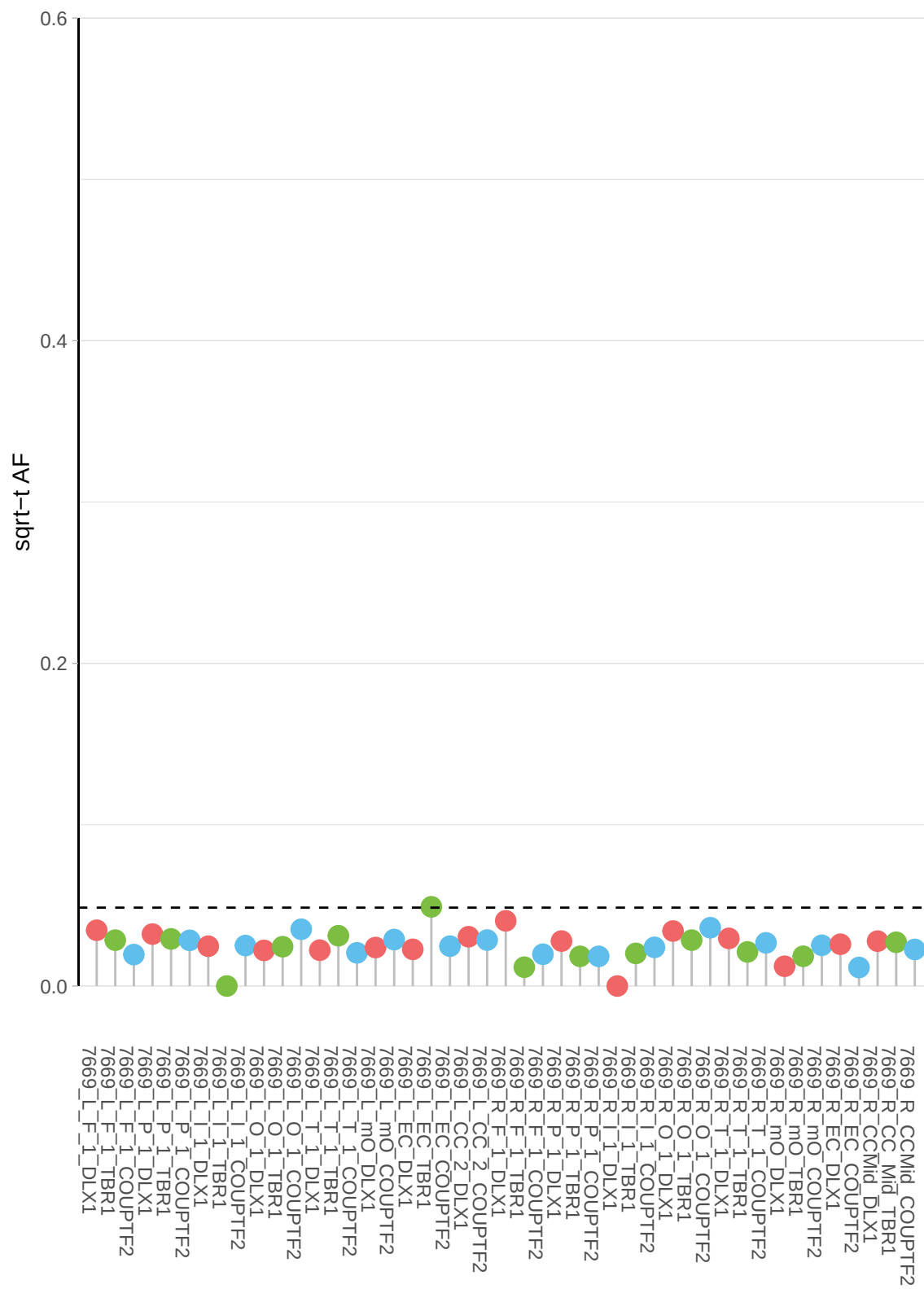

**b**

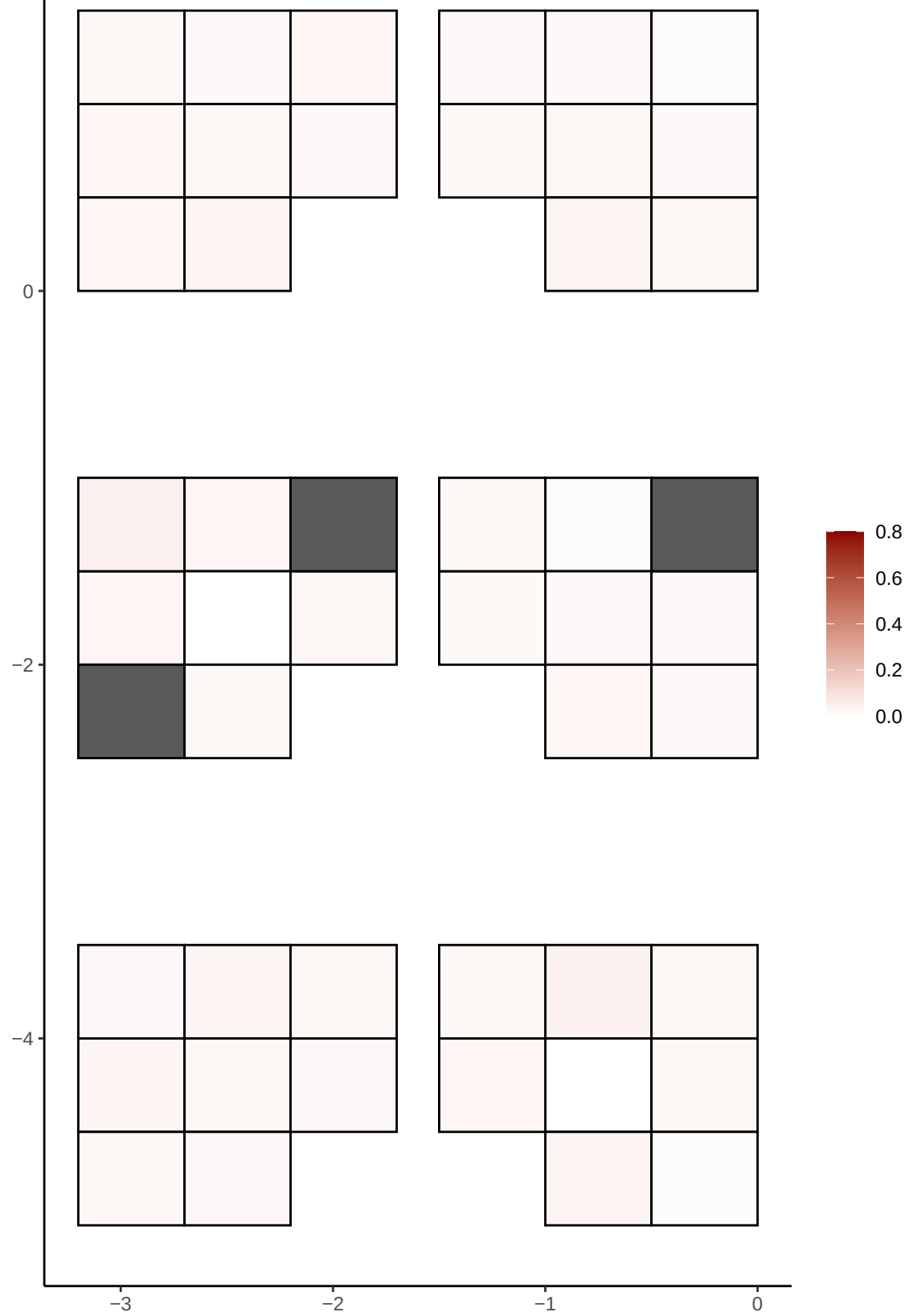

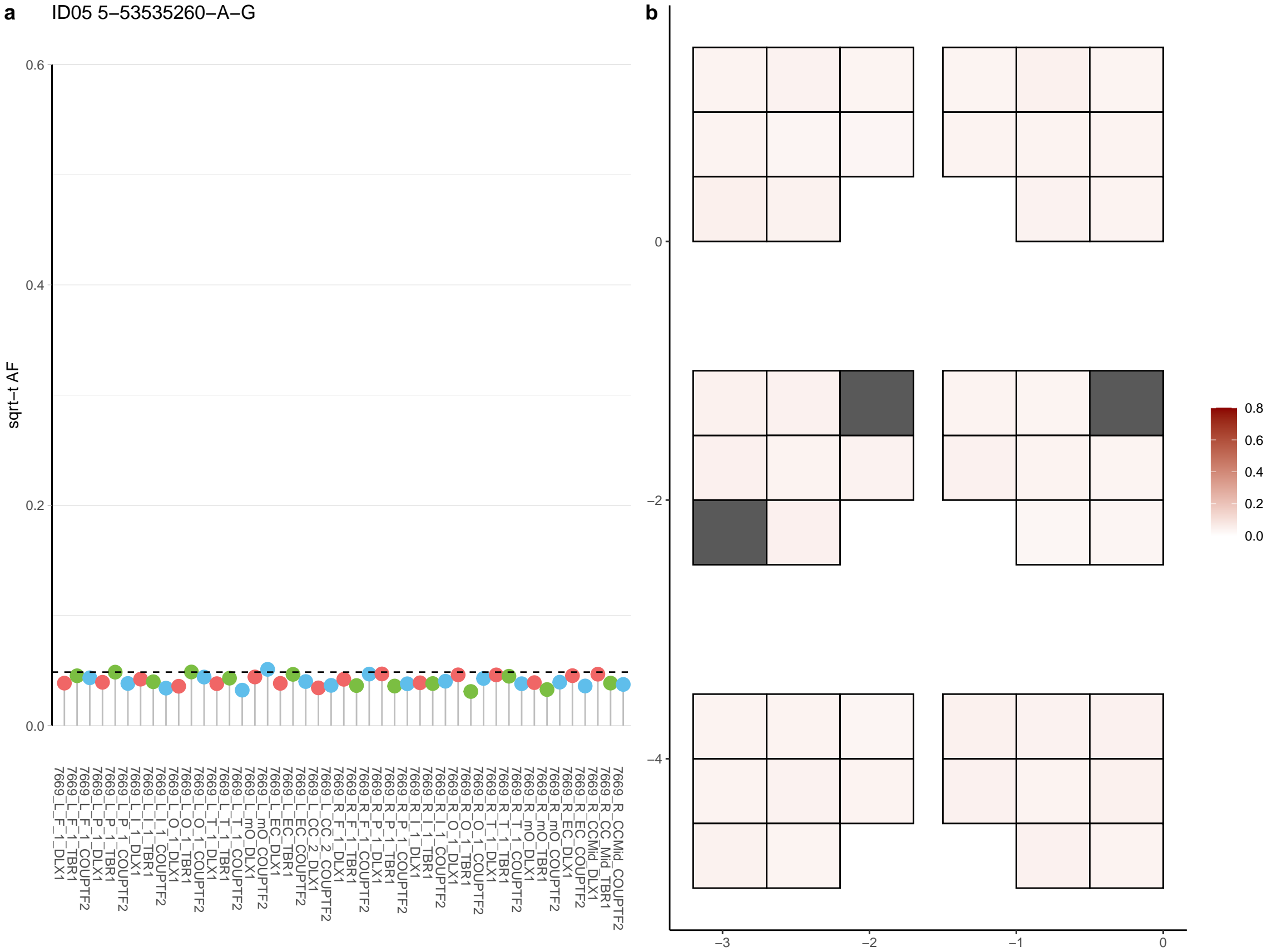

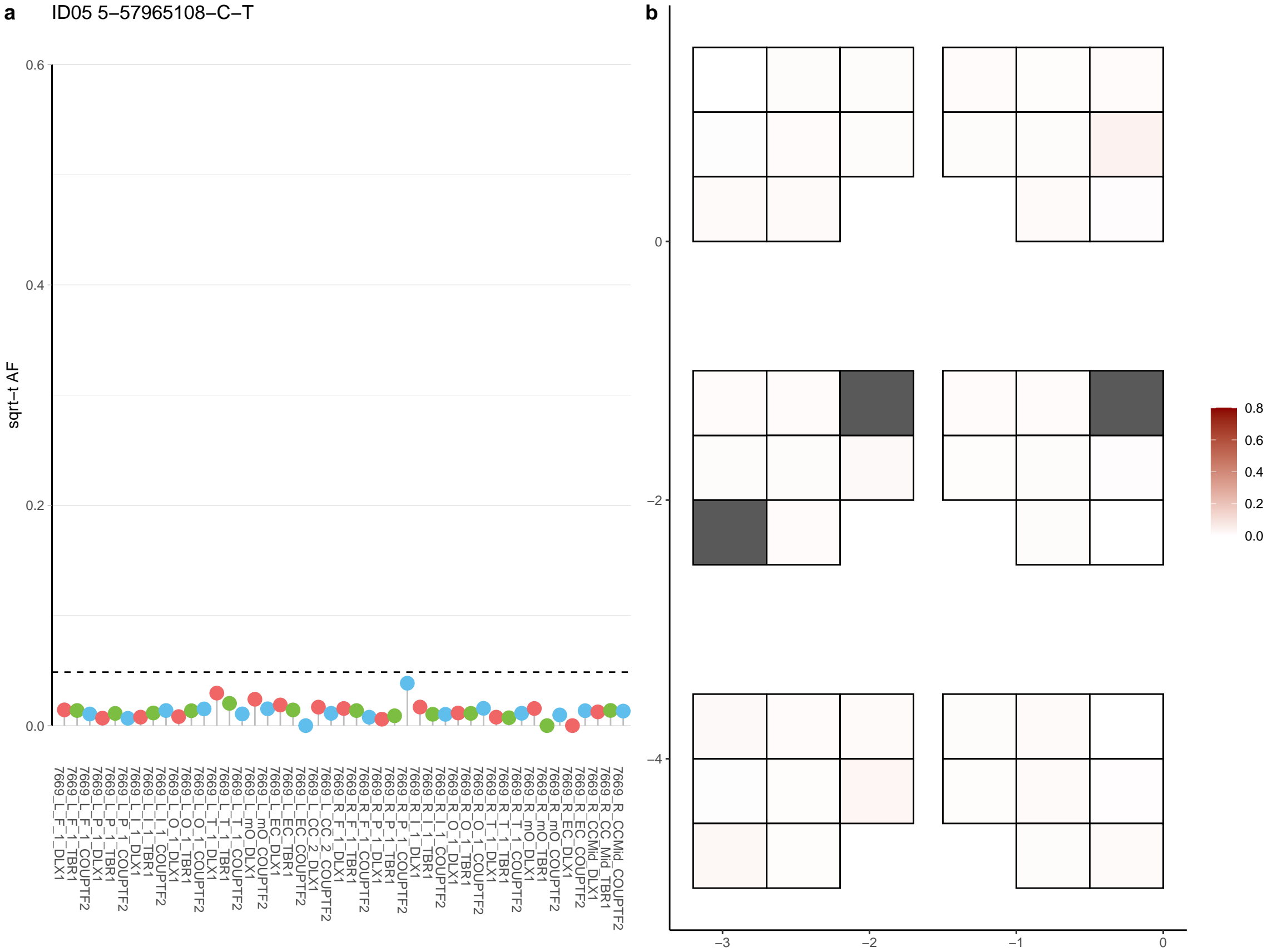

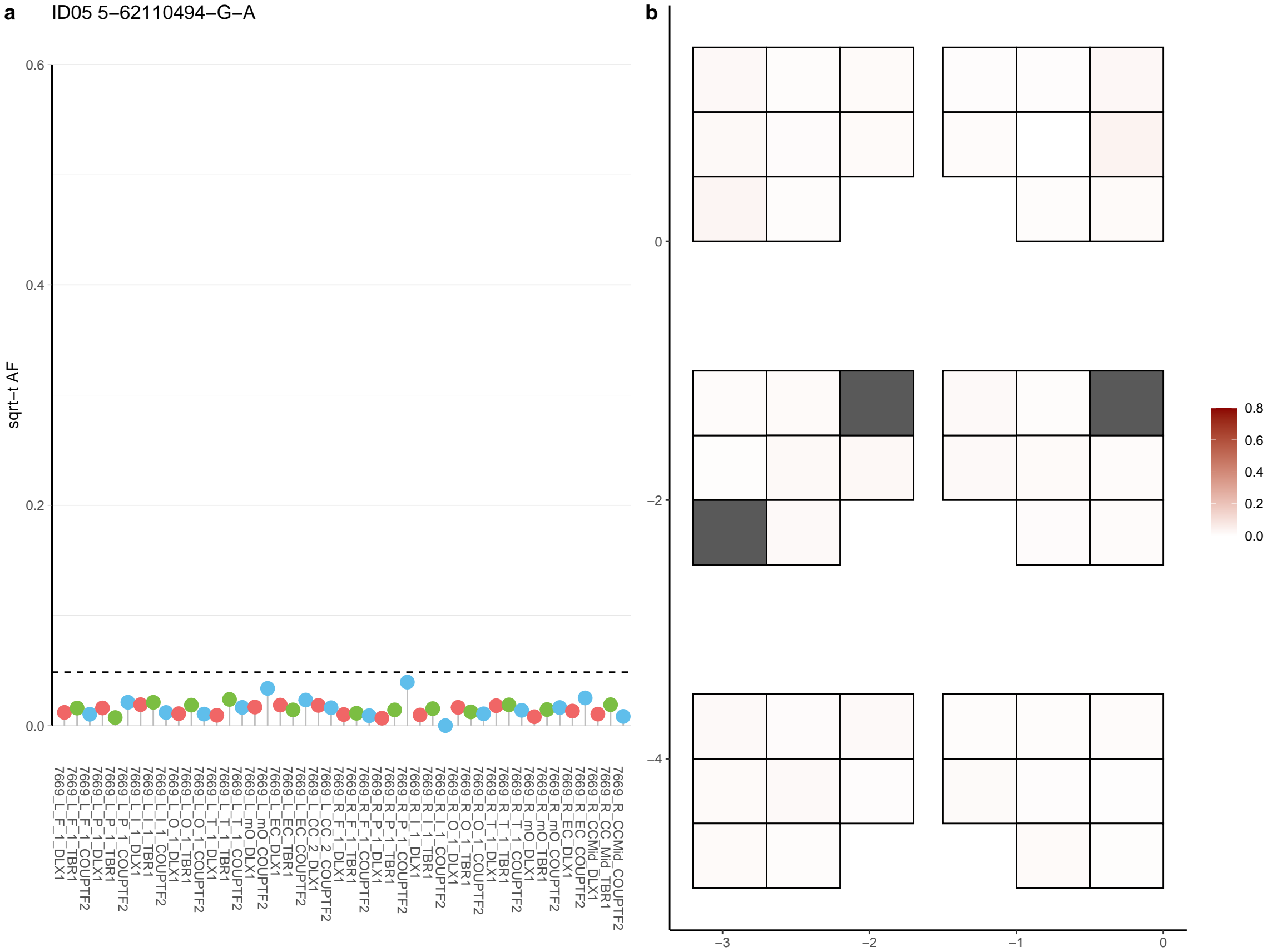

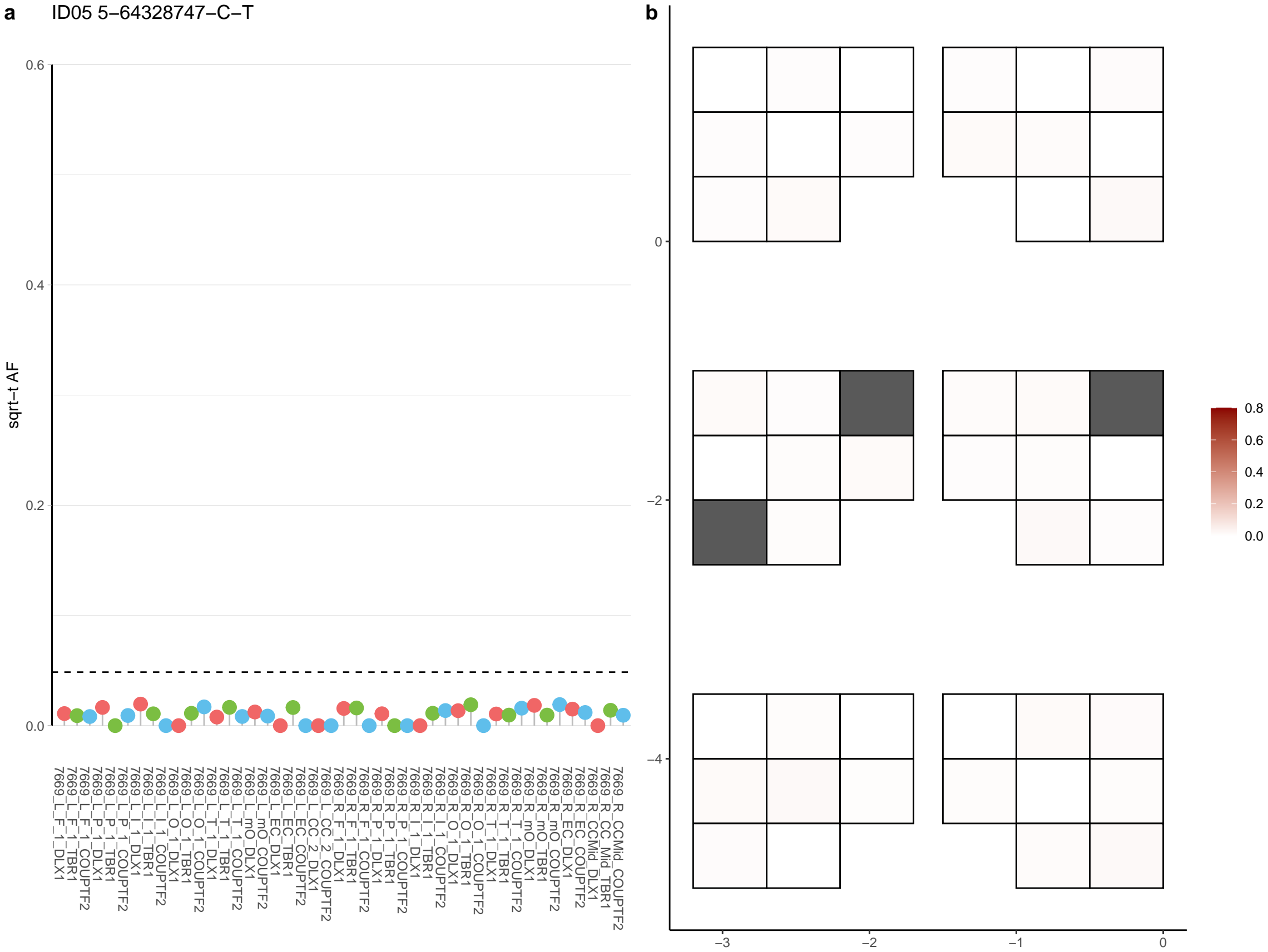

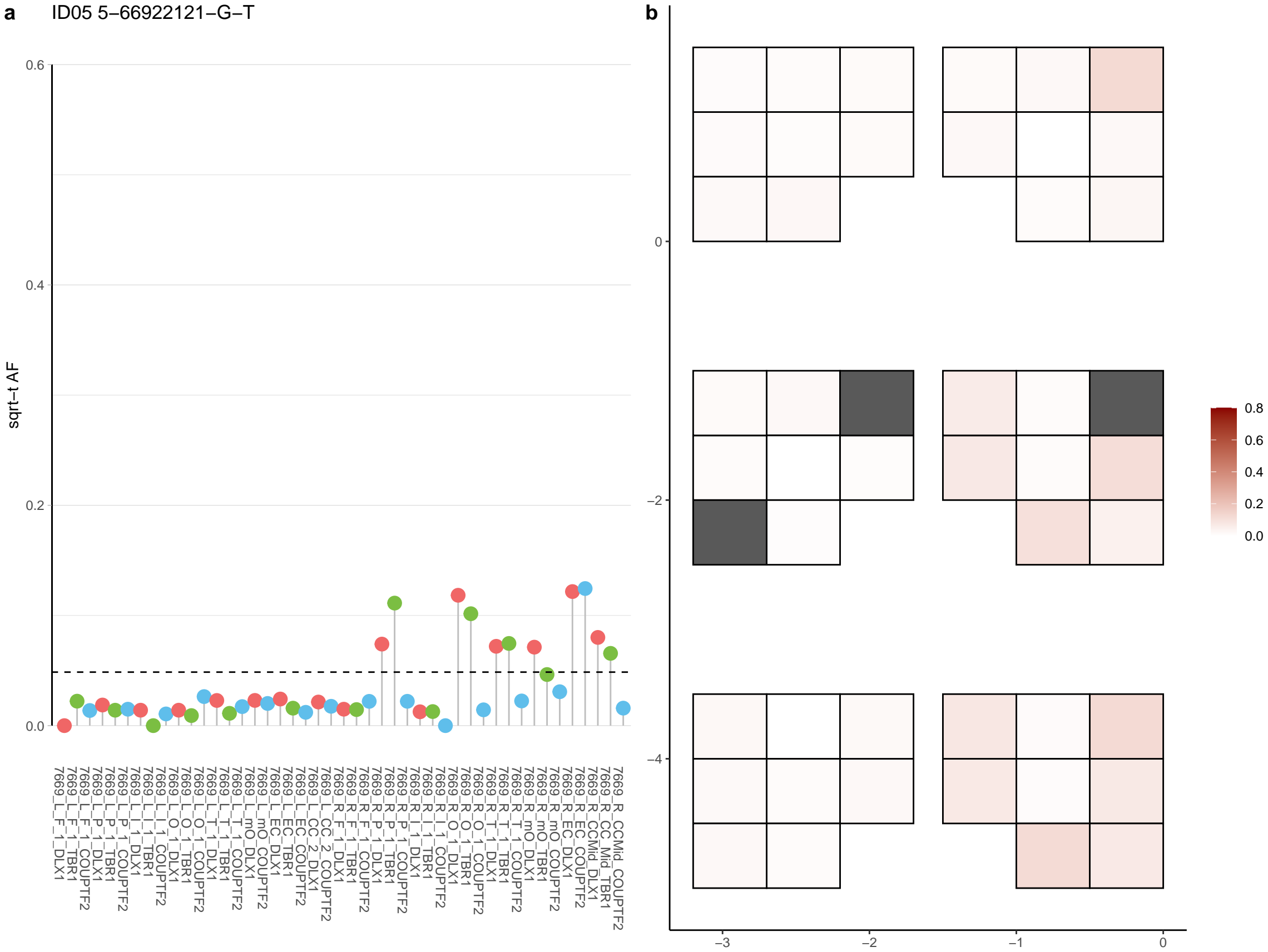

**a** ID05 5-67667744-C-T

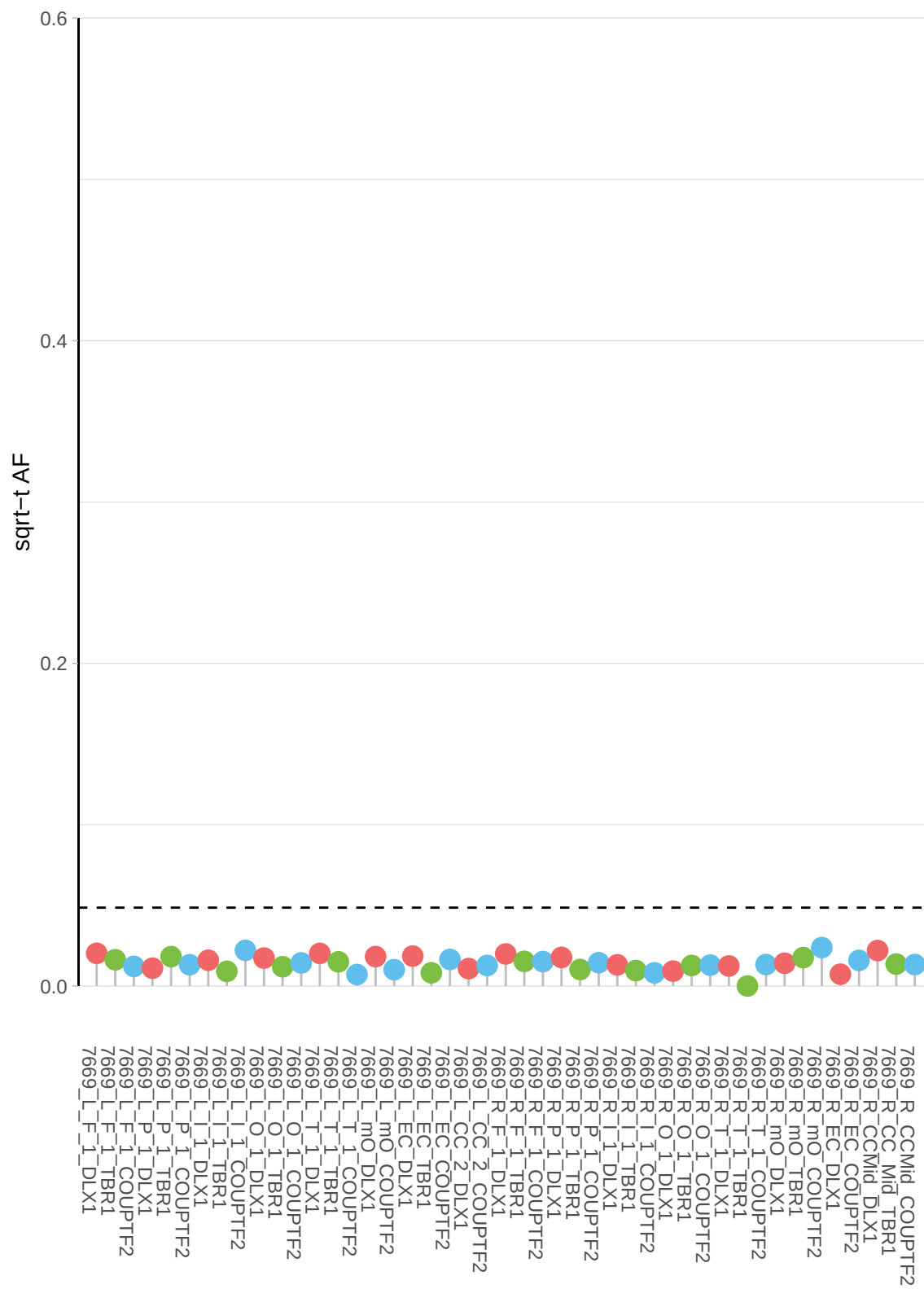

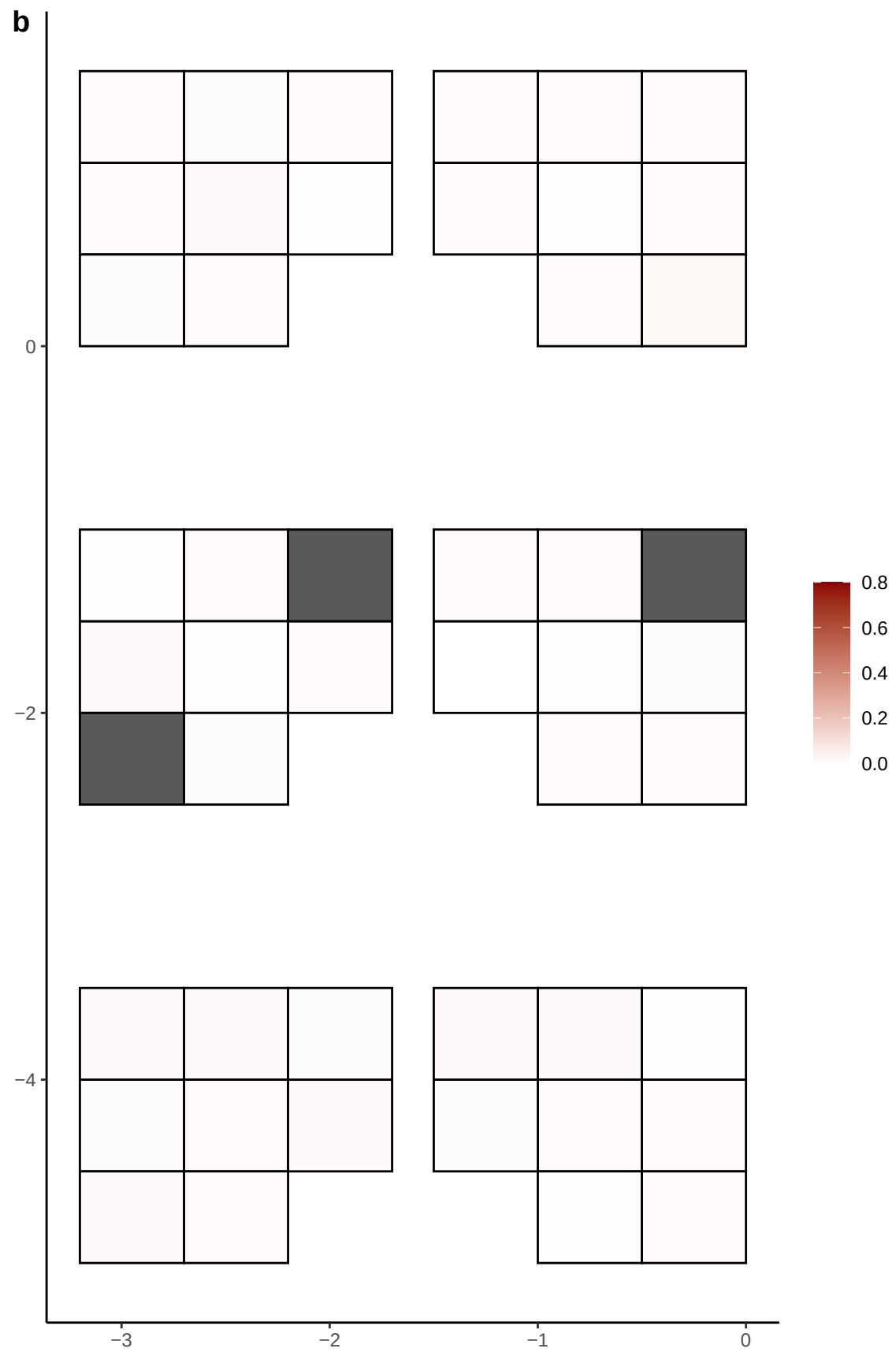

**a** ID05 5-7018116-G-A

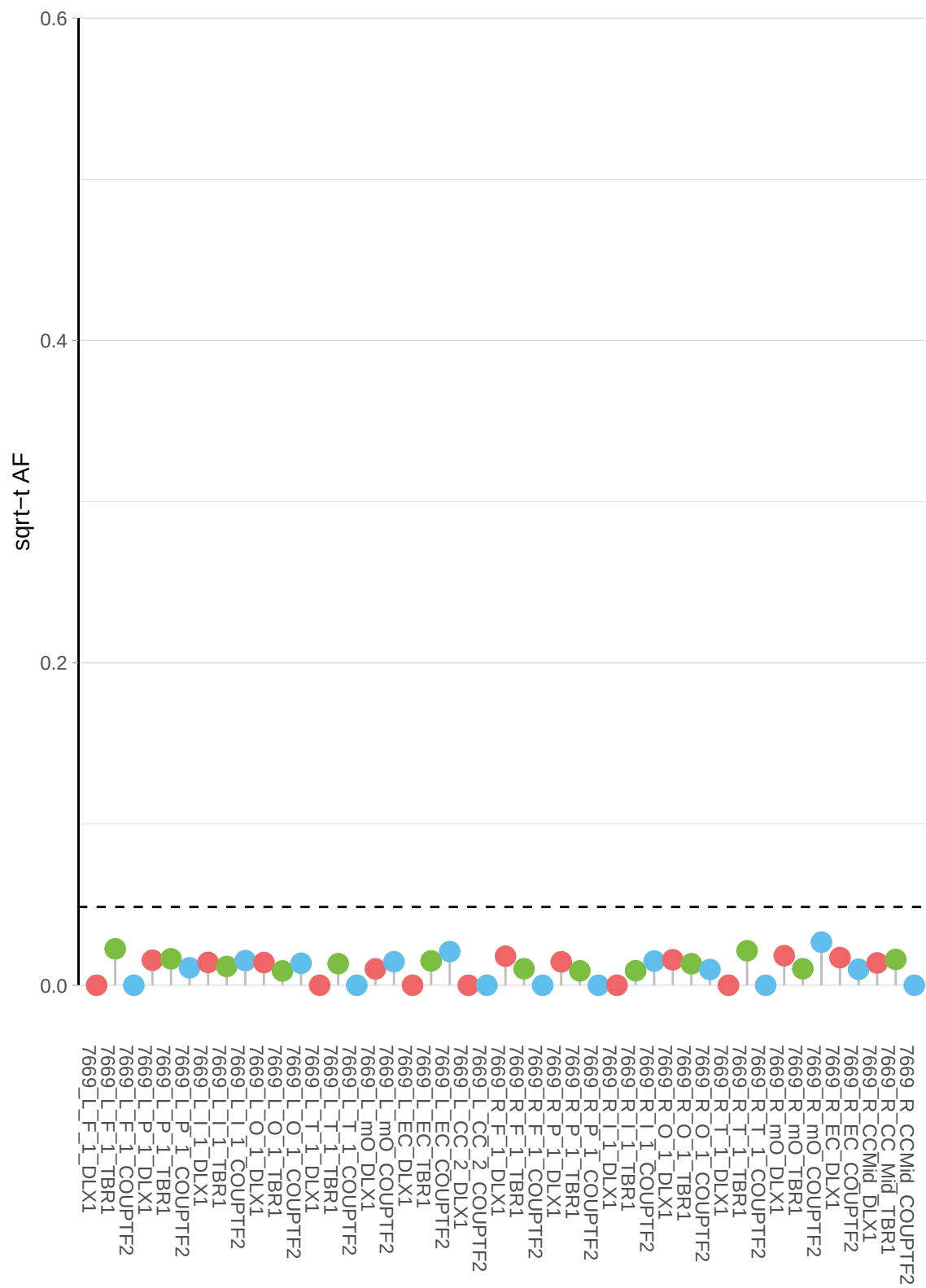

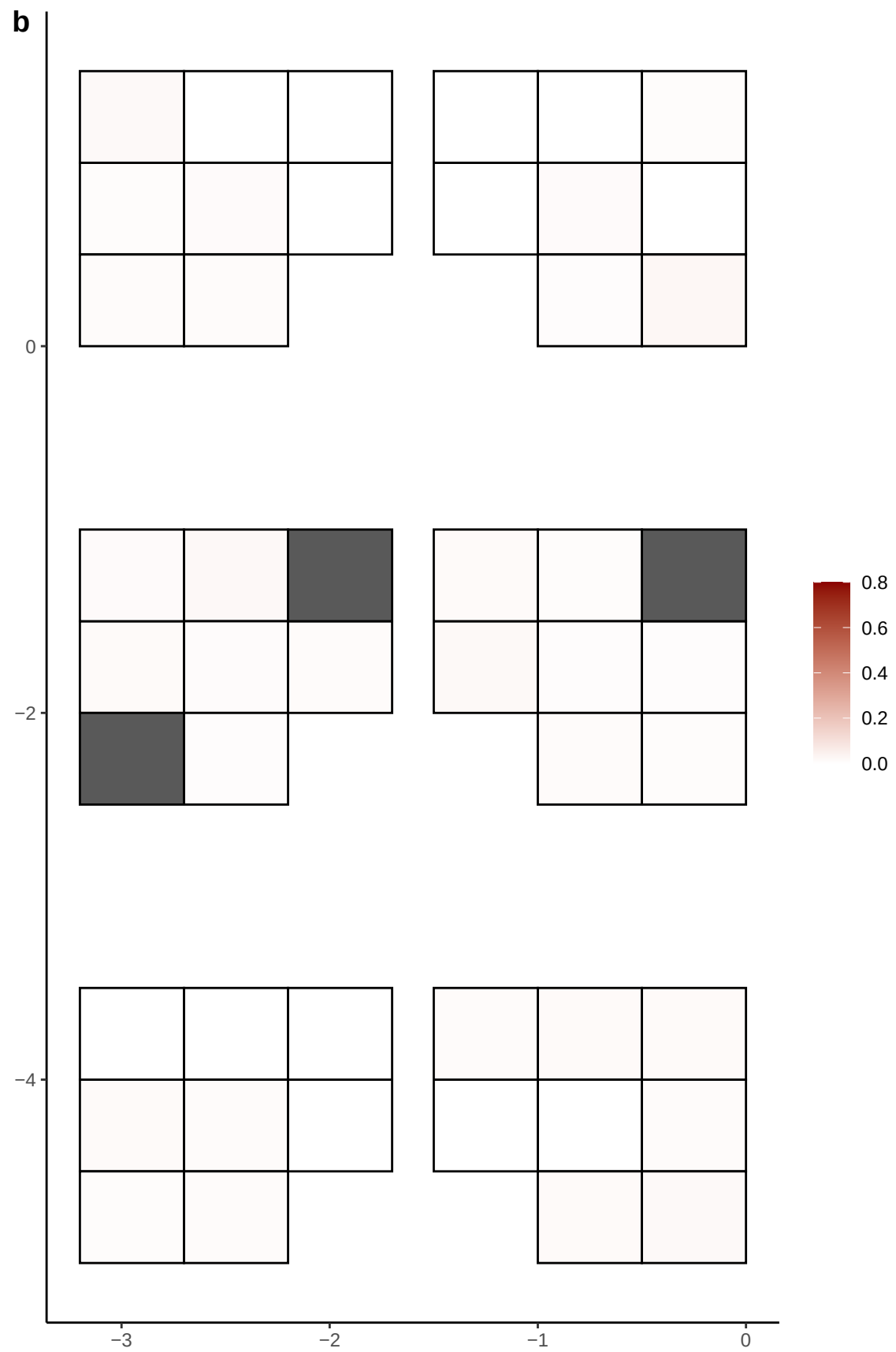

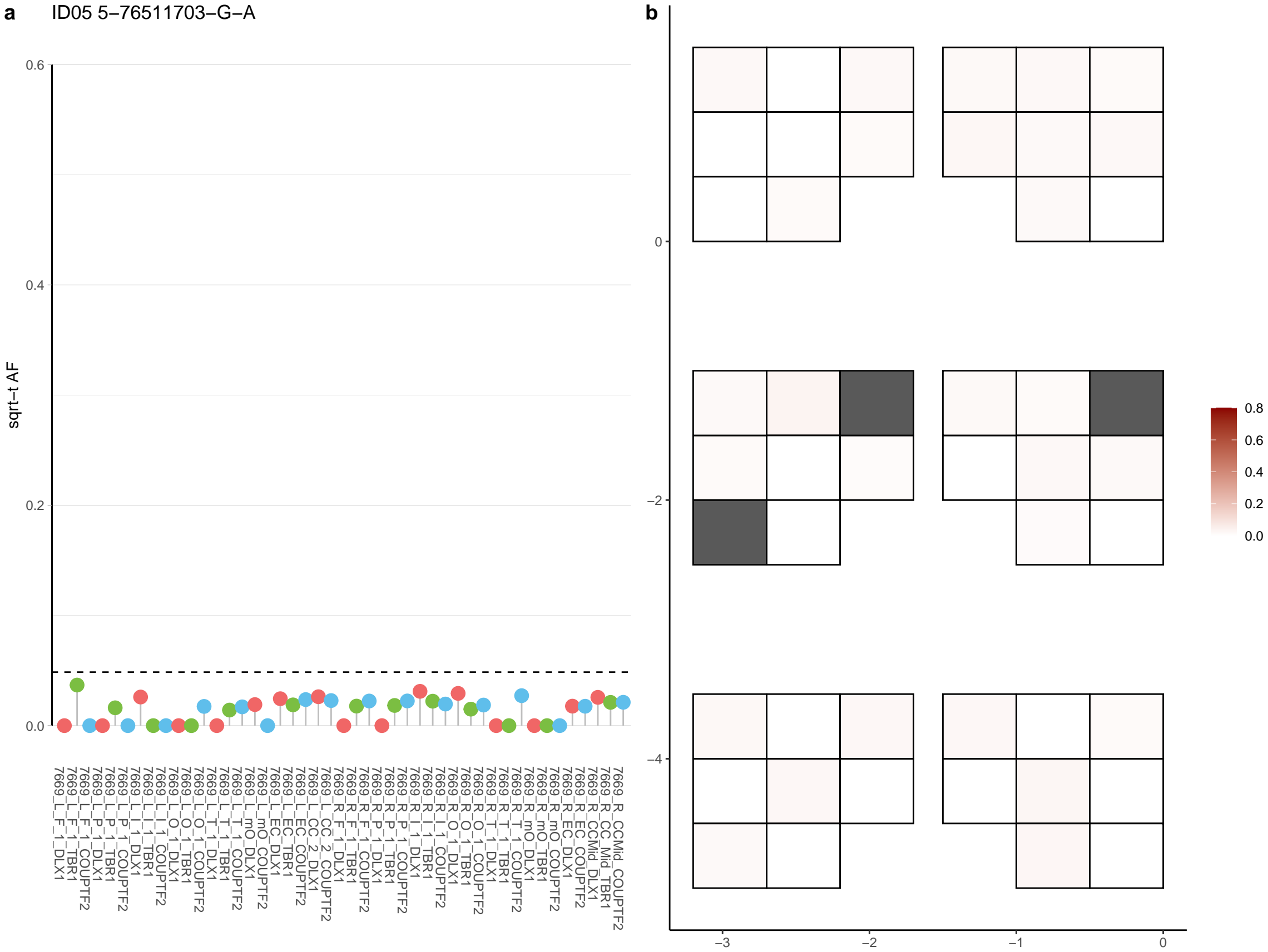

**a** ID05 5-7667409-C-G

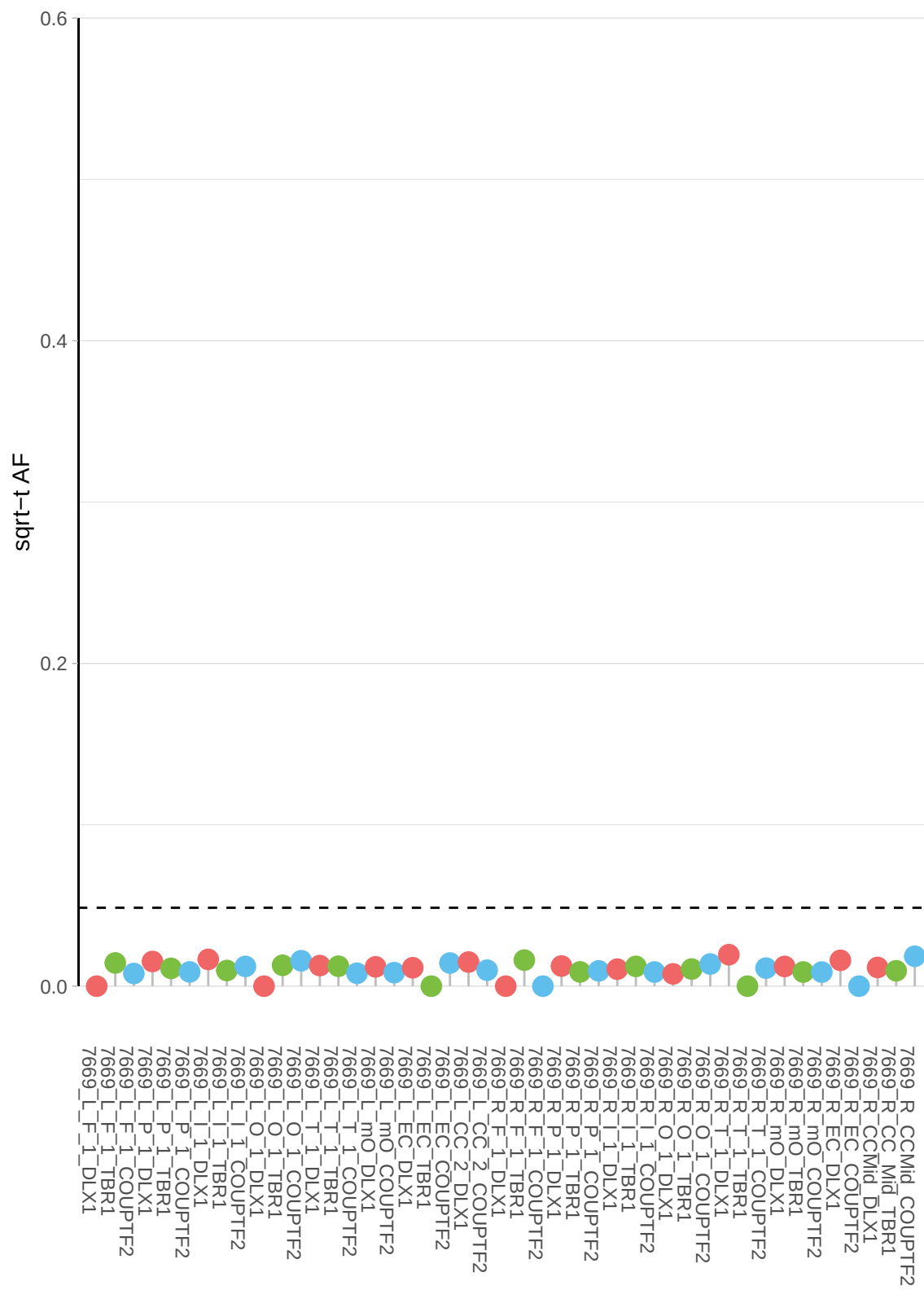

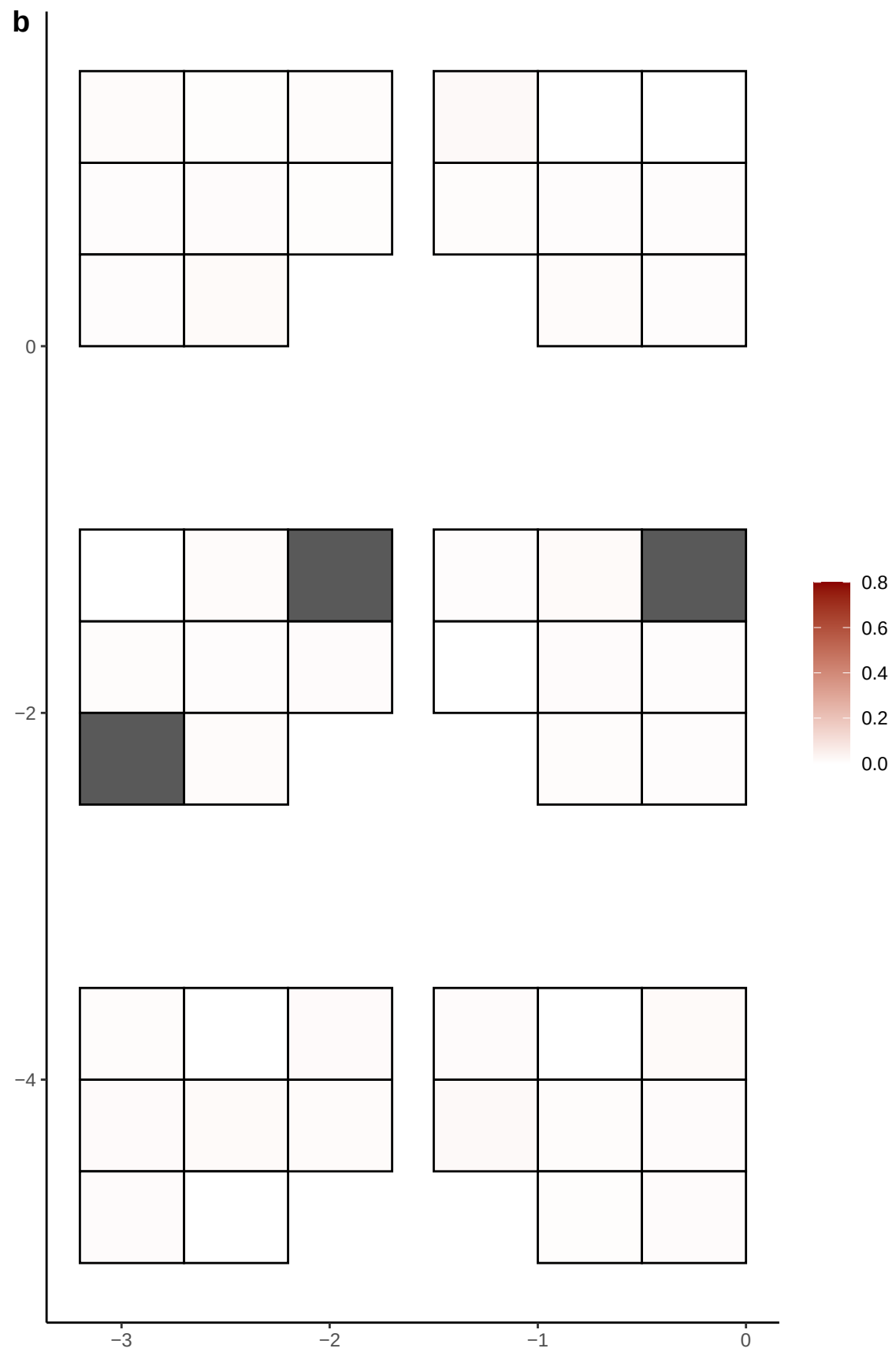

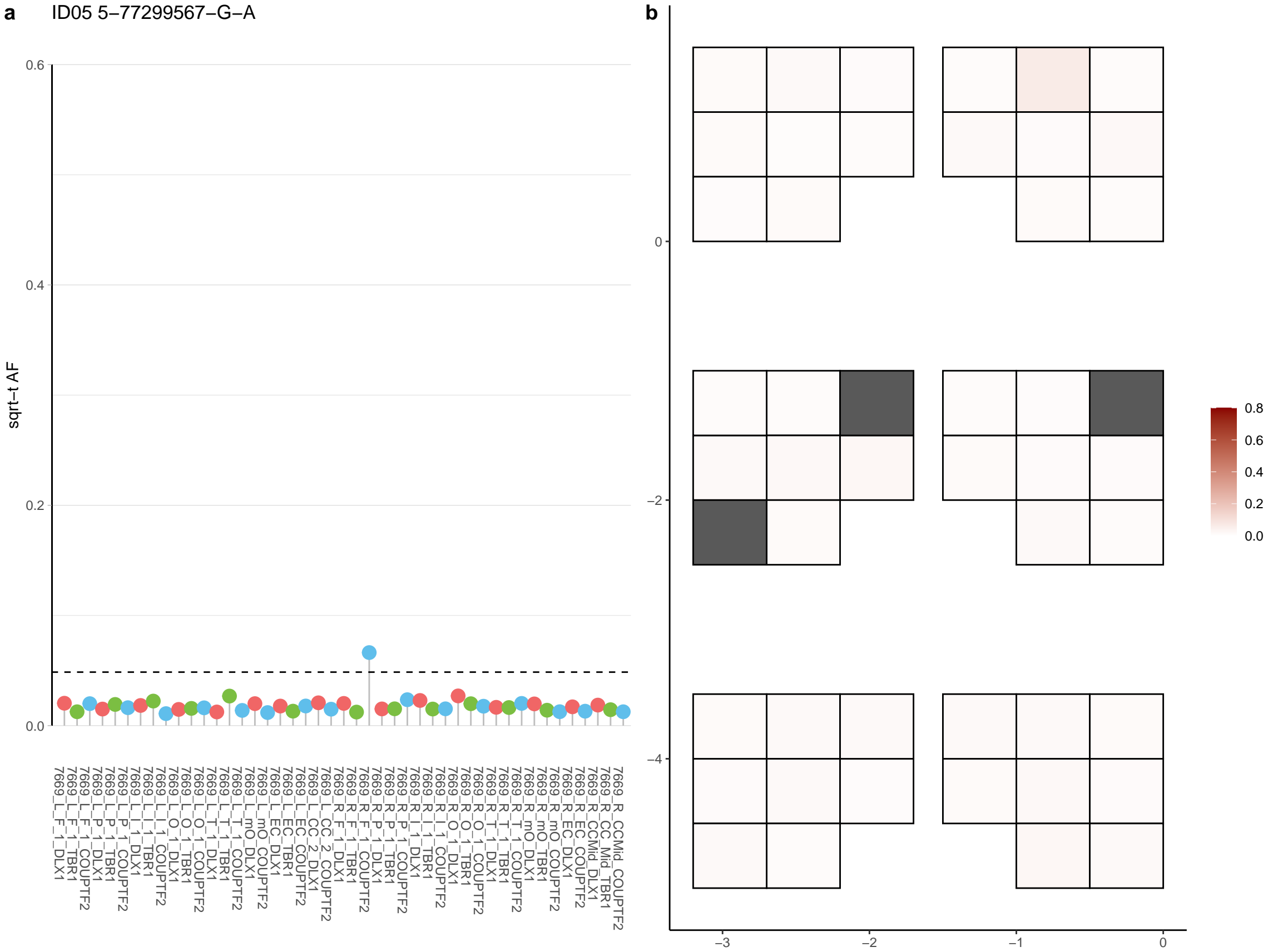

**a** ID05 5-7771752-C-T

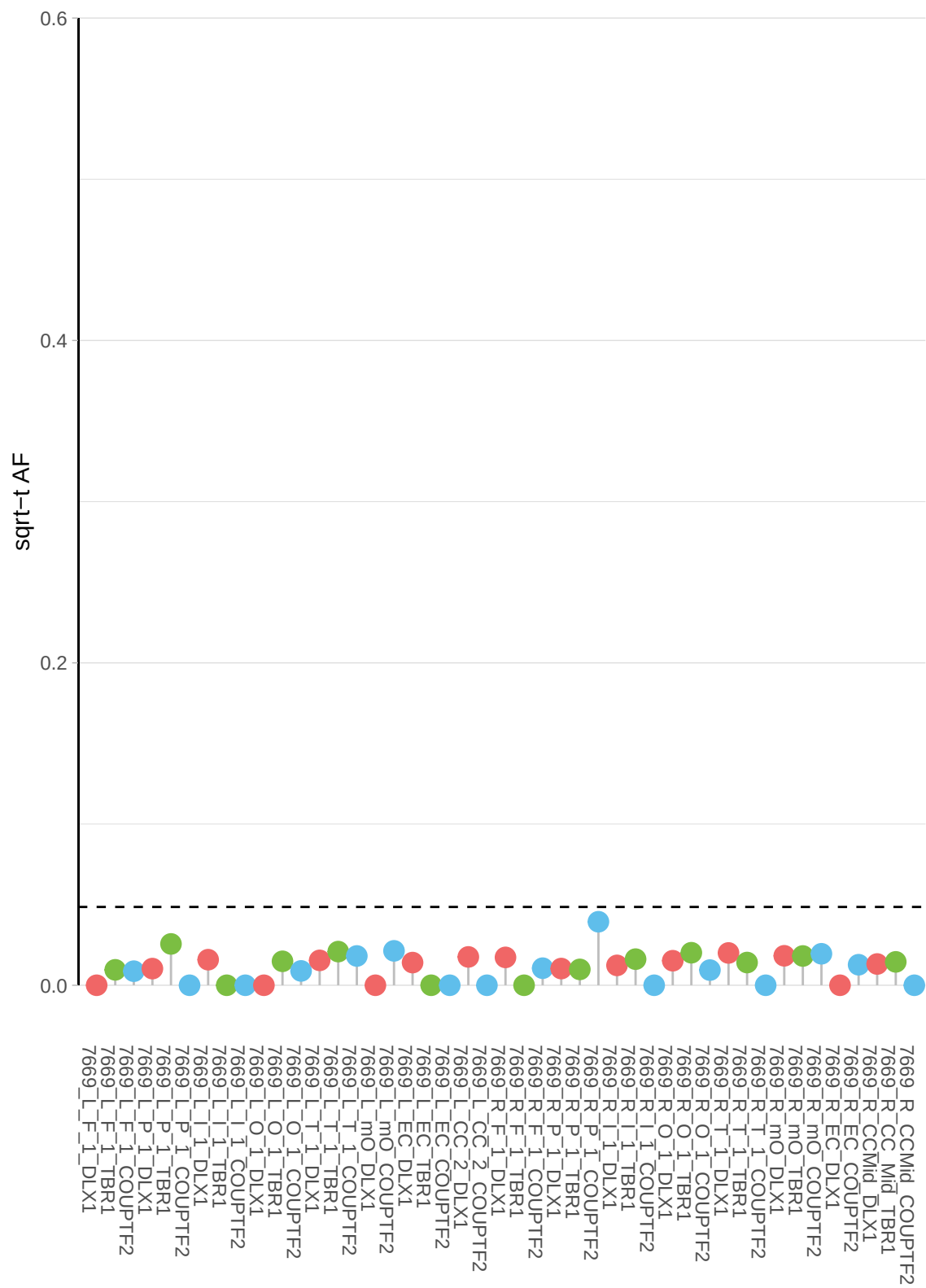

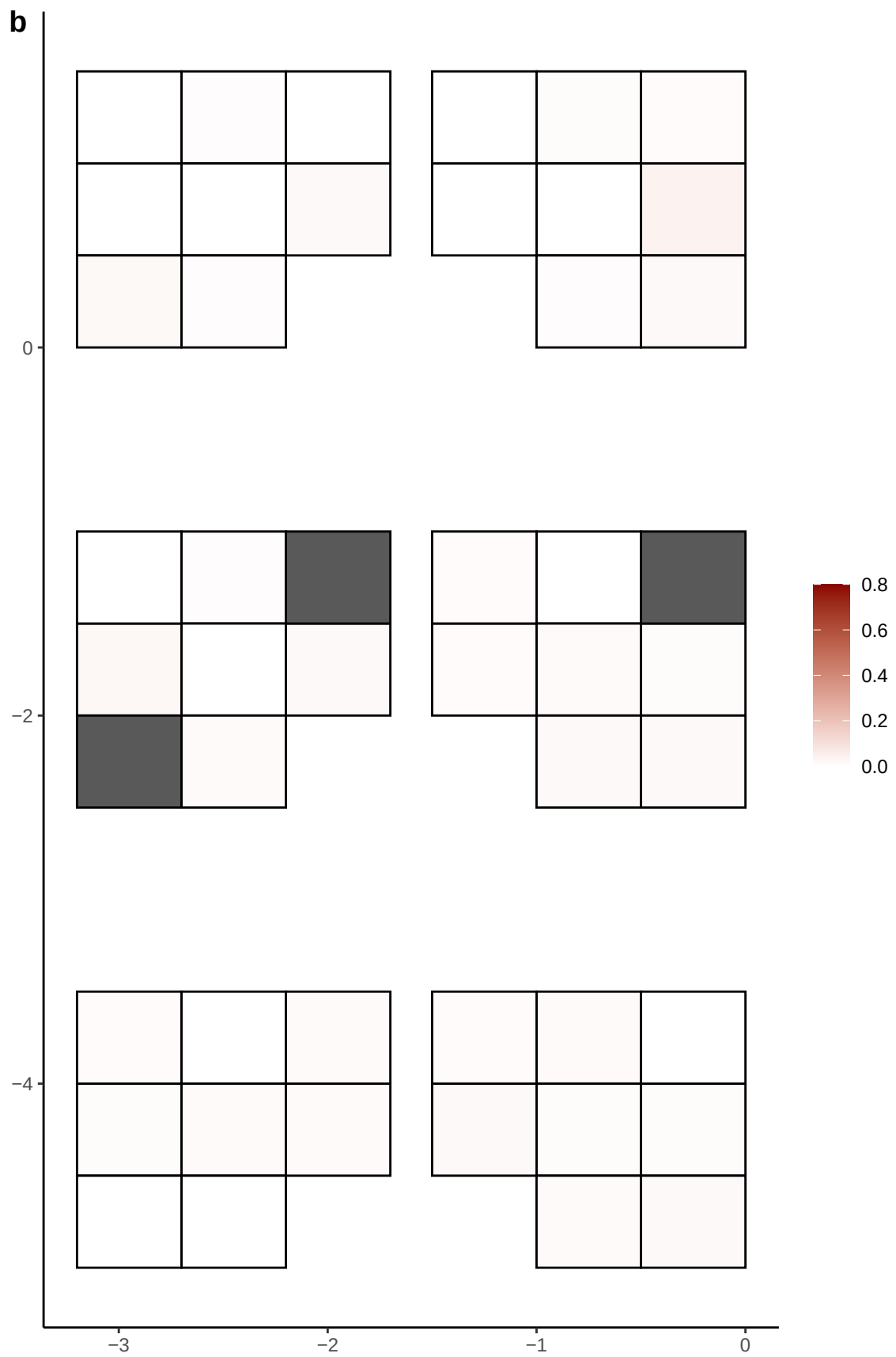

**a** ID05 5-96456832-G-A

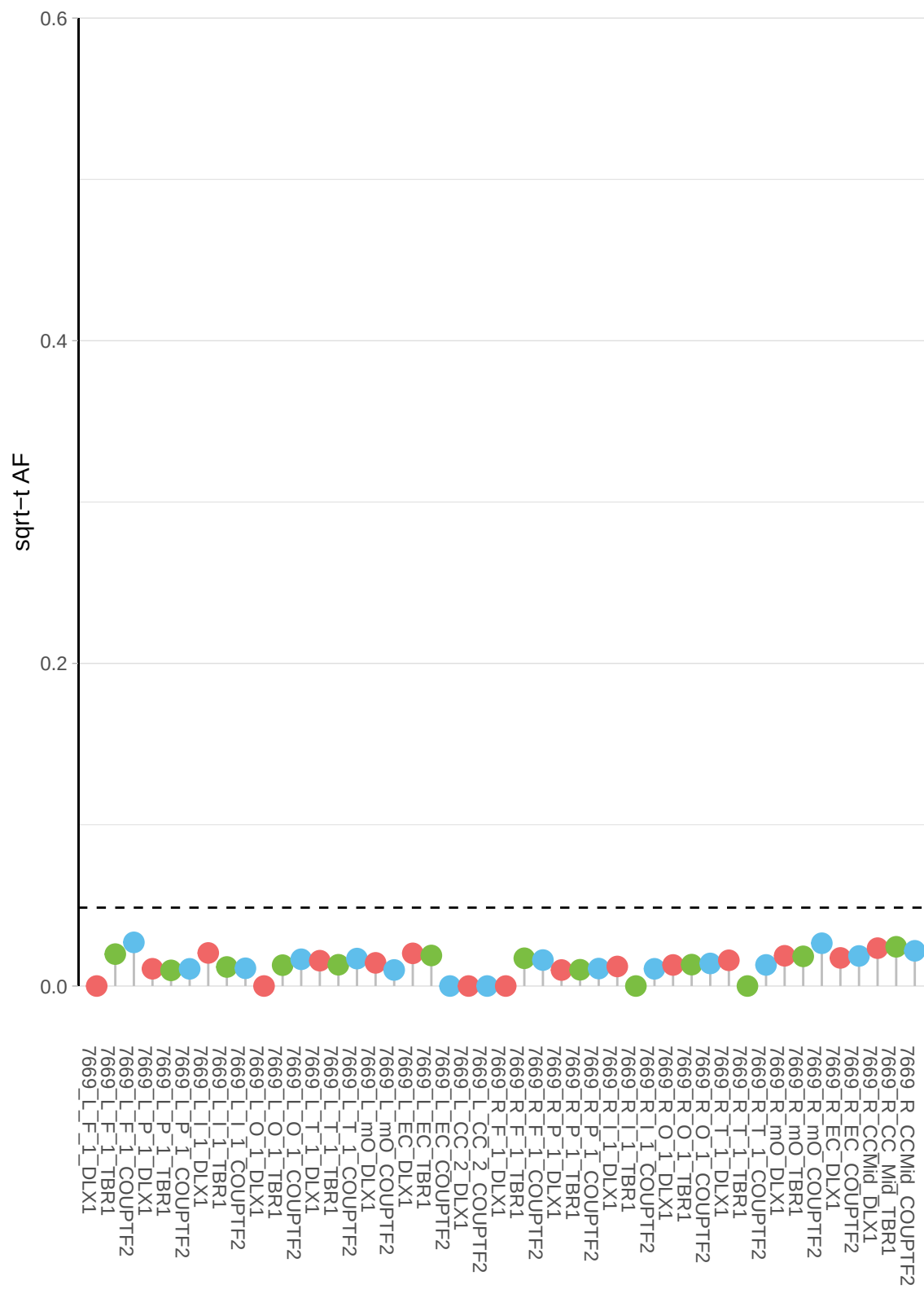

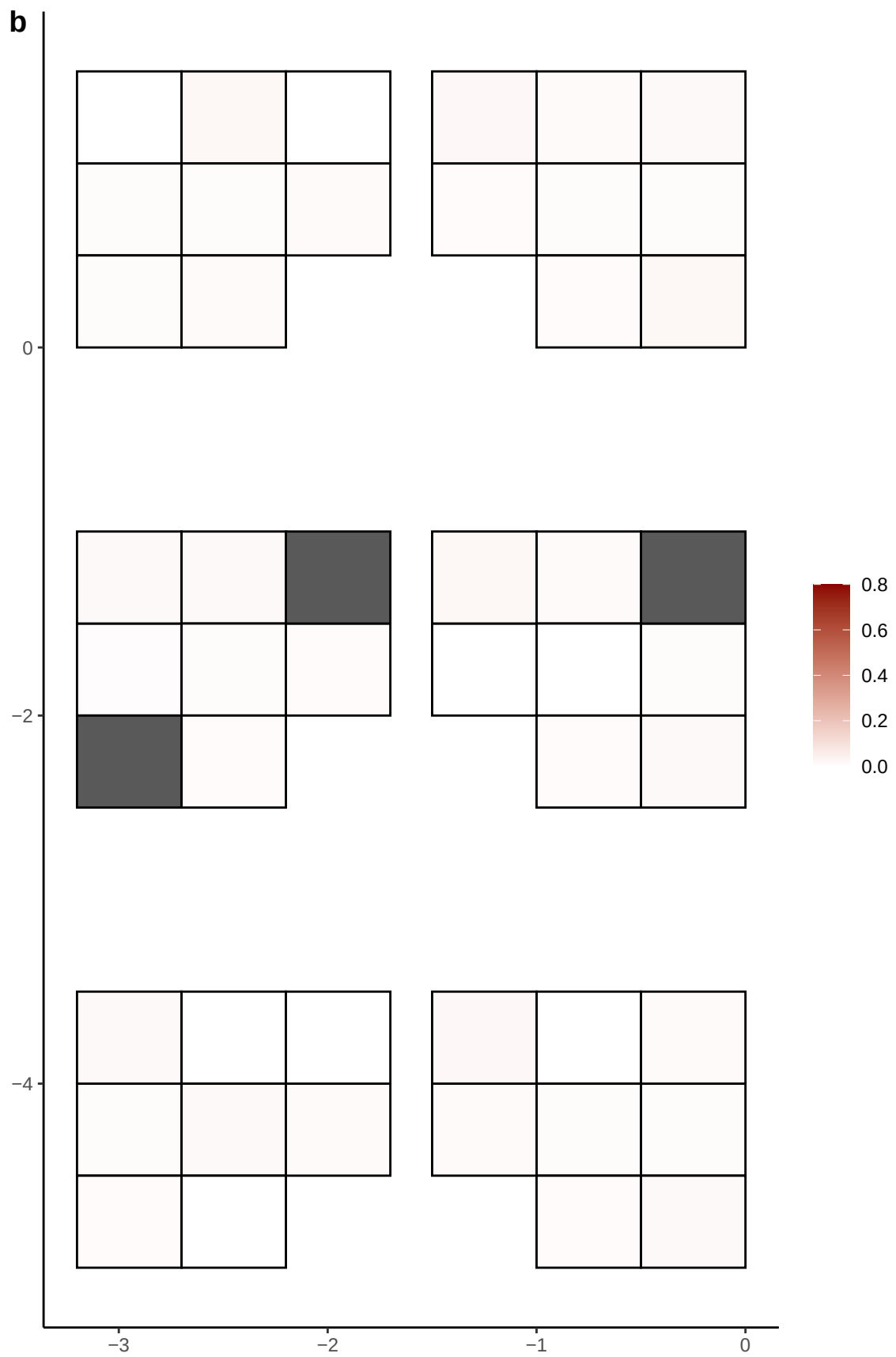

**a** ID05 5-99372353-G-A

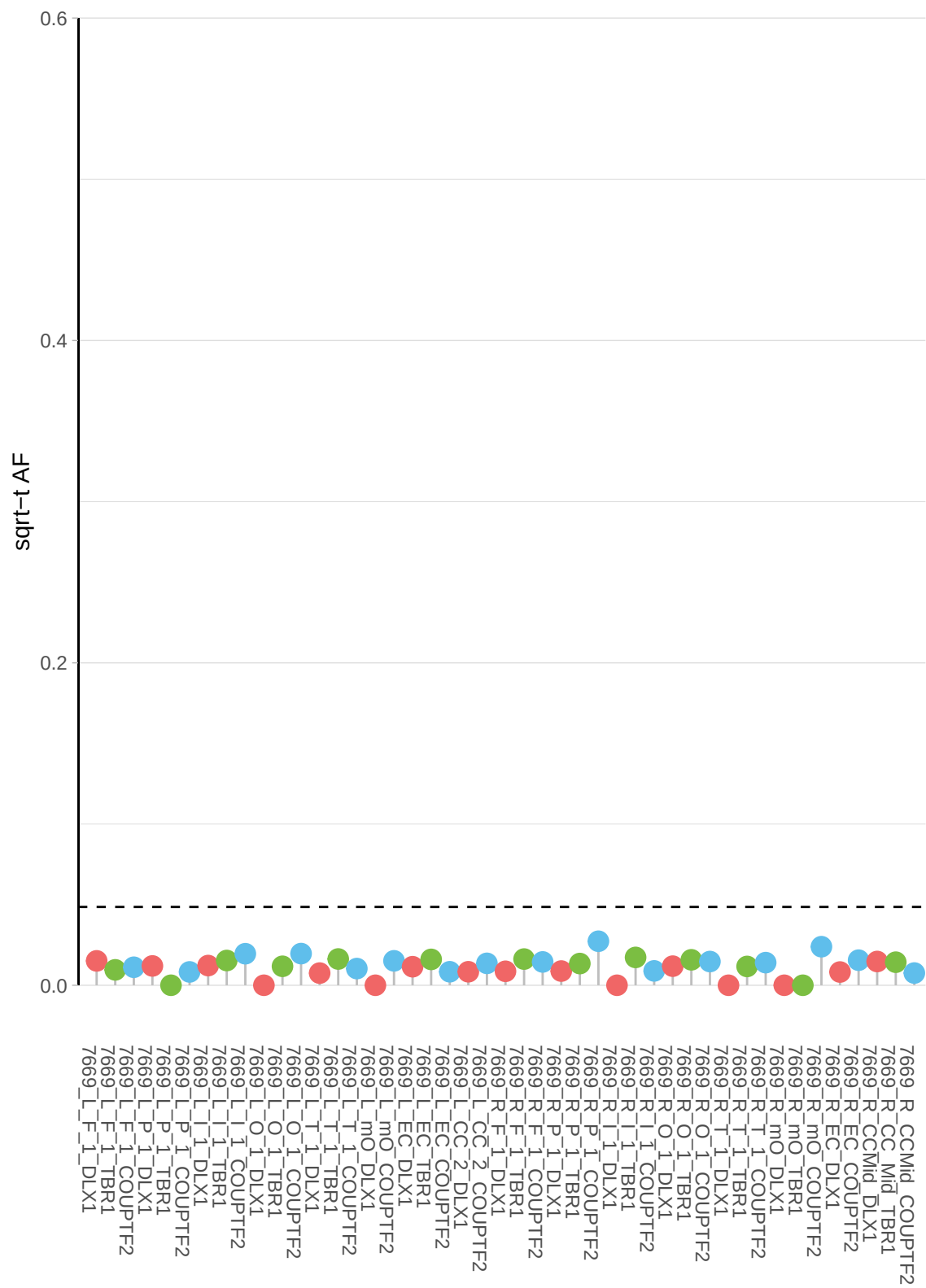

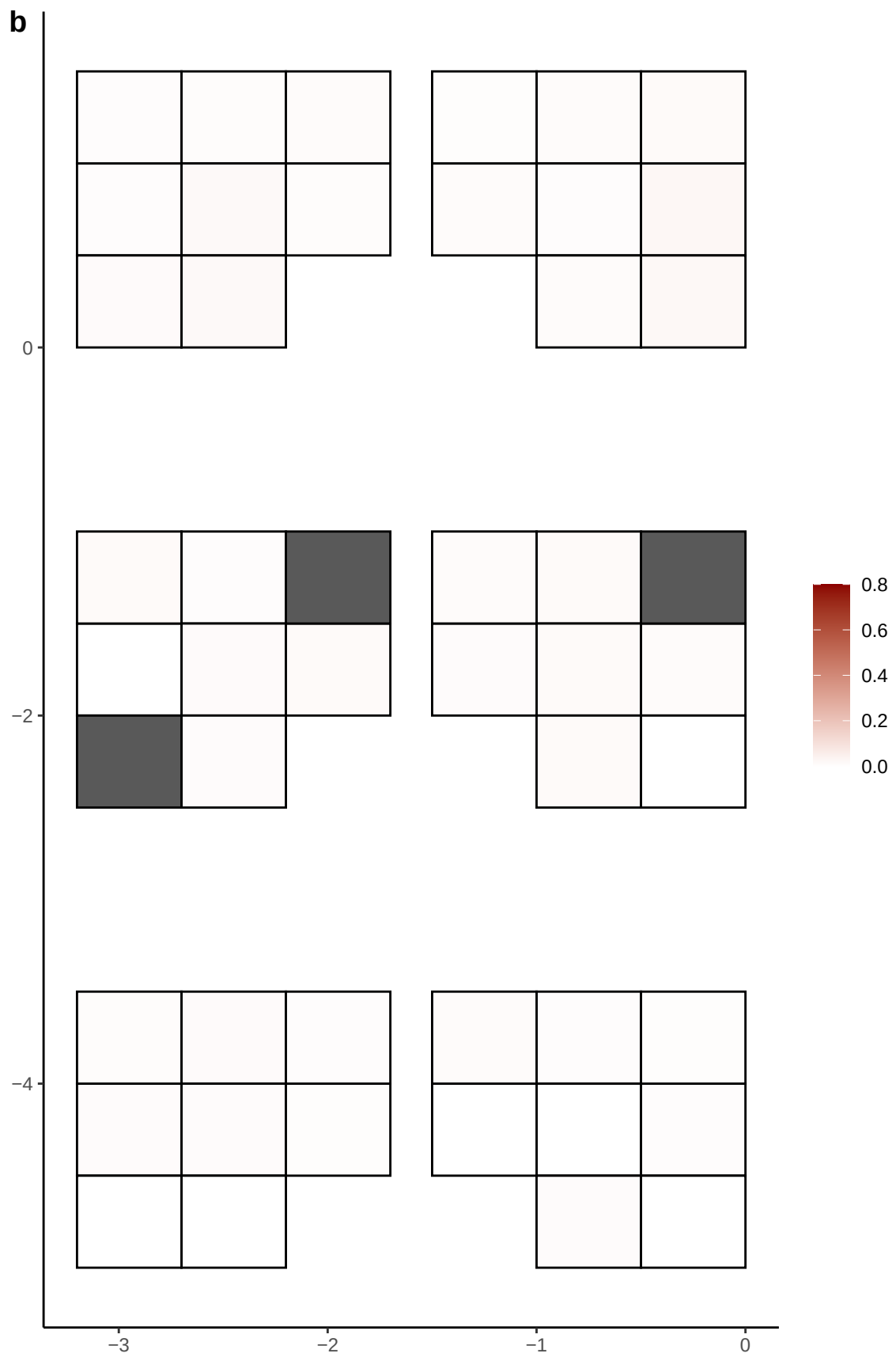

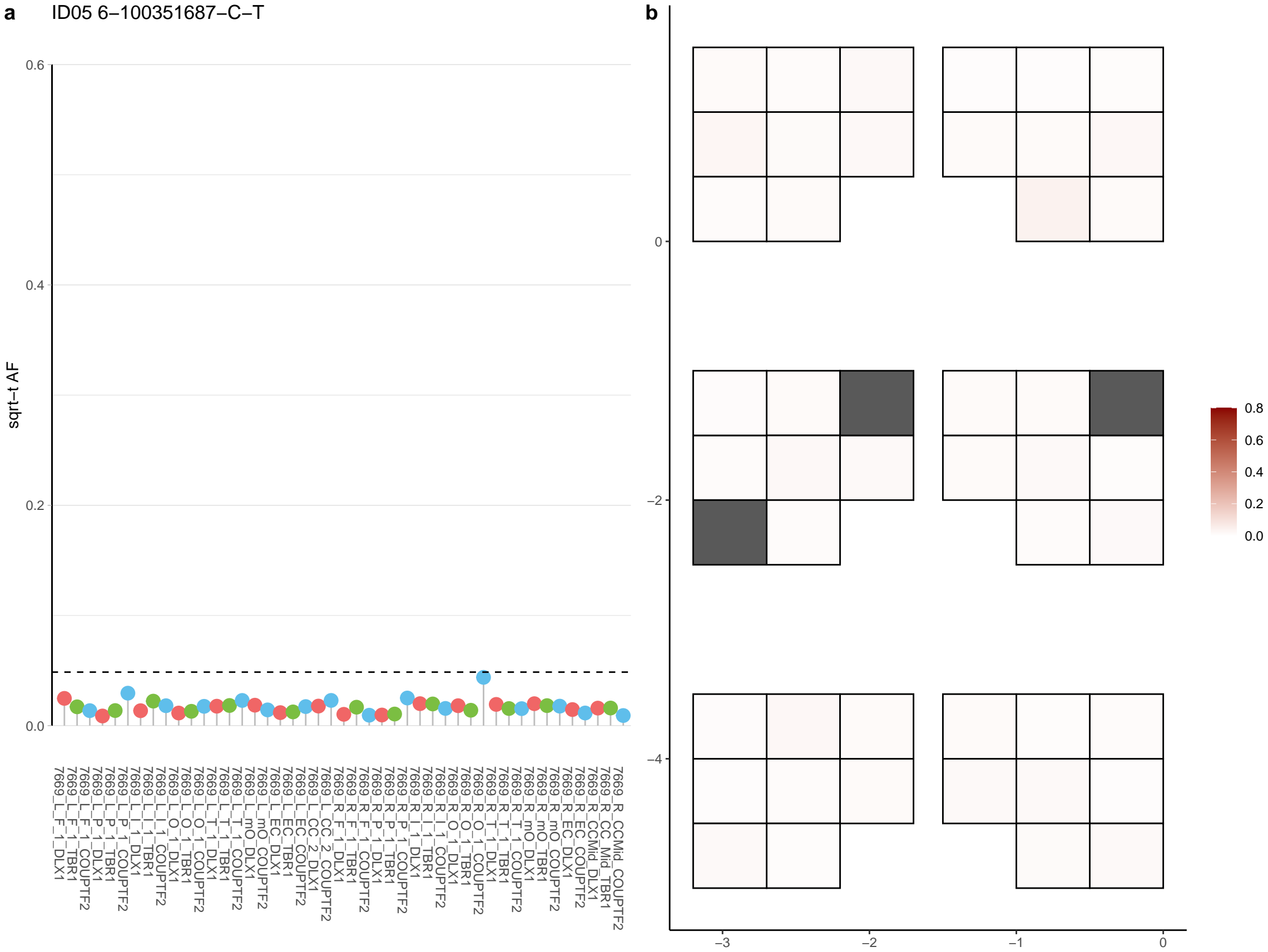

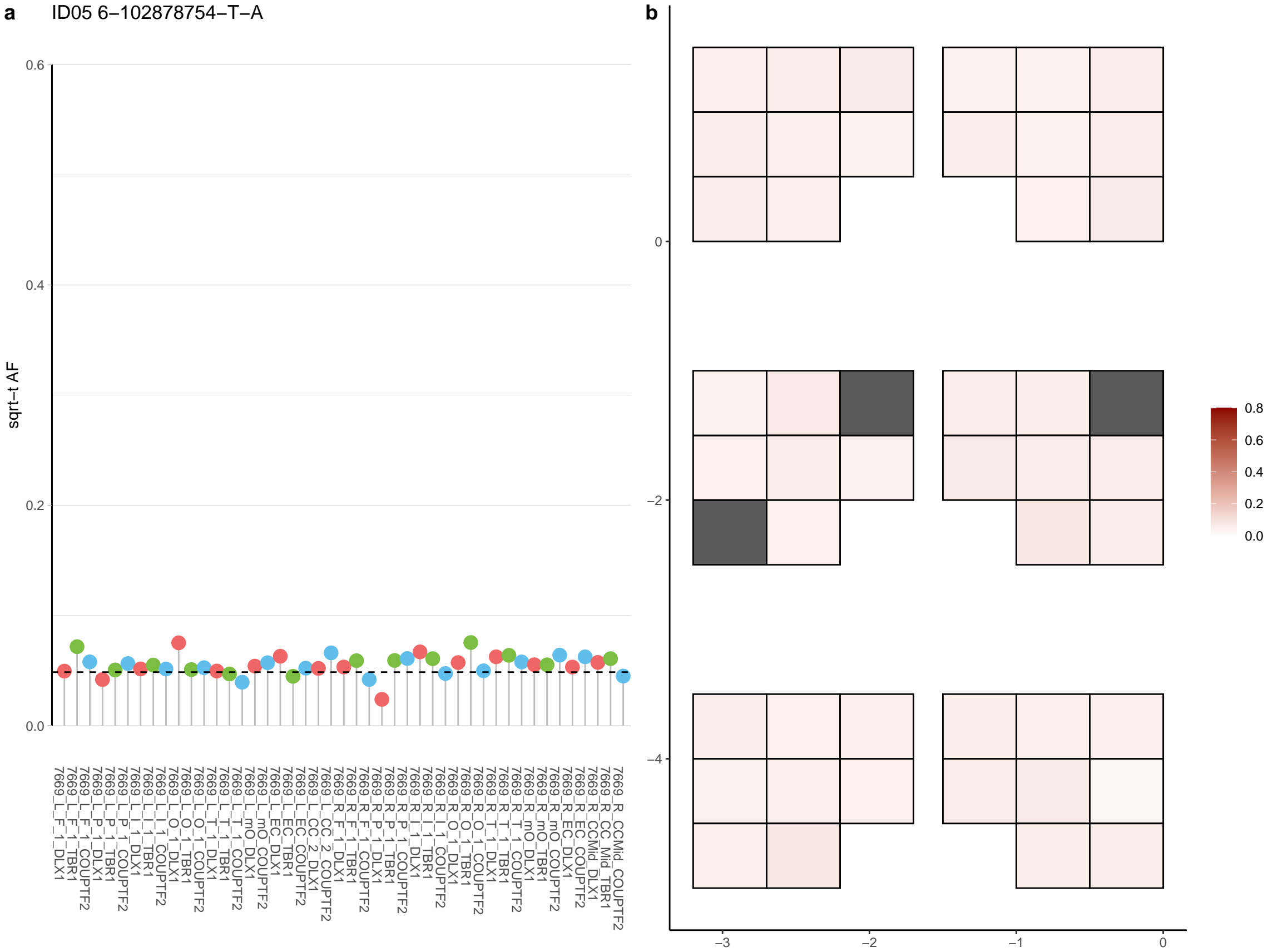

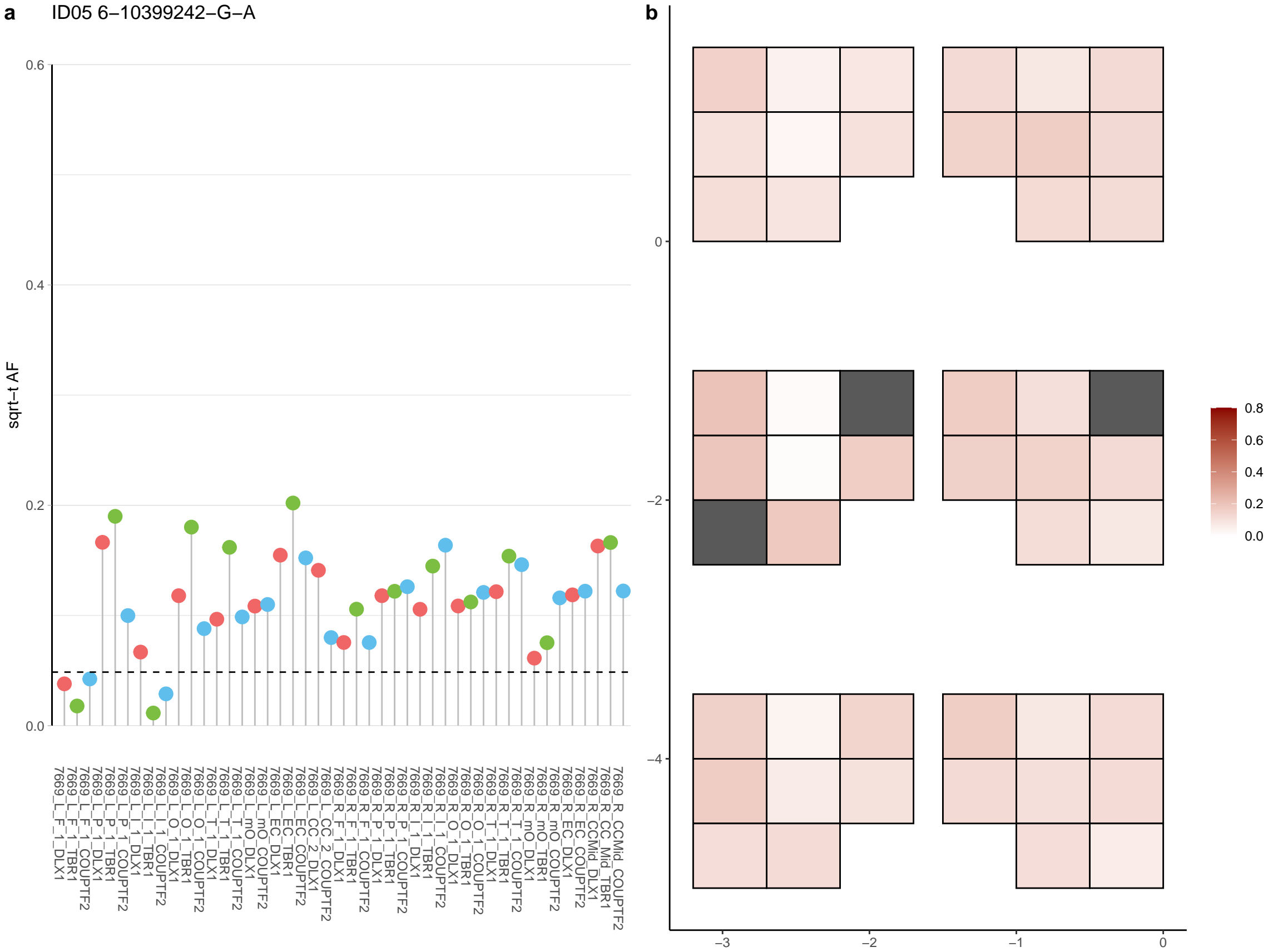

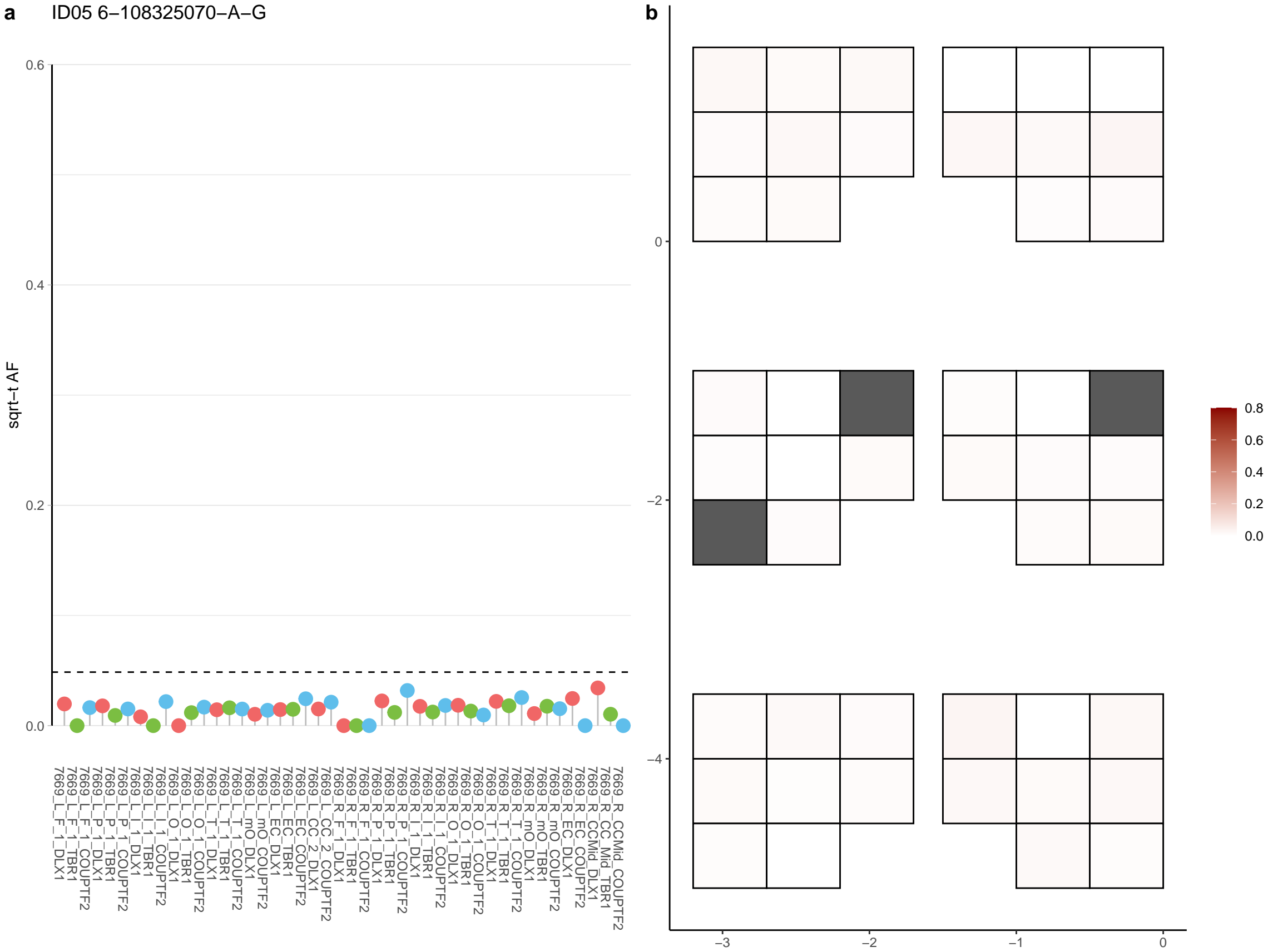

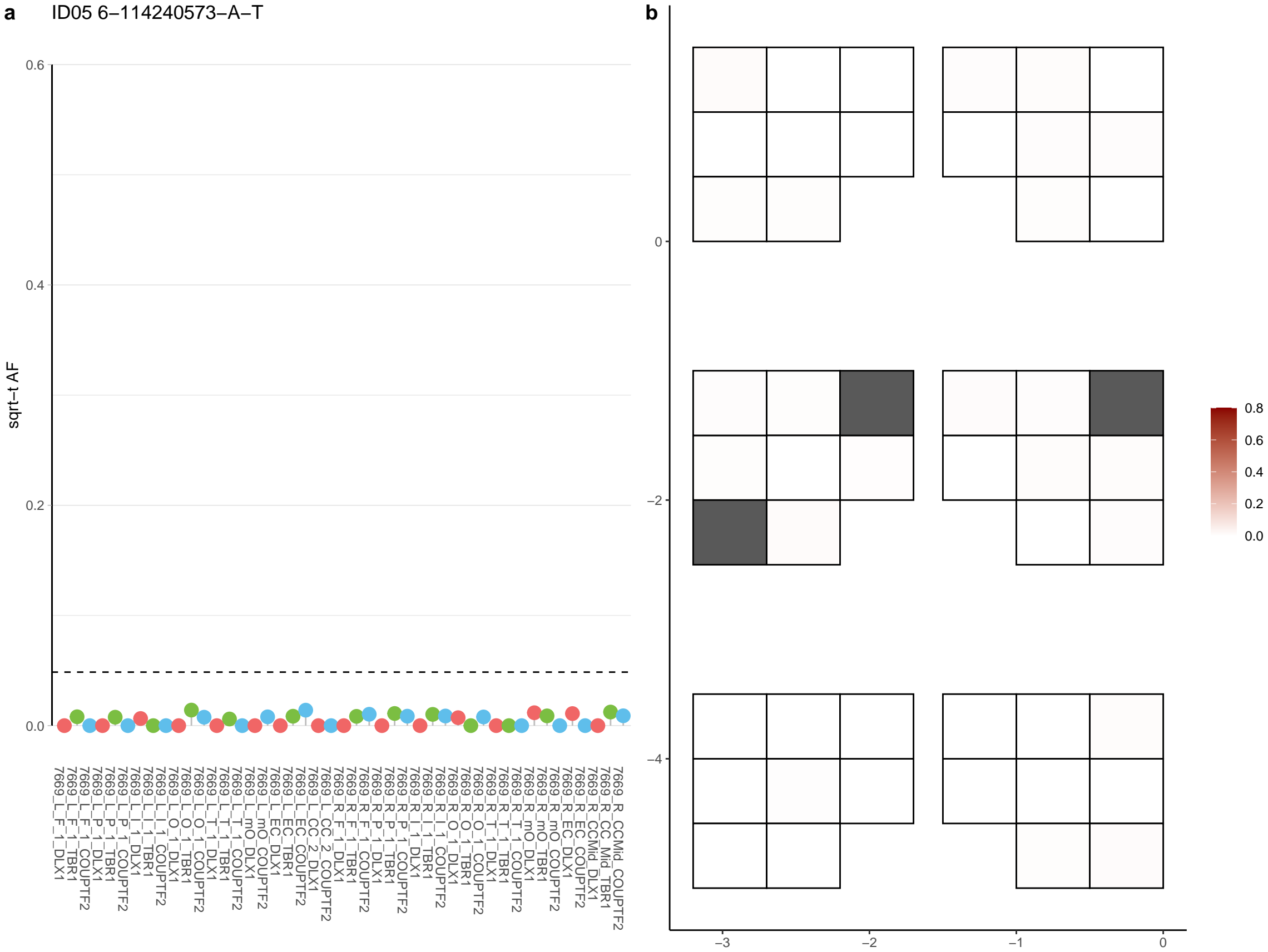

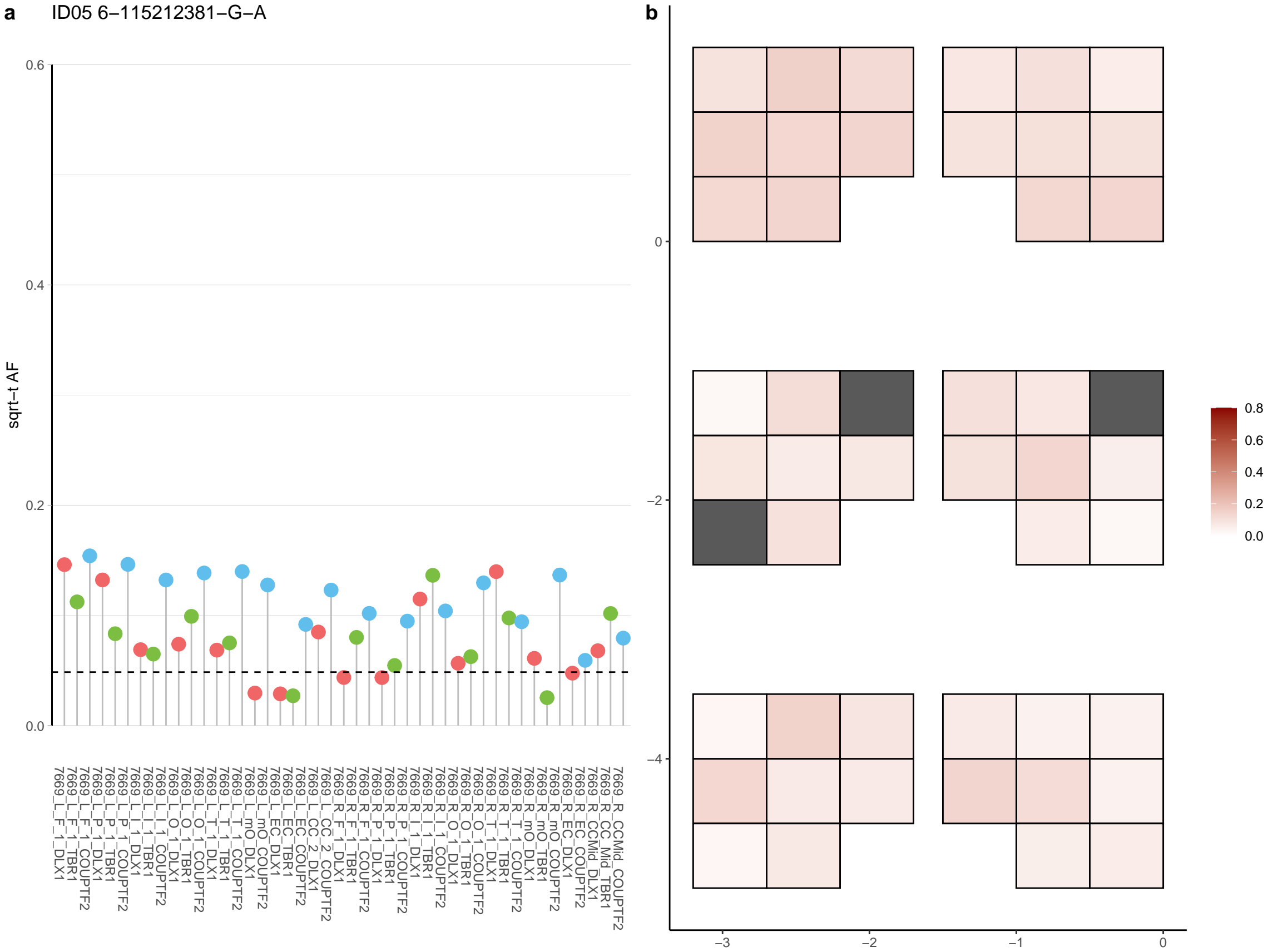

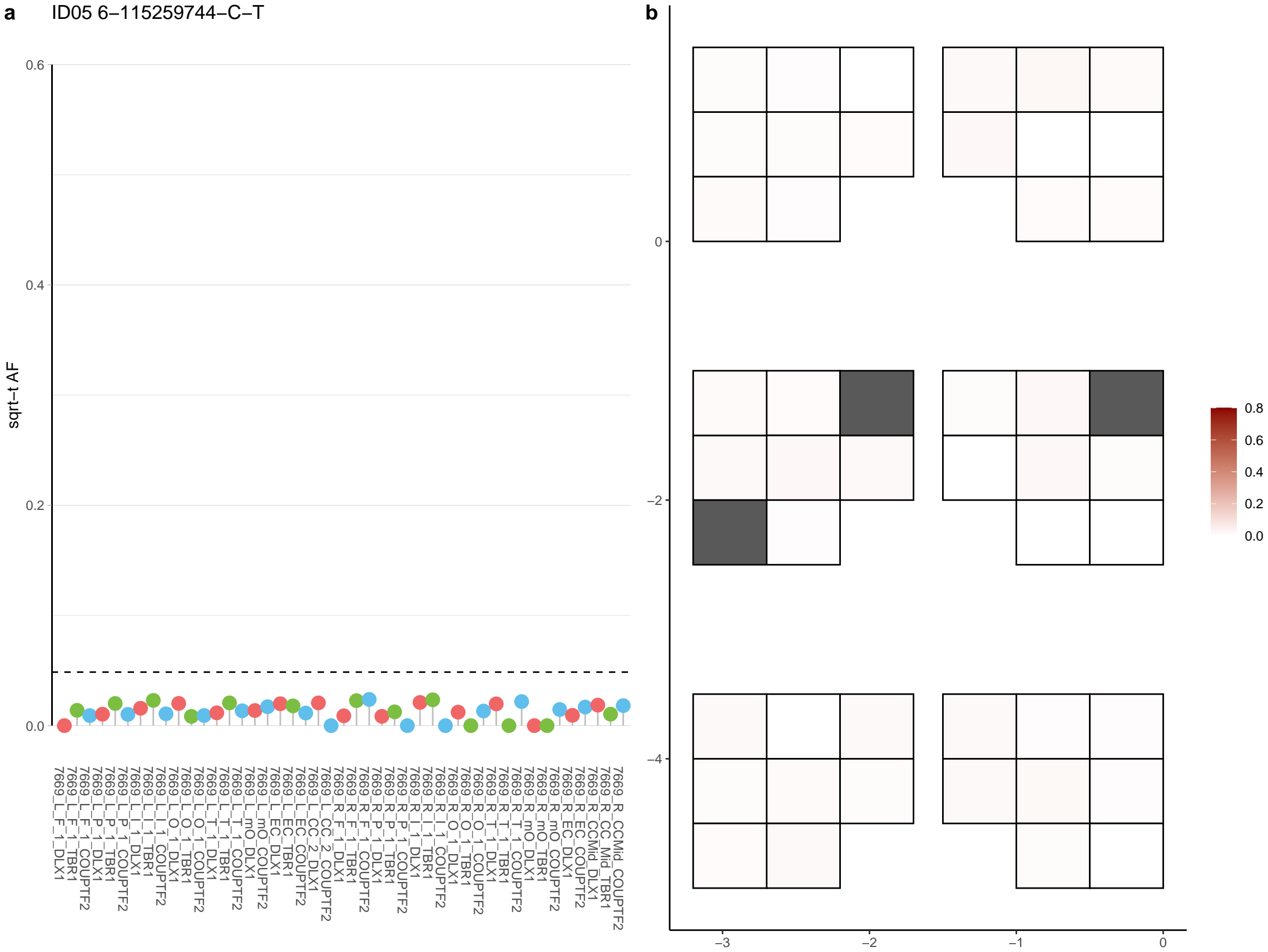

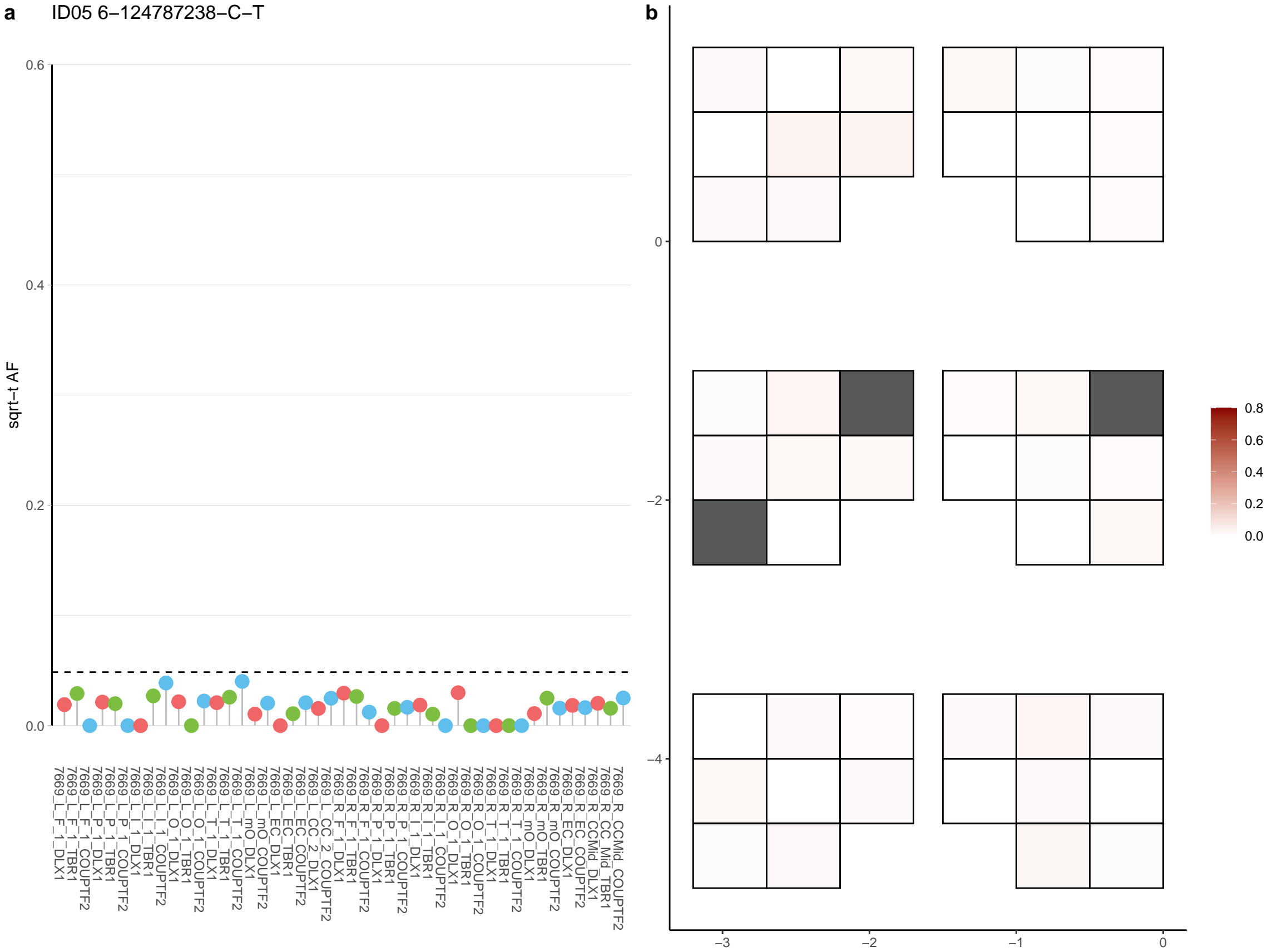

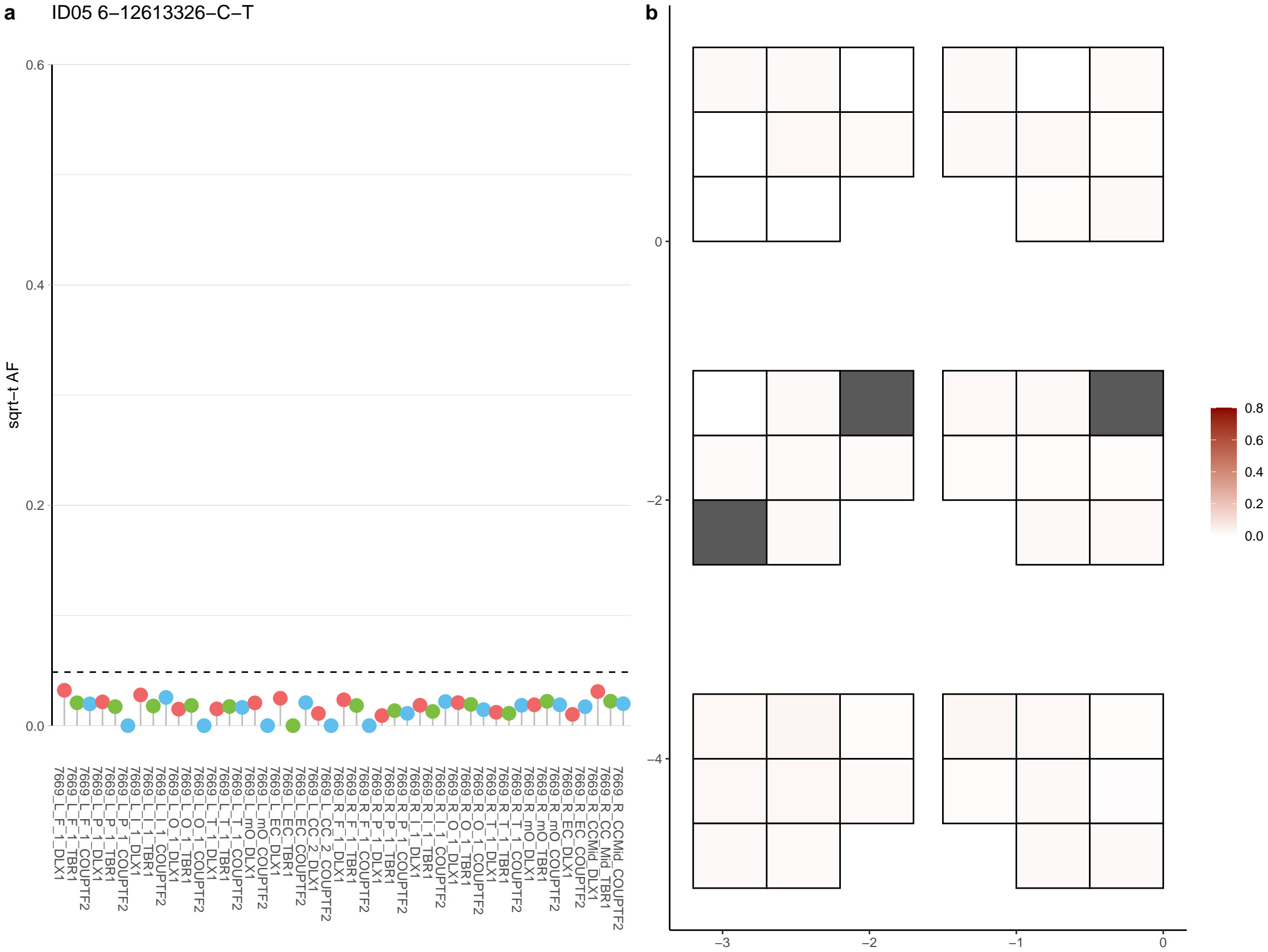

**a** ID05 6-132170864-G-A

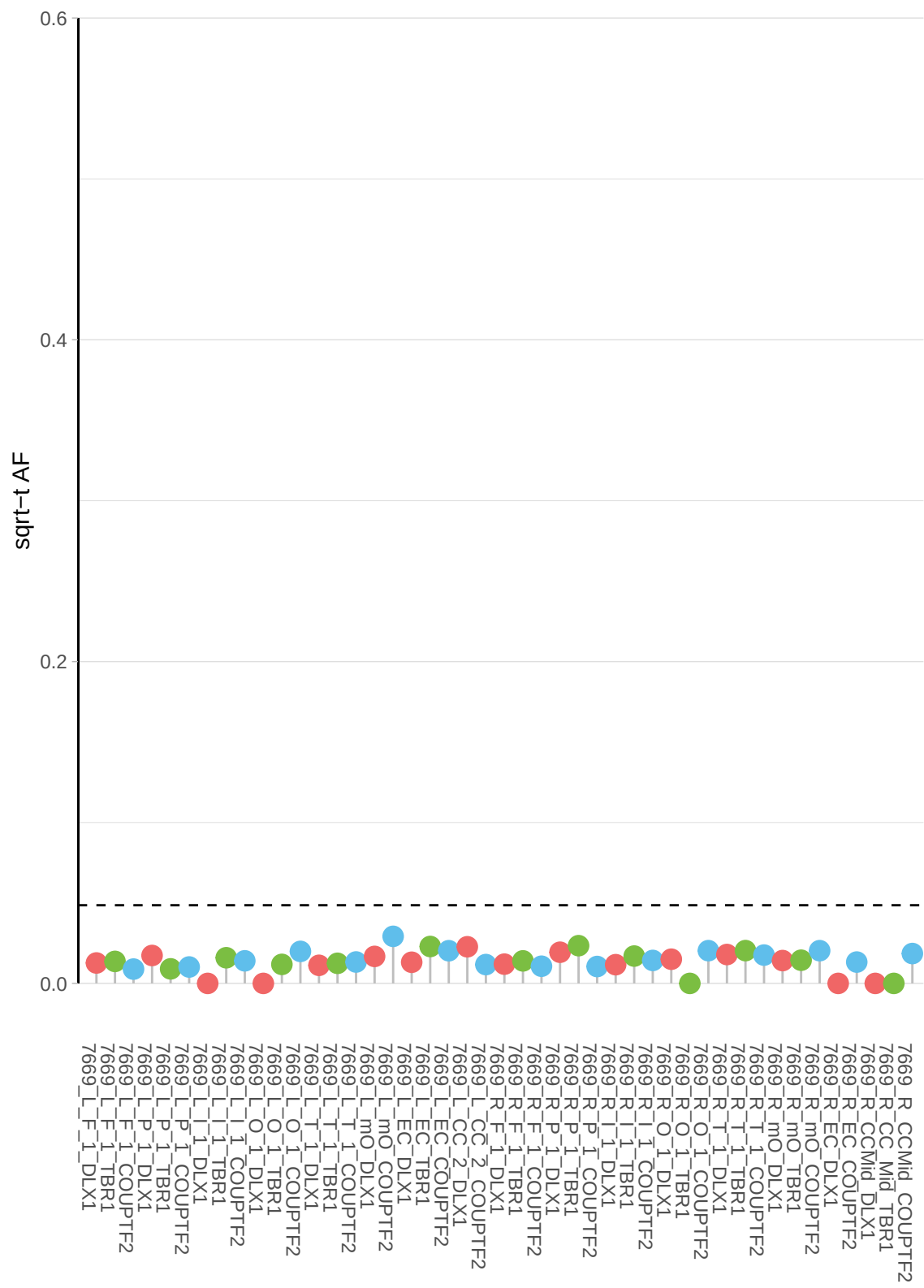

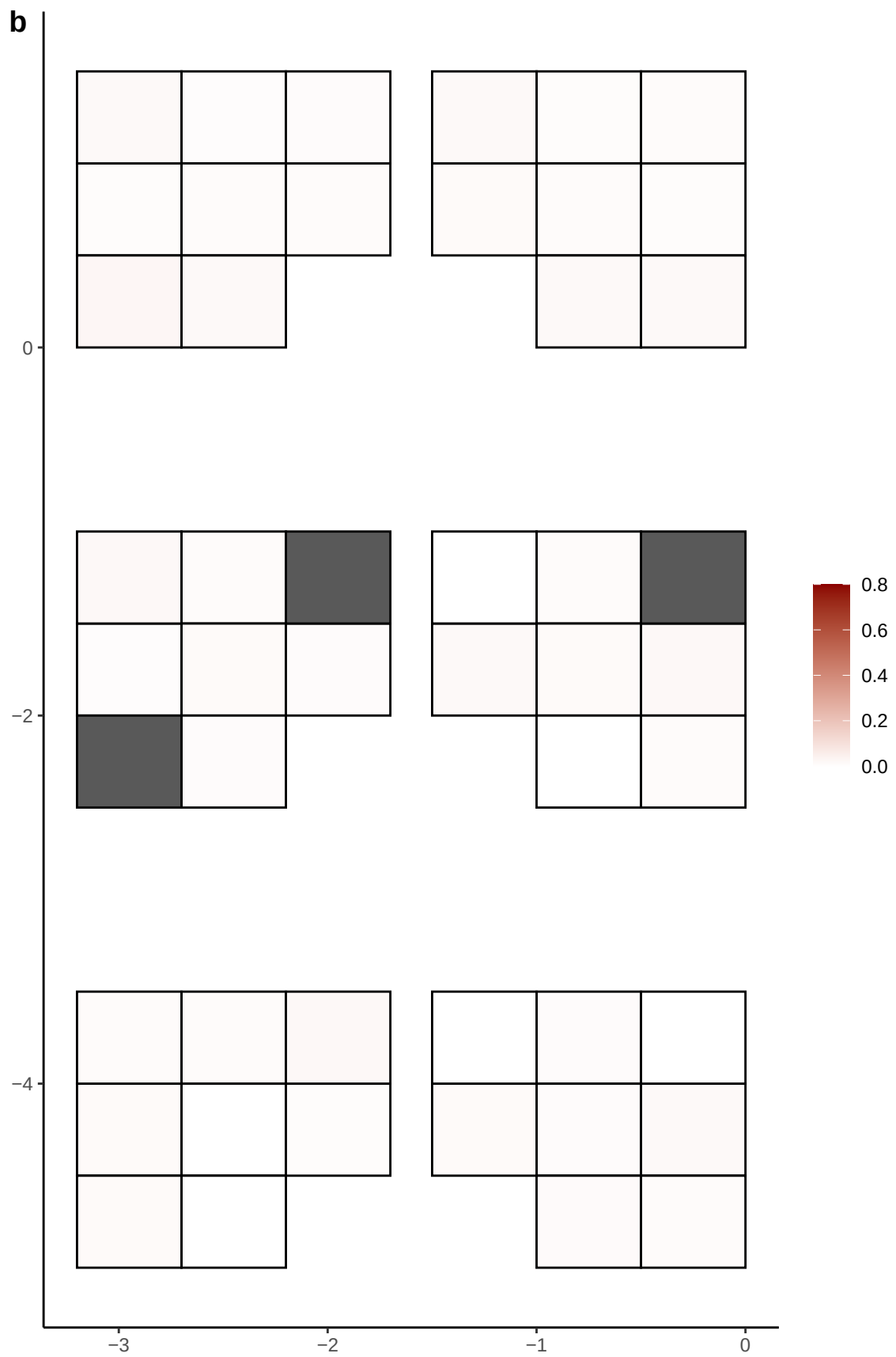

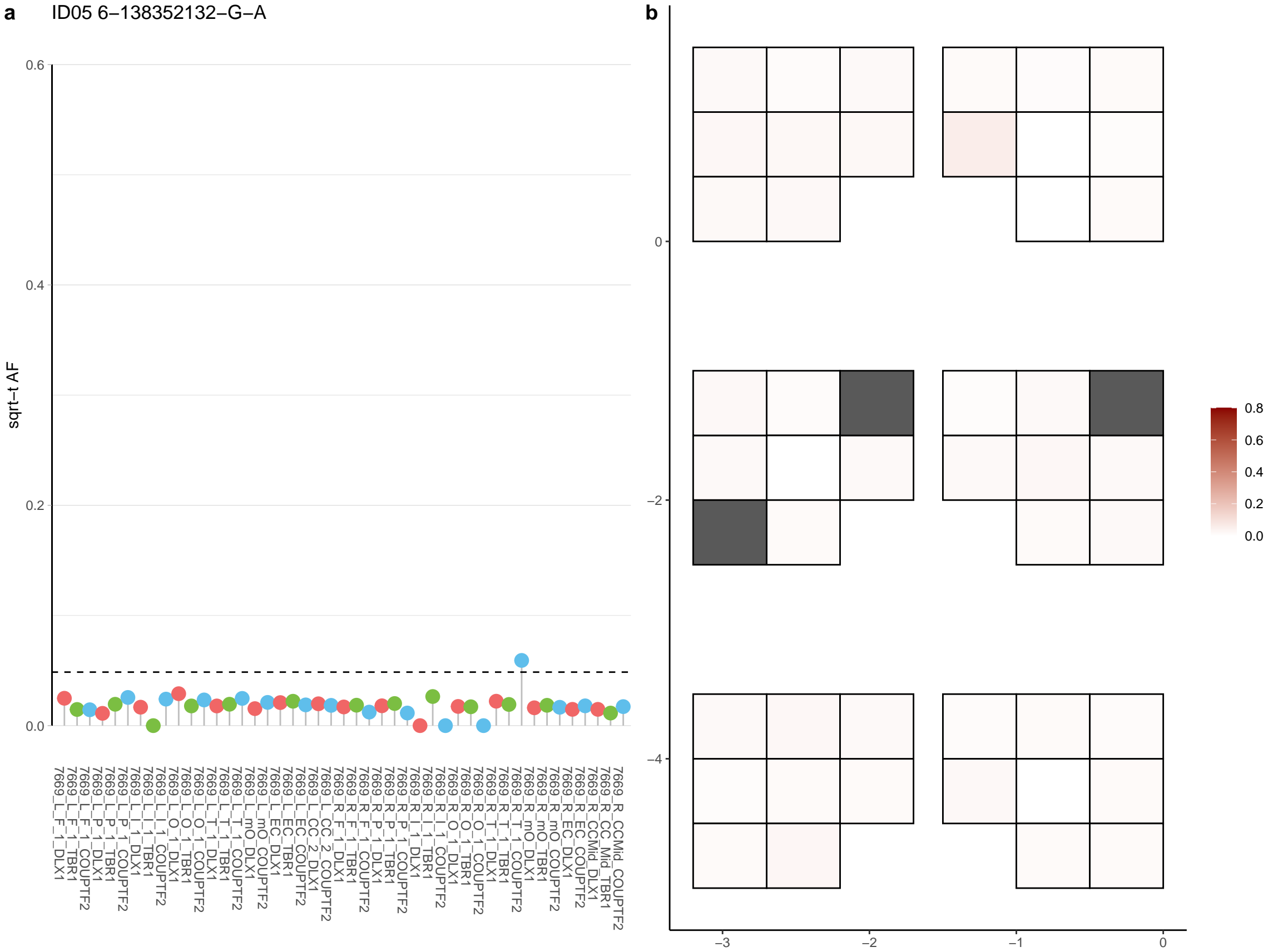

**a** ID05 6-139194275-C-T

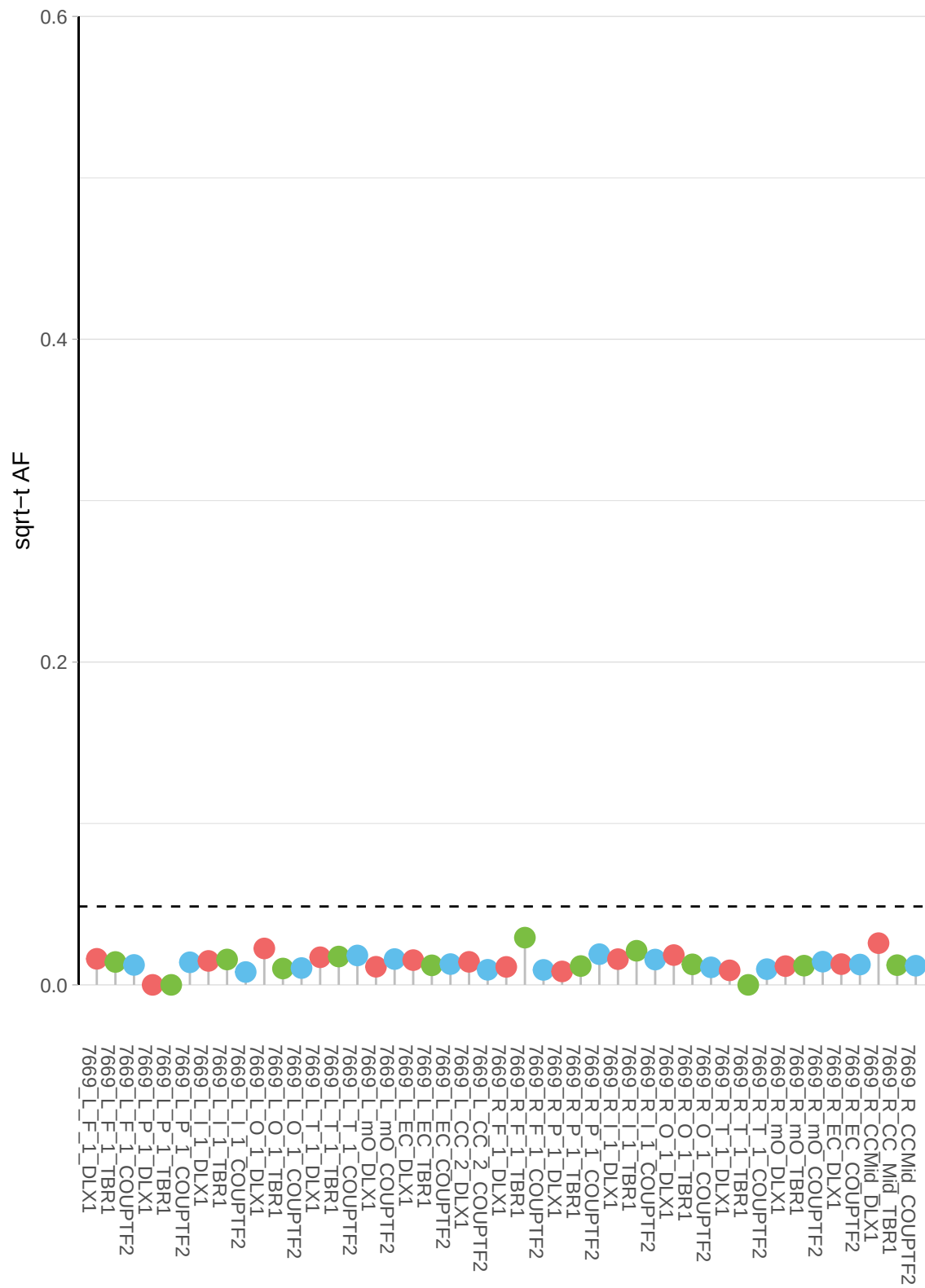

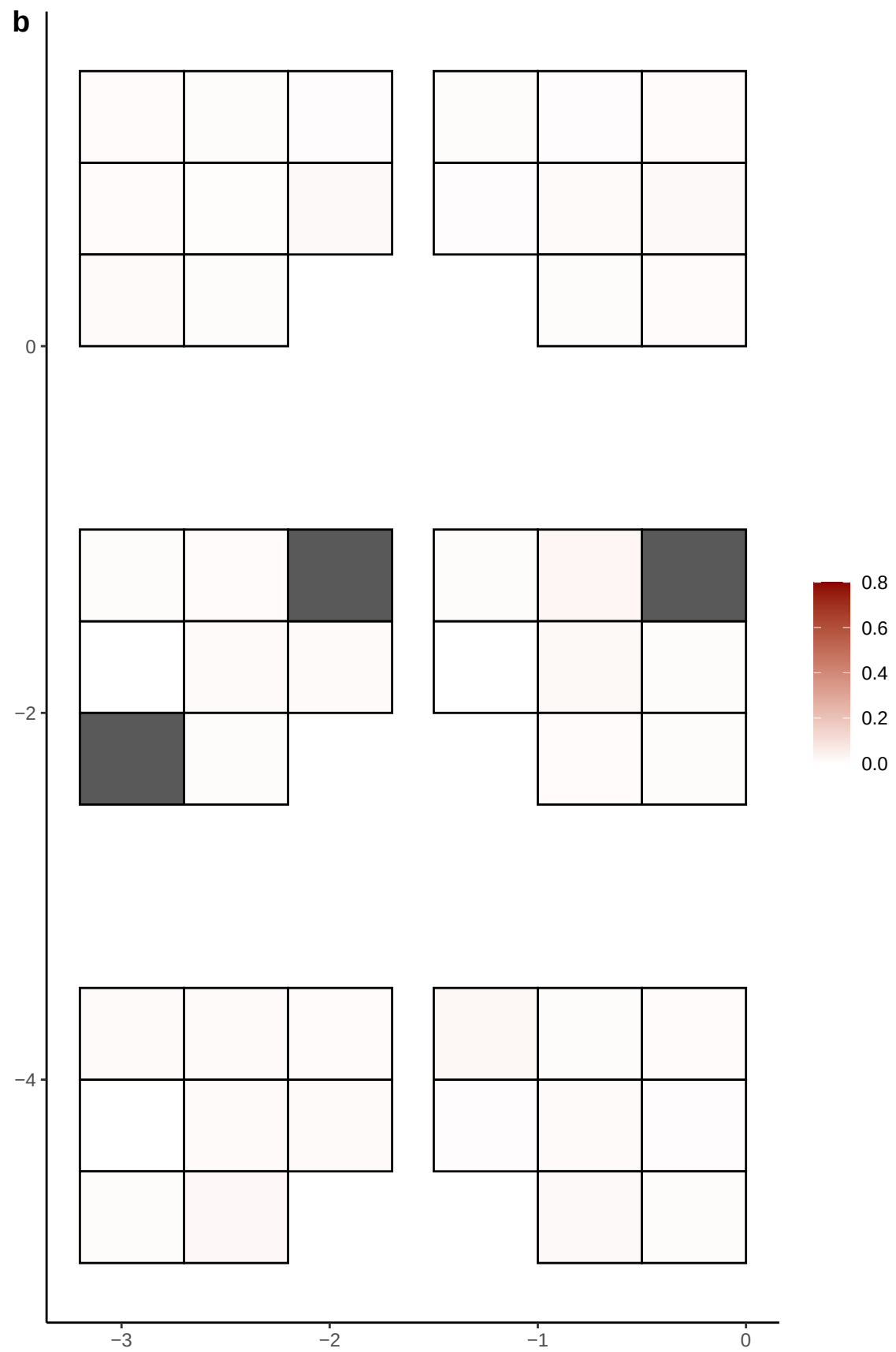

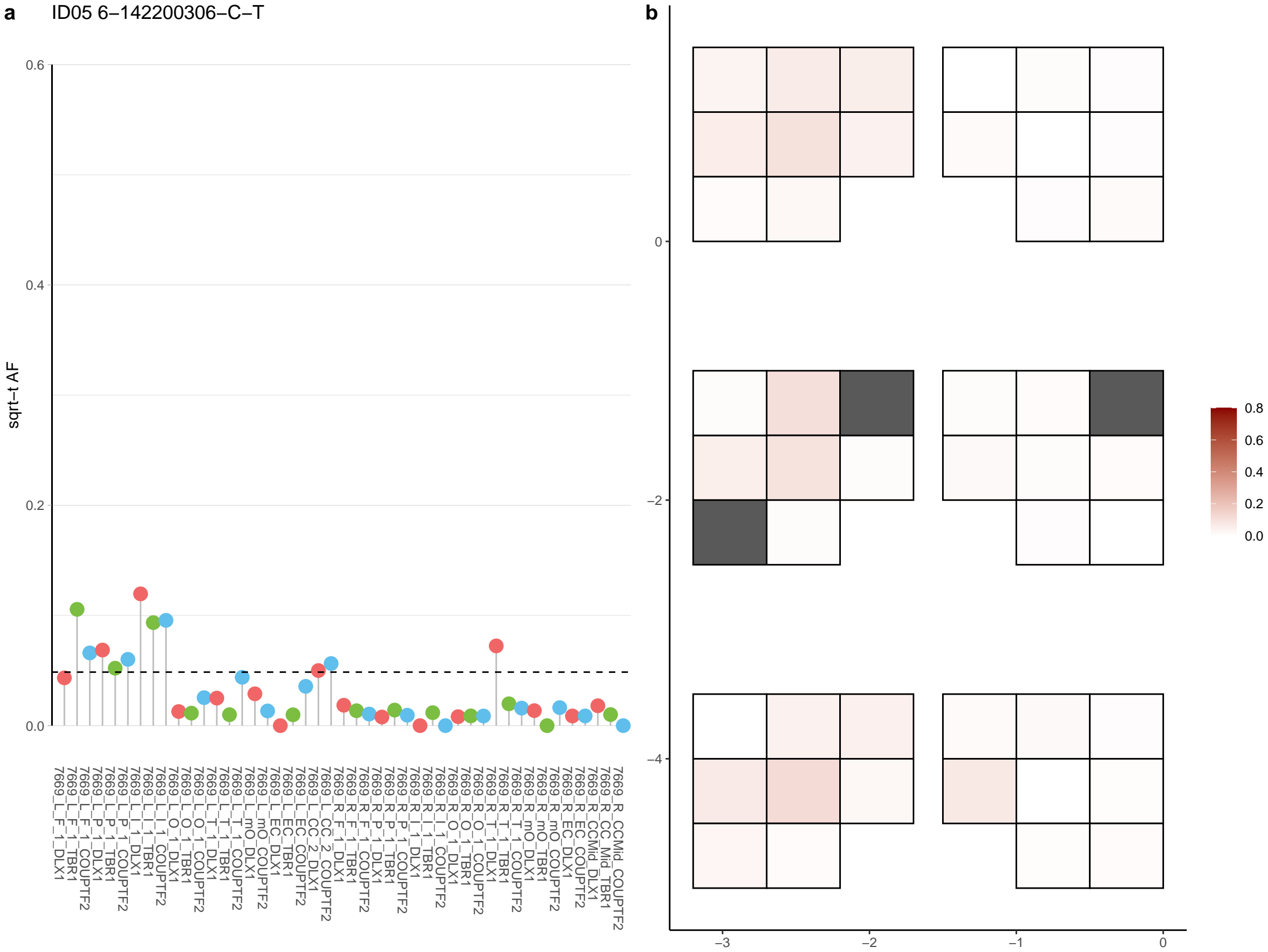

**a** ID05 6-144775505-C-T

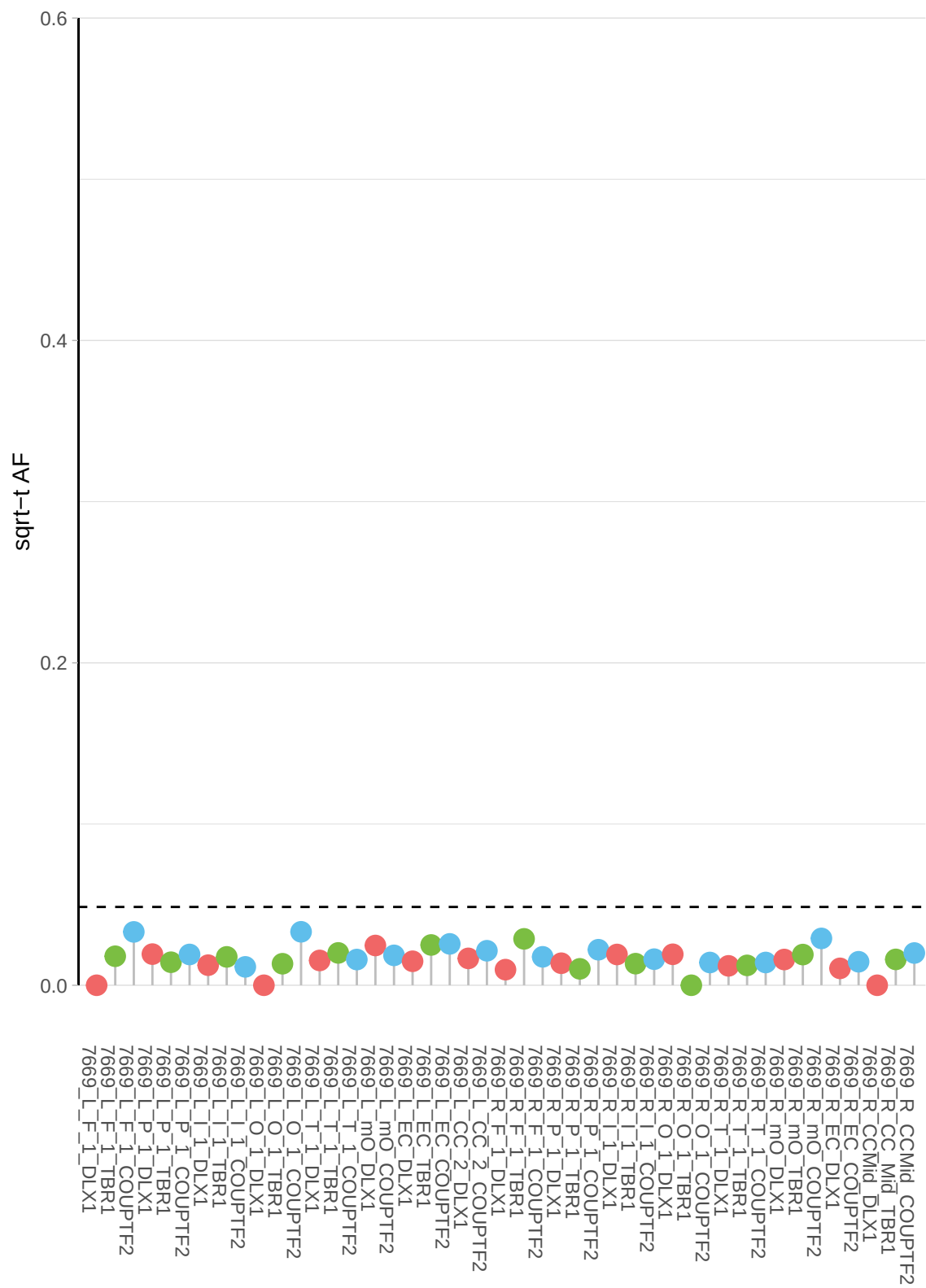

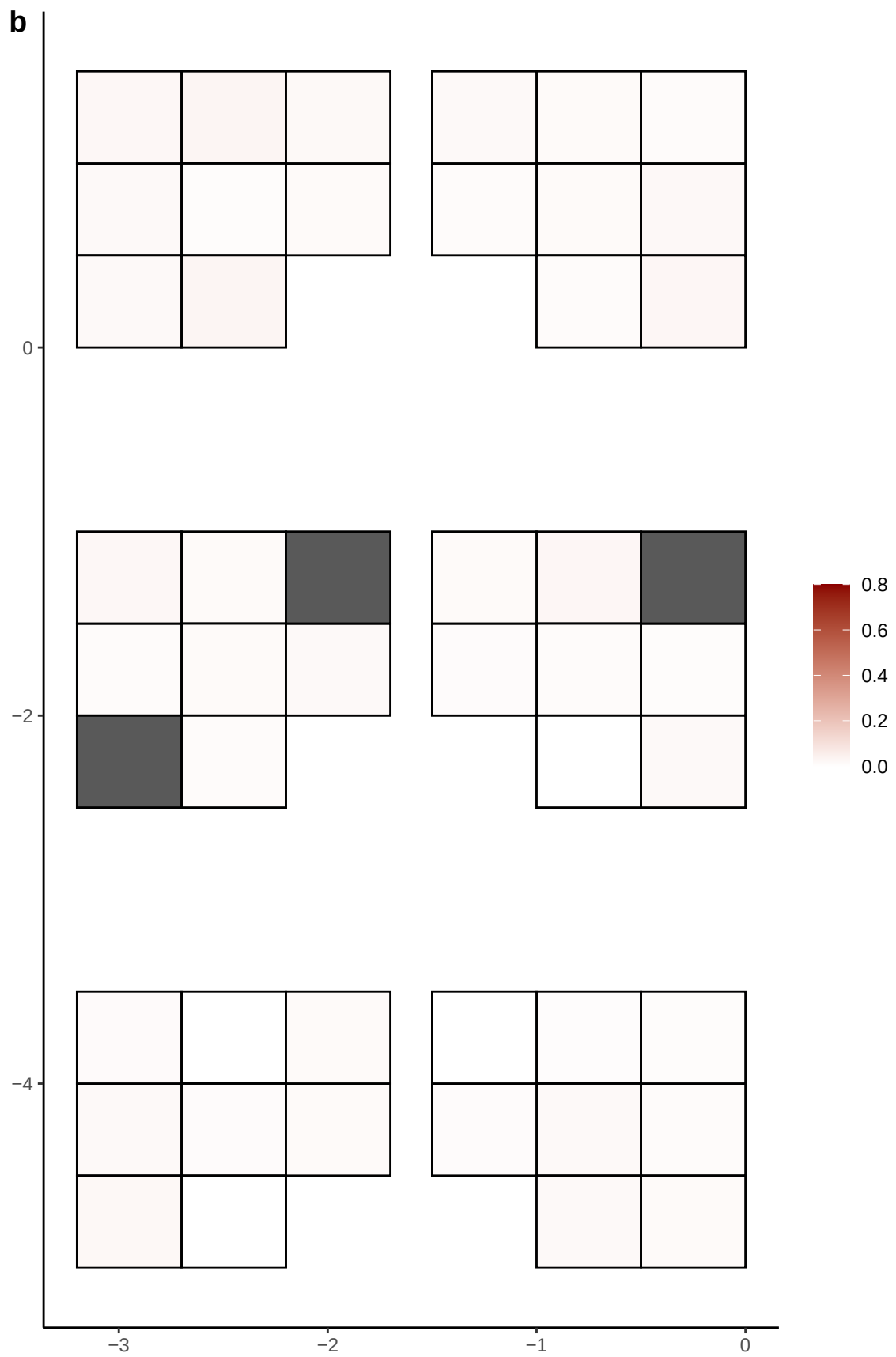

**a**

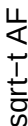

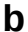

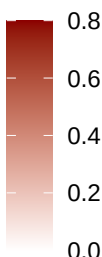

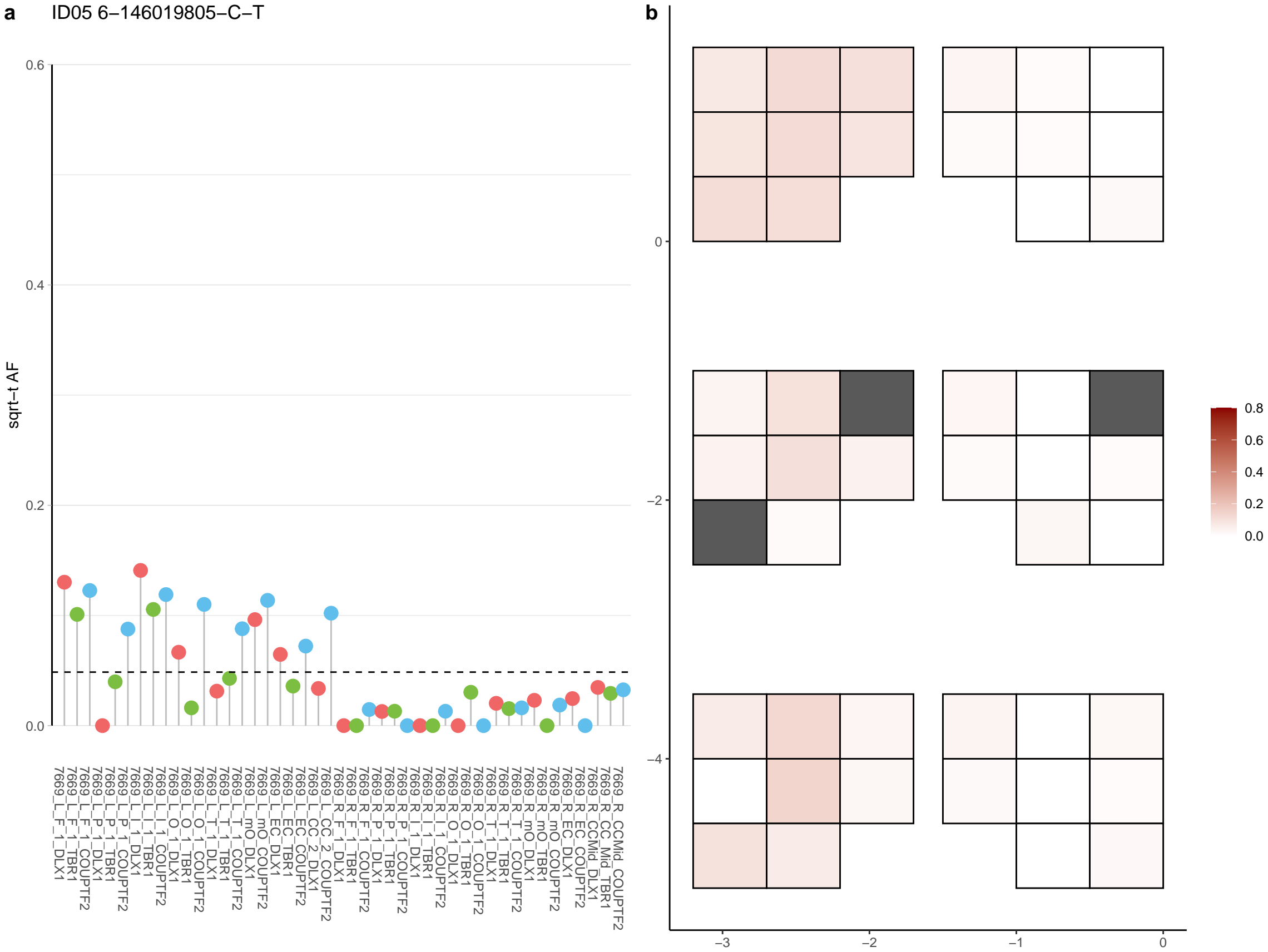

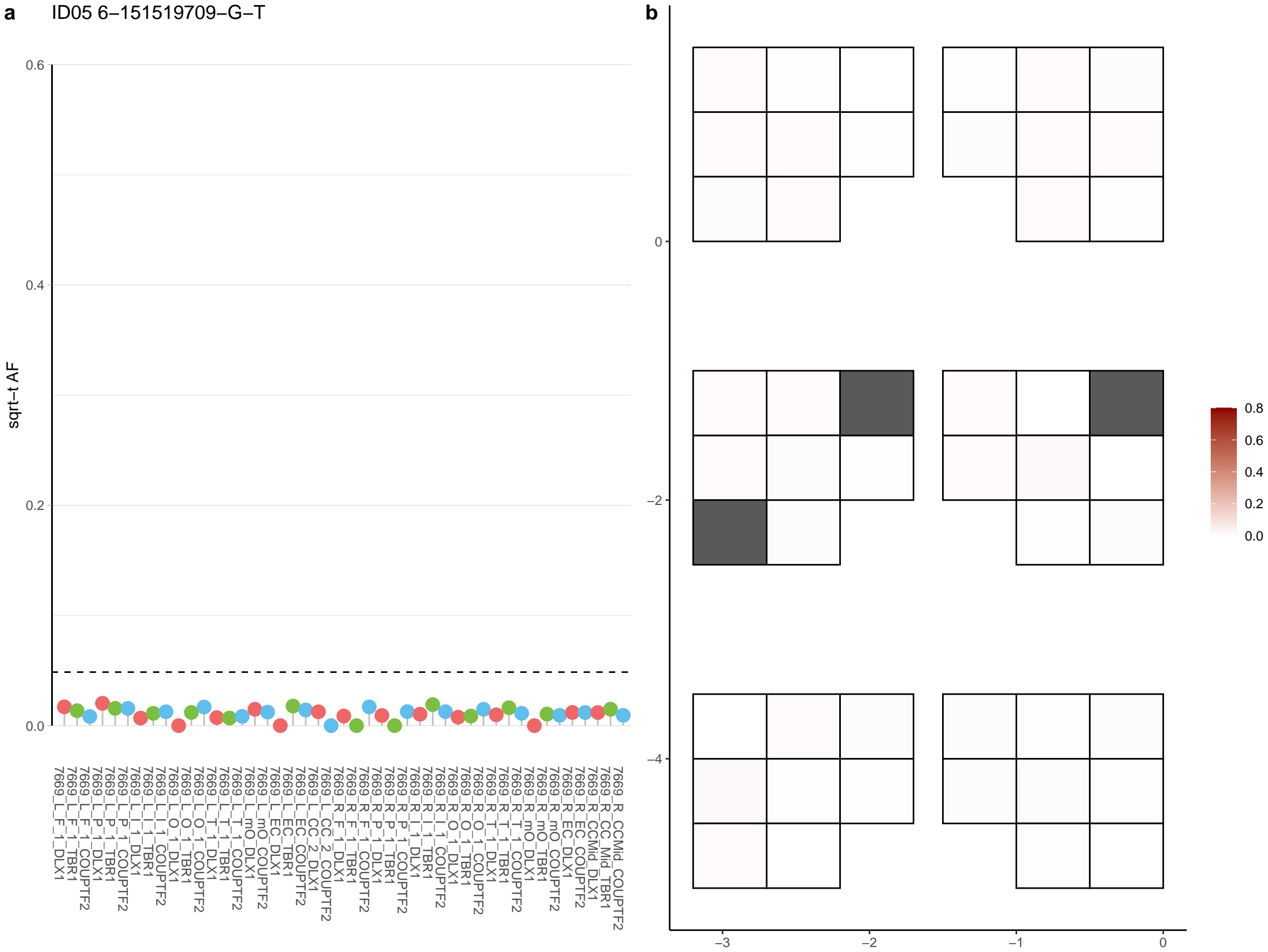

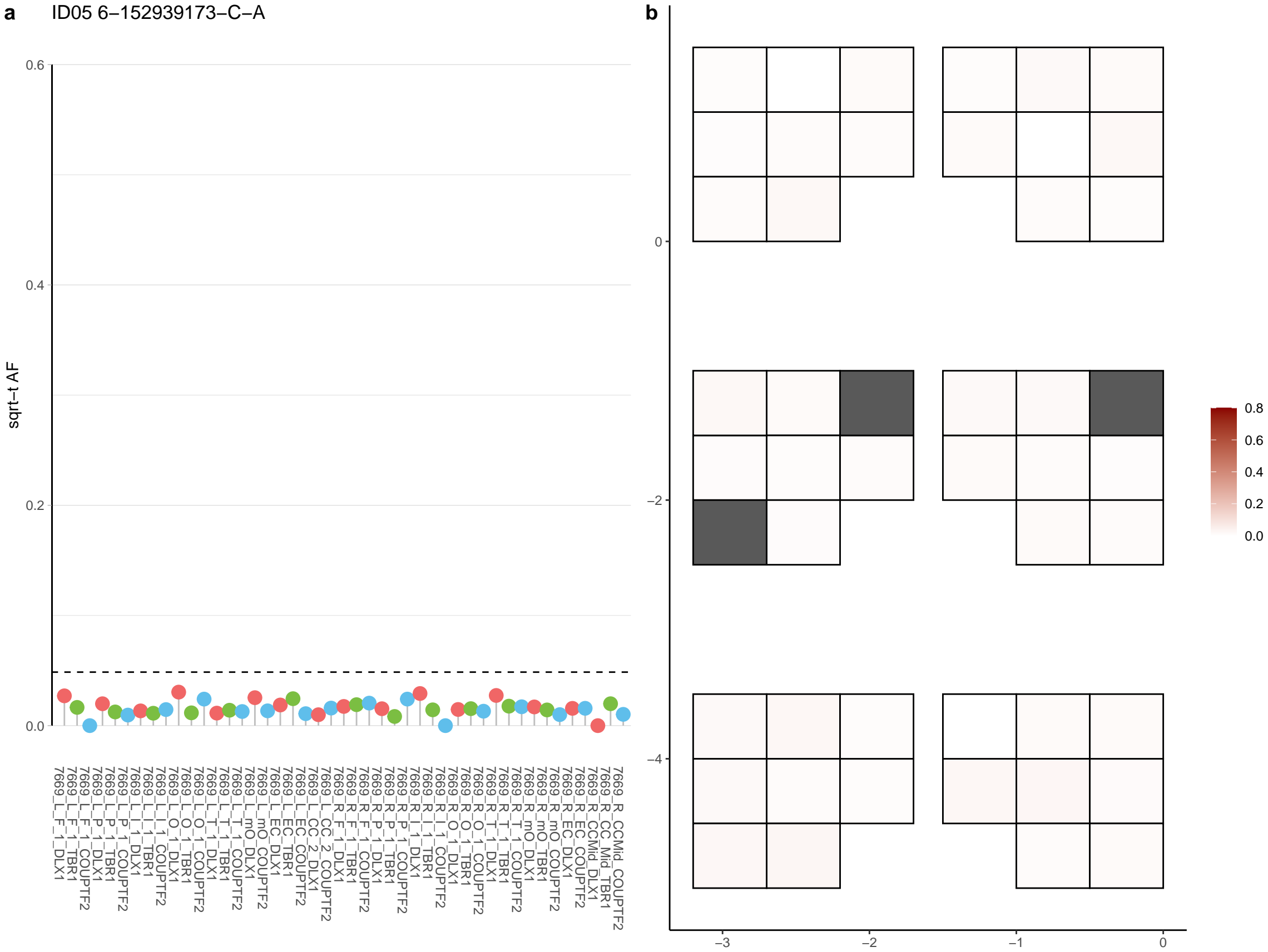

**a** ID05 6-153112948-G-T

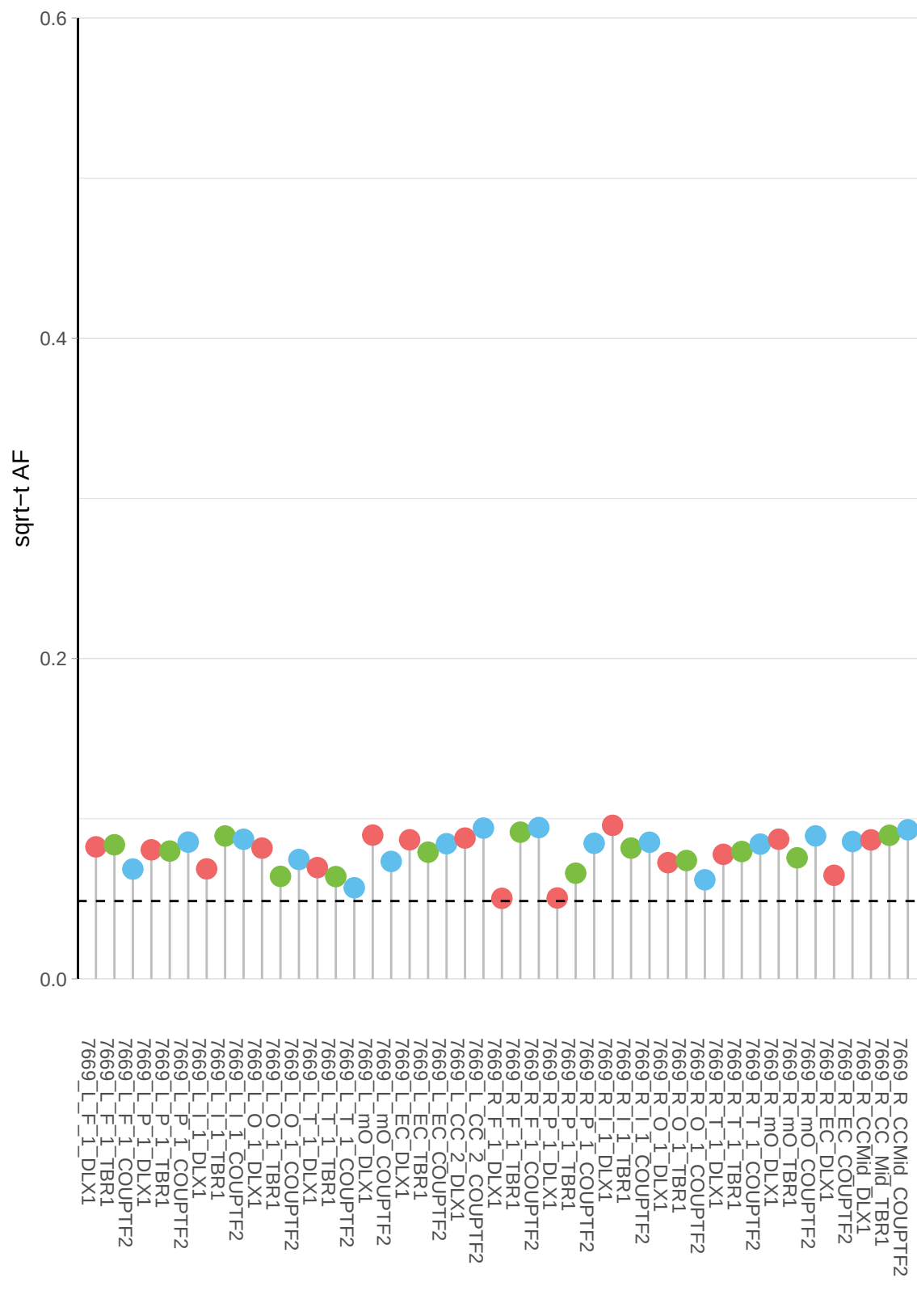

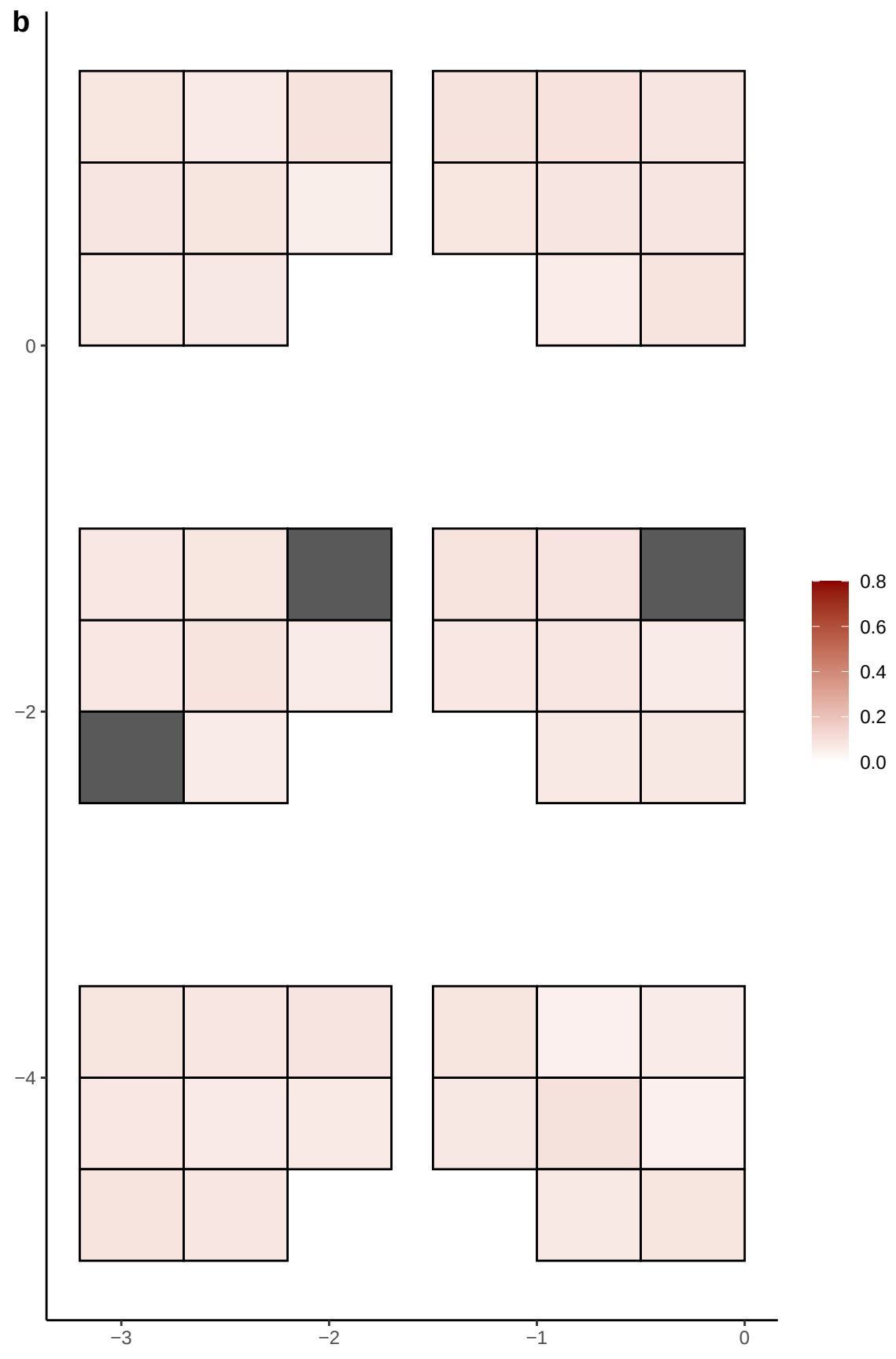

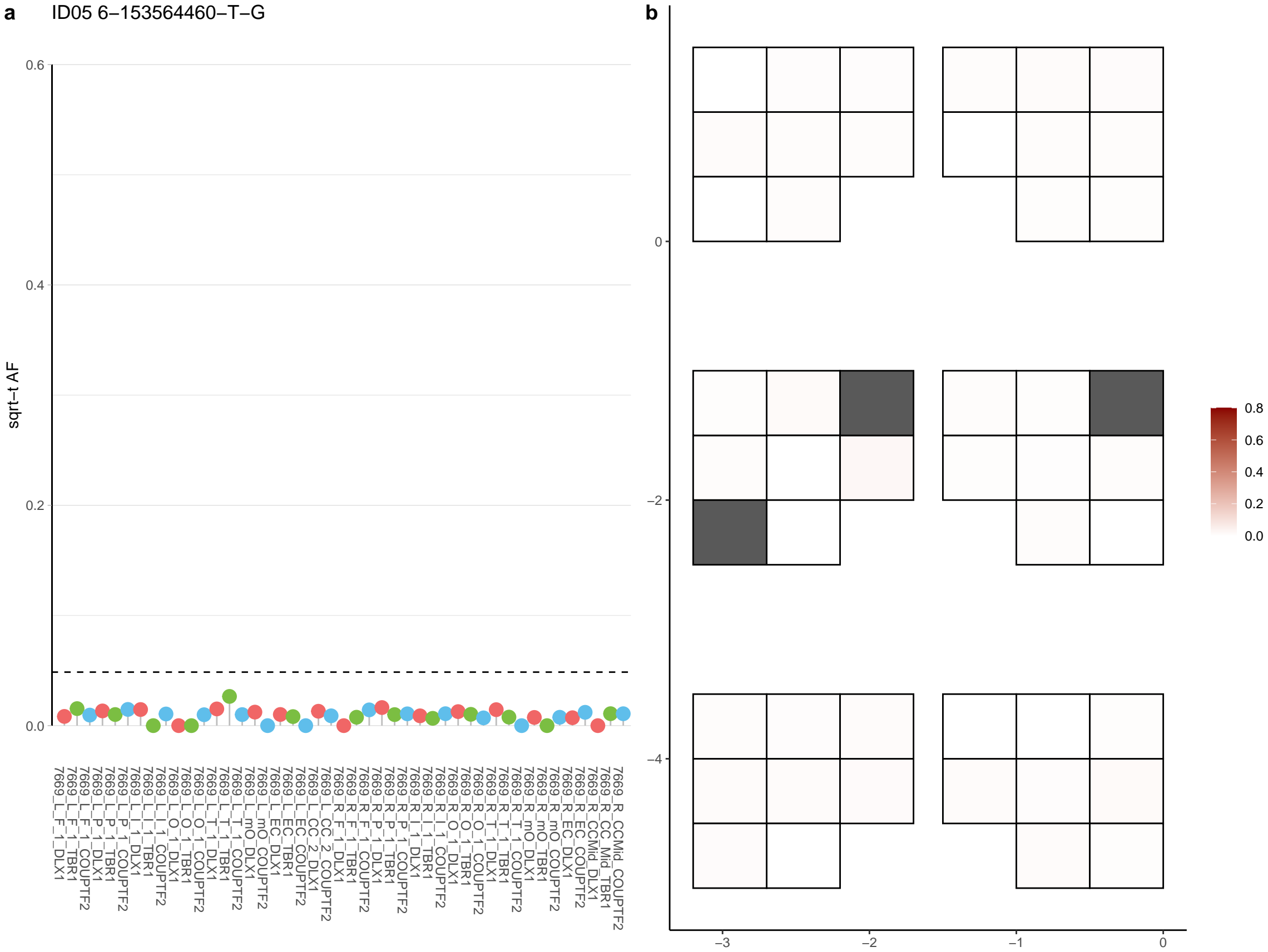

**a** ID05 6-155986031-C-T

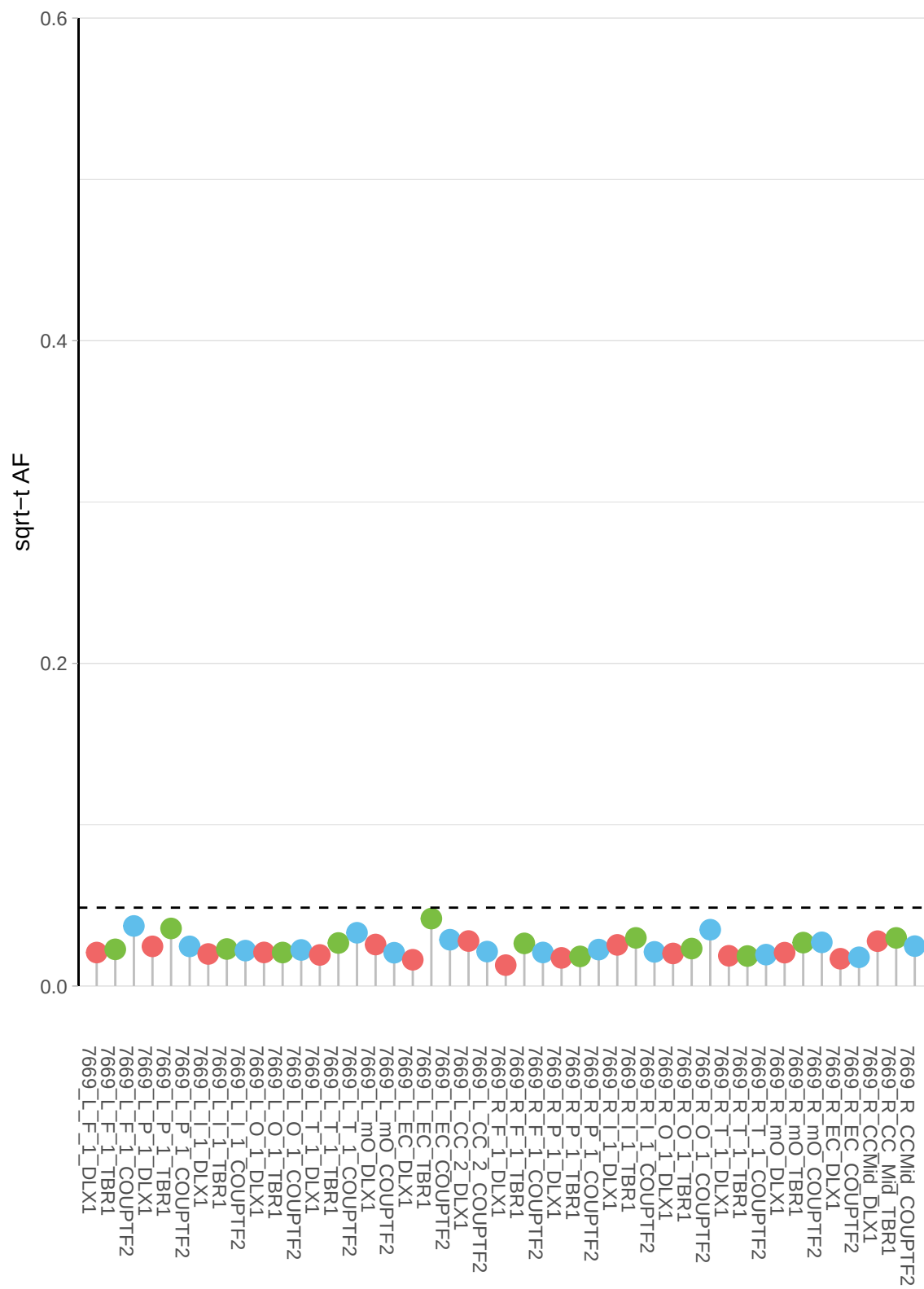

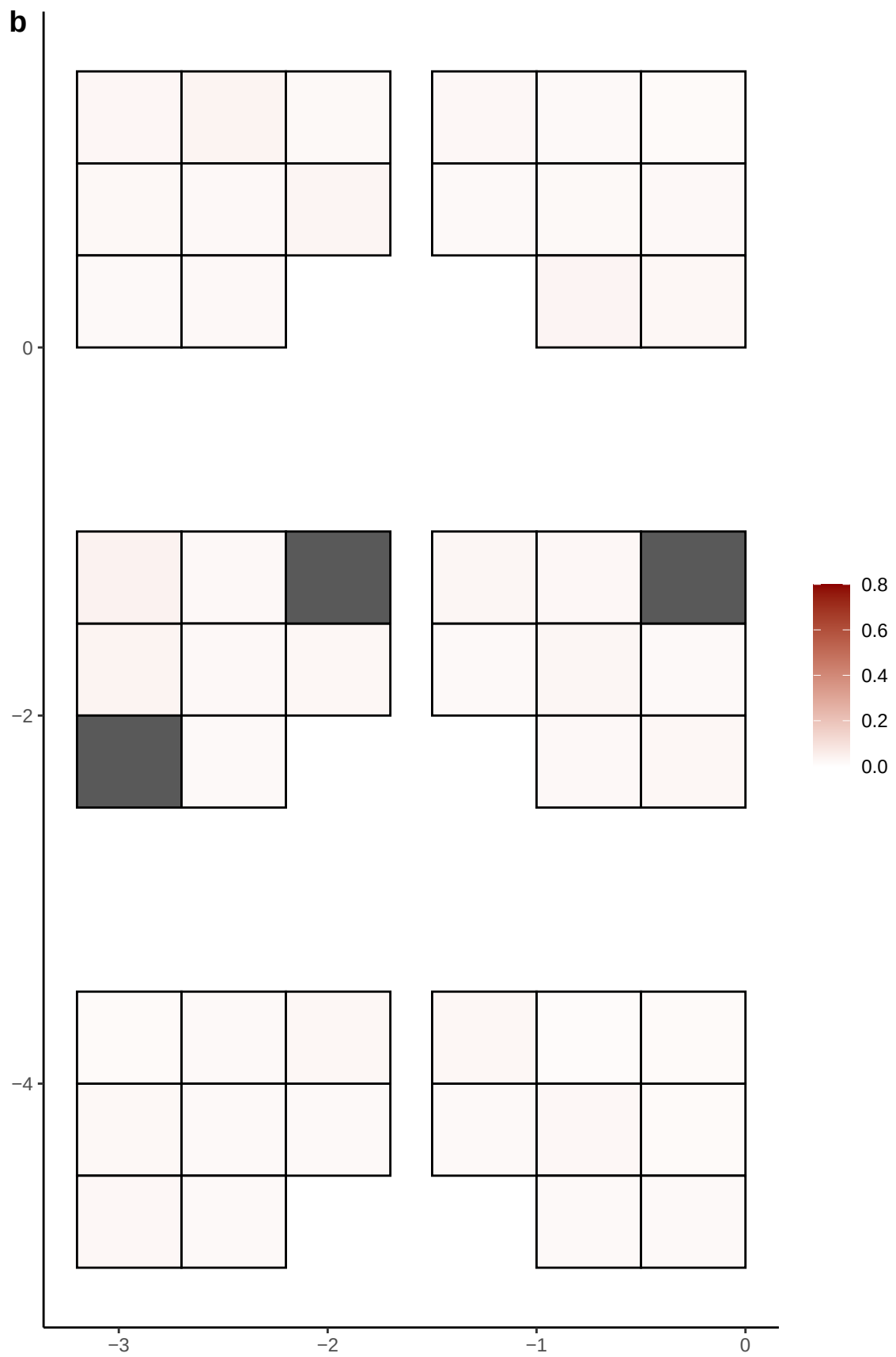

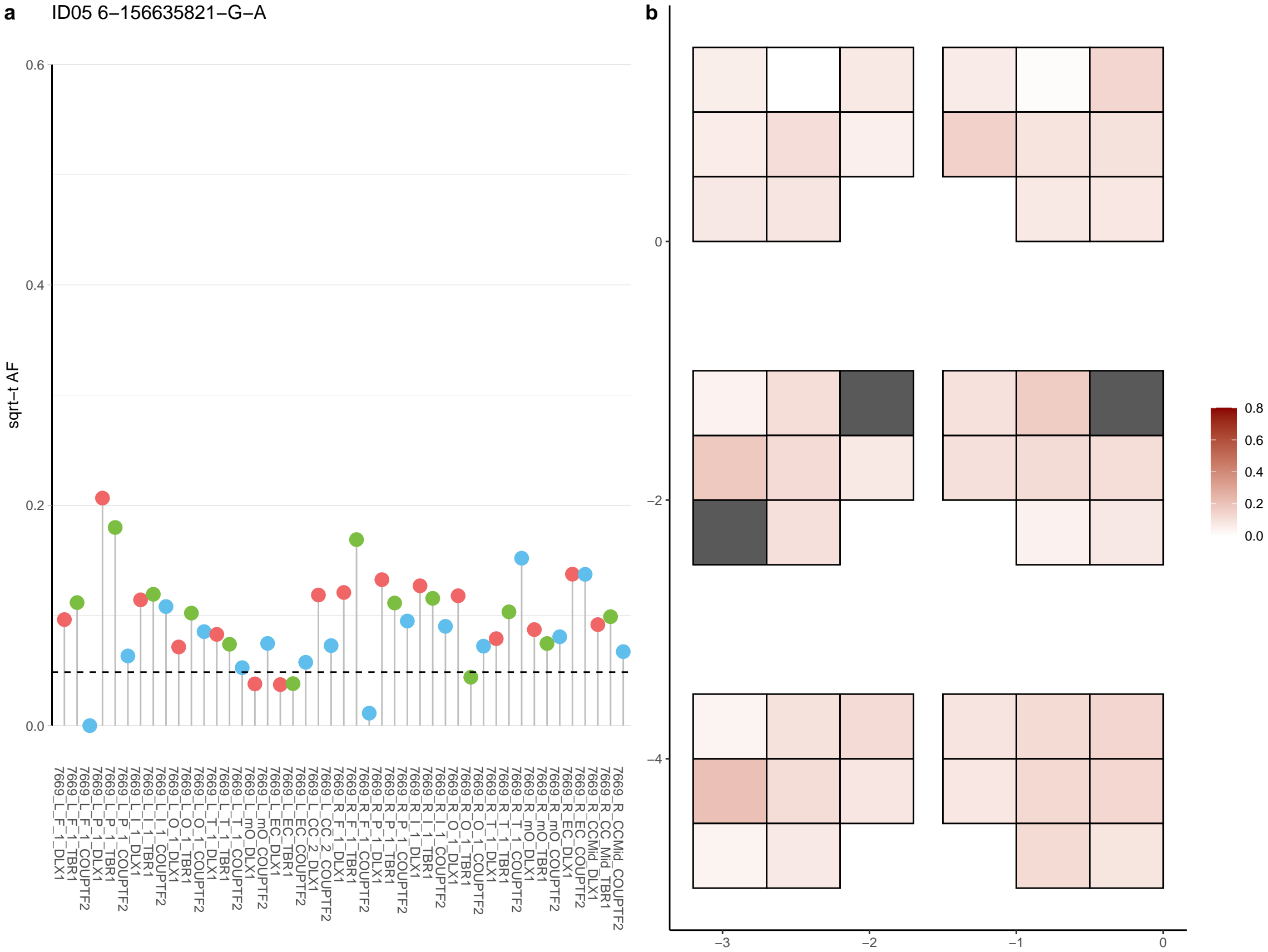

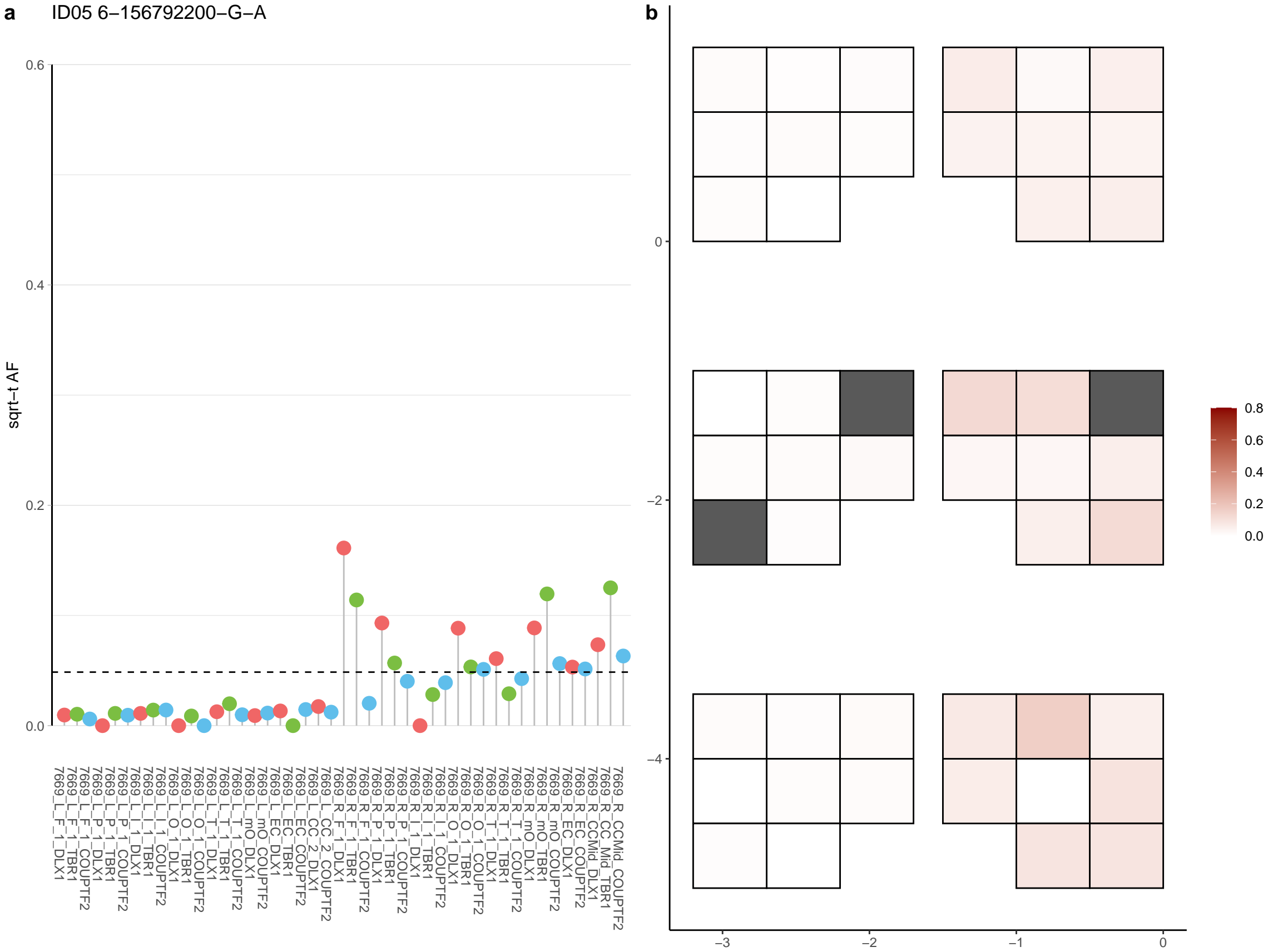

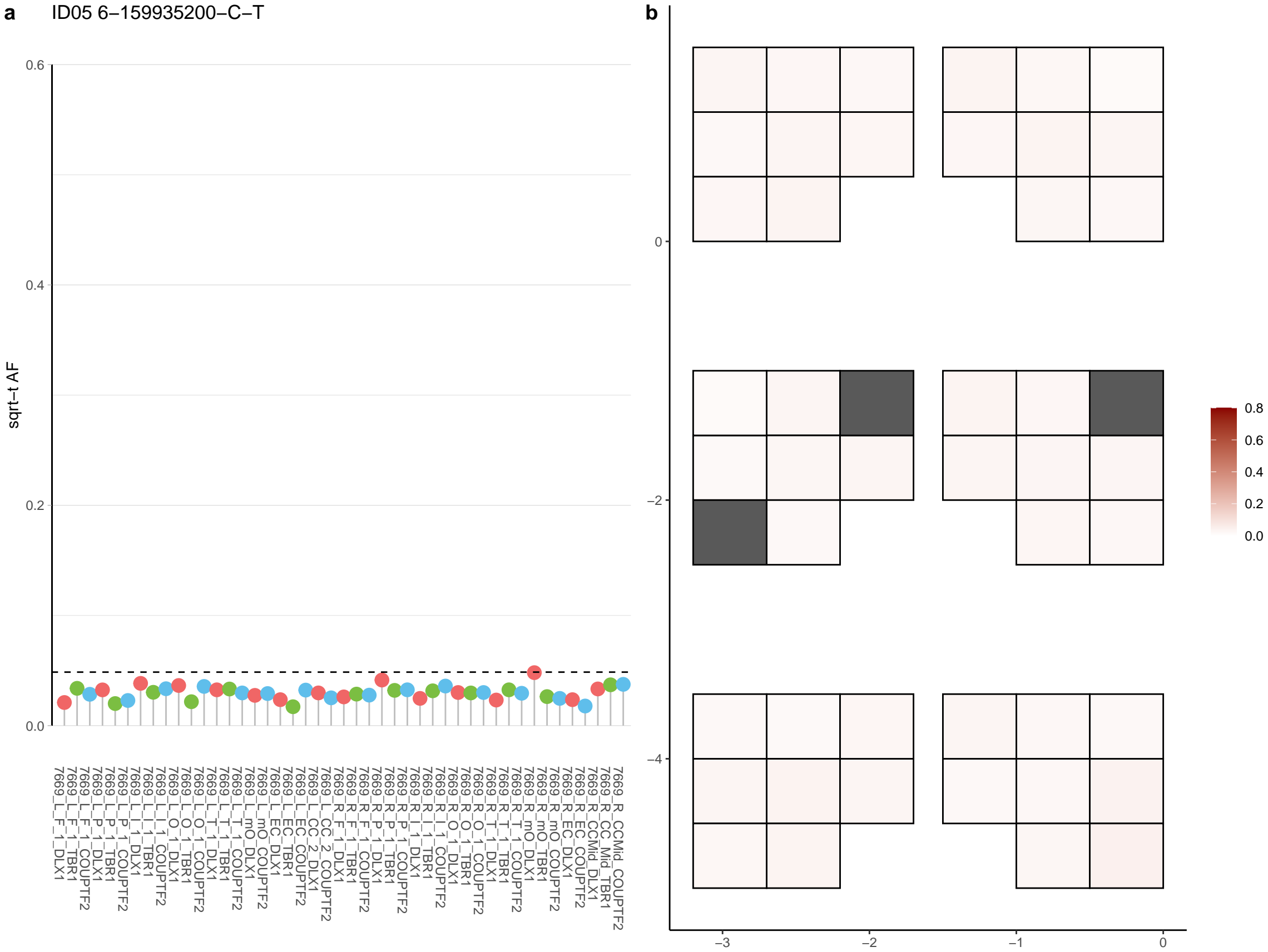

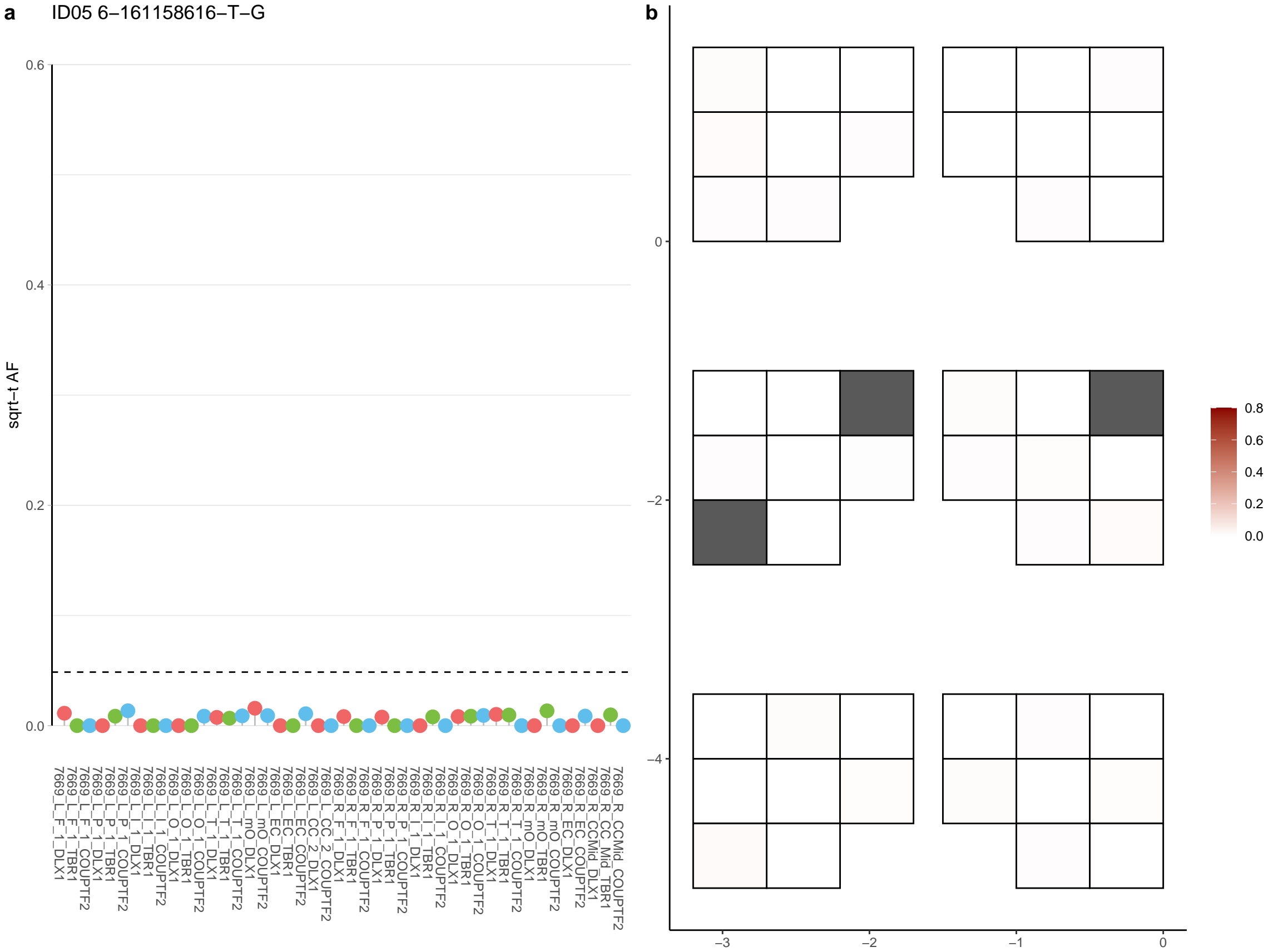

**a** ID05 6-162806617-C-T

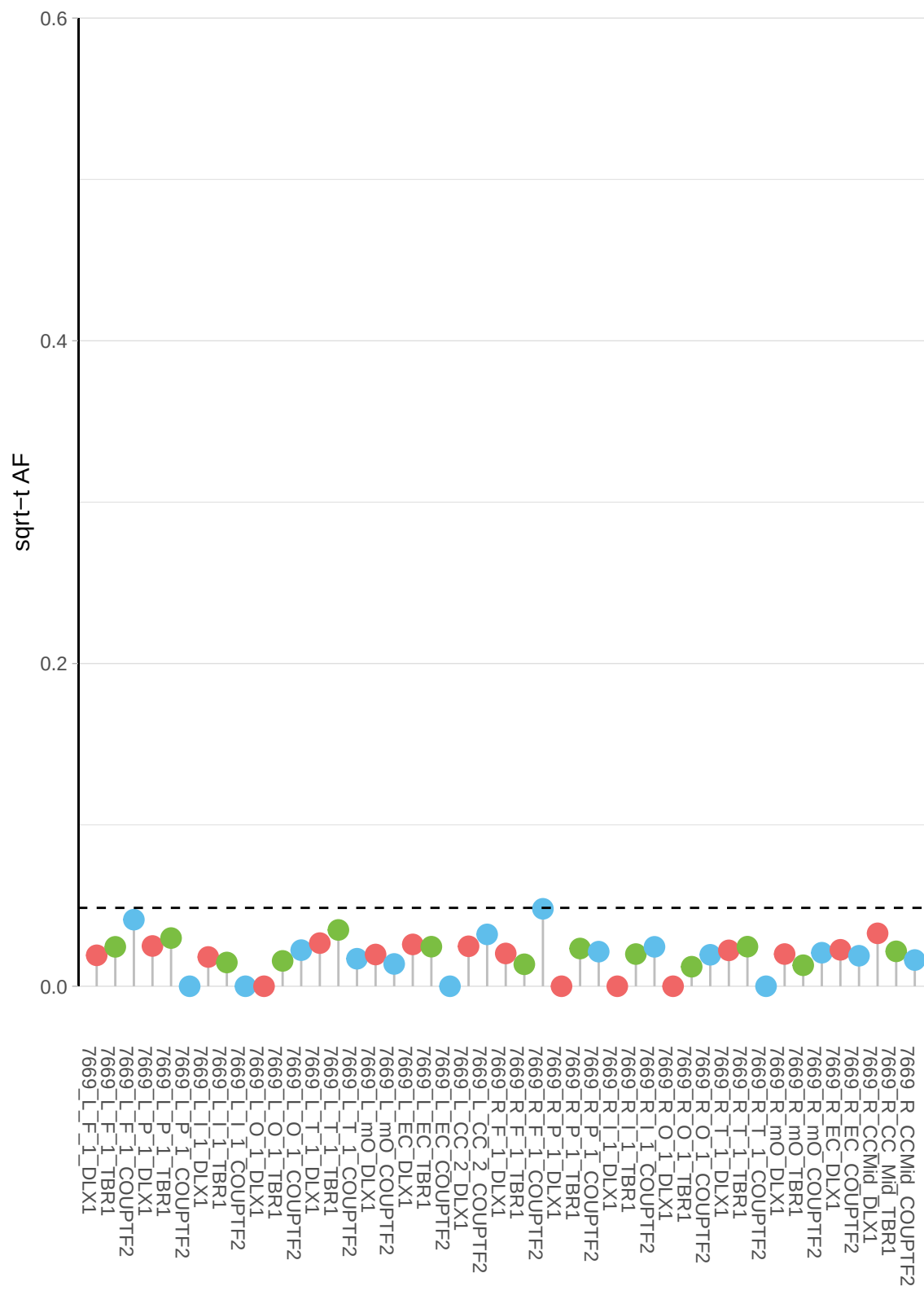

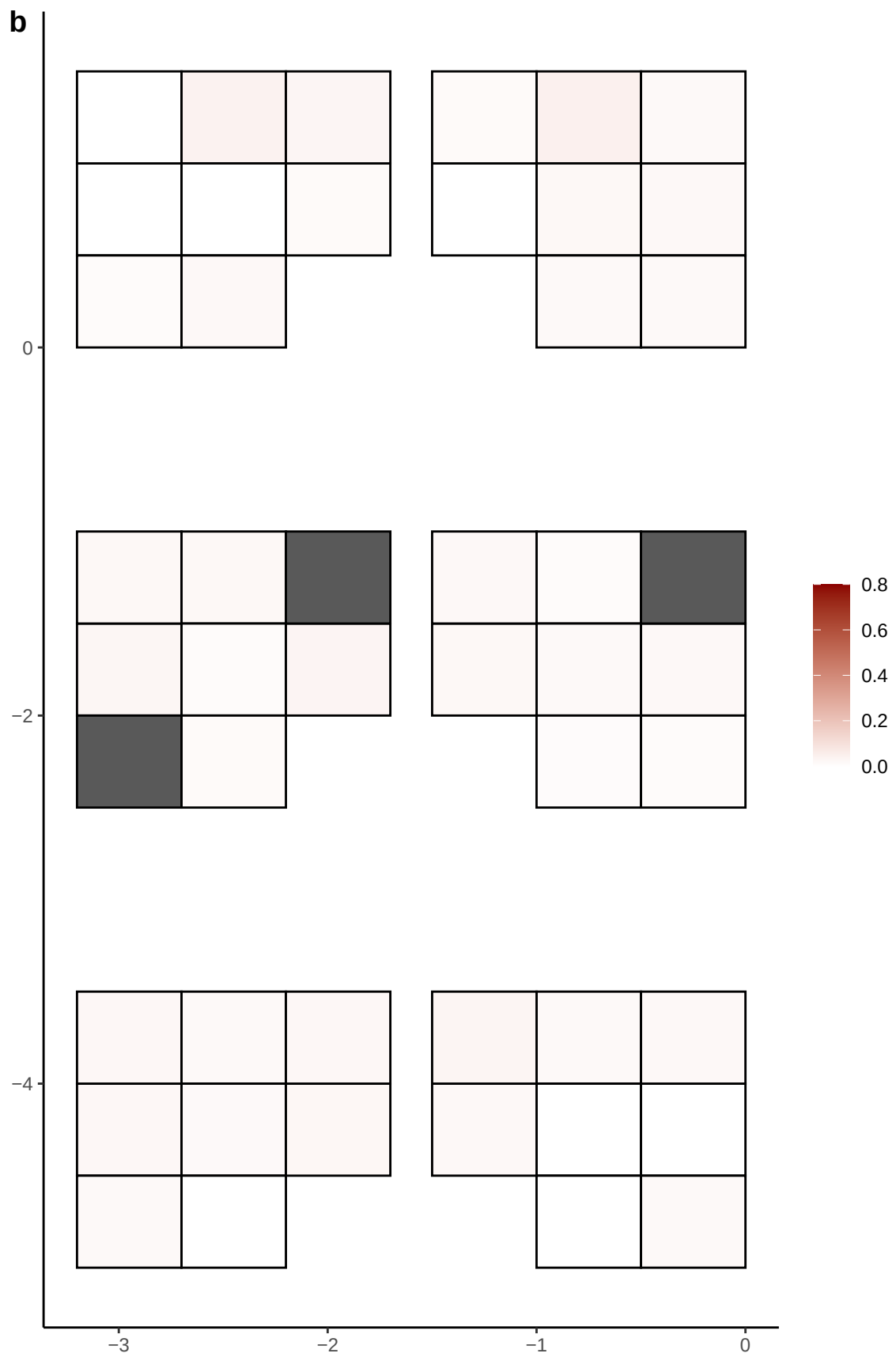

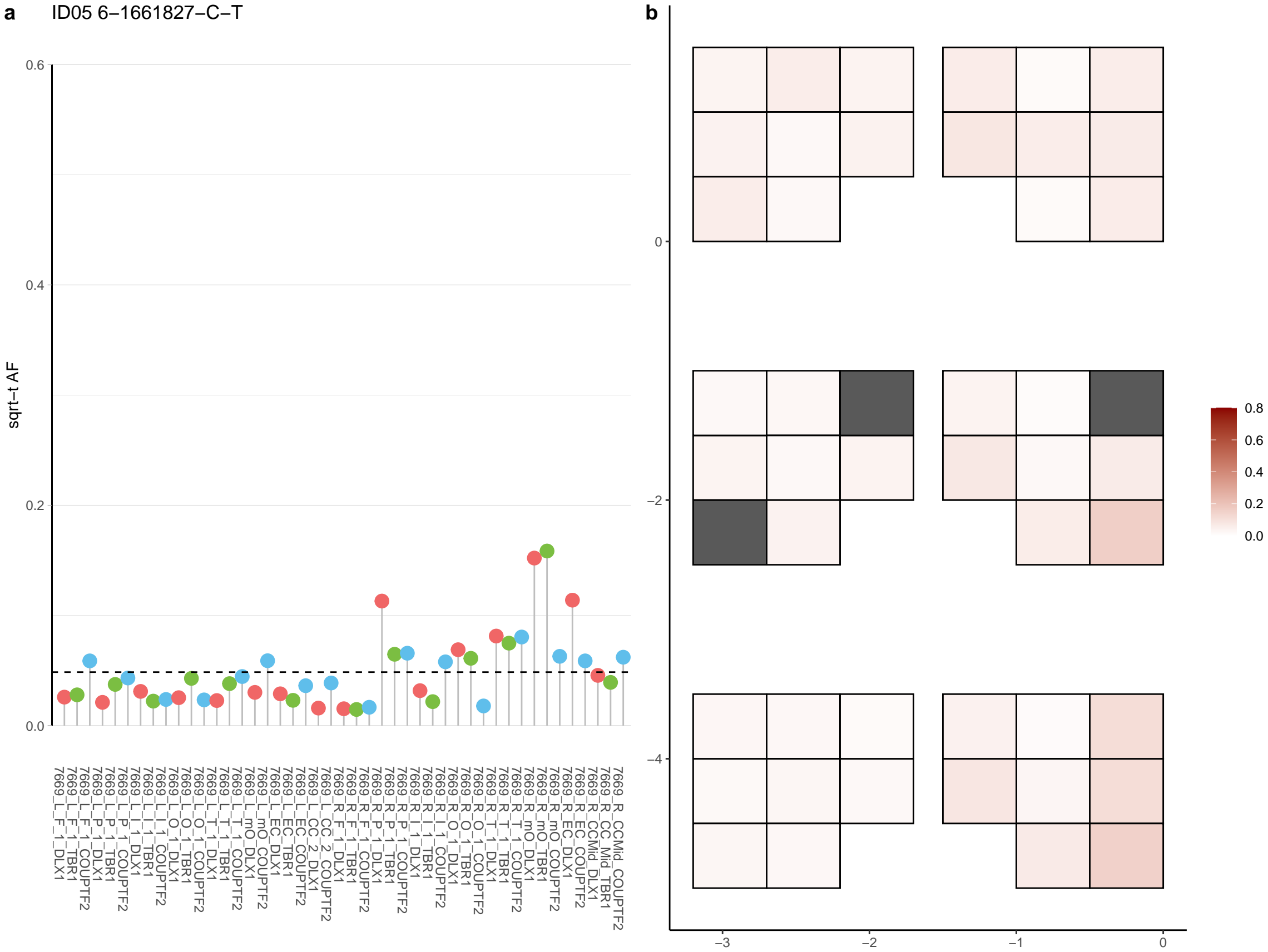

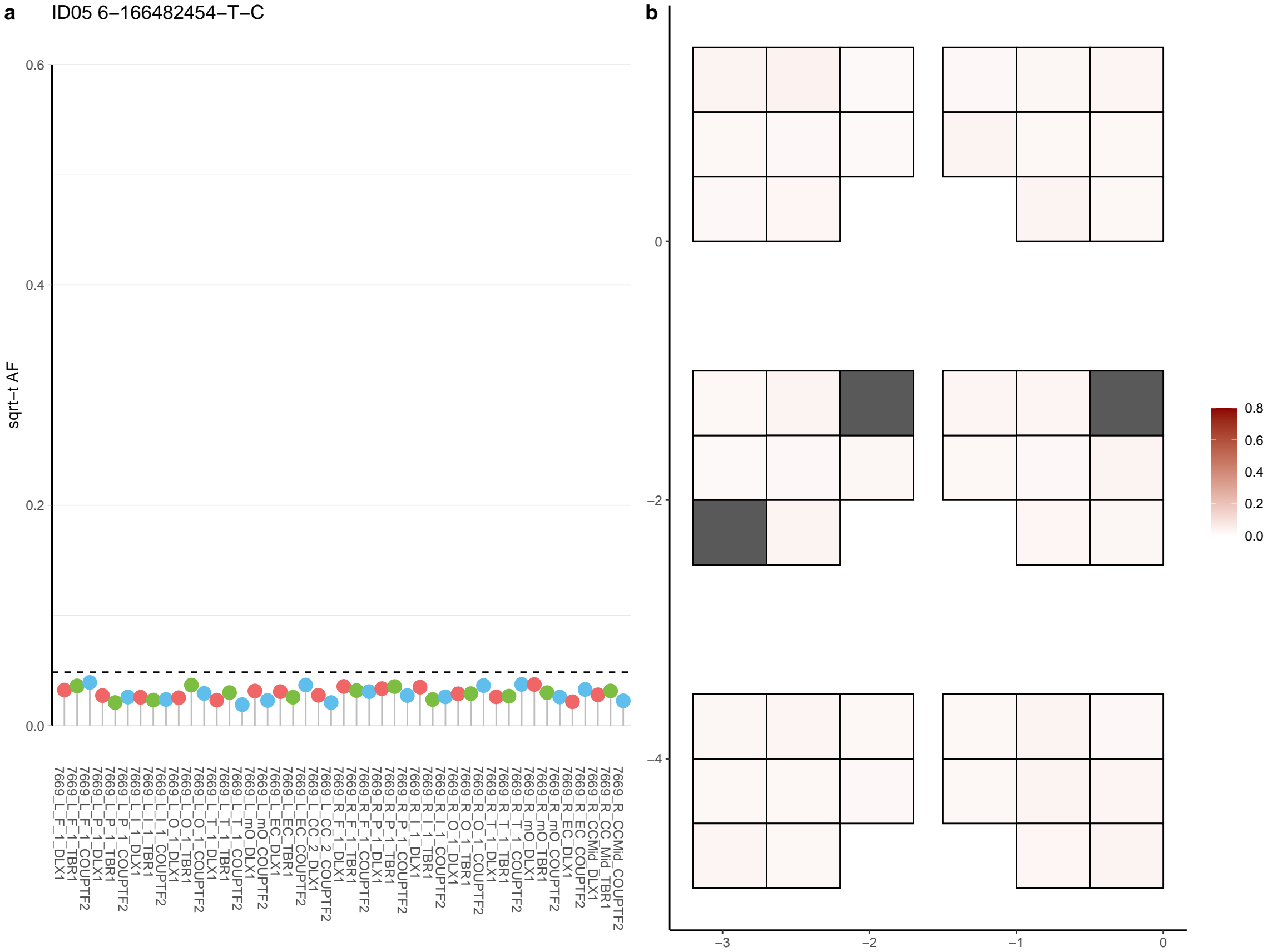

**a** ID05 6-169577791-G-A

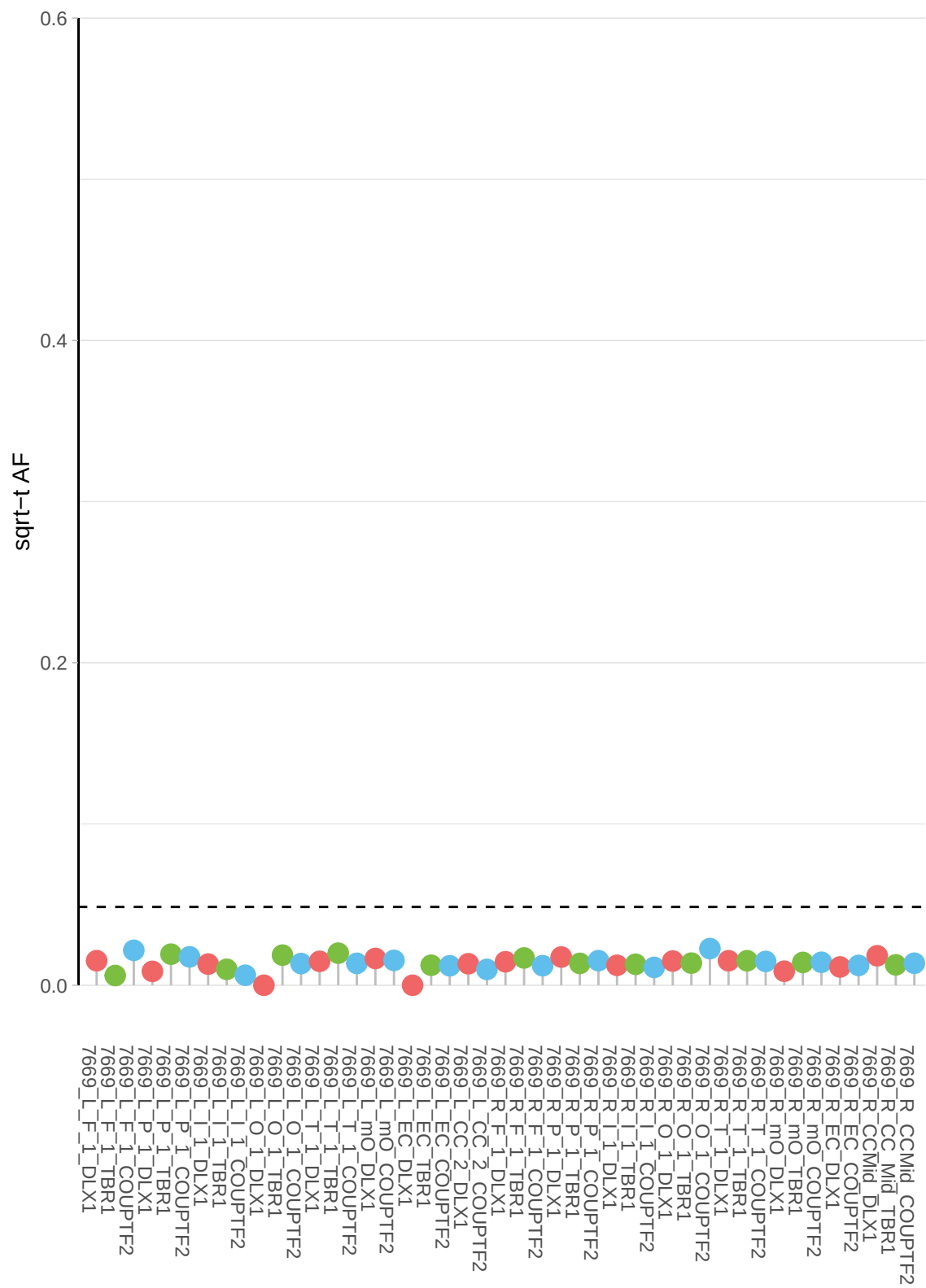

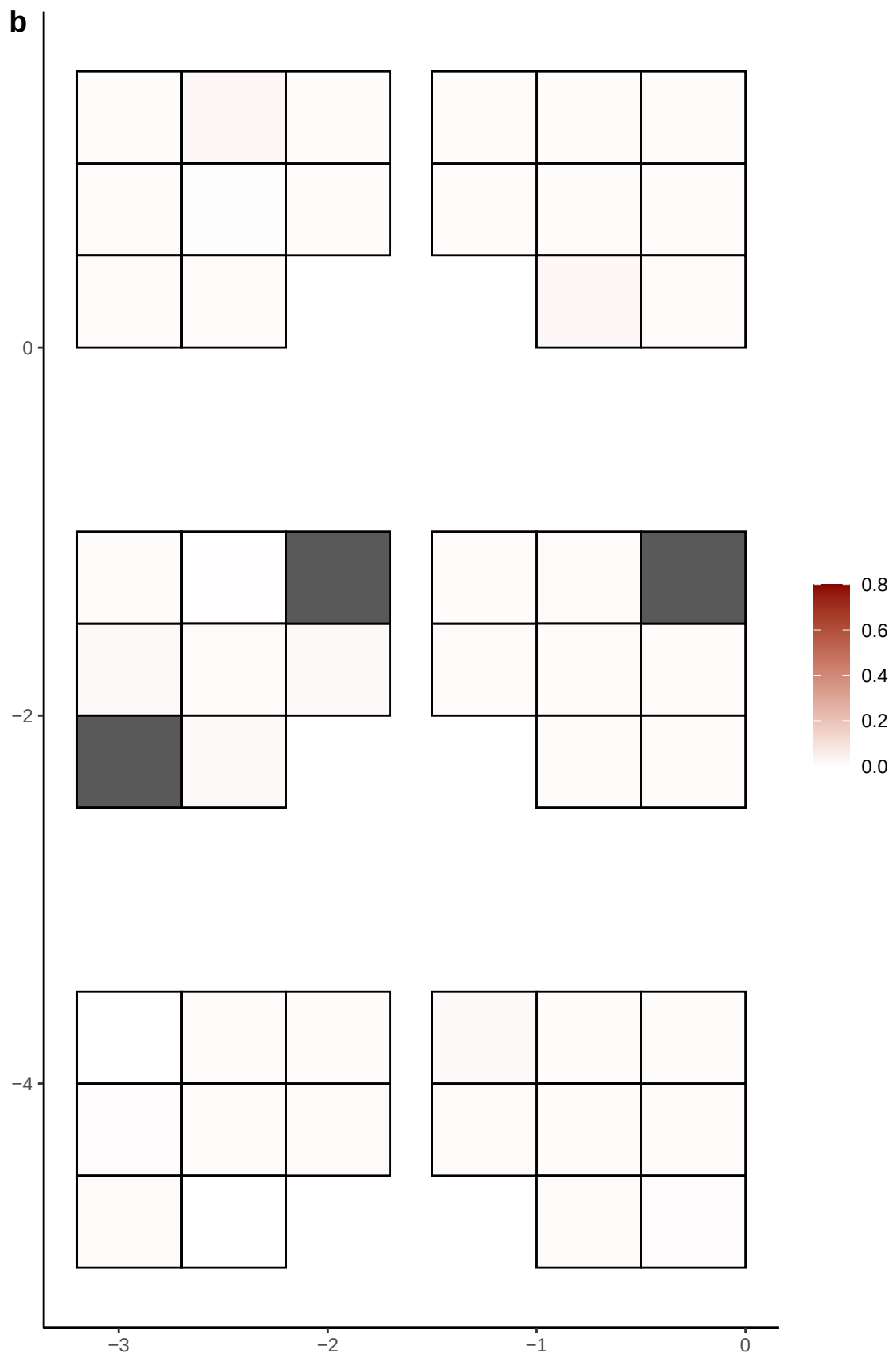

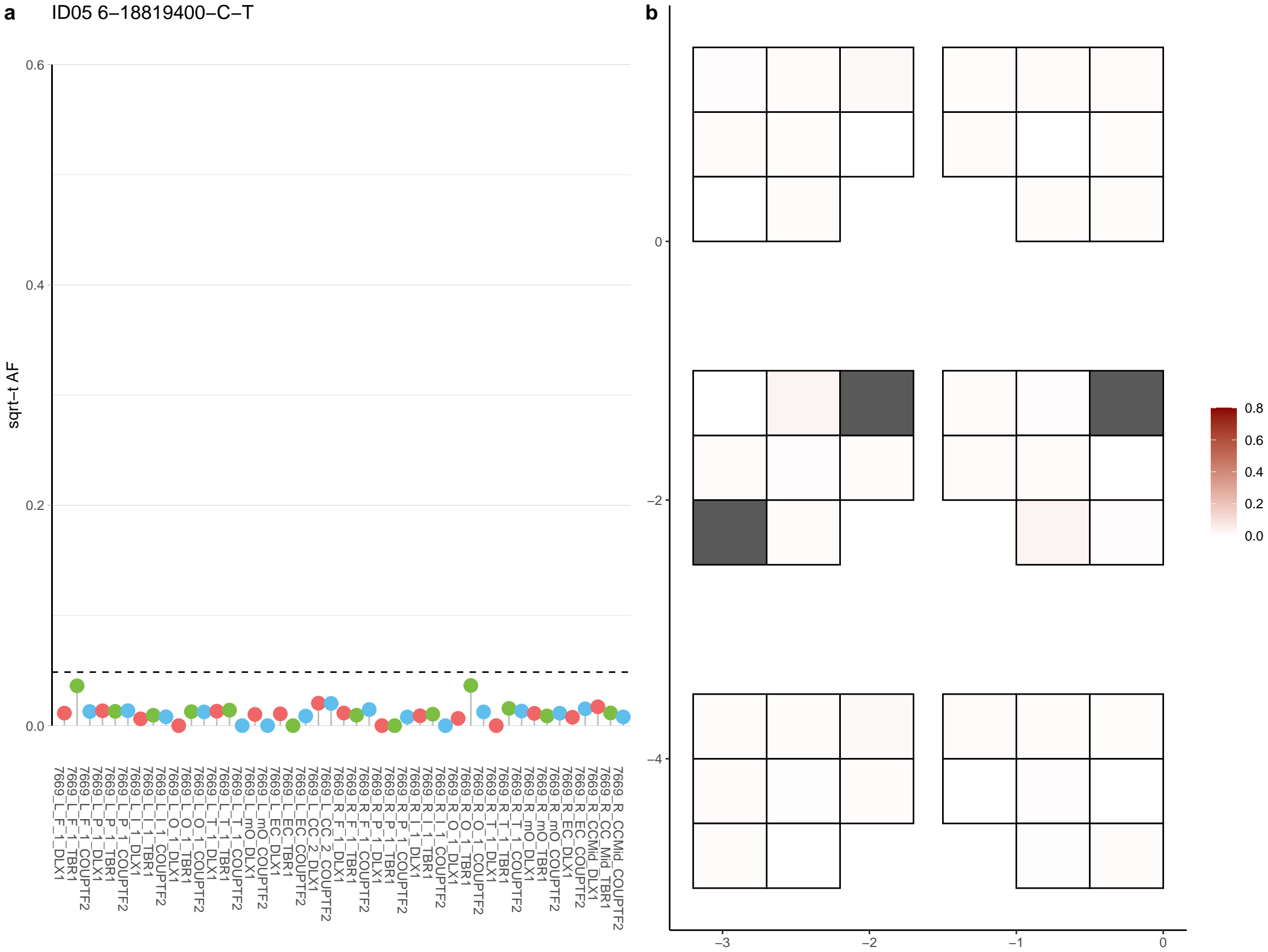

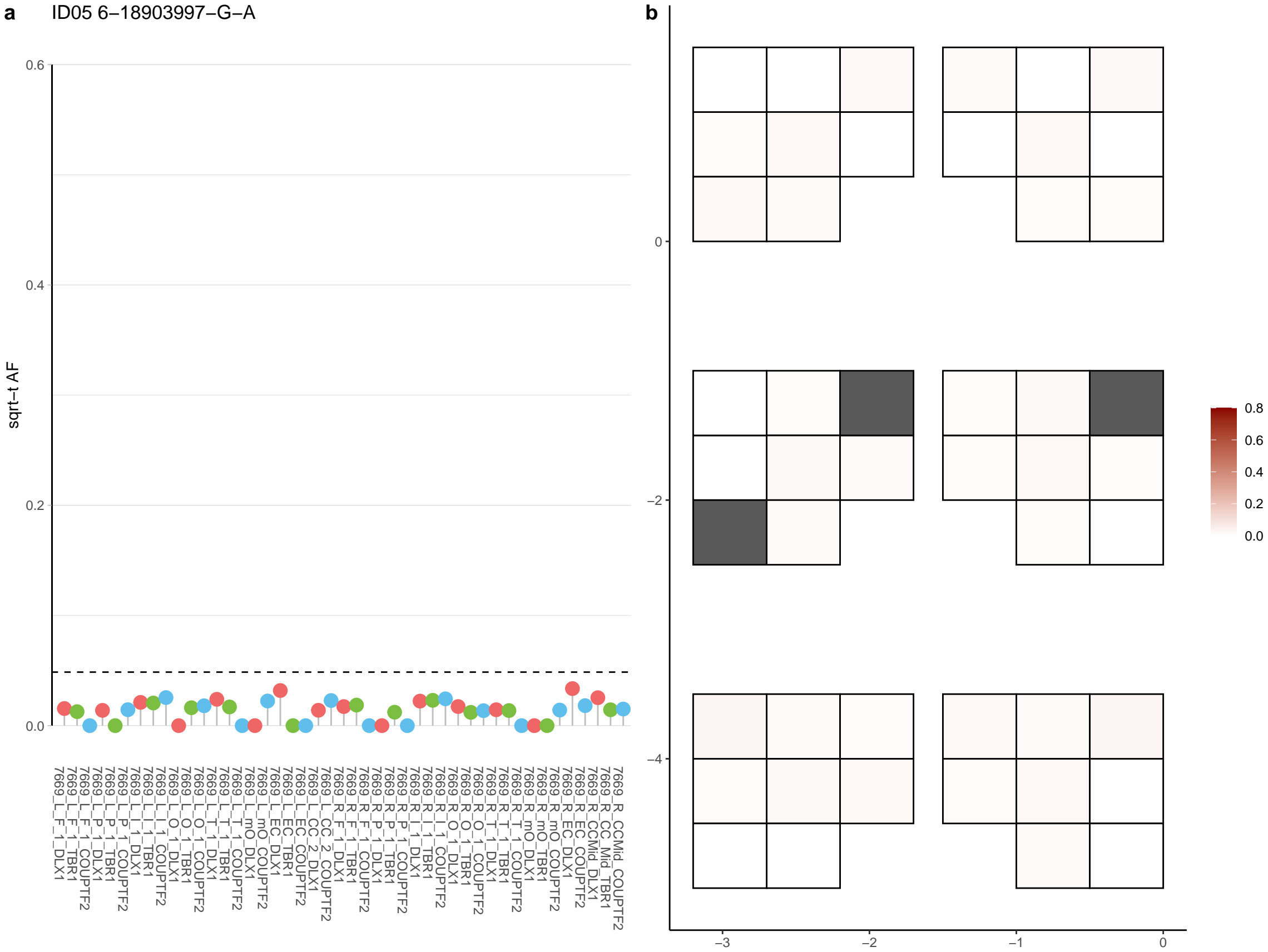

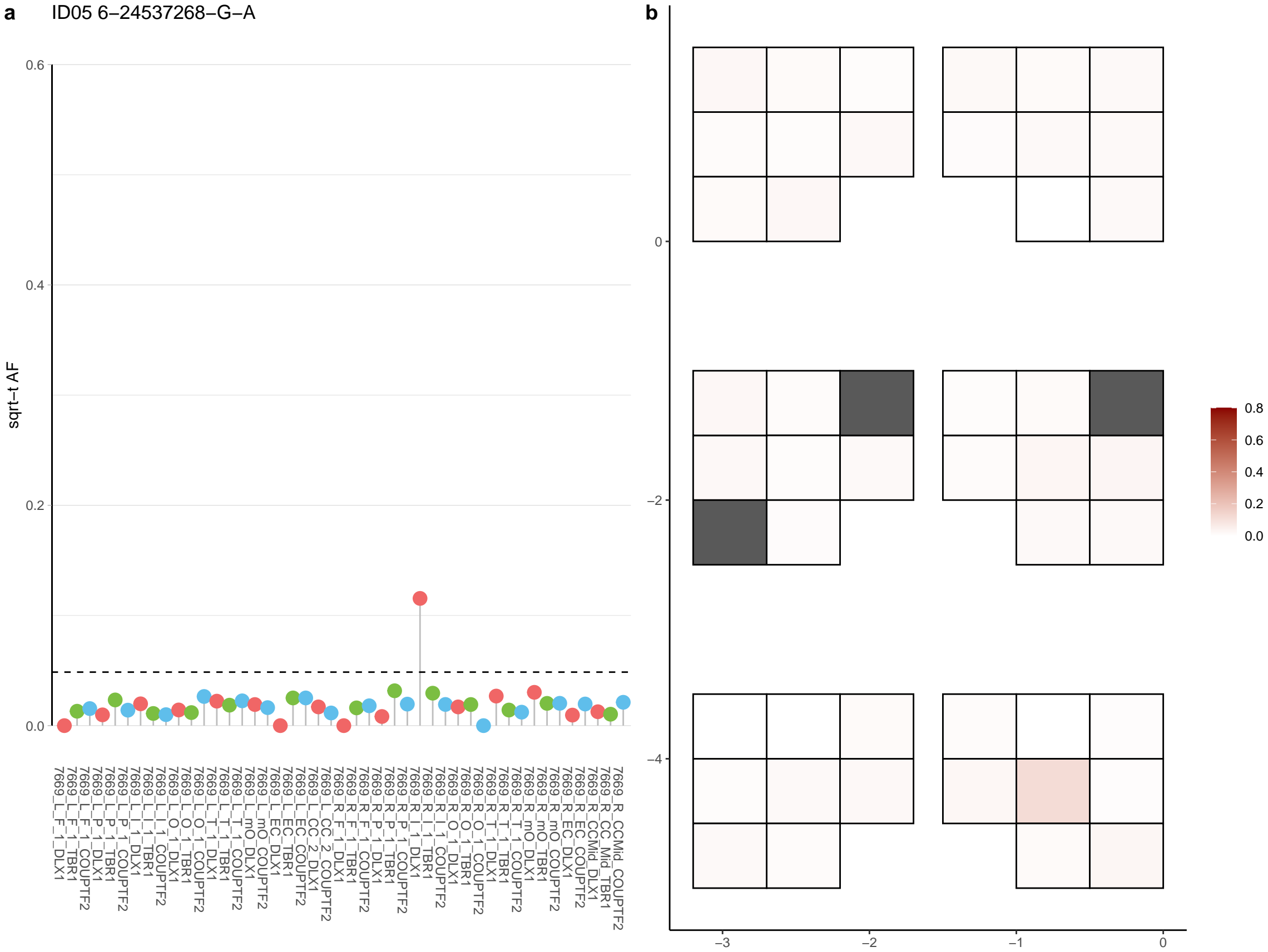

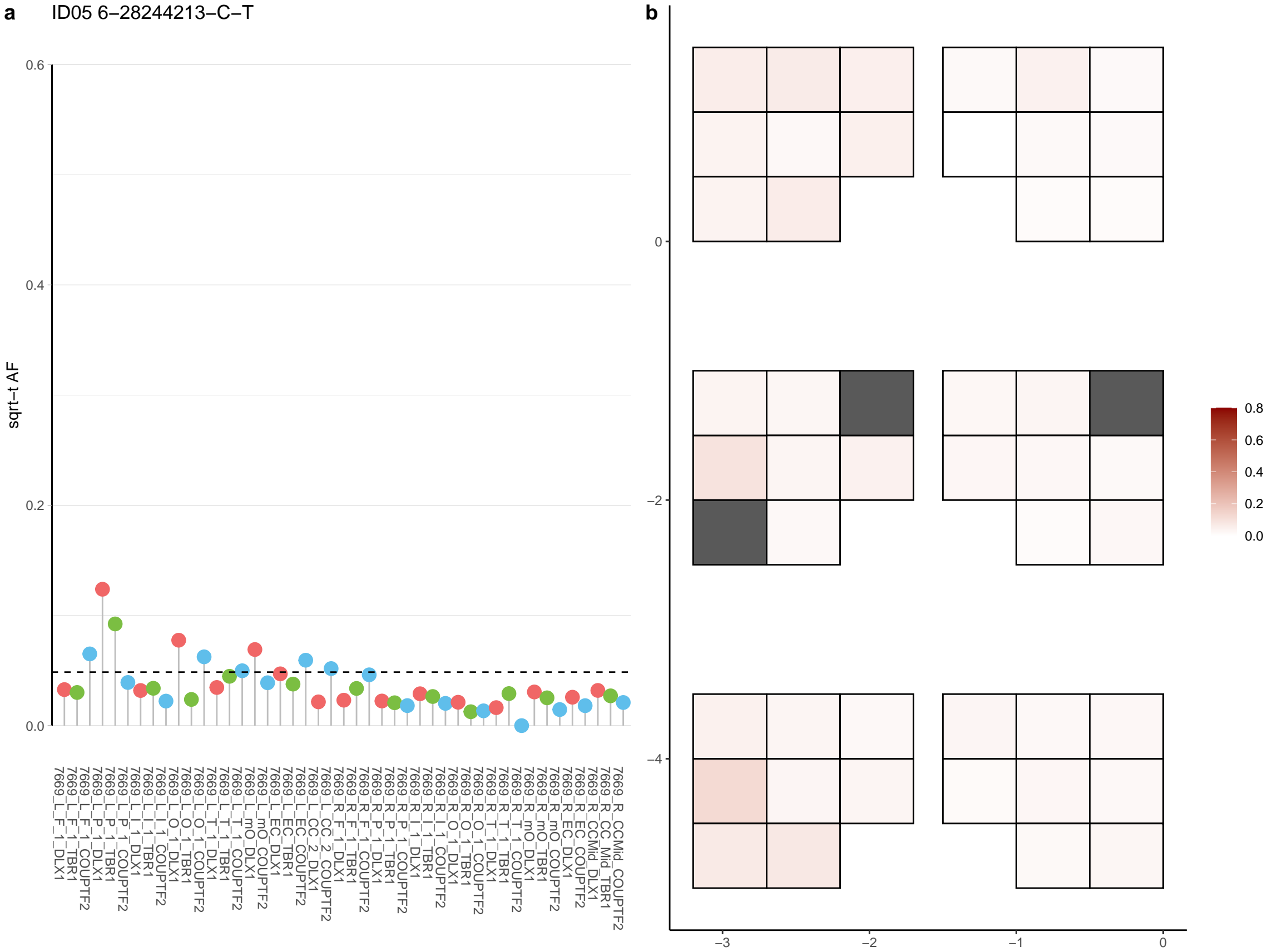

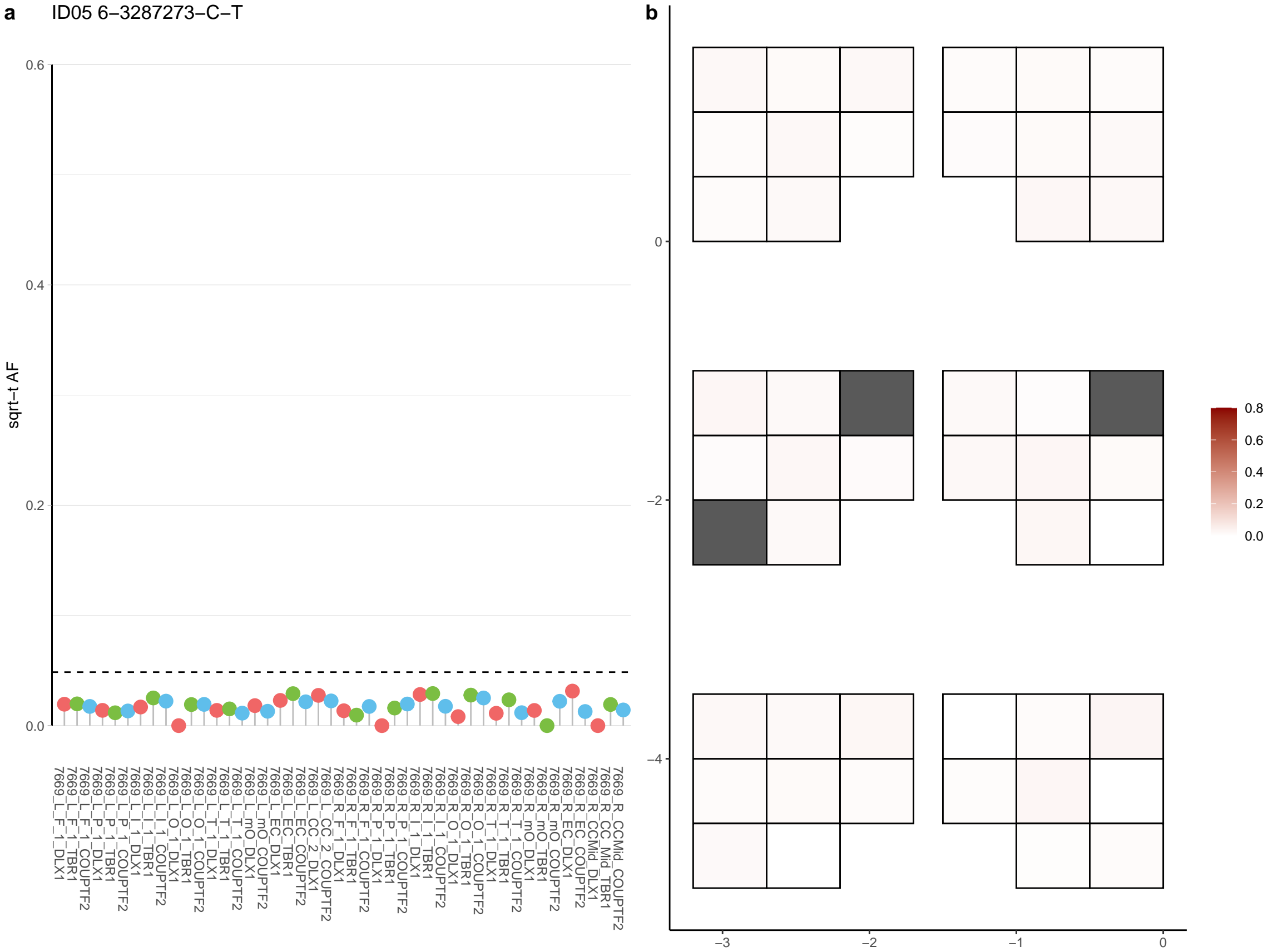

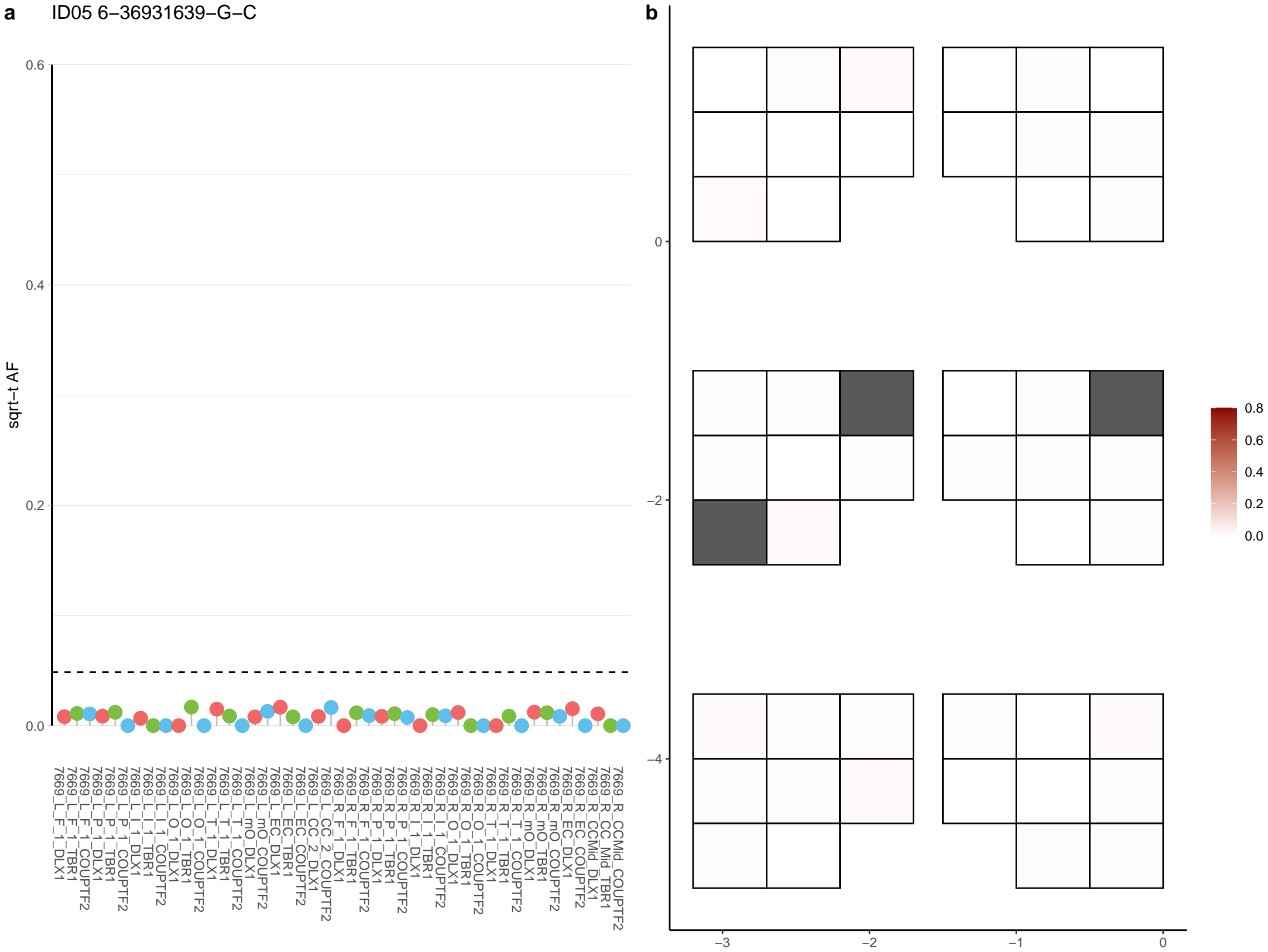

**a** ID05 6-41864337-C-T

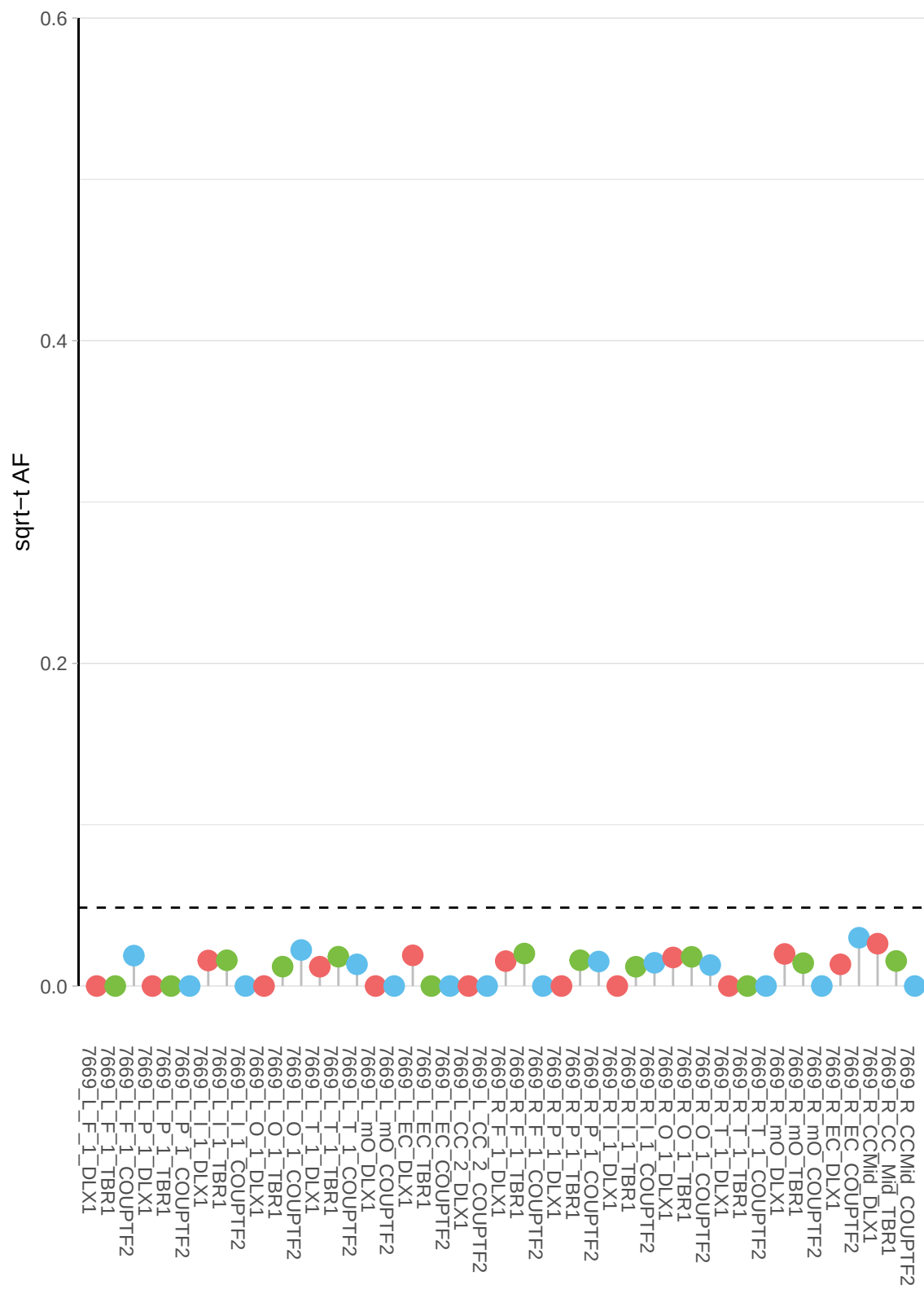

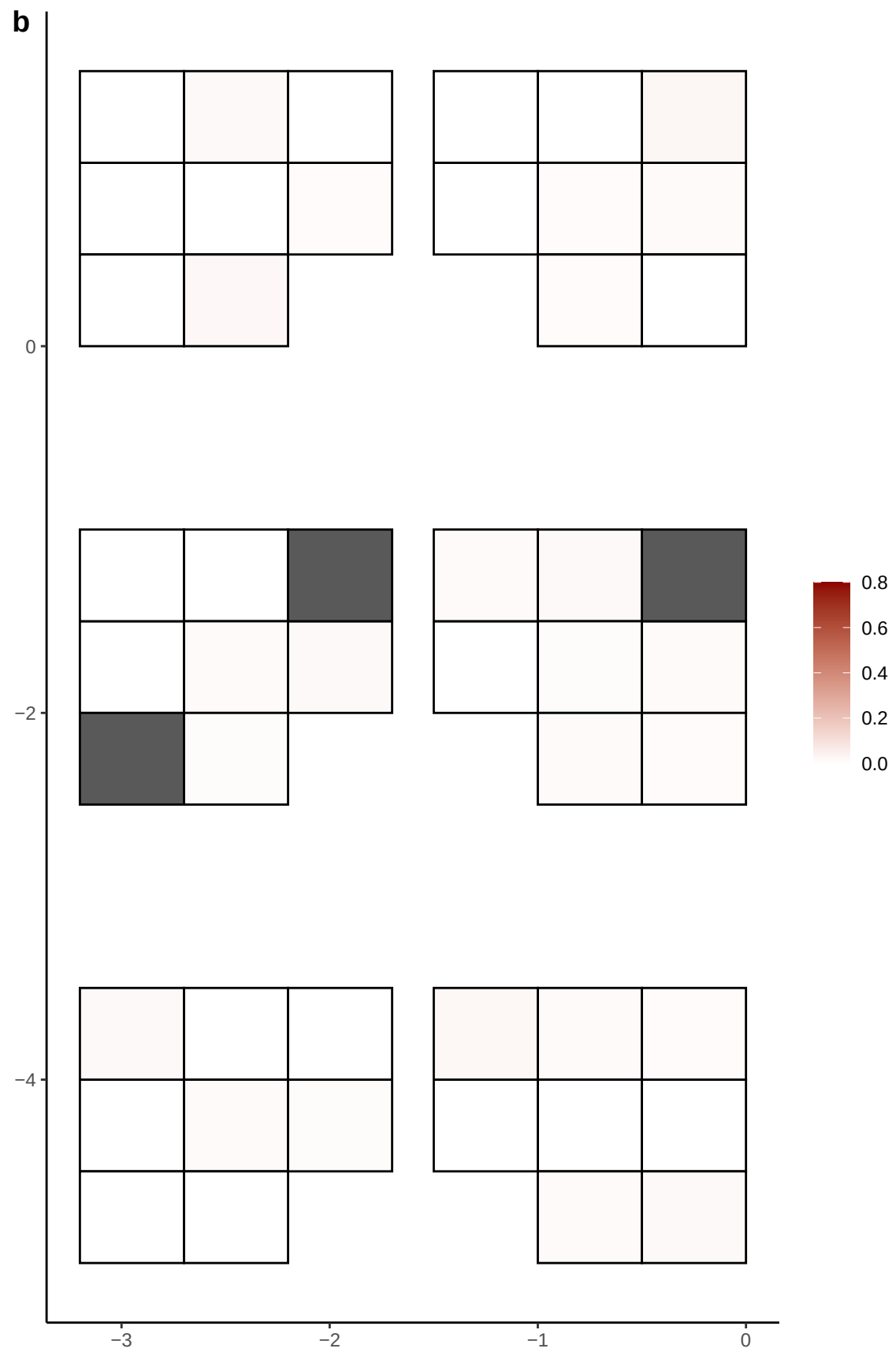

**a** ID05 6-47367037-G-A

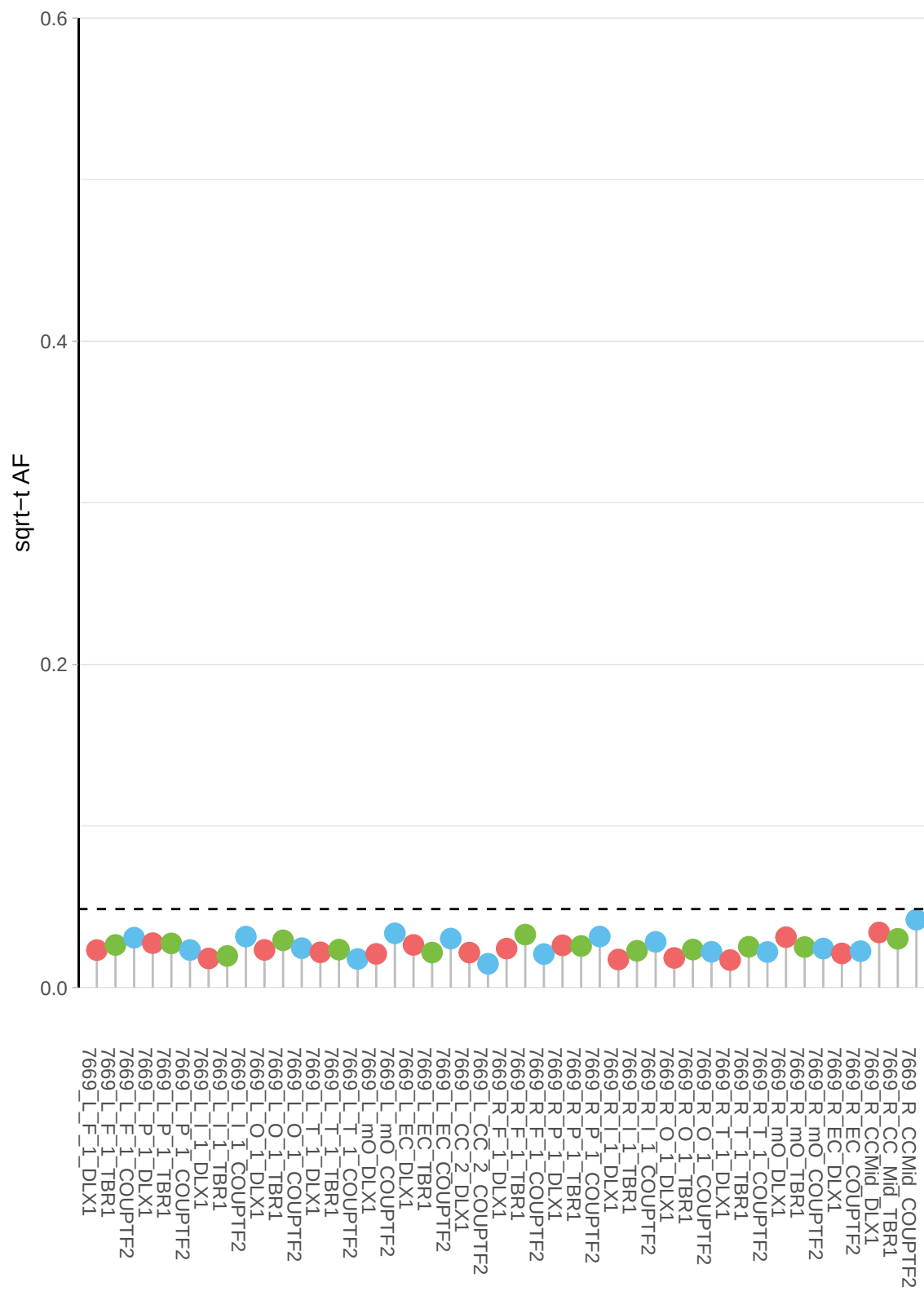

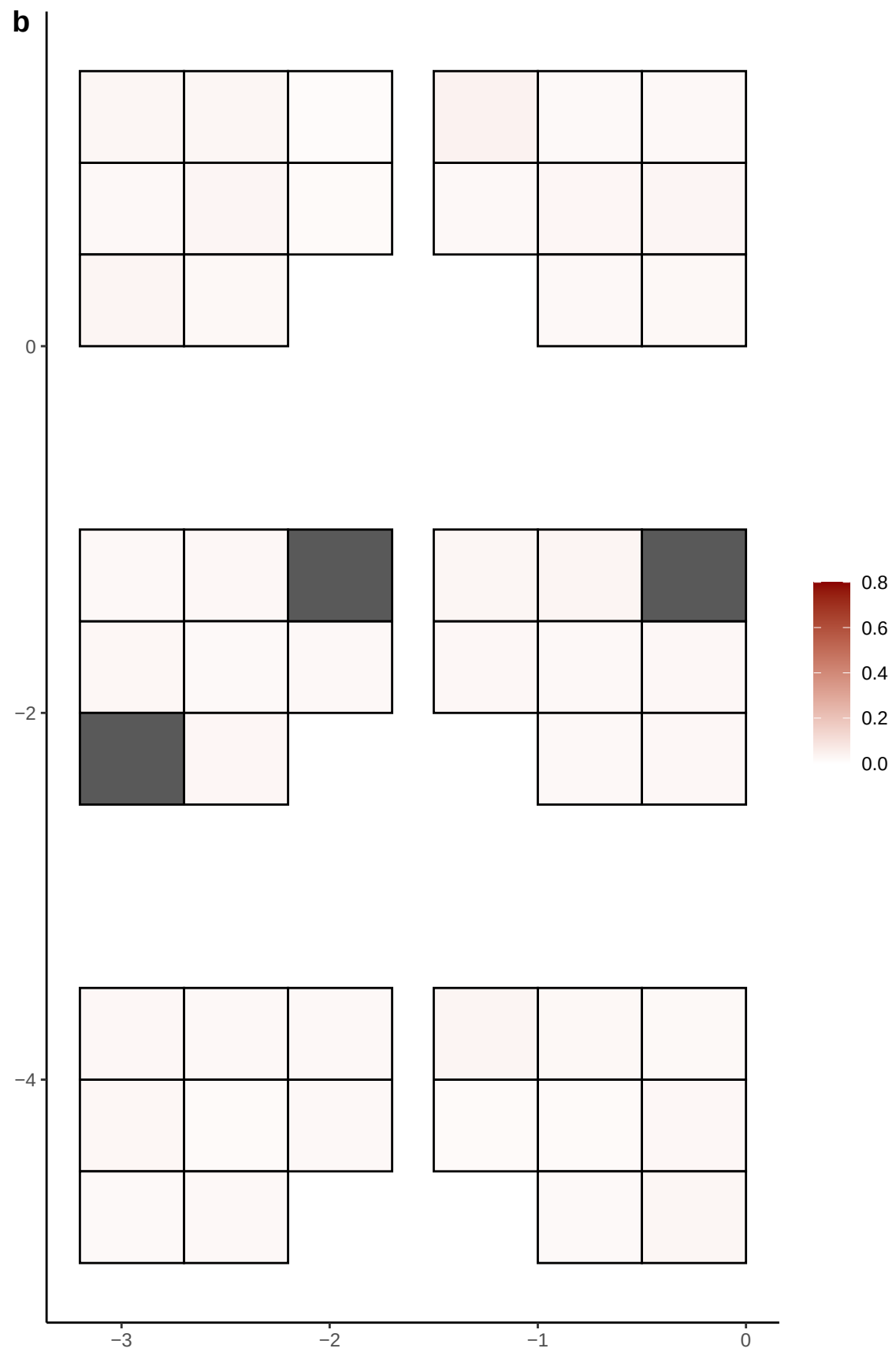

**a** ID05 6-49937325-C-G

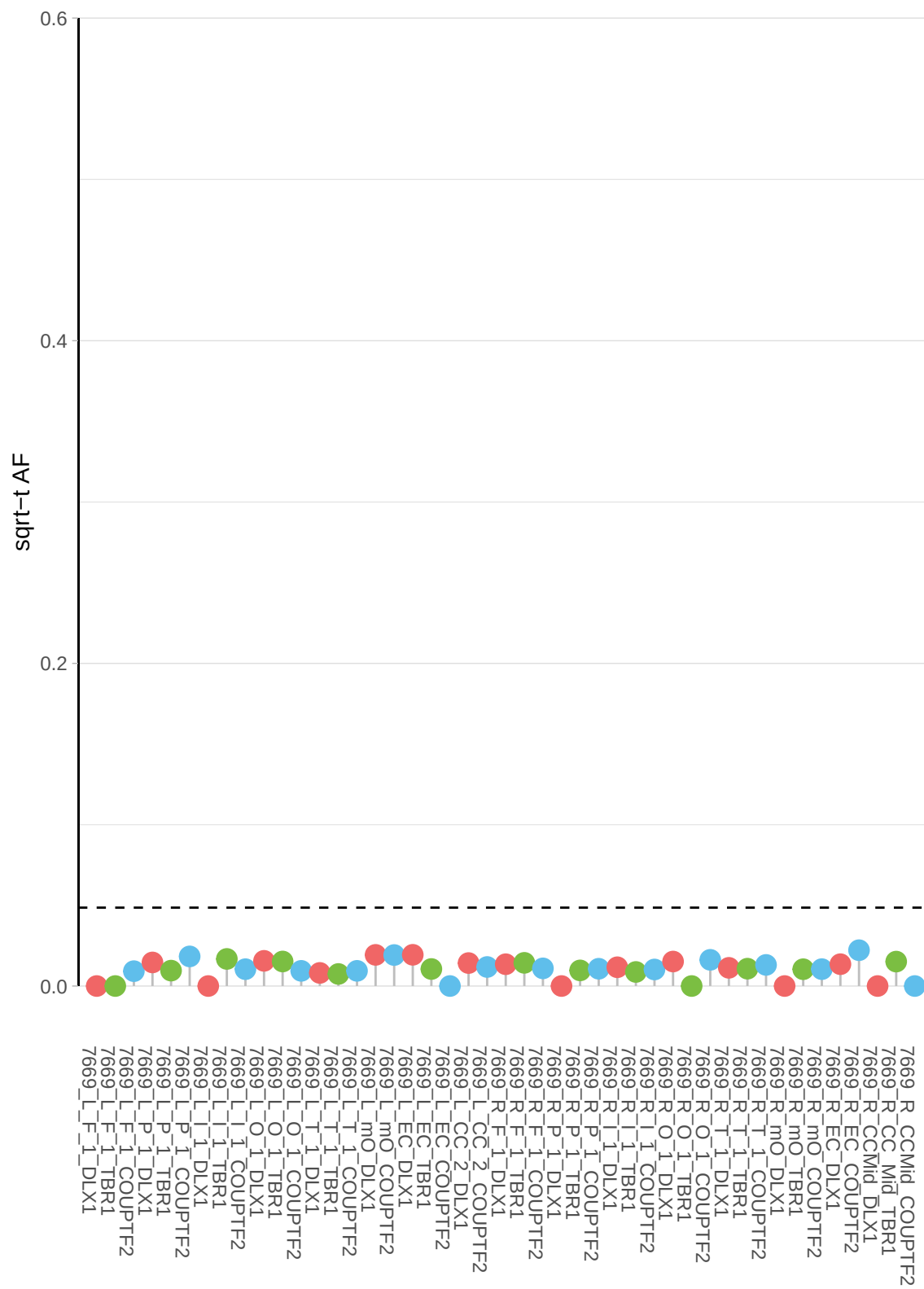

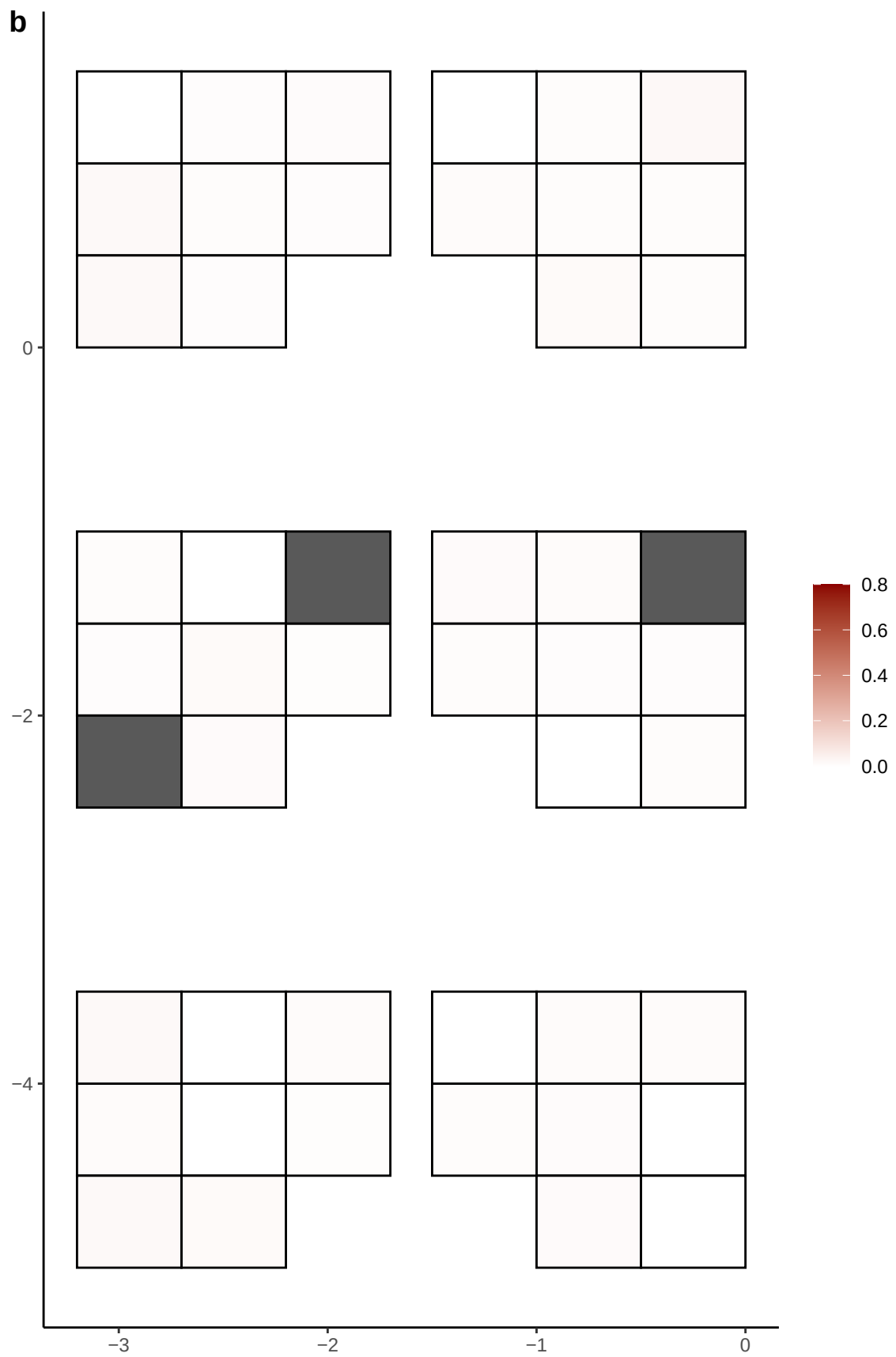

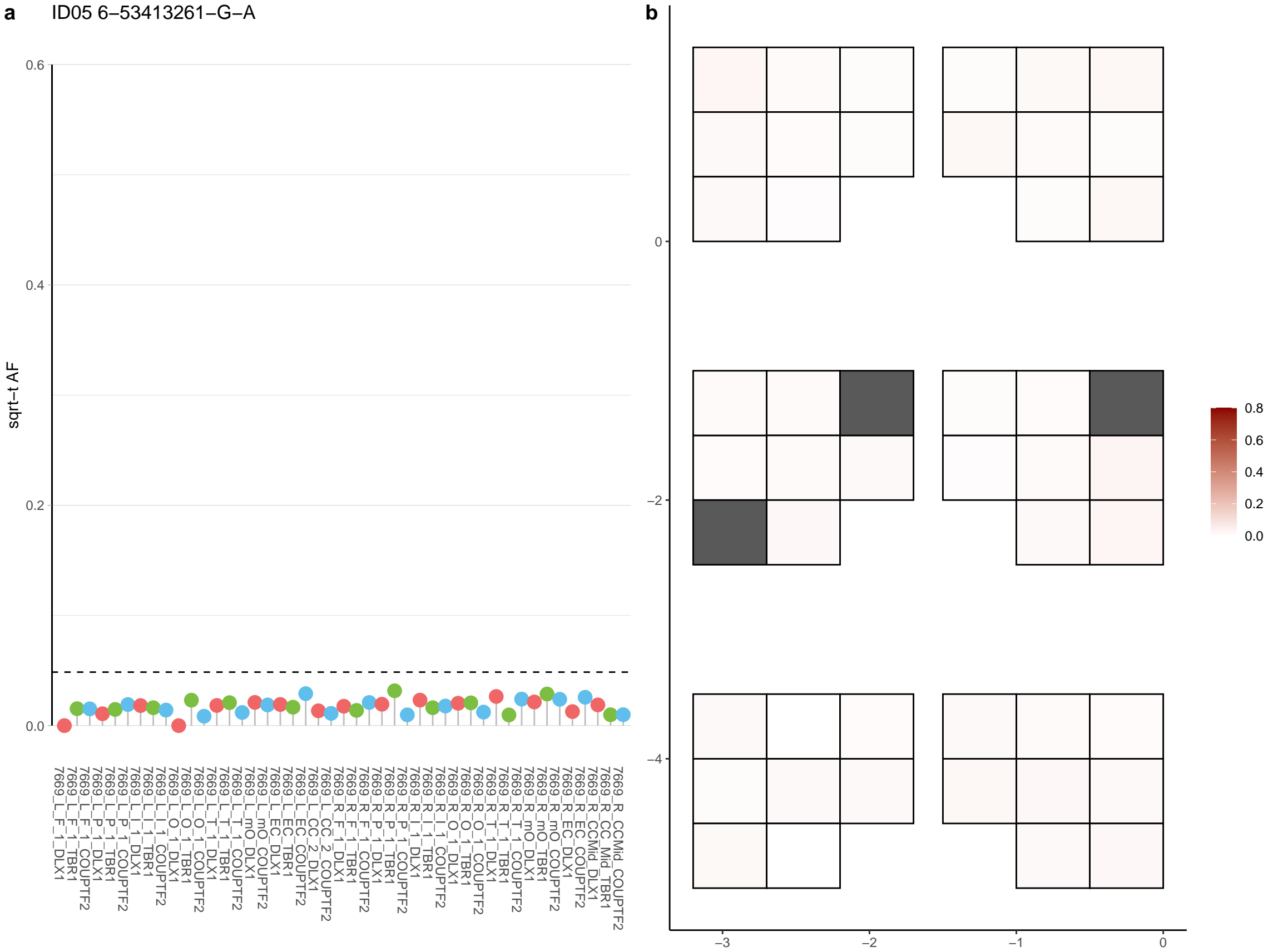

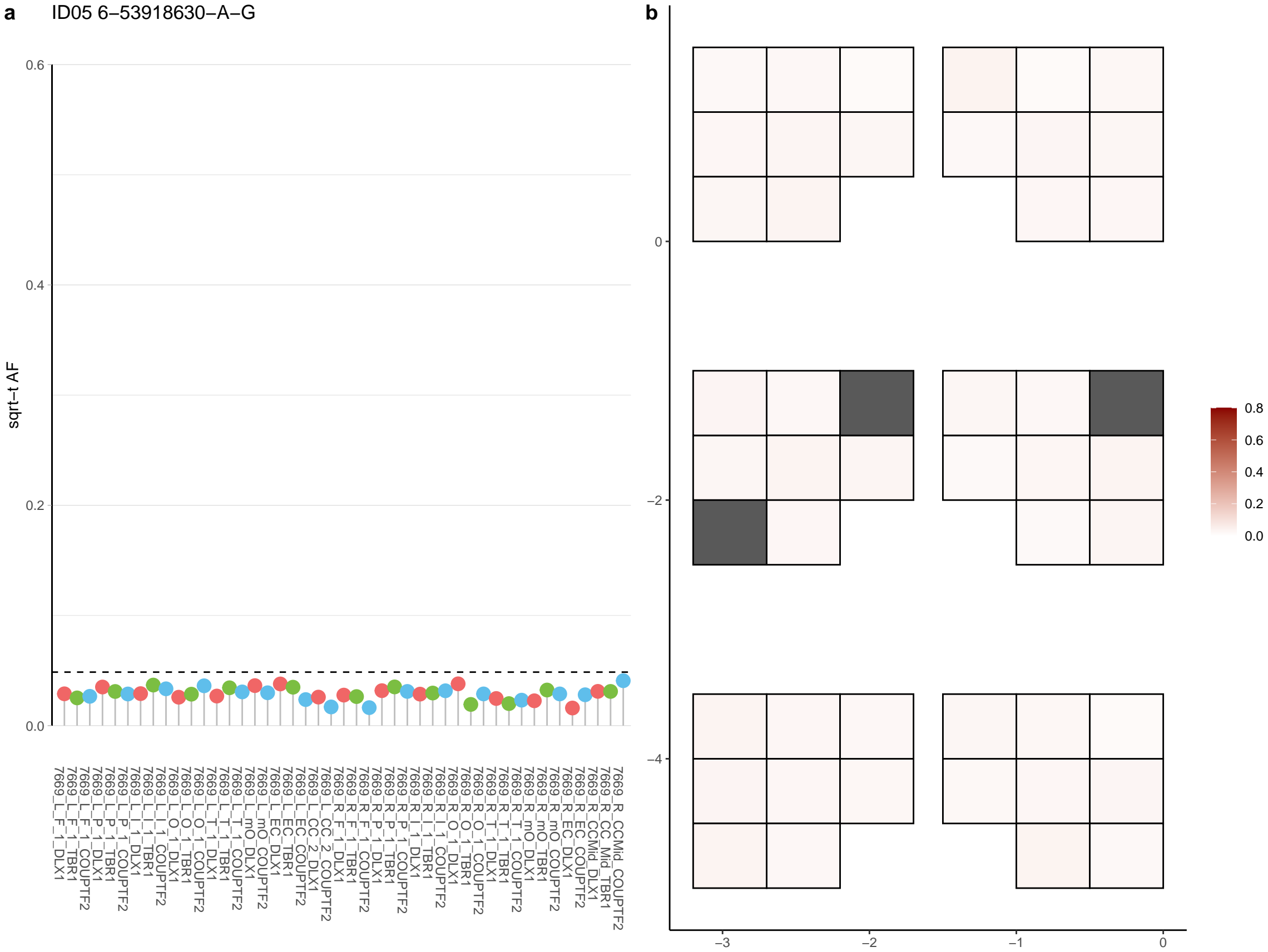

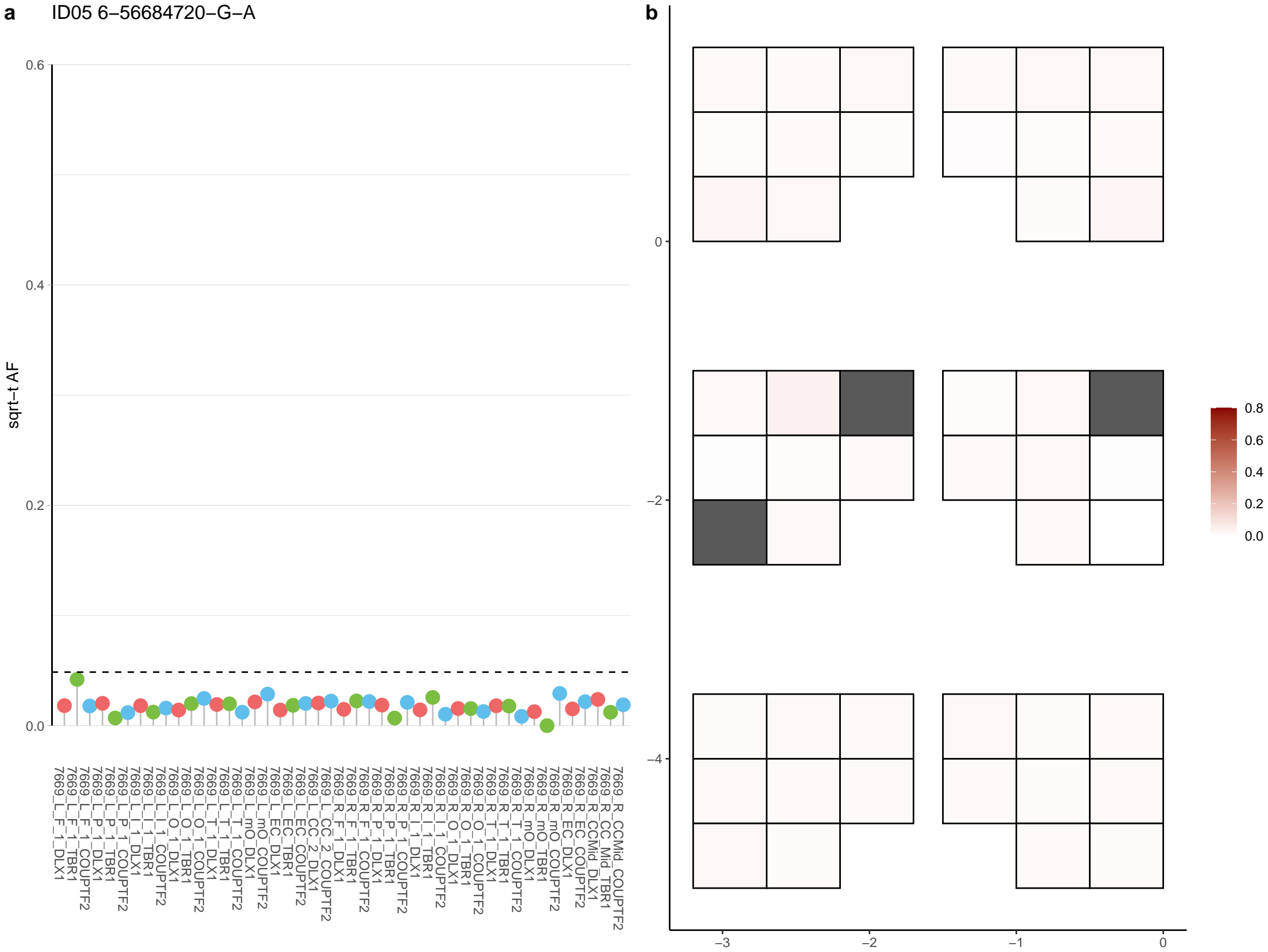

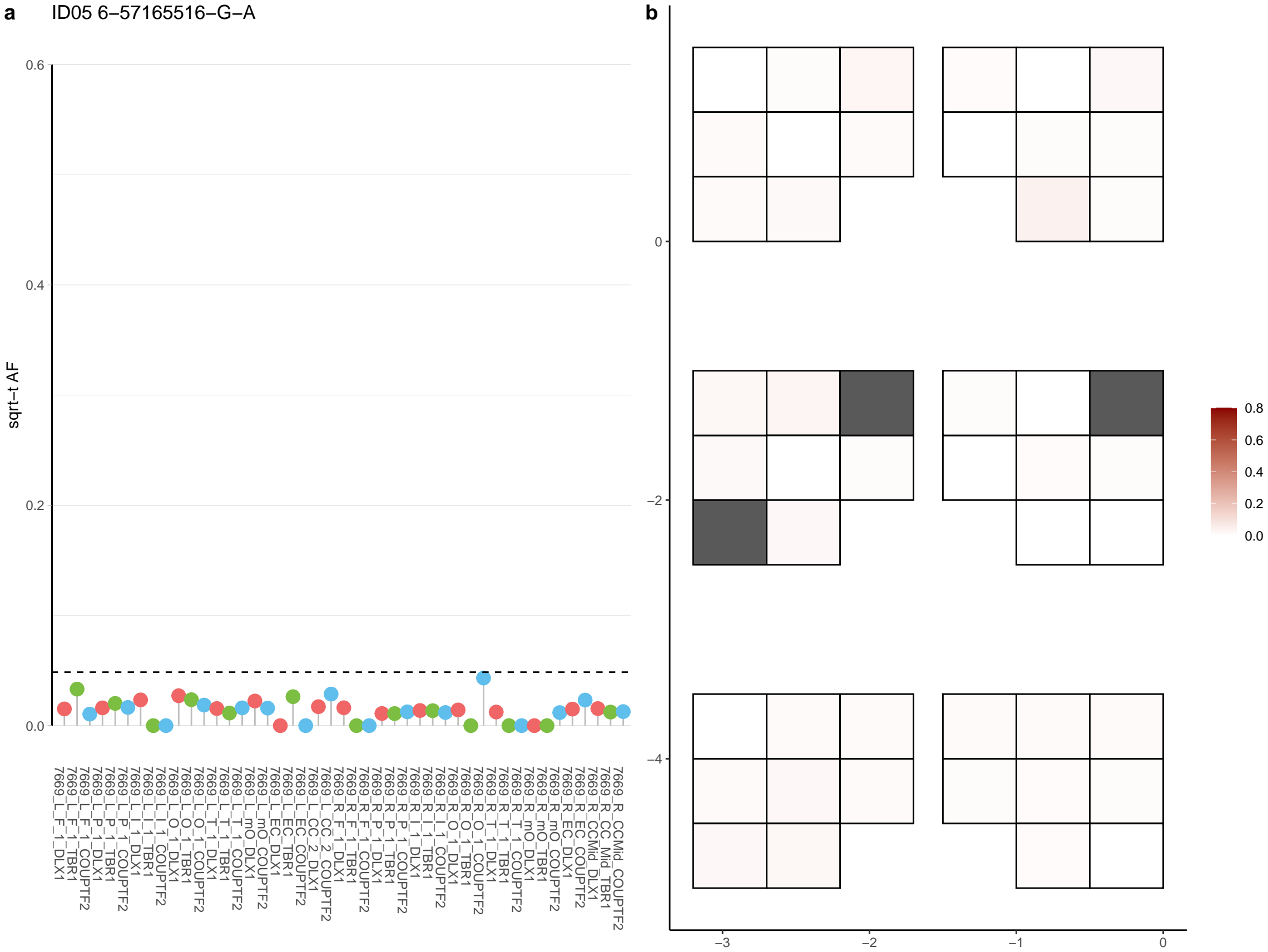

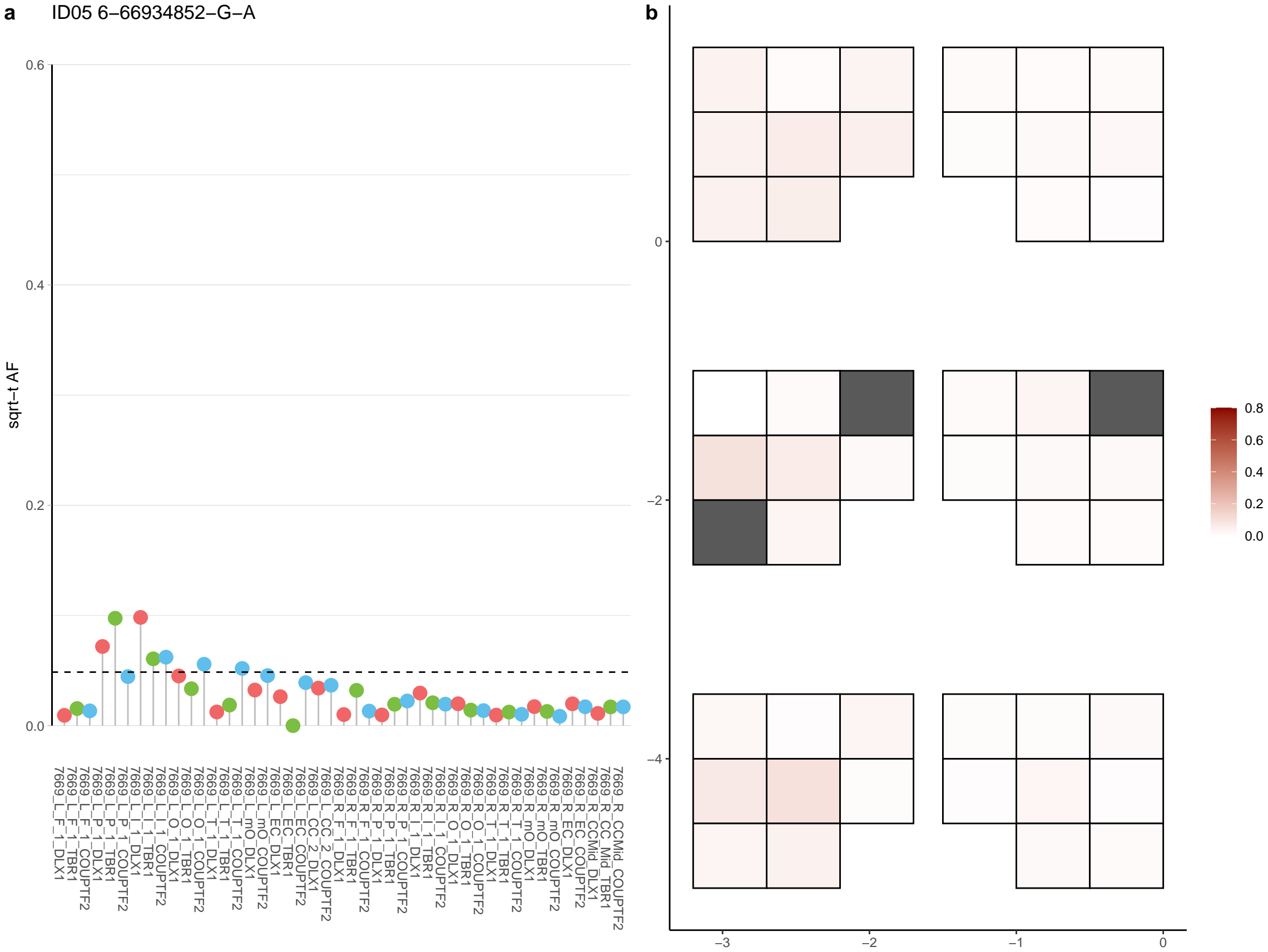

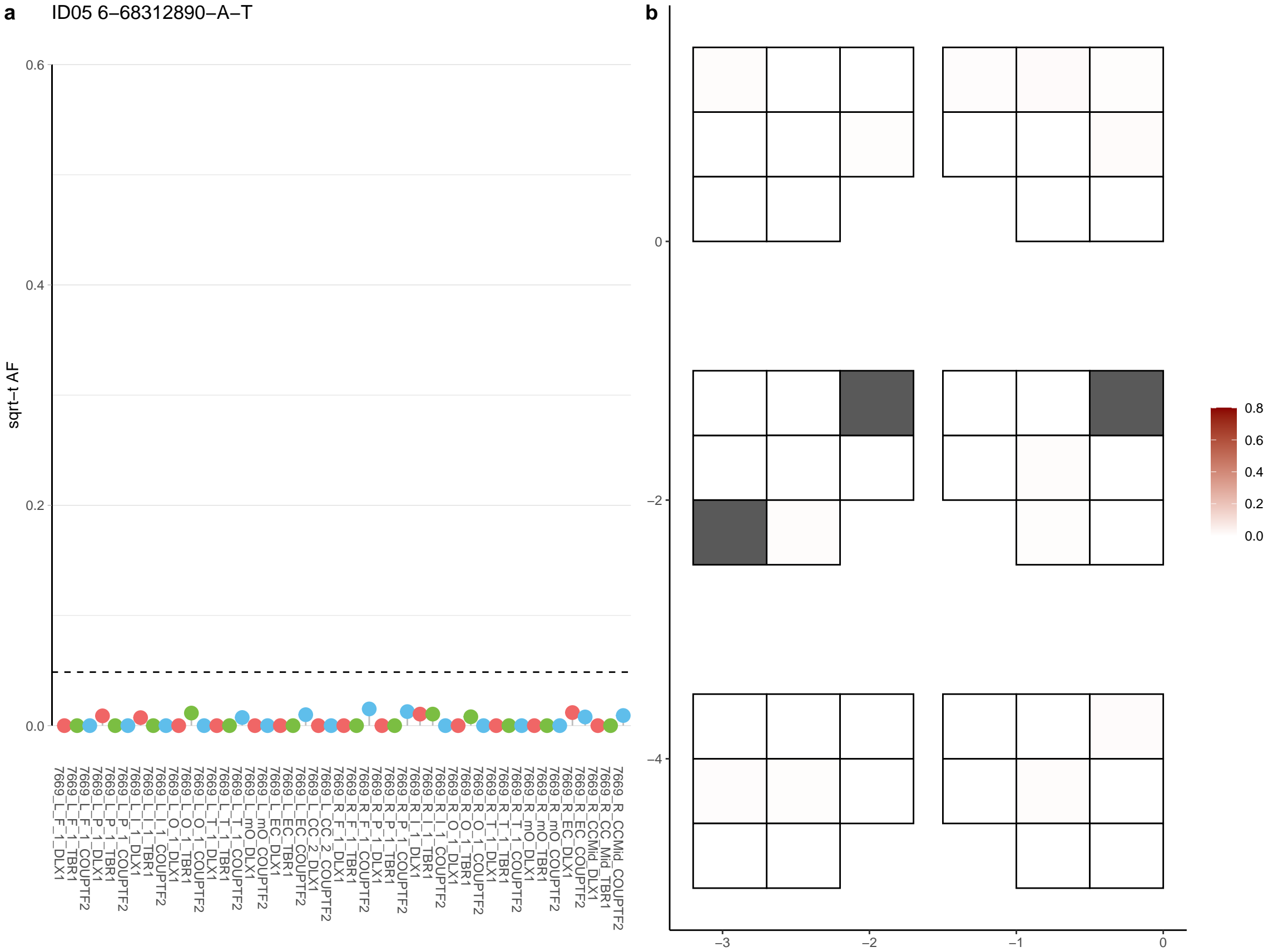

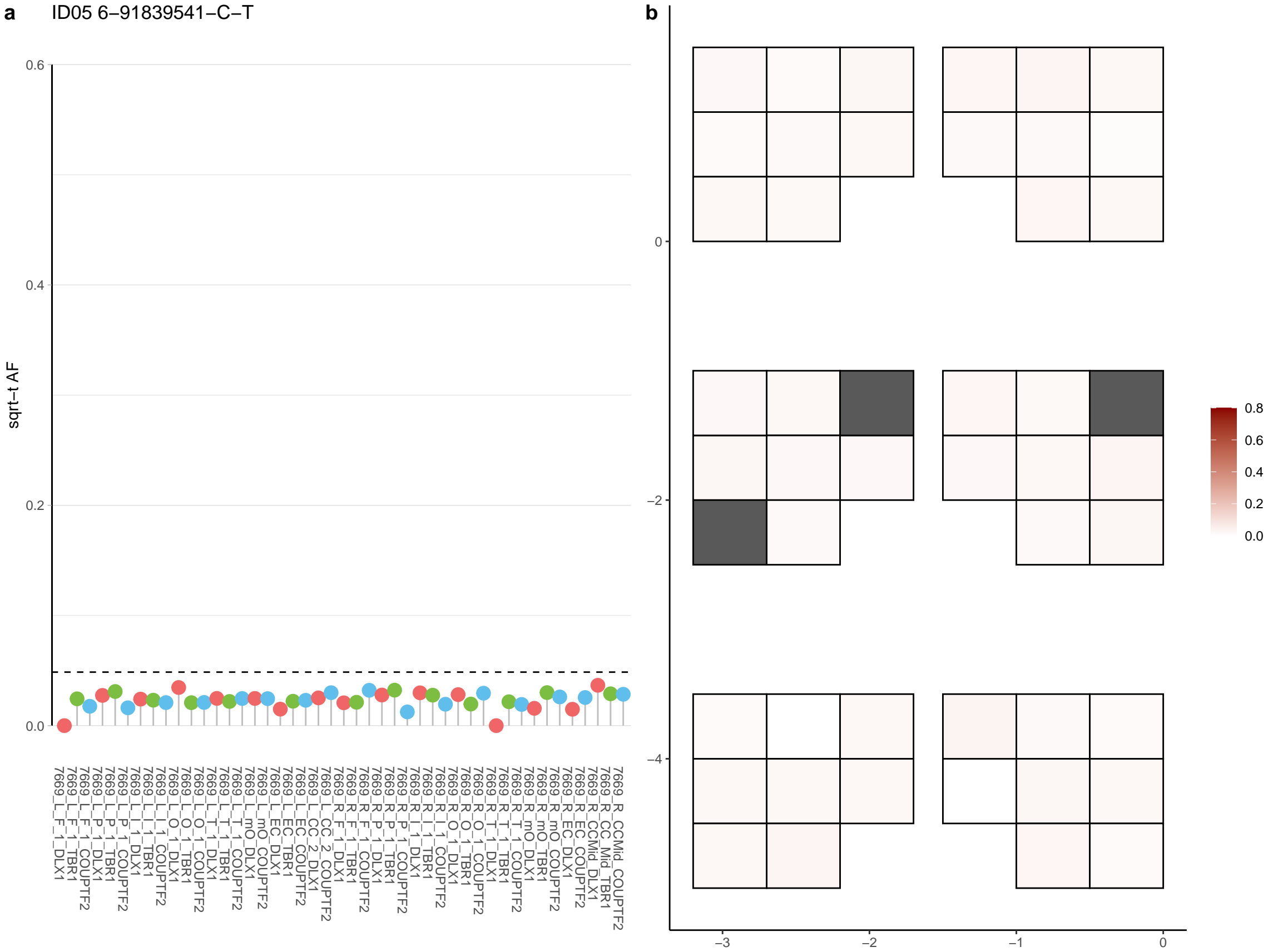

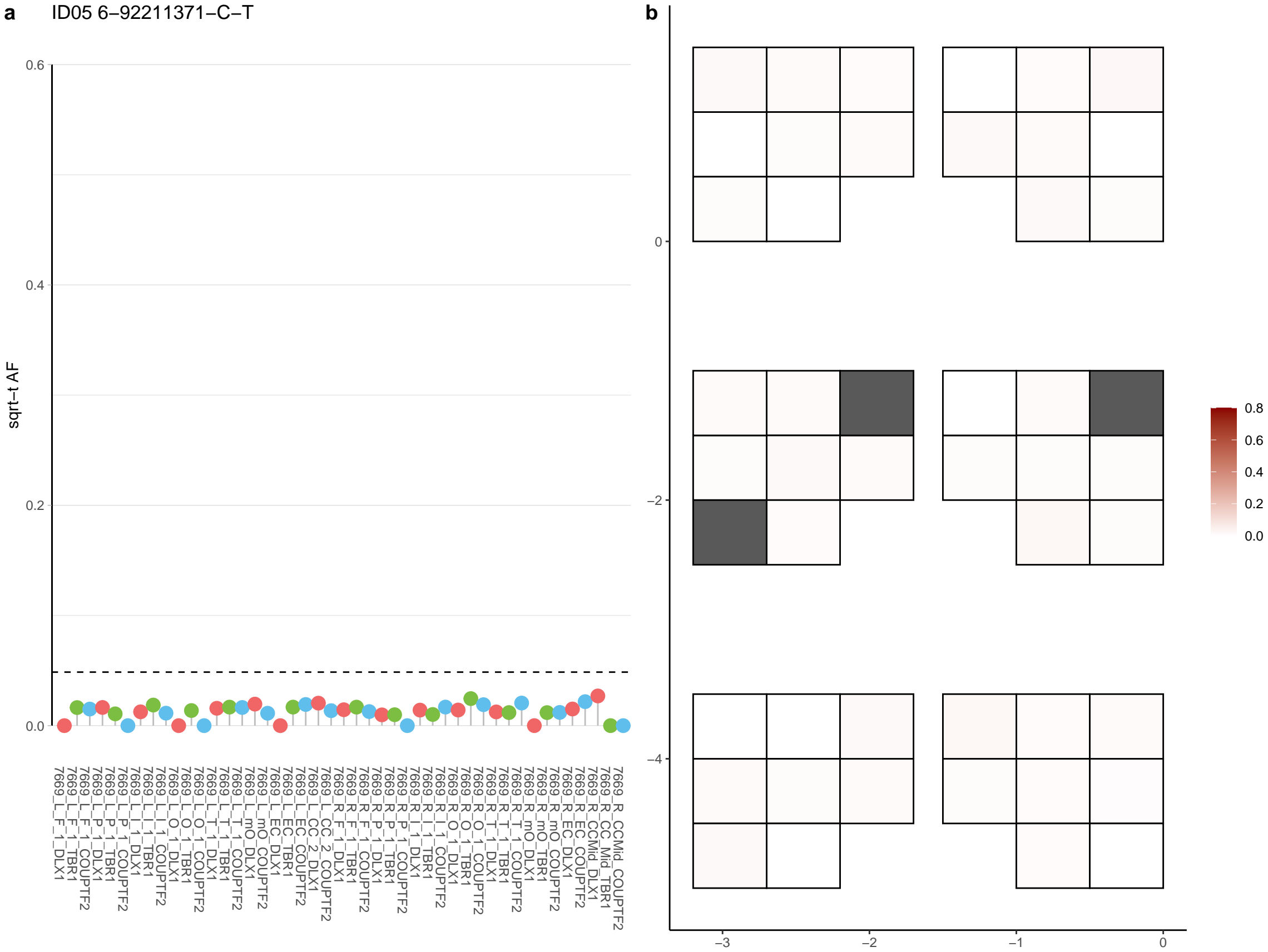

**a** ID05 6-94855972-G-A

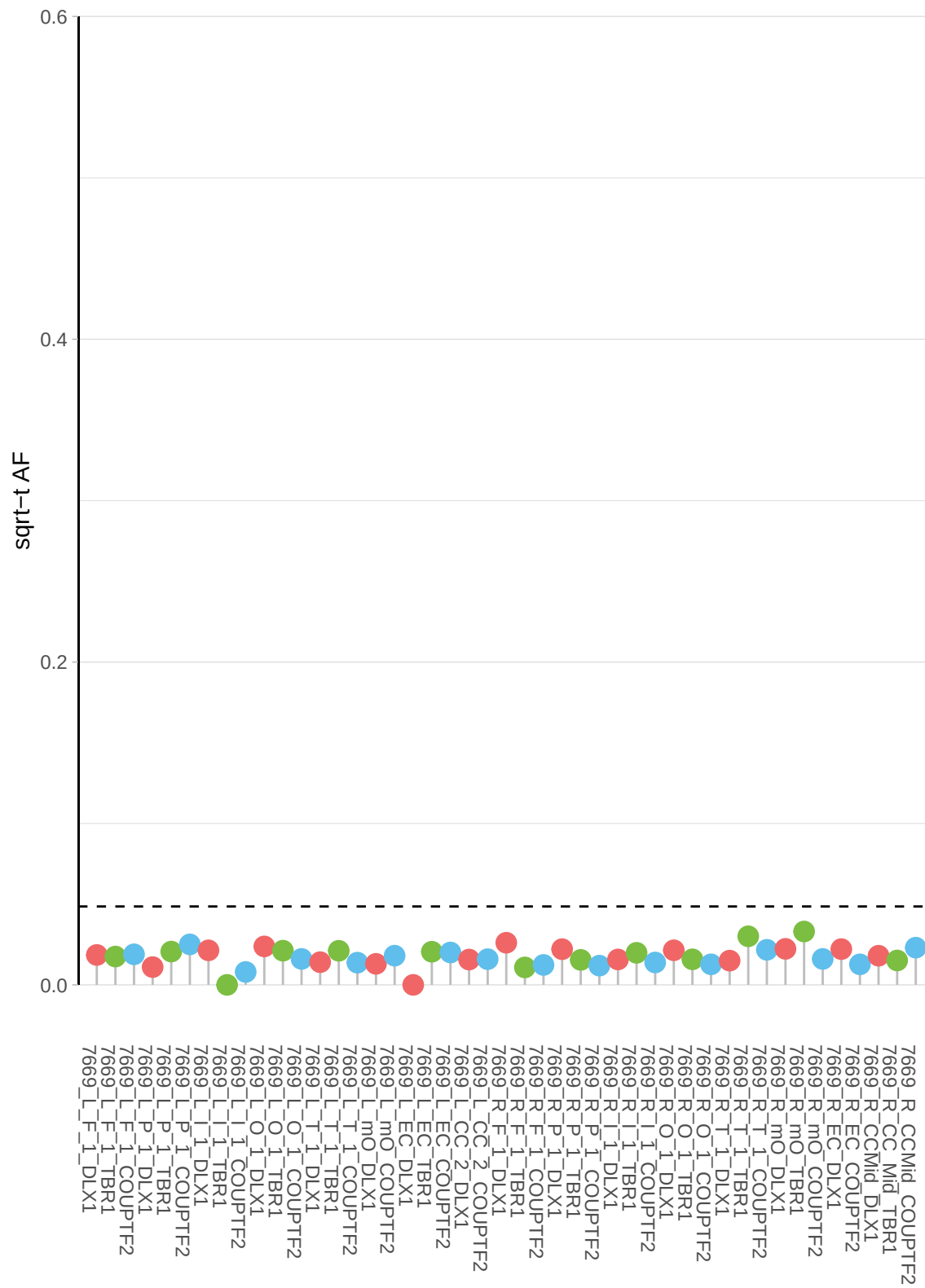

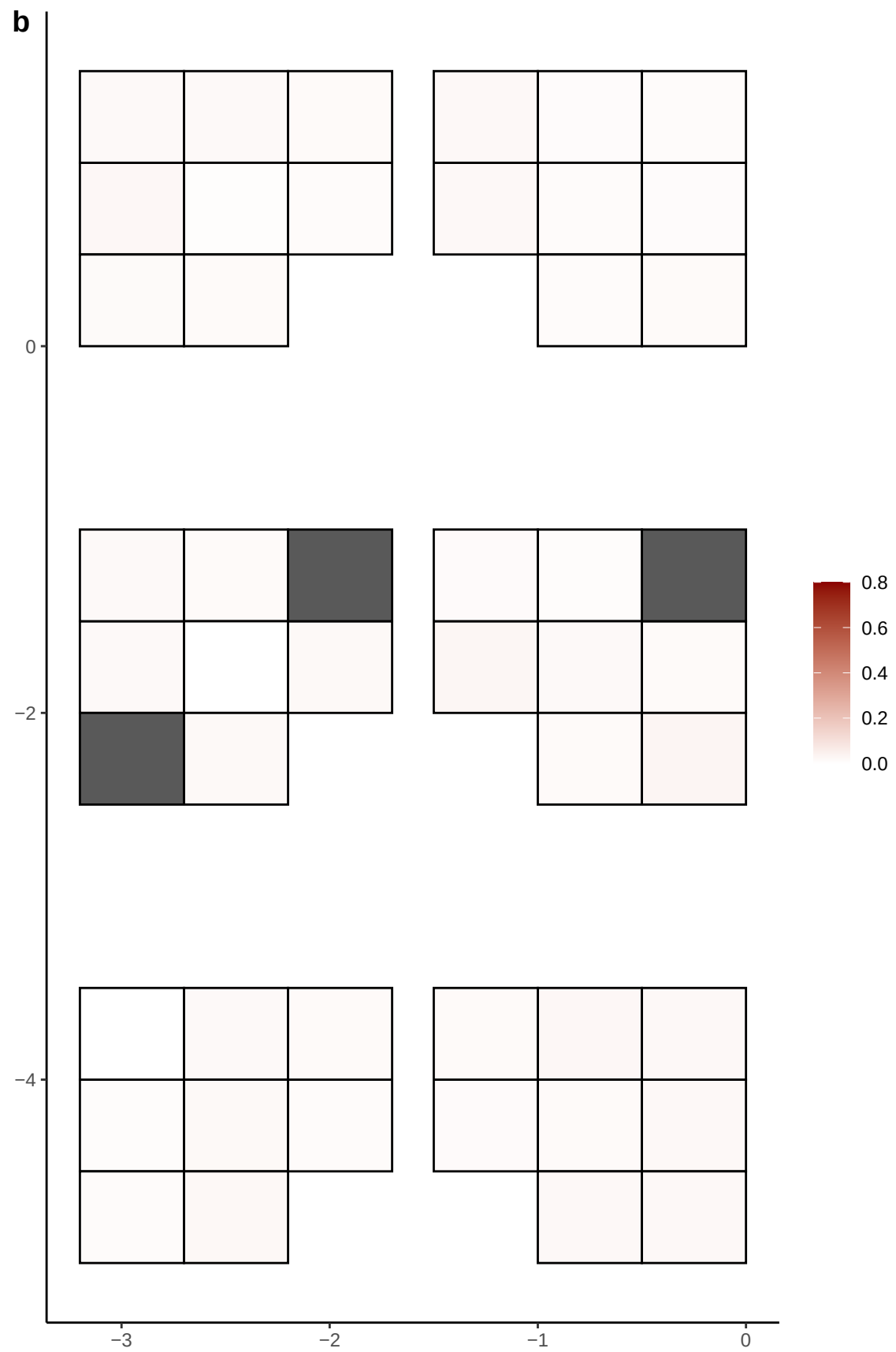

**a** ID05 6-96176402-T-C

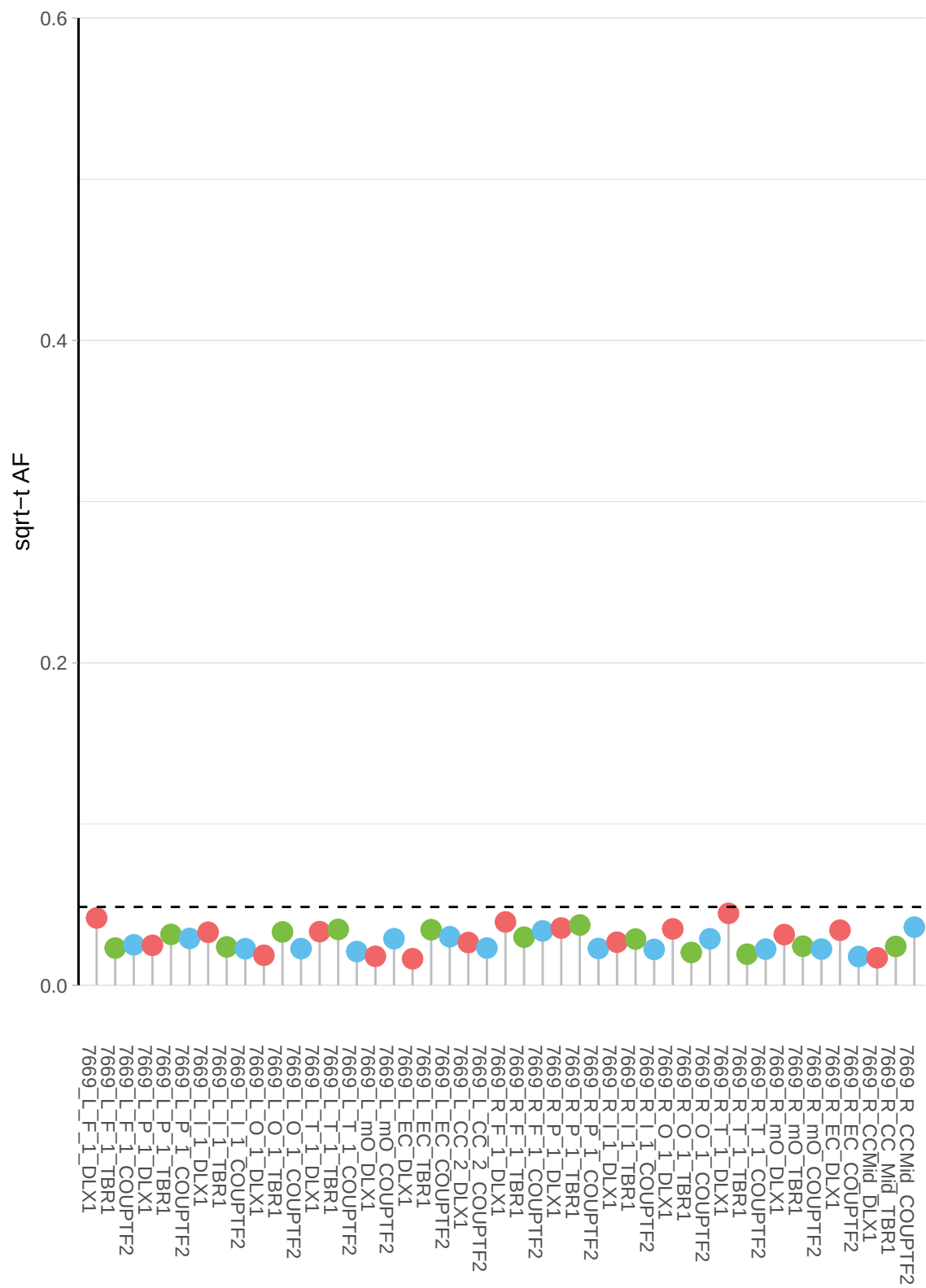

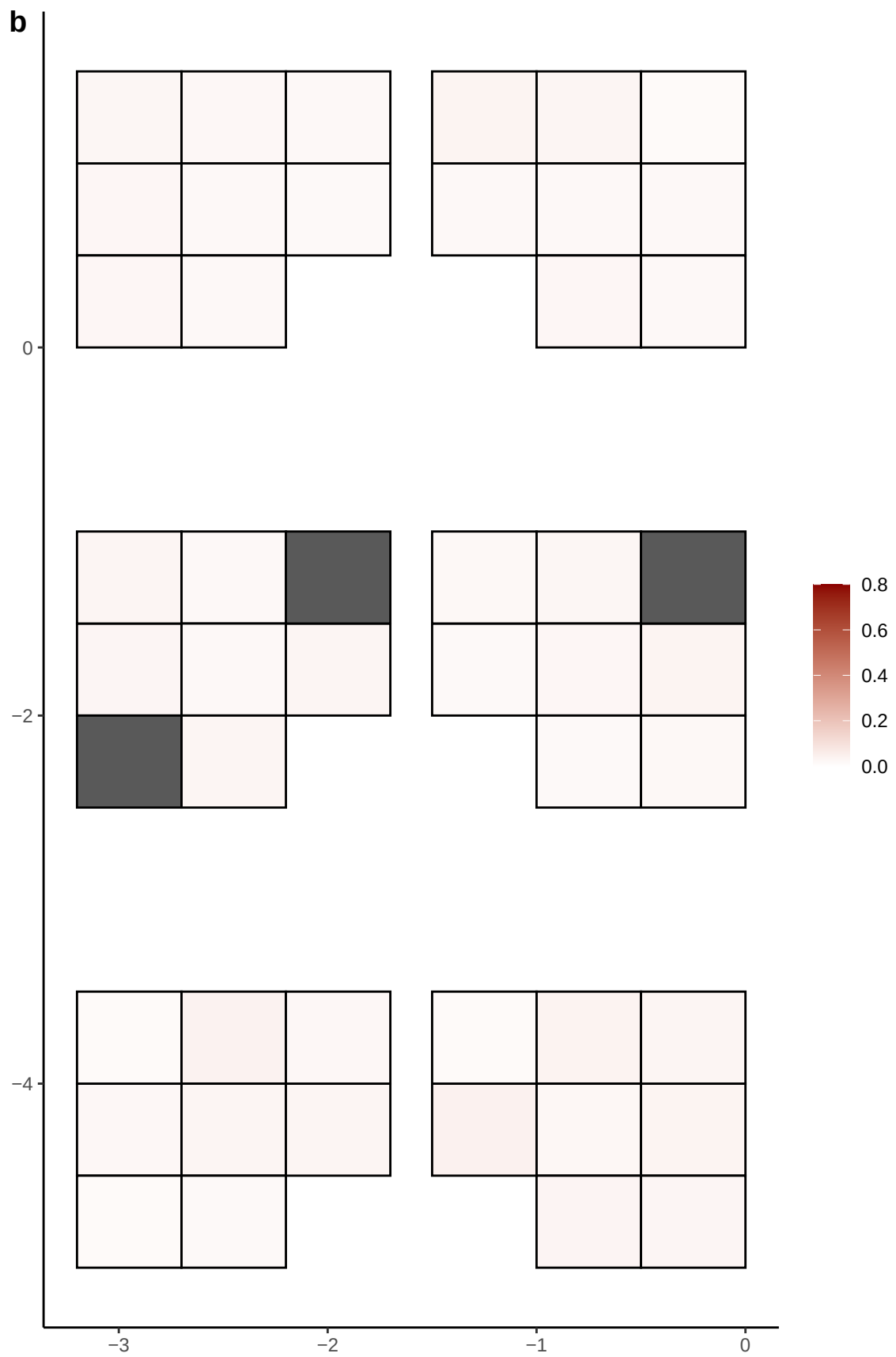

**a** ID05 6-98372583-C-T

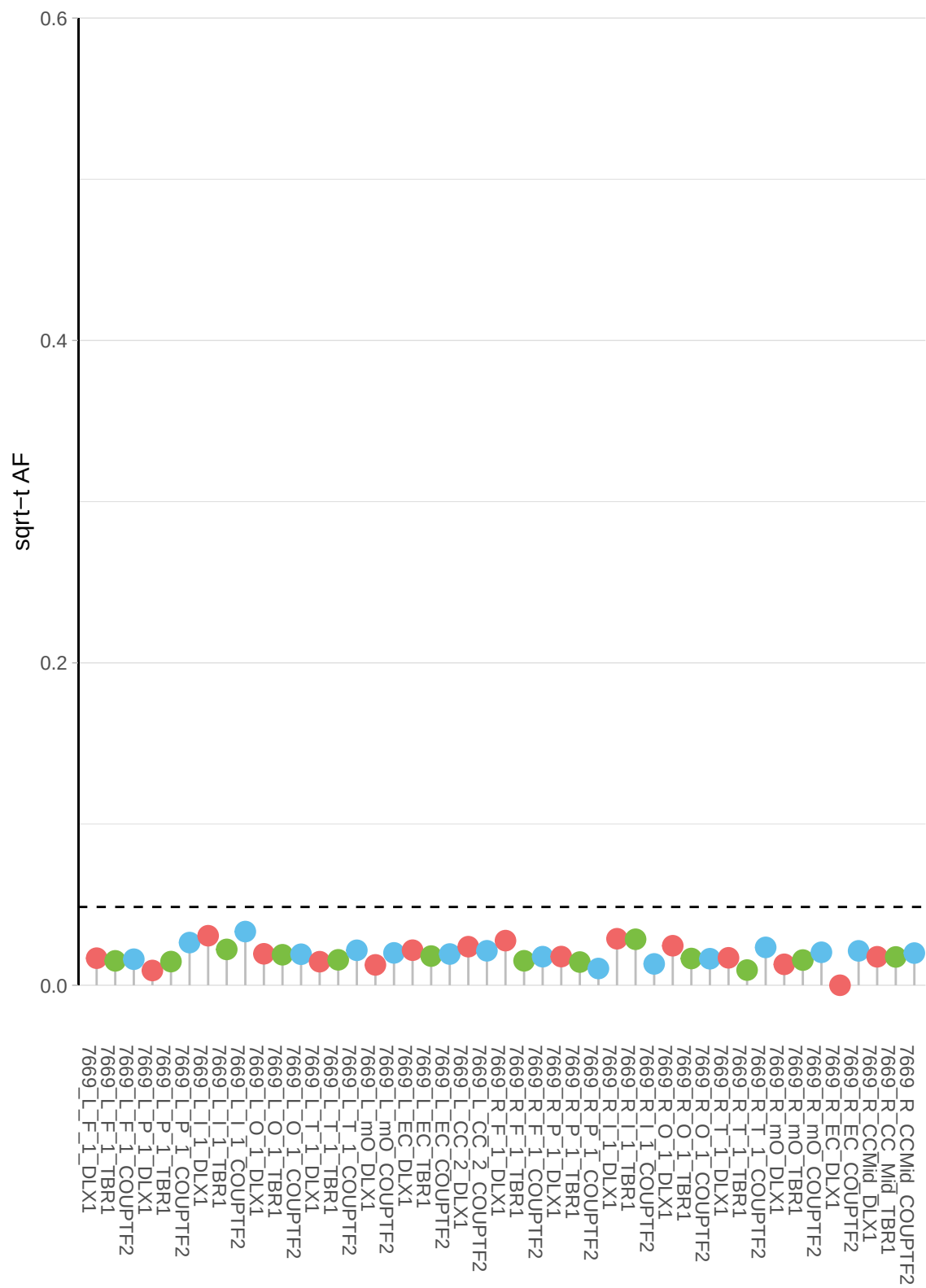

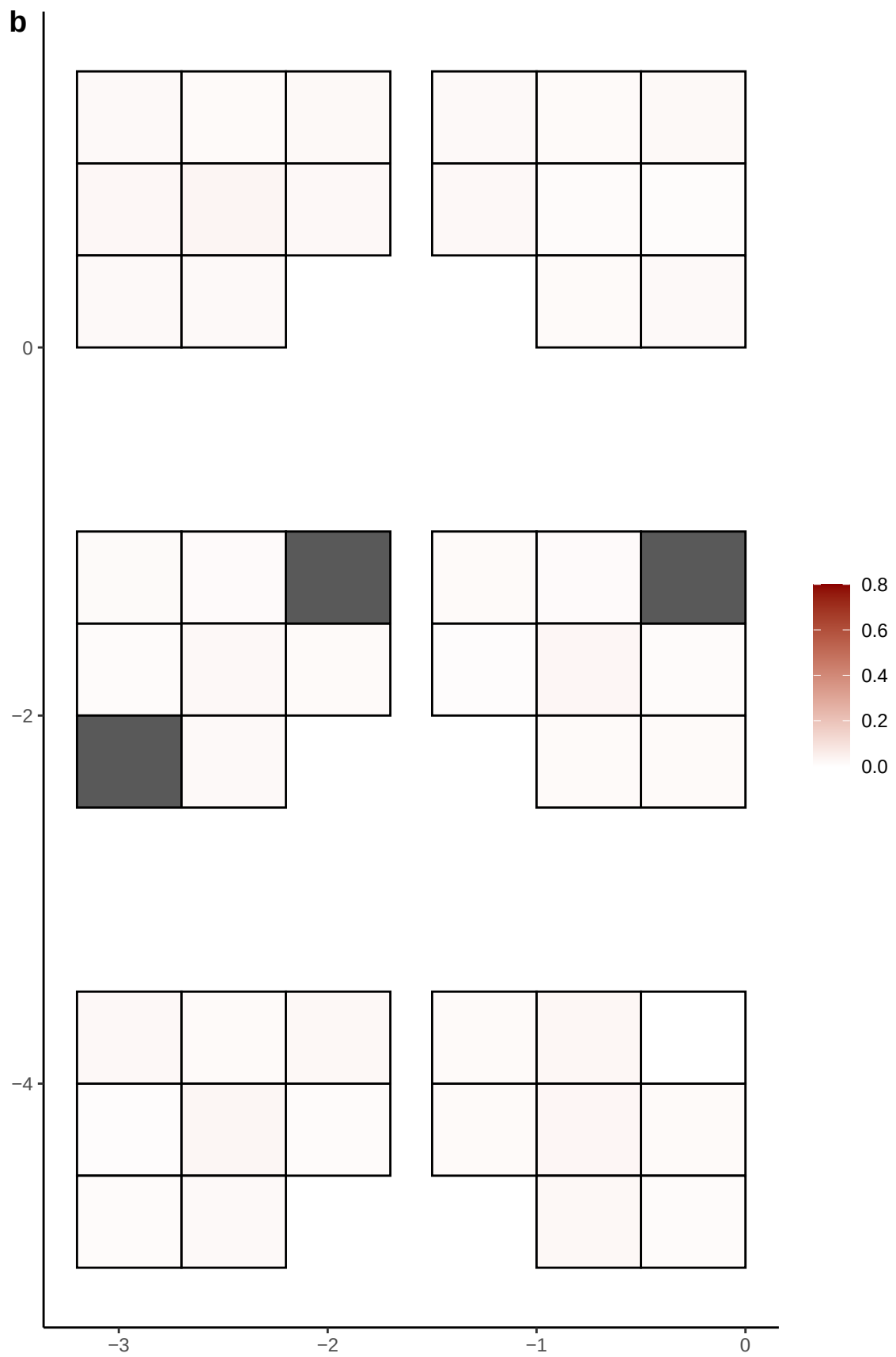

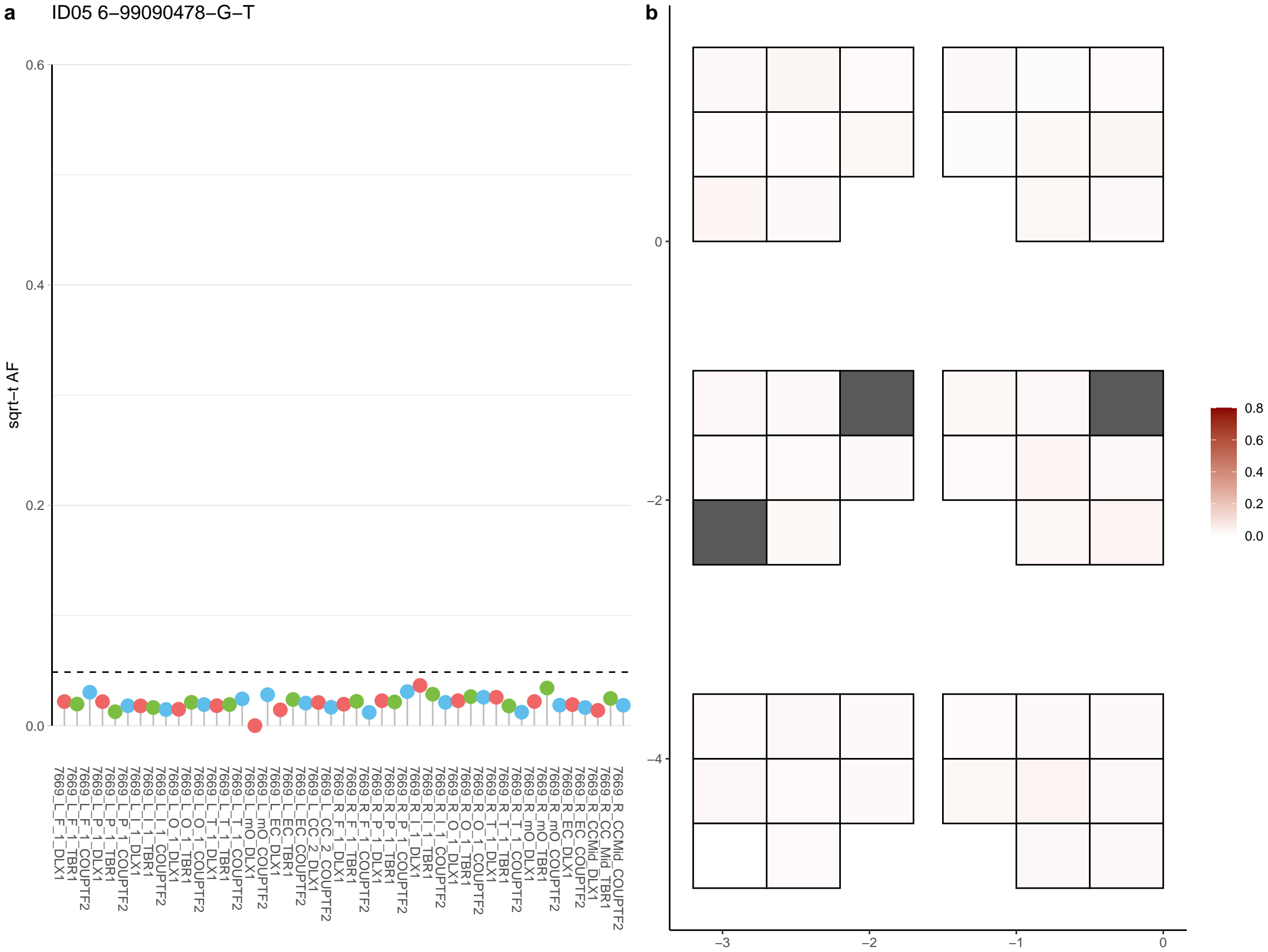

**a** ID05 6-99303058-CCT-C

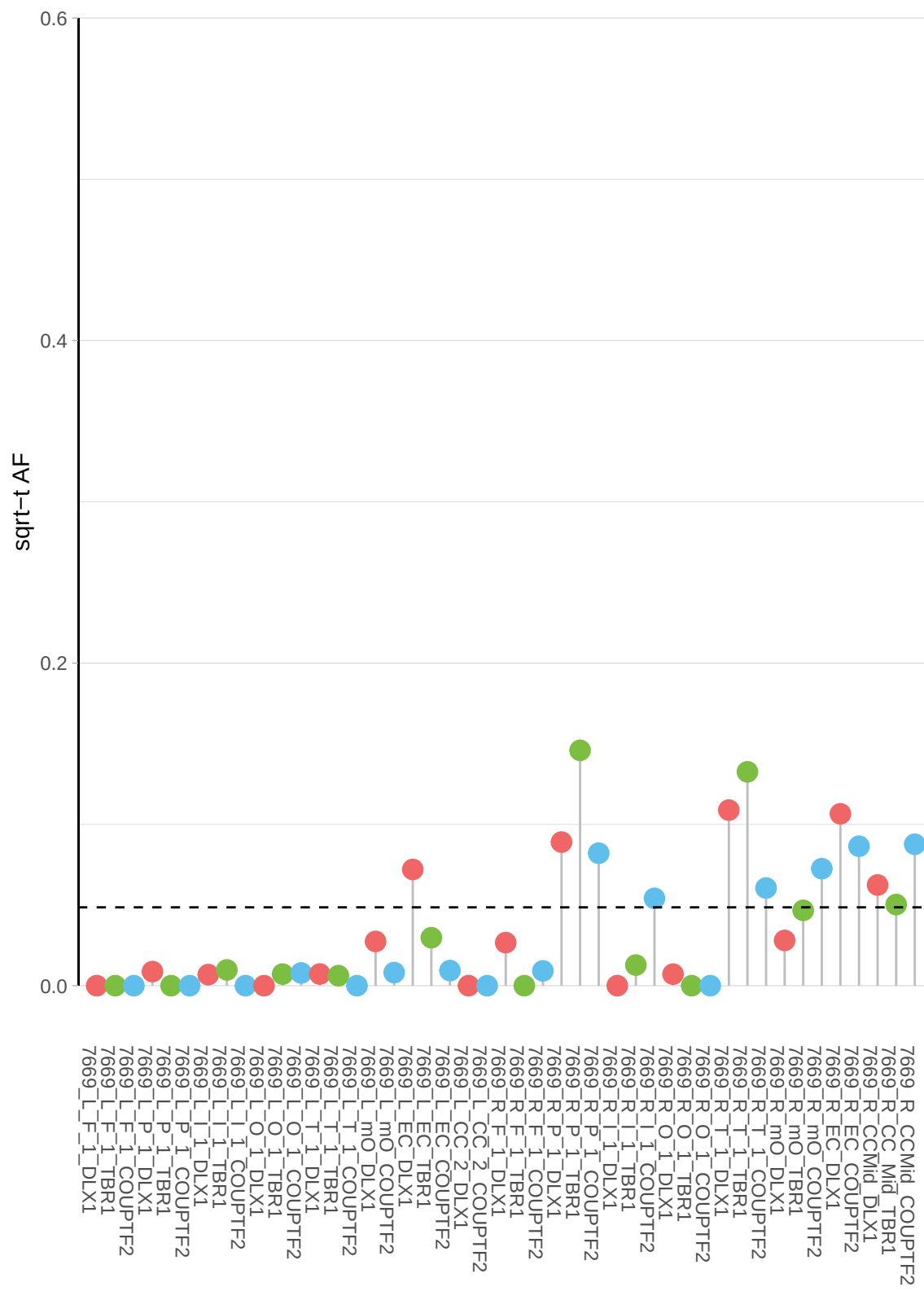

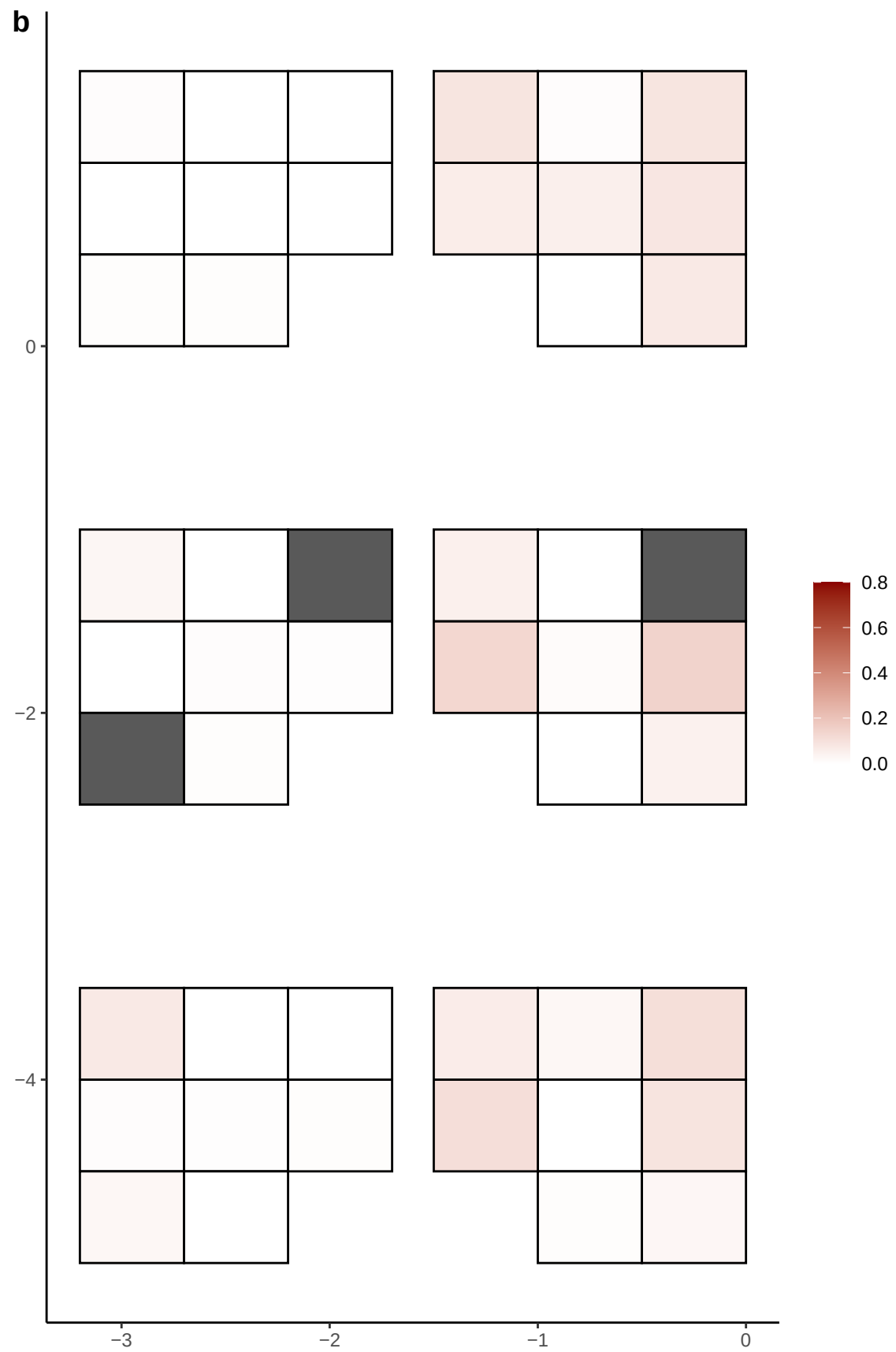

**a** ID05 7-10285800-T-C

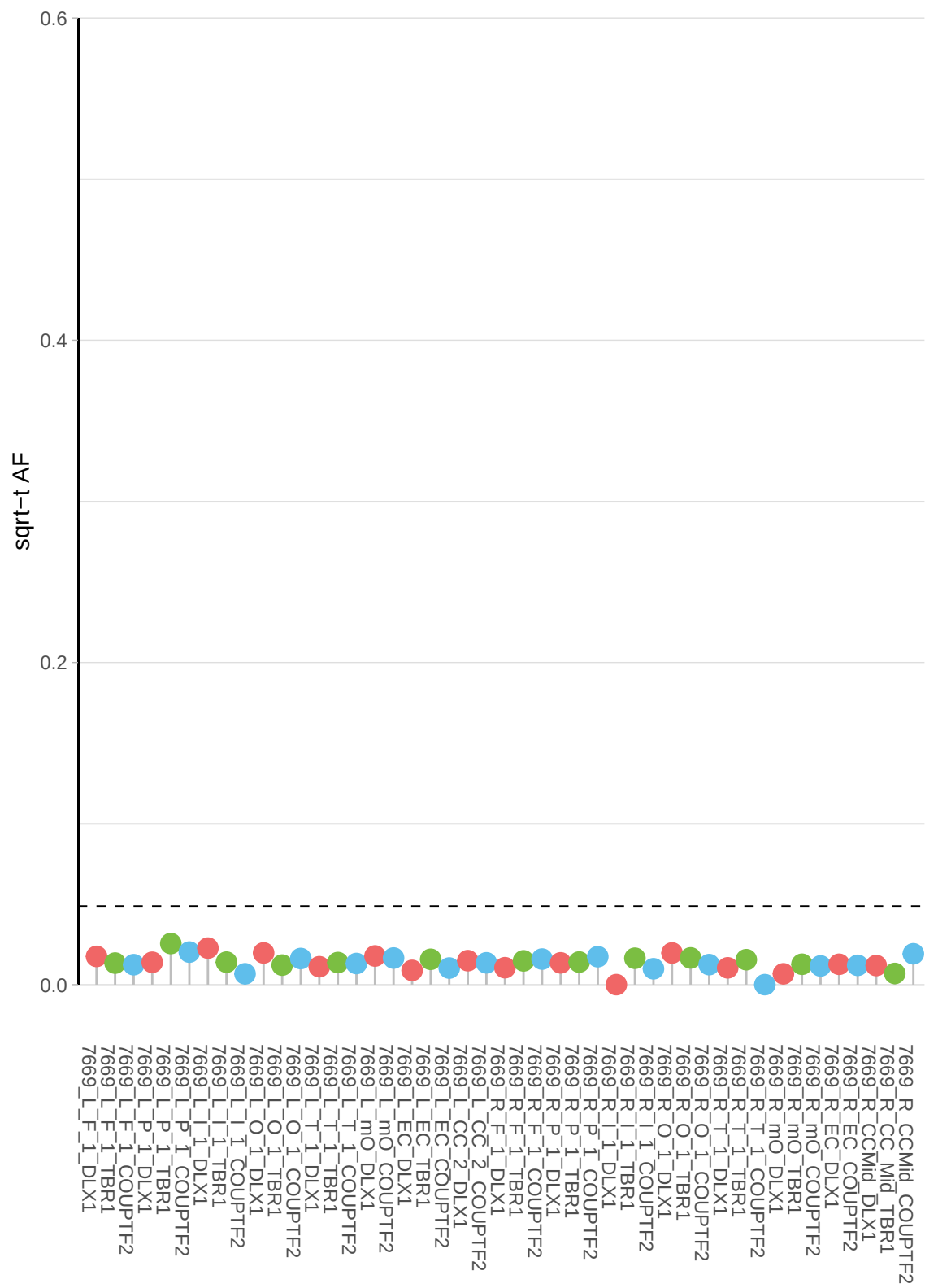

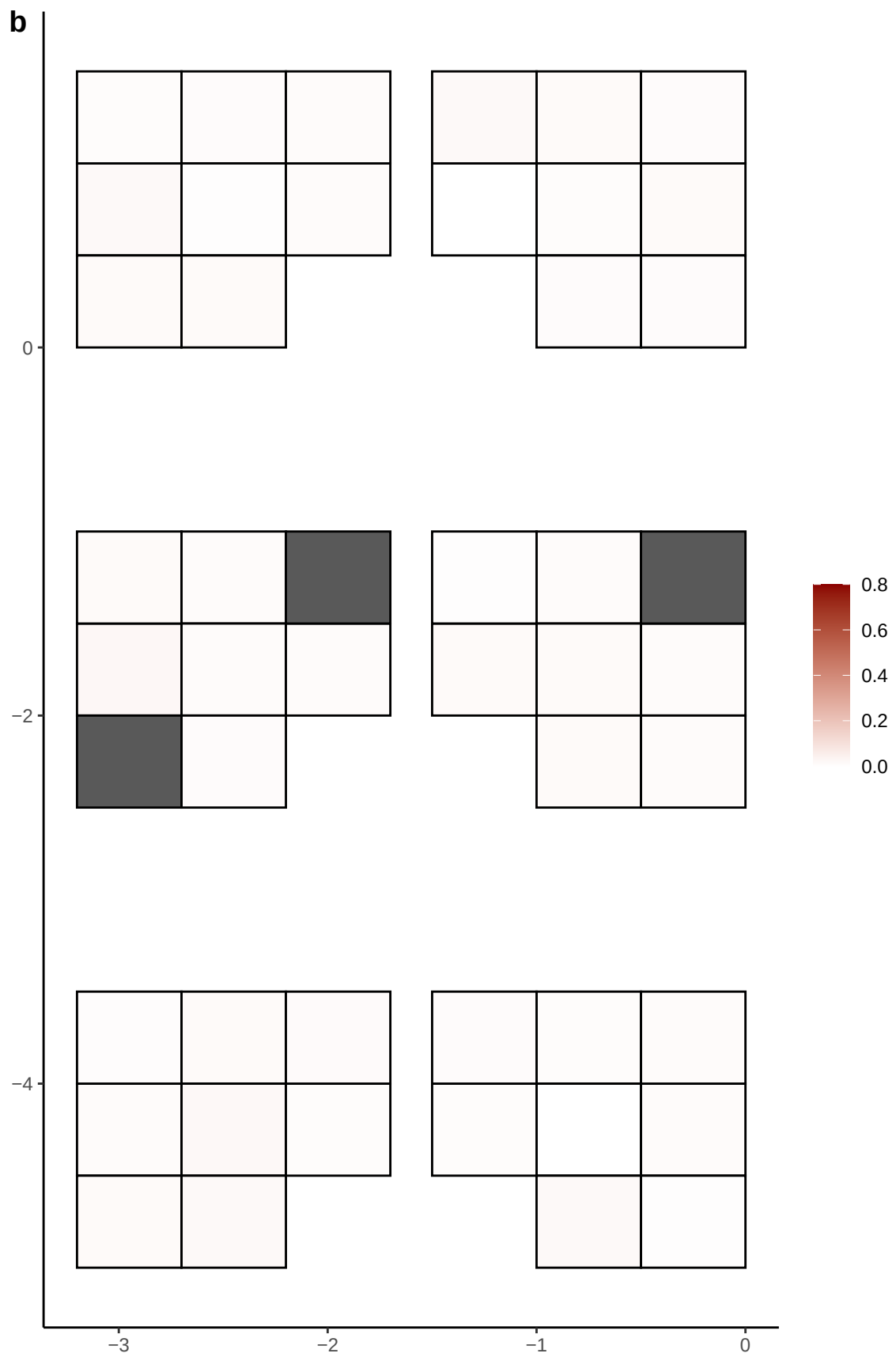

**a** ID05 7-109736410-A-T

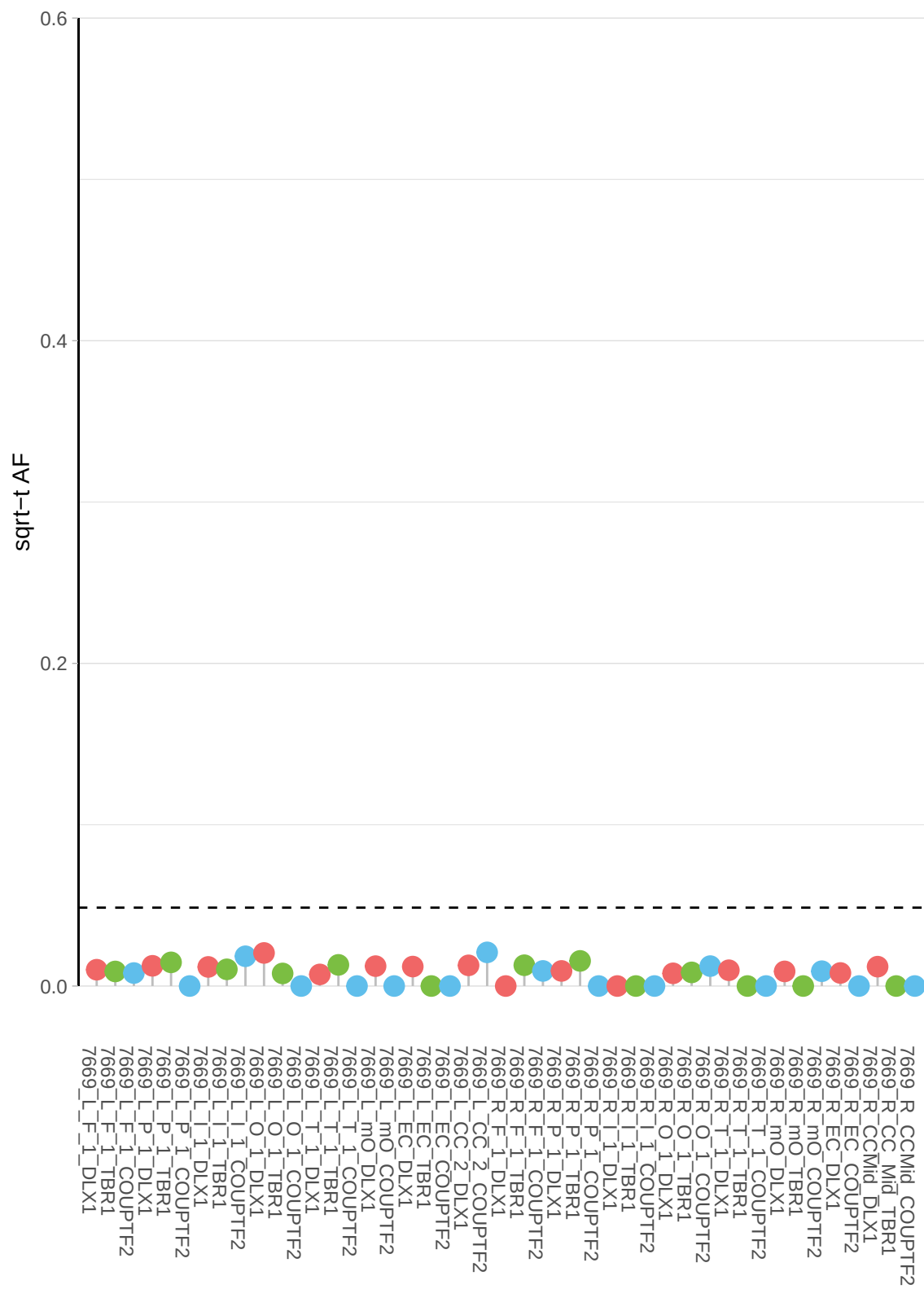

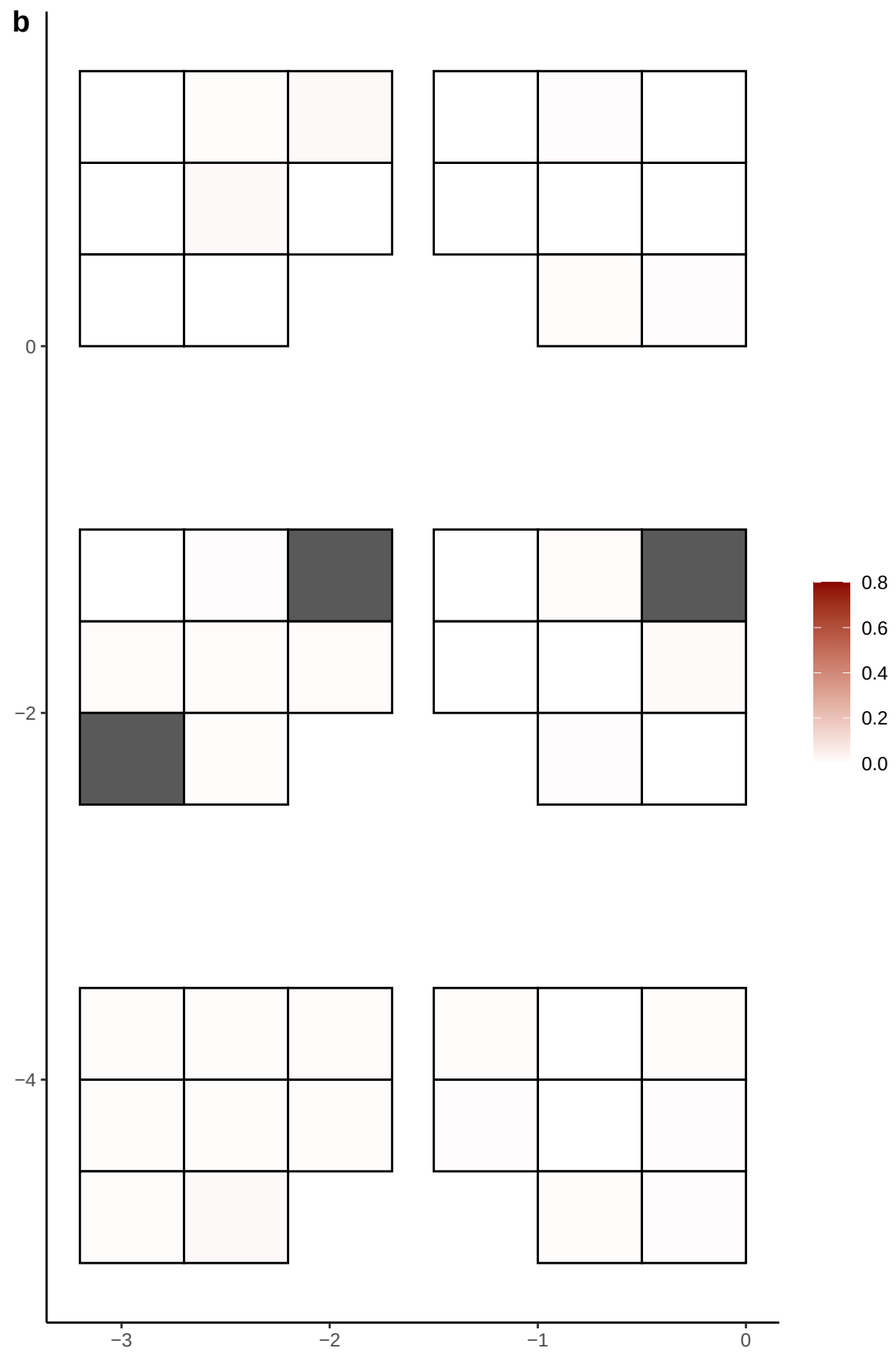

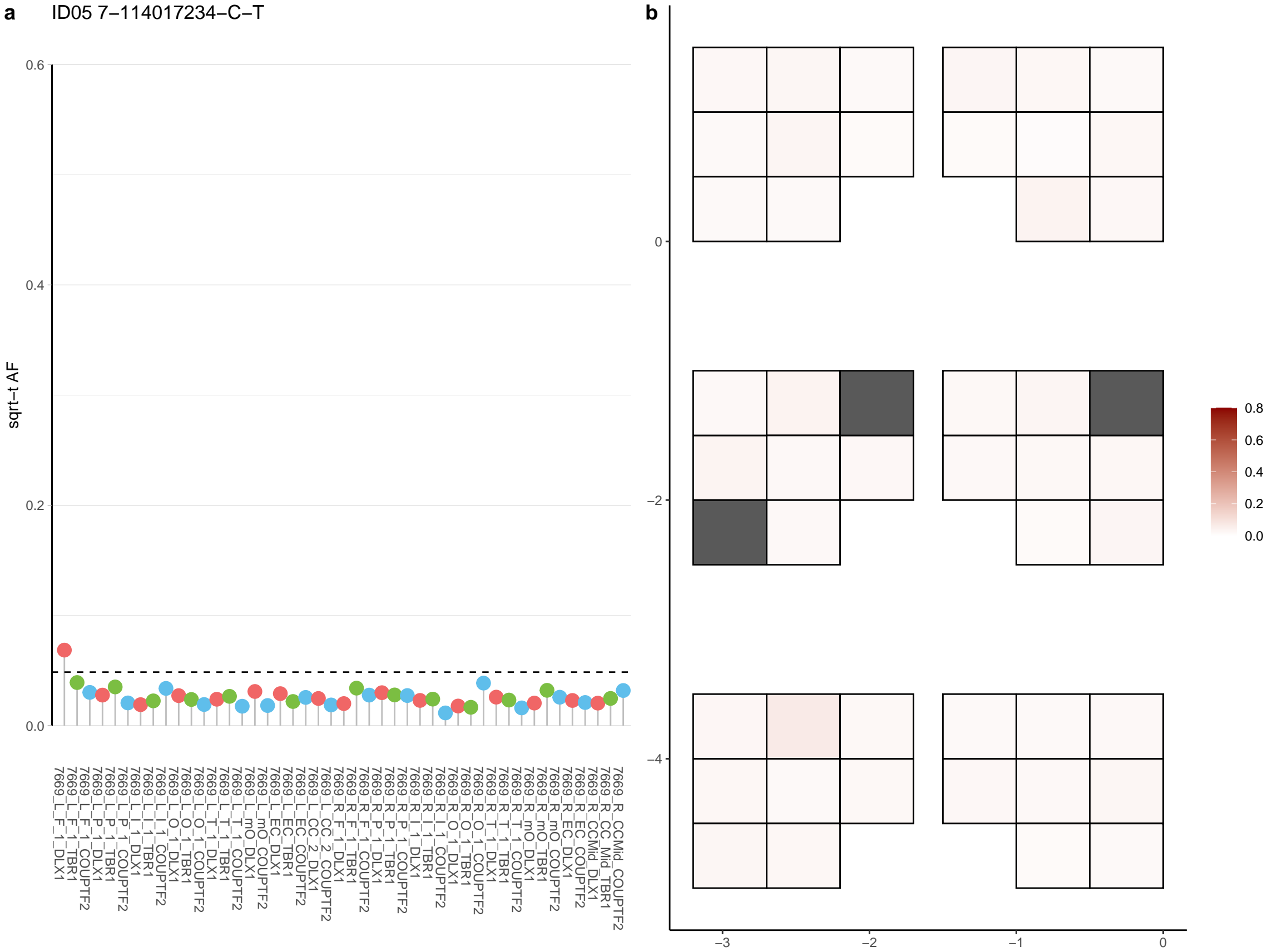

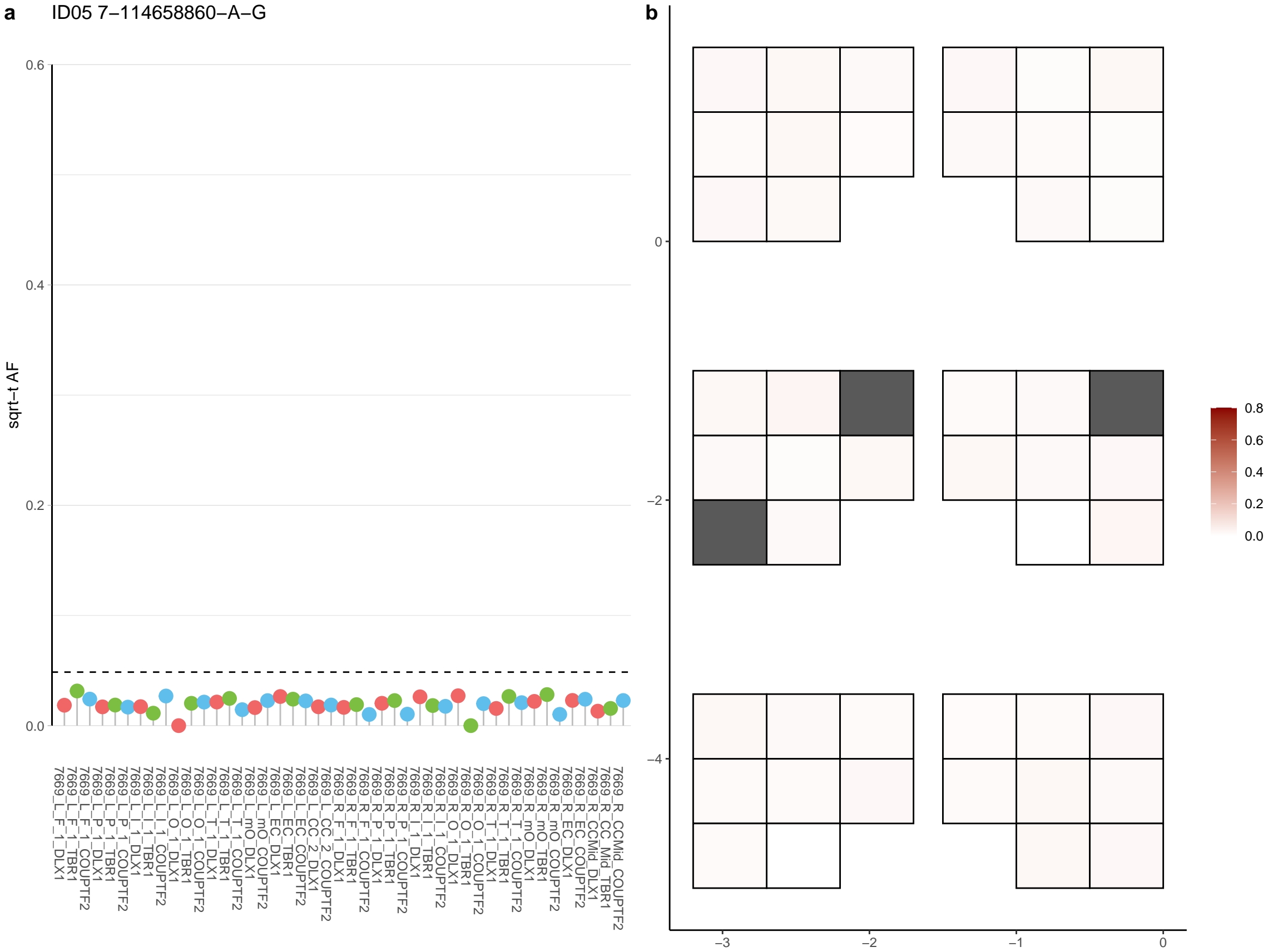

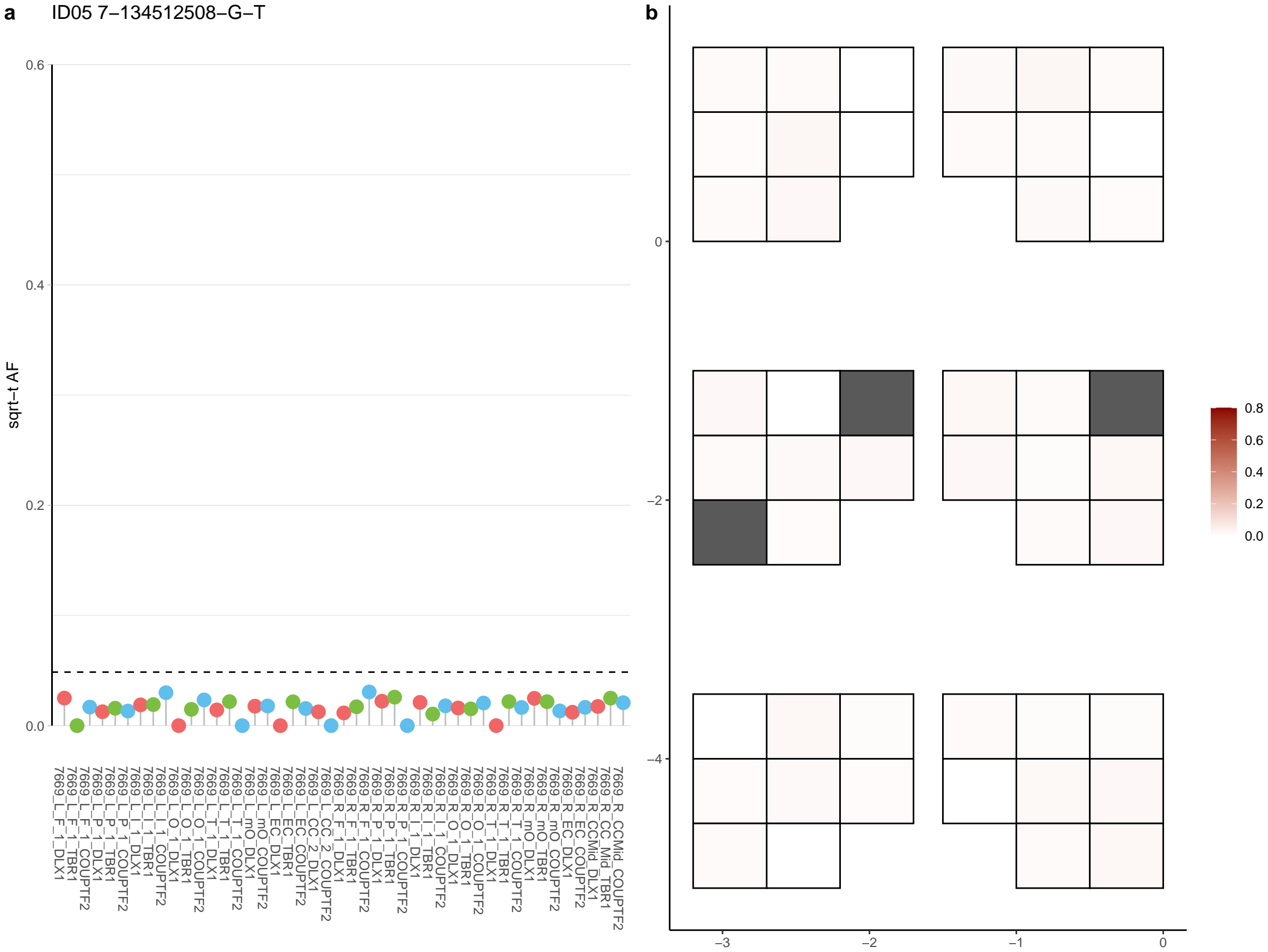

**a** ID05 7-137220703-G-A

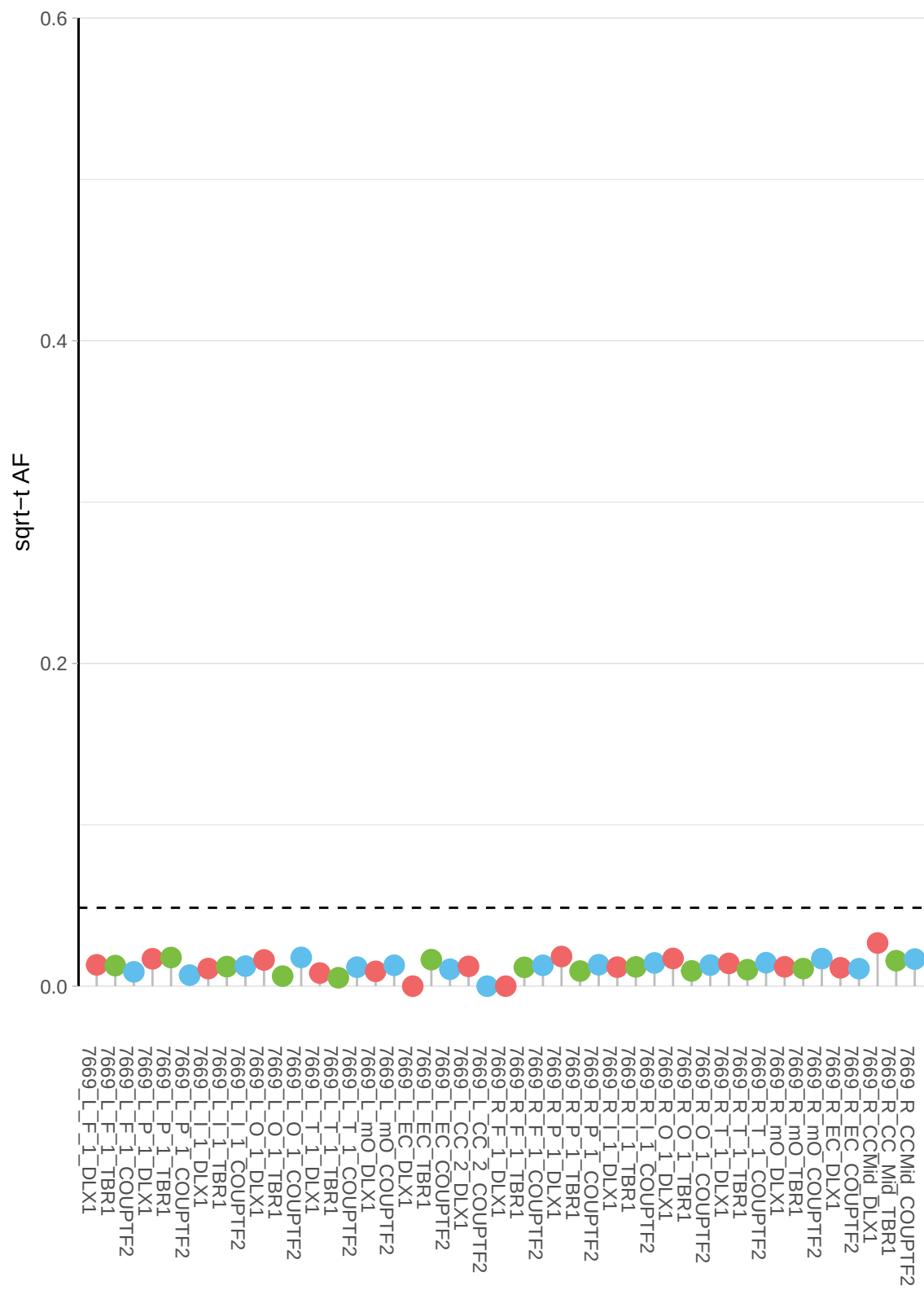

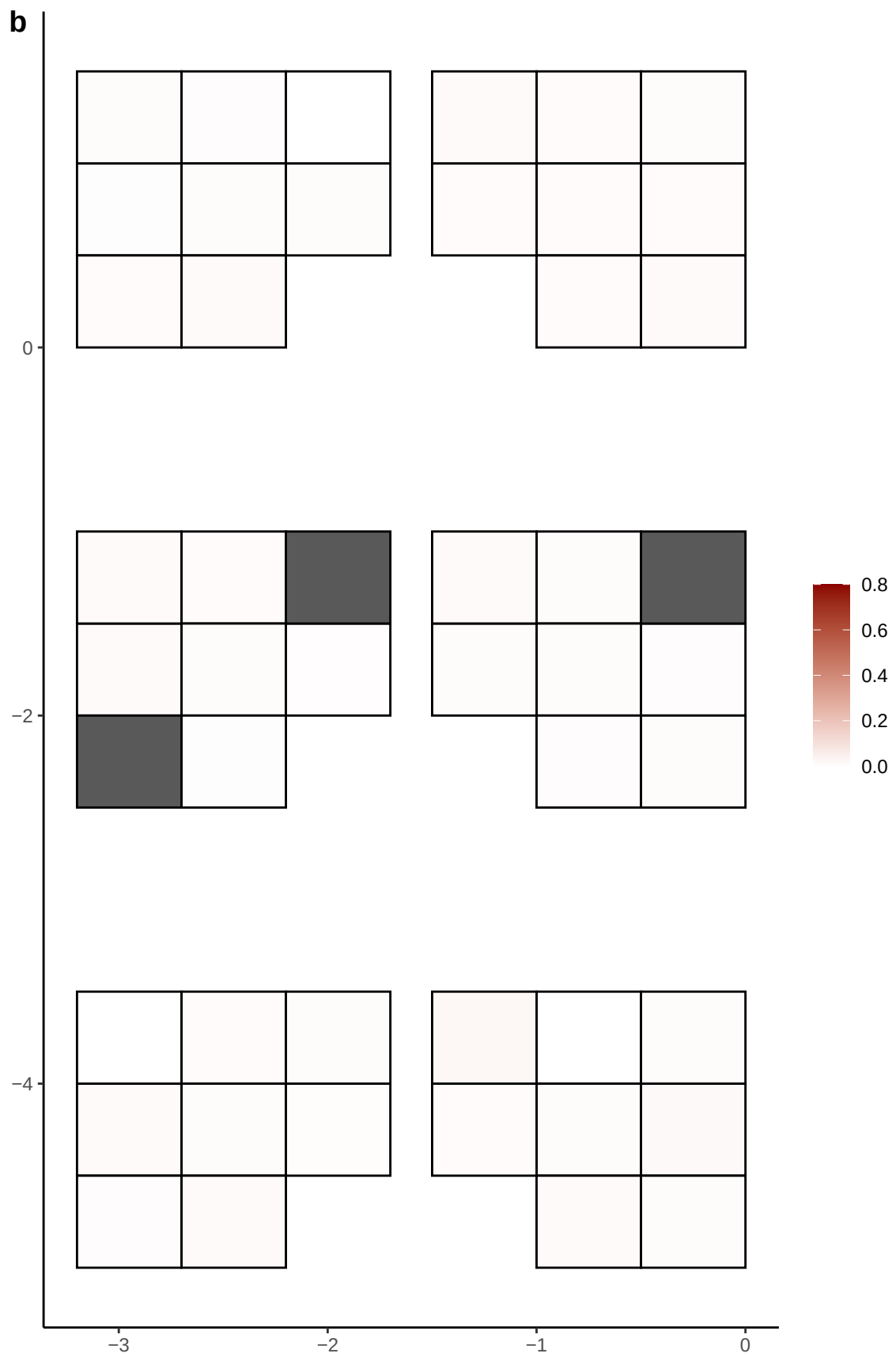

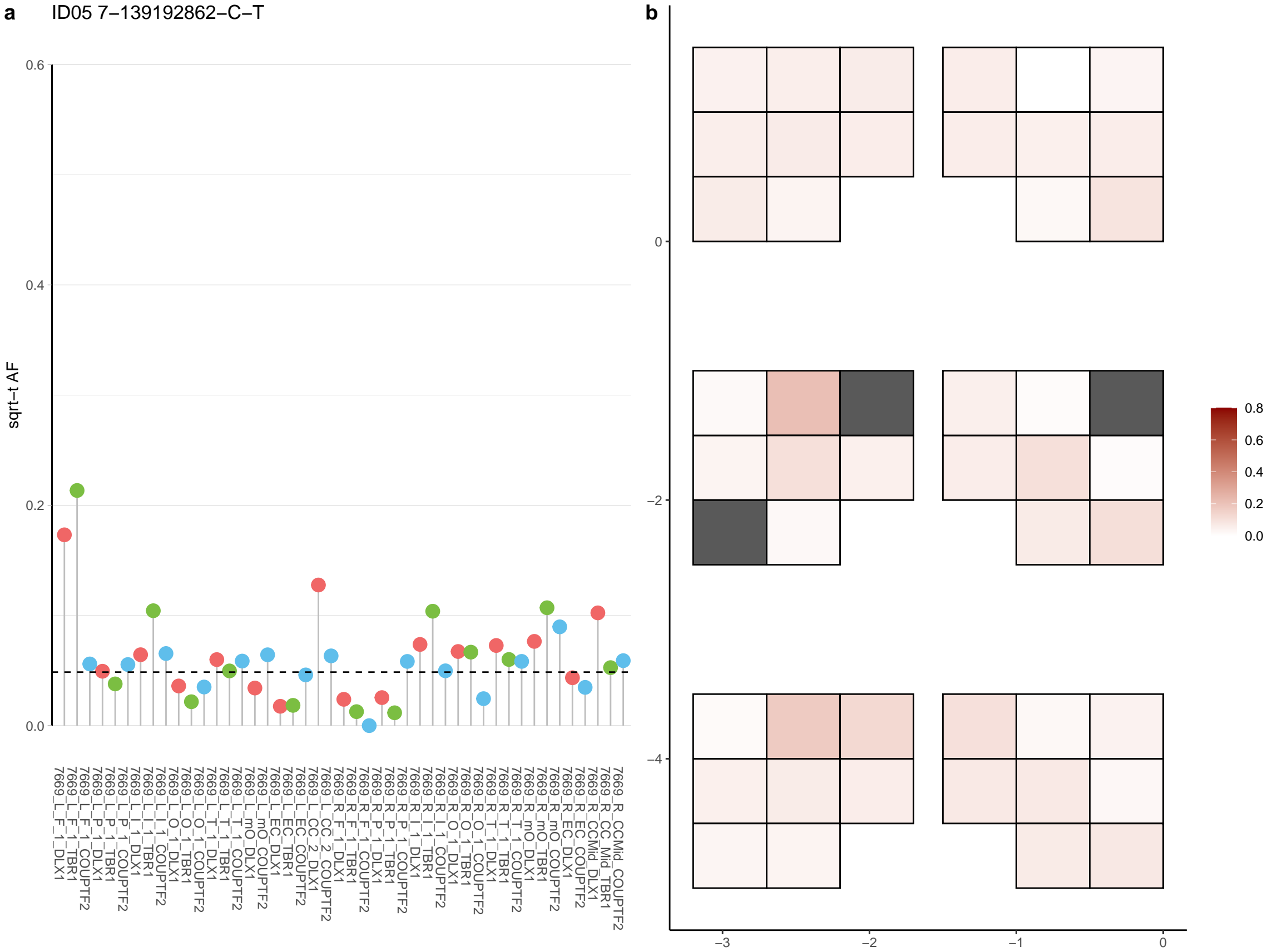

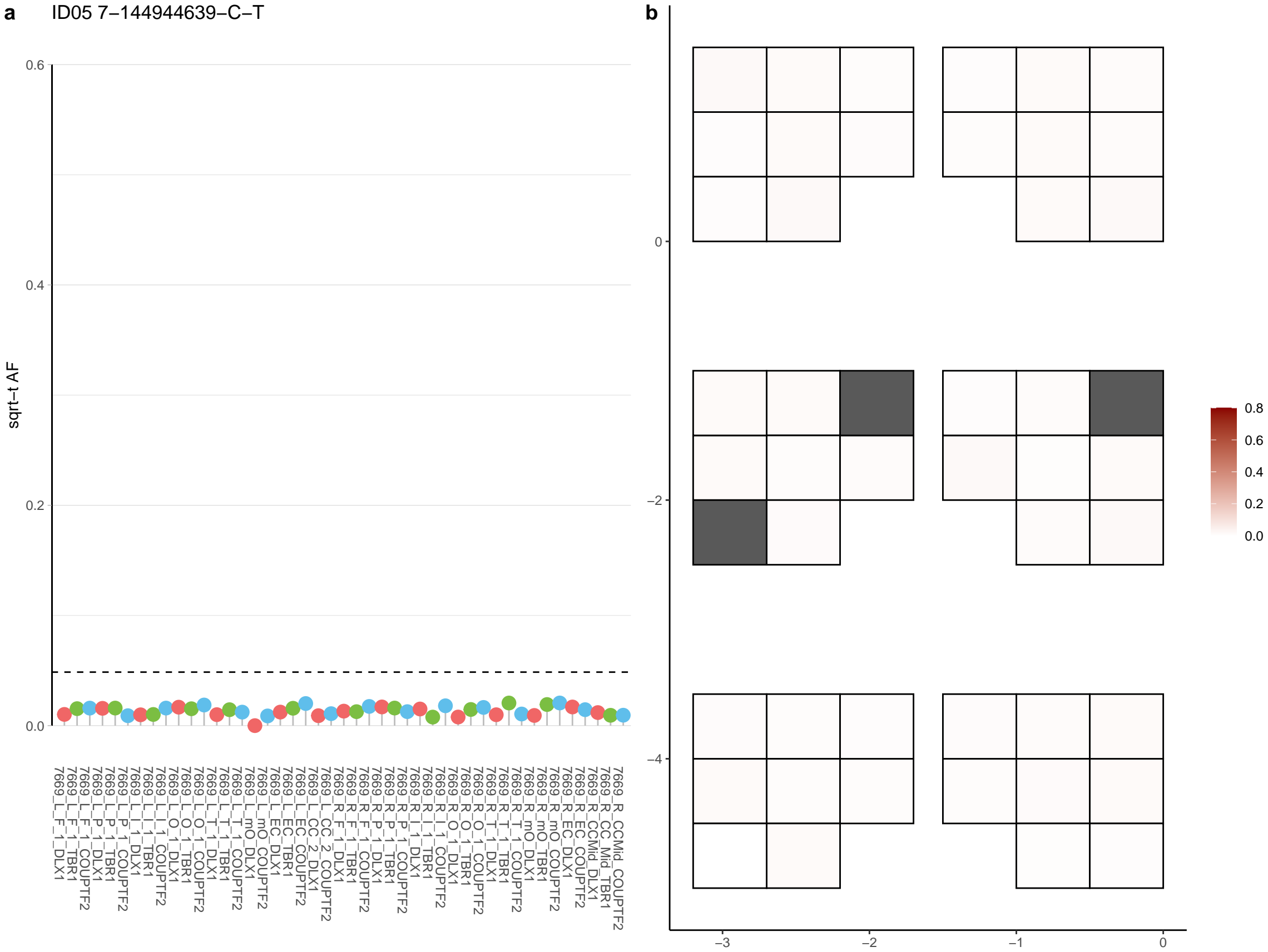

**a** ID05 7-146307789-T-A

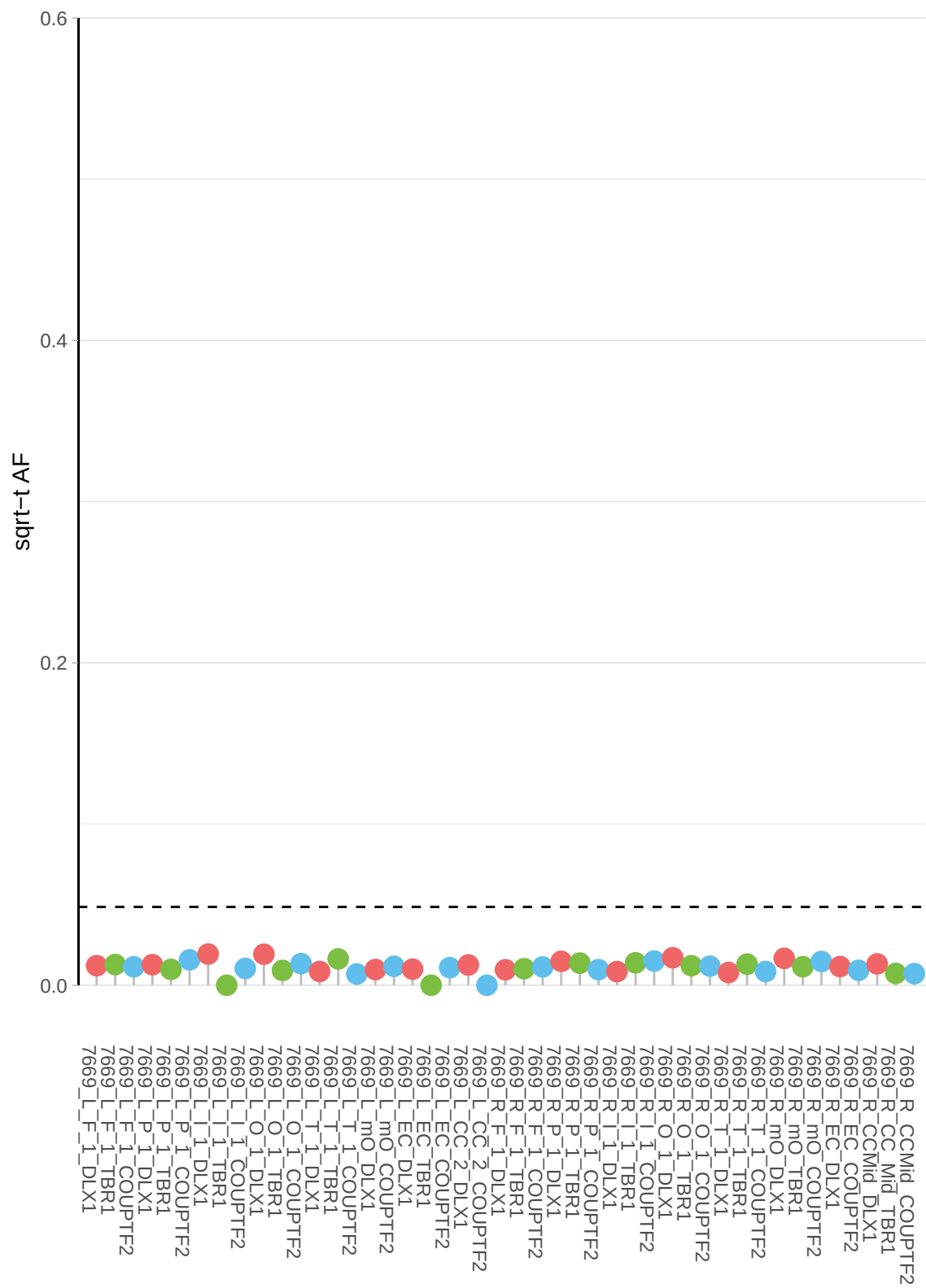

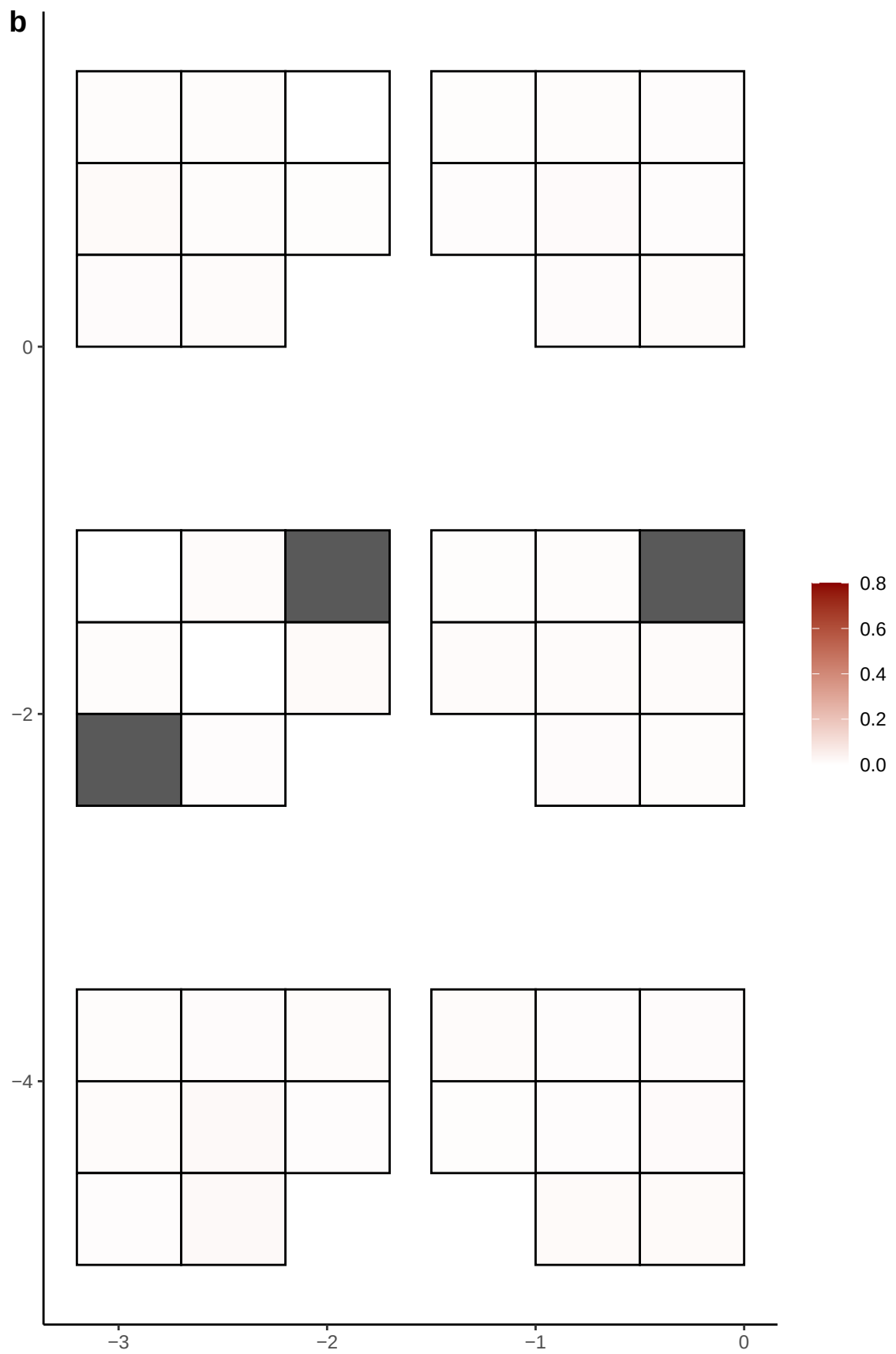

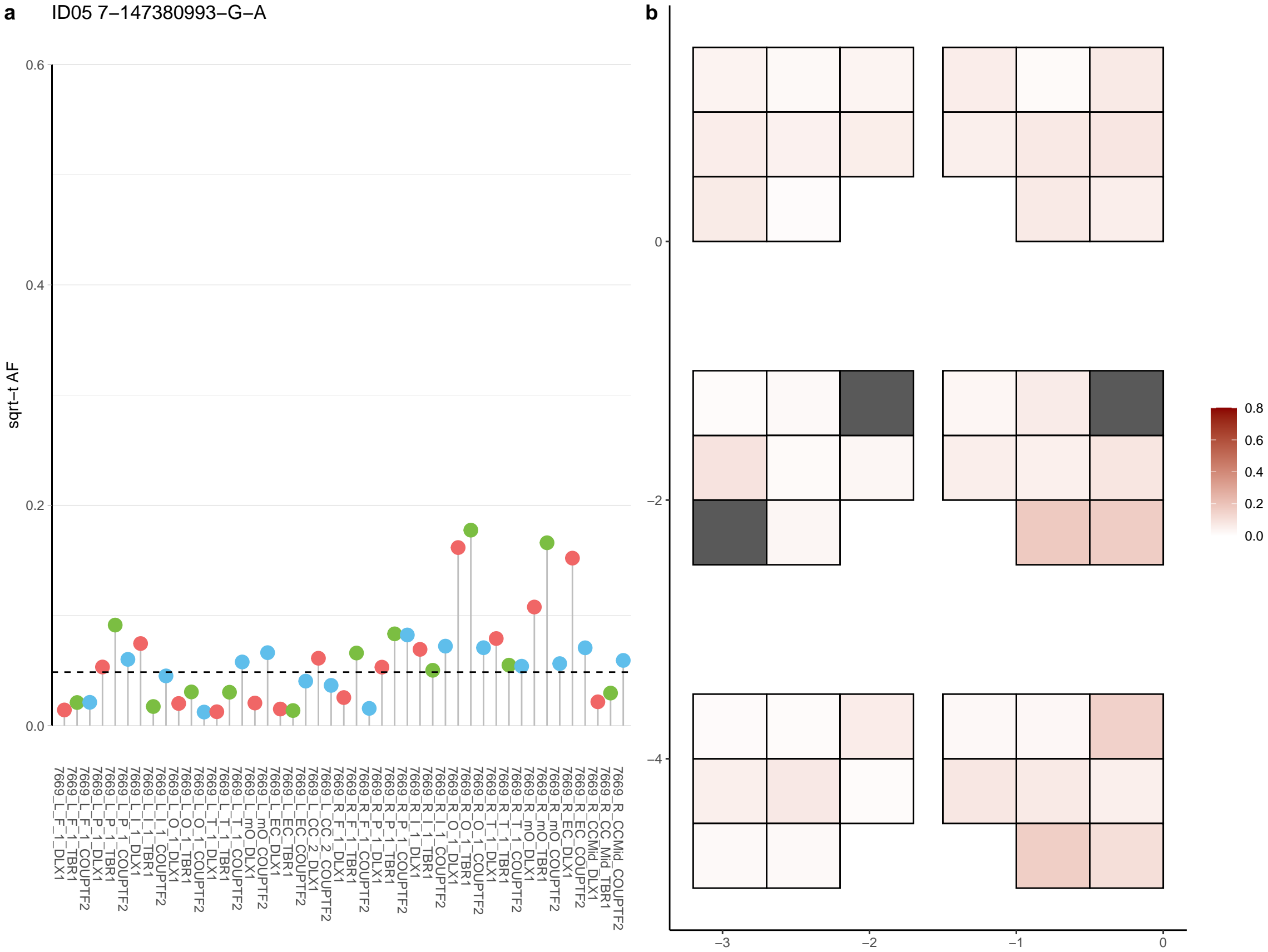

**a** ID05 7-151491278-C-T

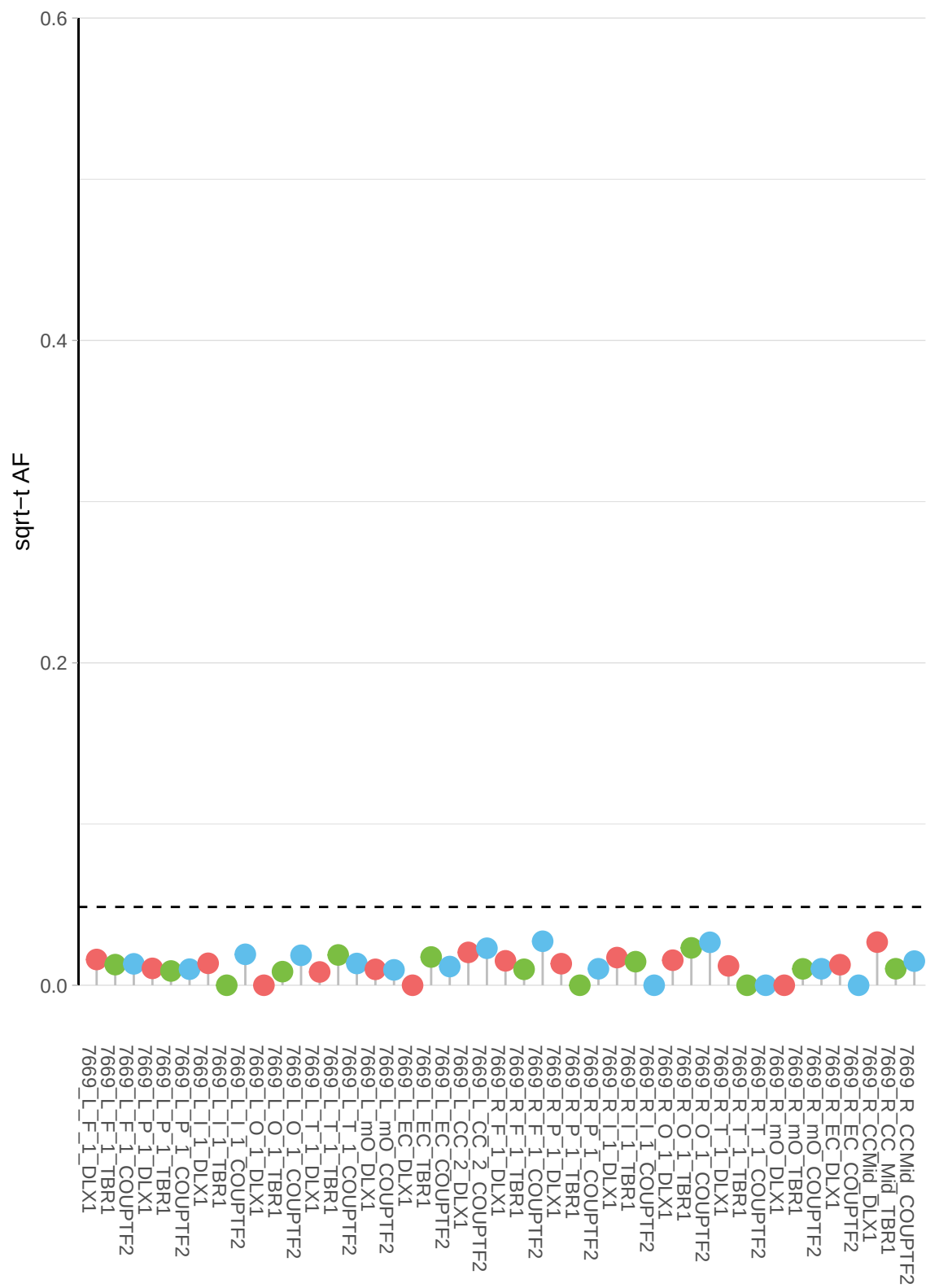

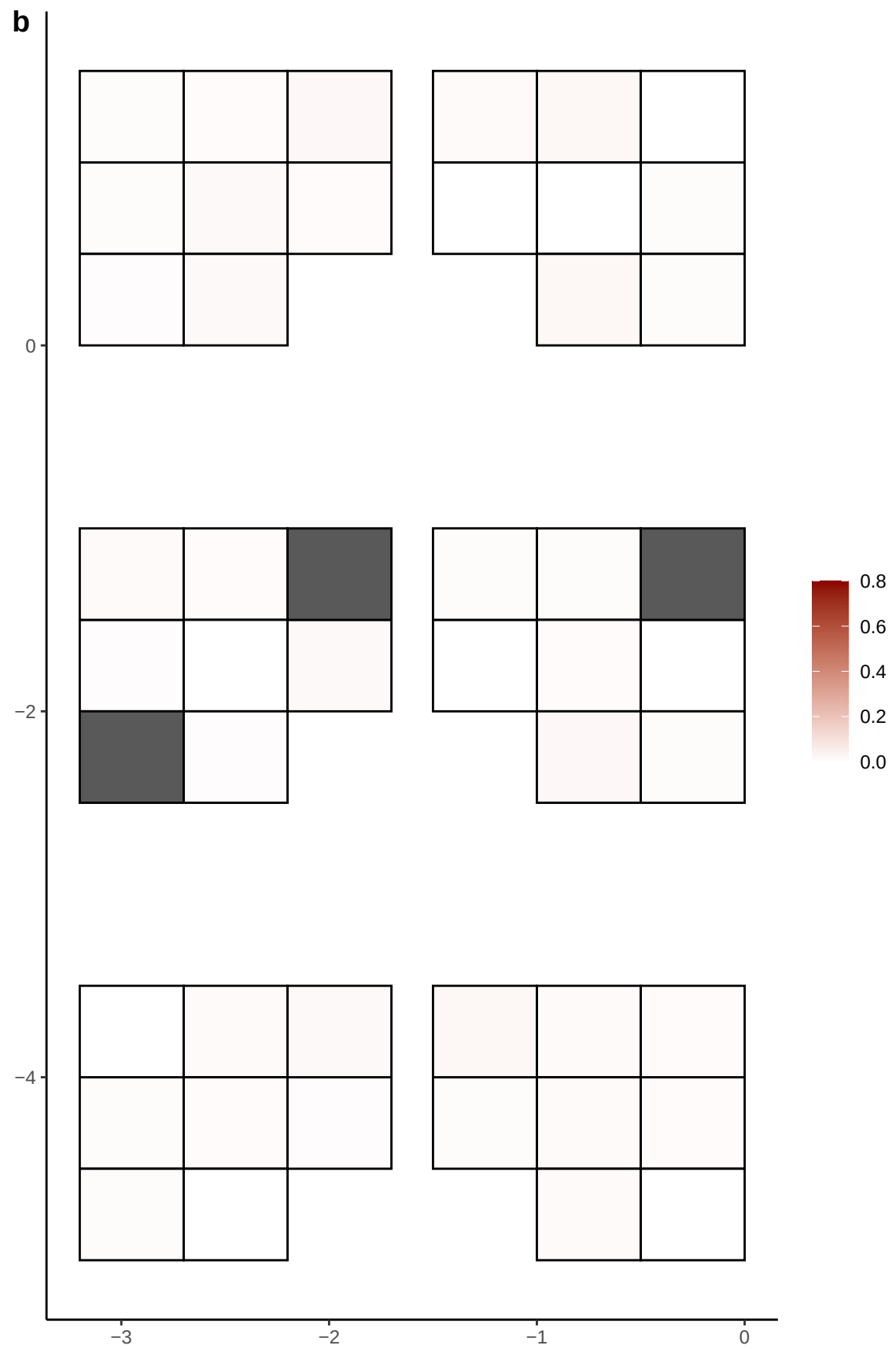

**a** ID05 7-154551172-G-A

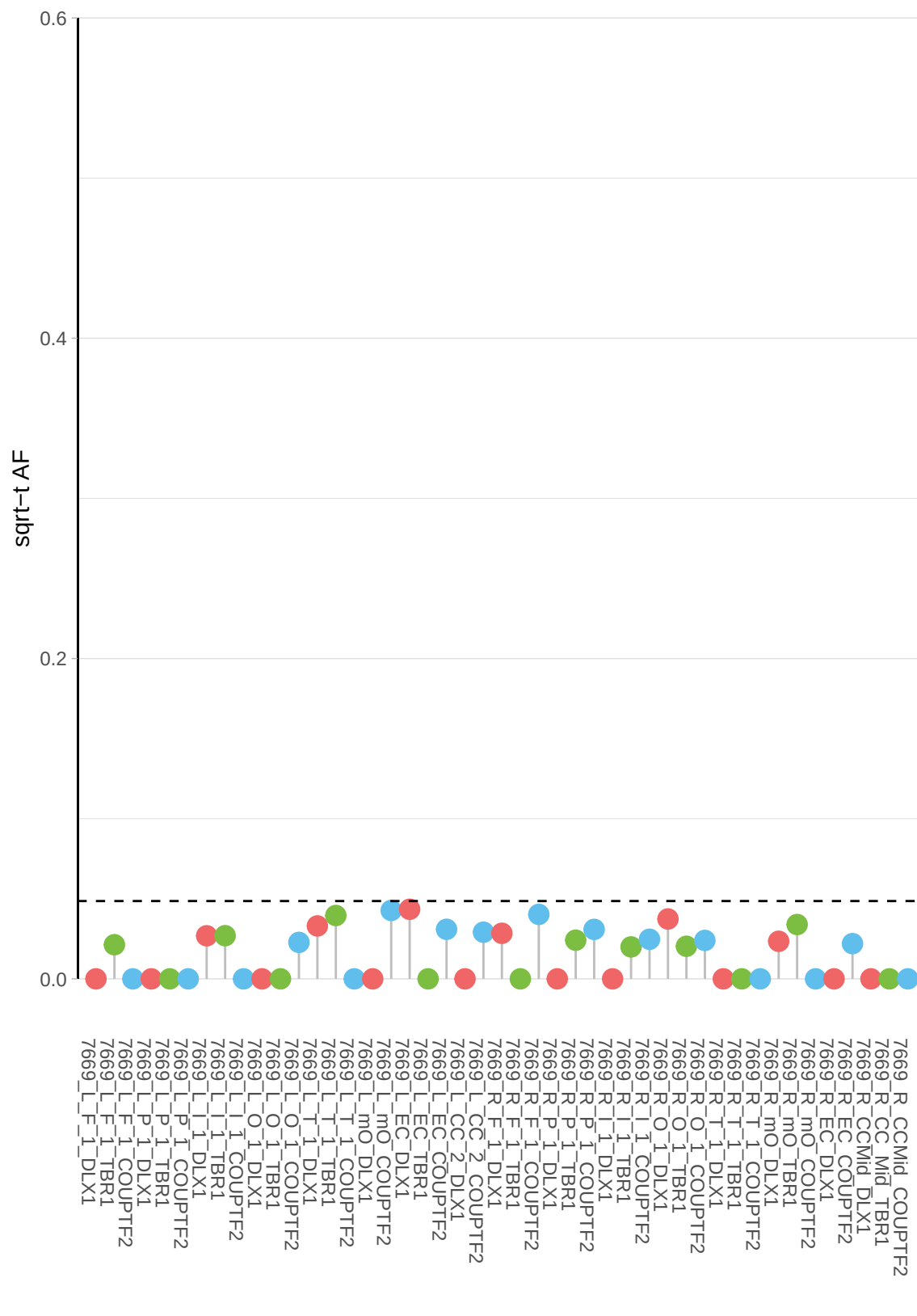

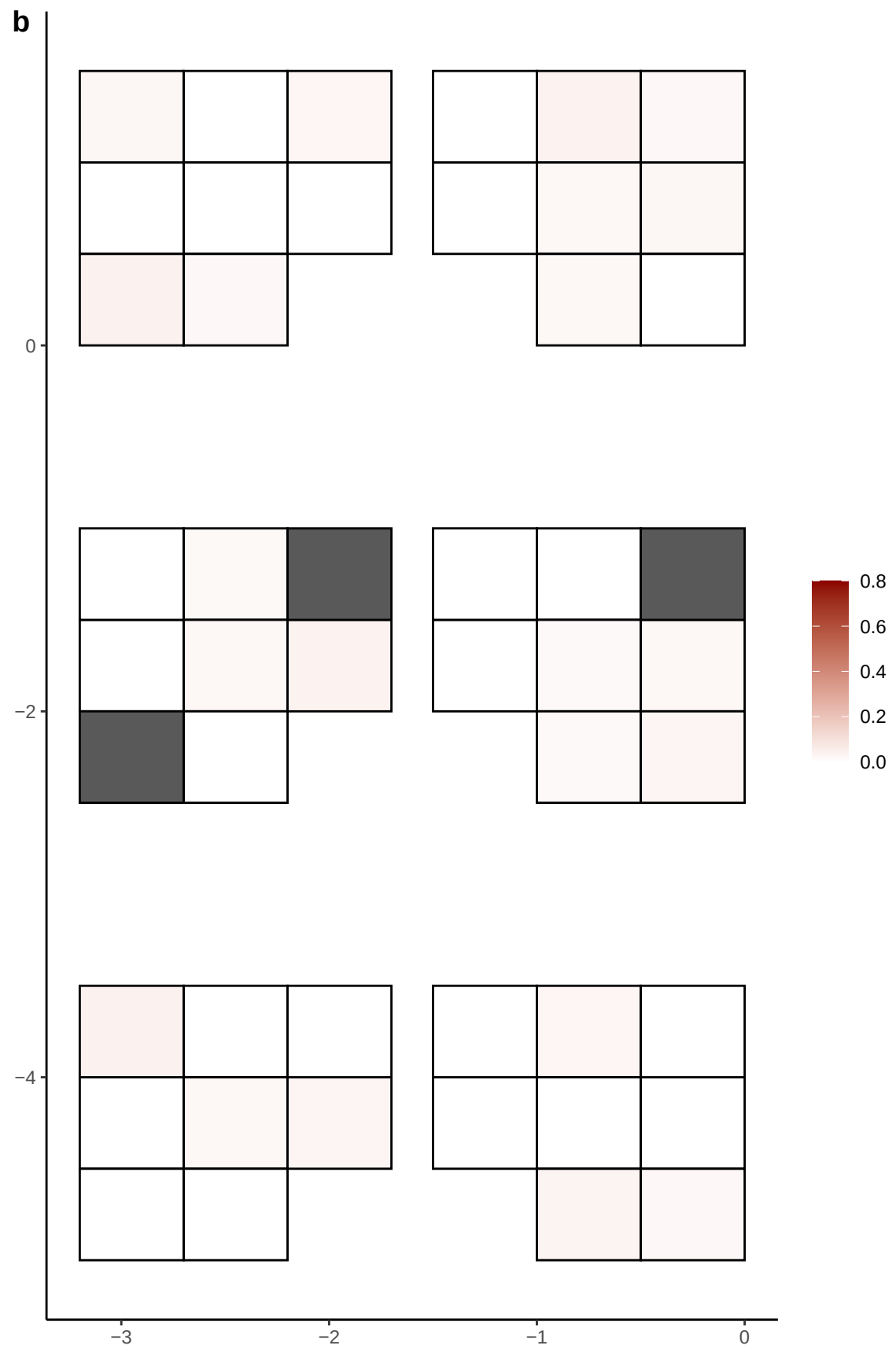

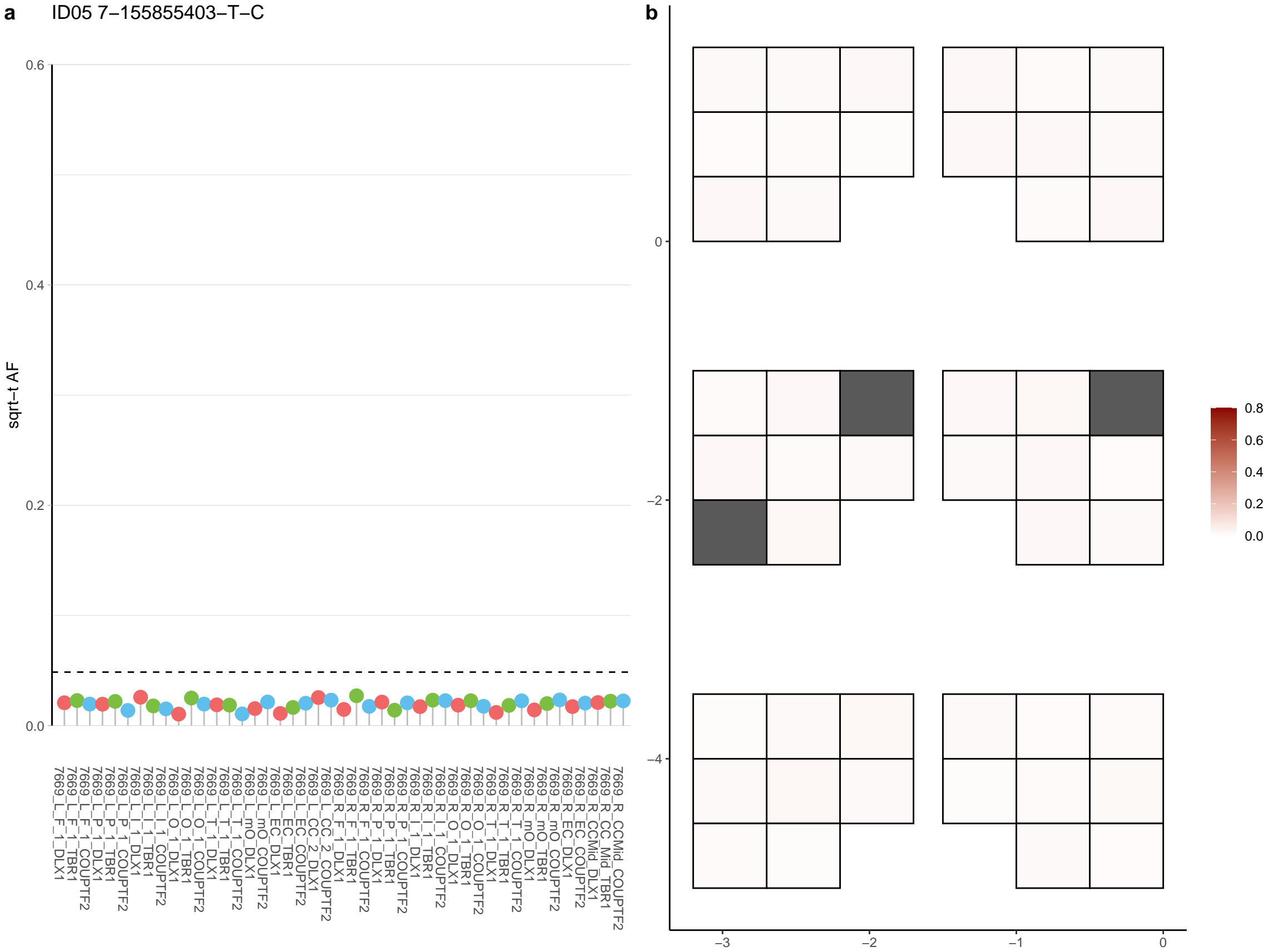

**a** ID05 7-16742061-C-T

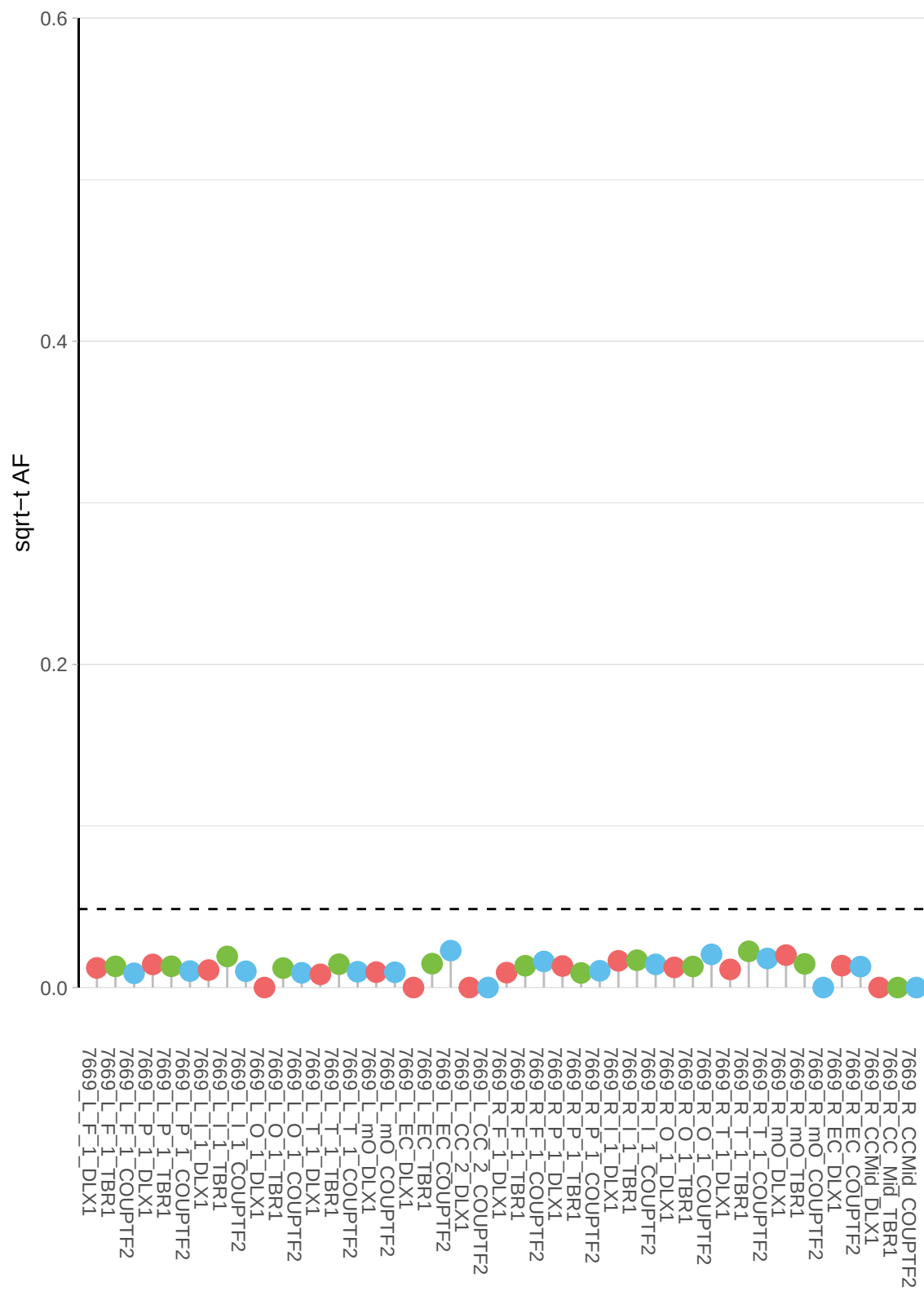

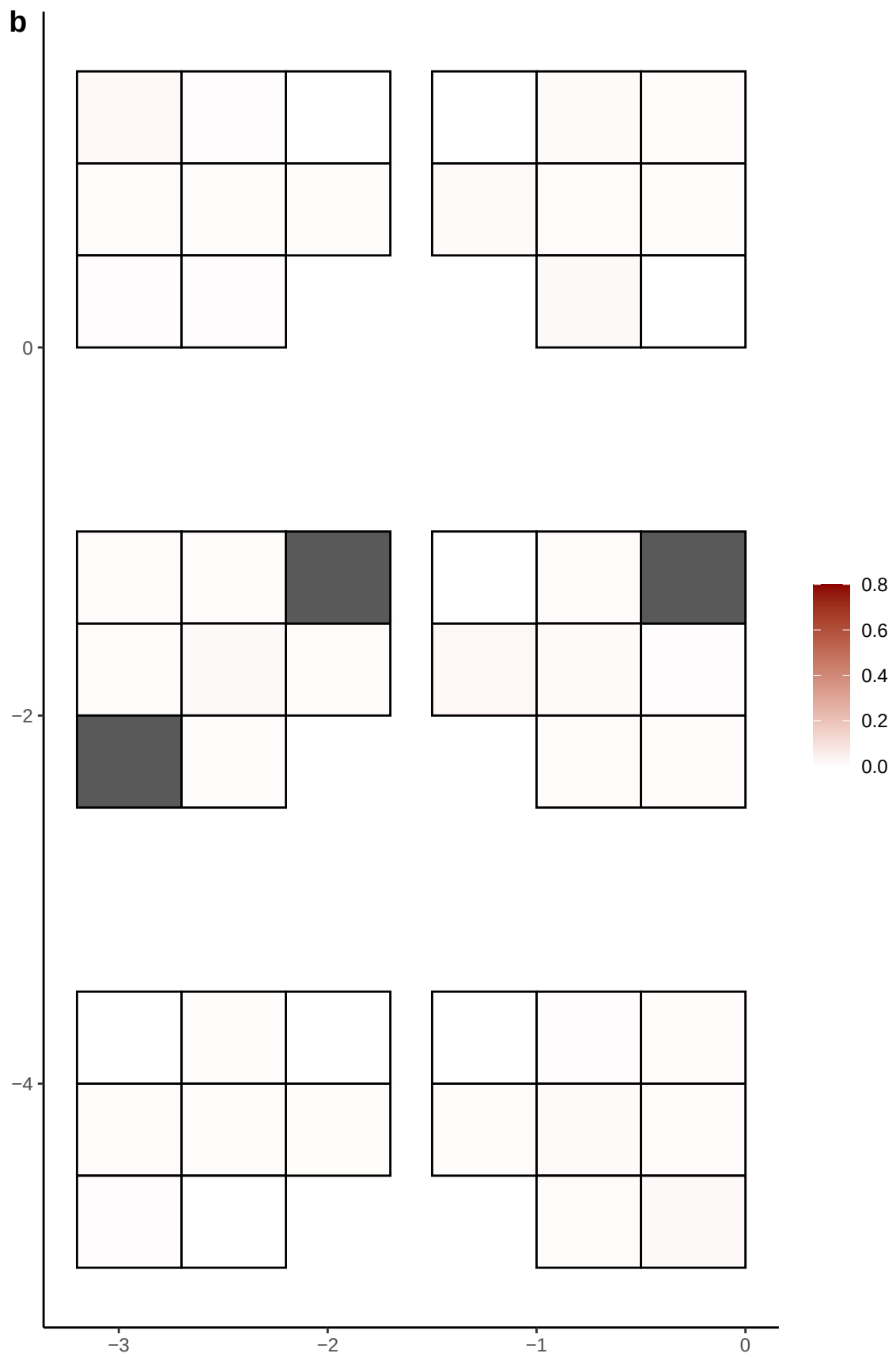

**a** ID05 7-19648848-A-G

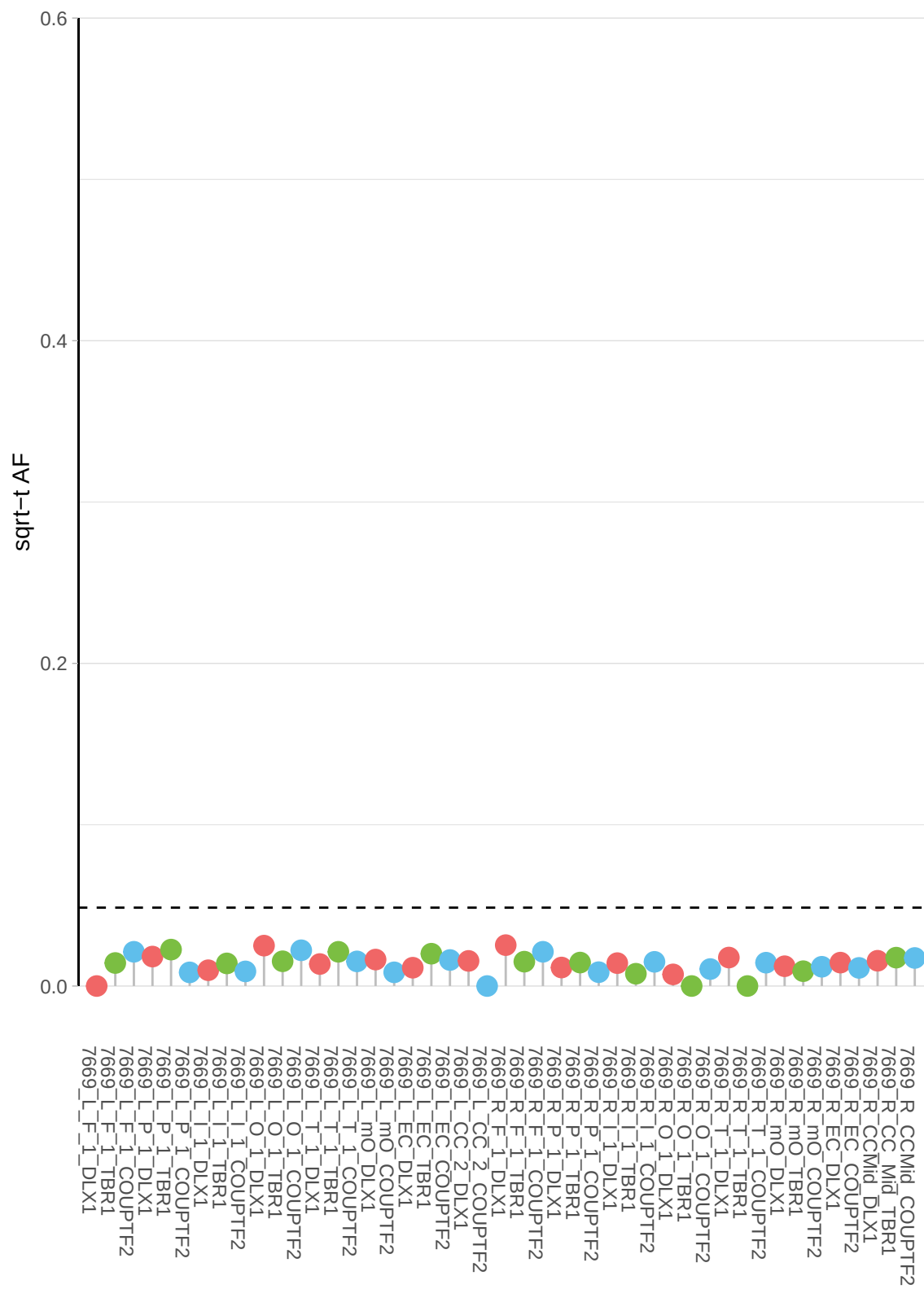

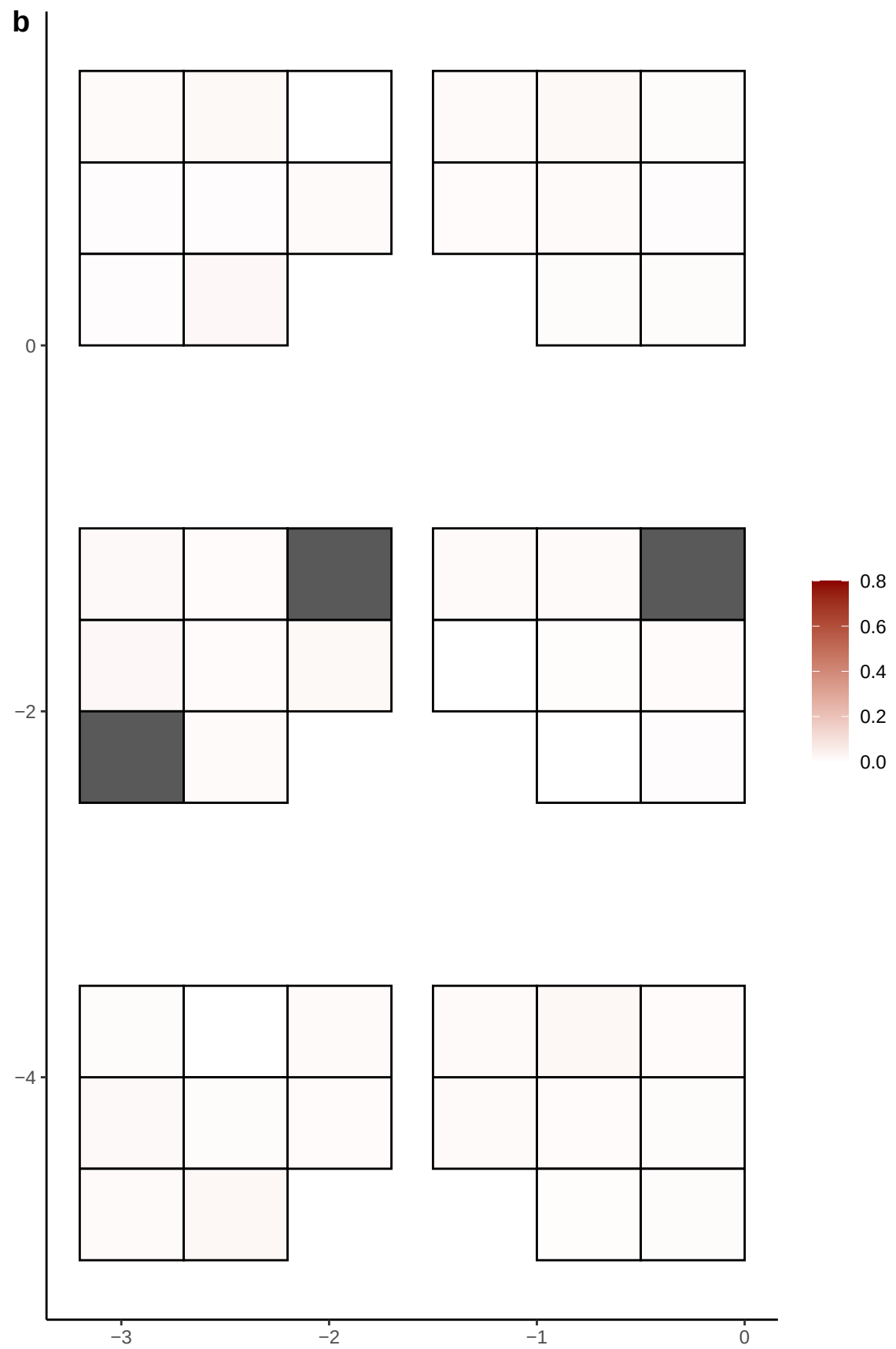

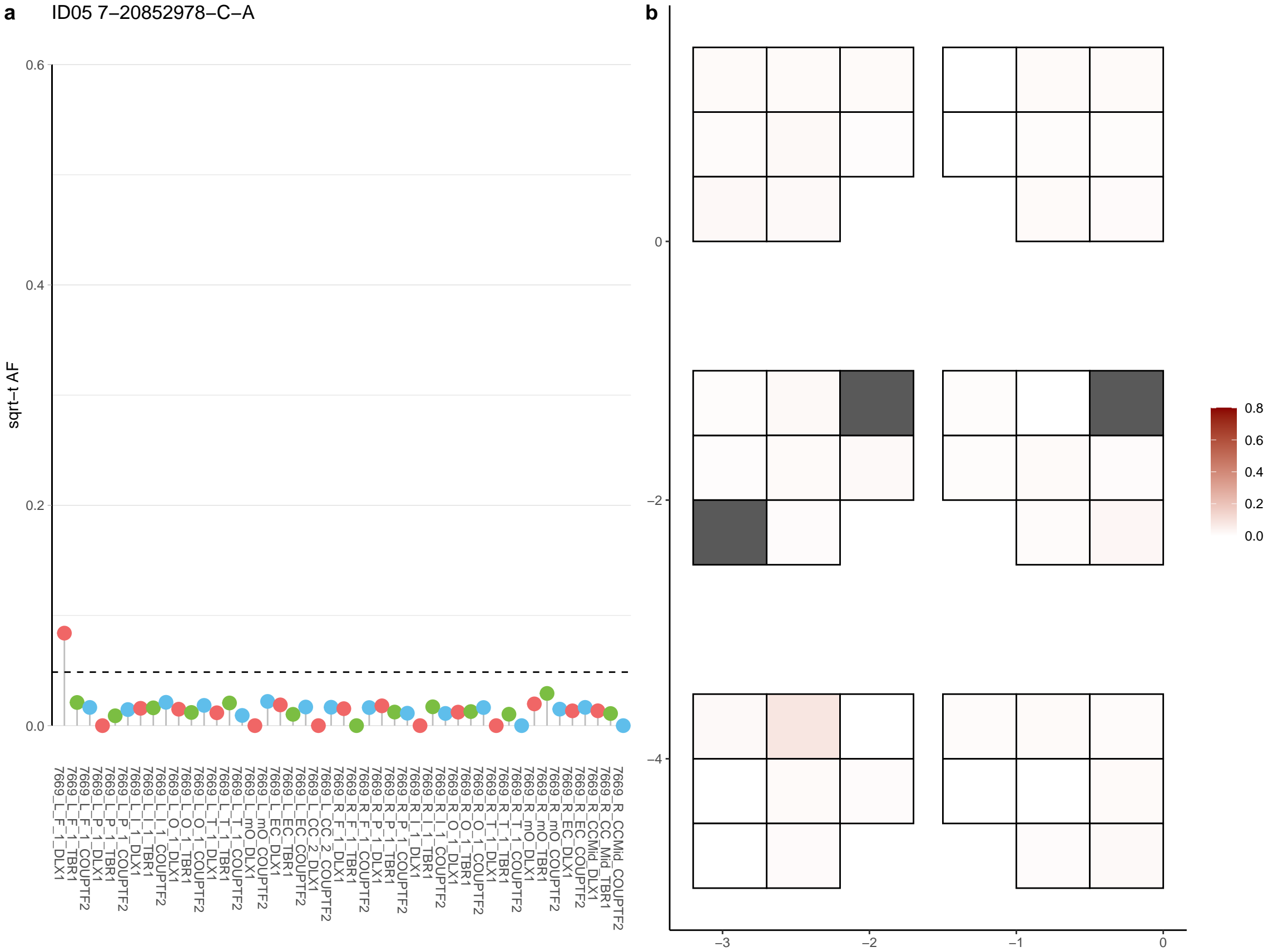

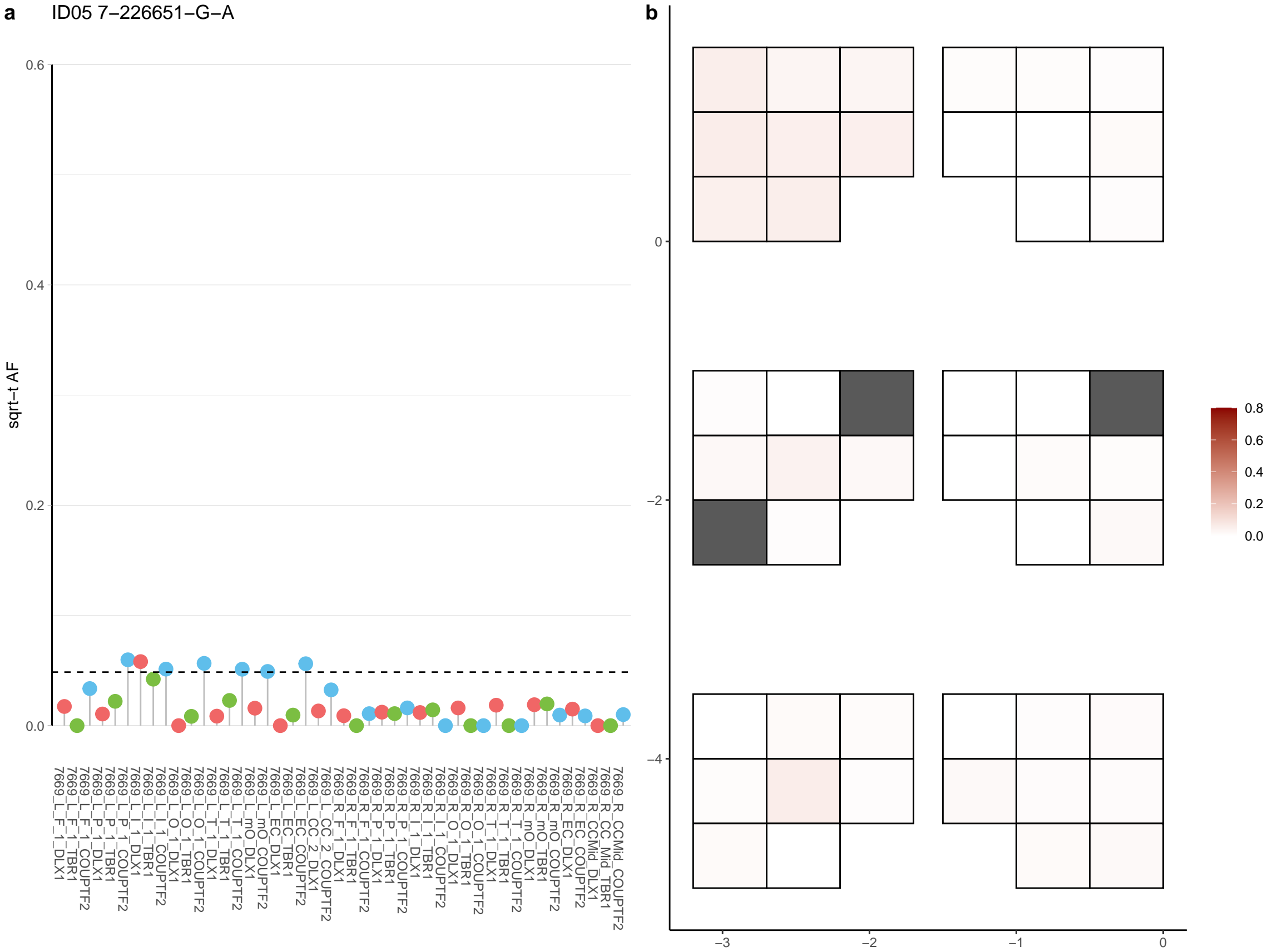

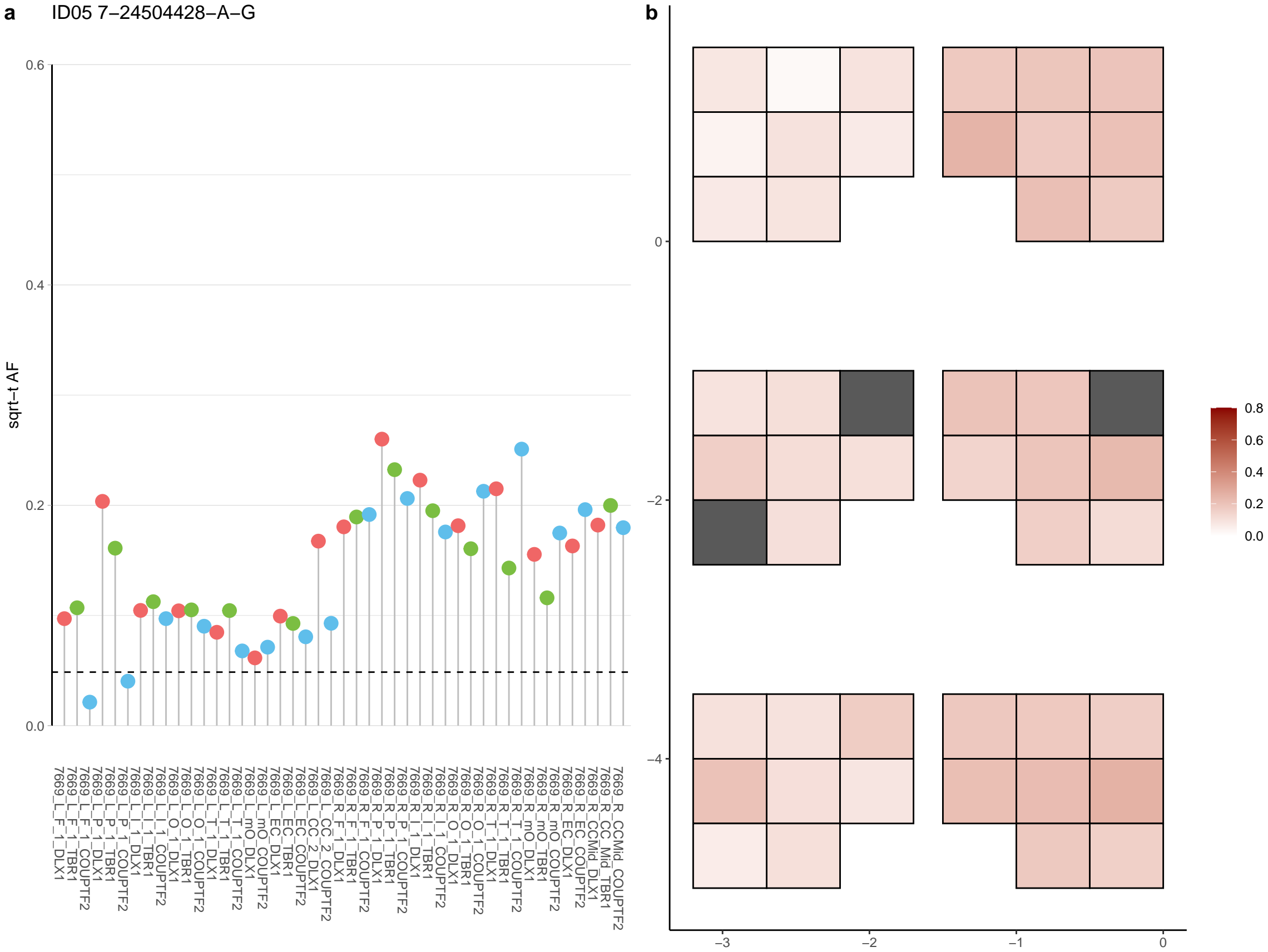

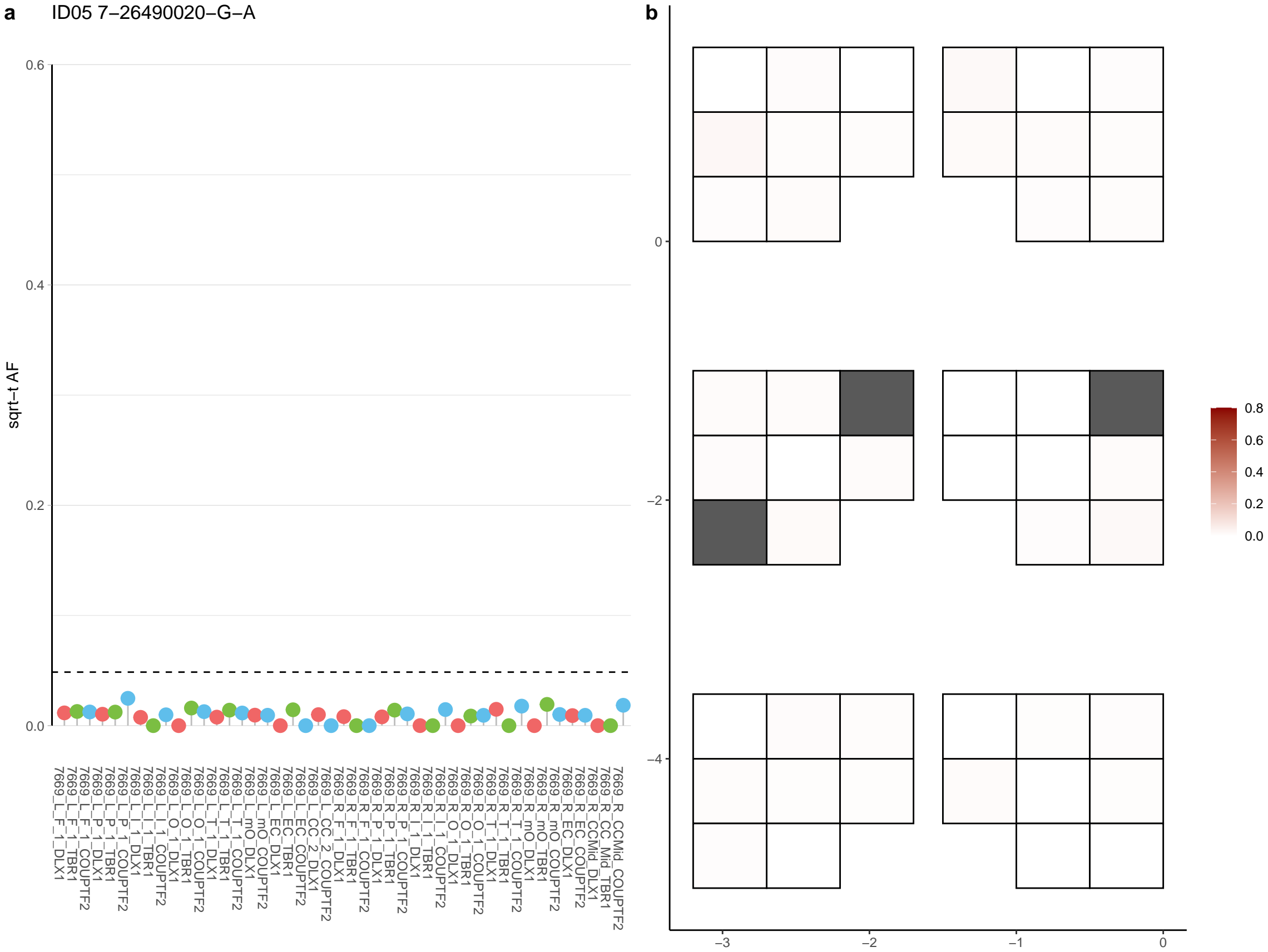

**a** ID05 7-30962886-G-A

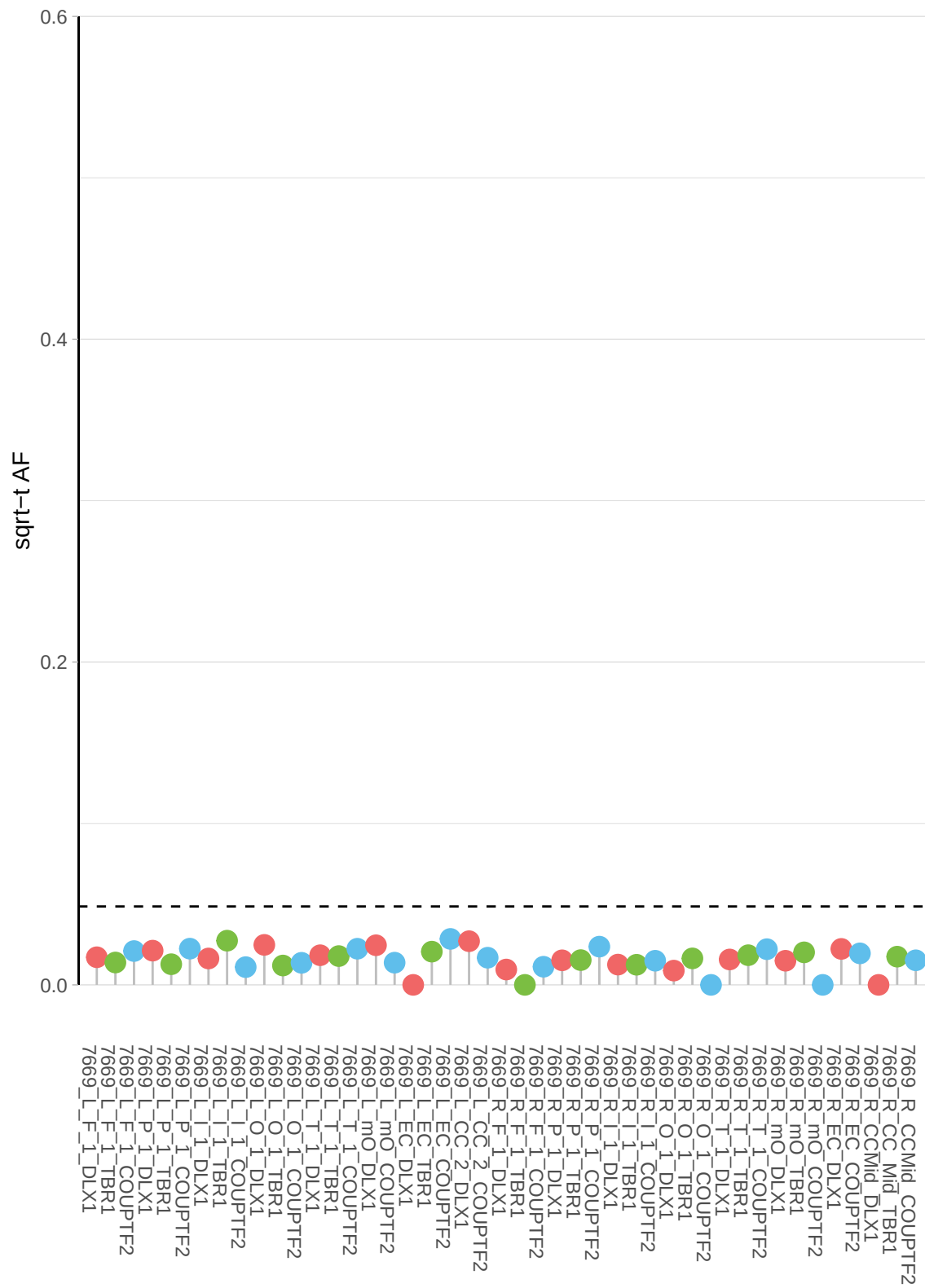

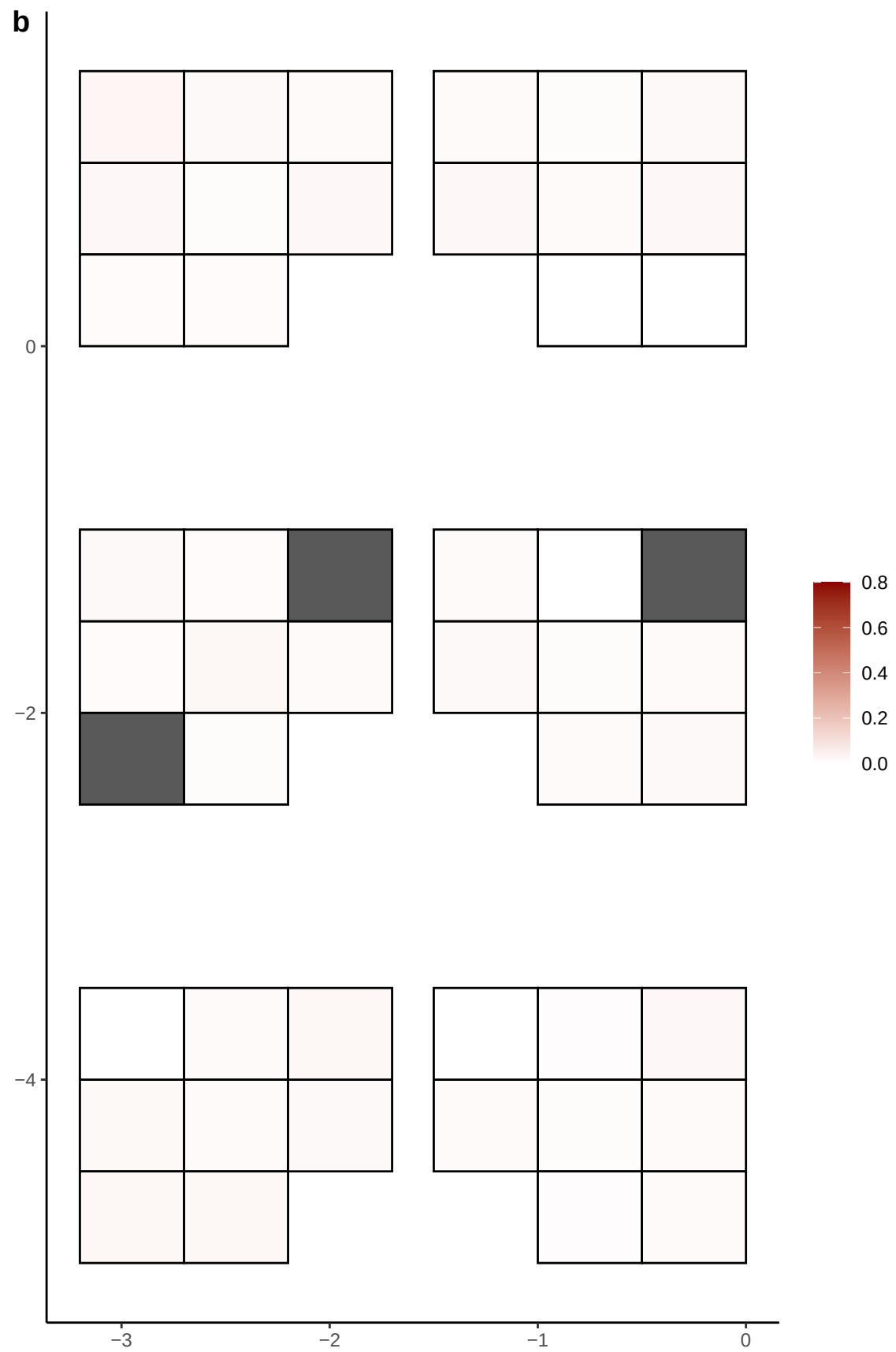

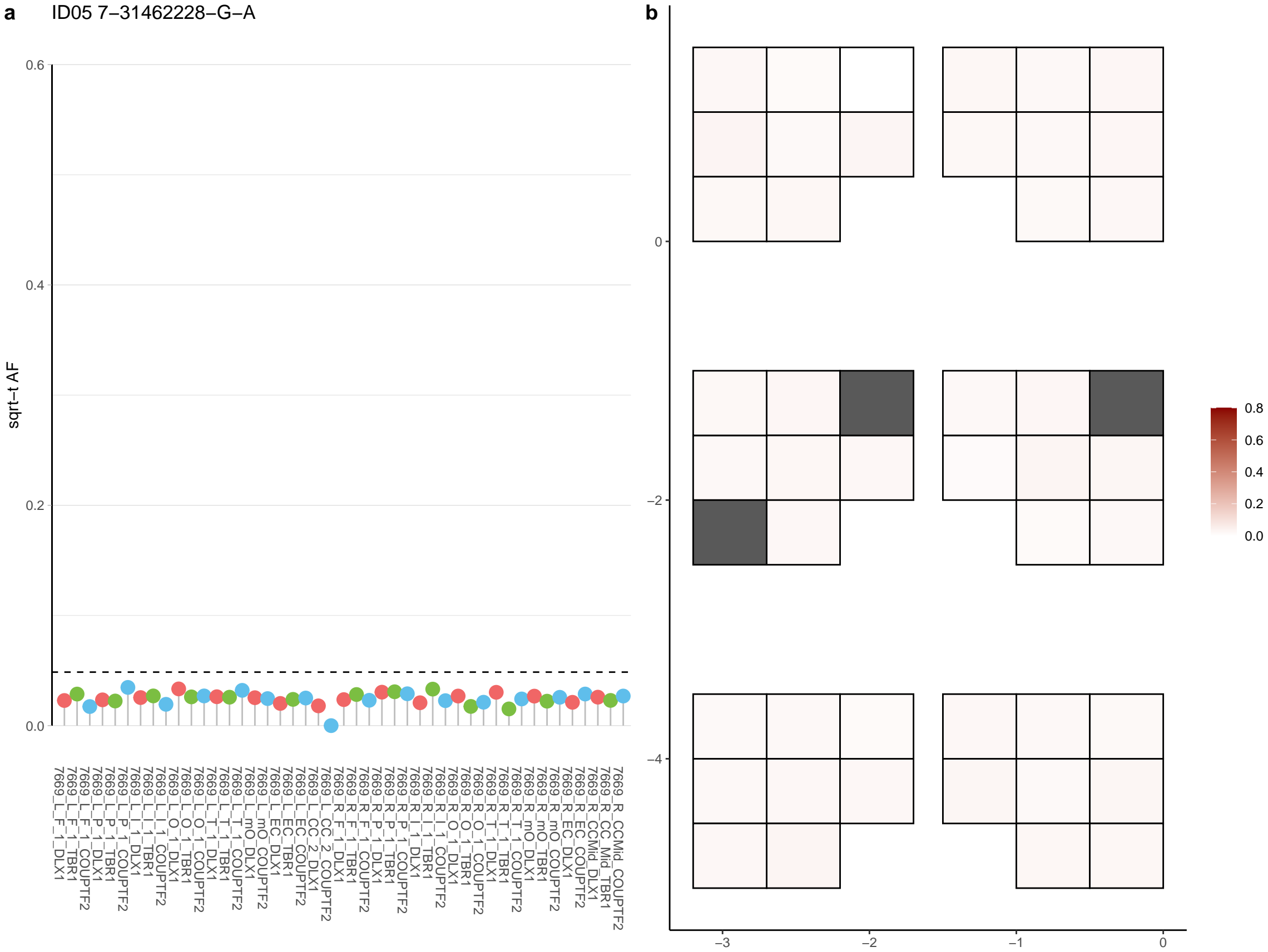

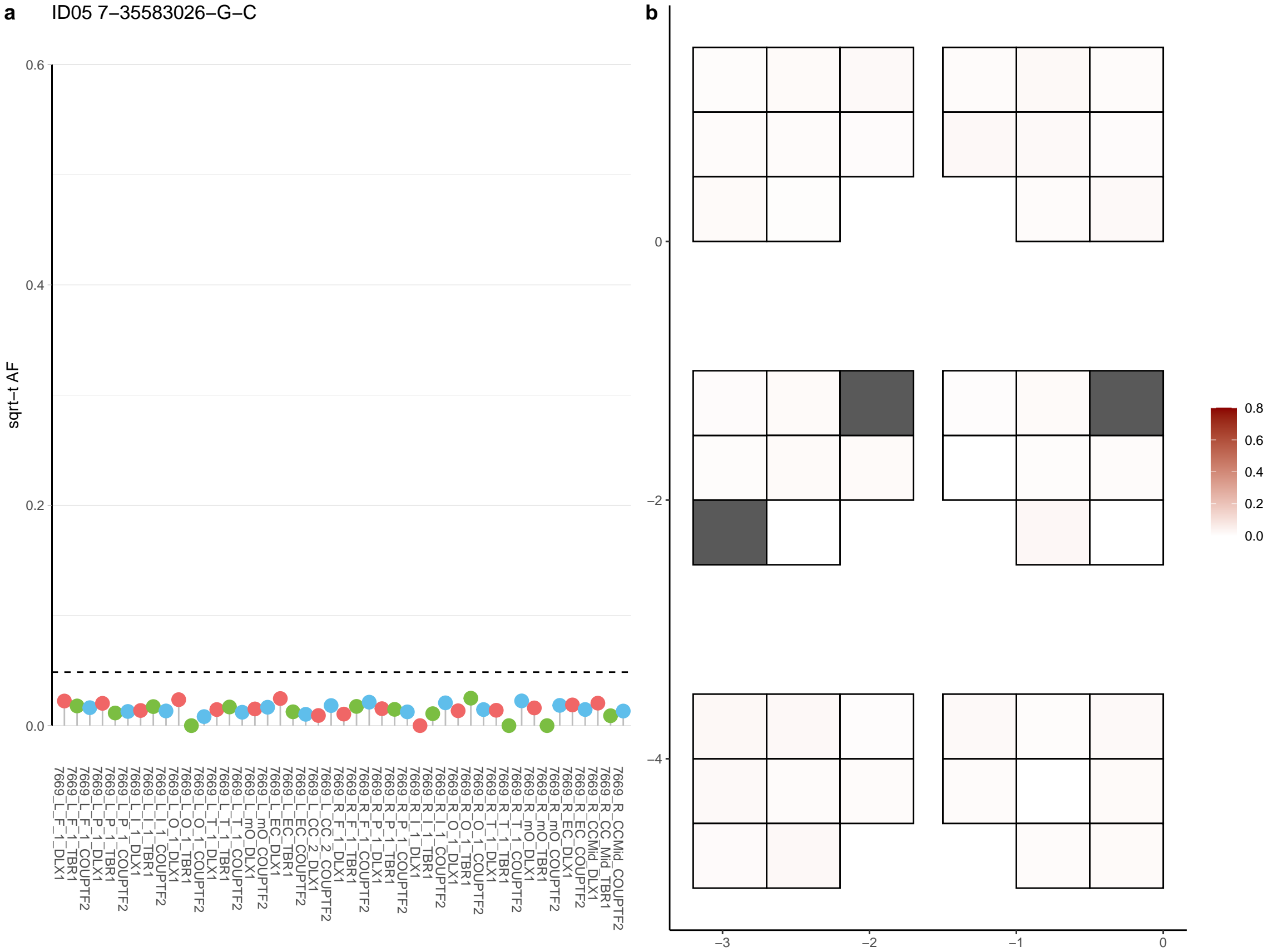

**a** ID05 7-41735414-A-G

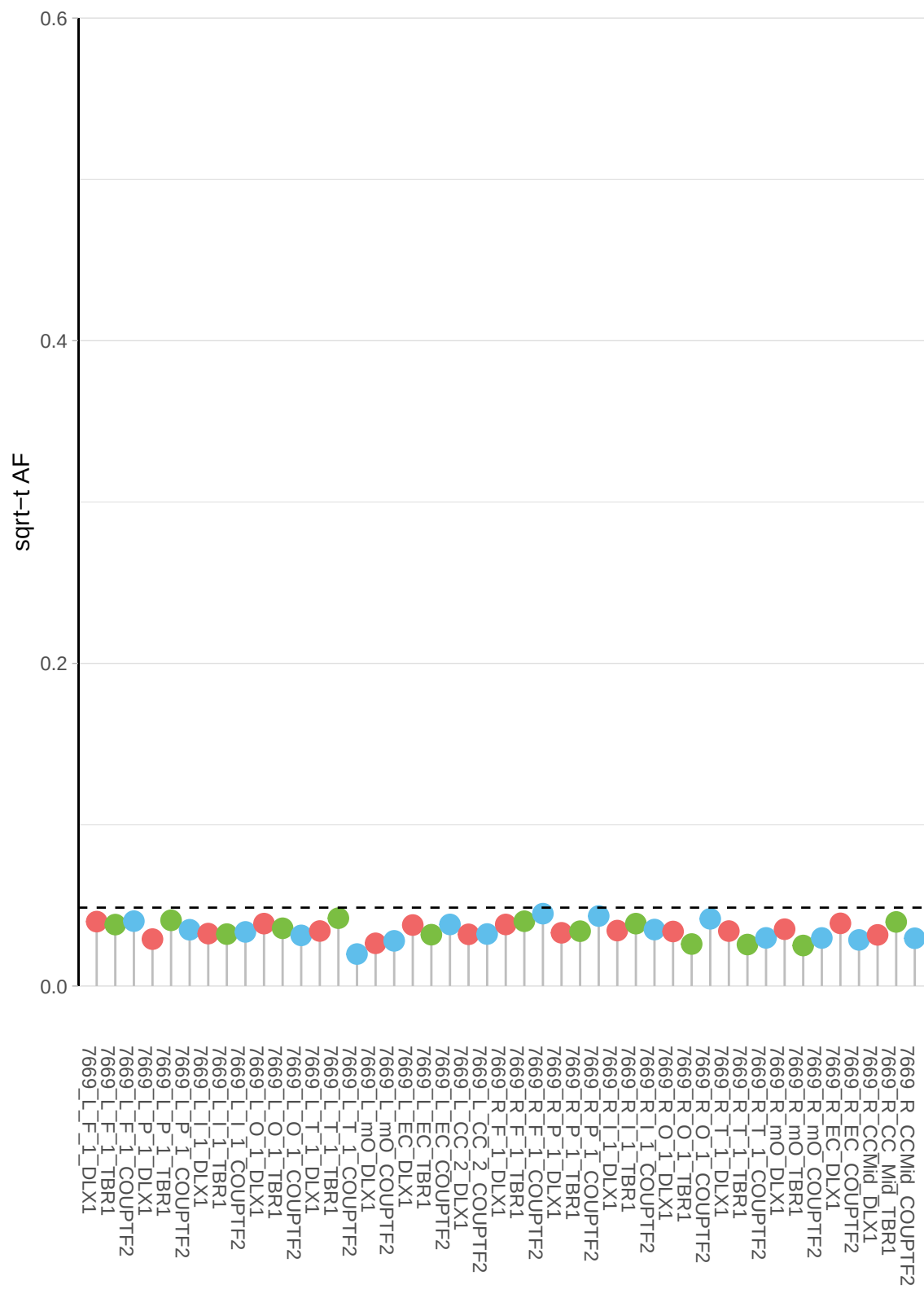

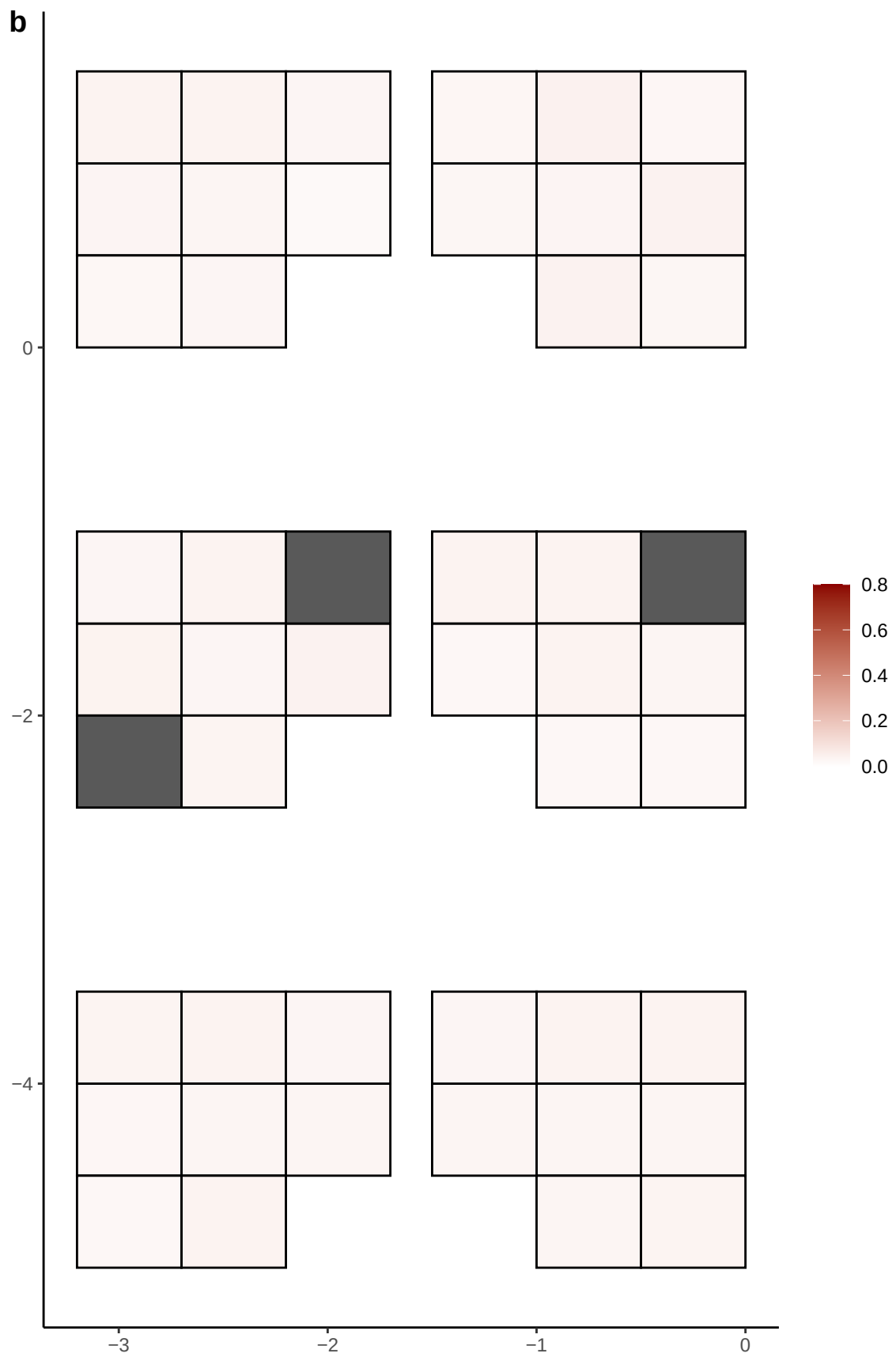

**a** ID05 7-46623362-G-C

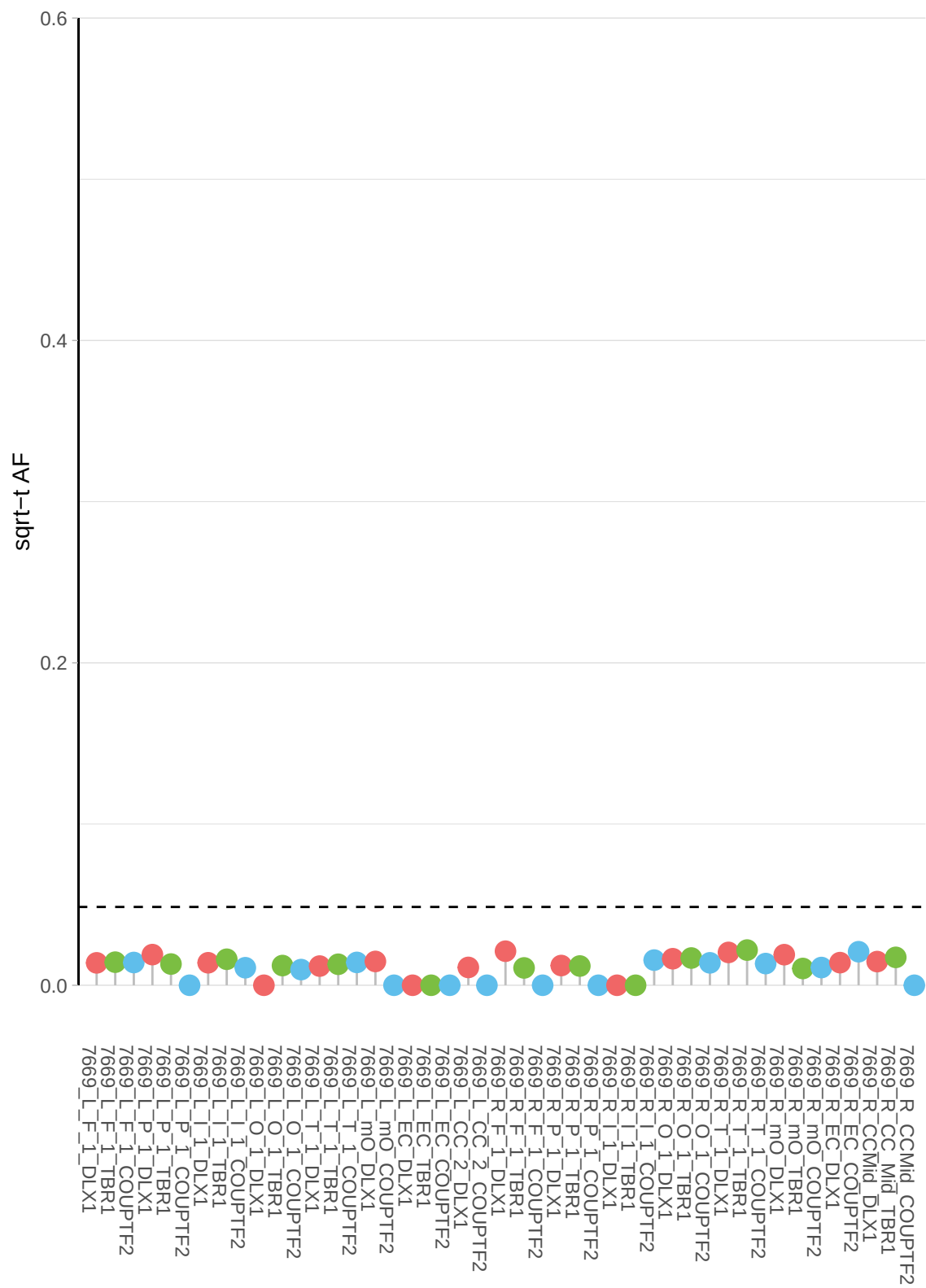

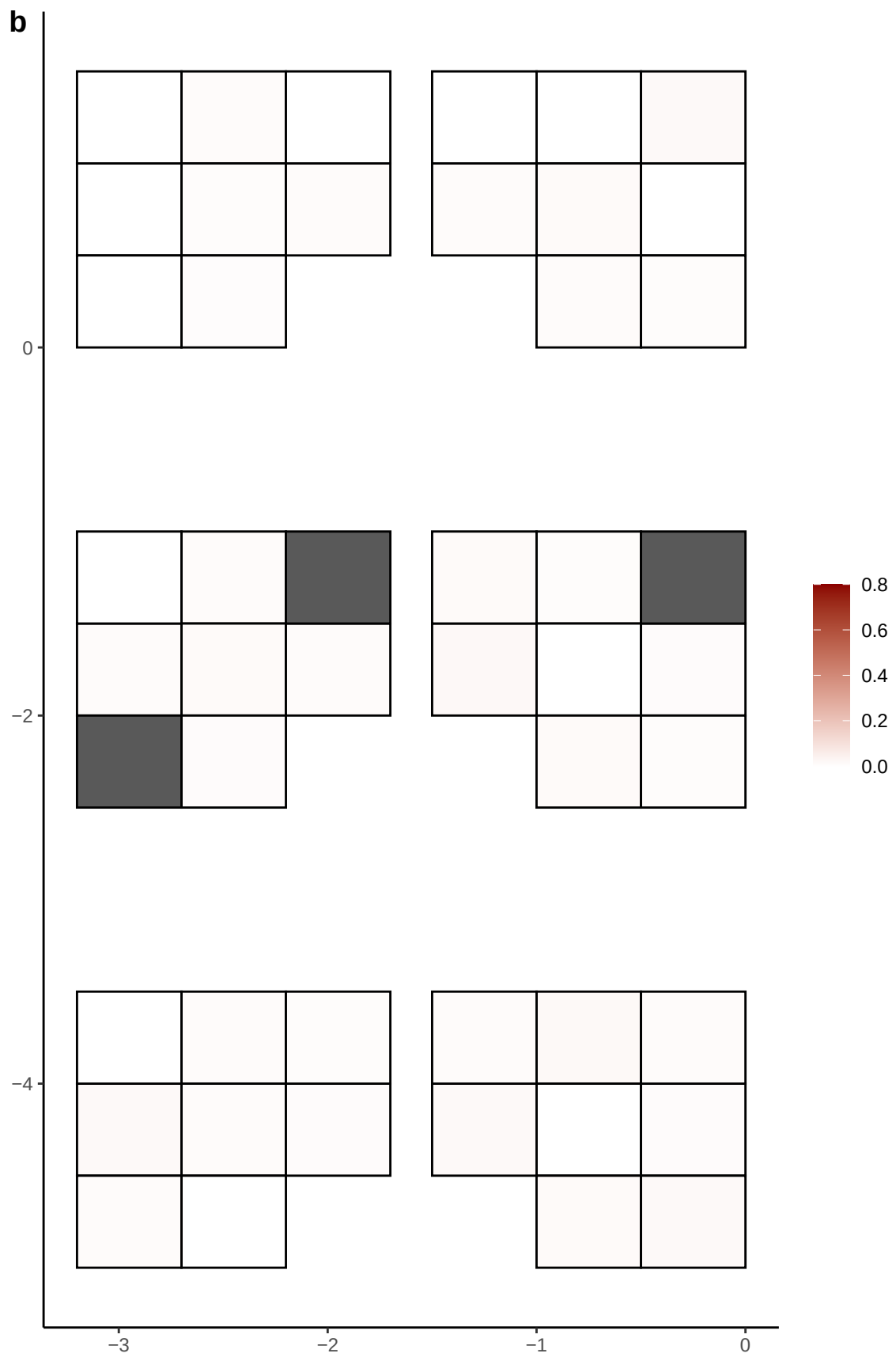

**a** ID05 7-54130903-T-G

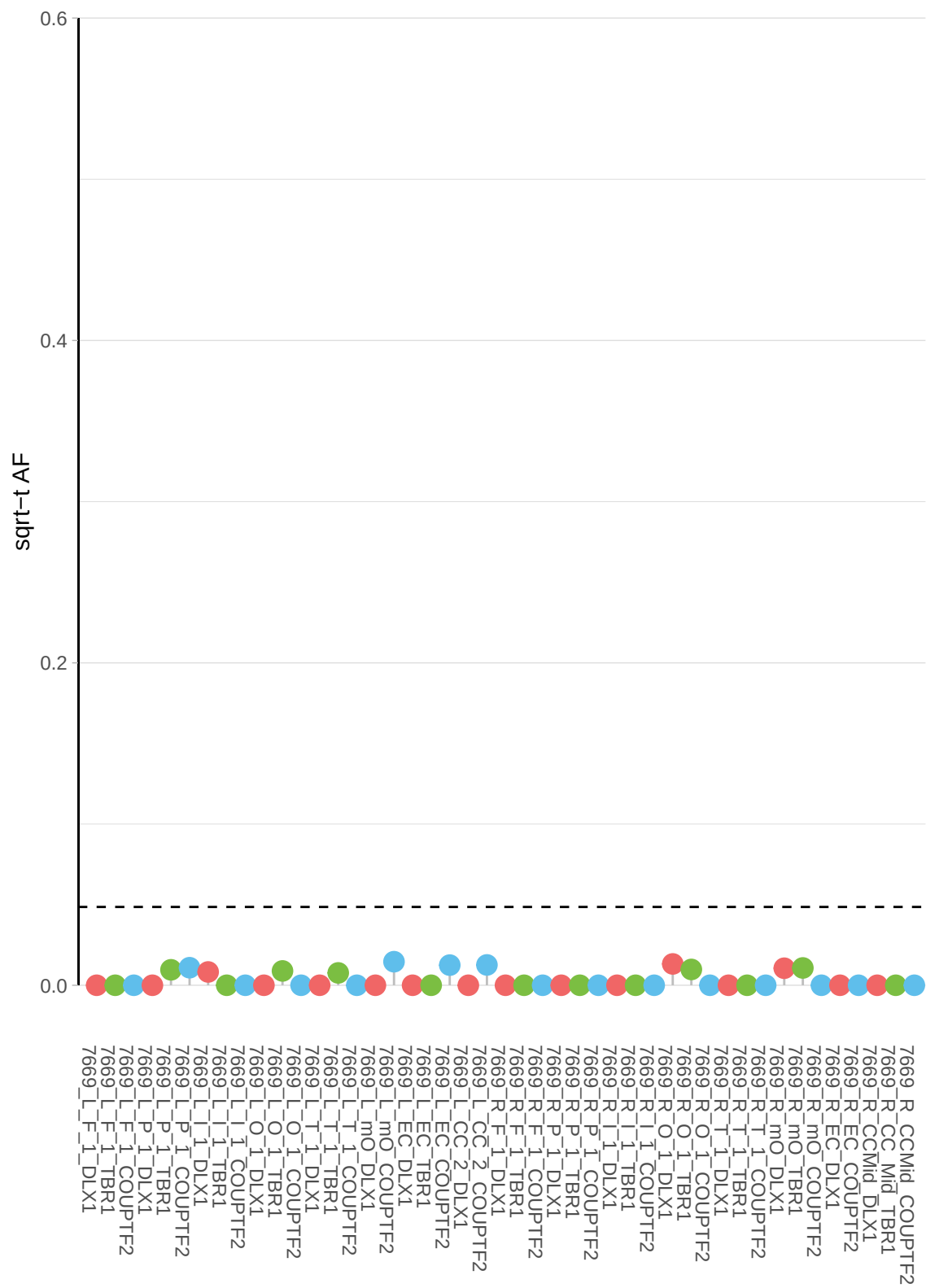

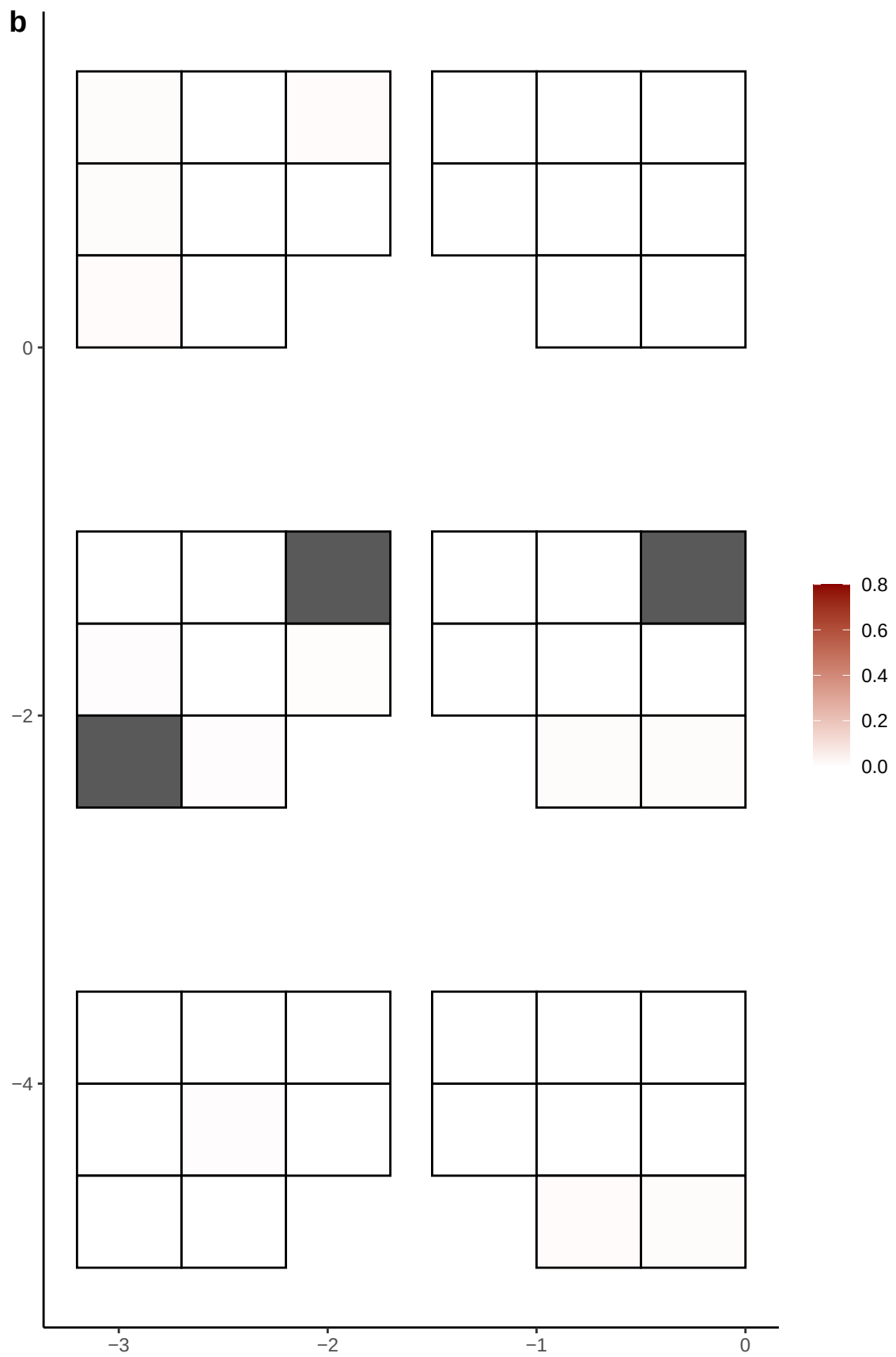

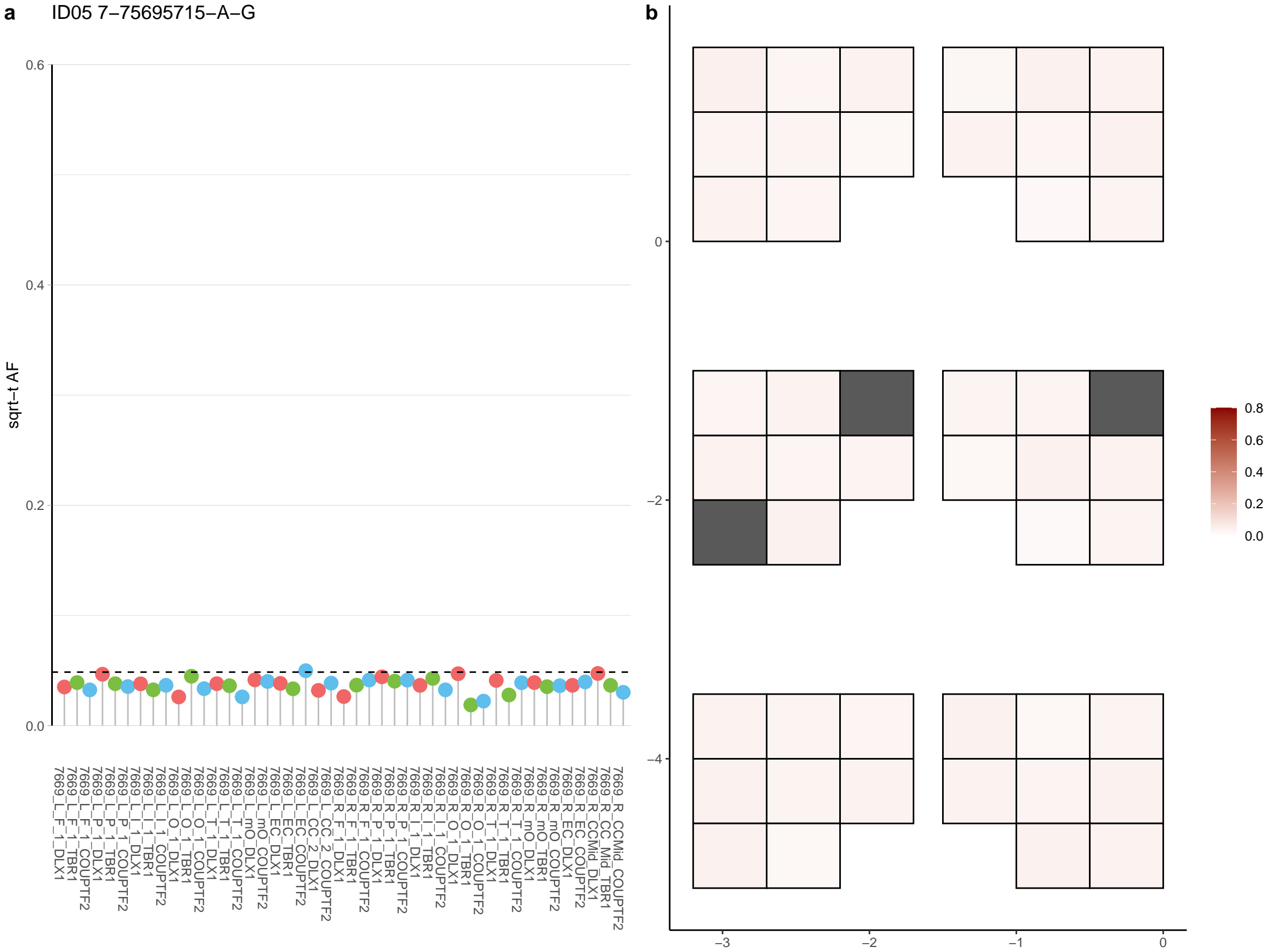

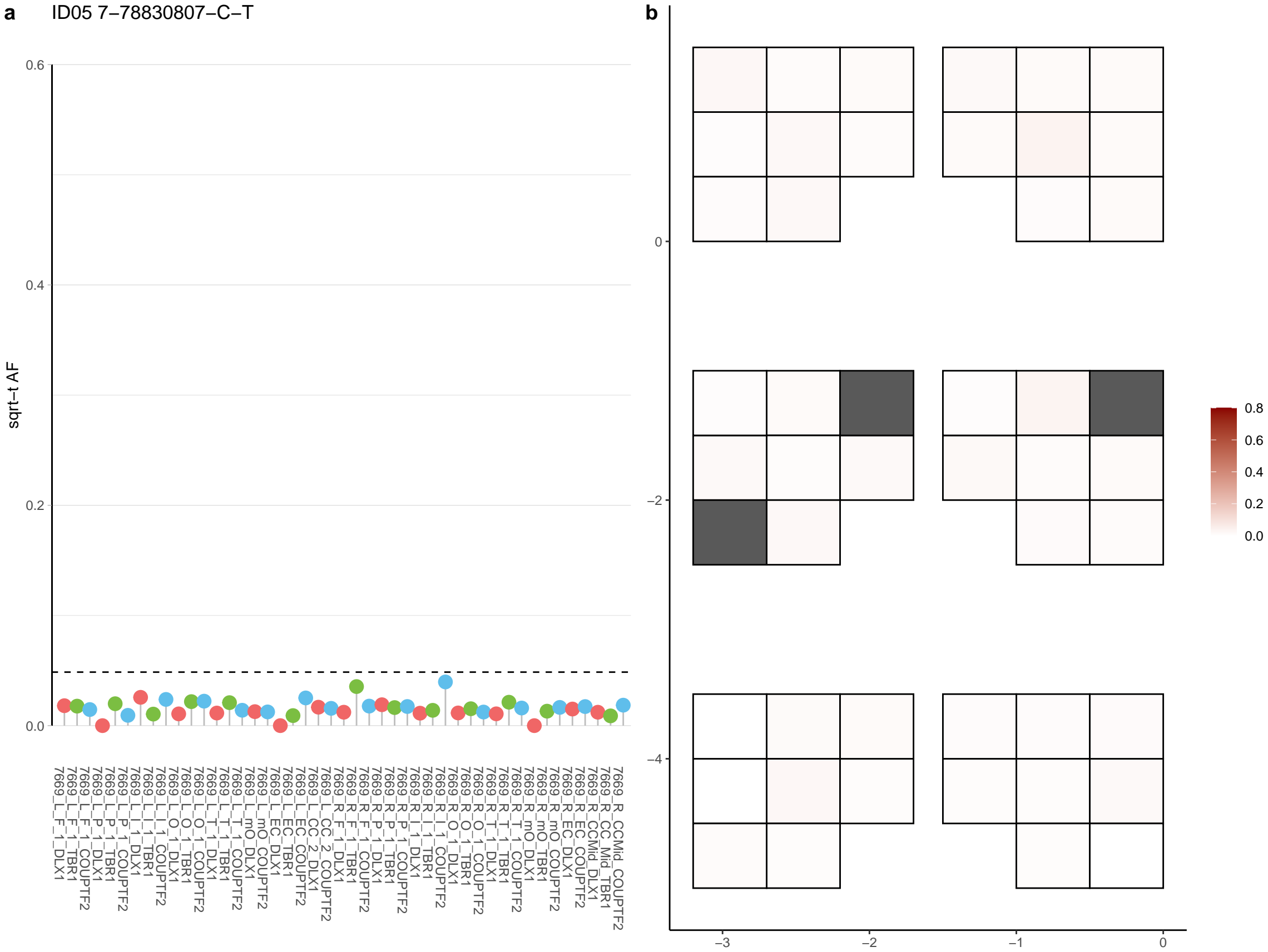

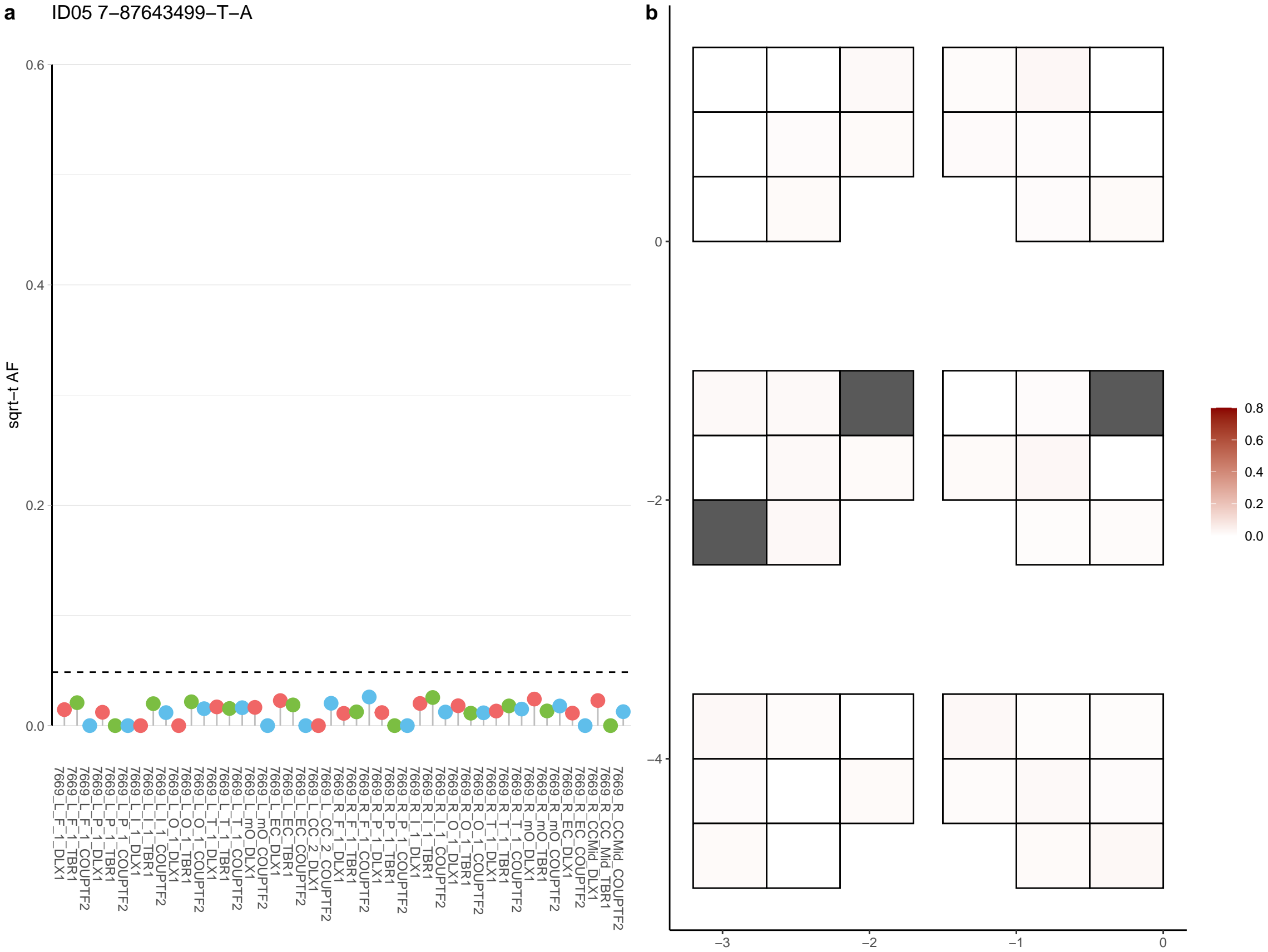

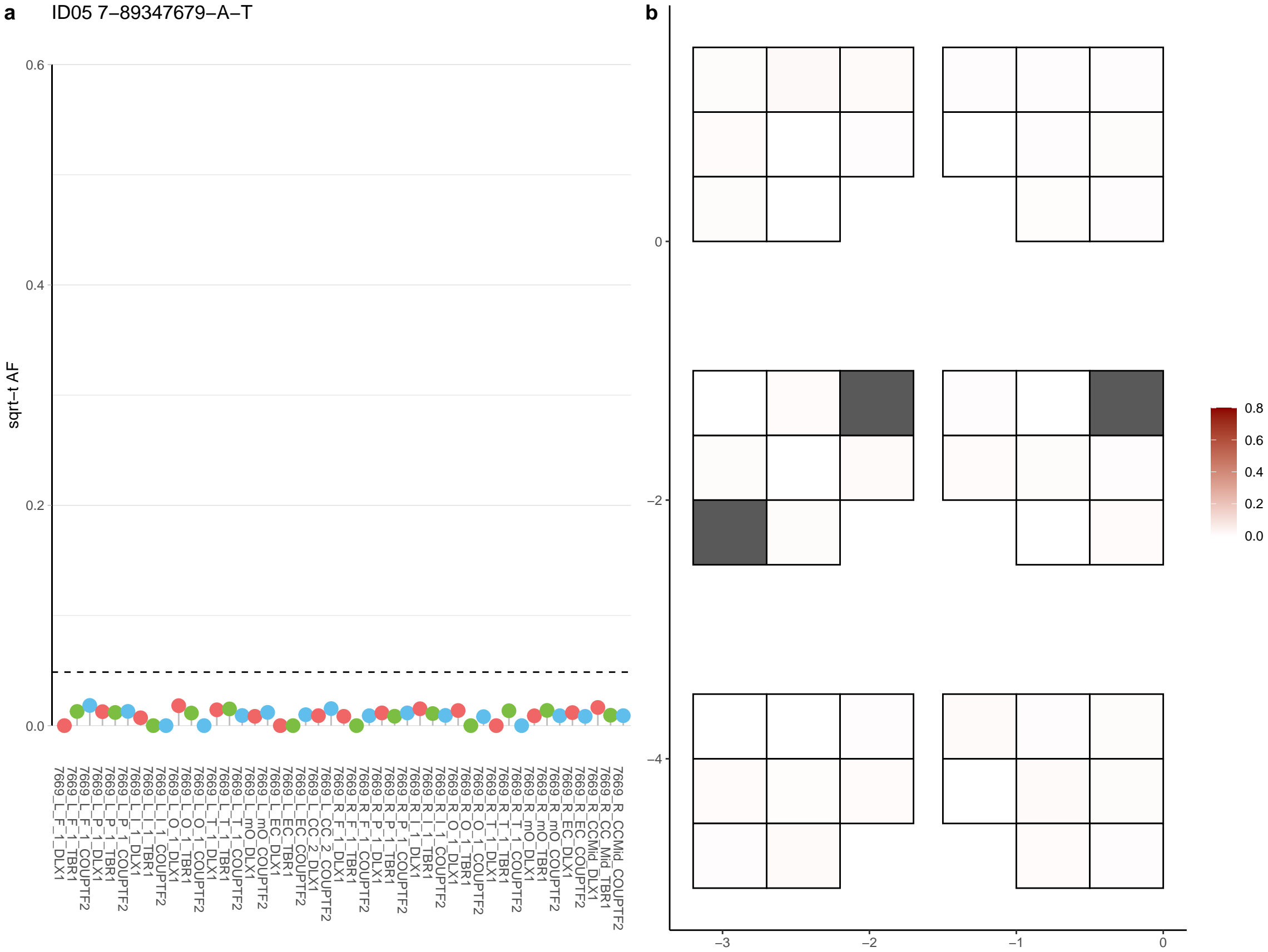

**a** ID05 7-9192631-C-T

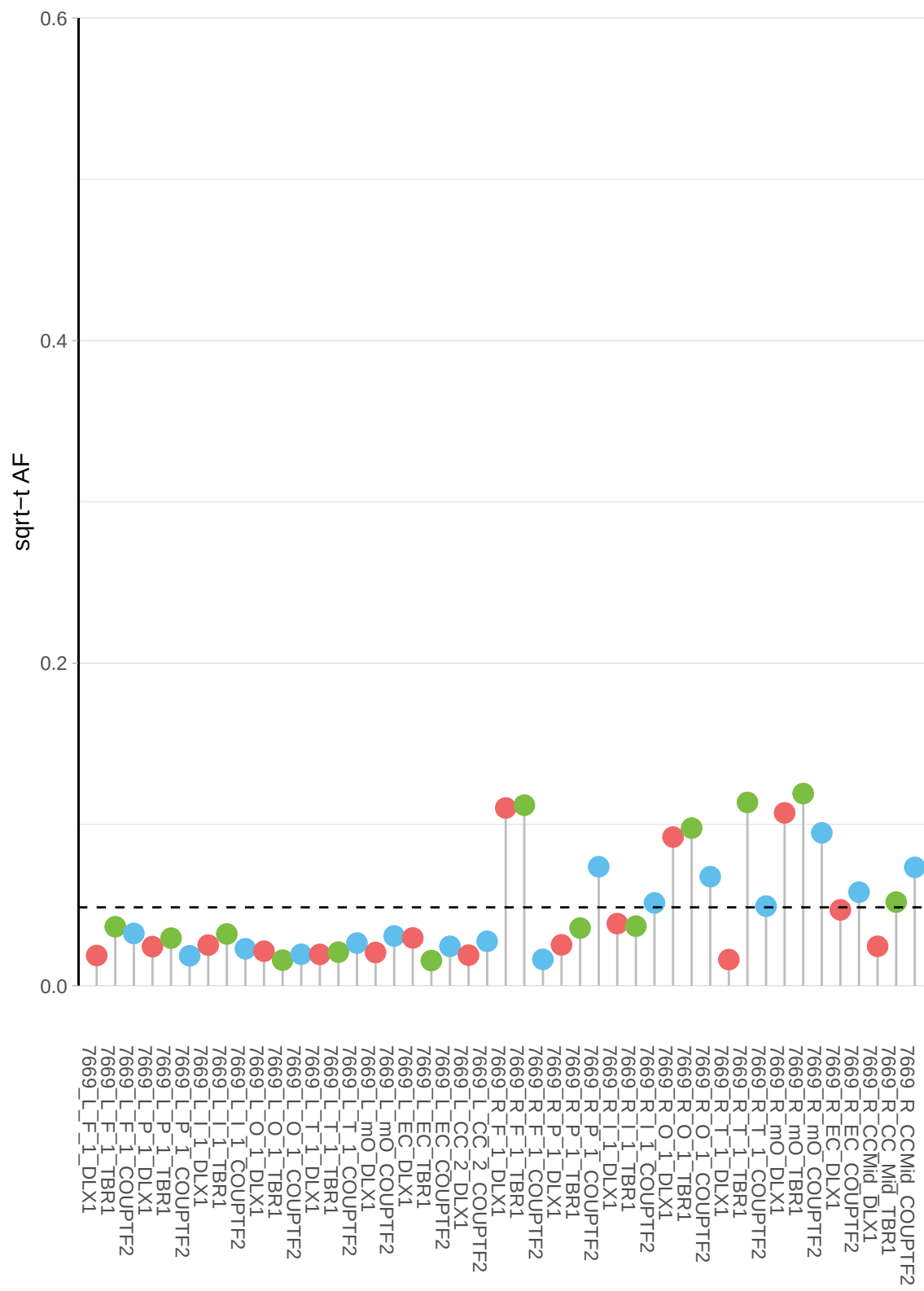

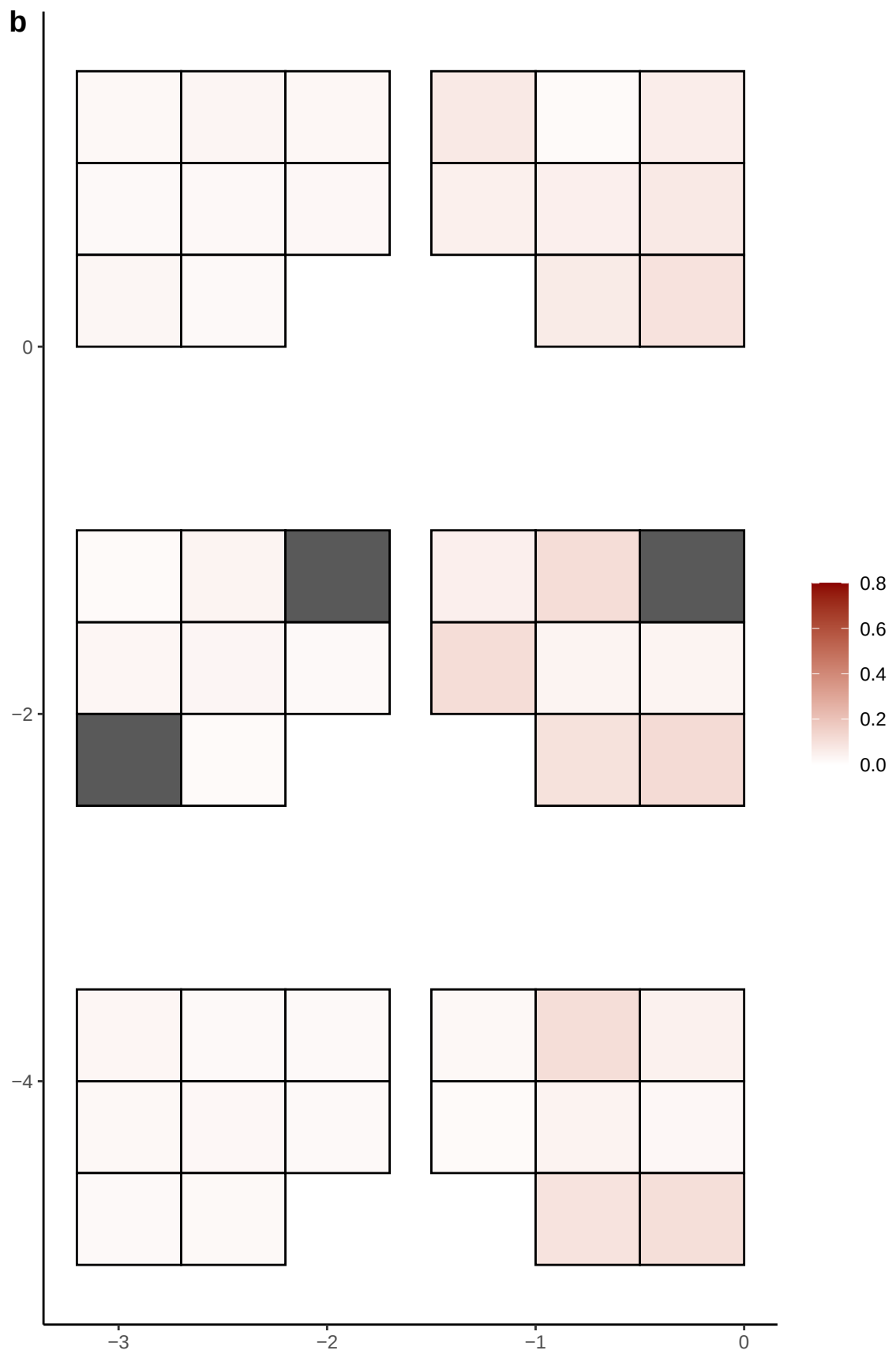

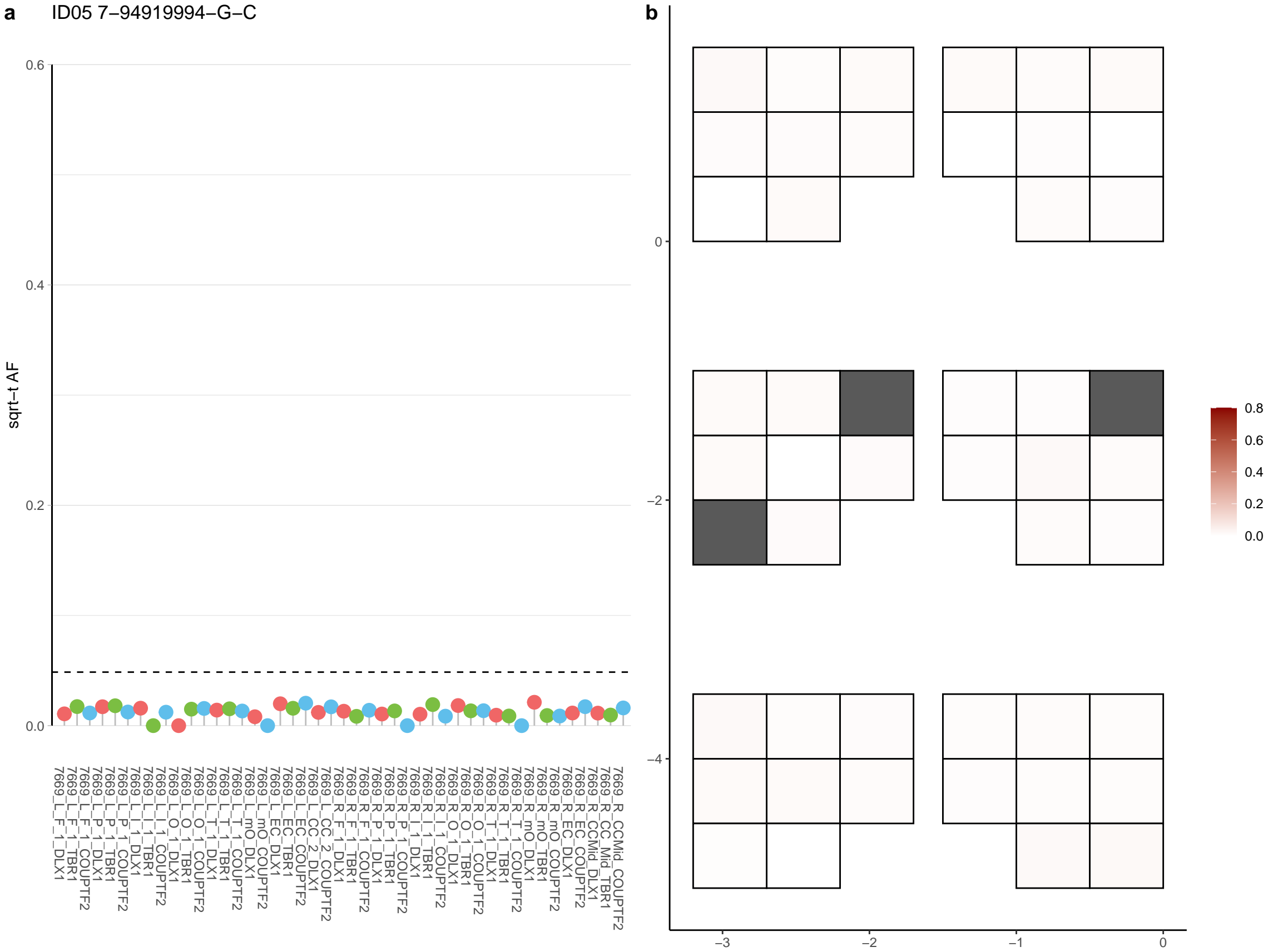

**a** ID05 7-98311420-C-T

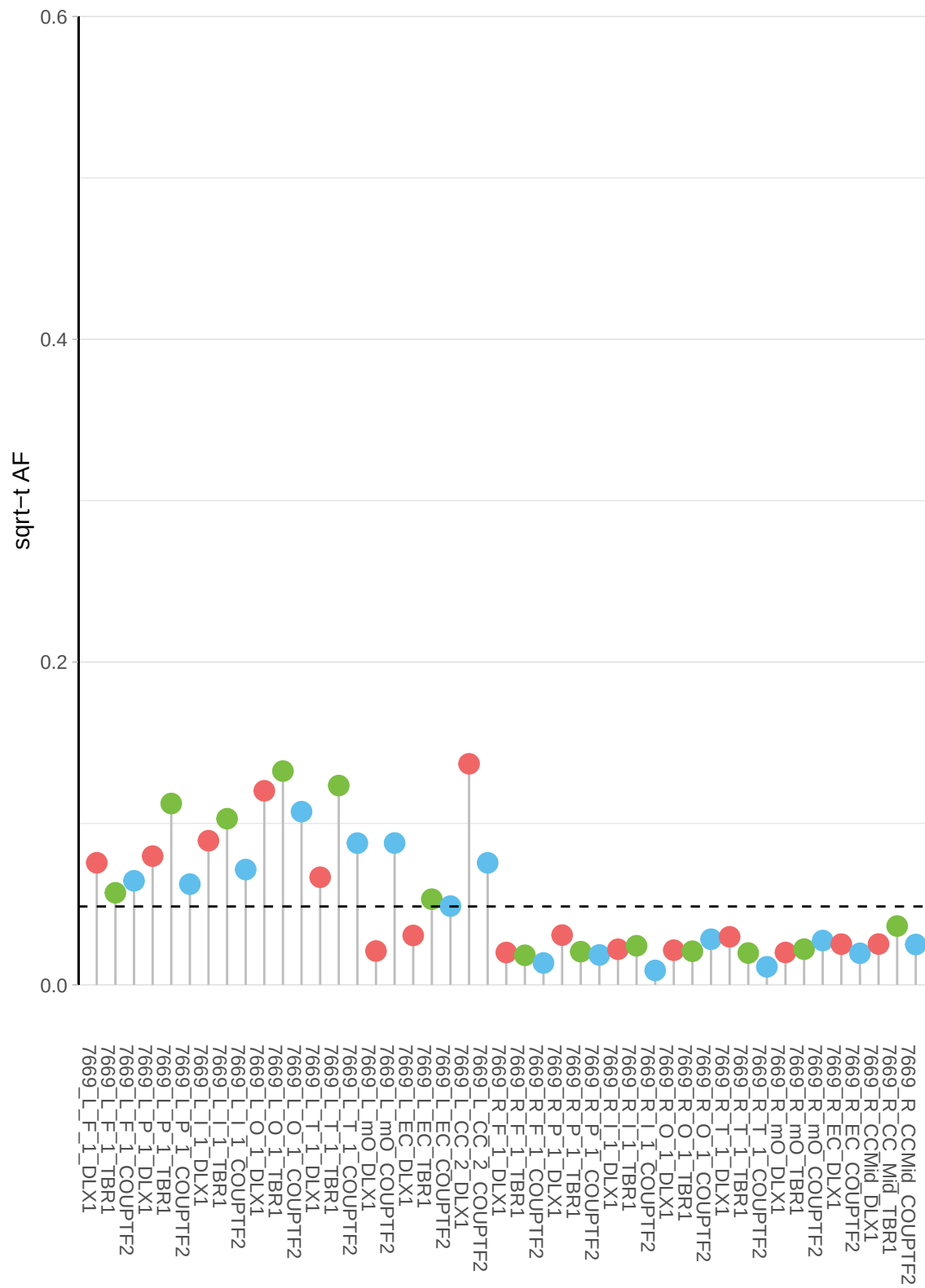

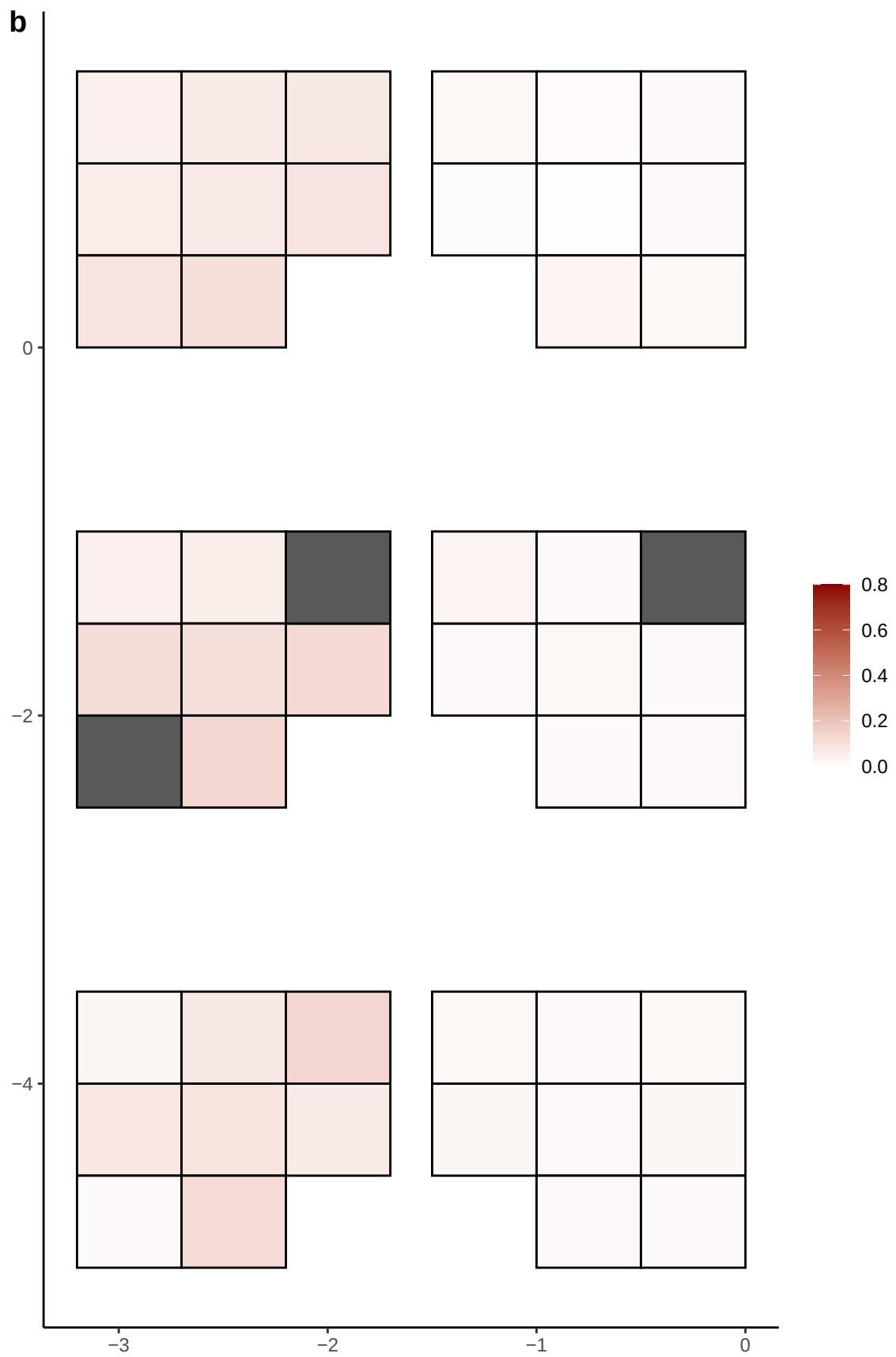

**a** ID05 8-101369438-G-A

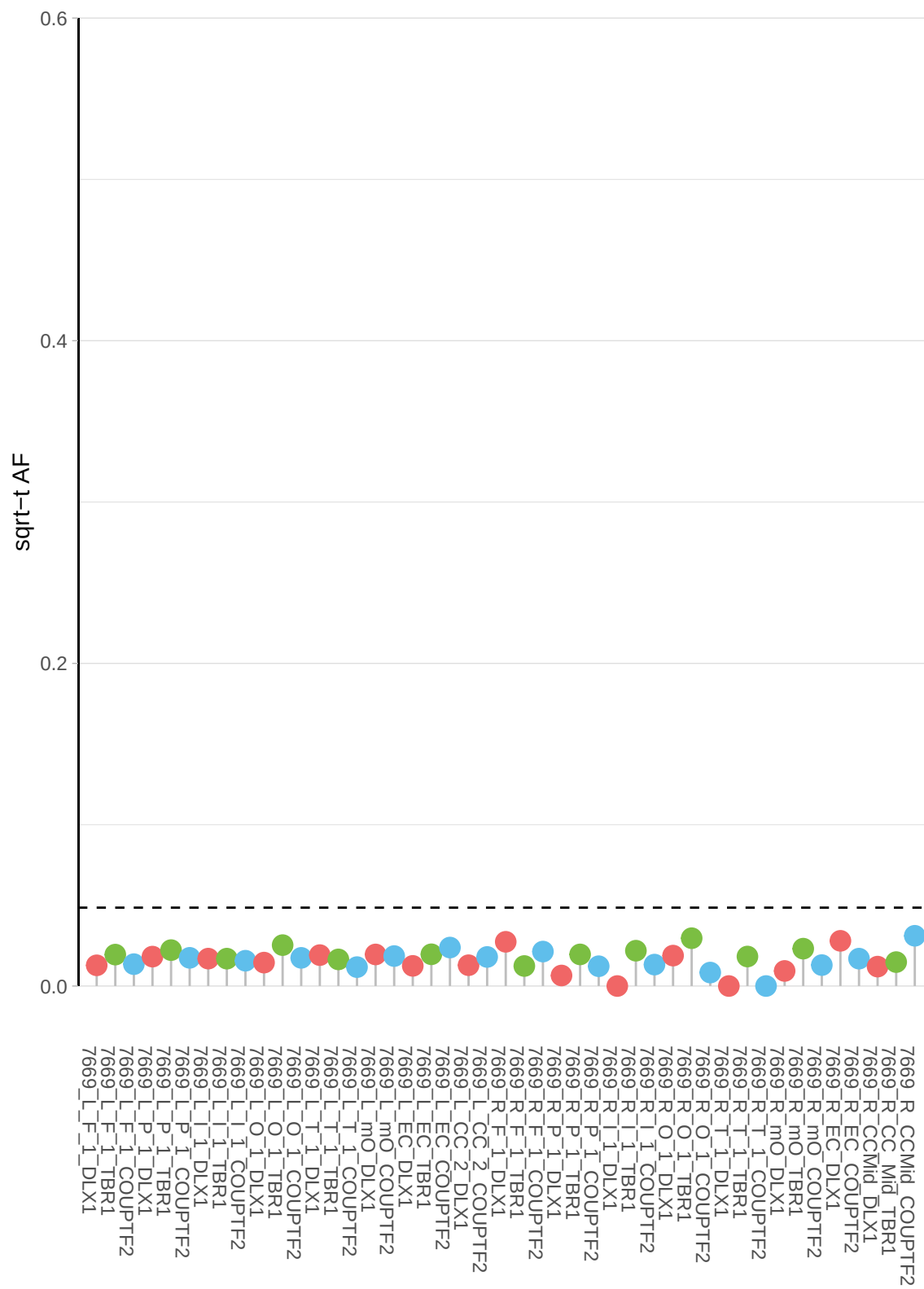

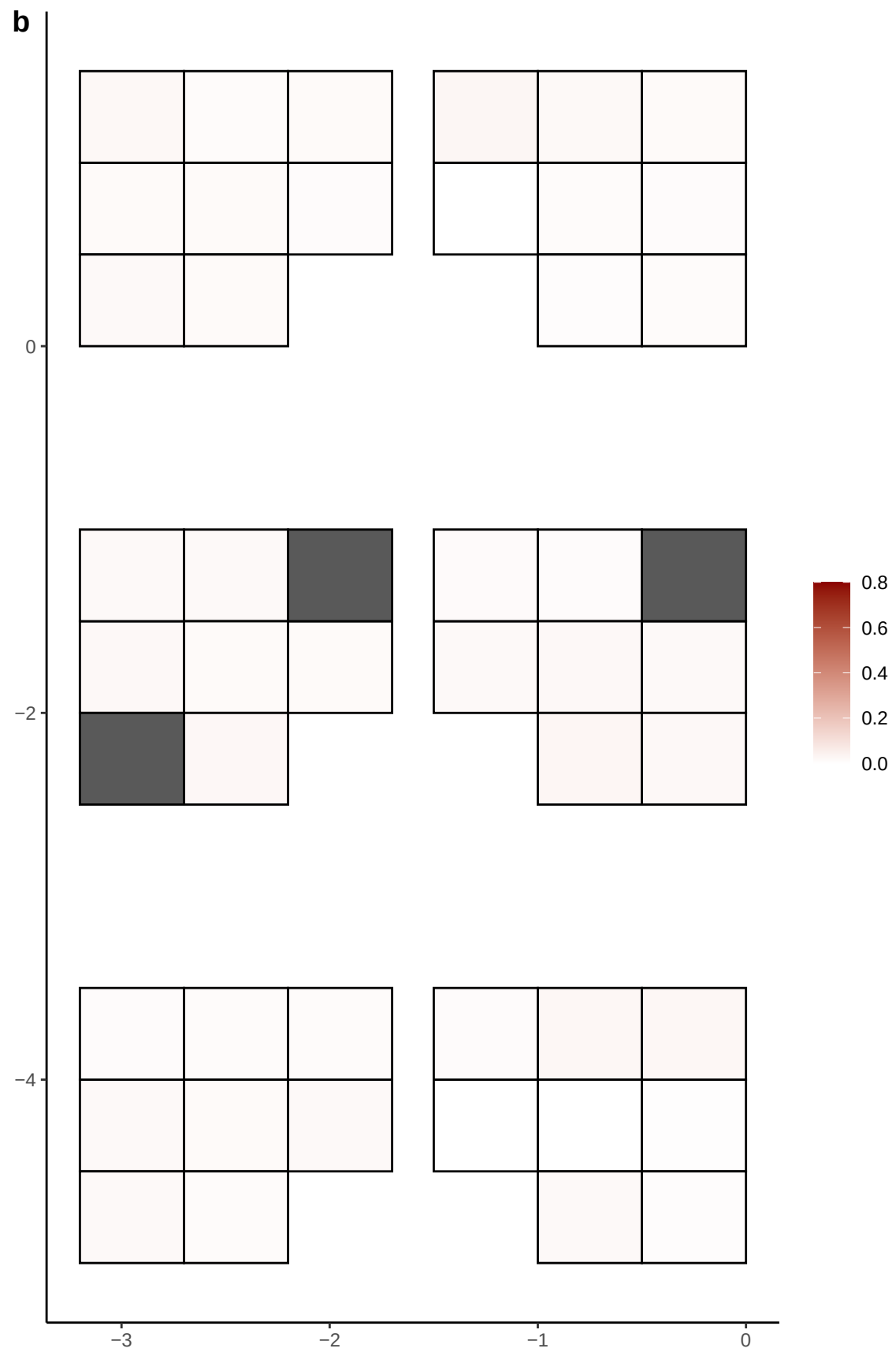

**a** ID05 8-101774660-C-A

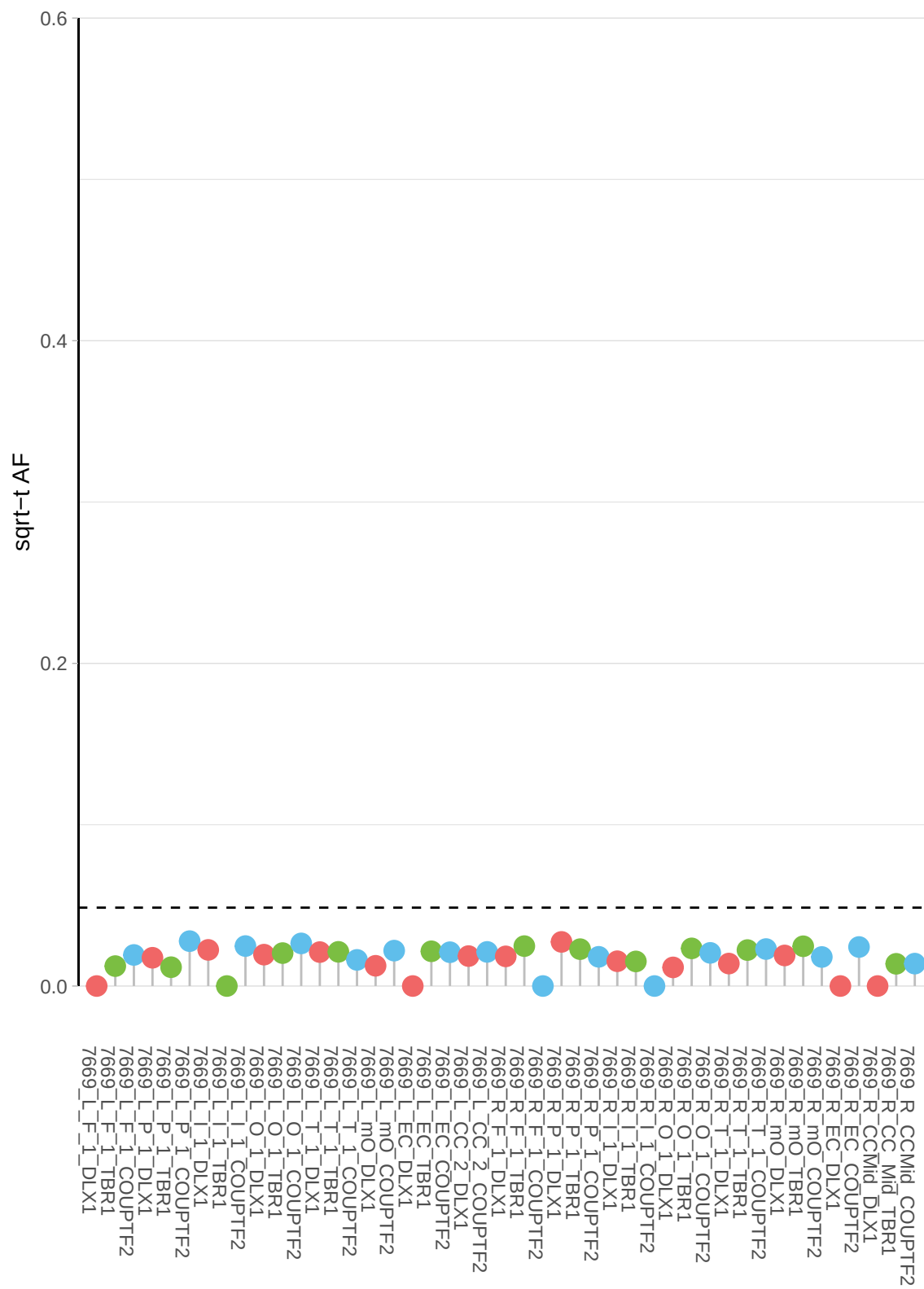

**b**

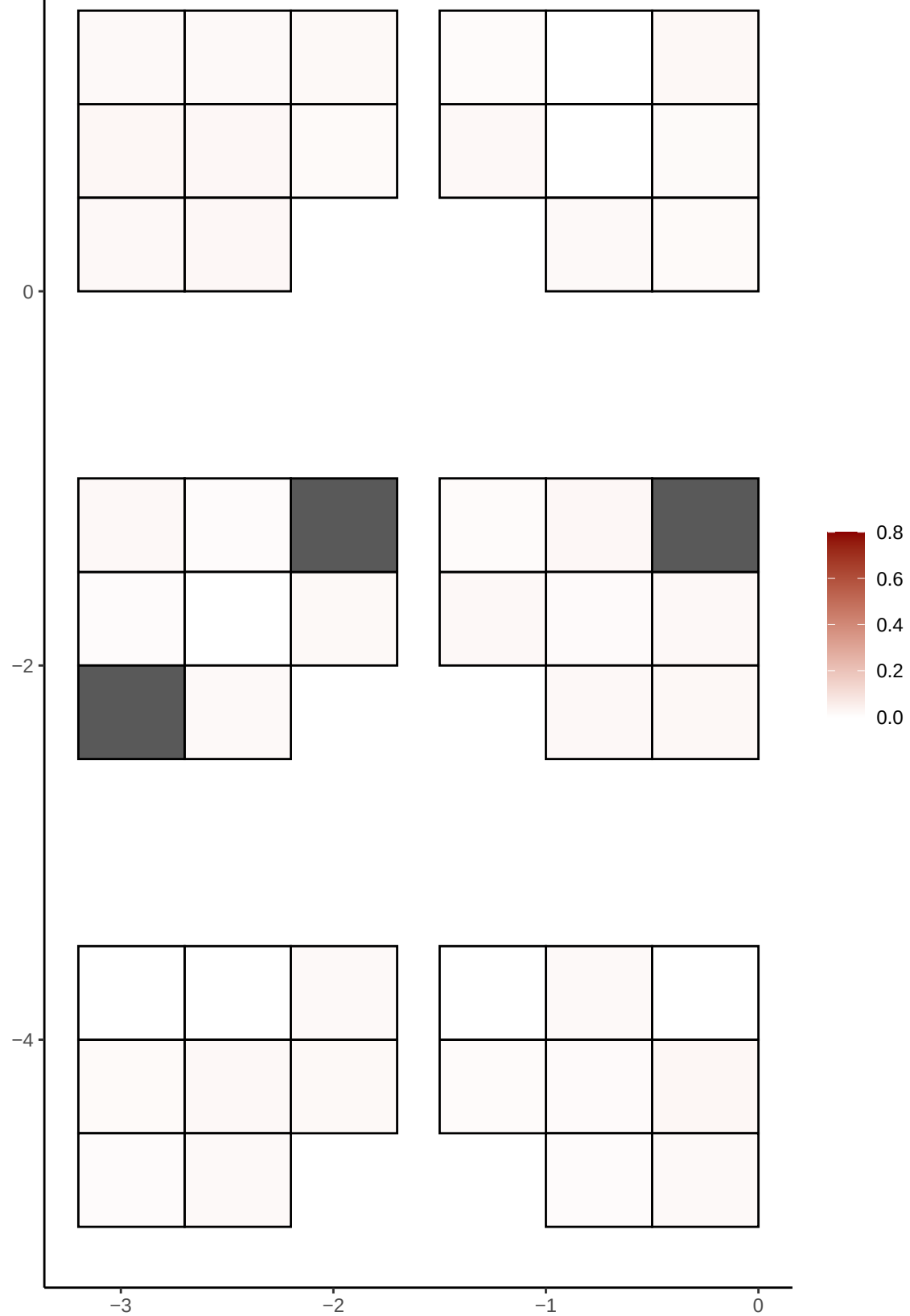

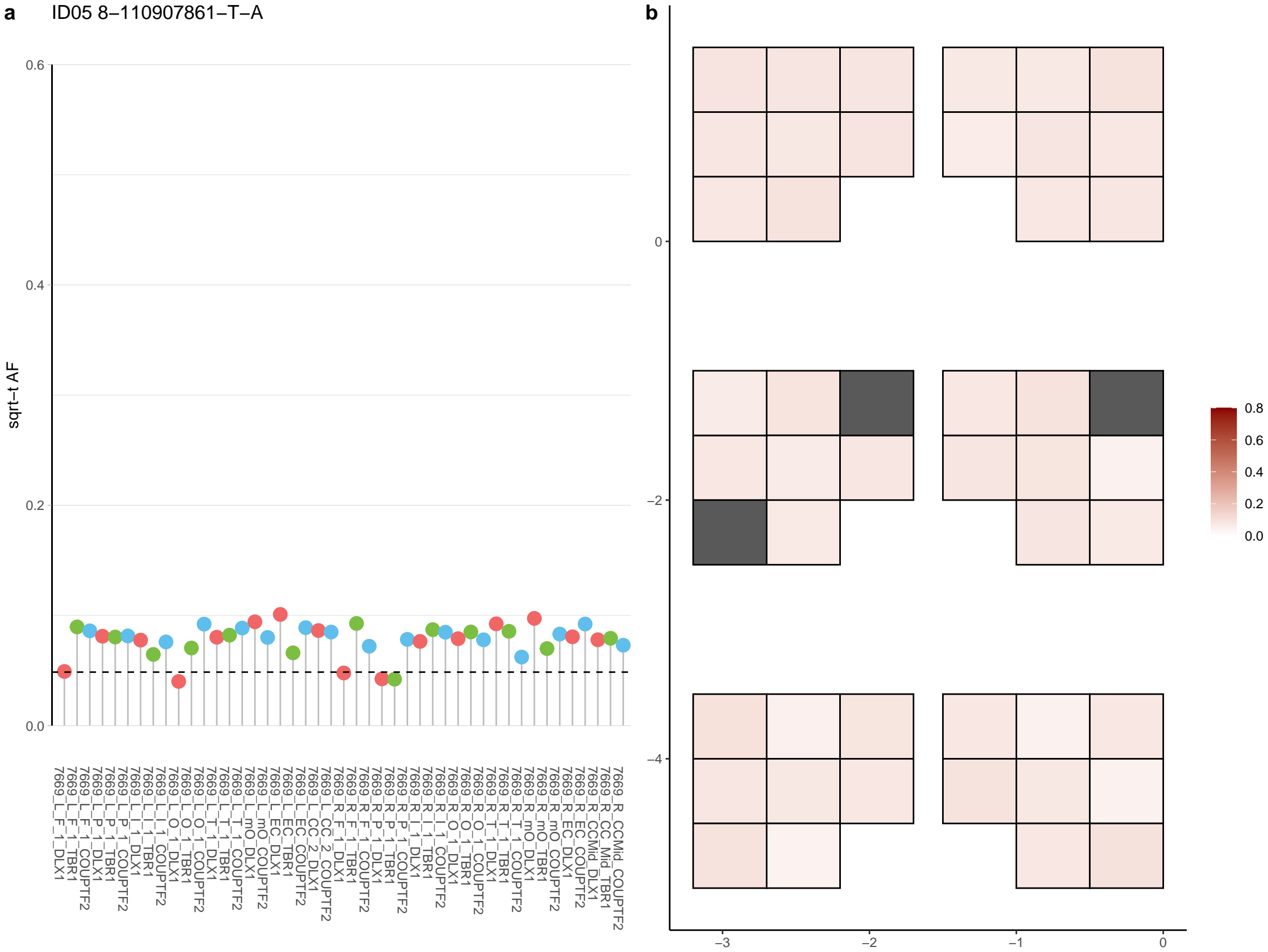

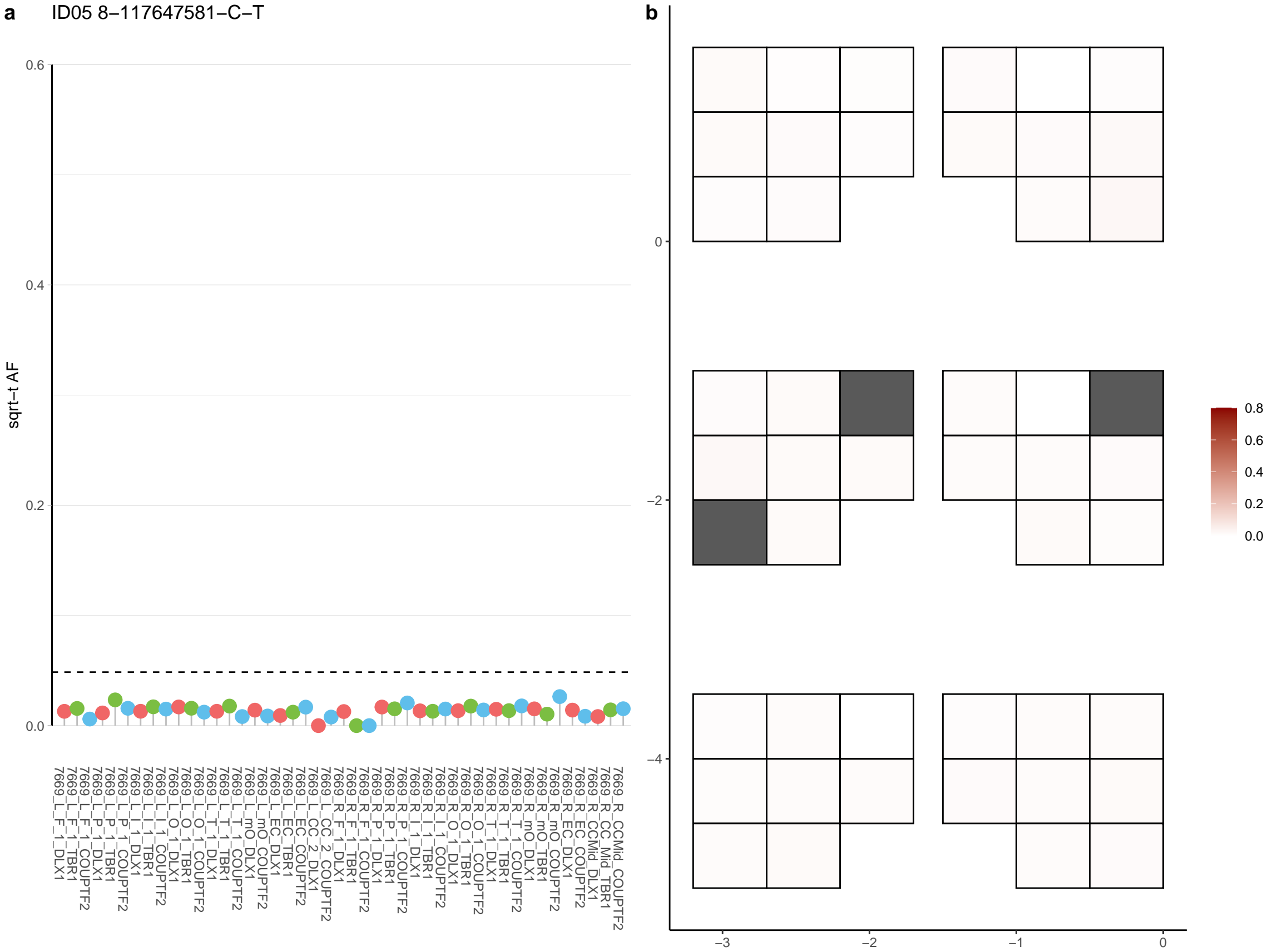

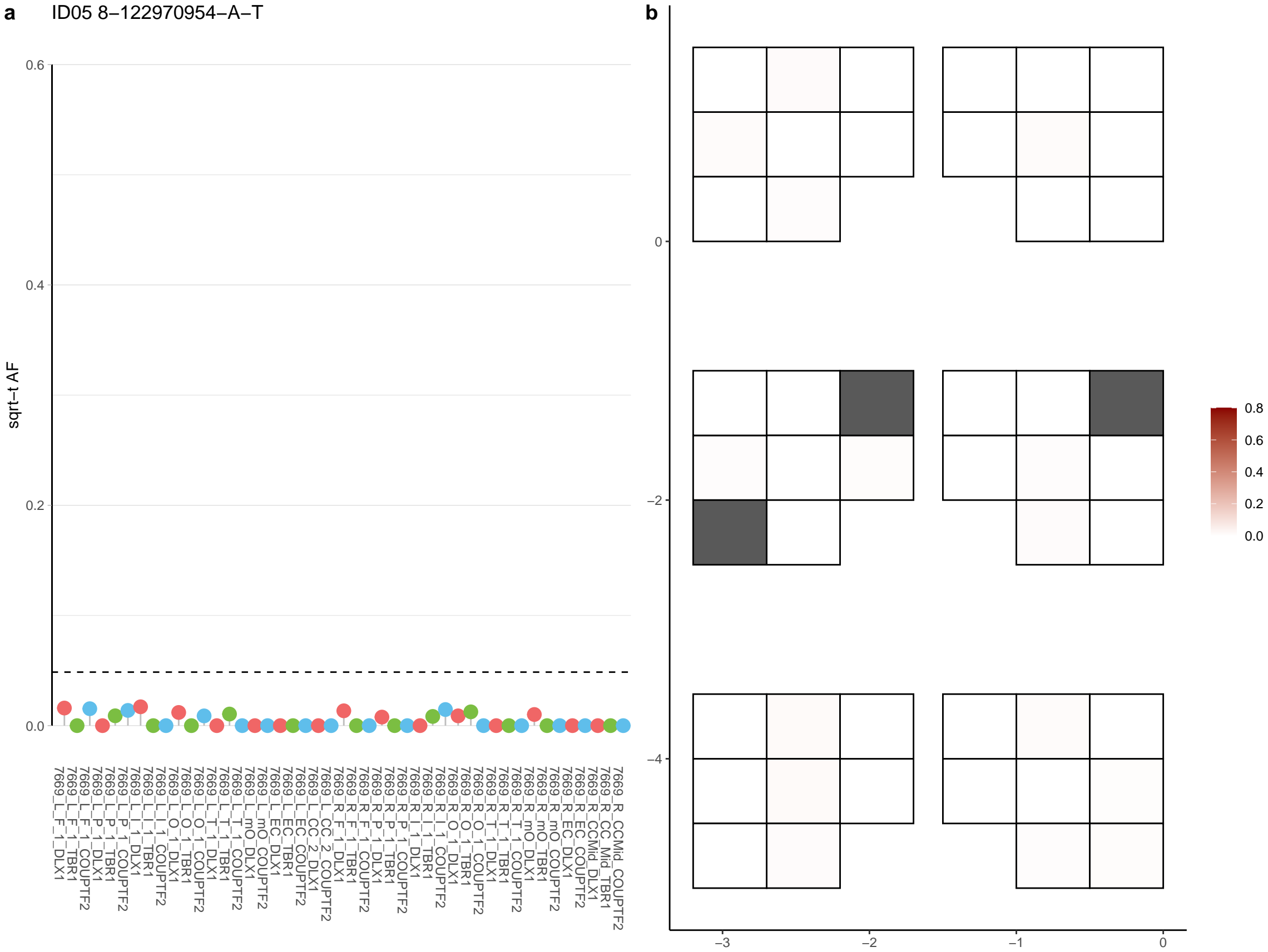

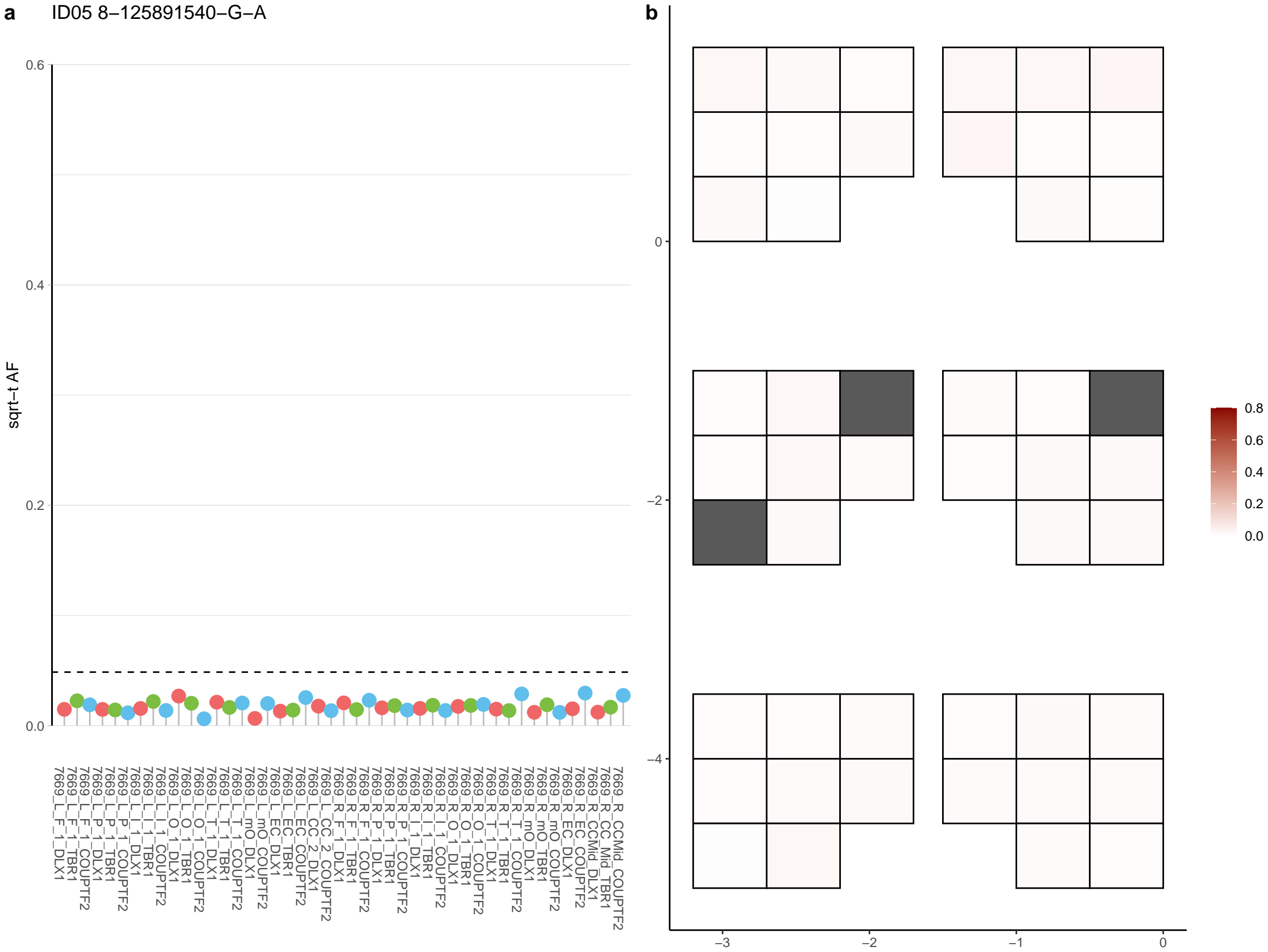

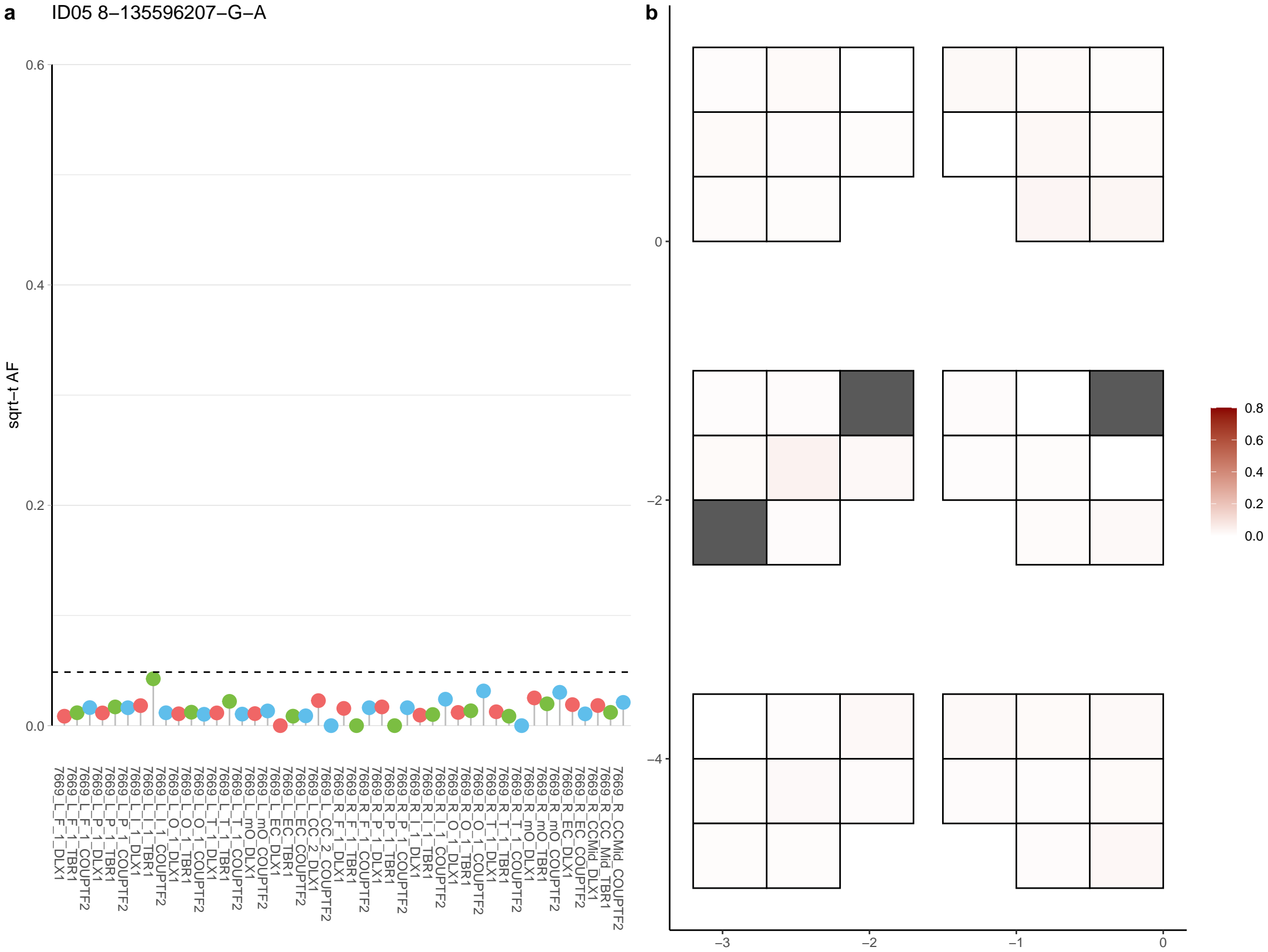

**a** ID05 8-140637848-C-A

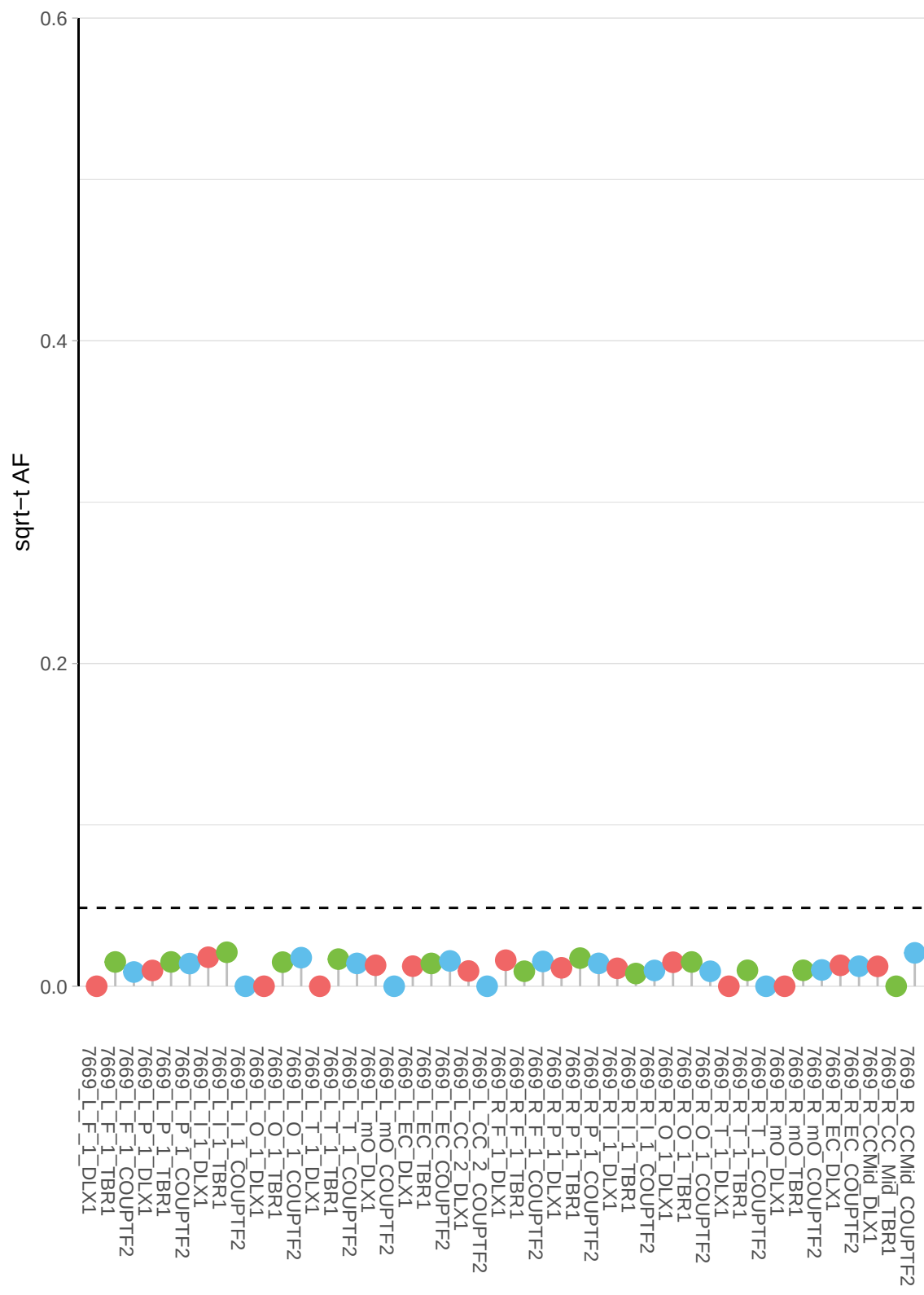

**b**

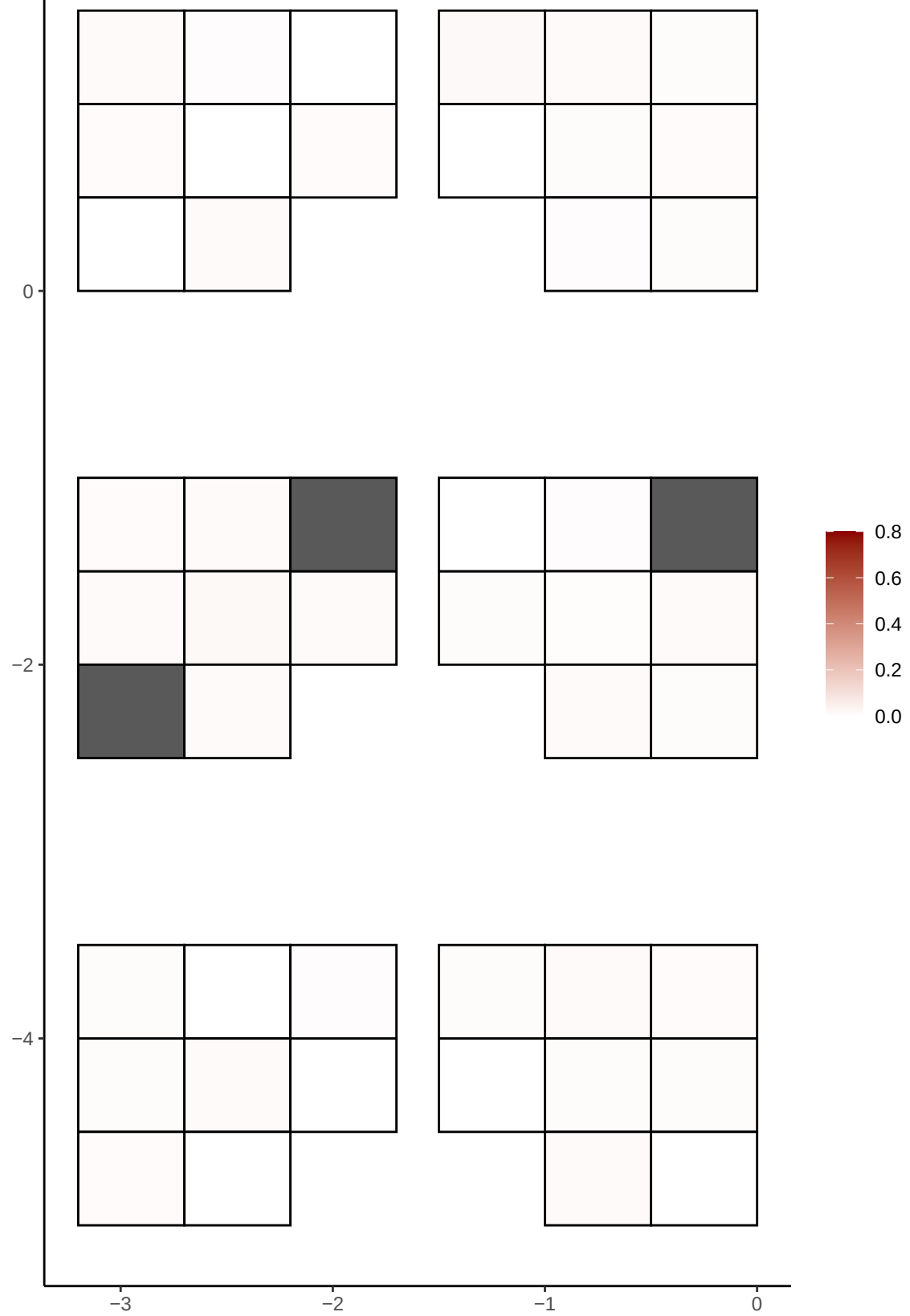

**a** ID05 8-20681102-G-A

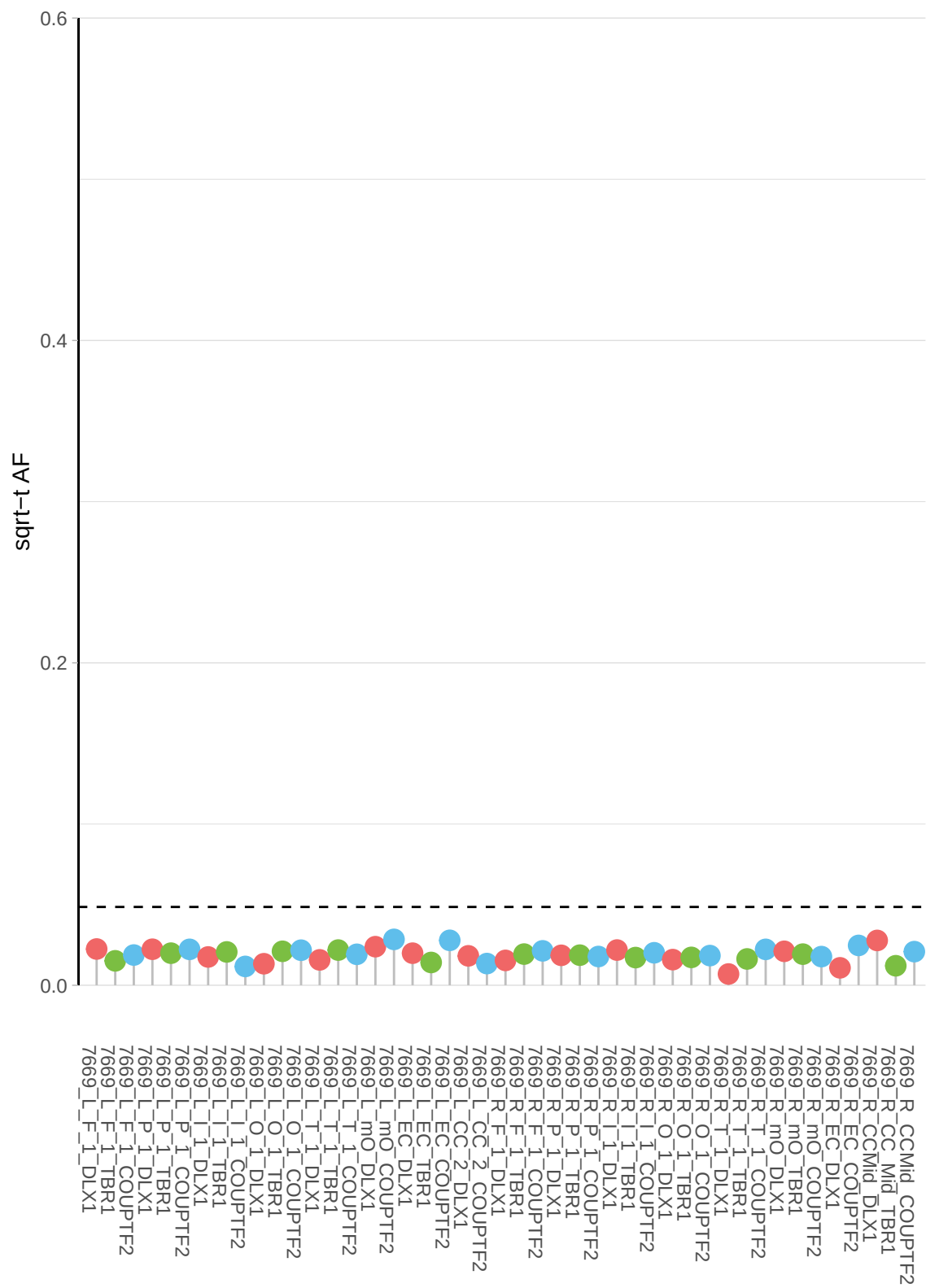

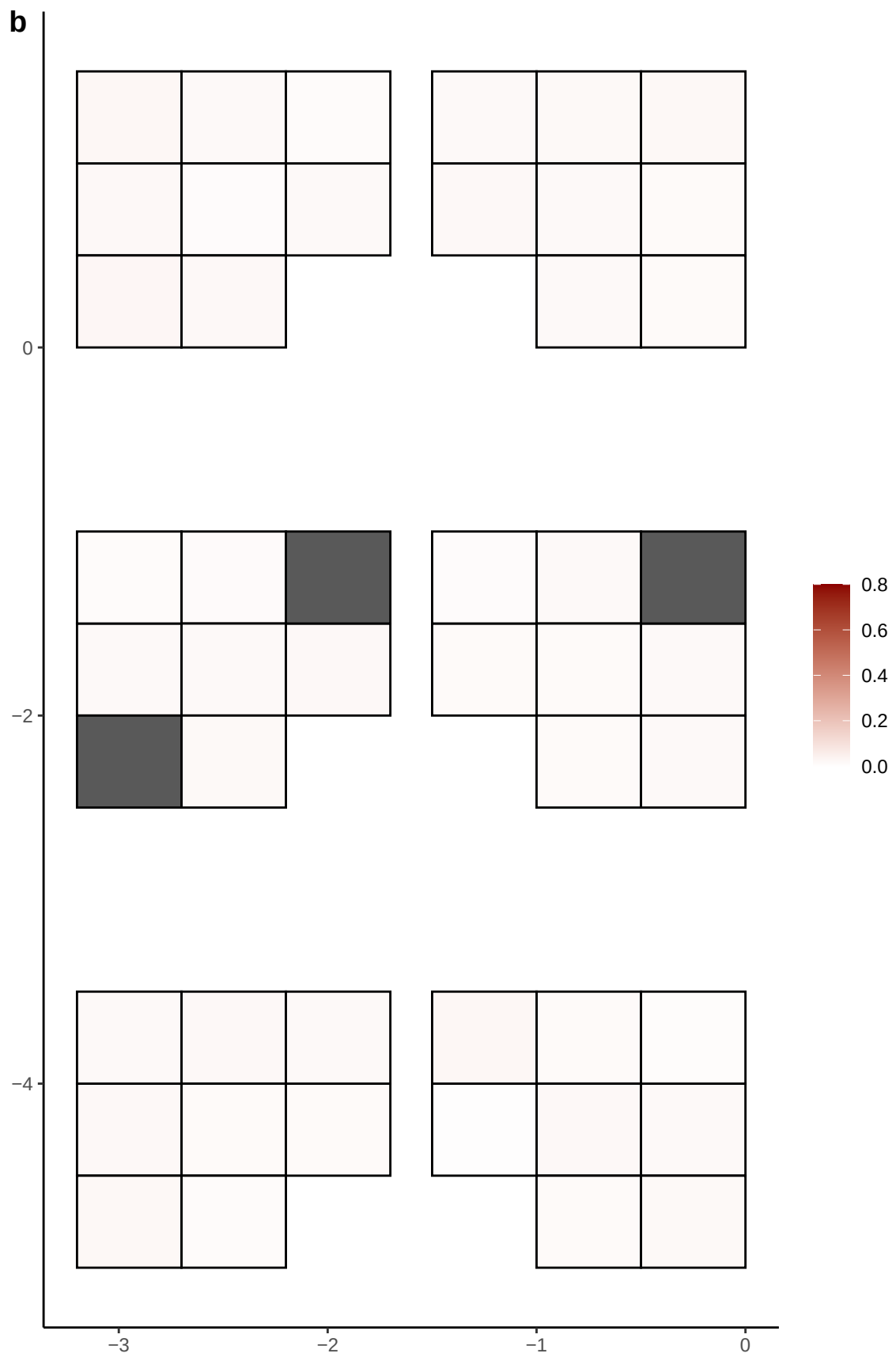

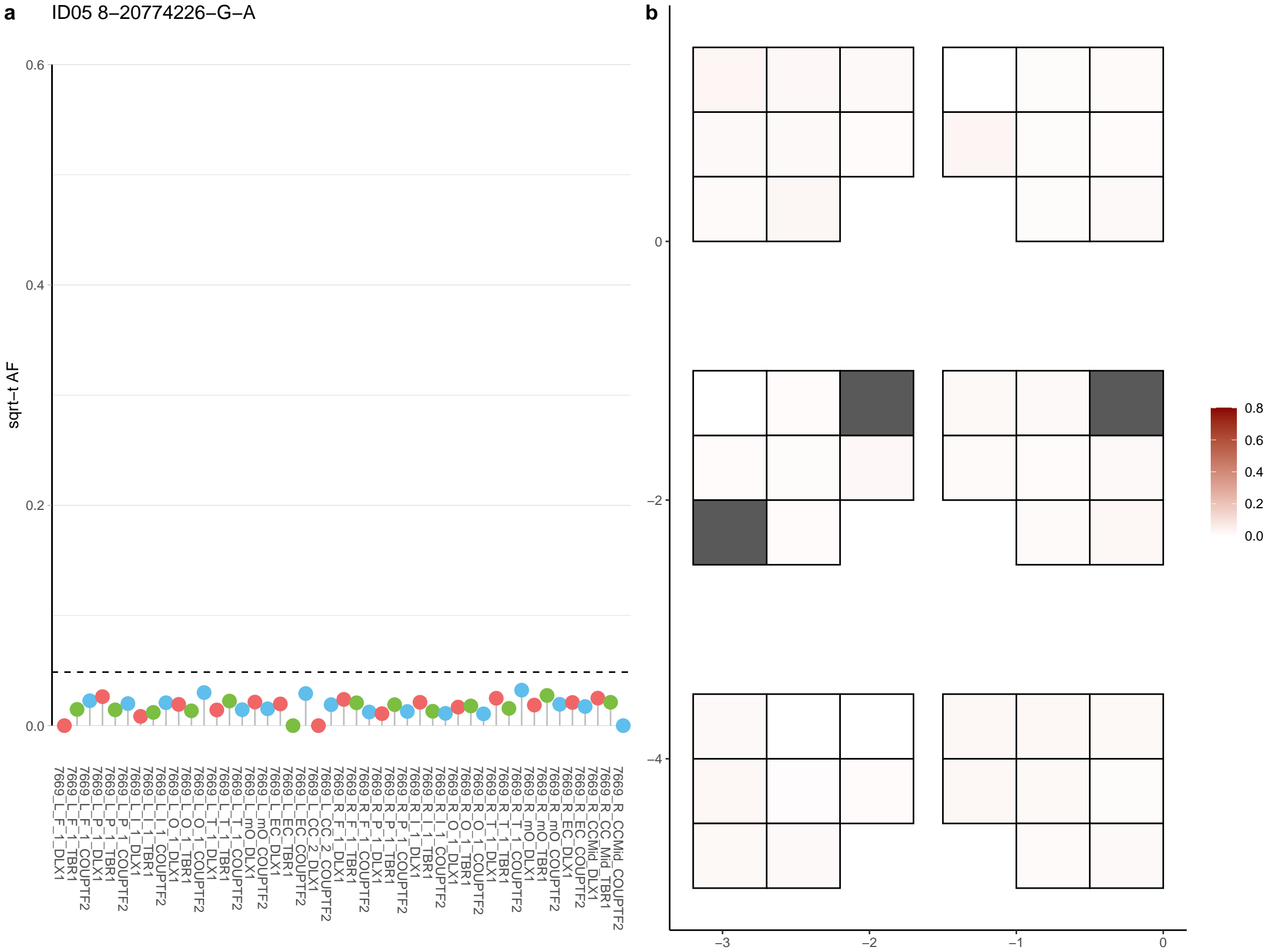

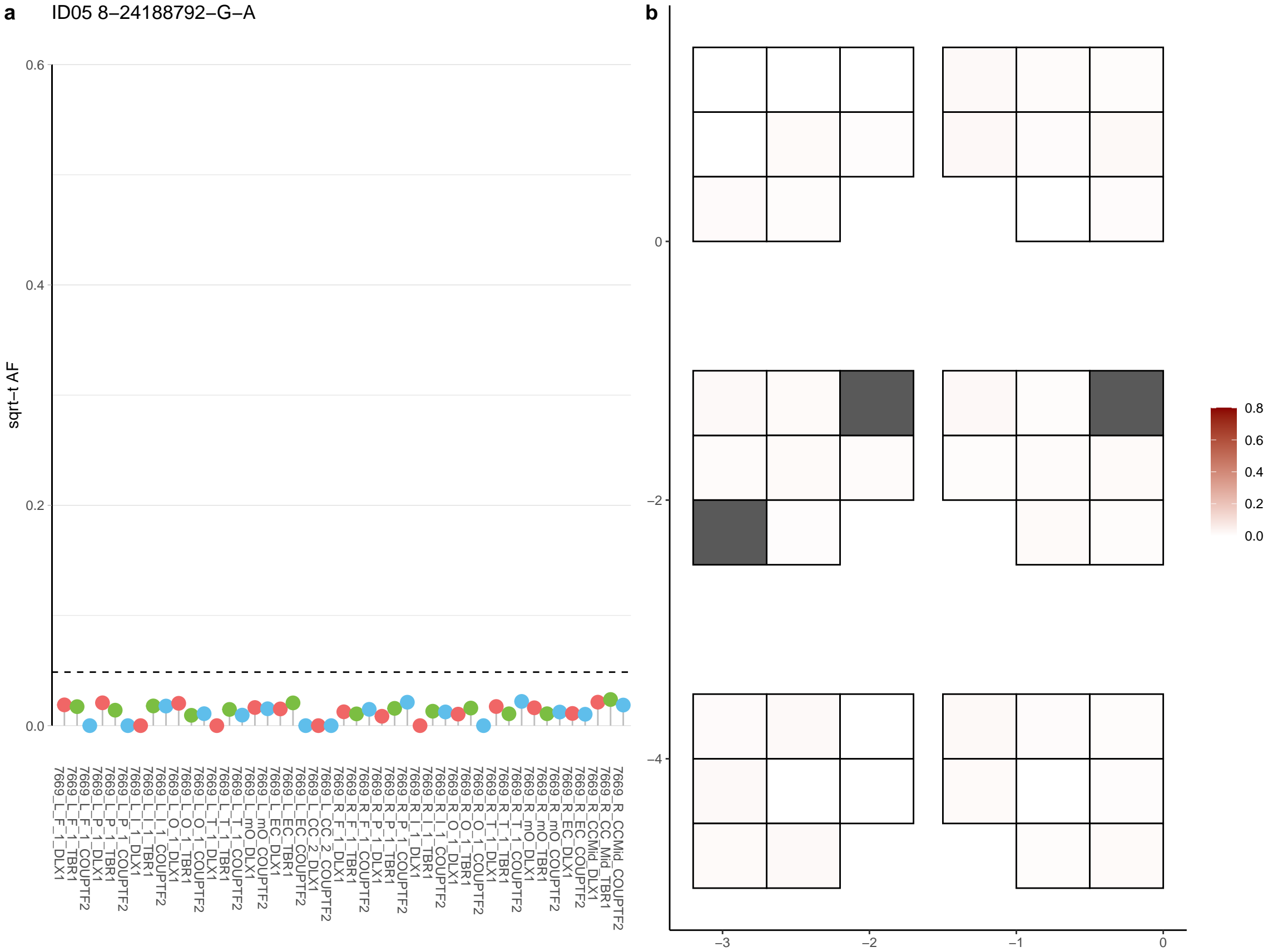

**a** ID05 8-29866666-C-T

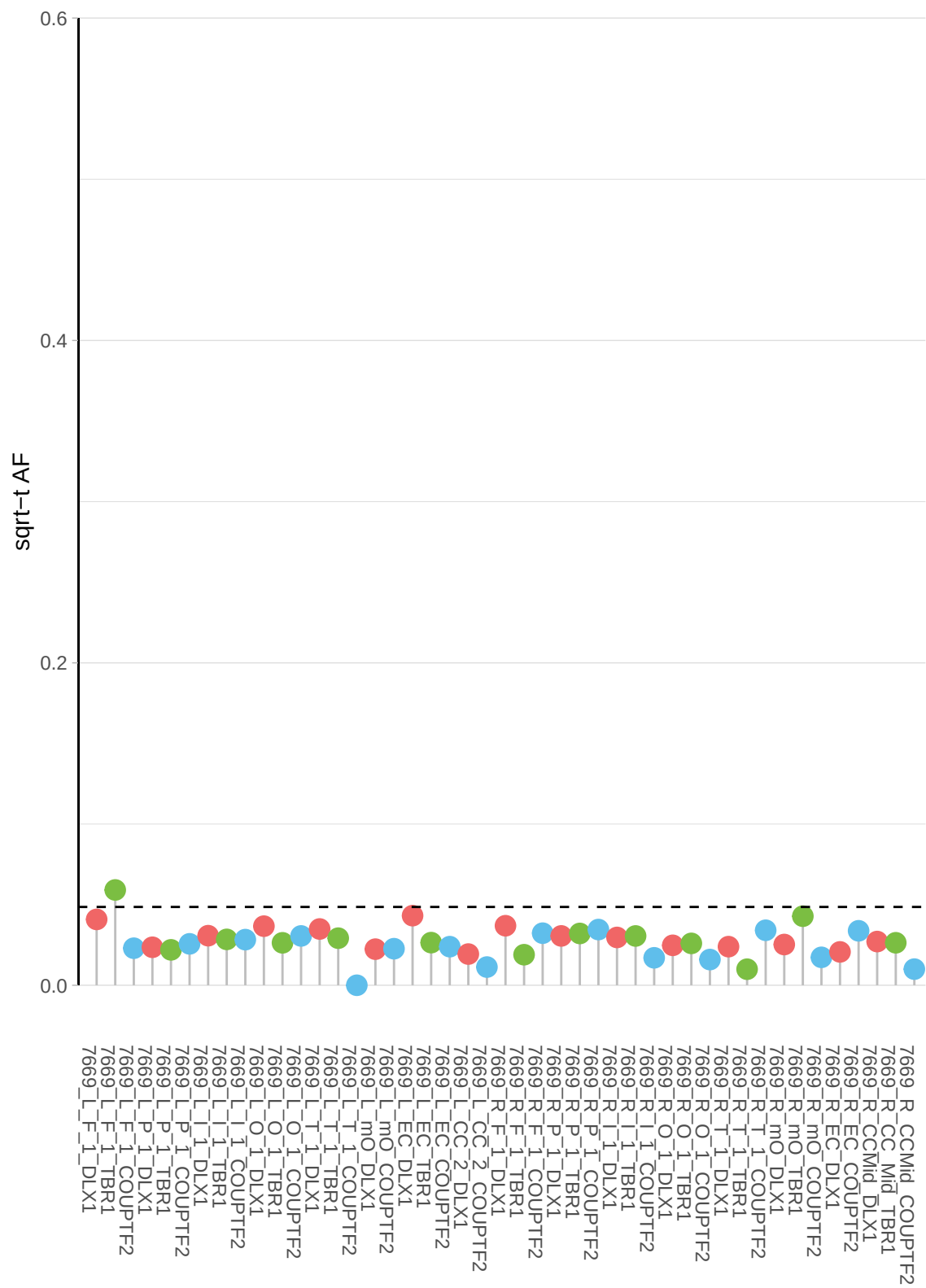

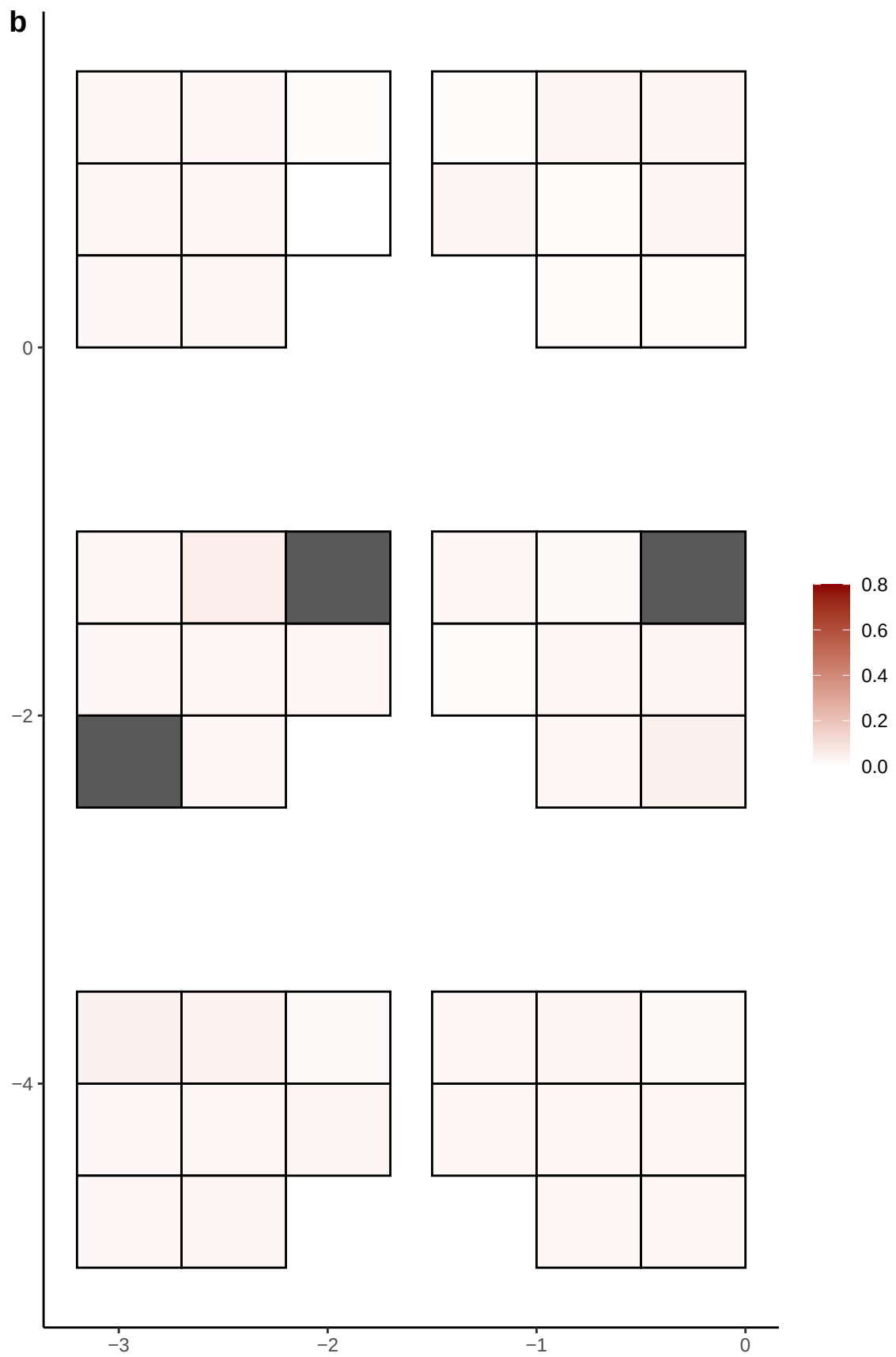

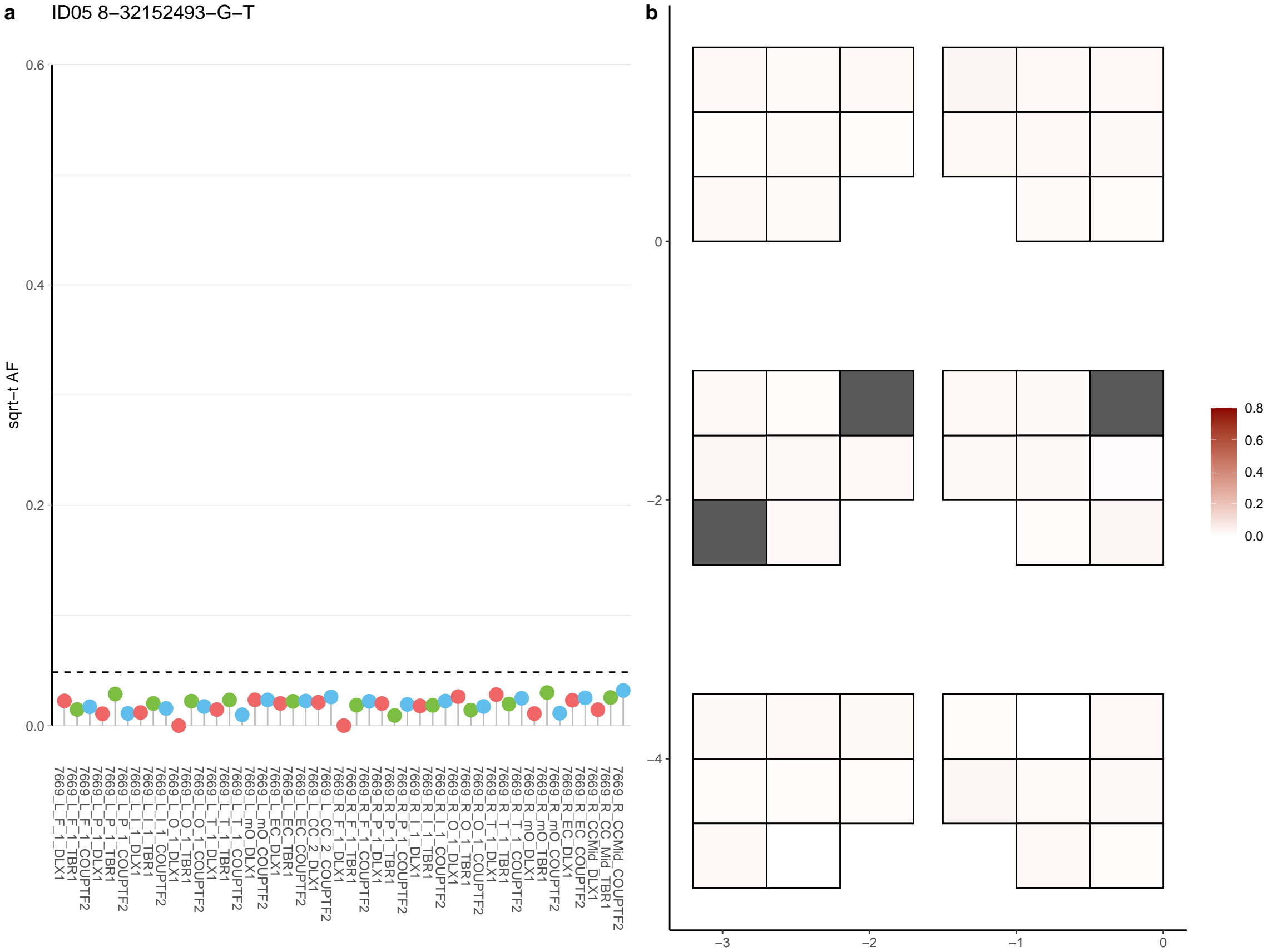

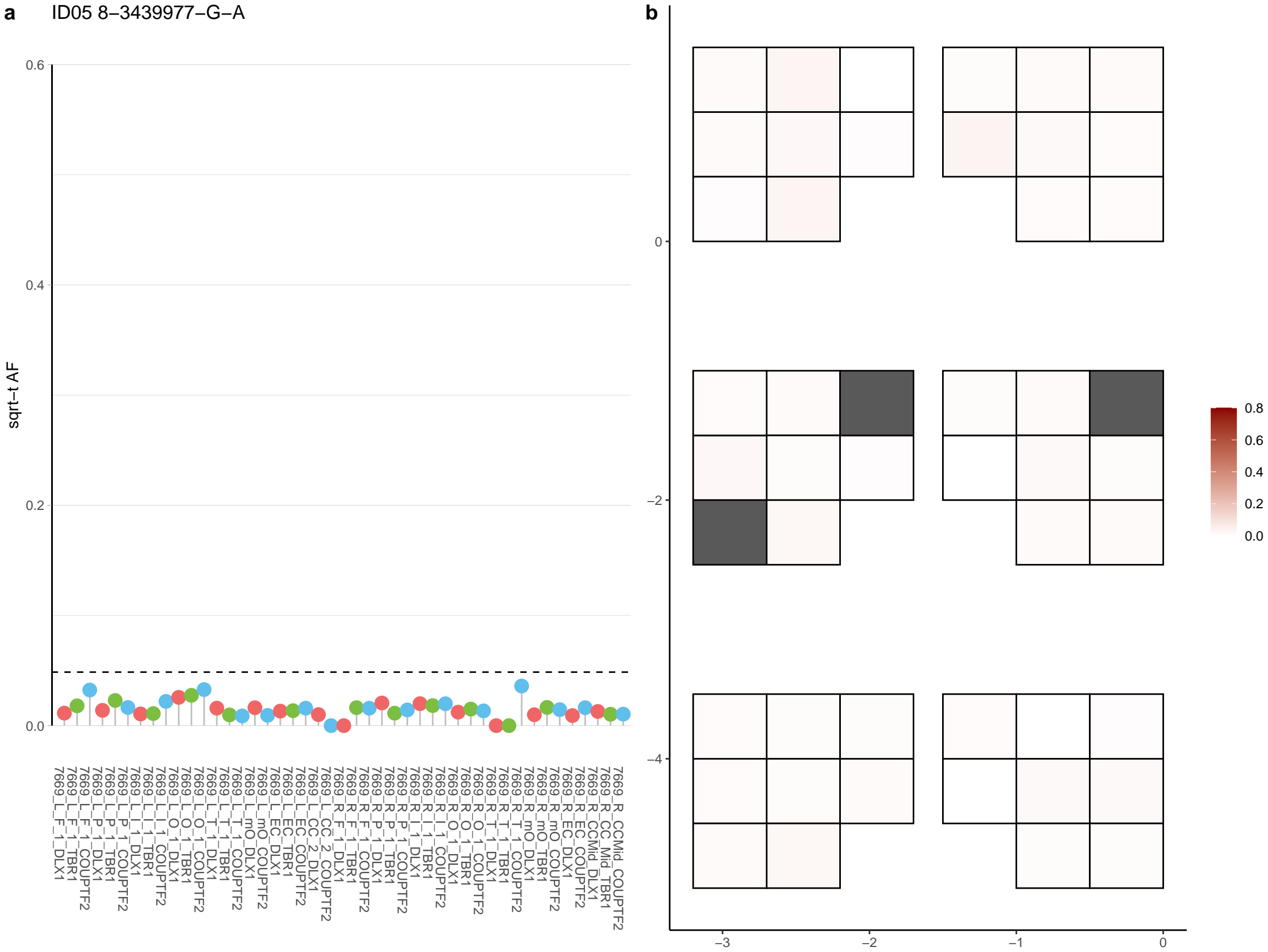

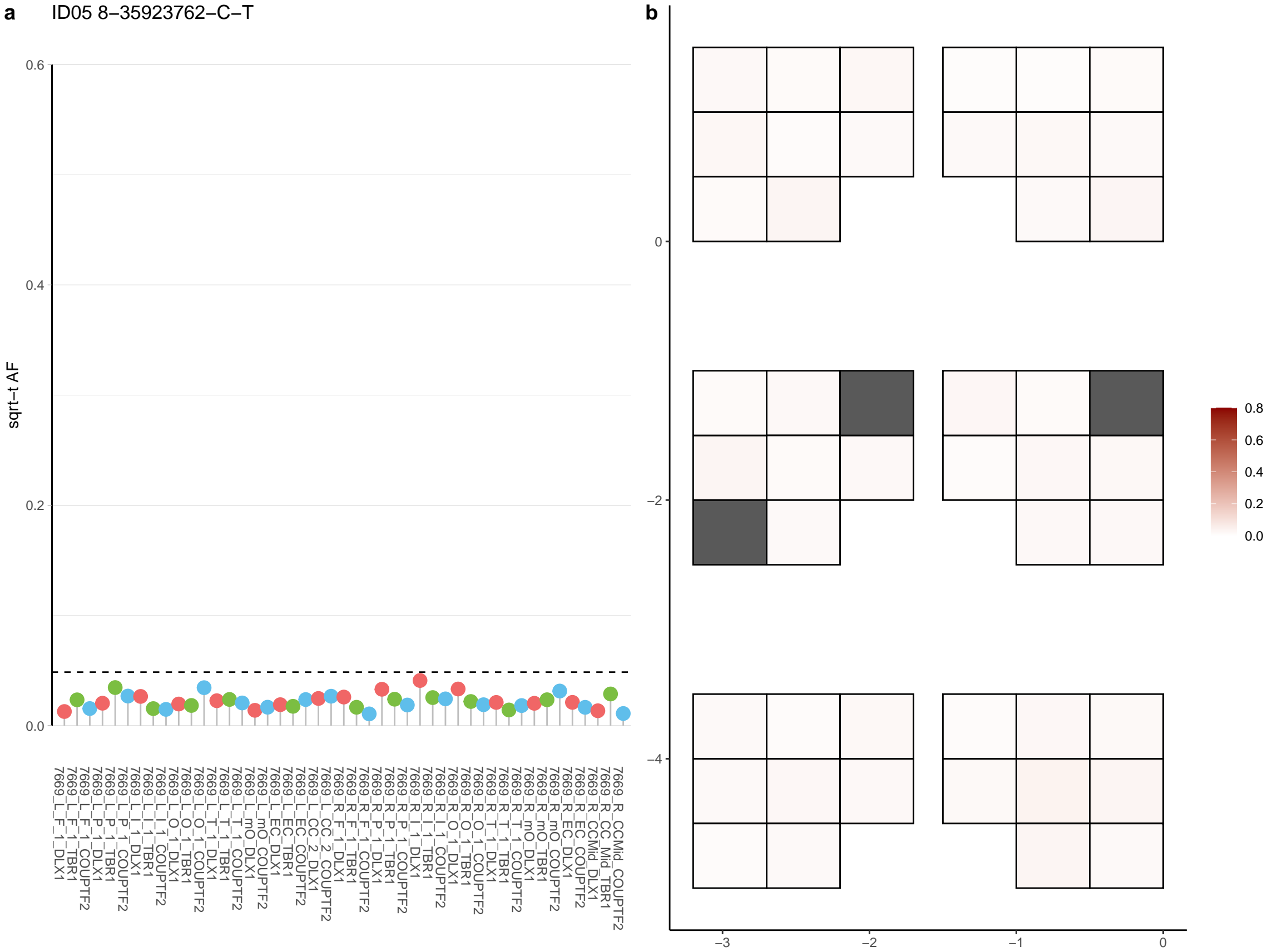

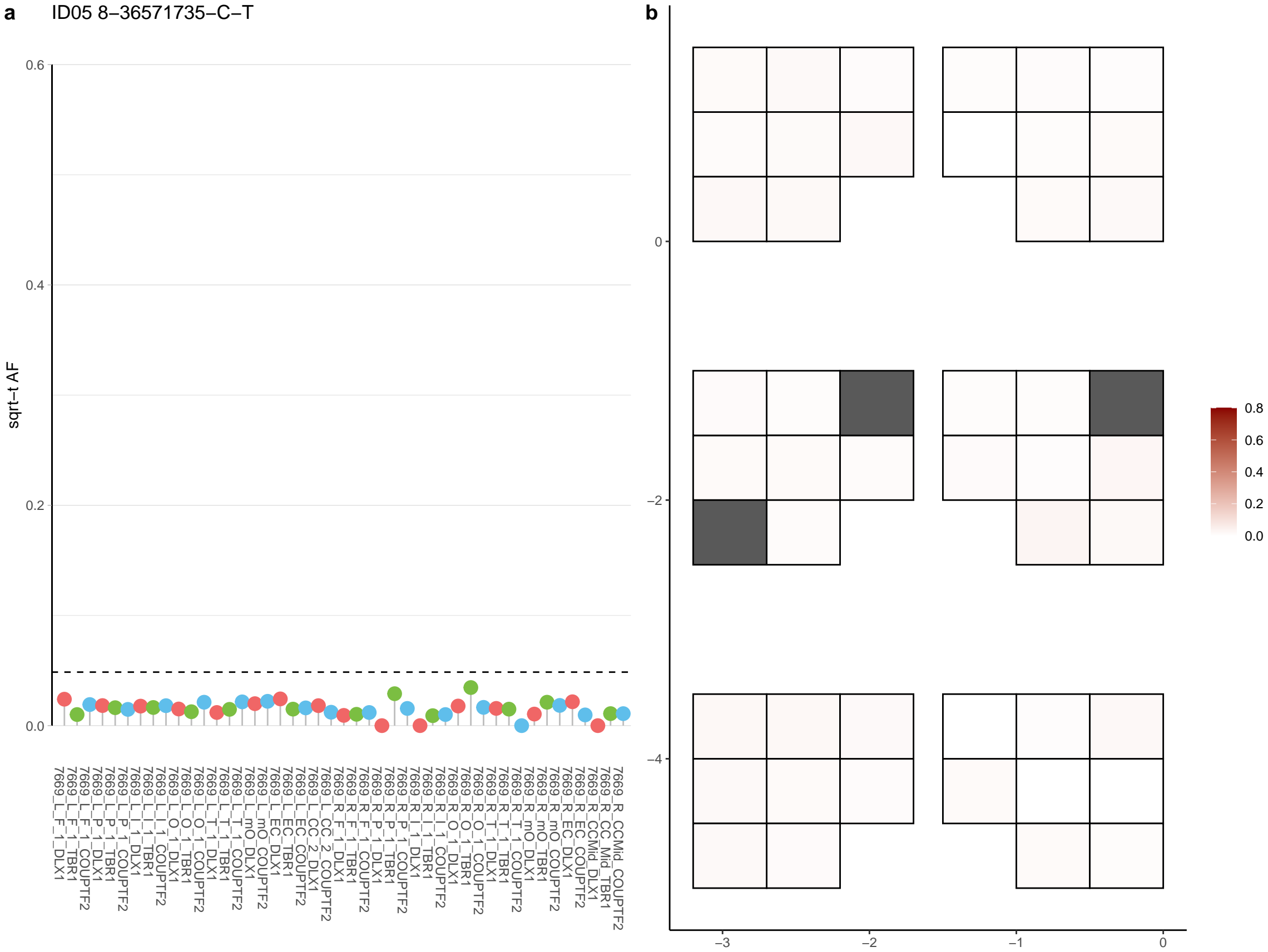

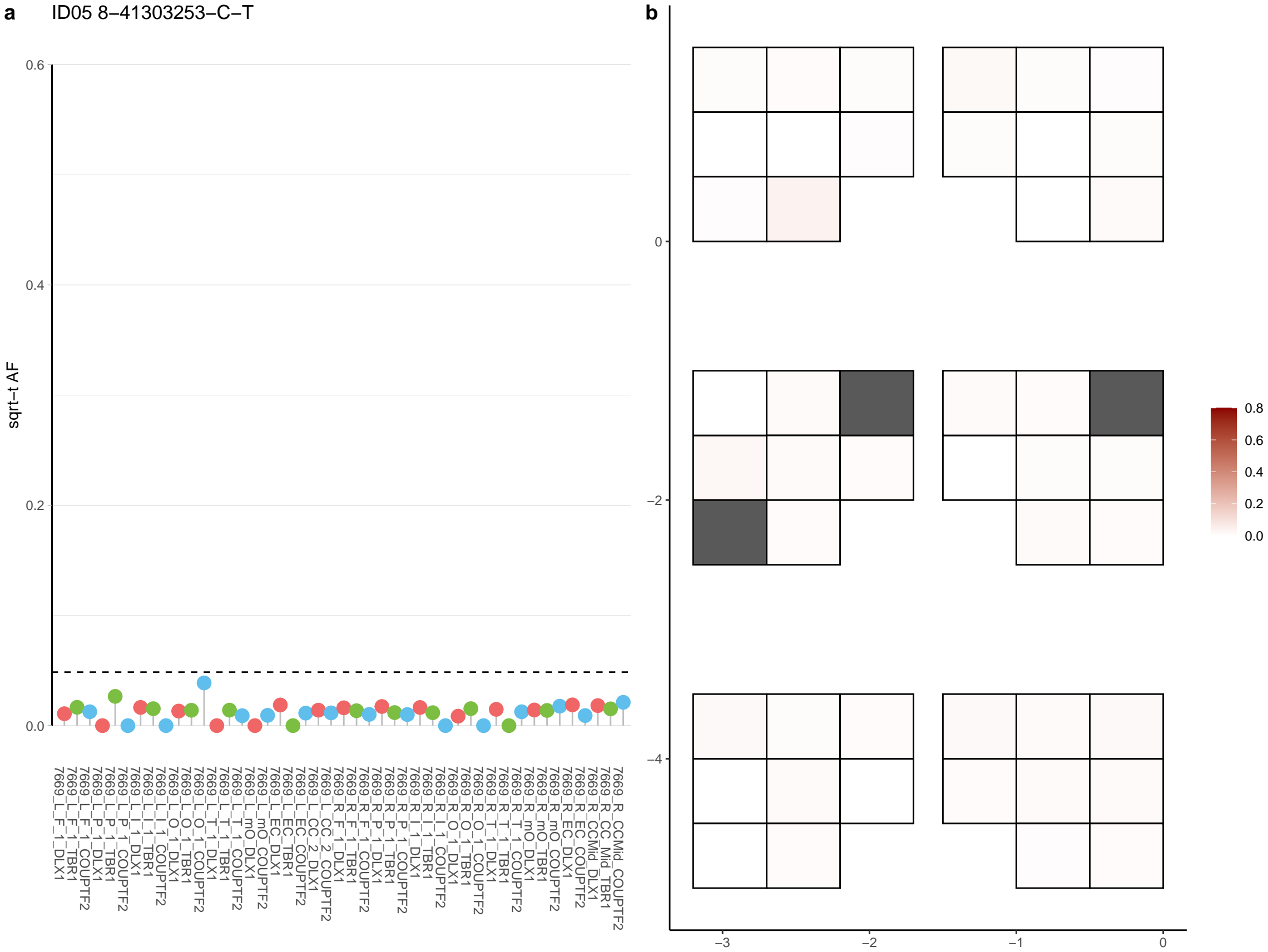

**a** ID05 8-4601722-C-G

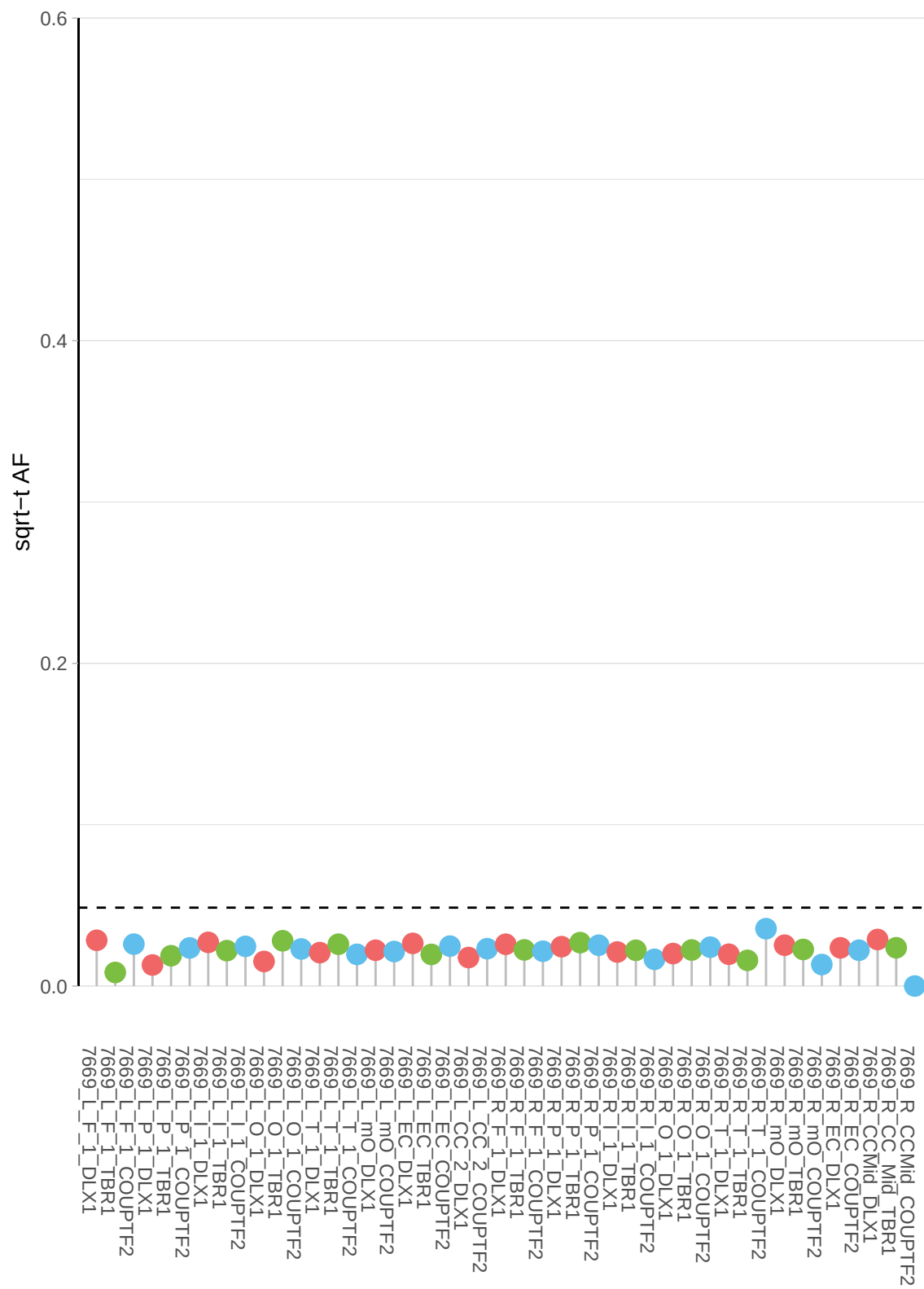

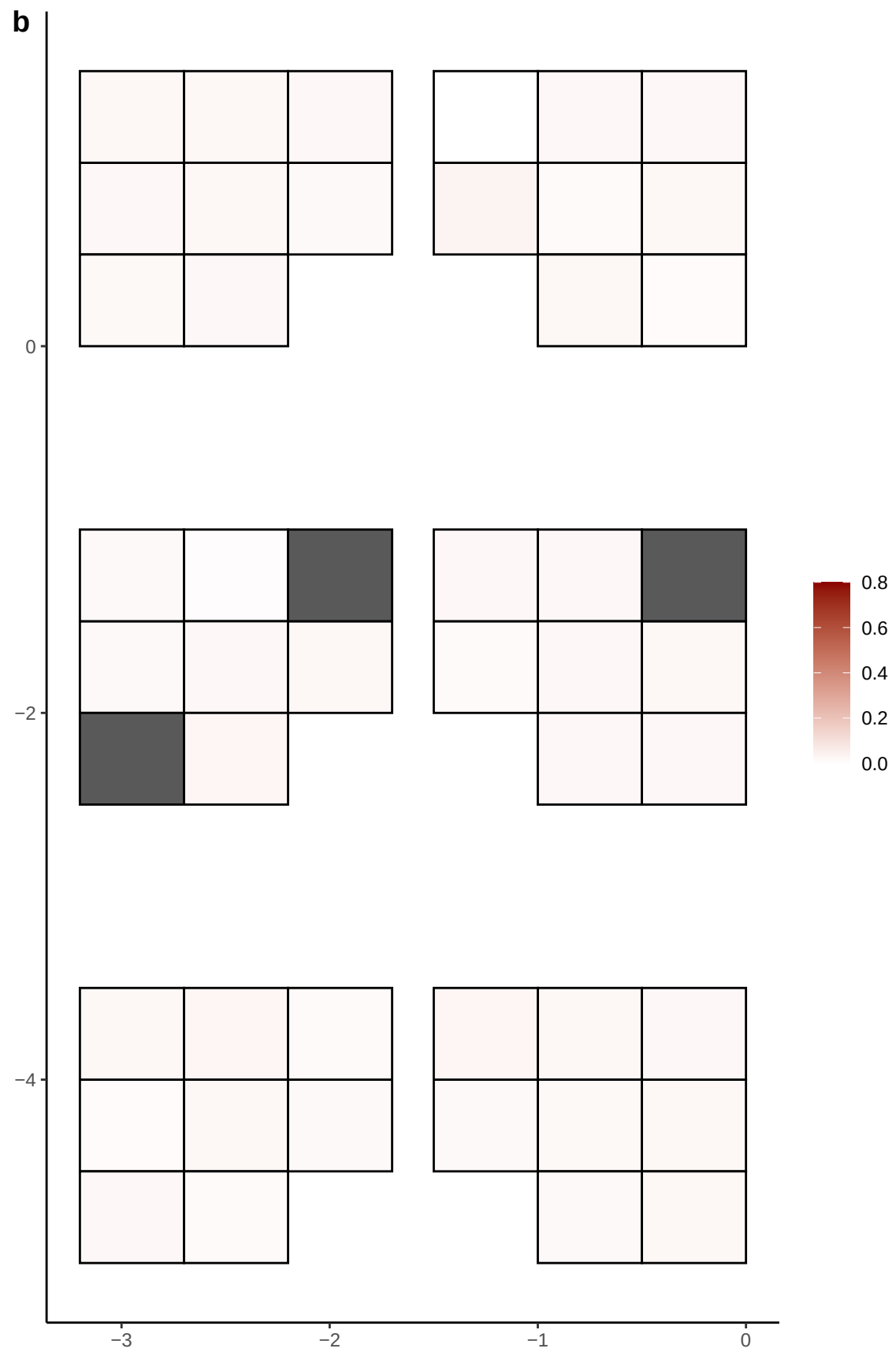

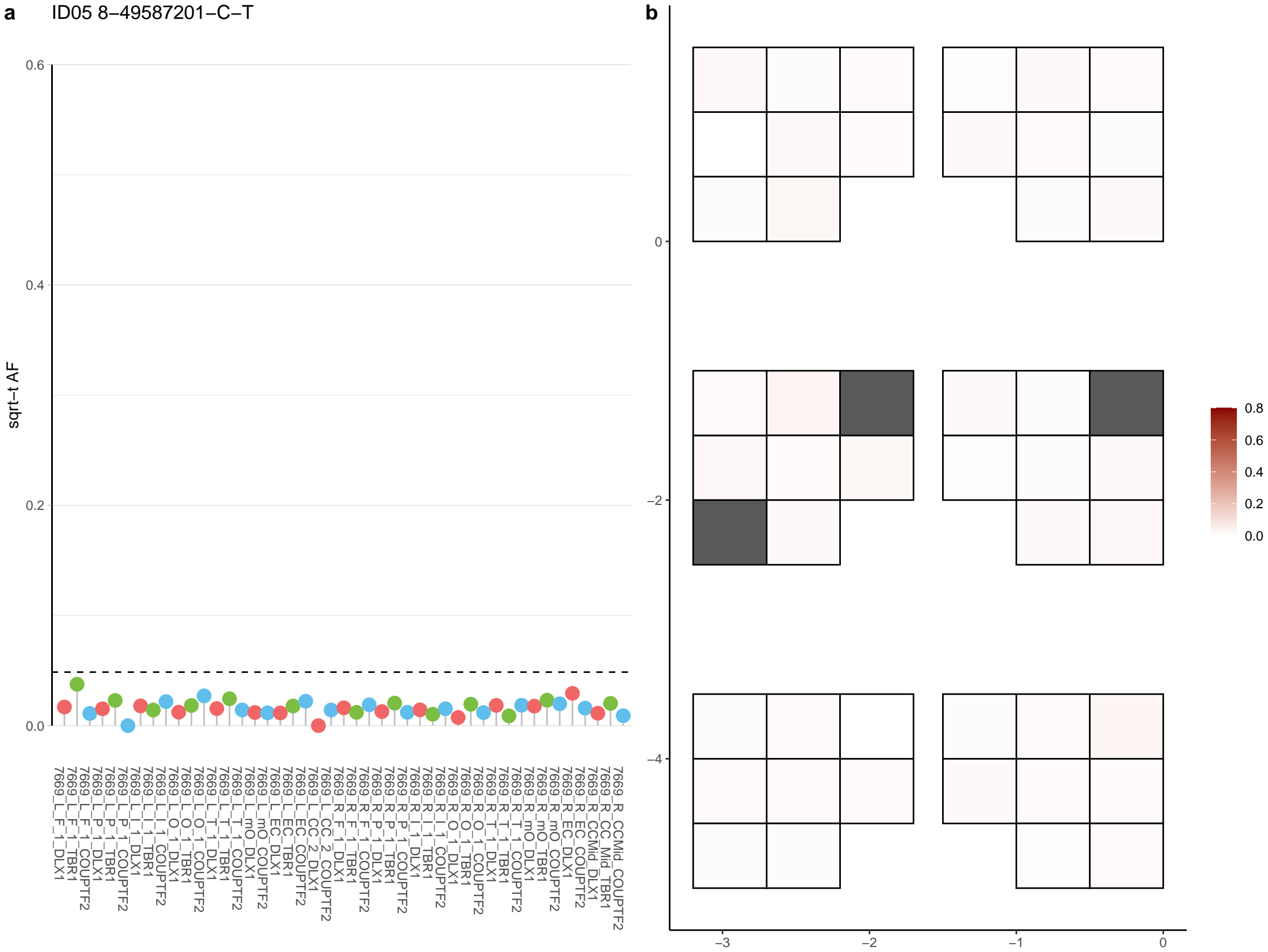

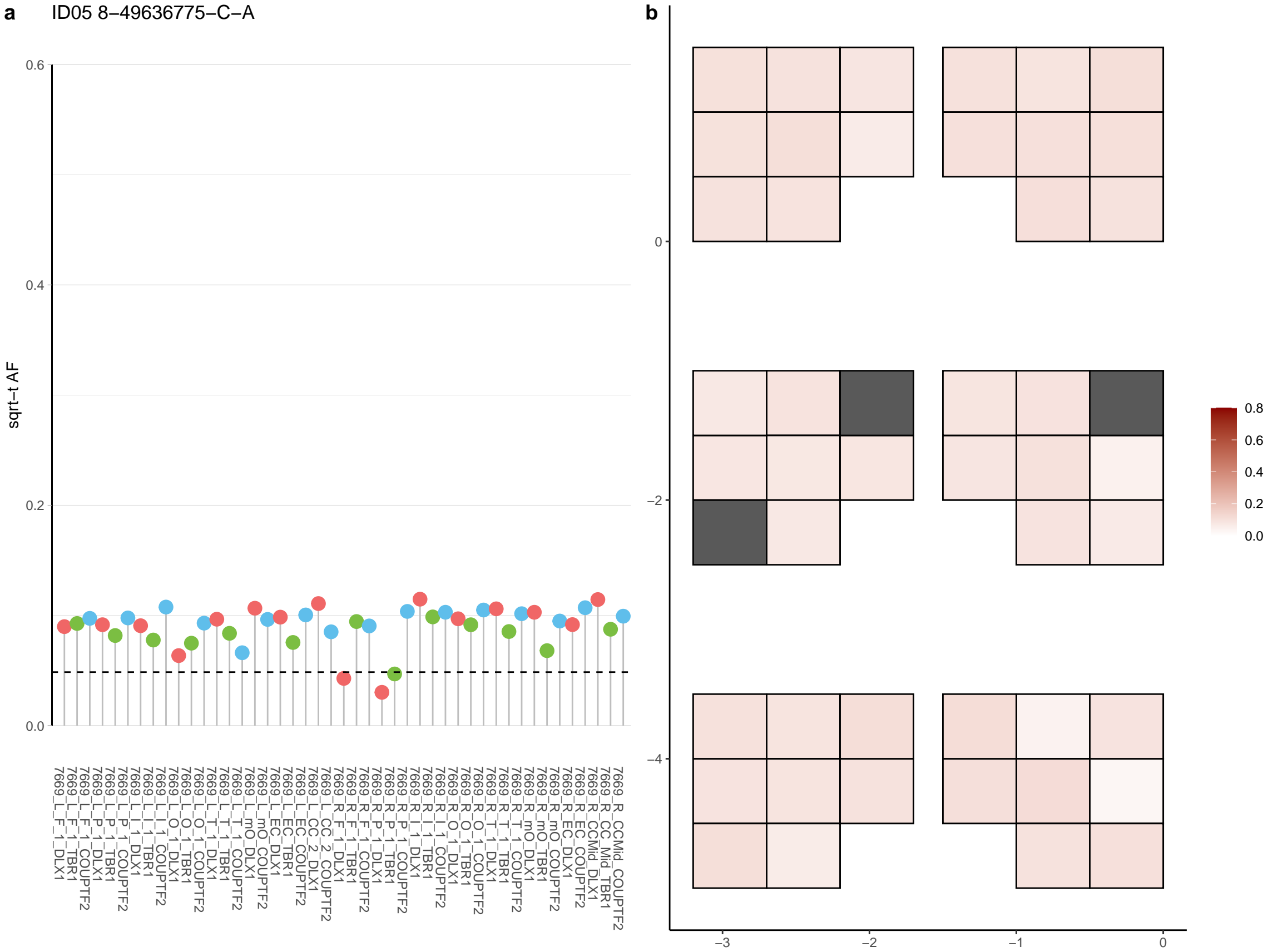

**a** ID05 8-50752909-G-C

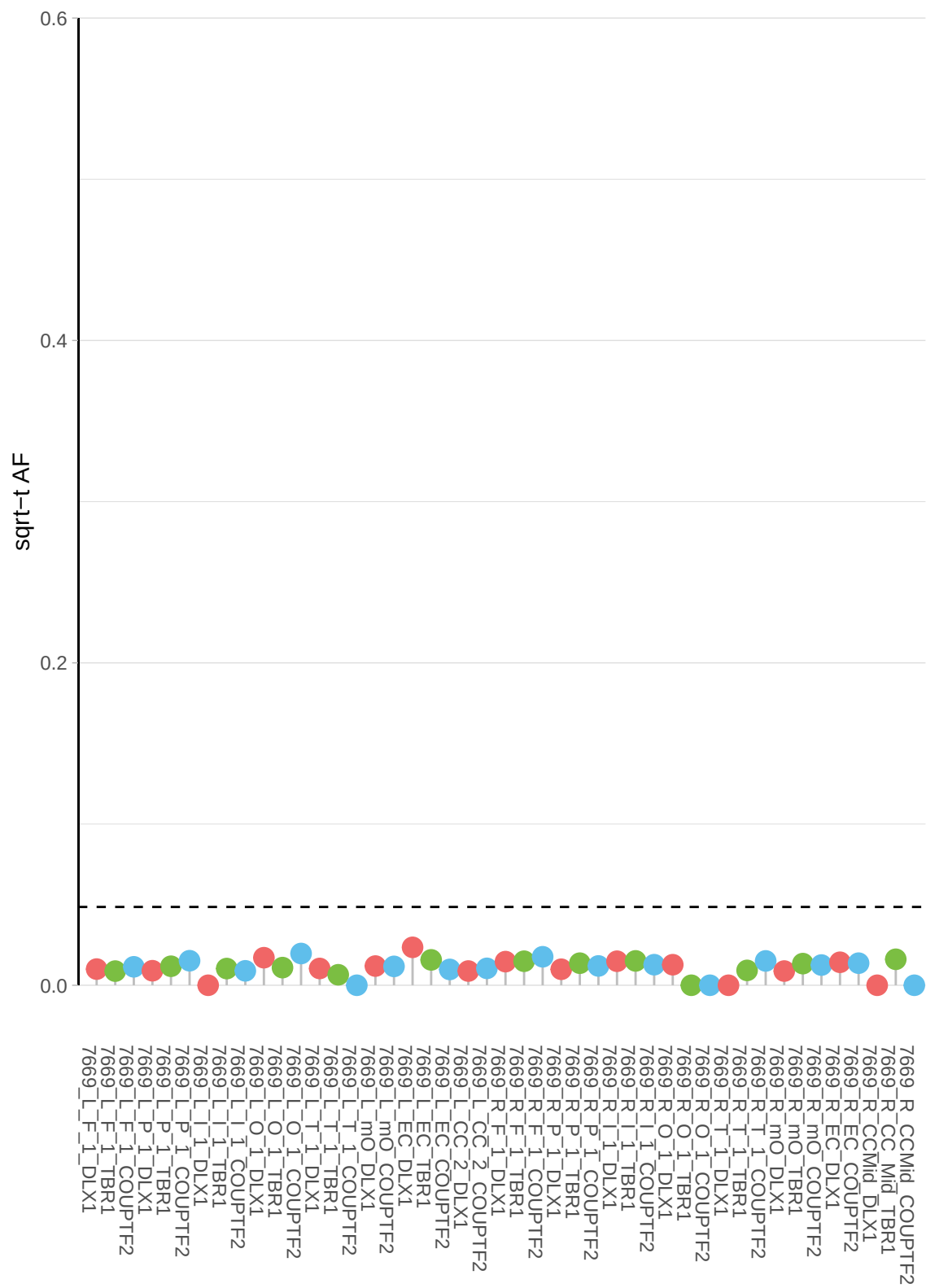

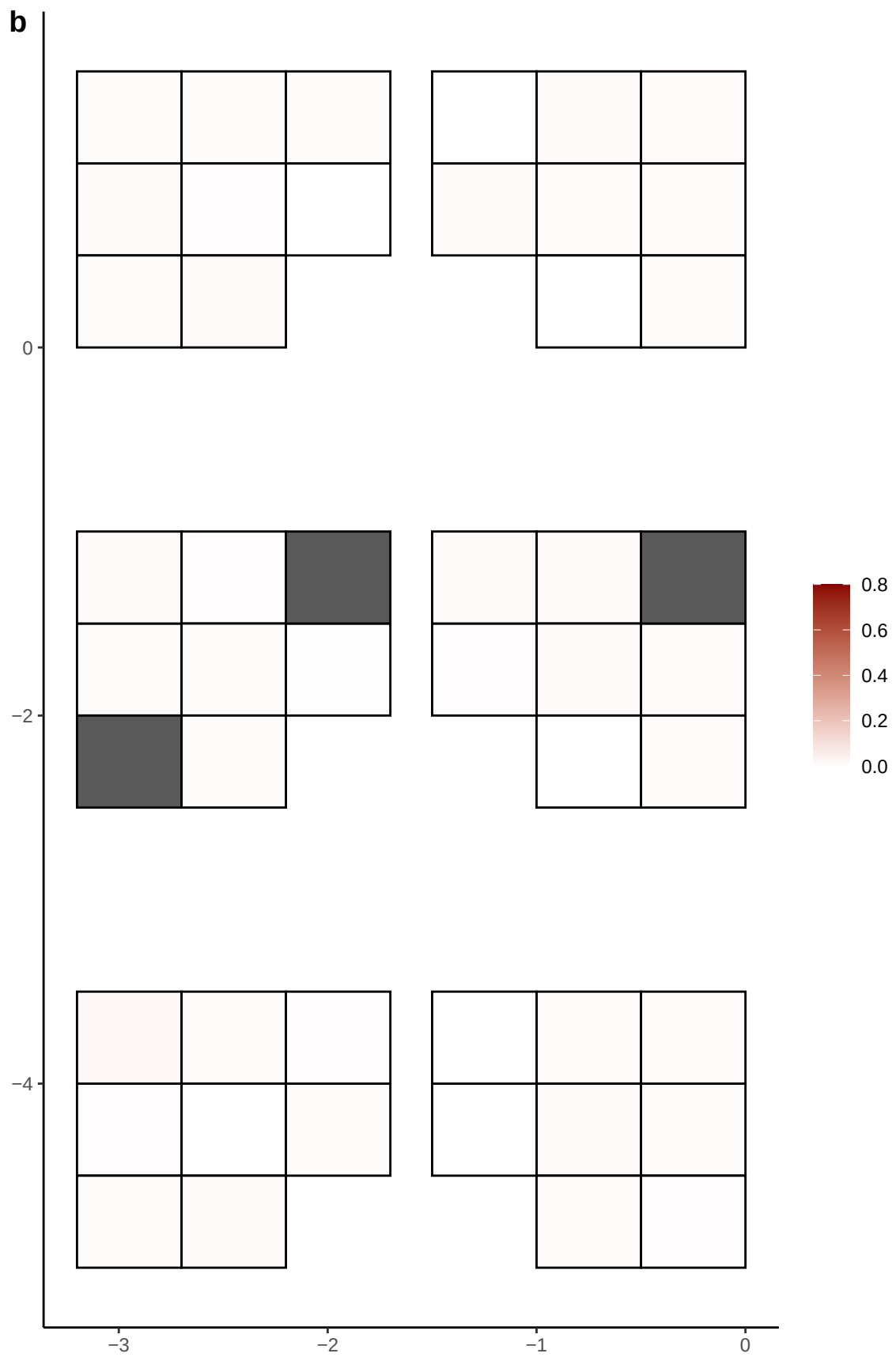

**a** ID05 8-56154213-C-T

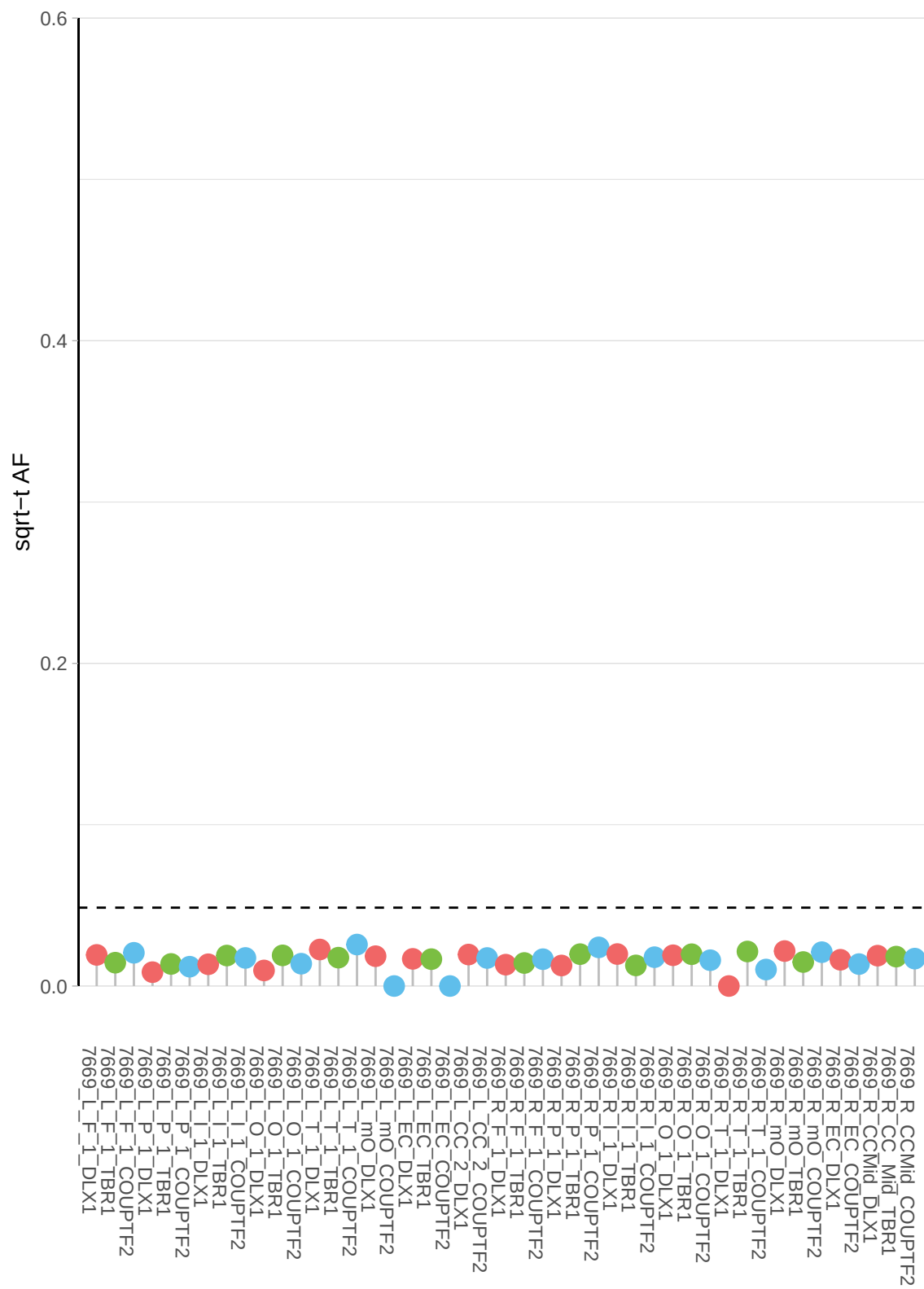

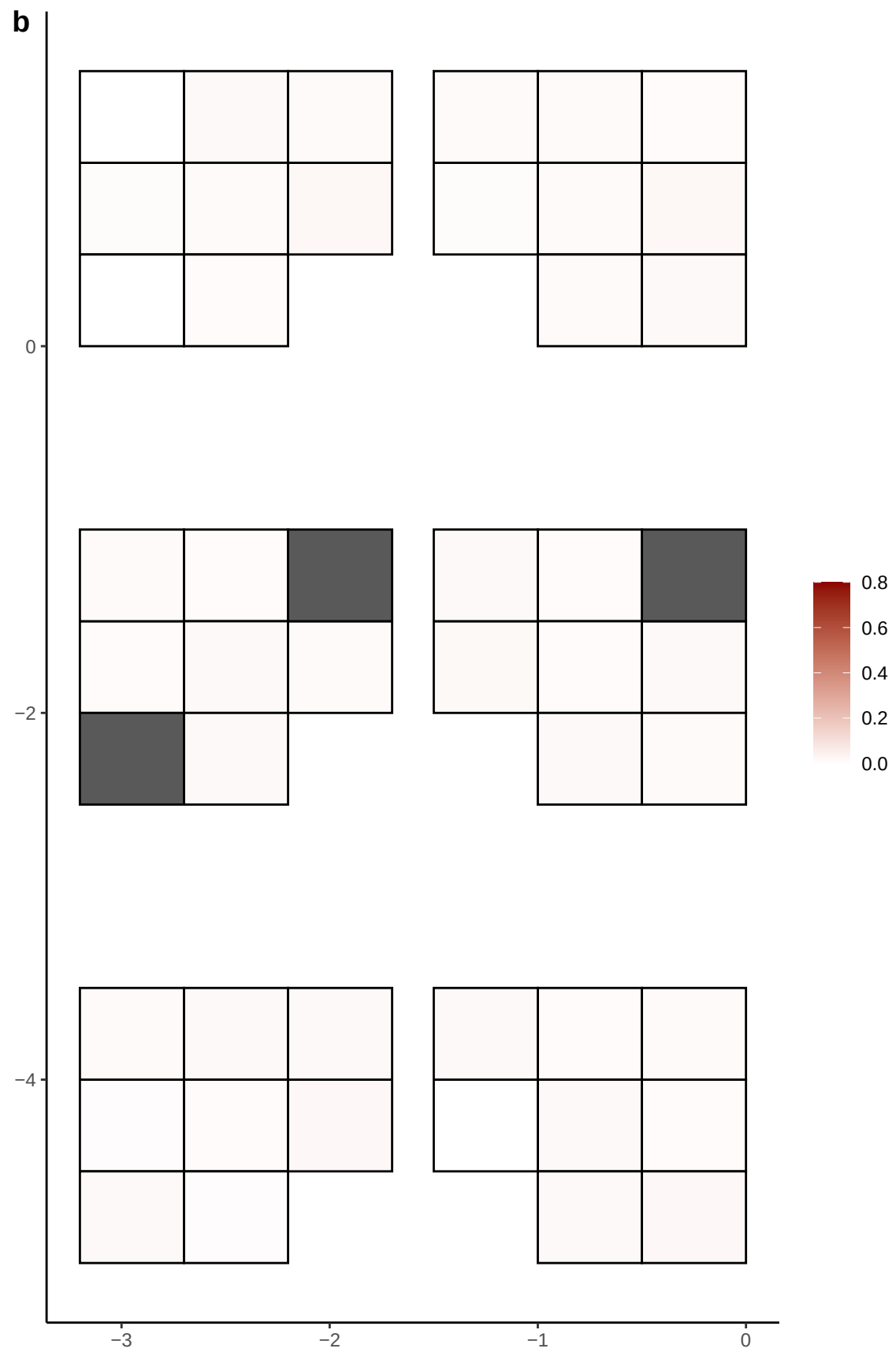

**a** ID05 8-57499201-TG-T

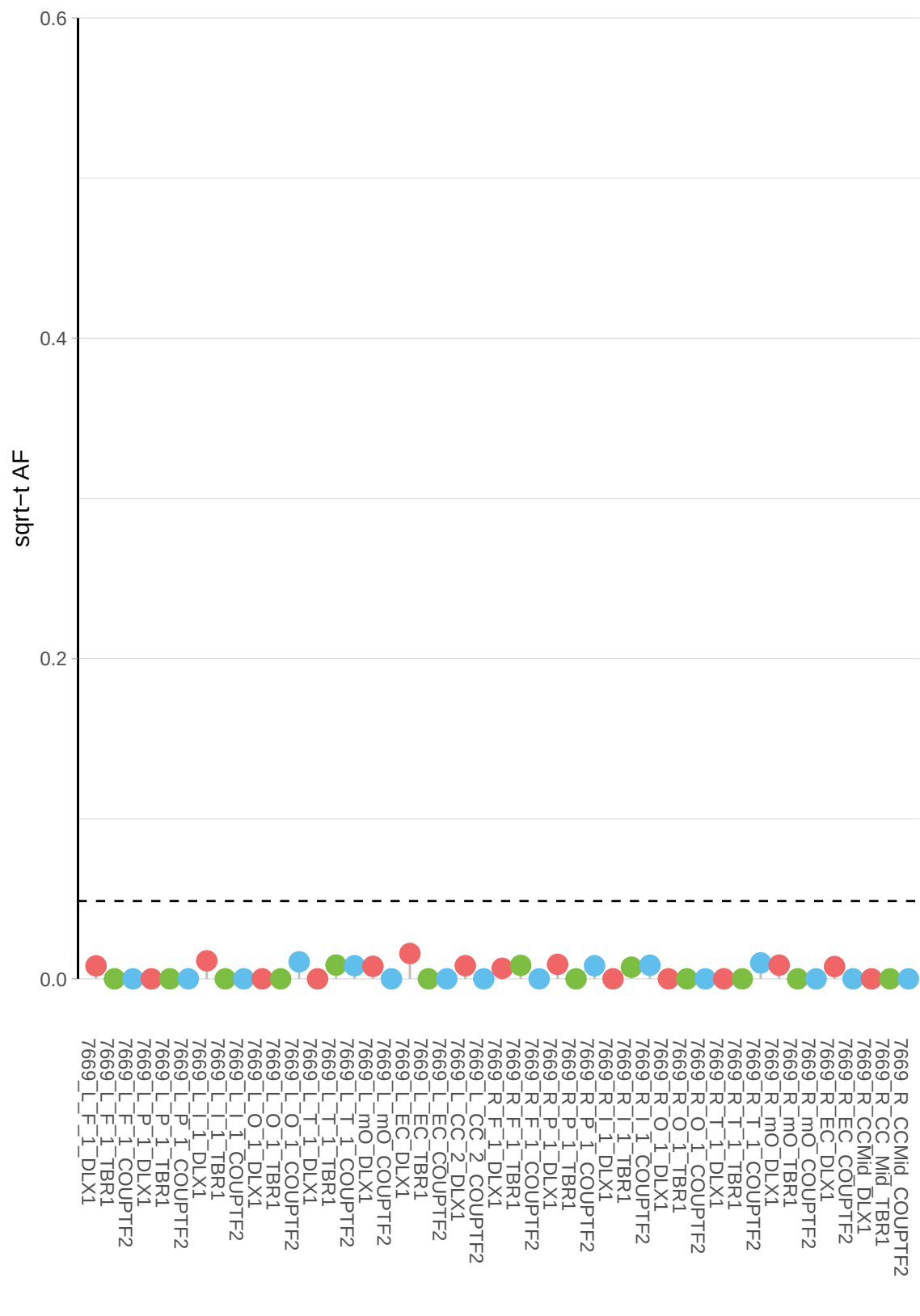

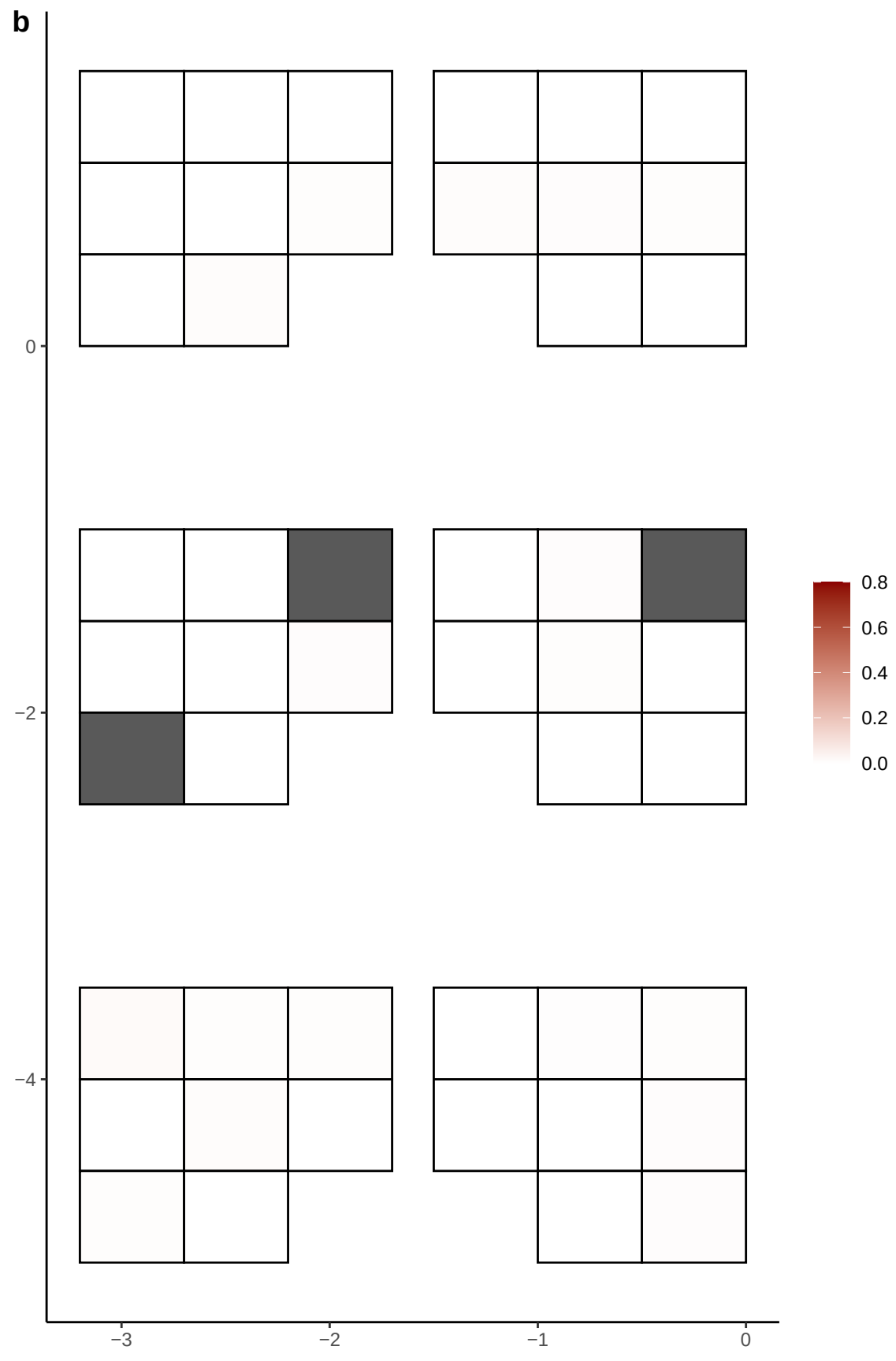

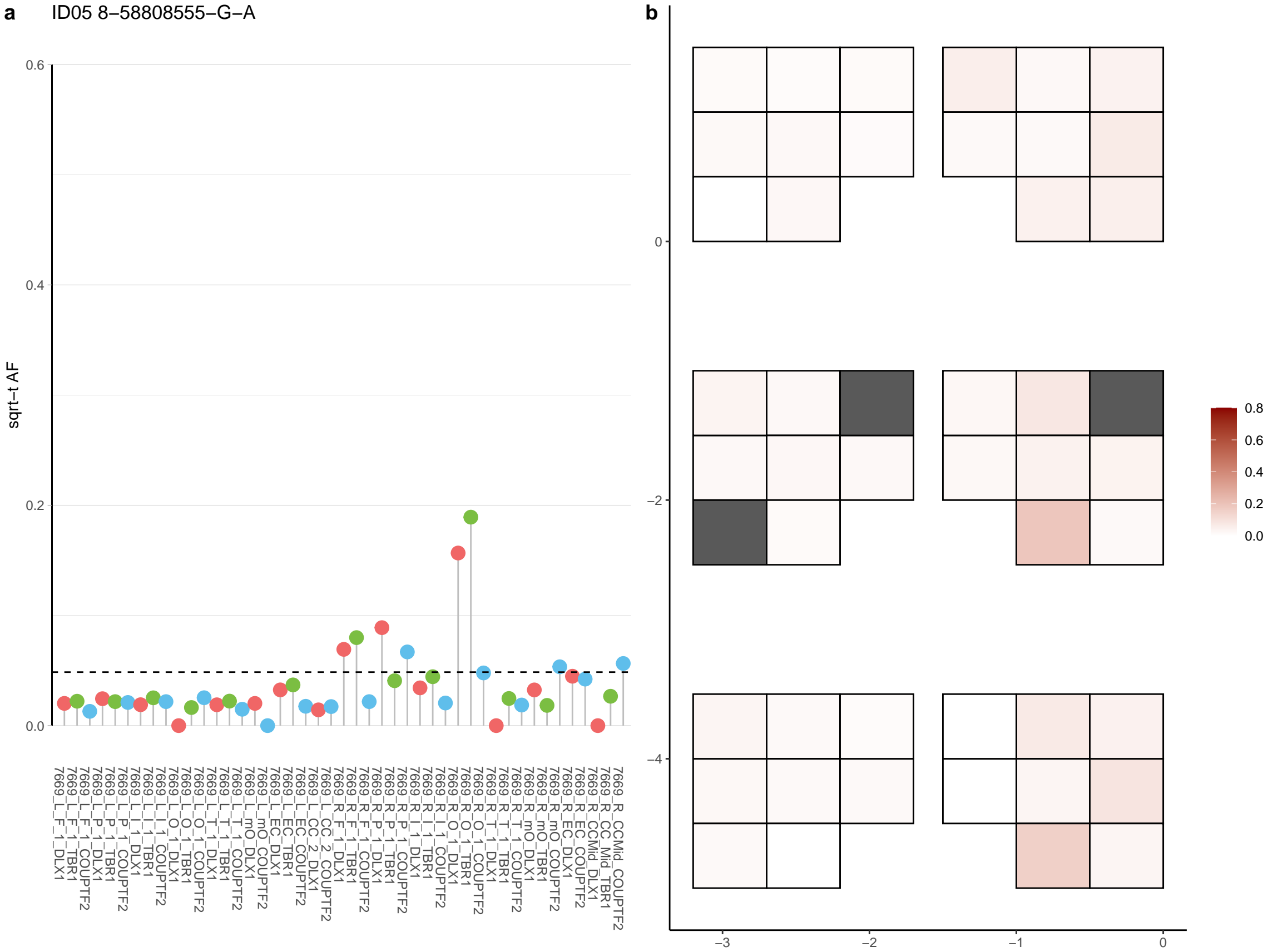

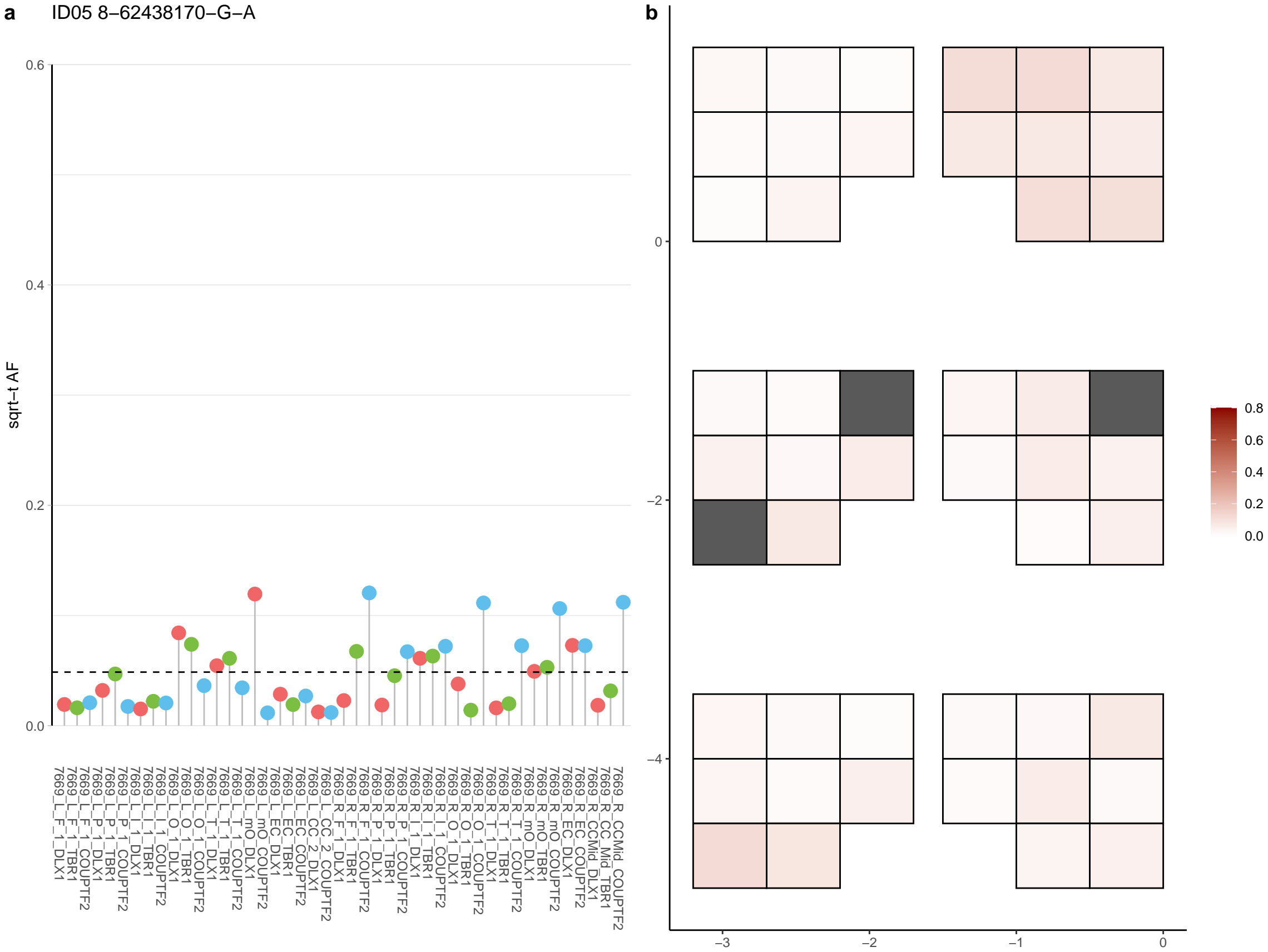

**a** ID05 8-65149924-C-T

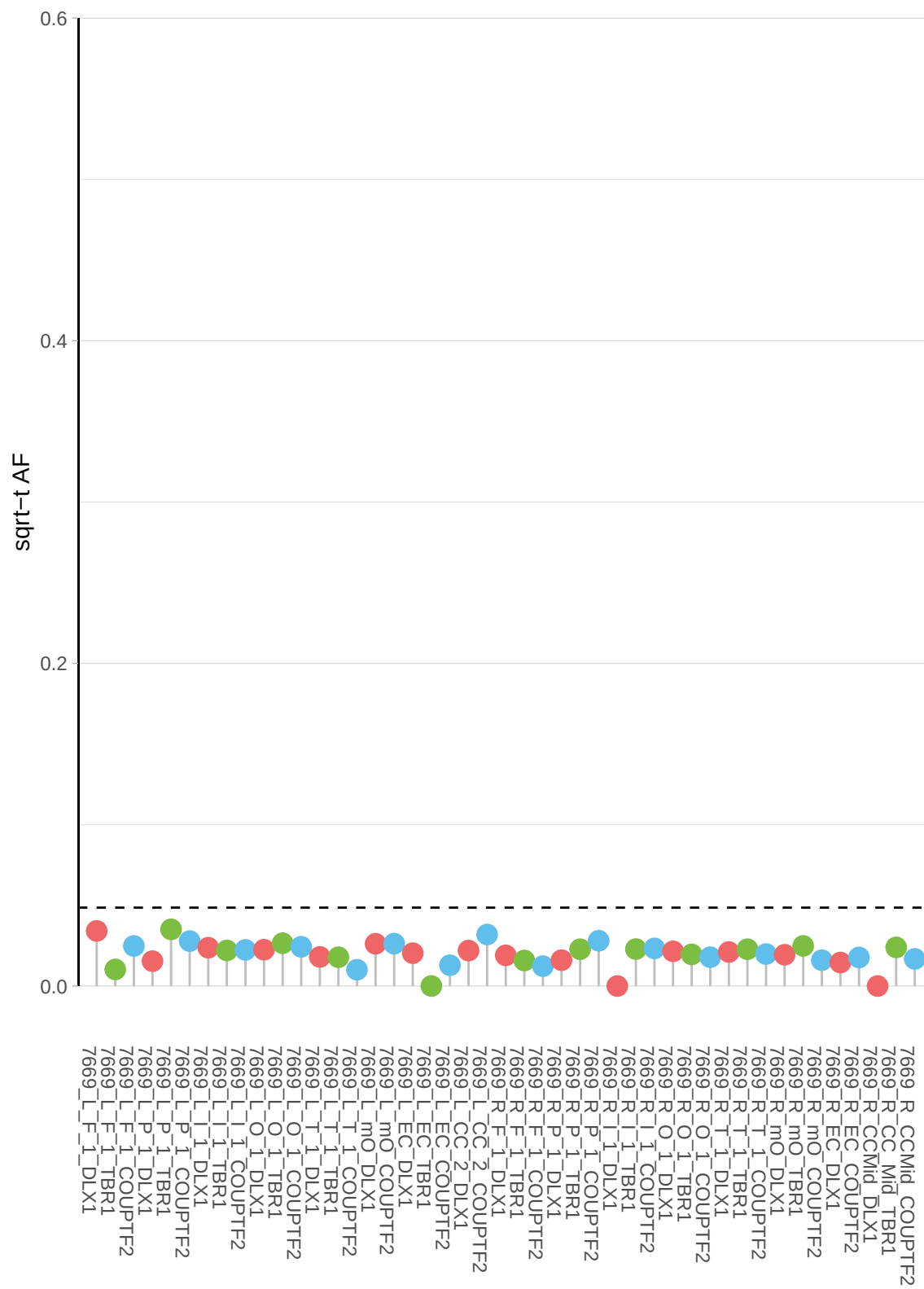

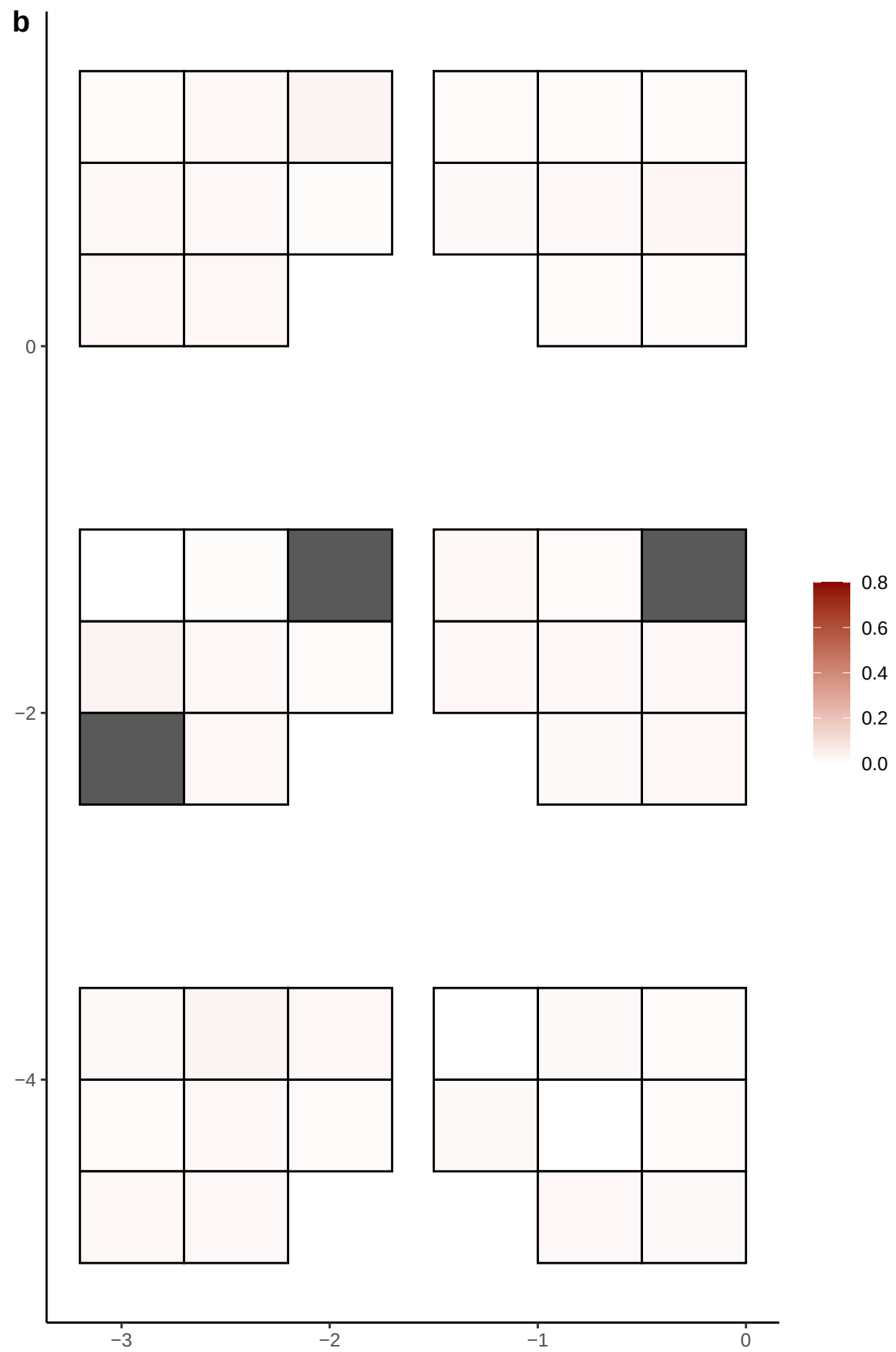

**a** ID05 8-66303042-C-T

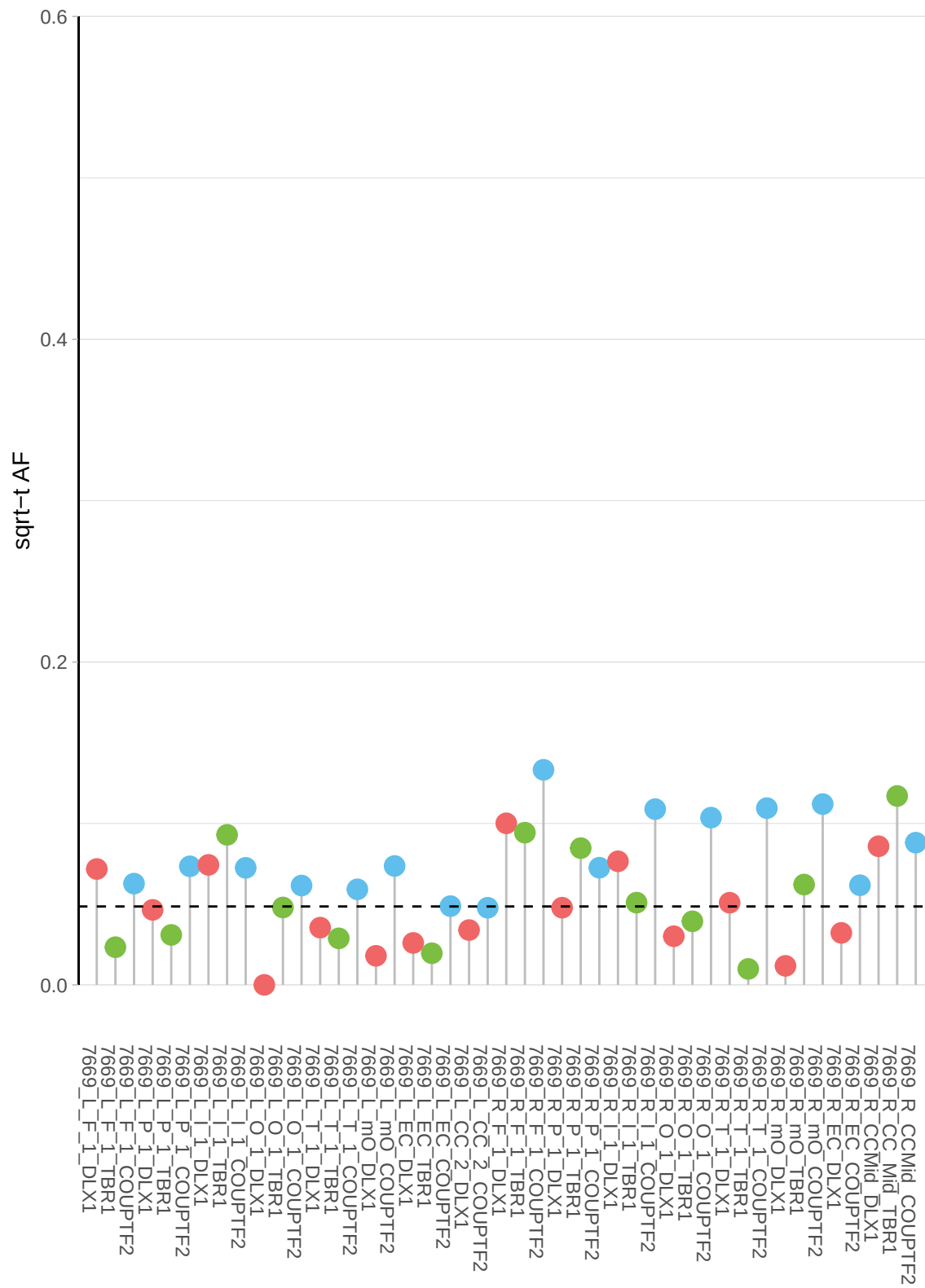

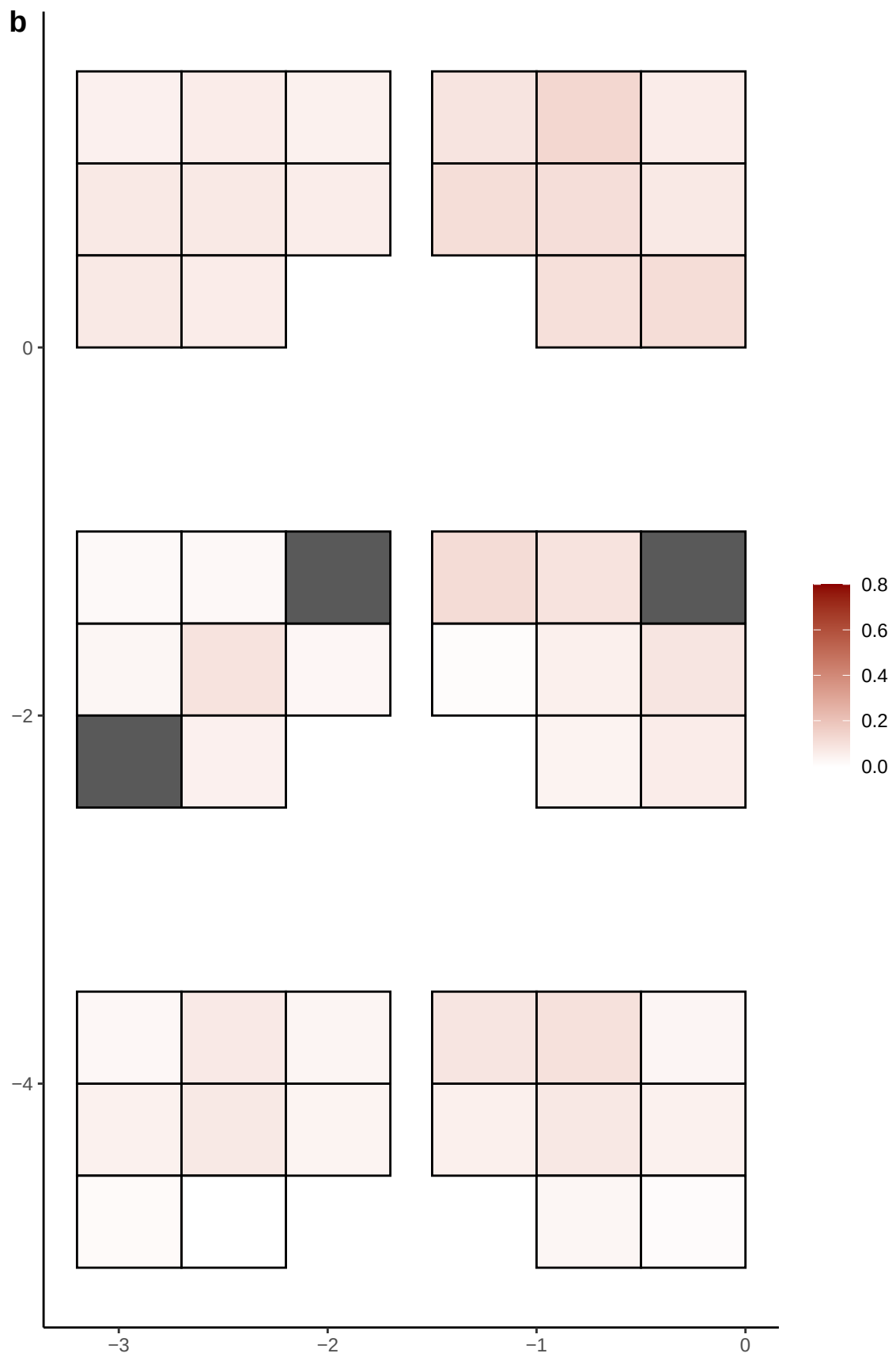

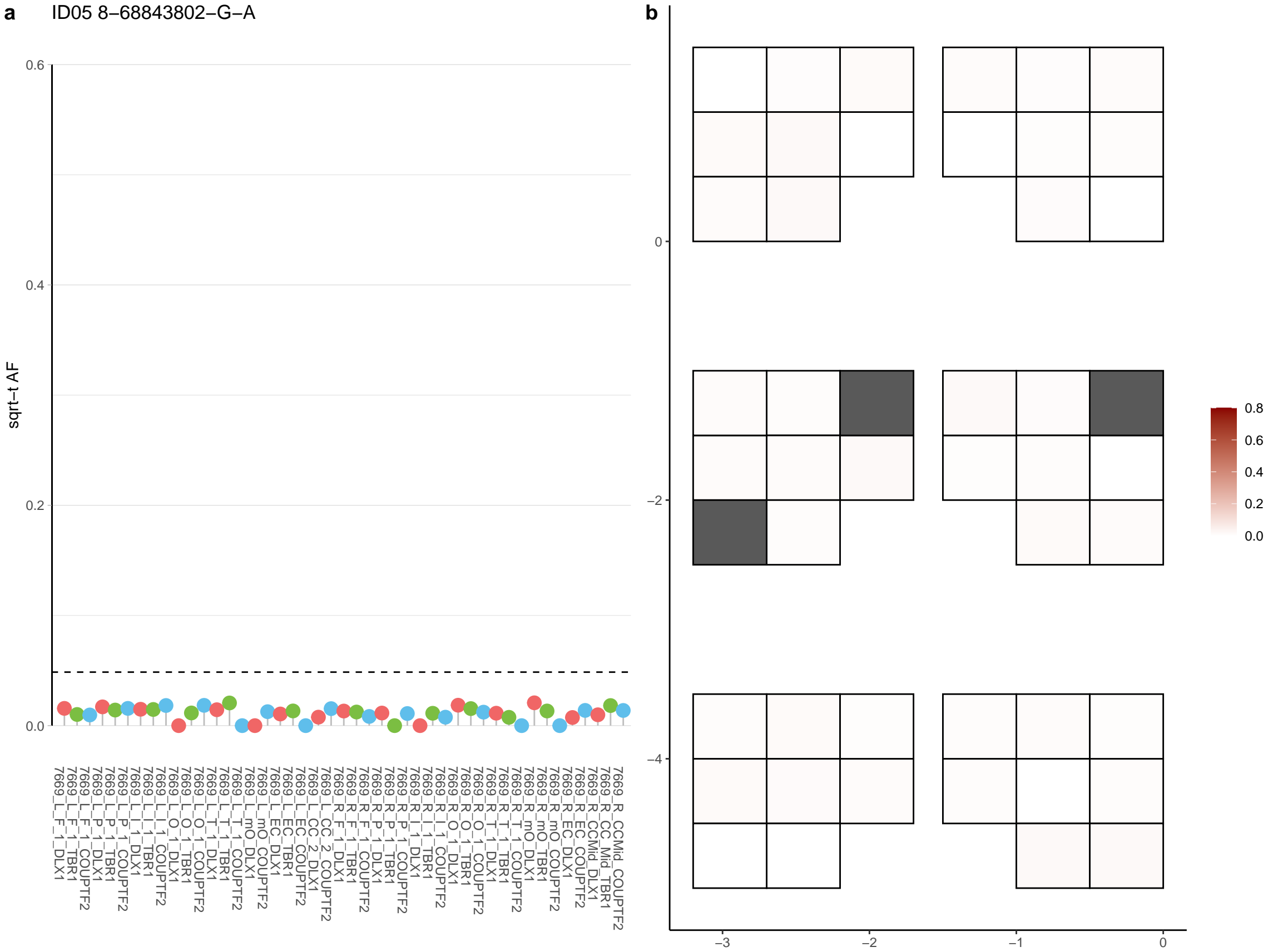

**a** ID05 8-72347819-C-T

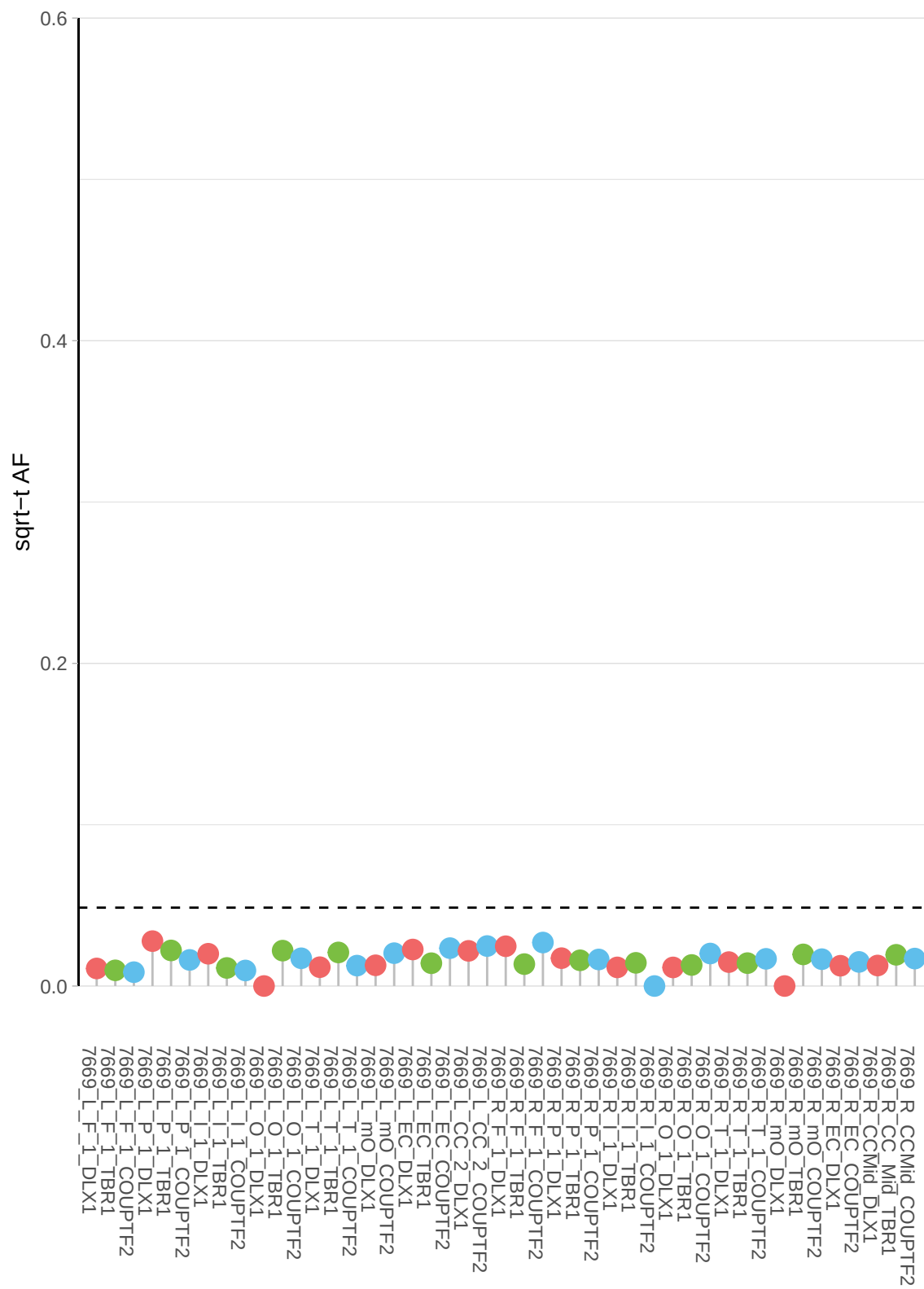

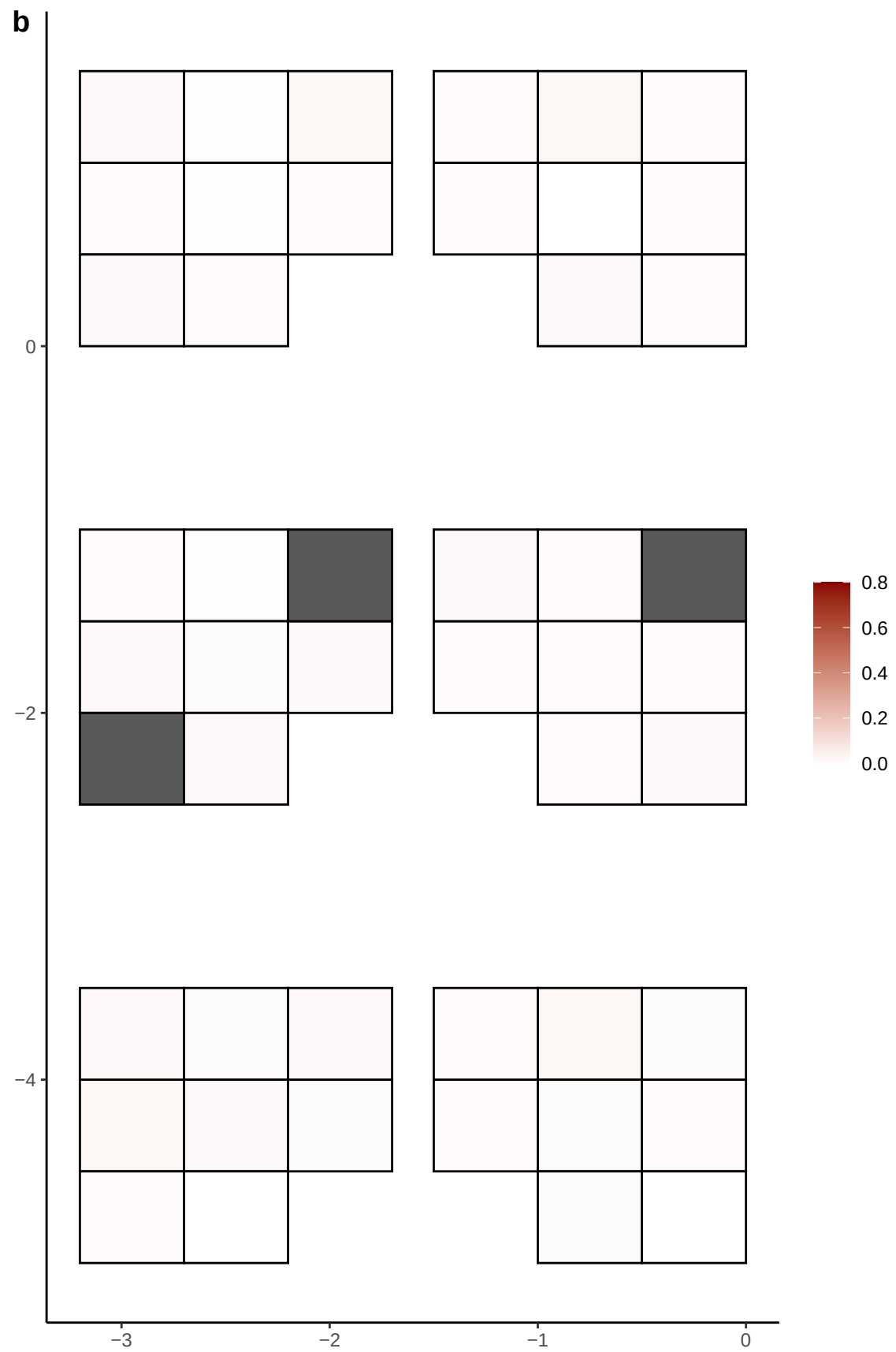

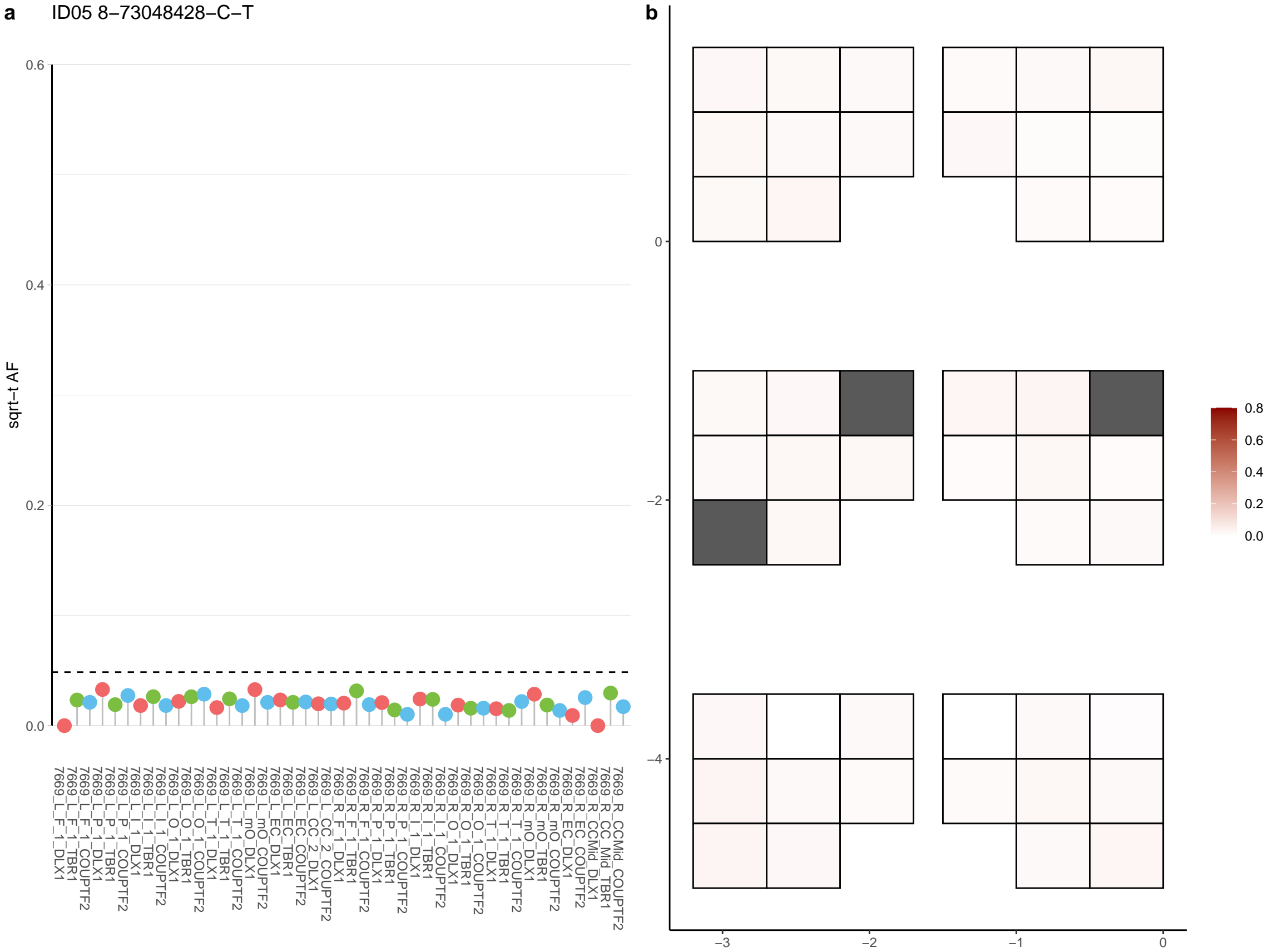

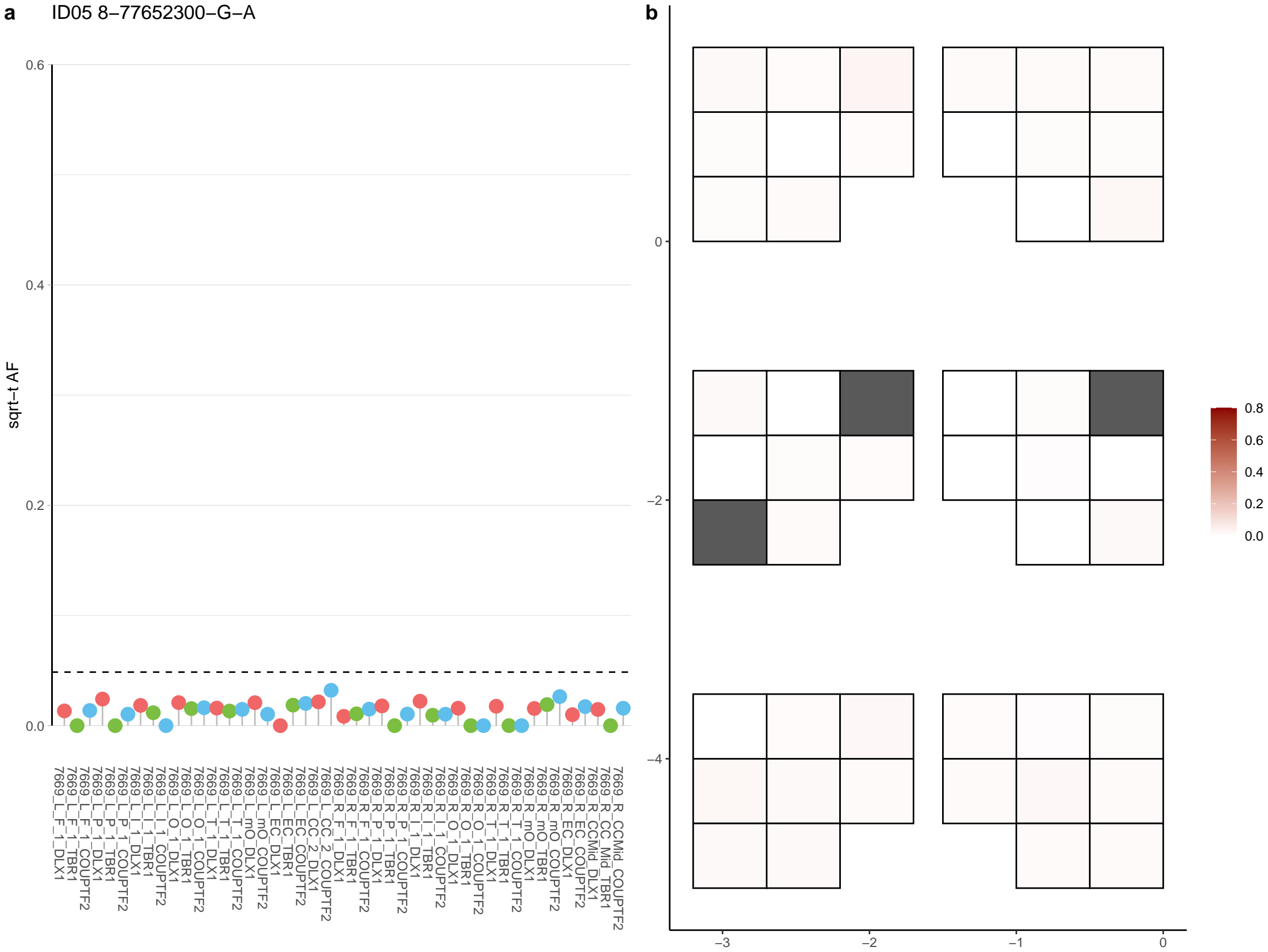

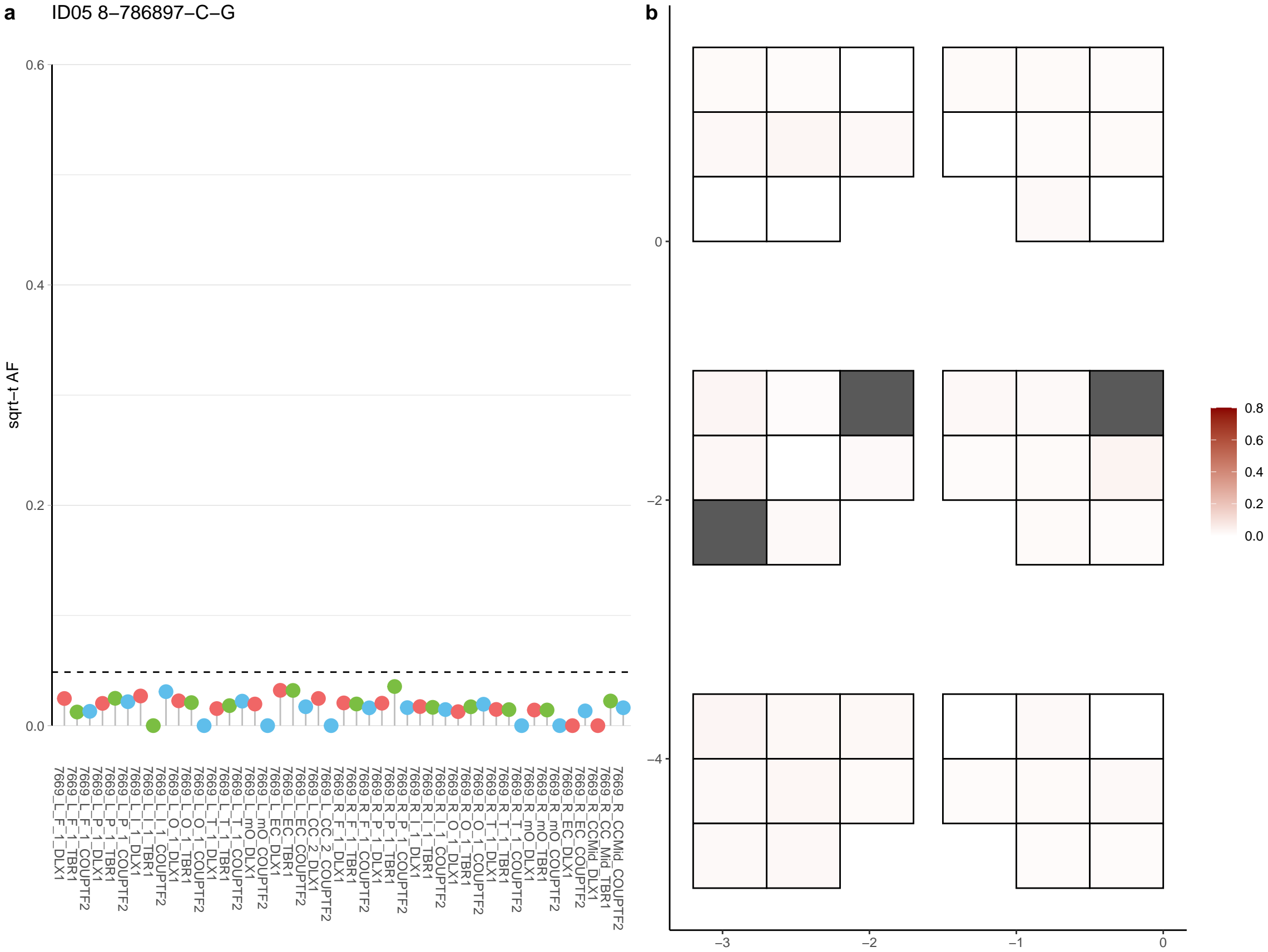

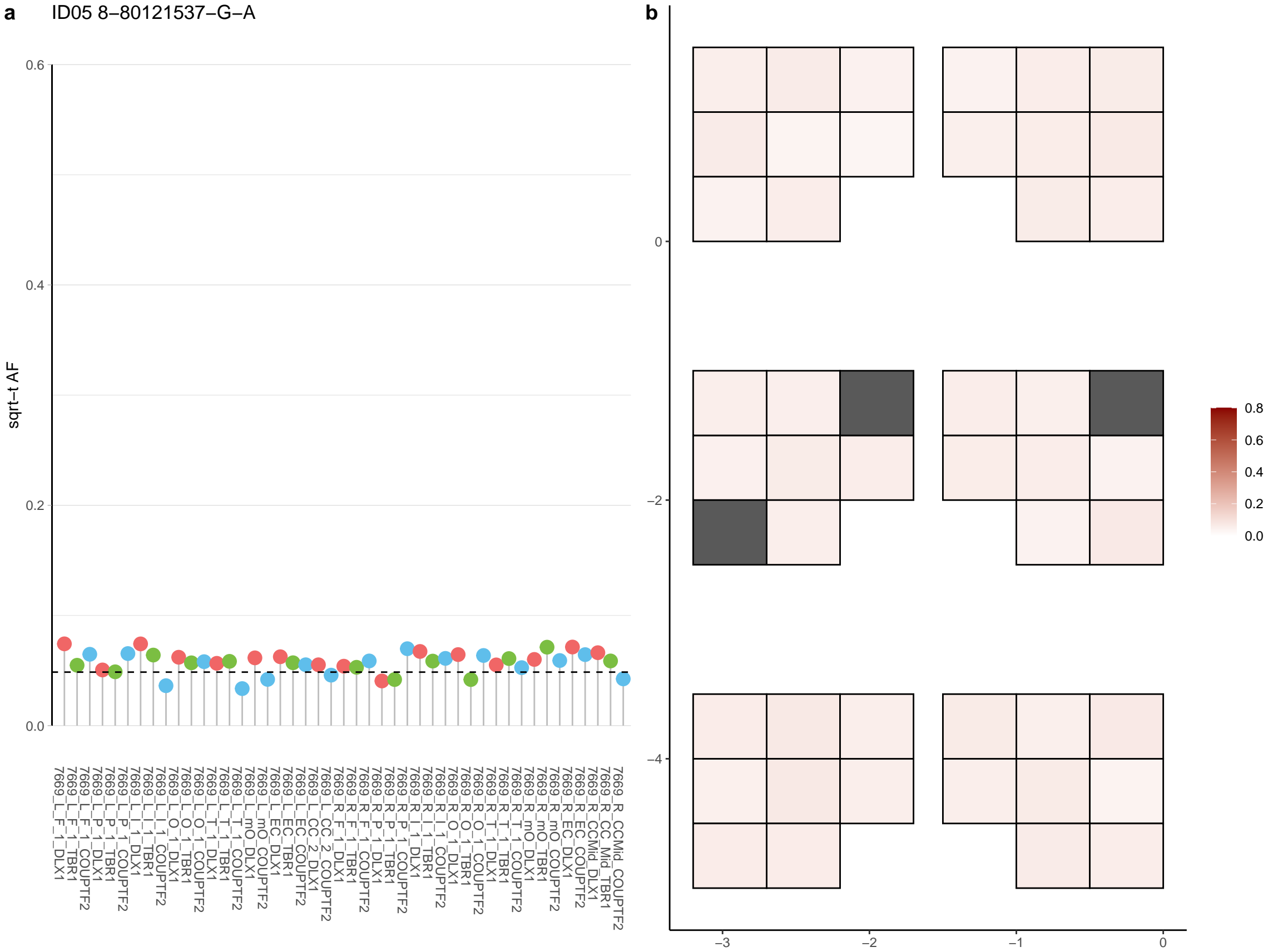

**a** ID05 8-80121538-T-C

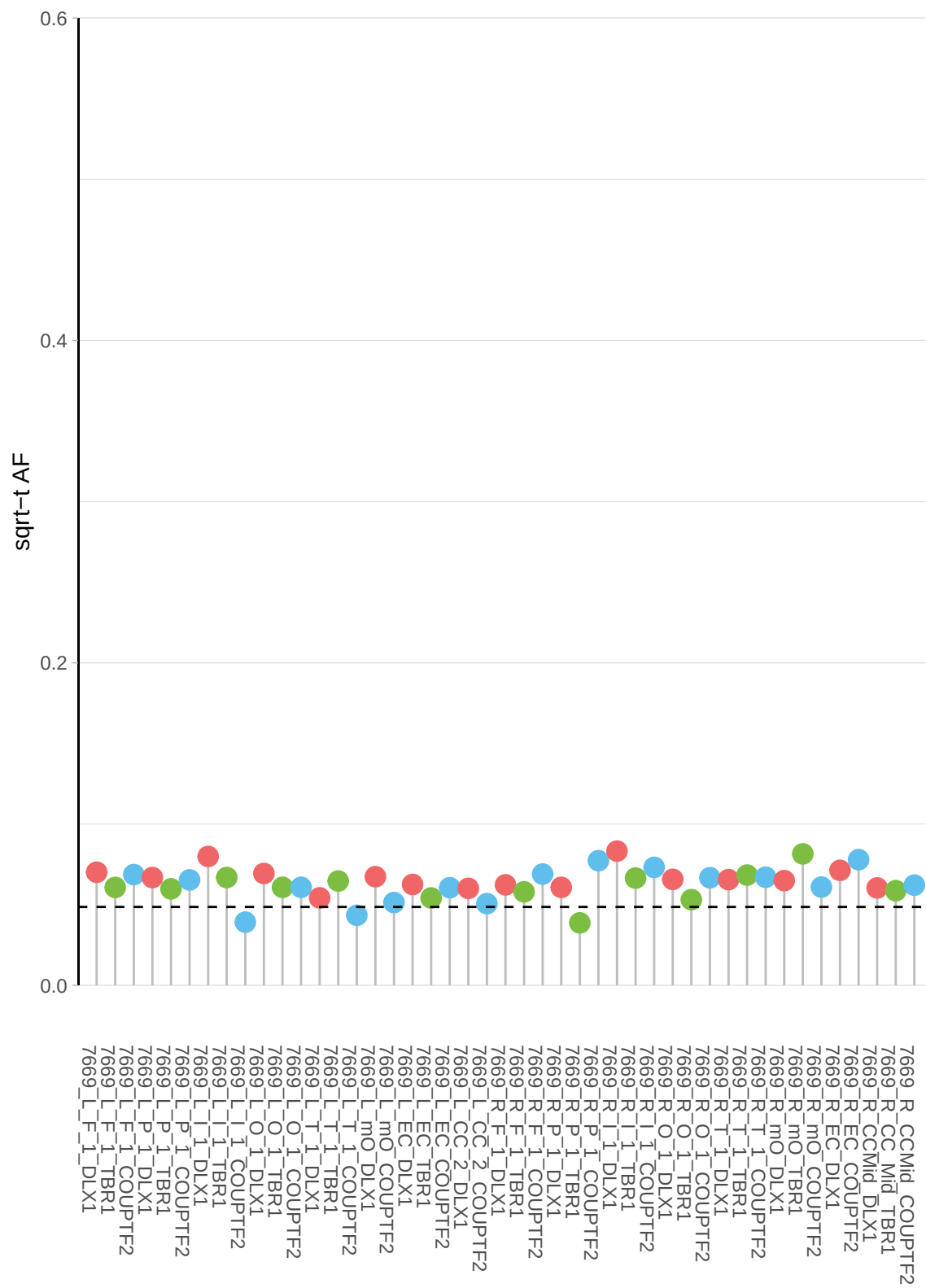

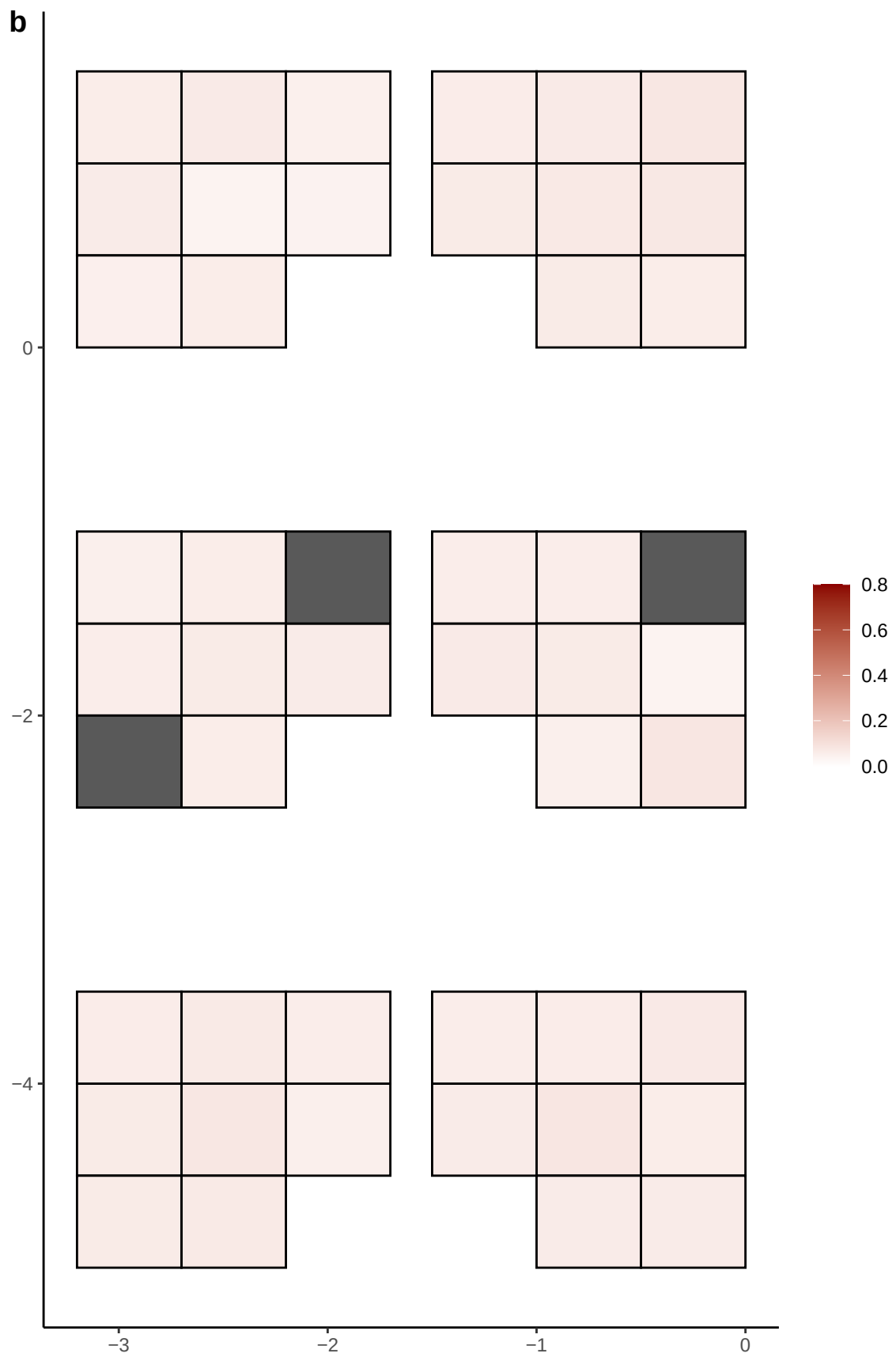

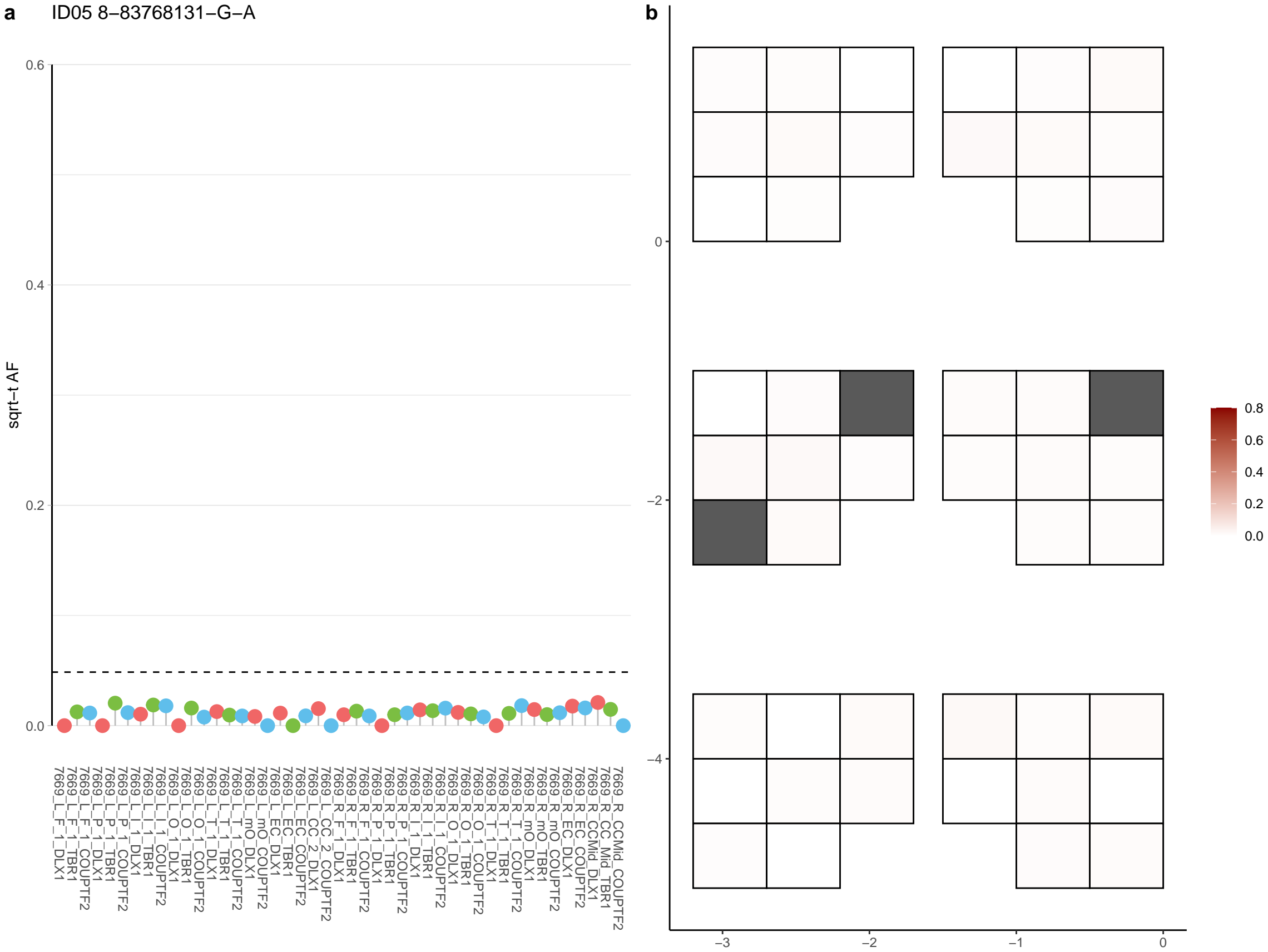

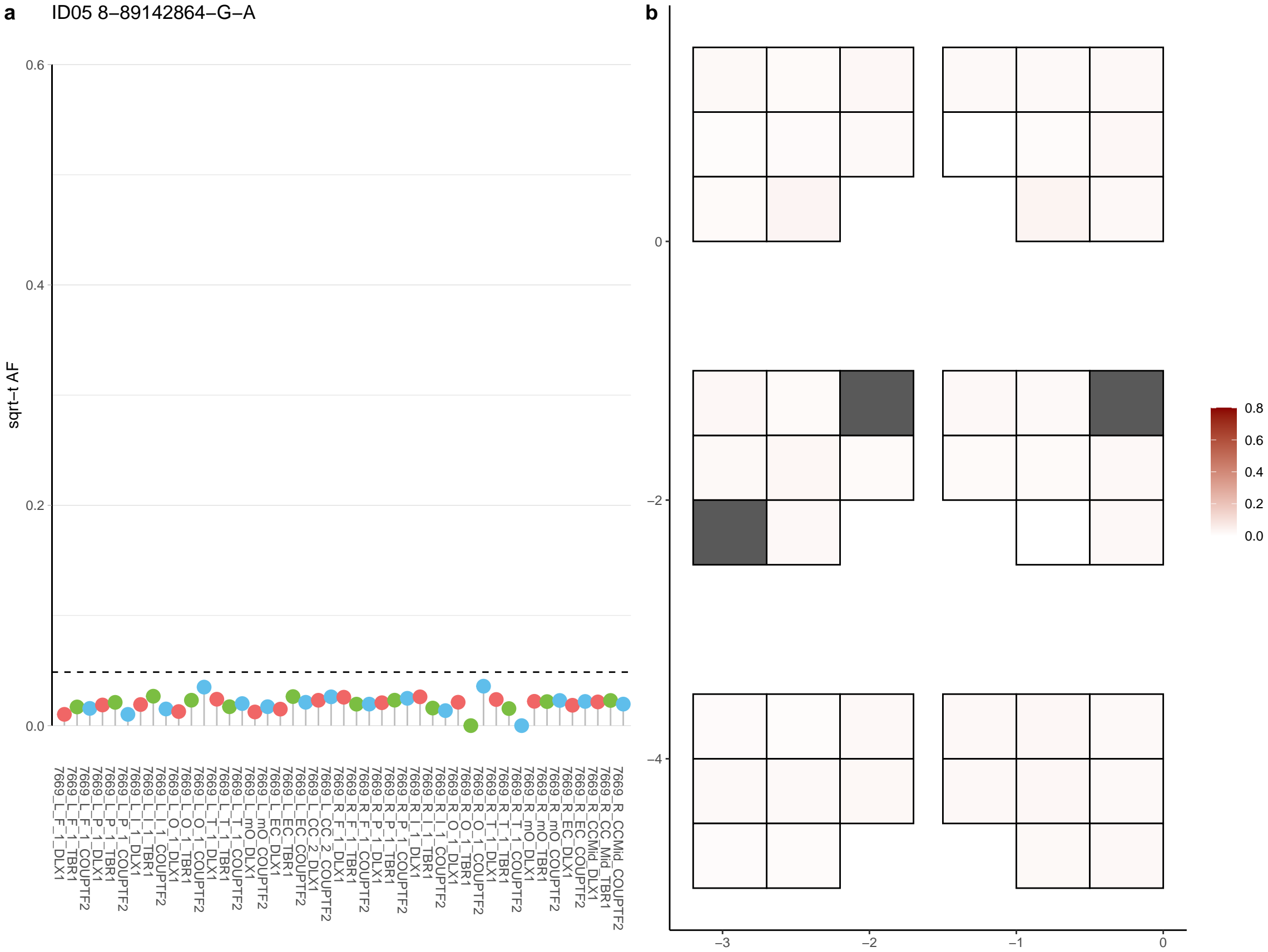

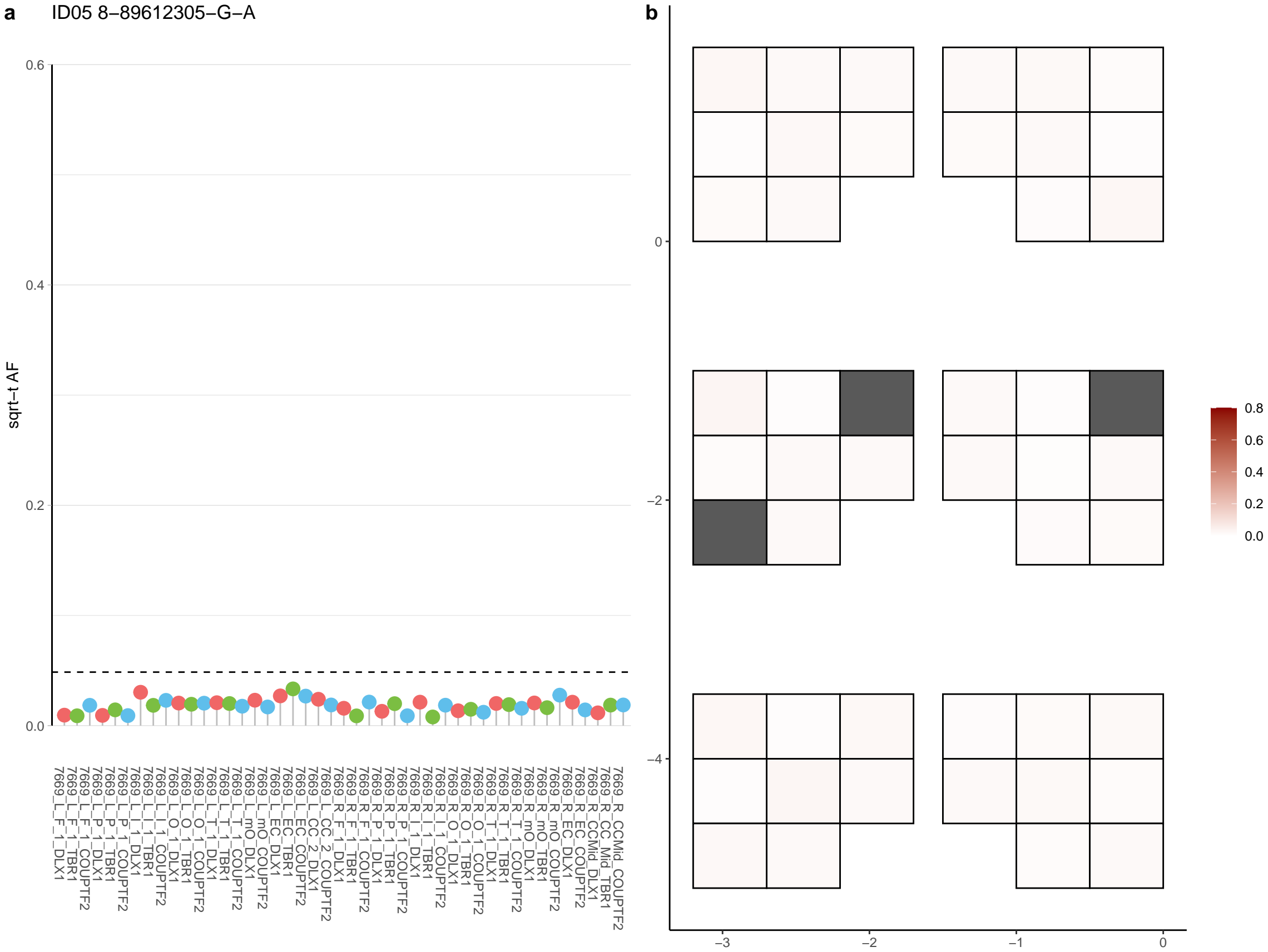

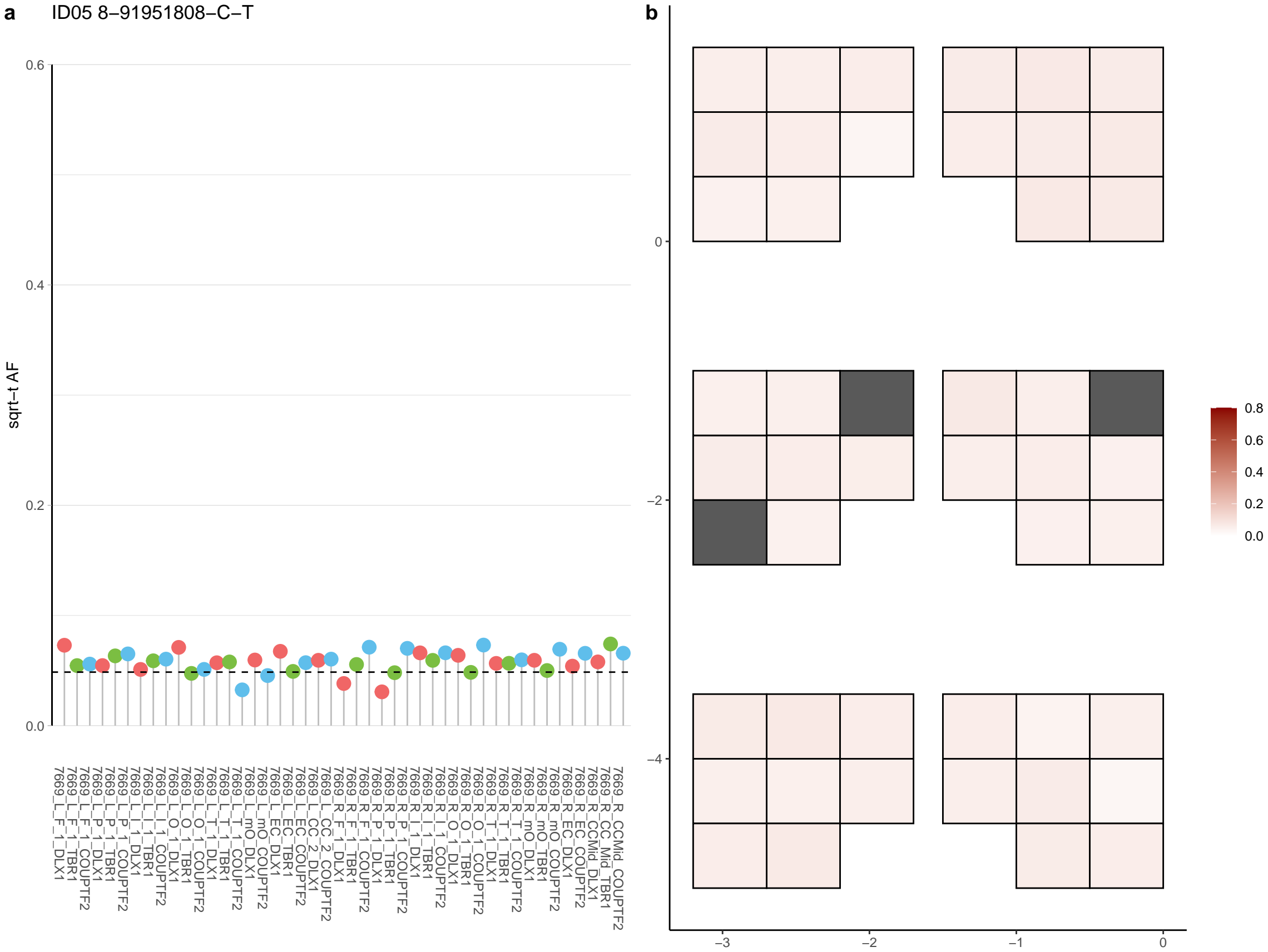

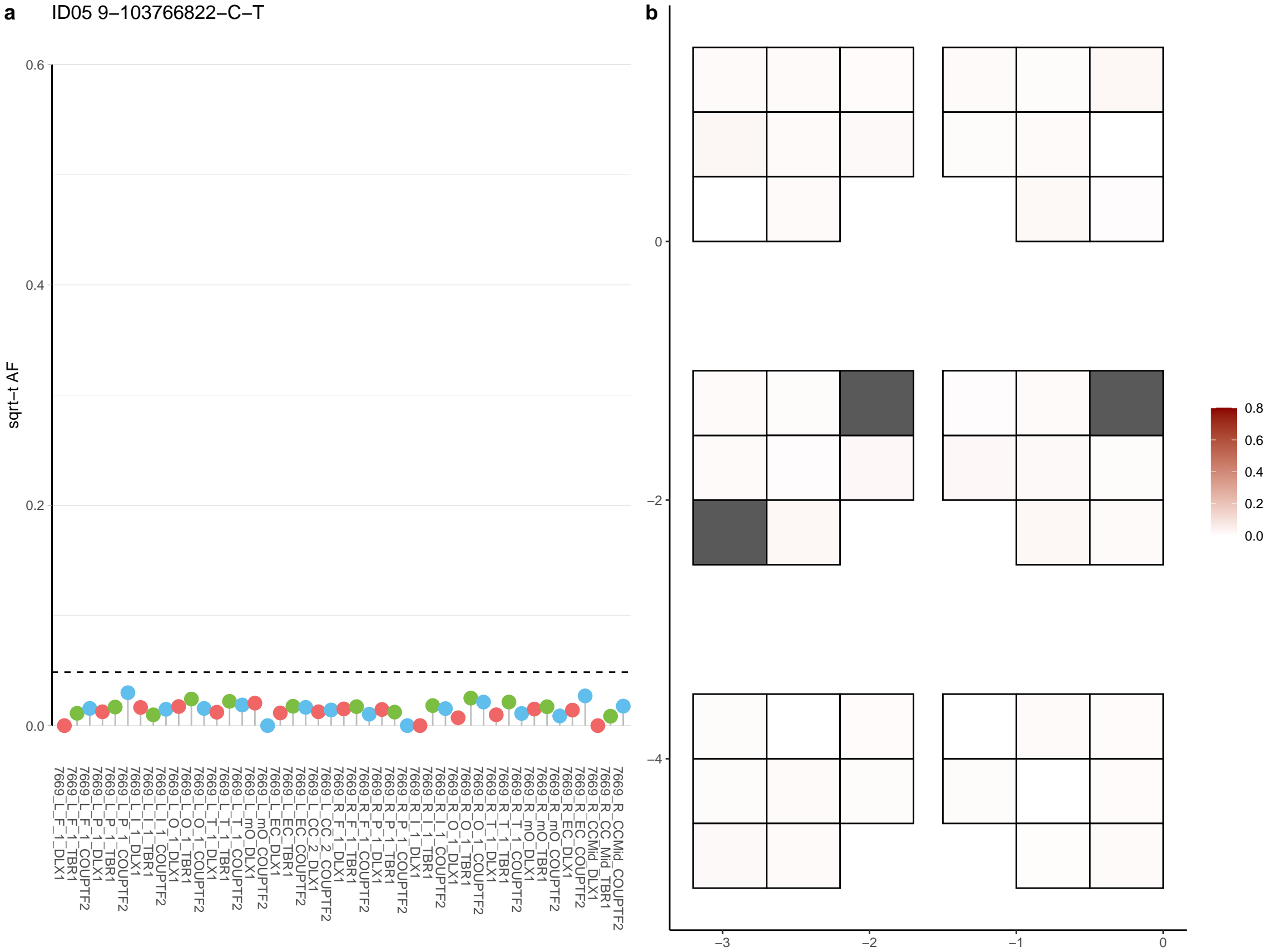

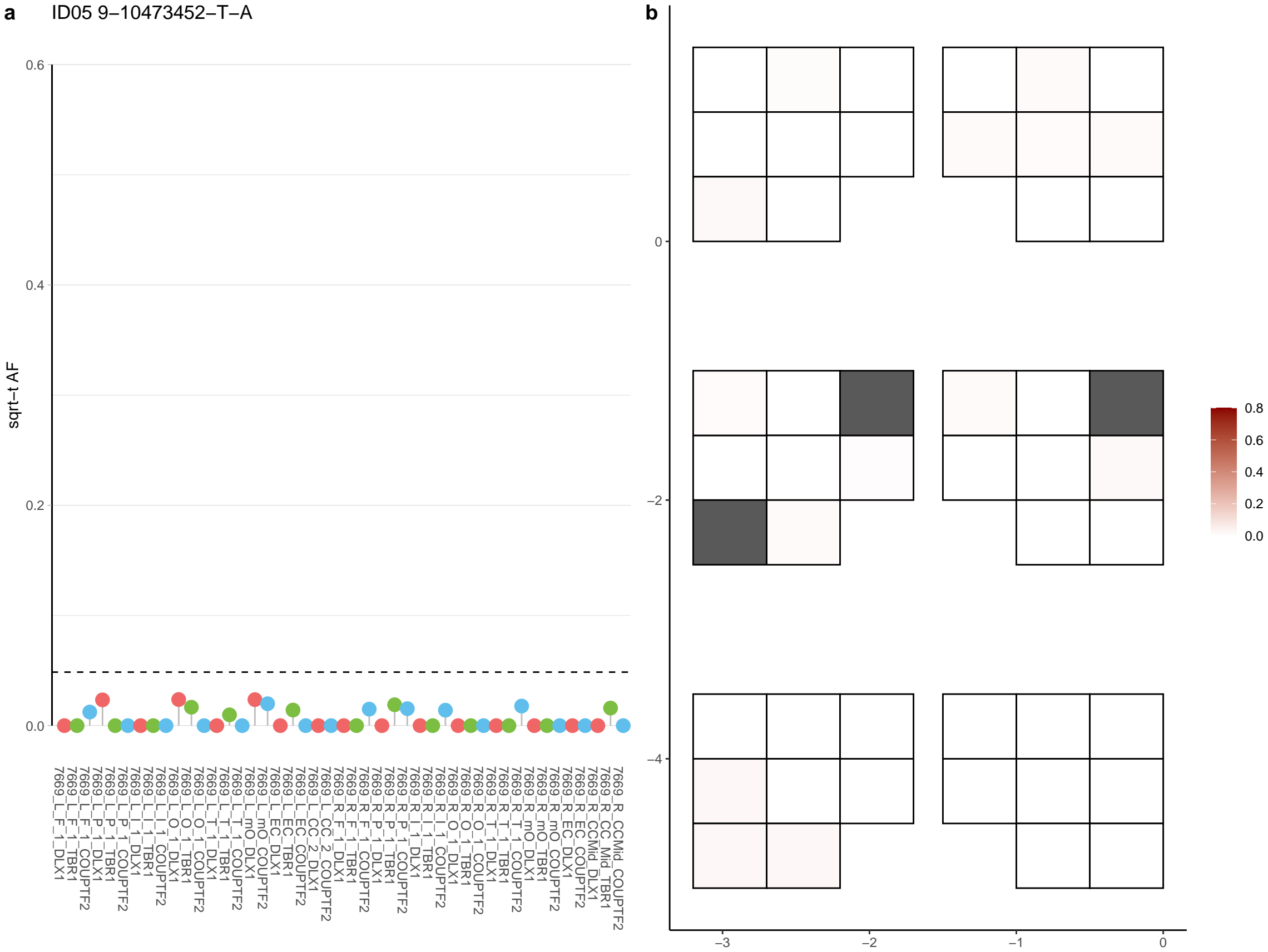

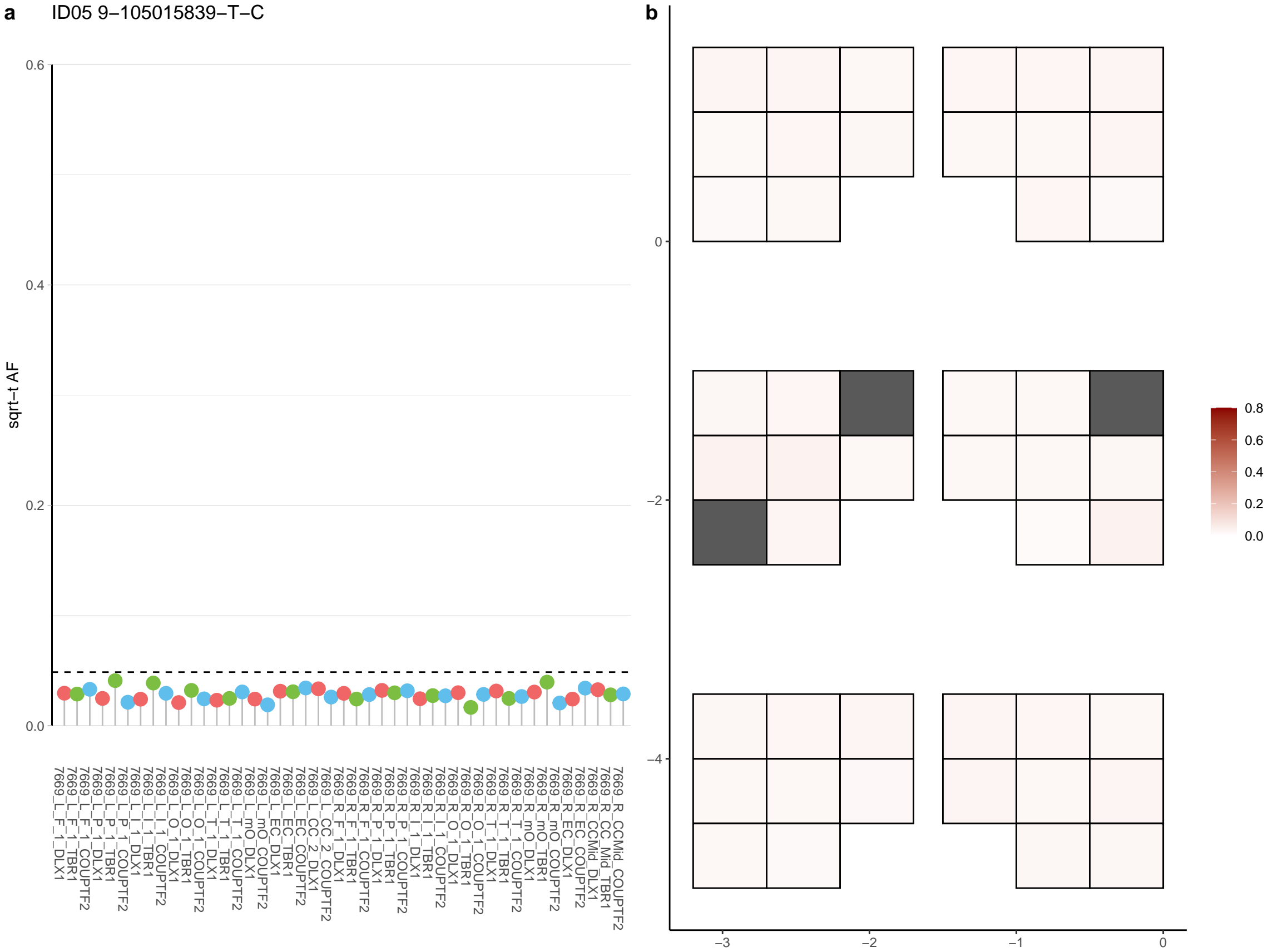

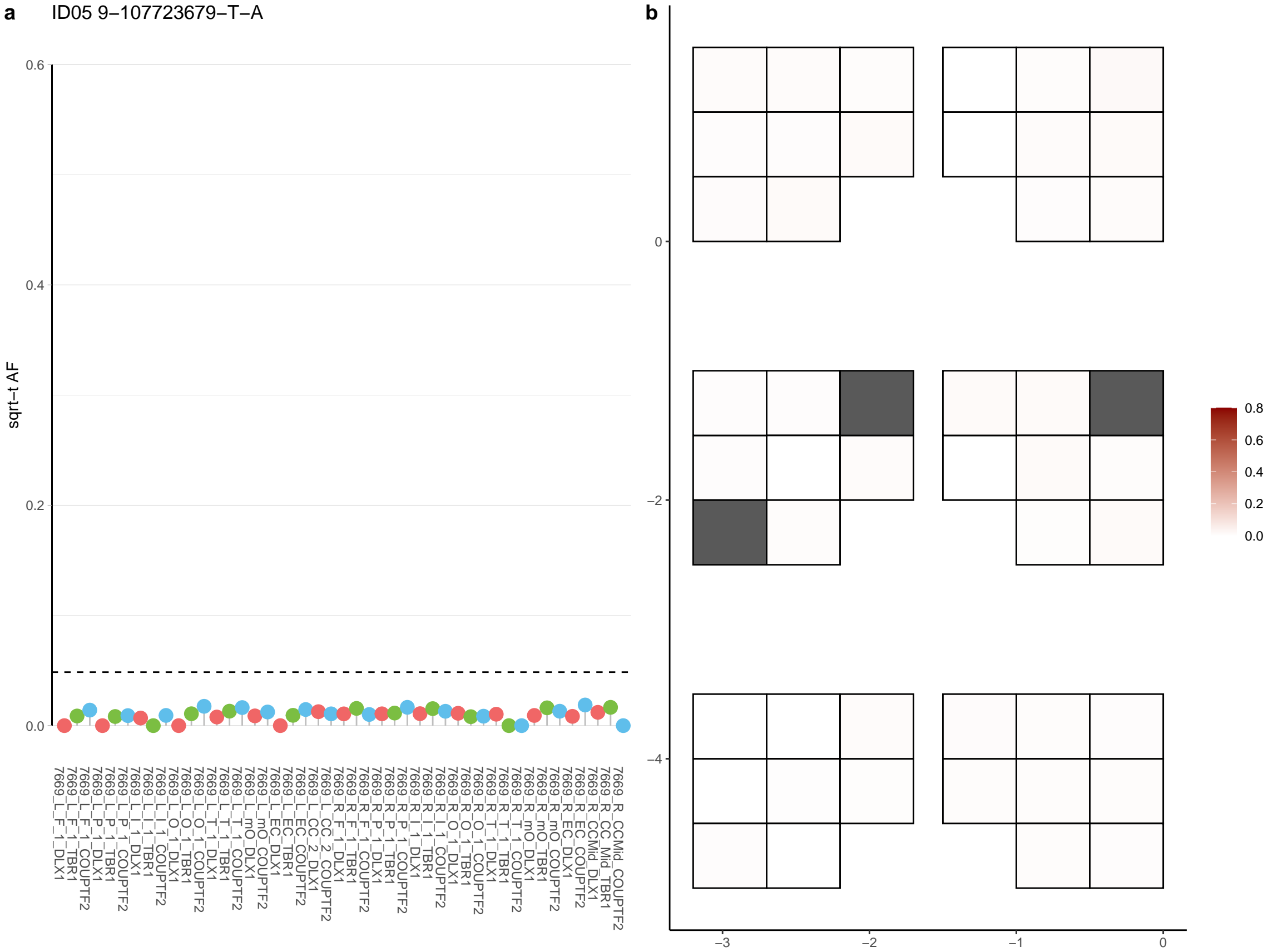

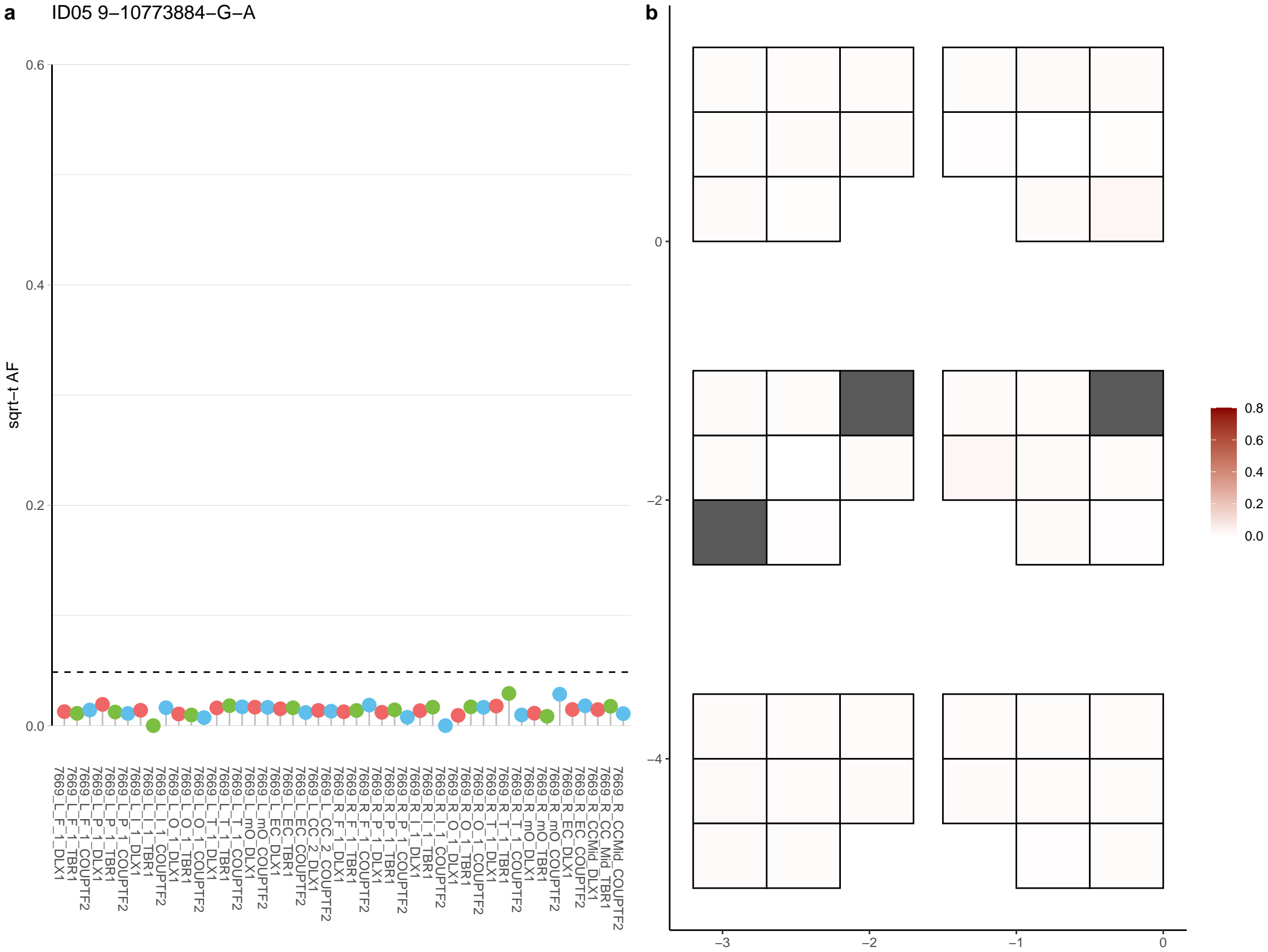

**a** ID05 9-119721731-C-T

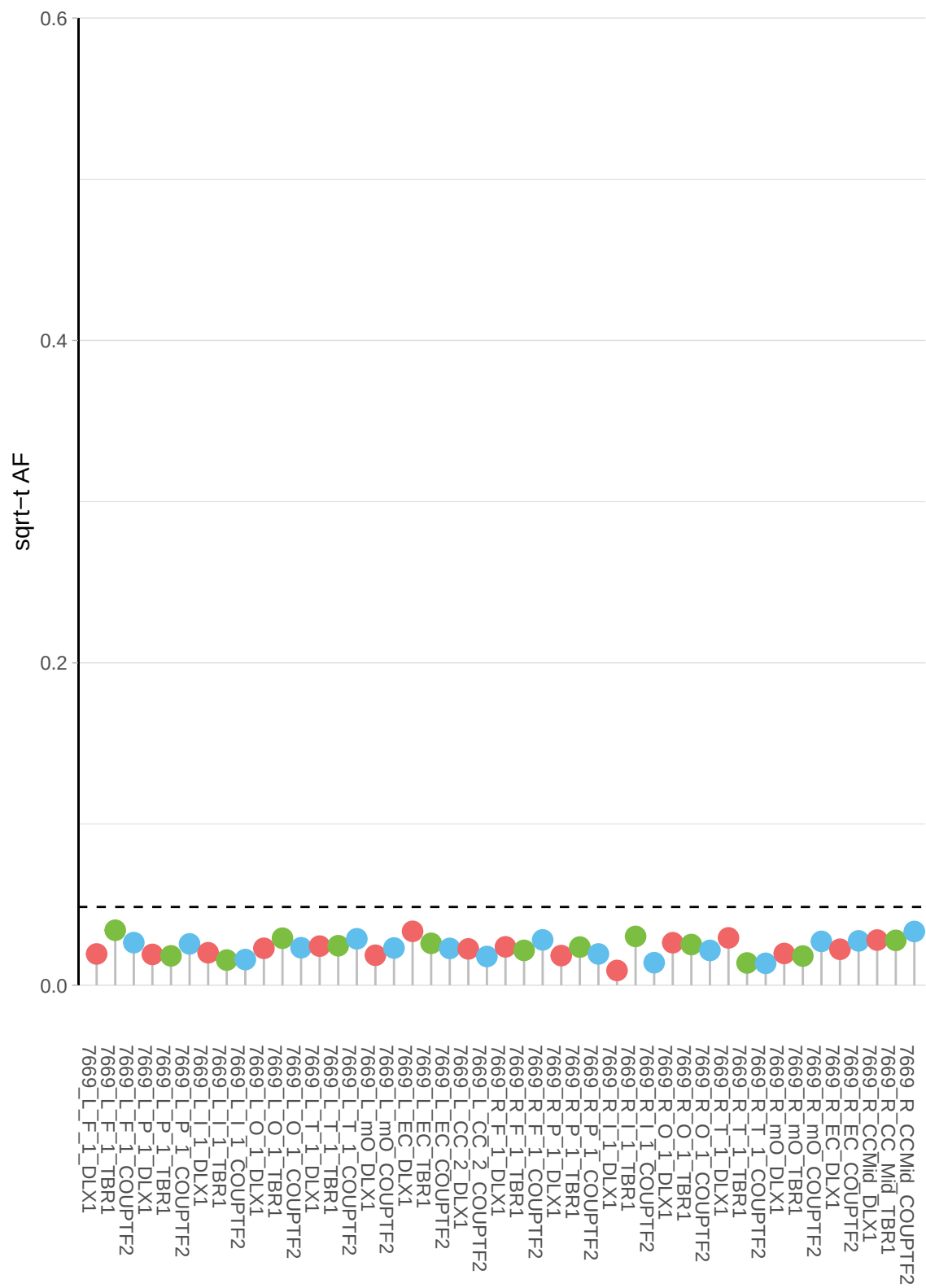

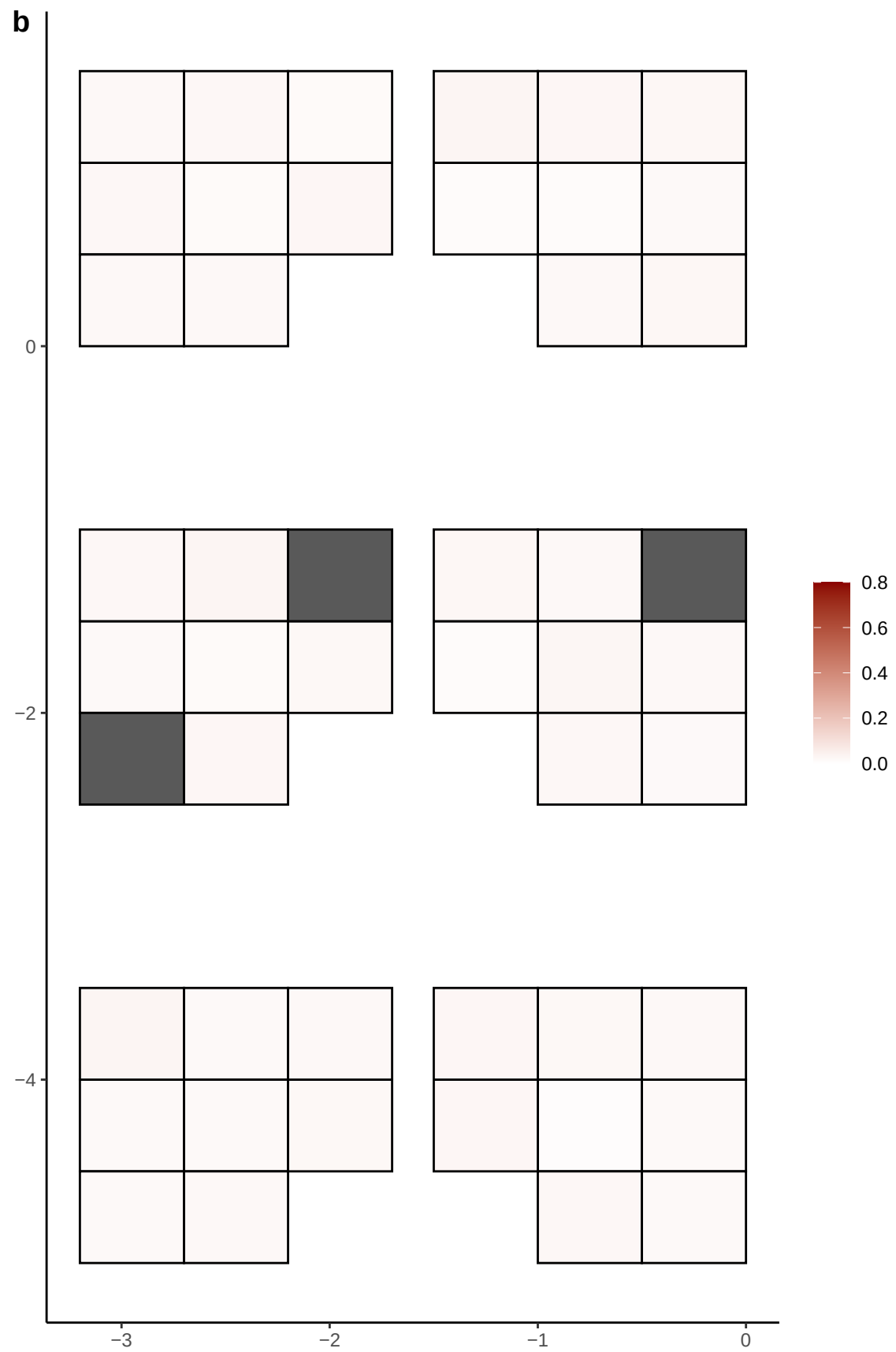

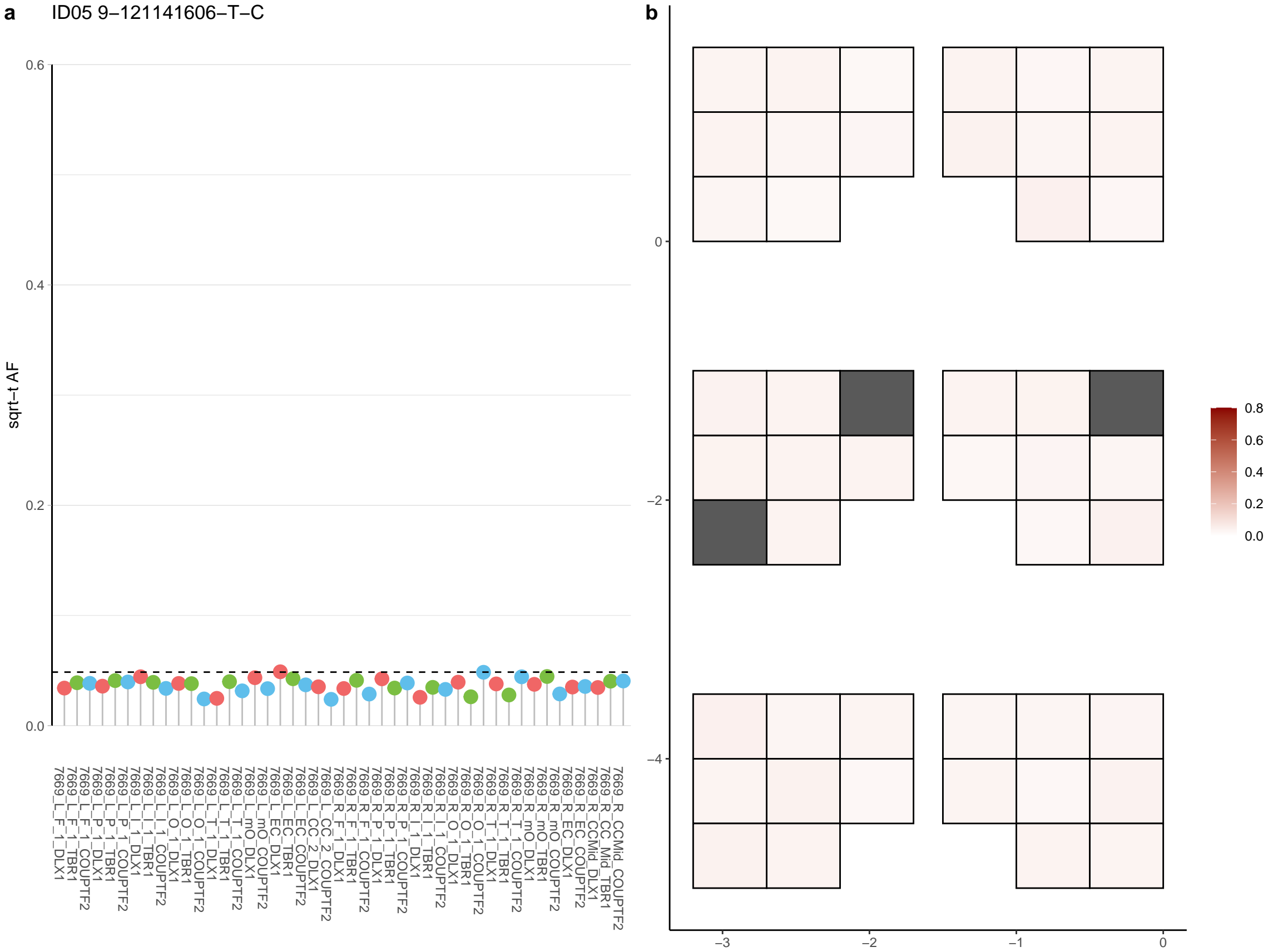

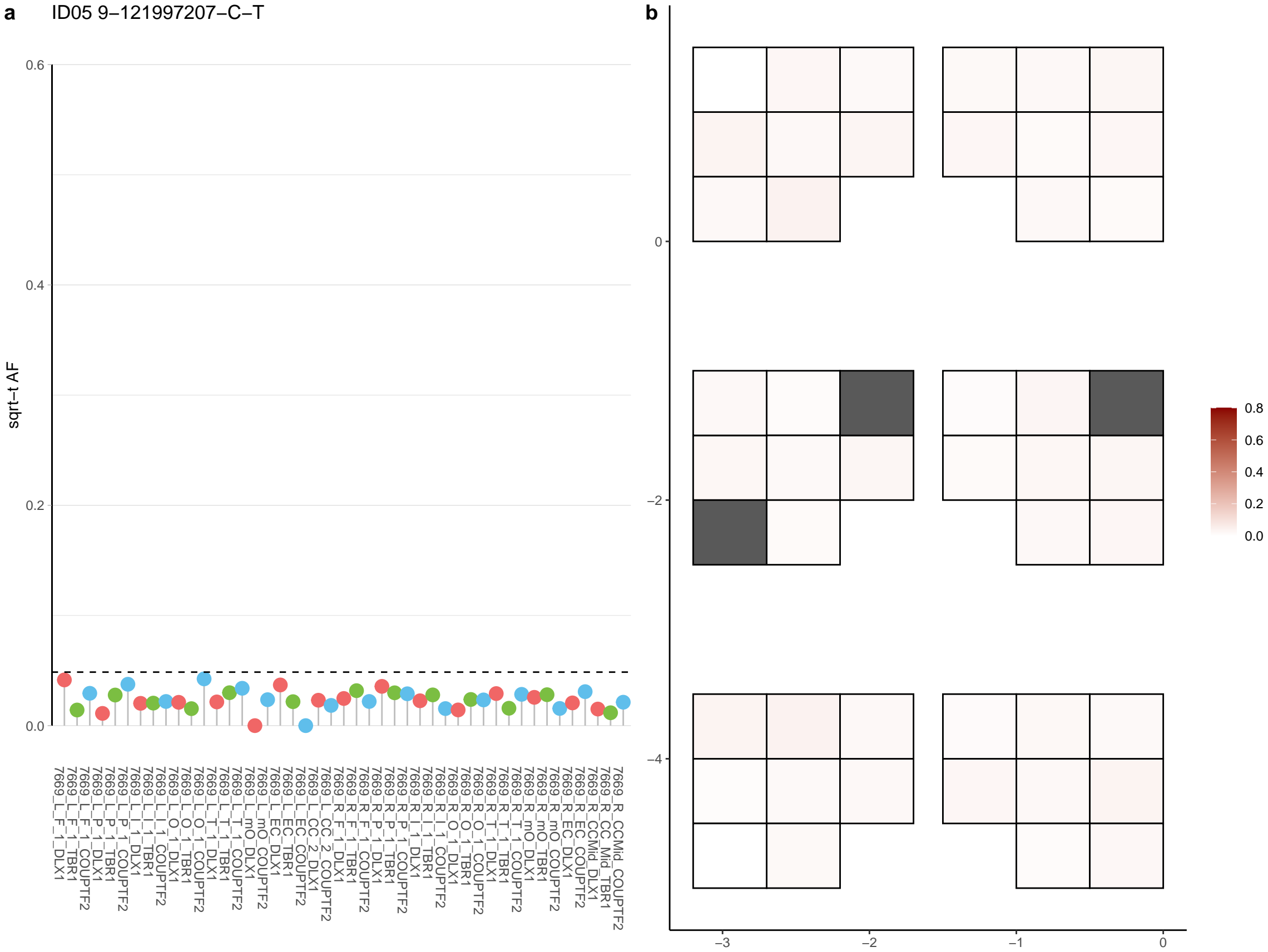

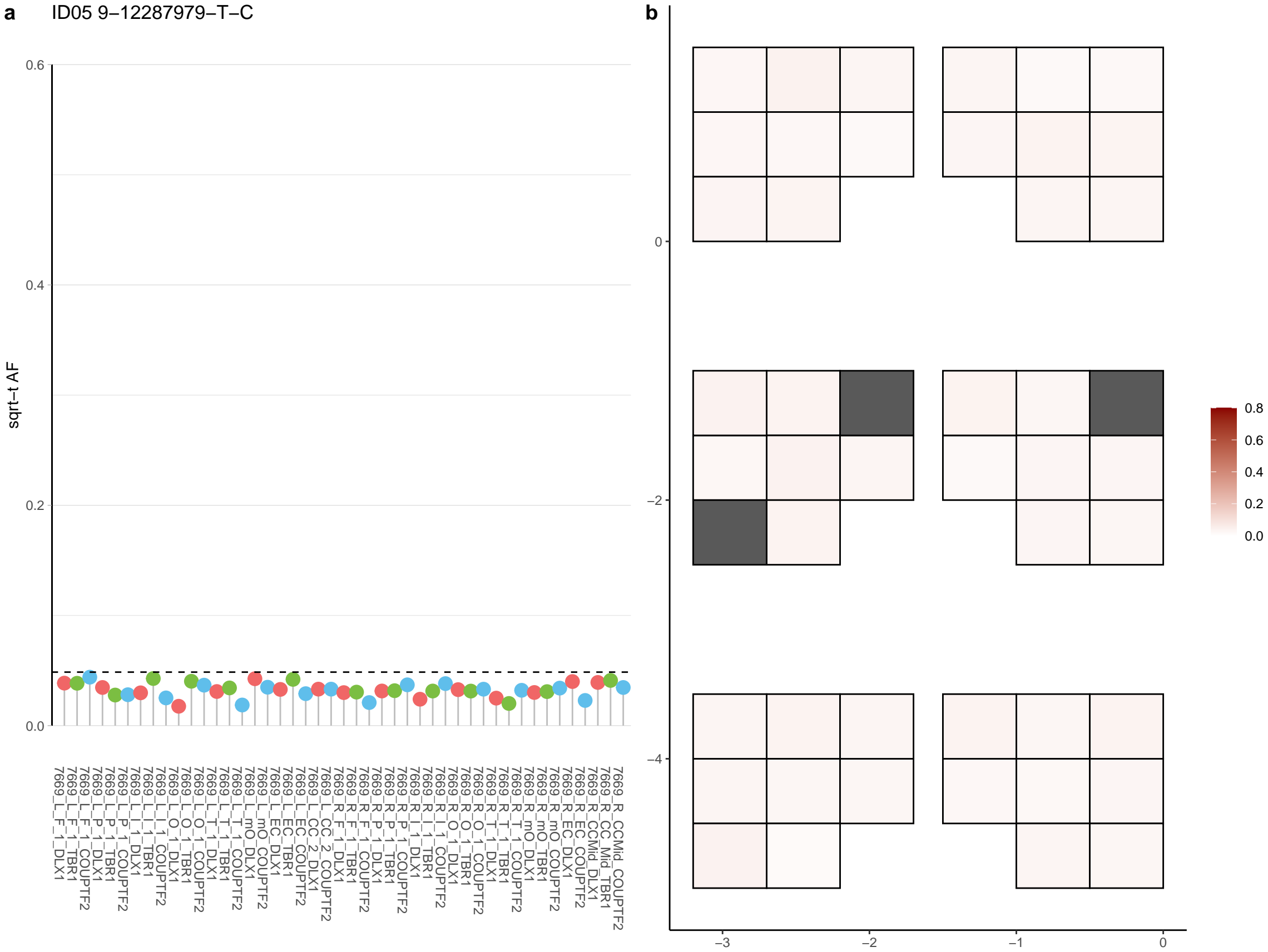

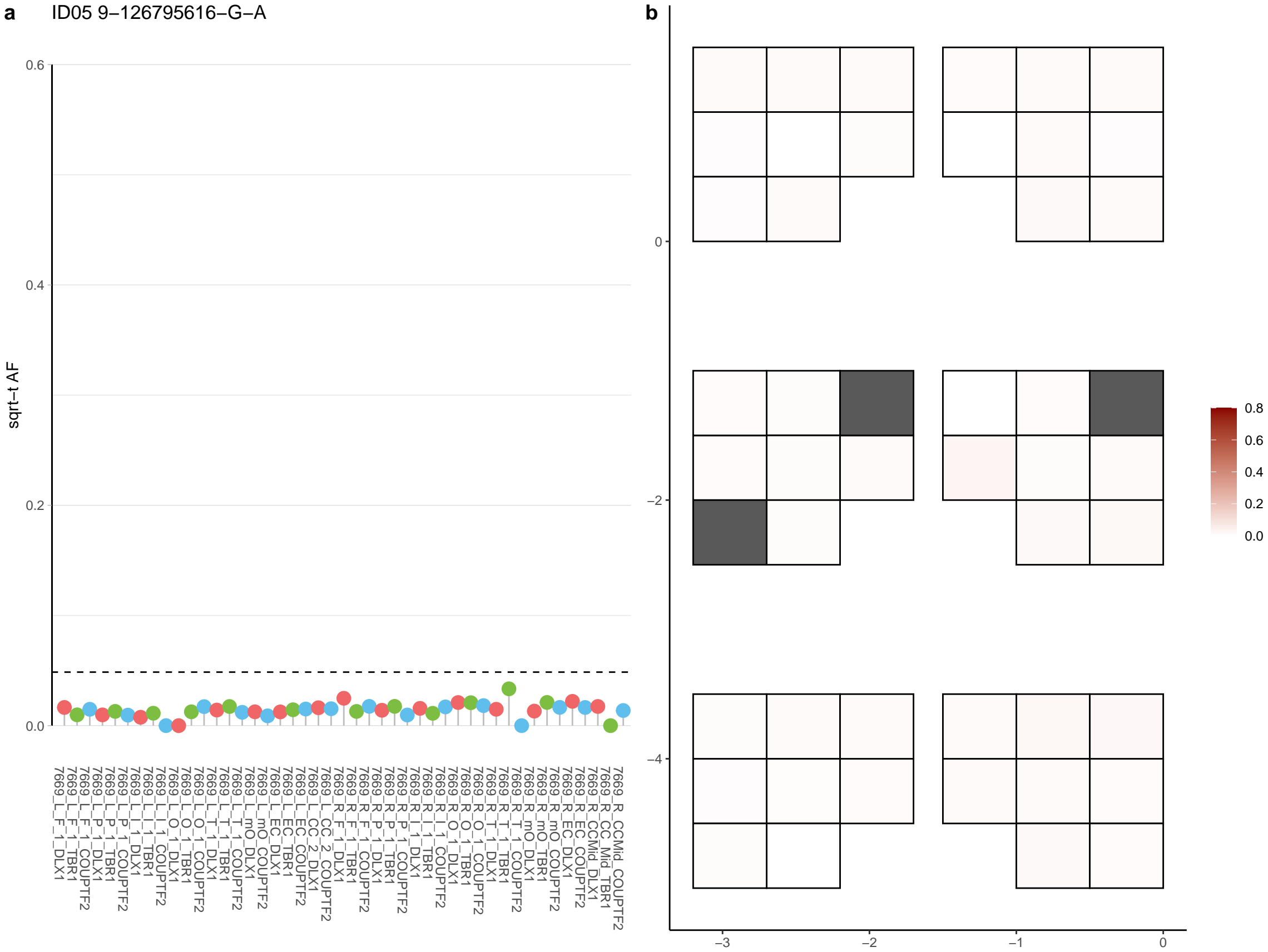

**a**

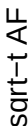

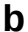

**a** ID05 9-13509007-G-A

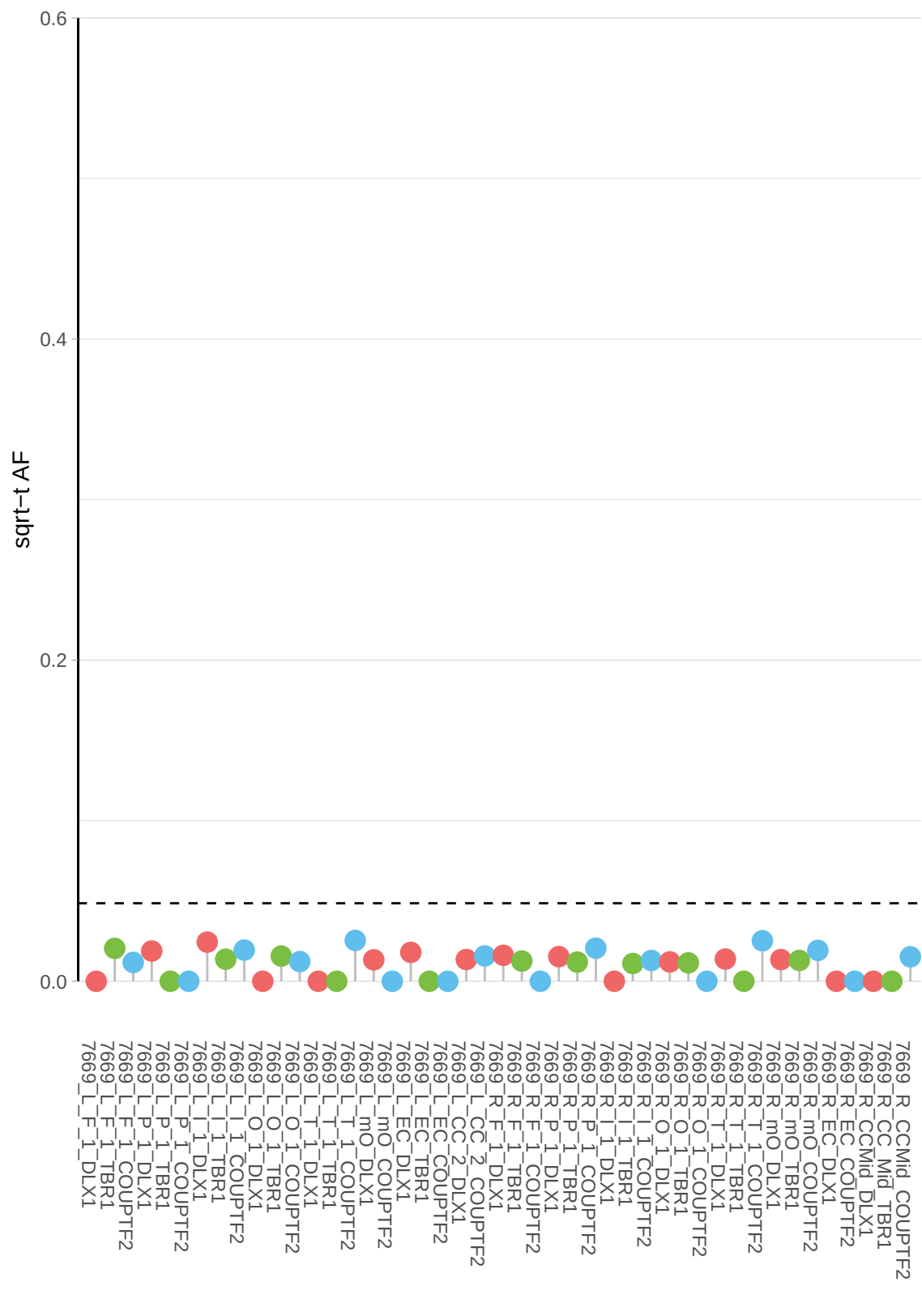

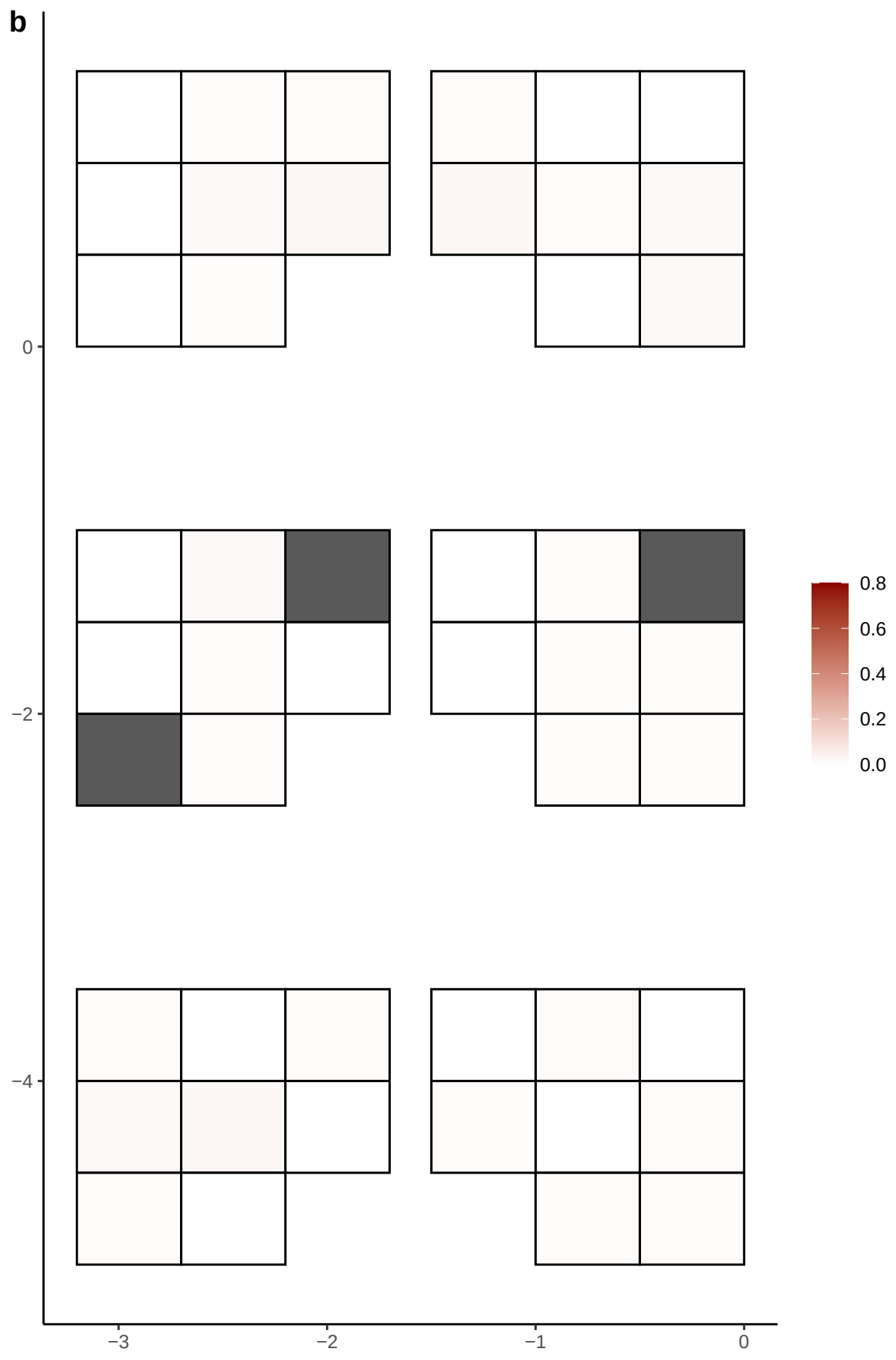

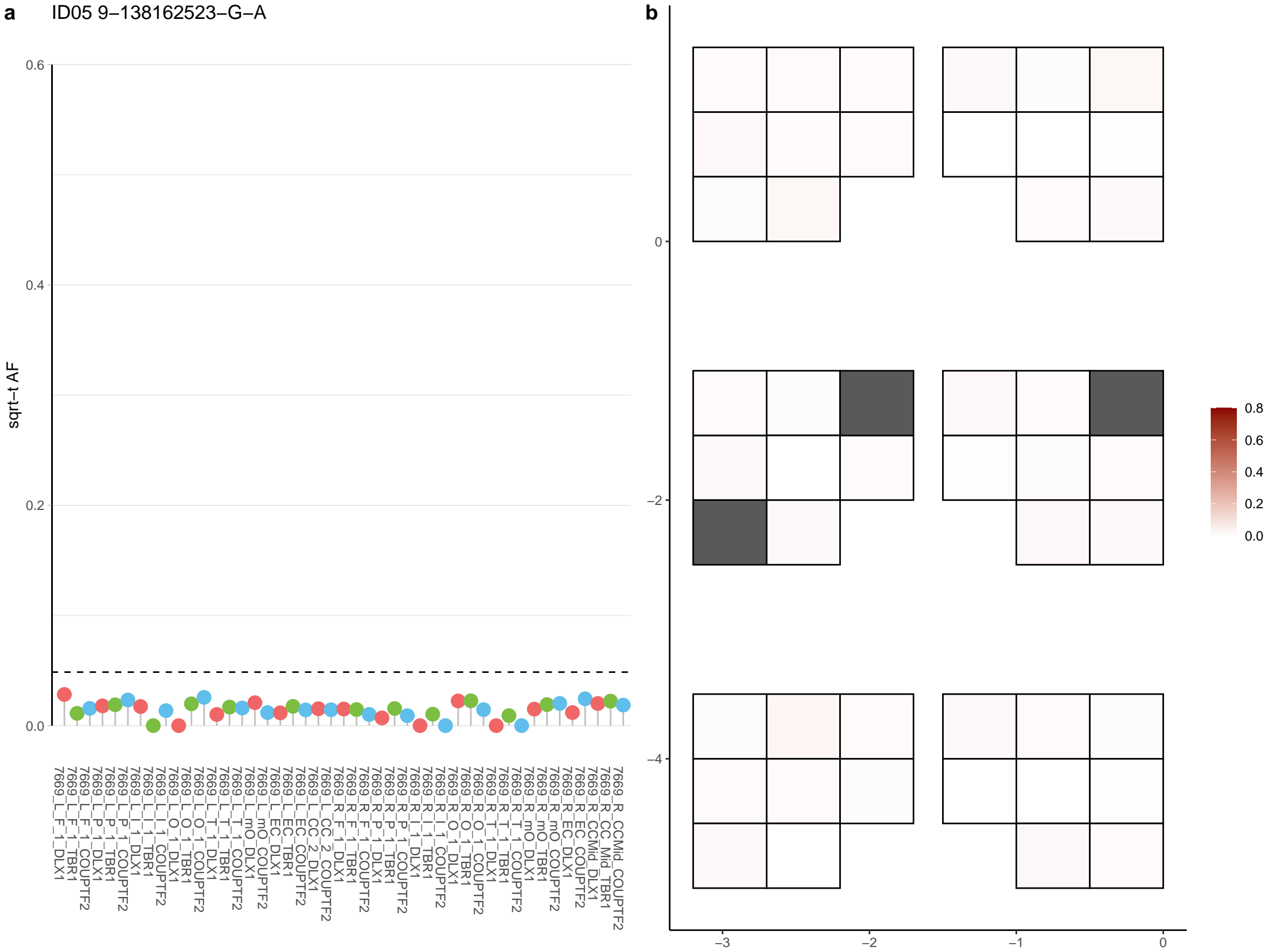

**a** ID05 9-14510306-G-A

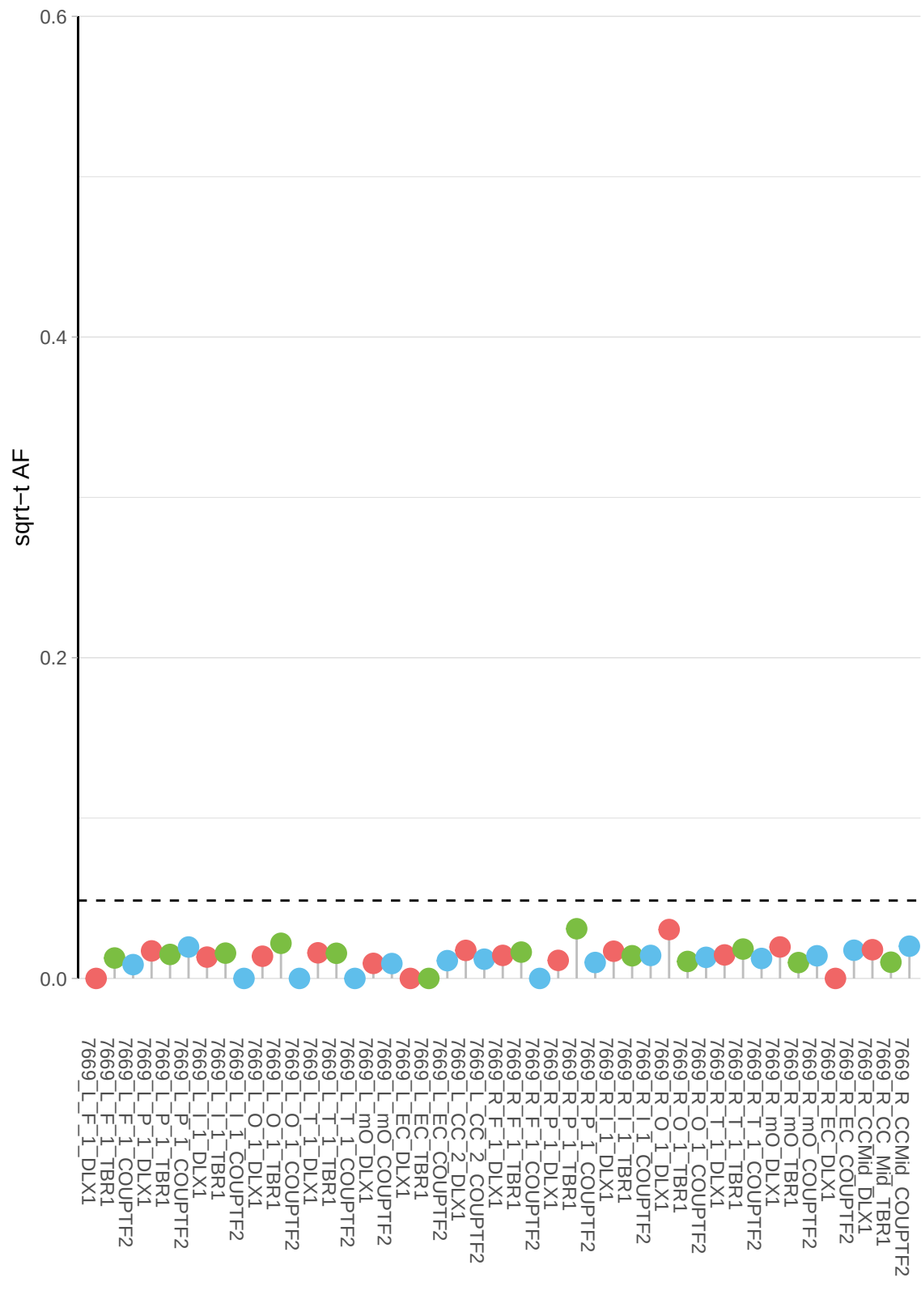

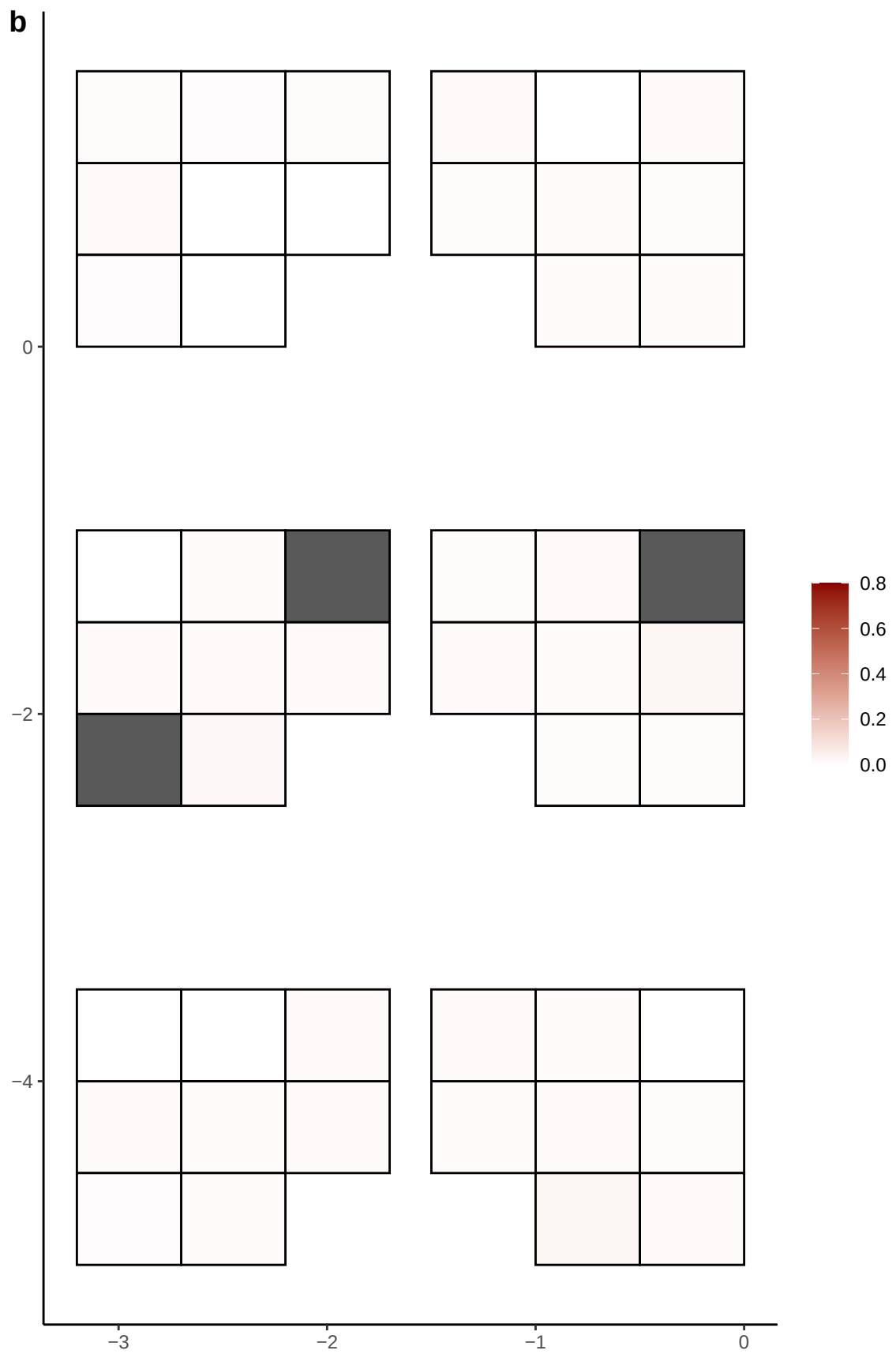

**a** ID05 9-19957121-C-T

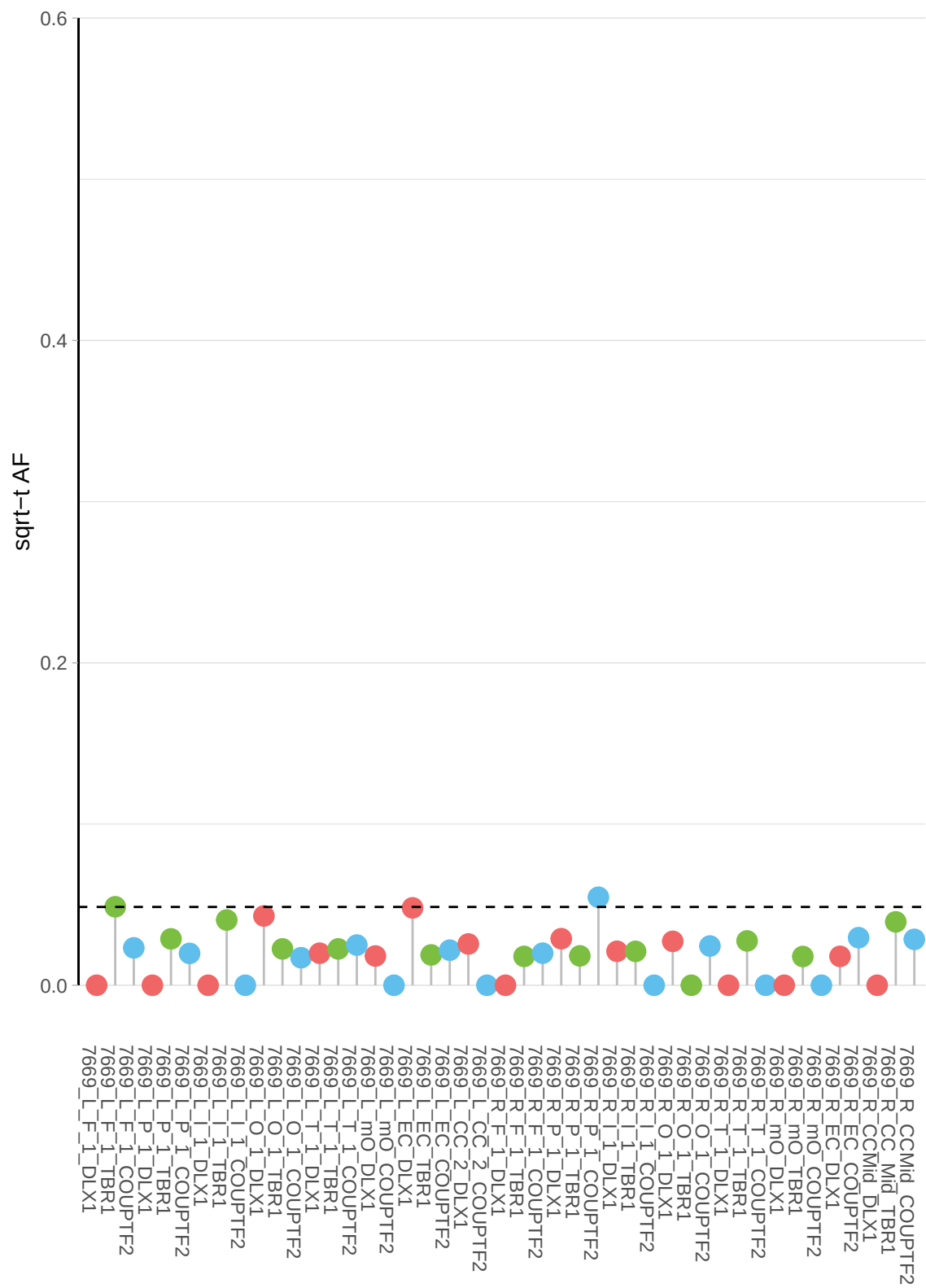

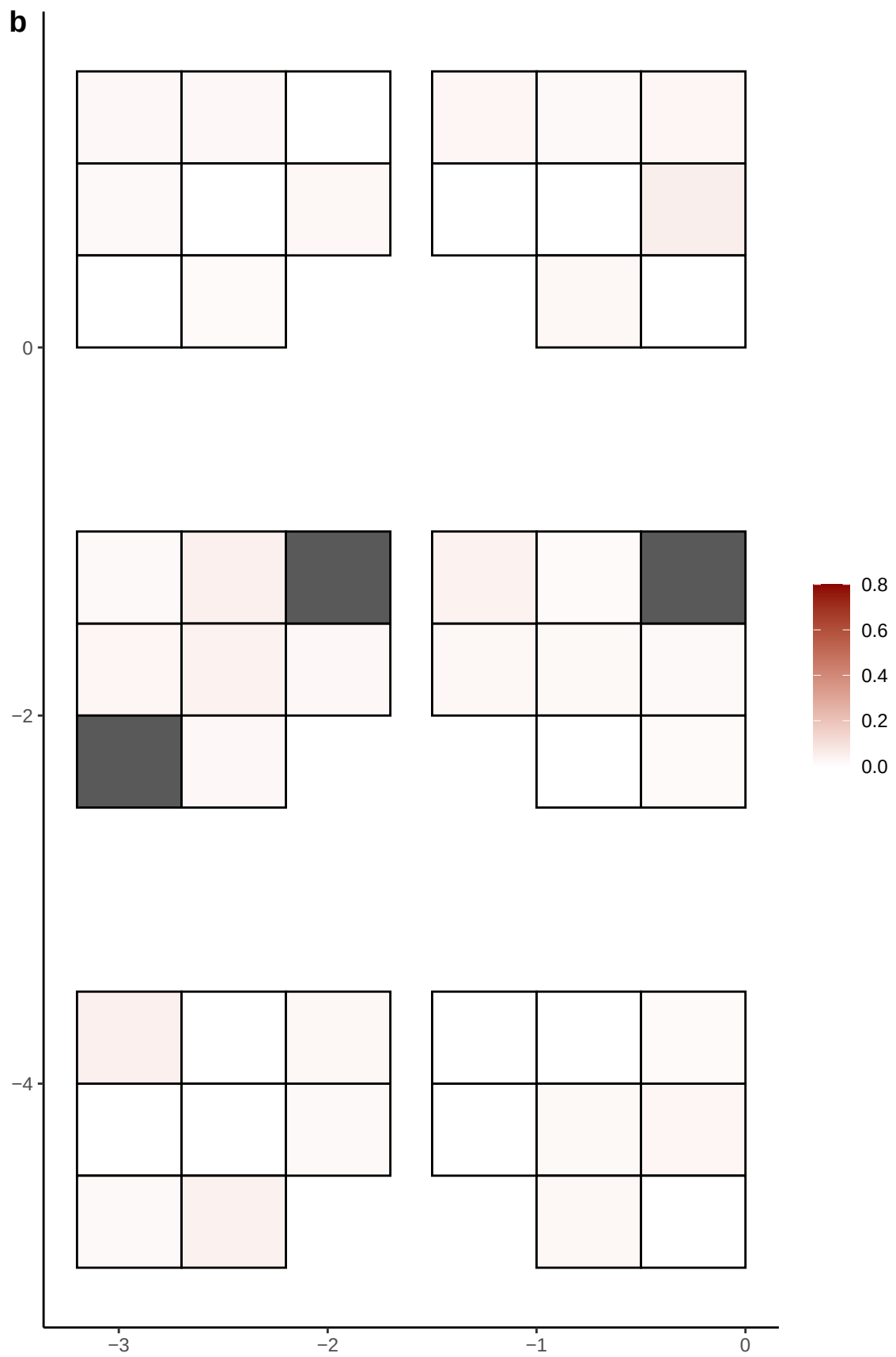

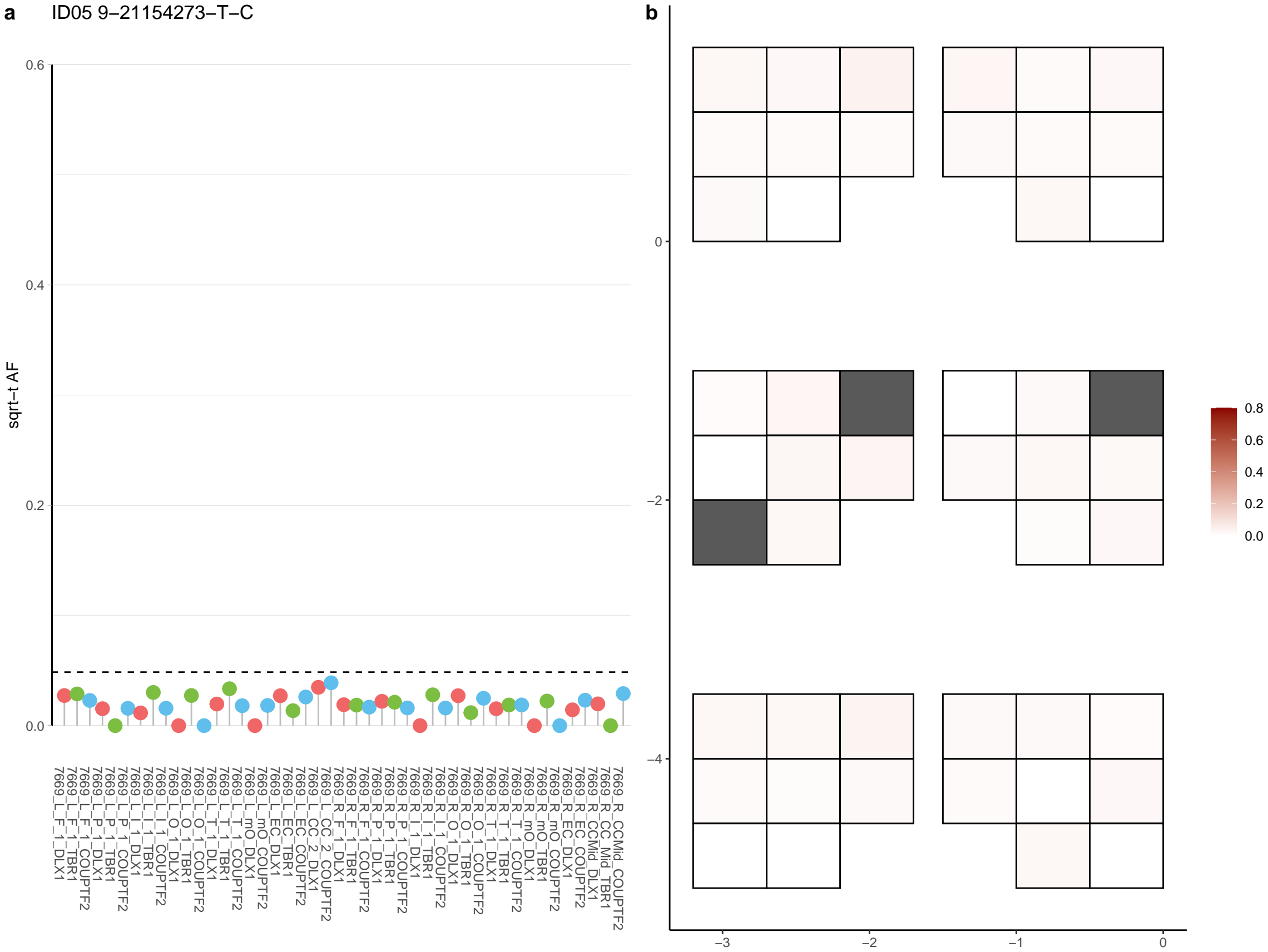

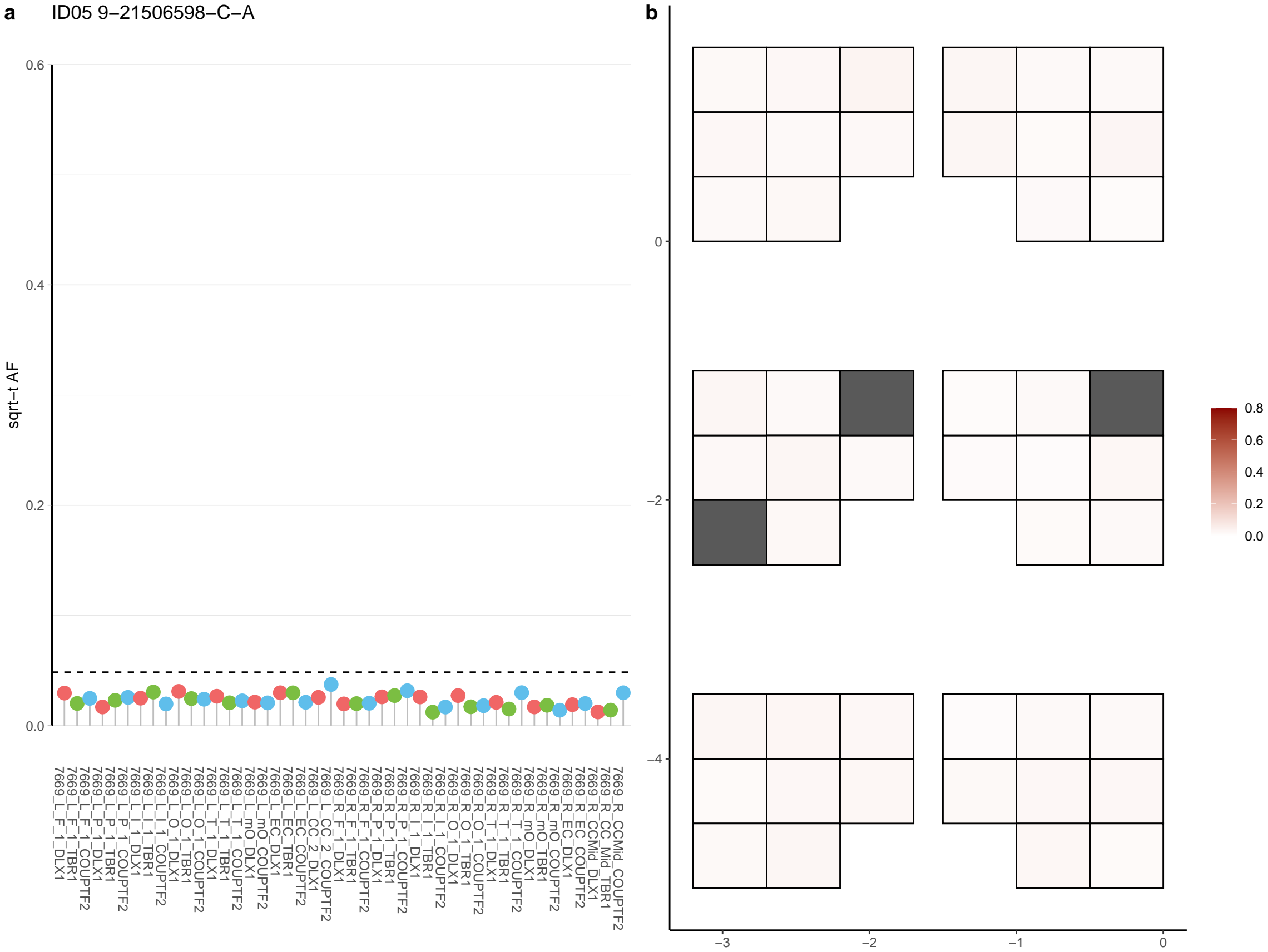

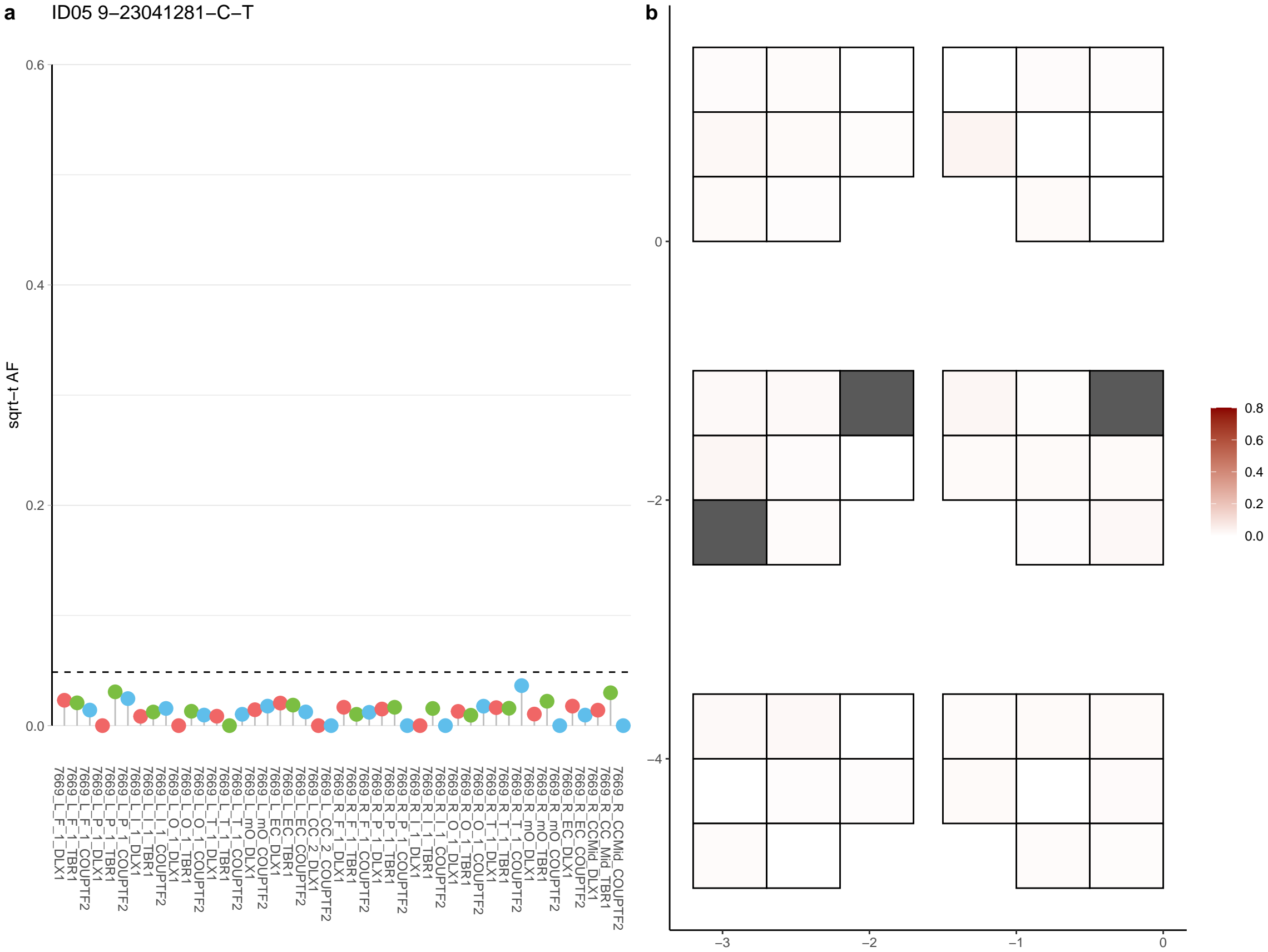

**a** ID05 9-28561999-C-T

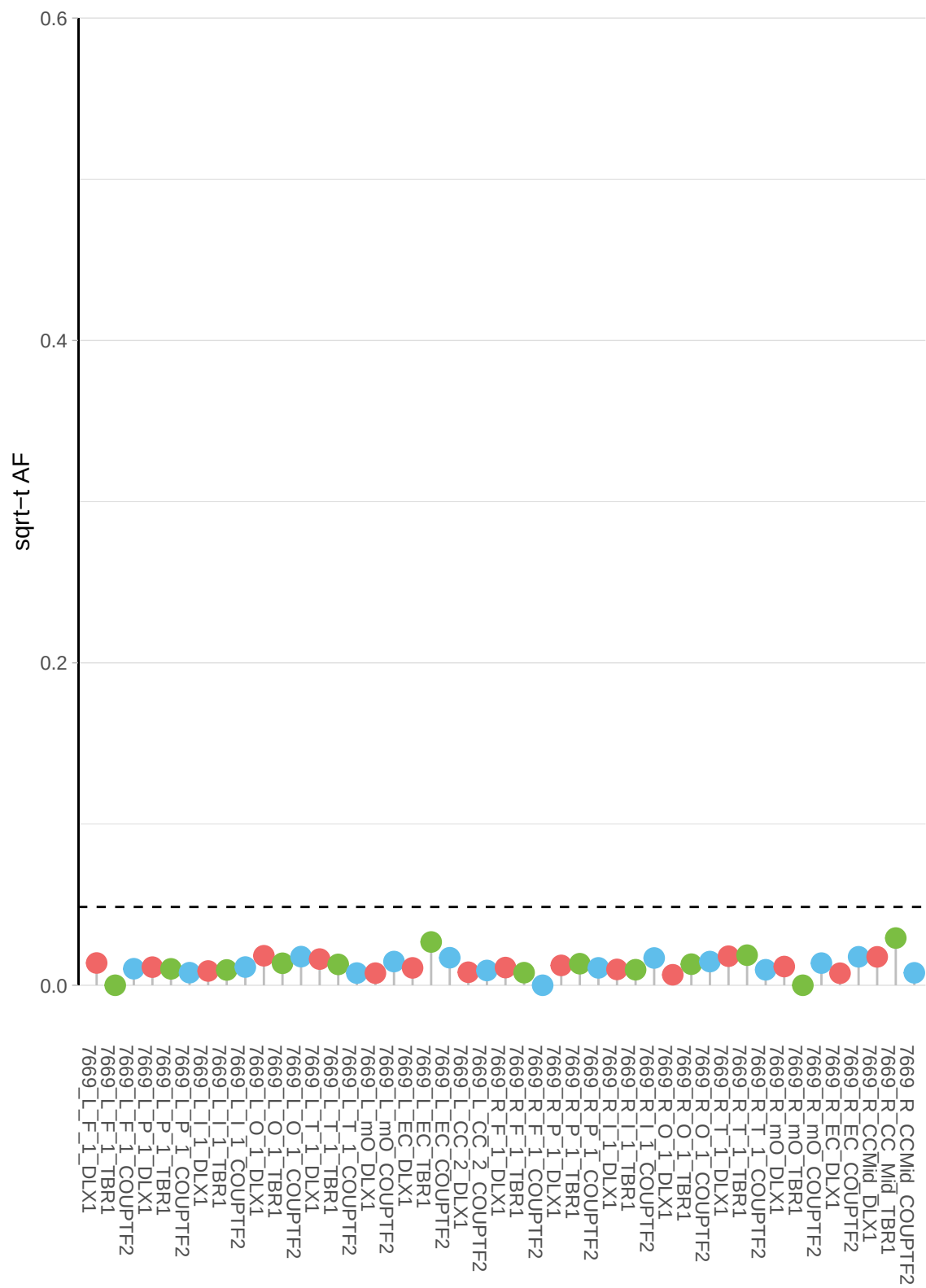

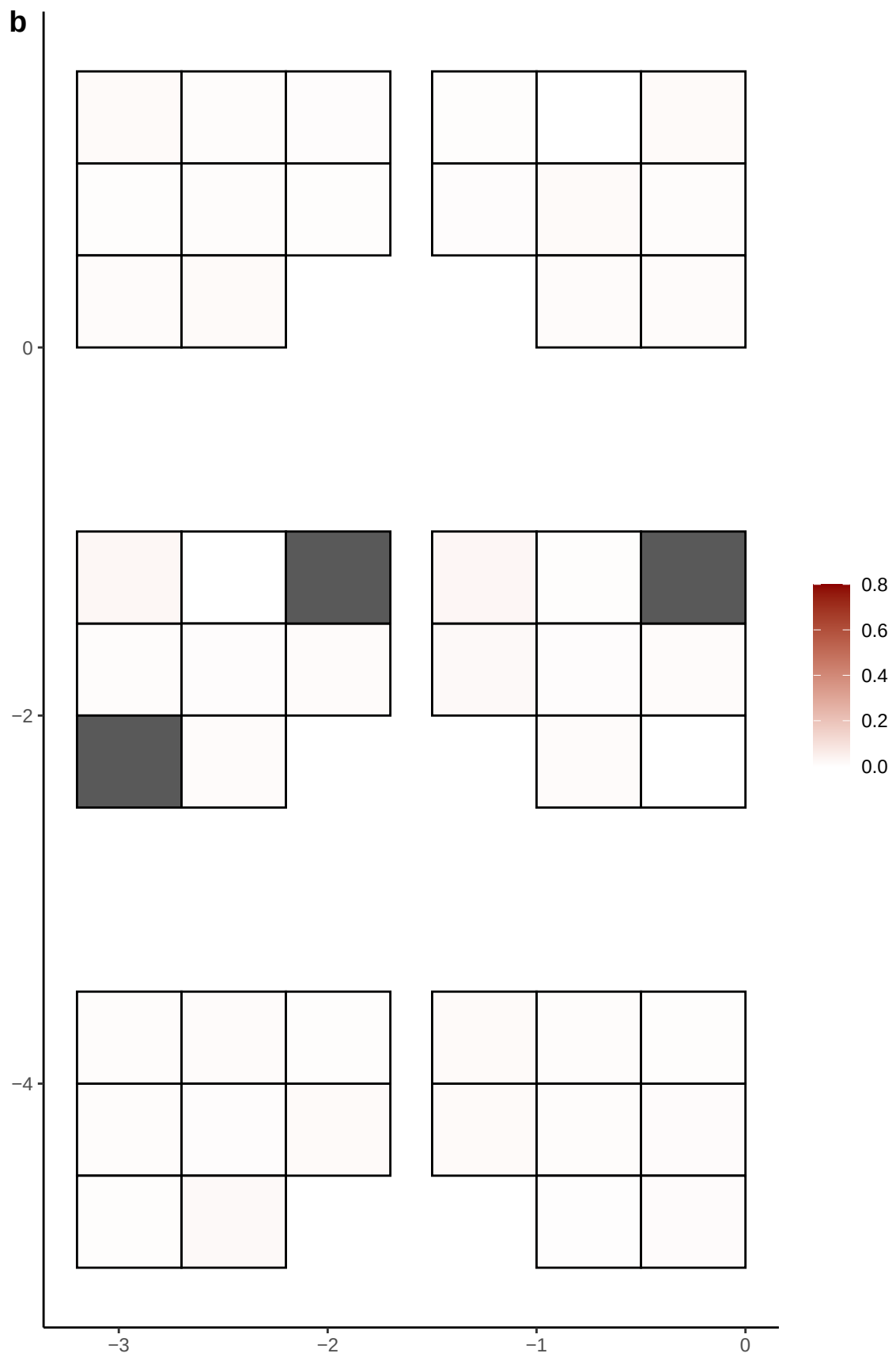

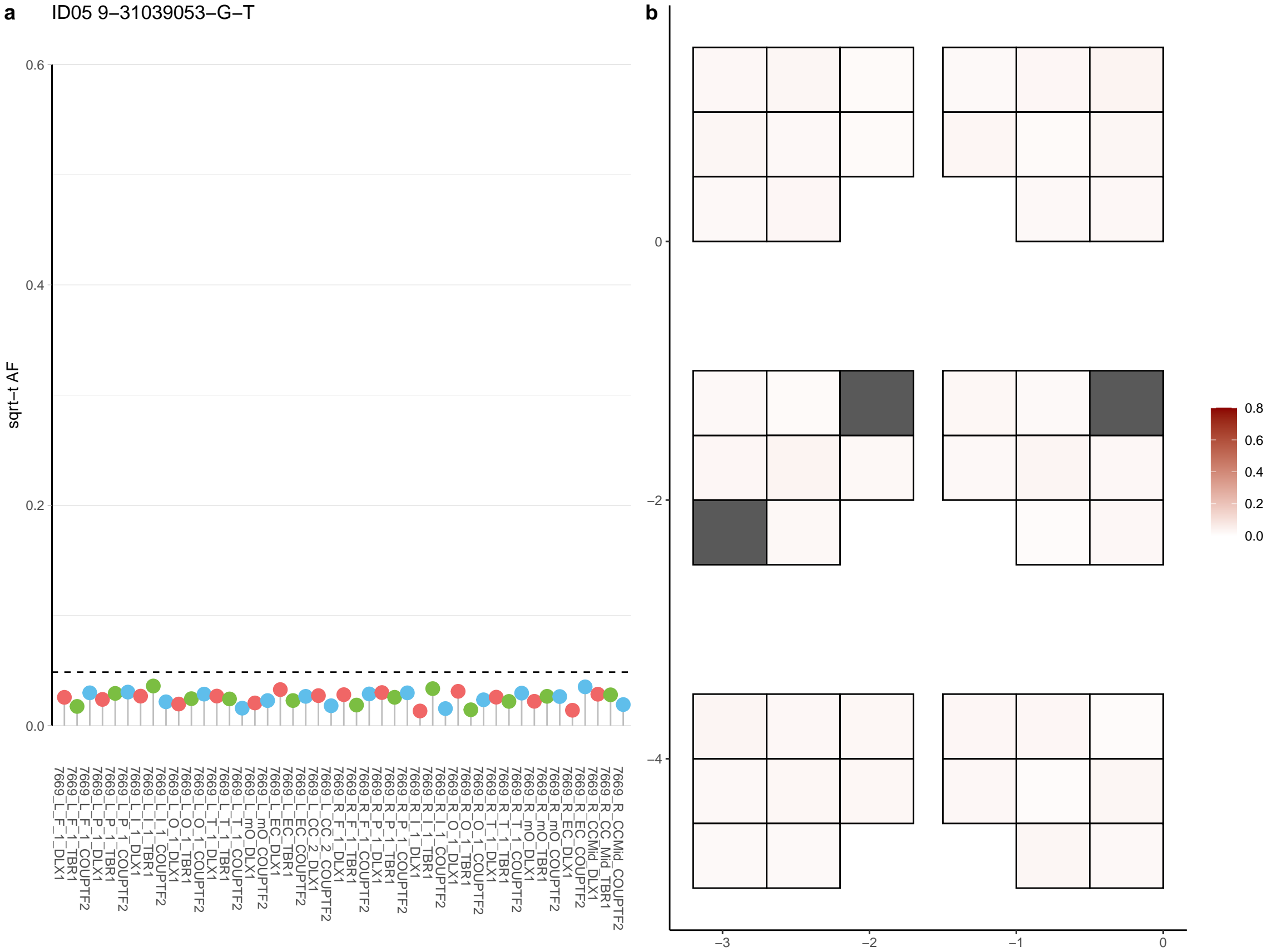

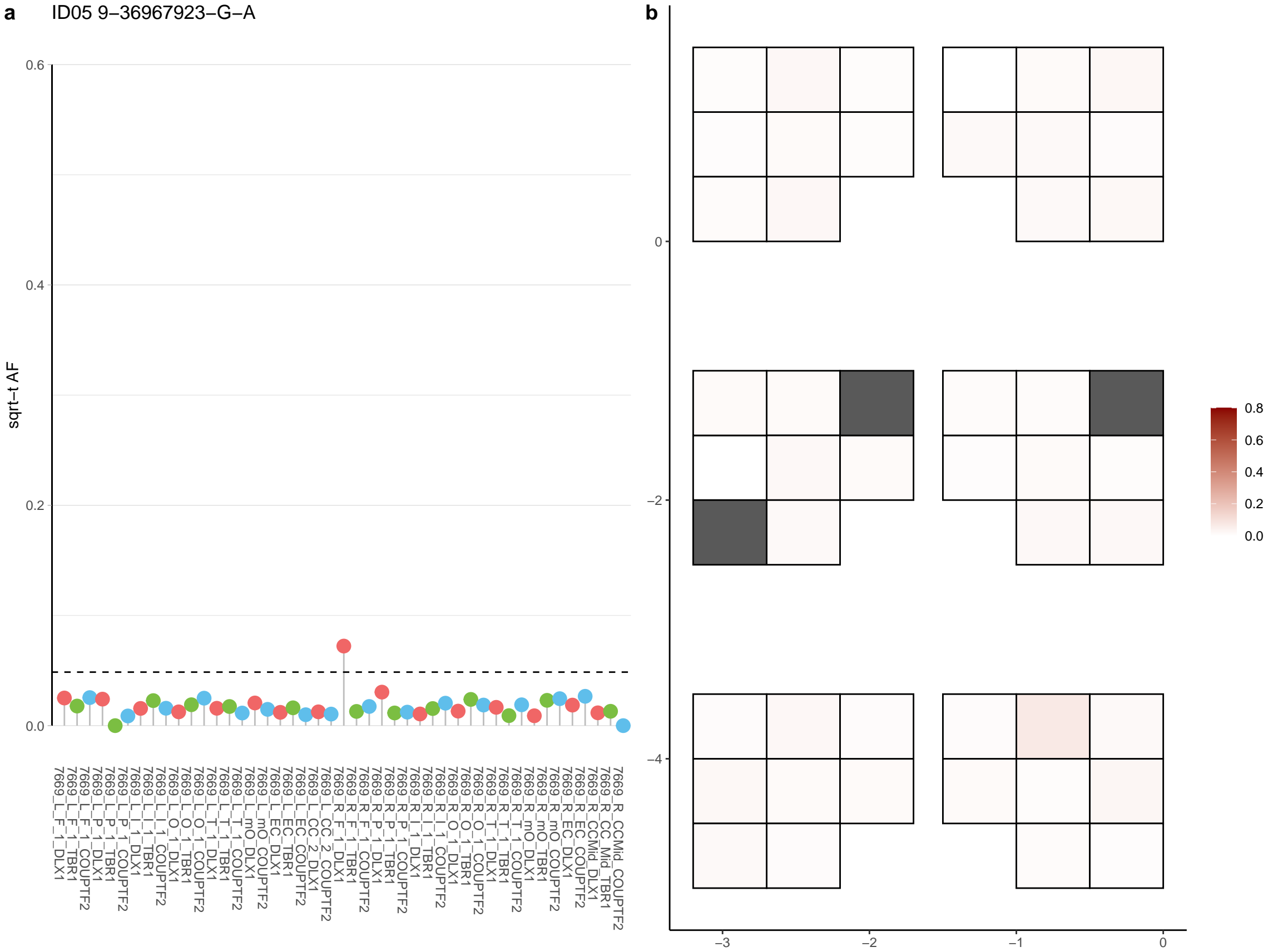

a ID05 9-559661-C-T

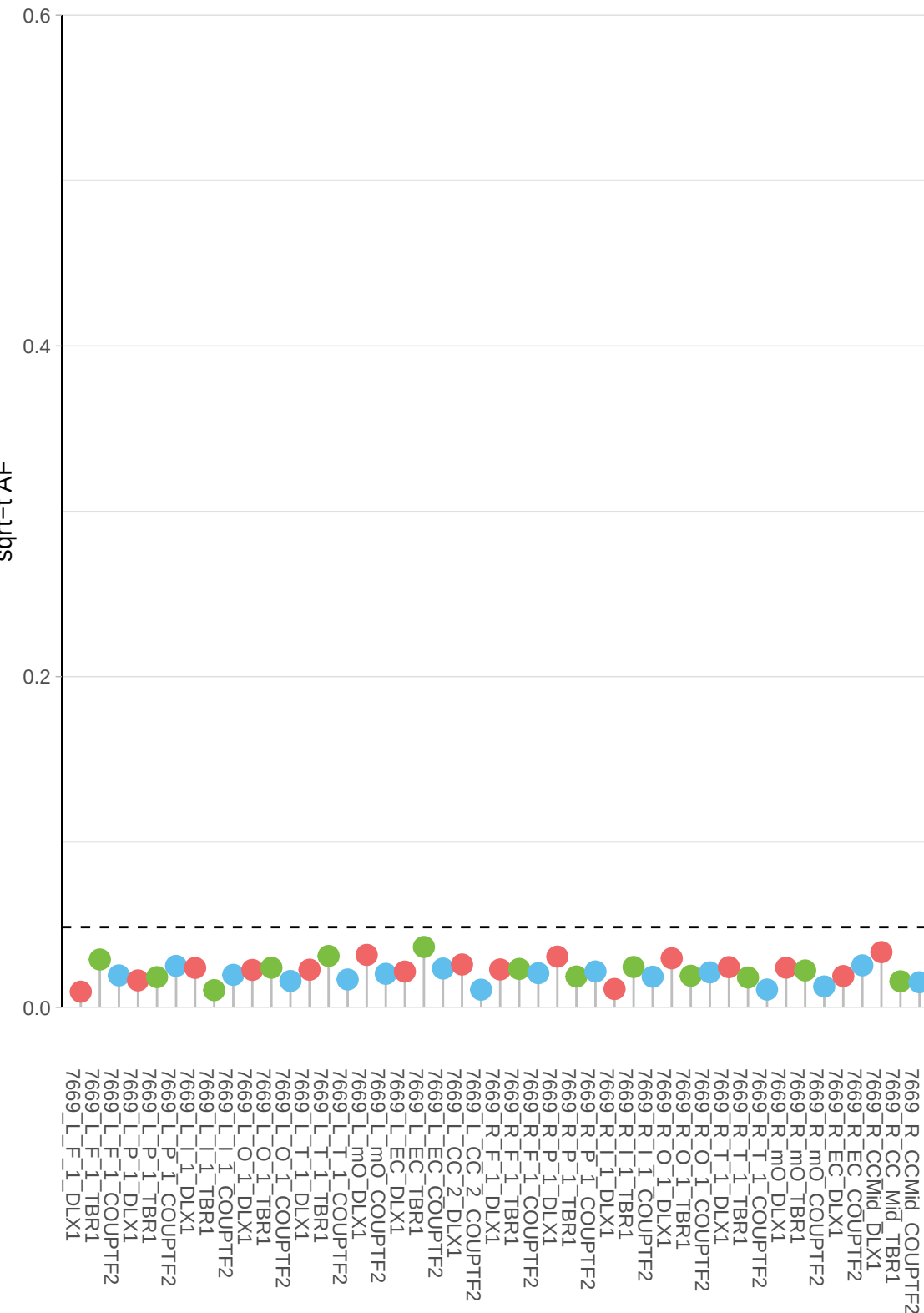

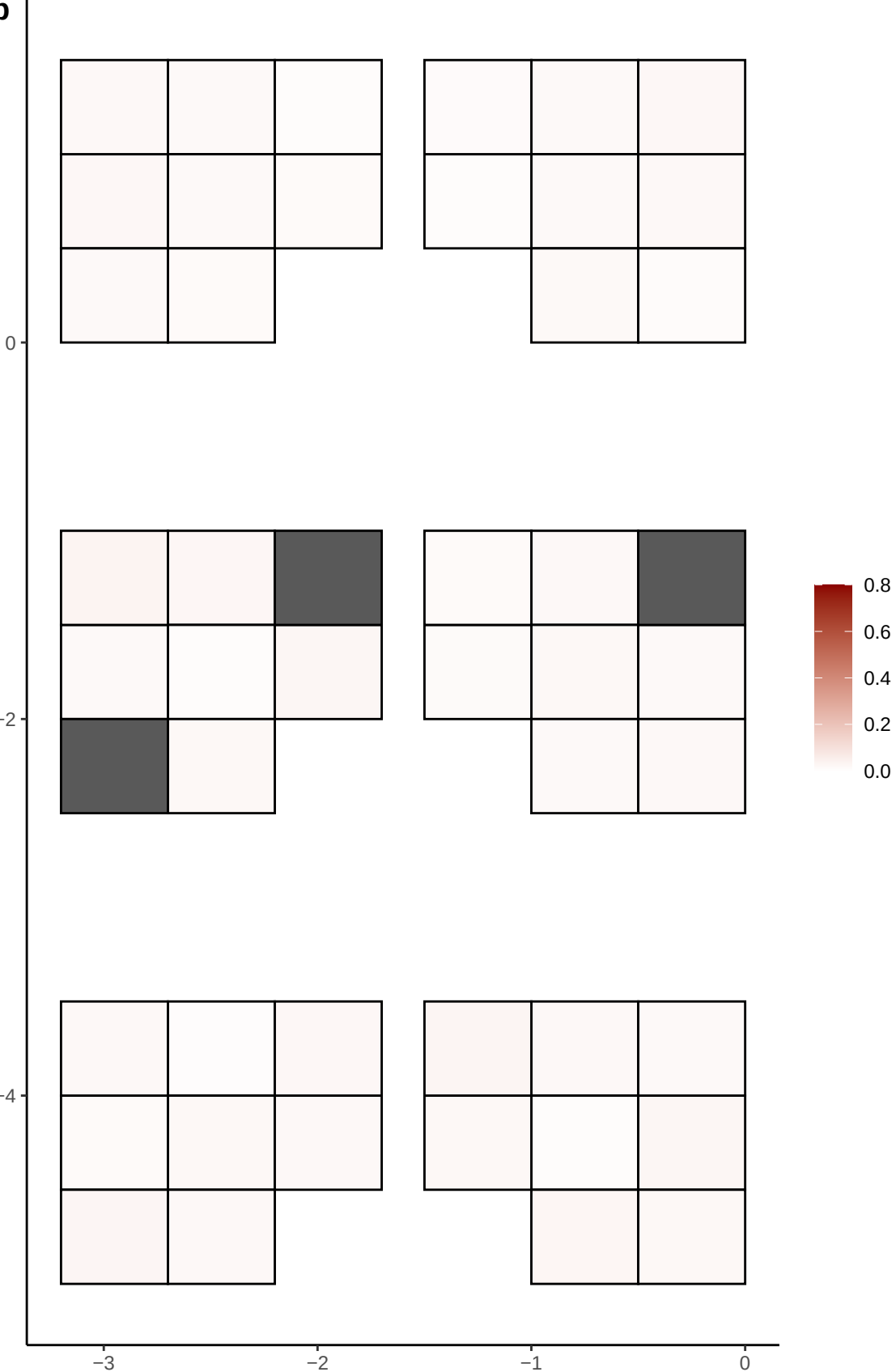

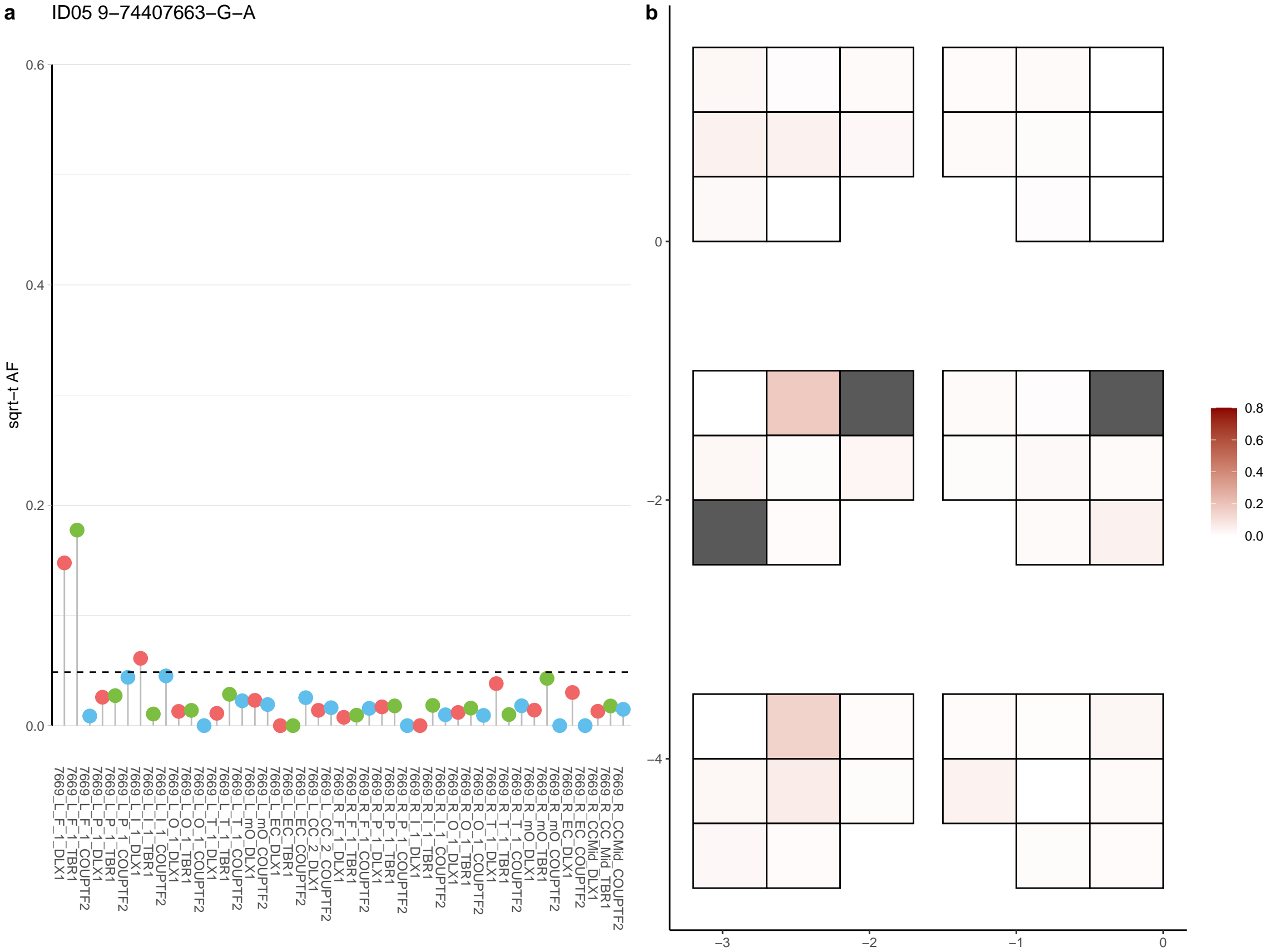

**a** ID05 9-76169299-T-C

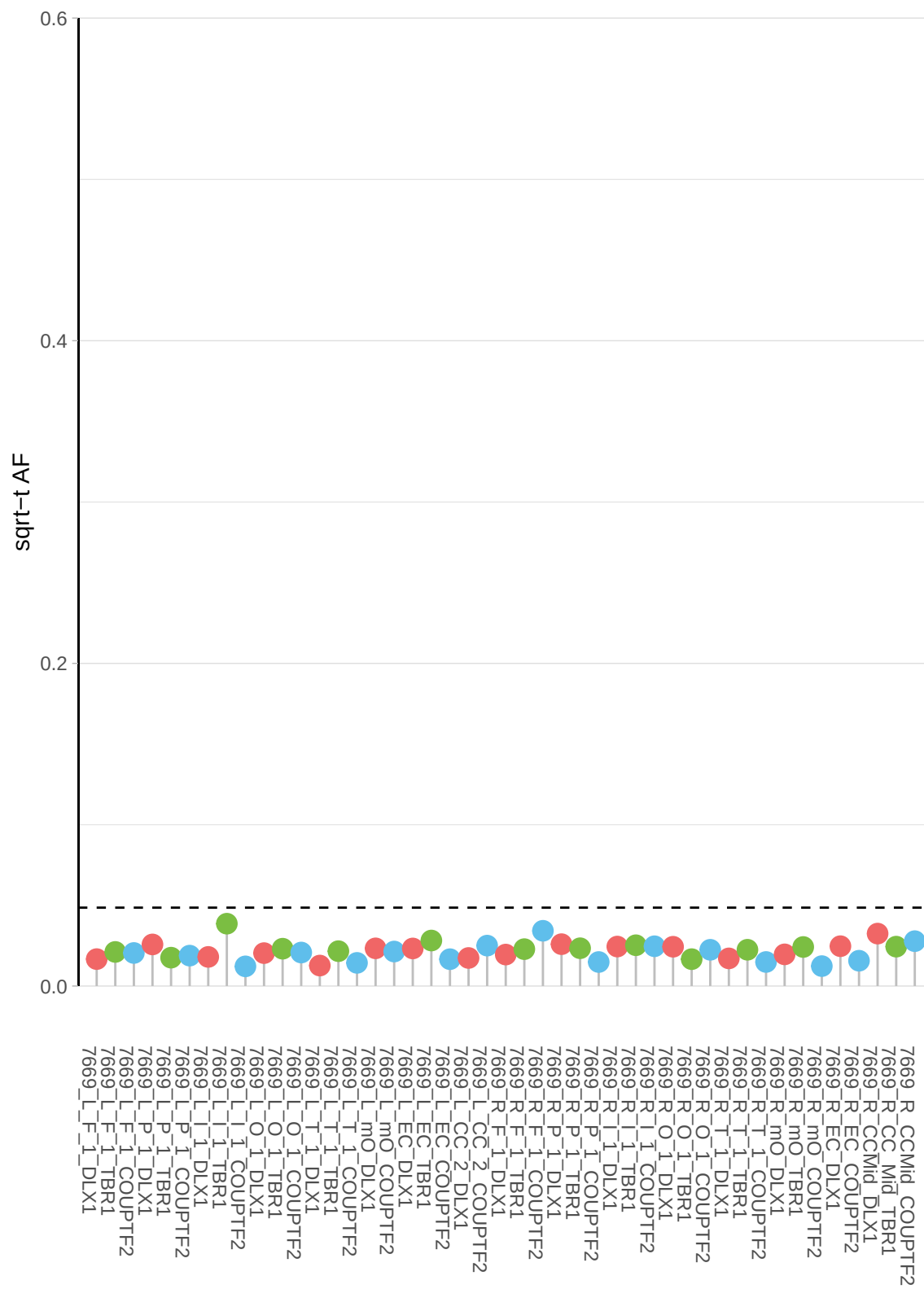

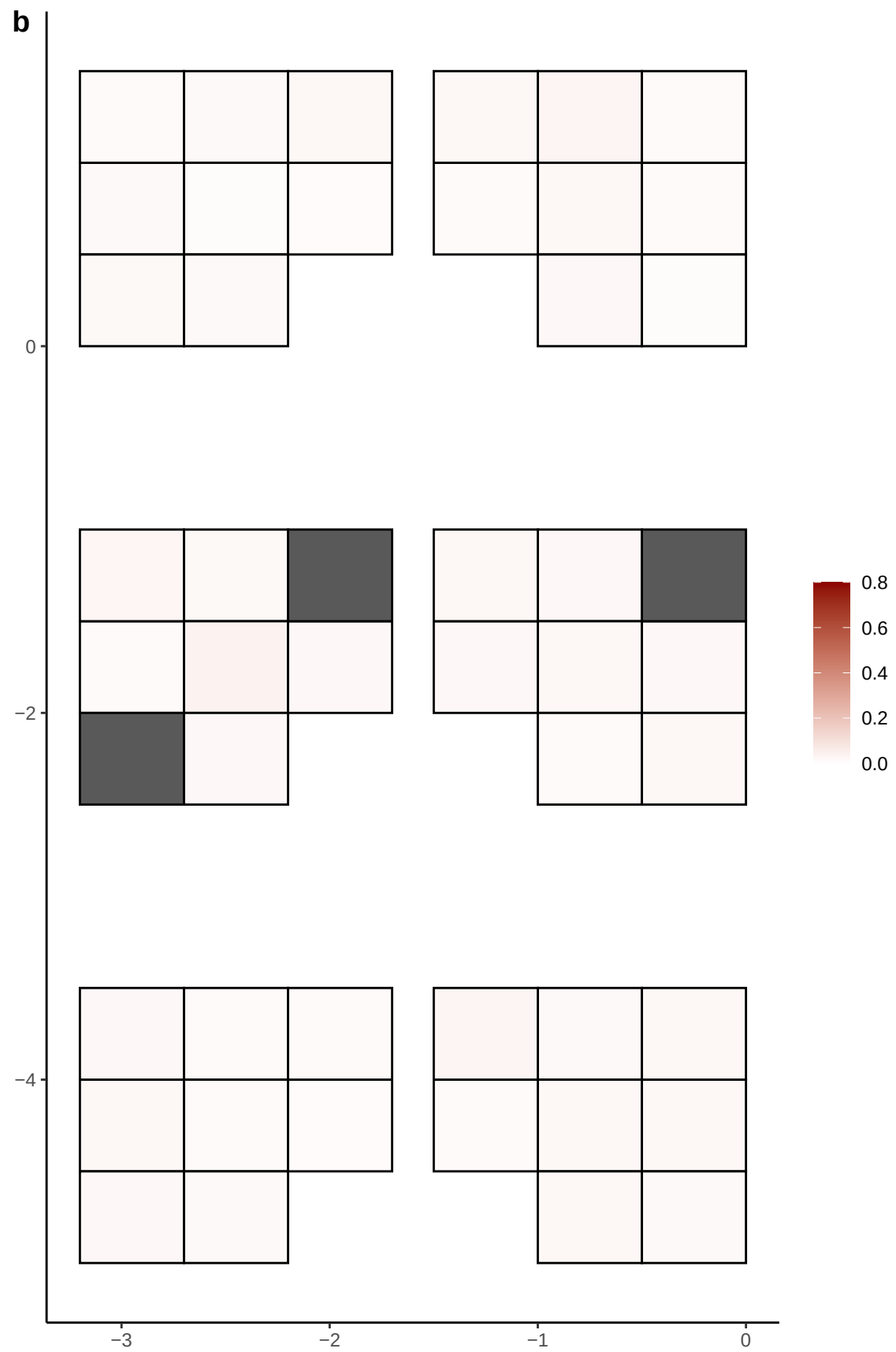

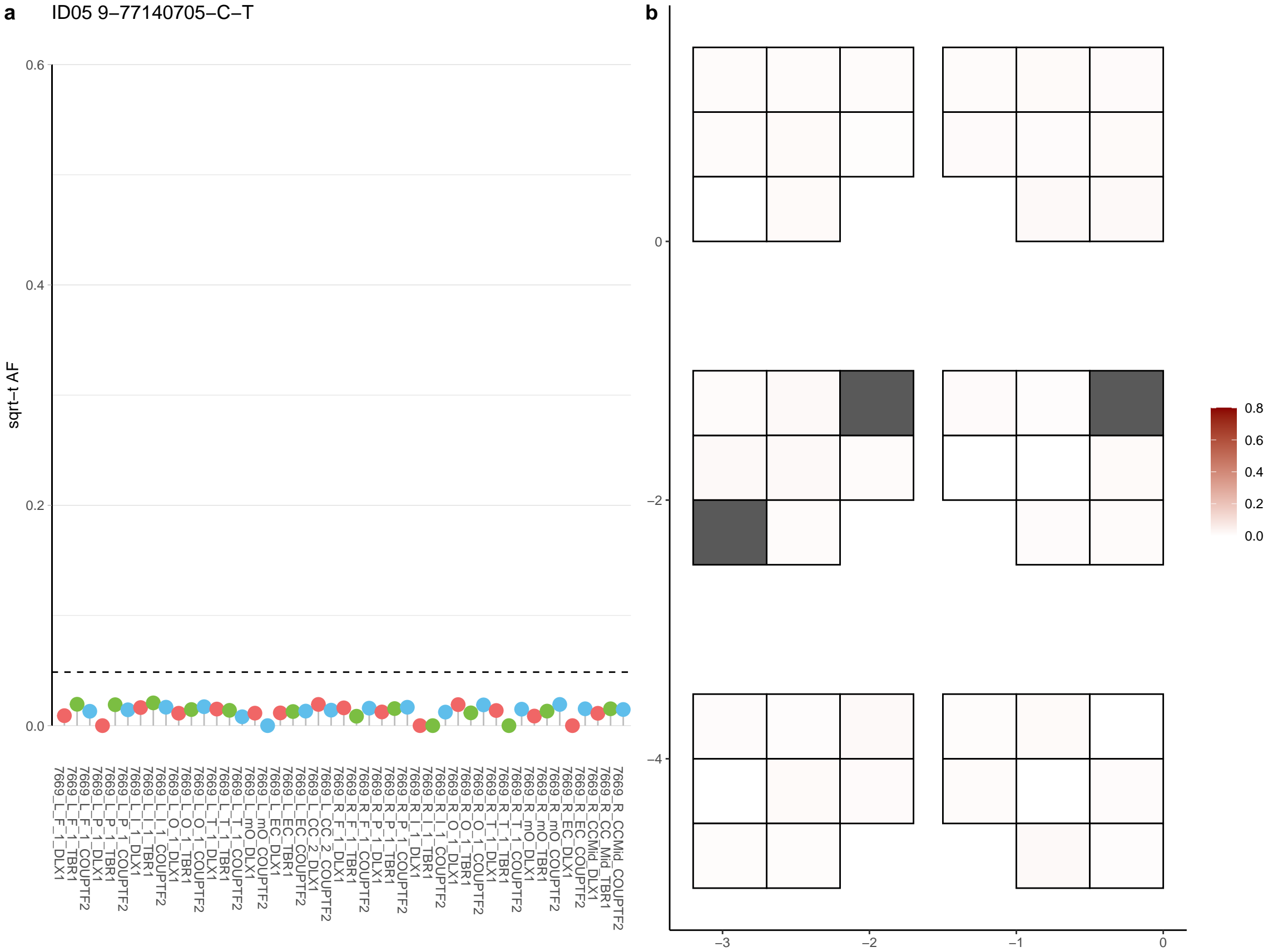

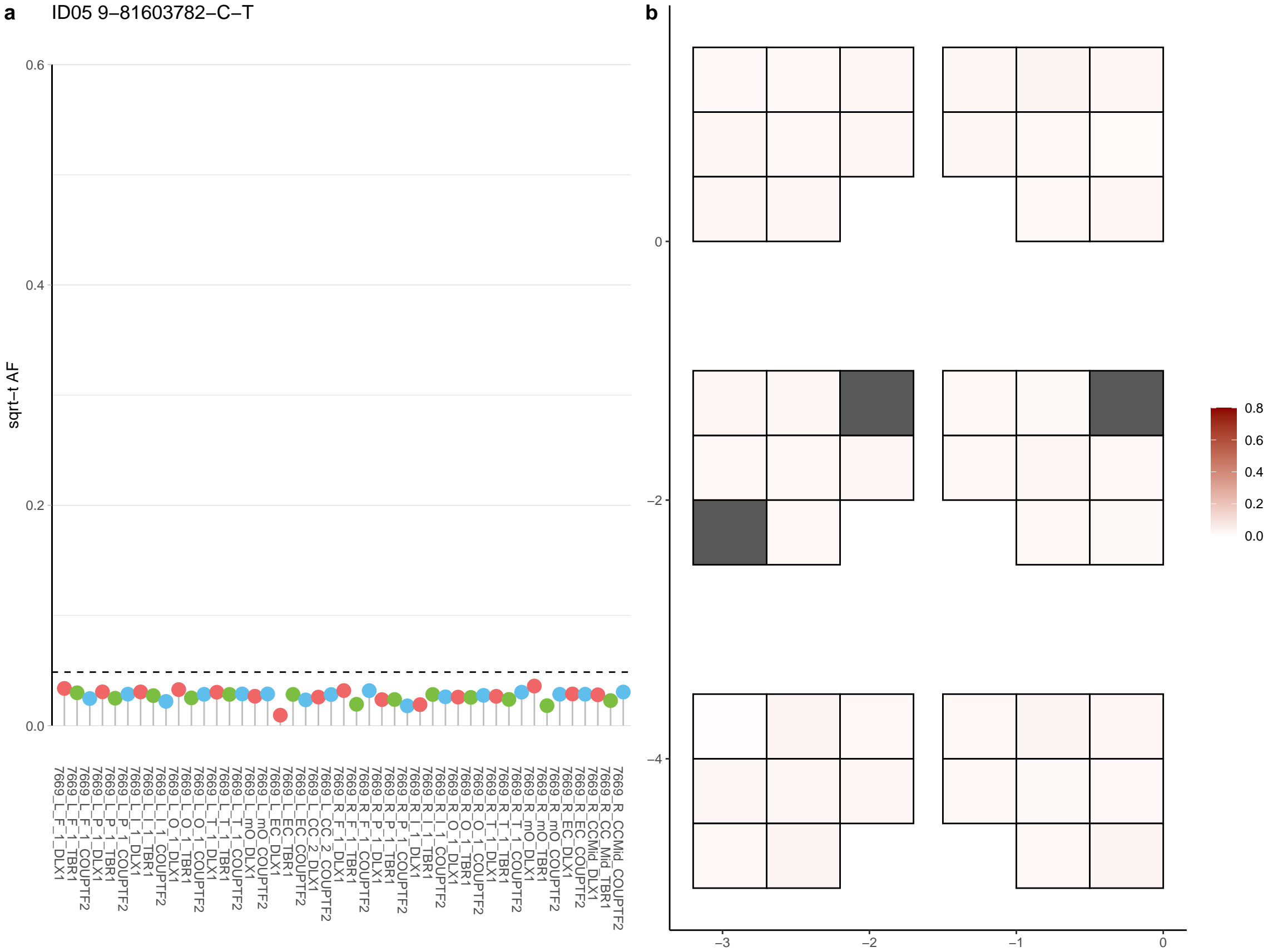

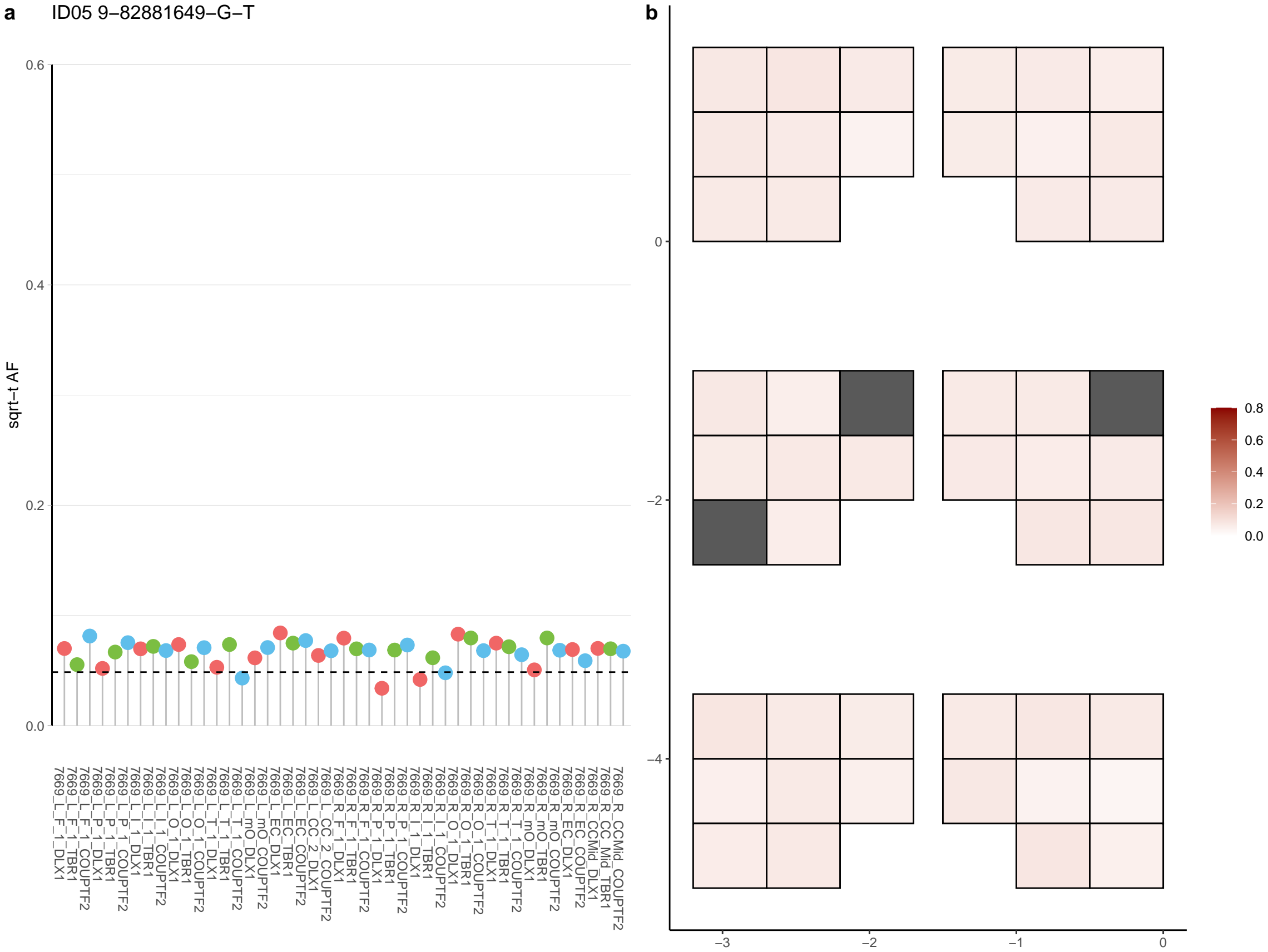

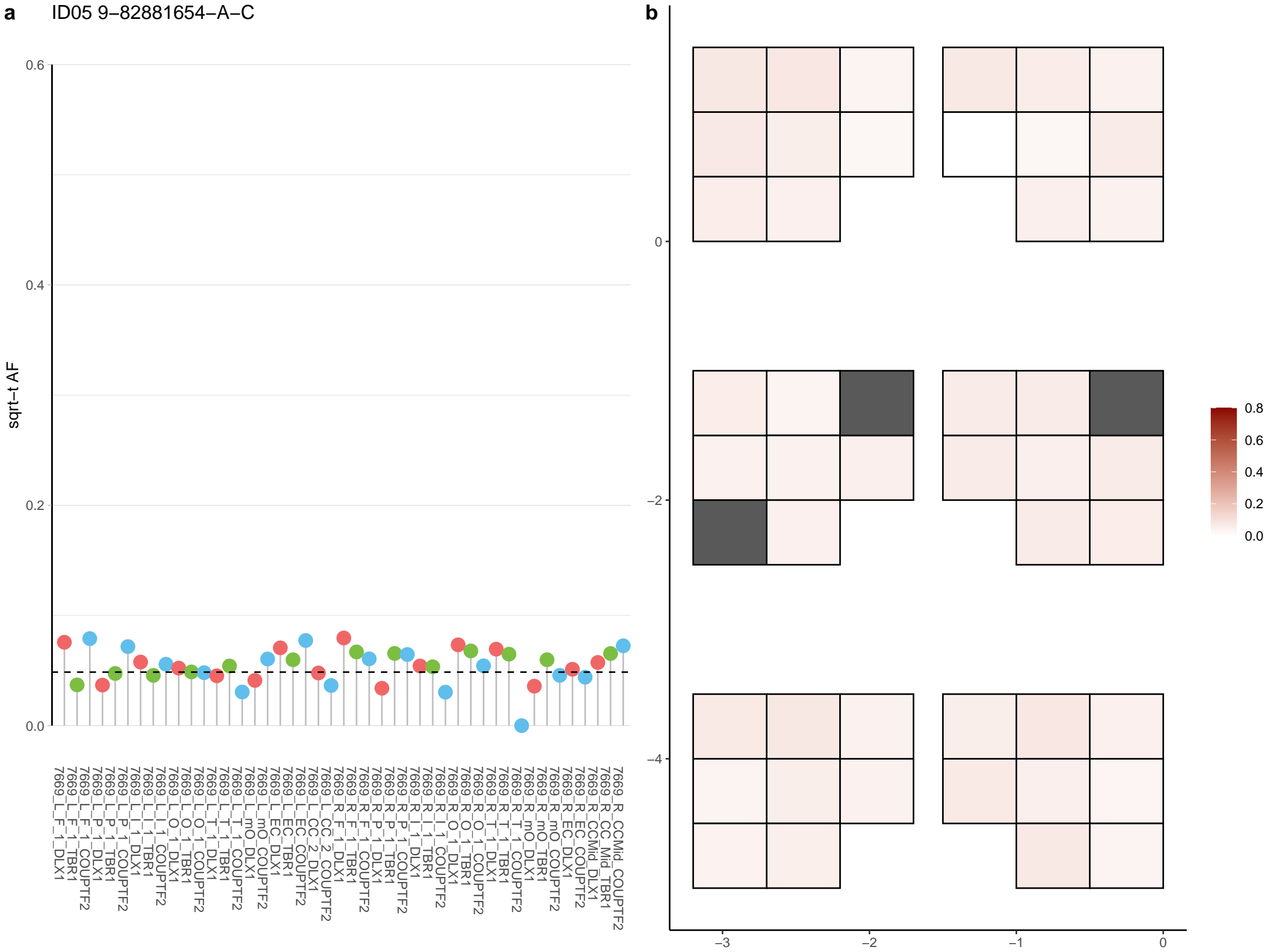

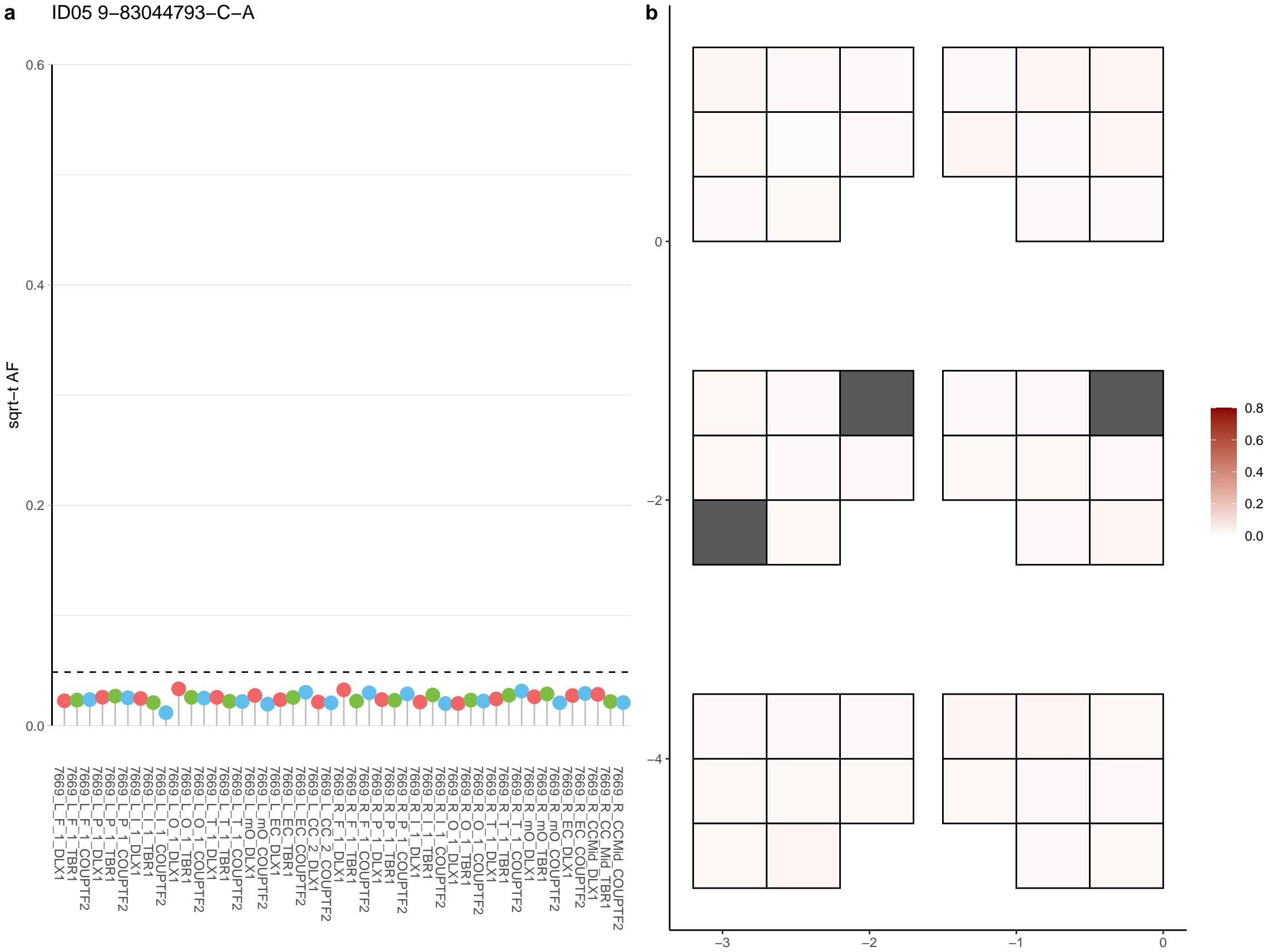

**a** ID05 9-83538985-A-G

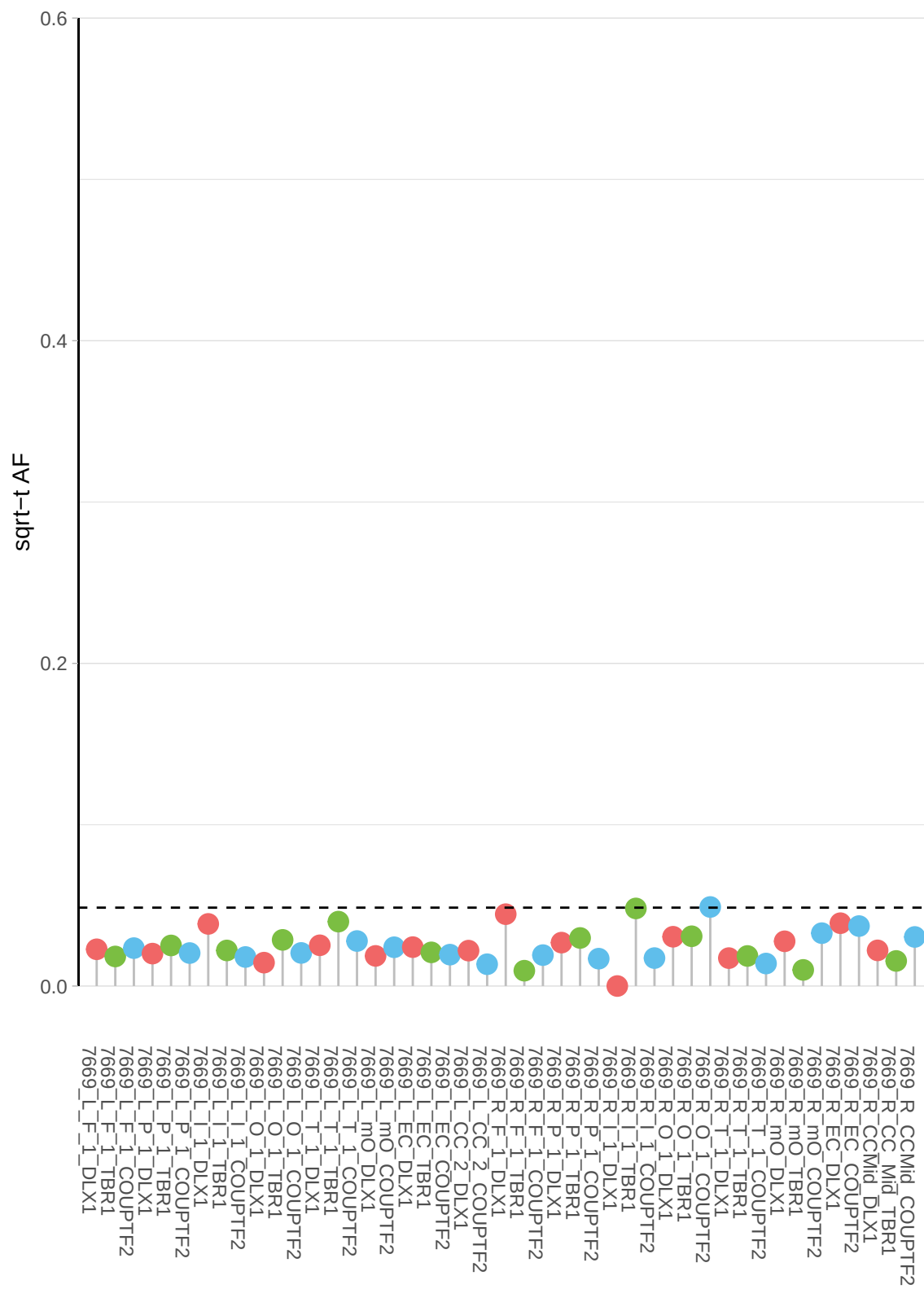

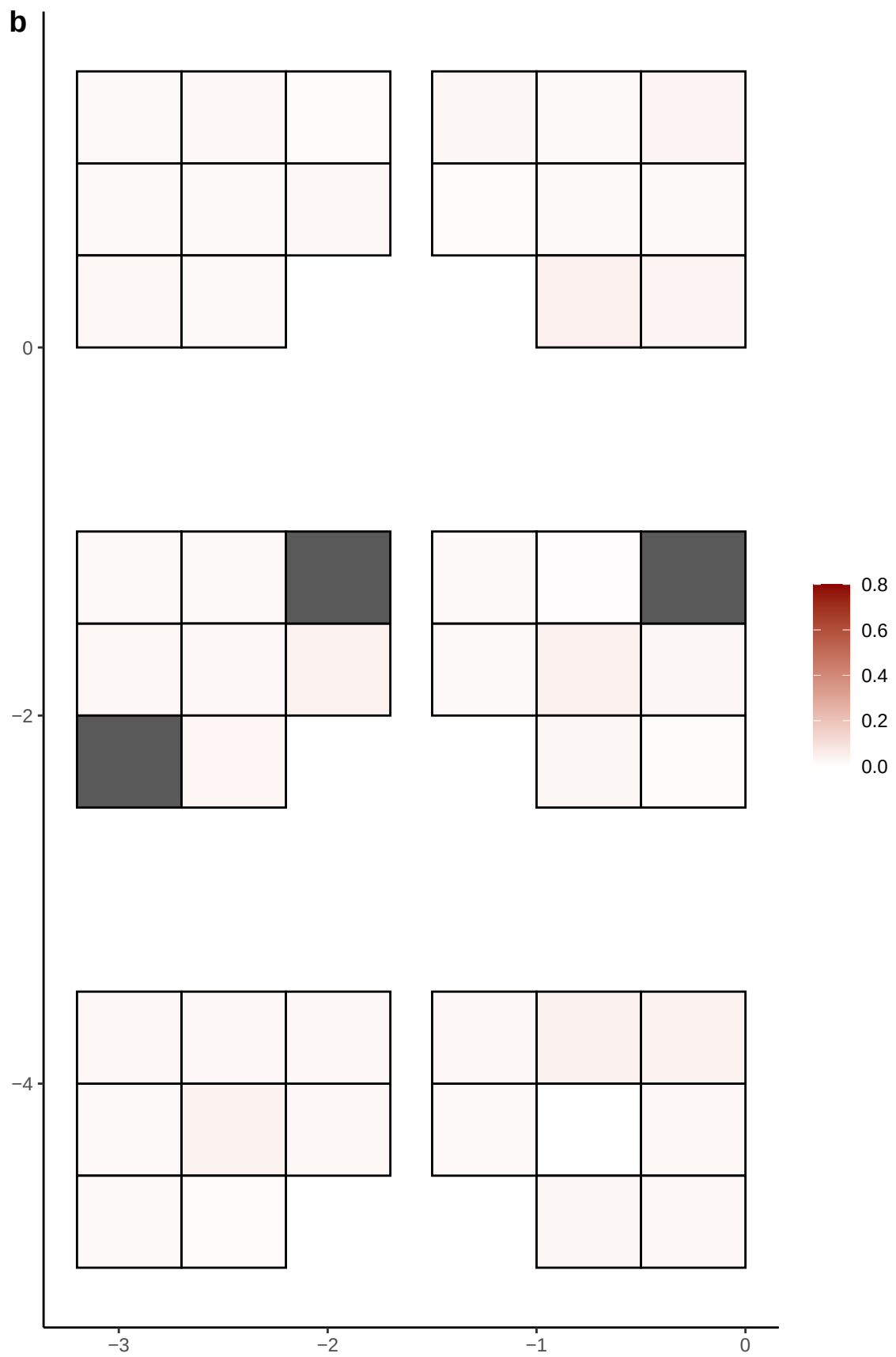

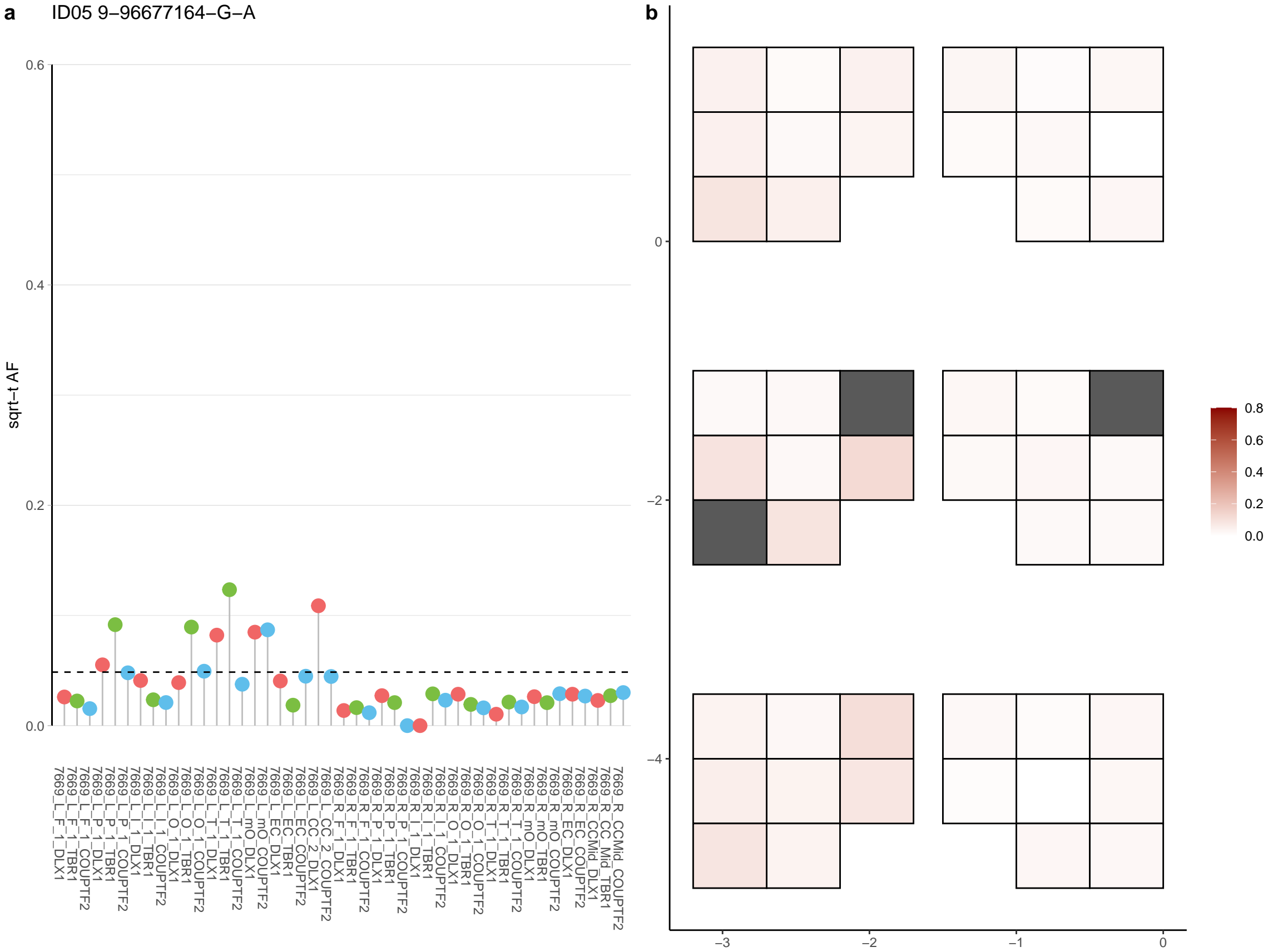

**a** ID05 9-9971800-C-T

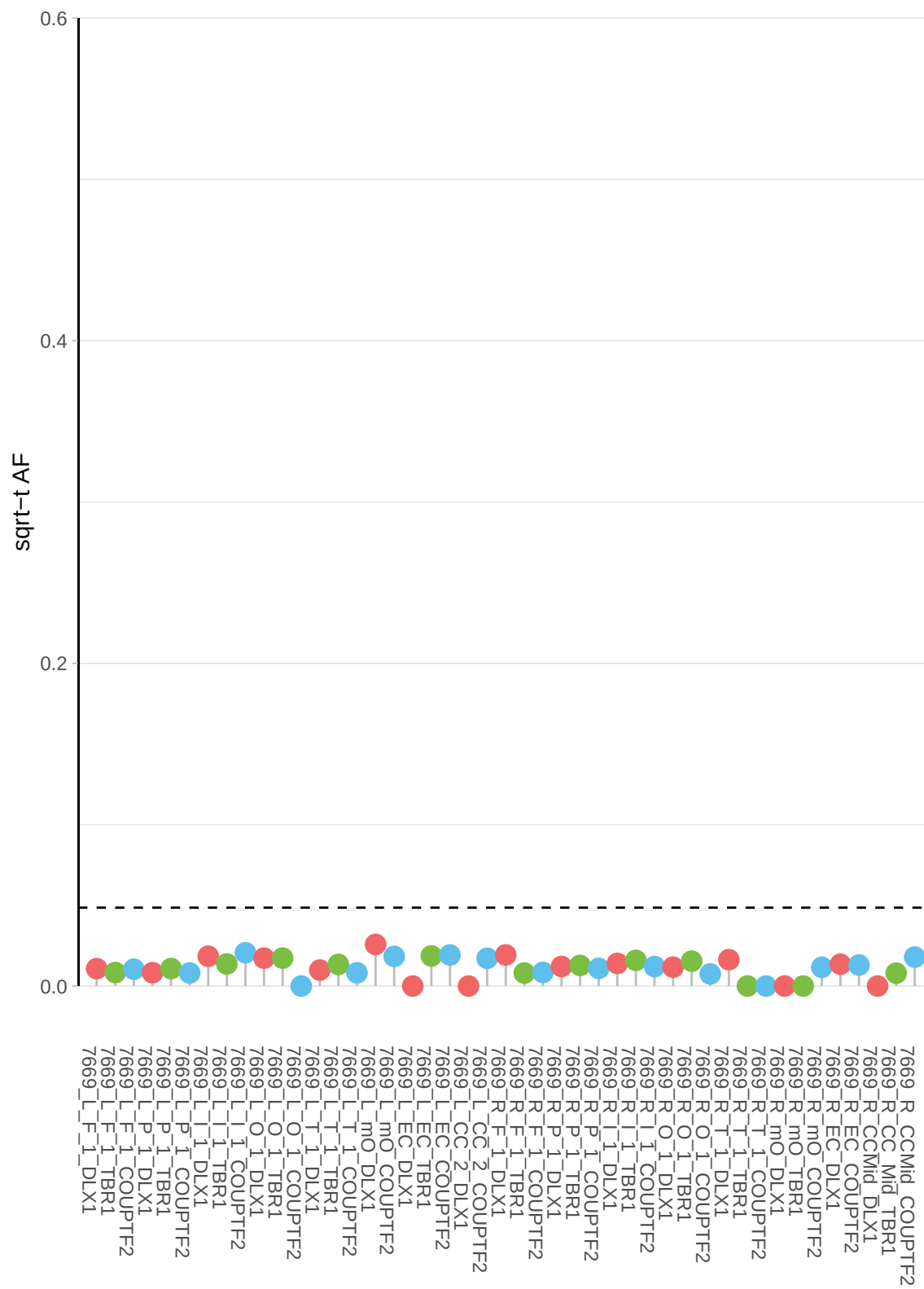

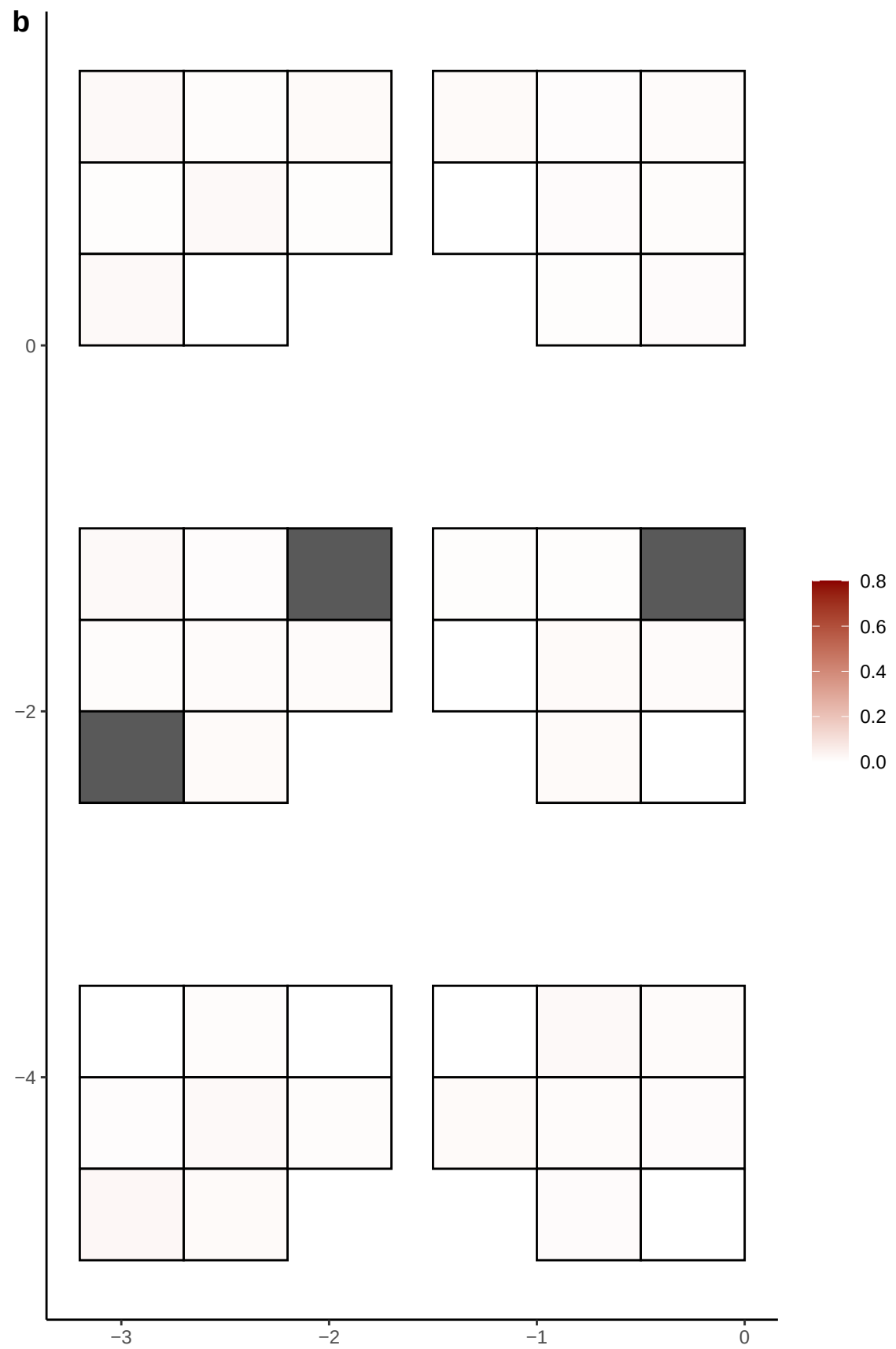

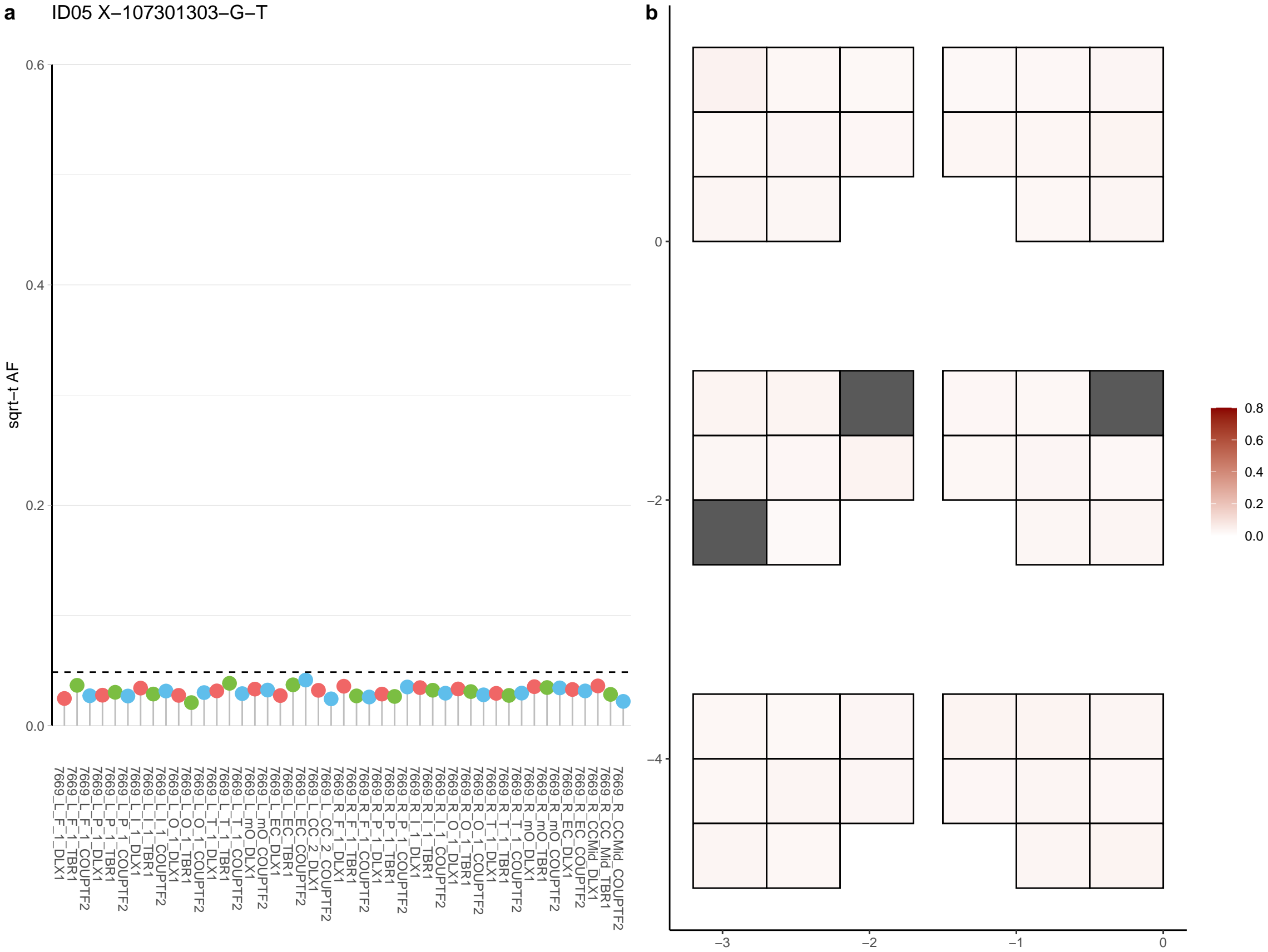

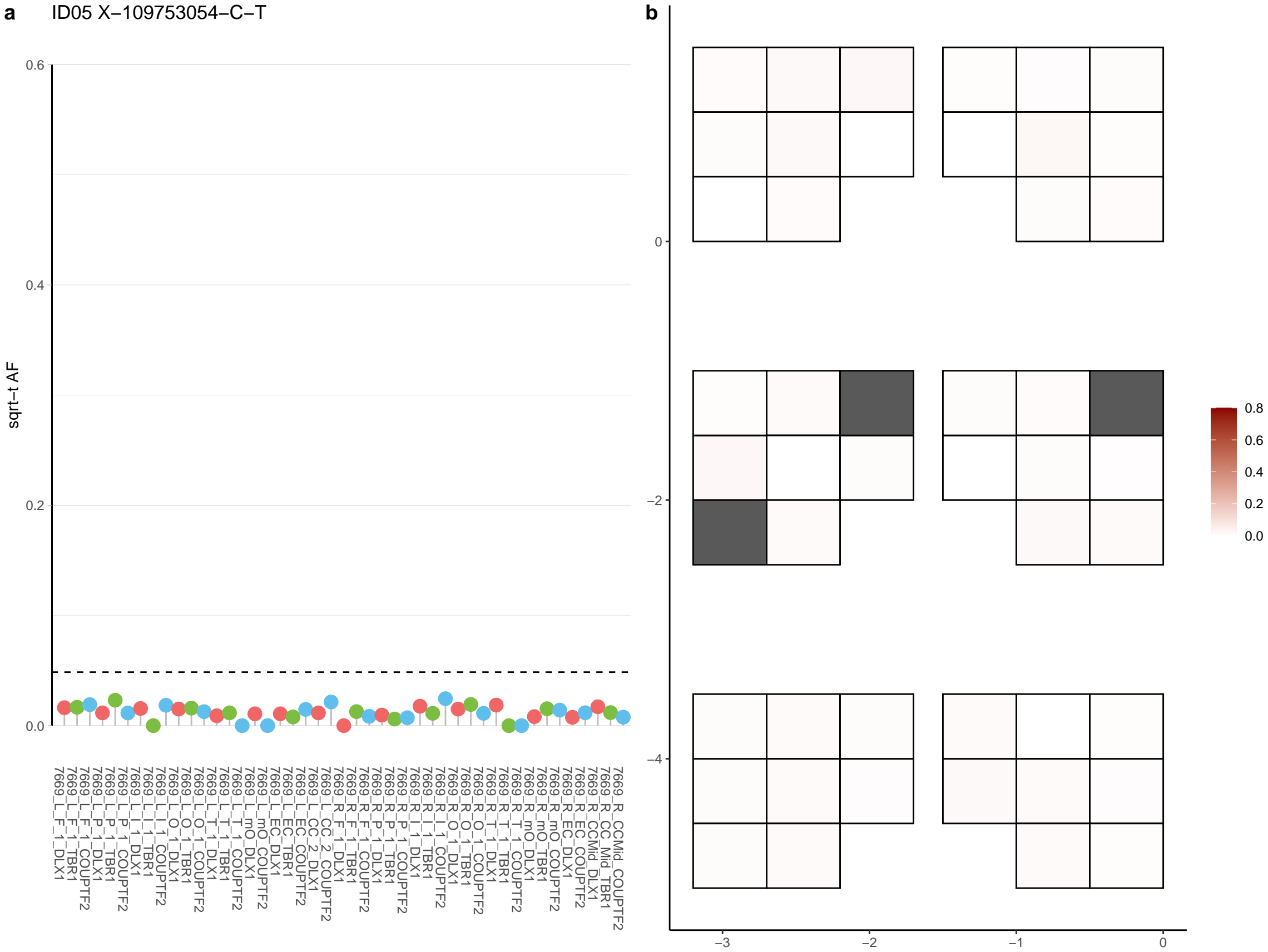

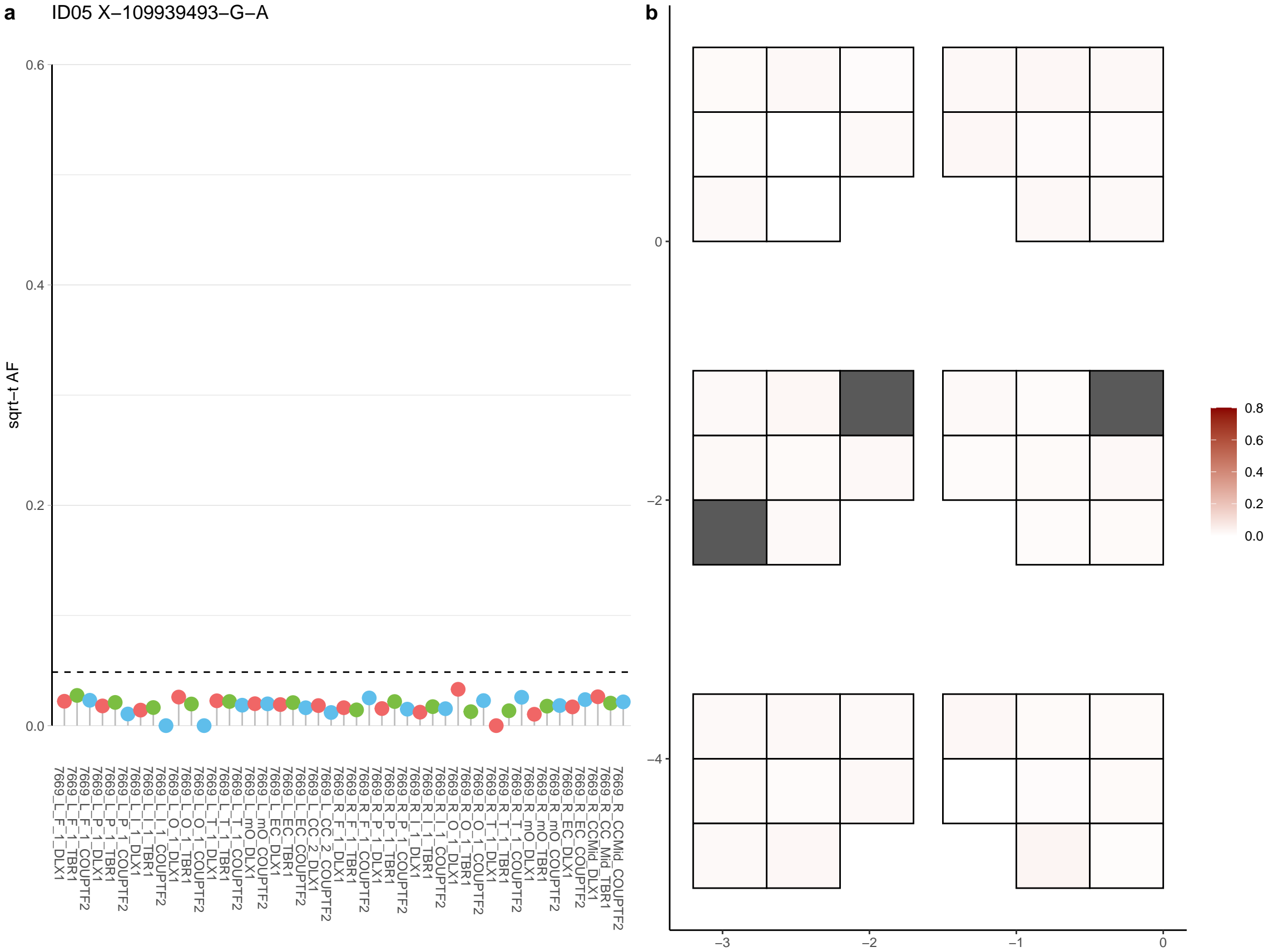

**a** ID05 X-117240525-A-C

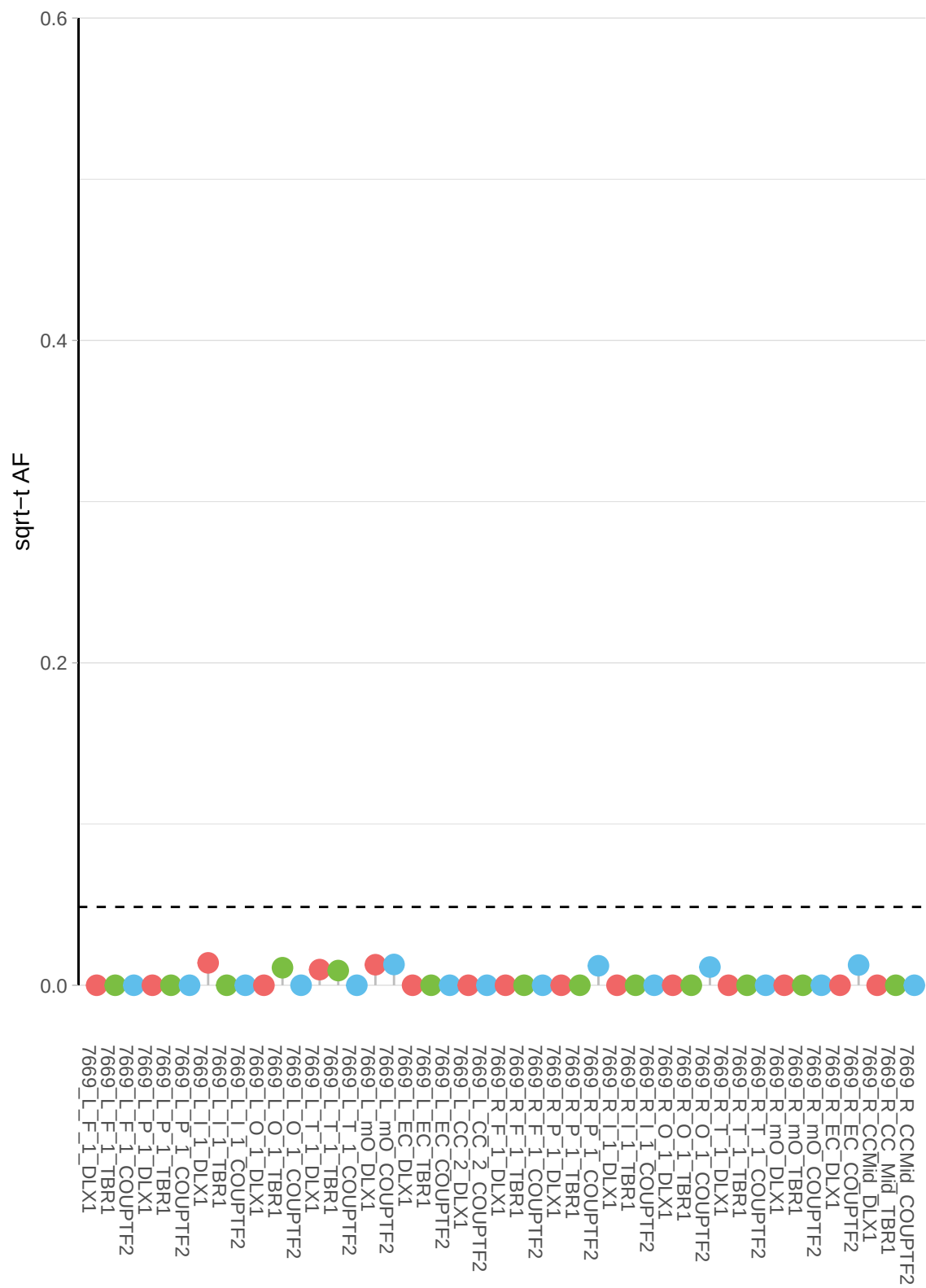

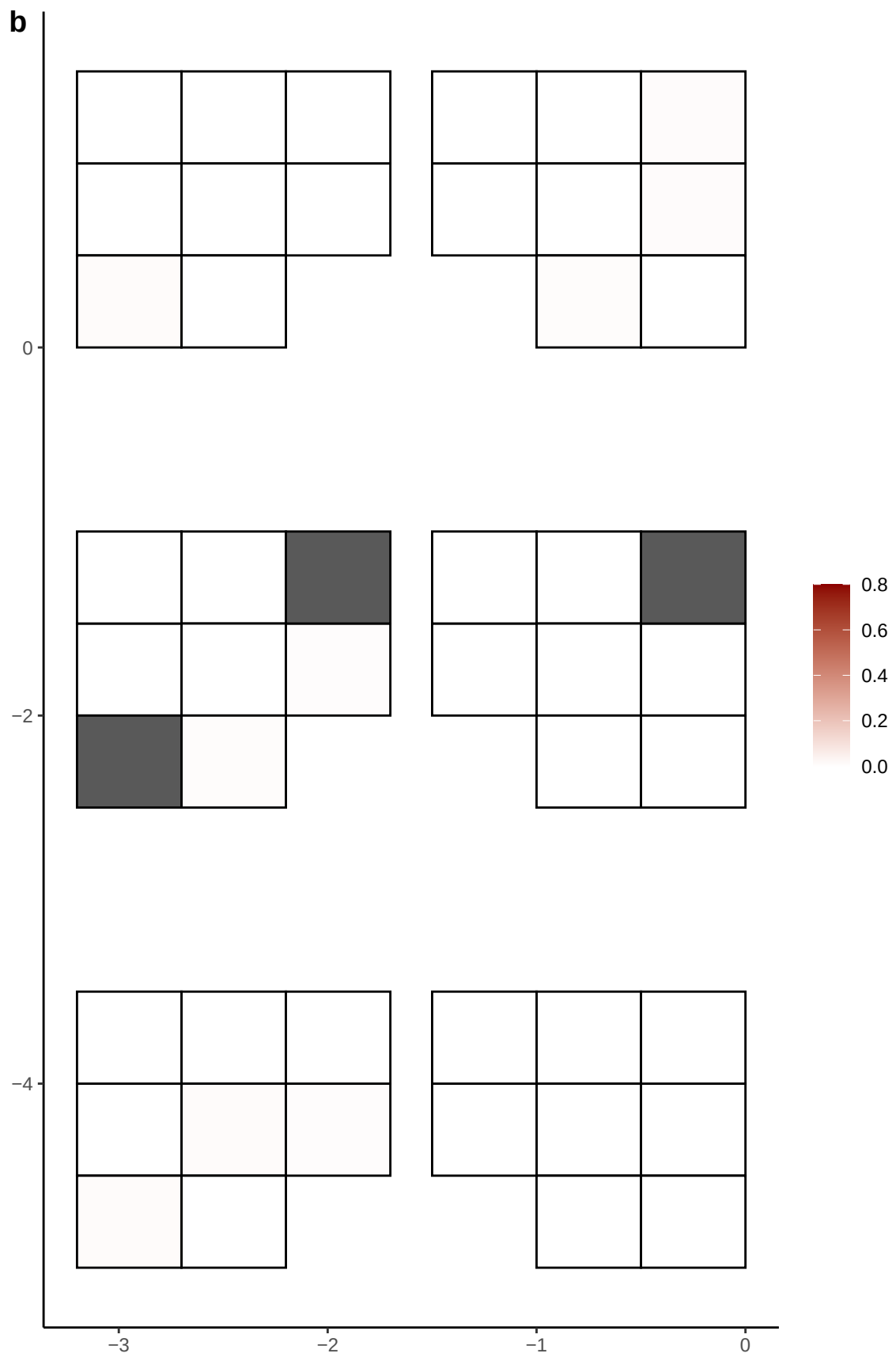

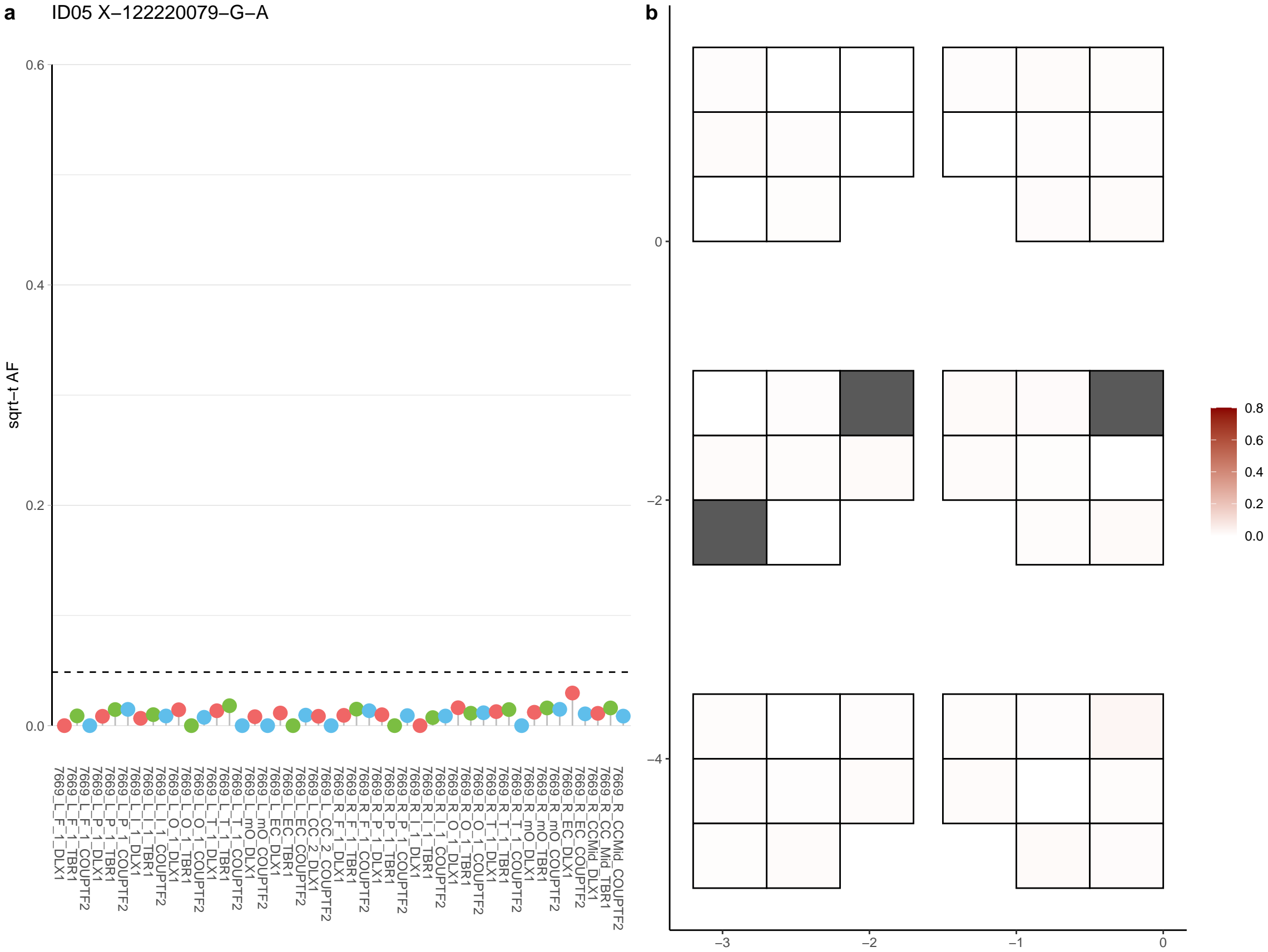

**a** ID05 X-125464024-G-A

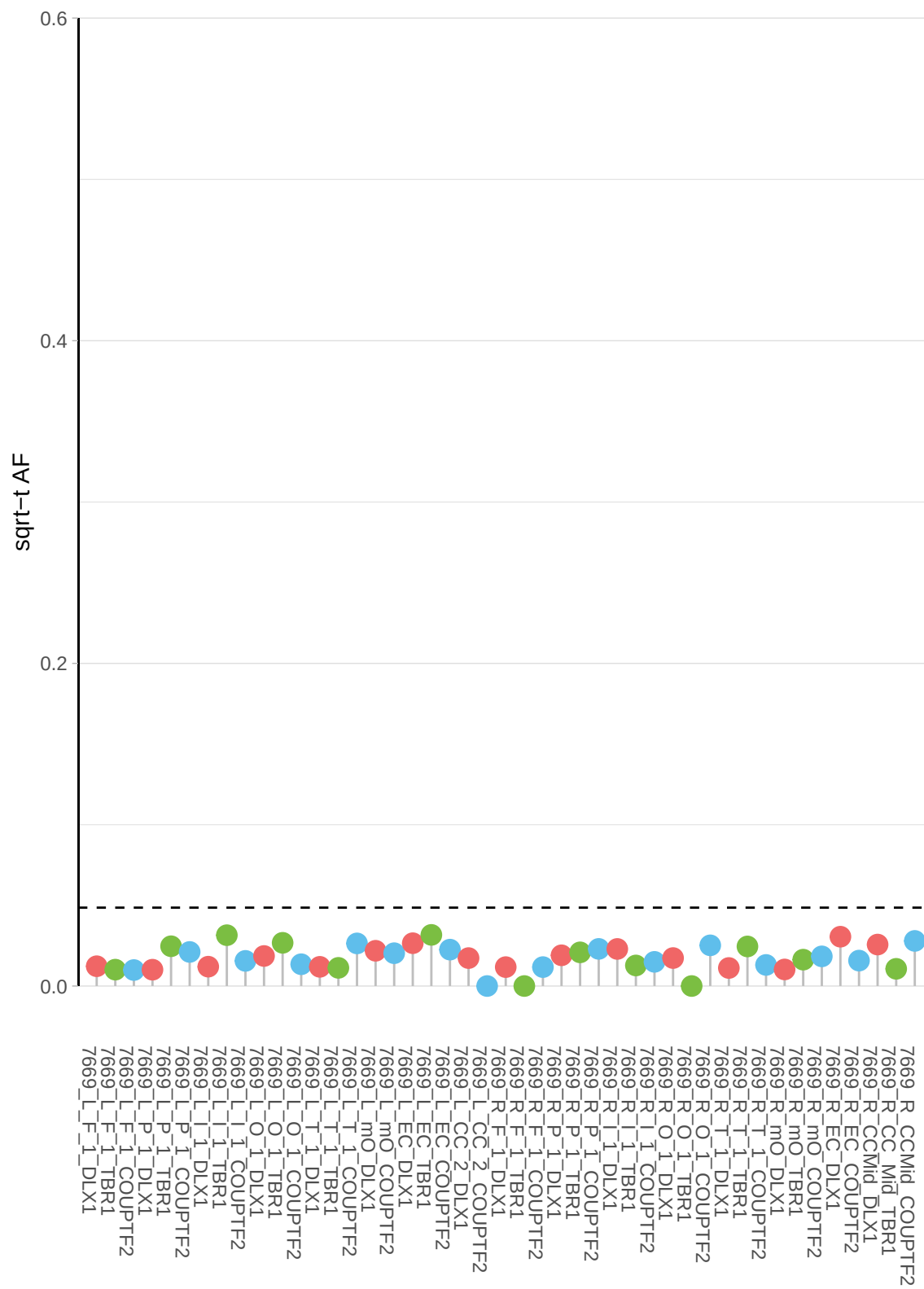

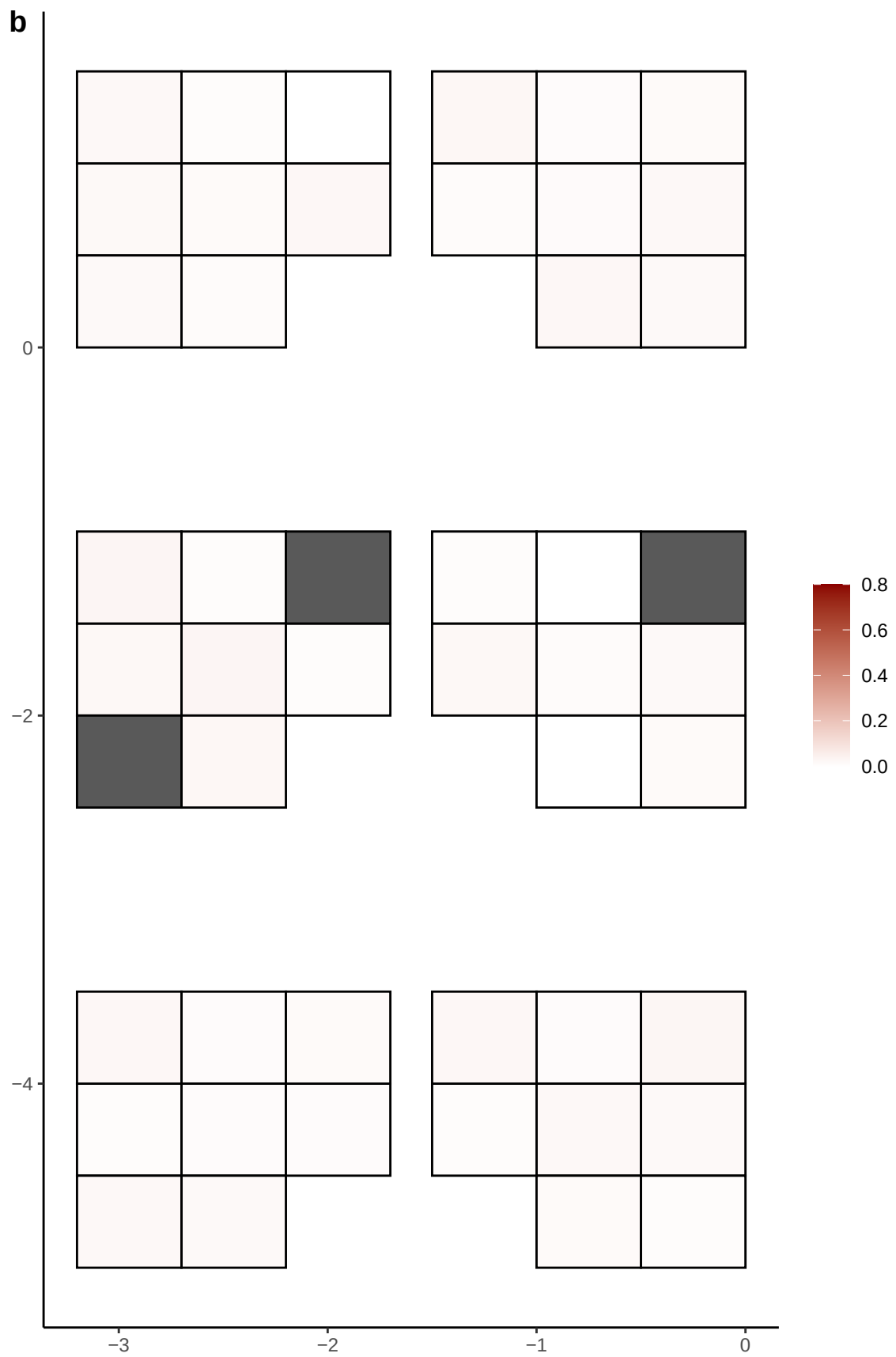

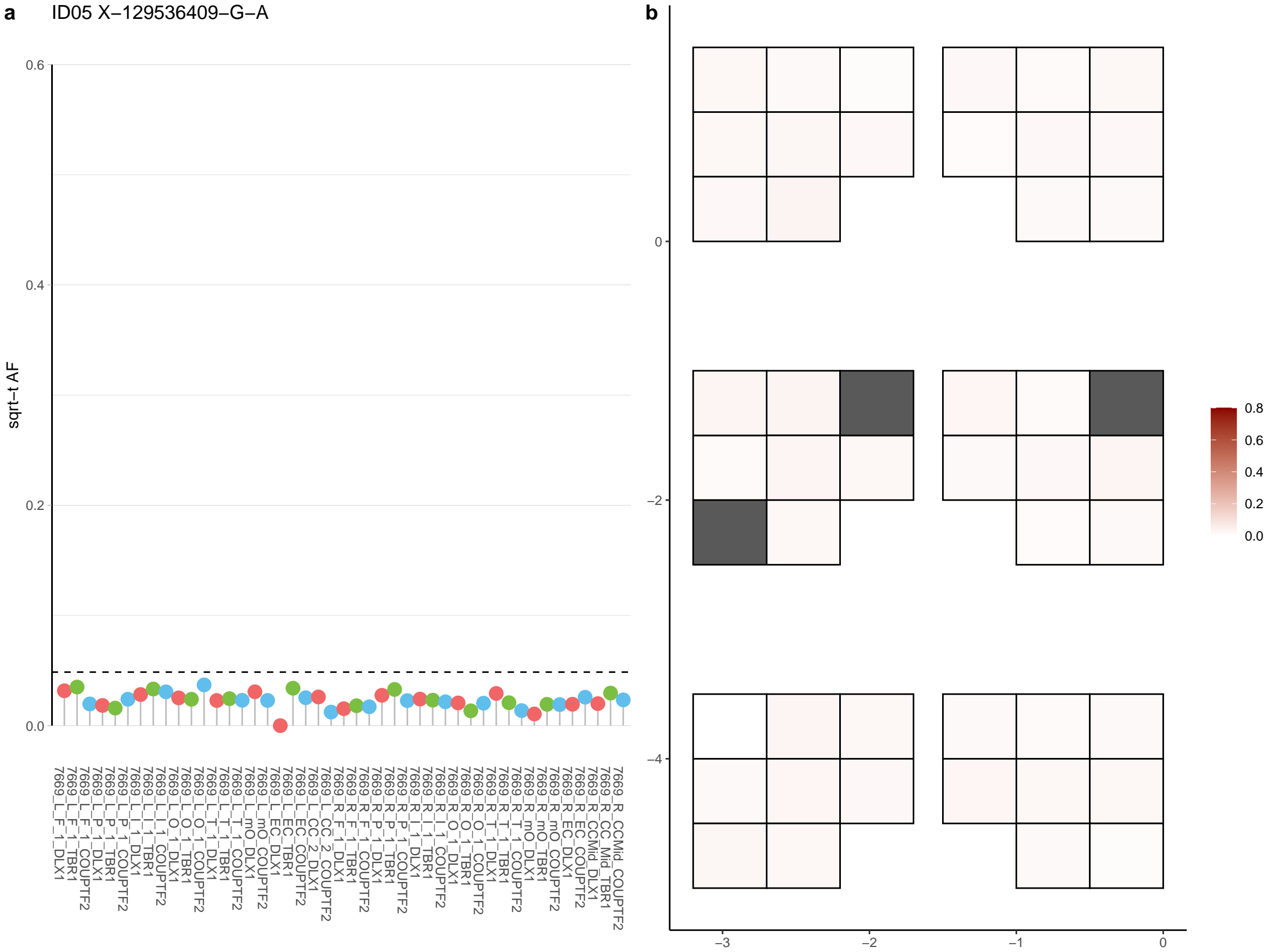

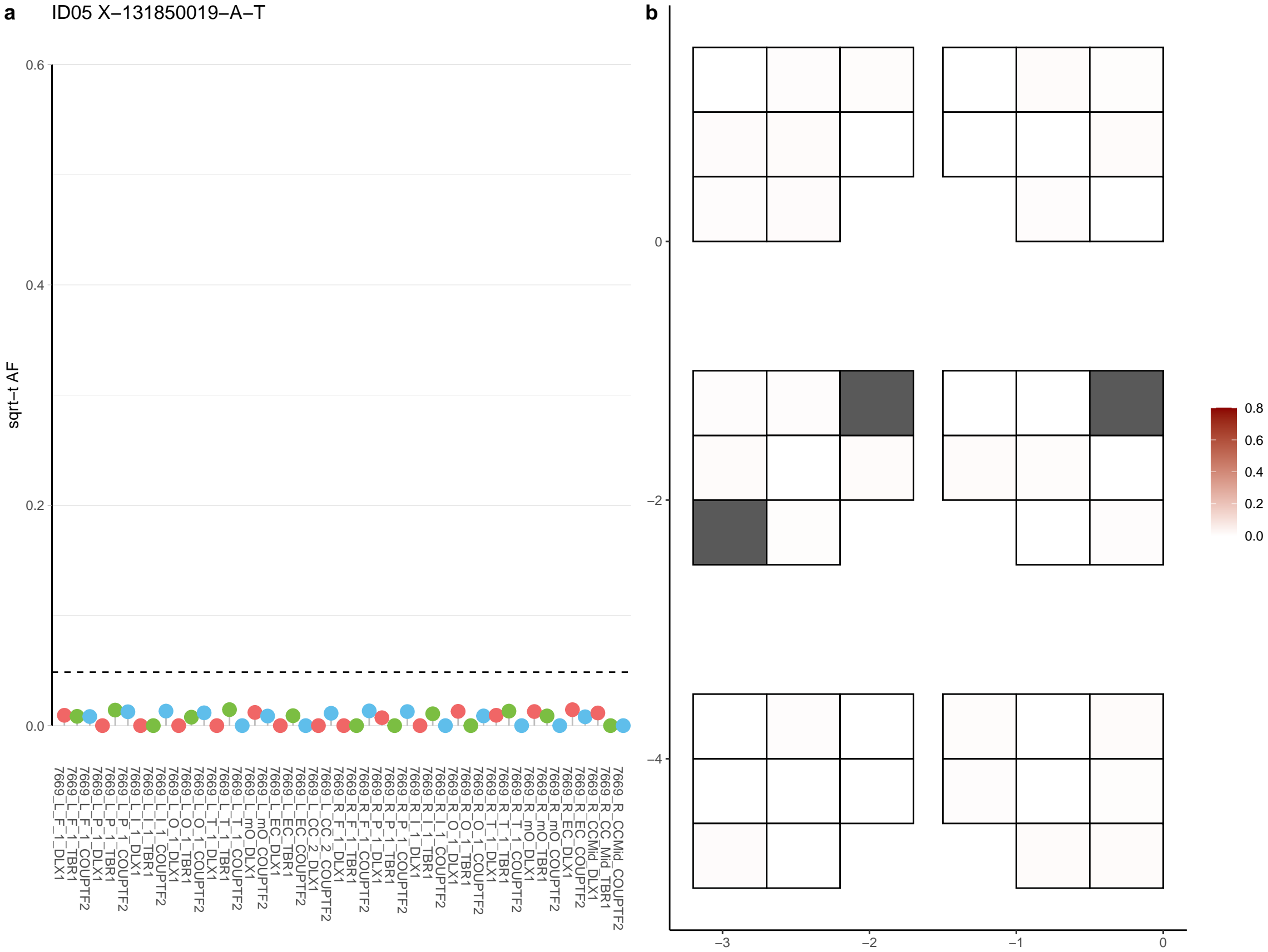

**a** ID05 X-138211912-G-A

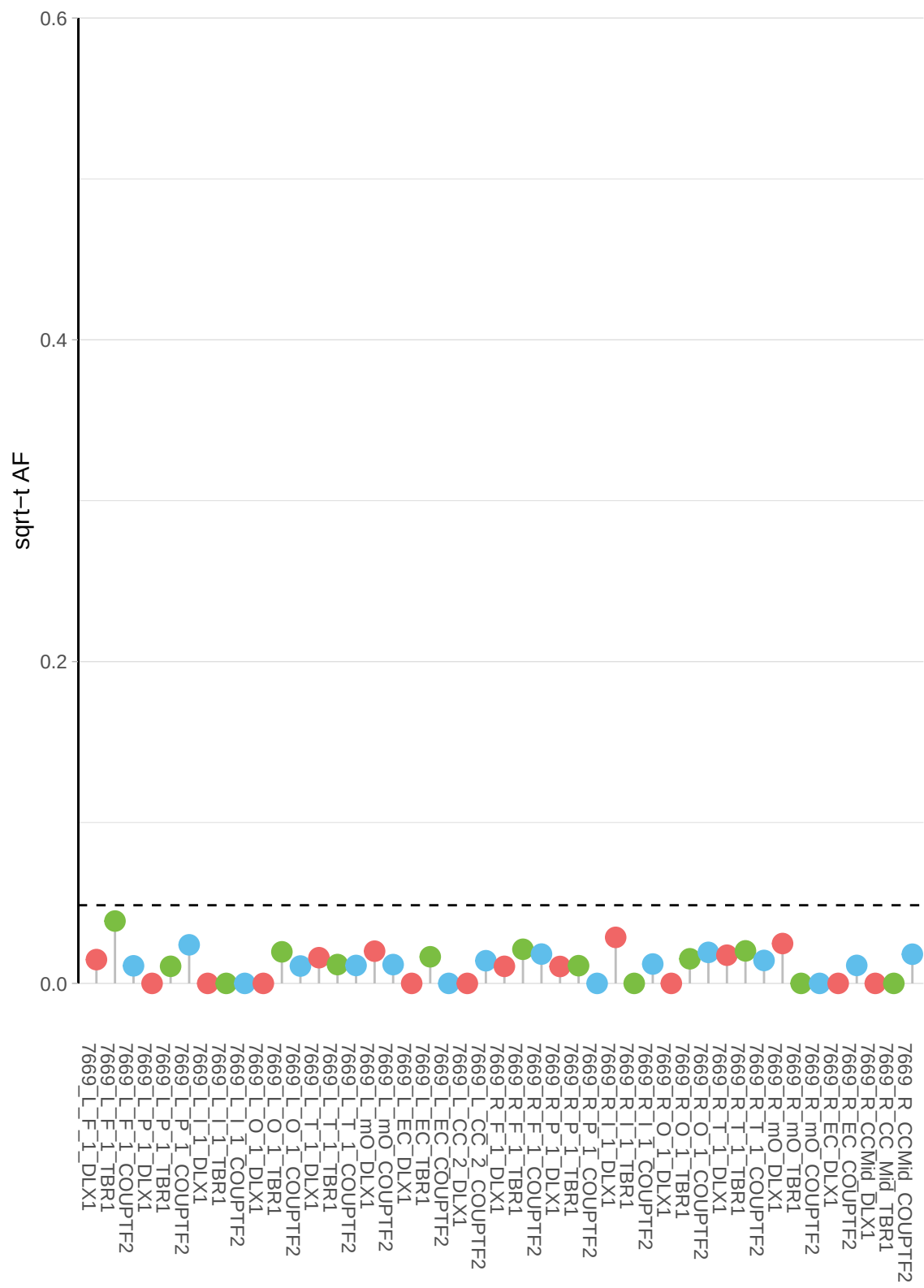

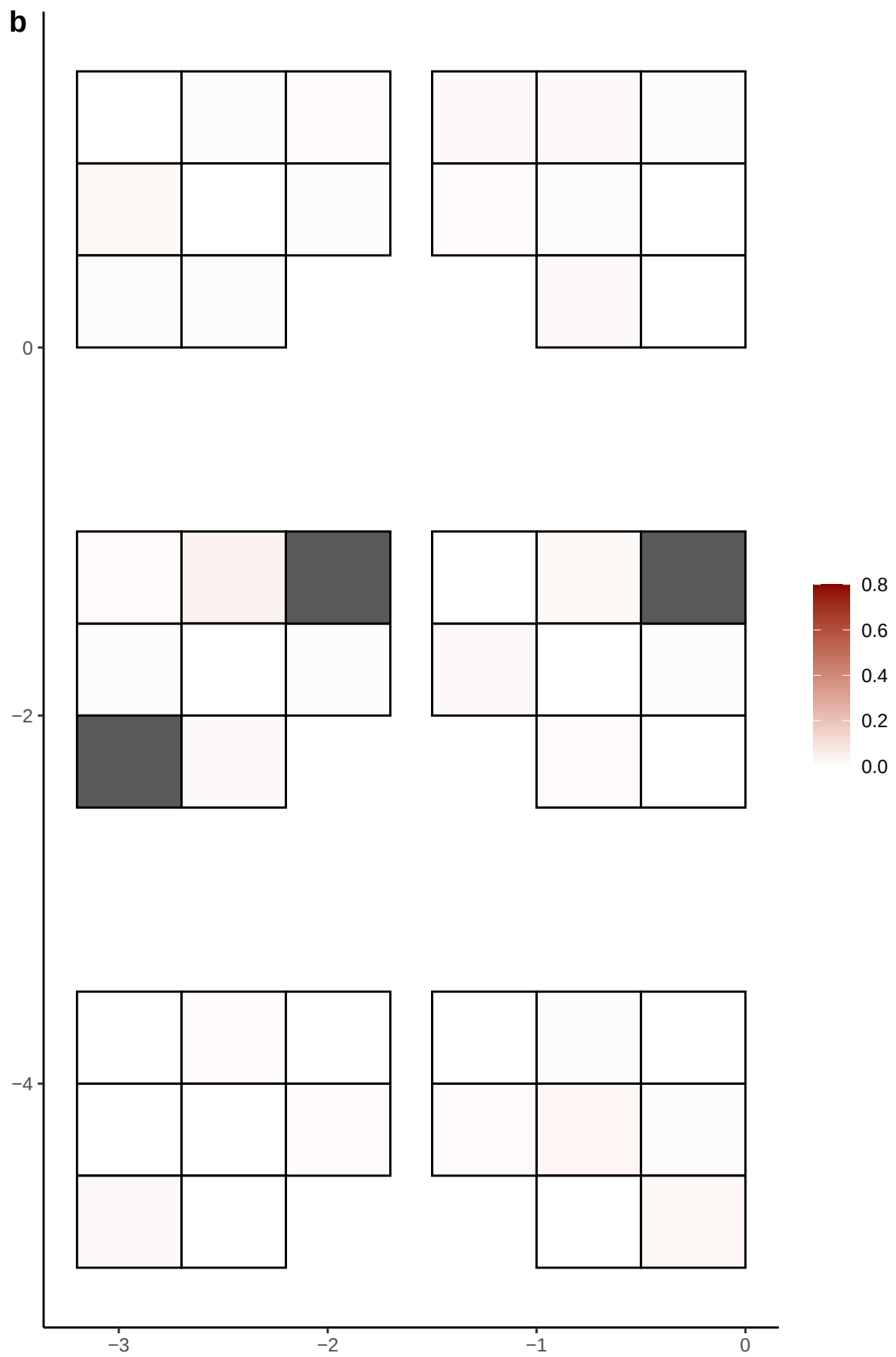

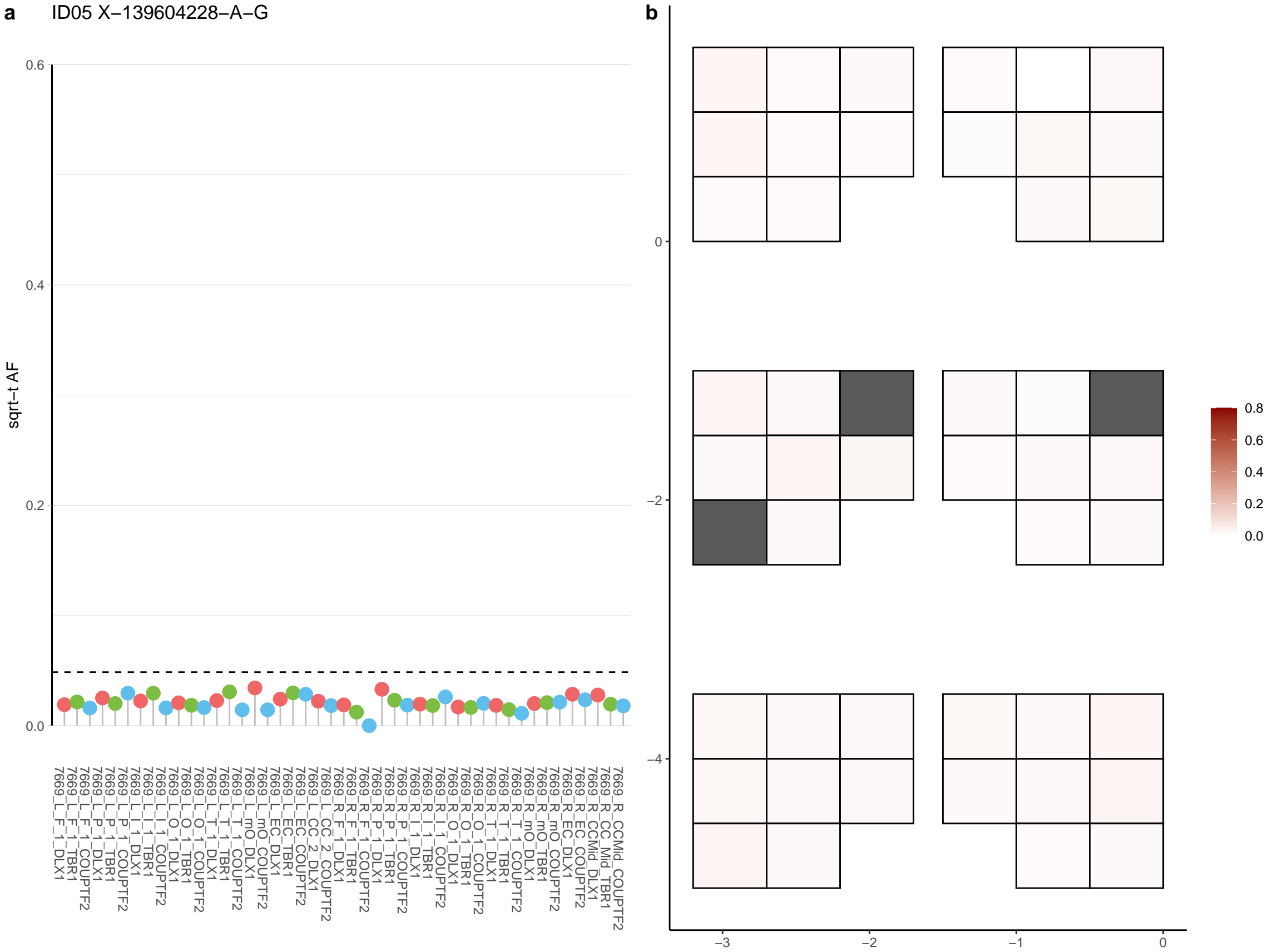

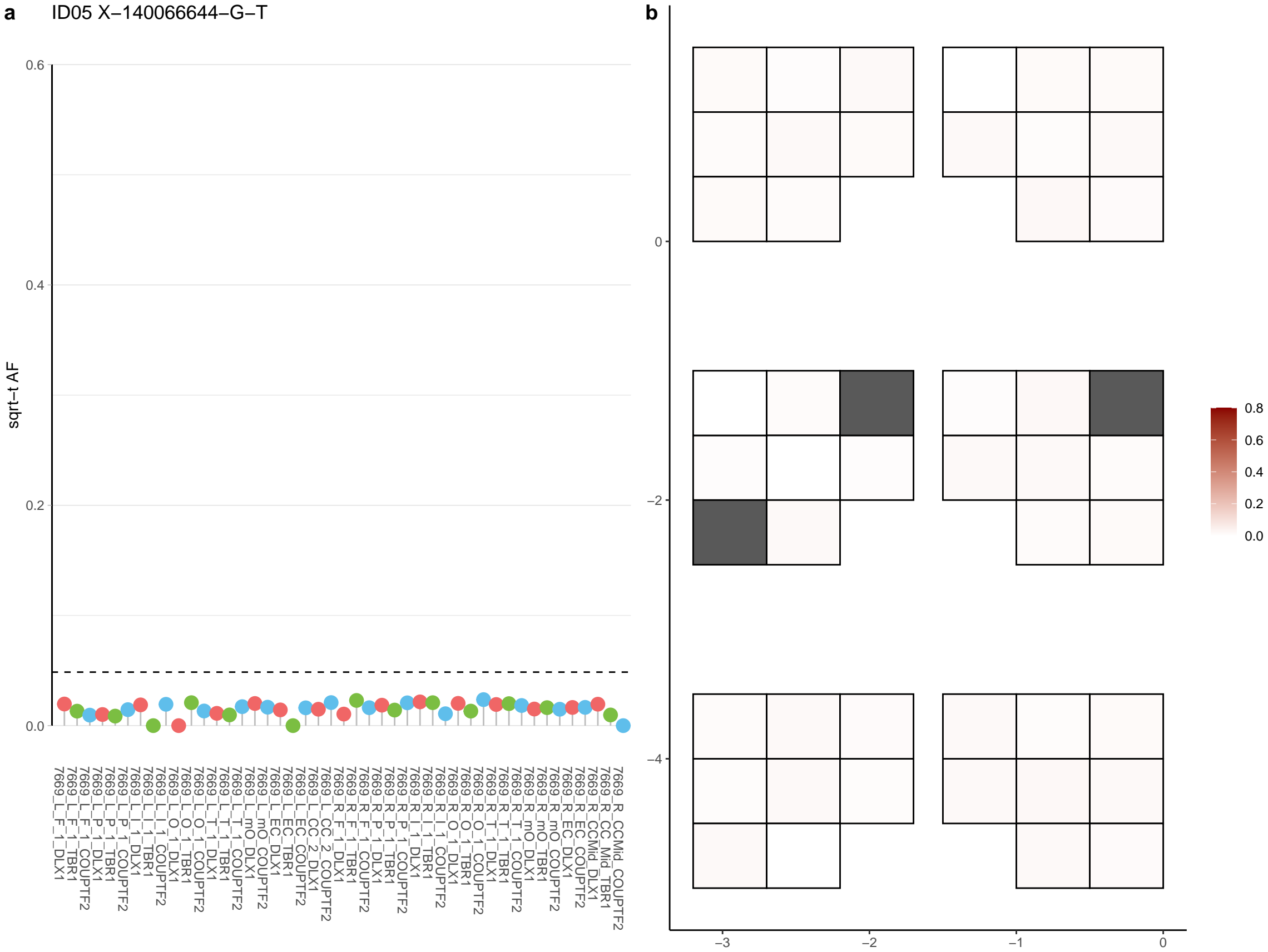

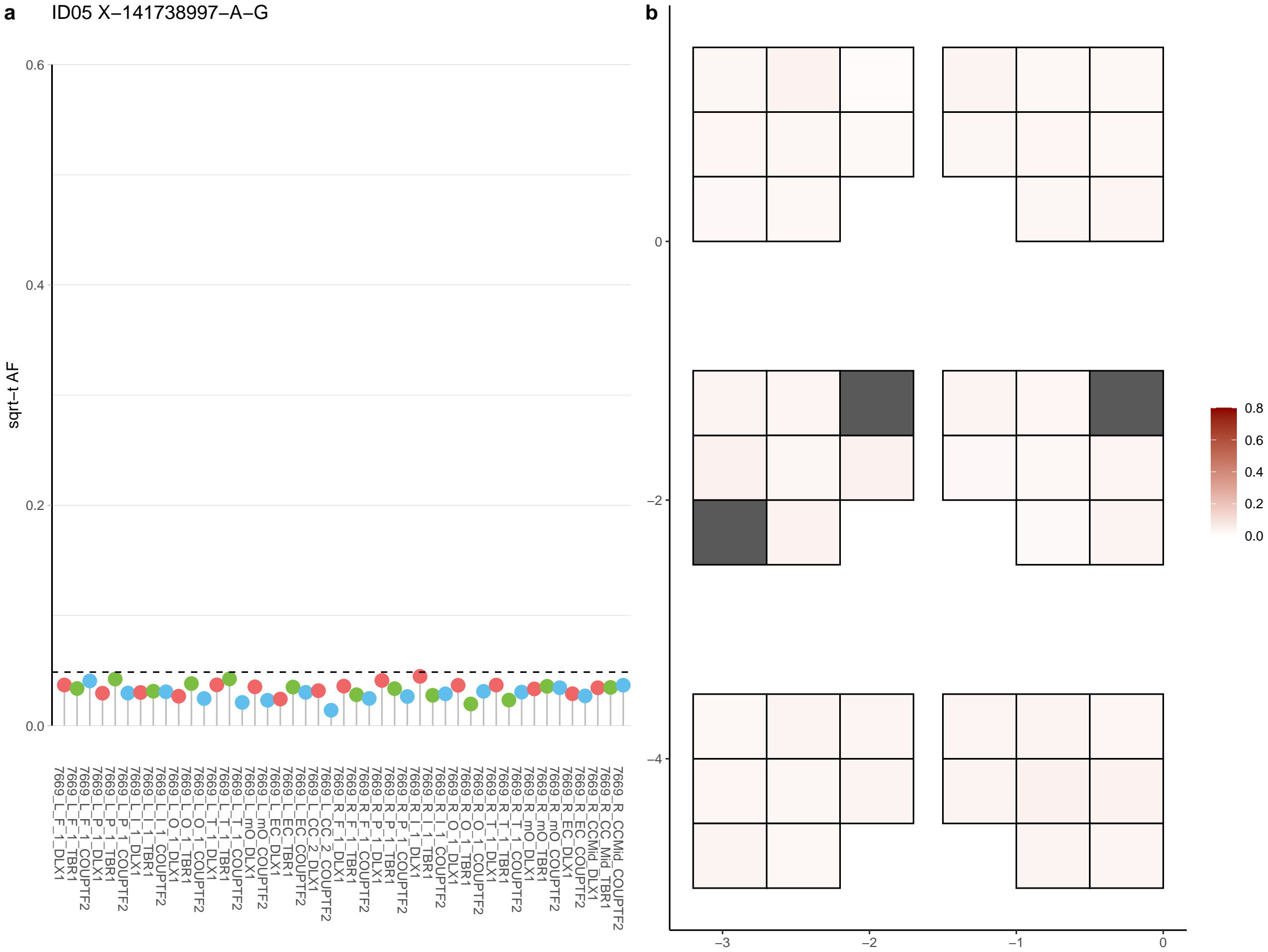

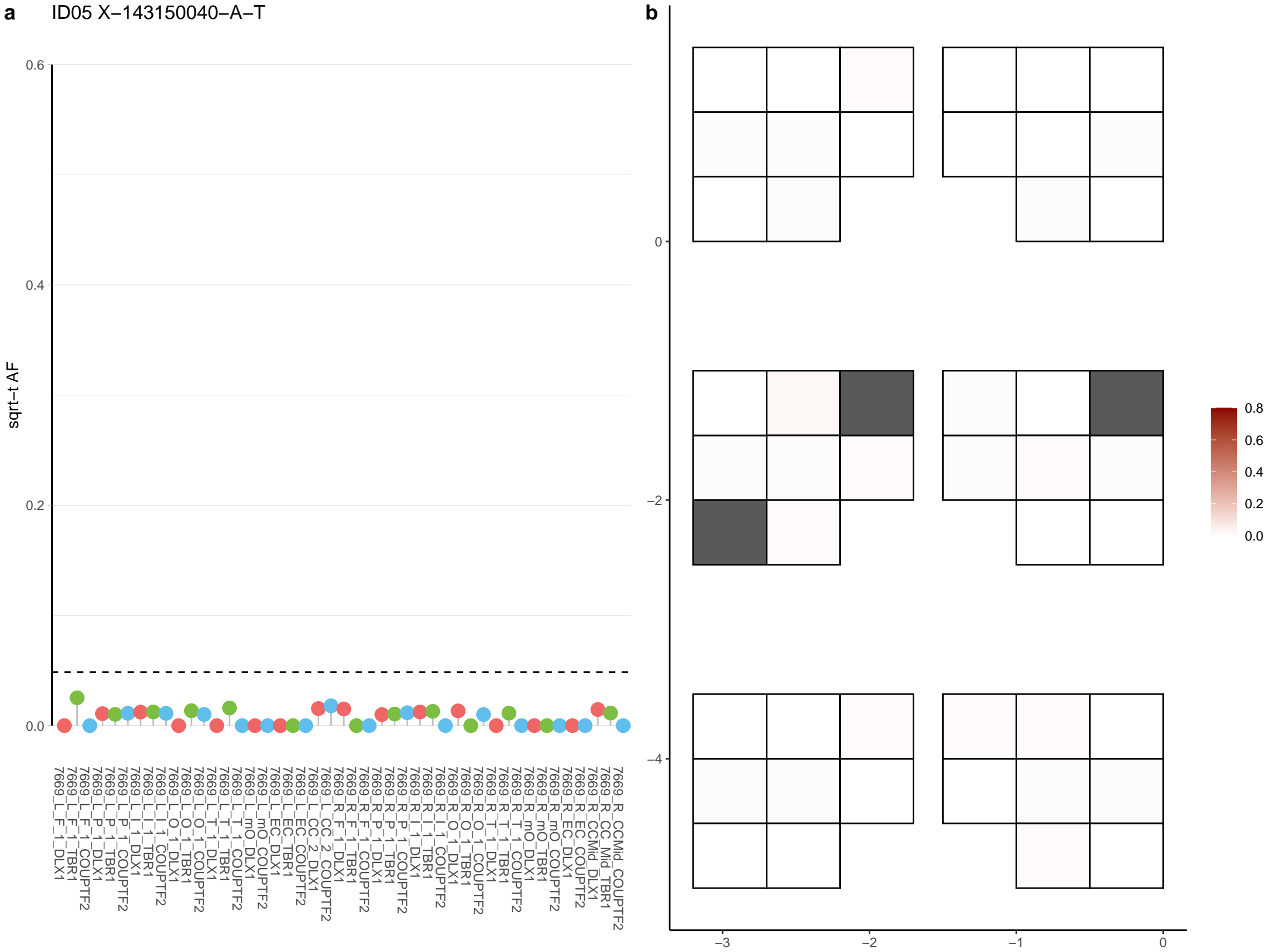

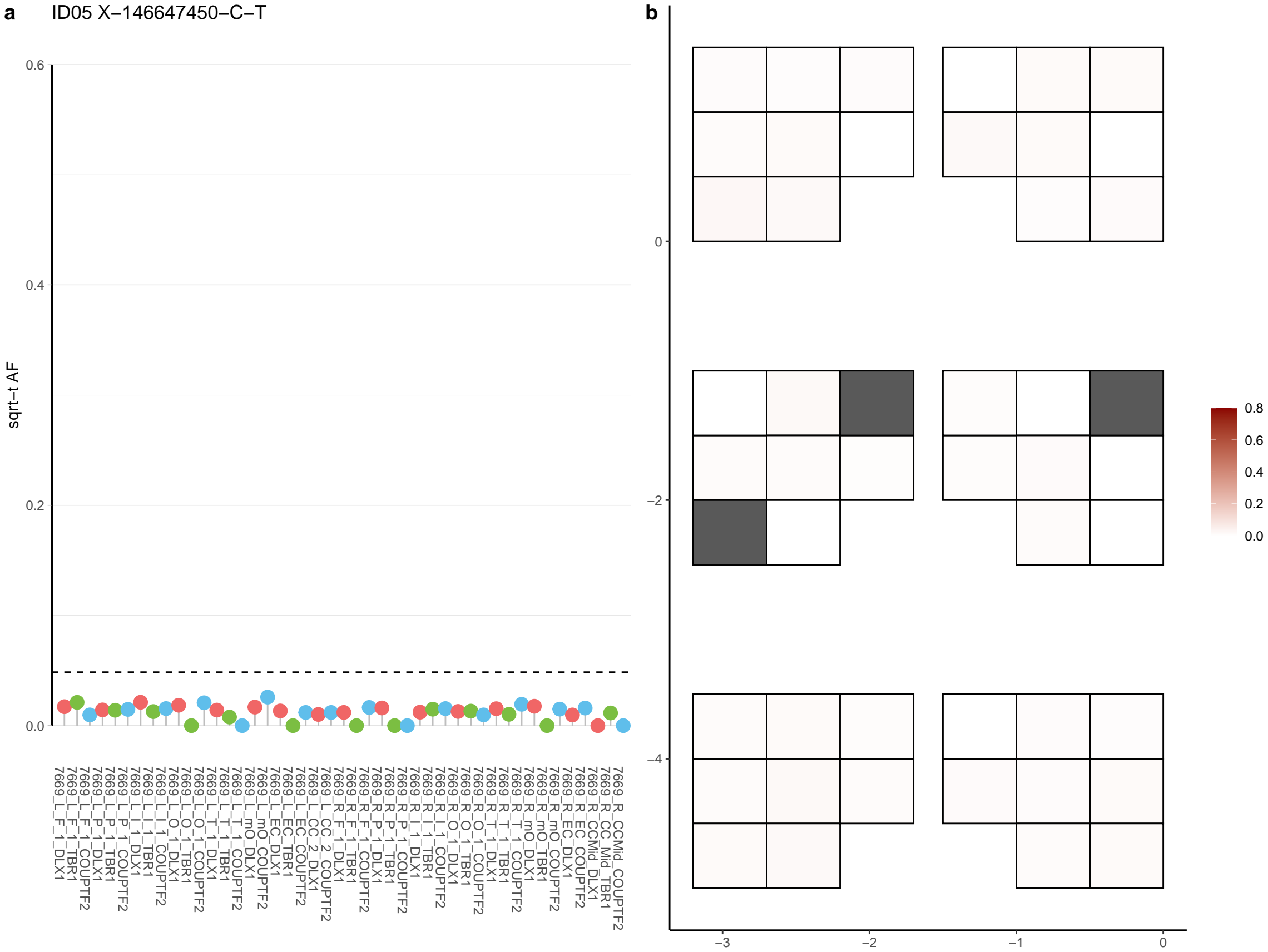

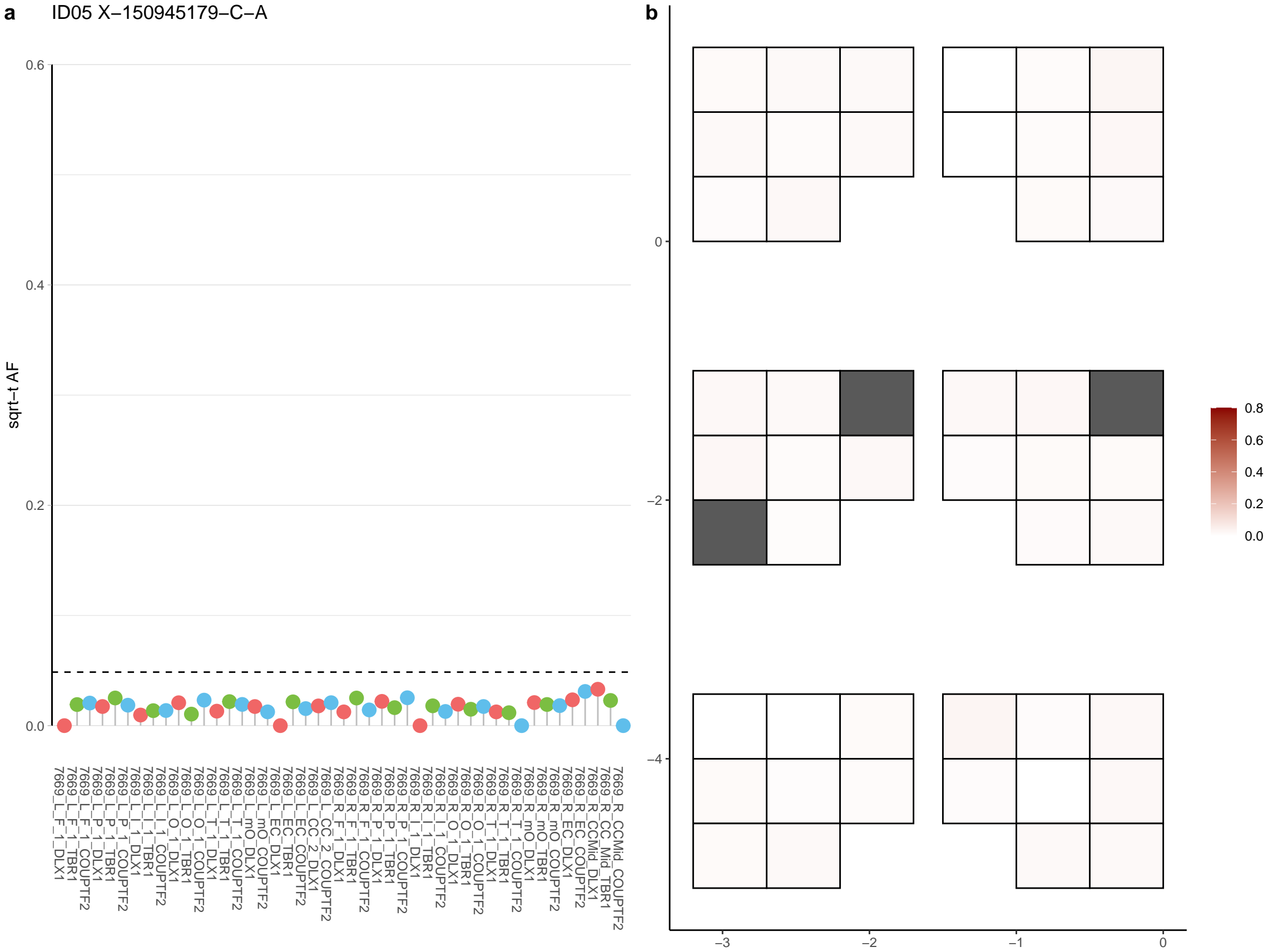

**a** ID05 X-151132909-G-A

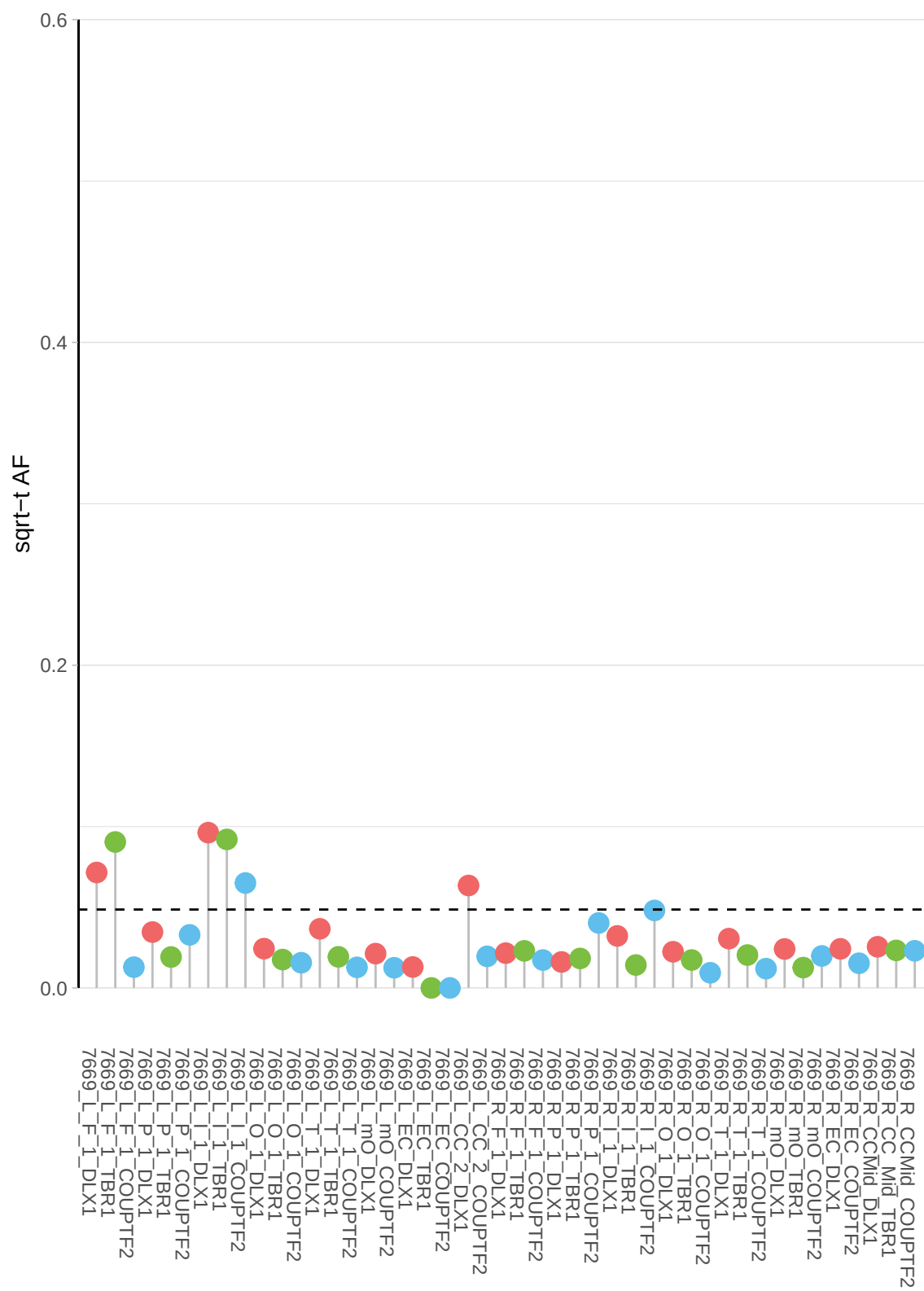

**b**

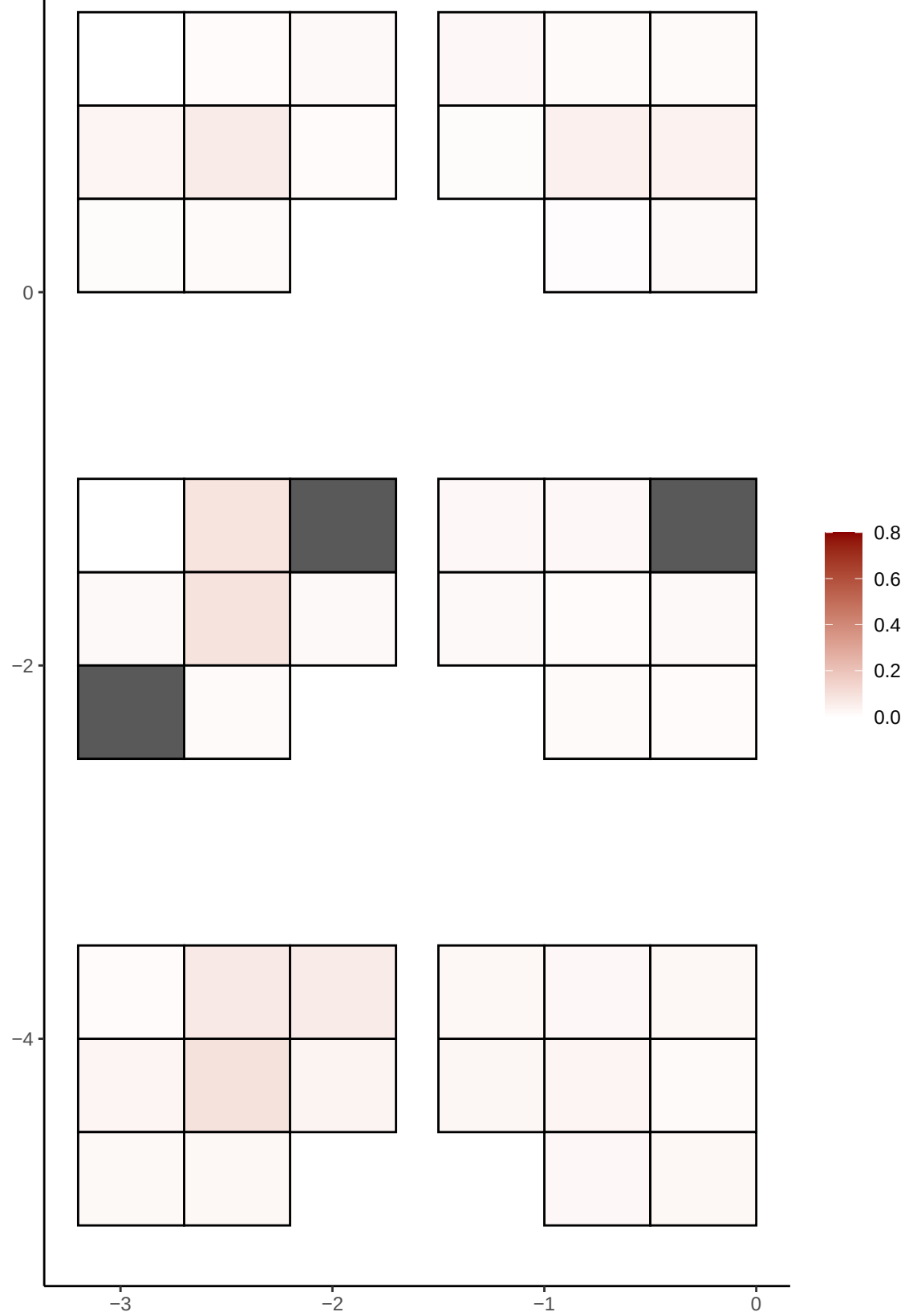

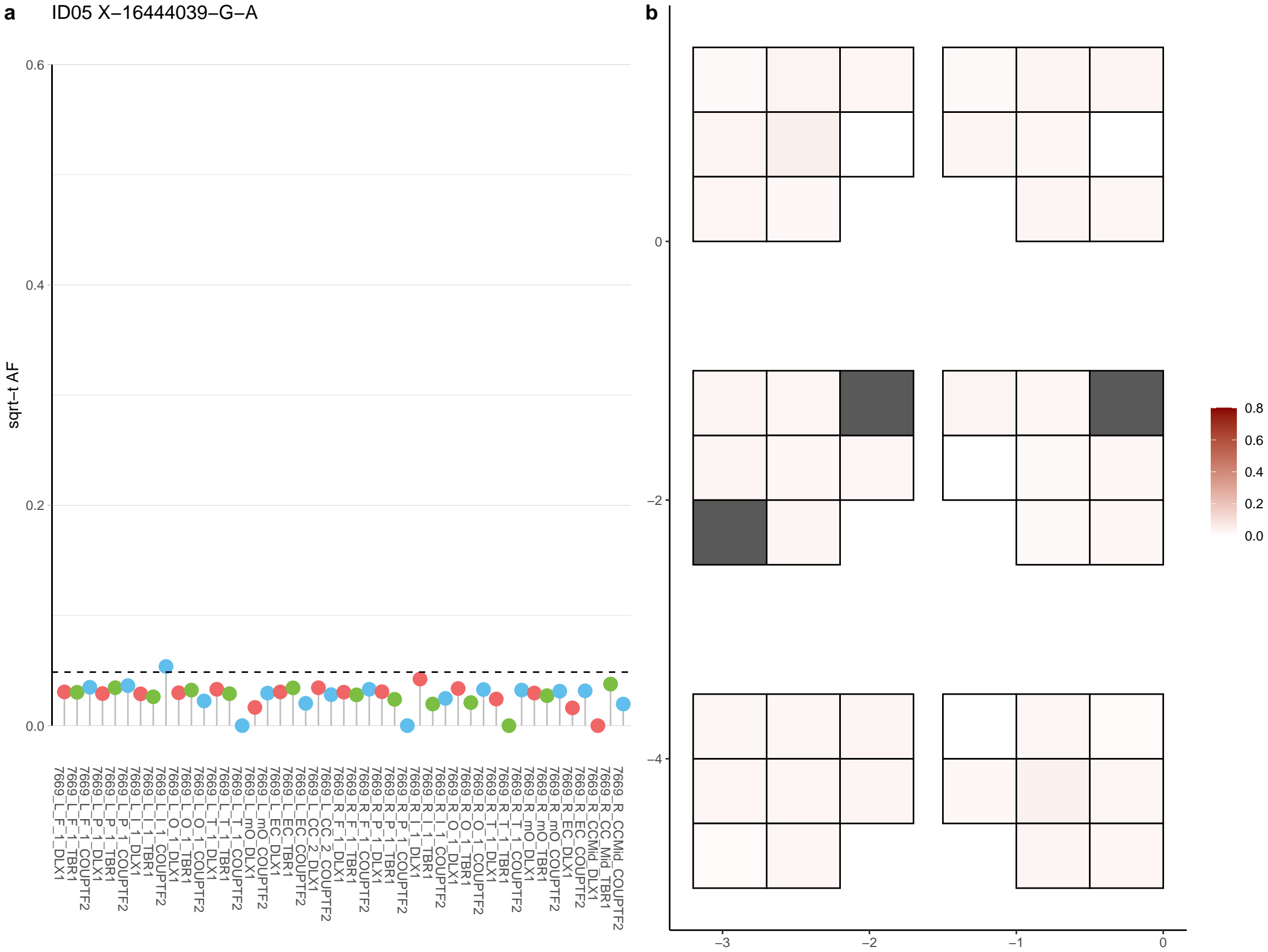

**a** ID05 X-17021591-G-A

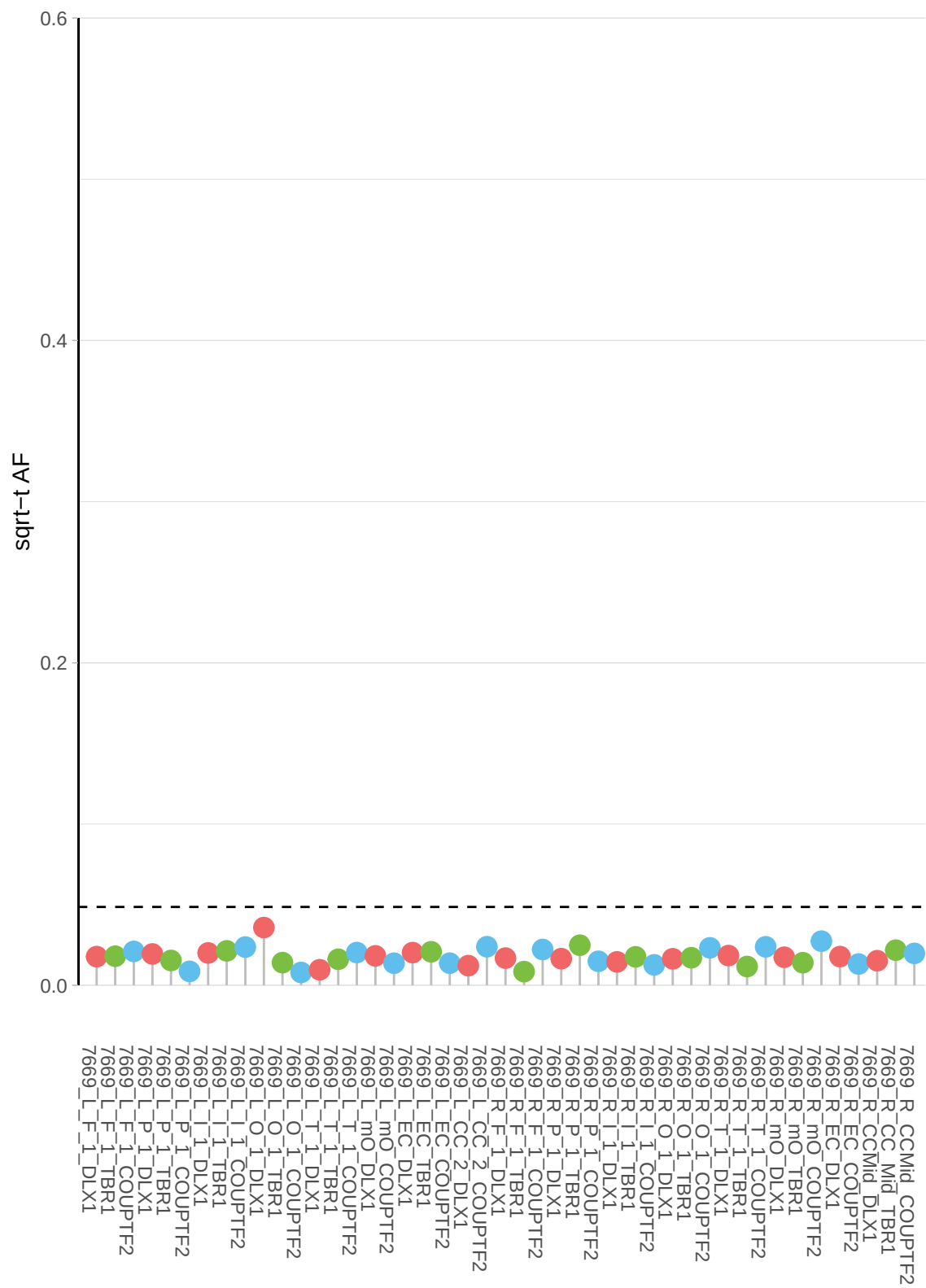

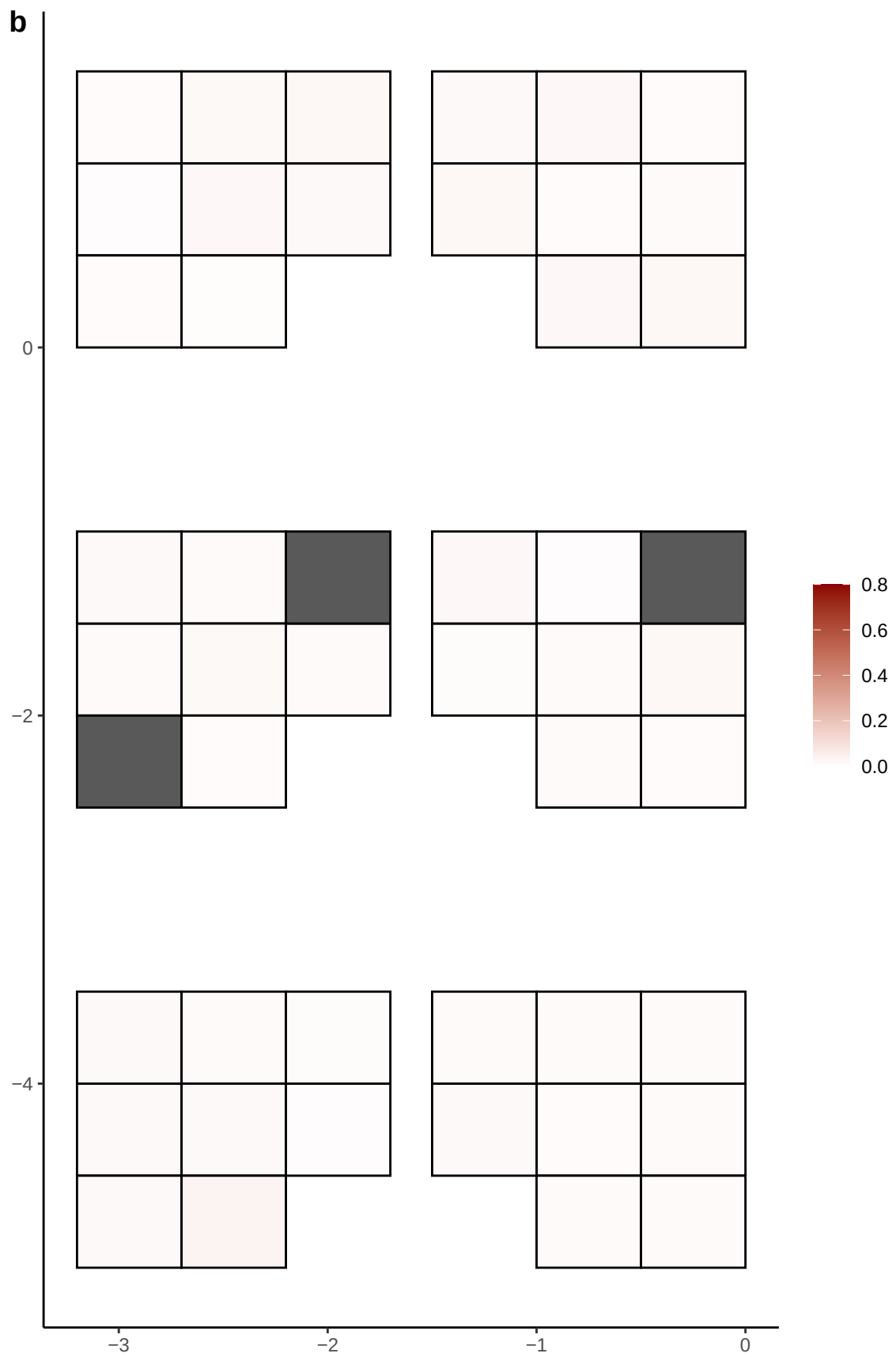

**a** ID05 X-17091941-T-G

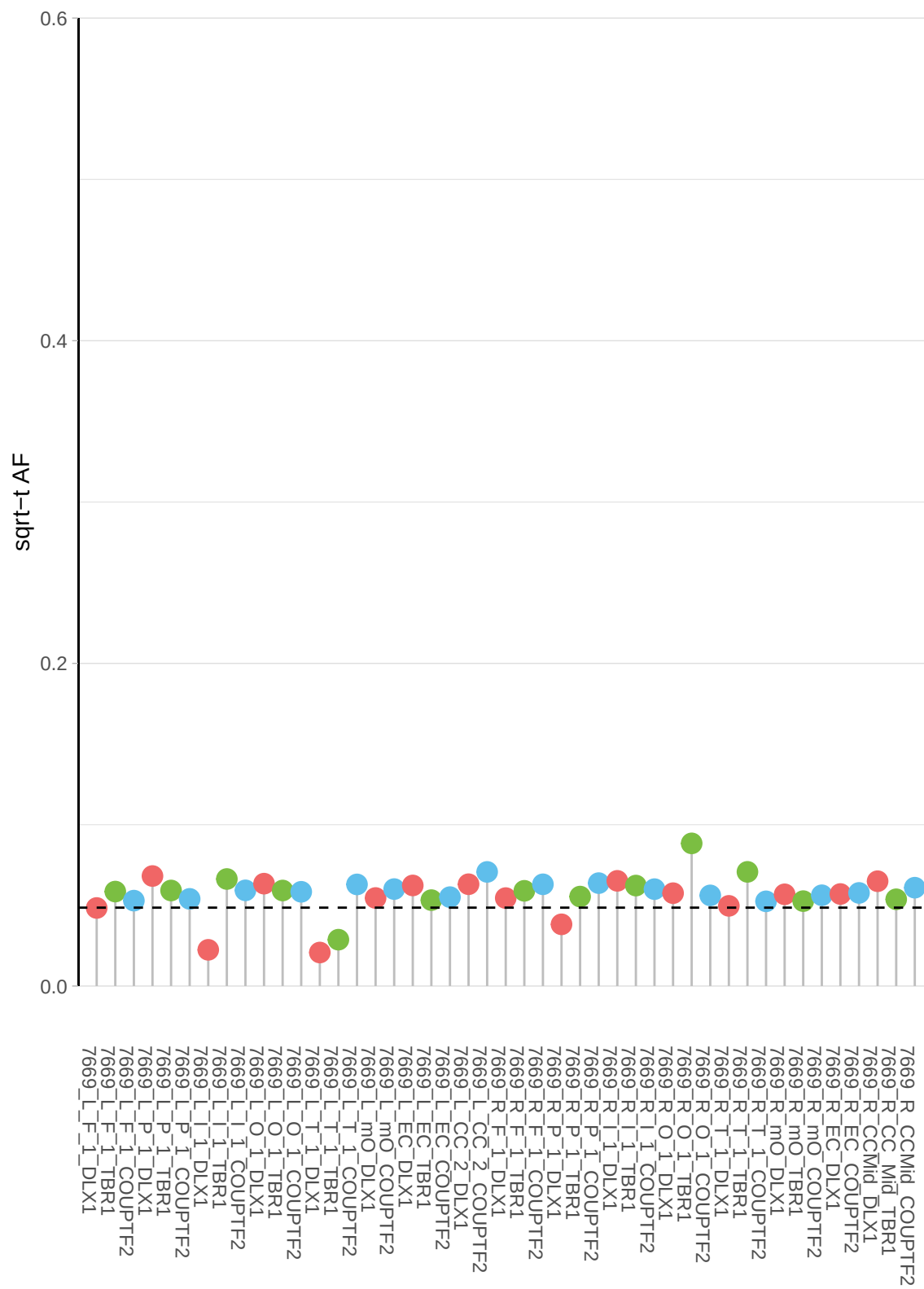

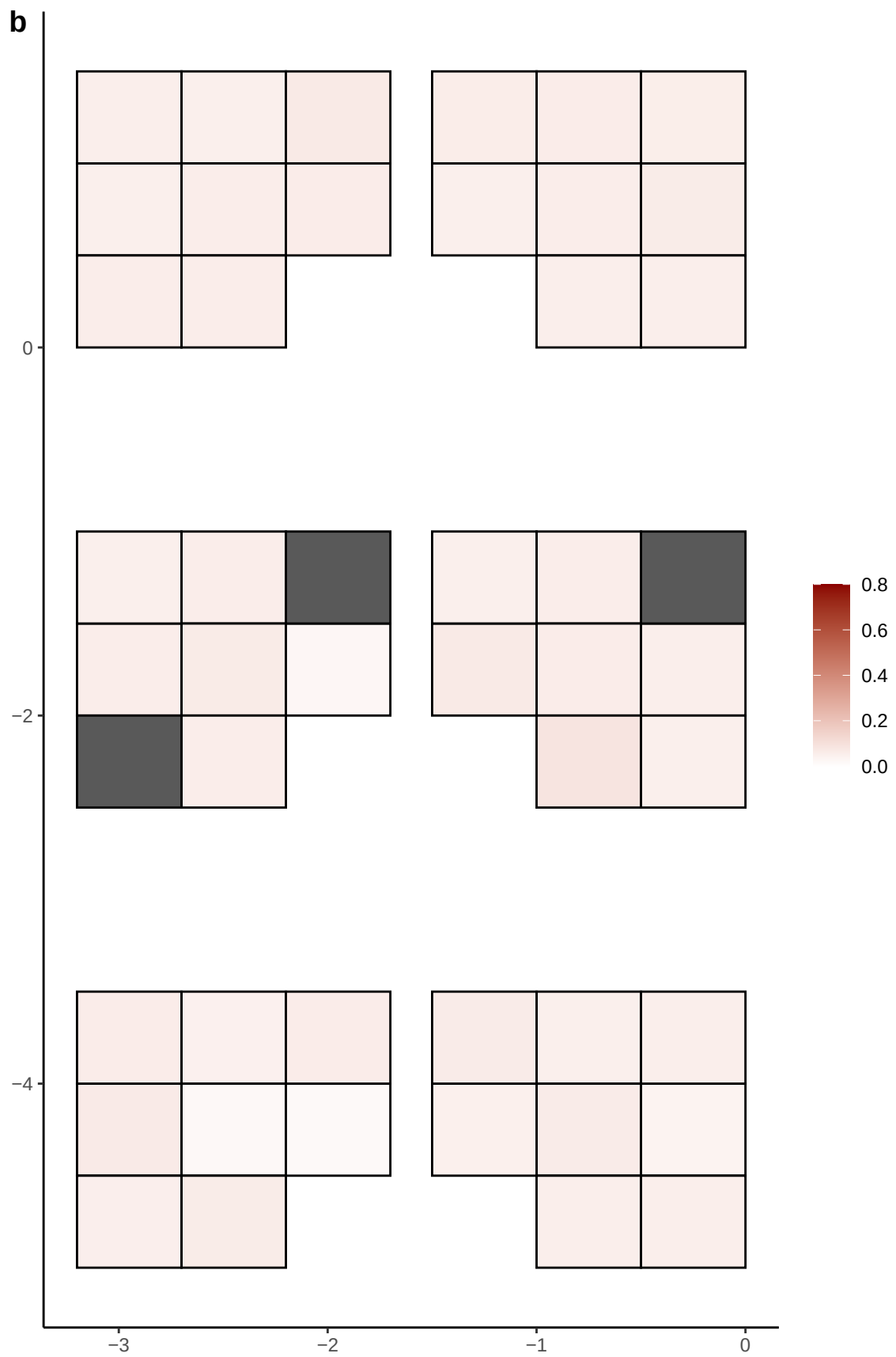

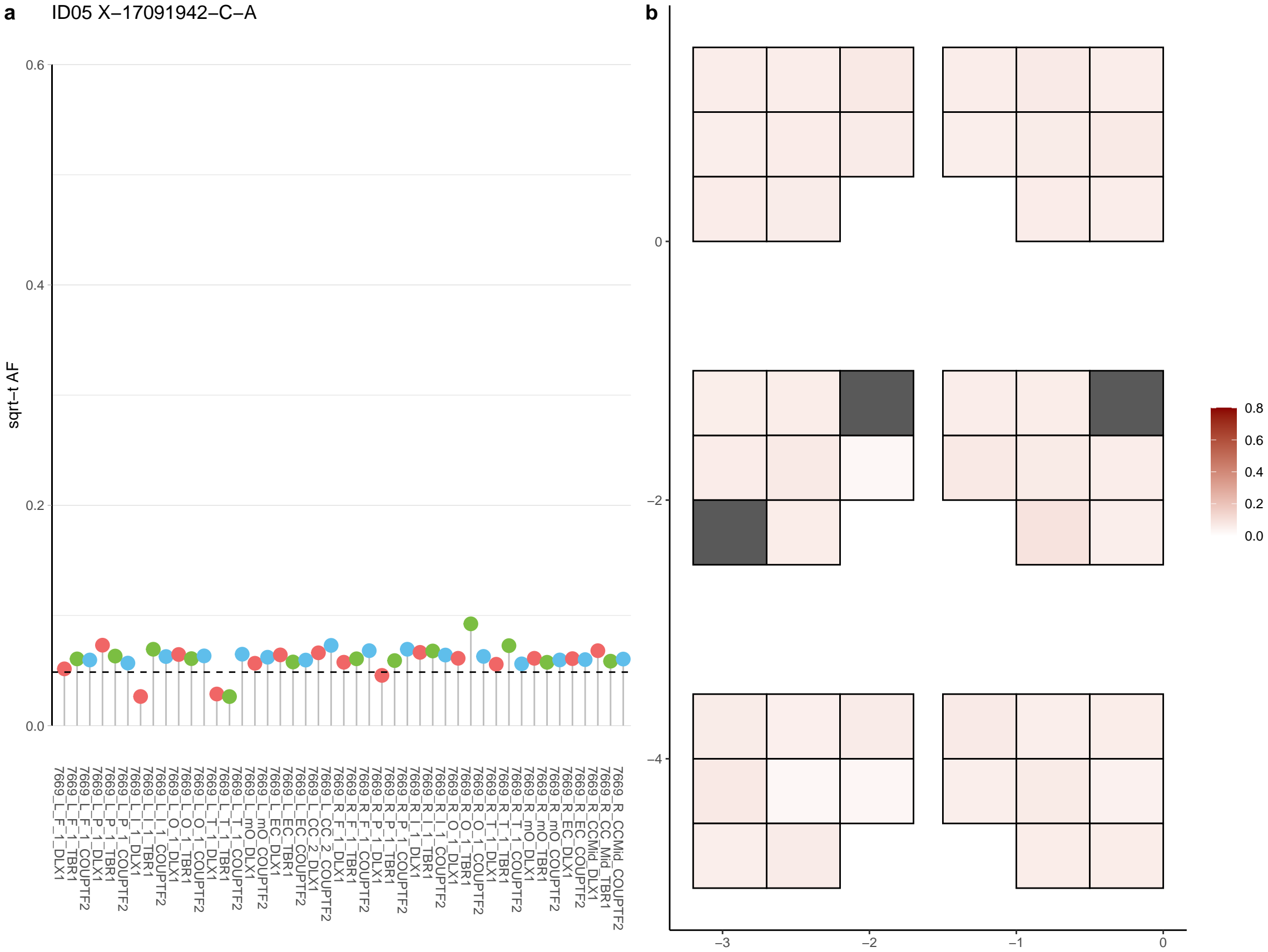

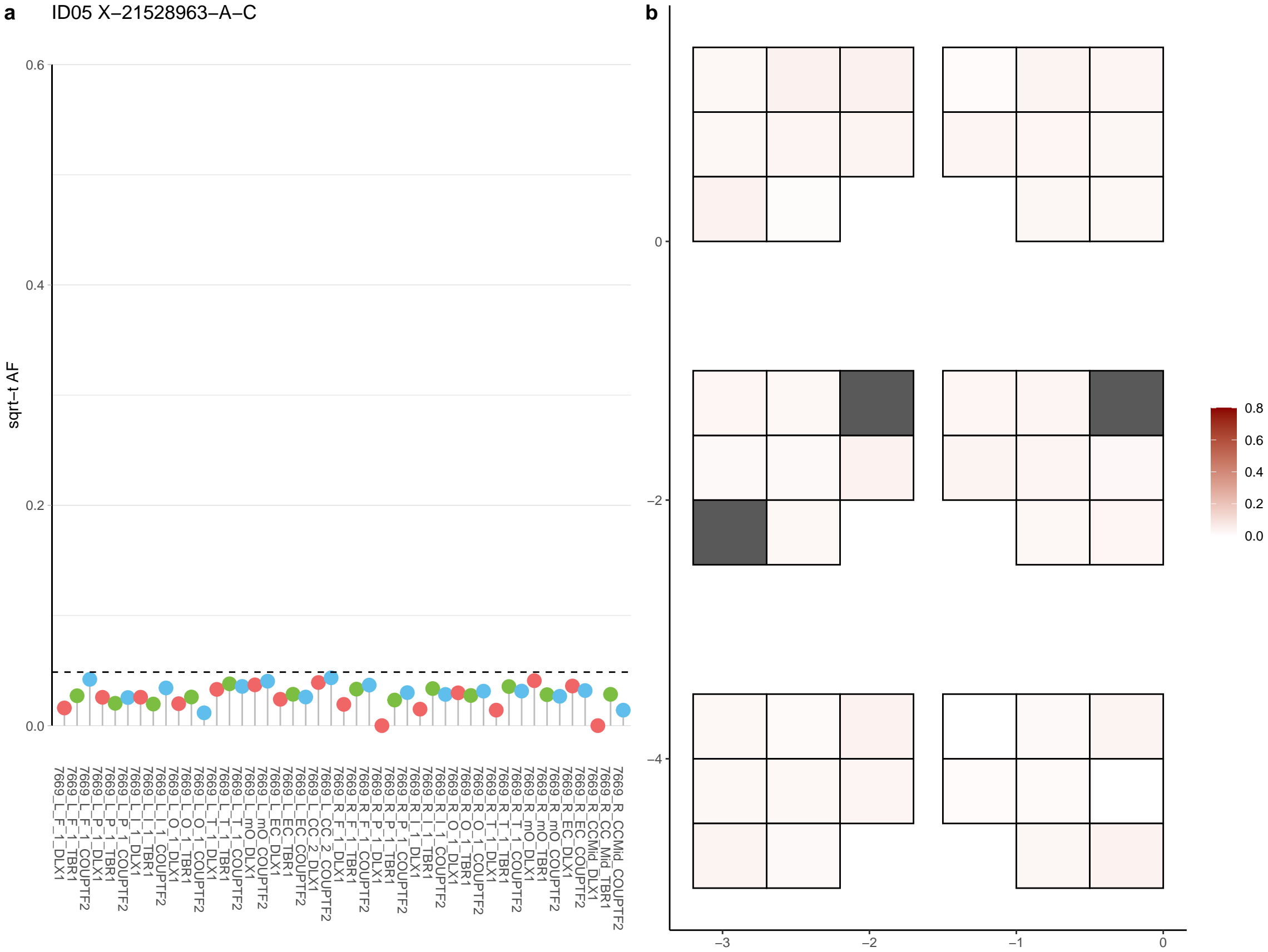

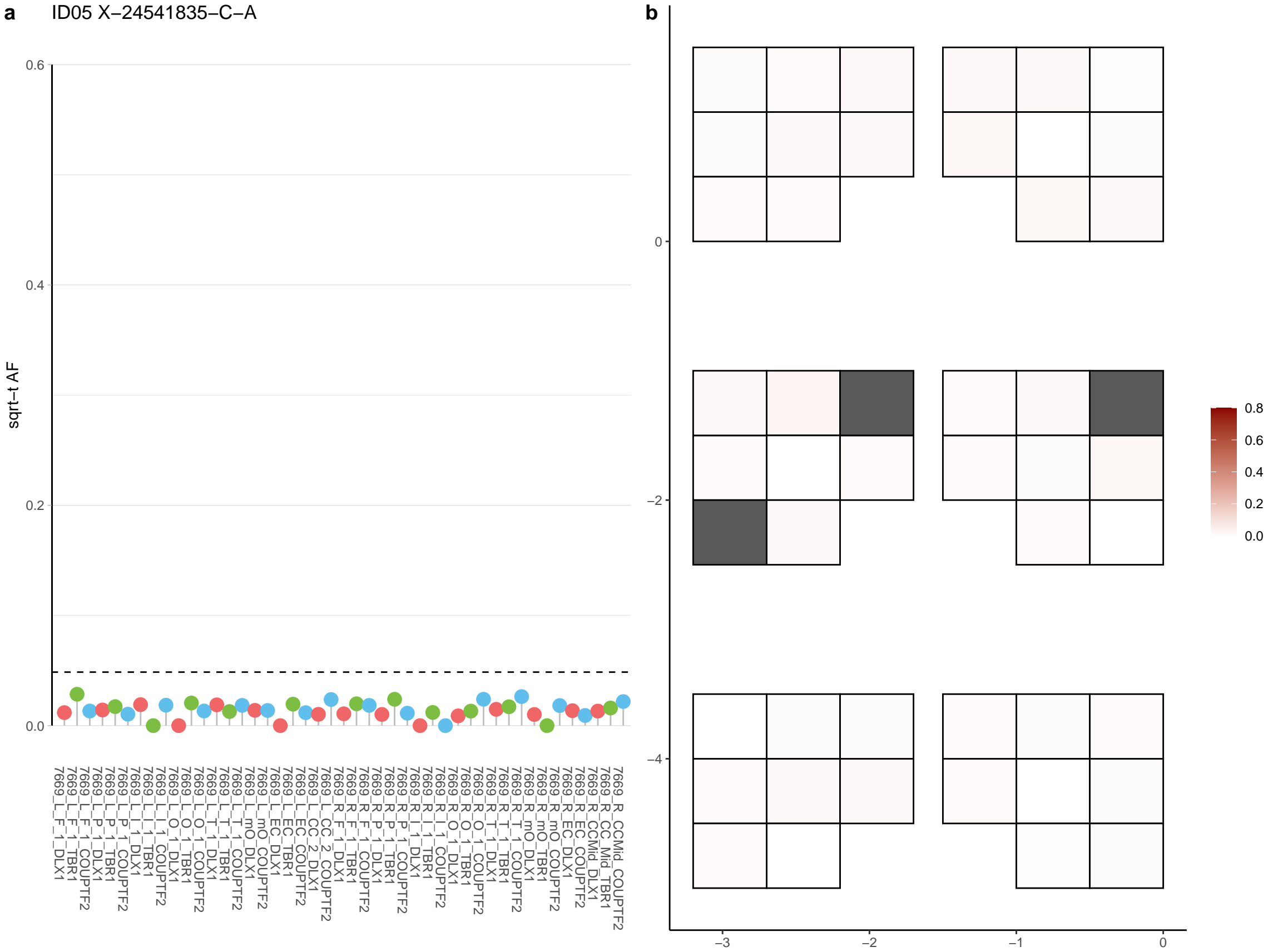

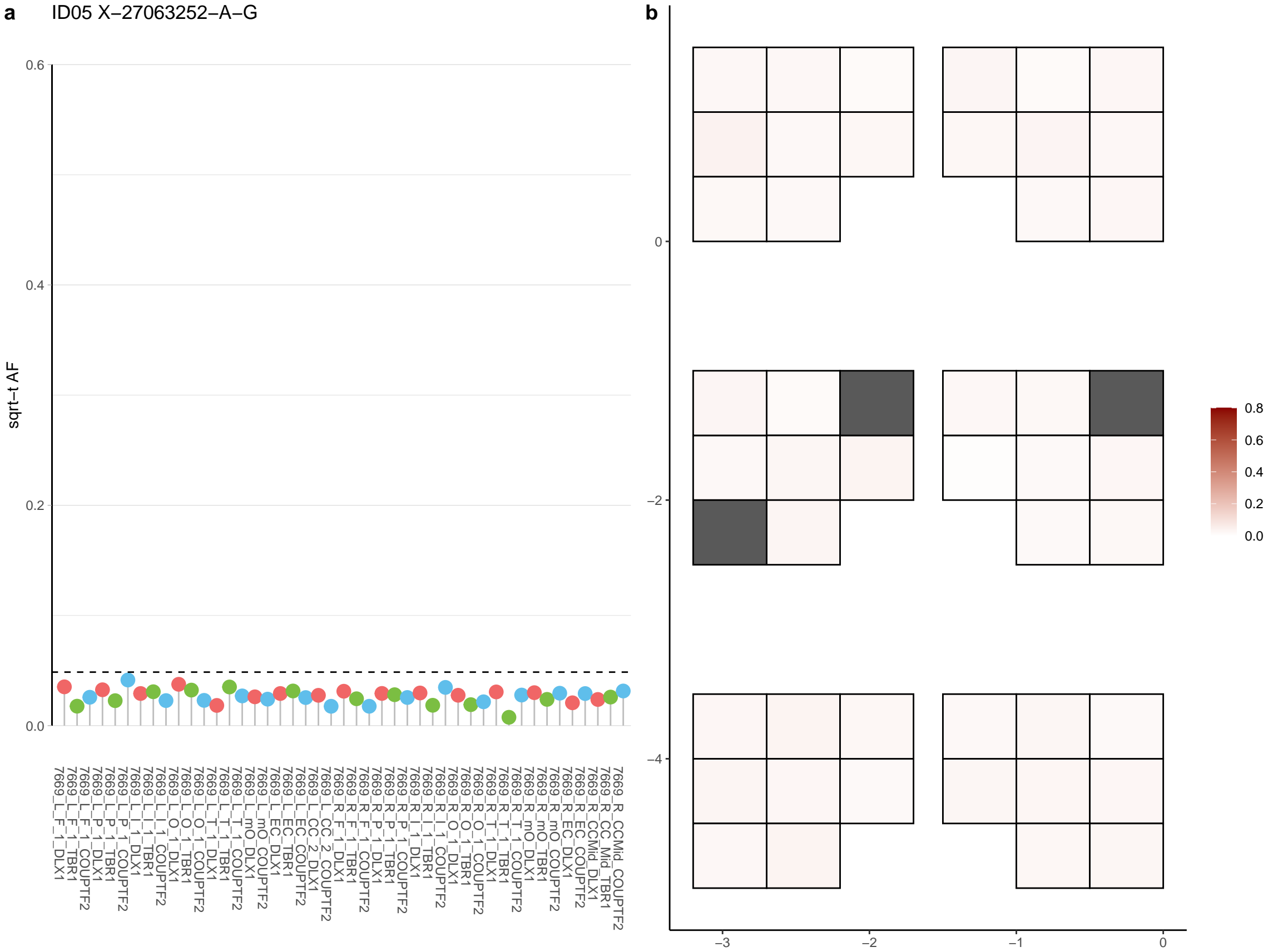

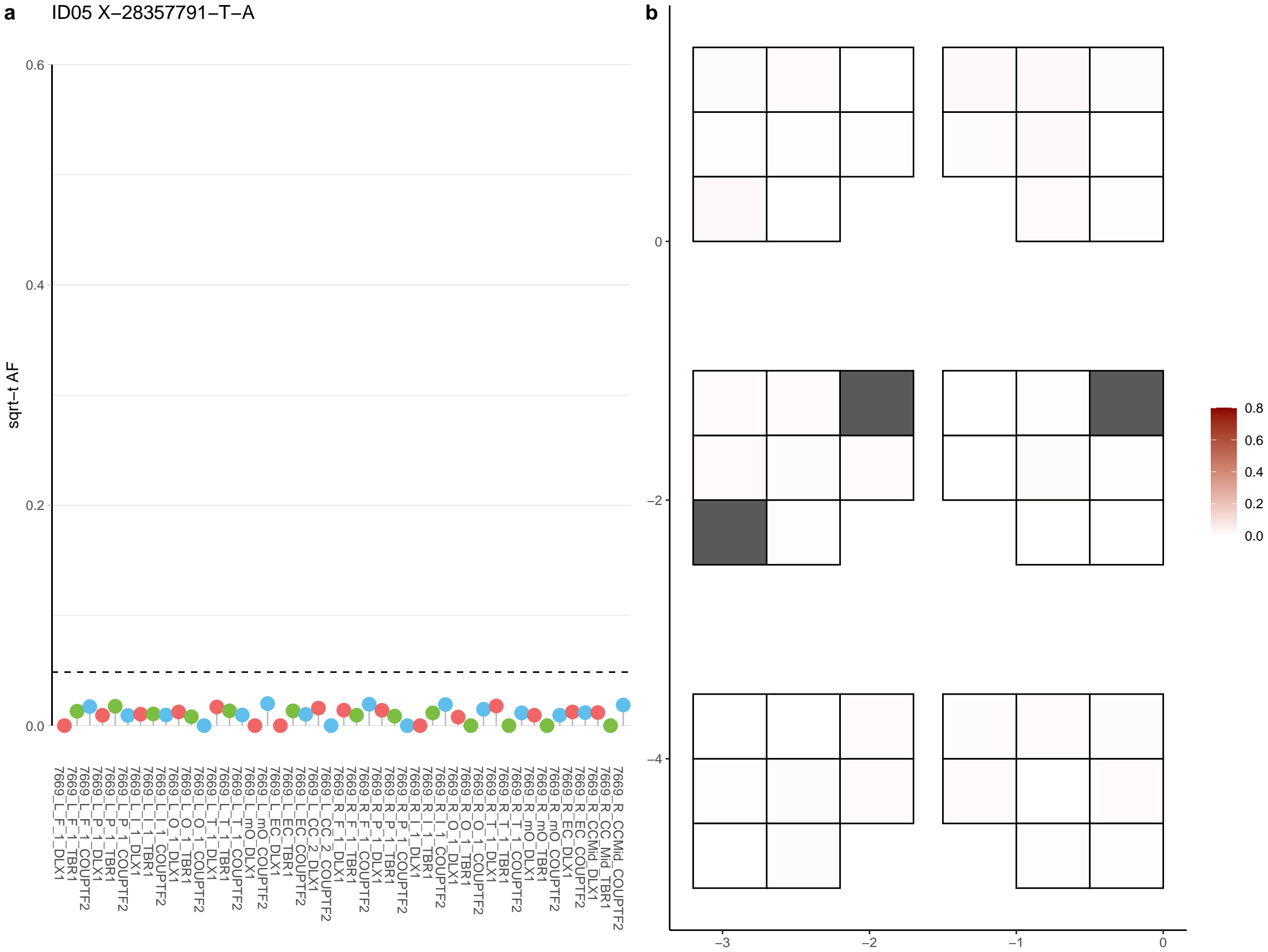

**a** ID05 X-32465017-G-A

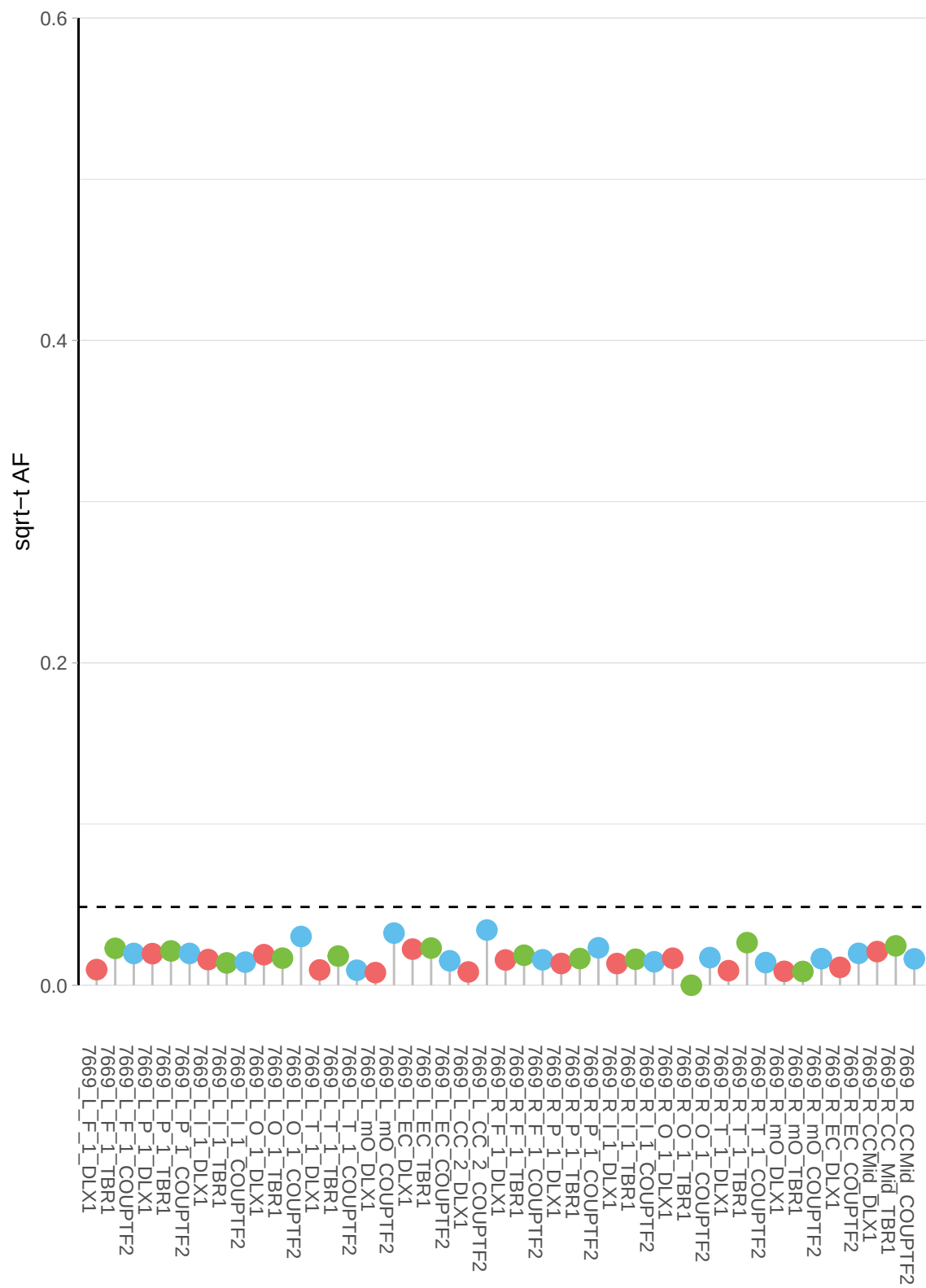

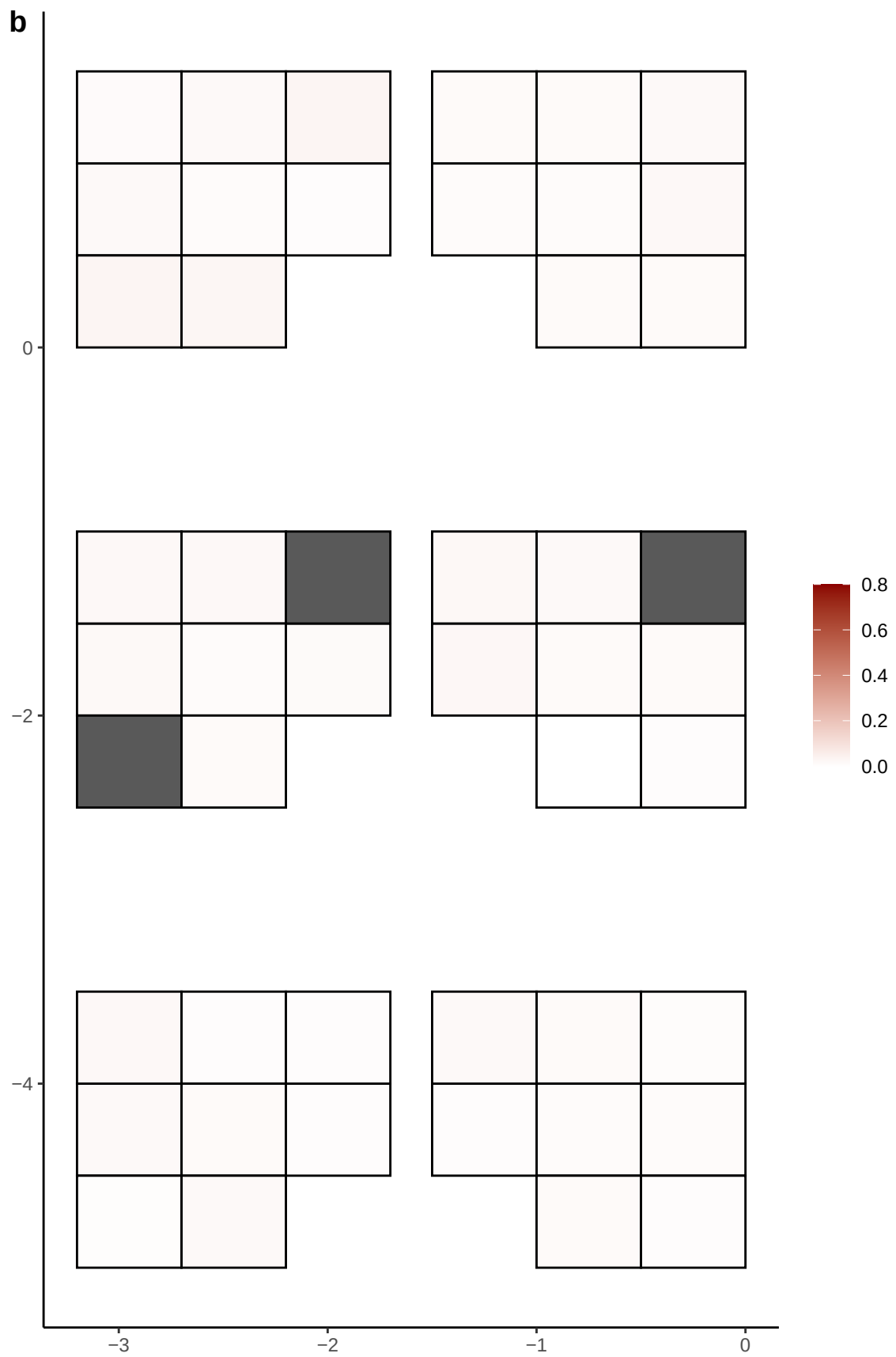

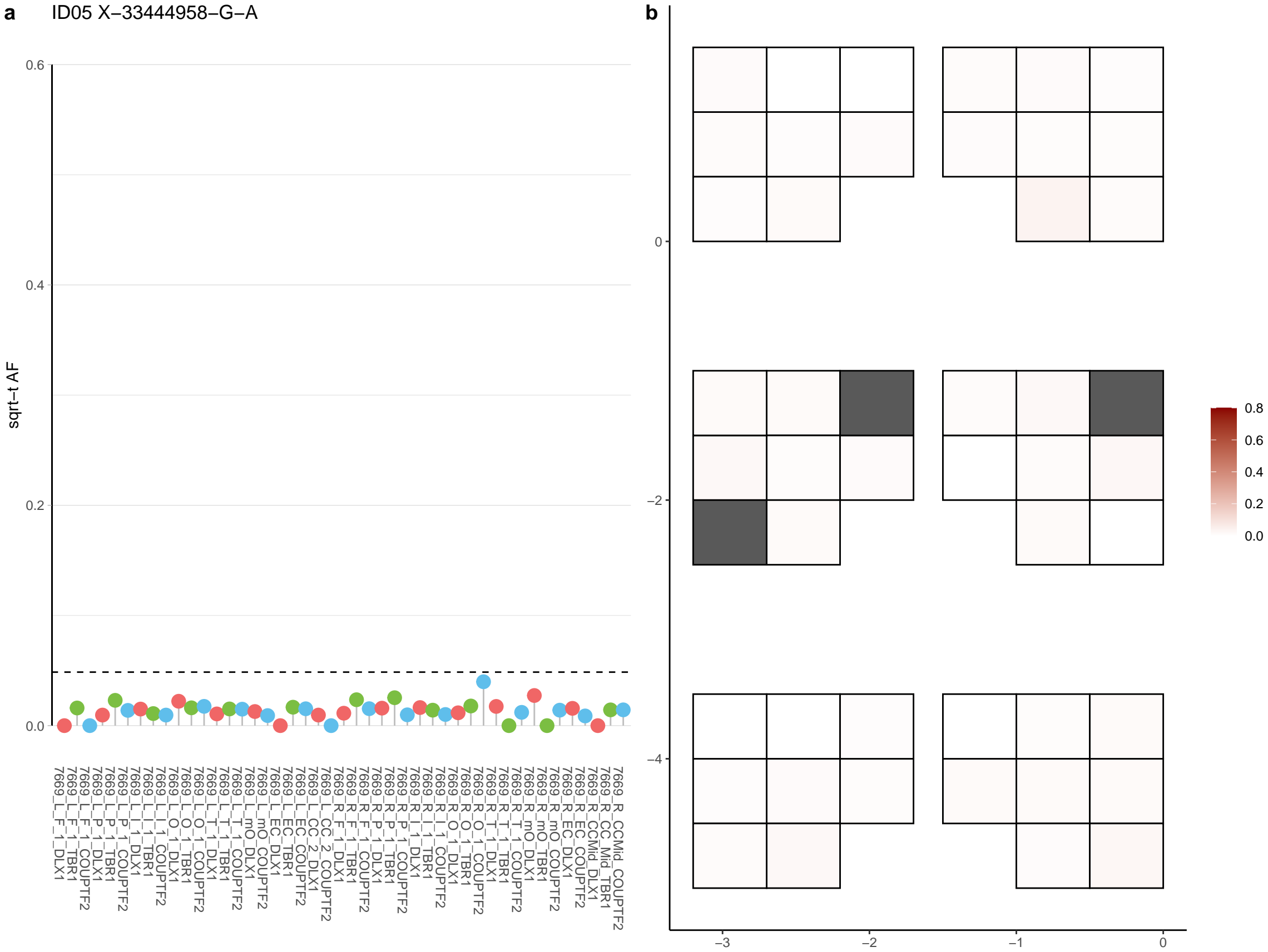

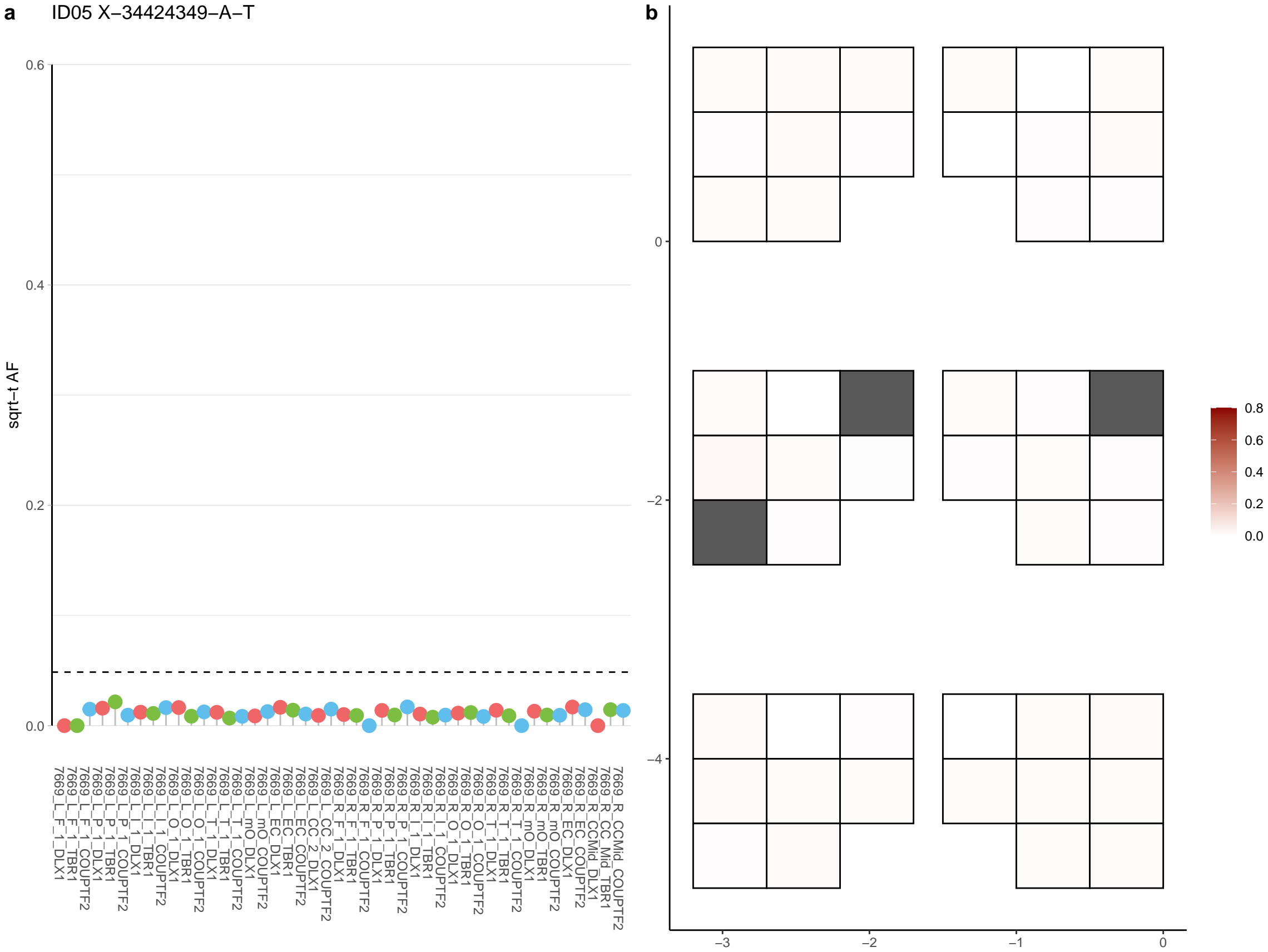

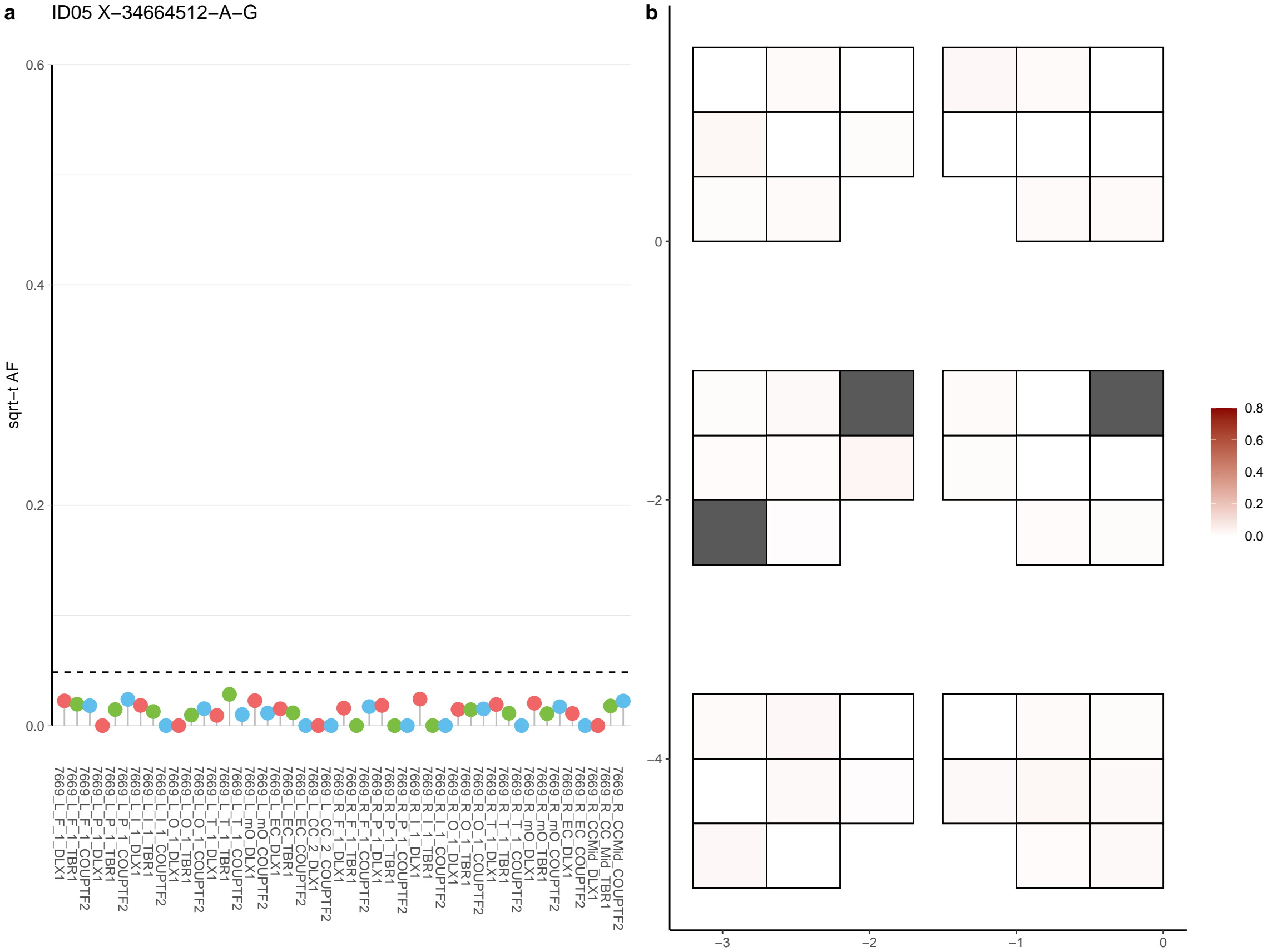

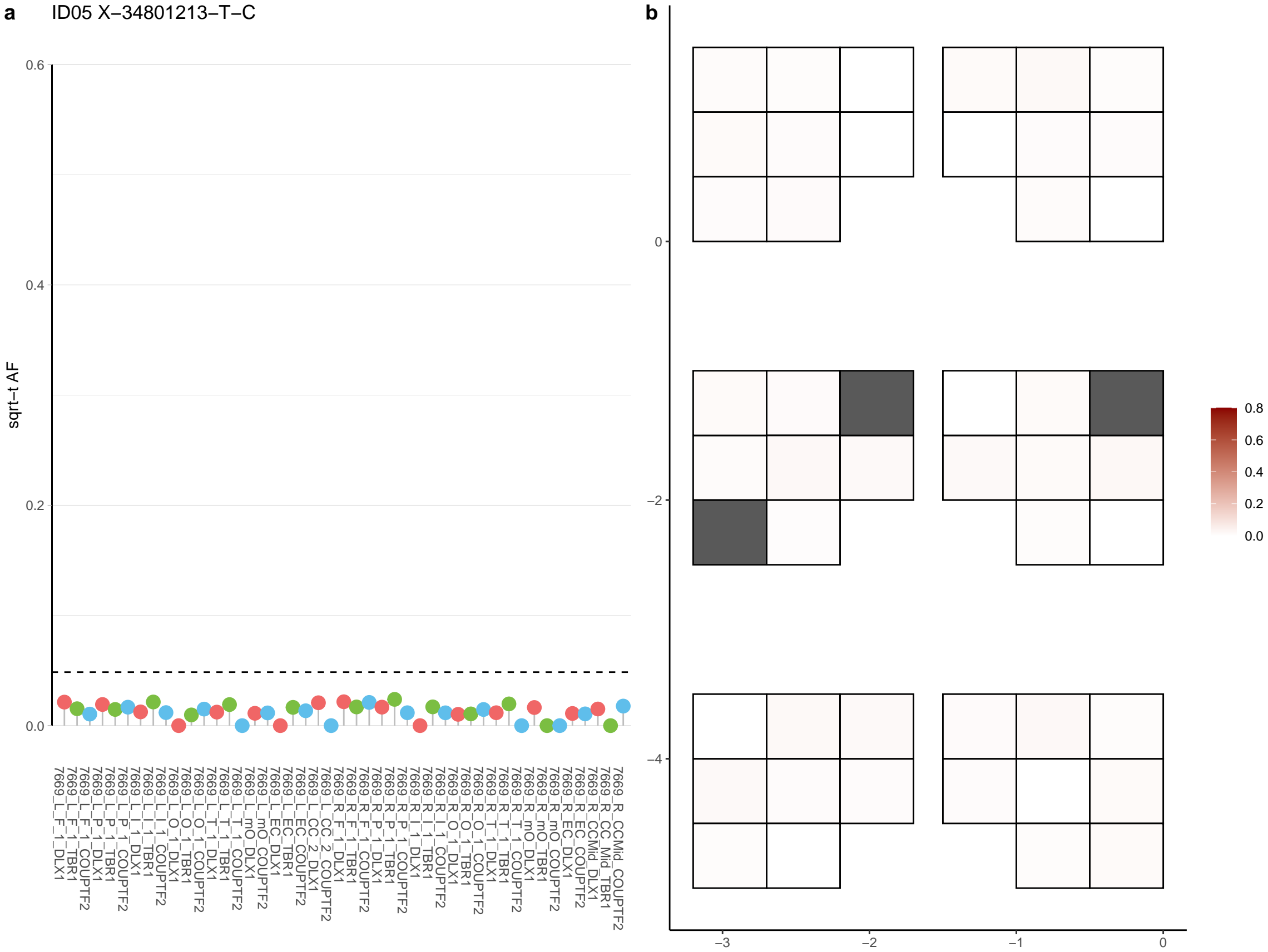

**a** ID05 X-35362338-C-T

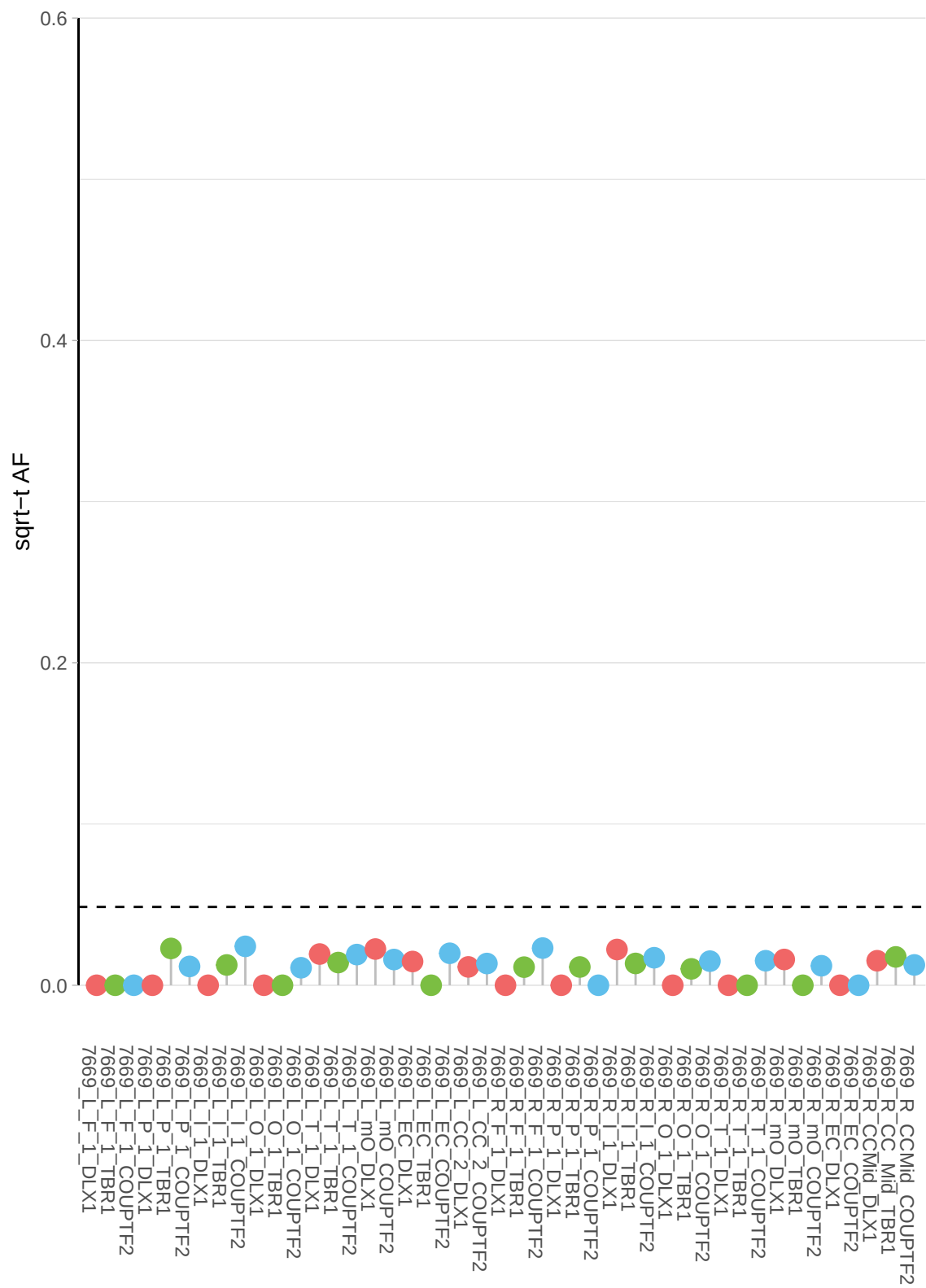

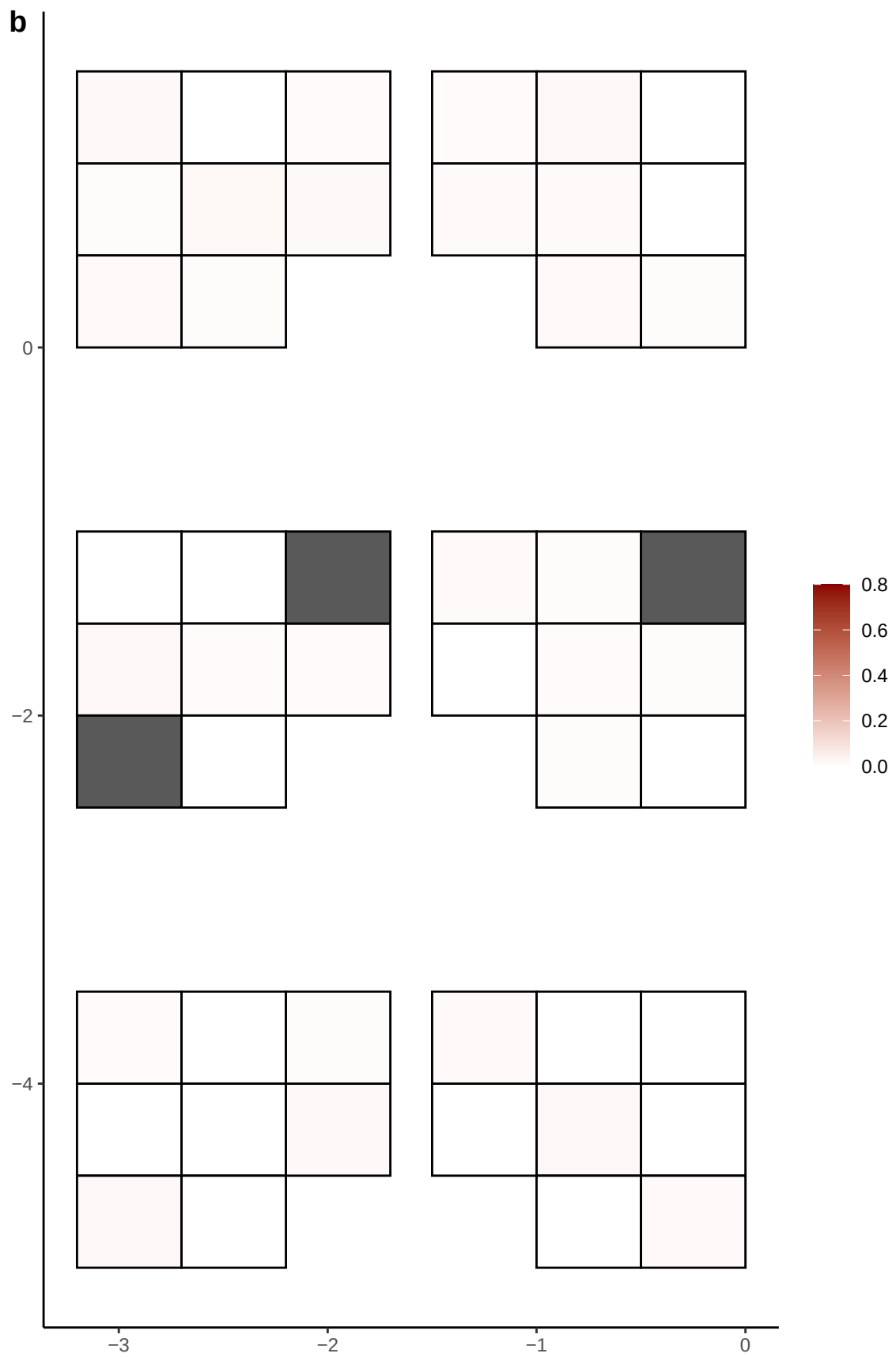

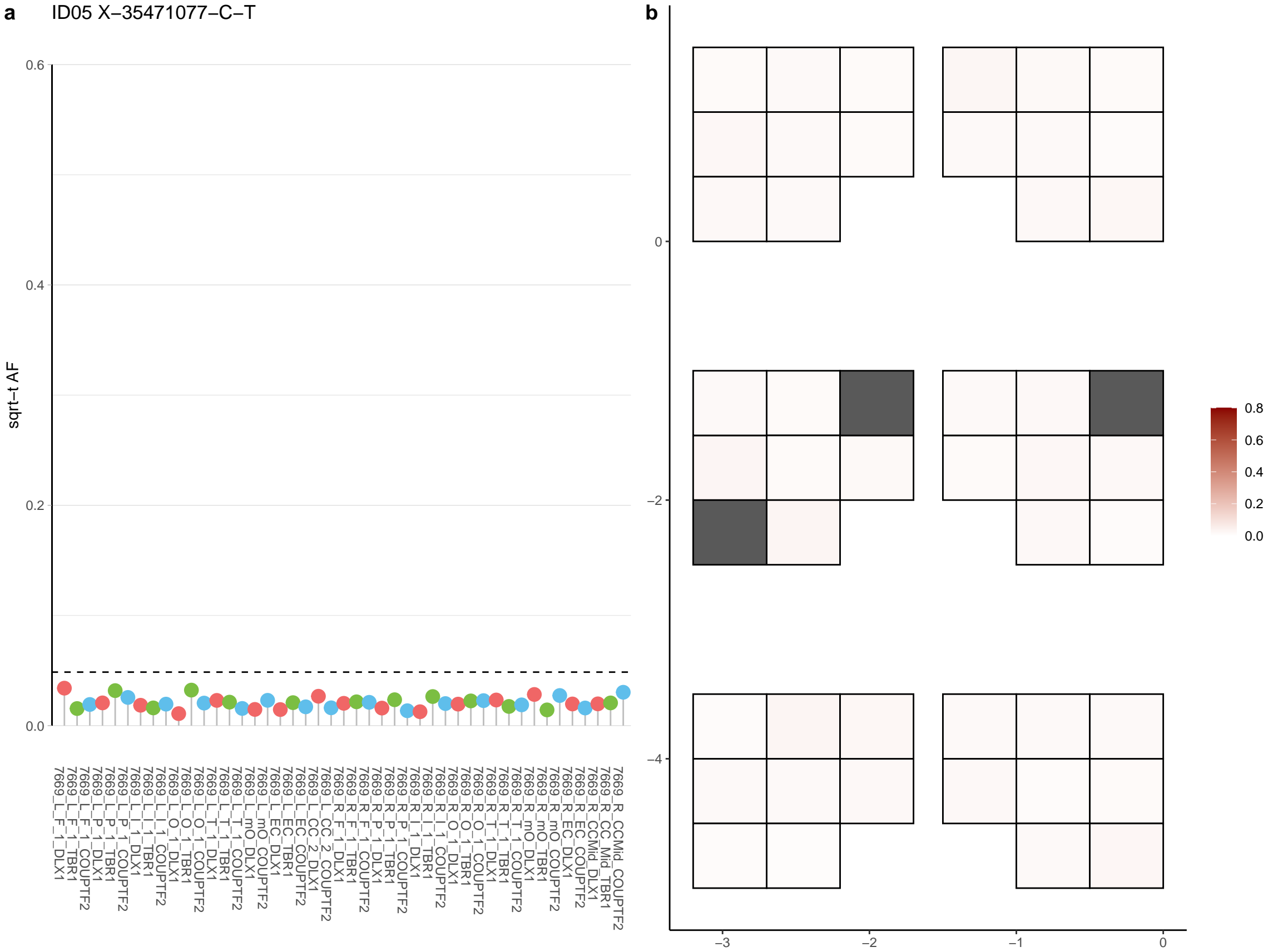

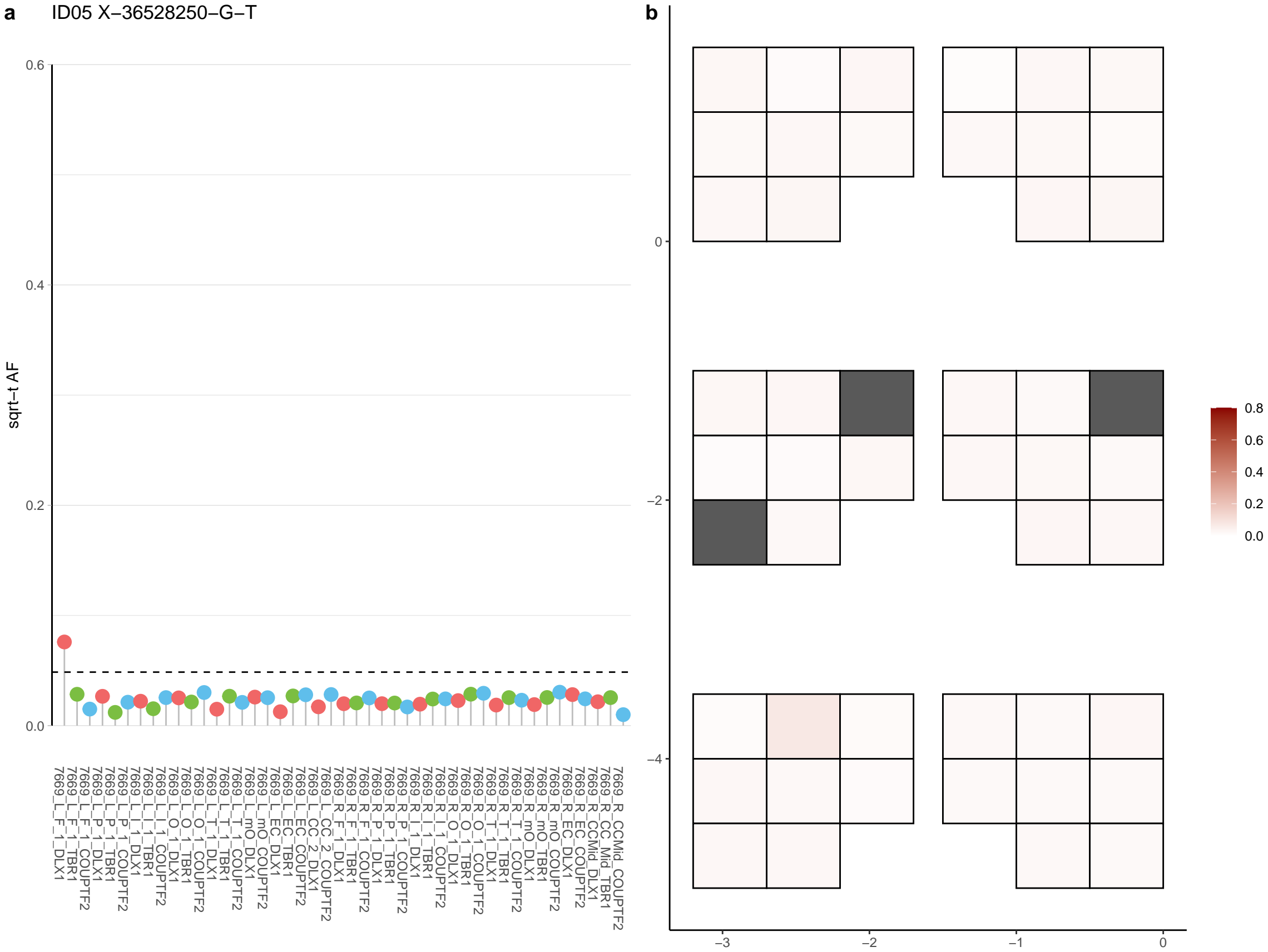

**a** ID05 X-39832087-T-A

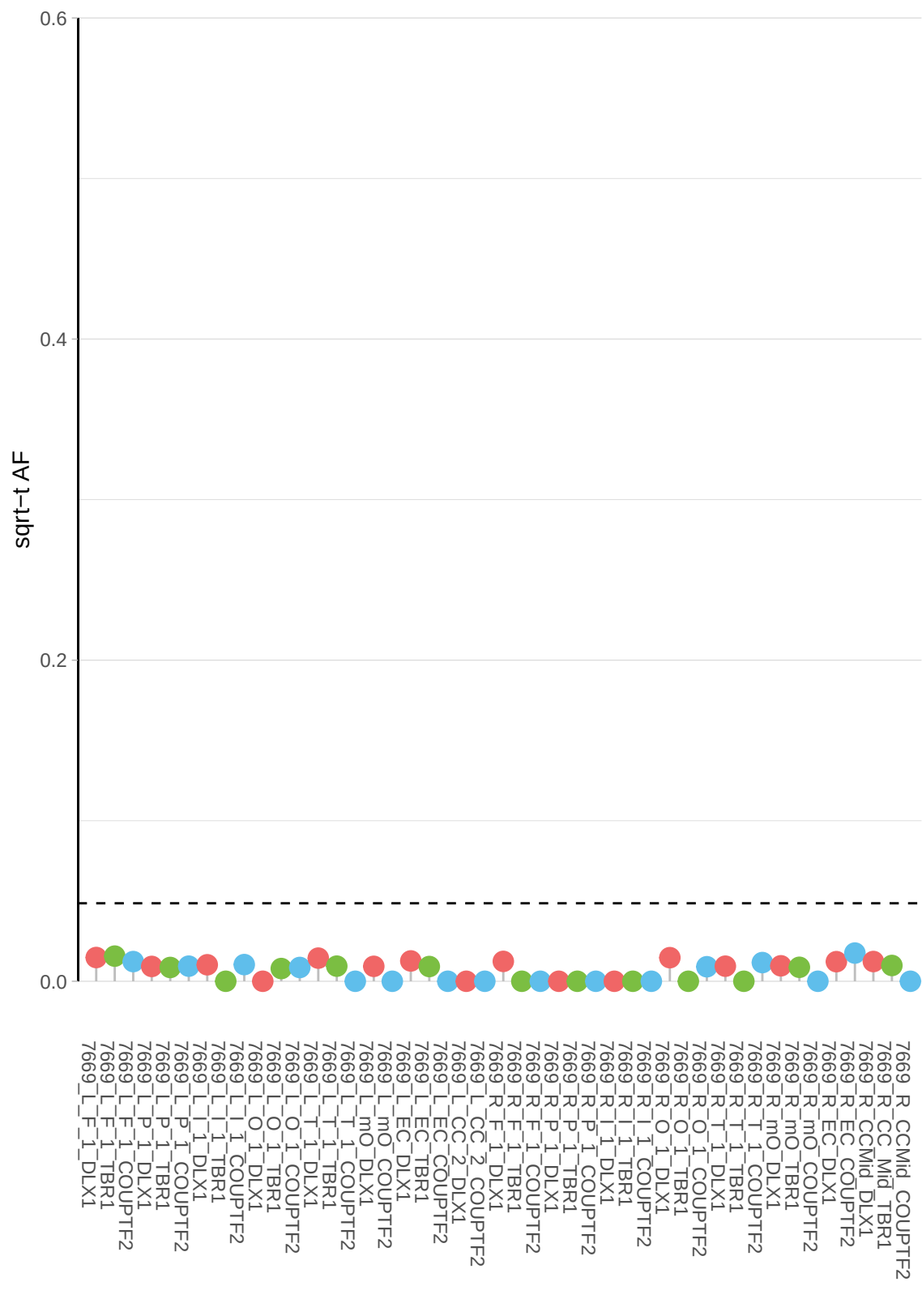

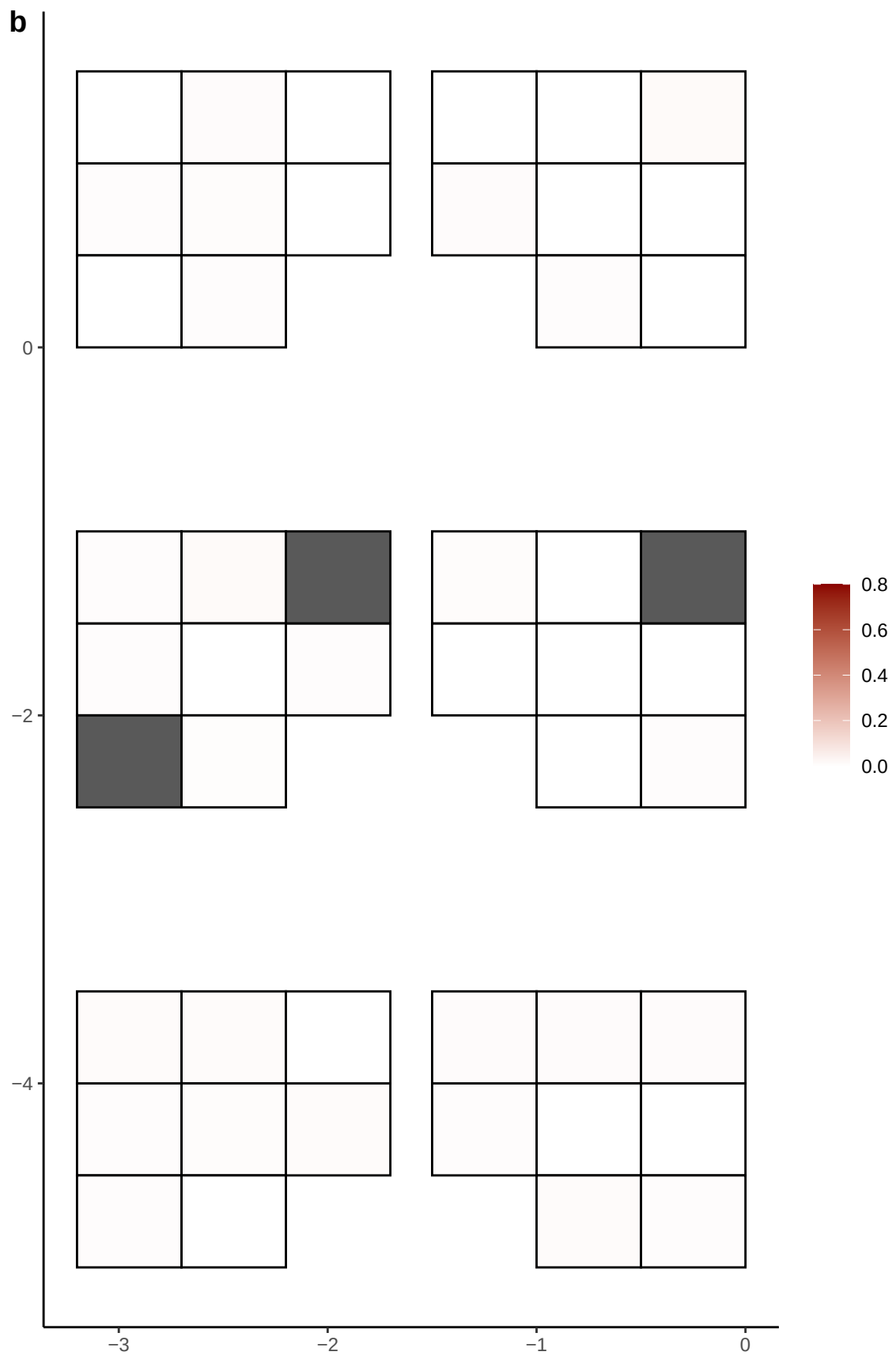

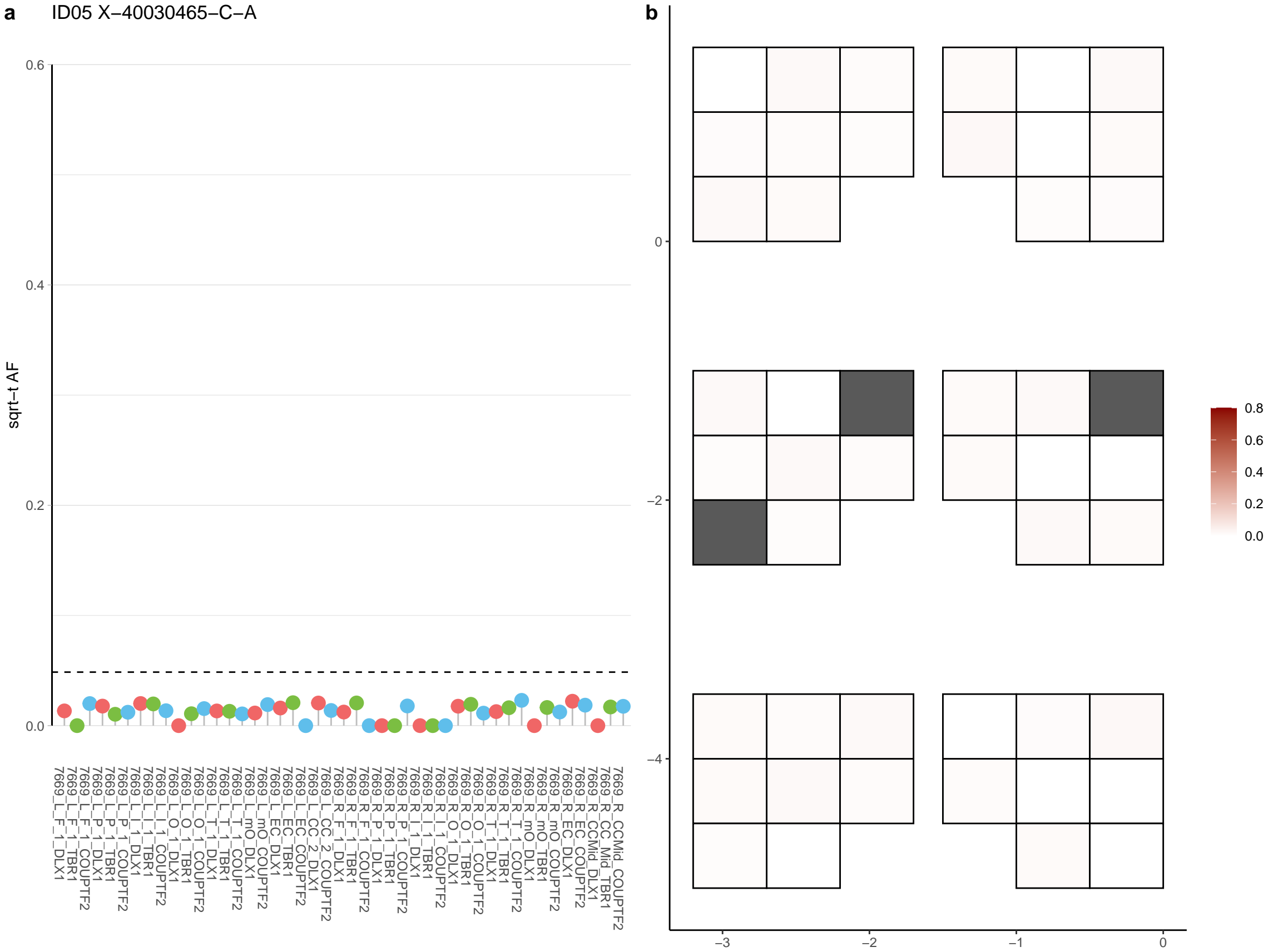

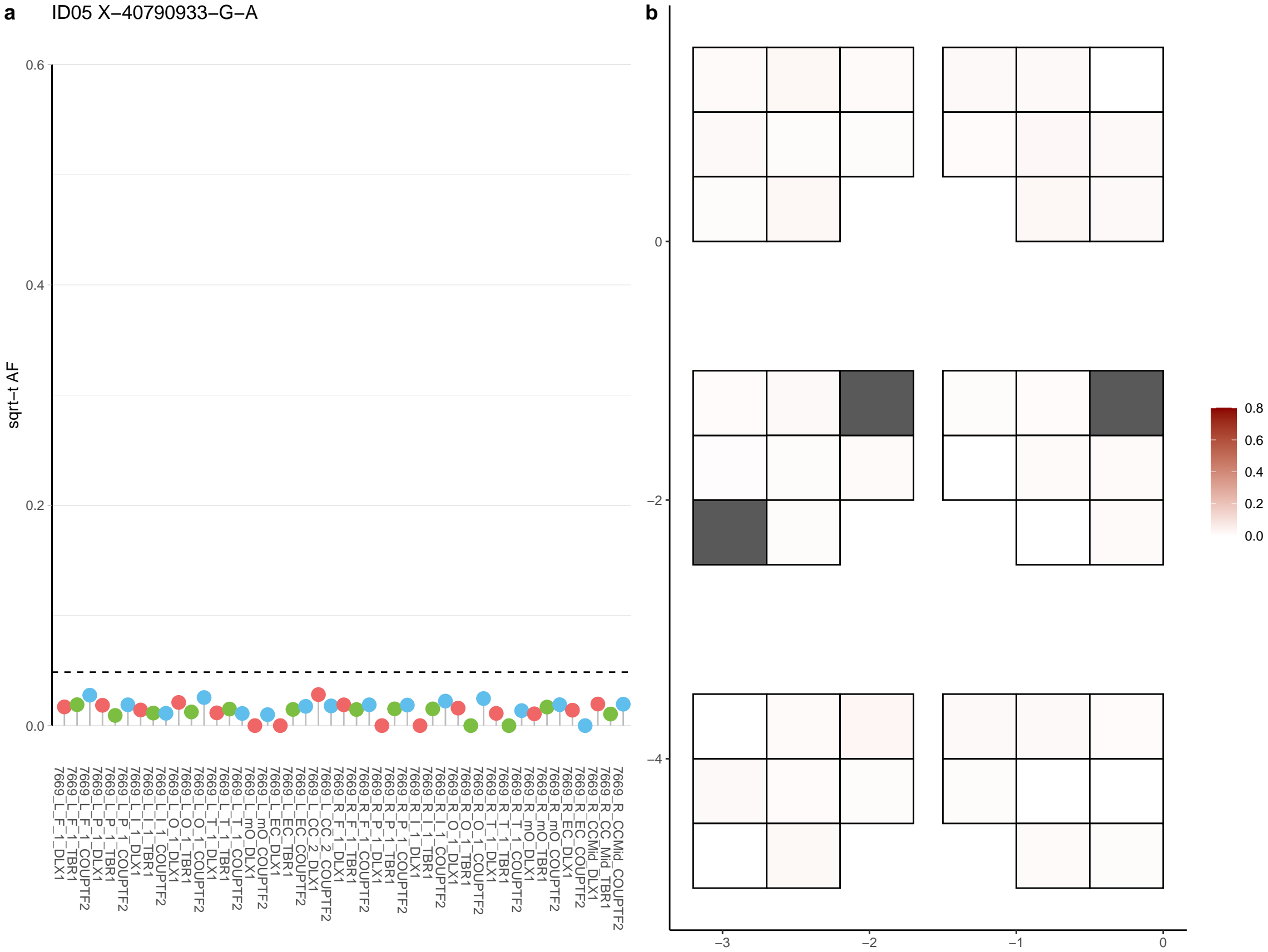

**a** ID05 X-41029320-G-A

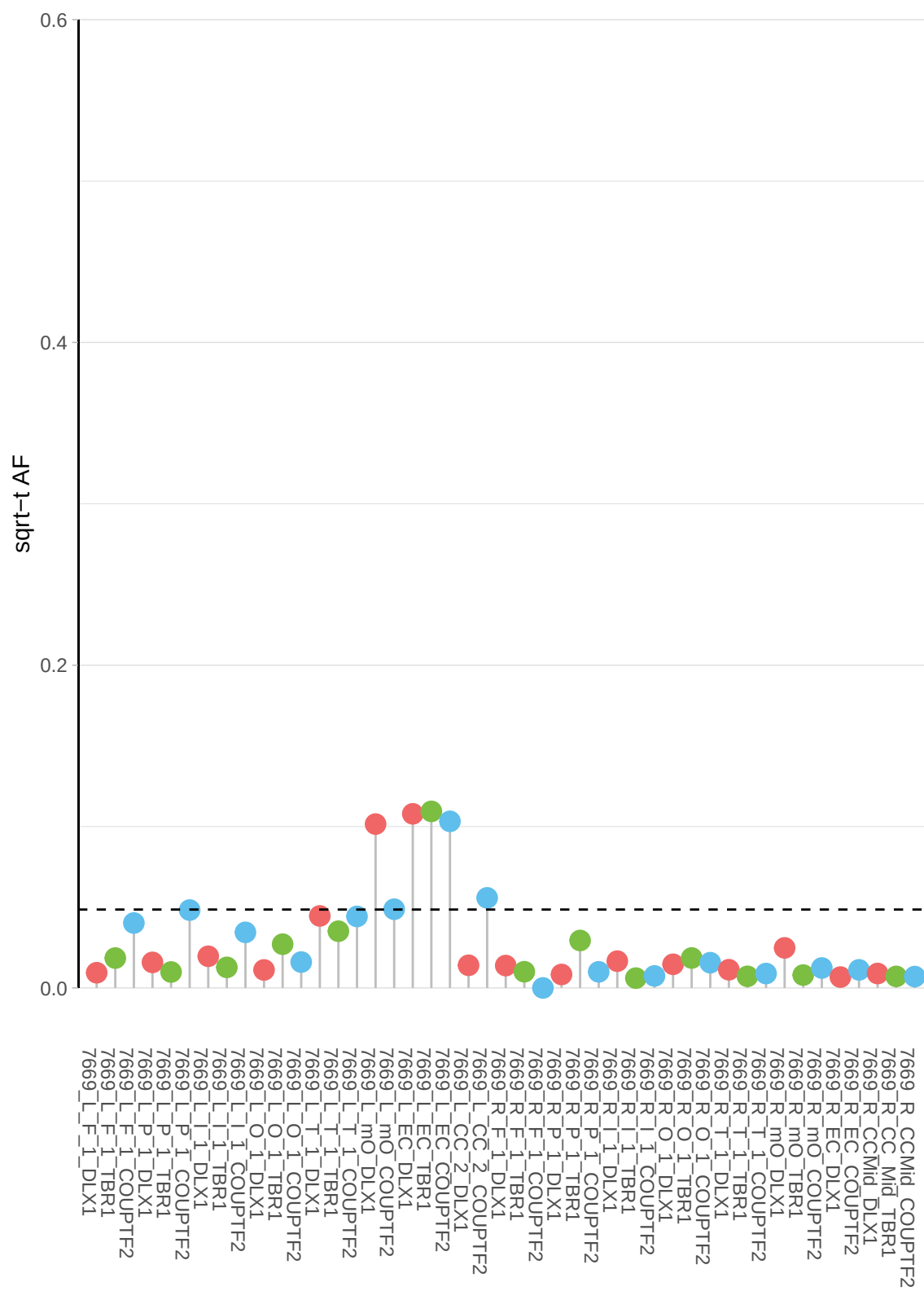

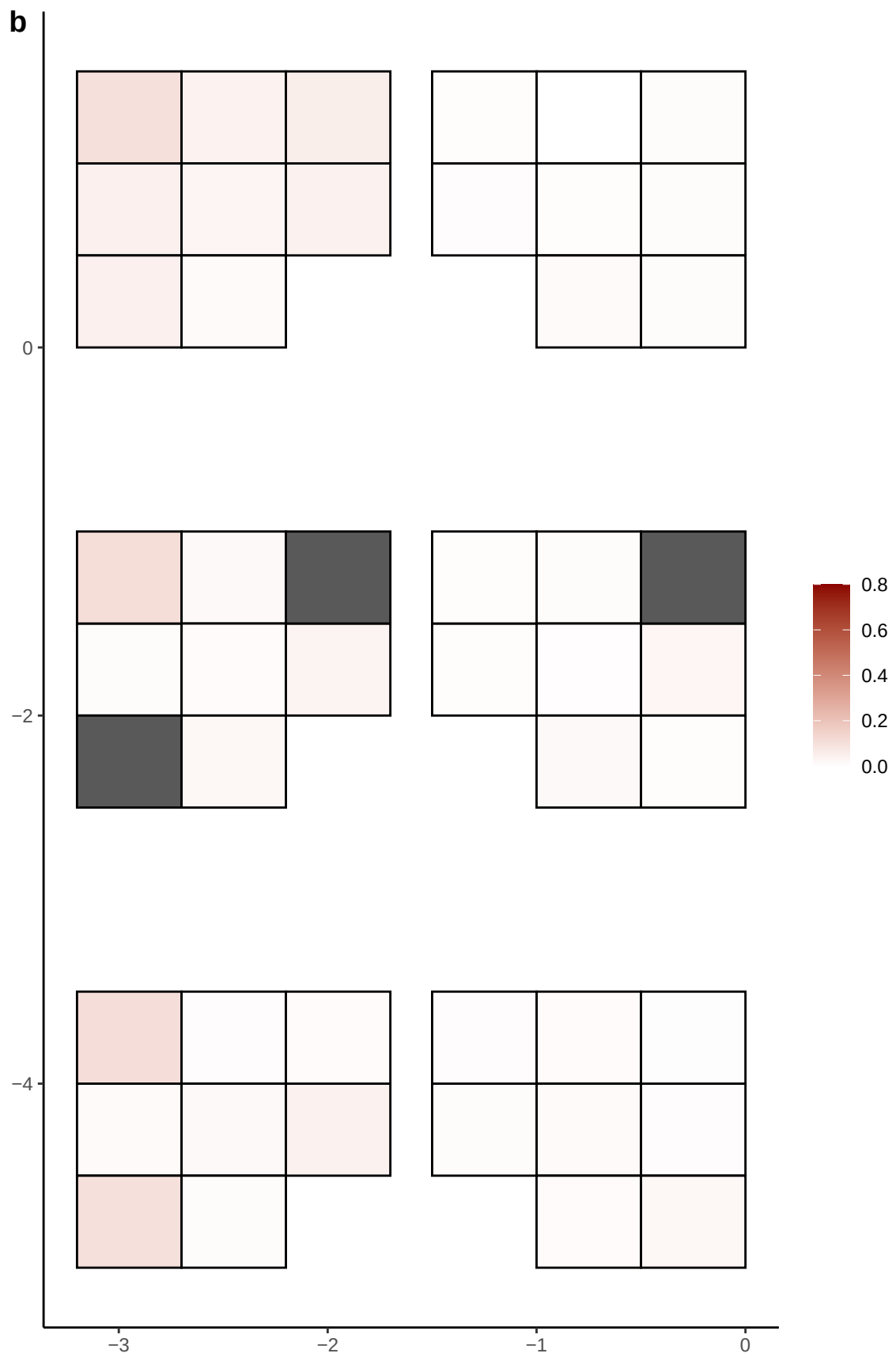

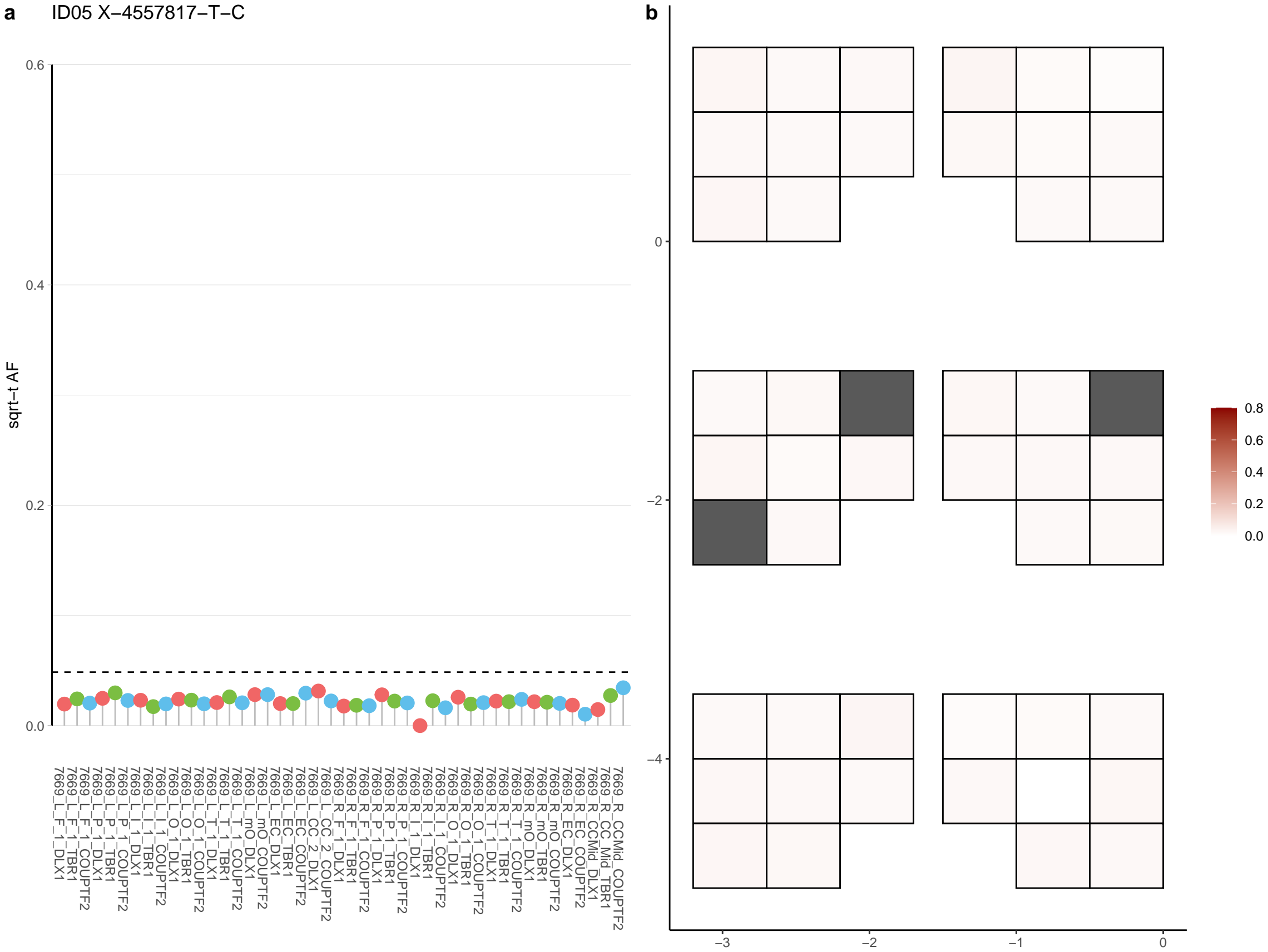

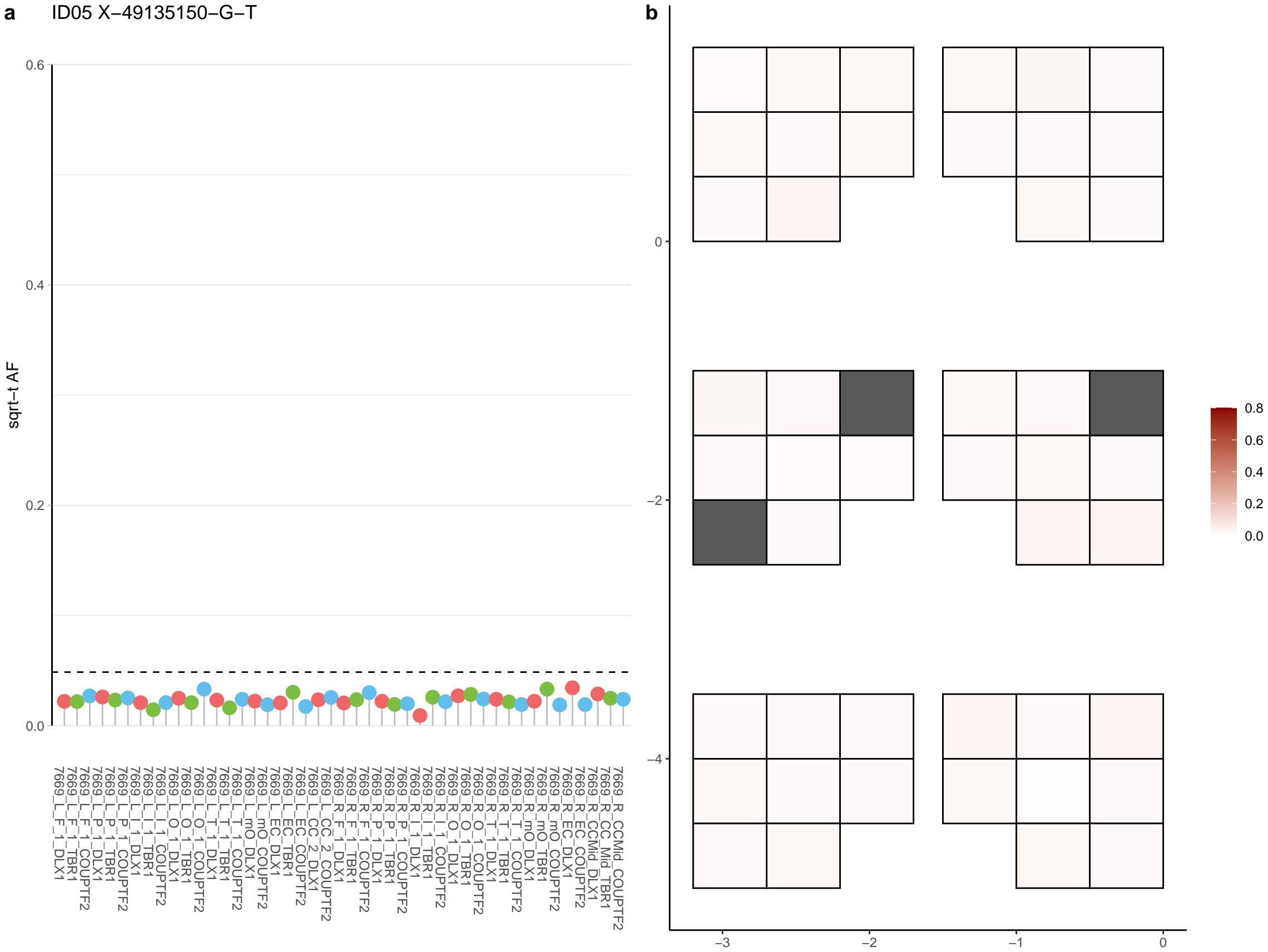

**a** ID05 X-53927483-G-A

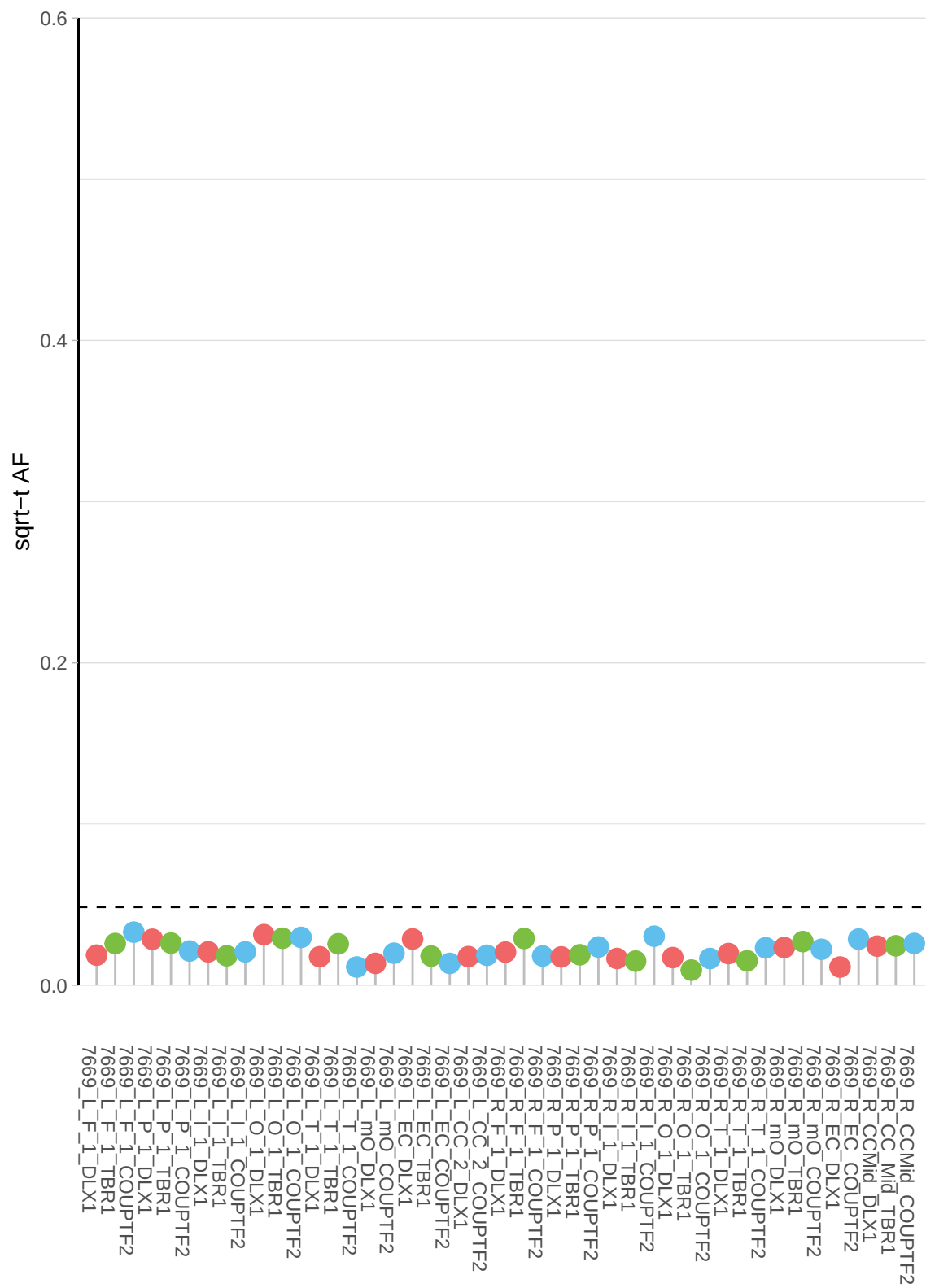

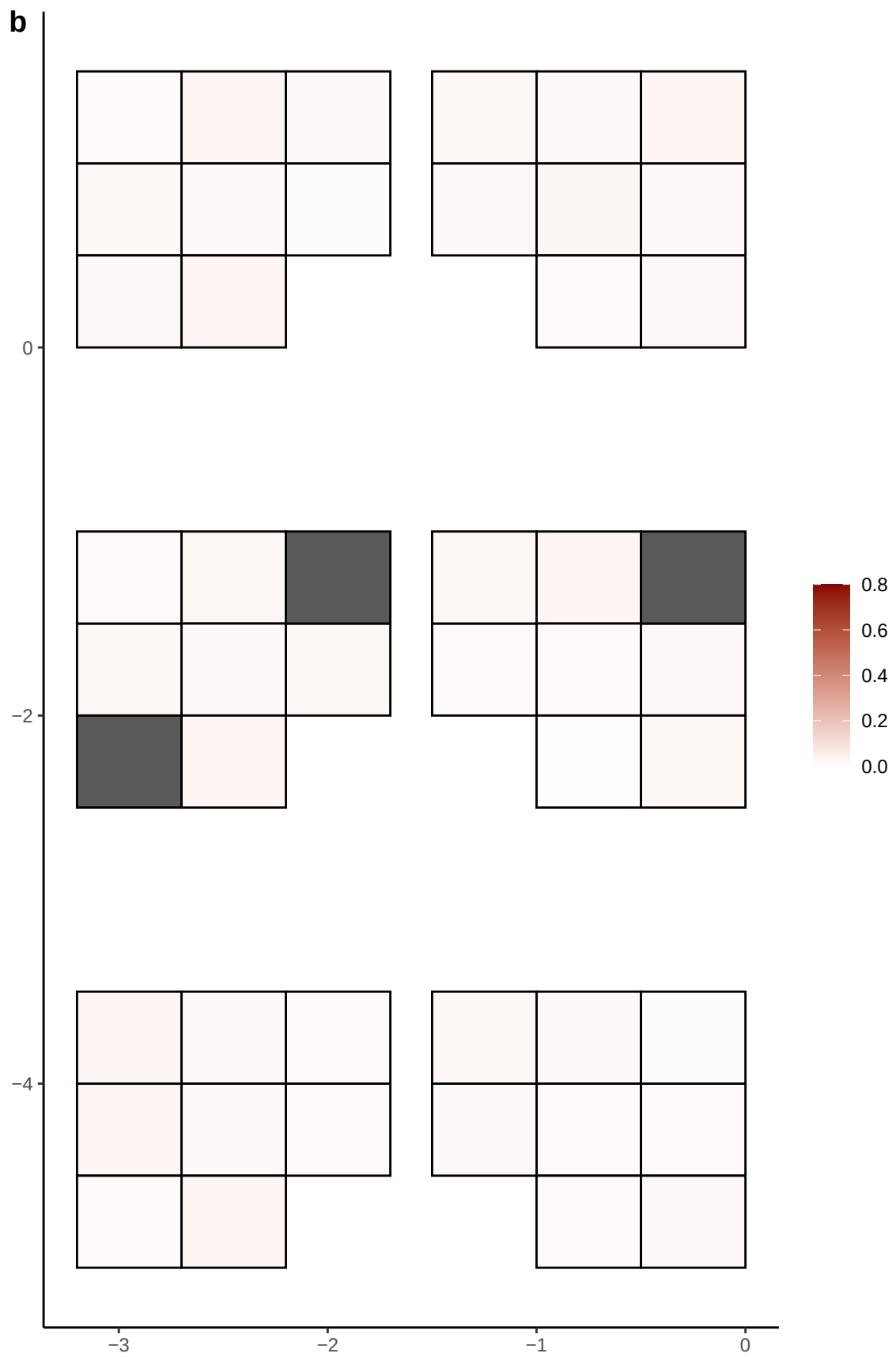

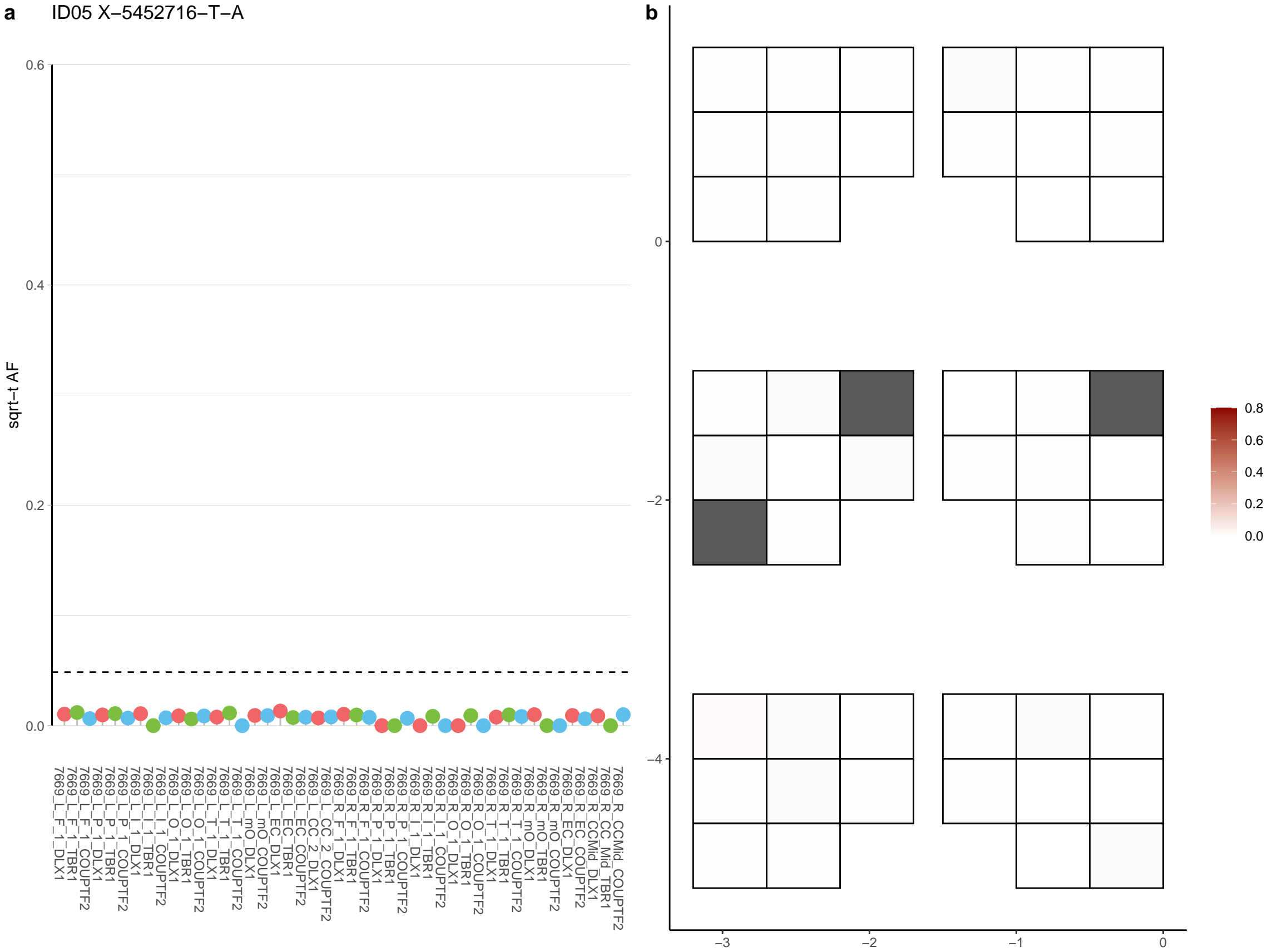

**a** ID05 X-6657783-T-A

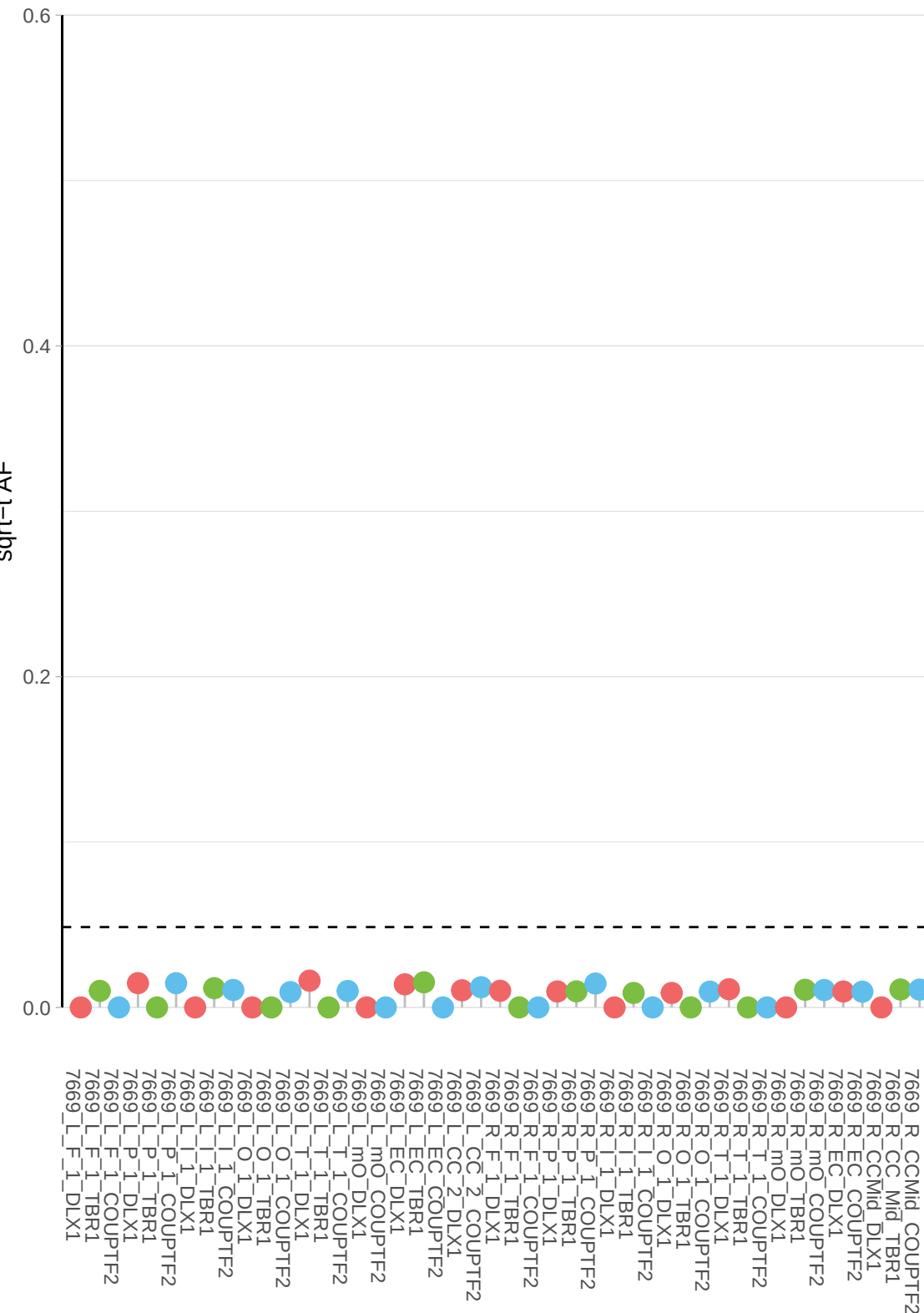

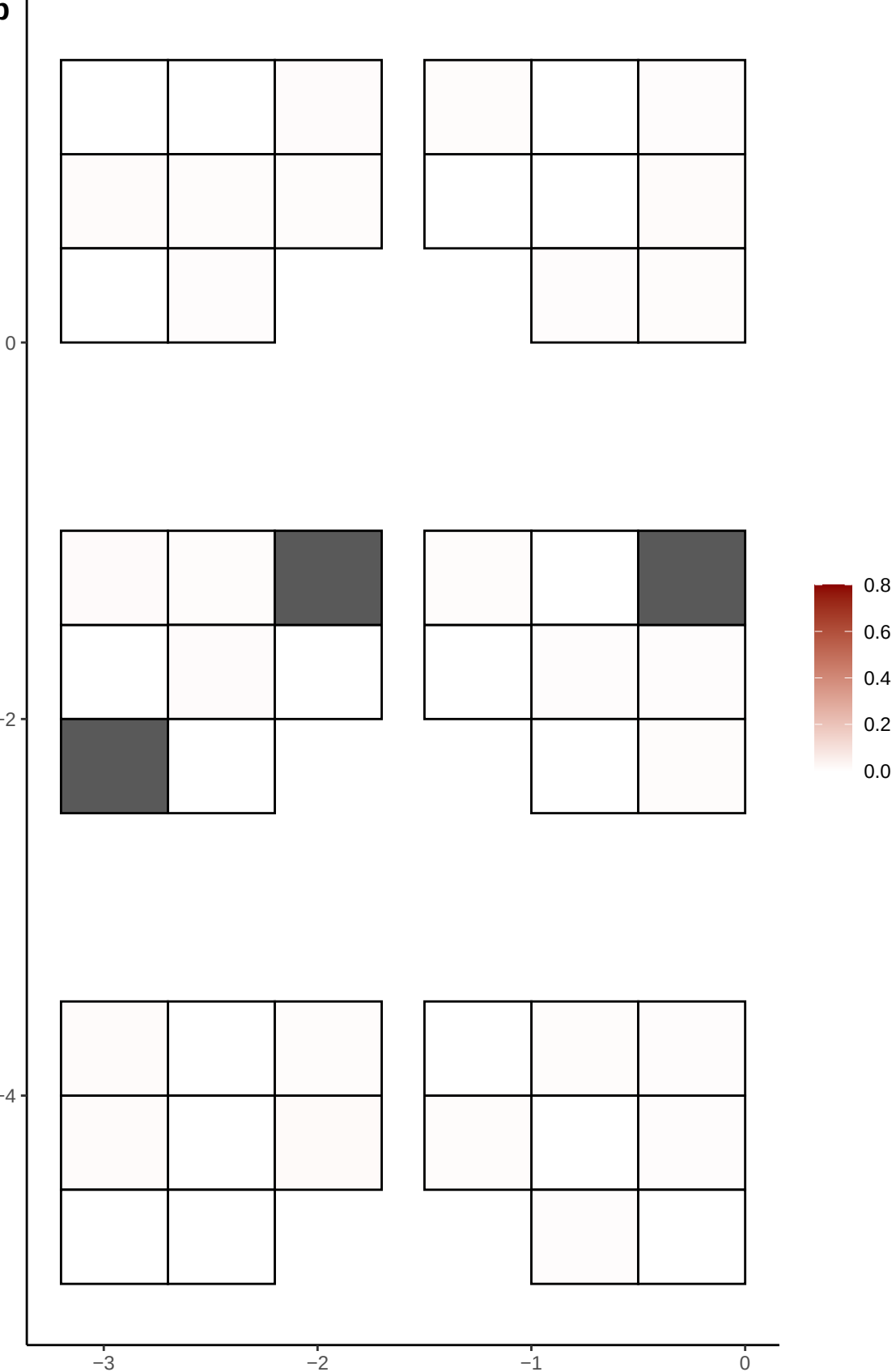

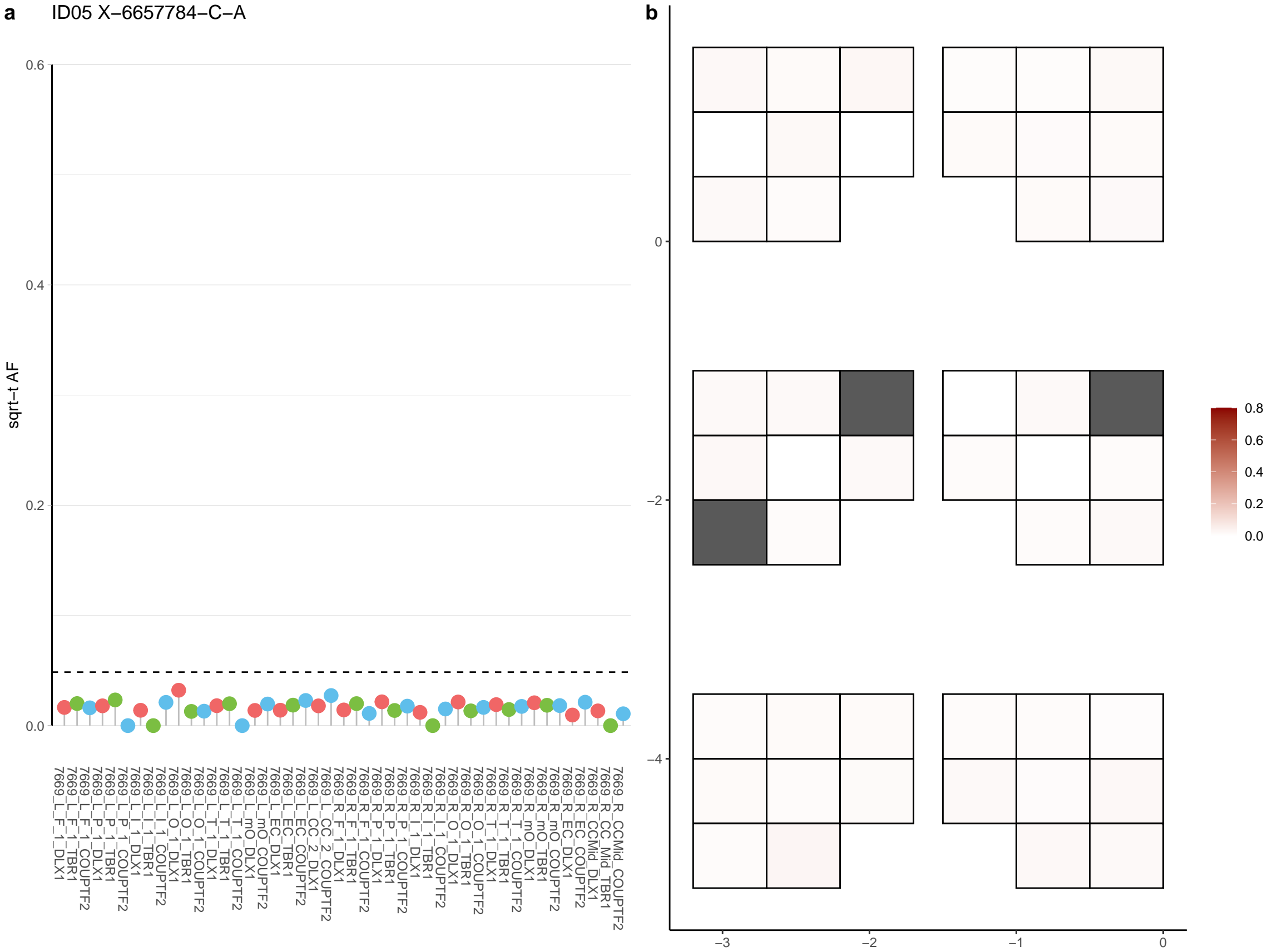

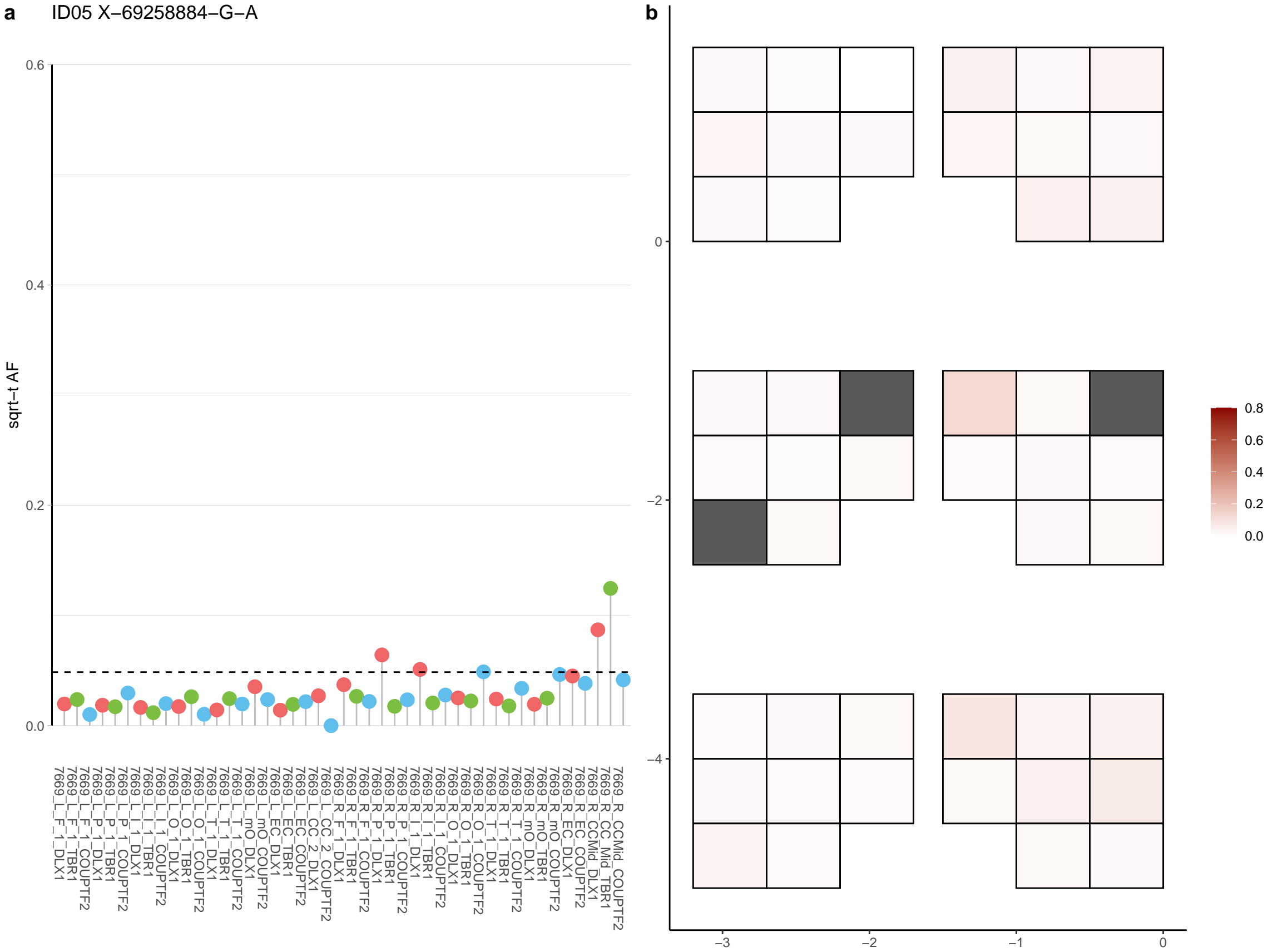

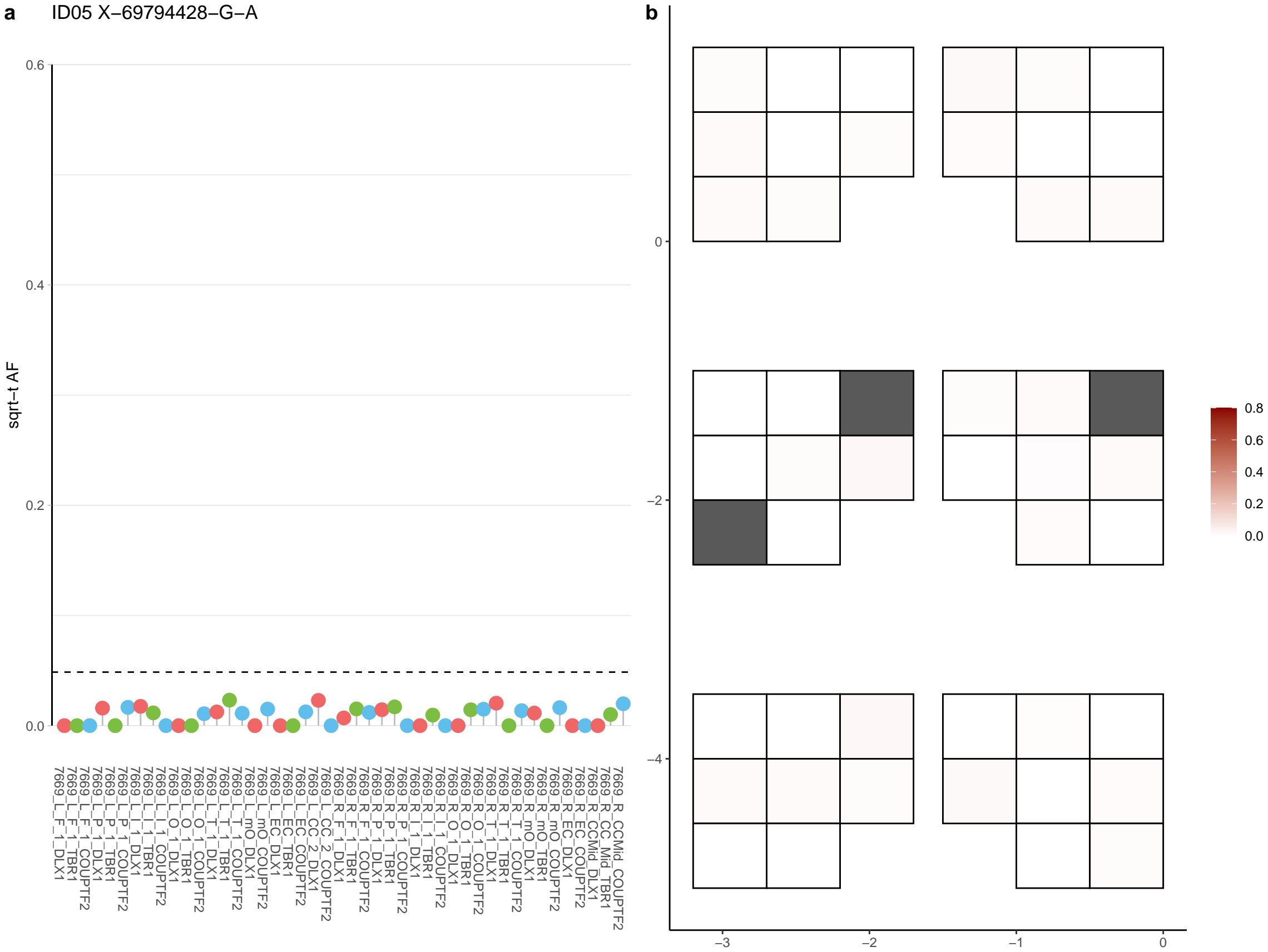

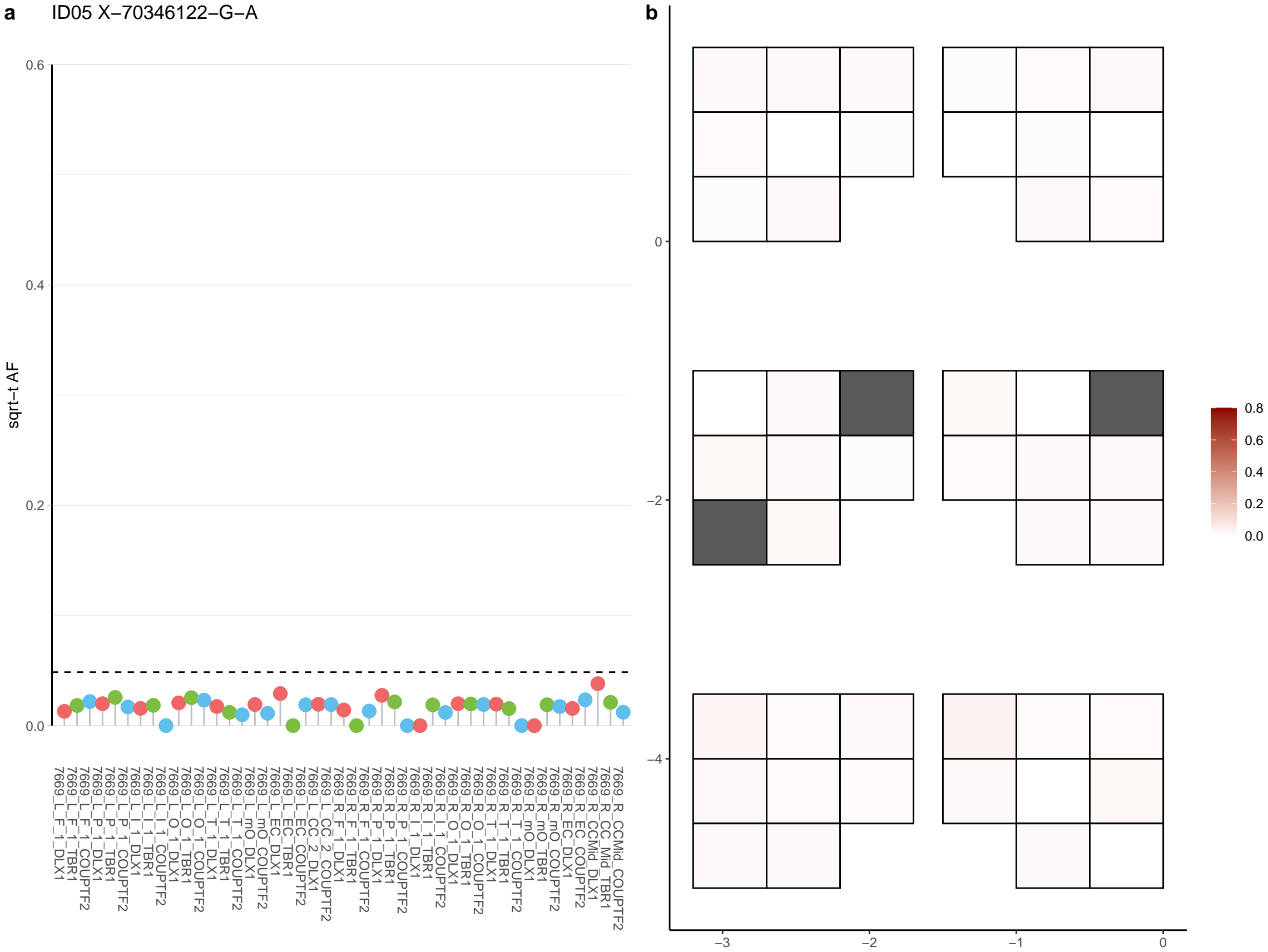

**a** ID05 X-72495149-C-A

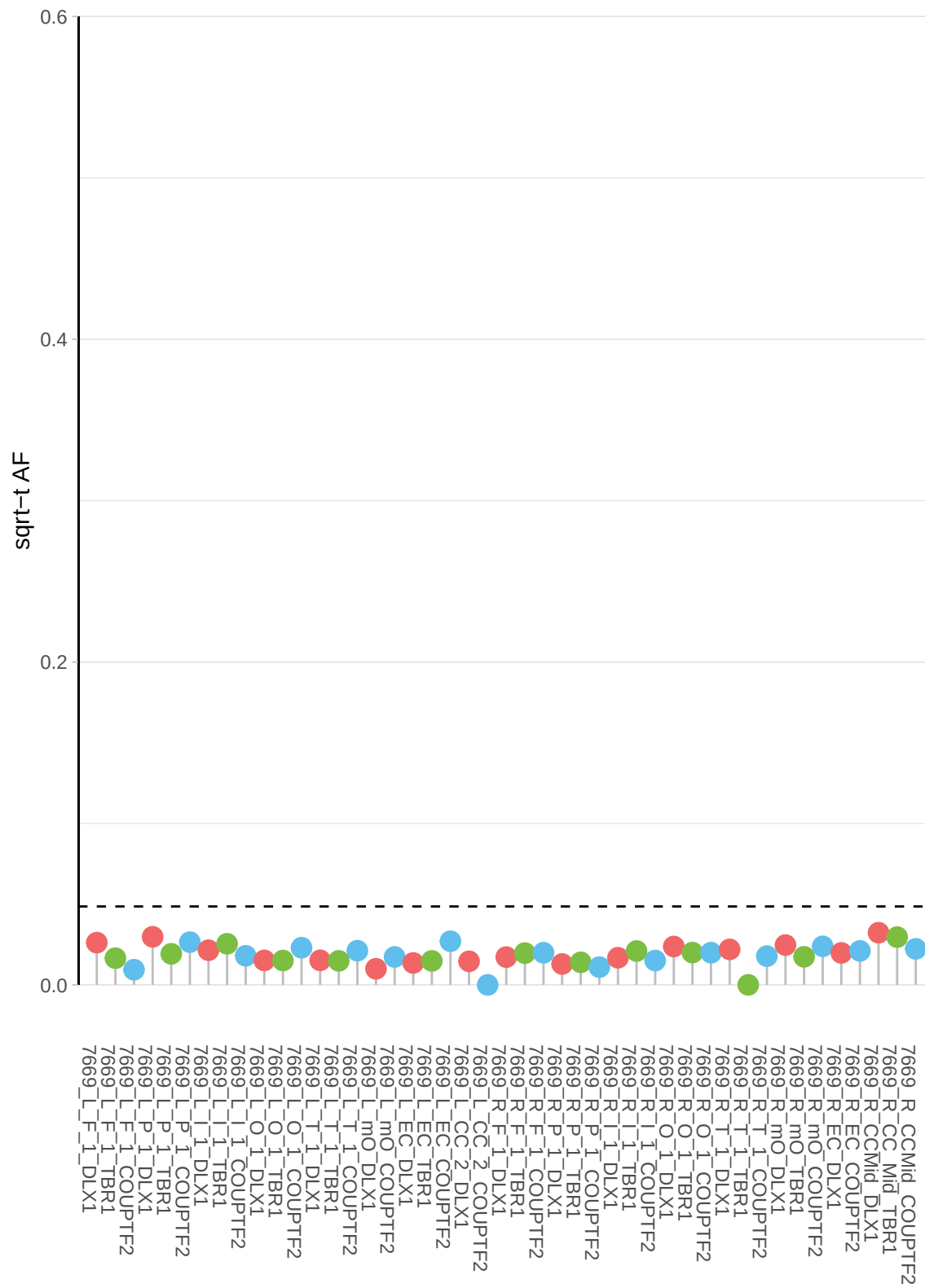

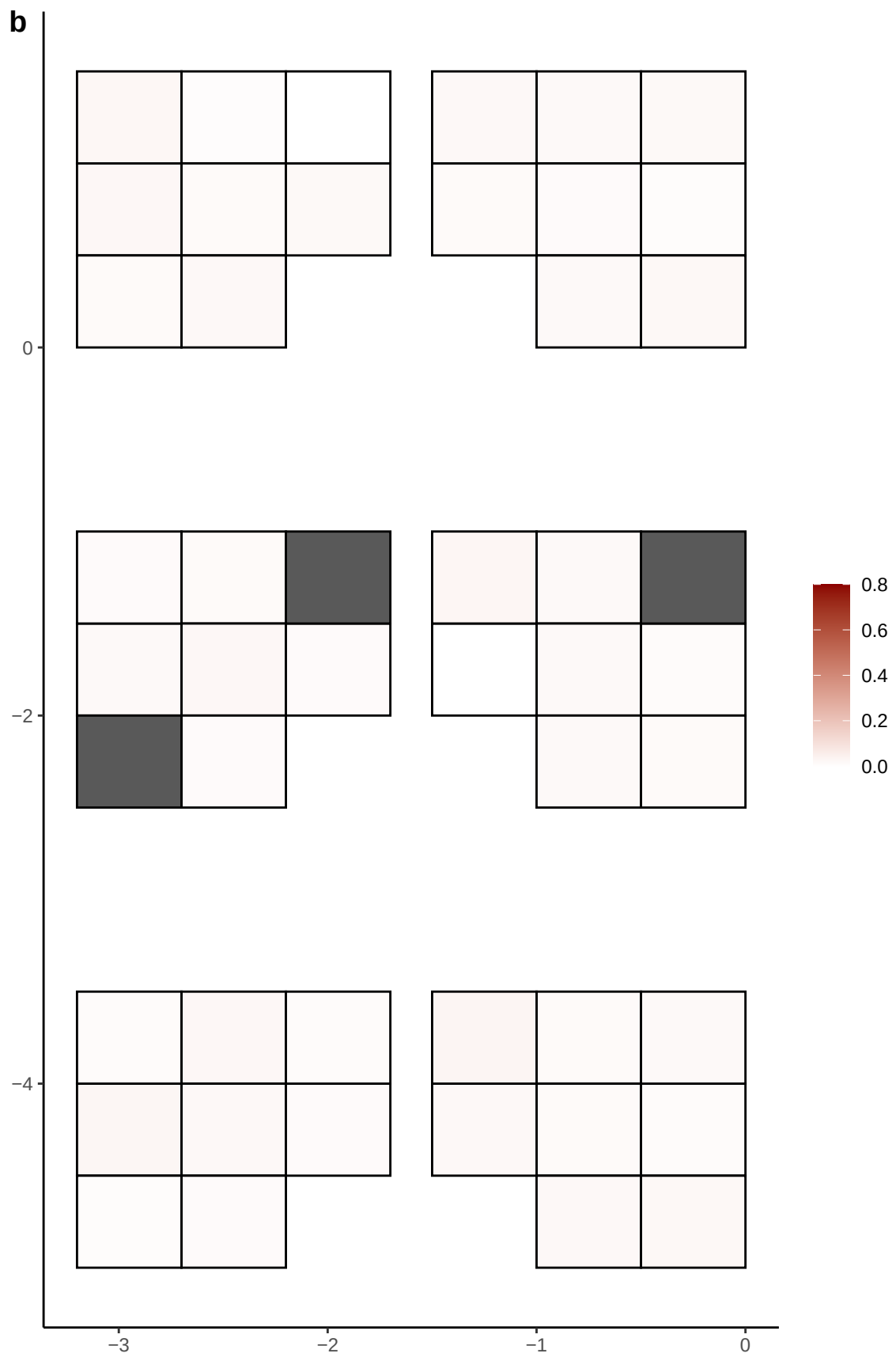

**a** ID05 X-7883974-C-T

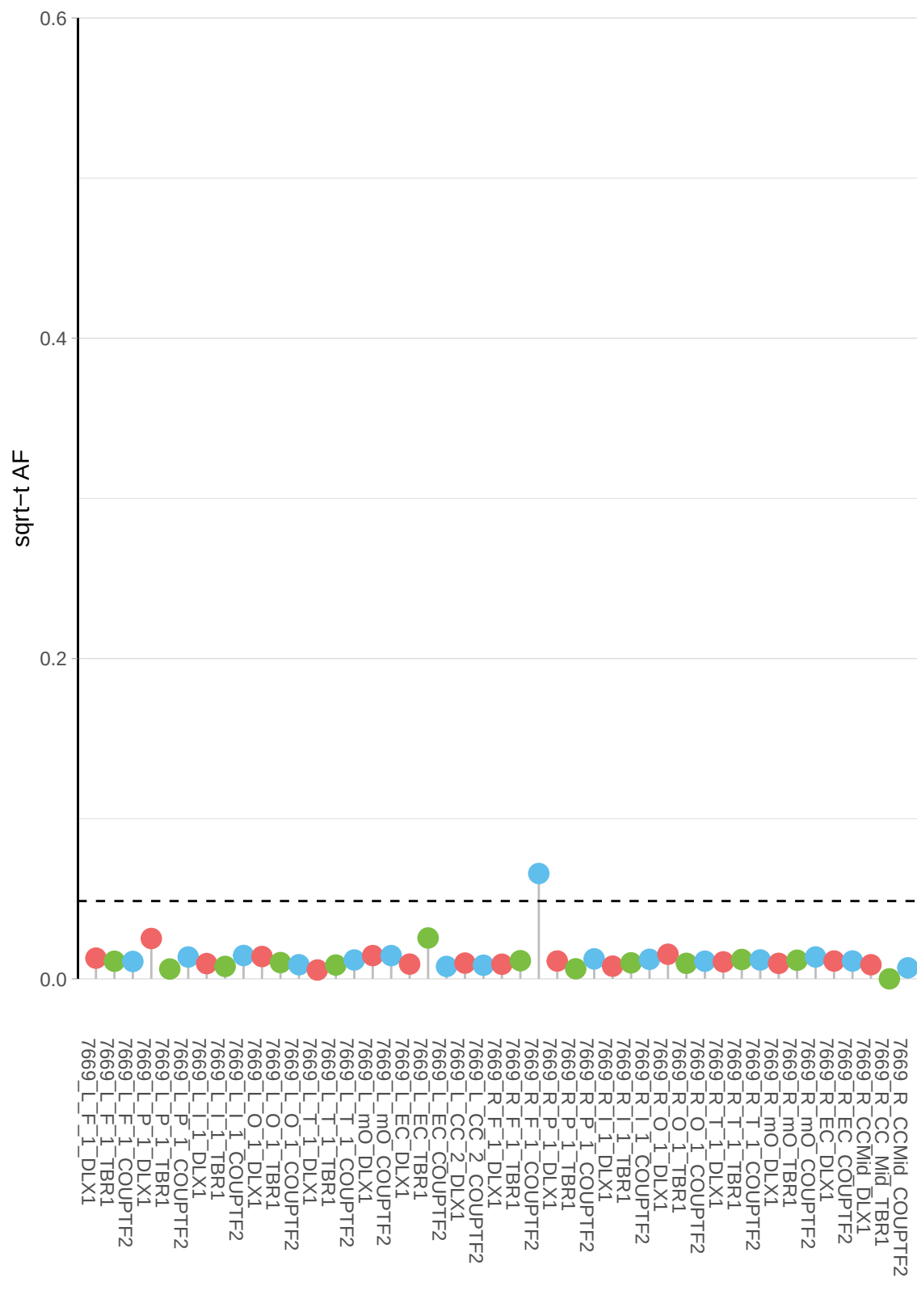

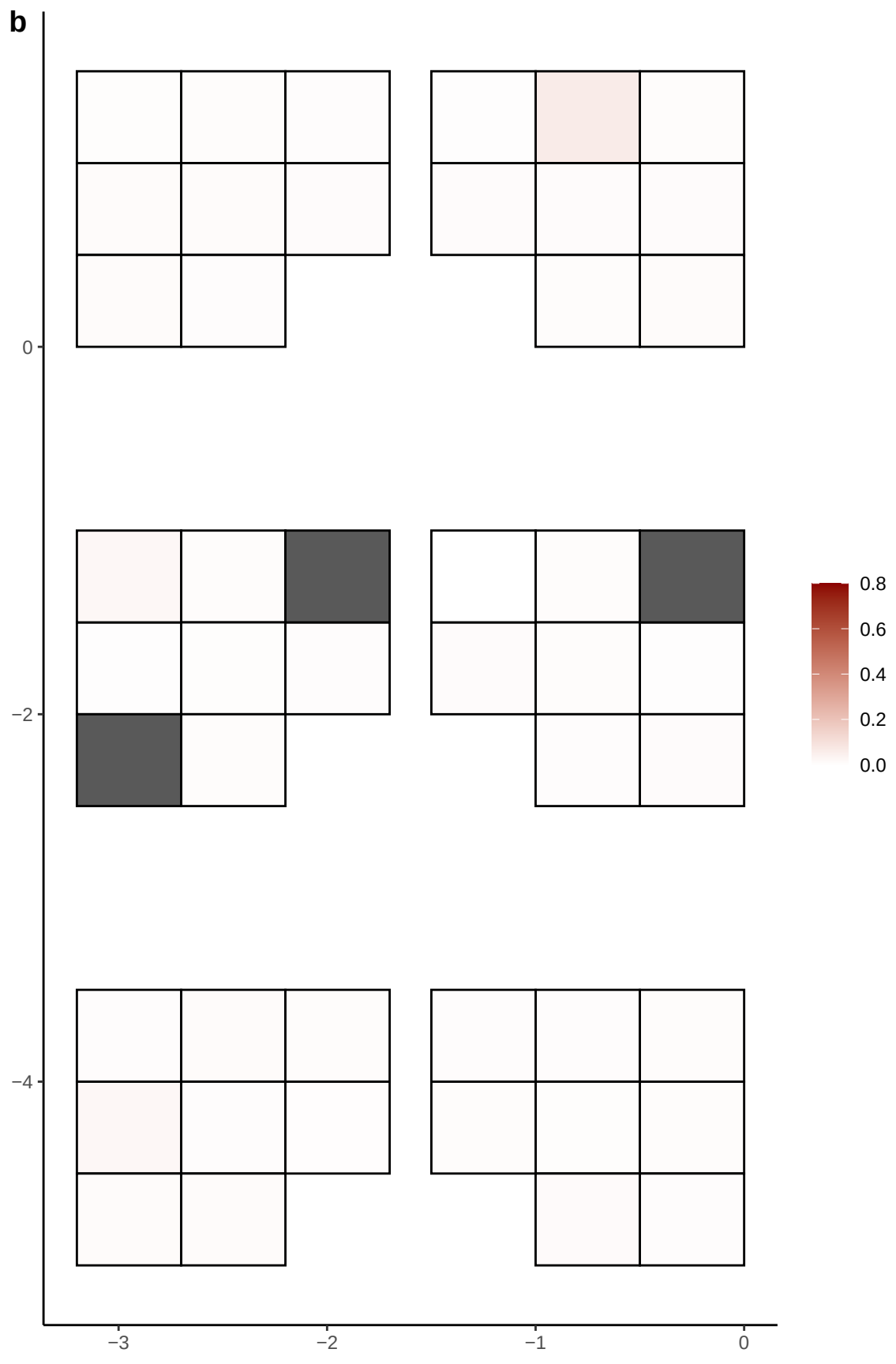

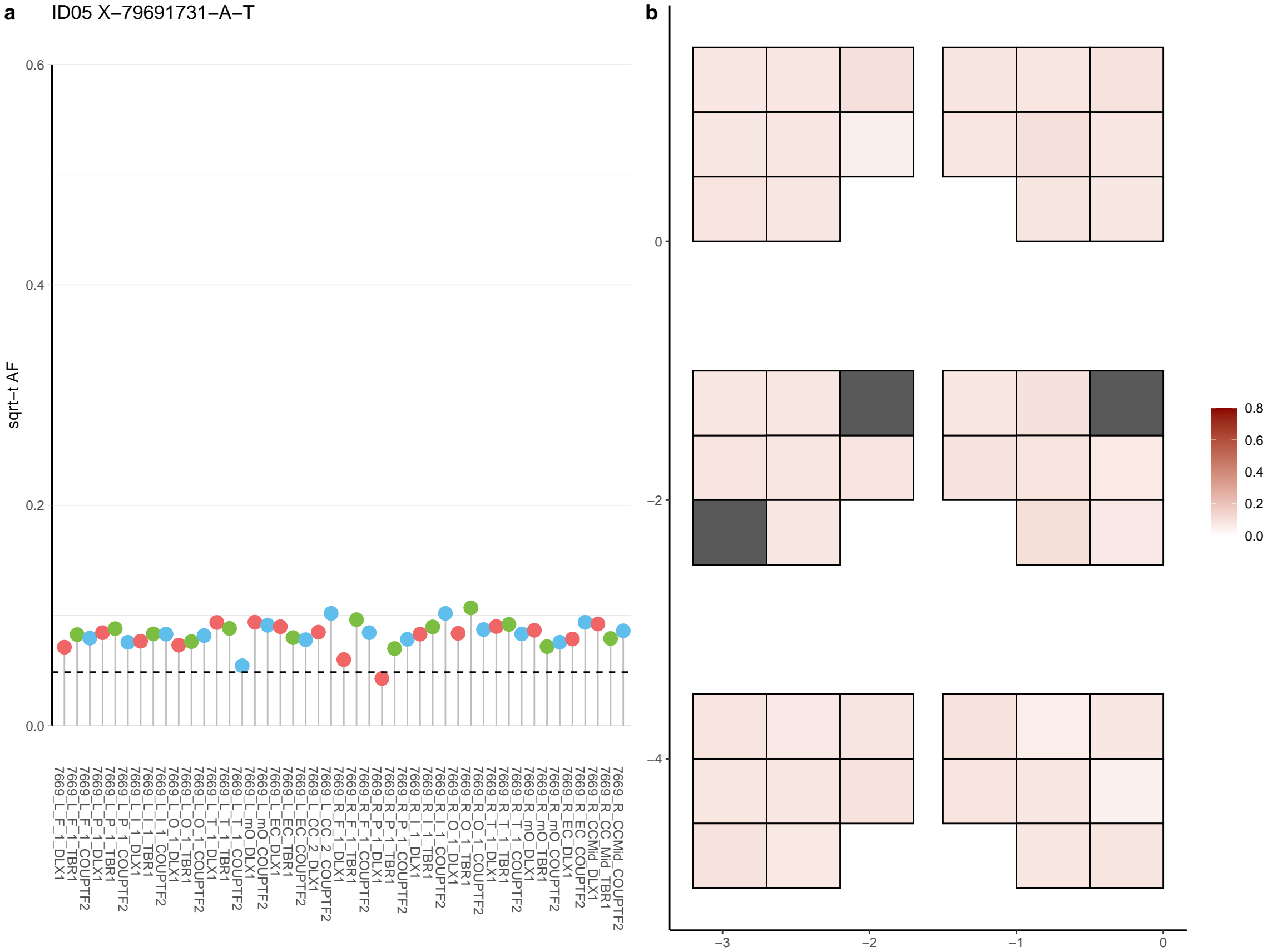

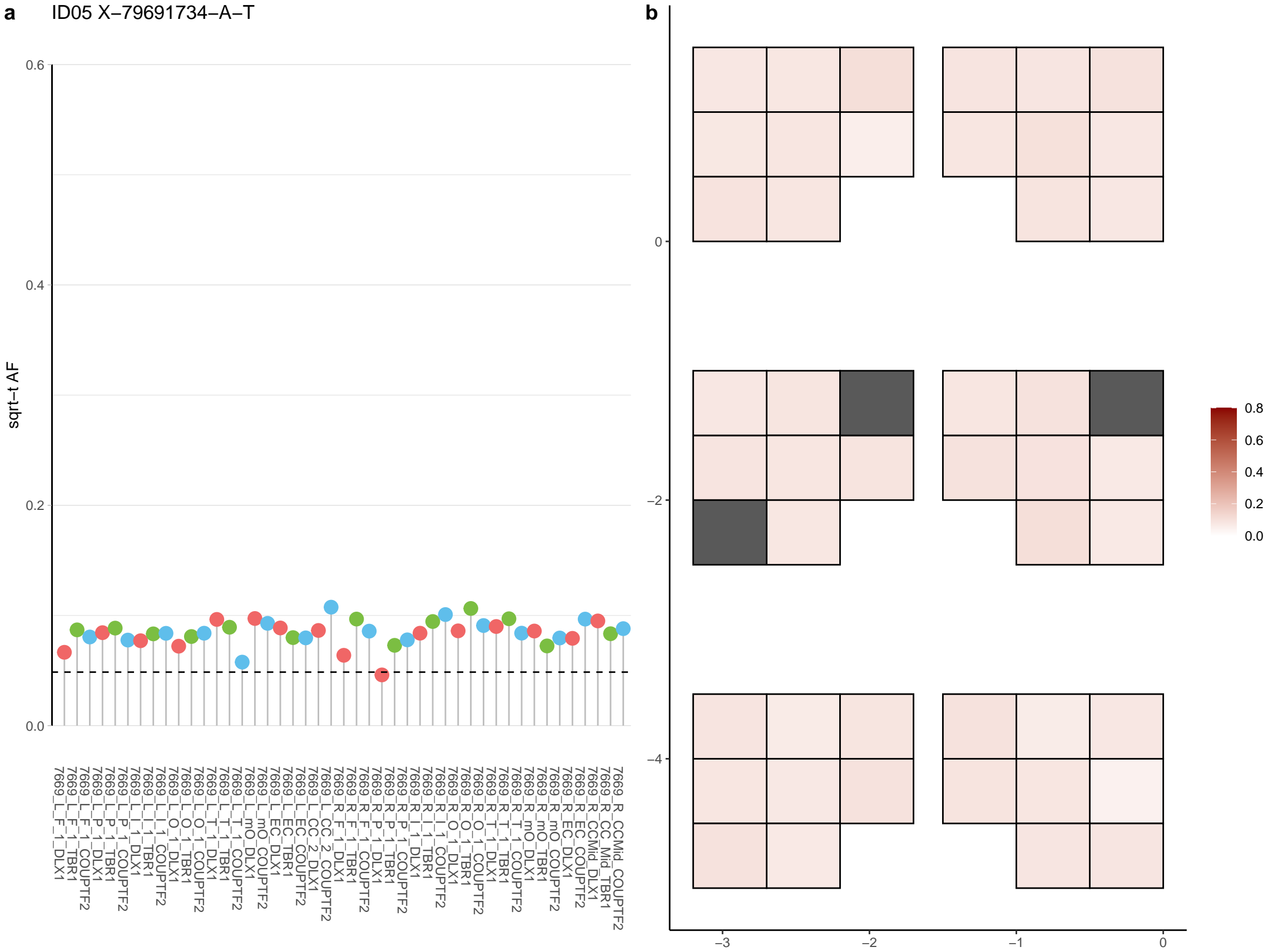

**a** ID05 X-83599149-G-A

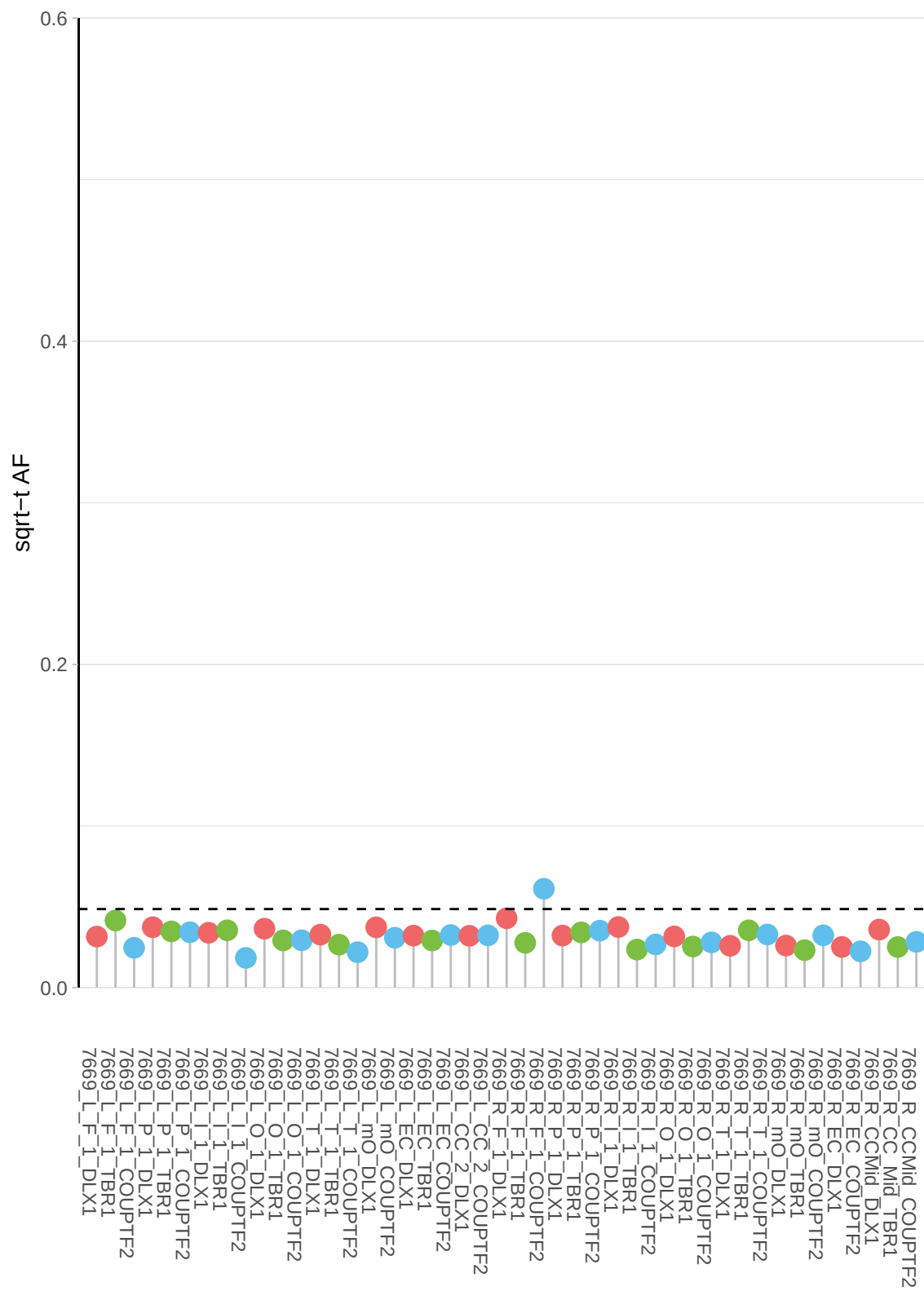

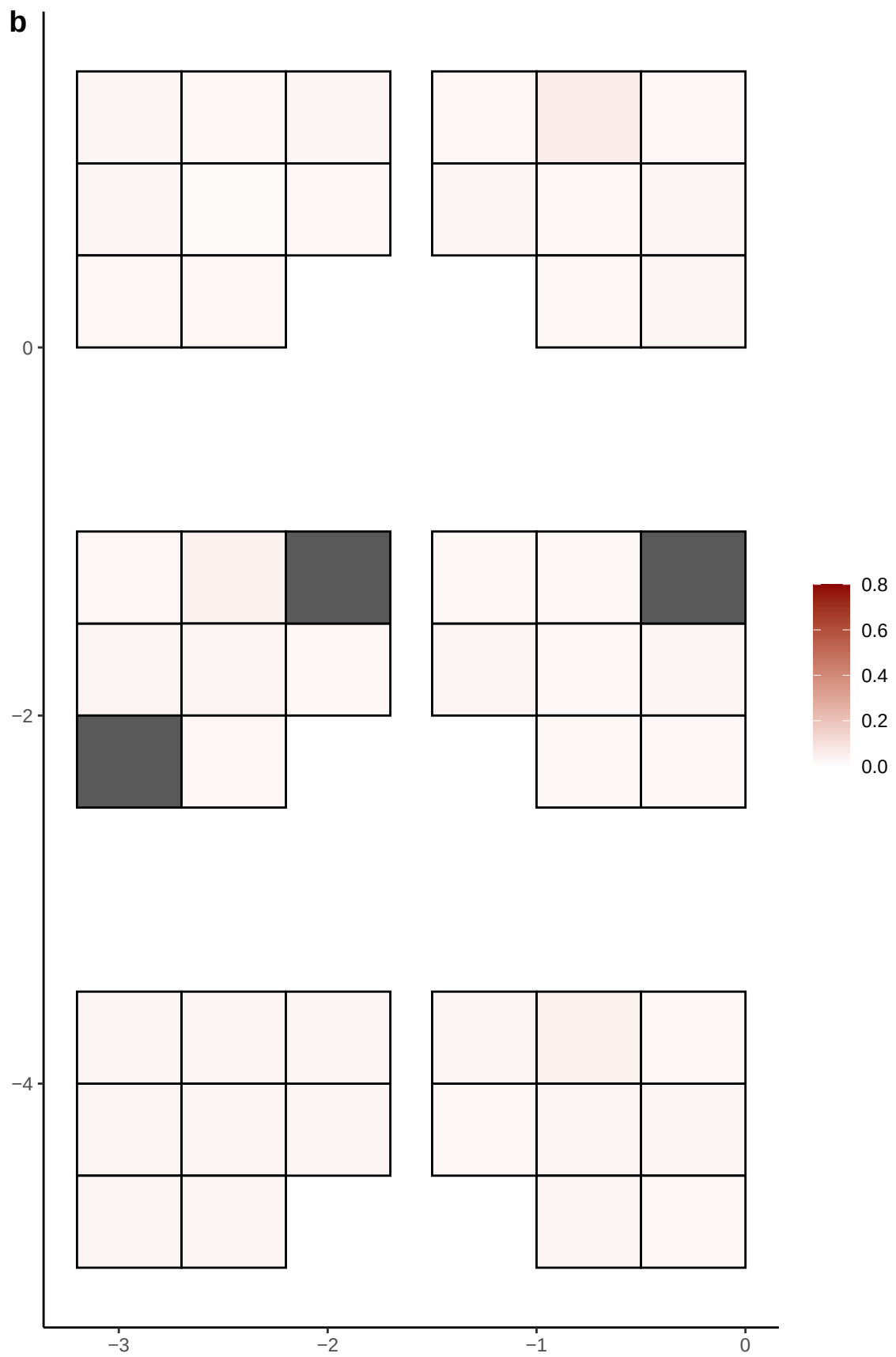

**a** ID05 X-88323908-T-G

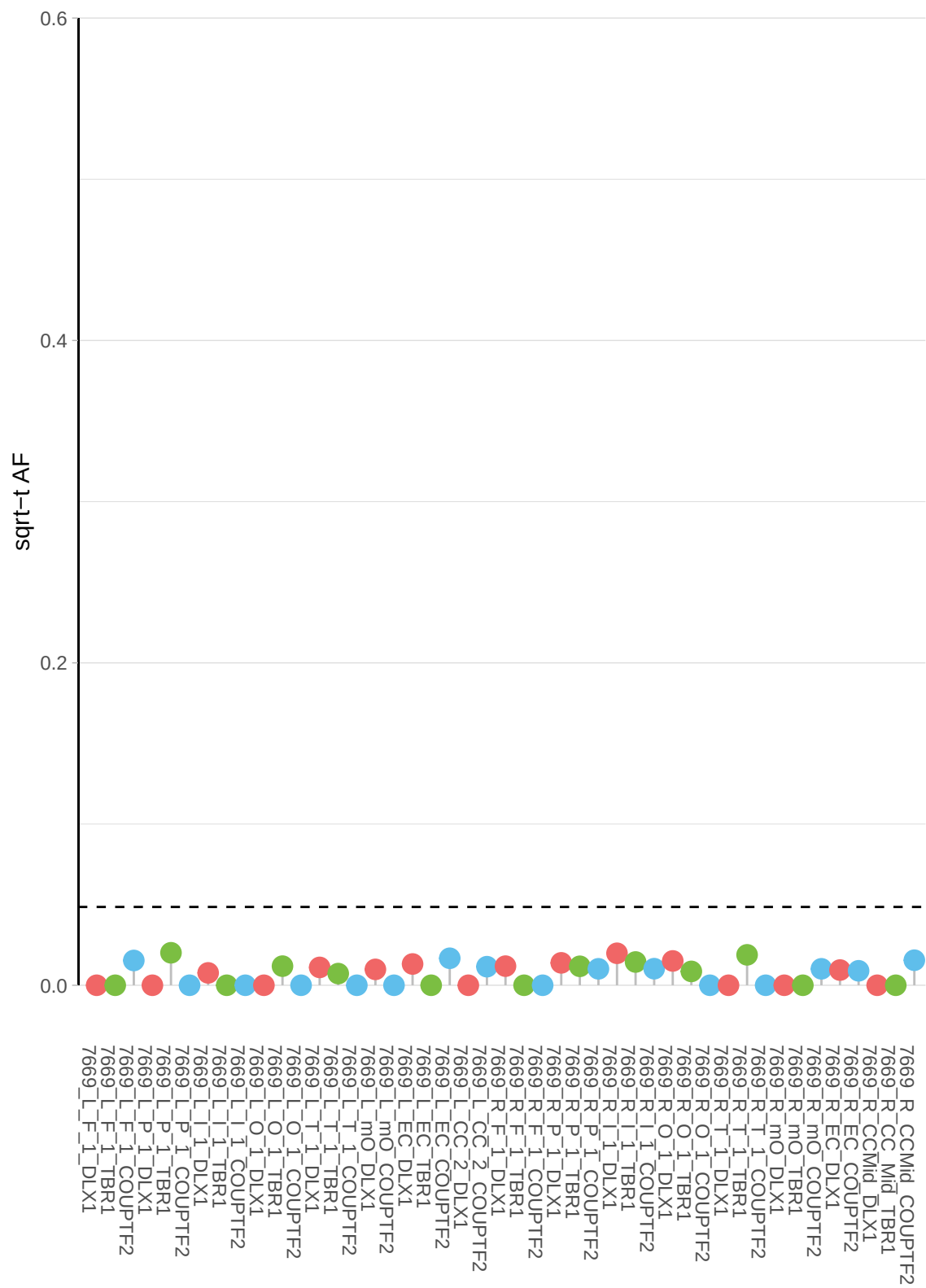

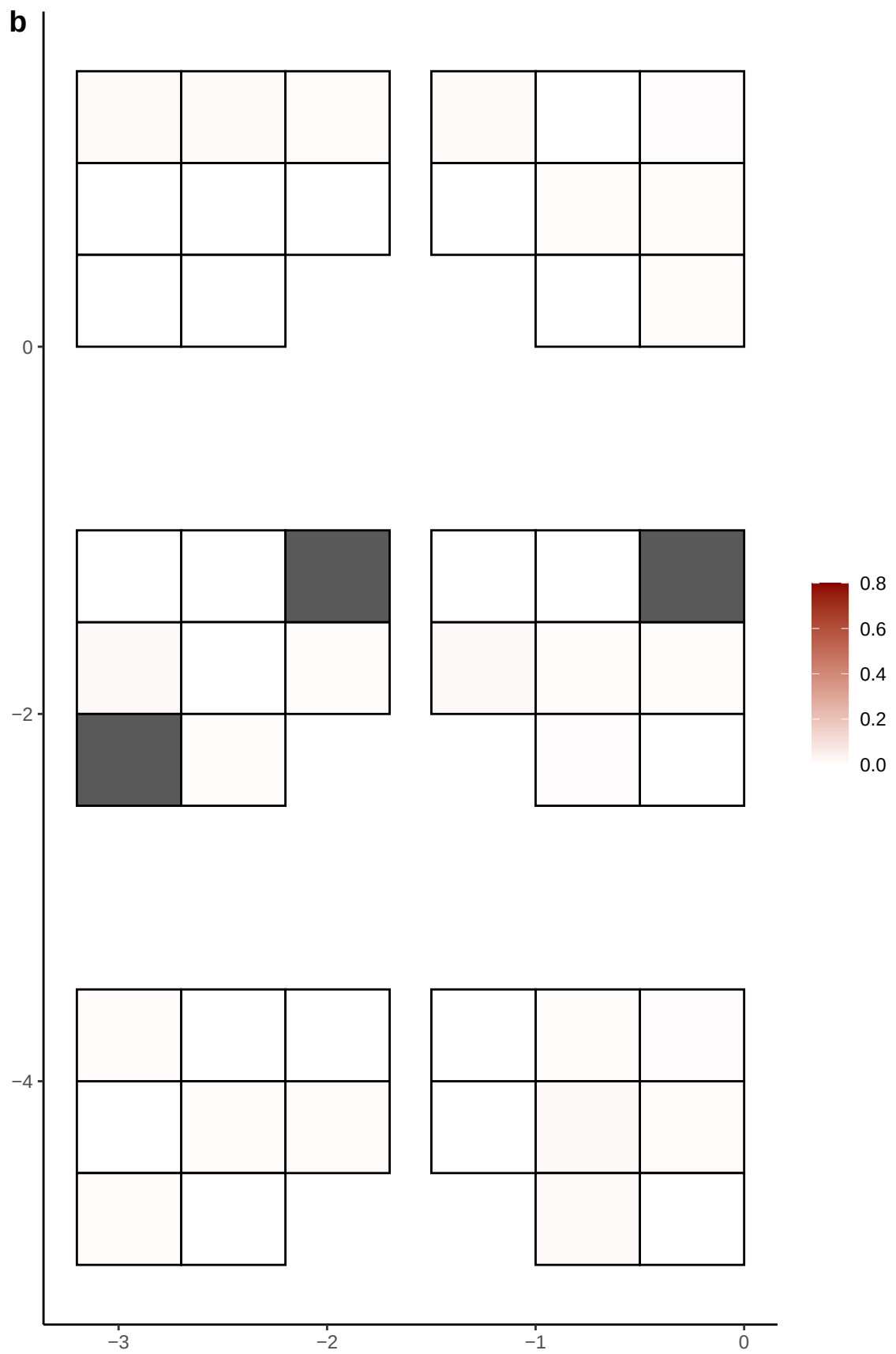

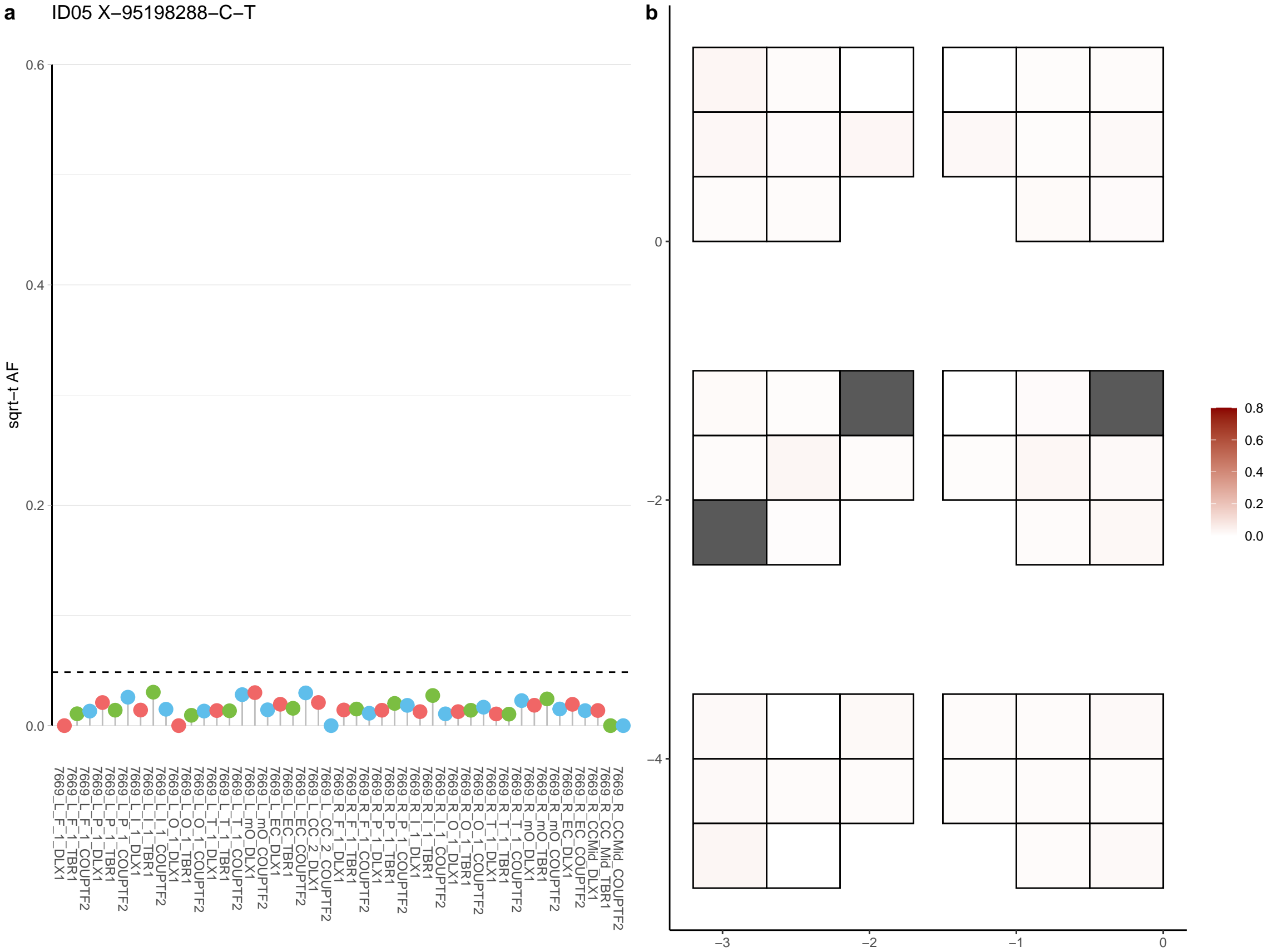

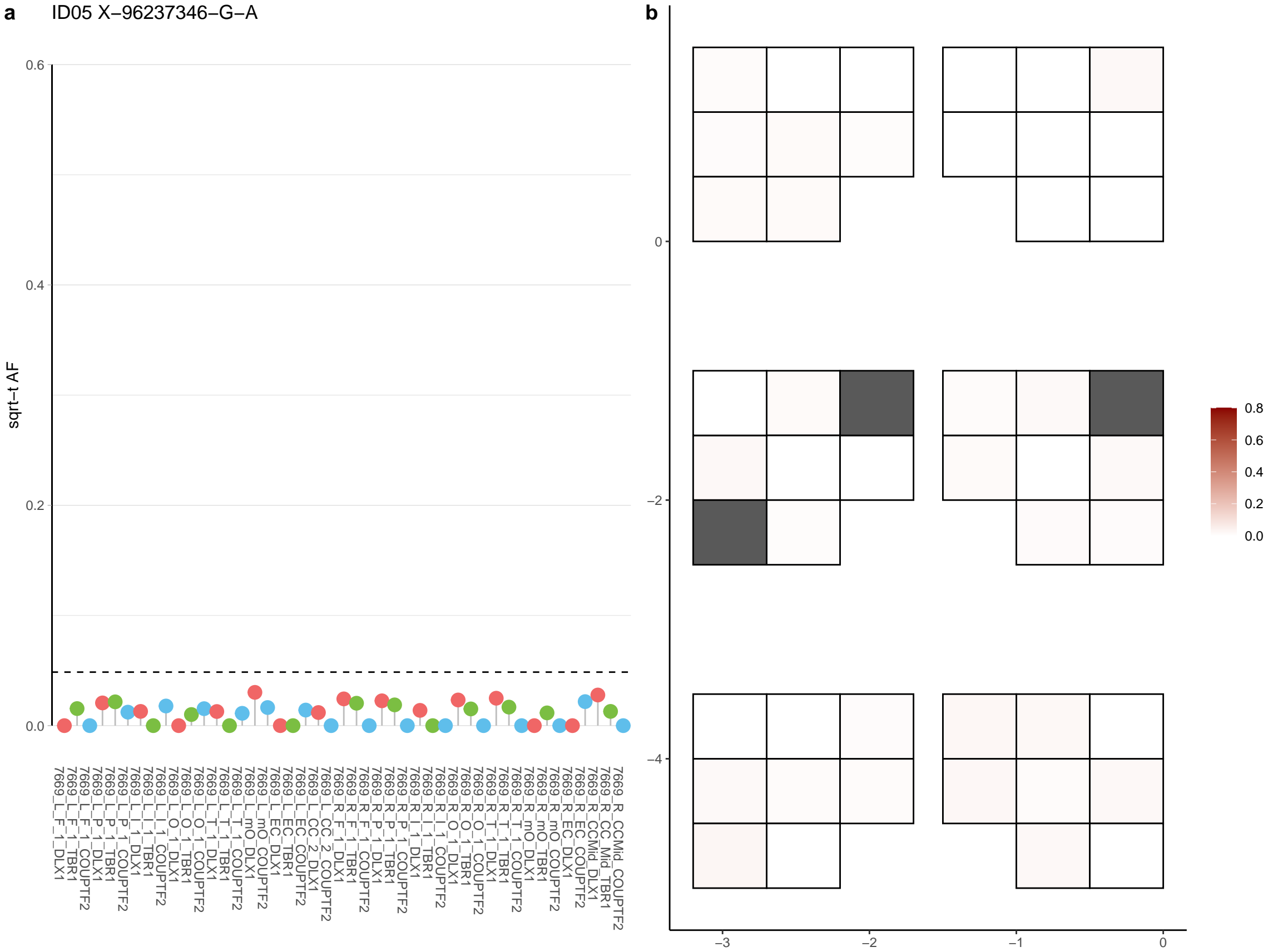

**a** ID05 X-98261573-C-T

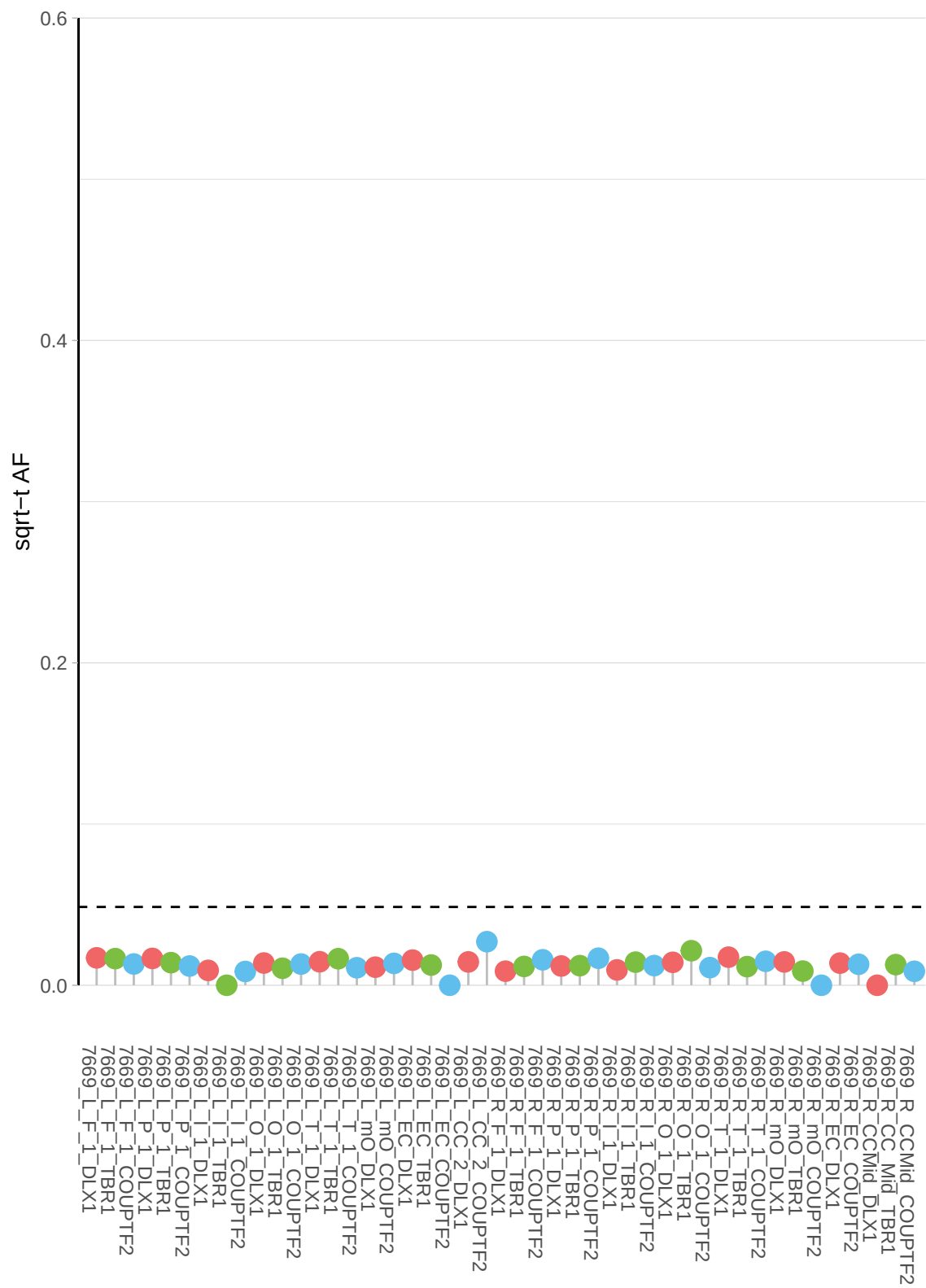

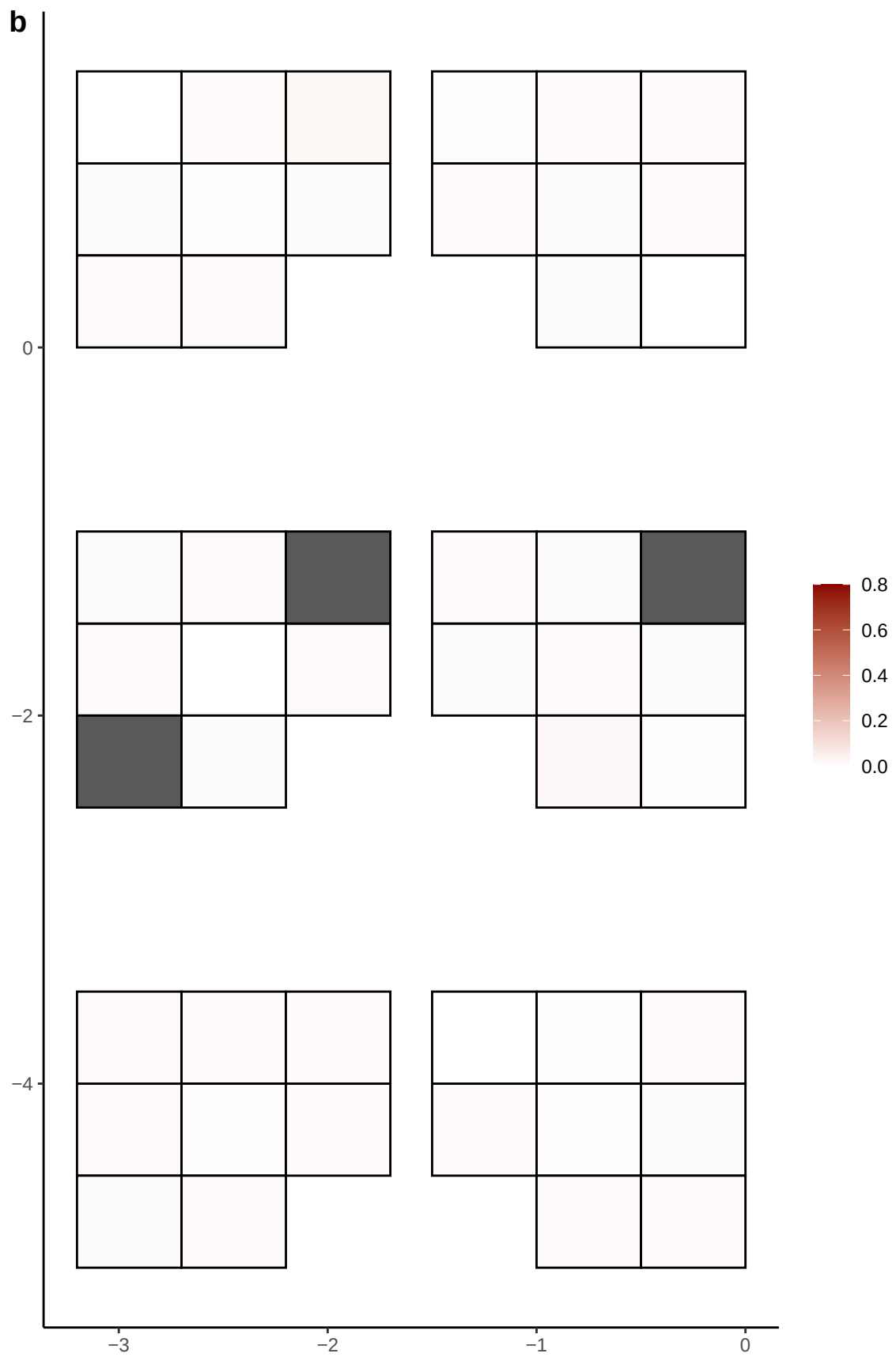

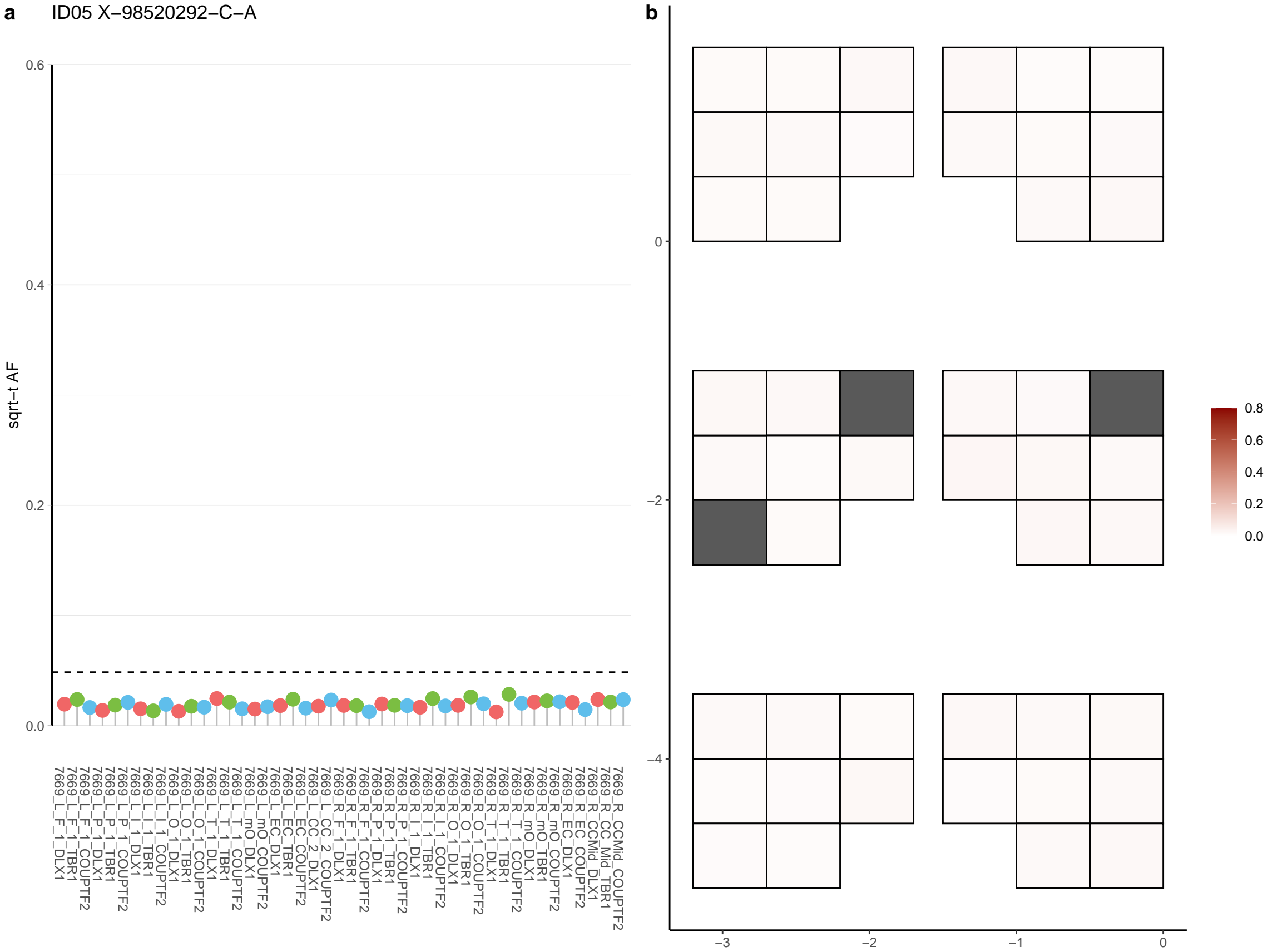

**a** ID05 X-98527468-AC-A

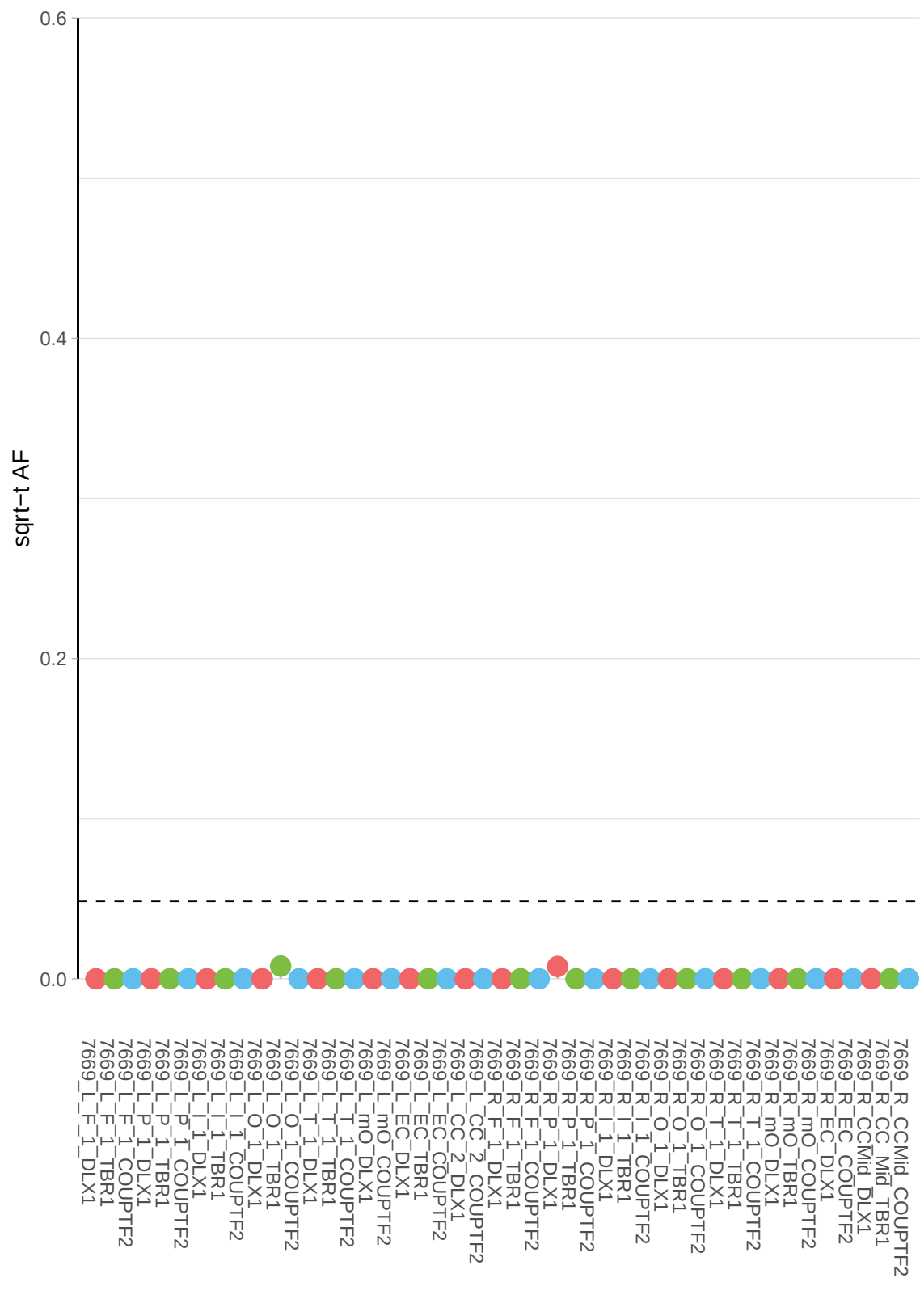

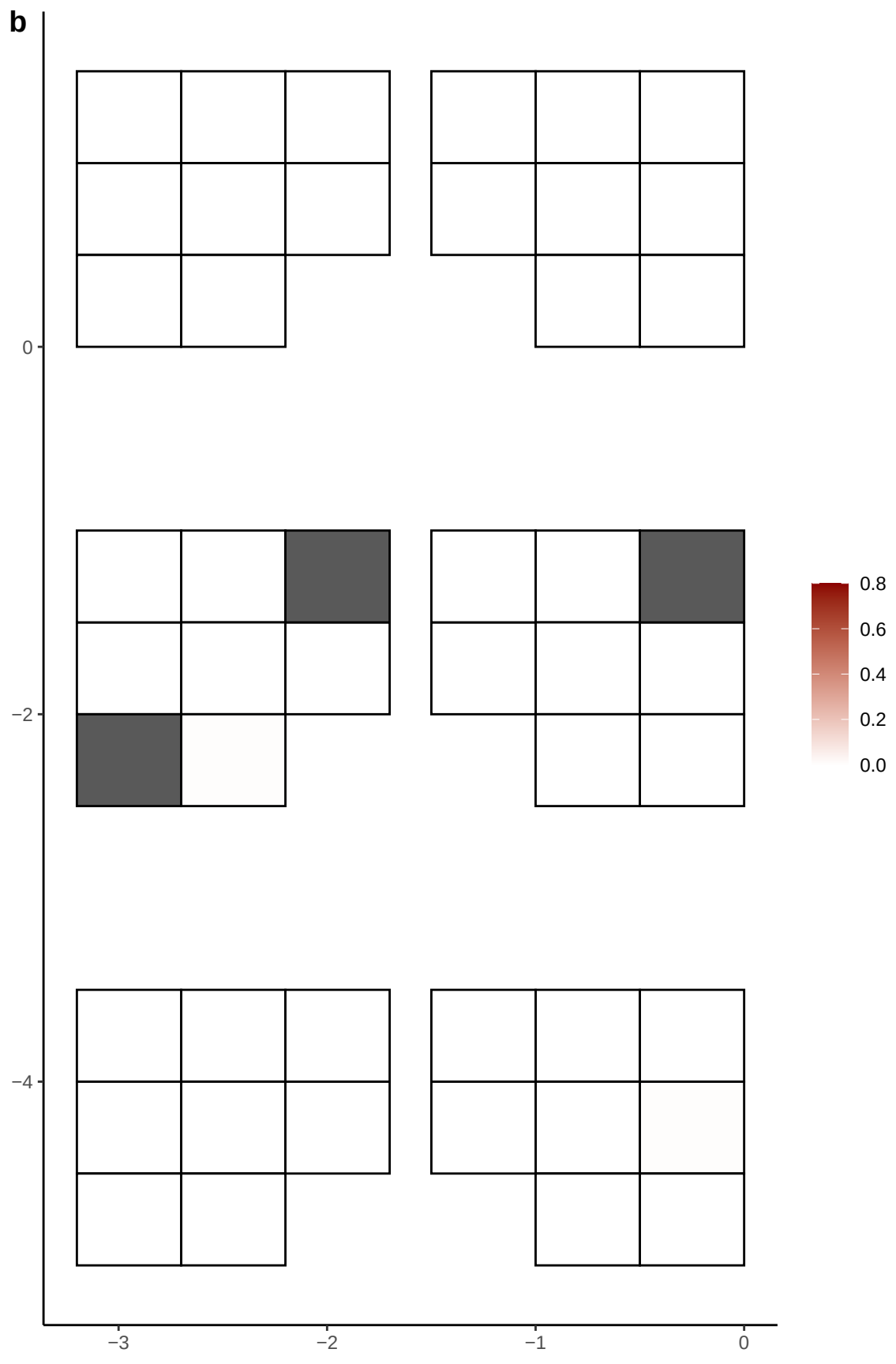

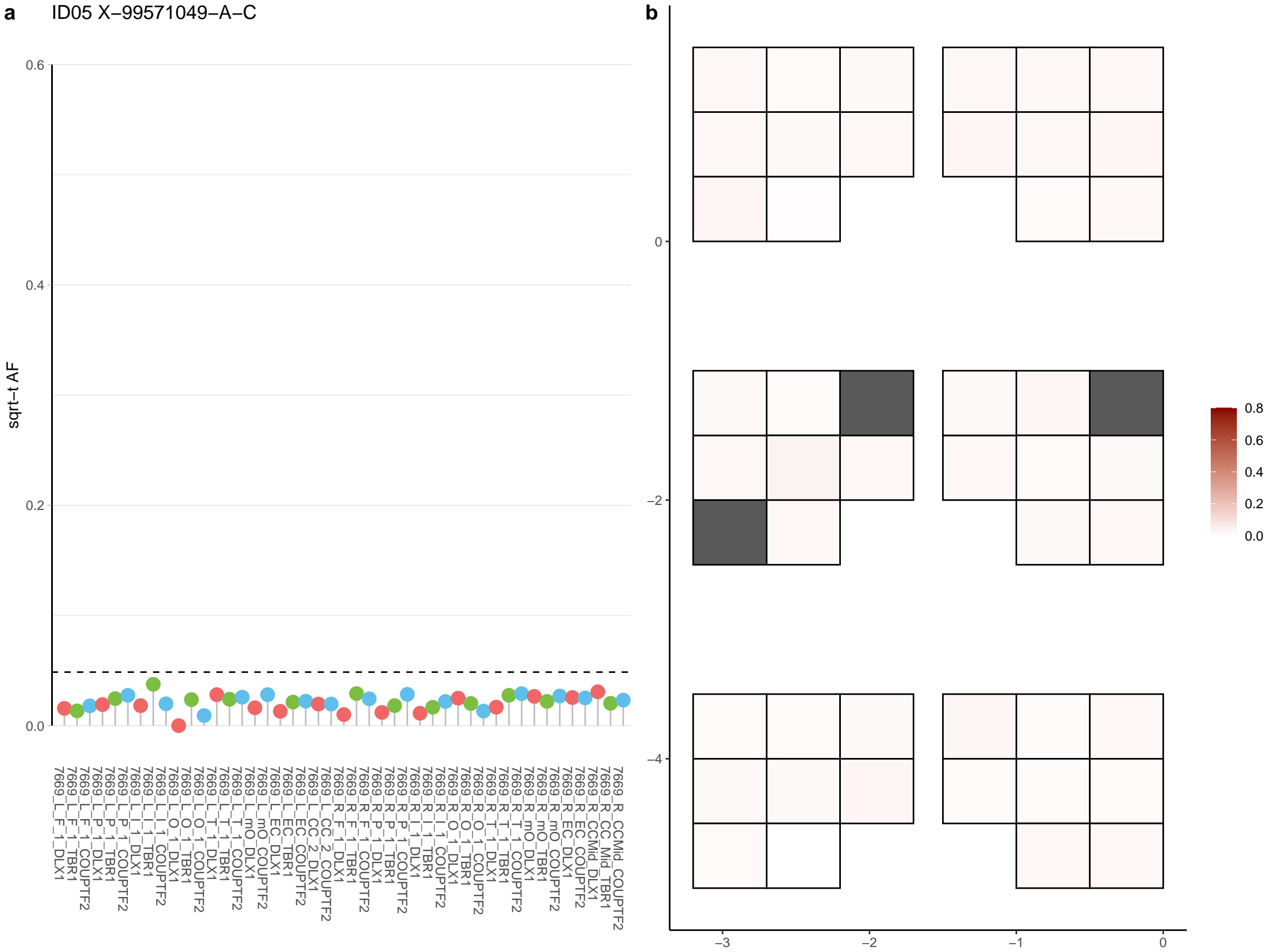

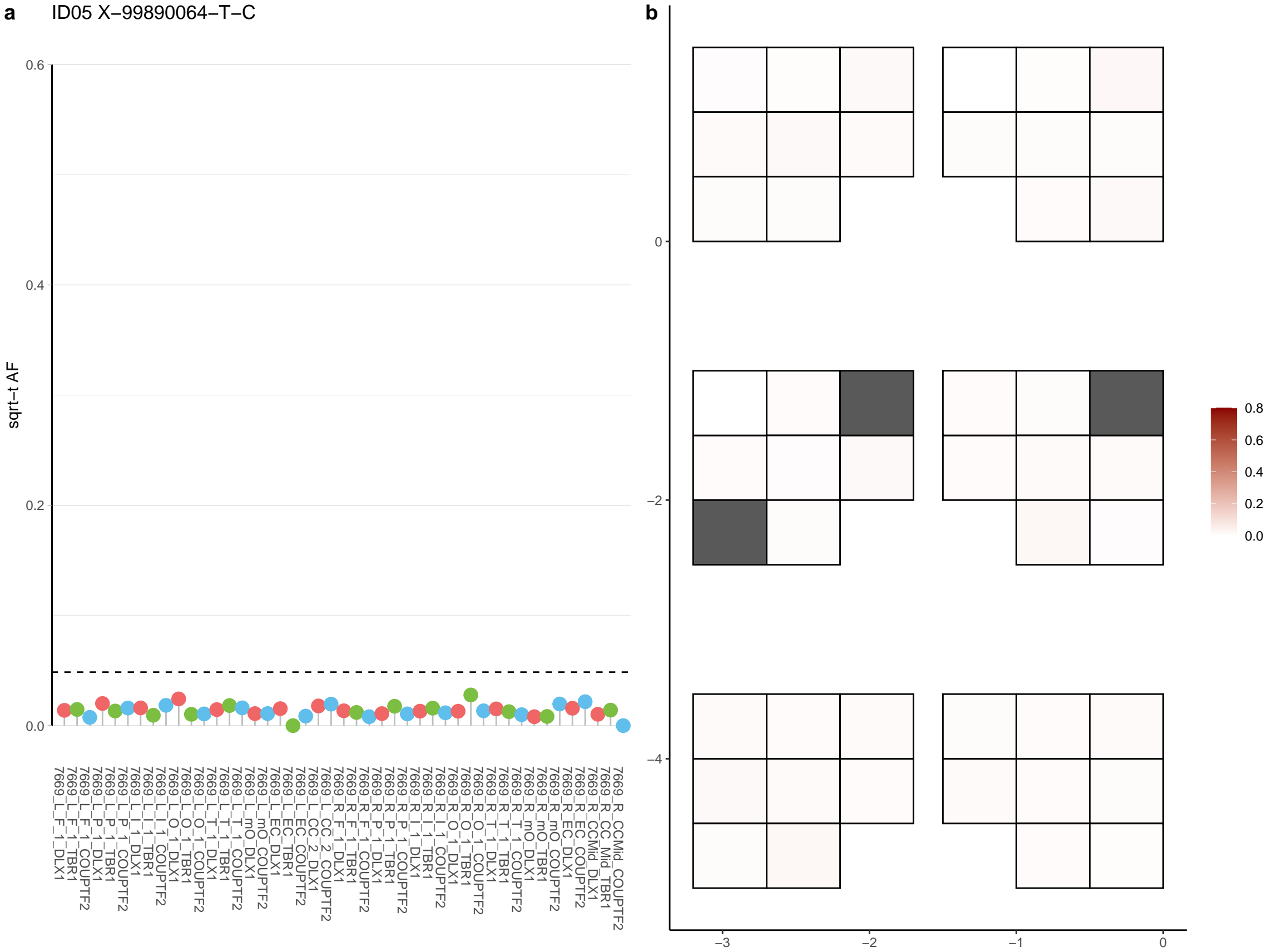
