## Supplementary Data 7 for "Cell-type-resolved somatic mosaicism reveals clonal dynamics of the human forebrain"

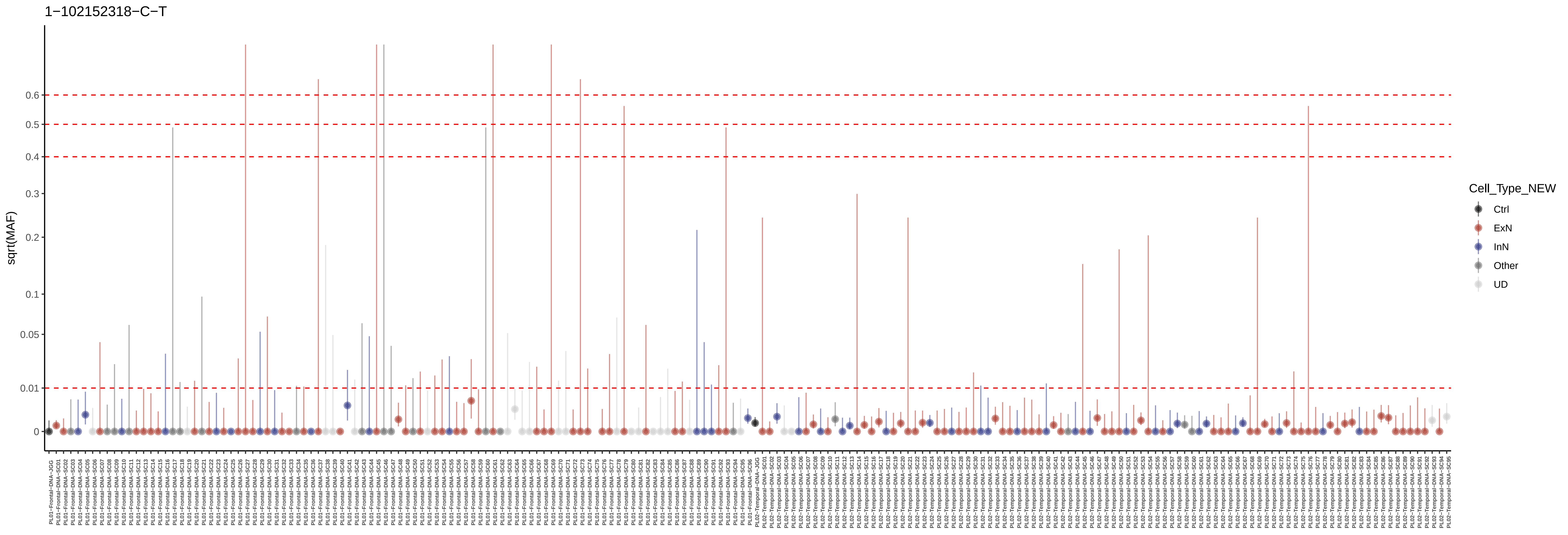
1−102152318−C−T
sqrt(MAF)
0.6
0.5
0.4
0.3
0.2
0.1
0.05
0.01
0
Cell_Type_NEW
Ctrl
ExN
InN
Other
UD

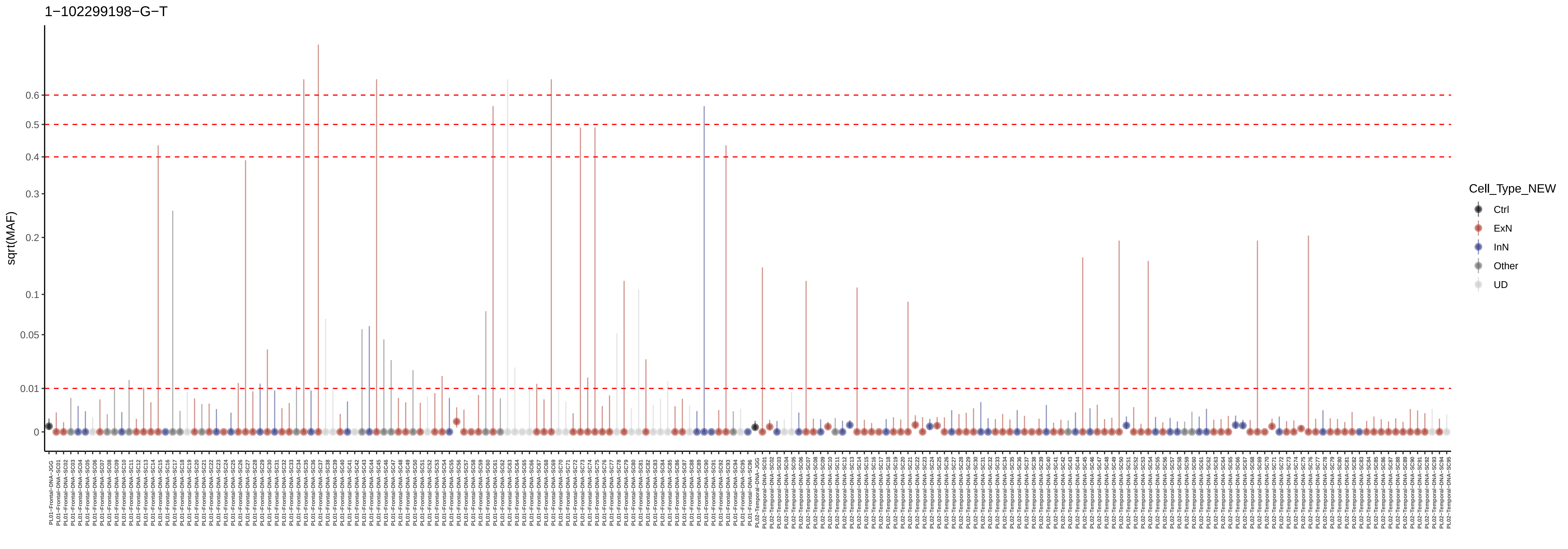
1−102299198−G−T
sqrt(MAF)
0.6
0.5
0.4
0.3
0.2
0.1
0.05
0.01
0
Cell_Type_NEW
Ctrl
ExN
InN
Other
UD

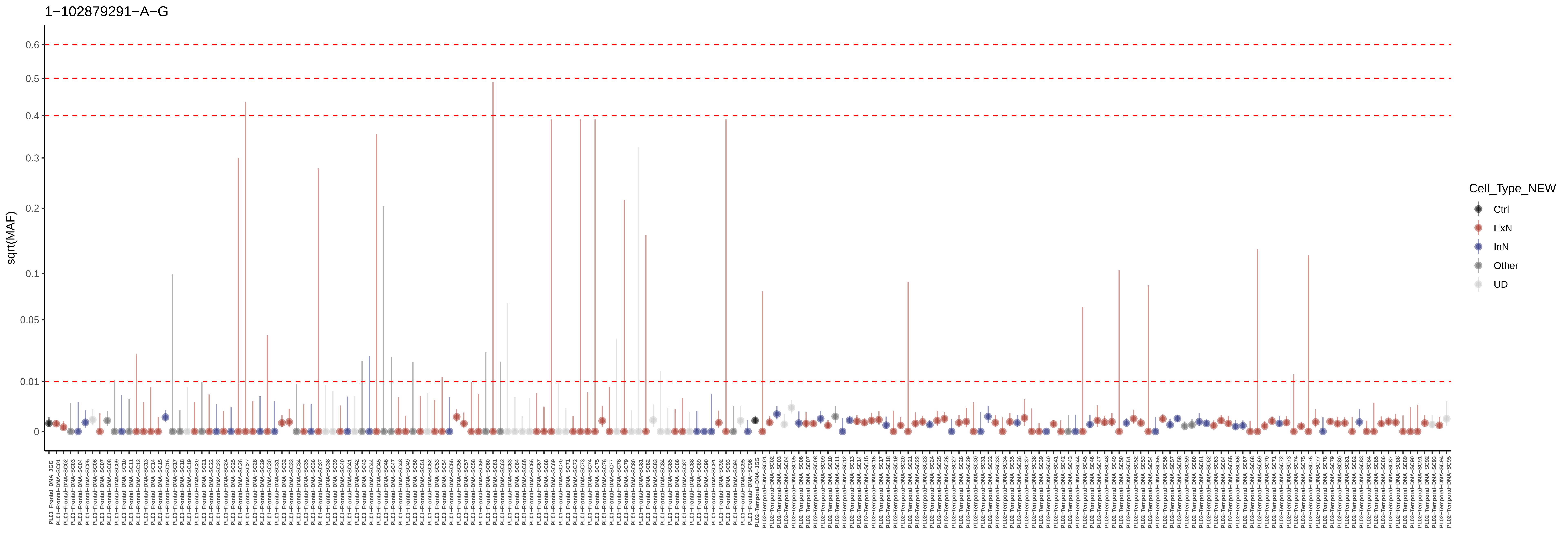
1−102879291−A−G
sqrt(MAF)
0.6
0.5
0.4
0.3
0.2
0.1
0.05
0.01
0
Cell_Type_NEW
Ctrl
ExN
InN
Other
UD

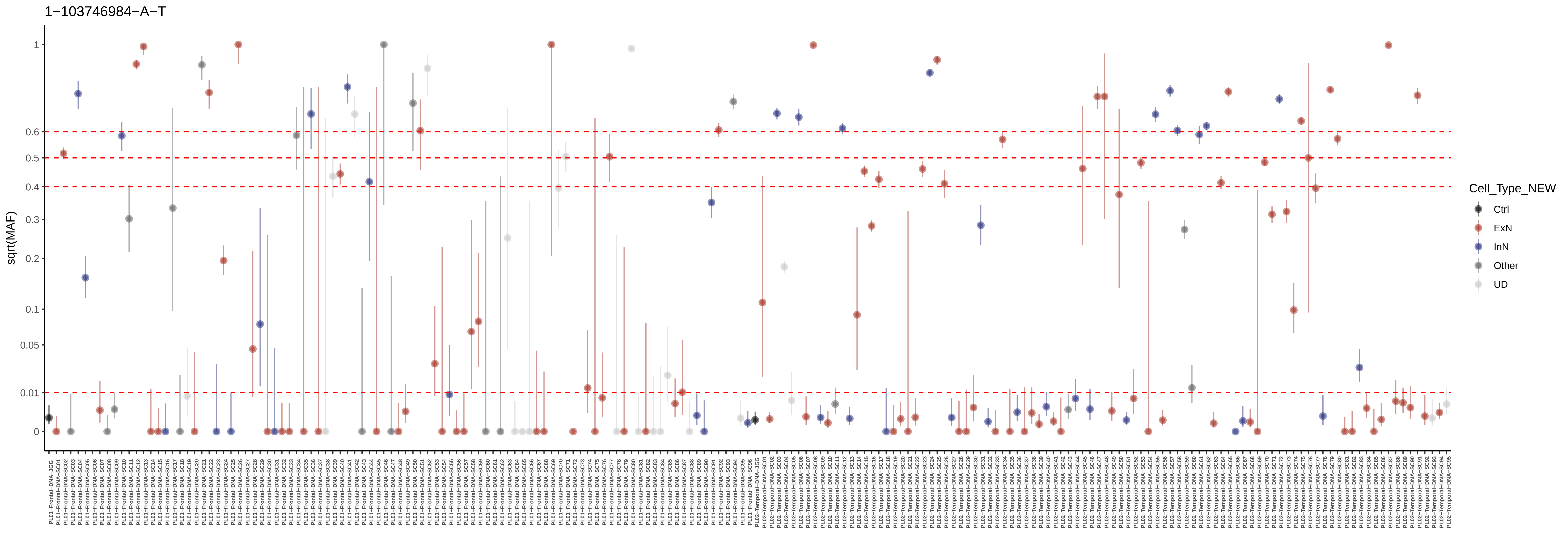
1−103746984−A−T
sqrt(MAF)
1
0.6
0.5
0.4
0.3
0.2
0.1
0.05
0.01
0
Cell_Type_NEW
Ctrl
ExN
InN
Other
UD

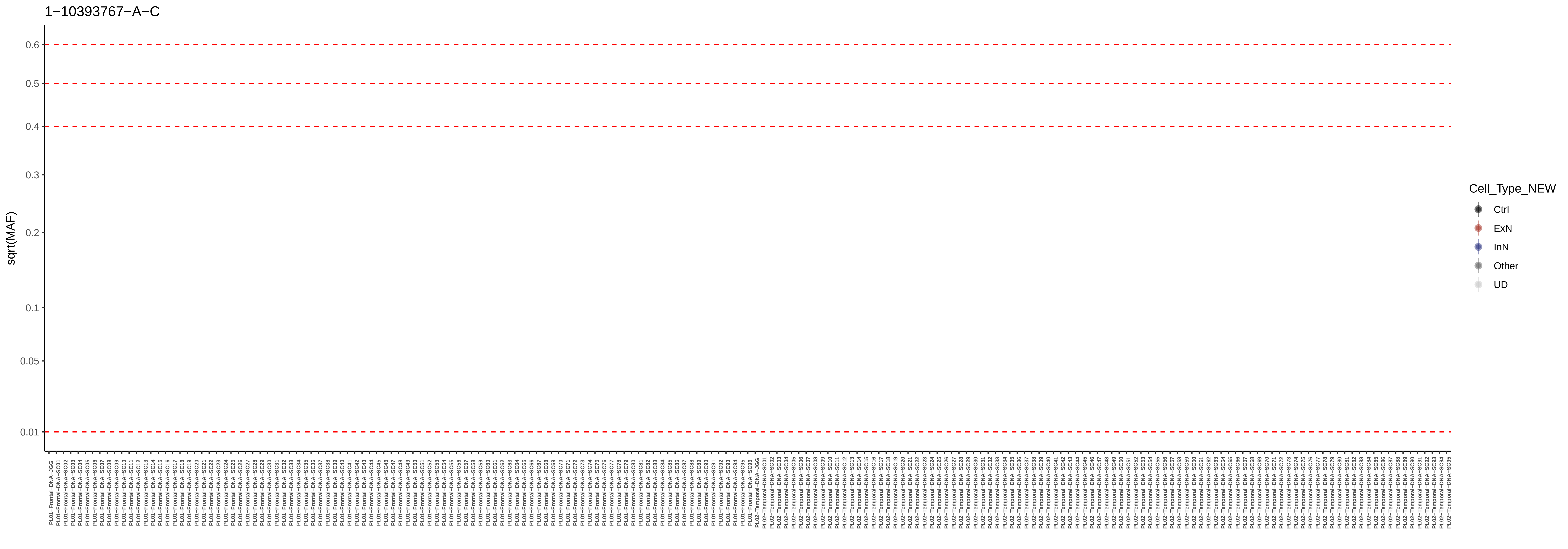
1−10393767−A−C
sqrt(MAF)
0.6
0.5
0.4
0.3
0.2
0.1
0.05
0.01
Cell_Type_NEW
Ctrl
ExN
InN
Other
UD

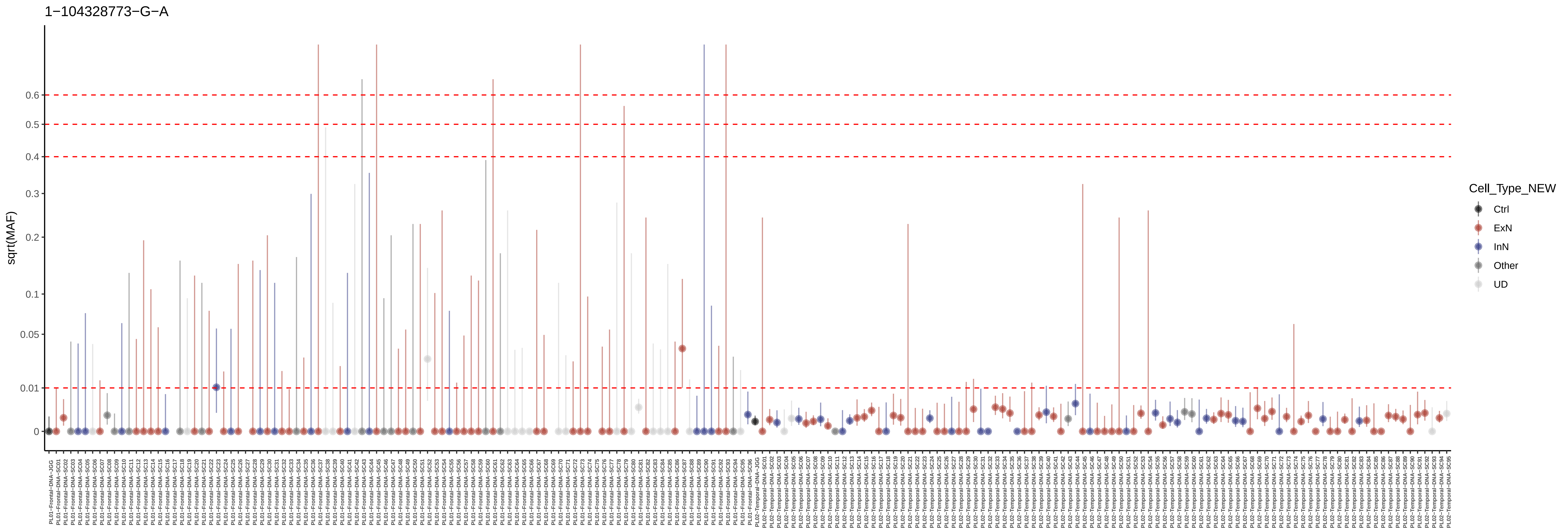
1−104328773−G−A
sqrt(MAF)
0.6
0.5
0.4
0.3
0.2
0.1
0.05
0.01
0
Cell_Type_NEW
Ctrl
ExN
InN
Other
UD

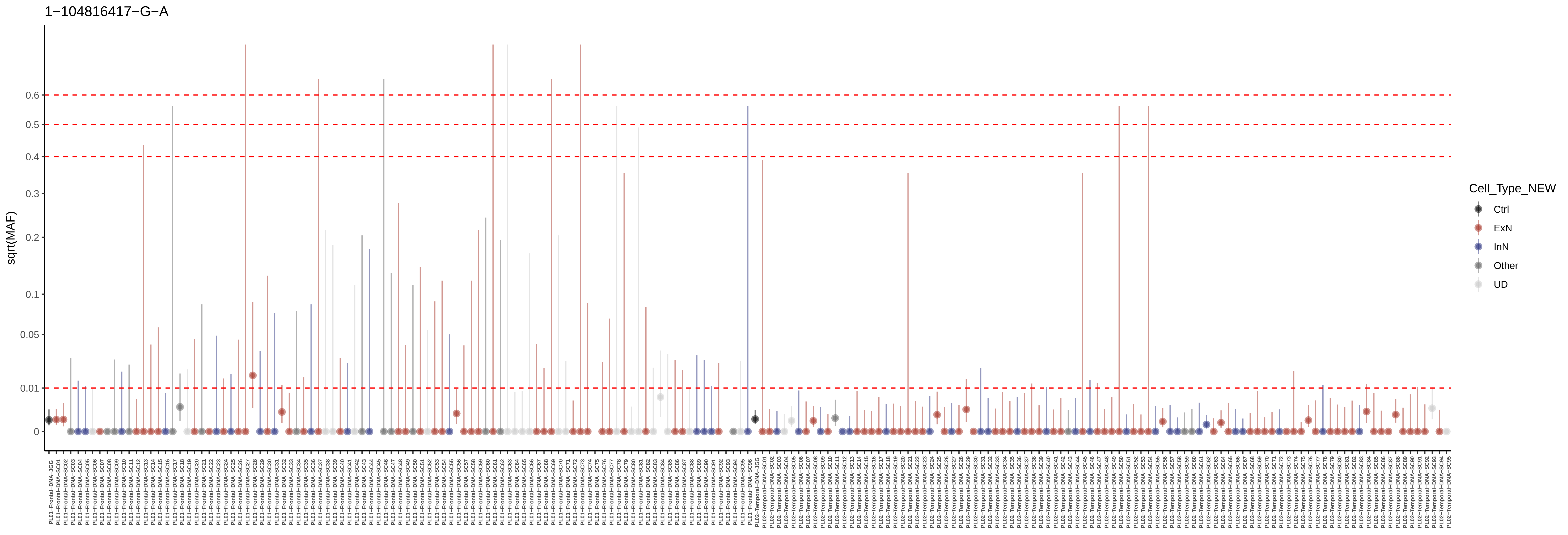
1−104816417−G−A
sqrt(MAF)
0.6
0.5
0.4
0.3
0.2
0.1
0.05
0.01
0
Cell_Type_NEW
Ctrl
ExN
InN
Other
UD

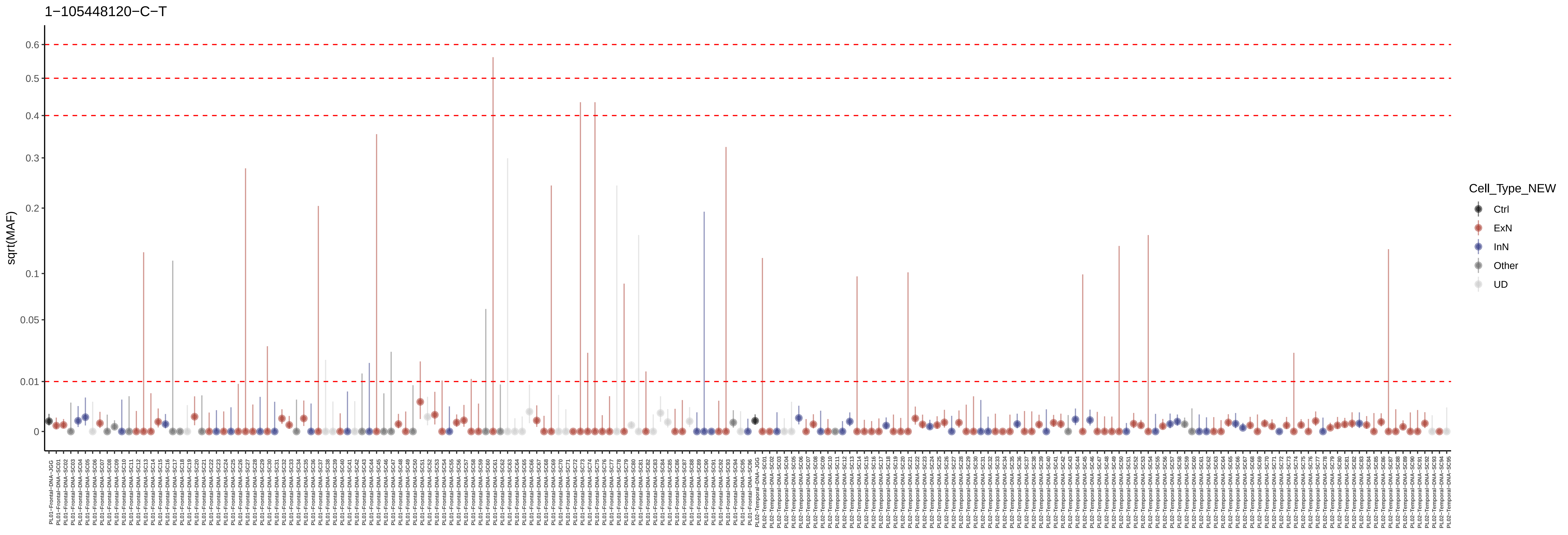
1−105448120−C−T
sqrt(MAF)
0.6
0.5
0.4
0.3
0.2
0.1
0.05
0.01
0
Cell_Type_NEW
Ctrl
ExN
InN
Other
UD

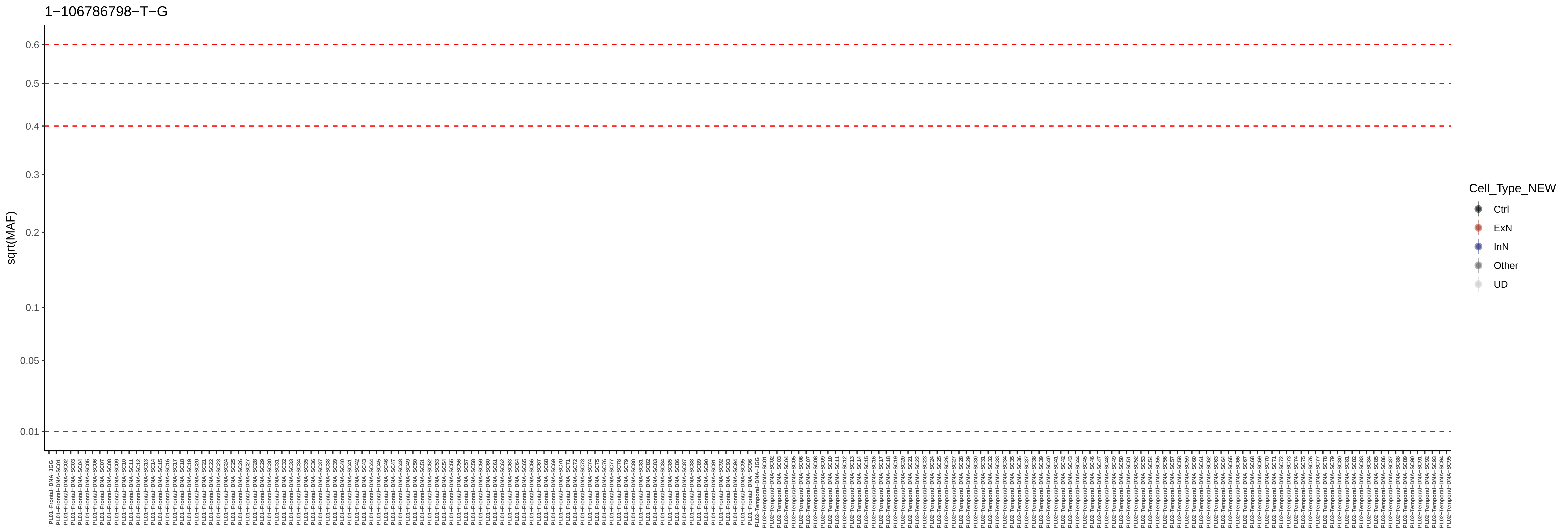
1−106786798−T−G
sqrt(MAF)
0.6
0.5
0.4
0.3
0.2
0.1
0.05
0.01
Cell_Type_NEW
Ctrl
ExN
InN
Other
UD

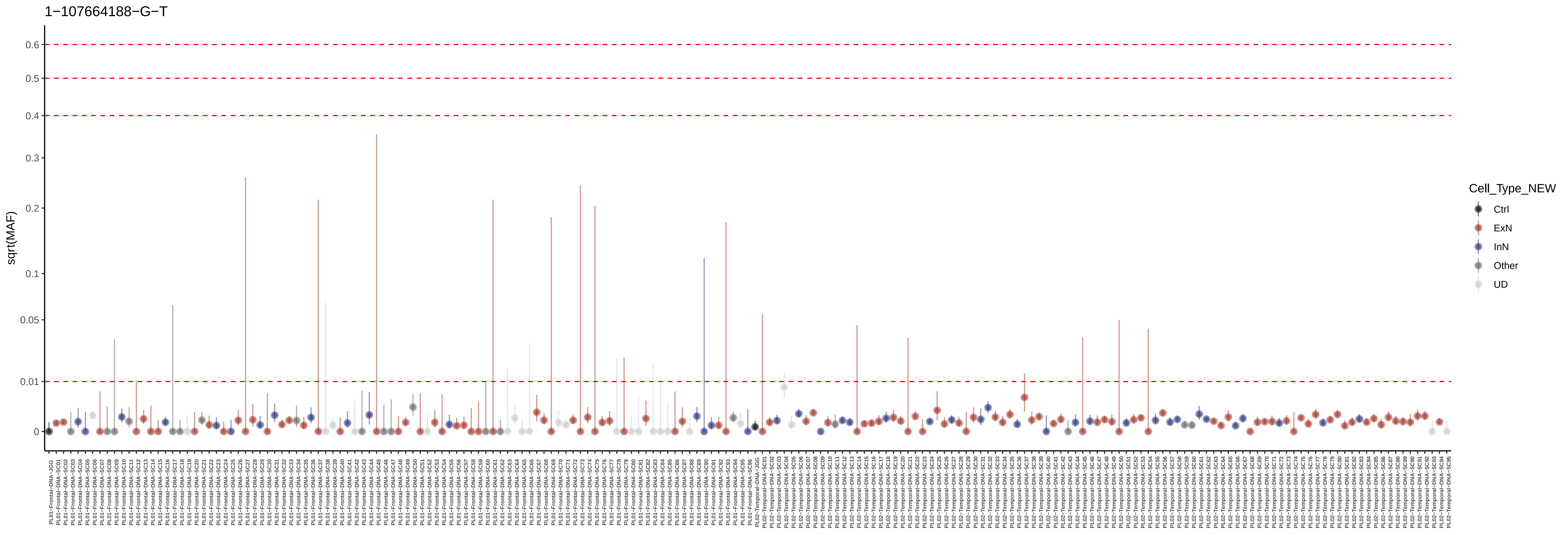

1−107664188−G−T
sqrt(MAF)
0.6
0.5
0.4
0.3
0.2
0.1
0.05
0.01
0
Cell_Type_NEW
Ctrl
ExN
InN
Other
UD

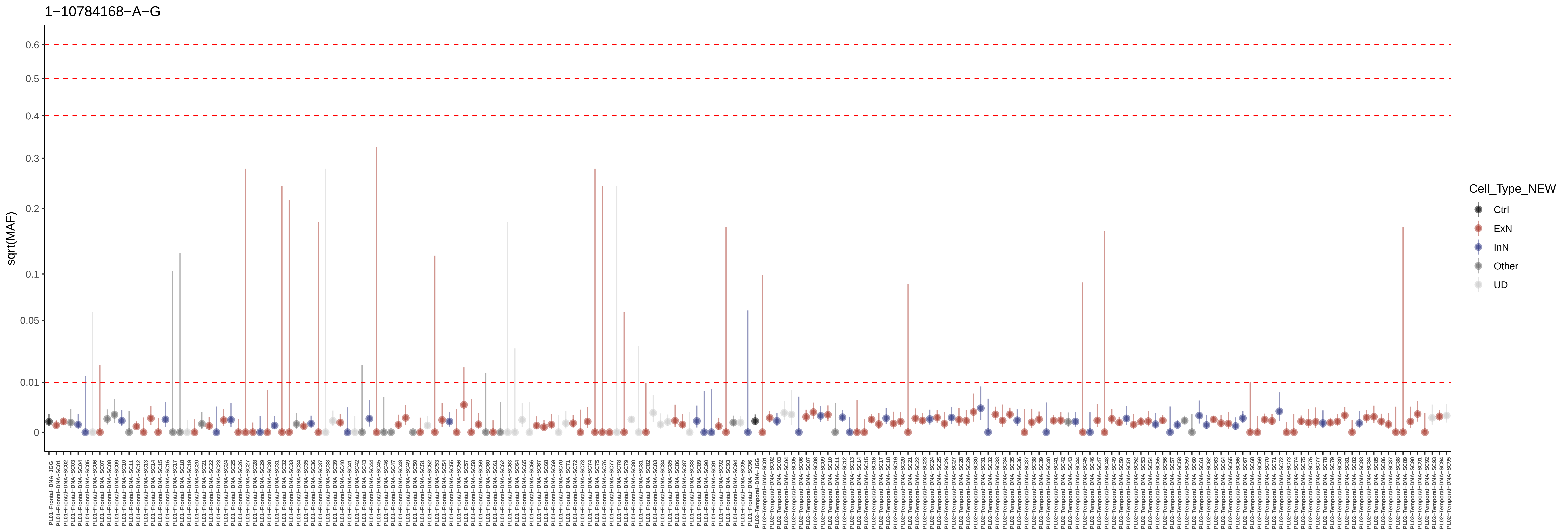
1−10784168−A−G
sqrt(MAF)
0.6
0.5
0.4
0.3
0.2
0.1
0.05
0.01
0
Cell_Type_NEW
Ctrl
ExN
InN
Other
UD

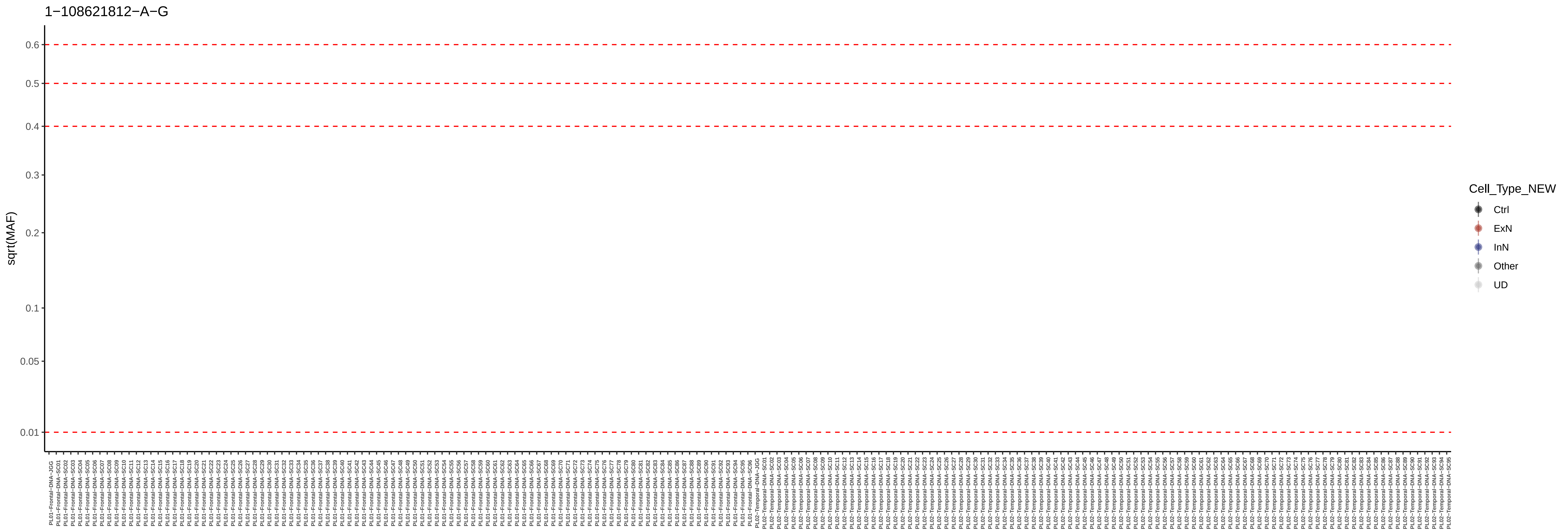
1−108621812−A−G
sqrt(MAF)
0.6
0.5
0.4
0.3
0.2
0.1
0.05
0.01
Cell_Type_NEW
Ctrl
ExN
InN
Other
UD

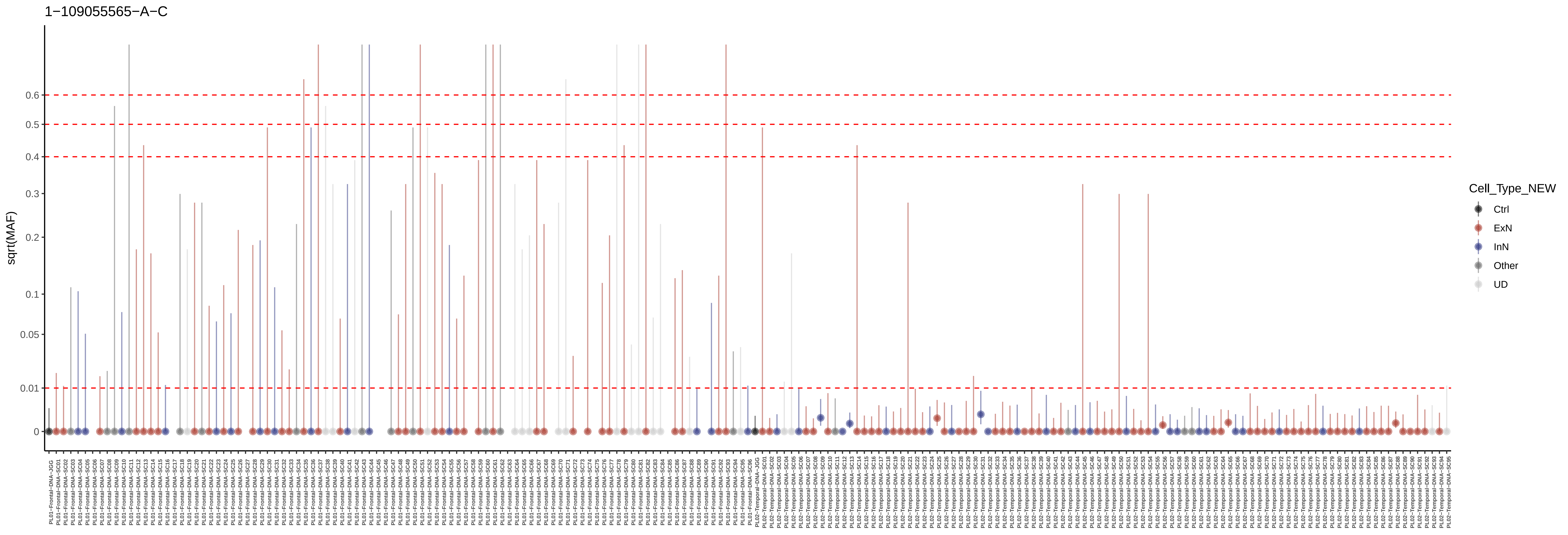
1−109055565−A−C
sqrt(MAF)
0.6
0.5
0.4
0.3
0.2
0.1
0.05
0.01
0
Cell_Type_NEW
Ctrl
ExN
InN
Other
UD

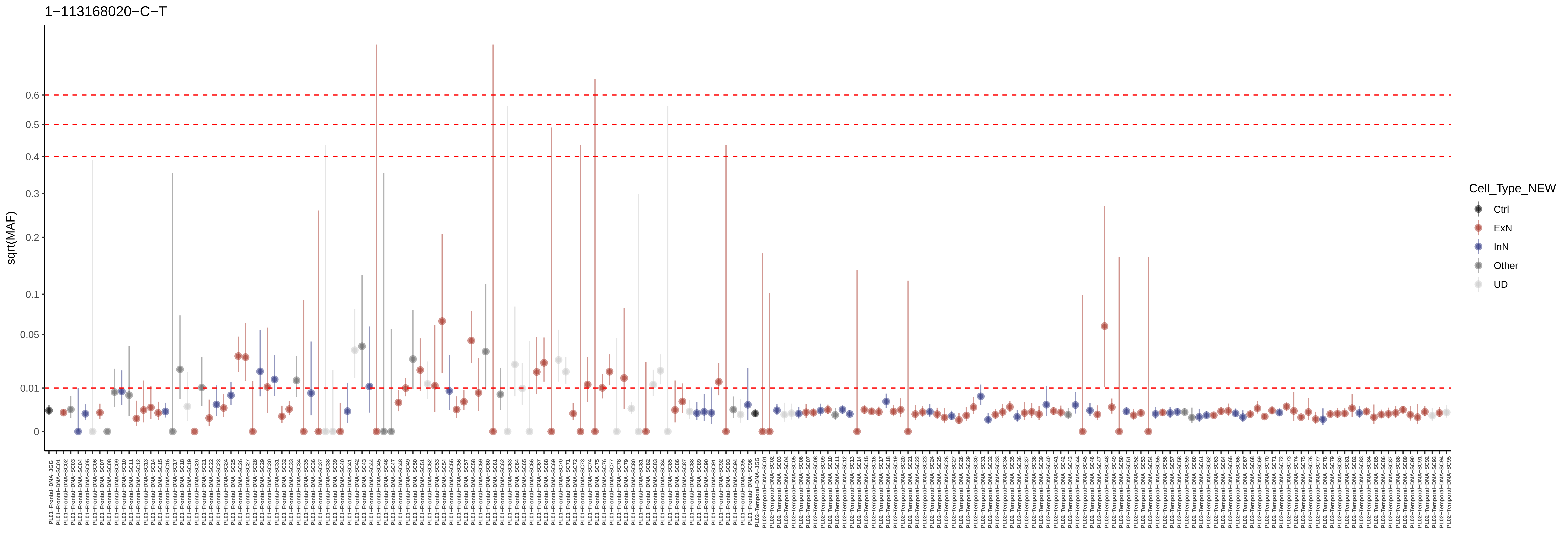

1−113168020−C−T
sqrt(MAF)
Cell_Type_NEW
Ctrl
ExN
InN
Other
UD

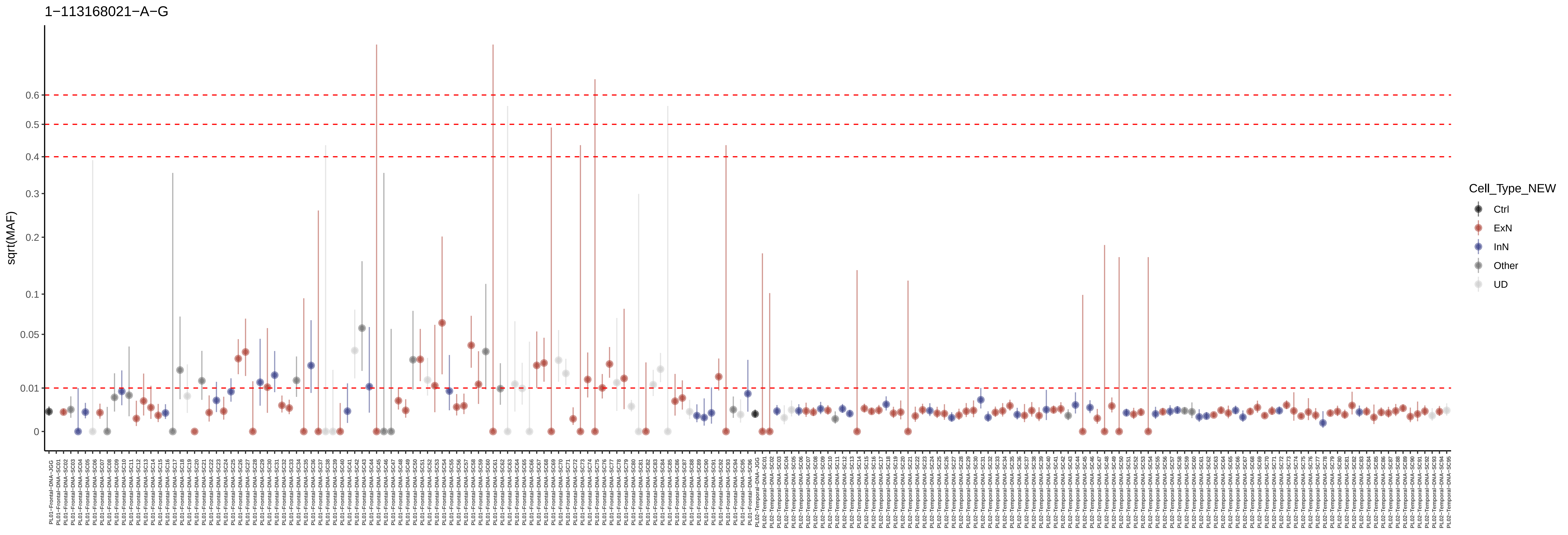
1−113168021−A−G
sqrt(MAF)
0.6
0.5
0.4
0.3
0.2
0.1
0.05
0.01
0
Cell_Type_NEW
Ctrl
ExN
InN
Other
UD

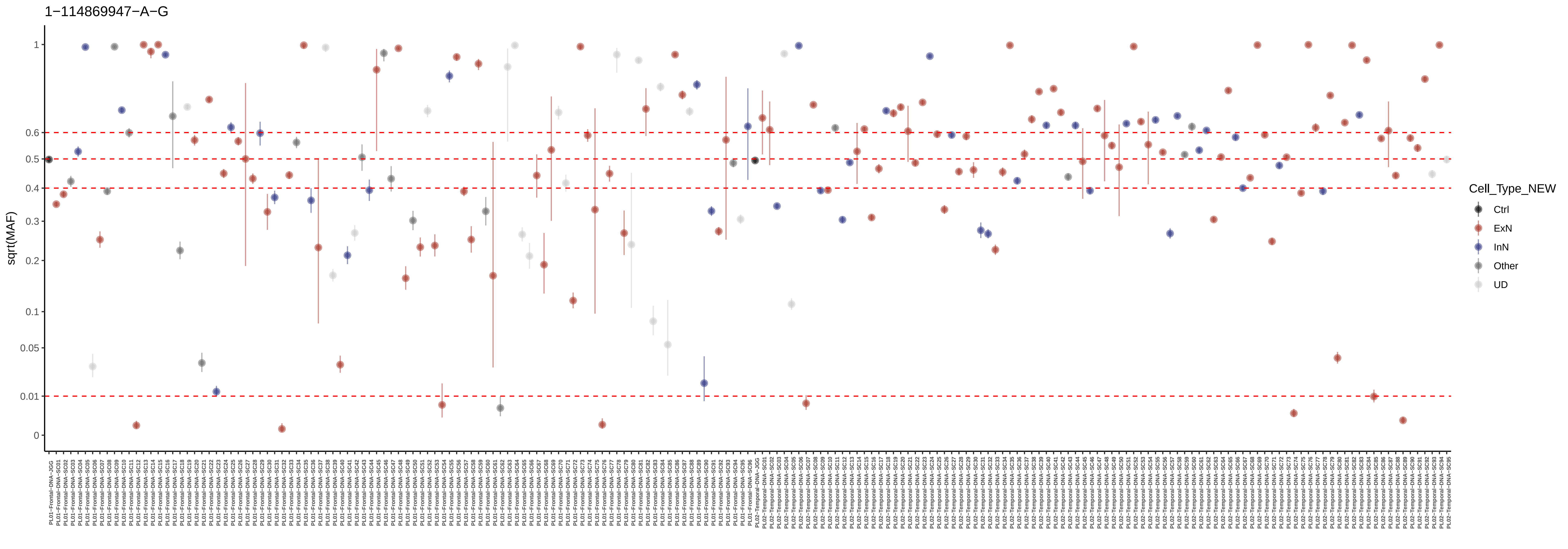

1−114869947−A−G
sqrt(MAF)
1
0.6
0.5
0.4
0.3
0.2
0.1
0.05
0.01
0
Cell_Type_NEW
Ctrl
ExN
InN
Other
UD

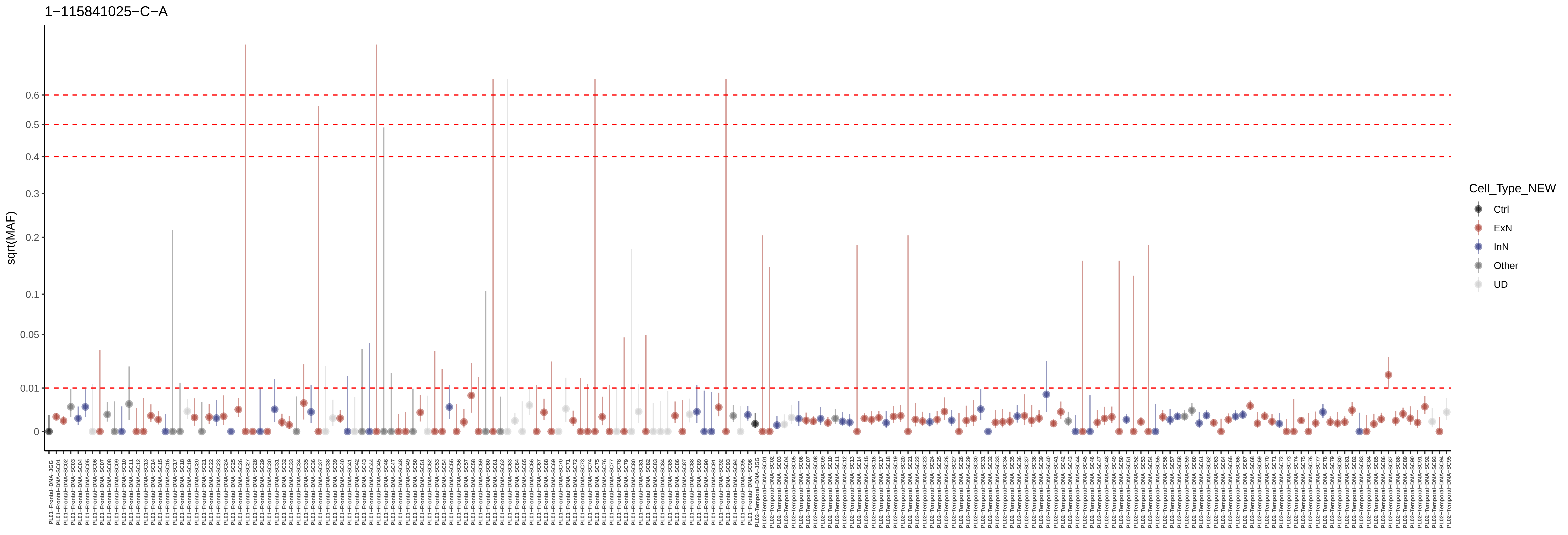
1−115841025−C−A
sqrt(MAF)
0.6
0.5
0.4
0.3
0.2
0.1
0.05
0.01
0
Cell_Type_NEW
Ctrl
ExN
InN
Other
UD

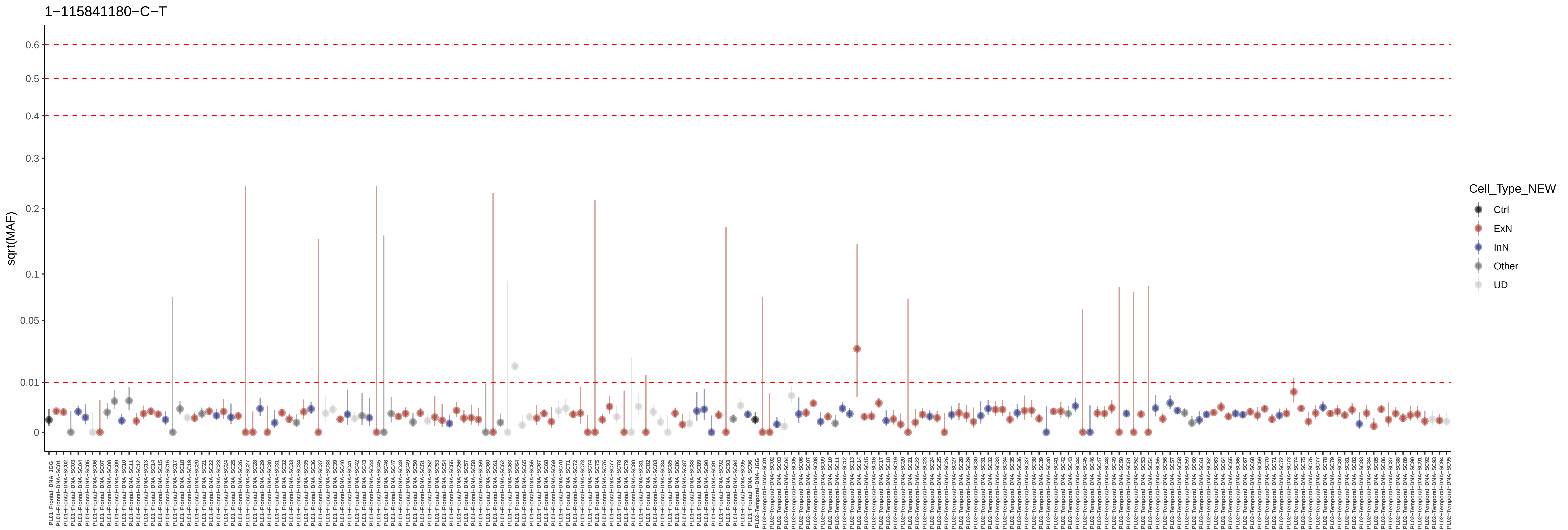
1−115841180−C−T
sqrt(MAF)
0.6
0.5
0.4
0.3
0.2
0.1
0.05
0.01
0
Cell_Type_NEW
Ctrl
ExN
InN
Other
UD

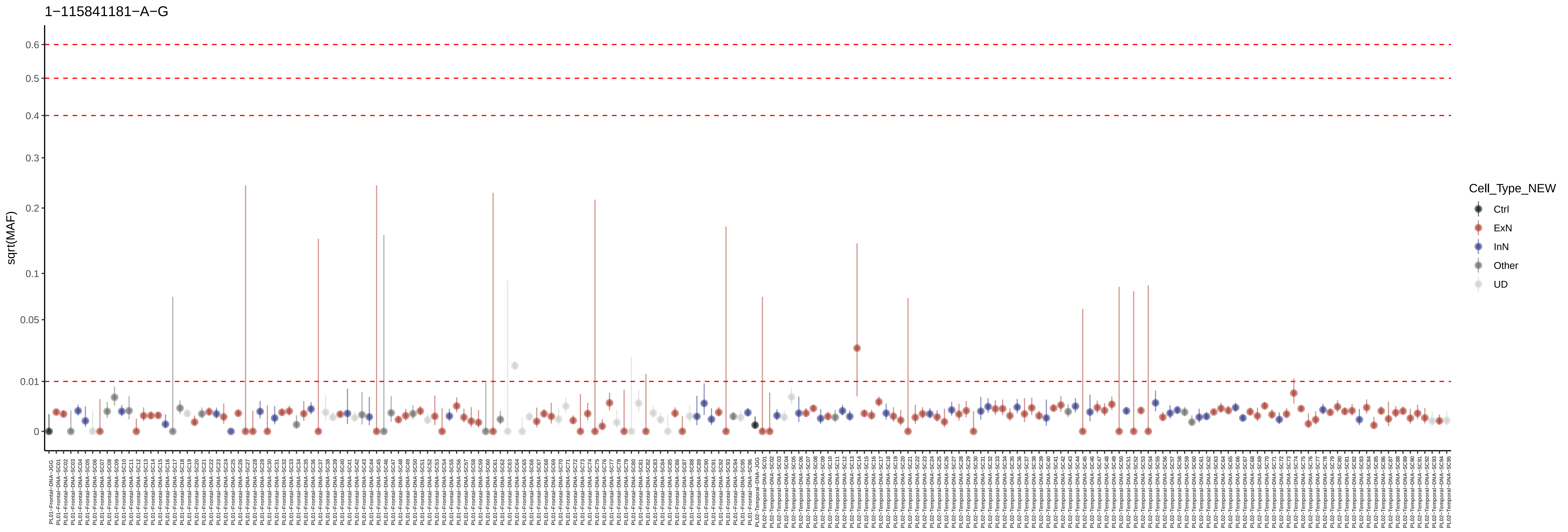
1−115841181−A−G
sqrt(MAF)
0.6
0.5
0.4
0.3
0.2
0.1
0.05
0.01
0
Cell_Type_NEW
Ctrl
ExN
InN
Other
UD

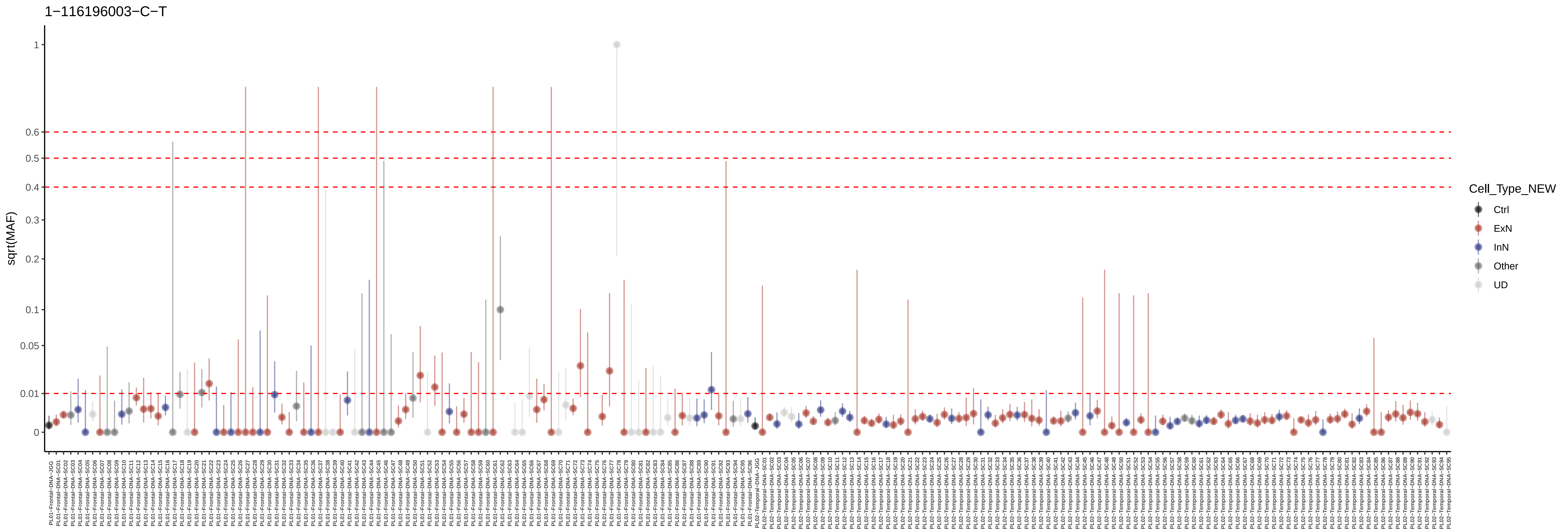
1−116196003−C−T
sqrt(MAF)
1
0.6
0.5
0.4
0.3
0.2
0.1
0.05
0.01
0
Cell_Type_NEW
Ctrl
ExN
InN
Other
UD

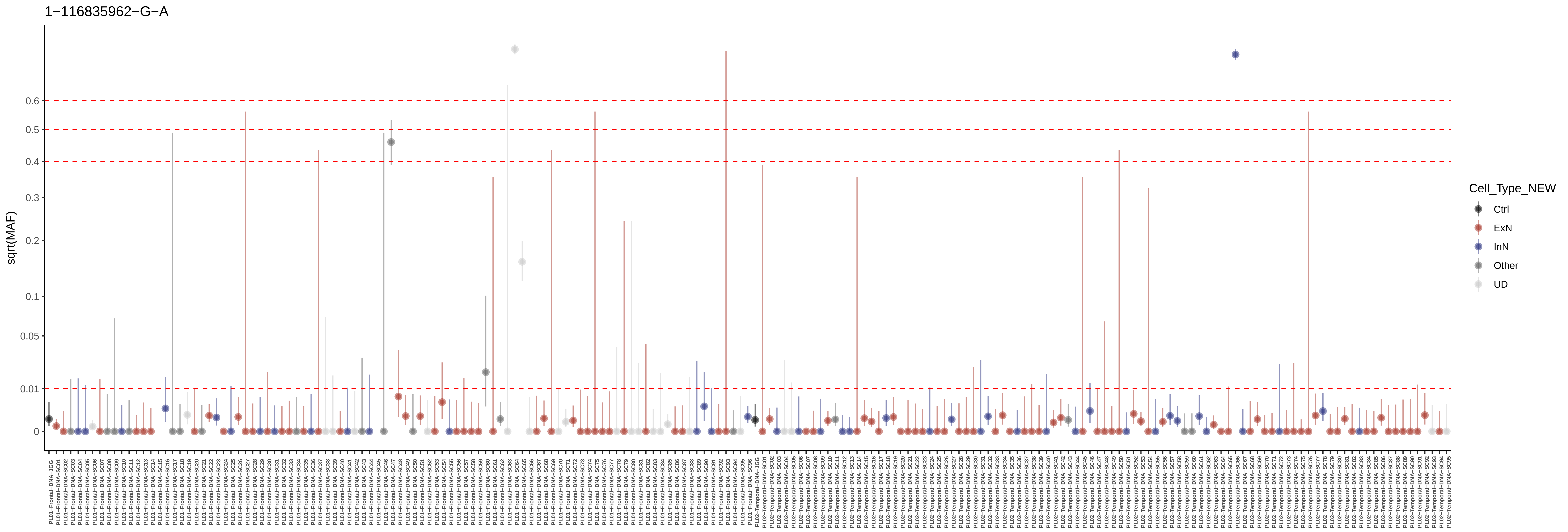
1−116835962−G−A
sqrt(MAF)
0.6
0.5
0.4
0.3
0.2
0.1
0.05
0.01
0
Cell_Type_NEW
Ctrl
ExN
InN
Other
UD

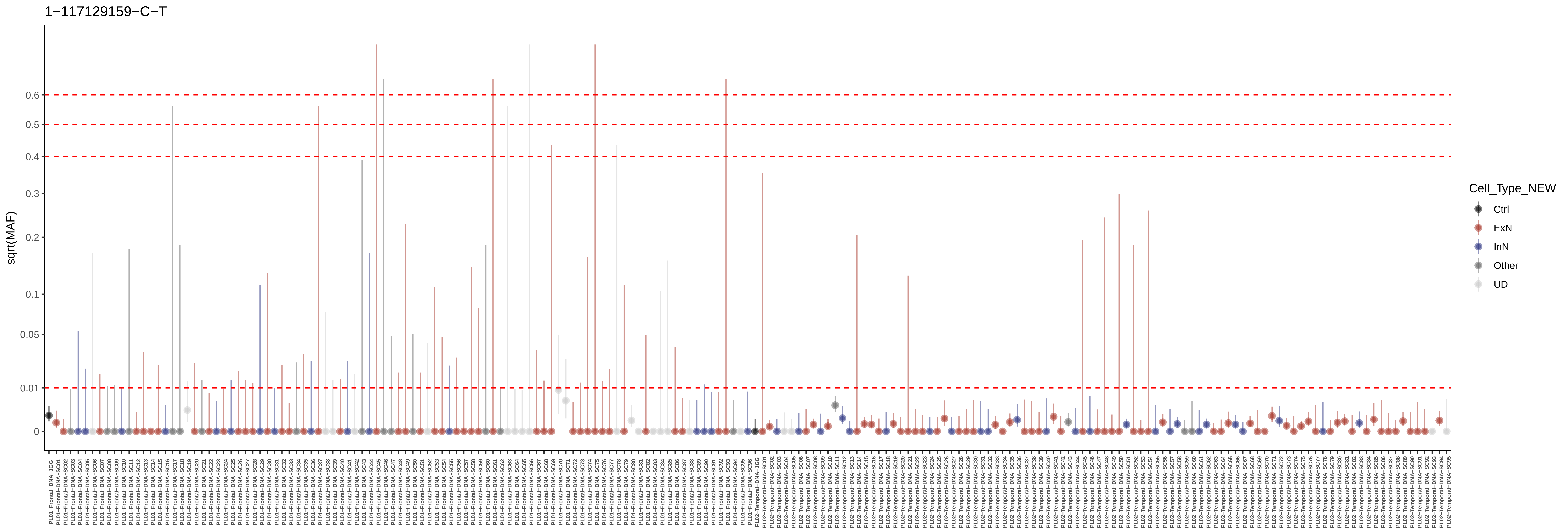
1−117129159−C−T
sqrt(MAF)
0.6
0.5
0.4
0.3
0.2
0.1
0.05
0.01
0
Cell_Type_NEW
Ctrl
ExN
InN
Other
UD

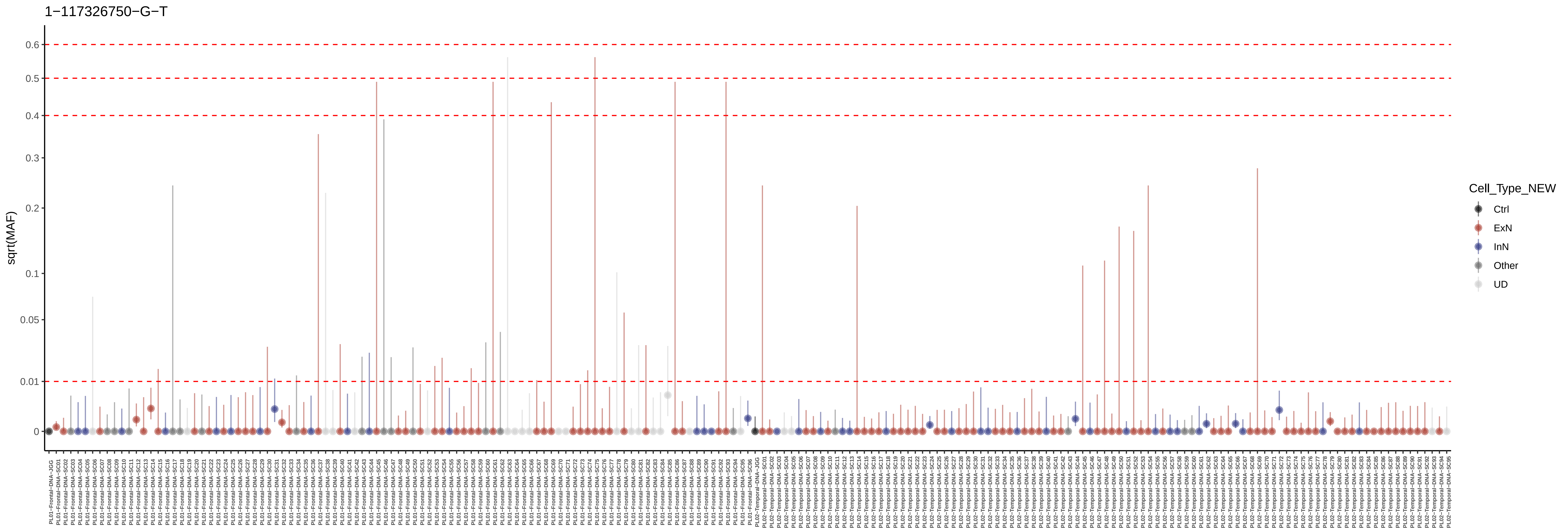
1−117326750−G−T
sqrt(MAF)
0.6
0.5
0.4
0.3
0.2
0.1
0.05
0.01
0
Cell_Type_NEW
Ctrl
ExN
InN
Other
UD

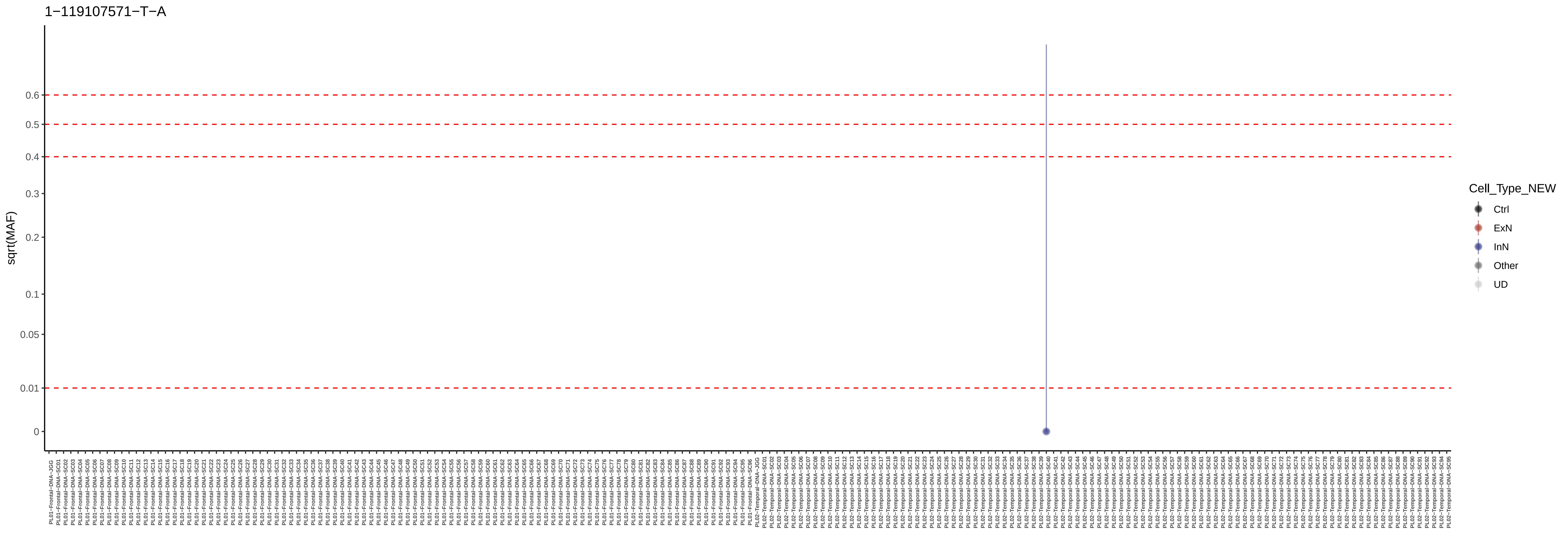
1−119107571−T−A
sqrt(MAF)
0.6
0.5
0.4
0.3
0.2
0.1
0.05
0.01
0
Cell_Type_NEW
Ctrl
ExN
InN
Other
UD

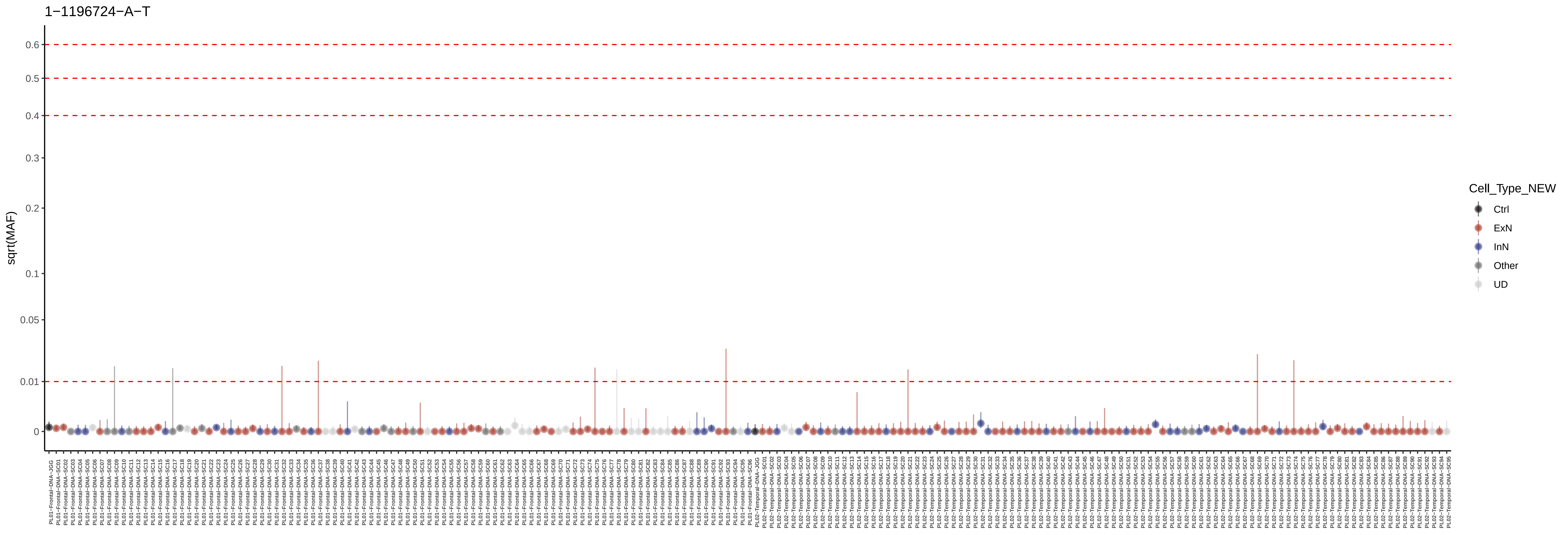
1−1196724−A−T
sqrt(MAF)
Cell_Type_NEW
Ctrl
ExN
InN
Other
UD

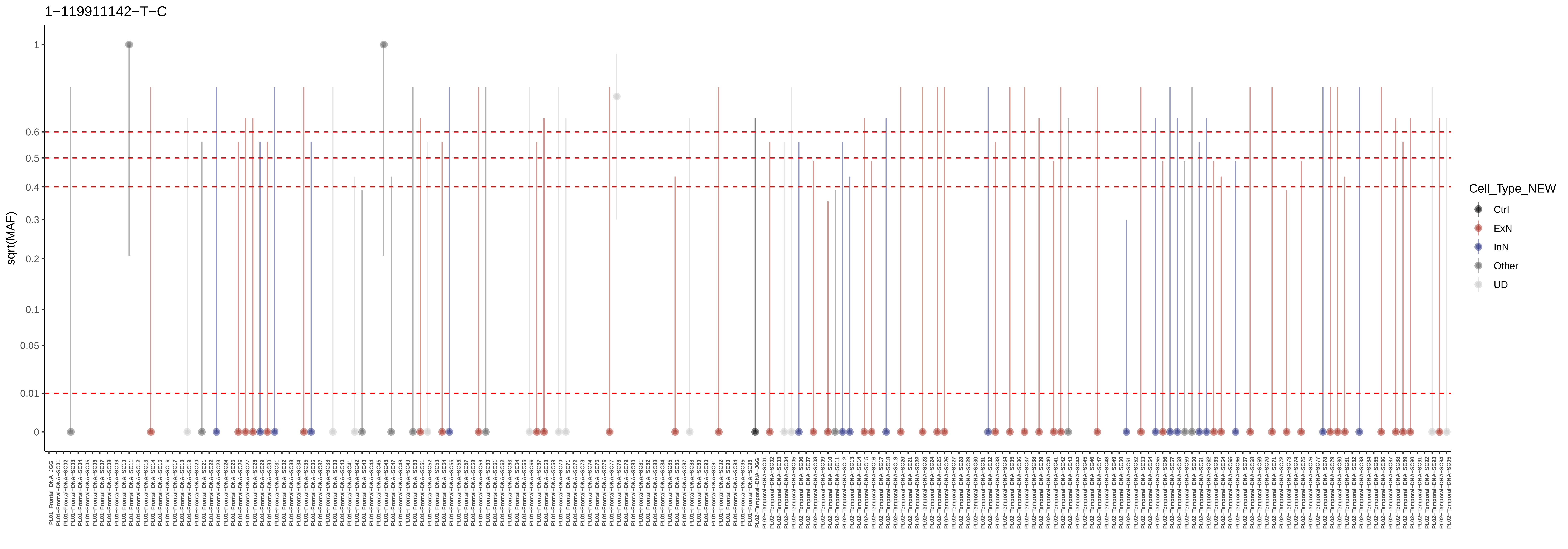
1−119911142−T−C
sqrt(MAF)
1
0.6
0.5
0.4
0.3
0.2
0.1
0.05
0.01
0
Cell_Type_NEW
Ctrl
ExN
InN
Other
UD

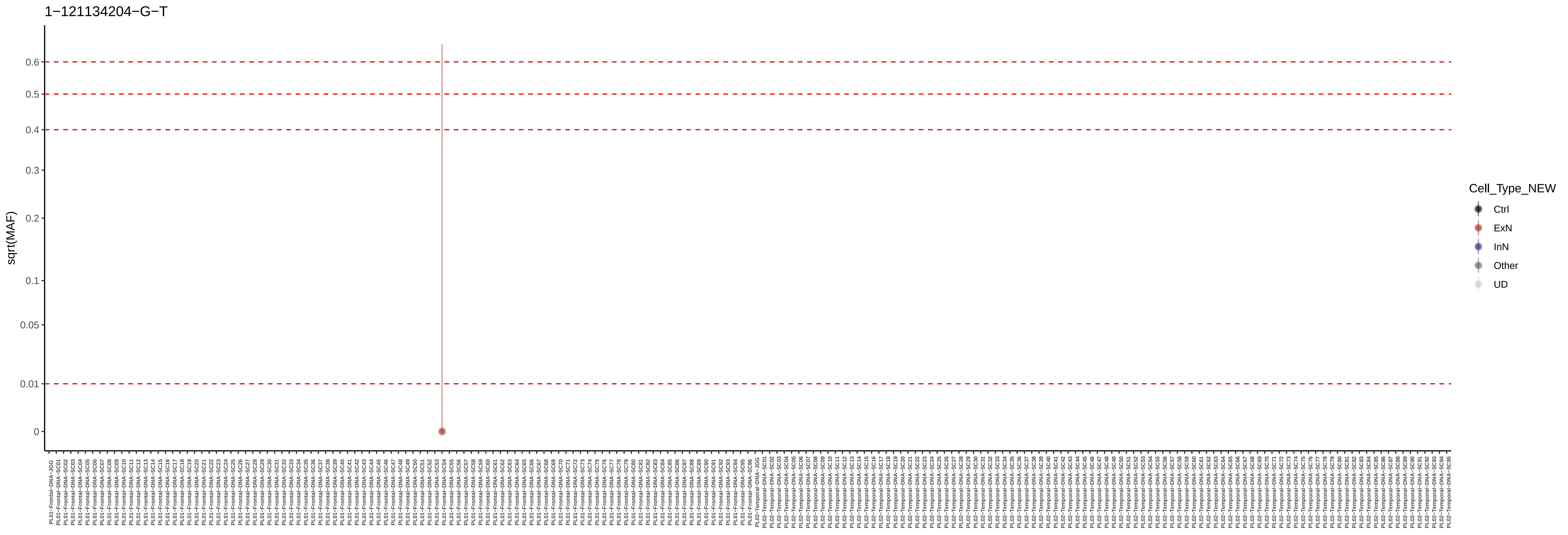
1−121134204−G−T
sqrt(MAF)
0.6
0.5
0.4
0.3
0.2
0.1
0.05
0.01
0
Cell_Type_NEW
Ctrl
ExN
InN
Other
UD

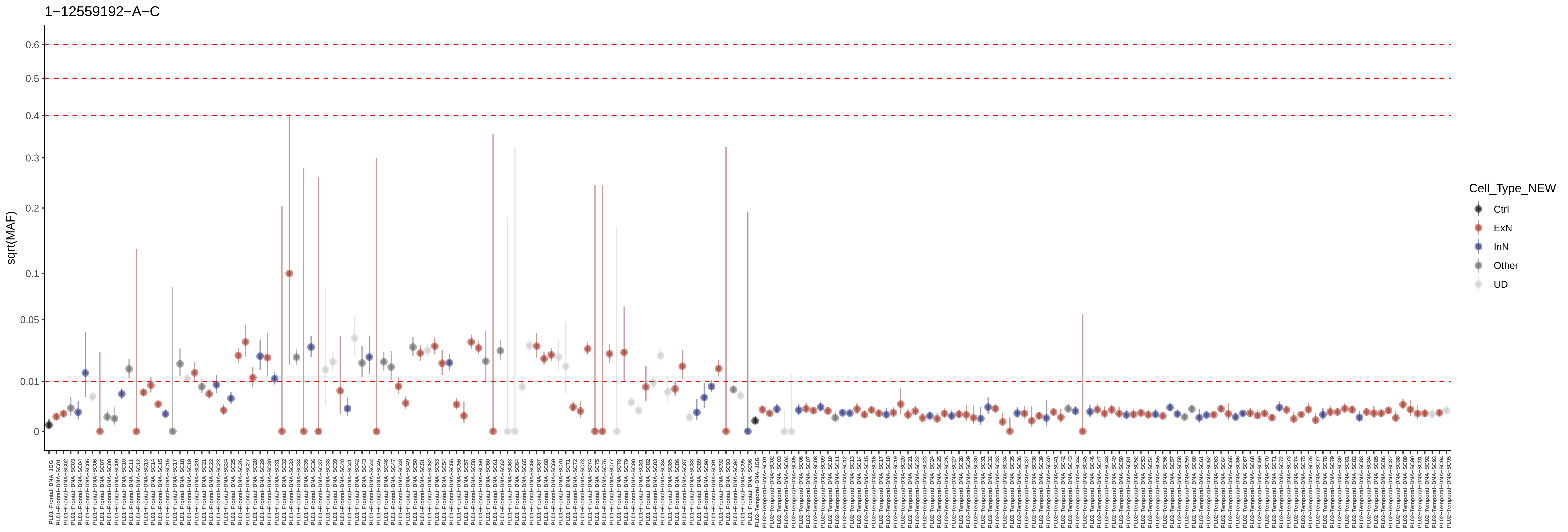
1−12559192−A−C
sqrt(MAF)
0.6
0.5
0.4
0.3
0.2
0.1
0.05
0.01
0
Cell_Type_NEW
Ctrl
ExN
InN
Other
UD

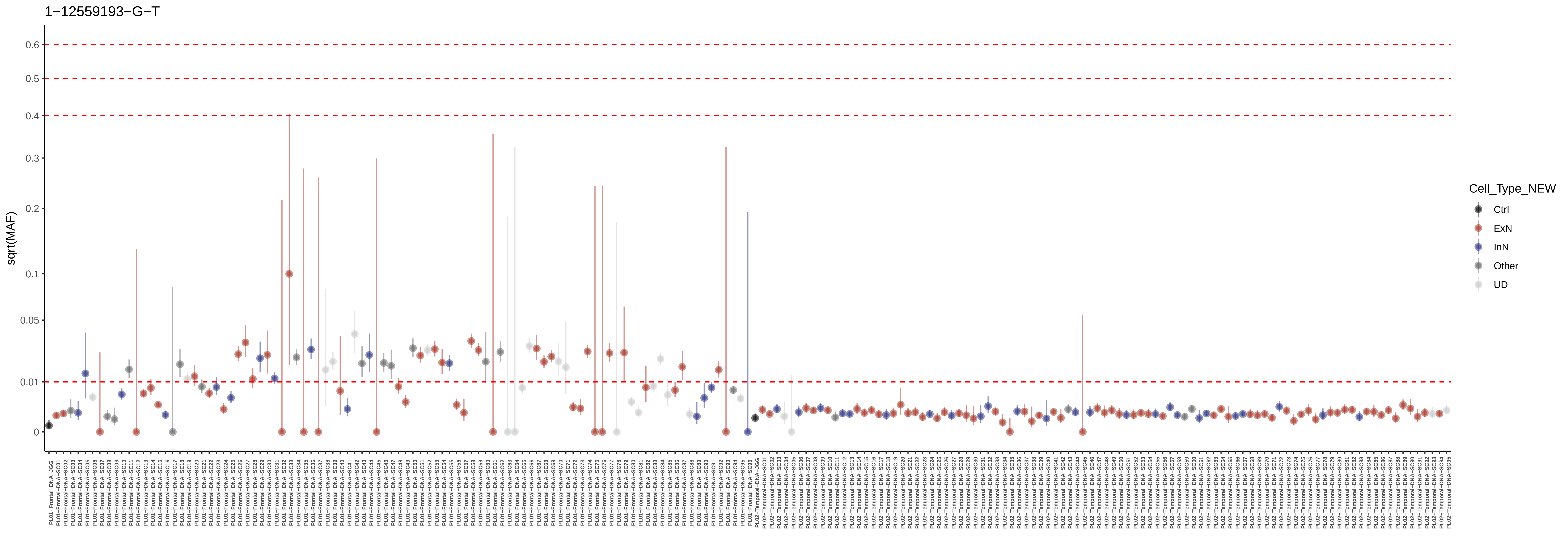
1−12559193−G−T
sqrt(MAF)
0.6
0.5
0.4
0.3
0.2
0.1
0.05
0.01
0
Cell_Type_NEW
Ctrl
ExN
InN
Other
UD

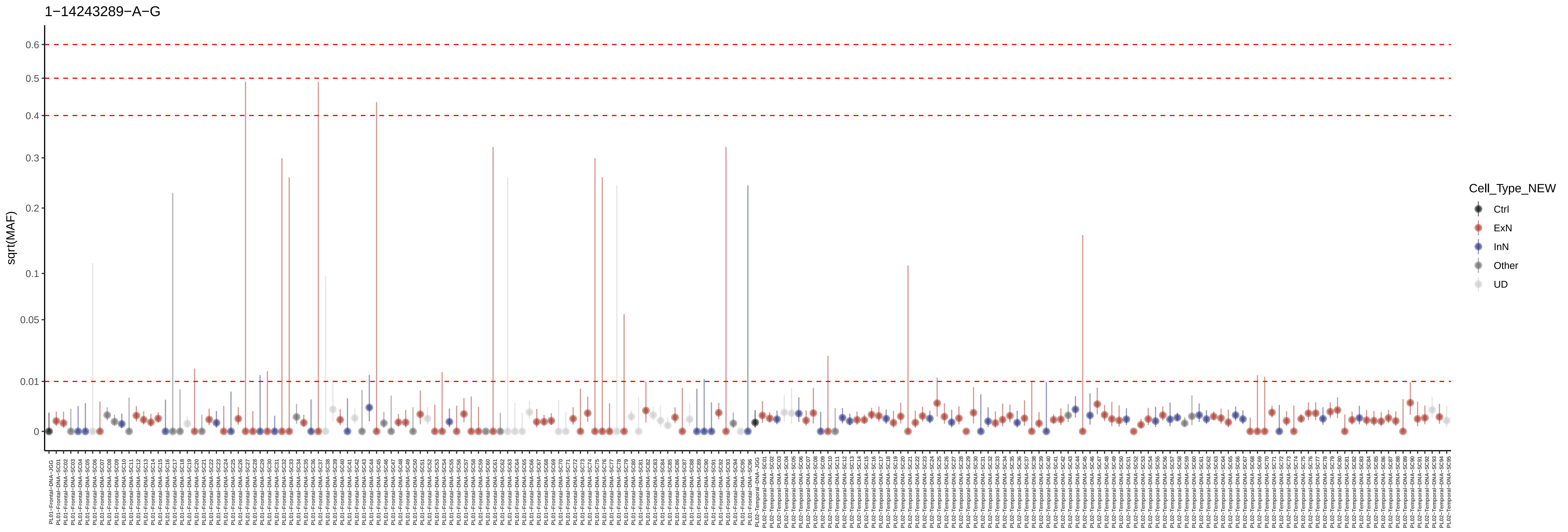
1−14243289−A−G
sqrt(MAF)
0.6
0.5
0.4
0.3
0.2
0.1
0.05
0.01
0
Cell_Type_NEW
Ctrl
ExN
InN
Other
UD

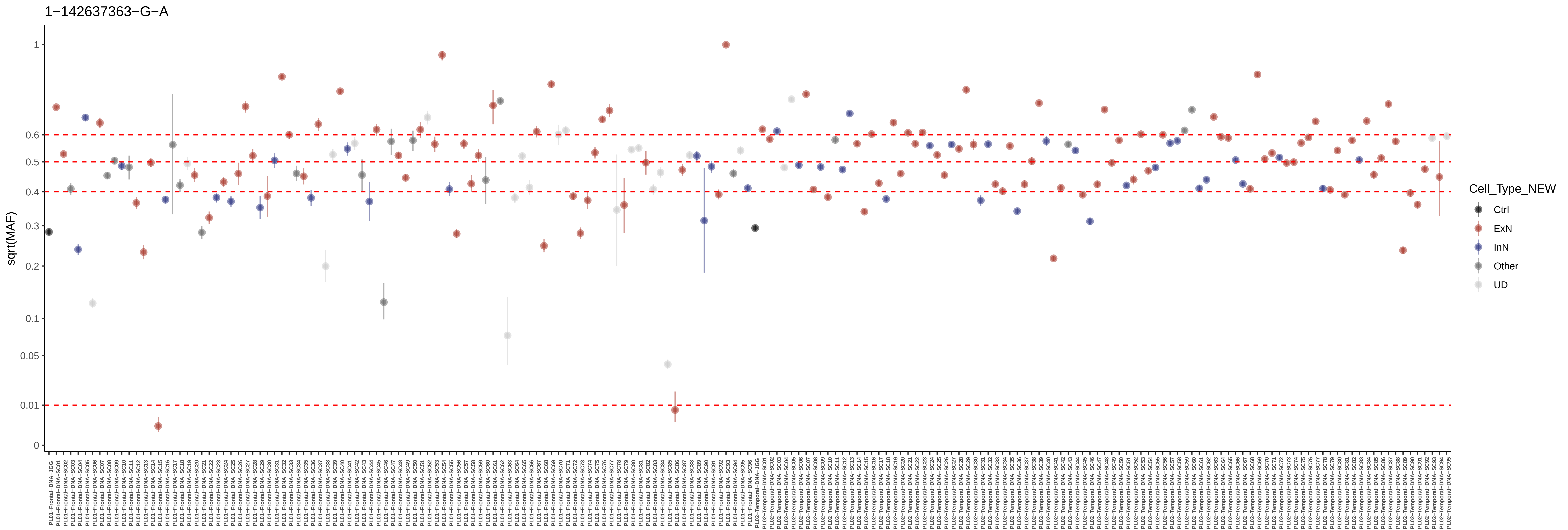

1−142637363−G−A
sqrt(MAF)
1
0.6
0.5
0.4
0.3
0.2
0.1
0.05
0.01
0
Cell_Type_NEW
Ctrl
ExN
InN
Other
UD

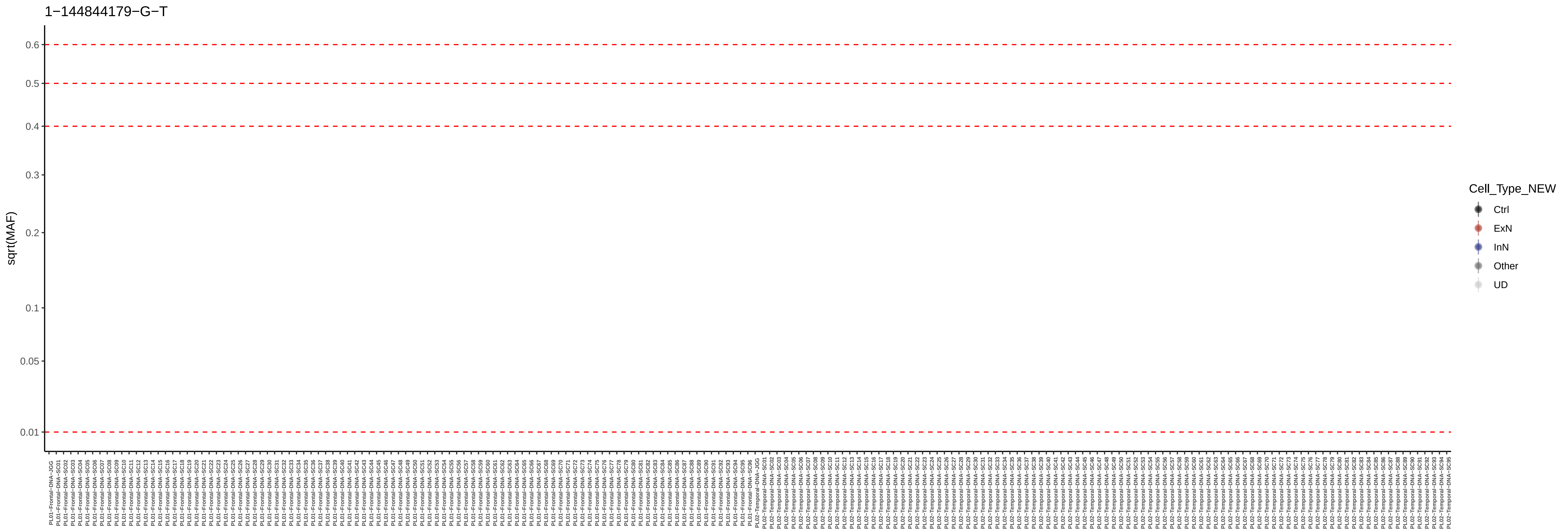

1−144844179−G−T
sqrt(MAF)
0.6
0.5
0.4
0.3
0.2
0.1
0.05
0.01
Cell_Type_NEW
Ctrl
ExN
InN
Other
UD

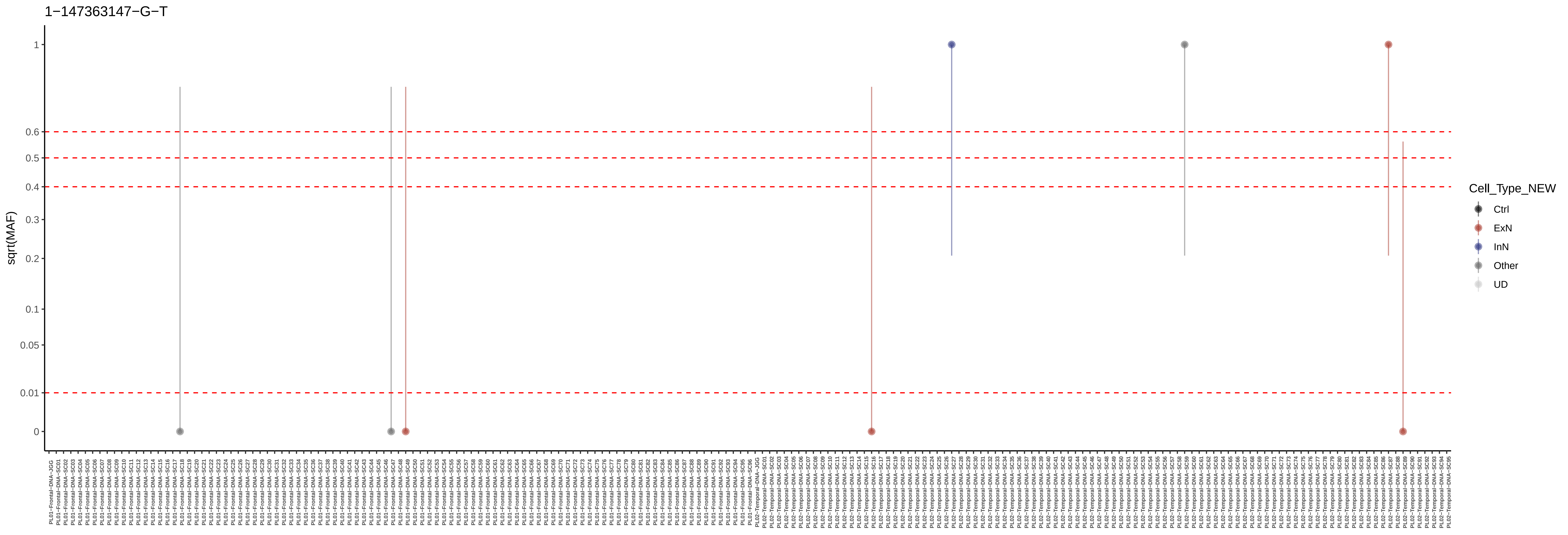

1−147363147−G−T
sqrt(MAF)
1
0.6
0.5
0.4
0.3
0.2
0.1
0.05
0.01
0
Cell_Type_NEW
Ctrl
ExN
InN
Other
UD

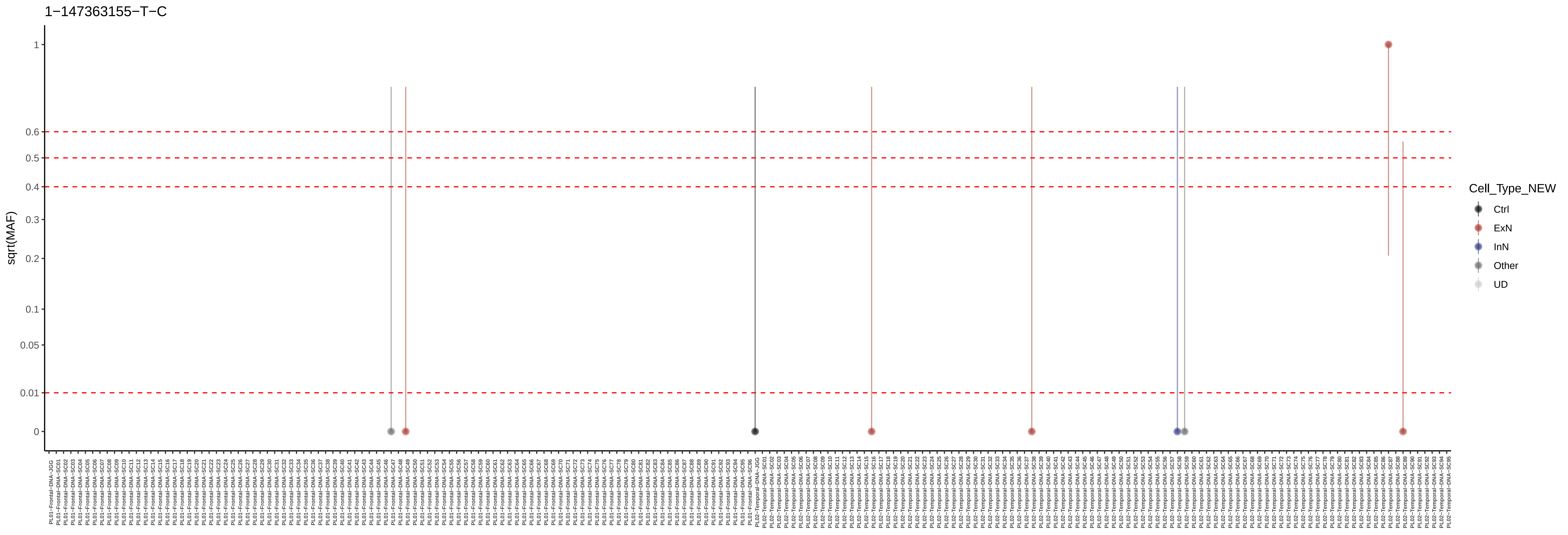
1−147363155−T−C
sqrt(MAF)
1
0.6
0.5
0.4
0.3
0.2
0.1
0.05
0.01
0
Cell_Type_NEW
Ctrl
ExN
InN
Other
UD

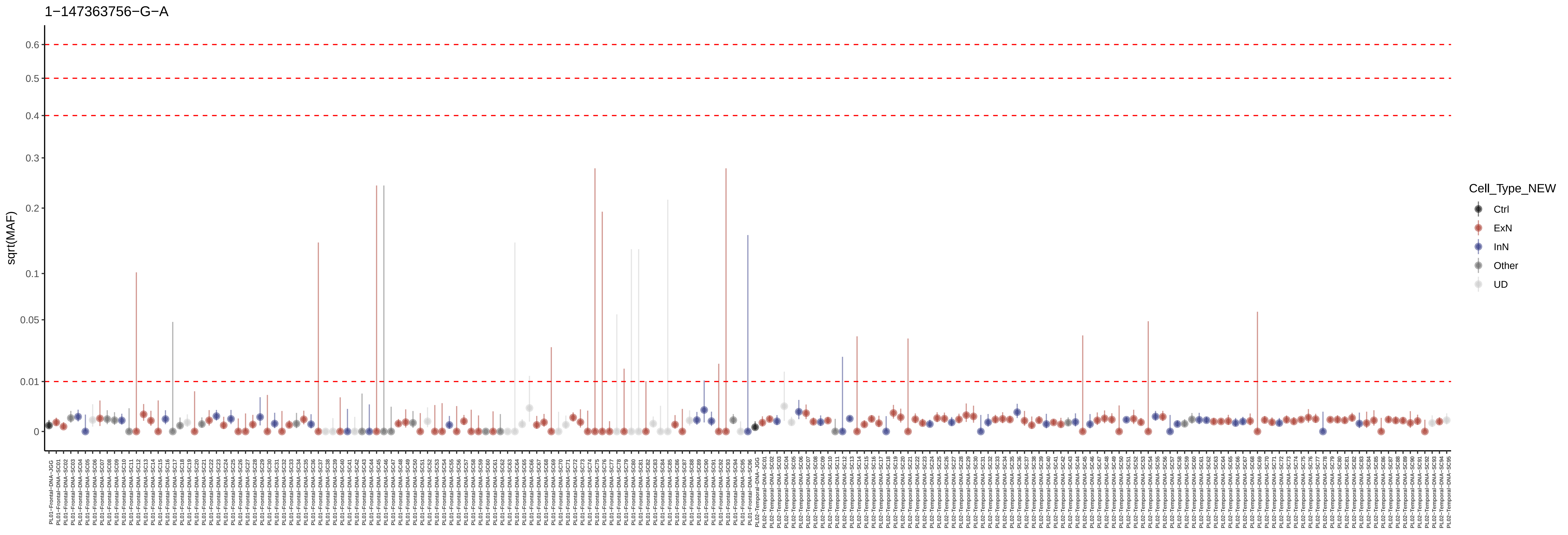
1−147363756−G−A
sqrt(MAF)
0.6
0.5
0.4
0.3
0.2
0.1
0.05
0.01
0
Cell_Type_NEW
Ctrl
ExN
InN
Other
UD

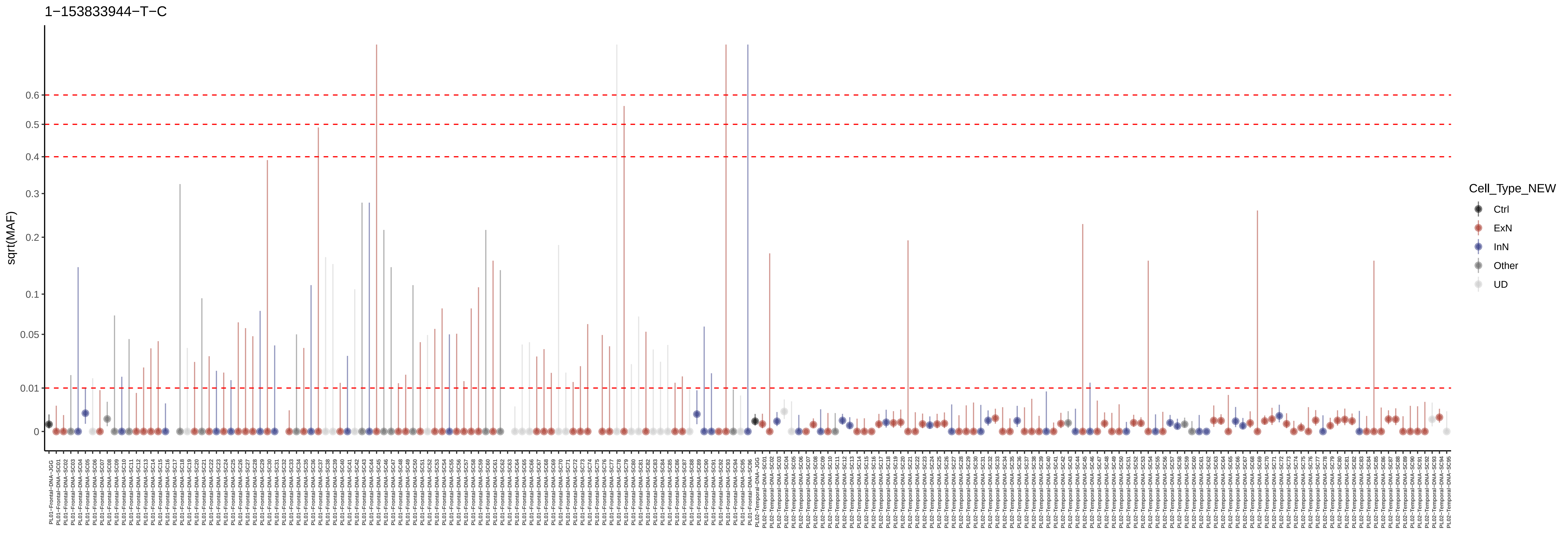
1−153833944−T−C
sqrt(MAF)
0.6
0.5
0.4
0.3
0.2
0.1
0.05
0.01
0
Cell_Type_NEW
Ctrl
ExN
InN
Other
UD

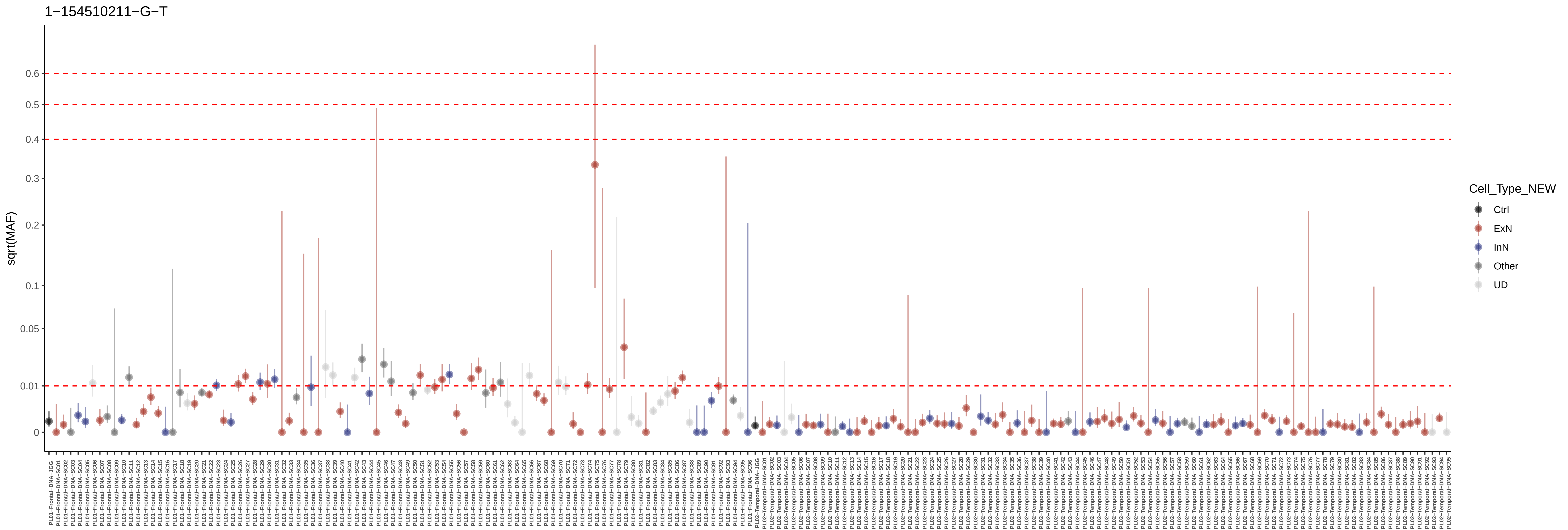
1−154510211−G−T
sqrt(MAF)
0.6
0.5
0.4
0.3
0.2
0.1
0.05
0.01
0
Cell_Type_NEW
Ctrl
ExN
InN
Other
UD

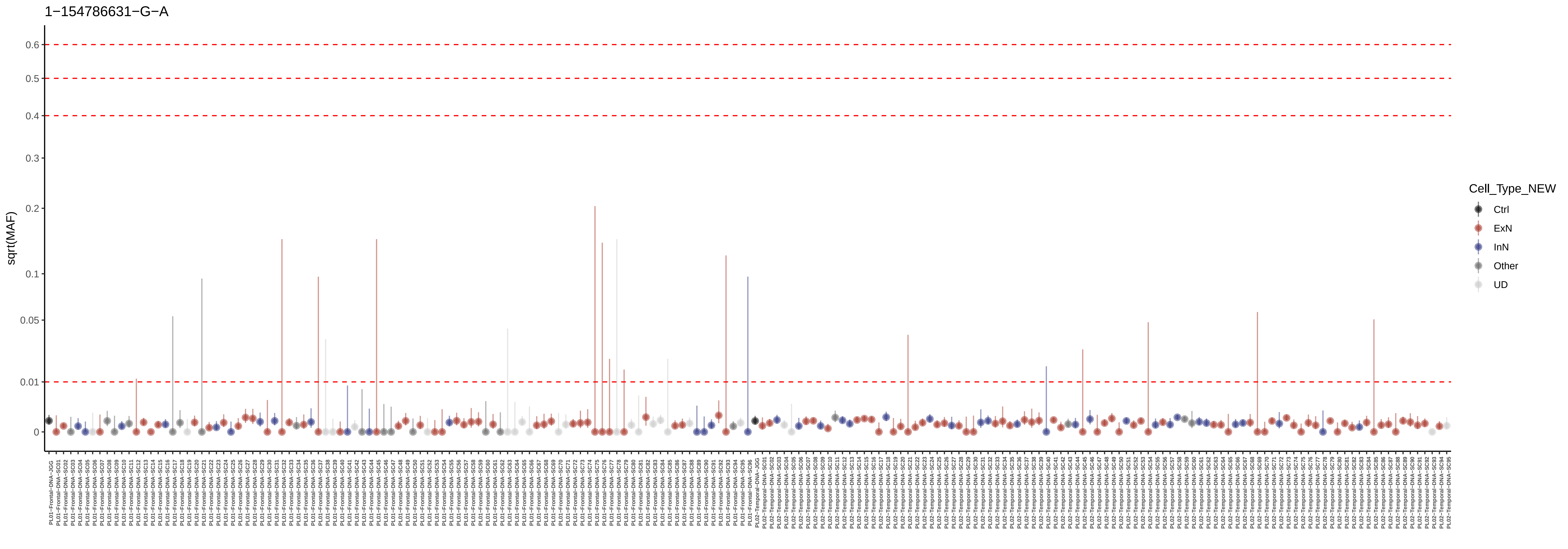
1−154786631−G−A
sqrt(MAF)
0.6
0.5
0.4
0.3
0.2
0.1
0.05
0.01
0
Cell_Type_NEW
Ctrl
ExN
InN
Other
UD

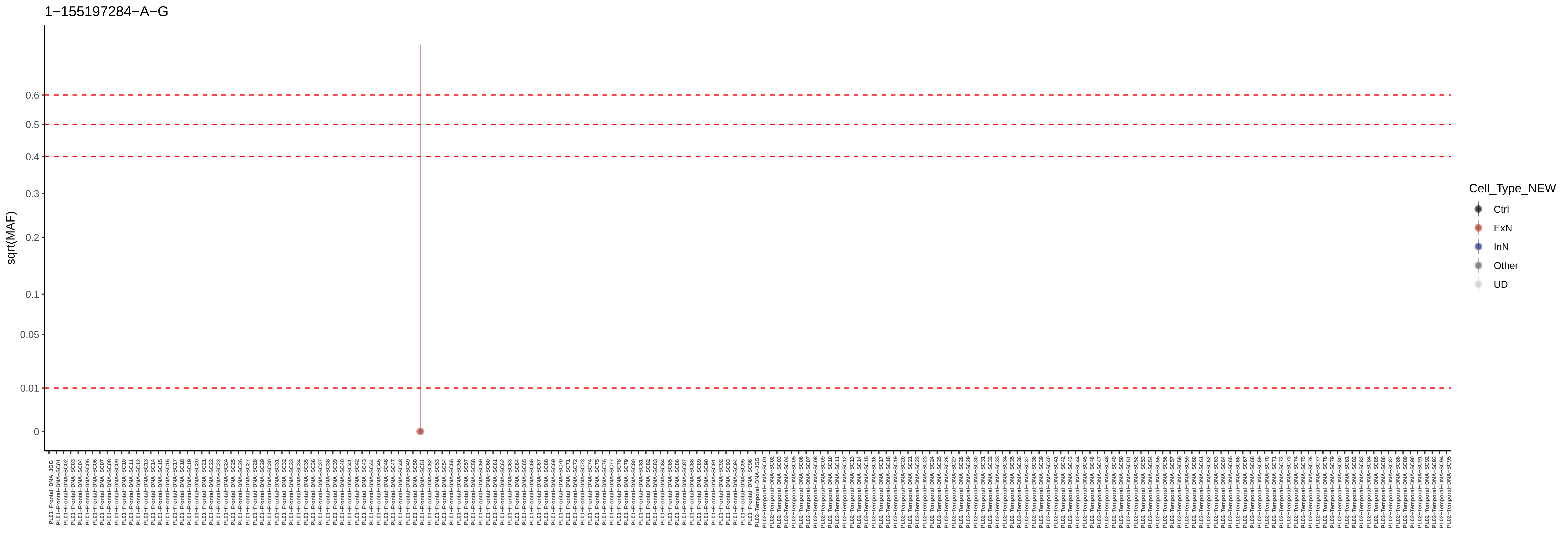

1−155197284−A−G
sqrt(MAF)
0.6
0.5
0.4
0.3
0.2
0.1
0.05
0.01
0
Cell_Type_NEW
Ctrl
ExN
InN
Other
UD

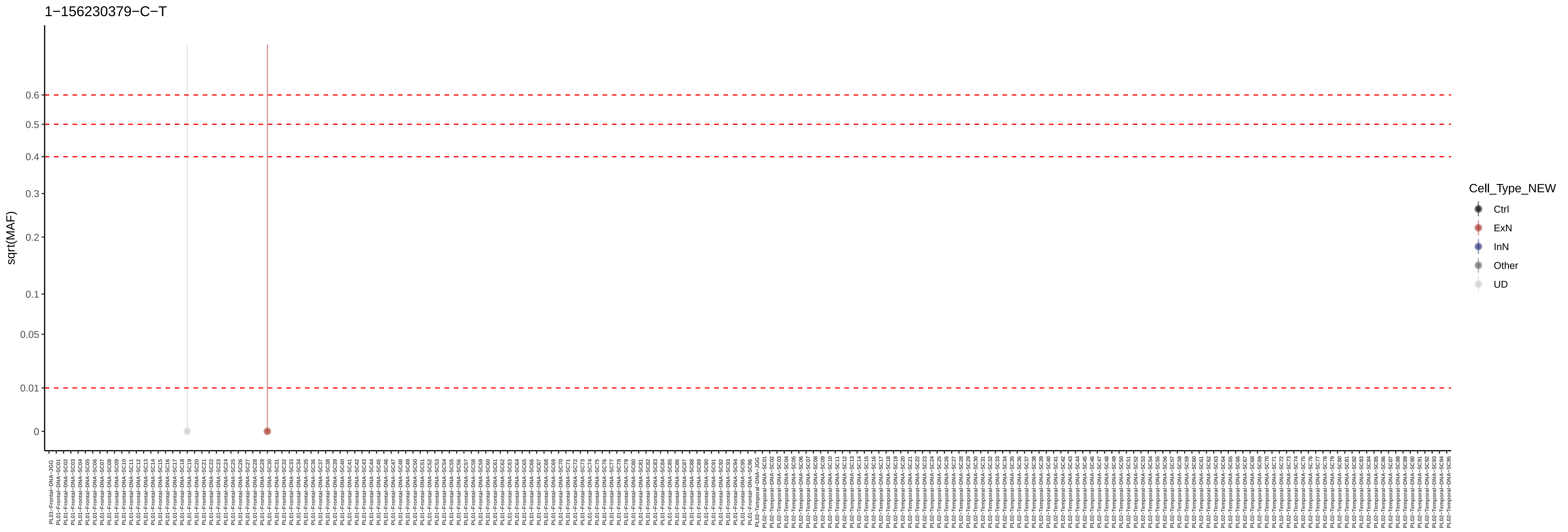
1−156230379−C−T
sqrt(MAF)
0.6
0.5
0.4
0.3
0.2
0.1
0.05
0.01
0
Cell_Type_NEW
Ctrl
ExN
InN
Other
UD

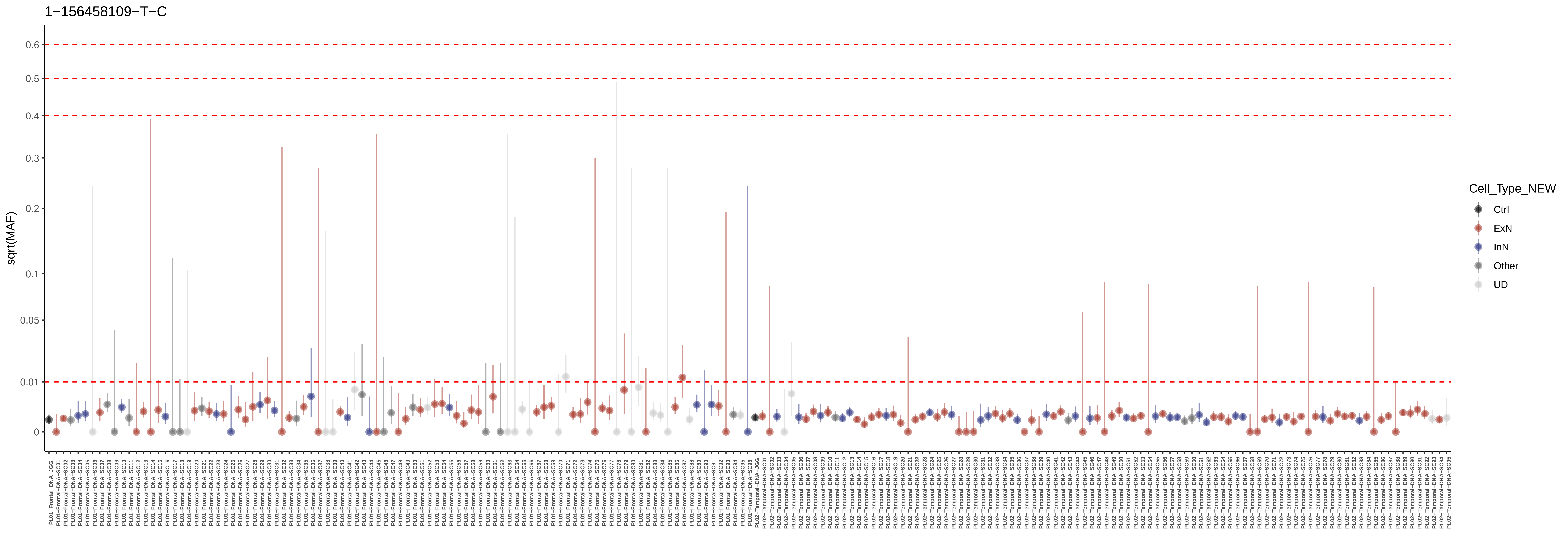
1−156458109−T−C
sqrt(MAF)
0.6
0.5
0.4
0.3
0.2
0.1
0.05
0.01
0
Cell_Type_NEW
Ctrl
ExN
InN
Other
UD

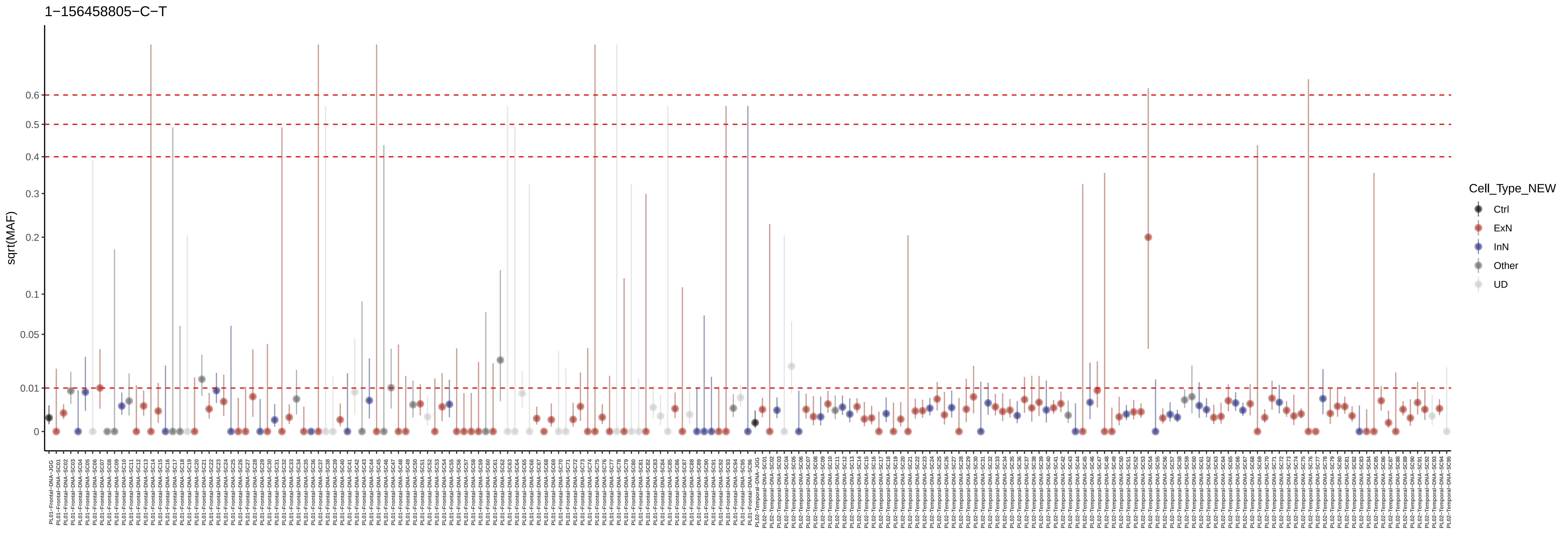
1−156458805−C−T
sqrt(MAF)
0.6
0.5
0.4
0.3
0.2
0.1
0.05
0.01
0
Cell_Type_NEW
Ctrl
ExN
InN
Other
UD

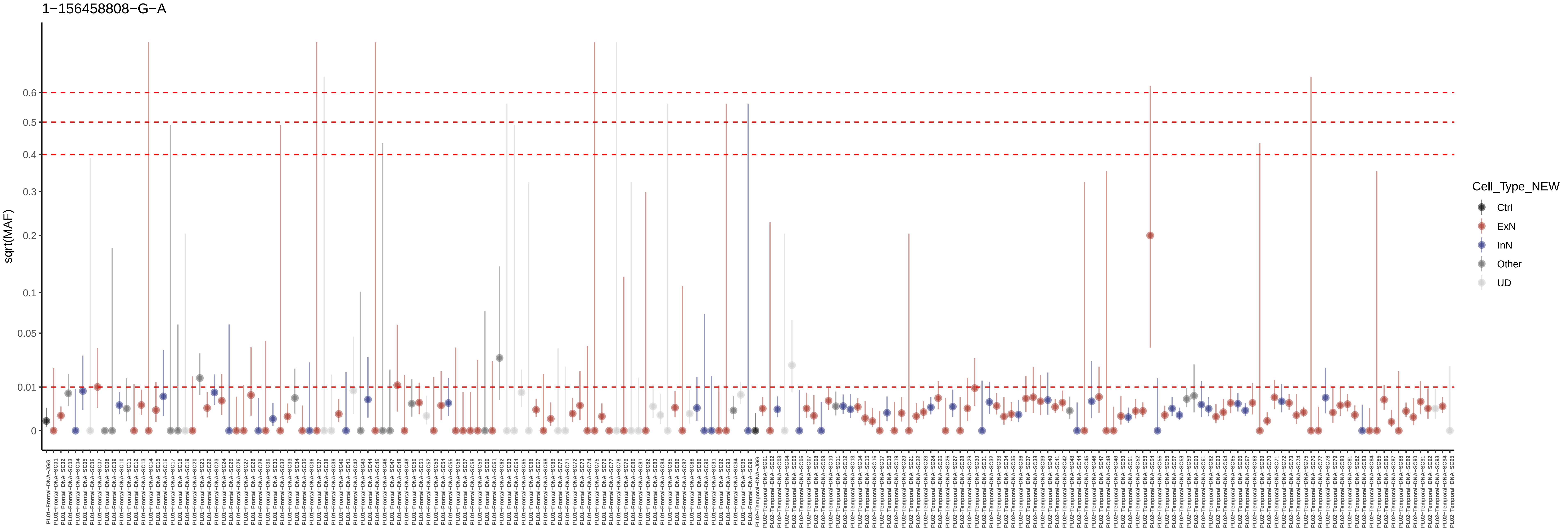
1−156458808−G−A
sqrt(MAF)
0.6
0.5
0.4
0.3
0.2
0.1
0.05
0.01
0
Cell_Type_NEW
Ctrl
ExN
InN
Other
UD

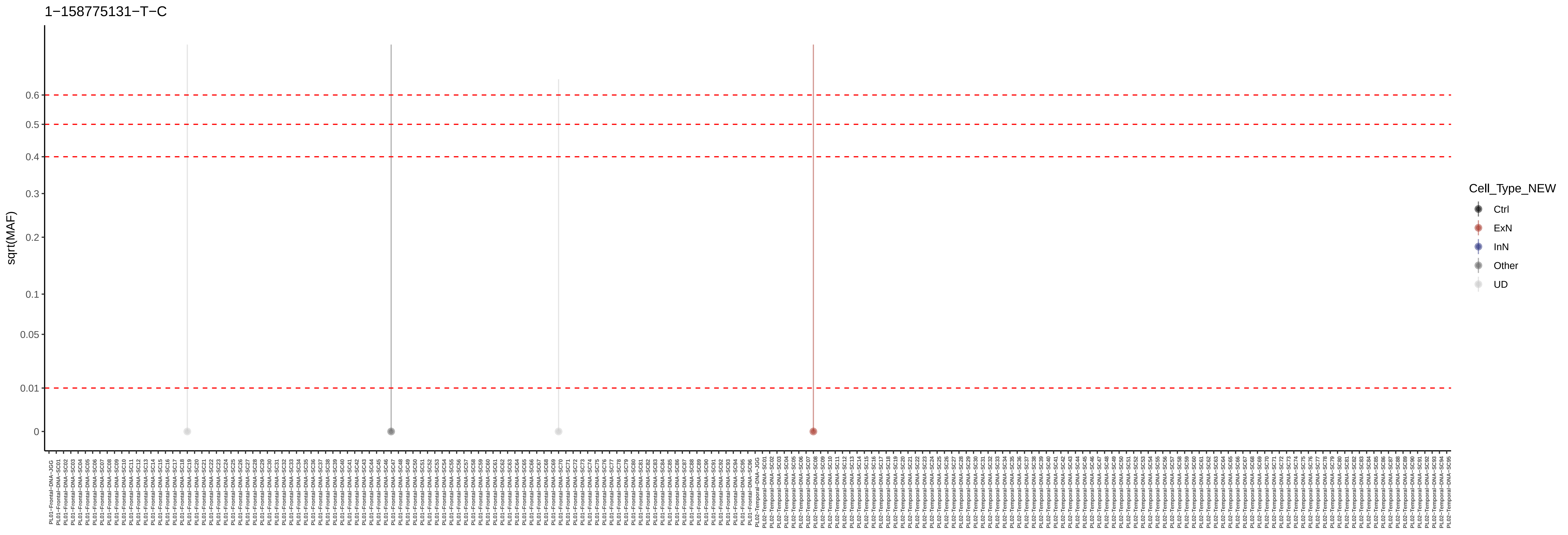
1−158775131−T−C
sqrt(MAF)
0.6
0.5
0.4
0.3
0.2
0.1
0.05
0.01
0
Cell_Type_NEW
Ctrl
ExN
InN
Other
UD

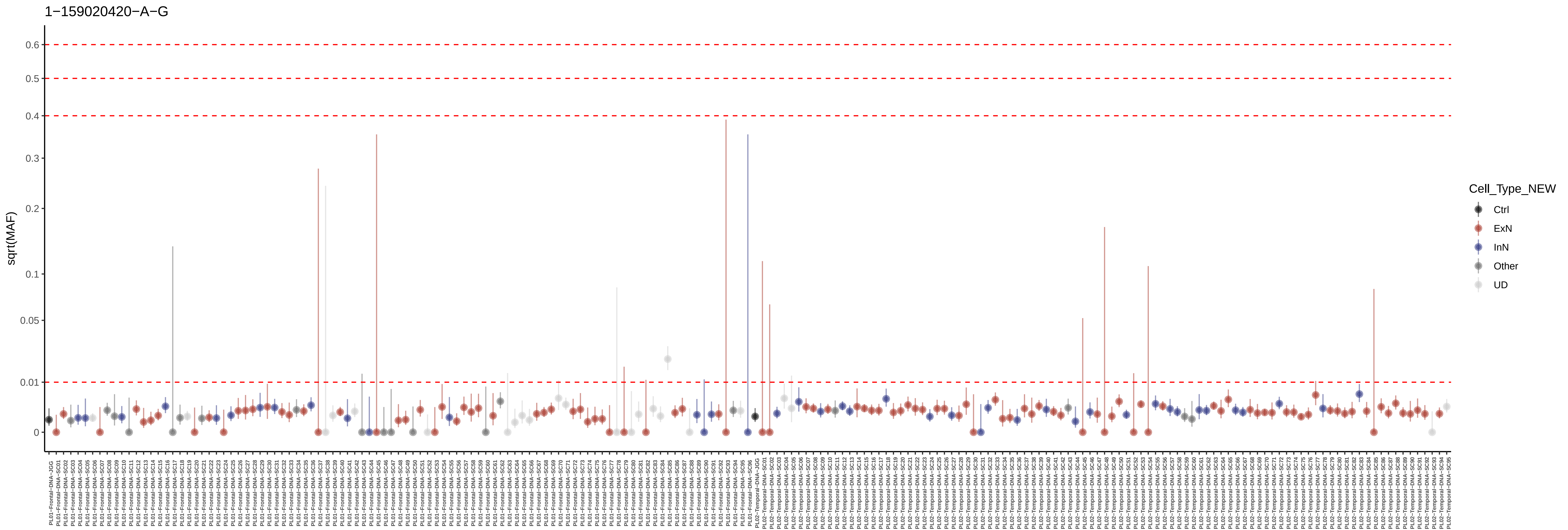

1−159020420−A−G
sqrt(MAF)
0.6
0.5
0.4
0.3
0.2
0.1
0.05
0.01
0
Cell_Type_NEW
Ctrl
ExN
InN
Other
UD

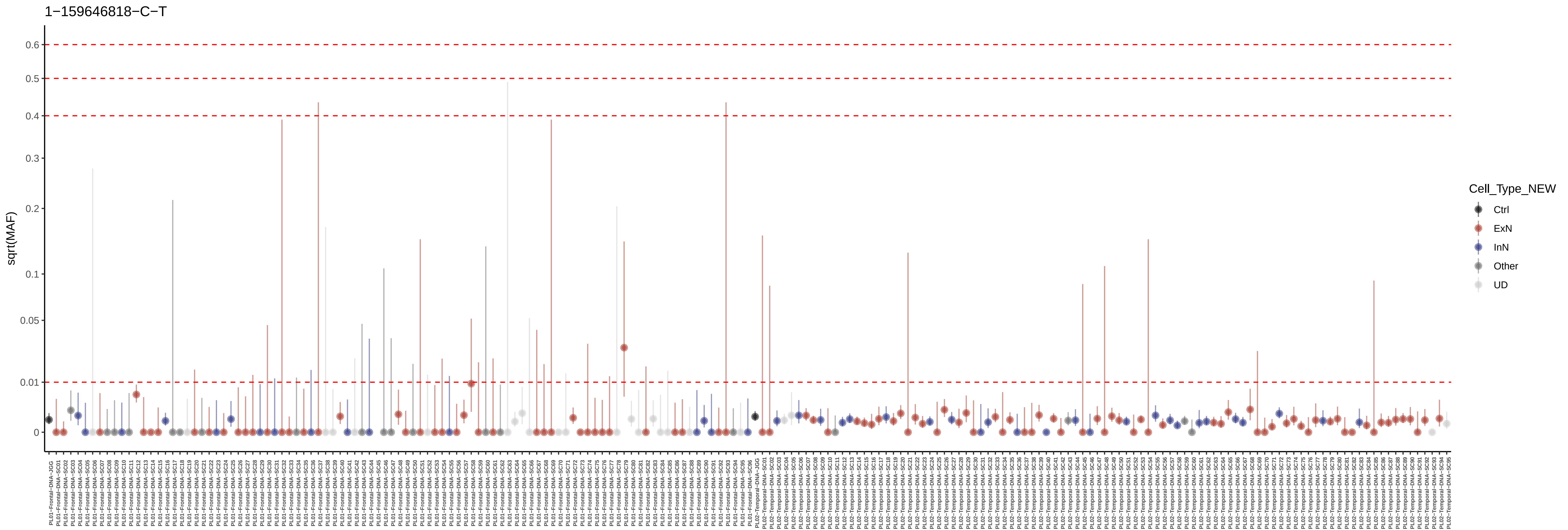

1−159646818−C−T
sqrt(MAF)
0.6
0.5
0.4
0.3
0.2
0.1
0.05
0.01
0
Cell_Type_NEW
Ctrl
ExN
InN
Other
UD

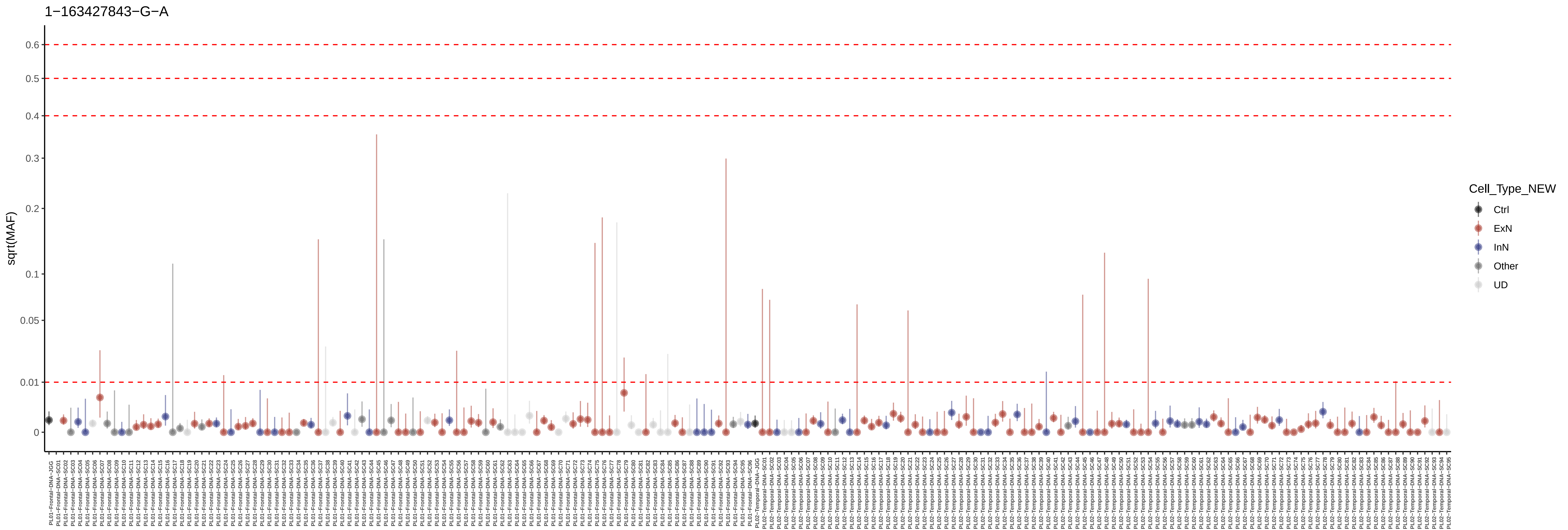

1−163427843−G−A
sqrt(MAF)
0.6
0.5
0.4
0.3
0.2
0.1
0.05
0.01
0
Cell_Type_NEW
Ctrl
ExN
InN
Other
UD

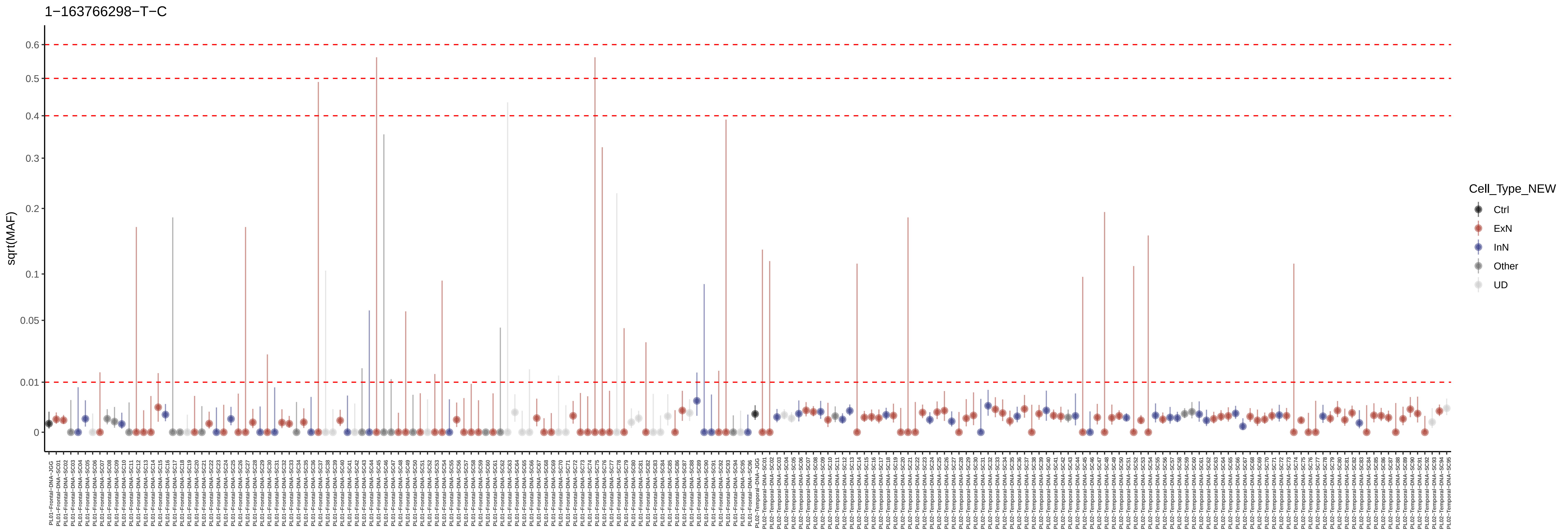
1−163766298−T−C
sqrt(MAF)
0.6
0.5
0.4
0.3
0.2
0.1
0.05
0.01
0
Cell_Type_NEW
Ctrl
ExN
InN
Other
UD

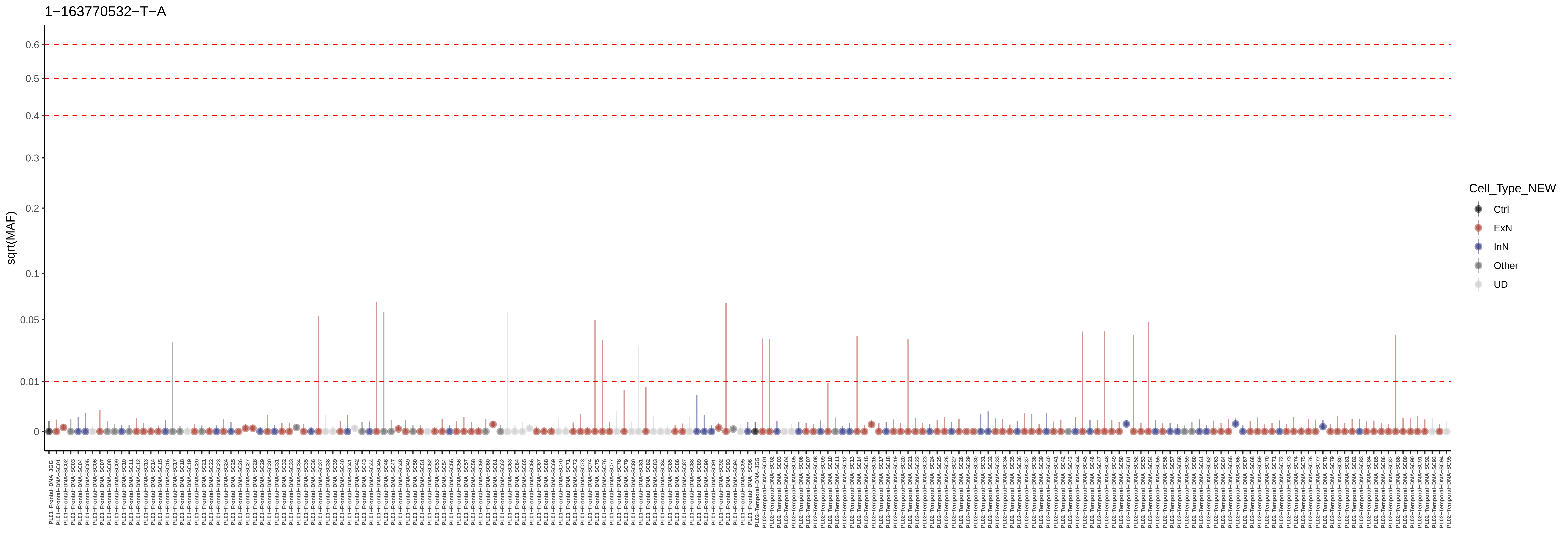
1−163770532−T−A
sqrt(MAF)
0.6
0.5
0.4
0.3
0.2
0.1
0.05
0.01
0
Cell_Type_NEW
Ctrl
ExN
InN
Other
UD

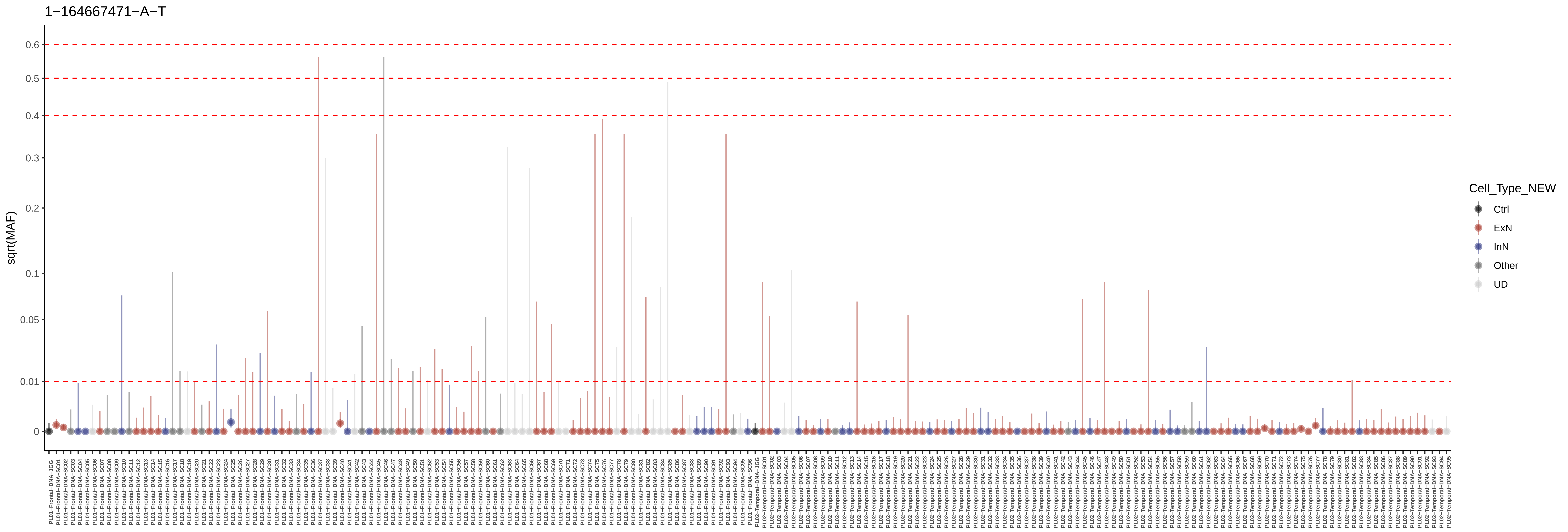
1−164667471−A−T
sqrt(MAF)
0.6
0.5
0.4
0.3
0.2
0.1
0.05
0.01
0
Cell_Type_NEW
Ctrl
ExN
InN
Other
UD

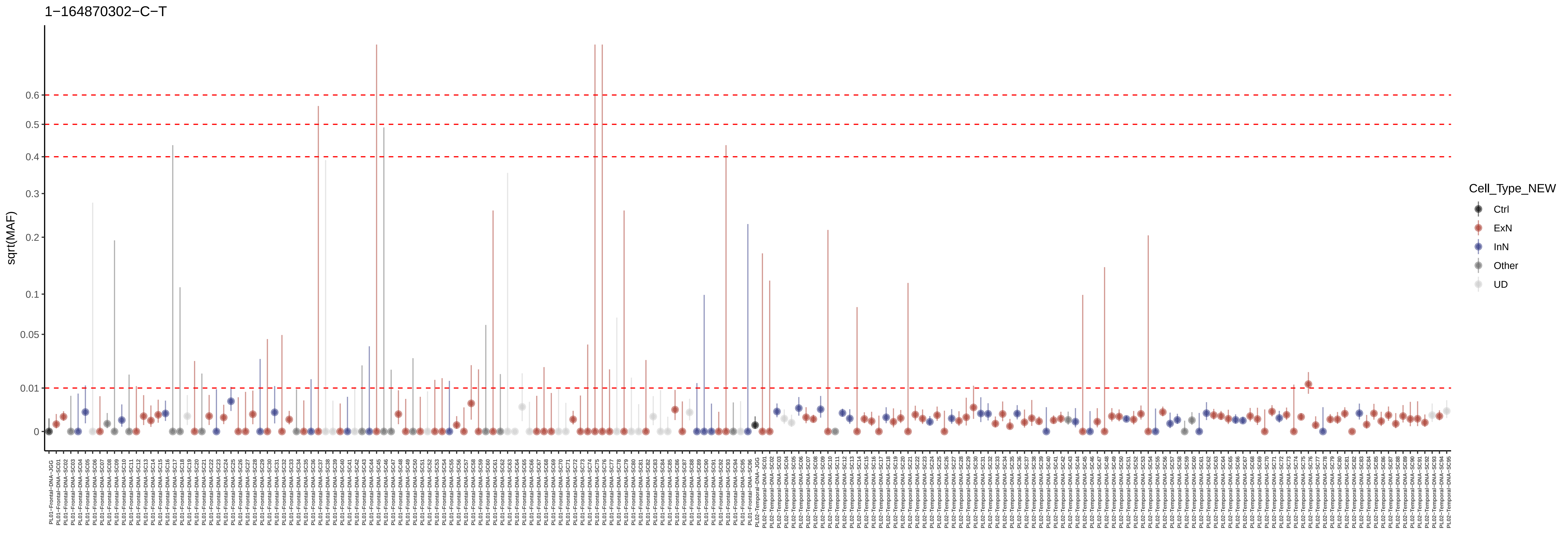

1−164870302−C−T
sqrt(MAF)
0.6
0.5
0.4
0.3
0.2
0.1
0.05
0.01
0
Cell_Type_NEW
Ctrl
ExN
InN
Other
UD

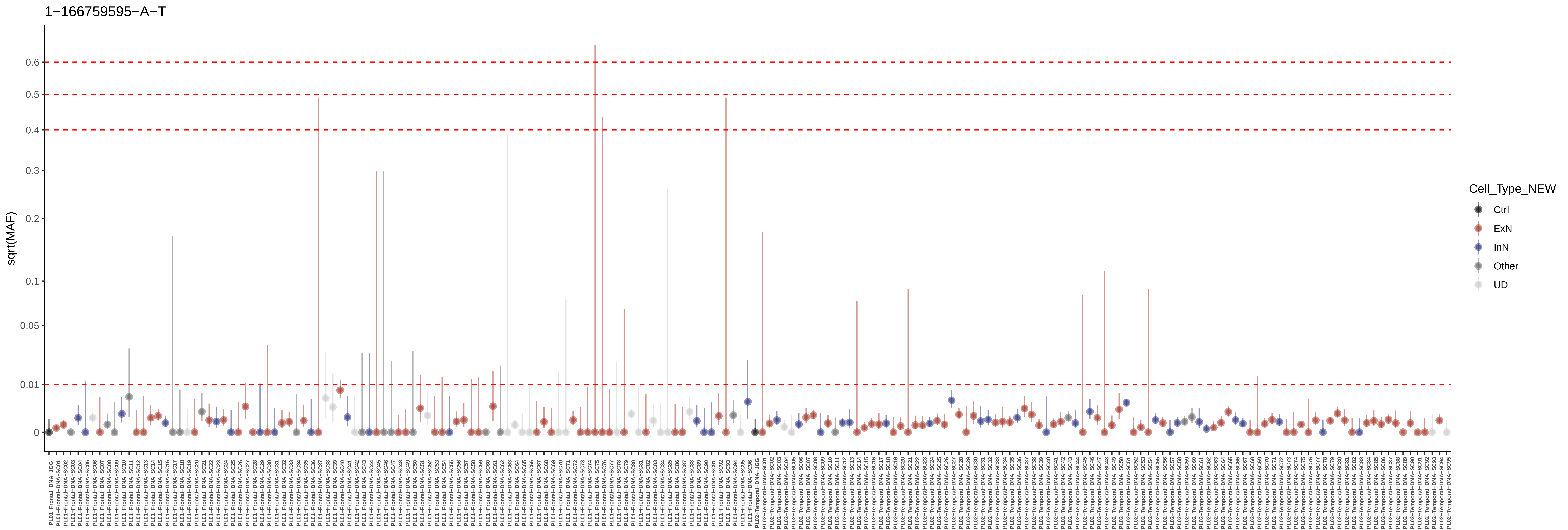
1−166759595−A−T
sqrt(MAF)
0.6
0.5
0.4
0.3
0.2
0.1
0.05
0.01
0
Cell_Type_NEW
Ctrl
ExN
InN
Other
UD

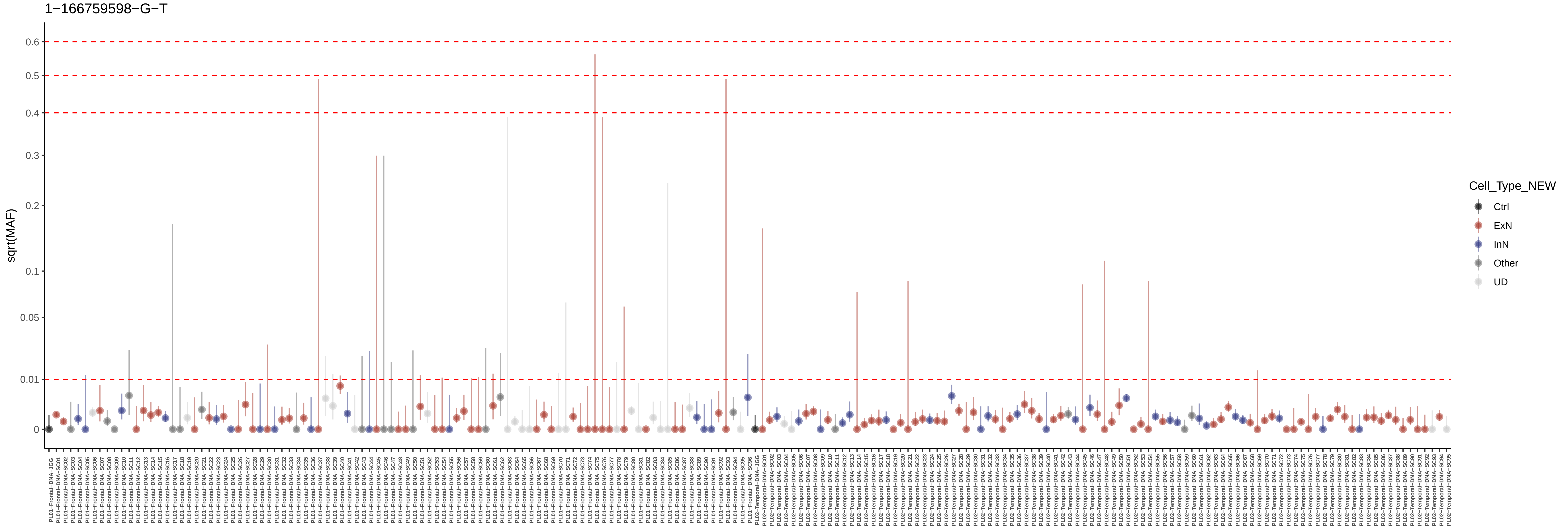

1−166759598−G−T
sqrt(MAF)
0.6
0.5
0.4
0.3
0.2
0.1
0.05
0.01
0
Cell_Type_NEW
Ctrl
ExN
InN
Other
UD

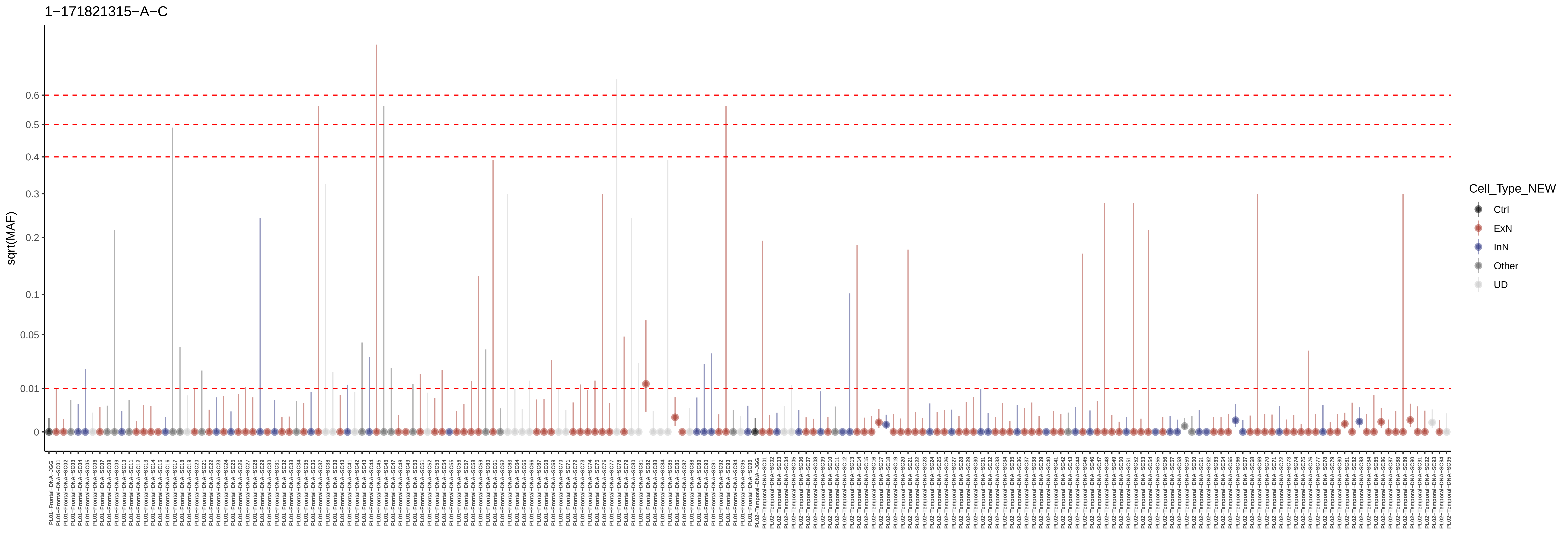
1−171821315−A−C
sqrt(MAF)
0.6
0.5
0.4
0.3
0.2
0.1
0.05
0.01
0
Cell_Type_NEW
Ctrl
ExN
InN
Other
UD

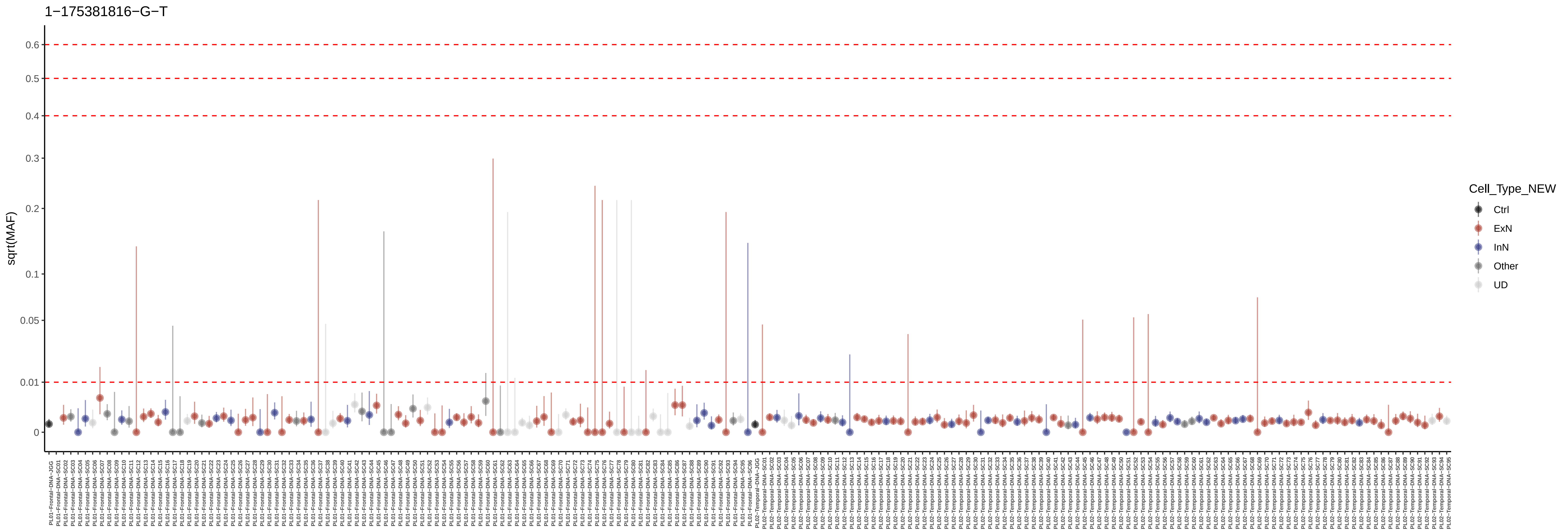
1−175381816−G−T
sqrt(MAF)
0.6
0.5
0.4
0.3
0.2
0.1
0.05
0.01
0
Cell_Type_NEW
Ctrl
ExN
InN
Other
UD

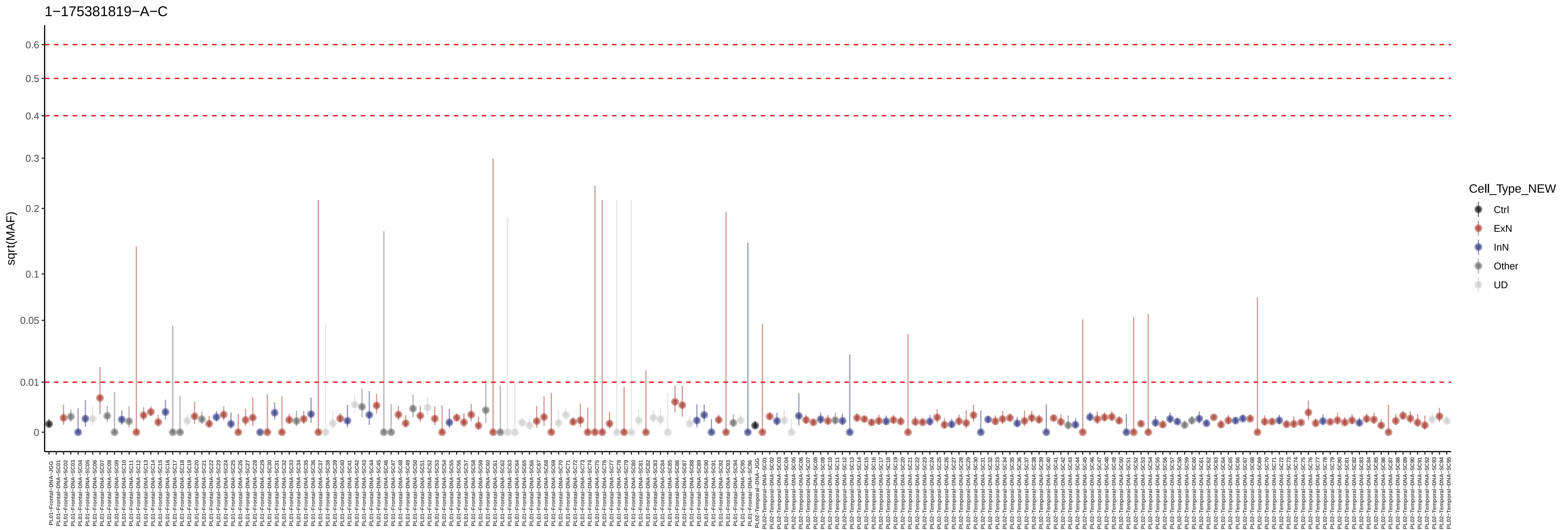
1−175381819−A−C
sqrt(MAF)
0.6
0.5
0.4
0.3
0.2
0.1
0.05
0.01
0
Cell_Type_NEW
Ctrl
ExN
InN
Other
UD

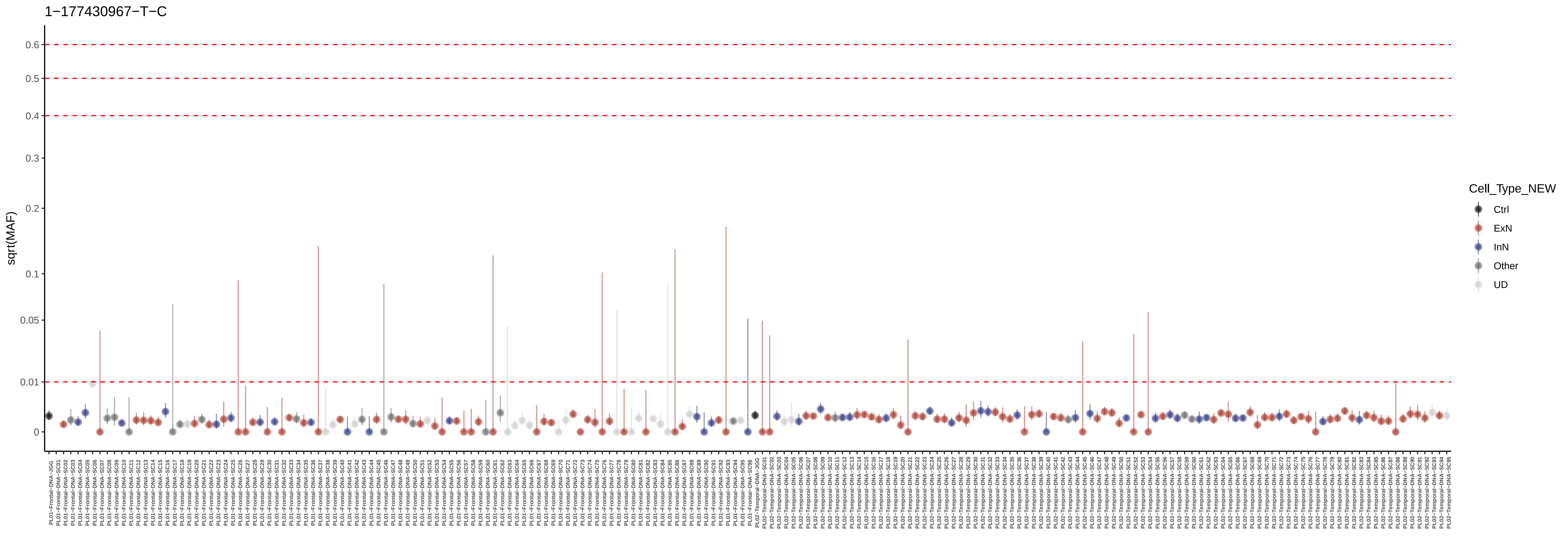
1−177430967−T−C
sqrt(MAF)
0.6
0.5
0.4
0.3
0.2
0.1
0.05
0.01
0
Cell_Type_NEW
Ctrl
ExN
InN
Other
UD

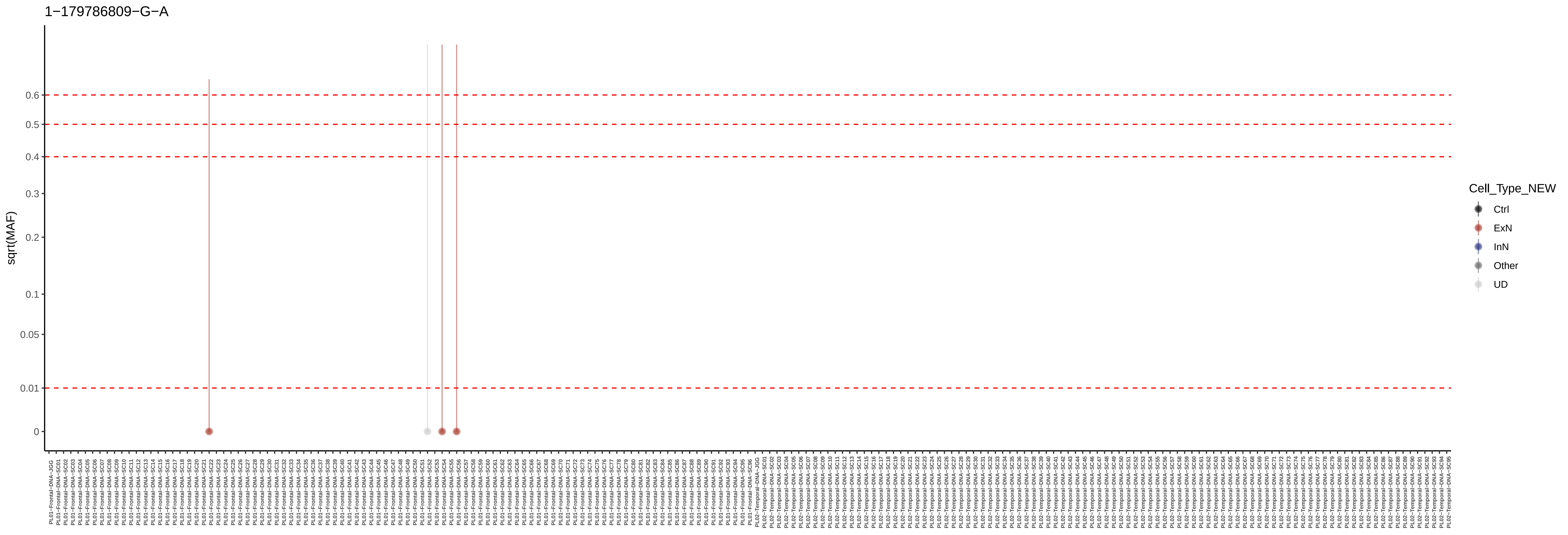
1−179786809−G−A
sqrt(MAF)
0.6
0.5
0.4
0.3
0.2
0.1
0.05
0.01
0
Cell_Type_NEW
Ctrl
ExN
InN
Other
UD

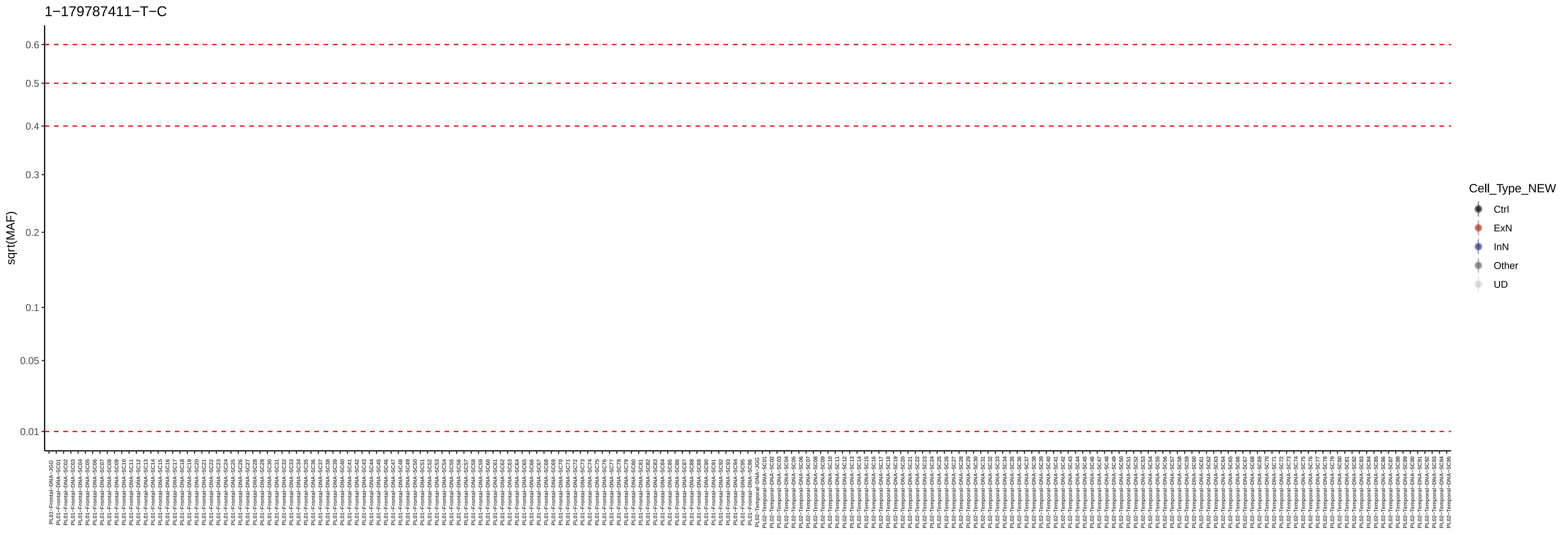
1−179787411−T−C
sqrt(MAF)
0.6
0.5
0.4
0.3
0.2
0.1
0.05
0.01
Cell_Type_NEW
Ctrl
ExN
InN
Other
UD

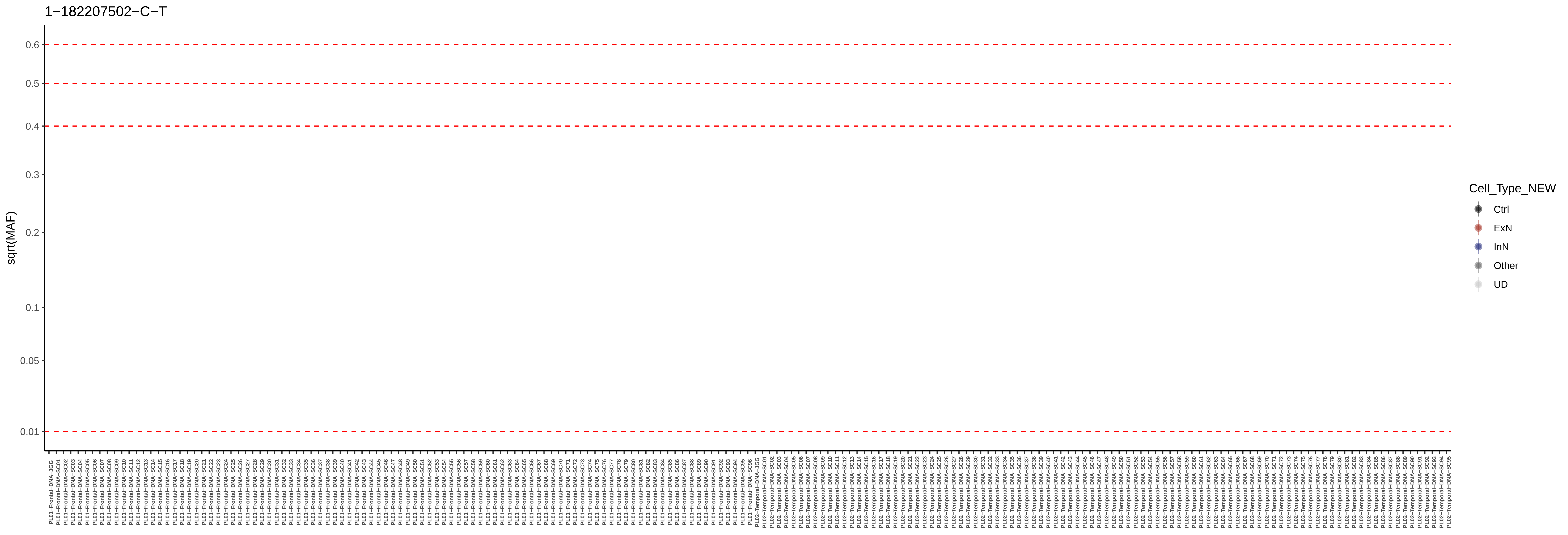
1−182207502−C−T
sqrt(MAF)
0.6
0.5
0.4
0.3
0.2
0.1
0.05
0.01
Cell_Type_NEW
Ctrl
ExN
InN
Other
UD

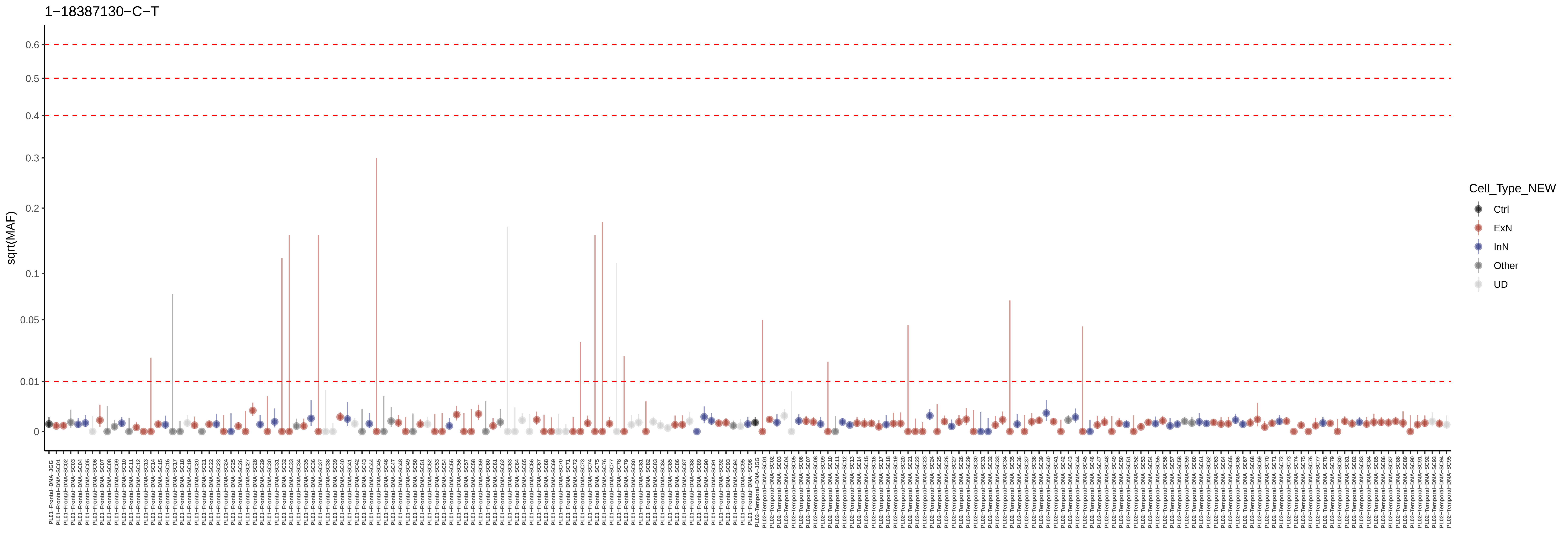
1−18387130−C−T
sqrt(MAF)
0.6
0.5
0.4
0.3
0.2
0.1
0.05
0.01
0
Cell_Type_NEW
Ctrl
ExN
InN
Other
UD

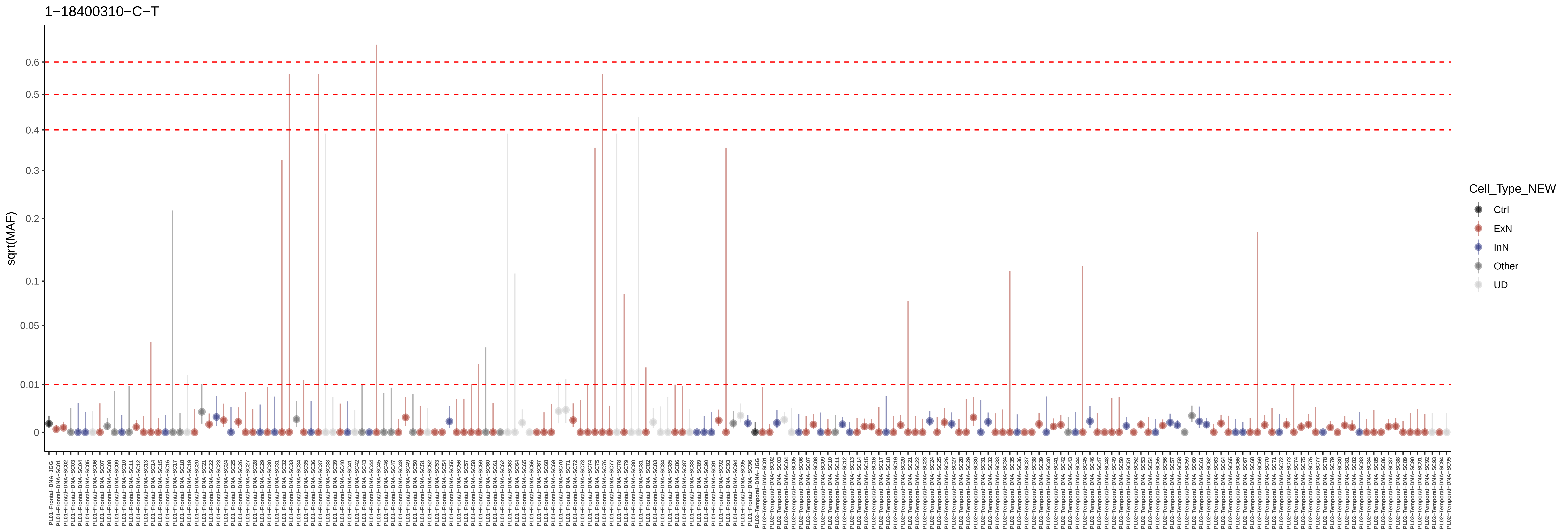
1−18400310−C−T
sqrt(MAF)
0.6
0.5
0.4
0.3
0.2
0.1
0.05
0.01
0
Cell_Type_NEW
Ctrl
ExN
InN
Other
UD

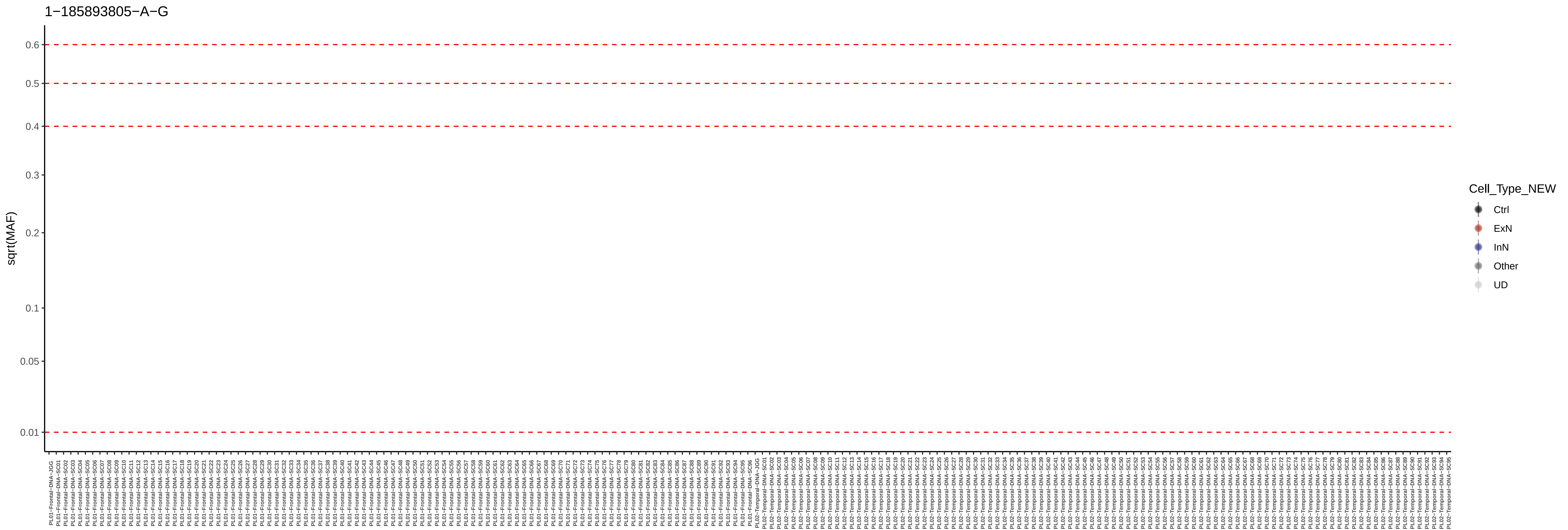
1−185893805−A−G
sqrt(MAF)
0.6
0.5
0.4
0.3
0.2
0.1
0.05
0.01
Cell_Type_NEW
Ctrl
ExN
InN
Other
UD

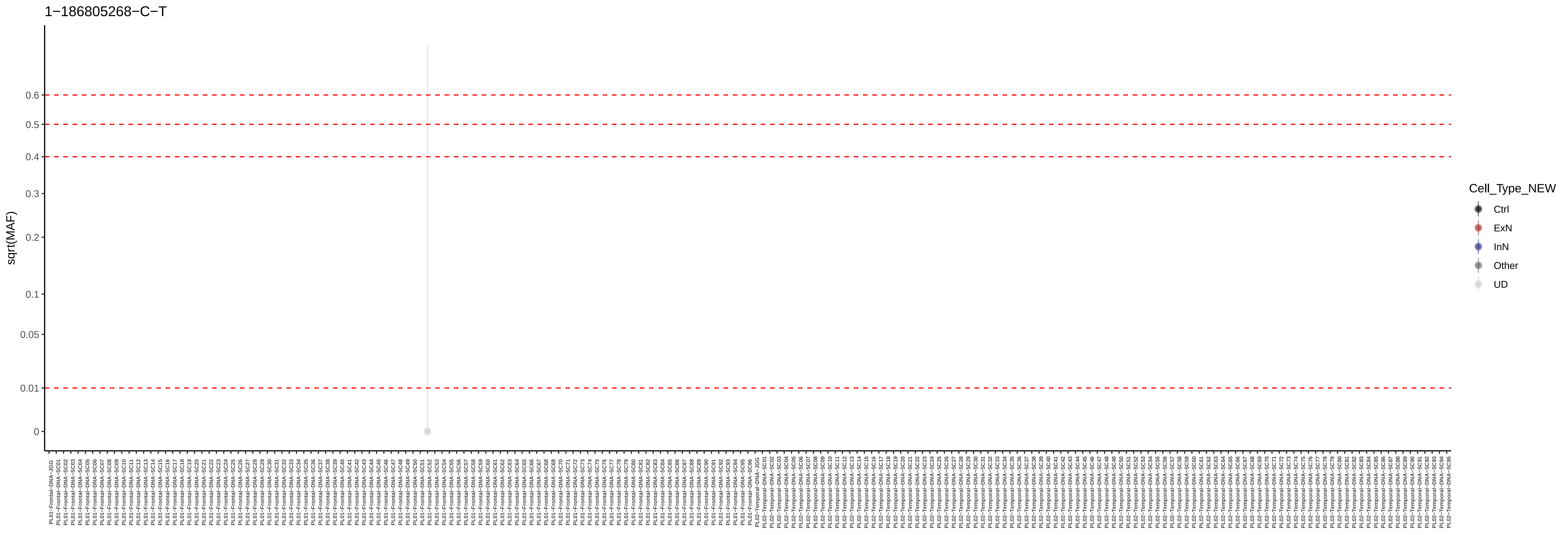
1−186805268−C−T
sqrt(MAF)
0.6
0.5
0.4
0.3
0.2
0.1
0.05
0.01
0
Cell_Type_NEW
Ctrl
ExN
InN
Other
UD

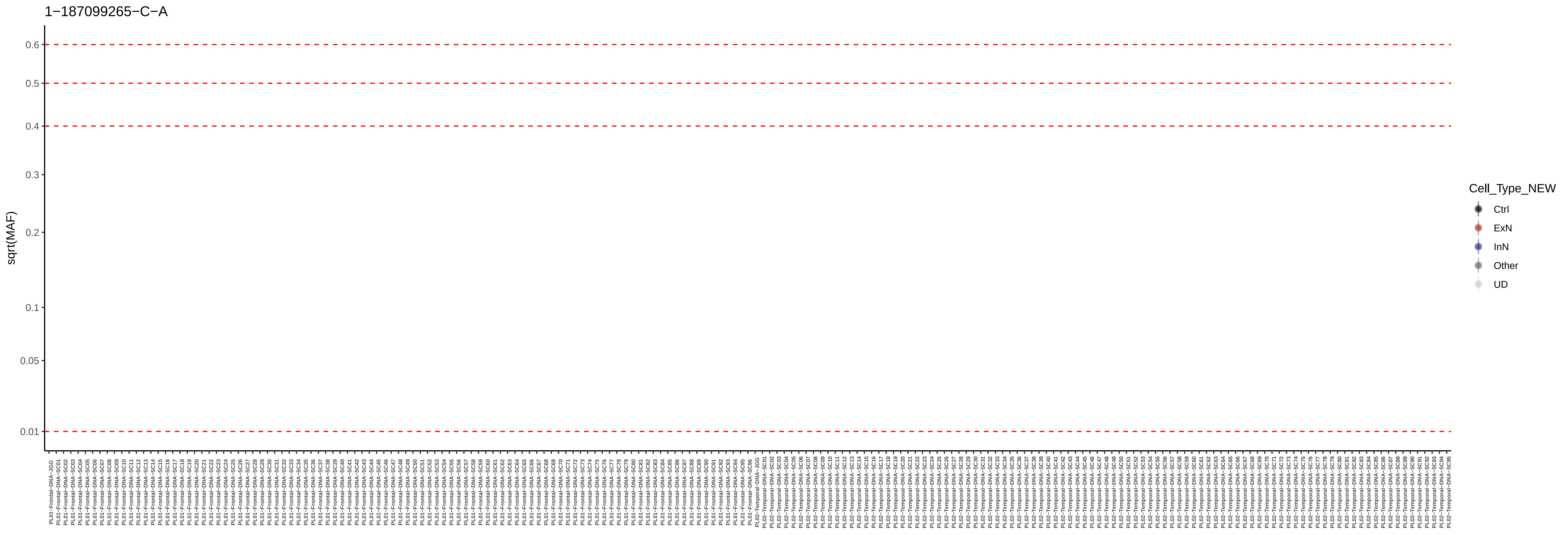

1−187099265−C−A
sqrt(MAF)
0.6
0.5
0.4
0.3
0.2
0.1
0.05
0.01
Cell_Type_NEW
Ctrl
ExN
InN
Other
UD

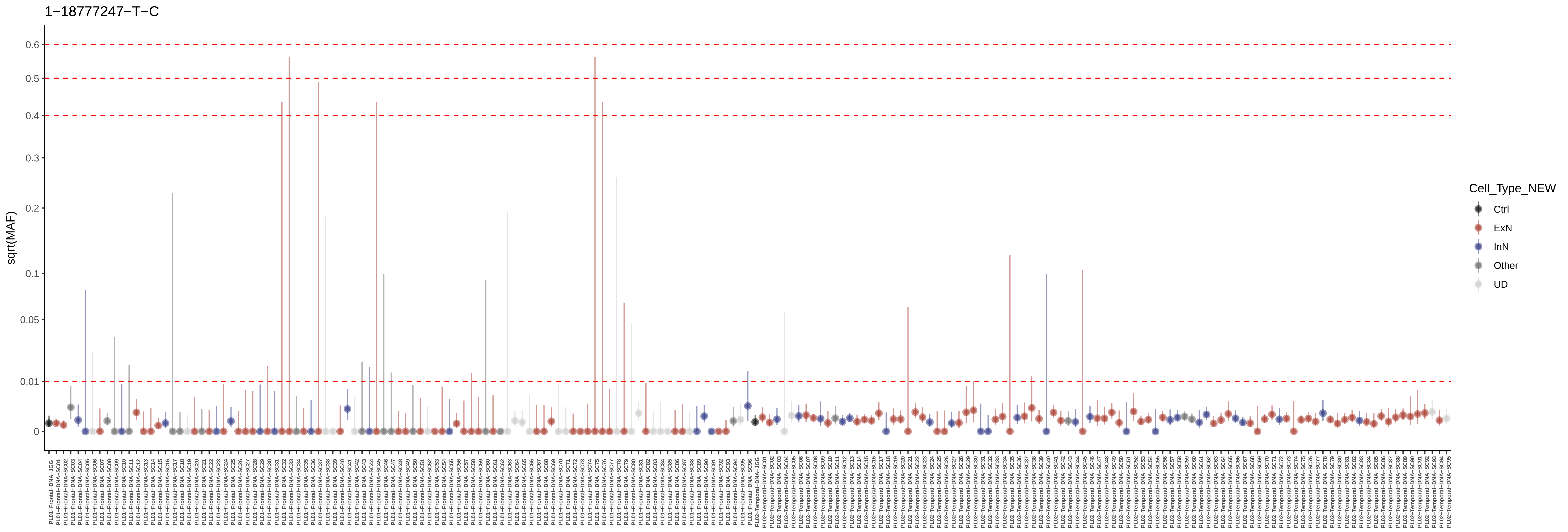
1−18777247−T−C
sqrt(MAF)
0.6
0.5
0.4
0.3
0.2
0.1
0.05
0.01
0
Cell_Type_NEW
Ctrl
ExN
InN
Other
UD

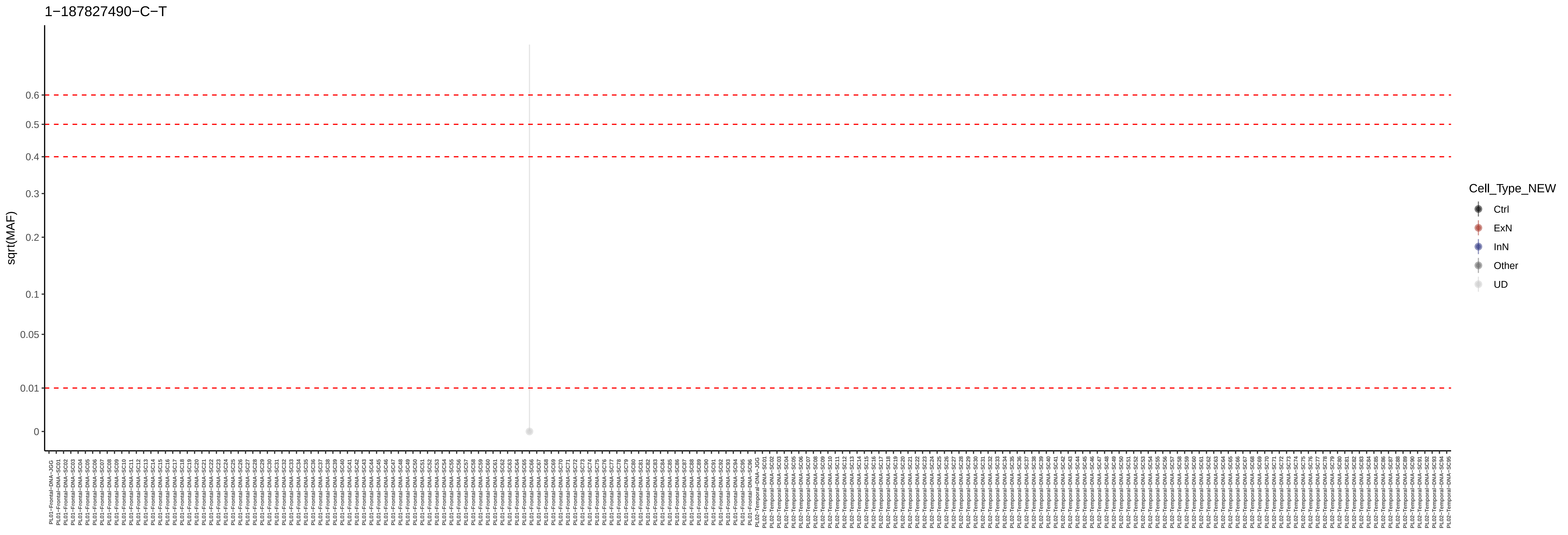
1−187827490−C−T
sqrt(MAF)
0.6
0.5
0.4
0.3
0.2
0.1
0.05
0.01
0
Cell_Type_NEW
Ctrl
ExN
InN
Other
UD

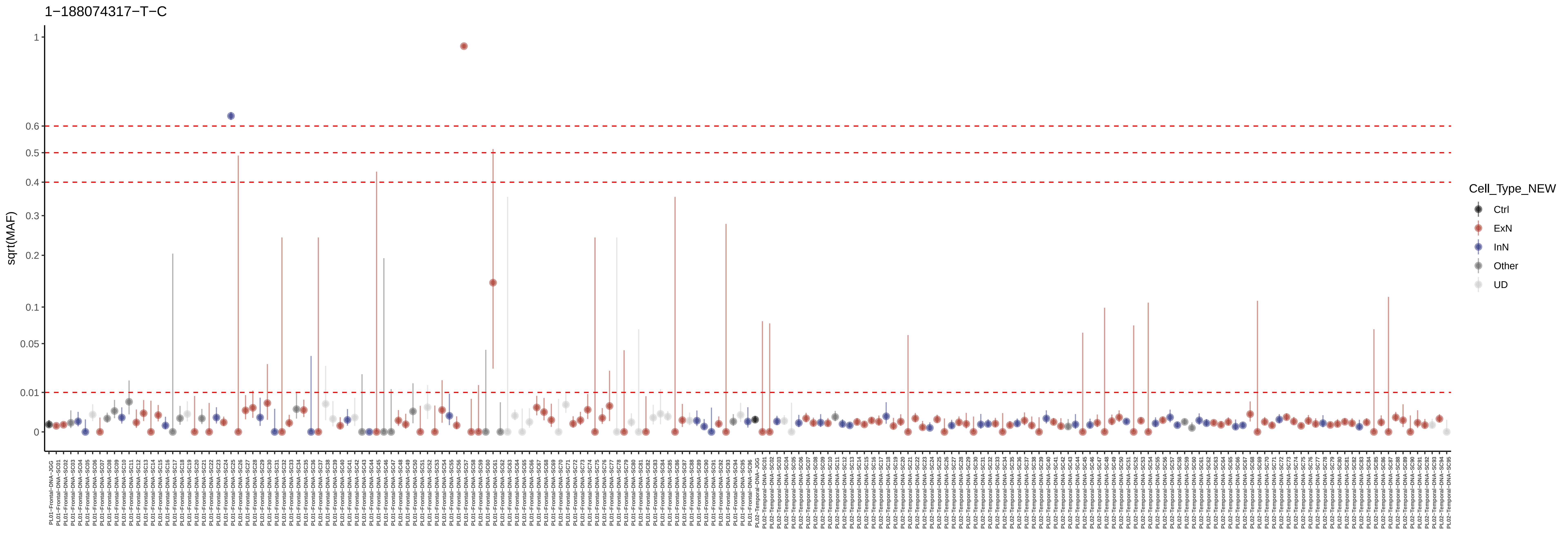
1−188074317−T−C
sqrt(MAF)
1
0.6
0.5
0.4
0.3
0.2
0.1
0.05
0.01
0
Cell_Type_NEW
Ctrl
ExN
InN
Other
UD

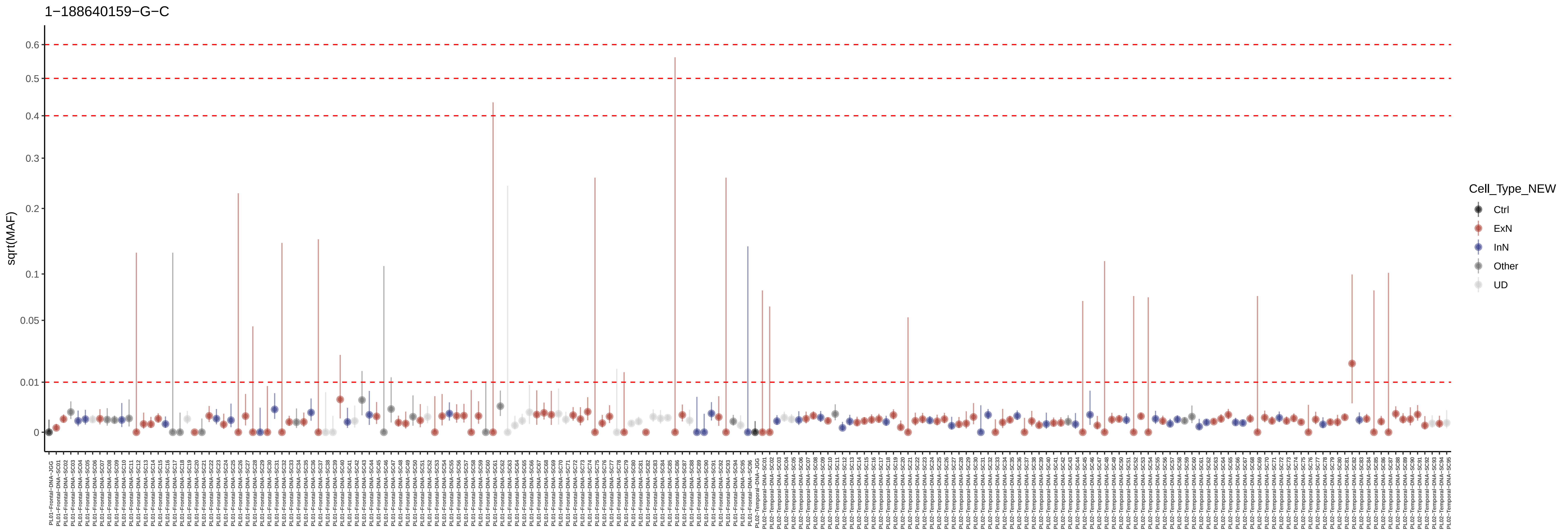
1−188640159−G−C
sqrt(MAF)
0.6
0.5
0.4
0.3
0.2
0.1
0.05
0.01
0
Cell_Type_NEW
Ctrl
ExN
InN
Other
UD

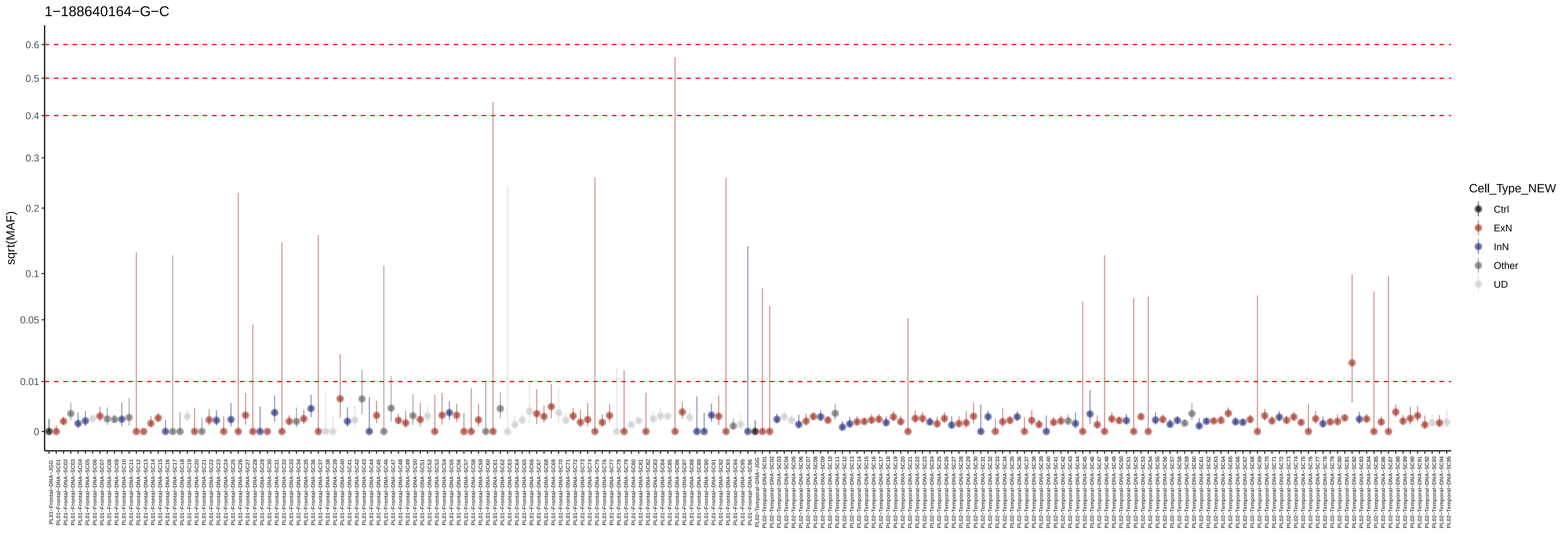
1−188640164−G−C
sqrt(MAF)
0.6
0.5
0.4
0.3
0.2
0.1
0.05
0.01
0
Cell_Type_NEW
Ctrl
ExN
InN
Other
UD

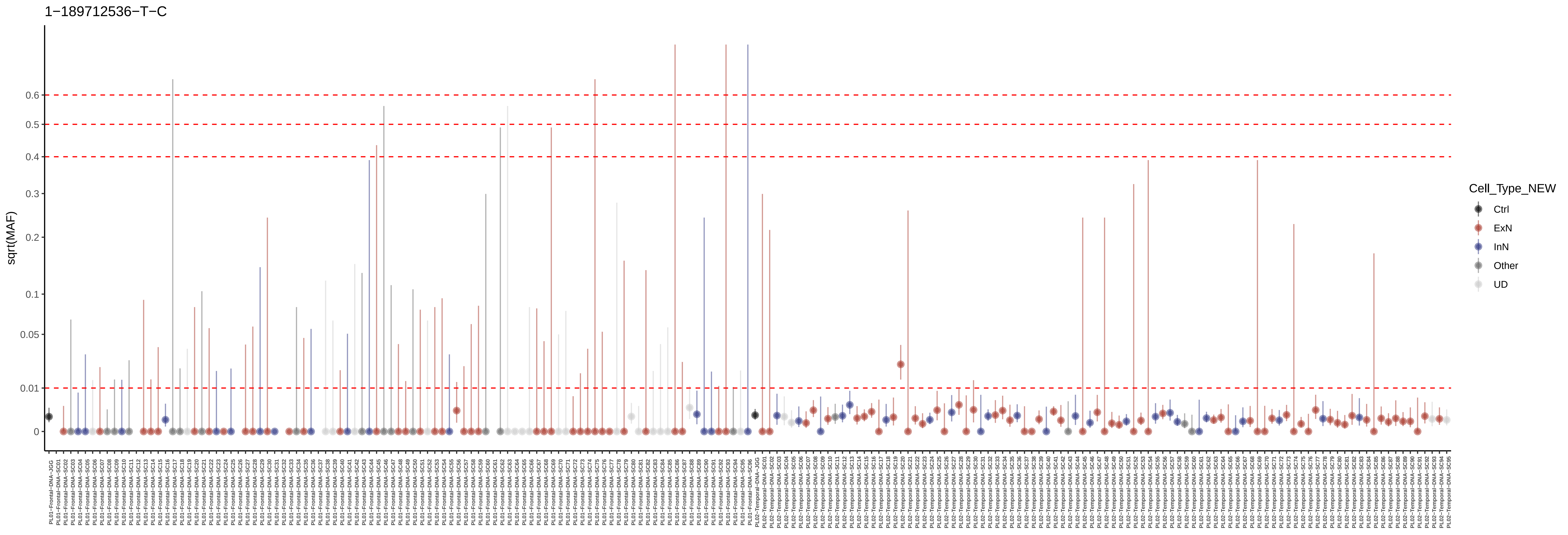
1−189712536−T−C
sqrt(MAF)
0.6
0.5
0.4
0.3
0.2
0.1
0.05
0.01
0
Cell_Type_NEW
Ctrl
ExN
InN
Other
UD

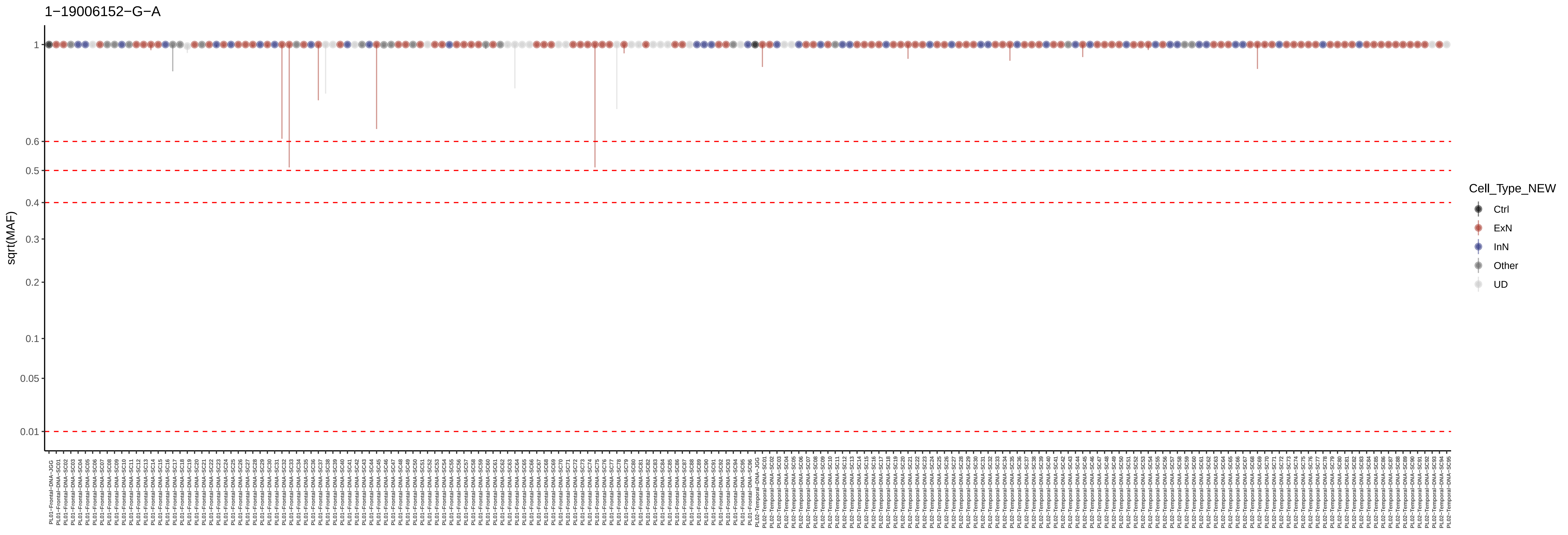
1−19006152−G−A
sqrt(MAF)
1
0.6
0.5
0.4
0.3
0.2
0.1
0.05
0.01
Cell_Type_NEW
Ctrl
ExN
InN
Other
UD
PL01−Frontal−DNA−JGG
PL02−Temporal−DNA−JGG

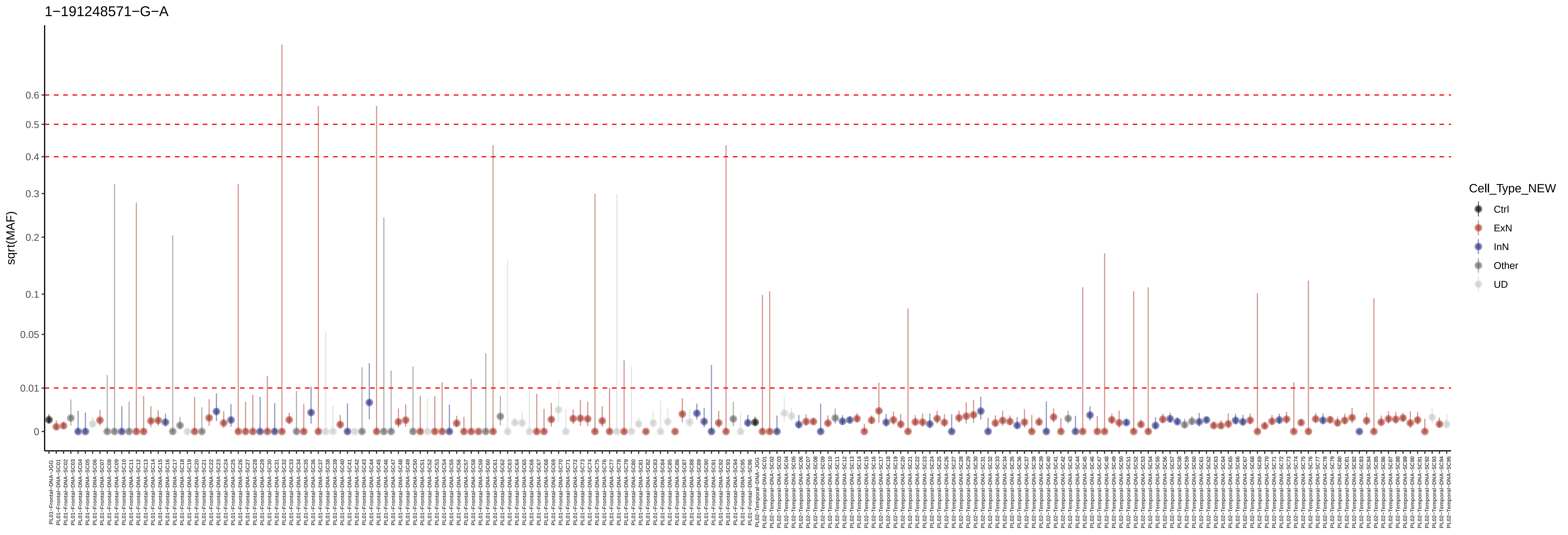

1−191248571−G−A
sqrt(MAF)
0.6
0.5
0.4
0.3
0.2
0.1
0.05
0.01
0
Cell_Type_NEW
Ctrl
ExN
InN
Other
UD

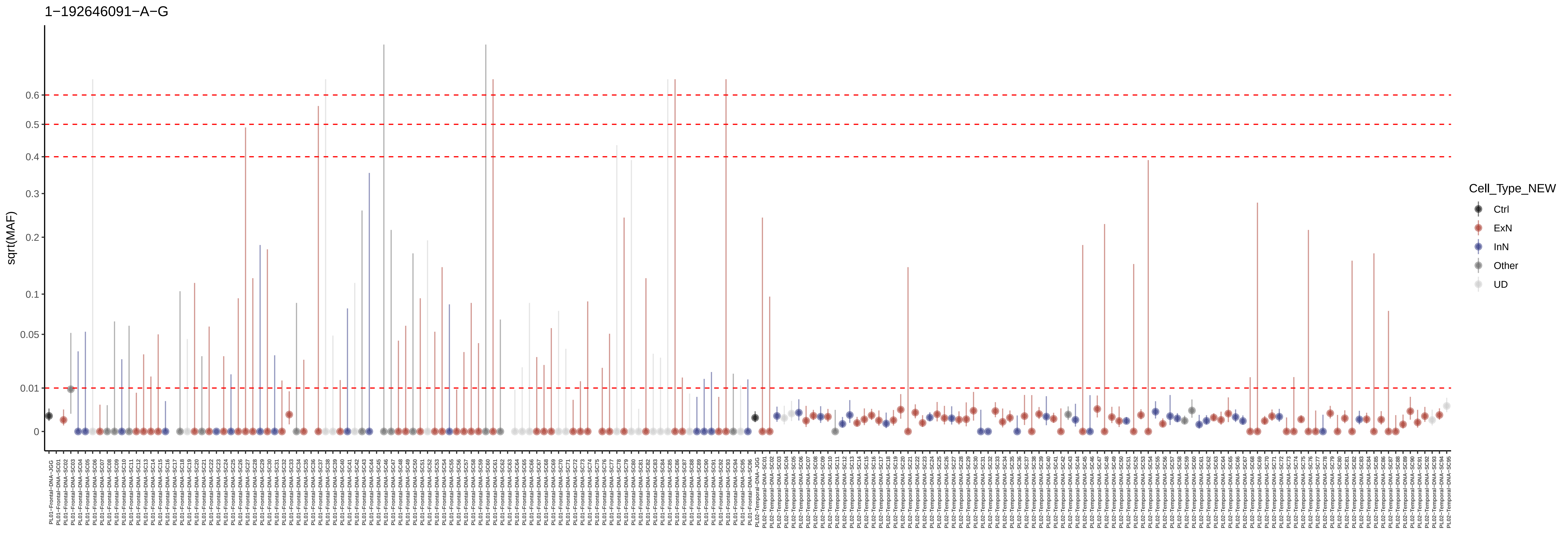
1−192646091−A−G
sqrt(MAF)
0.6
0.5
0.4
0.3
0.2
0.1
0.05
0.01
0
Cell_Type_NEW
Ctrl
ExN
InN
Other
UD

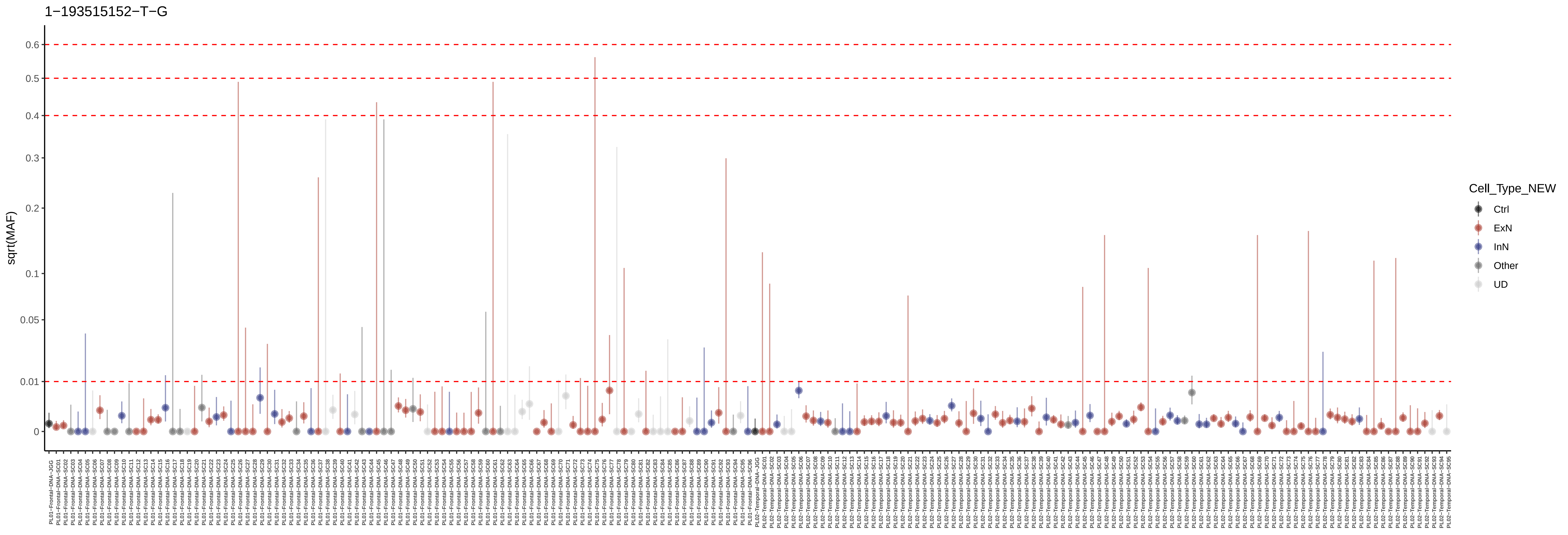
1−193515152−T−G
sqrt(MAF)
0.6
0.5
0.4
0.3
0.2
0.1
0.05
0.01
0
Cell_Type_NEW
Ctrl
ExN
InN
Other
UD

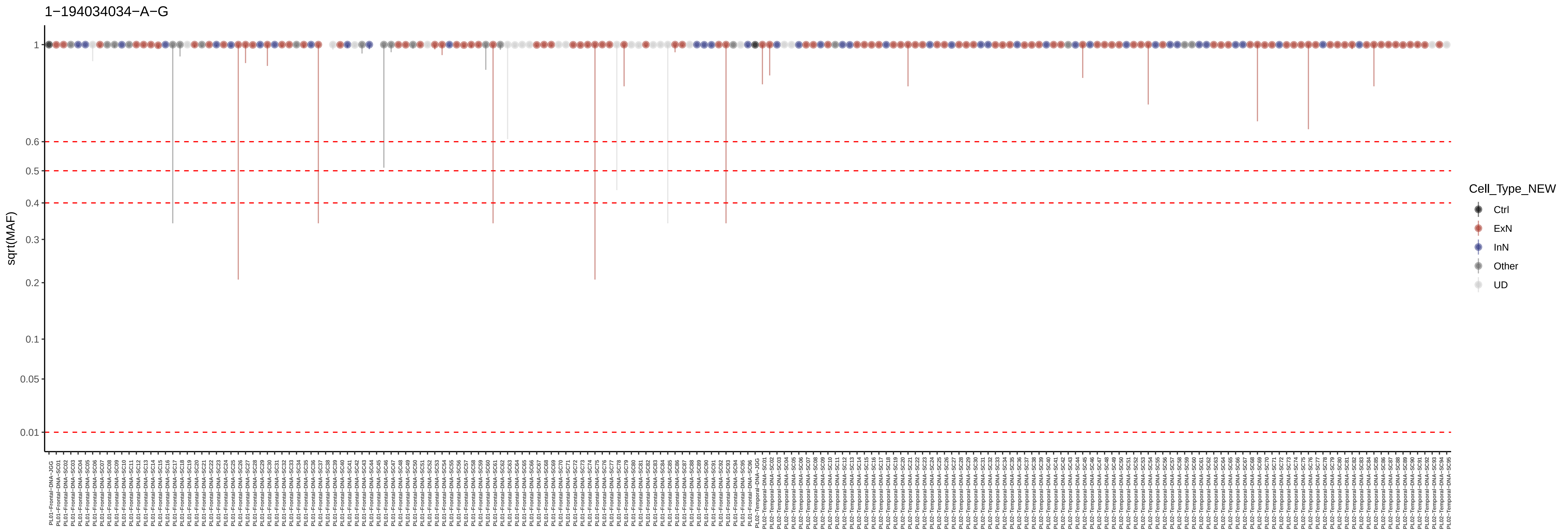

1−194034034−A−G
sqrt(MAF)
1
0.6
0.5
0.4
0.3
0.2
0.1
0.05
0.01
Cell_Type_NEW
Ctrl
ExN
InN
Other
UD

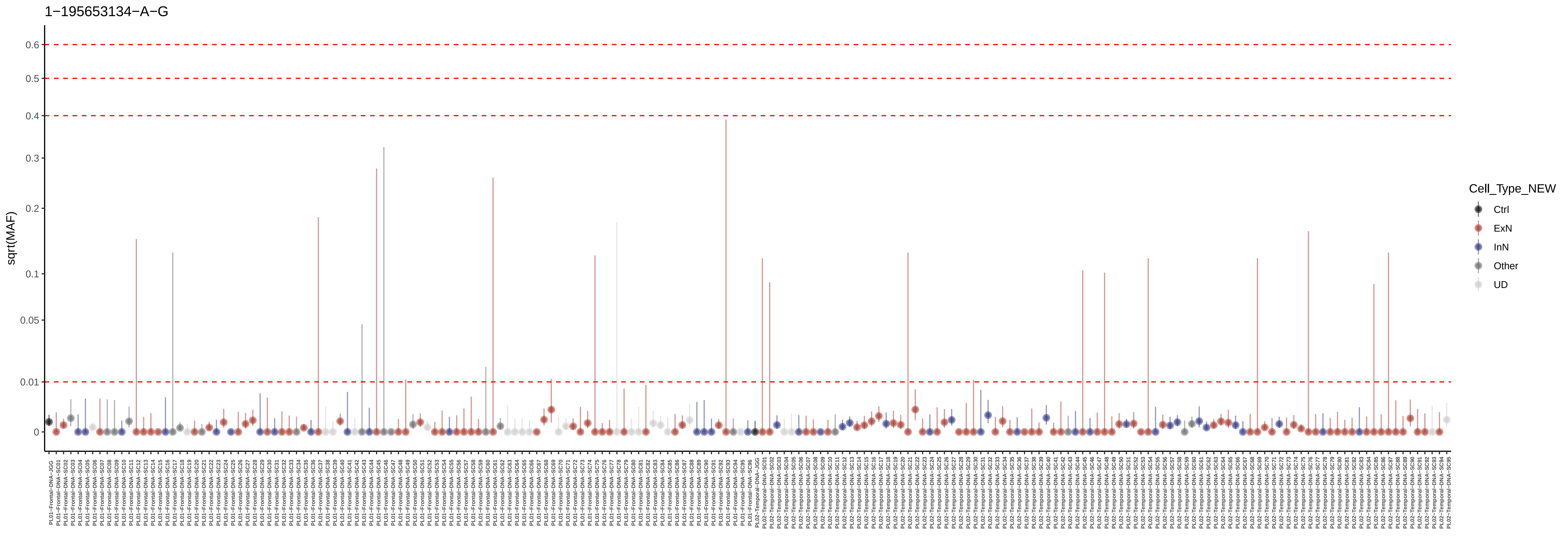

1−195653134−A−G
sqrt(MAF)
0.6
0.5
0.4
0.3
0.2
0.1
0.05
0.01
0
Cell_Type_NEW
Ctrl
ExN
InN
Other
UD

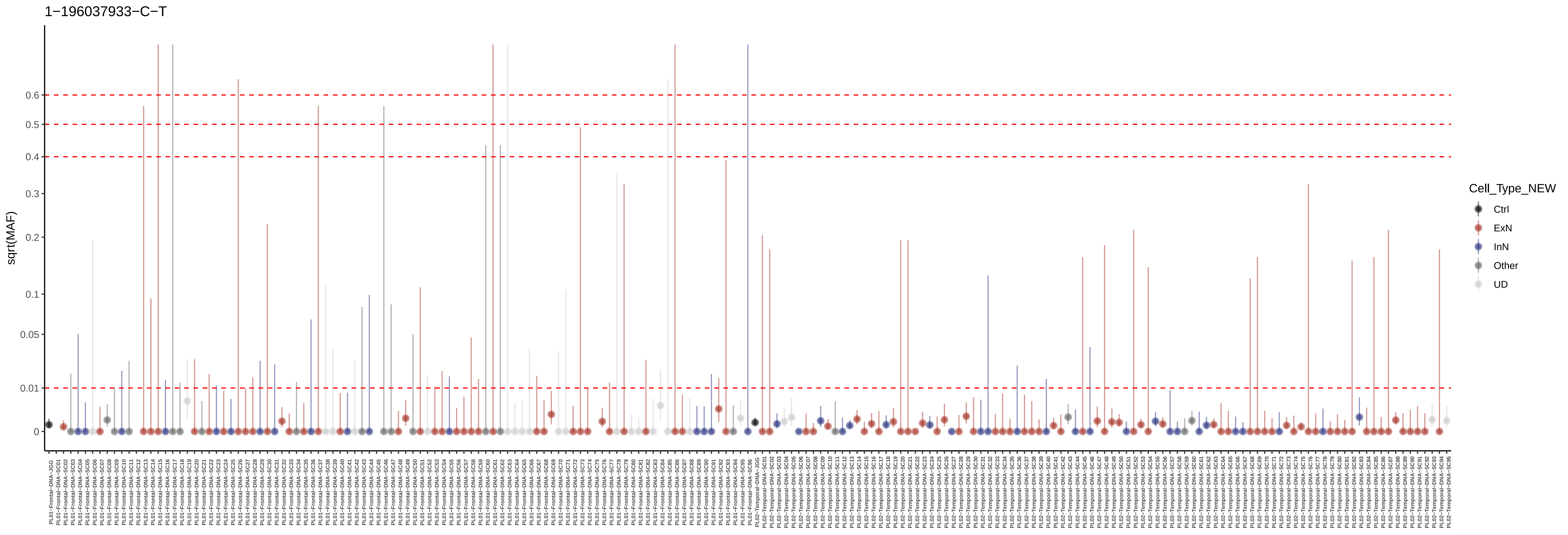
1−196037933−C−T
sqrt(MAF)
0
0.01
0.05
0.1
0.2
0.3
0.4
0.5
0.6
Cell_Type_NEW
Ctrl
ExN
InN
Other
UD

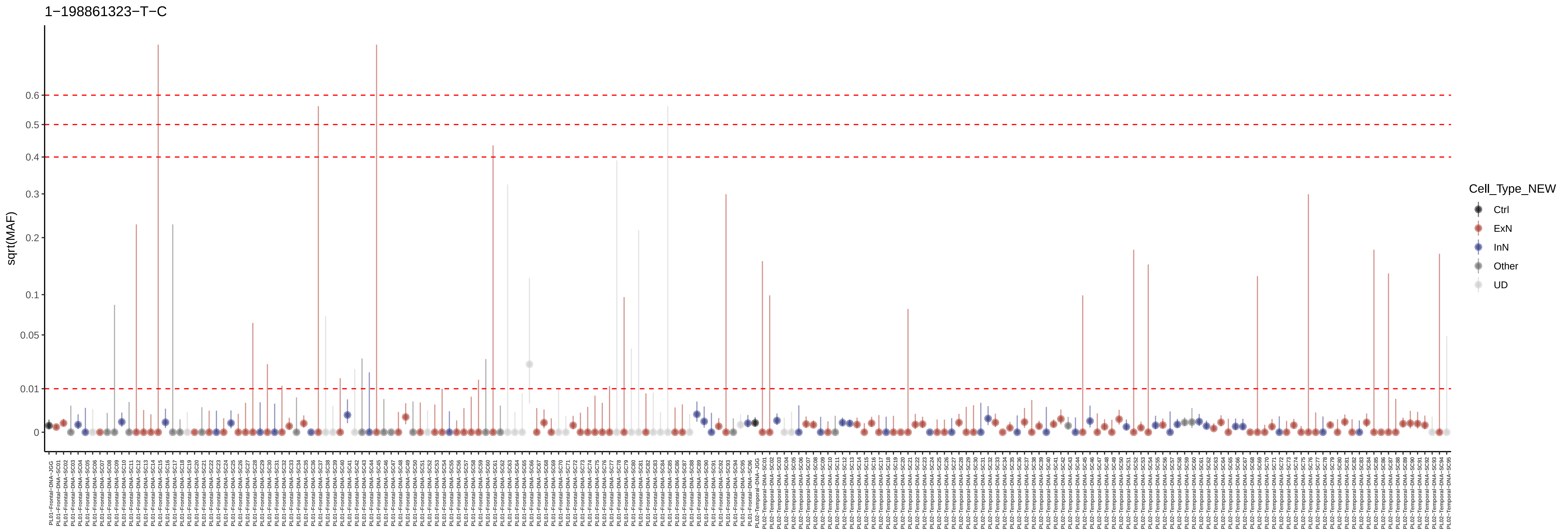
1−198861323−T−C
sqrt(MAF)
0.6
0.5
0.4
0.3
0.2
0.1
0.05
0.01
0
Cell_Type_NEW
Ctrl
ExN
InN
Other
UD

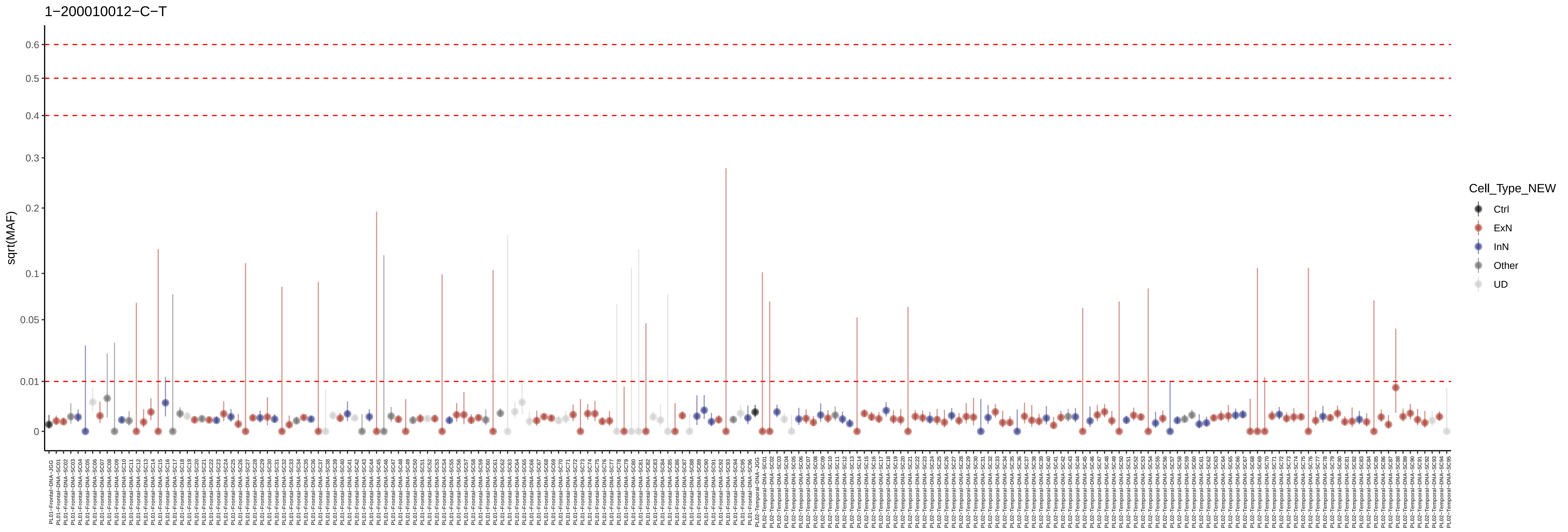
1−200010012−C−T
sqrt(MAF)
0.6
0.5
0.4
0.3
0.2
0.1
0.05
0.01
0
Cell_Type_NEW
Ctrl
ExN
InN
Other
UD

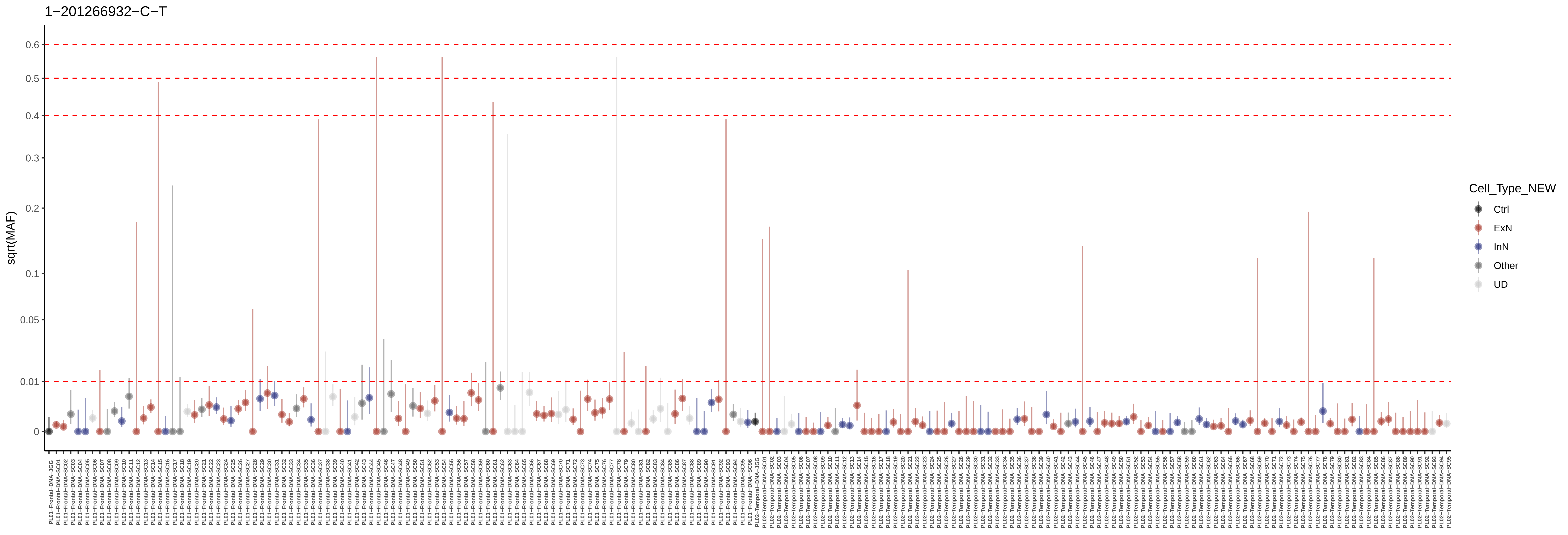
1−201266932−C−T
sqrt(MAF)
0.6
0.5
0.4
0.3
0.2
0.1
0.05
0.01
0
Cell_Type_NEW
Ctrl
ExN
InN
Other
UD

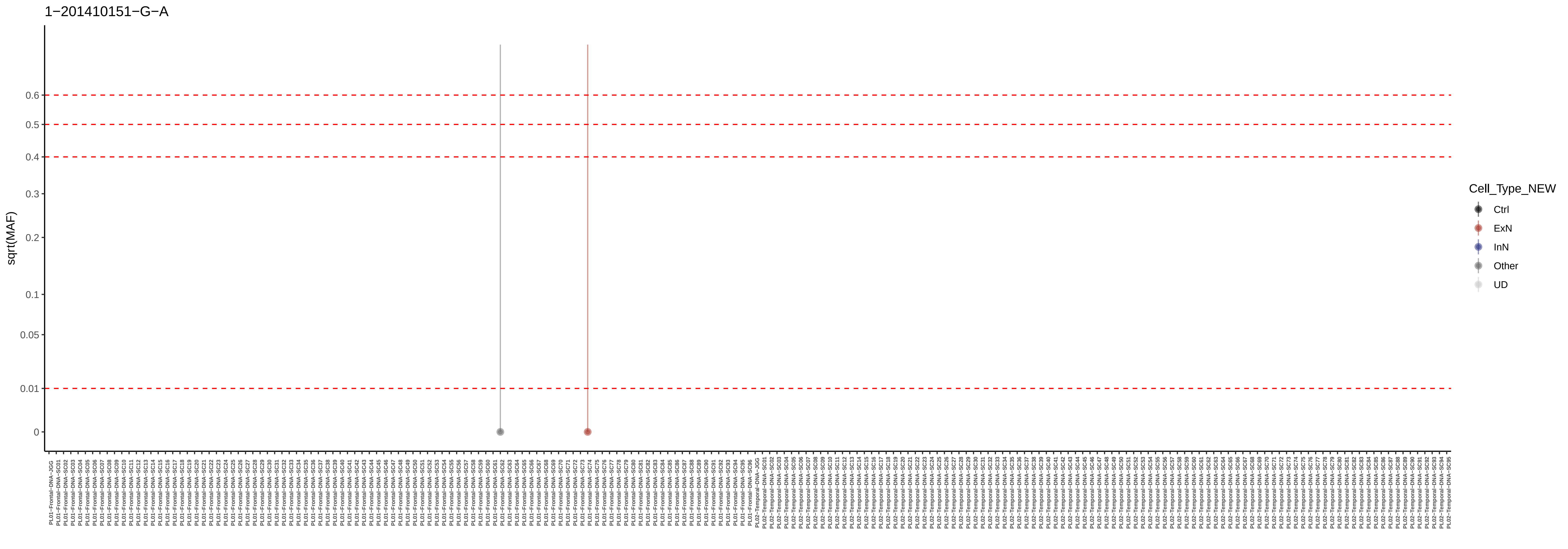

1−201410151−G−A
sqrt(MAF)
0
0.01
0.05
0.1
0.2
0.3
0.4
0.5
0.6
Cell_Type_NEW
Ctrl
ExN
InN
Other
UD

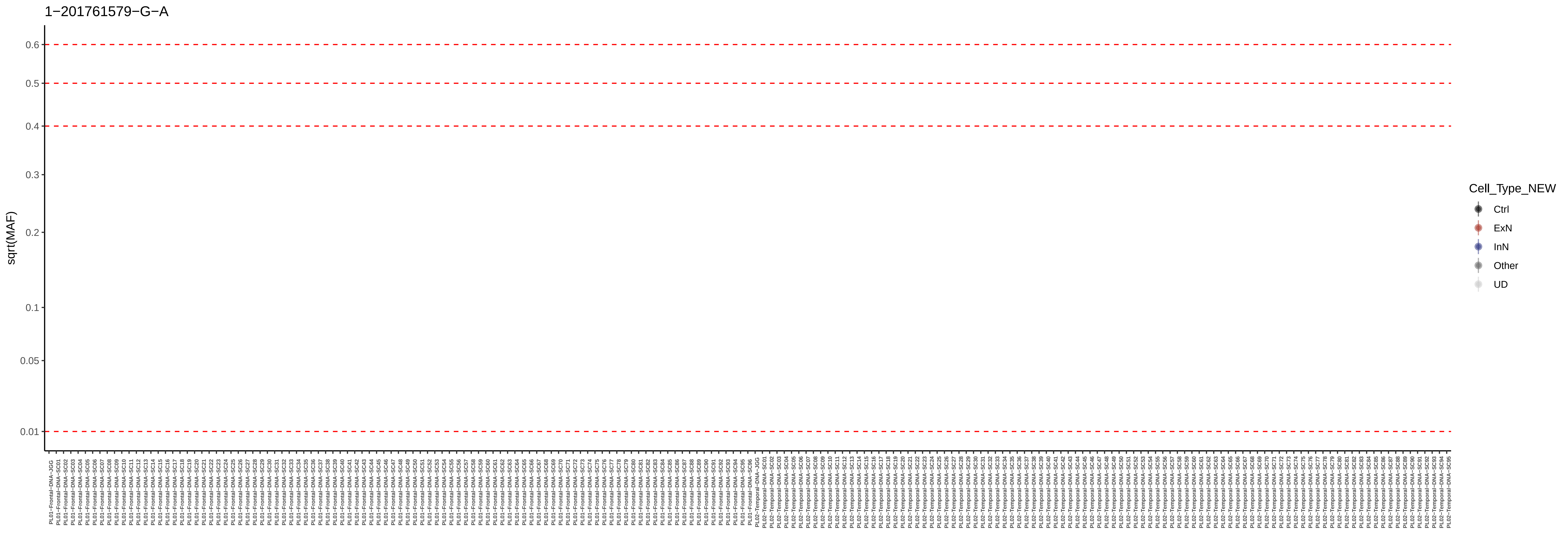
1−201761579−G−A
sqrt(MAF)
0.6
0.5
0.4
0.3
0.2
0.1
0.05
0.01
Cell_Type_NEW
Ctrl
ExN
InN
Other
UD

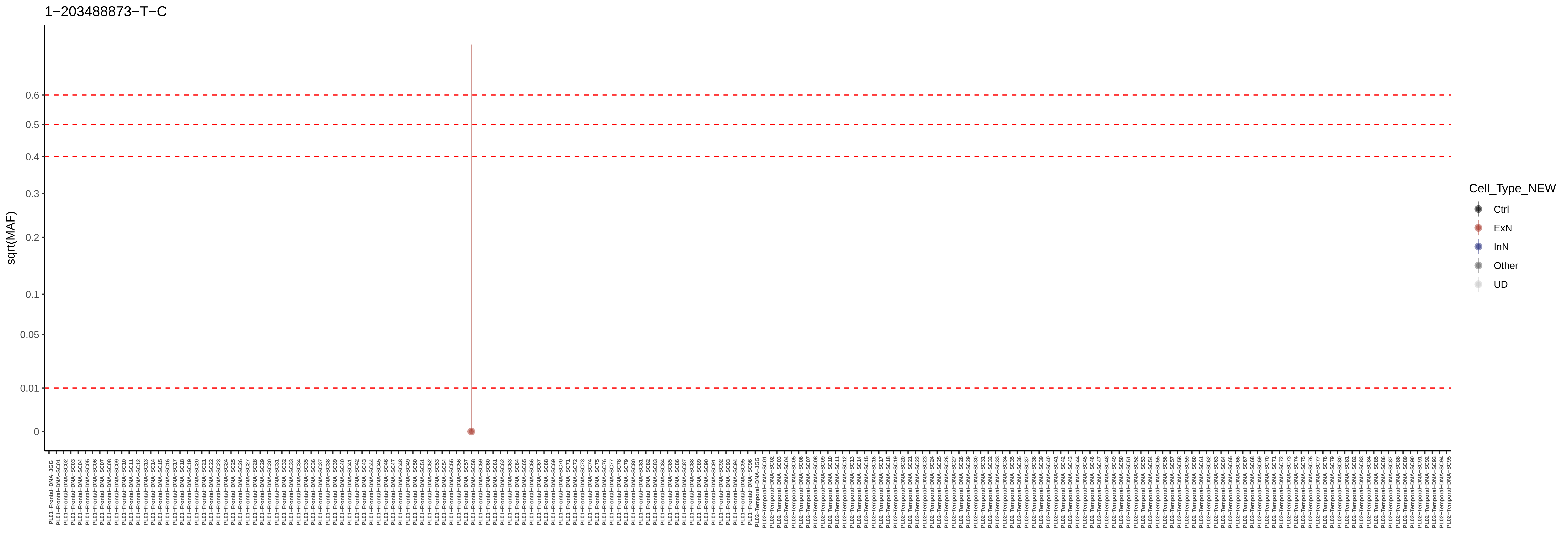
1−203488873−T−C
sqrt(MAF)
0.6
0.5
0.4
0.3
0.2
0.1
0.05
0.01
0
Cell_Type_NEW
Ctrl
ExN
InN
Other
UD

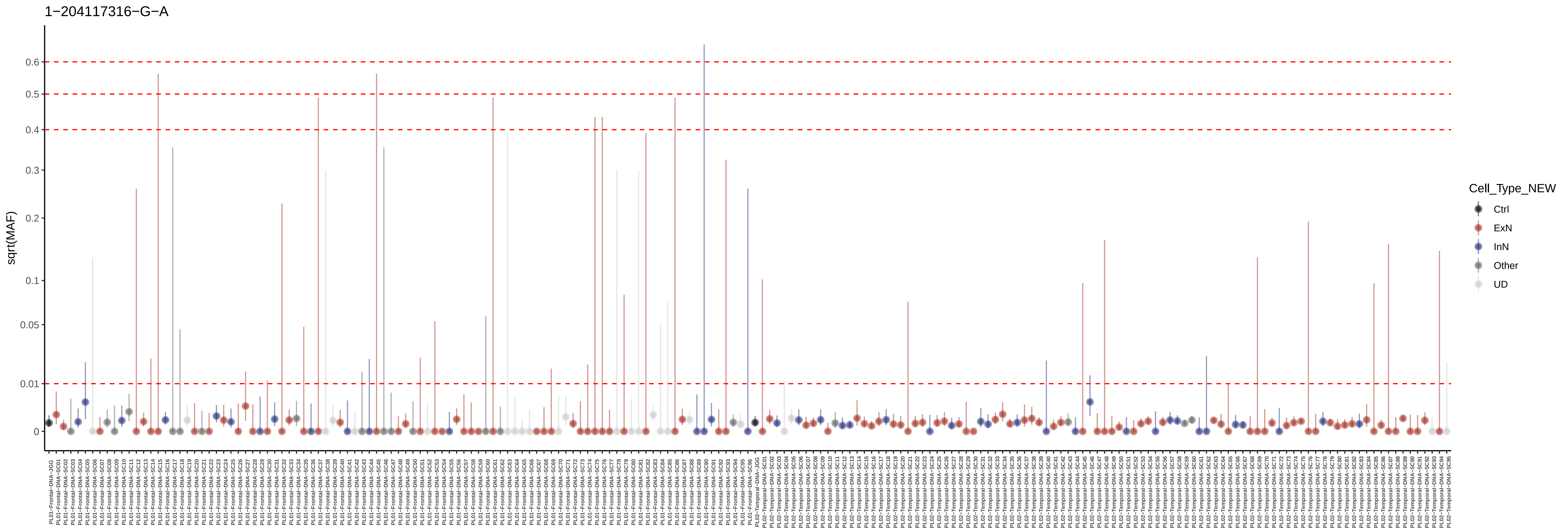
1−204117316−G−A
sqrt(MAF)
0.6
0.5
0.4
0.3
0.2
0.1
0.05
0.01
0
Cell_Type_NEW
Ctrl
ExN
InN
Other
UD

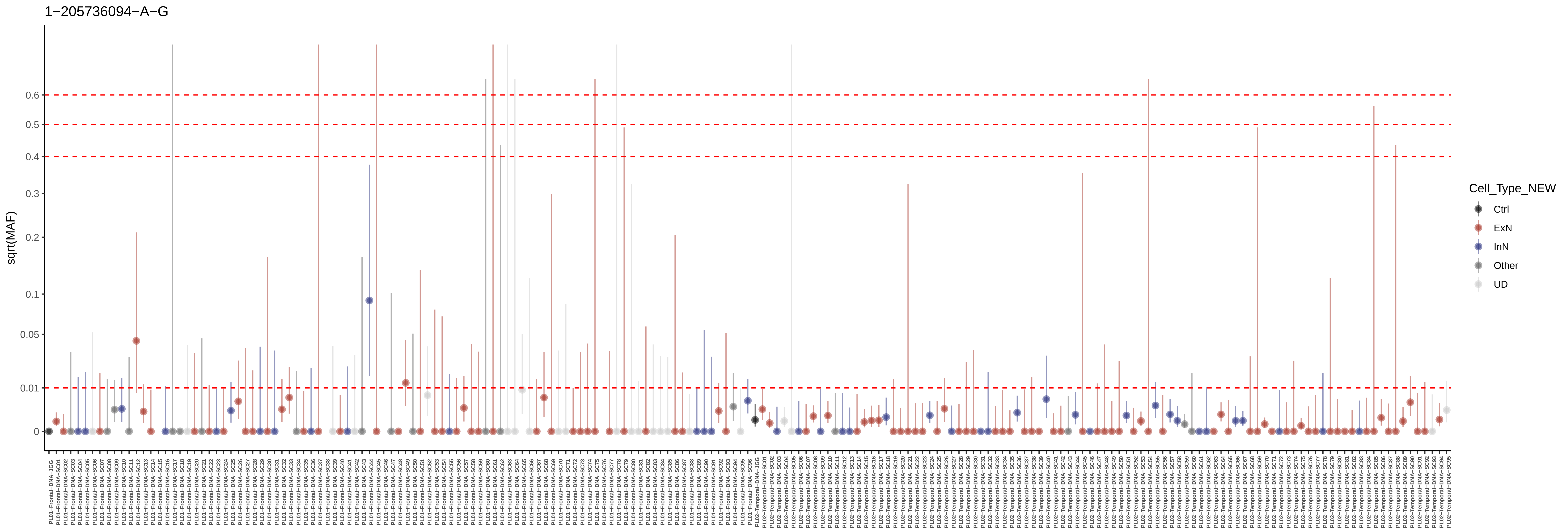
1−205736094−A−G
sqrt(MAF)
0.6
0.5
0.4
0.3
0.2
0.1
0.05
0.01
0
Cell_Type_NEW
Ctrl
ExN
InN
Other
UD

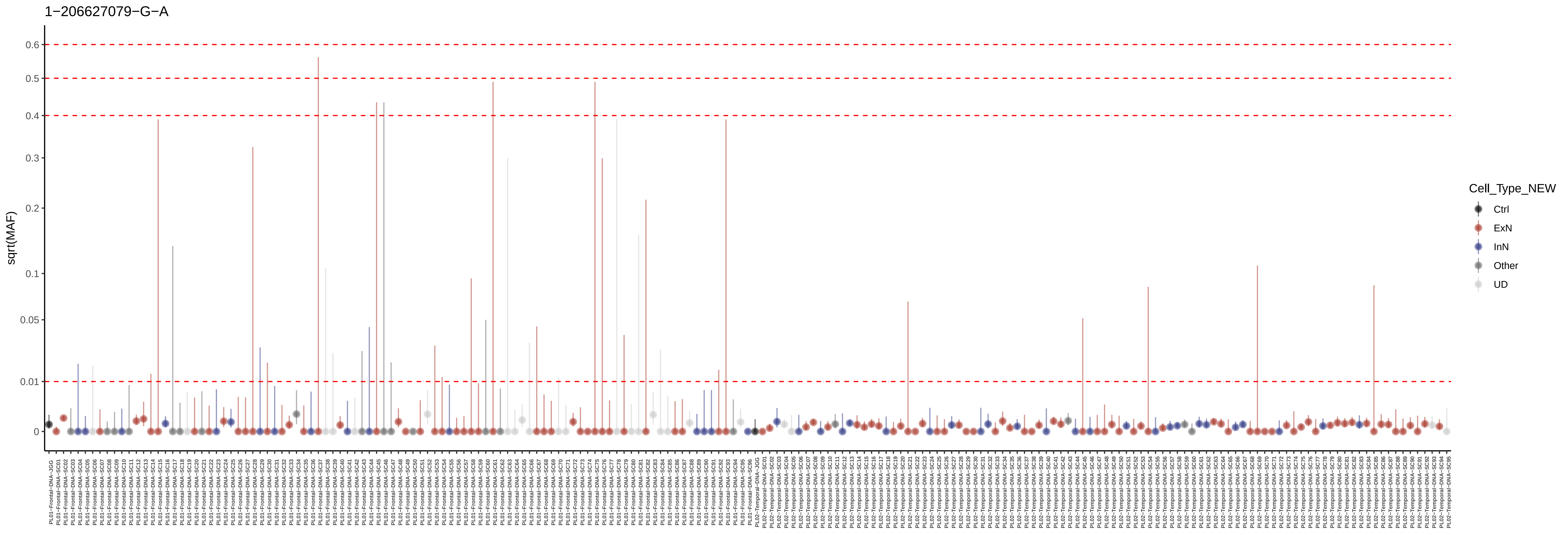
1−206627079−G−A
sqrt(MAF)
0.6
0.5
0.4
0.3
0.2
0.1
0.05
0.01
0
Cell_Type_NEW
Ctrl
ExN
InN
Other
UD

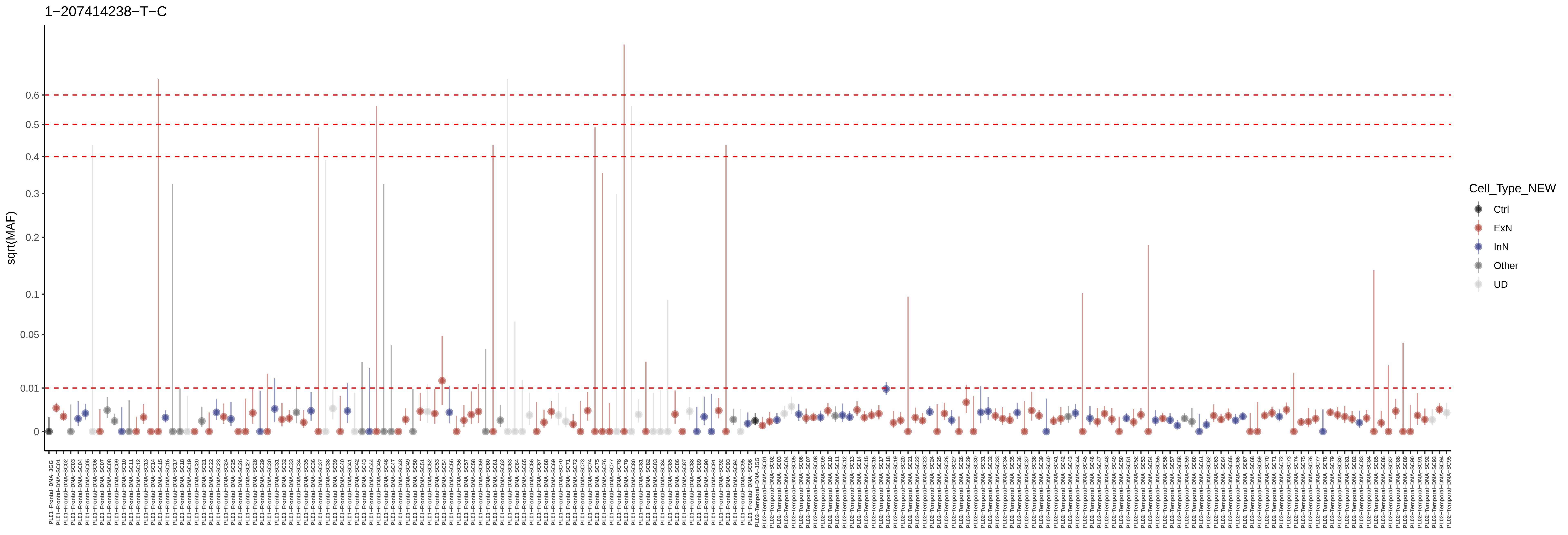
1−207414238−T−C
sqrt(MAF)
0.6
0.5
0.4
0.3
0.2
0.1
0.05
0.01
0
Cell_Type_NEW
Ctrl
ExN
InN
Other
UD

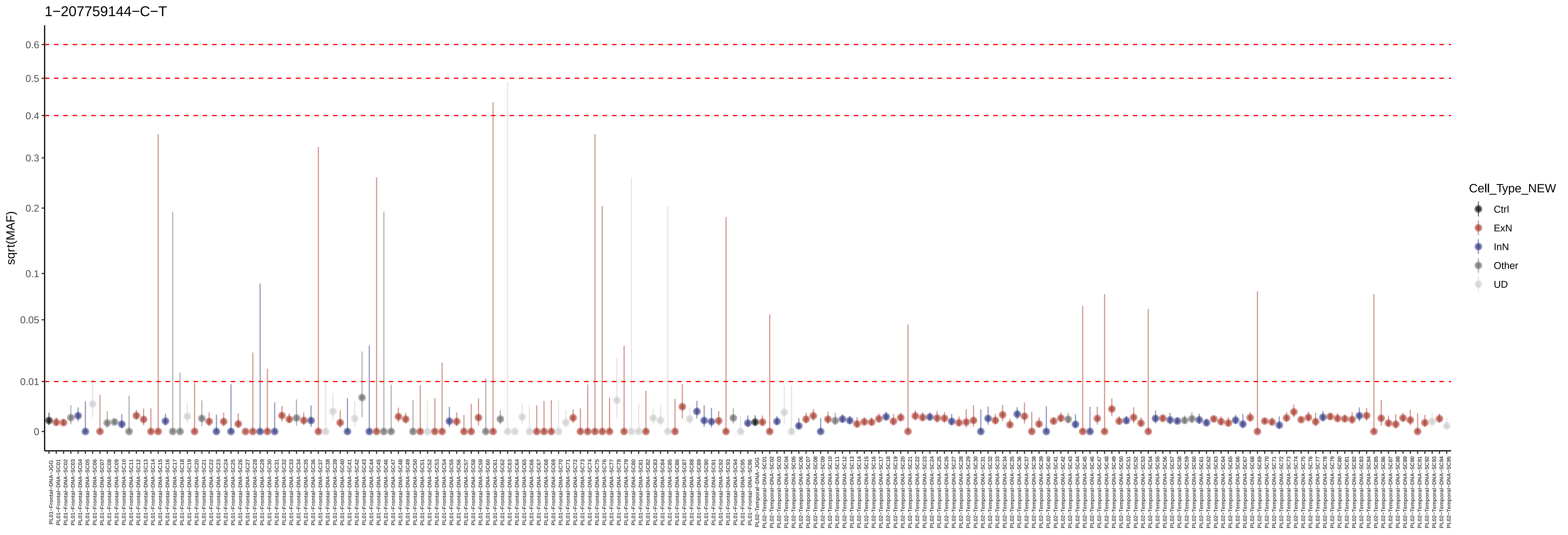
1−207759144−C−T
sqrt(MAF)
0.6
0.5
0.4
0.3
0.2
0.1
0.05
0.01
0
Cell_Type_NEW
Ctrl
ExN
InN
Other
UD

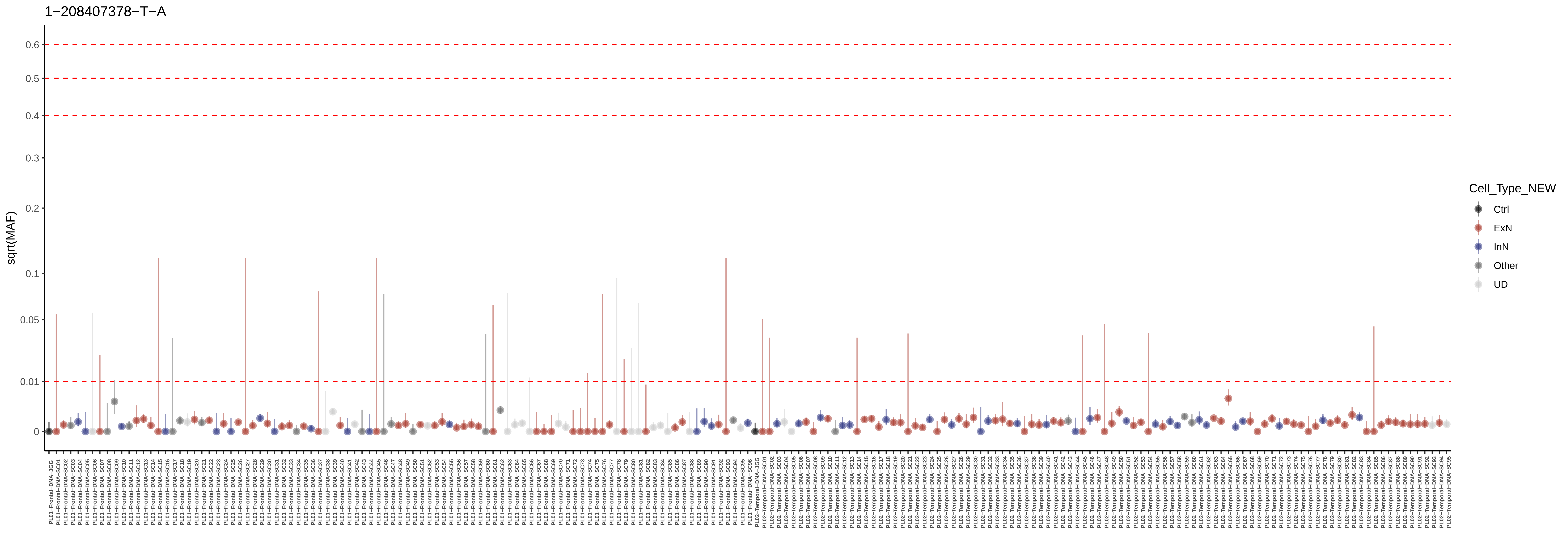
1−208407378−T−A
sqrt(MAF)
0.6
0.5
0.4
0.3
0.2
0.1
0.05
0.01
0
Cell_Type_NEW
Ctrl
ExN
InN
Other
UD

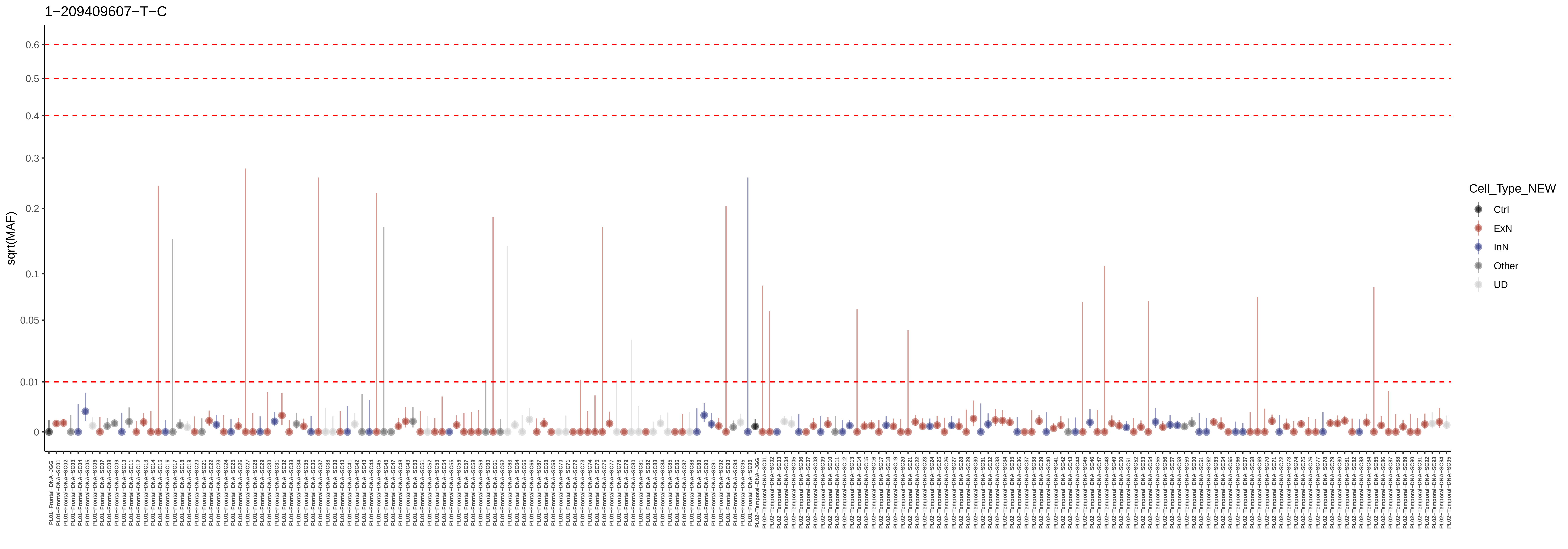
1−209409607−T−C
sqrt(MAF)
0.6
0.5
0.4
0.3
0.2
0.1
0.05
0.01
0
Cell_Type_NEW
Ctrl
ExN
InN
Other
UD

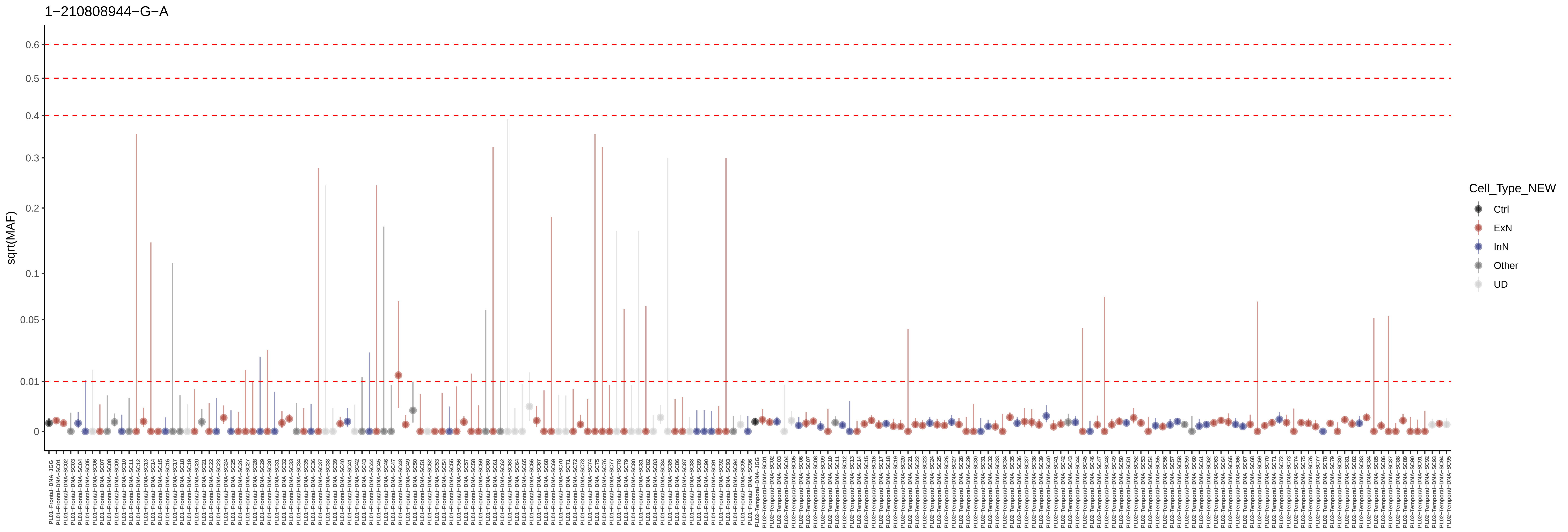
1−210808944−G−A
sqrt(MAF)
0.6
0.5
0.4
0.3
0.2
0.1
0.05
0.01
0
Cell_Type_NEW
Ctrl
ExN
InN
Other
UD

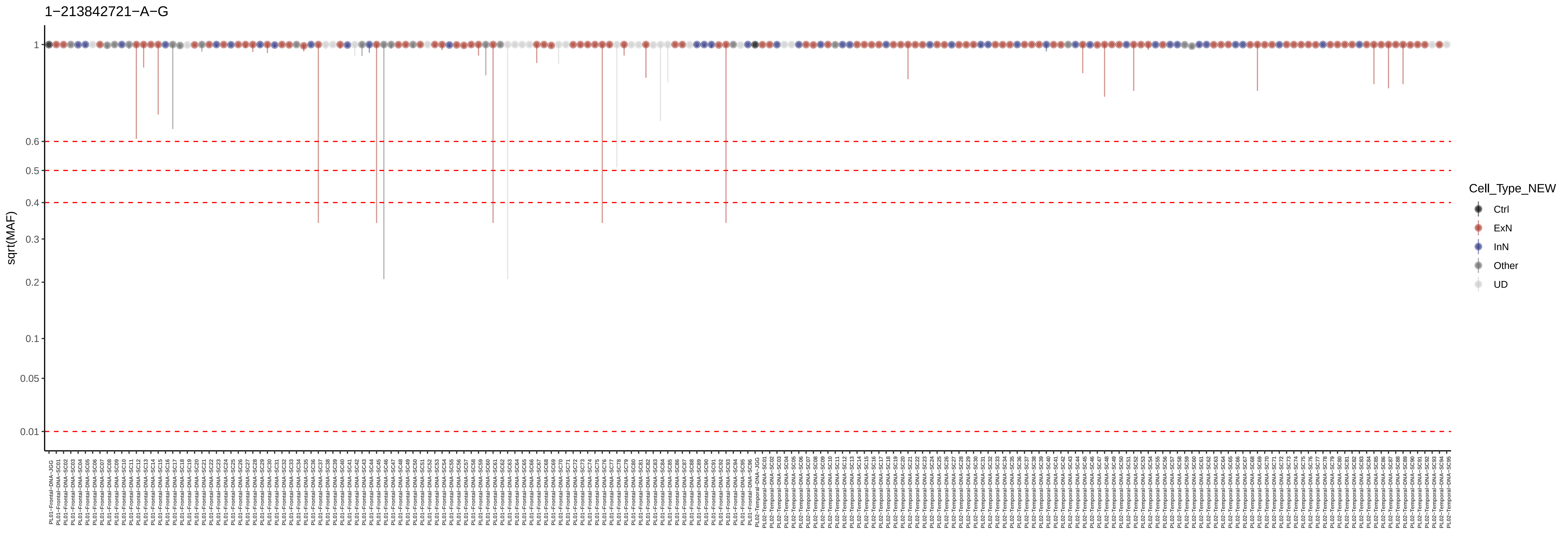
1−213842721−A−G
sqrt(MAF)
1
0.6
0.5
0.4
0.3
0.2
0.1
0.05
0.01
Cell_Type_NEW
Ctrl
ExN
InN
Other
UD

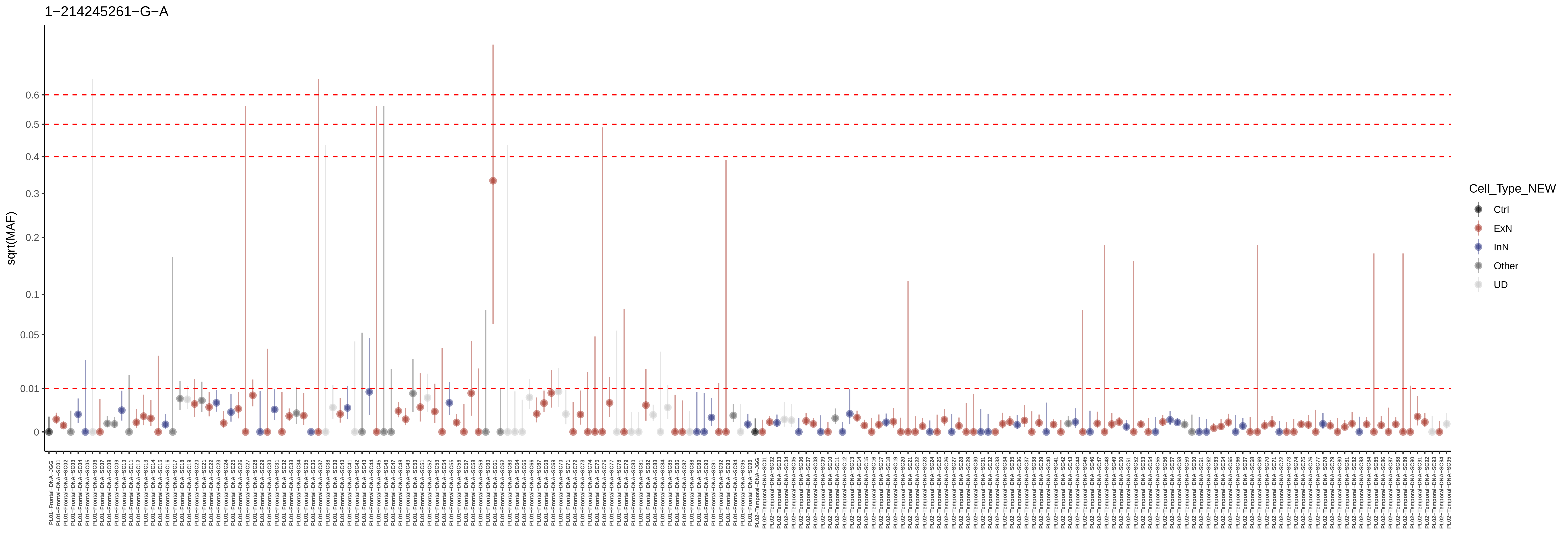

1−214245261−G−A
sqrt(MAF)
0.6
0.5
0.4
0.3
0.2
0.1
0.05
0.01
0
Cell_Type_NEW
Ctrl
ExN
InN
Other
UD

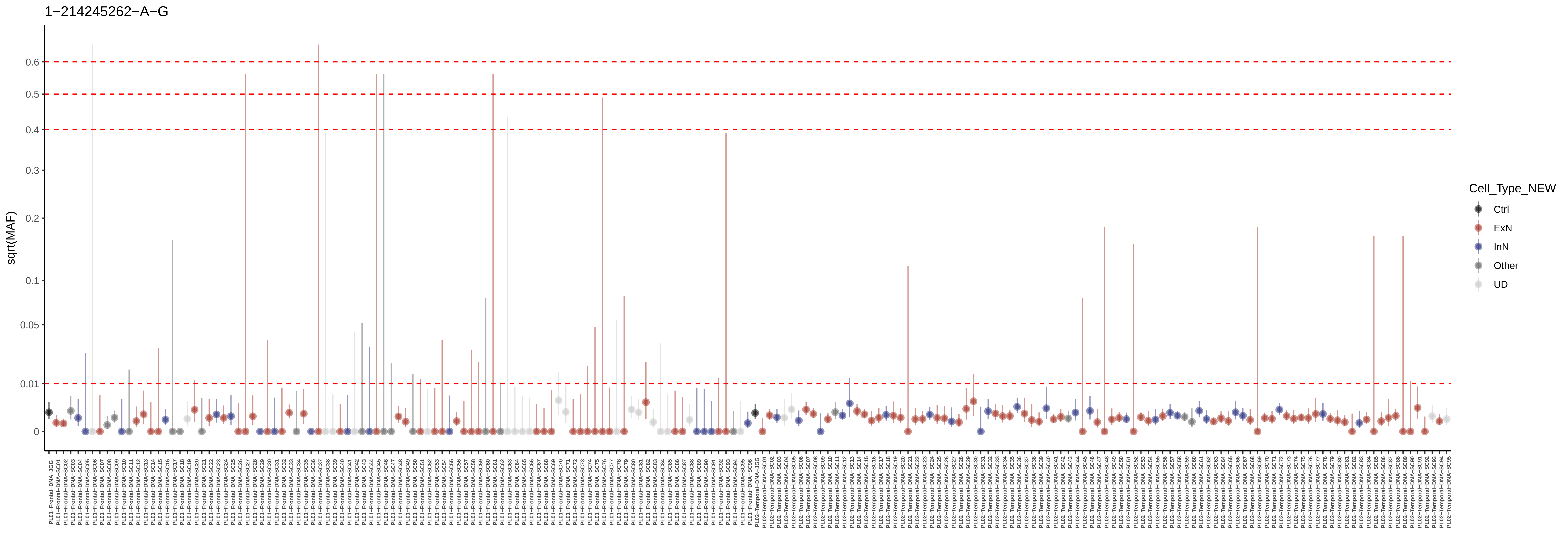

1−214245262−A−G
sqrt(MAF)
0.6
0.5
0.4
0.3
0.2
0.1
0.05
0.01
0
Cell_Type_NEW
Ctrl
ExN
InN
Other
UD

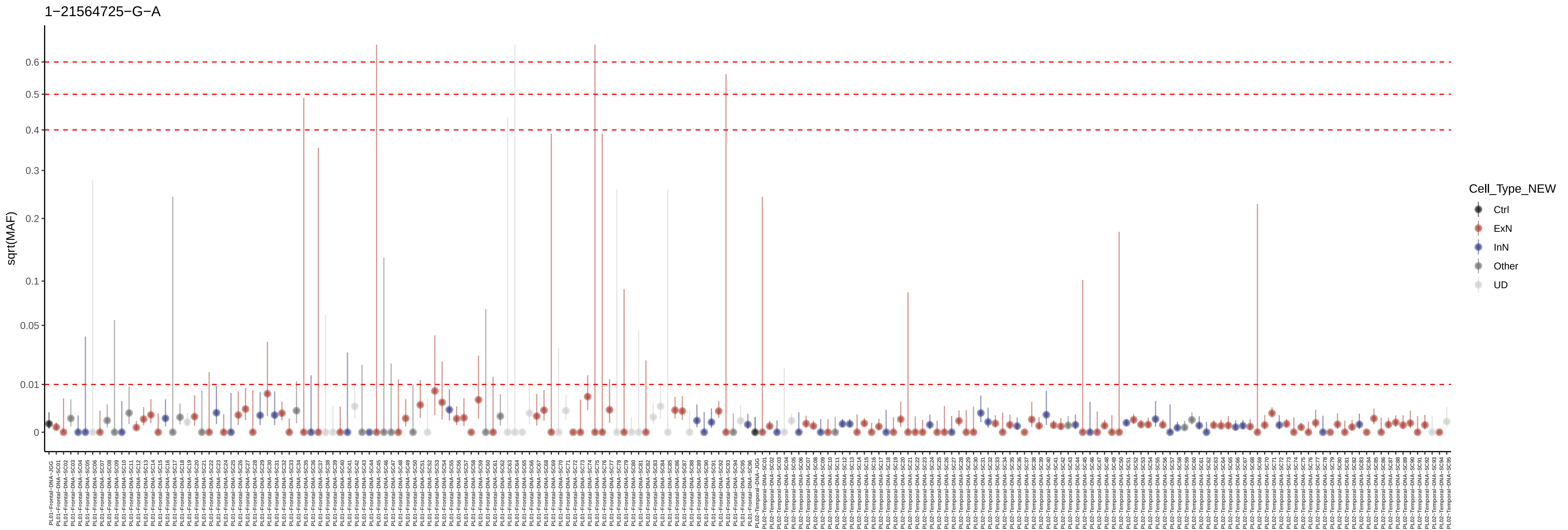

1−21564725−G−A
sqrt(MAF)
0
0.01
0.05
0.1
0.2
0.3
0.4
0.5
0.6
Cell_Type_NEW
Ctrl
ExN
InN
Other
UD

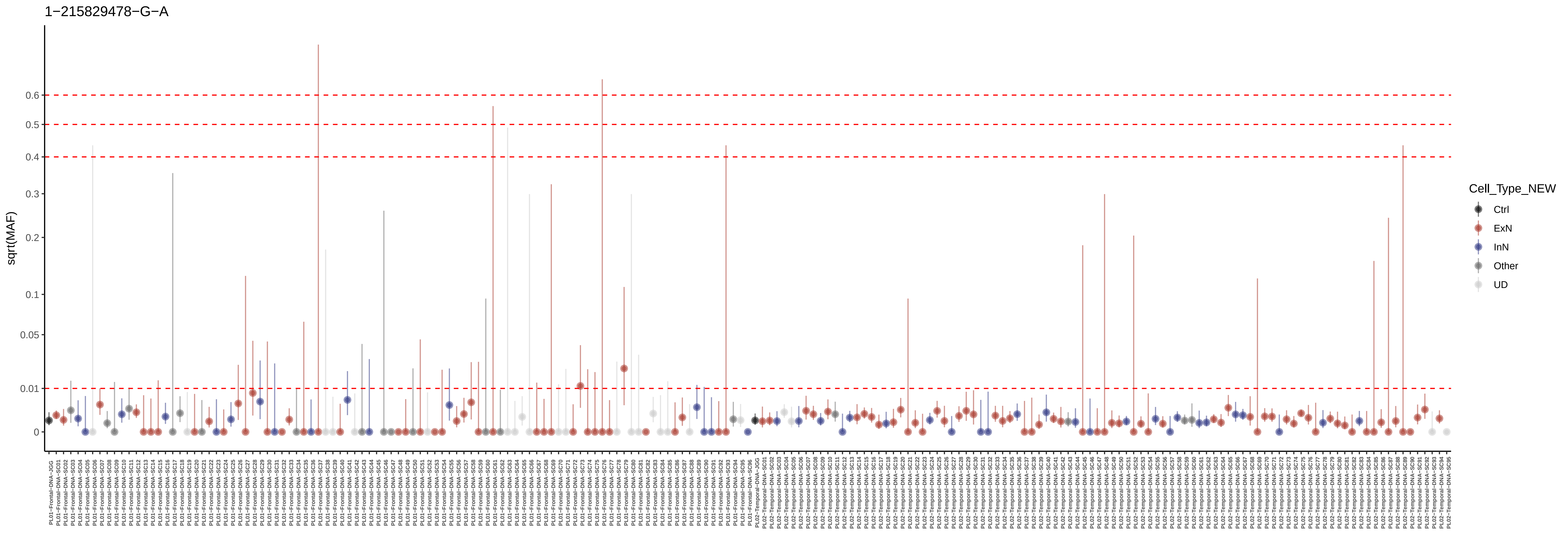

1−215829478−G−A
sqrt(MAF)
0.6
0.5
0.4
0.3
0.2
0.1
0.05
0.01
0
Cell_Type_NEW
Ctrl
ExN
InN
Other
UD

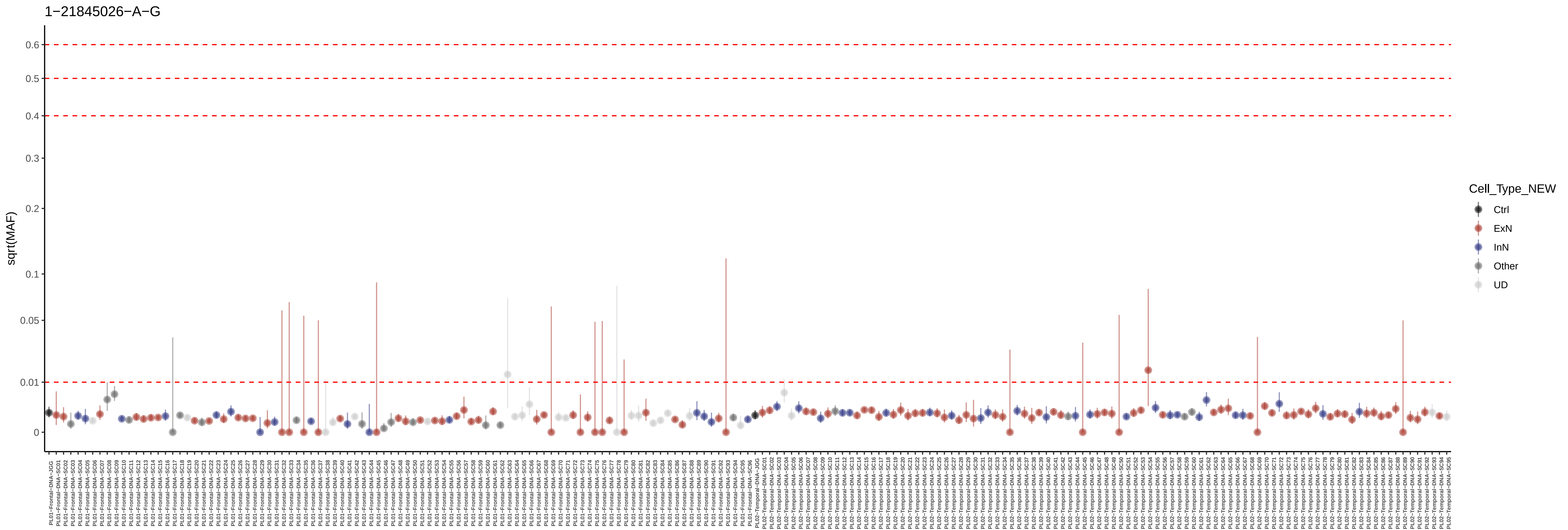
1−21845026−A−G
sqrt(MAF)
0.6
0.5
0.4
0.3
0.2
0.1
0.05
0.01
0
Cell_Type_NEW
Ctrl
ExN
InN
Other
UD

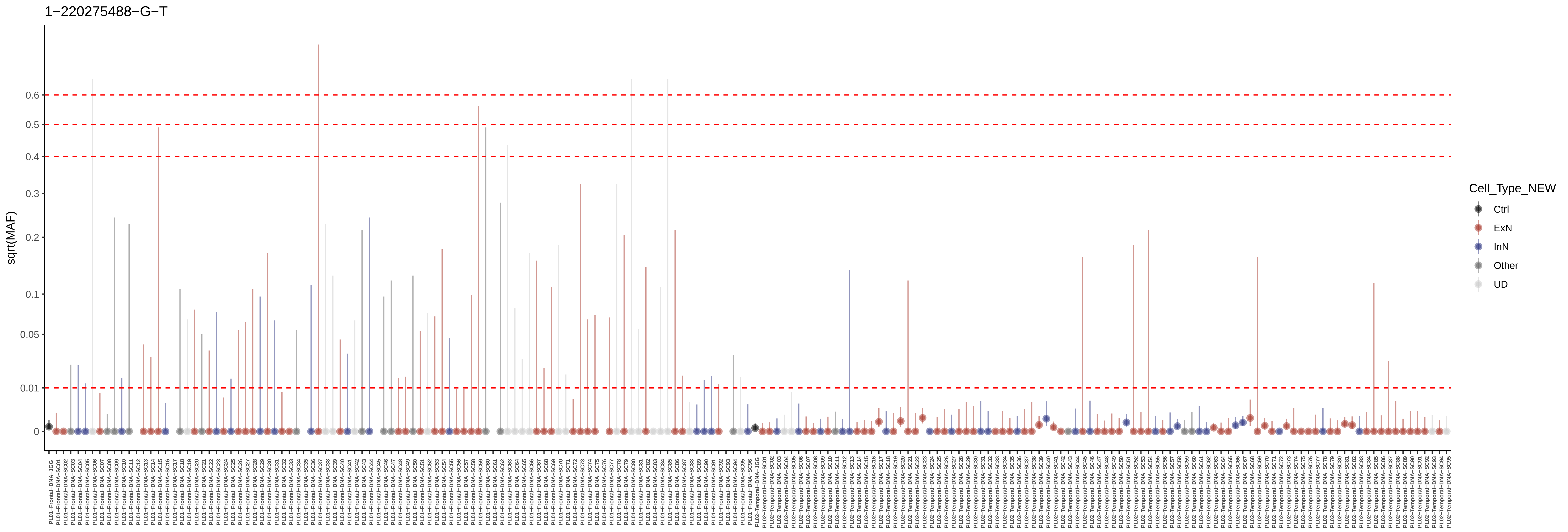

1−220275488−G−T
sqrt(MAF)
0.6
0.5
0.4
0.3
0.2
0.1
0.05
0.01
0
Cell_Type_NEW
Ctrl
ExN
InN
Other
UD

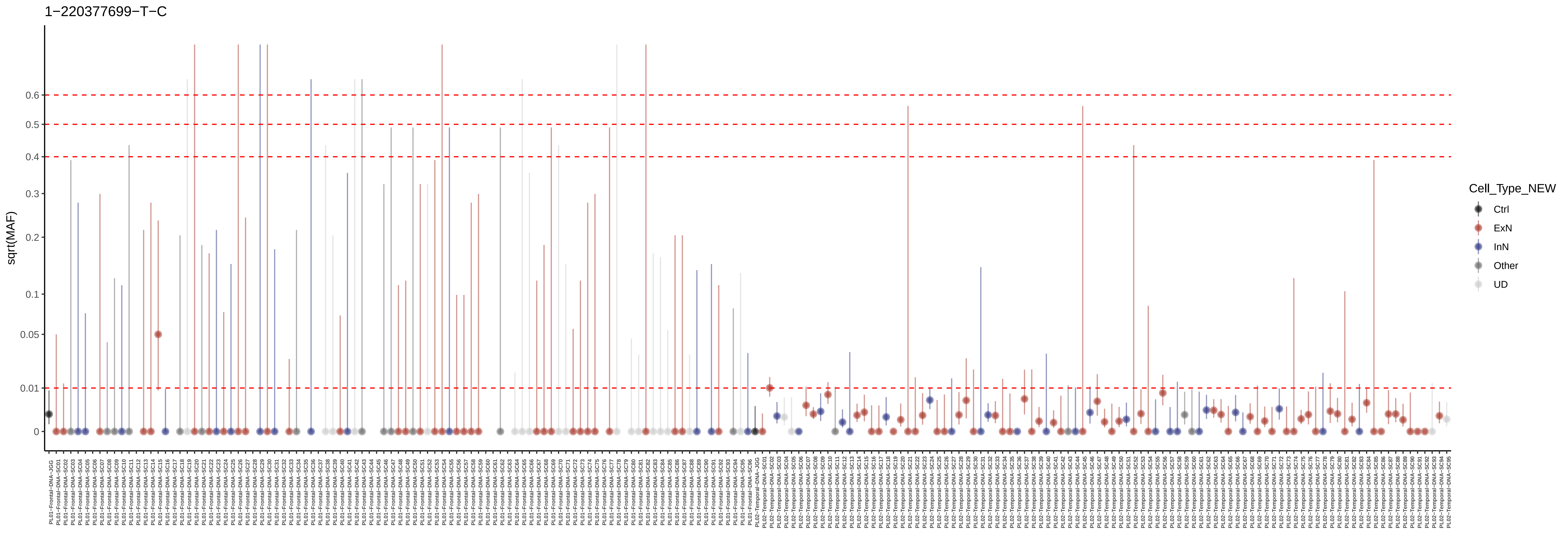
1−220377699−T−C
sqrt(MAF)
0.6
0.5
0.4
0.3
0.2
0.1
0.05
0.01
0
Cell_Type_NEW
Ctrl
ExN
InN
Other
UD

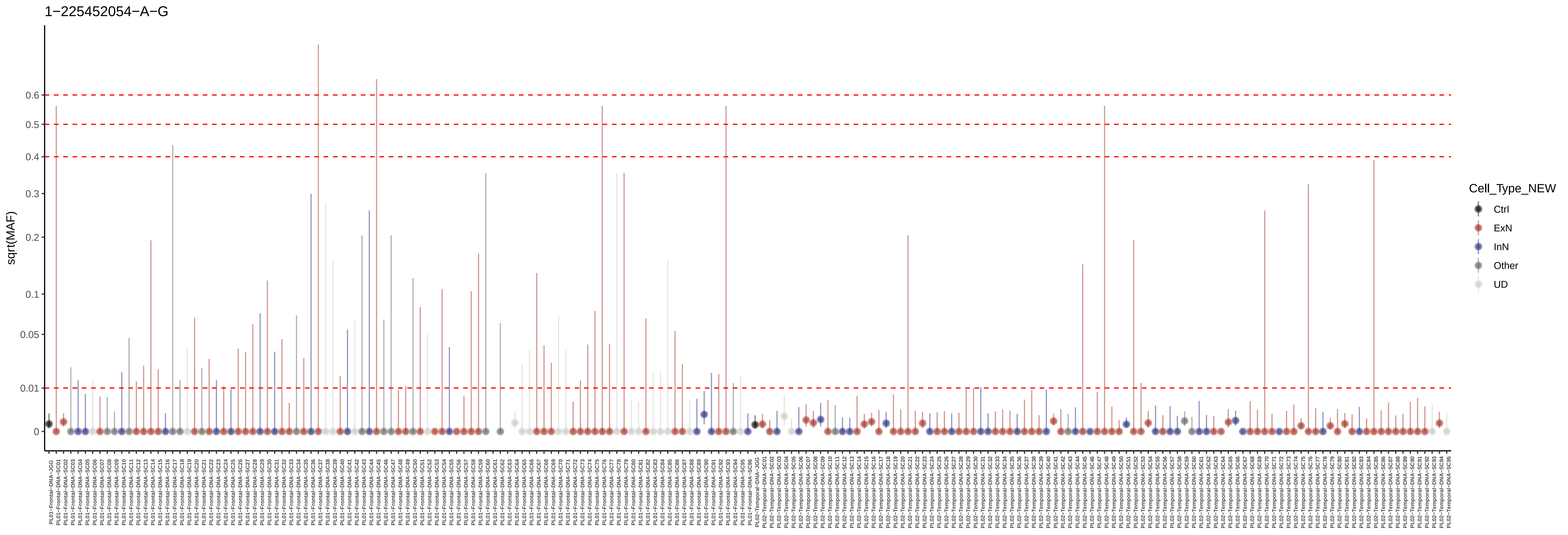
1−225452054−A−G
sqrt(MAF)
0.6
0.5
0.4
0.3
0.2
0.1
0.05
0.01
0
Cell_Type_NEW
Ctrl
ExN
InN
Other
UD

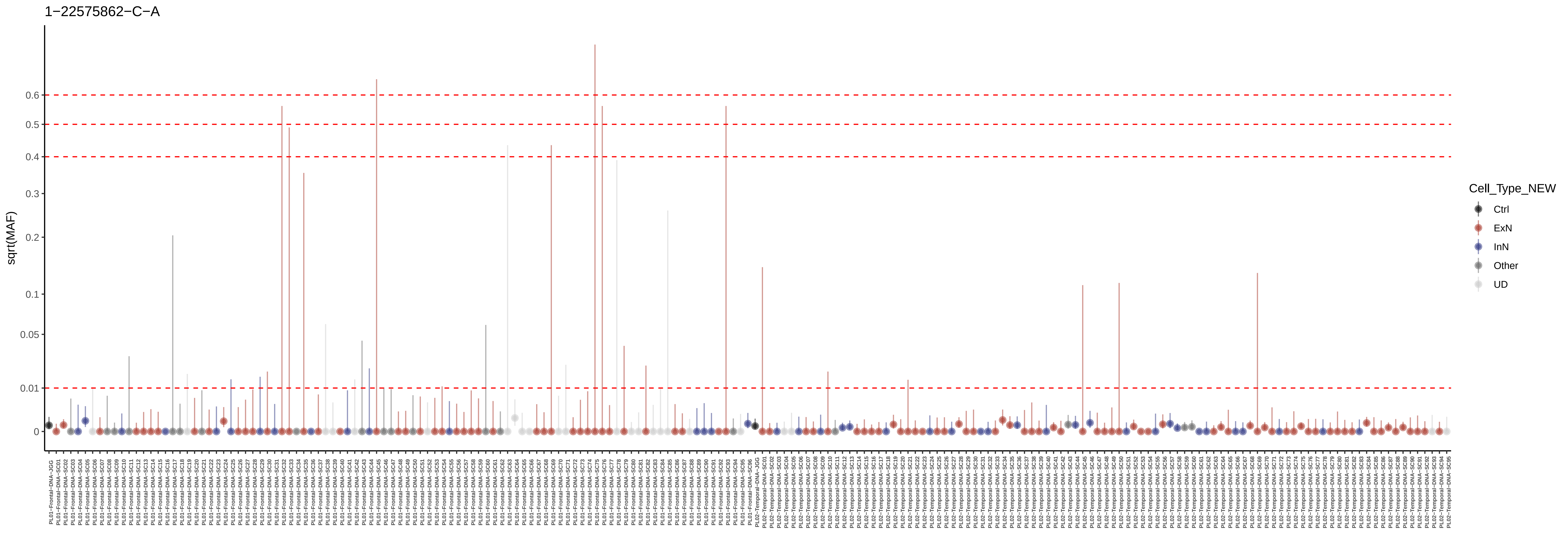
1−22575862−C−A
sqrt(MAF)
0.6
0.5
0.4
0.3
0.2
0.1
0.05
0.01
0
Cell_Type_NEW
Ctrl
ExN
InN
Other
UD

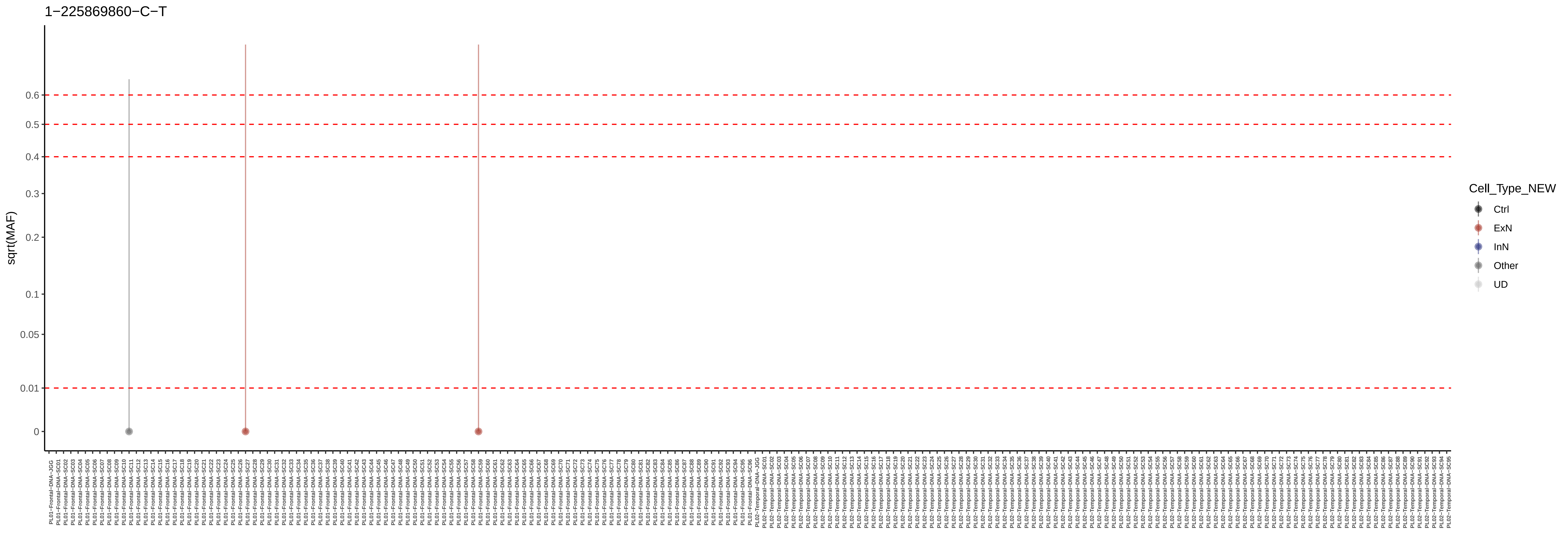
1−225869860−C−T
sqrt(MAF)
0.6
0.5
0.4
0.3
0.2
0.1
0.05
0.01
0
Cell_Type_NEW
Ctrl
ExN
InN
Other
UD

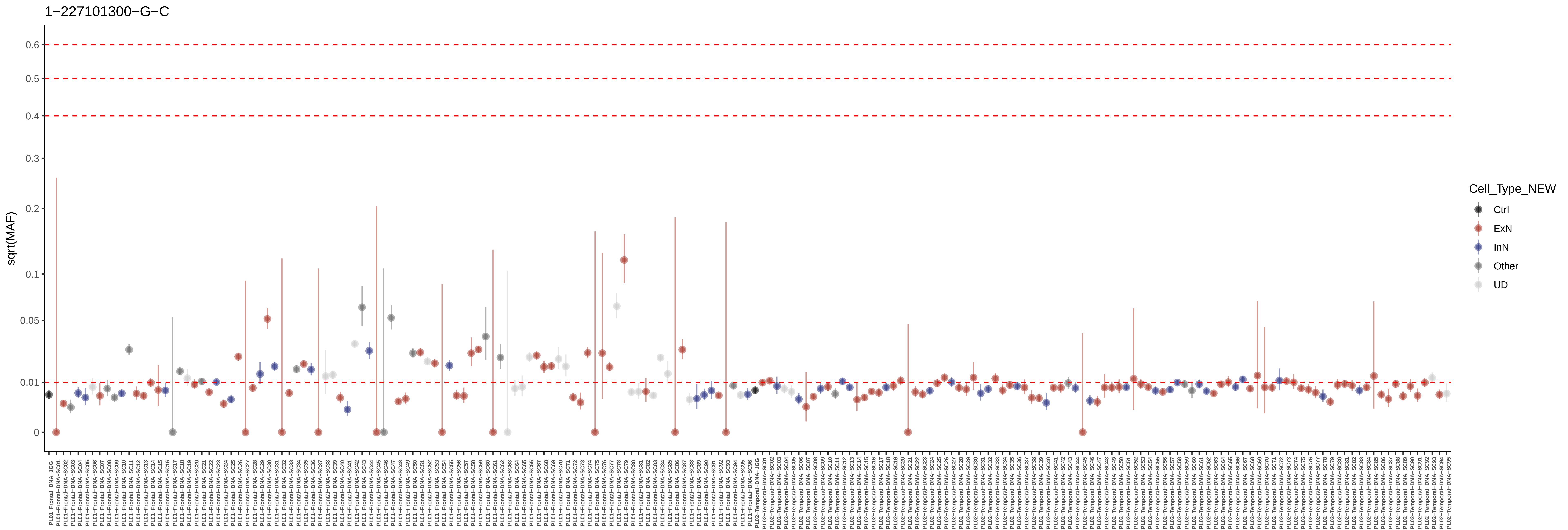
1−227101300−G−C
sqrt(MAF)
0.6
0.5
0.4
0.3
0.2
0.1
0.05
0.01
0
Cell_Type_NEW
Ctrl
ExN
InN
Other
UD

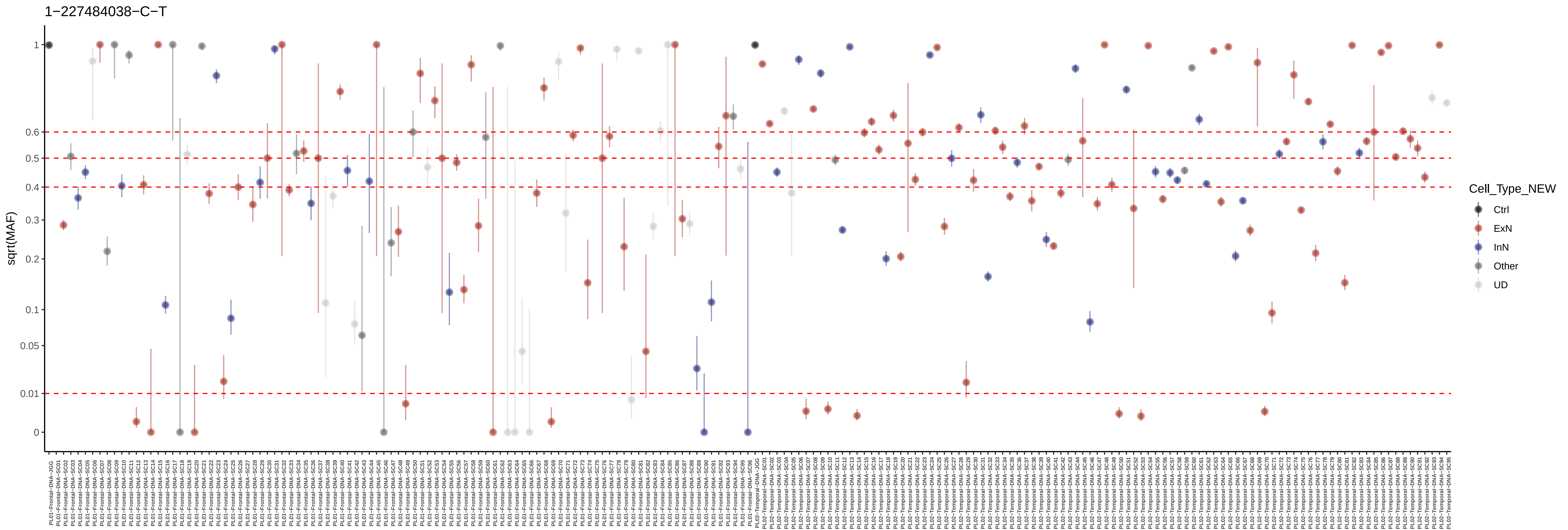
1−227484038−C−T
sqrt(MAF)
1
0.6
0.5
0.4
0.3
0.2
0.1
0.05
0.01
0
Cell_Type_NEW
Ctrl
ExN
InN
Other
UD

1−227887596−T−G

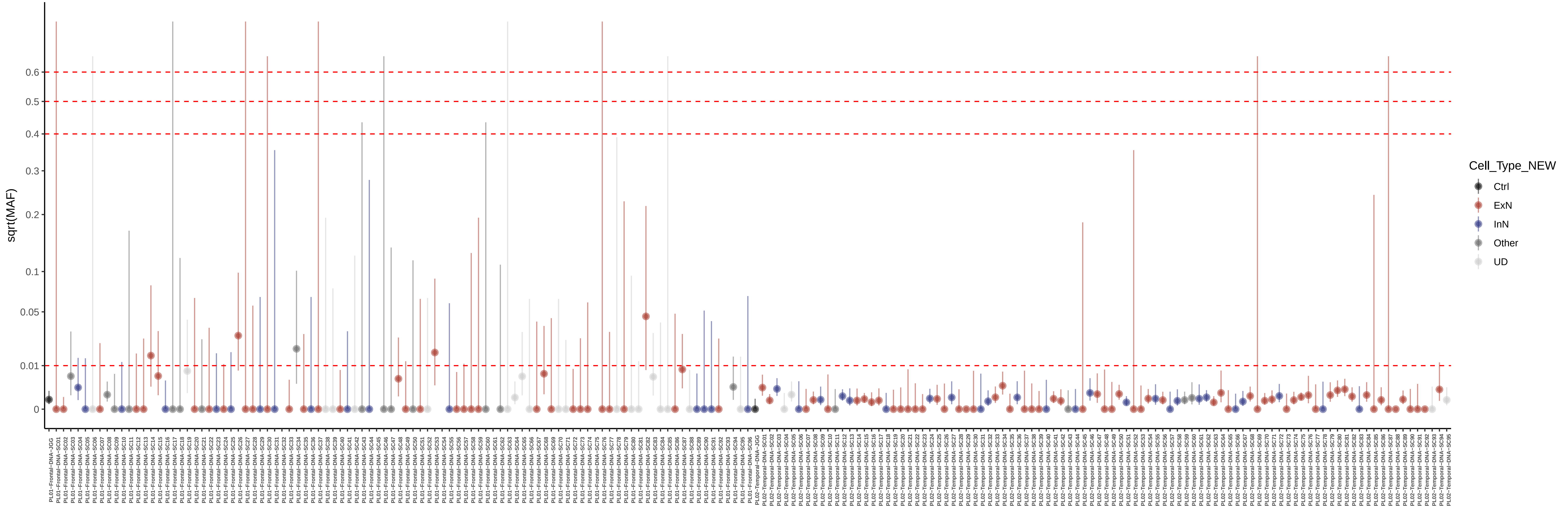

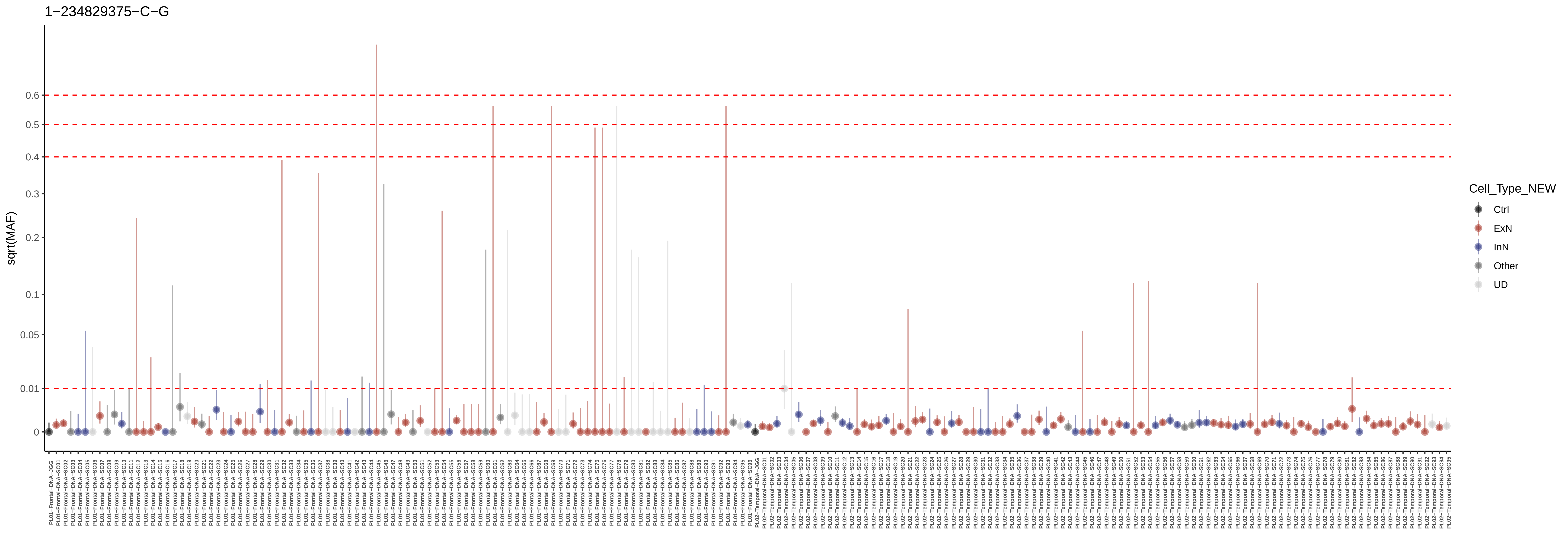

1−234829375−C−G
sqrt(MAF)
0.6
0.5
0.4
0.3
0.2
0.1
0.05
0.01
0
Cell_Type_NEW
Ctrl
ExN
InN
Other
UD

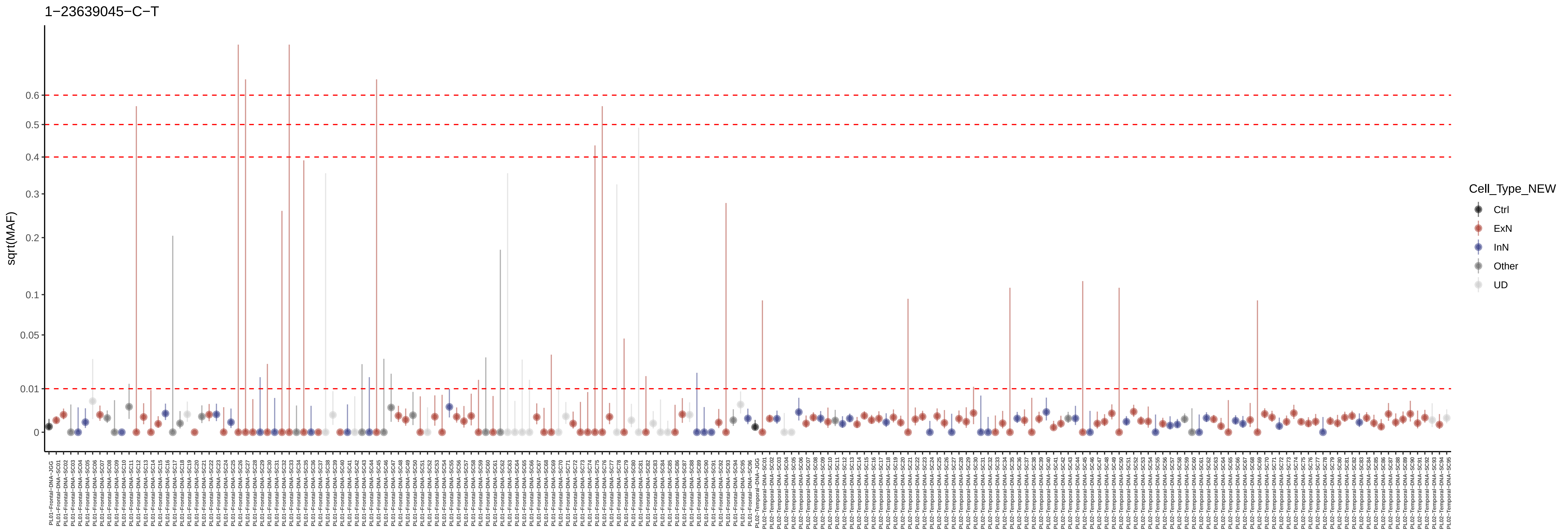
1−23639045−C−T
sqrt(MAF)
0.6
0.5
0.4
0.3
0.2
0.1
0.05
0.01
0
Cell_Type_NEW
Ctrl
ExN
InN
Other
UD

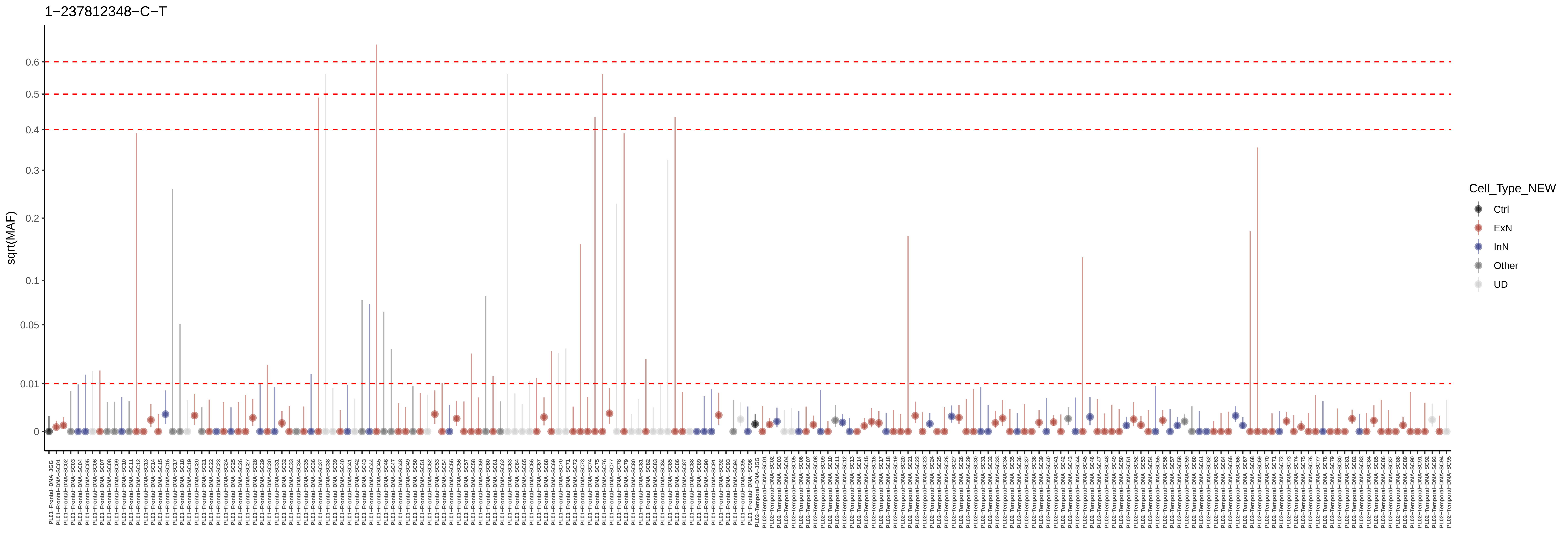
1−237812348−C−T
sqrt(MAF)
0.6
0.5
0.4
0.3
0.2
0.1
0.05
0.01
0
Cell_Type_NEW
Ctrl
ExN
InN
Other
UD

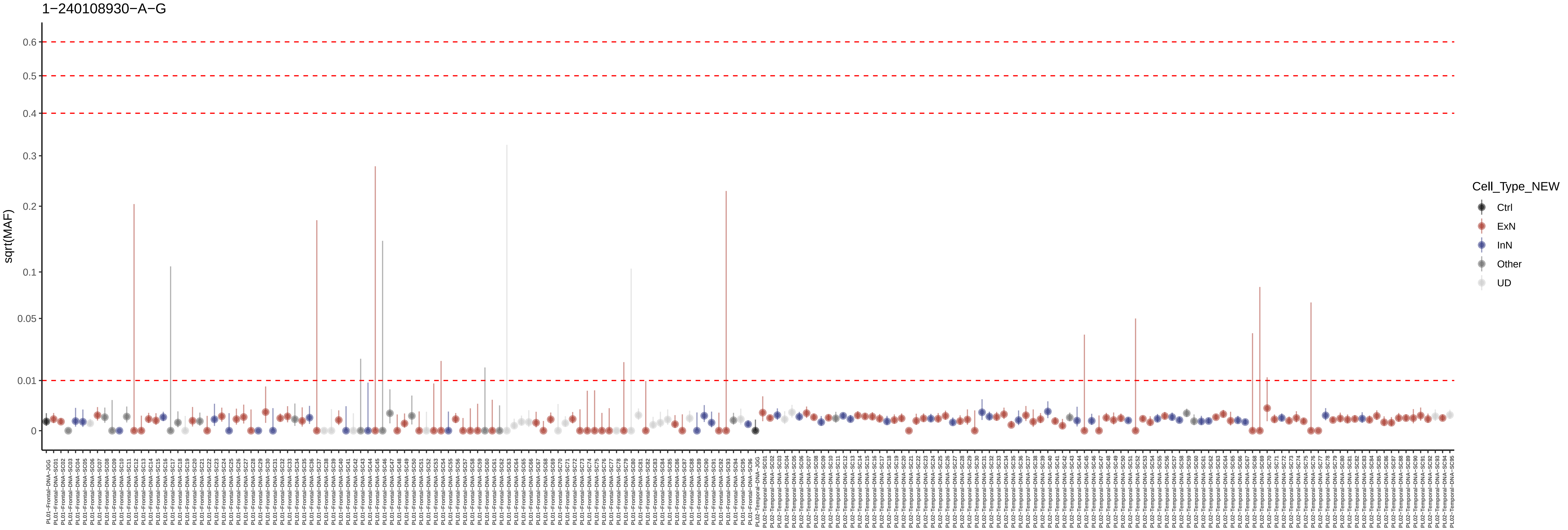
1−240108930−A−G
sqrt(MAF)
0.6
0.5
0.4
0.3
0.2
0.1
0.05
0.01
0
Cell_Type_NEW
Ctrl
ExN
InN
Other
UD

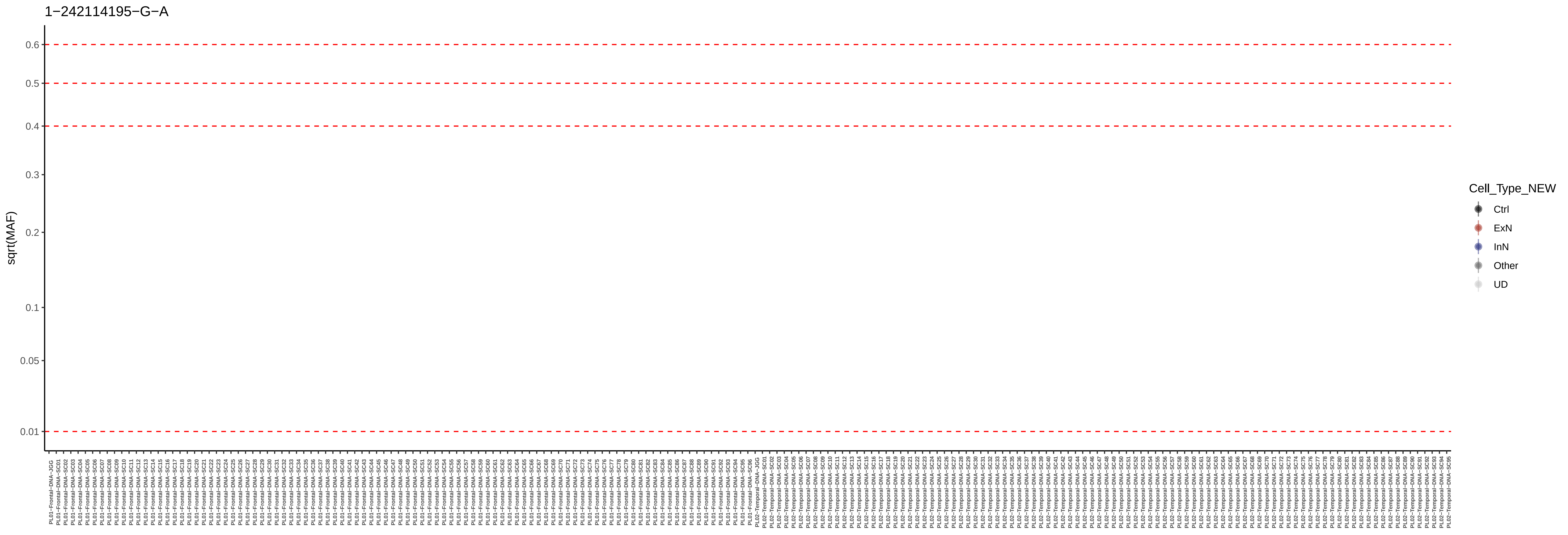
1−242114195−G−A
sqrt(MAF)
0.6
0.5
0.4
0.3
0.2
0.1
0.05
0.01
Cell_Type_NEW
Ctrl
ExN
InN
Other
UD

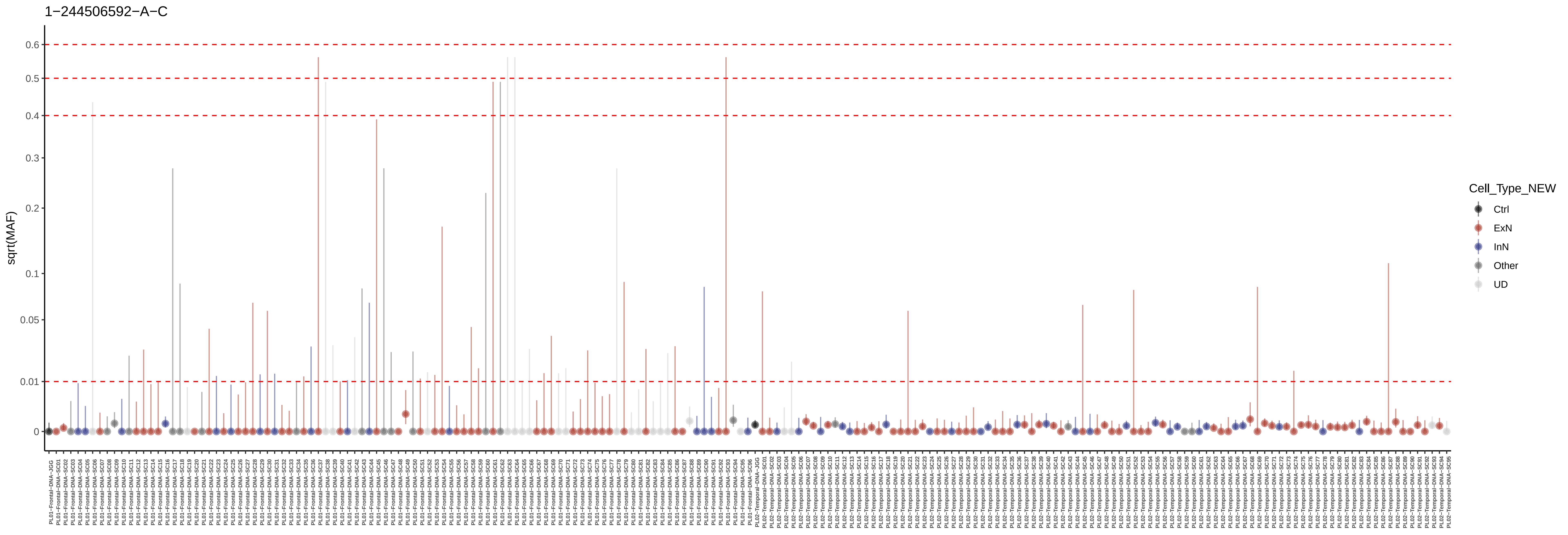
1−244506592−A−C
sqrt(MAF)
0.6
0.5
0.4
0.3
0.2
0.1
0.05
0.01
0
Cell_Type_NEW
Ctrl
ExN
InN
Other
UD

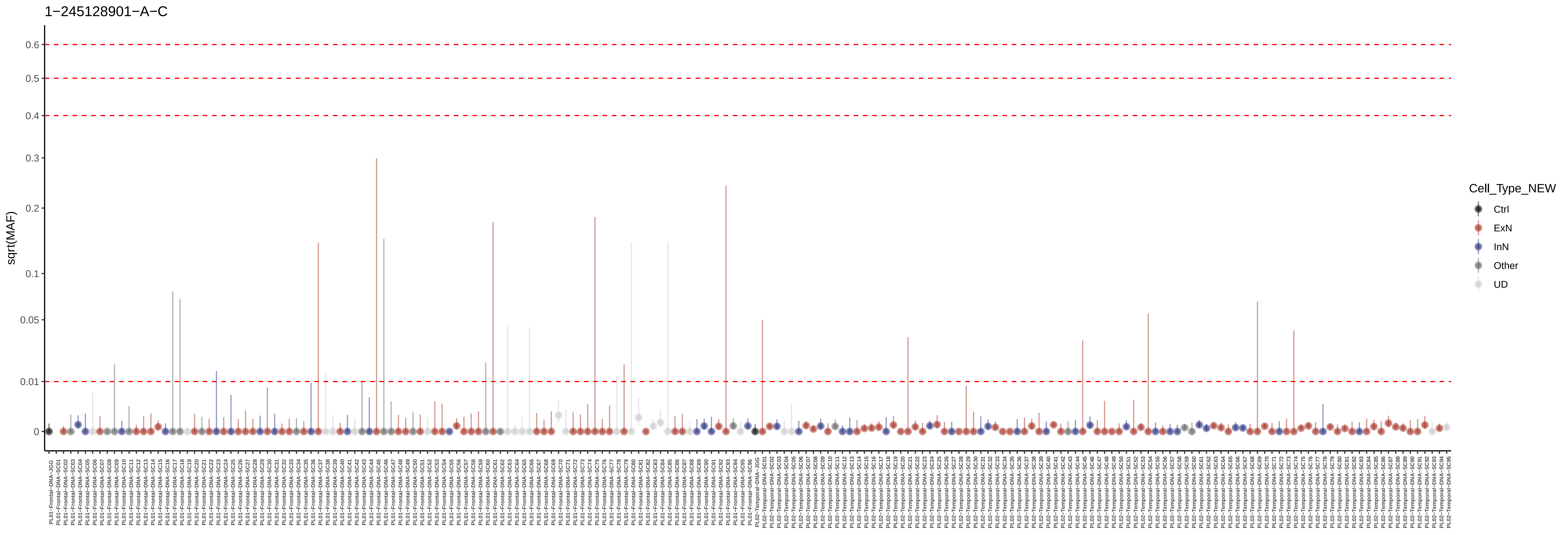
1−245128901−A−C
sqrt(MAF)
0.6
0.5
0.4
0.3
0.2
0.1
0.05
0.01
0
Cell_Type_NEW
Ctrl
ExN
InN
Other
UD

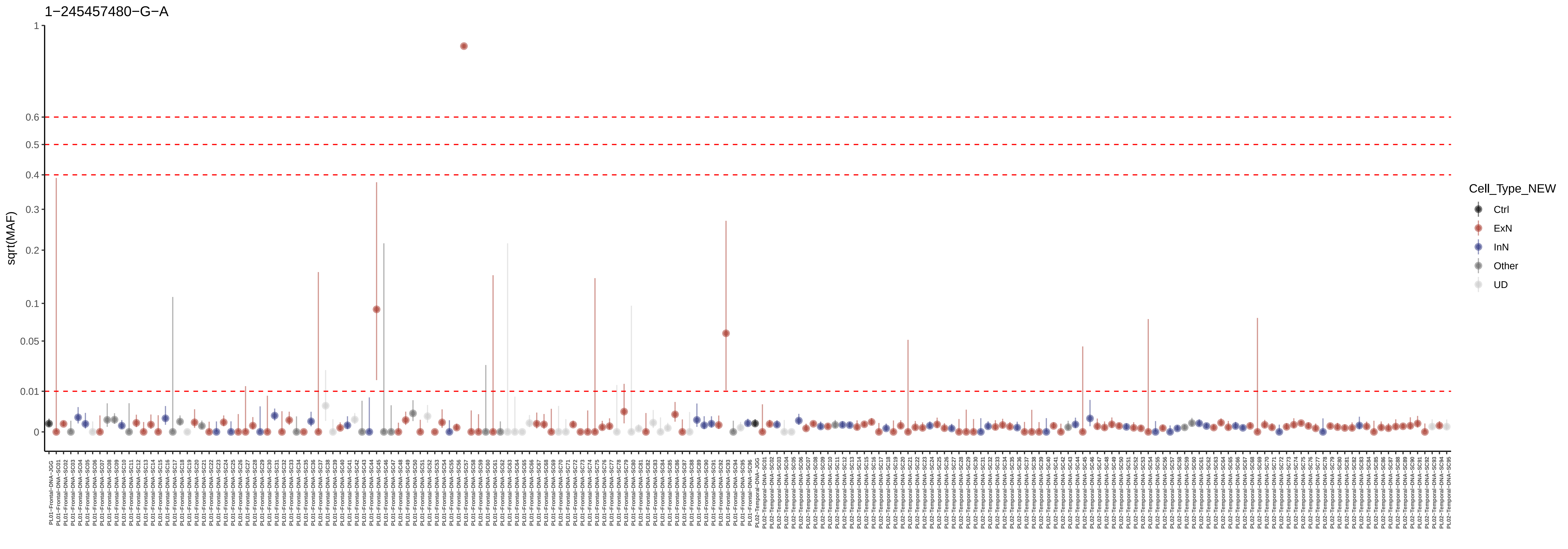
1−245457480−G−A
sqrt(MAF)
1
0.6
0.5
0.4
0.3
0.2
0.1
0.05
0.01
0
Cell_Type_NEW
Ctrl
ExN
InN
Other
UD

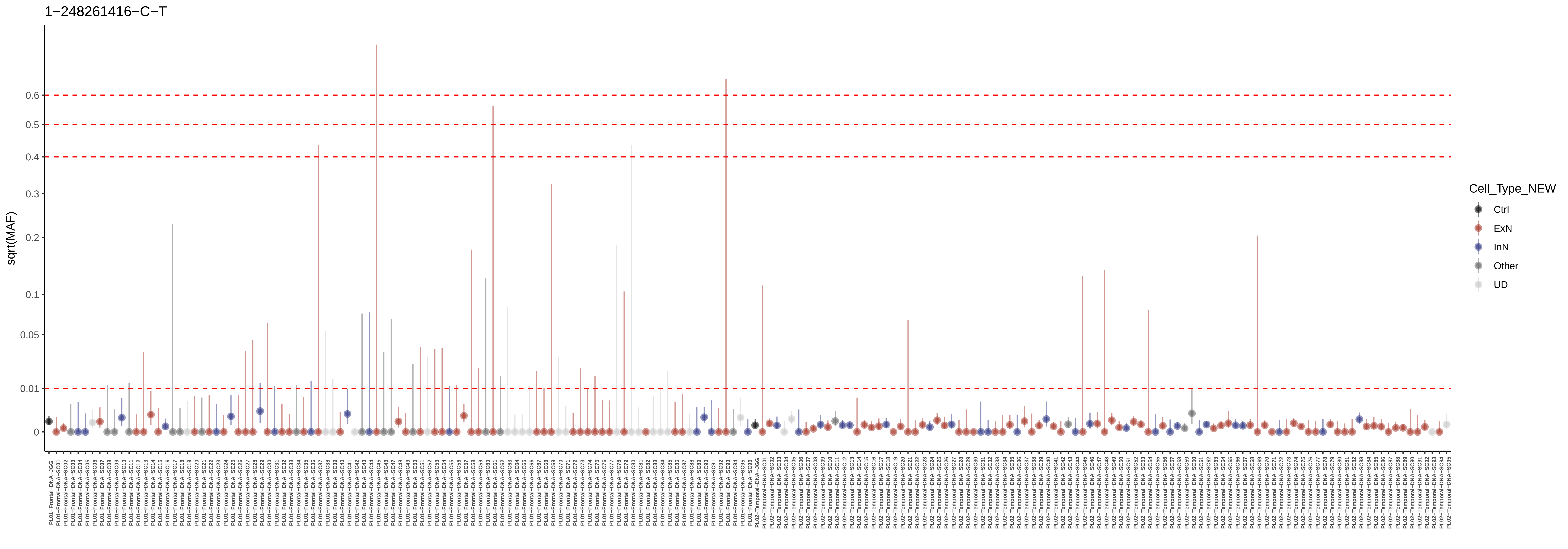
1−248261416−C−T
sqrt(MAF)
0.6
0.5
0.4
0.3
0.2
0.1
0.05
0.01
0
Cell_Type_NEW
Ctrl
ExN
InN
Other
UD

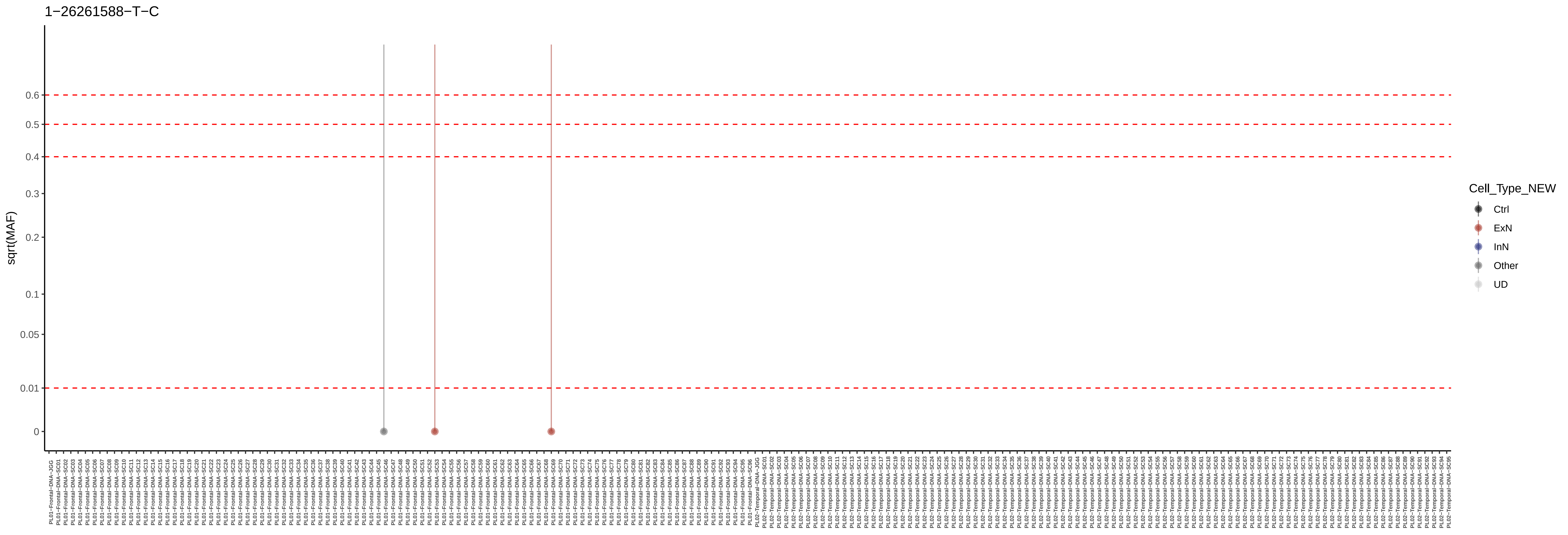
1−26261588−T−C
sqrt(MAF)
0.6
0.5
0.4
0.3
0.2
0.1
0.05
0.01
0
Cell_Type_NEW
Ctrl
ExN
InN
Other
UD

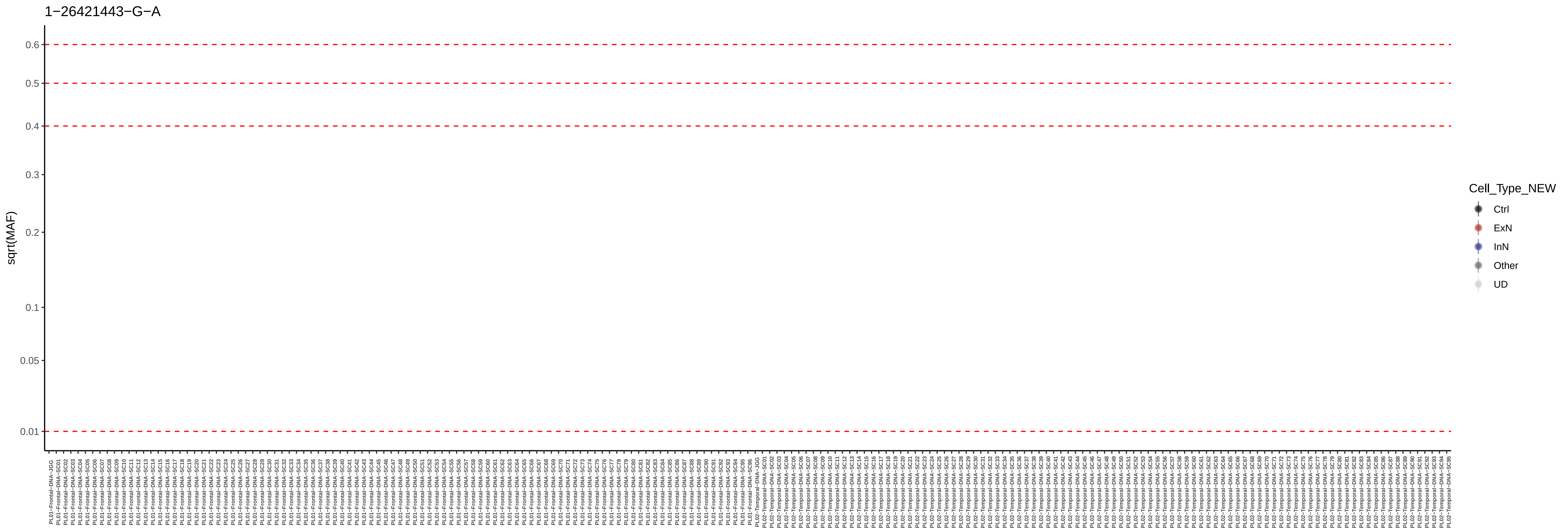

1−26421443−G−A
sqrt(MAF)
0.6
0.5
0.4
0.3
0.2
0.1
0.05
0.01
Cell_Type_NEW
Ctrl
ExN
InN
Other
UD

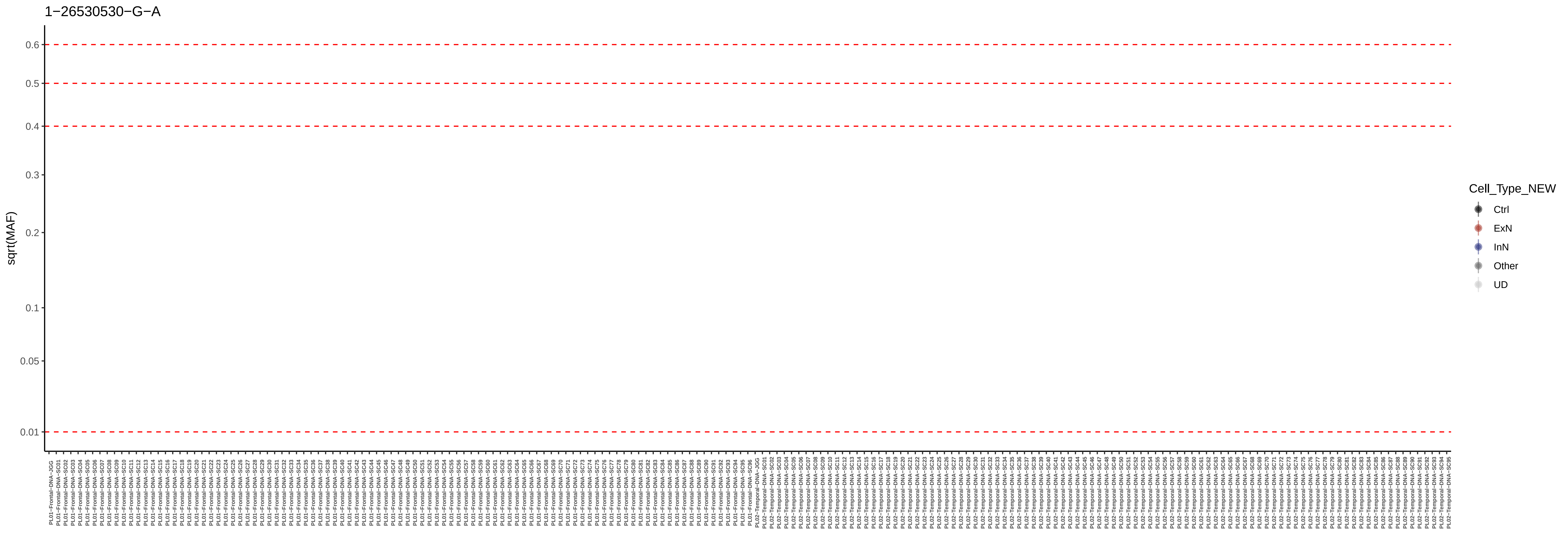
1−26530530−G−A
sqrt(MAF)
0.6
0.5
0.4
0.3
0.2
0.1
0.05
0.01
Cell_Type_NEW
Ctrl
ExN
InN
Other
UD

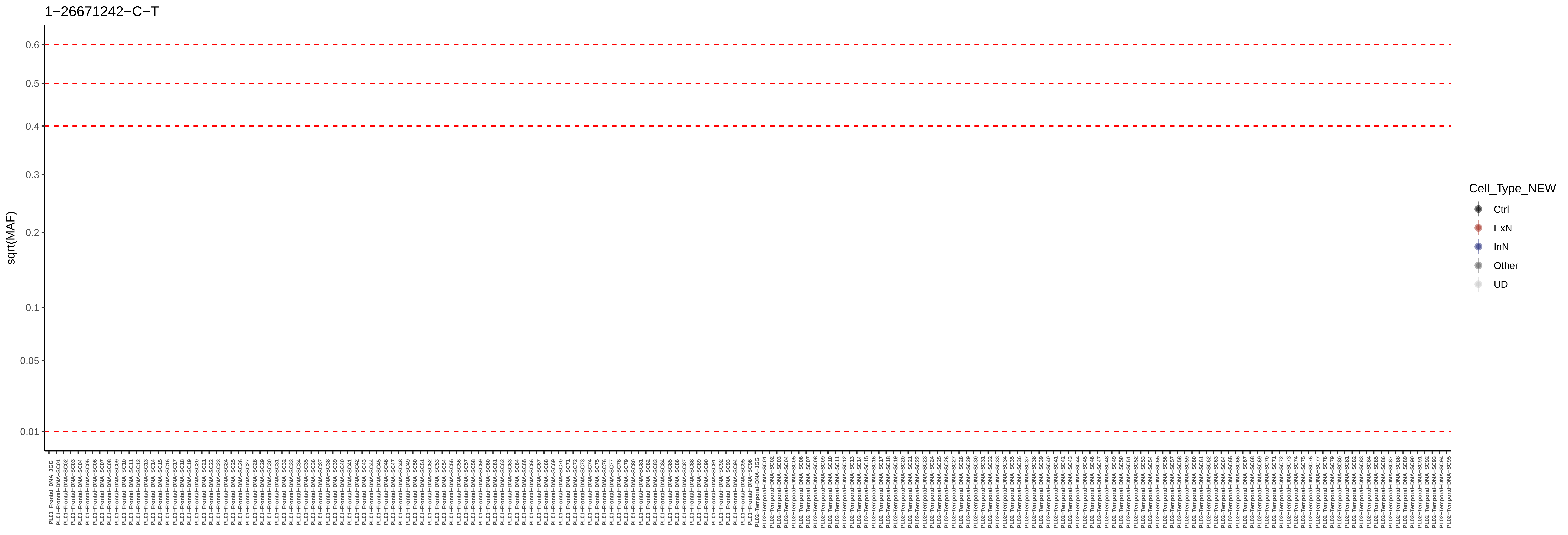
1−26671242−C−T
sqrt(MAF)
0.6
0.5
0.4
0.3
0.2
0.1
0.05
0.01
Cell_Type_NEW
Ctrl
ExN
InN
Other
UD

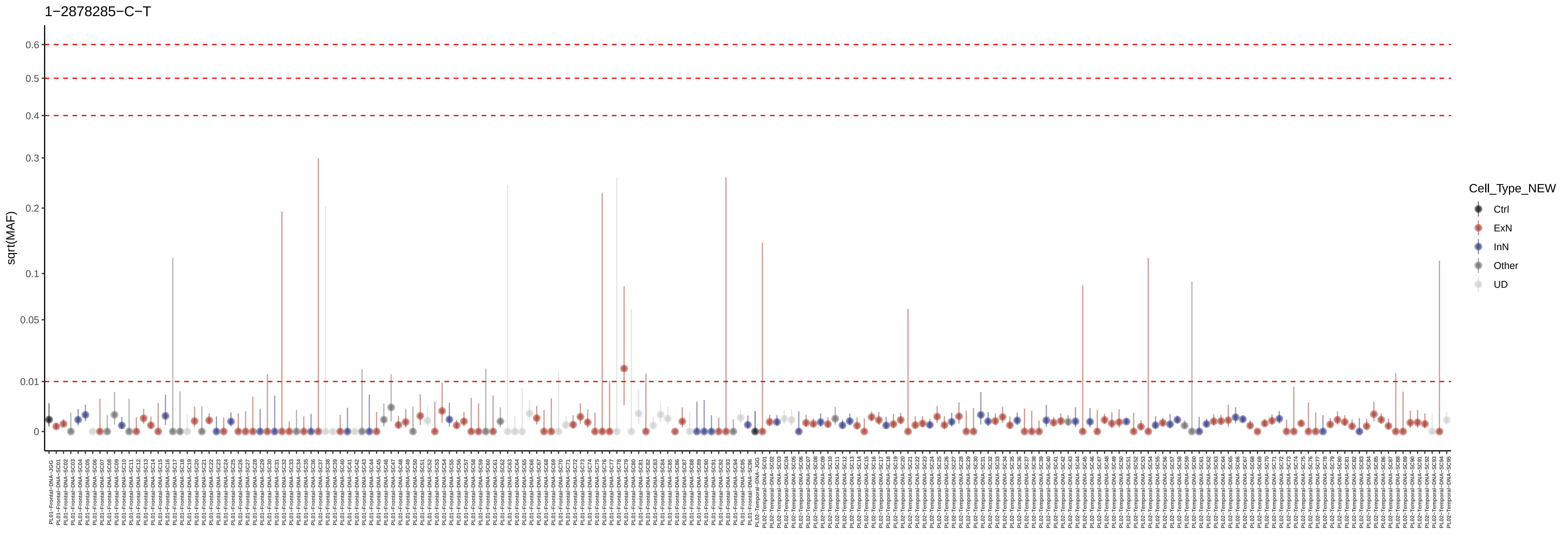
1−2878285−C−T
sqrt(MAF)
0.6
0.5
0.4
0.3
0.2
0.1
0.05
0.01
0
Cell_Type_NEW
Ctrl
ExN
InN
Other
UD

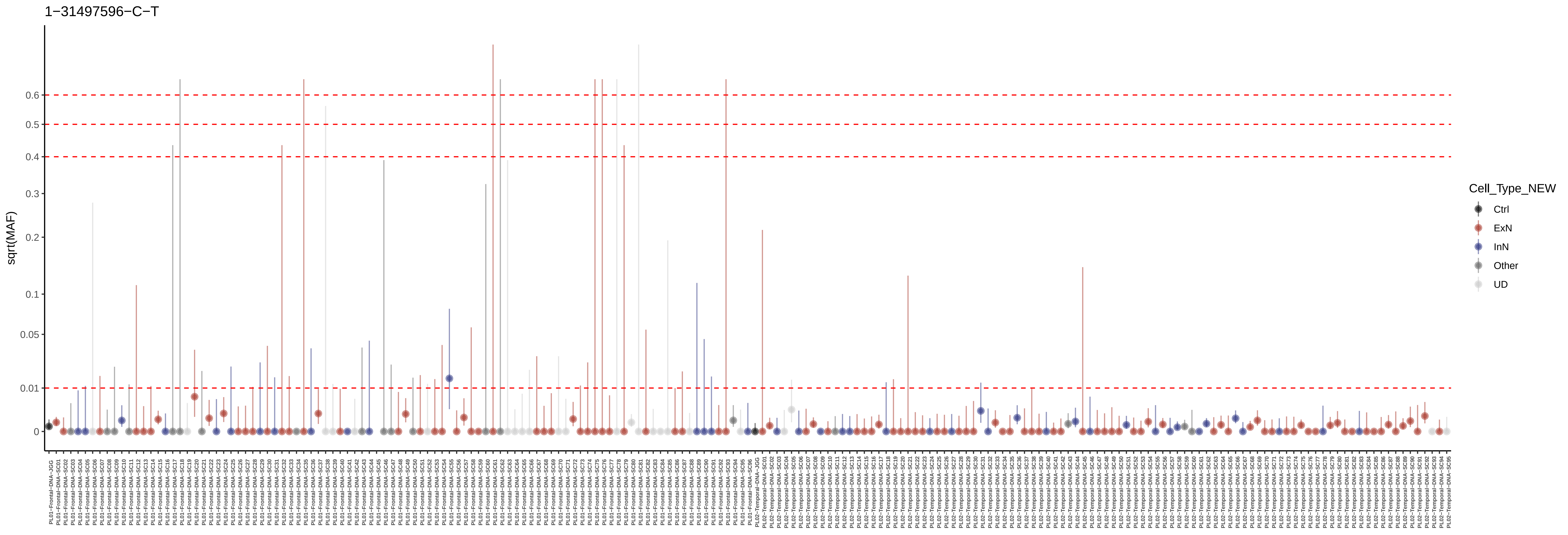
1−31497596−C−T
sqrt(MAF)
0.6
0.5
0.4
0.3
0.2
0.1
0.05
0.01
0
Cell_Type_NEW
Ctrl
ExN
InN
Other
UD

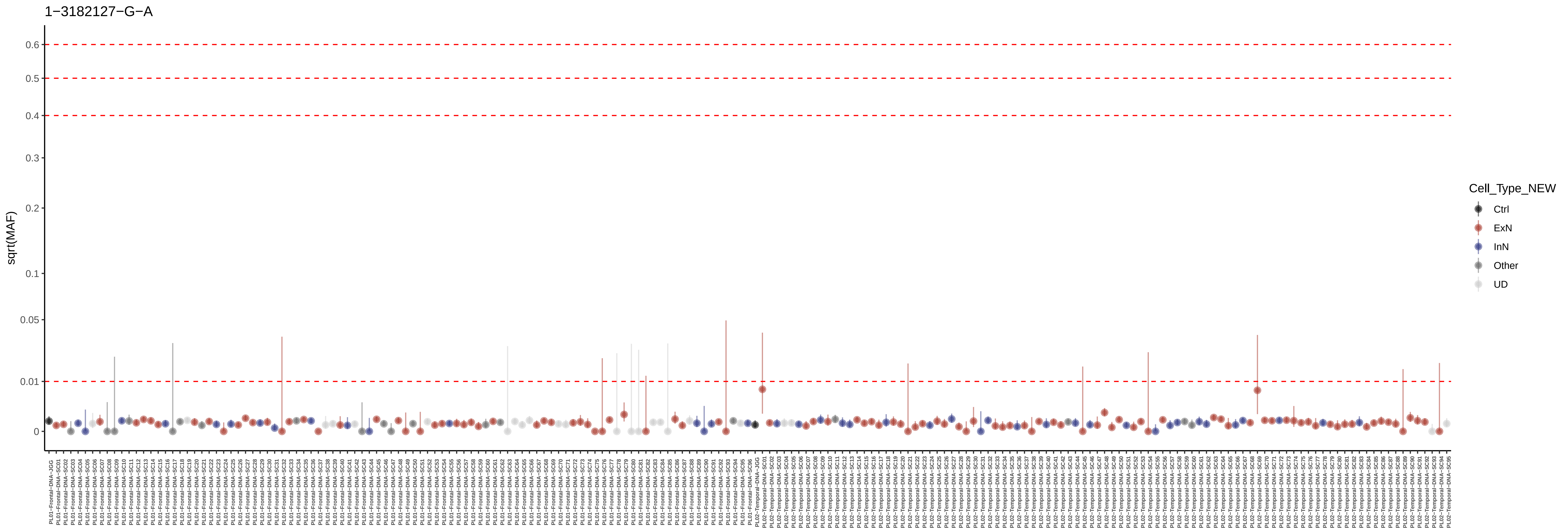
1−3182127−G−A
sqrt(MAF)
0.6
0.5
0.4
0.3
0.2
0.1
0.05
0.01
0
Cell_Type_NEW
Ctrl
ExN
InN
Other
UD

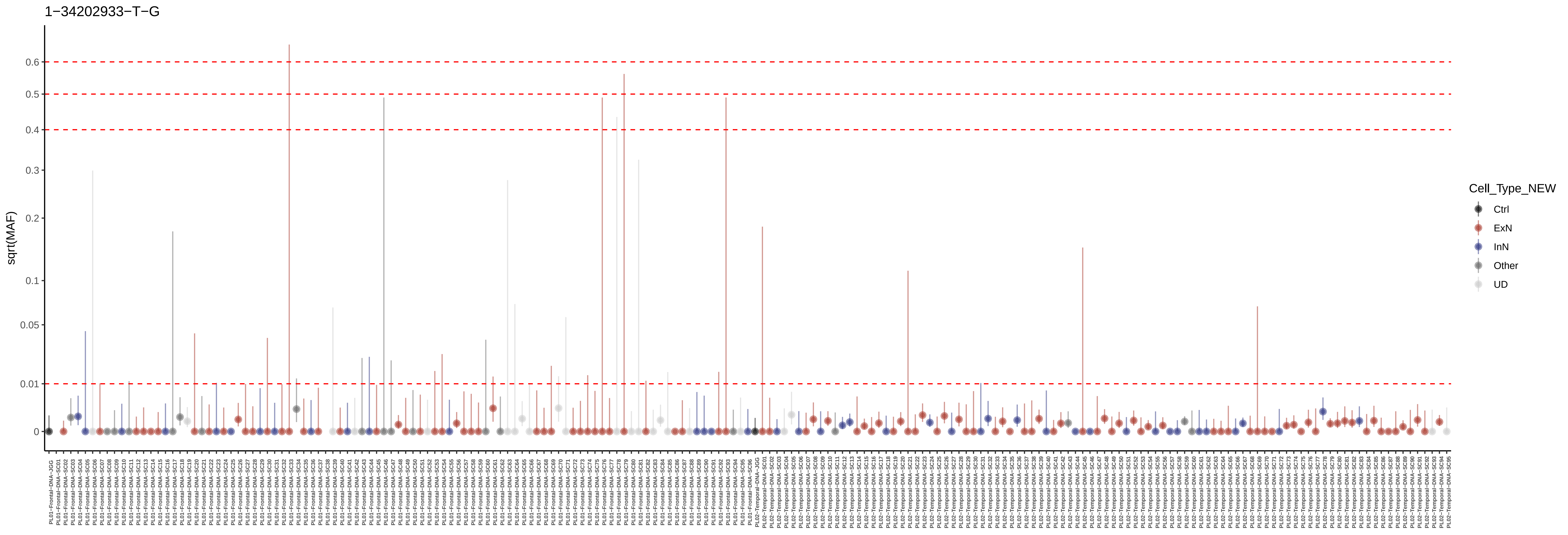

1−34202933−T−G
sqrt(MAF)
0.6
0.5
0.4
0.3
0.2
0.1
0.05
0.01
0
Cell_Type_NEW
Ctrl
ExN
InN
Other
UD

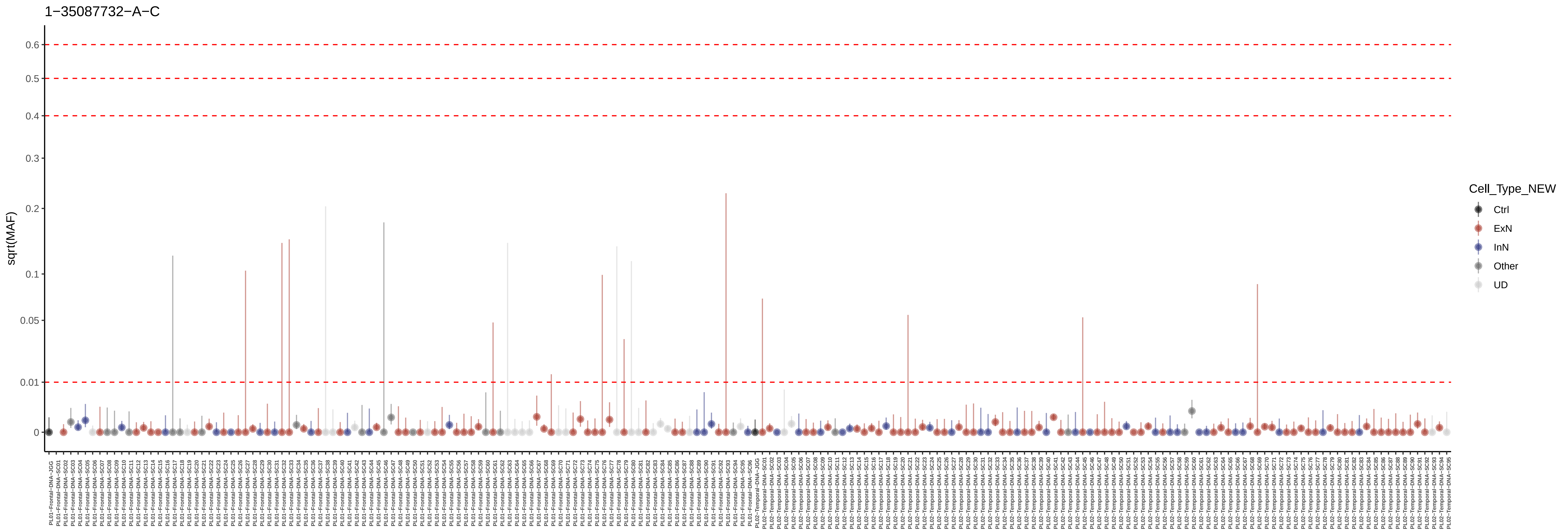
1−35087732−A−C
sqrt(MAF)
0.6
0.5
0.4
0.3
0.2
0.1
0.05
0.01
0
Cell_Type_NEW
Ctrl
ExN
InN
Other
UD

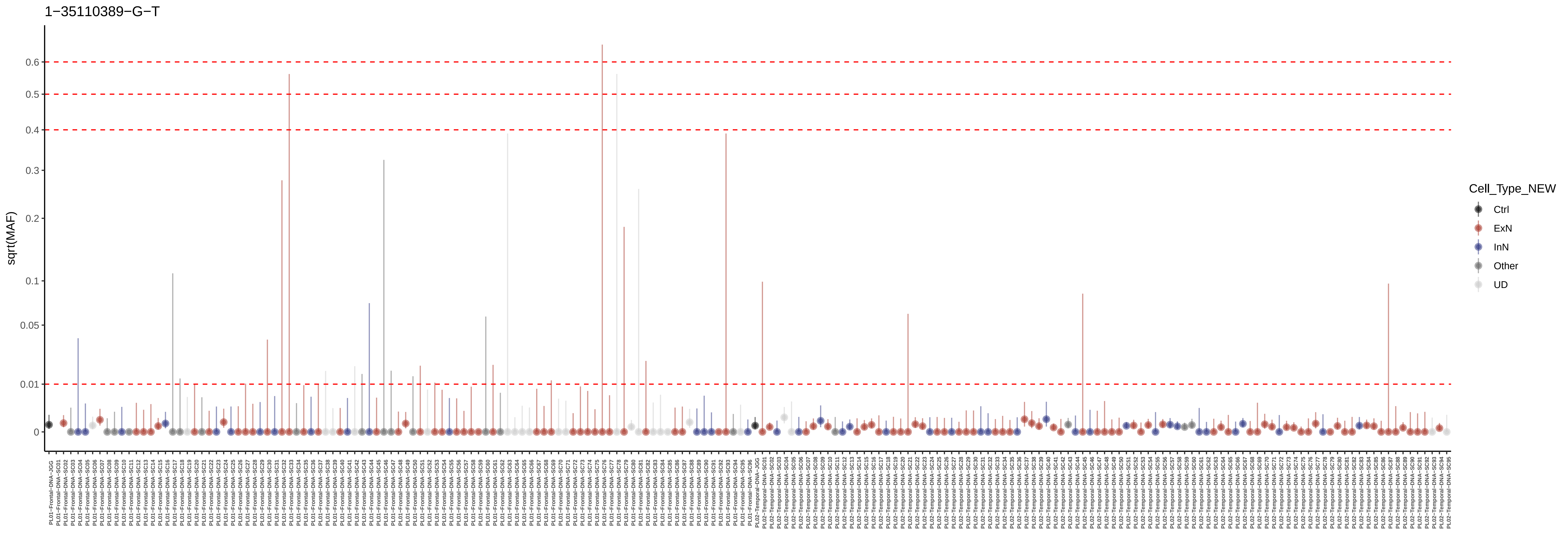
1−35110389−G−T
sqrt(MAF)
0.6
0.5
0.4
0.3
0.2
0.1
0.05
0.01
0
Cell_Type_NEW
Ctrl
ExN
InN
Other
UD

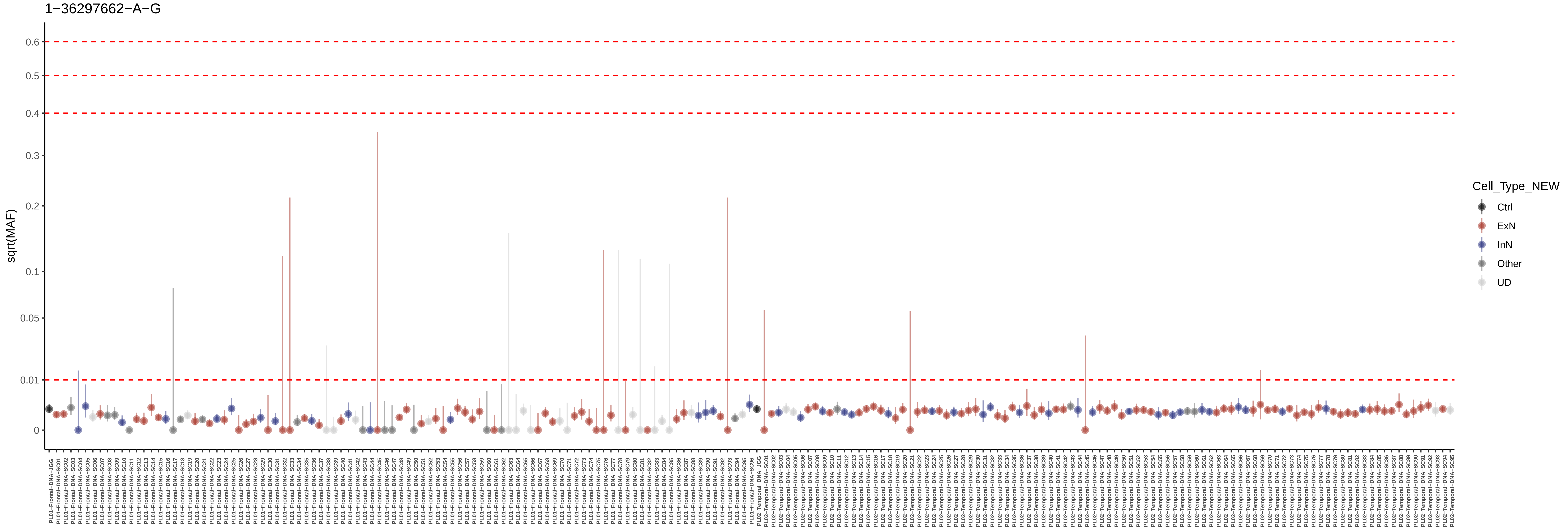
1−36297662−A−G
sqrt(MAF)
0.6
0.5
0.4
0.3
0.2
0.1
0.05
0.01
0
Cell_Type_NEW
Ctrl
ExN
InN
Other
UD

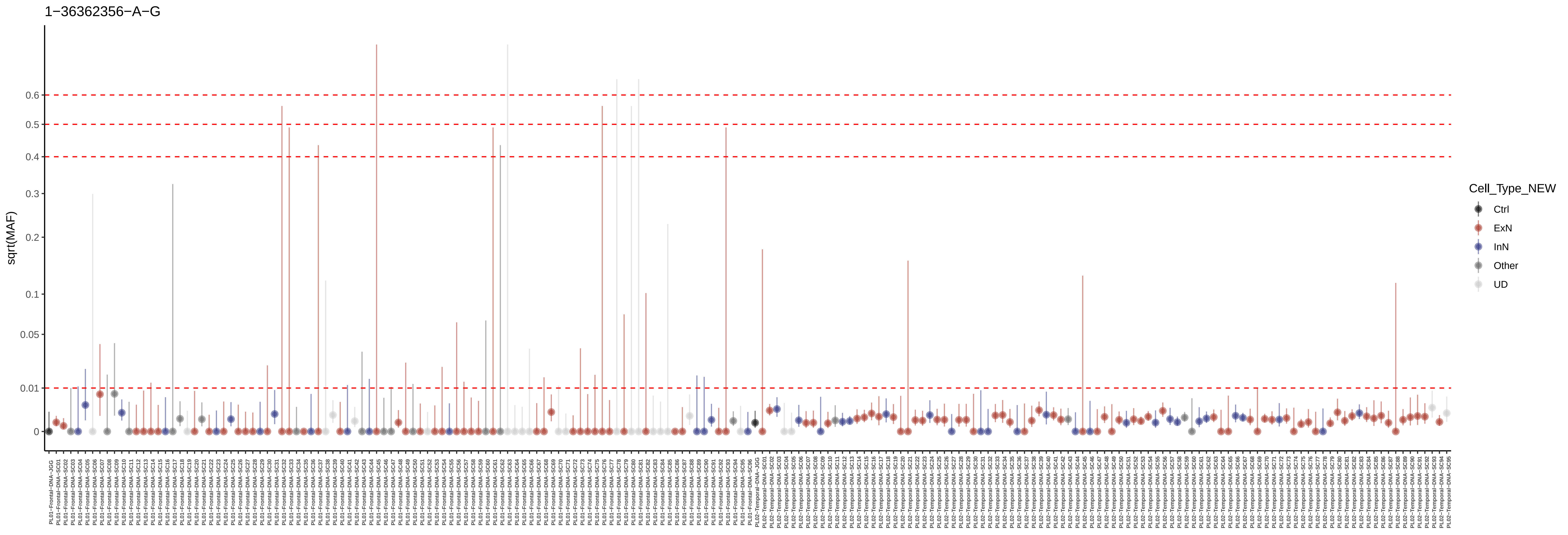
1−36362356−A−G
sqrt(MAF)
0.6
0.5
0.4
0.3
0.2
0.1
0.05
0.01
0
Cell_Type_NEW
Ctrl
ExN
InN
Other
UD

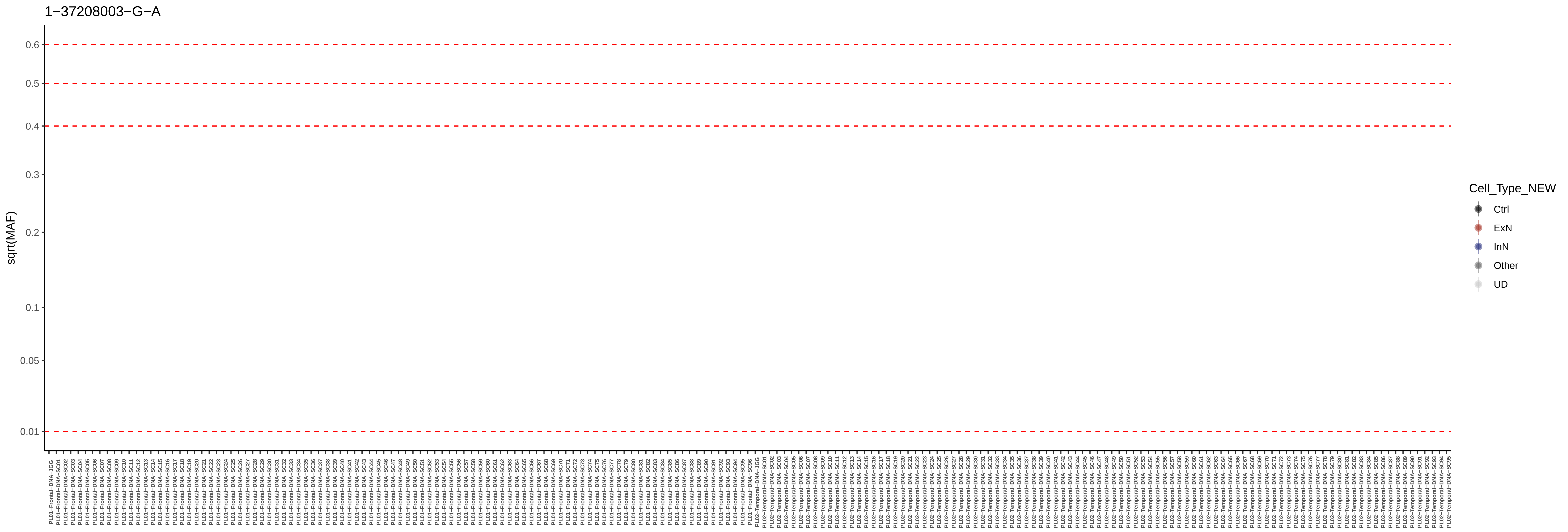
1−37208003−G−A
sqrt(MAF)
0.6
0.5
0.4
0.3
0.2
0.1
0.05
0.01
Cell_Type_NEW
Ctrl
ExN
InN
Other
UD

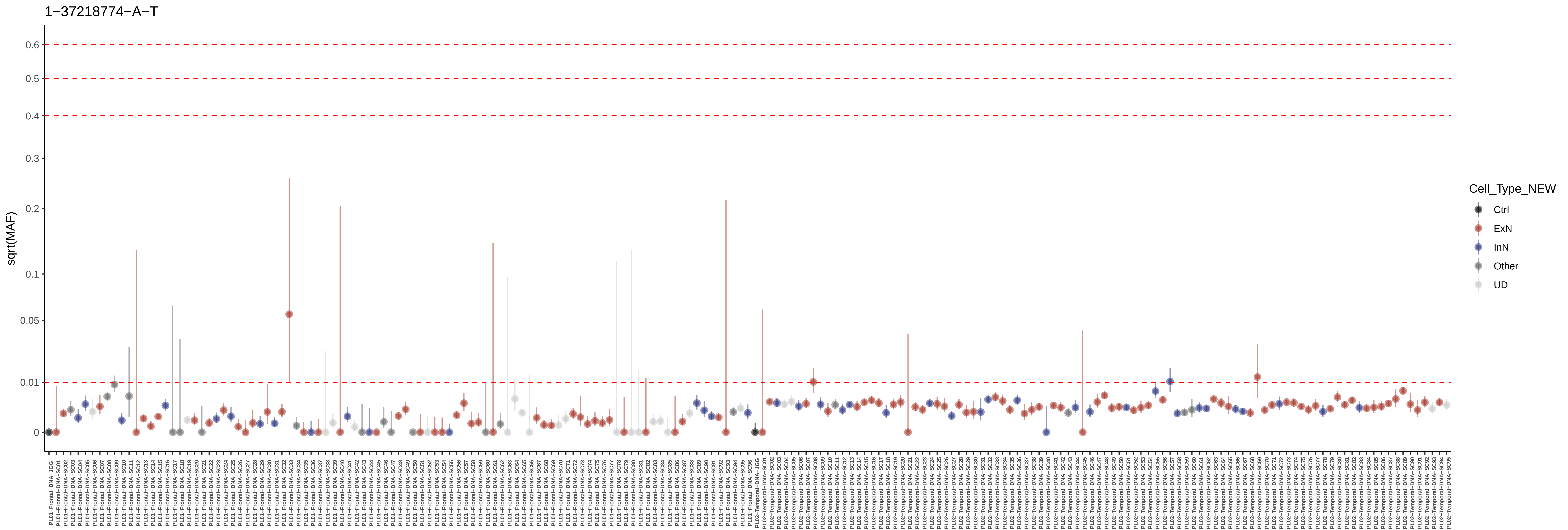
1−37218774−A−T
sqrt(MAF)
0.6
0.5
0.4
0.3
0.2
0.1
0.05
0.01
0
Cell_Type_NEW
Ctrl
ExN
InN
Other
UD

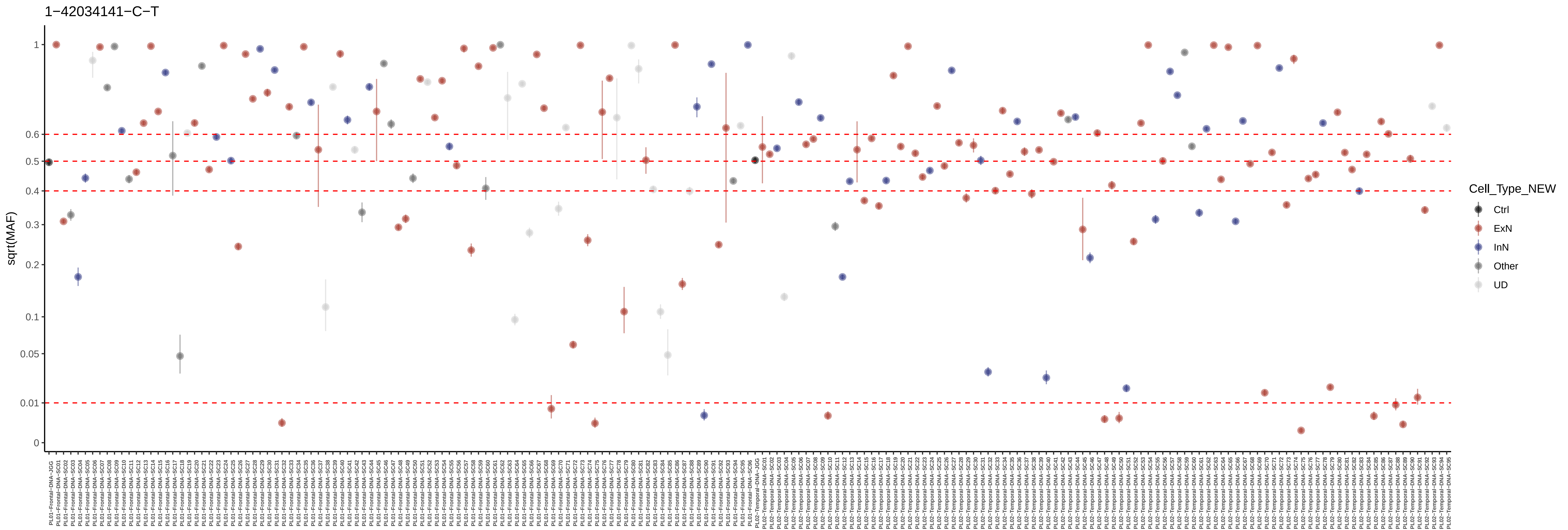
1−42034141−C−T
sqrt(MAF)
1
0.6
0.5
0.4
0.3
0.2
0.1
0.05
0.01
0
Cell_Type_NEW
Ctrl
ExN
InN
Other
UD

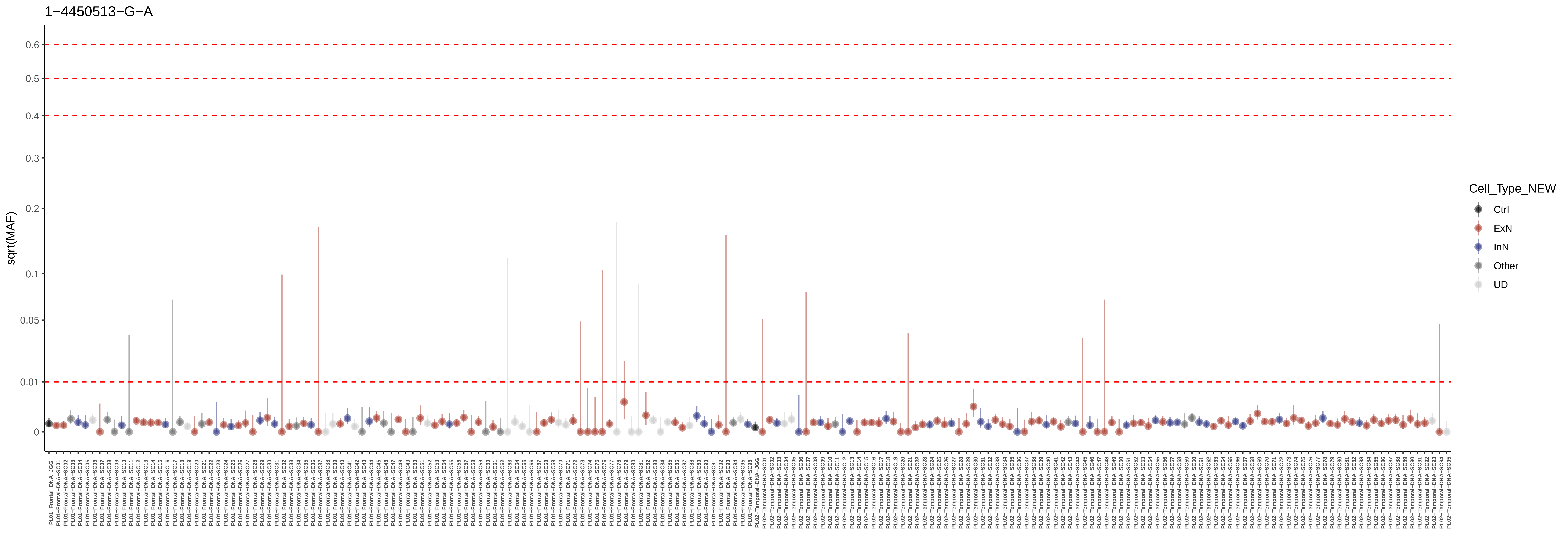
1−4450513−G−A
sqrt(MAF)
0.6
0.5
0.4
0.3
0.2
0.1
0.05
0.01
0
Cell_Type_NEW
Ctrl
ExN
InN
Other
UD

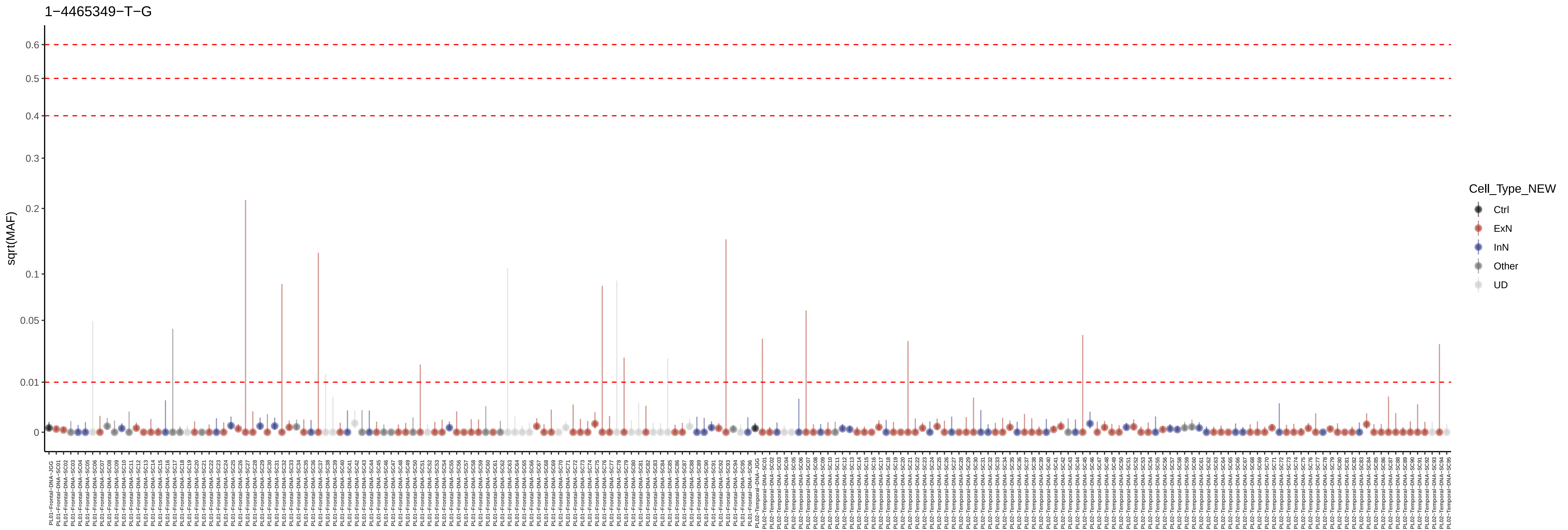

1−4465349−T−G
sqrt(MAF)
0.6
0.5
0.4
0.3
0.2
0.1
0.05
0.01
0
Cell_Type_NEW
Ctrl
ExN
InN
Other
UD

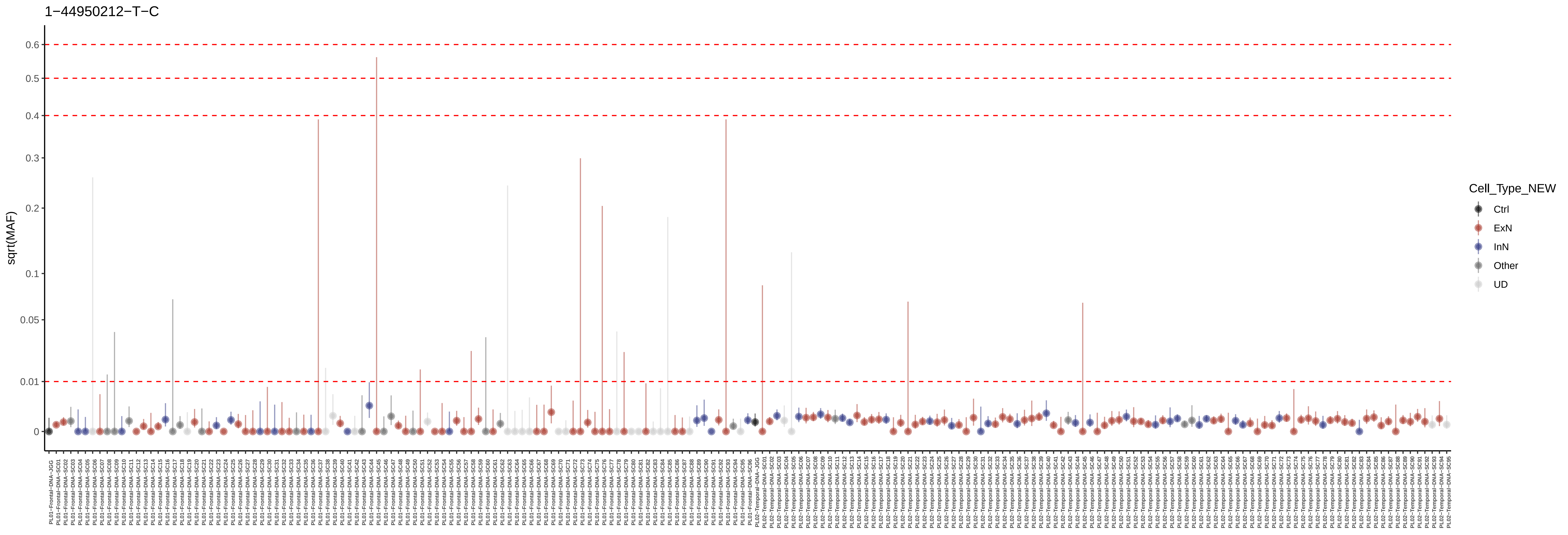

1−44950212−T−C
sqrt(MAF)
0.6
0.5
0.4
0.3
0.2
0.1
0.05
0.01
0
Cell_Type_NEW
Ctrl
ExN
InN
Other
UD

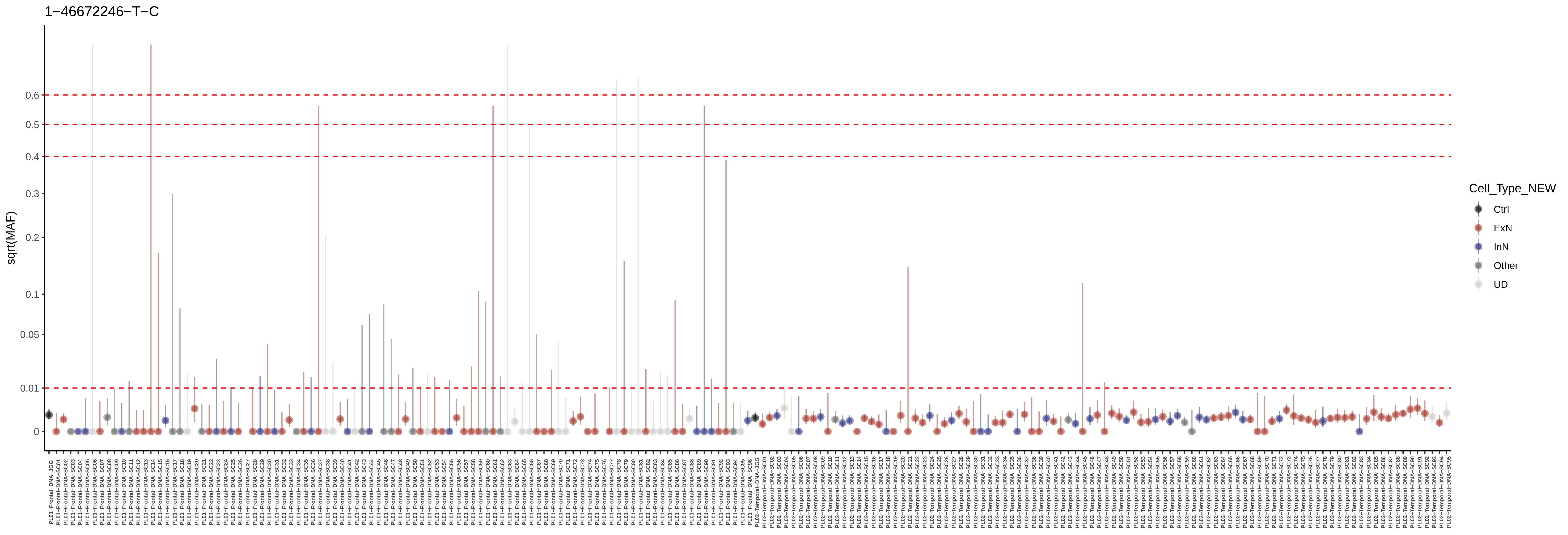

1−46672246−T−C
sqrt(MAF)
0.6
0.5
0.4
0.3
0.2
0.1
0.05
0.01
0
Cell_Type_NEW
Ctrl
ExN
InN
Other
UD

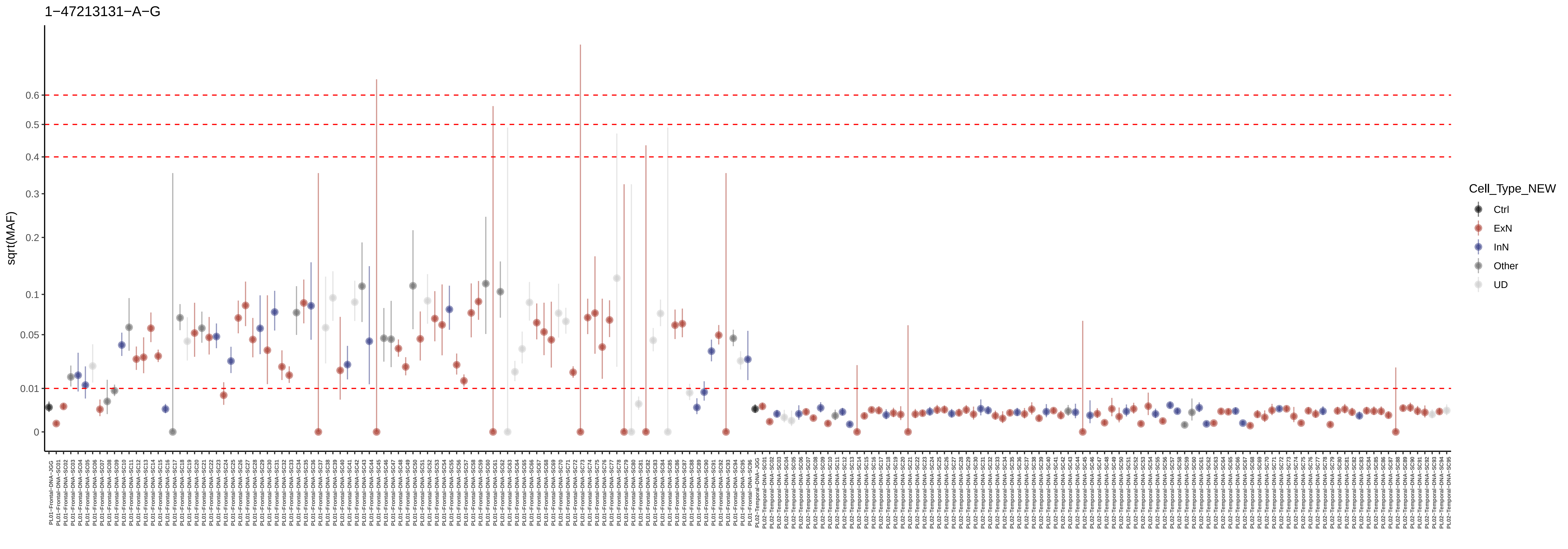
1−47213131−A−G
sqrt(MAF)
0.6
0.5
0.4
0.3
0.2
0.1
0.05
0.01
0
Cell_Type_NEW
Ctrl
ExN
InN
Other
UD

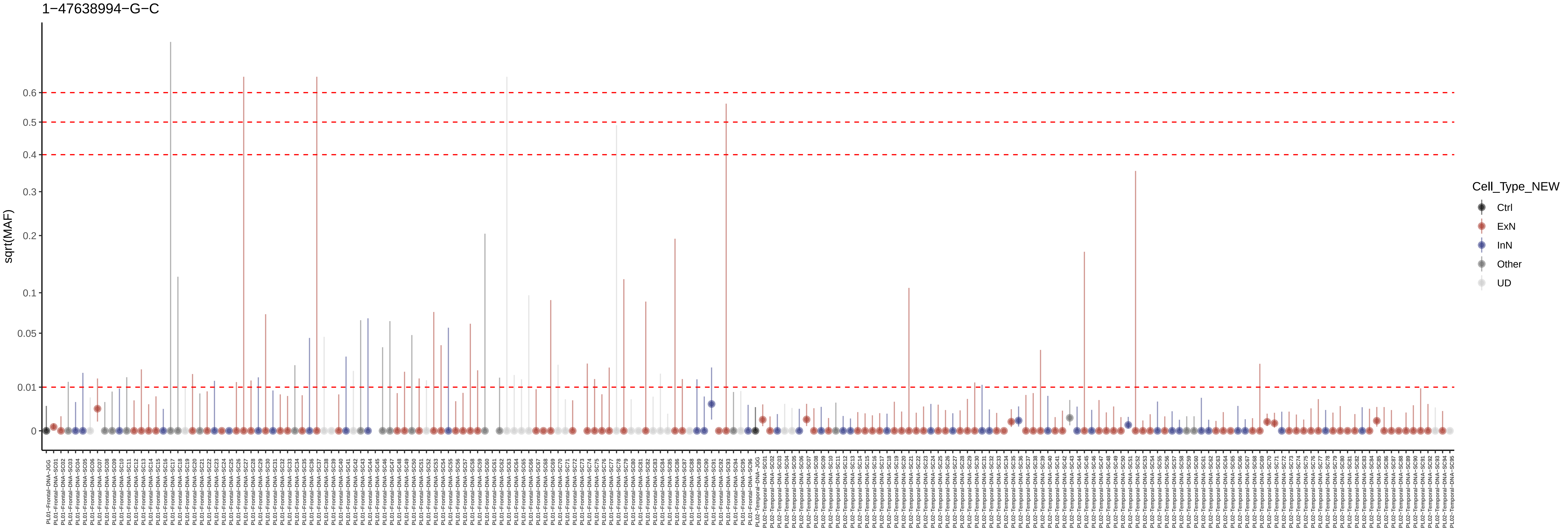
1−47638994−G−C
sqrt(MAF)
0.6
0.5
0.4
0.3
0.2
0.1
0.05
0.01
0
Cell_Type_NEW
Ctrl
ExN
InN
Other
UD

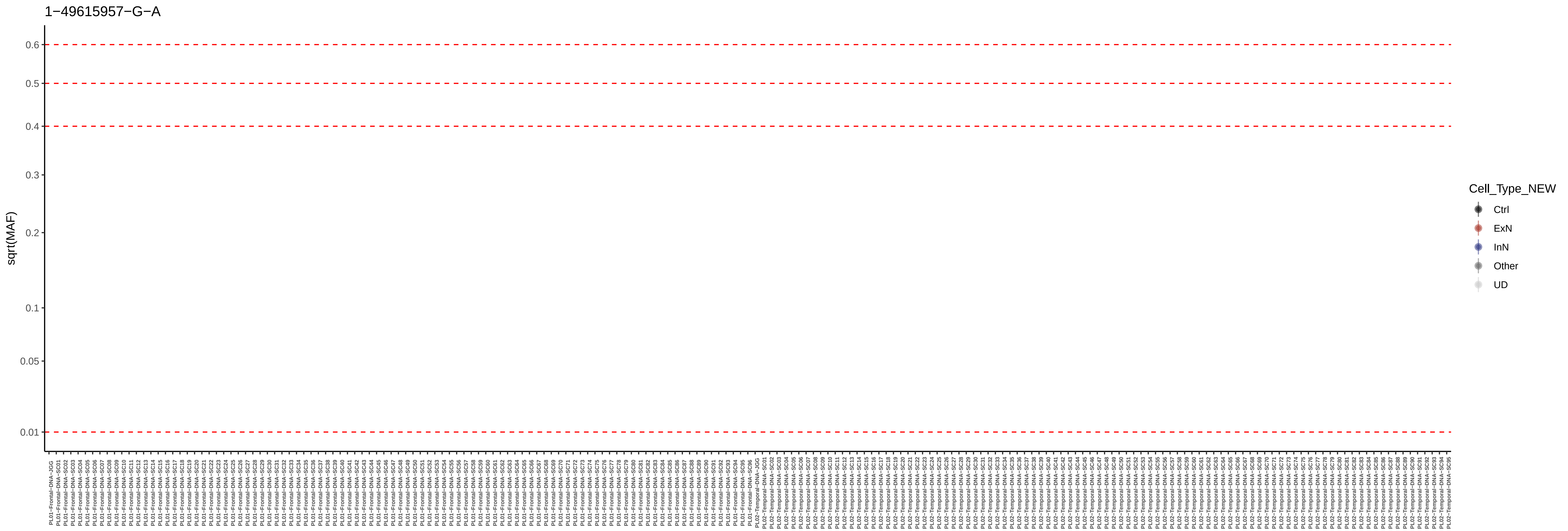
1−49615957−G−A
sqrt(MAF)
0.6
0.5
0.4
0.3
0.2
0.1
0.05
0.01
Cell_Type_NEW
Ctrl
ExN
InN
Other
UD

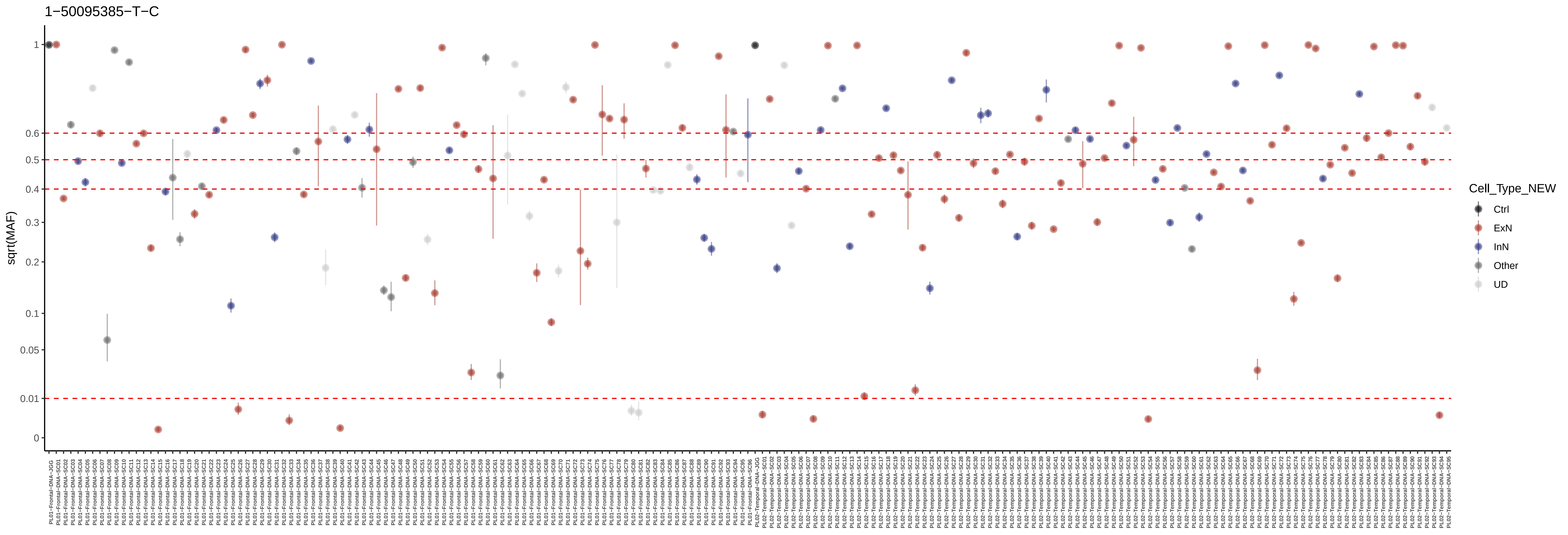

1−50095385−T−C
sqrt(MAF)
1
0.6
0.5
0.4
0.3
0.2
0.1
0.05
0.01
0
Cell_Type_NEW
Ctrl
ExN
InN
Other
UD

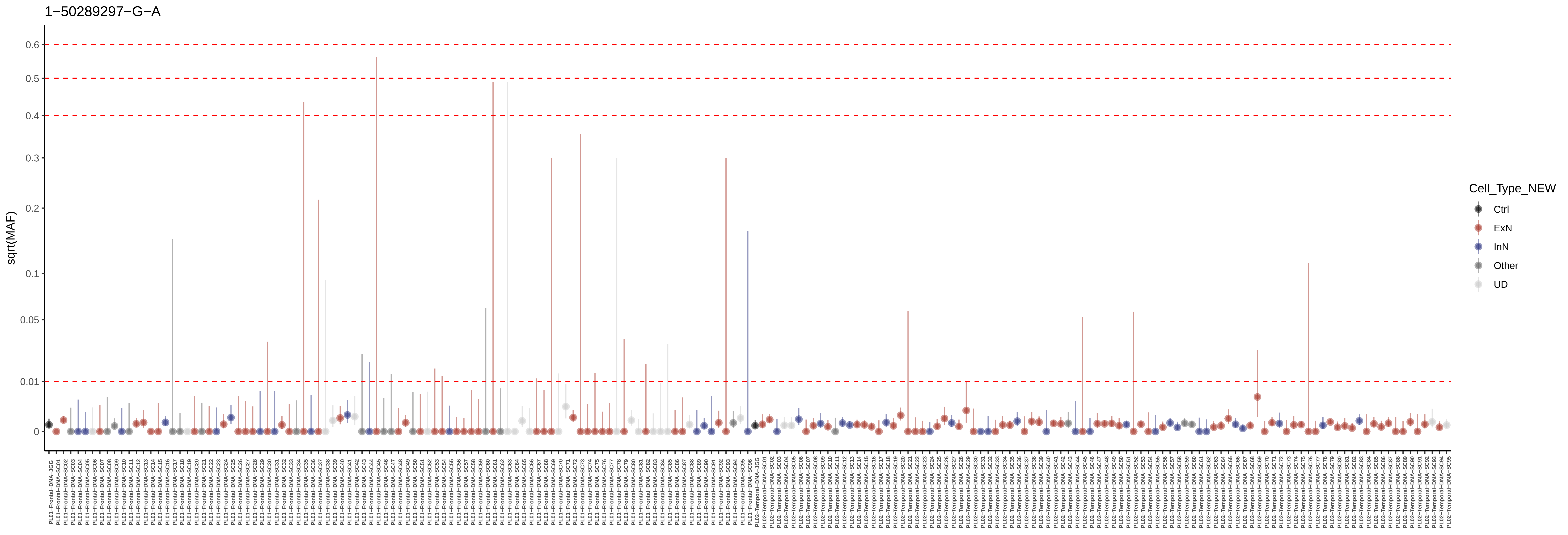
1−50289297−G−A
sqrt(MAF)
0.6
0.5
0.4
0.3
0.2
0.1
0.05
0.01
0
Cell_Type_NEW
Ctrl
ExN
InN
Other
UD

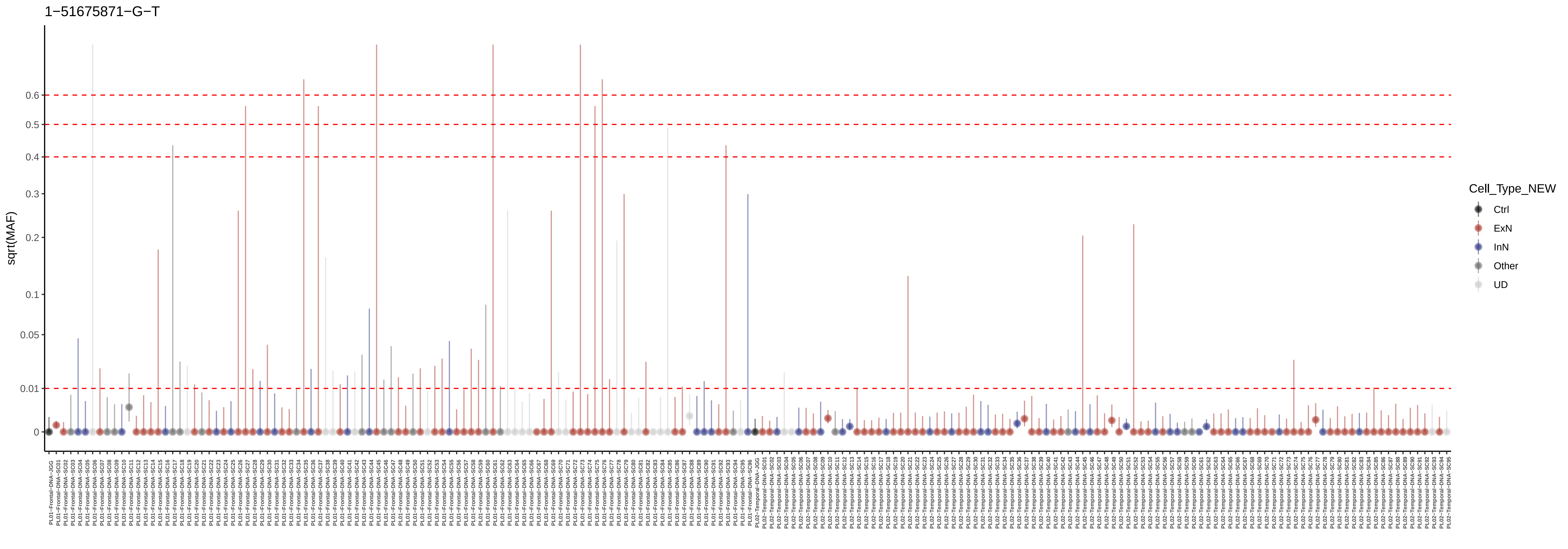
1−51675871−G−T
sqrt(MAF)
0.6
0.5
0.4
0.3
0.2
0.1
0.05
0.01
0
Cell_Type_NEW
Ctrl
ExN
InN
Other
UD

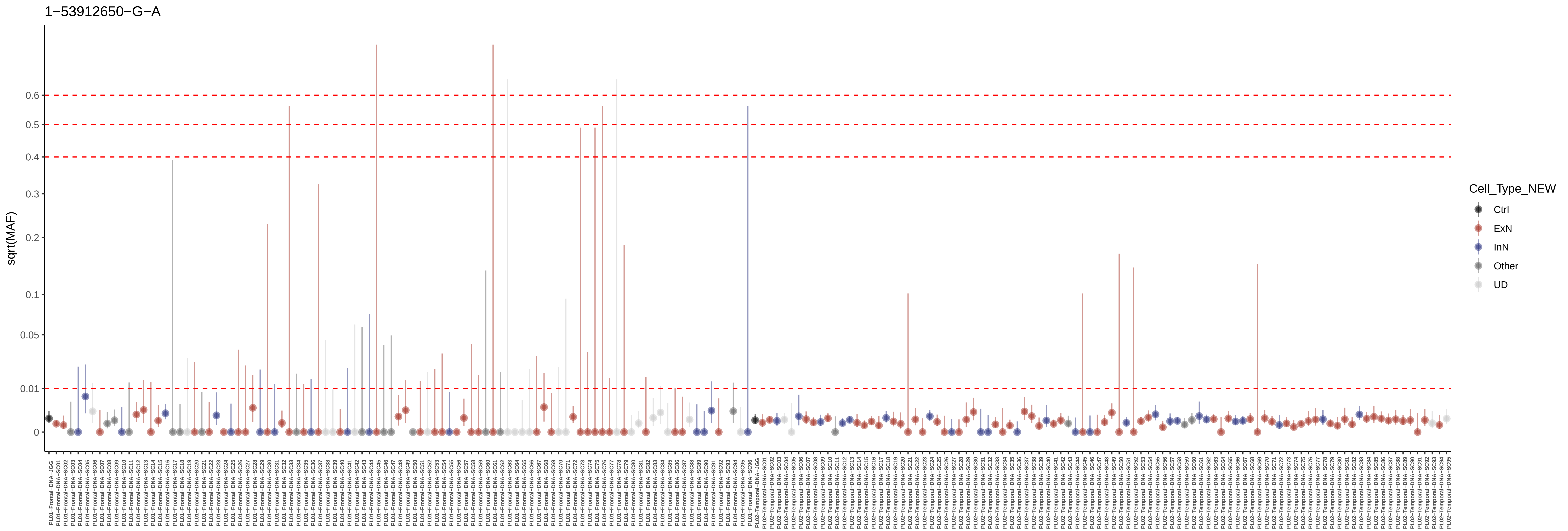
1−53912650−G−A
sqrt(MAF)
0.6
0.5
0.4
0.3
0.2
0.1
0.05
0.01
0
Cell_Type_NEW
Ctrl
ExN
InN
Other
UD

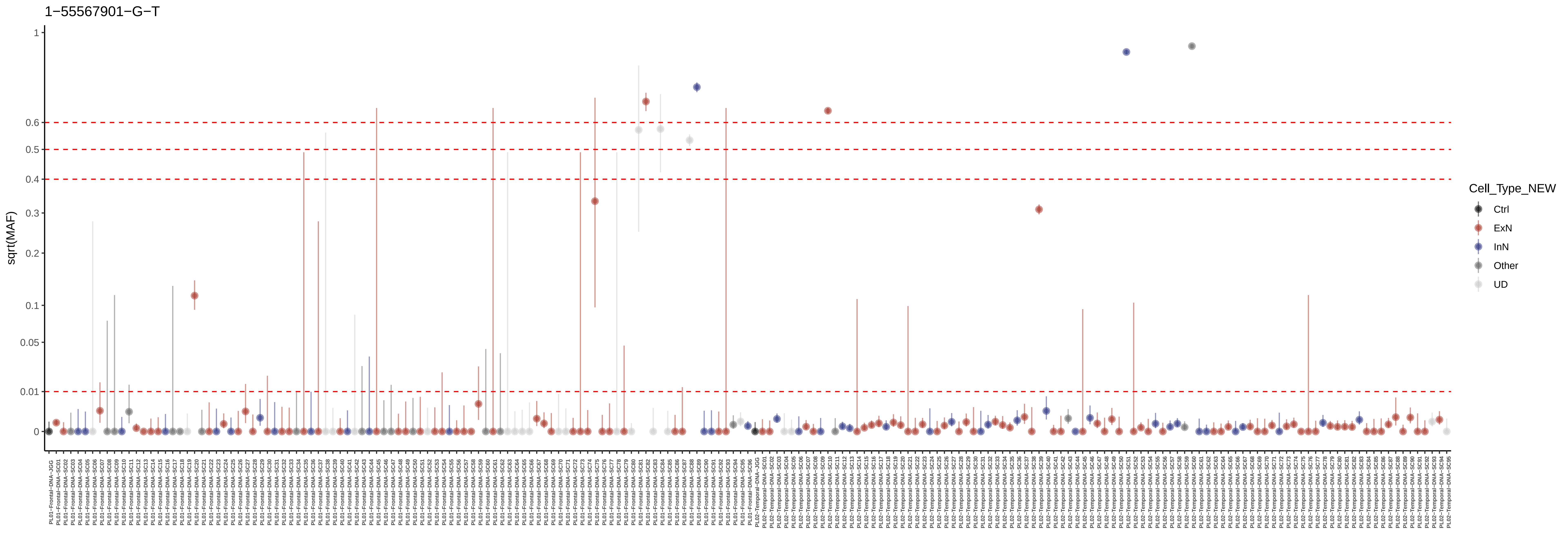
1−55567901−G−T
sqrt(MAF)
1
0.6
0.5
0.4
0.3
0.2
0.1
0.05
0.01
0
Cell_Type_NEW
Ctrl
ExN
InN
Other
UD

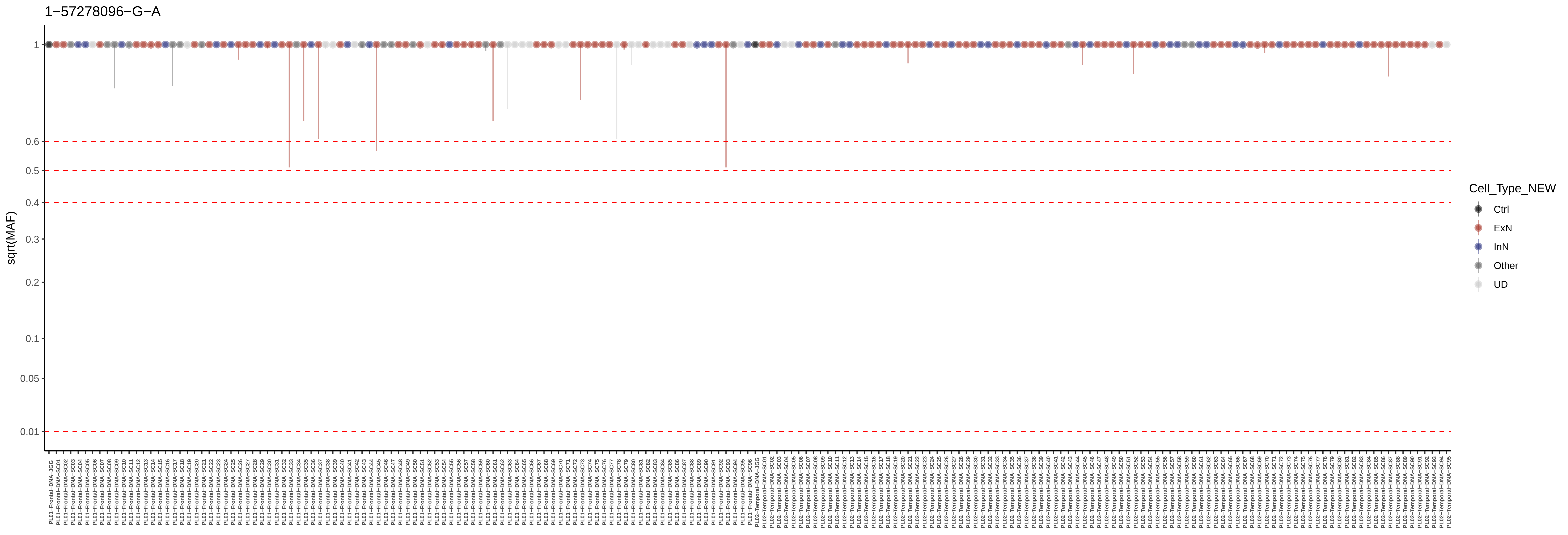
1−57278096−G−A
sqrt(MAF)
1
0.6
0.5
0.4
0.3
0.2
0.1
0.05
0.01
Cell_Type_NEW
Ctrl
ExN
InN
Other
UD

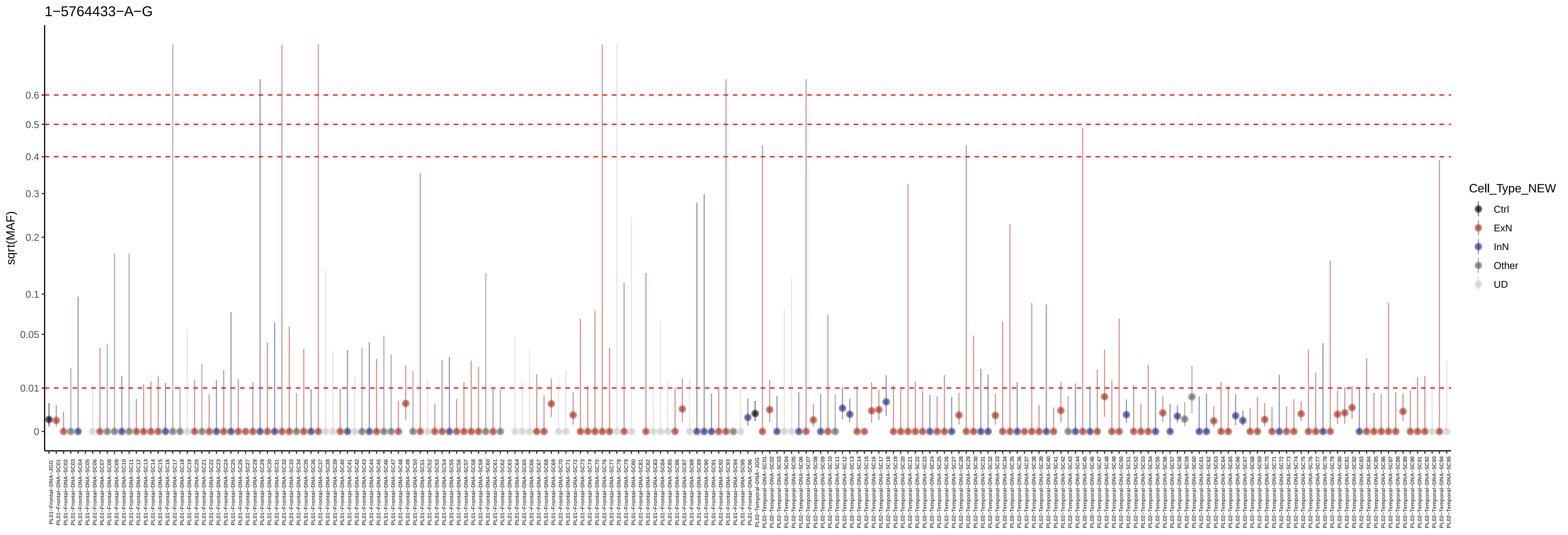
1−5764433−A−G
sqrt(MAF)
0.6
0.5
0.4
0.3
0.2
0.1
0.05
0.01
0
Cell_Type_NEW
Ctrl
ExN
InN
Other
UD

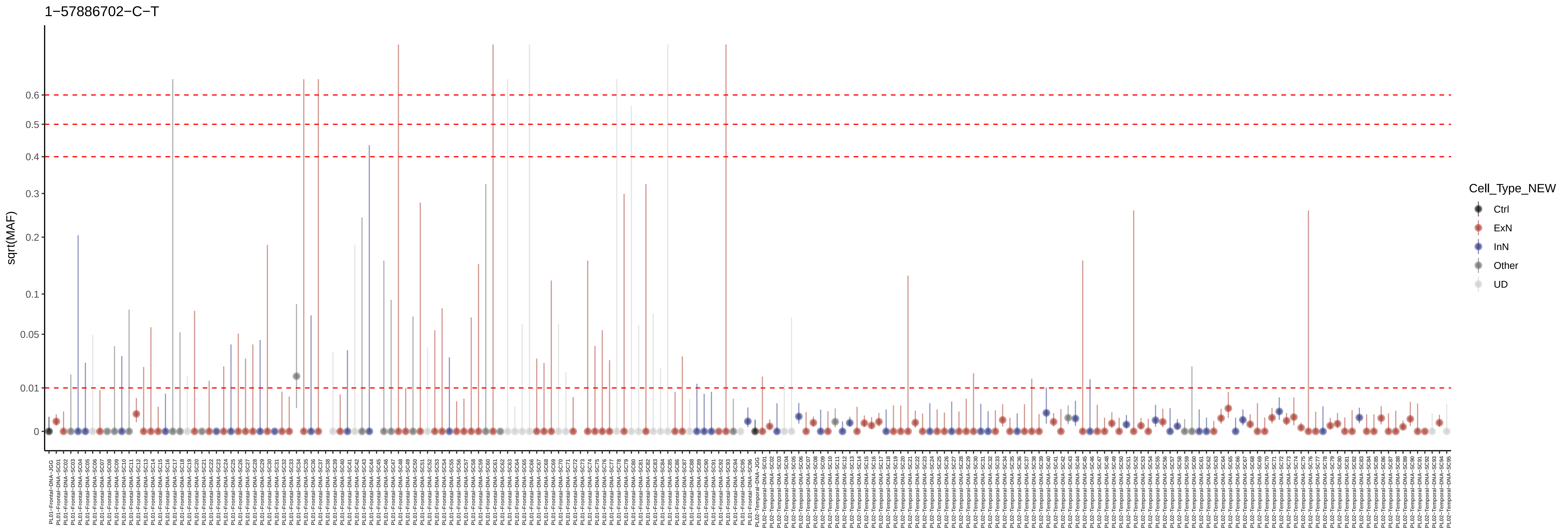
1−57886702−C−T
sqrt(MAF)
0.6
0.5
0.4
0.3
0.2
0.1
0.05
0.01
0
Cell_Type_NEW
Ctrl
ExN
InN
Other
UD

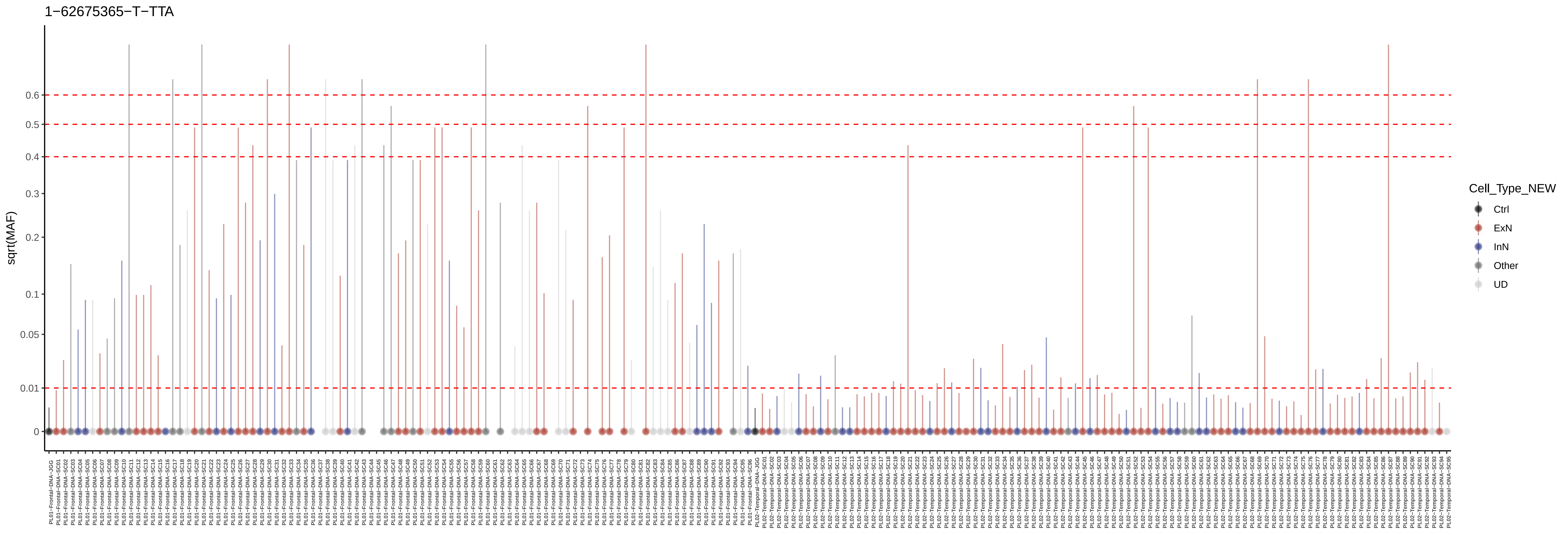
1−62675365−T−TTA
sqrt(MAF)
0.6
0.5
0.4
0.3
0.2
0.1
0.05
0.01
0
Cell_Type_NEW
Ctrl
ExN
InN
Other
UD

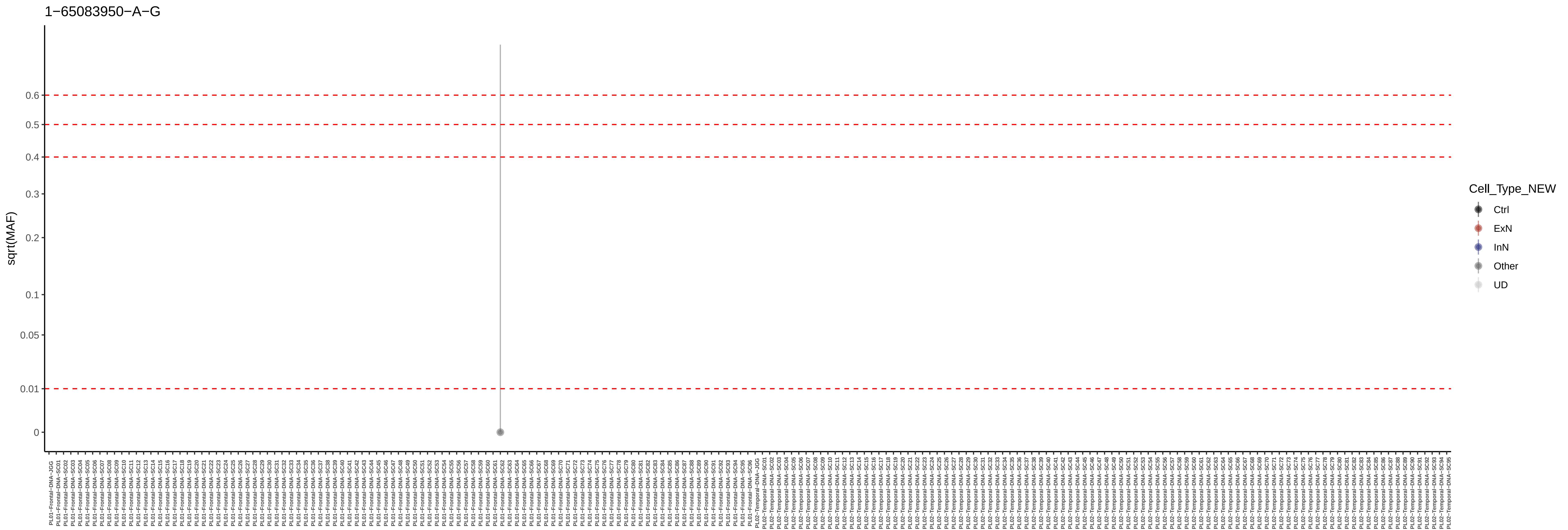
1−65083950−A−G
sqrt(MAF)
0.6
0.5
0.4
0.3
0.2
0.1
0.05
0.01
0
Cell_Type_NEW
Ctrl
ExN
InN
Other
UD

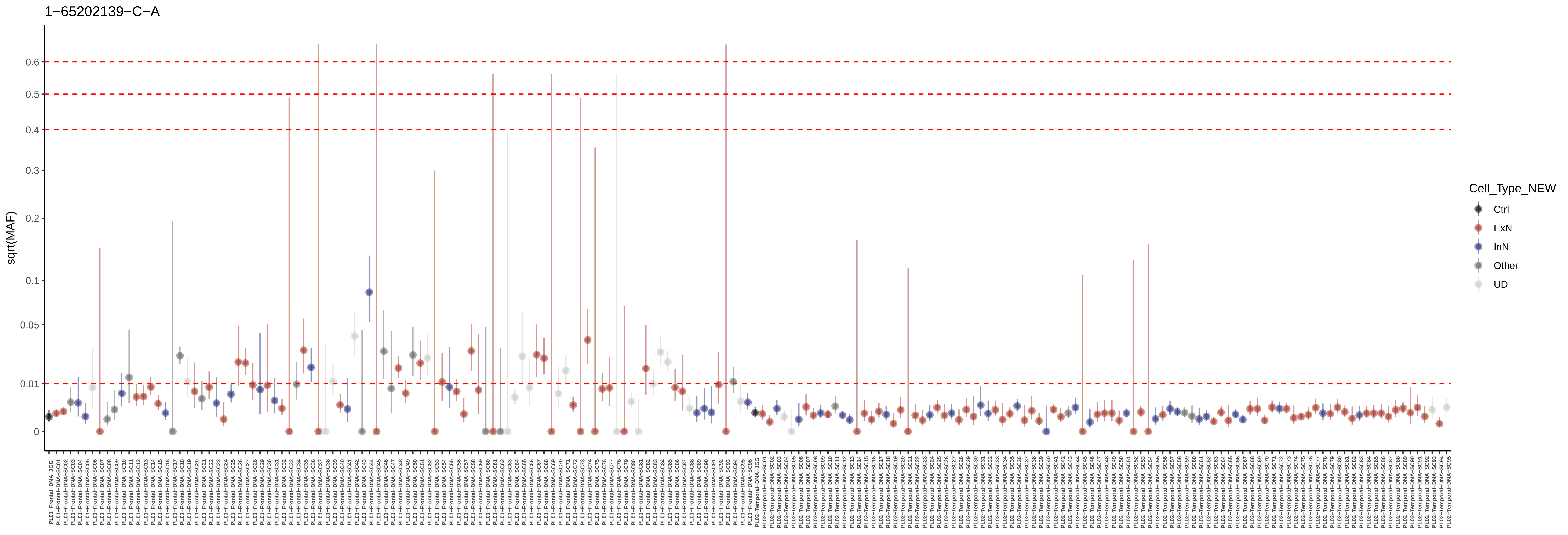
1−65202139−C−A
sqrt(MAF)
0.6
0.5
0.4
0.3
0.2
0.1
0.05
0.01
0
Cell_Type_NEW
Ctrl
ExN
InN
Other
UD

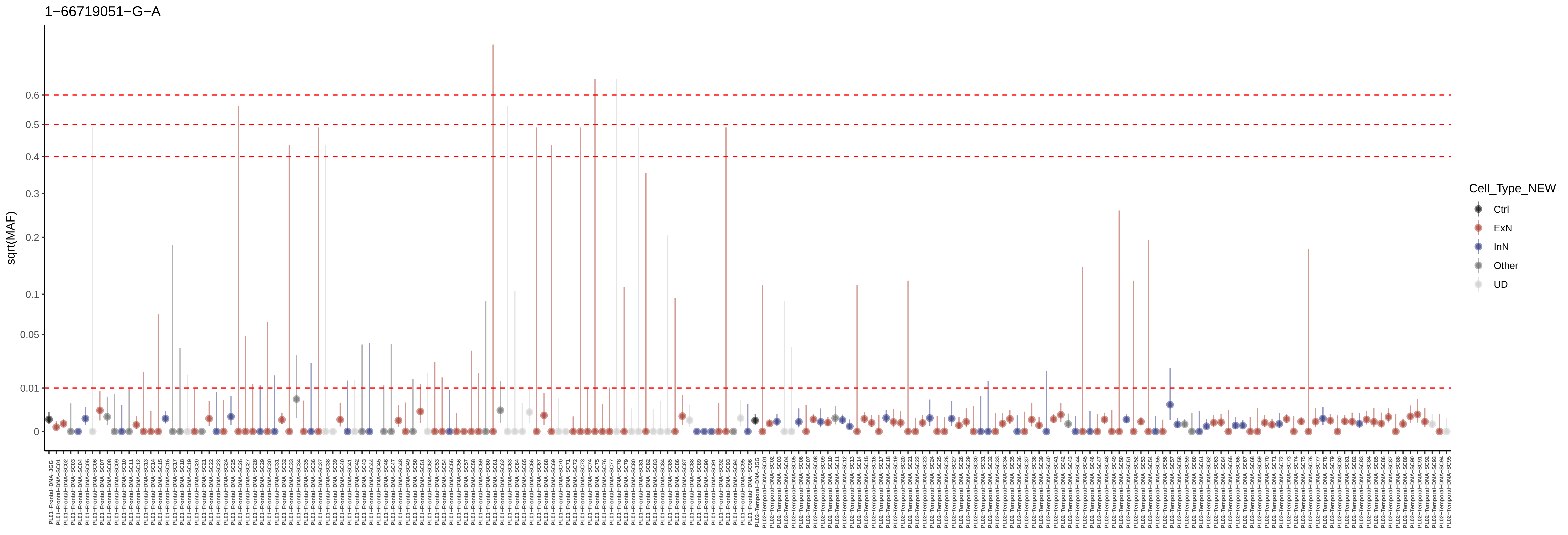
1−66719051−G−A
sqrt(MAF)
0.6
0.5
0.4
0.3
0.2
0.1
0.05
0.01
0
Cell_Type_NEW
Ctrl
ExN
InN
Other
UD

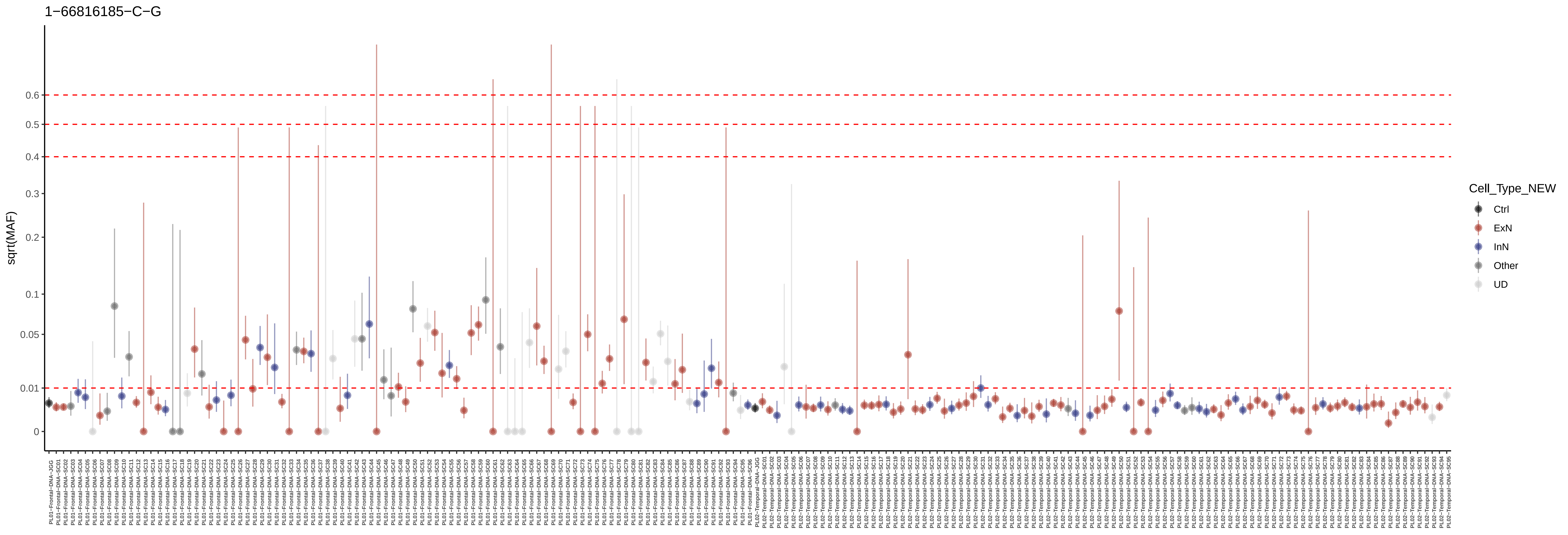
1−66816185−C−G
sqrt(MAF)
0.6
0.5
0.4
0.3
0.2
0.1
0.05
0.01
0
Cell_Type_NEW
Ctrl
ExN
InN
Other
UD

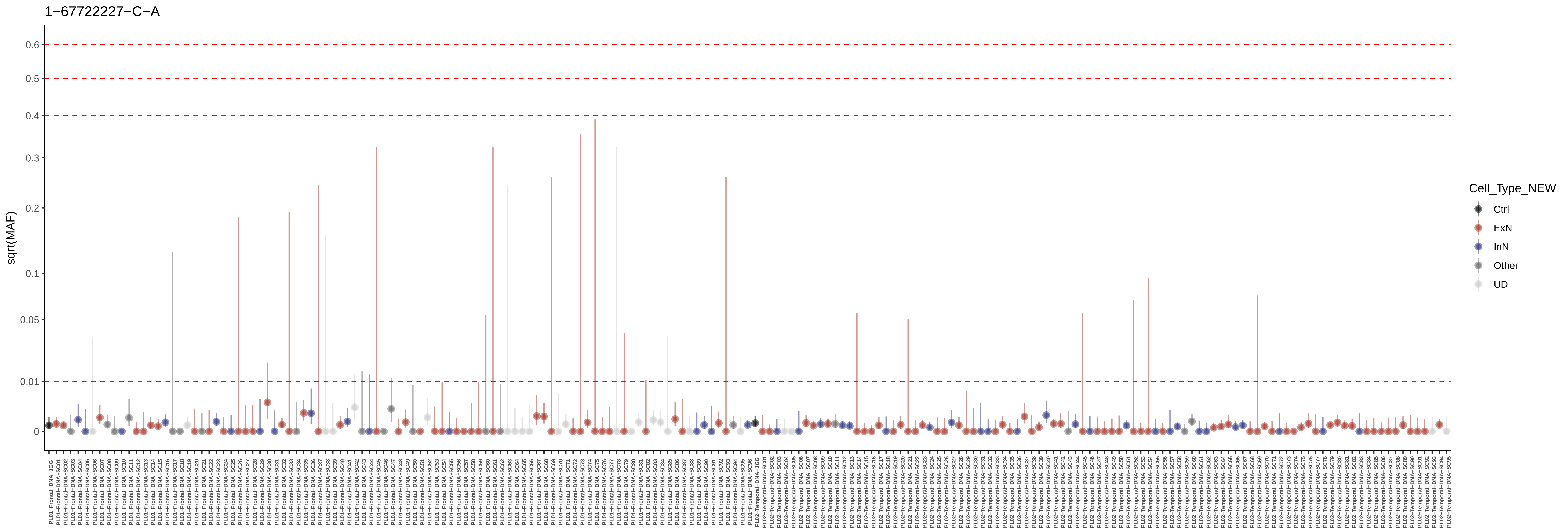

1−67722227−C−A
sqrt(MAF)
0.6
0.5
0.4
0.3
0.2
0.1
0.05
0.01
0
Cell_Type_NEW
Ctrl
ExN
InN
Other
UD

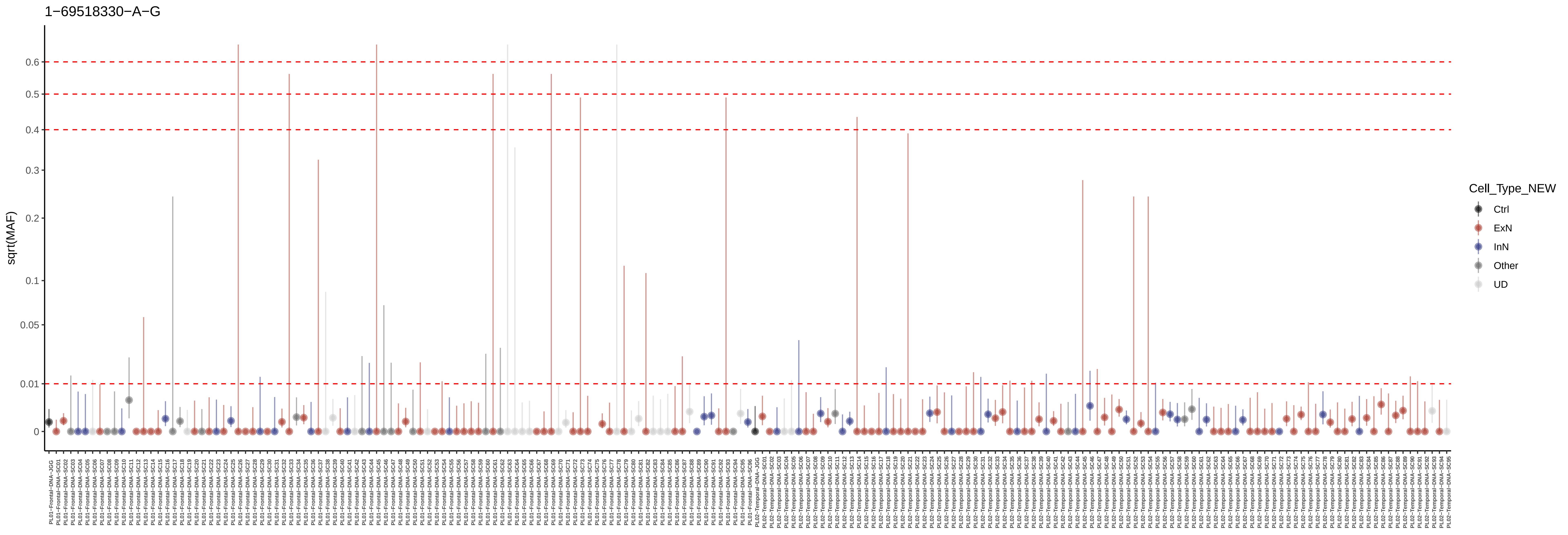
1−69518330−A−G
sqrt(MAF)
0.6
0.5
0.4
0.3
0.2
0.1
0.05
0.01
0
Cell_Type_NEW
Ctrl
ExN
InN
Other
UD

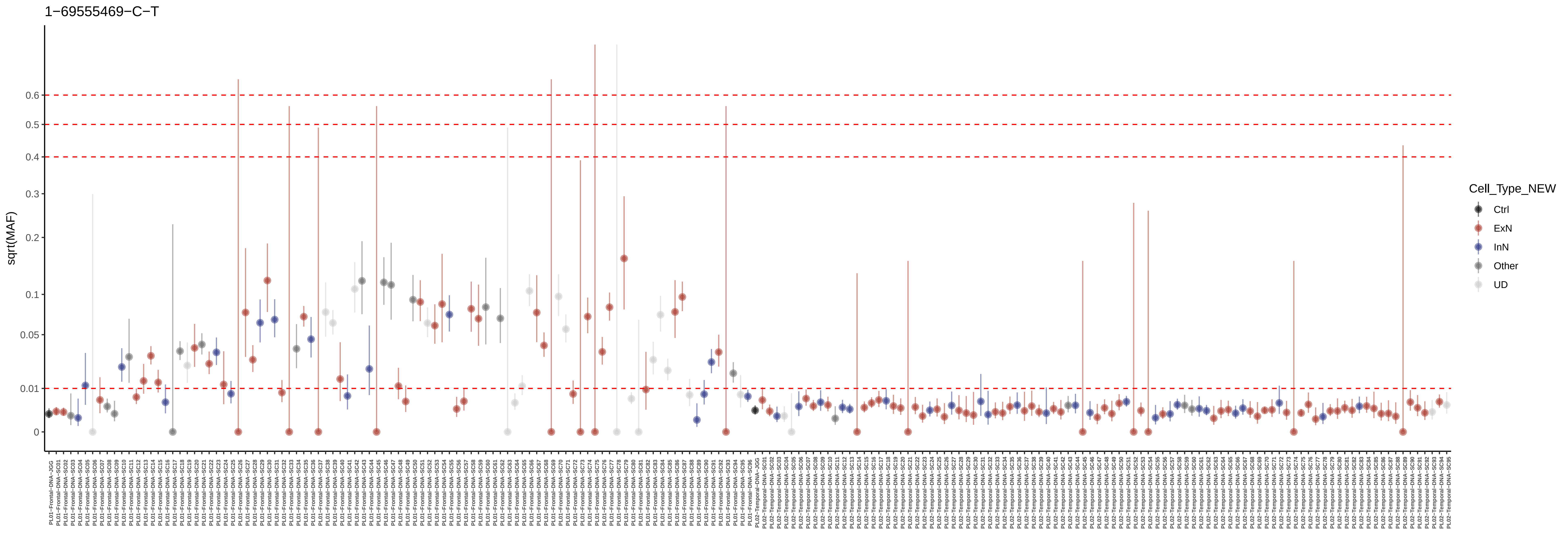

1−69555469−C−T
sqrt(MAF)
0.6
0.5
0.4
0.3
0.2
0.1
0.05
0.01
0
Cell_Type_NEW
Ctrl
ExN
InN
Other
UD

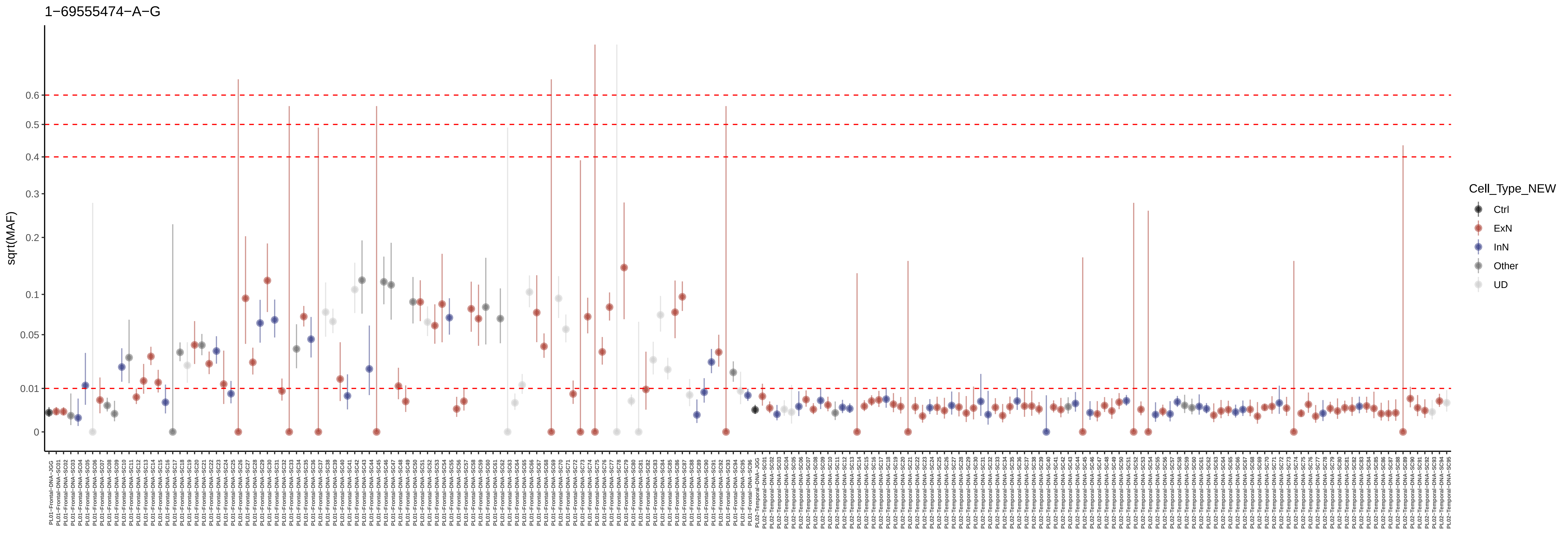
1−69555474−A−G
sqrt(MAF)
0.6
0.5
0.4
0.3
0.2
0.1
0.05
0.01
0
Cell_Type_NEW
Ctrl
ExN
InN
Other
UD

1−70091682−T−C

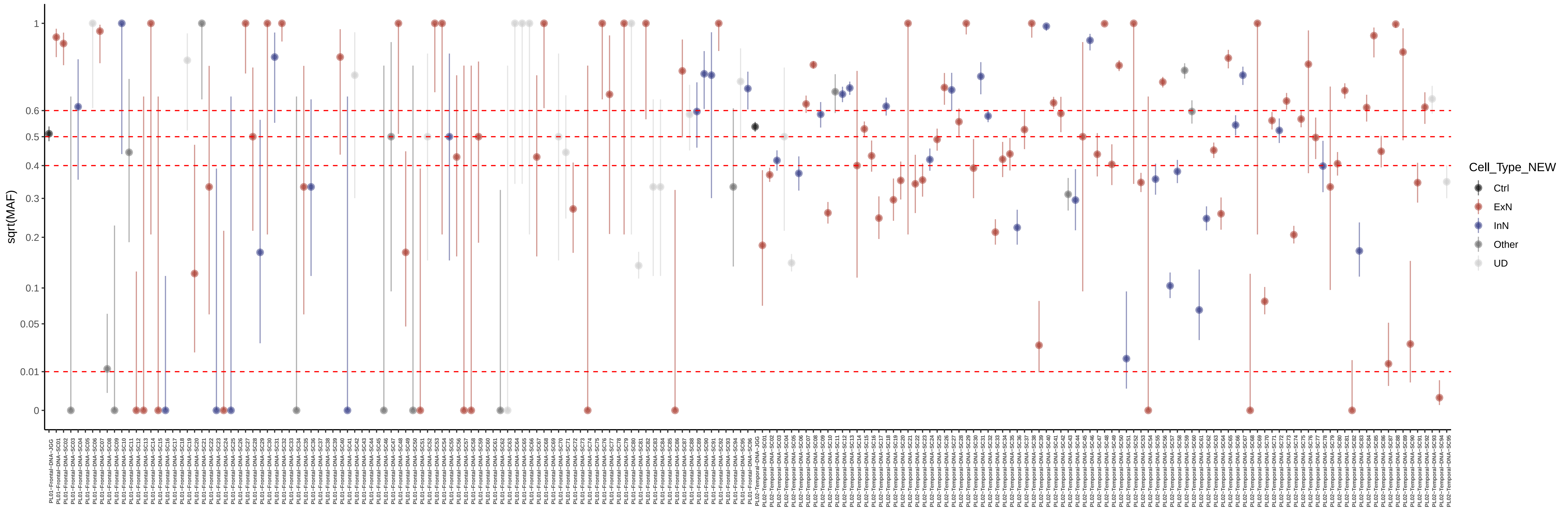

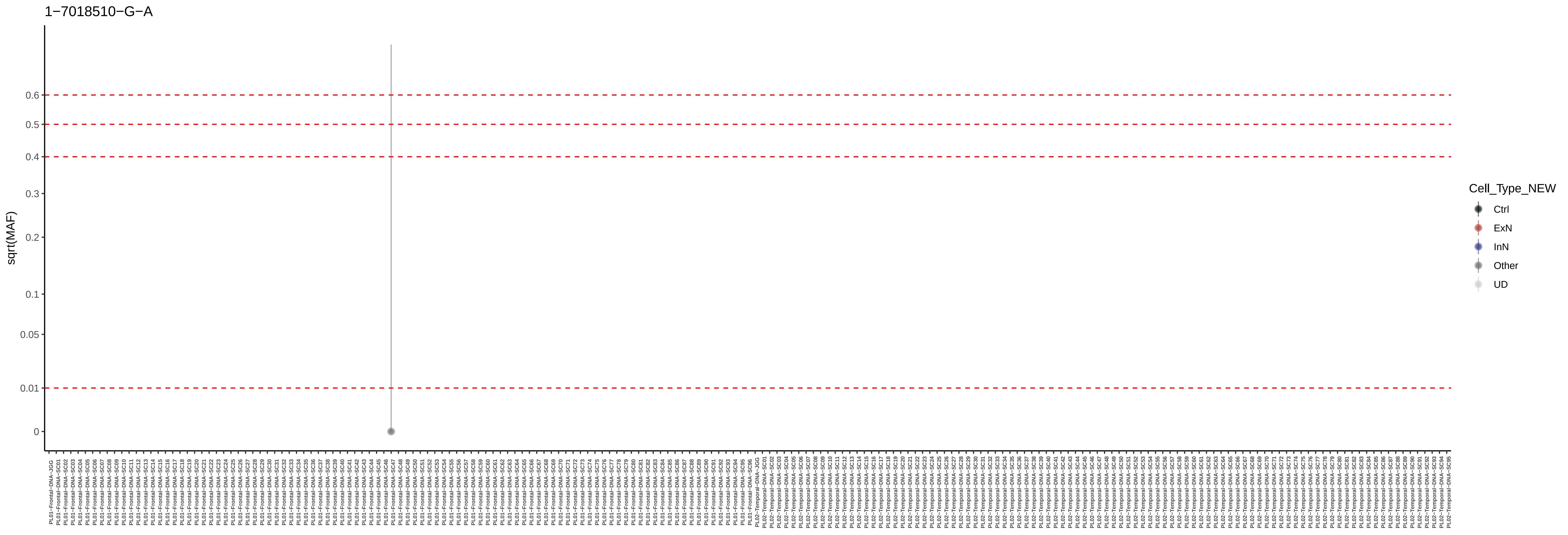
1−7018510−G−A
sqrt(MAF)
0.6
0.5
0.4
0.3
0.2
0.1
0.05
0.01
0
Cell_Type_NEW
Ctrl
ExN
InN
Other
UD

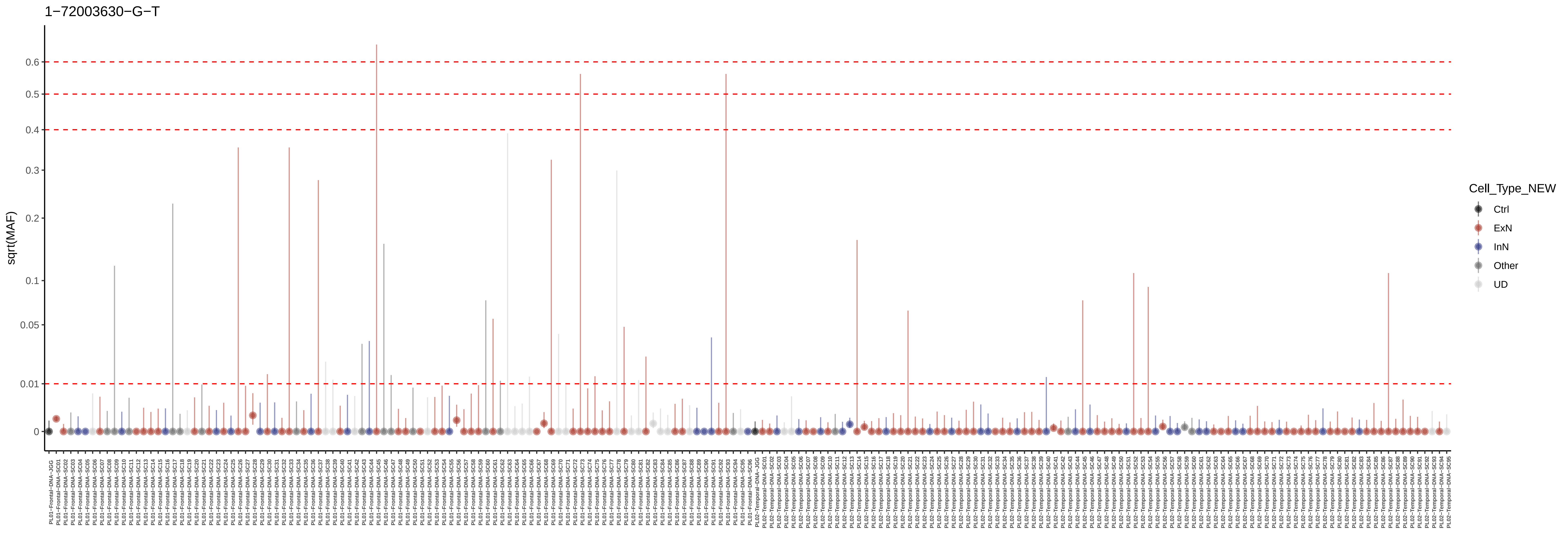
1−72003630−G−T
sqrt(MAF)
0.6
0.5
0.4
0.3
0.2
0.1
0.05
0.01
0
Cell_Type_NEW
Ctrl
ExN
InN
Other
UD

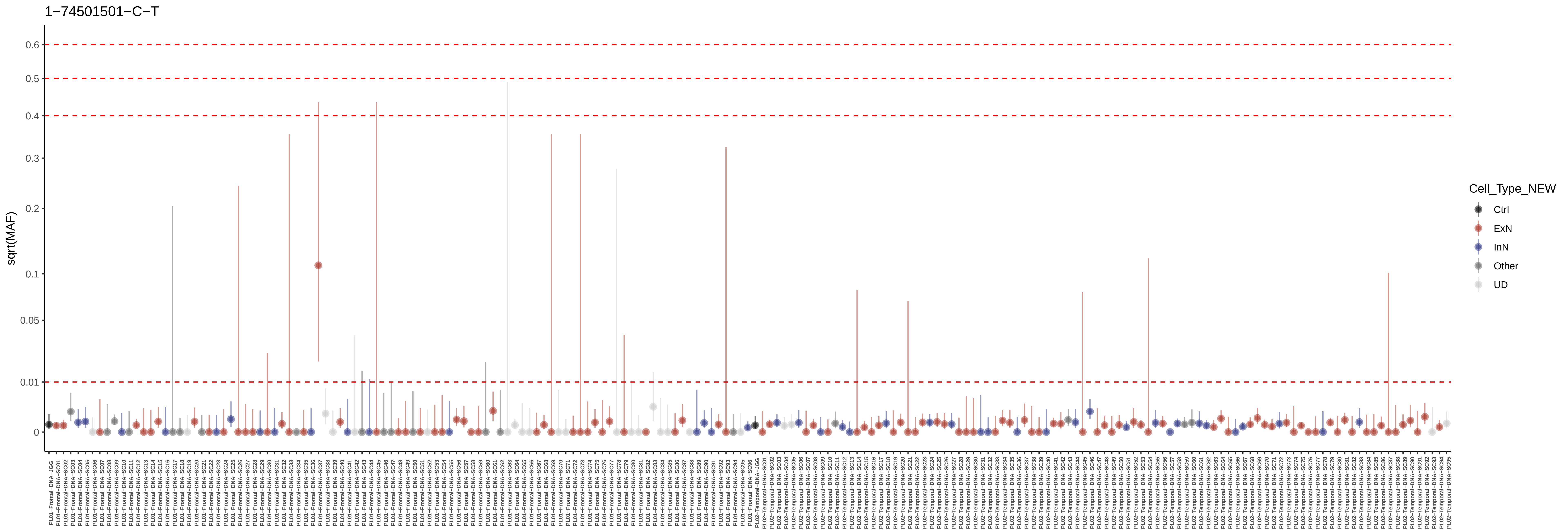
1−74501501−C−T
sqrt(MAF)
0.6
0.5
0.4
0.3
0.2
0.1
0.05
0.01
0
Cell_Type_NEW
Ctrl
ExN
InN
Other
UD

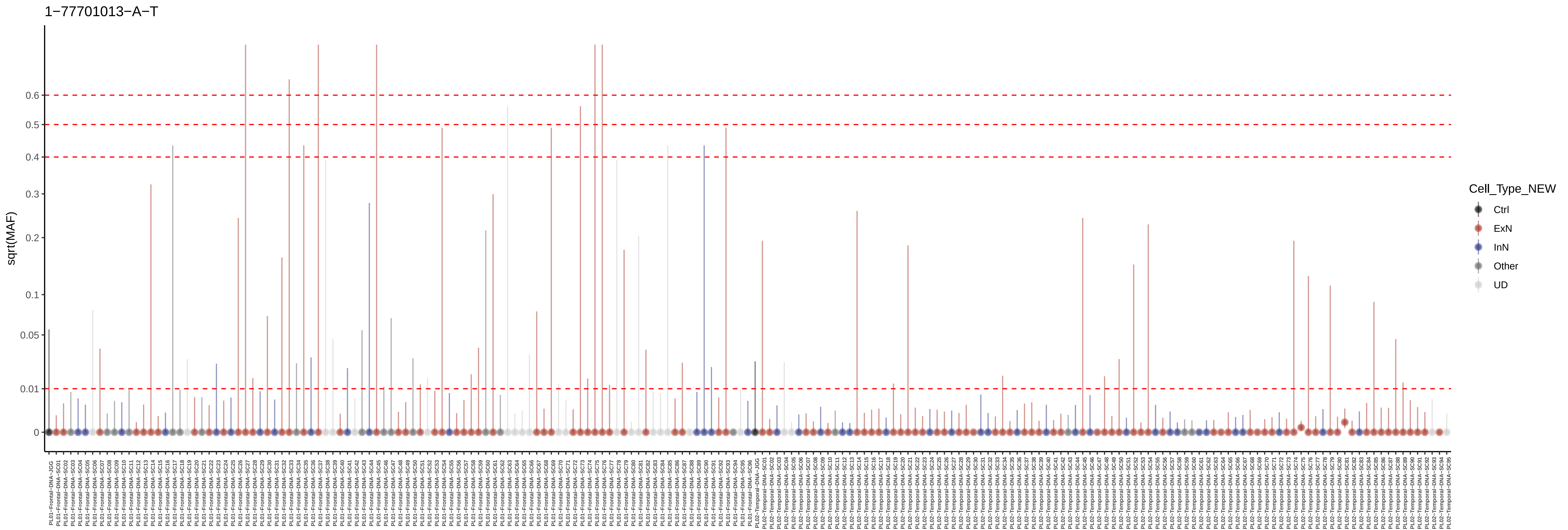
1−77701013−A−T
sqrt(MAF)
0.6
0.5
0.4
0.3
0.2
0.1
0.05
0.01
0
Cell_Type_NEW
Ctrl
ExN
InN
Other
UD

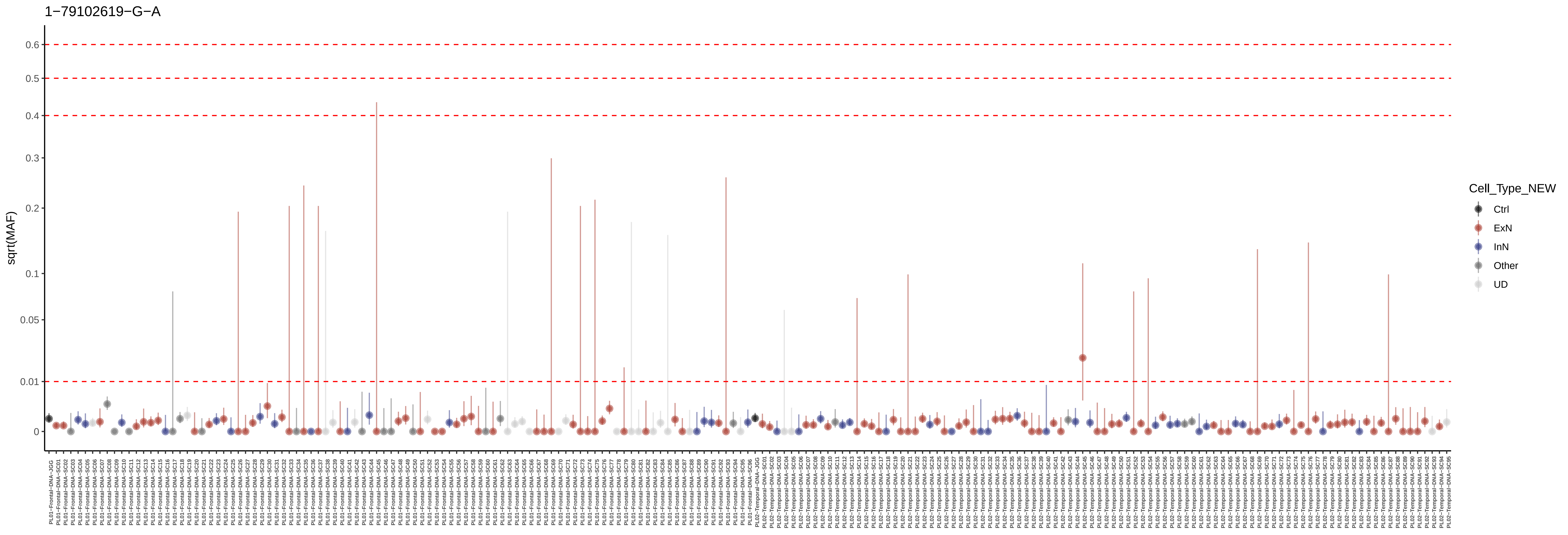

1−79102619−G−A
sqrt(MAF)
Cell_Type_NEW
Ctrl
ExN
InN
Other
UD

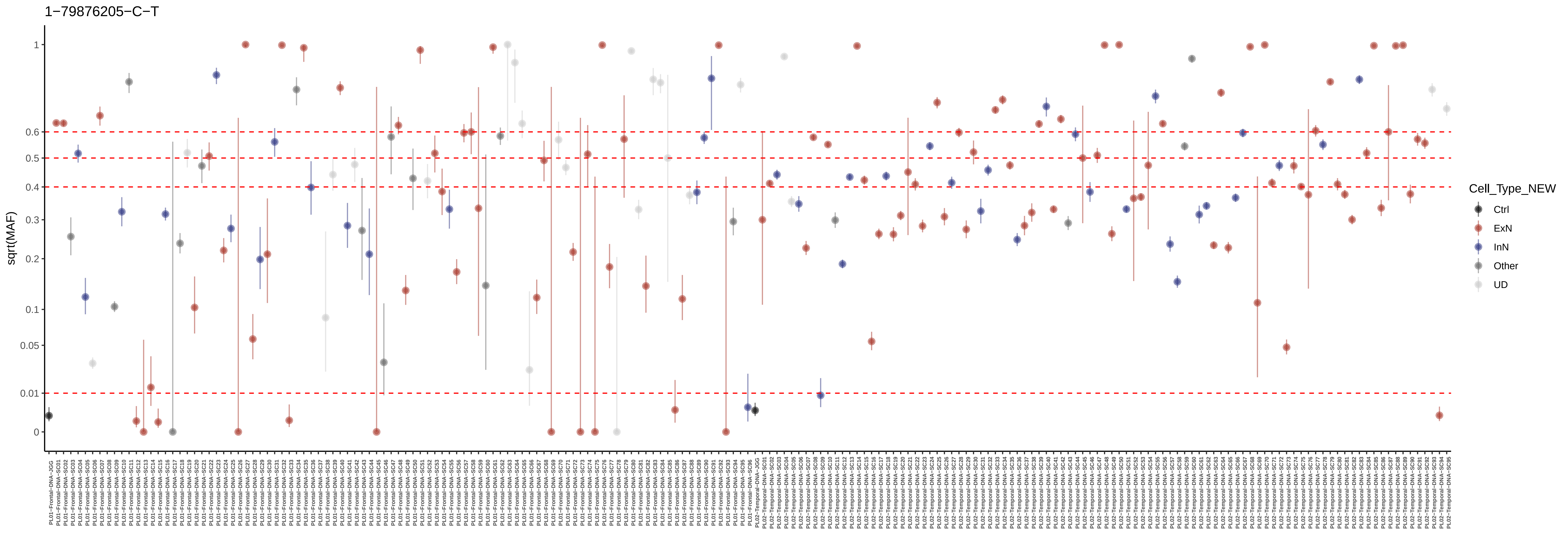

1−79876205−C−T
sqrt(MAF)
1
0.6
0.5
0.4
0.3
0.2
0.1
0.05
0.01
0
Cell_Type_NEW
Ctrl
ExN
InN
Other
UD

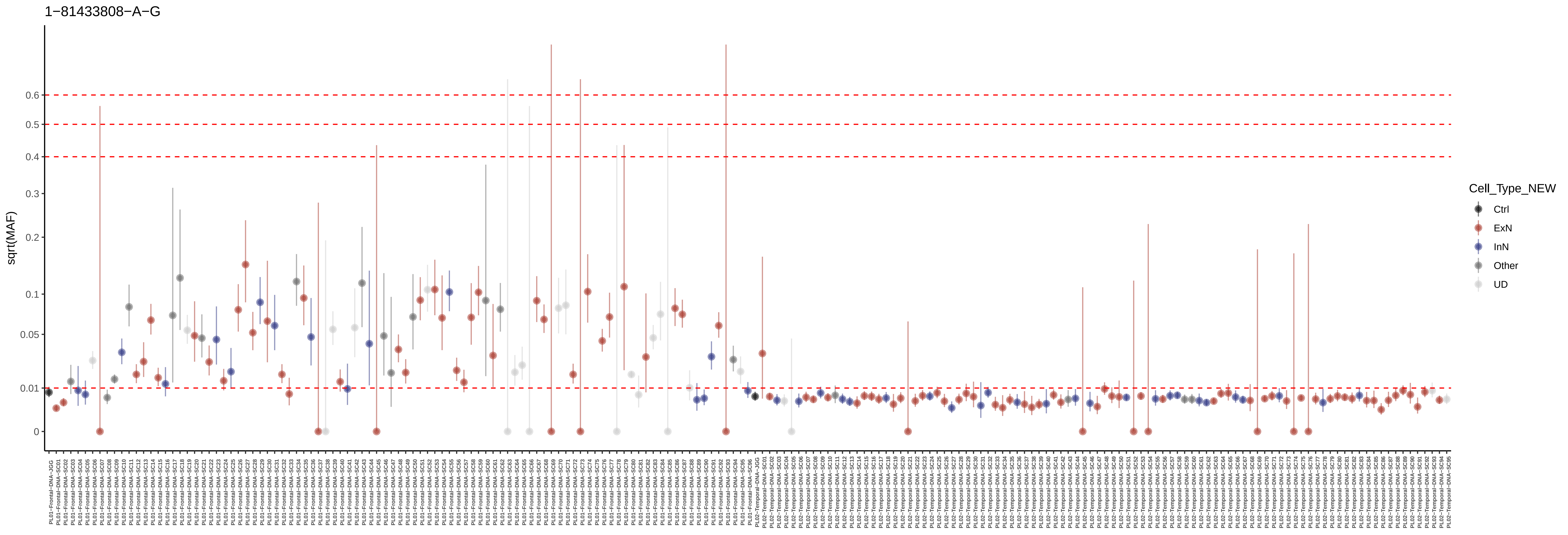

1−81433808−A−G
sqrt(MAF)
0.6
0.5
0.4
0.3
0.2
0.1
0.05
0.01
0
Cell_Type_NEW
Ctrl
ExN
InN
Other
UD

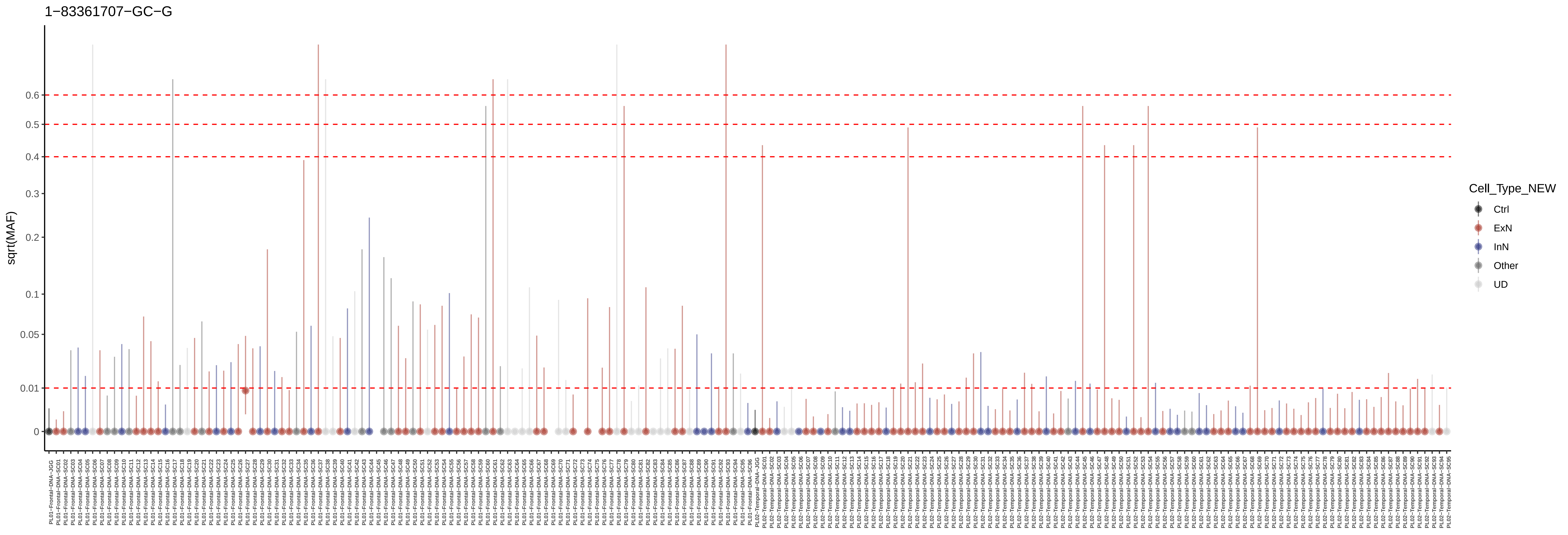
1−83361707−GC−G
sqrt(MAF)
0.6
0.5
0.4
0.3
0.2
0.1
0.05
0.01
0
Cell_Type_NEW
Ctrl
ExN
InN
Other
UD

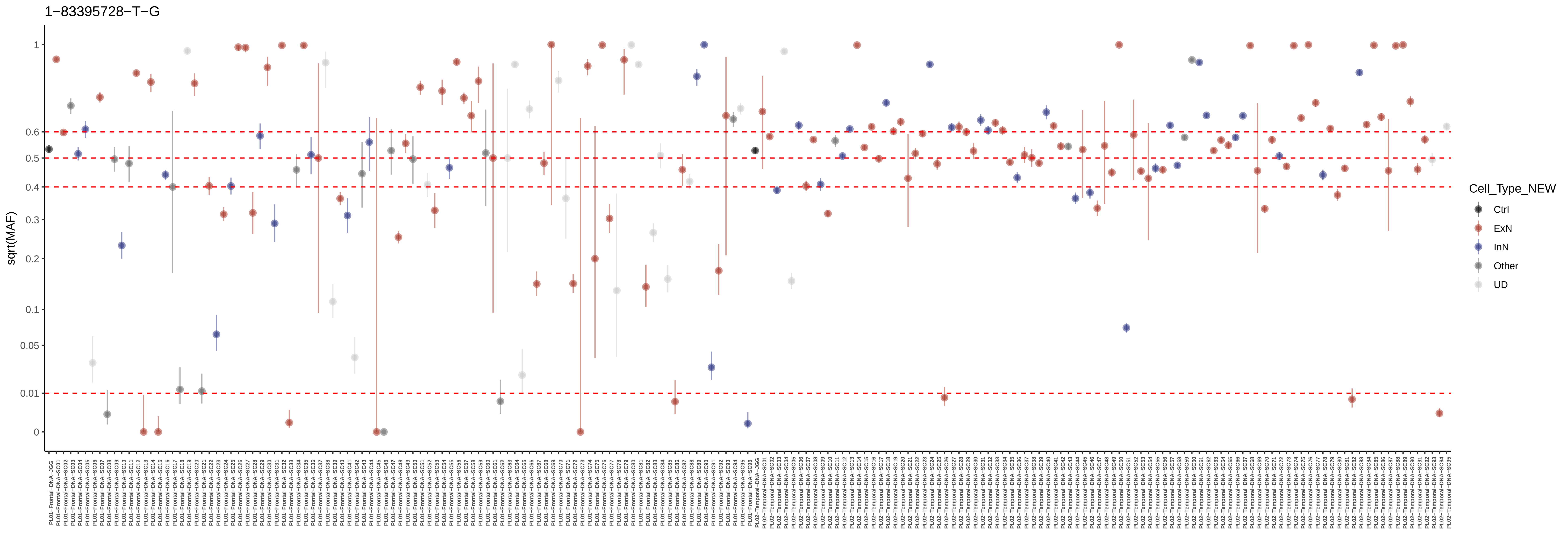

1−83395728−T−G
sqrt(MAF)
1
0.6
0.5
0.4
0.3
0.2
0.1
0.05
0.01
0
Cell_Type_NEW
Ctrl
ExN
InN
Other
UD

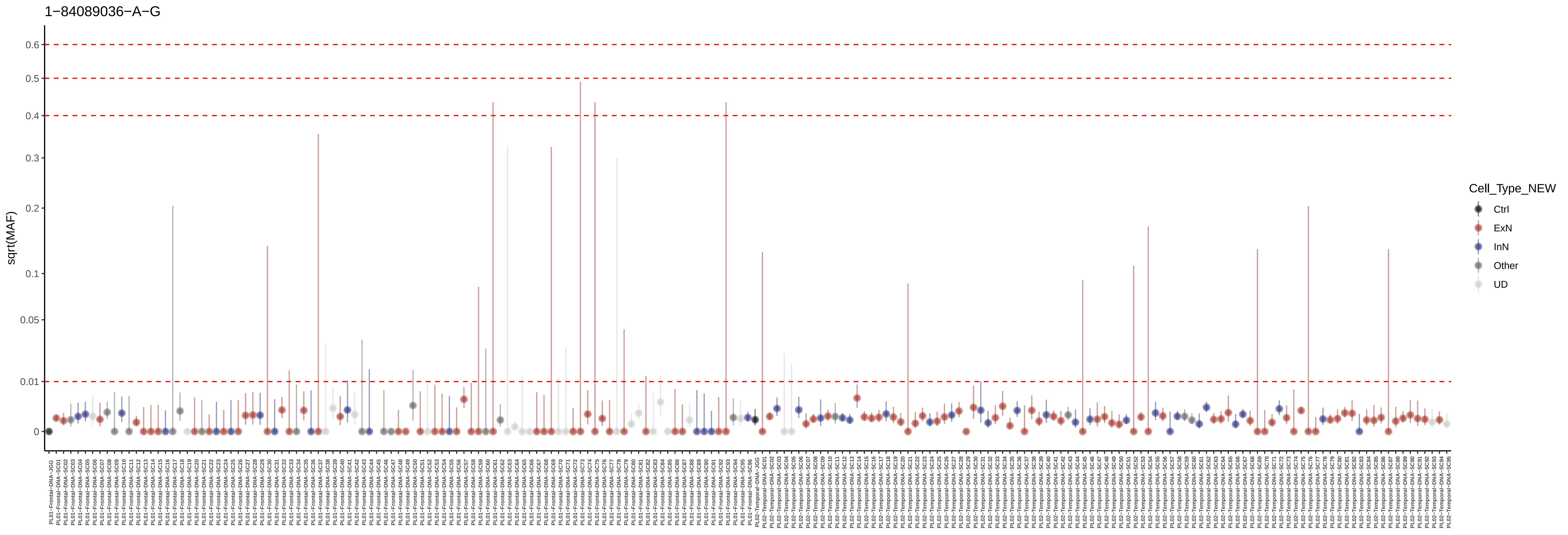
1−84089036−A−G
sqrt(MAF)
0.6
0.5
0.4
0.3
0.2
0.1
0.05
0.01
0
Cell_Type_NEW
Ctrl
ExN
InN
Other
UD

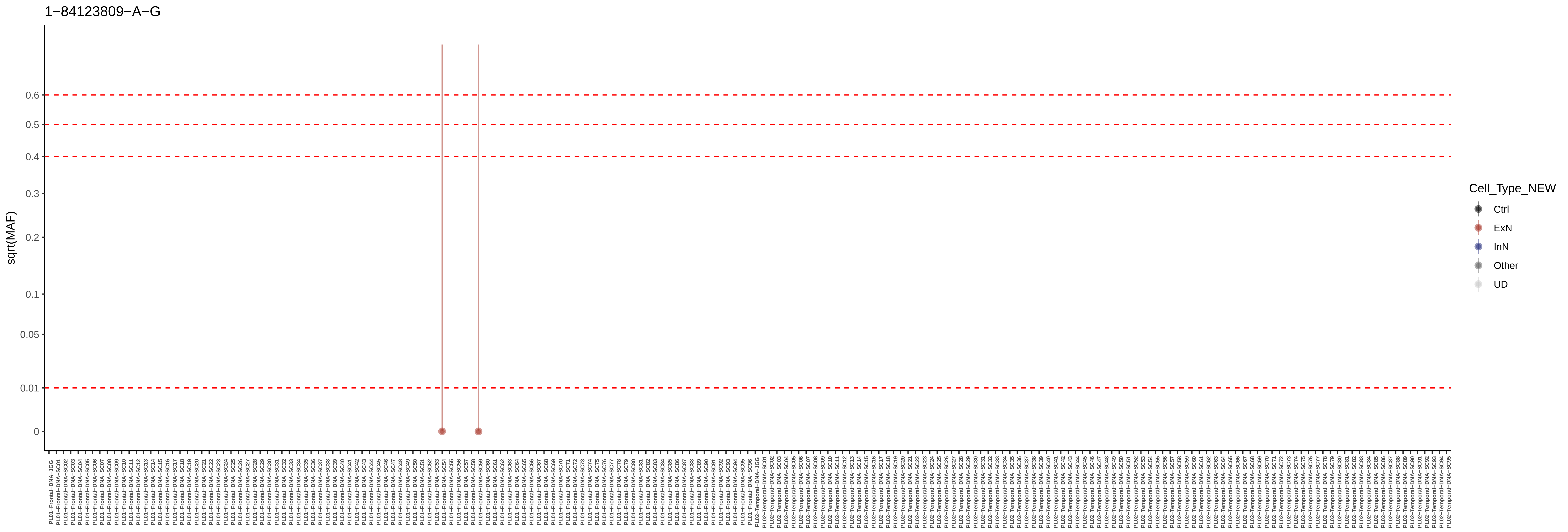
1−84123809−A−G
sqrt(MAF)
0.6
0.5
0.4
0.3
0.2
0.1
0.05
0.01
0
Cell_Type_NEW
Ctrl
ExN
InN
Other
UD

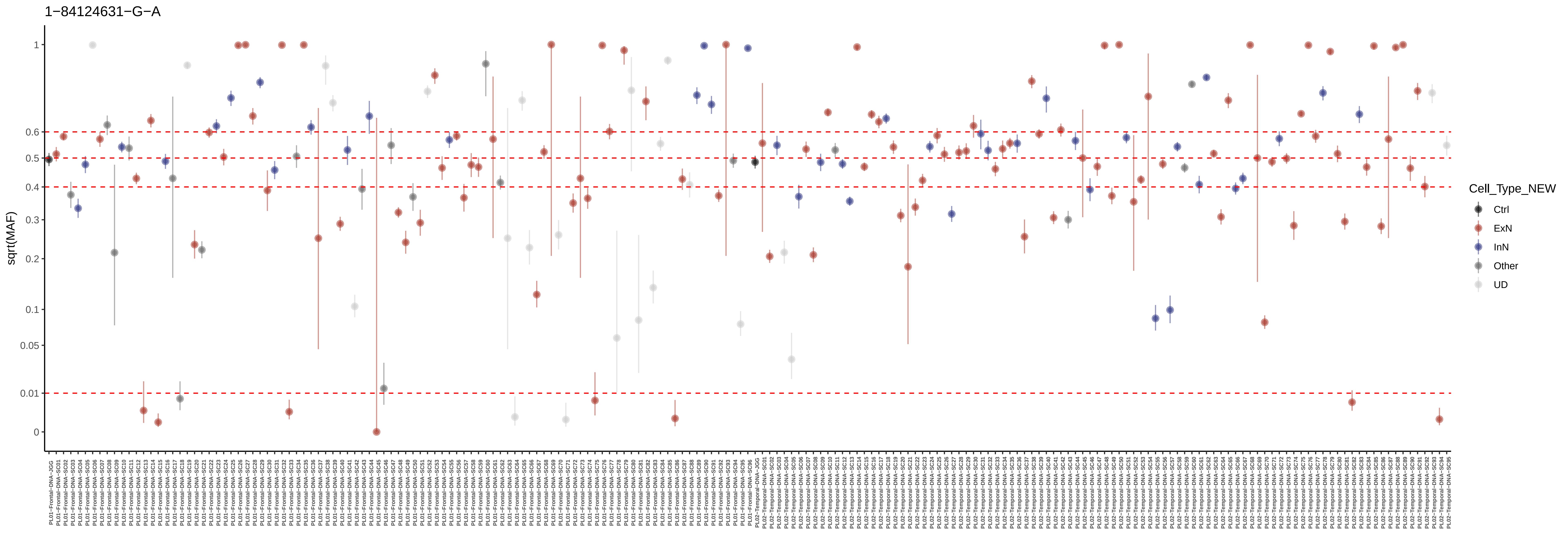
1−84124631−G−A
sqrt(MAF)
1
0.6
0.5
0.4
0.3
0.2
0.1
0.05
0.01
0
Cell_Type_NEW
Ctrl
ExN
InN
Other
UD

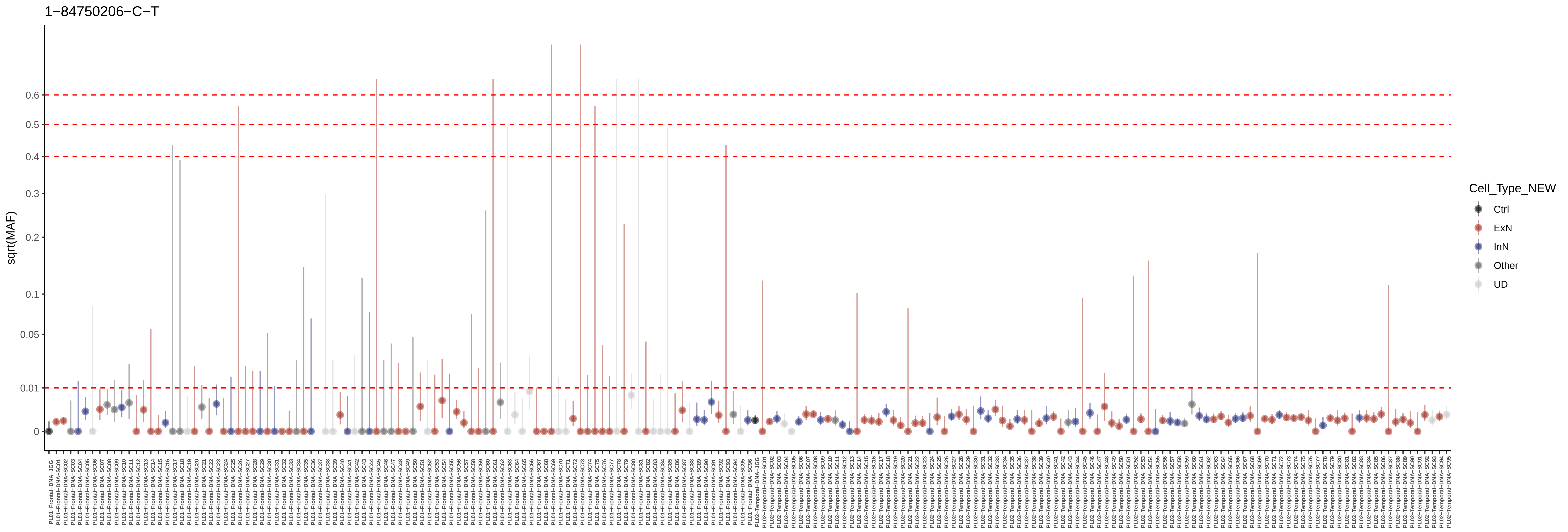
1−84750206−C−T
sqrt(MAF)
0.6
0.5
0.4
0.3
0.2
0.1
0.05
0.01
0
Cell_Type_NEW
Ctrl
ExN
InN
Other
UD

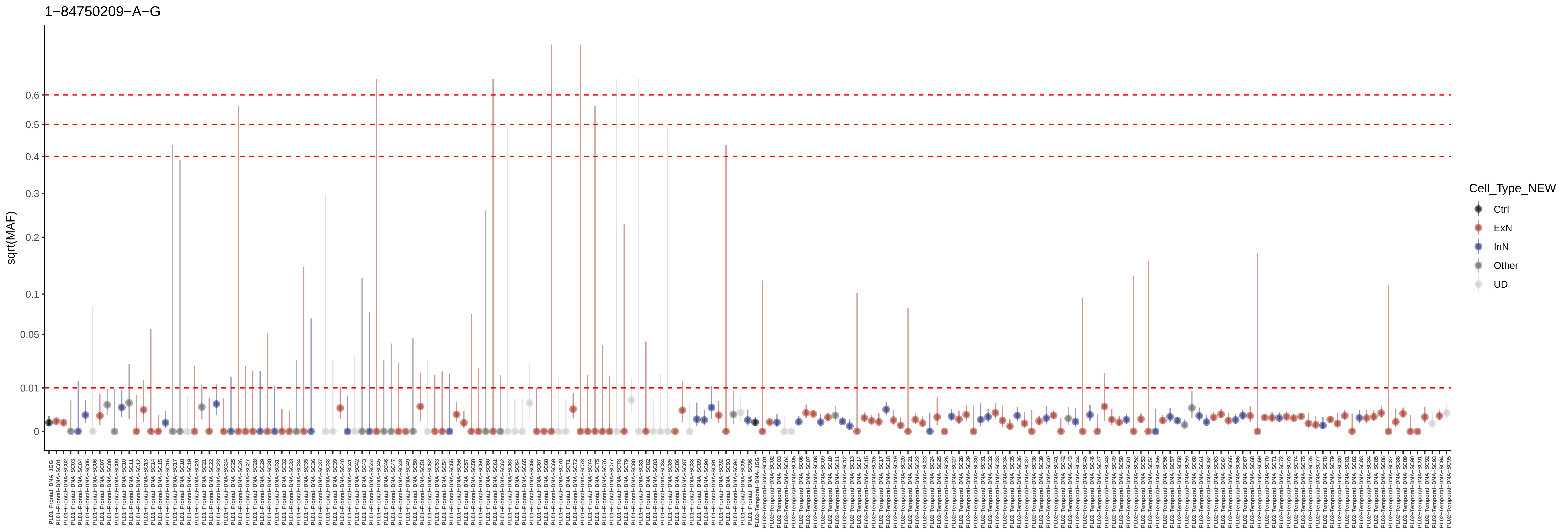
1−84750209−A−G
sqrt(MAF)
0.6
0.5
0.4
0.3
0.2
0.1
0.05
0.01
0
Cell_Type_NEW
Ctrl
ExN
InN
Other
UD

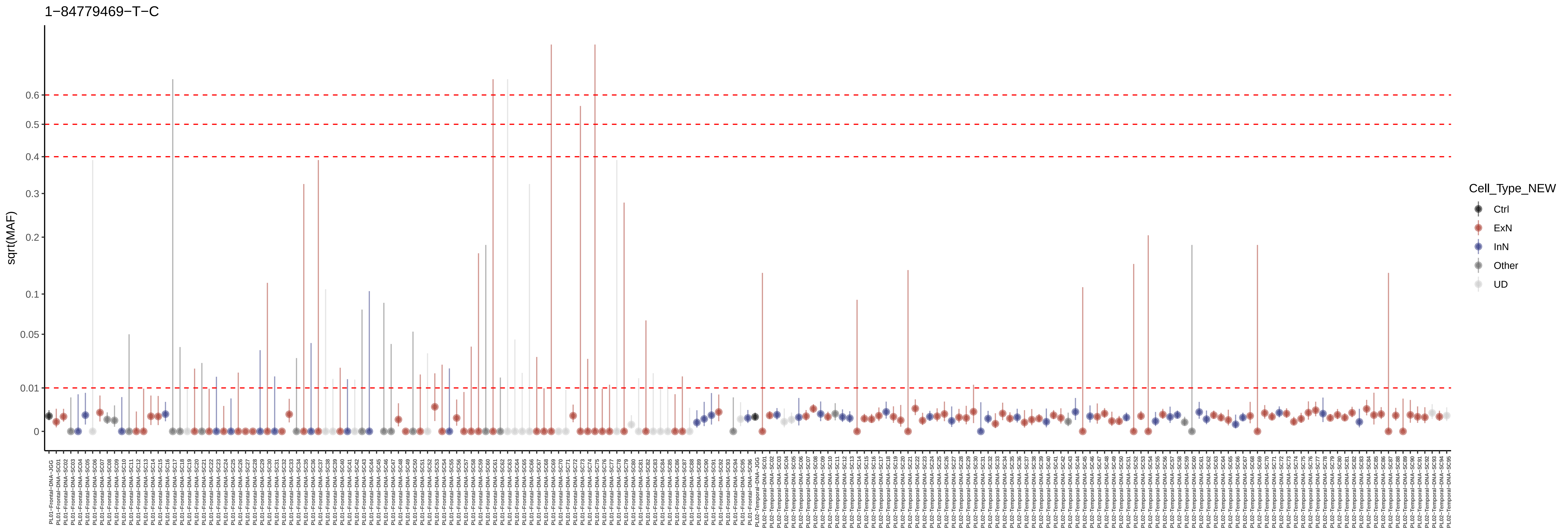
1−84779469−T−C
sqrt(MAF)
0.6
0.5
0.4
0.3
0.2
0.1
0.05
0.01
0
Cell_Type_NEW
Ctrl
ExN
InN
Other
UD

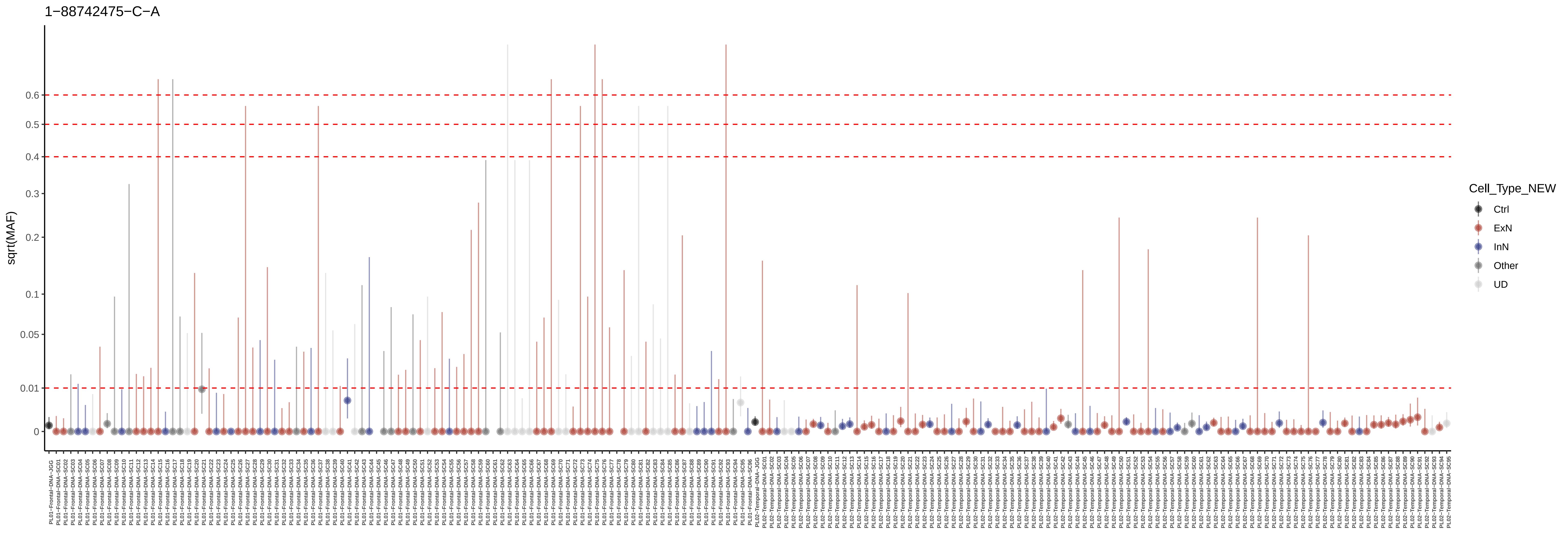
1−88742475−C−A
sqrt(MAF)
0.6
0.5
0.4
0.3
0.2
0.1
0.05
0.01
0
Cell_Type_NEW
Ctrl
ExN
InN
Other
UD

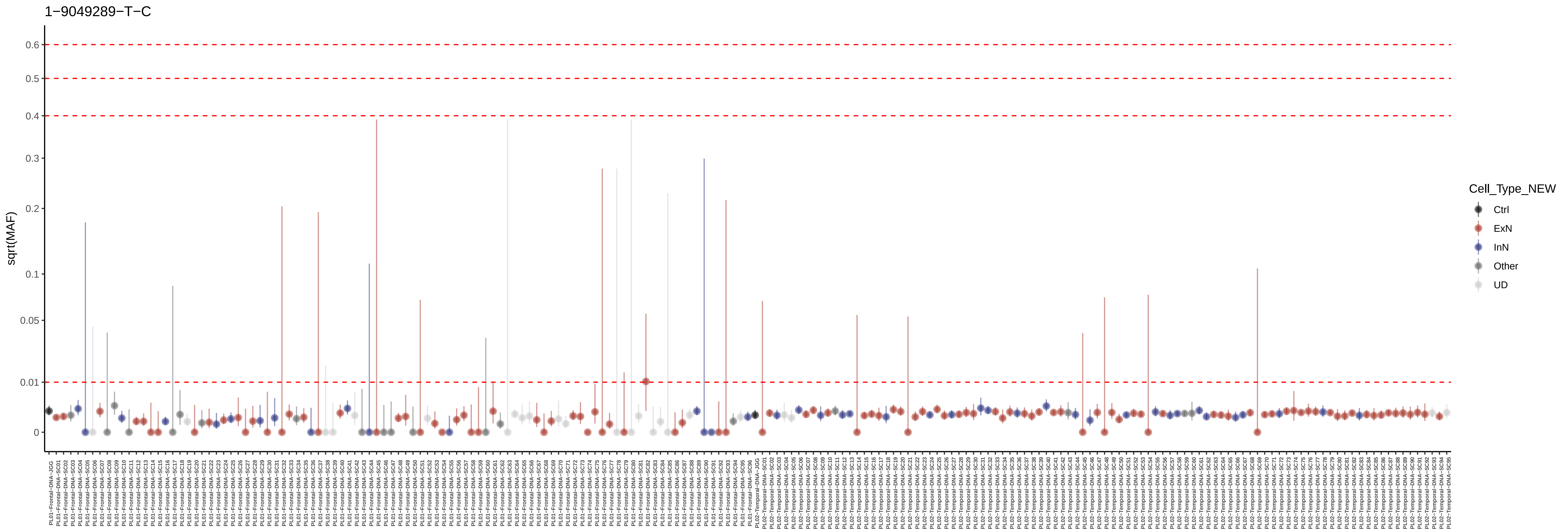
1−9049289−T−C
sqrt(MAF)
0.6
0.5
0.4
0.3
0.2
0.1
0.05
0.01
0
Cell_Type_NEW
Ctrl
ExN
InN
Other
UD

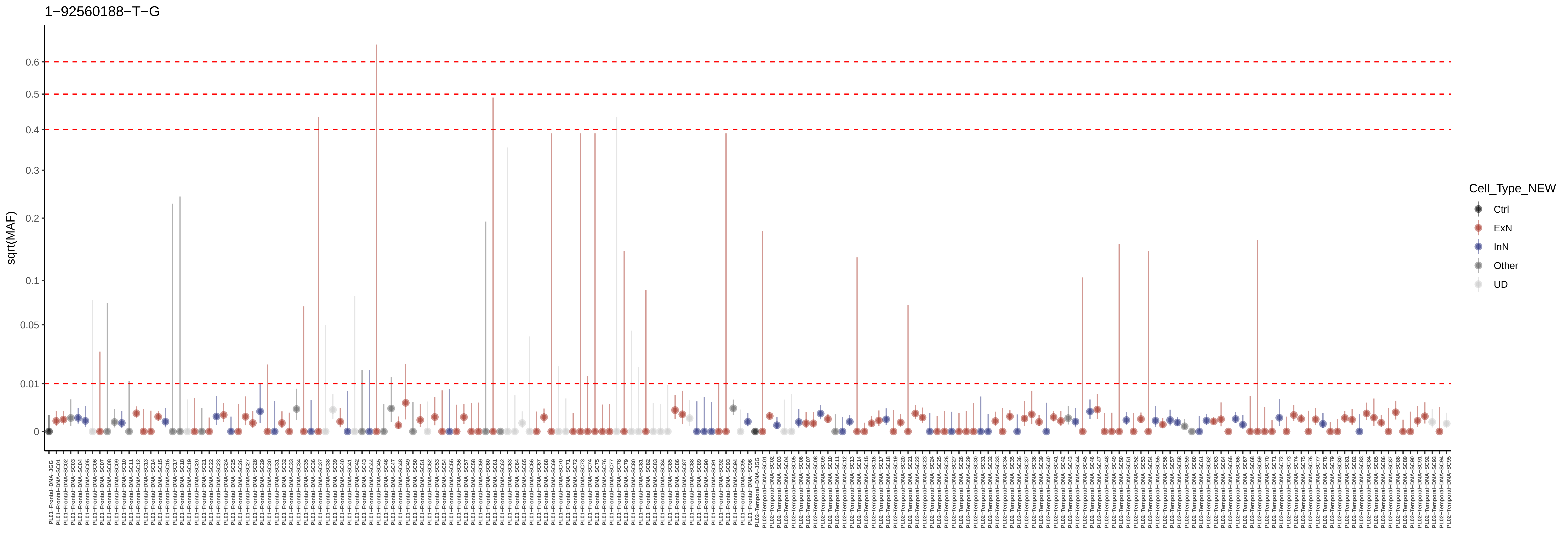
1−92560188−T−G
sqrt(MAF)
0.6
0.5
0.4
0.3
0.2
0.1
0.05
0.01
0
Cell_Type_NEW
Ctrl
ExN
InN
Other
UD

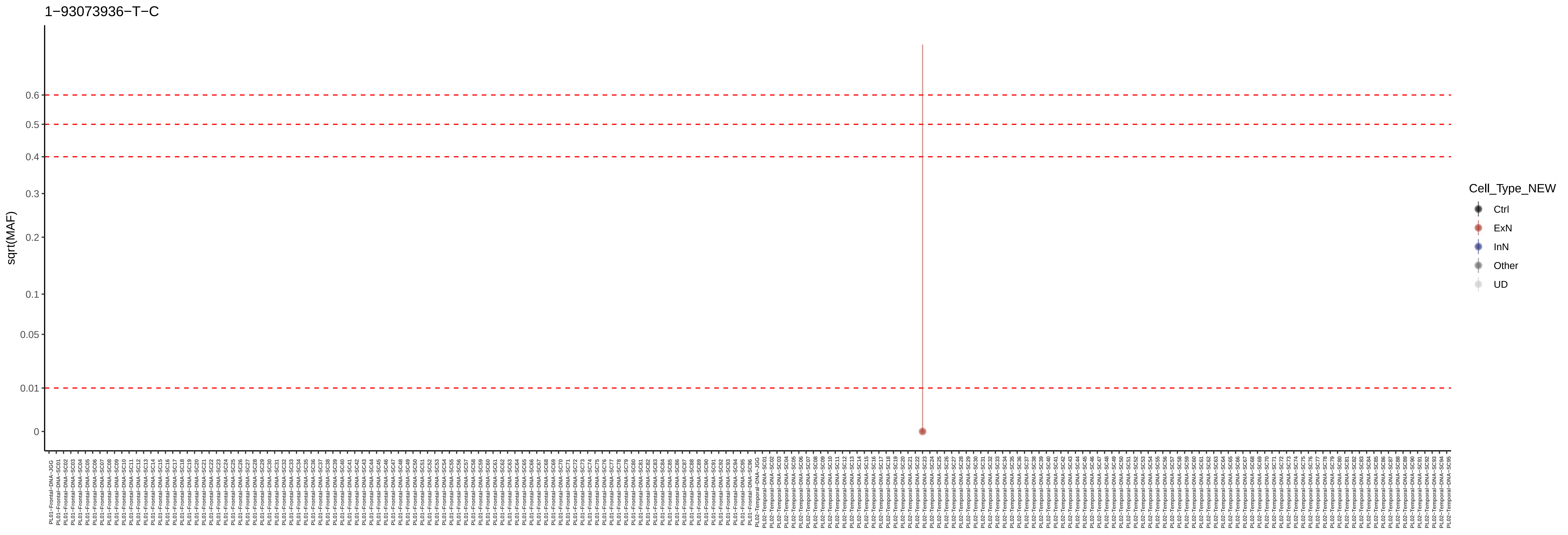
1−93073936−T−C
sqrt(MAF)
0.6
0.5
0.4
0.3
0.2
0.1
0.05
0.01
0
Cell_Type_NEW
Ctrl
ExN
InN
Other
UD

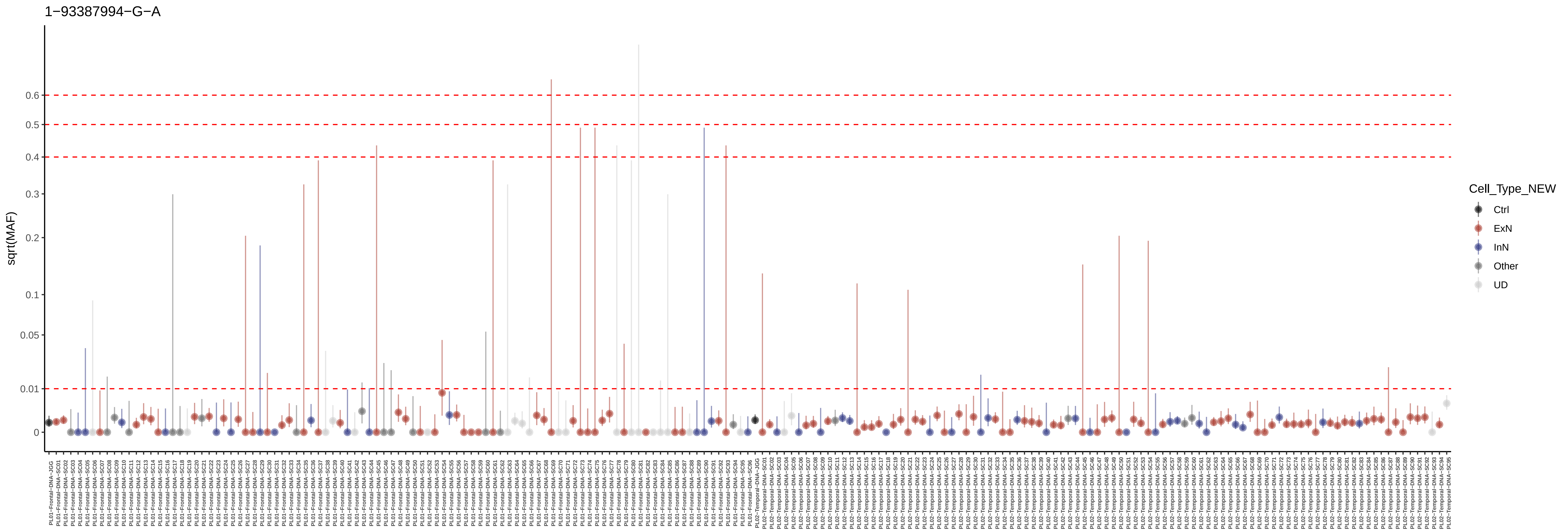
1−93387994−G−A
sqrt(MAF)
0.6
0.5
0.4
0.3
0.2
0.1
0.05
0.01
0
Cell_Type_NEW
Ctrl
ExN
InN
Other
UD

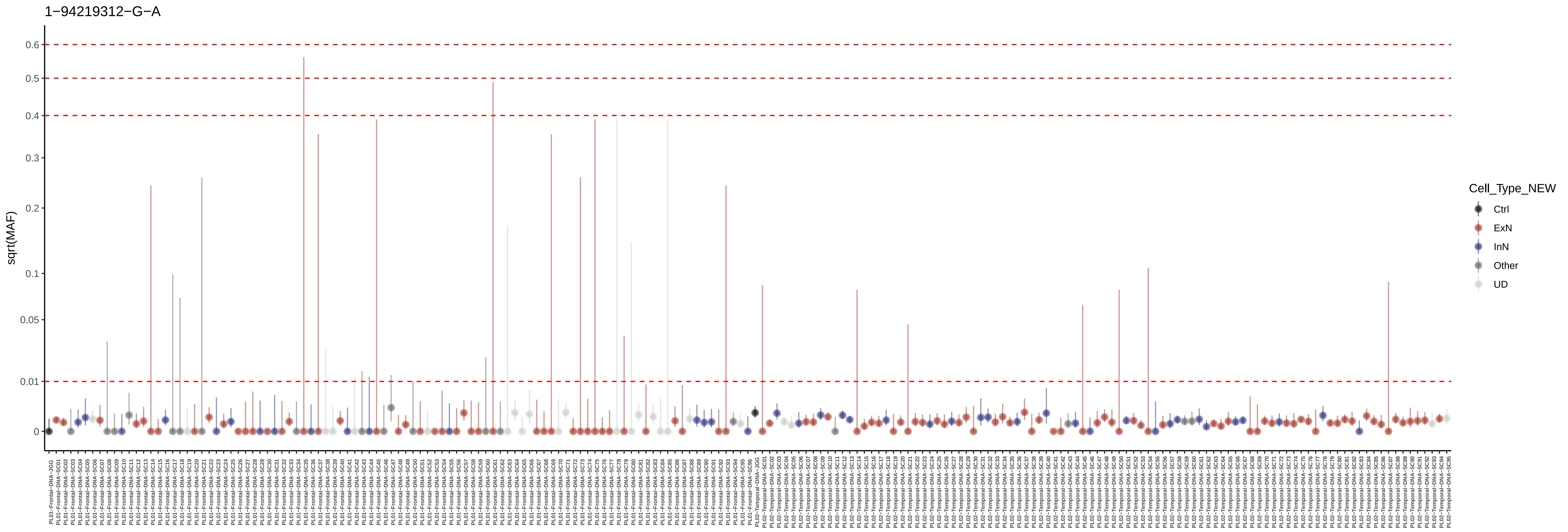
1−94219312−G−A
sqrt(MAF)
0.6
0.5
0.4
0.3
0.2
0.1
0.05
0.01
0
Cell_Type_NEW
Ctrl
ExN
InN
Other
UD

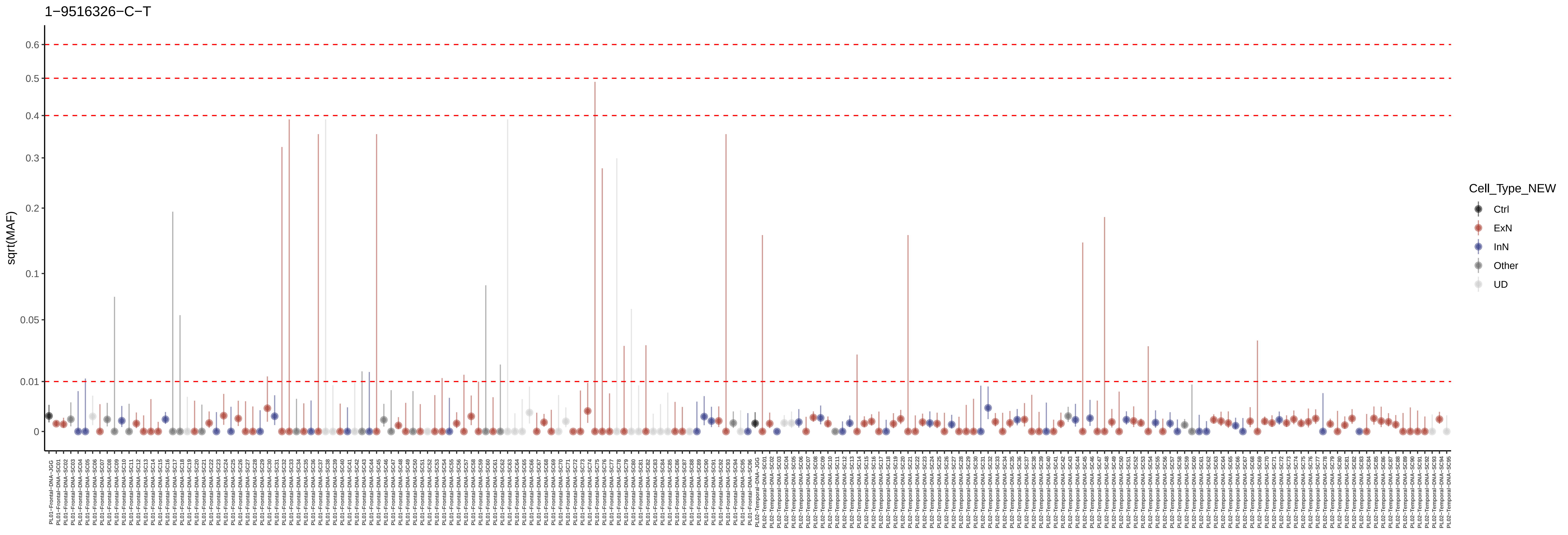
1−9516326−C−T
sqrt(MAF)
0.6
0.5
0.4
0.3
0.2
0.1
0.05
0.01
0
Cell_Type_NEW
Ctrl
ExN
InN
Other
UD

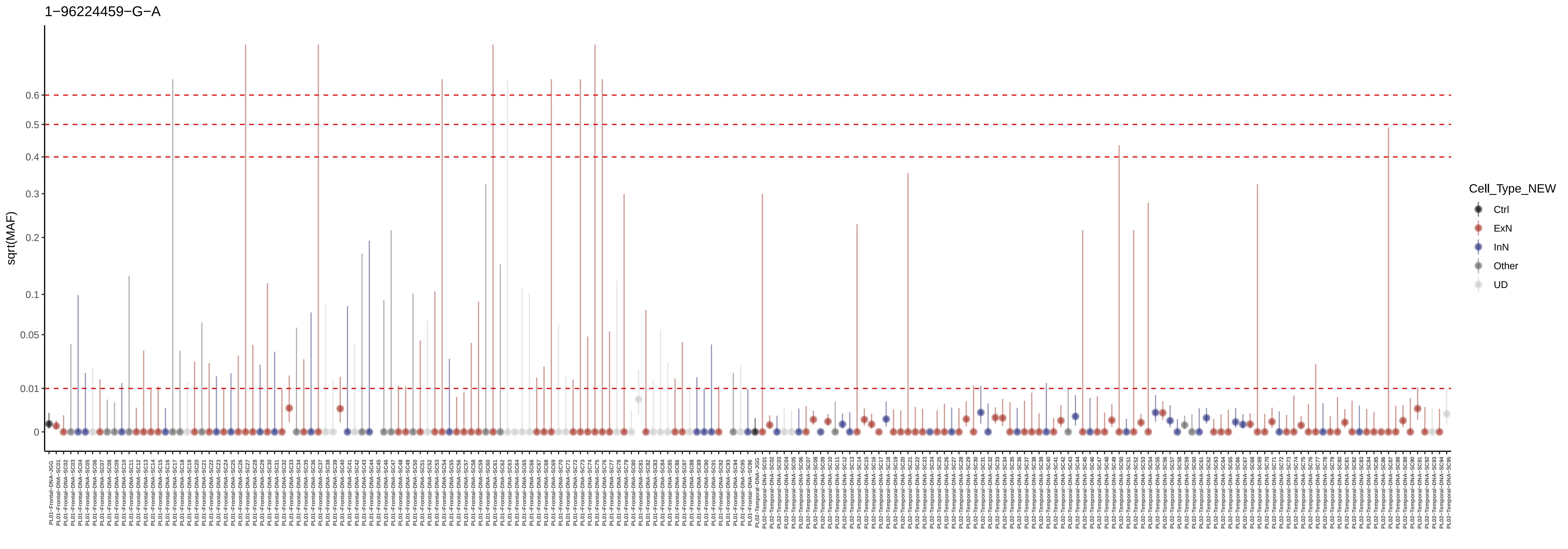
1−96224459−G−A
sqrt(MAF)
0.6
0.5
0.4
0.3
0.2
0.1
0.05
0.01
0
Cell_Type_NEW
Ctrl
ExN
InN
Other
UD

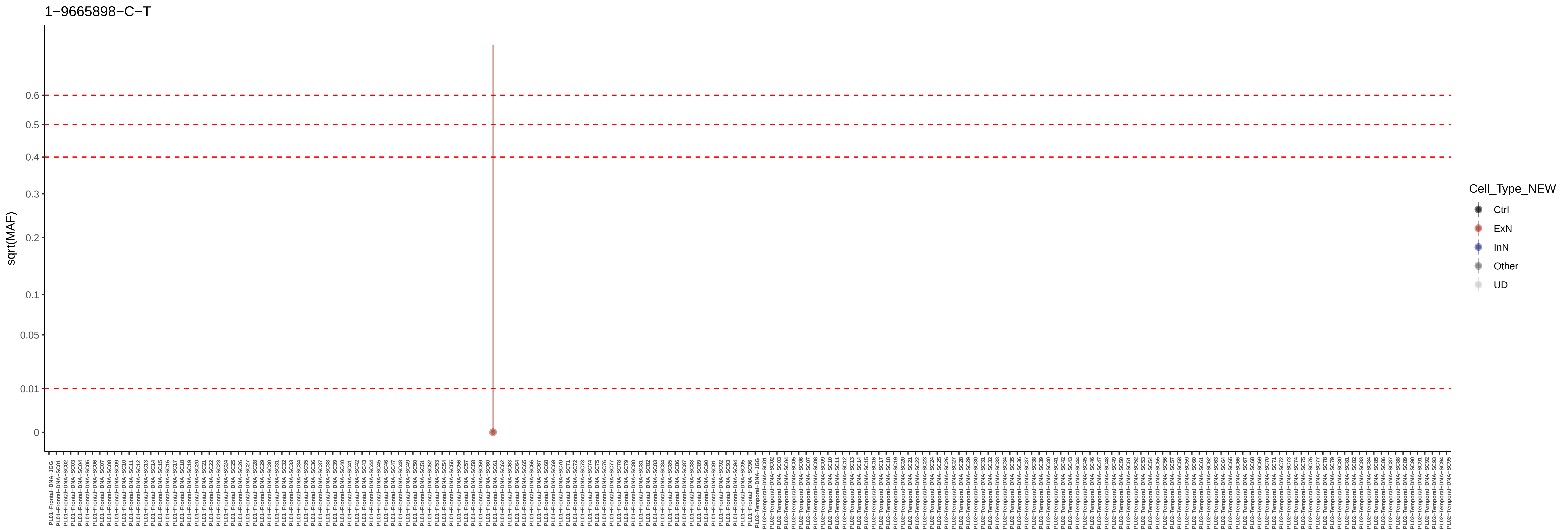

1−9665898−C−T
sqrt(MAF)
0.6
0.5
0.4
0.3
0.2
0.1
0.05
0.01
0
Cell_Type_NEW
Ctrl
ExN
InN
Other
UD

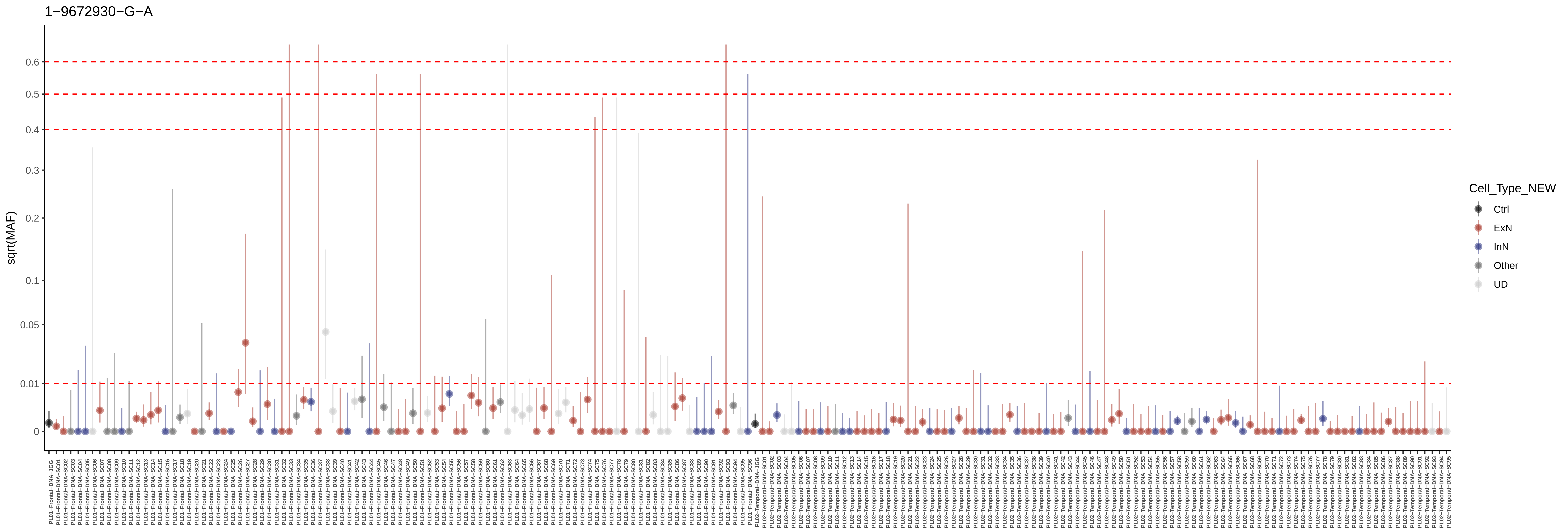
1−9672930−G−A
sqrt(MAF)
0.6
0.5
0.4
0.3
0.2
0.1
0.05
0.01
0
Cell_Type_NEW
Ctrl
ExN
InN
Other
UD

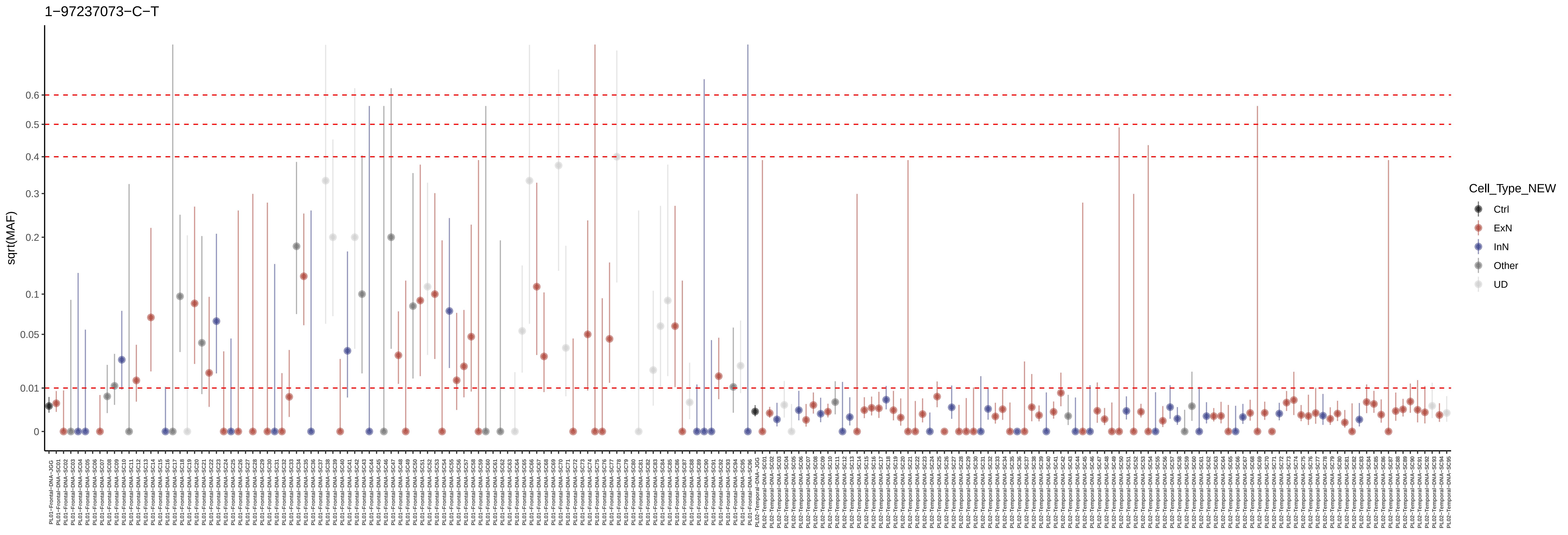
1−97237073−C−T
sqrt(MAF)
0.6
0.5
0.4
0.3
0.2
0.1
0.05
0.01
0
Cell_Type_NEW
Ctrl
ExN
InN
Other
UD

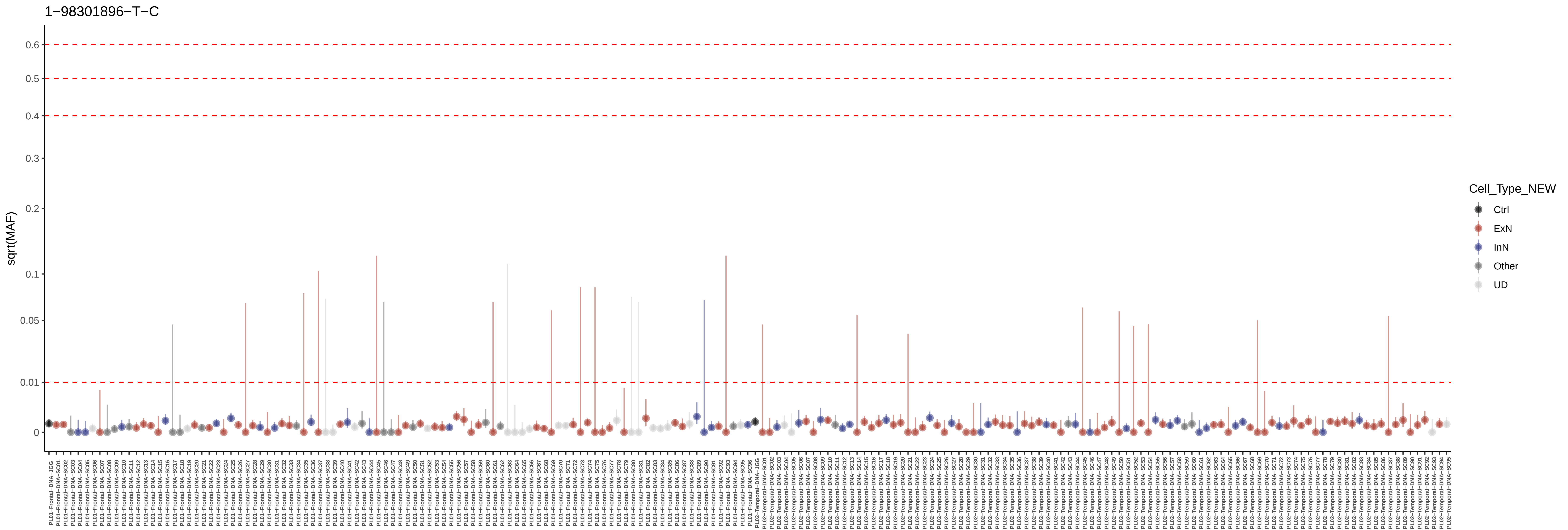
1−98301896−T−C
sqrt(MAF)
0.6
0.5
0.4
0.3
0.2
0.1
0.05
0.01
0
Cell_Type_NEW
Ctrl
ExN
InN
Other
UD

1−99606489−C−CT

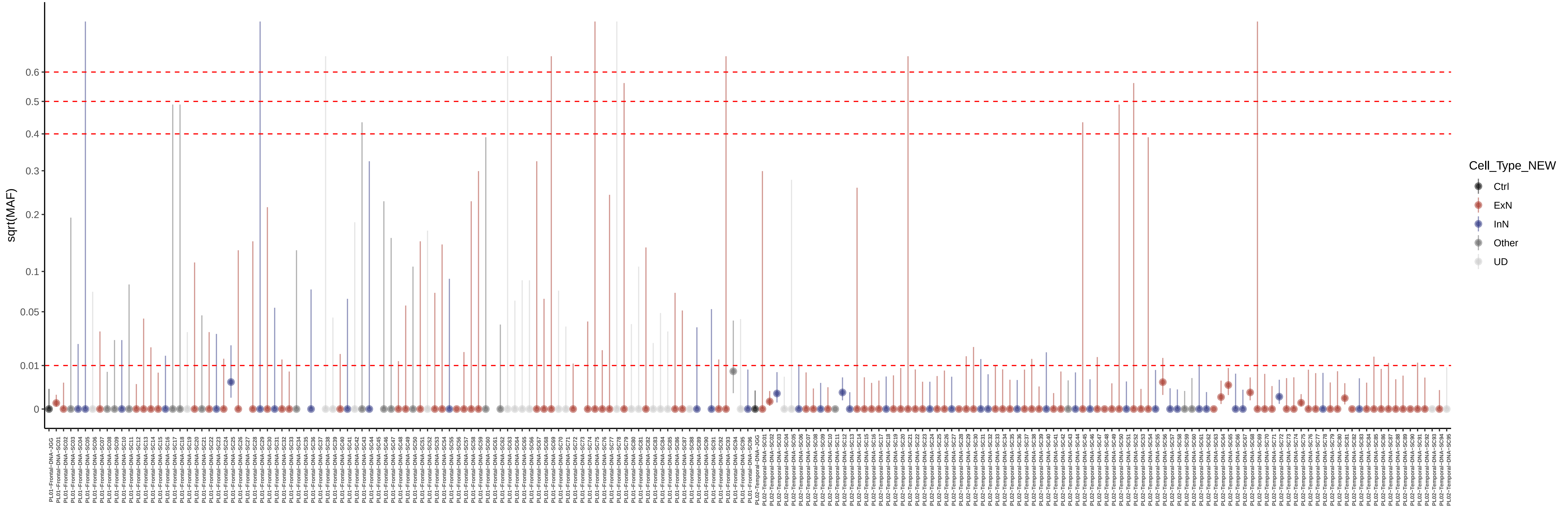

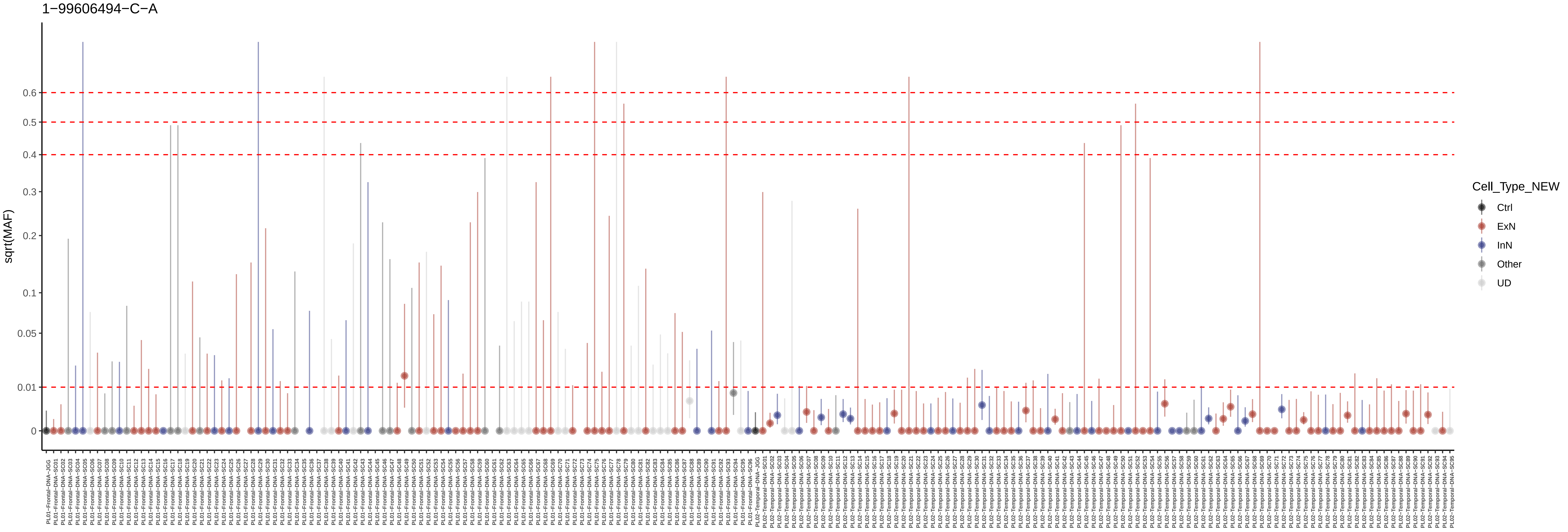
1−99606494−C−A
sqrt(MAF)
0.6
0.5
0.4
0.3
0.2
0.1
0.05
0.01
0
Cell_Type_NEW
Ctrl
ExN
InN
Other
UD

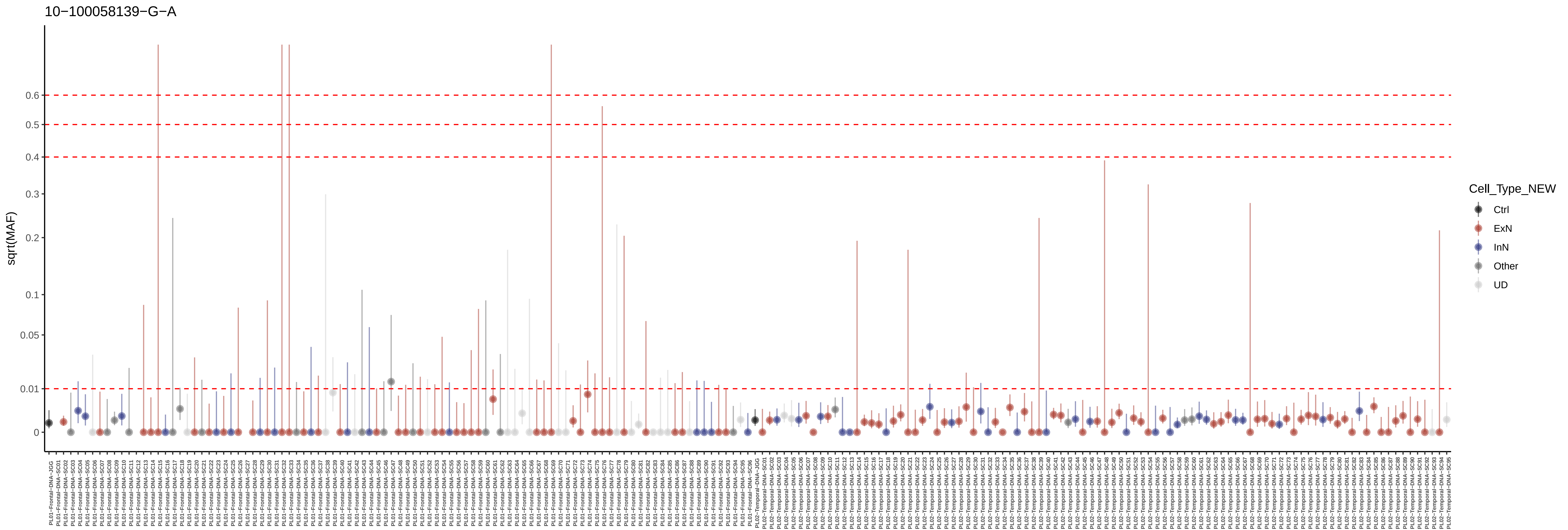
10−100058139−G−A
sqrt(MAF)
0.6
0.5
0.4
0.3
0.2
0.1
0.05
0.01
0
Cell_Type_NEW
Ctrl
ExN
InN
Other
UD

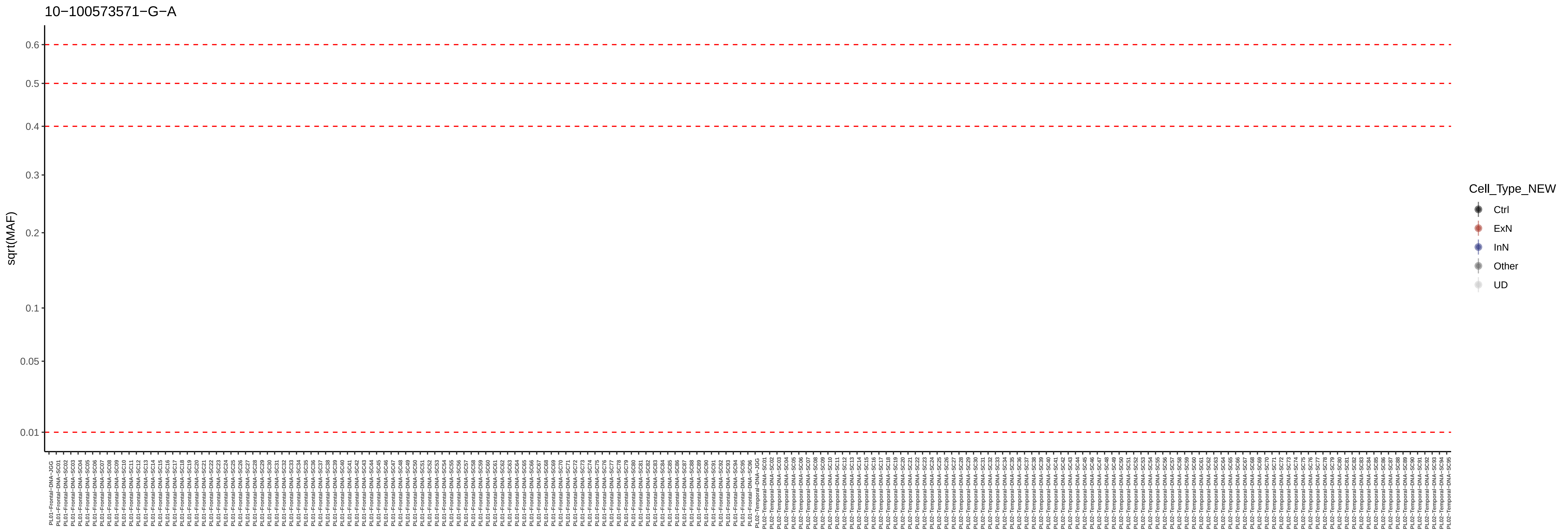

10−100573571−G−A
sqrt(MAF)
0.6
0.5
0.4
0.3
0.2
0.1
0.05
0.01
Cell_Type_NEW
Ctrl
ExN
InN
Other
UD

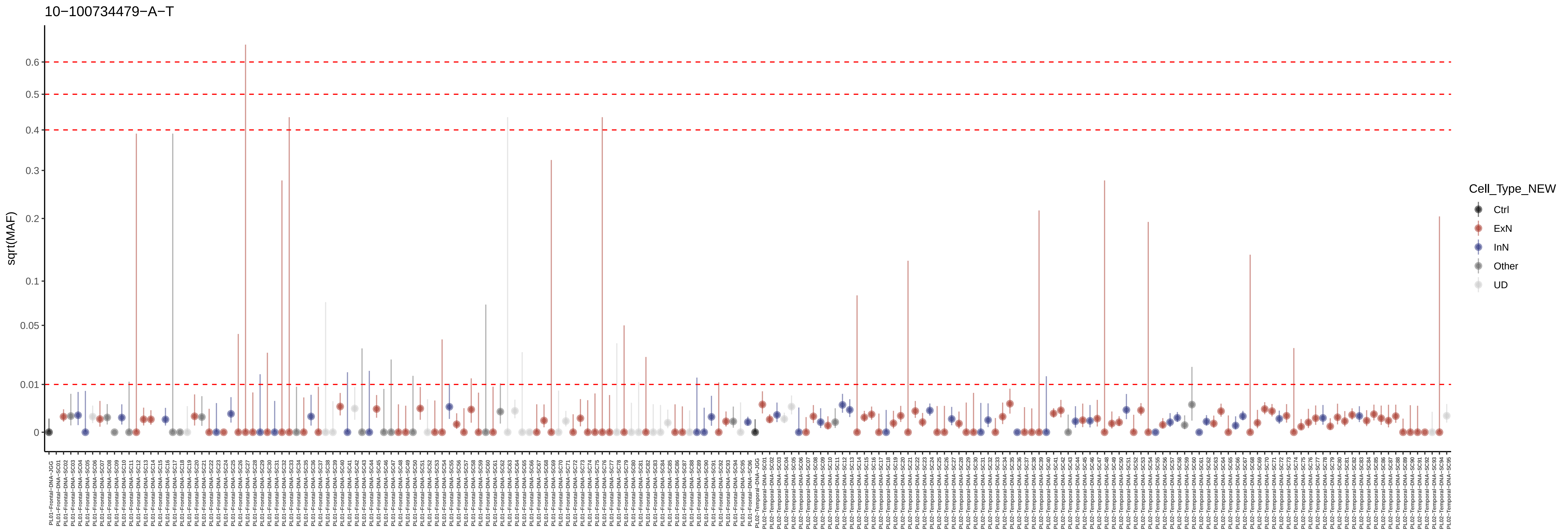

10−100734479−A−T
sqrt(MAF)
0.6
0.5
0.4
0.3
0.2
0.1
0.05
0.01
0
Cell_Type_NEW
Ctrl
ExN
InN
Other
UD

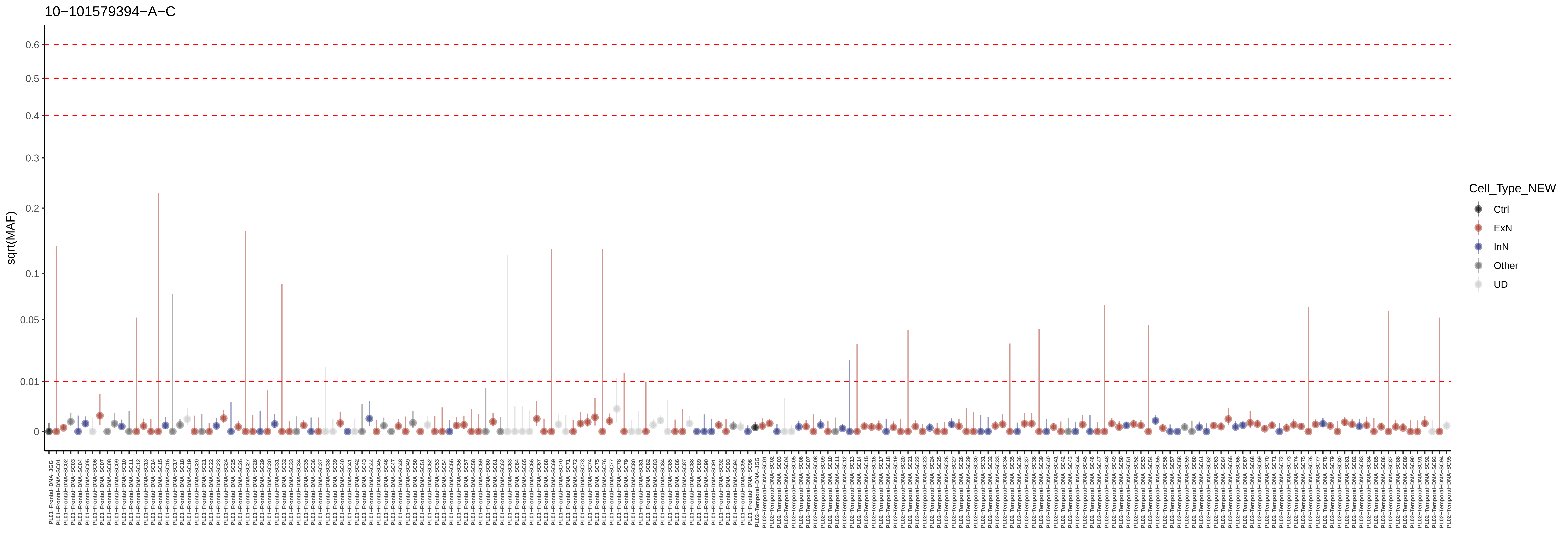

10−101579394−A−C
sqrt(MAF)
0.6
0.5
0.4
0.3
0.2
0.1
0.05
0.01
0
Cell_Type_NEW
Ctrl
ExN
InN
Other
UD

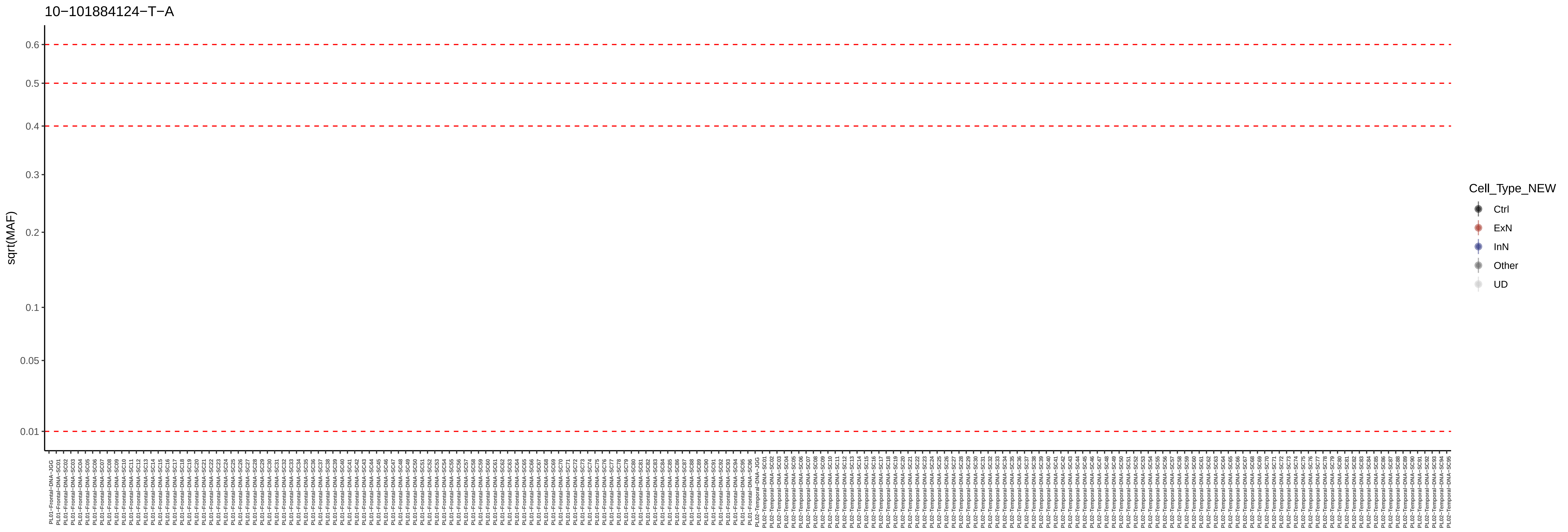
10−101884124−T−A
sqrt(MAF)
0.6
0.5
0.4
0.3
0.2
0.1
0.05
0.01
Cell_Type_NEW
Ctrl
ExN
InN
Other
UD

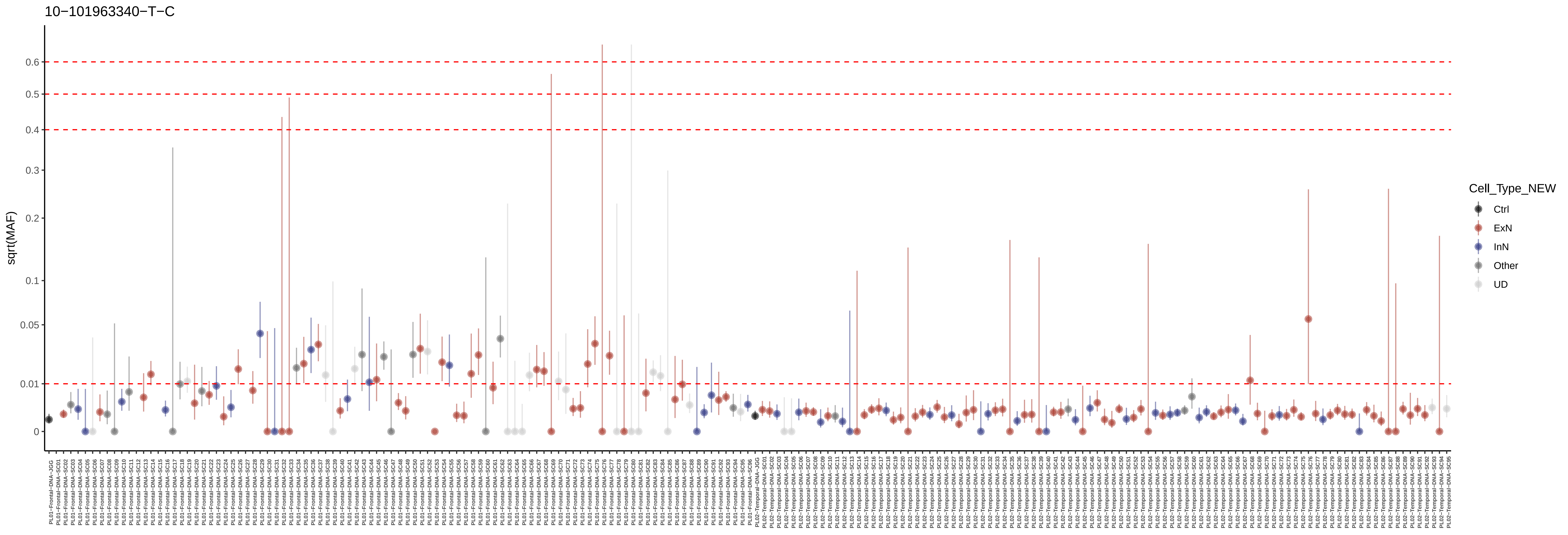
10−101963340−T−C
sqrt(MAF)
0.6
0.5
0.4
0.3
0.2
0.1
0.05
0.01
0
Cell_Type_NEW
Ctrl
ExN
InN
Other
UD

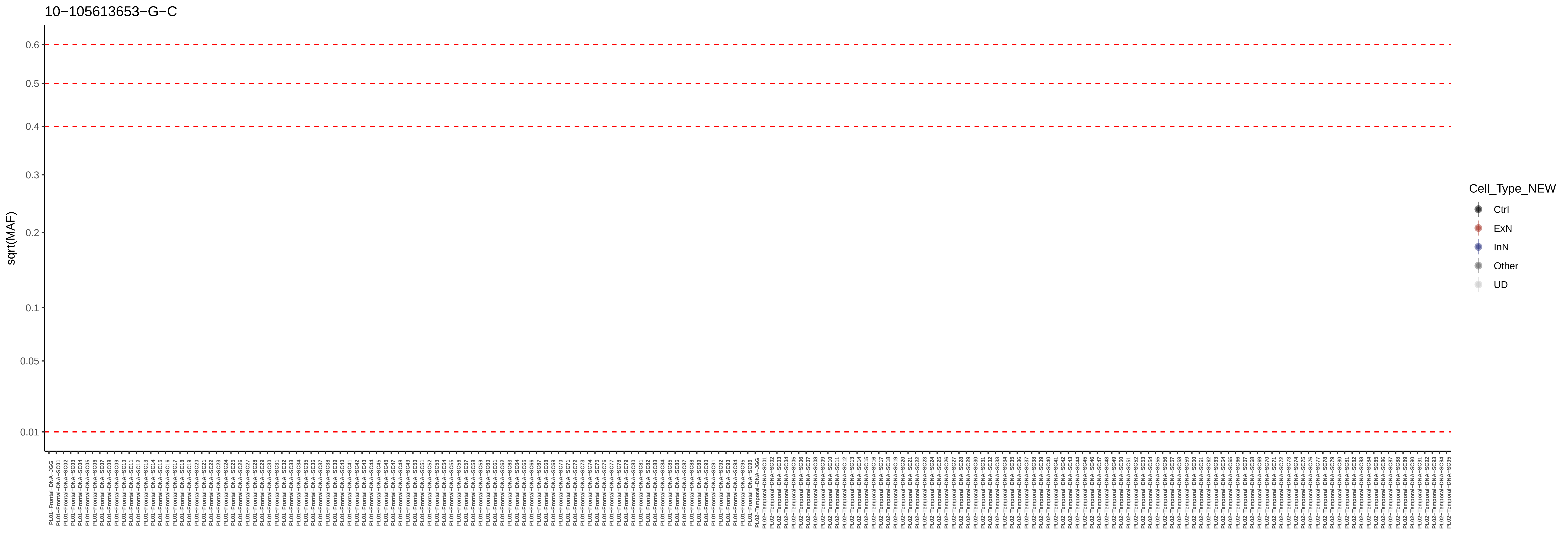

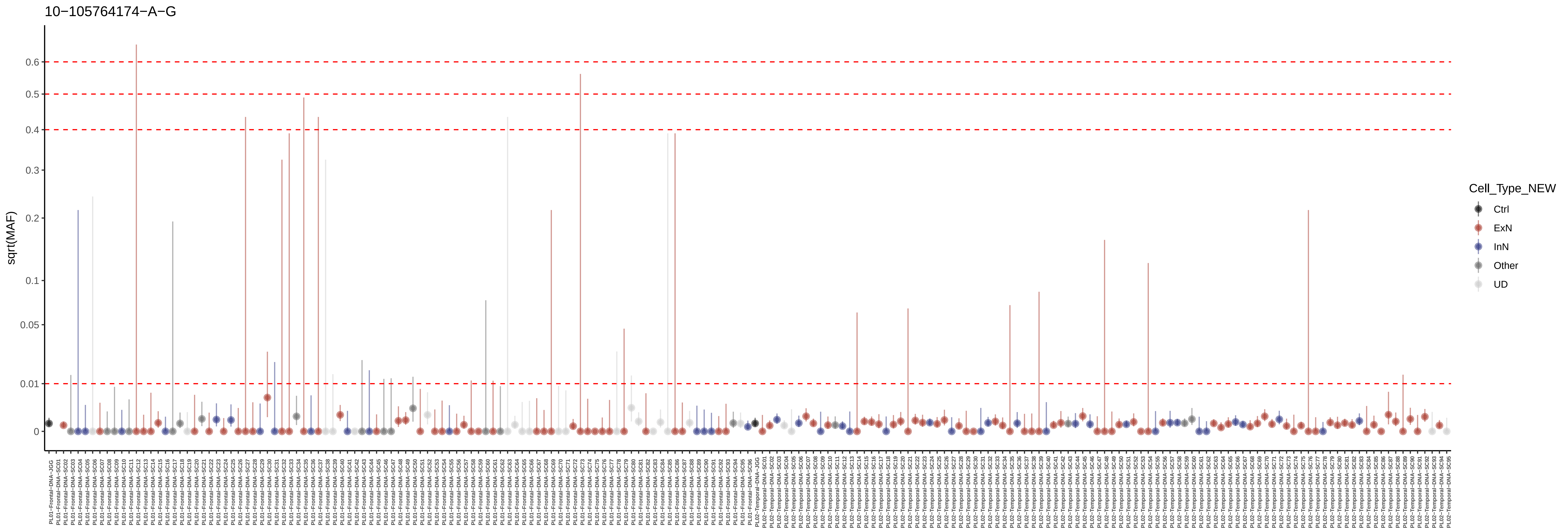
10−105764174−A−G
sqrt(MAF)
0.6
0.5
0.4
0.3
0.2
0.1
0.05
0.01
0
Cell_Type_NEW
Ctrl
ExN
InN
Other
UD

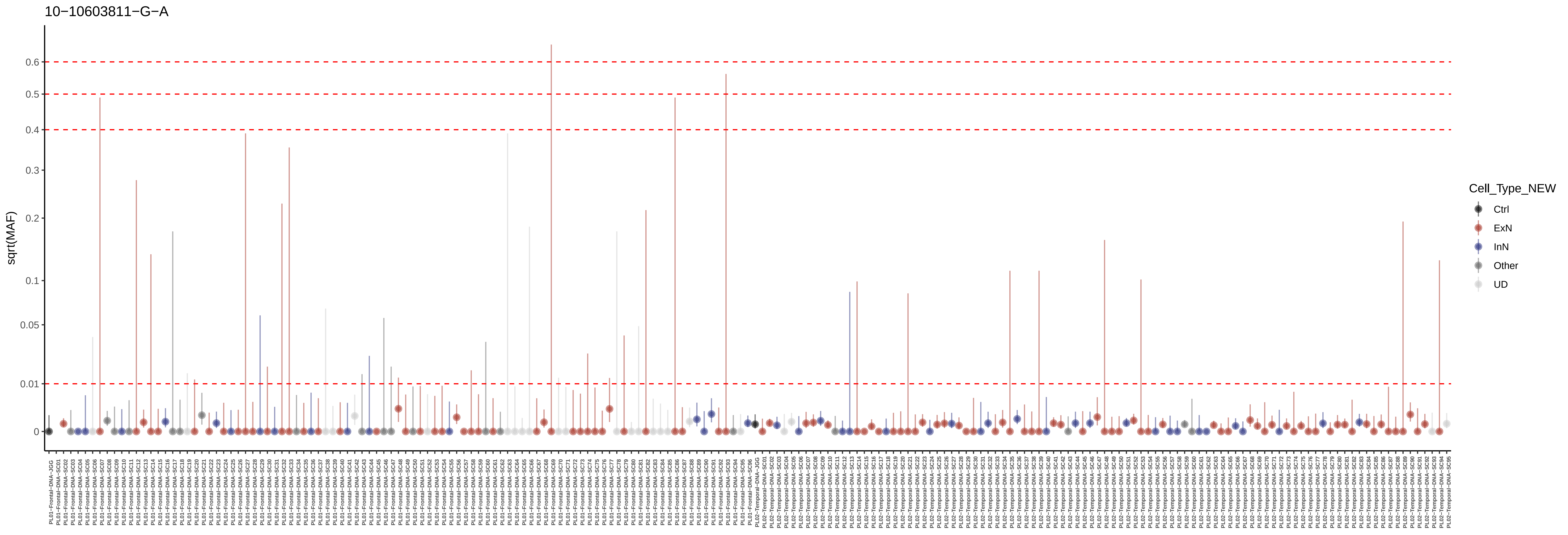

10−10603811−G−A
sqrt(MAF)
0.6
0.5
0.4
0.3
0.2
0.1
0.05
0.01
0
Cell_Type_NEW
Ctrl
ExN
InN
Other
UD

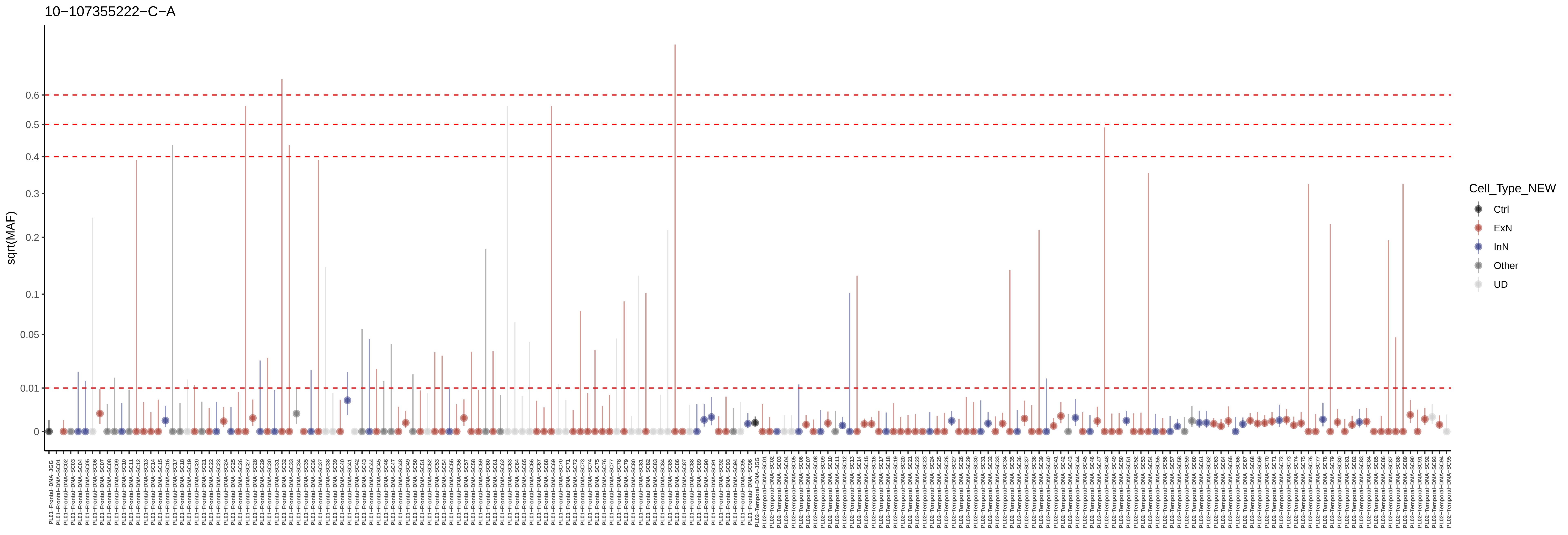
10−107355222−C−A
sqrt(MAF)
0.6
0.5
0.4
0.3
0.2
0.1
0.05
0.01
0
Cell_Type_NEW
Ctrl
ExN
InN
Other
UD

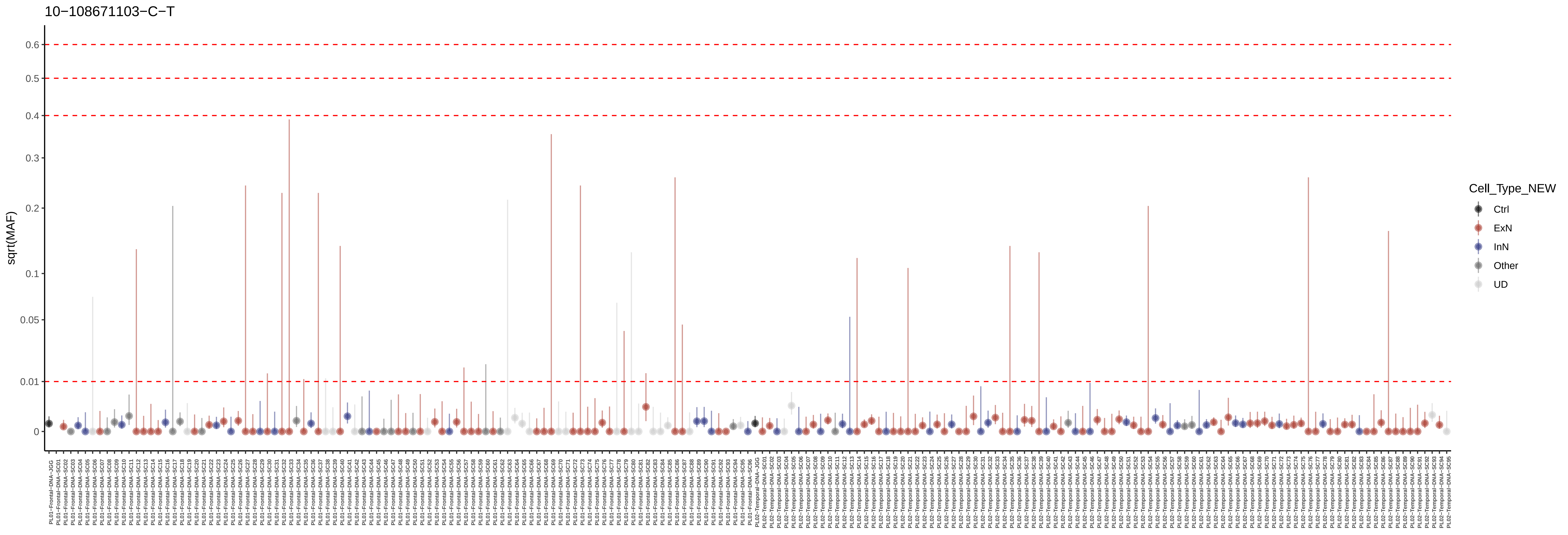

10−108671103−C−T
sqrt(MAF)
0.6
0.5
0.4
0.3
0.2
0.1
0.05
0.01
0
Cell_Type_NEW
Ctrl
ExN
InN
Other
UD

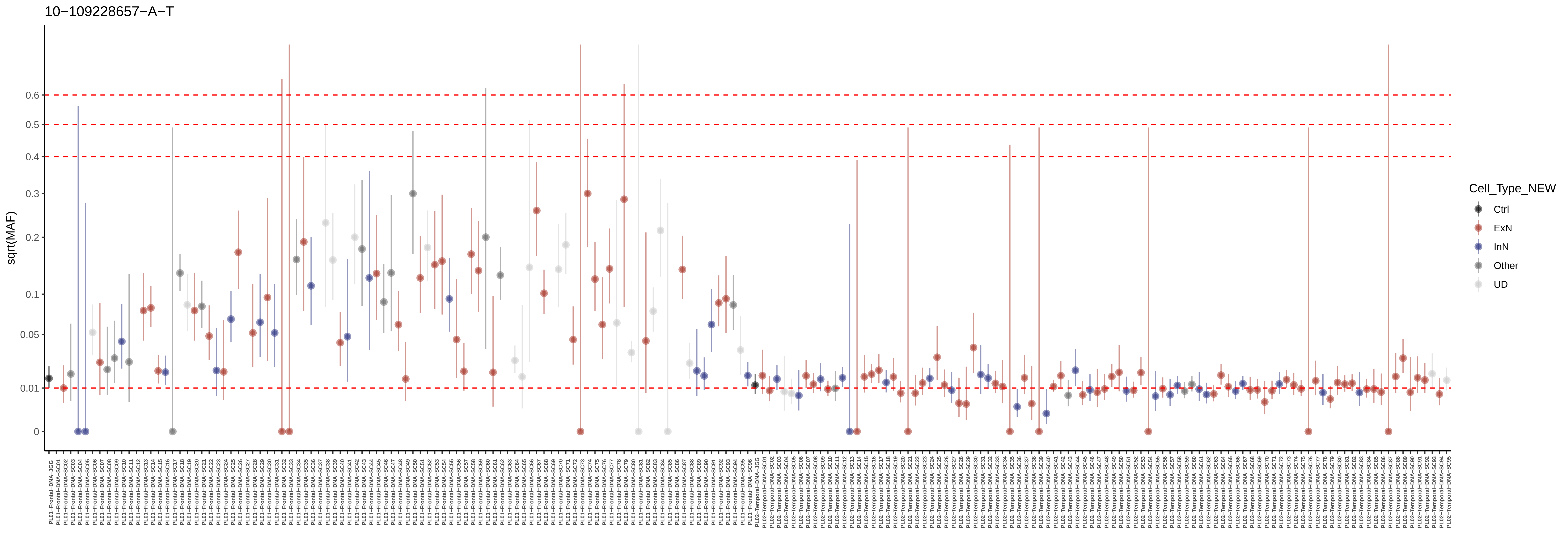
10−109228657−A−T
sqrt(MAF)
0.6
0.5
0.4
0.3
0.2
0.1
0.05
0.01
0
Cell_Type_NEW
Ctrl
ExN
InN
Other
UD

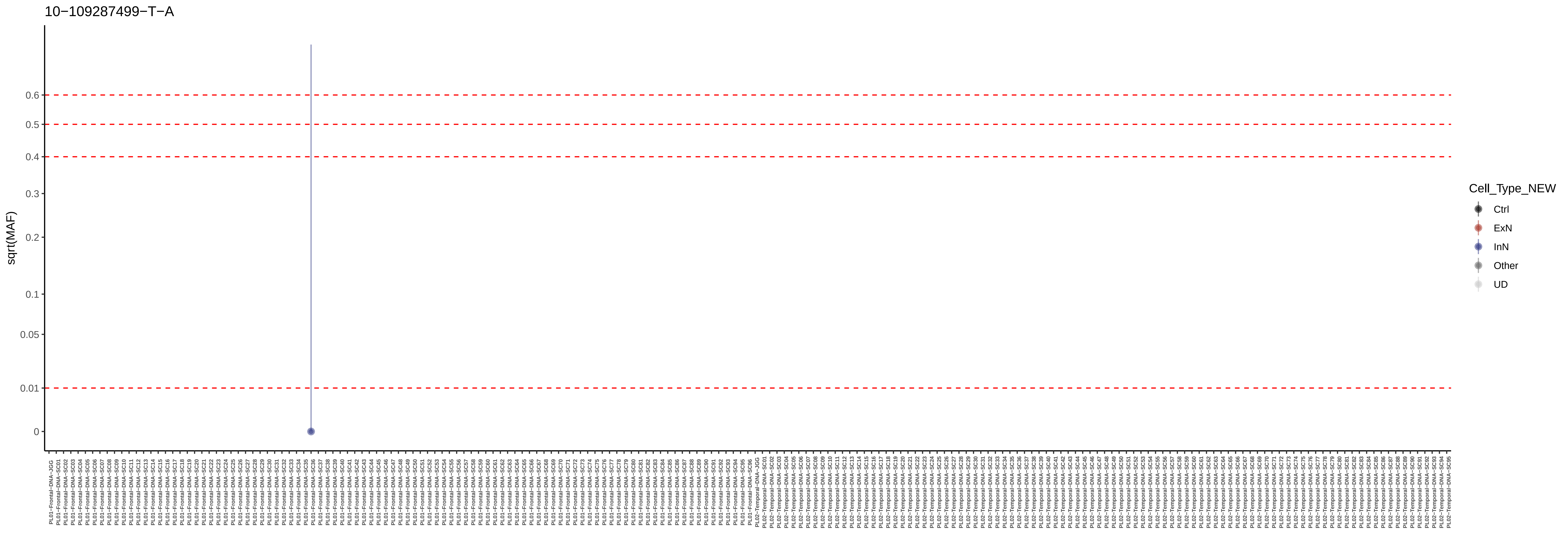
10−109287499−T−A
sqrt(MAF)
0.6
0.5
0.4
0.3
0.2
0.1
0.05
0.01
0
Cell_Type_NEW
Ctrl
ExN
InN
Other
UD

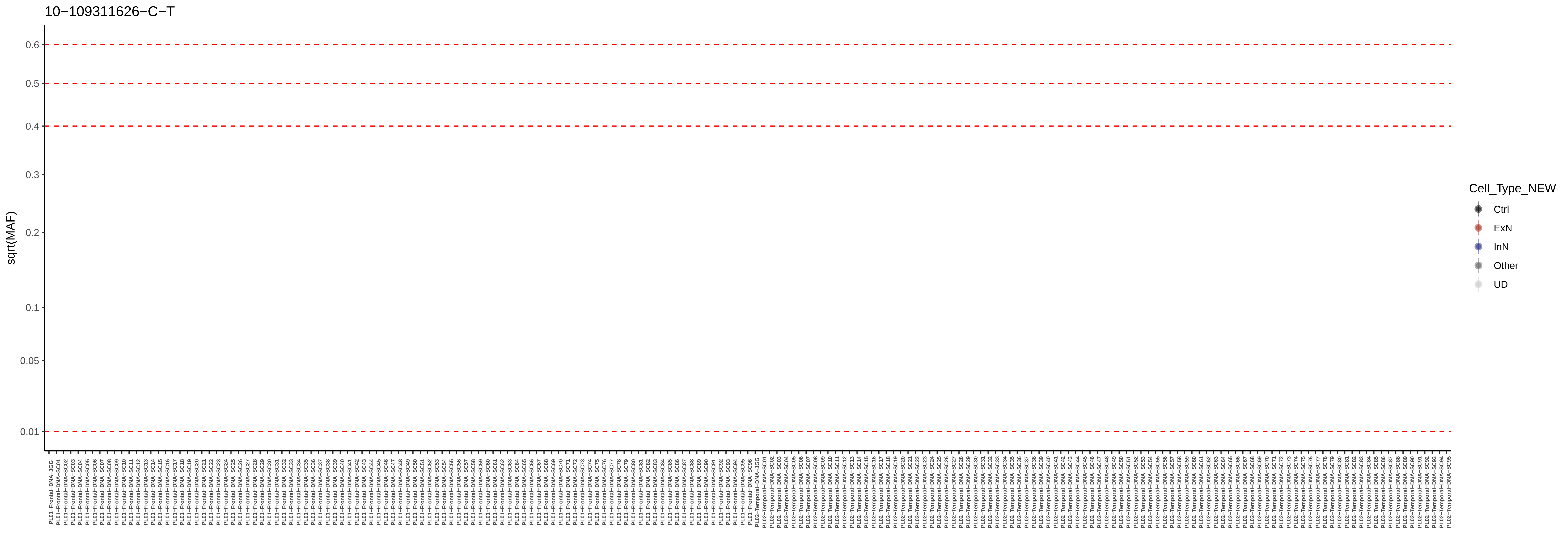
10−109311626−C−T
sqrt(MAF)
0.6
0.5
0.4
0.3
0.2
0.1
0.05
0.01
Cell_Type_NEW
Ctrl
ExN
InN
Other
UD

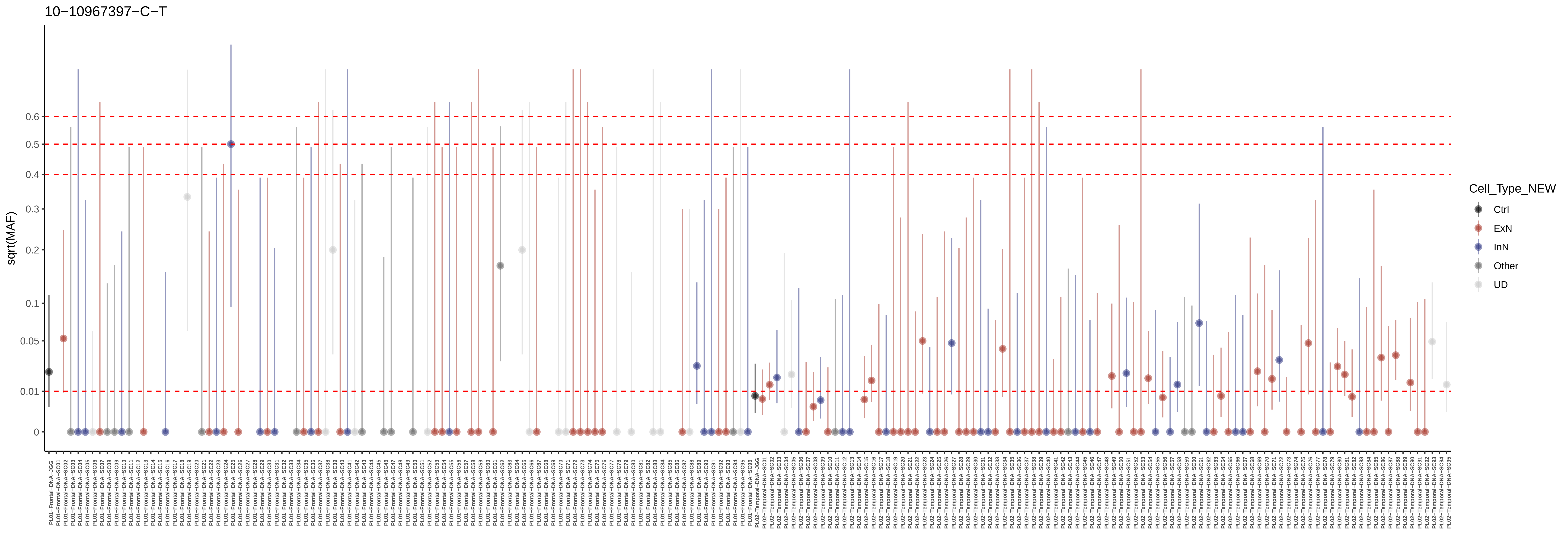
10−10967397−C−T
sqrt(MAF)
0.6
0.5
0.4
0.3
0.2
0.1
0.05
0.01
0
Cell_Type_NEW
Ctrl
ExN
InN
Other
UD

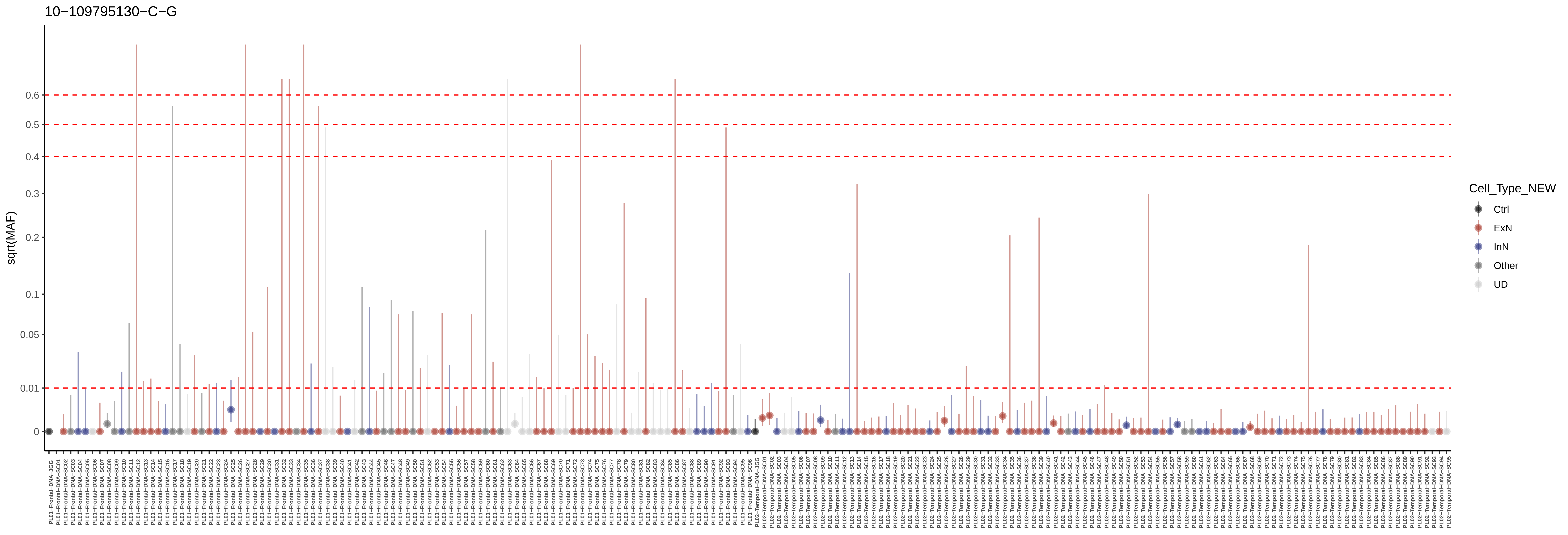
10−109795130−C−G
sqrt(MAF)
0.6
0.5
0.4
0.3
0.2
0.1
0.05
0.01
0
Cell_Type_NEW
Ctrl
ExN
InN
Other
UD

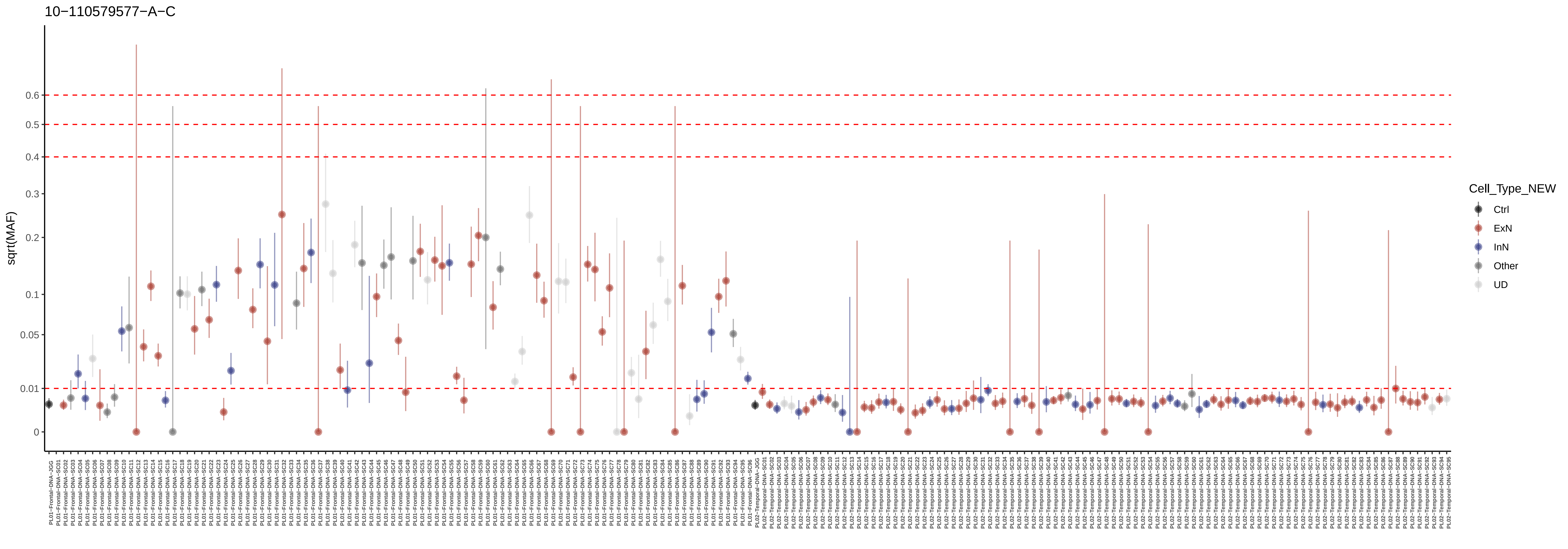
10−110579577−A−C
sqrt(MAF)
0.6
0.5
0.4
0.3
0.2
0.1
0.05
0.01
0
Cell_Type_NEW
Ctrl
ExN
InN
Other
UD

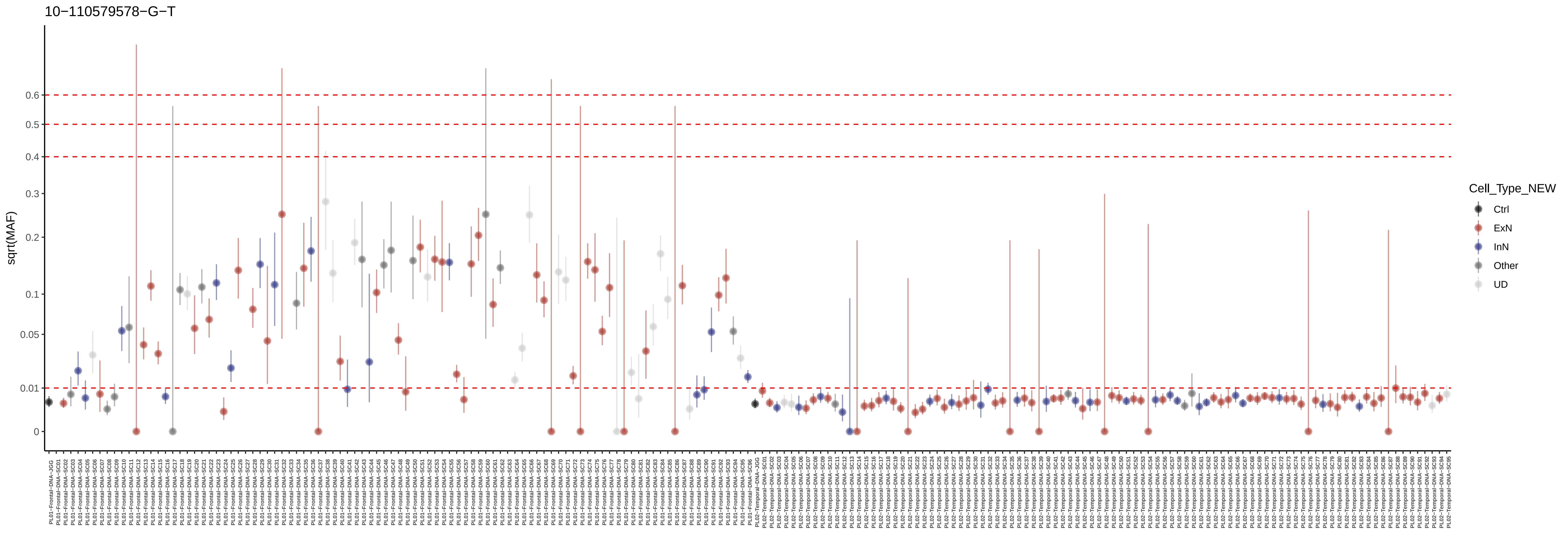

10−110579578−G−T
sqrt(MAF)
0.6
0.5
0.4
0.3
0.2
0.1
0.05
0.01
0
Cell_Type_NEW
Ctrl
ExN
InN
Other
UD

10−11281463−T−C

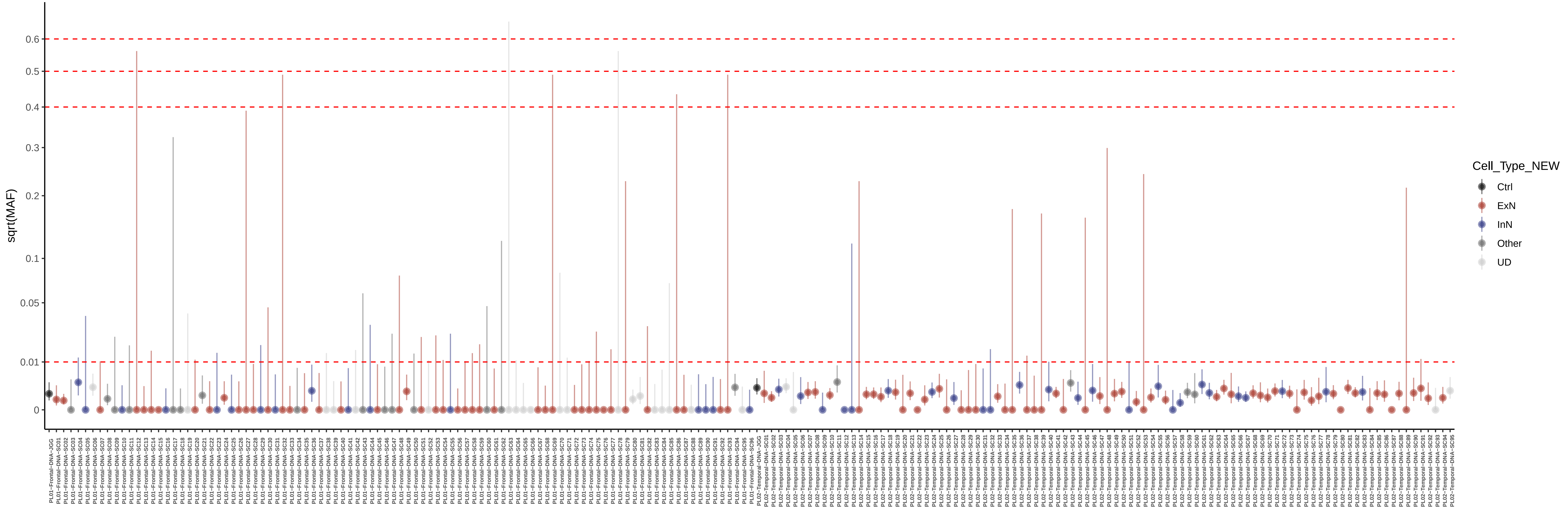

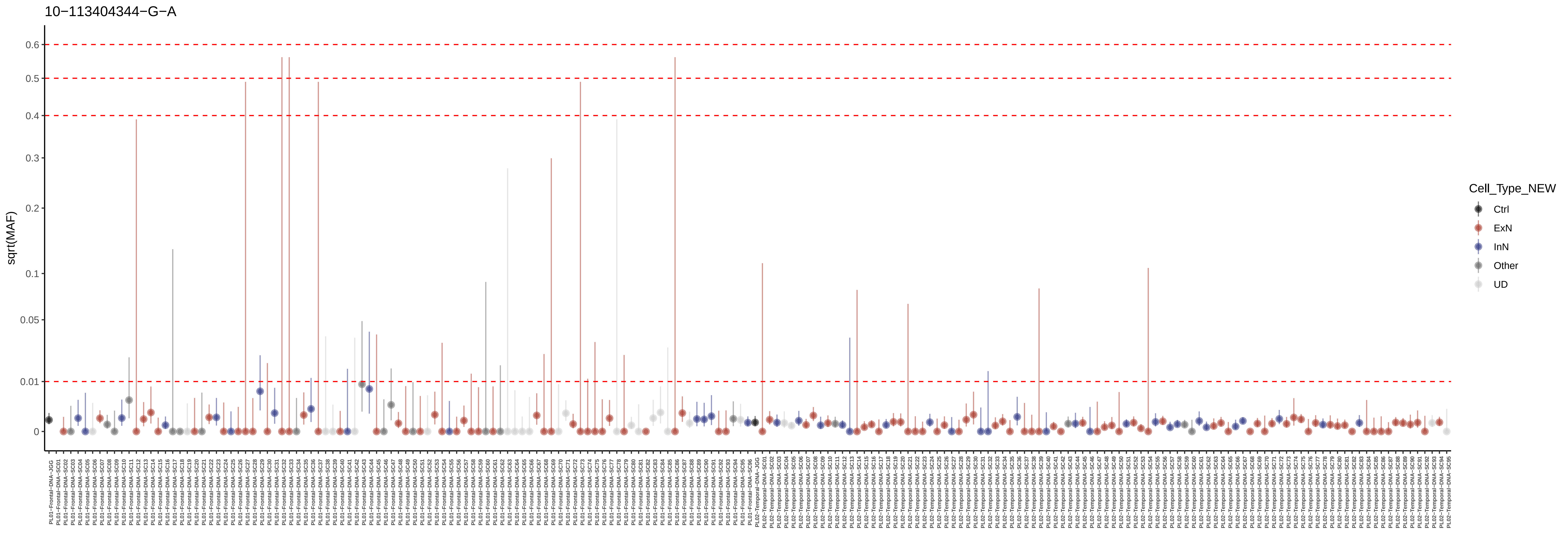
10−113404344−G−A
sqrt(MAF)
0.6
0.5
0.4
0.3
0.2
0.1
0.05
0.01
0
Cell_Type_NEW
Ctrl
ExN
InN
Other
UD

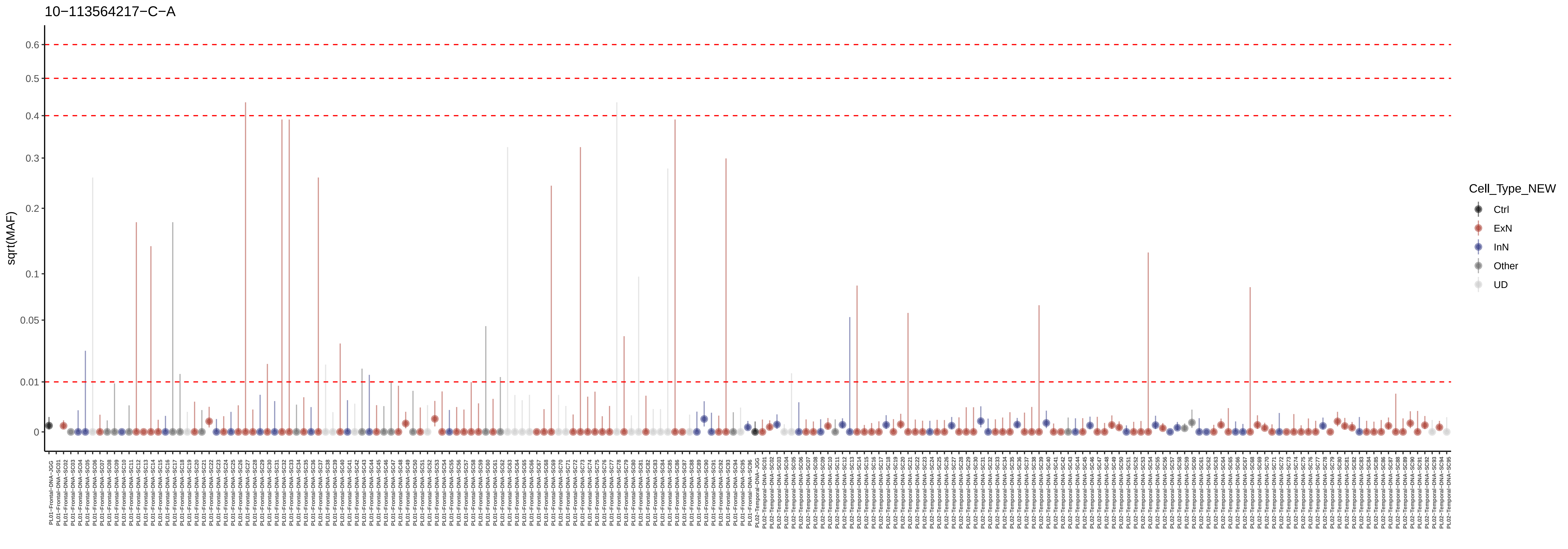
10−113564217−C−A
sqrt(MAF)
0.6
0.5
0.4
0.3
0.2
0.1
0.05
0.01
0
Cell_Type_NEW
Ctrl
ExN
InN
Other
UD

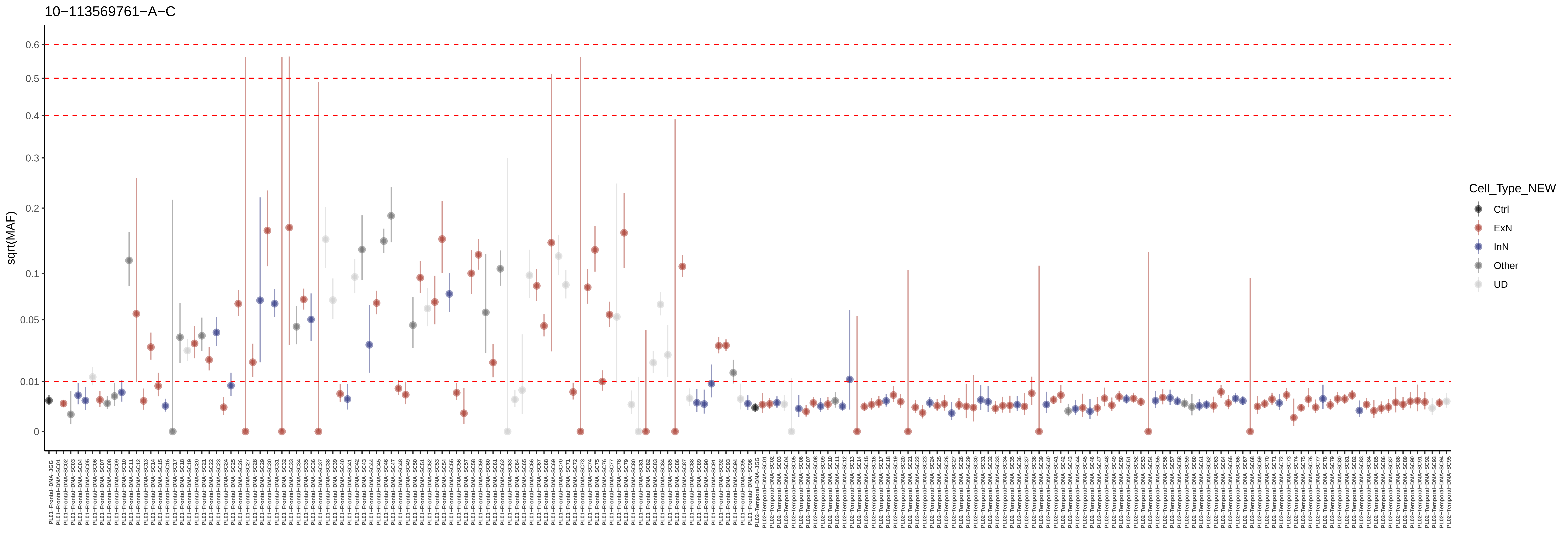

10−113569761−A−C
sqrt(MAF)
0.6
0.5
0.4
0.3
0.2
0.1
0.05
0.01
0
Cell_Type_NEW
Ctrl
ExN
InN
Other
UD

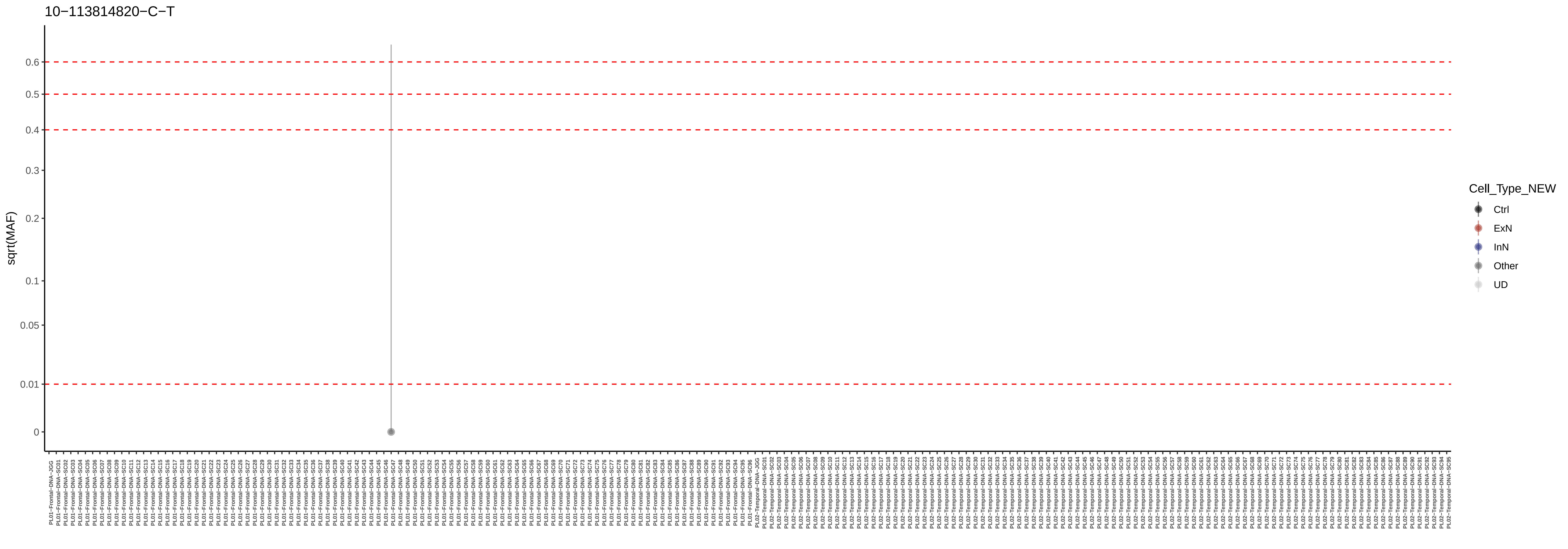
10−113814820−C−T
sqrt(MAF)
0.6
0.5
0.4
0.3
0.2
0.1
0.05
0.01
0
Cell_Type_NEW
Ctrl
ExN
InN
Other
UD

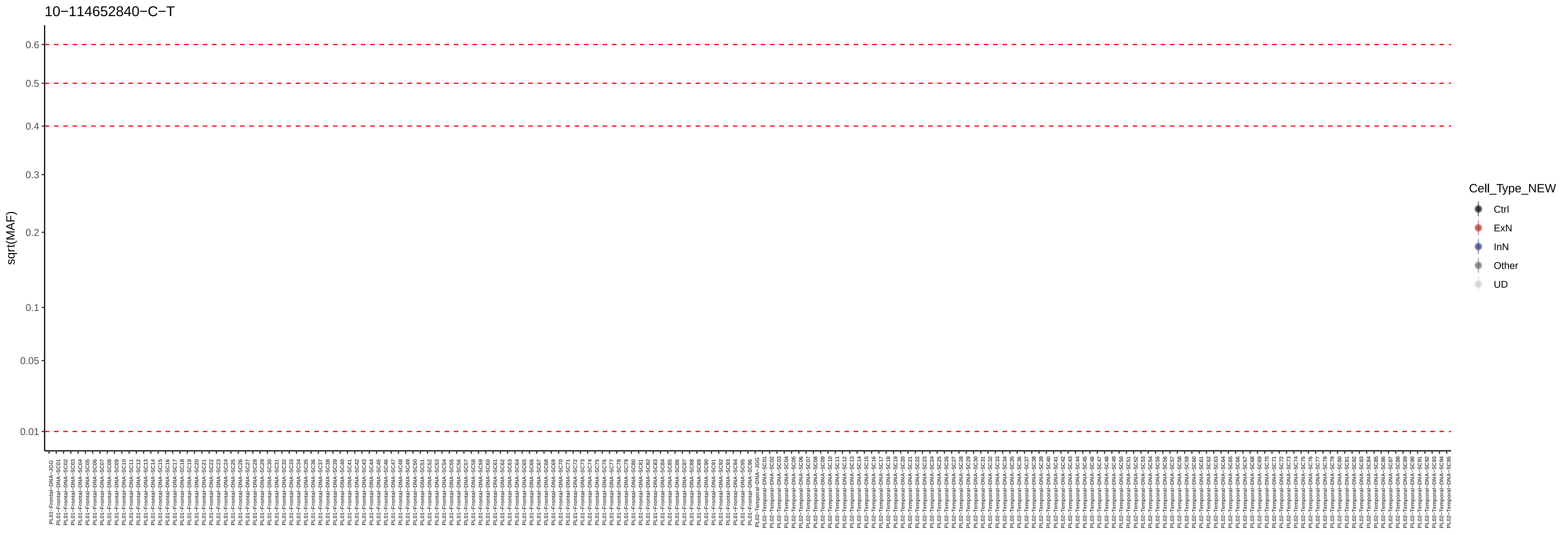
10−114652840−C−T
sqrt(MAF)
0.6
0.5
0.4
0.3
0.2
0.1
0.05
0.01
Cell_Type_NEW
Ctrl
ExN
InN
Other
UD

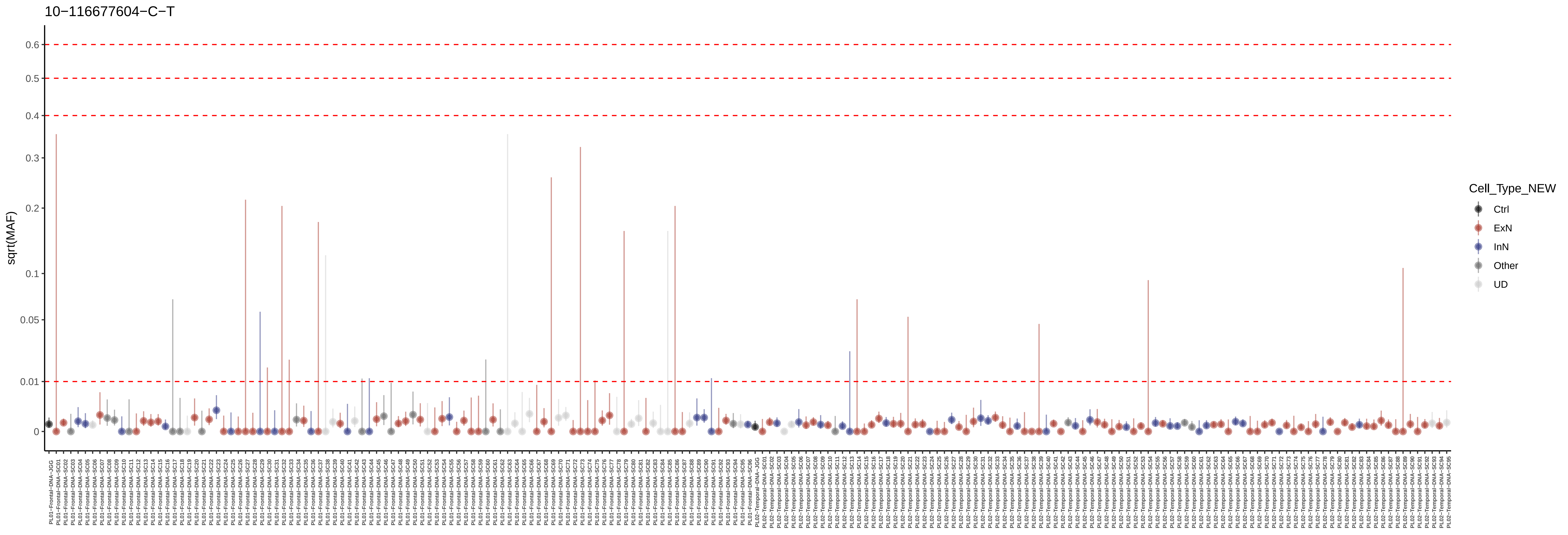
10−116677604−C−T
sqrt(MAF)
0.6
0.5
0.4
0.3
0.2
0.1
0.05
0.01
0
Cell_Type_NEW
Ctrl
ExN
InN
Other
UD

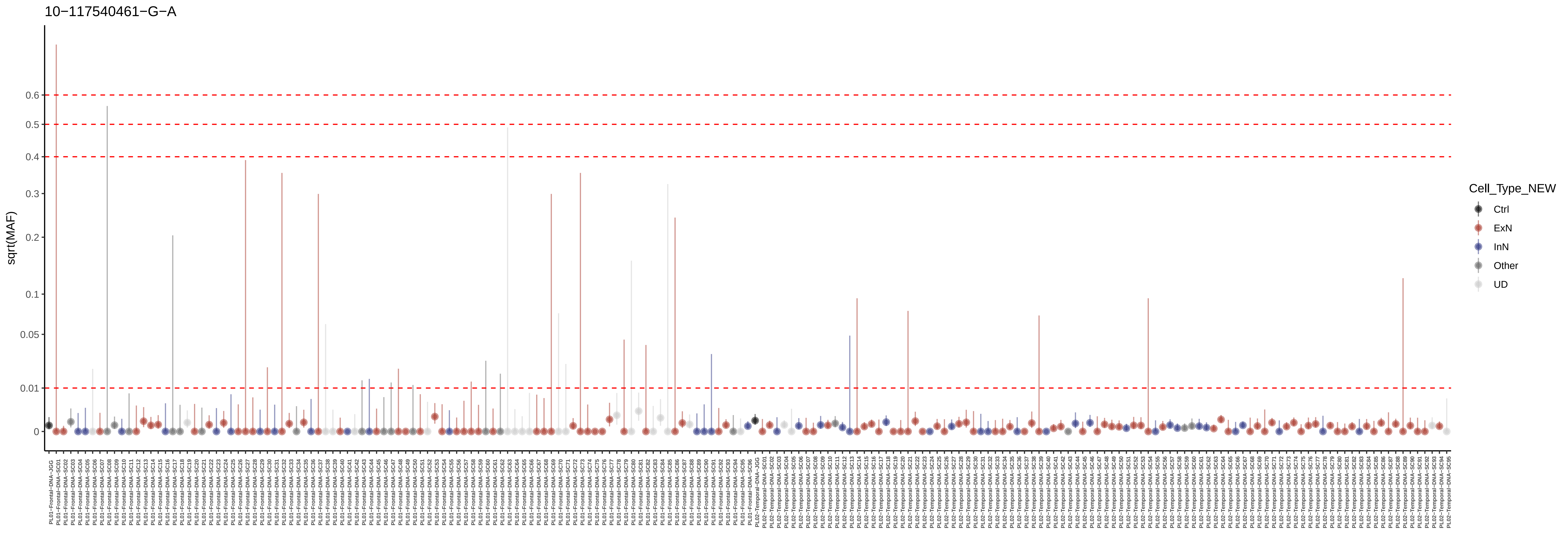
10−117540461−G−A
sqrt(MAF)
0.6
0.5
0.4
0.3
0.2
0.1
0.05
0.01
0
Cell_Type_NEW
Ctrl
ExN
InN
Other
UD

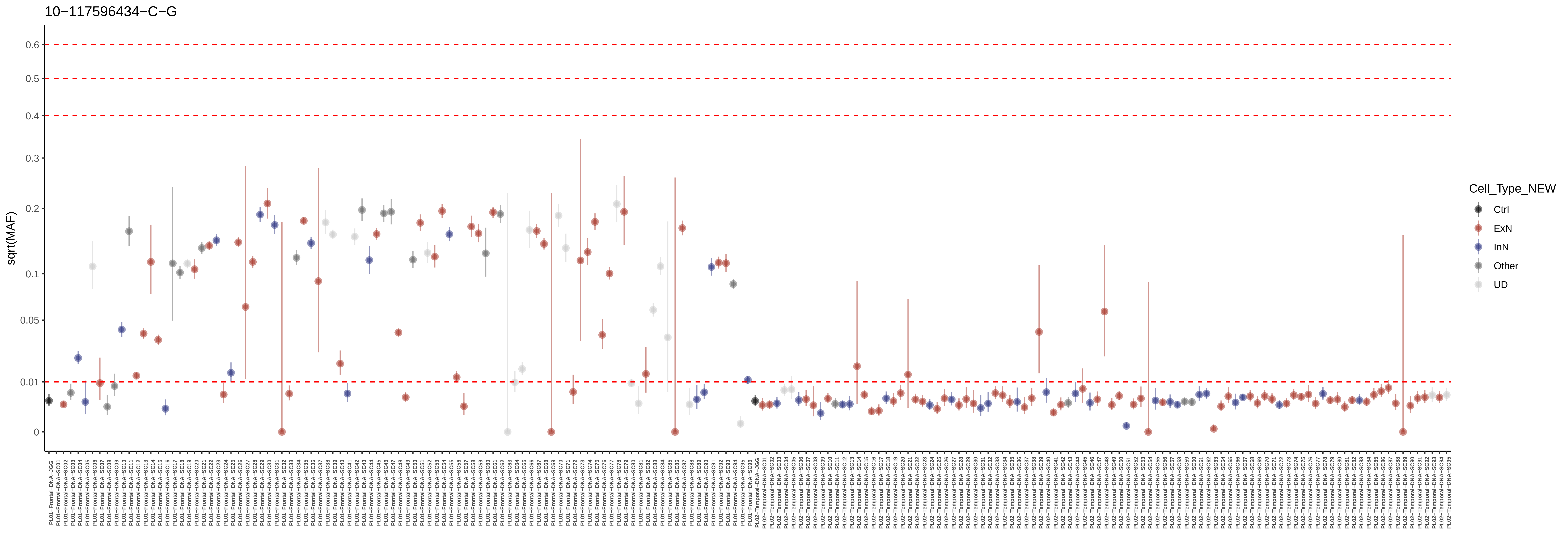
10−117596434−C−G
sqrt(MAF)
0.6
0.5
0.4
0.3
0.2
0.1
0.05
0.01
0
Cell_Type_NEW
Ctrl
ExN
InN
Other
UD

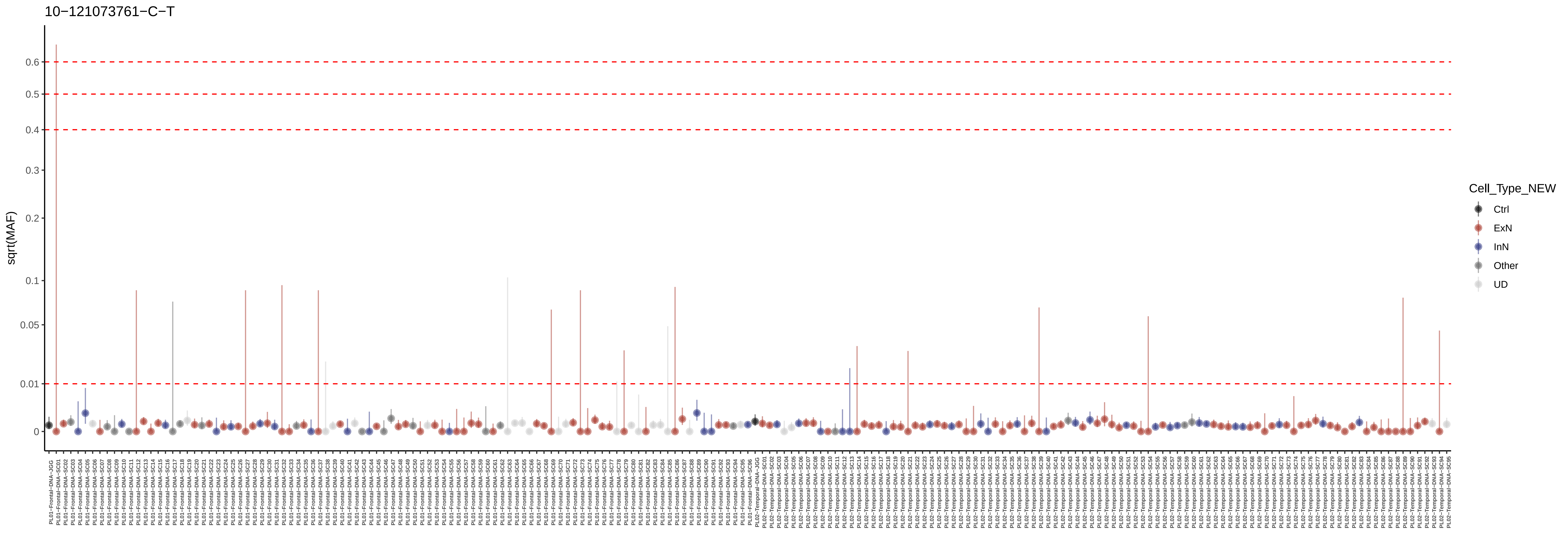
10−121073761−C−T
sqrt(MAF)
0.6
0.5
0.4
0.3
0.2
0.1
0.05
0.01
0
Cell_Type_NEW
Ctrl
ExN
InN
Other
UD

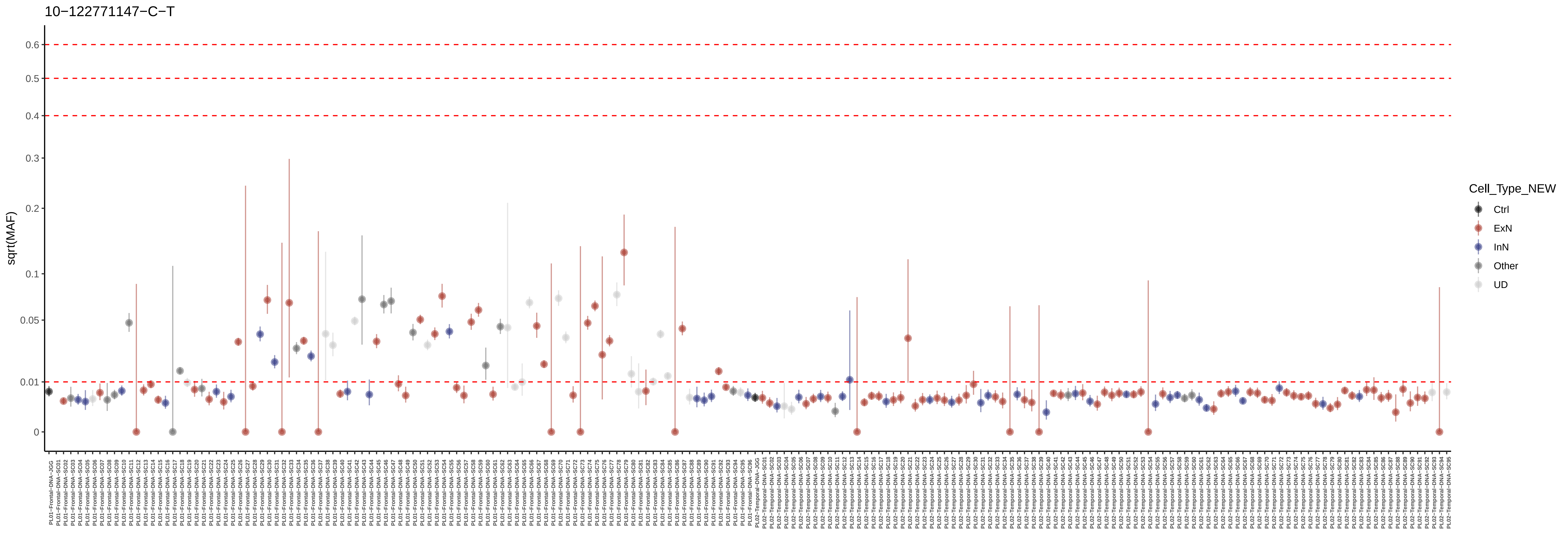
10−122771147−C−T
sqrt(MAF)
0.6
0.5
0.4
0.3
0.2
0.1
0.05
0.01
0
Cell_Type_NEW
Ctrl
ExN
InN
Other
UD

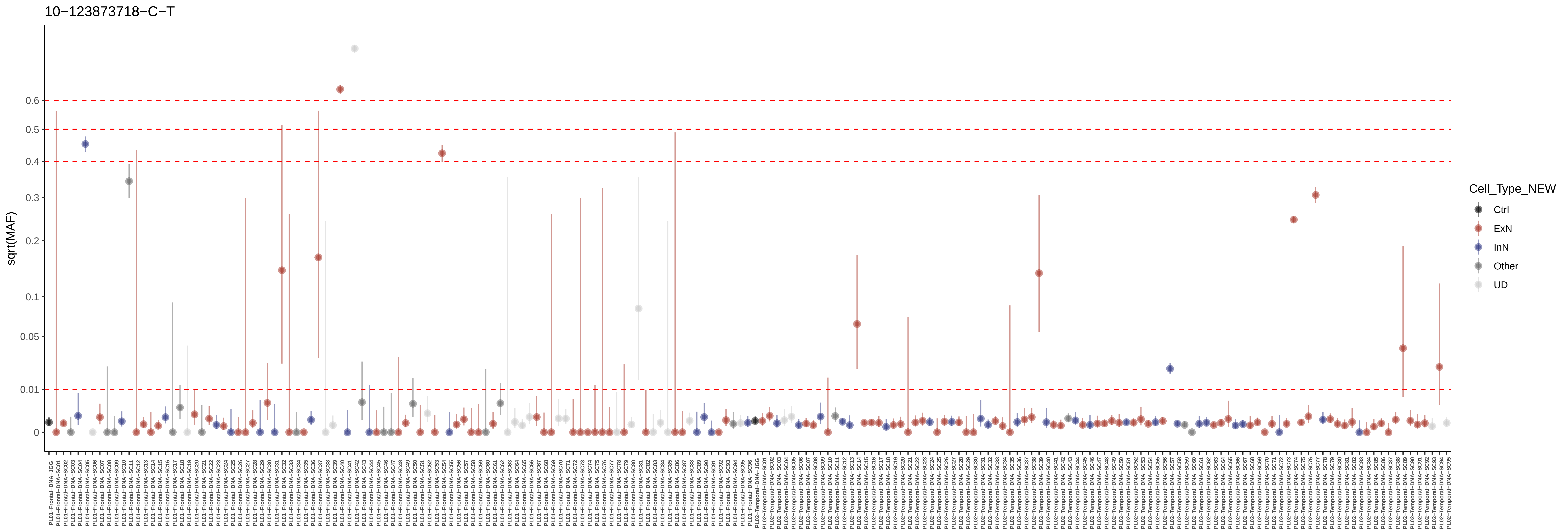
10−123873718−C−T
sqrt(MAF)
0.6
0.5
0.4
0.3
0.2
0.1
0.05
0.01
0
Cell_Type_NEW
Ctrl
ExN
InN
Other
UD

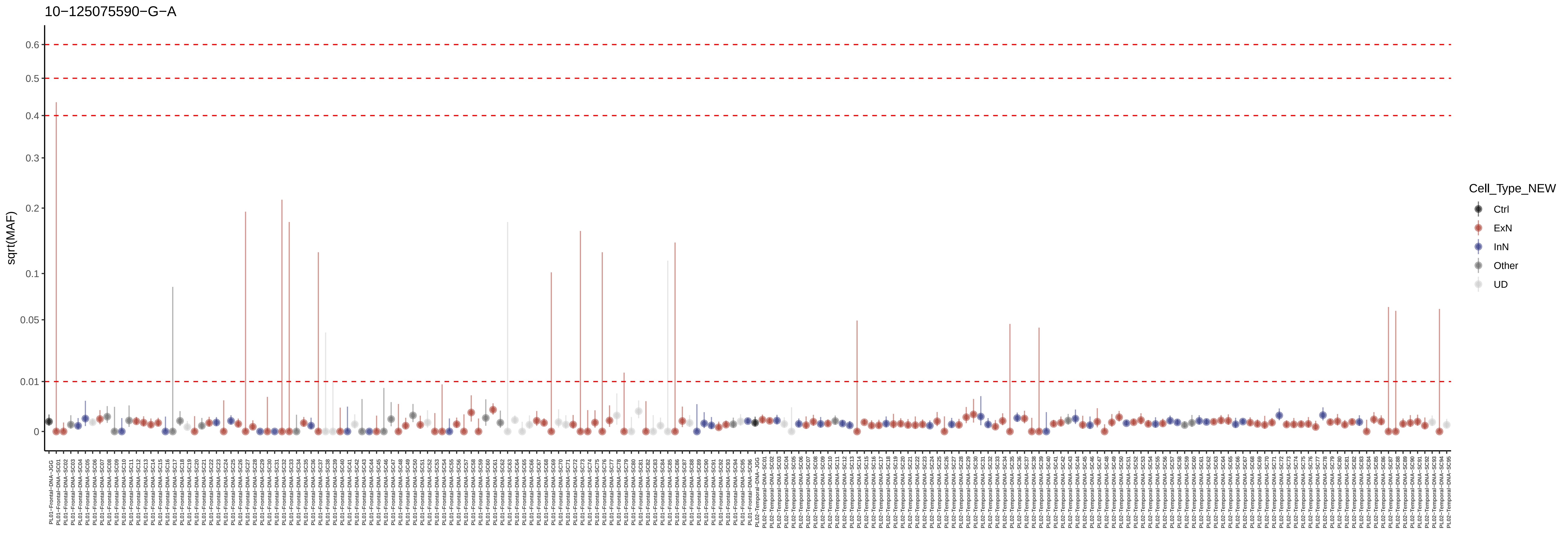
10−125075590−G−A
sqrt(MAF)
0.6
0.5
0.4
0.3
0.2
0.1
0.05
0.01
0
Cell_Type_NEW
Ctrl
ExN
InN
Other
UD

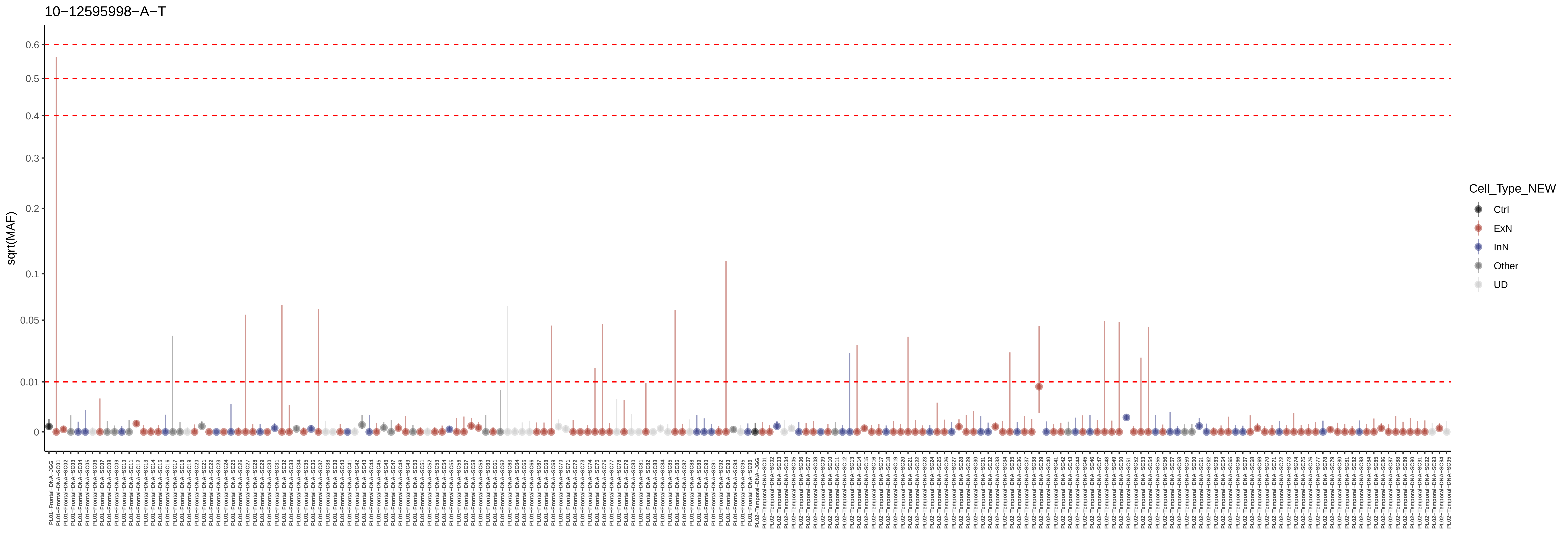
10−12595998−A−T
sqrt(MAF)
0.6
0.5
0.4
0.3
0.2
0.1
0.05
0.01
0
Cell_Type_NEW
Ctrl
ExN
InN
Other
UD

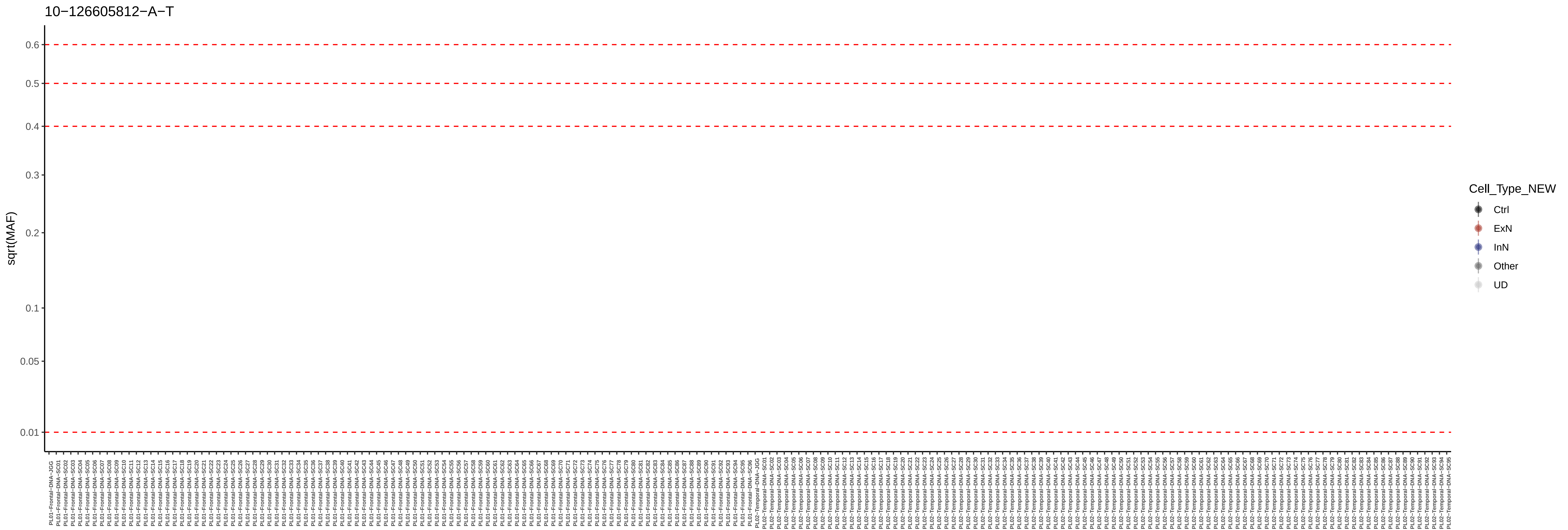
10−126605812−A−T
sqrt(MAF)
0.6
0.5
0.4
0.3
0.2
0.1
0.05
0.01
Cell_Type_NEW
Ctrl
ExN
InN
Other
UD

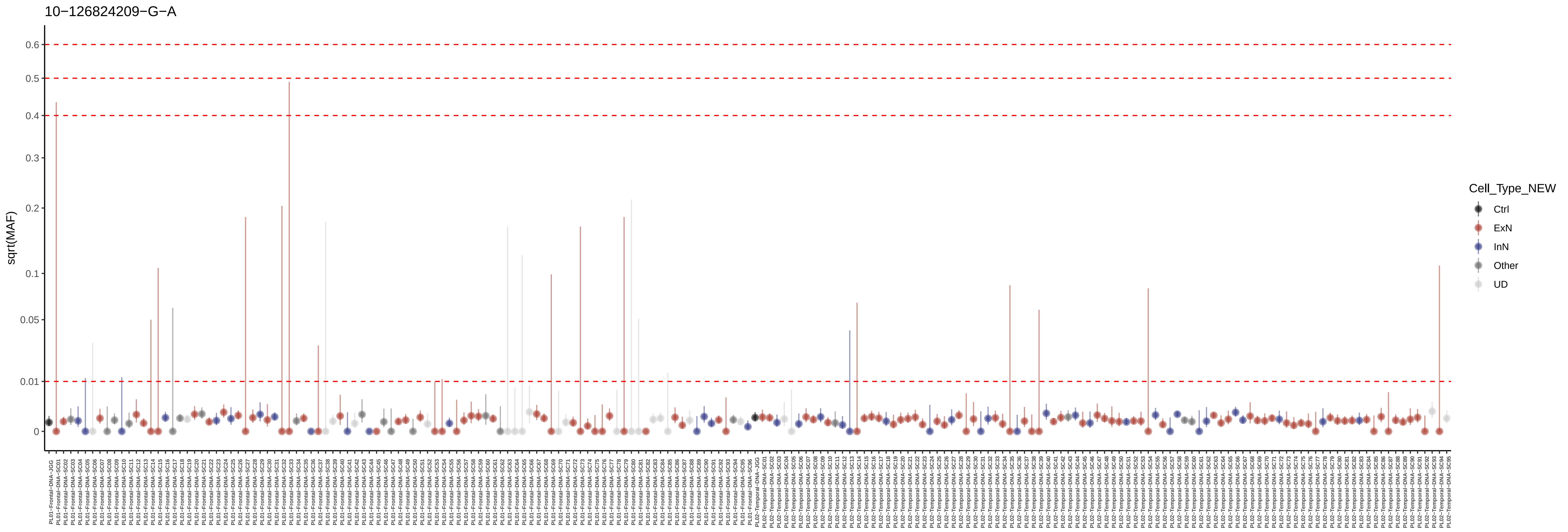
10−126824209−G−A
sqrt(MAF)
0.6
0.5
0.4
0.3
0.2
0.1
0.05
0.01
0
Cell_Type_NEW
Ctrl
ExN
InN
Other
UD

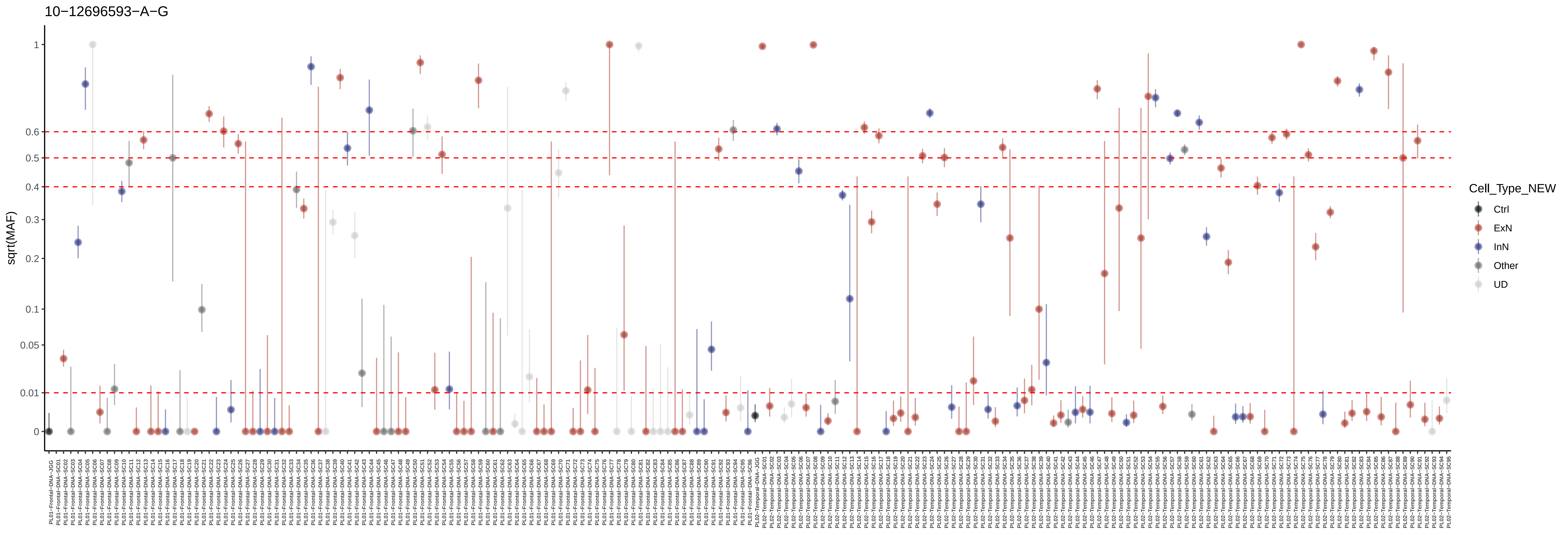
10−12696593−A−G
sqrt(MAF)
1
0.6
0.5
0.4
0.3
0.2
0.1
0.05
0.01
0
Cell_Type_NEW
Ctrl
ExN
InN
Other
UD

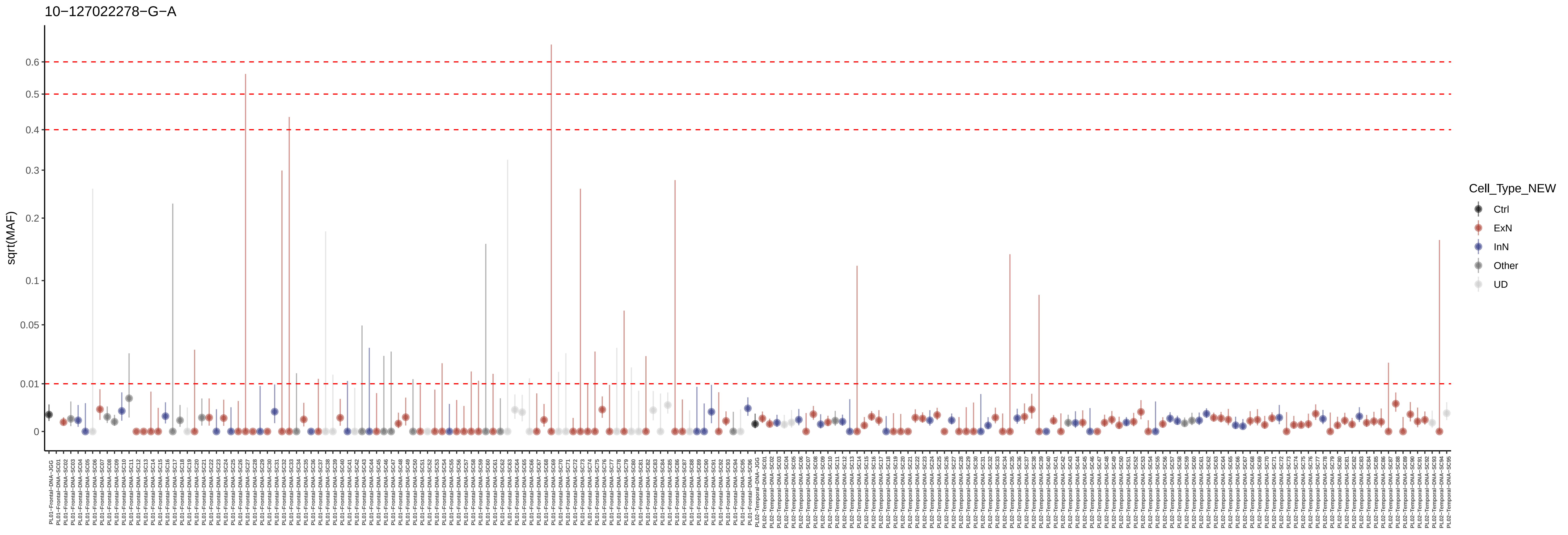
10−127022278−G−A
sqrt(MAF)
0.6
0.5
0.4
0.3
0.2
0.1
0.05
0.01
0
Cell_Type_NEW
Ctrl
ExN
InN
Other
UD

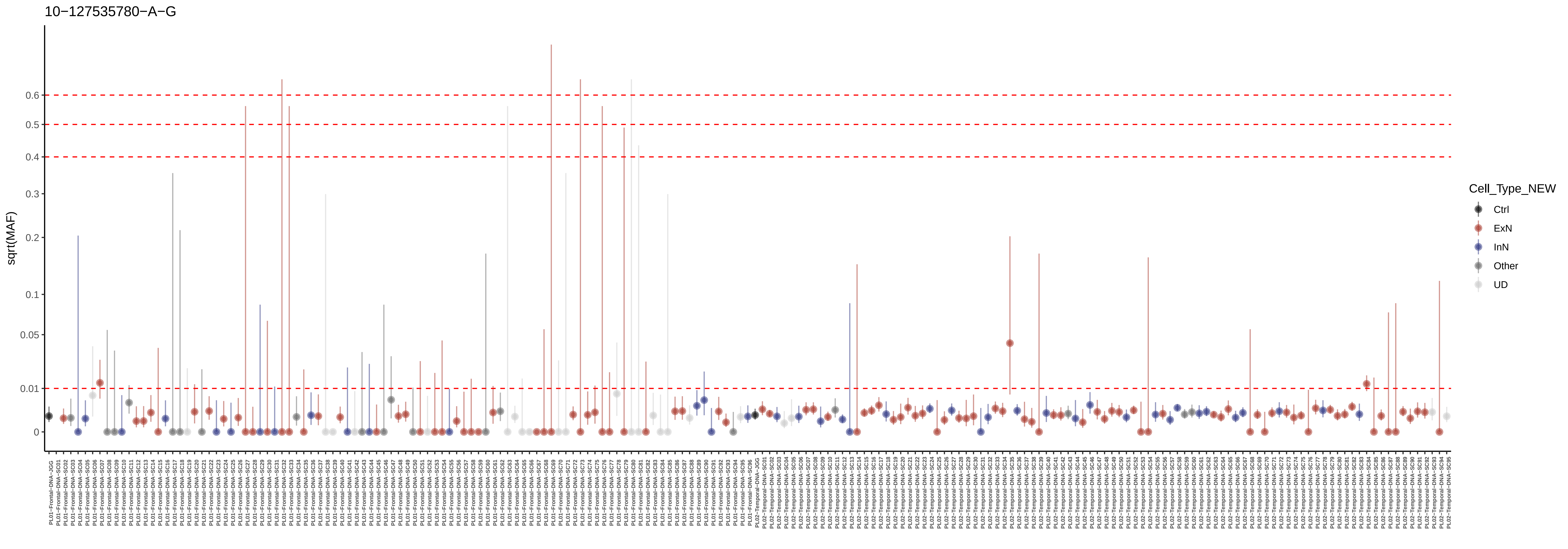
10−127535780−A−G
sqrt(MAF)
0.6
0.5
0.4
0.3
0.2
0.1
0.05
0.01
0
Cell_Type_NEW
Ctrl
ExN
InN
Other
UD

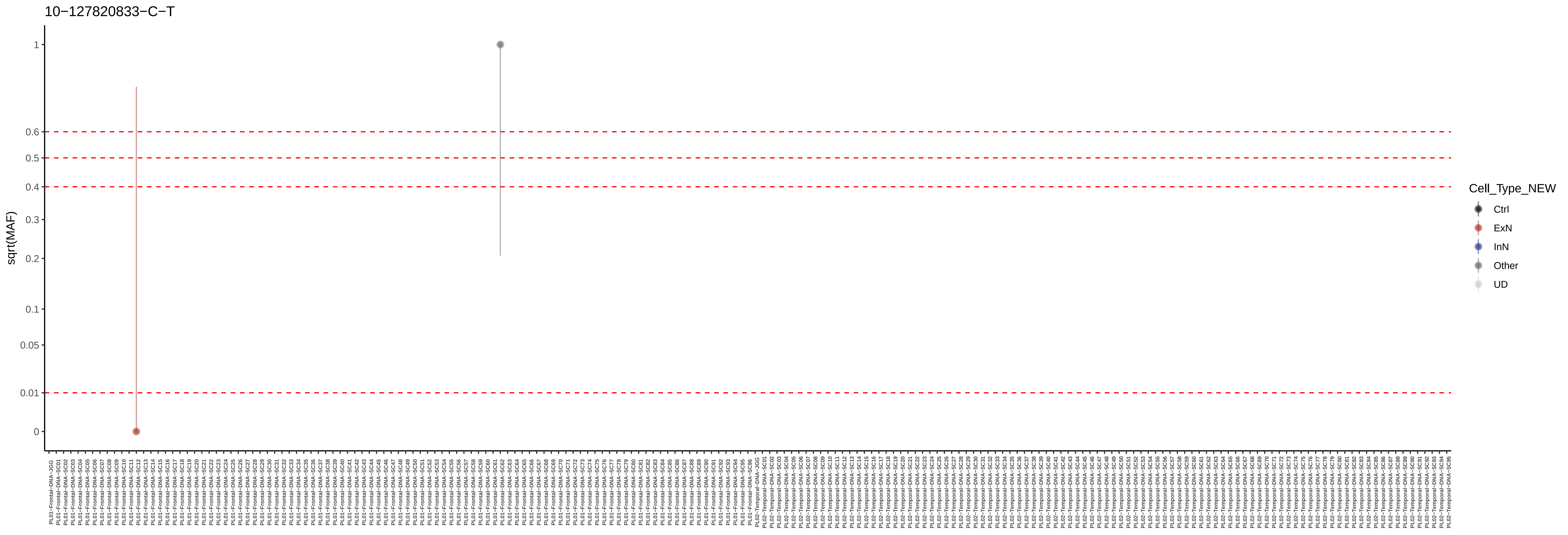

10−127820833−C−T
sqrt(MAF)
1
0.6
0.5
0.4
0.3
0.2
0.1
0.05
0.01
0
Cell_Type_NEW
Ctrl
ExN
InN
Other
UD

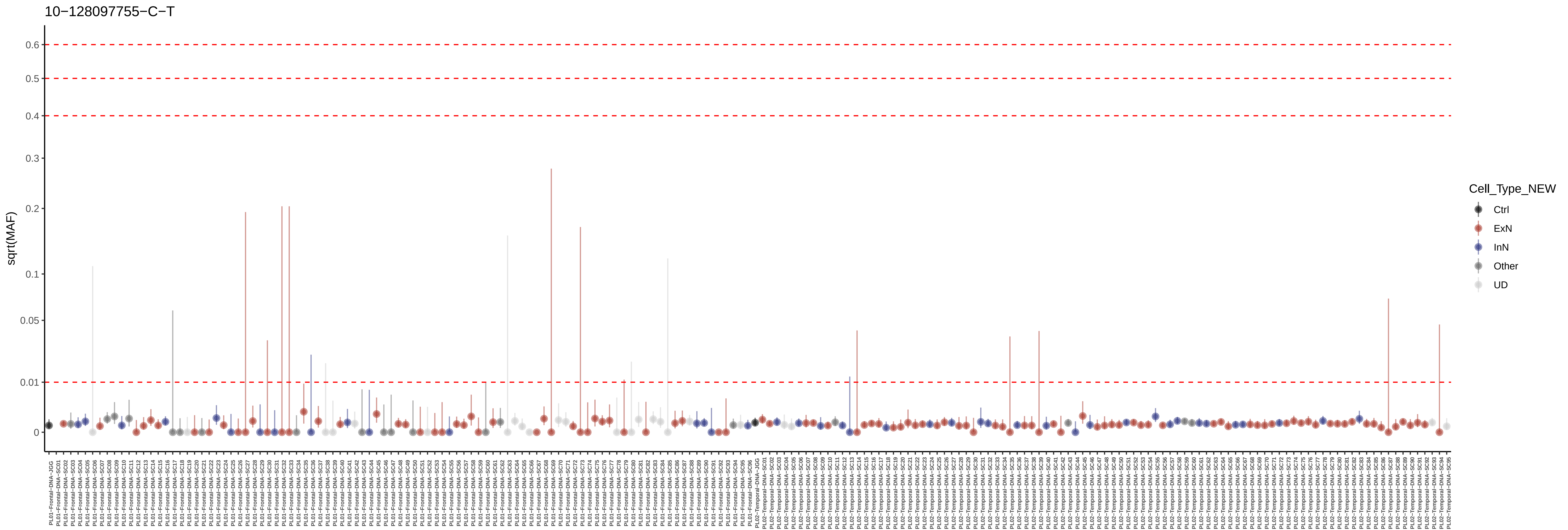
10−128097755−C−T
sqrt(MAF)
0.6
0.5
0.4
0.3
0.2
0.1
0.05
0.01
0
Cell_Type_NEW
Ctrl
ExN
InN
Other
UD

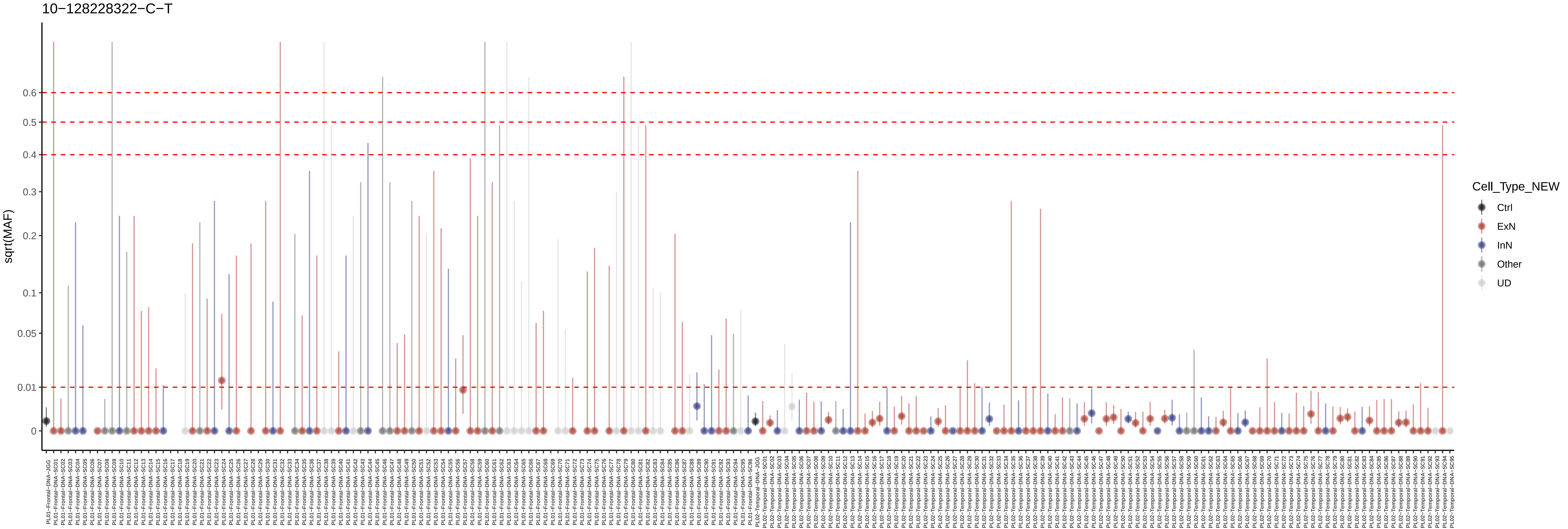
10−128228322−C−T
sqrt(MAF)
0.6
0.5
0.4
0.3
0.2
0.1
0.05
0.01
0
Cell_Type_NEW
Ctrl
ExN
InN
Other
UD

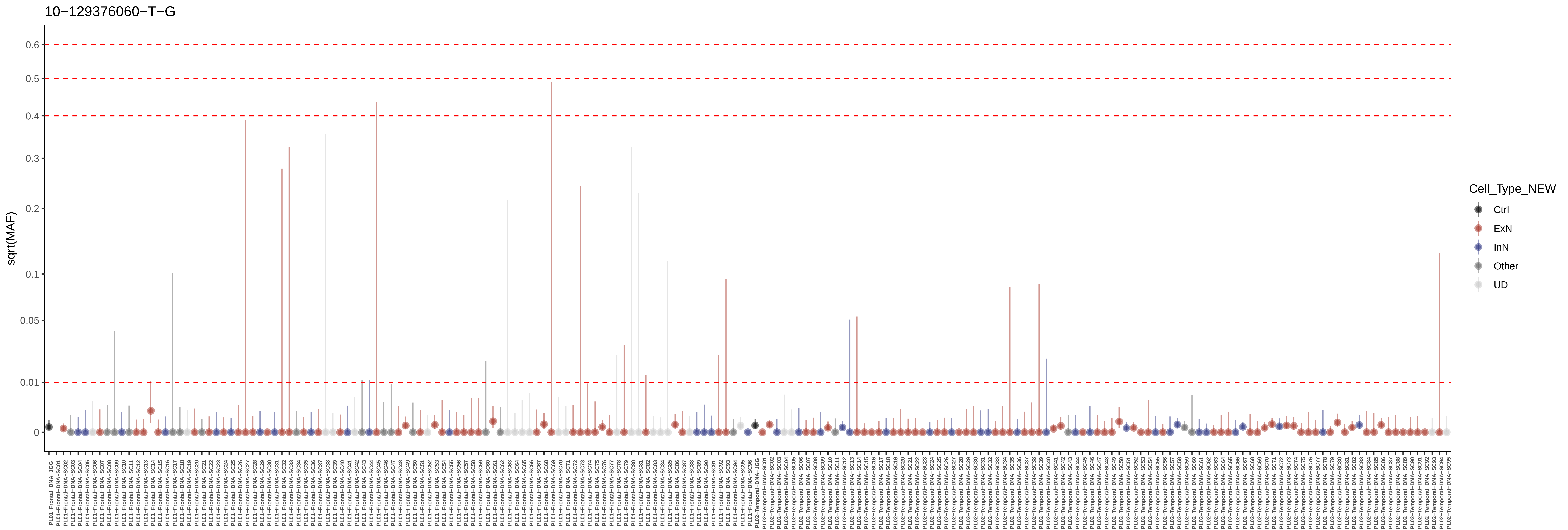
10−129376060−T−G
sqrt(MAF)
0.6
0.5
0.4
0.3
0.2
0.1
0.05
0.01
0
Cell_Type_NEW
Ctrl
ExN
InN
Other
UD

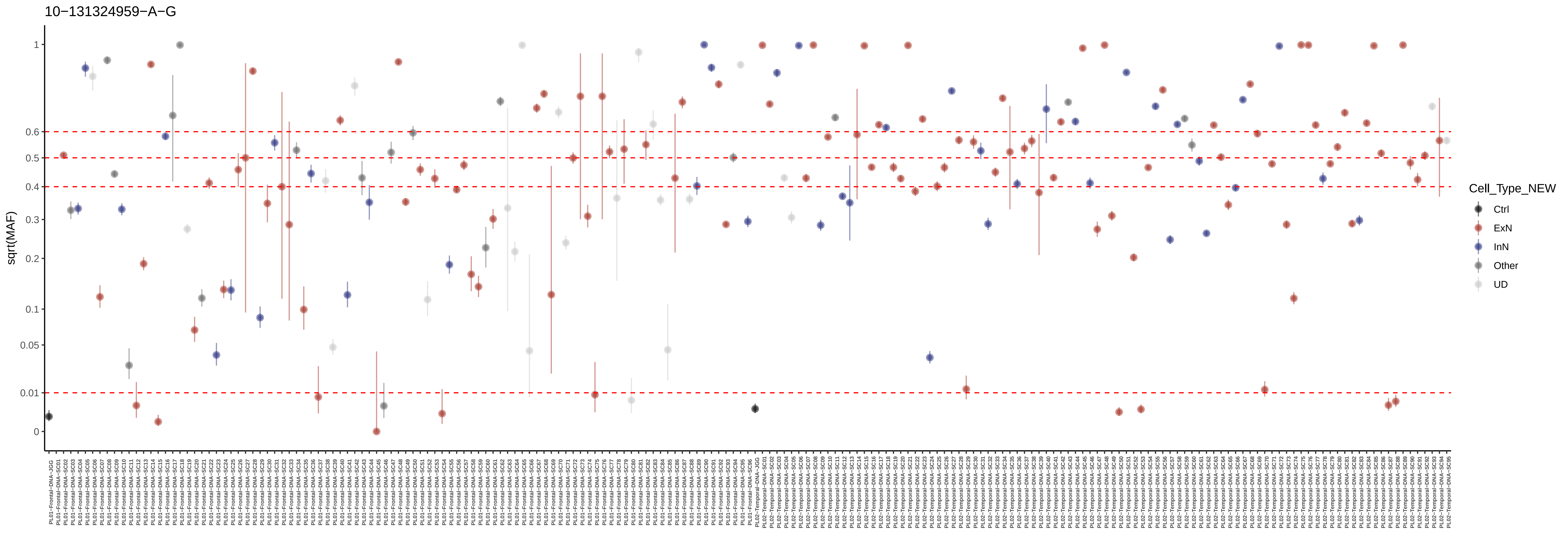
10−131324959−A−G
sqrt(MAF)
1
0.6
0.5
0.4
0.3
0.2
0.1
0.05
0.01
0
Cell_Type_NEW
Ctrl
ExN
InN
Other
UD

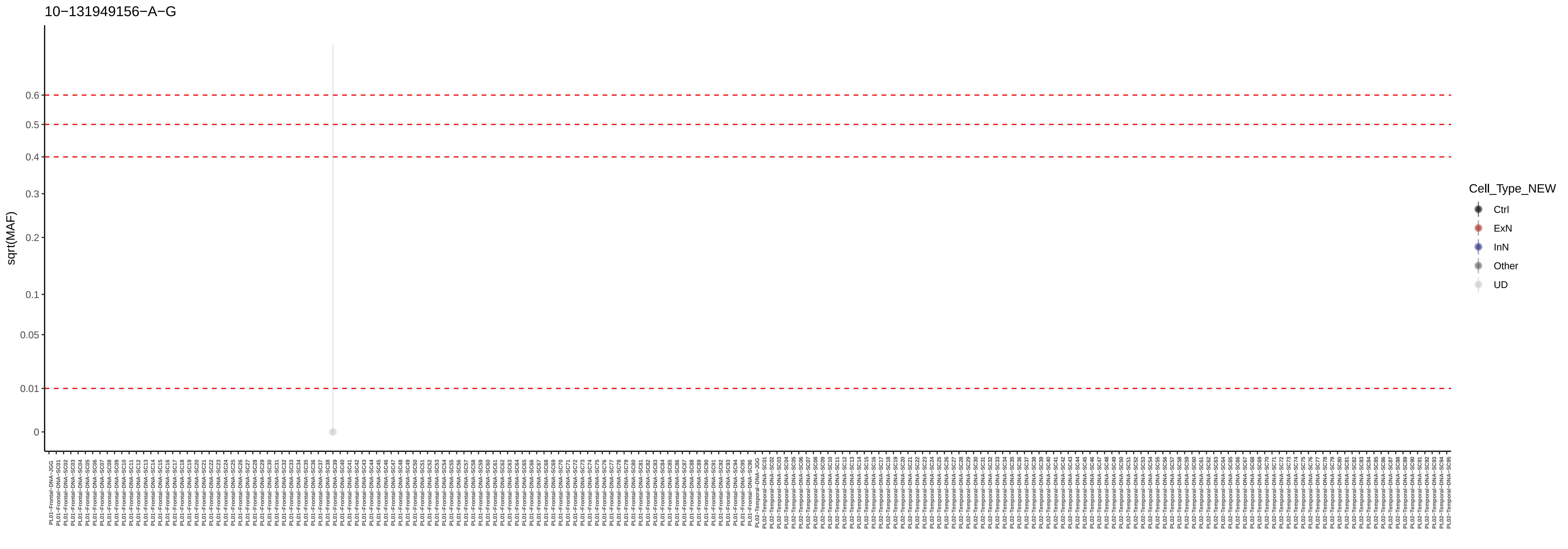
10−131949156−A−G
sqrt(MAF)
0.6
0.5
0.4
0.3
0.2
0.1
0.05
0.01
0
Cell_Type_NEW
Ctrl
ExN
InN
Other
UD

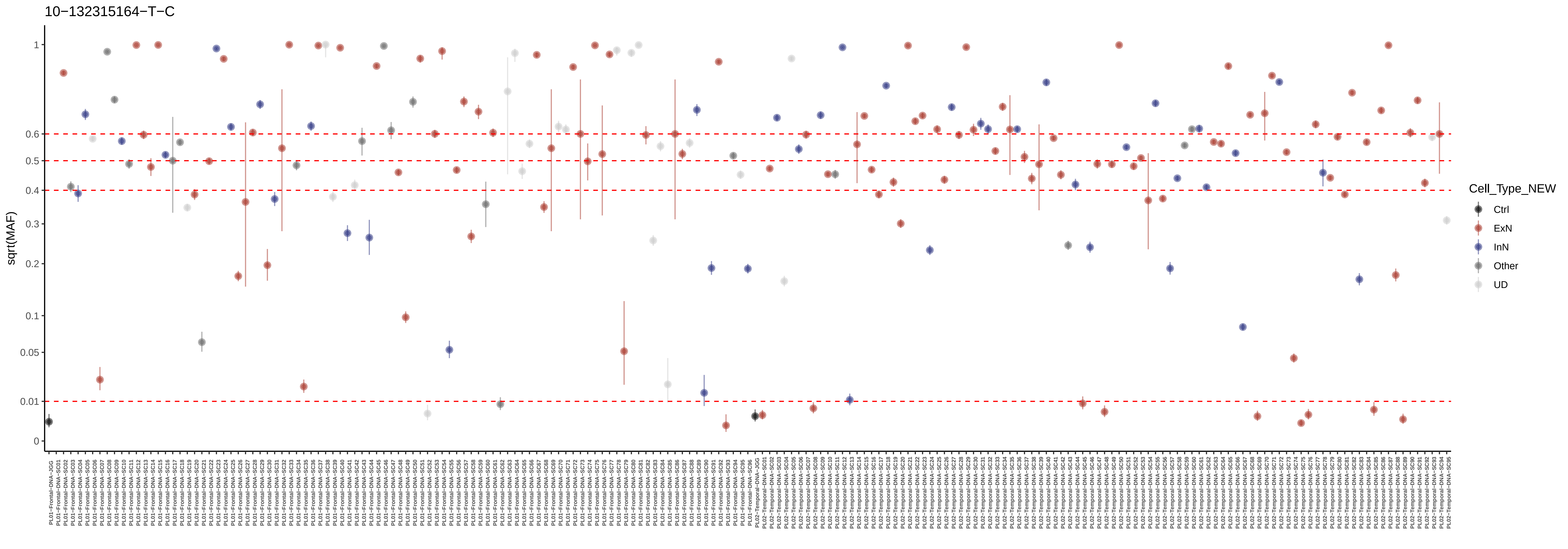
10−132315164−T−C
sqrt(MAF)
1
0.6
0.5
0.4
0.3
0.2
0.1
0.05
0.01
0
Cell_Type_NEW
Ctrl
ExN
InN
Other
UD

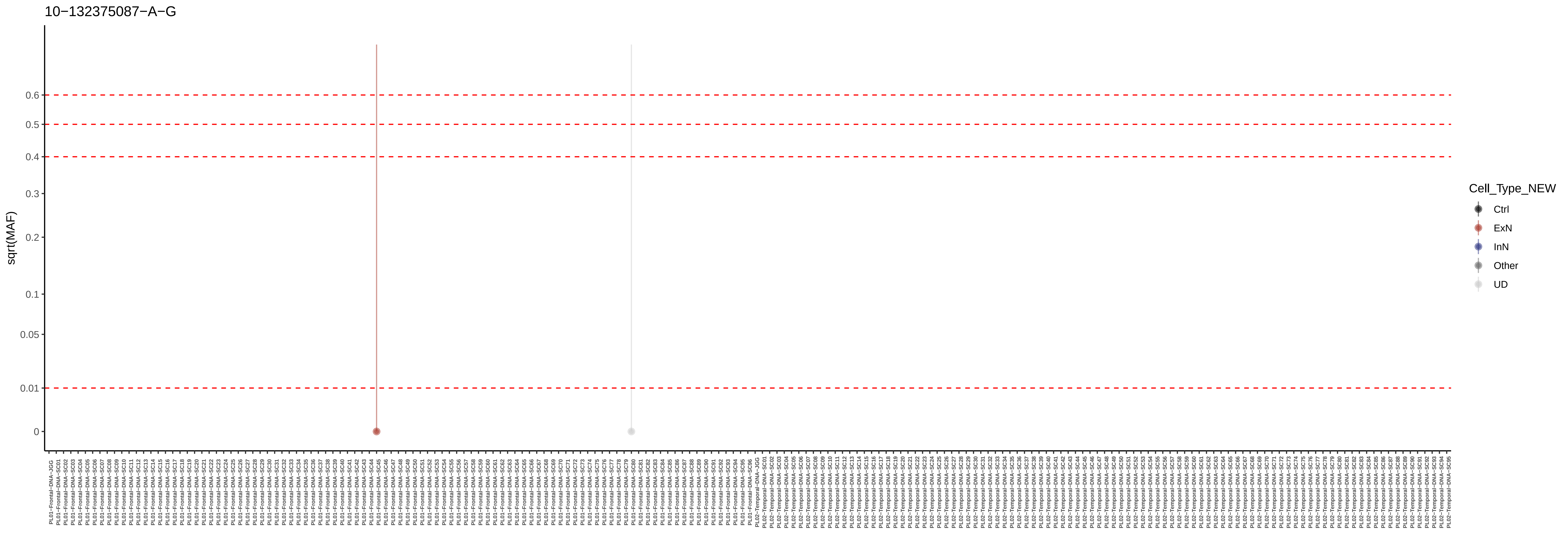
10−132375087−A−G
sqrt(MAF)
0.6
0.5
0.4
0.3
0.2
0.1
0.05
0.01
0
Cell_Type_NEW
Ctrl
ExN
InN
Other
UD

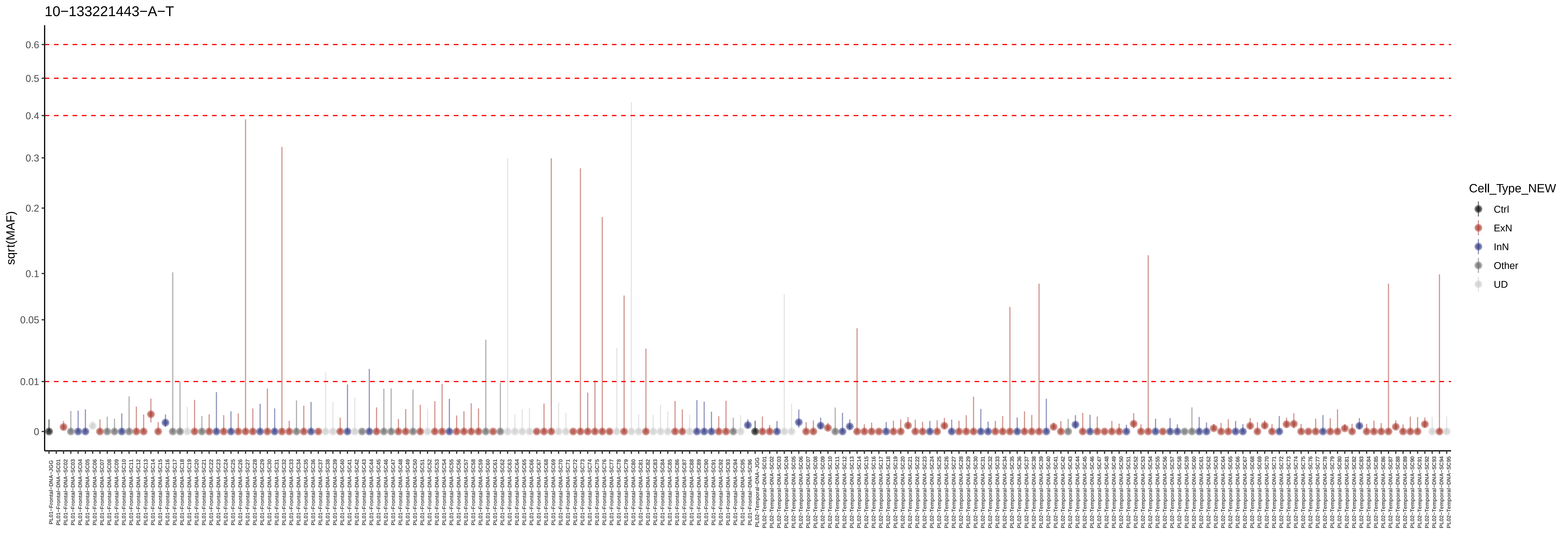
10−133221443−A−T
sqrt(MAF)
0.6
0.5
0.4
0.3
0.2
0.1
0.05
0.01
0
Cell_Type_NEW
Ctrl
ExN
InN
Other
UD

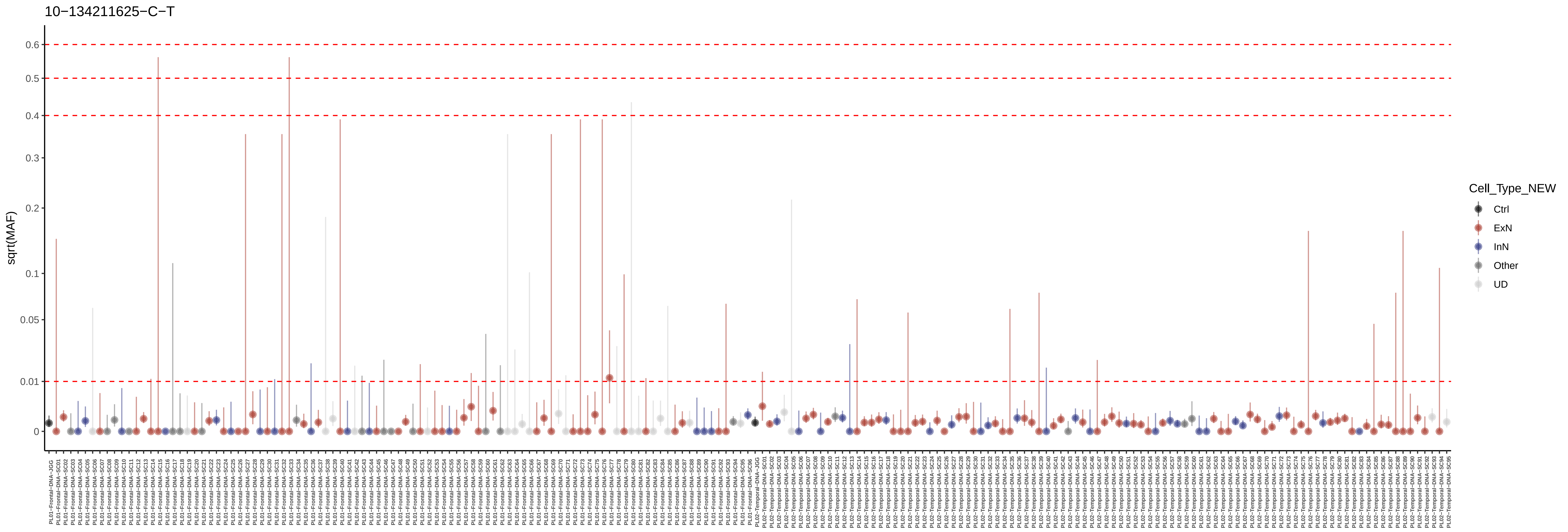
10−134211625−C−T
sqrt(MAF)
0.6
0.5
0.4
0.3
0.2
0.1
0.05
0.01
0
Cell_Type_NEW
Ctrl
ExN
InN
Other
UD

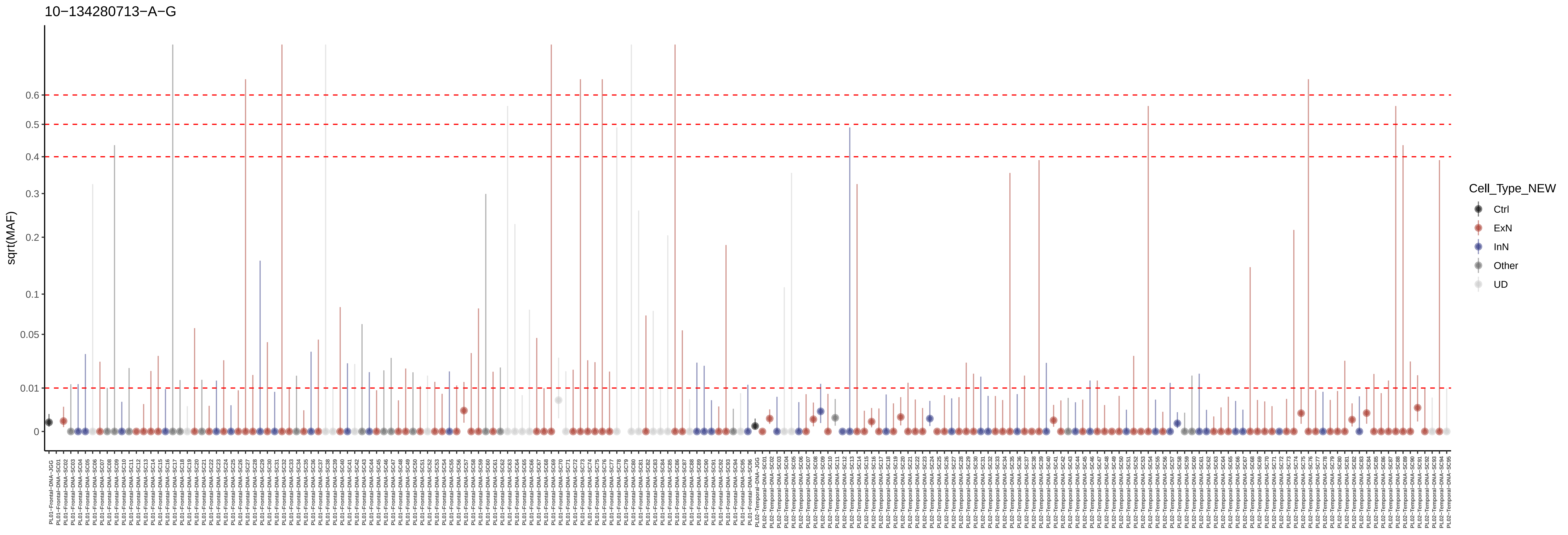
10−134280713−A−G
sqrt(MAF)
0.6
0.5
0.4
0.3
0.2
0.1
0.05
0.01
0
Cell_Type_NEW
Ctrl
ExN
InN
Other
UD

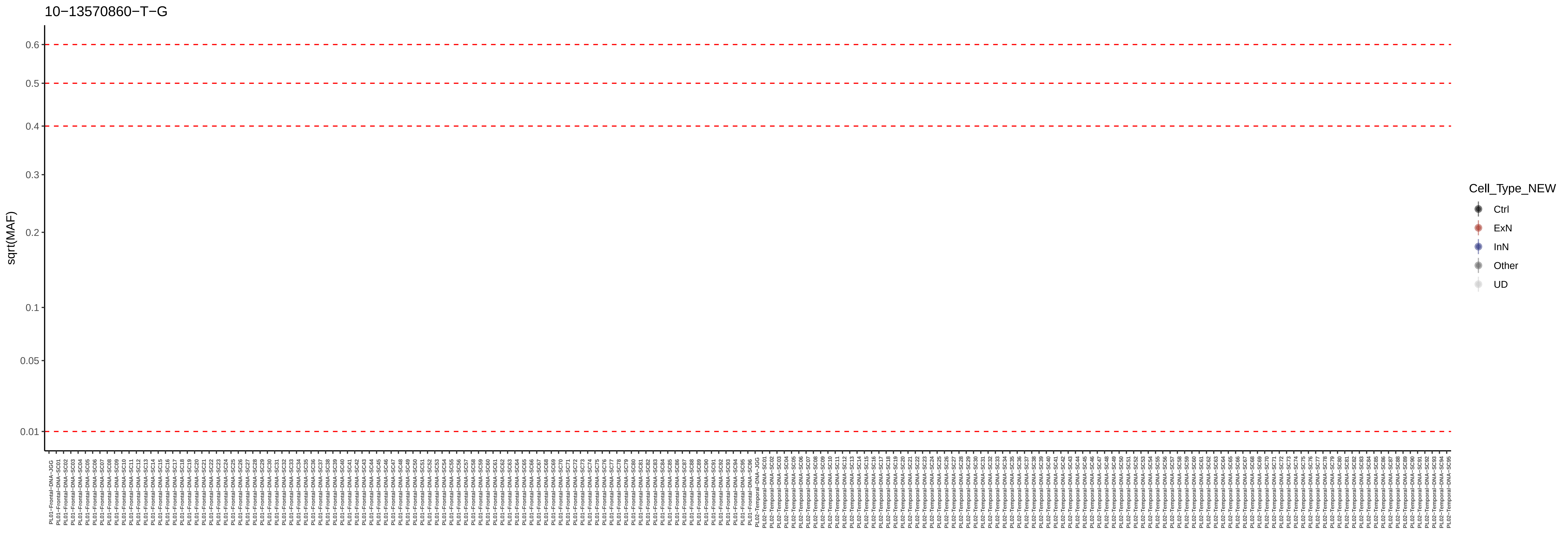
10−13570860−T−G
sqrt(MAF)
0.6
0.5
0.4
0.3
0.2
0.1
0.05
0.01
Cell_Type_NEW
Ctrl
ExN
InN
Other
UD

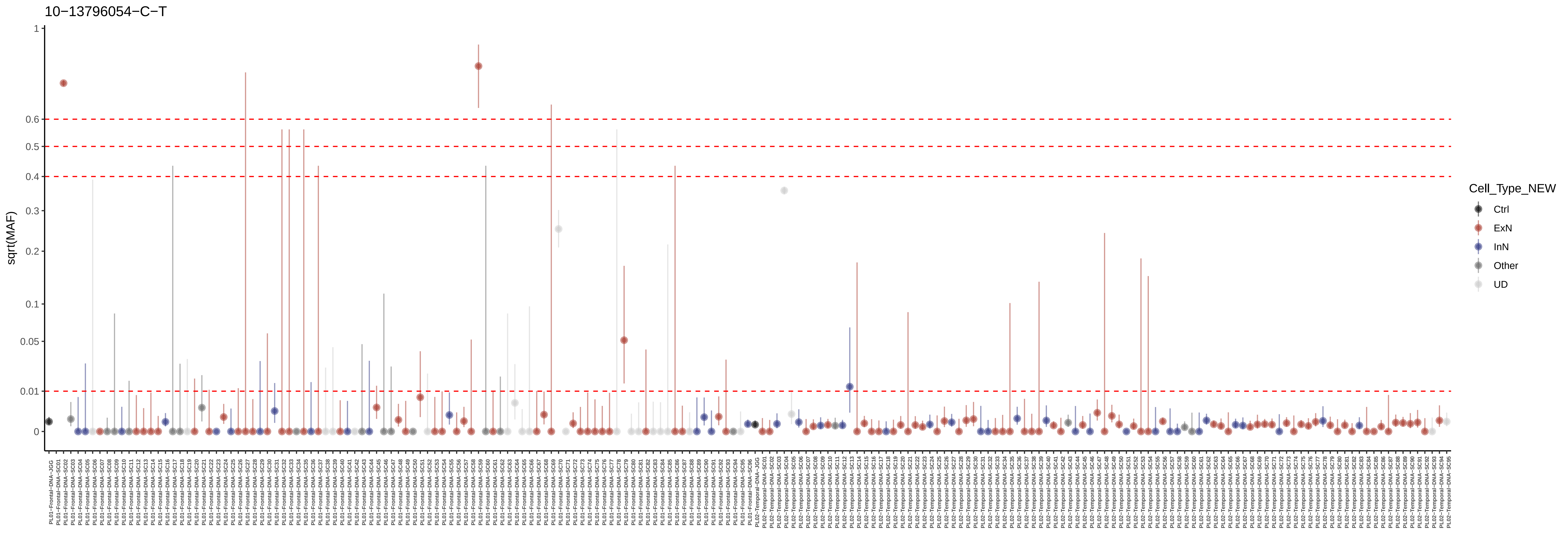
10−13796054−C−T
sqrt(MAF)
1
0.6
0.5
0.4
0.3
0.2
0.1
0.05
0.01
0
Cell_Type_NEW
Ctrl
ExN
InN
Other
UD

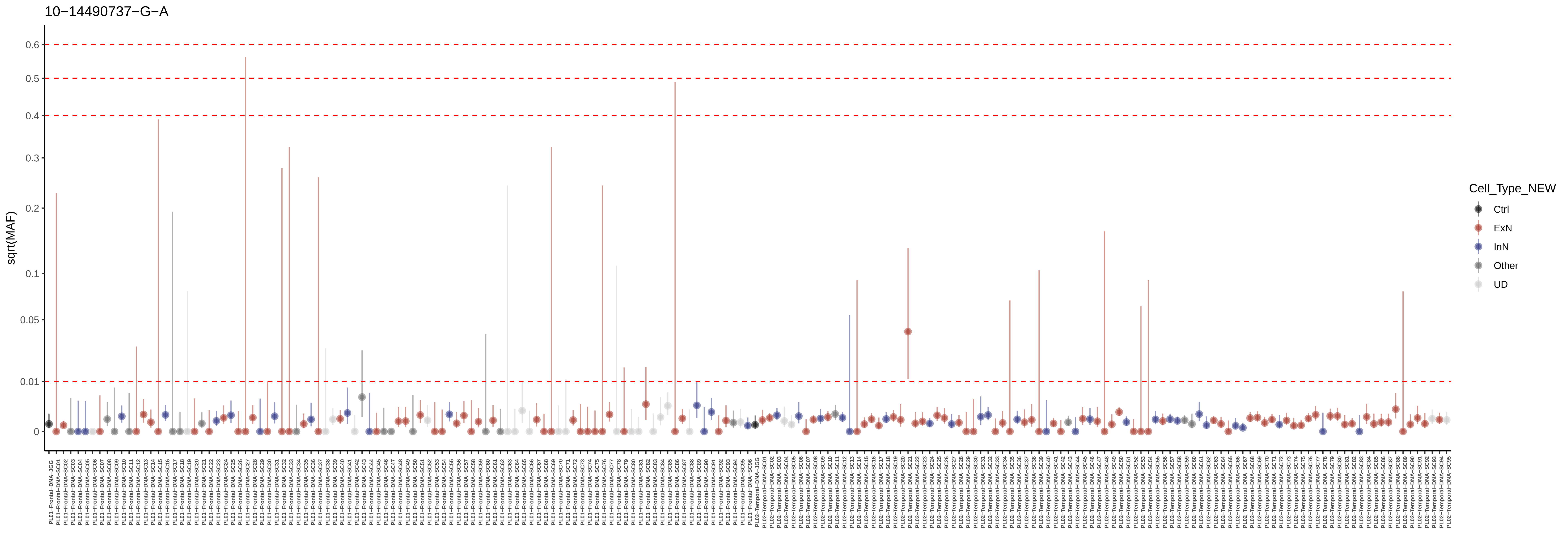
10−14490737−G−A
sqrt(MAF)
0.6
0.5
0.4
0.3
0.2
0.1
0.05
0.01
0
Cell_Type_NEW
Ctrl
ExN
InN
Other
UD

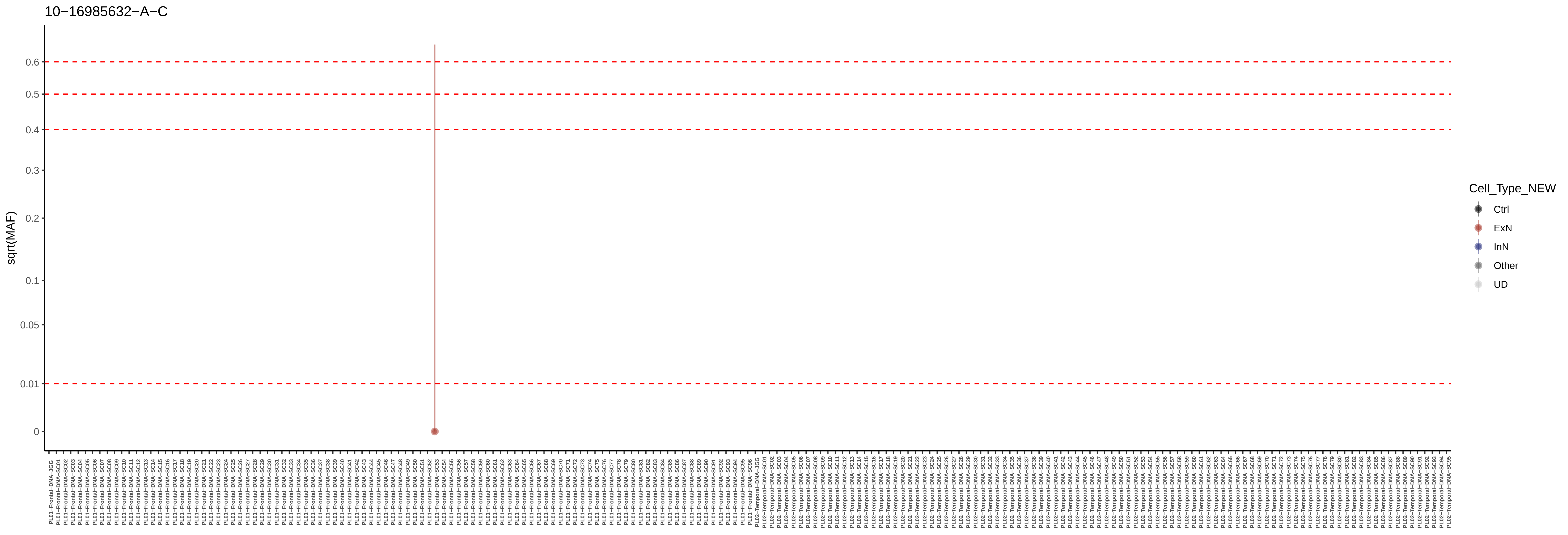
10−16985632−A−C
sqrt(MAF)
0.6
0.5
0.4
0.3
0.2
0.1
0.05
0.01
0
Cell_Type_NEW
Ctrl
ExN
InN
Other
UD

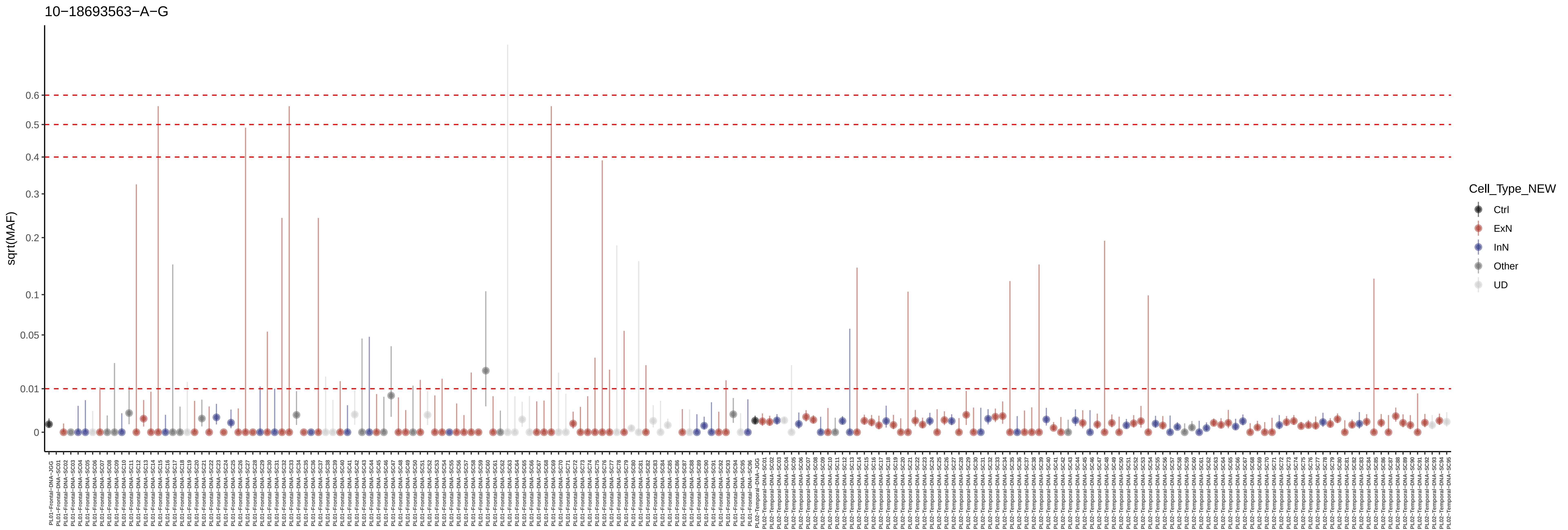

10−18693563−A−G
sqrt(MAF)
0.6
0.5
0.4
0.3
0.2
0.1
0.05
0.01
0
Cell_Type_NEW
Ctrl
ExN
InN
Other
UD

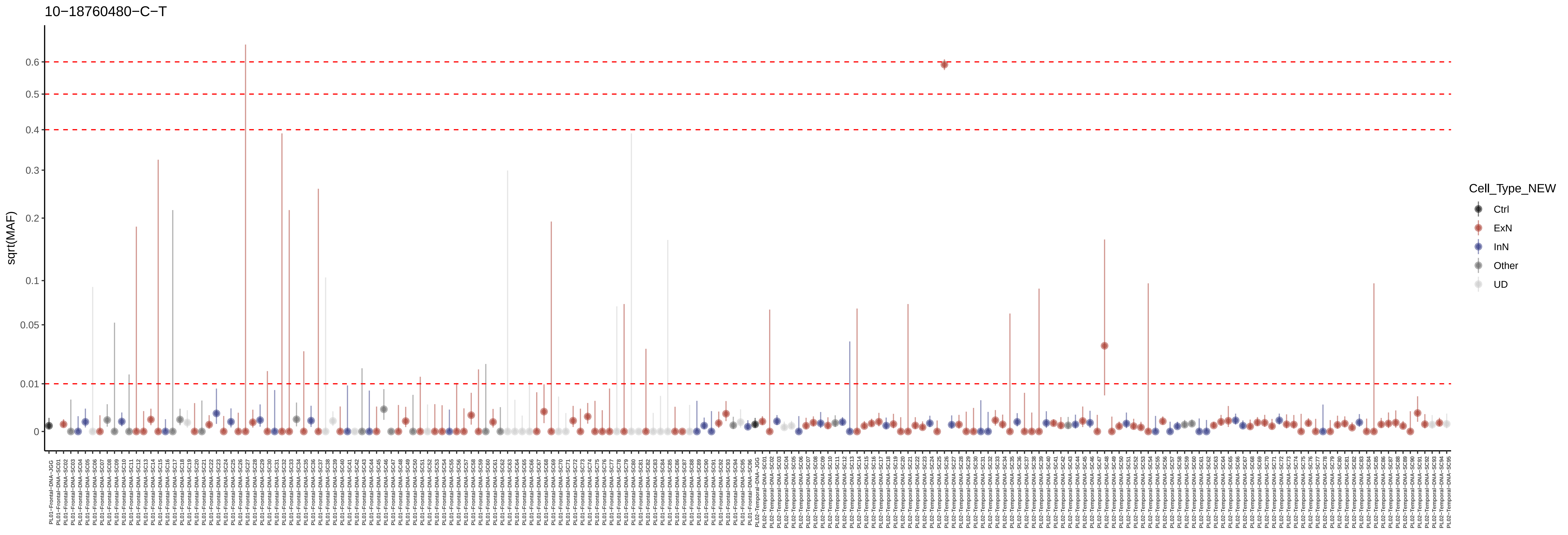
10−18760480−C−T
sqrt(MAF)
0.6
0.5
0.4
0.3
0.2
0.1
0.05
0.01
0
Cell_Type_NEW
Ctrl
ExN
InN
Other
UD

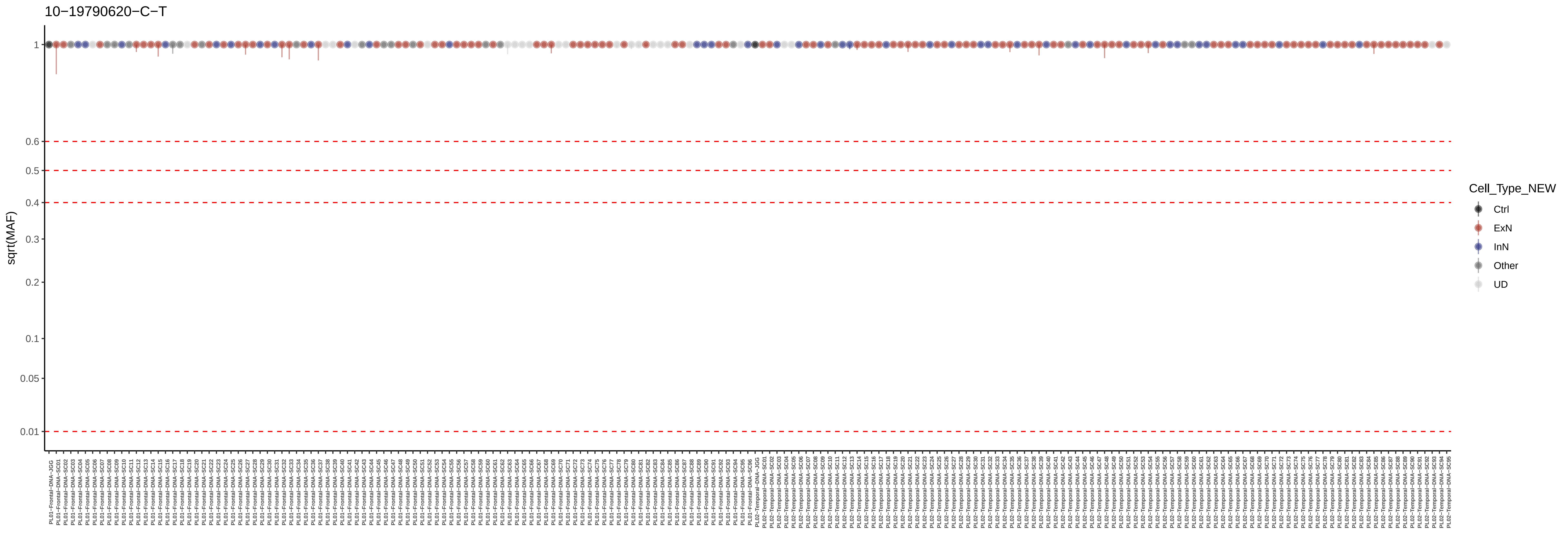

10−19790620−C−T
sqrt(MAF)
1
0.6
0.5
0.4
0.3
0.2
0.1
0.05
0.01
Cell_Type_NEW
Ctrl
ExN
InN
Other
UD

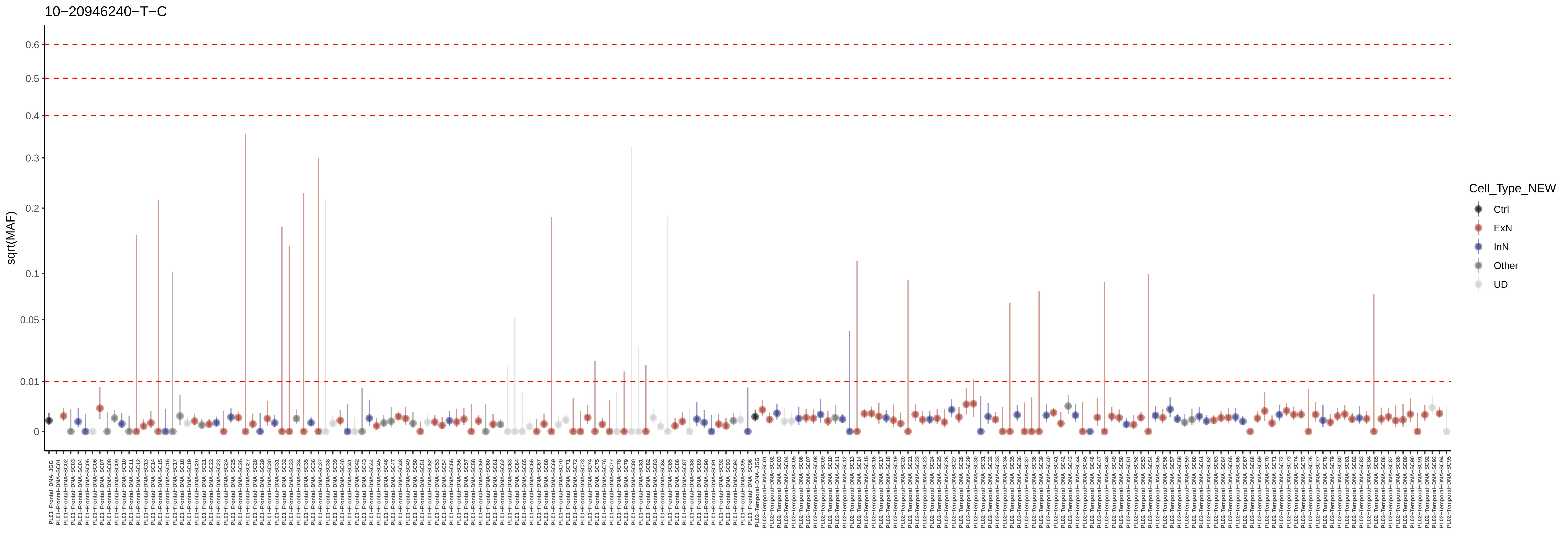
10−20946240−T−C
sqrt(MAF)
0.6
0.5
0.4
0.3
0.2
0.1
0.05
0.01
0
Cell_Type_NEW
Ctrl
ExN
InN
Other
UD

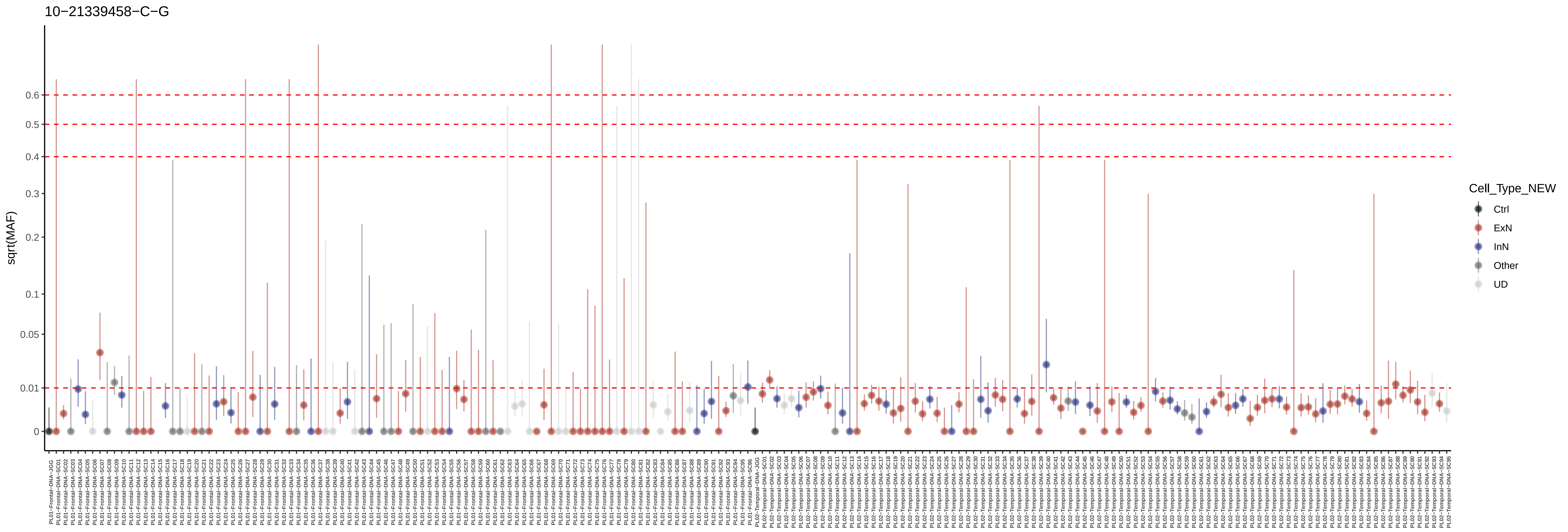
10−21339458−C−G
sqrt(MAF)
0.6
0.5
0.4
0.3
0.2
0.1
0.05
0.01
0
Cell_Type_NEW
Ctrl
ExN
InN
Other
UD

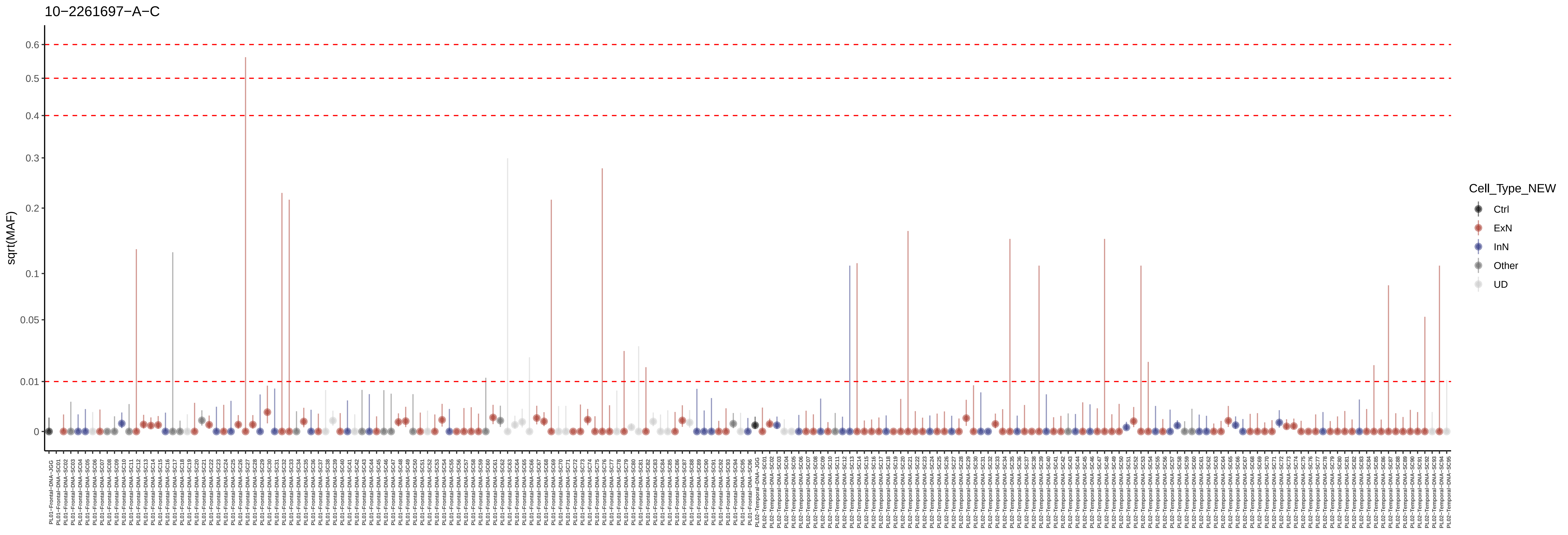
10−2261697−A−C
sqrt(MAF)
0.6
0.5
0.4
0.3
0.2
0.1
0.05
0.01
0
Cell_Type_NEW
Ctrl
ExN
InN
Other
UD
PL01−Frontal−DNA−JGG
PL02−Temporal−DNA−JGG

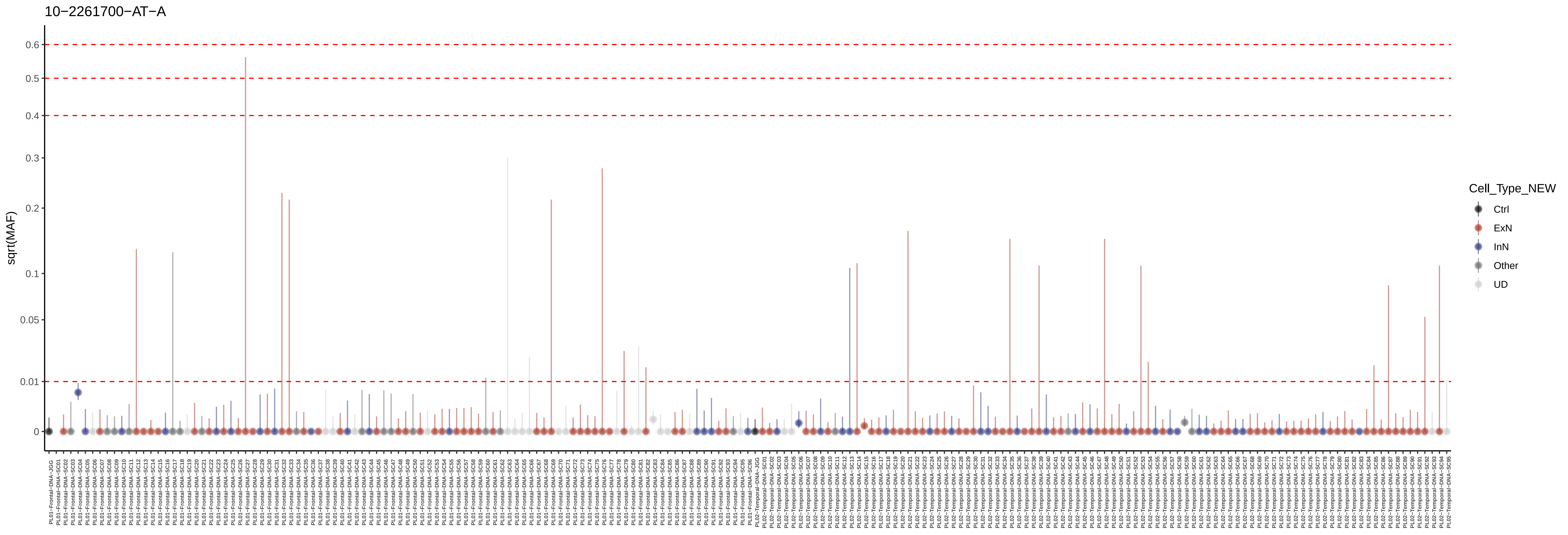
10−2261700−AT−A
sqrt(MAF)
0.6
0.5
0.4
0.3
0.2
0.1
0.05
0.01
0
Cell_Type_NEW
Ctrl
ExN
InN
Other
UD

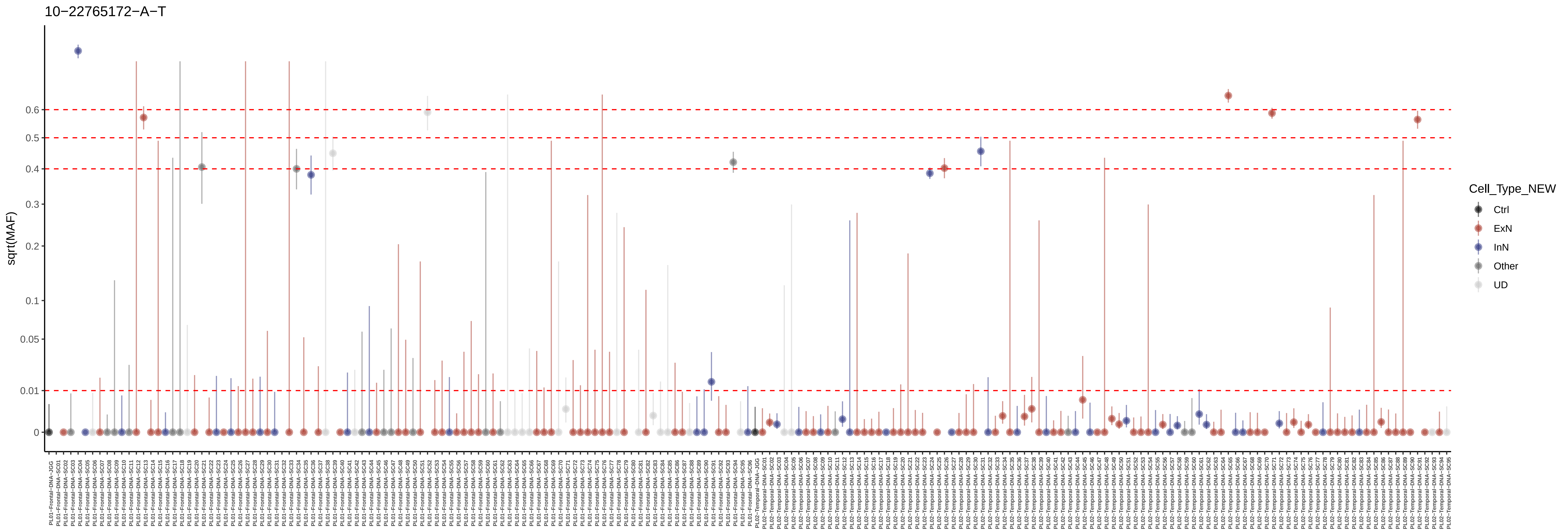
10−22765172−A−T
sqrt(MAF)
0.6
0.5
0.4
0.3
0.2
0.1
0.05
0.01
0
Cell_Type_NEW
Ctrl
ExN
InN
Other
UD

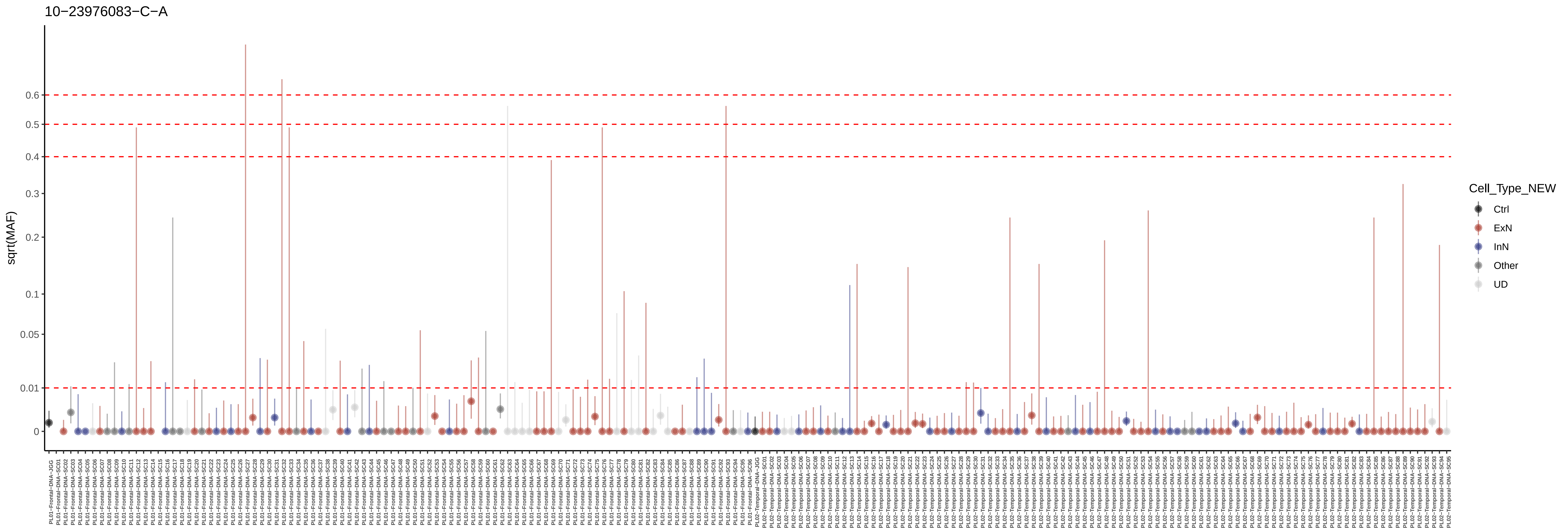
10−23976083−C−A
sqrt(MAF)
0.6
0.5
0.4
0.3
0.2
0.1
0.05
0.01
0
Cell_Type_NEW
Ctrl
ExN
InN
Other
UD

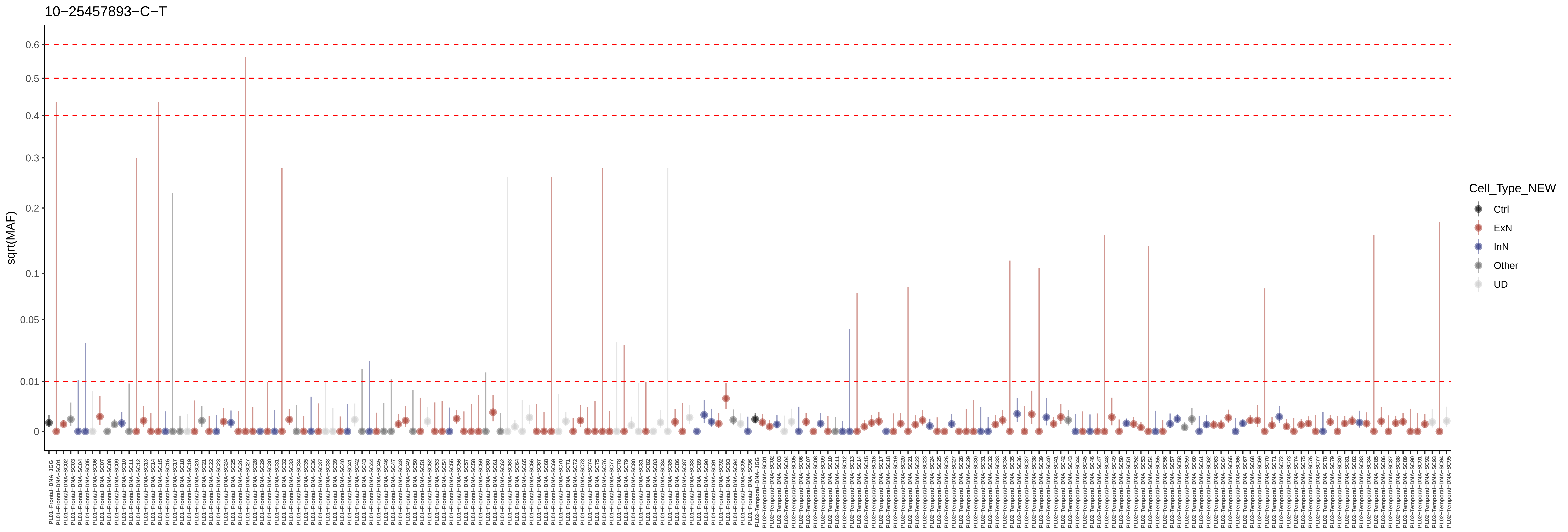
10−25457893−C−T
sqrt(MAF)
0.6
0.5
0.4
0.3
0.2
0.1
0.05
0.01
0
Cell_Type_NEW
Ctrl
ExN
InN
Other
UD

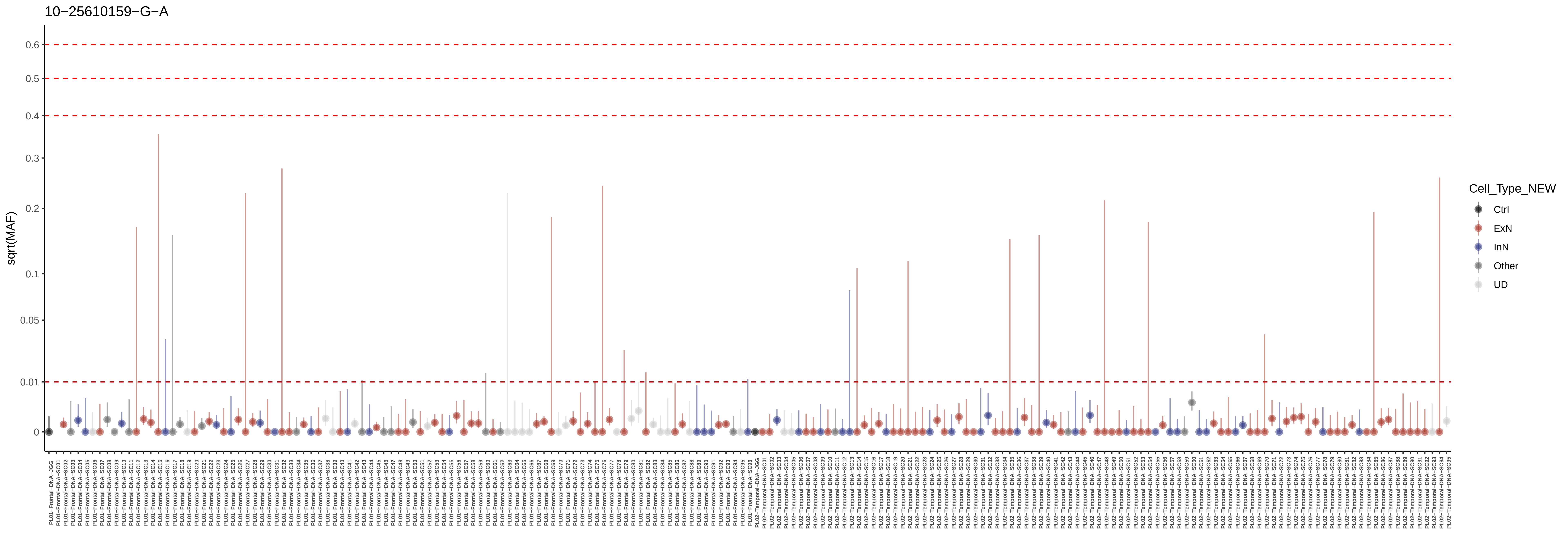

10−25610159−G−A
sqrt(MAF)
0.6
0.5
0.4
0.3
0.2
0.1
0.05
0.01
0
Cell_Type_NEW
Ctrl
ExN
InN
Other
UD

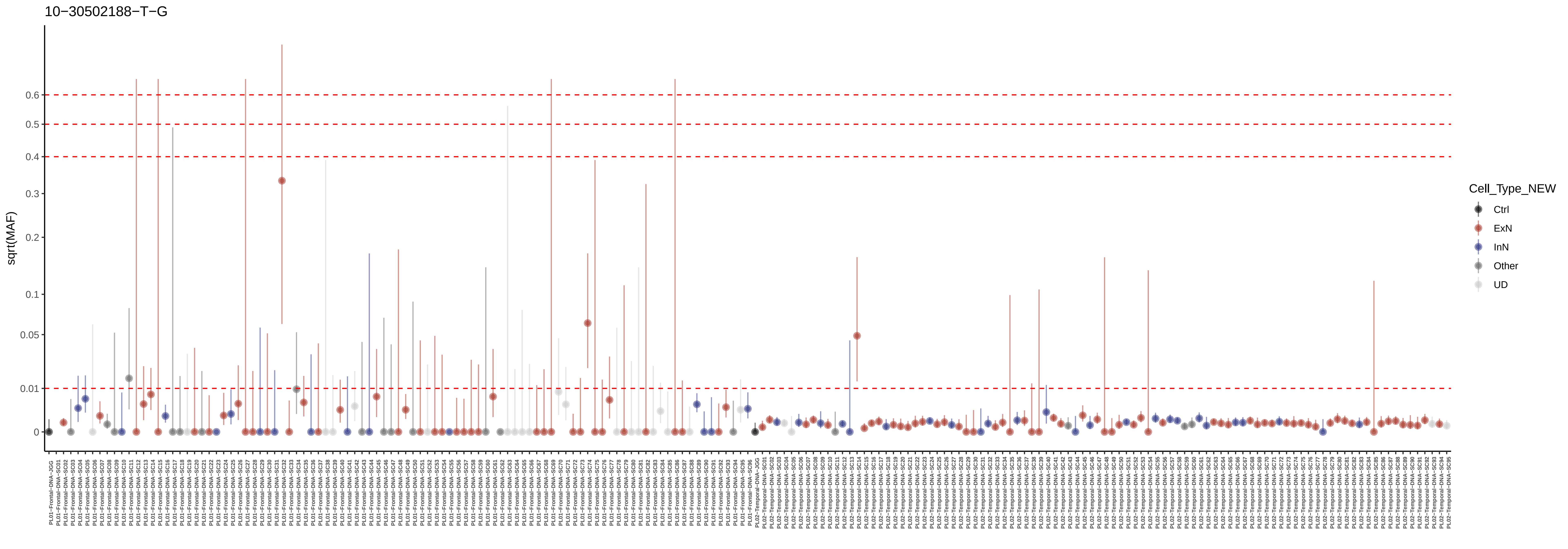
10−30502188−T−G
sqrt(MAF)
0.6
0.5
0.4
0.3
0.2
0.1
0.05
0.01
0
Cell_Type_NEW
Ctrl
ExN
InN
Other
UD

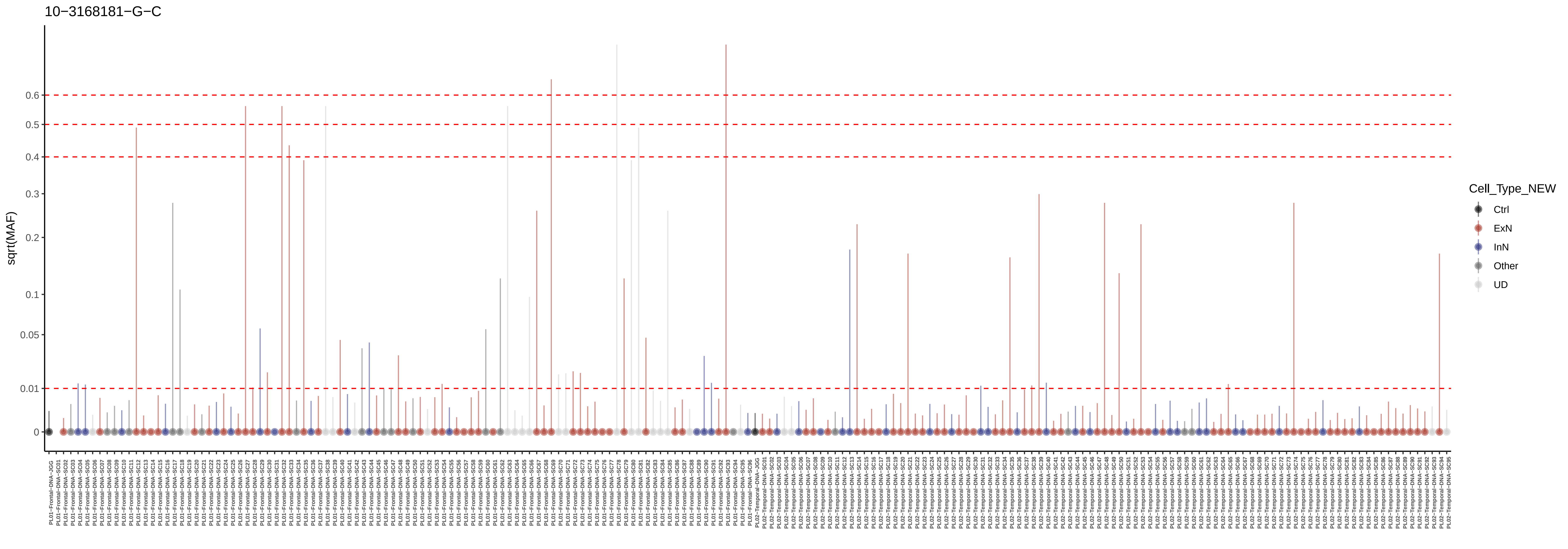
10−3168181−G−C
sqrt(MAF)
0.6
0.5
0.4
0.3
0.2
0.1
0.05
0.01
0
Cell_Type_NEW
Ctrl
ExN
InN
Other
UD

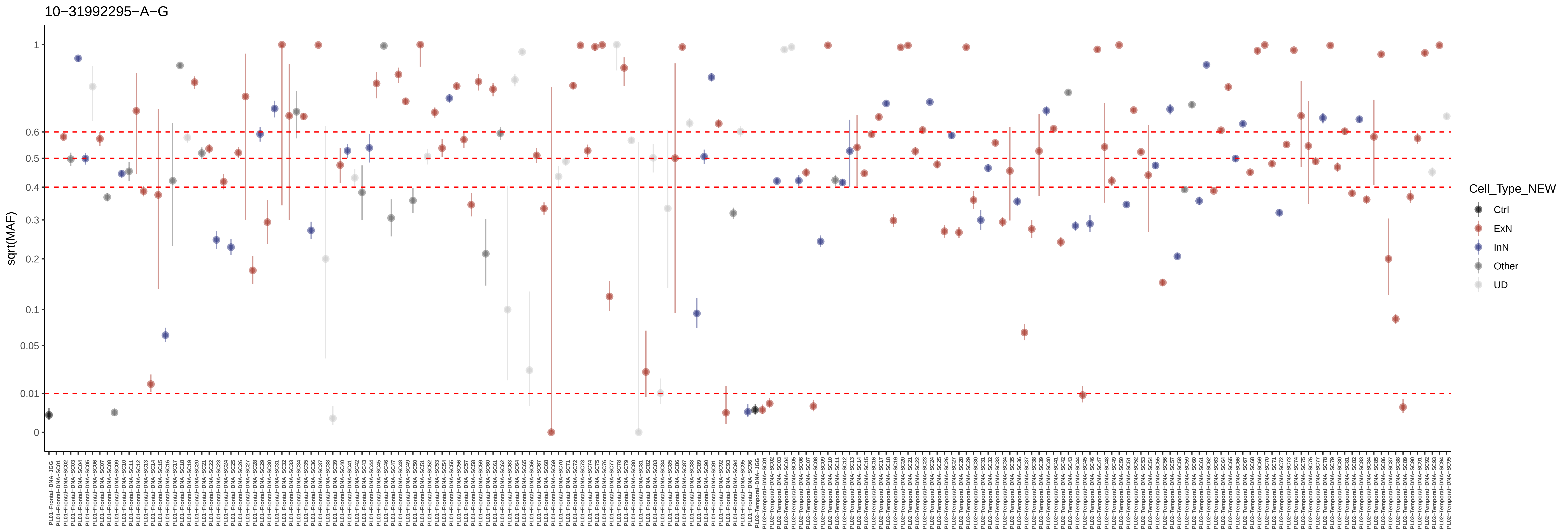
10−31992295−A−G
sqrt(MAF)
1
0.6
0.5
0.4
0.3
0.2
0.1
0.05
0.01
0
Cell_Type_NEW
Ctrl
ExN
InN
Other
UD

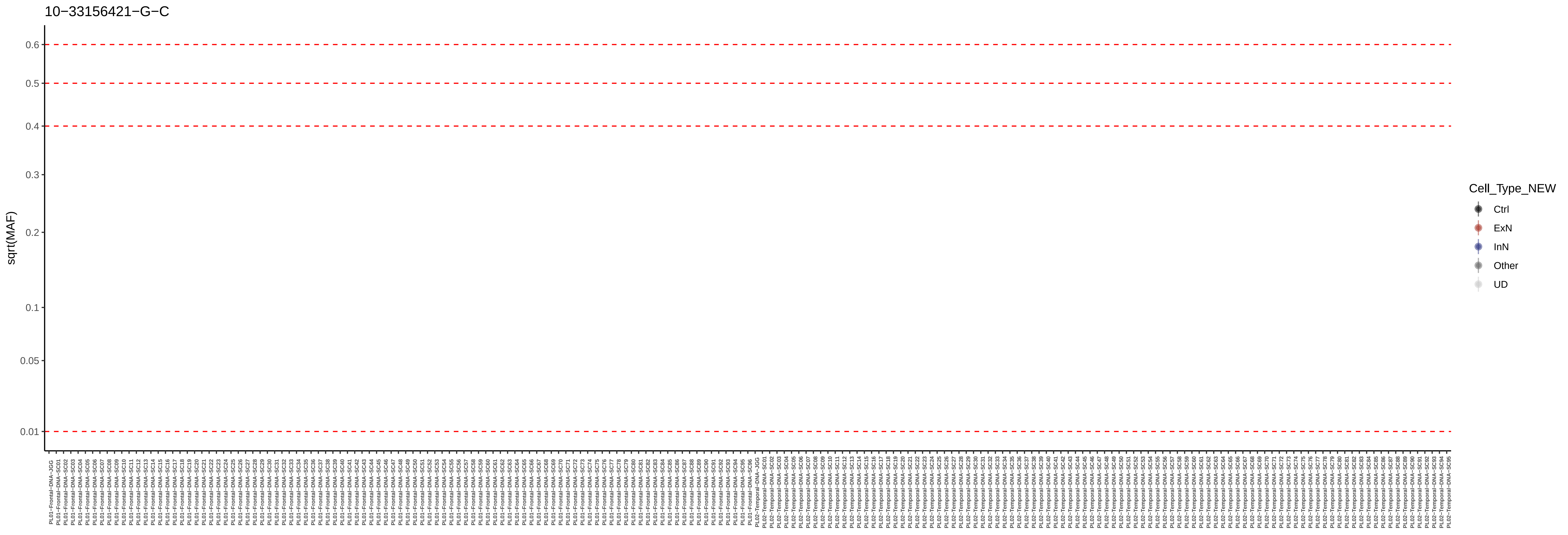

10−33156421−G−C
sqrt(MAF)
0.6
0.5
0.4
0.3
0.2
0.1
0.05
0.01
Cell_Type_NEW
Ctrl
ExN
InN
Other
UD

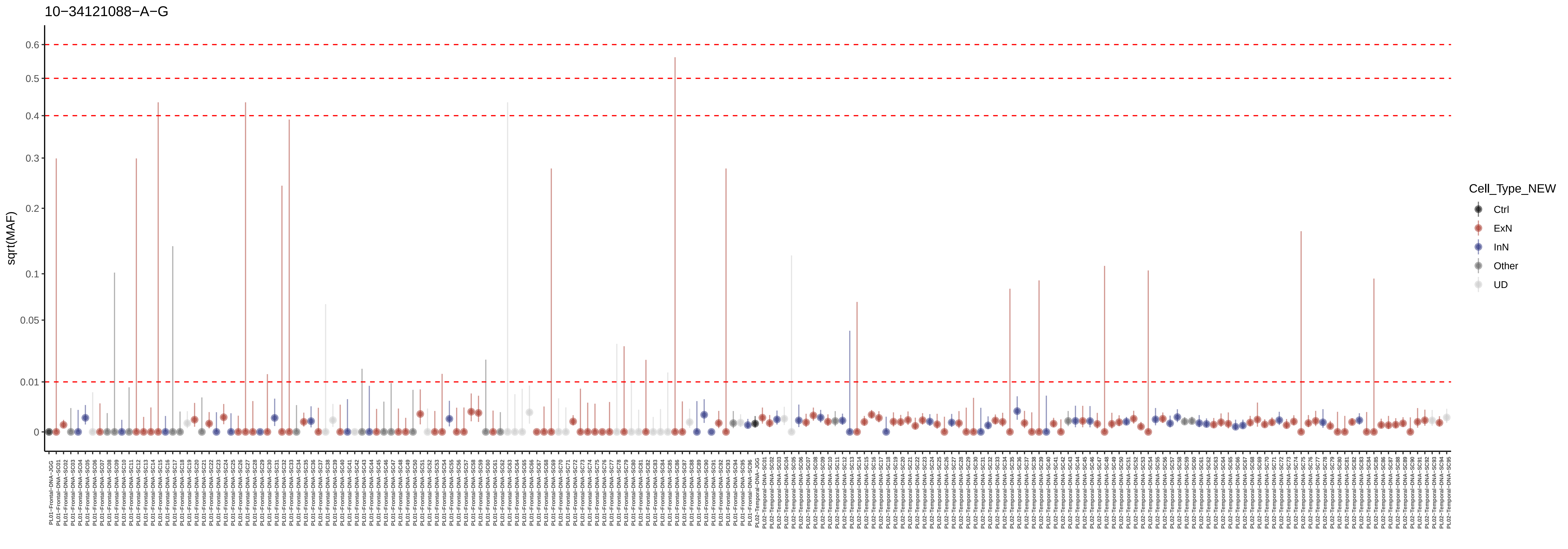
10−34121088−A−G
sqrt(MAF)
0.6
0.5
0.4
0.3
0.2
0.1
0.05
0.01
0
Cell_Type_NEW
Ctrl
ExN
InN
Other
UD

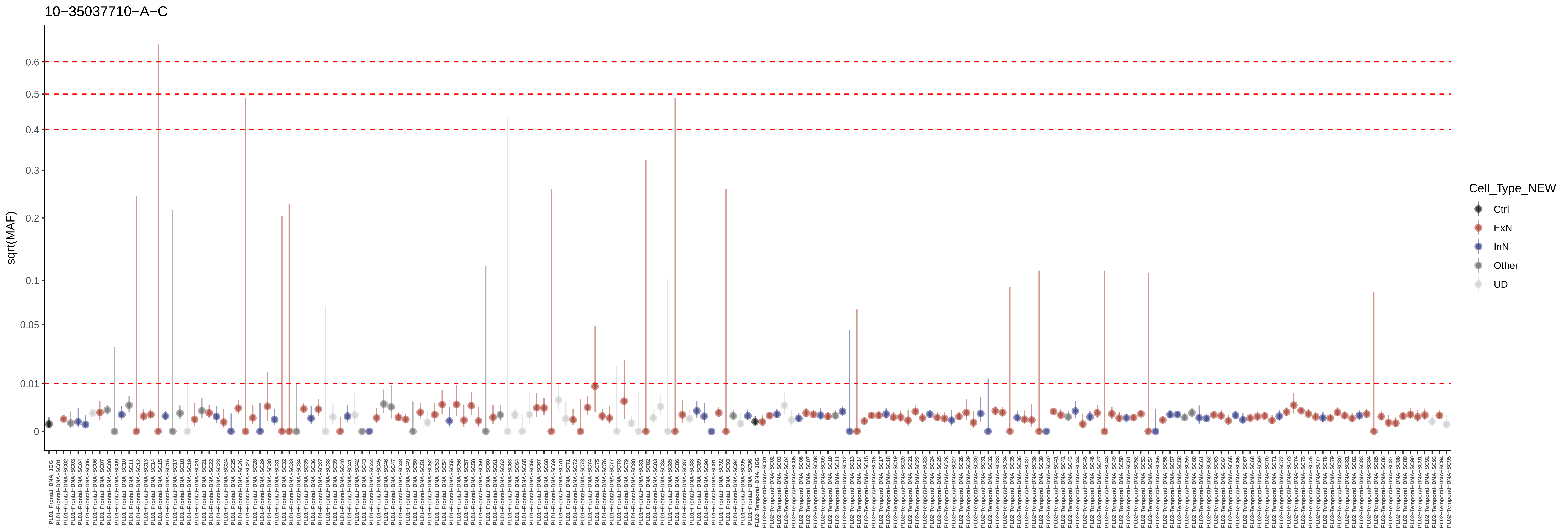
10−35037710−A−C
sqrt(MAF)
0.6
0.5
0.4
0.3
0.2
0.1
0.05
0.01
0
Cell_Type_NEW
Ctrl
ExN
InN
Other
UD

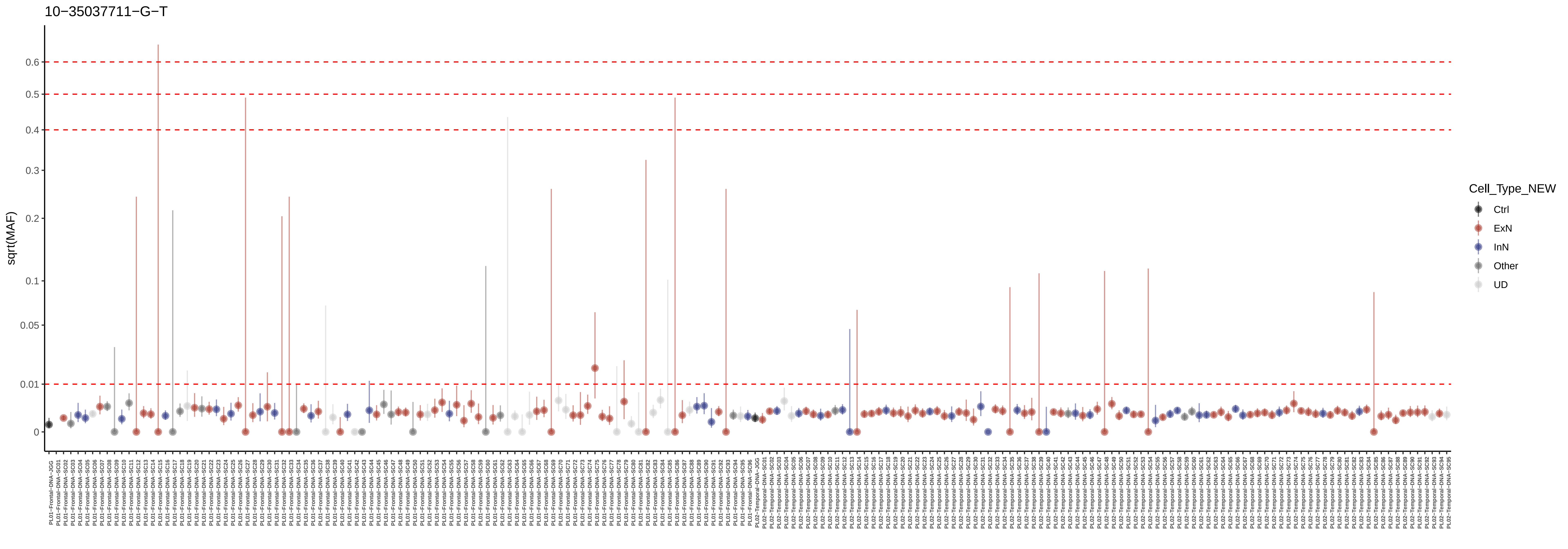
10−35037711−G−T
sqrt(MAF)
0.6
0.5
0.4
0.3
0.2
0.1
0.05
0.01
0
Cell_Type_NEW
Ctrl
ExN
InN
Other
UD

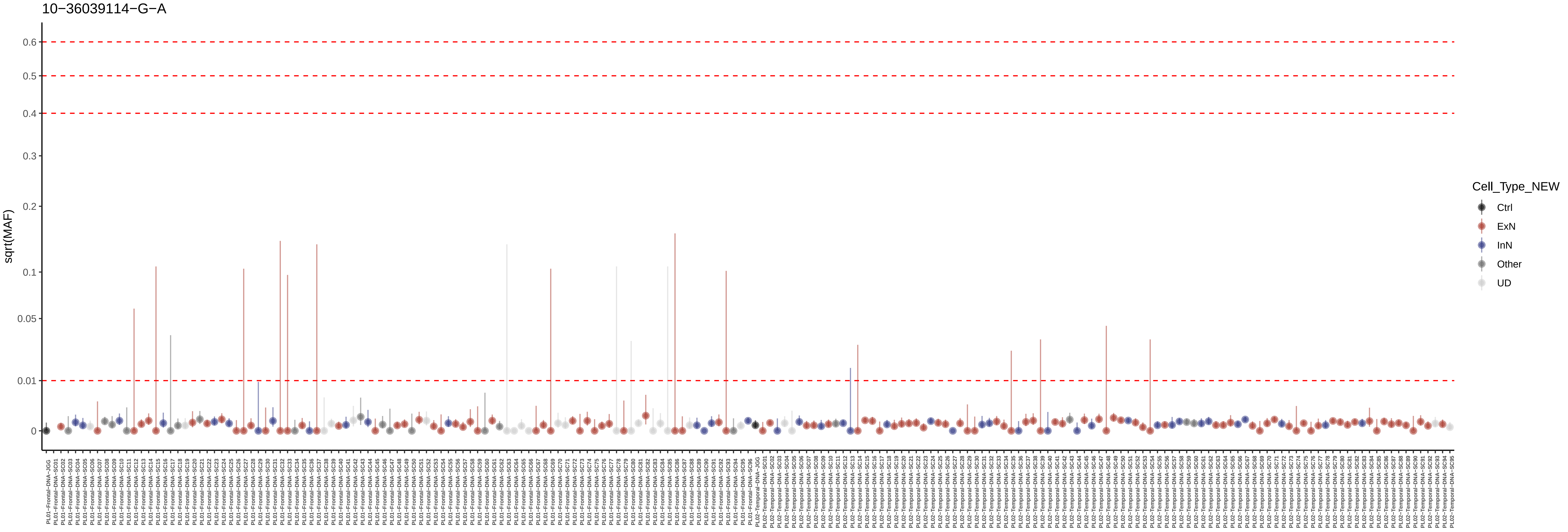

10−36039114−G−A
sqrt(MAF)
0.6
0.5
0.4
0.3
0.2
0.1
0.05
0.01
0
Cell_Type_NEW
Ctrl
ExN
InN
Other
UD

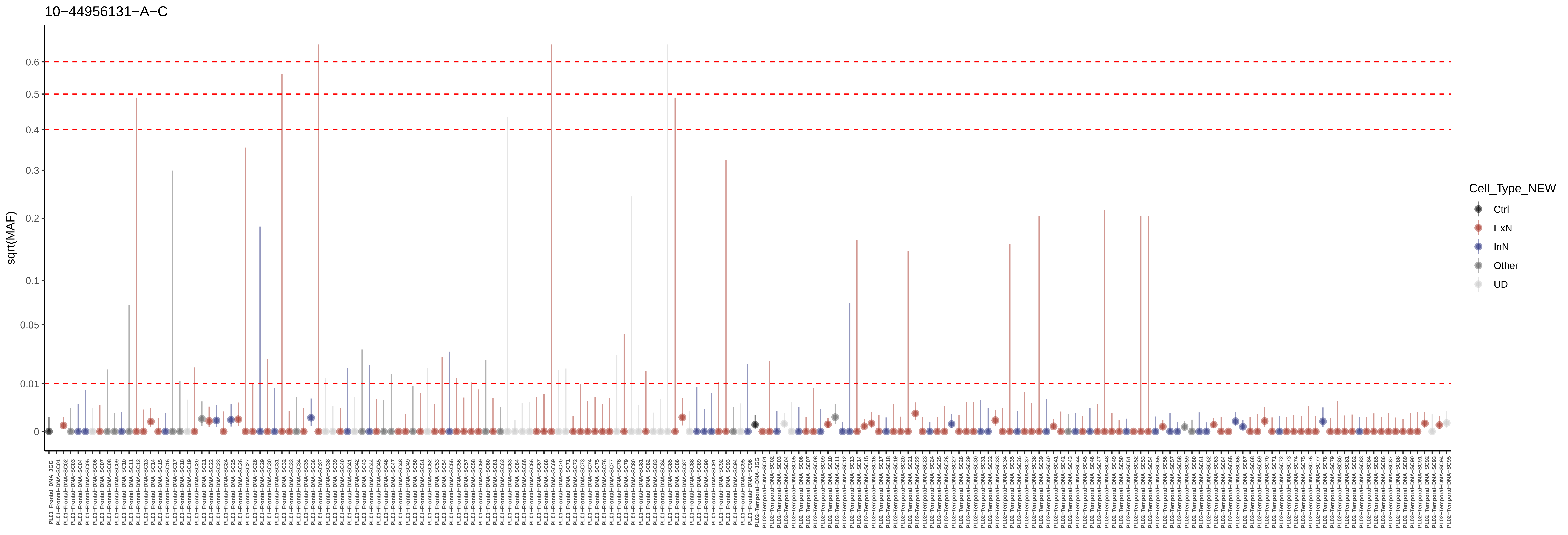

10−44956131−A−C
sqrt(MAF)
0.6
0.5
0.4
0.3
0.2
0.1
0.05
0.01
0
Cell_Type_NEW
Ctrl
ExN
InN
Other
UD

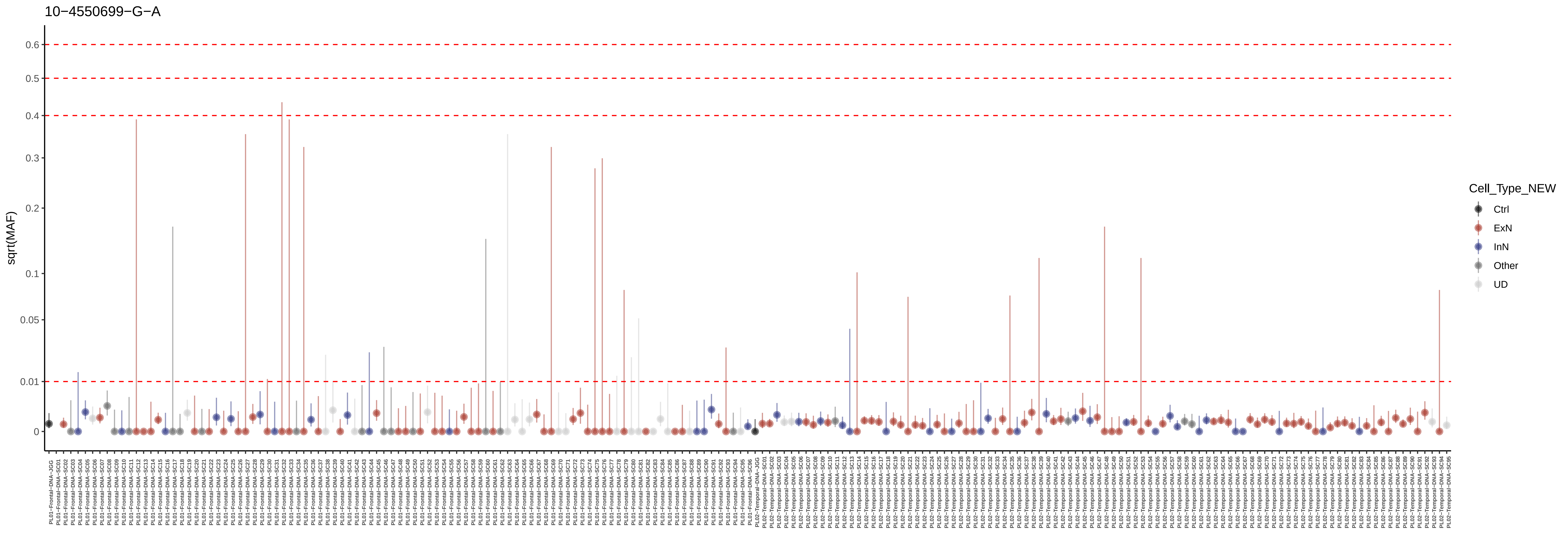
10−4550699−G−A
sqrt(MAF)
0.6
0.5
0.4
0.3
0.2
0.1
0.05
0.01
0
Cell_Type_NEW
Ctrl
ExN
InN
Other
UD

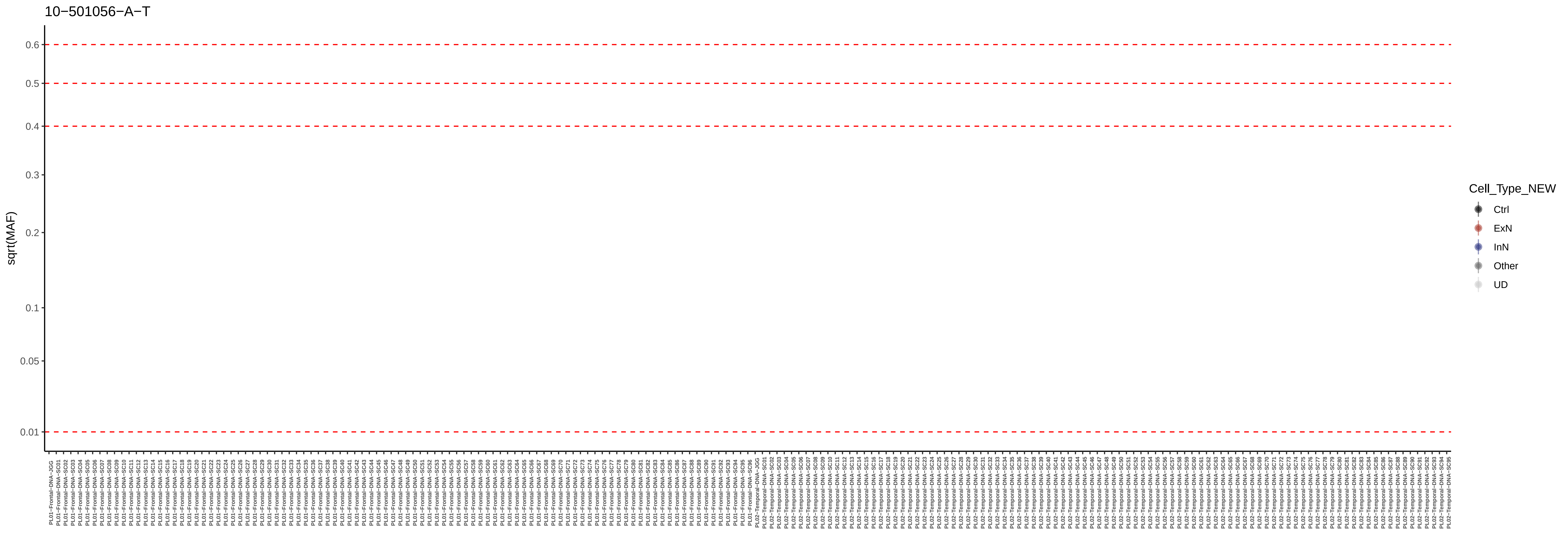
10−501056−A−T
sqrt(MAF)
0.6
0.5
0.4
0.3
0.2
0.1
0.05
0.01
Cell_Type_NEW
Ctrl
ExN
InN
Other
UD

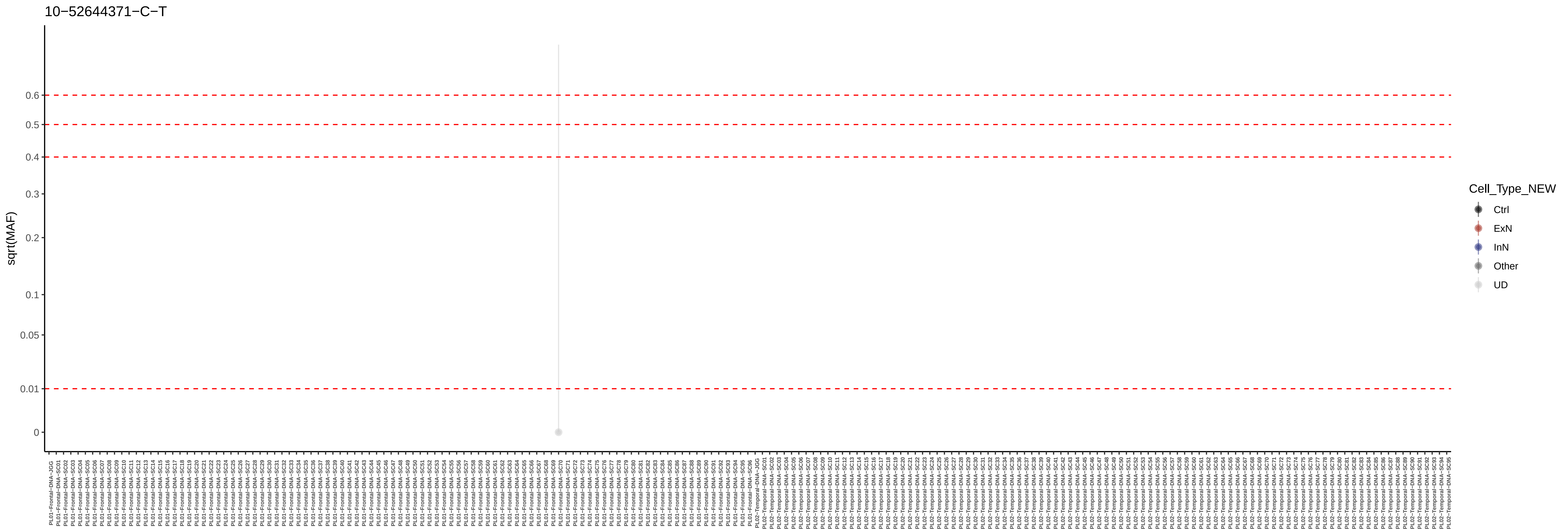

10−52644371−C−T
sqrt(MAF)
0.6
0.5
0.4
0.3
0.2
0.1
0.05
0.01
0
Cell_Type_NEW
Ctrl
ExN
InN
Other
UD

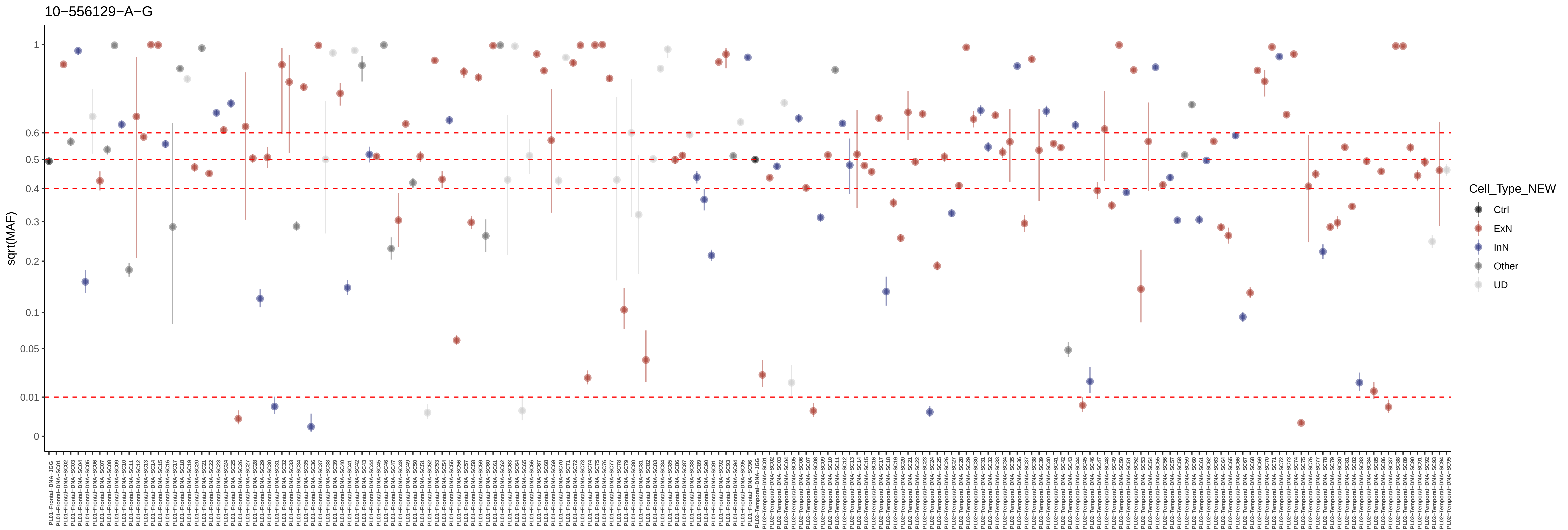
10−556129−A−G
sqrt(MAF)
1
0.6
0.5
0.4
0.3
0.2
0.1
0.05
0.01
0
Cell_Type_NEW
Ctrl
ExN
InN
Other
UD

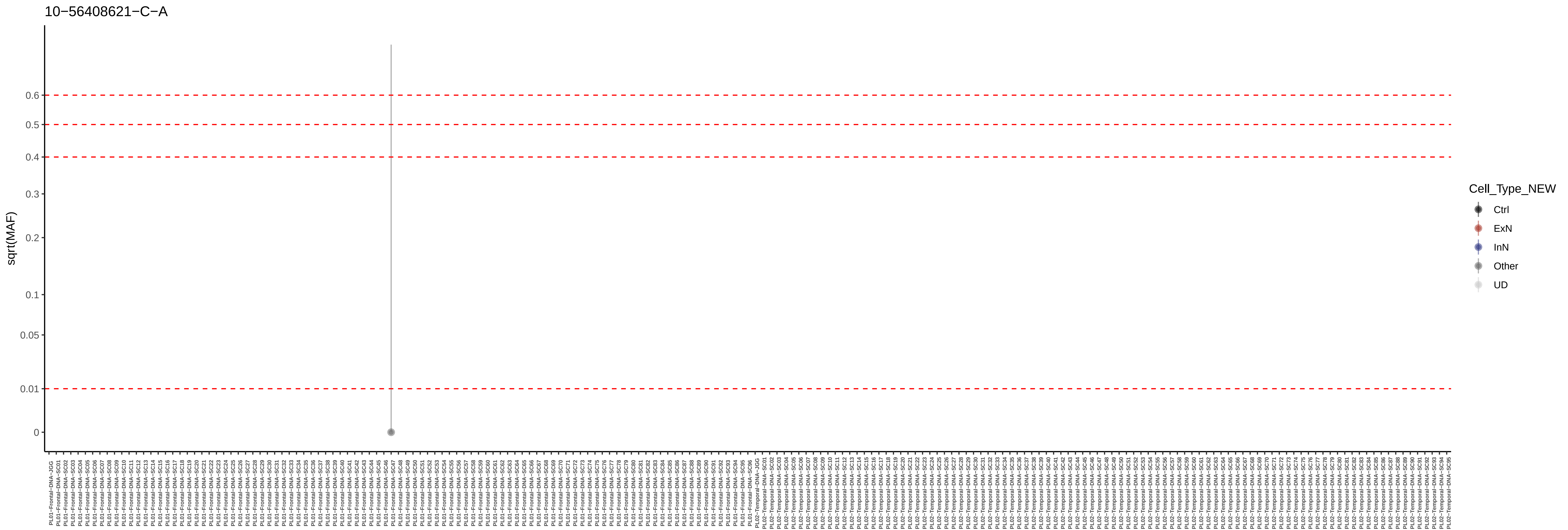
10−56408621−C−A
sqrt(MAF)
0.6
0.5
0.4
0.3
0.2
0.1
0.05
0.01
0
Cell_Type_NEW
Ctrl
ExN
InN
Other
UD

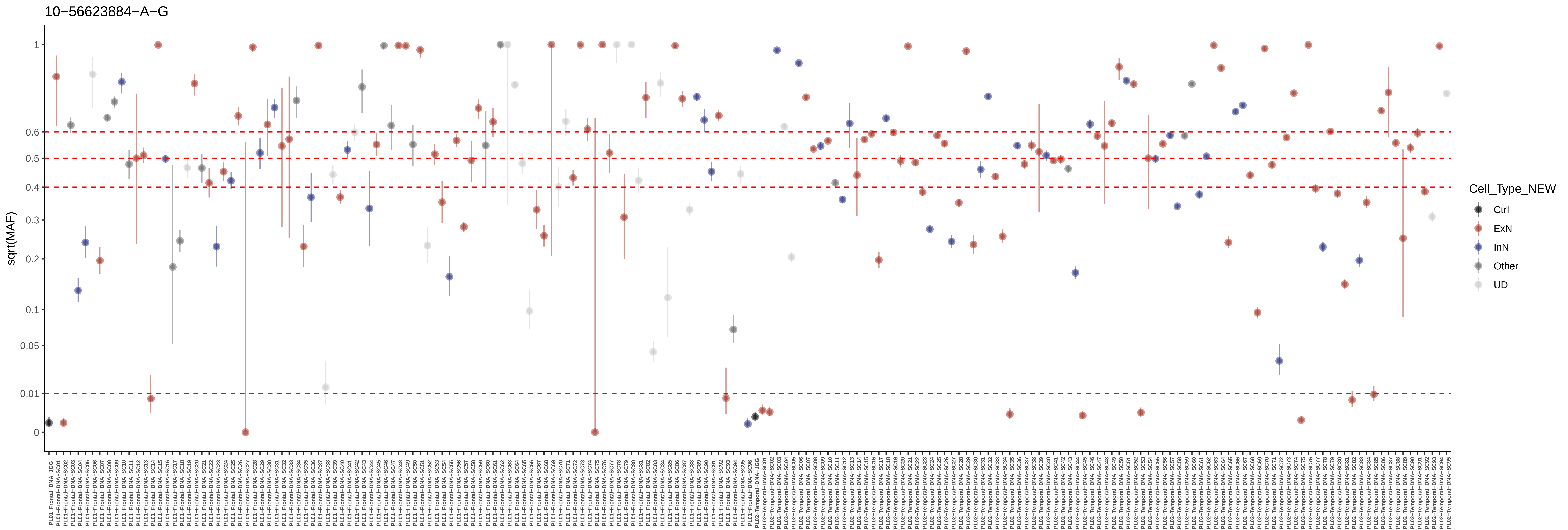

10−56623884−A−G
sqrt(MAF)
1
0.6
0.5
0.4
0.3
0.2
0.1
0.05
0.01
0
Cell_Type_NEW
Ctrl
ExN
InN
Other
UD

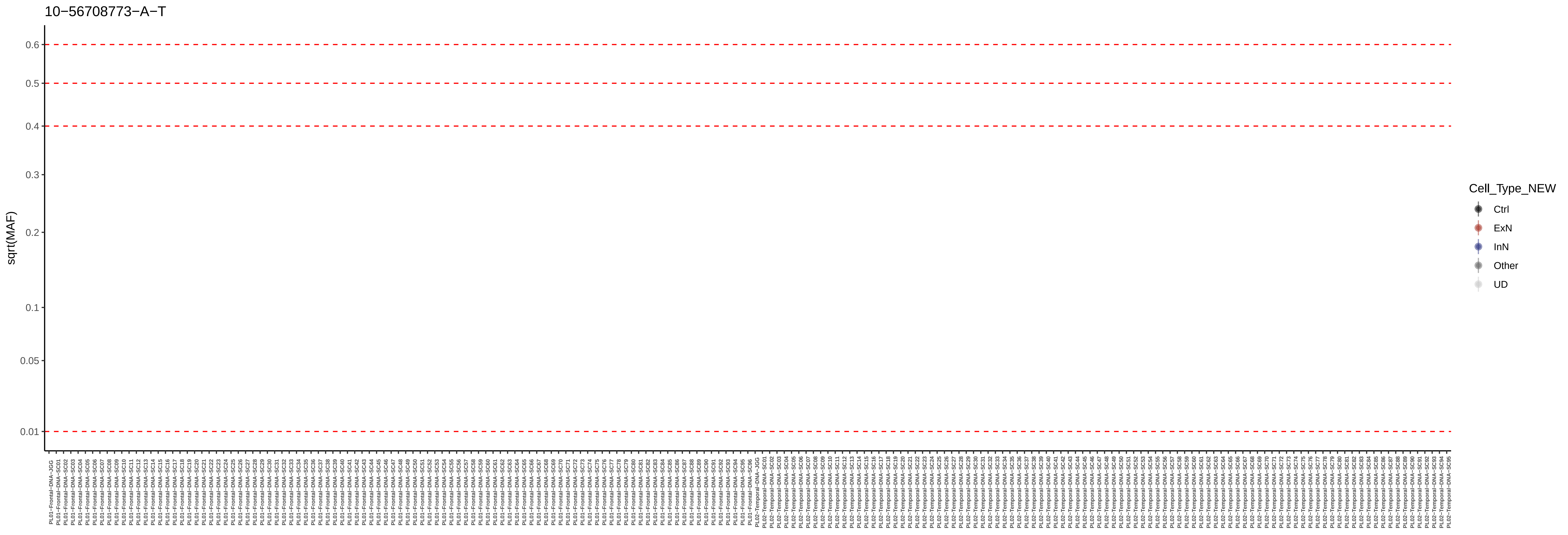
10−56708773−A−T
sqrt(MAF)
0.6
0.5
0.4
0.3
0.2
0.1
0.05
0.01
Cell_Type_NEW
Ctrl
ExN
InN
Other
UD

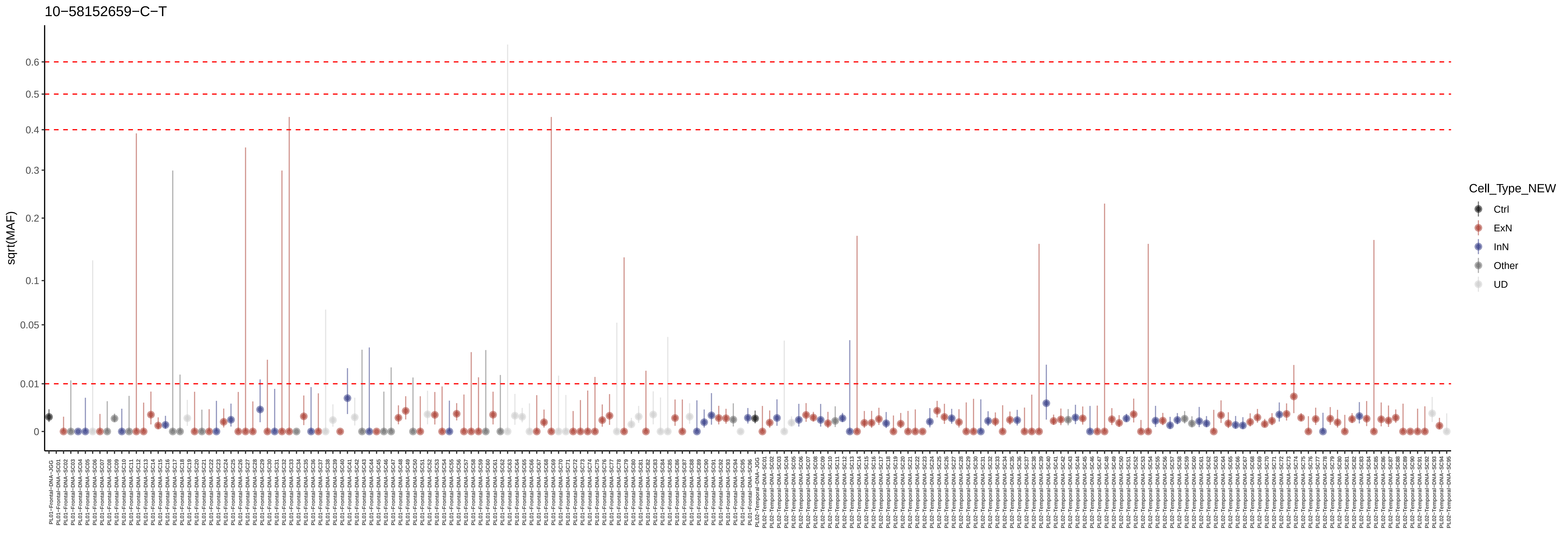
10−58152659−C−T
sqrt(MAF)
0.6
0.5
0.4
0.3
0.2
0.1
0.05
0.01
0
Cell_Type_NEW
Ctrl
ExN
InN
Other
UD

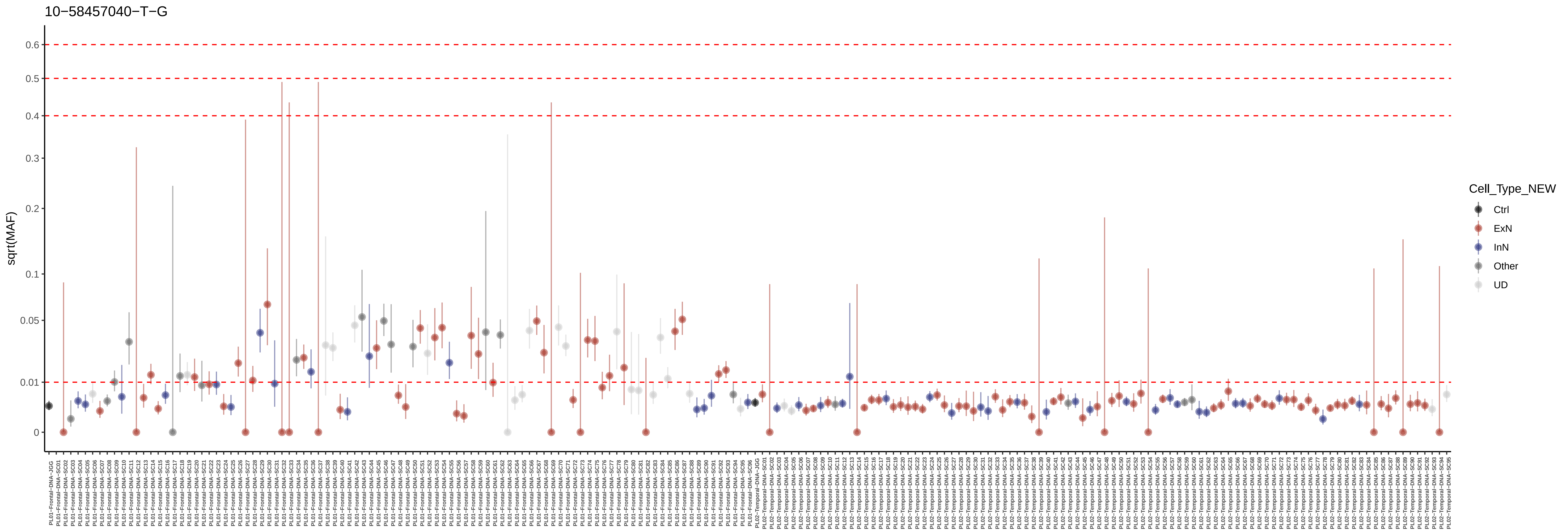
10−58457040−T−G
sqrt(MAF)
0.6
0.5
0.4
0.3
0.2
0.1
0.05
0.01
0
Cell_Type_NEW
Ctrl
ExN
InN
Other
UD

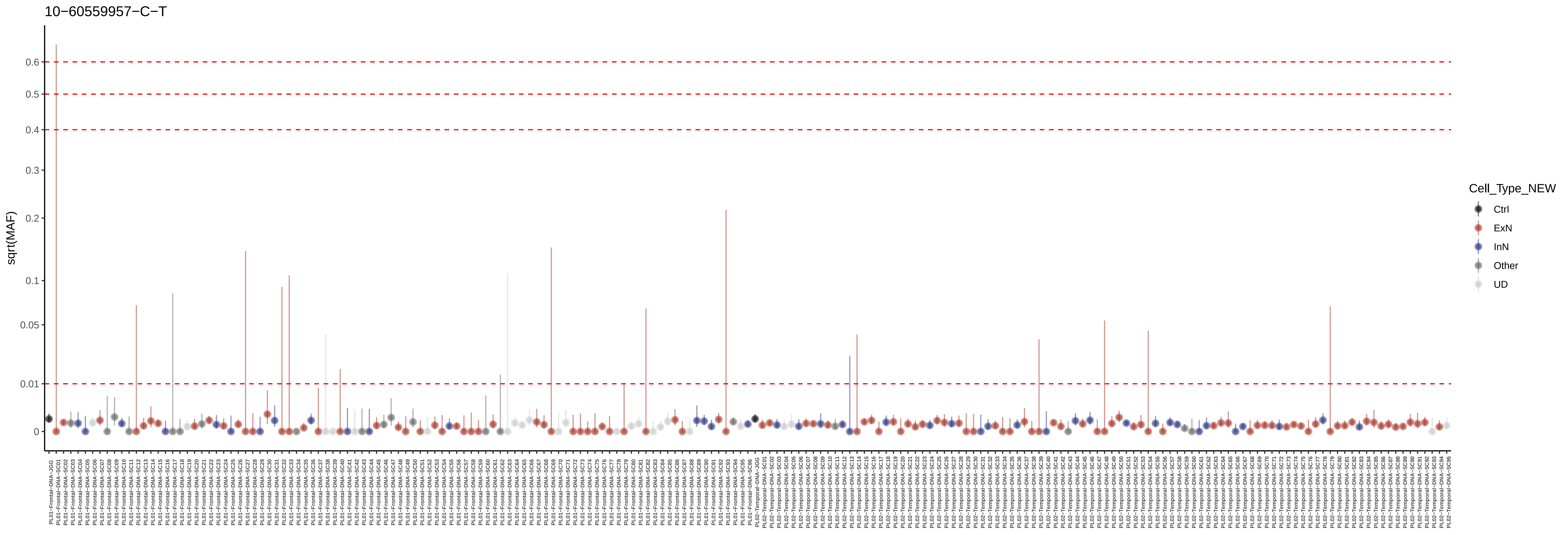

10−60559957−C−T
sqrt(MAF)
Cell_Type_NEW
Ctrl
ExN
InN
Other
UD

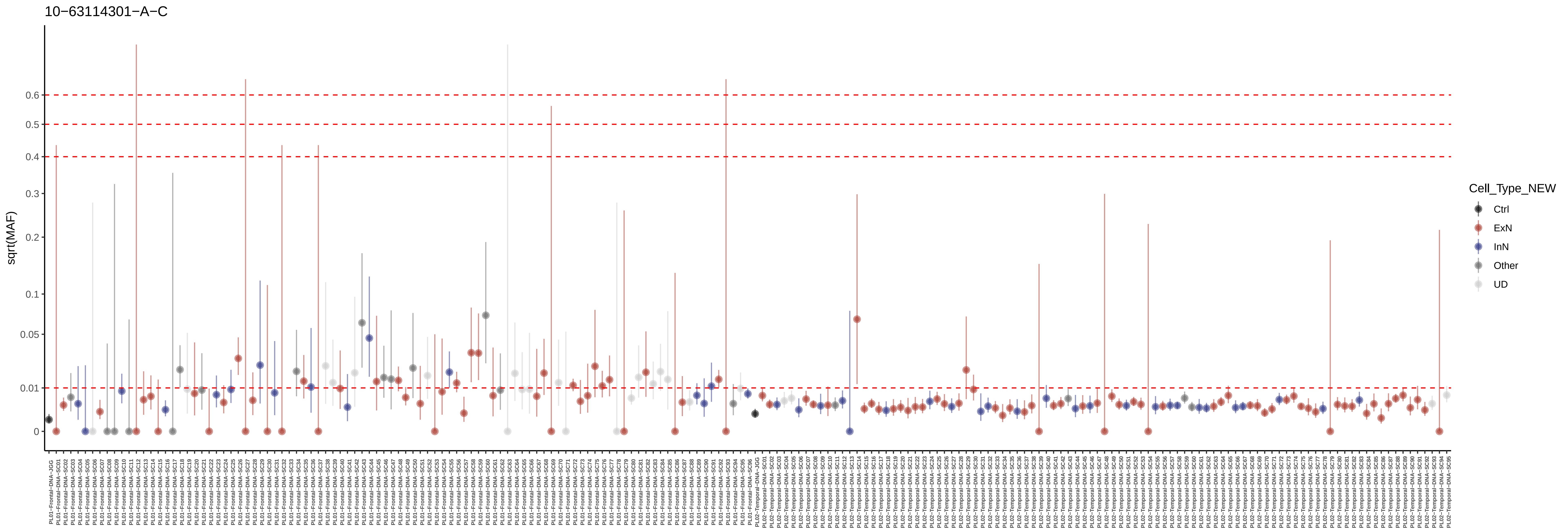
10−63114301−A−C
sqrt(MAF)
0.6
0.5
0.4
0.3
0.2
0.1
0.05
0.01
0
Cell_Type_NEW
Ctrl
ExN
InN
Other
UD

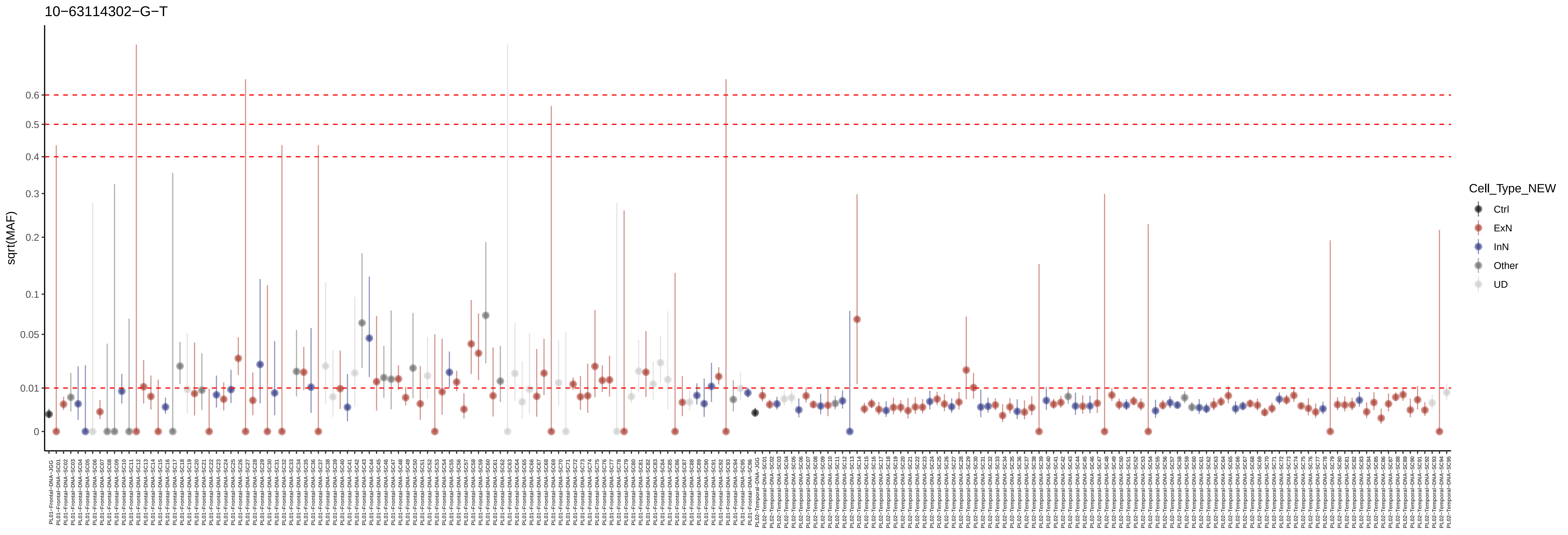
10−63114302−G−T
sqrt(MAF)
0.6
0.5
0.4
0.3
0.2
0.1
0.05
0.01
0
Cell_Type_NEW
Ctrl
ExN
InN
Other
UD

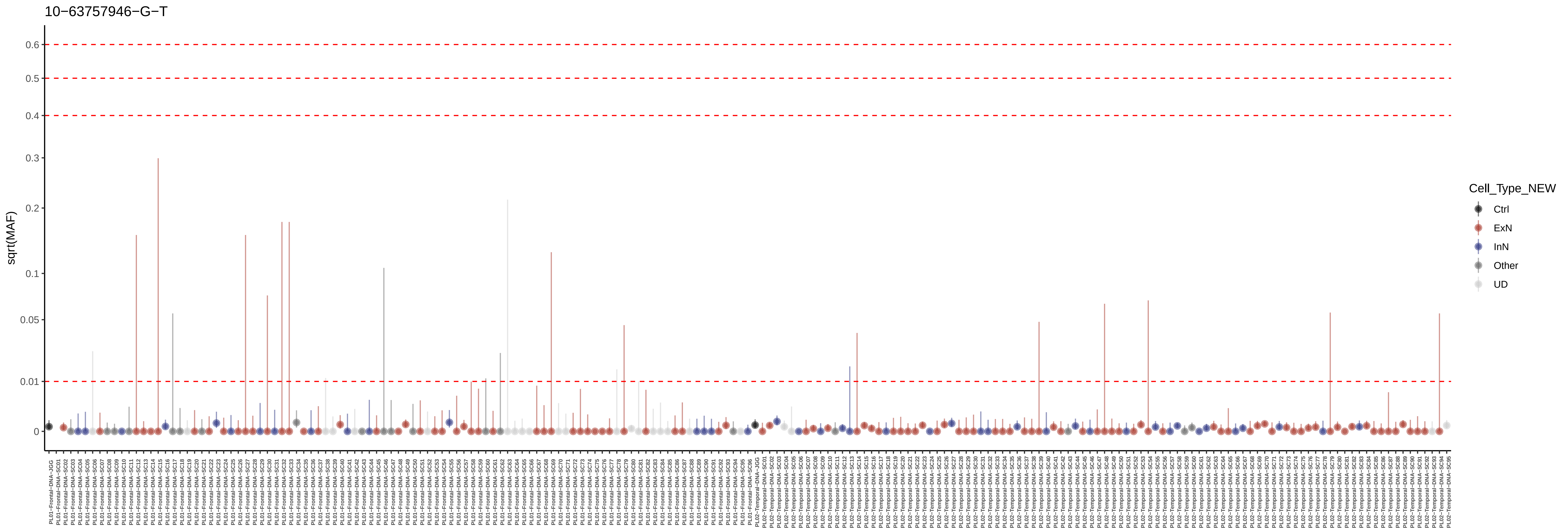

10−63757946−G−T
sqrt(MAF)
0.6
0.5
0.4
0.3
0.2
0.1
0.05
0.01
0
Cell_Type_NEW
Ctrl
ExN
InN
Other
UD

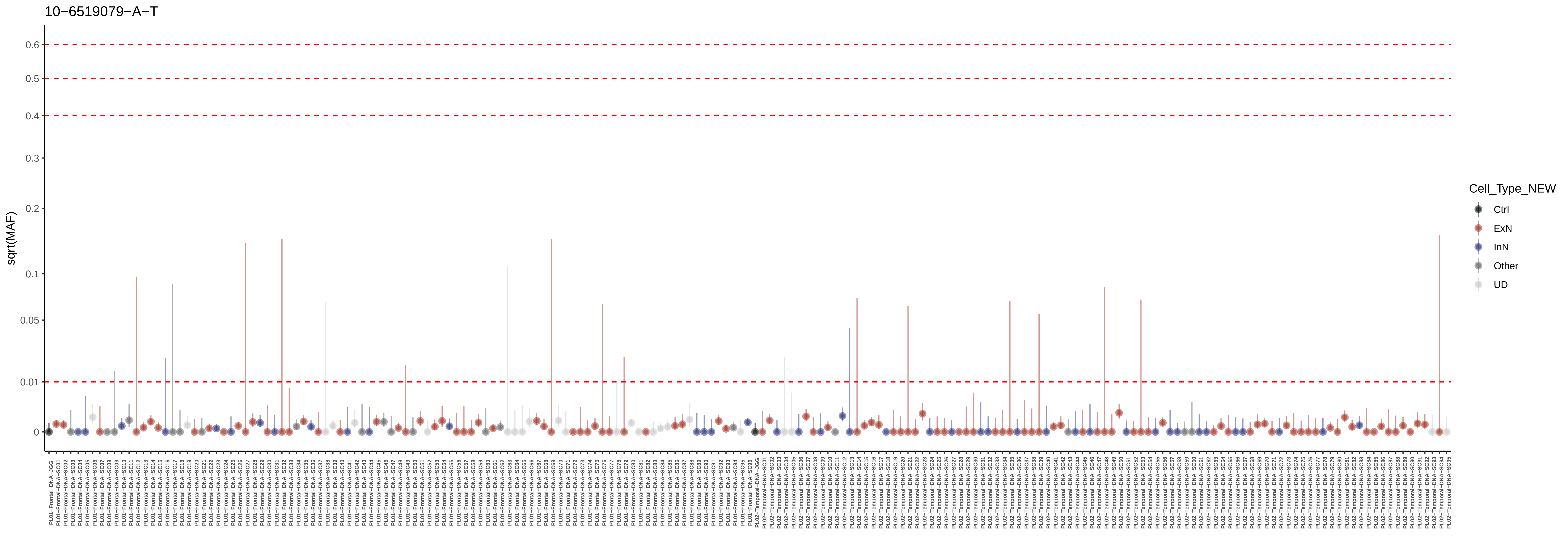
10−6519079−A−T
sqrt(MAF)
0.6
0.5
0.4
0.3
0.2
0.1
0.05
0.01
0
Cell_Type_NEW
Ctrl
ExN
InN
Other
UD

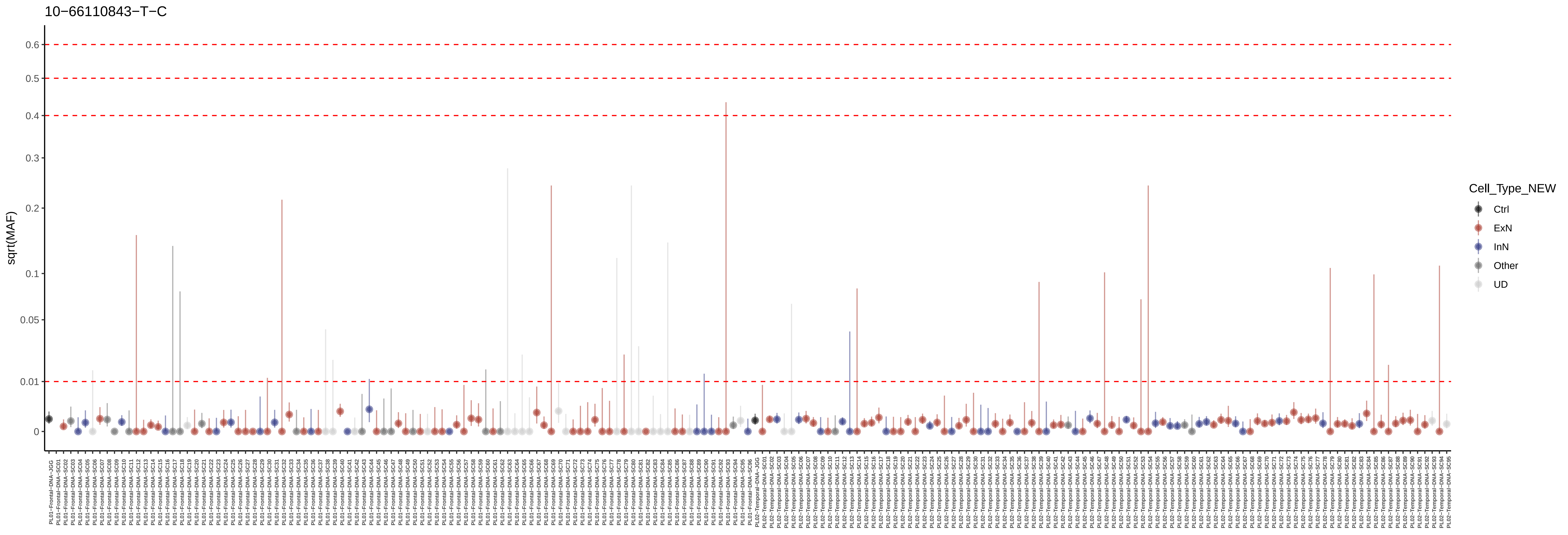
10−66110843−T−C
sqrt(MAF)
0.6
0.5
0.4
0.3
0.2
0.1
0.05
0.01
0
Cell_Type_NEW
Ctrl
ExN
InN
Other
UD

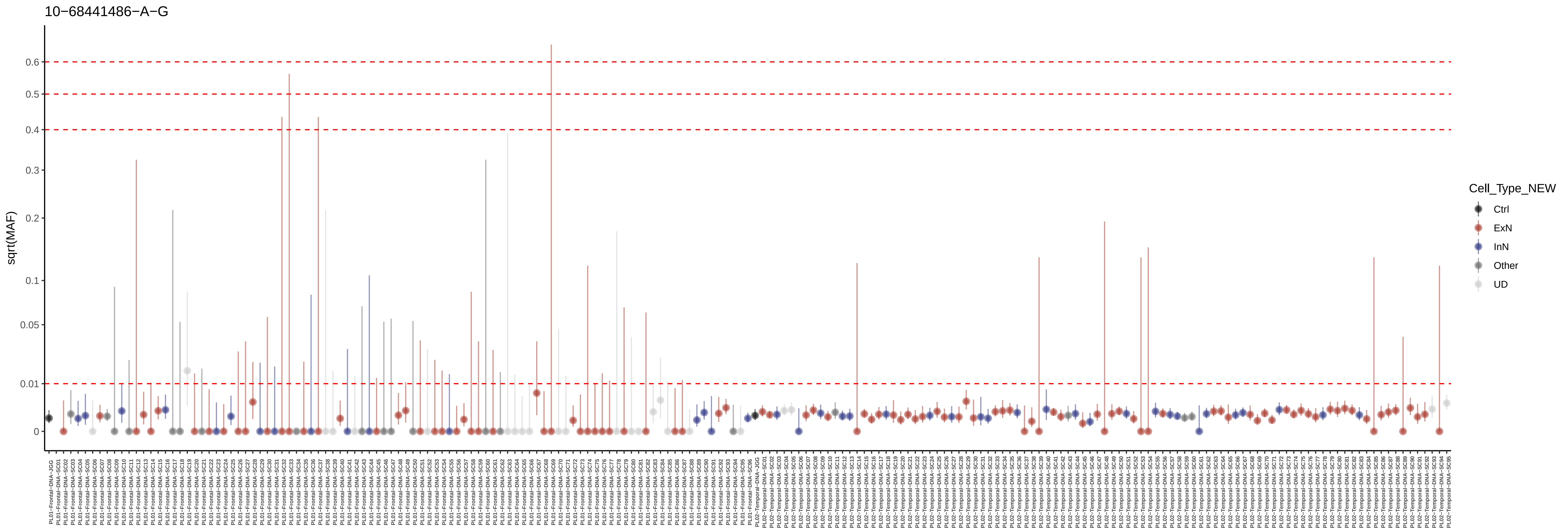
10−68441486−A−G
sqrt(MAF)
0.6
0.5
0.4
0.3
0.2
0.1
0.05
0.01
0
Cell_Type_NEW
Ctrl
ExN
InN
Other
UD

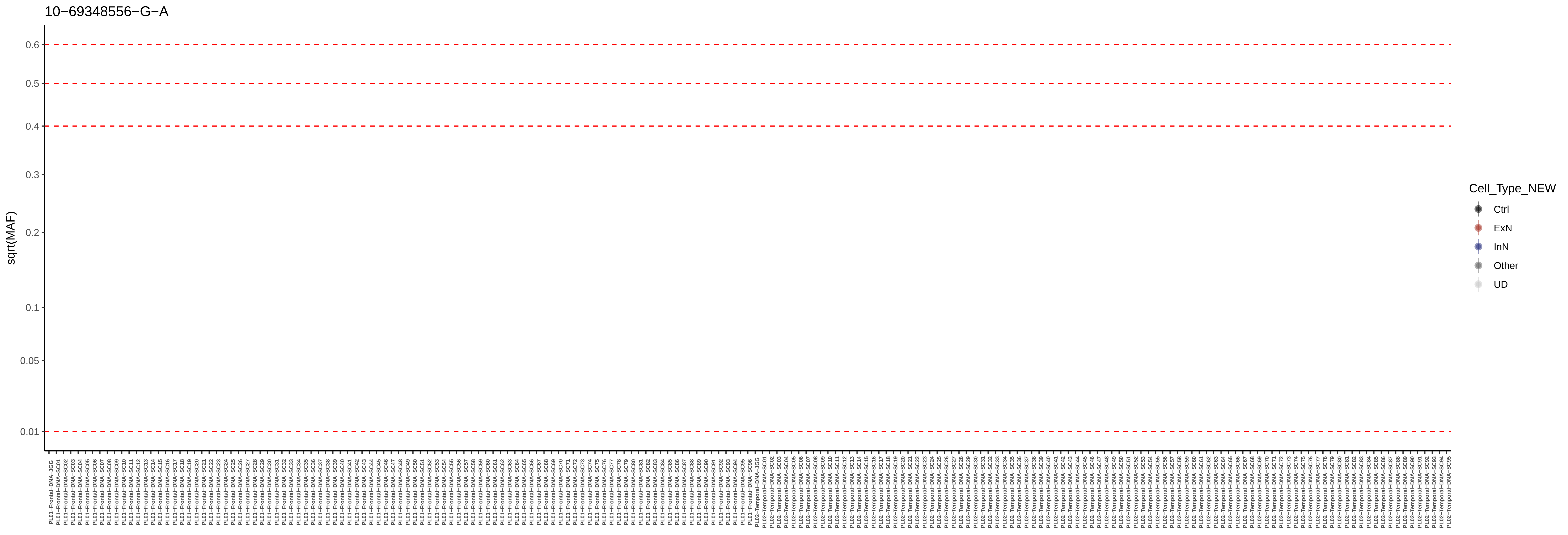
10−69348556−G−A
sqrt(MAF)
0.6
0.5
0.4
0.3
0.2
0.1
0.05
0.01
Cell_Type_NEW
Ctrl
ExN
InN
Other
UD

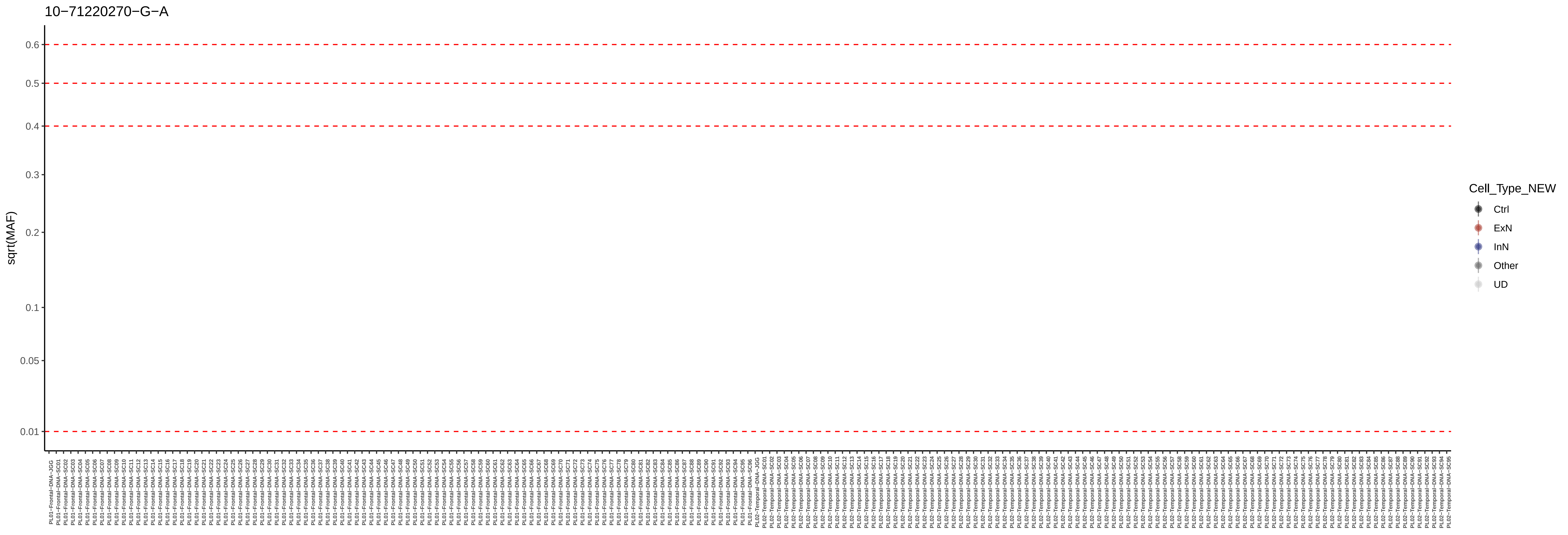
10−71220270−G−A
sqrt(MAF)
0.6
0.5
0.4
0.3
0.2
0.1
0.05
0.01
Cell_Type_NEW
Ctrl
ExN
InN
Other
UD

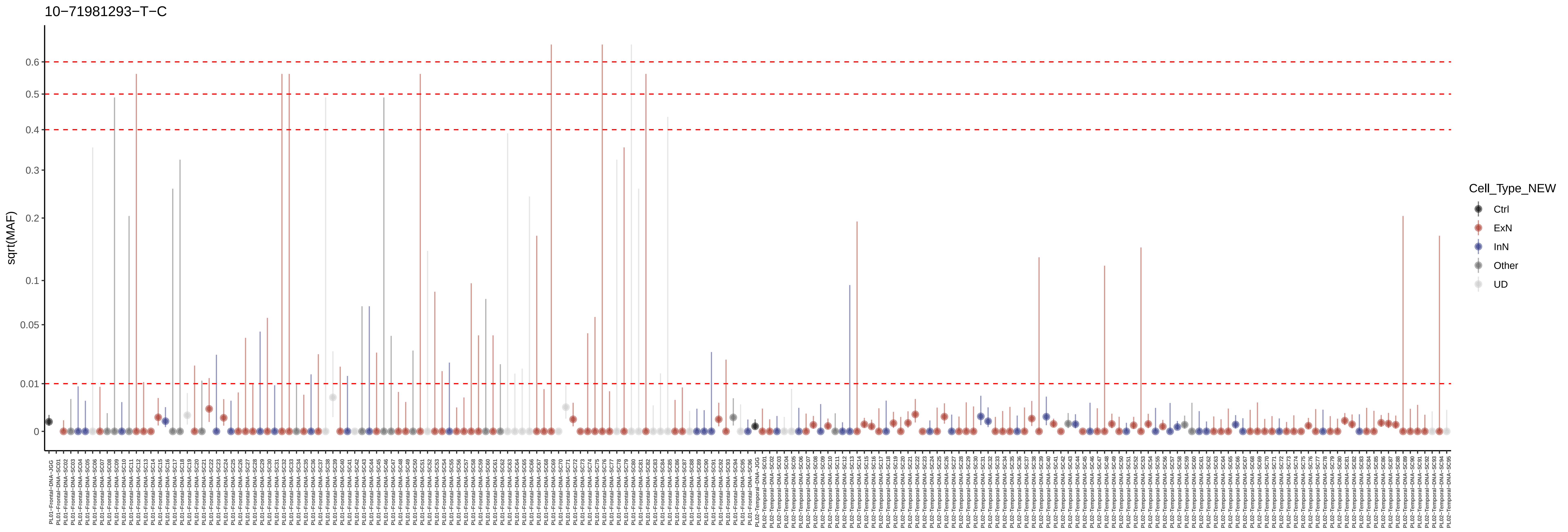
10−71981293−T−C
sqrt(MAF)
0.6
0.5
0.4
0.3
0.2
0.1
0.05
0.01
0
Cell_Type_NEW
Ctrl
ExN
InN
Other
UD

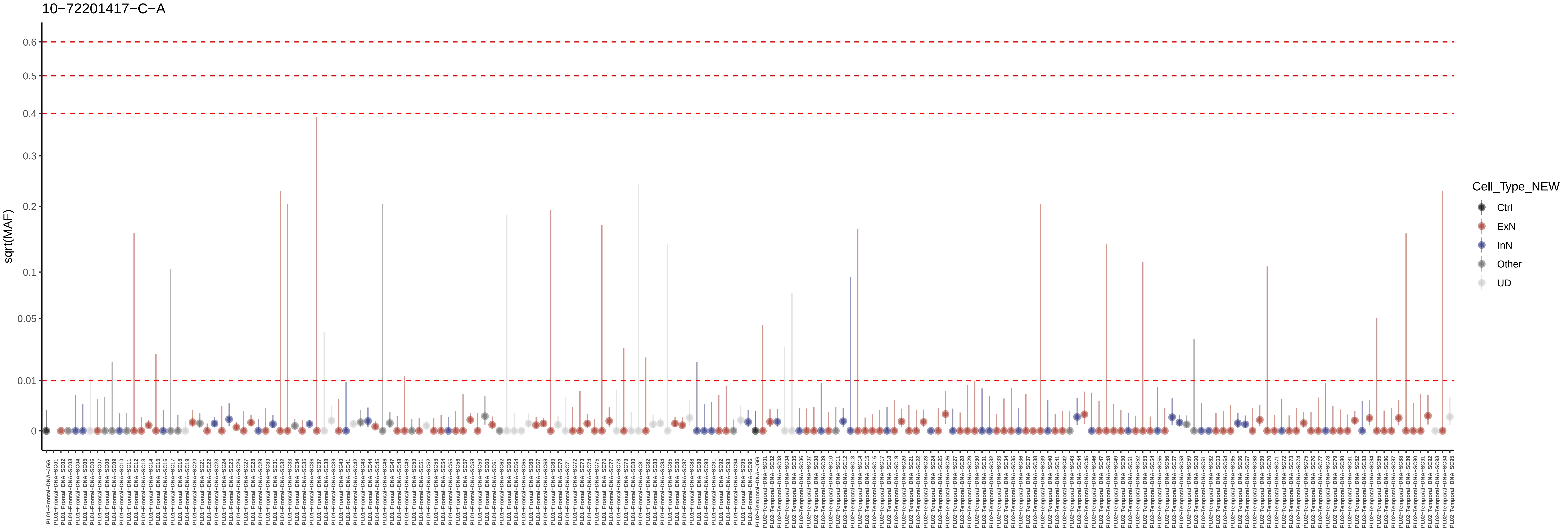
10−72201417−C−A
sqrt(MAF)
0.6
0.5
0.4
0.3
0.2
0.1
0.05
0.01
0
Cell_Type_NEW
Ctrl
ExN
InN
Other
UD

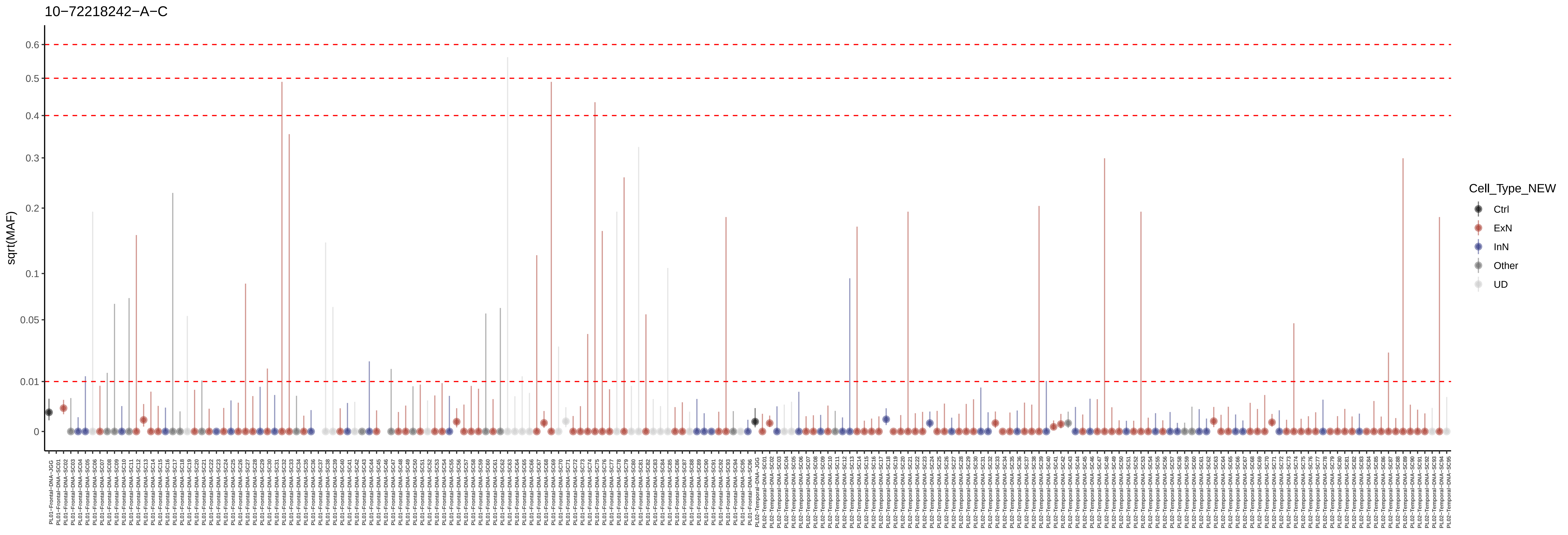
10−72218242−A−C
sqrt(MAF)
0.6
0.5
0.4
0.3
0.2
0.1
0.05
0.01
0
Cell_Type_NEW
Ctrl
ExN
InN
Other
UD

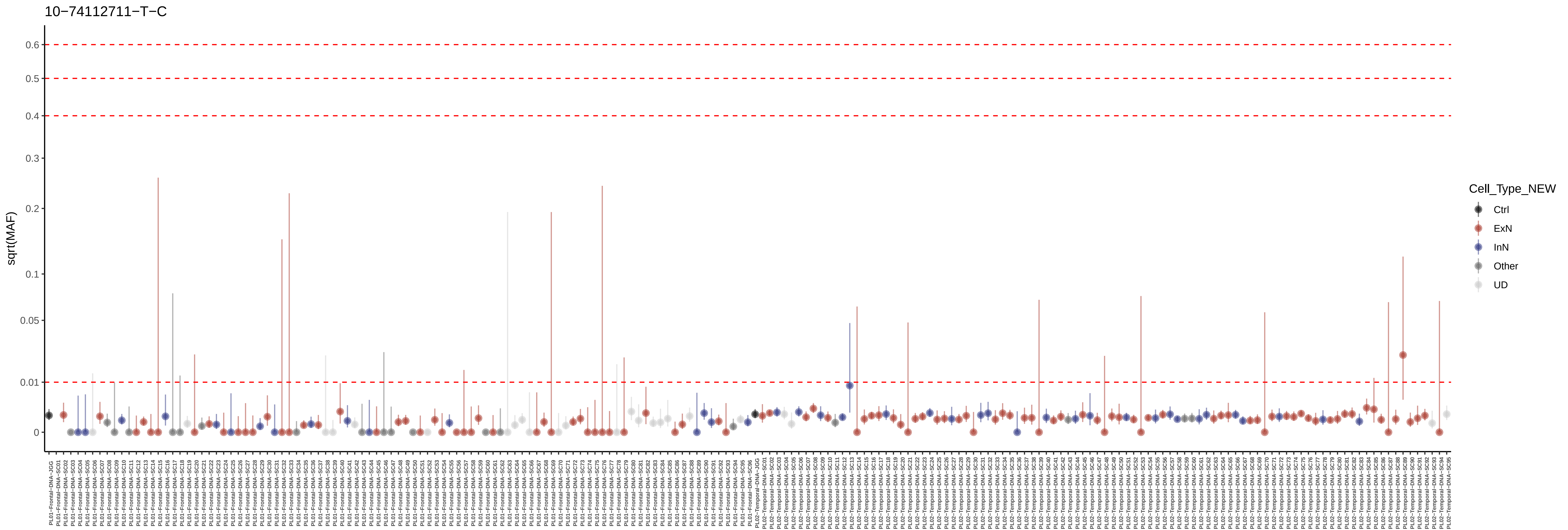

10−74112711−T−C
sqrt(MAF)
0.6
0.5
0.4
0.3
0.2
0.1
0.05
0.01
0
Cell_Type_NEW
Ctrl
ExN
InN
Other
UD

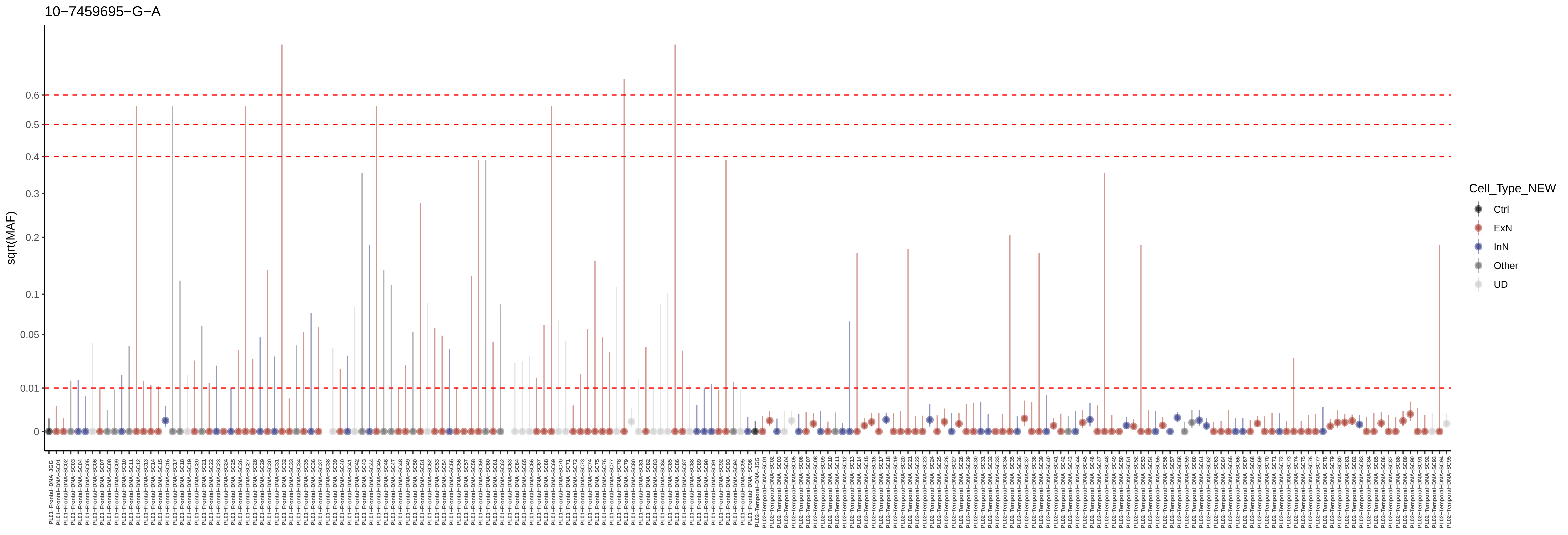

10−7459695−G−A
sqrt(MAF)
0.6
0.5
0.4
0.3
0.2
0.1
0.05
0.01
0
Cell_Type_NEW
Ctrl
ExN
InN
Other
UD

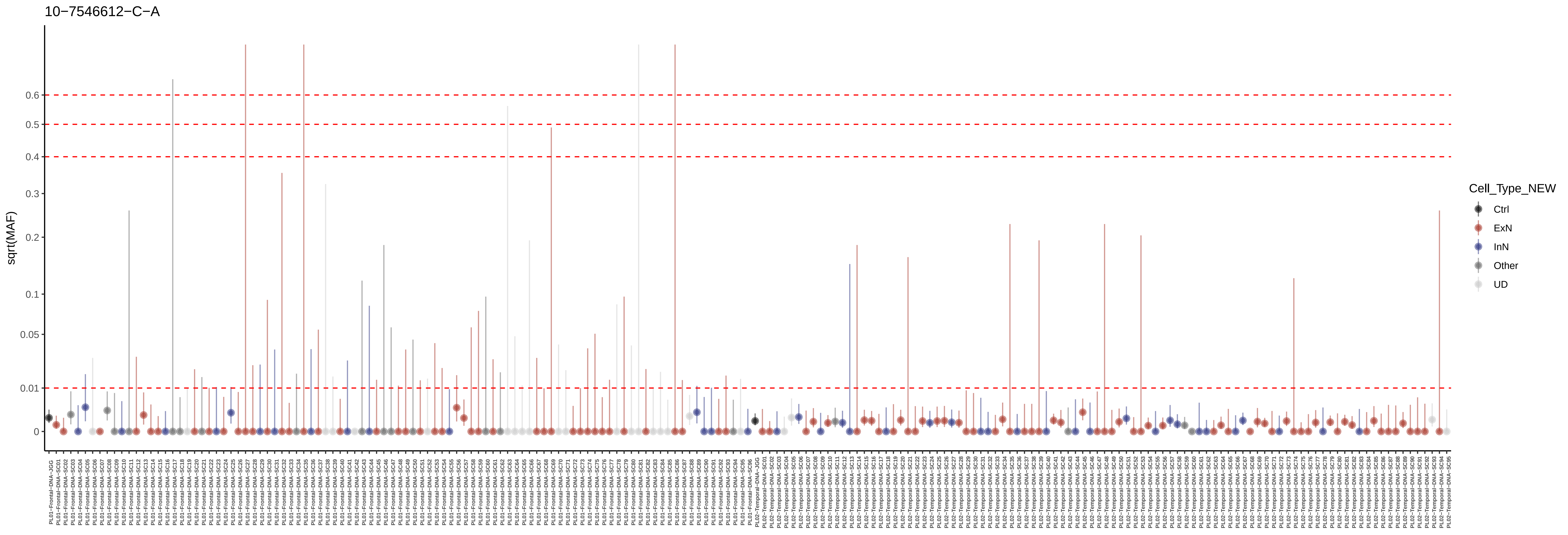
10−7546612−C−A
sqrt(MAF)
0.6
0.5
0.4
0.3
0.2
0.1
0.05
0.01
0
Cell_Type_NEW
Ctrl
ExN
InN
Other
UD

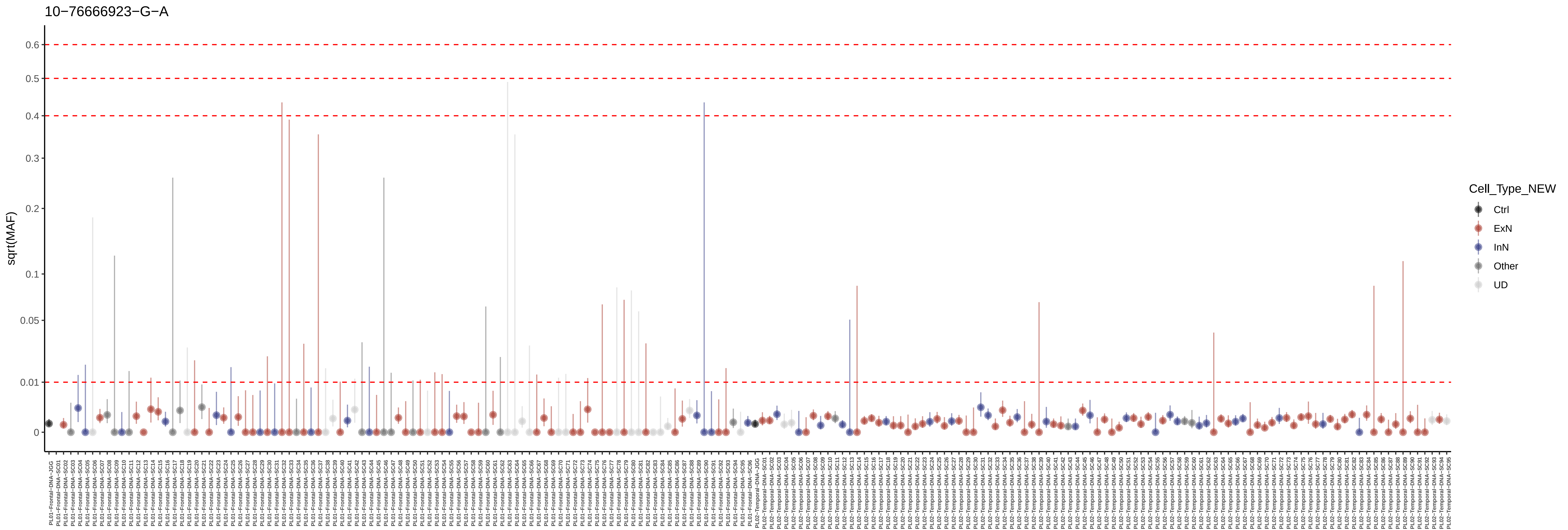
10−76666923−G−A
sqrt(MAF)
0.6
0.5
0.4
0.3
0.2
0.1
0.05
0.01
0
Cell_Type_NEW
Ctrl
ExN
InN
Other
UD

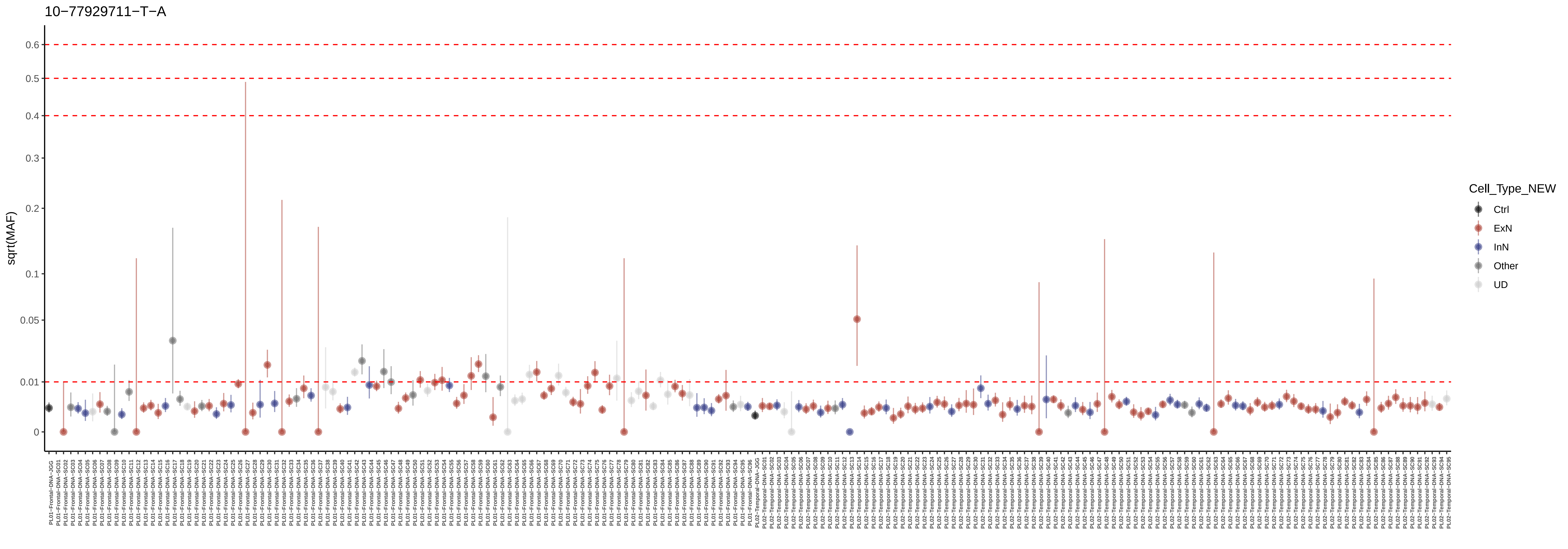

10−77929711−T−A
sqrt(MAF)
0.6
0.5
0.4
0.3
0.2
0.1
0.05
0.01
0
Cell_Type_NEW
Ctrl
ExN
InN
Other
UD

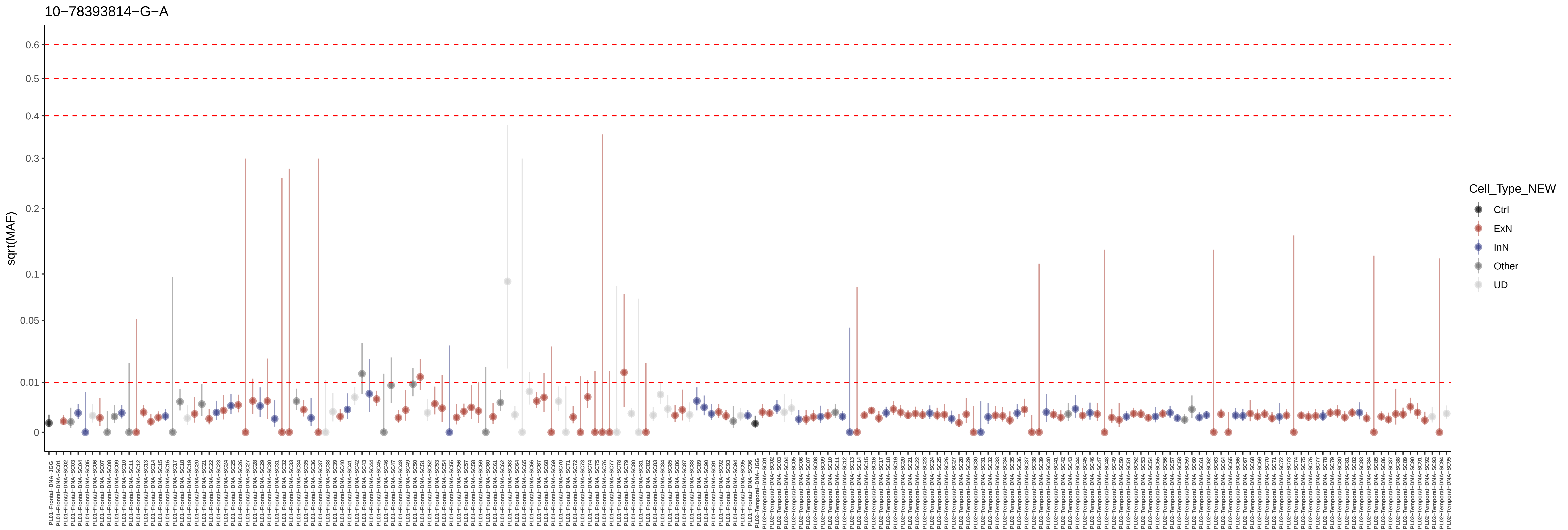
10−78393814−G−A
sqrt(MAF)
0.6
0.5
0.4
0.3
0.2
0.1
0.05
0.01
0
Cell_Type_NEW
Ctrl
ExN
InN
Other
UD

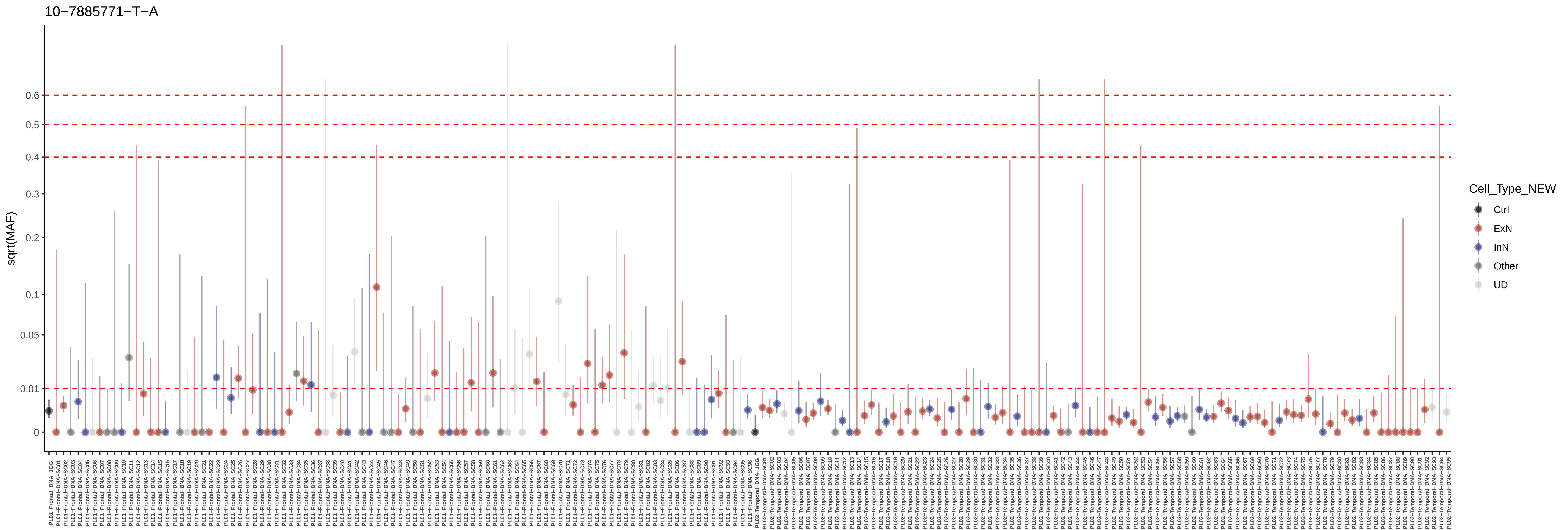

10−7885771−T−A
sqrt(MAF)
0.6
0.5
0.4
0.3
0.2
0.1
0.05
0.01
0
Cell_Type_NEW
Ctrl
ExN
InN
Other
UD

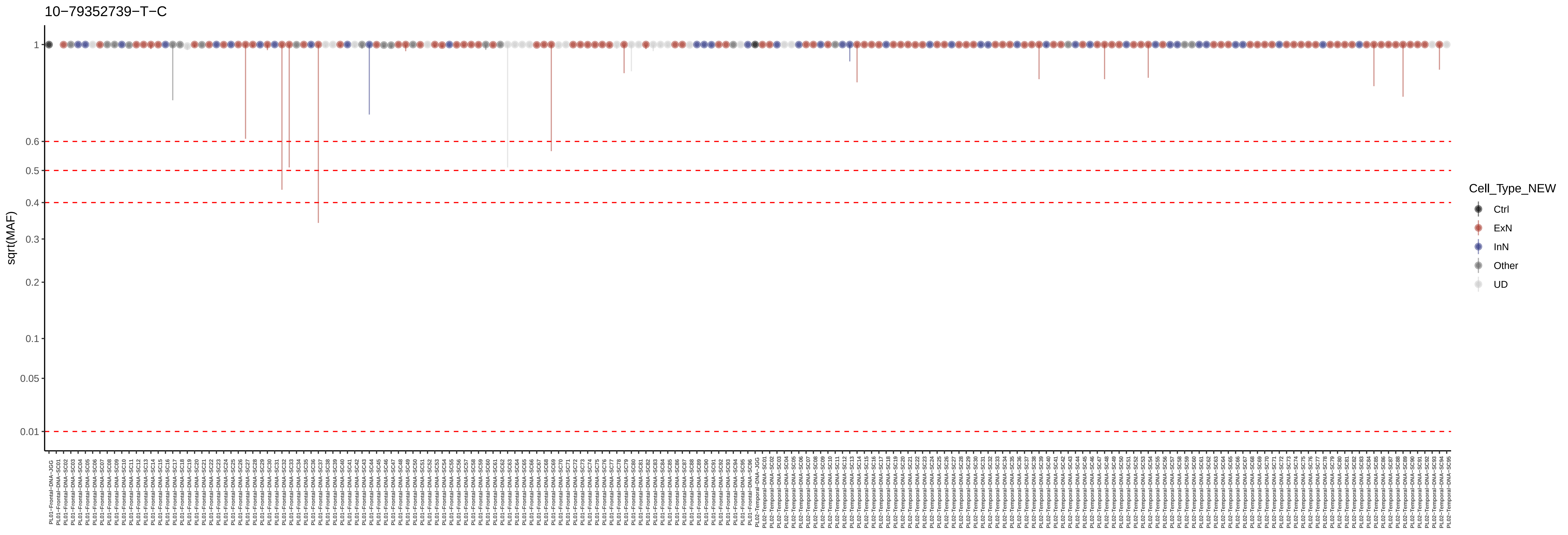
10−79352739−T−C
sqrt(MAF)
1
0.6
0.5
0.4
0.3
0.2
0.1
0.05
0.01
Cell_Type_NEW
Ctrl
ExN
InN
Other
UD

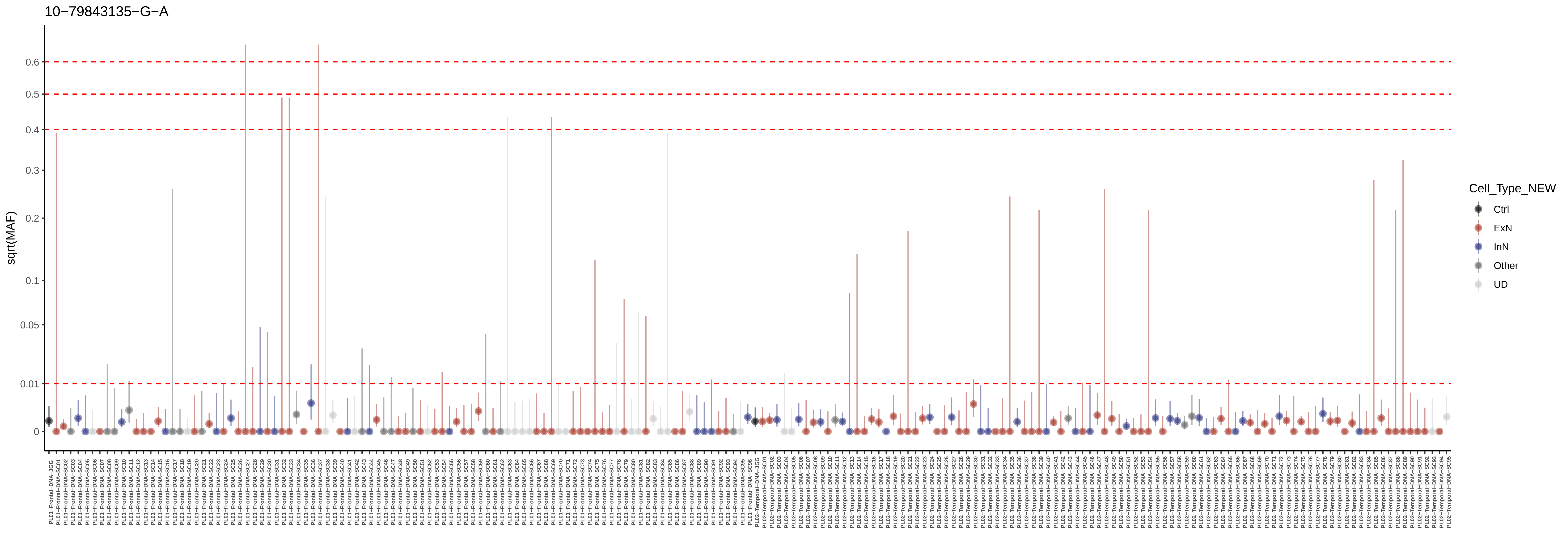

10−79843135−G−A
sqrt(MAF)
0.6
0.5
0.4
0.3
0.2
0.1
0.05
0.01
0
Cell_Type_NEW
Ctrl
ExN
InN
Other
UD

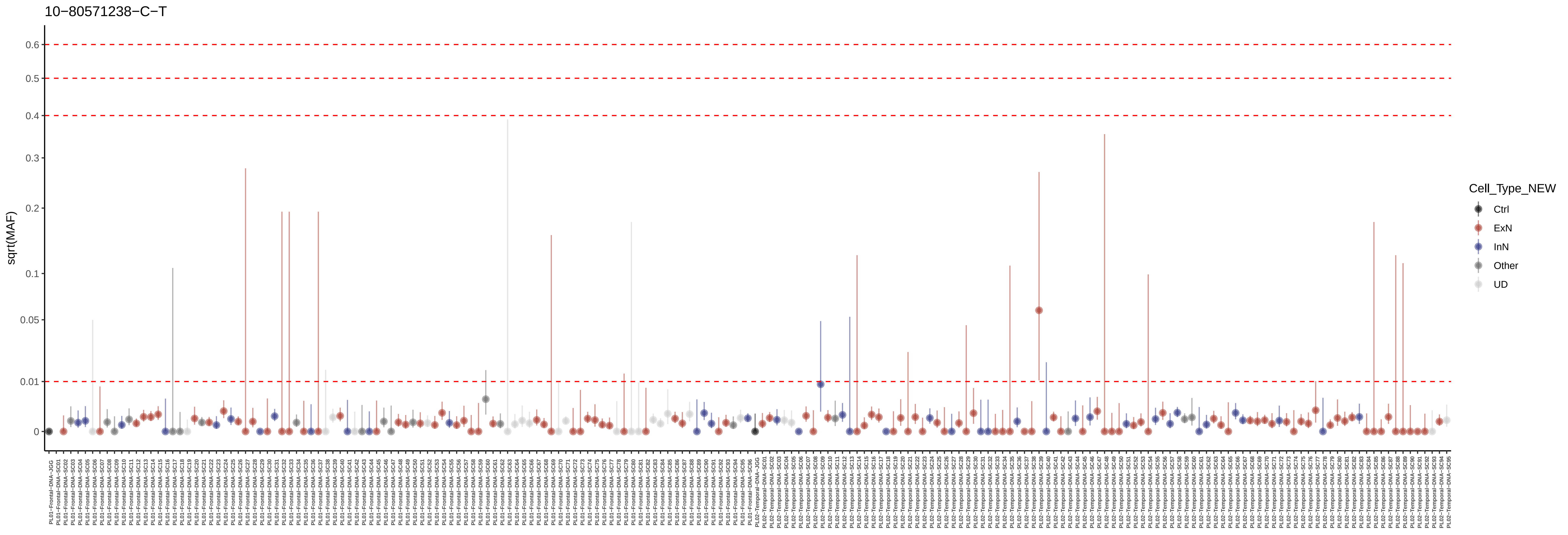
10−80571238−C−T
sqrt(MAF)
0.6
0.5
0.4
0.3
0.2
0.1
0.05
0.01
0
Cell_Type_NEW
Ctrl
ExN
InN
Other
UD

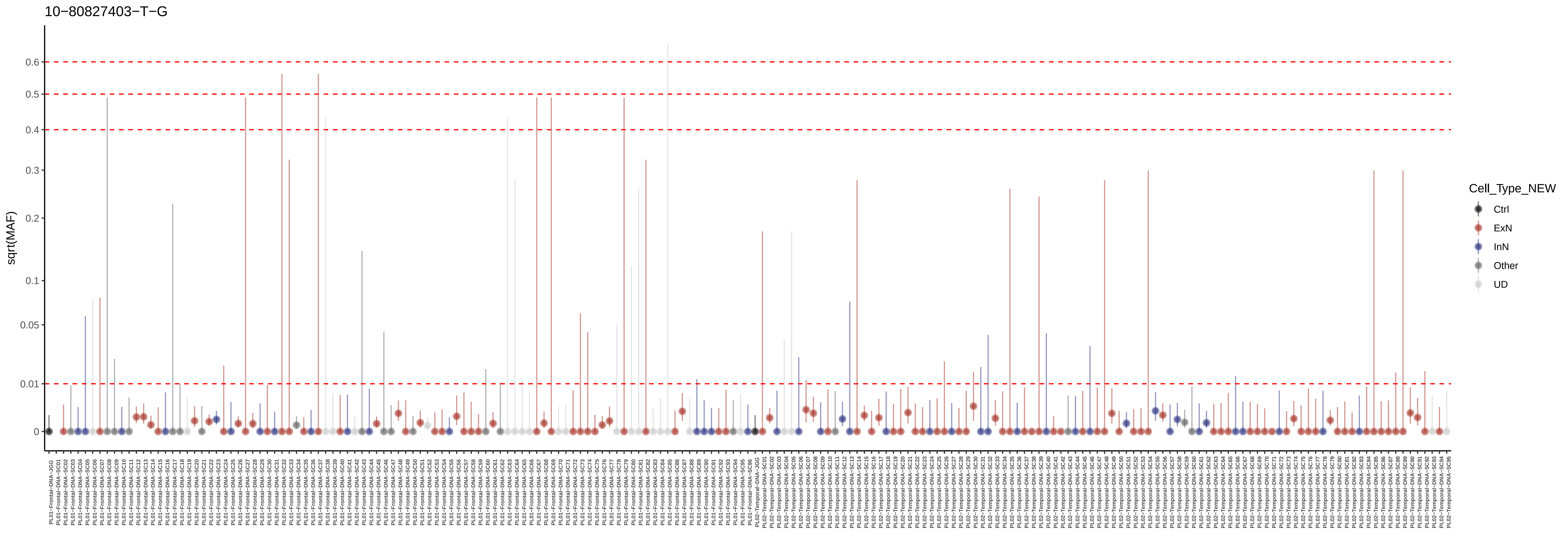
10−80827403−T−G
sqrt(MAF)
0.6
0.5
0.4
0.3
0.2
0.1
0.05
0.01
0
Cell_Type_NEW
Ctrl
ExN
InN
Other
UD

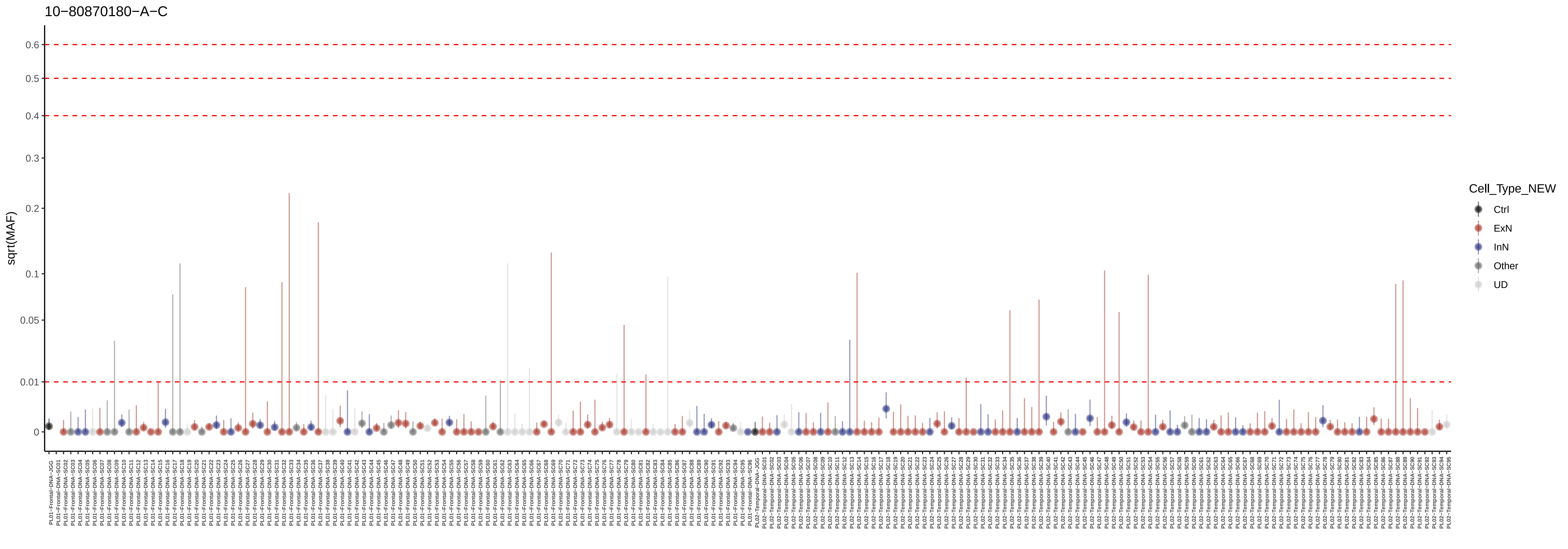
10−80870180−A−C
sqrt(MAF)
Cell_Type_NEW
Ctrl
ExN
InN
Other
UD

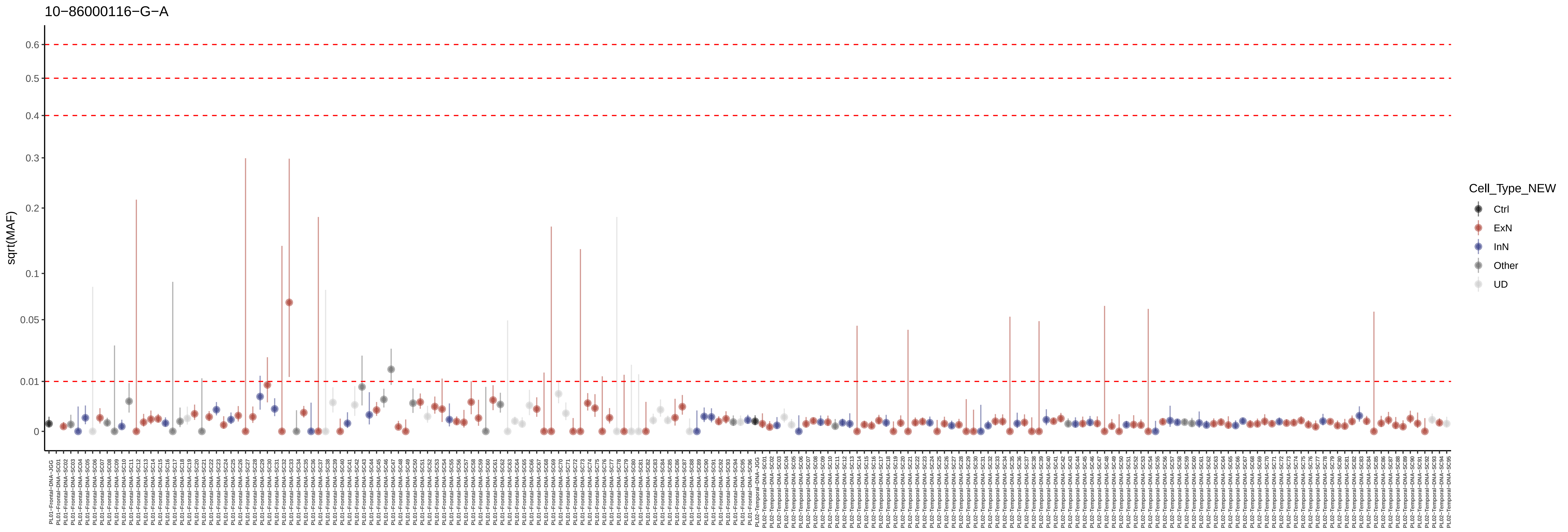
10−86000116−G−A
sqrt(MAF)
0.6
0.5
0.4
0.3
0.2
0.1
0.05
0.01
0
Cell_Type_NEW
Ctrl
ExN
InN
Other
UD

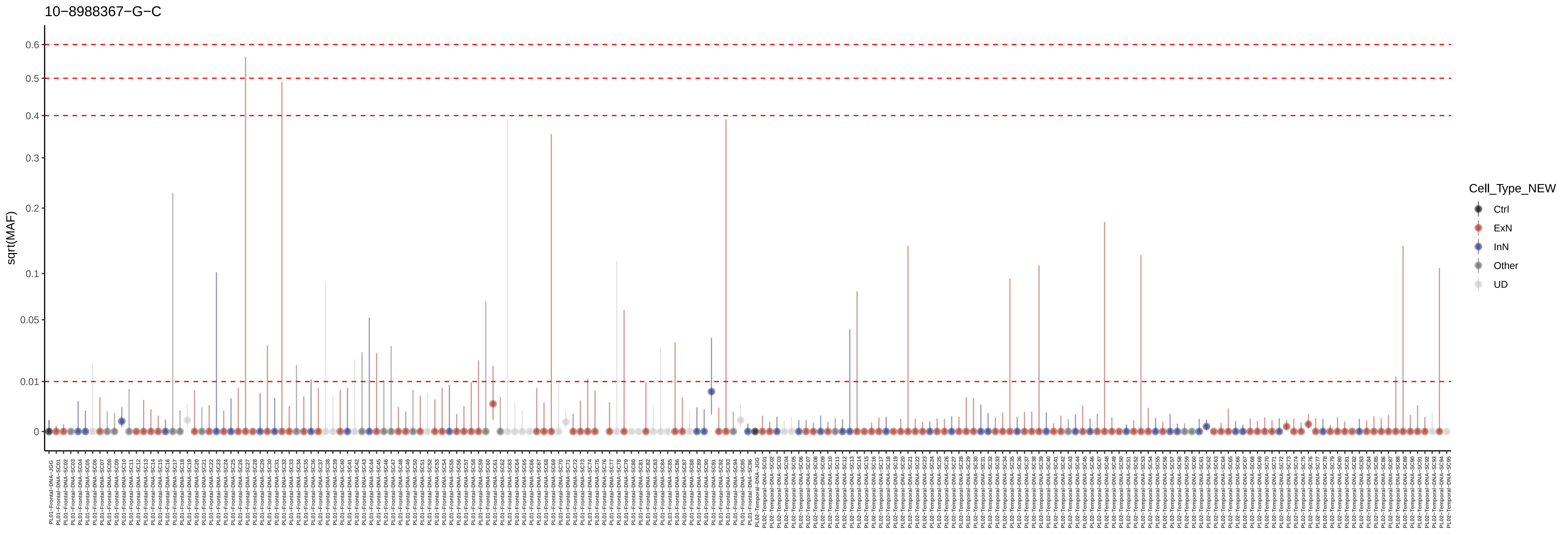
10−8988367−G−C
sqrt(MAF)
0.6
0.5
0.4
0.3
0.2
0.1
0.05
0.01
0
Cell_Type_NEW
Ctrl
ExN
InN
Other
UD

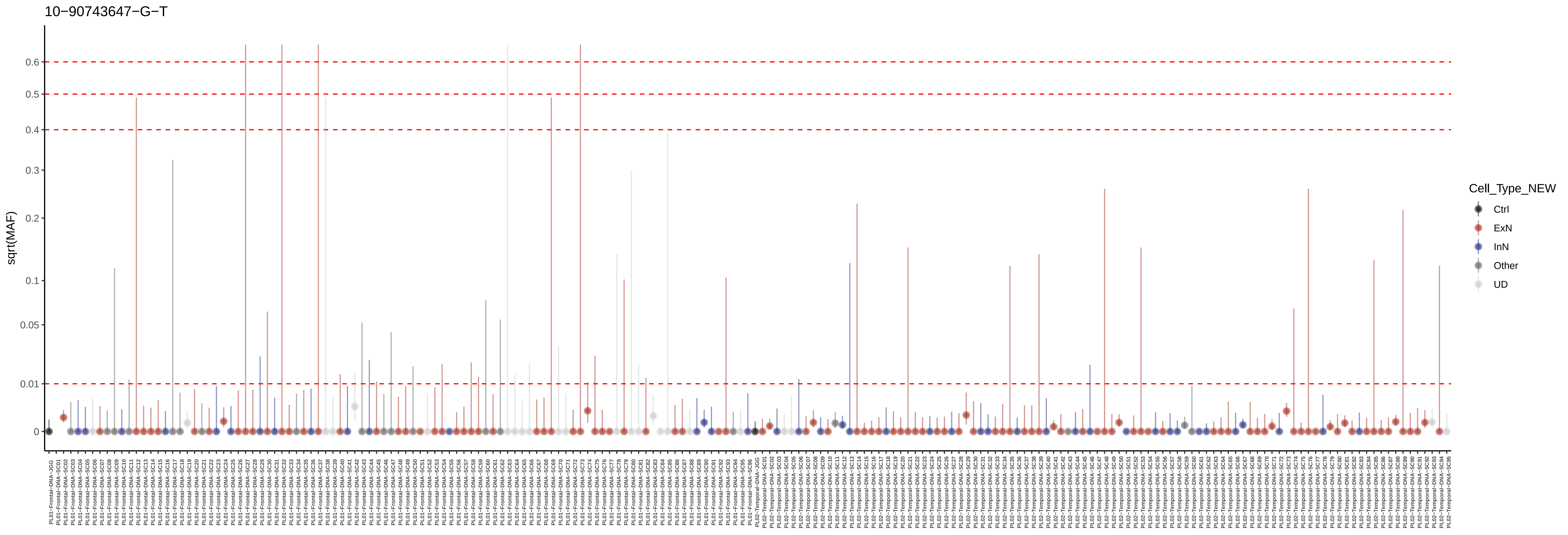
10−90743647−G−T
sqrt(MAF)
0.6
0.5
0.4
0.3
0.2
0.1
0.05
0.01
0
Cell_Type_NEW
Ctrl
ExN
InN
Other
UD

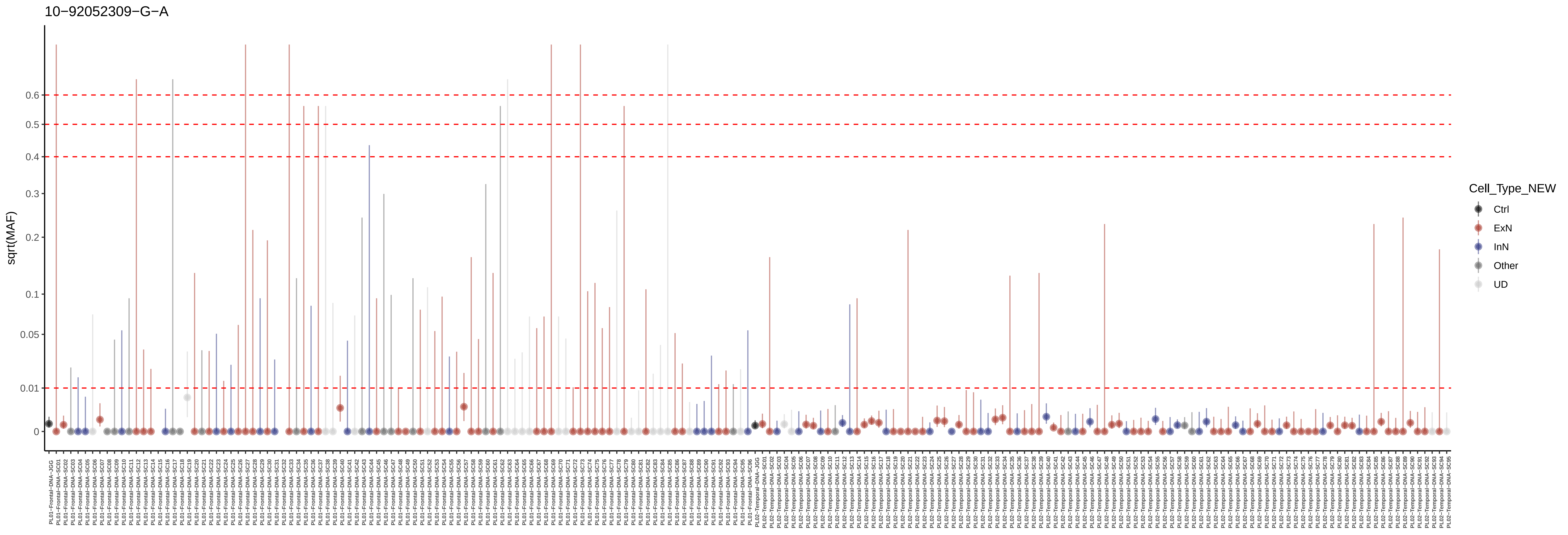
10−92052309−G−A
sqrt(MAF)
0.6
0.5
0.4
0.3
0.2
0.1
0.05
0.01
0
Cell_Type_NEW
Ctrl
ExN
InN
Other
UD

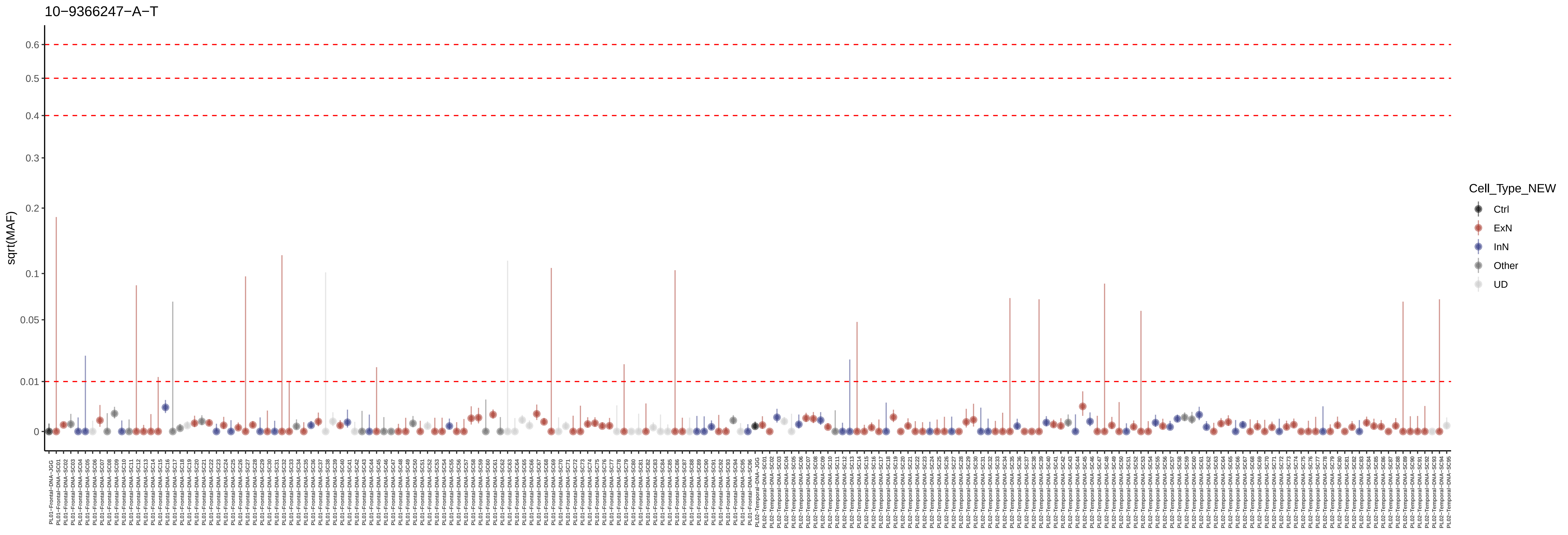

10−9366247−A−T
sqrt(MAF)
0.6
0.5
0.4
0.3
0.2
0.1
0.05
0.01
0
Cell_Type_NEW
Ctrl
ExN
InN
Other
UD

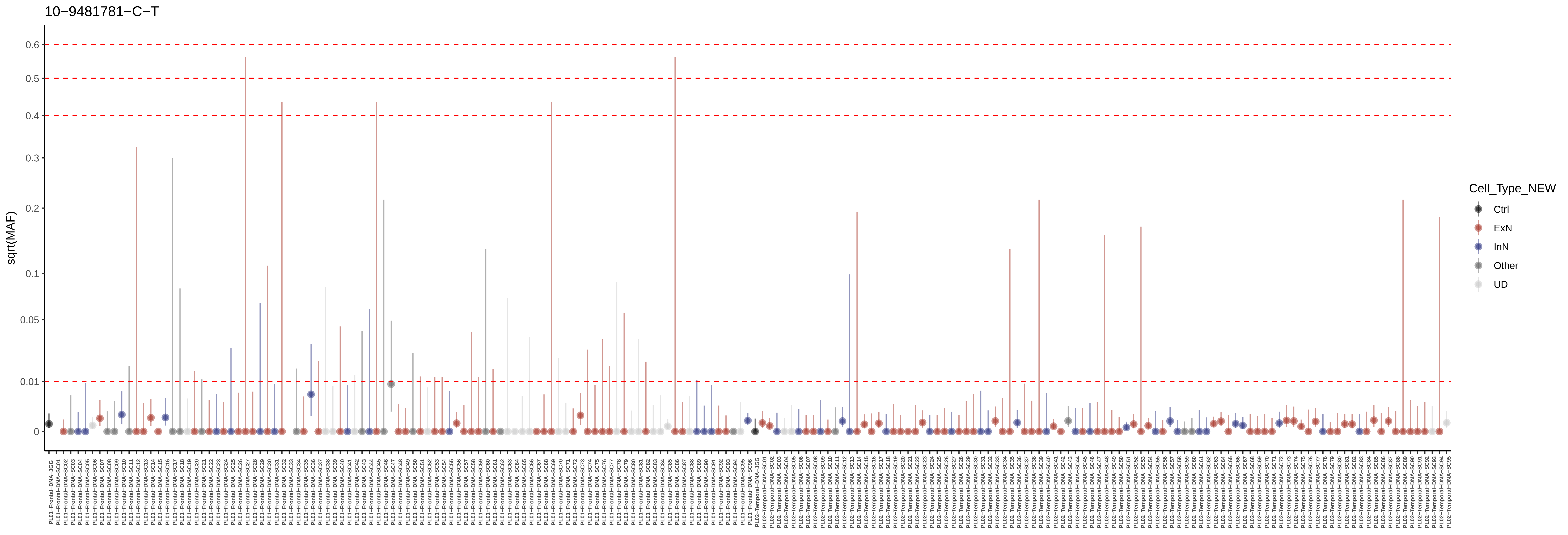

10−9481781−C−T
sqrt(MAF)
0.6
0.5
0.4
0.3
0.2
0.1
0.05
0.01
0
Cell_Type_NEW
Ctrl
ExN
InN
Other
UD

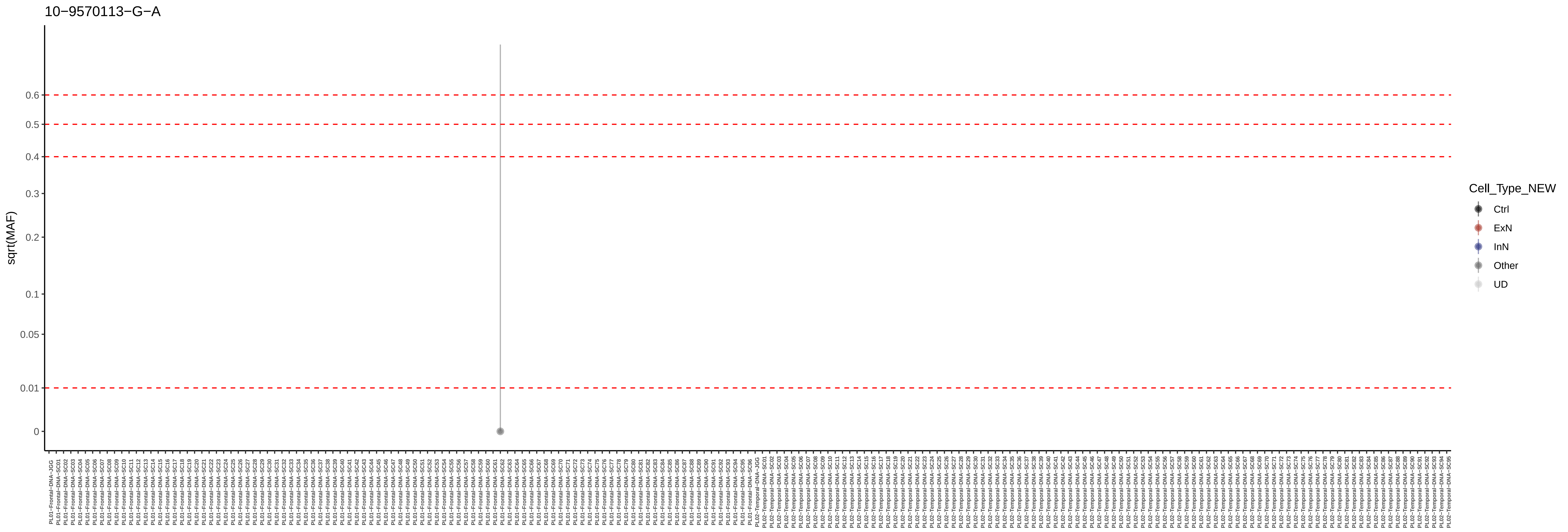
10−9570113−G−A
sqrt(MAF)
0.6
0.5
0.4
0.3
0.2
0.1
0.05
0.01
0
Cell_Type_NEW
Ctrl
ExN
InN
Other
UD

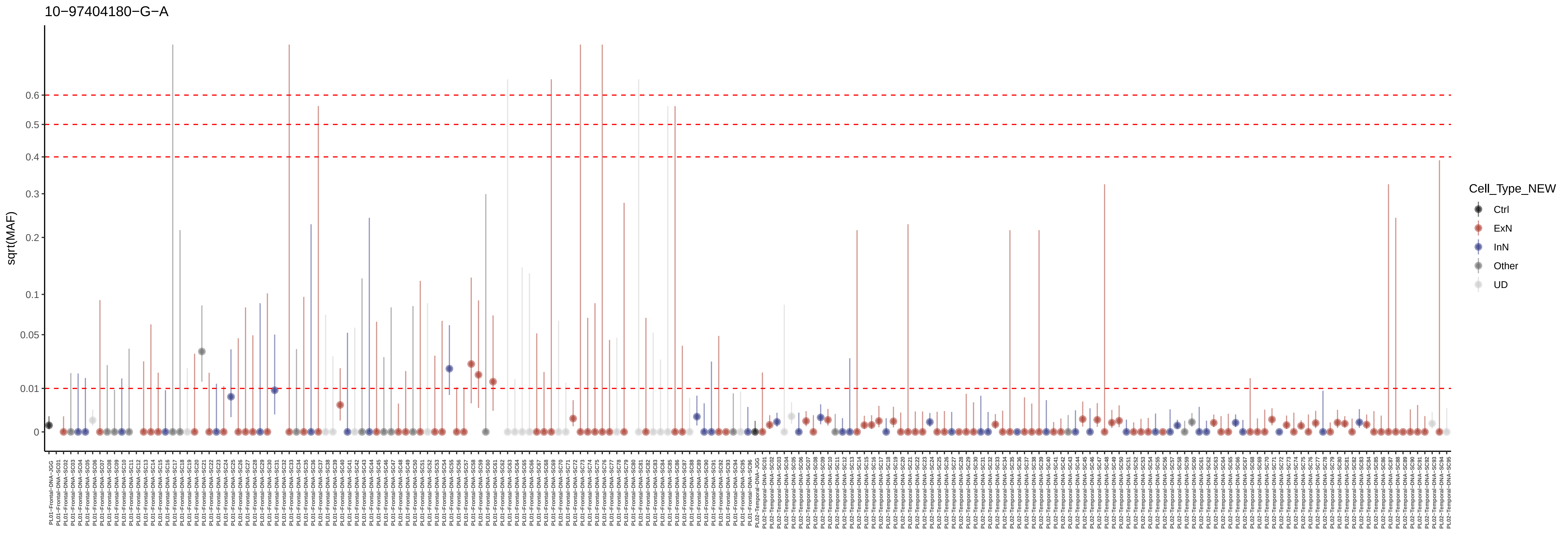

10−97404180−G−A
sqrt(MAF)
Cell_Type_NEW
Ctrl
ExN
InN
Other
UD

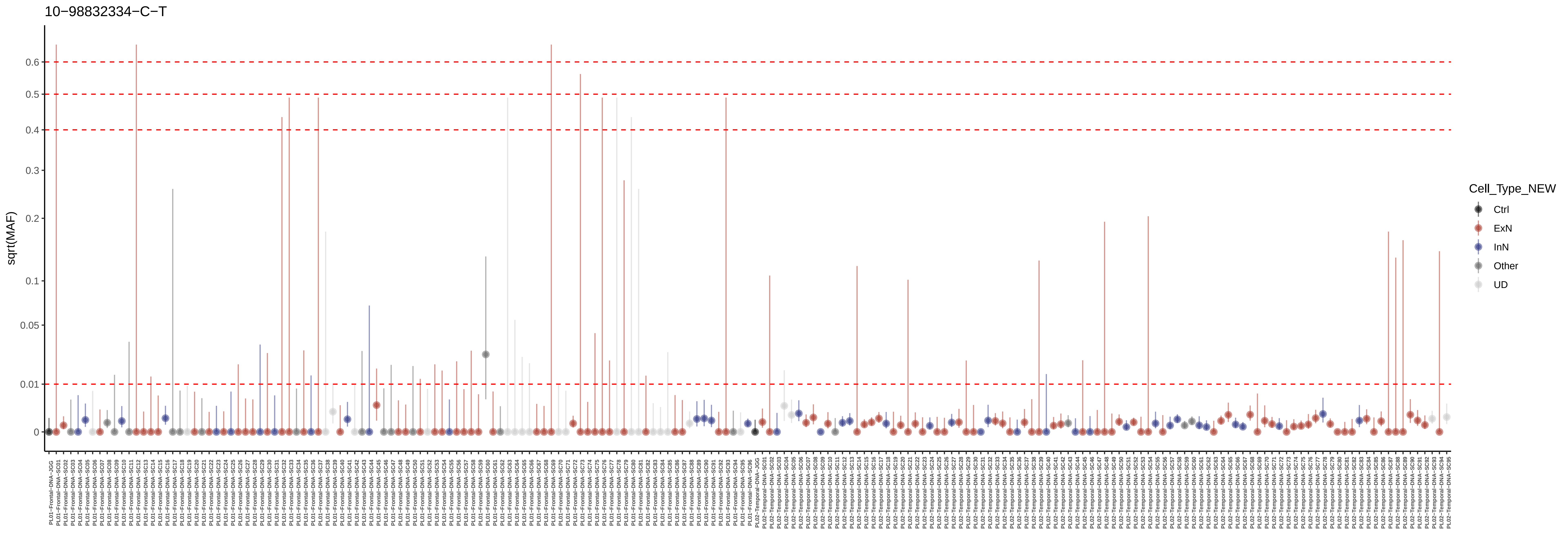

10−98832334−C−T
sqrt(MAF)
Cell_Type_NEW
Ctrl
ExN
InN
Other
UD

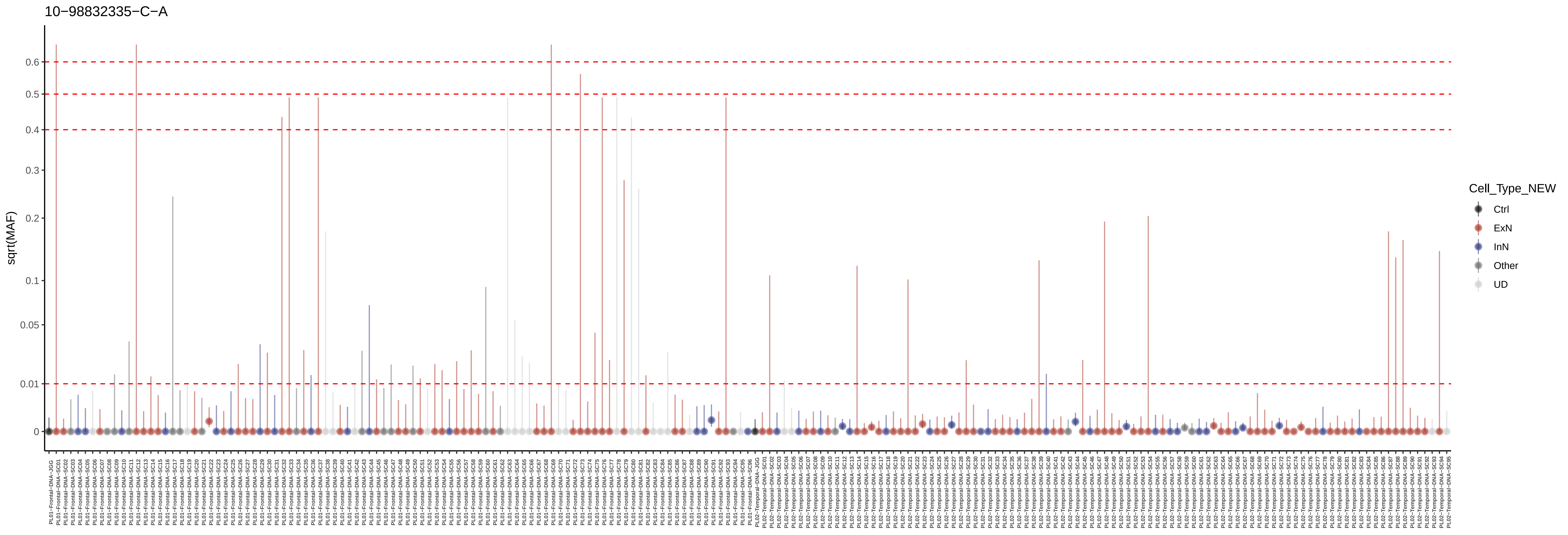
10−98832335−C−A
sqrt(MAF)
0.6
0.5
0.4
0.3
0.2
0.1
0.05
0.01
0
Cell_Type_NEW
Ctrl
ExN
InN
Other
UD

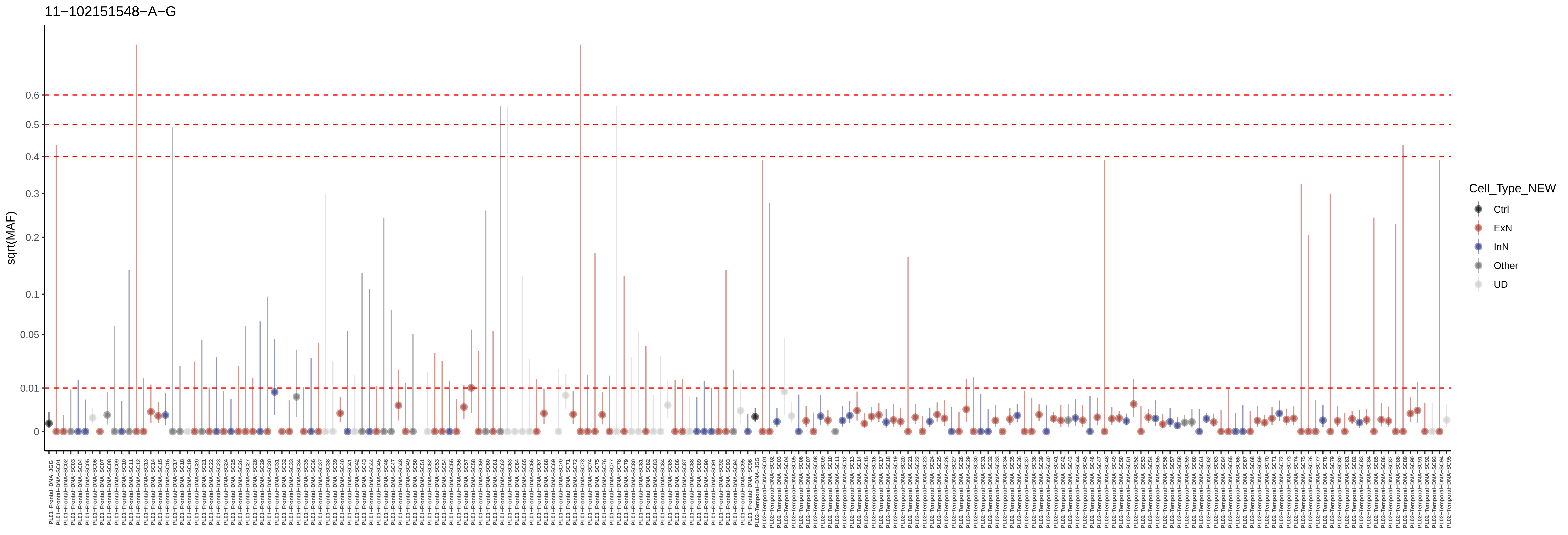
11−102151548−A−G
sqrt(MAF)
0.6
0.5
0.4
0.3
0.2
0.1
0.05
0.01
0
Cell_Type_NEW
Ctrl
ExN
InN
Other
UD

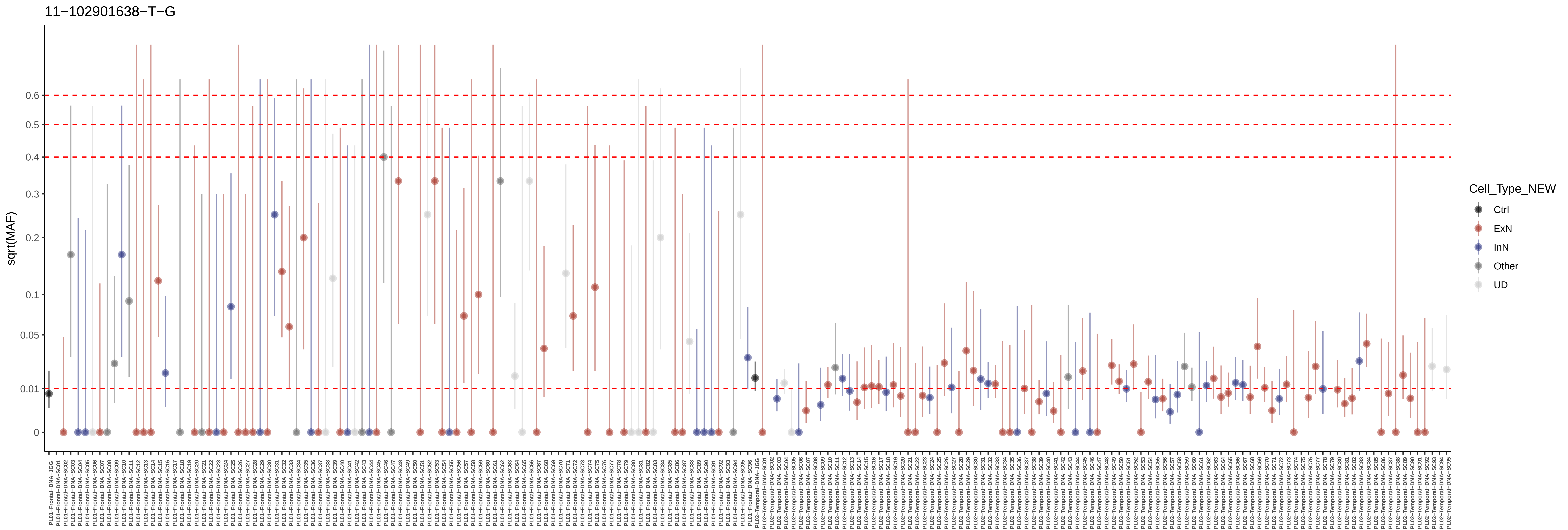
11−102901638−T−G
sqrt(MAF)
0.6
0.5
0.4
0.3
0.2
0.1
0.05
0.01
0
Cell_Type_NEW
Ctrl
ExN
InN
Other
UD

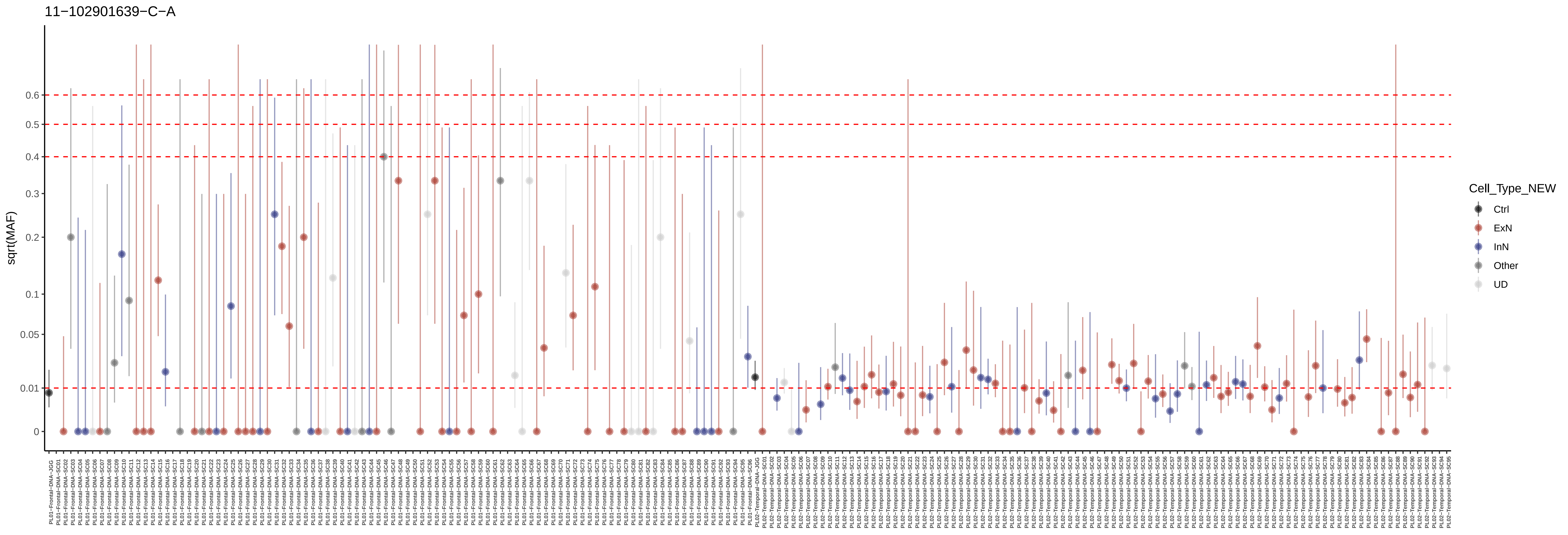
11−102901639−C−A
sqrt(MAF)
0.6
0.5
0.4
0.3
0.2
0.1
0.05
0.01
0
Cell_Type_NEW
Ctrl
ExN
InN
Other
UD

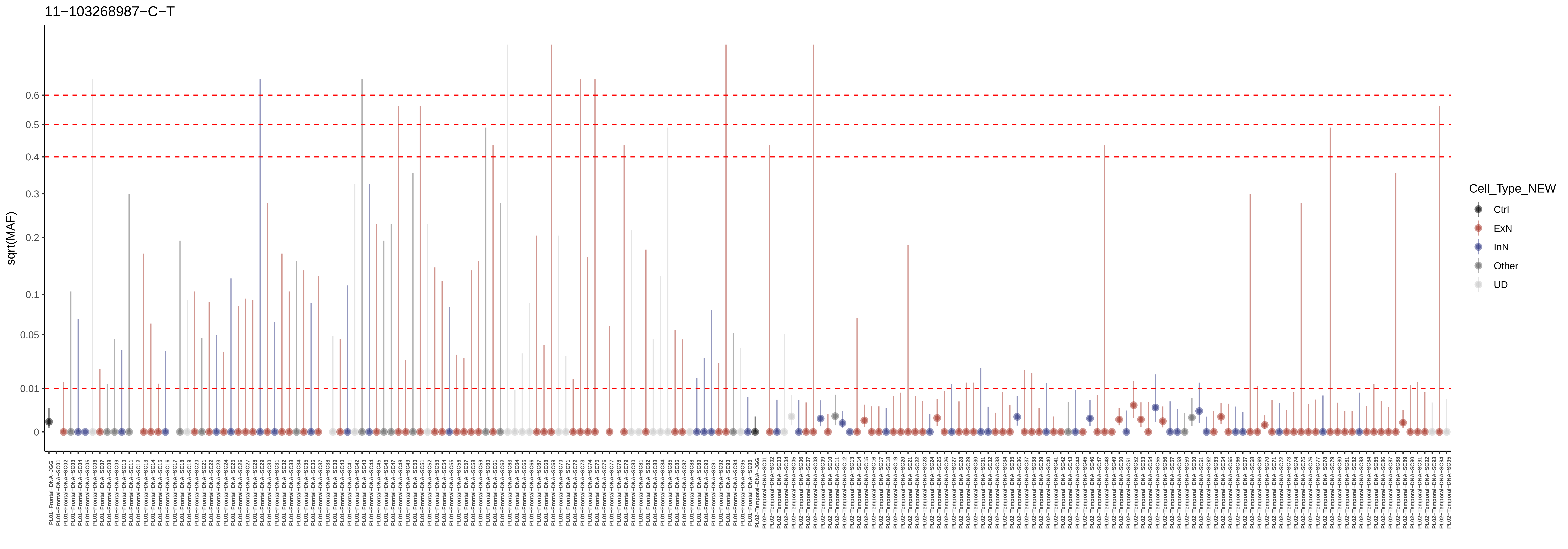
11−103268987−C−T
sqrt(MAF)
0.6
0.5
0.4
0.3
0.2
0.1
0.05
0.01
0
Cell_Type_NEW
Ctrl
ExN
InN
Other
UD

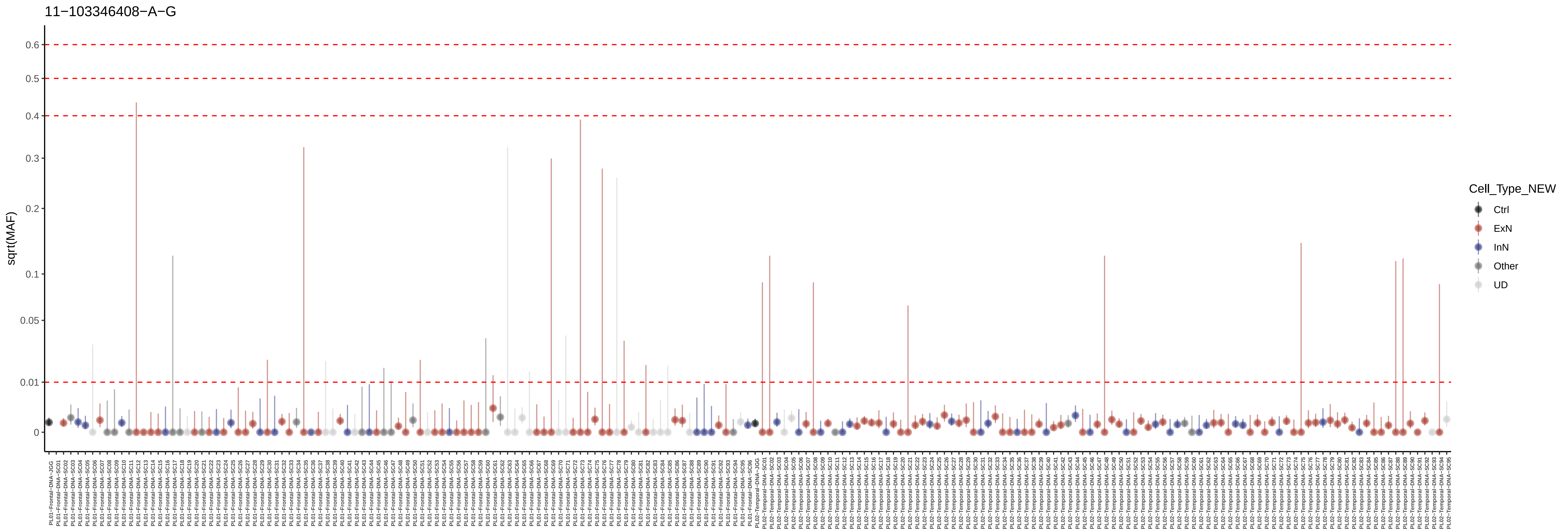

11−103346408−A−G
sqrt(MAF)
0.6
0.5
0.4
0.3
0.2
0.1
0.05
0.01
0
Cell_Type_NEW
Ctrl
ExN
InN
Other
UD

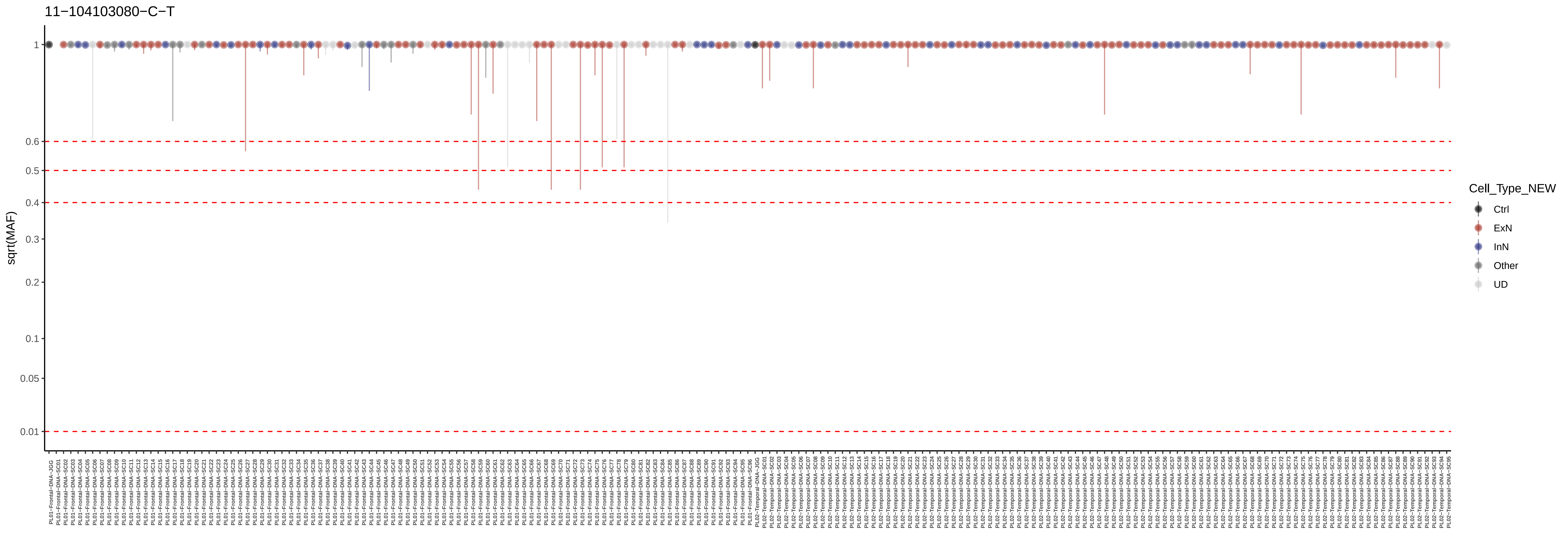
11−104103080−C−T
sqrt(MAF)
1
0.6
0.5
0.4
0.3
0.2
0.1
0.05
0.01
Cell_Type_NEW
Ctrl
ExN
InN
Other
UD

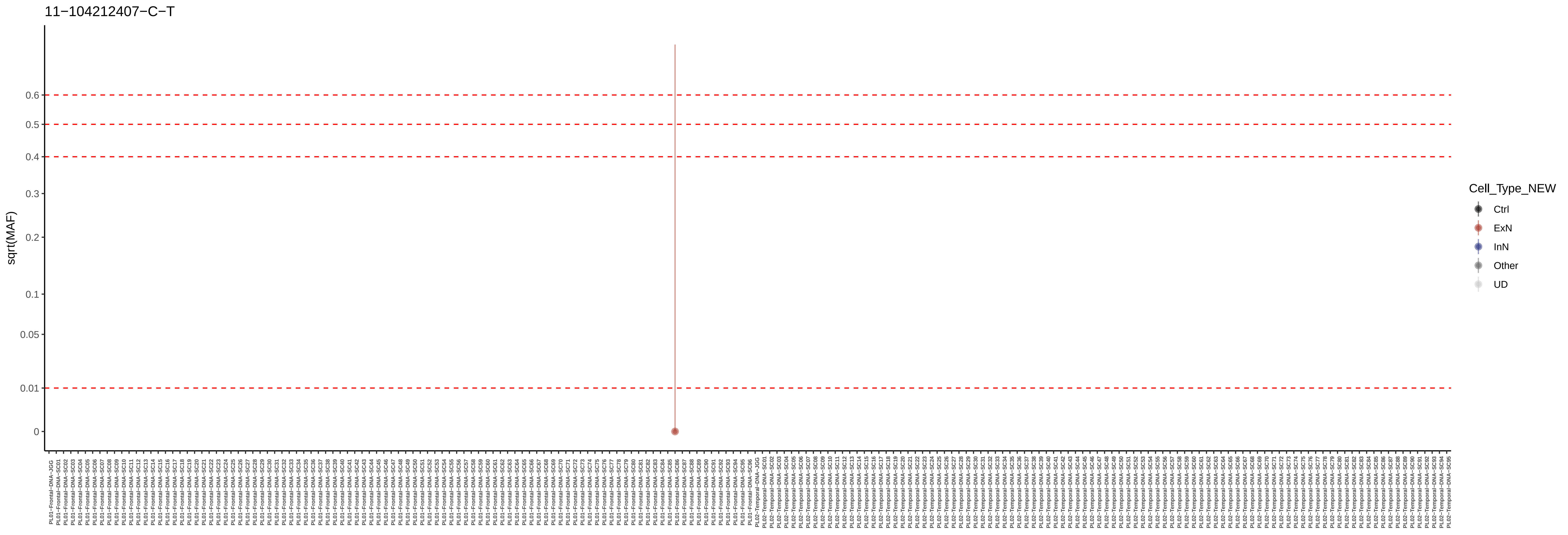

11−104212407−C−T
sqrt(MAF)
0.6
0.5
0.4
0.3
0.2
0.1
0.05
0.01
0
Cell_Type_NEW
Ctrl
ExN
InN
Other
UD

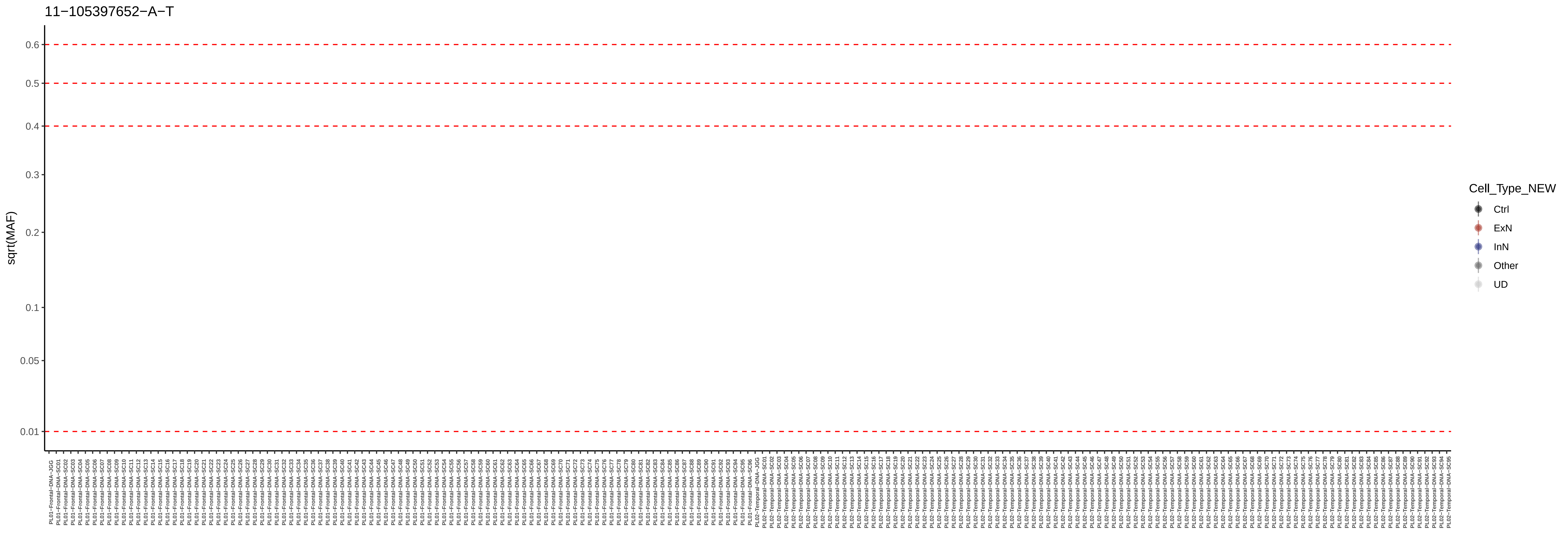
11−105397652−A−T
sqrt(MAF)
0.6
0.5
0.4
0.3
0.2
0.1
0.05
0.01
Cell_Type_NEW
Ctrl
ExN
InN
Other
UD

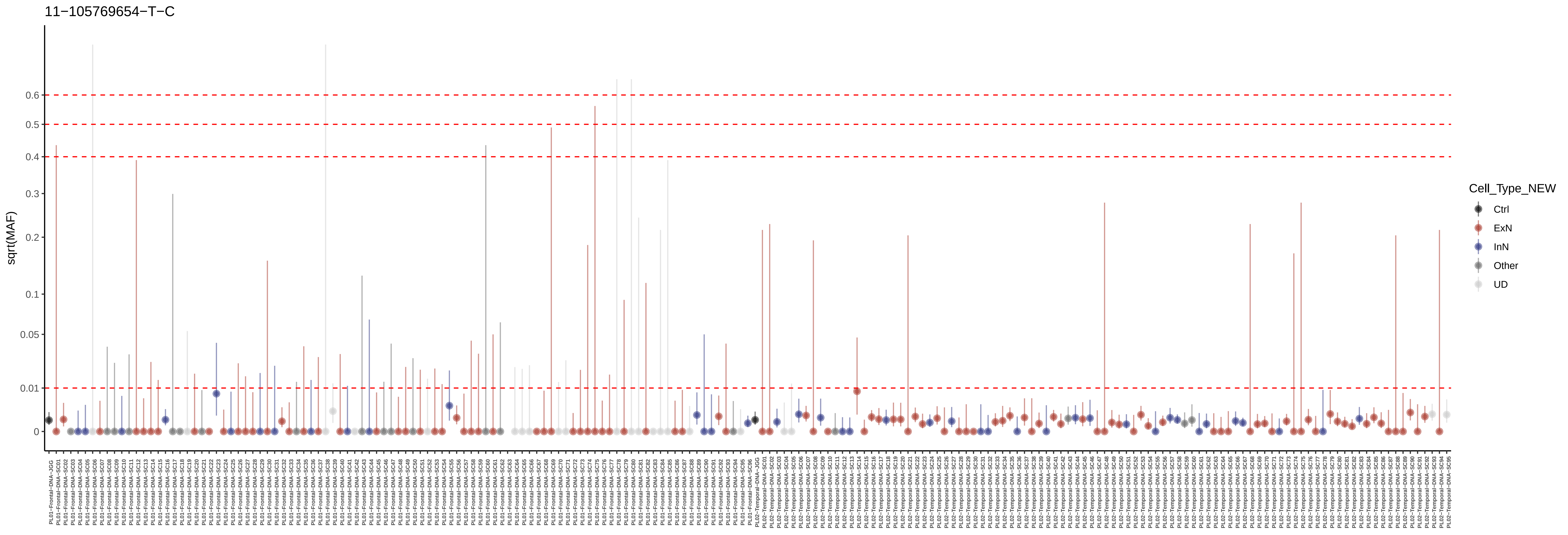

11−105769654−T−C
sqrt(MAF)
0.6
0.5
0.4
0.3
0.2
0.1
0.05
0.01
0
Cell_Type_NEW
Ctrl
ExN
InN
Other
UD

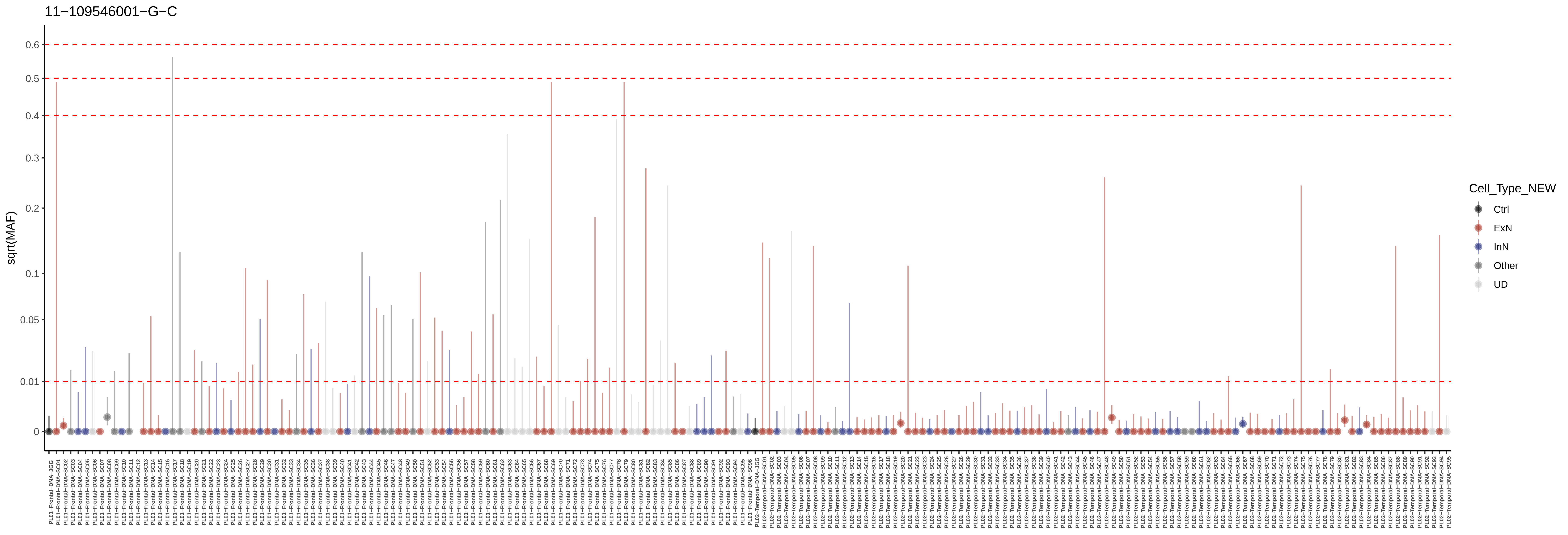
11−109546001−G−C
sqrt(MAF)
0.6
0.5
0.4
0.3
0.2
0.1
0.05
0.01
0
Cell_Type_NEW
Ctrl
ExN
InN
Other
UD

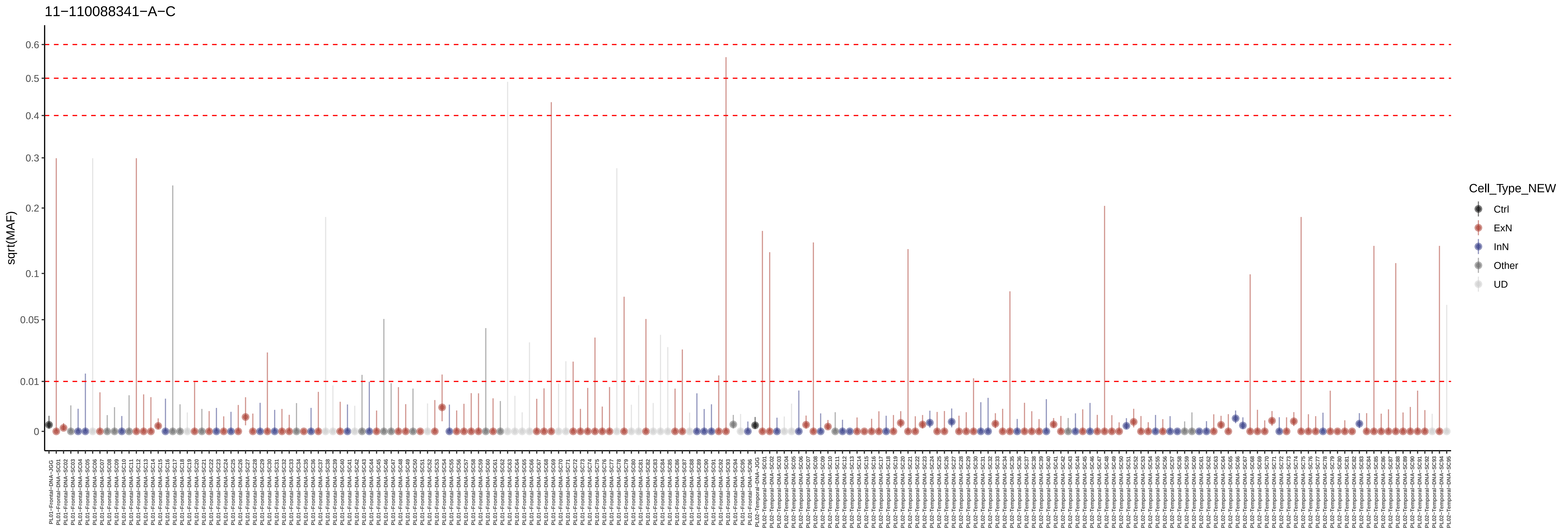
11−110088341−A−C
sqrt(MAF)
0.6
0.5
0.4
0.3
0.2
0.1
0.05
0.01
0
Cell_Type_NEW
Ctrl
ExN
InN
Other
UD

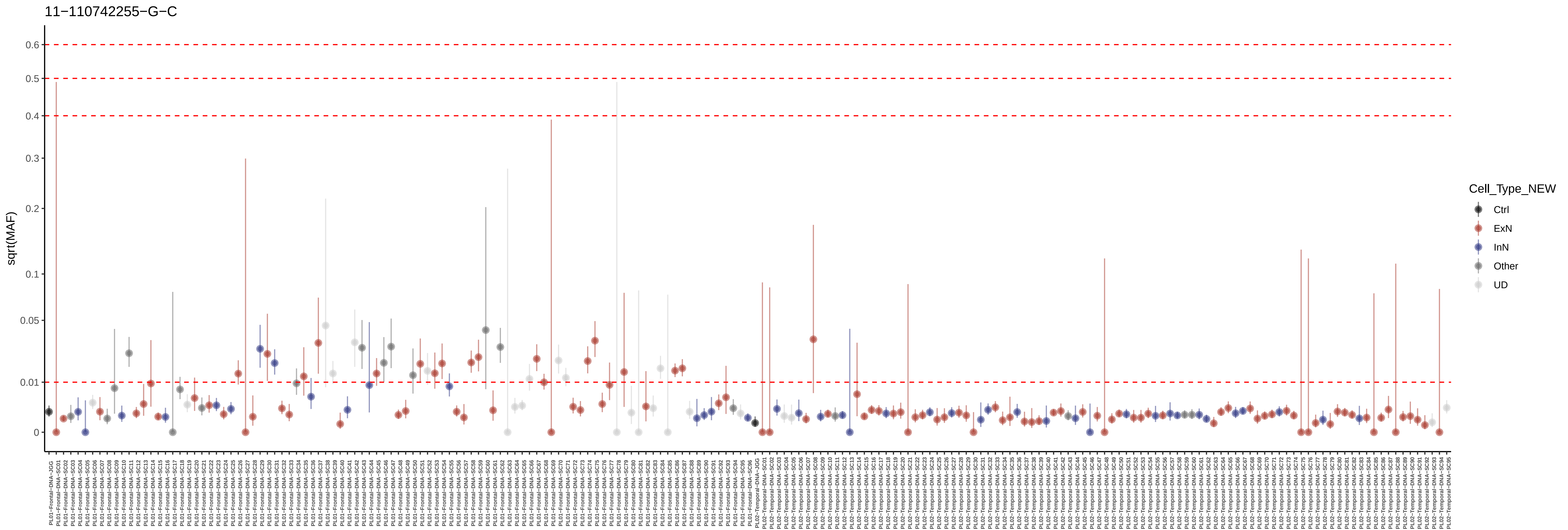

11−110742255−G−C
sqrt(MAF)
0.6
0.5
0.4
0.3
0.2
0.1
0.05
0.01
0
Cell_Type_NEW
Ctrl
ExN
InN
Other
UD

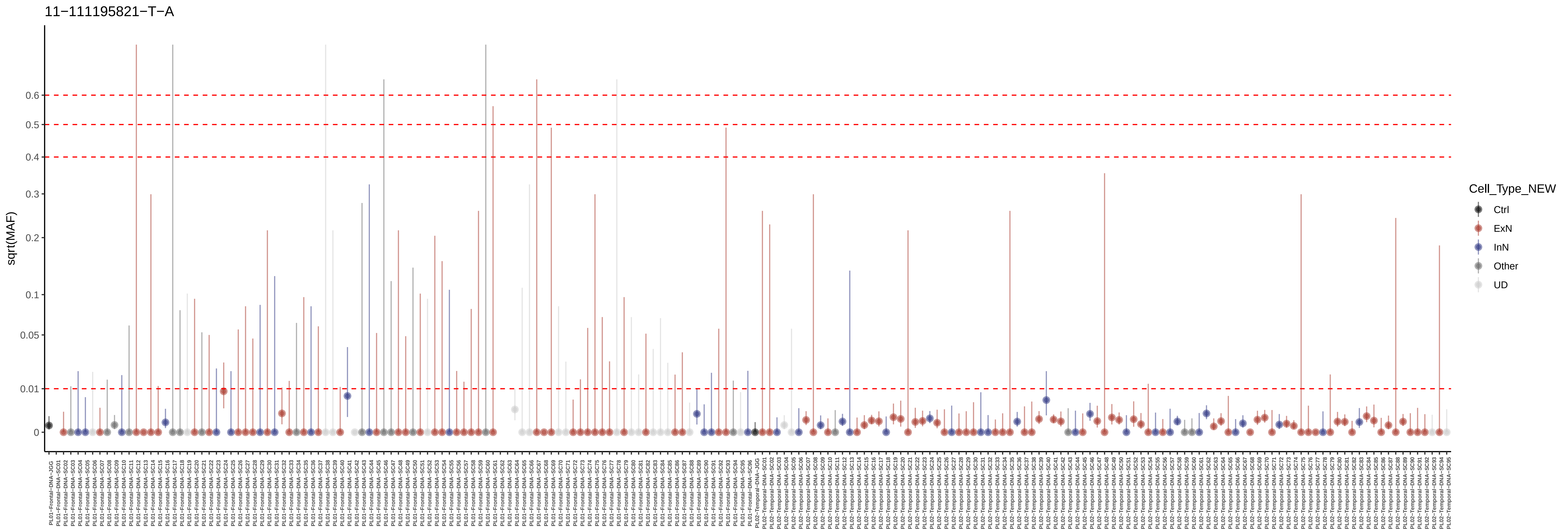
11−111195821−T−A
sqrt(MAF)
0.6
0.5
0.4
0.3
0.2
0.1
0.05
0.01
0
Cell_Type_NEW
Ctrl
ExN
InN
Other
UD

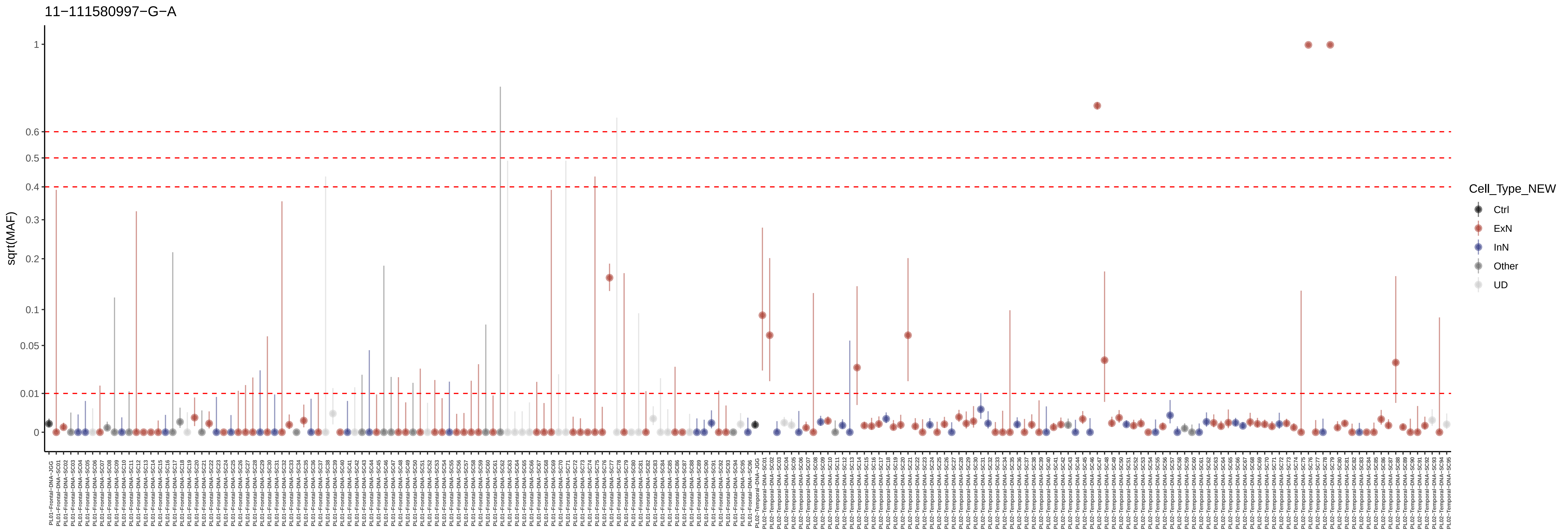

11−111580997−G−A
sqrt(MAF)
1
0.6
0.5
0.4
0.3
0.2
0.1
0.05
0.01
0
Cell_Type_NEW
Ctrl
ExN
InN
Other
UD

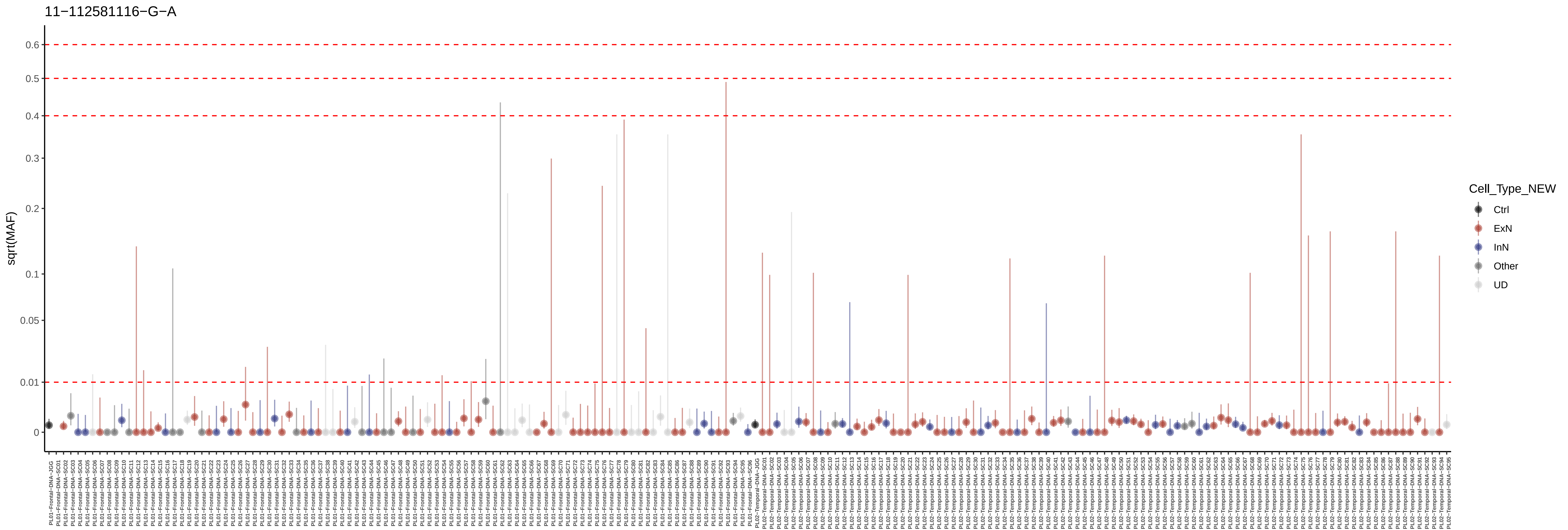
11−112581116−G−A
sqrt(MAF)
0.6
0.5
0.4
0.3
0.2
0.1
0.05
0.01
0
Cell_Type_NEW
Ctrl
ExN
InN
Other
UD

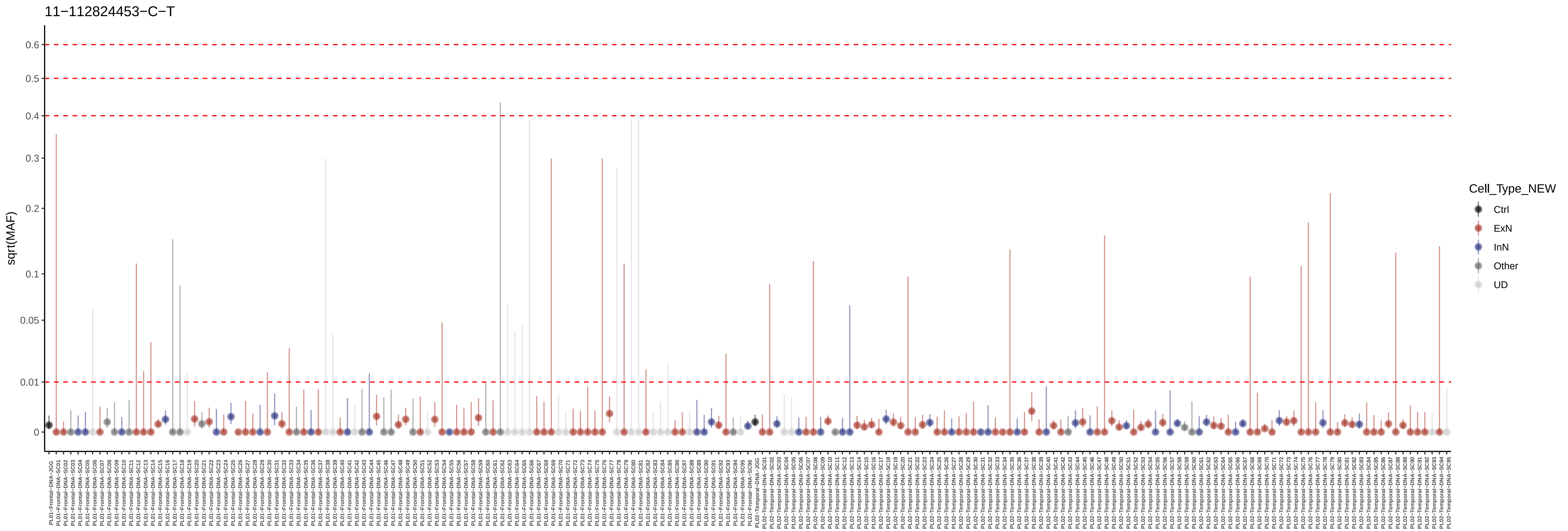
11−112824453−C−T
sqrt(MAF)
0.6
0.5
0.4
0.3
0.2
0.1
0.05
0.01
0
Cell_Type_NEW
Ctrl
ExN
InN
Other
UD

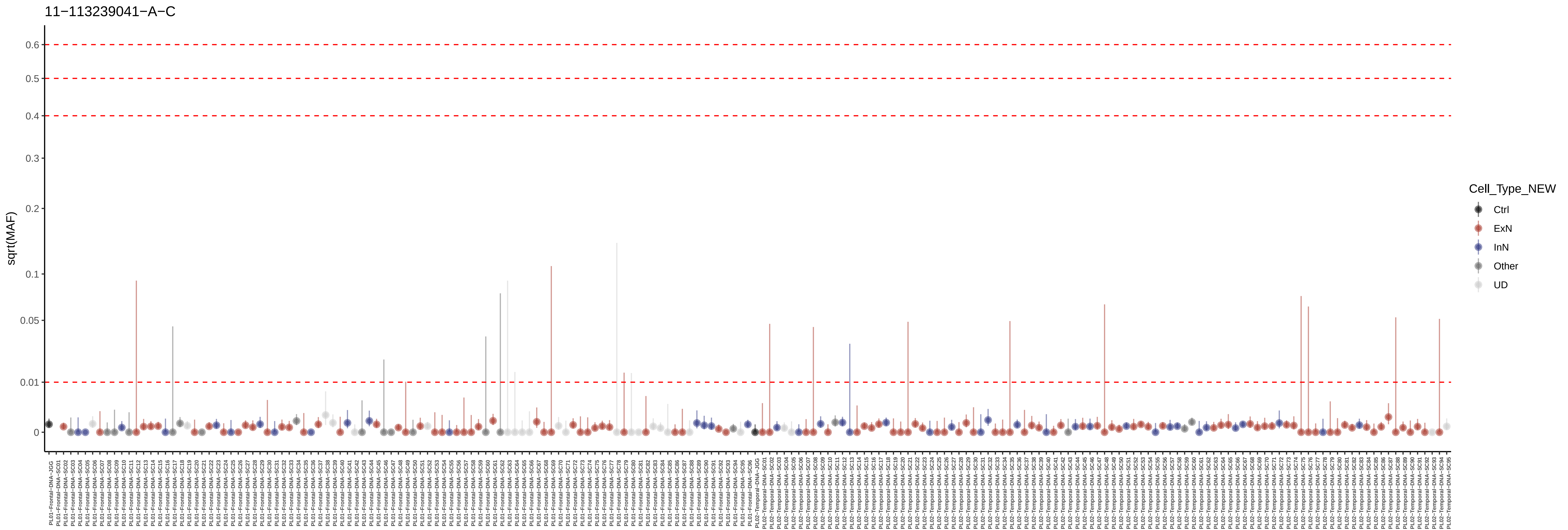
11−113239041−A−C
sqrt(MAF)
Cell_Type_NEW
Ctrl
ExN
InN
Other
UD

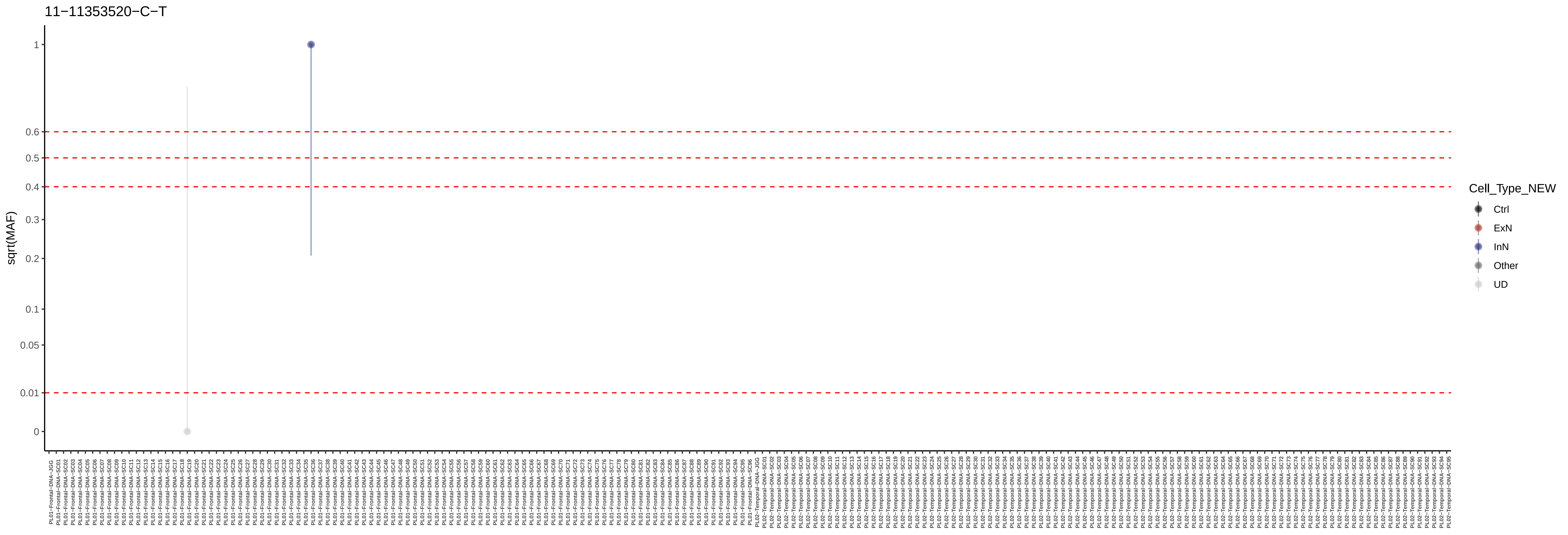
11−11353520−C−T
sqrt(MAF)
1
0.6
0.5
0.4
0.3
0.2
0.1
0.05
0.01
0
Cell_Type_NEW
Ctrl
ExN
InN
Other
UD

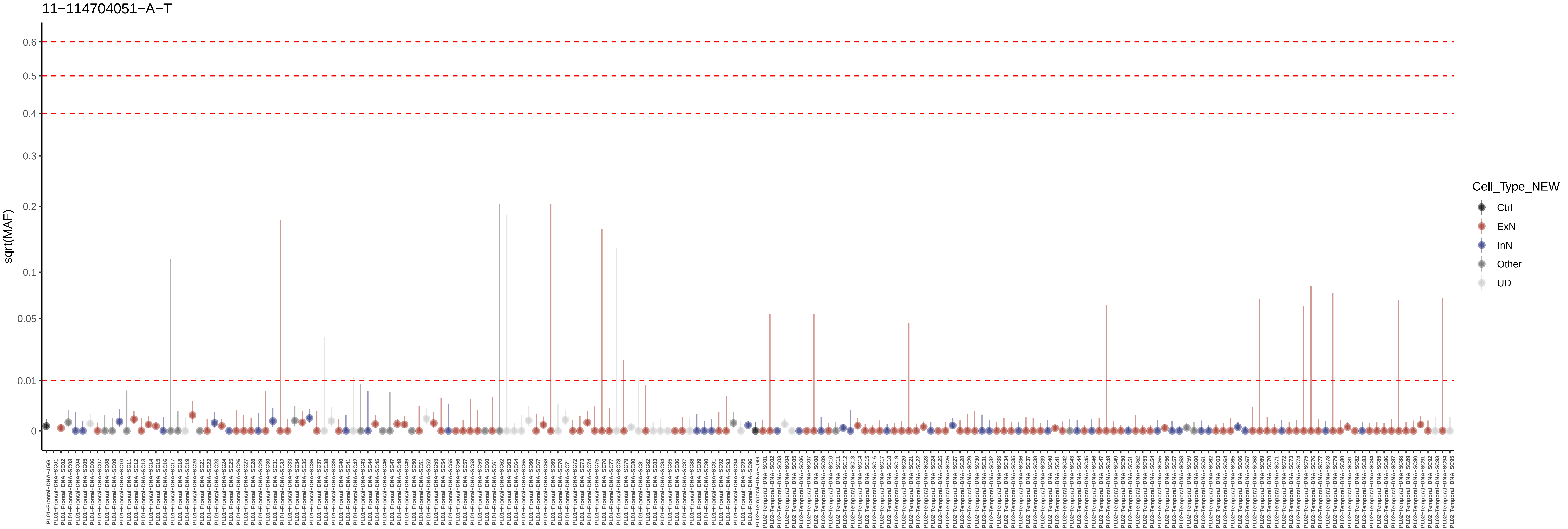
11−114704051−A−T
sqrt(MAF)
0.6
0.5
0.4
0.3
0.2
0.1
0.05
0.01
0
Cell_Type_NEW
Ctrl
ExN
InN
Other
UD

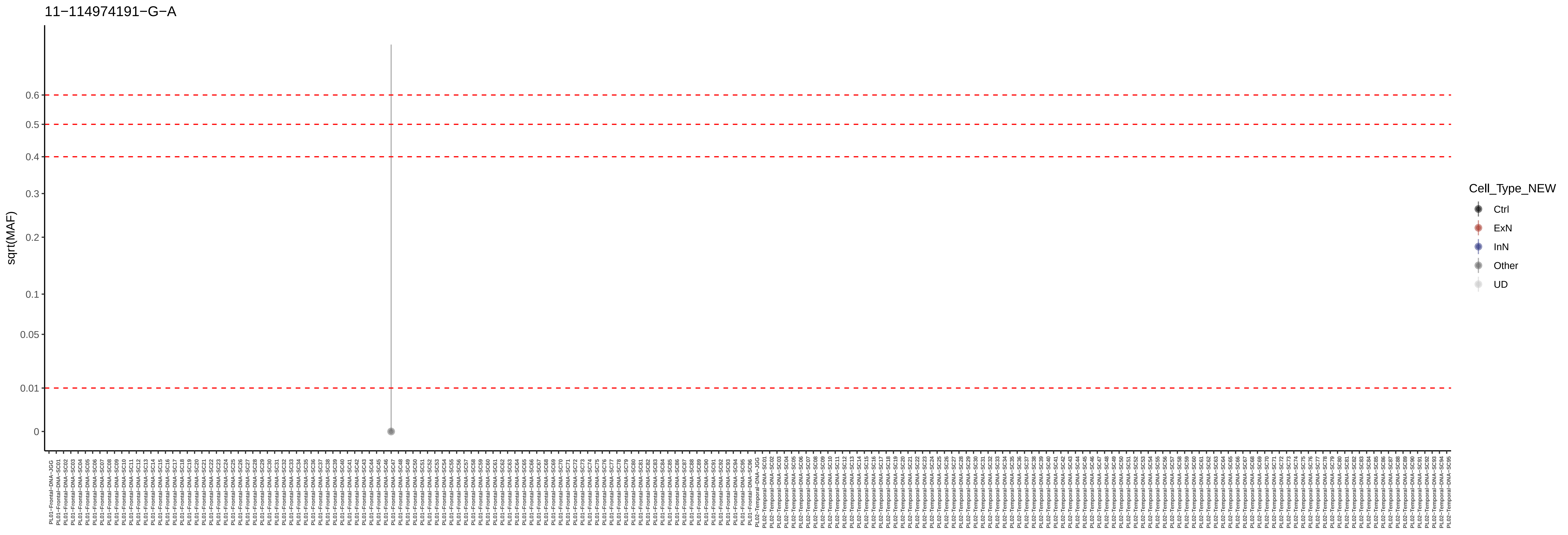

11−114974191−G−A
sqrt(MAF)
0.6
0.5
0.4
0.3
0.2
0.1
0.05
0.01
0
Cell_Type_NEW
Ctrl
ExN
InN
Other
UD

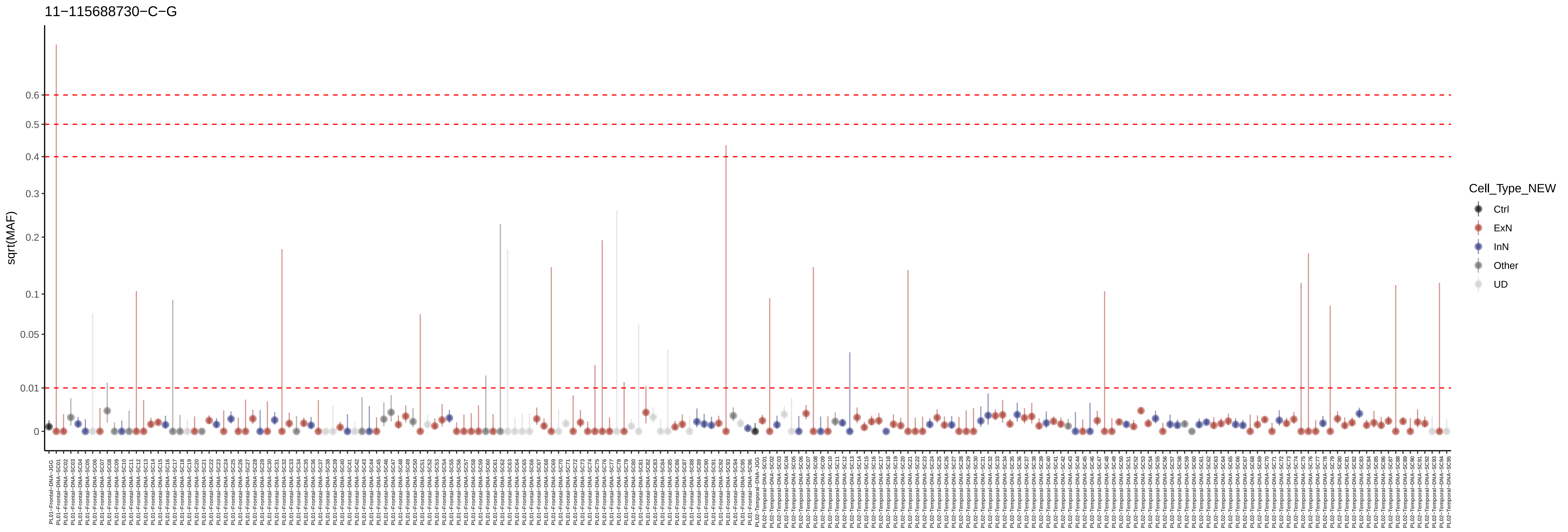
11−115688730−C−G
sqrt(MAF)
0.6
0.5
0.4
0.3
0.2
0.1
0.05
0.01
0
Cell_Type_NEW
Ctrl
ExN
InN
Other
UD

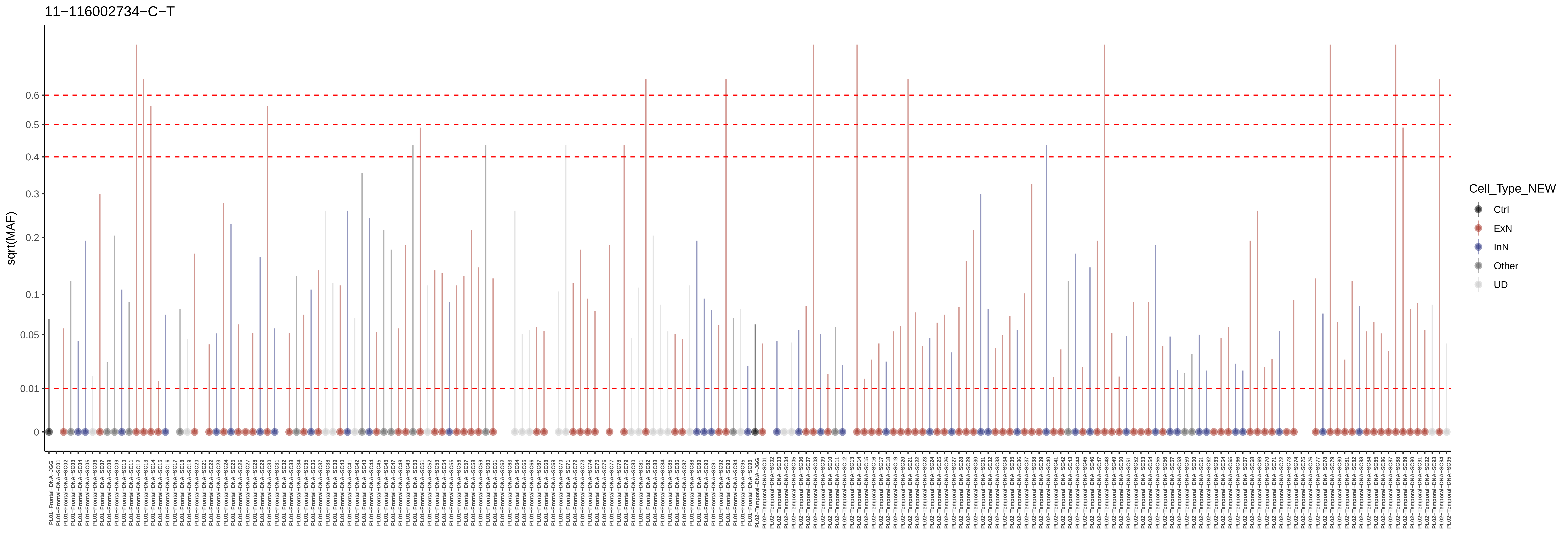
11−116002734−C−T
sqrt(MAF)
0.6
0.5
0.4
0.3
0.2
0.1
0.05
0.01
0
Cell_Type_NEW
Ctrl
ExN
InN
Other
UD

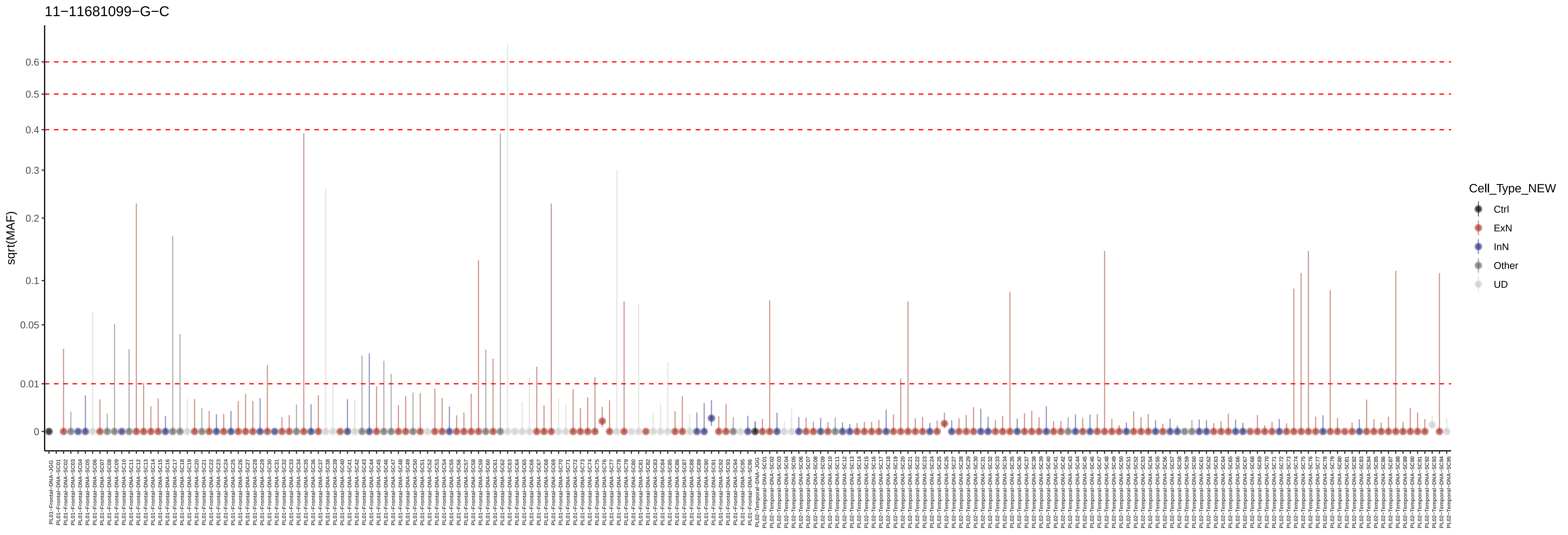
11−11681099−G−C
sqrt(MAF)
0.6
0.5
0.4
0.3
0.2
0.1
0.05
0.01
0
Cell_Type_NEW
Ctrl
ExN
InN
Other
UD

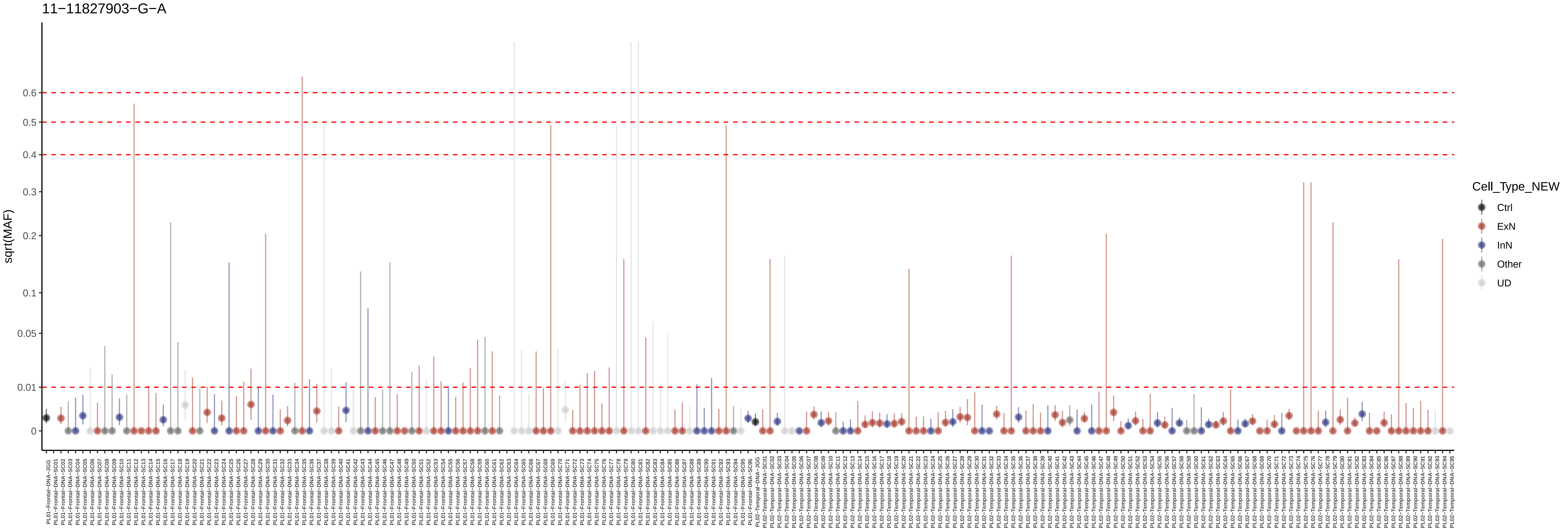
11−11827903−G−A
sqrt(MAF)
0.6
0.5
0.4
0.3
0.2
0.1
0.05
0.01
0
Cell_Type_NEW
Ctrl
ExN
InN
Other
UD

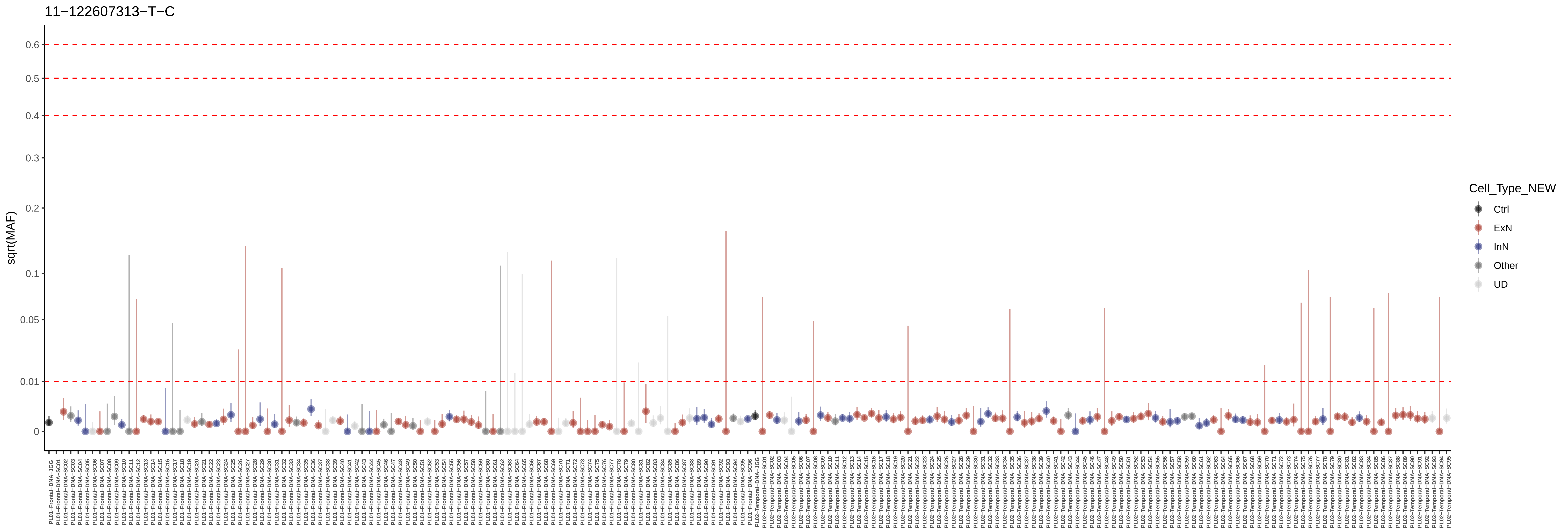
11−122607313−T−C
sqrt(MAF)
0.6
0.5
0.4
0.3
0.2
0.1
0.05
0.01
0
Cell_Type_NEW
Ctrl
ExN
InN
Other
UD

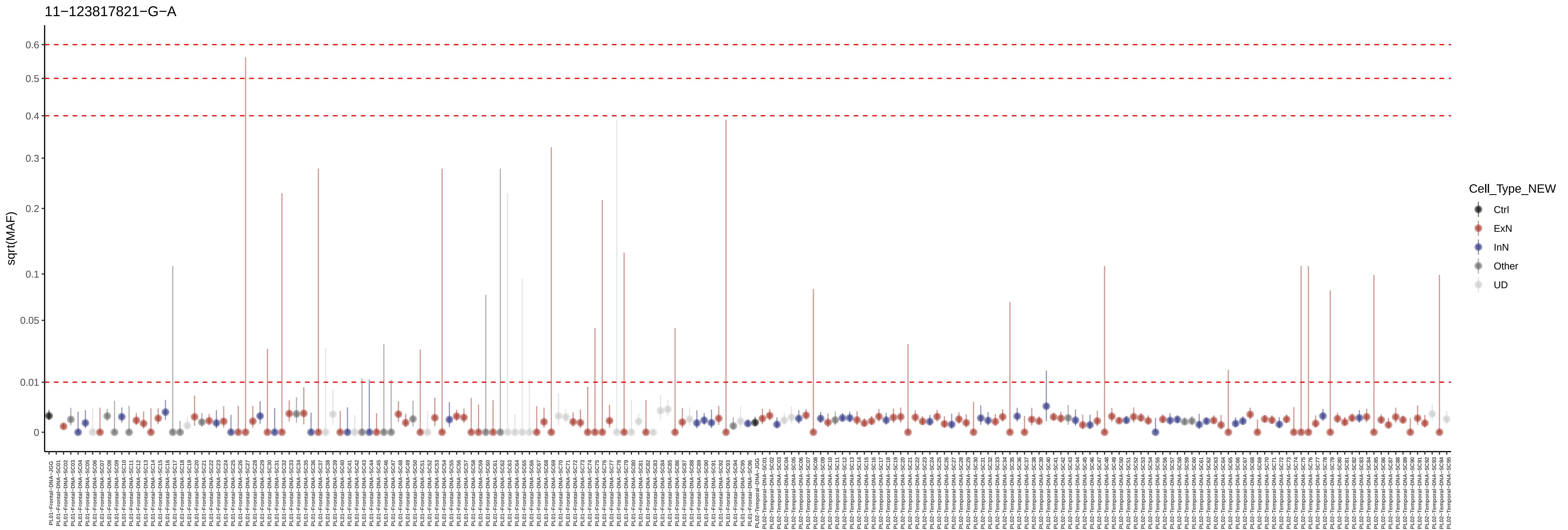
11−123817821−G−A
sqrt(MAF)
0.6
0.5
0.4
0.3
0.2
0.1
0.05
0.01
0
Cell_Type_NEW
Ctrl
ExN
InN
Other
UD

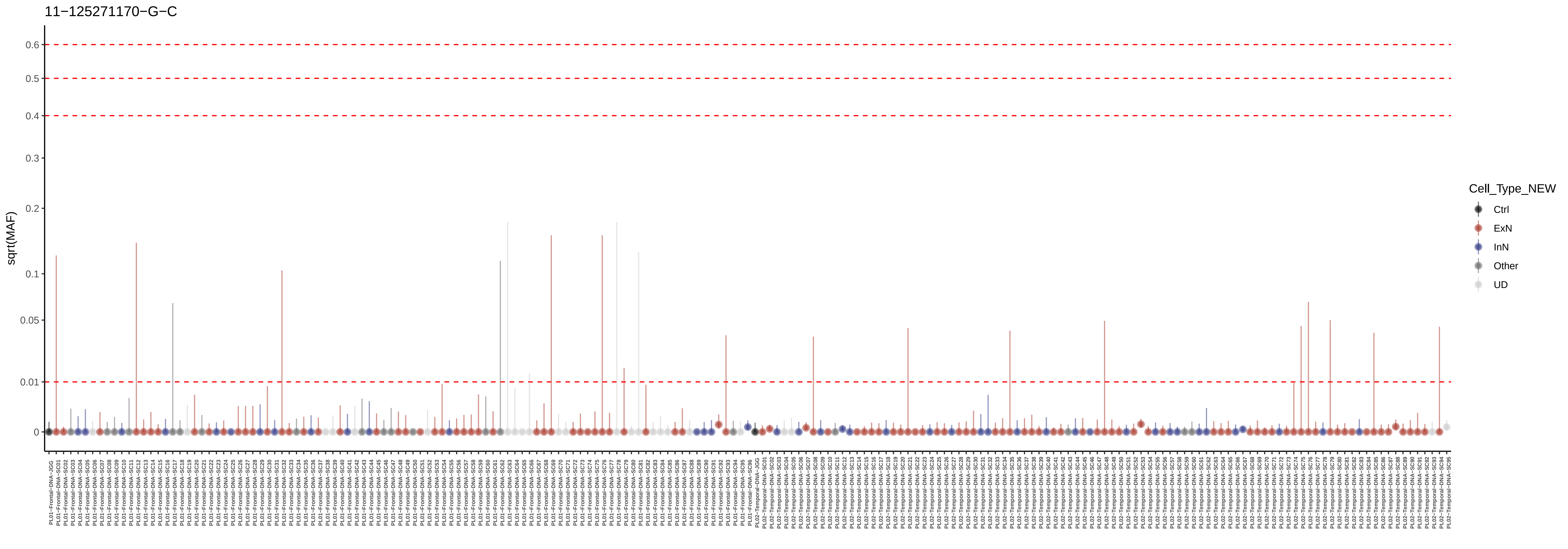
11−125271170−G−C
sqrt(MAF)
0.6
0.5
0.4
0.3
0.2
0.1
0.05
0.01
0
Cell_Type_NEW
Ctrl
ExN
InN
Other
UD

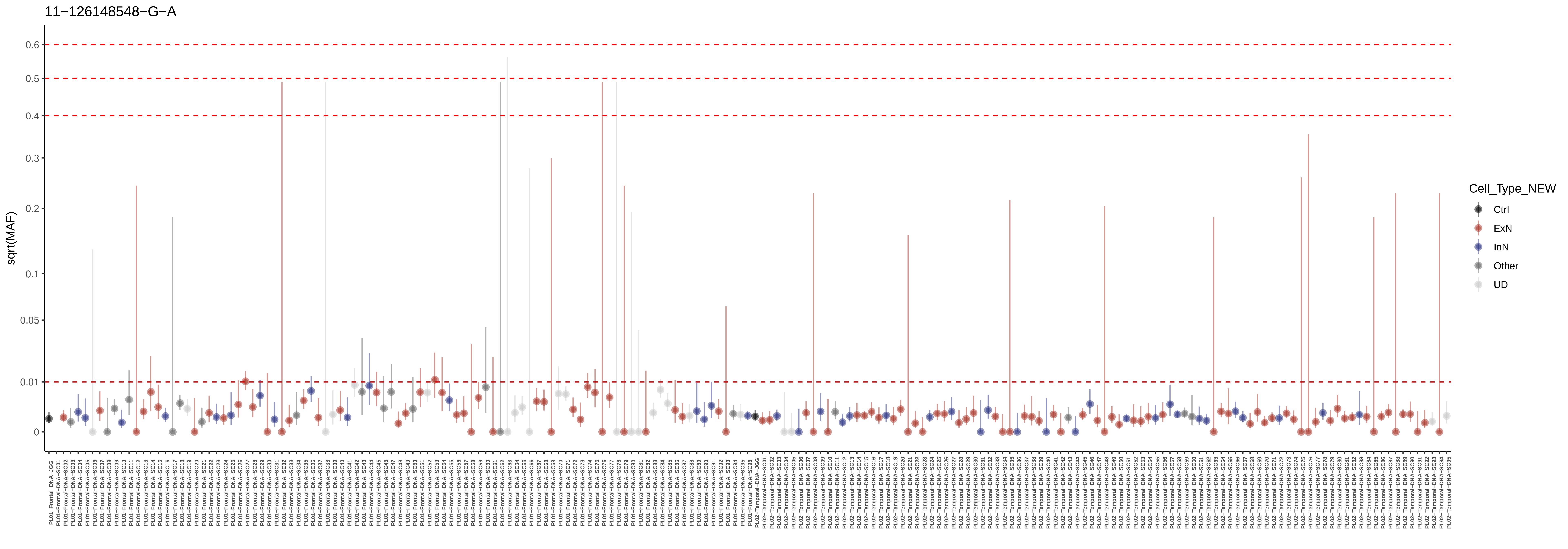
11−126148548−G−A
sqrt(MAF)
0.6
0.5
0.4
0.3
0.2
0.1
0.05
0.01
0
Cell_Type_NEW
Ctrl
ExN
InN
Other
UD

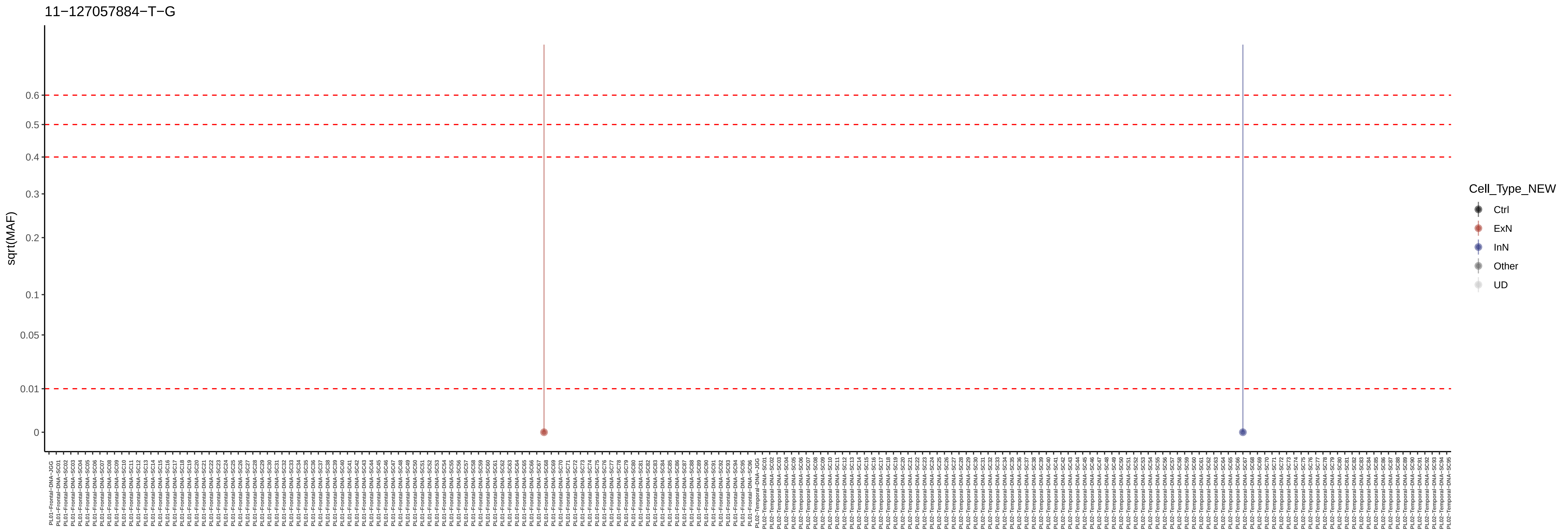
11−127057884−T−G
sqrt(MAF)
0.6
0.5
0.4
0.3
0.2
0.1
0.05
0.01
0
Cell_Type_NEW
Ctrl
ExN
InN
Other
UD

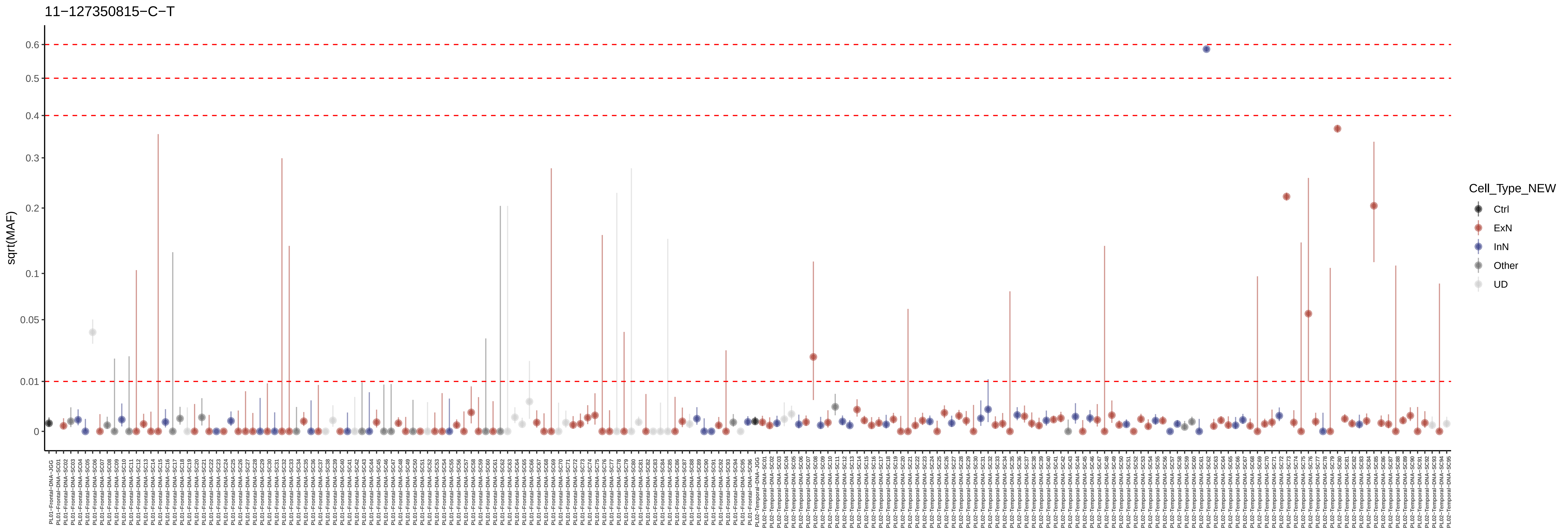
11−127350815−C−T
sqrt(MAF)
0.6
0.5
0.4
0.3
0.2
0.1
0.05
0.01
0
Cell_Type_NEW
Ctrl
ExN
InN
Other
UD

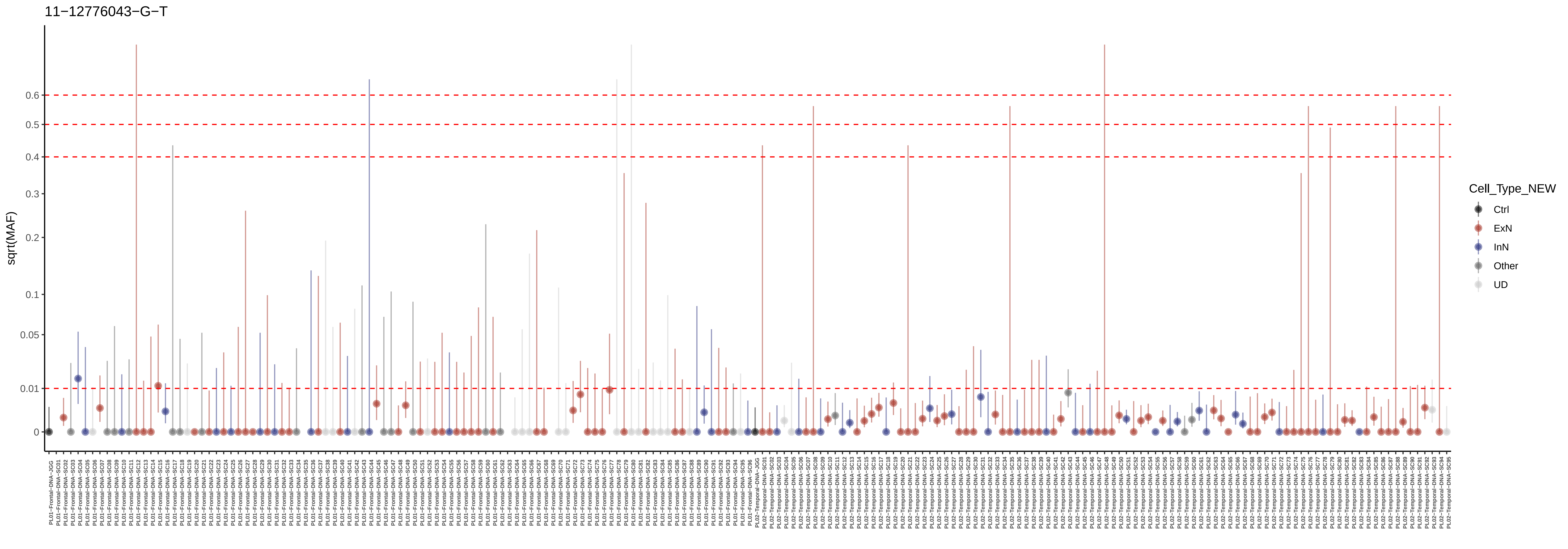

11−12776043−G−T
sqrt(MAF)
0.6
0.5
0.4
0.3
0.2
0.1
0.05
0.01
0
Cell_Type_NEW
Ctrl
ExN
InN
Other
UD

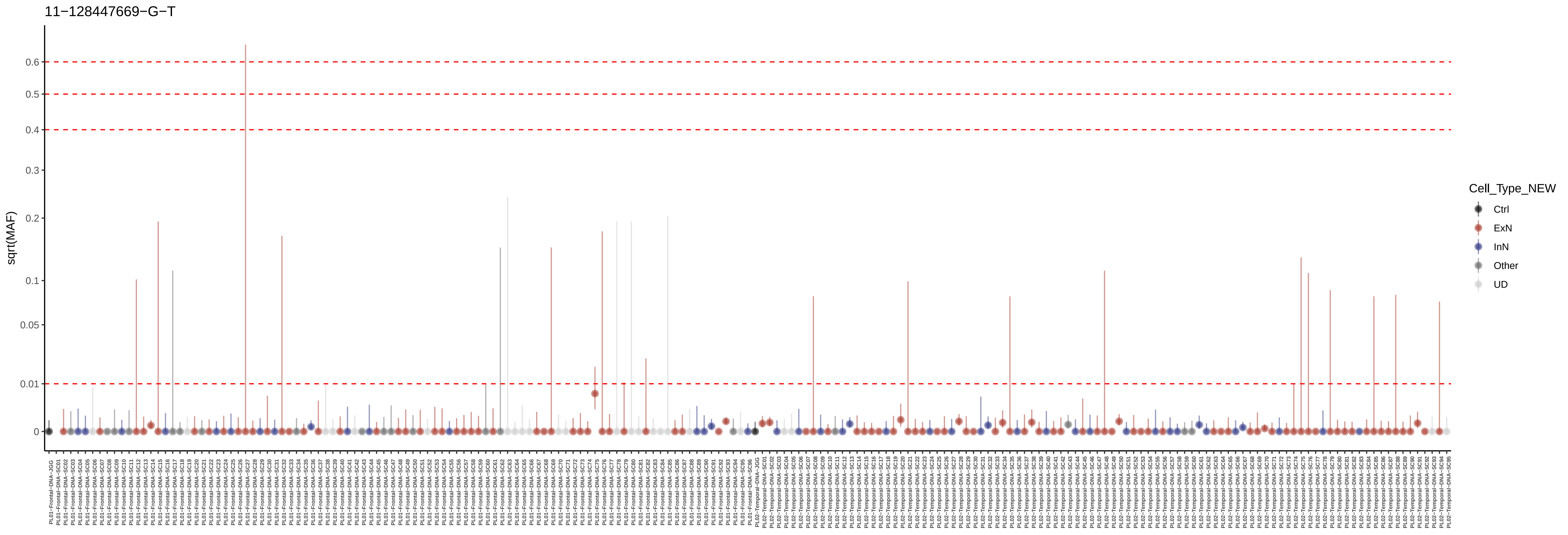
11−128447669−G−T
sqrt(MAF)
0.6
0.5
0.4
0.3
0.2
0.1
0.05
0.01
0
Cell_Type_NEW
Ctrl
ExN
InN
Other
UD

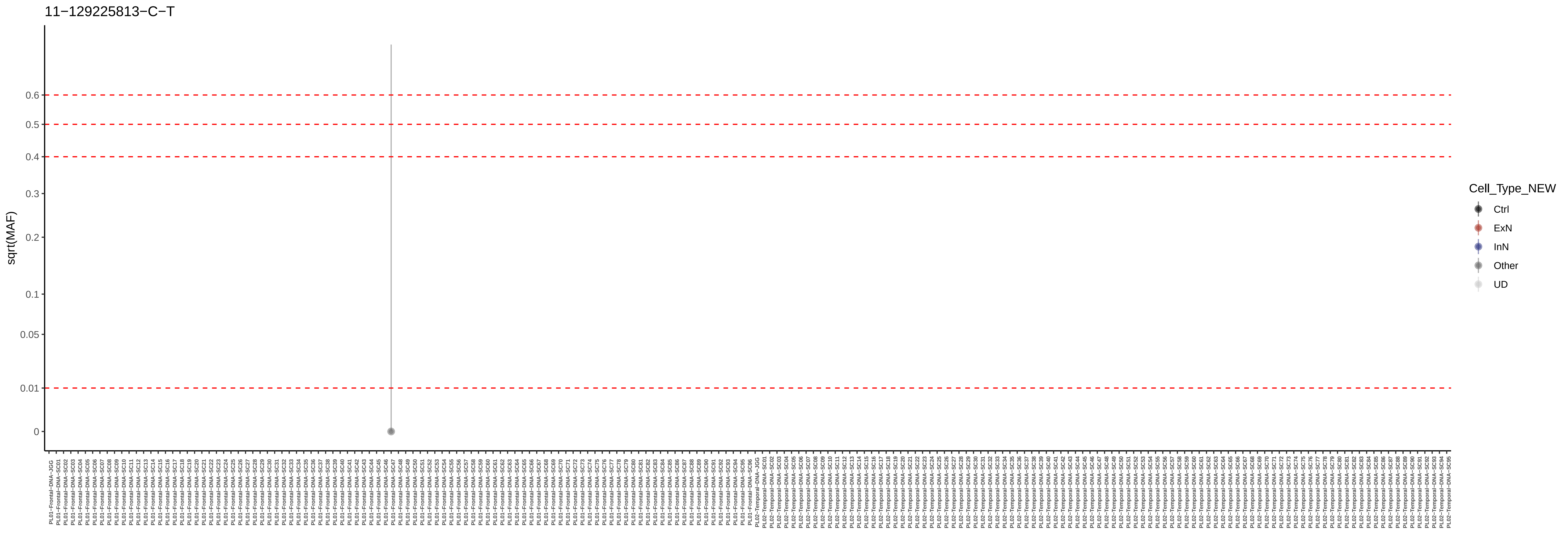

11−129225813−C−T
sqrt(MAF)
0
0.01
0.05
0.1
0.2
0.3
0.4
0.5
0.6
Cell_Type_NEW
Ctrl
ExN
InN
Other
UD

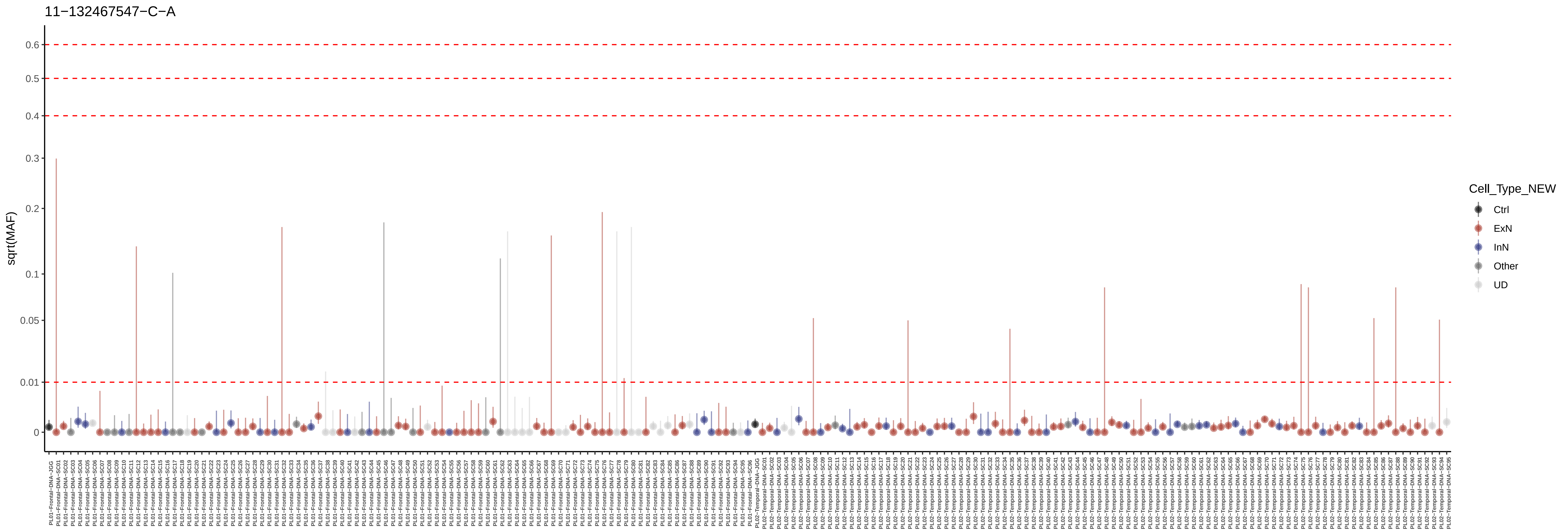
11−132467547−C−A
sqrt(MAF)
0.6
0.5
0.4
0.3
0.2
0.1
0.05
0.01
0
Cell_Type_NEW
Ctrl
ExN
InN
Other
UD

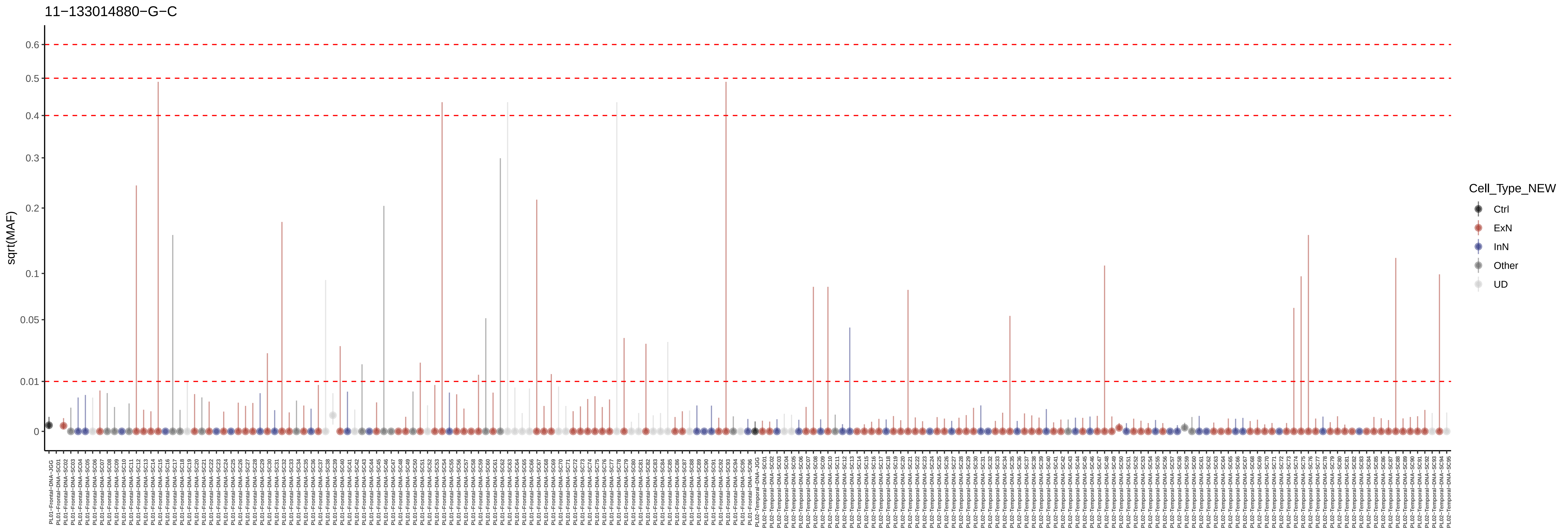
11−133014880−G−C
sqrt(MAF)
0.6
0.5
0.4
0.3
0.2
0.1
0.05
0.01
0
Cell_Type_NEW
Ctrl
ExN
InN
Other
UD

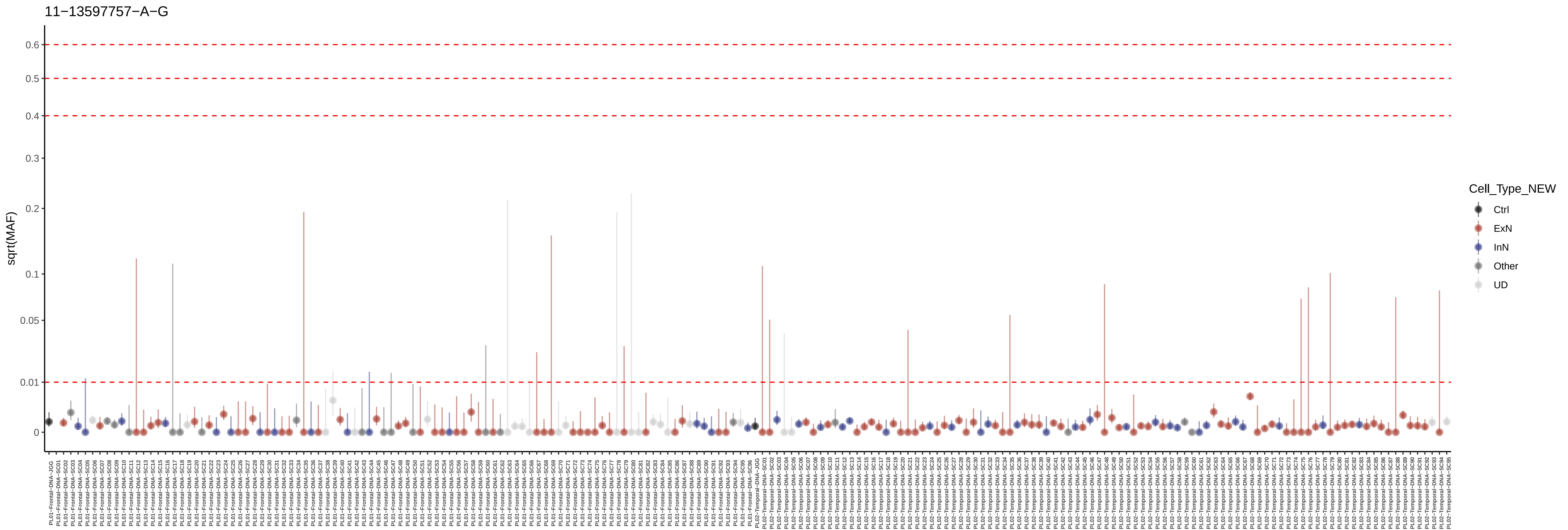
11−13597757−A−G
sqrt(MAF)
0.6
0.5
0.4
0.3
0.2
0.1
0.05
0.01
0
Cell_Type_NEW
Ctrl
ExN
InN
Other
UD

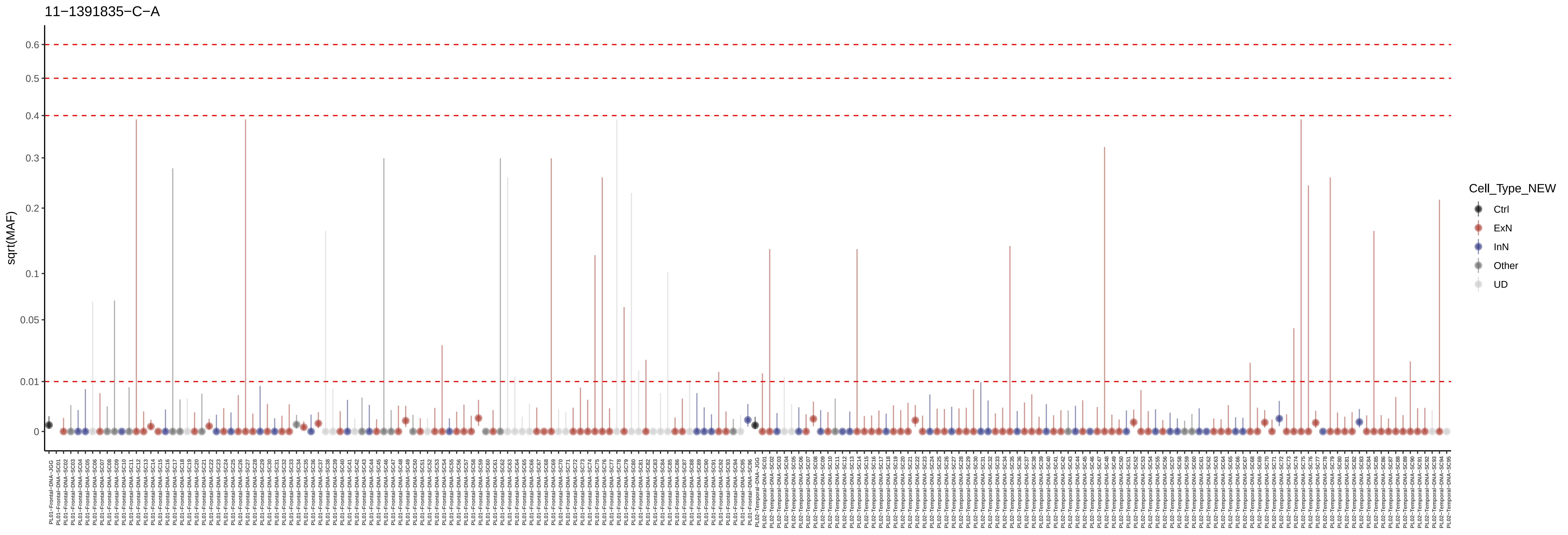

11−1391835−C−A
sqrt(MAF)
0.6
0.5
0.4
0.3
0.2
0.1
0.05
0.01
0
Cell_Type_NEW
Ctrl
ExN
InN
Other
UD

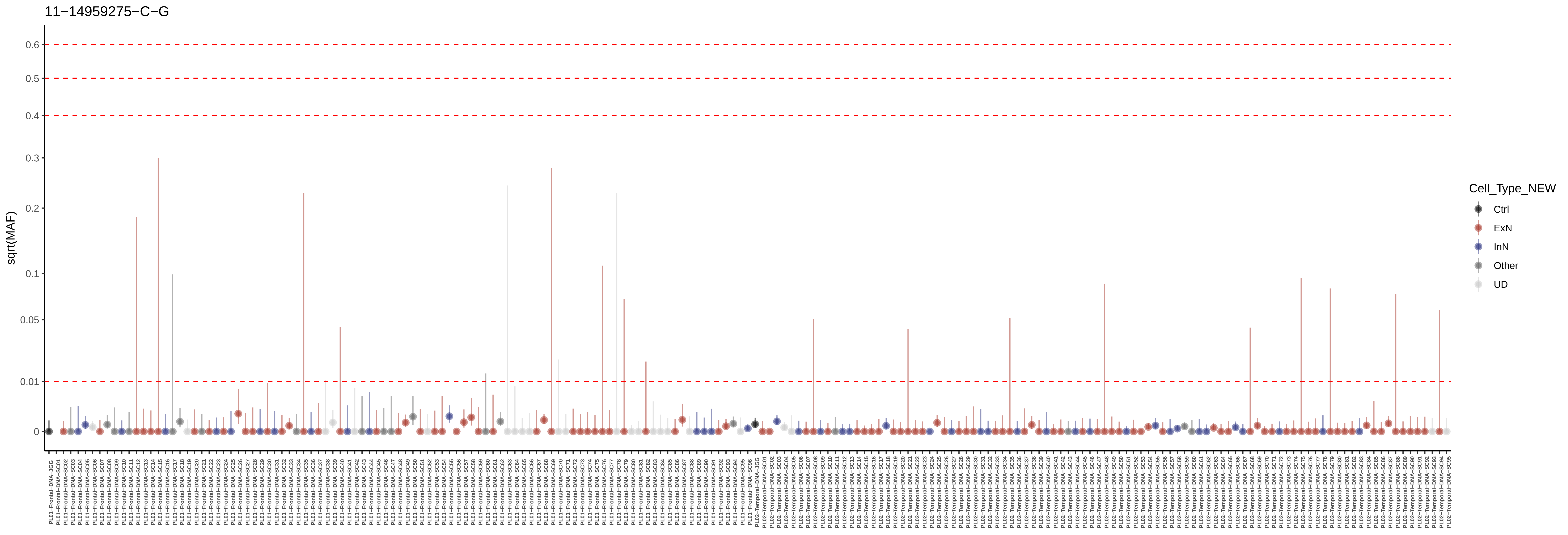
11−14959275−C−G
sqrt(MAF)
0.6
0.5
0.4
0.3
0.2
0.1
0.05
0.01
0
Cell_Type_NEW
Ctrl
ExN
InN
Other
UD

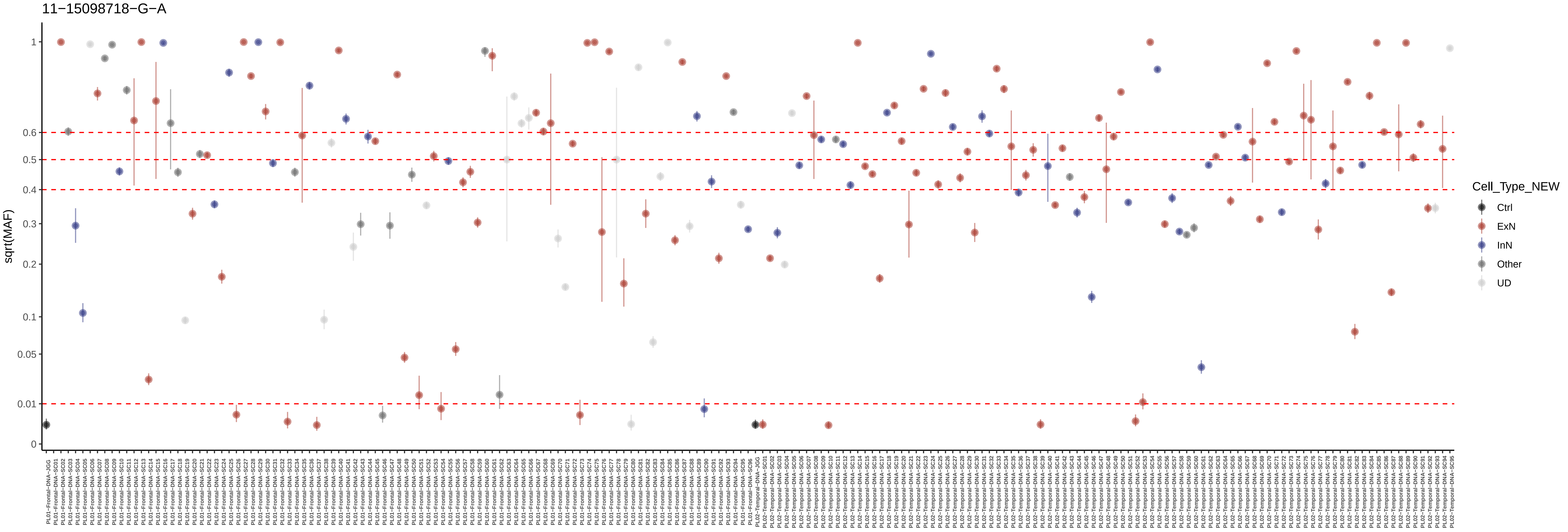
11−15098718−G−A
sqrt(MAF)
1
0.6
0.5
0.4
0.3
0.2
0.1
0.05
0.01
0
Cell_Type_NEW
Ctrl
ExN
InN
Other
UD

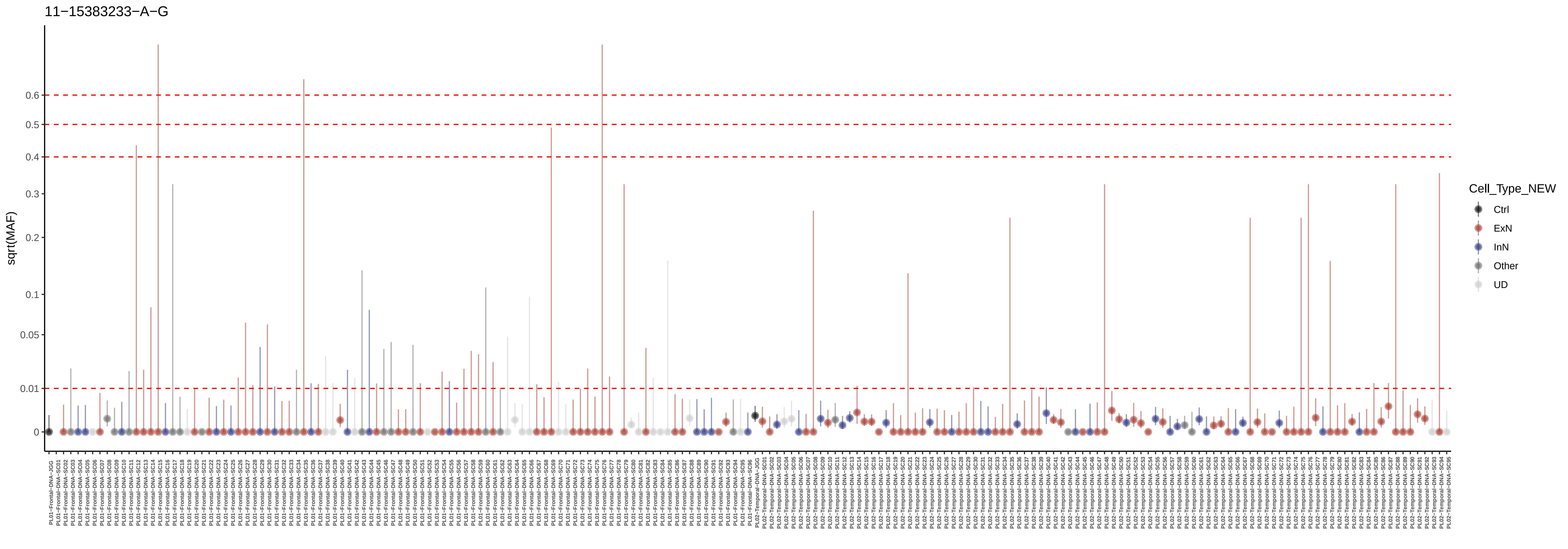

11−15383233−A−G
sqrt(MAF)
0.6
0.5
0.4
0.3
0.2
0.1
0.05
0.01
0
Cell_Type_NEW
Ctrl
ExN
InN
Other
UD

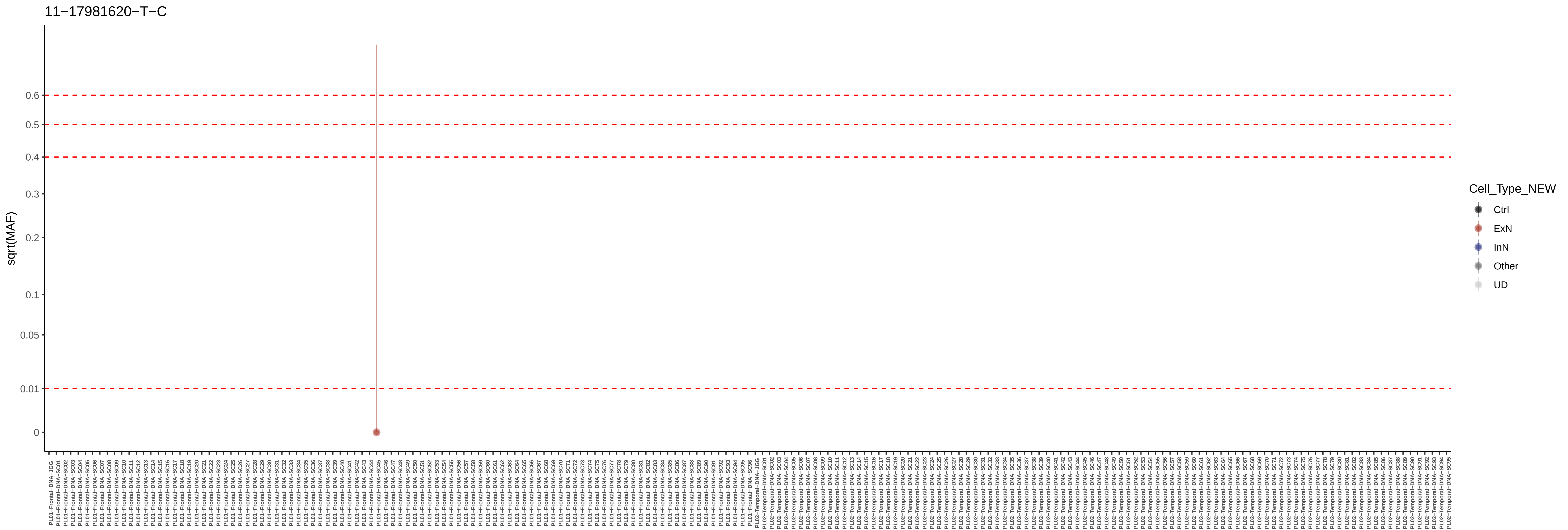
11−17981620−T−C
sqrt(MAF)
0.6
0.5
0.4
0.3
0.2
0.1
0.05
0.01
0
Cell_Type_NEW
Ctrl
ExN
InN
Other
UD

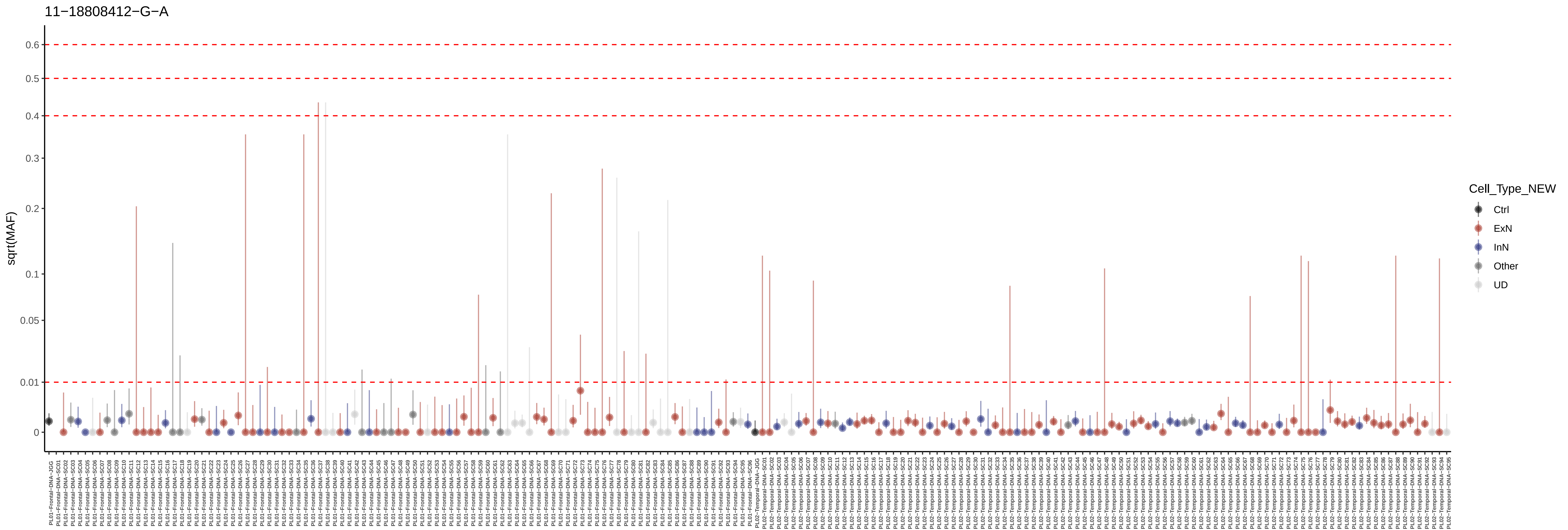
11−18808412−G−A
sqrt(MAF)
0.6
0.5
0.4
0.3
0.2
0.1
0.05
0.01
0
Cell_Type_NEW
Ctrl
ExN
InN
Other
UD

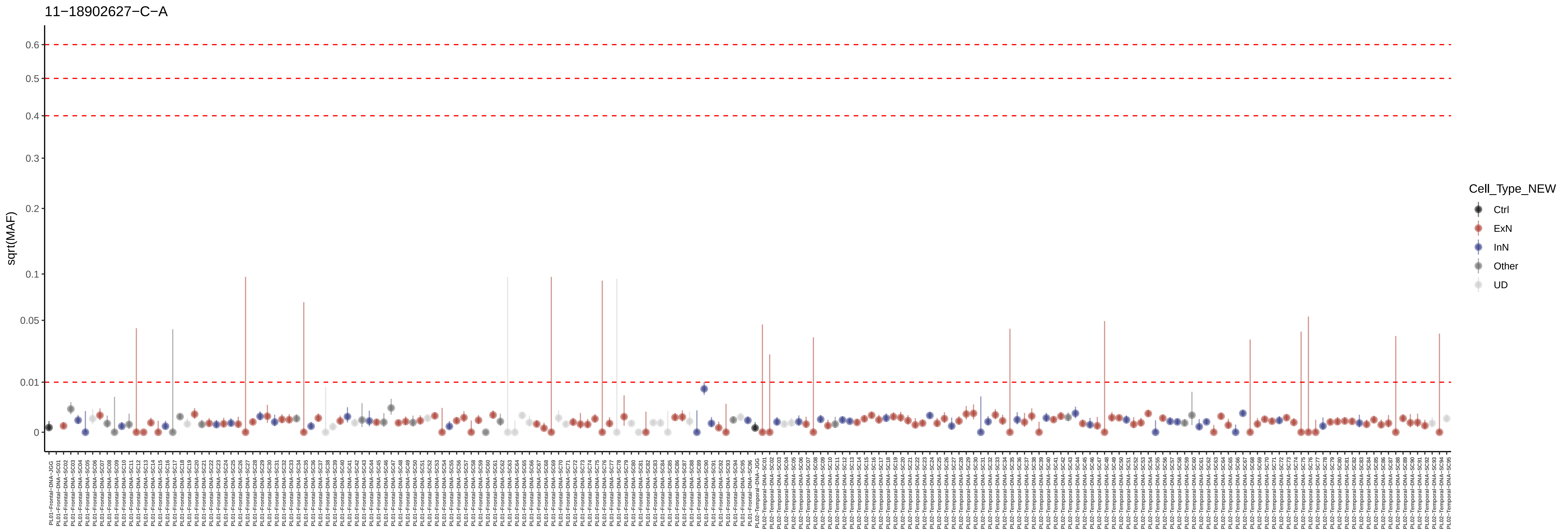

11−18902627−C−A
sqrt(MAF)
0.6
0.5
0.4
0.3
0.2
0.1
0.05
0.01
0
Cell_Type_NEW
Ctrl
ExN
InN
Other
UD

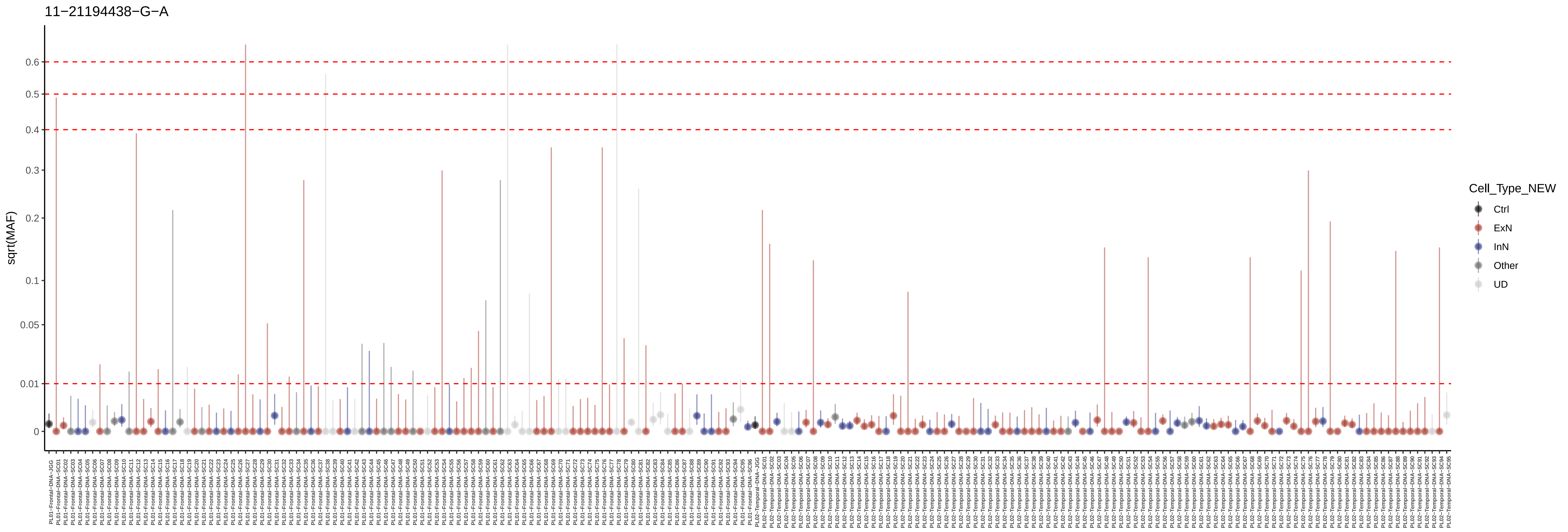
11−21194438−G−A
sqrt(MAF)
0.6
0.5
0.4
0.3
0.2
0.1
0.05
0.01
0
Cell_Type_NEW
Ctrl
ExN
InN
Other
UD

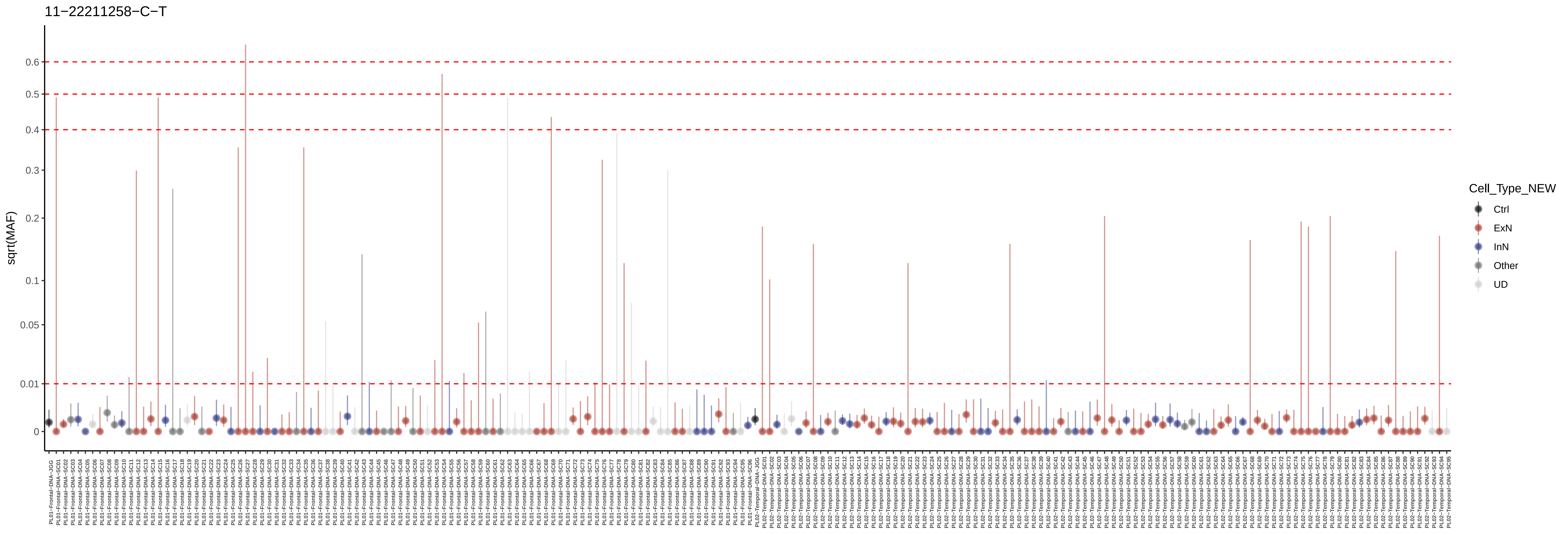

11−22211258−C−T
sqrt(MAF)
0.6
0.5
0.4
0.3
0.2
0.1
0.05
0.01
0
Cell_Type_NEW
Ctrl
ExN
InN
Other
UD

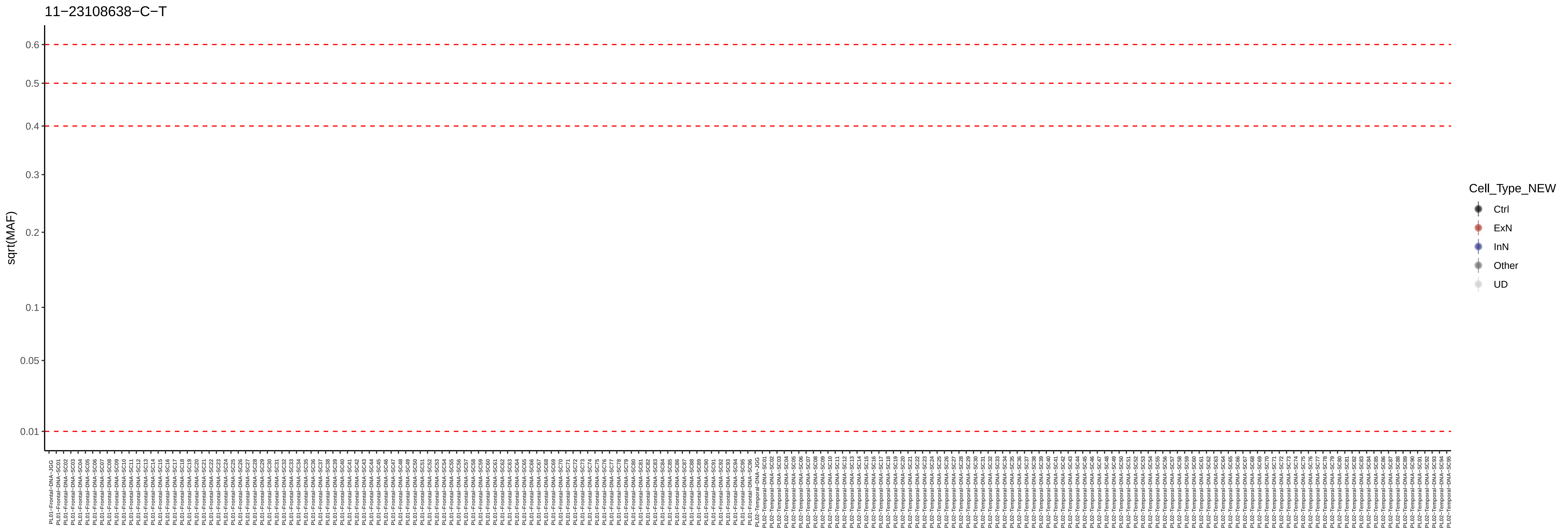
11−23108638−C−T
sqrt(MAF)
0.6
0.5
0.4
0.3
0.2
0.1
0.05
0.01
Cell_Type_NEW
Ctrl
ExN
InN
Other
UD

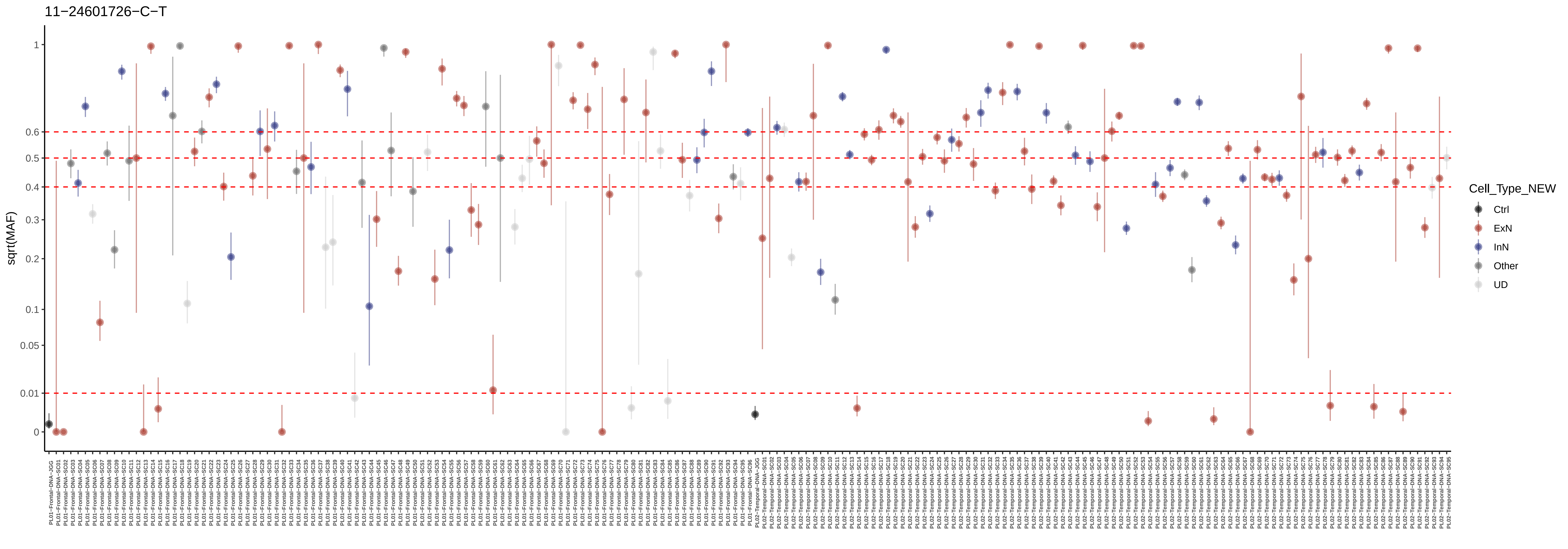

11−24601726−C−T
sqrt(MAF)
1
0.6
0.5
0.4
0.3
0.2
0.1
0.05
0.01
0
Cell_Type_NEW
Ctrl
ExN
InN
Other
UD

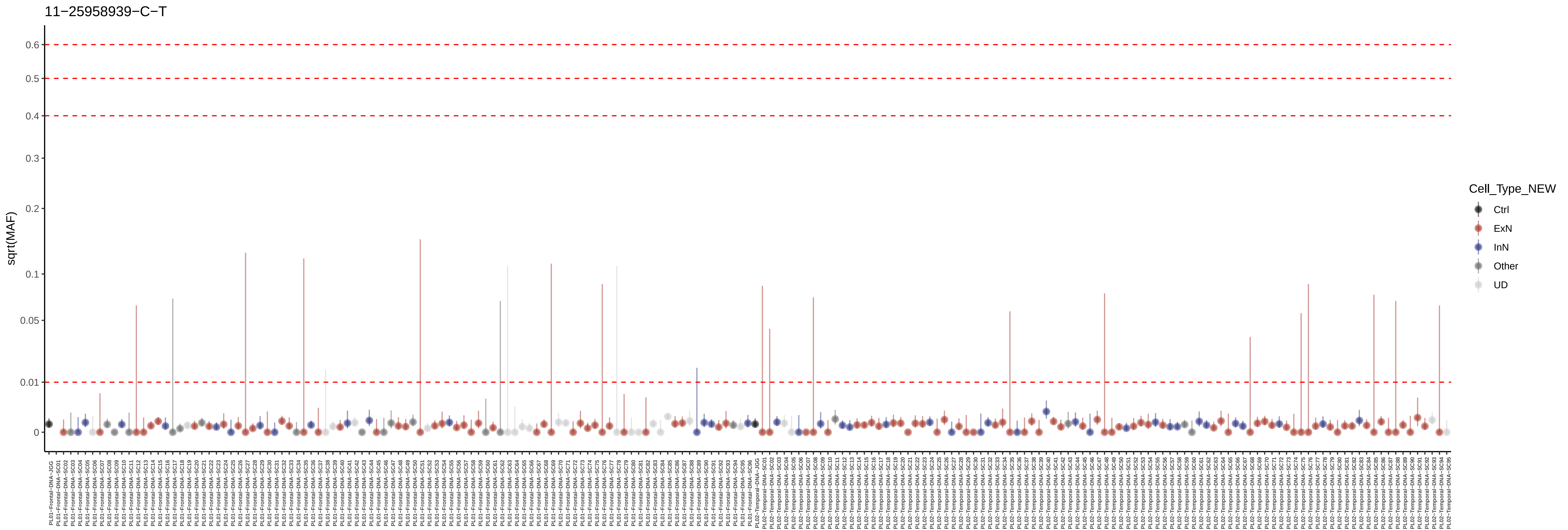
11−25958939−C−T
sqrt(MAF)
0.6
0.5
0.4
0.3
0.2
0.1
0.05
0.01
0
Cell_Type_NEW
Ctrl
ExN
InN
Other
UD

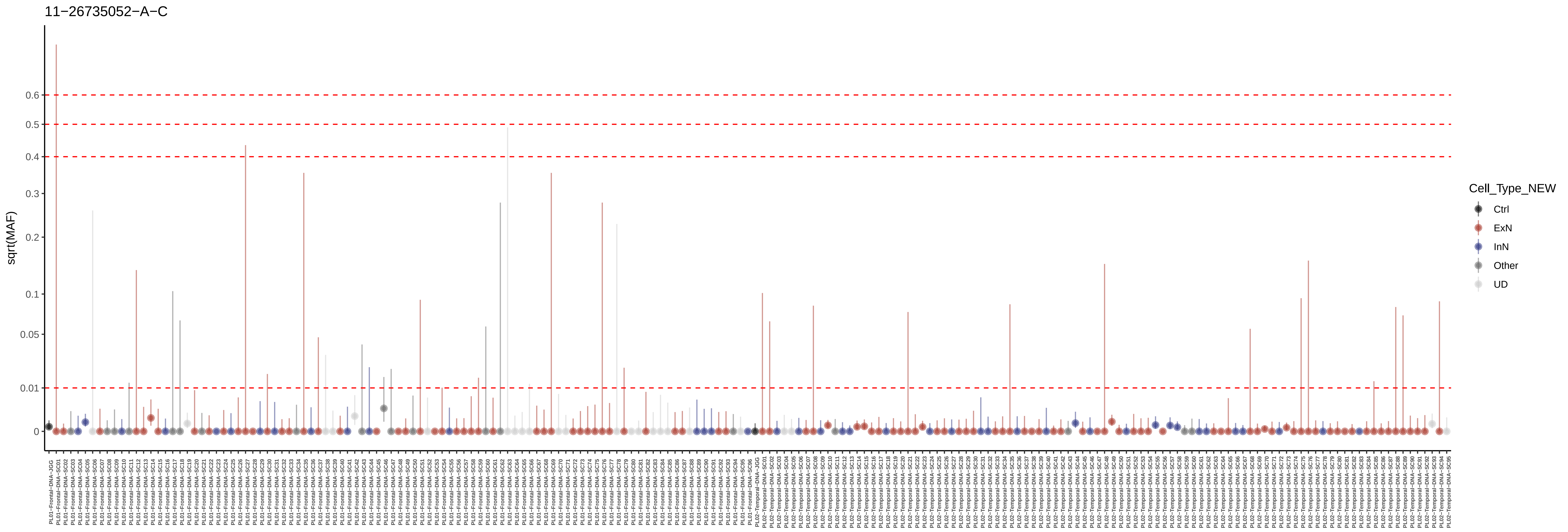

11−26735052−A−C
sqrt(MAF)
0.6
0.5
0.4
0.3
0.2
0.1
0.05
0.01
0
Cell_Type_NEW
Ctrl
ExN
InN
Other
UD

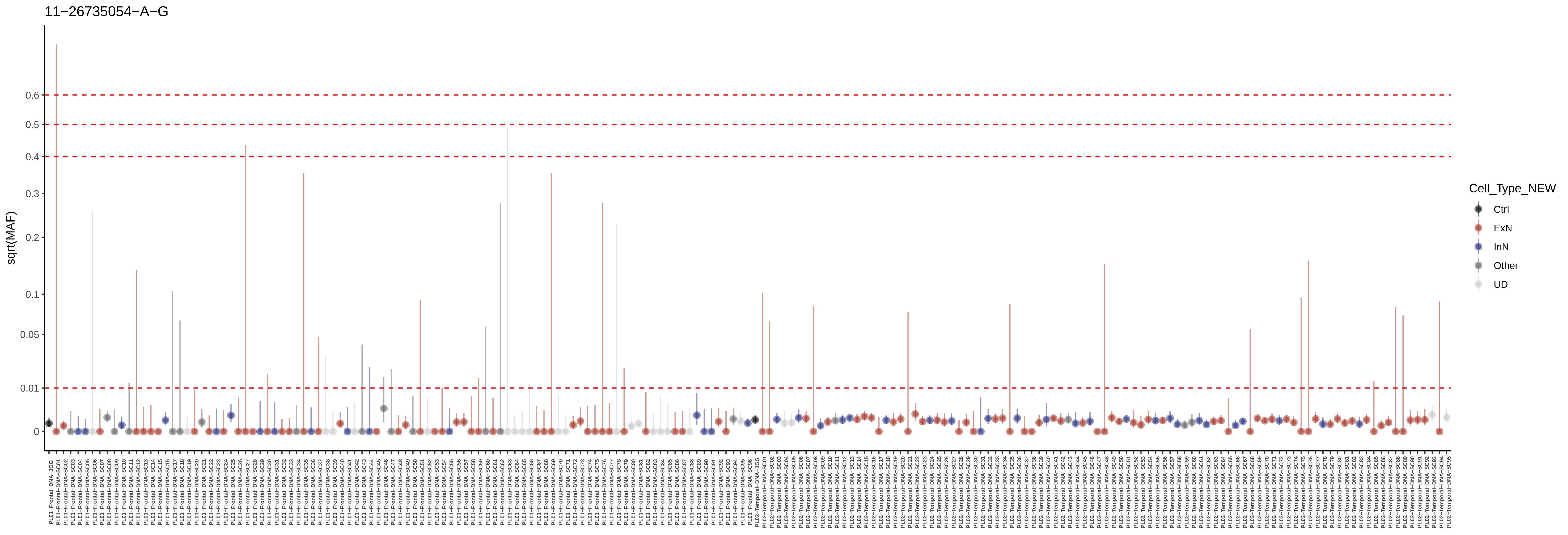
11−26735054−A−G
sqrt(MAF)
0.6
0.5
0.4
0.3
0.2
0.1
0.05
0.01
0
Cell_Type_NEW
Ctrl
ExN
InN
Other
UD

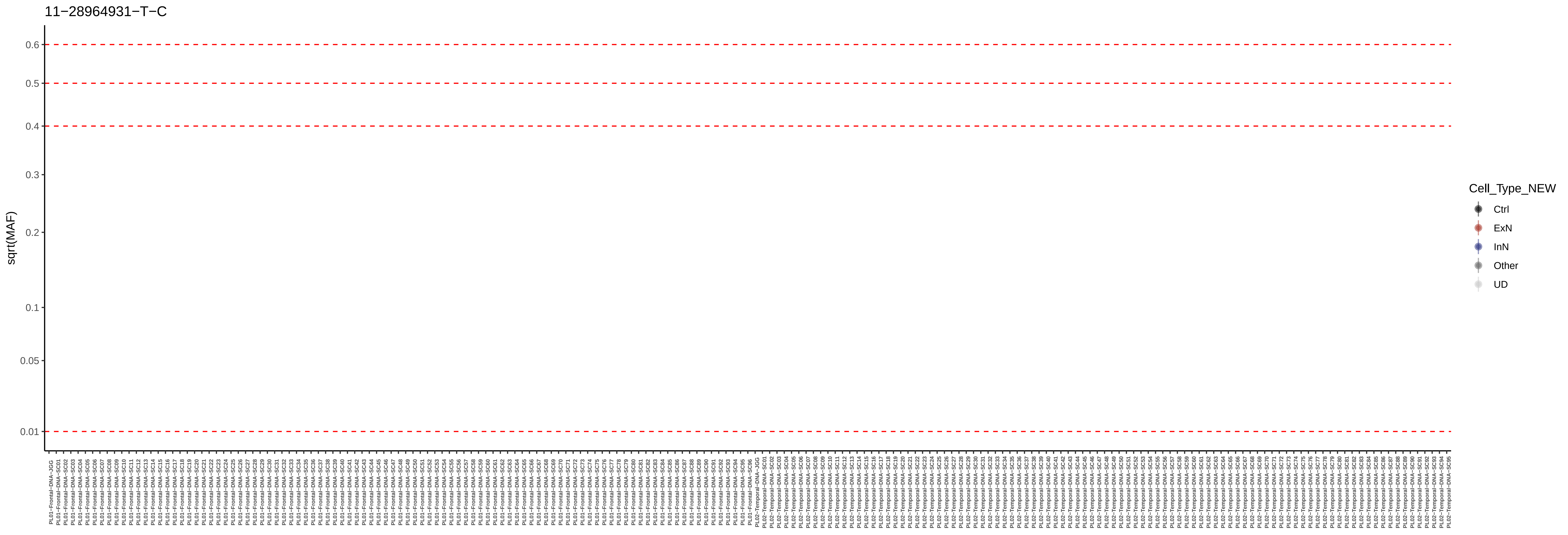
11−28964931−T−C
sqrt(MAF)
0.6
0.5
0.4
0.3
0.2
0.1
0.05
0.01
Cell_Type_NEW
Ctrl
ExN
InN
Other
UD

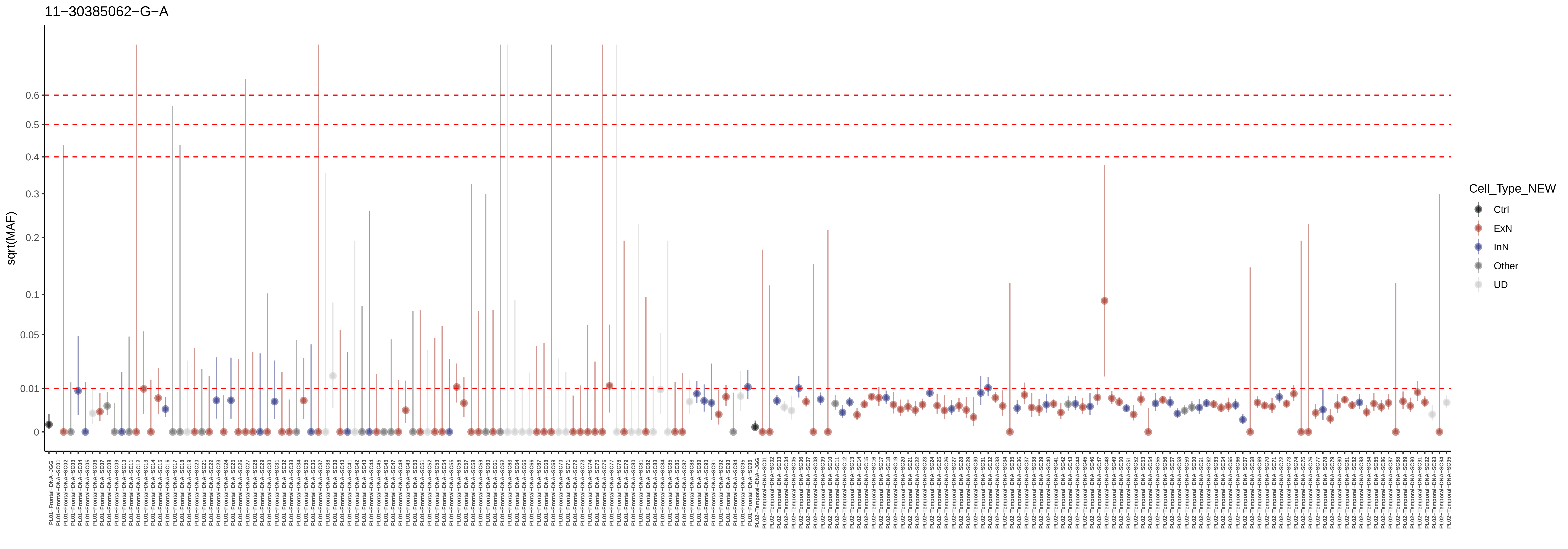

11−30385062−G−A
sqrt(MAF)
0.6
0.5
0.4
0.3
0.2
0.1
0.05
0.01
0
Cell_Type_NEW
Ctrl
ExN
InN
Other
UD

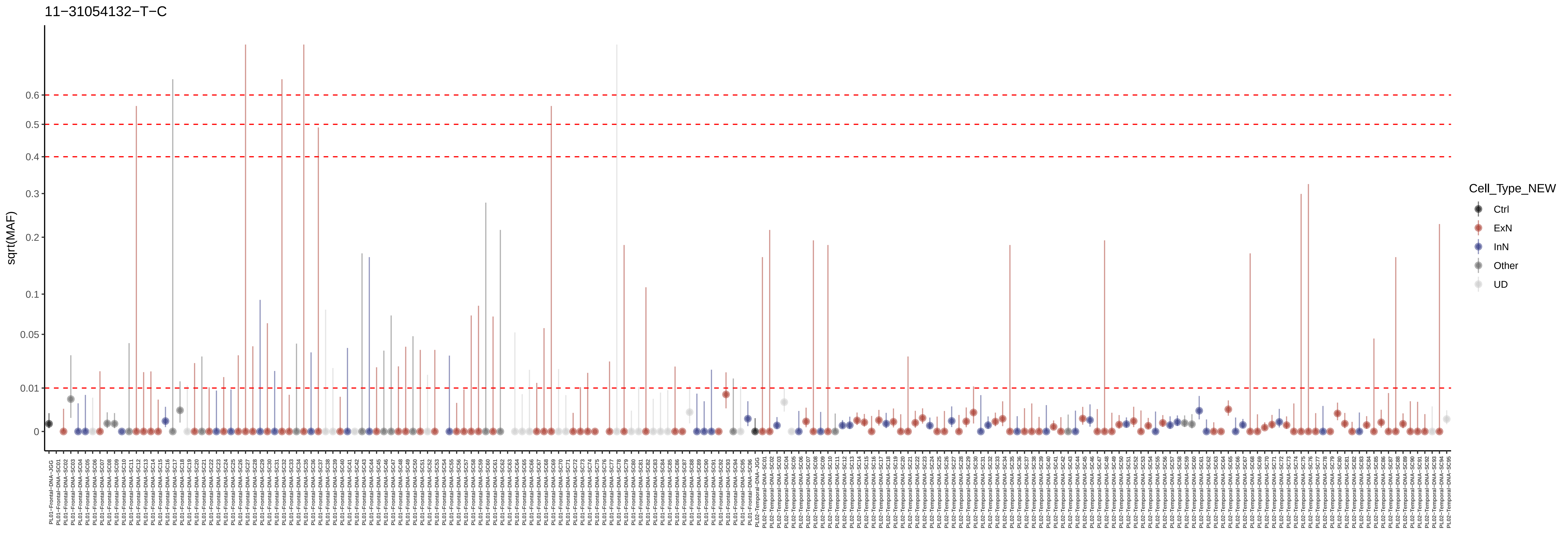

11−31054132−T−C
sqrt(MAF)
0.6
0.5
0.4
0.3
0.2
0.1
0.05
0.01
0
Cell_Type_NEW
Ctrl
ExN
InN
Other
UD

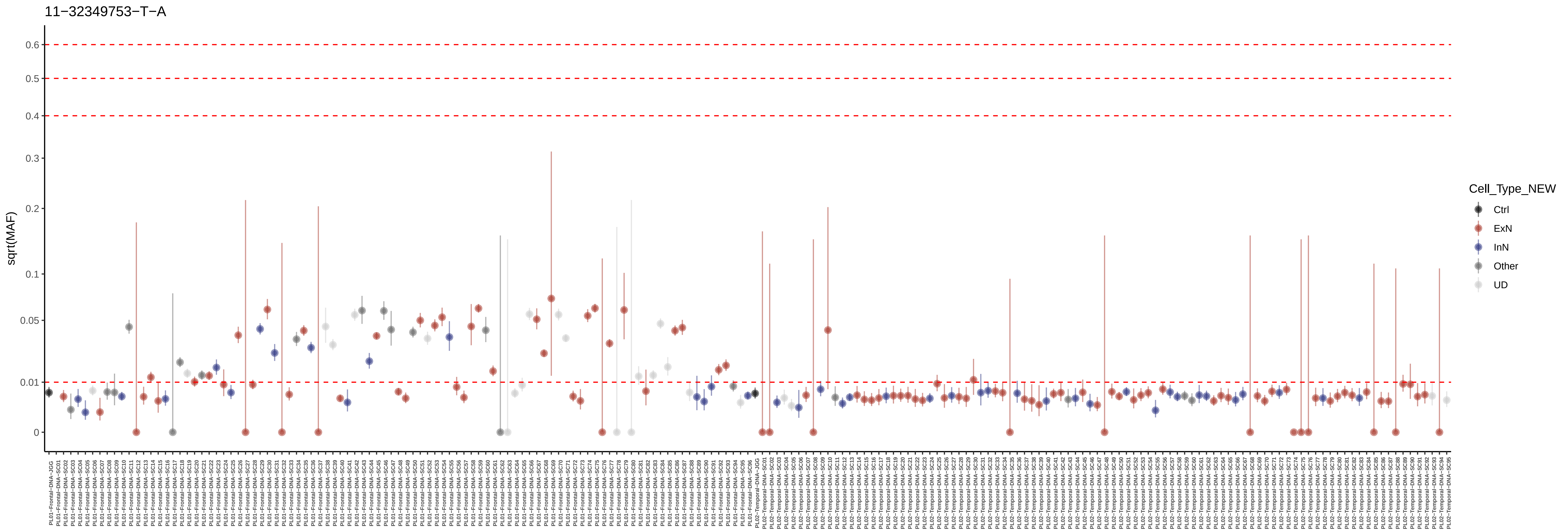

11−32349753−T−A
sqrt(MAF)
0.6
0.5
0.4
0.3
0.2
0.1
0.05
0.01
0
Cell_Type_NEW
Ctrl
ExN
InN
Other
UD

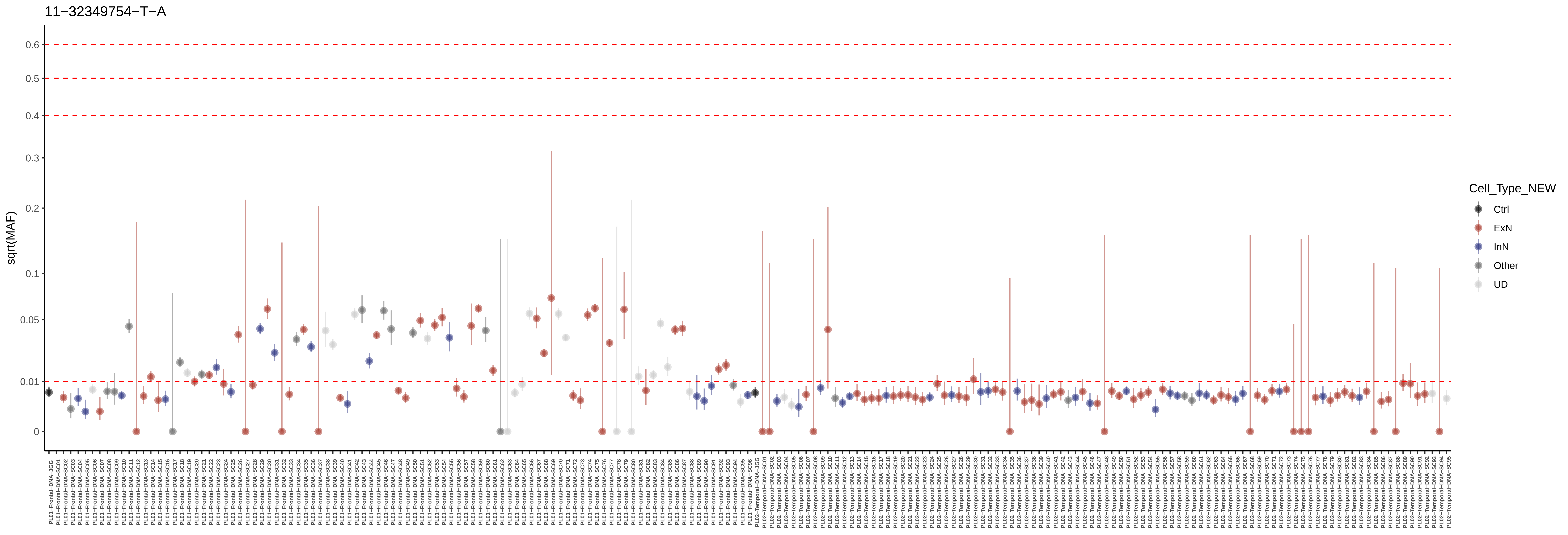

11−32349754−T−A
sqrt(MAF)
0.6
0.5
0.4
0.3
0.2
0.1
0.05
0.01
0
Cell_Type_NEW
Ctrl
ExN
InN
Other
UD

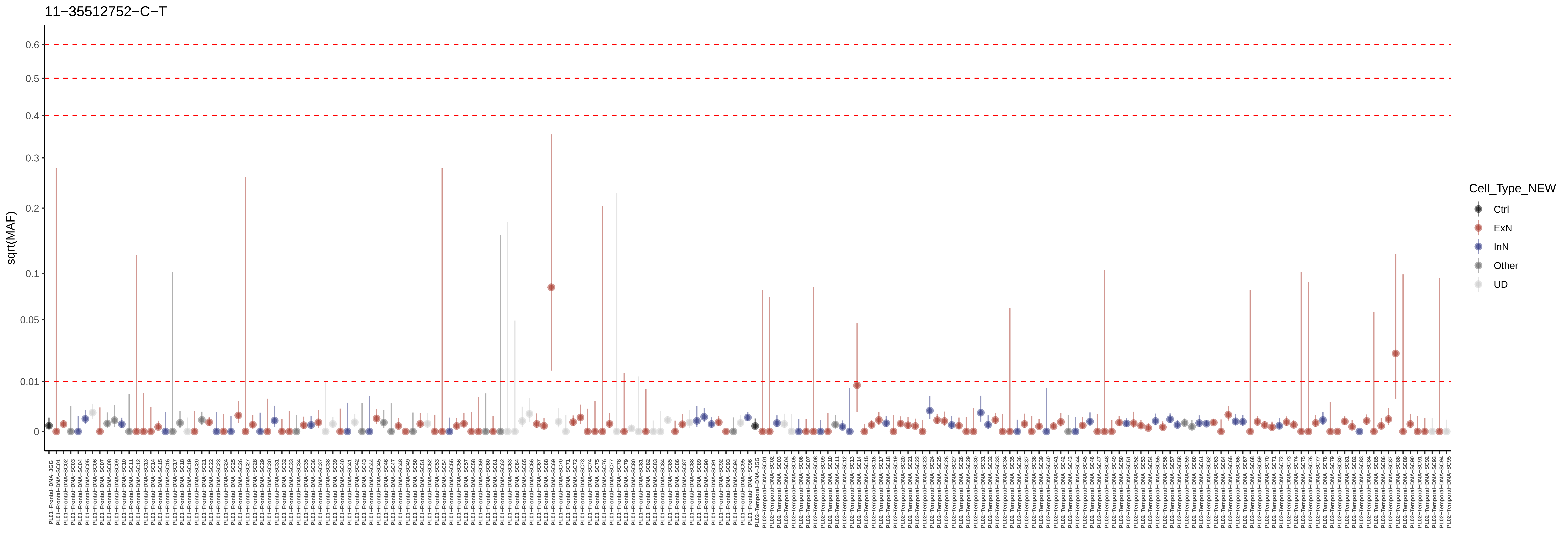
11−35512752−C−T
sqrt(MAF)
0.6
0.5
0.4
0.3
0.2
0.1
0.05
0.01
0
Cell_Type_NEW
Ctrl
ExN
InN
Other
UD

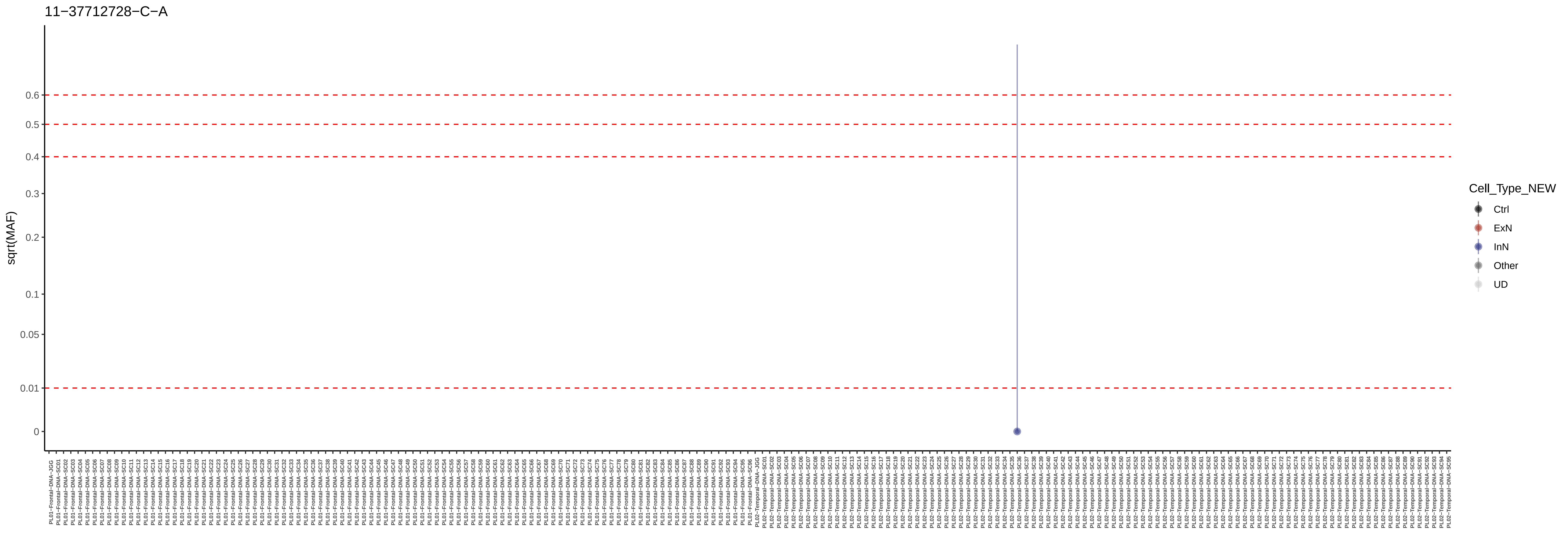

11−37712728−C−A
sqrt(MAF)
0.6
0.5
0.4
0.3
0.2
0.1
0.05
0.01
0
Cell_Type_NEW
Ctrl
ExN
InN
Other
UD

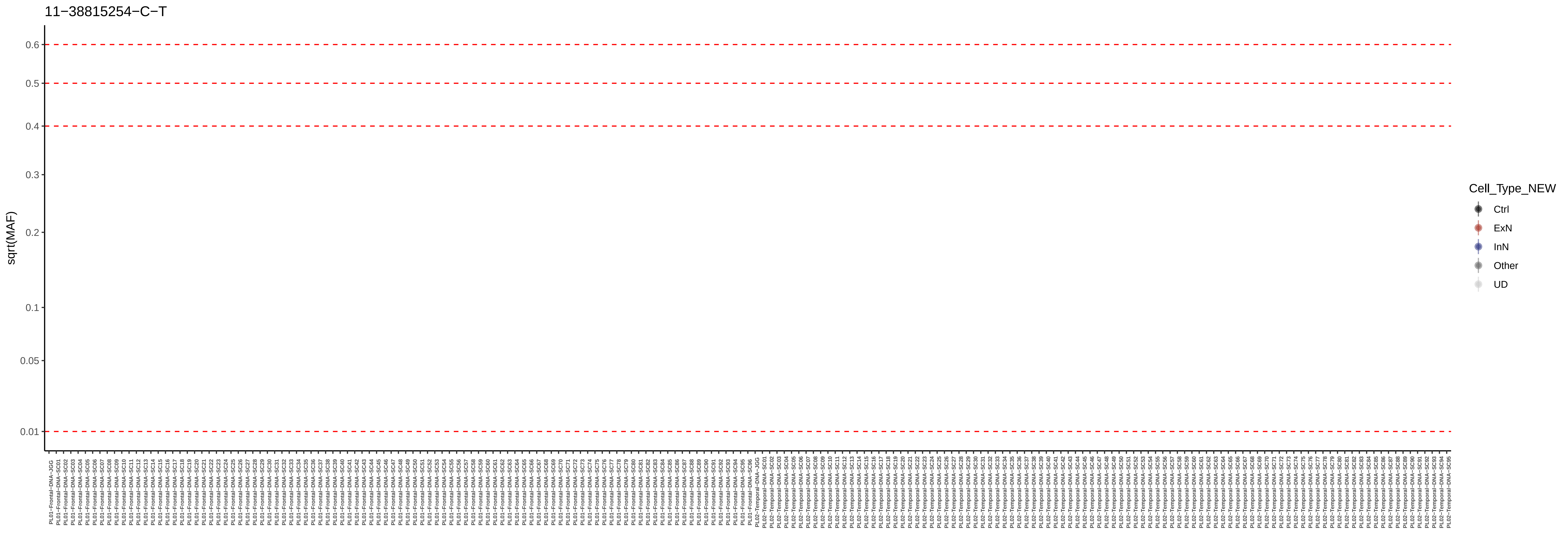

11−38815254−C−T
sqrt(MAF)
0.6
0.5
0.4
0.3
0.2
0.1
0.05
0.01
Cell_Type_NEW
Ctrl
ExN
InN
Other
UD

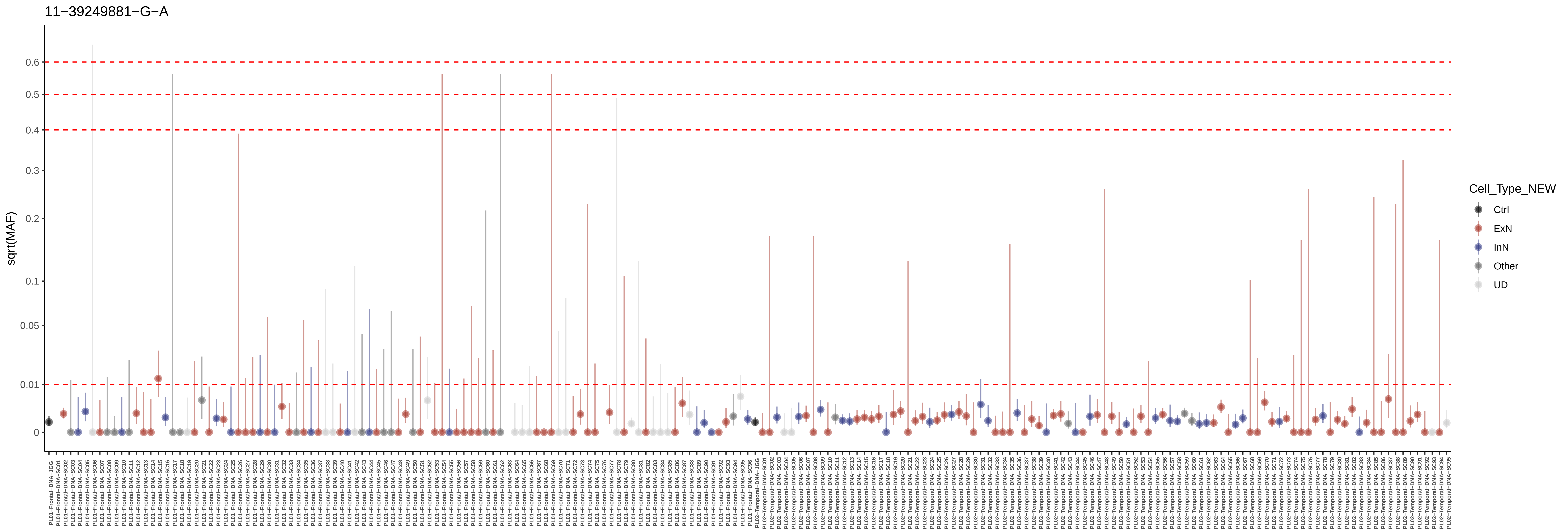
11−39249881−G−A
sqrt(MAF)
0.6
0.5
0.4
0.3
0.2
0.1
0.05
0.01
0
Cell_Type_NEW
Ctrl
ExN
InN
Other
UD

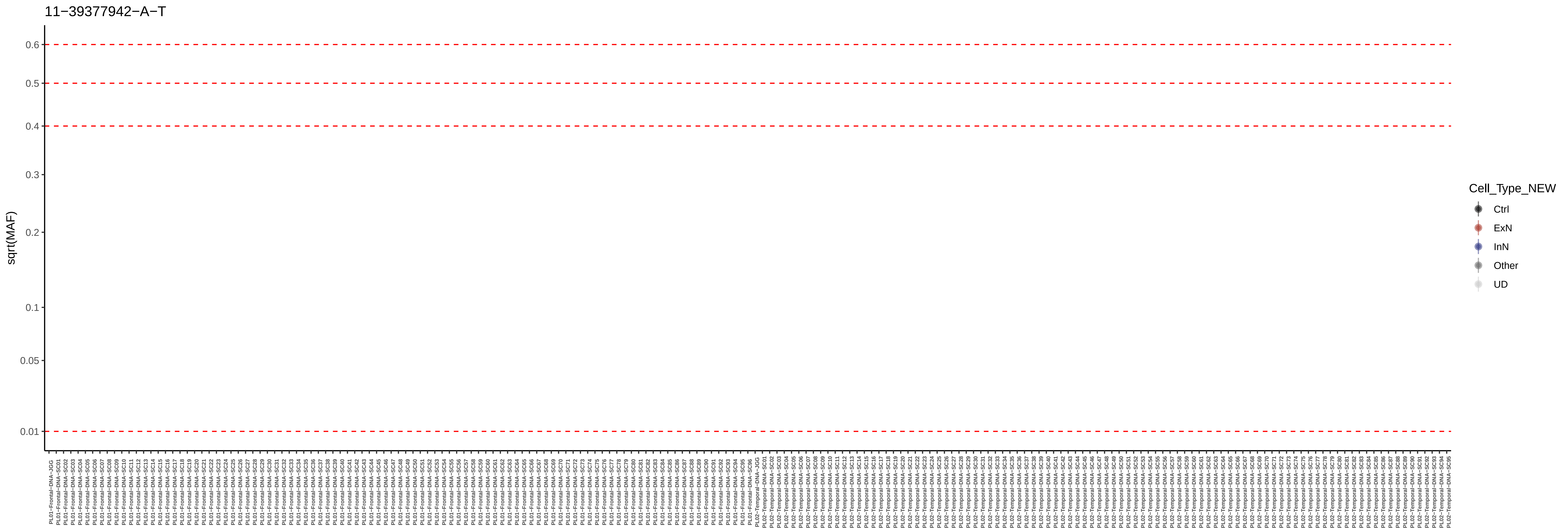

11−39377942−A−T
sqrt(MAF)
0.6
0.5
0.4
0.3
0.2
0.1
0.05
0.01
Cell_Type_NEW
Ctrl
ExN
InN
Other
UD

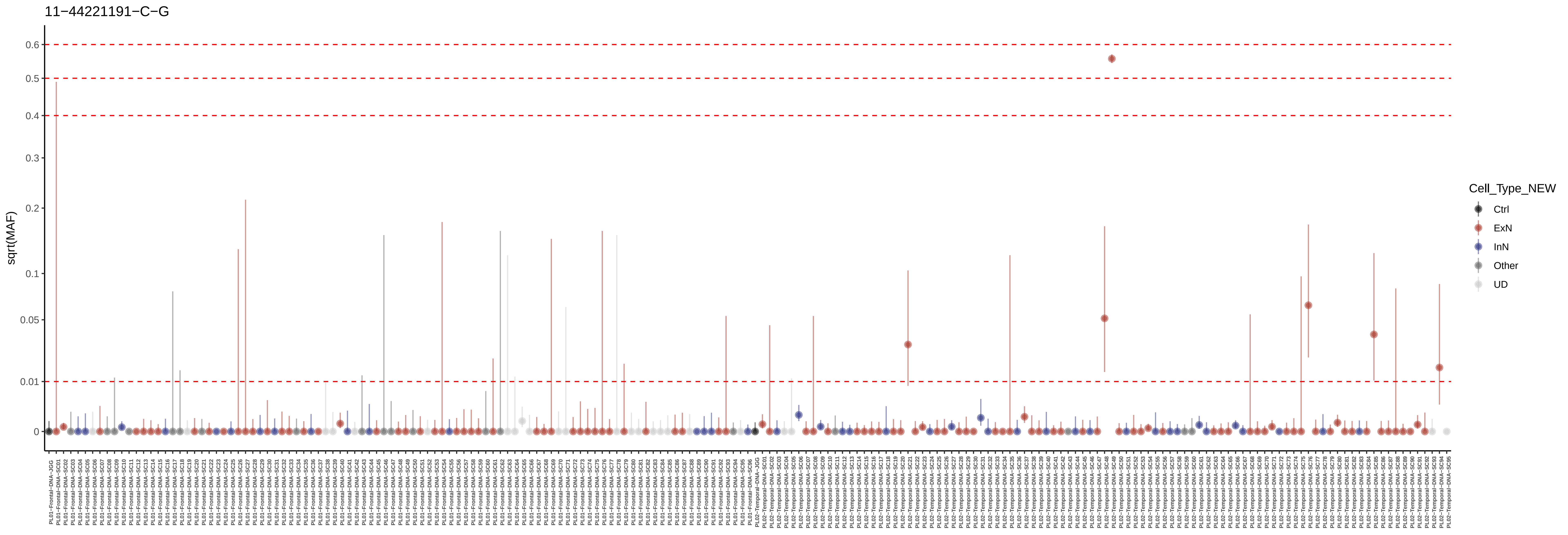
11−44221191−C−G
sqrt(MAF)
0.6
0.5
0.4
0.3
0.2
0.1
0.05
0.01
0
Cell_Type_NEW
Ctrl
ExN
InN
Other
UD

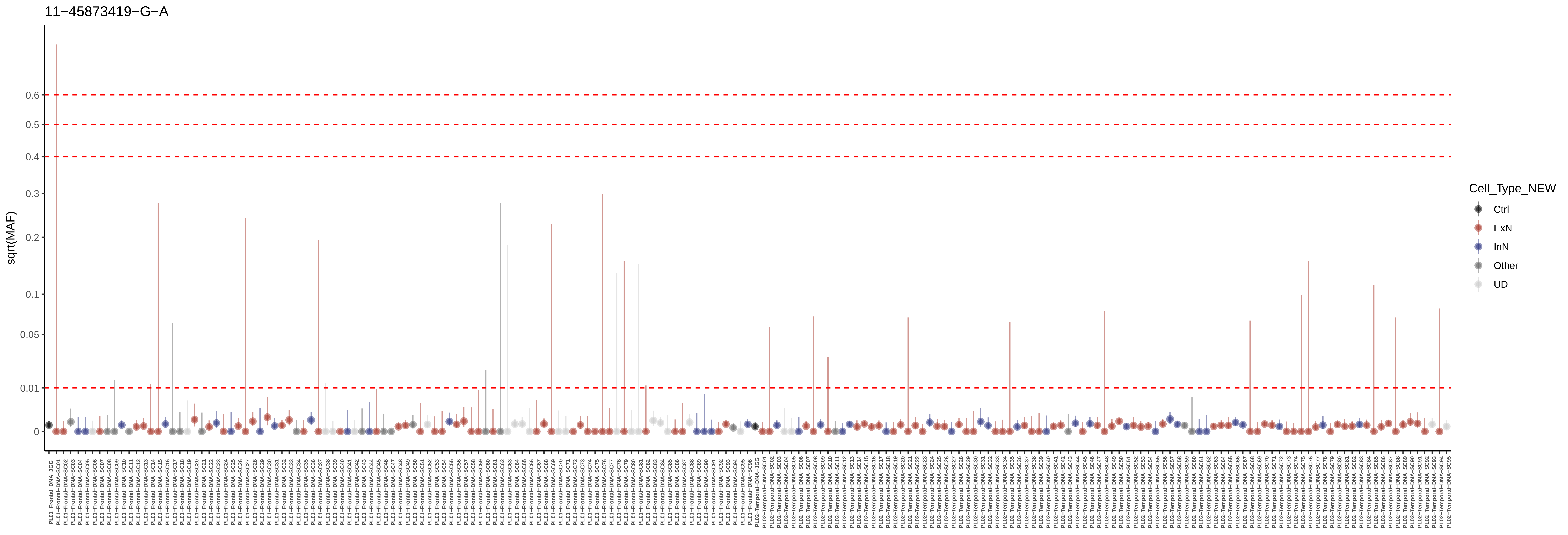
11−45873419−G−A
sqrt(MAF)
Cell_Type_NEW
Ctrl
ExN
InN
Other
UD

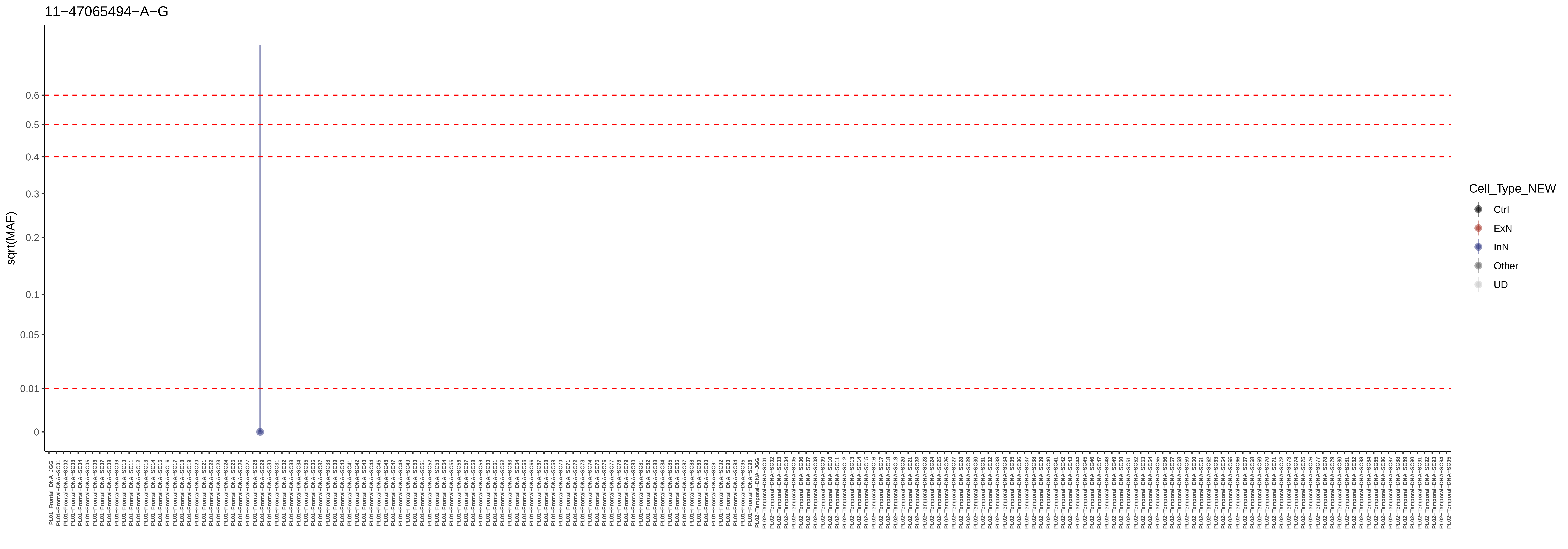
11−47065494−A−G
sqrt(MAF)
0.6
0.5
0.4
0.3
0.2
0.1
0.05
0.01
0
Cell_Type_NEW
Ctrl
ExN
InN
Other
UD

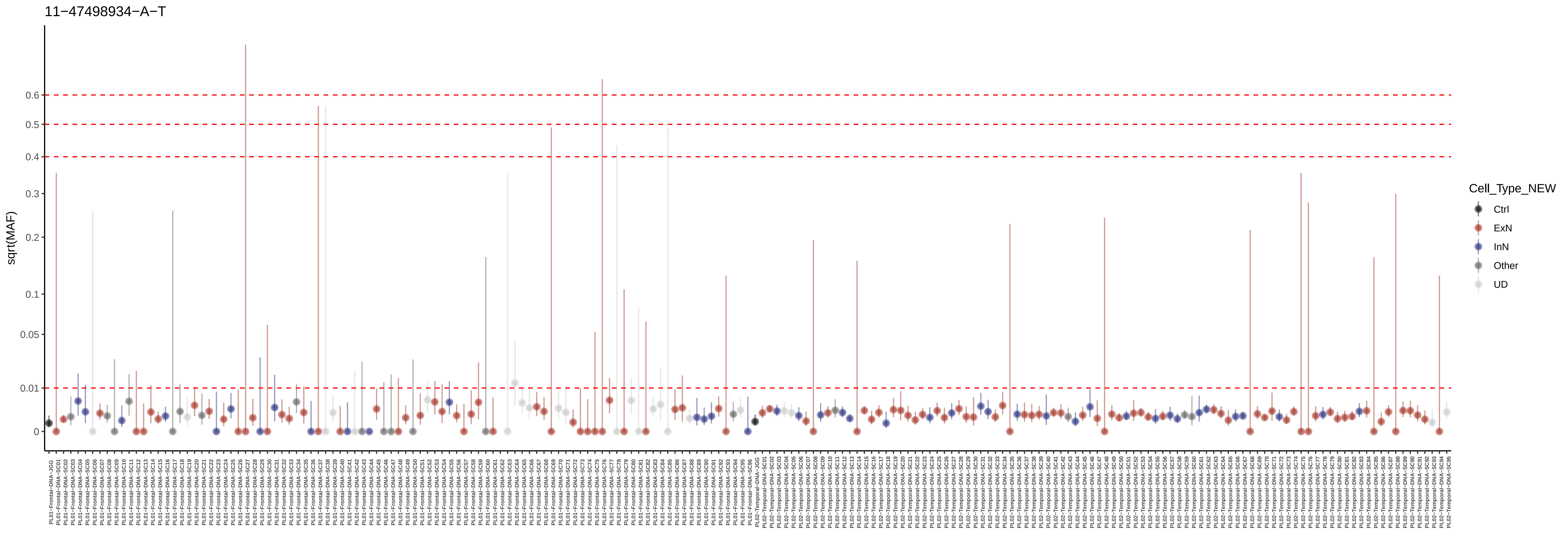
11−47498934−A−T
sqrt(MAF)
0.6
0.5
0.4
0.3
0.2
0.1
0.05
0.01
0
Cell_Type_NEW
Ctrl
ExN
InN
Other
UD

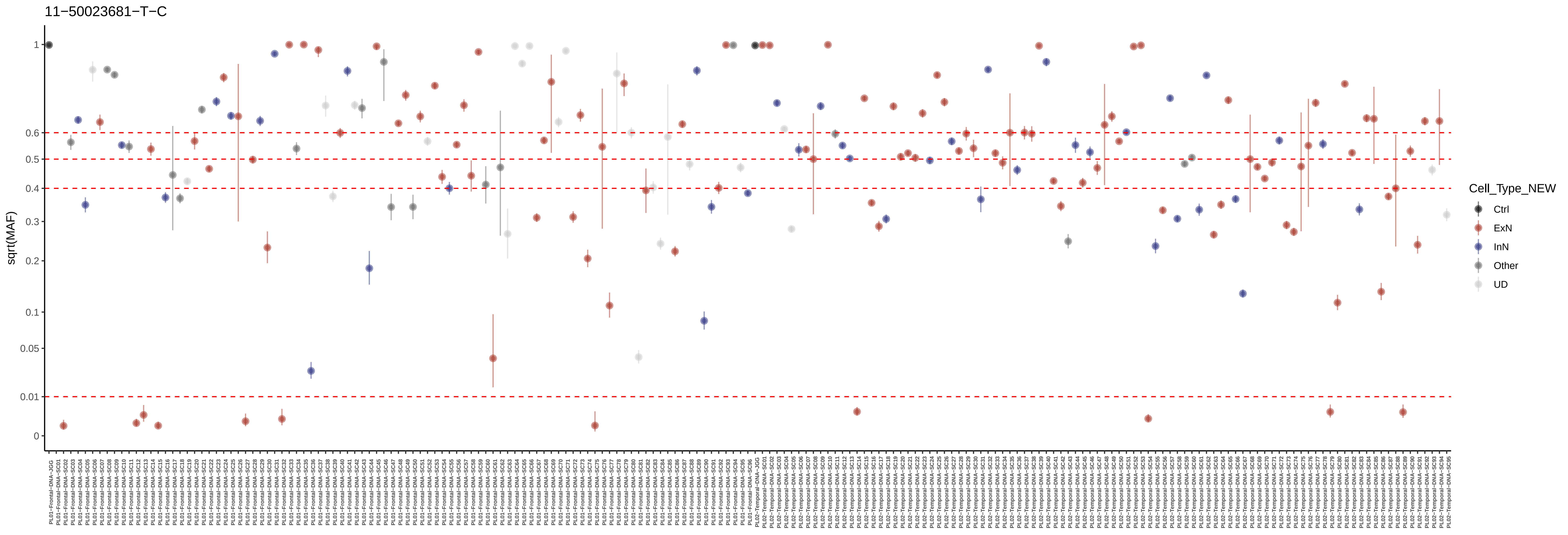
11−50023681−T−C
sqrt(MAF)
1
0.6
0.5
0.4
0.3
0.2
0.1
0.05
0.01
0
Cell_Type_NEW
Ctrl
ExN
InN
Other
UD

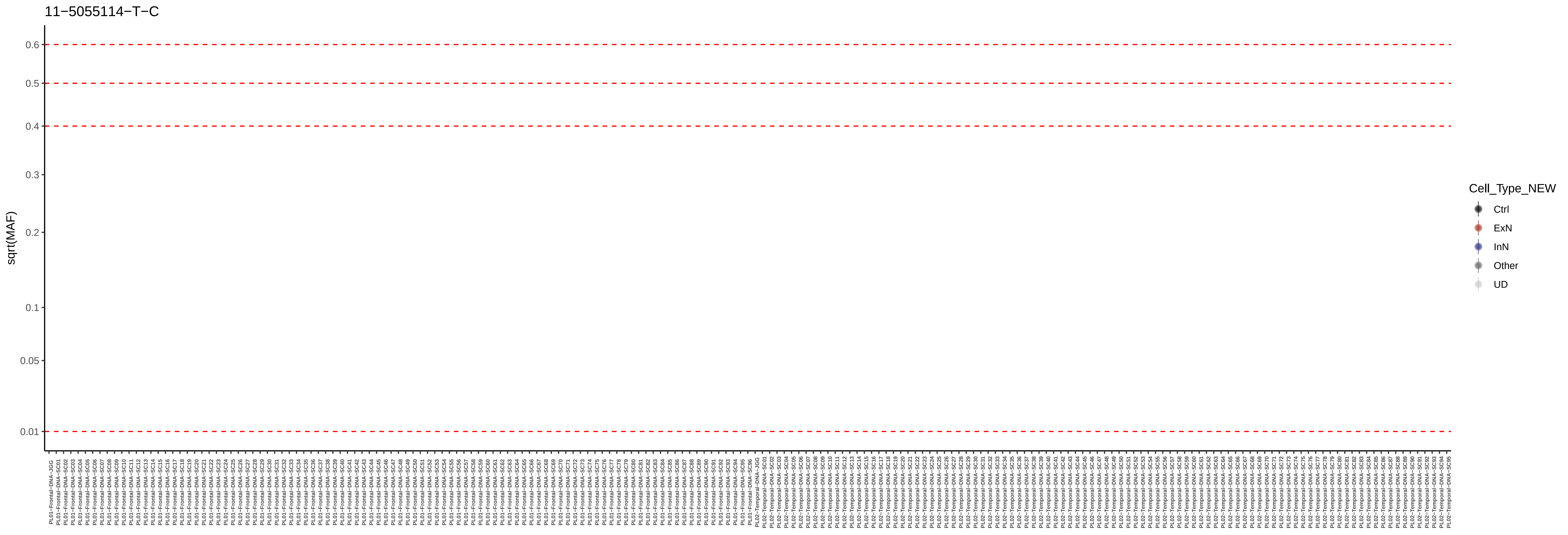
11−5055114−T−C
sqrt(MAF)
0.6
0.5
0.4
0.3
0.2
0.1
0.05
0.01
Cell_Type_NEW
Ctrl
ExN
InN
Other
UD

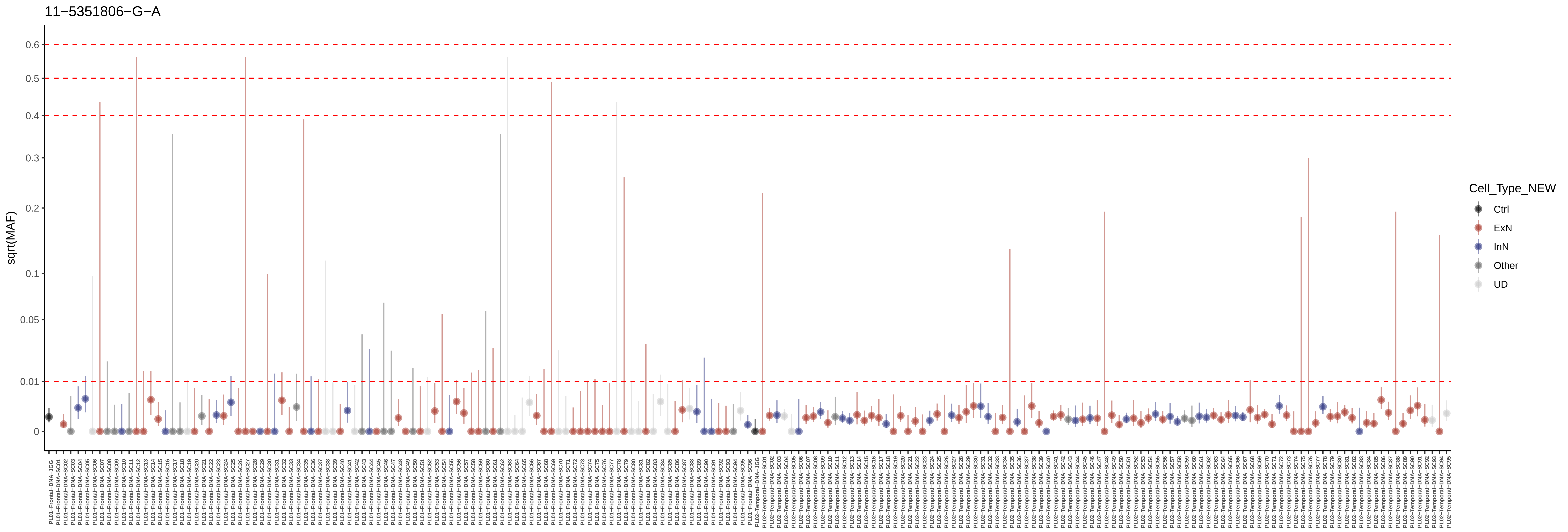
11−5351806−G−A
sqrt(MAF)
0.6
0.5
0.4
0.3
0.2
0.1
0.05
0.01
0
Cell_Type_NEW
Ctrl
ExN
InN
Other
UD

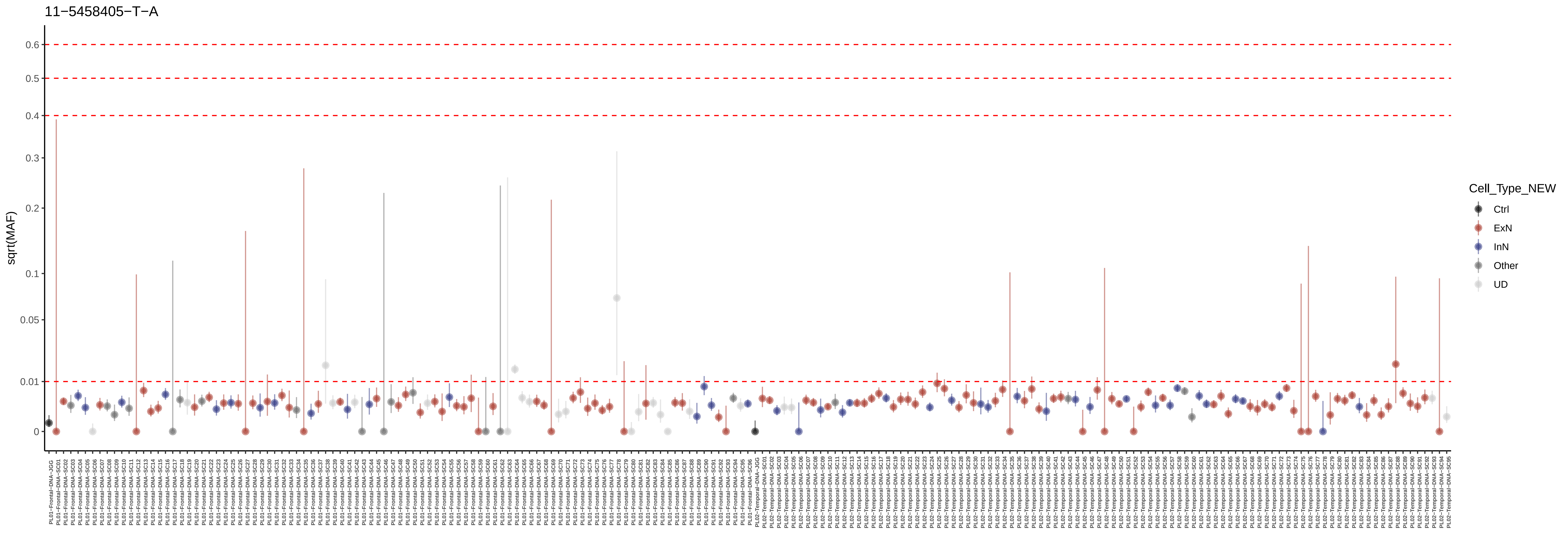
11−5458405−T−A
sqrt(MAF)
0.6
0.5
0.4
0.3
0.2
0.1
0.05
0.01
0
Cell_Type_NEW
Ctrl
ExN
InN
Other
UD

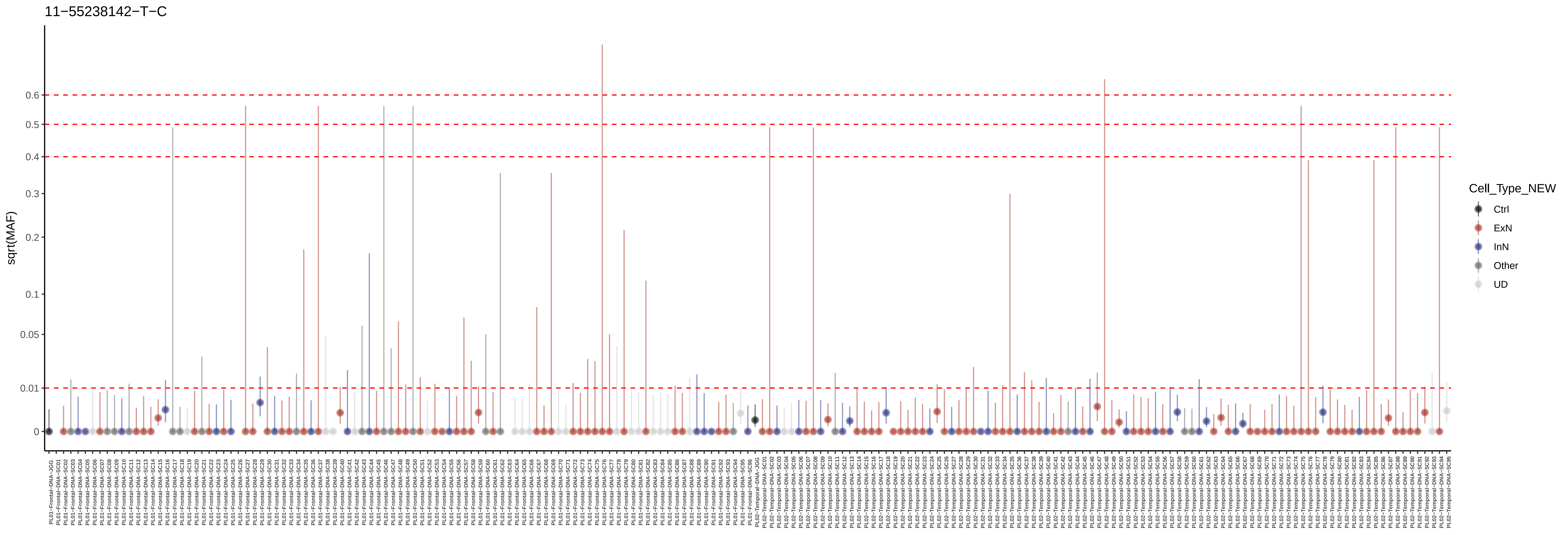

11−55238142−T−C
sqrt(MAF)
0.6
0.5
0.4
0.3
0.2
0.1
0.05
0.01
0
Cell_Type_NEW
Ctrl
ExN
InN
Other
UD

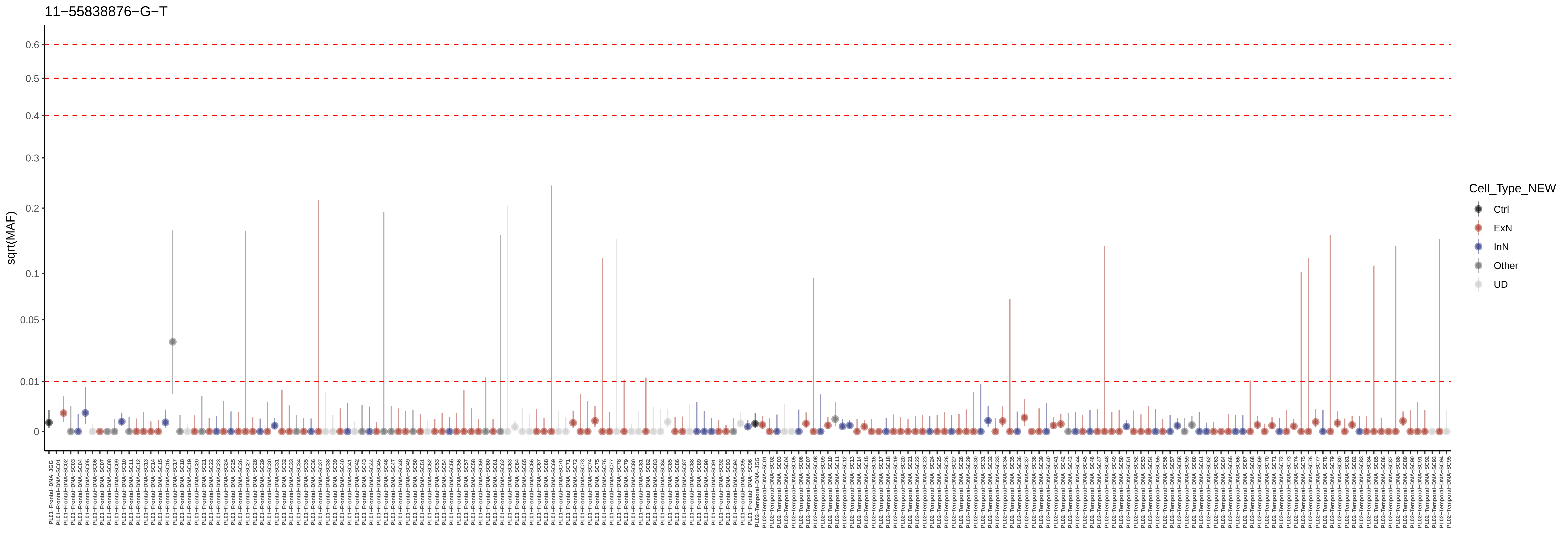

11−55838876−G−T
sqrt(MAF)
0.6
0.5
0.4
0.3
0.2
0.1
0.05
0.01
0
Cell_Type_NEW
Ctrl
ExN
InN
Other
UD

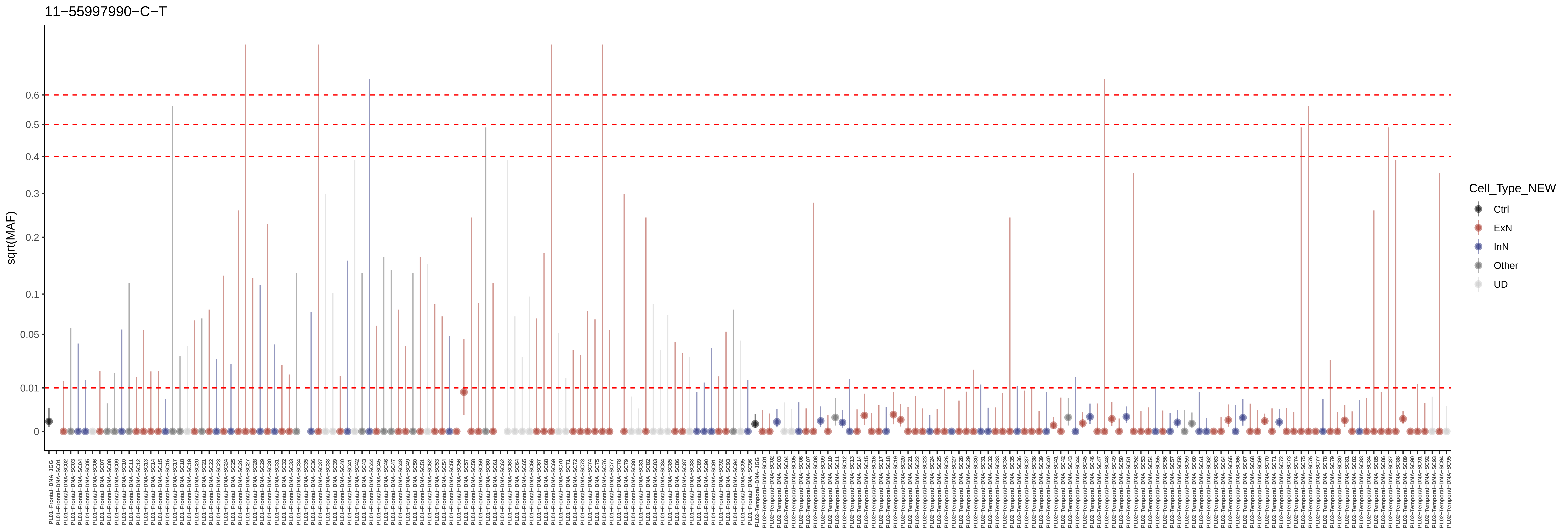

11−55997990−C−T
sqrt(MAF)
0.6
0.5
0.4
0.3
0.2
0.1
0.05
0.01
0
Cell_Type_NEW
Ctrl
ExN
InN
Other
UD

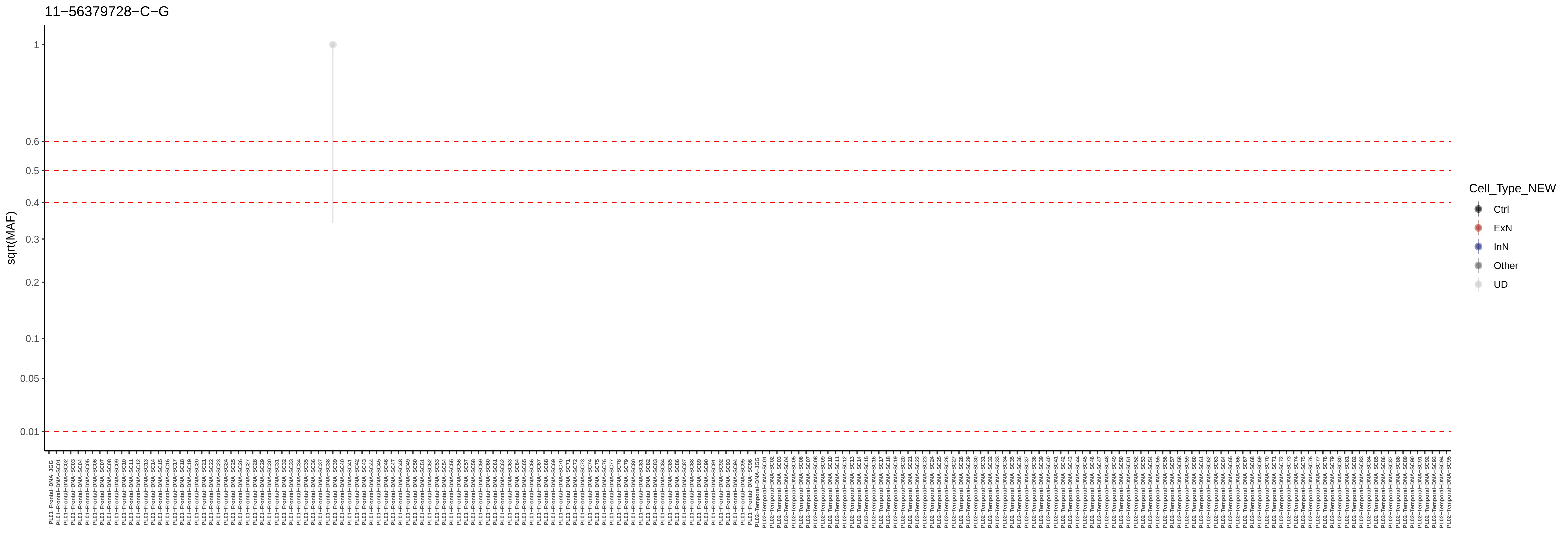
11−56379728−C−G
sqrt(MAF)
1
0.6
0.5
0.4
0.3
0.2
0.1
0.05
0.01
Cell_Type_NEW
Ctrl
ExN
InN
Other
UD

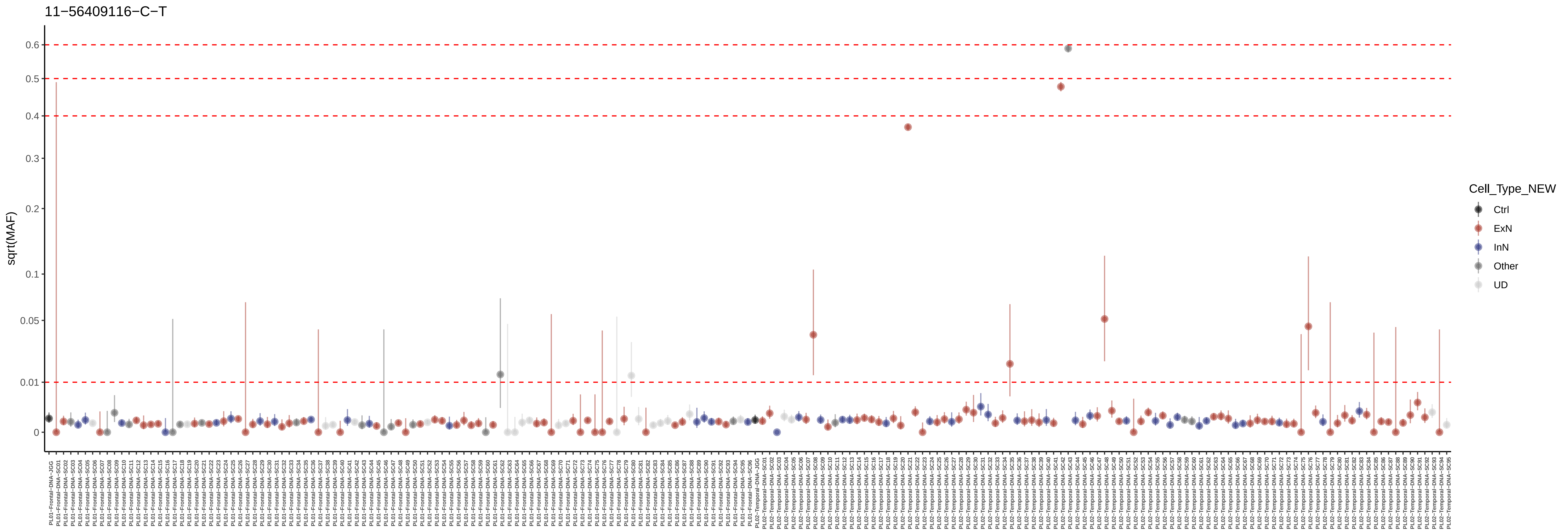
11−56409116−C−T
sqrt(MAF)
0.6
0.5
0.4
0.3
0.2
0.1
0.05
0.01
0
Cell_Type_NEW
Ctrl
ExN
InN
Other
UD

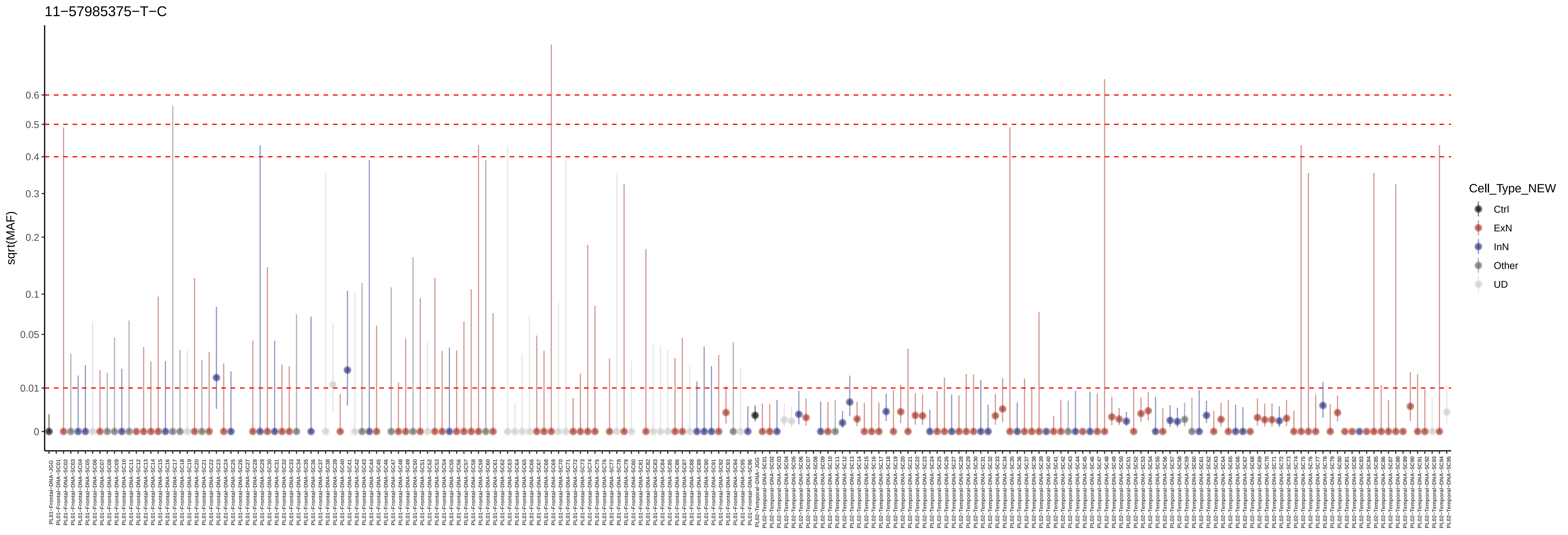
11−57985375−T−C
sqrt(MAF)
0.6
0.5
0.4
0.3
0.2
0.1
0.05
0.01
0
Cell_Type_NEW
Ctrl
ExN
InN
Other
UD

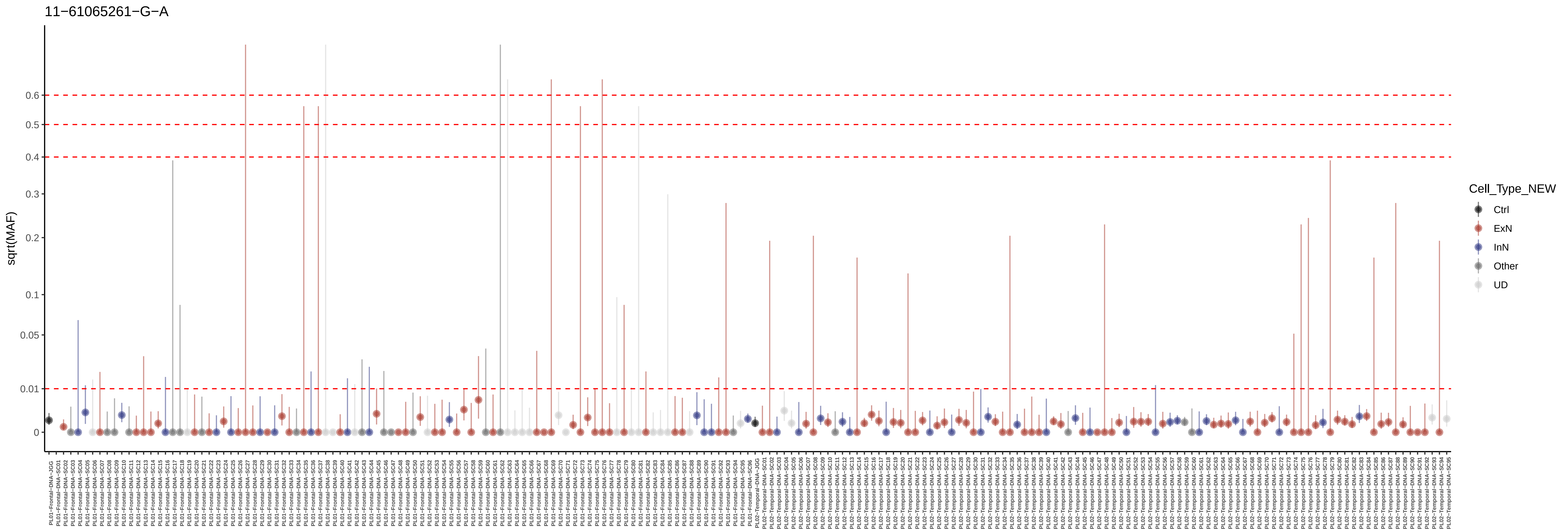
11−61065261−G−A
sqrt(MAF)
0.6
0.5
0.4
0.3
0.2
0.1
0.05
0.01
0
Cell_Type_NEW
Ctrl
ExN
InN
Other
UD

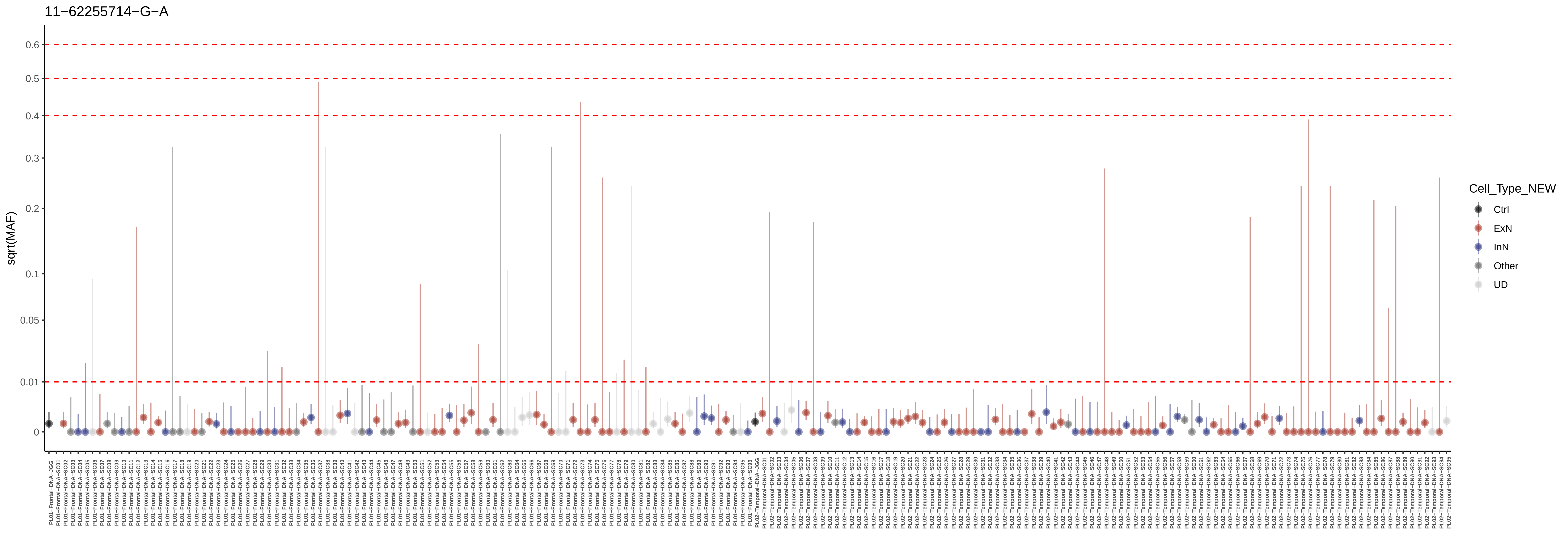
11−62255714−G−A
sqrt(MAF)
0.6
0.5
0.4
0.3
0.2
0.1
0.05
0.01
0
Cell_Type_NEW
Ctrl
ExN
InN
Other
UD

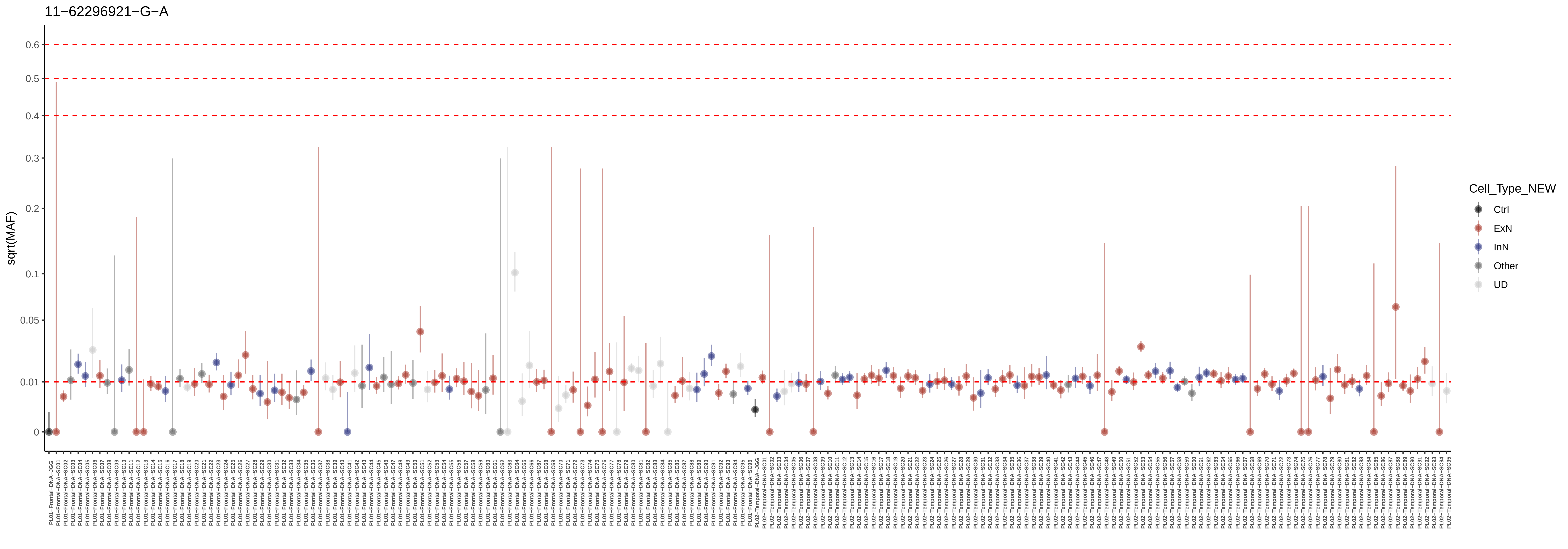

11−62296921−G−A
sqrt(MAF)
0.6
0.5
0.4
0.3
0.2
0.1
0.05
0.01
0
Cell_Type_NEW
Ctrl
ExN
InN
Other
UD

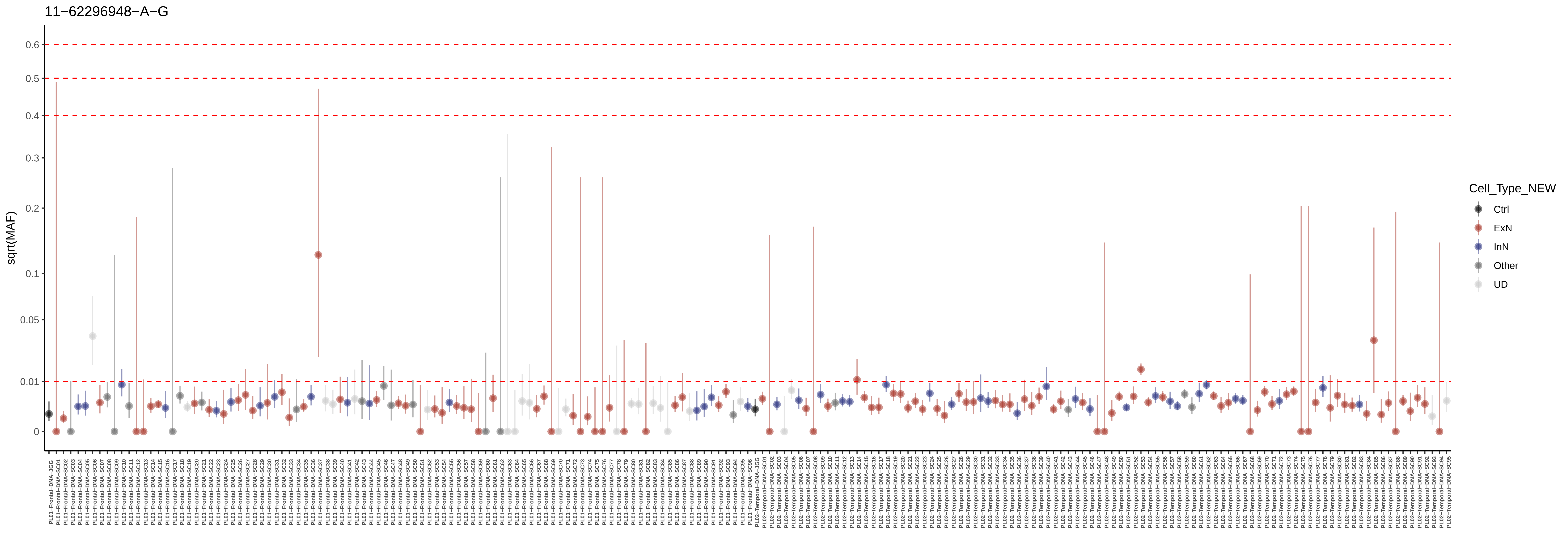
11−62296948−A−G
sqrt(MAF)
0.6
0.5
0.4
0.3
0.2
0.1
0.05
0.01
0
Cell_Type_NEW
Ctrl
ExN
InN
Other
UD

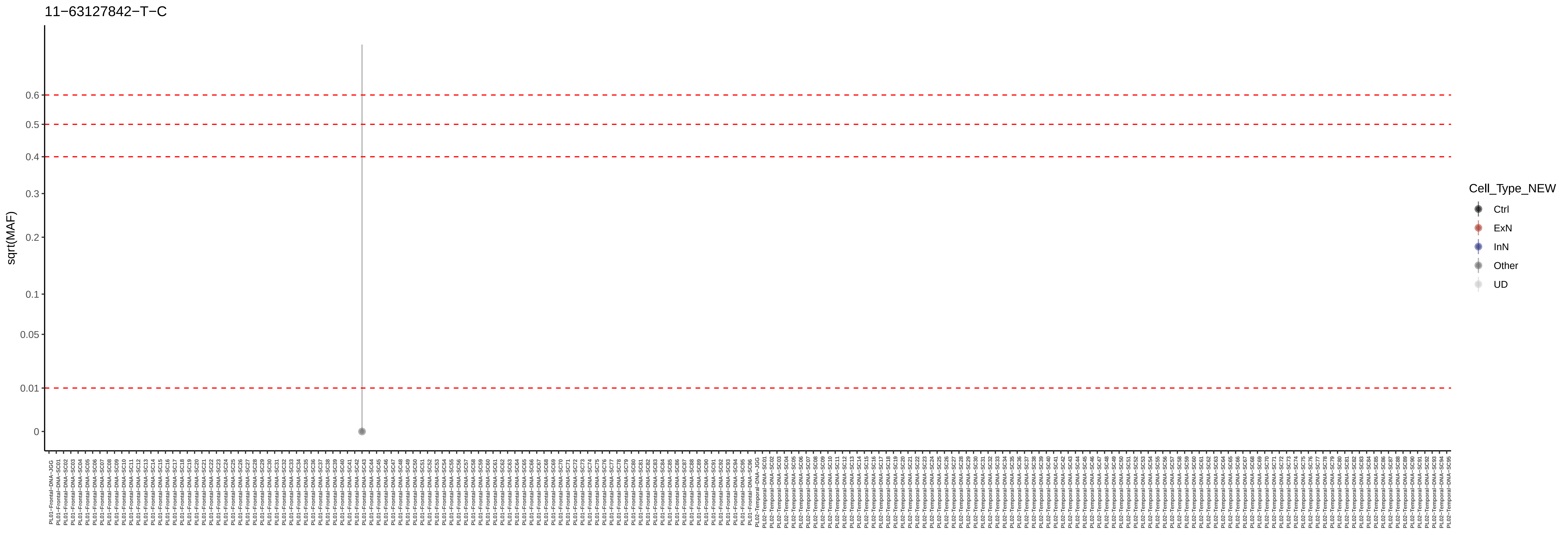
11−63127842−T−C
sqrt(MAF)
0.6
0.5
0.4
0.3
0.2
0.1
0.05
0.01
0
Cell_Type_NEW
Ctrl
ExN
InN
Other
UD

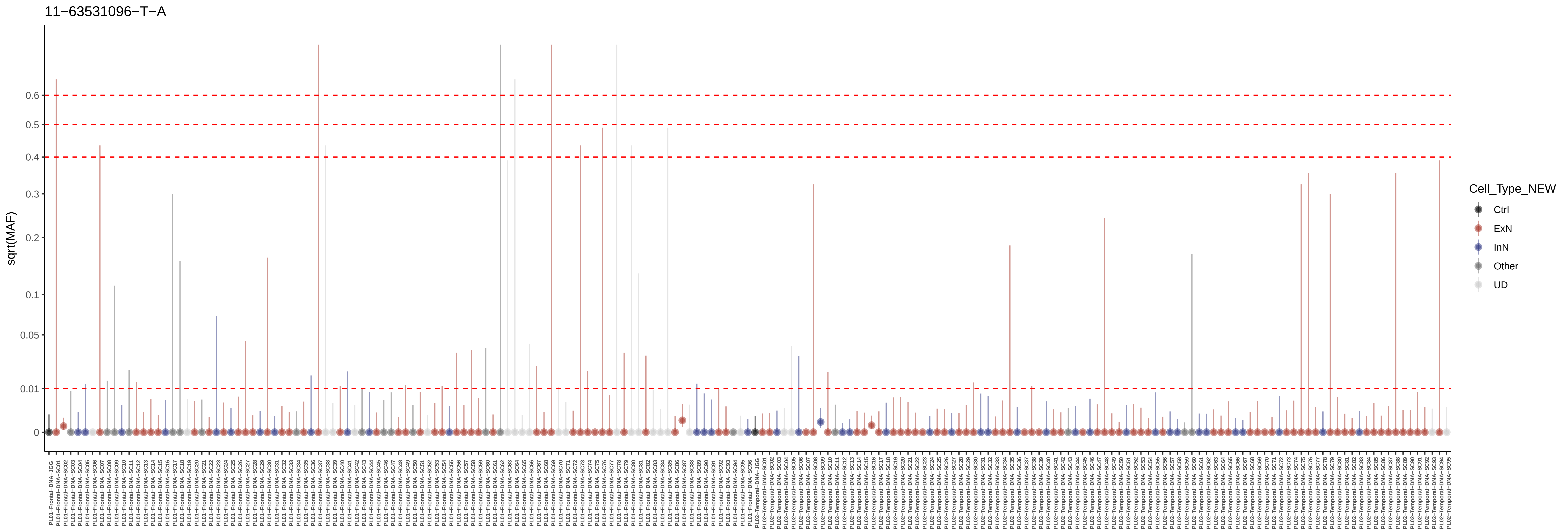

11−63531096−T−A
sqrt(MAF)
0.6
0.5
0.4
0.3
0.2
0.1
0.05
0.01
0
Cell_Type_NEW
Ctrl
ExN
InN
Other
UD

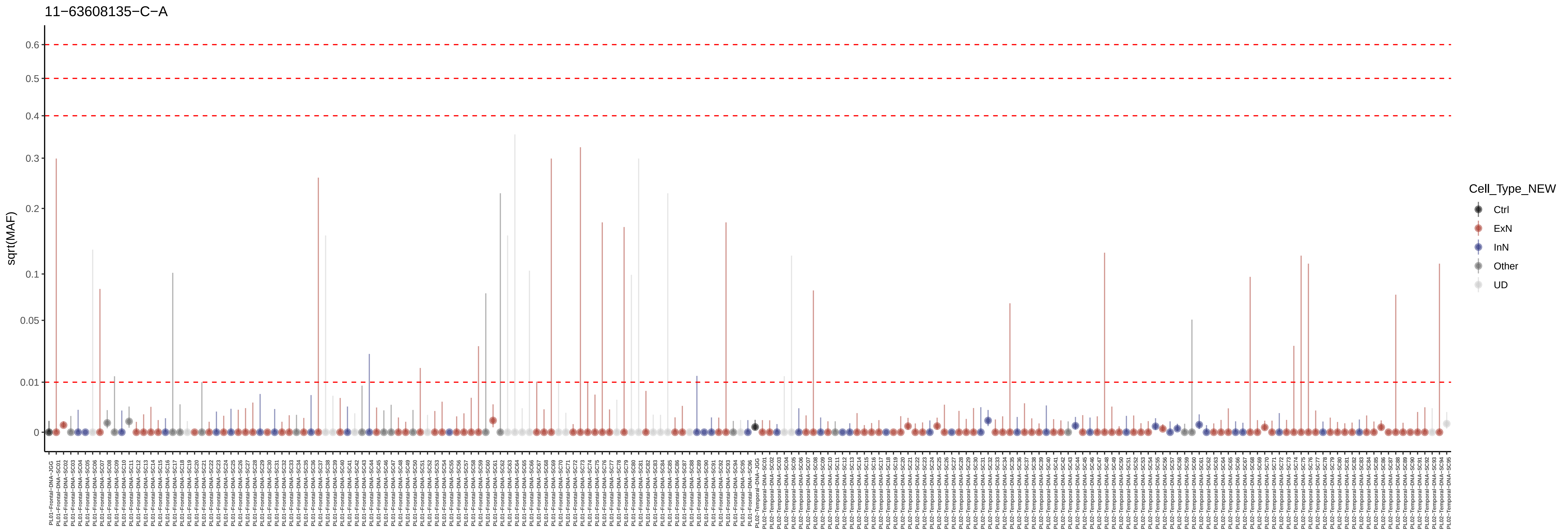
11−63608135−C−A
sqrt(MAF)
0.6
0.5
0.4
0.3
0.2
0.1
0.05
0.01
0
Cell_Type_NEW
Ctrl
ExN
InN
Other
UD

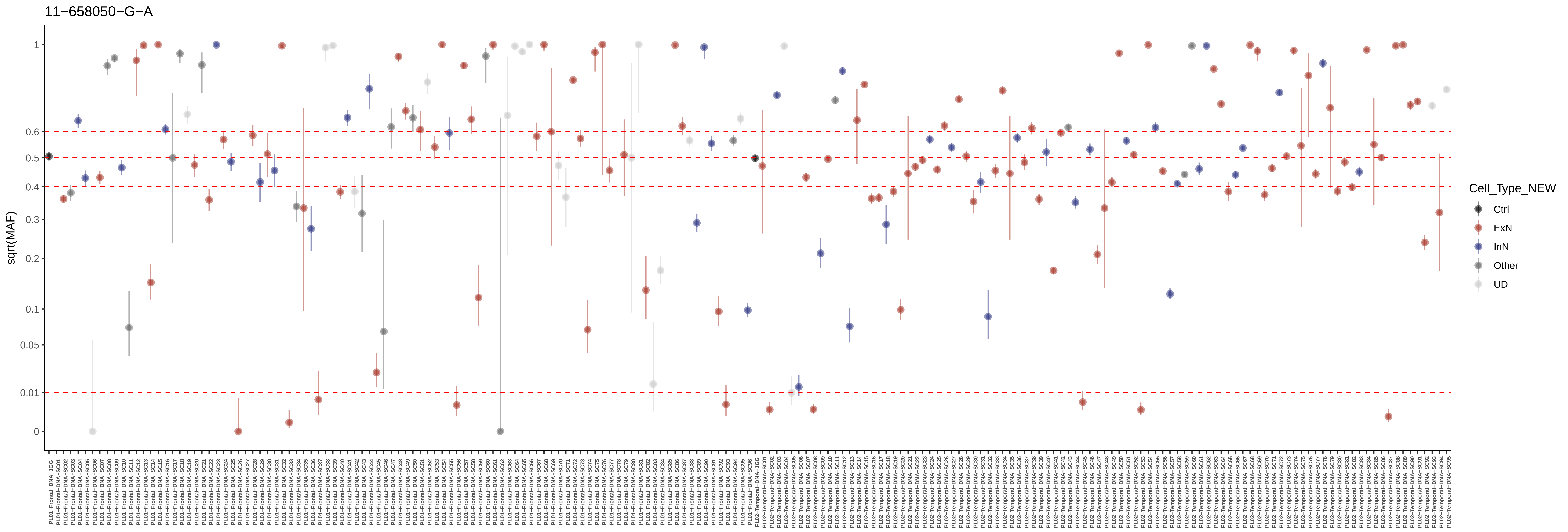
11−658050−G−A
sqrt(MAF)
1
0.6
0.5
0.4
0.3
0.2
0.1
0.05
0.01
0
Cell_Type_NEW
Ctrl
ExN
InN
Other
UD

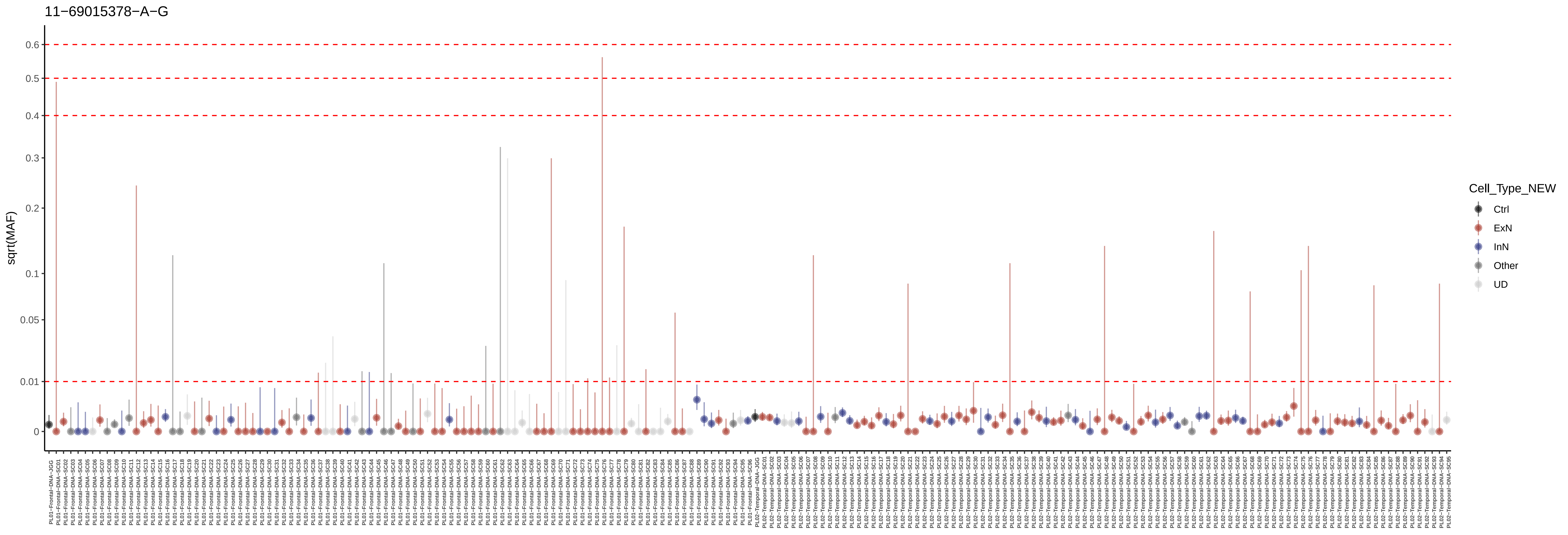

11−69015378−A−G
sqrt(MAF)
0.6
0.5
0.4
0.3
0.2
0.1
0.05
0.01
0
Cell_Type_NEW
Ctrl
ExN
InN
Other
UD

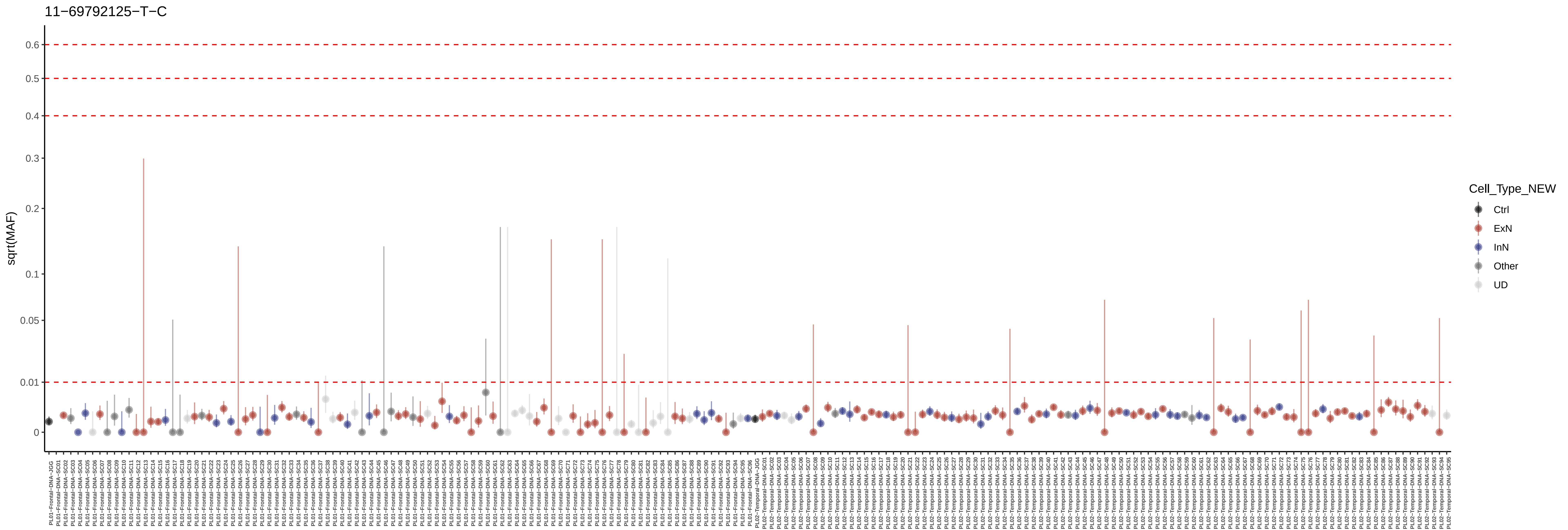
11−69792125−T−C
sqrt(MAF)
0.6
0.5
0.4
0.3
0.2
0.1
0.05
0.01
0
Cell_Type_NEW
Ctrl
ExN
InN
Other
UD

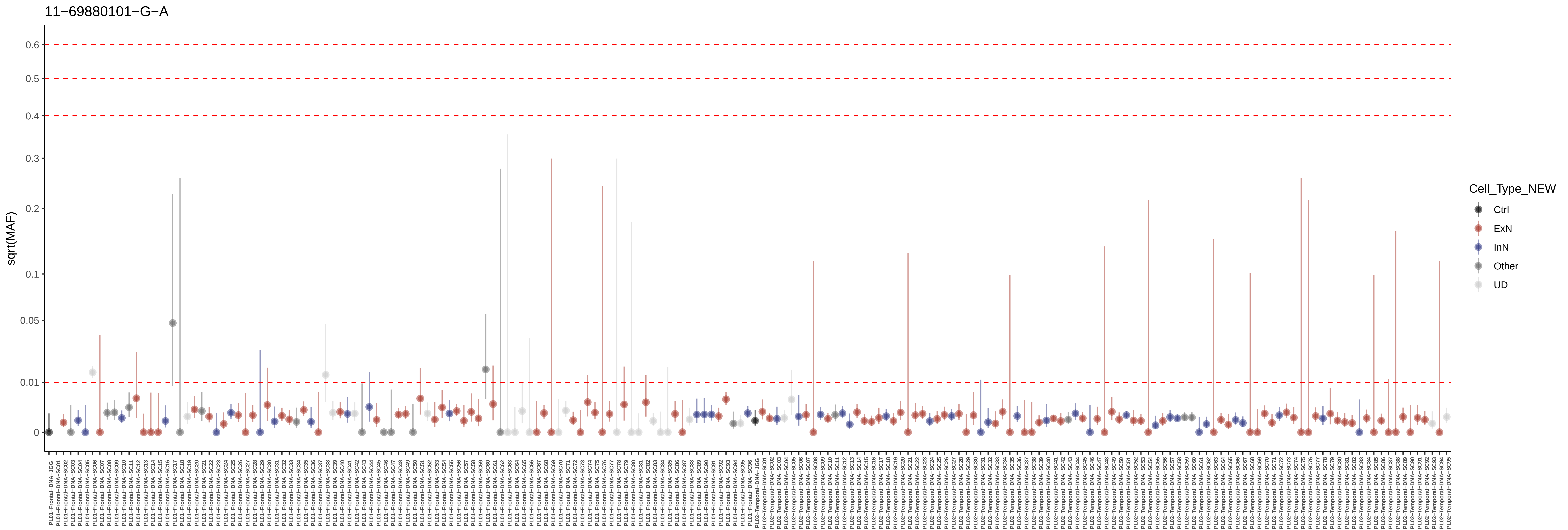

11−69880101−G−A
sqrt(MAF)
0.6
0.5
0.4
0.3
0.2
0.1
0.05
0.01
0
Cell_Type_NEW
Ctrl
ExN
InN
Other
UD

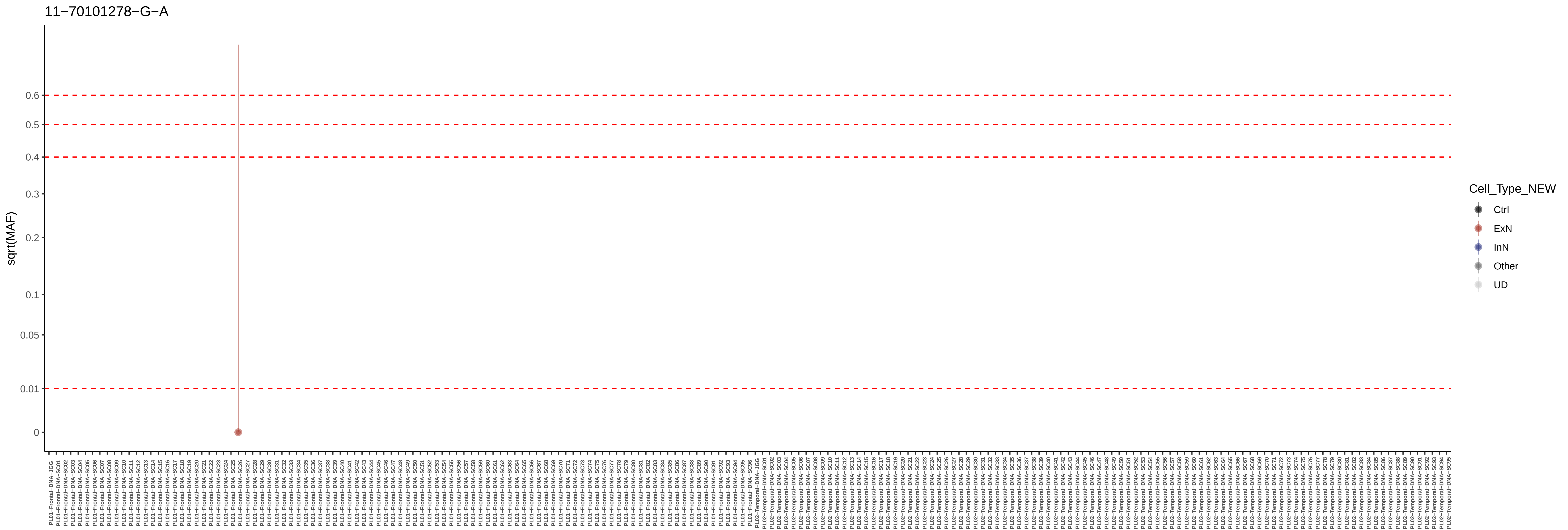
11−70101278−G−A
sqrt(MAF)
0.6
0.5
0.4
0.3
0.2
0.1
0.05
0.01
0
Cell_Type_NEW
Ctrl
ExN
InN
Other
UD

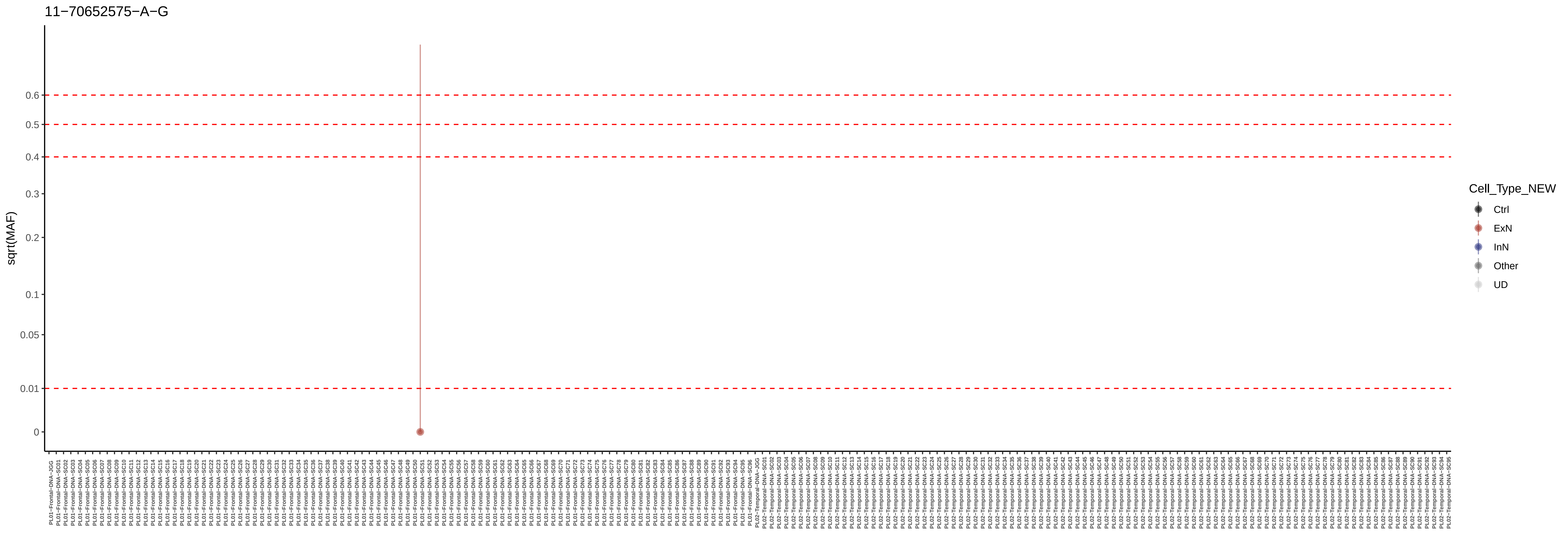
11−70652575−A−G
sqrt(MAF)
0.6
0.5
0.4
0.3
0.2
0.1
0.05
0.01
0
Cell_Type_NEW
Ctrl
ExN
InN
Other
UD

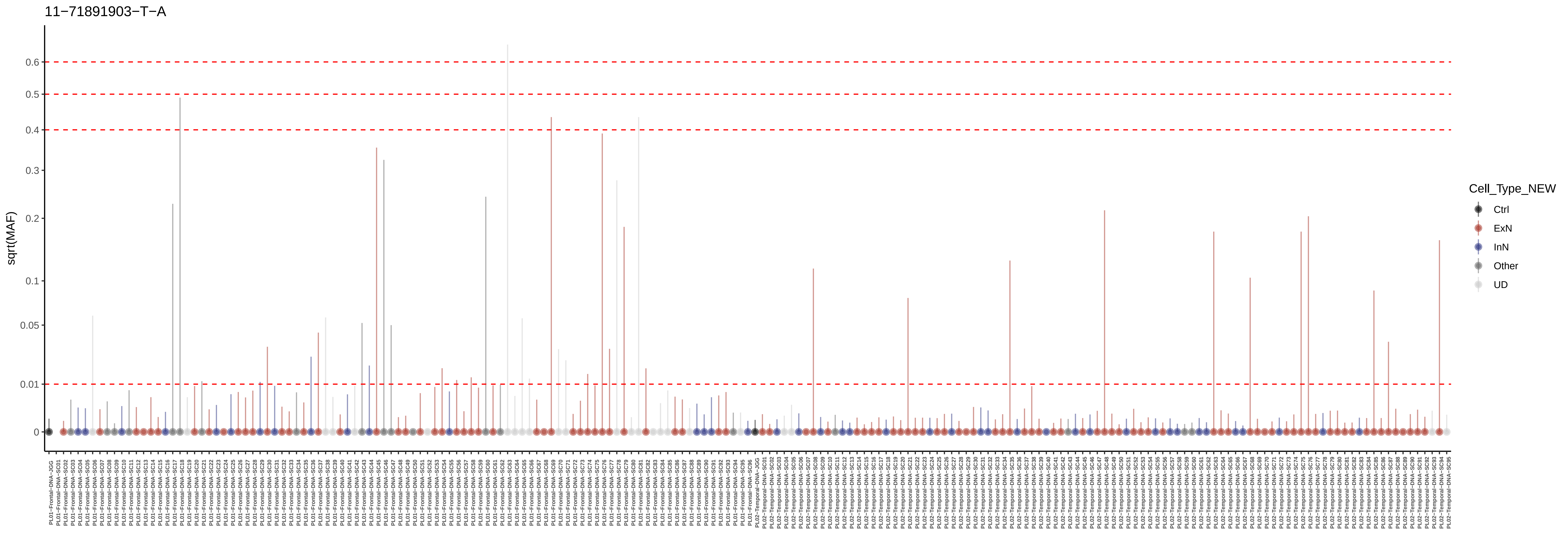
11−71891903−T−A
sqrt(MAF)
0.6
0.5
0.4
0.3
0.2
0.1
0.05
0.01
0
Cell_Type_NEW
Ctrl
ExN
InN
Other
UD

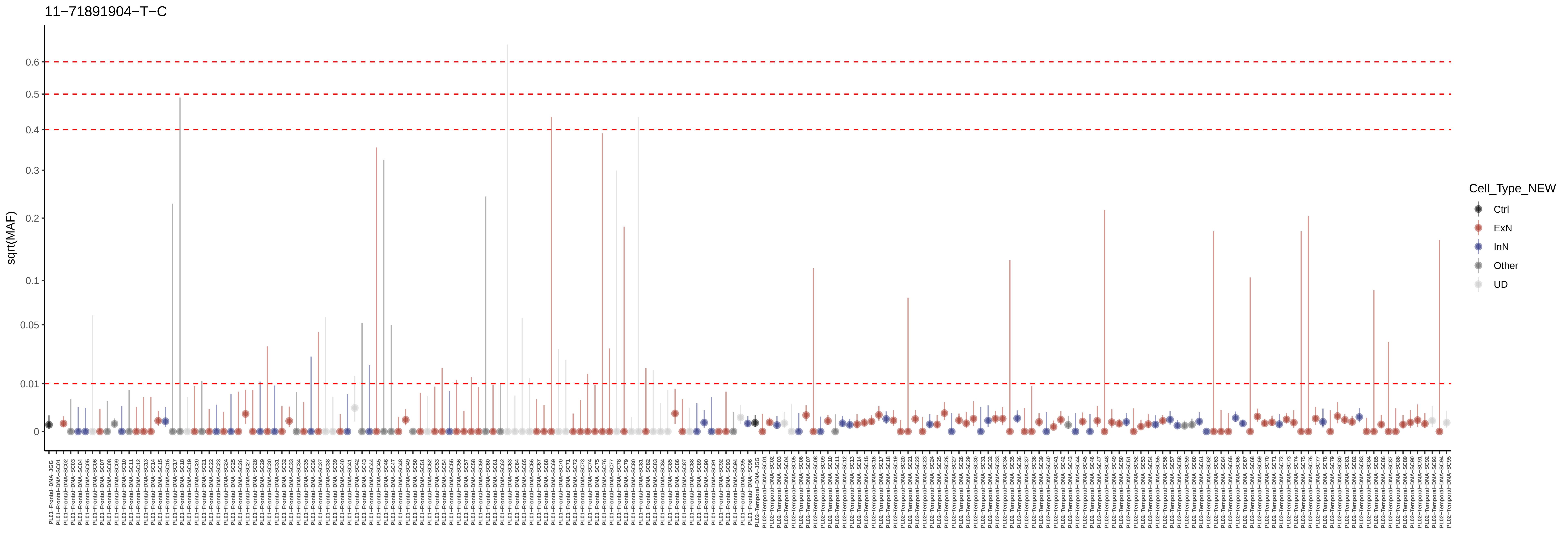
11−71891904−T−C
sqrt(MAF)
0.6
0.5
0.4
0.3
0.2
0.1
0.05
0.01
0
Cell_Type_NEW
Ctrl
ExN
InN
Other
UD

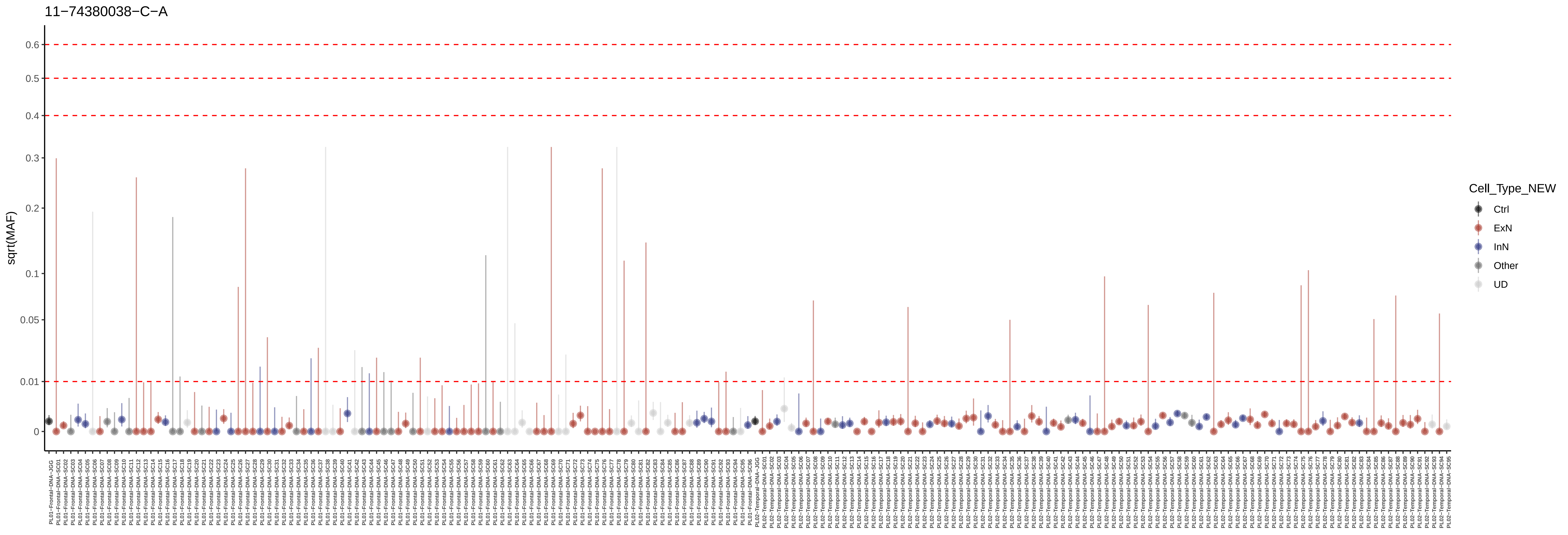
11−74380038−C−A
sqrt(MAF)
0.6
0.5
0.4
0.3
0.2
0.1
0.05
0.01
0
Cell_Type_NEW
Ctrl
ExN
InN
Other
UD

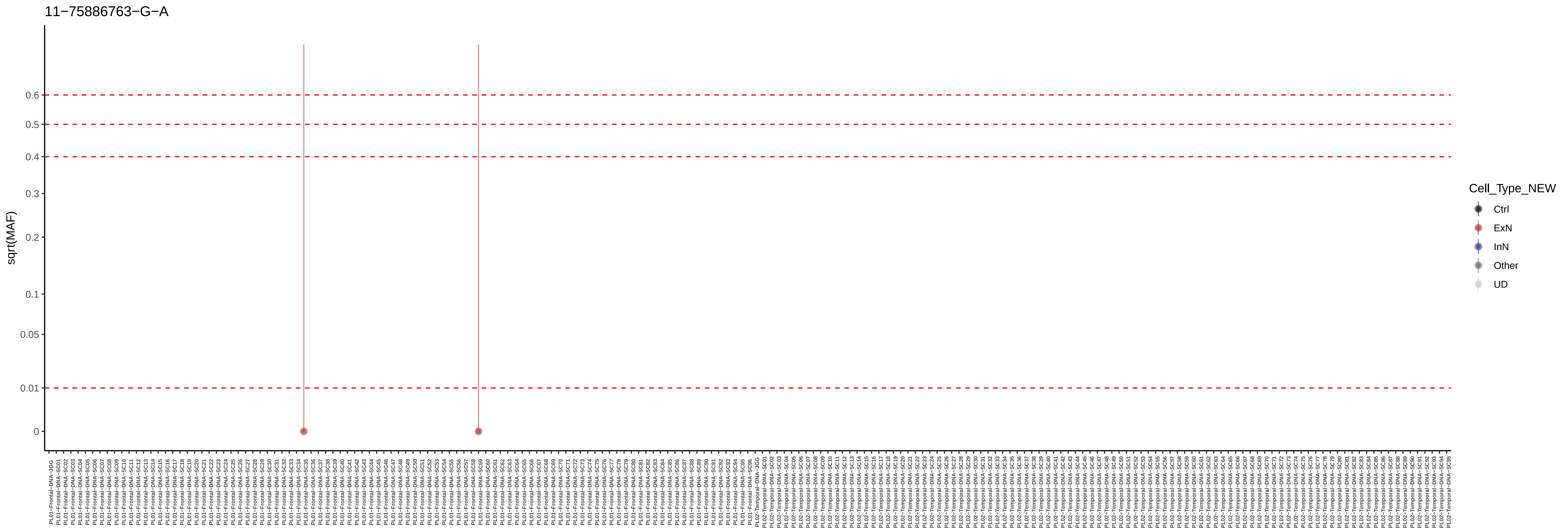
11−75886763−G−A
sqrt(MAF)
0.6
0.5
0.4
0.3
0.2
0.1
0.05
0.01
0
Cell_Type_NEW
Ctrl
ExN
InN
Other
UD

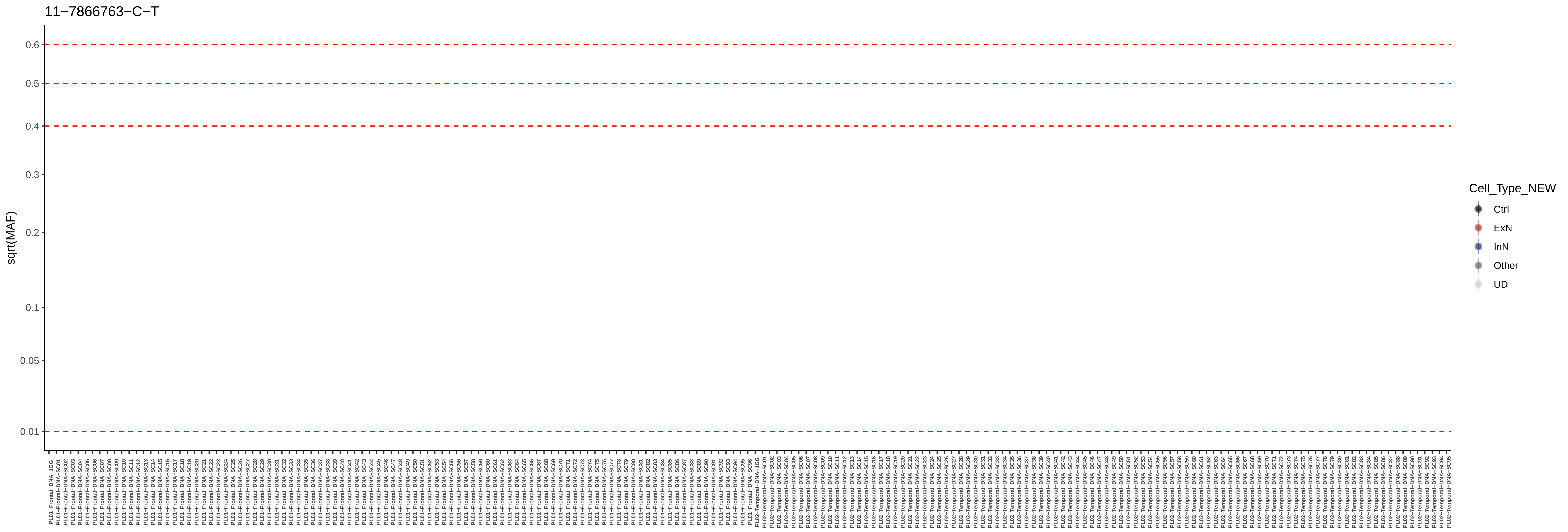
11−7866763−C−T
sqrt(MAF)
0.6
0.5
0.4
0.3
0.2
0.1
0.05
0.01
Cell_Type_NEW
Ctrl
ExN
InN
Other
UD

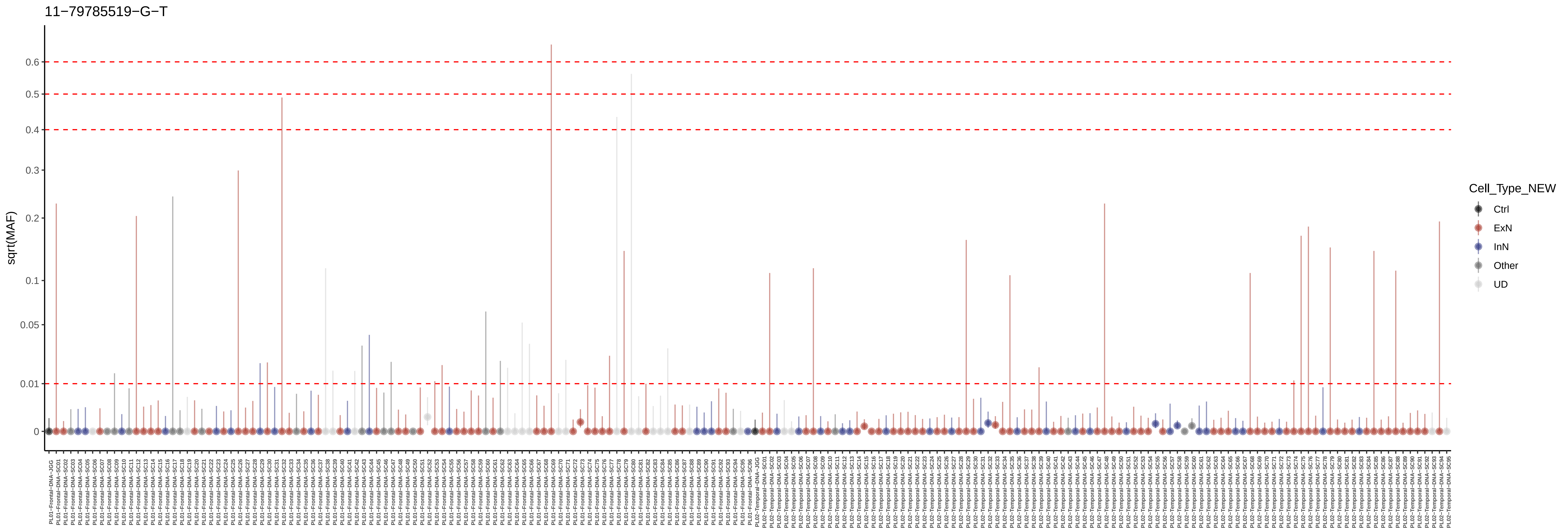
11−79785519−G−T
sqrt(MAF)
0.6
0.5
0.4
0.3
0.2
0.1
0.05
0.01
0
Cell_Type_NEW
Ctrl
ExN
InN
Other
UD

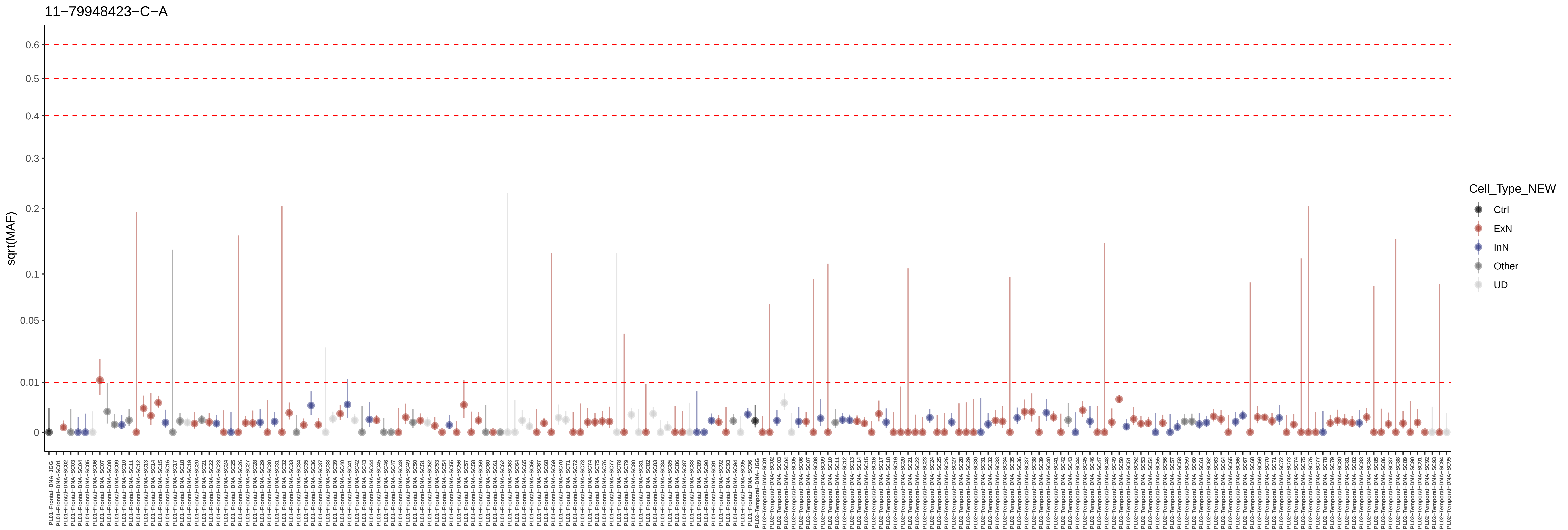
11−79948423−C−A
sqrt(MAF)
0.6
0.5
0.4
0.3
0.2
0.1
0.05
0.01
0
Cell_Type_NEW
Ctrl
ExN
InN
Other
UD

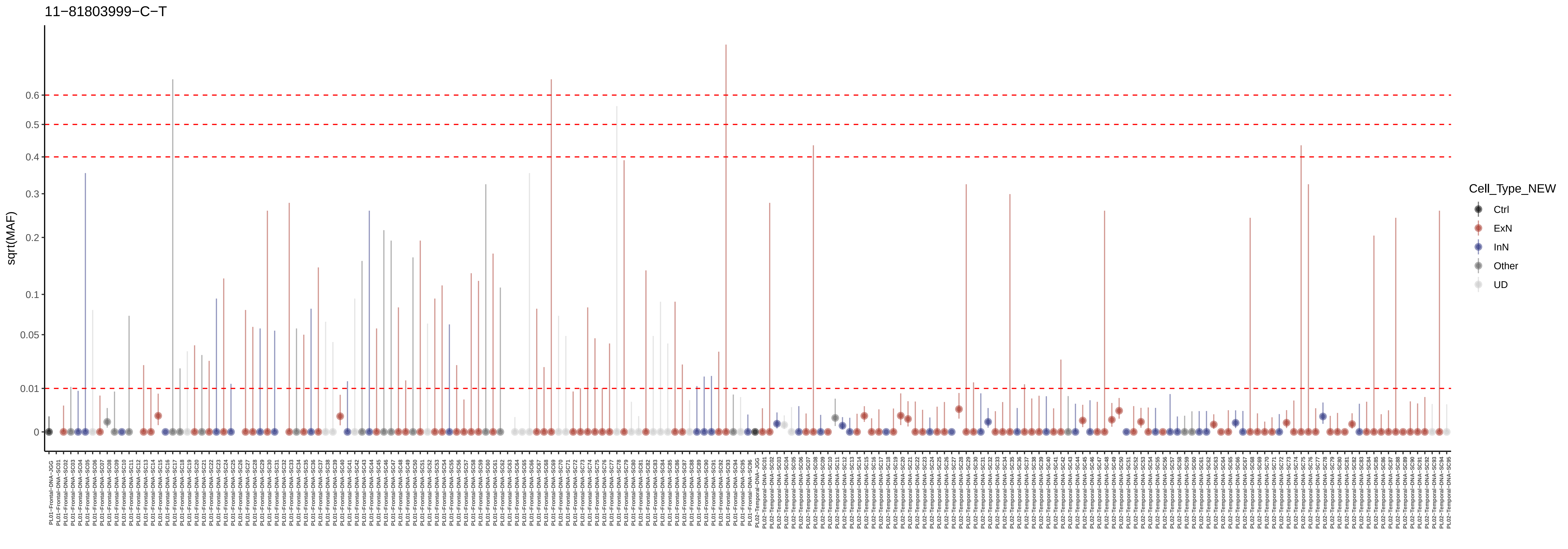
11−81803999−C−T
sqrt(MAF)
0.6
0.5
0.4
0.3
0.2
0.1
0.05
0.01
0
Cell_Type_NEW
Ctrl
ExN
InN
Other
UD

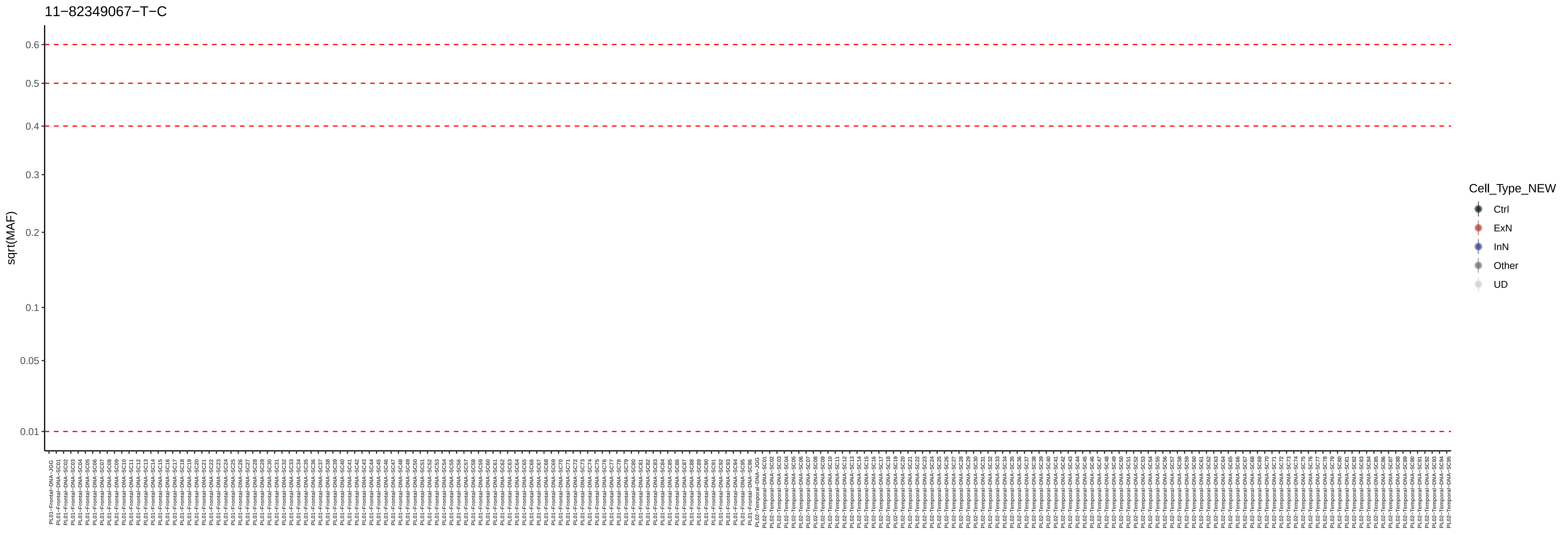
11−82349067−T−C
sqrt(MAF)
0.6
0.5
0.4
0.3
0.2
0.1
0.05
0.01
Cell_Type_NEW
Ctrl
ExN
InN
Other
UD

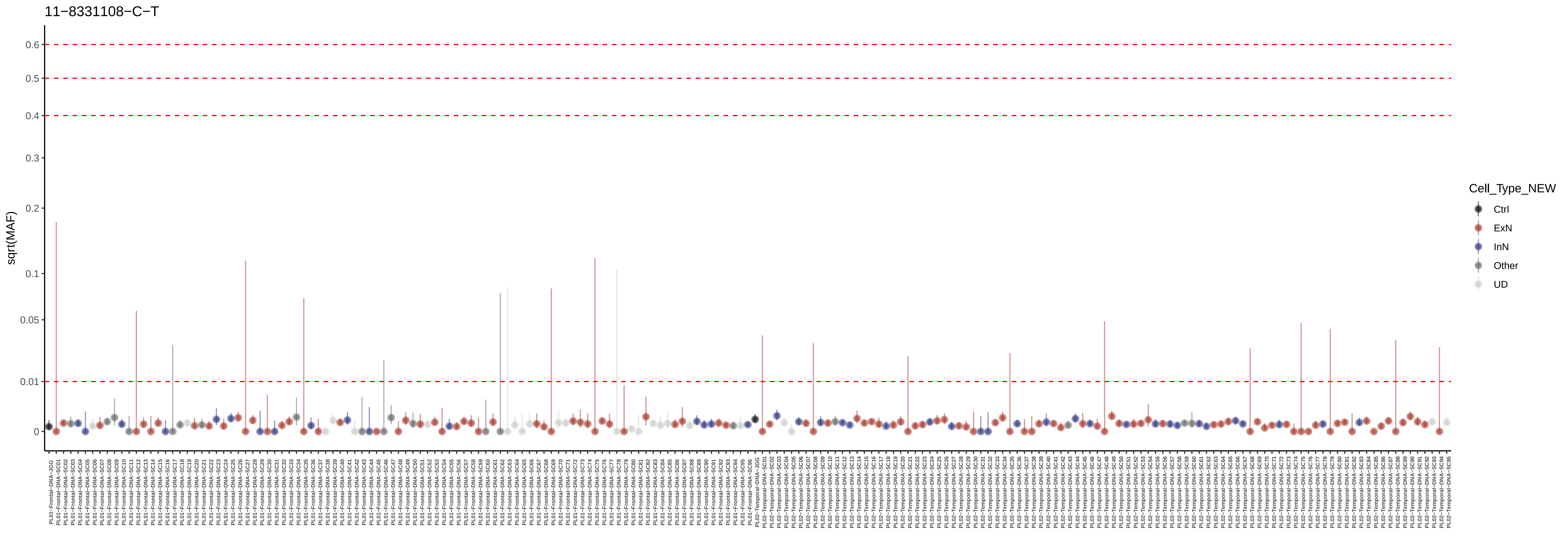
11−8331108−C−T
sqrt(MAF)
0.6
0.5
0.4
0.3
0.2
0.1
0.05
0.01
0
Cell_Type_NEW
Ctrl
ExN
InN
Other
UD

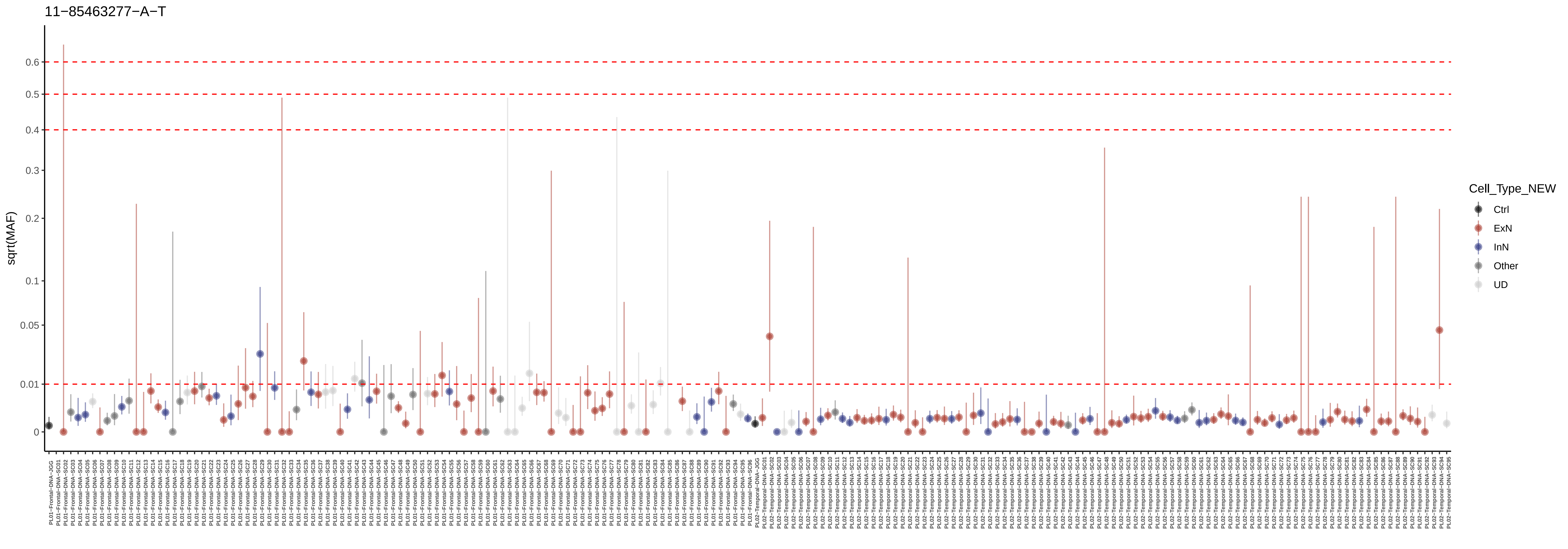

11−85463277−A−T
sqrt(MAF)
Cell_Type_NEW
Ctrl
ExN
InN
Other
UD

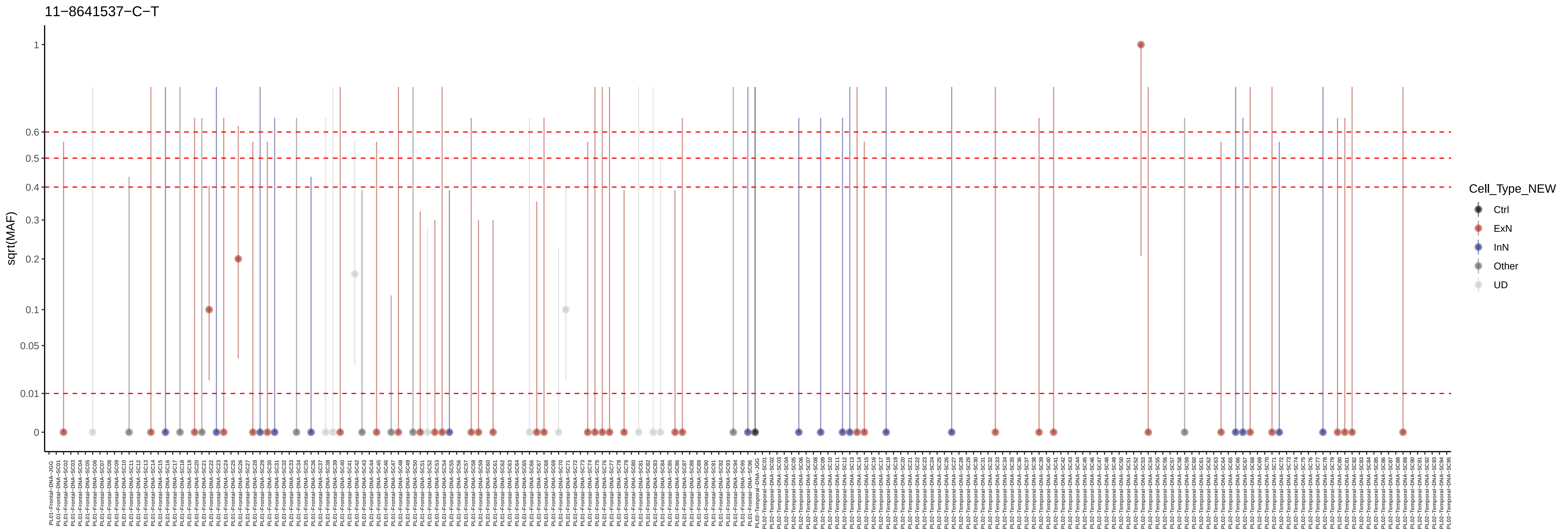
11−8641537−C−T
sqrt(MAF)
1
0.6
0.5
0.4
0.3
0.2
0.1
0.05
0.01
0
Cell_Type_NEW
Ctrl
ExN
InN
Other
UD

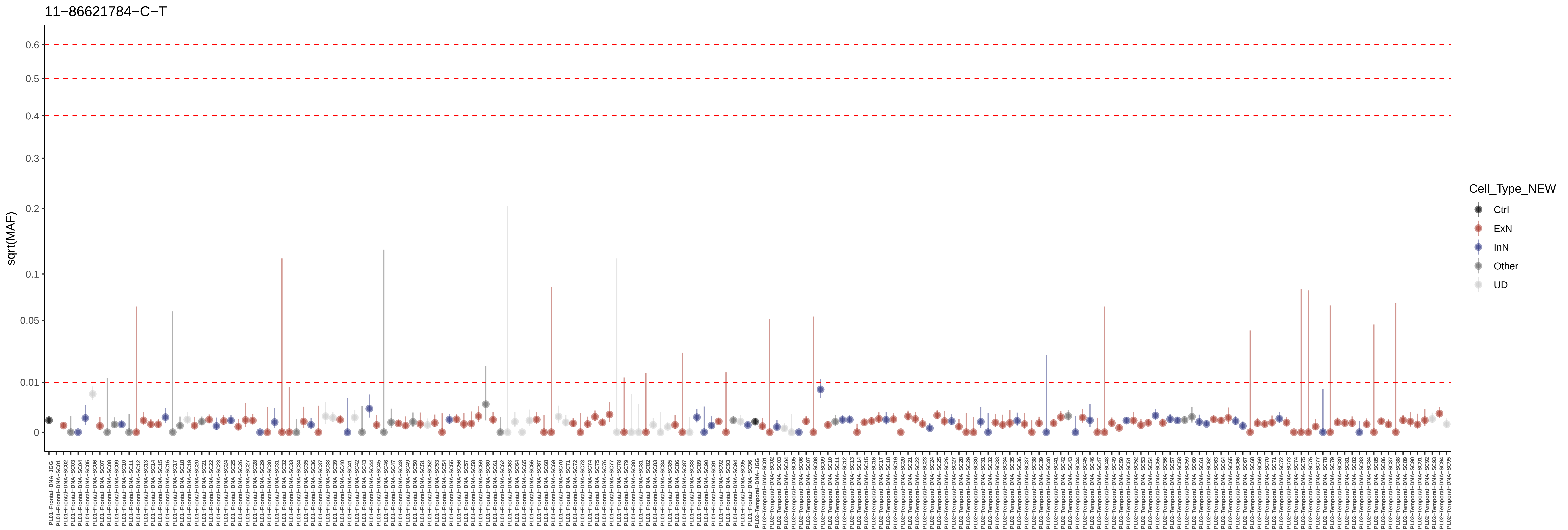
11−86621784−C−T
sqrt(MAF)
0.6
0.5
0.4
0.3
0.2
0.1
0.05
0.01
0
Cell_Type_NEW
Ctrl
ExN
InN
Other
UD

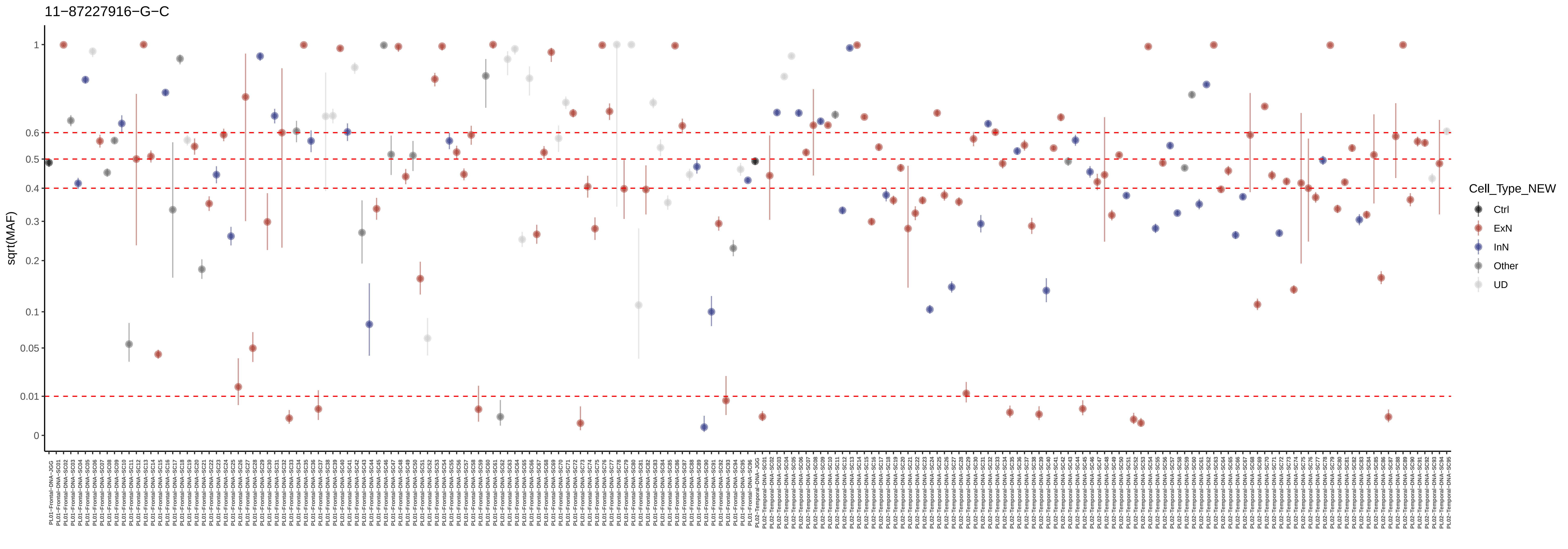

11−87227916−G−C
sqrt(MAF)
1
0.6
0.5
0.4
0.3
0.2
0.1
0.05
0.01
0
Cell_Type_NEW
Ctrl
ExN
InN
Other
UD

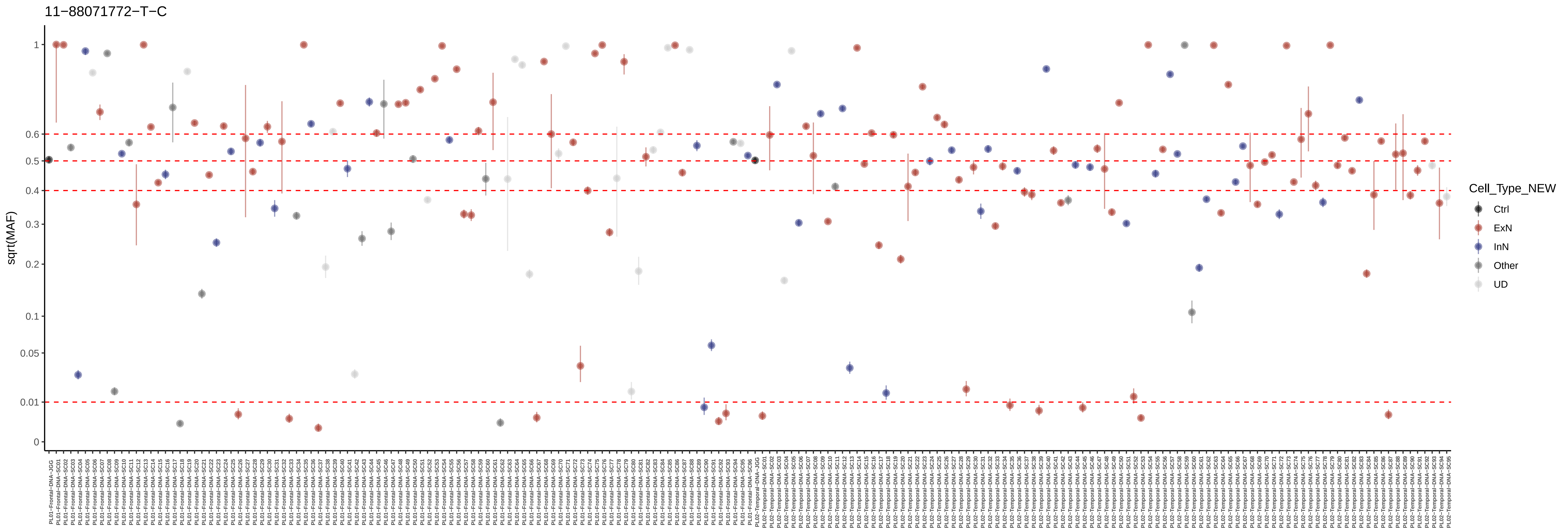

11−88071772−T−C
sqrt(MAF)
1
0.6
0.5
0.4
0.3
0.2
0.1
0.05
0.01
0
Cell_Type_NEW
Ctrl
ExN
InN
Other
UD

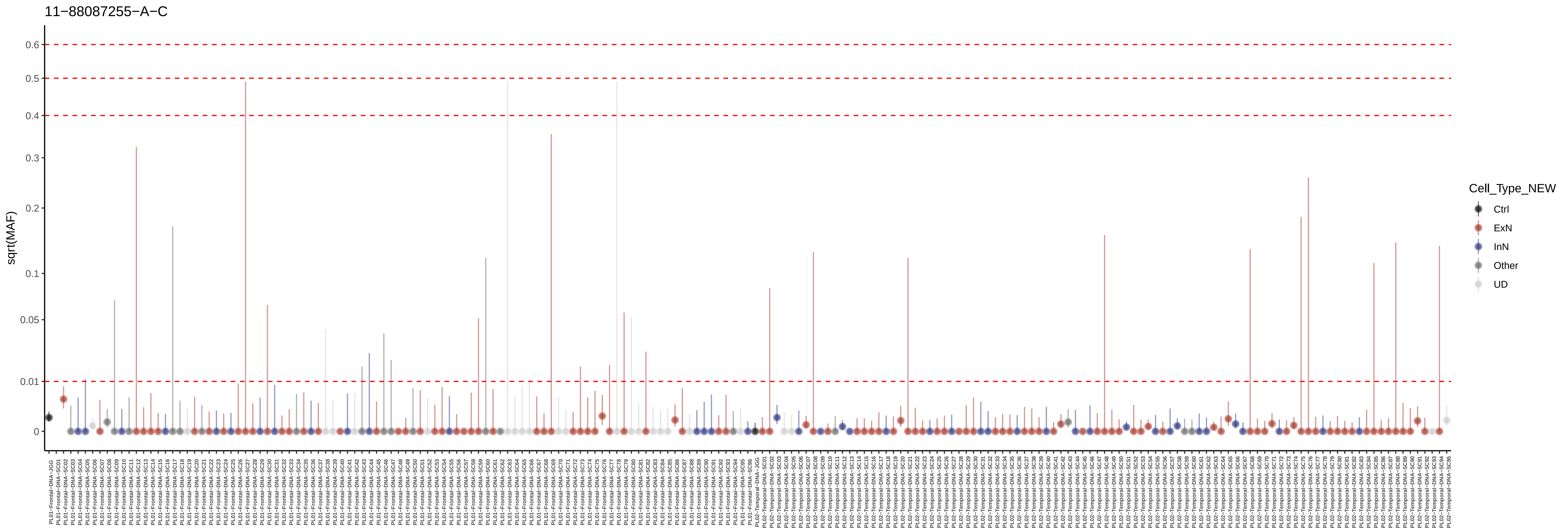
11−88087255−A−C
sqrt(MAF)
0.6
0.5
0.4
0.3
0.2
0.1
0.05
0.01
0
Cell_Type_NEW
Ctrl
ExN
InN
Other
UD

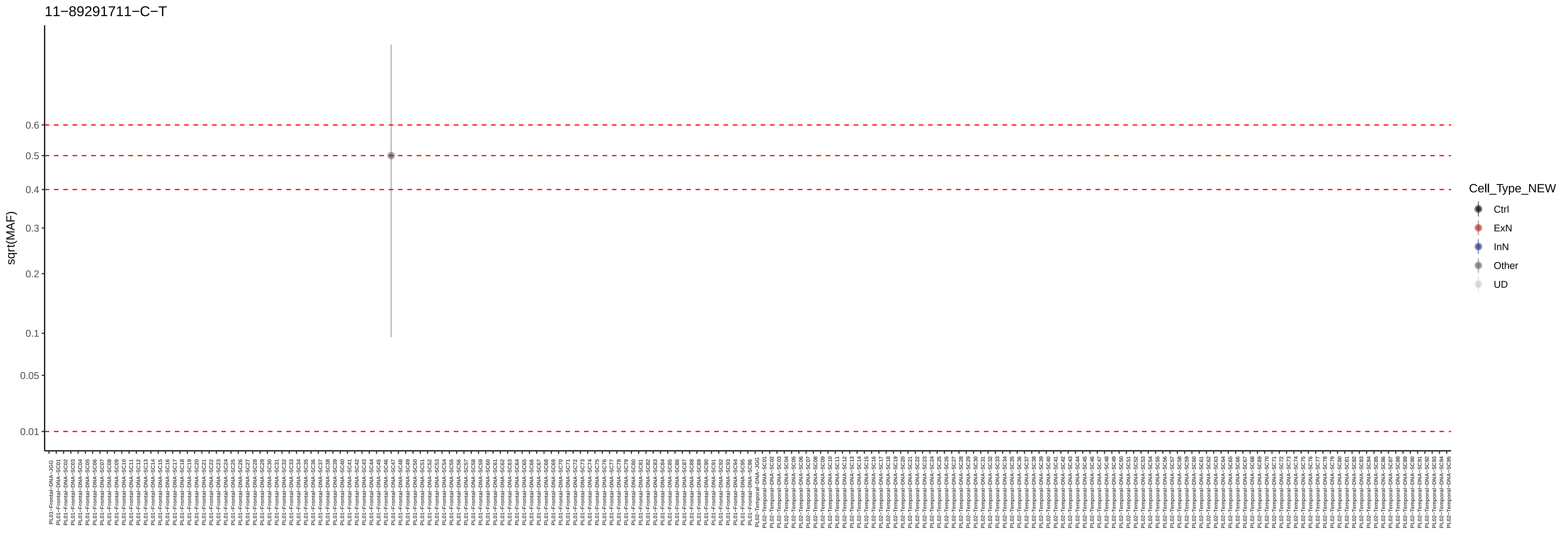

11−89291711−C−T
sqrt(MAF)
0.6
0.5
0.4
0.3
0.2
0.1
0.05
0.01
Cell_Type_NEW
Ctrl
ExN
InN
Other
UD

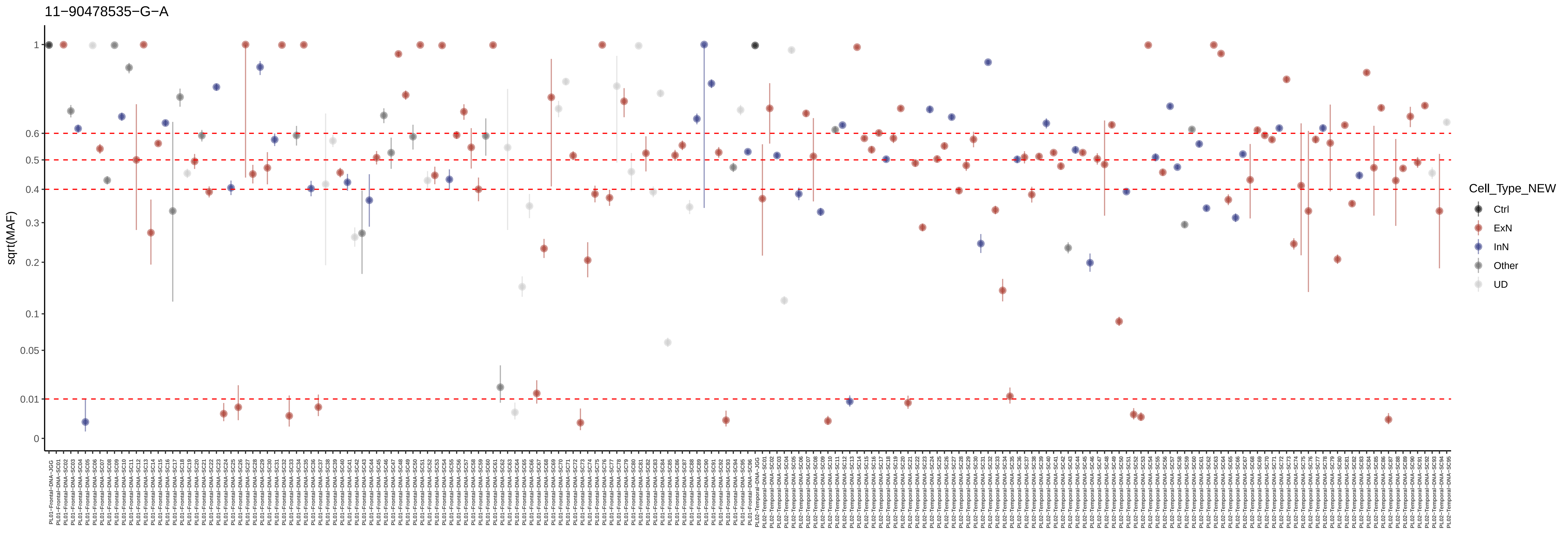

11−90478535−G−A
sqrt(MAF)
Cell_Type_NEW
Ctrl
ExN
InN
Other
UD

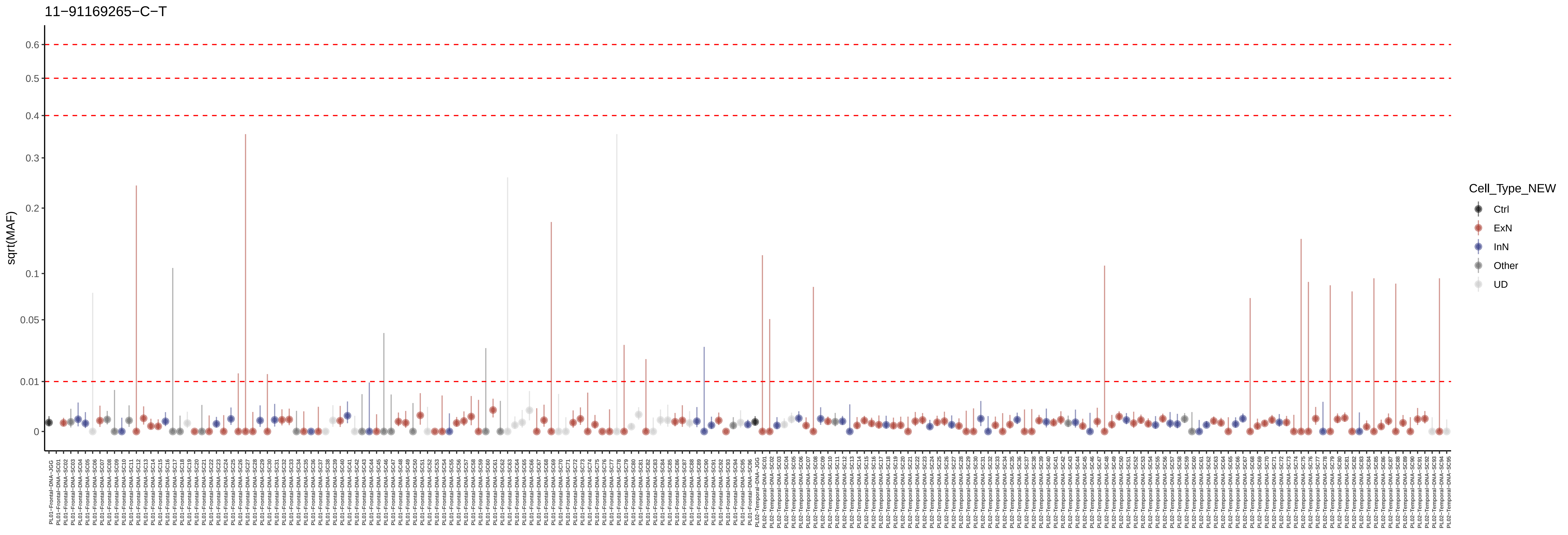
11−91169265−C−T
sqrt(MAF)
0.6
0.5
0.4
0.3
0.2
0.1
0.05
0.01
0
Cell_Type_NEW
Ctrl
ExN
InN
Other
UD

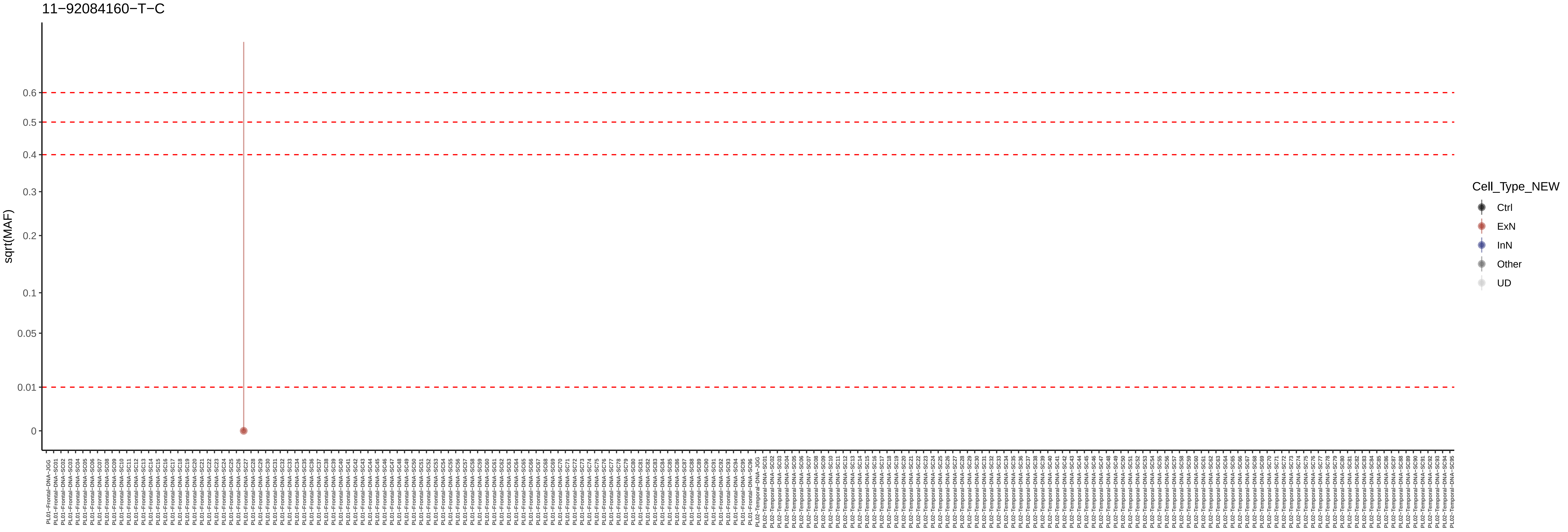
11−92084160−T−C
sqrt(MAF)
0.6
0.5
0.4
0.3
0.2
0.1
0.05
0.01
0
Cell_Type_NEW
Ctrl
ExN
InN
Other
UD

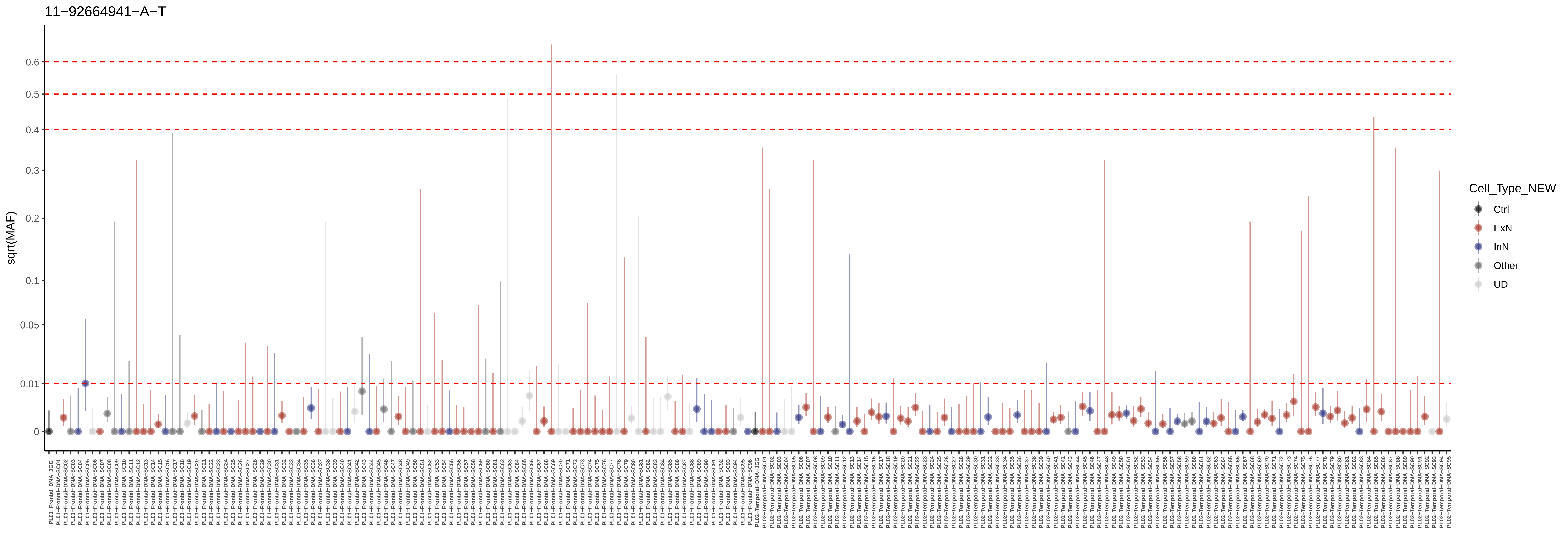
11−92664941−A−T
sqrt(MAF)
0.6
0.5
0.4
0.3
0.2
0.1
0.05
0.01
0
Cell_Type_NEW
Ctrl
ExN
InN
Other
UD

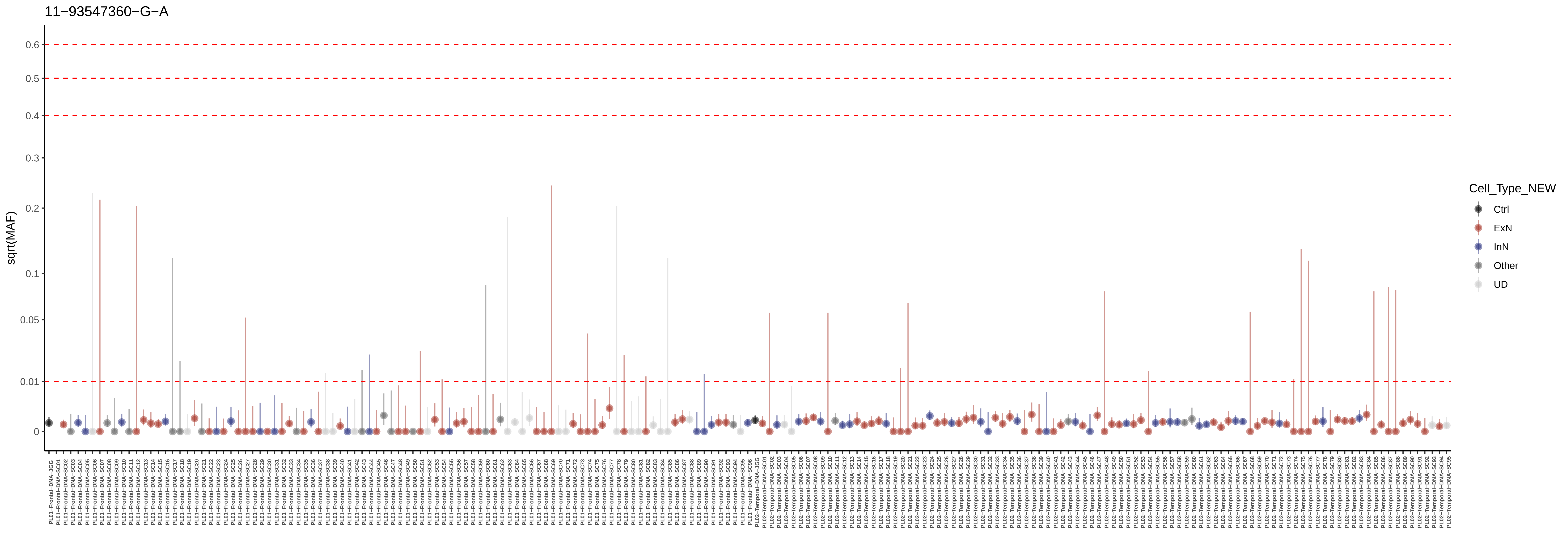
11−93547360−G−A
sqrt(MAF)
0.6
0.5
0.4
0.3
0.2
0.1
0.05
0.01
0
Cell_Type_NEW
Ctrl
ExN
InN
Other
UD

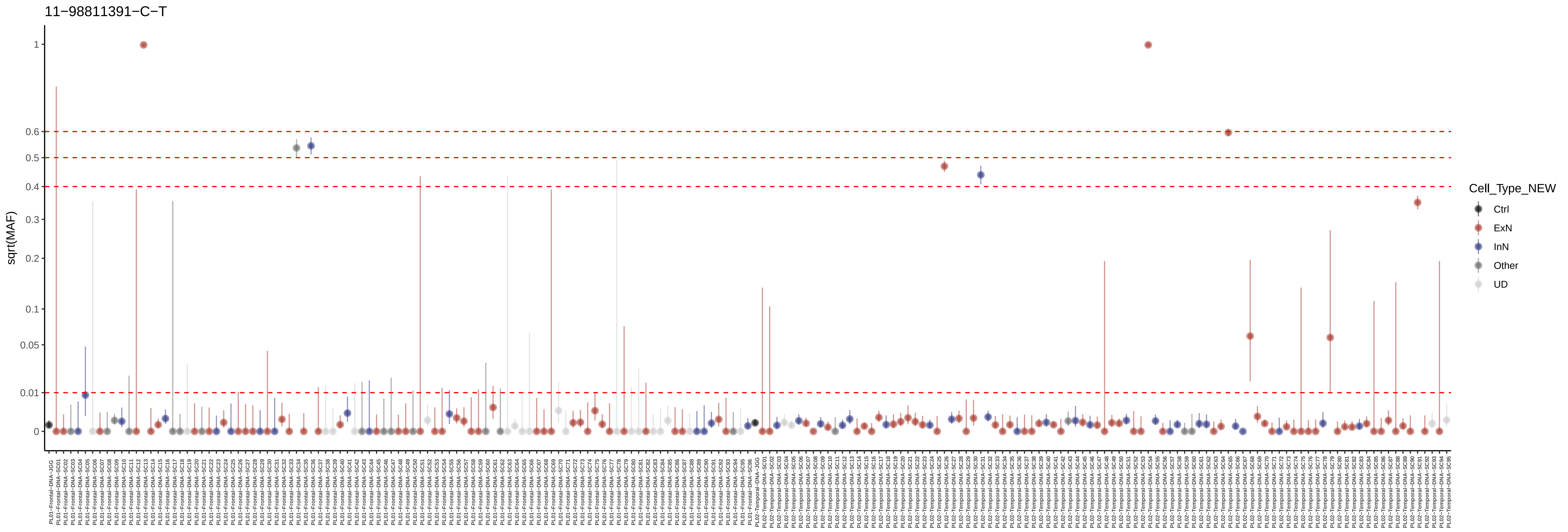

11−98811391−C−T
sqrt(MAF)
1
0.6
0.5
0.4
0.3
0.2
0.1
0.05
0.01
0
Cell_Type_NEW
Ctrl
ExN
InN
Other
UD

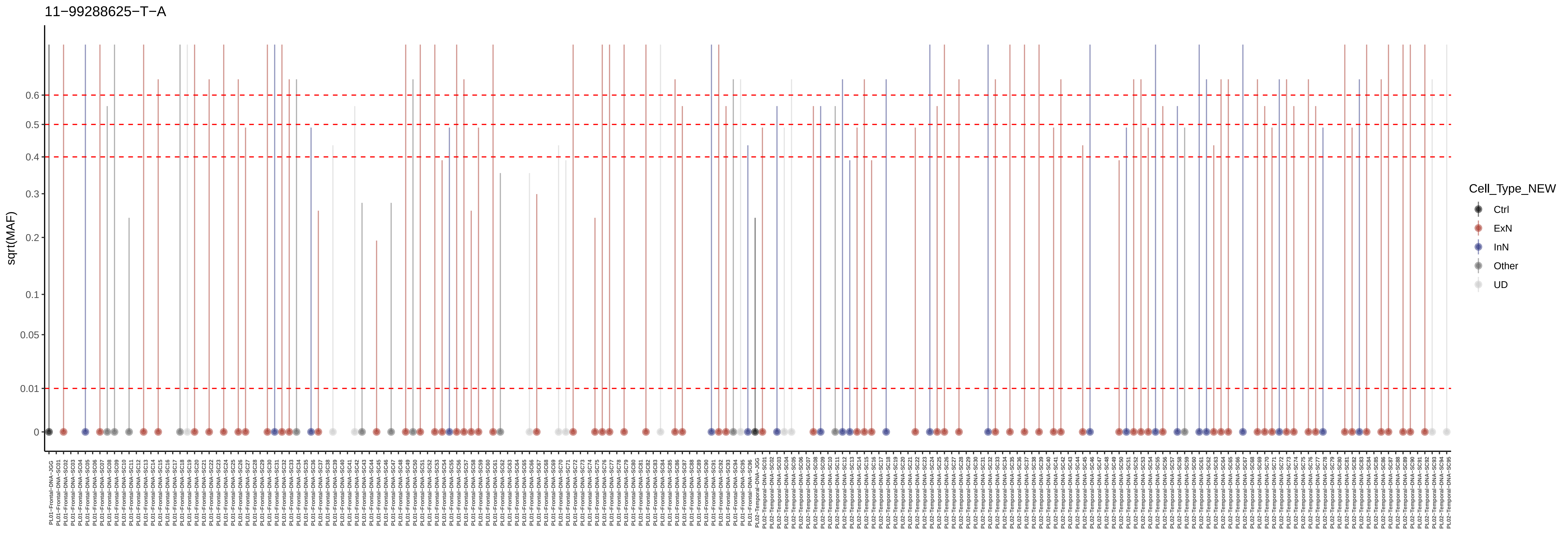
11−99288625−T−A
sqrt(MAF)
0.6
0.5
0.4
0.3
0.2
0.1
0.05
0.01
0
Cell_Type_NEW
Ctrl
ExN
InN
Other
UD

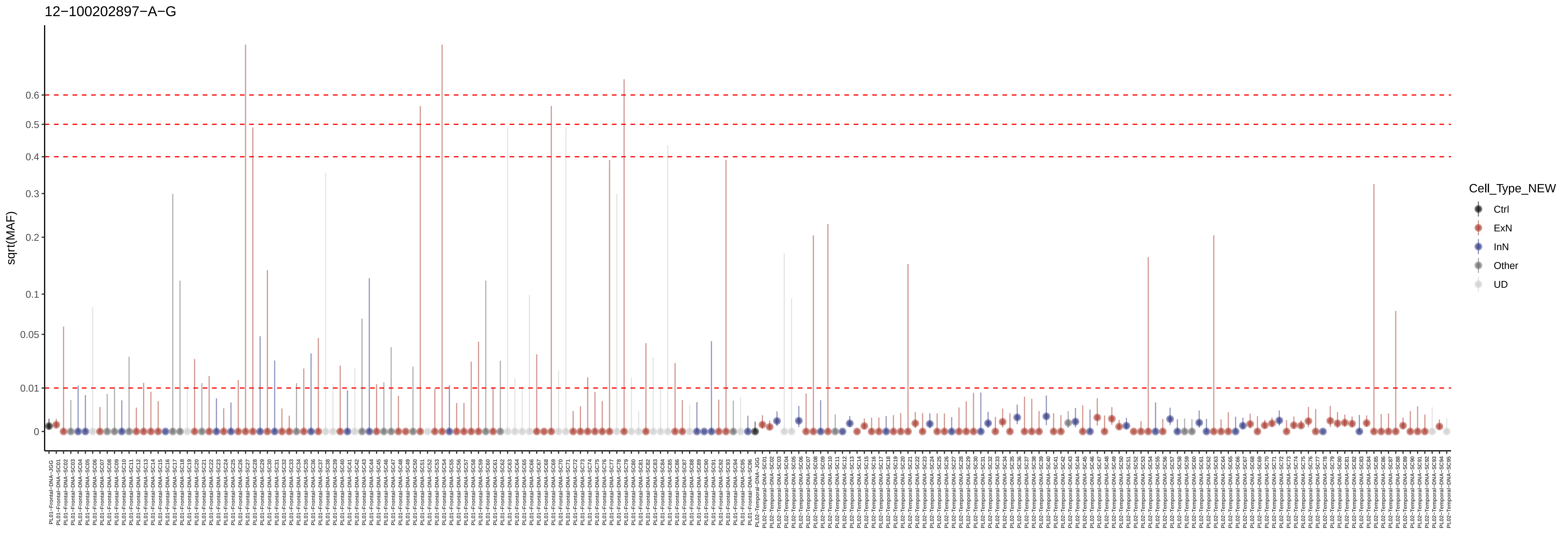
12−100202897−A−G
sqrt(MAF)
0.6
0.5
0.4
0.3
0.2
0.1
0.05
0.01
0
Cell_Type_NEW
Ctrl
ExN
InN
Other
UD

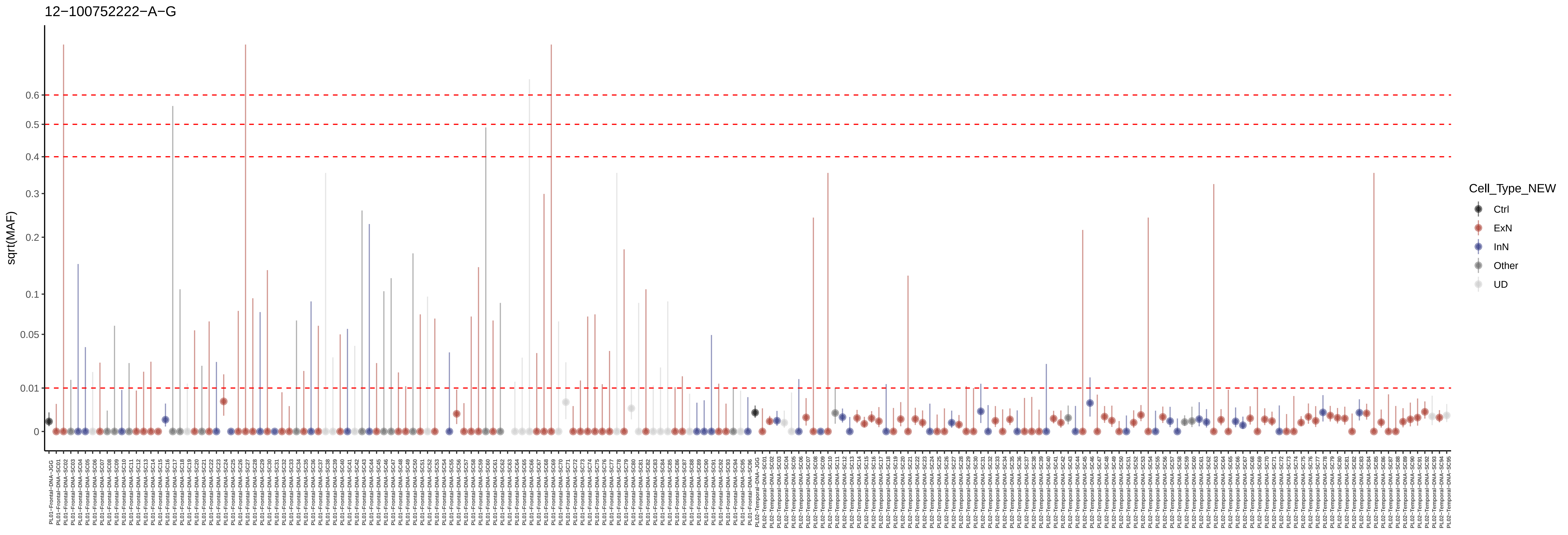
12−100752222−A−G
sqrt(MAF)
0.6
0.5
0.4
0.3
0.2
0.1
0.05
0.01
0
Cell_Type_NEW
Ctrl
ExN
InN
Other
UD

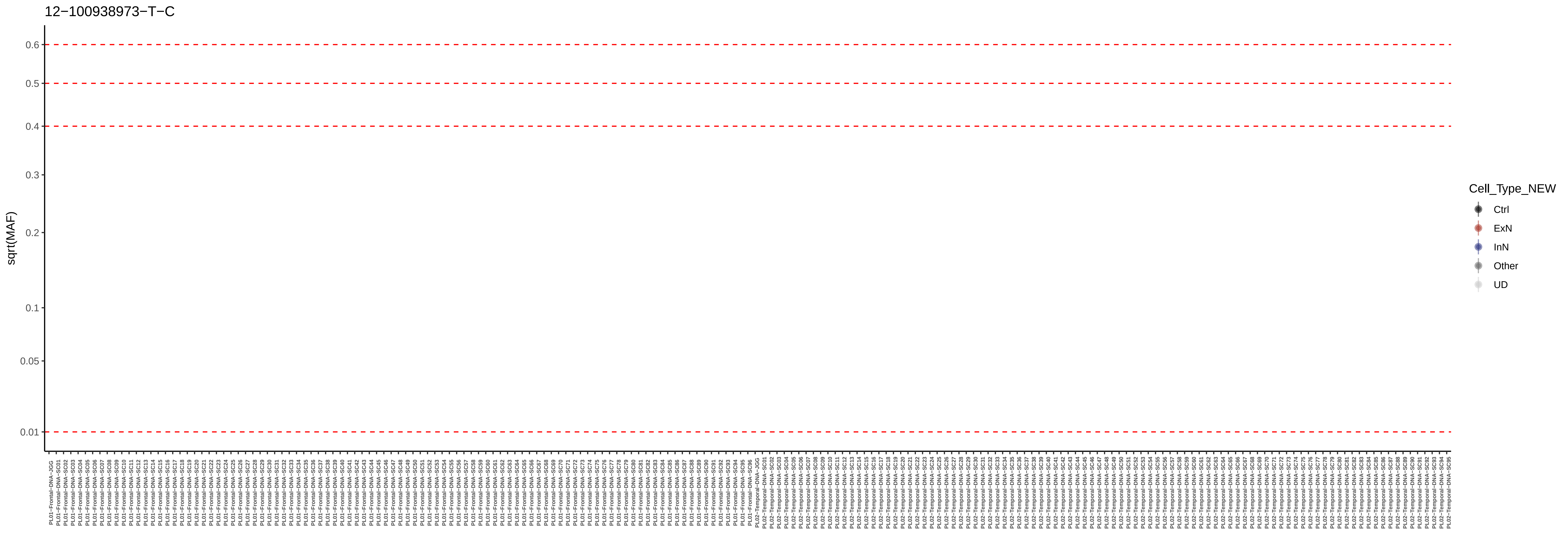
12−100938973−T−C
sqrt(MAF)
0.6
0.5
0.4
0.3
0.2
0.1
0.05
0.01
Cell_Type_NEW
Ctrl
ExN
InN
Other
UD
PL01−Frontal−DNA−JGG
PL01−Frontal−DNA−SC01 … PL01−Frontal−DNA−SC96
PL02−Temporal−DNA−JGG
PL02−Temporal−DNA−SC01 … PL02−Temporal−DNA−SC95

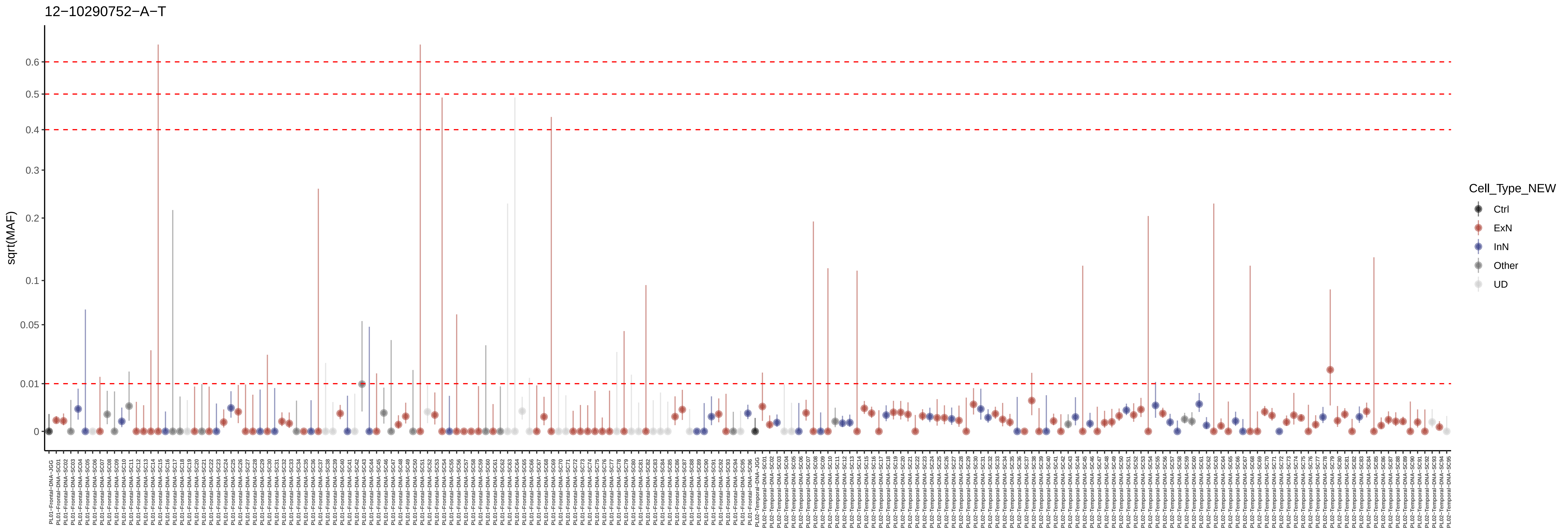
12−10290752−A−T
sqrt(MAF)
0.6
0.5
0.4
0.3
0.2
0.1
0.05
0.01
0
Cell_Type_NEW
Ctrl
ExN
InN
Other
UD

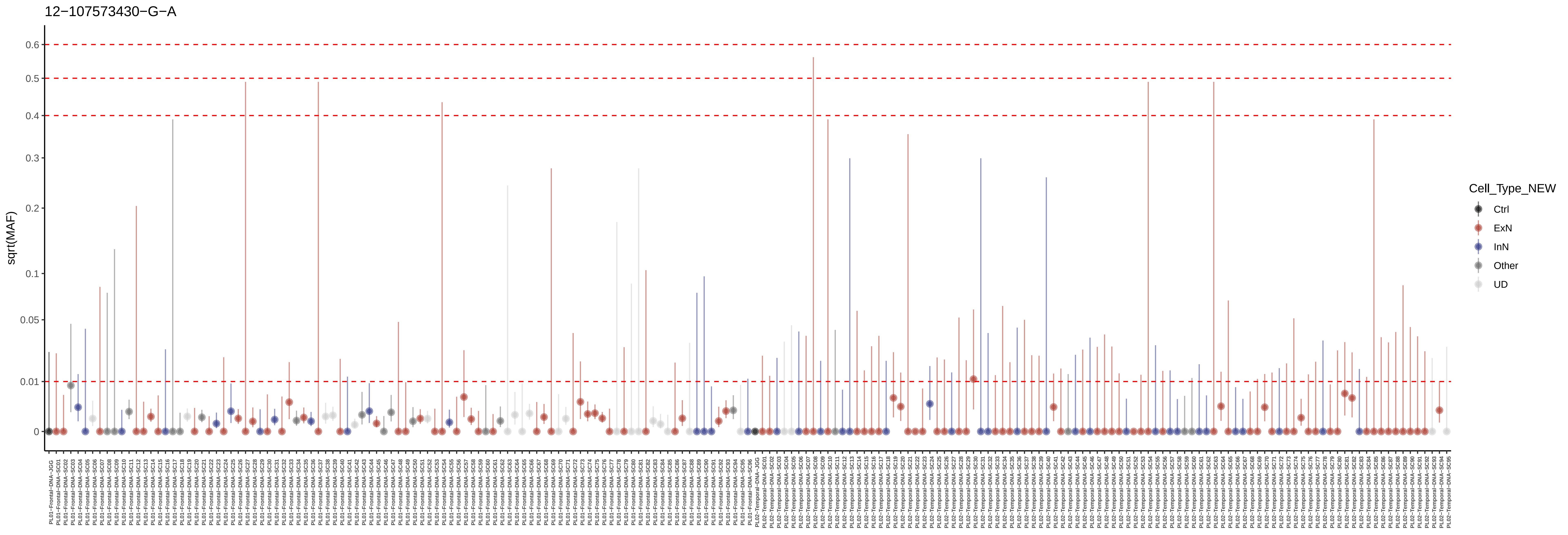
12−107573430−G−A
sqrt(MAF)
0.6
0.5
0.4
0.3
0.2
0.1
0.05
0.01
0
Cell_Type_NEW
Ctrl
ExN
InN
Other
UD

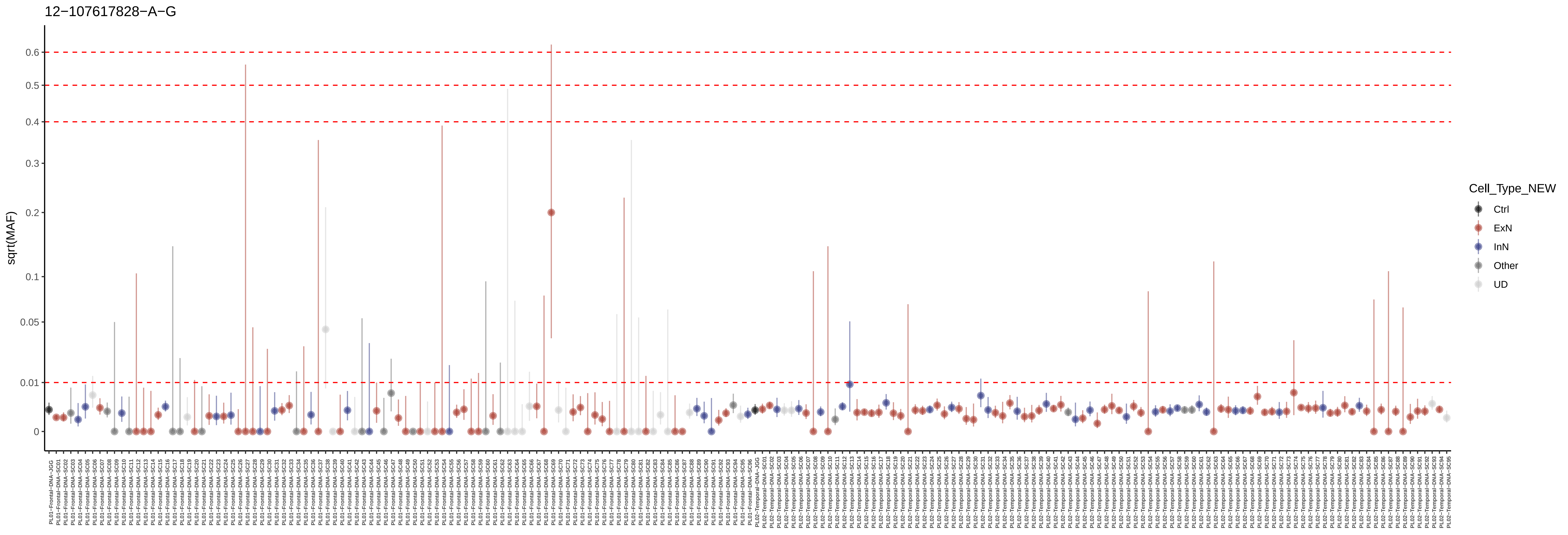
12−107617828−A−G
sqrt(MAF)
0.6
0.5
0.4
0.3
0.2
0.1
0.05
0.01
0
Cell_Type_NEW
Ctrl
ExN
InN
Other
UD

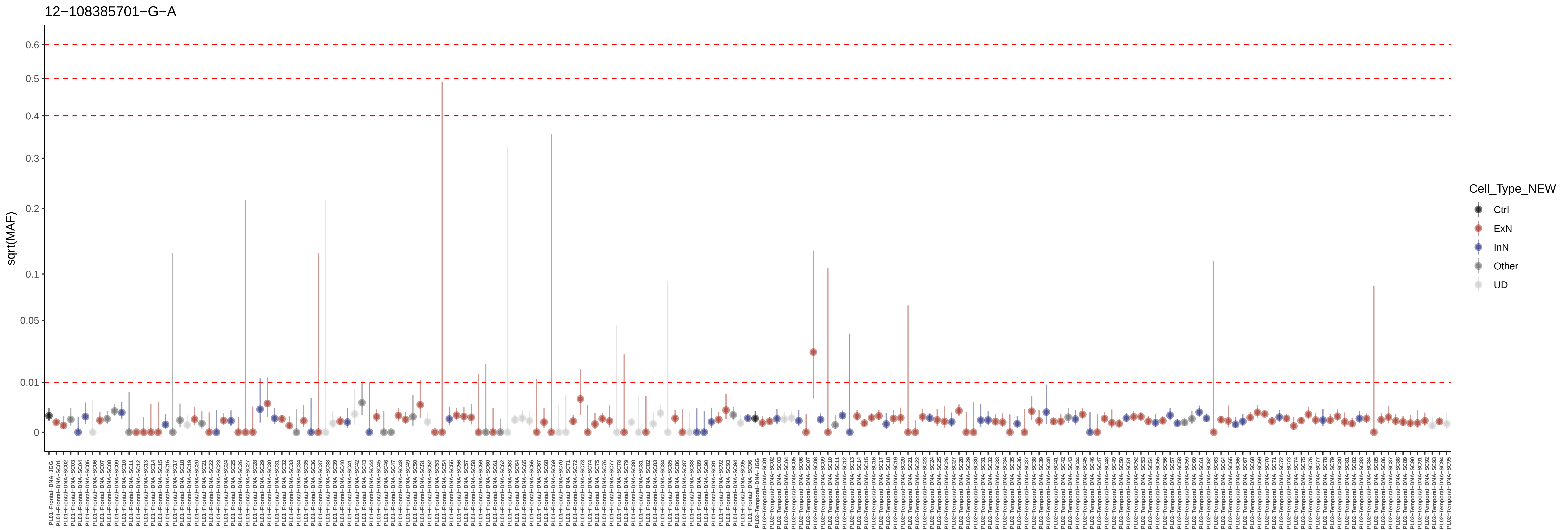
12−108385701−G−A
sqrt(MAF)
0.6
0.5
0.4
0.3
0.2
0.1
0.05
0.01
0
Cell_Type_NEW
Ctrl
ExN
InN
Other
UD

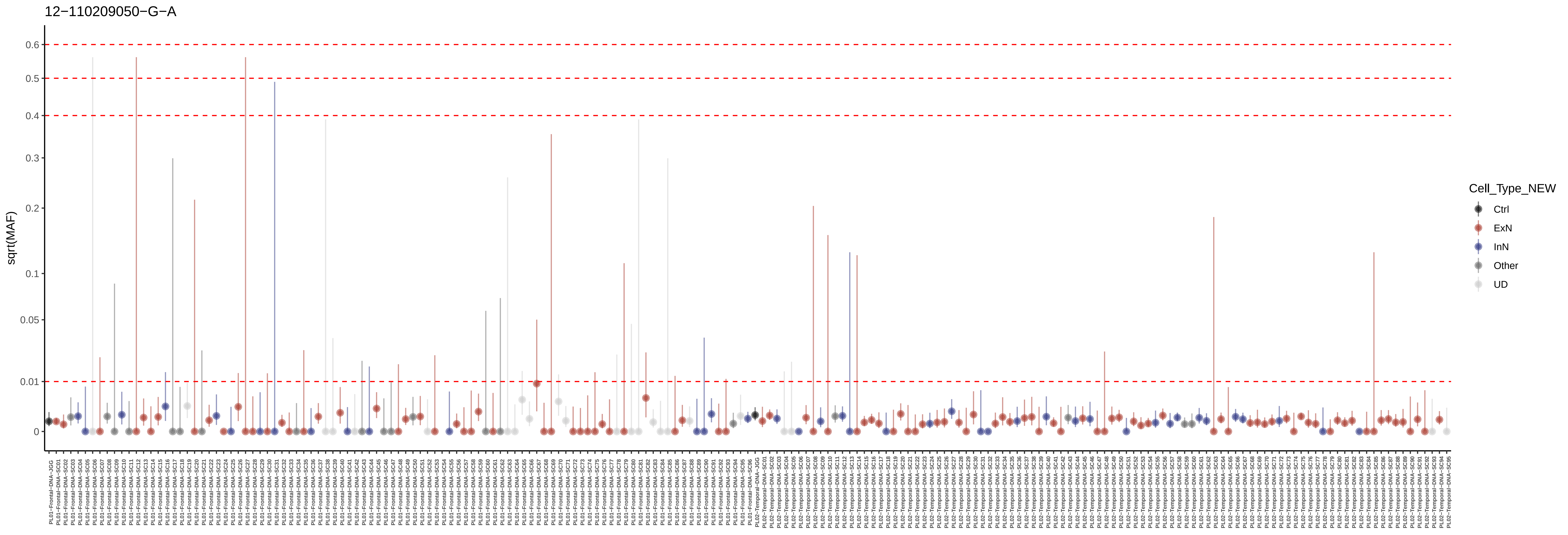
12−110209050−G−A
sqrt(MAF)
0.6
0.5
0.4
0.3
0.2
0.1
0.05
0.01
0
Cell_Type_NEW
Ctrl
ExN
InN
Other
UD

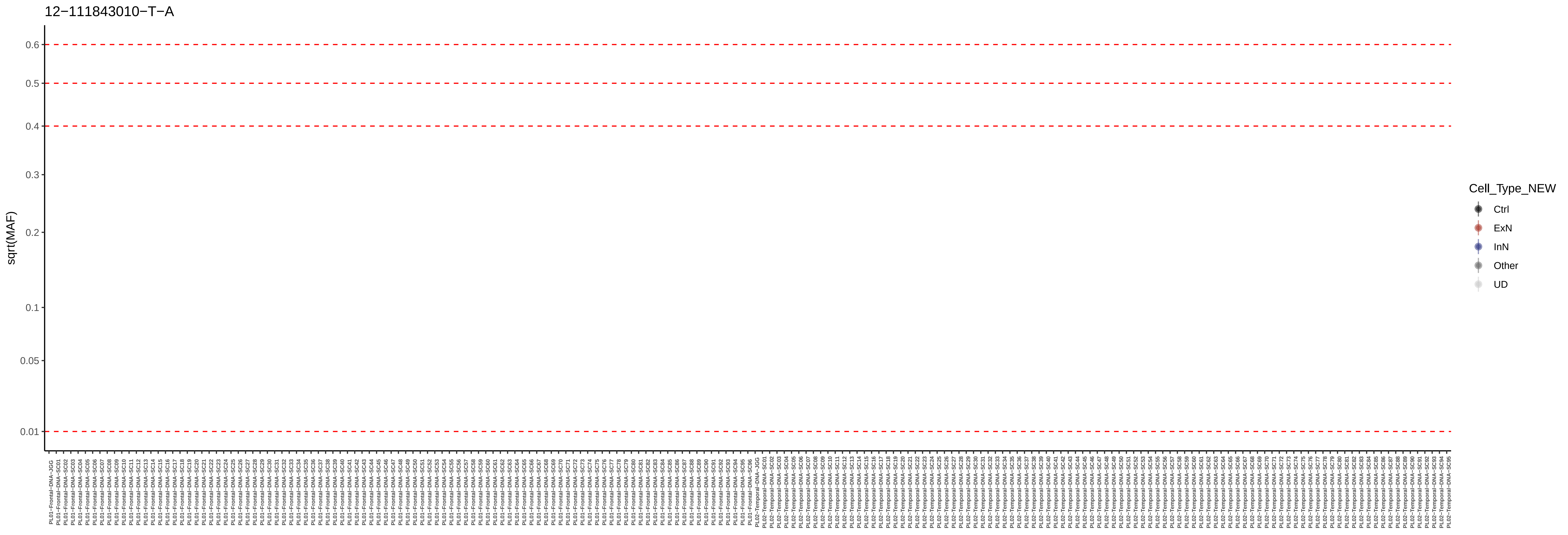
12−111843010−T−A
sqrt(MAF)
0.6
0.5
0.4
0.3
0.2
0.1
0.05
0.01
Cell_Type_NEW
Ctrl
ExN
InN
Other
UD

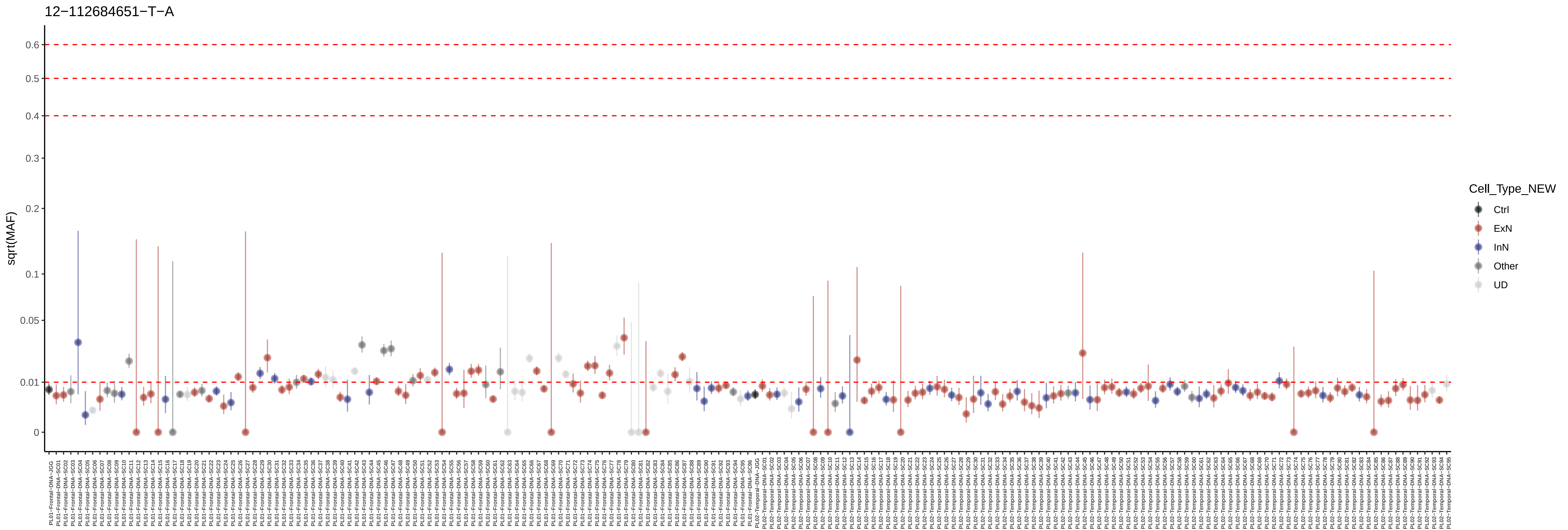

12−112684651−T−A
sqrt(MAF)
0.6
0.5
0.4
0.3
0.2
0.1
0.05
0.01
0
Cell_Type_NEW
Ctrl
ExN
InN
Other
UD

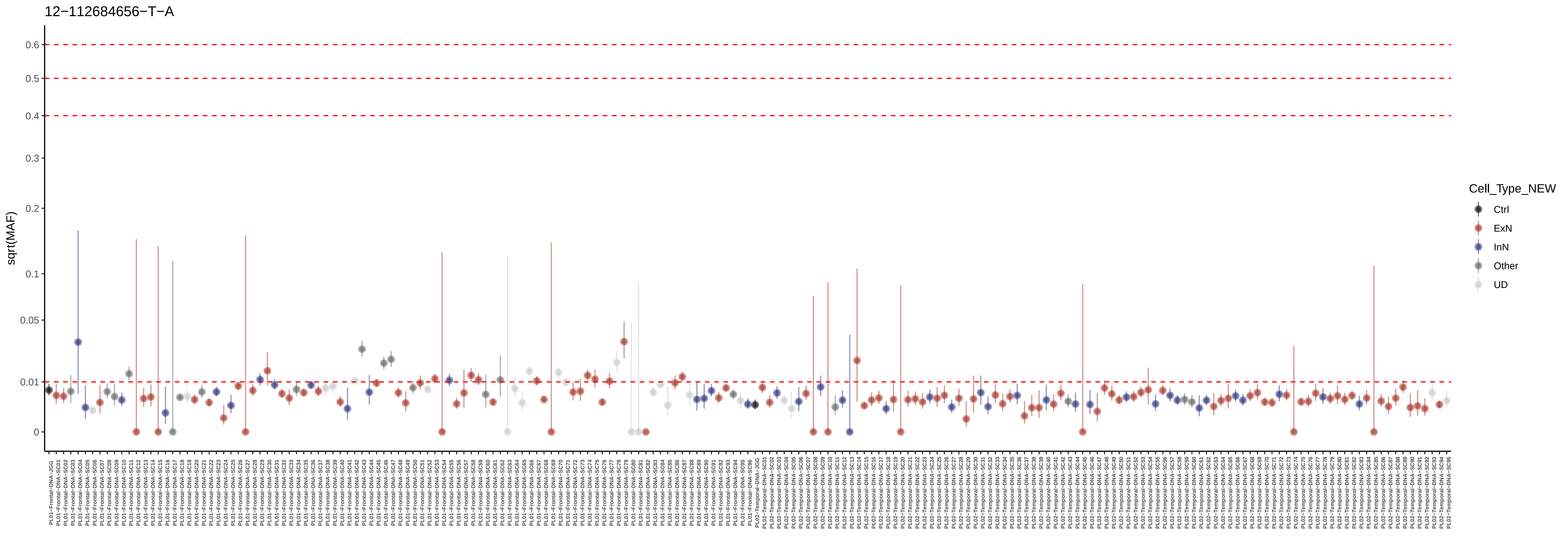

12−112684656−T−A
sqrt(MAF)
0.6
0.5
0.4
0.3
0.2
0.1
0.05
0.01
0
Cell_Type_NEW
Ctrl
ExN
InN
Other
UD

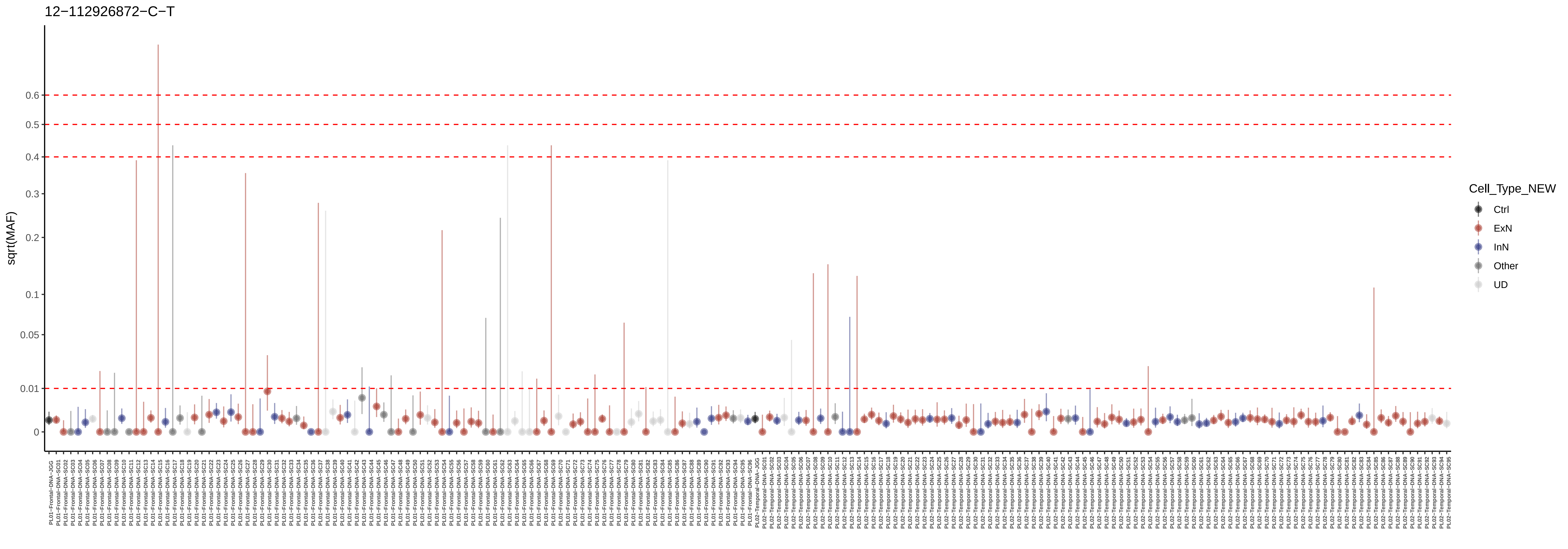
12−112926872−C−T
sqrt(MAF)
0.6
0.5
0.4
0.3
0.2
0.1
0.05
0.01
0
Cell_Type_NEW
Ctrl
ExN
InN
Other
UD

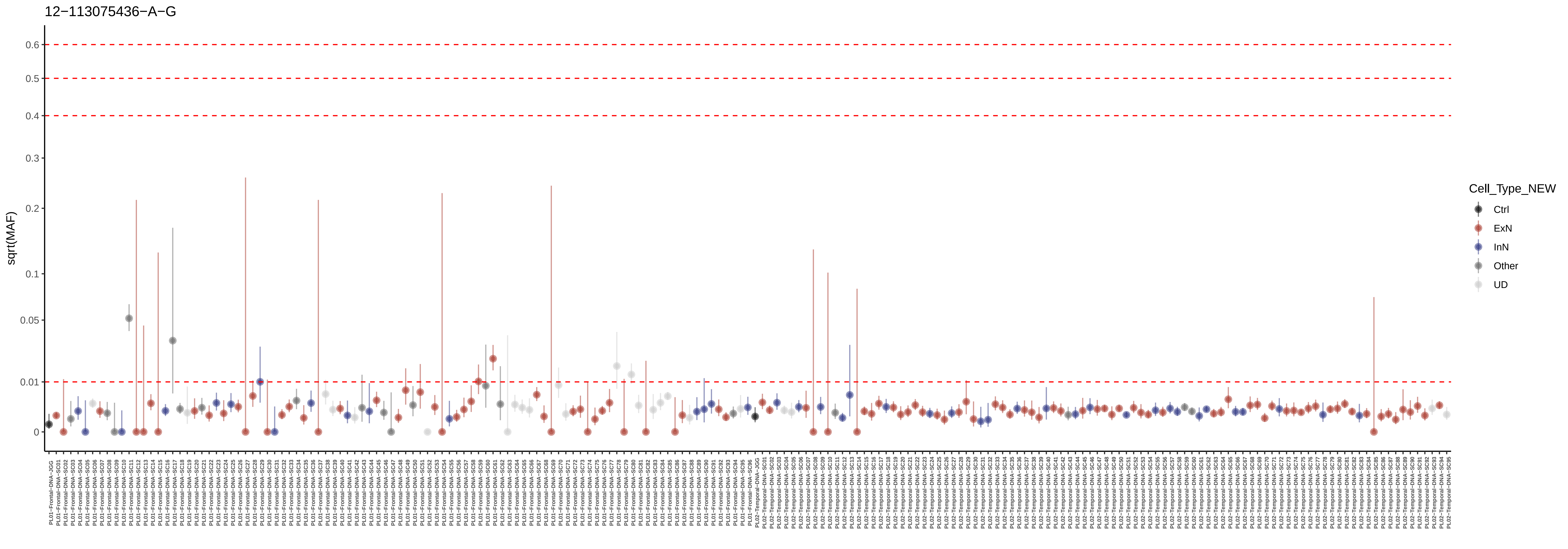

12−113075436−A−G
sqrt(MAF)
0.6
0.5
0.4
0.3
0.2
0.1
0.05
0.01
0
Cell_Type_NEW
Ctrl
ExN
InN
Other
UD

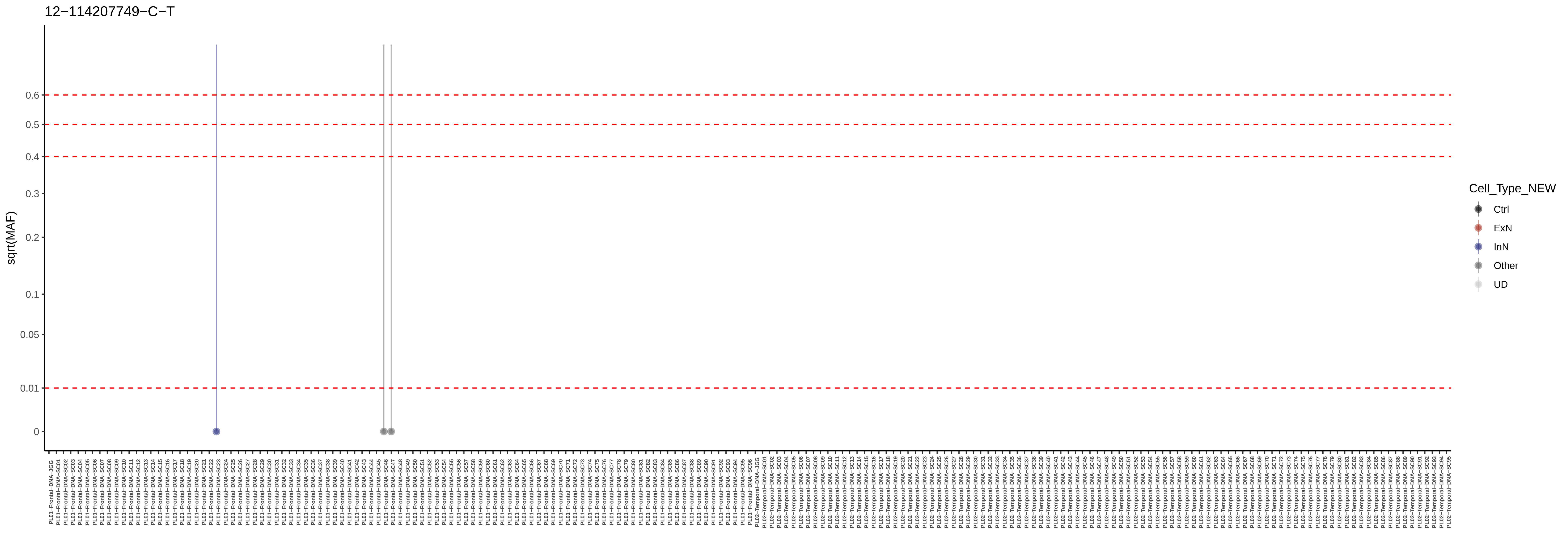
12−114207749−C−T
sqrt(MAF)
Cell_Type_NEW
Ctrl
ExN
InN
Other
UD

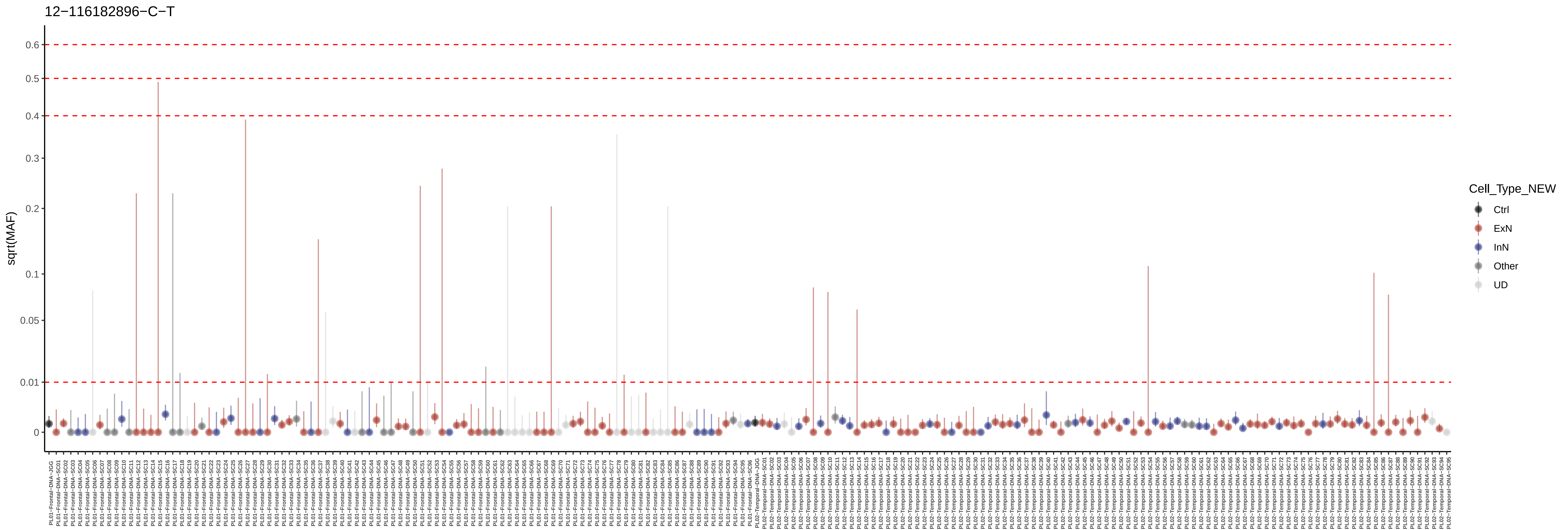
12−116182896−C−T
sqrt(MAF)
0.6
0.5
0.4
0.3
0.2
0.1
0.05
0.01
0
Cell_Type_NEW
Ctrl
ExN
InN
Other
UD

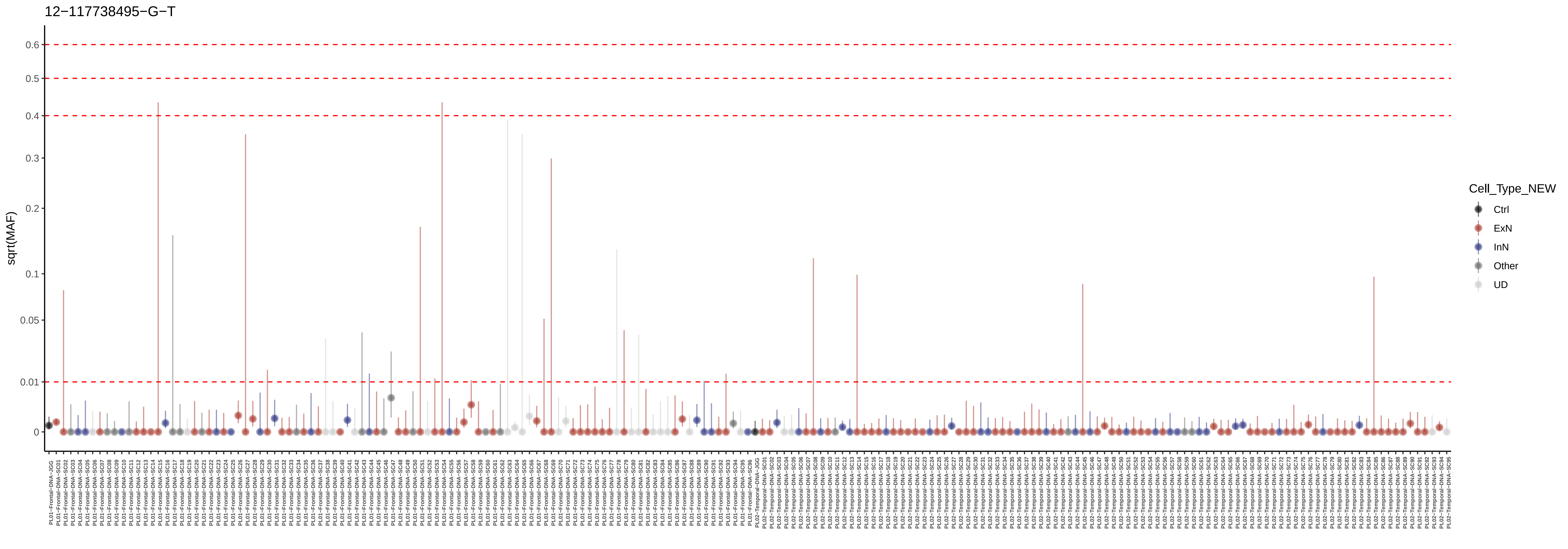
12−117738495−G−T
sqrt(MAF)
0.6
0.5
0.4
0.3
0.2
0.1
0.05
0.01
0
Cell_Type_NEW
Ctrl
ExN
InN
Other
UD

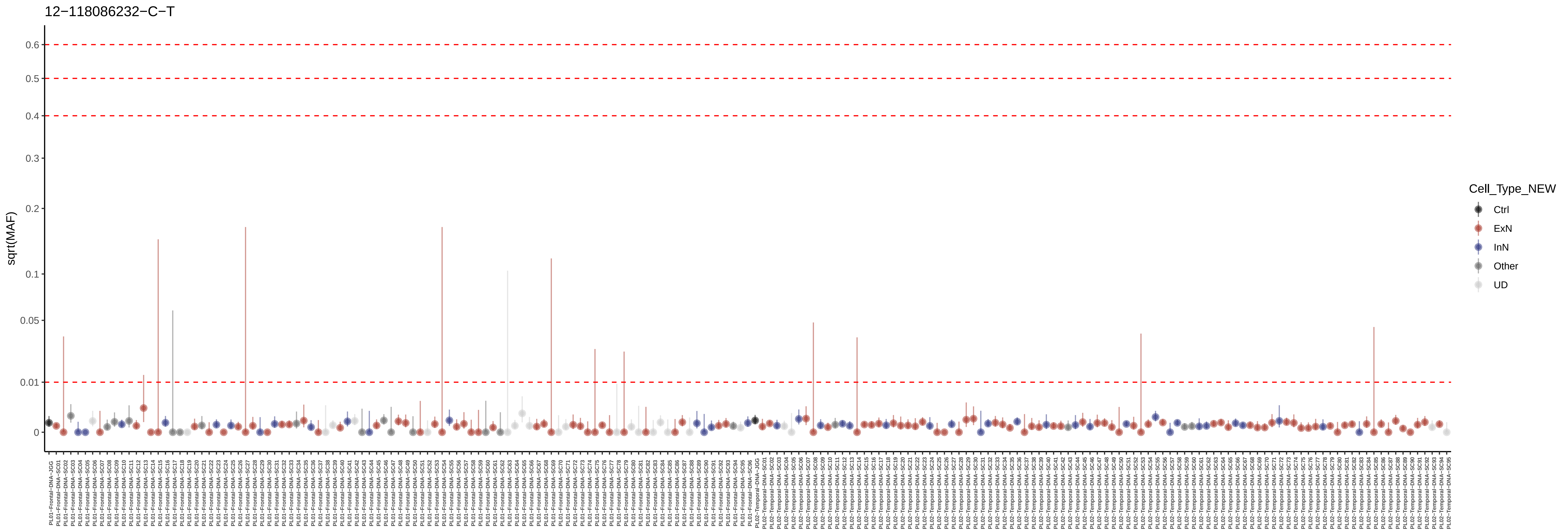

12−118086232−C−T
sqrt(MAF)
0.6
0.5
0.4
0.3
0.2
0.1
0.05
0.01
0
Cell_Type_NEW
Ctrl
ExN
InN
Other
UD

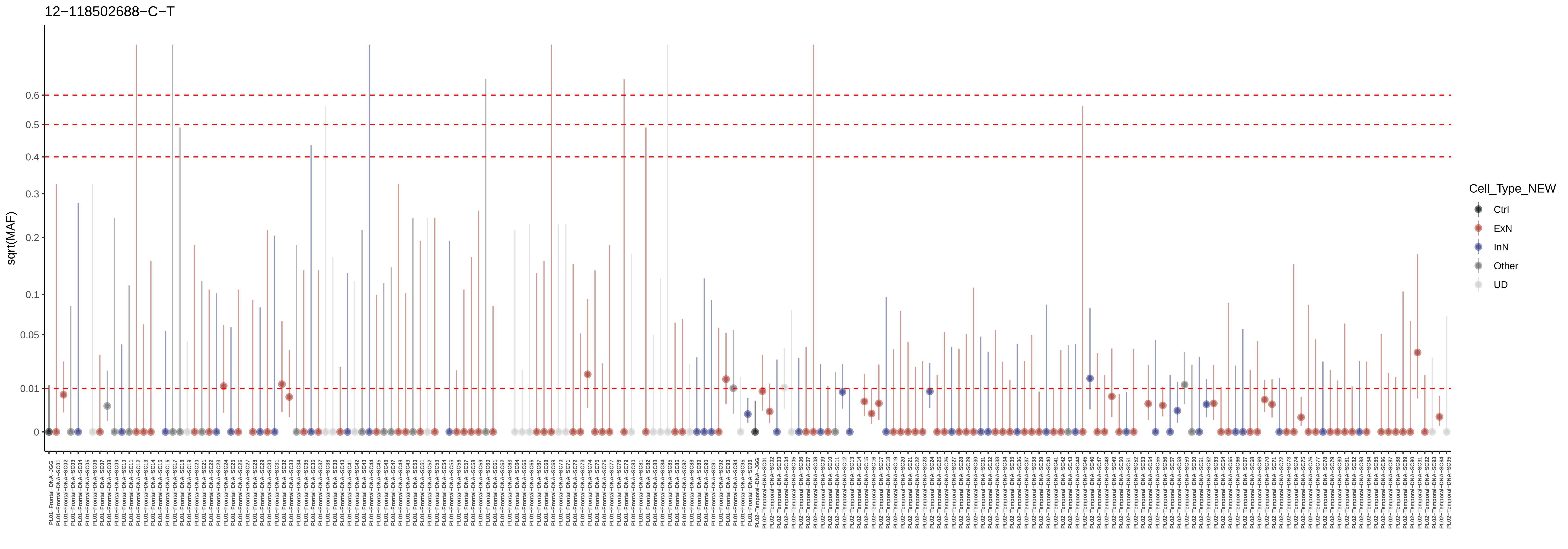
12−118502688−C−T
sqrt(MAF)
0
0.01
0.05
0.1
0.2
0.3
0.4
0.5
0.6
Cell_Type_NEW
Ctrl
ExN
InN
Other
UD

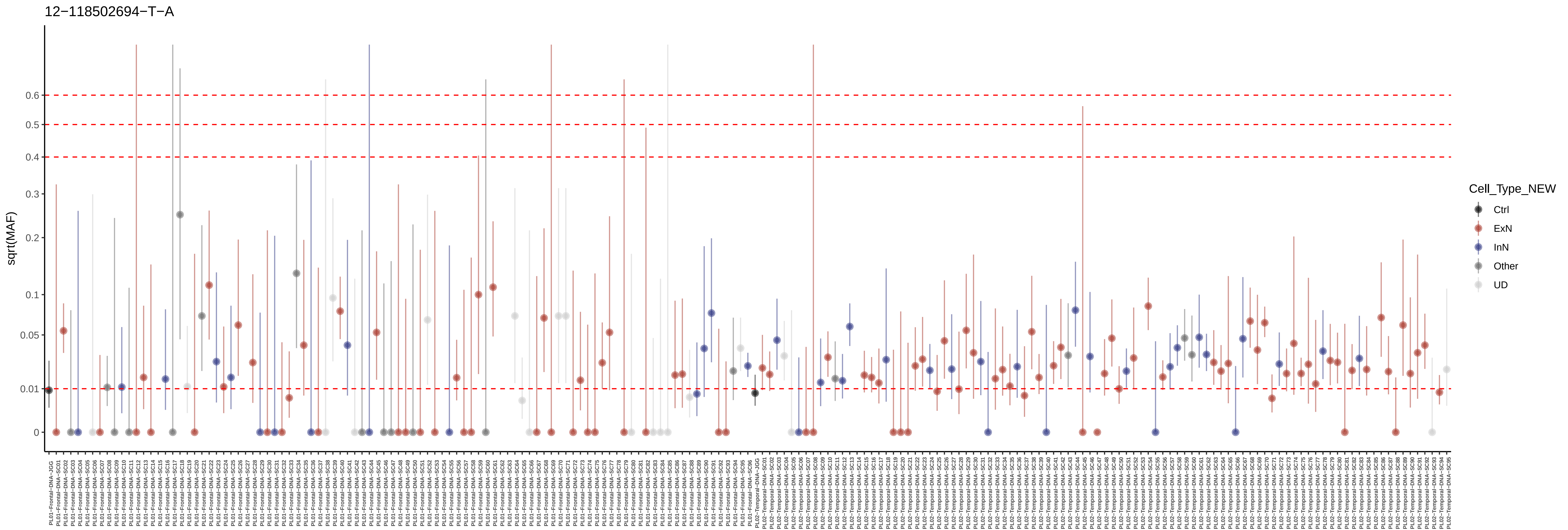

12−118502694−T−A
sqrt(MAF)
0.6
0.5
0.4
0.3
0.2
0.1
0.05
0.01
0
Cell_Type_NEW
Ctrl
ExN
InN
Other
UD

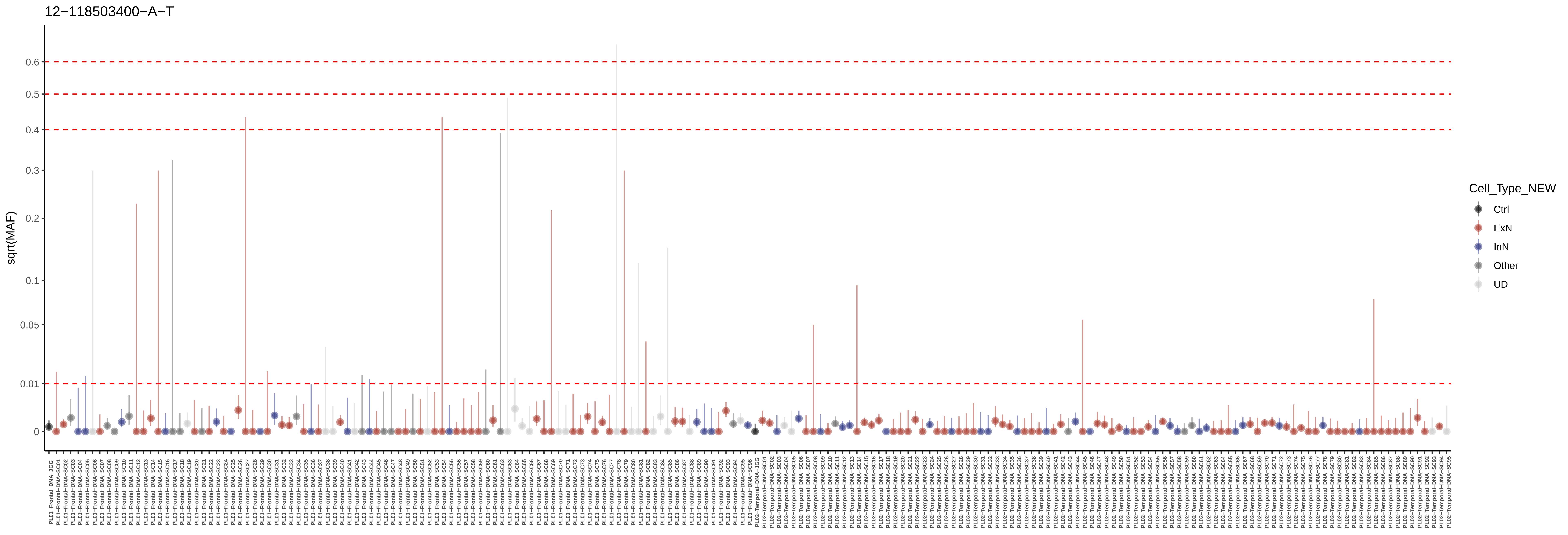
12−118503400−A−T
sqrt(MAF)
0.6
0.5
0.4
0.3
0.2
0.1
0.05
0.01
0
Cell_Type_NEW
Ctrl
ExN
InN
Other
UD

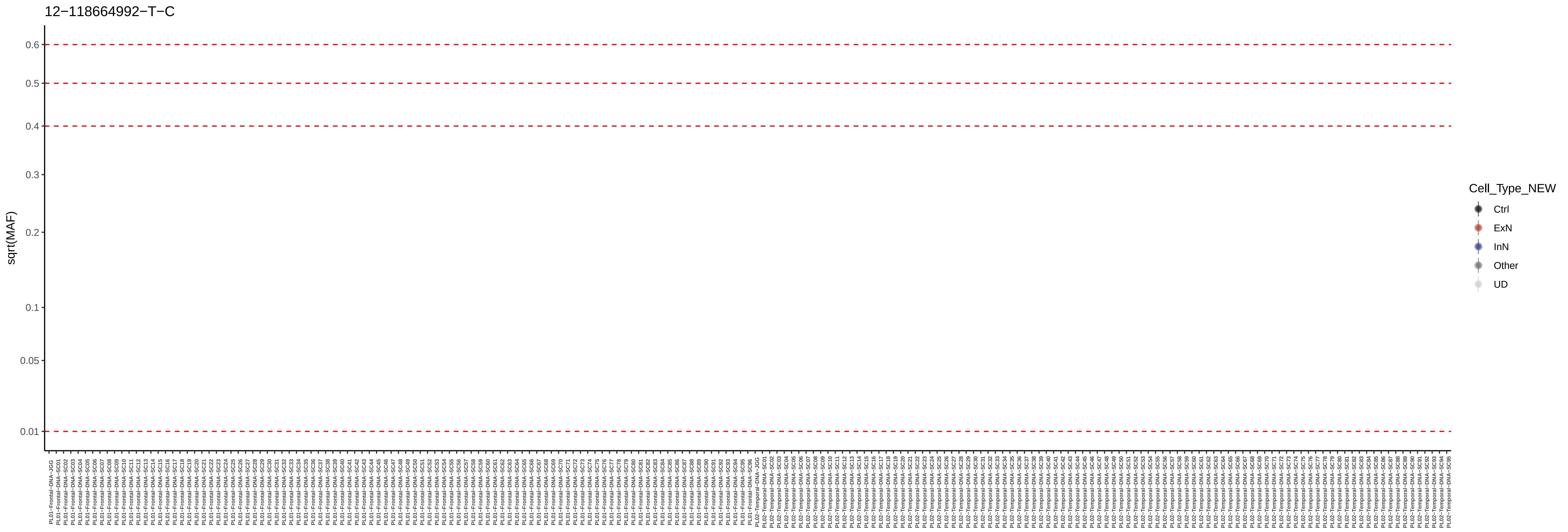
12−118664992−T−C
sqrt(MAF)
0.6
0.5
0.4
0.3
0.2
0.1
0.05
0.01
Cell_Type_NEW
Ctrl
ExN
InN
Other
UD

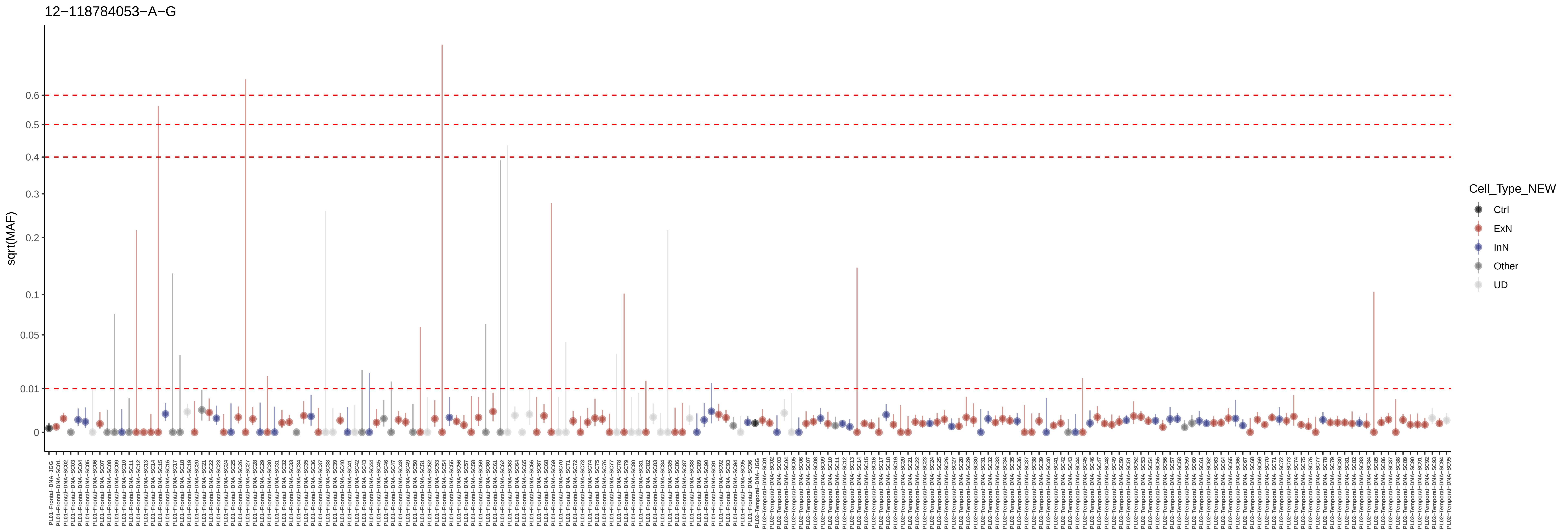
12−118784053−A−G
sqrt(MAF)
0.6
0.5
0.4
0.3
0.2
0.1
0.05
0.01
0
Cell_Type_NEW
Ctrl
ExN
InN
Other
UD

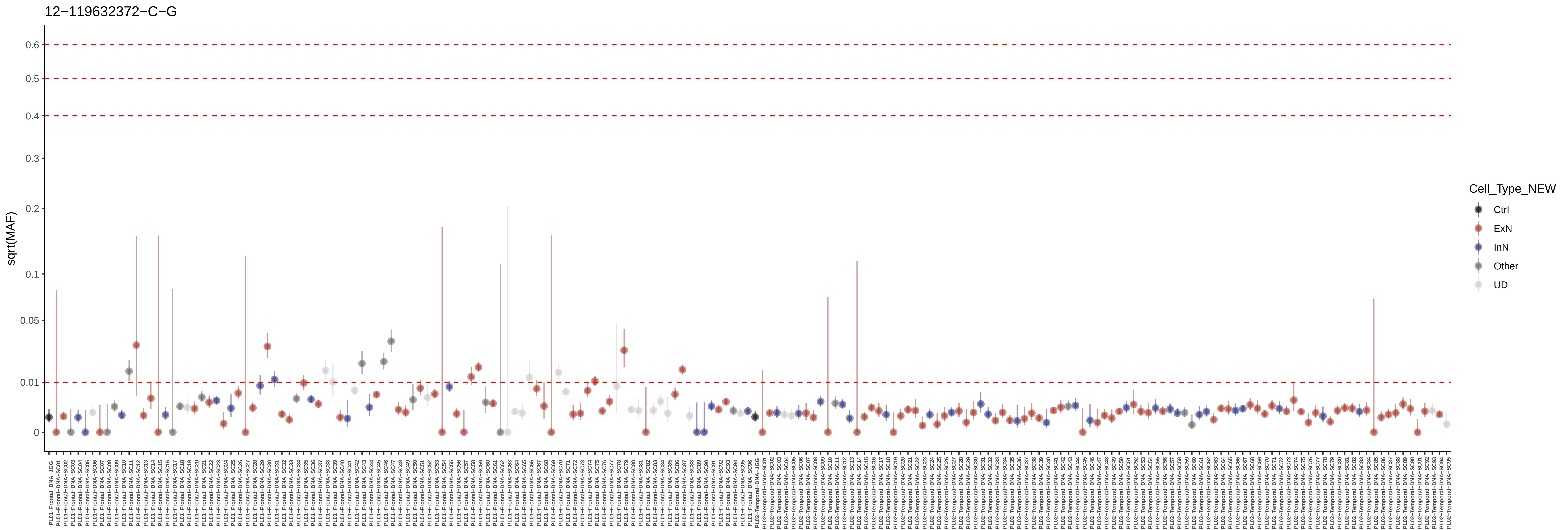

12−119632372−C−G
sqrt(MAF)
0.6
0.5
0.4
0.3
0.2
0.1
0.05
0.01
0
Cell_Type_NEW
Ctrl
ExN
InN
Other
UD

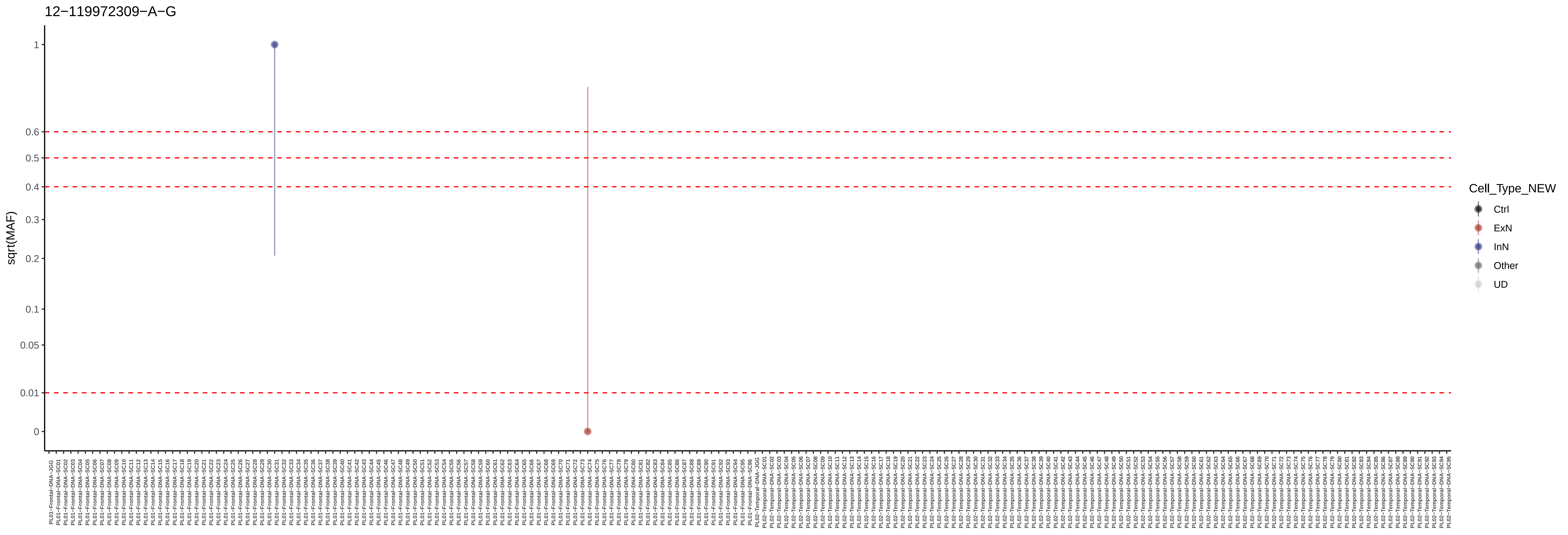
12−119972309−A−G
sqrt(MAF)
1
0.6
0.5
0.4
0.3
0.2
0.1
0.05
0.01
0
Cell_Type_NEW
Ctrl
ExN
InN
Other
UD

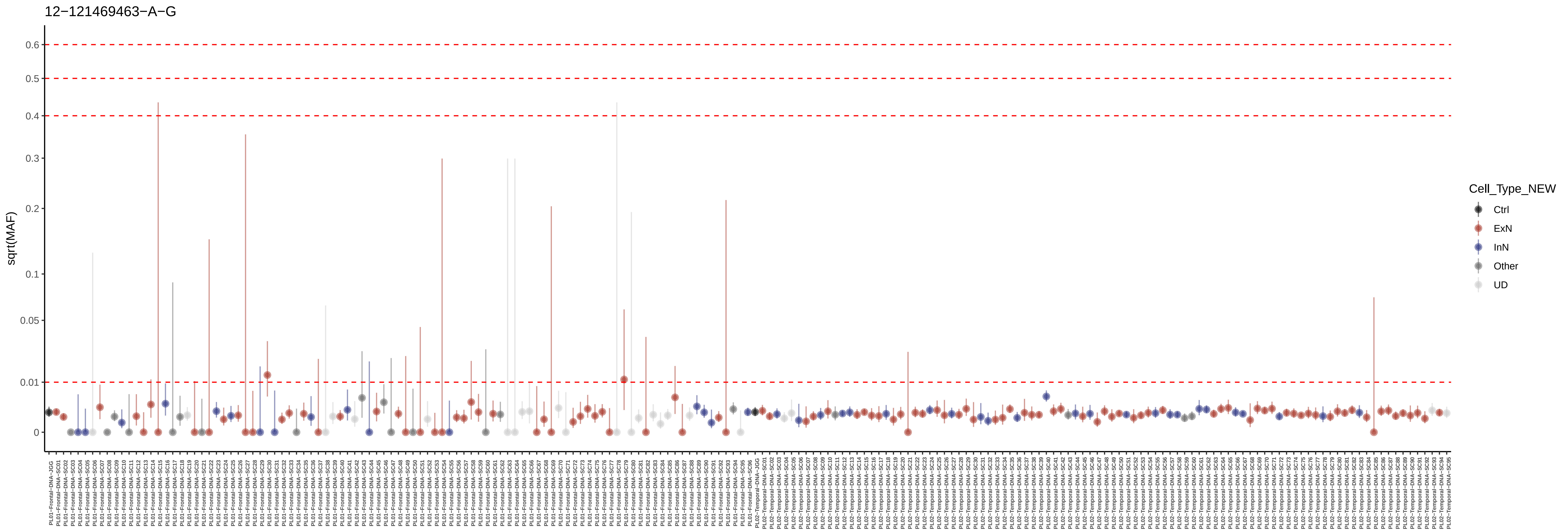
12−121469463−A−G
sqrt(MAF)
0.6
0.5
0.4
0.3
0.2
0.1
0.05
0.01
0
Cell_Type_NEW
Ctrl
ExN
InN
Other
UD

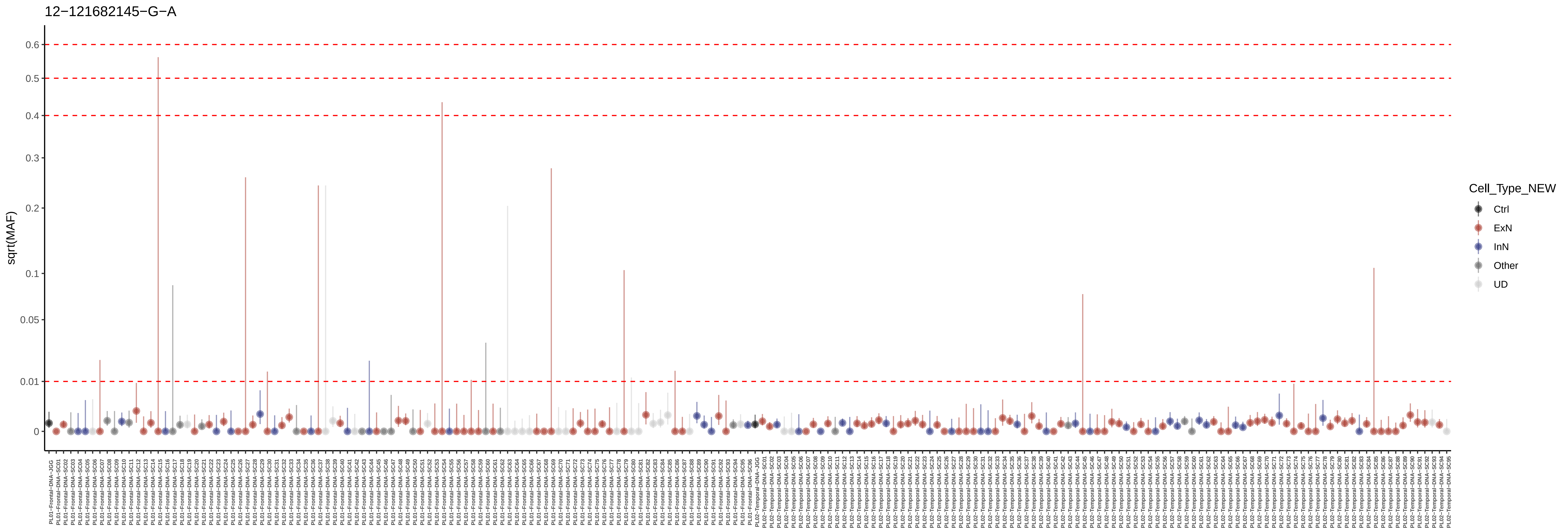
12−121682145−G−A
sqrt(MAF)
0.6
0.5
0.4
0.3
0.2
0.1
0.05
0.01
0
Cell_Type_NEW
Ctrl
ExN
InN
Other
UD

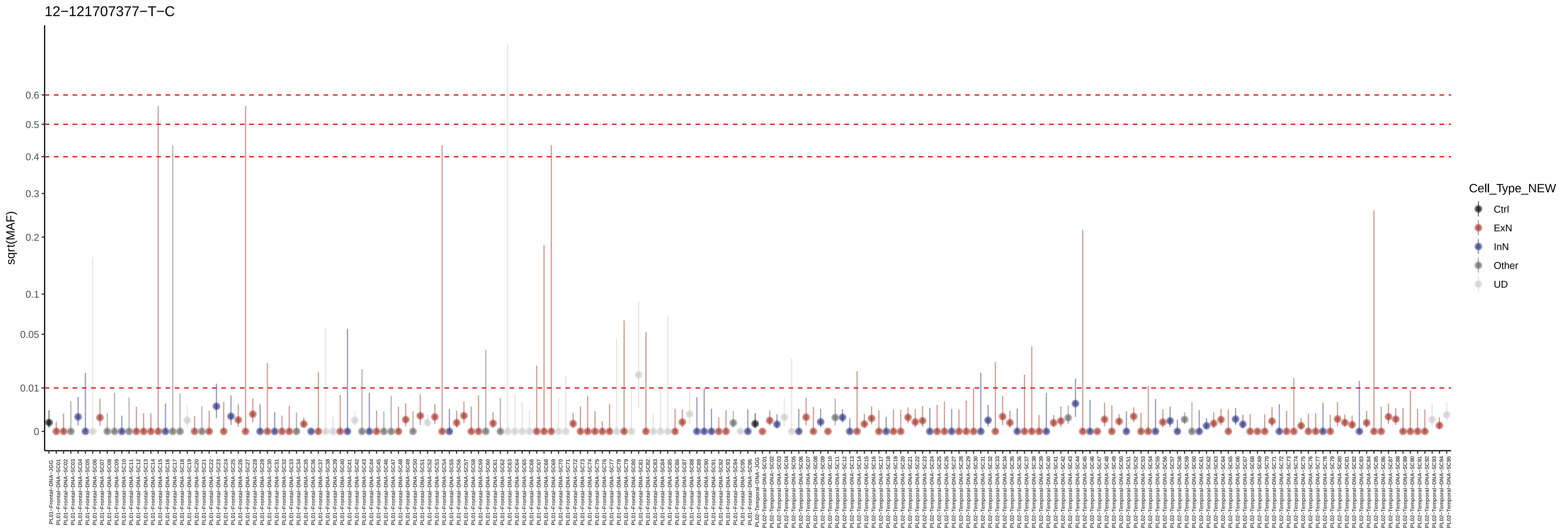
12−121707377−T−C
sqrt(MAF)
0.6
0.5
0.4
0.3
0.2
0.1
0.05
0.01
0
Cell_Type_NEW
Ctrl
ExN
InN
Other
UD

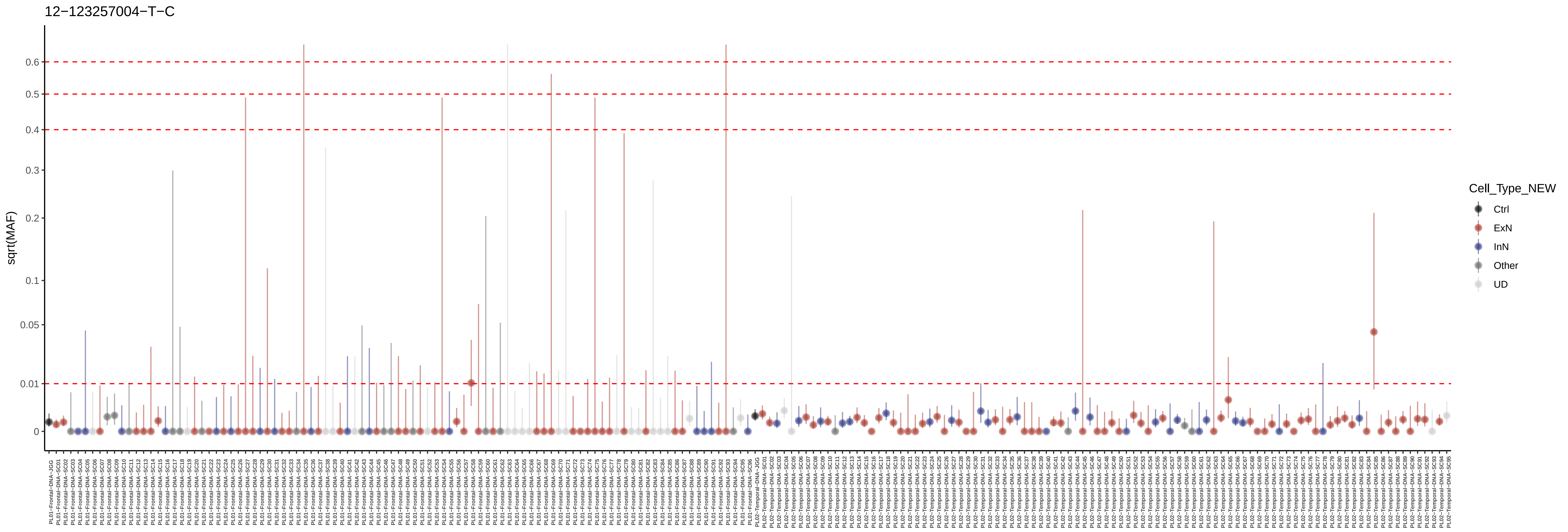

12−123257004−T−C
sqrt(MAF)
0.6
0.5
0.4
0.3
0.2
0.1
0.05
0.01
0
Cell_Type_NEW
Ctrl
ExN
InN
Other
UD

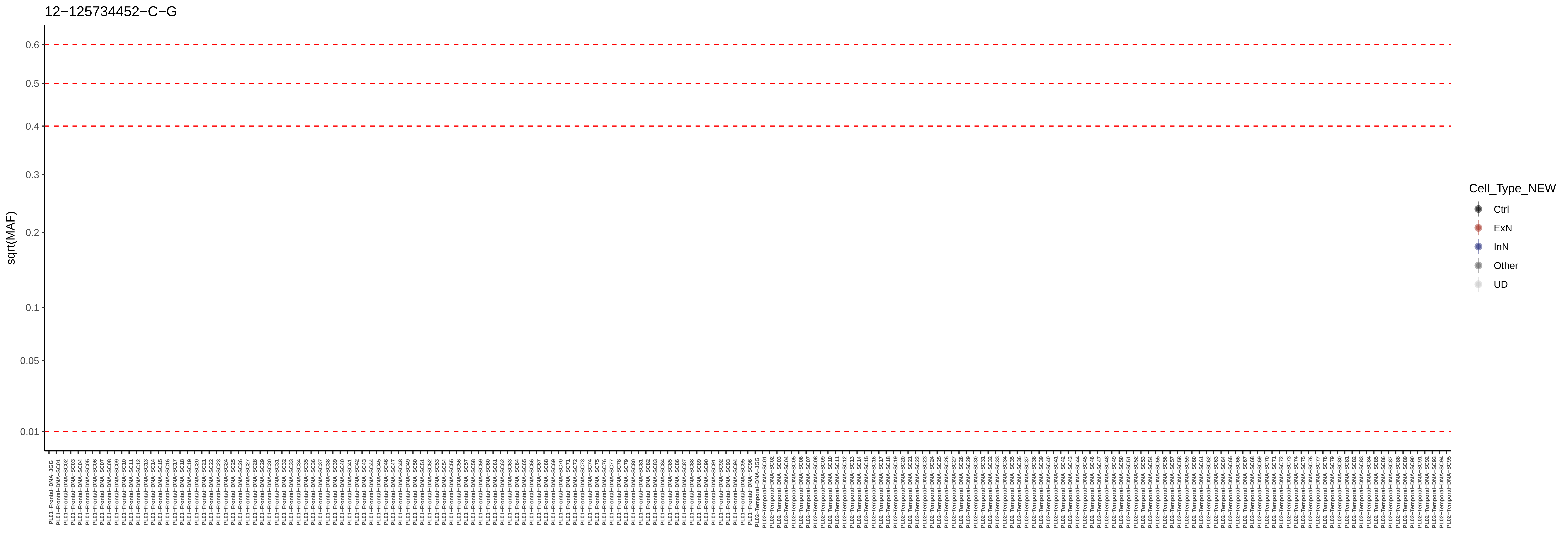
12−125734452−C−G
sqrt(MAF)
0.6
0.5
0.4
0.3
0.2
0.1
0.05
0.01
Cell_Type_NEW
Ctrl
ExN
InN
Other
UD

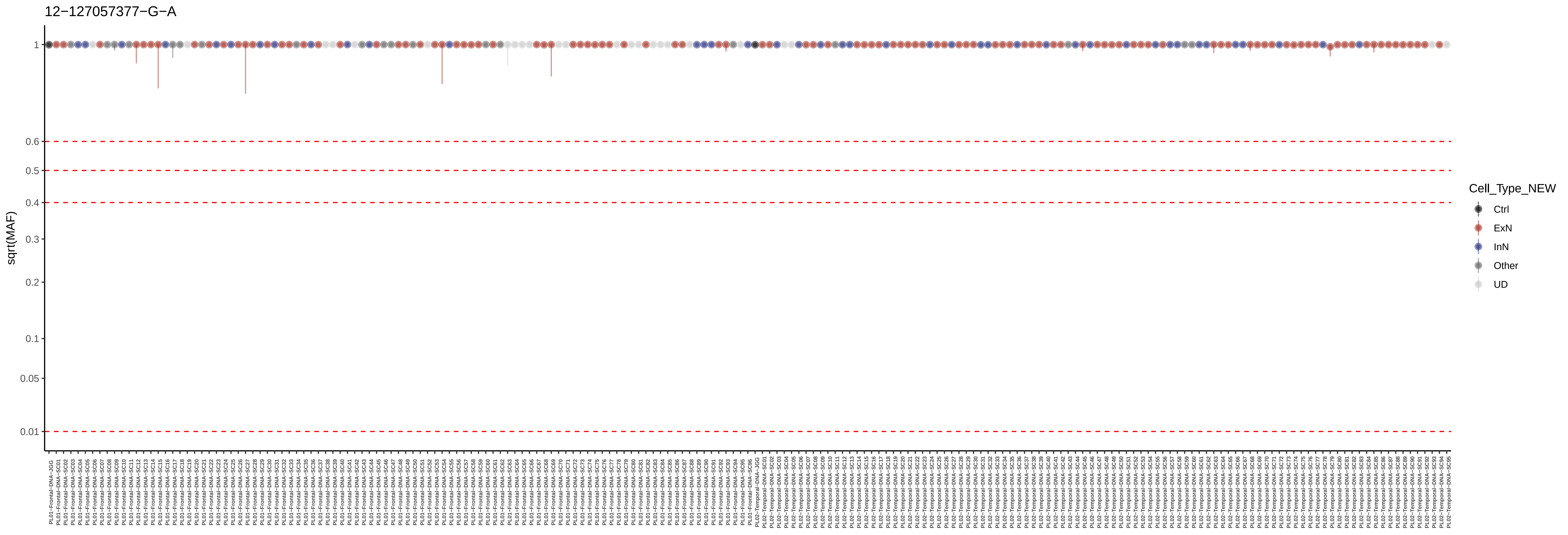
12−127057377−G−A
sqrt(MAF)
1
0.6
0.5
0.4
0.3
0.2
0.1
0.05
0.01
Cell_Type_NEW
Ctrl
ExN
InN
Other
UD

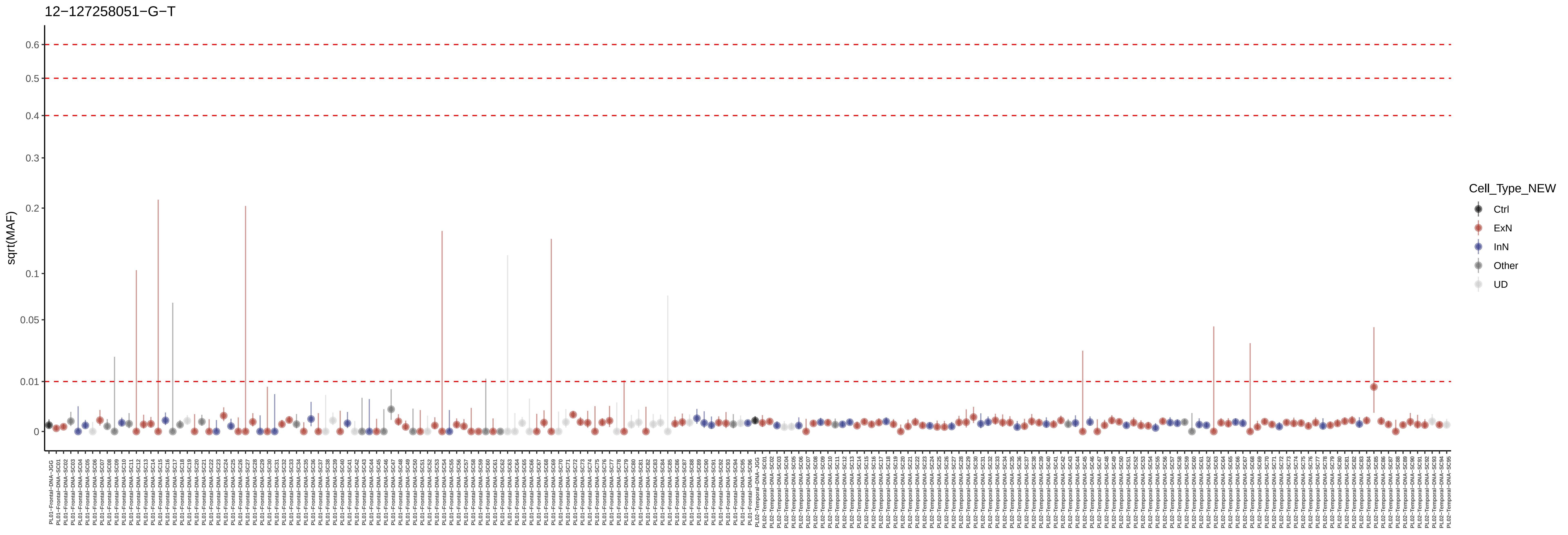
12−127258051−G−T
sqrt(MAF)
0.6
0.5
0.4
0.3
0.2
0.1
0.05
0.01
0
Cell_Type_NEW
Ctrl
ExN
InN
Other
UD

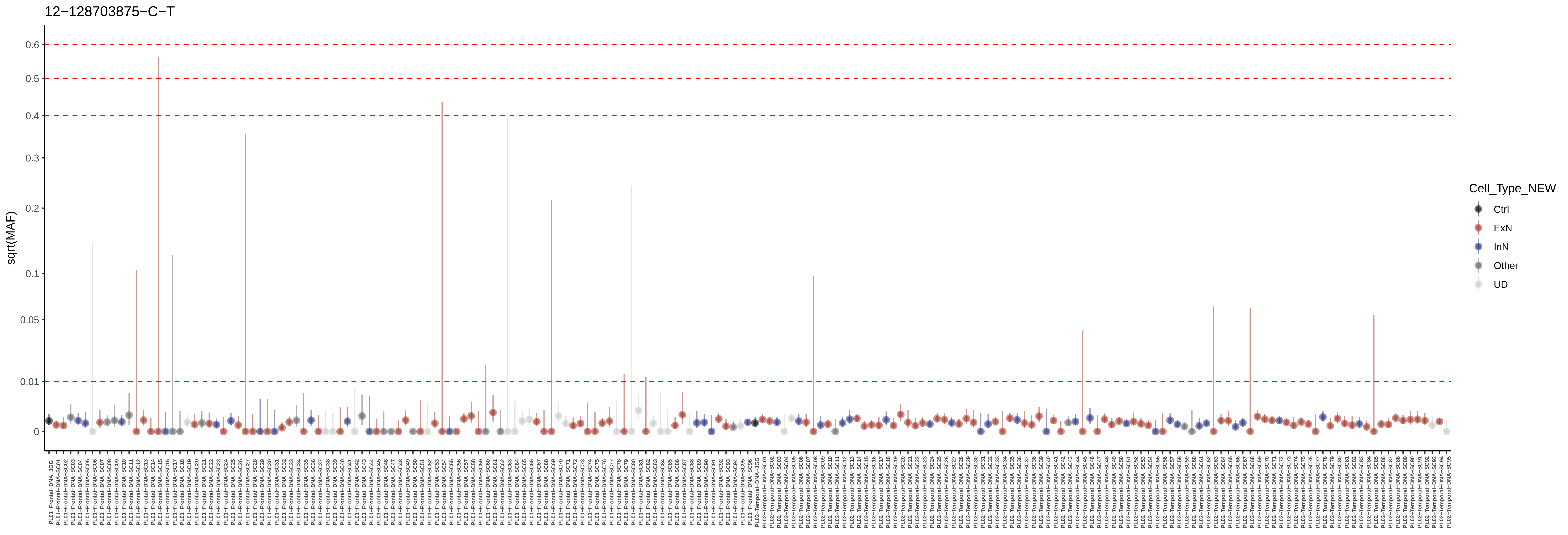
12−128703875−C−T
sqrt(MAF)
0.6
0.5
0.4
0.3
0.2
0.1
0.05
0.01
0
Cell_Type_NEW
Ctrl
ExN
InN
Other
UD

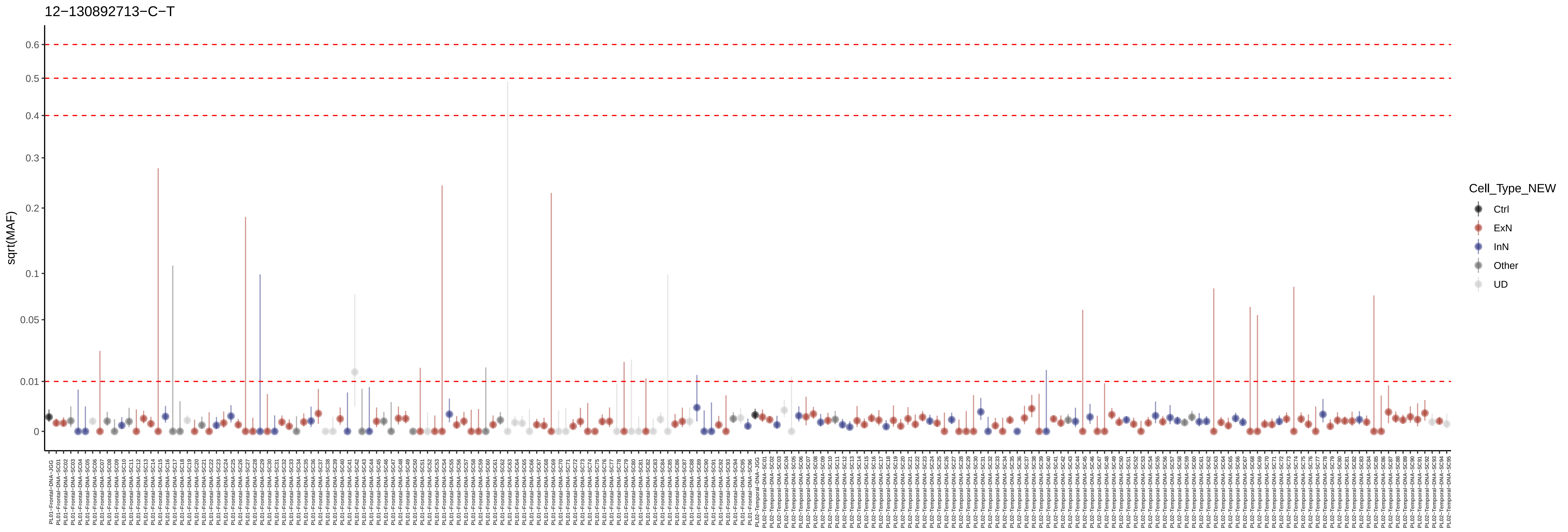

12−130892713−C−T
sqrt(MAF)
0.6
0.5
0.4
0.3
0.2
0.1
0.05
0.01
0
Cell_Type_NEW
Ctrl
ExN
InN
Other
UD

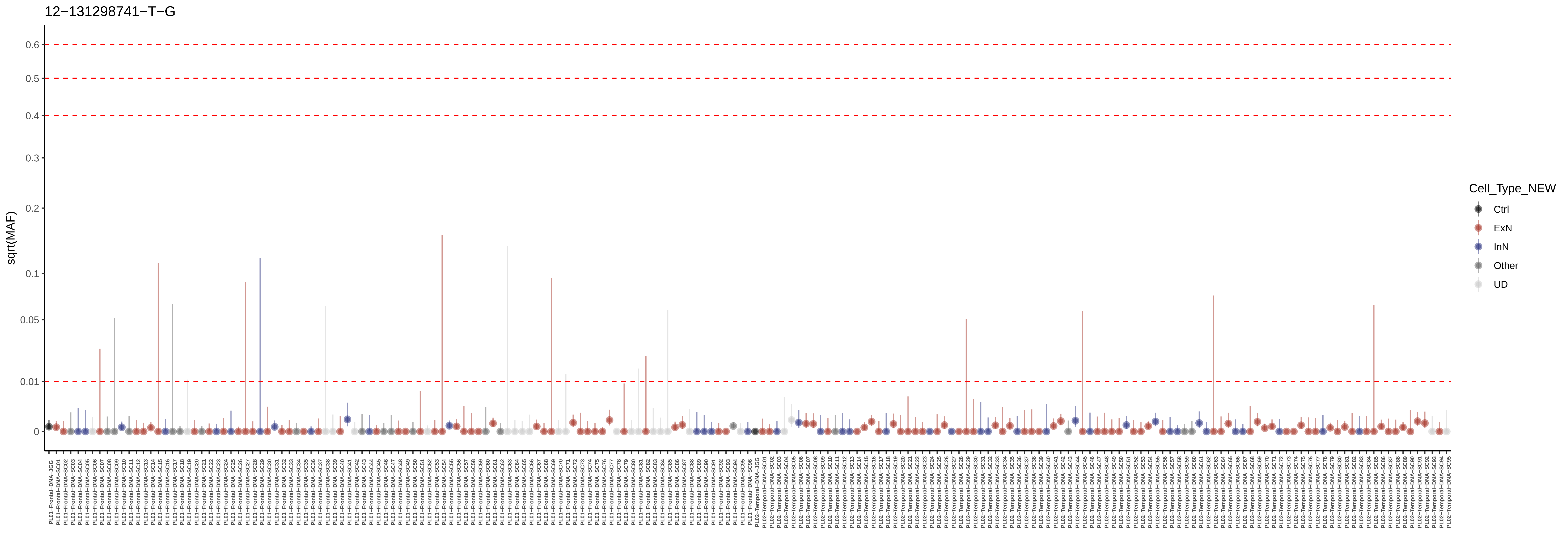
12−131298741−T−G
sqrt(MAF)
0.6
0.5
0.4
0.3
0.2
0.1
0.05
0.01
0
Cell_Type_NEW
Ctrl
ExN
InN
Other
UD

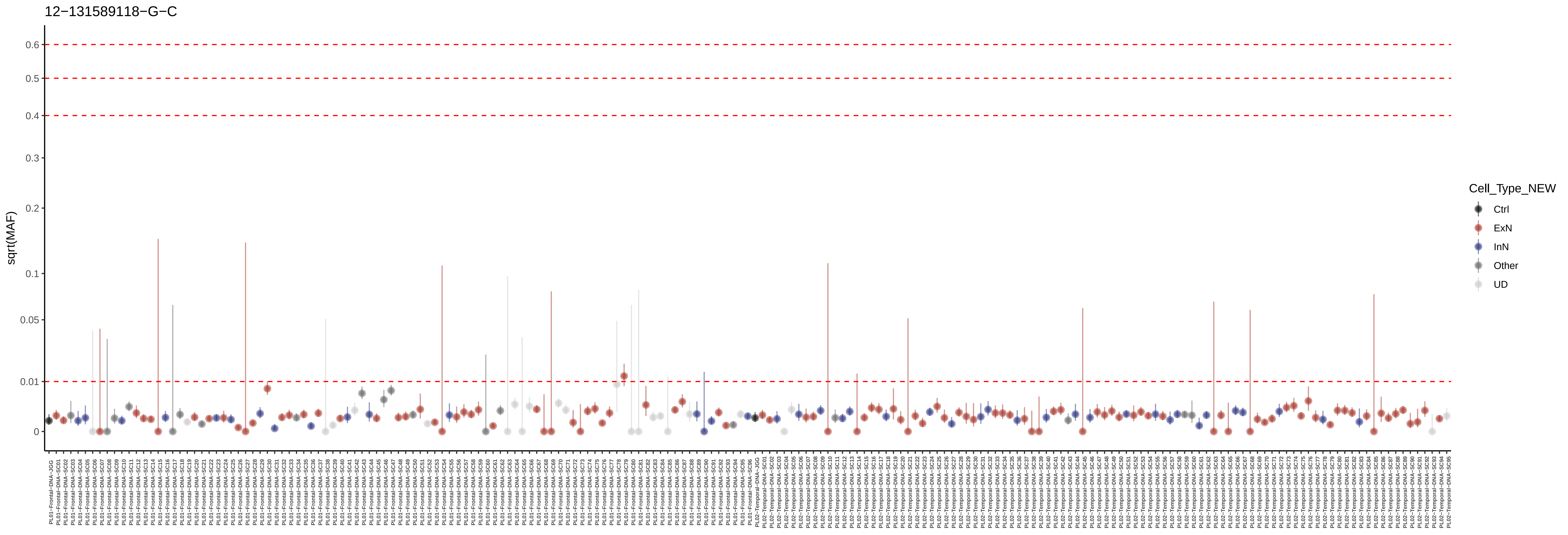

12−131589118−G−C
sqrt(MAF)
0.6
0.5
0.4
0.3
0.2
0.1
0.05
0.01
0
Cell_Type_NEW
Ctrl
ExN
InN
Other
UD

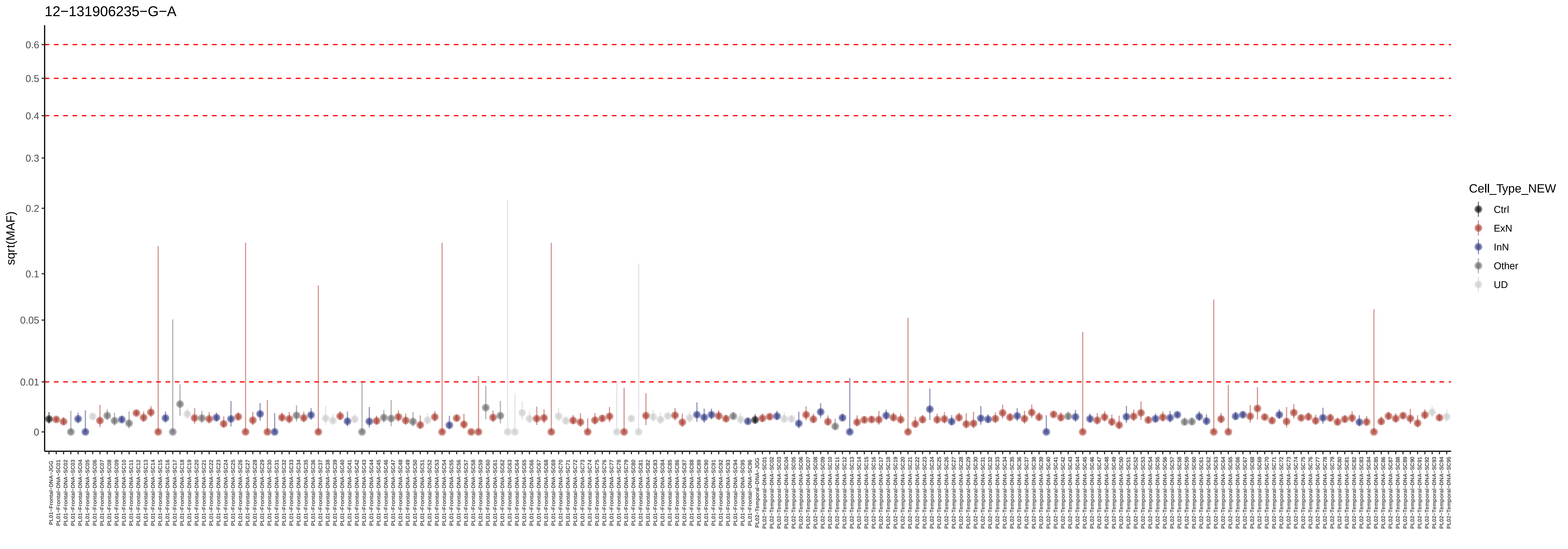
12−131906235−G−A
sqrt(MAF)
0.6
0.5
0.4
0.3
0.2
0.1
0.05
0.01
0
Cell_Type_NEW
Ctrl
ExN
InN
Other
UD

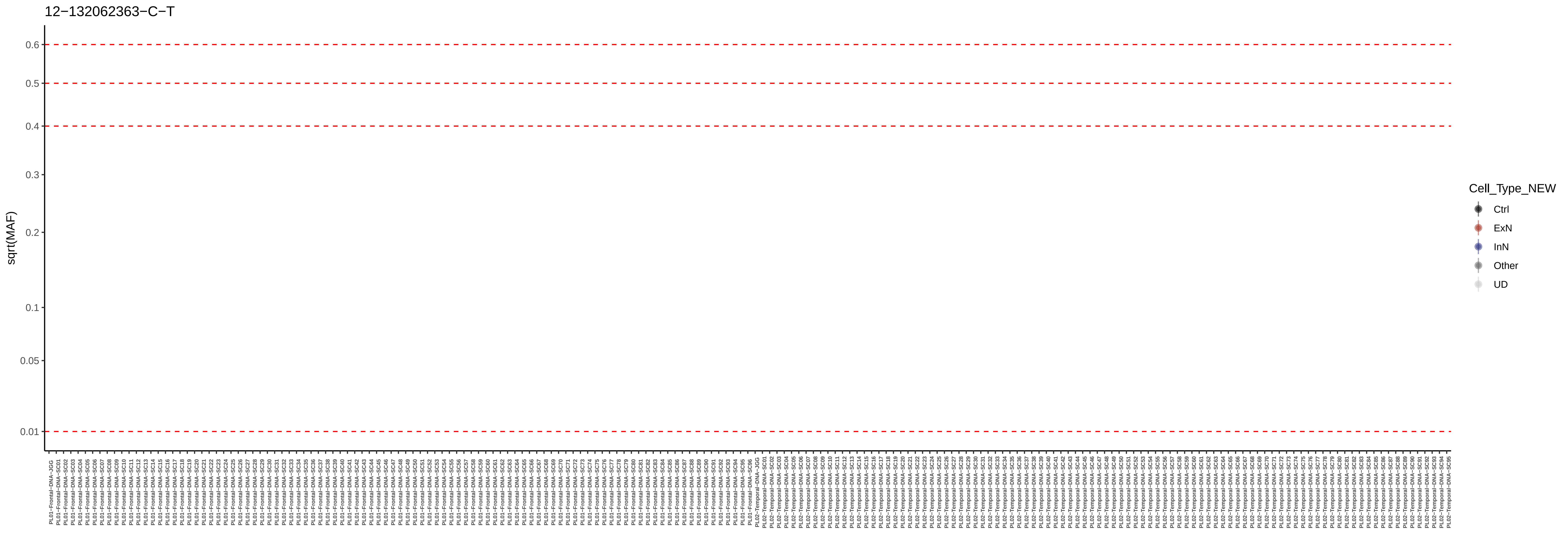

12−132062363−C−T
sqrt(MAF)
Cell_Type_NEW
Ctrl
ExN
InN
Other
UD

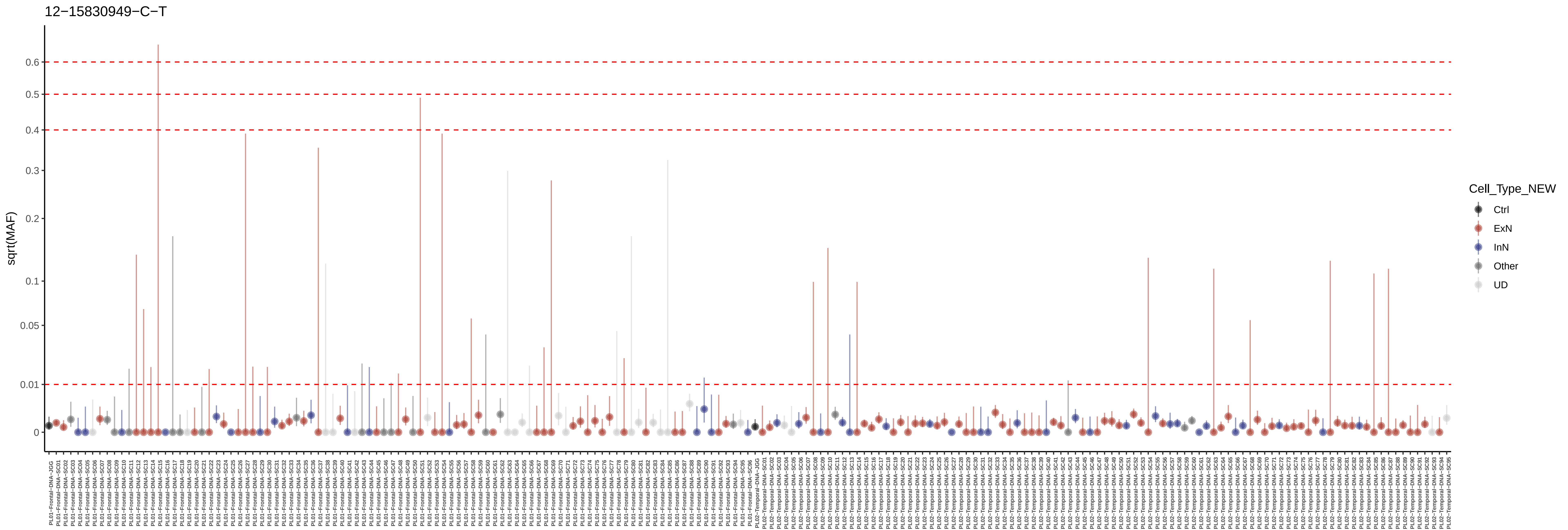

12−15830949−C−T
sqrt(MAF)
0.6
0.5
0.4
0.3
0.2
0.1
0.05
0.01
0
Cell_Type_NEW
Ctrl
ExN
InN
Other
UD

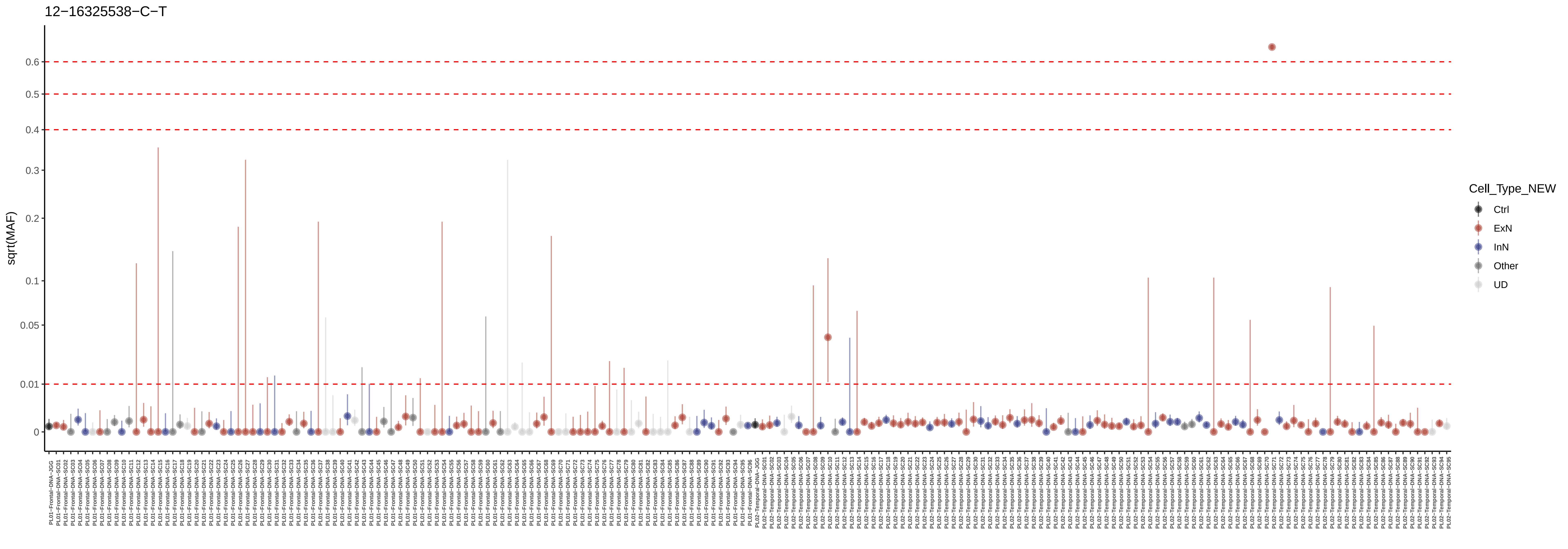

12−16325538−C−T
sqrt(MAF)
Cell_Type_NEW
Ctrl
ExN
InN
Other
UD

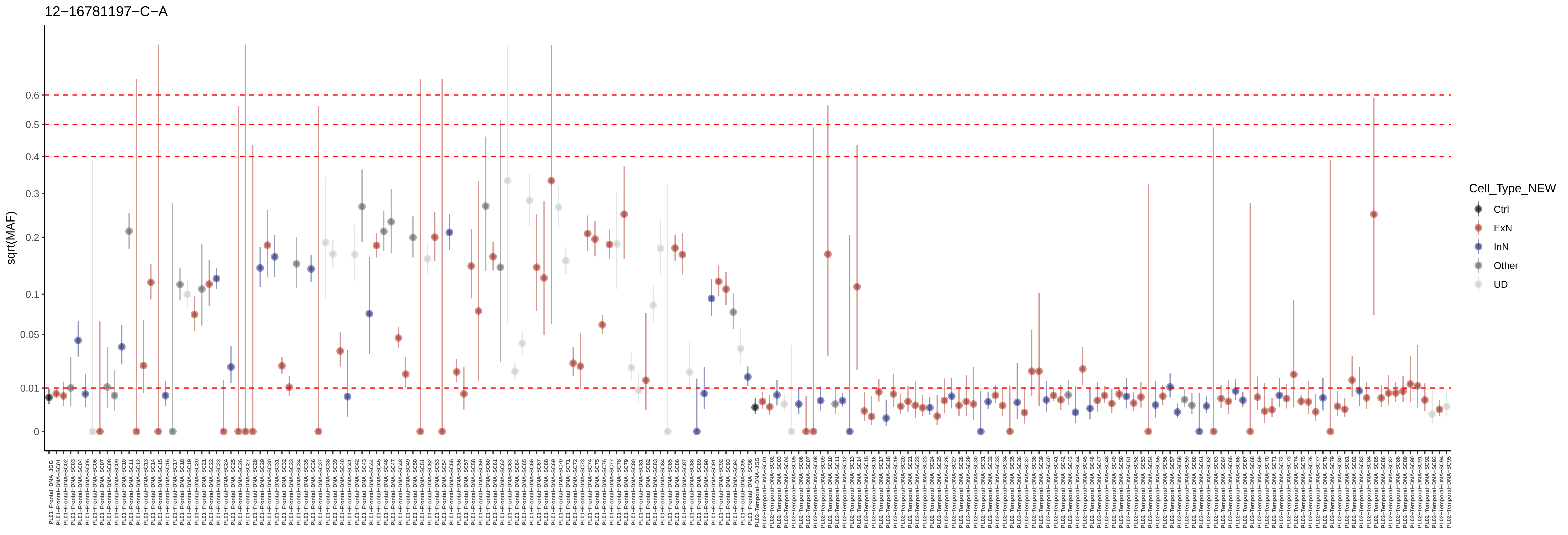

12−16781197−C−A
sqrt(MAF)
0.6
0.5
0.4
0.3
0.2
0.1
0.05
0.01
0
Cell_Type_NEW
Ctrl
ExN
InN
Other
UD

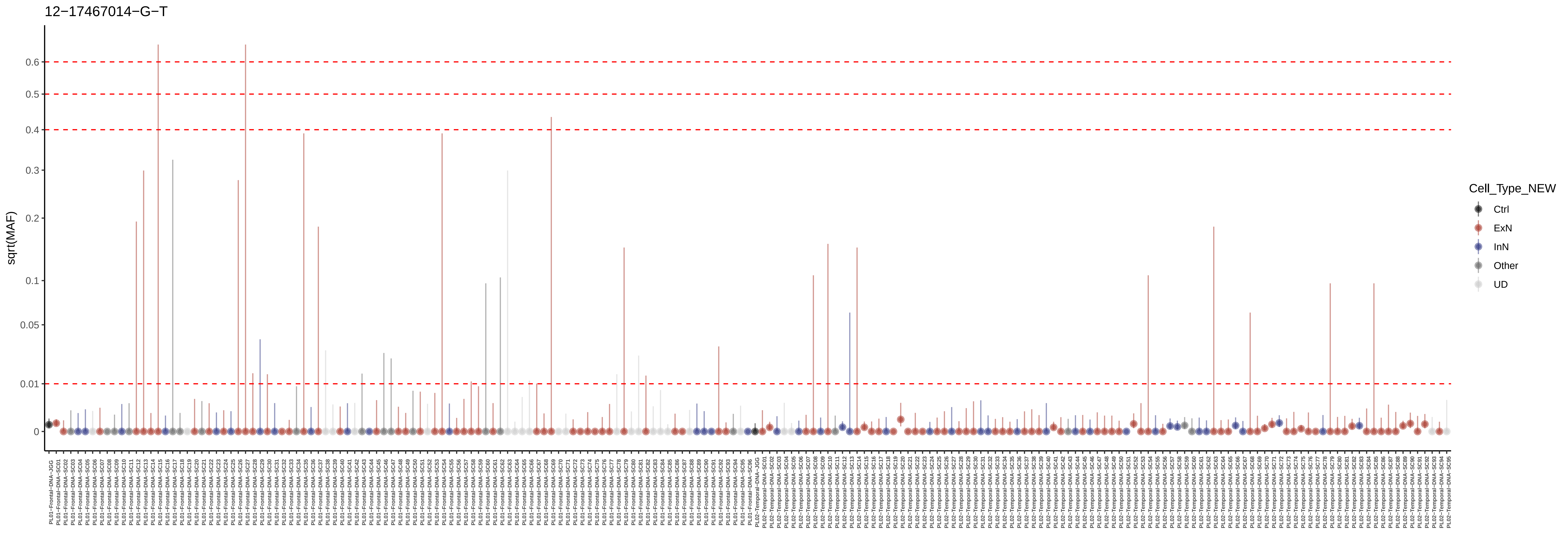
12−17467014−G−T
sqrt(MAF)
0.6
0.5
0.4
0.3
0.2
0.1
0.05
0.01
0
Cell_Type_NEW
Ctrl
ExN
InN
Other
UD

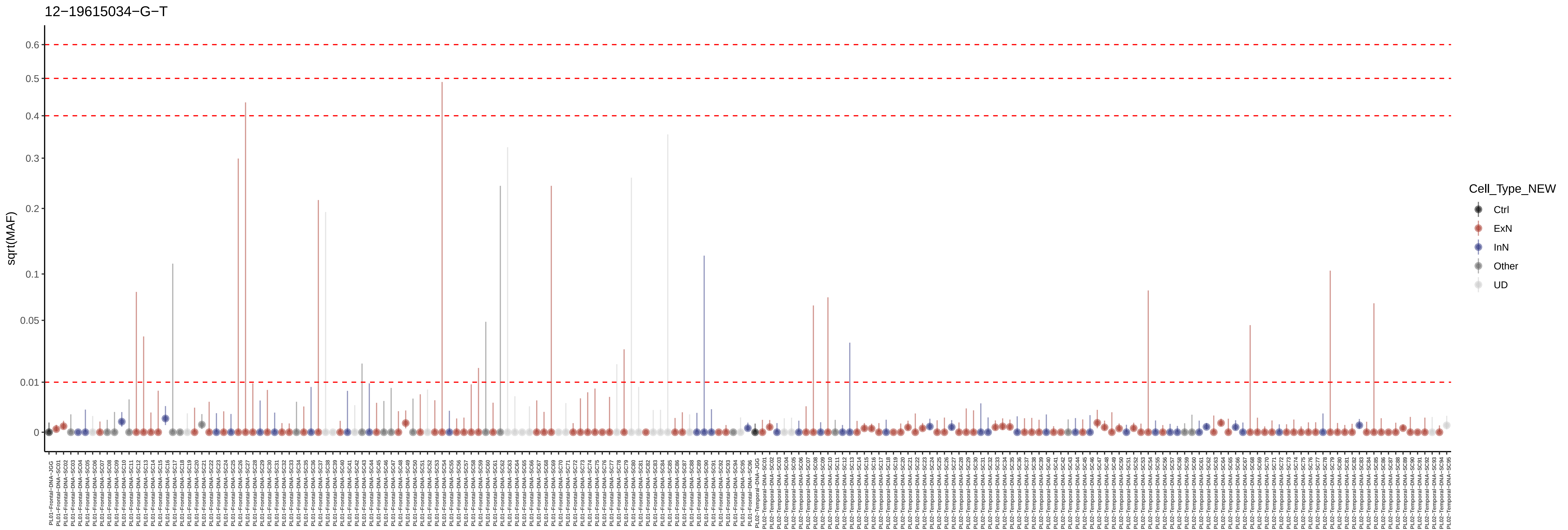

12−19615034−G−T
sqrt(MAF)
0.6
0.5
0.4
0.3
0.2
0.1
0.05
0.01
0
Cell_Type_NEW
Ctrl
ExN
InN
Other
UD

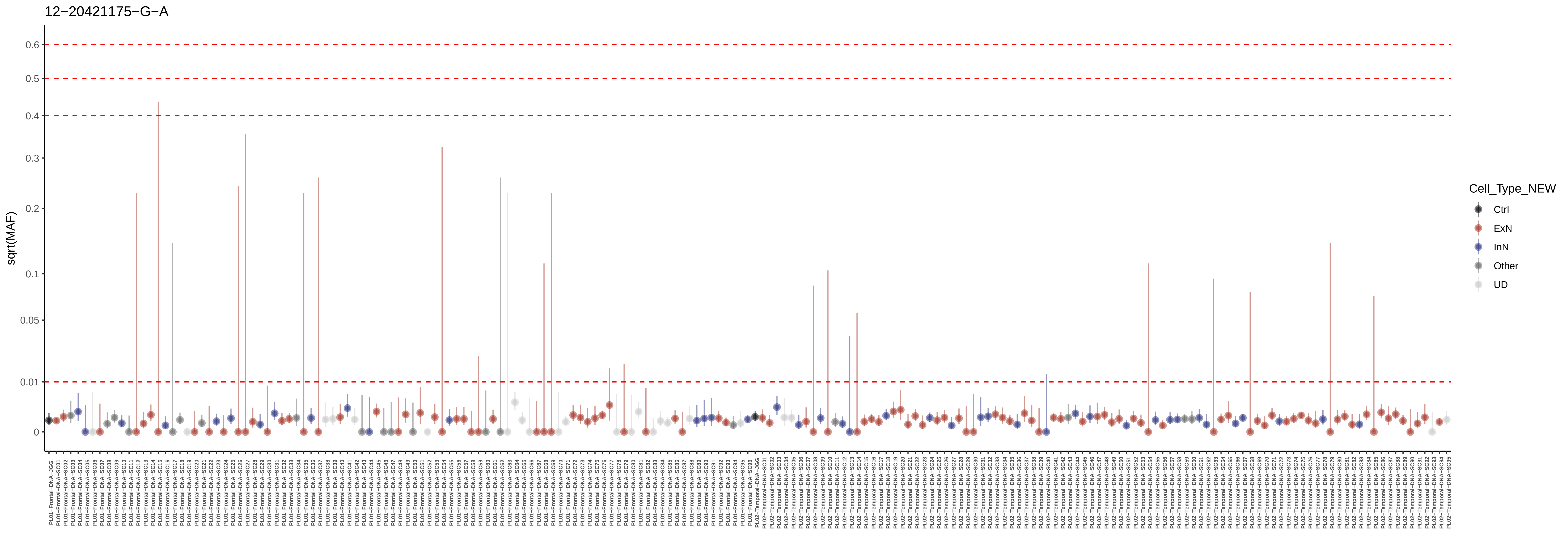

12−20421175−G−A
sqrt(MAF)
0.6
0.5
0.4
0.3
0.2
0.1
0.05
0.01
0
Cell_Type_NEW
Ctrl
ExN
InN
Other
UD

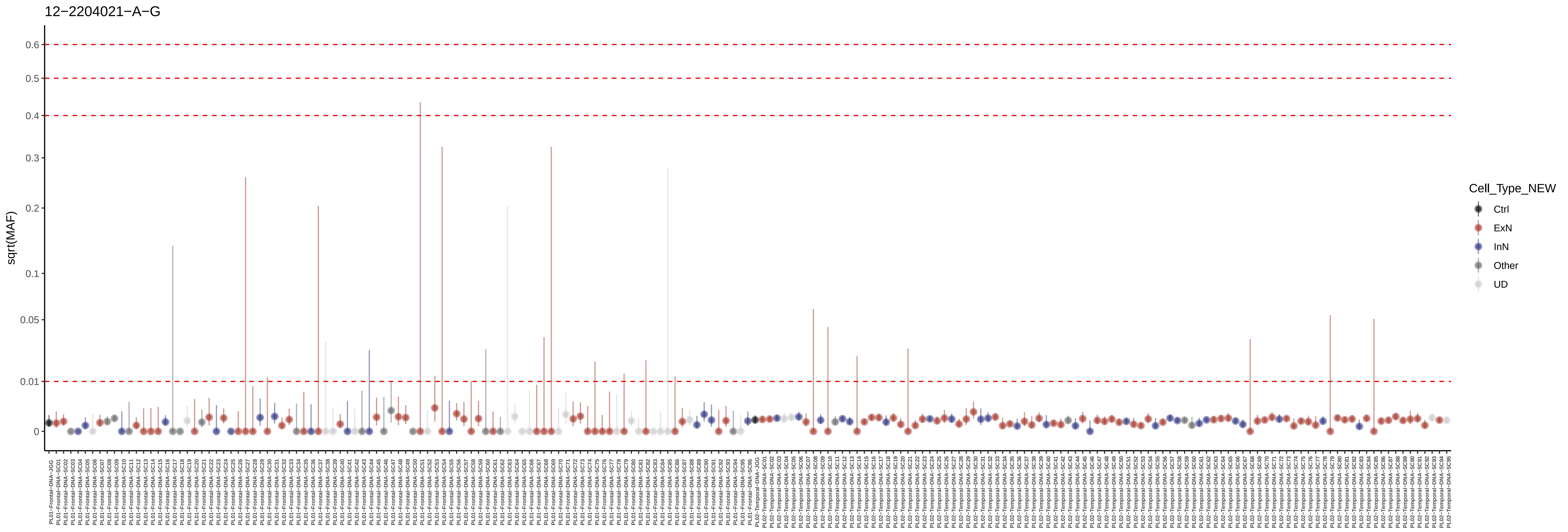
12−2204021−A−G
sqrt(MAF)
0.6
0.5
0.4
0.3
0.2
0.1
0.05
0.01
0
Cell_Type_NEW
Ctrl
ExN
InN
Other
UD

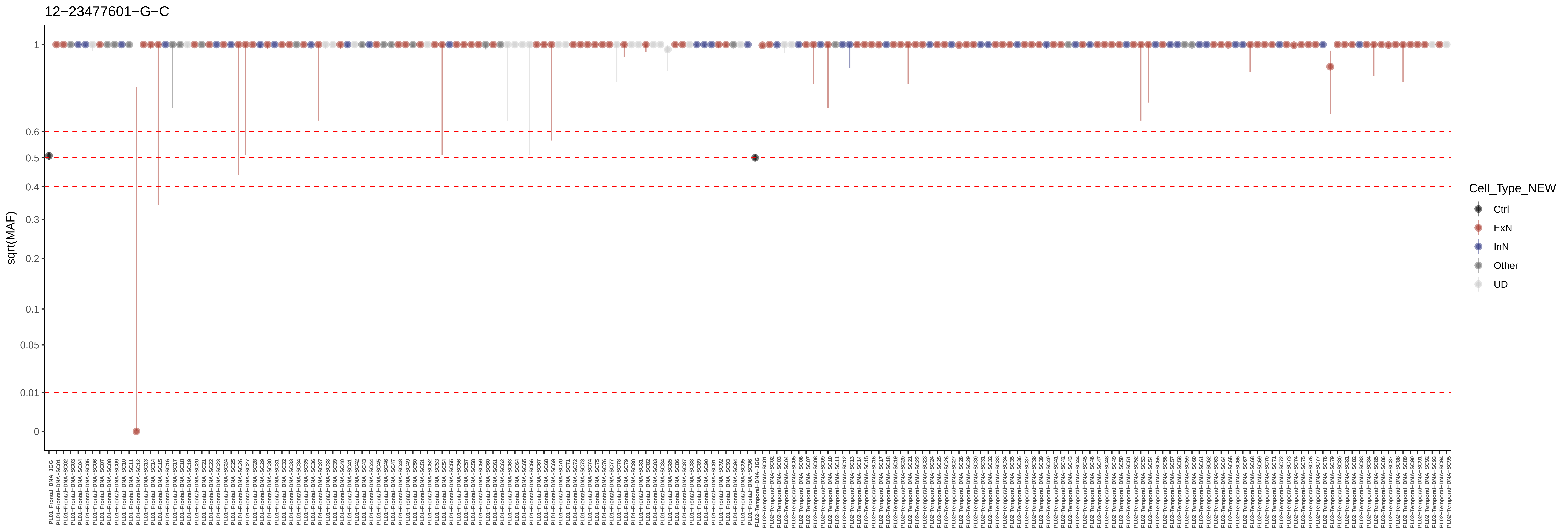
12−23477601−G−C
sqrt(MAF)
1
0.6
0.5
0.4
0.3
0.2
0.1
0.05
0.01
0
Cell_Type_NEW
Ctrl
ExN
InN
Other
UD

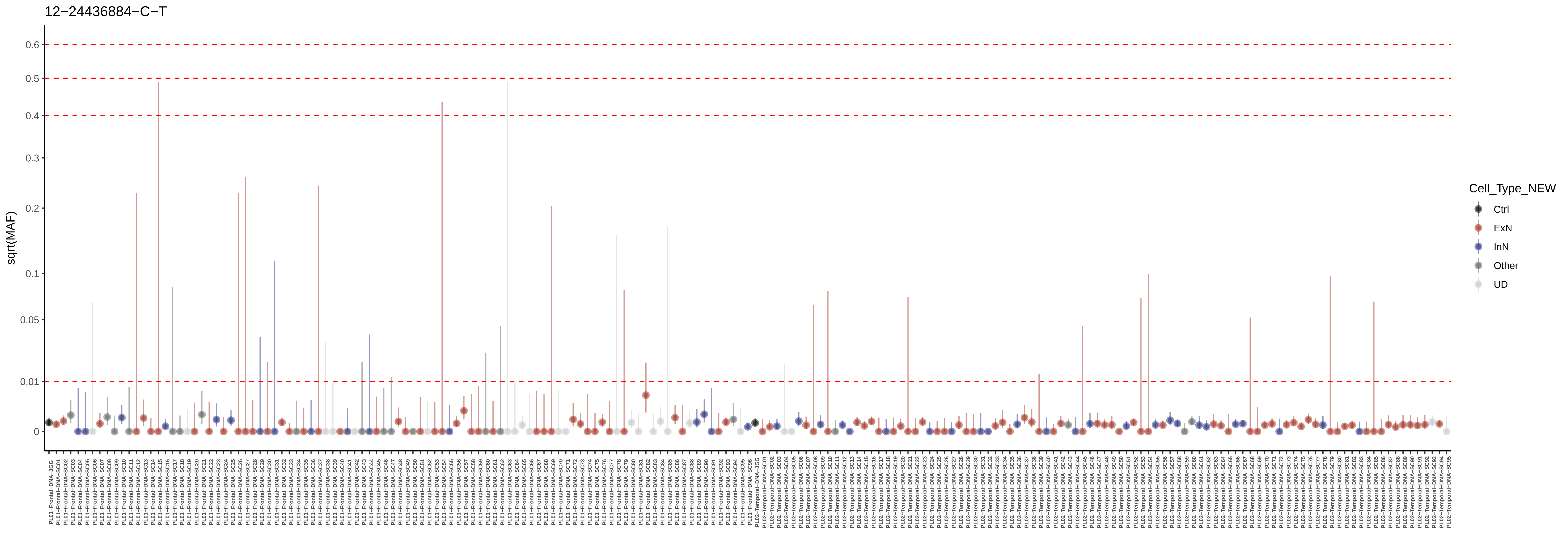
12−24436884−C−T
sqrt(MAF)
0.6
0.5
0.4
0.3
0.2
0.1
0.05
0.01
0
Cell_Type_NEW
Ctrl
ExN
InN
Other
UD

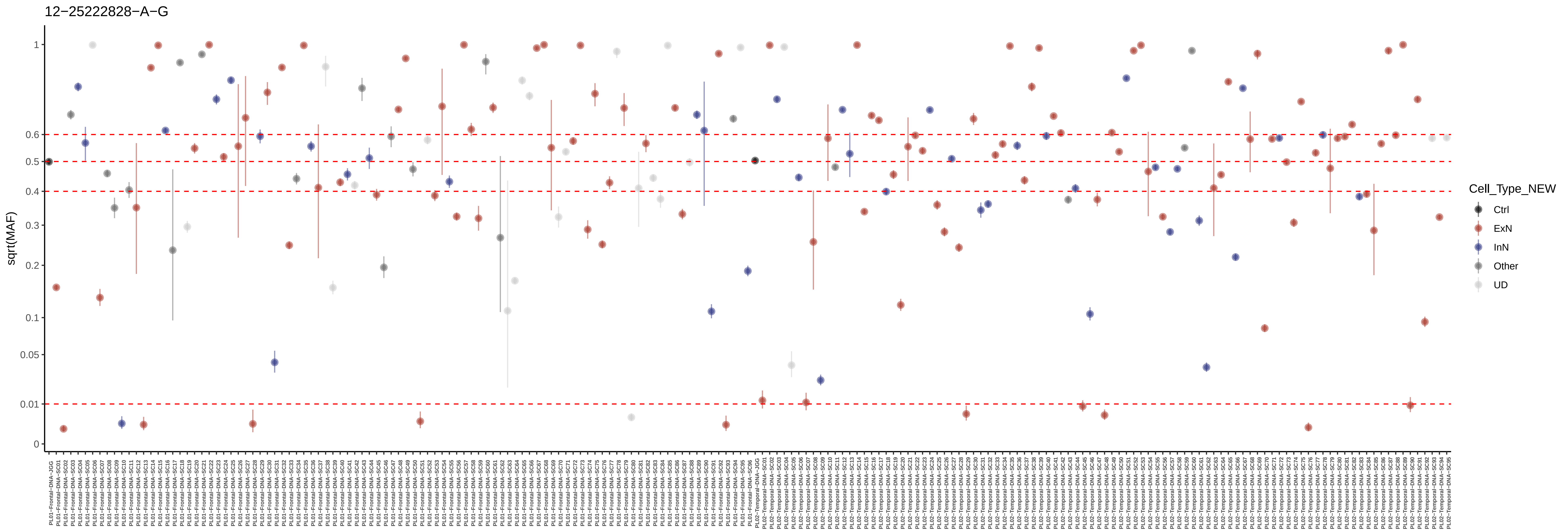

12−25222828−A−G
sqrt(MAF)
1
0.6
0.5
0.4
0.3
0.2
0.1
0.05
0.01
0
Cell_Type_NEW
Ctrl
ExN
InN
Other
UD

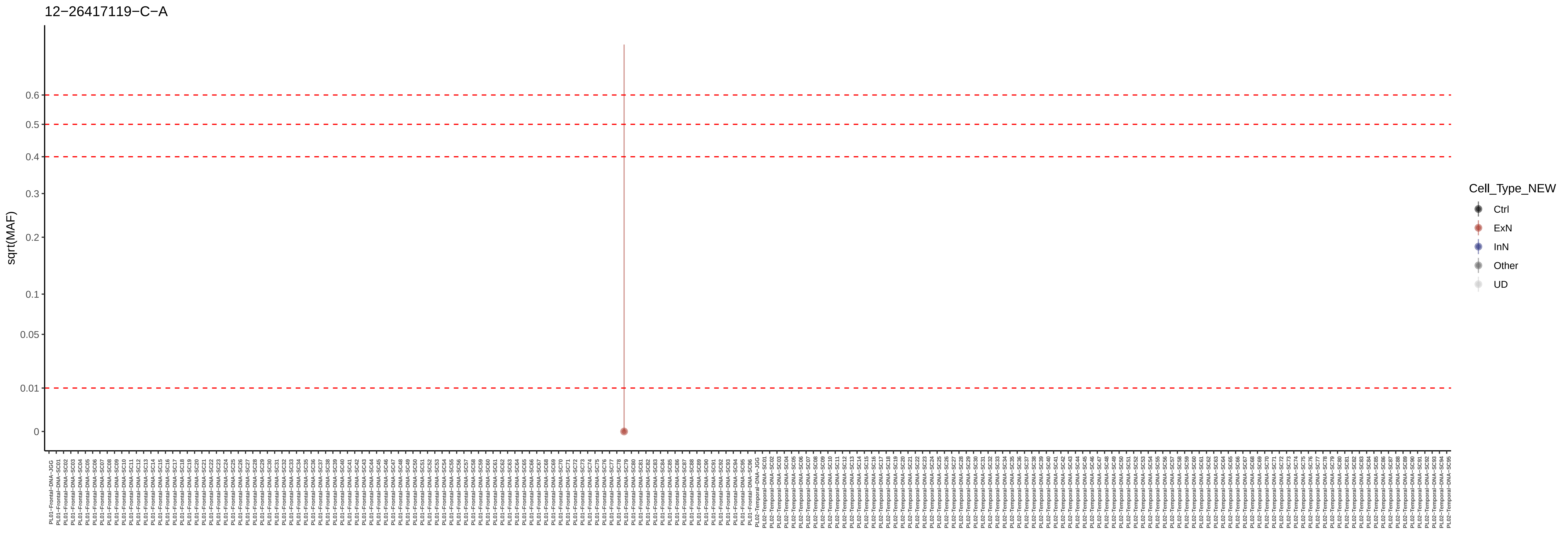
12−26417119−C−A
sqrt(MAF)
0.6
0.5
0.4
0.3
0.2
0.1
0.05
0.01
0
Cell_Type_NEW
Ctrl
ExN
InN
Other
UD

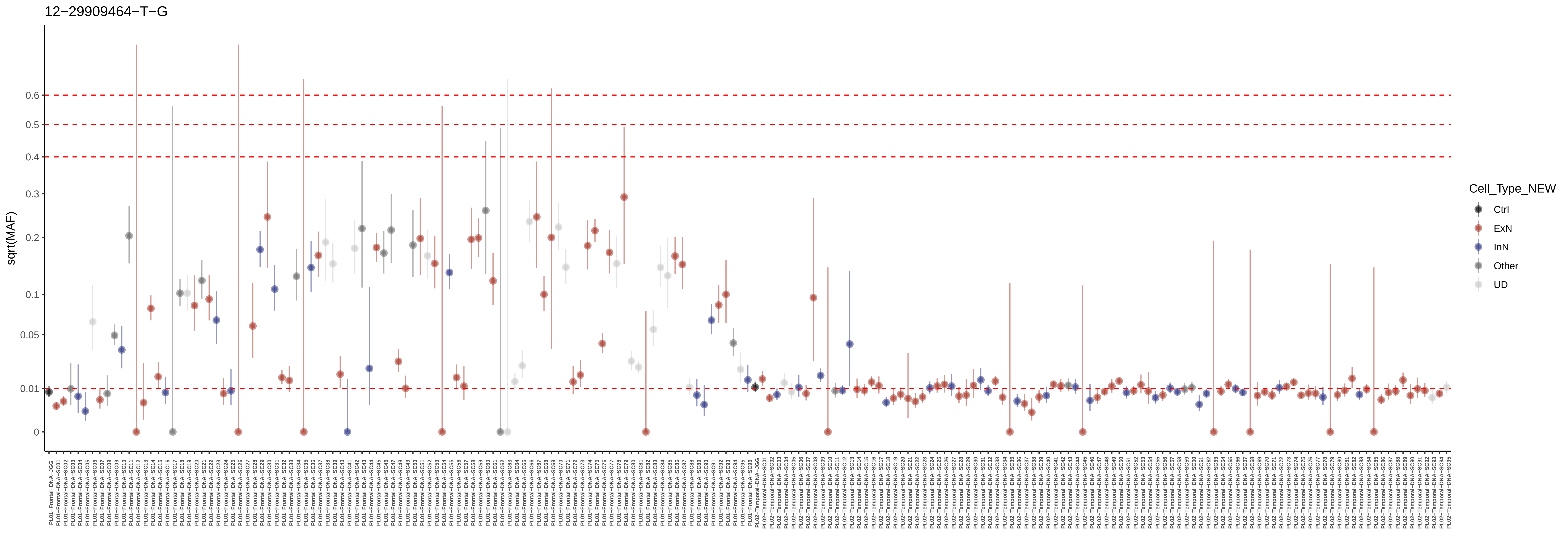

12−29909464−T−G
sqrt(MAF)
0.6
0.5
0.4
0.3
0.2
0.1
0.05
0.01
0
Cell_Type_NEW
Ctrl
ExN
InN
Other
UD

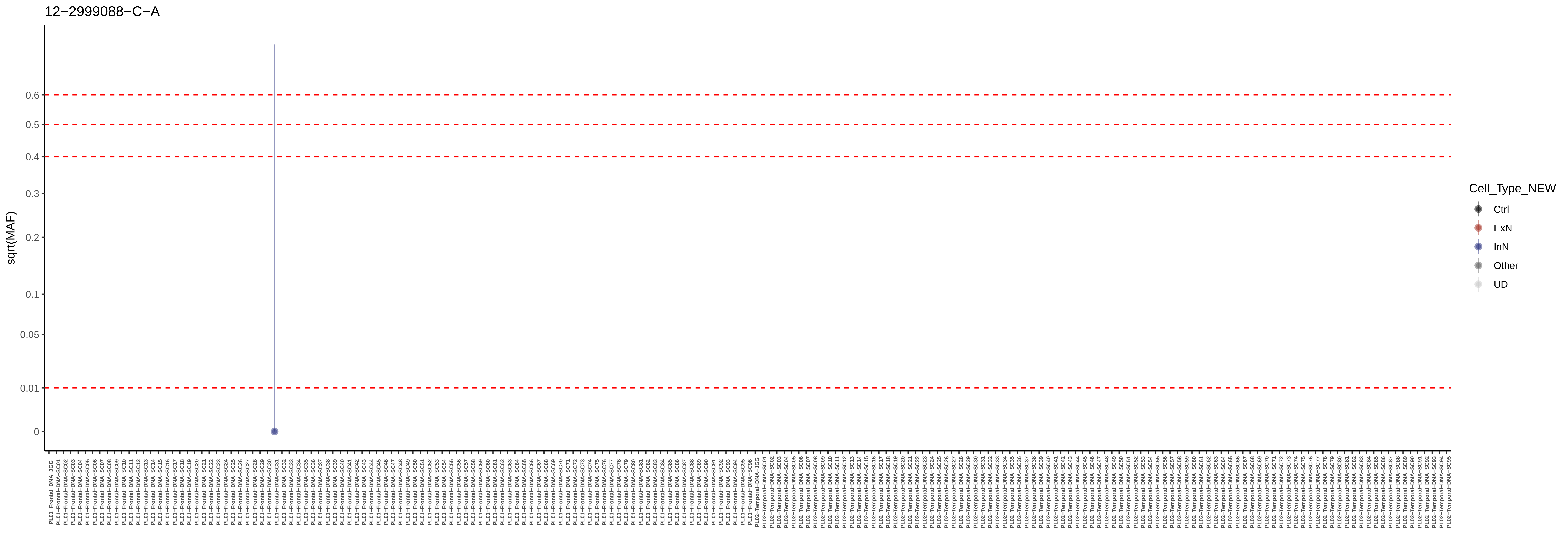
12−2999088−C−A
sqrt(MAF)
0.6
0.5
0.4
0.3
0.2
0.1
0.05
0.01
0
Cell_Type_NEW
Ctrl
ExN
InN
Other
UD

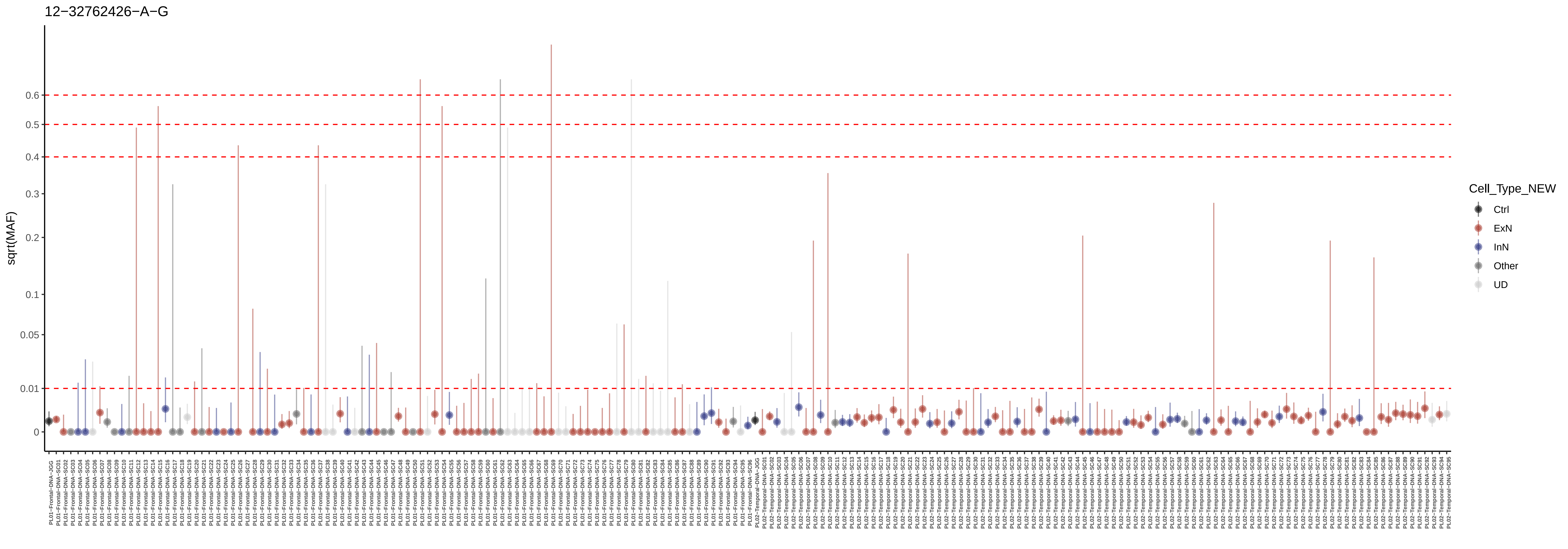

12−32762426−A−G
sqrt(MAF)
Cell_Type_NEW
Ctrl
ExN
InN
Other
UD

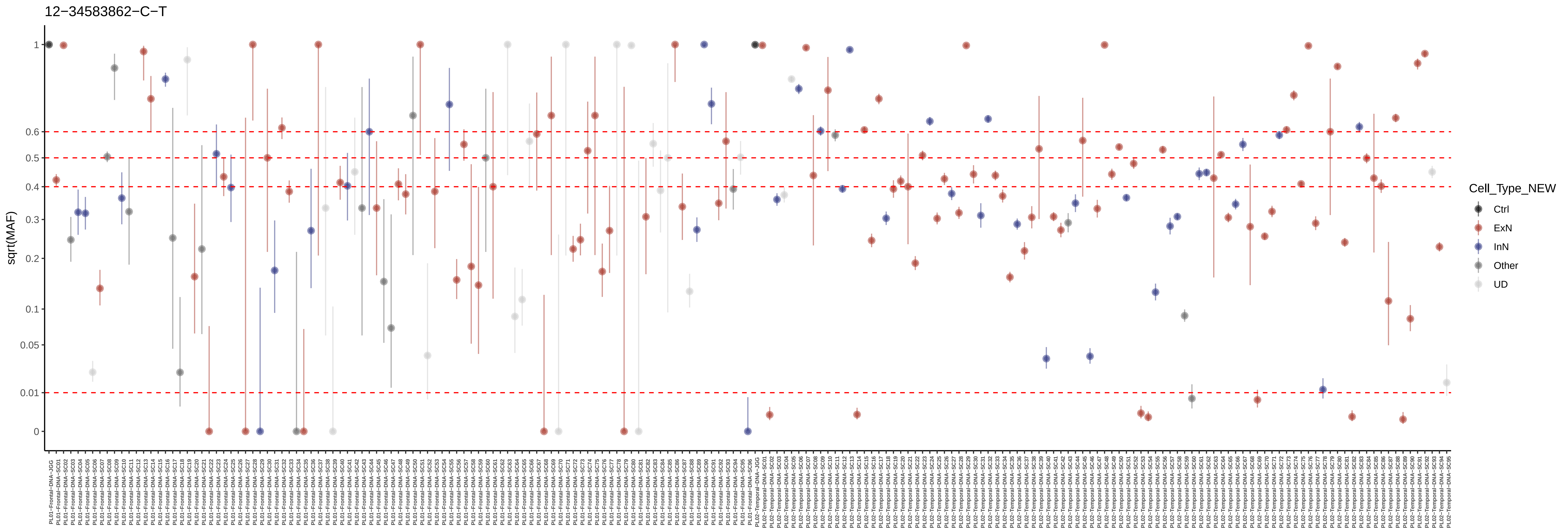

12−34583862−C−T
sqrt(MAF)
1
0.6
0.5
0.4
0.3
0.2
0.1
0.05
0.01
0
Cell_Type_NEW
Ctrl
ExN
InN
Other
UD

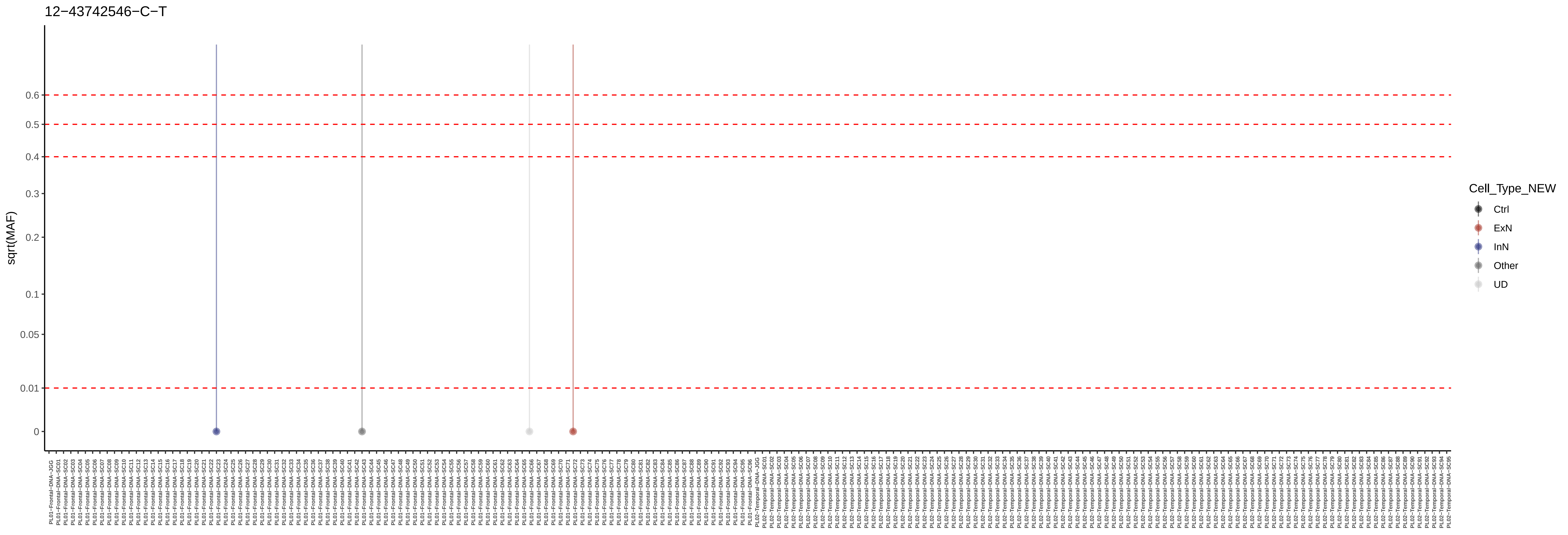

12−43742546−C−T
sqrt(MAF)
0.6
0.5
0.4
0.3
0.2
0.1
0.05
0.01
0
Cell_Type_NEW
Ctrl
ExN
InN
Other
UD

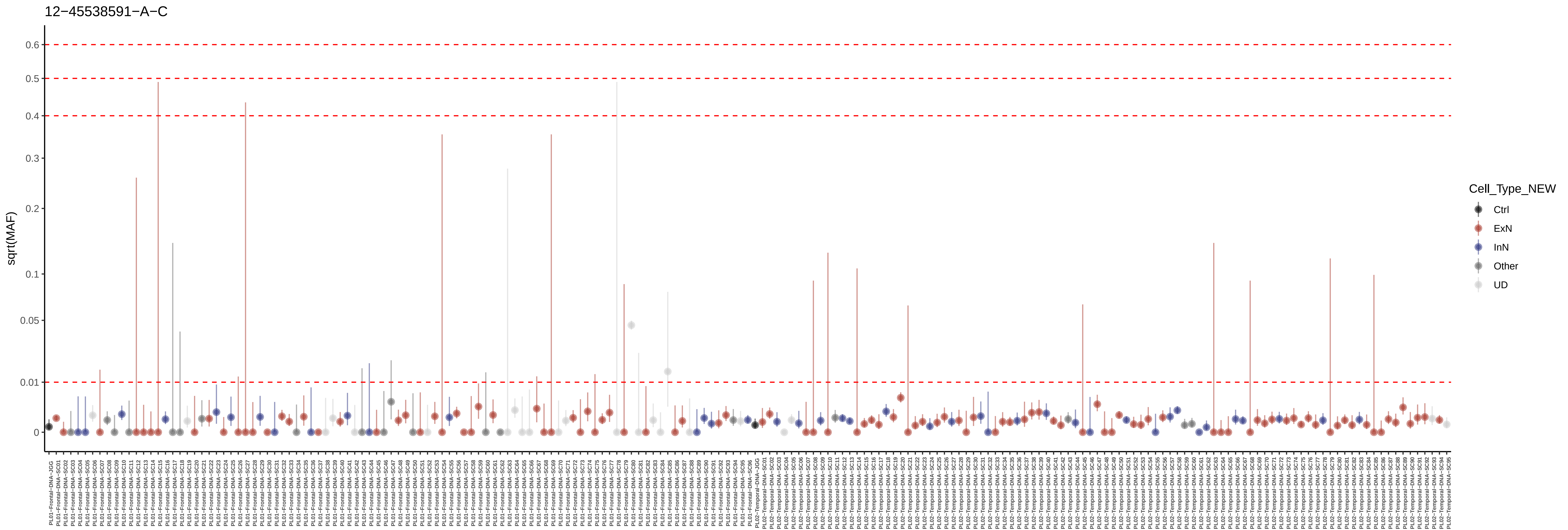
12−45538591−A−C
sqrt(MAF)
0.6
0.5
0.4
0.3
0.2
0.1
0.05
0.01
0
Cell_Type_NEW
Ctrl
ExN
InN
Other
UD

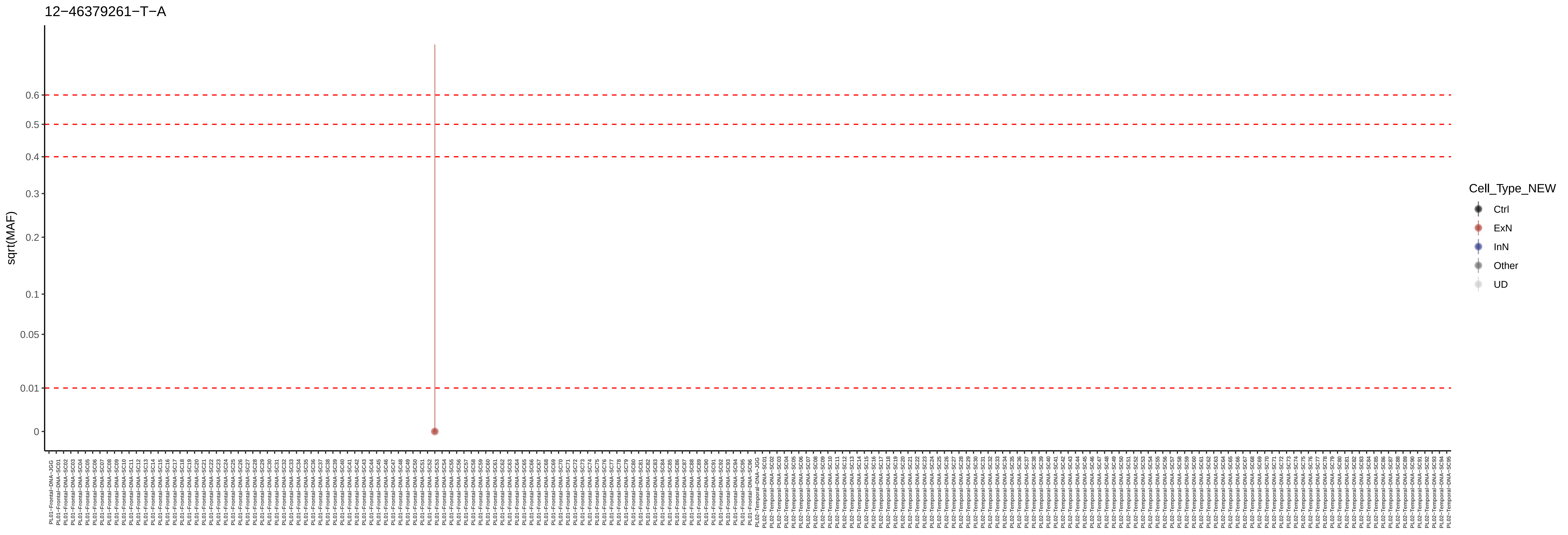
12−46379261−T−A
sqrt(MAF)
0.6
0.5
0.4
0.3
0.2
0.1
0.05
0.01
0
Cell_Type_NEW
Ctrl
ExN
InN
Other
UD

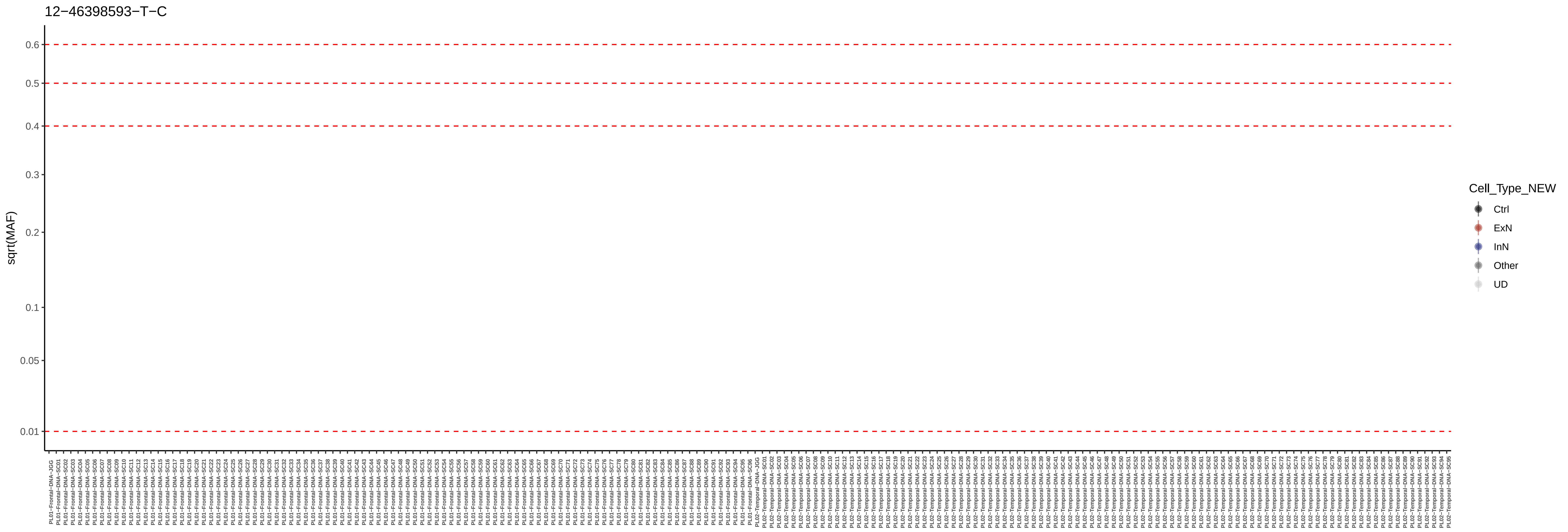
12−46398593−T−C
sqrt(MAF)
0.6
0.5
0.4
0.3
0.2
0.1
0.05
0.01
Cell_Type_NEW
Ctrl
ExN
InN
Other
UD

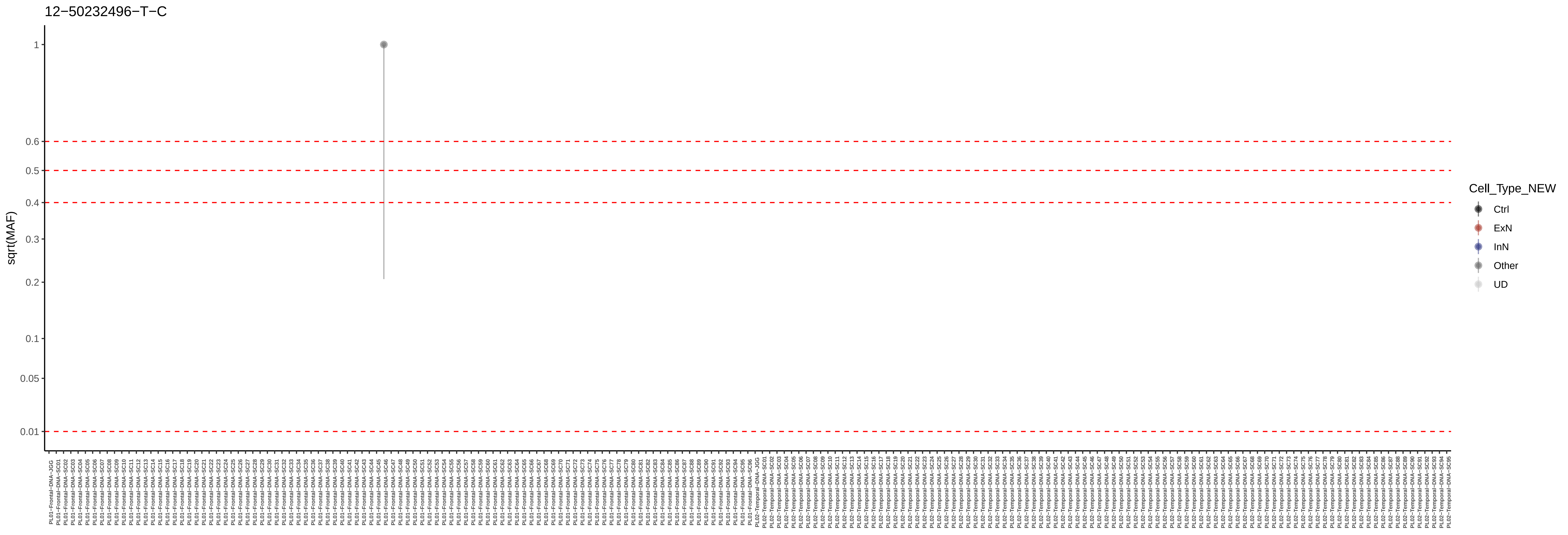
12−50232496−T−C
sqrt(MAF)
1
0.6
0.5
0.4
0.3
0.2
0.1
0.05
0.01
Cell_Type_NEW
Ctrl
ExN
InN
Other
UD

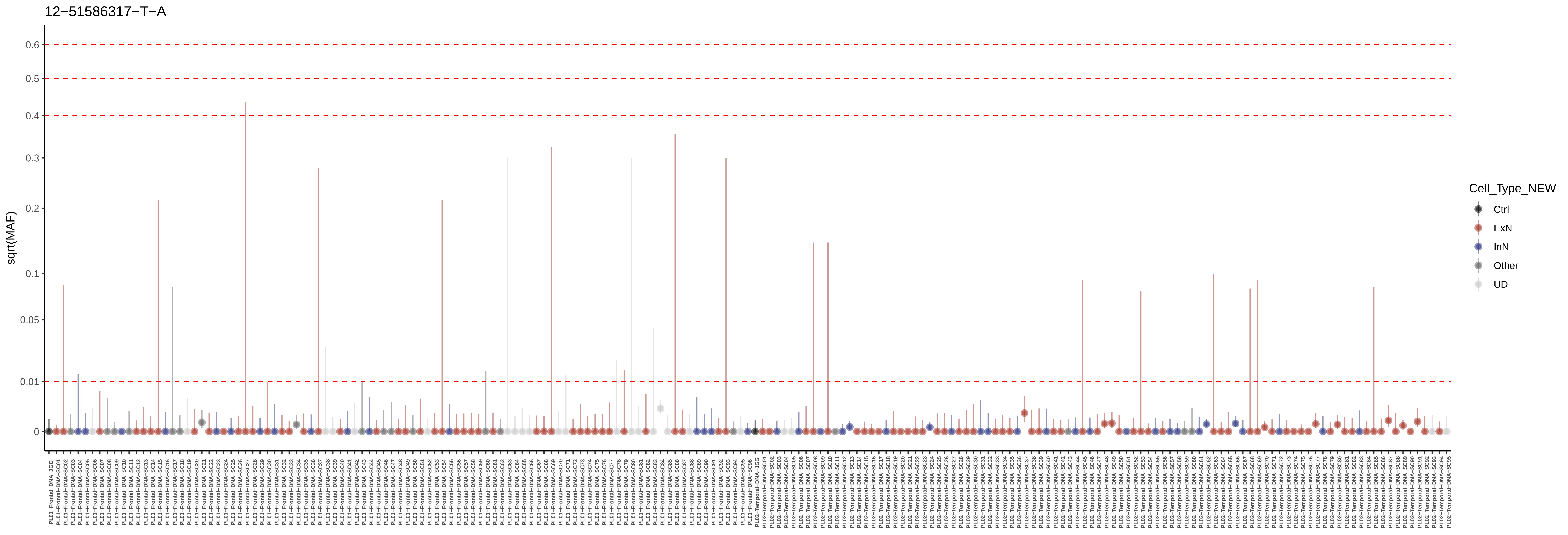

12−51586317−T−A
sqrt(MAF)
0.6
0.5
0.4
0.3
0.2
0.1
0.05
0.01
0
Cell_Type_NEW
Ctrl
ExN
InN
Other
UD

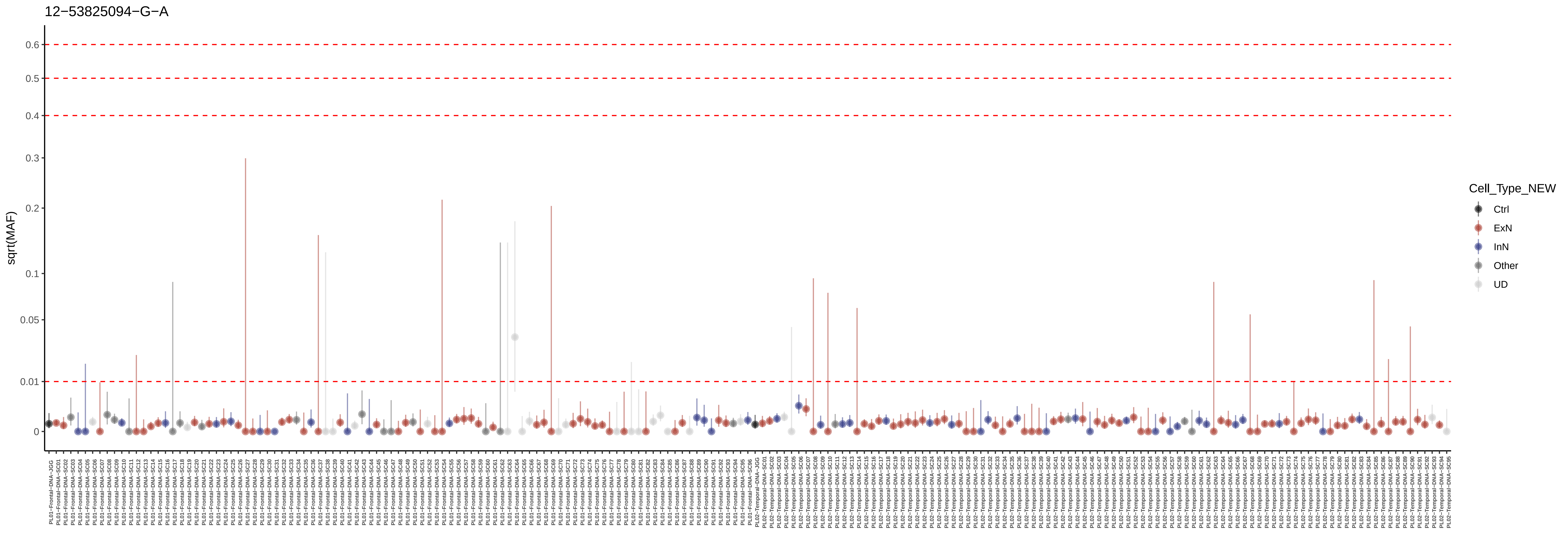
12−53825094−G−A
sqrt(MAF)
0.6
0.5
0.4
0.3
0.2
0.1
0.05
0.01
0
Cell_Type_NEW
Ctrl
ExN
InN
Other
UD

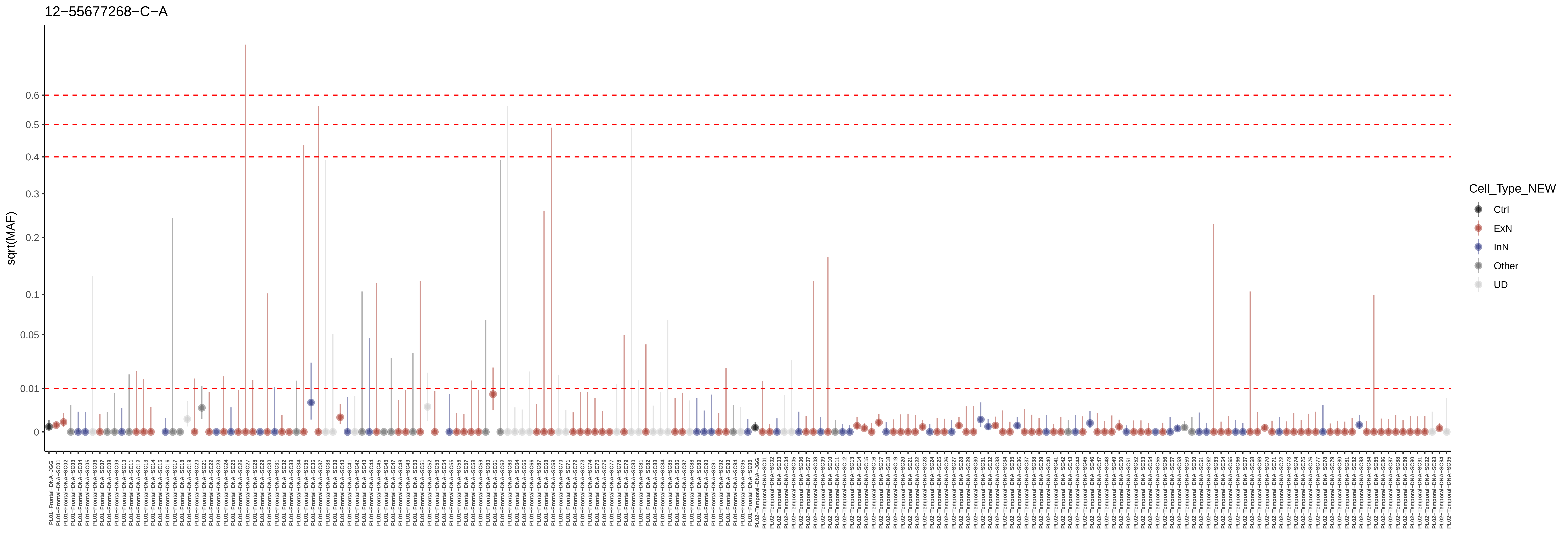
12−55677268−C−A
sqrt(MAF)
0.6
0.5
0.4
0.3
0.2
0.1
0.05
0.01
0
Cell_Type_NEW
Ctrl
ExN
InN
Other
UD

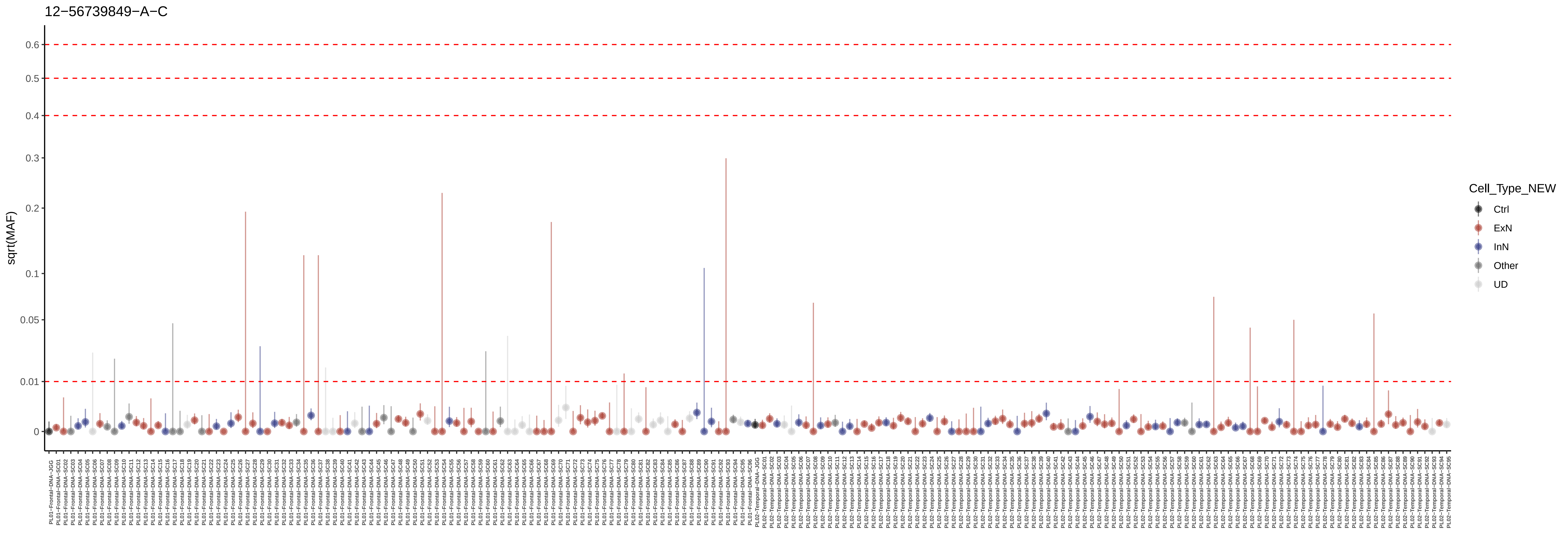

12−56739849−A−C
sqrt(MAF)
0.6
0.5
0.4
0.3
0.2
0.1
0.05
0.01
0
Cell_Type_NEW
Ctrl
ExN
InN
Other
UD

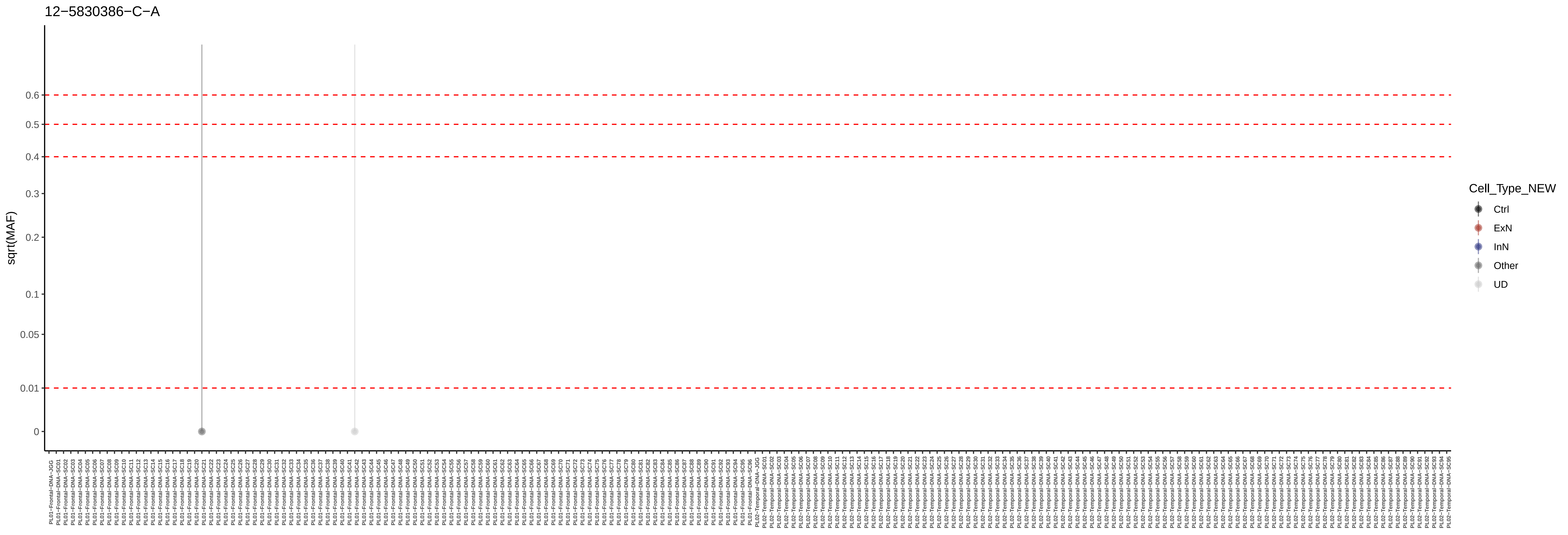
12−5830386−C−A
sqrt(MAF)
0.6
0.5
0.4
0.3
0.2
0.1
0.05
0.01
0
Cell_Type_NEW
Ctrl
ExN
InN
Other
UD

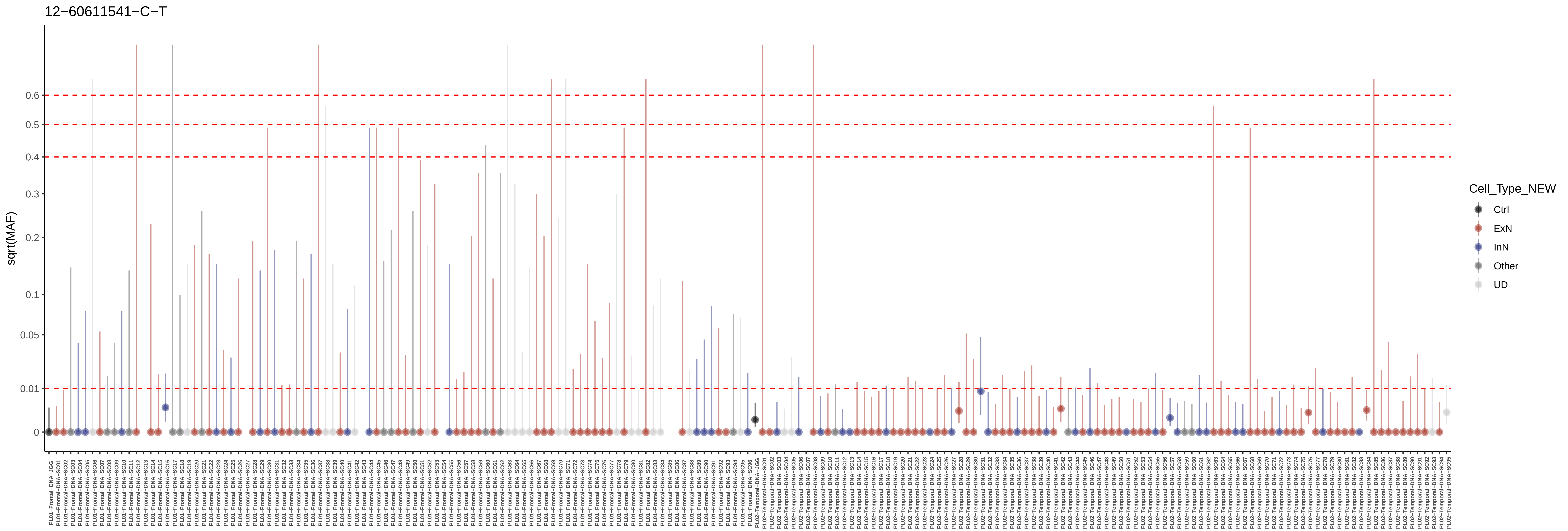
12−60611541−C−T
sqrt(MAF)
0.6
0.5
0.4
0.3
0.2
0.1
0.05
0.01
0
Cell_Type_NEW
Ctrl
ExN
InN
Other
UD

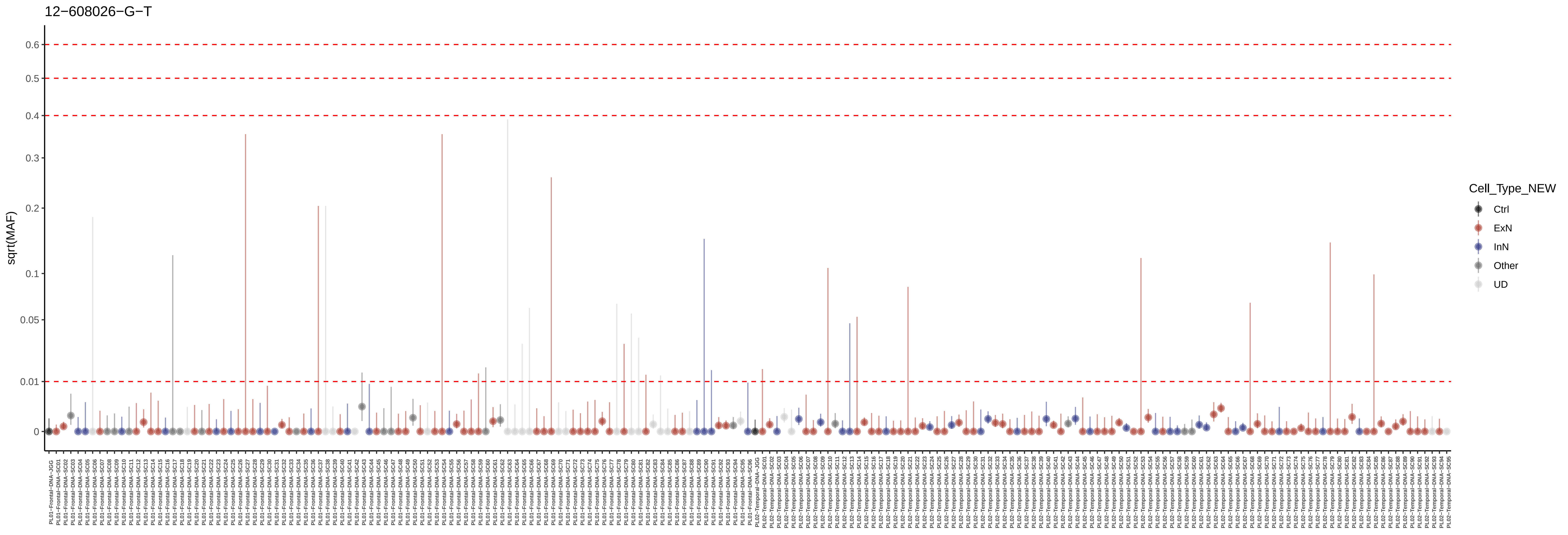

12−608026−G−T
sqrt(MAF)
0.6
0.5
0.4
0.3
0.2
0.1
0.05
0.01
0
Cell_Type_NEW
Ctrl
ExN
InN
Other
UD

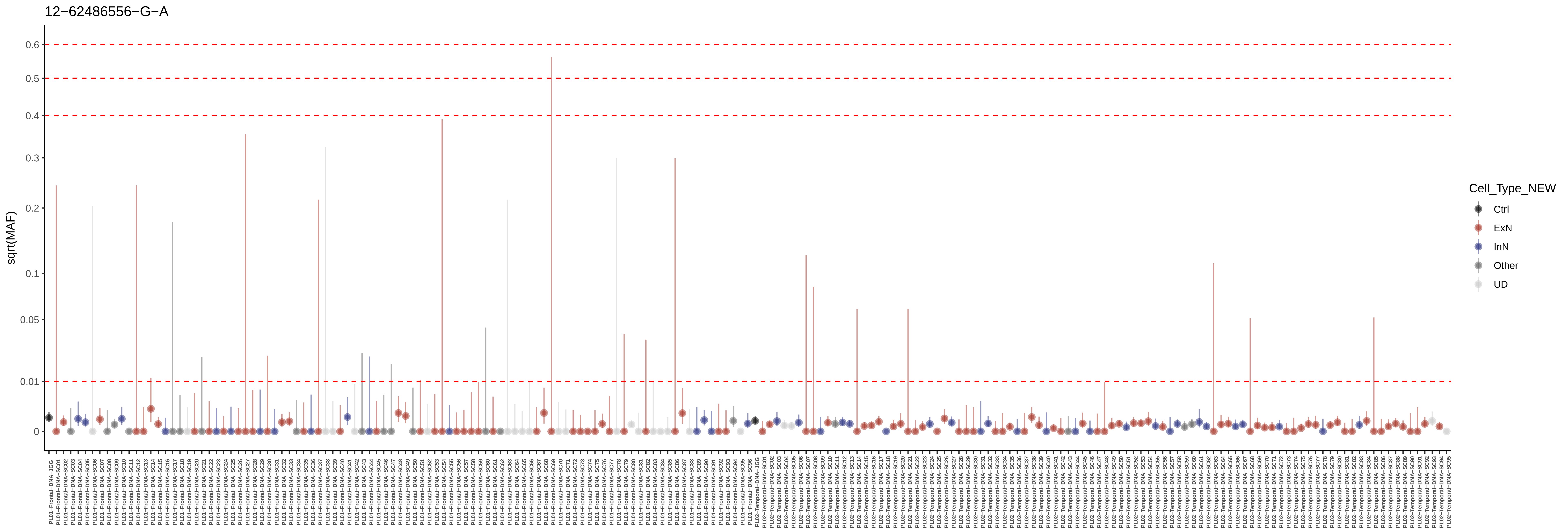
12−62486556−G−A
sqrt(MAF)
0.6
0.5
0.4
0.3
0.2
0.1
0.05
0.01
0
Cell_Type_NEW
Ctrl
ExN
InN
Other
UD

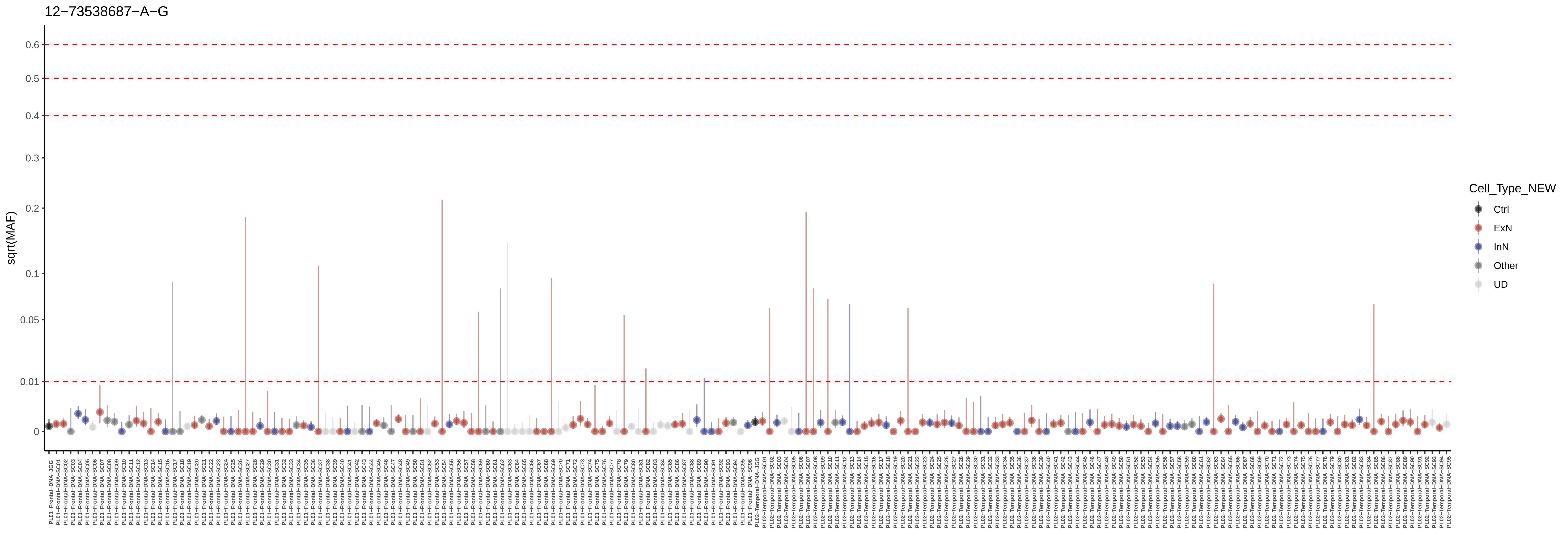
12−73538687−A−G
sqrt(MAF)
0.6
0.5
0.4
0.3
0.2
0.1
0.05
0.01
0
Cell_Type_NEW
Ctrl
ExN
InN
Other
UD

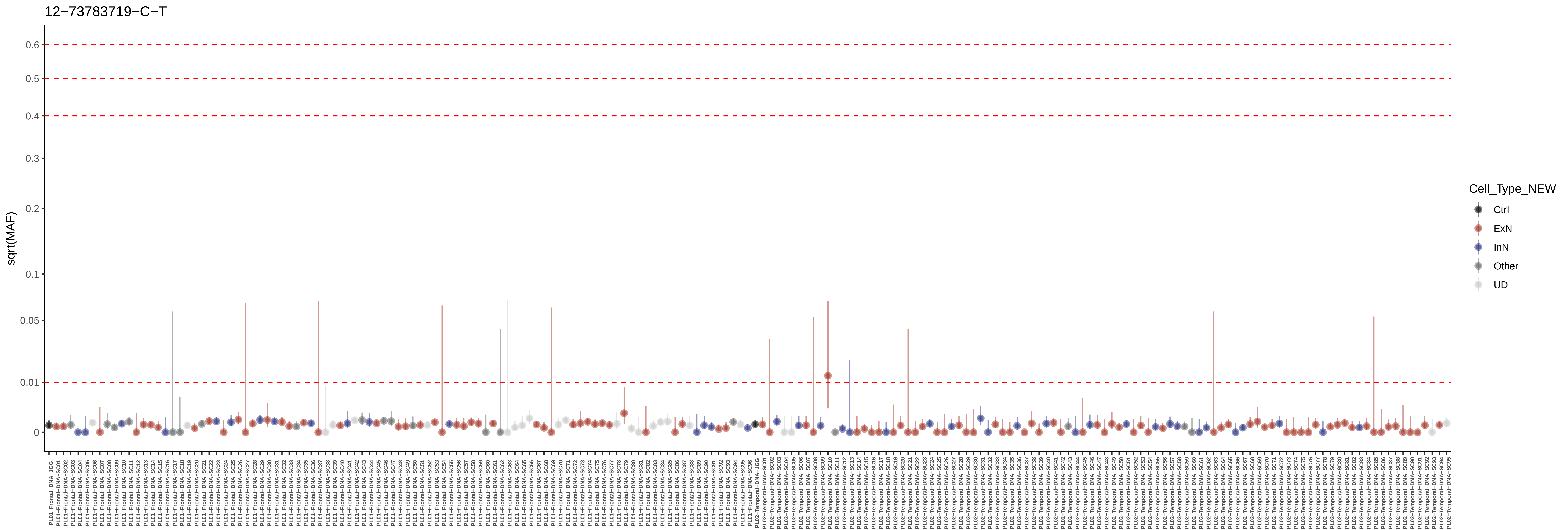
12−73783719−C−T
sqrt(MAF)
0.6
0.5
0.4
0.3
0.2
0.1
0.05
0.01
0
Cell_Type_NEW
Ctrl
ExN
InN
Other
UD

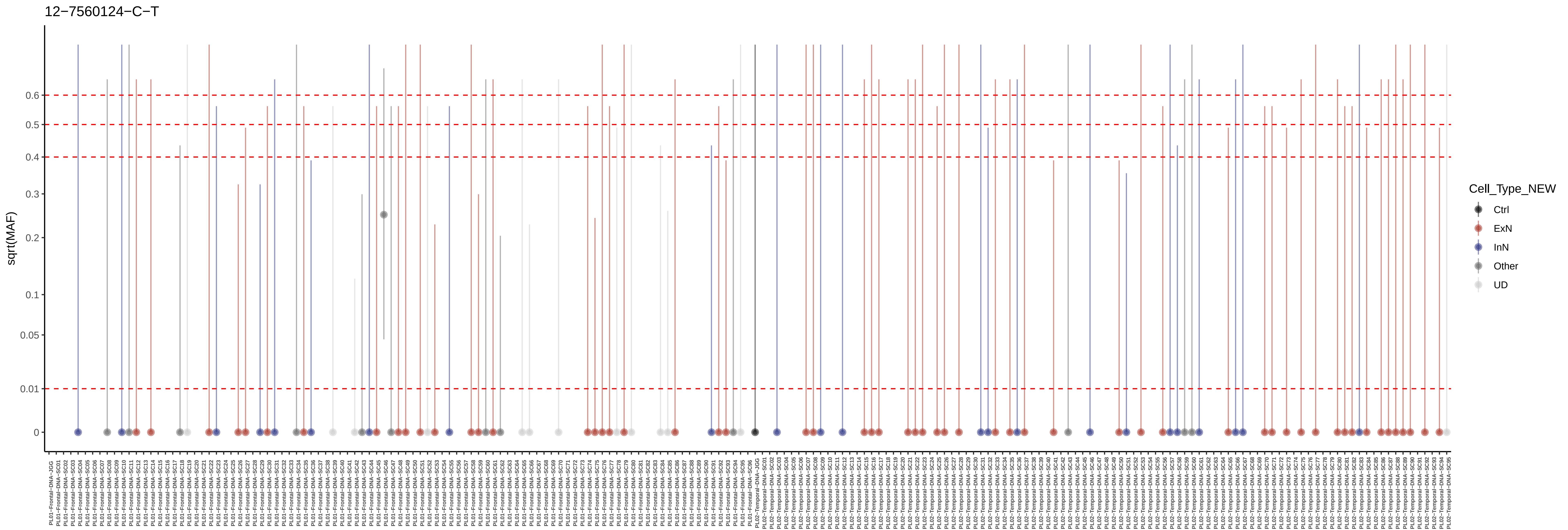
12−7560124−C−T
sqrt(MAF)
0.6
0.5
0.4
0.3
0.2
0.1
0.05
0.01
0
Cell_Type_NEW
Ctrl
ExN
InN
Other
UD

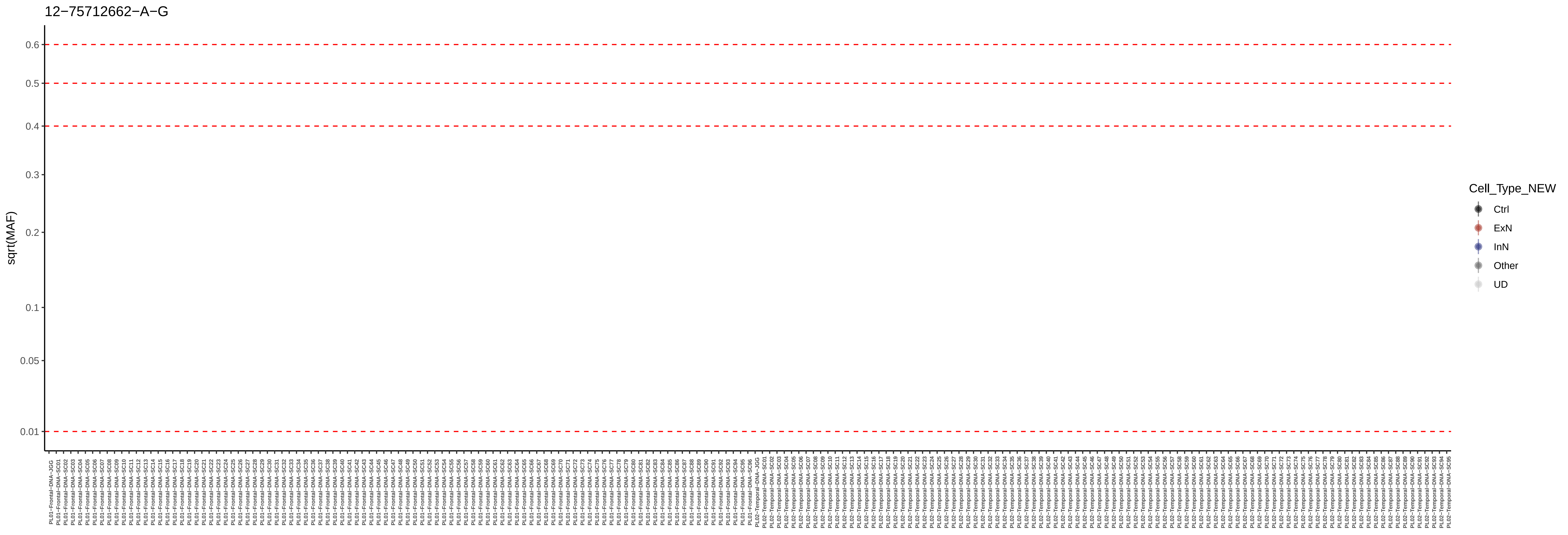
12−75712662−A−G
sqrt(MAF)
0.6
0.5
0.4
0.3
0.2
0.1
0.05
0.01
Cell_Type_NEW
Ctrl
ExN
InN
Other
UD

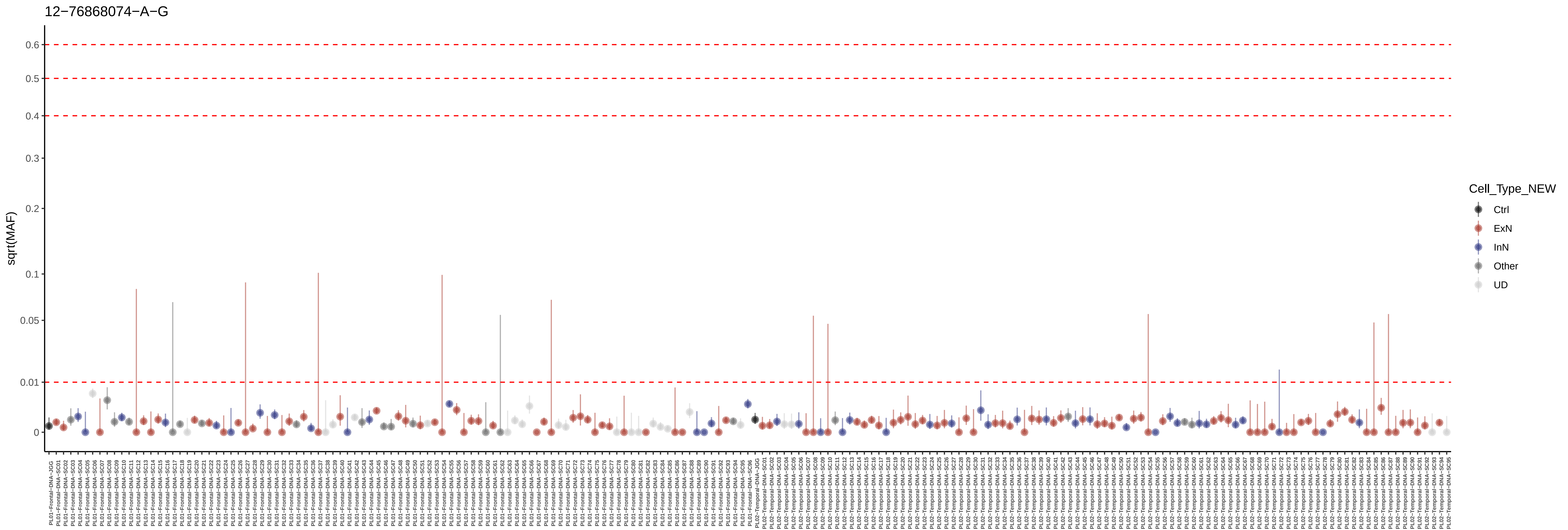

12−76868074−A−G
sqrt(MAF)
0.6
0.5
0.4
0.3
0.2
0.1
0.05
0.01
0
Cell_Type_NEW
Ctrl
ExN
InN
Other
UD

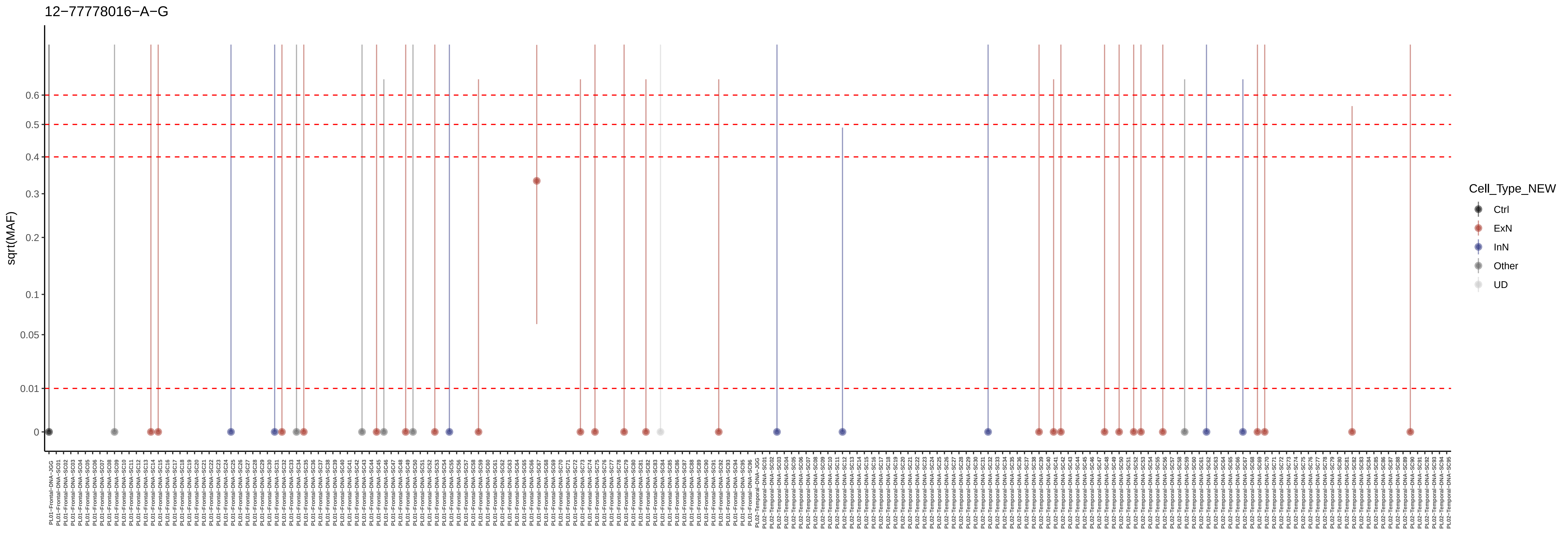
12−77778016−A−G
sqrt(MAF)
0.6
0.5
0.4
0.3
0.2
0.1
0.05
0.01
0
Cell_Type_NEW
Ctrl
ExN
InN
Other
UD

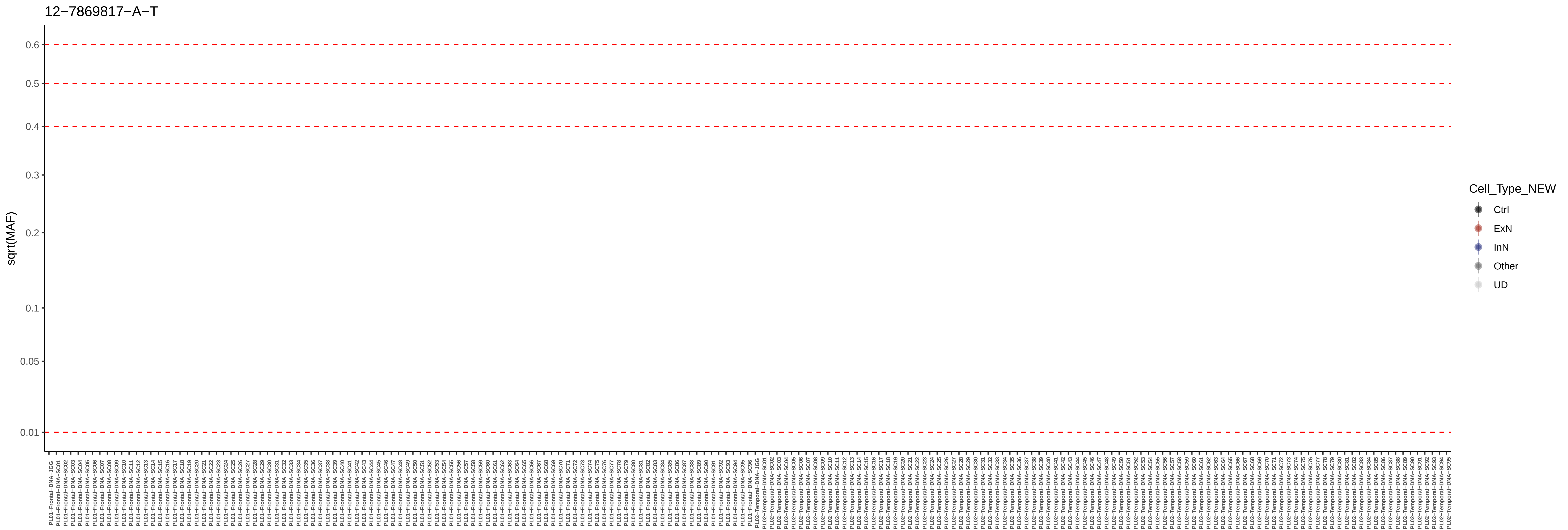

12−7869817−A−T
sqrt(MAF)
0.6
0.5
0.4
0.3
0.2
0.1
0.05
0.01
Cell_Type_NEW
Ctrl
ExN
InN
Other
UD

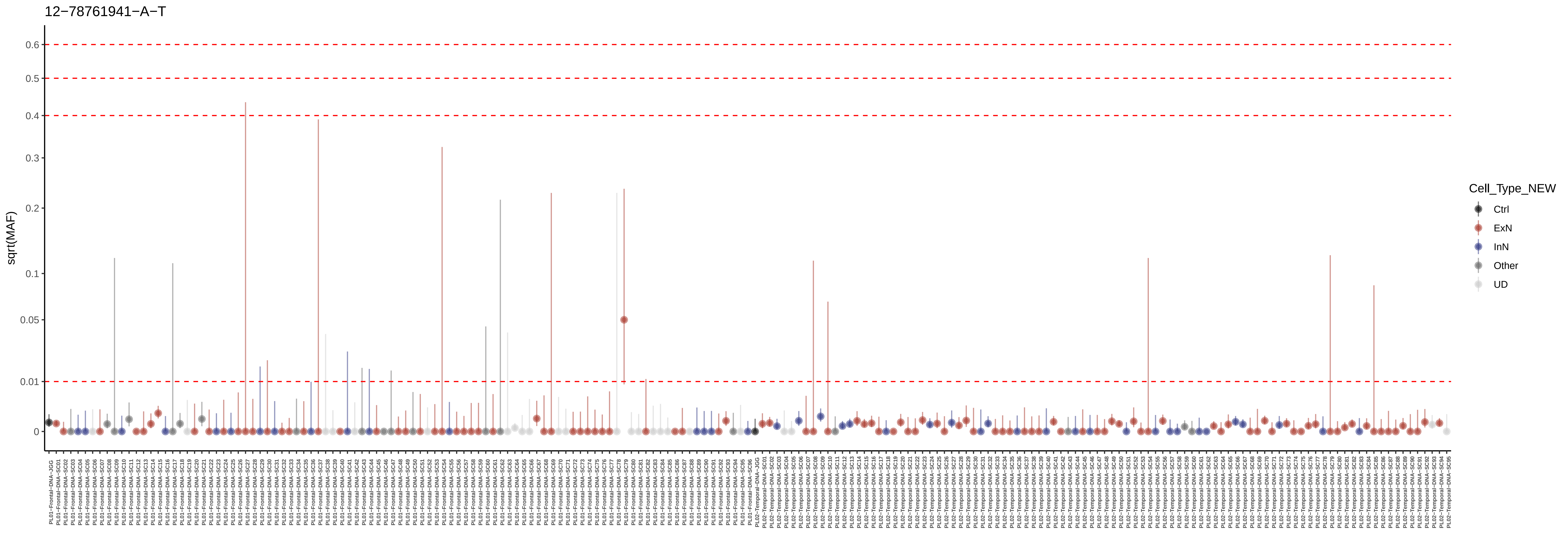

12−78761941−A−T
sqrt(MAF)
0.6
0.5
0.4
0.3
0.2
0.1
0.05
0.01
0
Cell_Type_NEW
Ctrl
ExN
InN
Other
UD

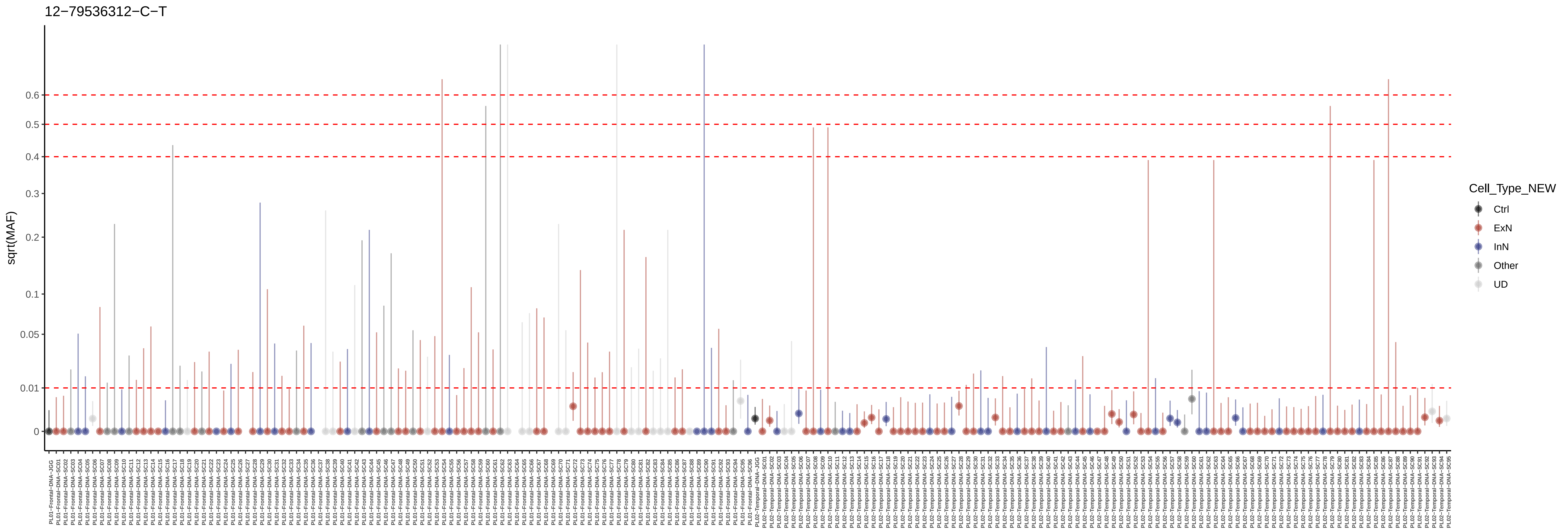
12−79536312−C−T
sqrt(MAF)
0.6
0.5
0.4
0.3
0.2
0.1
0.05
0.01
0
Cell_Type_NEW
Ctrl
ExN
InN
Other
UD

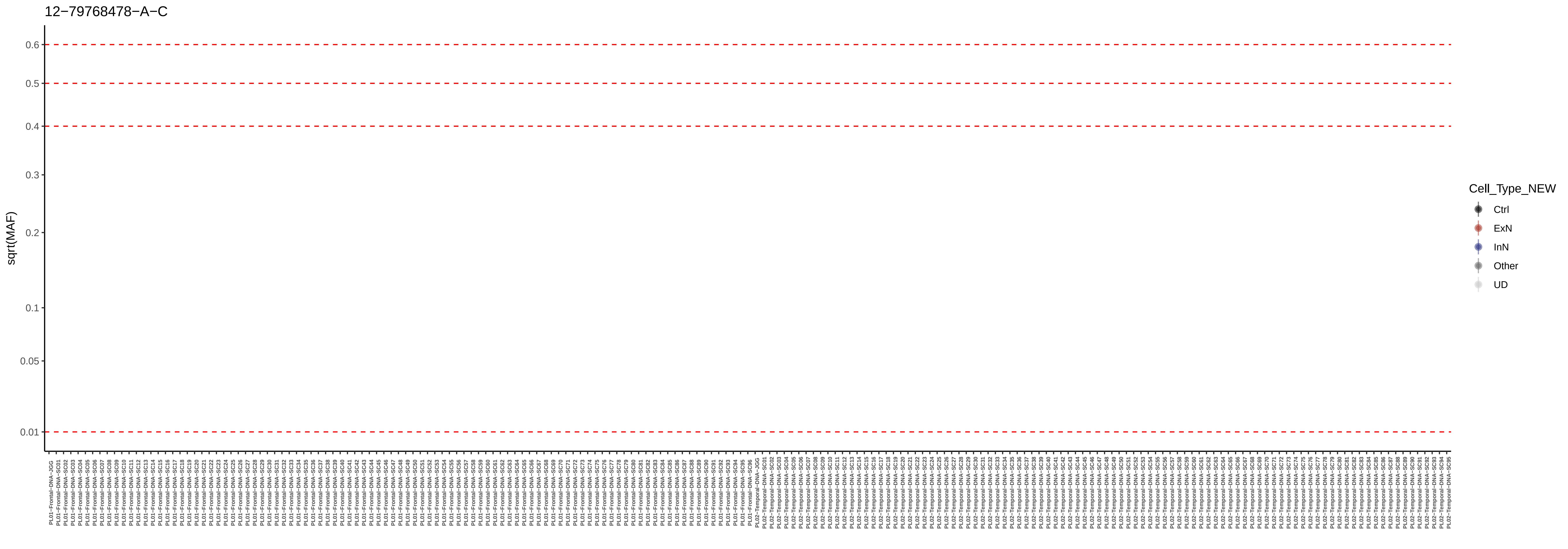
12−79768478−A−C
sqrt(MAF)
0.6
0.5
0.4
0.3
0.2
0.1
0.05
0.01
Cell_Type_NEW
Ctrl
ExN
InN
Other
UD

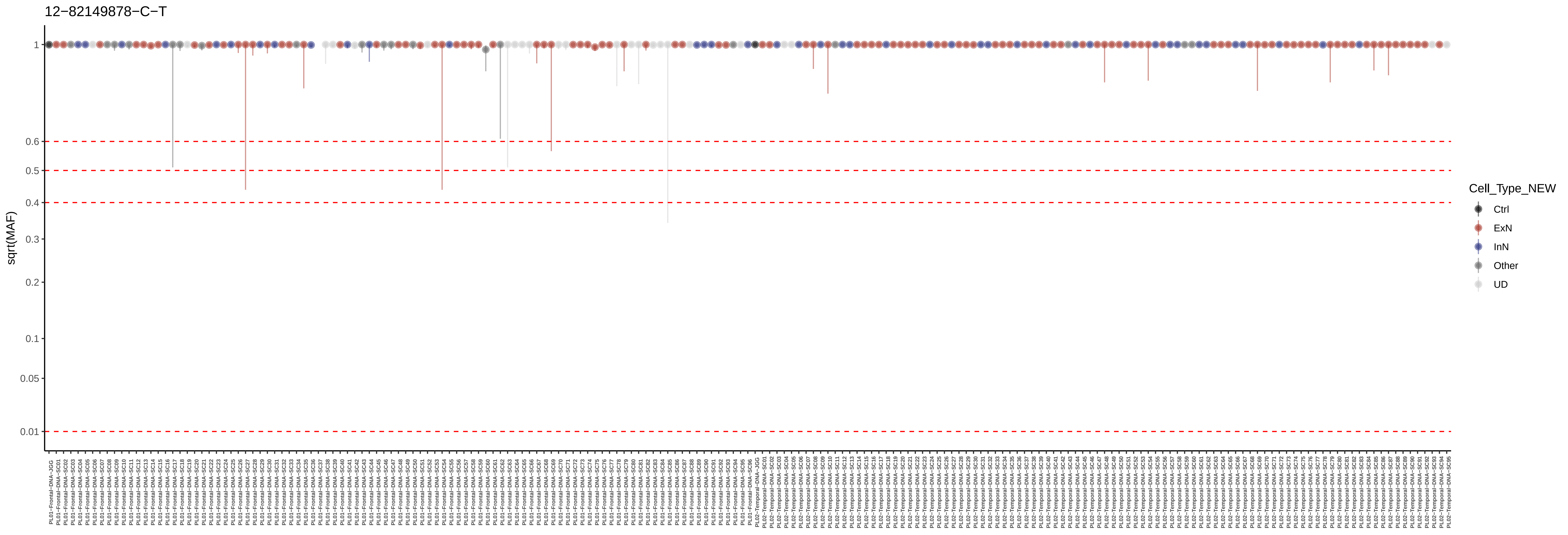
12−82149878−C−T
sqrt(MAF)
1
0.6
0.5
0.4
0.3
0.2
0.1
0.05
0.01
Cell_Type_NEW
Ctrl
ExN
InN
Other
UD

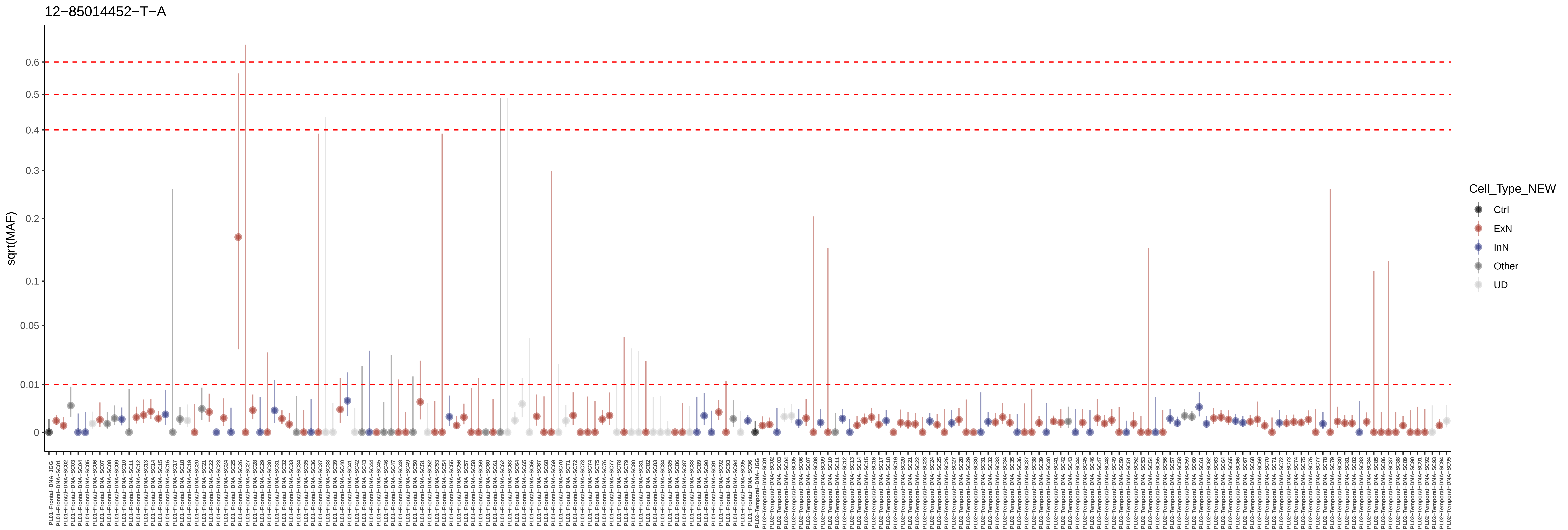
12−85014452−T−A
sqrt(MAF)
0.6
0.5
0.4
0.3
0.2
0.1
0.05
0.01
0
Cell_Type_NEW
Ctrl
ExN
InN
Other
UD

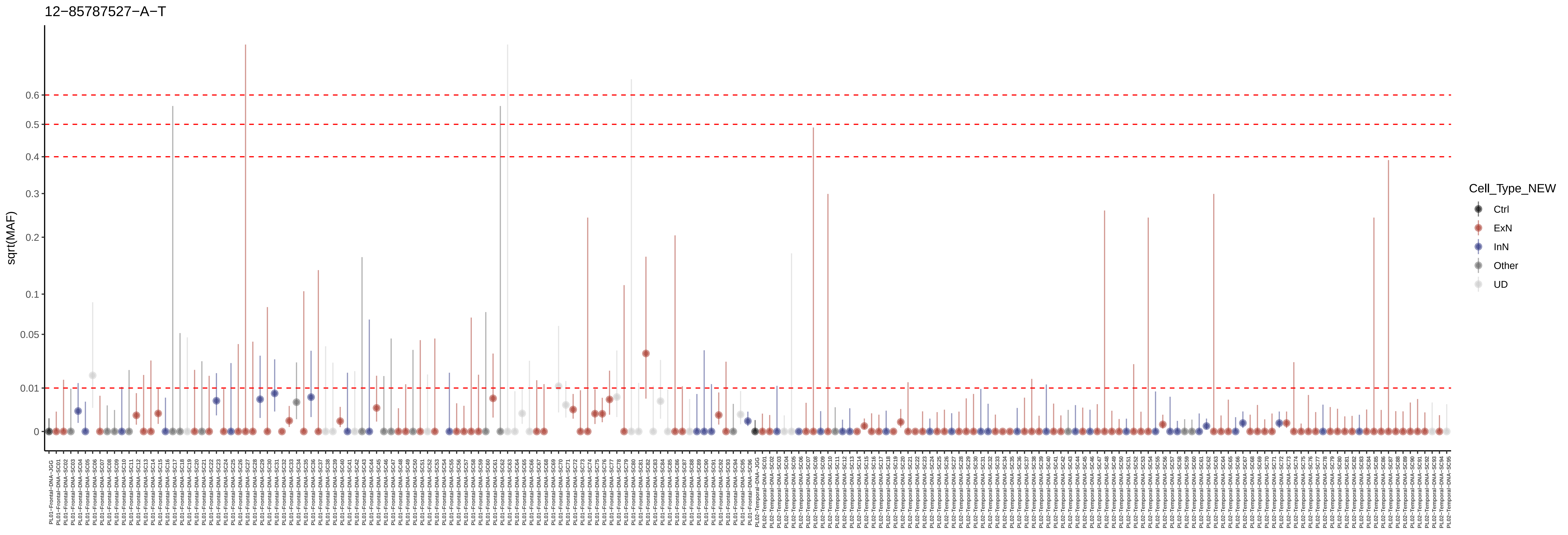

12−85787527−A−T
sqrt(MAF)
0.6
0.5
0.4
0.3
0.2
0.1
0.05
0.01
0
Cell_Type_NEW
Ctrl
ExN
InN
Other
UD

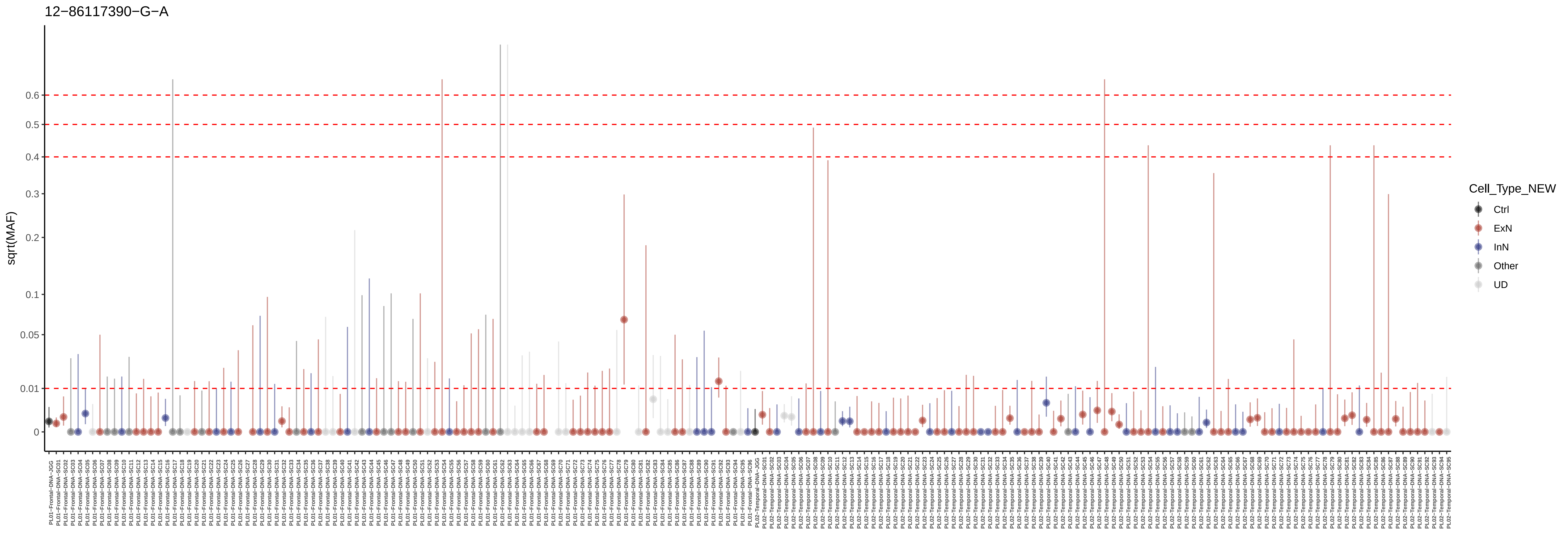

12−86117390−G−A
sqrt(MAF)
0.6
0.5
0.4
0.3
0.2
0.1
0.05
0.01
0
Cell_Type_NEW
Ctrl
ExN
InN
Other
UD

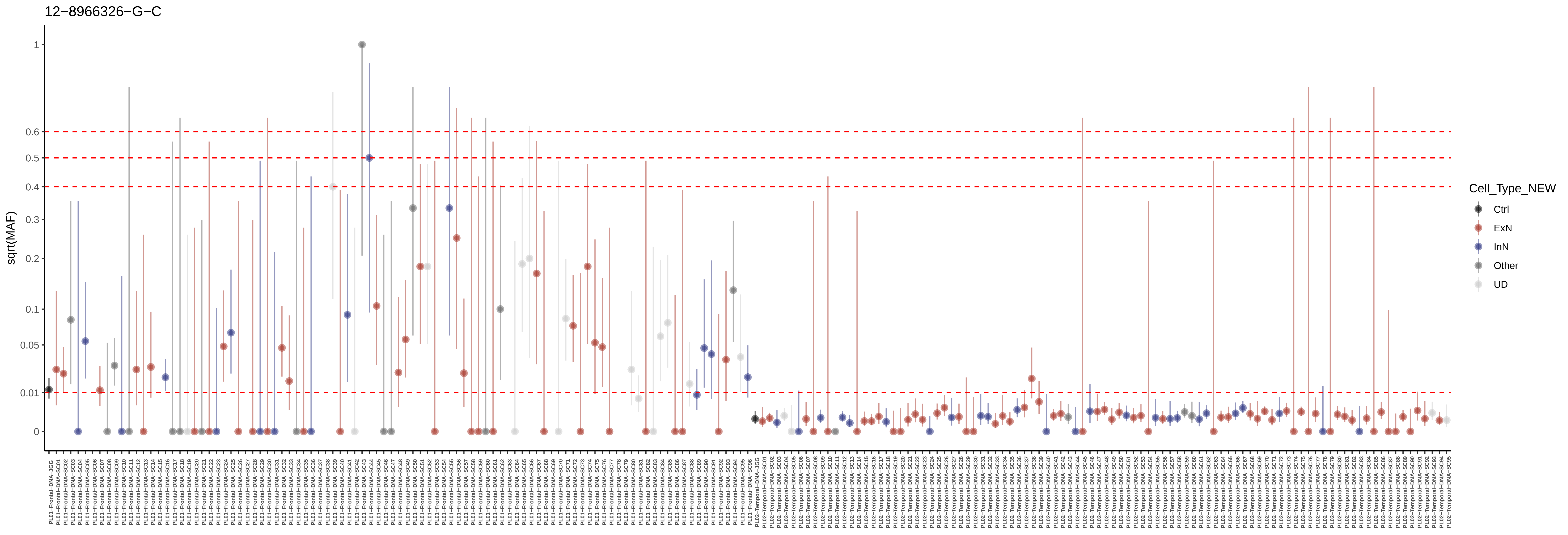
12−8966326−G−C
sqrt(MAF)
1
0.6
0.5
0.4
0.3
0.2
0.1
0.05
0.01
0
Cell_Type_NEW
Ctrl
ExN
InN
Other
UD

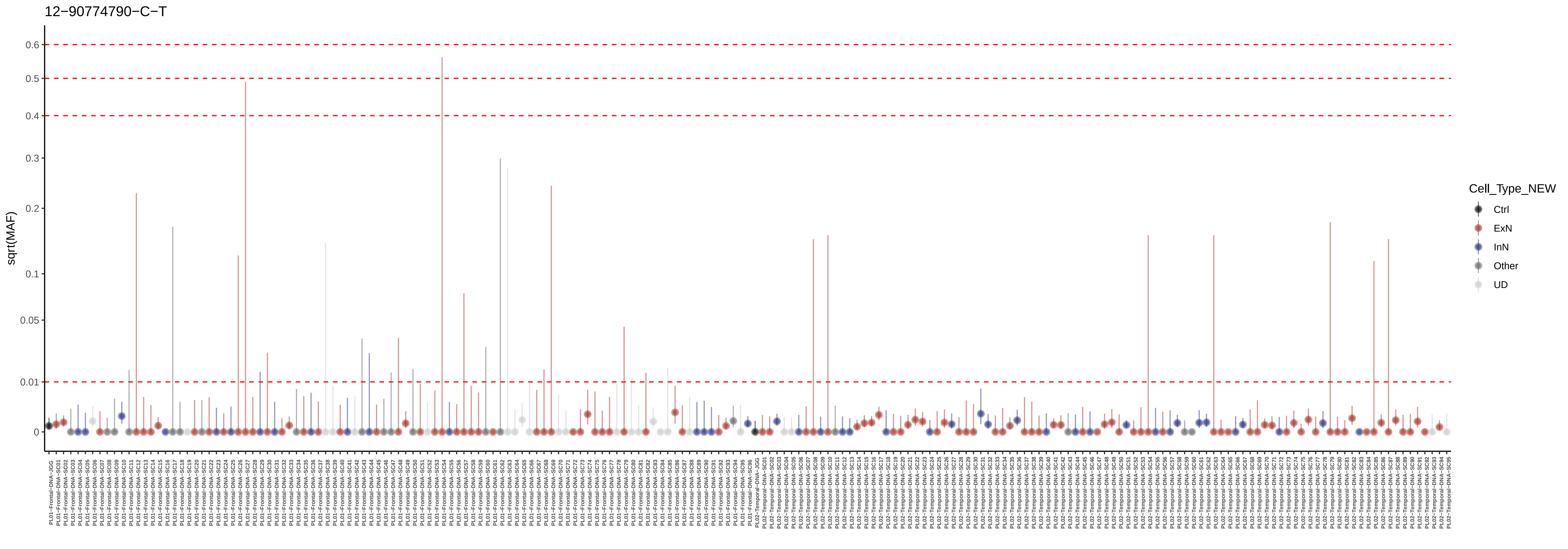
12−90774790−C−T
sqrt(MAF)
0.6
0.5
0.4
0.3
0.2
0.1
0.05
0.01
0
Cell_Type_NEW
Ctrl
ExN
InN
Other
UD

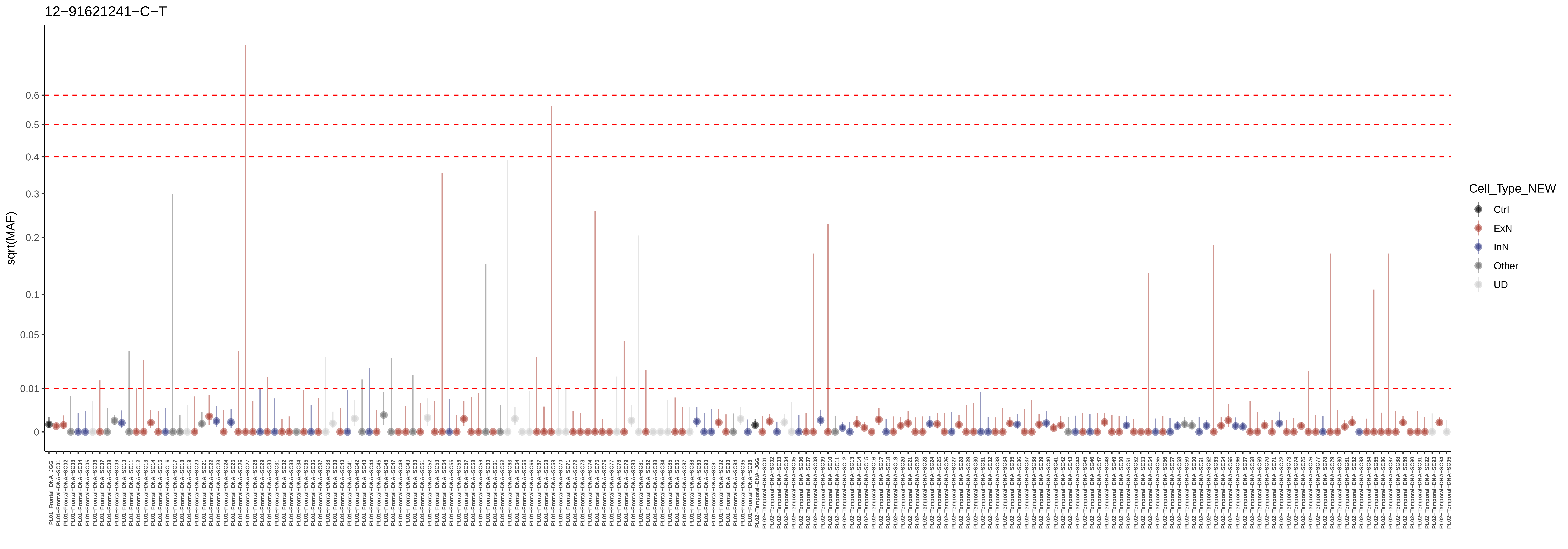
12−91621241−C−T
sqrt(MAF)
0.6
0.5
0.4
0.3
0.2
0.1
0.05
0.01
0
Cell_Type_NEW
Ctrl
ExN
InN
Other
UD

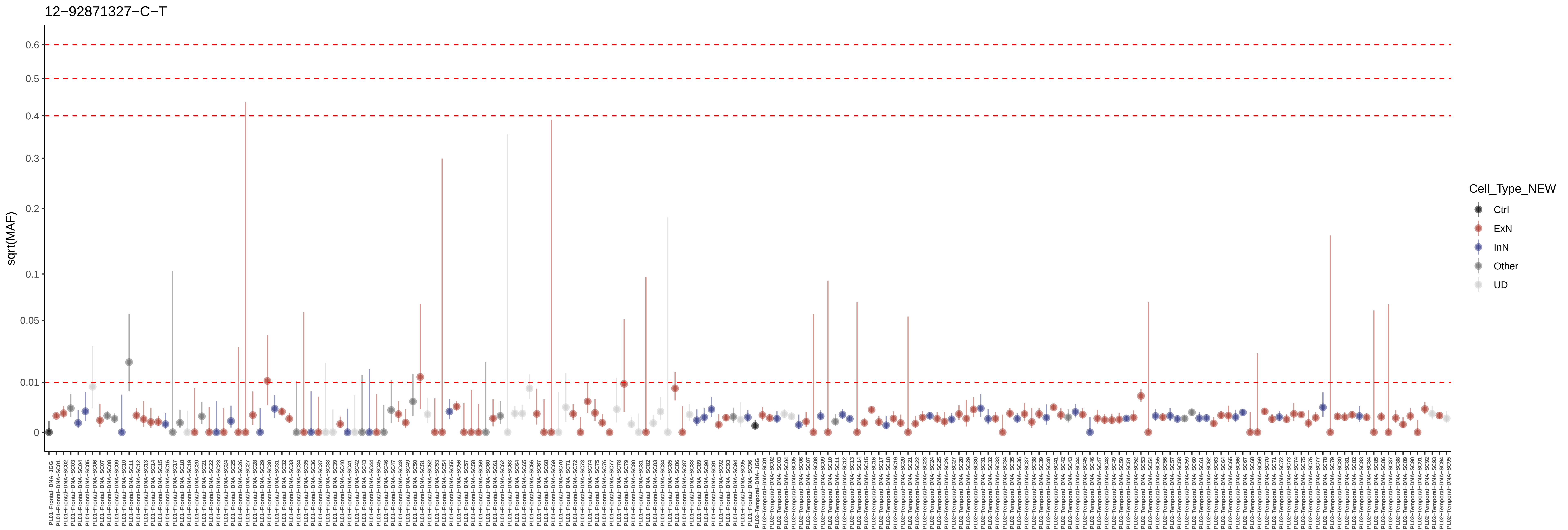

12−92871327−C−T
sqrt(MAF)
0.6
0.5
0.4
0.3
0.2
0.1
0.05
0.01
0
Cell_Type_NEW
Ctrl
ExN
InN
Other
UD

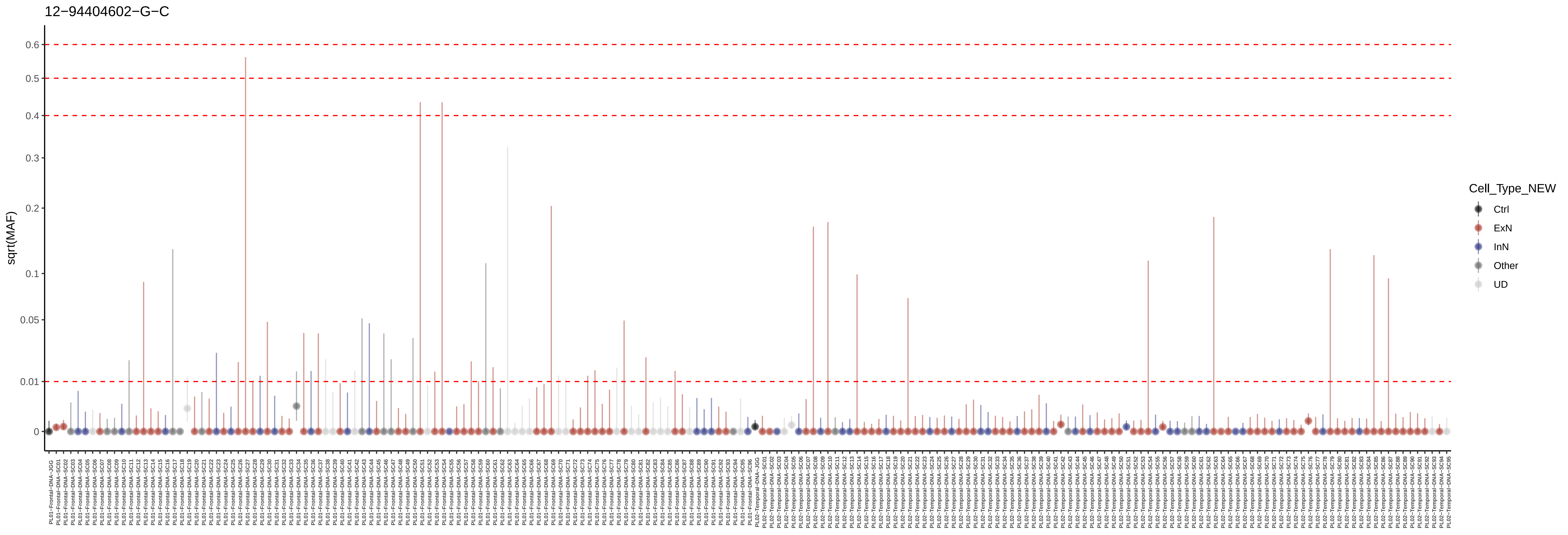
12−94404602−G−C
sqrt(MAF)
0.6
0.5
0.4
0.3
0.2
0.1
0.05
0.01
0
Cell_Type_NEW
Ctrl
ExN
InN
Other
UD

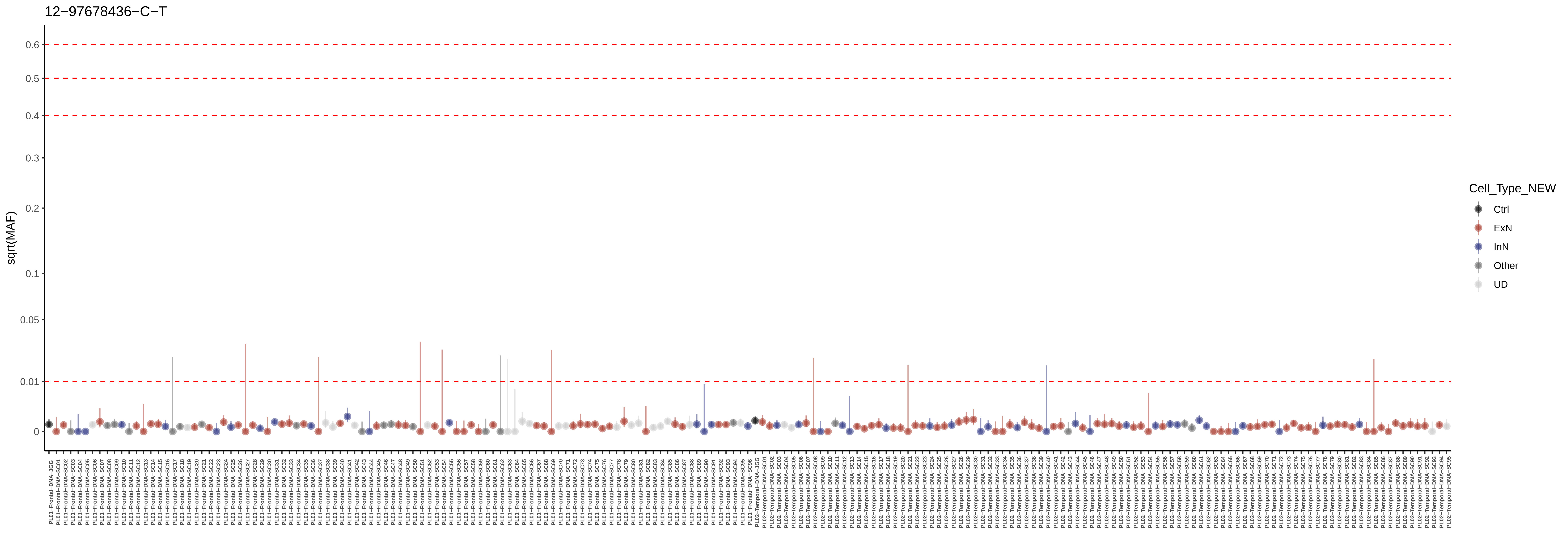

12−97678436−C−T
sqrt(MAF)
0.6
0.5
0.4
0.3
0.2
0.1
0.05
0.01
0
Cell_Type_NEW
Ctrl
ExN
InN
Other
UD

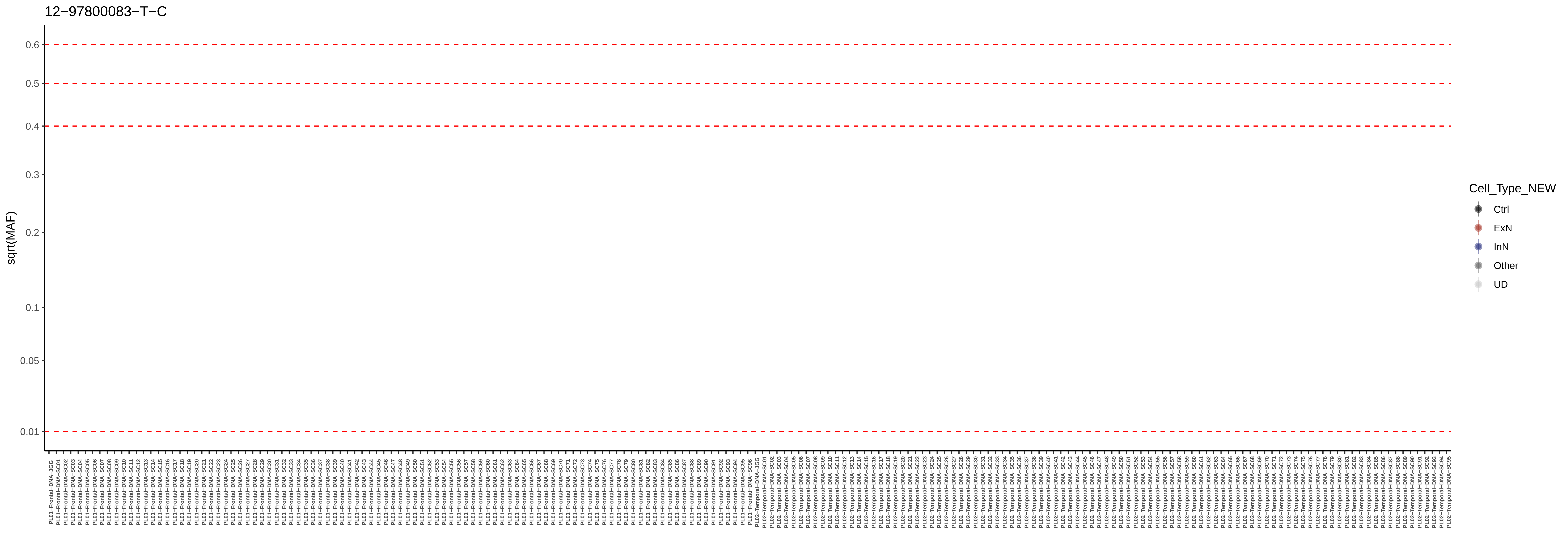
12−97800083−T−C
sqrt(MAF)
0.6
0.5
0.4
0.3
0.2
0.1
0.05
0.01
Cell_Type_NEW
Ctrl
ExN
InN
Other
UD

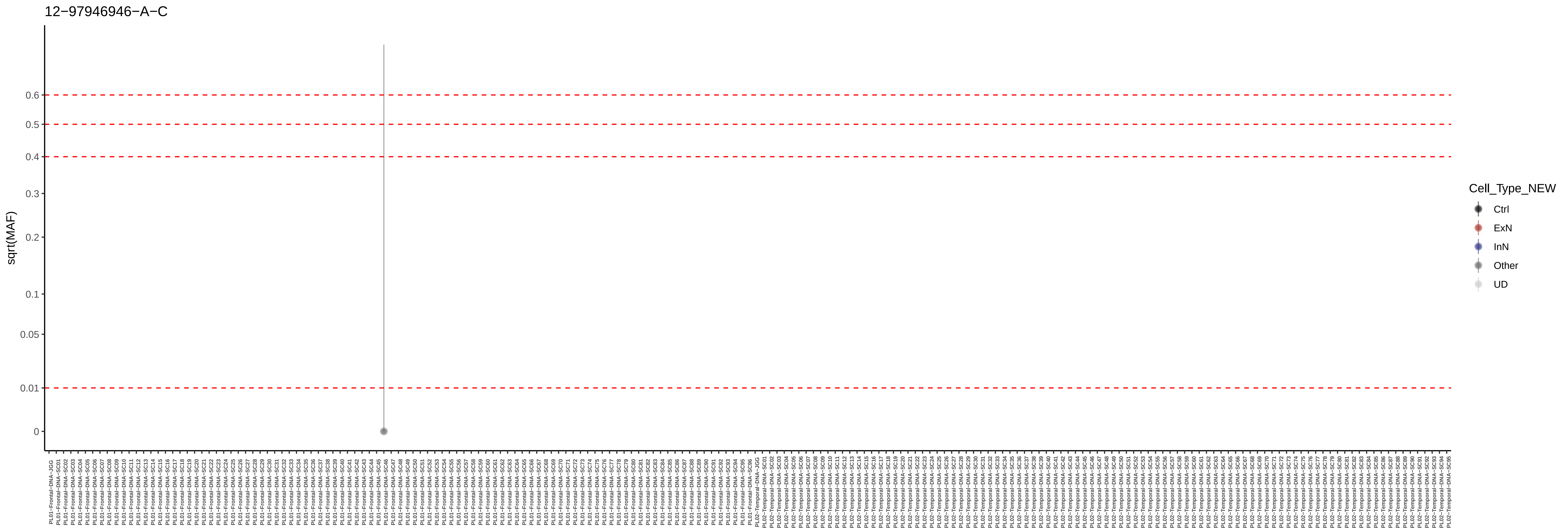
12−97946946−A−C
sqrt(MAF)
0.6
0.5
0.4
0.3
0.2
0.1
0.05
0.01
0
Cell_Type_NEW
Ctrl
ExN
InN
Other
UD

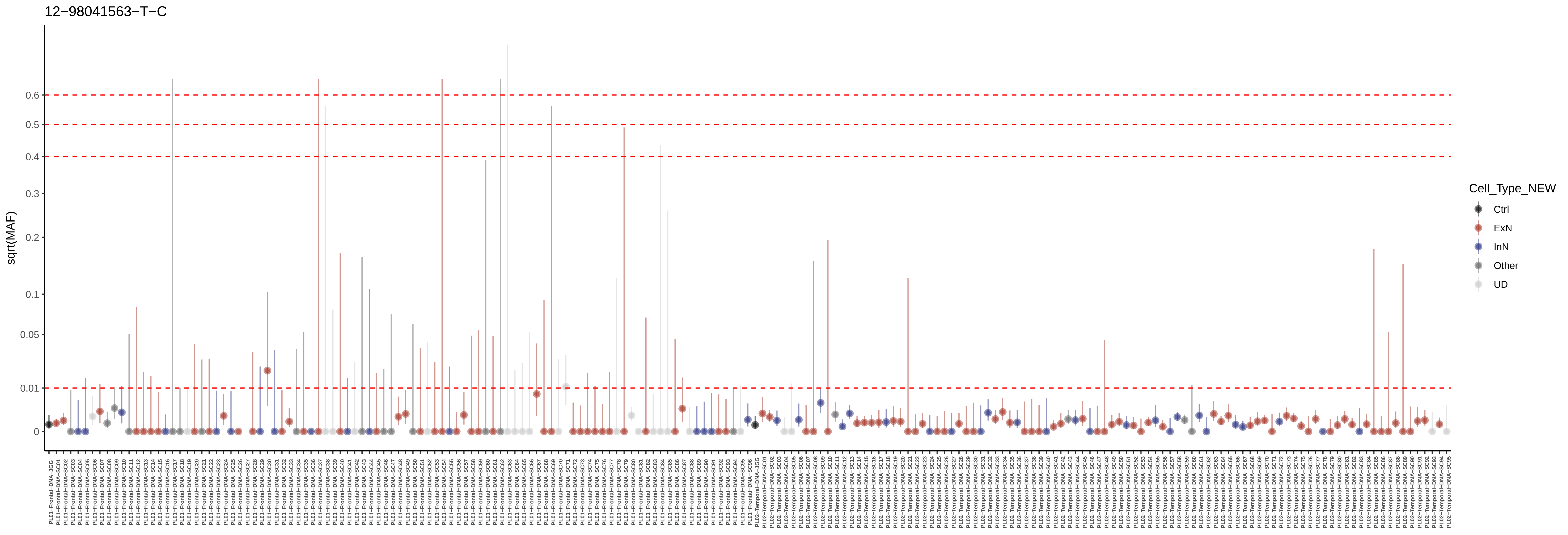
12−98041563−T−C
sqrt(MAF)
0.6
0.5
0.4
0.3
0.2
0.1
0.05
0.01
0
Cell_Type_NEW
Ctrl
ExN
InN
Other
UD

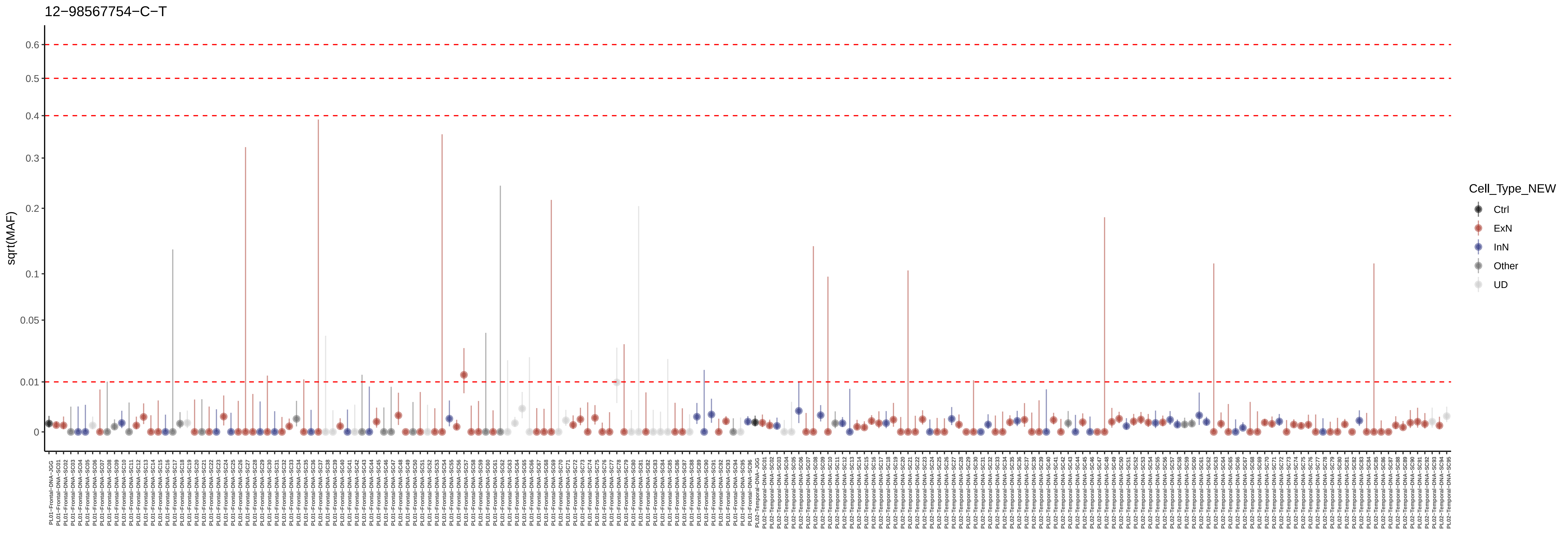
12−98567754−C−T
sqrt(MAF)
0.6
0.5
0.4
0.3
0.2
0.1
0.05
0.01
0
Cell_Type_NEW
Ctrl
ExN
InN
Other
UD

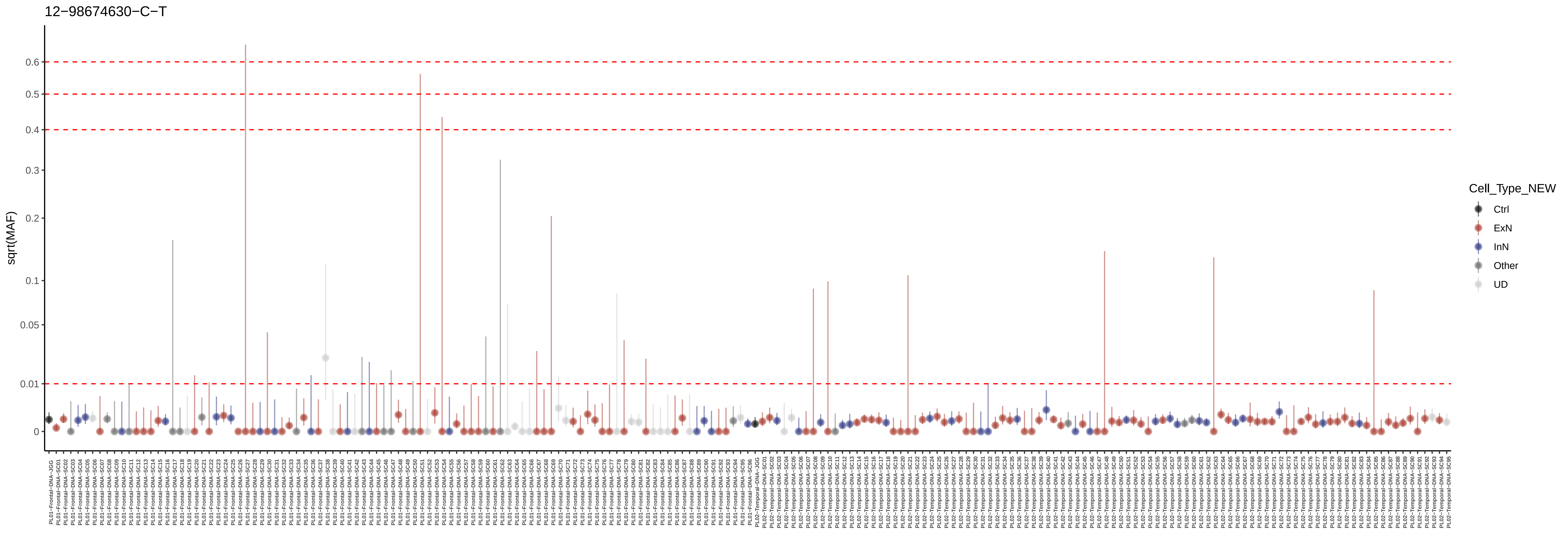
12−98674630−C−T
sqrt(MAF)
0.6
0.5
0.4
0.3
0.2
0.1
0.05
0.01
0
Cell_Type_NEW
Ctrl
ExN
InN
Other
UD

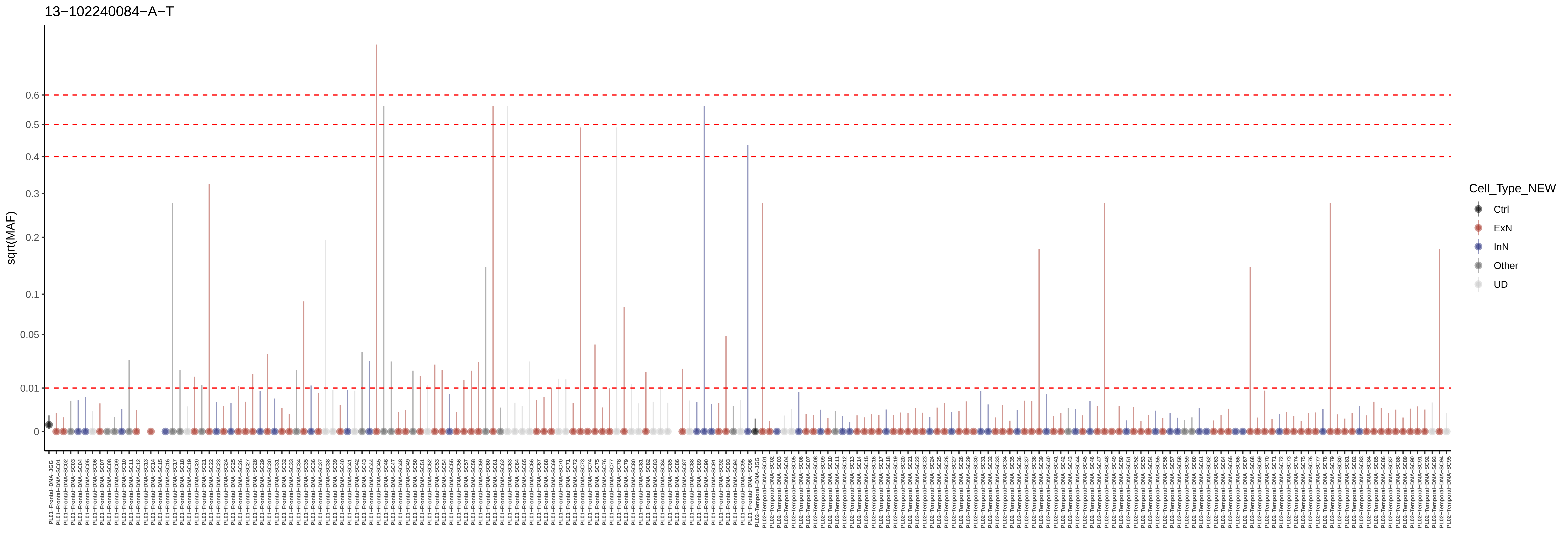

13−102240084−A−T
sqrt(MAF)
0.6
0.5
0.4
0.3
0.2
0.1
0.05
0.01
0
Cell_Type_NEW
Ctrl
ExN
InN
Other
UD

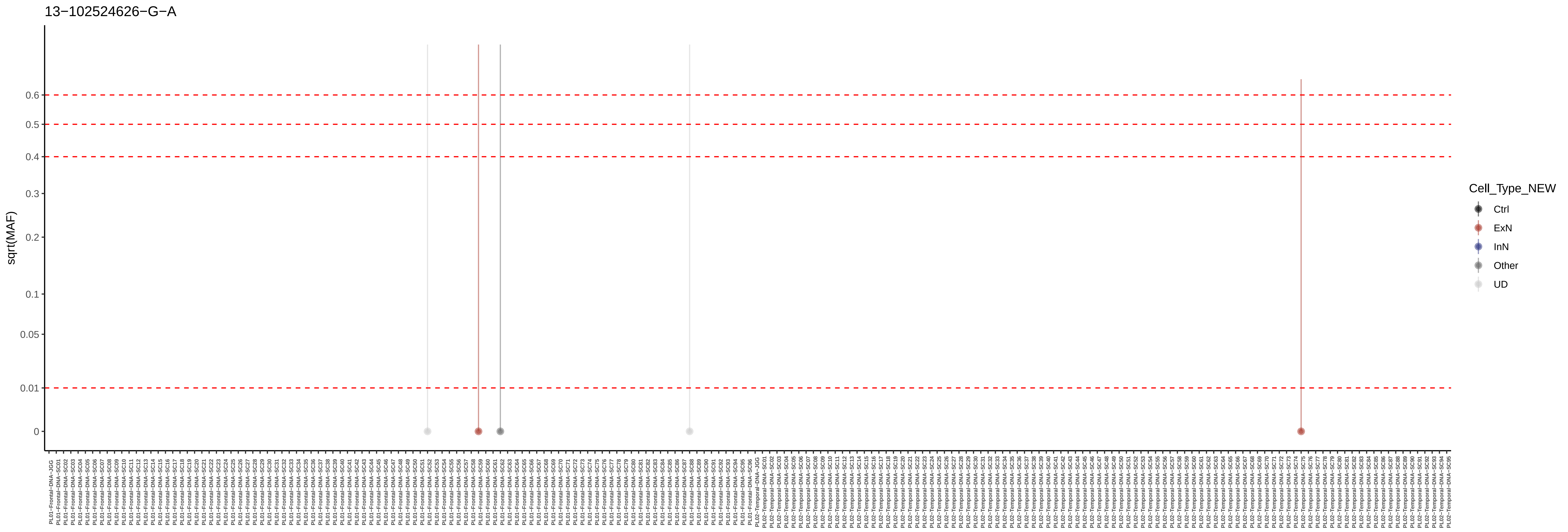
13−102524626−G−A
sqrt(MAF)
0.6
0.5
0.4
0.3
0.2
0.1
0.05
0.01
0
Cell_Type_NEW
Ctrl
ExN
InN
Other
UD

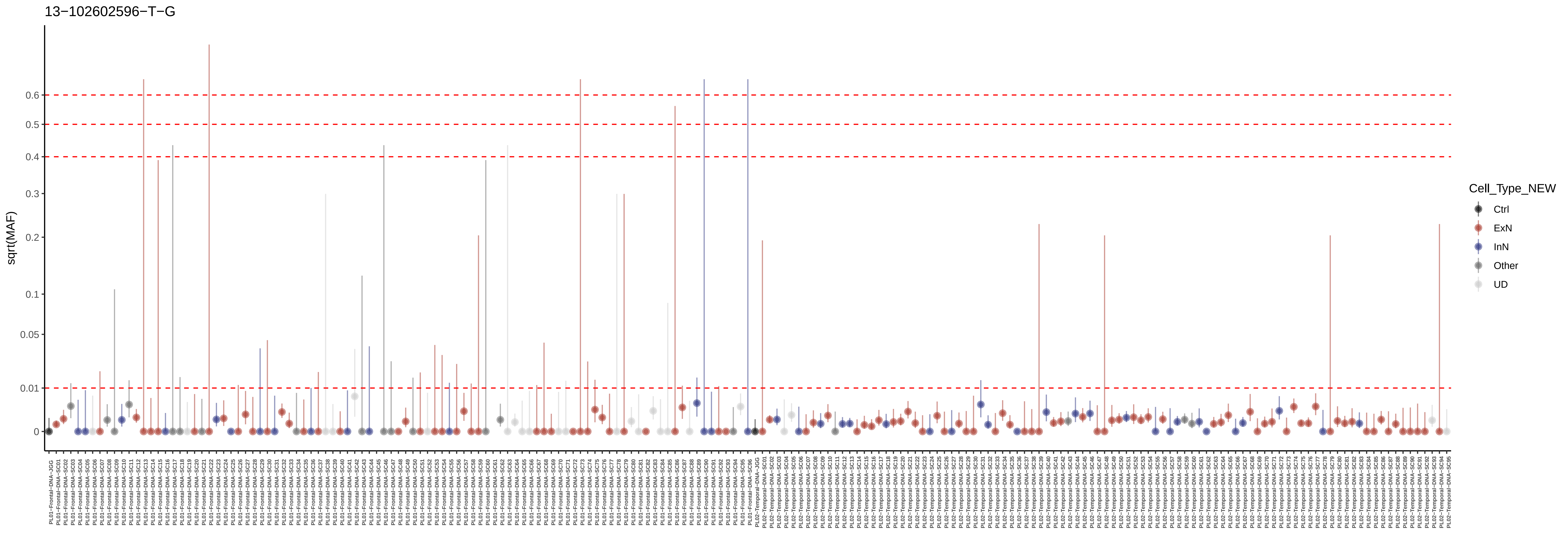
13−102602596−T−G
sqrt(MAF)
0.6
0.5
0.4
0.3
0.2
0.1
0.05
0.01
0
Cell_Type_NEW
Ctrl
ExN
InN
Other
UD

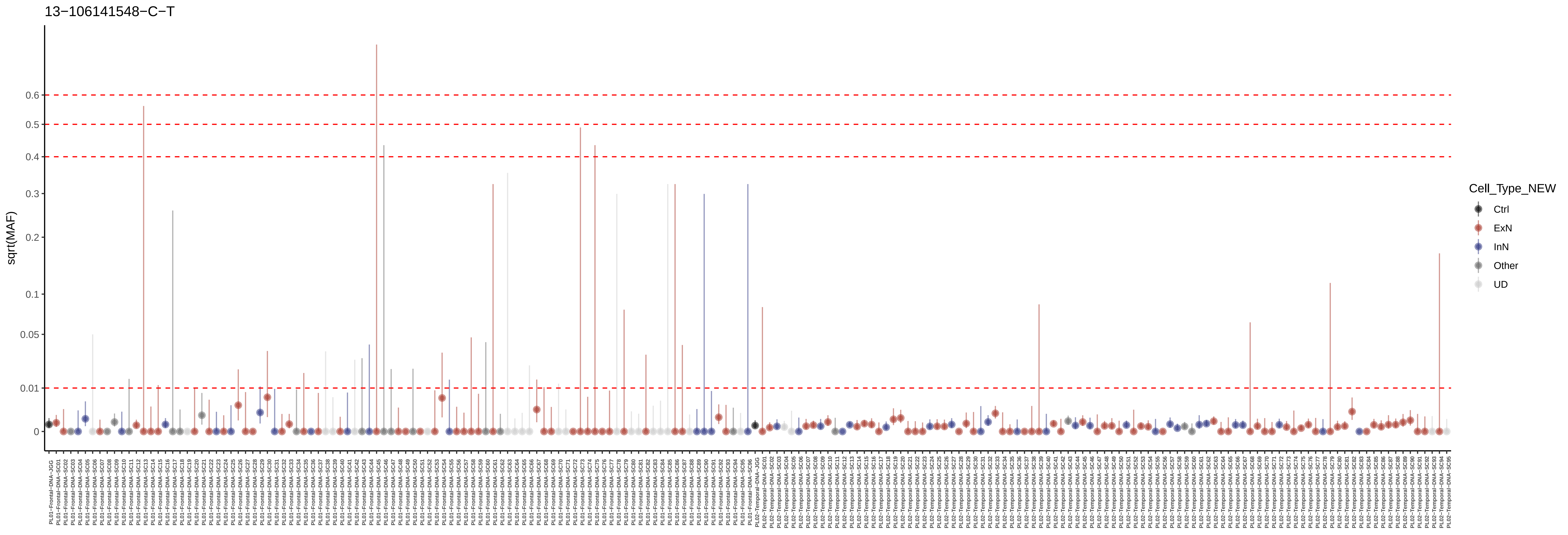

13−106141548−C−T
sqrt(MAF)
0
0.01
0.05
0.1
0.2
0.3
0.4
0.5
0.6
Cell_Type_NEW
Ctrl
ExN
InN
Other
UD

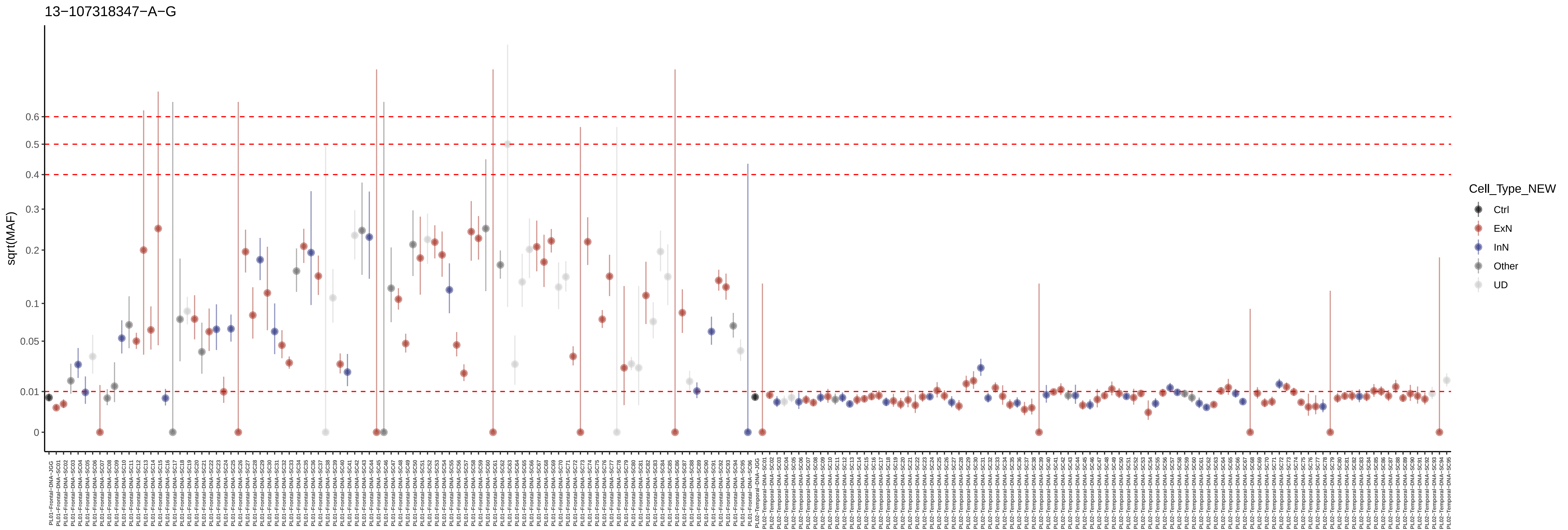

13−107318347−A−G
sqrt(MAF)
0.6
0.5
0.4
0.3
0.2
0.1
0.05
0.01
0
Cell_Type_NEW
Ctrl
ExN
InN
Other
UD

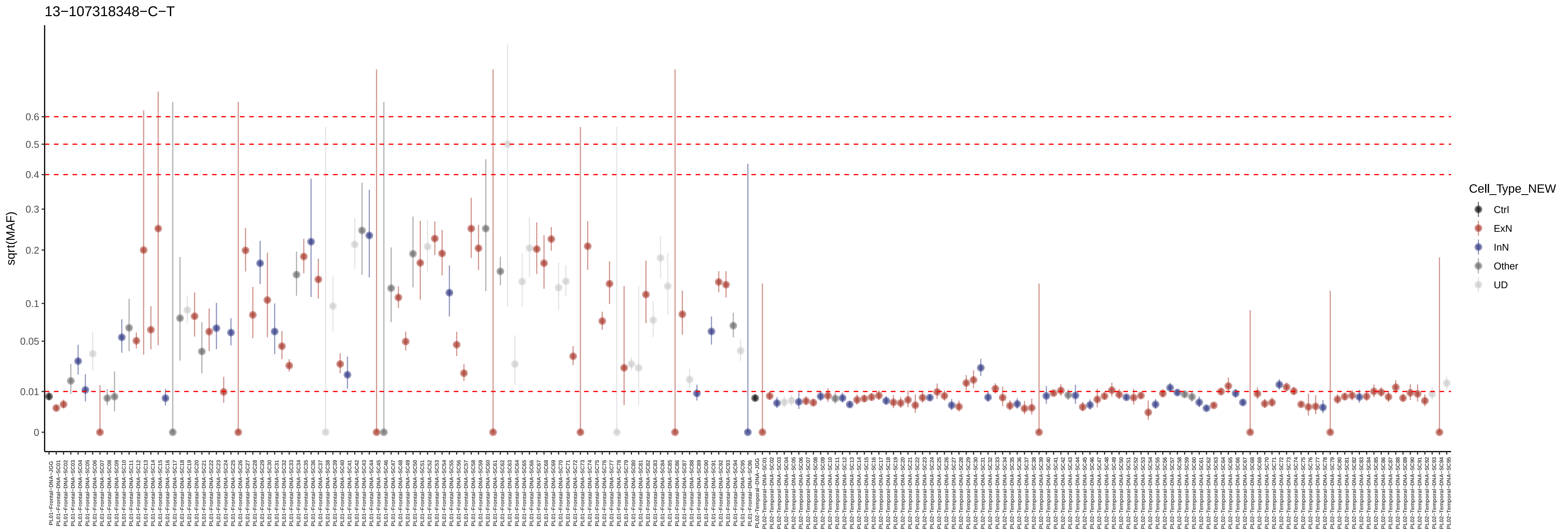

13−107318348−C−T
sqrt(MAF)
0.6
0.5
0.4
0.3
0.2
0.1
0.05
0.01
0
Cell_Type_NEW
Ctrl
ExN
InN
Other
UD

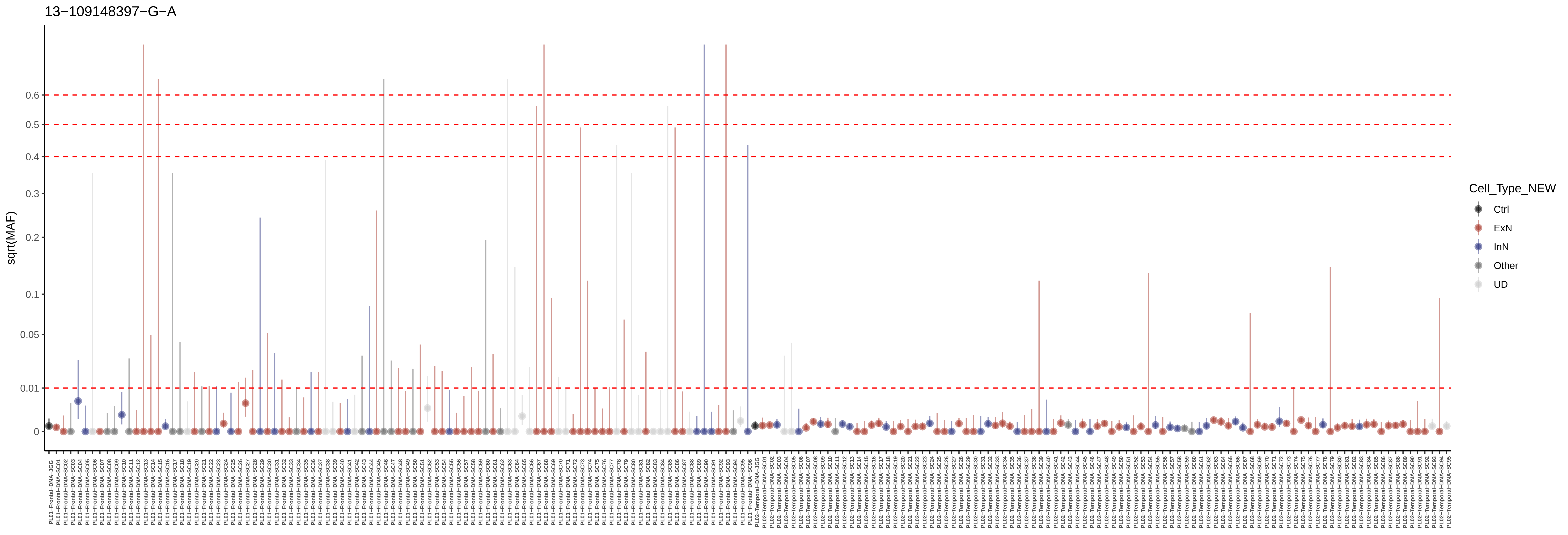

13−109148397−G−A
sqrt(MAF)
0.6
0.5
0.4
0.3
0.2
0.1
0.05
0.01
0
Cell_Type_NEW
Ctrl
ExN
InN
Other
UD

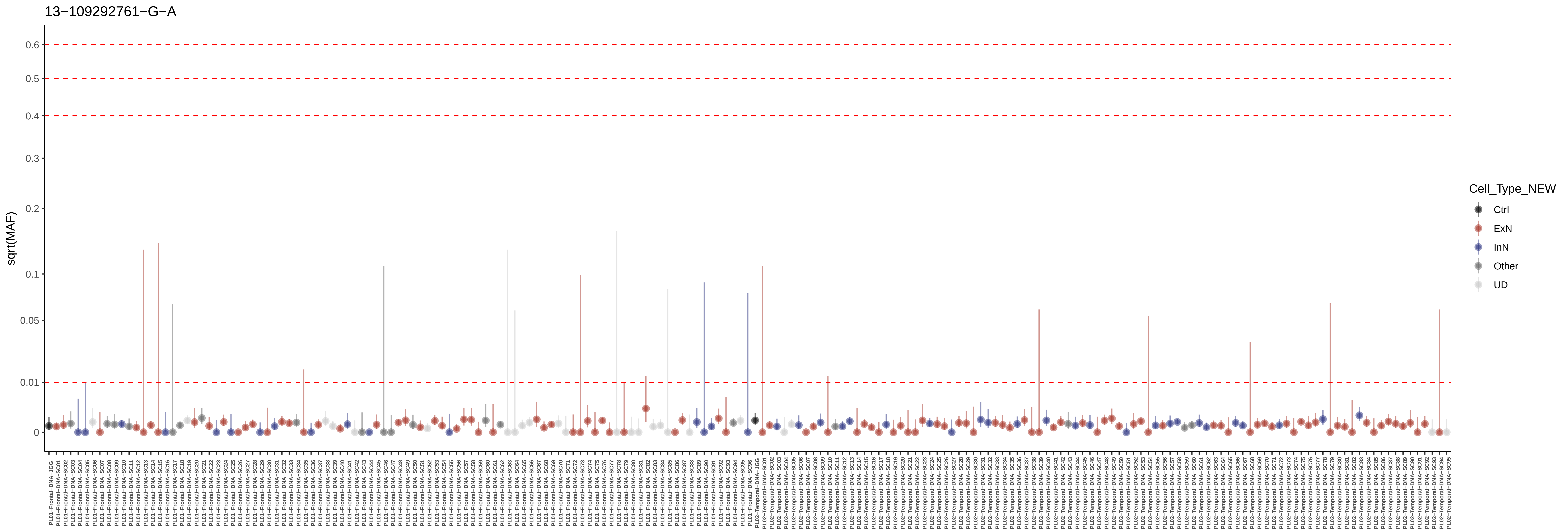

13−109292761−G−A
sqrt(MAF)
0.6
0.5
0.4
0.3
0.2
0.1
0.05
0.01
0
Cell_Type_NEW
Ctrl
ExN
InN
Other
UD

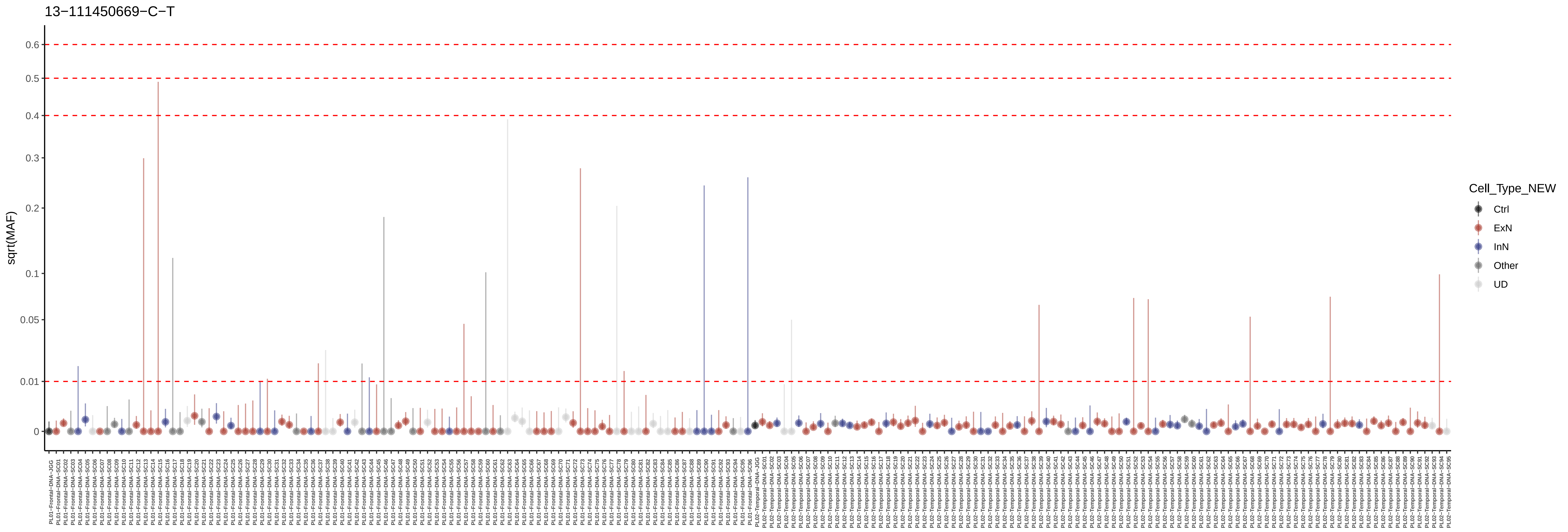

13−111450669−C−T
sqrt(MAF)
0.6
0.5
0.4
0.3
0.2
0.1
0.05
0.01
0
Cell_Type_NEW
Ctrl
ExN
InN
Other
UD

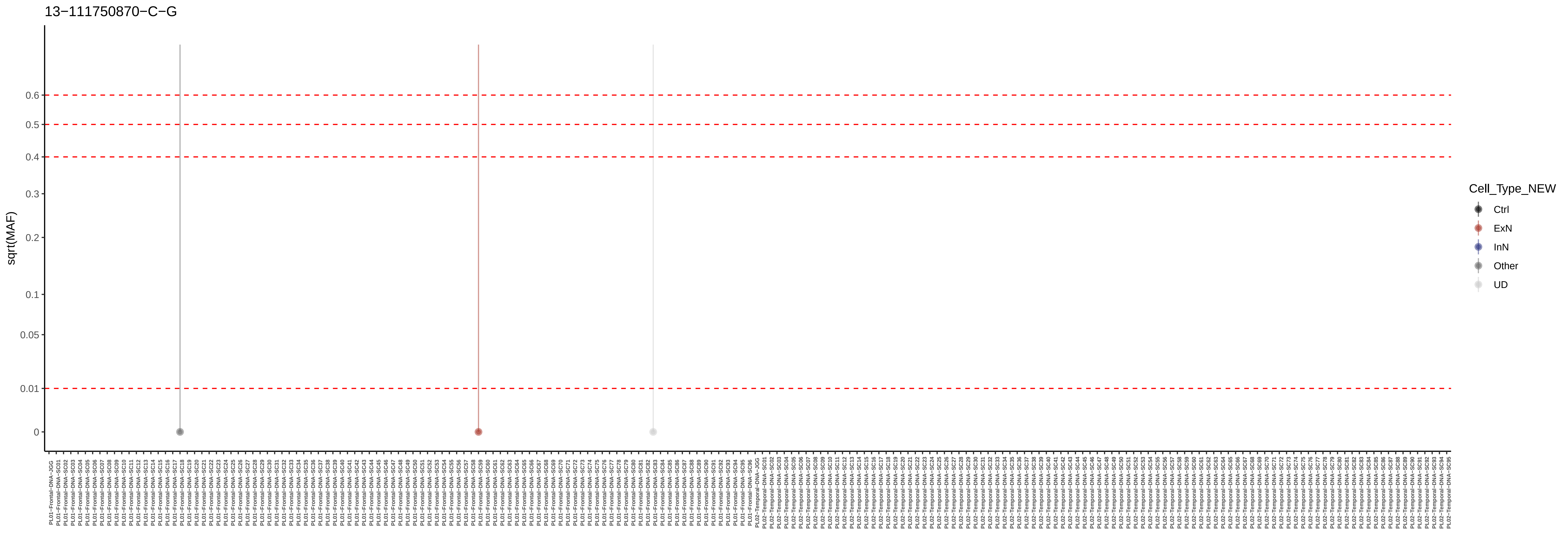

13−111750870−C−G
sqrt(MAF)
0.6
0.5
0.4
0.3
0.2
0.1
0.05
0.01
0
Cell_Type_NEW
Ctrl
ExN
InN
Other
UD

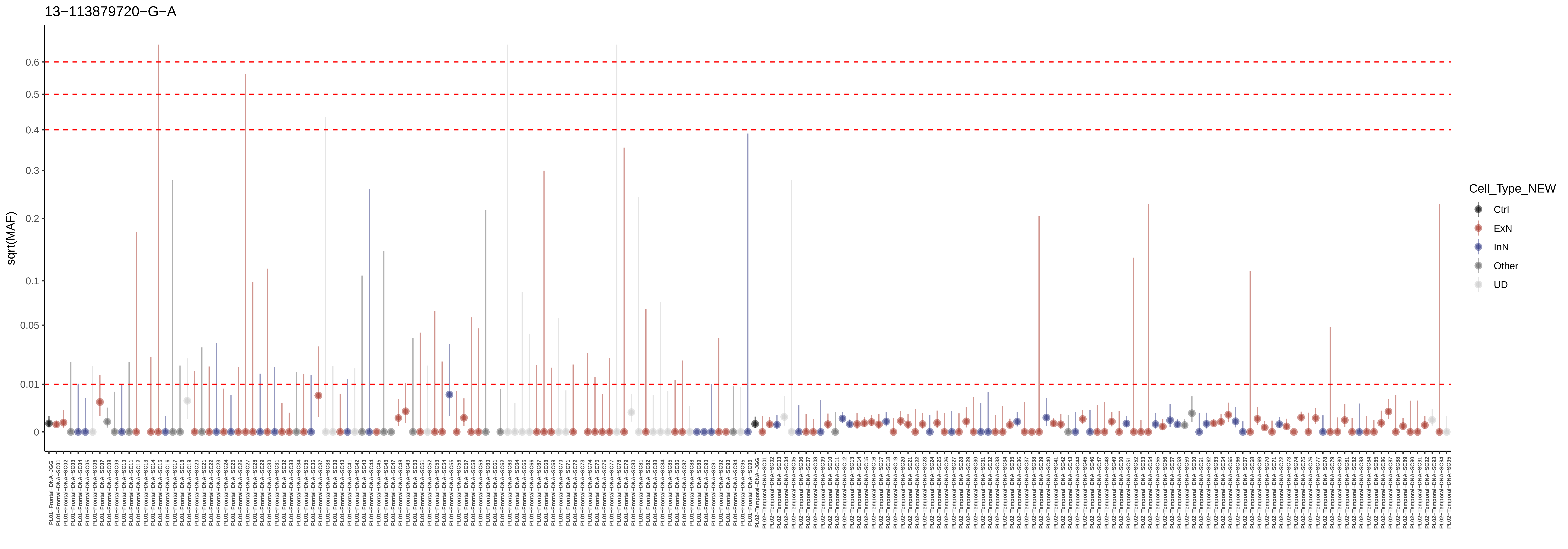

13−113879720−G−A
sqrt(MAF)
0
0.01
0.05
0.1
0.2
0.3
0.4
0.5
0.6
Cell_Type_NEW
Ctrl
ExN
InN
Other
UD

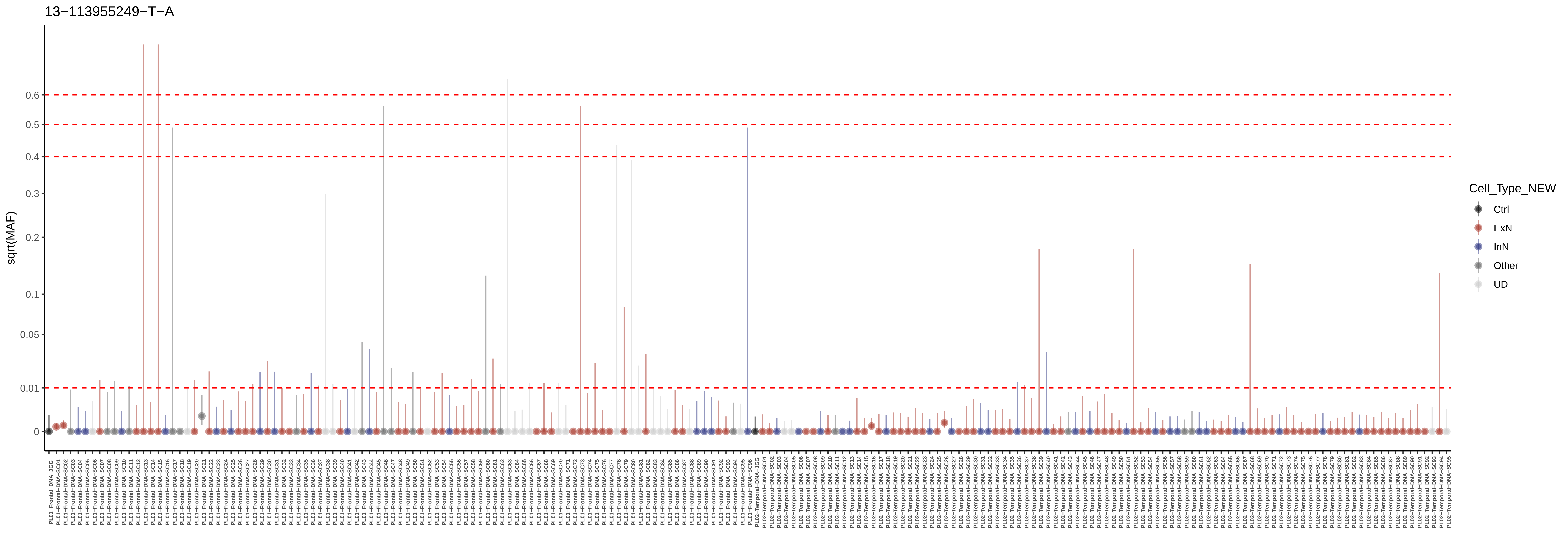
13−113955249−T−A
sqrt(MAF)
0.6
0.5
0.4
0.3
0.2
0.1
0.05
0.01
0
Cell_Type_NEW
Ctrl
ExN
InN
Other
UD

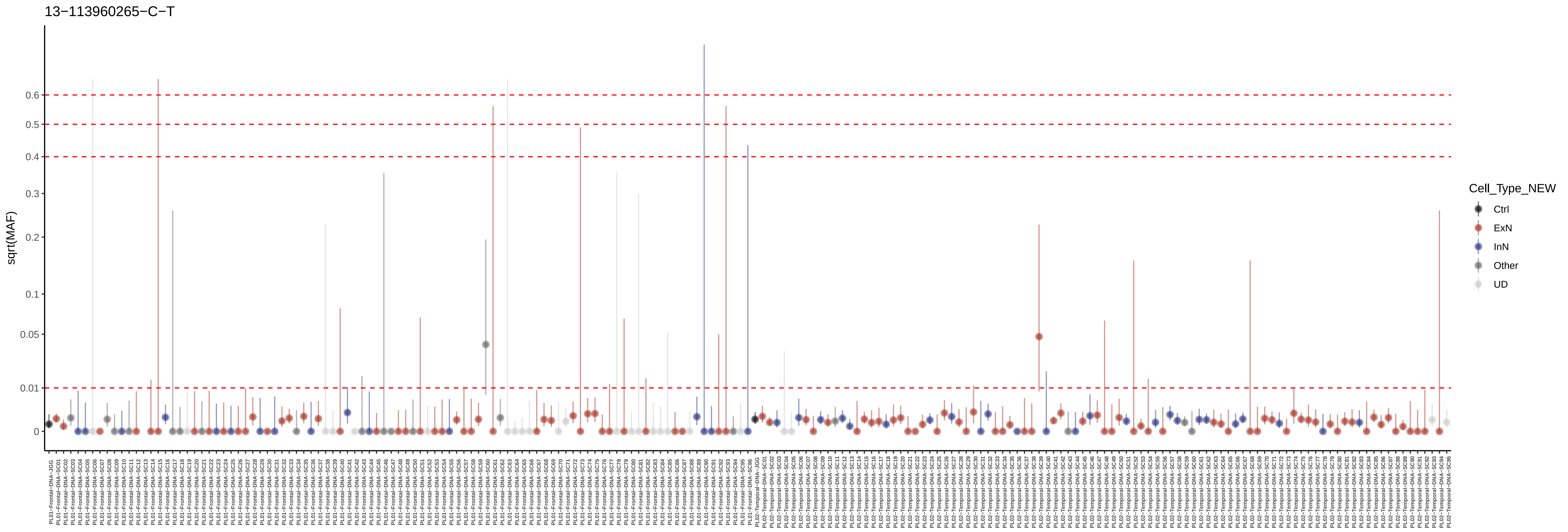
13−113960265−C−T
sqrt(MAF)
0.6
0.5
0.4
0.3
0.2
0.1
0.05
0.01
0
Cell_Type_NEW
Ctrl
ExN
InN
Other
UD

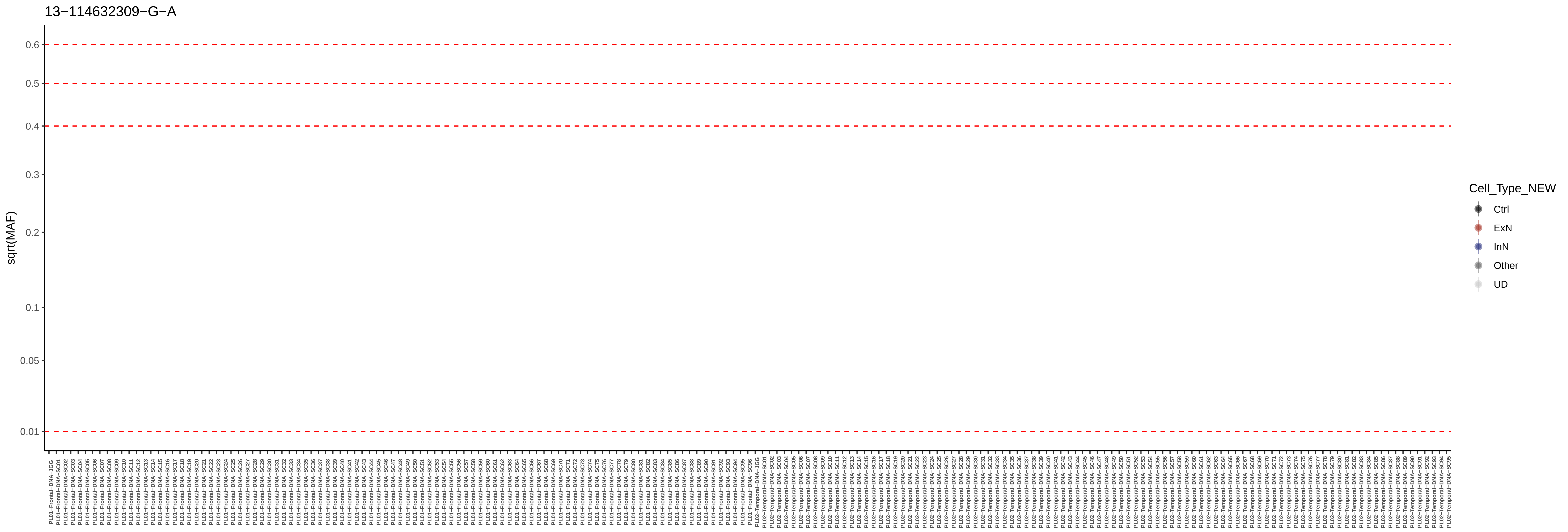
13−114632309−G−A
sqrt(MAF)
0.6
0.5
0.4
0.3
0.2
0.1
0.05
0.01
Cell_Type_NEW
Ctrl
ExN
InN
Other
UD

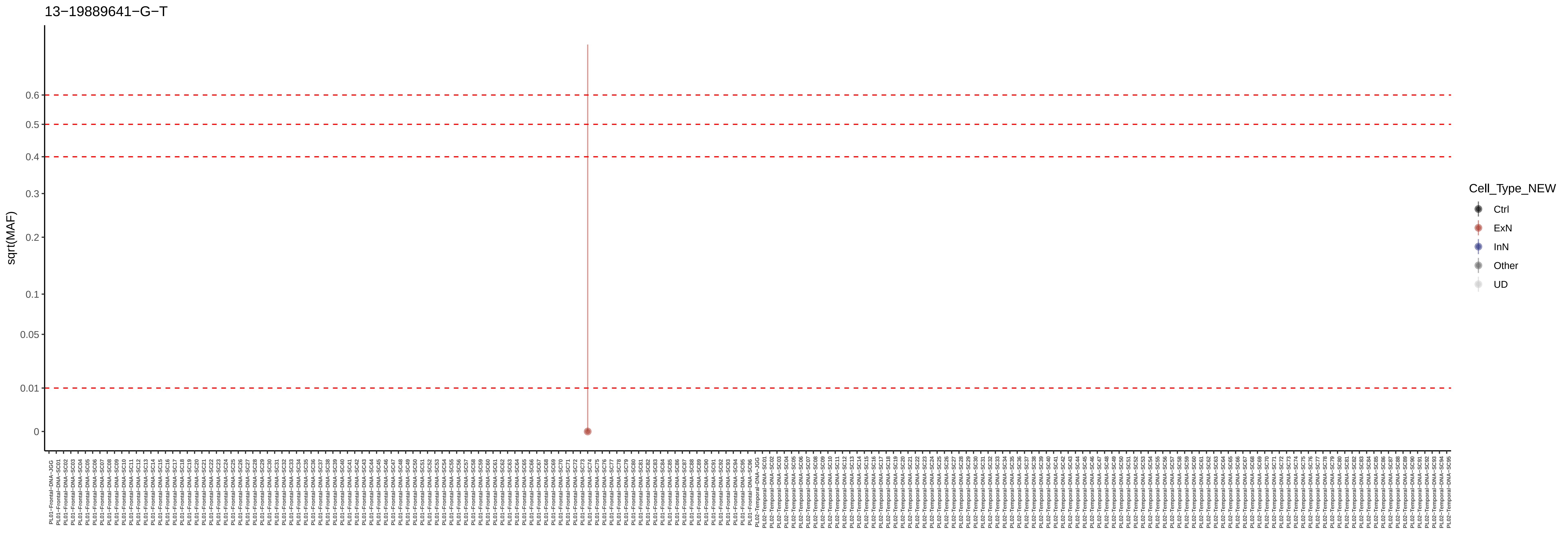
13−19889641−G−T
sqrt(MAF)
0.6
0.5
0.4
0.3
0.2
0.1
0.05
0.01
0
Cell_Type_NEW
Ctrl
ExN
InN
Other
UD

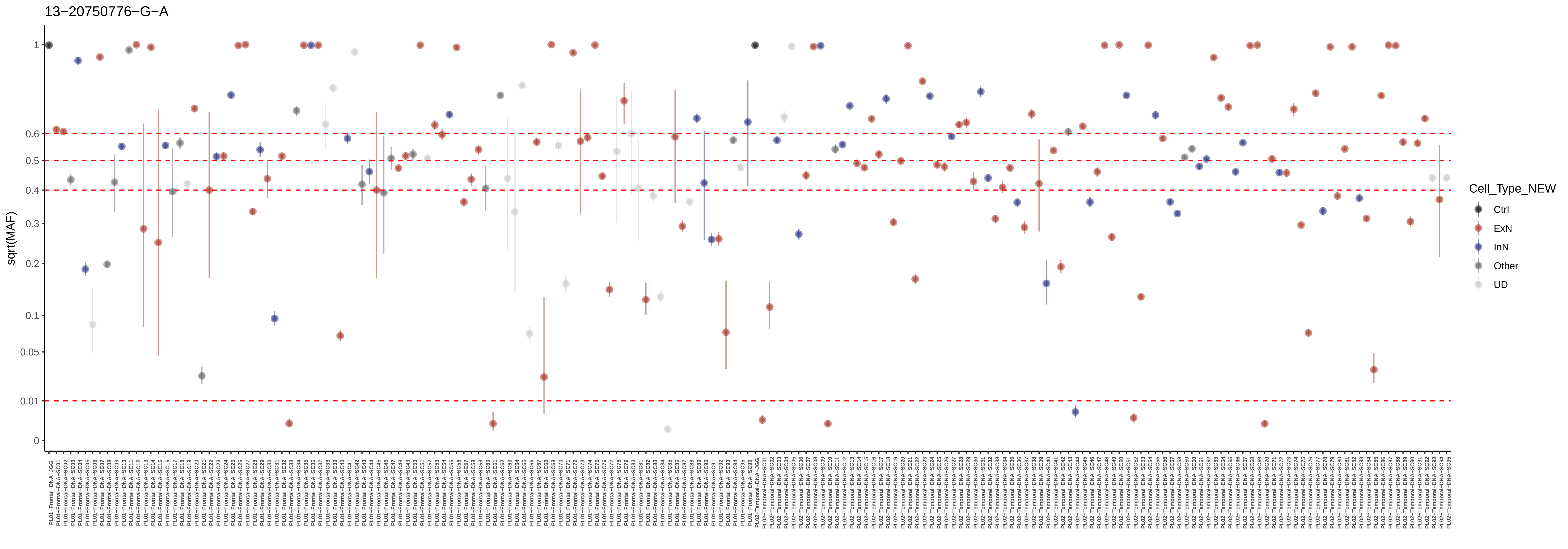

13−20750776−G−A
sqrt(MAF)
1
0.6
0.5
0.4
0.3
0.2
0.1
0.05
0.01
0
Cell_Type_NEW
Ctrl
ExN
InN
Other
UD

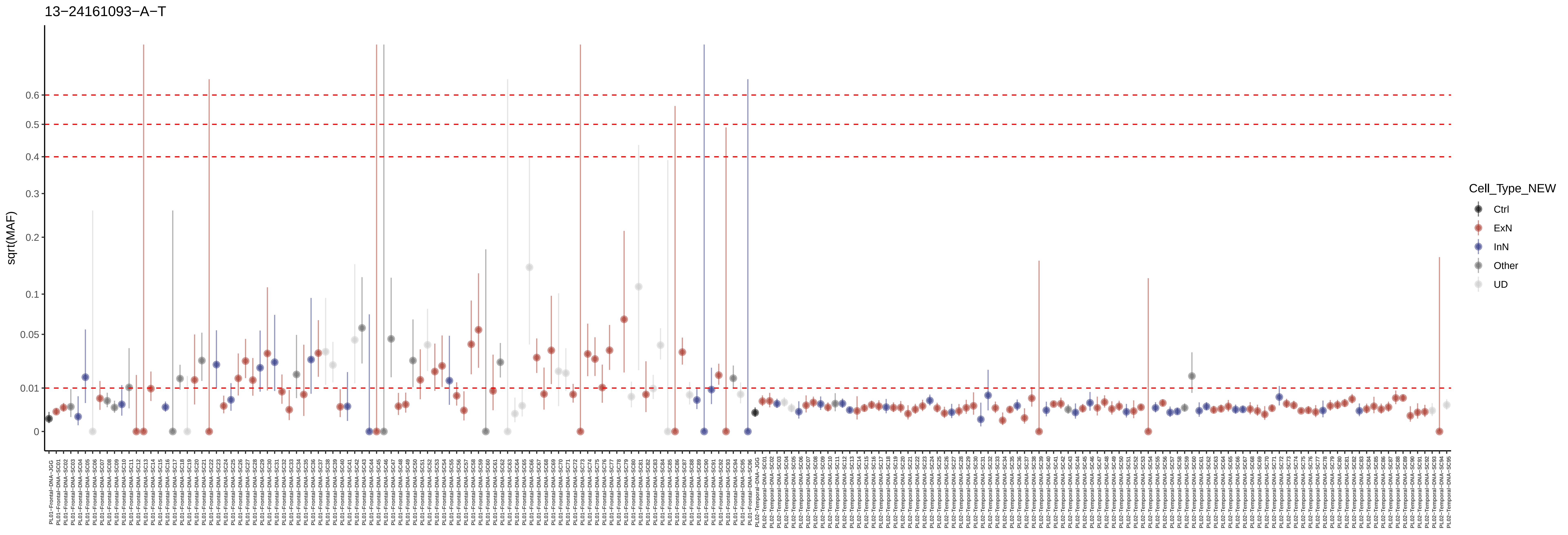

13−24161093−A−T
sqrt(MAF)
0.6
0.5
0.4
0.3
0.2
0.1
0.05
0.01
0
Cell_Type_NEW
Ctrl
ExN
InN
Other
UD

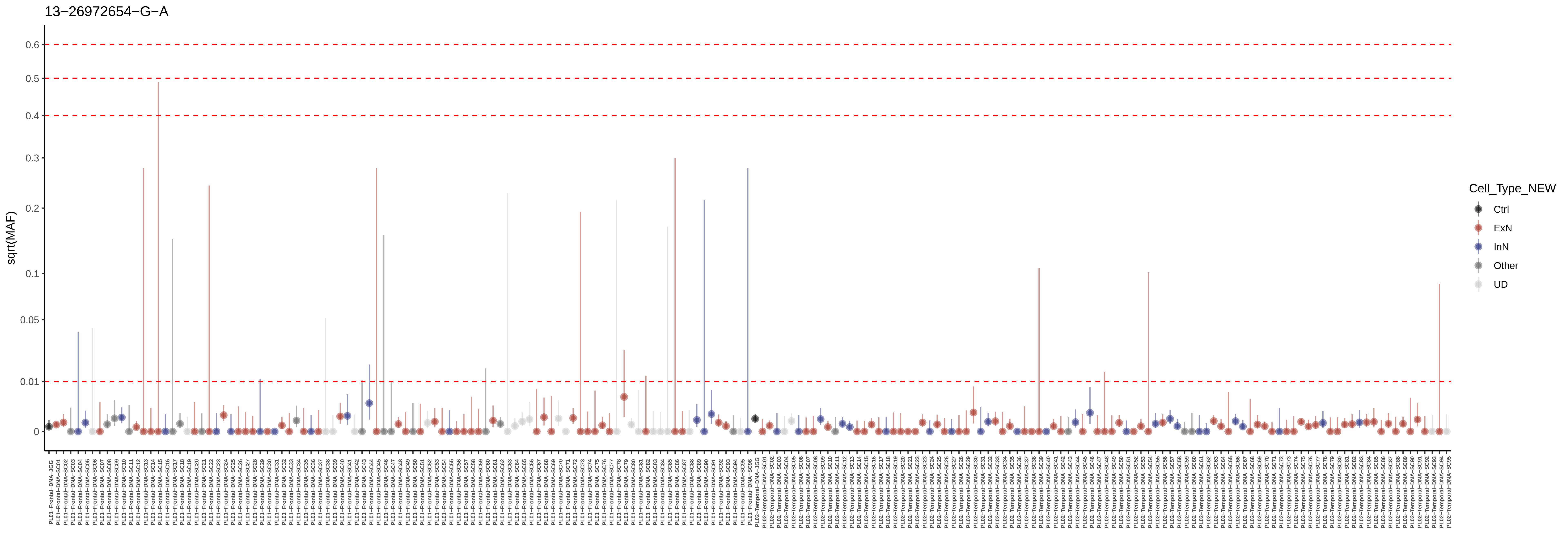
13−26972654−G−A
sqrt(MAF)
0.6
0.5
0.4
0.3
0.2
0.1
0.05
0.01
0
Cell_Type_NEW
Ctrl
ExN
InN
Other
UD

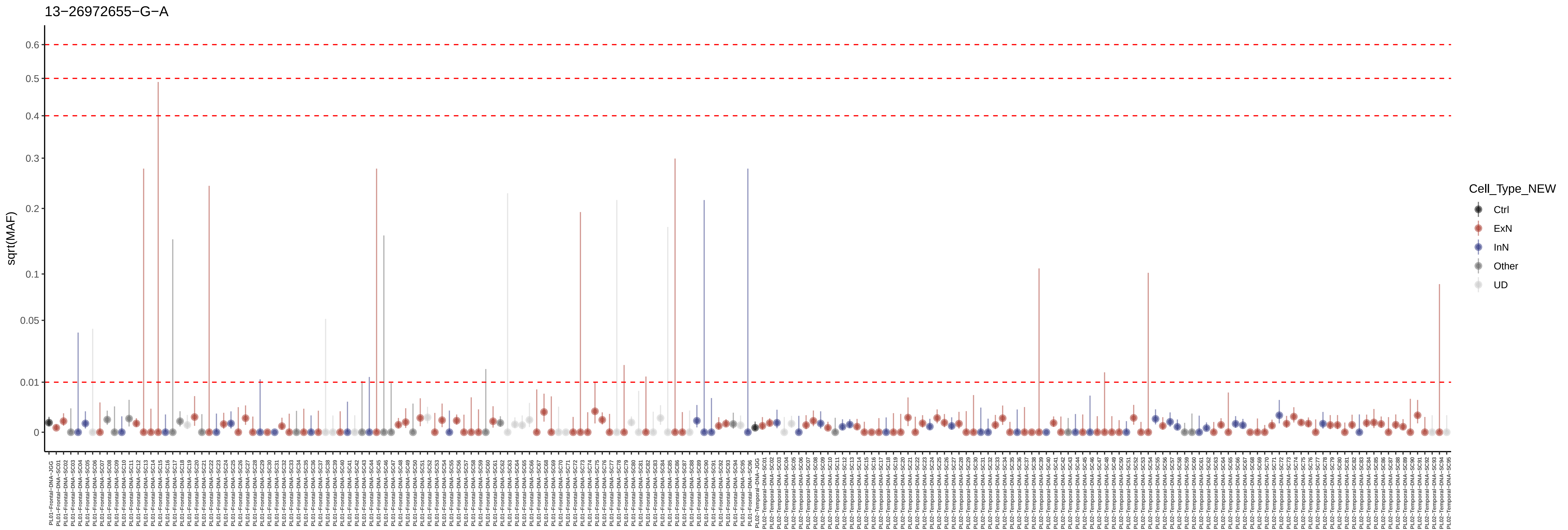

13−26972655−G−A
sqrt(MAF)
0.6
0.5
0.4
0.3
0.2
0.1
0.05
0.01
0
Cell_Type_NEW
Ctrl
ExN
InN
Other
UD

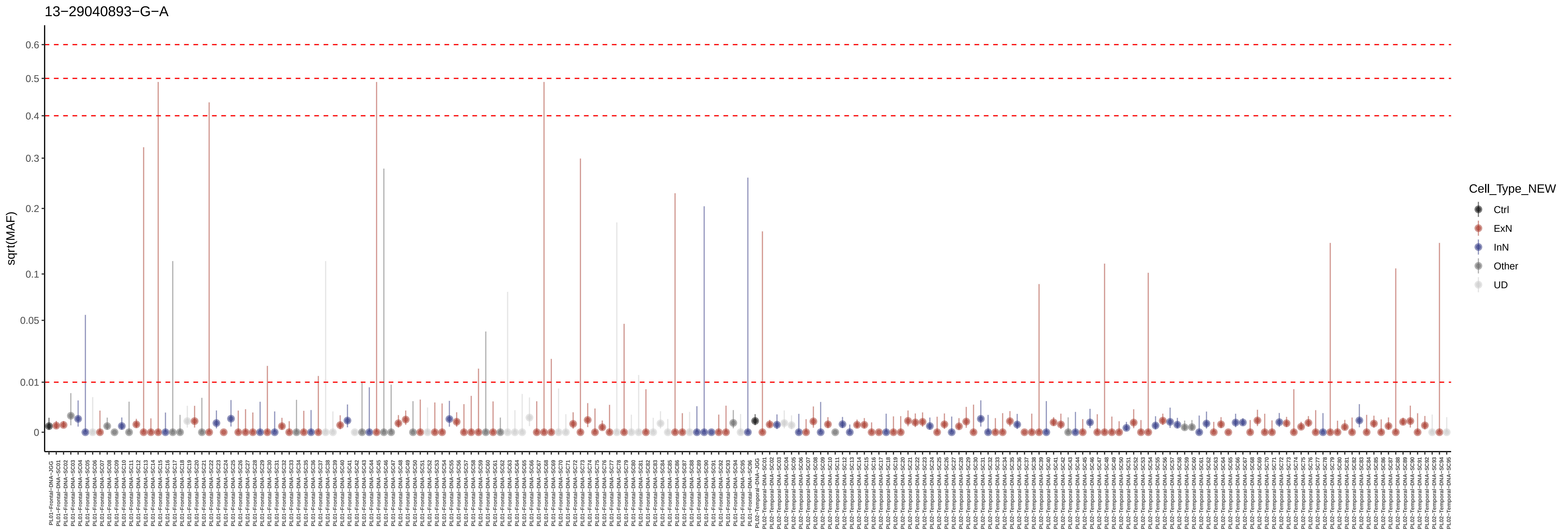
13−29040893−G−A
sqrt(MAF)
0.6
0.5
0.4
0.3
0.2
0.1
0.05
0.01
0
Cell_Type_NEW
Ctrl
ExN
InN
Other
UD

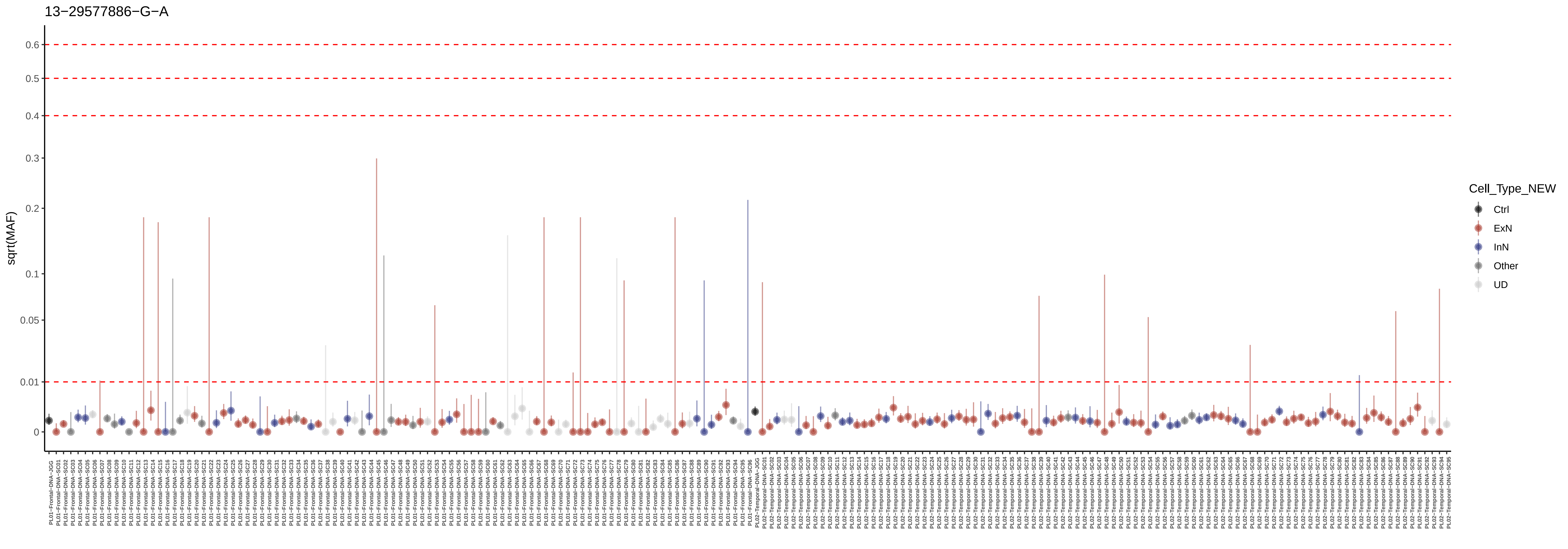
13−29577886−G−A
sqrt(MAF)
0.6
0.5
0.4
0.3
0.2
0.1
0.05
0.01
0
Cell_Type_NEW
Ctrl
ExN
InN
Other
UD

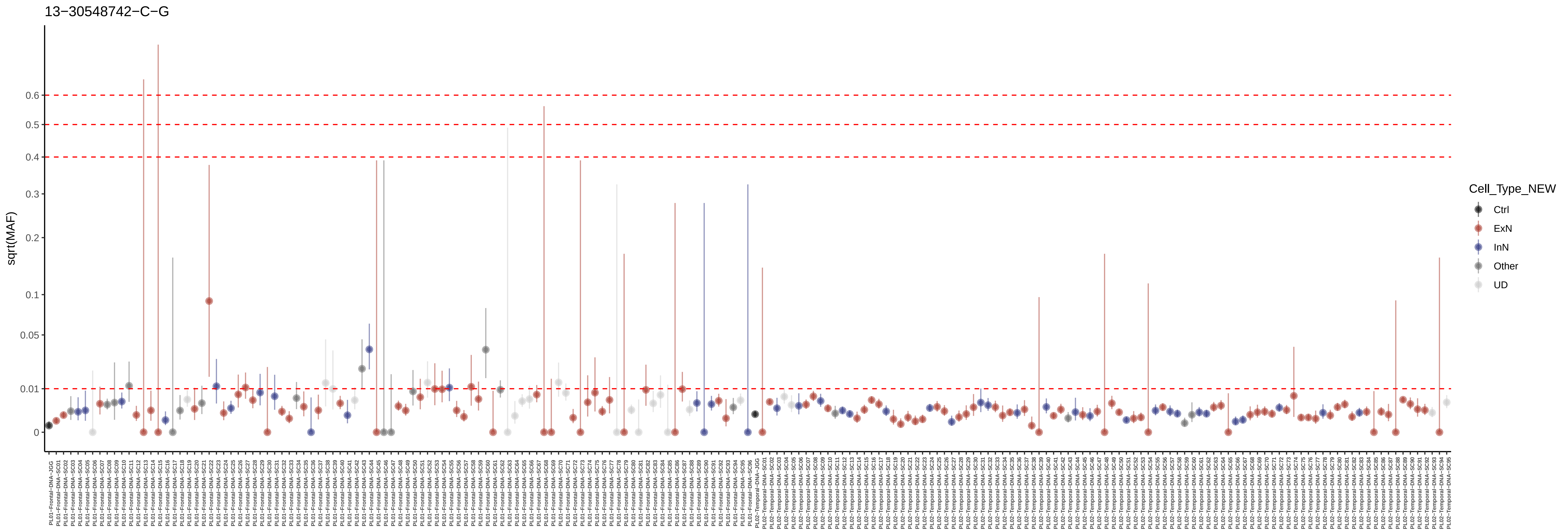
13−30548742−C−G
sqrt(MAF)
0.6
0.5
0.4
0.3
0.2
0.1
0.05
0.01
0
Cell_Type_NEW
Ctrl
ExN
InN
Other
UD

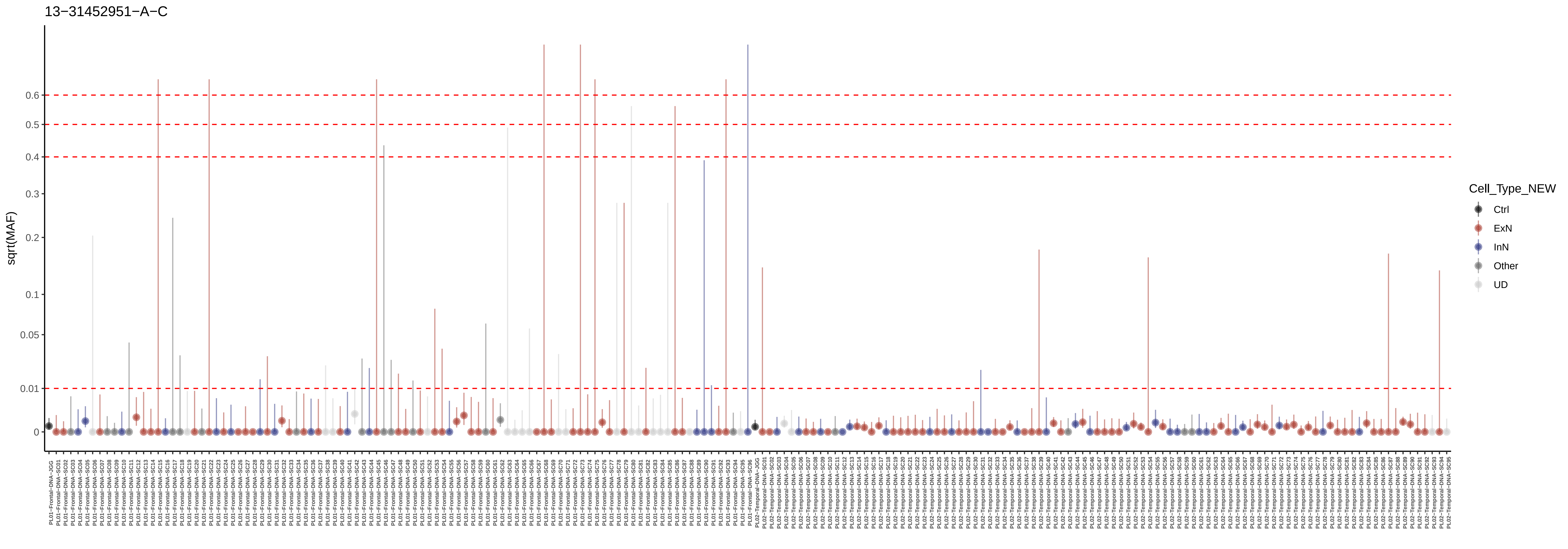
13−31452951−A−C
sqrt(MAF)
0.6
0.5
0.4
0.3
0.2
0.1
0.05
0.01
0
Cell_Type_NEW
Ctrl
ExN
InN
Other
UD

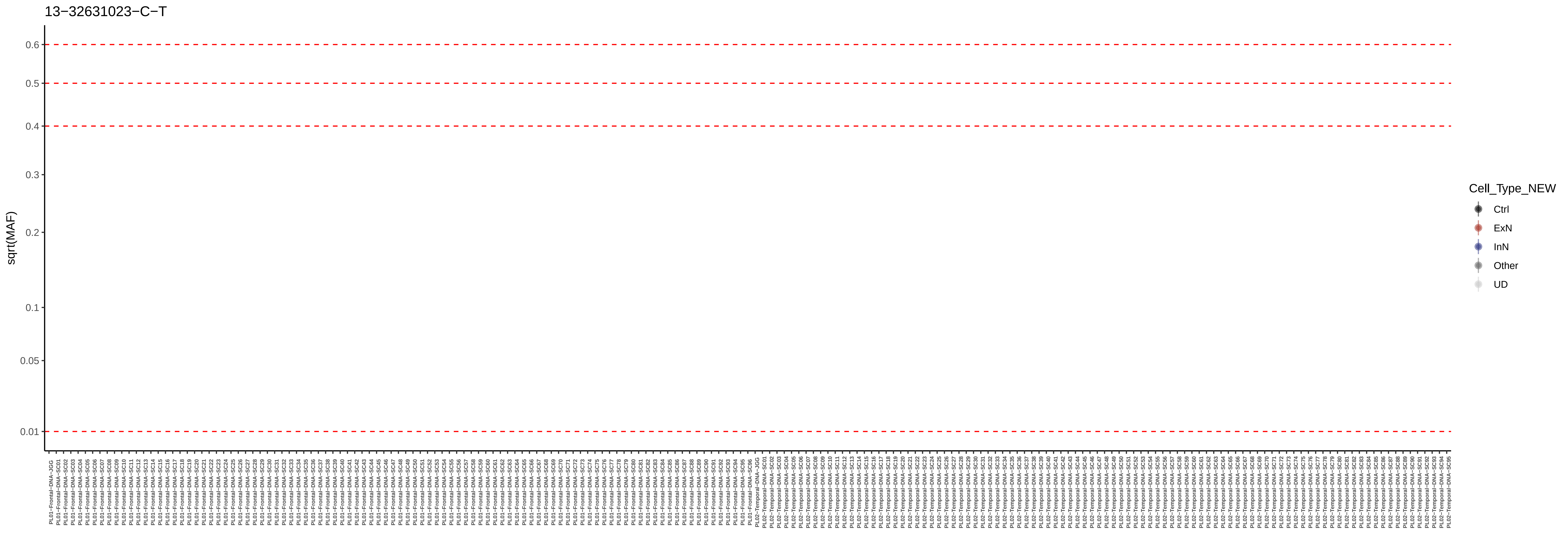
13−32631023−C−T
sqrt(MAF)
0.6
0.5
0.4
0.3
0.2
0.1
0.05
0.01
Cell_Type_NEW
Ctrl
ExN
InN
Other
UD

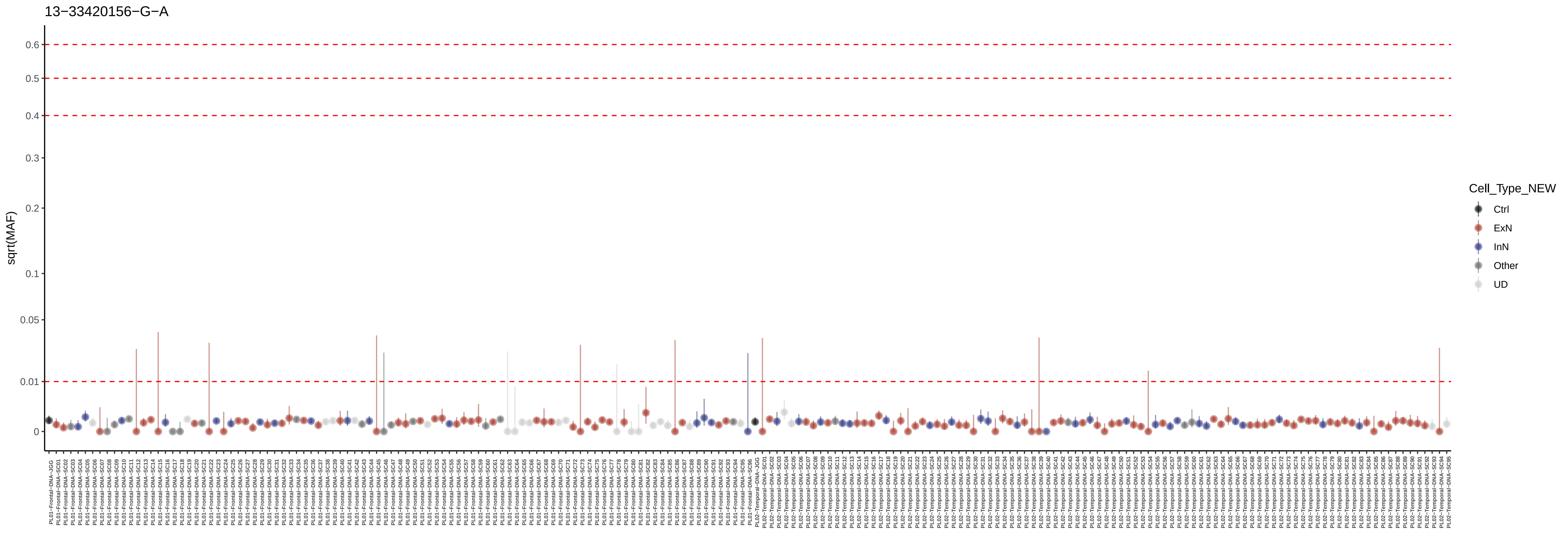

13−33420156−G−A
sqrt(MAF)
0.6
0.5
0.4
0.3
0.2
0.1
0.05
0.01
0
Cell_Type_NEW
Ctrl
ExN
InN
Other
UD

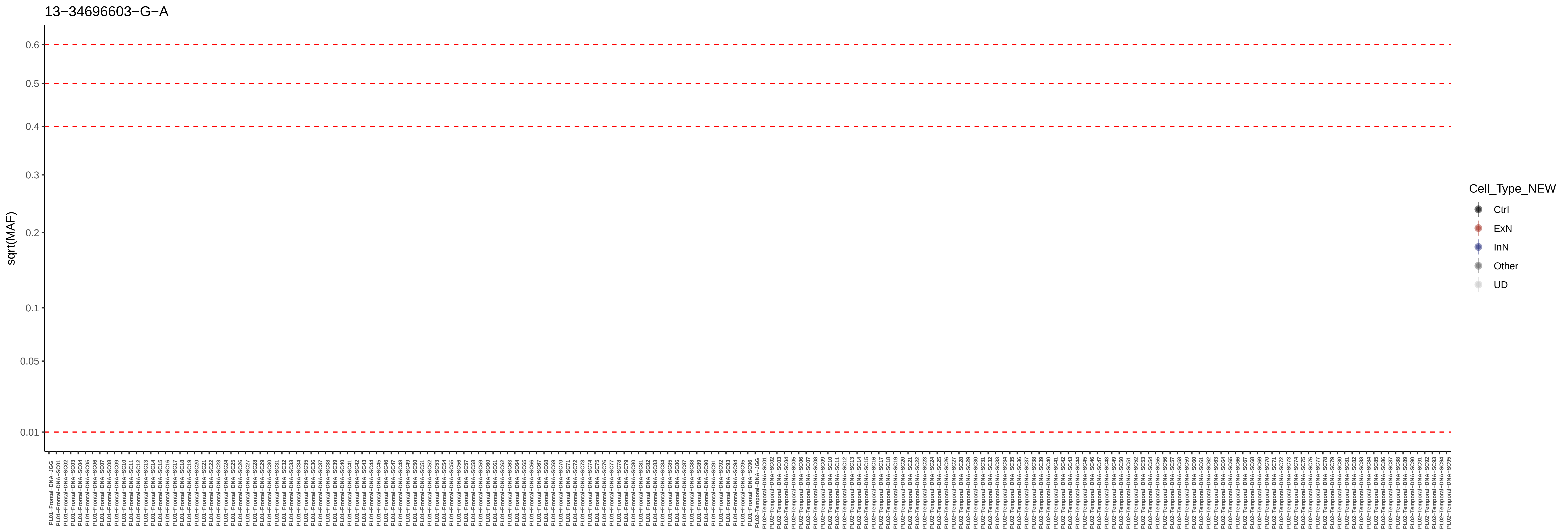
13−34696603−G−A
sqrt(MAF)
0.6
0.5
0.4
0.3
0.2
0.1
0.05
0.01
Cell_Type_NEW
Ctrl
ExN
InN
Other
UD

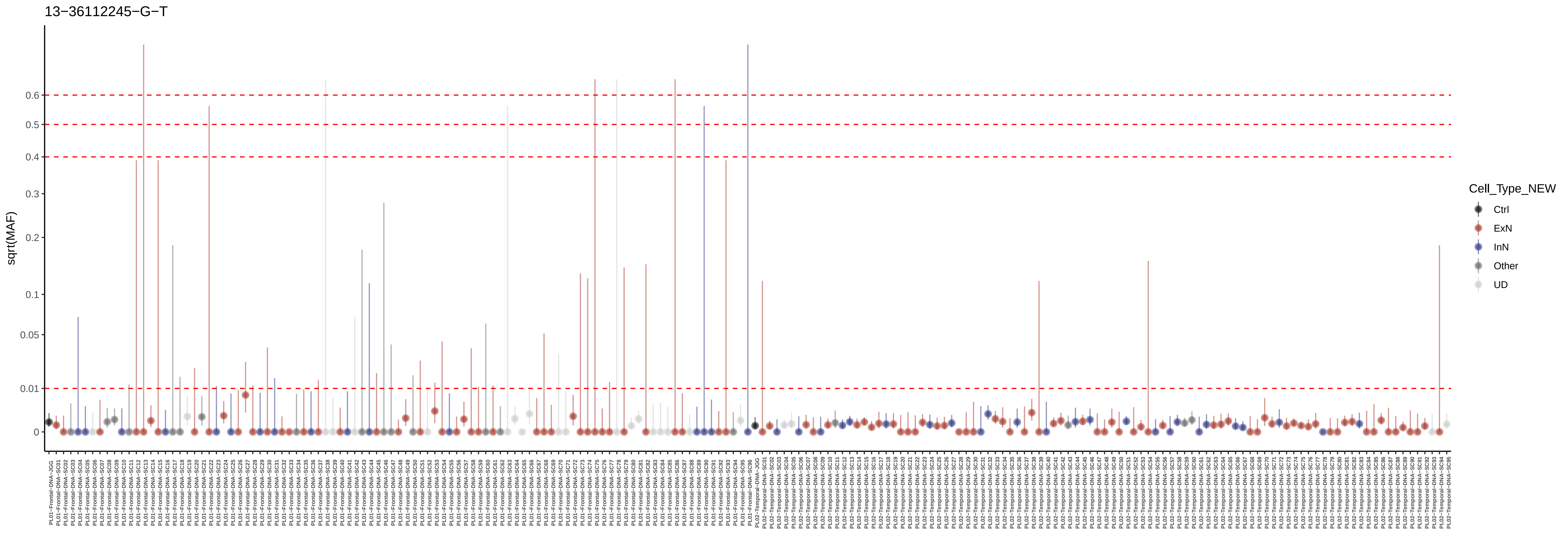

13−36112245−G−T
sqrt(MAF)
Cell_Type_NEW
Ctrl
ExN
InN
Other
UD

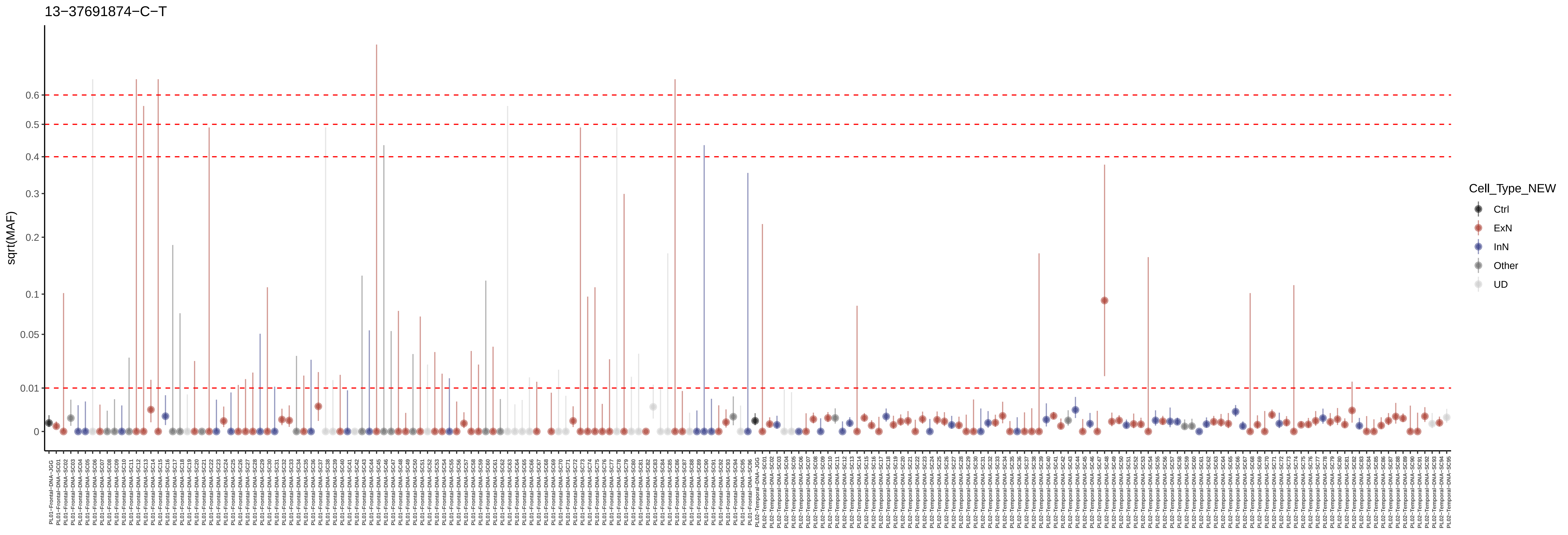
13−37691874−C−T
sqrt(MAF)
0.6
0.5
0.4
0.3
0.2
0.1
0.05
0.01
0
Cell_Type_NEW
Ctrl
ExN
InN
Other
UD

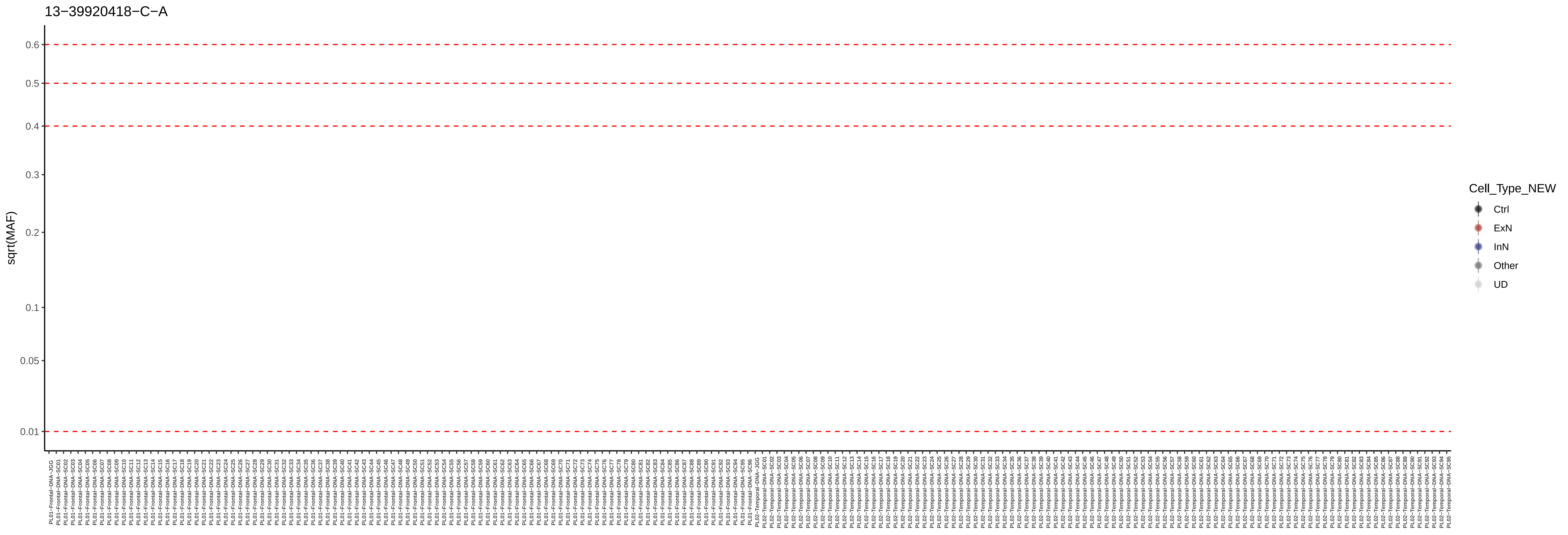
13−39920418−C−A
sqrt(MAF)
0.6
0.5
0.4
0.3
0.2
0.1
0.05
0.01
Cell_Type_NEW
Ctrl
ExN
InN
Other
UD

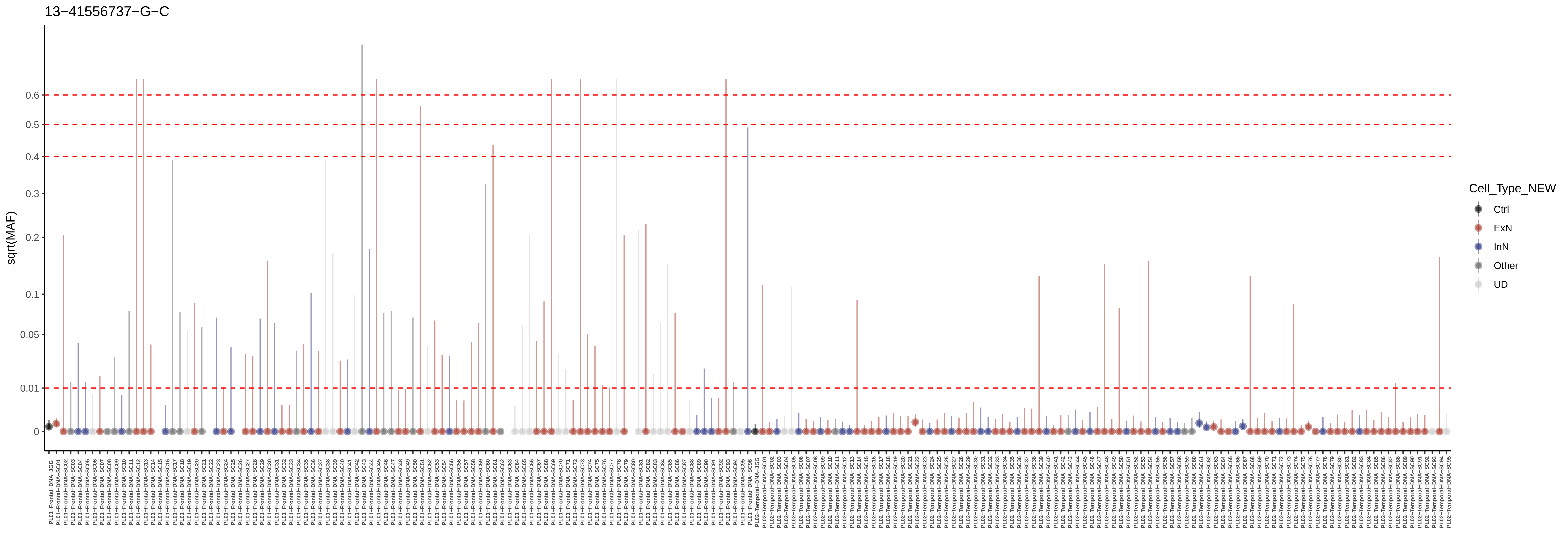
13−41556737−G−C
sqrt(MAF)
0.6
0.5
0.4
0.3
0.2
0.1
0.05
0.01
0
Cell_Type_NEW
Ctrl
ExN
InN
Other
UD

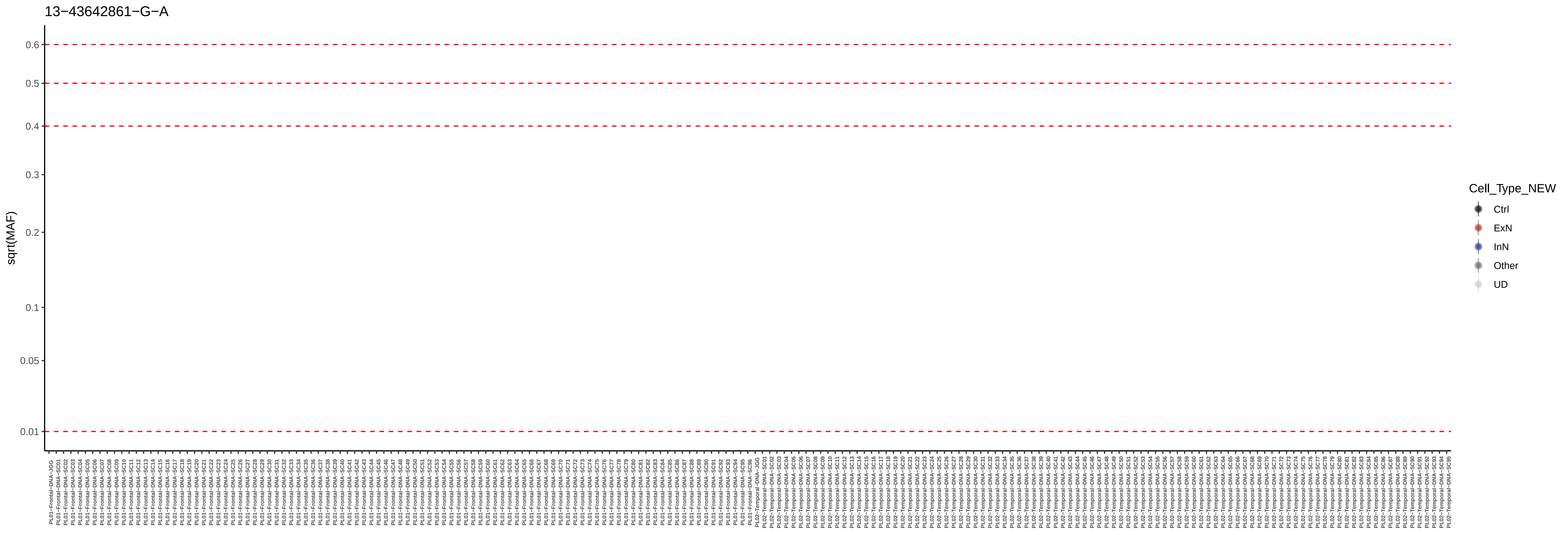
13−43642861−G−A
sqrt(MAF)
0.6
0.5
0.4
0.3
0.2
0.1
0.05
0.01
Cell_Type_NEW
Ctrl
ExN
InN
Other
UD

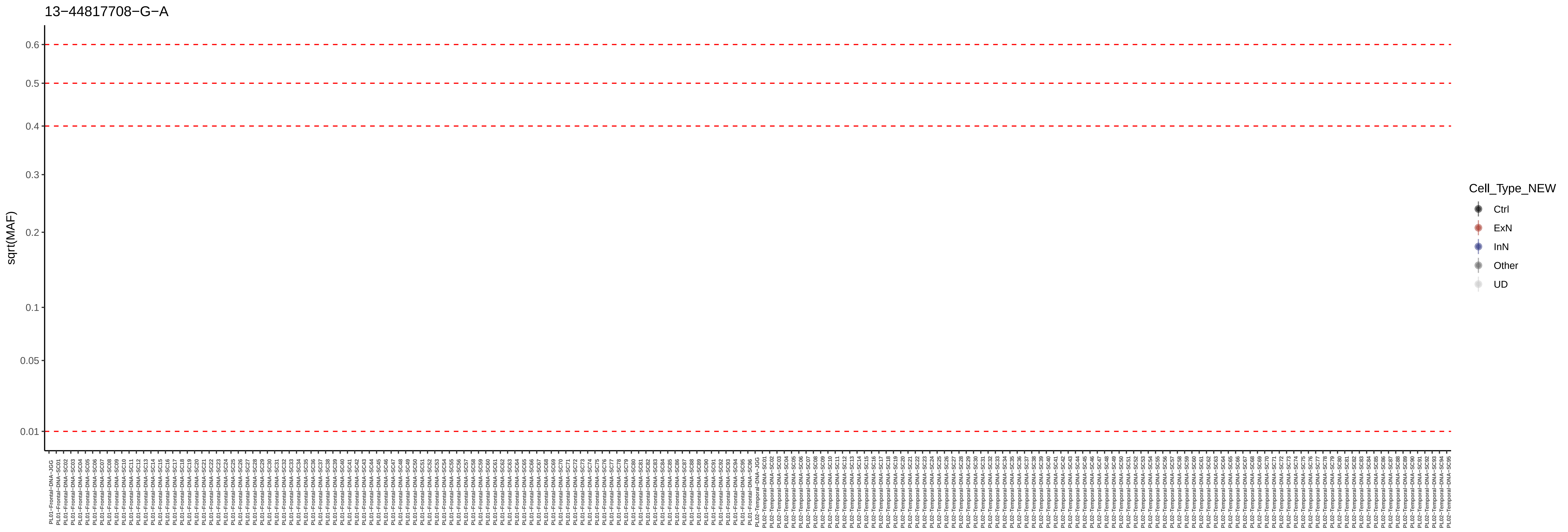
13−44817708−G−A
sqrt(MAF)
0.6
0.5
0.4
0.3
0.2
0.1
0.05
0.01
Cell_Type_NEW
Ctrl
ExN
InN
Other
UD

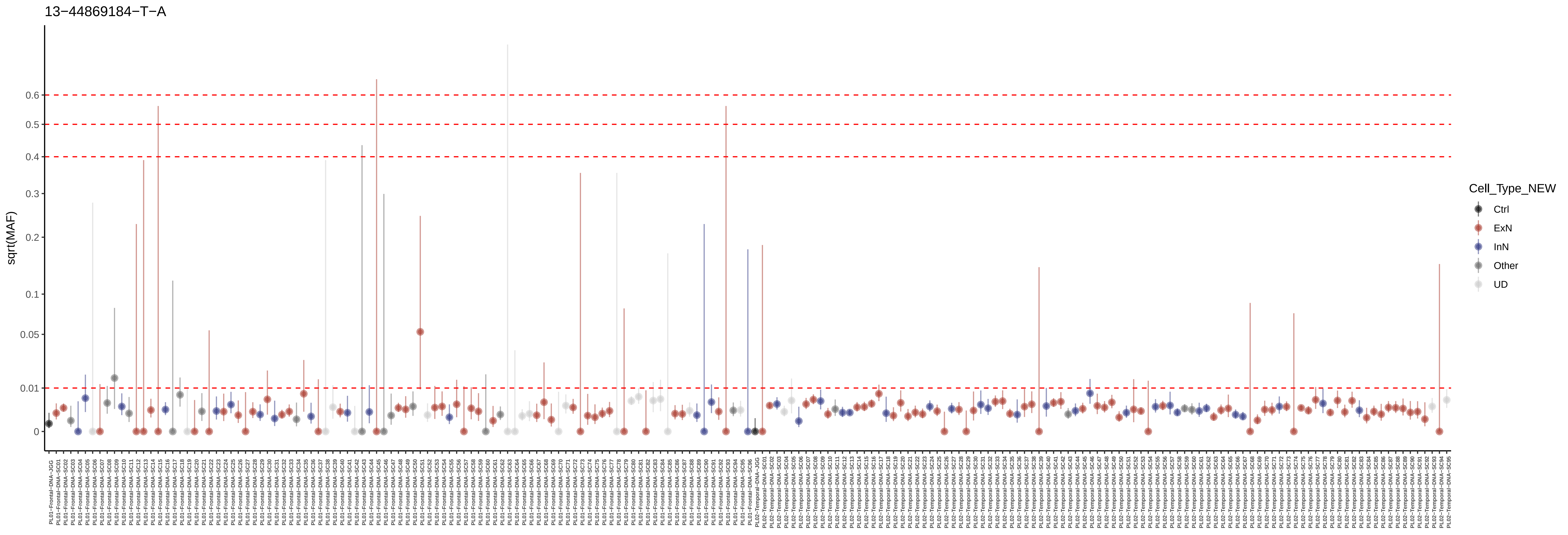
13−44869184−T−A
sqrt(MAF)
0.6
0.5
0.4
0.3
0.2
0.1
0.05
0.01
0
Cell_Type_NEW
Ctrl
ExN
InN
Other
UD

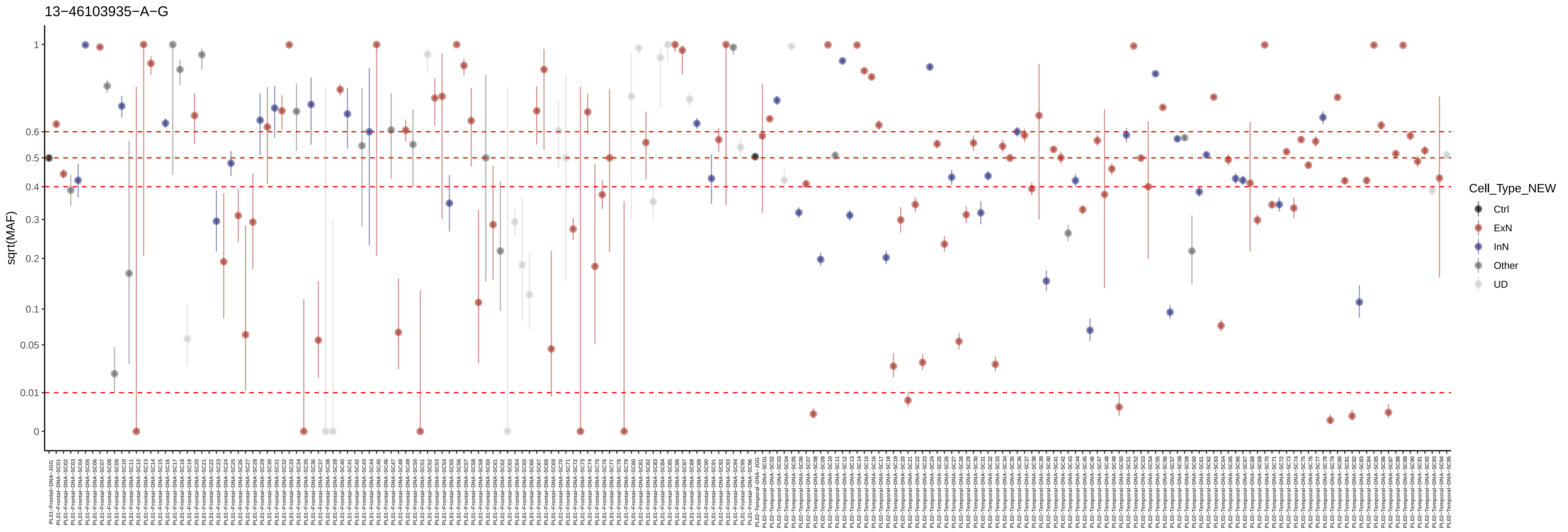
13−46103935−A−G
sqrt(MAF)
1
0.6
0.5
0.4
0.3
0.2
0.1
0.05
0.01
0
Cell_Type_NEW
Ctrl
ExN
InN
Other
UD

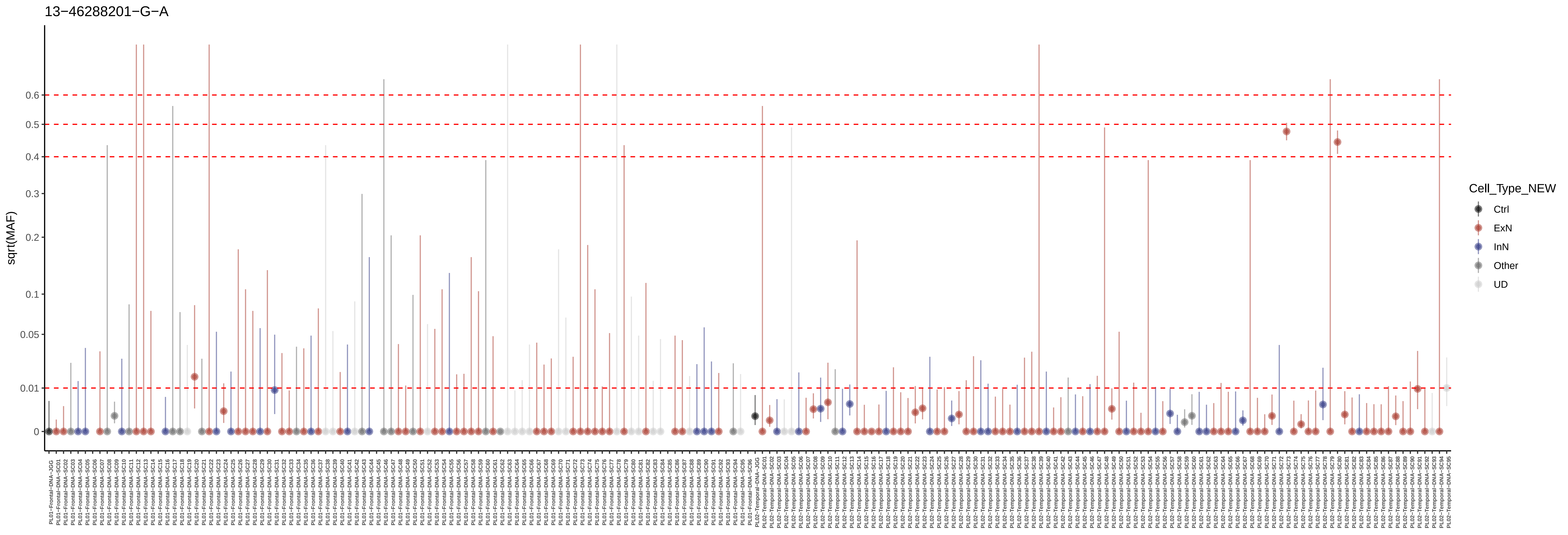
13−46288201−G−A
sqrt(MAF)
0
0.01
0.05
0.1
0.2
0.3
0.4
0.5
0.6
Cell_Type_NEW
Ctrl
ExN
InN
Other
UD

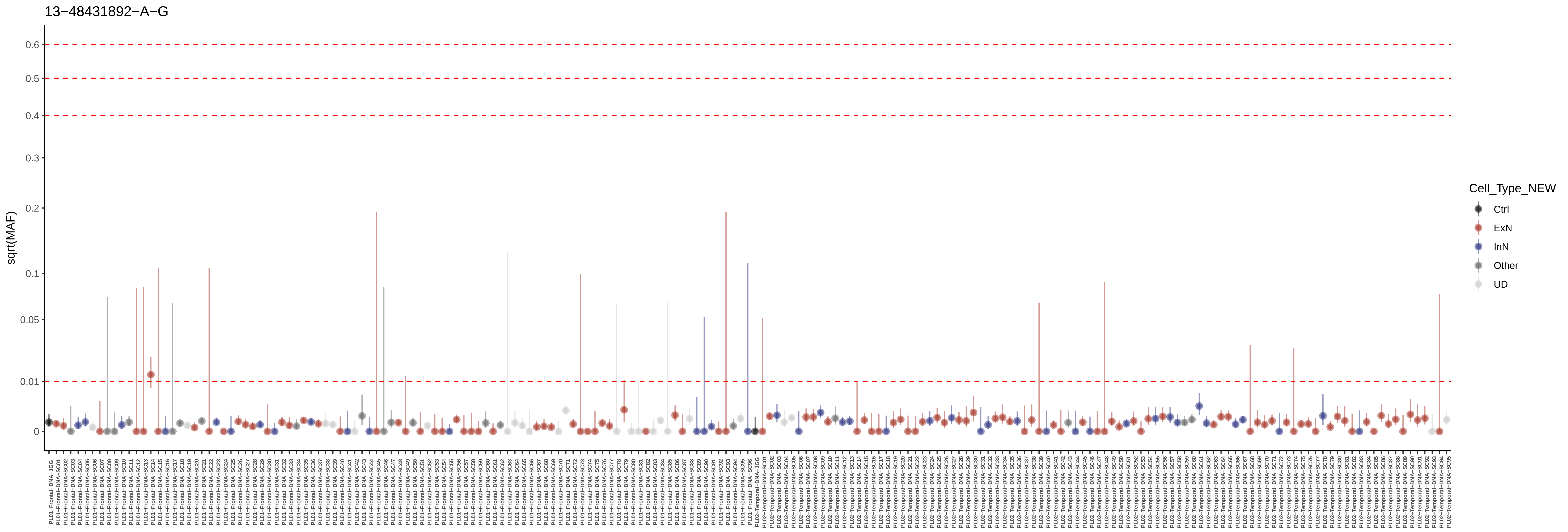
13−48431892−A−G
sqrt(MAF)
0.6
0.5
0.4
0.3
0.2
0.1
0.05
0.01
0
Cell_Type_NEW
Ctrl
ExN
InN
Other
UD

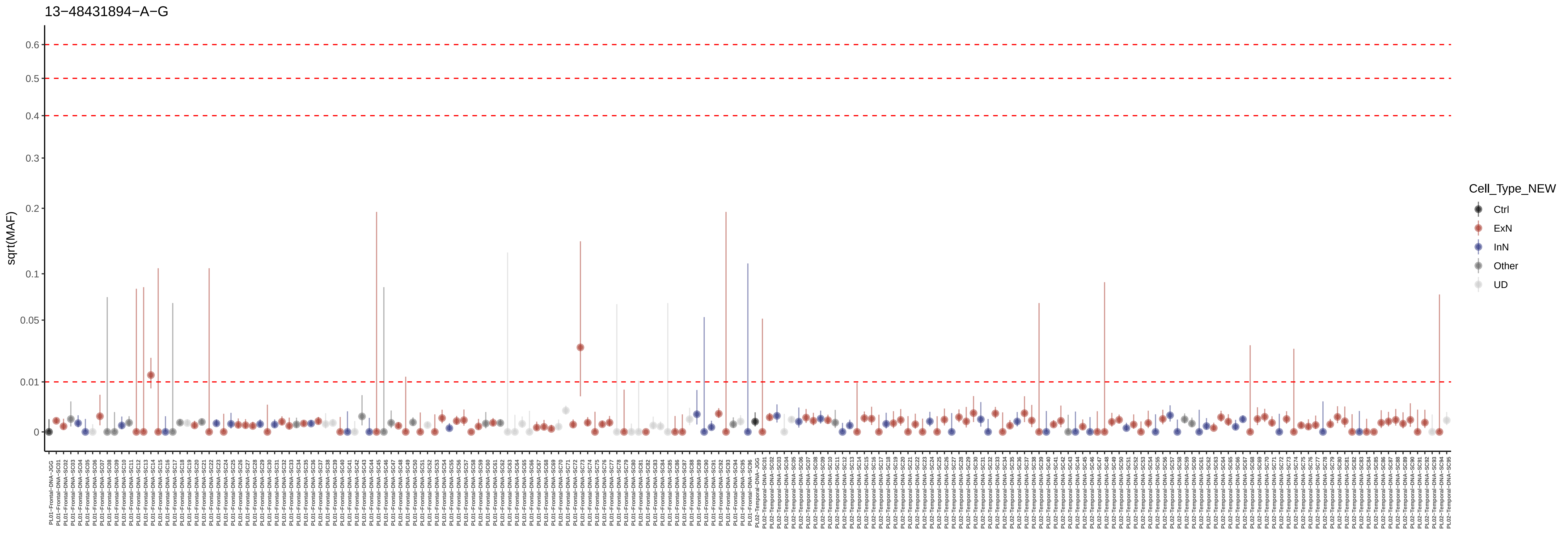
13−48431894−A−G
sqrt(MAF)
0.6
0.5
0.4
0.3
0.2
0.1
0.05
0.01
0
Cell_Type_NEW
Ctrl
ExN
InN
Other
UD

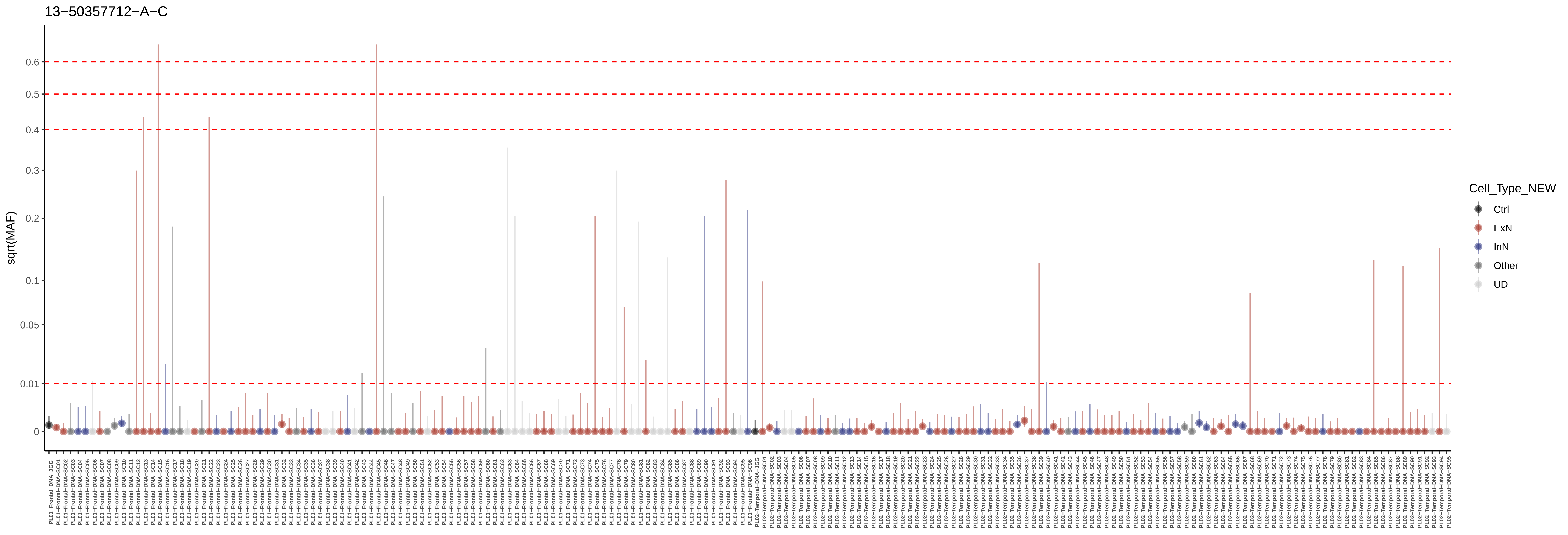
13−50357712−A−C
sqrt(MAF)
0.6
0.5
0.4
0.3
0.2
0.1
0.05
0.01
0
Cell_Type_NEW
Ctrl
ExN
InN
Other
UD

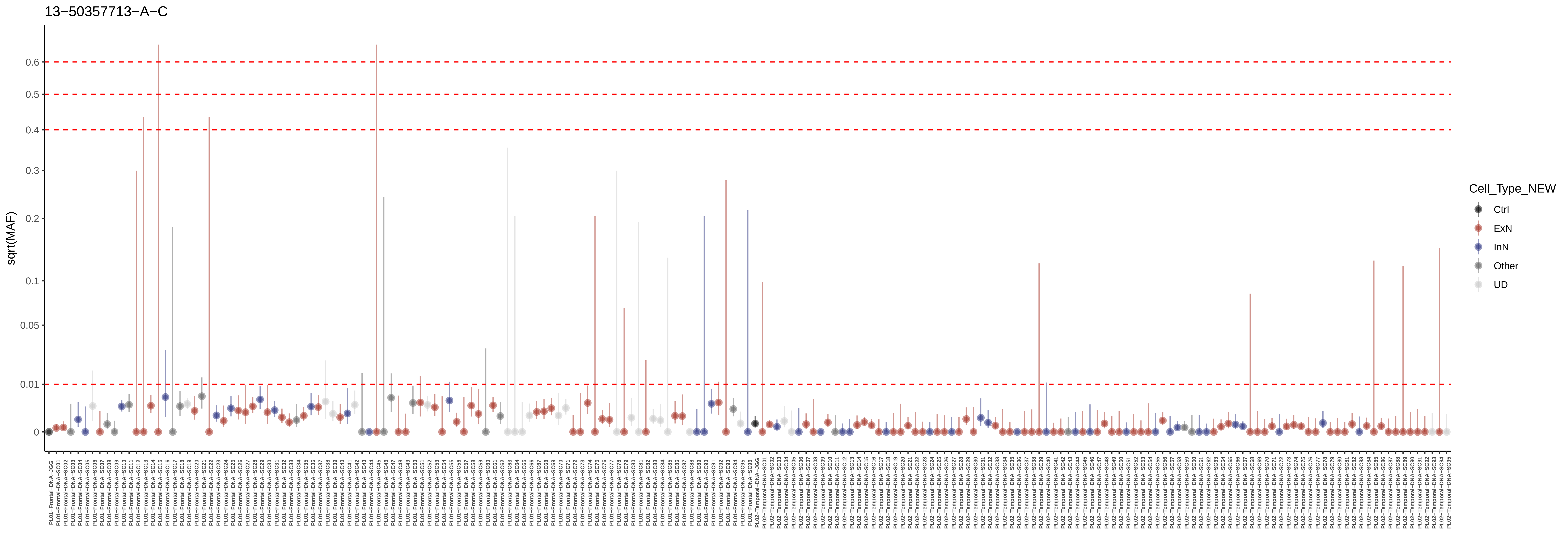

13−50357713−A−C
sqrt(MAF)
0.6
0.5
0.4
0.3
0.2
0.1
0.05
0.01
0
Cell_Type_NEW
Ctrl
ExN
InN
Other
UD

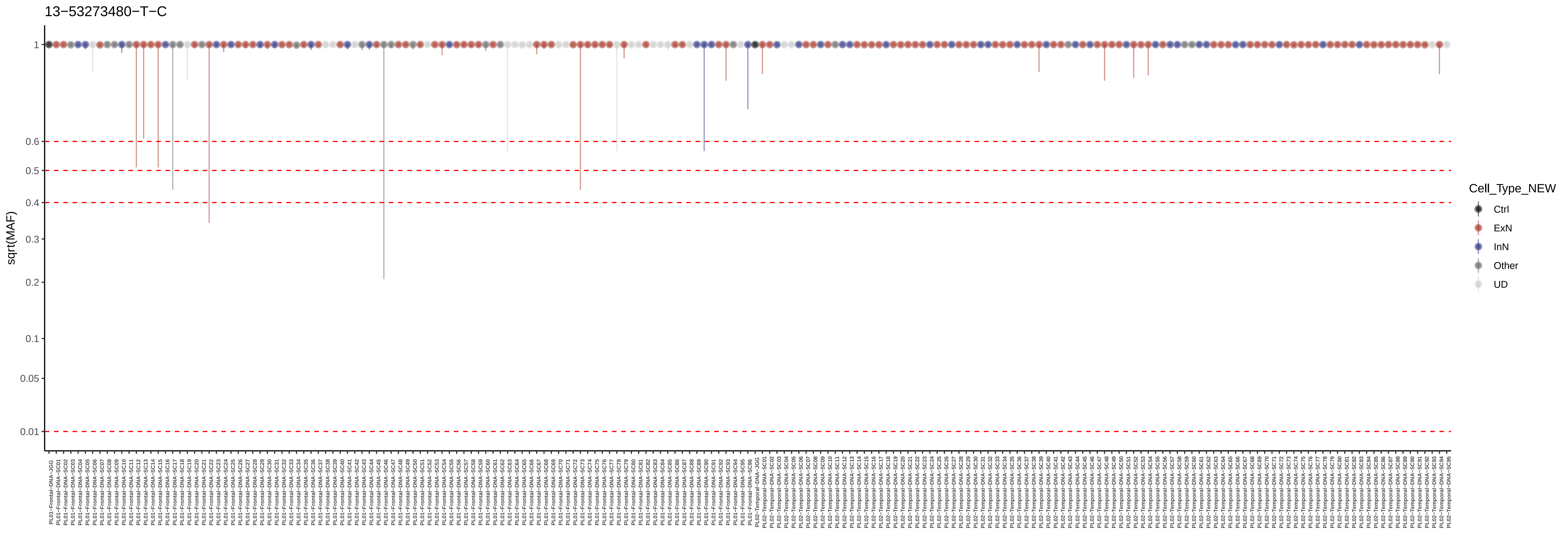

13−53273480−T−C
sqrt(MAF)
1
0.6
0.5
0.4
0.3
0.2
0.1
0.05
0.01
Cell_Type_NEW
Ctrl
ExN
InN
Other
UD

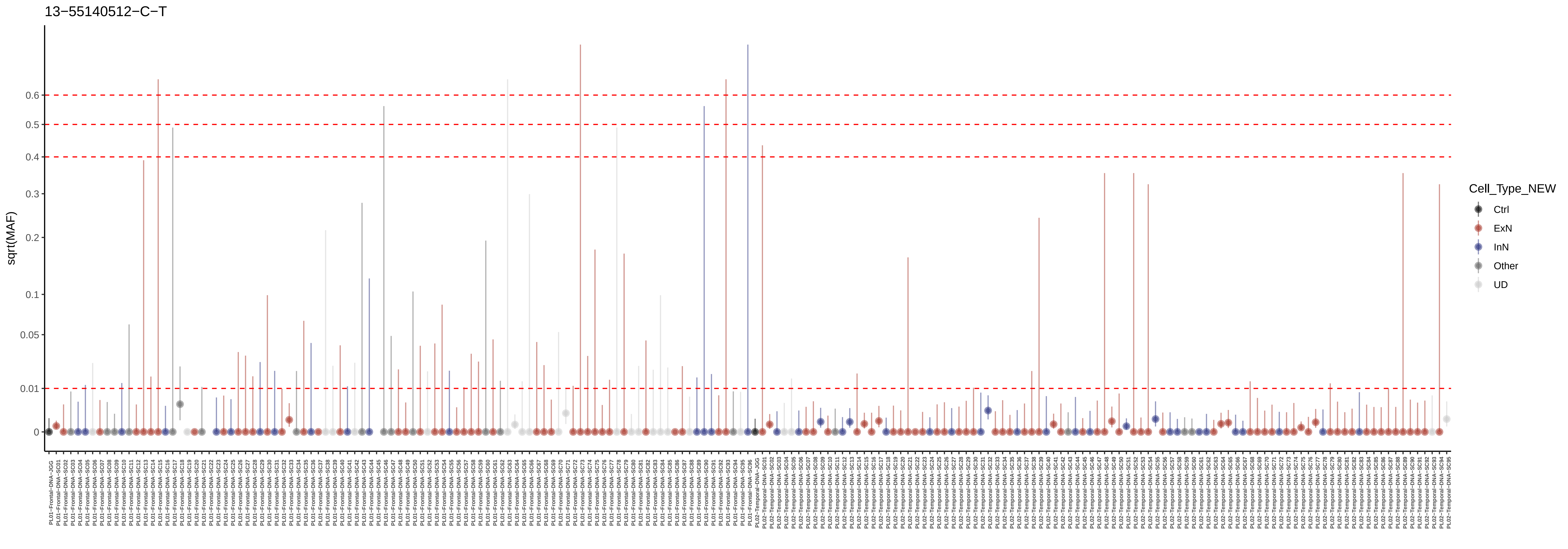

13−55140512−C−T
sqrt(MAF)
0.6
0.5
0.4
0.3
0.2
0.1
0.05
0.01
0
Cell_Type_NEW
Ctrl
ExN
InN
Other
UD

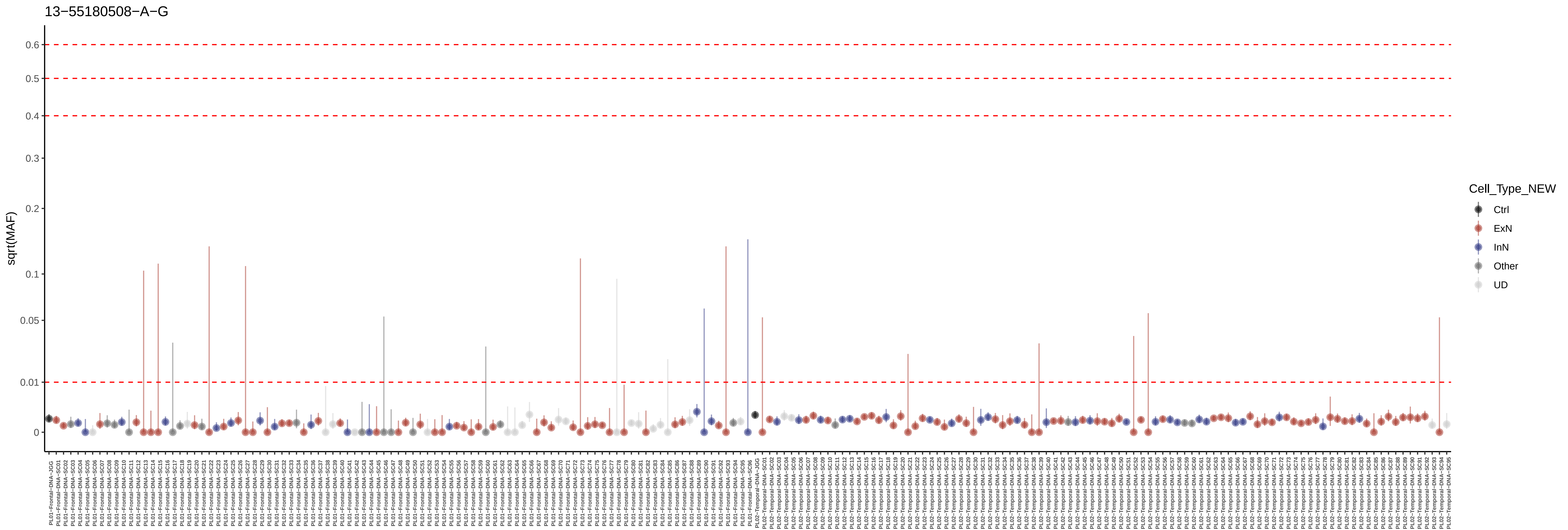
13−55180508−A−G
sqrt(MAF)
0.6
0.5
0.4
0.3
0.2
0.1
0.05
0.01
0
Cell_Type_NEW
Ctrl
ExN
InN
Other
UD

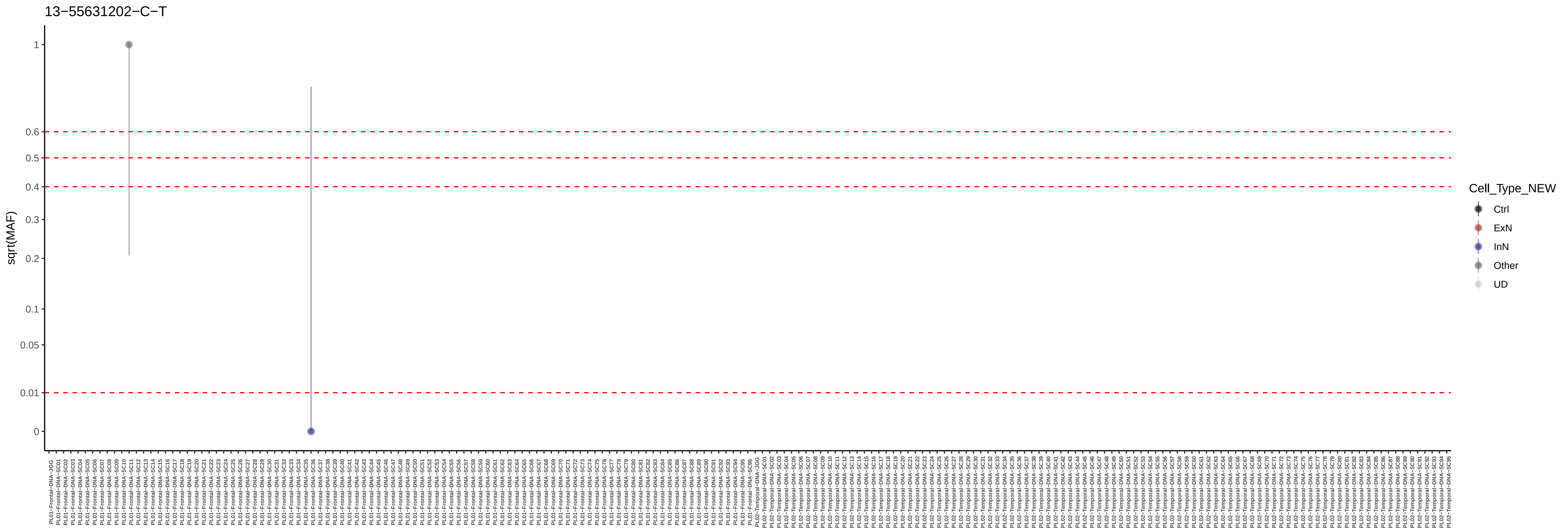
13−55631202−C−T
sqrt(MAF)
1
0.6
0.5
0.4
0.3
0.2
0.1
0.05
0.01
0
Cell_Type_NEW
Ctrl
ExN
InN
Other
UD

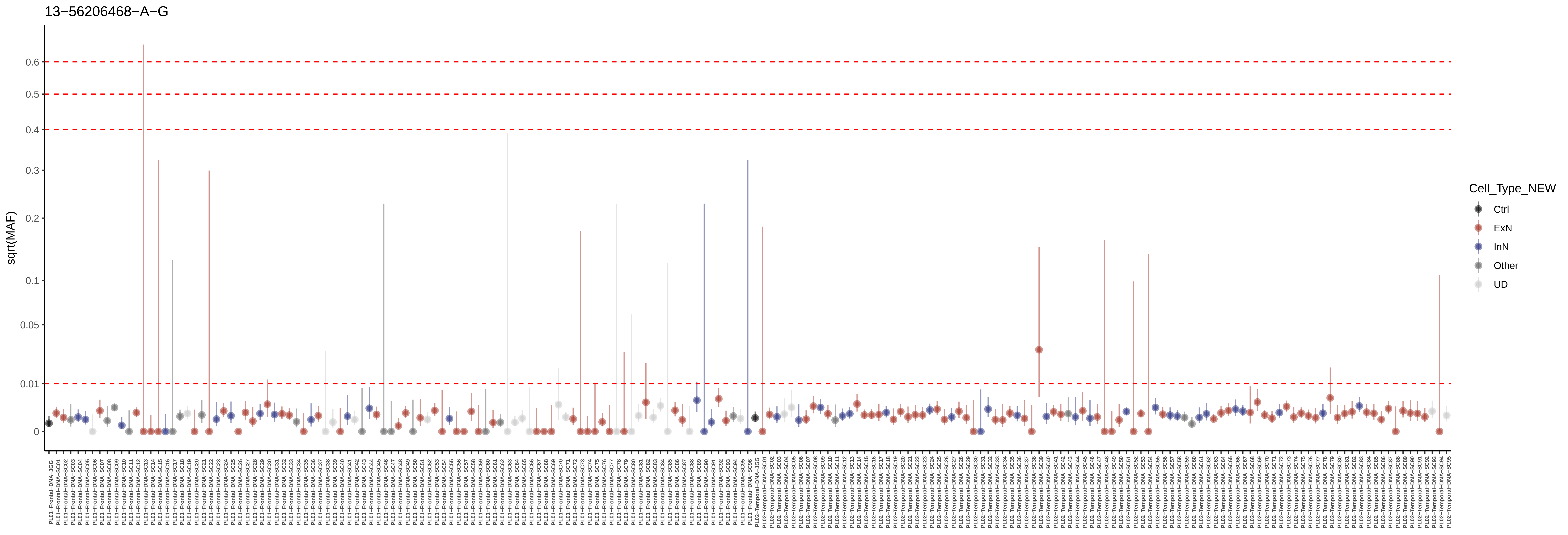
13−56206468−A−G
sqrt(MAF)
0.6
0.5
0.4
0.3
0.2
0.1
0.05
0.01
0
Cell_Type_NEW
Ctrl
ExN
InN
Other
UD

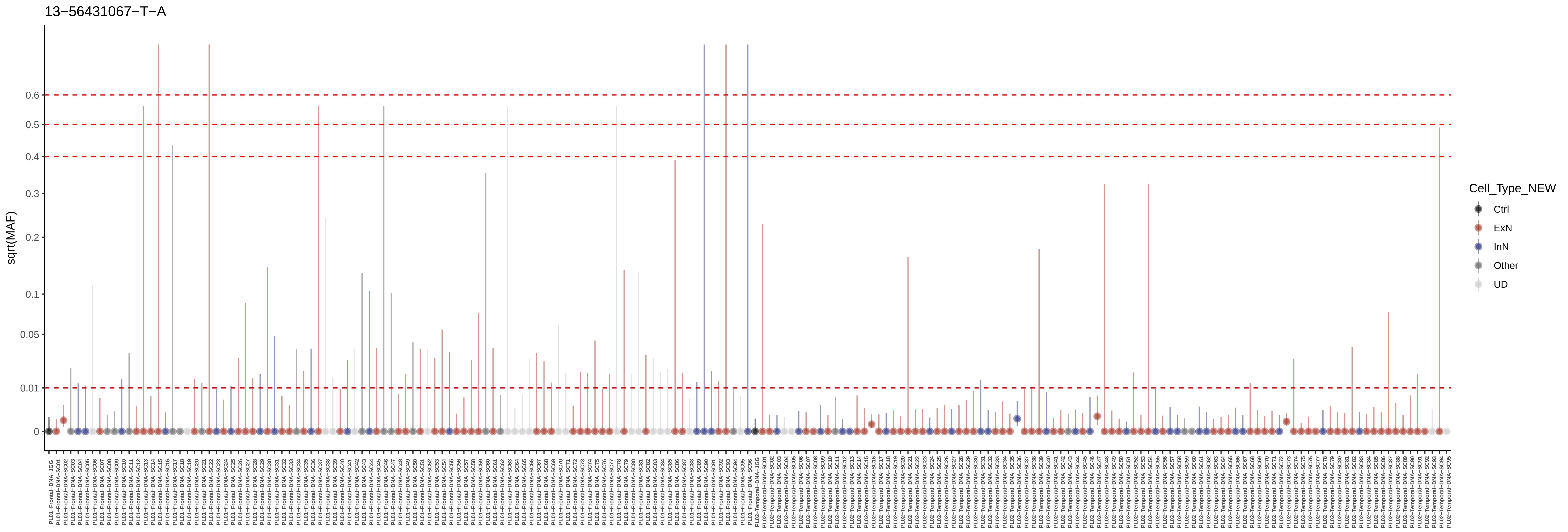
13−56431067−T−A
sqrt(MAF)
0.6
0.5
0.4
0.3
0.2
0.1
0.05
0.01
0
Cell_Type_NEW
Ctrl
ExN
InN
Other
UD

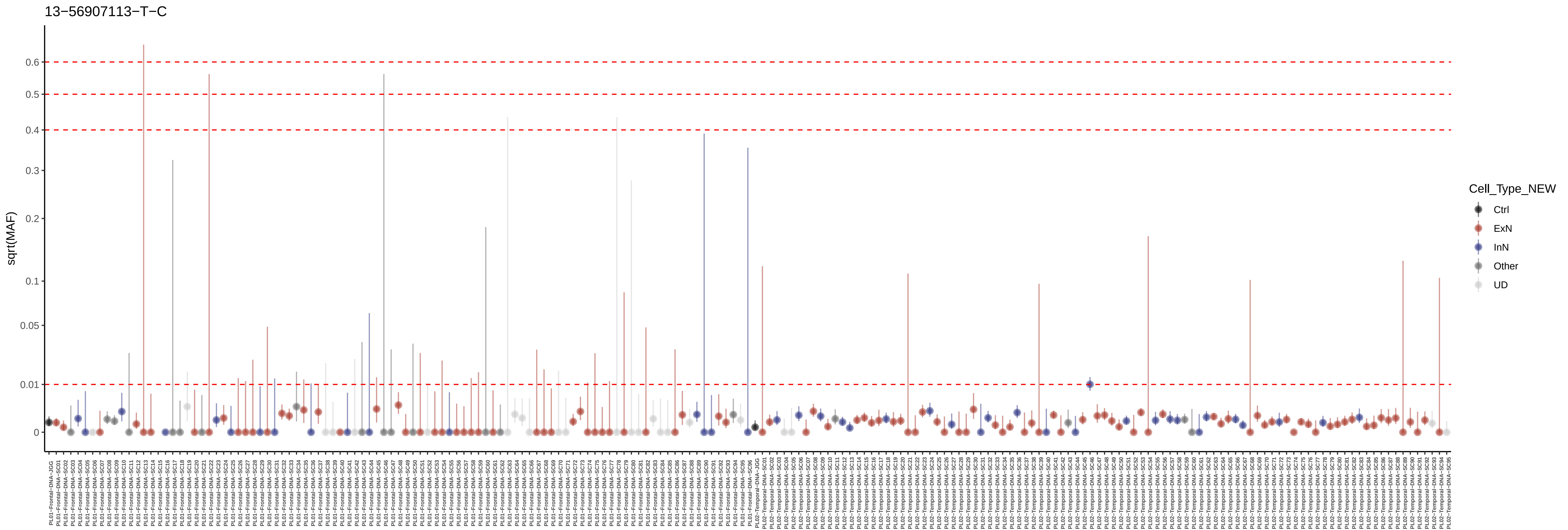

13−56907113−T−C
sqrt(MAF)
0.6
0.5
0.4
0.3
0.2
0.1
0.05
0.01
0
Cell_Type_NEW
Ctrl
ExN
InN
Other
UD

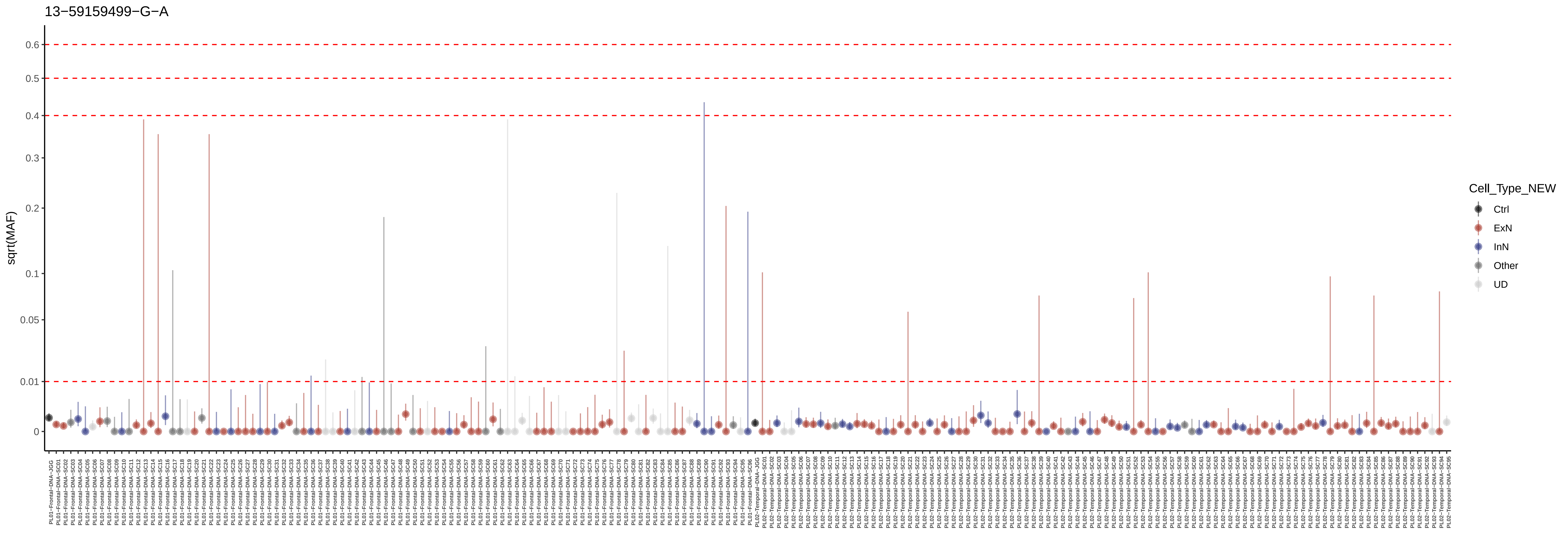
13−59159499−G−A
sqrt(MAF)
0.6
0.5
0.4
0.3
0.2
0.1
0.05
0.01
0
Cell_Type_NEW
Ctrl
ExN
InN
Other
UD

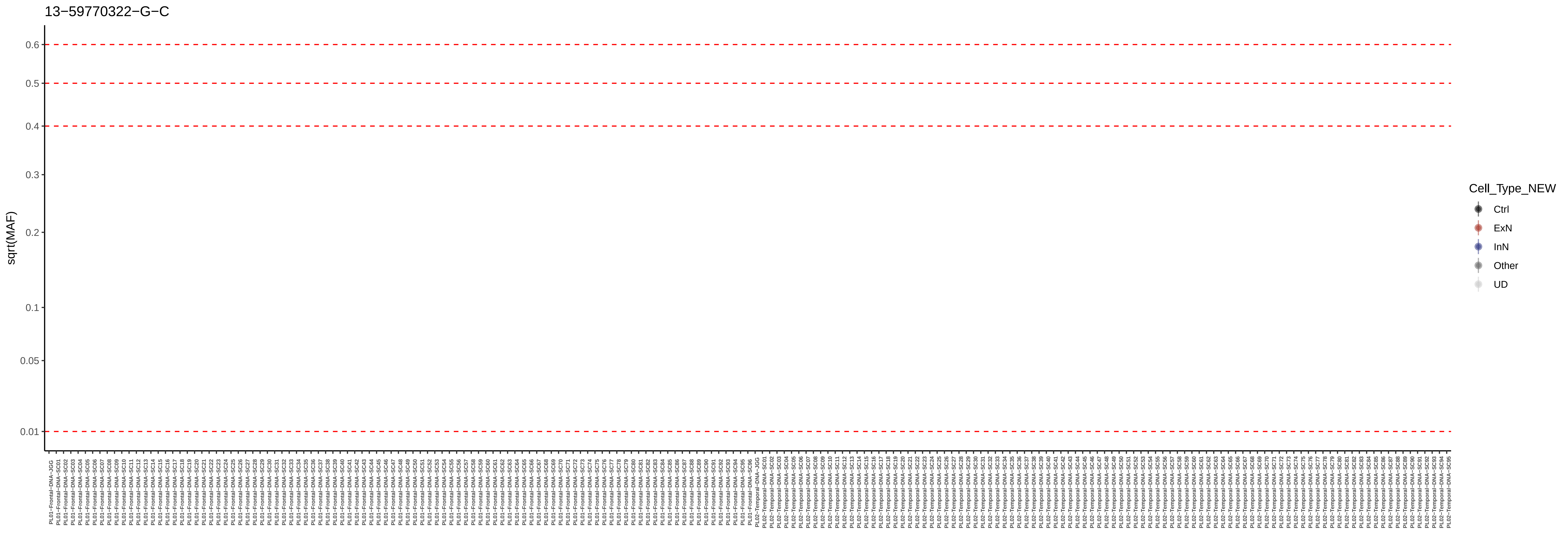
13−59770322−G−C
sqrt(MAF)
0.6
0.5
0.4
0.3
0.2
0.1
0.05
0.01
Cell_Type_NEW
Ctrl
ExN
InN
Other
UD

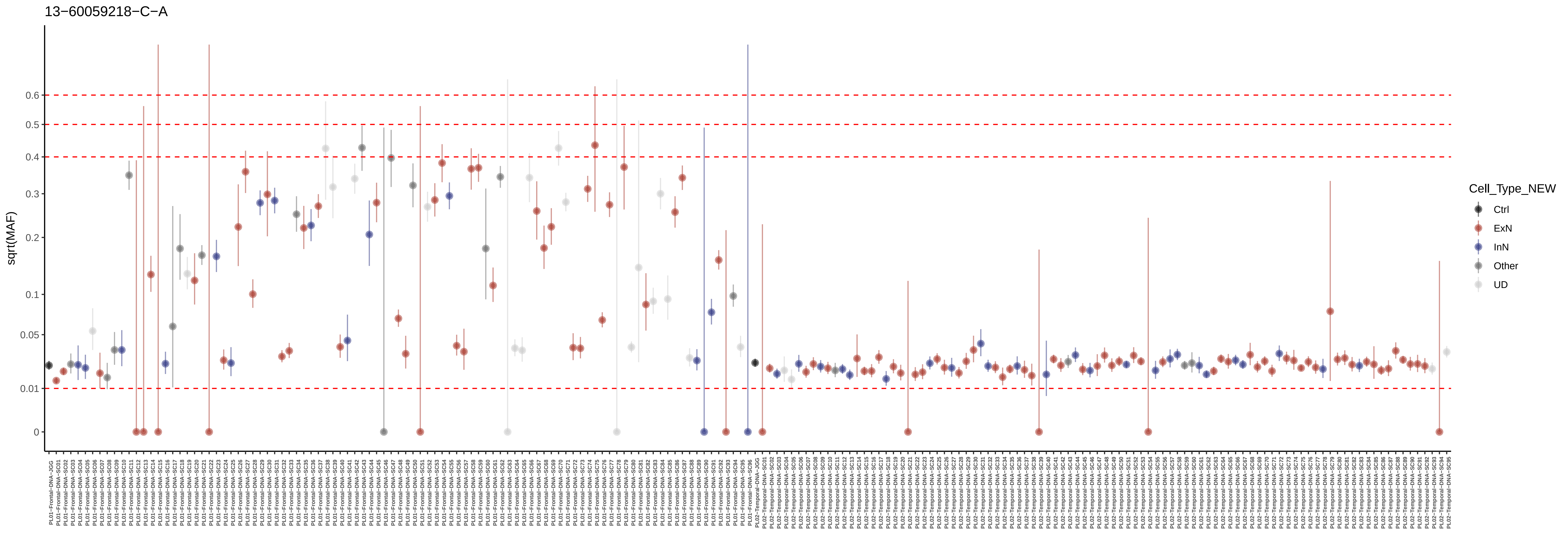

13−60059218−C−A
sqrt(MAF)
0.6
0.5
0.4
0.3
0.2
0.1
0.05
0.01
0
Cell_Type_NEW
Ctrl
ExN
InN
Other
UD

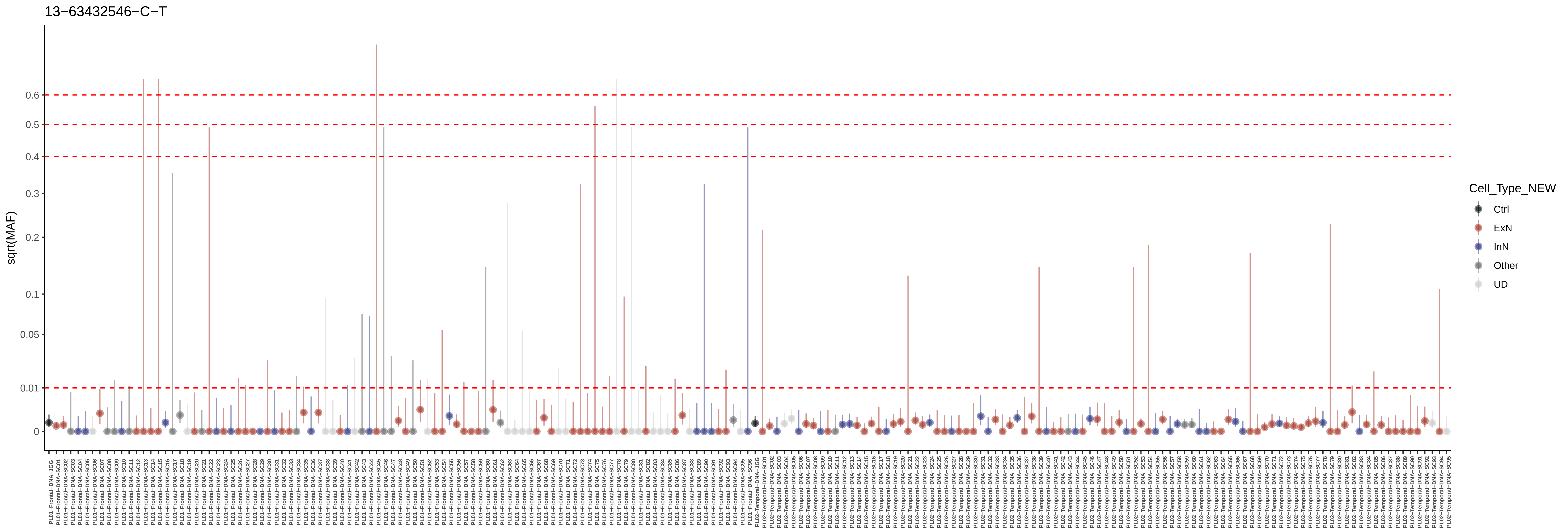
13−63432546−C−T
sqrt(MAF)
0.6
0.5
0.4
0.3
0.2
0.1
0.05
0.01
0
Cell_Type_NEW
Ctrl
ExN
InN
Other
UD

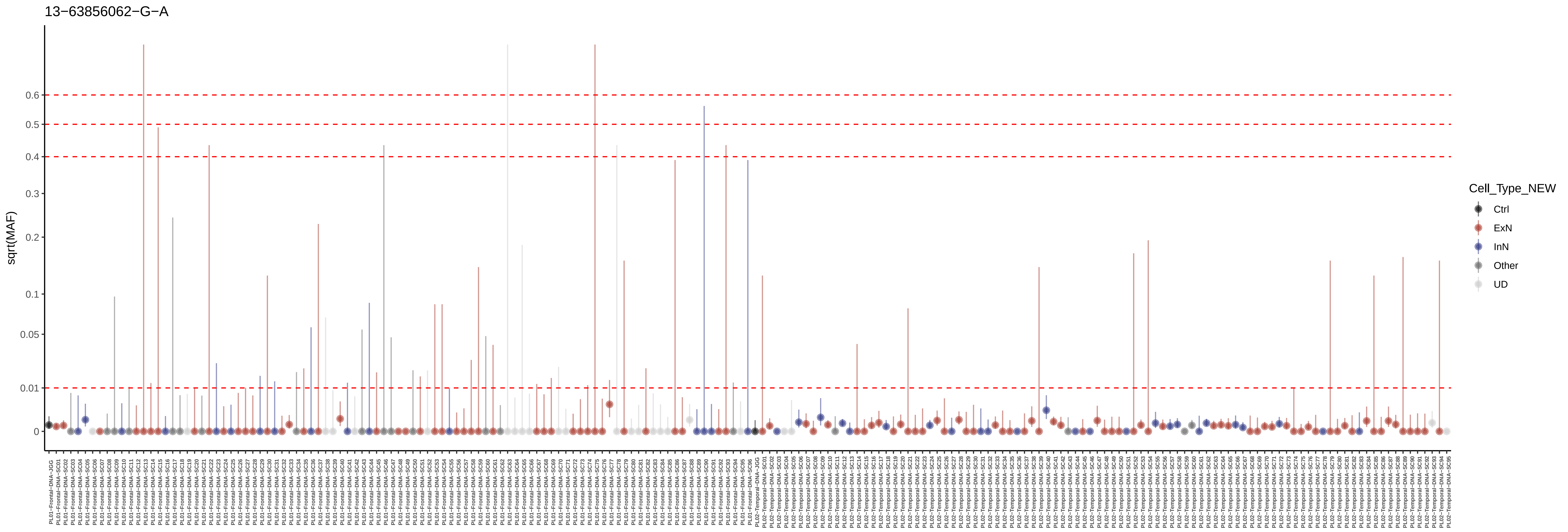
13−63856062−G−A
sqrt(MAF)
0.6
0.5
0.4
0.3
0.2
0.1
0.05
0.01
0
Cell_Type_NEW
Ctrl
ExN
InN
Other
UD

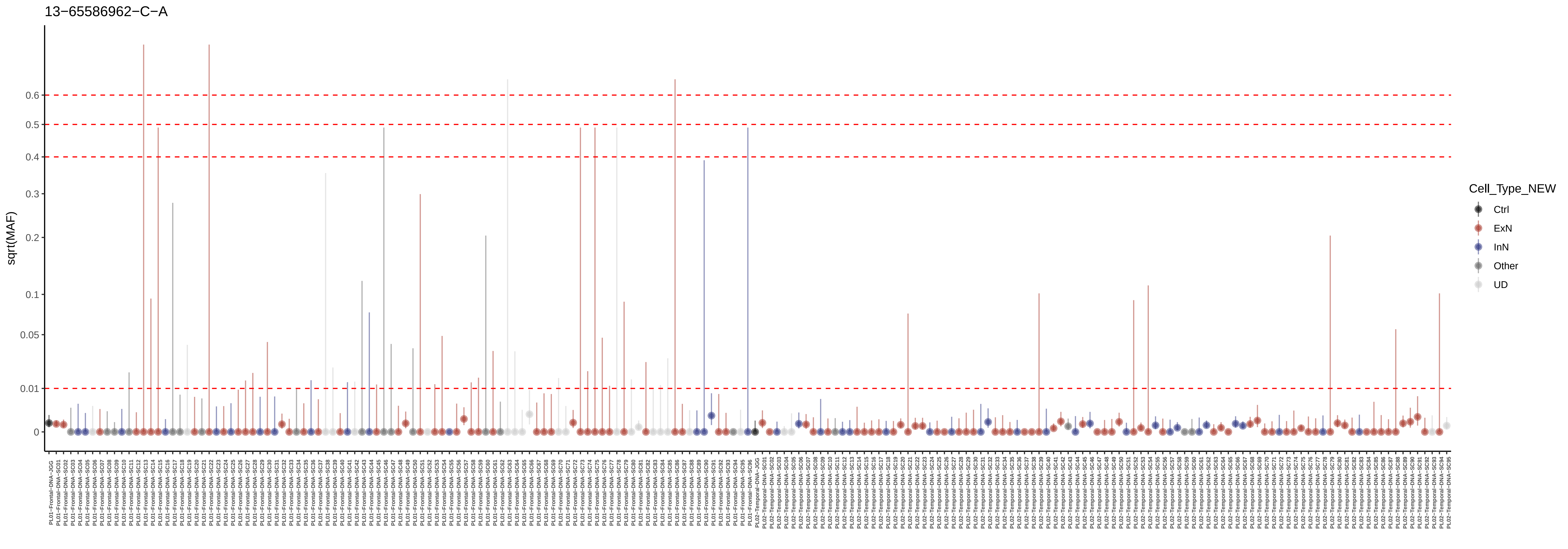
13−65586962−C−A
sqrt(MAF)
0.6
0.5
0.4
0.3
0.2
0.1
0.05
0.01
0
Cell_Type_NEW
Ctrl
ExN
InN
Other
UD

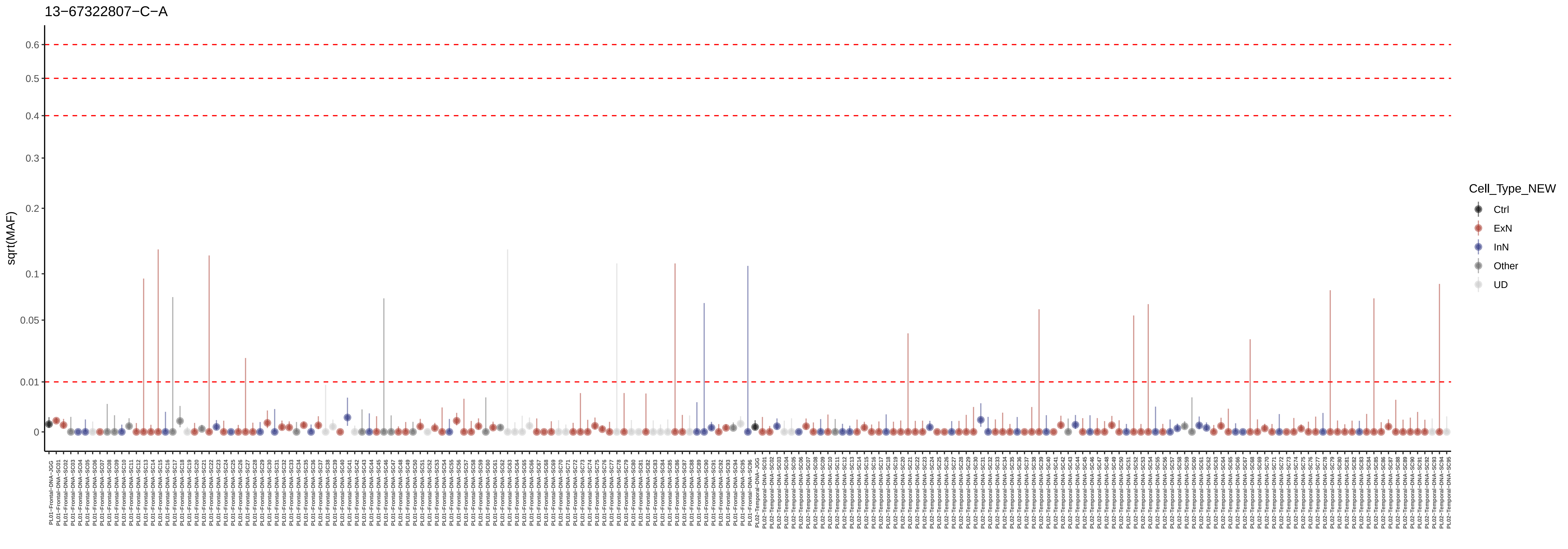
13−67322807−C−A
sqrt(MAF)
0.6
0.5
0.4
0.3
0.2
0.1
0.05
0.01
0
Cell_Type_NEW
Ctrl
ExN
InN
Other
UD

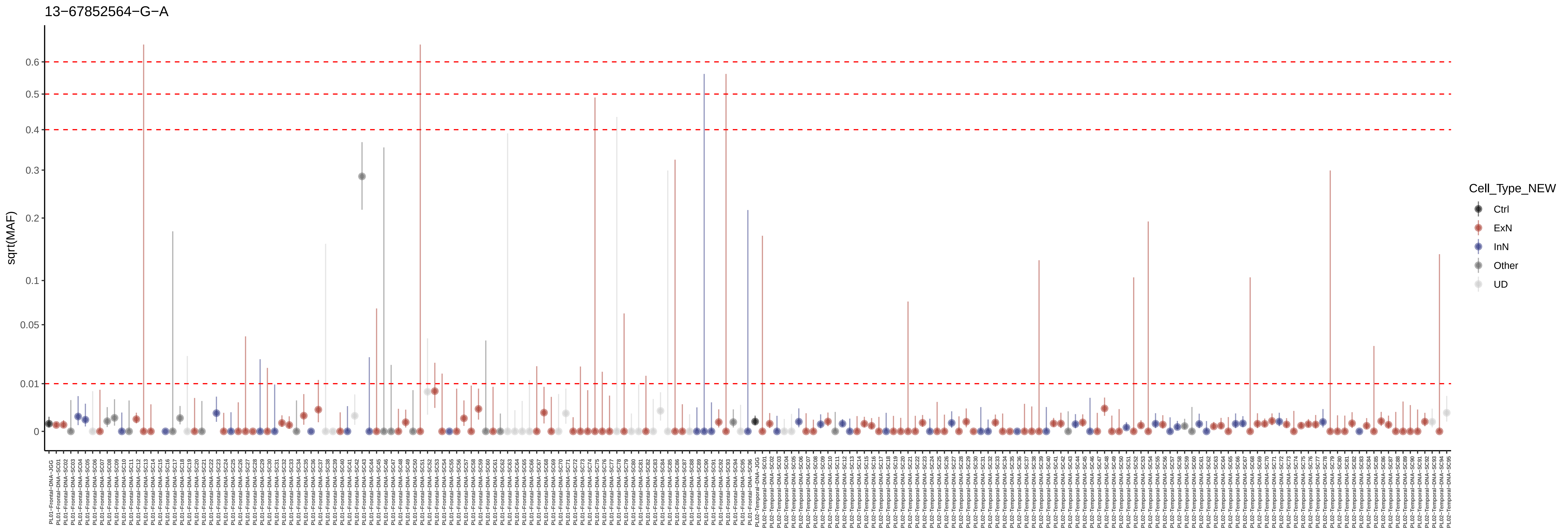
13−67852564−G−A
sqrt(MAF)
0.6
0.5
0.4
0.3
0.2
0.1
0.05
0.01
0
Cell_Type_NEW
Ctrl
ExN
InN
Other
UD

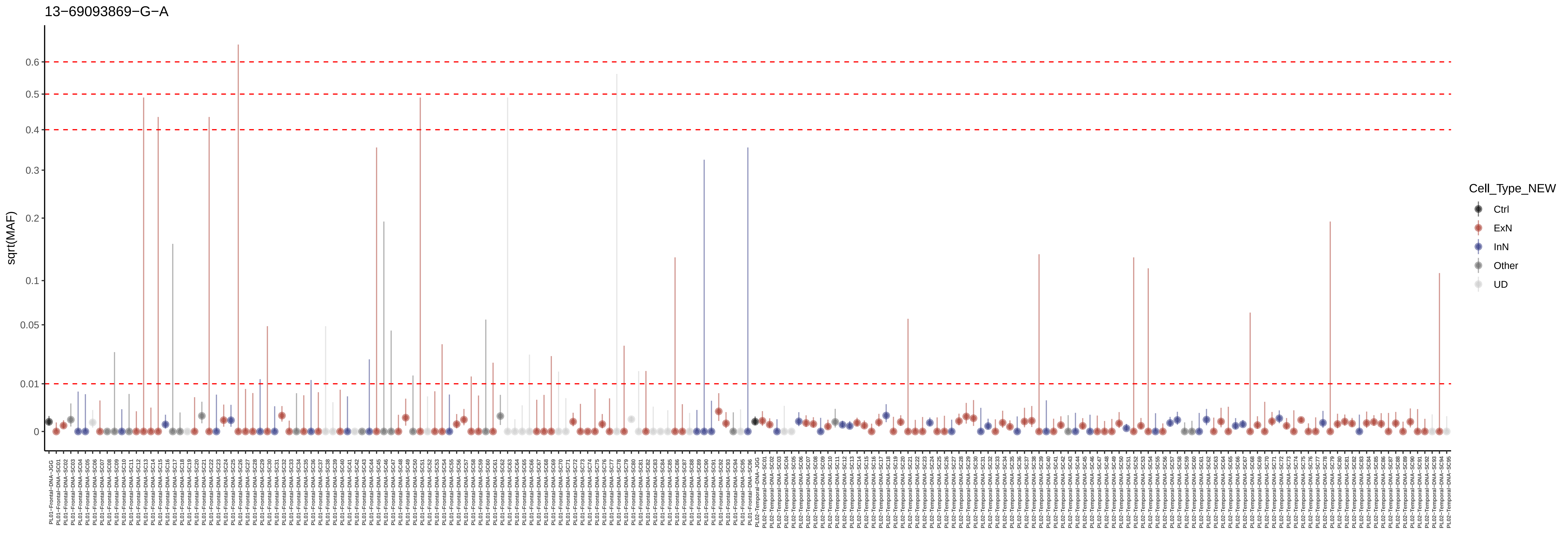
13−69093869−G−A
sqrt(MAF)
0.6
0.5
0.4
0.3
0.2
0.1
0.05
0.01
0
Cell_Type_NEW
Ctrl
ExN
InN
Other
UD

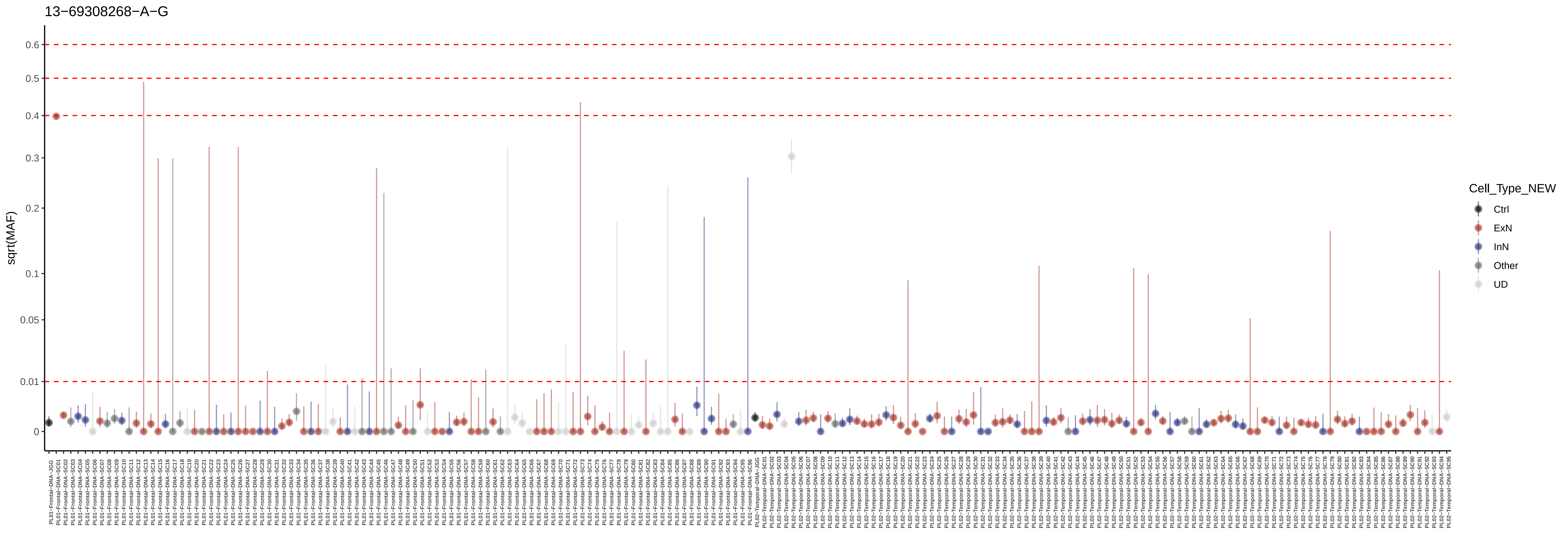
13−69308268−A−G
sqrt(MAF)
0.6
0.5
0.4
0.3
0.2
0.1
0.05
0.01
0
Cell_Type_NEW
Ctrl
ExN
InN
Other
UD

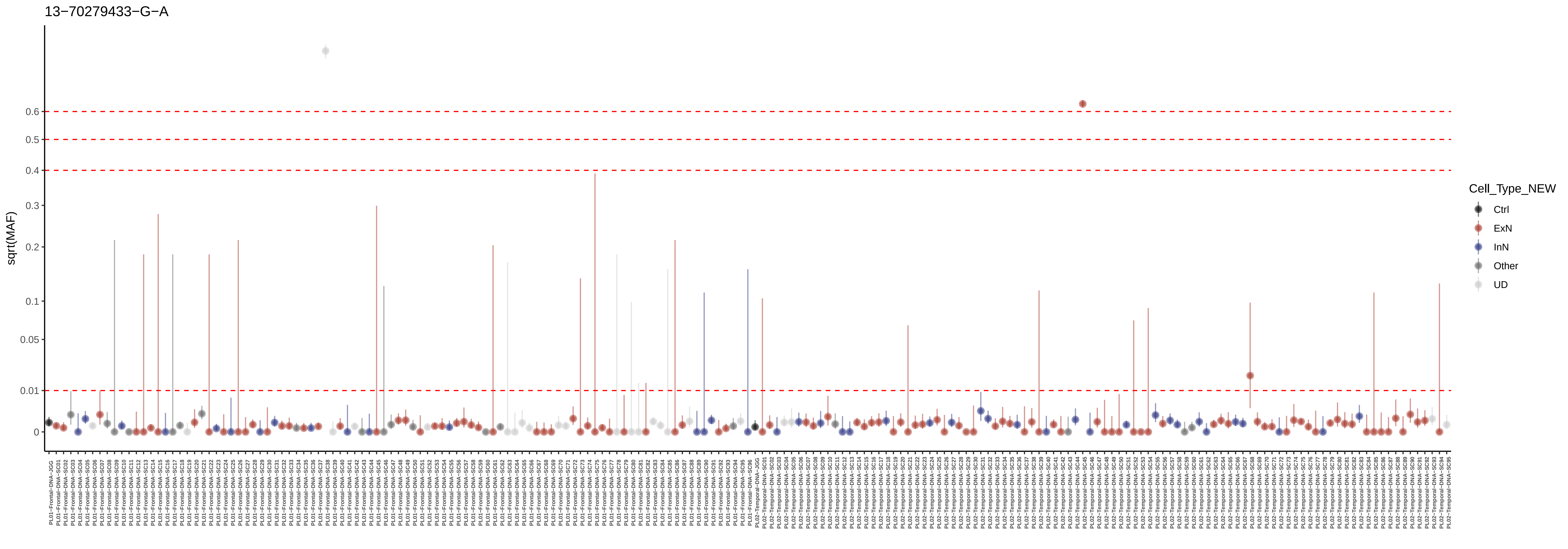

13−70279433−G−A
sqrt(MAF)
0.6
0.5
0.4
0.3
0.2
0.1
0.05
0.01
0
Cell_Type_NEW
Ctrl
ExN
InN
Other
UD

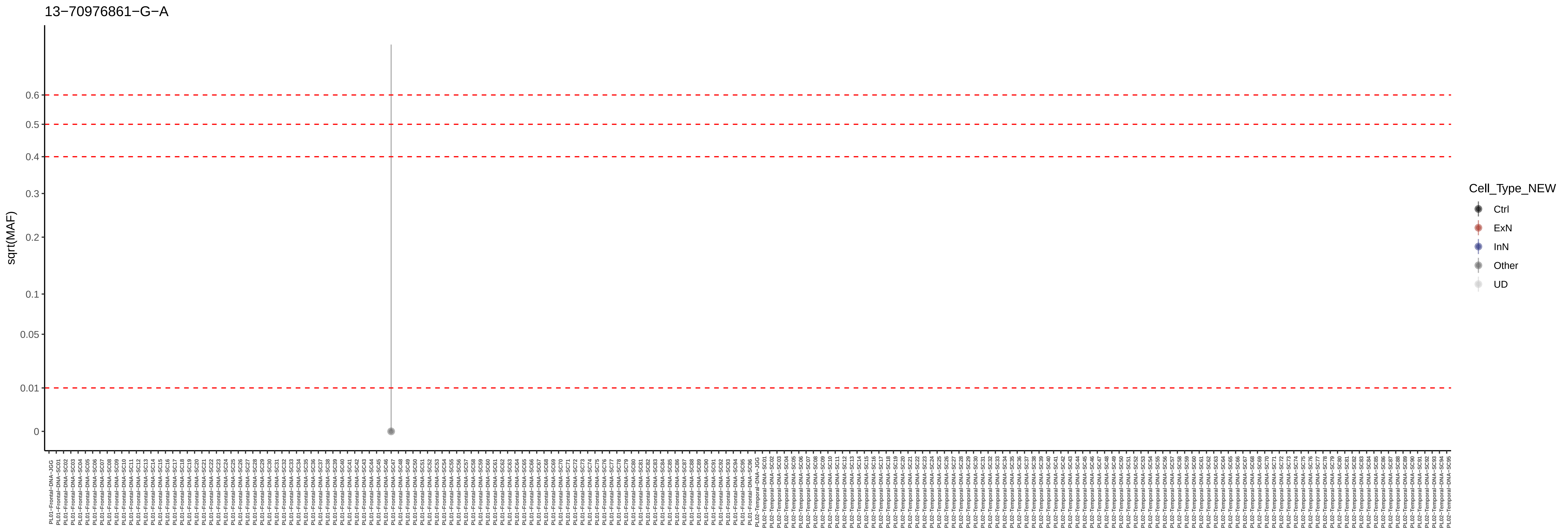

13−70976861−G−A
sqrt(MAF)
0.6
0.5
0.4
0.3
0.2
0.1
0.05
0.01
0
Cell_Type_NEW
Ctrl
ExN
InN
Other
UD

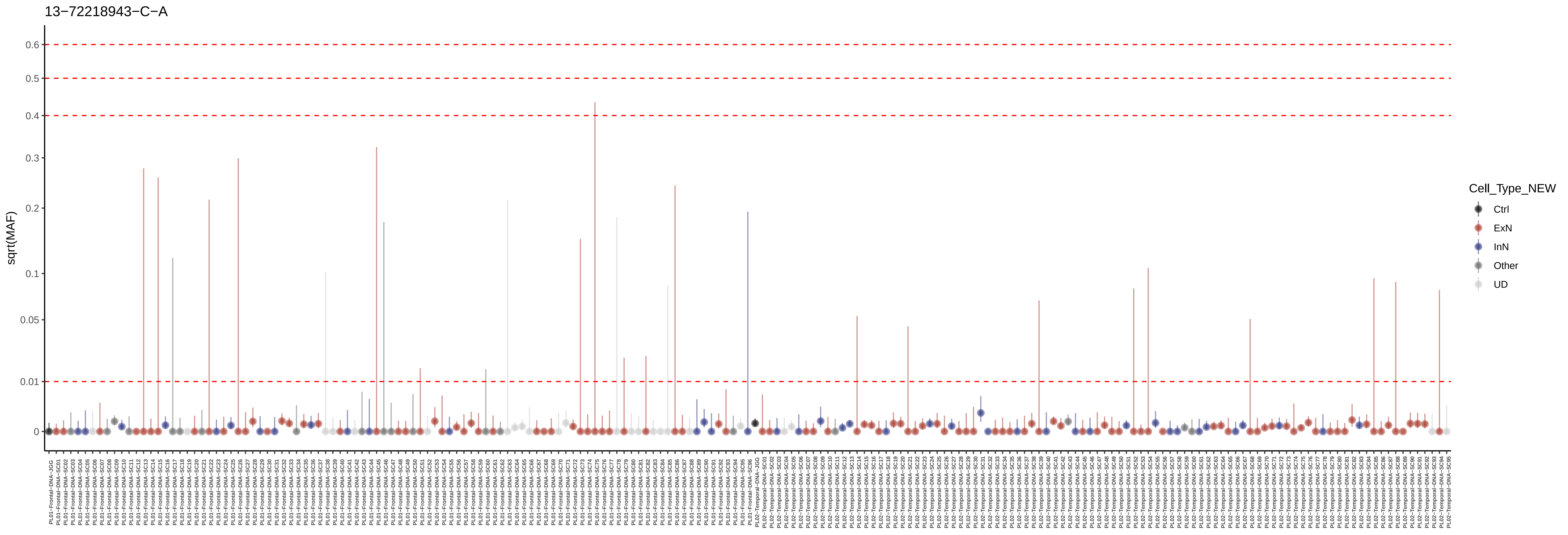

13−72218943−C−A
sqrt(MAF)
0.6
0.5
0.4
0.3
0.2
0.1
0.05
0.01
0
Cell_Type_NEW
Ctrl
ExN
InN
Other
UD

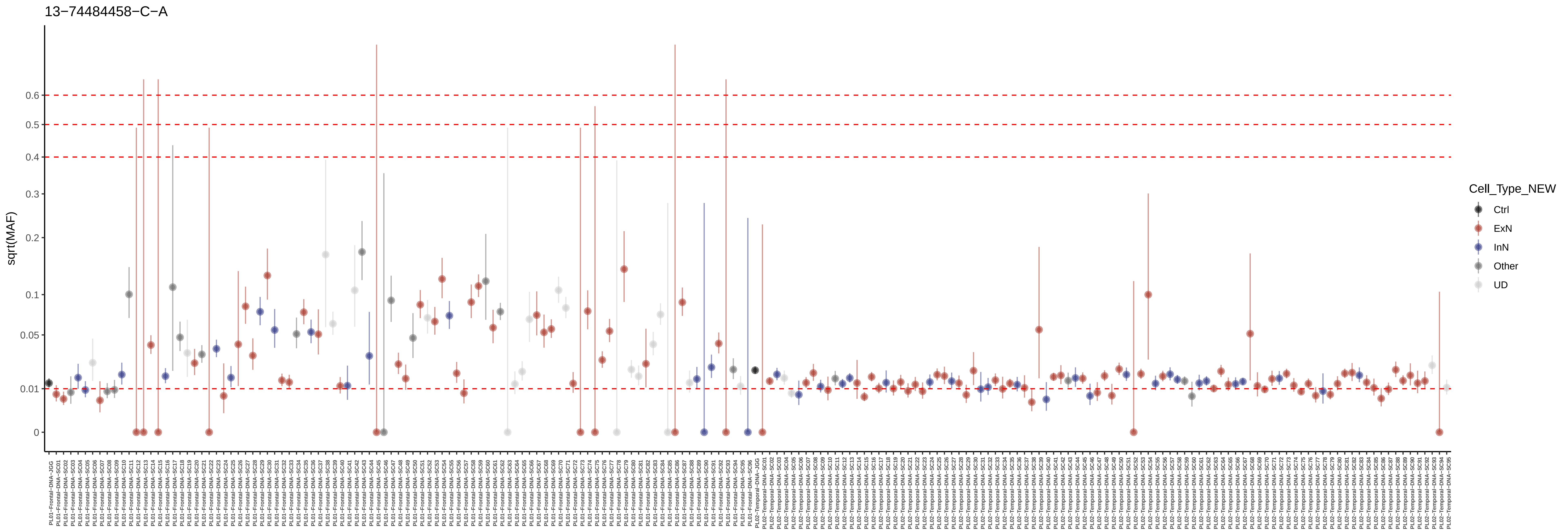
13−74484458−C−A
sqrt(MAF)
0.6
0.5
0.4
0.3
0.2
0.1
0.05
0.01
0
Cell_Type_NEW
Ctrl
ExN
InN
Other
UD

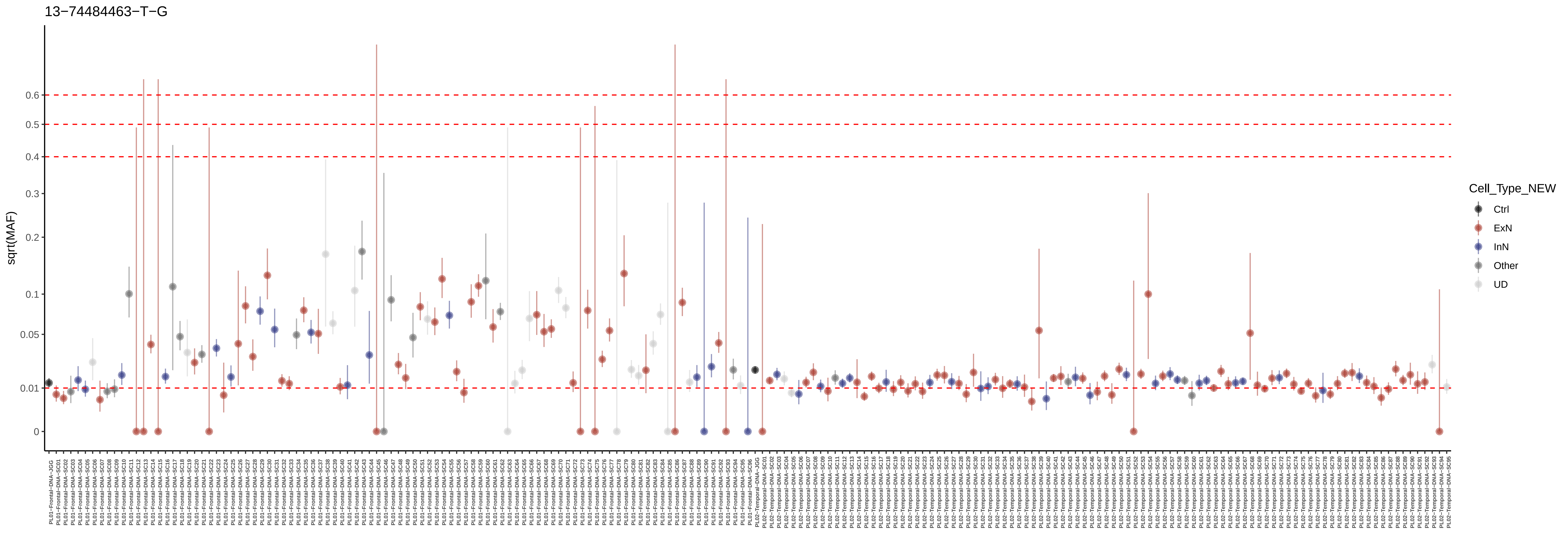
13−74484463−T−G
sqrt(MAF)
0.6
0.5
0.4
0.3
0.2
0.1
0.05
0.01
0
Cell_Type_NEW
Ctrl
ExN
InN
Other
UD

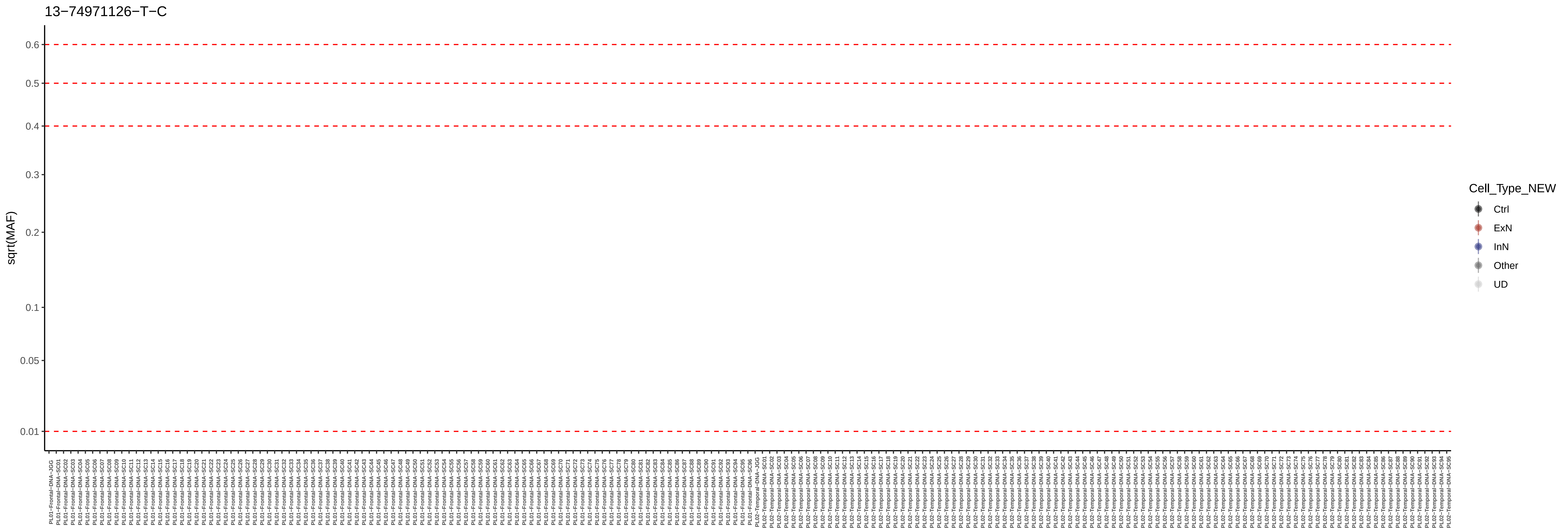
13−74971126−T−C
sqrt(MAF)
0.6
0.5
0.4
0.3
0.2
0.1
0.05
0.01
Cell_Type_NEW
Ctrl
ExN
InN
Other
UD

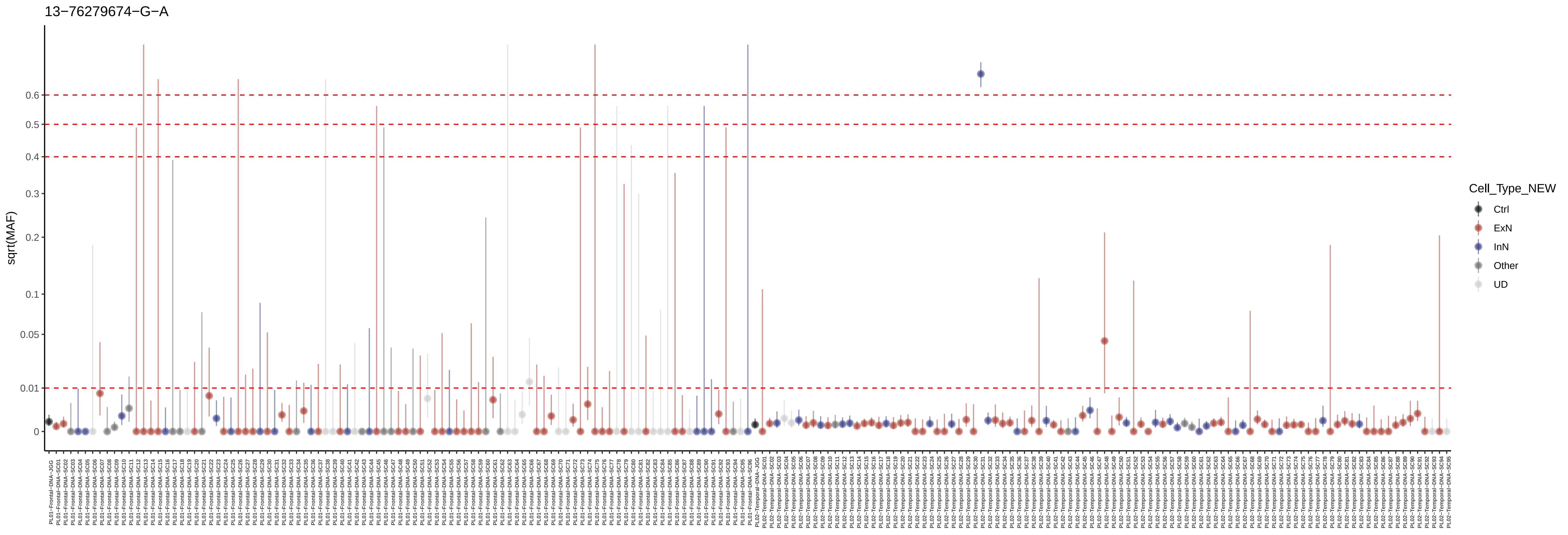
13−76279674−G−A
sqrt(MAF)
0.6
0.5
0.4
0.3
0.2
0.1
0.05
0.01
0
Cell_Type_NEW
Ctrl
ExN
InN
Other
UD

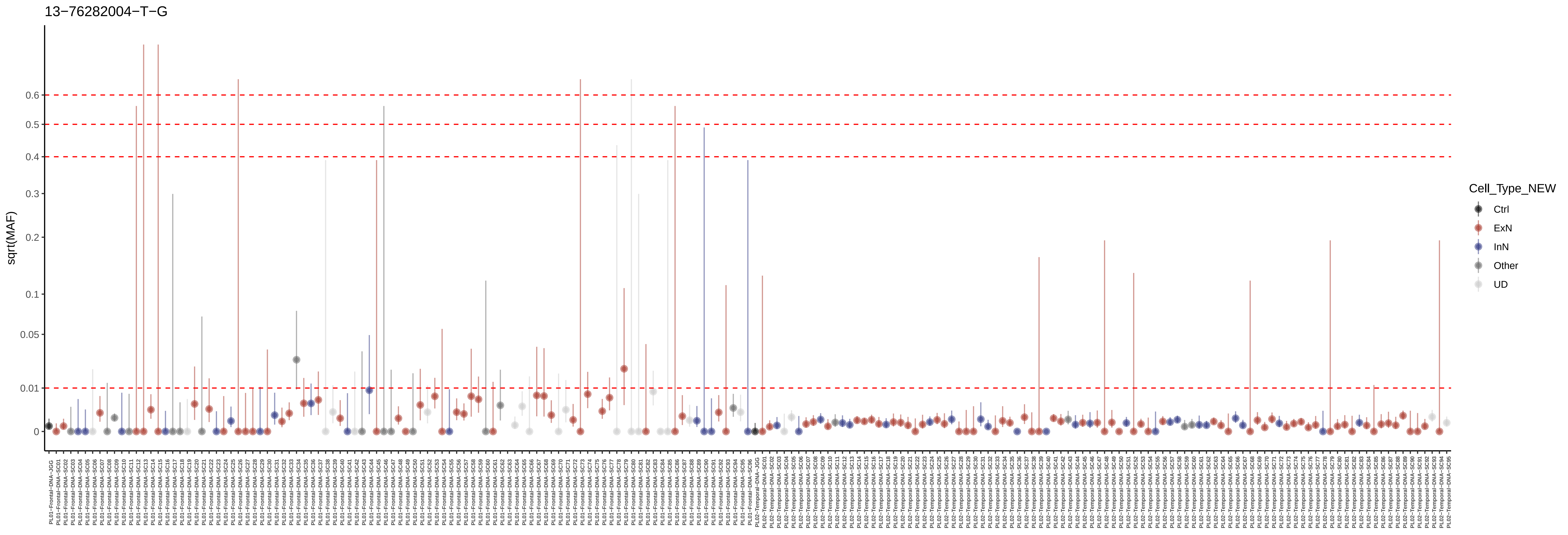
13−76282004−T−G
sqrt(MAF)
0.6
0.5
0.4
0.3
0.2
0.1
0.05
0.01
0
Cell_Type_NEW
Ctrl
ExN
InN
Other
UD

13−76874198−C−T

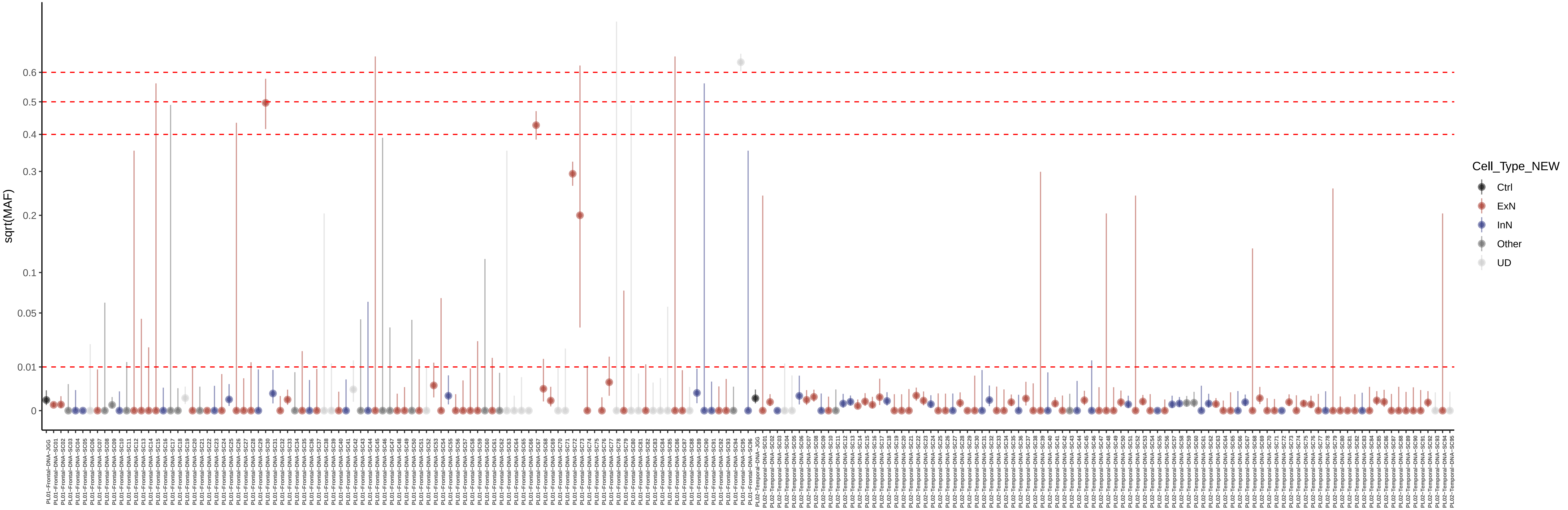

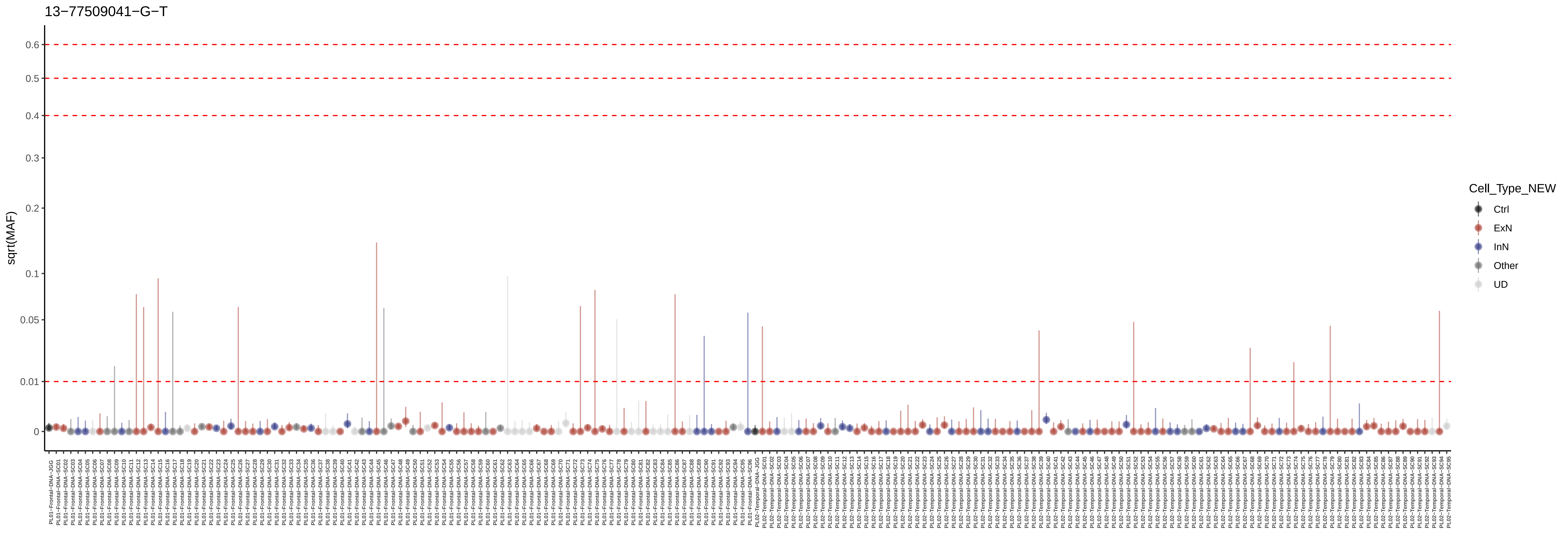
13−77509041−G−T
sqrt(MAF)
0.6
0.5
0.4
0.3
0.2
0.1
0.05
0.01
0
Cell_Type_NEW
Ctrl
ExN
InN
Other
UD

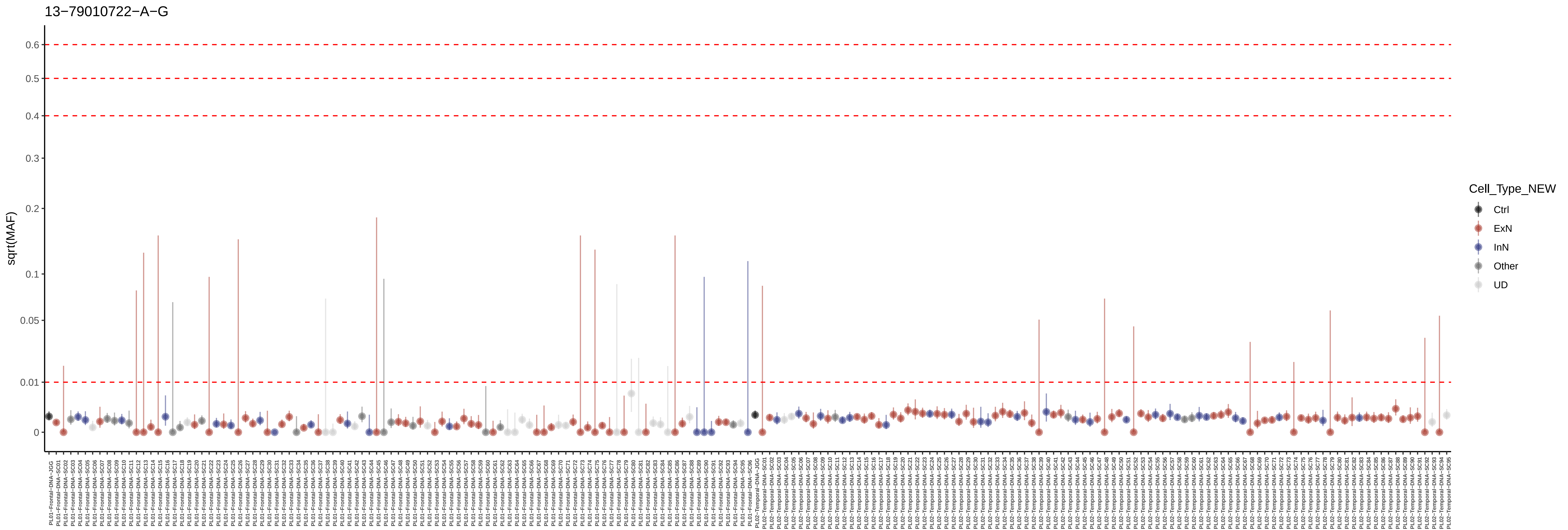
13−79010722−A−G
sqrt(MAF)
0.6
0.5
0.4
0.3
0.2
0.1
0.05
0.01
0
Cell_Type_NEW
Ctrl
ExN
InN
Other
UD

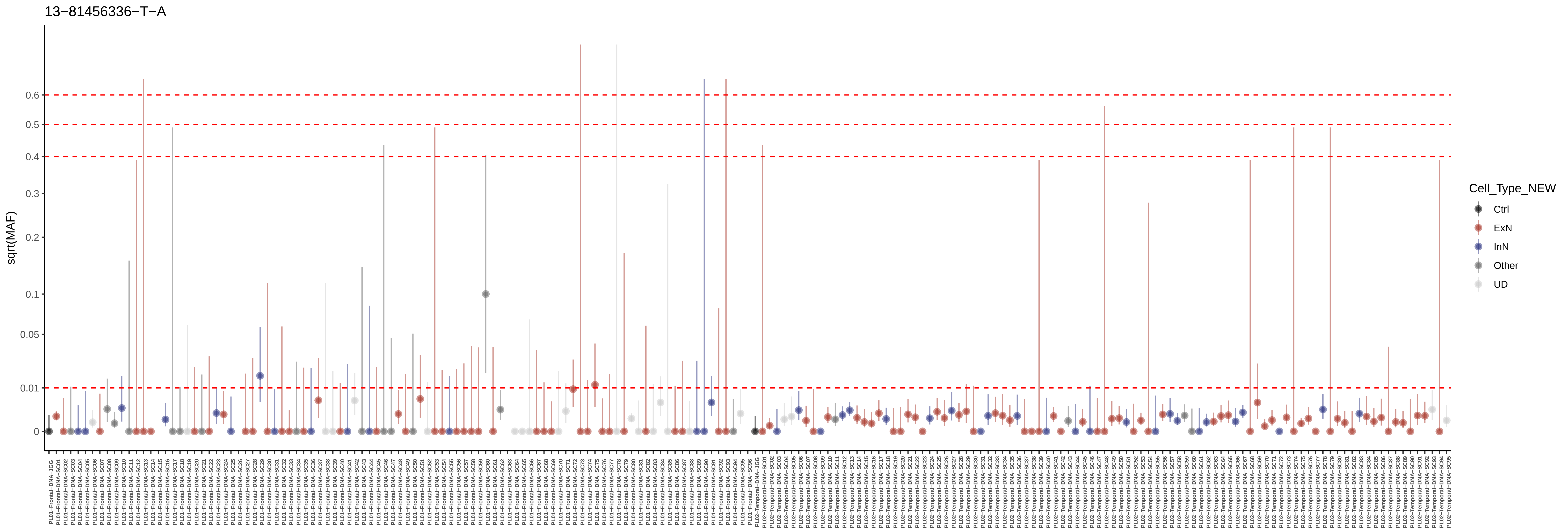
13−81456336−T−A
sqrt(MAF)
0.6
0.5
0.4
0.3
0.2
0.1
0.05
0.01
0
Cell_Type_NEW
Ctrl
ExN
InN
Other
UD

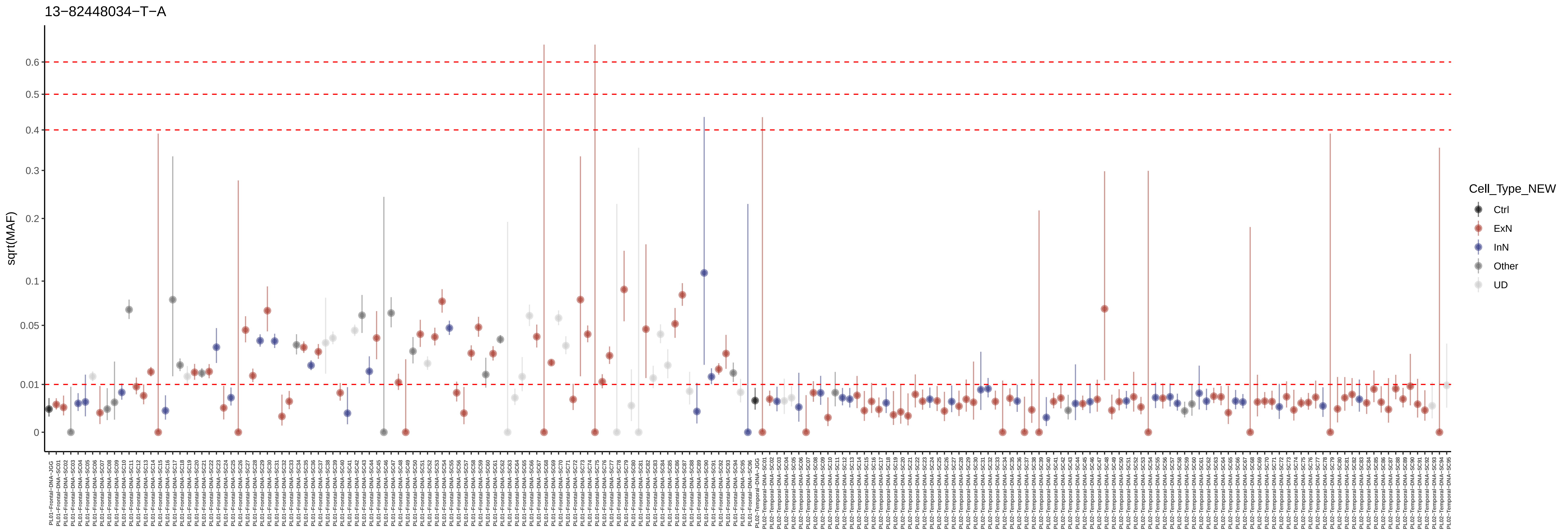

13−82448034−T−A
sqrt(MAF)
0.6
0.5
0.4
0.3
0.2
0.1
0.05
0.01
0
Cell_Type_NEW
Ctrl
ExN
InN
Other
UD

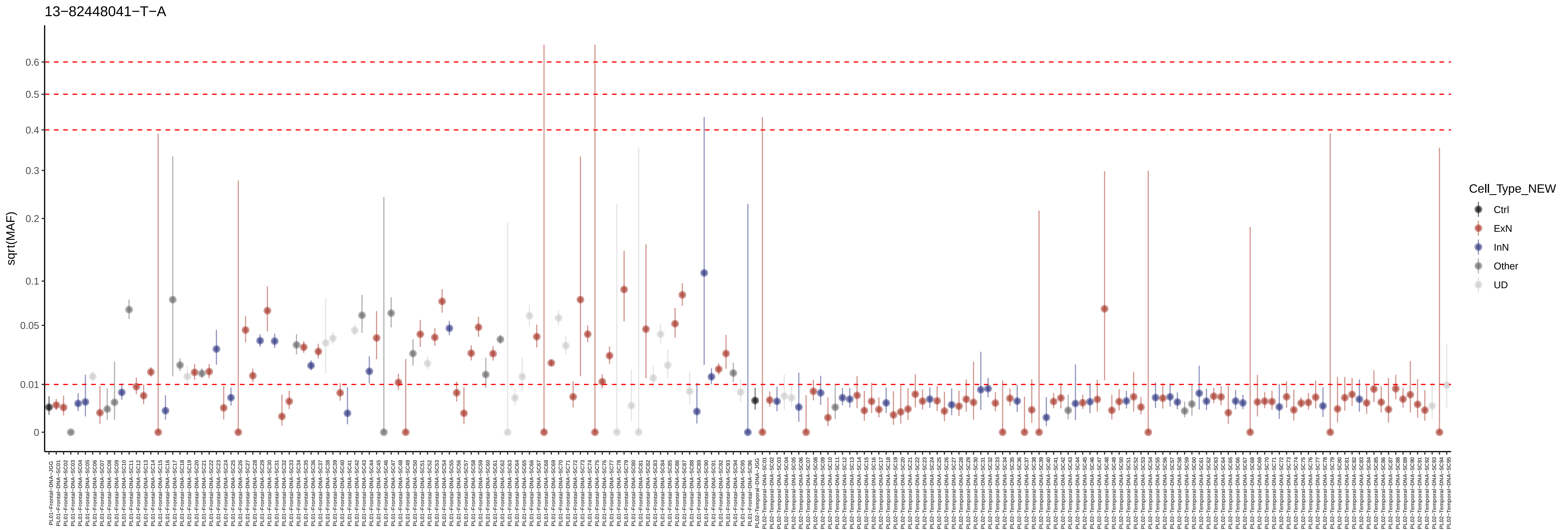
13−82448041−T−A
sqrt(MAF)
0.6
0.5
0.4
0.3
0.2
0.1
0.05
0.01
0
Cell_Type_NEW
Ctrl
ExN
InN
Other
UD

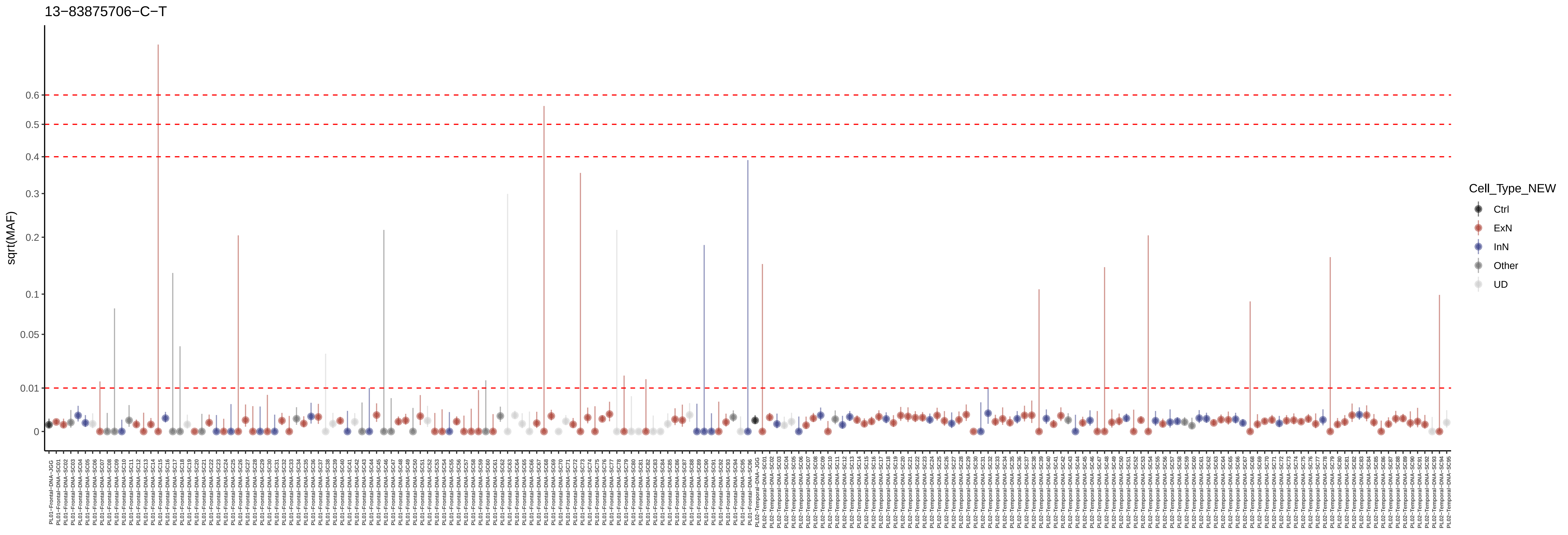
13−83875706−C−T
sqrt(MAF)
0.6
0.5
0.4
0.3
0.2
0.1
0.05
0.01
0
Cell_Type_NEW
Ctrl
ExN
InN
Other
UD

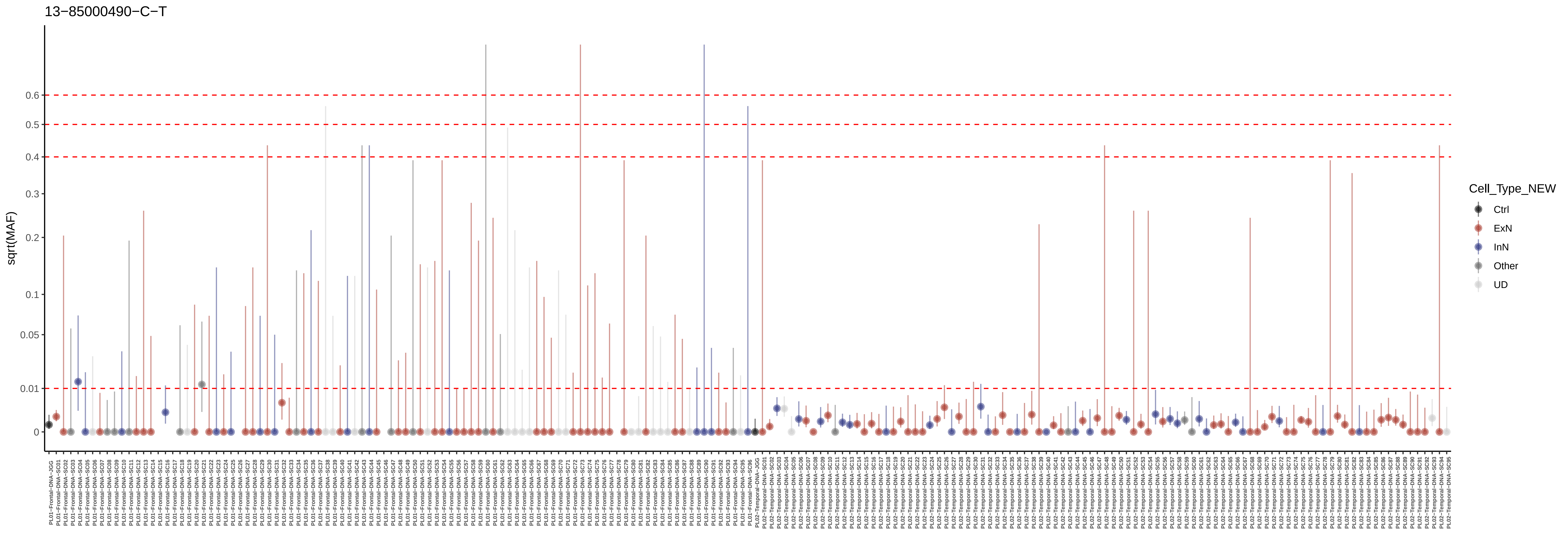
13−85000490−C−T
sqrt(MAF)
0.6
0.5
0.4
0.3
0.2
0.1
0.05
0.01
0
Cell_Type_NEW
Ctrl
ExN
InN
Other
UD

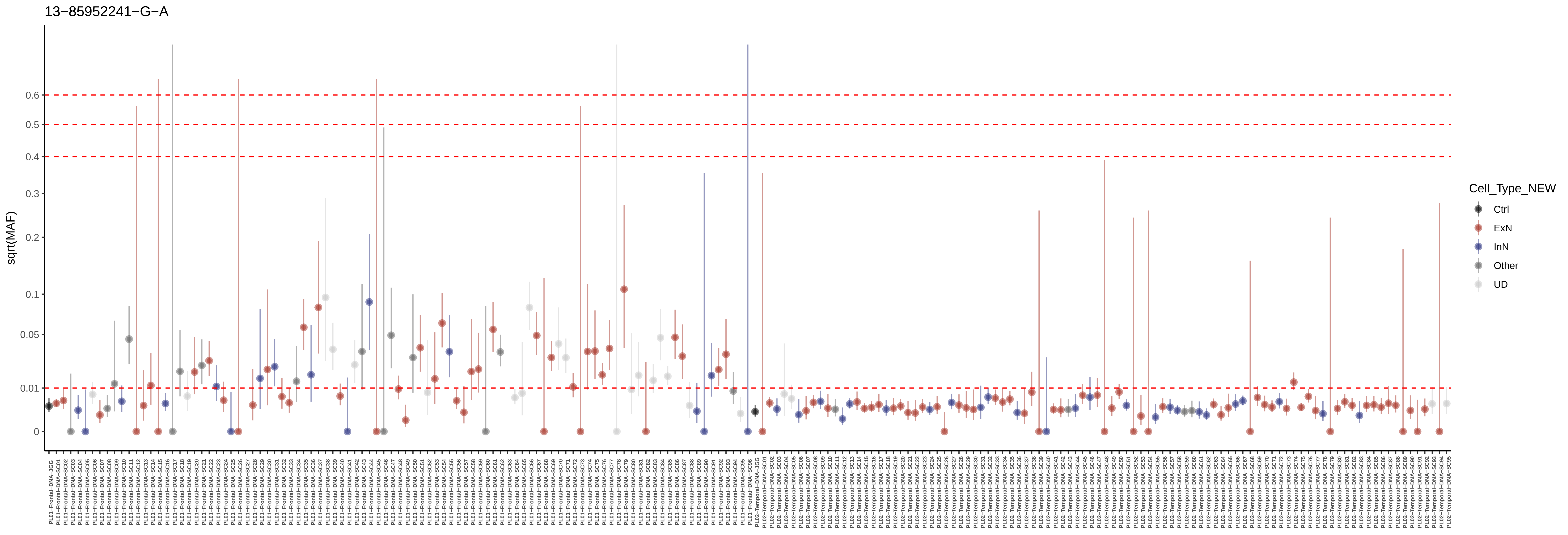

13−85952241−G−A
sqrt(MAF)
0.6
0.5
0.4
0.3
0.2
0.1
0.05
0.01
0
Cell_Type_NEW
Ctrl
ExN
InN
Other
UD

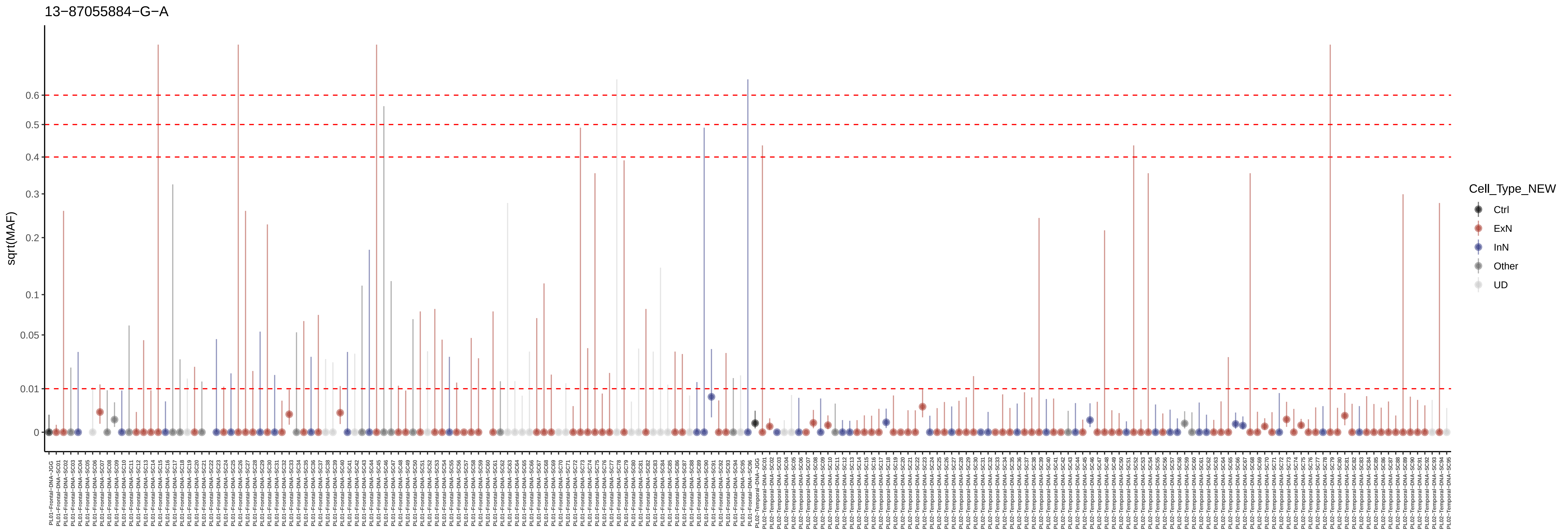

13−87055884−G−A
sqrt(MAF)
0.6
0.5
0.4
0.3
0.2
0.1
0.05
0.01
0
Cell_Type_NEW
Ctrl
ExN
InN
Other
UD

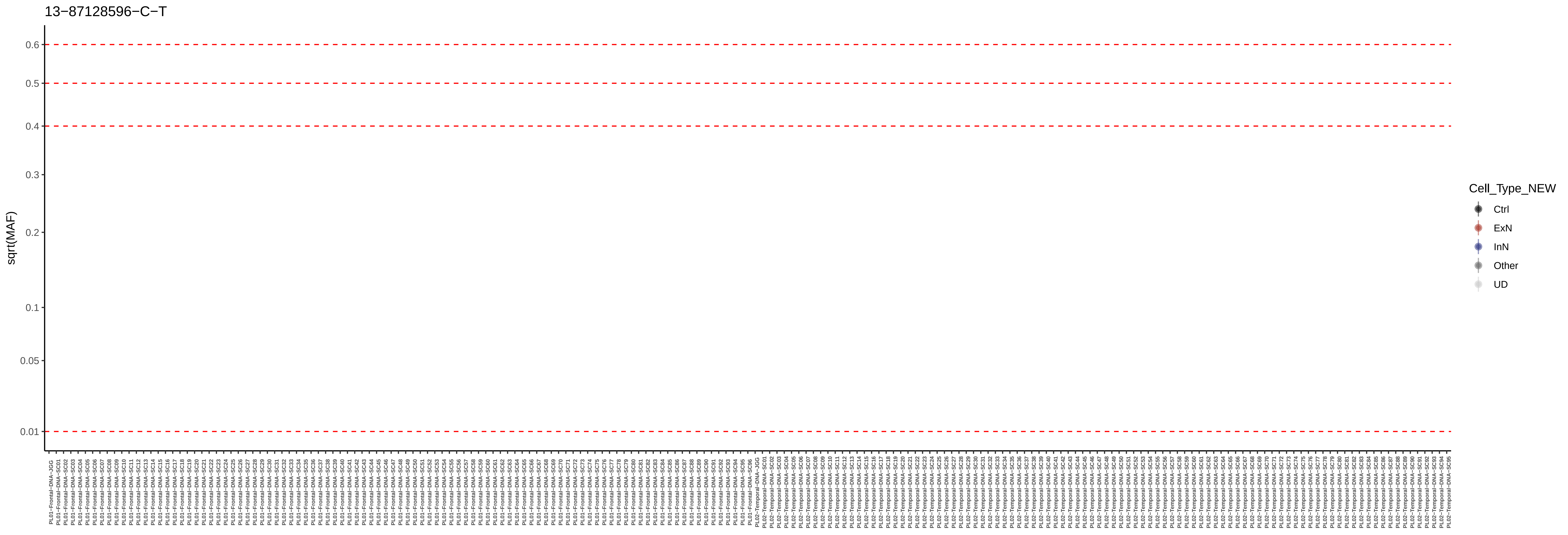
13−87128596−C−T
sqrt(MAF)
0.6
0.5
0.4
0.3
0.2
0.1
0.05
0.01
Cell_Type_NEW
Ctrl
ExN
InN
Other
UD

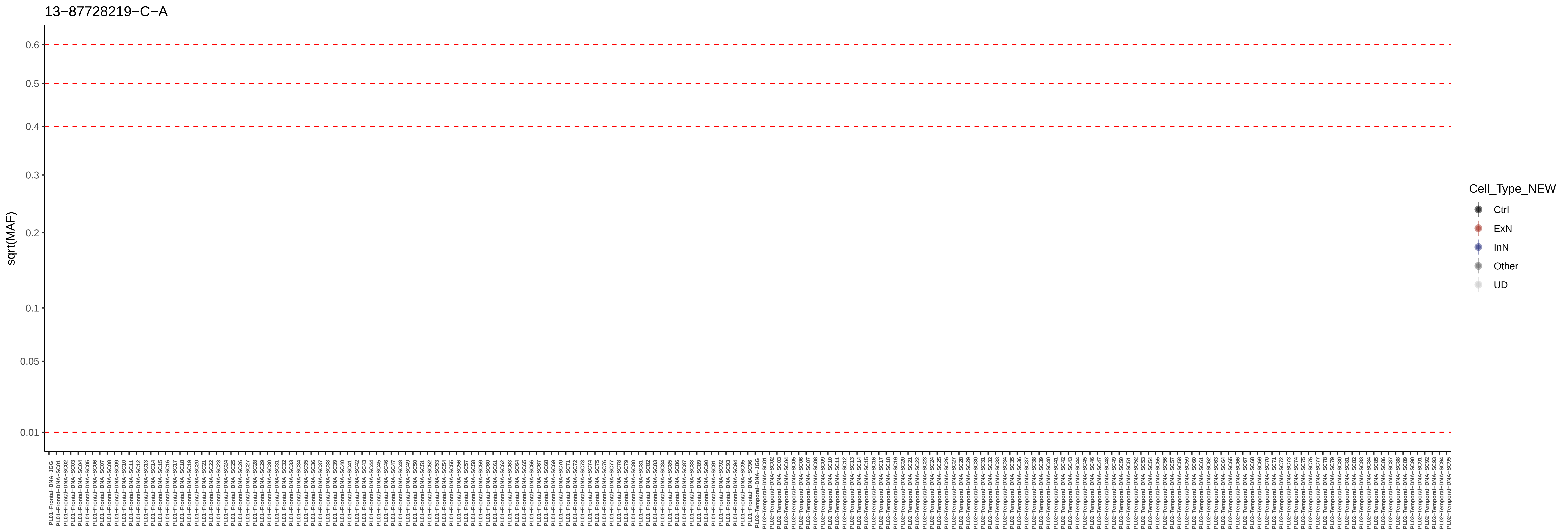
13−87728219−C−A
sqrt(MAF)
0.6
0.5
0.4
0.3
0.2
0.1
0.05
0.01
Cell_Type_NEW
Ctrl
ExN
InN
Other
UD

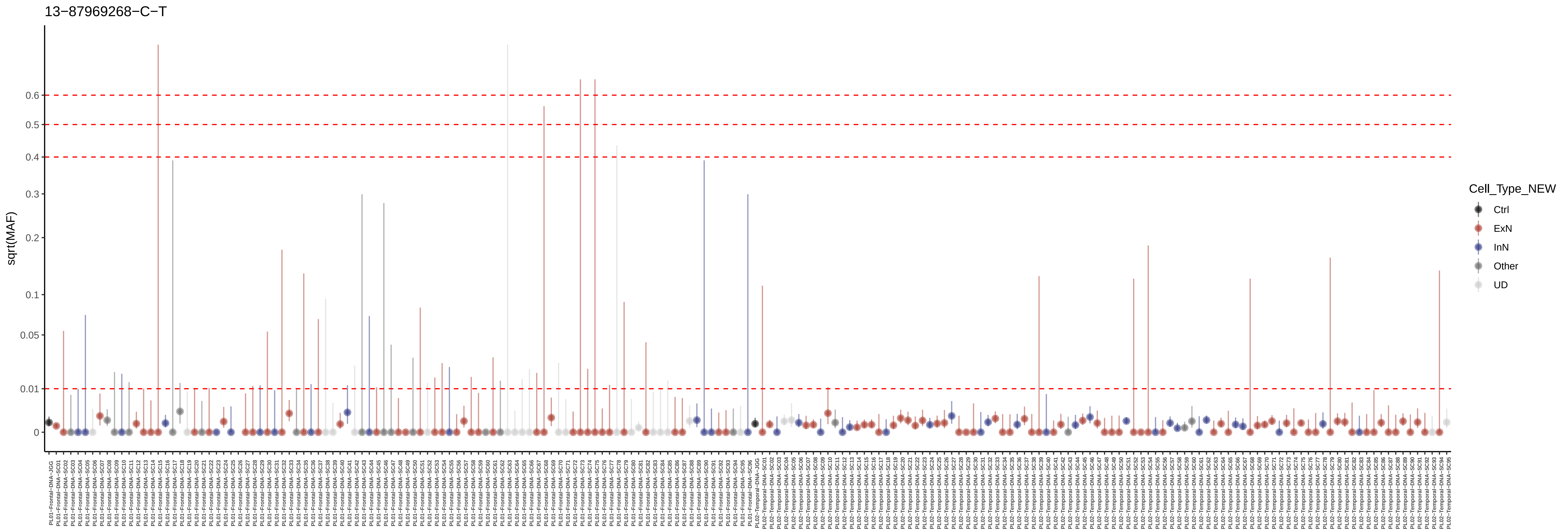
13−87969268−C−T
sqrt(MAF)
0.6
0.5
0.4
0.3
0.2
0.1
0.05
0.01
0
Cell_Type_NEW
Ctrl
ExN
InN
Other
UD

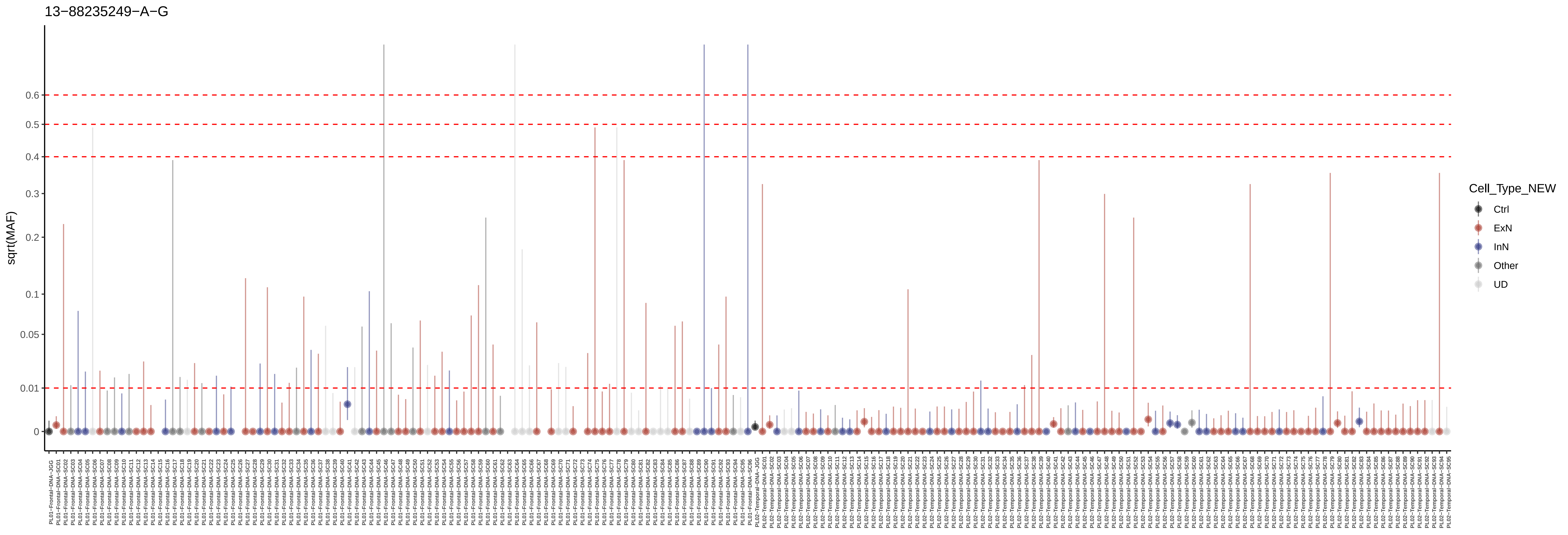
13−88235249−A−G
sqrt(MAF)
0.6
0.5
0.4
0.3
0.2
0.1
0.05
0.01
0
Cell_Type_NEW
Ctrl
ExN
InN
Other
UD

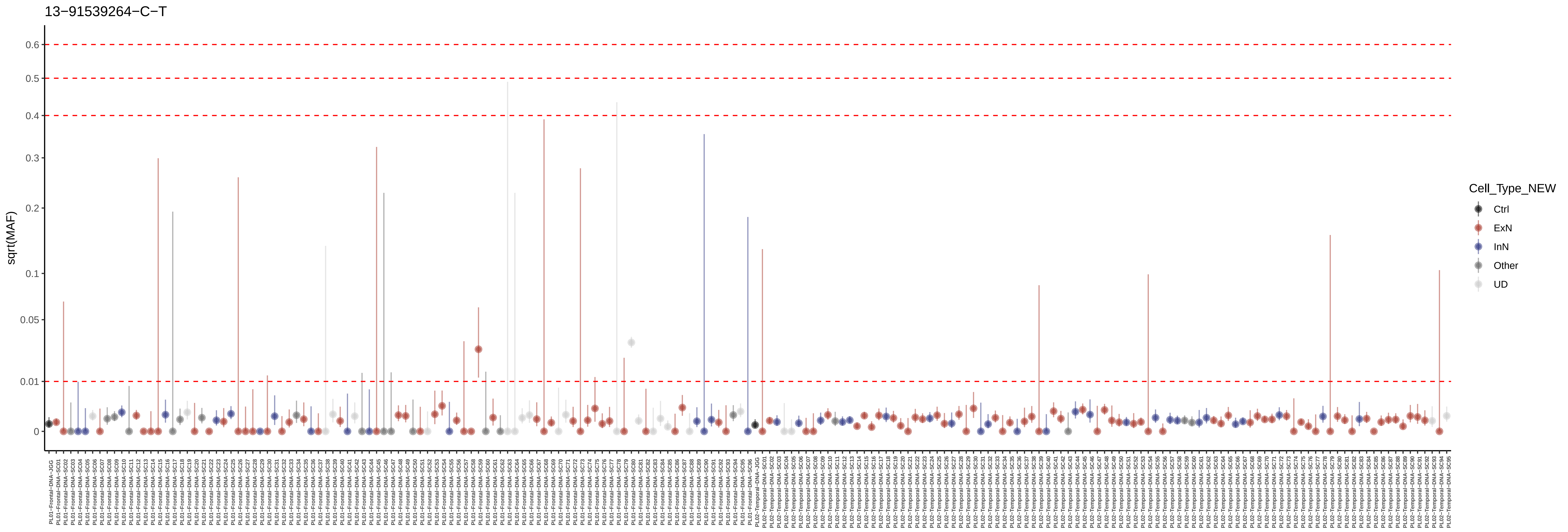
13−91539264−C−T
sqrt(MAF)
0.6
0.5
0.4
0.3
0.2
0.1
0.05
0.01
0
Cell_Type_NEW
Ctrl
ExN
InN
Other
UD

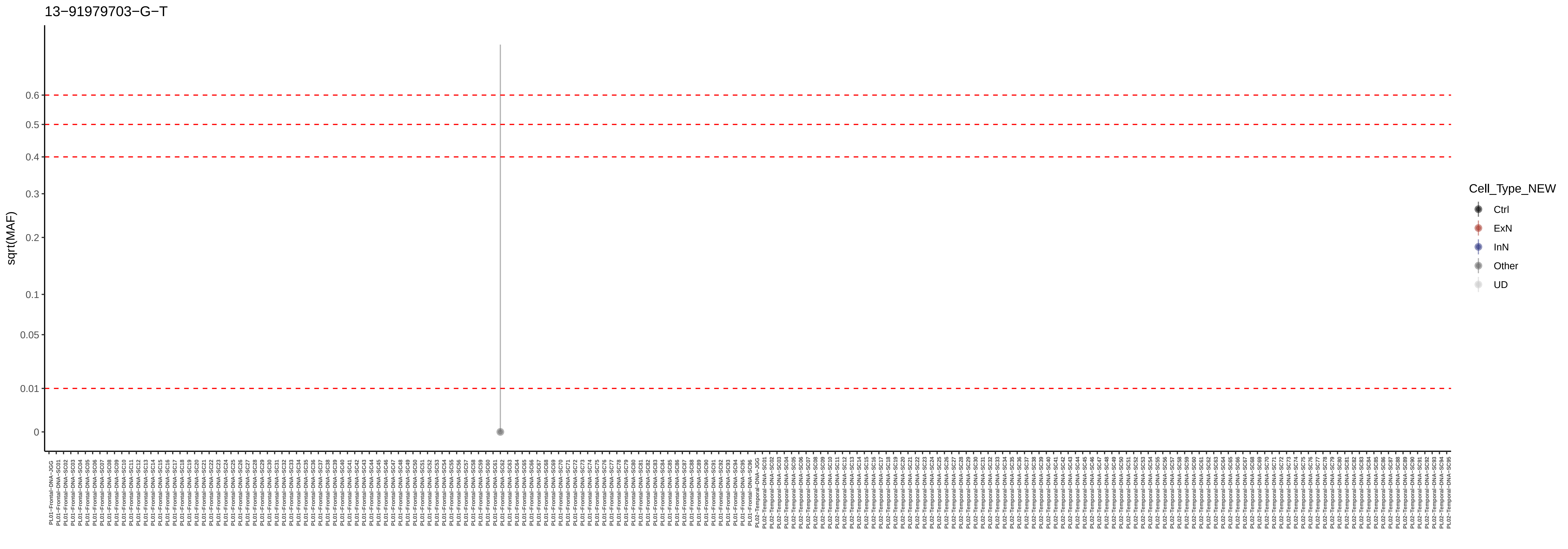
13−91979703−G−T
sqrt(MAF)
0.6
0.5
0.4
0.3
0.2
0.1
0.05
0.01
0
Cell_Type_NEW
Ctrl
ExN
InN
Other
UD

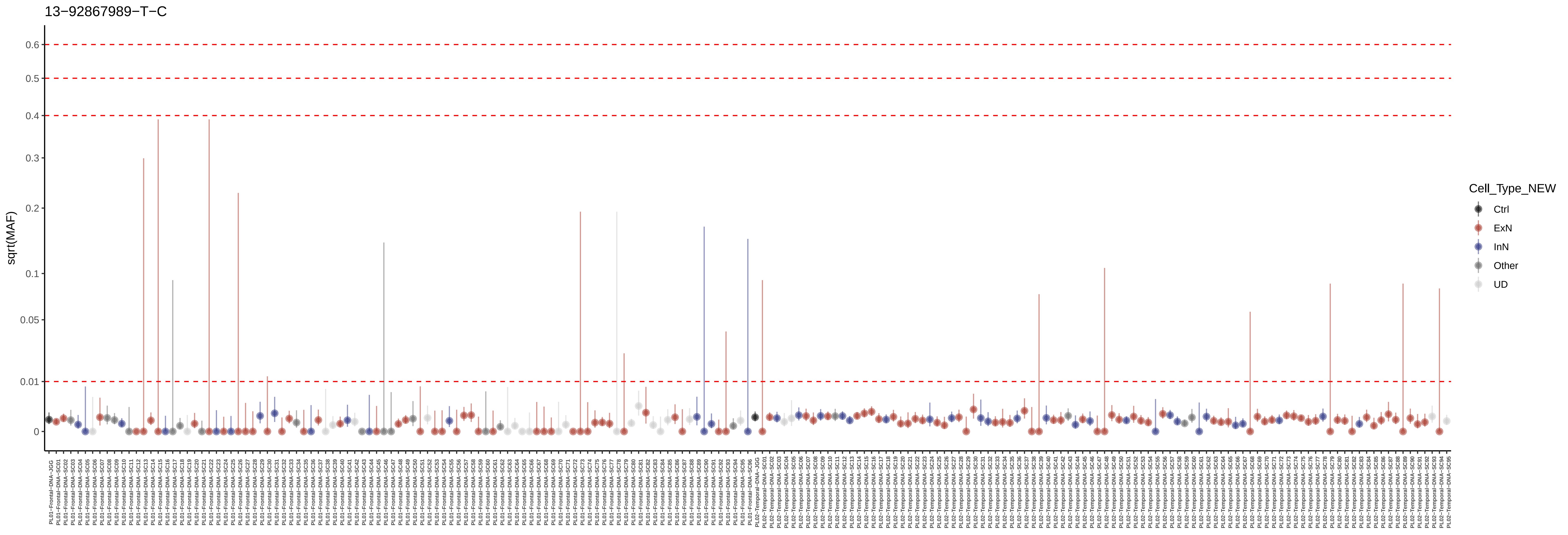

13−92867989−T−C
sqrt(MAF)
0.6
0.5
0.4
0.3
0.2
0.1
0.05
0.01
0
Cell_Type_NEW
Ctrl
ExN
InN
Other
UD

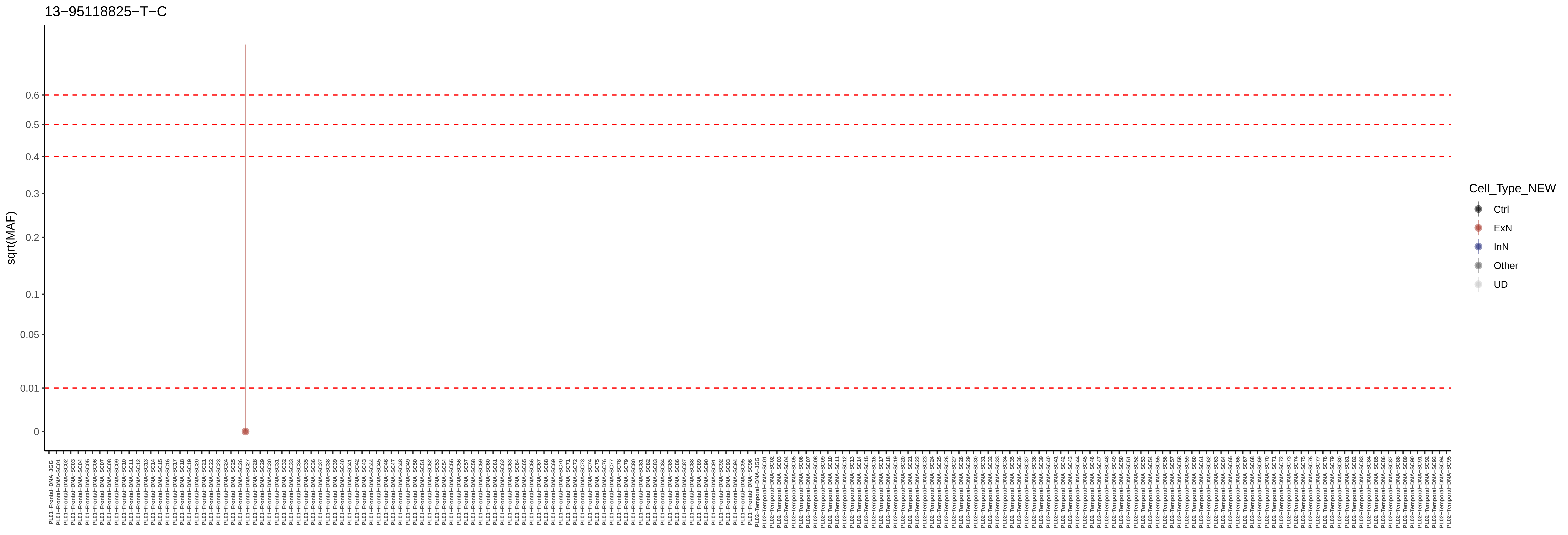
13−95118825−T−C
sqrt(MAF)
0.6
0.5
0.4
0.3
0.2
0.1
0.05
0.01
0
Cell_Type_NEW
Ctrl
ExN
InN
Other
UD

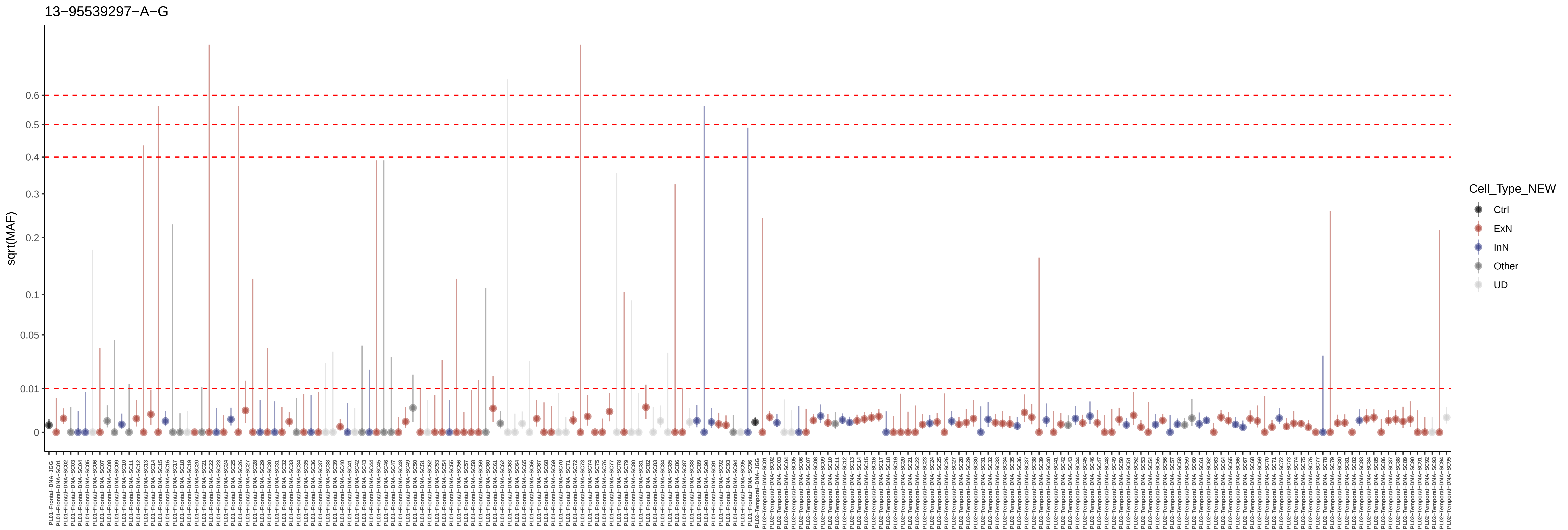

13−95539297−A−G
sqrt(MAF)
0.6
0.5
0.4
0.3
0.2
0.1
0.05
0.01
0
Cell_Type_NEW
Ctrl
ExN
InN
Other
UD

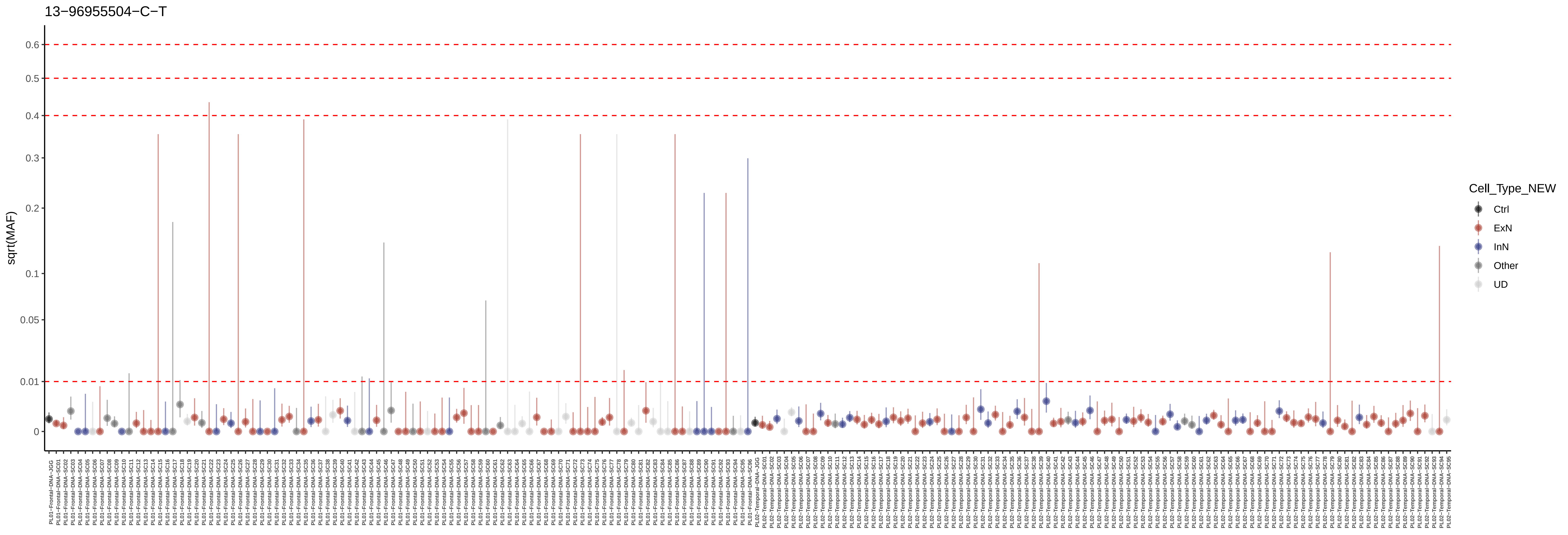
13−96955504−C−T
sqrt(MAF)
0.6
0.5
0.4
0.3
0.2
0.1
0.05
0.01
0
Cell_Type_NEW
Ctrl
ExN
InN
Other
UD

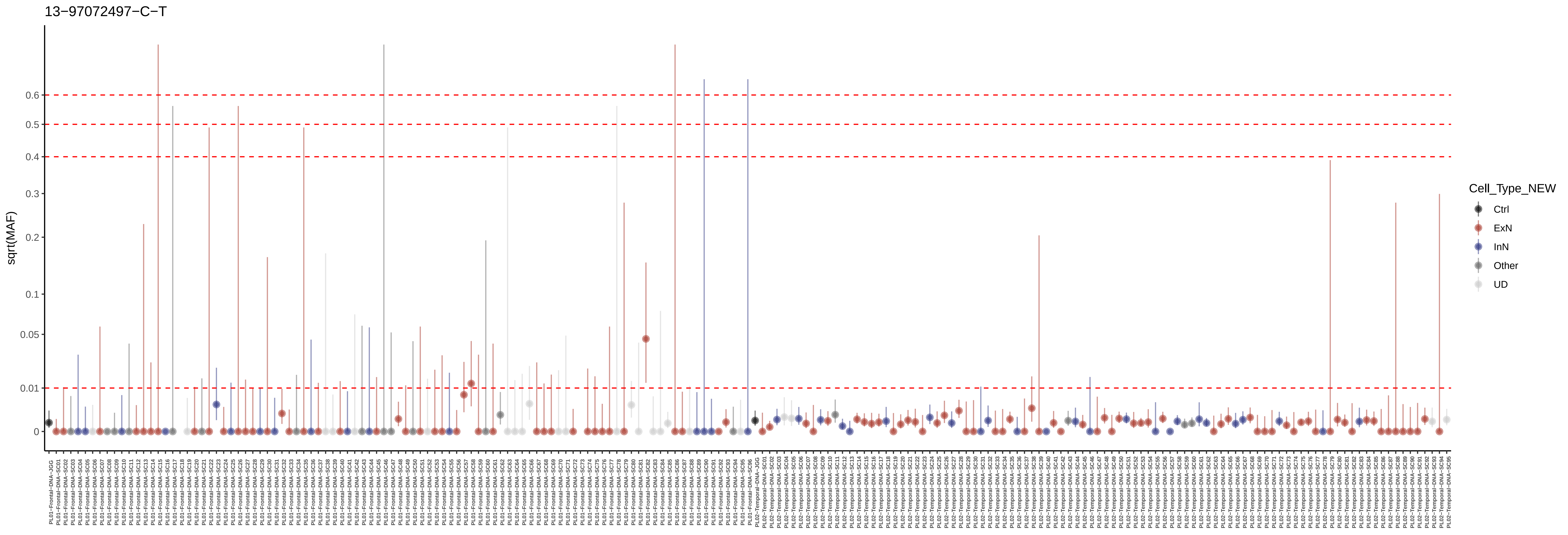

13−97072497−C−T
sqrt(MAF)
0
0.01
0.05
0.1
0.2
0.3
0.4
0.5
0.6
Cell_Type_NEW
Ctrl
ExN
InN
Other
UD

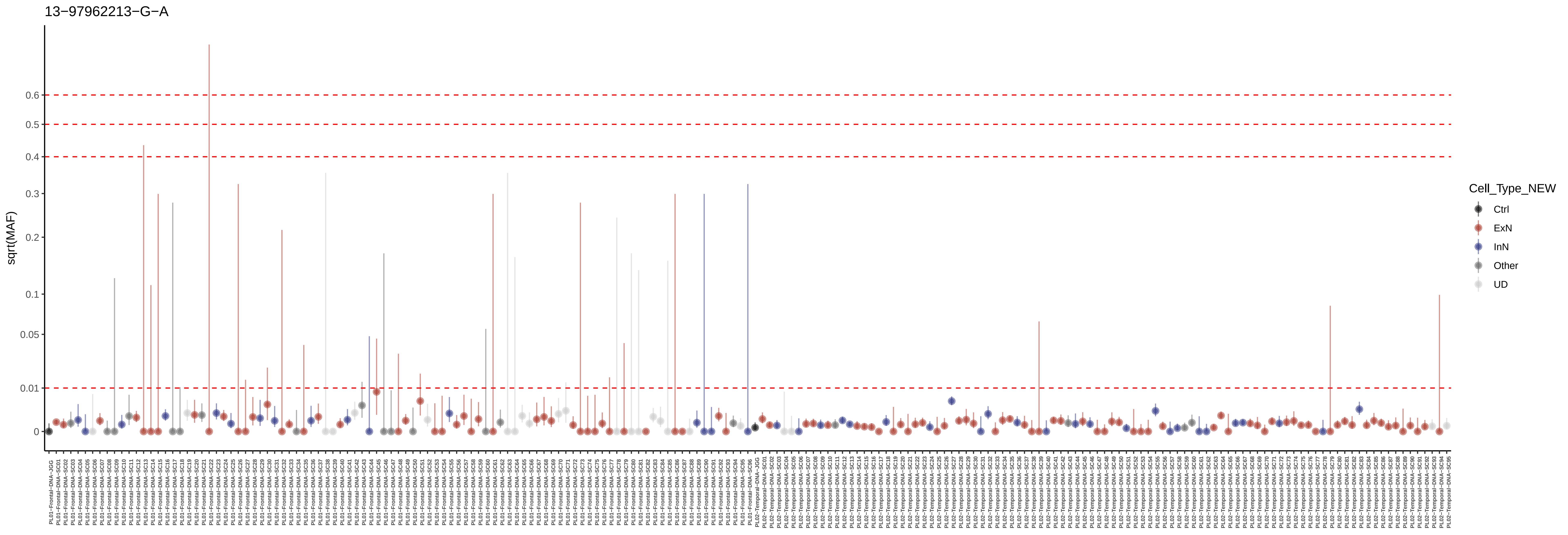
13−97962213−G−A
sqrt(MAF)
0.6
0.5
0.4
0.3
0.2
0.1
0.05
0.01
0
Cell_Type_NEW
Ctrl
ExN
InN
Other
UD

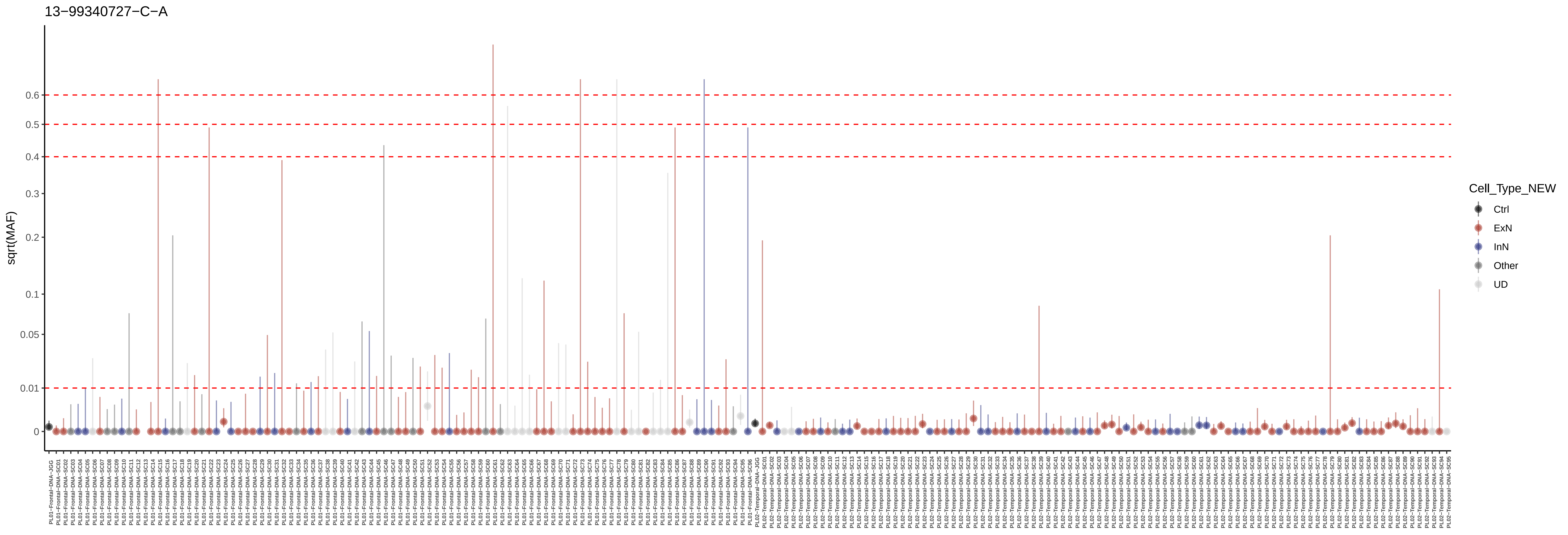
13−99340727−C−A
sqrt(MAF)
0.6
0.5
0.4
0.3
0.2
0.1
0.05
0.01
0
Cell_Type_NEW
Ctrl
ExN
InN
Other
UD

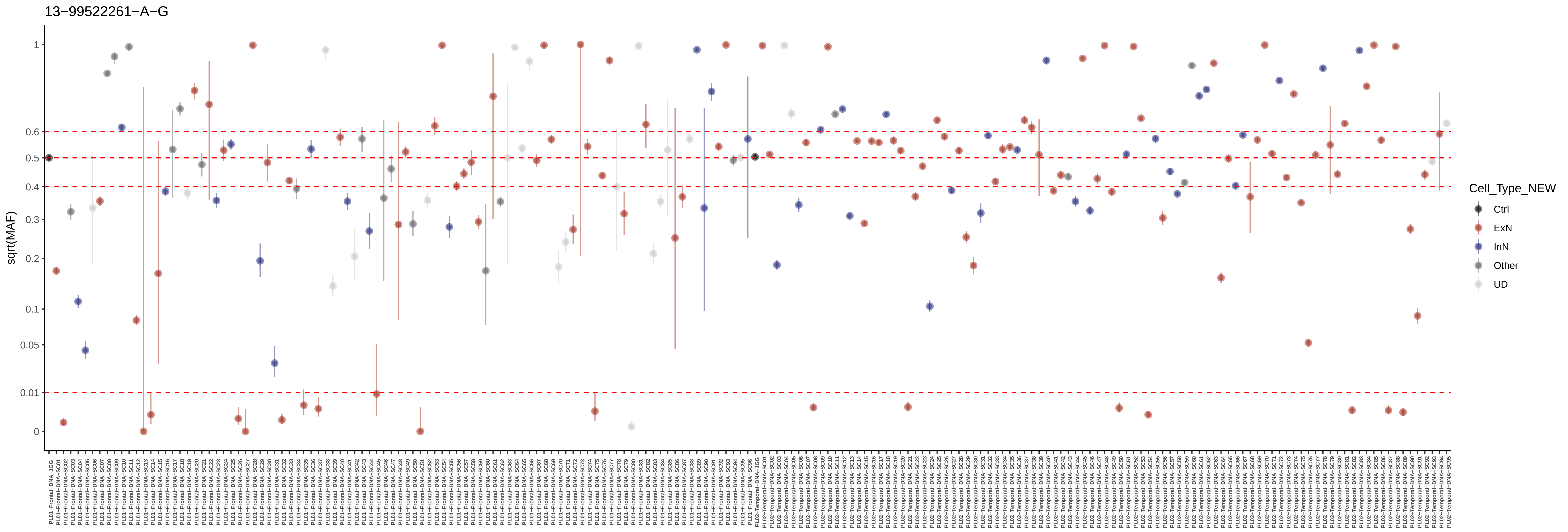
13−99522261−A−G
sqrt(MAF)
1
0.6
0.5
0.4
0.3
0.2
0.1
0.05
0.01
0
Cell_Type_NEW
Ctrl
ExN
InN
Other
UD

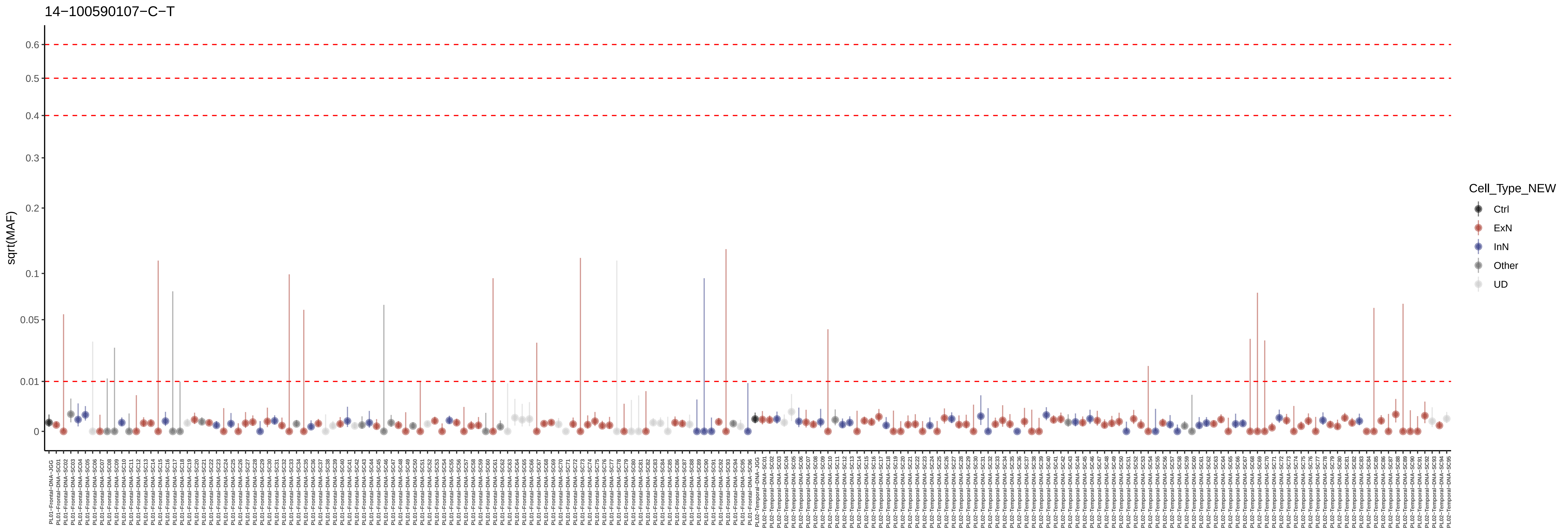
14−100590107−C−T
sqrt(MAF)
0.6
0.5
0.4
0.3
0.2
0.1
0.05
0.01
0
Cell_Type_NEW
Ctrl
ExN
InN
Other
UD

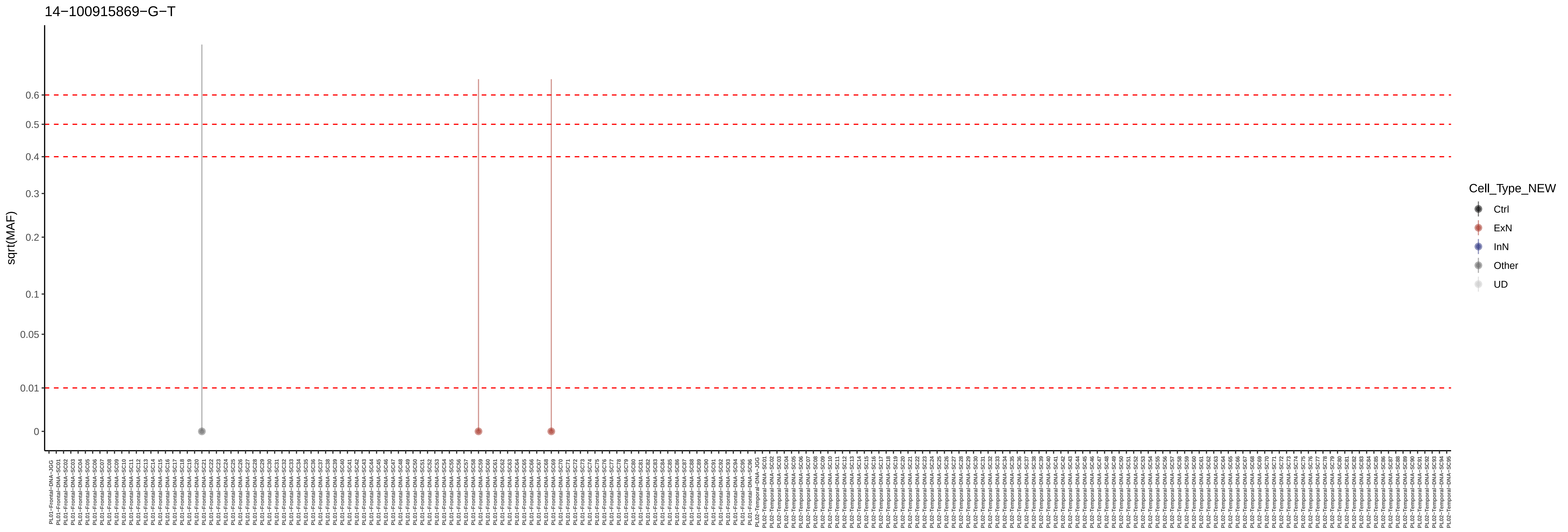
14−100915869−G−T
sqrt(MAF)
0.6
0.5
0.4
0.3
0.2
0.1
0.05
0.01
0
Cell_Type_NEW
Ctrl
ExN
InN
Other
UD

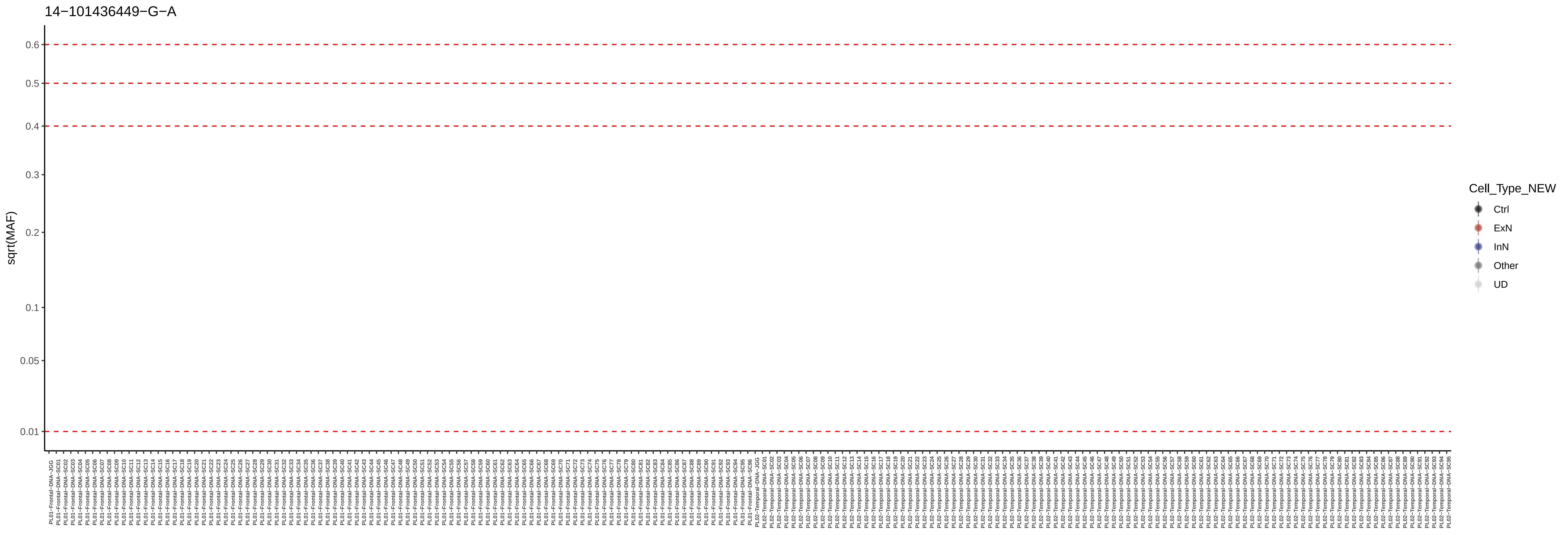
14−101436449−G−A
sqrt(MAF)
0.6
0.5
0.4
0.3
0.2
0.1
0.05
0.01
Cell_Type_NEW
Ctrl
ExN
InN
Other
UD

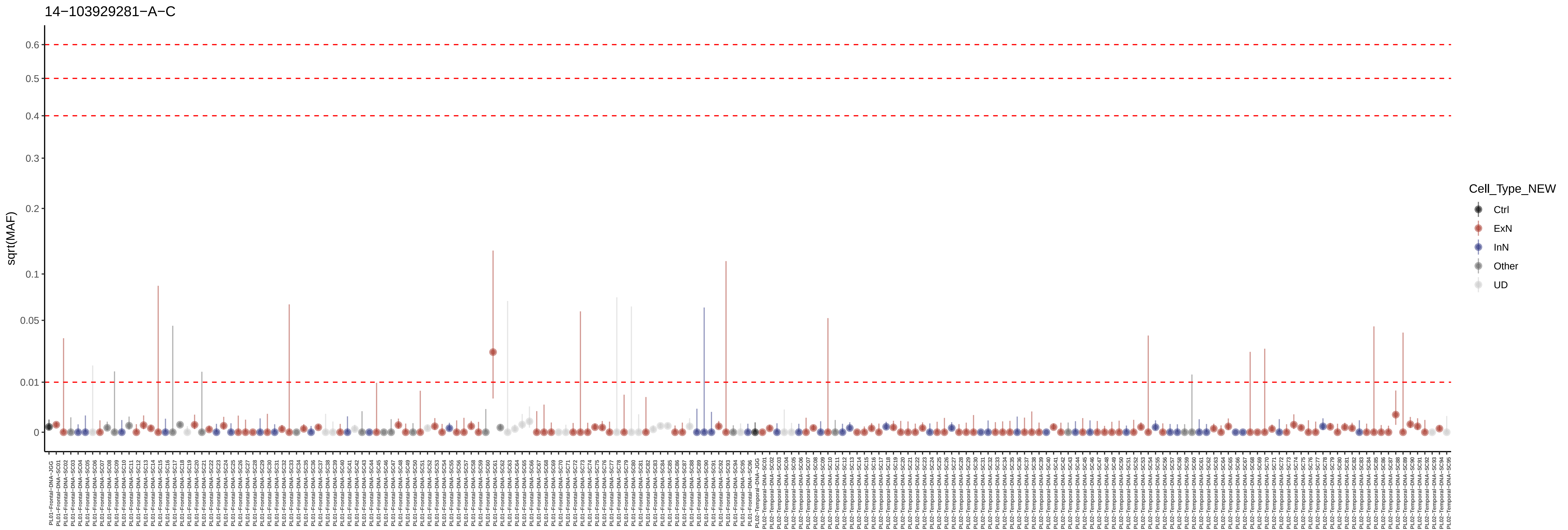
14−103929281−A−C
sqrt(MAF)
0.6
0.5
0.4
0.3
0.2
0.1
0.05
0.01
0
Cell_Type_NEW
Ctrl
ExN
InN
Other
UD

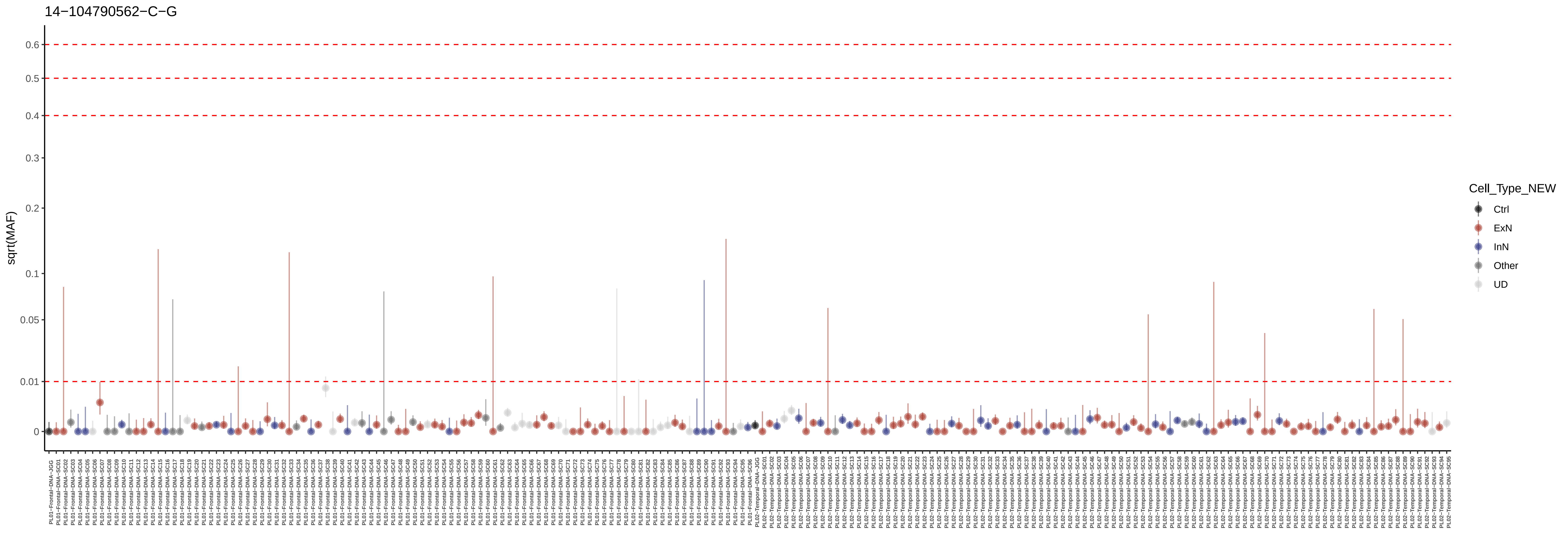
14−104790562−C−G
sqrt(MAF)
0.6
0.5
0.4
0.3
0.2
0.1
0.05
0.01
0
Cell_Type_NEW
Ctrl
ExN
InN
Other
UD

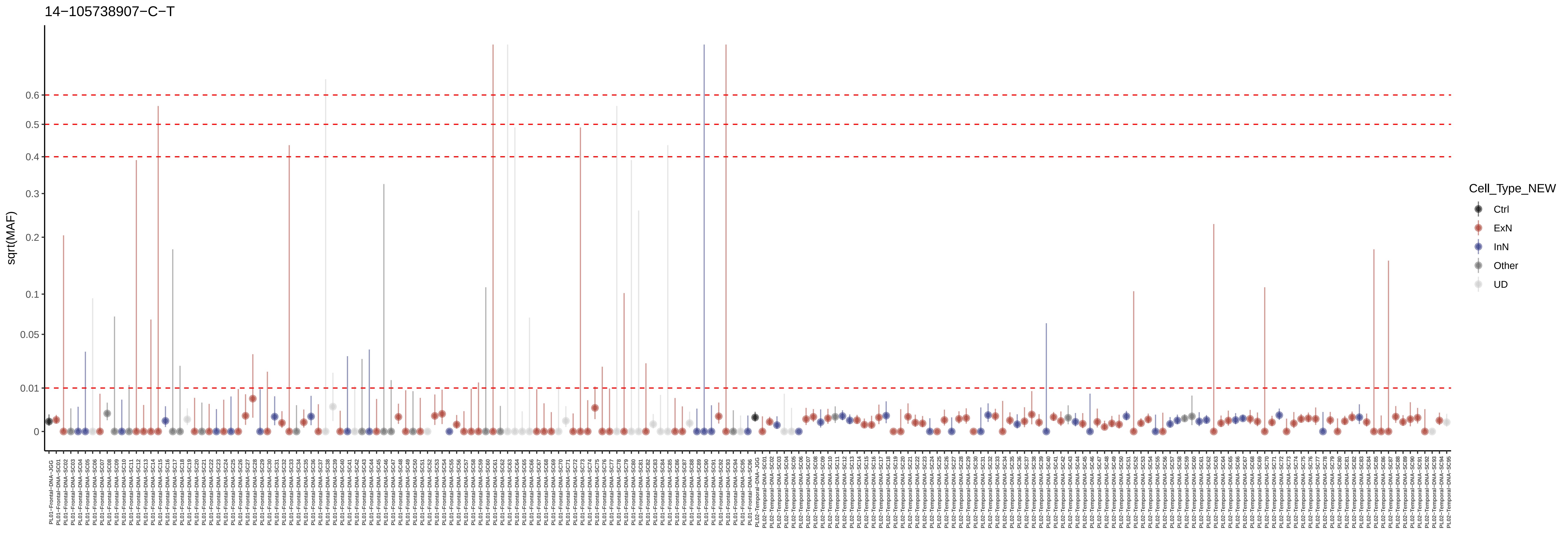

14−105738907−C−T
sqrt(MAF)
0.6
0.5
0.4
0.3
0.2
0.1
0.05
0.01
0
Cell_Type_NEW
Ctrl
ExN
InN
Other
UD

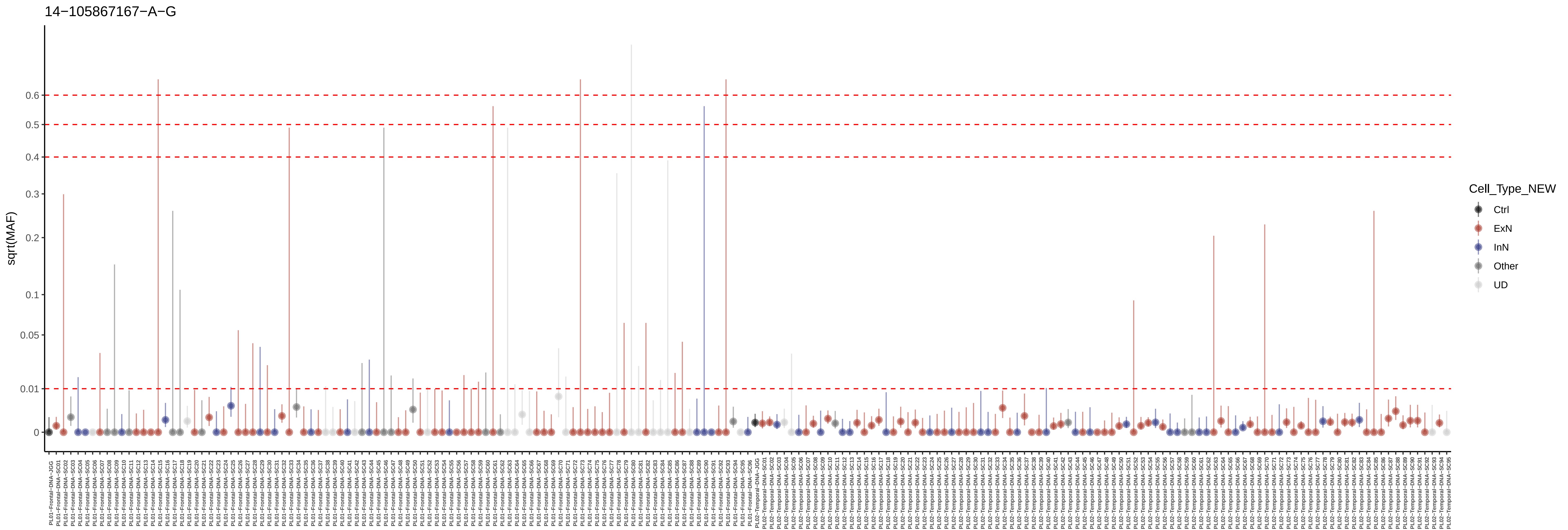
14−105867167−A−G
sqrt(MAF)
0.6
0.5
0.4
0.3
0.2
0.1
0.05
0.01
0
Cell_Type_NEW
Ctrl
ExN
InN
Other
UD

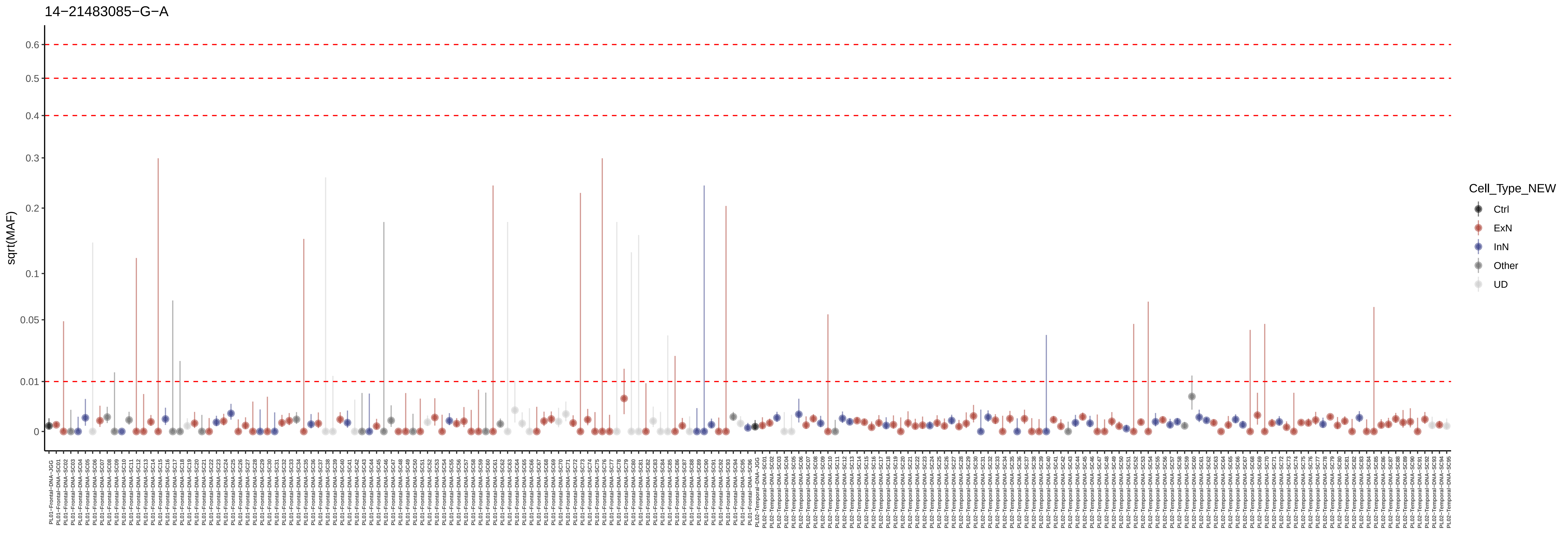
14−21483085−G−A
sqrt(MAF)
0.6
0.5
0.4
0.3
0.2
0.1
0.05
0.01
0
Cell_Type_NEW
Ctrl
ExN
InN
Other
UD

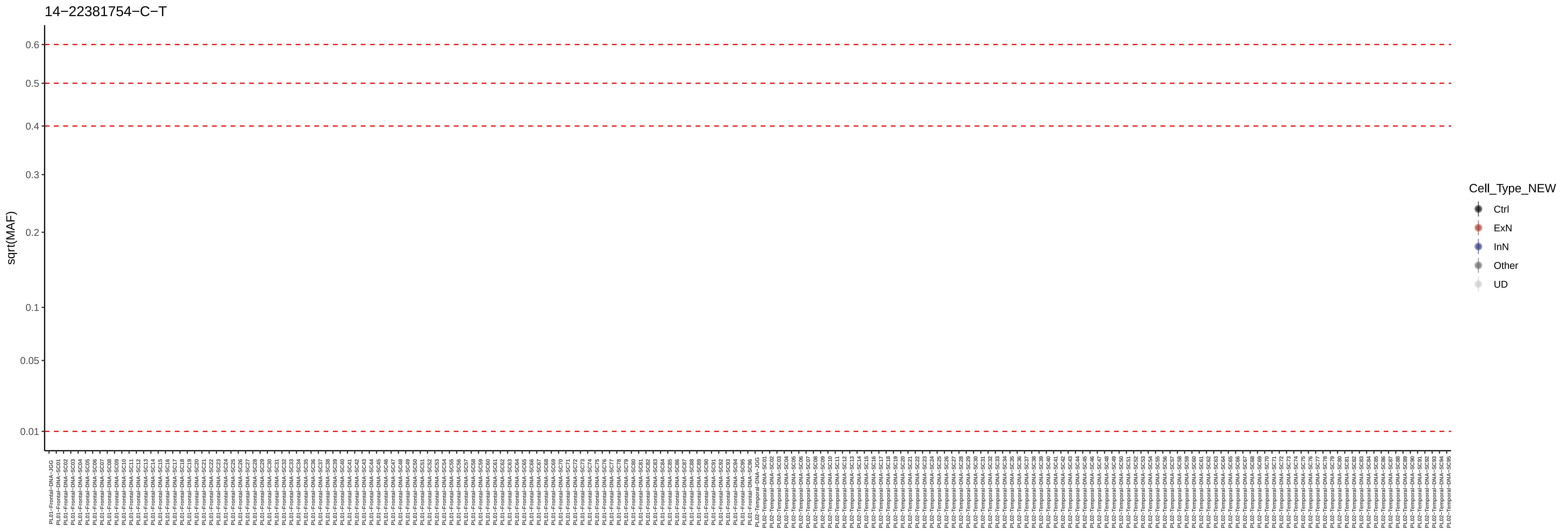
14−22381754−C−T
sqrt(MAF)
0.6
0.5
0.4
0.3
0.2
0.1
0.05
0.01
Cell_Type_NEW
Ctrl
ExN
InN
Other
UD

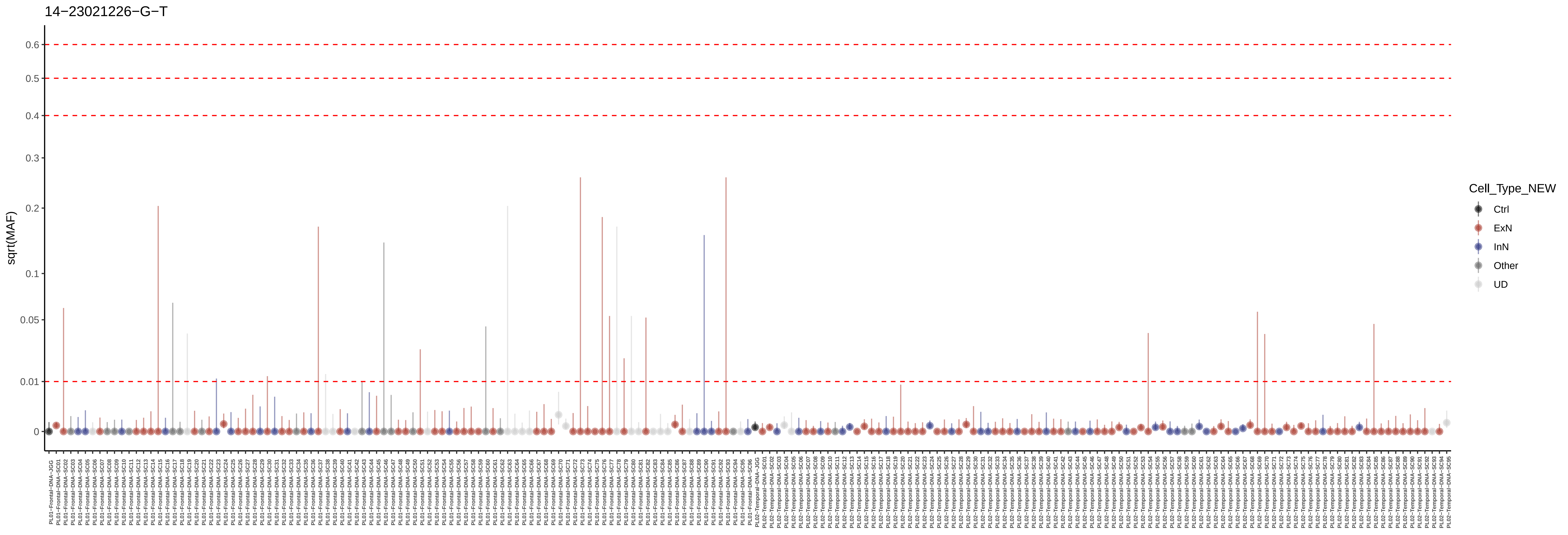
14−23021226−G−T
sqrt(MAF)
0.6
0.5
0.4
0.3
0.2
0.1
0.05
0.01
0
Cell_Type_NEW
Ctrl
ExN
InN
Other
UD

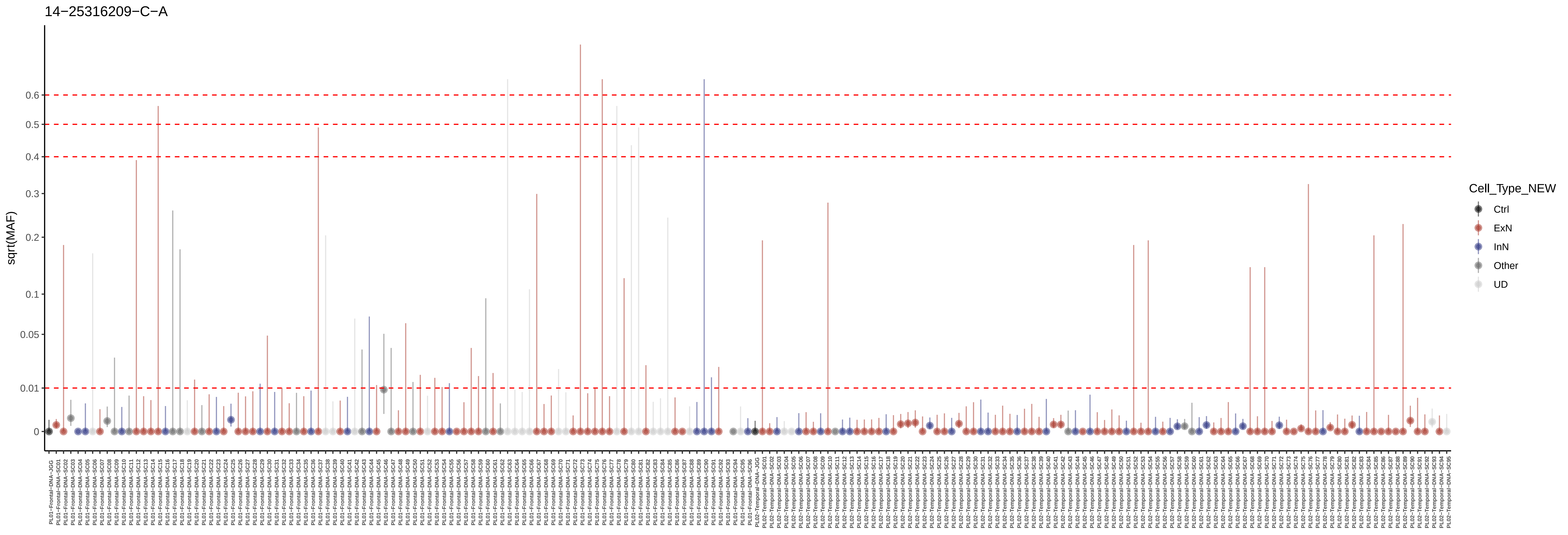

14−25316209−C−A
sqrt(MAF)
0.6
0.5
0.4
0.3
0.2
0.1
0.05
0.01
0
Cell_Type_NEW
Ctrl
ExN
InN
Other
UD

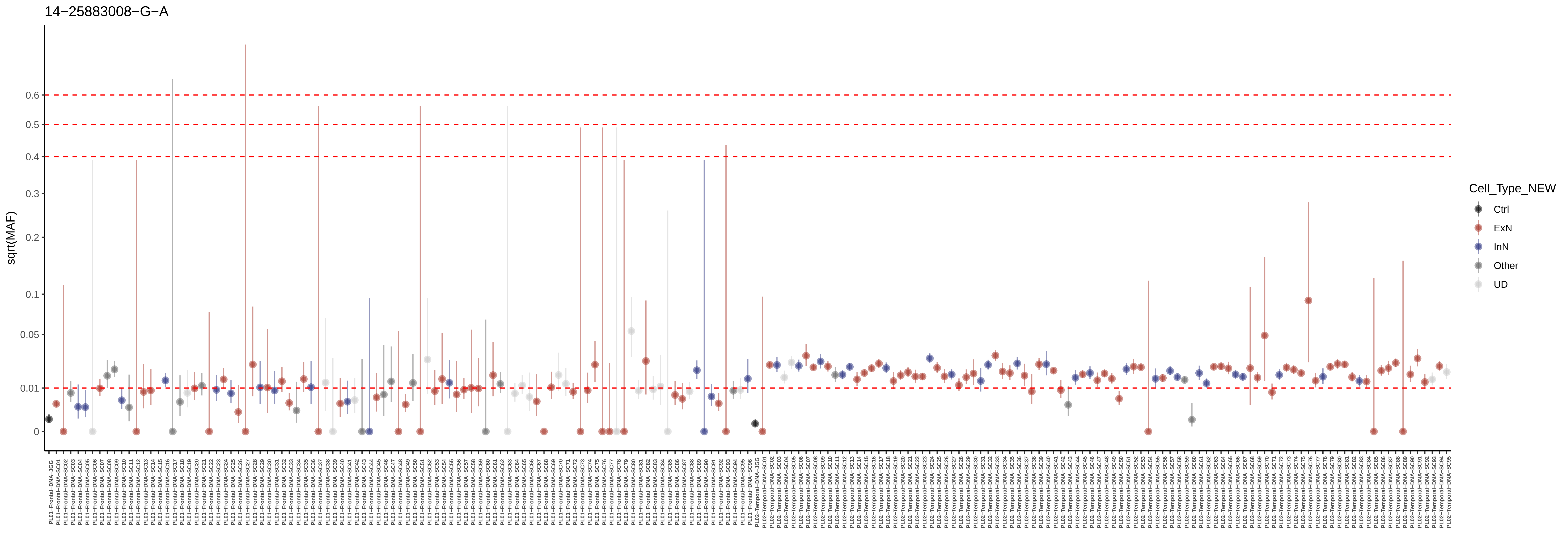
14−25883008−G−A
sqrt(MAF)
0.6
0.5
0.4
0.3
0.2
0.1
0.05
0.01
0
Cell_Type_NEW
Ctrl
ExN
InN
Other
UD

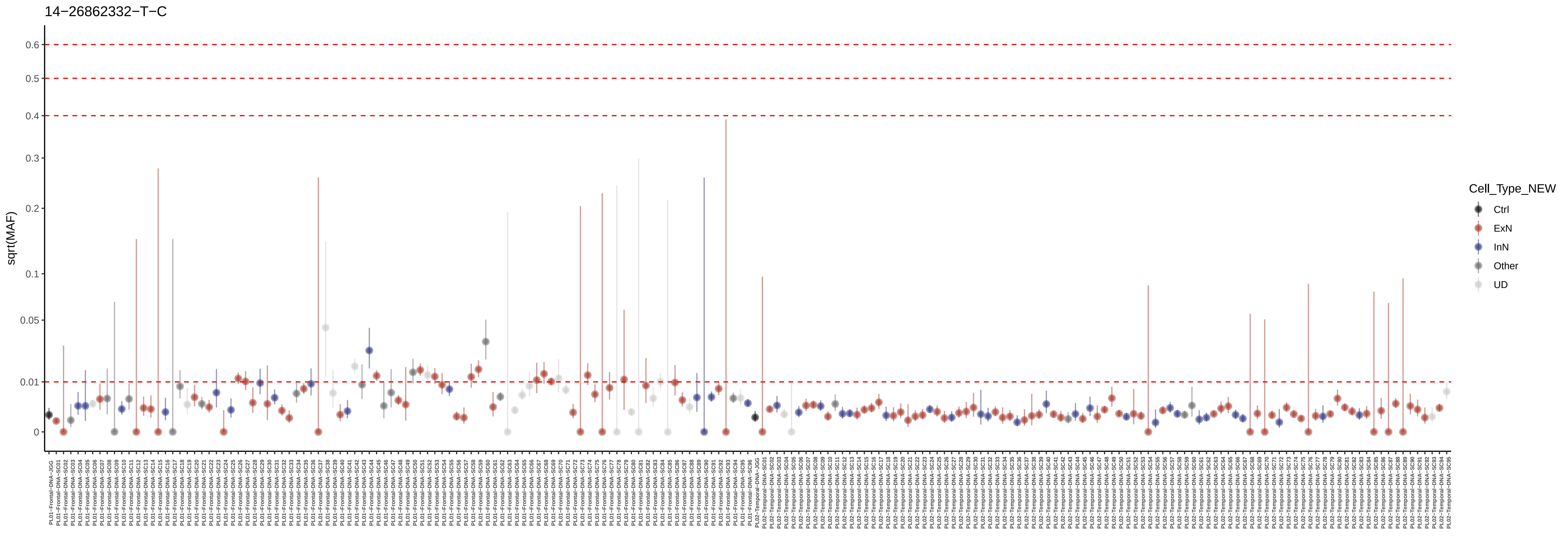
14−26862332−T−C
sqrt(MAF)
0.6
0.5
0.4
0.3
0.2
0.1
0.05
0.01
0
Cell_Type_NEW
Ctrl
ExN
InN
Other
UD

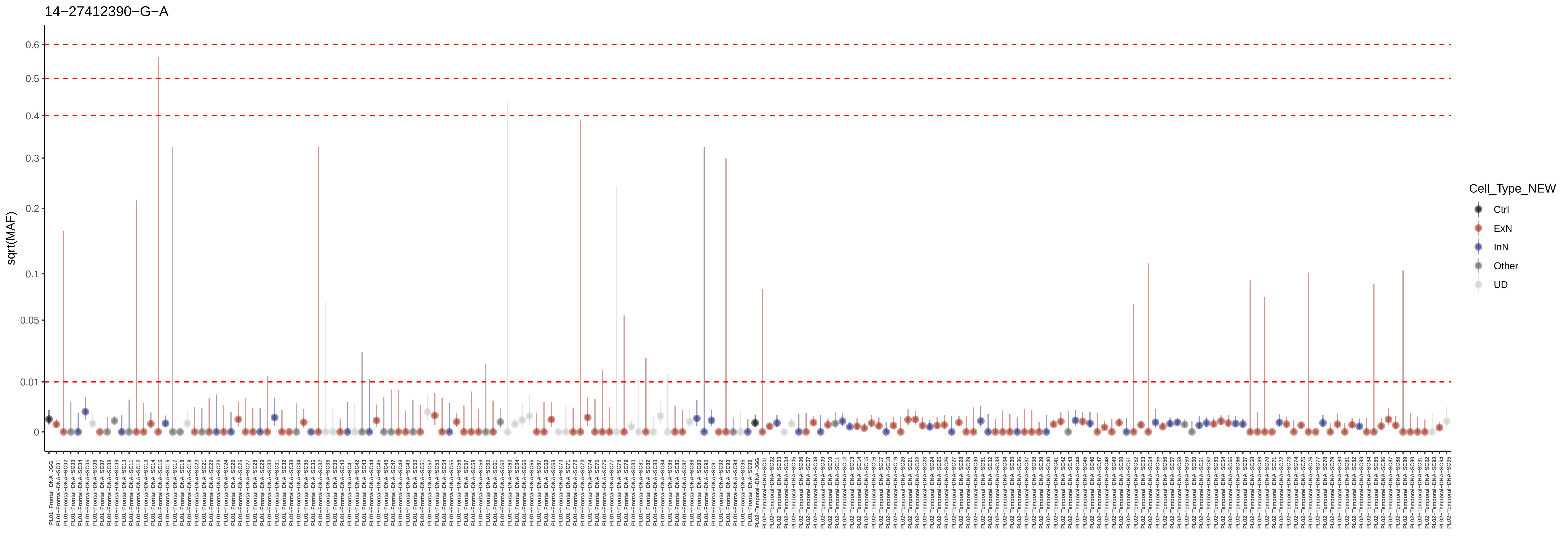
14−27412390−G−A
sqrt(MAF)
0.6
0.5
0.4
0.3
0.2
0.1
0.05
0.01
0
Cell_Type_NEW
Ctrl
ExN
InN
Other
UD

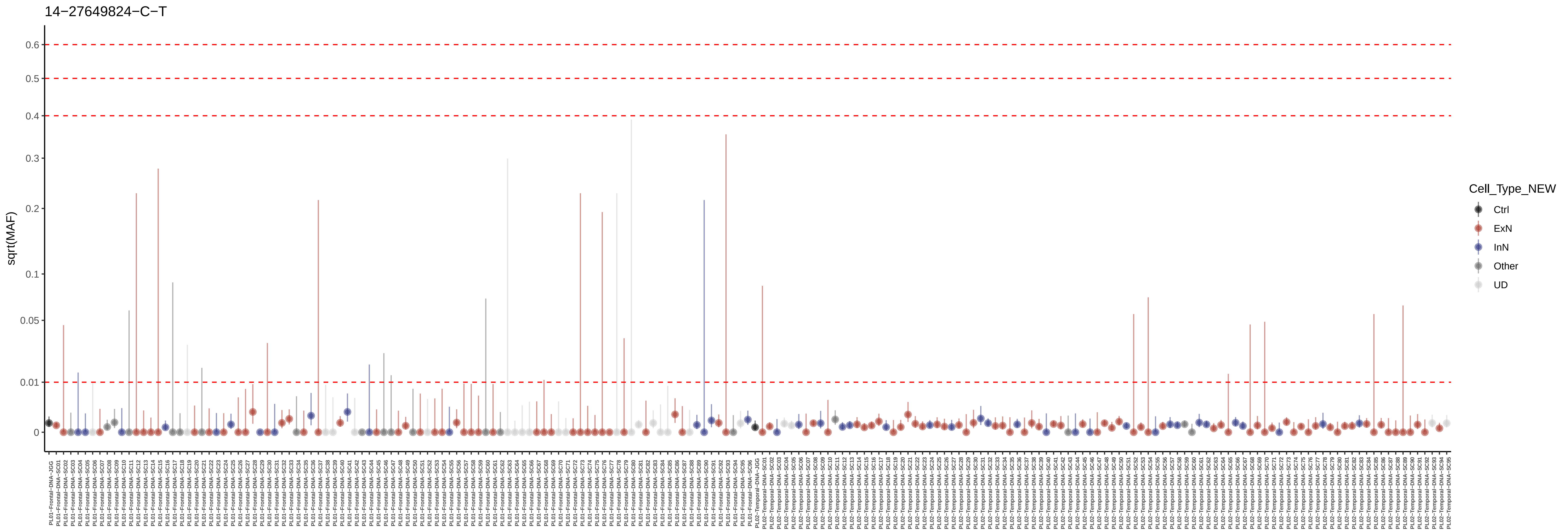

14−27649824−C−T
sqrt(MAF)
0.6
0.5
0.4
0.3
0.2
0.1
0.05
0.01
0
Cell_Type_NEW
Ctrl
ExN
InN
Other
UD

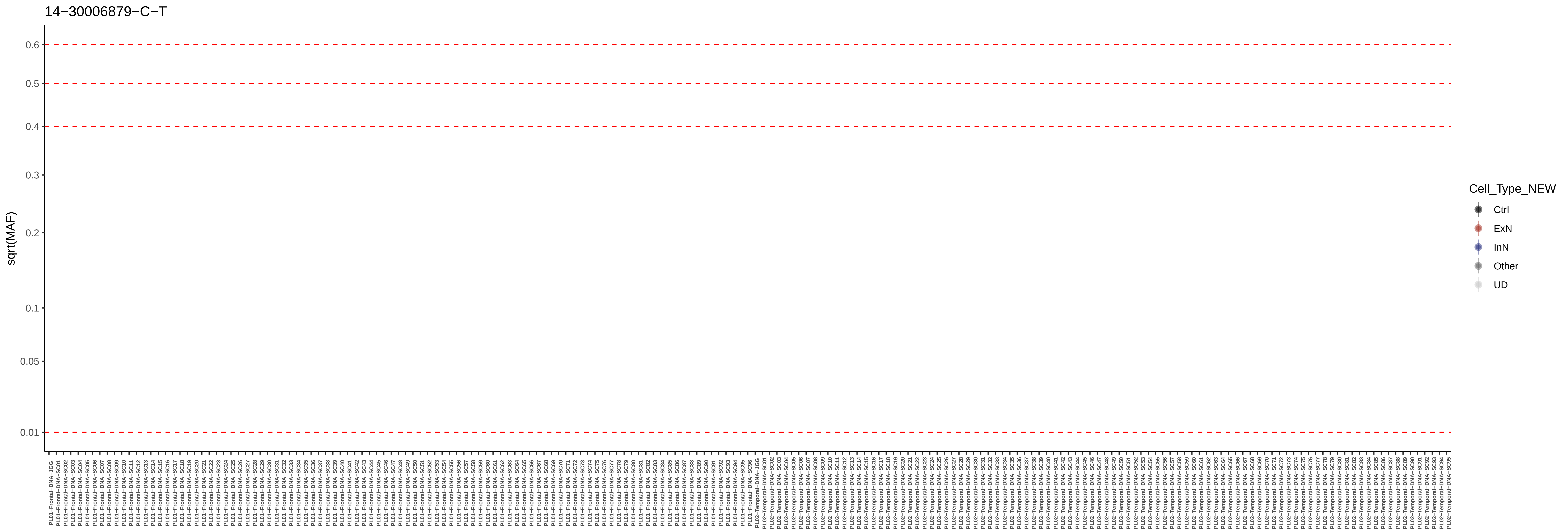
14−30006879−C−T
sqrt(MAF)
0.6
0.5
0.4
0.3
0.2
0.1
0.05
0.01
Cell_Type_NEW
Ctrl
ExN
InN
Other
UD

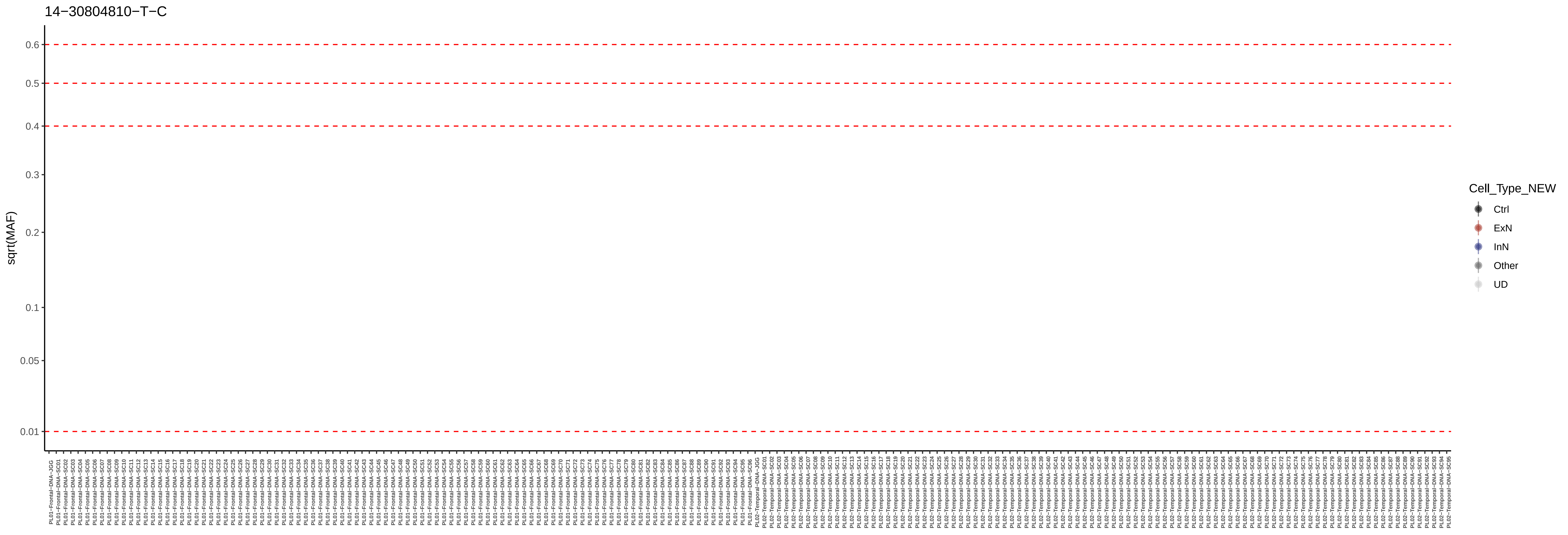
14−30804810−T−C
sqrt(MAF)
0.6
0.5
0.4
0.3
0.2
0.1
0.05
0.01
Cell_Type_NEW
Ctrl
ExN
InN
Other
UD

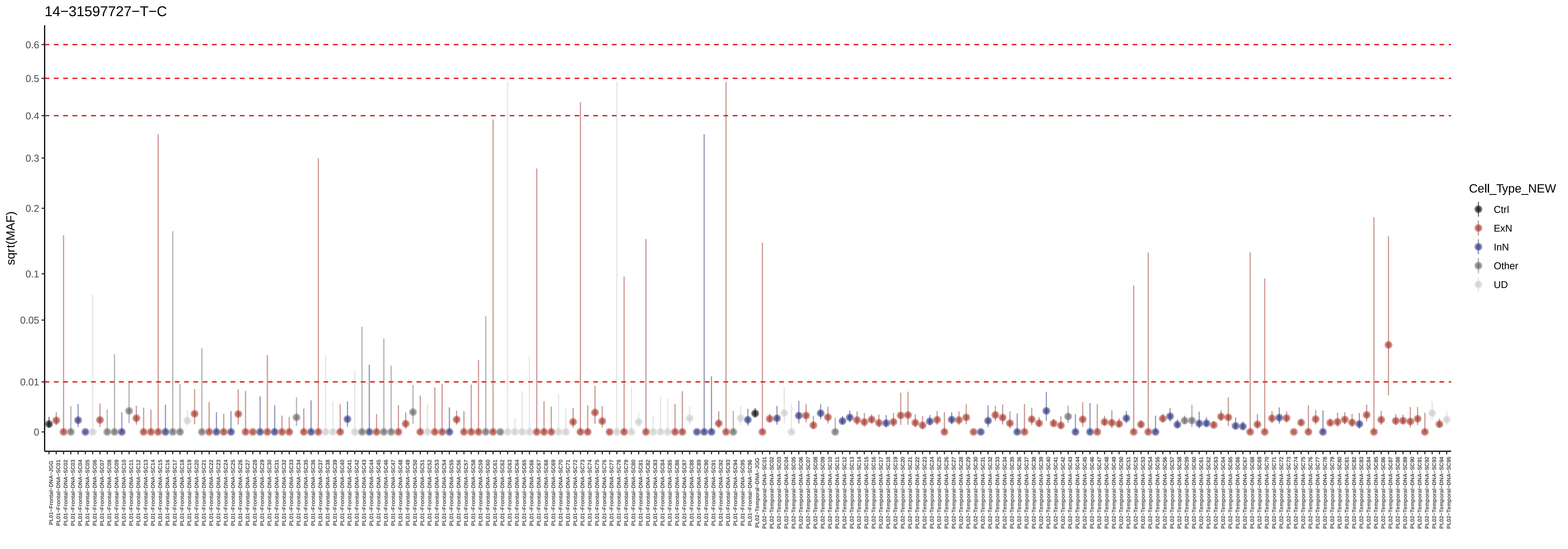
14−31597727−T−C
sqrt(MAF)
0.6
0.5
0.4
0.3
0.2
0.1
0.05
0.01
0
Cell_Type_NEW
Ctrl
ExN
InN
Other
UD

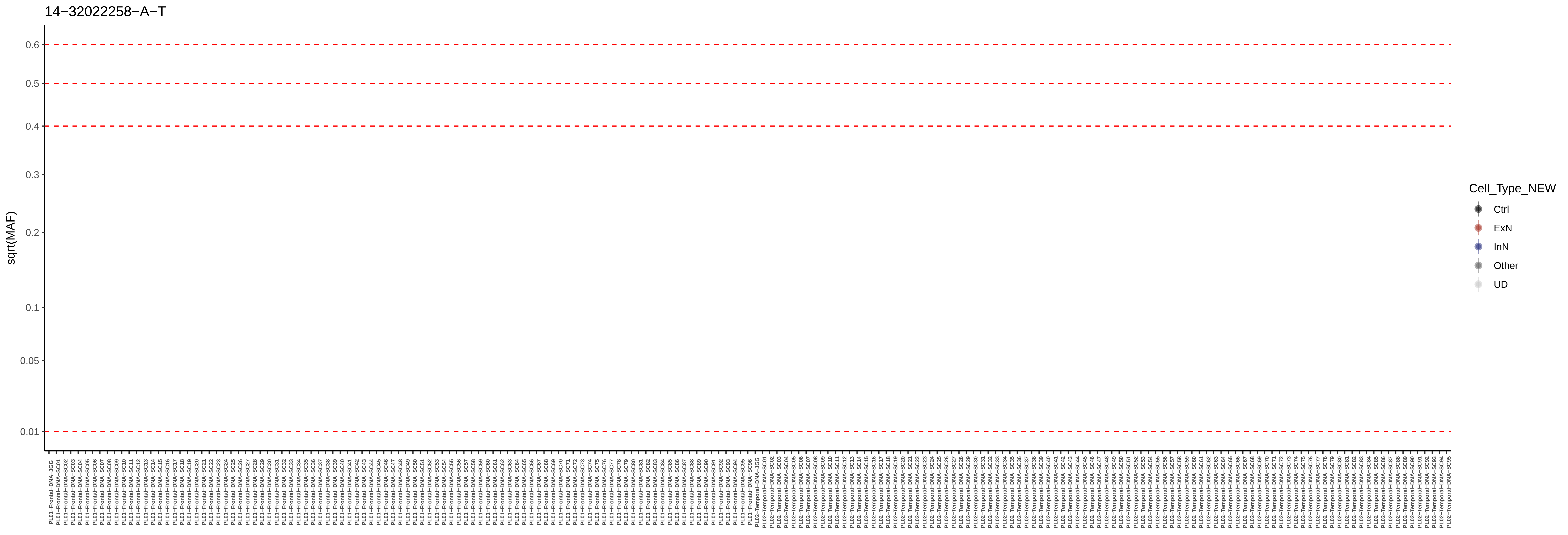
14−32022258−A−T
sqrt(MAF)
0.6
0.5
0.4
0.3
0.2
0.1
0.05
0.01
Cell_Type_NEW
Ctrl
ExN
InN
Other
UD

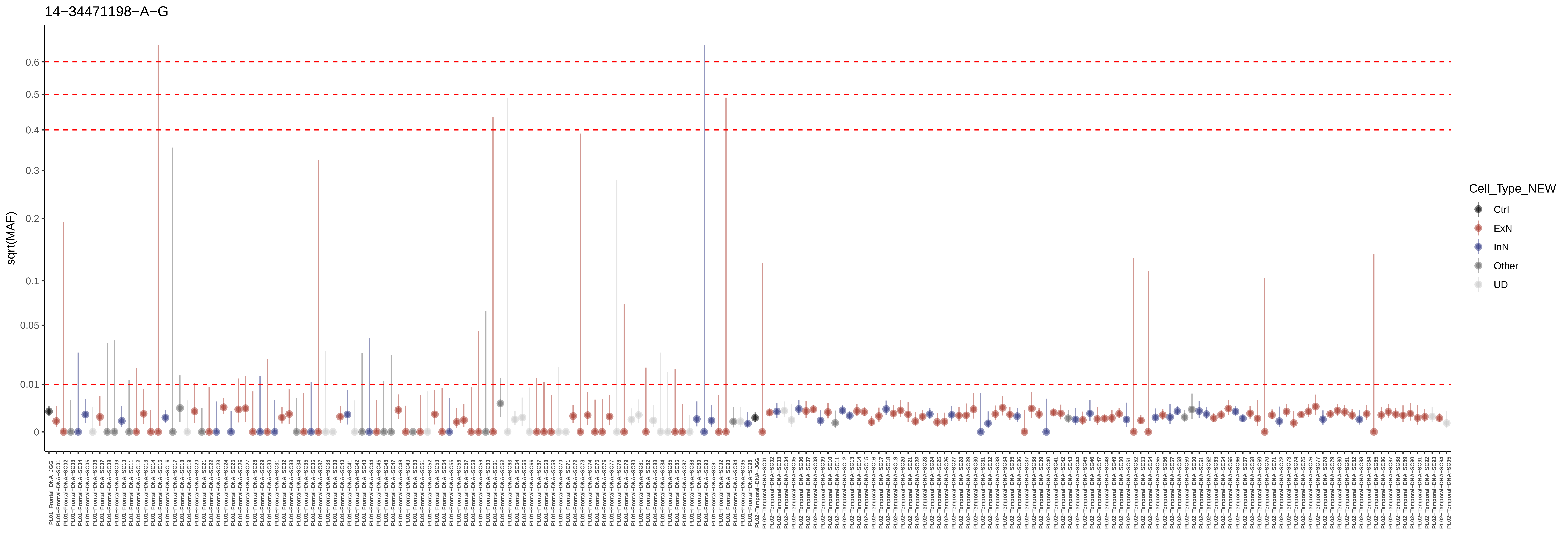

14−34471198−A−G
sqrt(MAF)
Cell_Type_NEW
Ctrl
ExN
InN
Other
UD

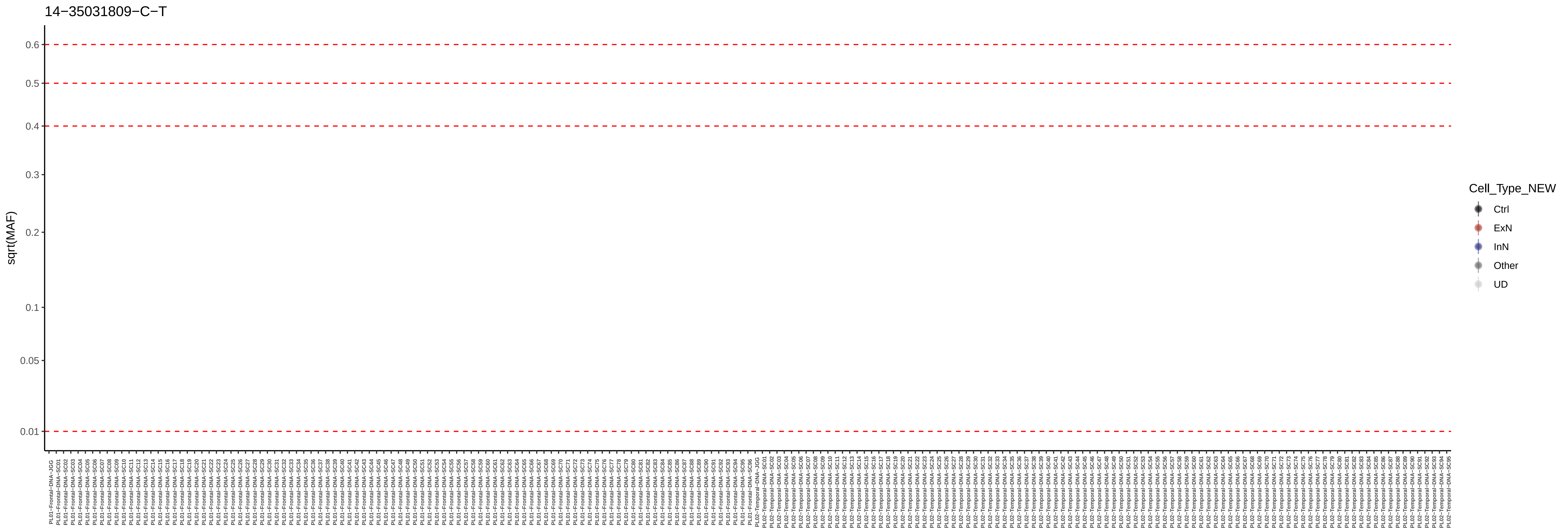
14−35031809−C−T
sqrt(MAF)
0.6
0.5
0.4
0.3
0.2
0.1
0.05
0.01
Cell_Type_NEW
Ctrl
ExN
InN
Other
UD

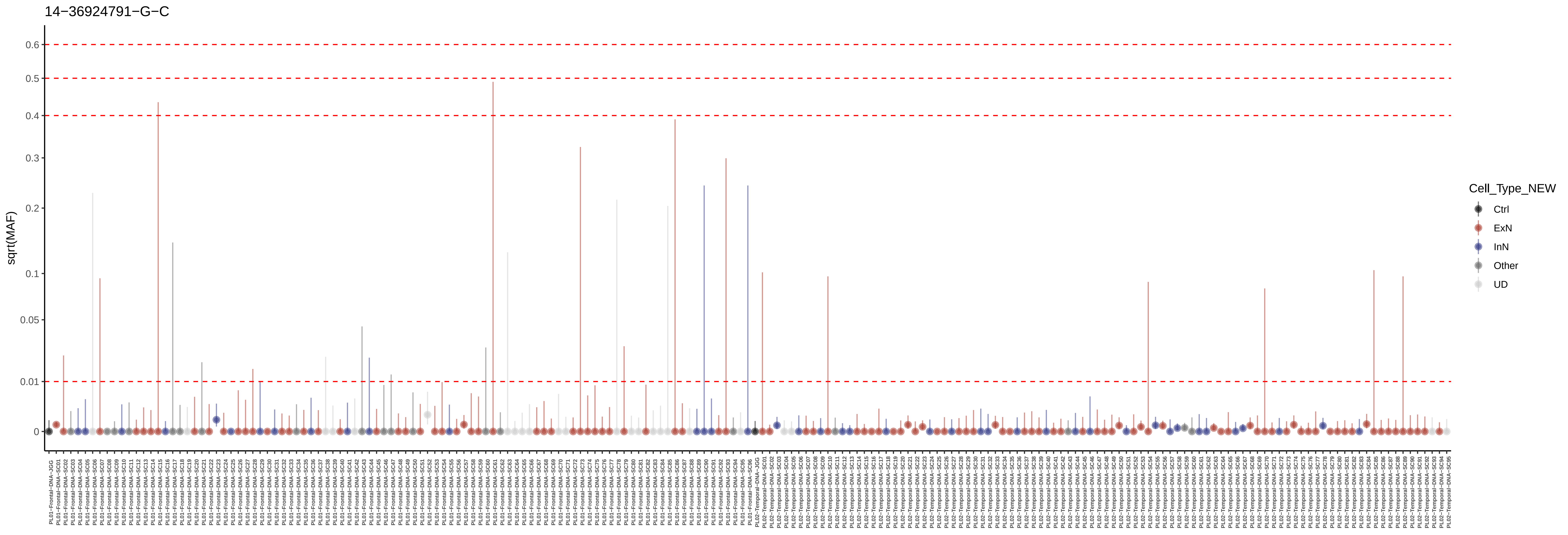
14−36924791−G−C
sqrt(MAF)
0.6
0.5
0.4
0.3
0.2
0.1
0.05
0.01
0
Cell_Type_NEW
Ctrl
ExN
InN
Other
UD

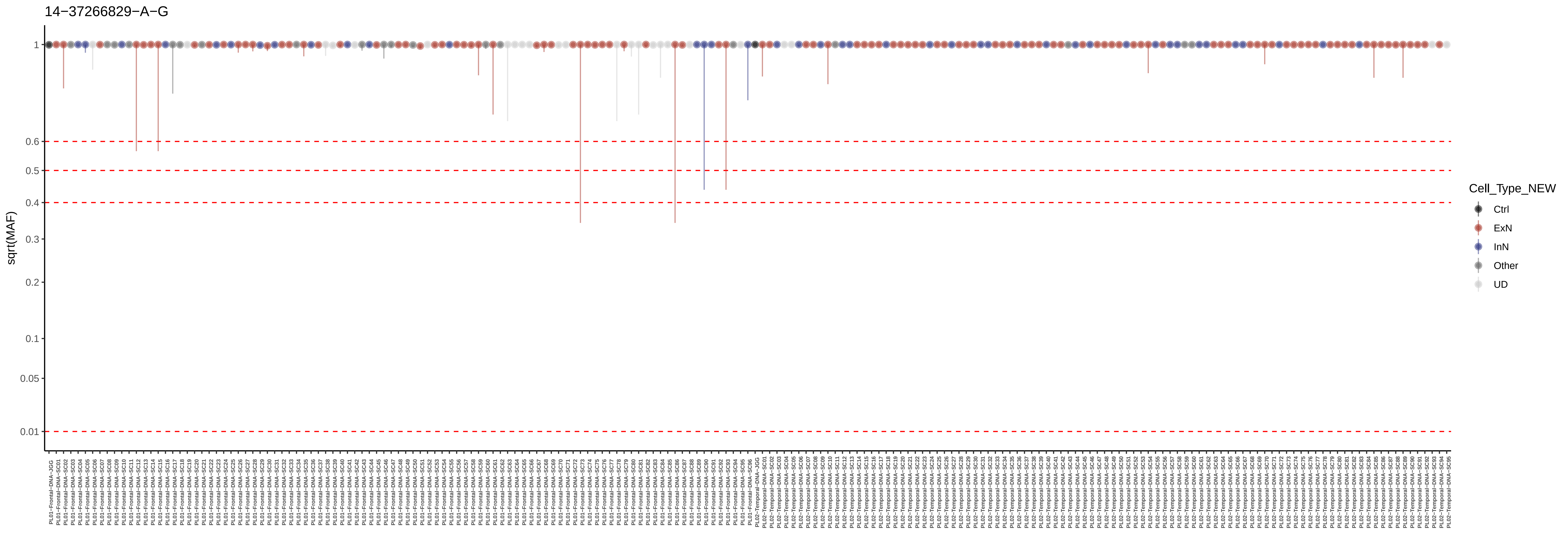
14−37266829−A−G
sqrt(MAF)
1
0.6
0.5
0.4
0.3
0.2
0.1
0.05
0.01
Cell_Type_NEW
Ctrl
ExN
InN
Other
UD

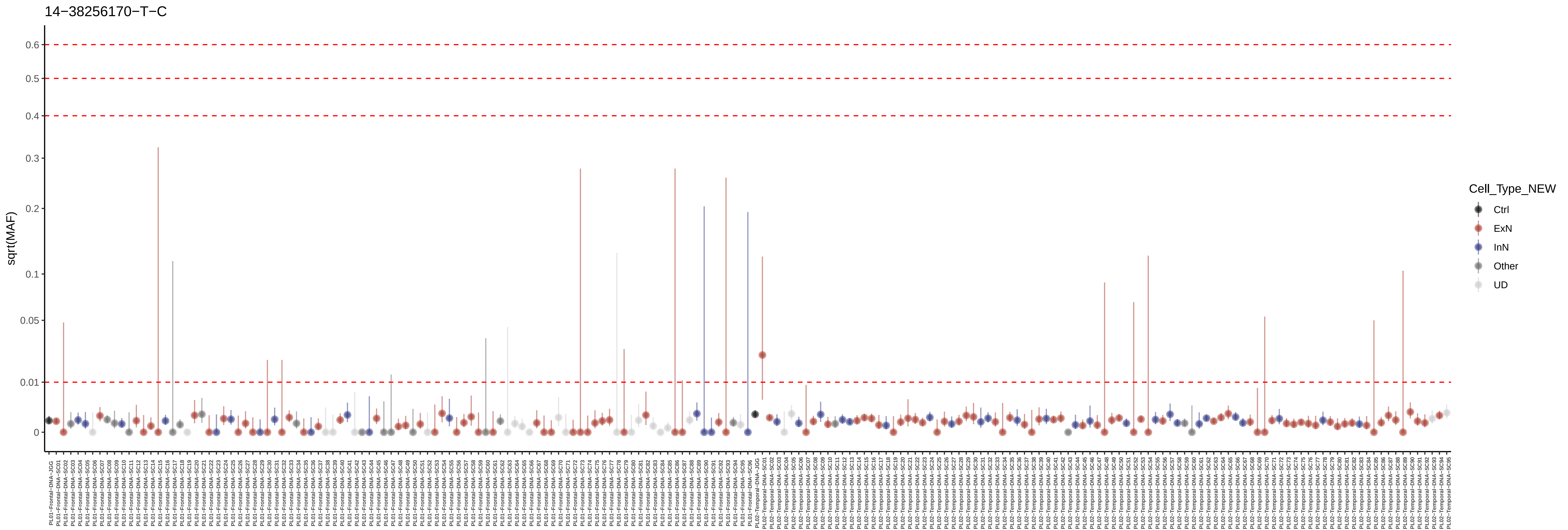
14−38256170−T−C
sqrt(MAF)
0.6
0.5
0.4
0.3
0.2
0.1
0.05
0.01
0
Cell_Type_NEW
Ctrl
ExN
InN
Other
UD

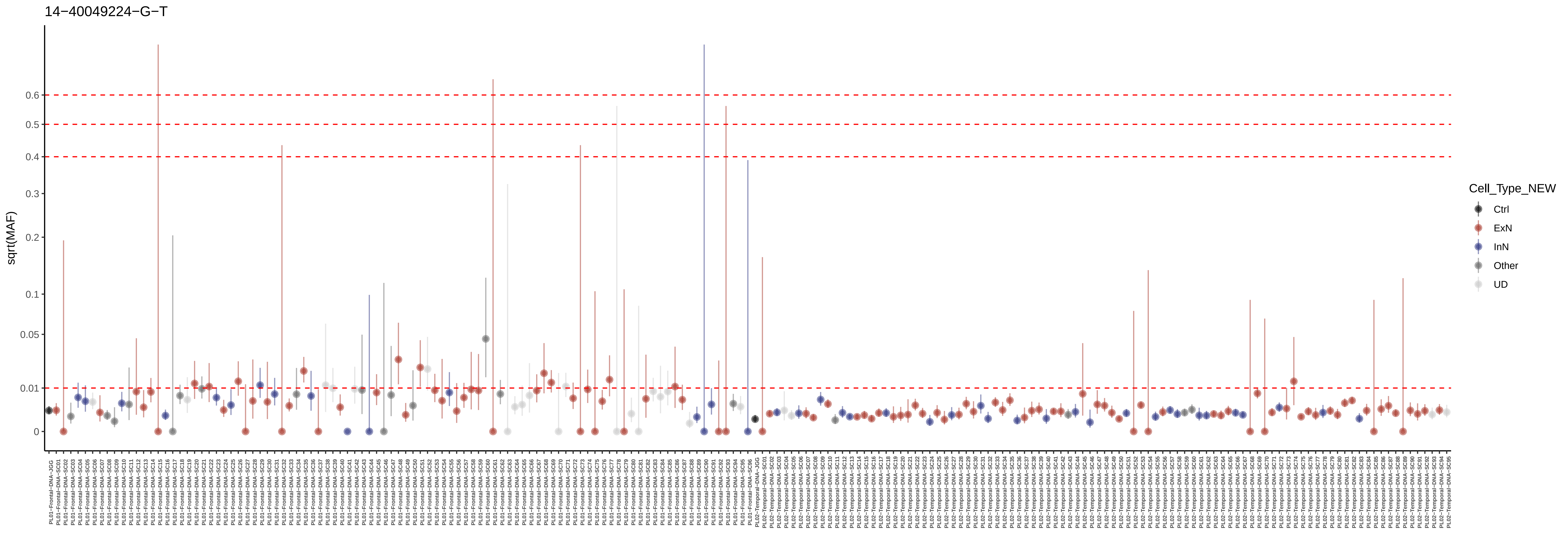
14−40049224−G−T
sqrt(MAF)
0.6
0.5
0.4
0.3
0.2
0.1
0.05
0.01
0
Cell_Type_NEW
Ctrl
ExN
InN
Other
UD

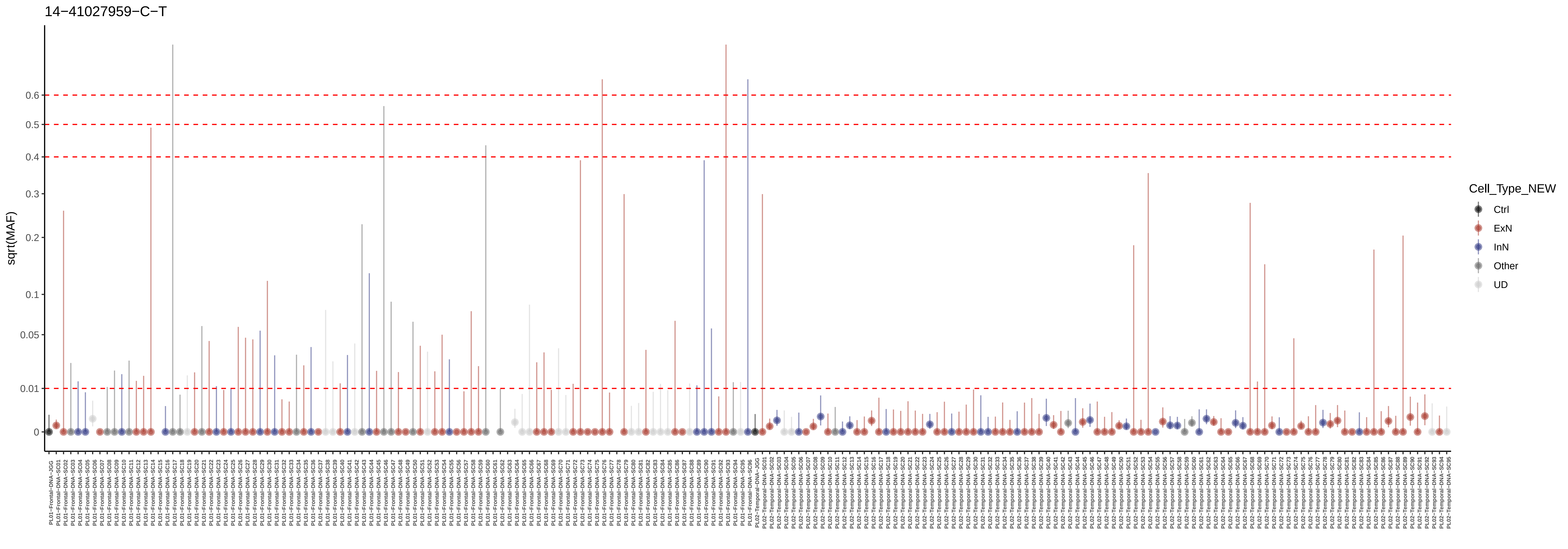
14−41027959−C−T
sqrt(MAF)
0.6
0.5
0.4
0.3
0.2
0.1
0.05
0.01
0
Cell_Type_NEW
Ctrl
ExN
InN
Other
UD

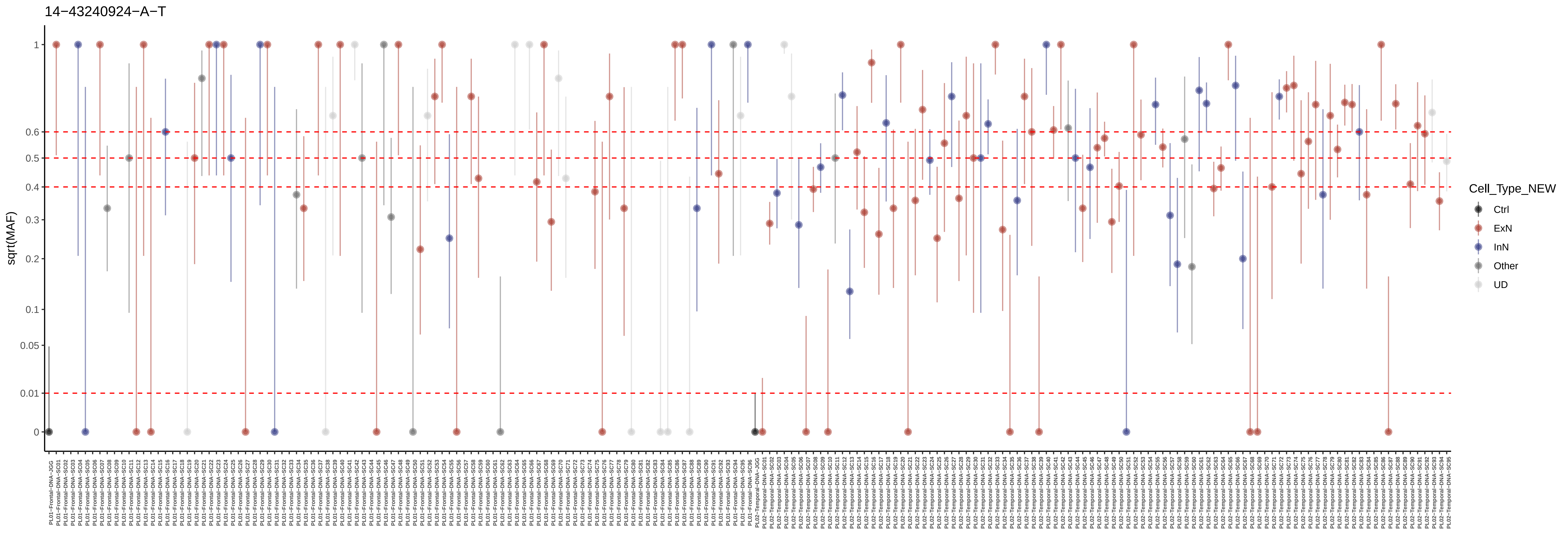
14-43240924-A-T
sqrt(MAF)
1
0.6
0.5
0.4
0.3
0.2
0.1
0.05
0.01
0
Cell_Type_NEW
Ctrl
ExN
InN
Other
UD

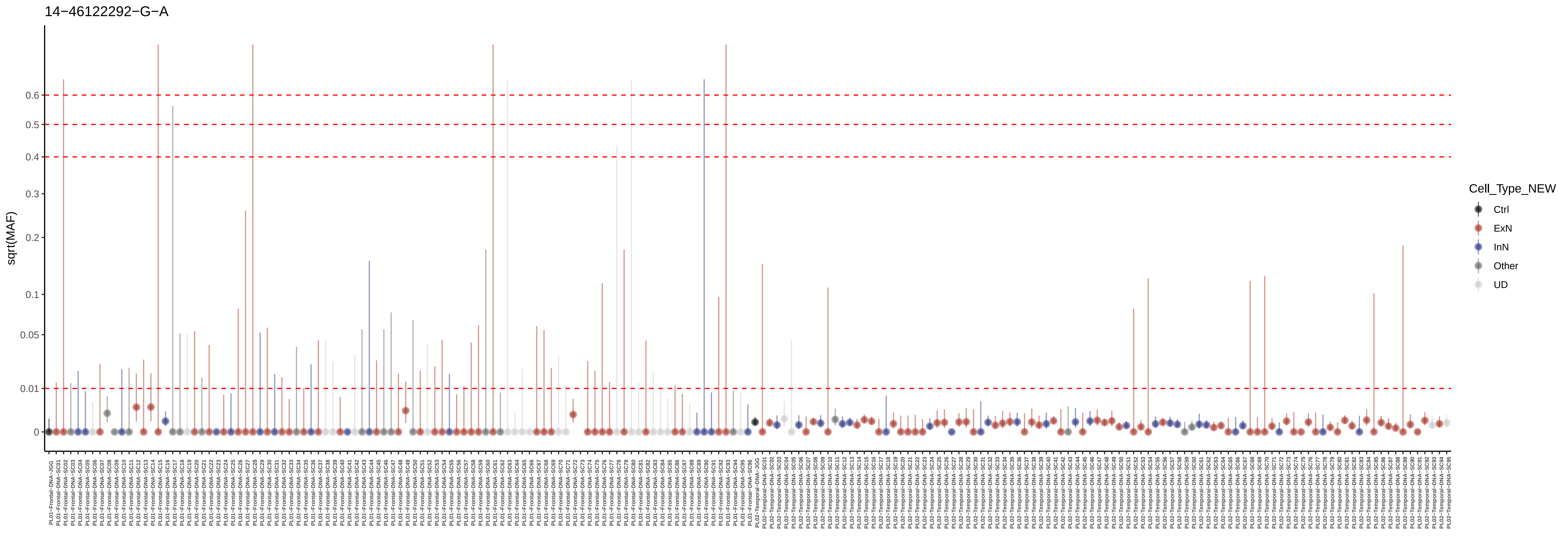
14−46122292−G−A
sqrt(MAF)
0
0.01
0.05
0.1
0.2
0.3
0.4
0.5
0.6
Cell_Type_NEW
Ctrl
ExN
InN
Other
UD

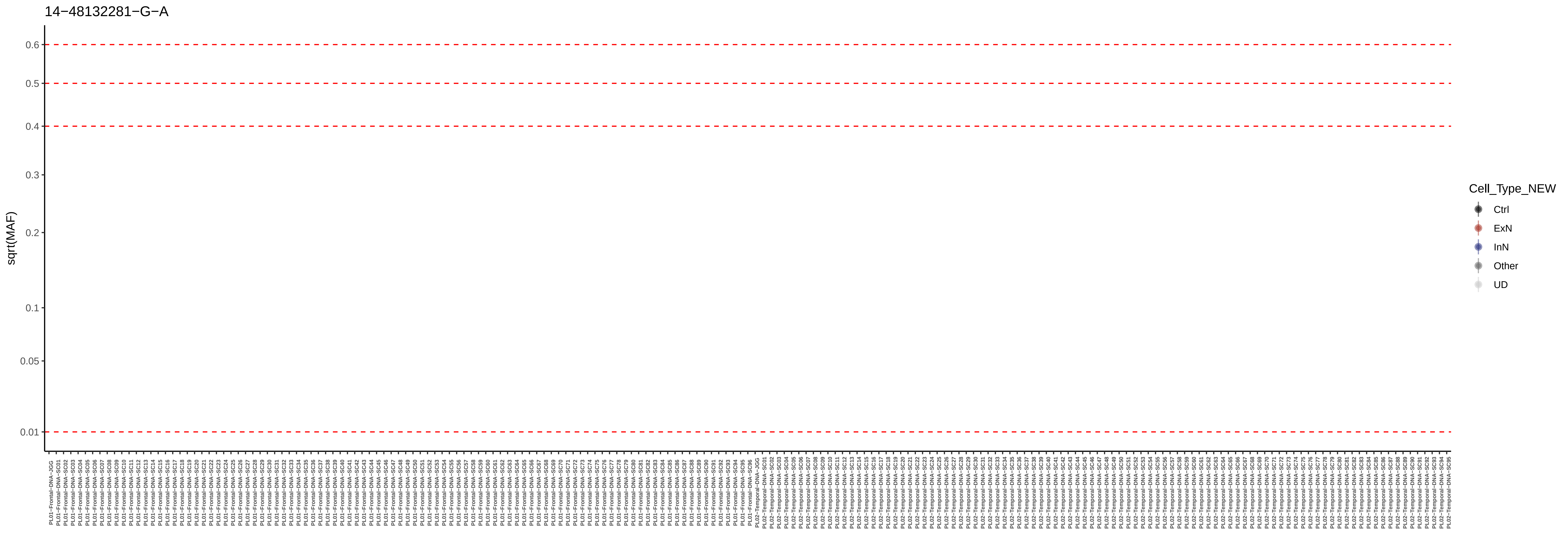
14−48132281−G−A
sqrt(MAF)
0.6
0.5
0.4
0.3
0.2
0.1
0.05
0.01
Cell_Type_NEW
Ctrl
ExN
InN
Other
UD

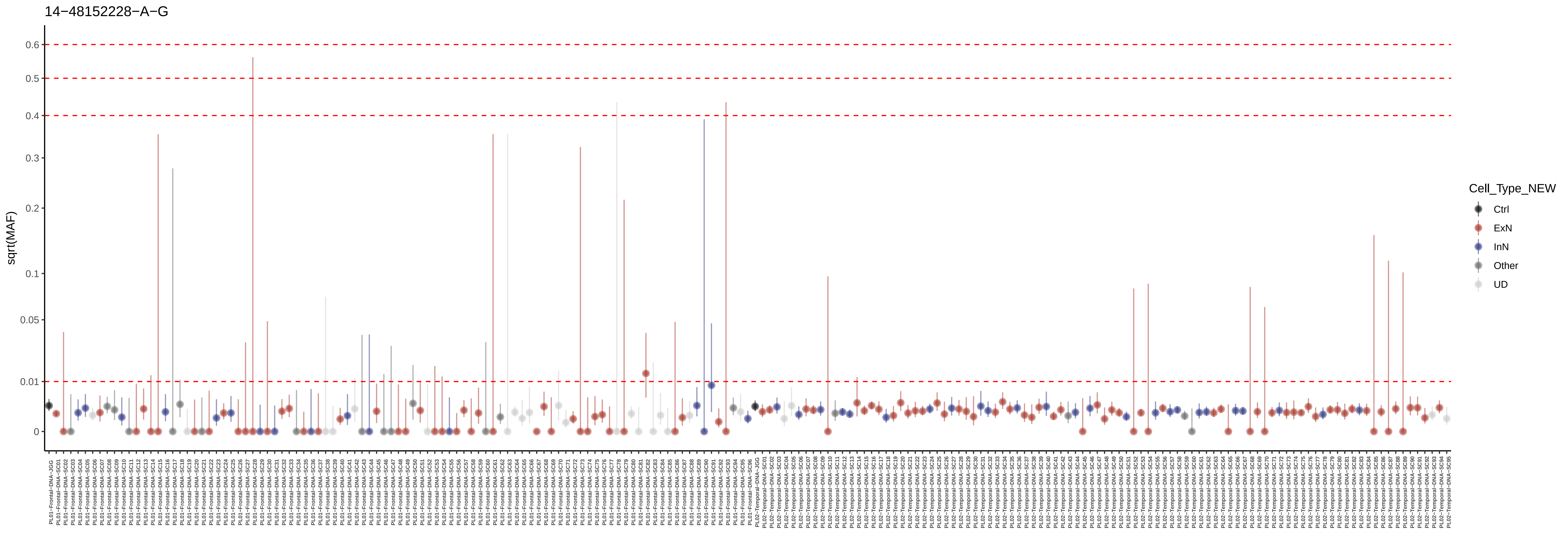

14−48152228−A−G
sqrt(MAF)
0.6
0.5
0.4
0.3
0.2
0.1
0.05
0.01
0
Cell_Type_NEW
Ctrl
ExN
InN
Other
UD

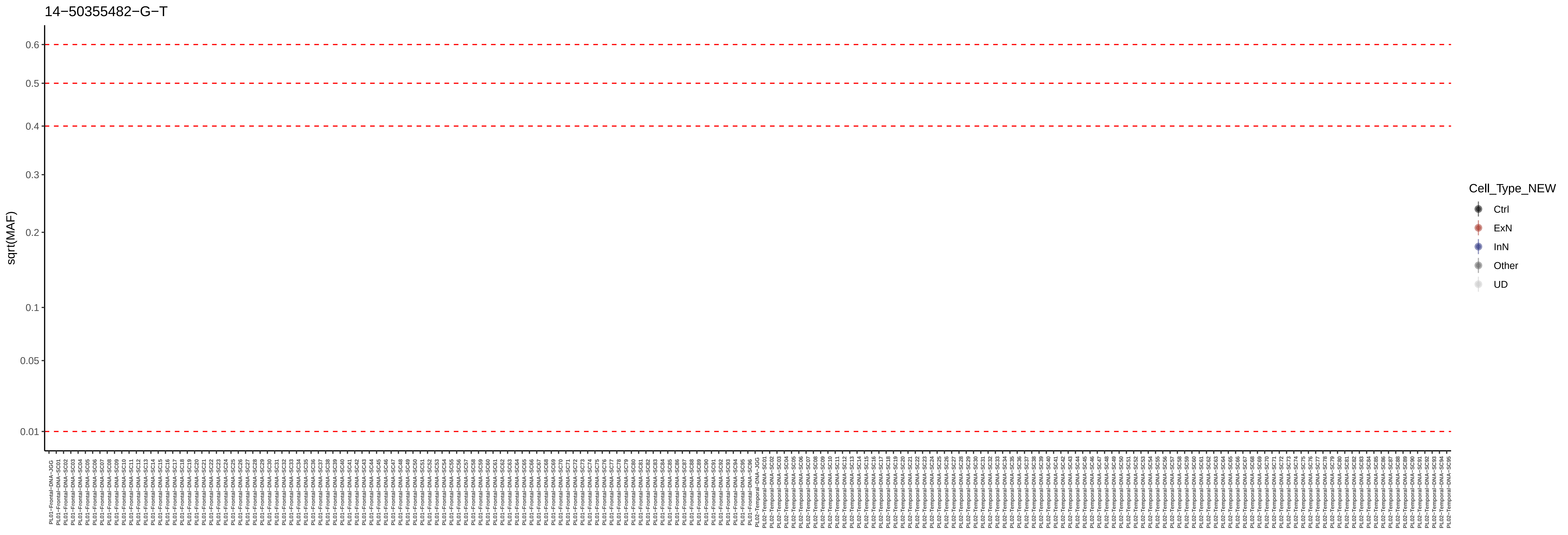

14−50355482−G−T
sqrt(MAF)
0.6
0.5
0.4
0.3
0.2
0.1
0.05
0.01
Cell_Type_NEW
Ctrl
ExN
InN
Other
UD

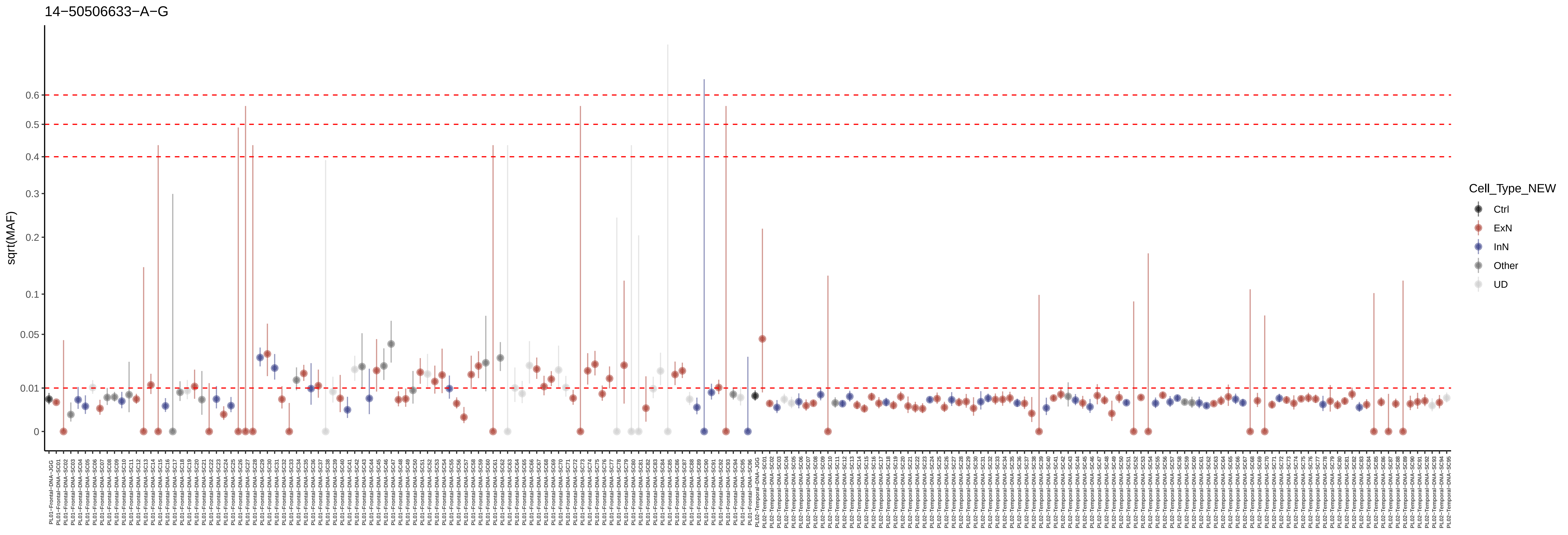
14−50506633−A−G
sqrt(MAF)
0.6
0.5
0.4
0.3
0.2
0.1
0.05
0.01
0
Cell_Type_NEW
Ctrl
ExN
InN
Other
UD

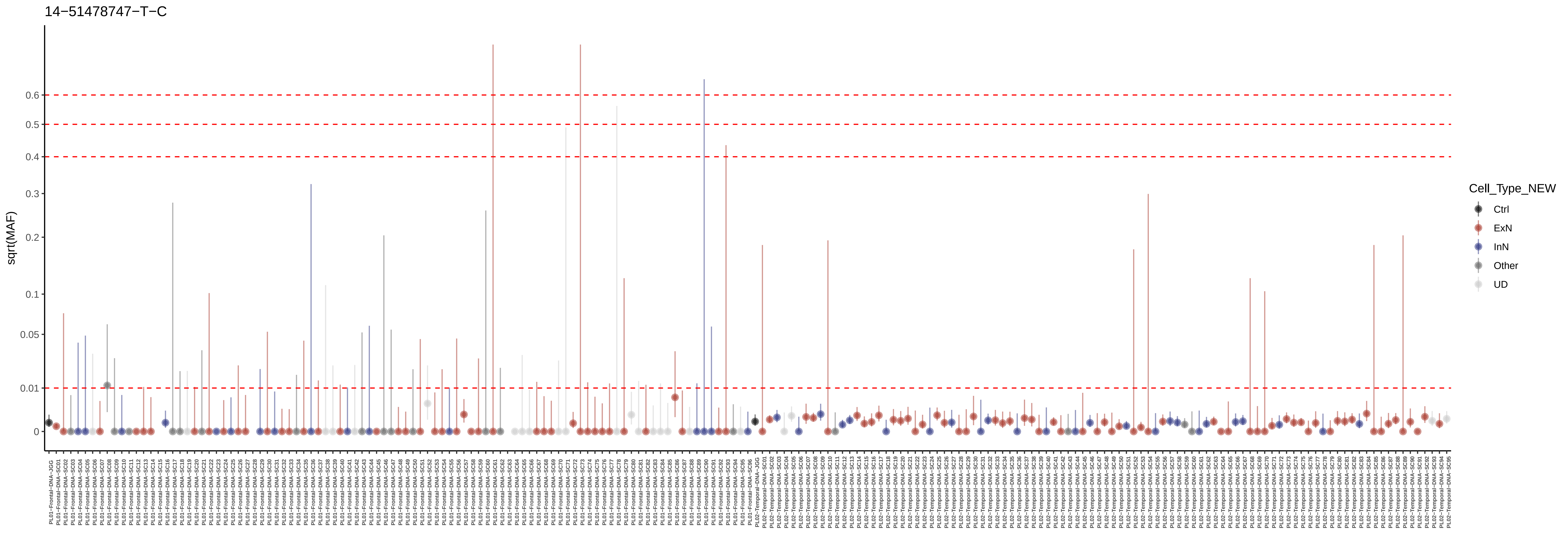
14−51478747−T−C
sqrt(MAF)
0.6
0.5
0.4
0.3
0.2
0.1
0.05
0.01
0
Cell_Type_NEW
Ctrl
ExN
InN
Other
UD

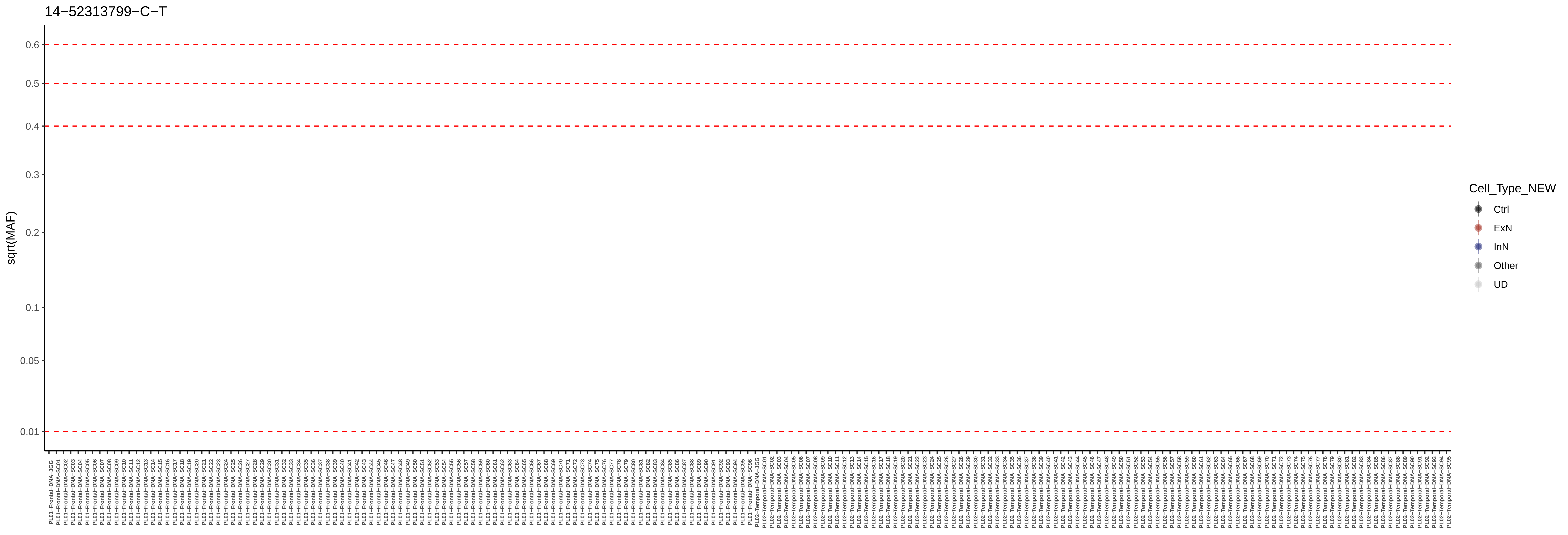
14−52313799−C−T
sqrt(MAF)
0.6
0.5
0.4
0.3
0.2
0.1
0.05
0.01
Cell_Type_NEW
Ctrl
ExN
InN
Other
UD

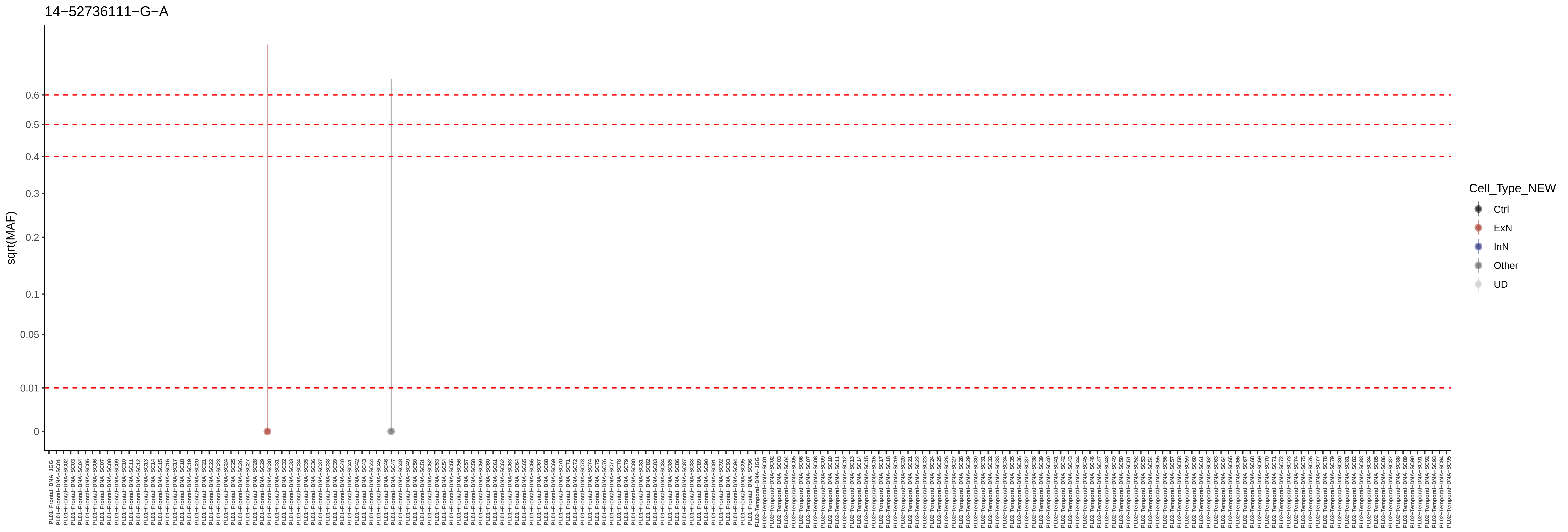

14−52736111−G−A
sqrt(MAF)
0.6
0.5
0.4
0.3
0.2
0.1
0.05
0.01
0
Cell_Type_NEW
Ctrl
ExN
InN
Other
UD

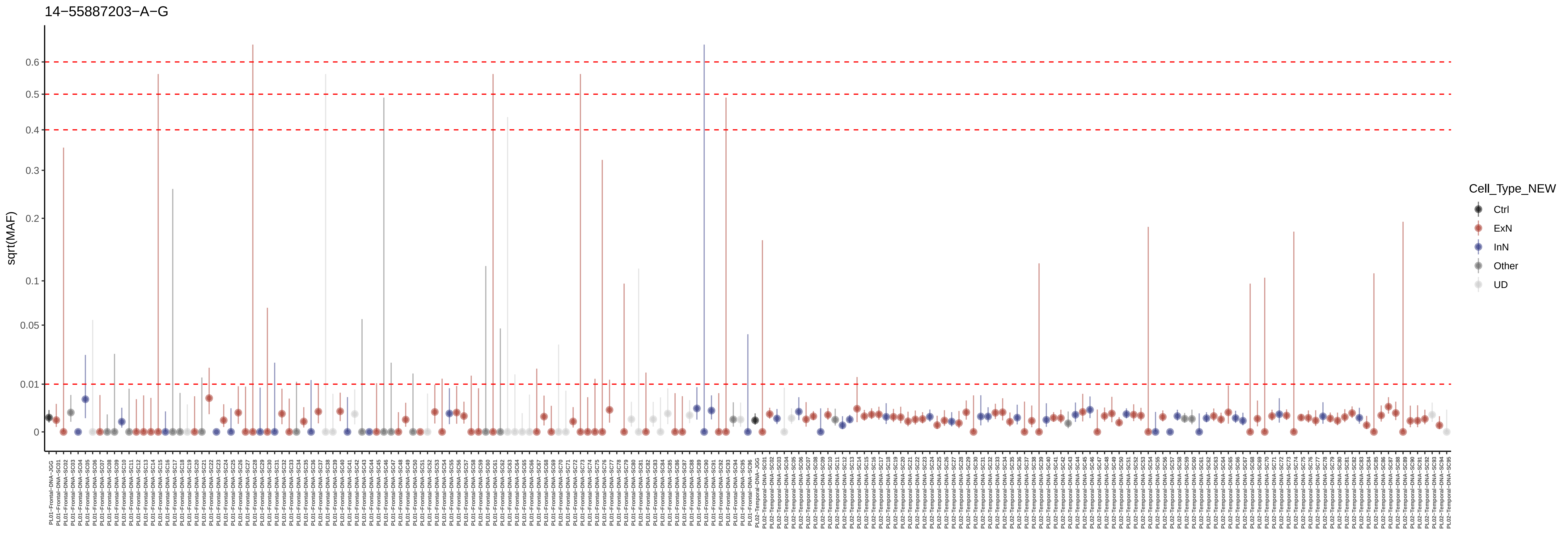
14−55887203−A−G
sqrt(MAF)
0.6
0.5
0.4
0.3
0.2
0.1
0.05
0.01
0
Cell_Type_NEW
Ctrl
ExN
InN
Other
UD

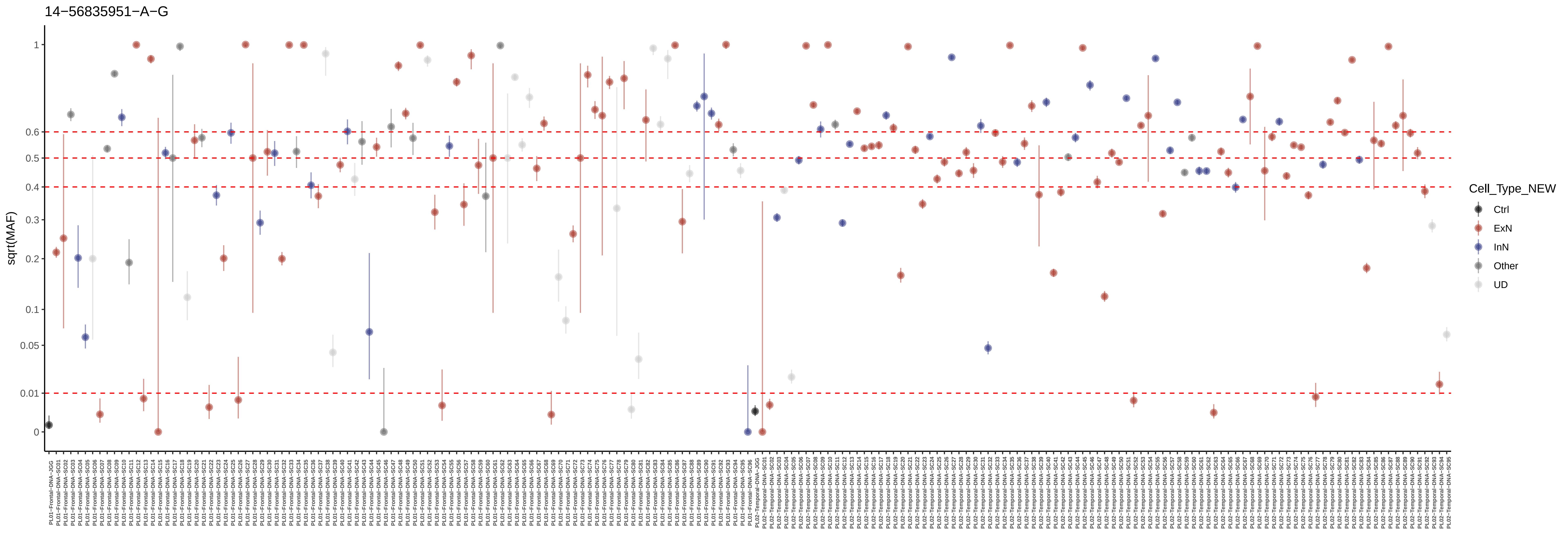
14−56835951−A−G
sqrt(MAF)
1
0.6
0.5
0.4
0.3
0.2
0.1
0.05
0.01
0
Cell_Type_NEW
Ctrl
ExN
InN
Other
UD

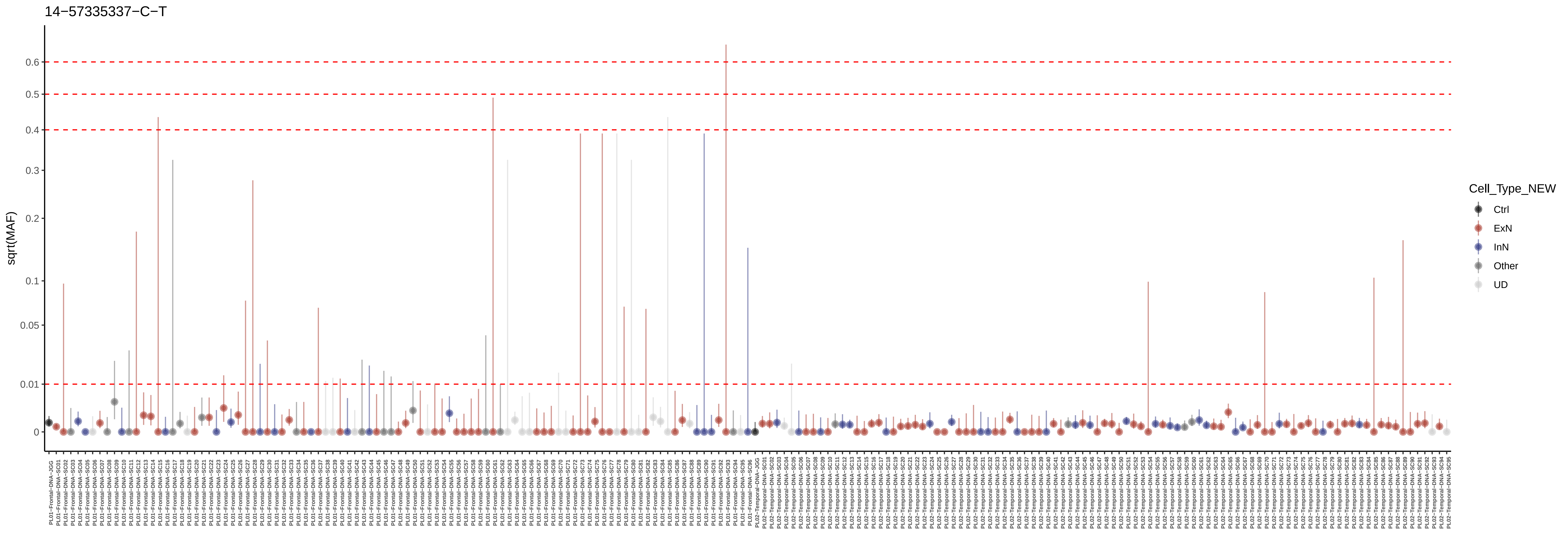
14−57335337−C−T
sqrt(MAF)
0.6
0.5
0.4
0.3
0.2
0.1
0.05
0.01
0
Cell_Type_NEW
Ctrl
ExN
InN
Other
UD

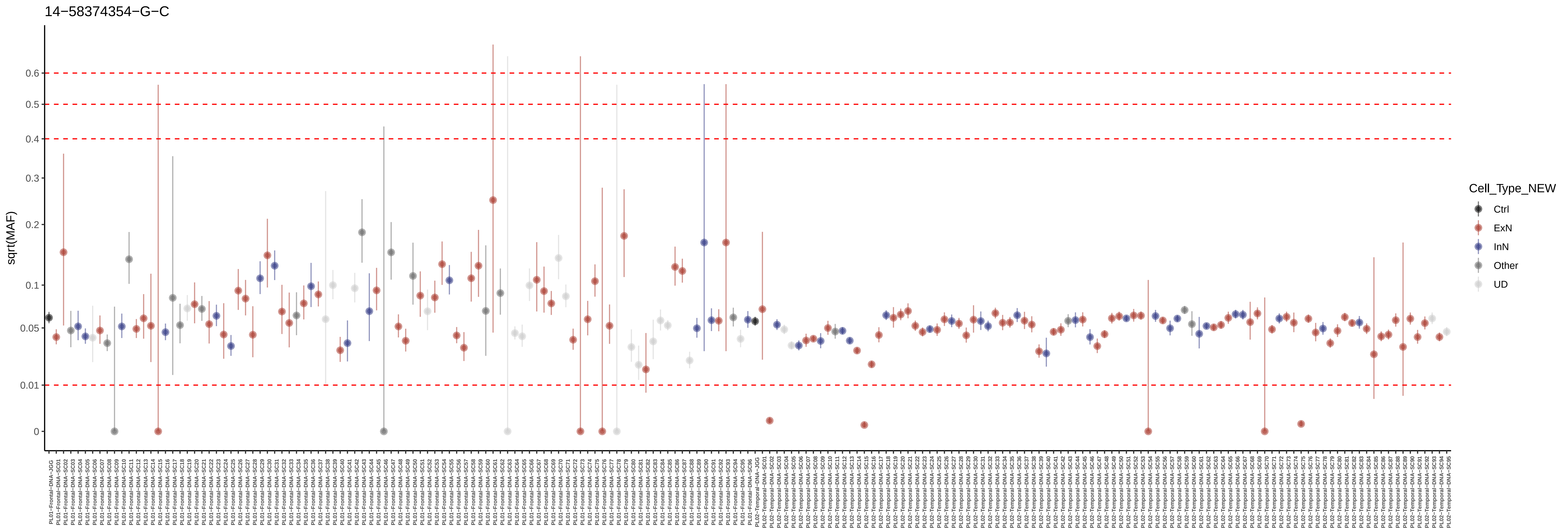
14−58374354−G−C
sqrt(MAF)
0.6
0.5
0.4
0.3
0.2
0.1
0.05
0.01
0
Cell_Type_NEW
Ctrl
ExN
InN
Other
UD

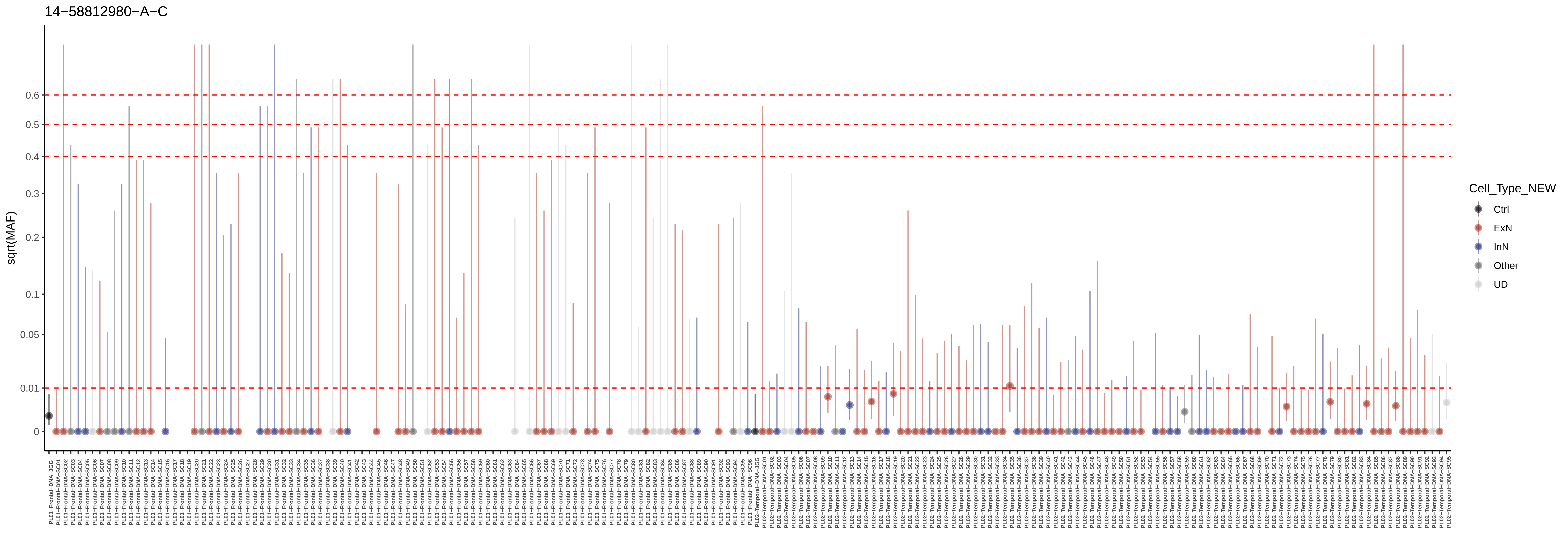
14−58812980−A−C
sqrt(MAF)
0.6
0.5
0.4
0.3
0.2
0.1
0.05
0.01
0
Cell_Type_NEW
Ctrl
ExN
InN
Other
UD

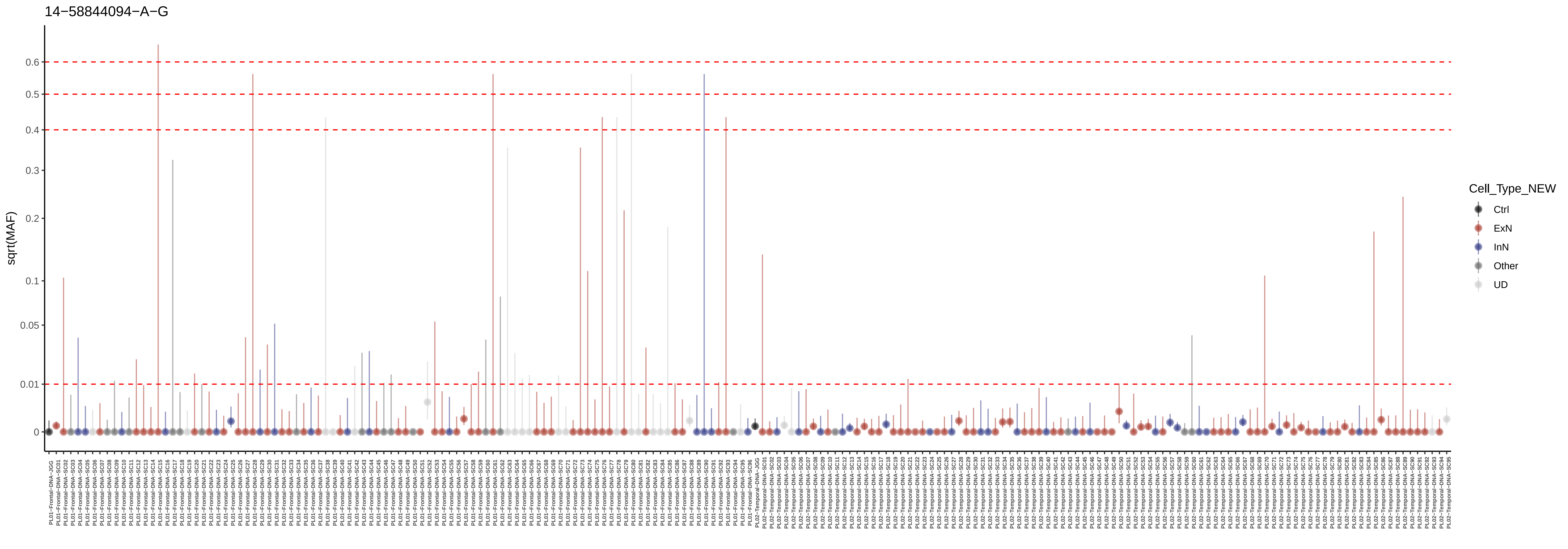

14−58844094−A−G
sqrt(MAF)
0.6
0.5
0.4
0.3
0.2
0.1
0.05
0.01
0
Cell_Type_NEW
Ctrl
ExN
InN
Other
UD

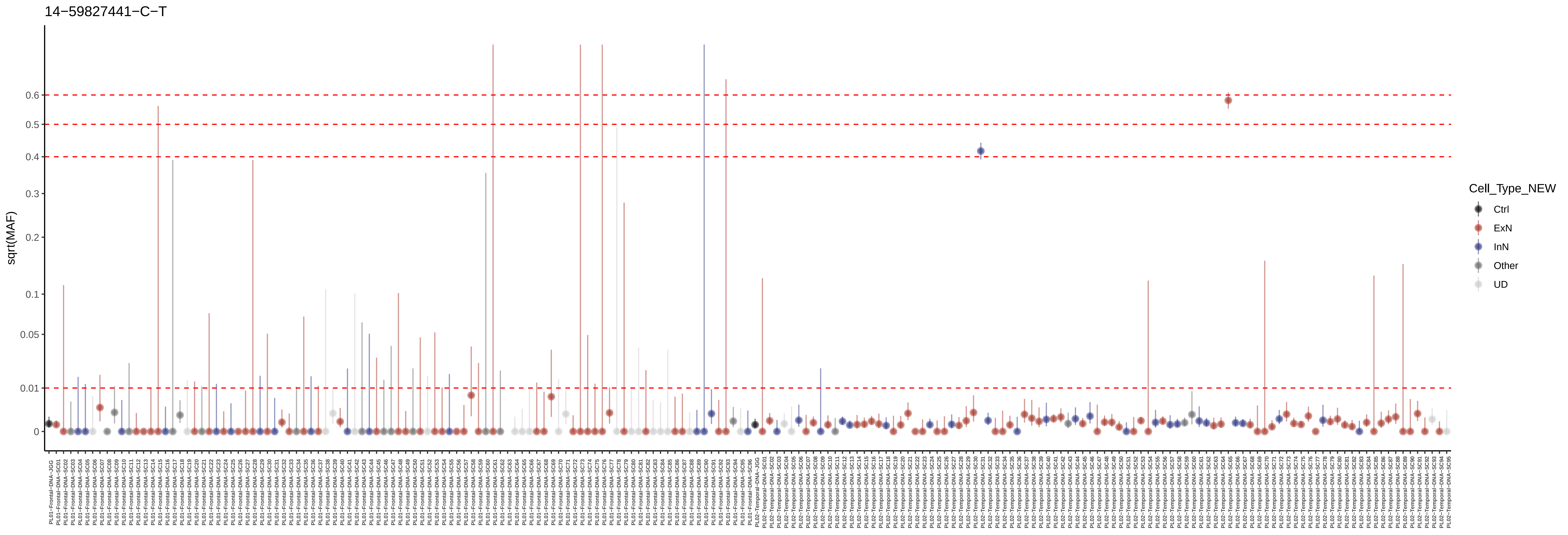

14−59827441−C−T
sqrt(MAF)
0.6
0.5
0.4
0.3
0.2
0.1
0.05
0.01
0
Cell_Type_NEW
Ctrl
ExN
InN
Other
UD

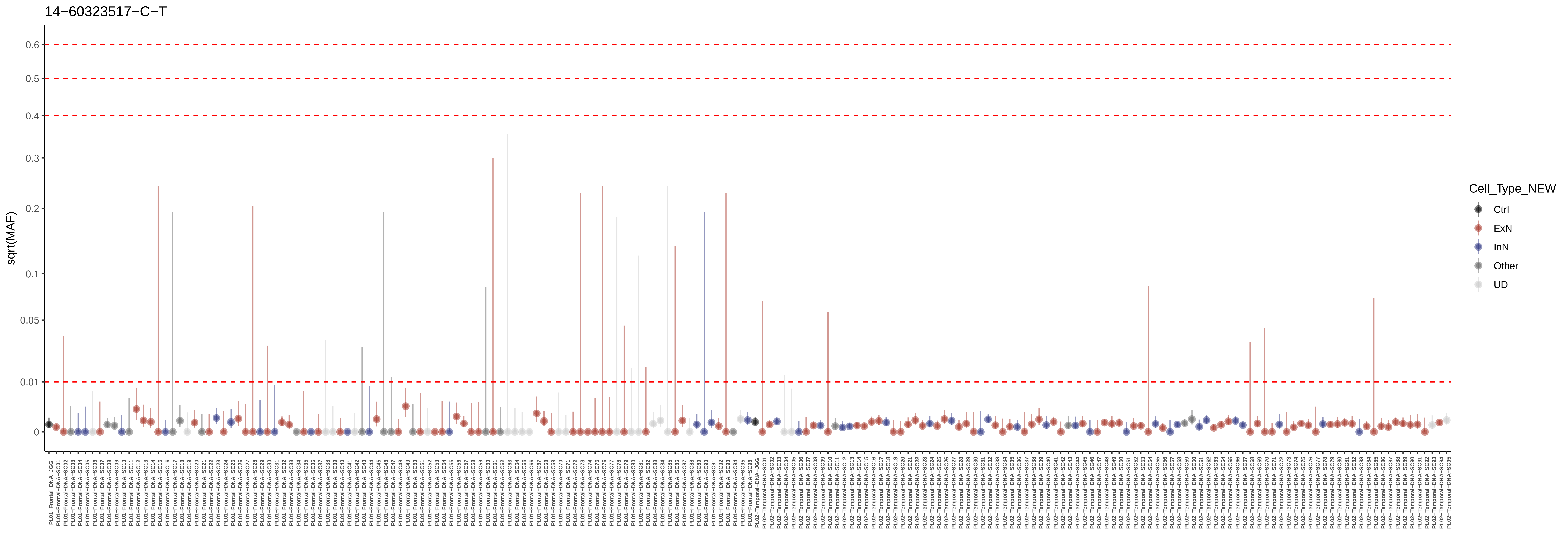

14−60323517−C−T
sqrt(MAF)
0.6
0.5
0.4
0.3
0.2
0.1
0.05
0.01
0
Cell_Type_NEW
Ctrl
ExN
InN
Other
UD

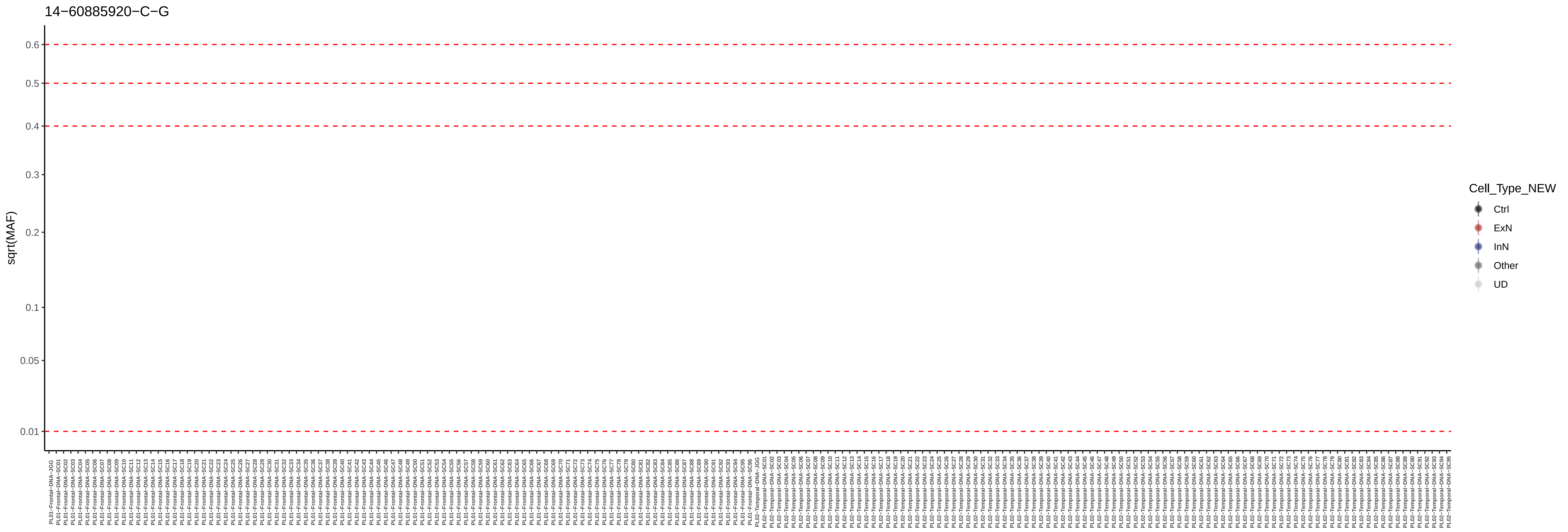
14−60885920−C−G
sqrt(MAF)
0.6
0.5
0.4
0.3
0.2
0.1
0.05
0.01
Cell_Type_NEW
Ctrl
ExN
InN
Other
UD

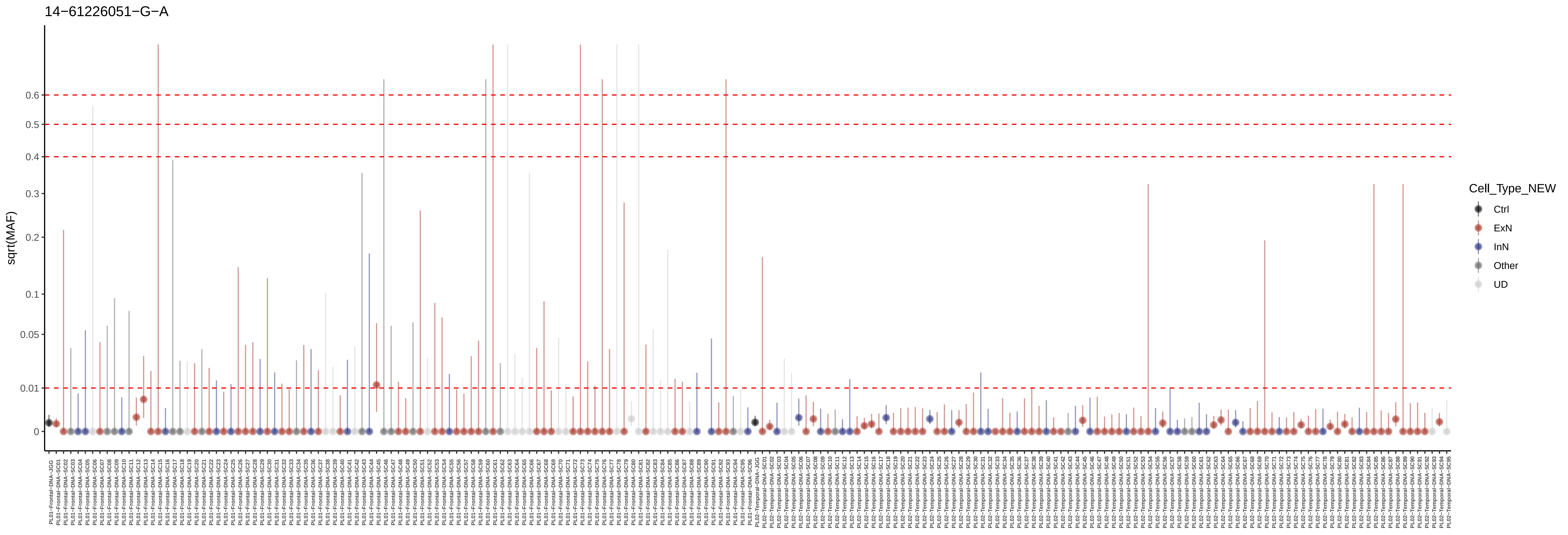
14−61226051−G−A
sqrt(MAF)
0.6
0.5
0.4
0.3
0.2
0.1
0.05
0.01
0
Cell_Type_NEW
Ctrl
ExN
InN
Other
UD

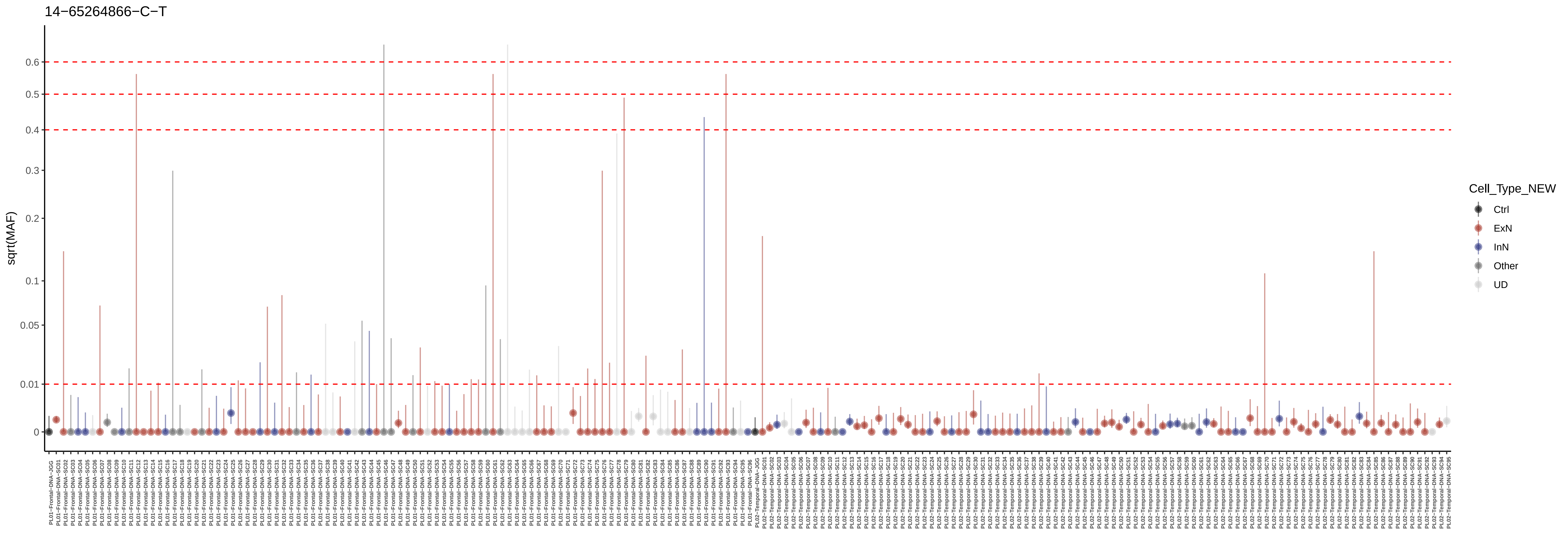
14−65264866−C−T
sqrt(MAF)
0.6
0.5
0.4
0.3
0.2
0.1
0.05
0.01
0
Cell_Type_NEW
Ctrl
ExN
InN
Other
UD

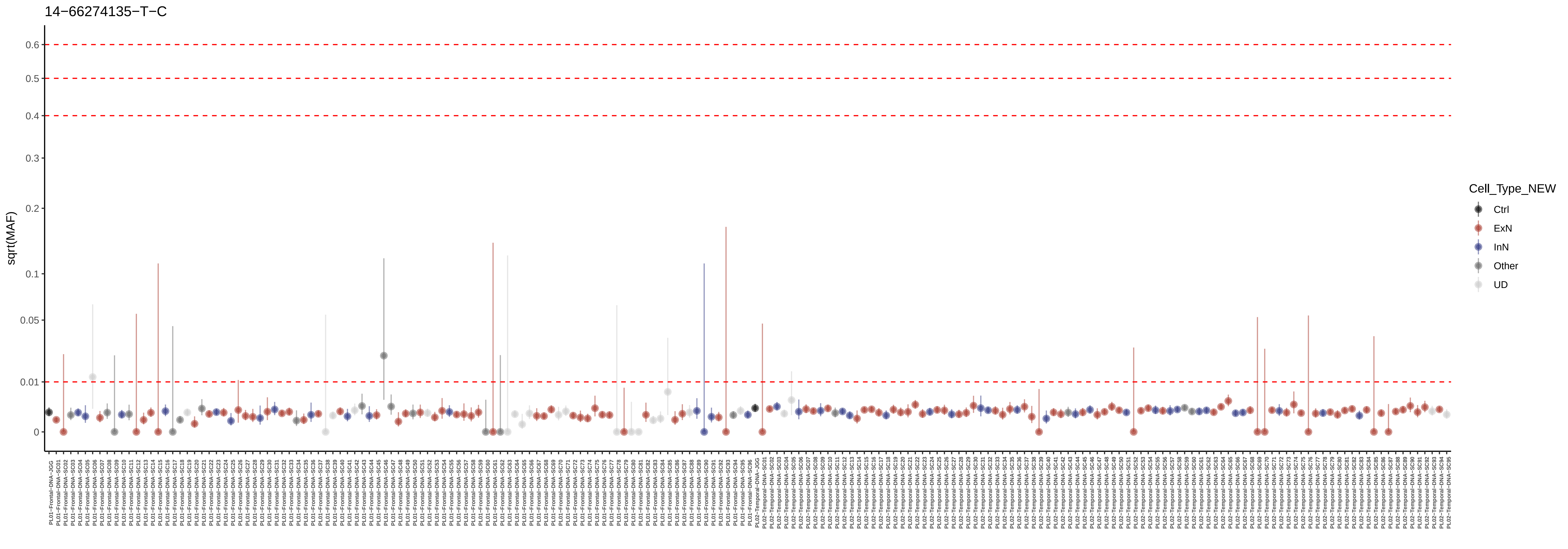

14−66274135−T−C
sqrt(MAF)
0.6
0.5
0.4
0.3
0.2
0.1
0.05
0.01
0
Cell_Type_NEW
Ctrl
ExN
InN
Other
UD

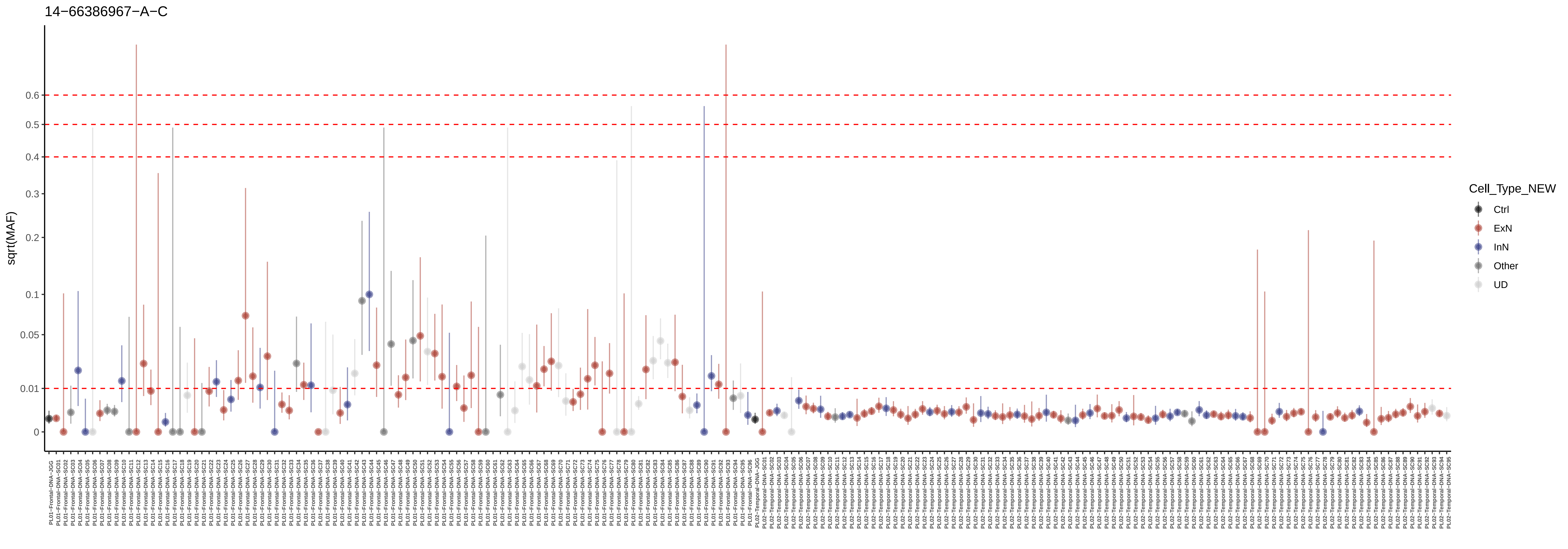
14−66386967−A−C
sqrt(MAF)
0.6
0.5
0.4
0.3
0.2
0.1
0.05
0.01
0
Cell_Type_NEW
Ctrl
ExN
InN
Other
UD

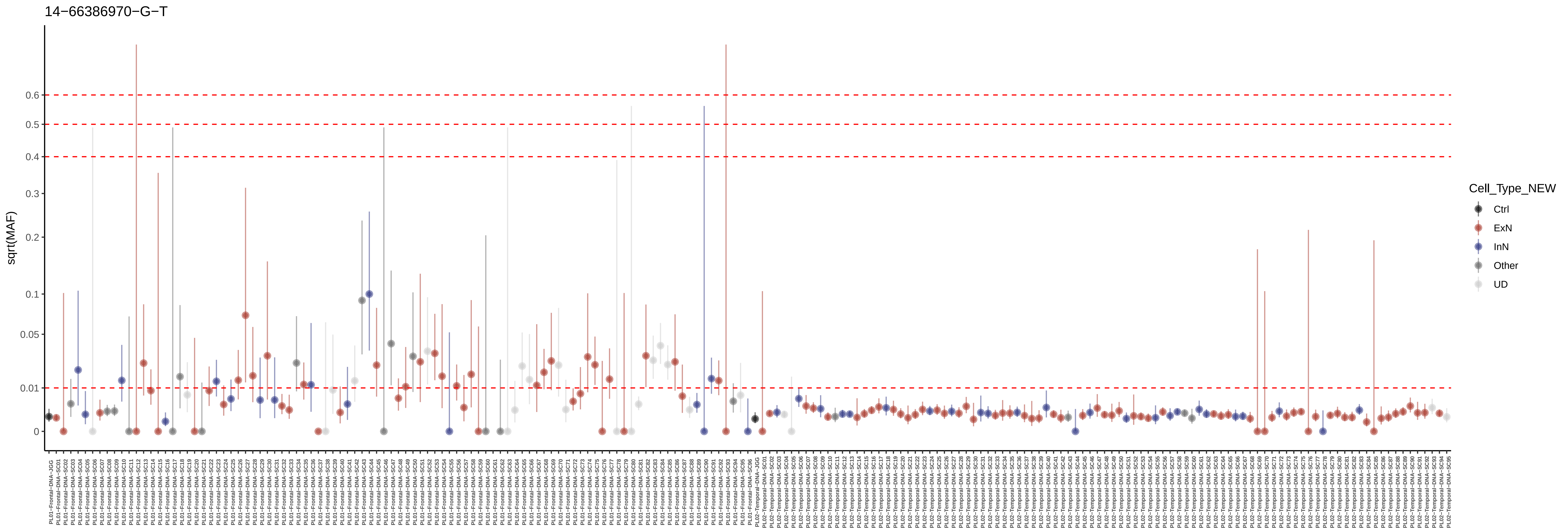

14−66386970−G−T
sqrt(MAF)
0.6
0.5
0.4
0.3
0.2
0.1
0.05
0.01
0
Cell_Type_NEW
Ctrl
ExN
InN
Other
UD

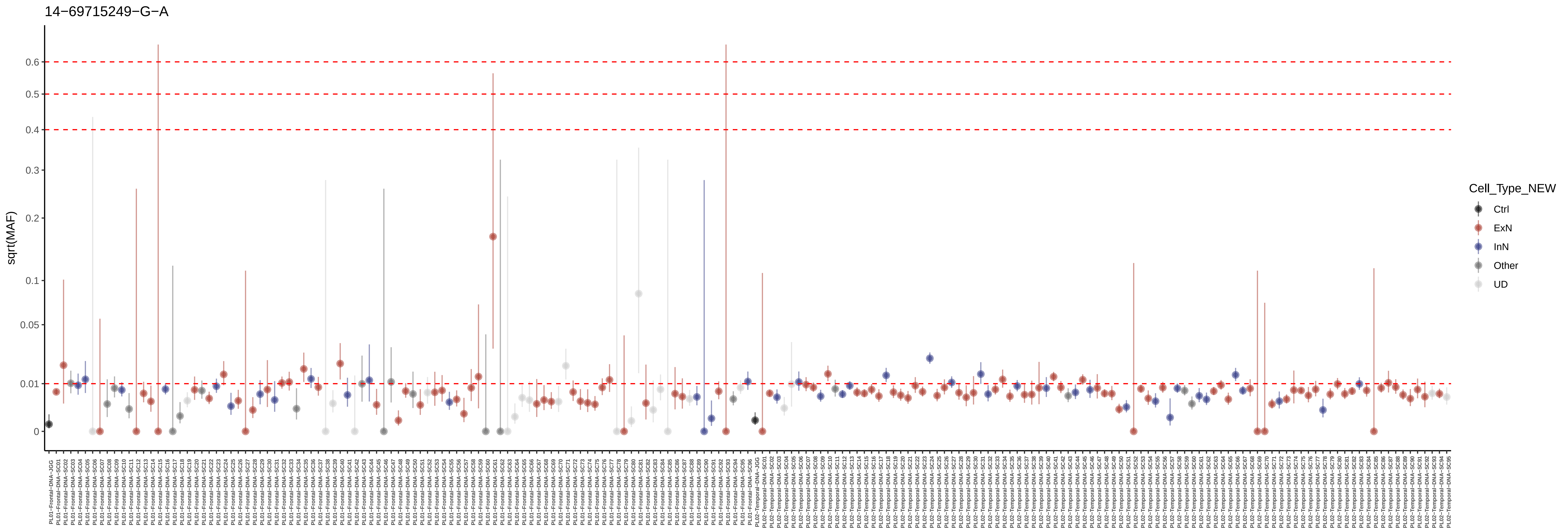
14−69715249−G−A
sqrt(MAF)
0.6
0.5
0.4
0.3
0.2
0.1
0.05
0.01
0
Cell_Type_NEW
Ctrl
ExN
InN
Other
UD

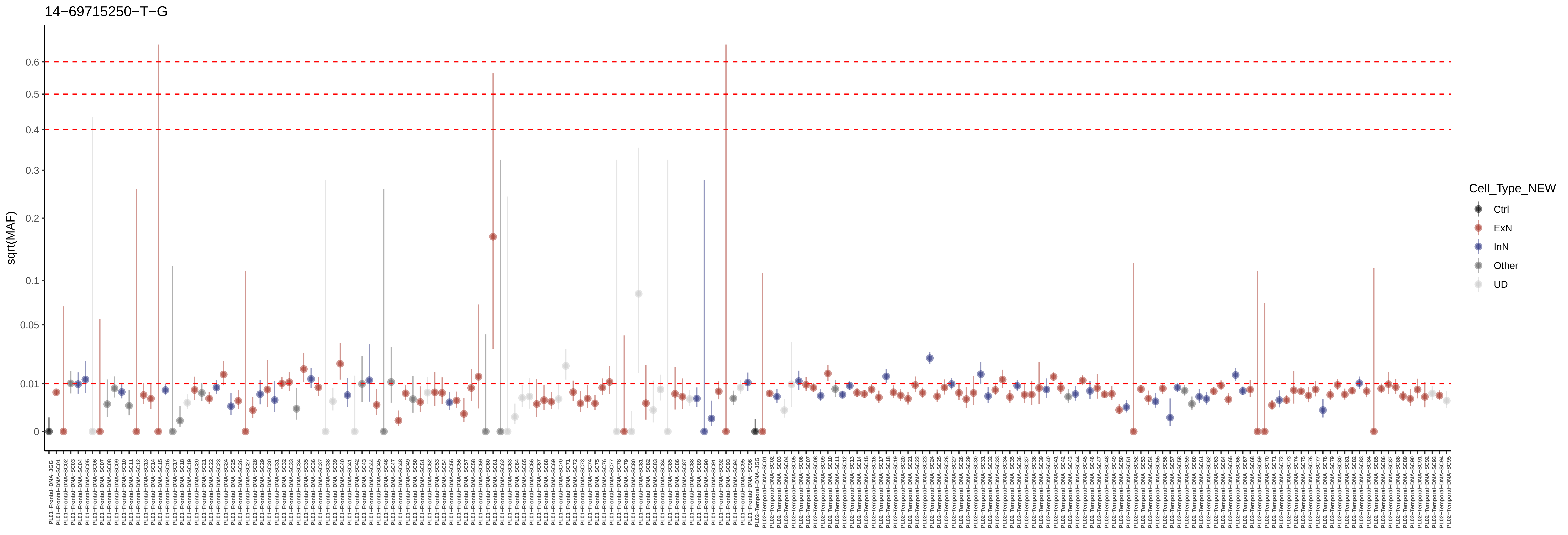
14−69715250−T−G
sqrt(MAF)
0.6
0.5
0.4
0.3
0.2
0.1
0.05
0.01
0
Cell_Type_NEW
Ctrl
ExN
InN
Other
UD

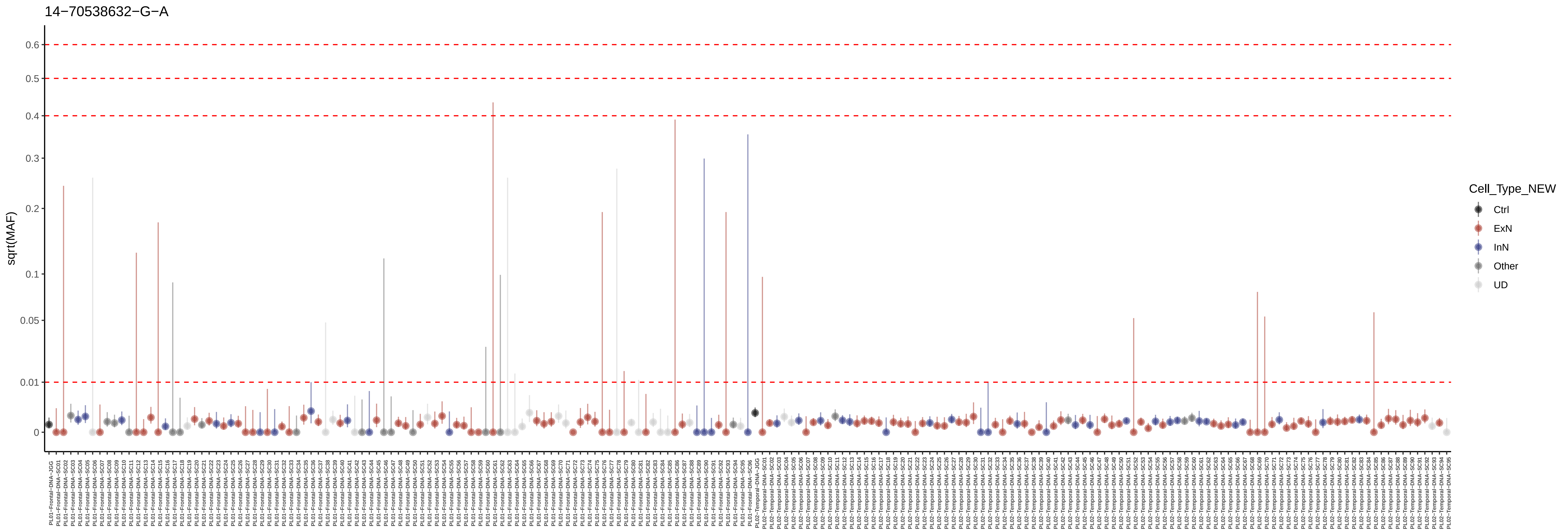
14−70538632−G−A
sqrt(MAF)
0.6
0.5
0.4
0.3
0.2
0.1
0.05
0.01
0
Cell_Type_NEW
Ctrl
ExN
InN
Other
UD

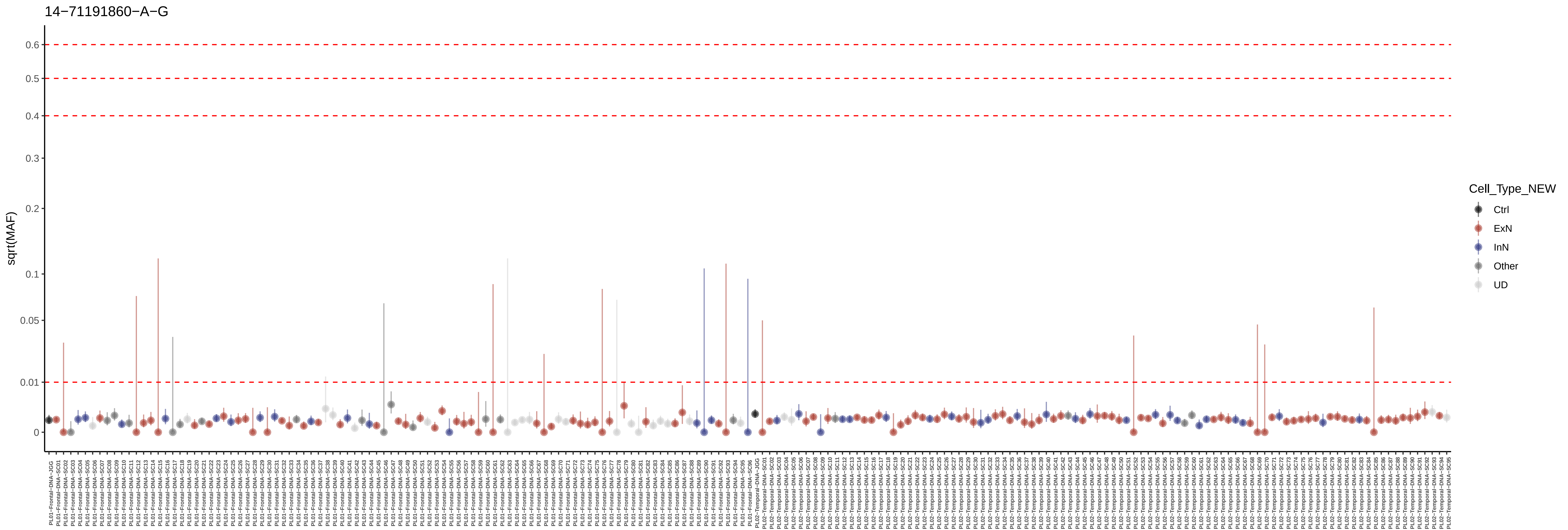

14−71191860−A−G
sqrt(MAF)
0.6
0.5
0.4
0.3
0.2
0.1
0.05
0.01
0
Cell_Type_NEW
Ctrl
ExN
InN
Other
UD

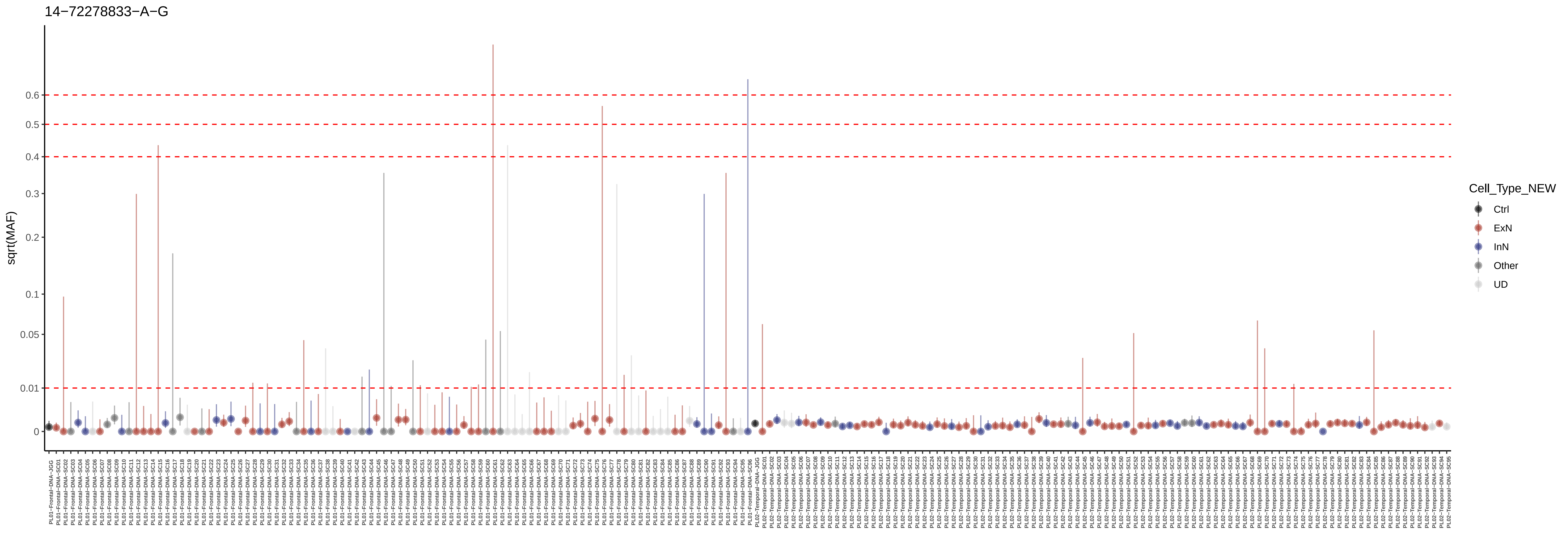
14−72278833−A−G
sqrt(MAF)
0.6
0.5
0.4
0.3
0.2
0.1
0.05
0.01
0
Cell_Type_NEW
Ctrl
ExN
InN
Other
UD

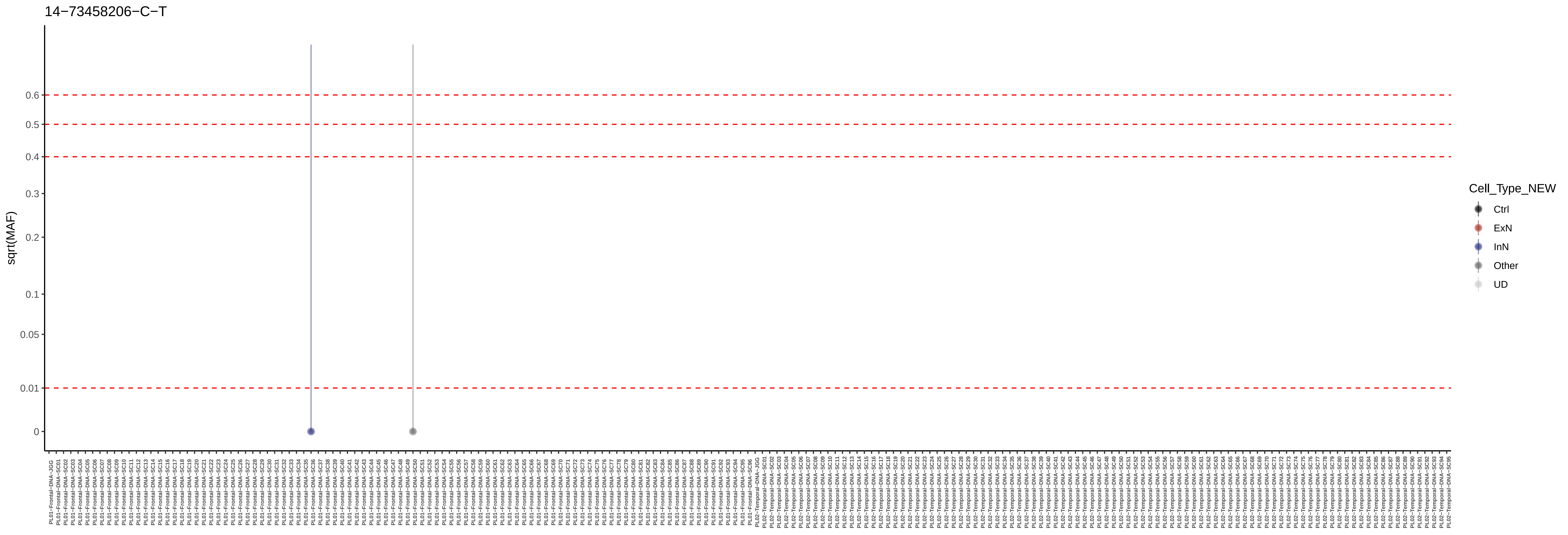

14−73458206−C−T
sqrt(MAF)
0.6
0.5
0.4
0.3
0.2
0.1
0.05
0.01
0
Cell_Type_NEW
Ctrl
ExN
InN
Other
UD

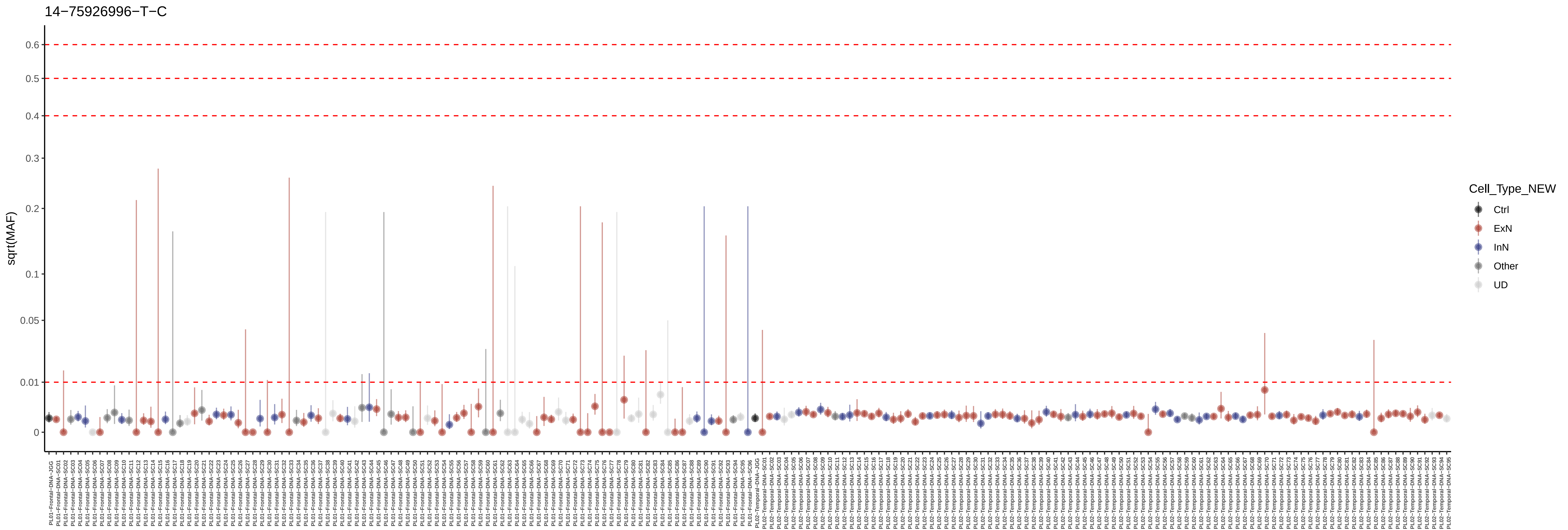
14−75926996−T−C
sqrt(MAF)
0.6
0.5
0.4
0.3
0.2
0.1
0.05
0.01
0
Cell_Type_NEW
Ctrl
ExN
InN
Other
UD

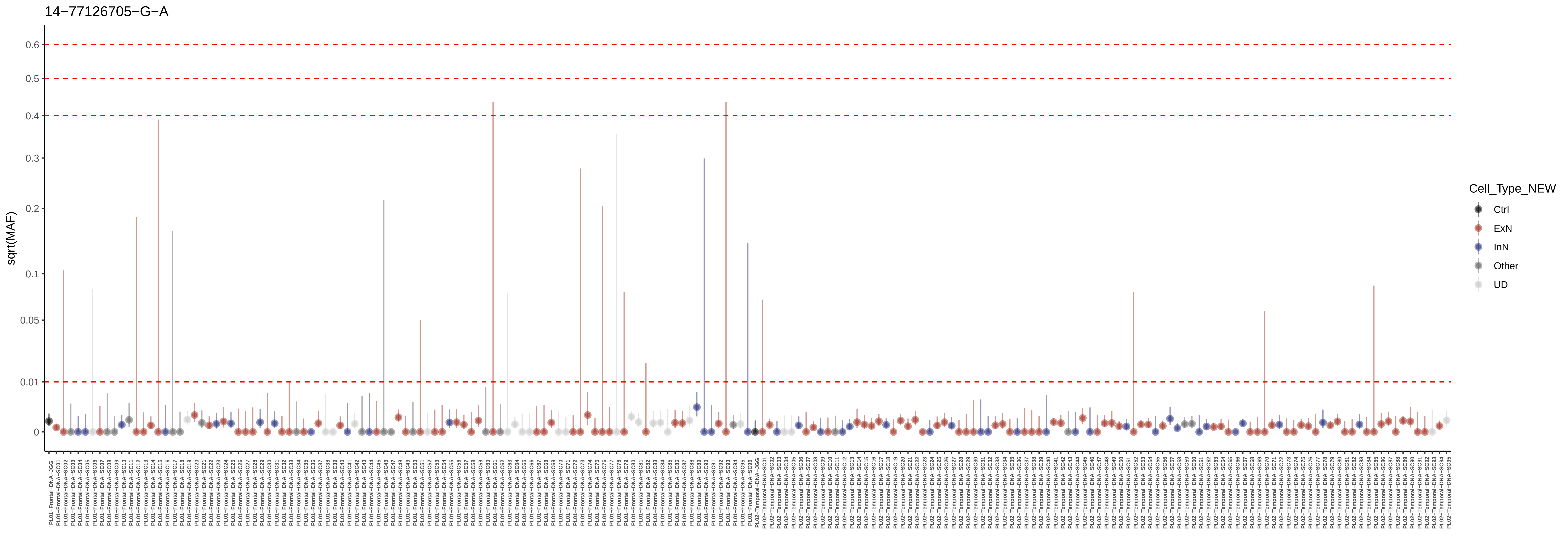

14−77126705−G−A
sqrt(MAF)
Cell_Type_NEW
Ctrl
ExN
InN
Other
UD

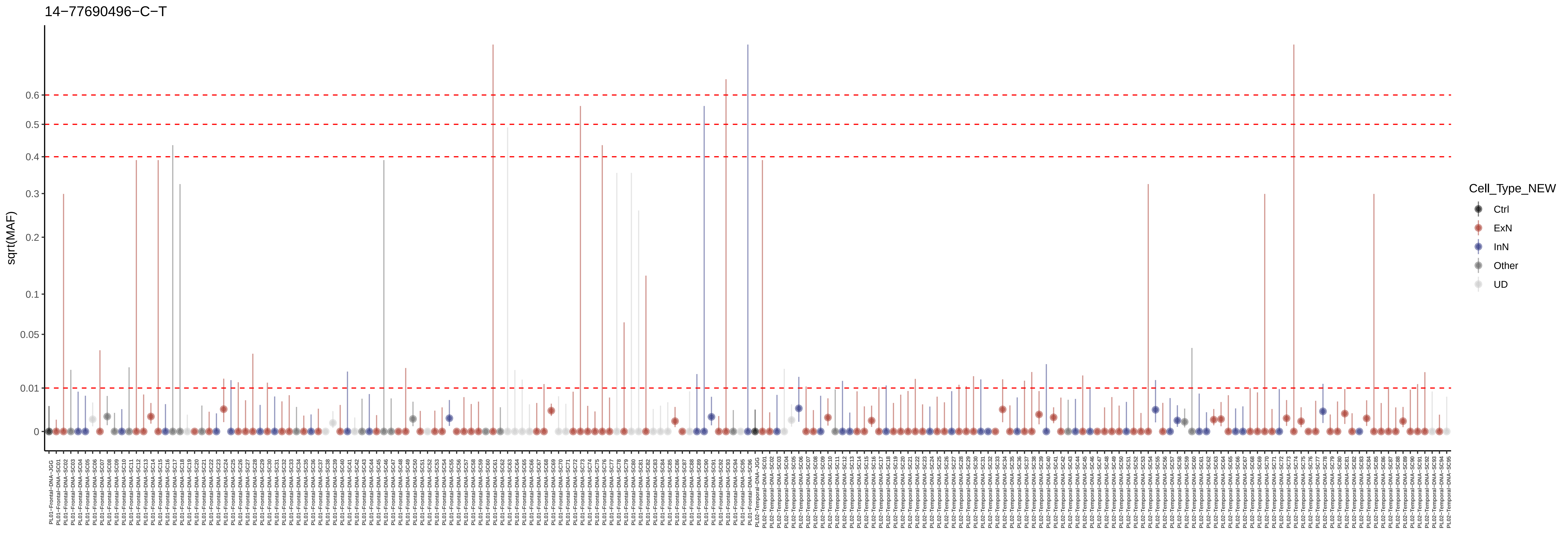
14−77690496−C−T
sqrt(MAF)
0.6
0.5
0.4
0.3
0.2
0.1
0.05
0.01
0
Cell_Type_NEW
Ctrl
ExN
InN
Other
UD

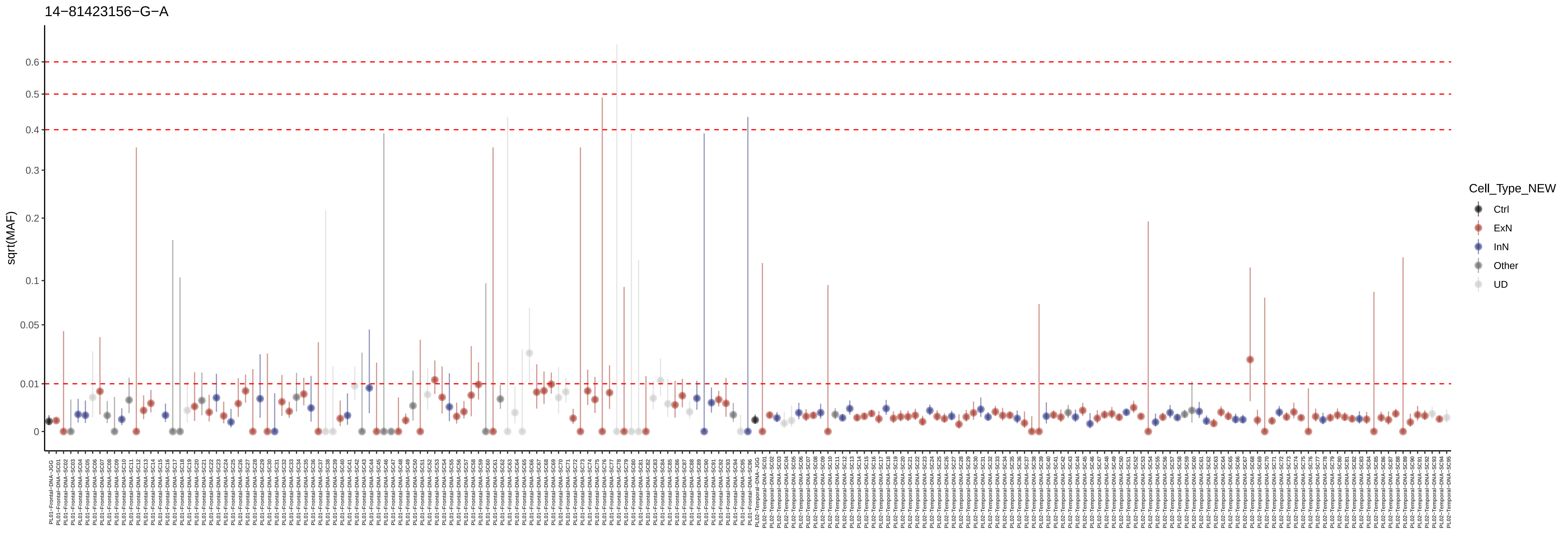
14−81423156−G−A
sqrt(MAF)
0.6
0.5
0.4
0.3
0.2
0.1
0.05
0.01
0
Cell_Type_NEW
Ctrl
ExN
InN
Other
UD

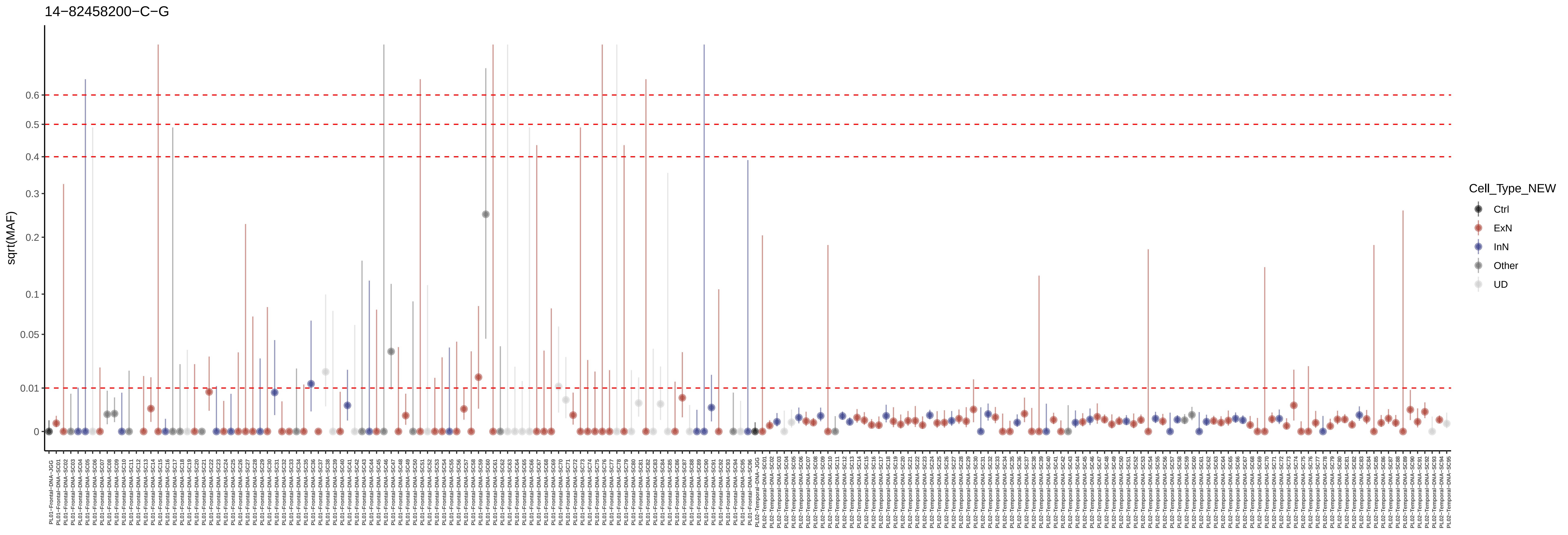

14−82458200−C−G
sqrt(MAF)
0.6
0.5
0.4
0.3
0.2
0.1
0.05
0.01
0
Cell_Type_NEW
Ctrl
ExN
InN
Other
UD

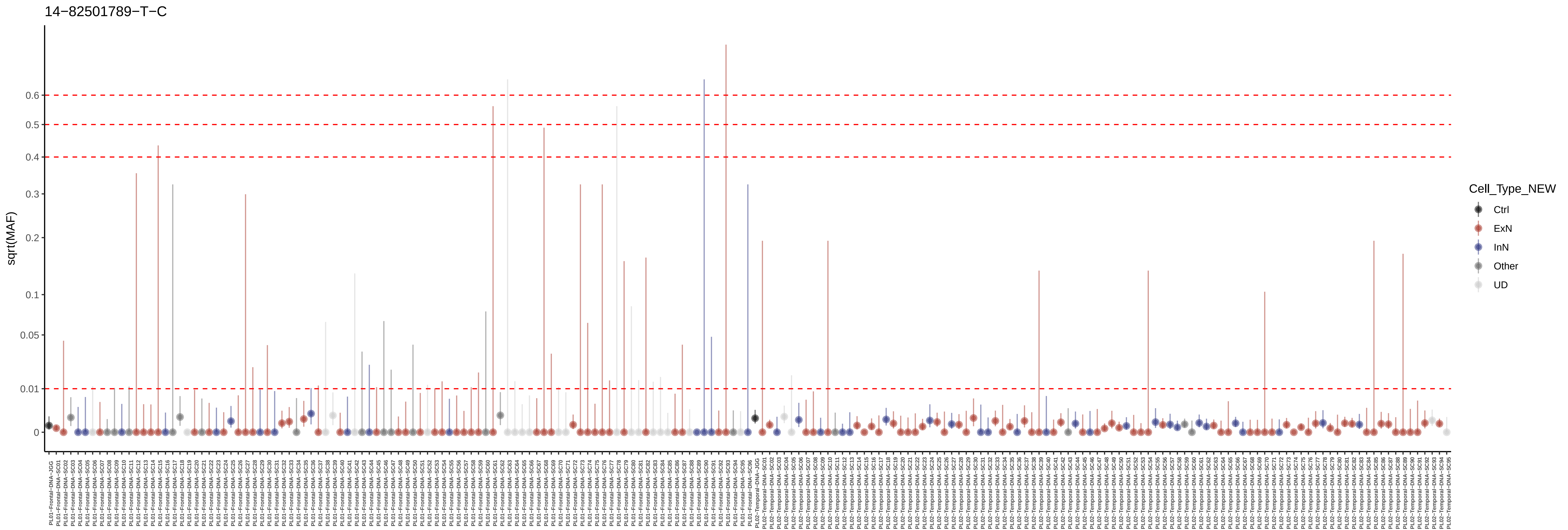
14−82501789−T−C
sqrt(MAF)
0.6
0.5
0.4
0.3
0.2
0.1
0.05
0.01
0
Cell_Type_NEW
Ctrl
ExN
InN
Other
UD

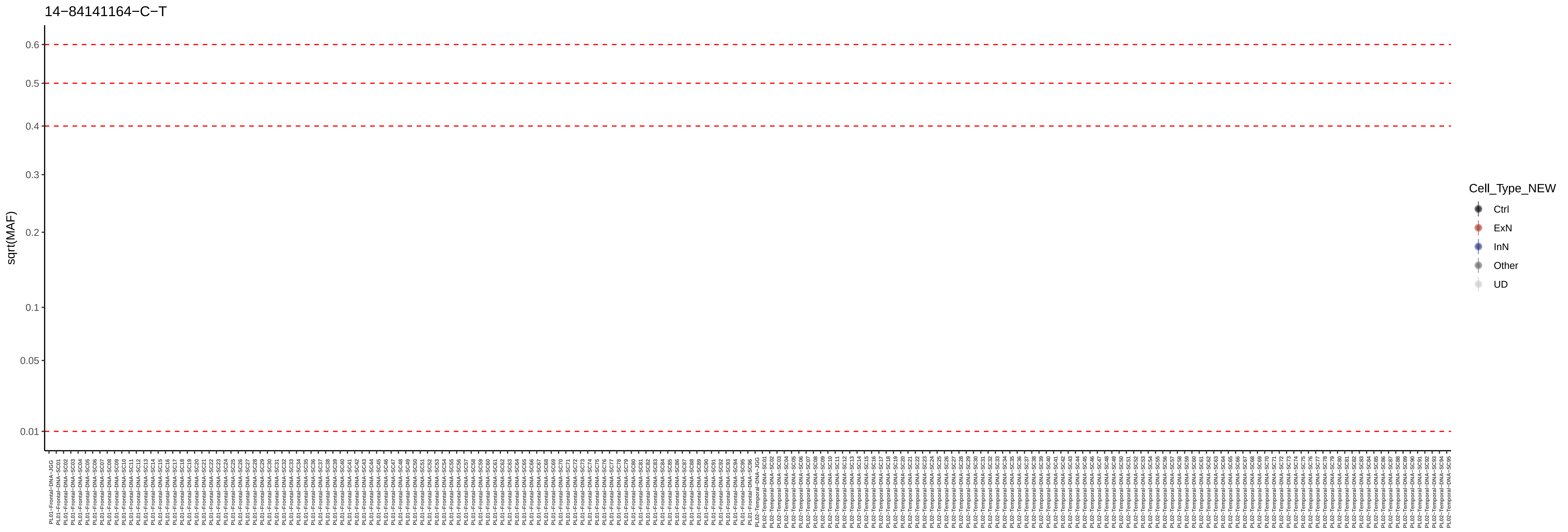
14−84141164−C−T
sqrt(MAF)
0.6
0.5
0.4
0.3
0.2
0.1
0.05
0.01
Cell_Type_NEW
Ctrl
ExN
InN
Other
UD

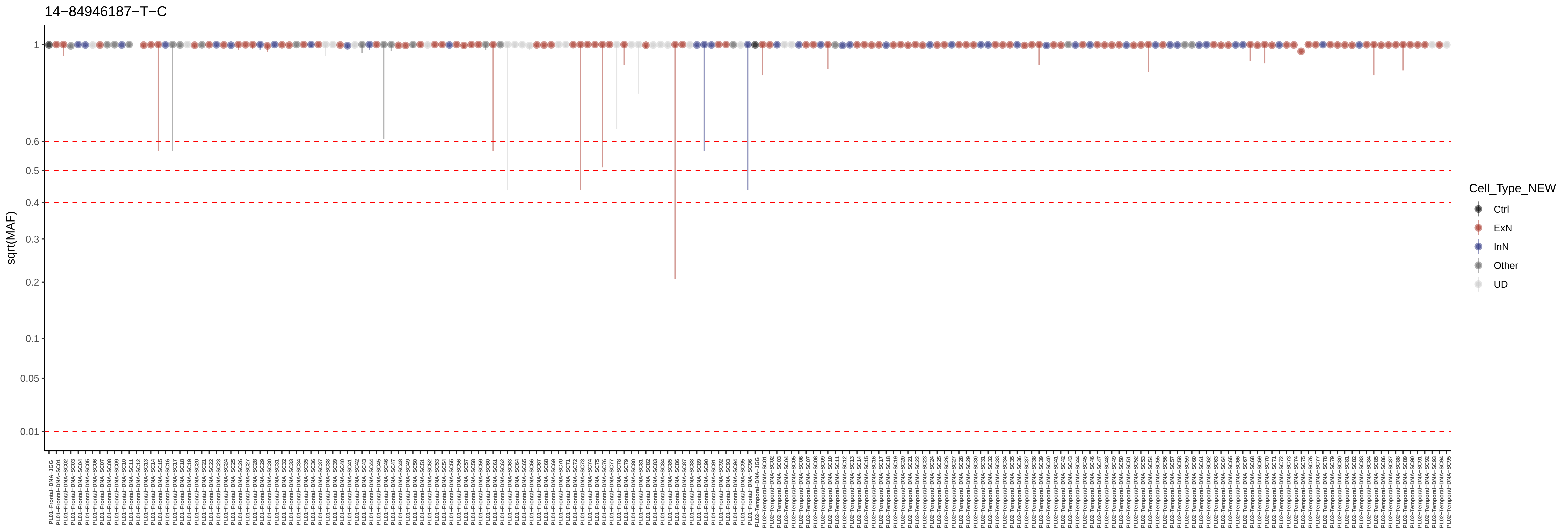
14−84946187−T−C
sqrt(MAF)
1
0.6
0.5
0.4
0.3
0.2
0.1
0.05
0.01
Cell_Type_NEW
Ctrl
ExN
InN
Other
UD

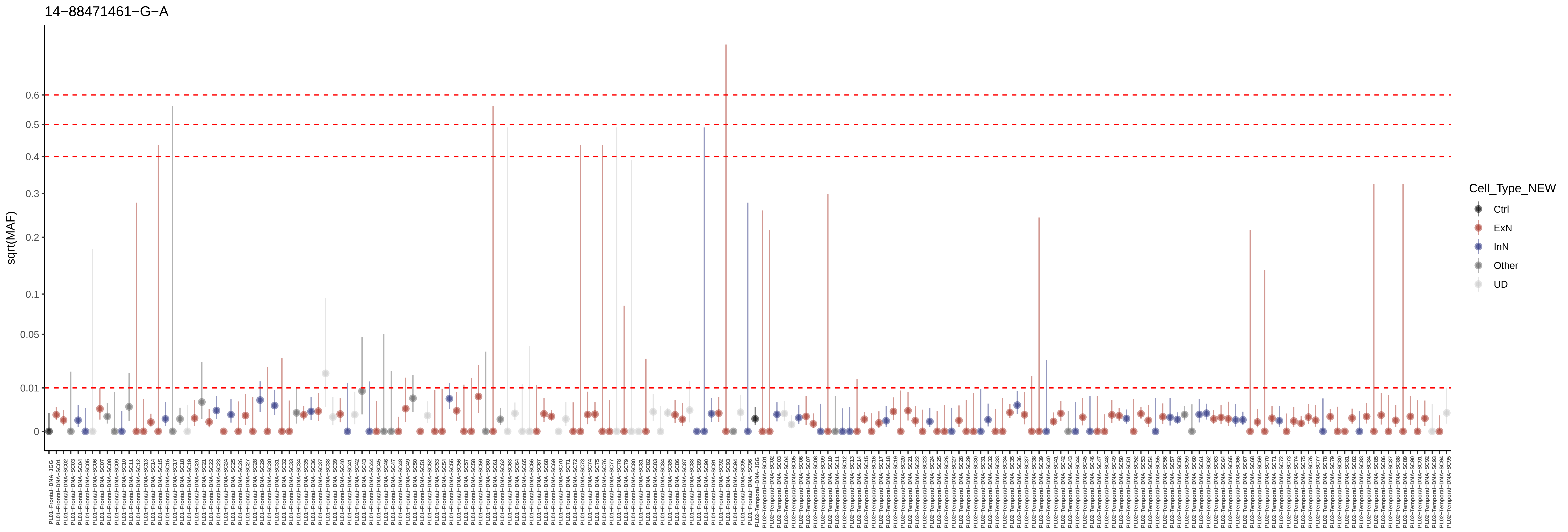
14−88471461−G−A
sqrt(MAF)
0.6
0.5
0.4
0.3
0.2
0.1
0.05
0.01
0
Cell_Type_NEW
Ctrl
ExN
InN
Other
UD

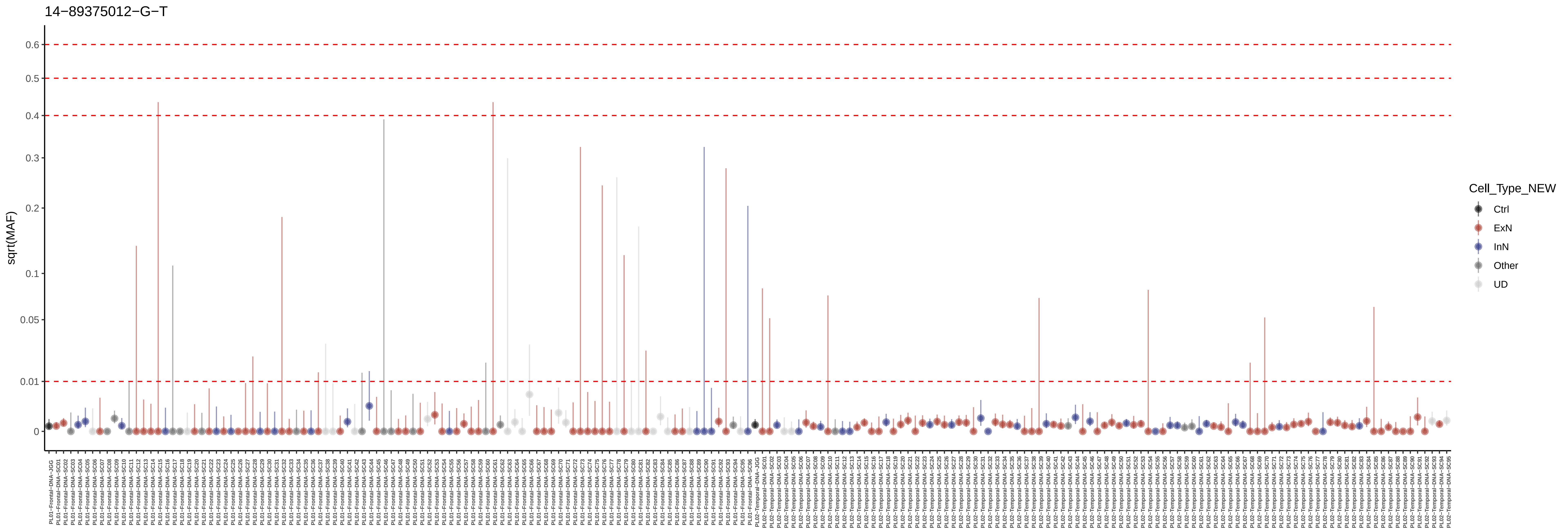
14−89375012−G−T
sqrt(MAF)
0.6
0.5
0.4
0.3
0.2
0.1
0.05
0.01
0
Cell_Type_NEW
Ctrl
ExN
InN
Other
UD

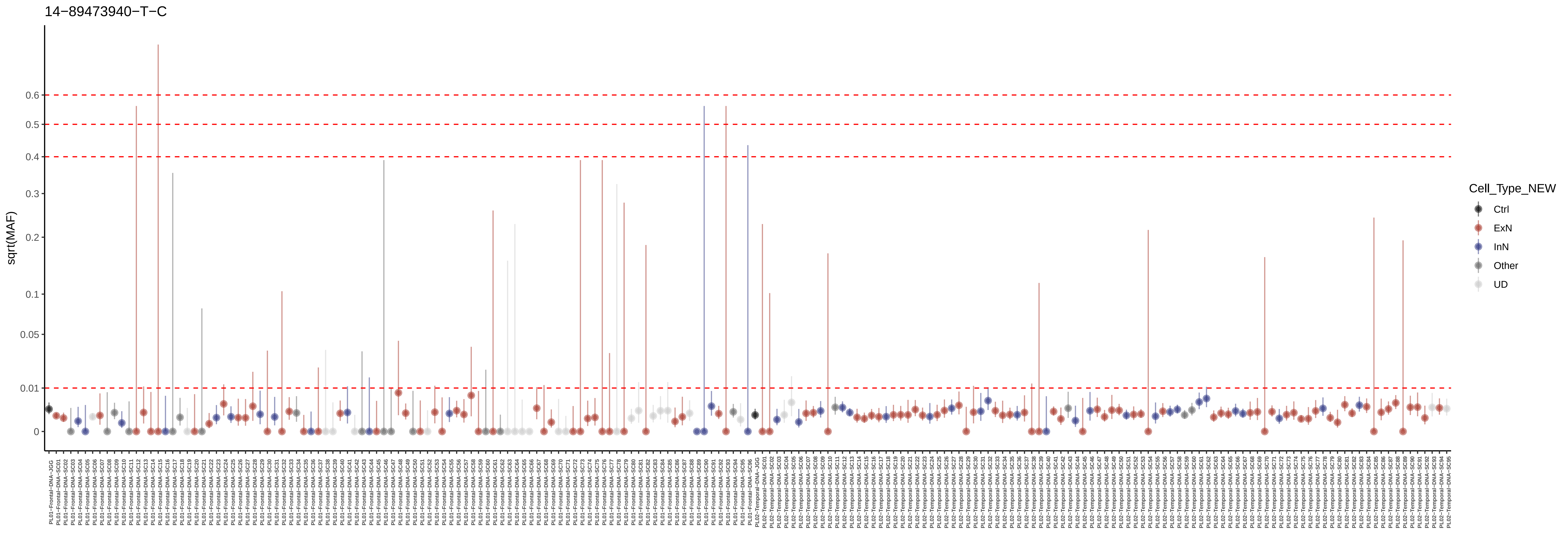
14−89473940−T−C
sqrt(MAF)
0.6
0.5
0.4
0.3
0.2
0.1
0.05
0.01
0
Cell_Type_NEW
Ctrl
ExN
InN
Other
UD

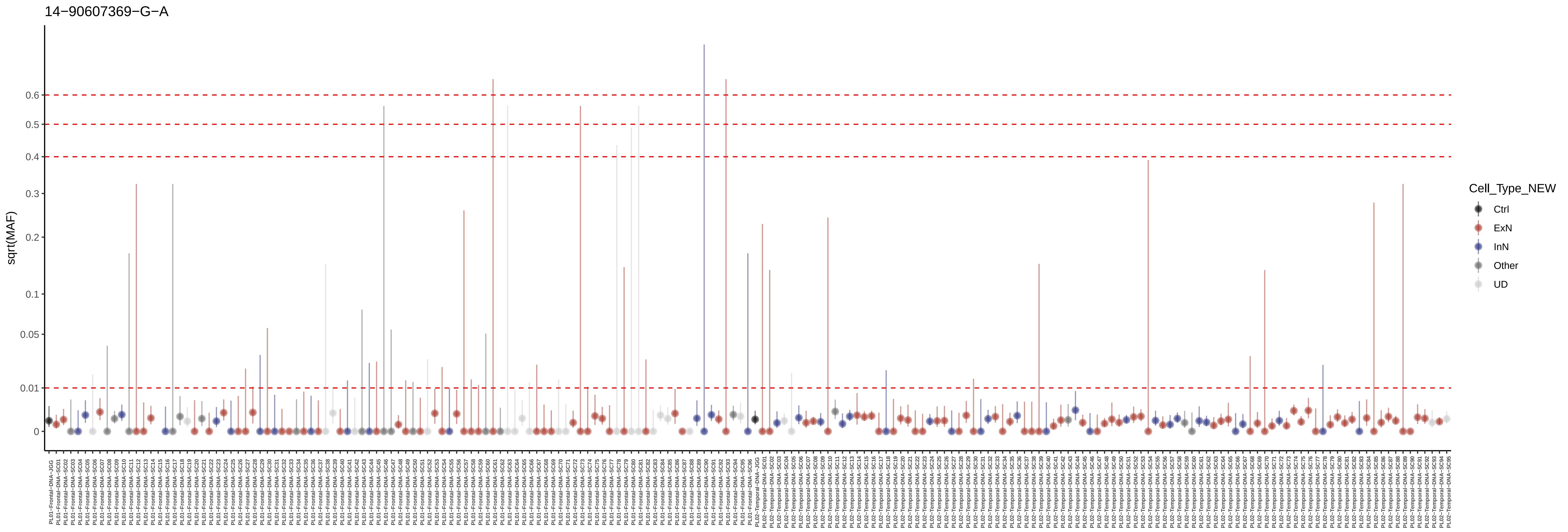

14−90607369−G−A
sqrt(MAF)
0.6
0.5
0.4
0.3
0.2
0.1
0.05
0.01
0
Cell_Type_NEW
Ctrl
ExN
InN
Other
UD

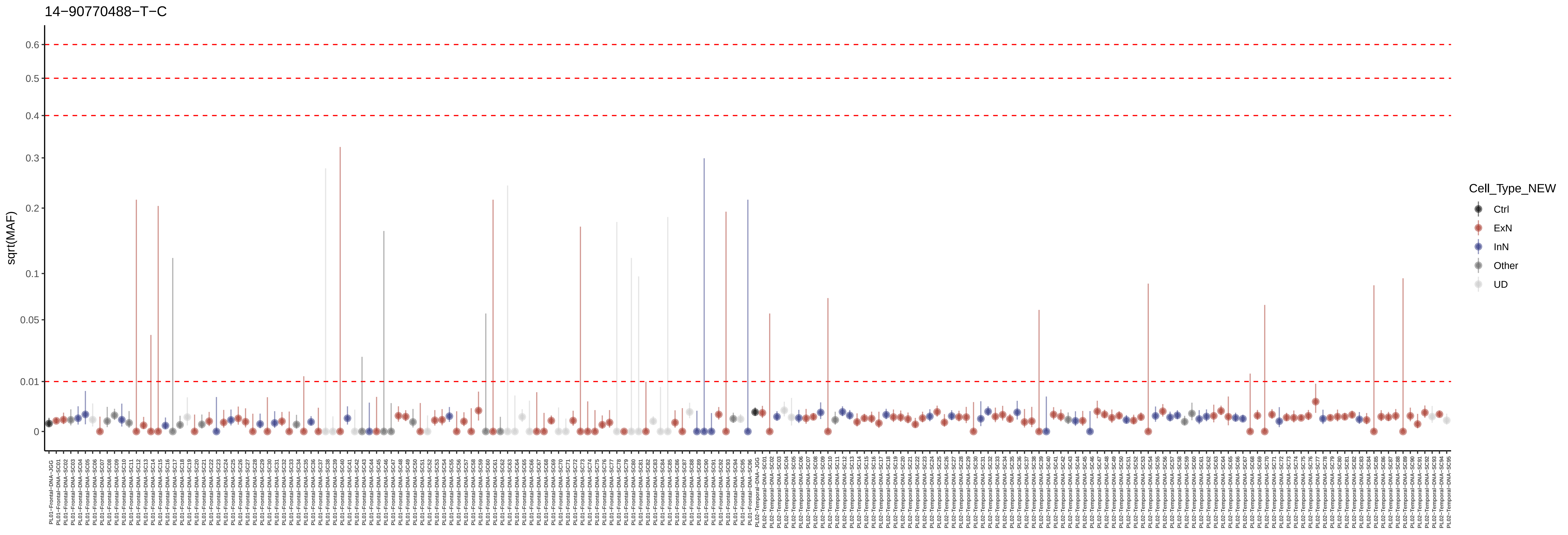

14−90770488−T−C
sqrt(MAF)
0.6
0.5
0.4
0.3
0.2
0.1
0.05
0.01
0
Cell_Type_NEW
Ctrl
ExN
InN
Other
UD

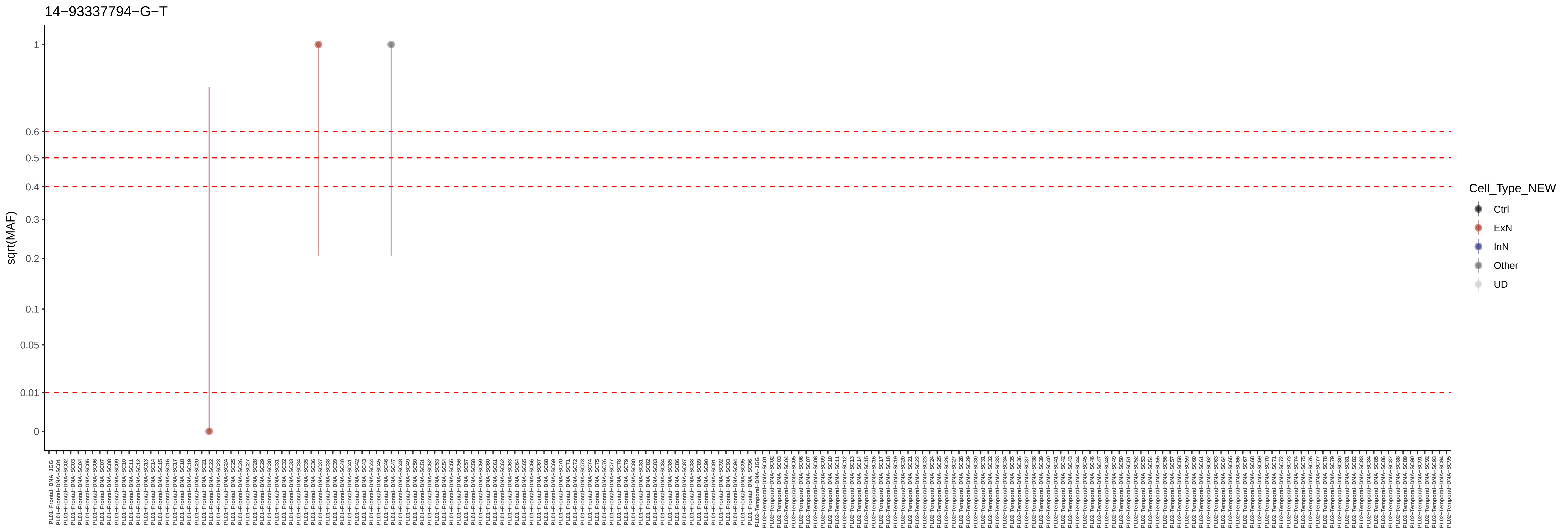
14−93337794−G−T
sqrt(MAF)
1
0.6
0.5
0.4
0.3
0.2
0.1
0.05
0.01
0
Cell_Type_NEW
Ctrl
ExN
InN
Other
UD

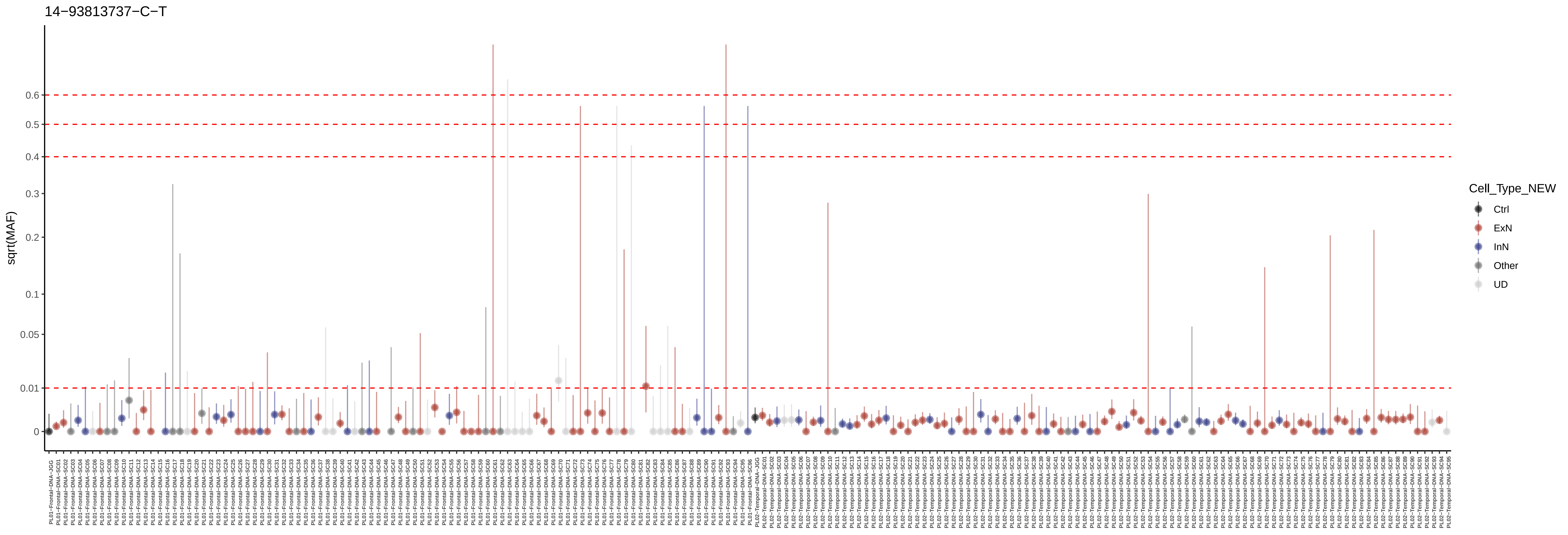
14−93813737−C−T
sqrt(MAF)
0.6
0.5
0.4
0.3
0.2
0.1
0.05
0.01
0
Cell_Type_NEW
Ctrl
ExN
InN
Other
UD

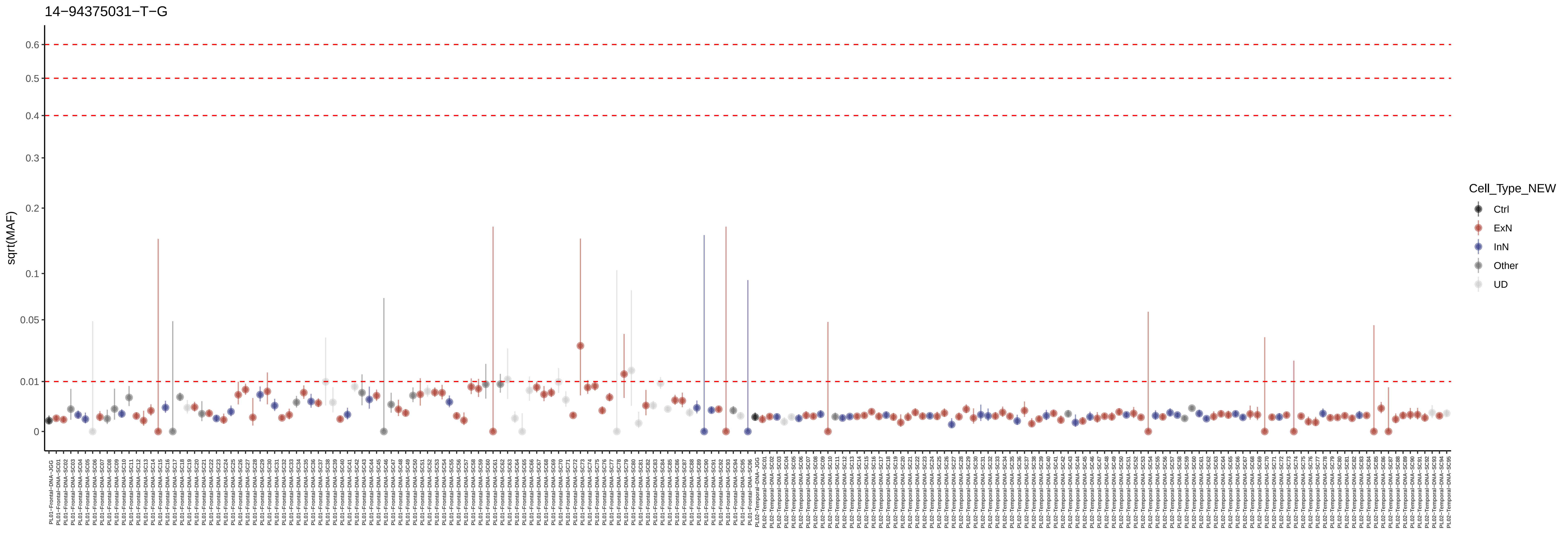

14−94375031−T−G
sqrt(MAF)
0.6
0.5
0.4
0.3
0.2
0.1
0.05
0.01
0
Cell_Type_NEW
Ctrl
ExN
InN
Other
UD

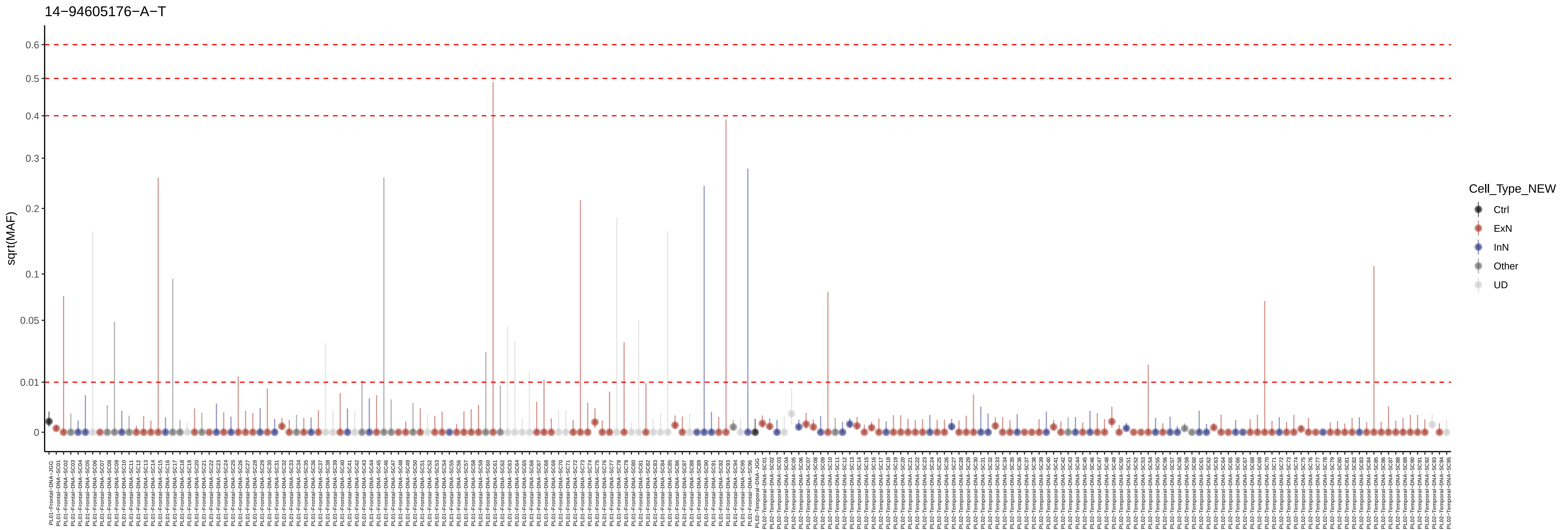

14−94605176−A−T
sqrt(MAF)
0.6
0.5
0.4
0.3
0.2
0.1
0.05
0.01
0
Cell_Type_NEW
Ctrl
ExN
InN
Other
UD

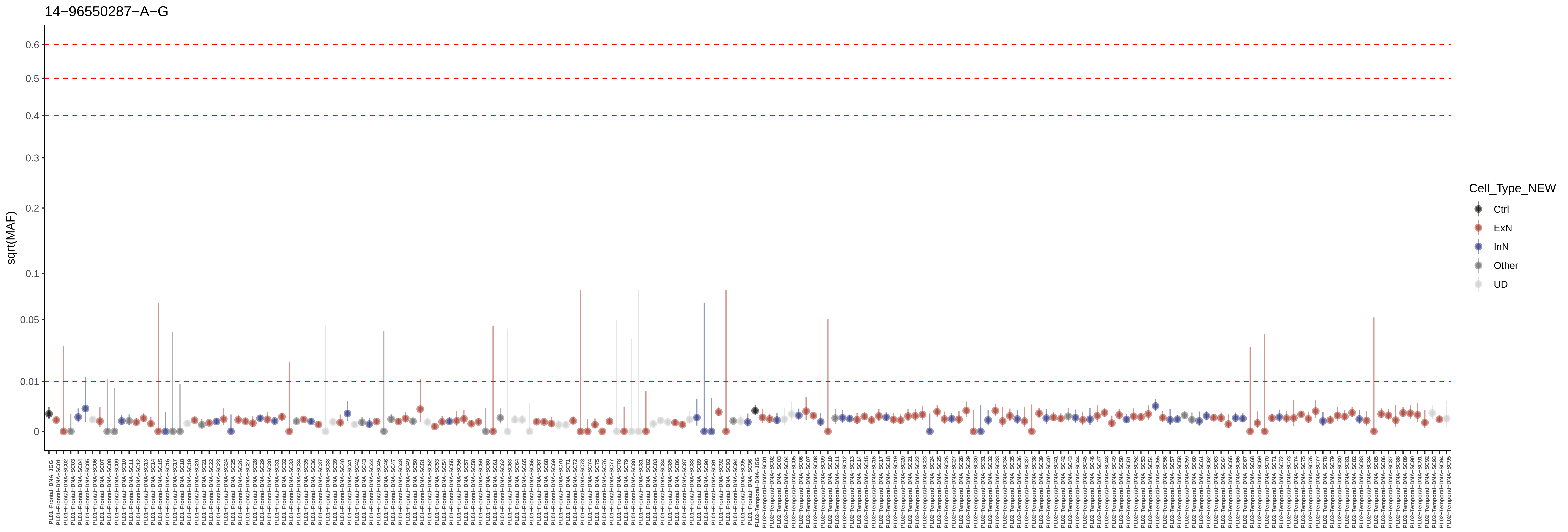

14−96550287−A−G
sqrt(MAF)
0.6
0.5
0.4
0.3
0.2
0.1
0.05
0.01
0
Cell_Type_NEW
Ctrl
ExN
InN
Other
UD

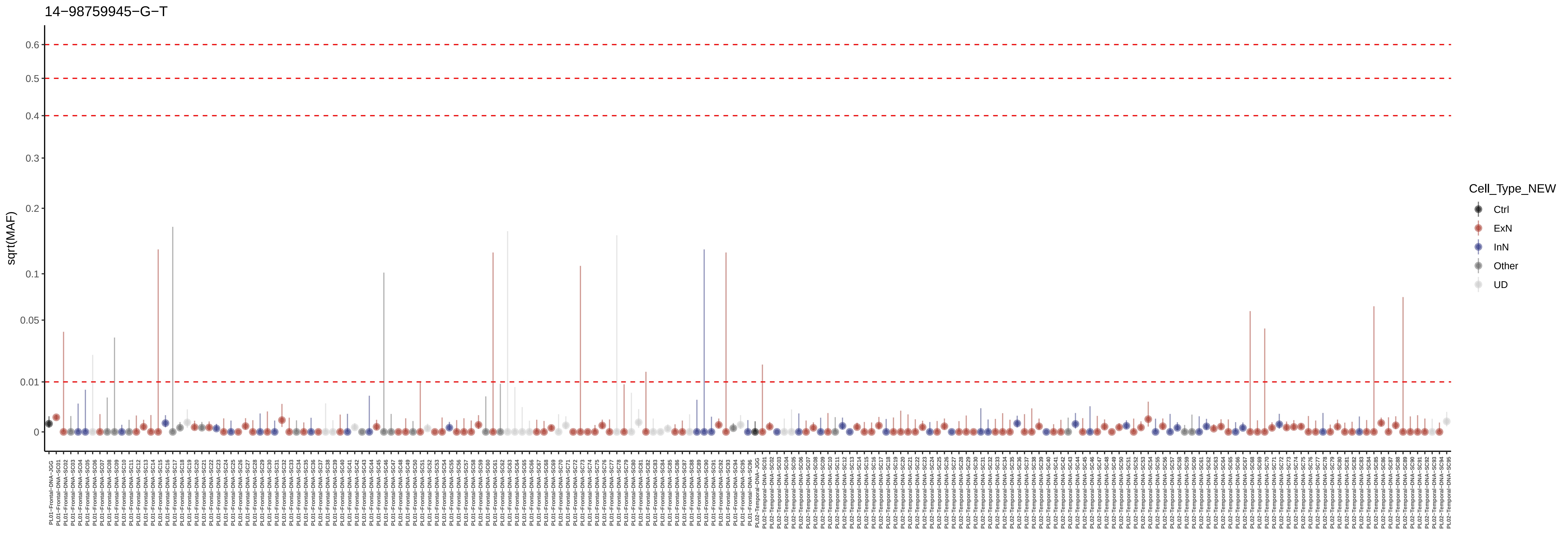

14−98759945−G−T
sqrt(MAF)
0.6
0.5
0.4
0.3
0.2
0.1
0.05
0.01
0
Cell_Type_NEW
Ctrl
ExN
InN
Other
UD

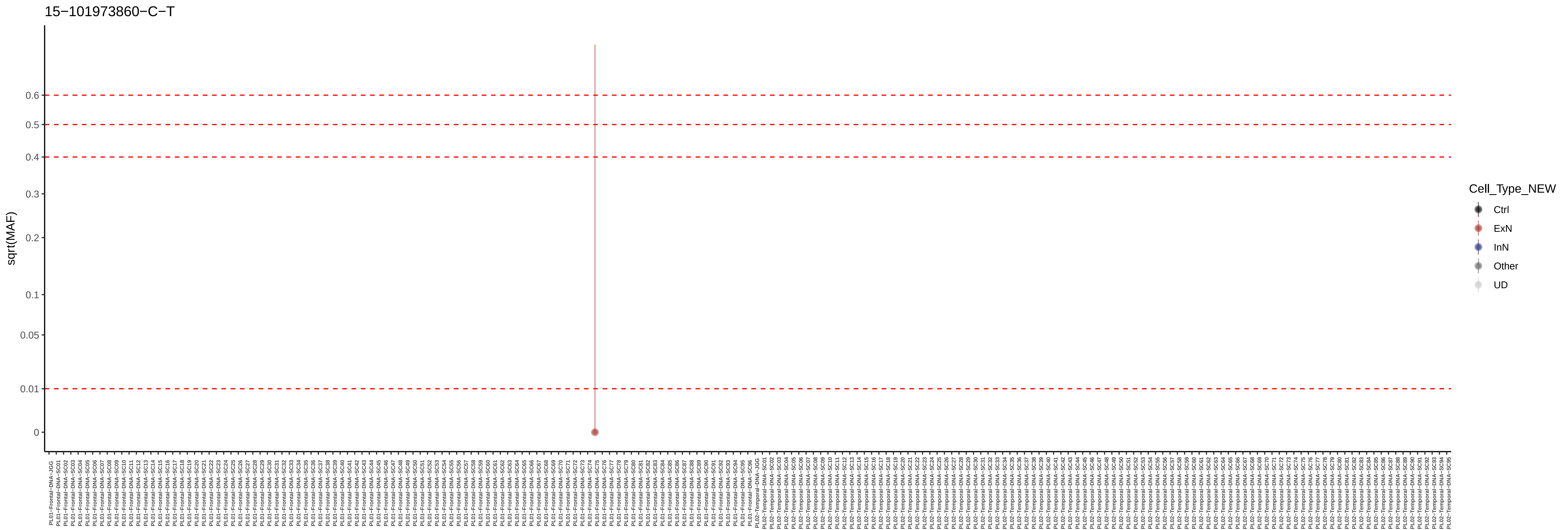

15−101973860−C−T
sqrt(MAF)
0.6
0.5
0.4
0.3
0.2
0.1
0.05
0.01
0
Cell_Type_NEW
Ctrl
ExN
InN
Other
UD

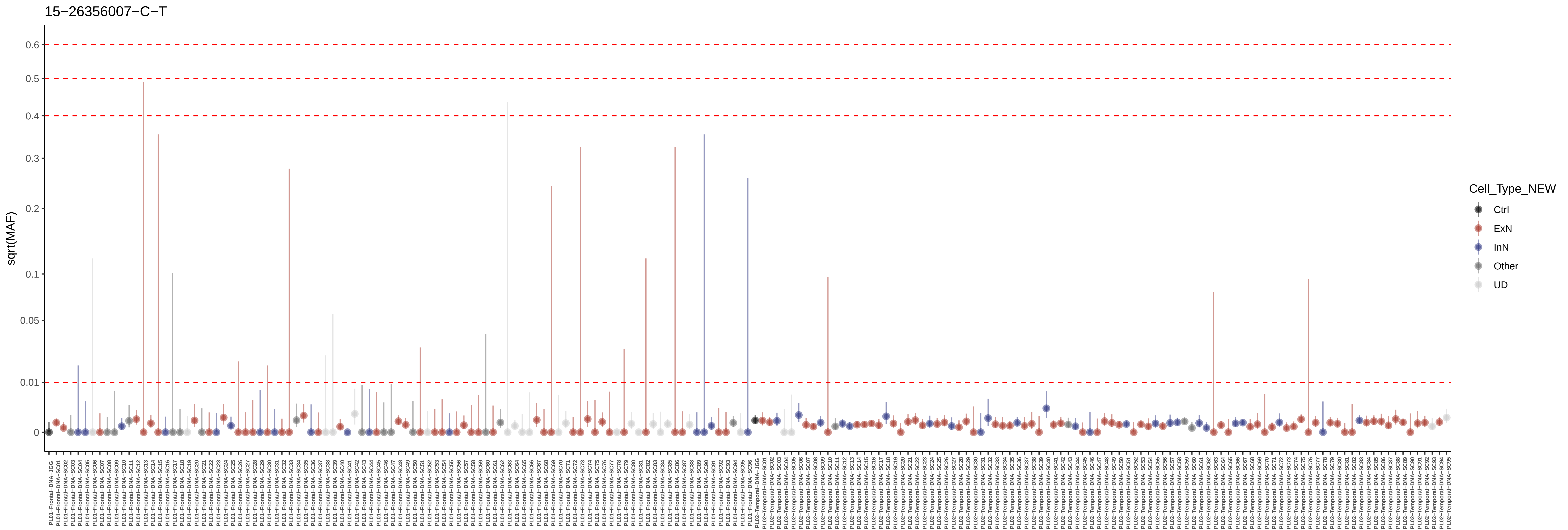
15−26356007−C−T
sqrt(MAF)
0.6
0.5
0.4
0.3
0.2
0.1
0.05
0.01
0
Cell_Type_NEW
Ctrl
ExN
InN
Other
UD

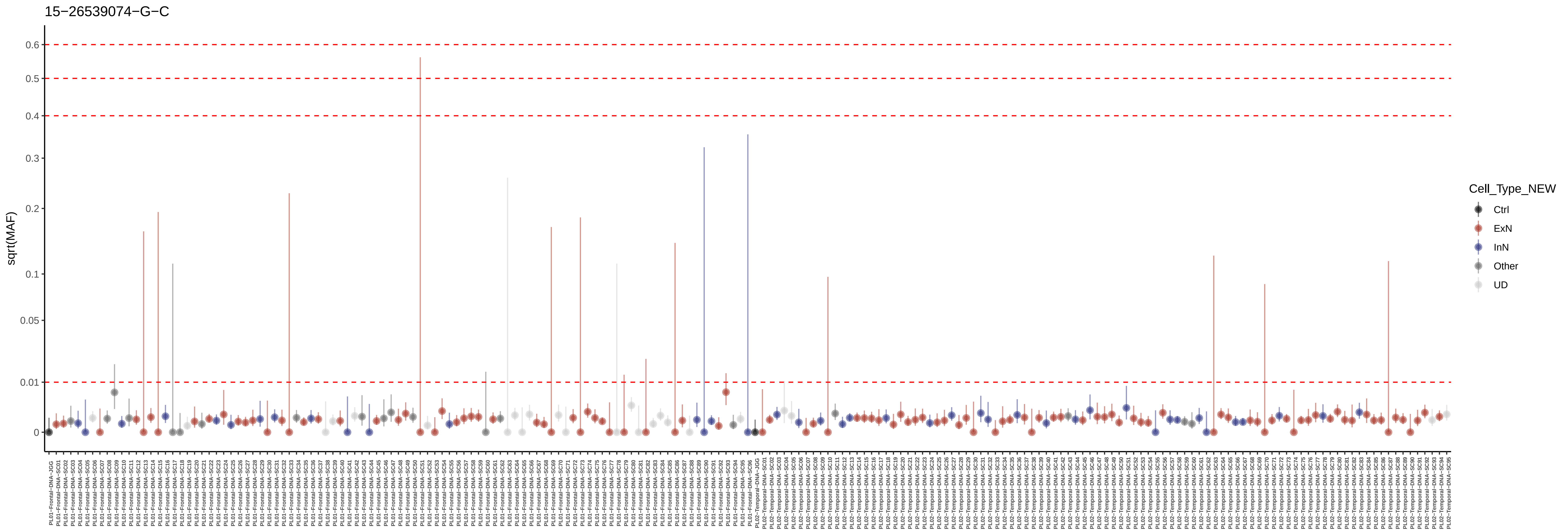

15−26539074−G−C
sqrt(MAF)
0.6
0.5
0.4
0.3
0.2
0.1
0.05
0.01
0
Cell_Type_NEW
Ctrl
ExN
InN
Other
UD

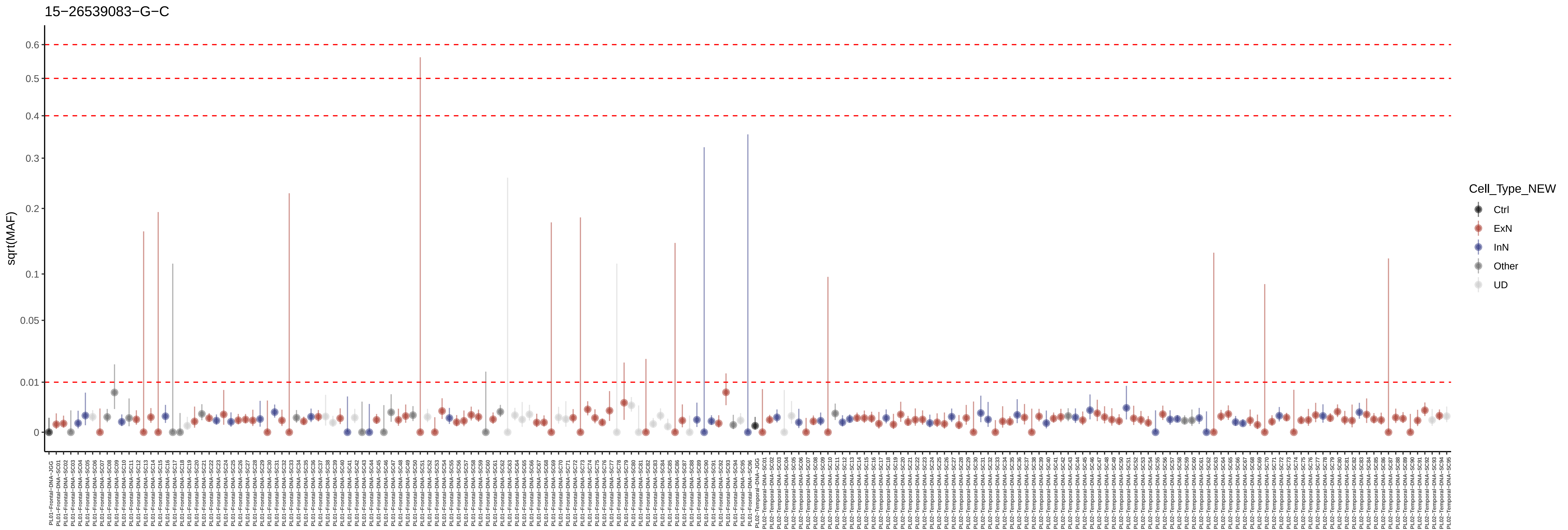
15−26539083−G−C
sqrt(MAF)
0.6
0.5
0.4
0.3
0.2
0.1
0.05
0.01
0
Cell_Type_NEW
Ctrl
ExN
InN
Other
UD

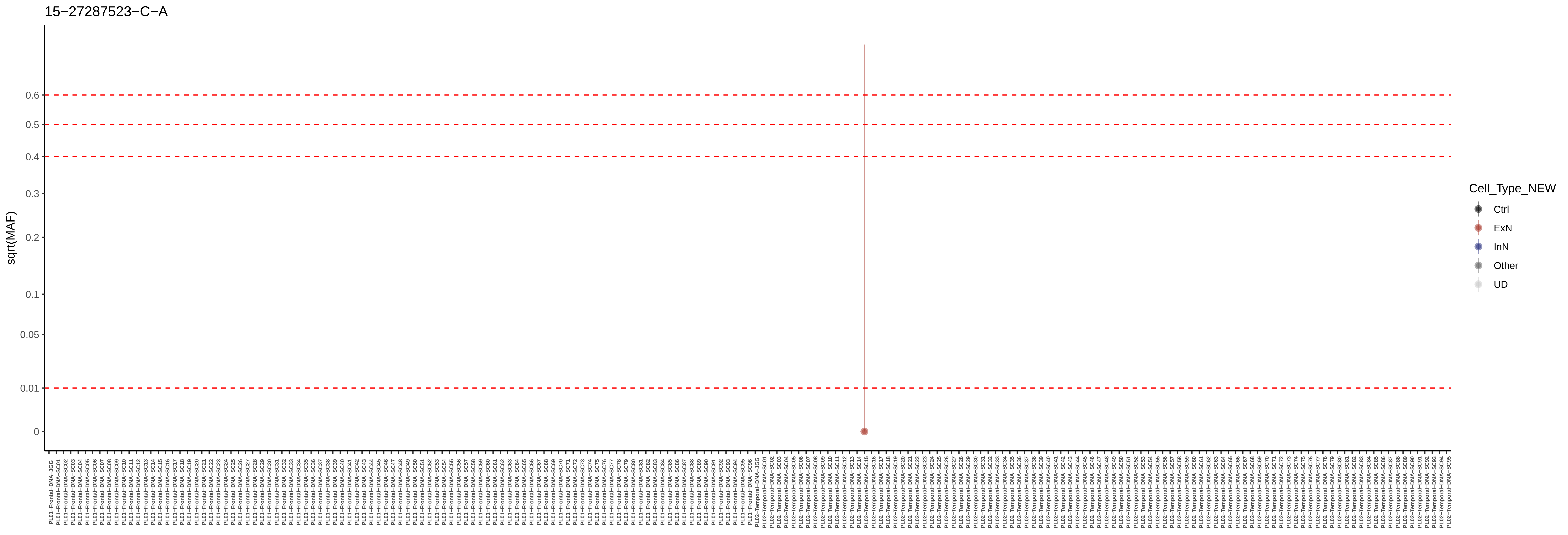

15−27287523−C−A
sqrt(MAF)
0.6
0.5
0.4
0.3
0.2
0.1
0.05
0.01
0
Cell_Type_NEW
Ctrl
ExN
InN
Other
UD

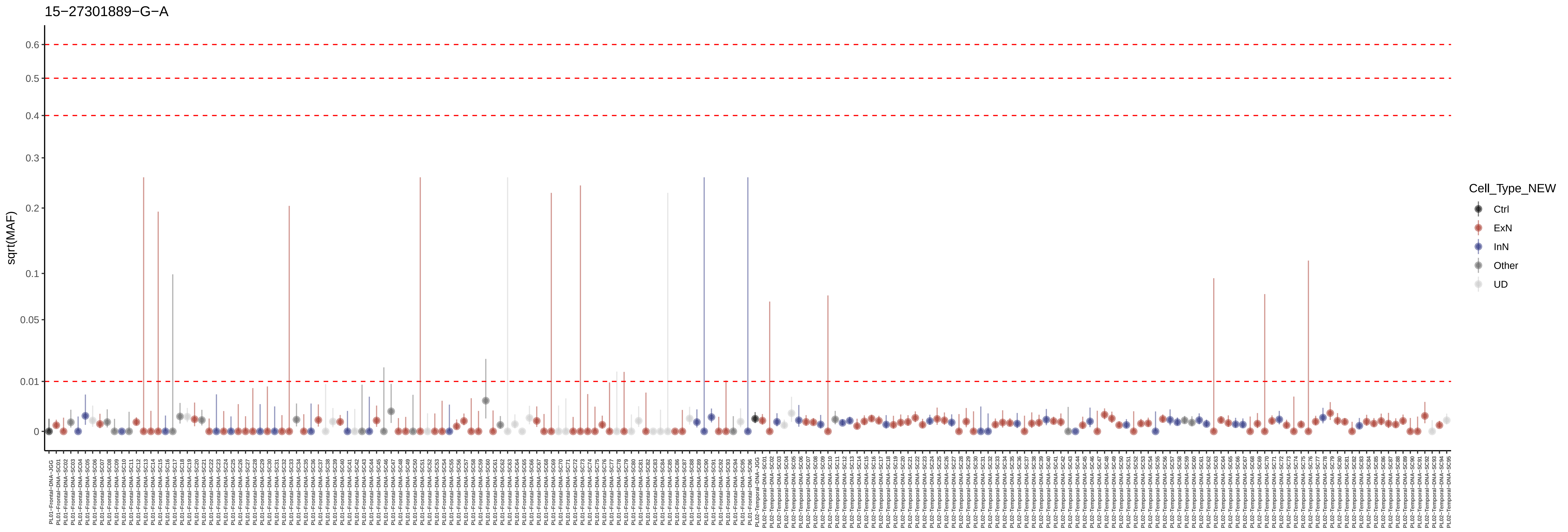
15−27301889−G−A
sqrt(MAF)
0.6
0.5
0.4
0.3
0.2
0.1
0.05
0.01
0
Cell_Type_NEW
Ctrl
ExN
InN
Other
UD

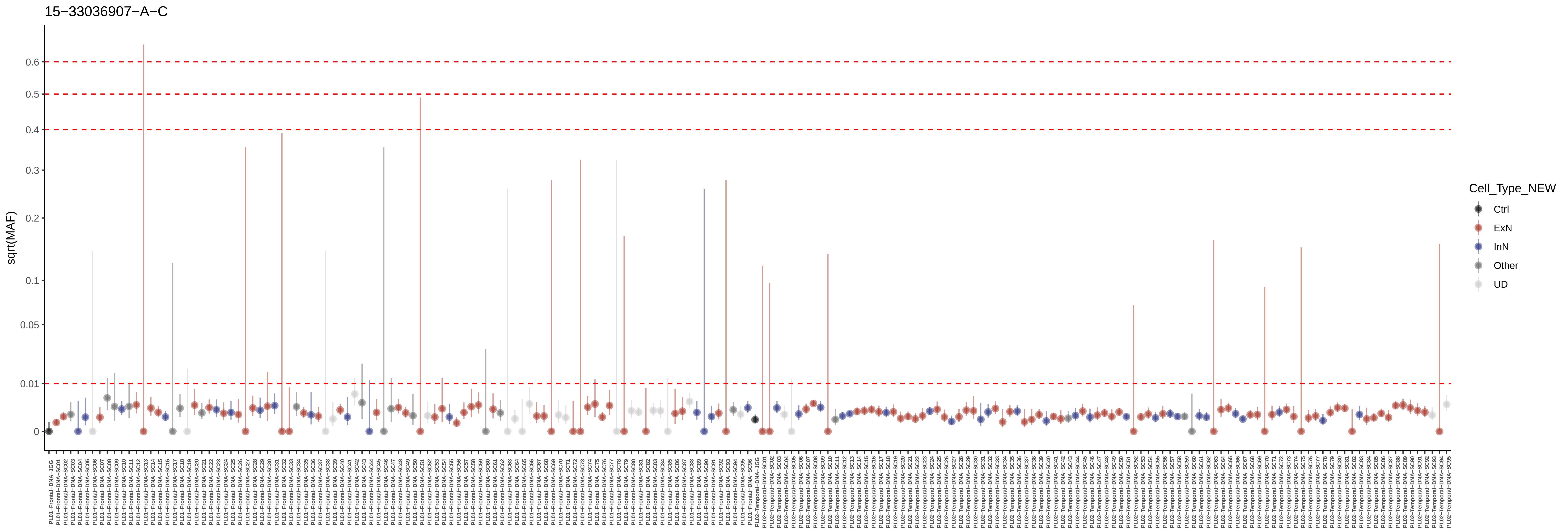
15−33036907−A−C
sqrt(MAF)
0.6
0.5
0.4
0.3
0.2
0.1
0.05
0.01
0
Cell_Type_NEW
Ctrl
ExN
InN
Other
UD

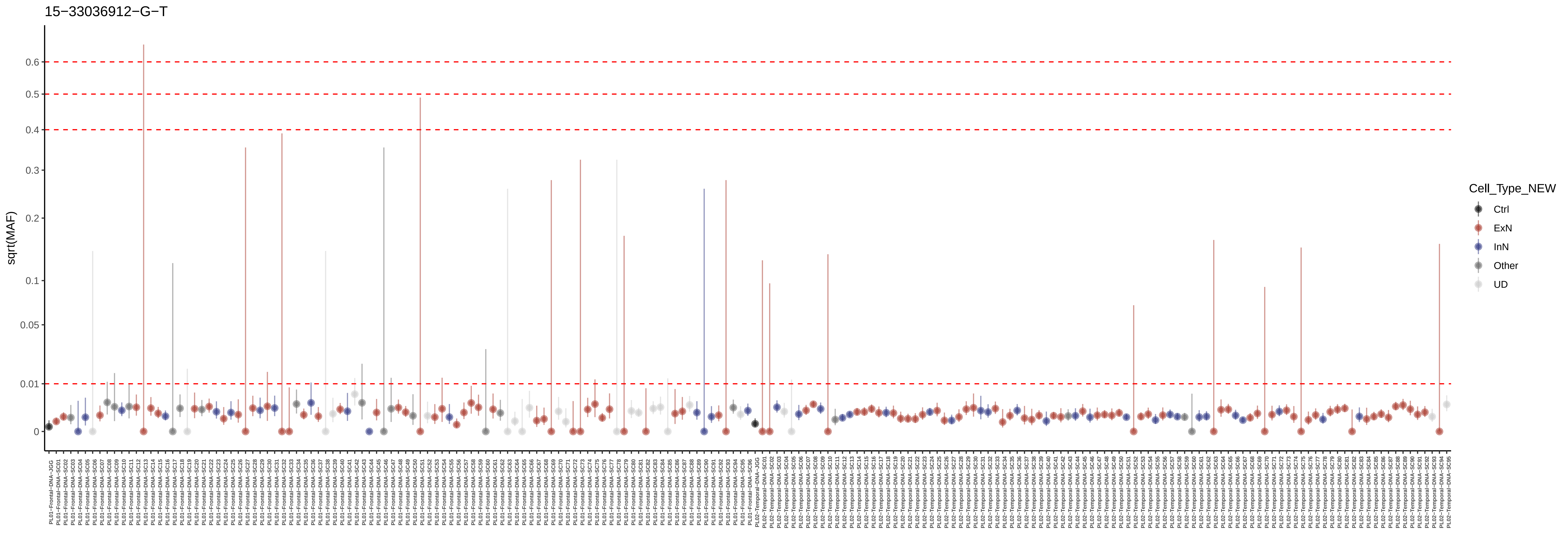

15−33036912−G−T
sqrt(MAF)
0.6
0.5
0.4
0.3
0.2
0.1
0.05
0.01
0
Cell_Type_NEW
Ctrl
ExN
InN
Other
UD

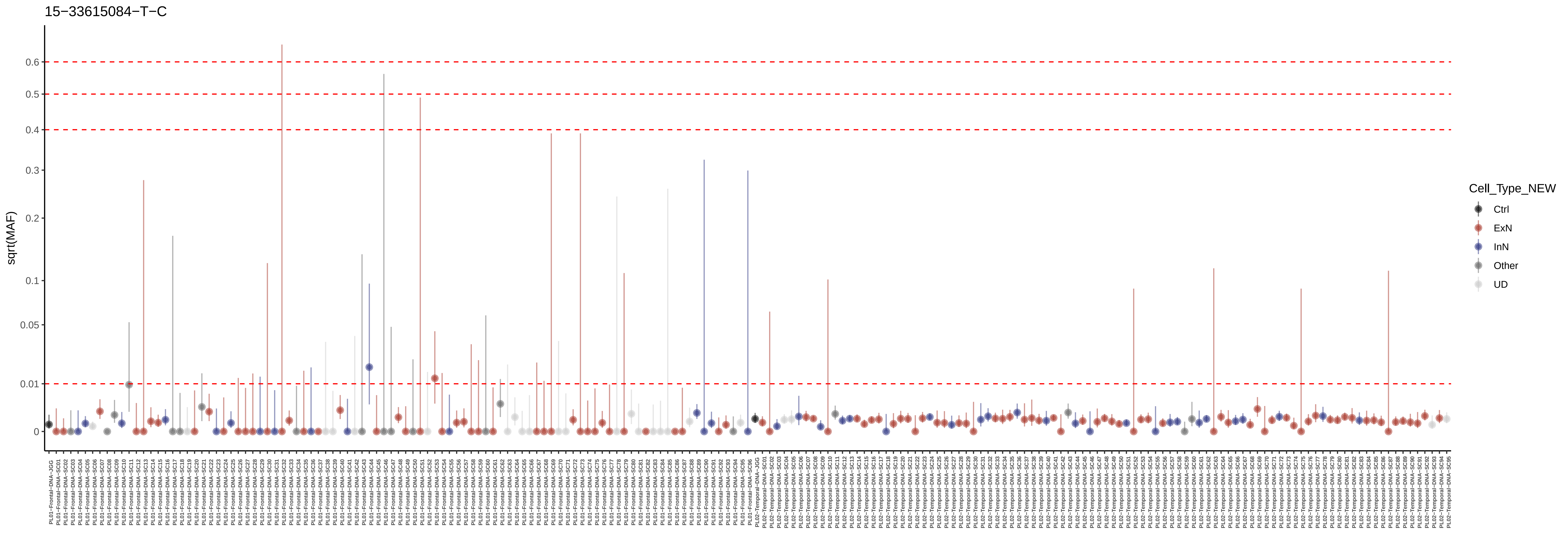
15−33615084−T−C
sqrt(MAF)
0.6
0.5
0.4
0.3
0.2
0.1
0.05
0.01
0
Cell_Type_NEW
Ctrl
ExN
InN
Other
UD

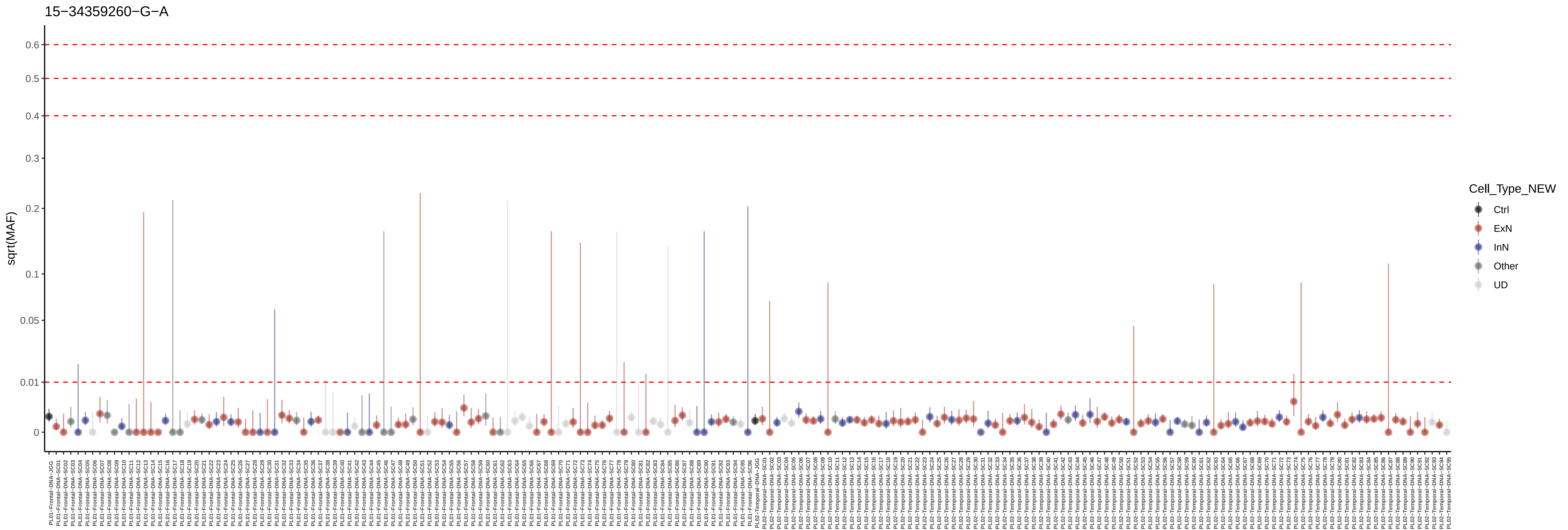
15−34359260−G−A
sqrt(MAF)
0.6
0.5
0.4
0.3
0.2
0.1
0.05
0.01
0
Cell_Type_NEW
Ctrl
ExN
InN
Other
UD

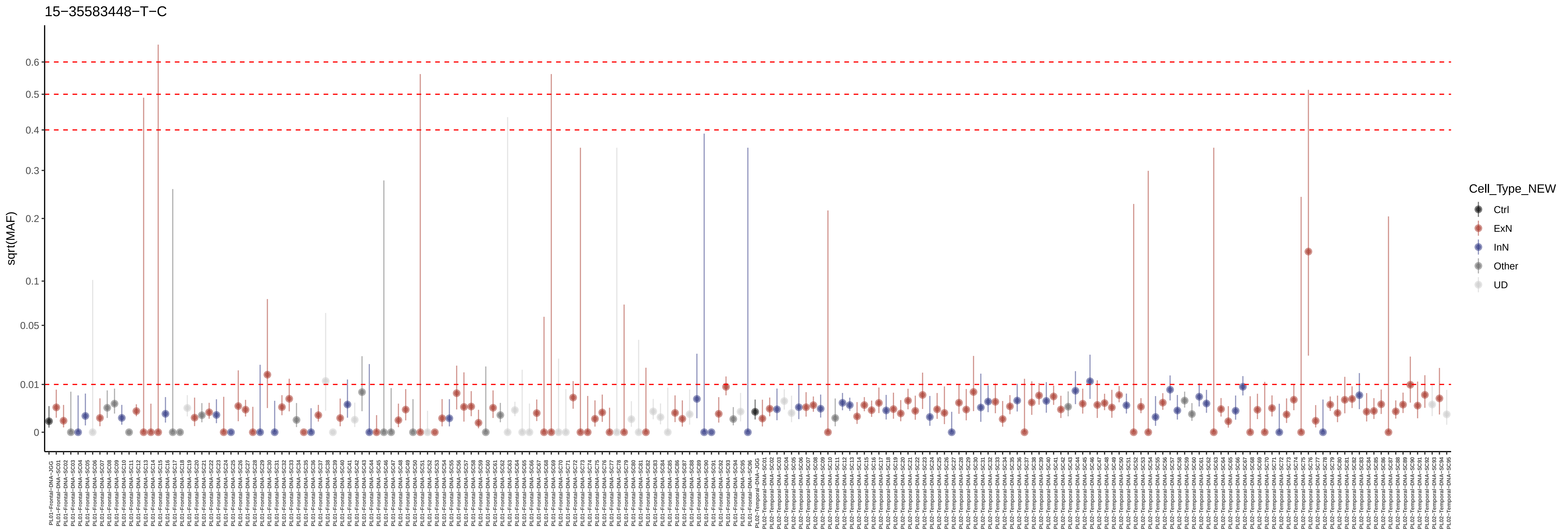

15−35583448−T−C
sqrt(MAF)
0.6
0.5
0.4
0.3
0.2
0.1
0.05
0.01
0
Cell_Type_NEW
Ctrl
ExN
InN
Other
UD

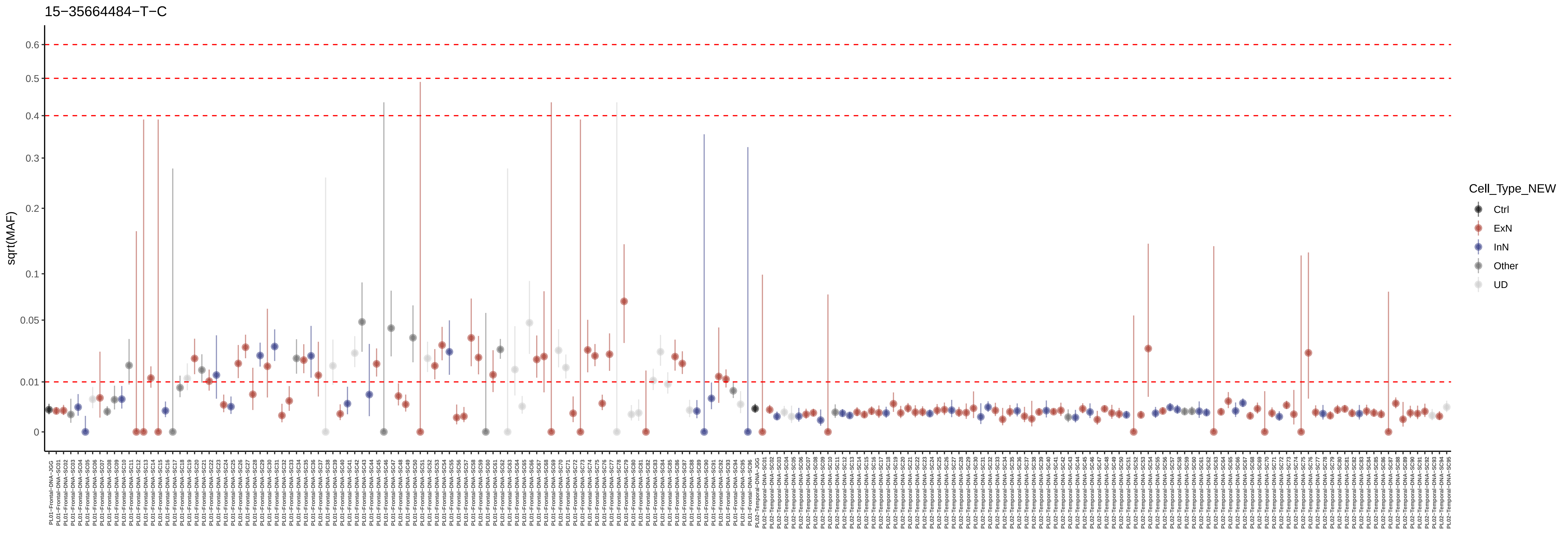

15−35664484−T−C
sqrt(MAF)
0.6
0.5
0.4
0.3
0.2
0.1
0.05
0.01
0
Cell_Type_NEW
Ctrl
ExN
InN
Other
UD

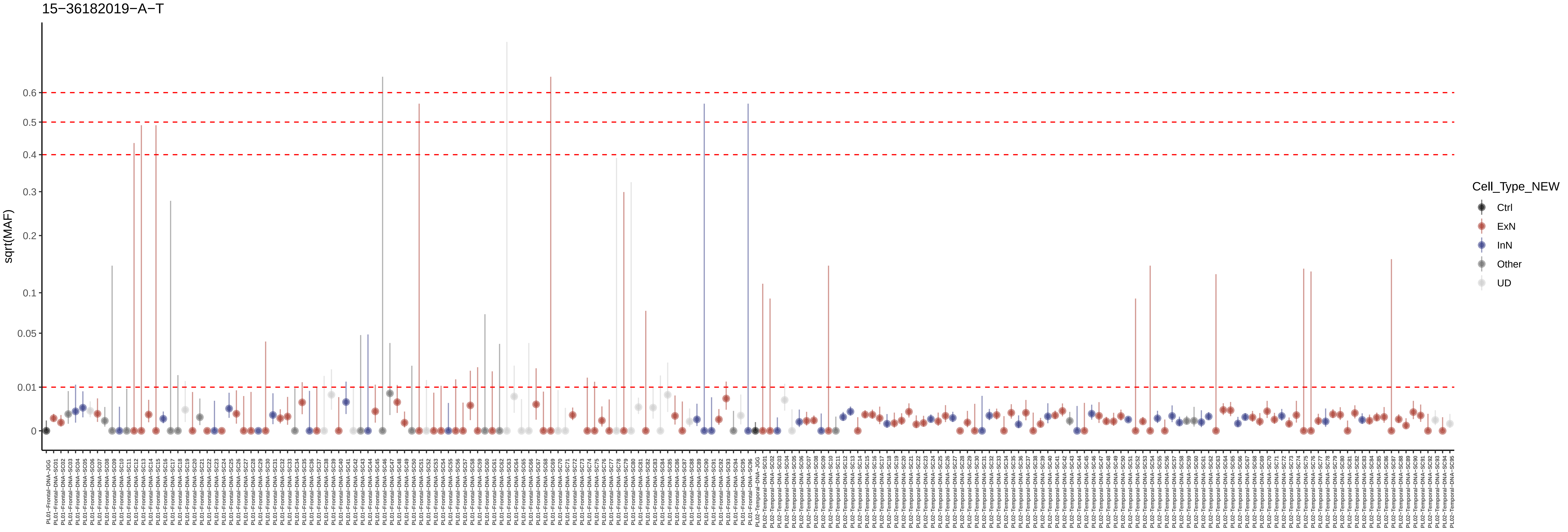
15−36182019−A−T
sqrt(MAF)
0.6
0.5
0.4
0.3
0.2
0.1
0.05
0.01
0
Cell_Type_NEW
Ctrl
ExN
InN
Other
UD

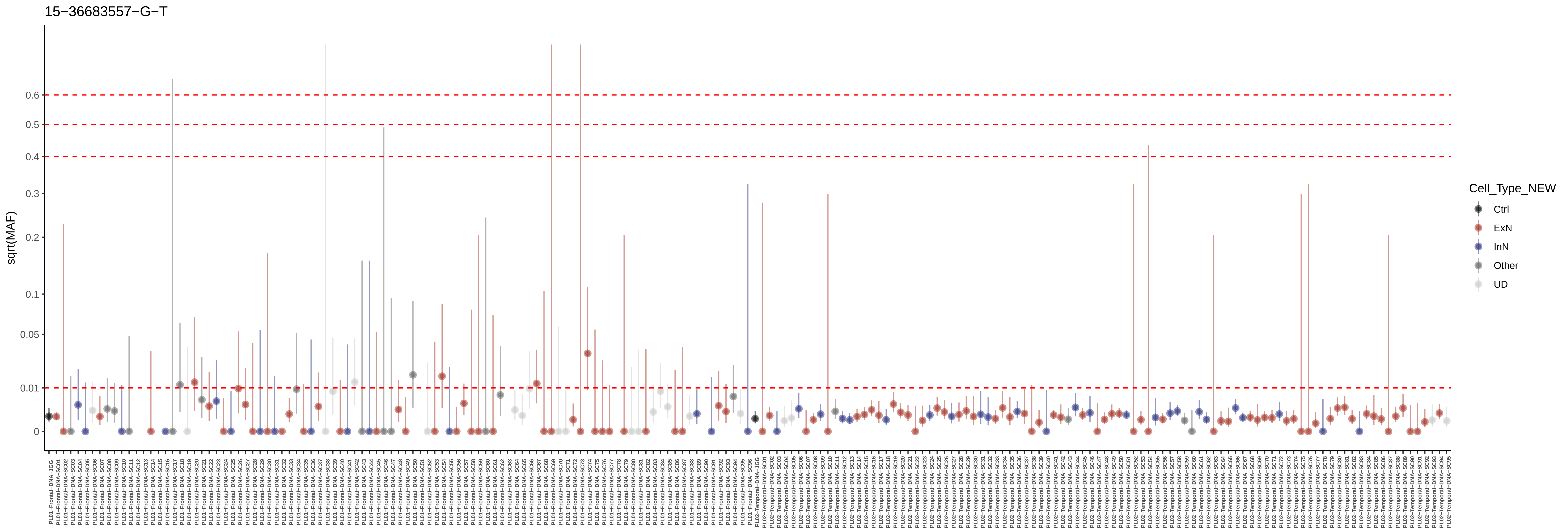

15−36683557−G−T
sqrt(MAF)
0.6
0.5
0.4
0.3
0.2
0.1
0.05
0.01
0
Cell_Type_NEW
Ctrl
ExN
InN
Other
UD

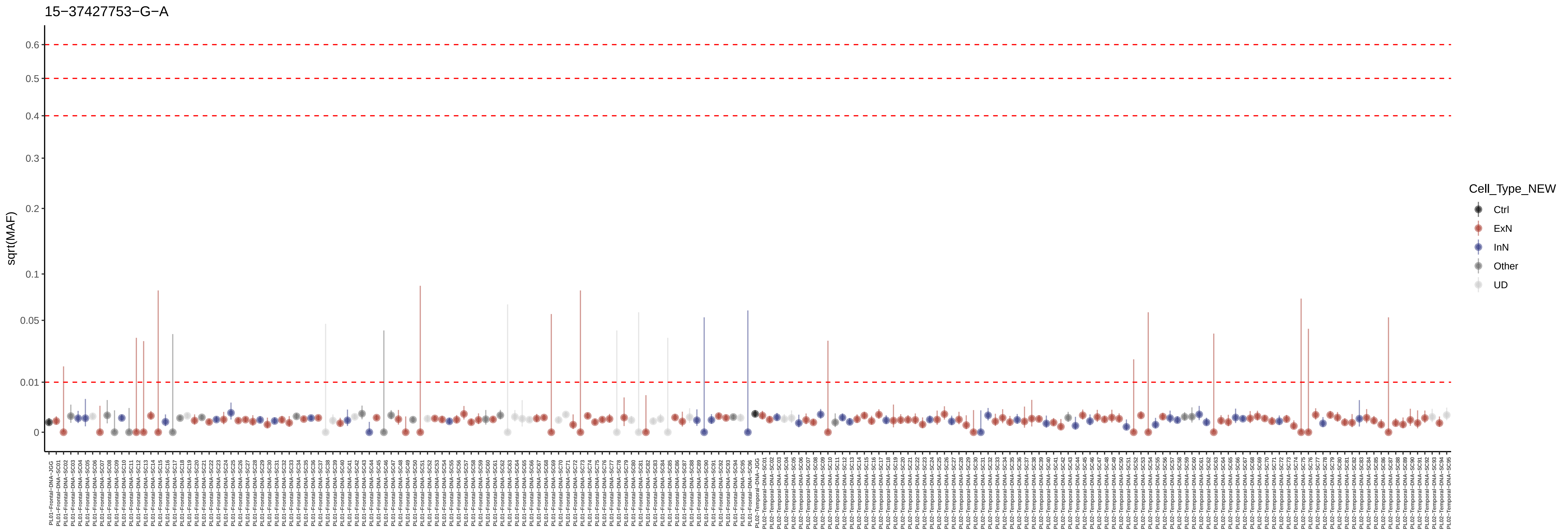

15−37427753−G−A
sqrt(MAF)
0.6
0.5
0.4
0.3
0.2
0.1
0.05
0.01
0
Cell_Type_NEW
Ctrl
ExN
InN
Other
UD

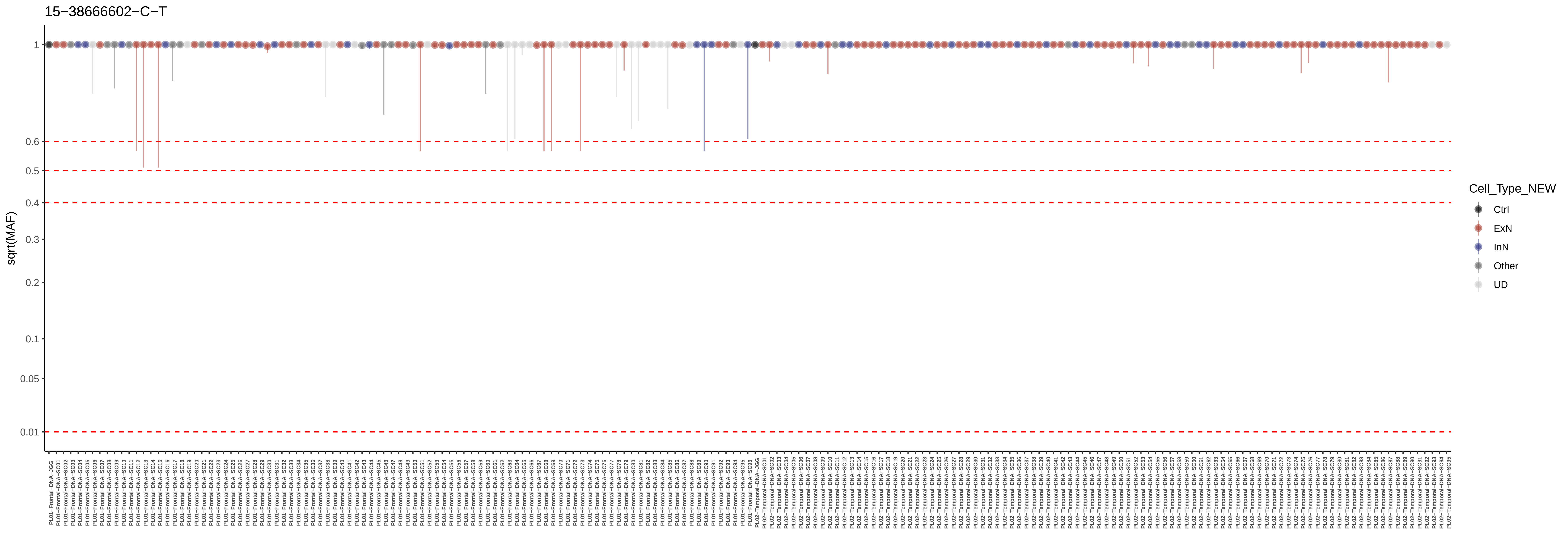
15−38666602−C−T
sqrt(MAF)
1
0.6
0.5
0.4
0.3
0.2
0.1
0.05
0.01
Cell_Type_NEW
Ctrl
ExN
InN
Other
UD

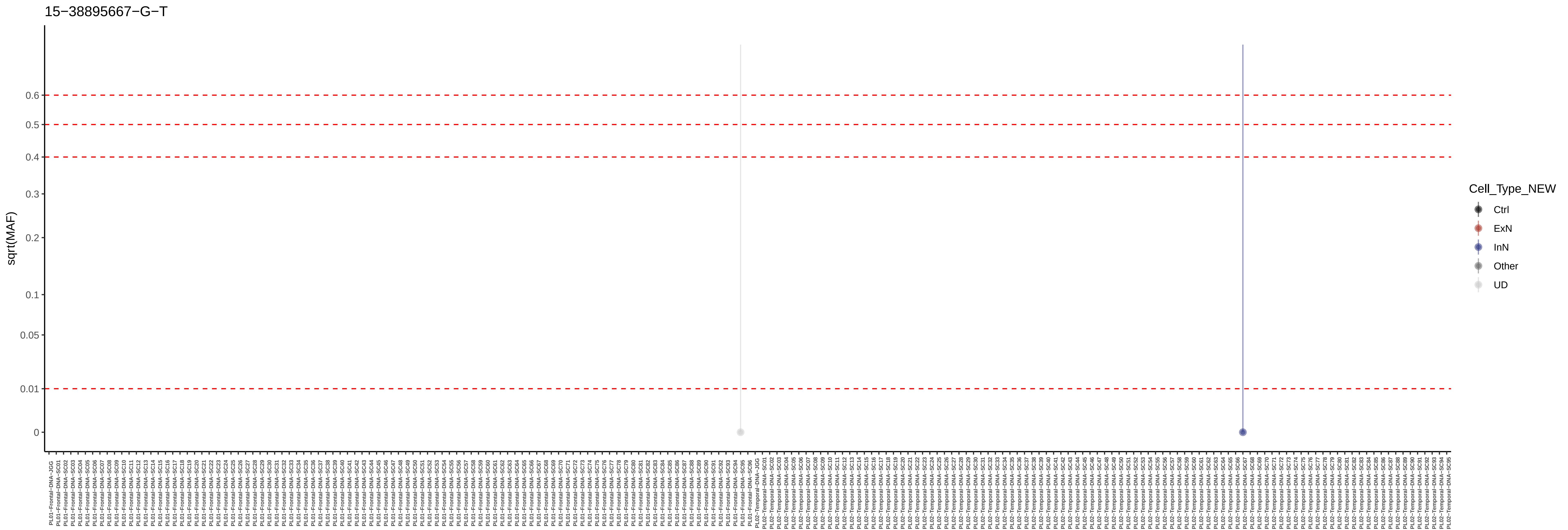
15−38895667−G−T
sqrt(MAF)
0.6
0.5
0.4
0.3
0.2
0.1
0.05
0.01
0
Cell_Type_NEW
Ctrl
ExN
InN
Other
UD

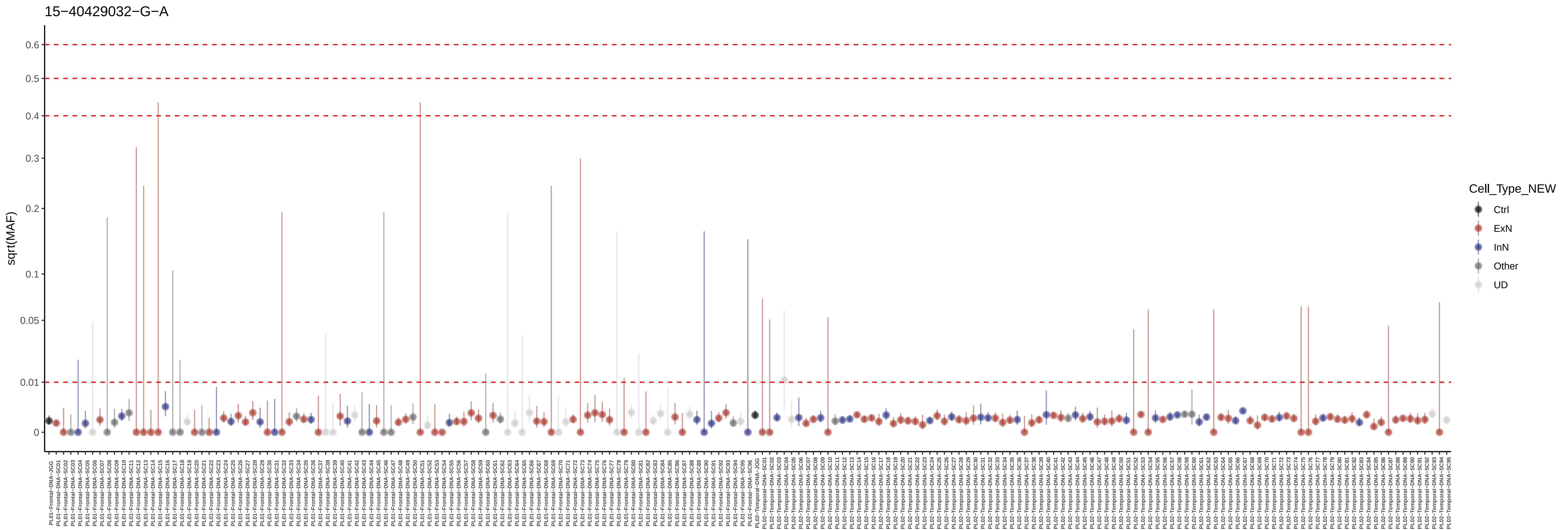
15−40429032−G−A
sqrt(MAF)
0.6
0.5
0.4
0.3
0.2
0.1
0.05
0.01
0
Cell_Type_NEW
Ctrl
ExN
InN
Other
UD

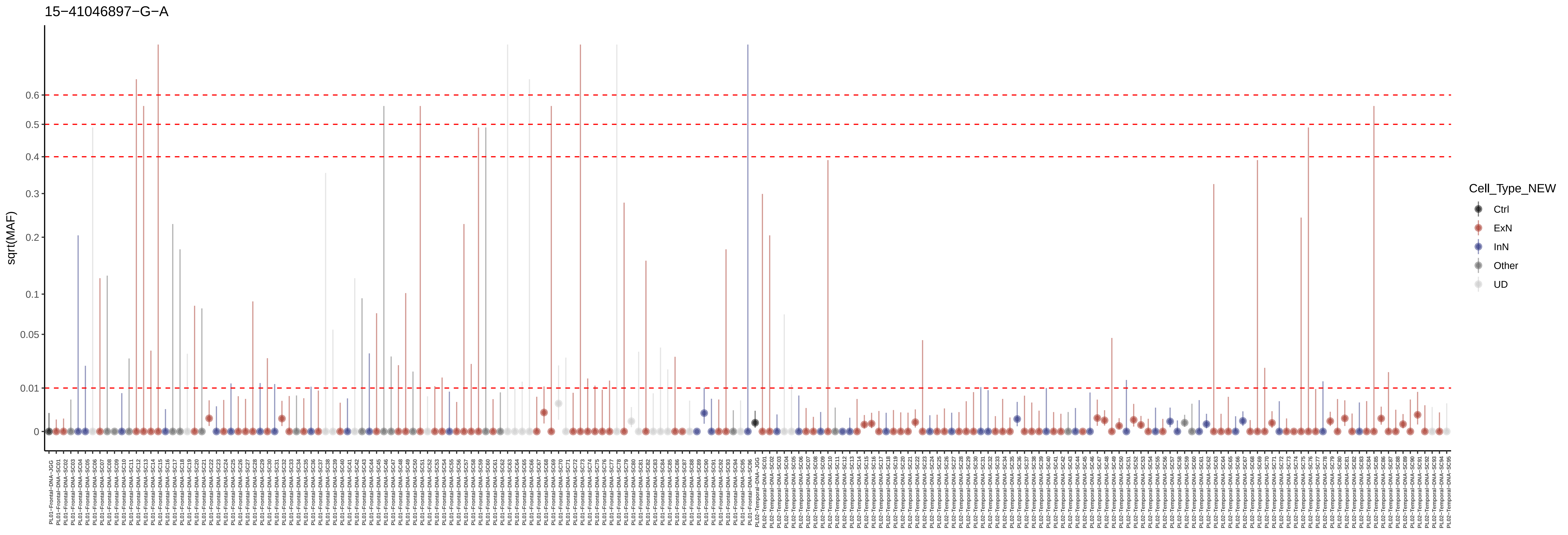
15−41046897−G−A
sqrt(MAF)
0.6
0.5
0.4
0.3
0.2
0.1
0.05
0.01
0
Cell_Type_NEW
Ctrl
ExN
InN
Other
UD

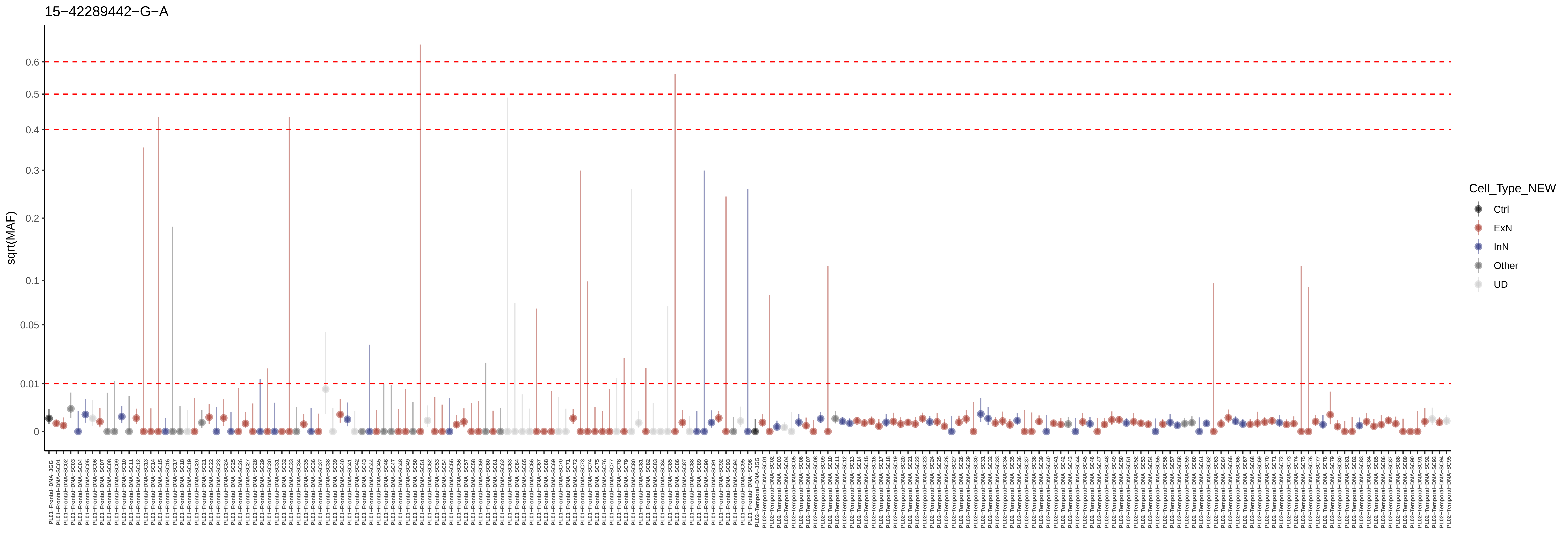

15−42289442−G−A
sqrt(MAF)
Cell_Type_NEW
Ctrl
ExN
InN
Other
UD

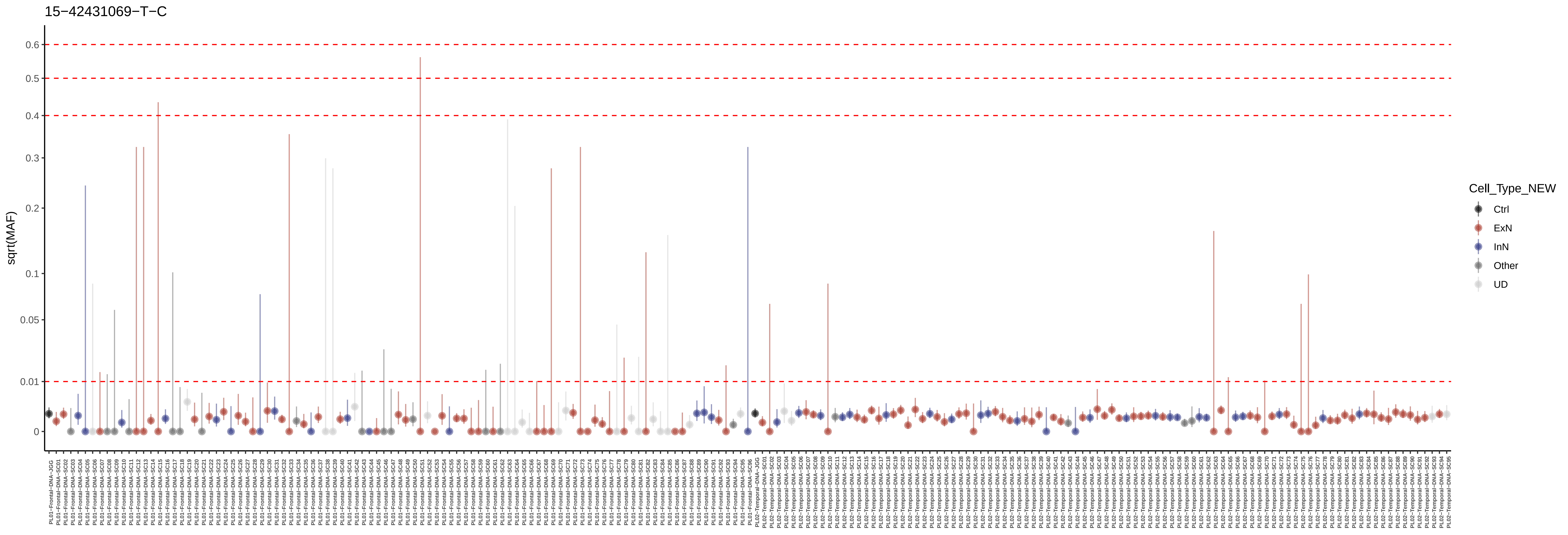
15−42431069−T−C
sqrt(MAF)
0.6
0.5
0.4
0.3
0.2
0.1
0.05
0.01
0
Cell_Type_NEW
Ctrl
ExN
InN
Other
UD

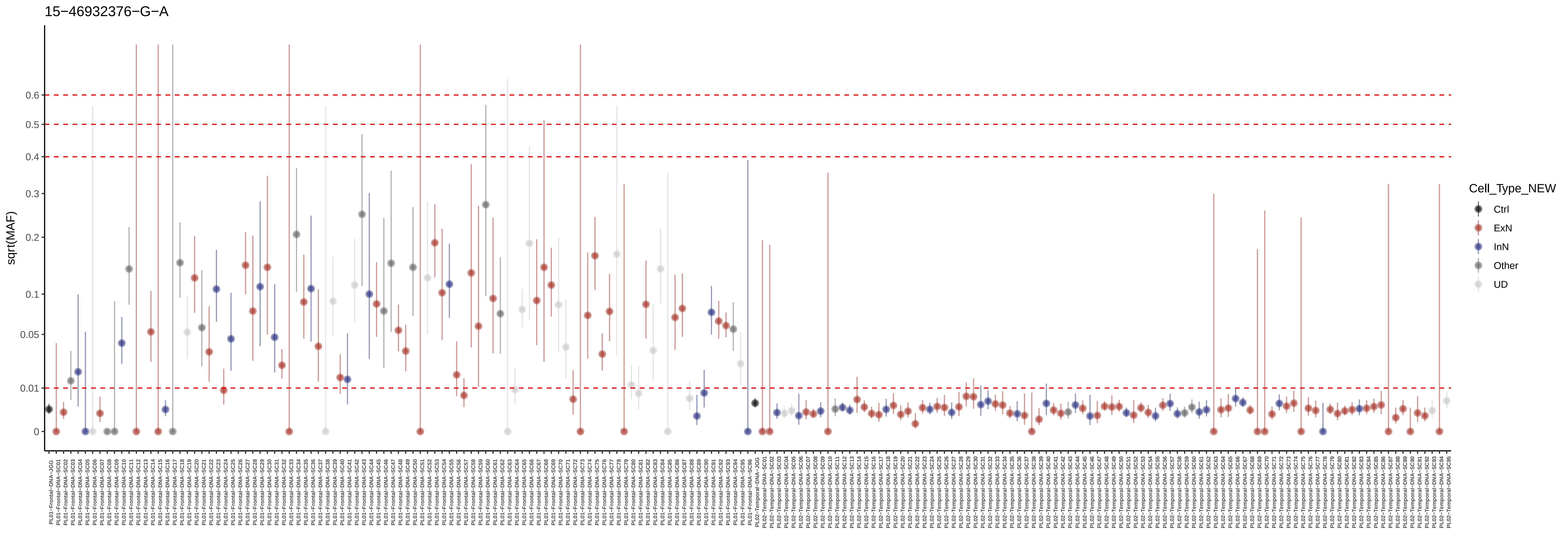
15−46932376−G−A
sqrt(MAF)
0.6
0.5
0.4
0.3
0.2
0.1
0.05
0.01
0
Cell_Type_NEW
Ctrl
ExN
InN
Other
UD

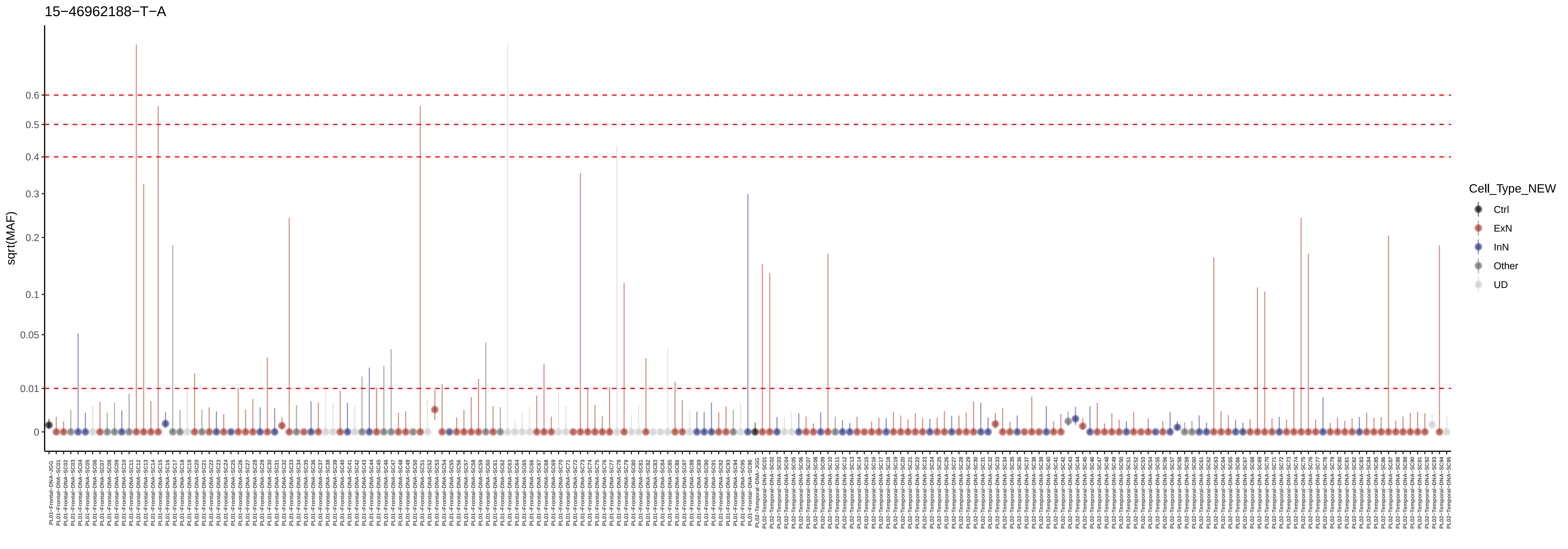
15−46962188−T−A
sqrt(MAF)
0.6
0.5
0.4
0.3
0.2
0.1
0.05
0.01
0
Cell_Type_NEW
Ctrl
ExN
InN
Other
UD

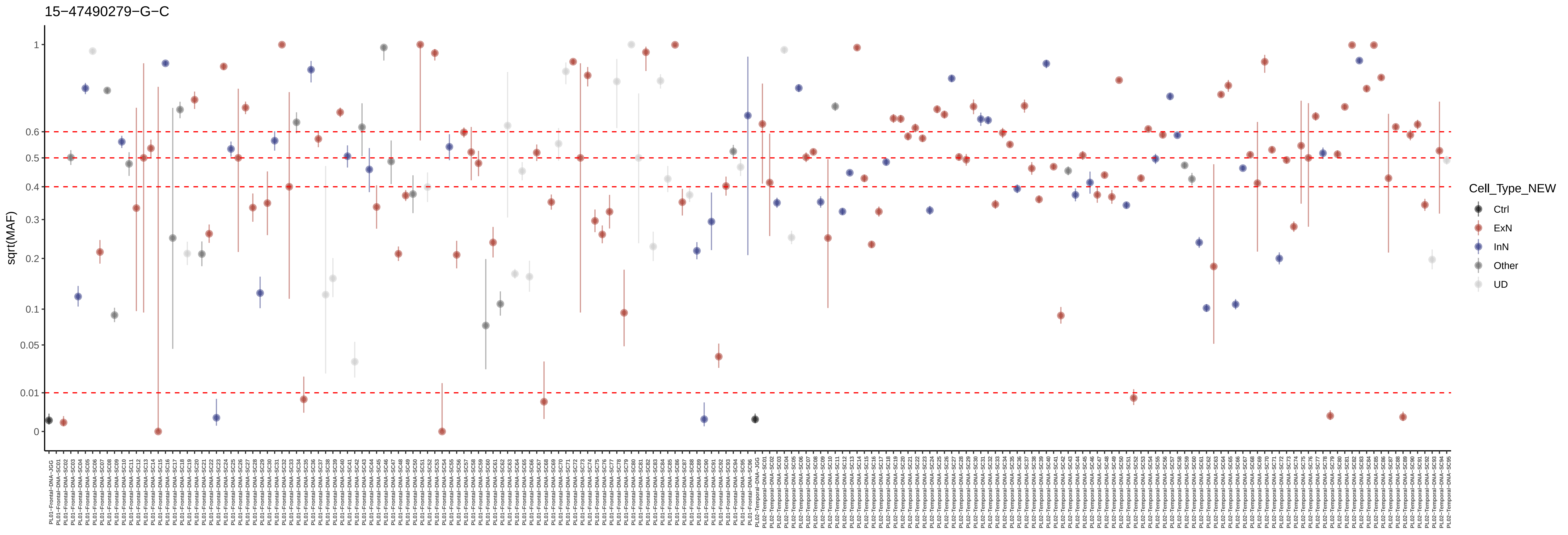

15−47490279−G−C
sqrt(MAF)
Cell_Type_NEW
Ctrl
ExN
InN
Other
UD

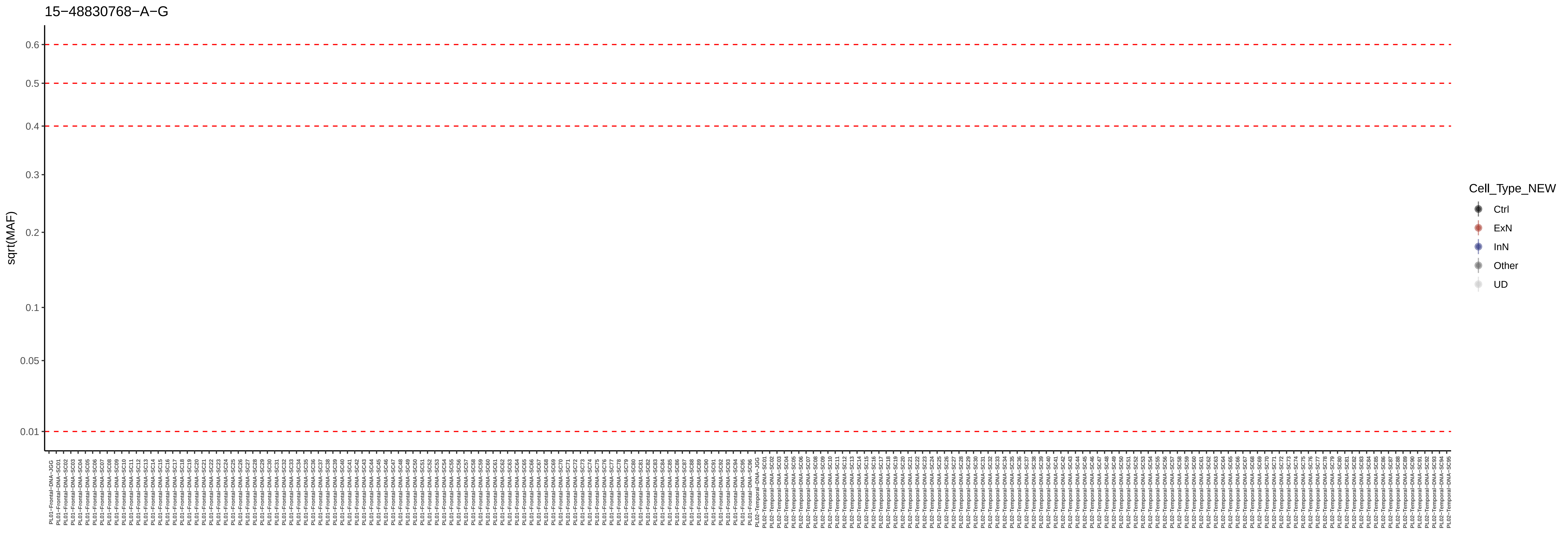
15−48830768−A−G
sqrt(MAF)
0.6
0.5
0.4
0.3
0.2
0.1
0.05
0.01
Cell_Type_NEW
Ctrl
ExN
InN
Other
UD

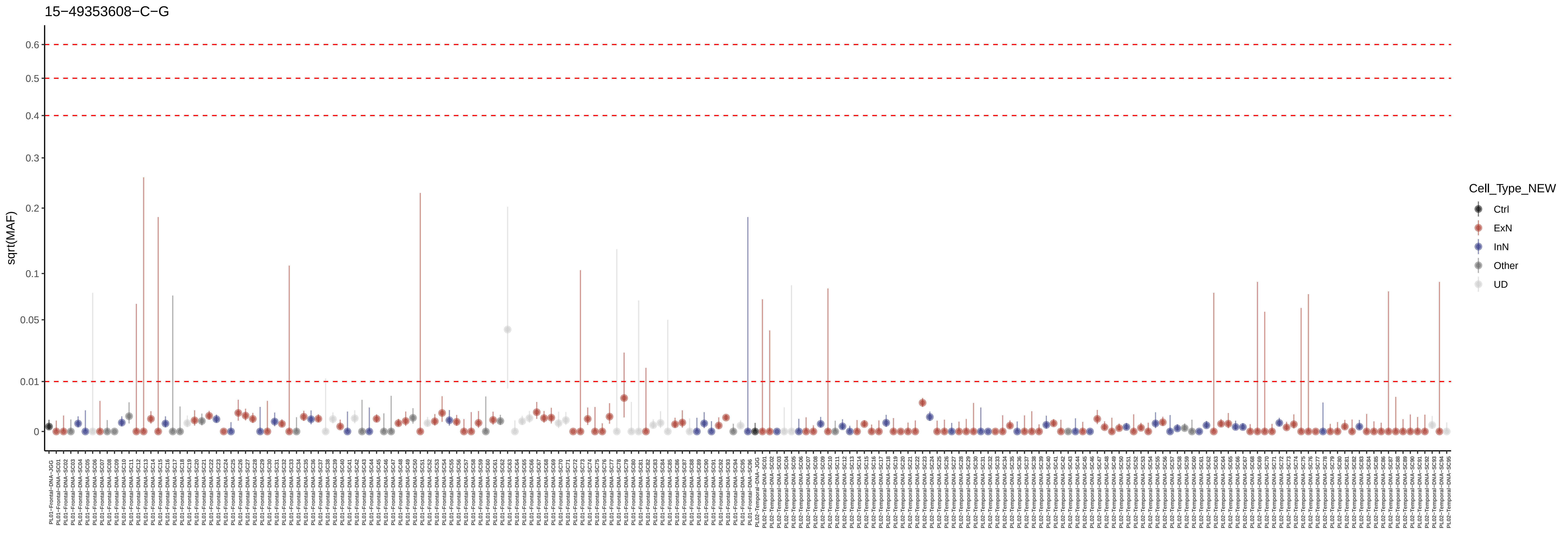
15−49353608−C−G
sqrt(MAF)
0.6
0.5
0.4
0.3
0.2
0.1
0.05
0.01
0
Cell_Type_NEW
Ctrl
ExN
InN
Other
UD

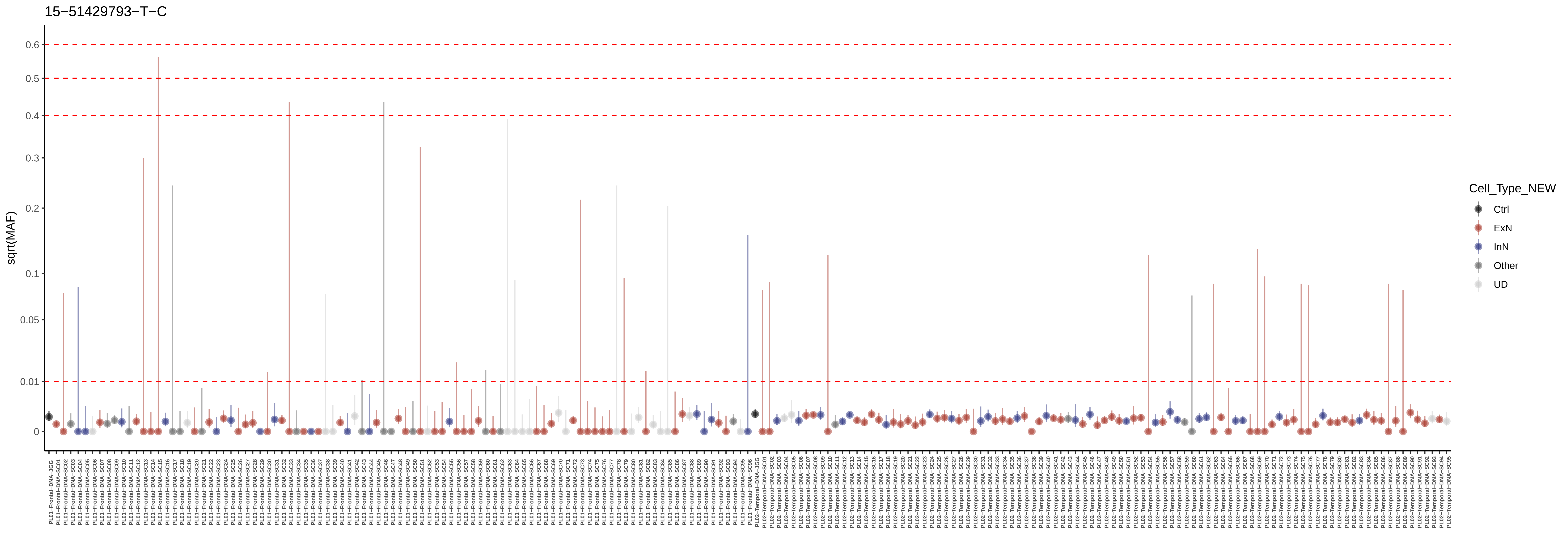

15−51429793−T−C
sqrt(MAF)
0.6
0.5
0.4
0.3
0.2
0.1
0.05
0.01
0
Cell_Type_NEW
Ctrl
ExN
InN
Other
UD

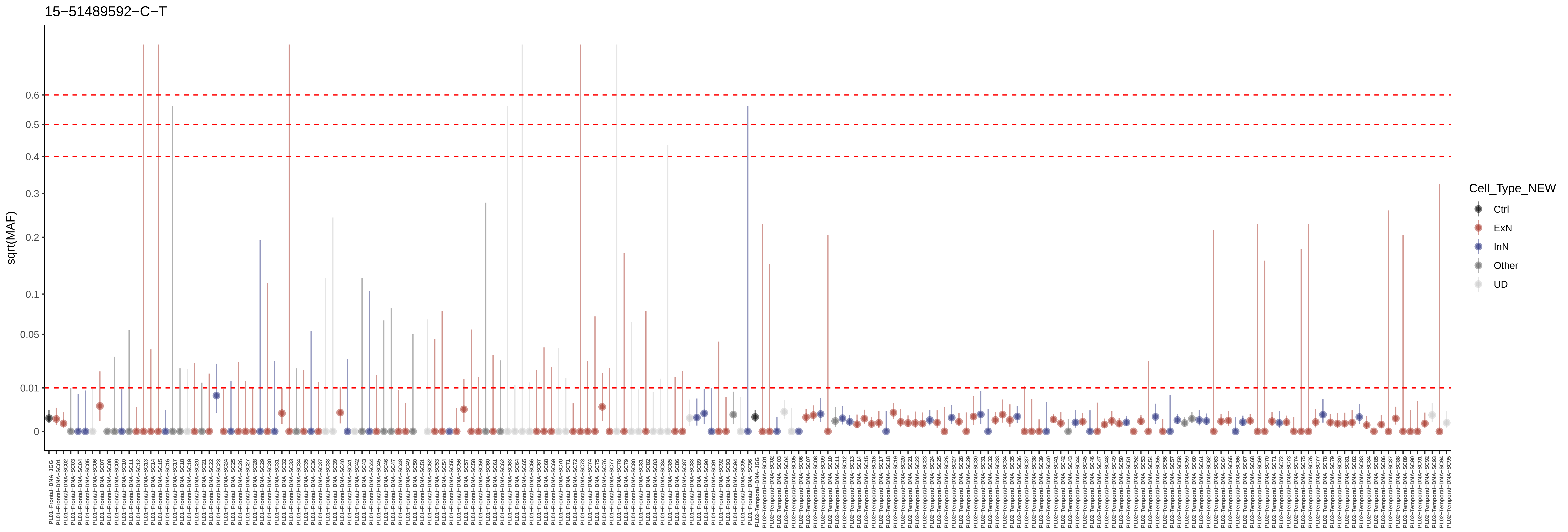

15−51489592−C−T
sqrt(MAF)
0.6
0.5
0.4
0.3
0.2
0.1
0.05
0.01
0
Cell_Type_NEW
Ctrl
ExN
InN
Other
UD

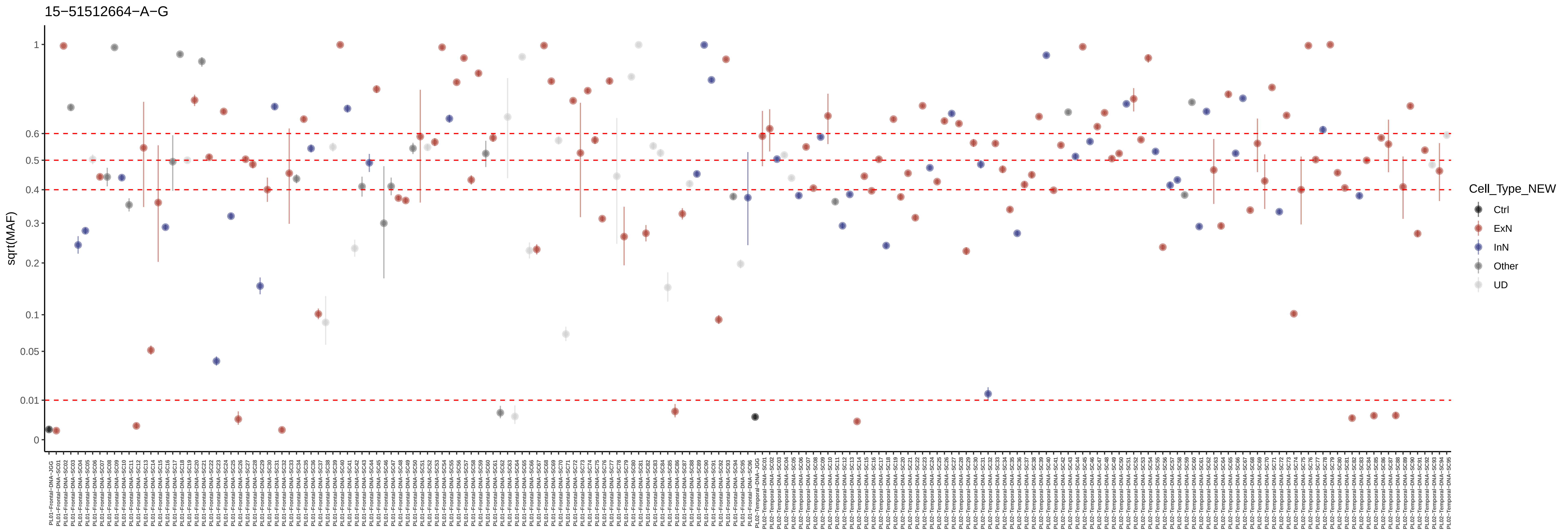

15−51512664−A−G
sqrt(MAF)
1
0.6
0.5
0.4
0.3
0.2
0.1
0.05
0.01
0
Cell_Type_NEW
Ctrl
ExN
InN
Other
UD

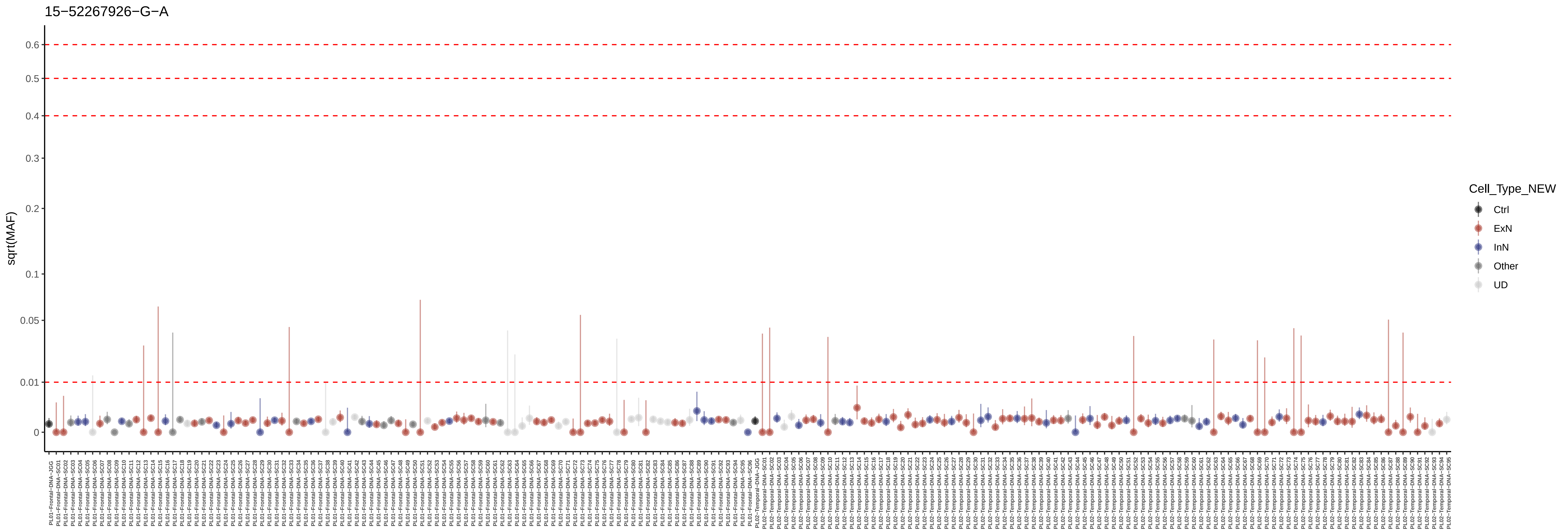

15−52267926−G−A
sqrt(MAF)
0.6
0.5
0.4
0.3
0.2
0.1
0.05
0.01
0
Cell_Type_NEW
Ctrl
ExN
InN
Other
UD

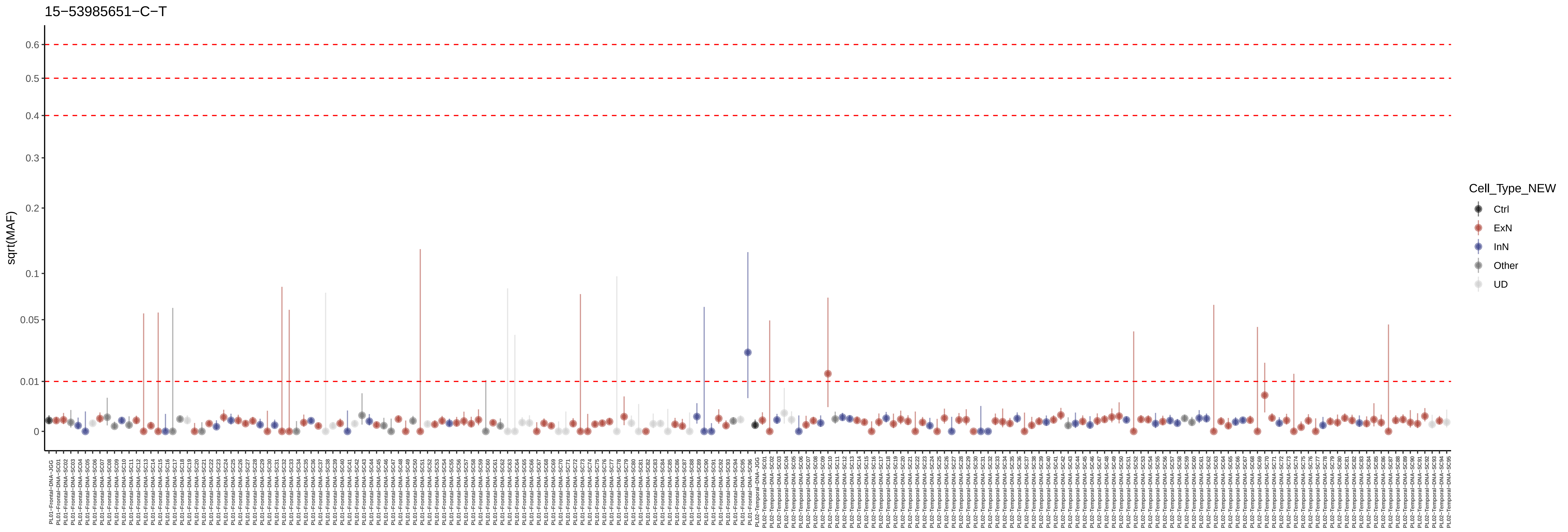
15−53985651−C−T
sqrt(MAF)
0.6
0.5
0.4
0.3
0.2
0.1
0.05
0.01
0
Cell_Type_NEW
Ctrl
ExN
InN
Other
UD

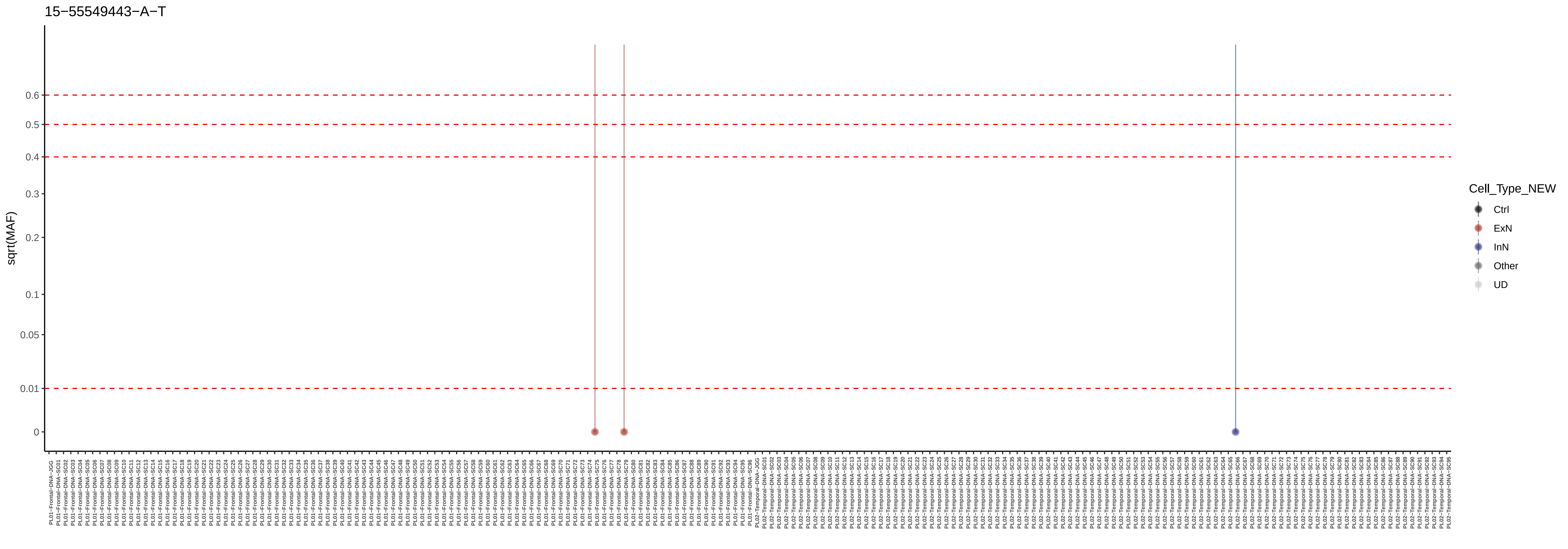

15−55549443−A−T
sqrt(MAF)
0.6
0.5
0.4
0.3
0.2
0.1
0.05
0.01
0
Cell_Type_NEW
Ctrl
ExN
InN
Other
UD

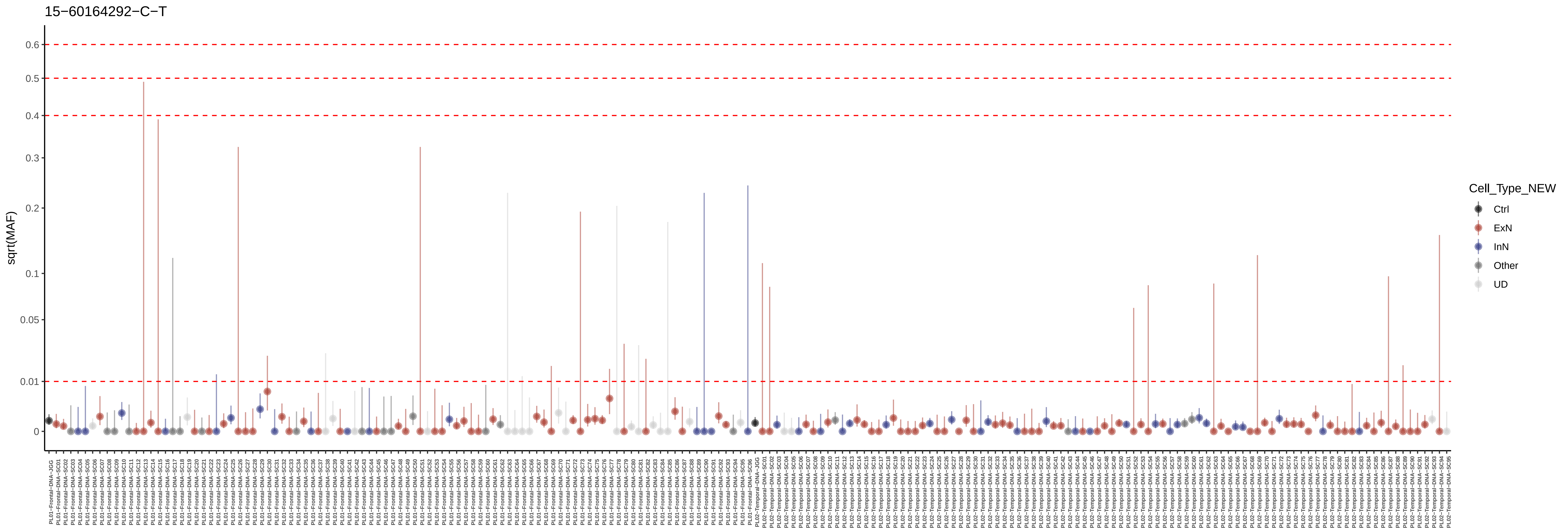
15−60164292−C−T
sqrt(MAF)
0.6
0.5
0.4
0.3
0.2
0.1
0.05
0.01
0
Cell_Type_NEW
Ctrl
ExN
InN
Other
UD

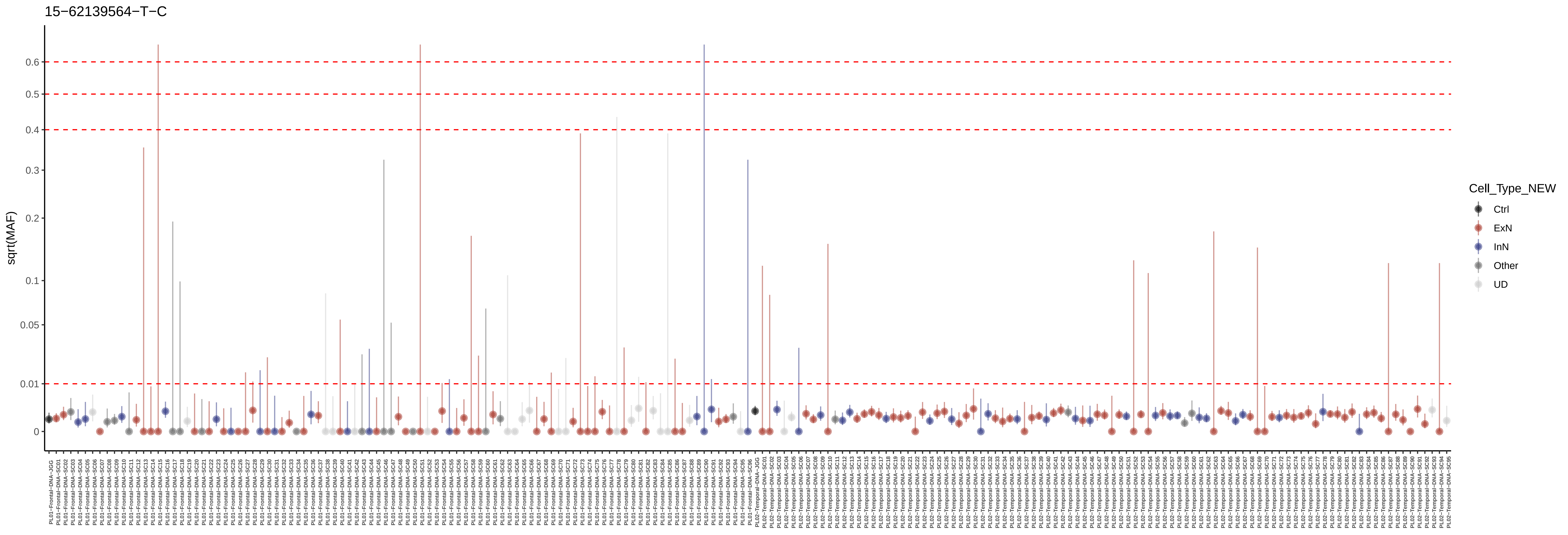
15−62139564−T−C
sqrt(MAF)
0.6
0.5
0.4
0.3
0.2
0.1
0.05
0.01
0
Cell_Type_NEW
Ctrl
ExN
InN
Other
UD

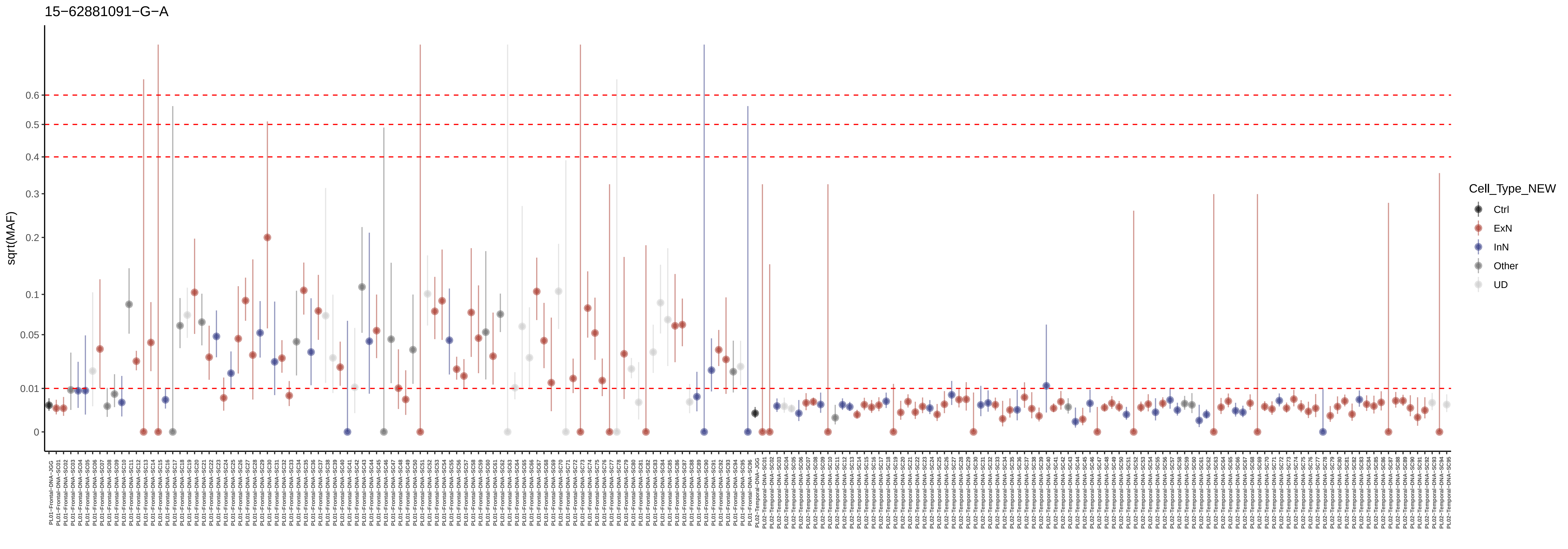
15−62881091−G−A
sqrt(MAF)
0.6
0.5
0.4
0.3
0.2
0.1
0.05
0.01
0
Cell_Type_NEW
Ctrl
ExN
InN
Other
UD

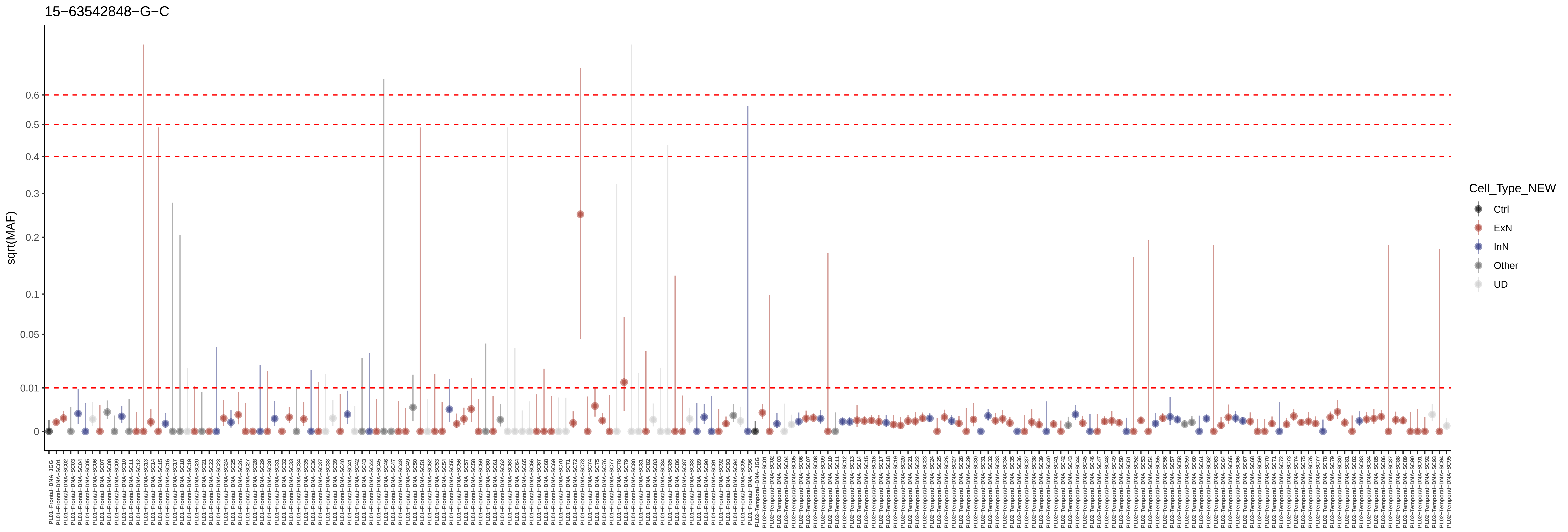
15−63542848−G−C
sqrt(MAF)
0.6
0.5
0.4
0.3
0.2
0.1
0.05
0.01
0
Cell_Type_NEW
Ctrl
ExN
InN
Other
UD

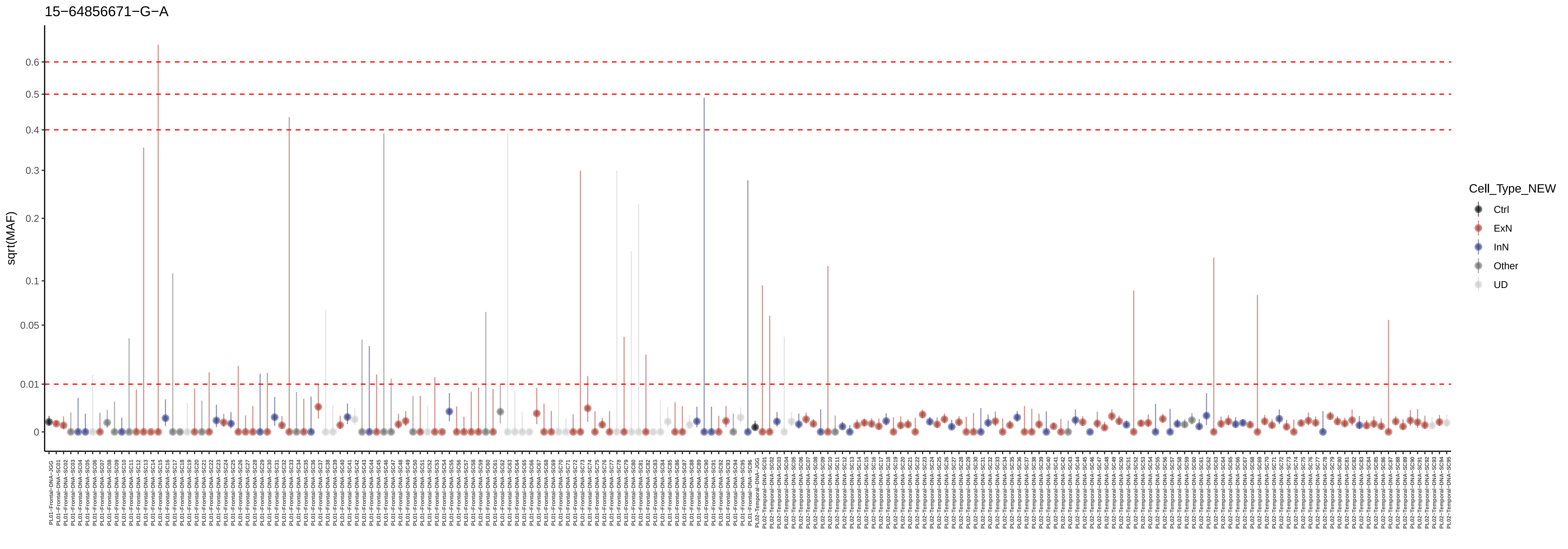
15−64856671−G−A
sqrt(MAF)
0.6
0.5
0.4
0.3
0.2
0.1
0.05
0.01
0
Cell_Type_NEW
Ctrl
ExN
InN
Other
UD

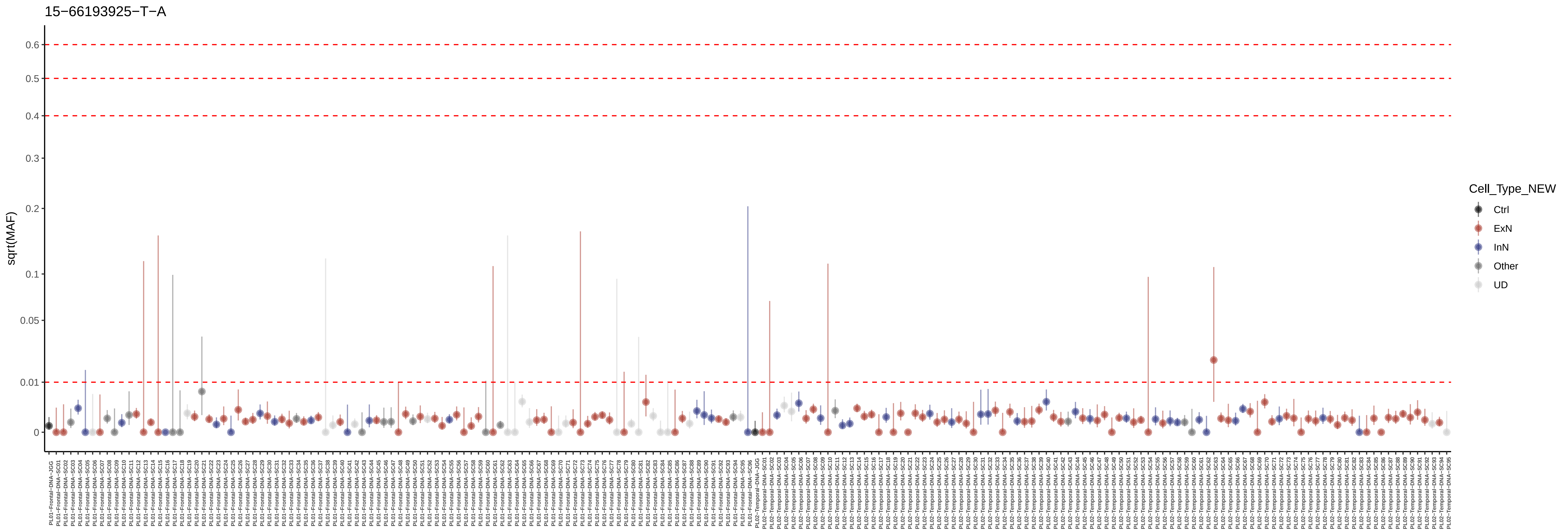

15−66193925−T−A
sqrt(MAF)
0.6
0.5
0.4
0.3
0.2
0.1
0.05
0.01
0
Cell_Type_NEW
Ctrl
ExN
InN
Other
UD

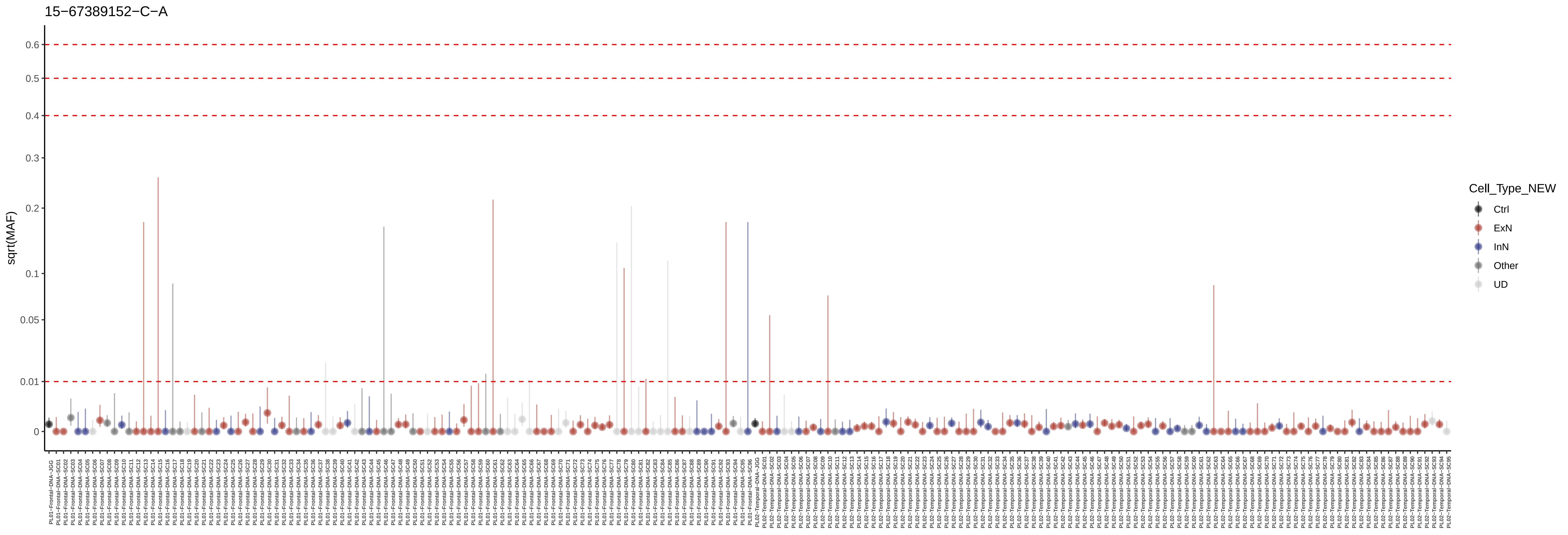
15−67389152−C−A
sqrt(MAF)
0.6
0.5
0.4
0.3
0.2
0.1
0.05
0.01
0
Cell_Type_NEW
Ctrl
ExN
InN
Other
UD

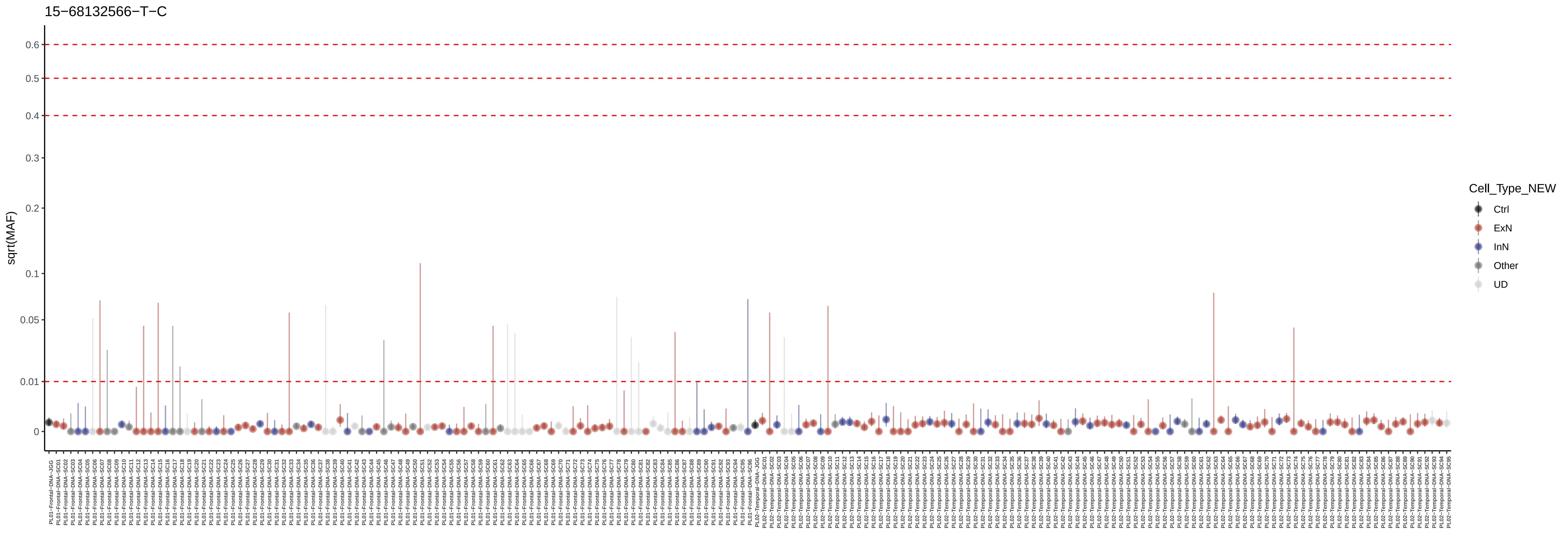
15−68132566−T−C
sqrt(MAF)
0.6
0.5
0.4
0.3
0.2
0.1
0.05
0.01
0
Cell_Type_NEW
Ctrl
ExN
InN
Other
UD

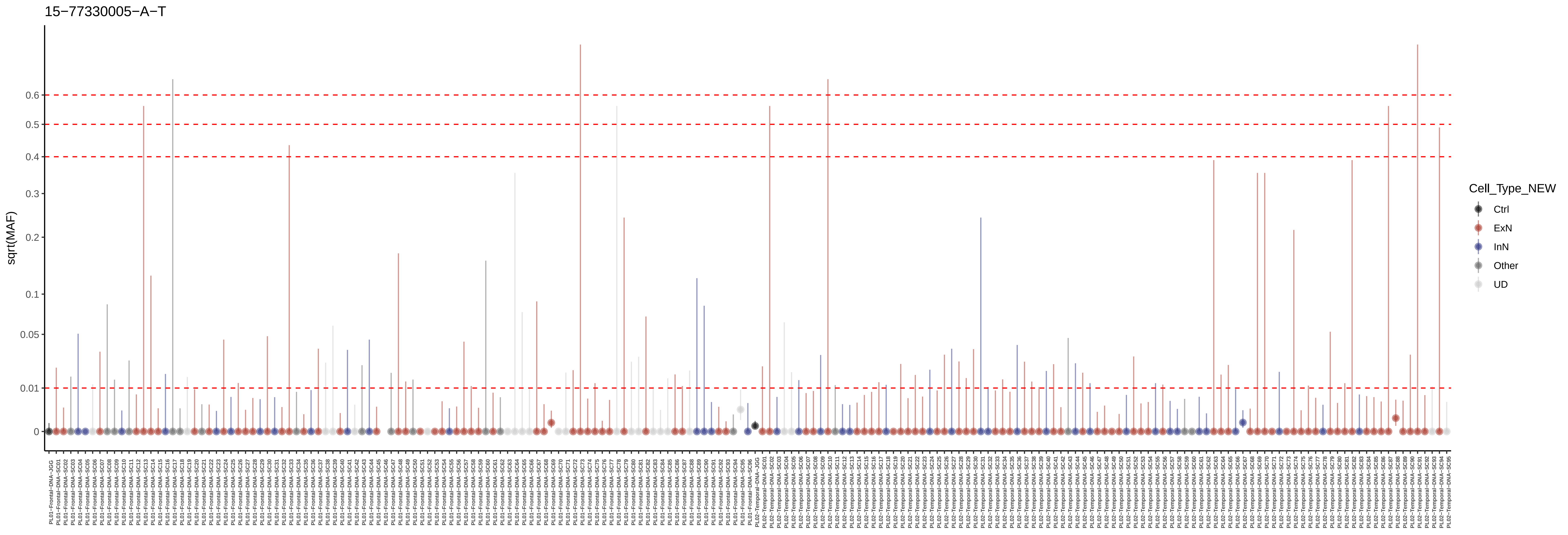
15−77330005−A−T
sqrt(MAF)
0.6
0.5
0.4
0.3
0.2
0.1
0.05
0.01
0
Cell_Type_NEW
Ctrl
ExN
InN
Other
UD

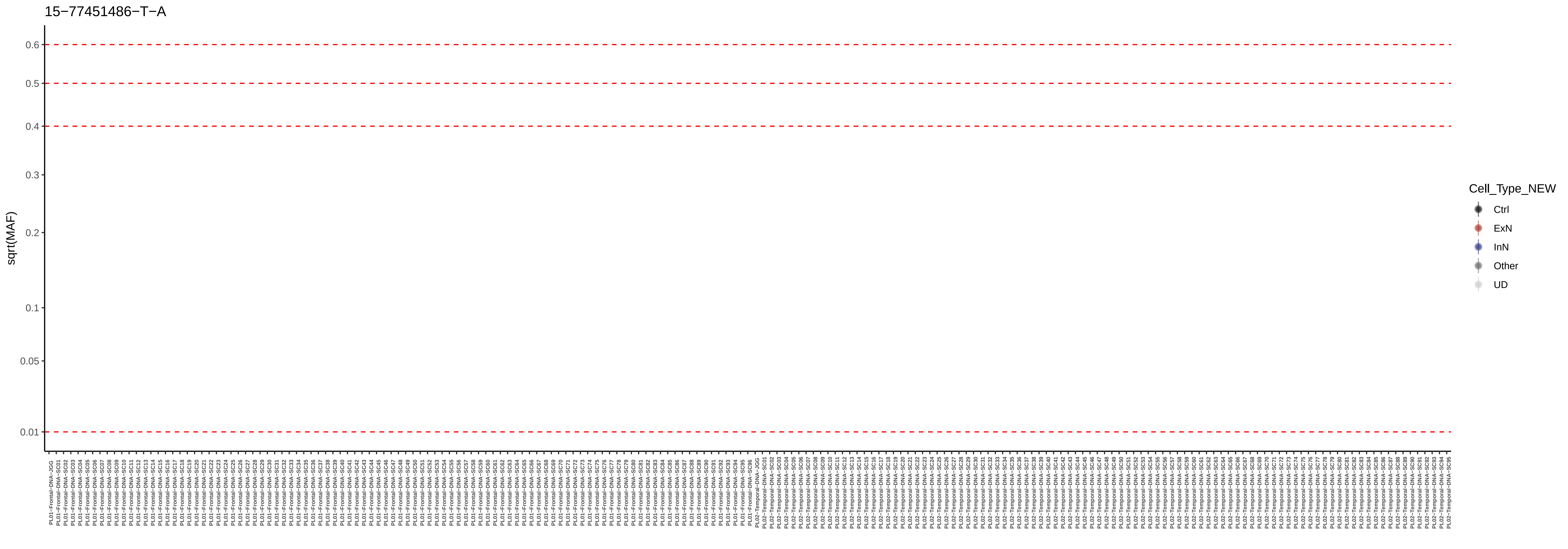

15−77451486−T−A
sqrt(MAF)
Cell_Type_NEW
Ctrl
ExN
InN
Other
UD

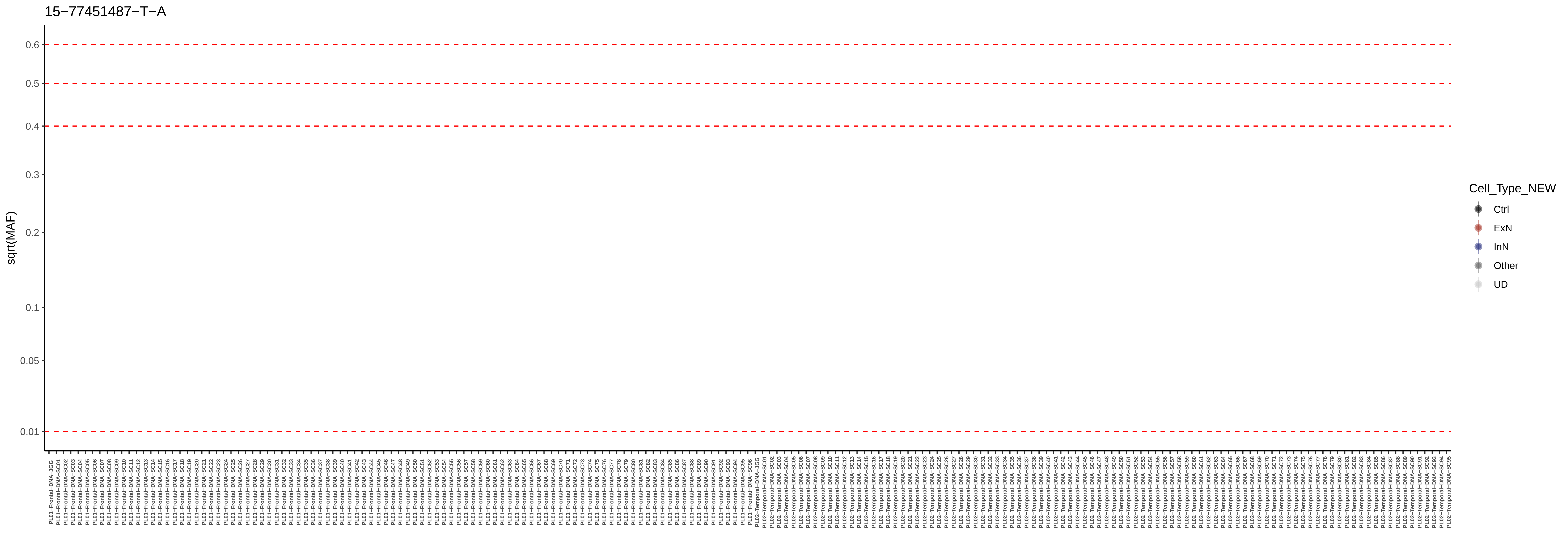
15−77451487−T−A
sqrt(MAF)
0.6
0.5
0.4
0.3
0.2
0.1
0.05
0.01
Cell_Type_NEW
Ctrl
ExN
InN
Other
UD

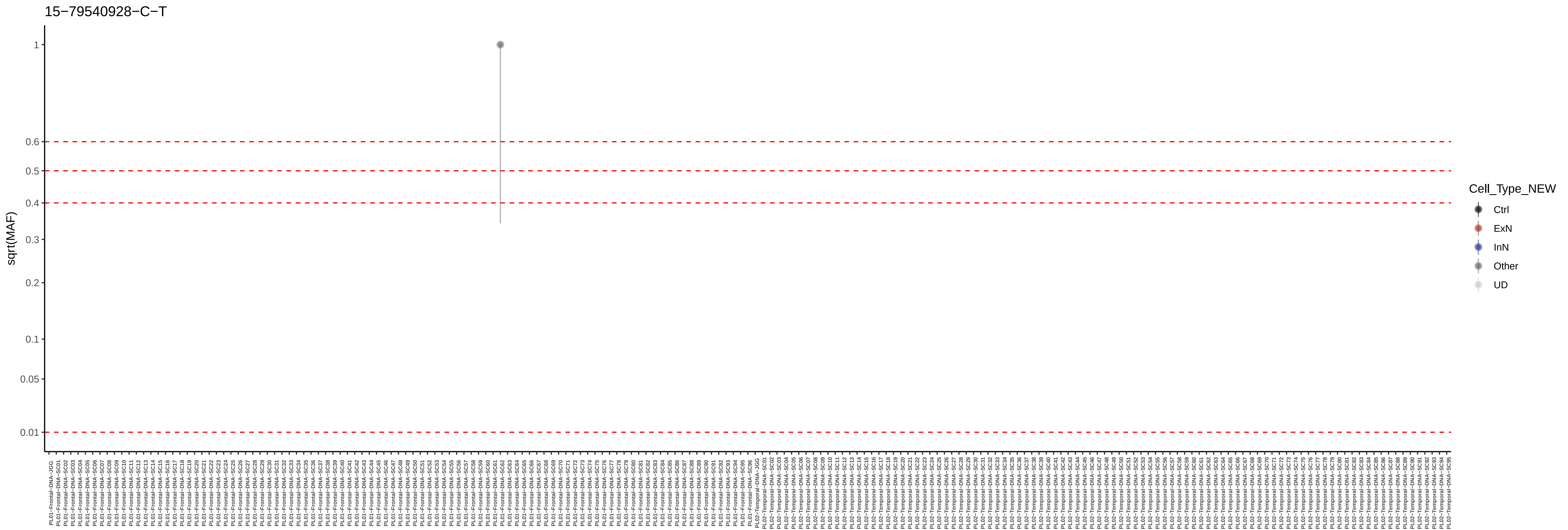

15−79540928−C−T
sqrt(MAF)
1
0.6
0.5
0.4
0.3
0.2
0.1
0.05
0.01
Cell_Type_NEW
Ctrl
ExN
InN
Other
UD

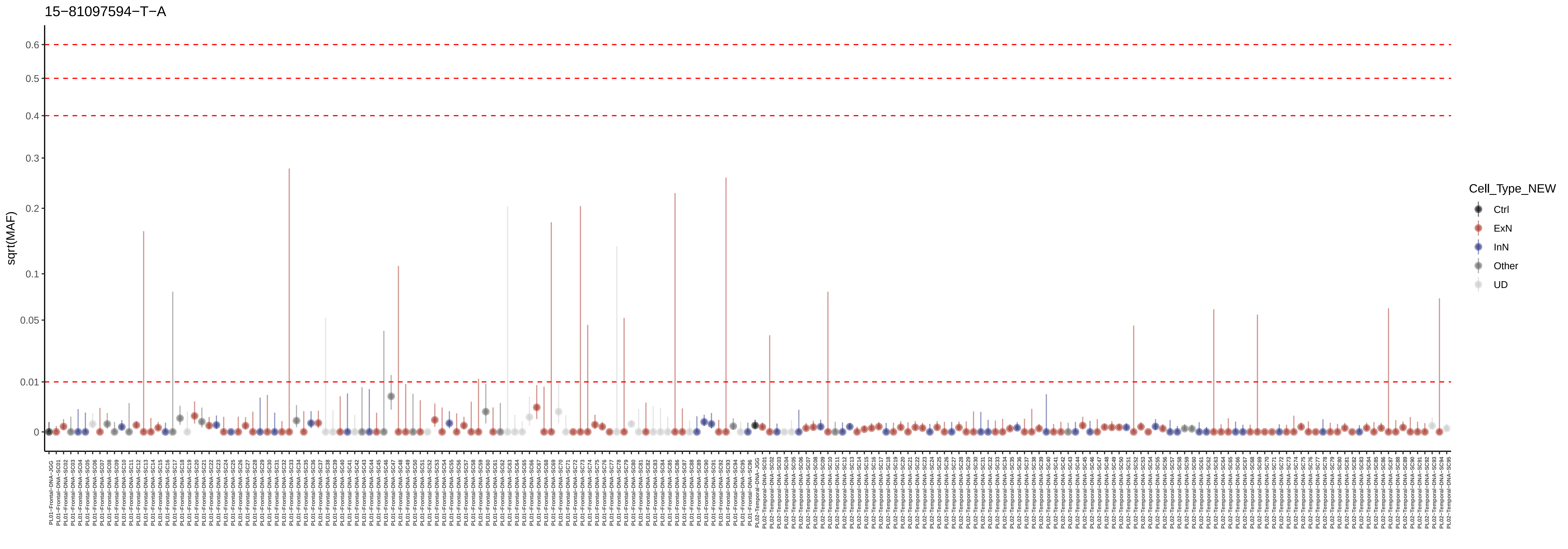

15−81097594−T−A
sqrt(MAF)
0.6
0.5
0.4
0.3
0.2
0.1
0.05
0.01
0
Cell_Type_NEW
Ctrl
ExN
InN
Other
UD

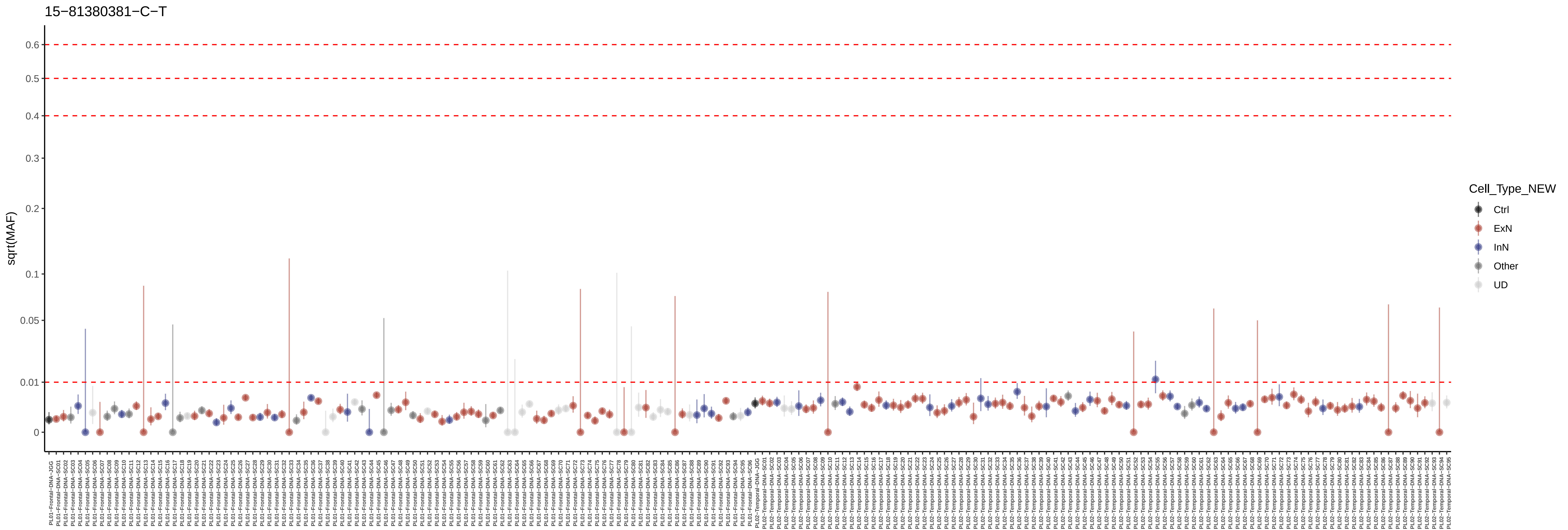
15−81380381−C−T
sqrt(MAF)
0.6
0.5
0.4
0.3
0.2
0.1
0.05
0.01
0
Cell_Type_NEW
Ctrl
ExN
InN
Other
UD

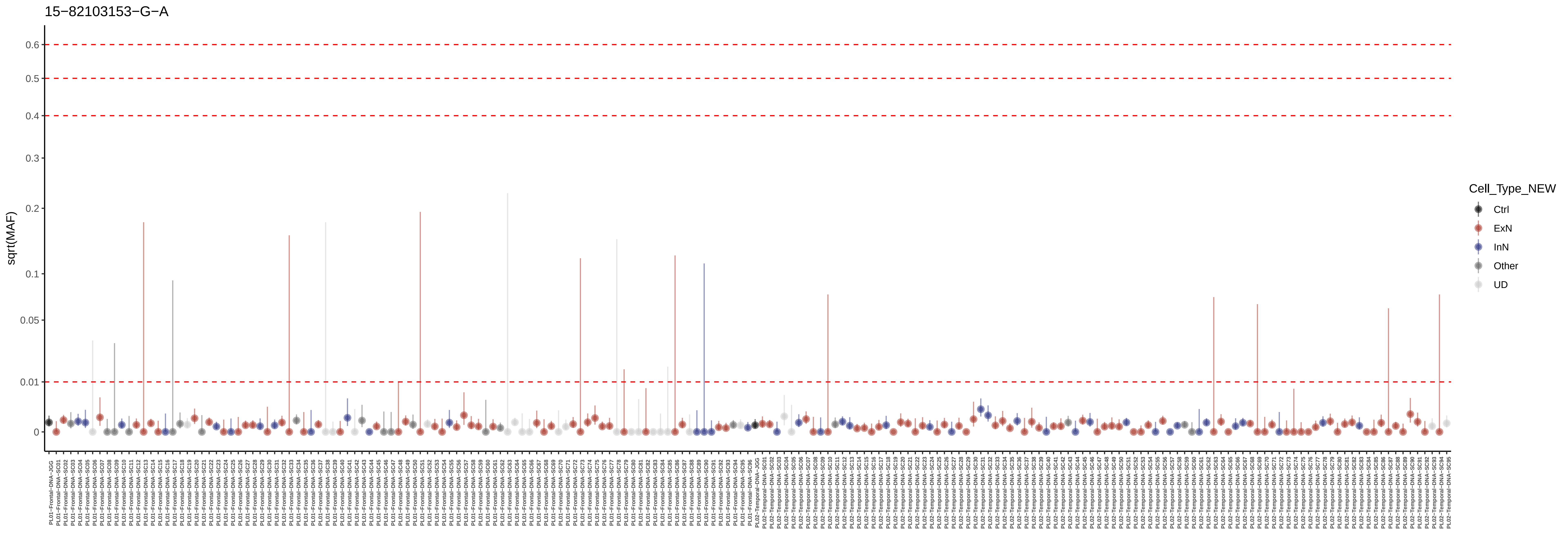

15−82103153−G−A
sqrt(MAF)
0.6
0.5
0.4
0.3
0.2
0.1
0.05
0.01
0
Cell_Type_NEW
Ctrl
ExN
InN
Other
UD

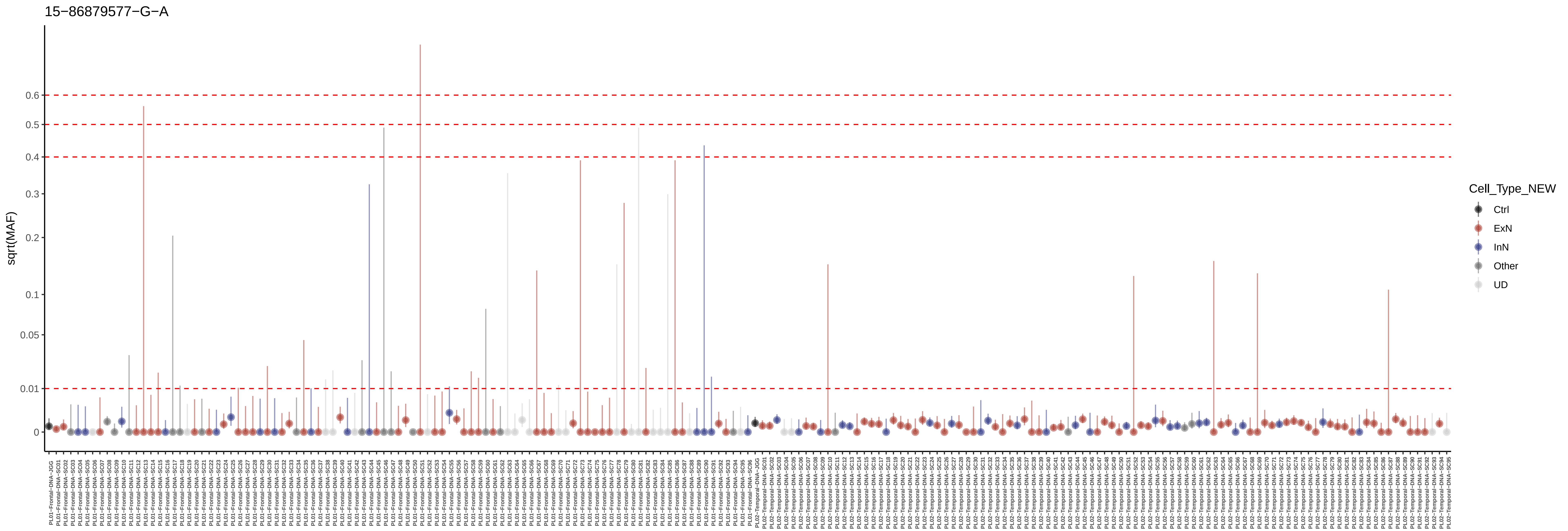

15−86879577−G−A
sqrt(MAF)
0.6
0.5
0.4
0.3
0.2
0.1
0.05
0.01
0
Cell_Type_NEW
Ctrl
ExN
InN
Other
UD

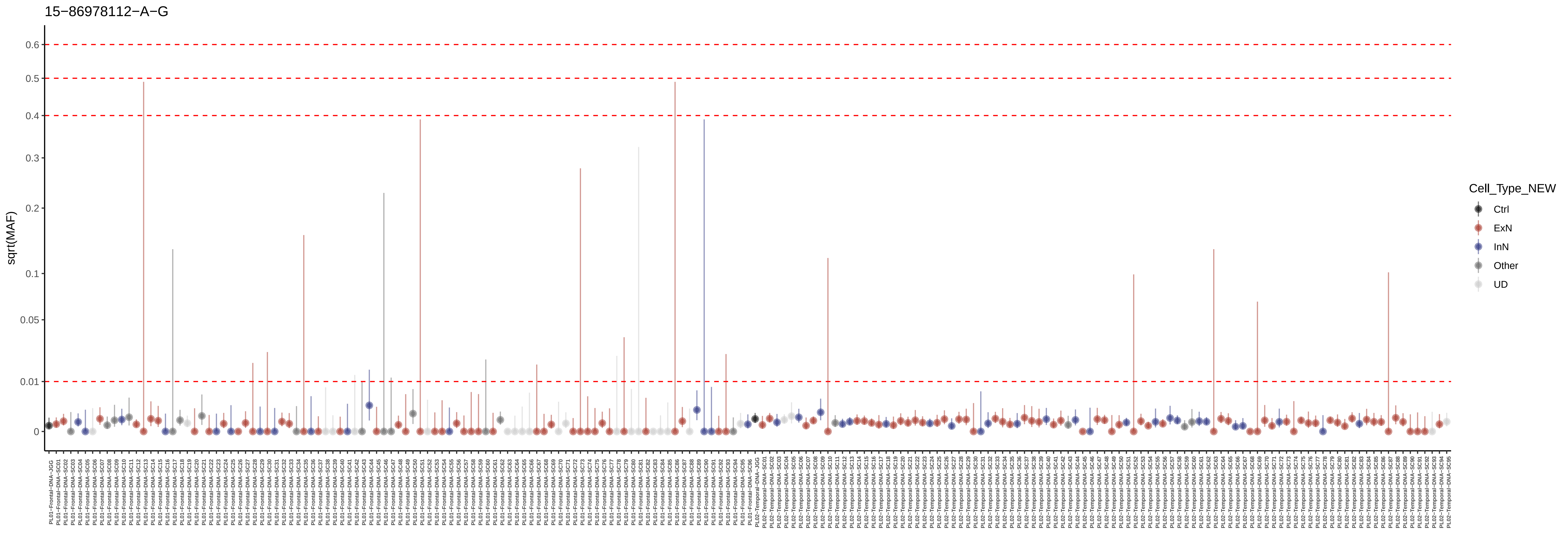
15−86978112−A−G
sqrt(MAF)
0.6
0.5
0.4
0.3
0.2
0.1
0.05
0.01
0
Cell_Type_NEW
Ctrl
ExN
InN
Other
UD

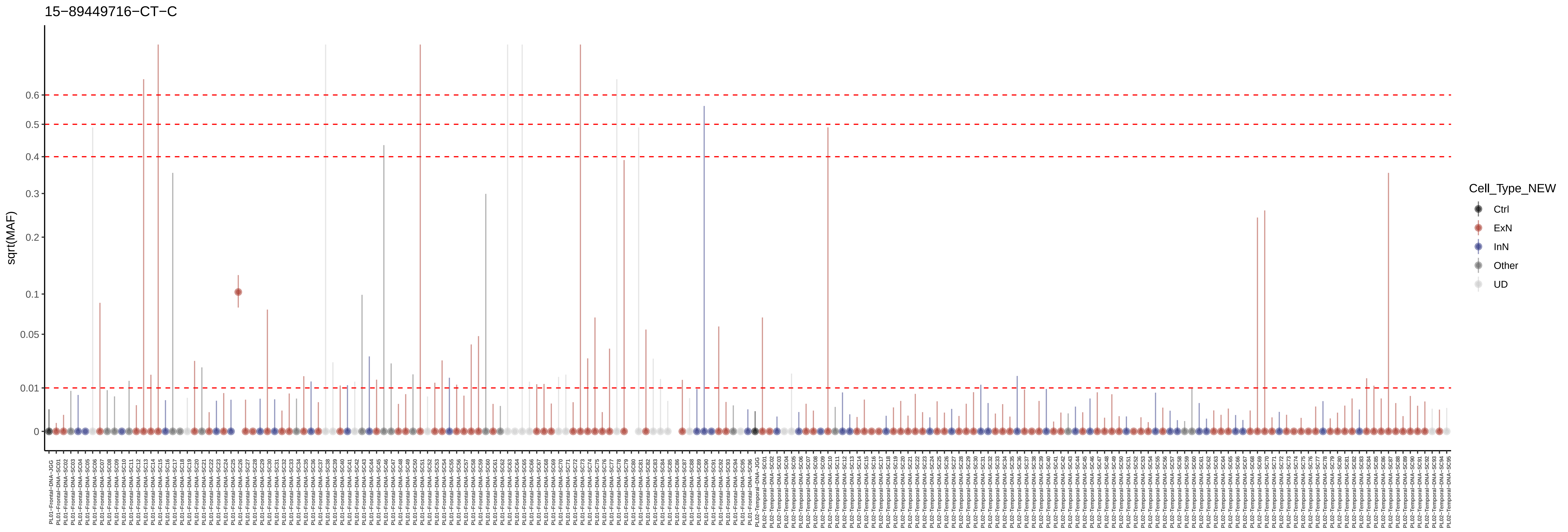
15−89449716−CT−C
sqrt(MAF)
0.6
0.5
0.4
0.3
0.2
0.1
0.05
0.01
0
Cell_Type_NEW
Ctrl
ExN
InN
Other
UD

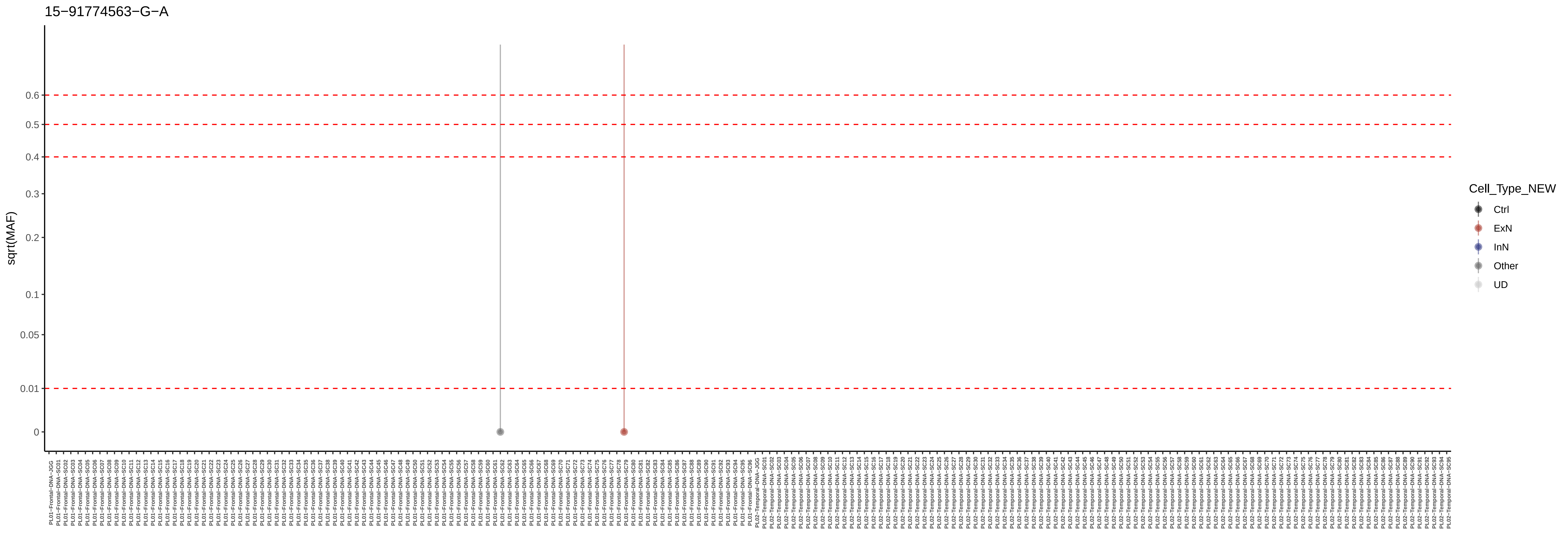

15−91774563−G−A
sqrt(MAF)
0.6
0.5
0.4
0.3
0.2
0.1
0.05
0.01
0
Cell_Type_NEW
Ctrl
ExN
InN
Other
UD

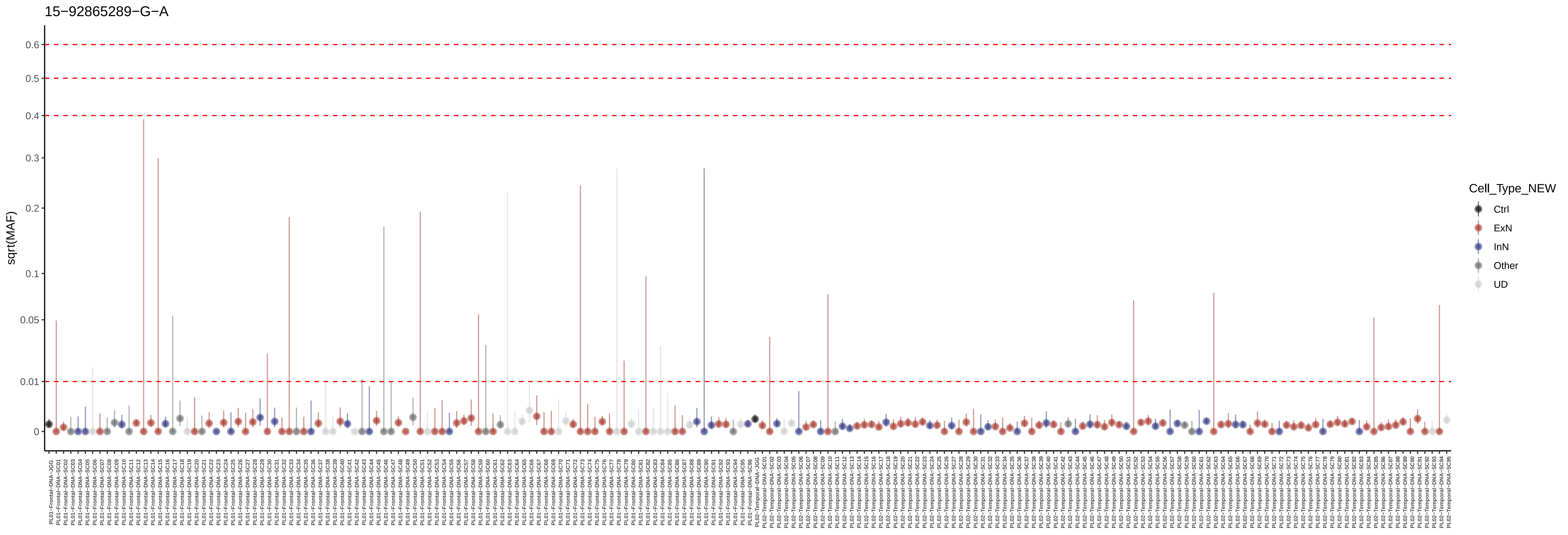
15−92865289−G−A
sqrt(MAF)
0.6
0.5
0.4
0.3
0.2
0.1
0.05
0.01
0
Cell_Type_NEW
Ctrl
ExN
InN
Other
UD

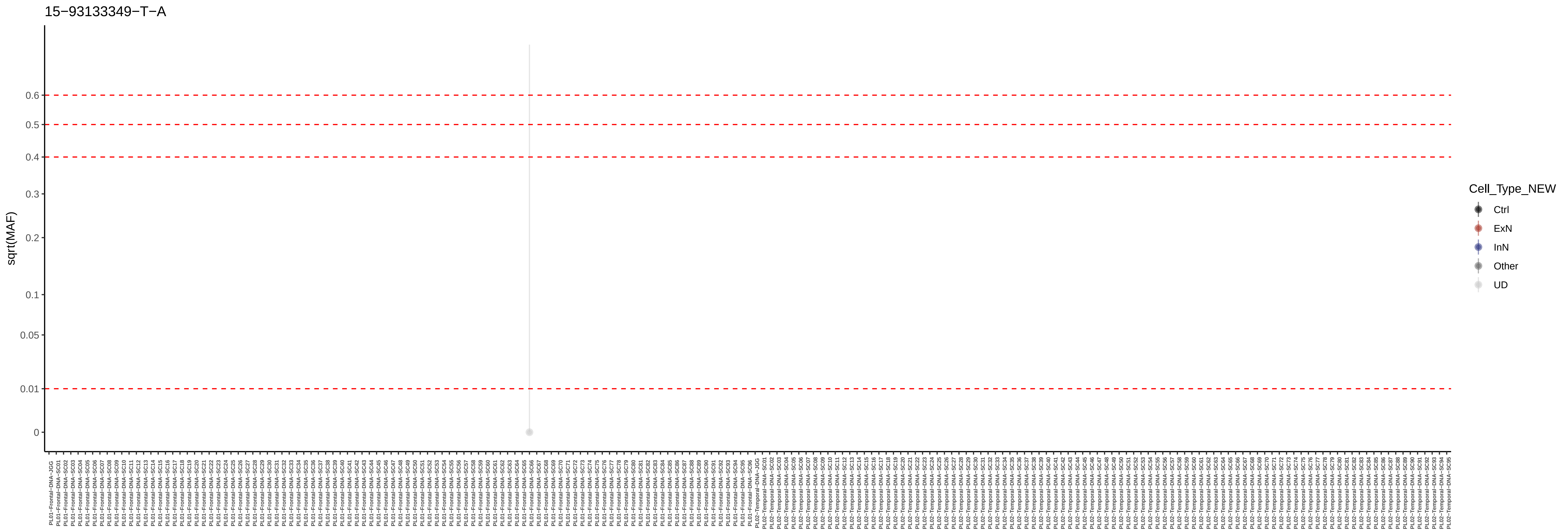
15−93133349−T−A
sqrt(MAF)
0.6
0.5
0.4
0.3
0.2
0.1
0.05
0.01
0
Cell_Type_NEW
Ctrl
ExN
InN
Other
UD

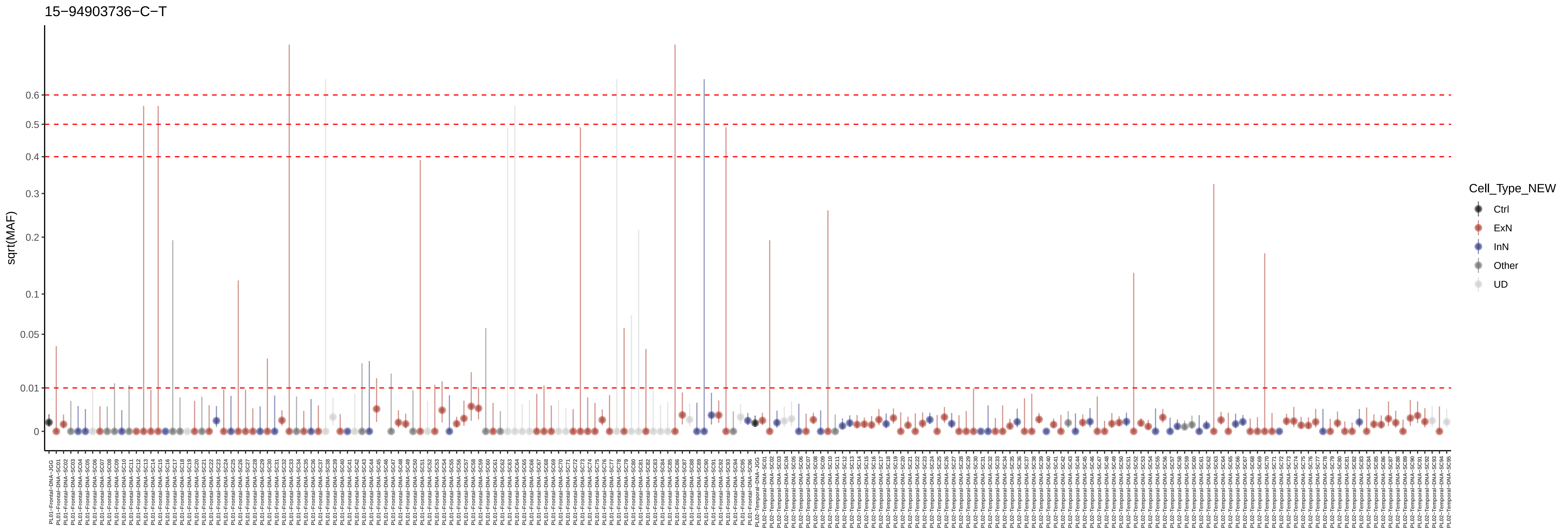
15−94903736−C−T
sqrt(MAF)
0.6
0.5
0.4
0.3
0.2
0.1
0.05
0.01
0
Cell_Type_NEW
Ctrl
ExN
InN
Other
UD

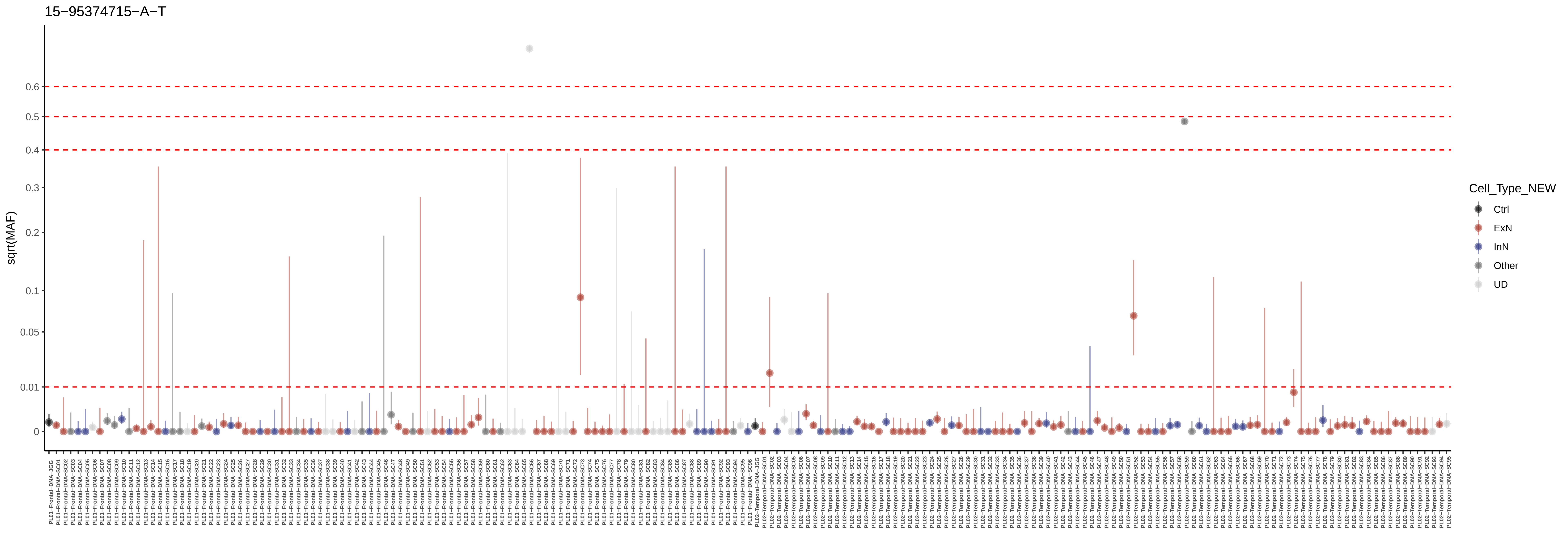
15−95374715−A−T
sqrt(MAF)
0.6
0.5
0.4
0.3
0.2
0.1
0.05
0.01
0
Cell_Type_NEW
Ctrl
ExN
InN
Other
UD

15−95789506−C−T

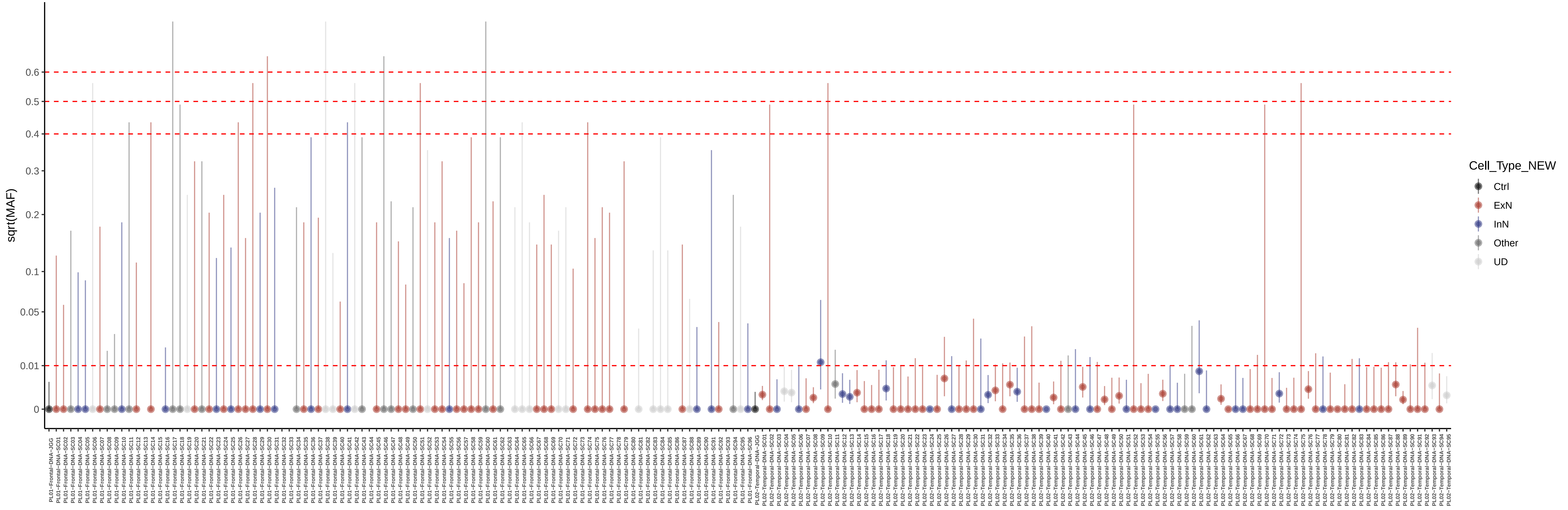

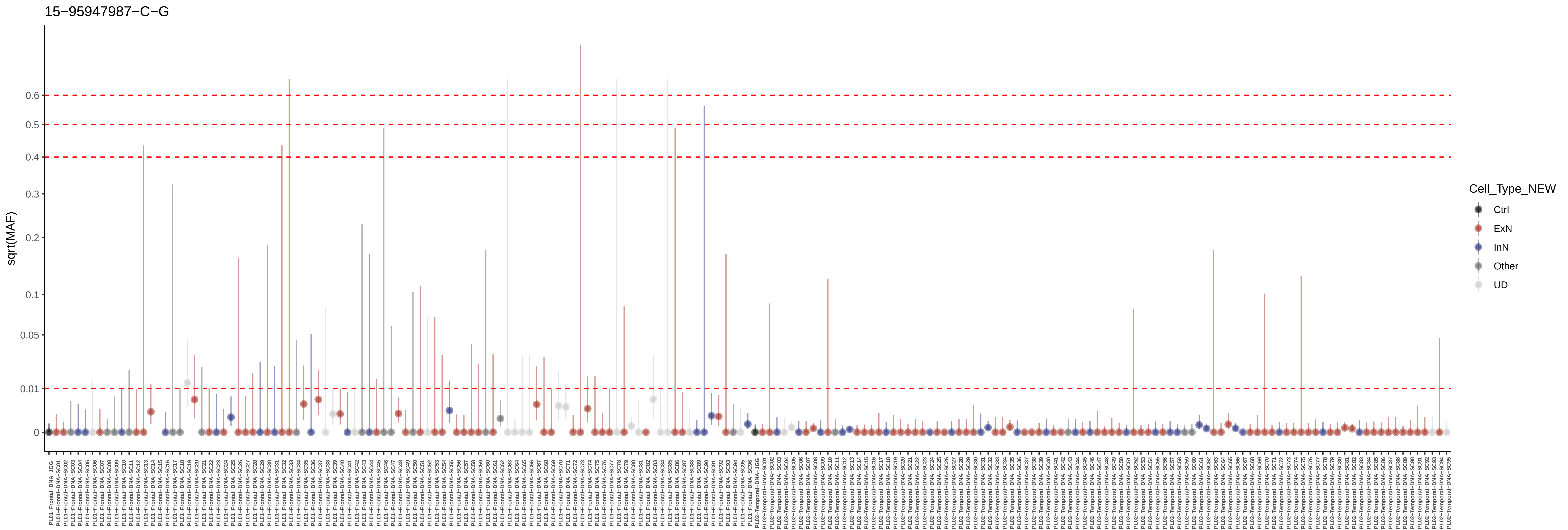

15−95947987−C−G
sqrt(MAF)
0.6
0.5
0.4
0.3
0.2
0.1
0.05
0.01
0
Cell_Type_NEW
Ctrl
ExN
InN
Other
UD

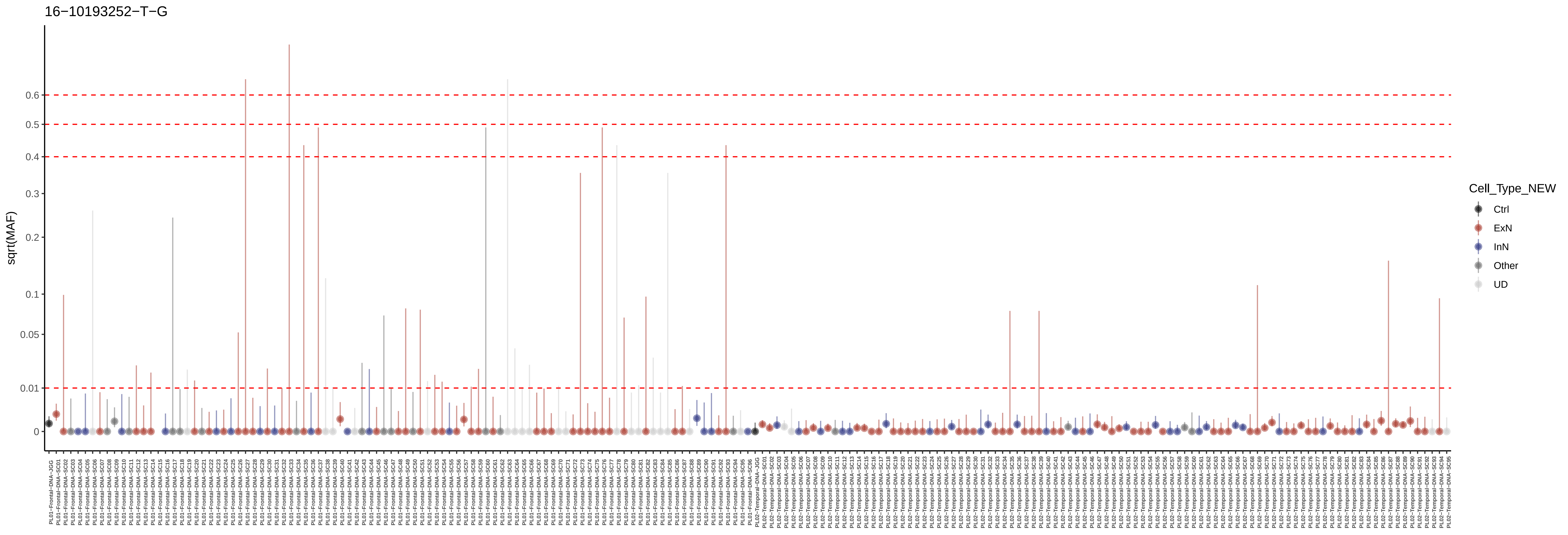

16−10193252−T−G
sqrt(MAF)
0.6
0.5
0.4
0.3
0.2
0.1
0.05
0.01
0
Cell_Type_NEW
Ctrl
ExN
InN
Other
UD

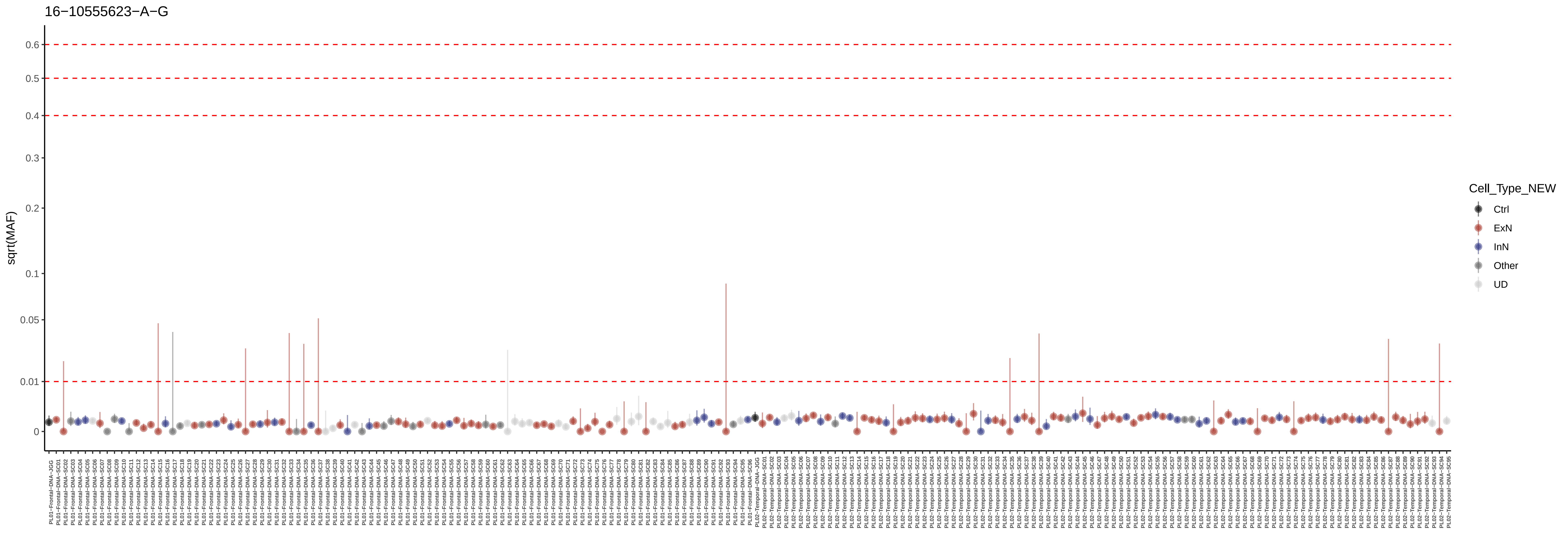
16−10555623−A−G
sqrt(MAF)
0.6
0.5
0.4
0.3
0.2
0.1
0.05
0.01
0
Cell_Type_NEW
Ctrl
ExN
InN
Other
UD

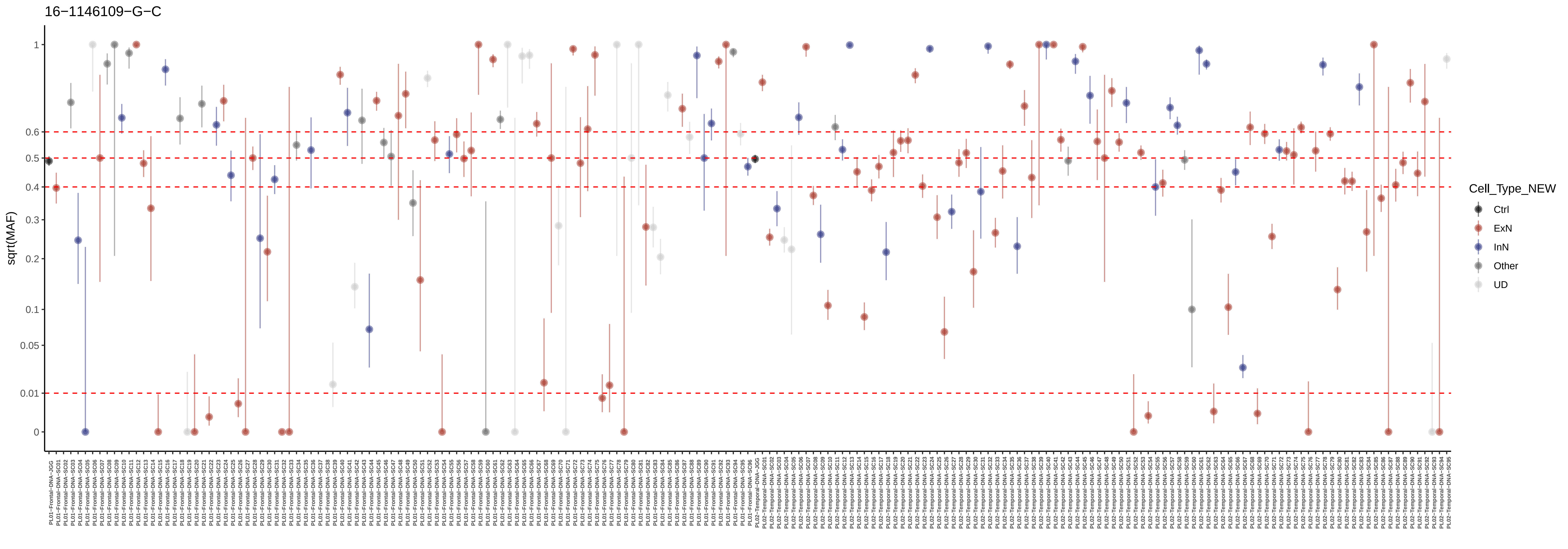
16−1146109−G−C
sqrt(MAF)
1
0.6
0.5
0.4
0.3
0.2
0.1
0.05
0.01
0
Cell_Type_NEW
Ctrl
ExN
InN
Other
UD

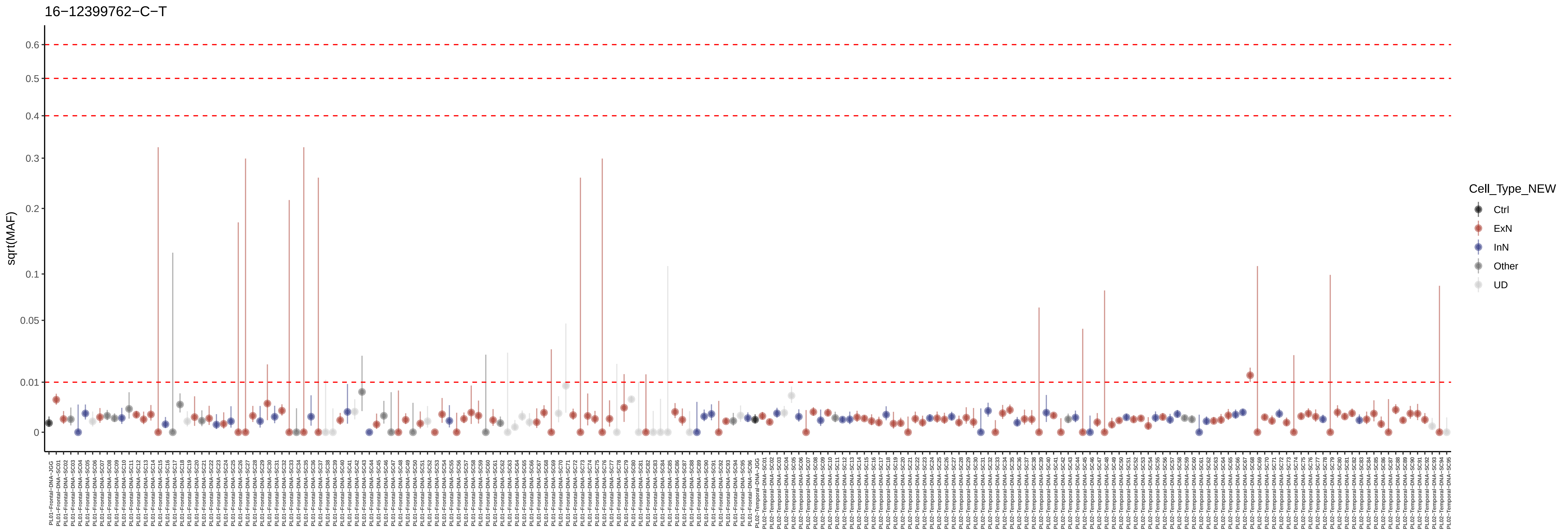
16−12399762−C−T
sqrt(MAF)
0.6
0.5
0.4
0.3
0.2
0.1
0.05
0.01
0
Cell_Type_NEW
Ctrl
ExN
InN
Other
UD

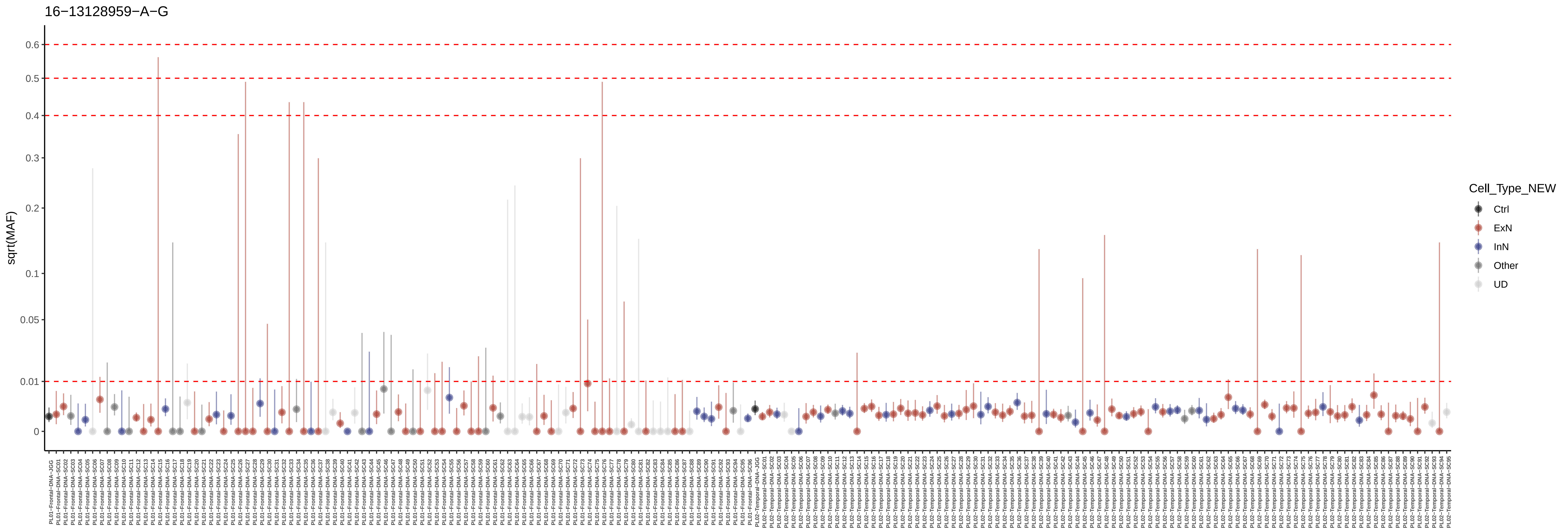

16−13128959−A−G
sqrt(MAF)
0.6
0.5
0.4
0.3
0.2
0.1
0.05
0.01
0
Cell_Type_NEW
Ctrl
ExN
InN
Other
UD

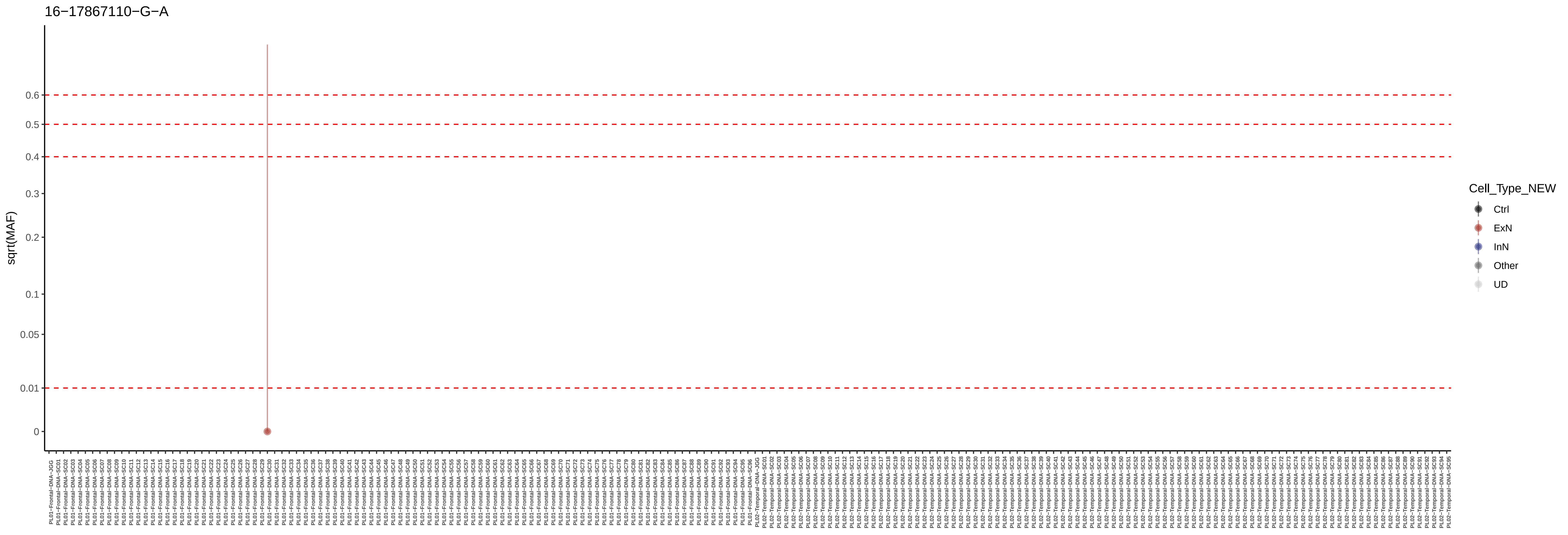
16−17867110−G−A
sqrt(MAF)
0.6
0.5
0.4
0.3
0.2
0.1
0.05
0.01
0
Cell_Type_NEW
Ctrl
ExN
InN
Other
UD

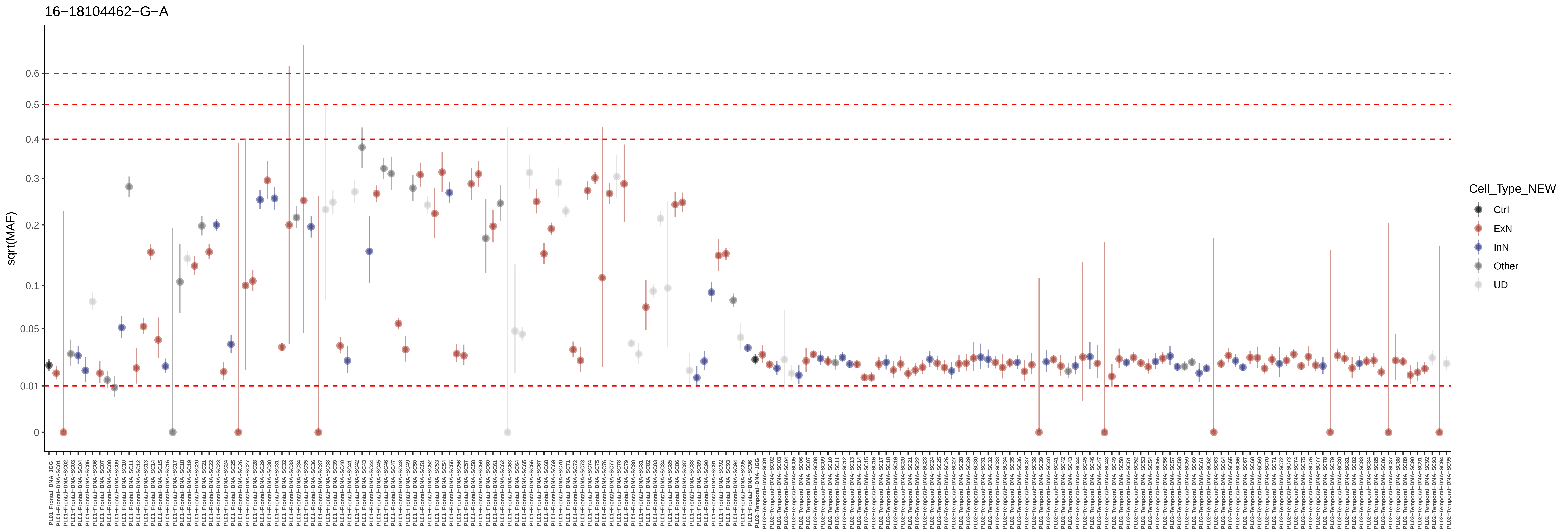
16−18104462−G−A
sqrt(MAF)
0.6
0.5
0.4
0.3
0.2
0.1
0.05
0.01
0
Cell_Type_NEW
Ctrl
ExN
InN
Other
UD

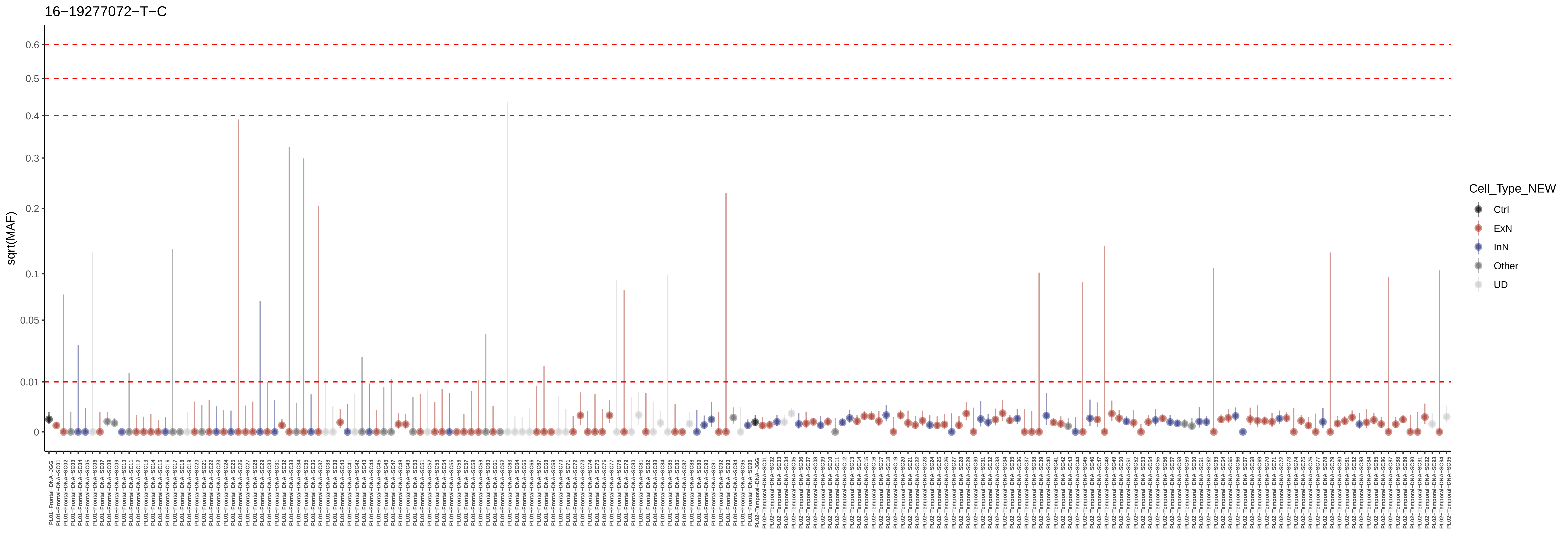
16−19277072−T−C
sqrt(MAF)
0.6
0.5
0.4
0.3
0.2
0.1
0.05
0.01
0
Cell_Type_NEW
Ctrl
ExN
InN
Other
UD

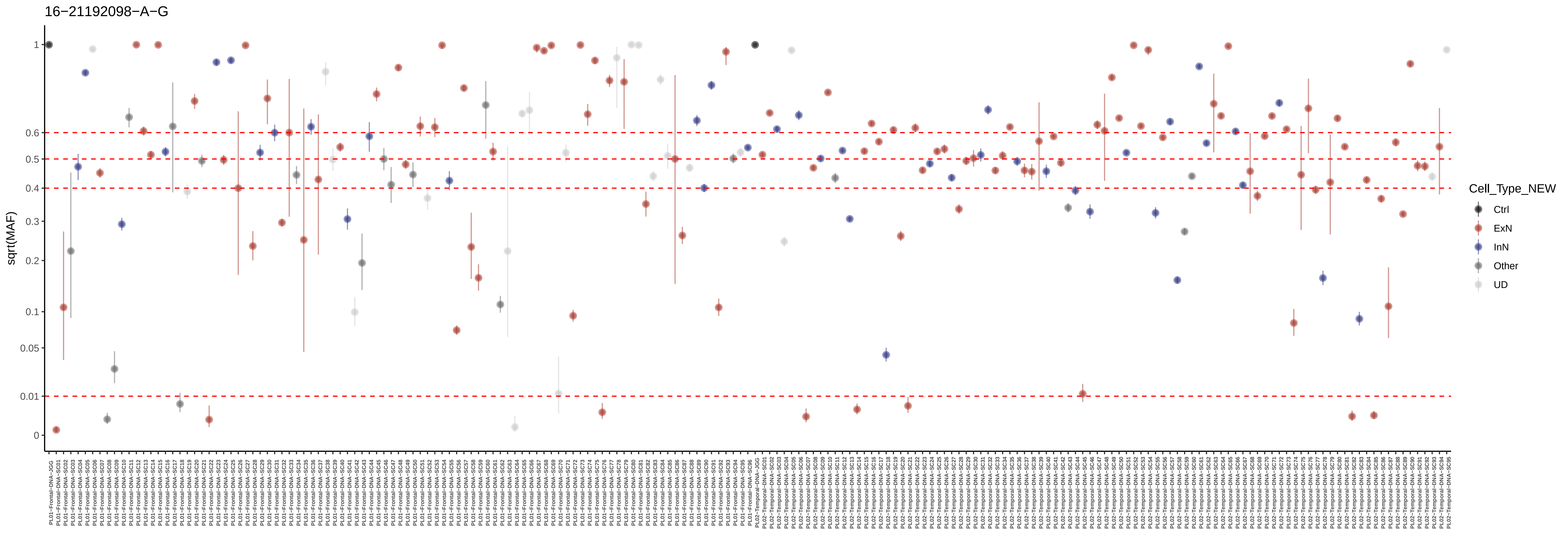
16−21192098−A−G
sqrt(MAF)
1
0.6
0.5
0.4
0.3
0.2
0.1
0.05
0.01
0
Cell_Type_NEW
Ctrl
ExN
InN
Other
UD

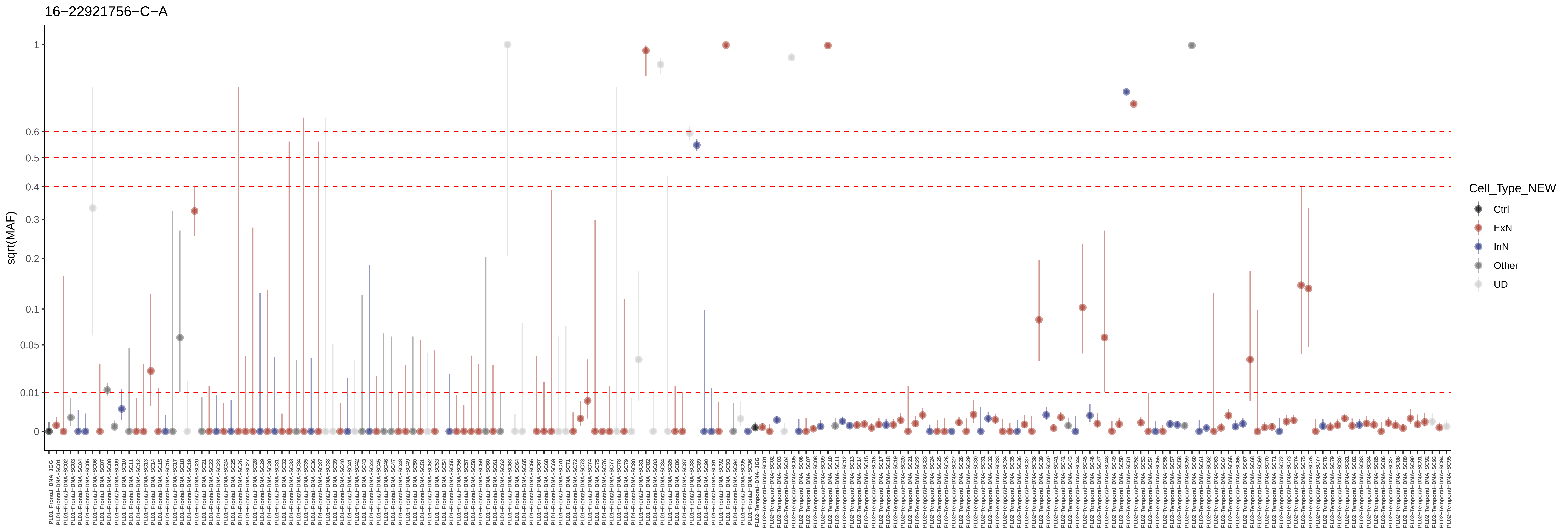
16−22921756−C−A
sqrt(MAF)
1
0.6
0.5
0.4
0.3
0.2
0.1
0.05
0.01
0
Cell_Type_NEW
Ctrl
ExN
InN
Other
UD

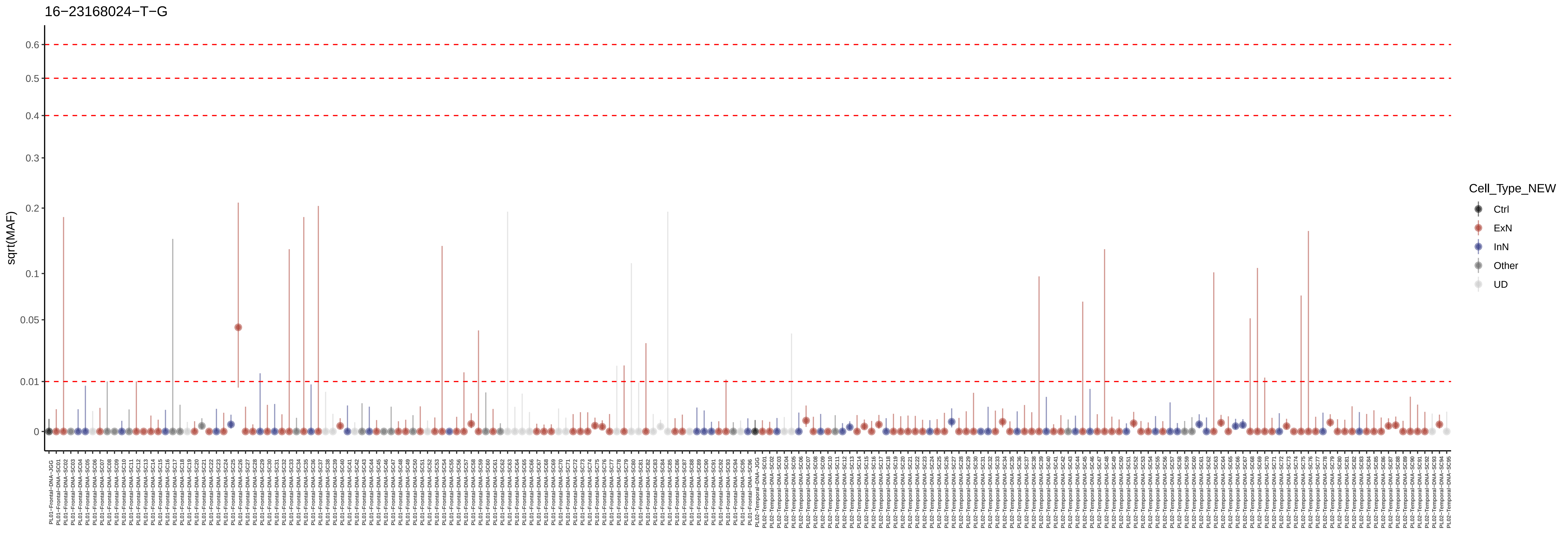

16−23168024−T−G
sqrt(MAF)
0.6
0.5
0.4
0.3
0.2
0.1
0.05
0.01
0
Cell_Type_NEW
Ctrl
ExN
InN
Other
UD

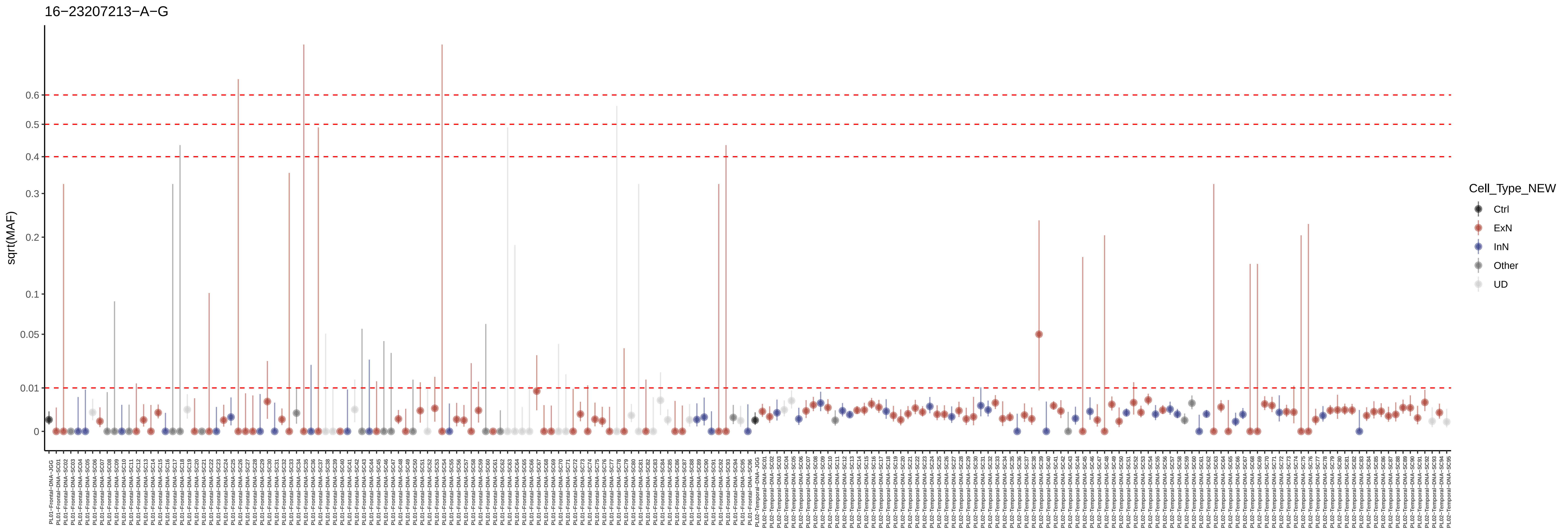
16−23207213−A−G
sqrt(MAF)
0.6
0.5
0.4
0.3
0.2
0.1
0.05
0.01
0
Cell_Type_NEW
Ctrl
ExN
InN
Other
UD

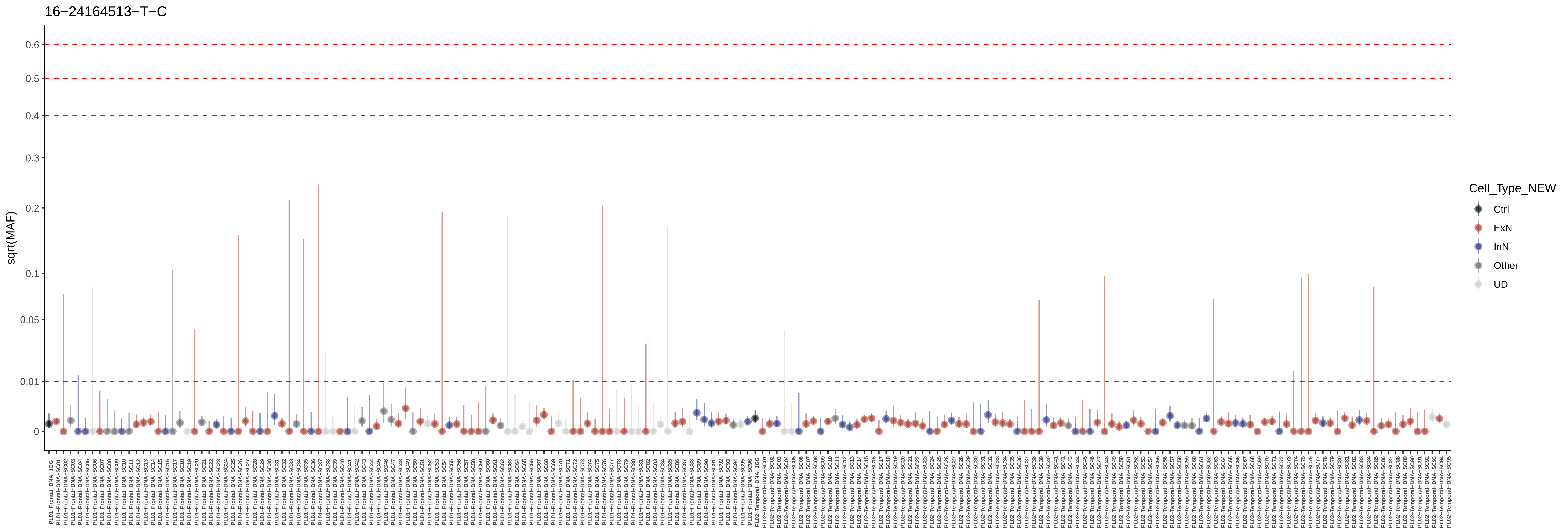
16−24164513−T−C
sqrt(MAF)
0.6
0.5
0.4
0.3
0.2
0.1
0.05
0.01
0
Cell_Type_NEW
Ctrl
ExN
InN
Other
UD

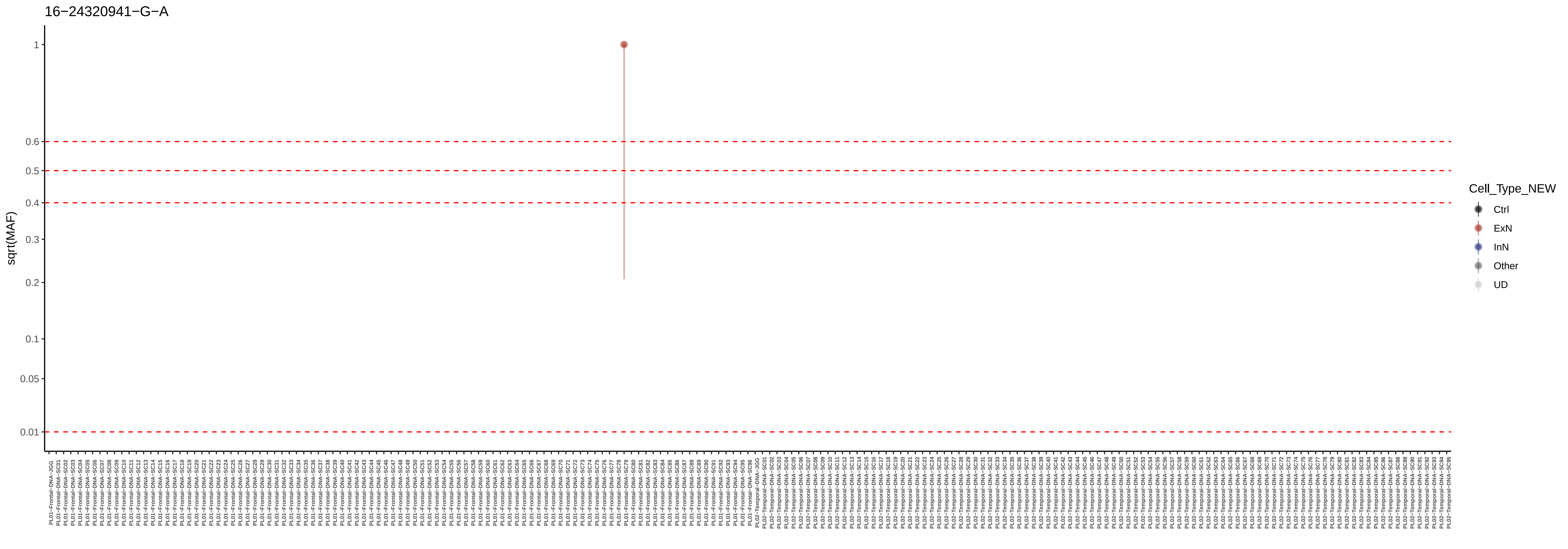
16−24320941−G−A
sqrt(MAF)
1
0.6
0.5
0.4
0.3
0.2
0.1
0.05
0.01
Cell_Type_NEW
Ctrl
ExN
InN
Other
UD

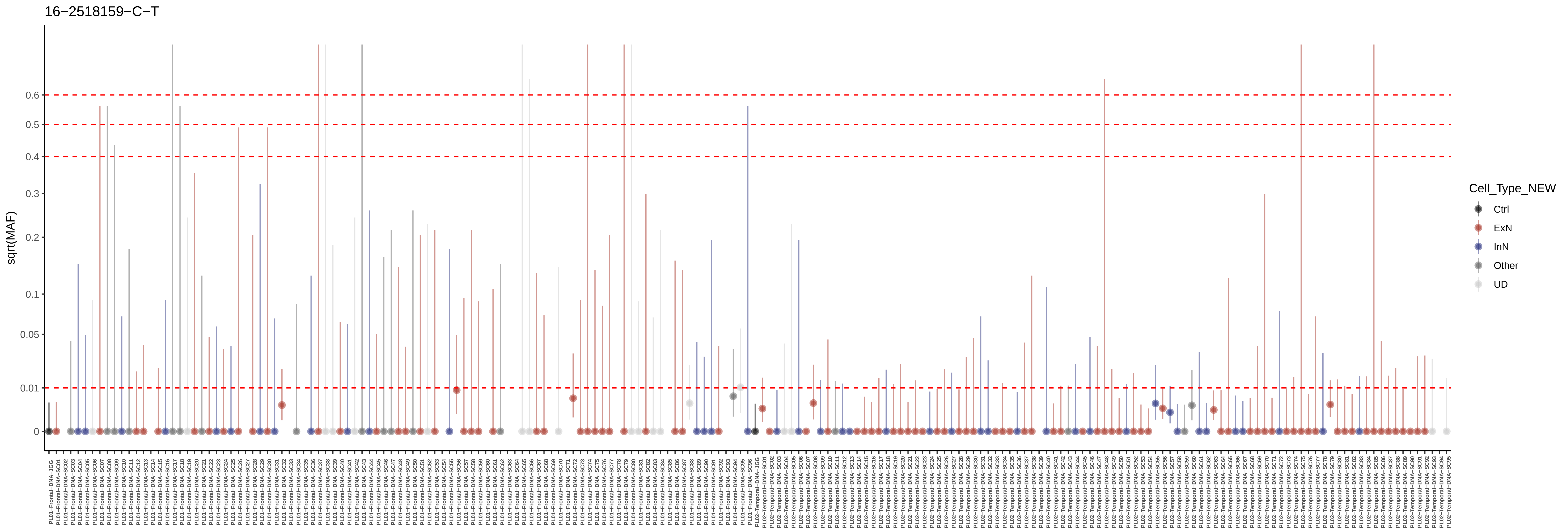
16−2518159−C−T
sqrt(MAF)
0.6
0.5
0.4
0.3
0.2
0.1
0.05
0.01
0
Cell_Type_NEW
Ctrl
ExN
InN
Other
UD

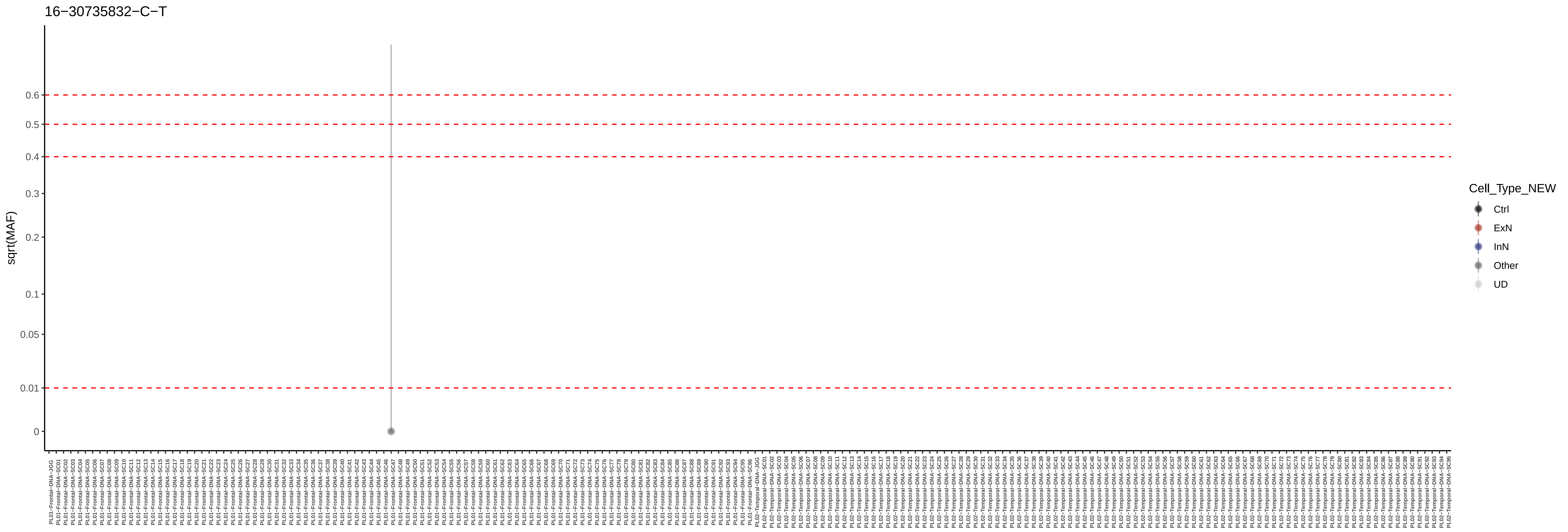
16−30735832−C−T
sqrt(MAF)
0.6
0.5
0.4
0.3
0.2
0.1
0.05
0.01
0
Cell_Type_NEW
Ctrl
ExN
InN
Other
UD

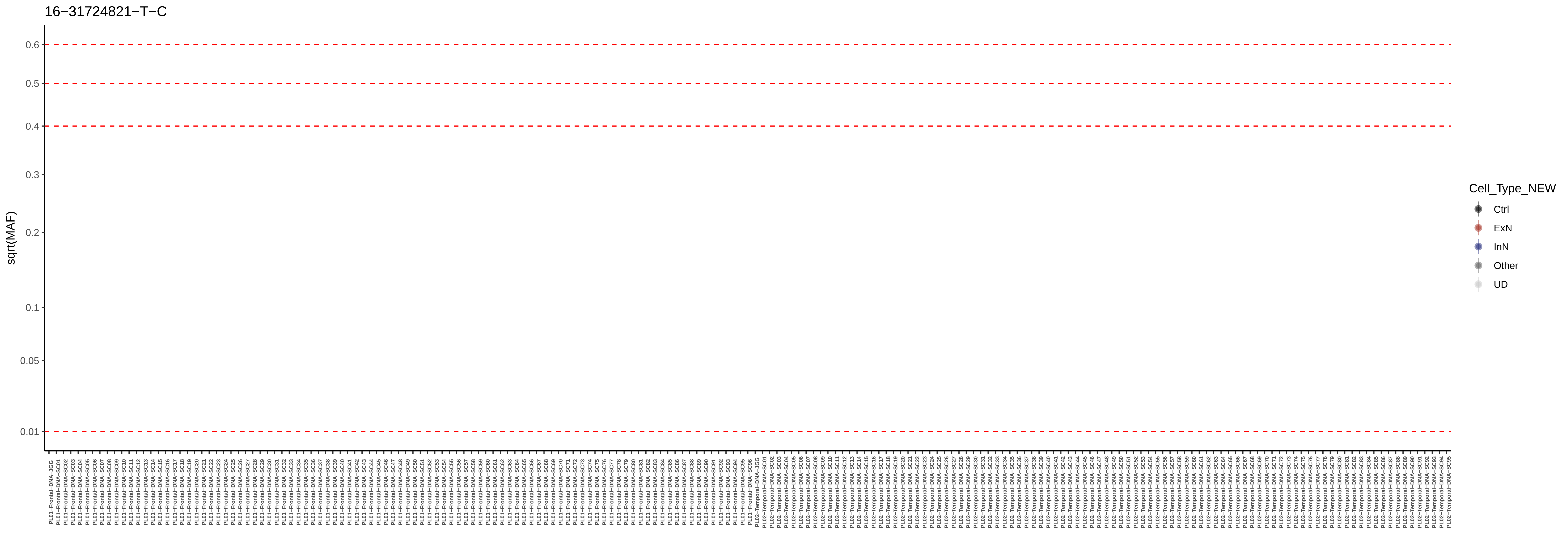
16−31724821−T−C
sqrt(MAF)
0.6
0.5
0.4
0.3
0.2
0.1
0.05
0.01
Cell_Type_NEW
Ctrl
ExN
InN
Other
UD

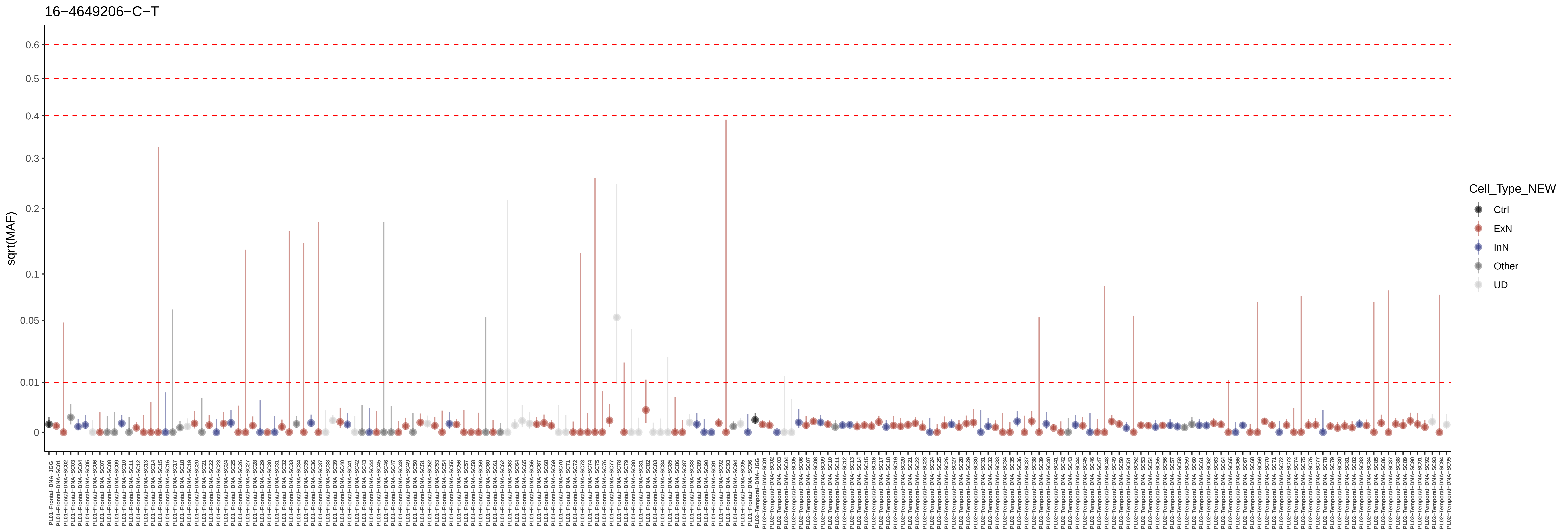

16−4649206−C−T
sqrt(MAF)
0.6
0.5
0.4
0.3
0.2
0.1
0.05
0.01
0
Cell_Type_NEW
Ctrl
ExN
InN
Other
UD

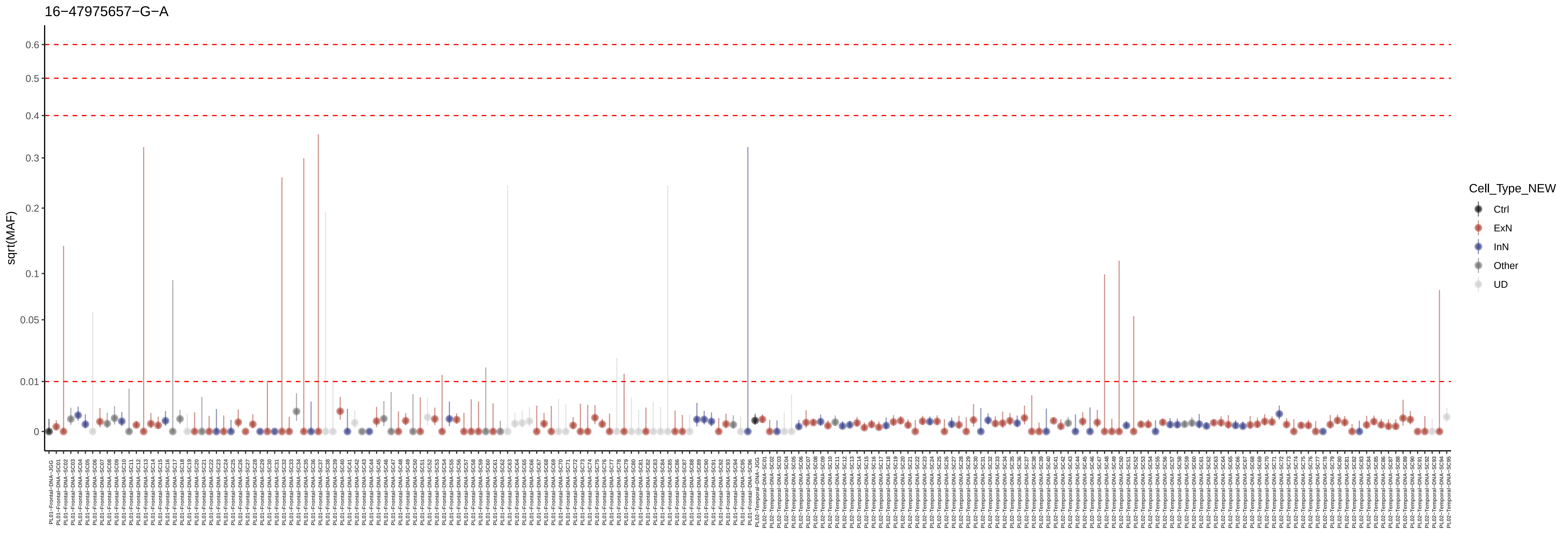
16−47975657−G−A
sqrt(MAF)
0.6
0.5
0.4
0.3
0.2
0.1
0.05
0.01
0
Cell_Type_NEW
Ctrl
ExN
InN
Other
UD

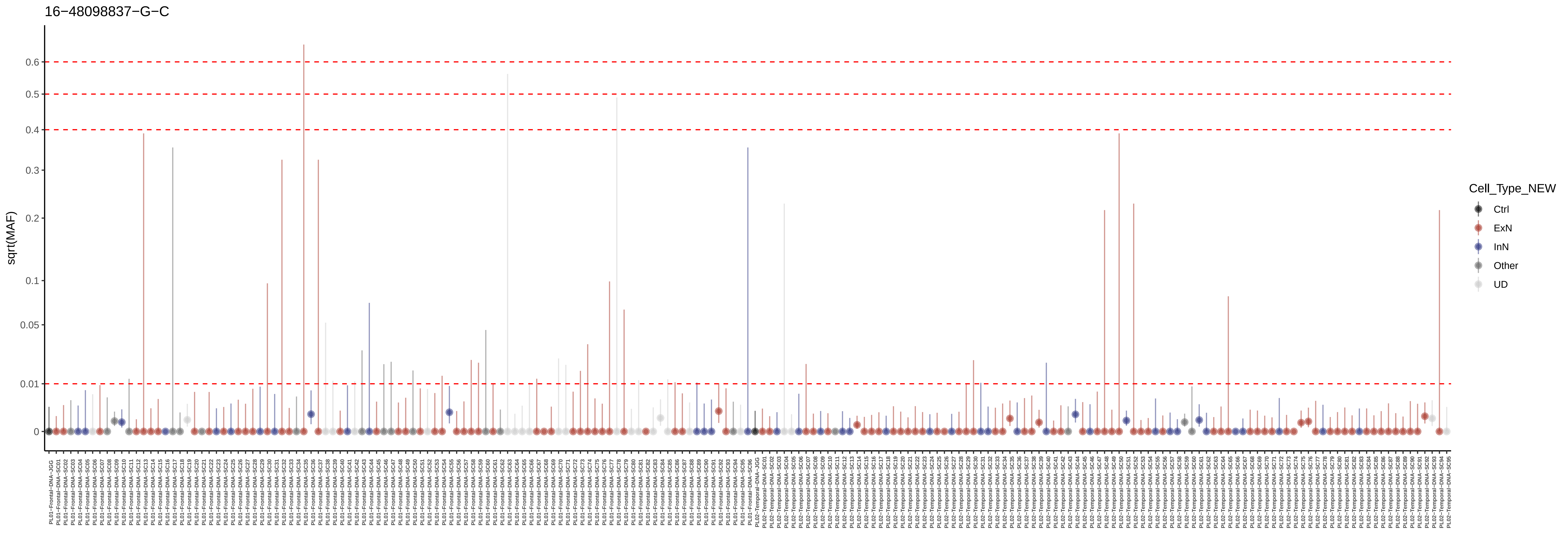
16−48098837−G−C
sqrt(MAF)
0.6
0.5
0.4
0.3
0.2
0.1
0.05
0.01
0
Cell_Type_NEW
Ctrl
ExN
InN
Other
UD

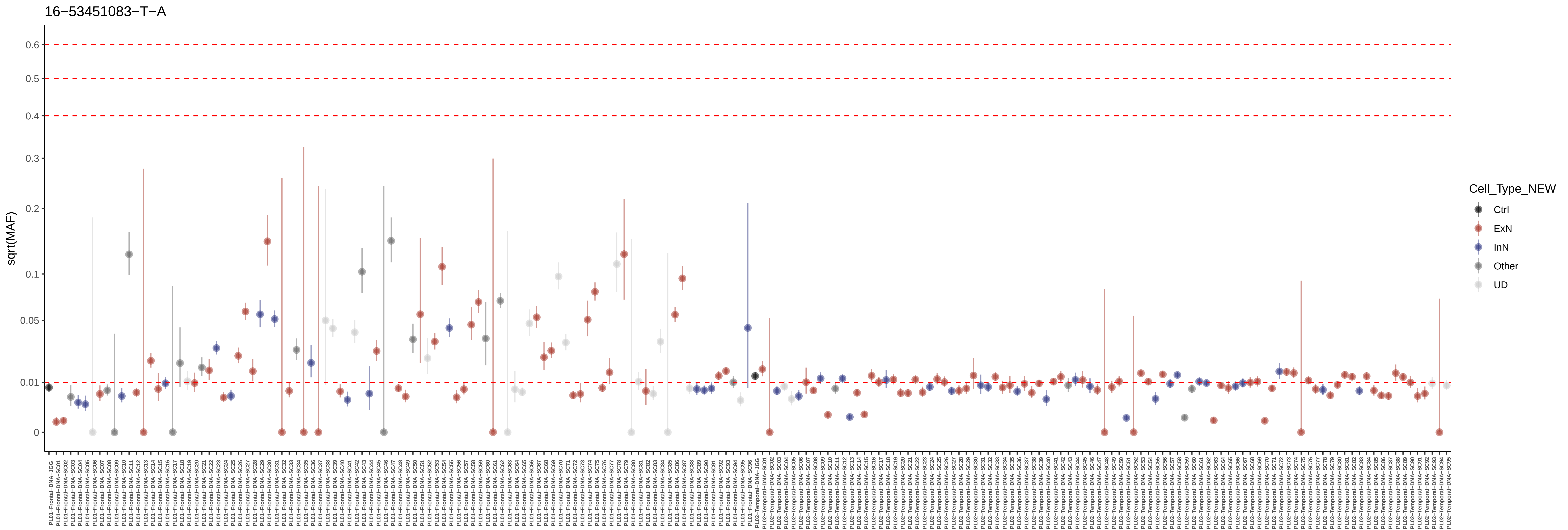
16−53451083−T−A
sqrt(MAF)
0.6
0.5
0.4
0.3
0.2
0.1
0.05
0.01
0
Cell_Type_NEW
Ctrl
ExN
InN
Other
UD

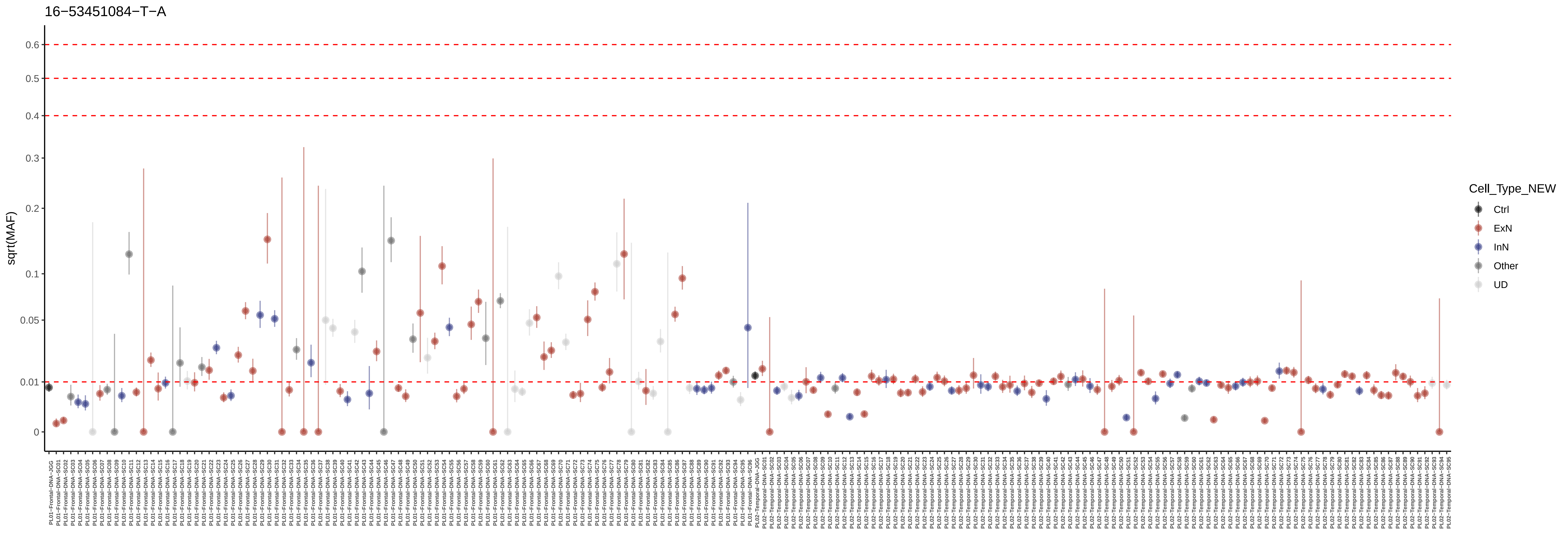

16−53451084−T−A
sqrt(MAF)
0.6
0.5
0.4
0.3
0.2
0.1
0.05
0.01
0
Cell_Type_NEW
Ctrl
ExN
InN
Other
UD

16−5483913−T−A

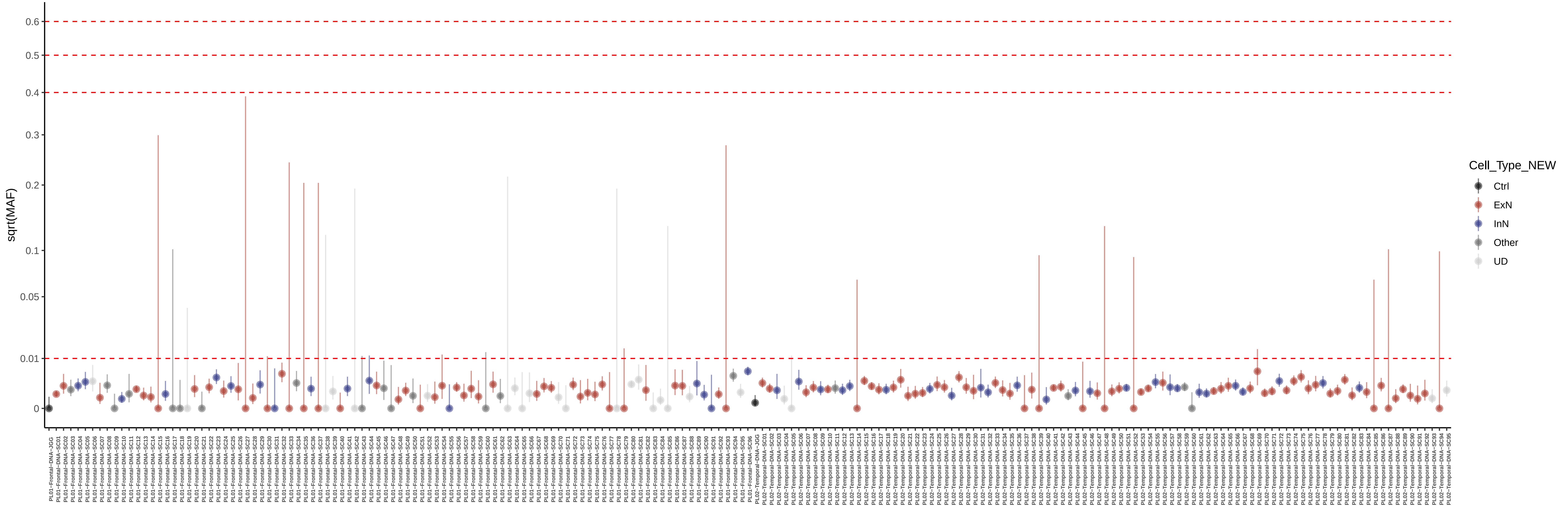

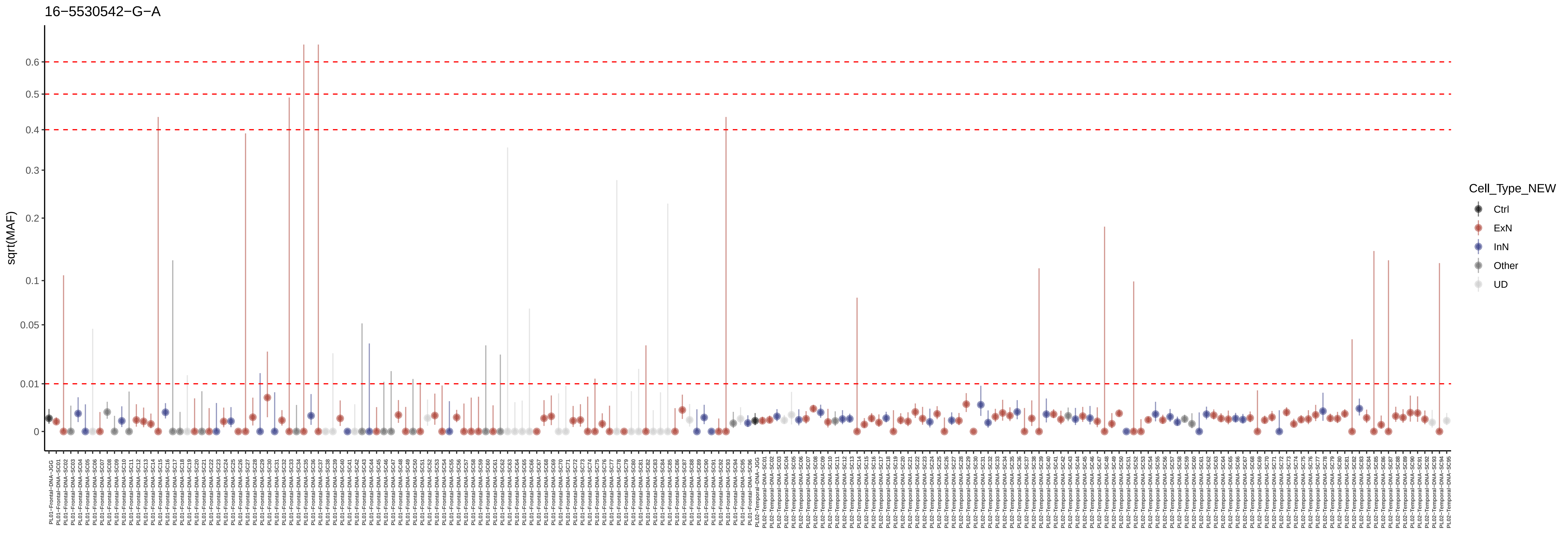
16−5530542−G−A
sqrt(MAF)
0.6
0.5
0.4
0.3
0.2
0.1
0.05
0.01
0
Cell_Type_NEW
Ctrl
ExN
InN
Other
UD

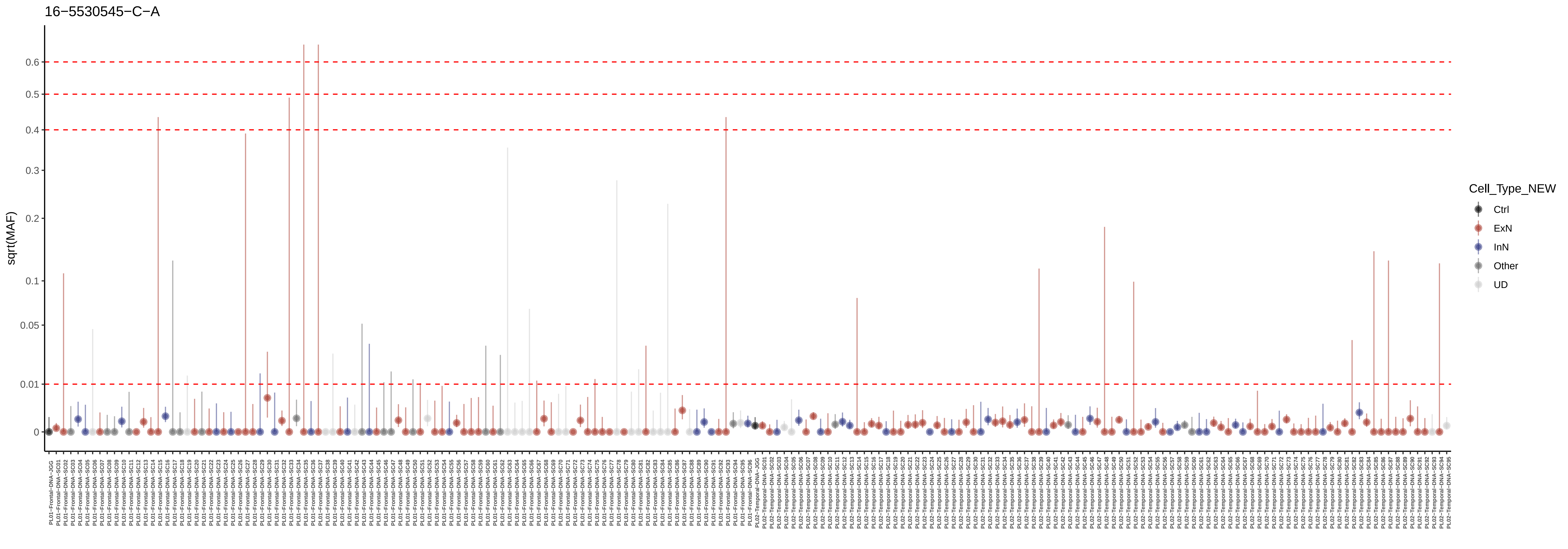
16−5530545−C−A
sqrt(MAF)
0.6
0.5
0.4
0.3
0.2
0.1
0.05
0.01
0
Cell_Type_NEW
Ctrl
ExN
InN
Other
UD

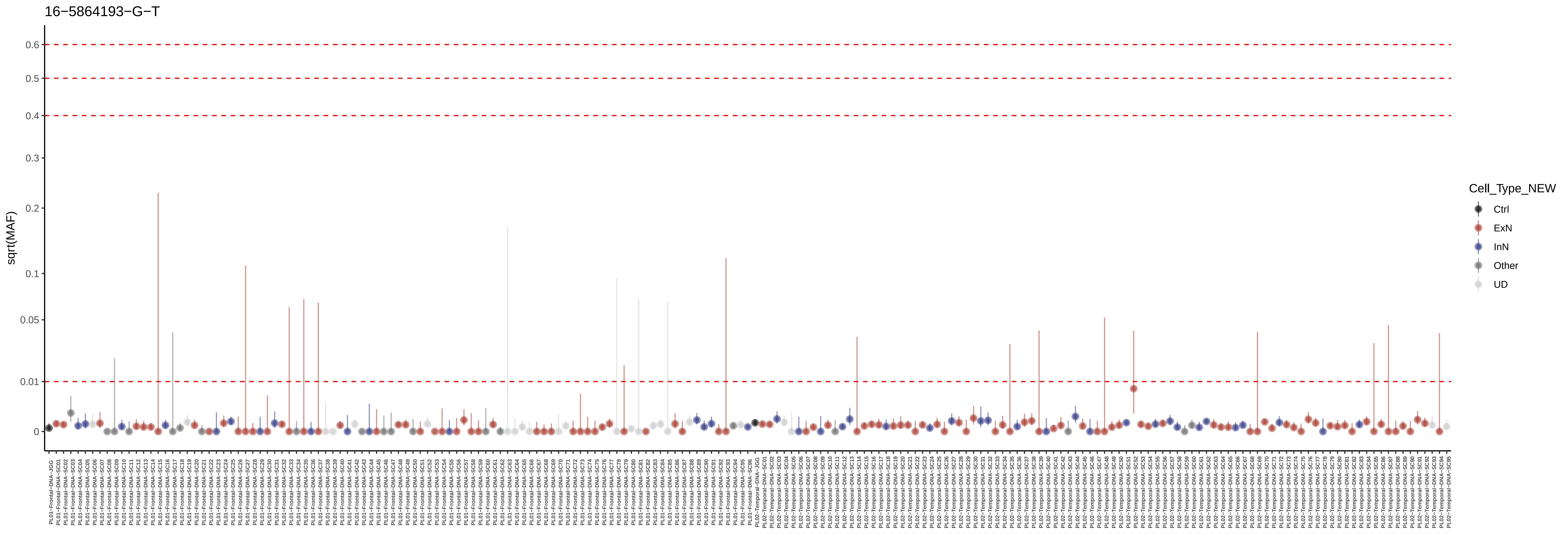
16−5864193−G−T
sqrt(MAF)
0.6
0.5
0.4
0.3
0.2
0.1
0.05
0.01
0
Cell_Type_NEW
Ctrl
ExN
InN
Other
UD

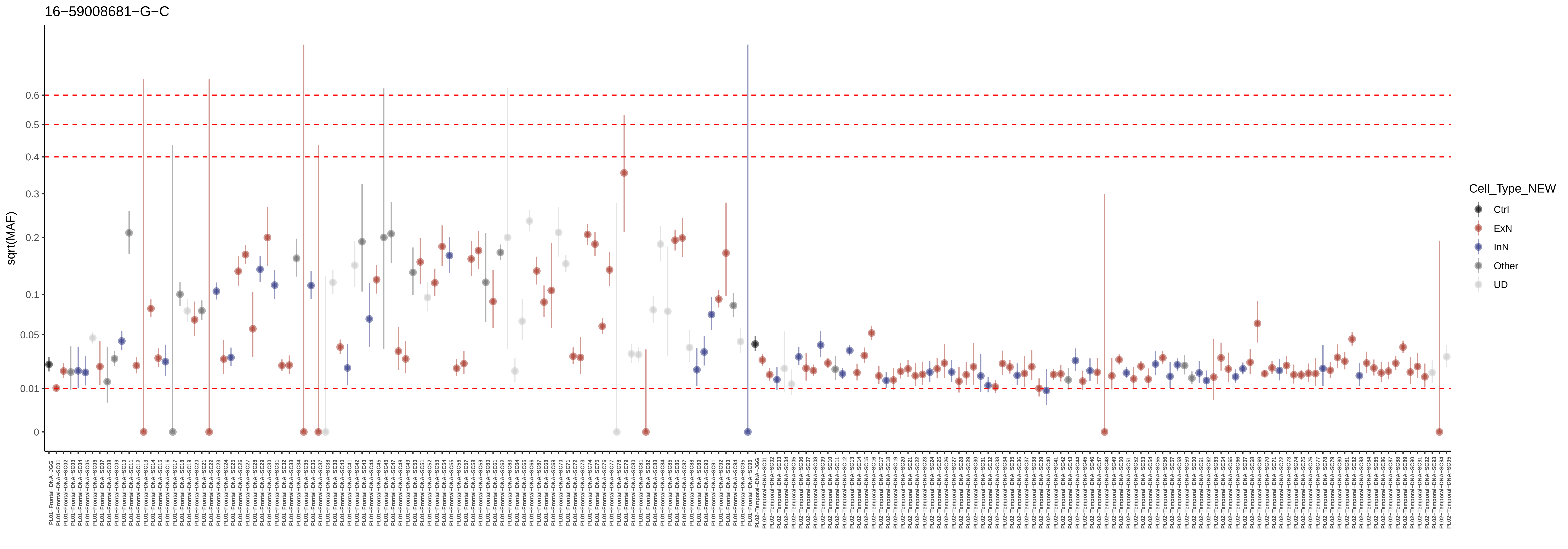
16−59008681−G−C
sqrt(MAF)
0.6
0.5
0.4
0.3
0.2
0.1
0.05
0.01
0
Cell_Type_NEW
Ctrl
ExN
InN
Other
UD

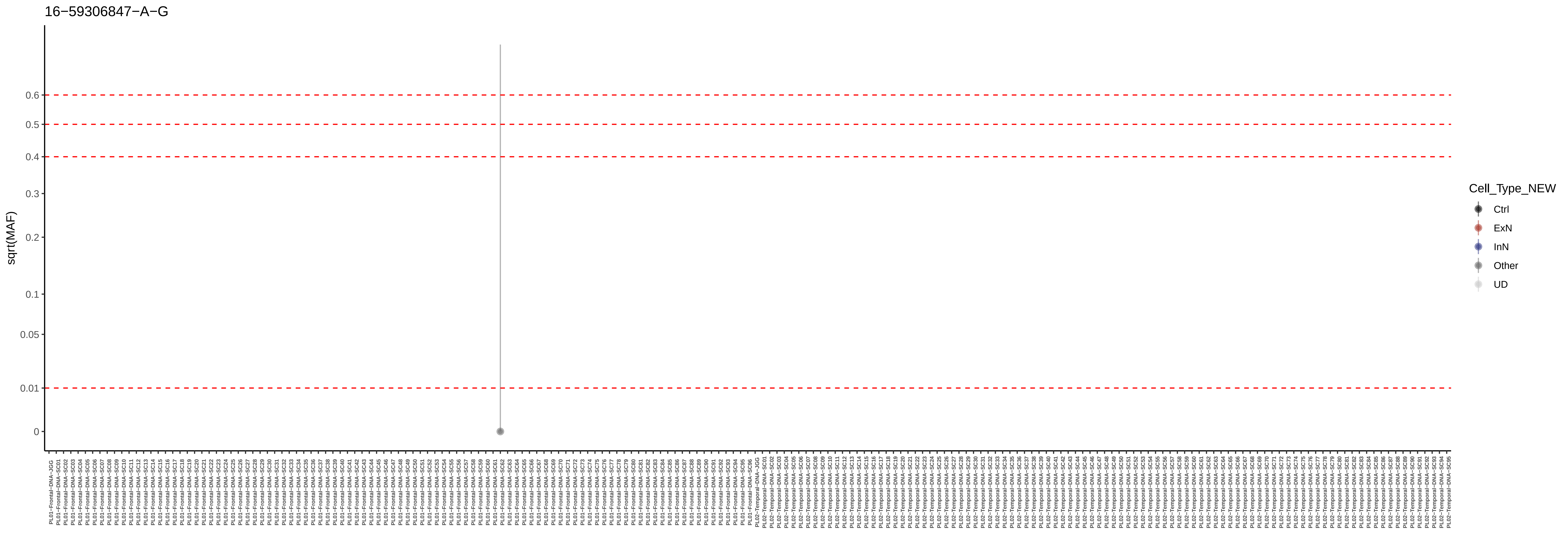
16−59306847−A−G
sqrt(MAF)
0.6
0.5
0.4
0.3
0.2
0.1
0.05
0.01
0
Cell_Type_NEW
Ctrl
ExN
InN
Other
UD

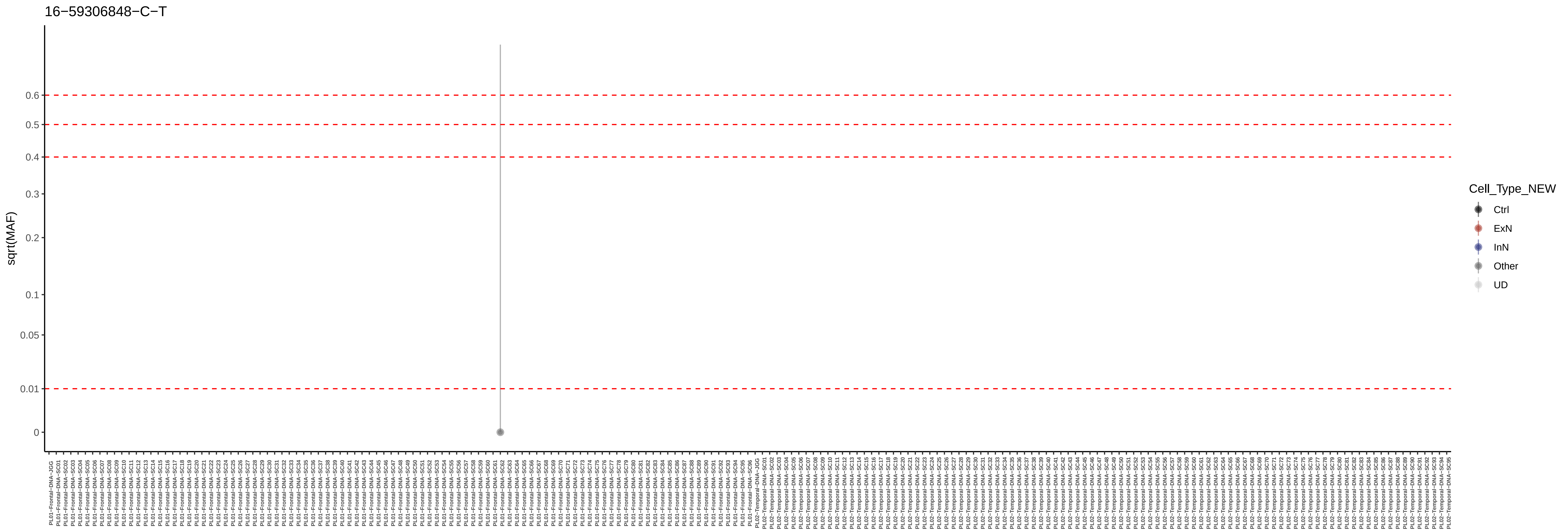

16−59306848−C−T
sqrt(MAF)
0.6
0.5
0.4
0.3
0.2
0.1
0.05
0.01
0
Cell_Type_NEW
Ctrl
ExN
InN
Other
UD

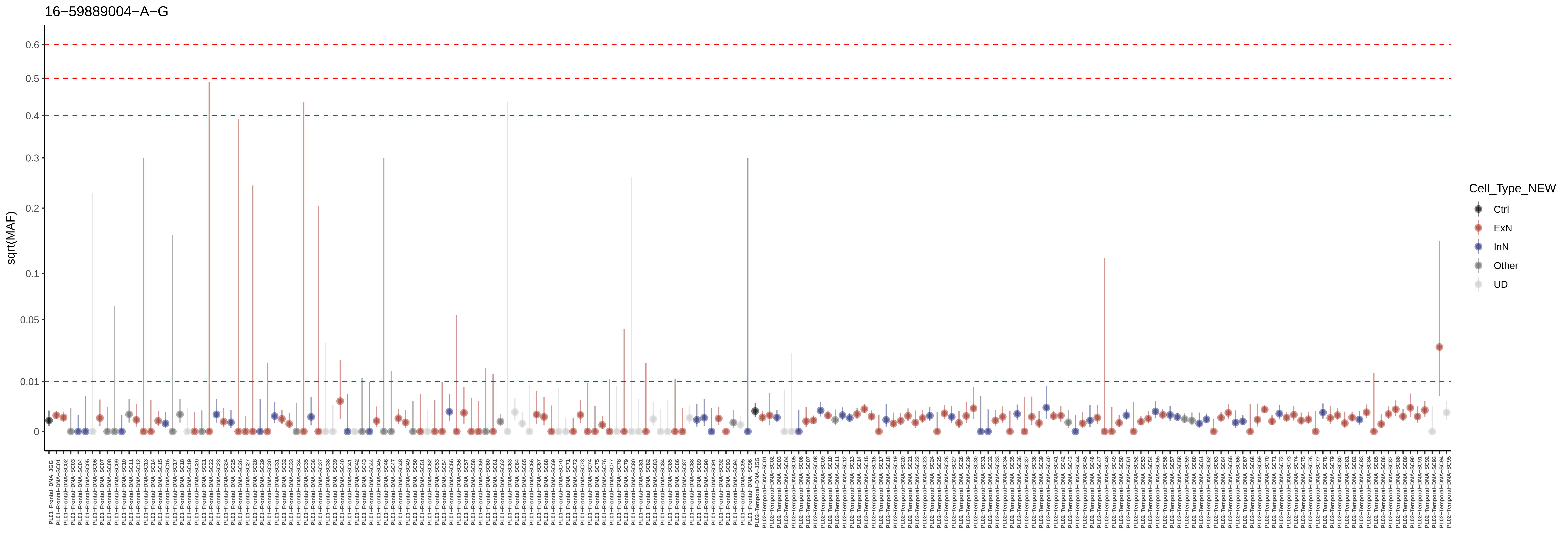
16−59889004−A−G
sqrt(MAF)
0.6
0.5
0.4
0.3
0.2
0.1
0.05
0.01
0
Cell_Type_NEW
Ctrl
ExN
InN
Other
UD

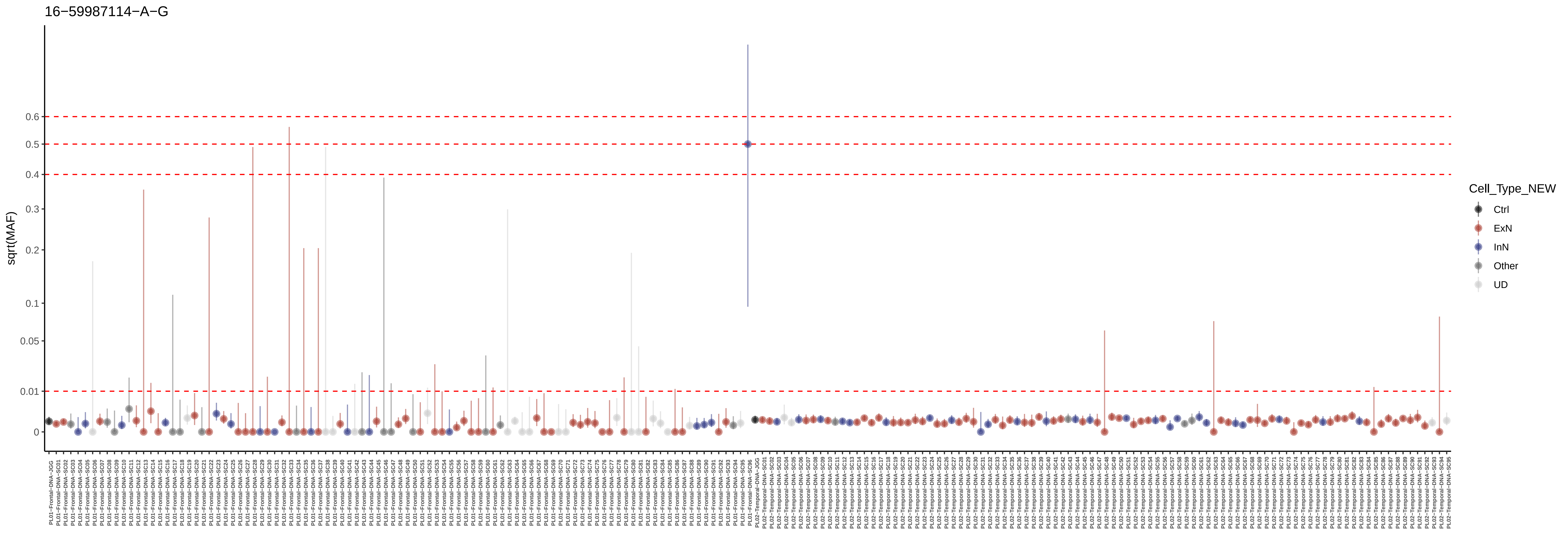
16−59987114−A−G
sqrt(MAF)
0.6
0.5
0.4
0.3
0.2
0.1
0.05
0.01
0
Cell_Type_NEW
Ctrl
ExN
InN
Other
UD

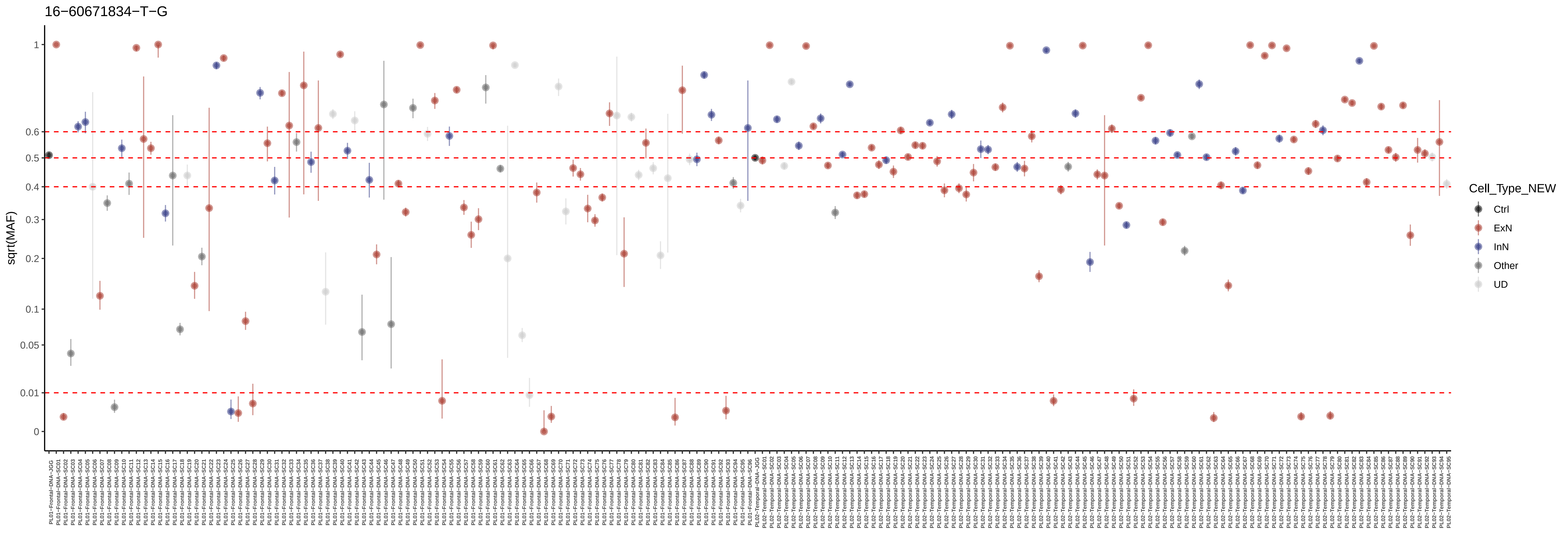
16−60671834−T−G
sqrt(MAF)
1
0.6
0.5
0.4
0.3
0.2
0.1
0.05
0.01
0
Cell_Type_NEW
Ctrl
ExN
InN
Other
UD

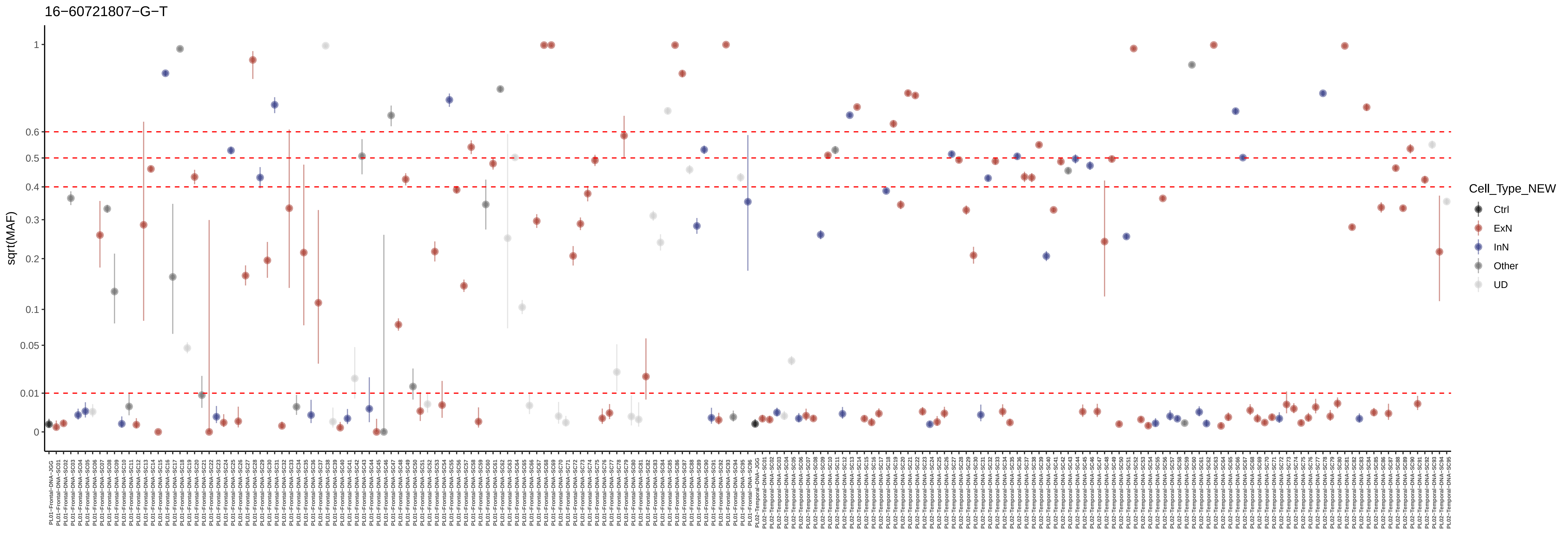
16−60721807−G−T
sqrt(MAF)
1
0.6
0.5
0.4
0.3
0.2
0.1
0.05
0.01
0
Cell_Type_NEW
Ctrl
ExN
InN
Other
UD

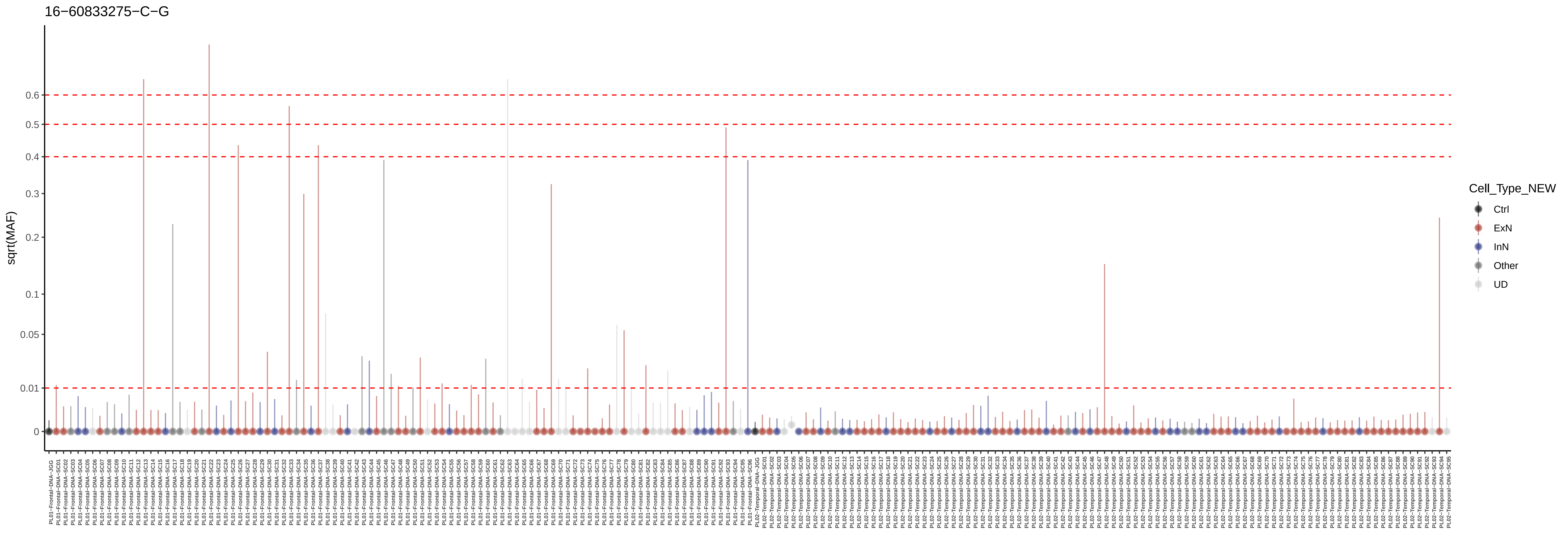
16−60833275−C−G
sqrt(MAF)
0.6
0.5
0.4
0.3
0.2
0.1
0.05
0.01
0
Cell_Type_NEW
Ctrl
ExN
InN
Other
UD

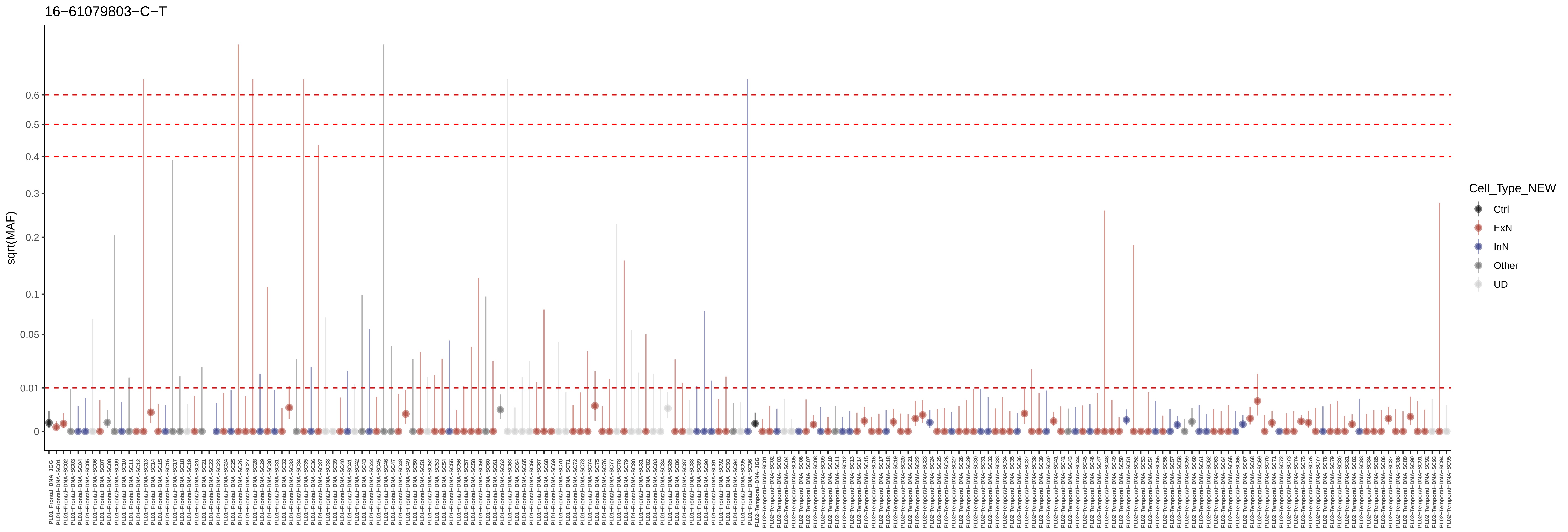
16−61079803−C−T
sqrt(MAF)
0.6
0.5
0.4
0.3
0.2
0.1
0.05
0.01
0
Cell_Type_NEW
Ctrl
ExN
InN
Other
UD

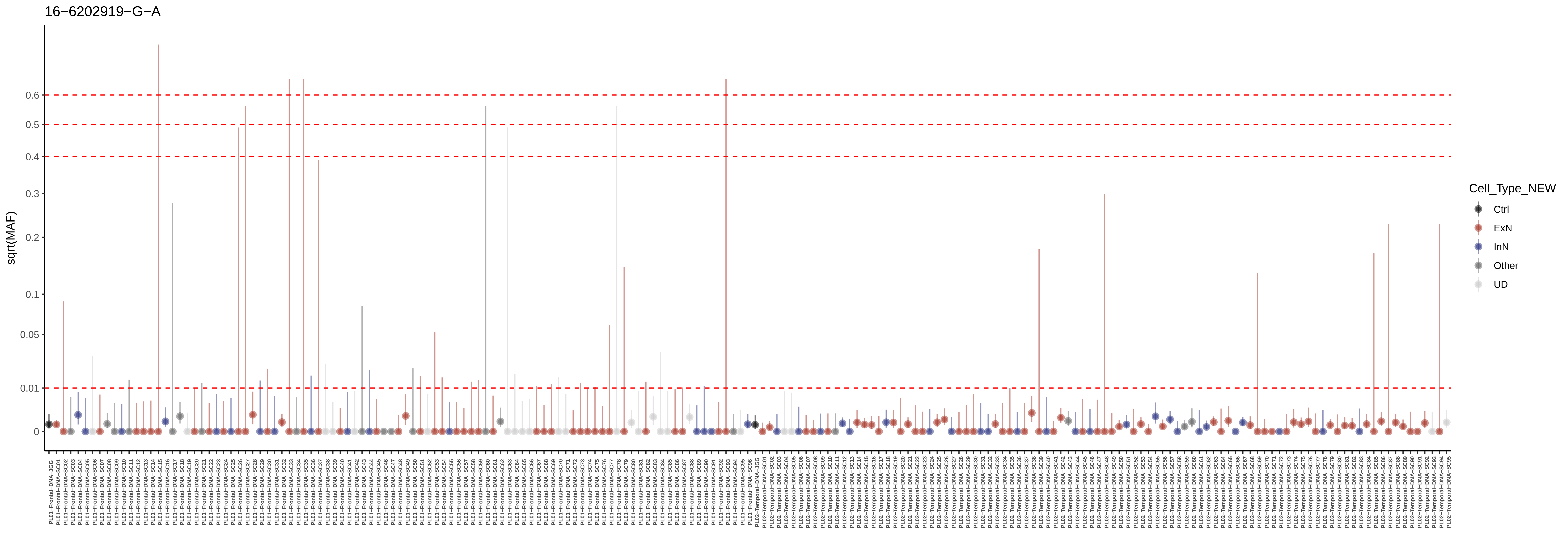
16−6202919−G−A
sqrt(MAF)
0.6
0.5
0.4
0.3
0.2
0.1
0.05
0.01
0
Cell_Type_NEW
Ctrl
ExN
InN
Other
UD

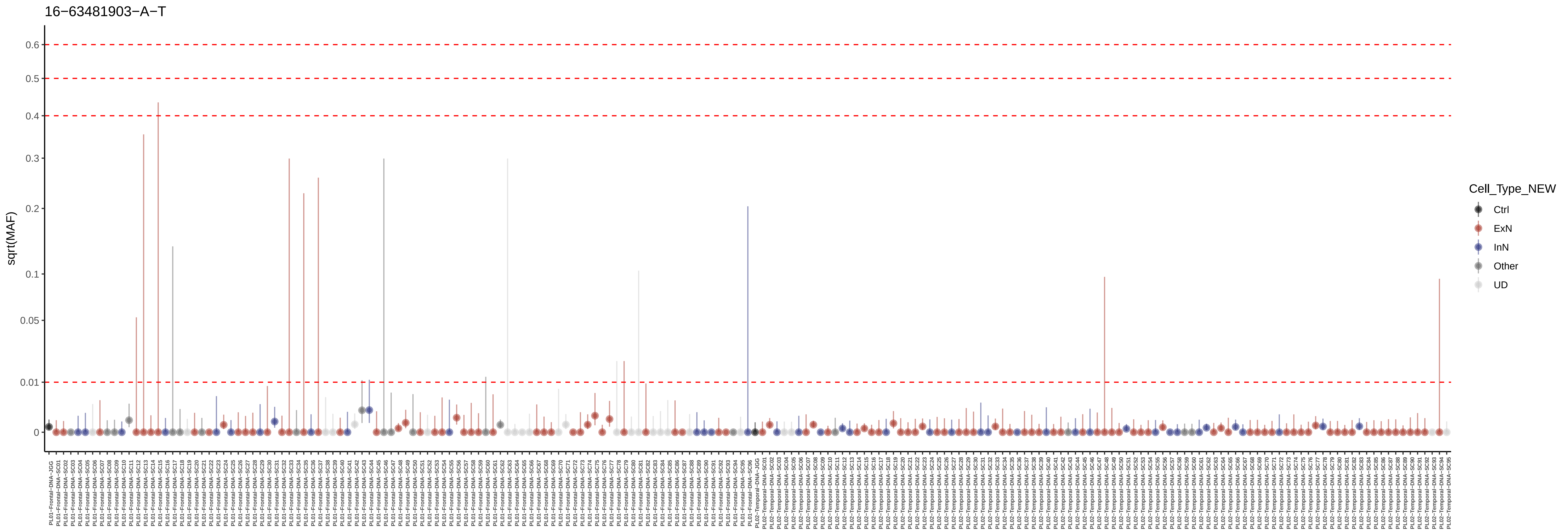

16−63481903−A−T
sqrt(MAF)
0.6
0.5
0.4
0.3
0.2
0.1
0.05
0.01
0
Cell_Type_NEW
Ctrl
ExN
InN
Other
UD

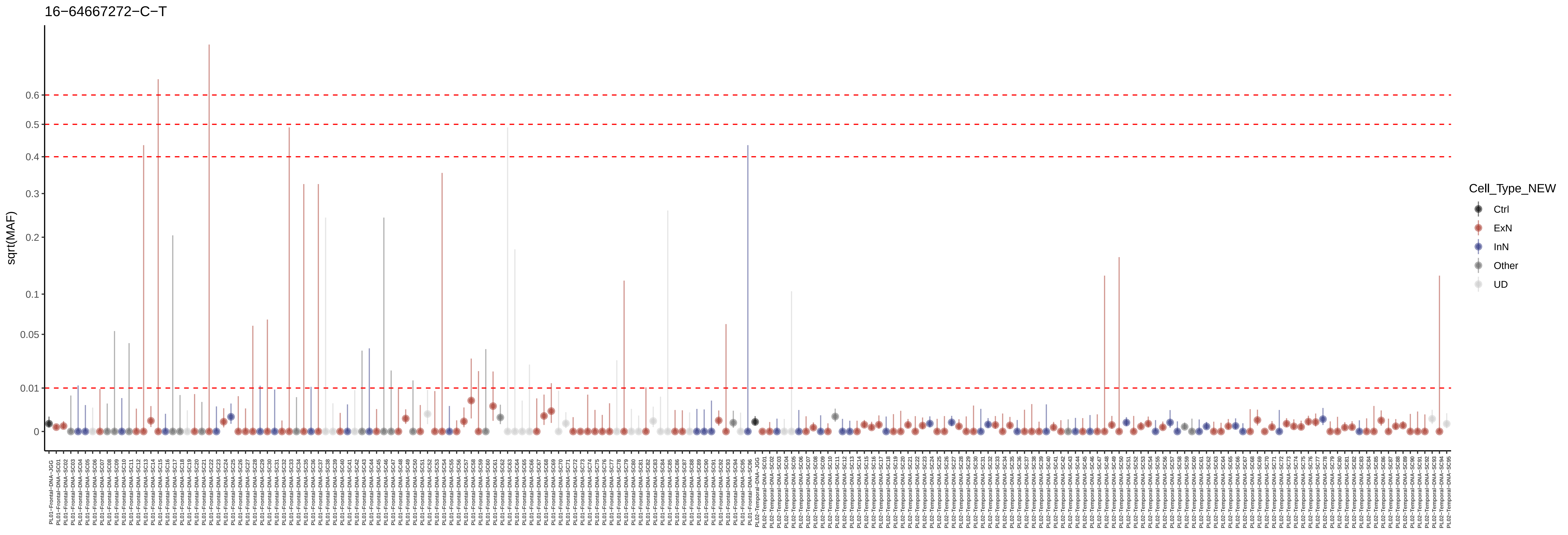

16−64667272−C−T
sqrt(MAF)
0.6
0.5
0.4
0.3
0.2
0.1
0.05
0.01
0
Cell_Type_NEW
Ctrl
ExN
InN
Other
UD

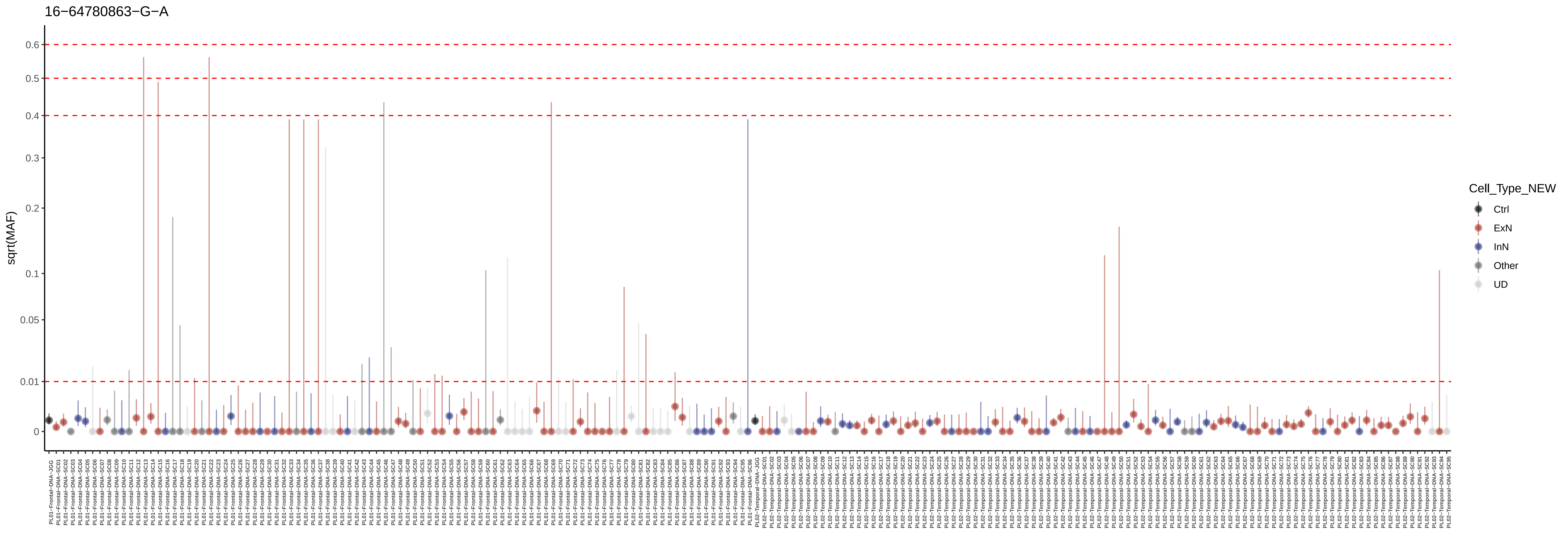
16−64780863−G−A
sqrt(MAF)
0.6
0.5
0.4
0.3
0.2
0.1
0.05
0.01
0
Cell_Type_NEW
Ctrl
ExN
InN
Other
UD

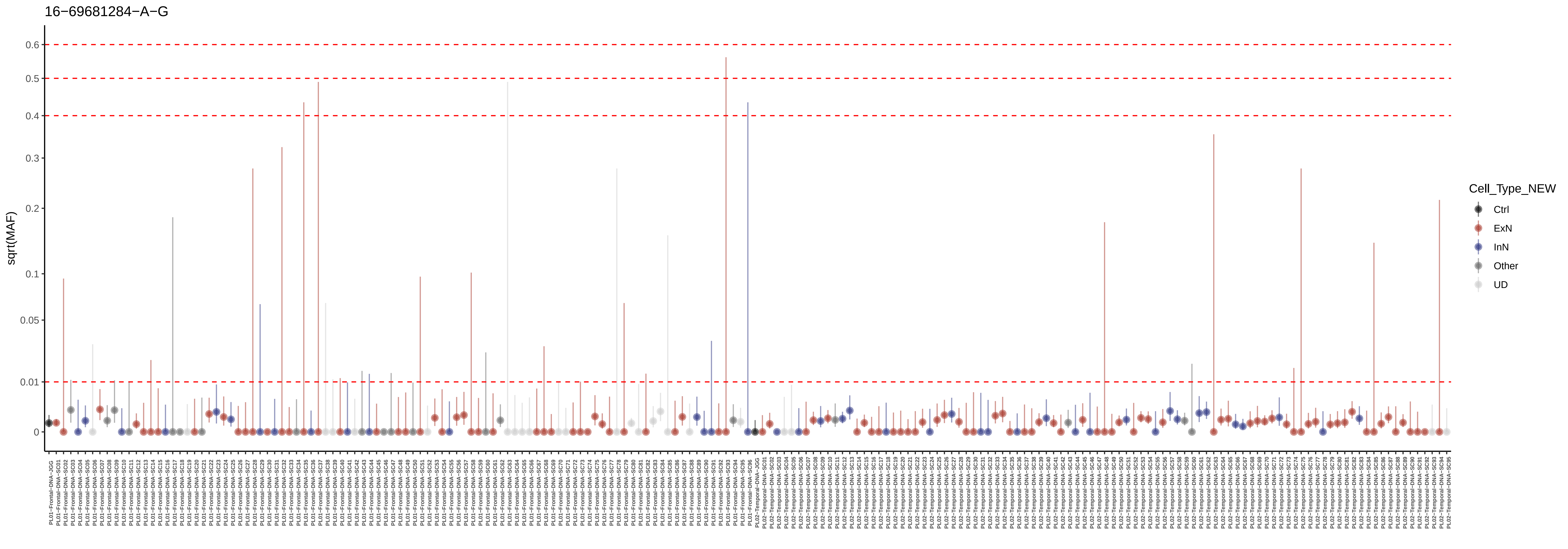
16−69681284−A−G
sqrt(MAF)
0.6
0.5
0.4
0.3
0.2
0.1
0.05
0.01
0
Cell_Type_NEW
Ctrl
ExN
InN
Other
UD

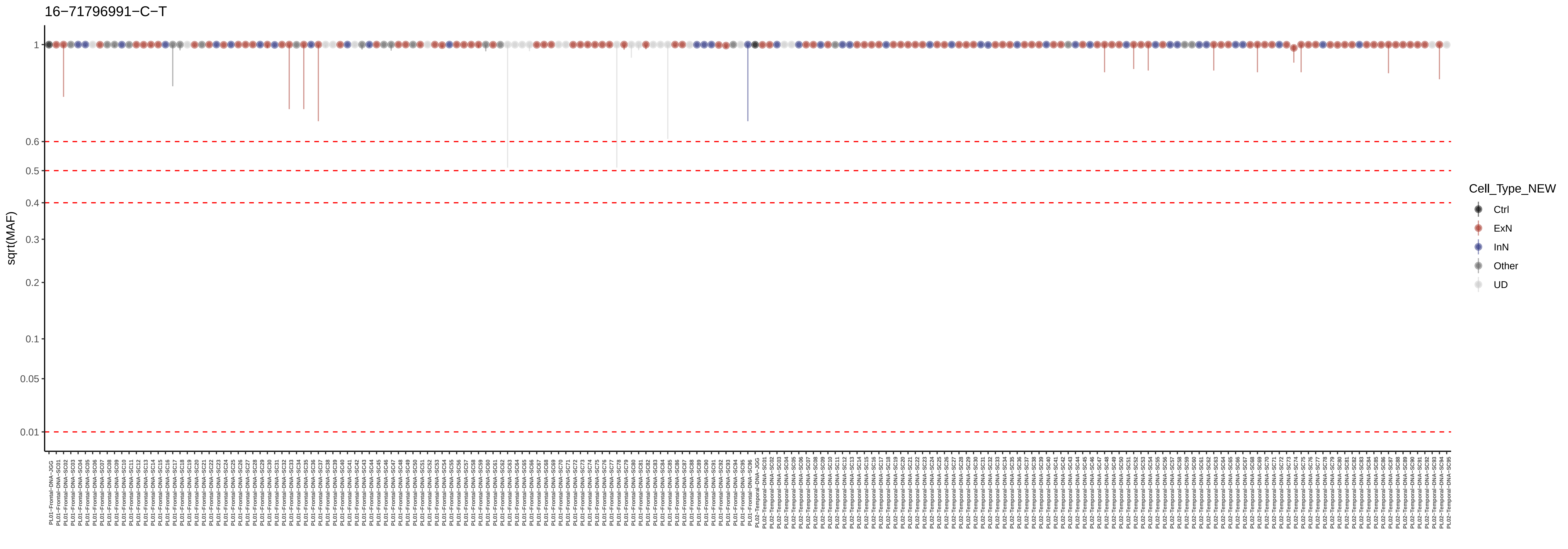
16−71796991−C−T
sqrt(MAF)
1
0.6
0.5
0.4
0.3
0.2
0.1
0.05
0.01
Cell_Type_NEW
Ctrl
ExN
InN
Other
UD

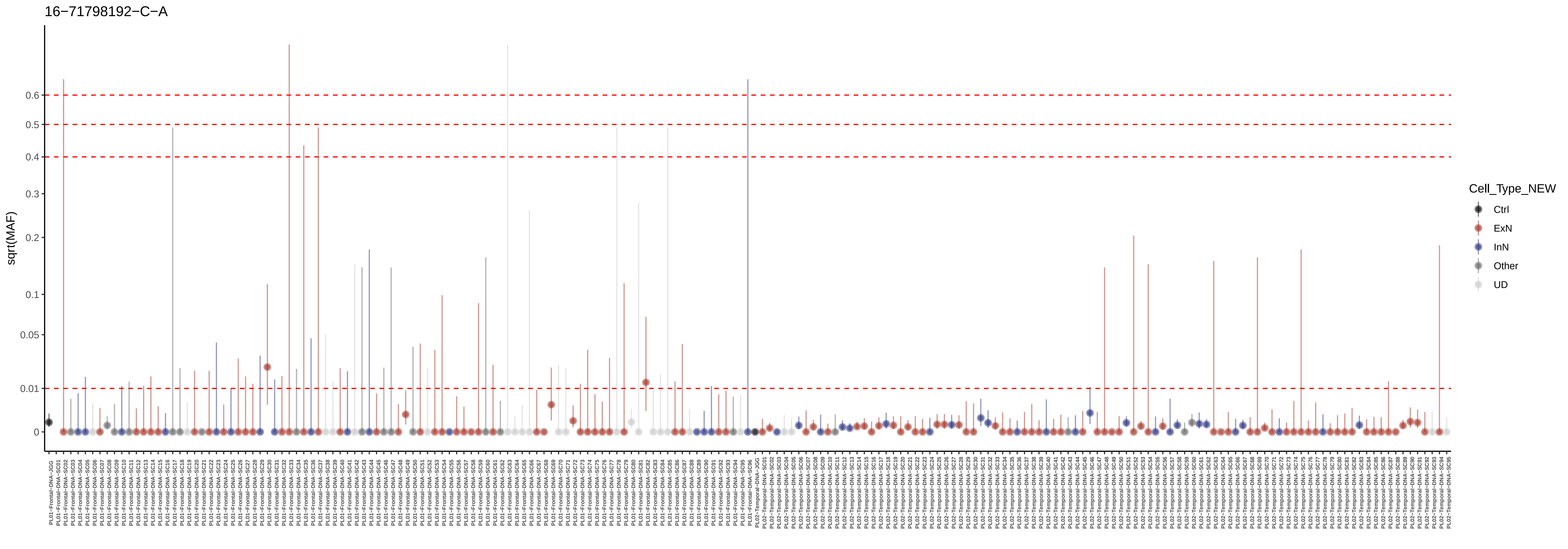
16−71798192−C−A
sqrt(MAF)
0.6
0.5
0.4
0.3
0.2
0.1
0.05
0.01
0
Cell_Type_NEW
Ctrl
ExN
InN
Other
UD

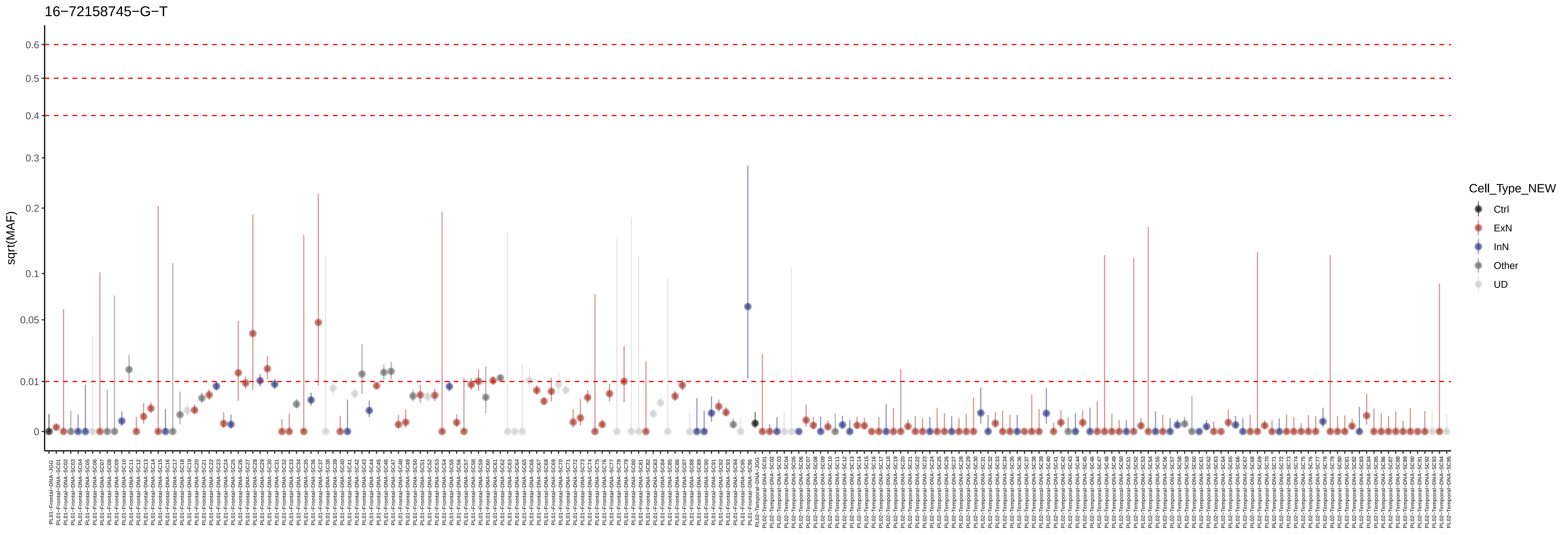
16−72158745−G−T
sqrt(MAF)
0.6
0.5
0.4
0.3
0.2
0.1
0.05
0.01
0
Cell_Type_NEW
Ctrl
ExN
InN
Other
UD

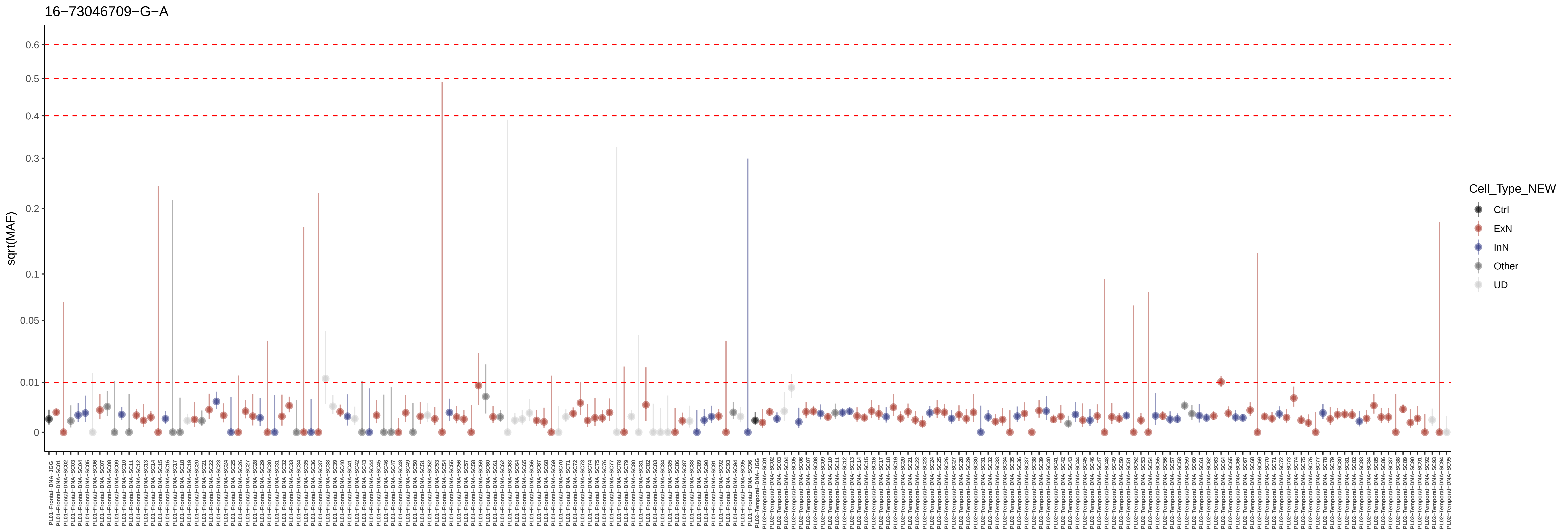
16−73046709−G−A
sqrt(MAF)
0.6
0.5
0.4
0.3
0.2
0.1
0.05
0.01
0
Cell_Type_NEW
Ctrl
ExN
InN
Other
UD

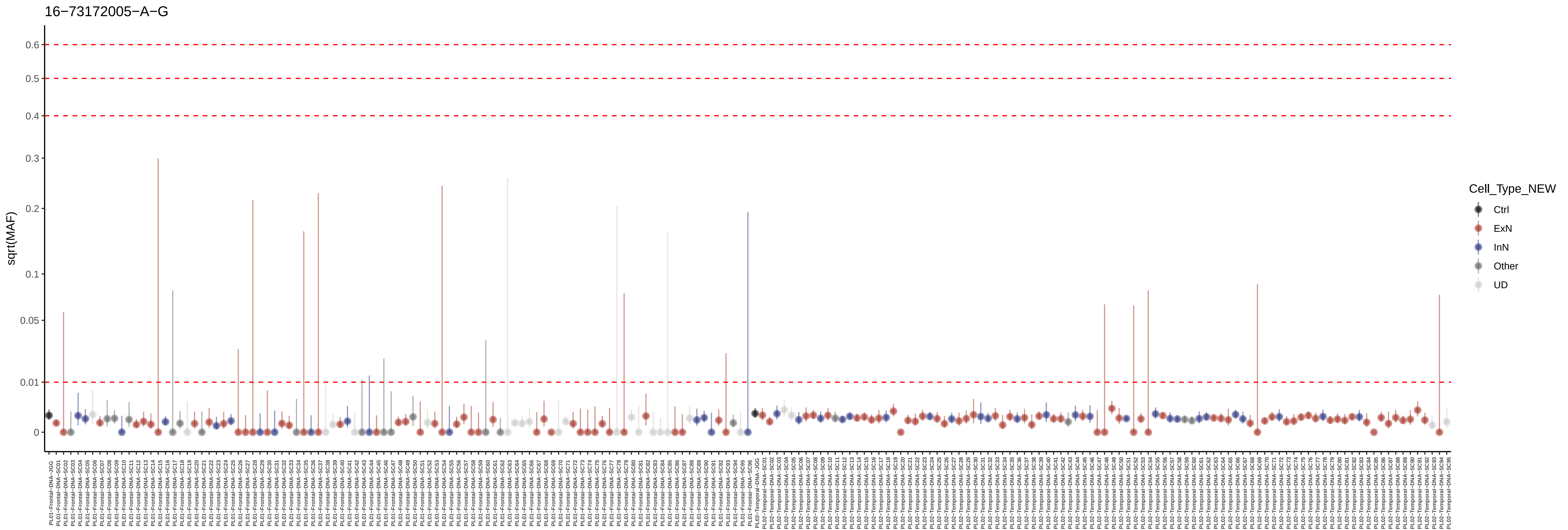
16−73172005−A−G
sqrt(MAF)
0.6
0.5
0.4
0.3
0.2
0.1
0.05
0.01
0
Cell_Type_NEW
Ctrl
ExN
InN
Other
UD

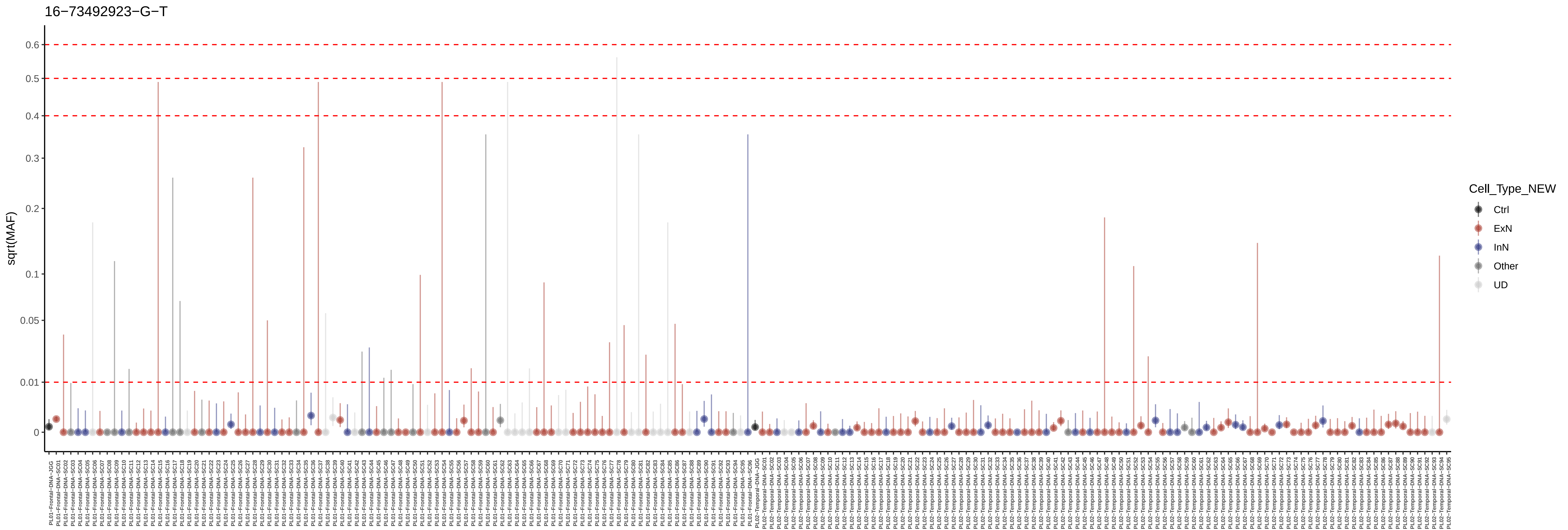
16−73492923−G−T
sqrt(MAF)
0.6
0.5
0.4
0.3
0.2
0.1
0.05
0.01
0
Cell_Type_NEW
Ctrl
ExN
InN
Other
UD

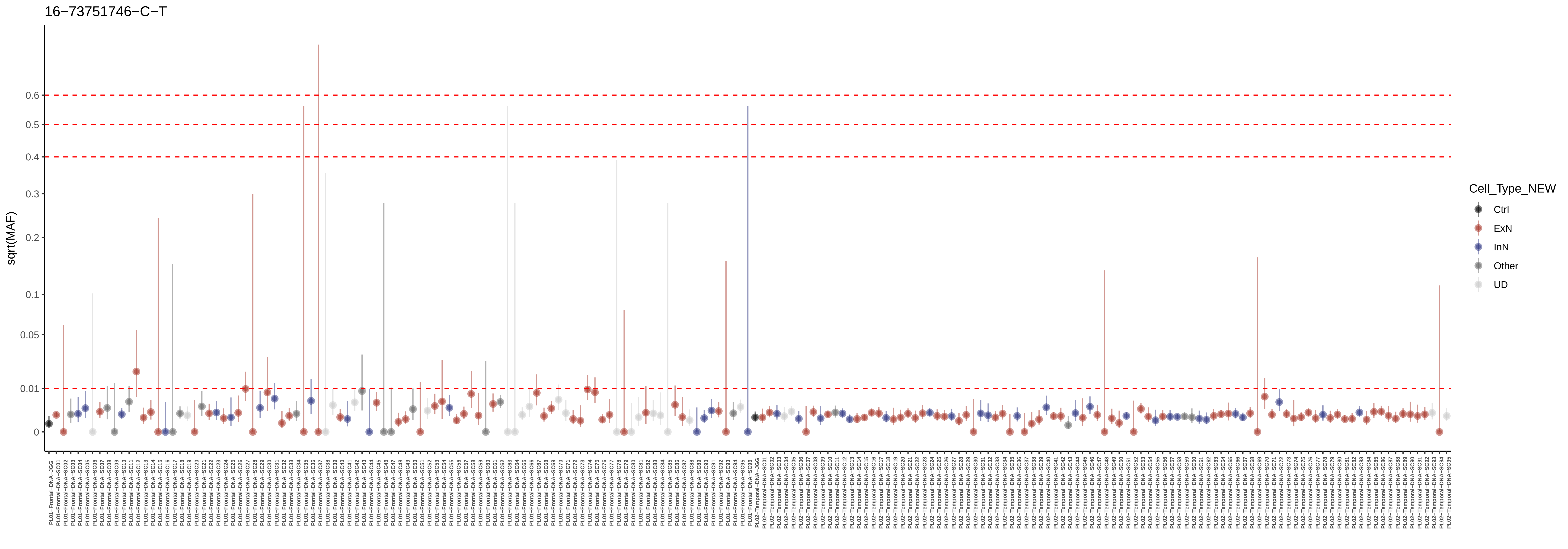

16−73751746−C−T
sqrt(MAF)
0.6
0.5
0.4
0.3
0.2
0.1
0.05
0.01
0
Cell_Type_NEW
Ctrl
ExN
InN
Other
UD

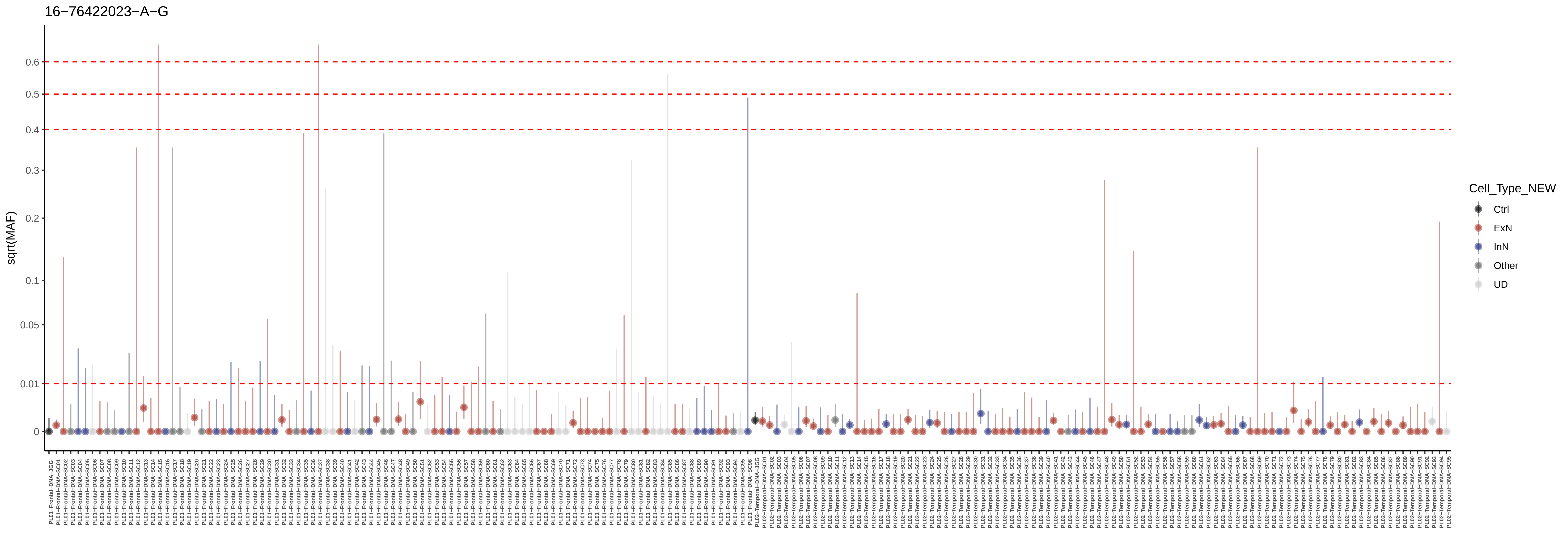
16−76422023−A−G
sqrt(MAF)
0.6
0.5
0.4
0.3
0.2
0.1
0.05
0.01
0
Cell_Type_NEW
Ctrl
ExN
InN
Other
UD

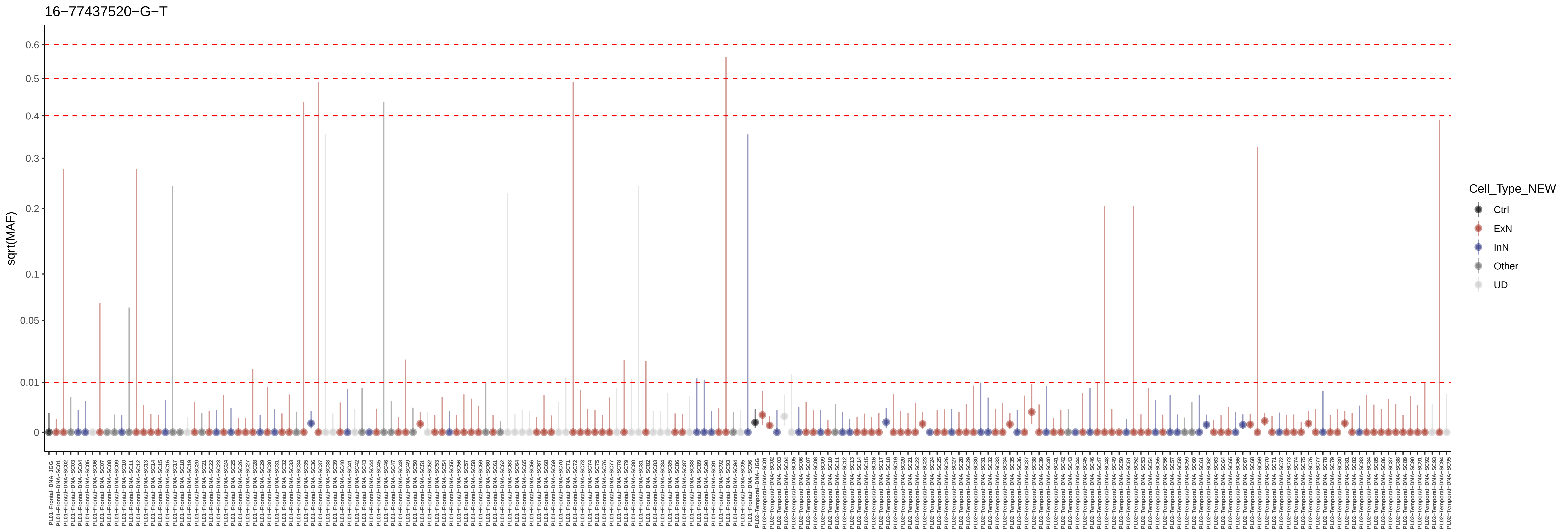
16−77437520−G−T
sqrt(MAF)
0.6
0.5
0.4
0.3
0.2
0.1
0.05
0.01
0
Cell_Type_NEW
Ctrl
ExN
InN
Other
UD

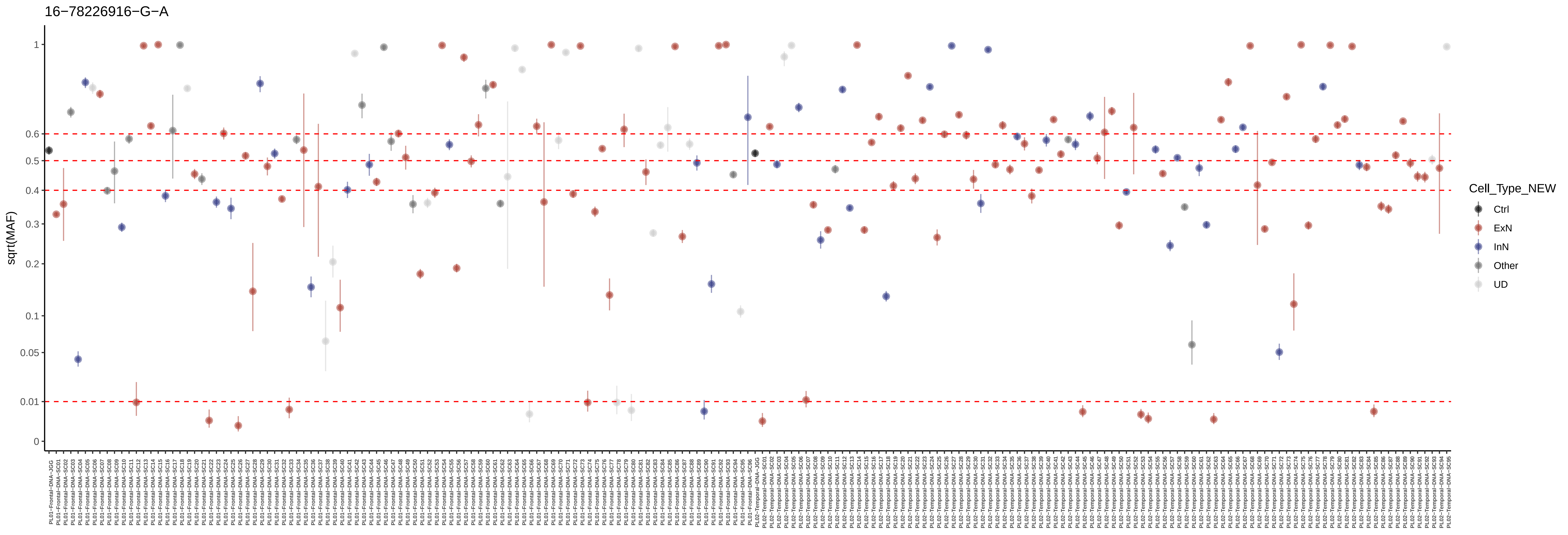
16−78226916−G−A
sqrt(MAF)
1
0.6
0.5
0.4
0.3
0.2
0.1
0.05
0.01
0
Cell_Type_NEW
Ctrl
ExN
InN
Other
UD

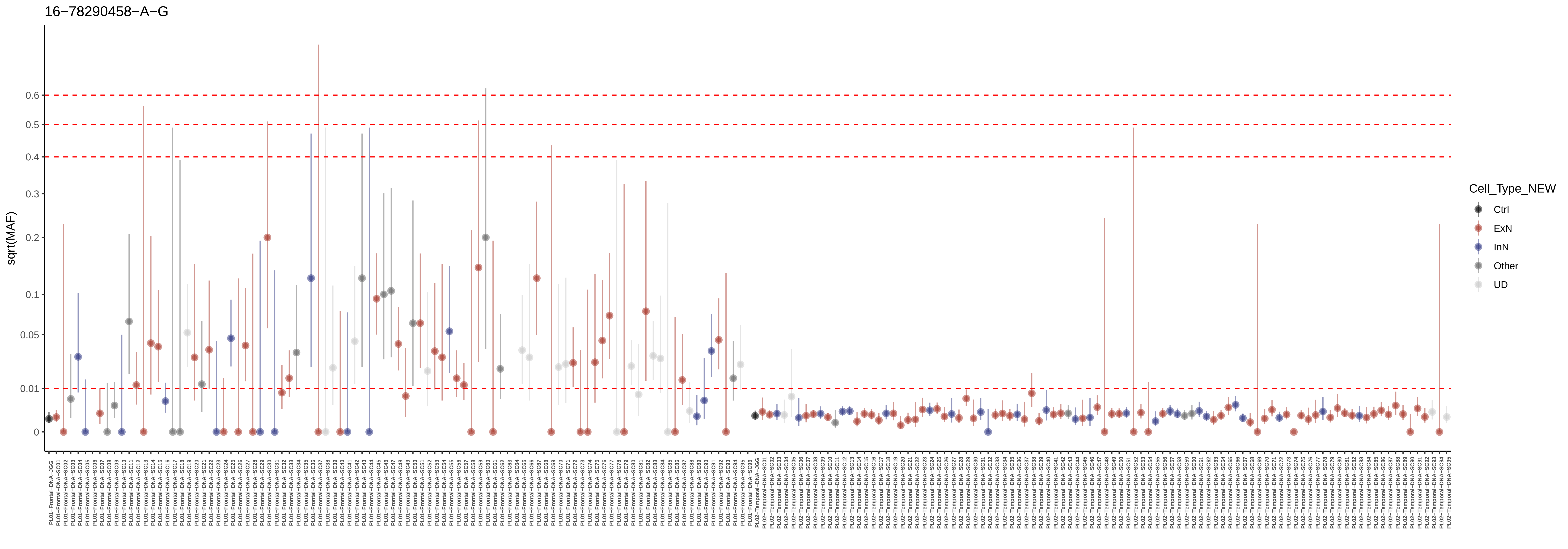
16−78290458−A−G
sqrt(MAF)
0.6
0.5
0.4
0.3
0.2
0.1
0.05
0.01
0
Cell_Type_NEW
Ctrl
ExN
InN
Other
UD

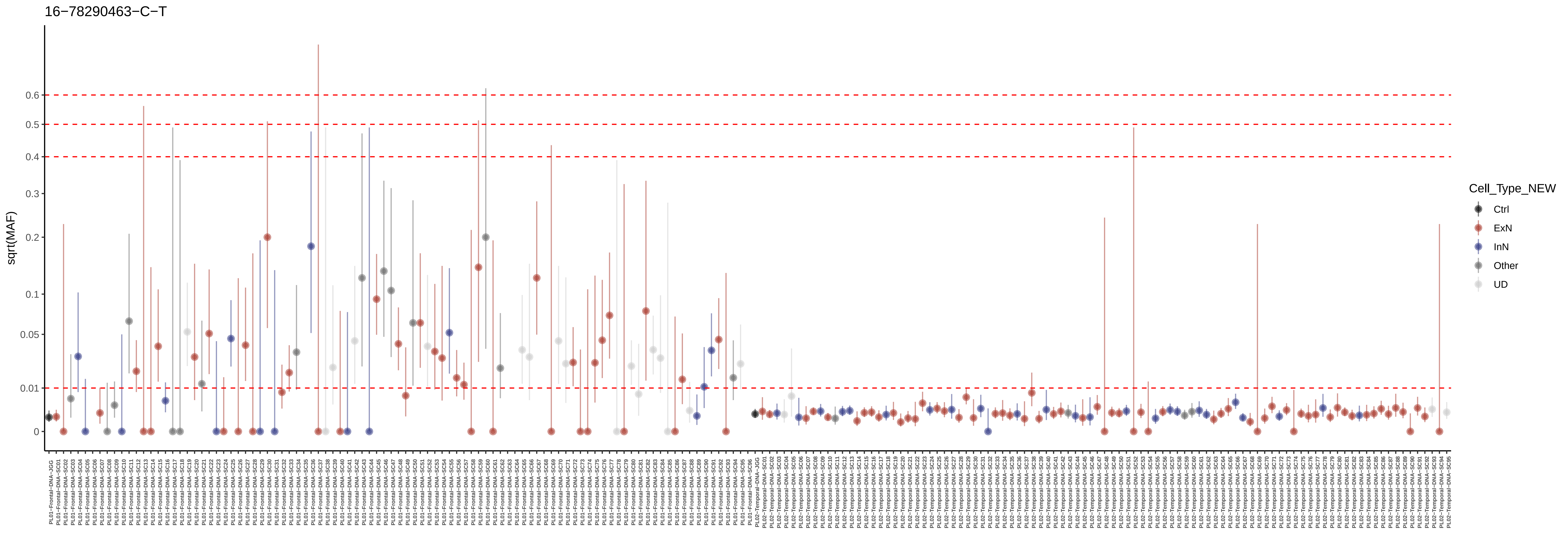

16−78290463−C−T
sqrt(MAF)
0.6
0.5
0.4
0.3
0.2
0.1
0.05
0.01
0
Cell_Type_NEW
Ctrl
ExN
InN
Other
UD

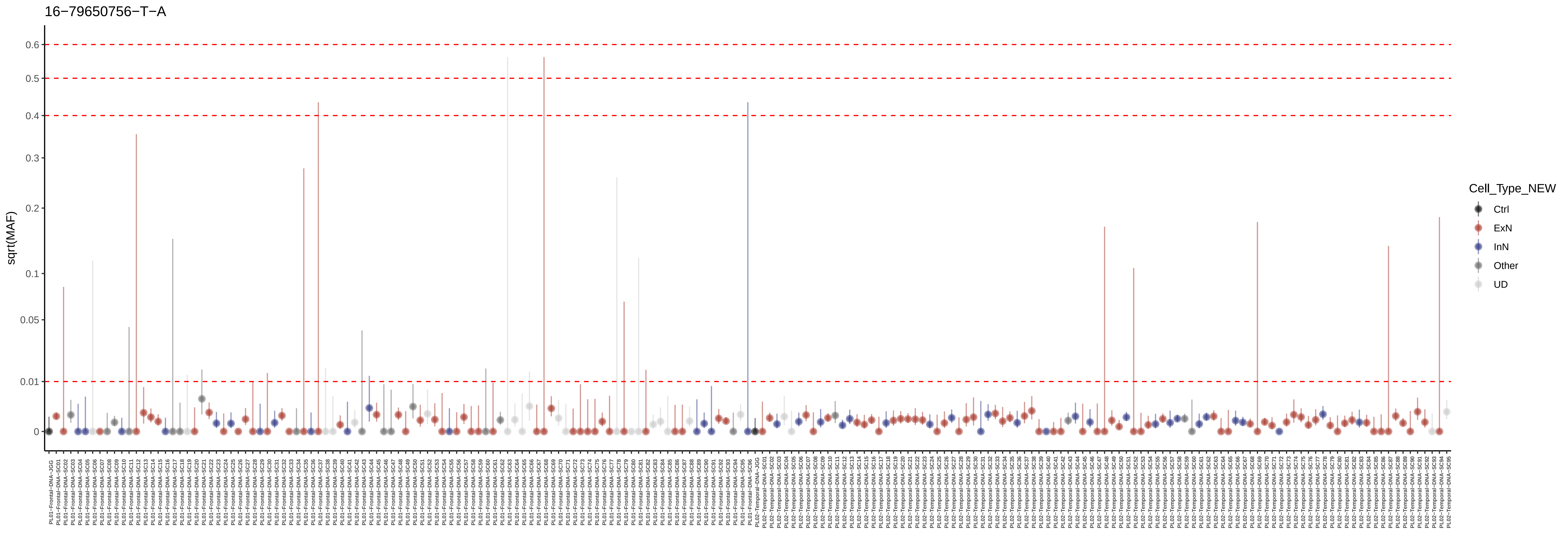

16−79650756−T−A
sqrt(MAF)
0.6
0.5
0.4
0.3
0.2
0.1
0.05
0.01
0
Cell_Type_NEW
Ctrl
ExN
InN
Other
UD

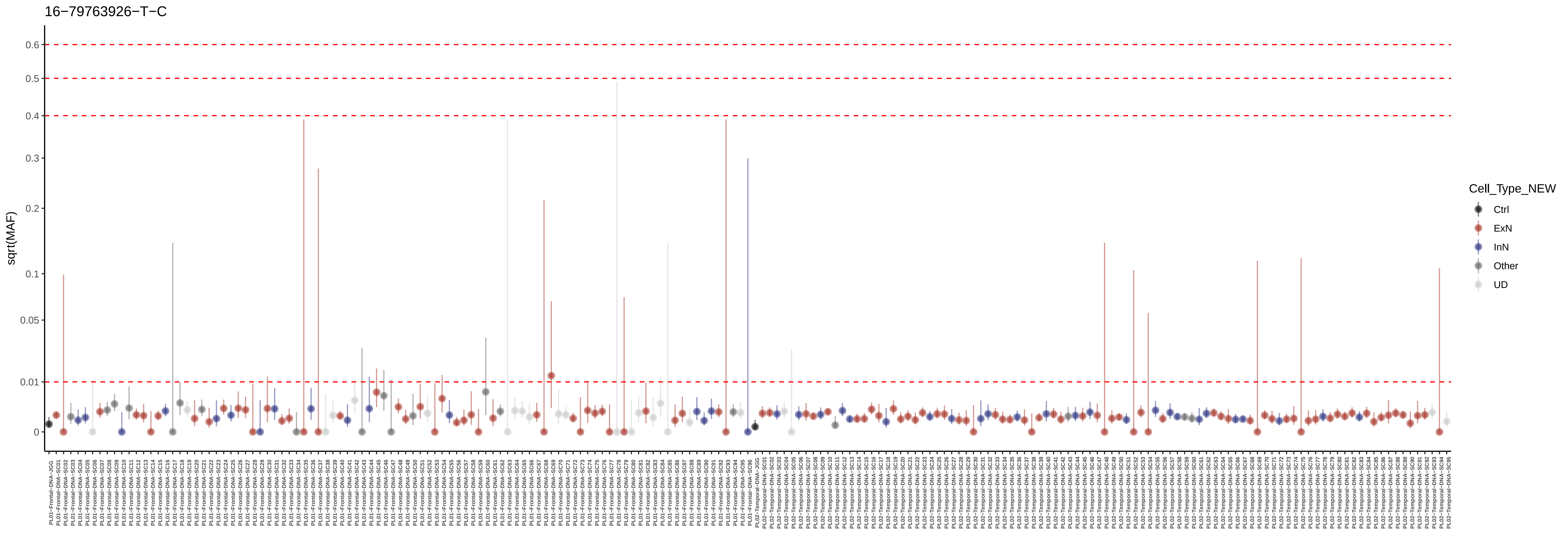
16−79763926−T−C
sqrt(MAF)
0.6
0.5
0.4
0.3
0.2
0.1
0.05
0.01
0
Cell_Type_NEW
Ctrl
ExN
InN
Other
UD

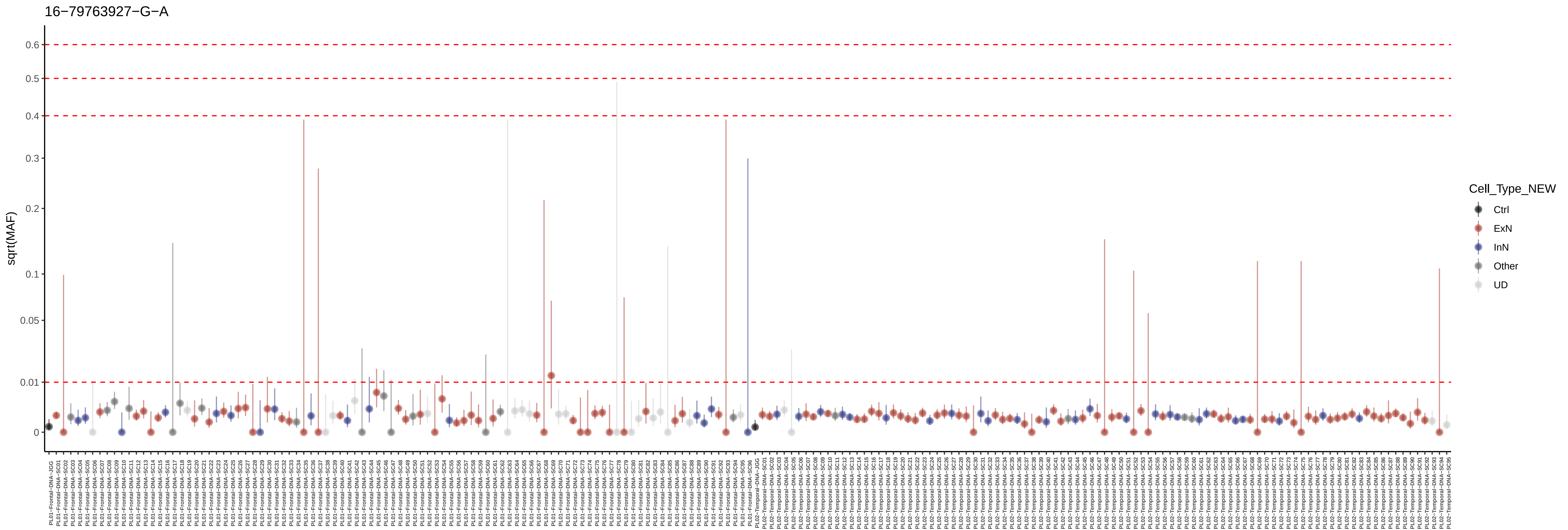
16−79763927−G−A
sqrt(MAF)
0.6
0.5
0.4
0.3
0.2
0.1
0.05
0.01
0
Cell_Type_NEW
Ctrl
ExN
InN
Other
UD

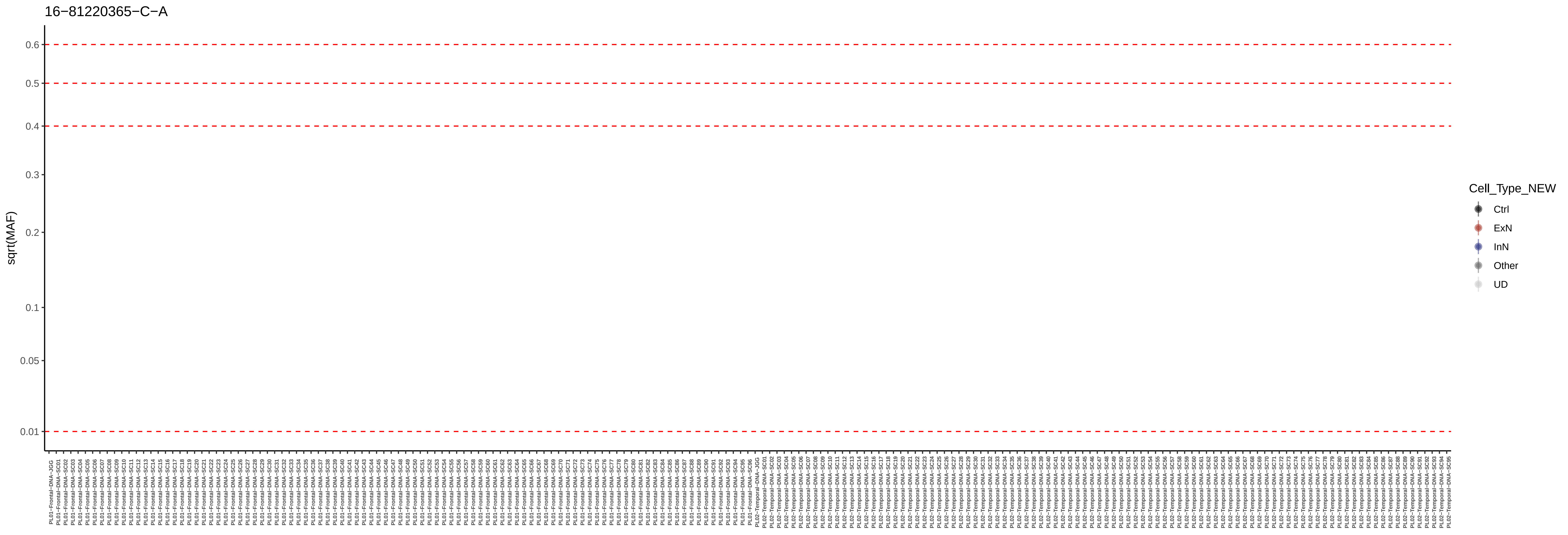
16−81220365−C−A
sqrt(MAF)
0.6
0.5
0.4
0.3
0.2
0.1
0.05
0.01
Cell_Type_NEW
Ctrl
ExN
InN
Other
UD

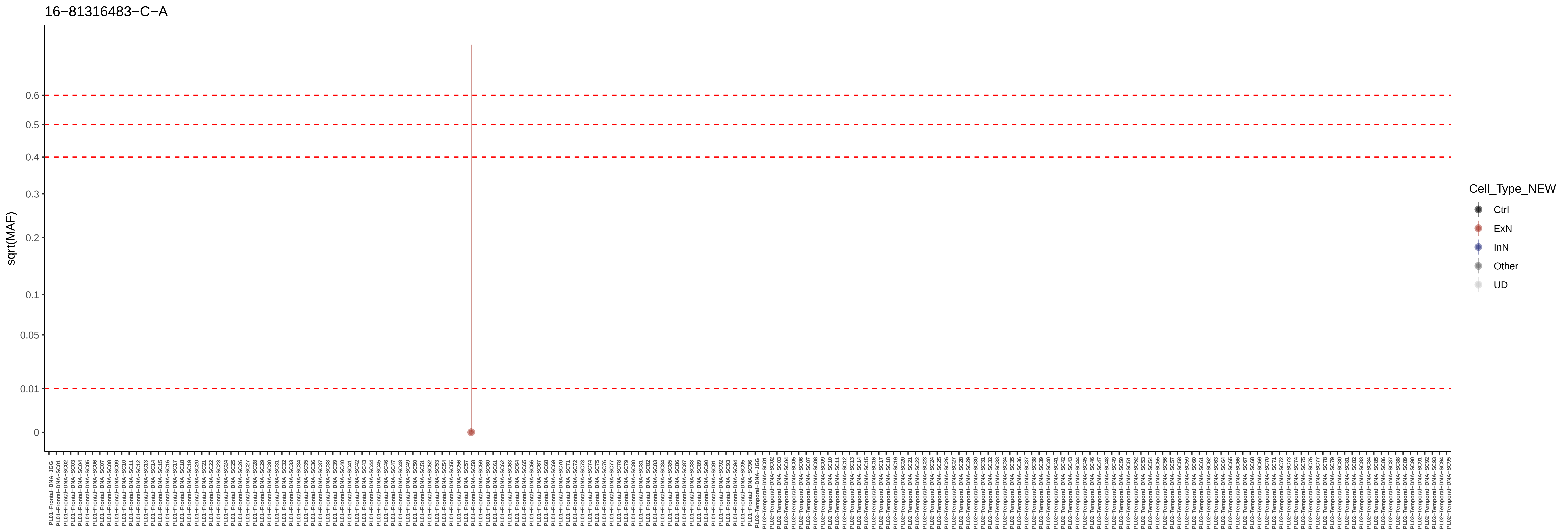

16−81316483−C−A
sqrt(MAF)
0.6
0.5
0.4
0.3
0.2
0.1
0.05
0.01
0
Cell_Type_NEW
Ctrl
ExN
InN
Other
UD

16−82026404−A−T

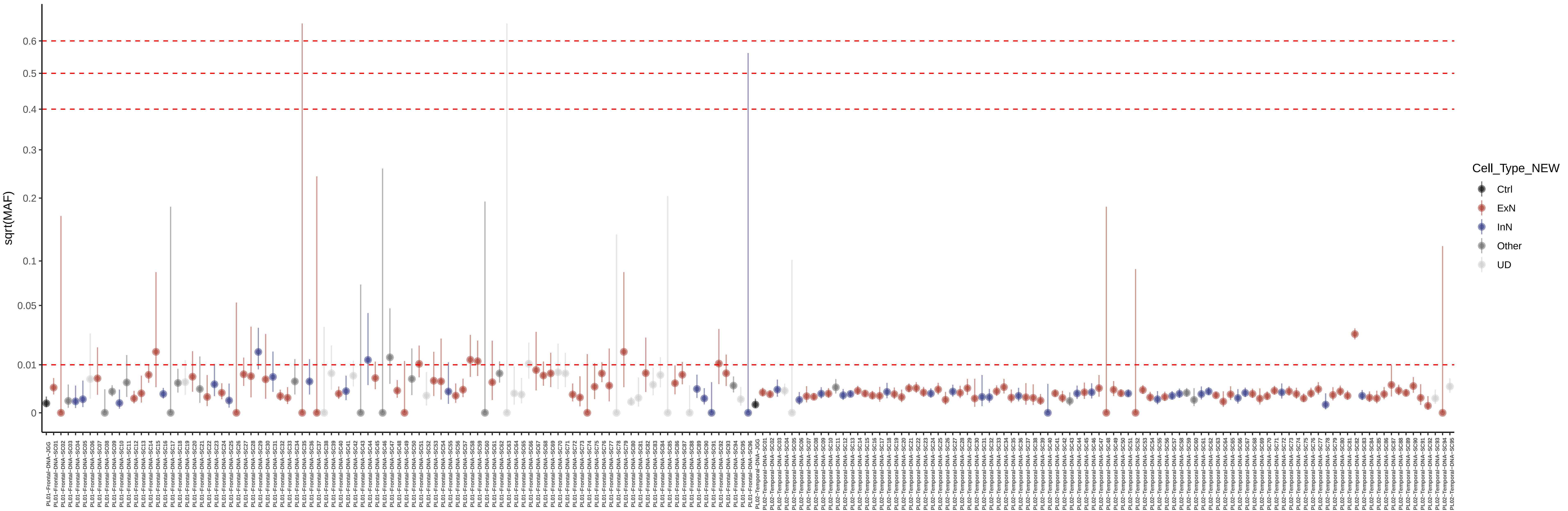

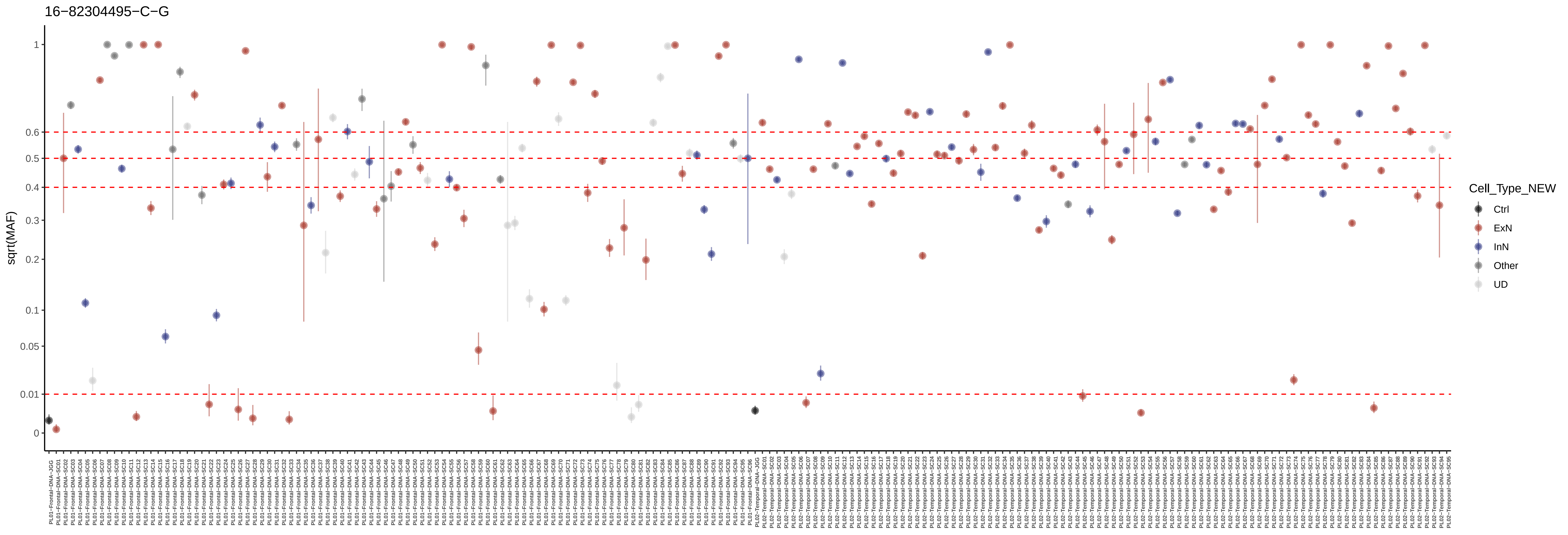

16−82304495−C−G
sqrt(MAF)
1
0.6
0.5
0.4
0.3
0.2
0.1
0.05
0.01
0
Cell_Type_NEW
Ctrl
ExN
InN
Other
UD

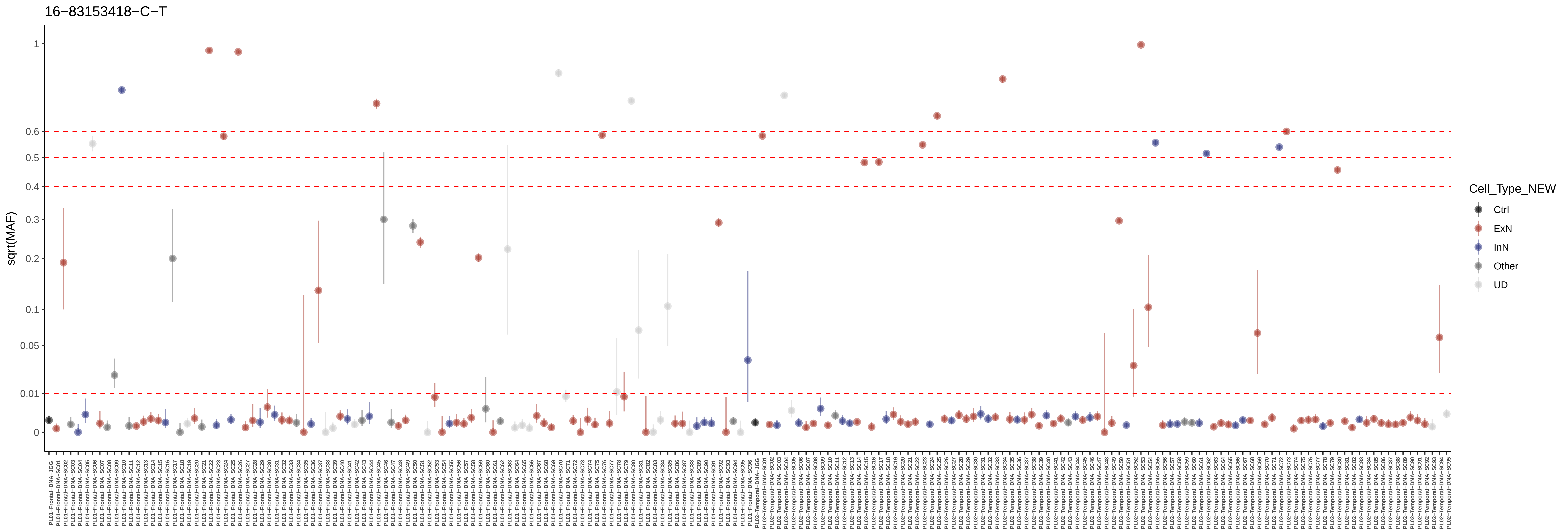
16−83153418−C−T
sqrt(MAF)
1
0.6
0.5
0.4
0.3
0.2
0.1
0.05
0.01
0
Cell_Type_NEW
Ctrl
ExN
InN
Other
UD

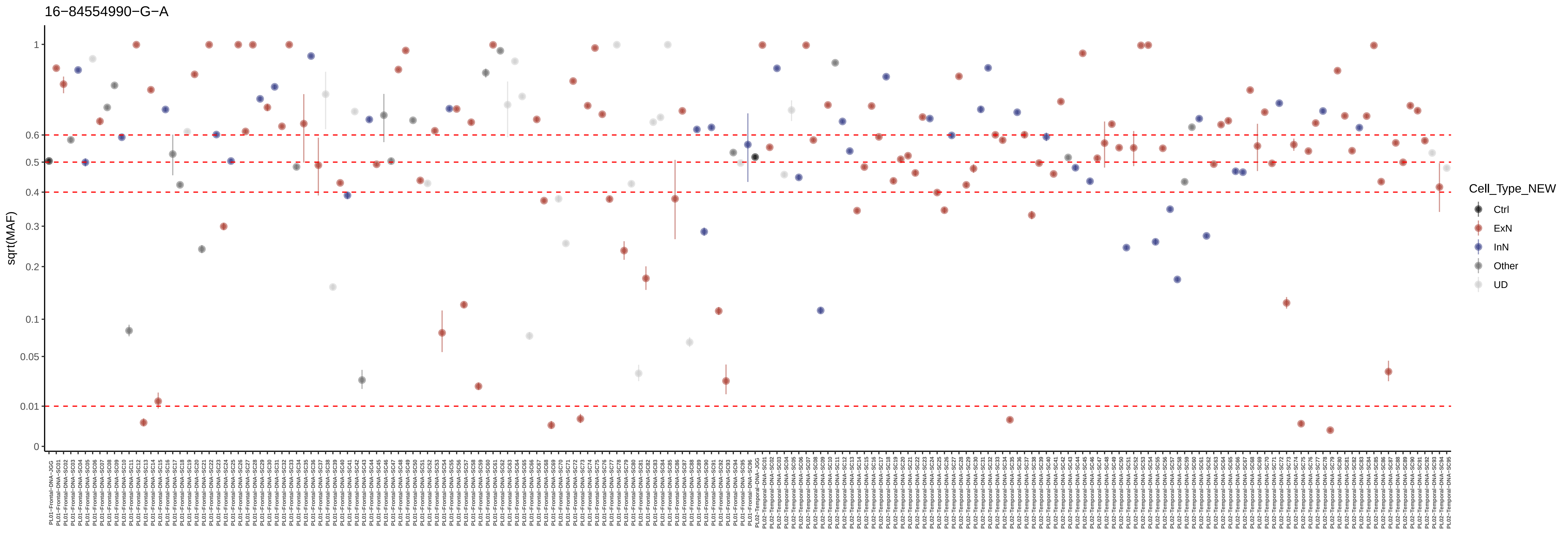
16−84554990−G−A
sqrt(MAF)
1
0.6
0.5
0.4
0.3
0.2
0.1
0.05
0.01
0
Cell_Type_NEW
Ctrl
ExN
InN
Other
UD

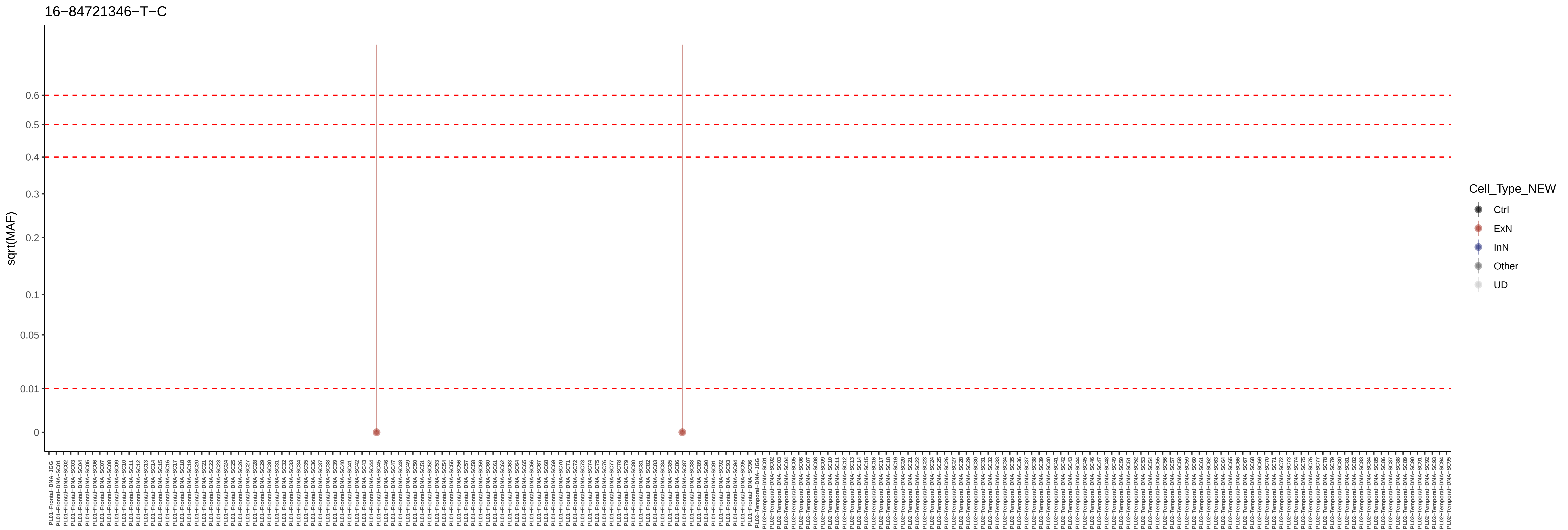
16−84721346−T−C
sqrt(MAF)
0.6
0.5
0.4
0.3
0.2
0.1
0.05
0.01
0
Cell_Type_NEW
Ctrl
ExN
InN
Other
UD

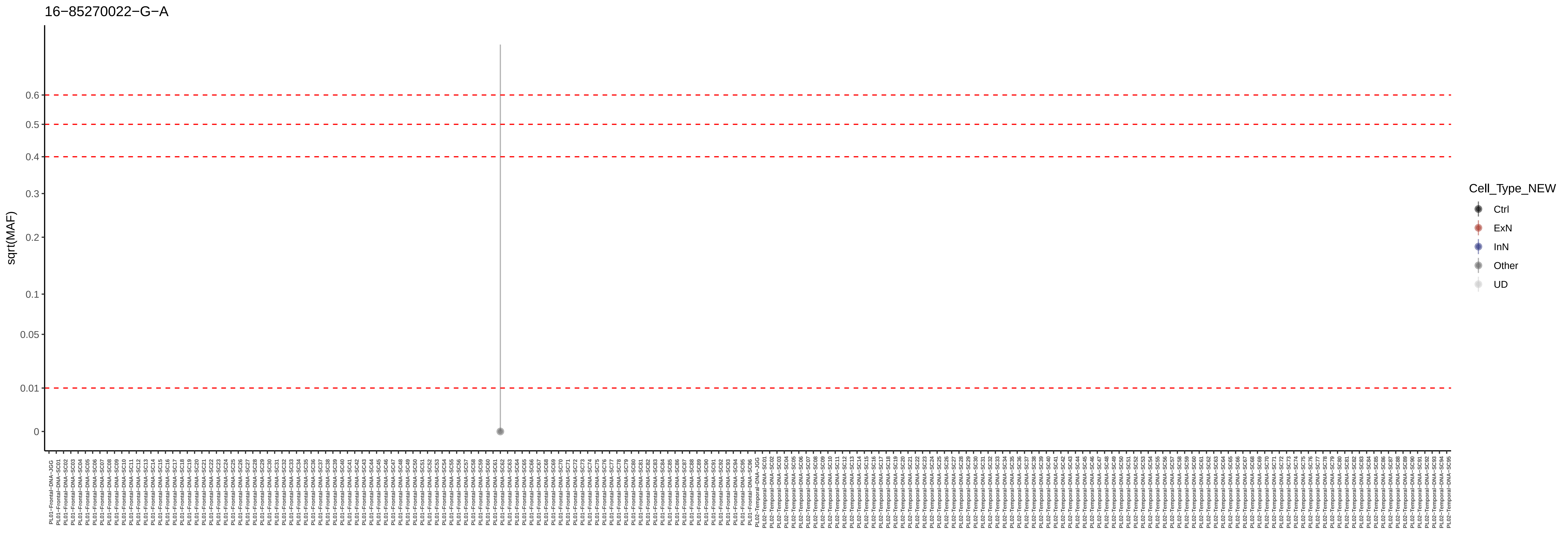
16−85270022−G−A
sqrt(MAF)
0.6
0.5
0.4
0.3
0.2
0.1
0.05
0.01
0
Cell_Type_NEW
Ctrl
ExN
InN
Other
UD

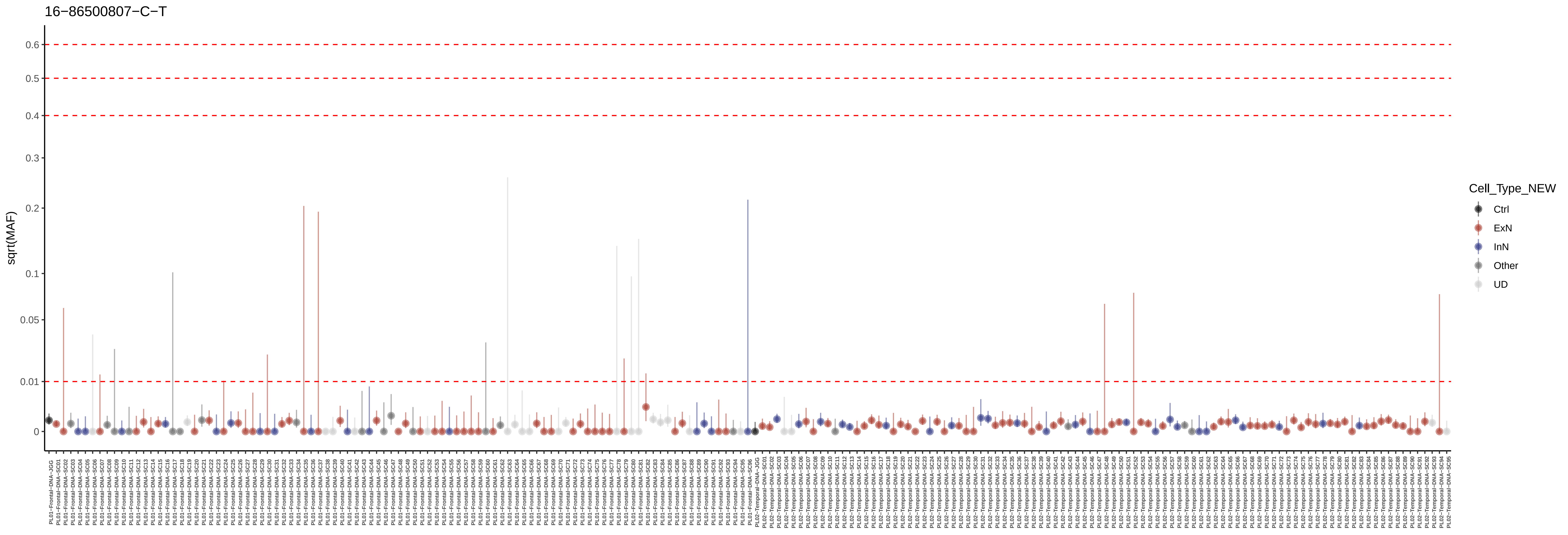
16−86500807−C−T
sqrt(MAF)
0.6
0.5
0.4
0.3
0.2
0.1
0.05
0.01
0
Cell_Type_NEW
Ctrl
ExN
InN
Other
UD

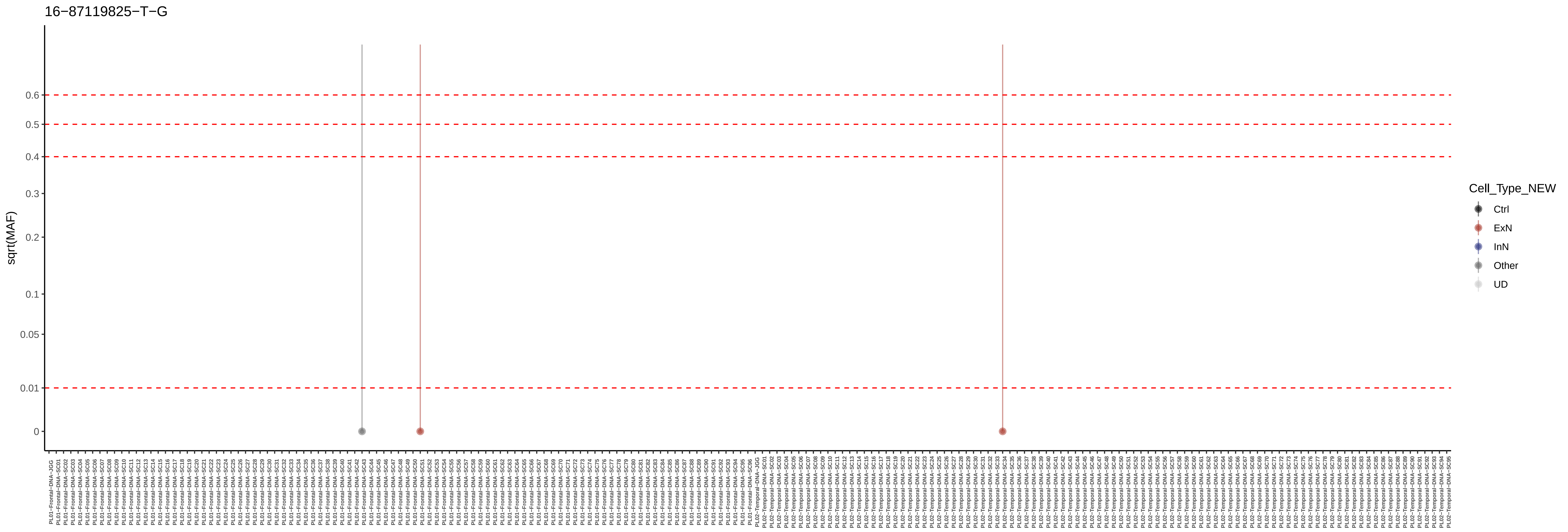

16−87119825−T−G
sqrt(MAF)
Cell_Type_NEW
Ctrl
ExN
InN
Other
UD

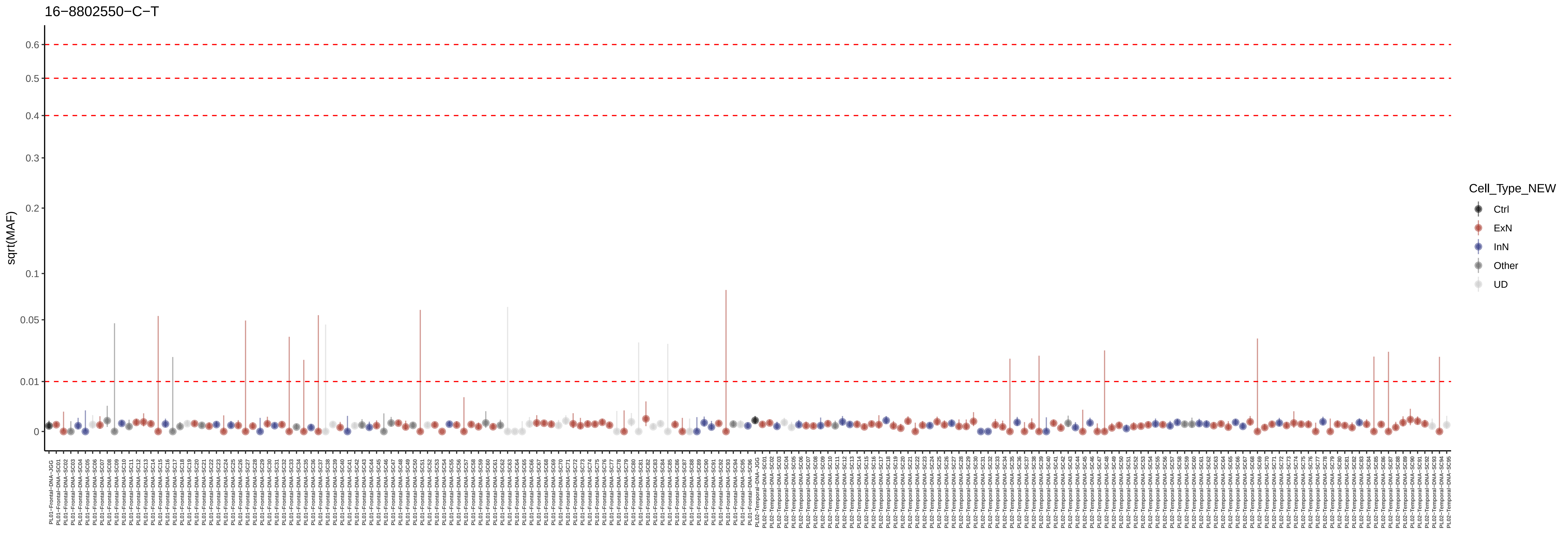
16−8802550−C−T
sqrt(MAF)
0.6
0.5
0.4
0.3
0.2
0.1
0.05
0.01
0
Cell_Type_NEW
Ctrl
ExN
InN
Other
UD

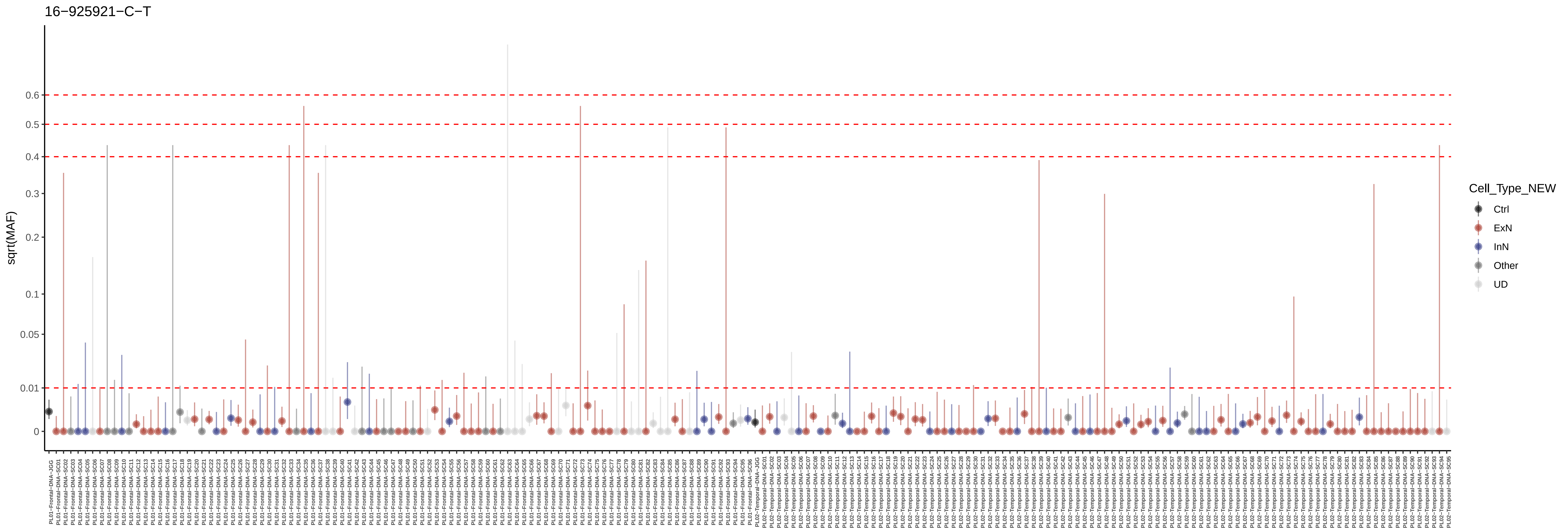
16−925921−C−T
sqrt(MAF)
0.6
0.5
0.4
0.3
0.2
0.1
0.05
0.01
0
Cell_Type_NEW
Ctrl
ExN
InN
Other
UD

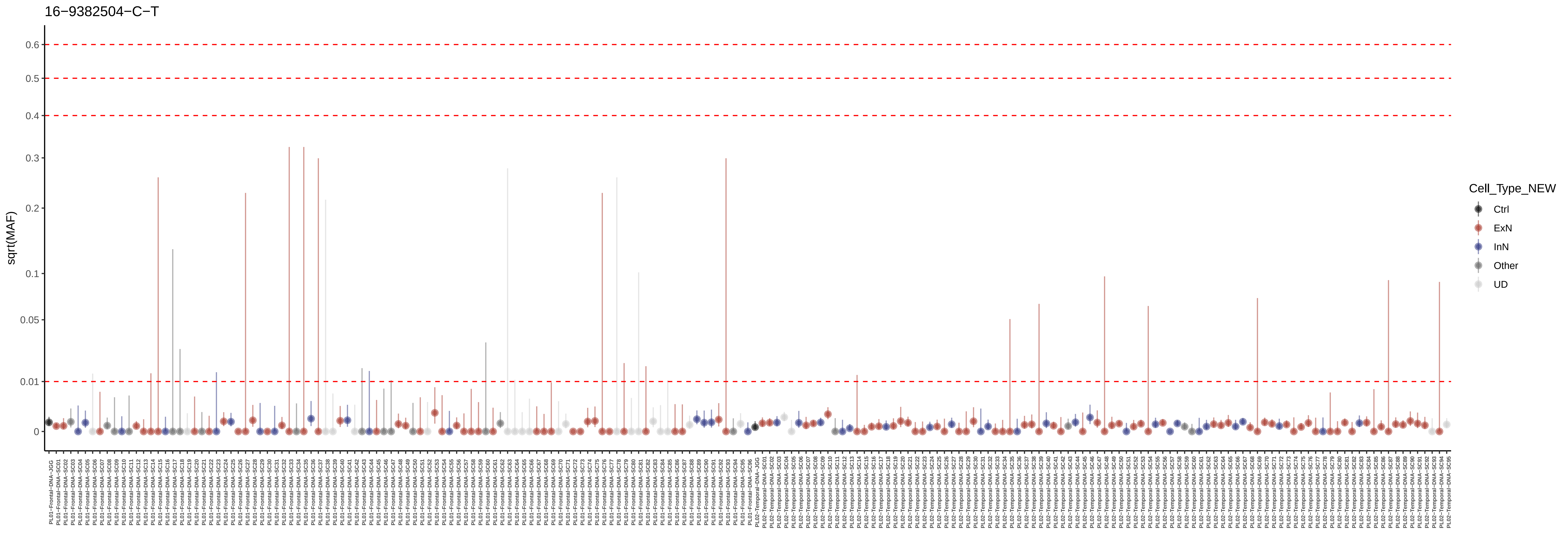
16−9382504−C−T
sqrt(MAF)
0.6
0.5
0.4
0.3
0.2
0.1
0.05
0.01
0
Cell_Type_NEW
Ctrl
ExN
InN
Other
UD

17−10257817−T−A

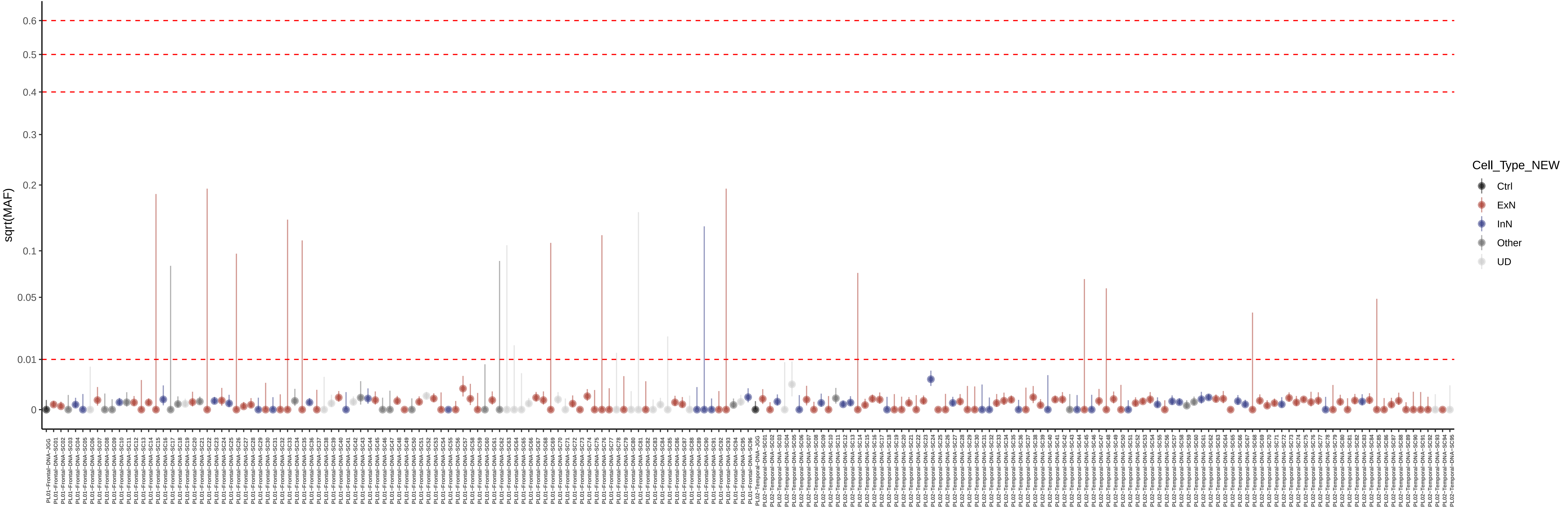

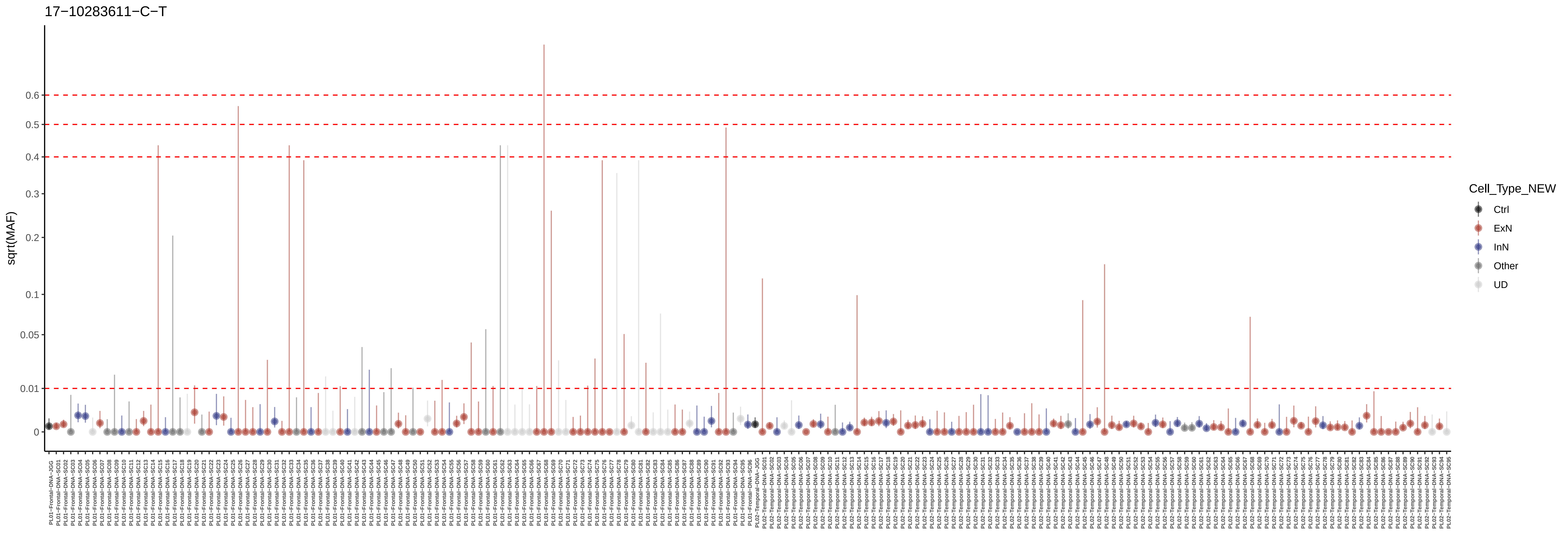
17−10283611−C−T
sqrt(MAF)
0.6
0.5
0.4
0.3
0.2
0.1
0.05
0.01
0
Cell_Type_NEW
Ctrl
ExN
InN
Other
UD

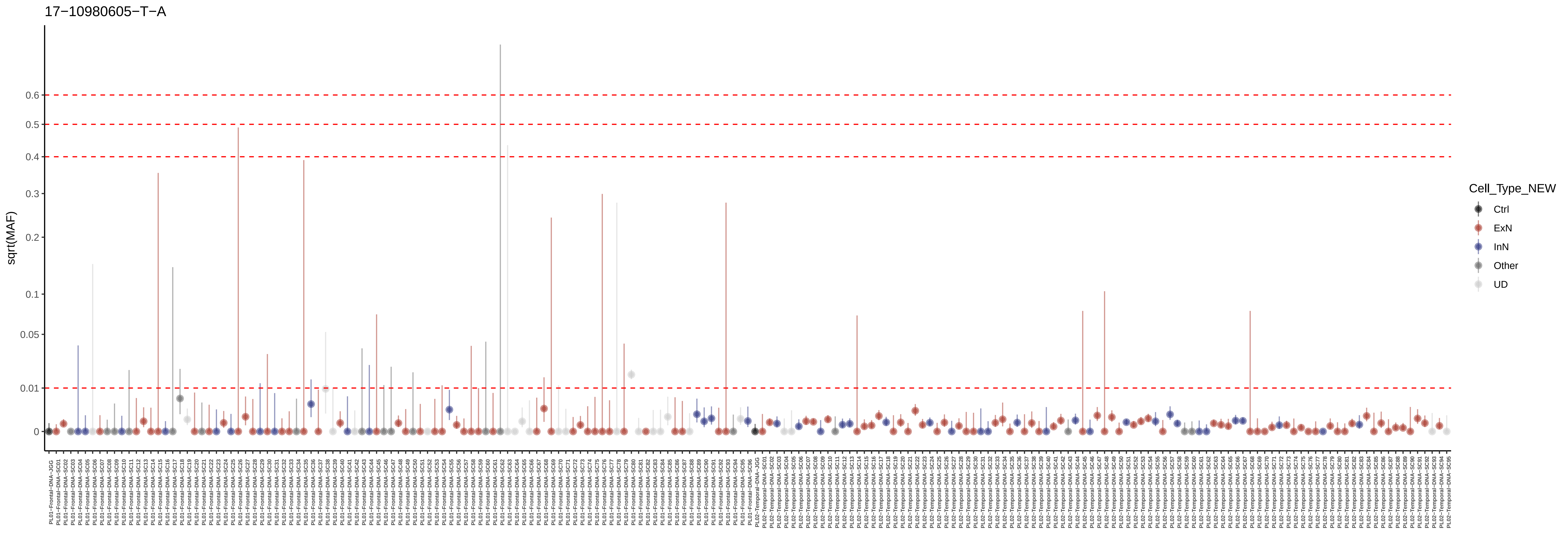
17−10980605−T−A
sqrt(MAF)
0.6
0.5
0.4
0.3
0.2
0.1
0.05
0.01
0
Cell_Type_NEW
Ctrl
ExN
InN
Other
UD

17−12450661−G−A

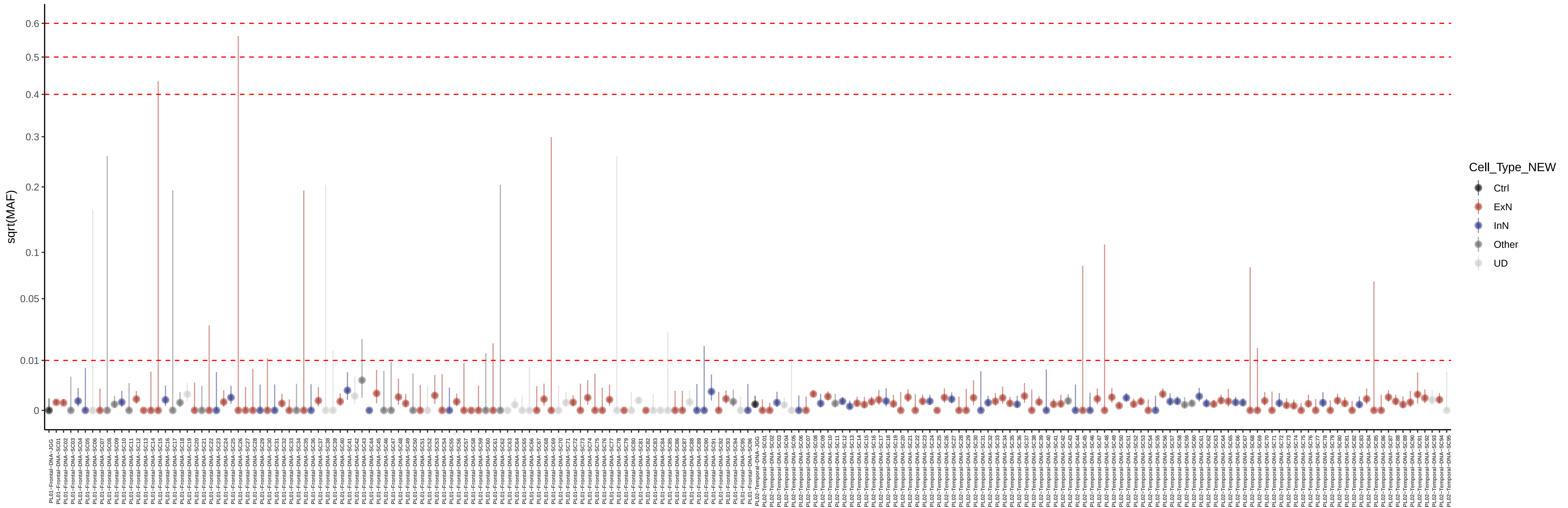

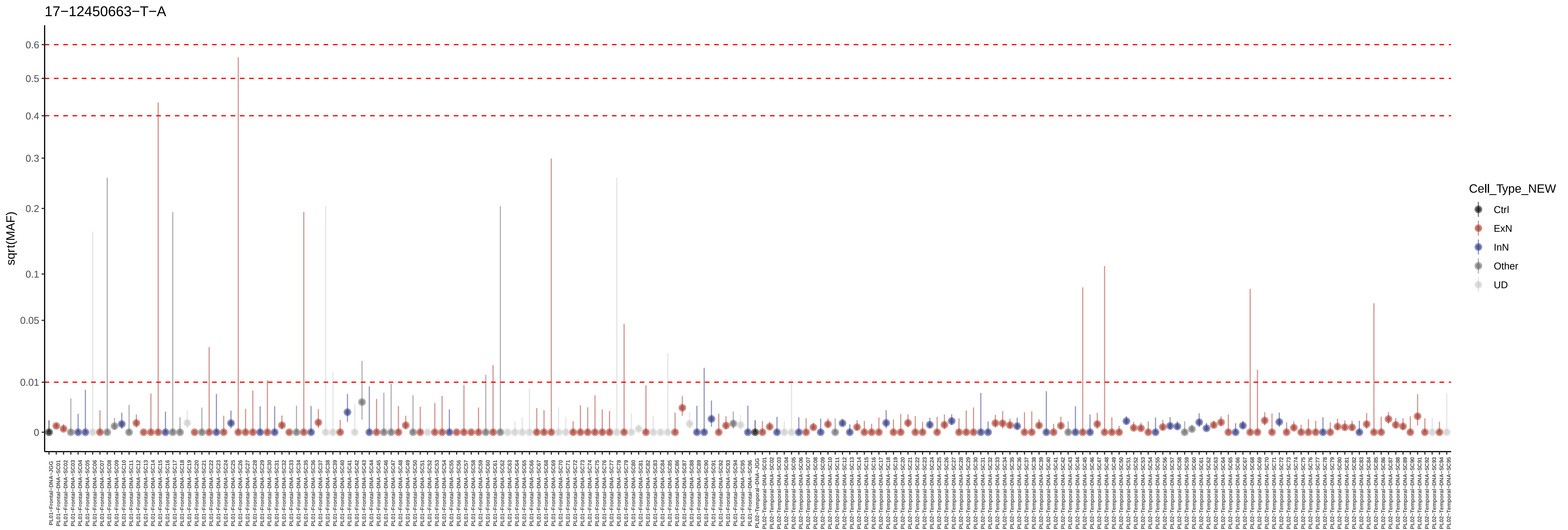

17−12450663−T−A
sqrt(MAF)
0.6
0.5
0.4
0.3
0.2
0.1
0.05
0.01
0
Cell_Type_NEW
Ctrl
ExN
InN
Other
UD

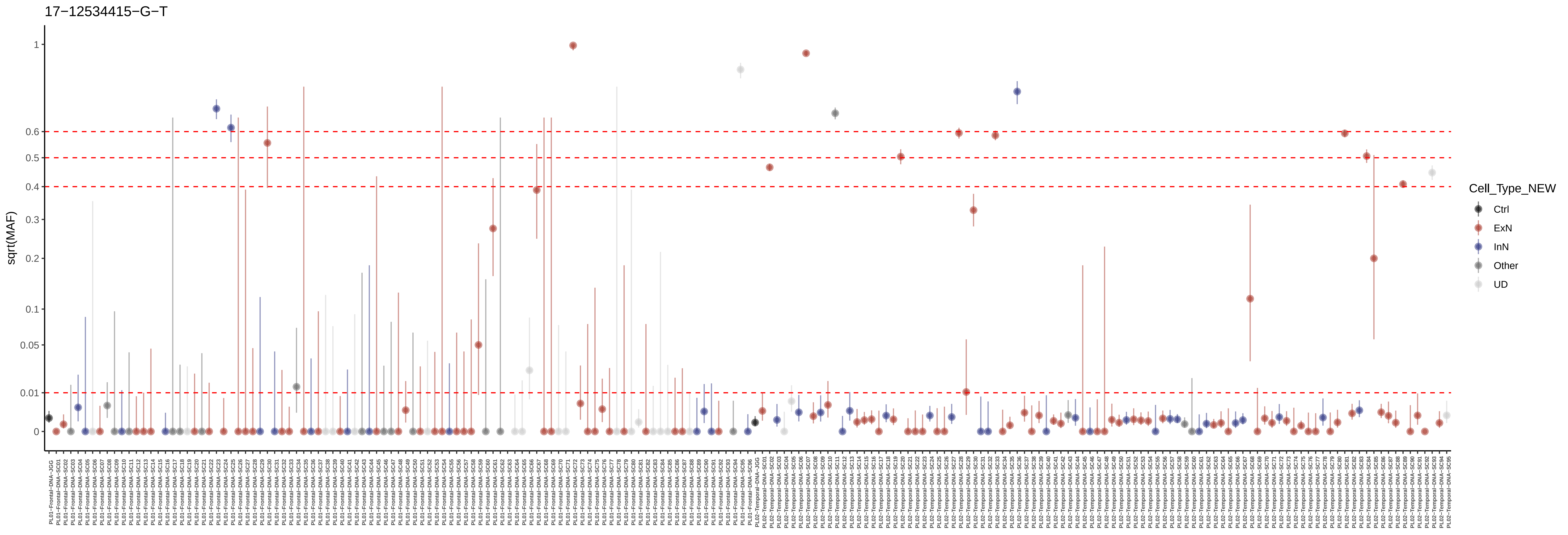
17−12534415−G−T
sqrt(MAF)
1
0.6
0.5
0.4
0.3
0.2
0.1
0.05
0.01
0
Cell_Type_NEW
Ctrl
ExN
InN
Other
UD

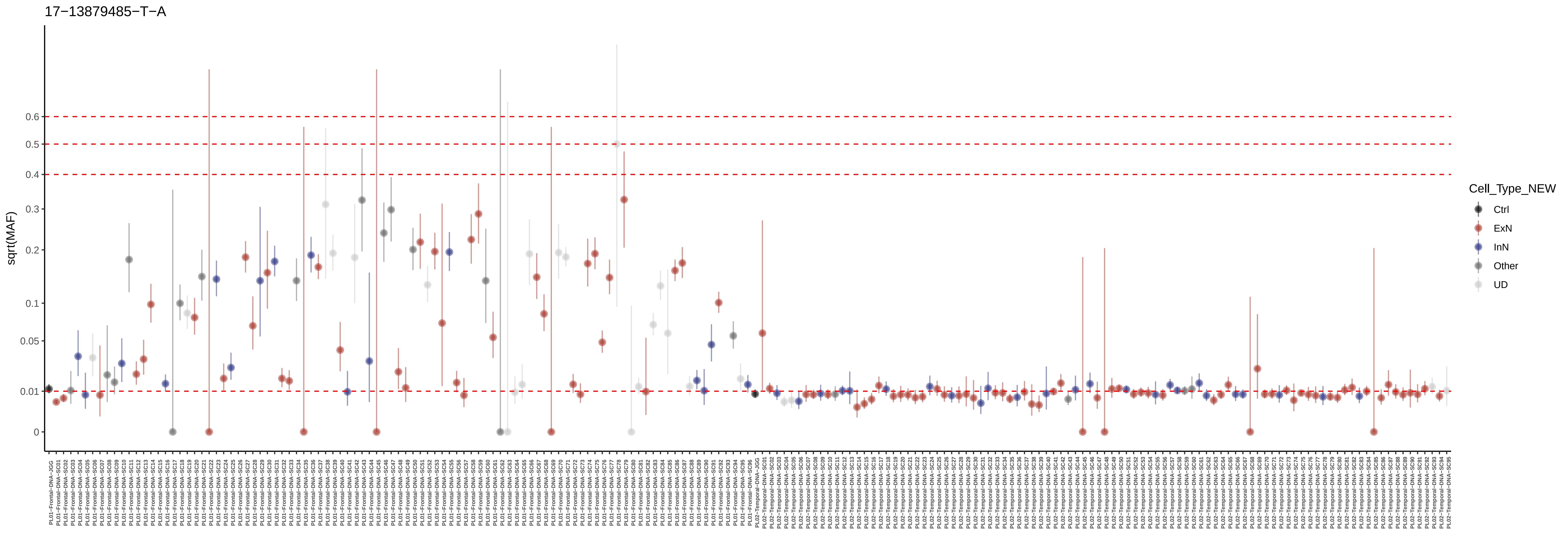
17−13879485−T−A
sqrt(MAF)
0.6
0.5
0.4
0.3
0.2
0.1
0.05
0.01
0
Cell_Type_NEW
Ctrl
ExN
InN
Other
UD

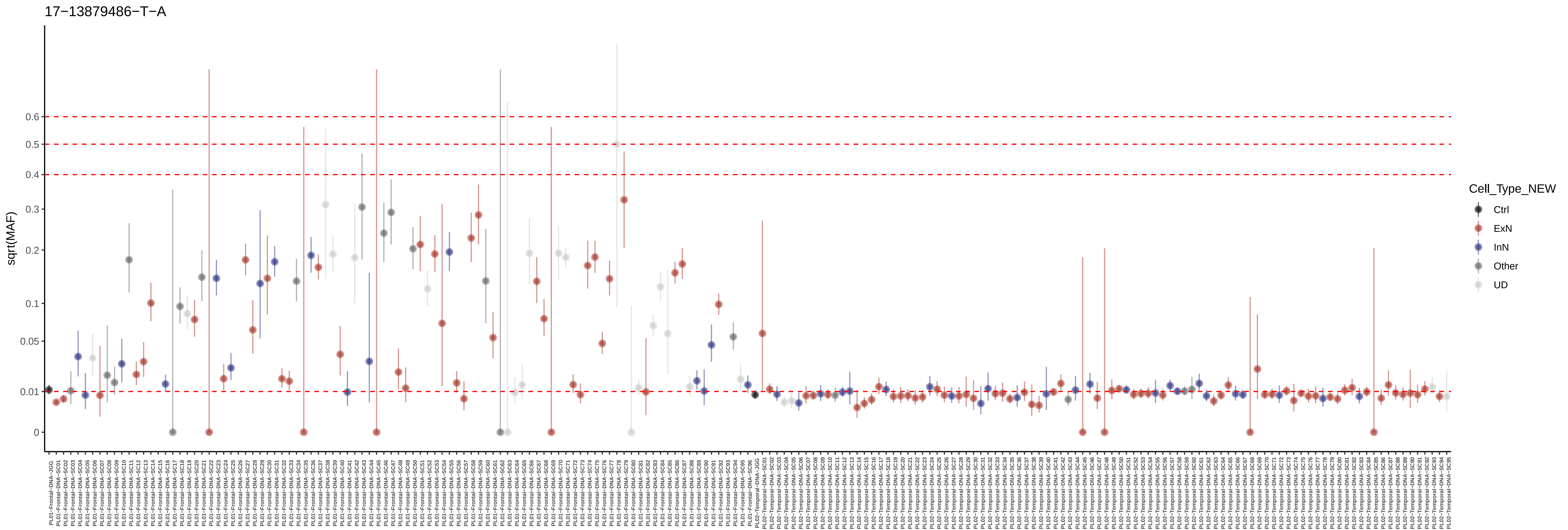
17−13879486−T−A
sqrt(MAF)
0.6
0.5
0.4
0.3
0.2
0.1
0.05
0.01
0
Cell_Type_NEW
Ctrl
ExN
InN
Other
UD

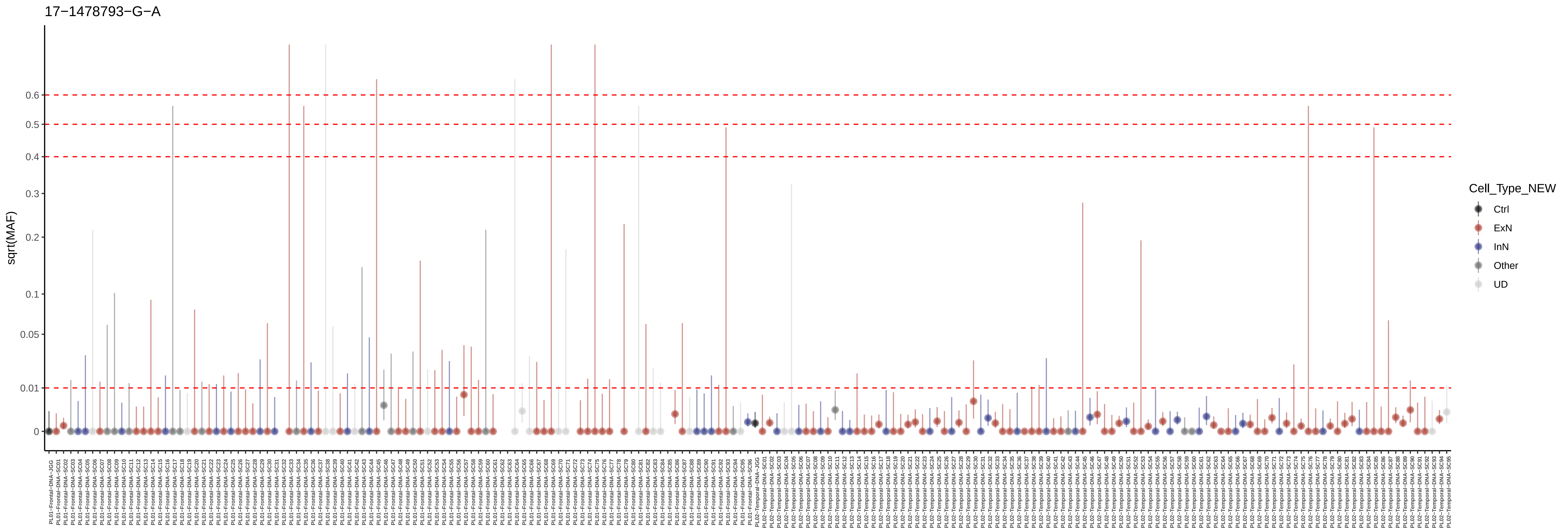
17−1478793−G−A
sqrt(MAF)
0.6
0.5
0.4
0.3
0.2
0.1
0.05
0.01
0
Cell_Type_NEW
Ctrl
ExN
InN
Other
UD

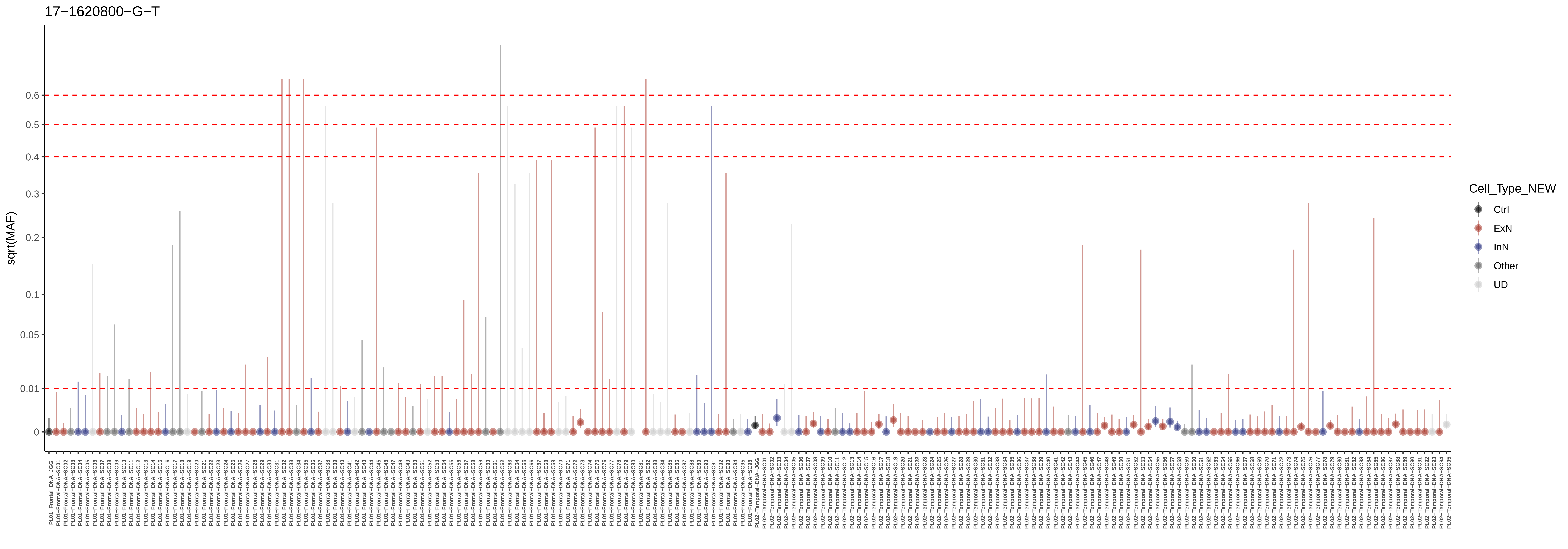
17−1620800−G−T
sqrt(MAF)
Cell_Type_NEW
Ctrl
ExN
InN
Other
UD

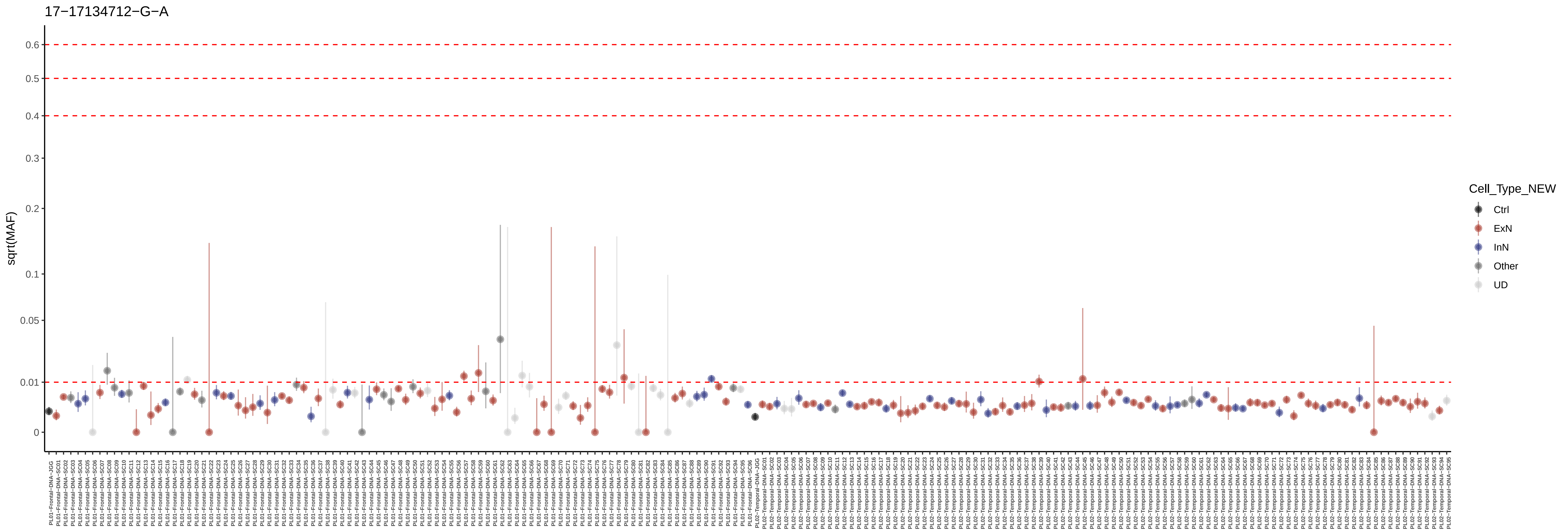
17−17134712−G−A
sqrt(MAF)
0.6
0.5
0.4
0.3
0.2
0.1
0.05
0.01
0
Cell_Type_NEW
Ctrl
ExN
InN
Other
UD

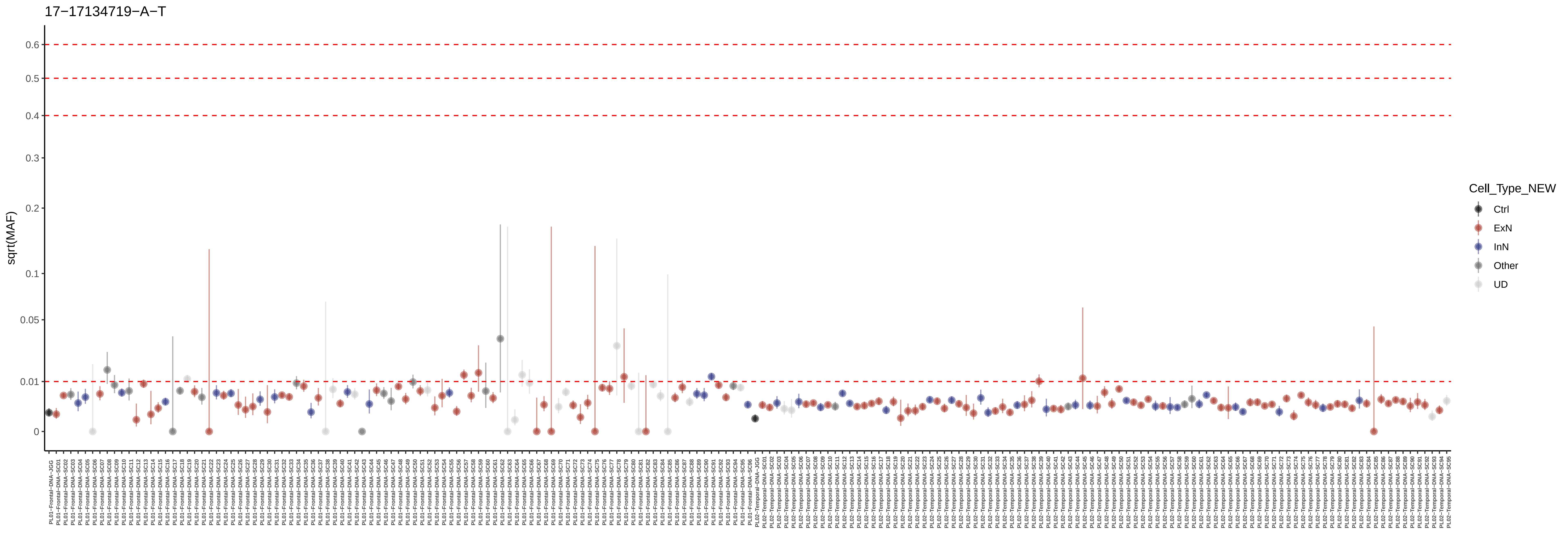
17−17134719−A−T
sqrt(MAF)
0.6
0.5
0.4
0.3
0.2
0.1
0.05
0.01
0
Cell_Type_NEW
Ctrl
ExN
InN
Other
UD

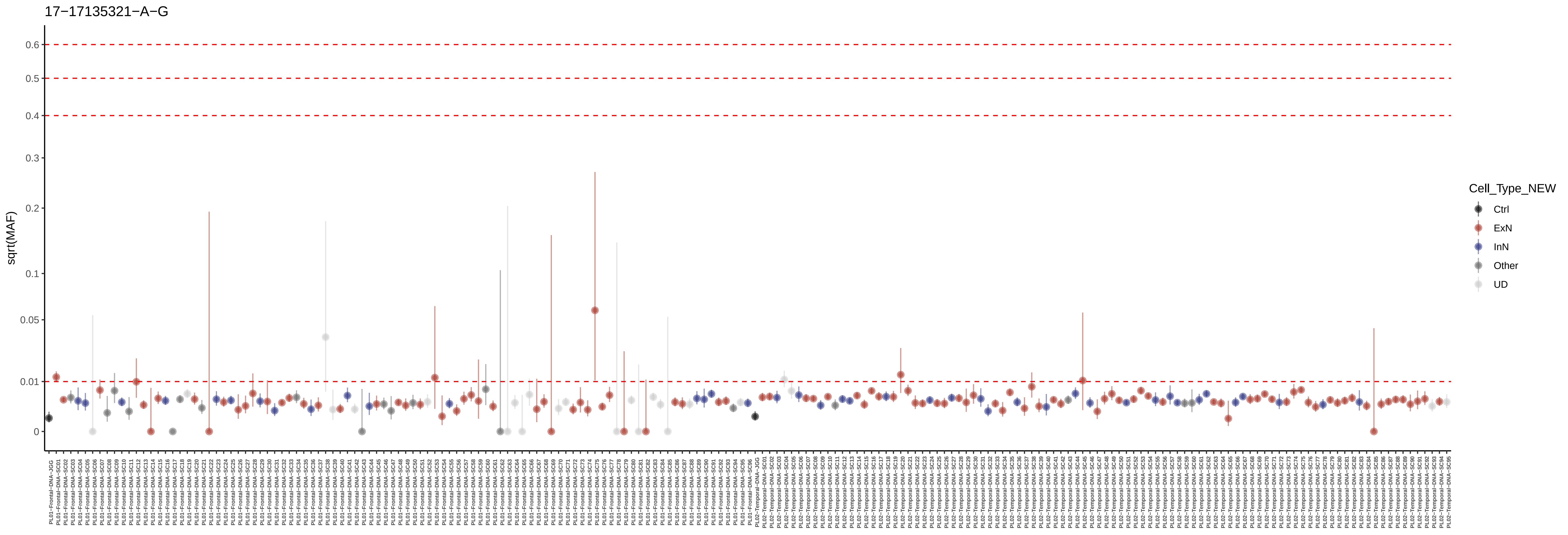
17−17135321−A−G
sqrt(MAF)
0.6
0.5
0.4
0.3
0.2
0.1
0.05
0.01
0
Cell_Type_NEW
Ctrl
ExN
InN
Other
UD

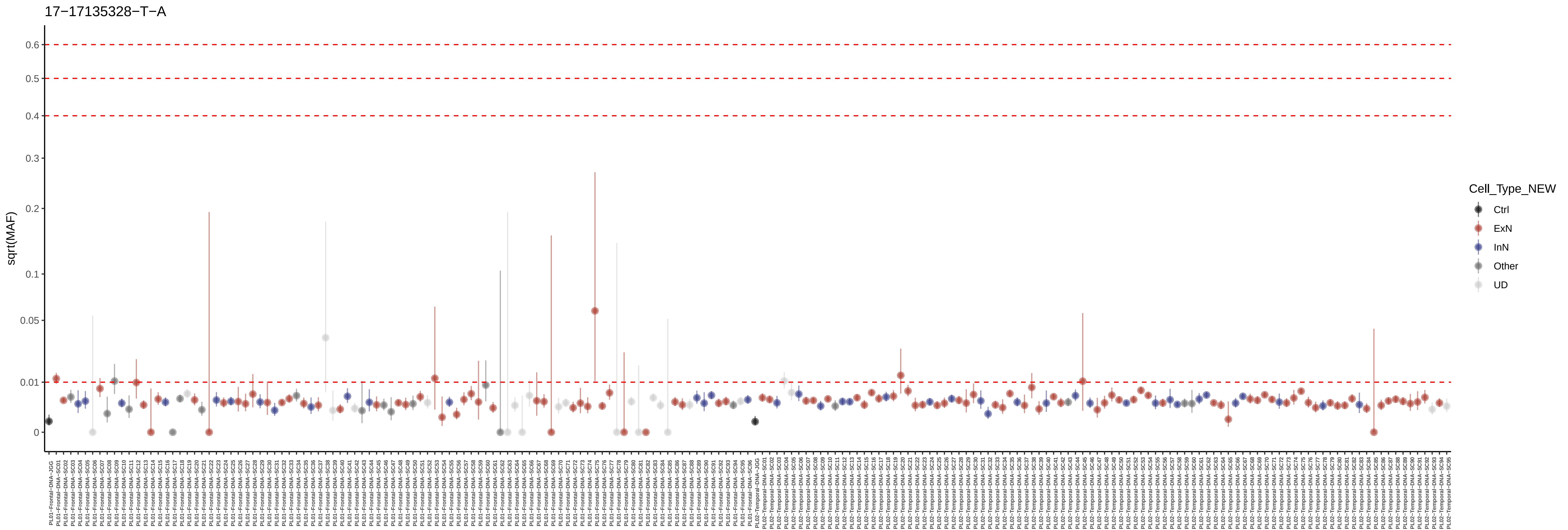

17−17135328−T−A
sqrt(MAF)
0.6
0.5
0.4
0.3
0.2
0.1
0.05
0.01
0
Cell_Type_NEW
Ctrl
ExN
InN
Other
UD

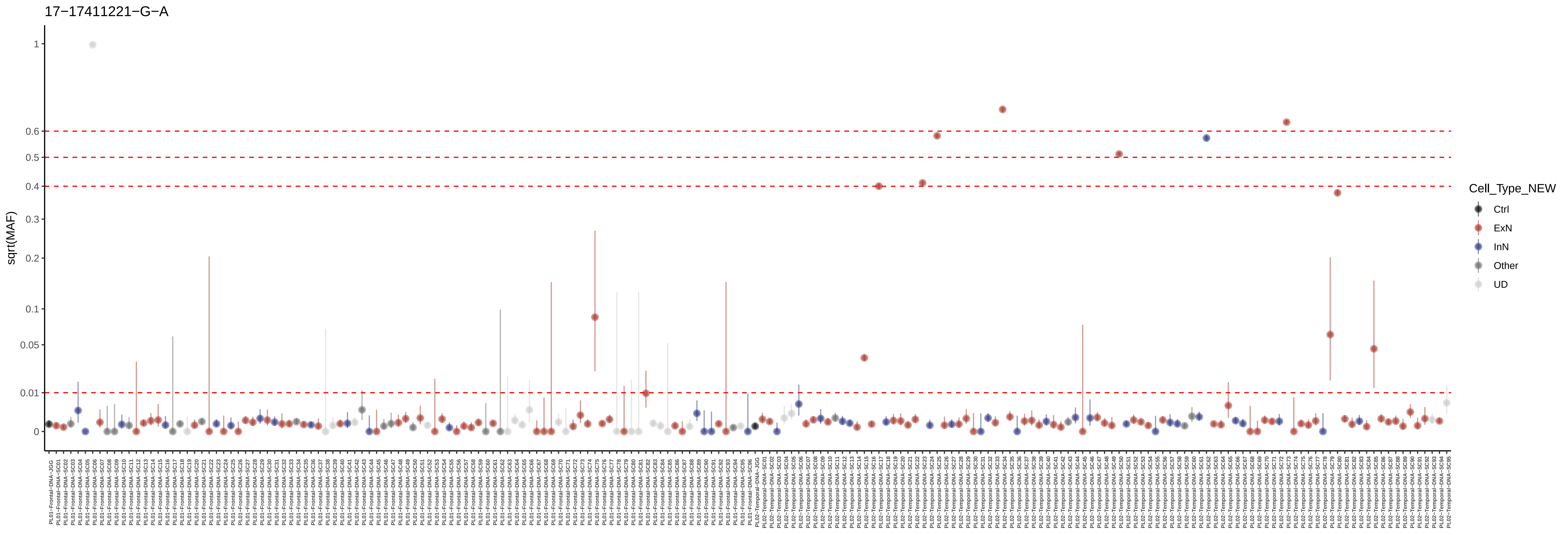

17−17411221−G−A
sqrt(MAF)
1
0.6
0.5
0.4
0.3
0.2
0.1
0.05
0.01
0
Cell_Type_NEW
Ctrl
ExN
InN
Other
UD

17−184881−G−A

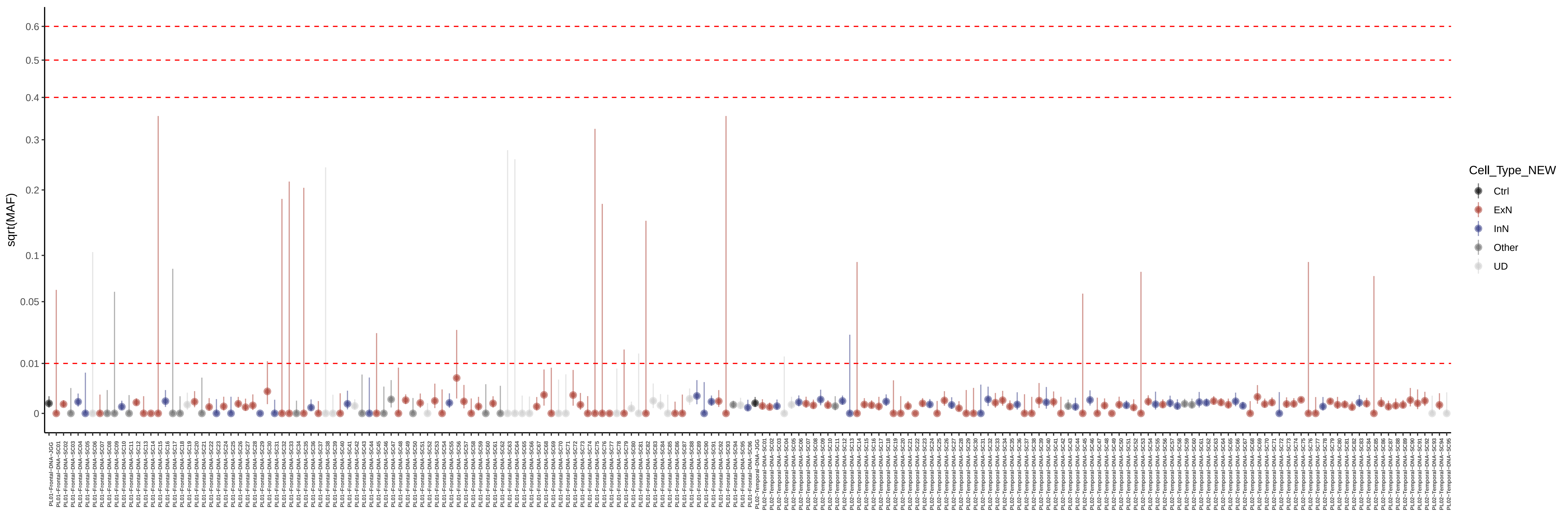

sqrt(MAF)
0.6
0.5
0.4
0.3
0.2
0.1
0.05
0.01
0
Cell_Type_NEW
Ctrl
ExN
InN
Other
UD

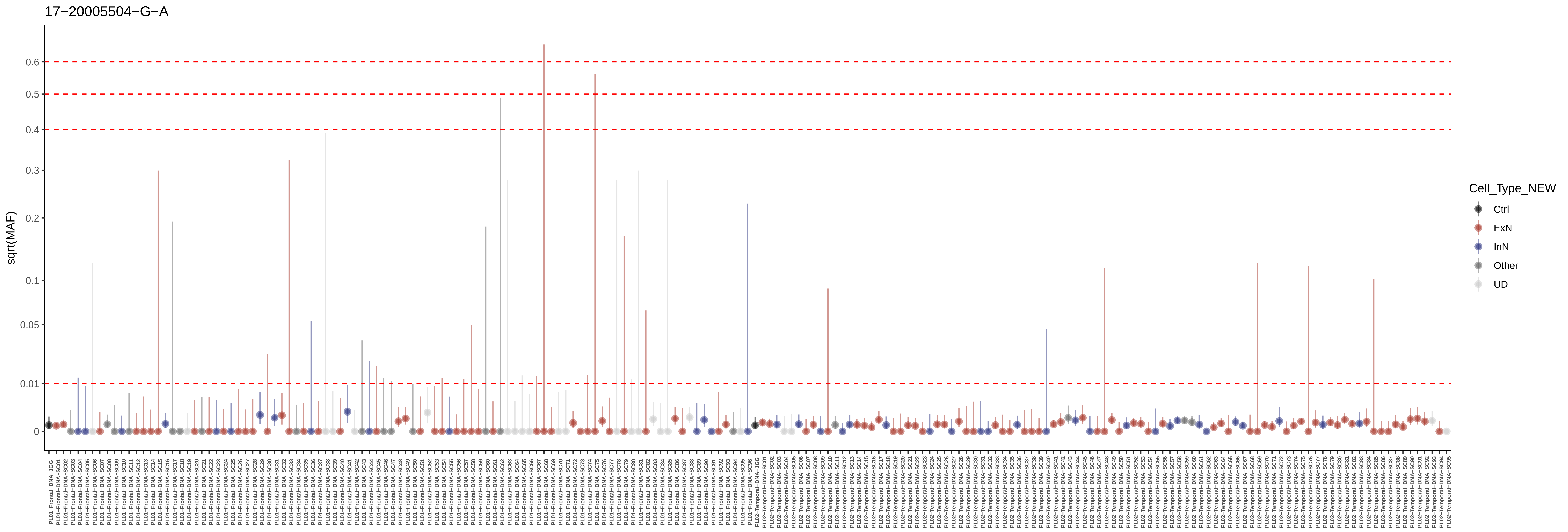
17−20005504−G−A
sqrt(MAF)
0.6
0.5
0.4
0.3
0.2
0.1
0.05
0.01
0
Cell_Type_NEW
Ctrl
ExN
InN
Other
UD

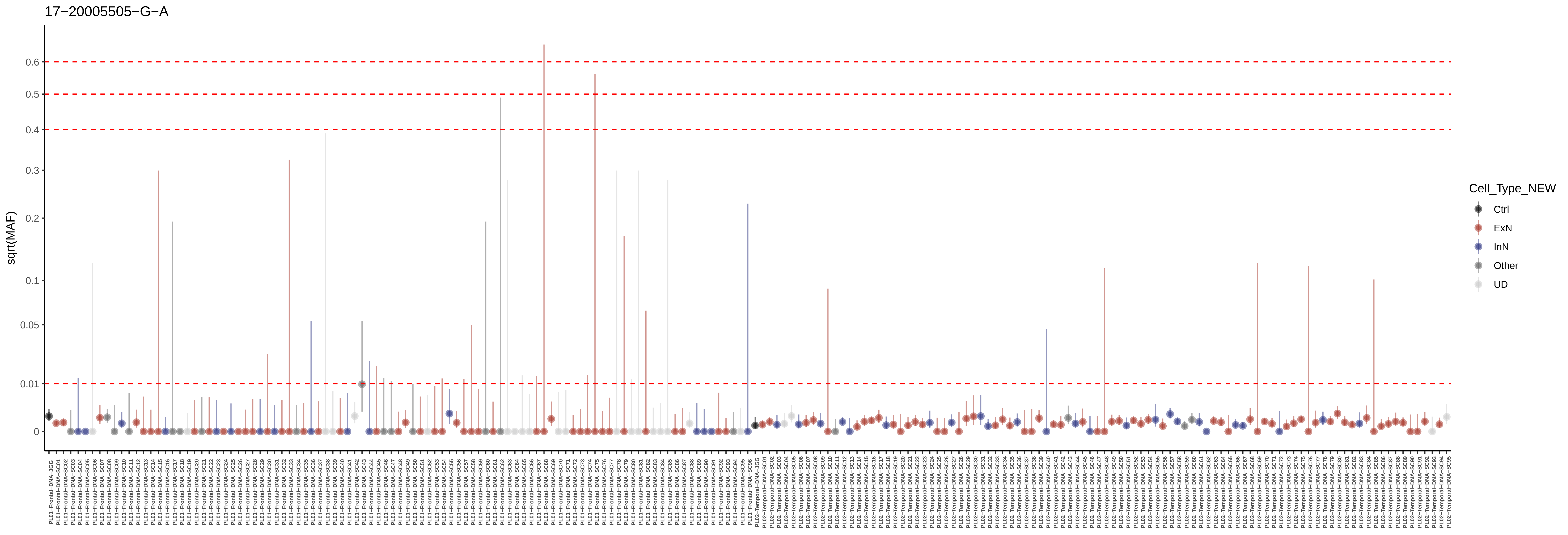

17−20005505−G−A
sqrt(MAF)
0.6
0.5
0.4
0.3
0.2
0.1
0.05
0.01
0
Cell_Type_NEW
Ctrl
ExN
InN
Other
UD

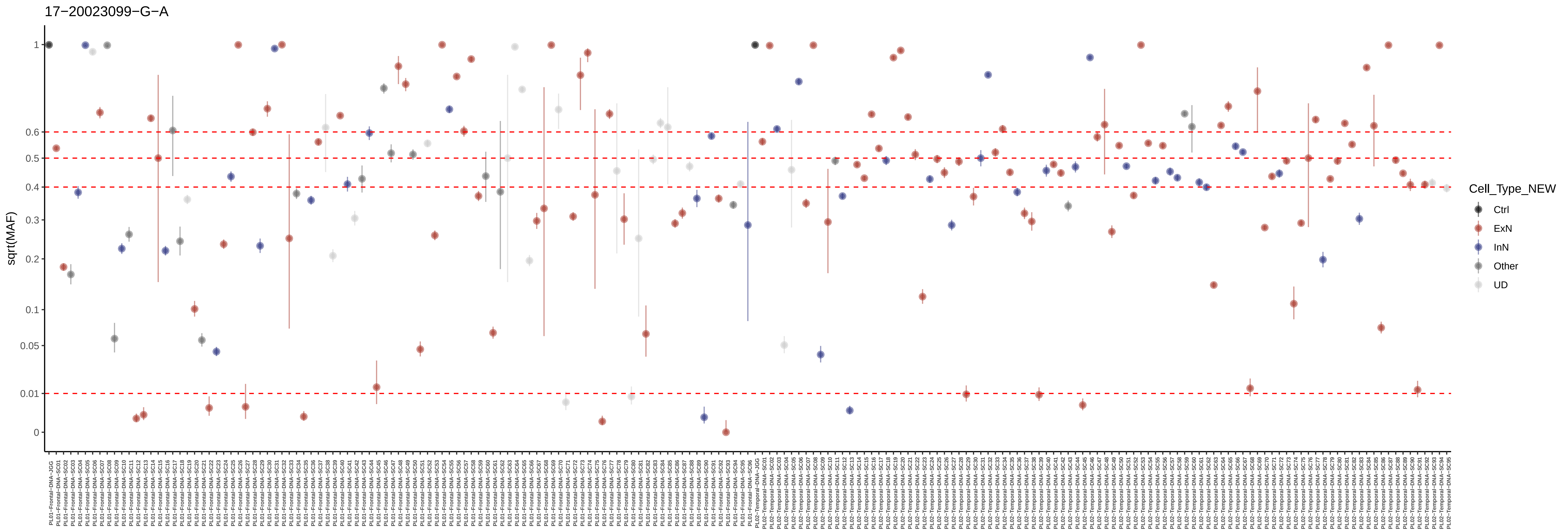
17−20023099−G−A
sqrt(MAF)
1
0.6
0.5
0.4
0.3
0.2
0.1
0.05
0.01
0
Cell_Type_NEW
Ctrl
ExN
InN
Other
UD

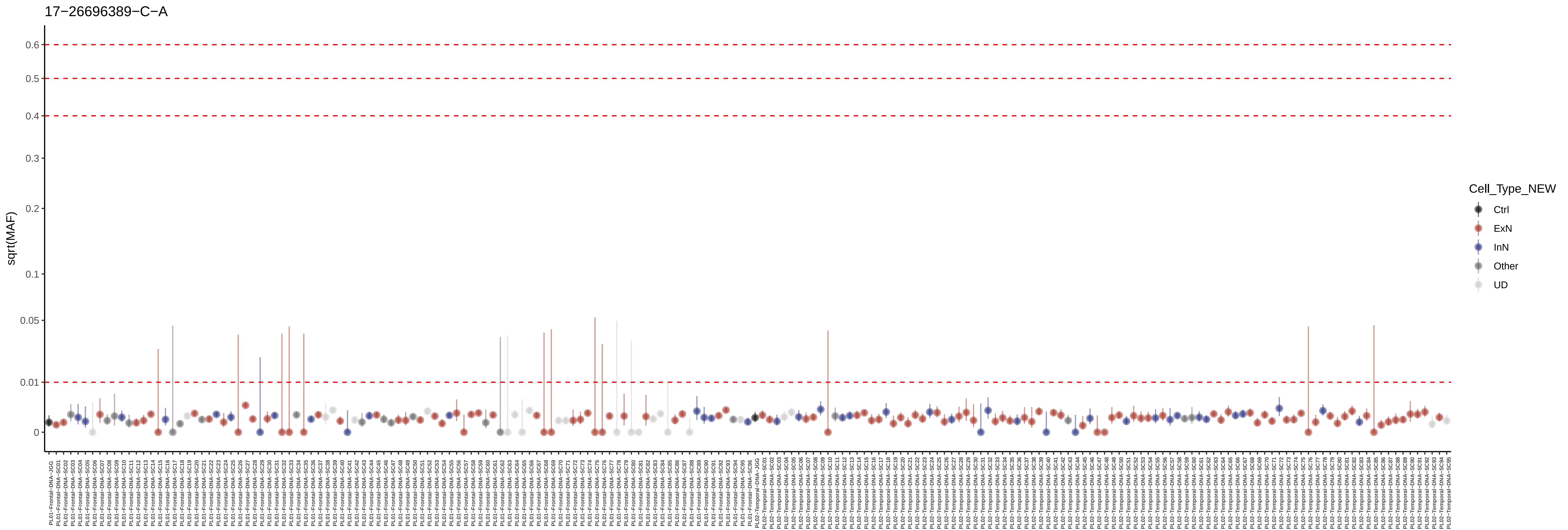

17−26696389−C−A
sqrt(MAF)
0.6
0.5
0.4
0.3
0.2
0.1
0.05
0.01
0
Cell_Type_NEW
Ctrl
ExN
InN
Other
UD

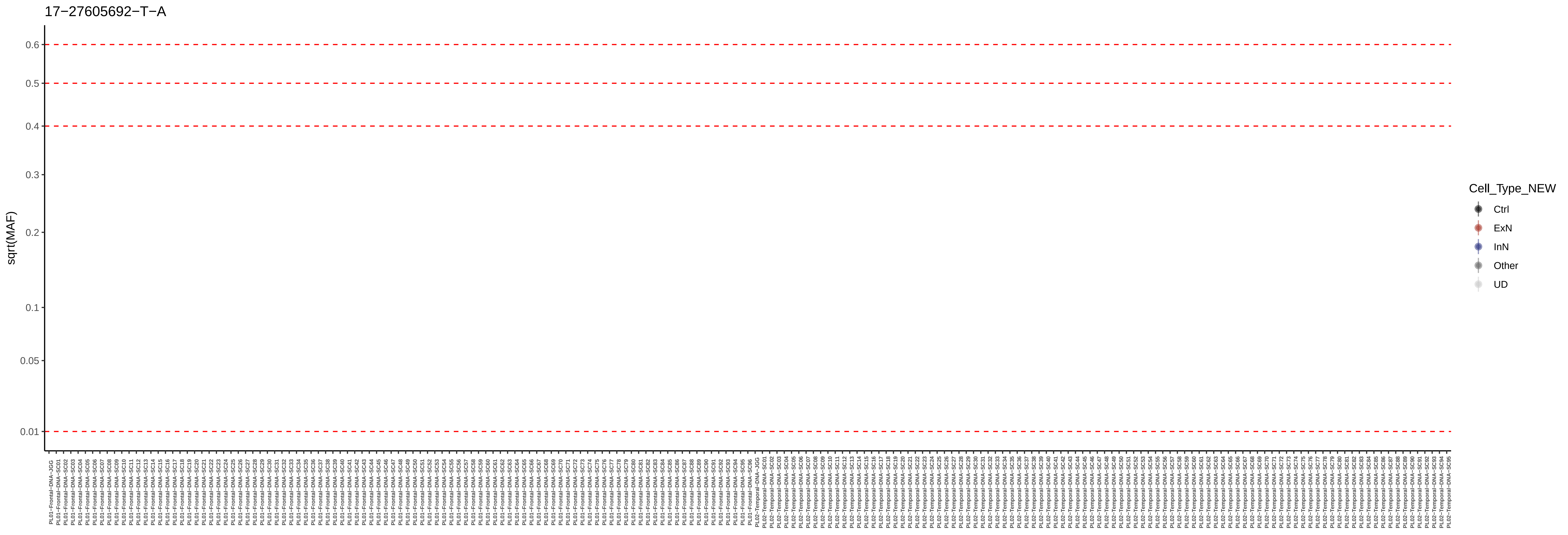
17−27605692−T−A
sqrt(MAF)
0.6
0.5
0.4
0.3
0.2
0.1
0.05
0.01
Cell_Type_NEW
Ctrl
ExN
InN
Other
UD

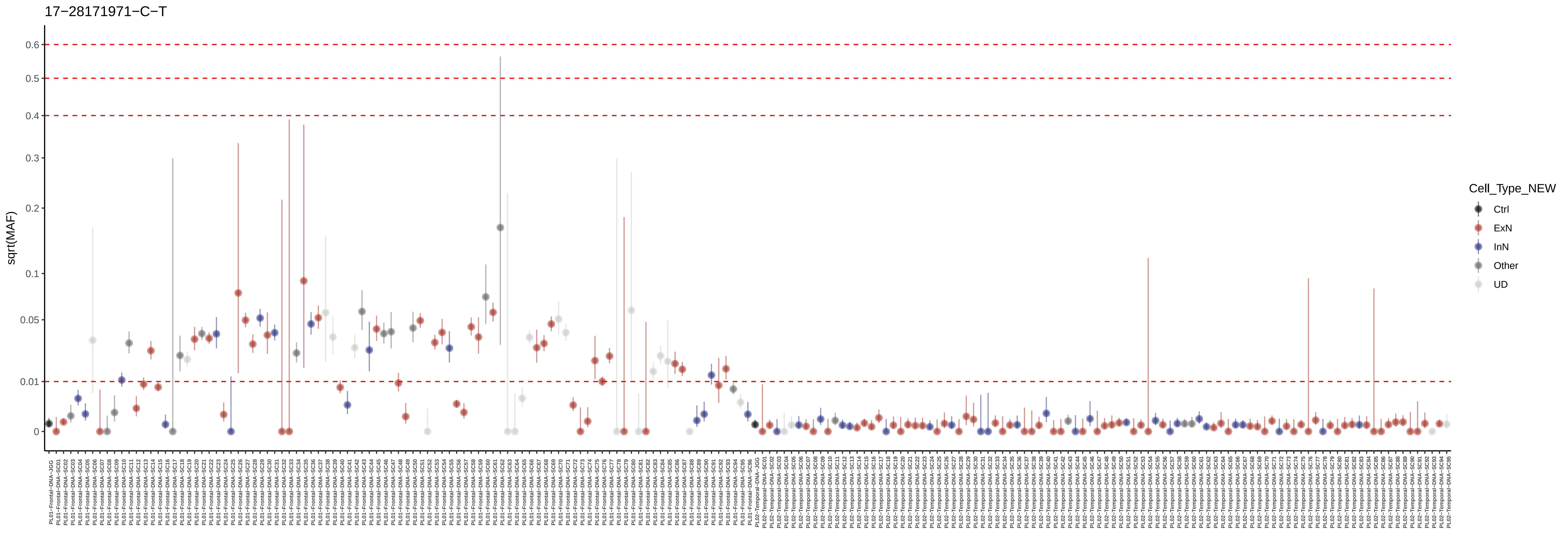
17−28171971−C−T
sqrt(MAF)
0.6
0.5
0.4
0.3
0.2
0.1
0.05
0.01
0
Cell_Type_NEW
Ctrl
ExN
InN
Other
UD

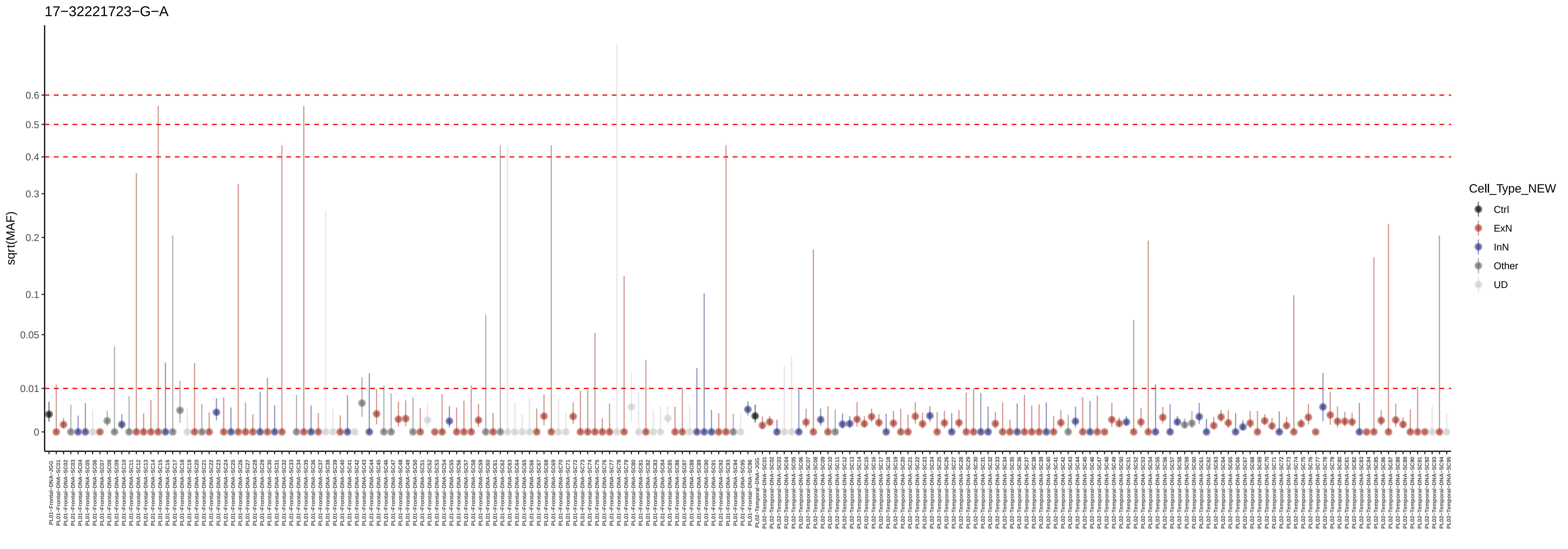

17−32221723−G−A
sqrt(MAF)
0.6
0.5
0.4
0.3
0.2
0.1
0.05
0.01
0
Cell_Type_NEW
Ctrl
ExN
InN
Other
UD

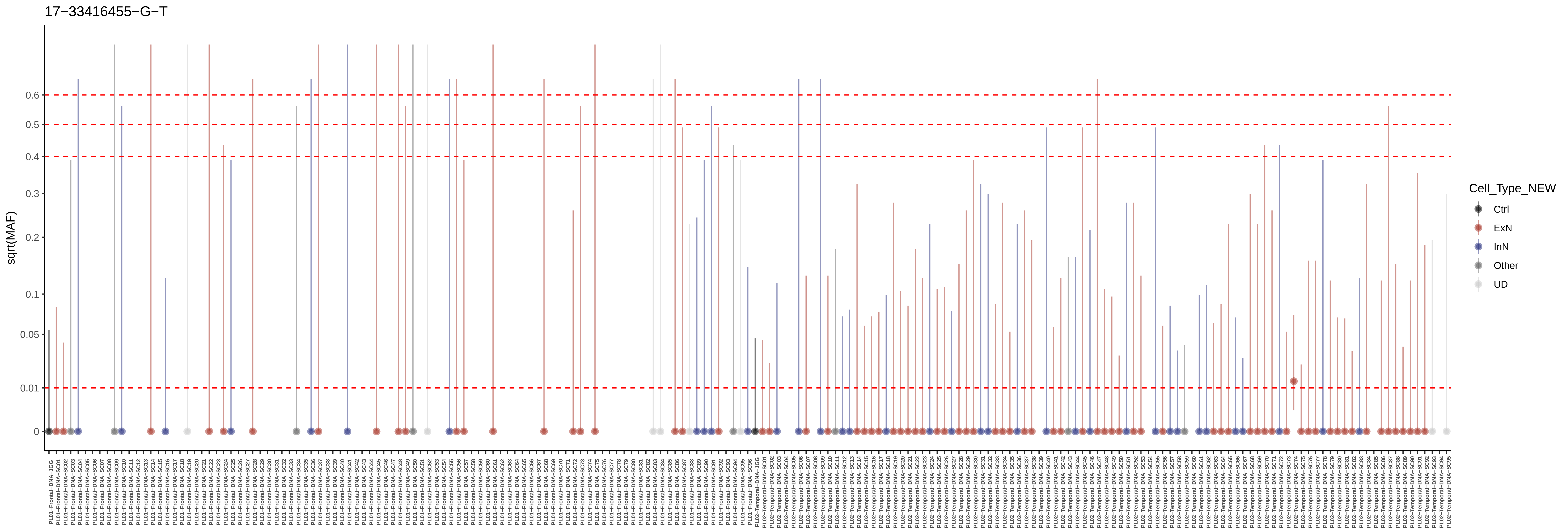

17−33416455−G−T
sqrt(MAF)
0.6
0.5
0.4
0.3
0.2
0.1
0.05
0.01
0
Cell_Type_NEW
Ctrl
ExN
InN
Other
UD

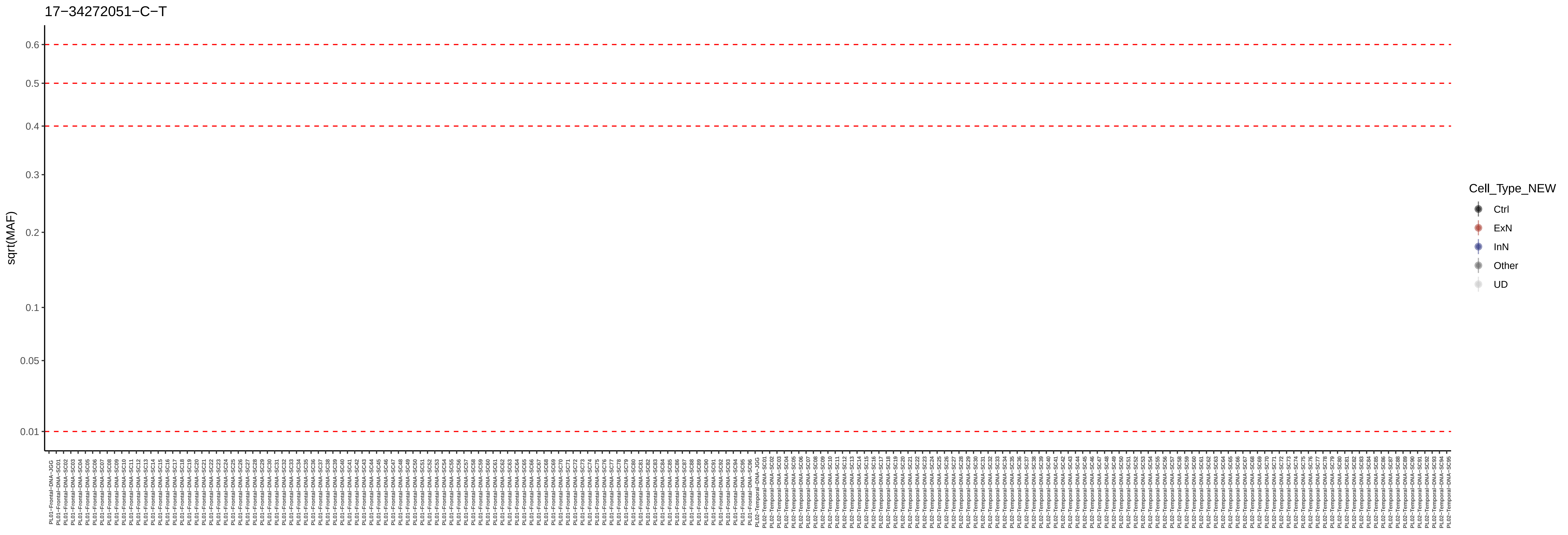
17−34272051−C−T
sqrt(MAF)
0.6
0.5
0.4
0.3
0.2
0.1
0.05
0.01
Cell_Type_NEW
Ctrl
ExN
InN
Other
UD

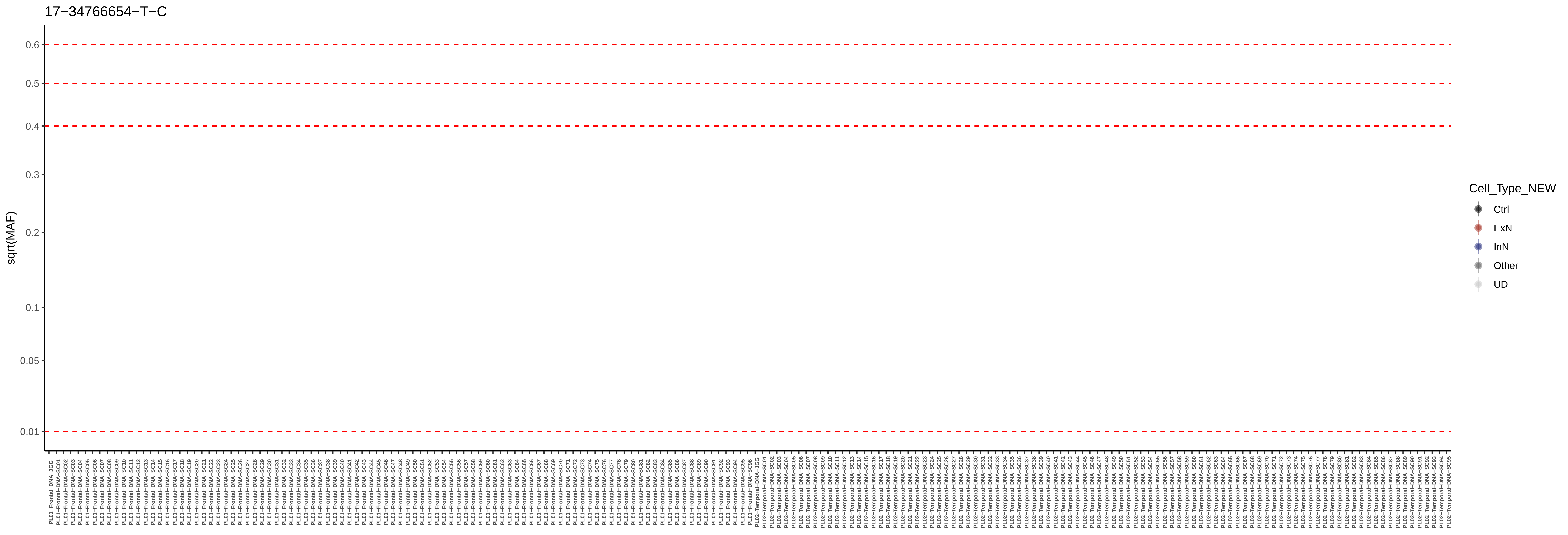

17−34766654−T−C
sqrt(MAF)
0.6
0.5
0.4
0.3
0.2
0.1
0.05
0.01
Cell_Type_NEW
Ctrl
ExN
InN
Other
UD

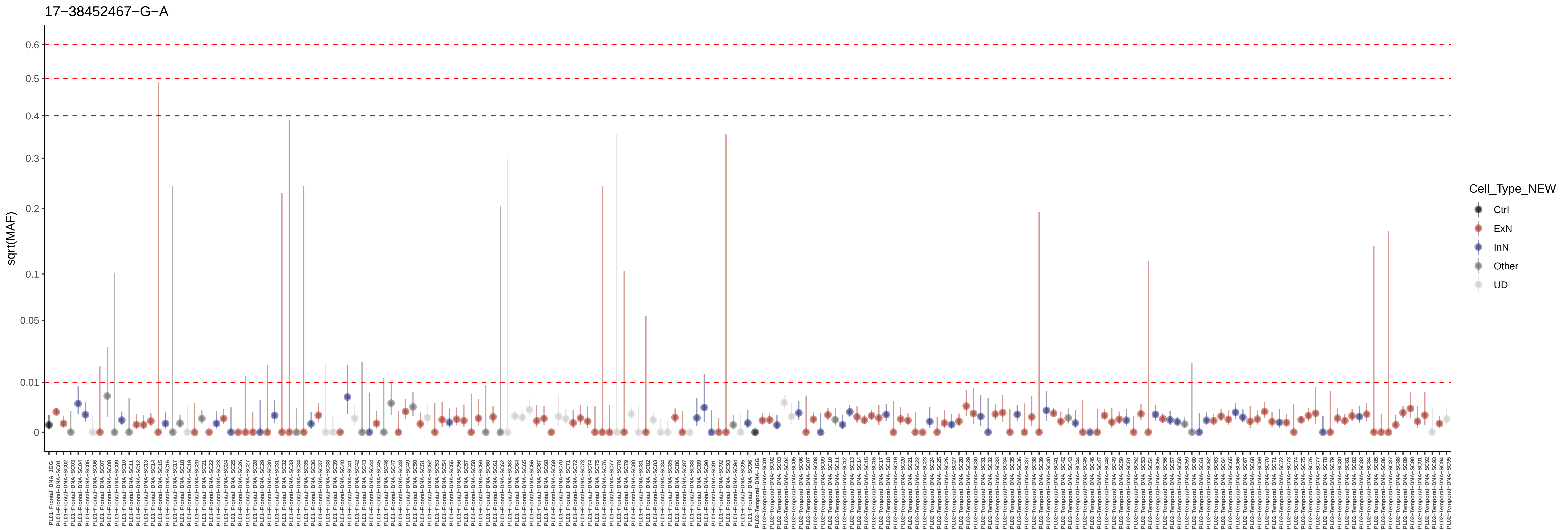

17−38452467−G−A
sqrt(MAF)
0.6
0.5
0.4
0.3
0.2
0.1
0.05
0.01
0
Cell_Type_NEW
Ctrl
ExN
InN
Other
UD

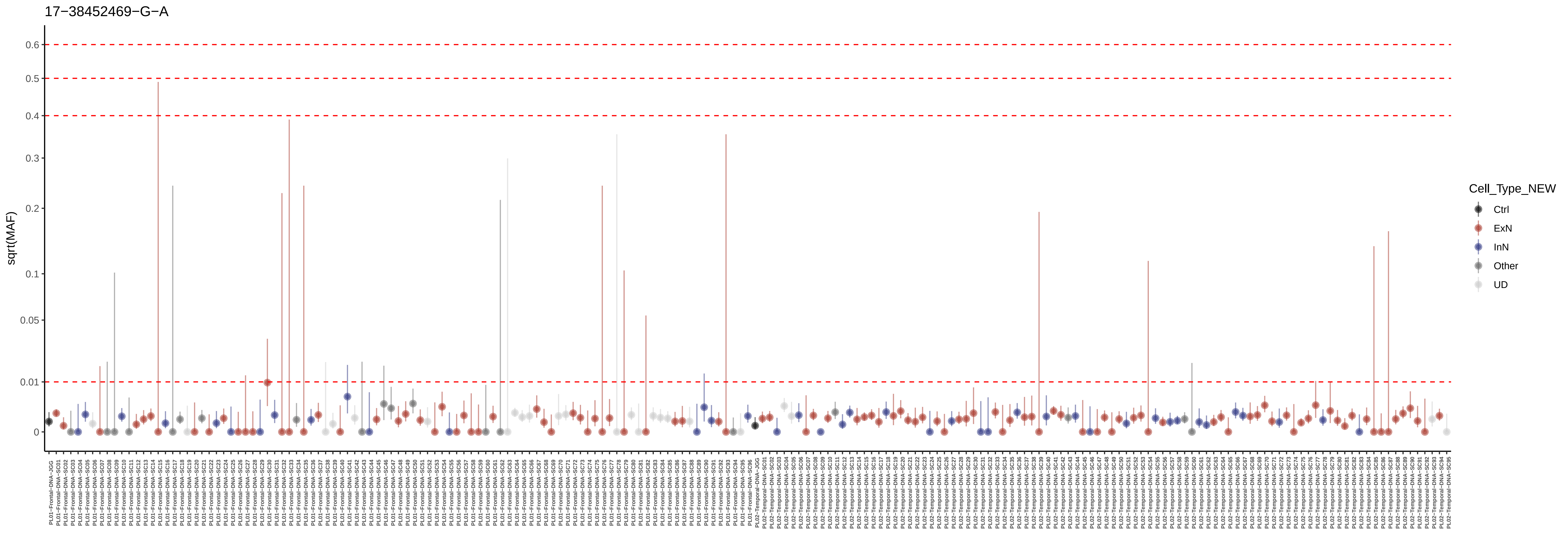
17−38452469−G−A
sqrt(MAF)
0.6
0.5
0.4
0.3
0.2
0.1
0.05
0.01
0
Cell_Type_NEW
Ctrl
ExN
InN
Other
UD

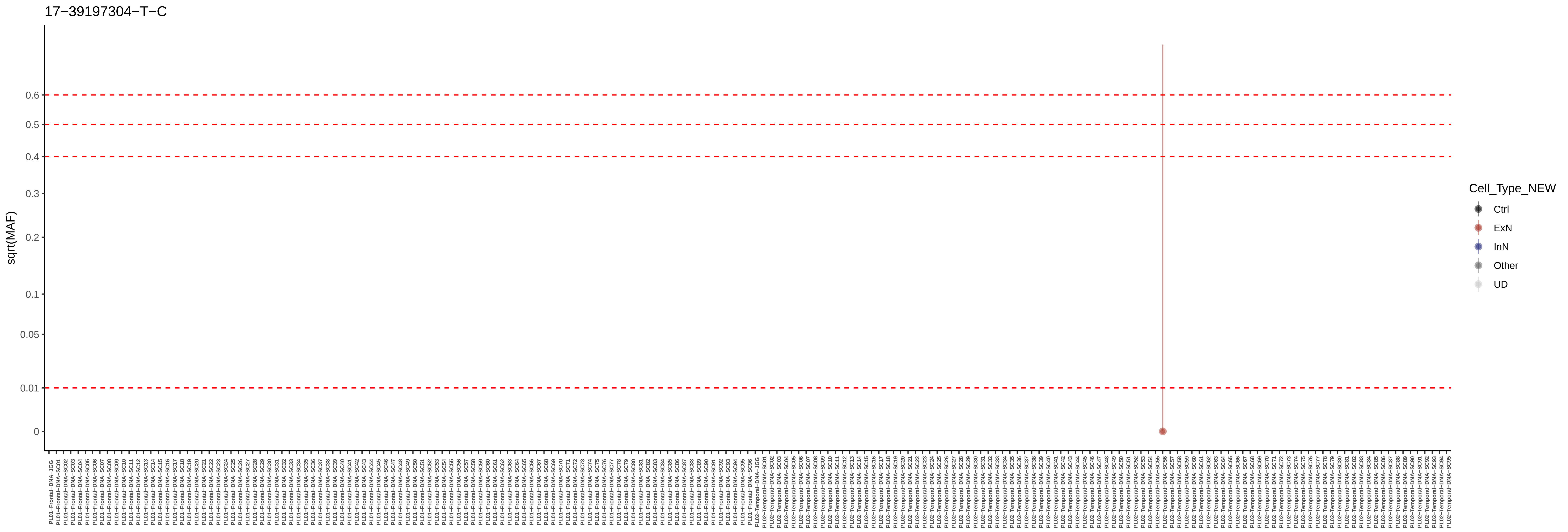
17−39197304−T−C
sqrt(MAF)
0.6
0.5
0.4
0.3
0.2
0.1
0.05
0.01
0
Cell_Type_NEW
Ctrl
ExN
InN
Other
UD

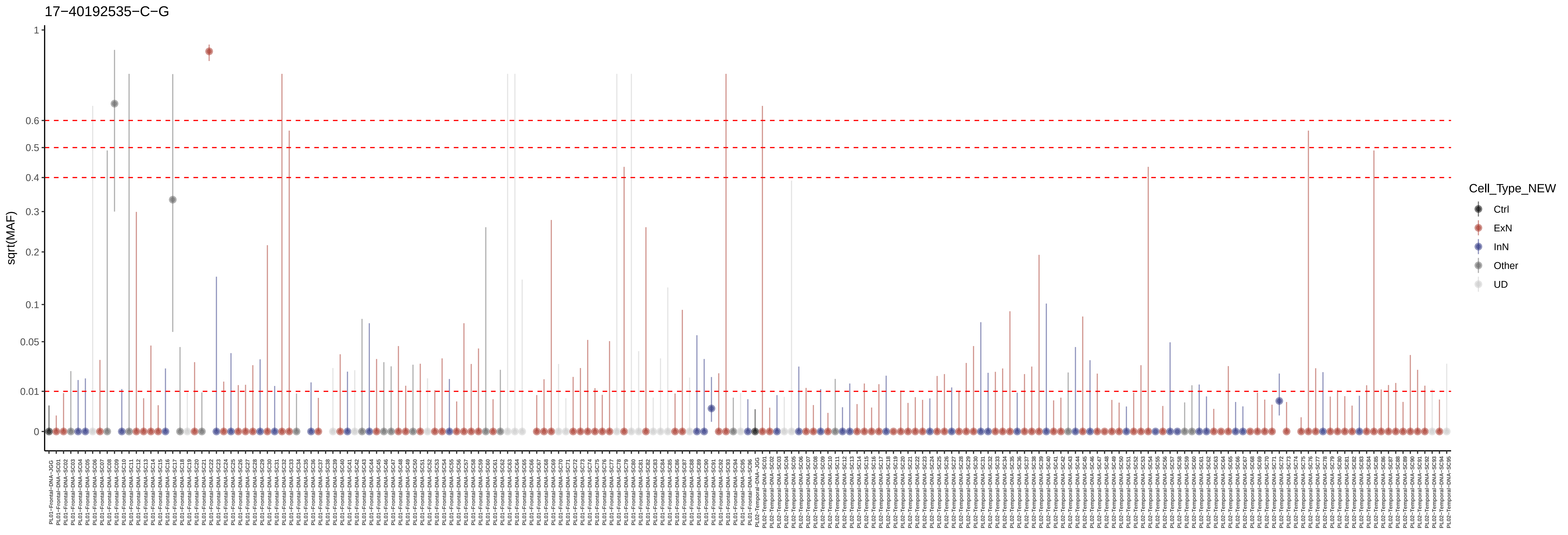
17−40192535−C−G
sqrt(MAF)
1
0.6
0.5
0.4
0.3
0.2
0.1
0.05
0.01
0
Cell_Type_NEW
Ctrl
ExN
InN
Other
UD

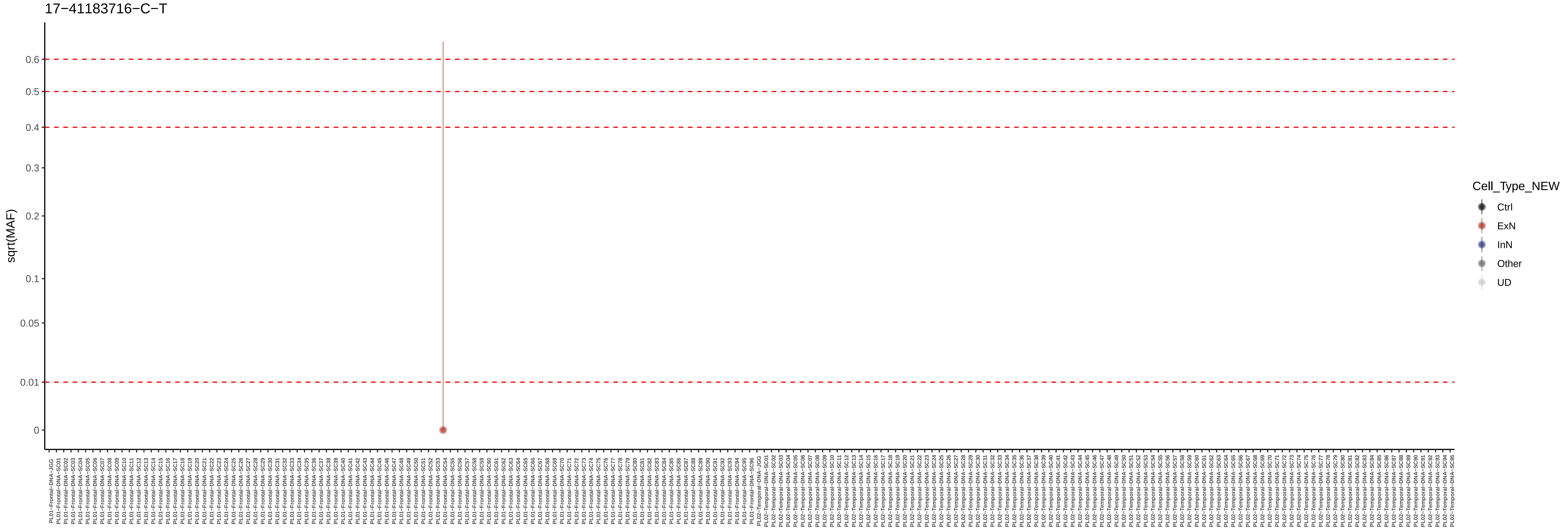
17−41183716−C−T
sqrt(MAF)
0.6
0.5
0.4
0.3
0.2
0.1
0.05
0.01
0
Cell_Type_NEW
Ctrl
ExN
InN
Other
UD

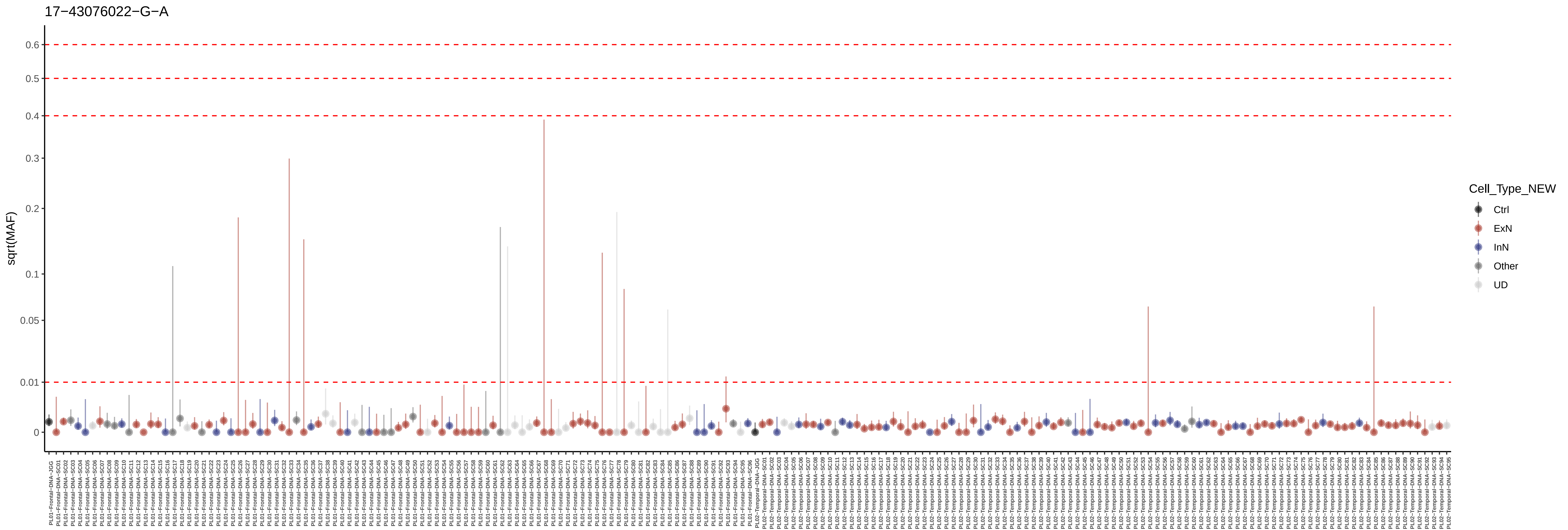
17−43076022−G−A
sqrt(MAF)
0.6
0.5
0.4
0.3
0.2
0.1
0.05
0.01
0
Cell_Type_NEW
Ctrl
ExN
InN
Other
UD

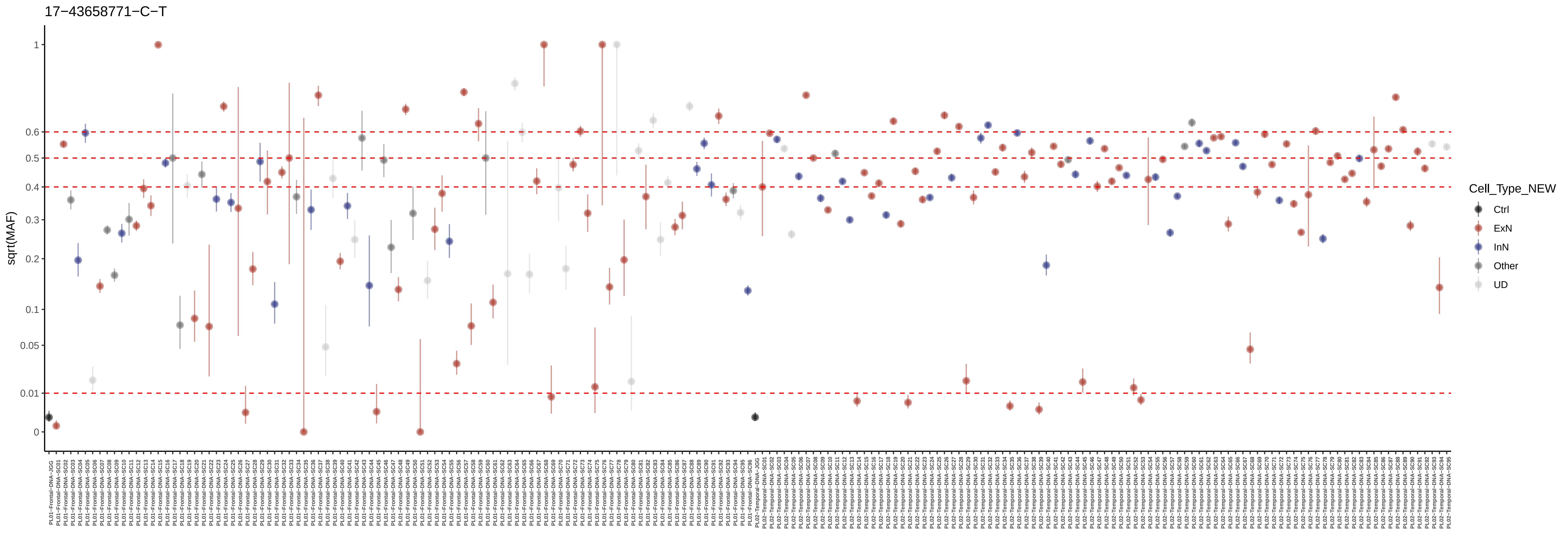
17−43658771−C−T
sqrt(MAF)
1
0.6
0.5
0.4
0.3
0.2
0.1
0.05
0.01
0
Cell_Type_NEW
Ctrl
ExN
InN
Other
UD

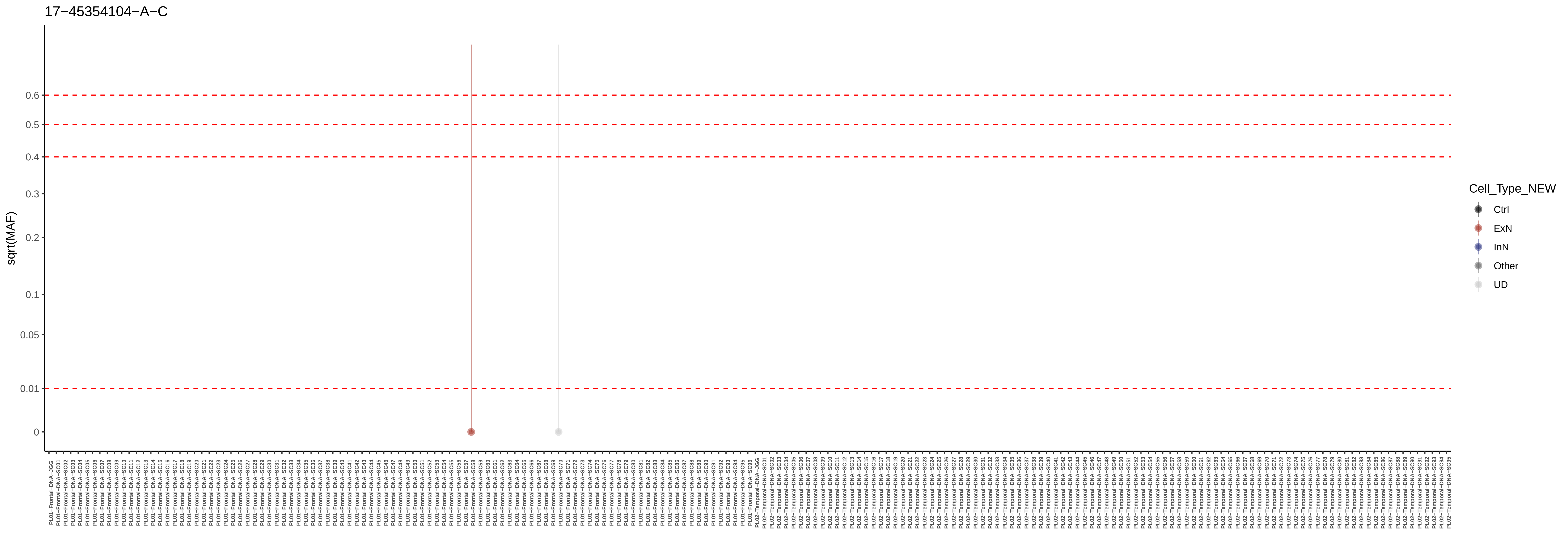
17−45354104−A−C
sqrt(MAF)
0.6
0.5
0.4
0.3
0.2
0.1
0.05
0.01
0
Cell_Type_NEW
Ctrl
ExN
InN
Other
UD

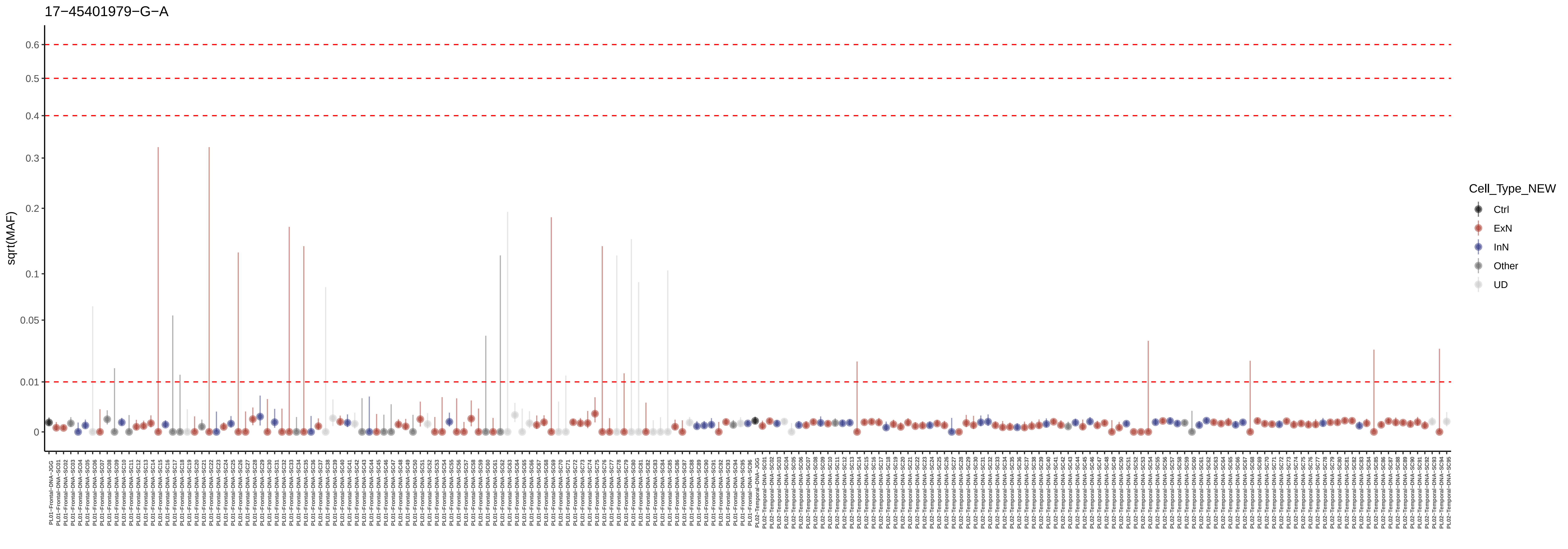
17−45401979−G−A
sqrt(MAF)
0.6
0.5
0.4
0.3
0.2
0.1
0.05
0.01
0
Cell_Type_NEW
Ctrl
ExN
InN
Other
UD

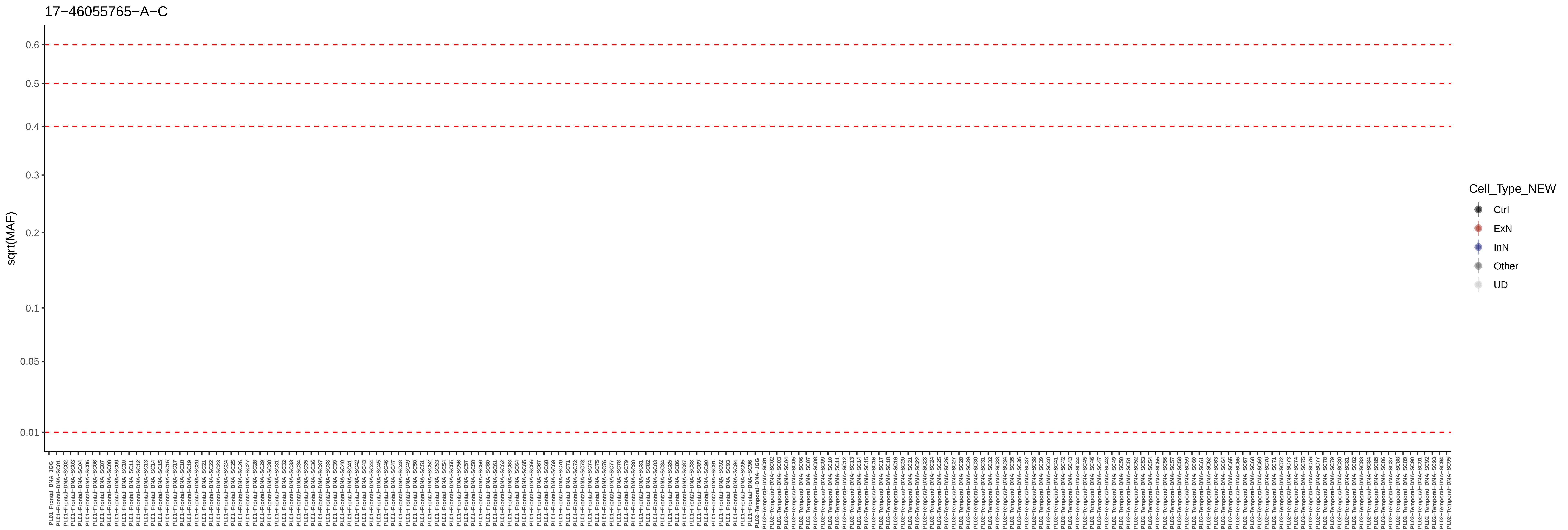
17−46055765−A−C
sqrt(MAF)
0.6
0.5
0.4
0.3
0.2
0.1
0.05
0.01
Cell_Type_NEW
Ctrl
ExN
InN
Other
UD

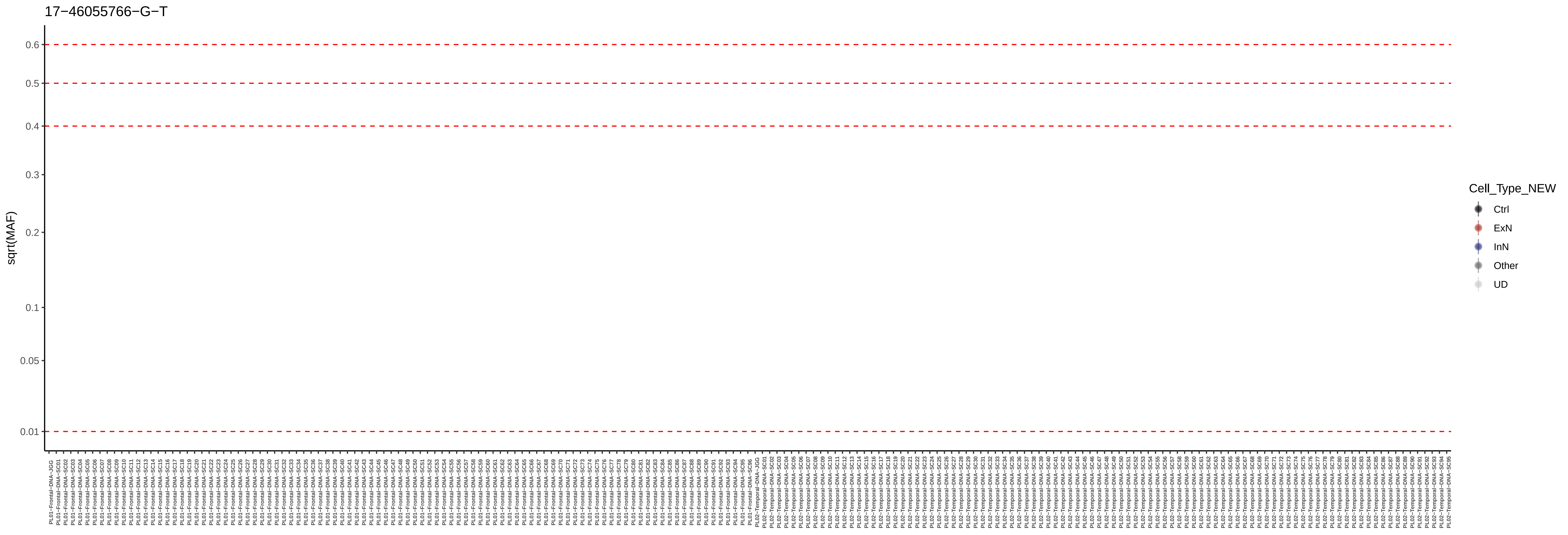

17−46055766−G−T
sqrt(MAF)
0.6
0.5
0.4
0.3
0.2
0.1
0.05
0.01
Cell_Type_NEW
Ctrl
ExN
InN
Other
UD

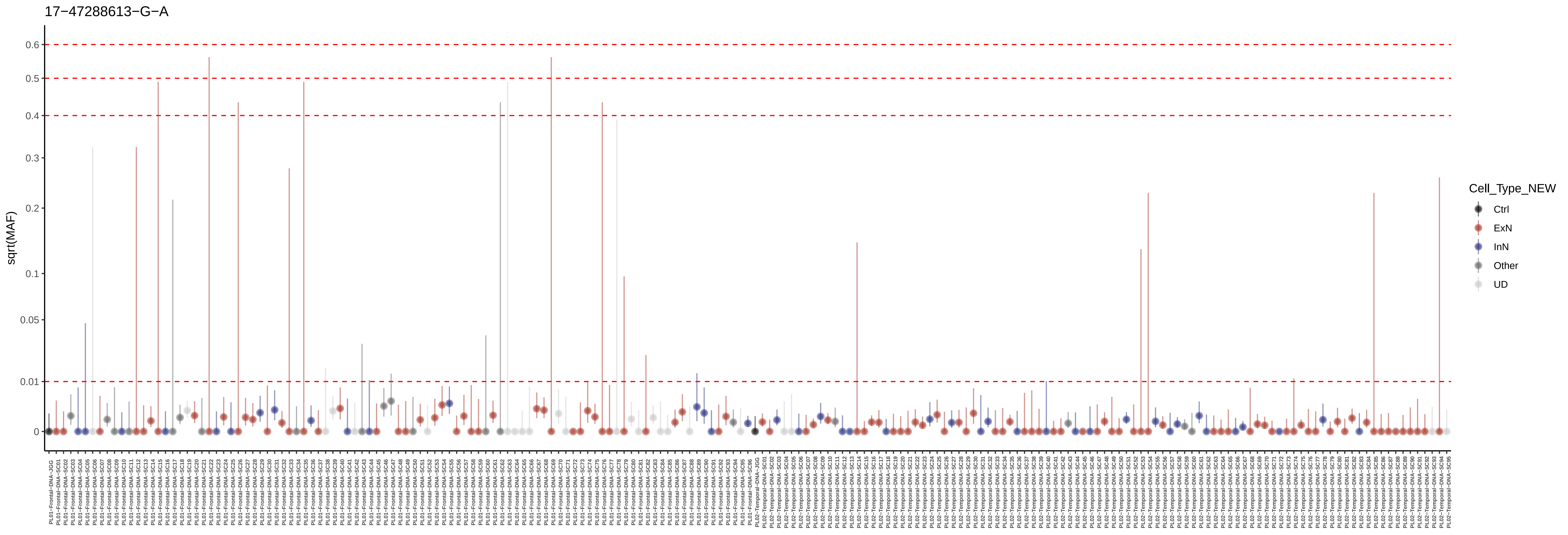
17−47288613−G−A
sqrt(MAF)
0.6
0.5
0.4
0.3
0.2
0.1
0.05
0.01
0
Cell_Type_NEW
Ctrl
ExN
InN
Other
UD

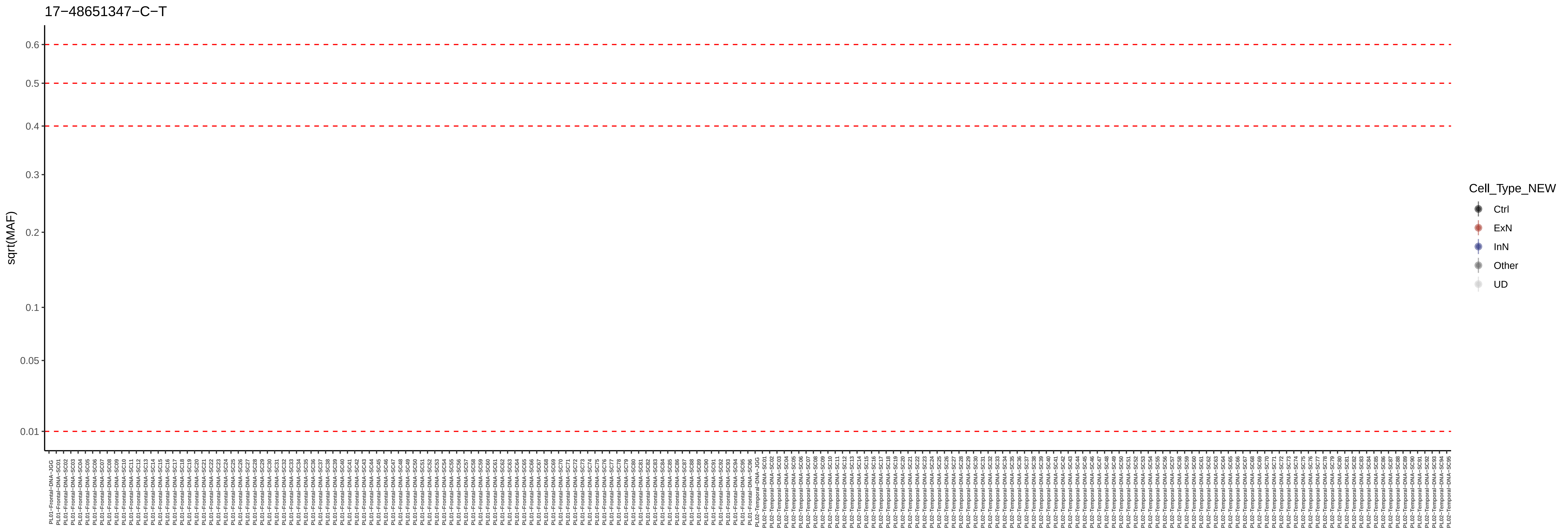
17−48651347−C−T
sqrt(MAF)
0.6
0.5
0.4
0.3
0.2
0.1
0.05
0.01
Cell_Type_NEW
Ctrl
ExN
InN
Other
UD

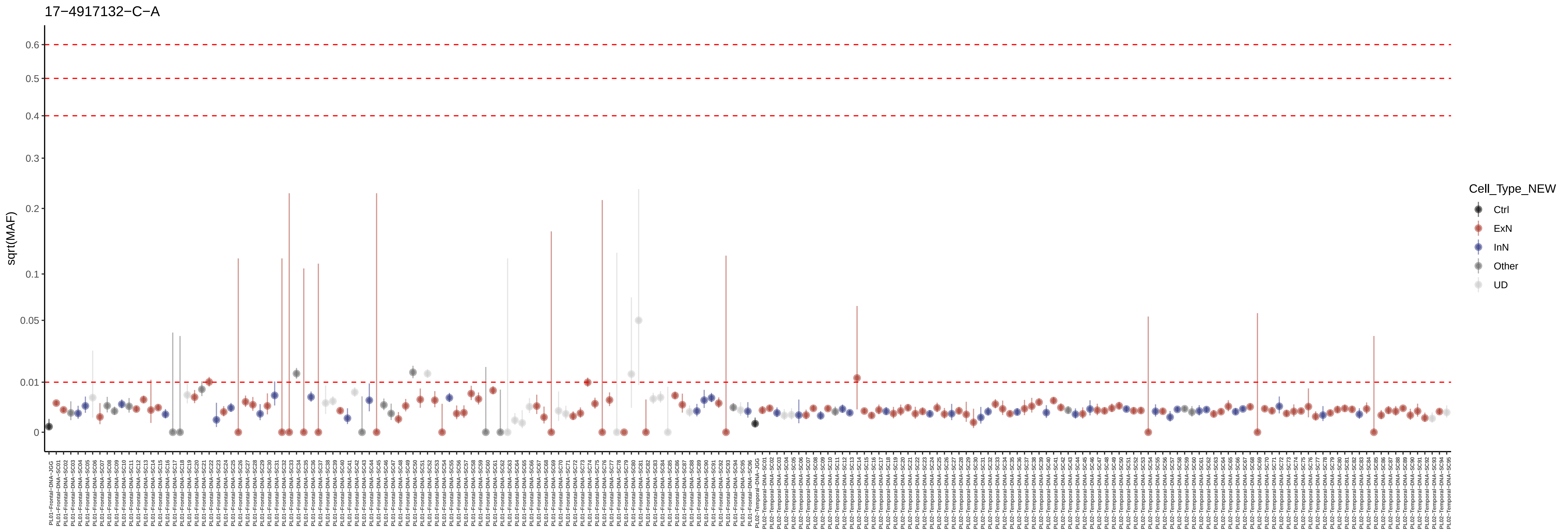

17−4917132−C−A
sqrt(MAF)
0.6
0.5
0.4
0.3
0.2
0.1
0.05
0.01
0
Cell_Type_NEW
Ctrl
ExN
InN
Other
UD

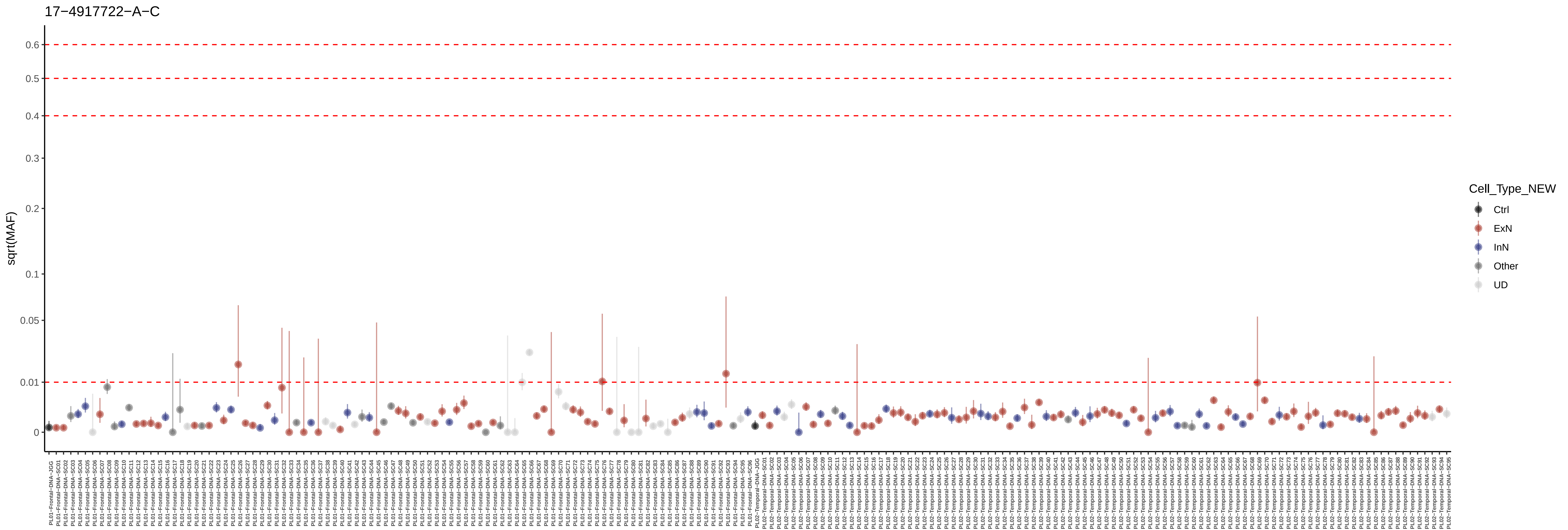

17−4917722−A−C
sqrt(MAF)
0.6
0.5
0.4
0.3
0.2
0.1
0.05
0.01
0
Cell_Type_NEW
Ctrl
ExN
InN
Other
UD

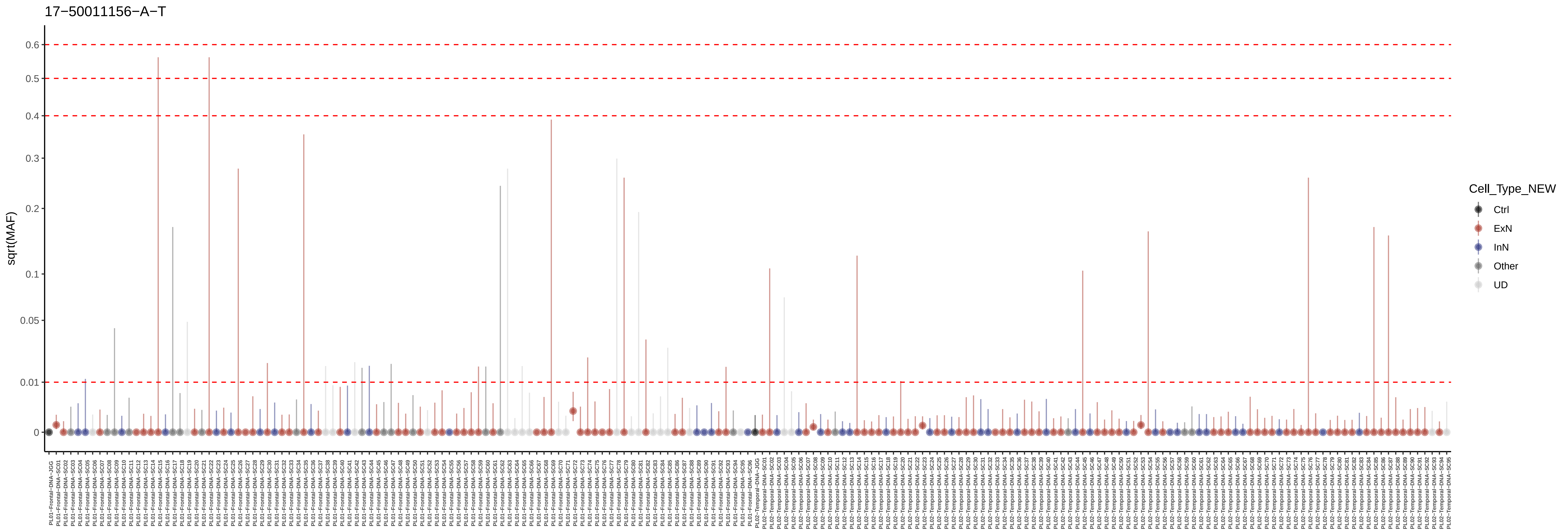
17−50011156−A−T
sqrt(MAF)
0.6
0.5
0.4
0.3
0.2
0.1
0.05
0.01
0
Cell_Type_NEW
Ctrl
ExN
InN
Other
UD

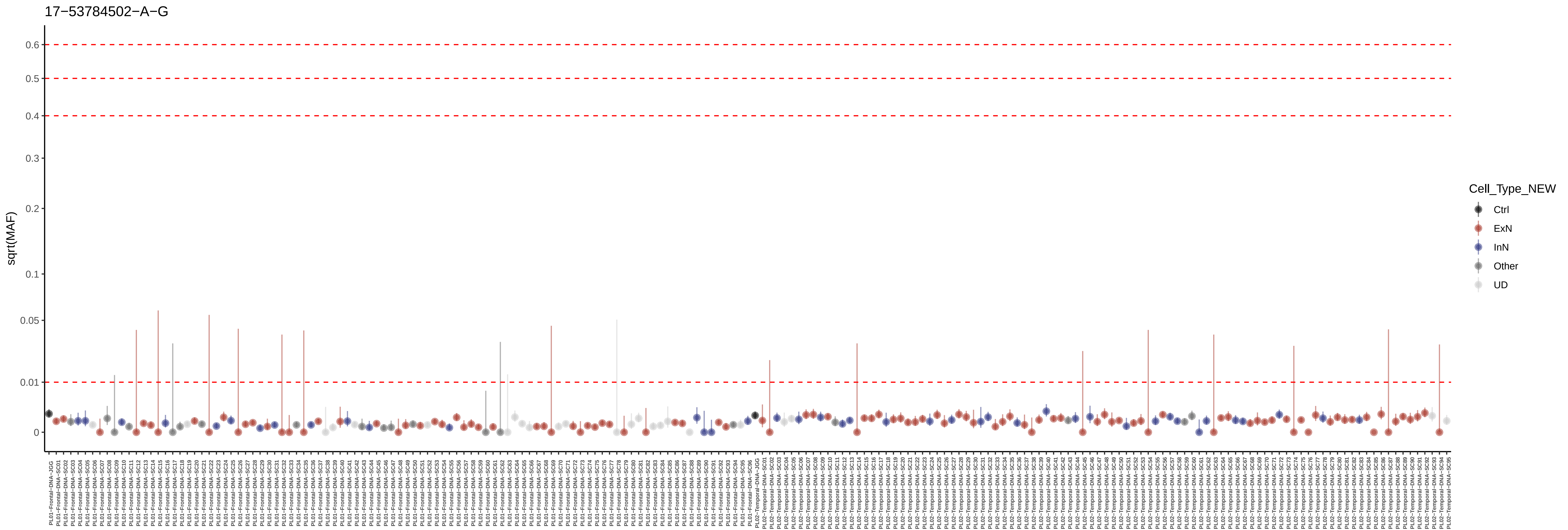

17−53784502−A−G
sqrt(MAF)
0.6
0.5
0.4
0.3
0.2
0.1
0.05
0.01
0
Cell_Type_NEW
Ctrl
ExN
InN
Other
UD

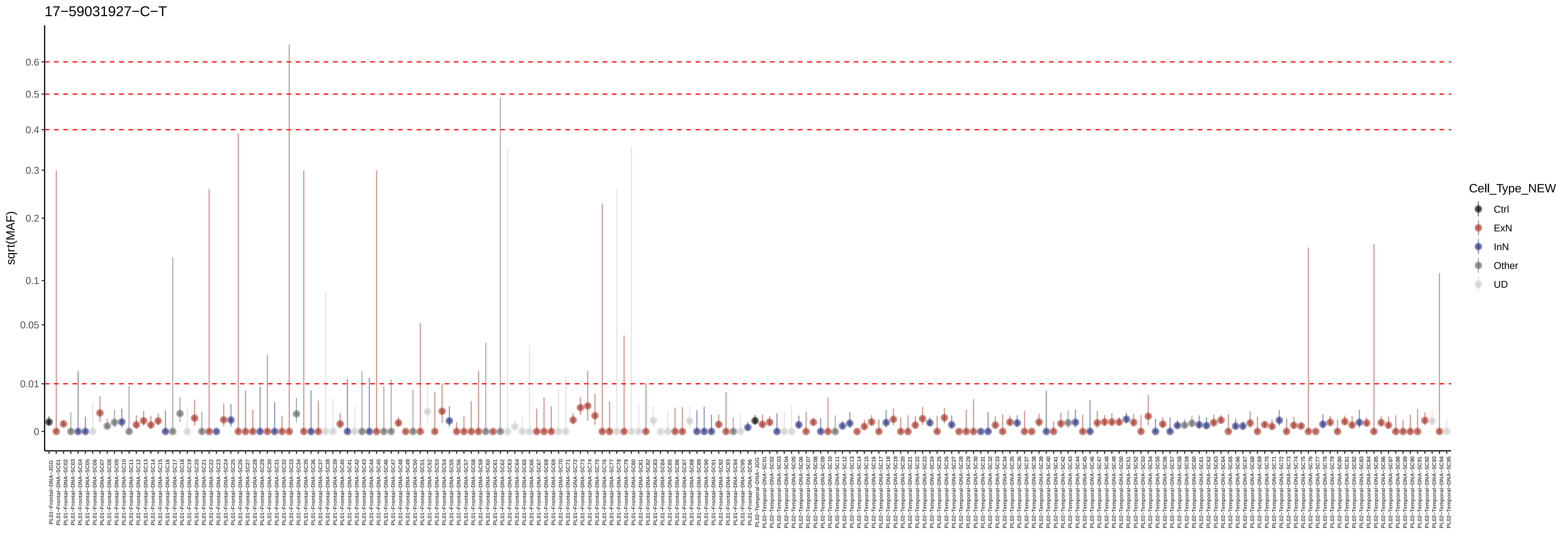
17−59031927−C−T
sqrt(MAF)
0.6
0.5
0.4
0.3
0.2
0.1
0.05
0.01
0
Cell_Type_NEW
Ctrl
ExN
InN
Other
UD

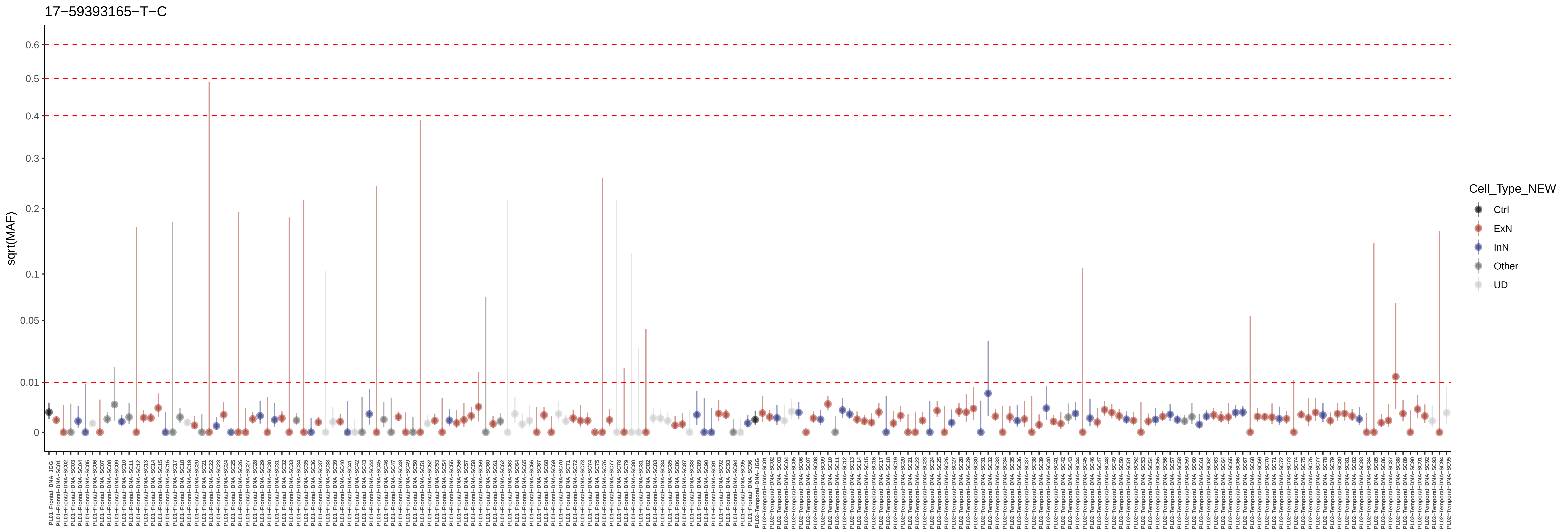

17−59393165−T−C
sqrt(MAF)
0.6
0.5
0.4
0.3
0.2
0.1
0.05
0.01
0
Cell_Type_NEW
Ctrl
ExN
InN
Other
UD

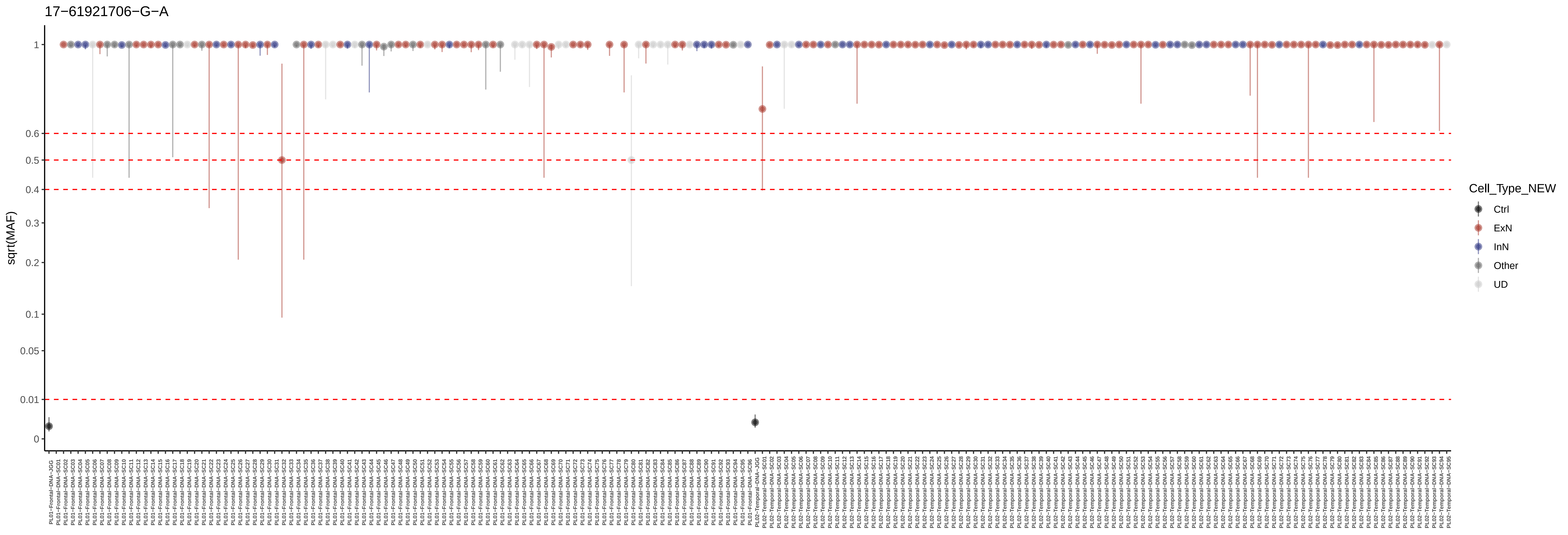
17−61921706−G−A
sqrt(MAF)
1
0.6
0.5
0.4
0.3
0.2
0.1
0.05
0.01
0
Cell_Type_NEW
Ctrl
ExN
InN
Other
UD

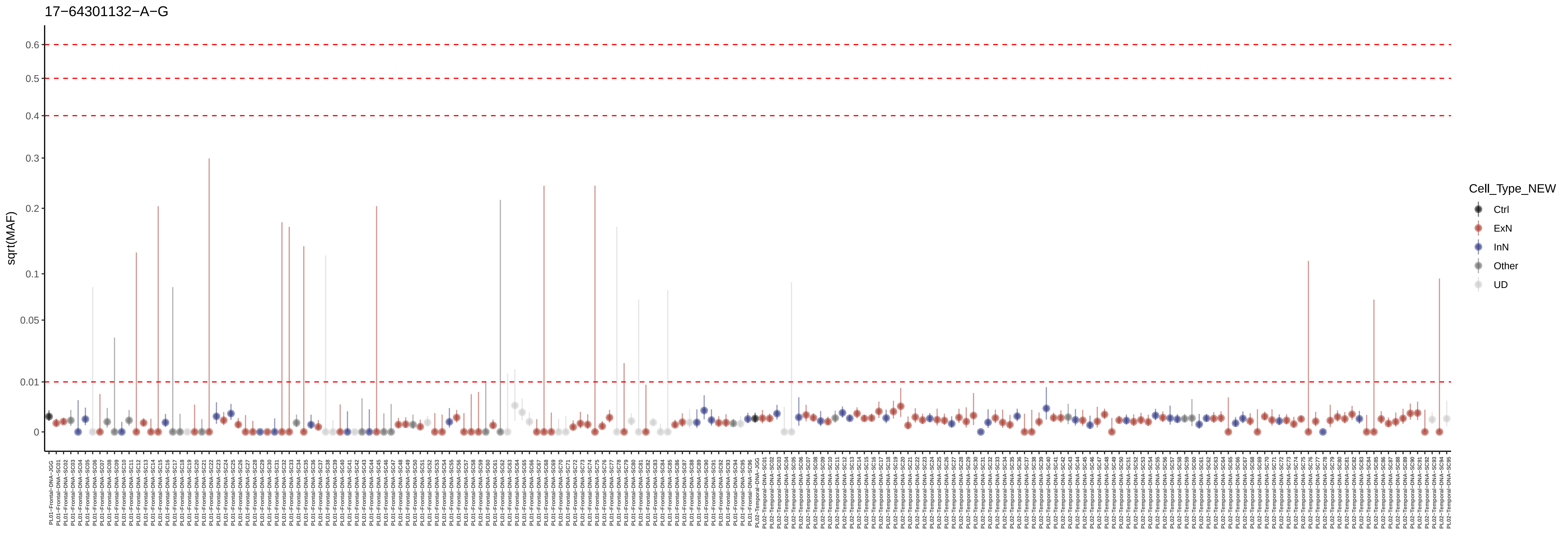

17−64301132−A−G
sqrt(MAF)
0.6
0.5
0.4
0.3
0.2
0.1
0.05
0.01
0
Cell_Type_NEW
Ctrl
ExN
InN
Other
UD

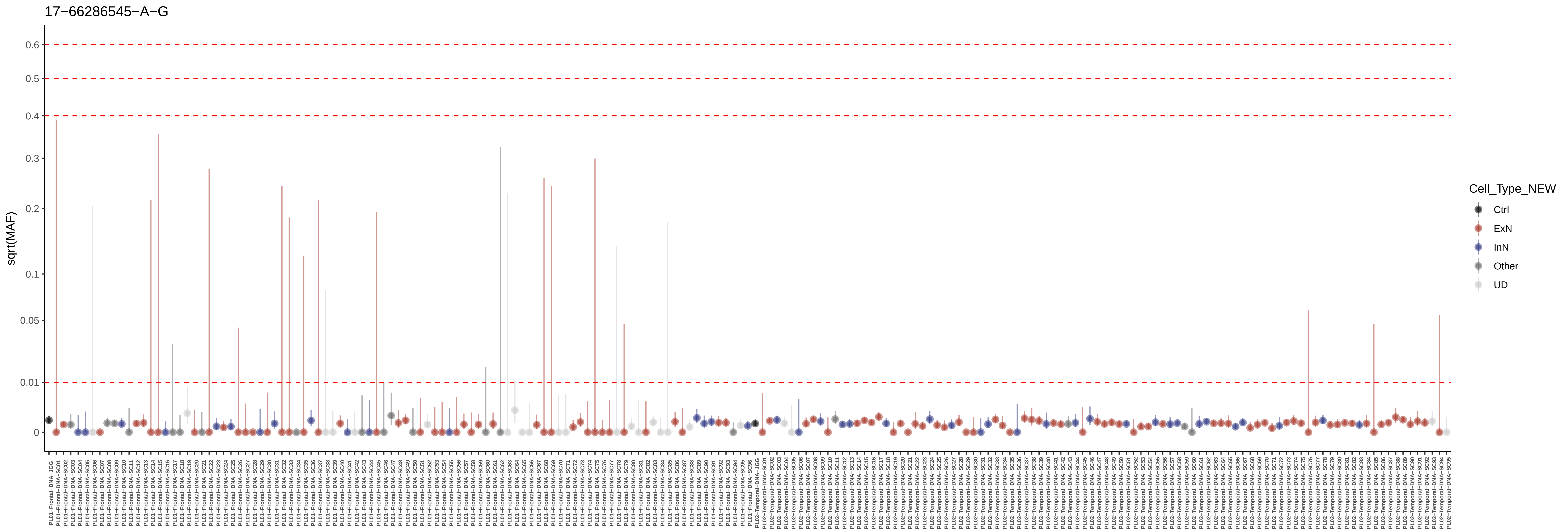
17−66286545−A−G
sqrt(MAF)
0.6
0.5
0.4
0.3
0.2
0.1
0.05
0.01
0
Cell_Type_NEW
Ctrl
ExN
InN
Other
UD

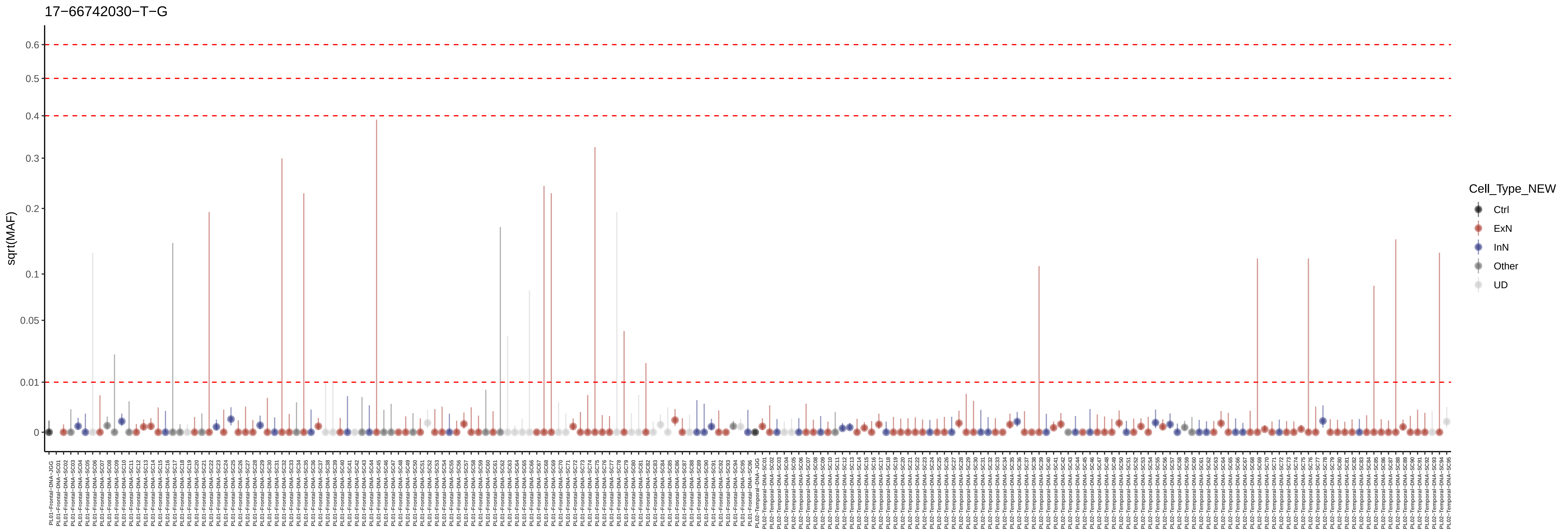
17−66742030−T−G
sqrt(MAF)
0.6
0.5
0.4
0.3
0.2
0.1
0.05
0.01
0
Cell_Type_NEW
Ctrl
ExN
InN
Other
UD

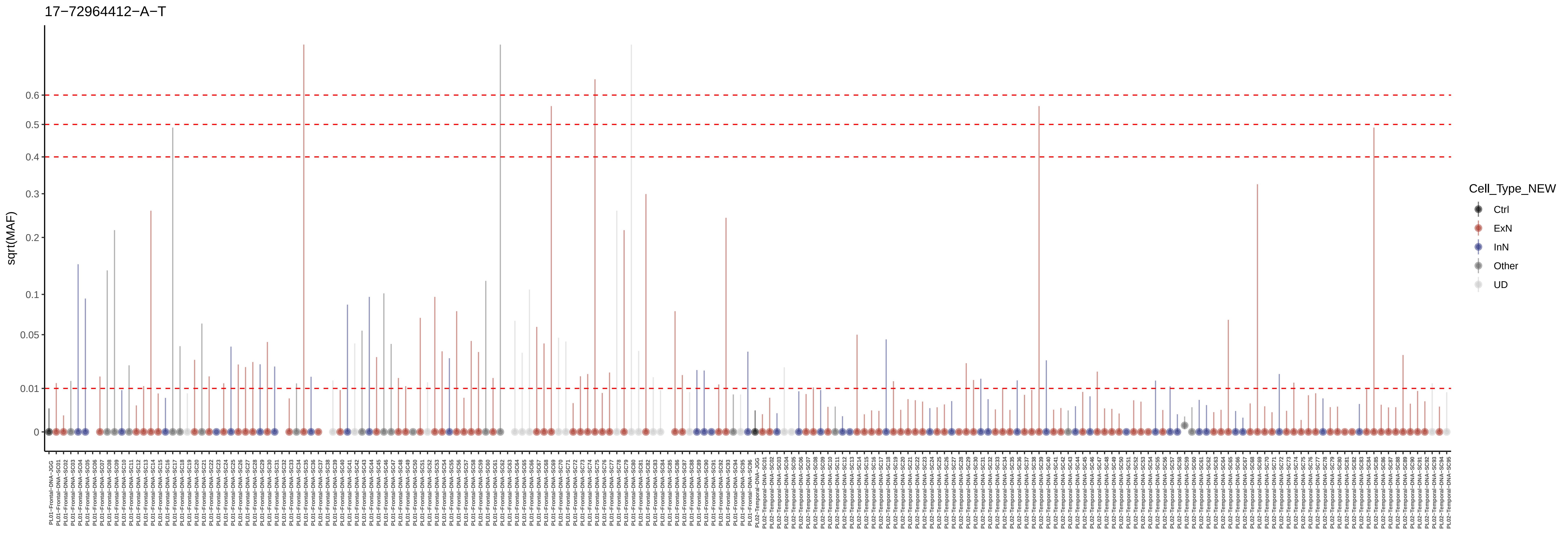
17−72964412−A−T
sqrt(MAF)
0.6
0.5
0.4
0.3
0.2
0.1
0.05
0.01
0
Cell_Type_NEW
Ctrl
ExN
InN
Other
UD

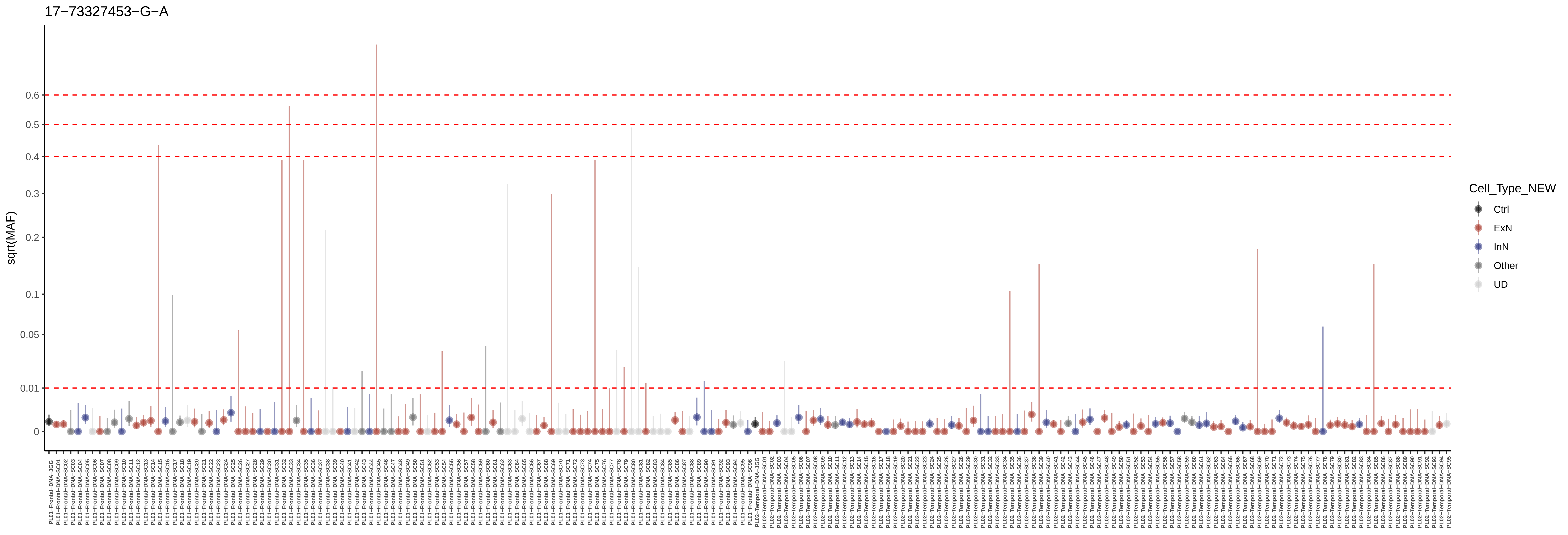

17−73327453−G−A
sqrt(MAF)
0.6
0.5
0.4
0.3
0.2
0.1
0.05
0.01
0
Cell_Type_NEW
Ctrl
ExN
InN
Other
UD

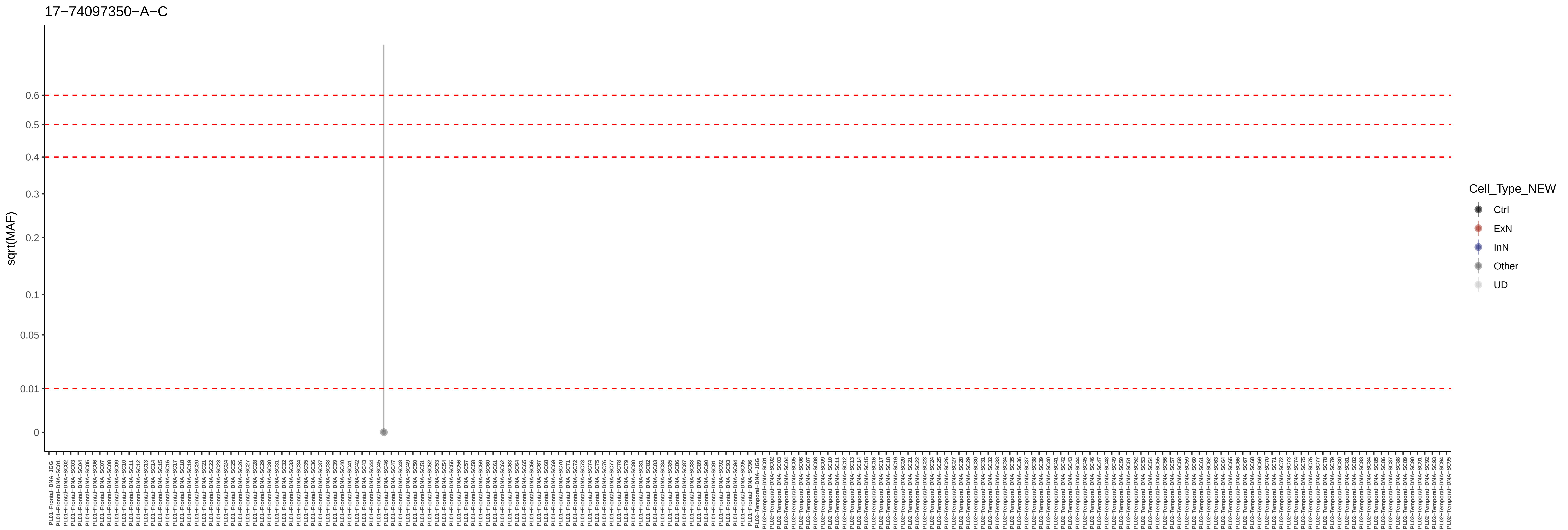

17−74097350−A−C
sqrt(MAF)
0.6
0.5
0.4
0.3
0.2
0.1
0.05
0.01
0
Cell_Type_NEW
Ctrl
ExN
InN
Other
UD

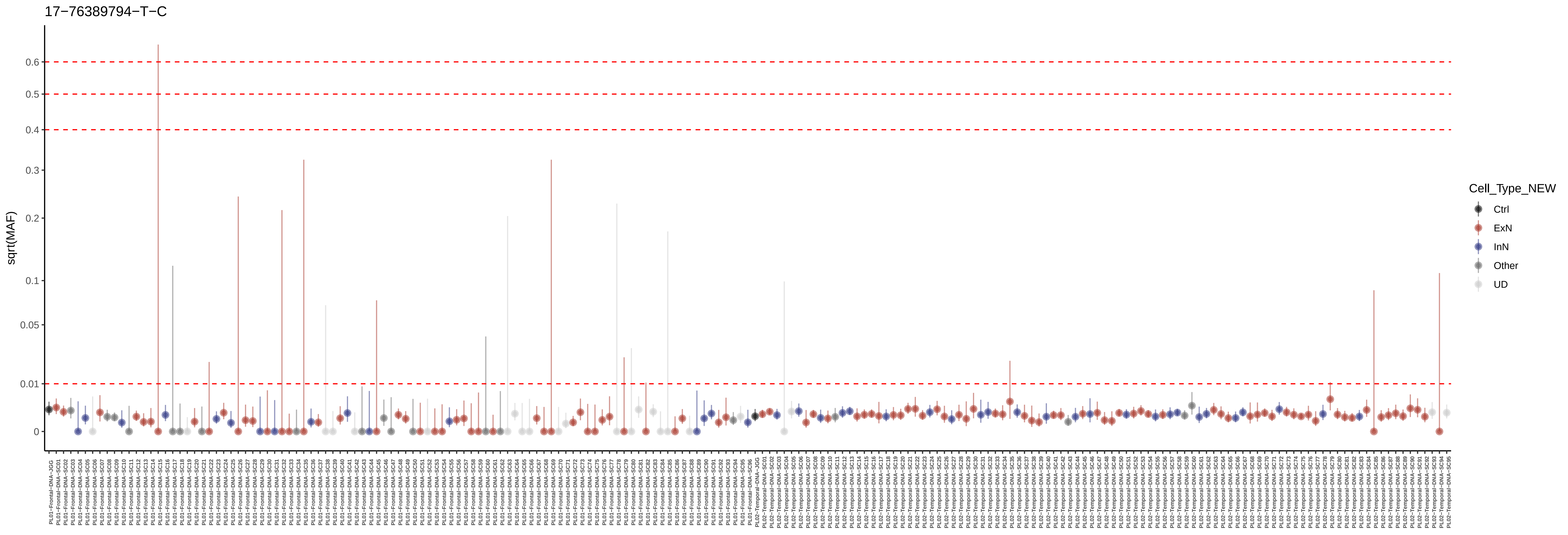
17−76389794−T−C
sqrt(MAF)
0.6
0.5
0.4
0.3
0.2
0.1
0.05
0.01
0
Cell_Type_NEW
Ctrl
ExN
InN
Other
UD

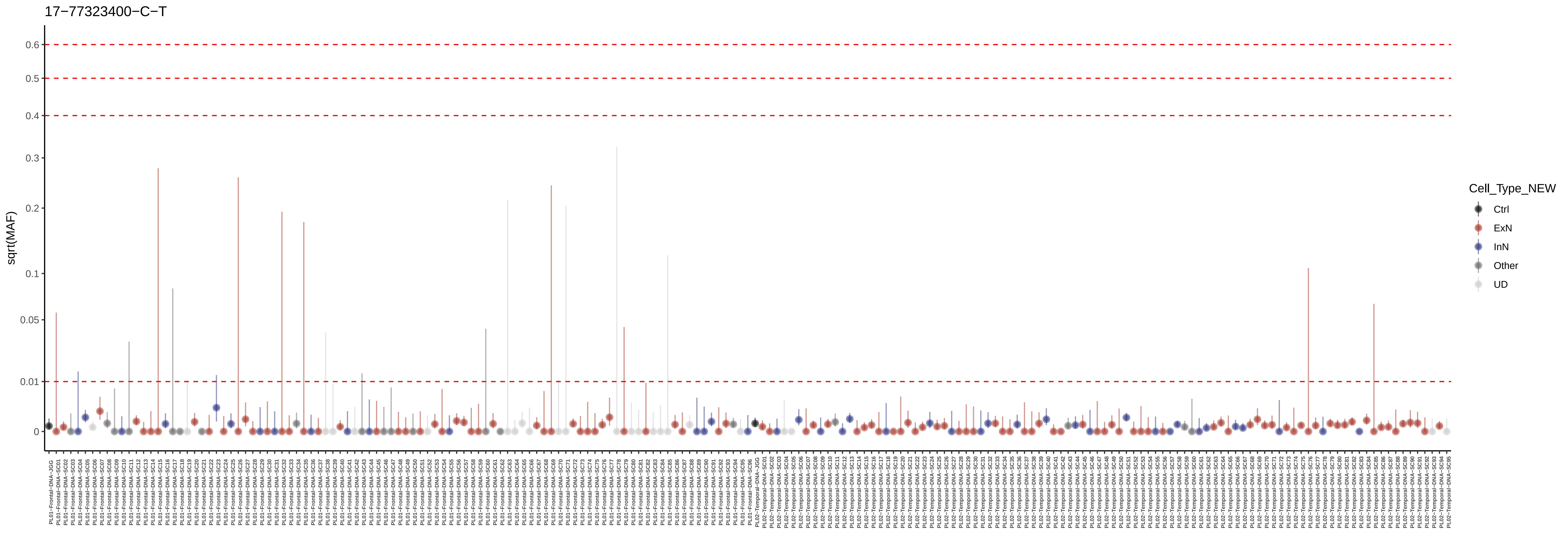
17−77323400−C−T
sqrt(MAF)
0.6
0.5
0.4
0.3
0.2
0.1
0.05
0.01
0
Cell_Type_NEW
Ctrl
ExN
InN
Other
UD

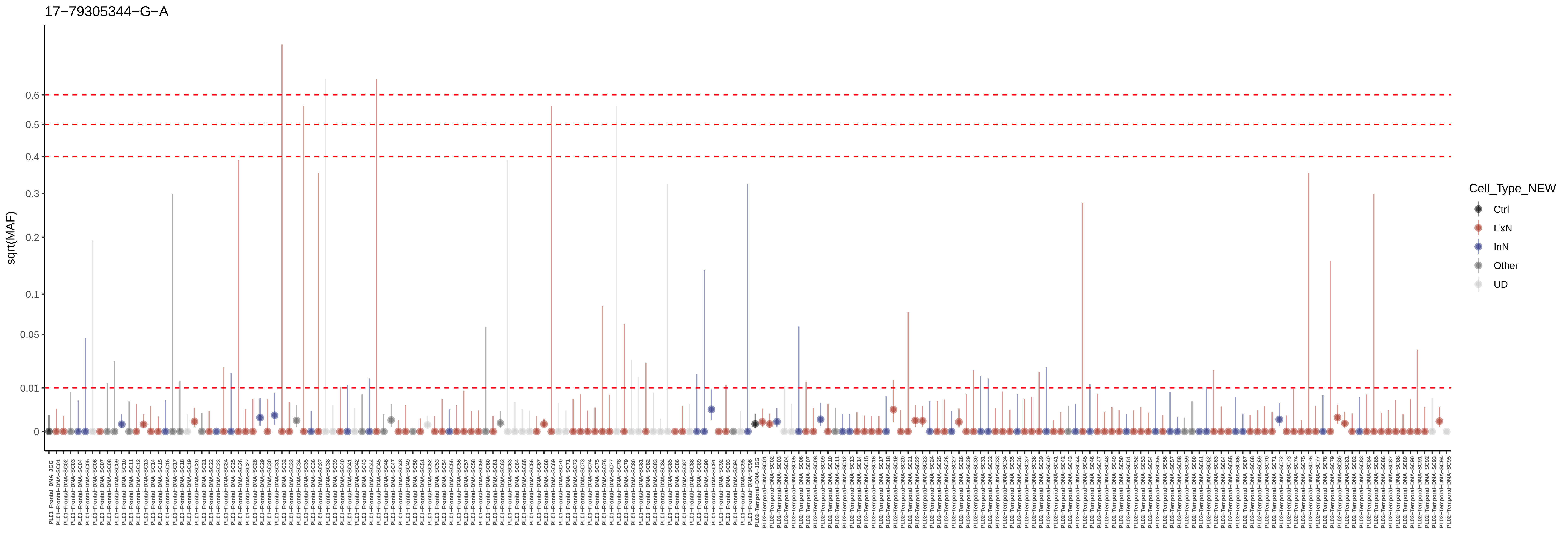
17−79305344−G−A
sqrt(MAF)
0.6
0.5
0.4
0.3
0.2
0.1
0.05
0.01
0
Cell_Type_NEW
Ctrl
ExN
InN
Other
UD

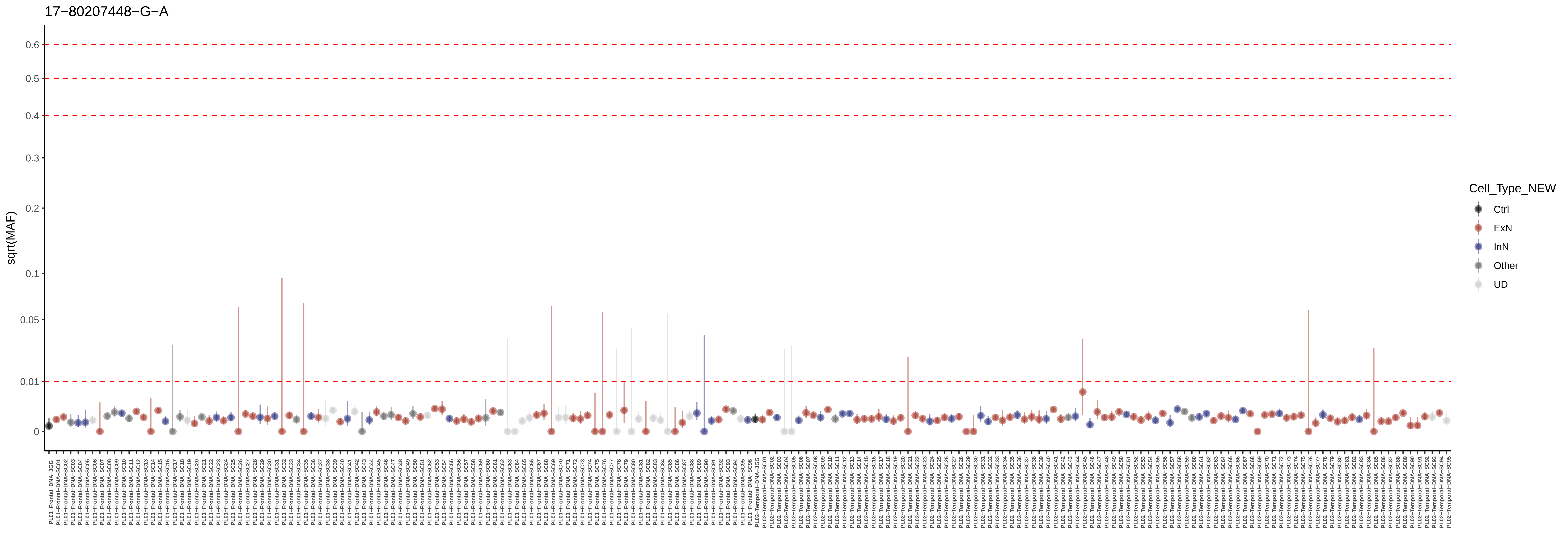
17−80207448−G−A
sqrt(MAF)
0
0.01
0.05
0.1
0.2
0.3
0.4
0.5
0.6
Cell_Type_NEW
Ctrl
ExN
InN
Other
UD

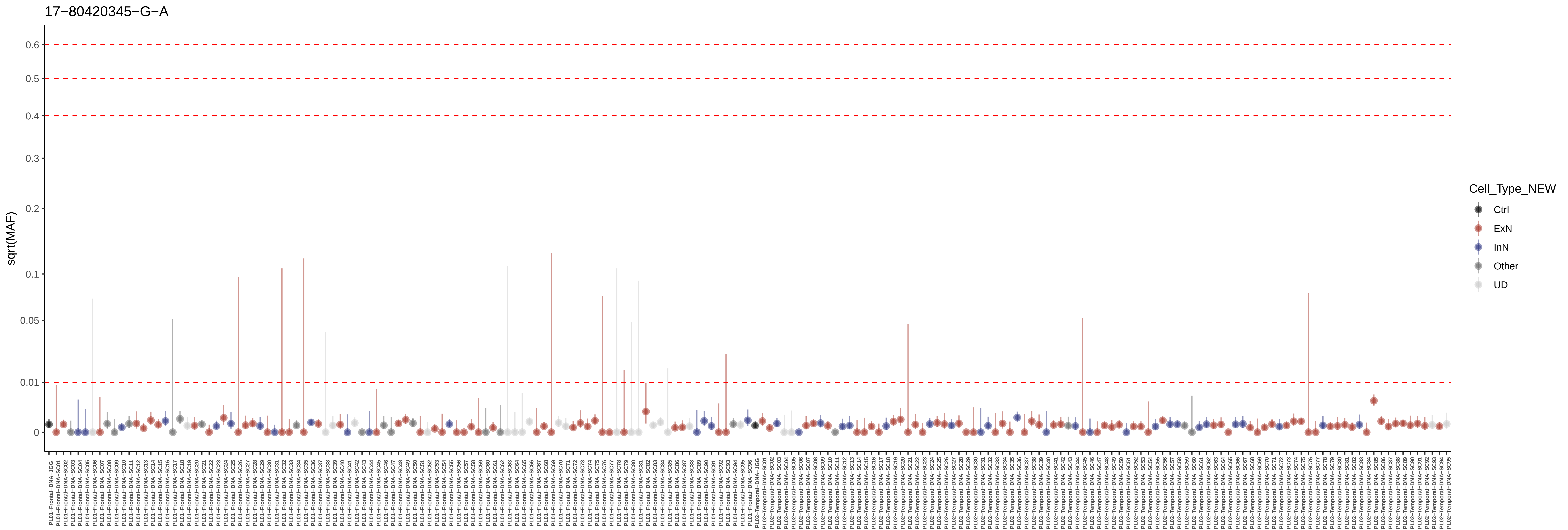
17−80420345−G−A
sqrt(MAF)
0.6
0.5
0.4
0.3
0.2
0.1
0.05
0.01
0
Cell_Type_NEW
Ctrl
ExN
InN
Other
UD

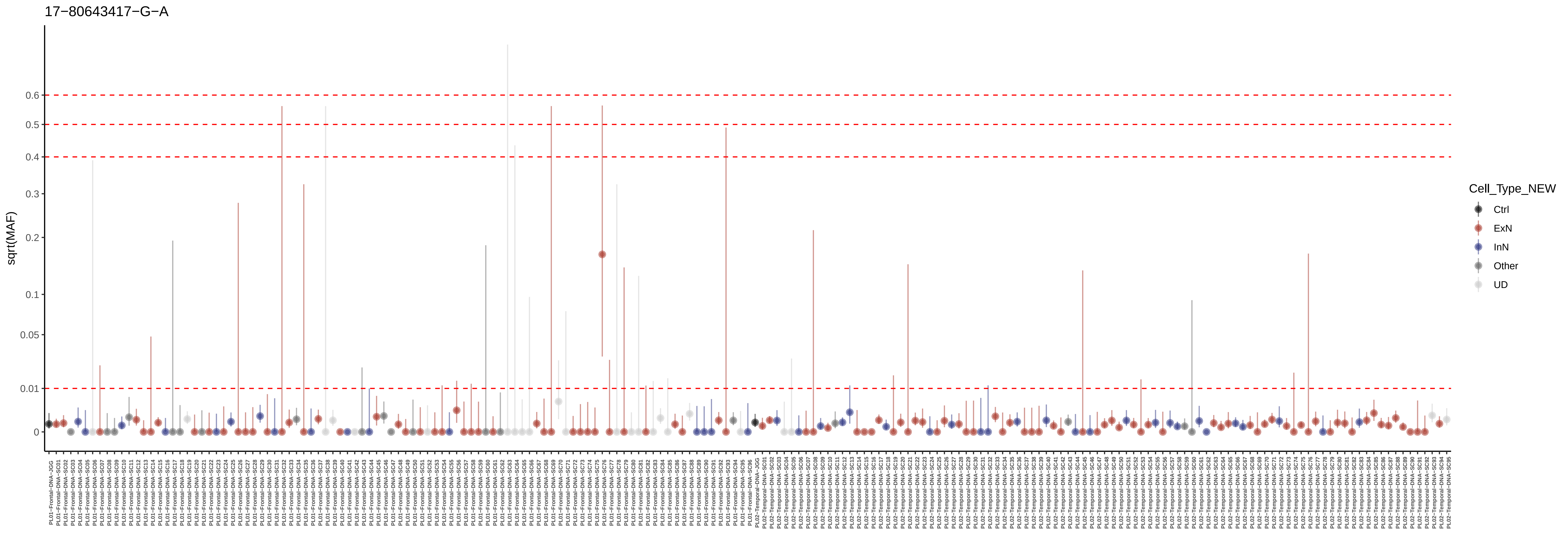

17−80643417−G−A
sqrt(MAF)
0.6
0.5
0.4
0.3
0.2
0.1
0.05
0.01
0
Cell_Type_NEW
Ctrl
ExN
InN
Other
UD

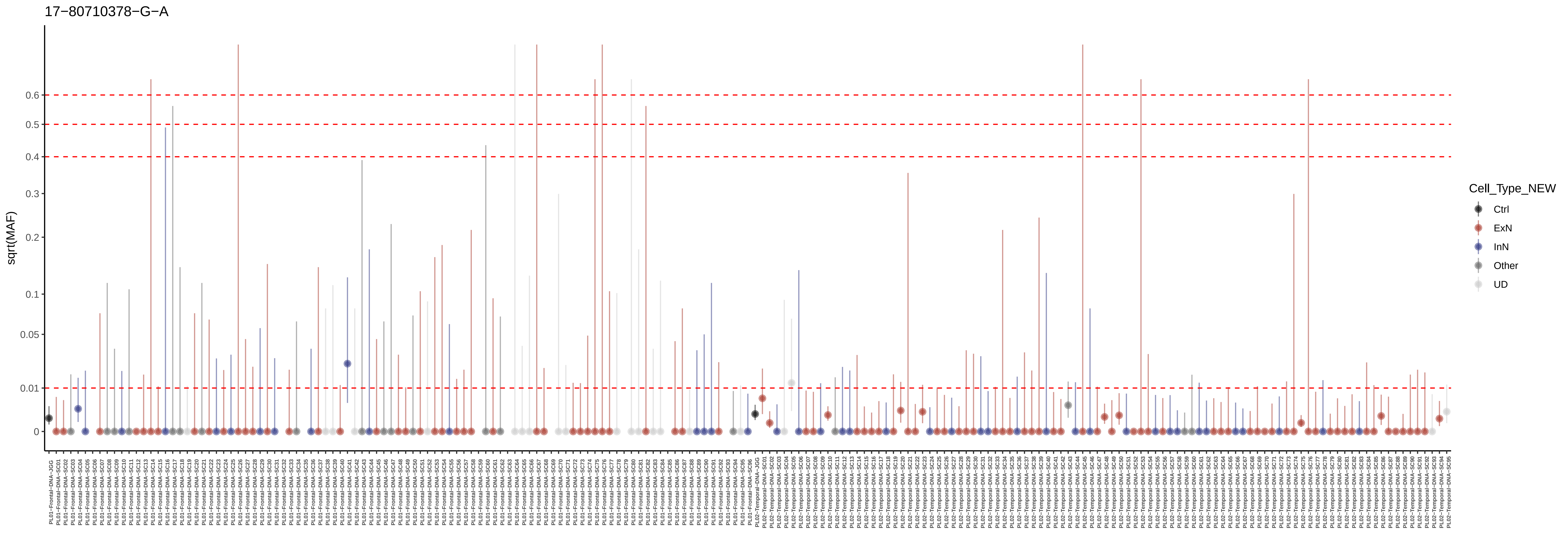
17−80710378−G−A
sqrt(MAF)
0.6
0.5
0.4
0.3
0.2
0.1
0.05
0.01
0
Cell_Type_NEW
Ctrl
ExN
InN
Other
UD

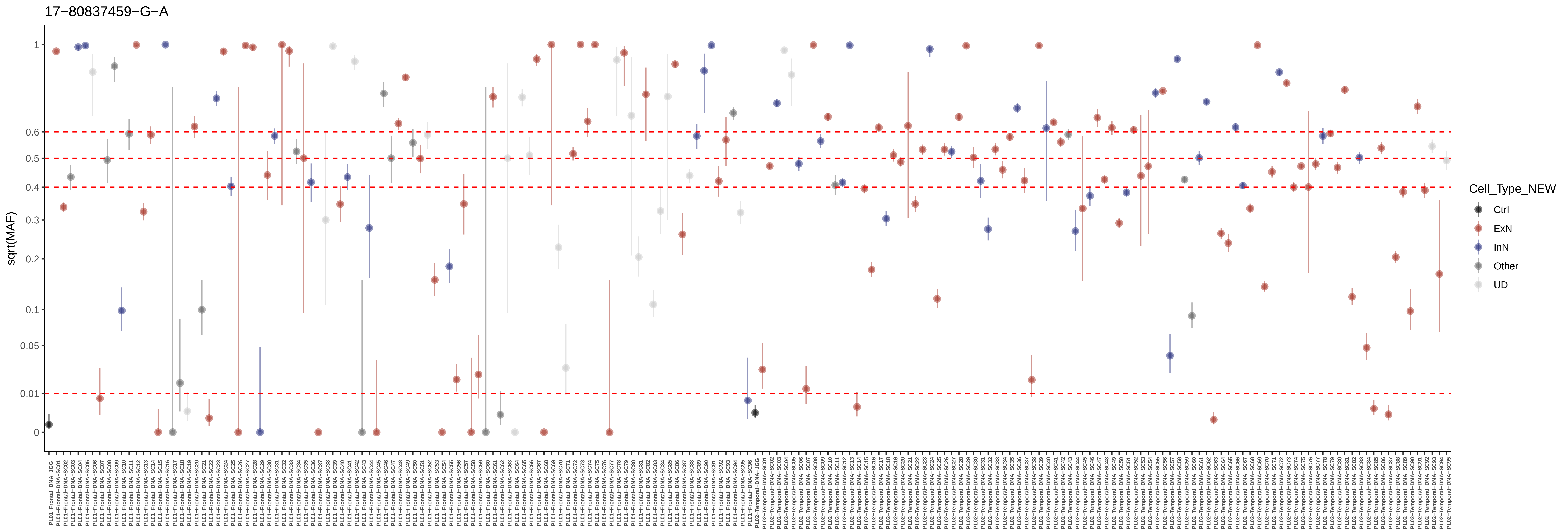

17−80837459−G−A
sqrt(MAF)
1
0.6
0.5
0.4
0.3
0.2
0.1
0.05
0.01
0
Cell_Type_NEW
Ctrl
ExN
InN
Other
UD

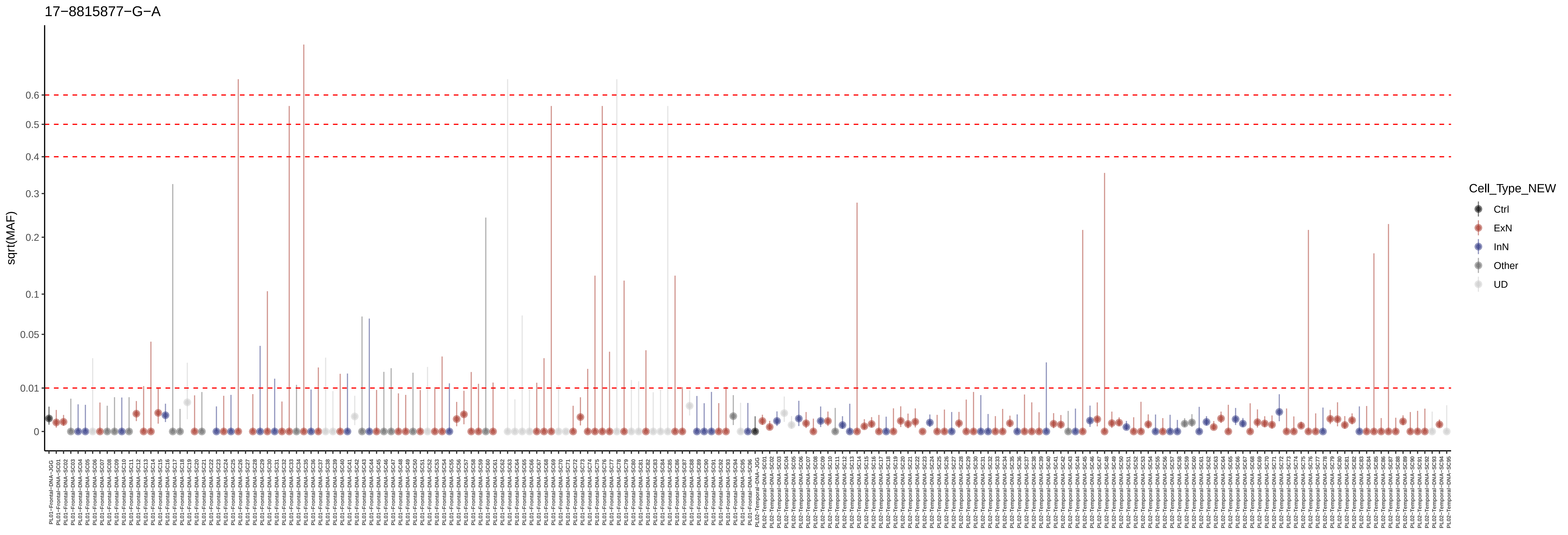
17−8815877−G−A
sqrt(MAF)
0.6
0.5
0.4
0.3
0.2
0.1
0.05
0.01
0
Cell_Type_NEW
Ctrl
ExN
InN
Other
UD

17−902278−A−C

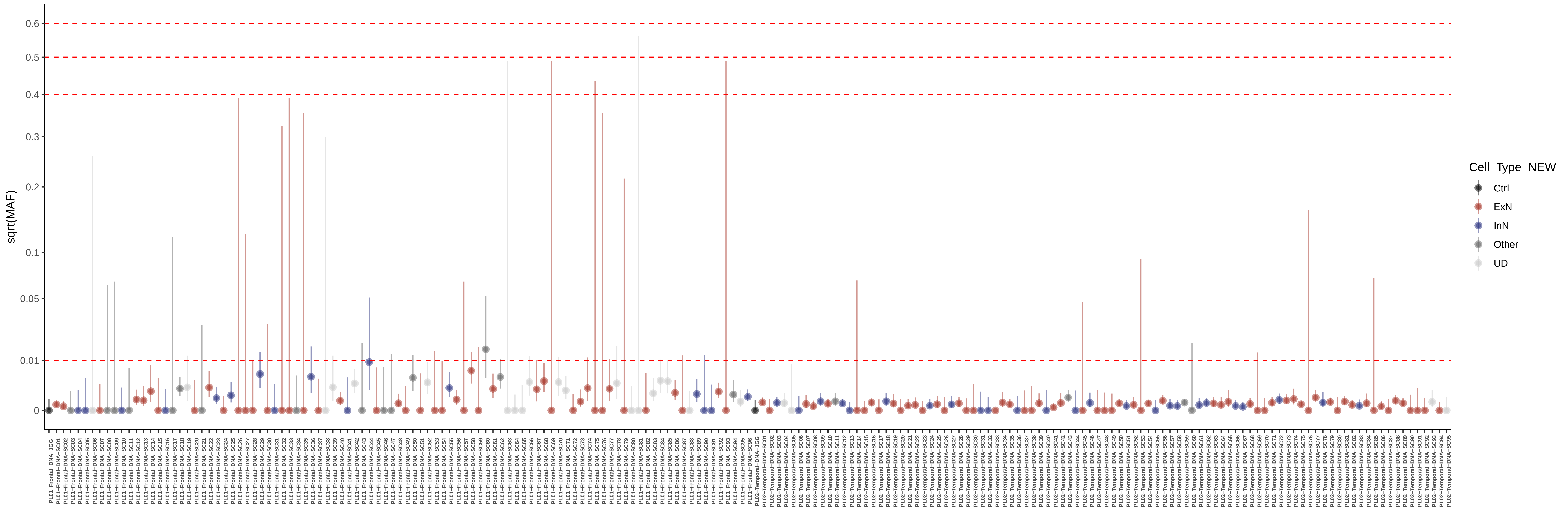
sqrt(MAF)
0.6
0.5
0.4
0.3
0.2
0.1
0.05
0.01
0
Cell_Type_NEW
Ctrl
ExN
InN
Other
UD

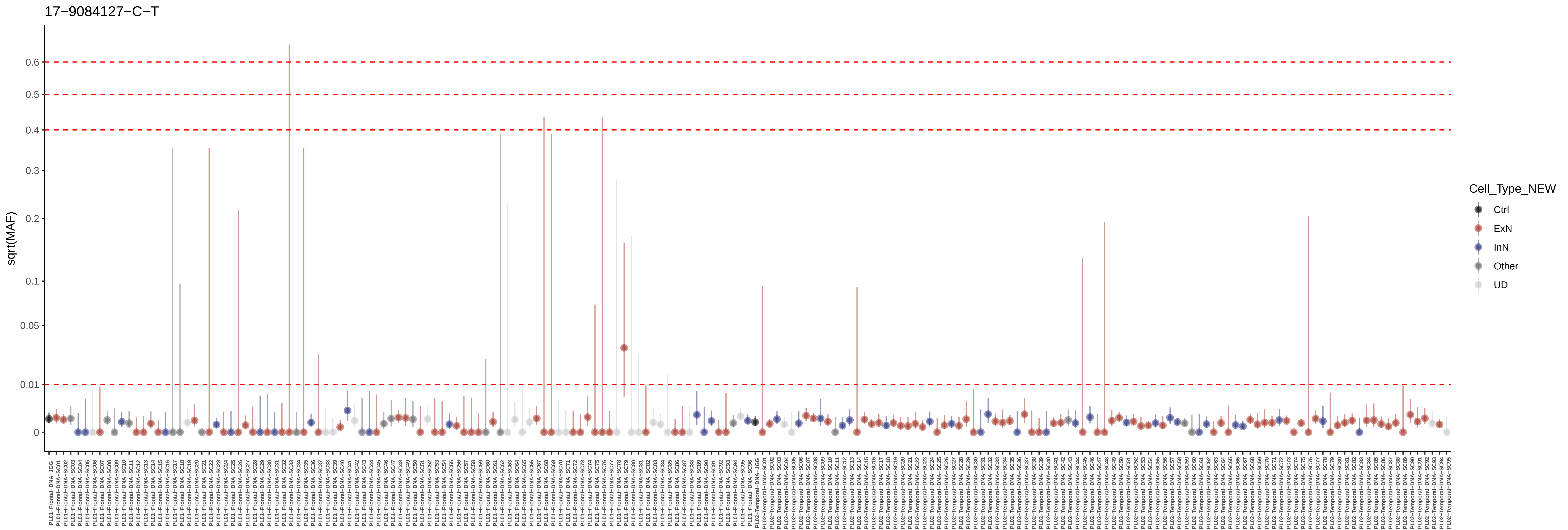
17−9084127−C−T
sqrt(MAF)
0.6
0.5
0.4
0.3
0.2
0.1
0.05
0.01
0
Cell_Type_NEW
Ctrl
ExN
InN
Other
UD

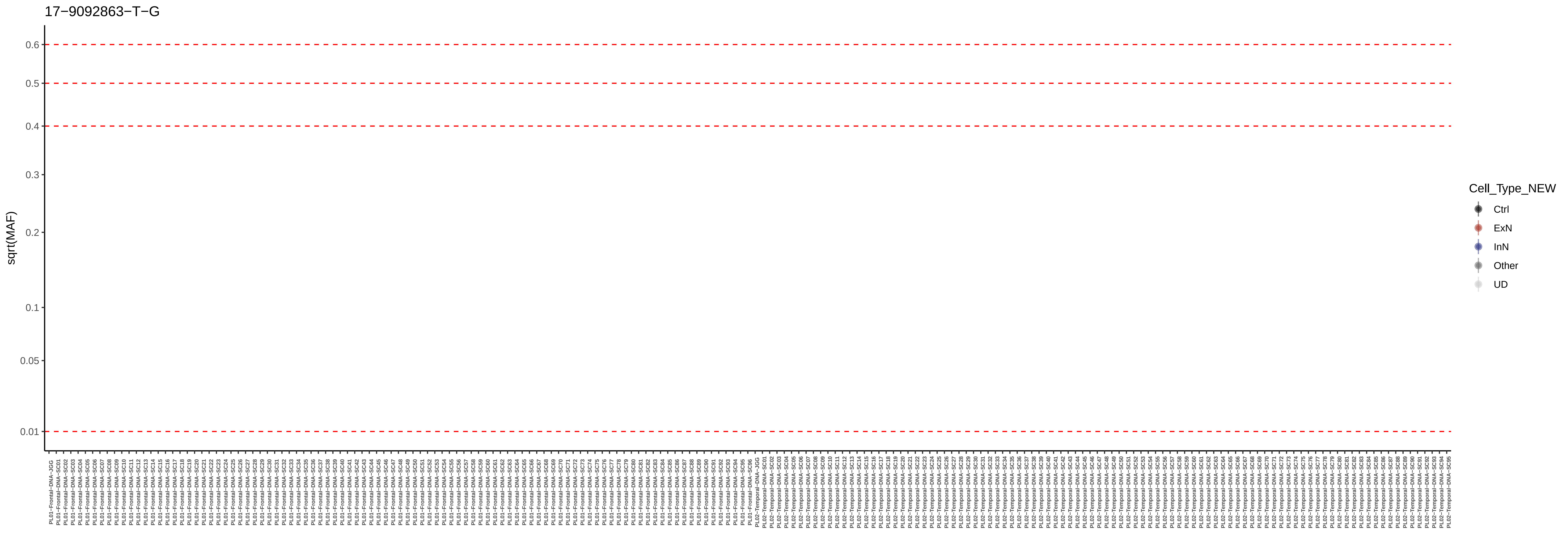
17−9092863−T−G
sqrt(MAF)
0.6
0.5
0.4
0.3
0.2
0.1
0.05
0.01
Cell_Type_NEW
Ctrl
ExN
InN
Other
UD

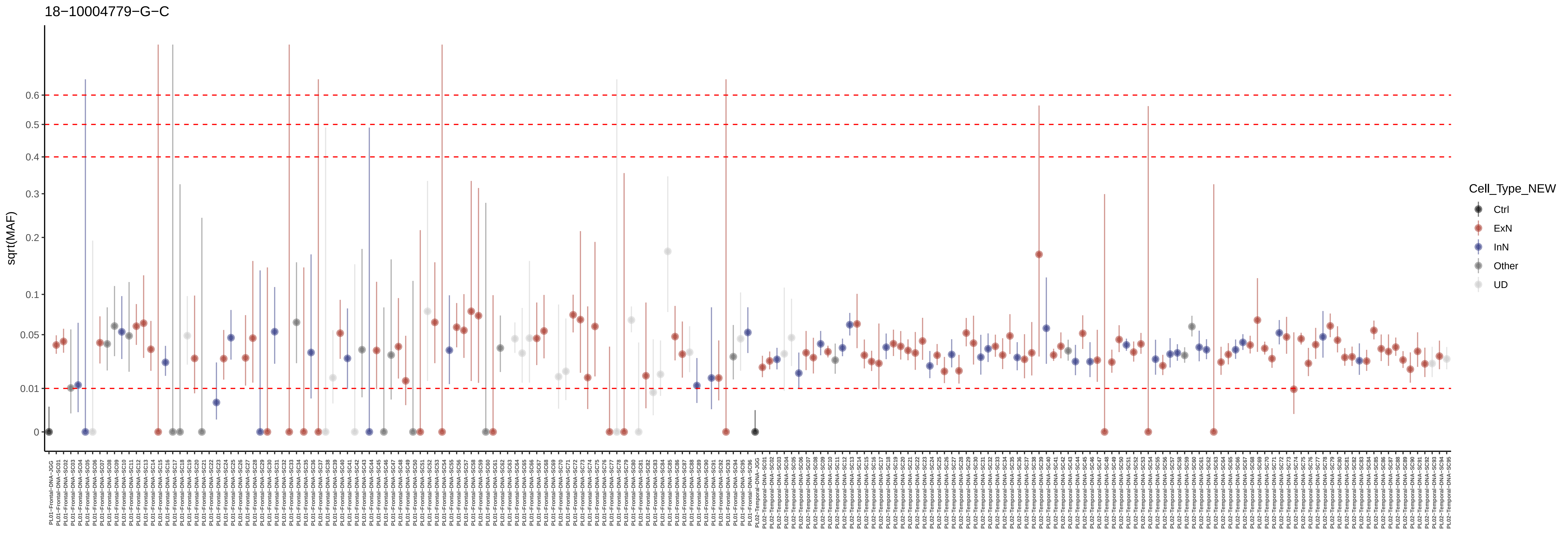

18−10004779−G−C
sqrt(MAF)
0.6
0.5
0.4
0.3
0.2
0.1
0.05
0.01
0
Cell_Type_NEW
Ctrl
ExN
InN
Other
UD

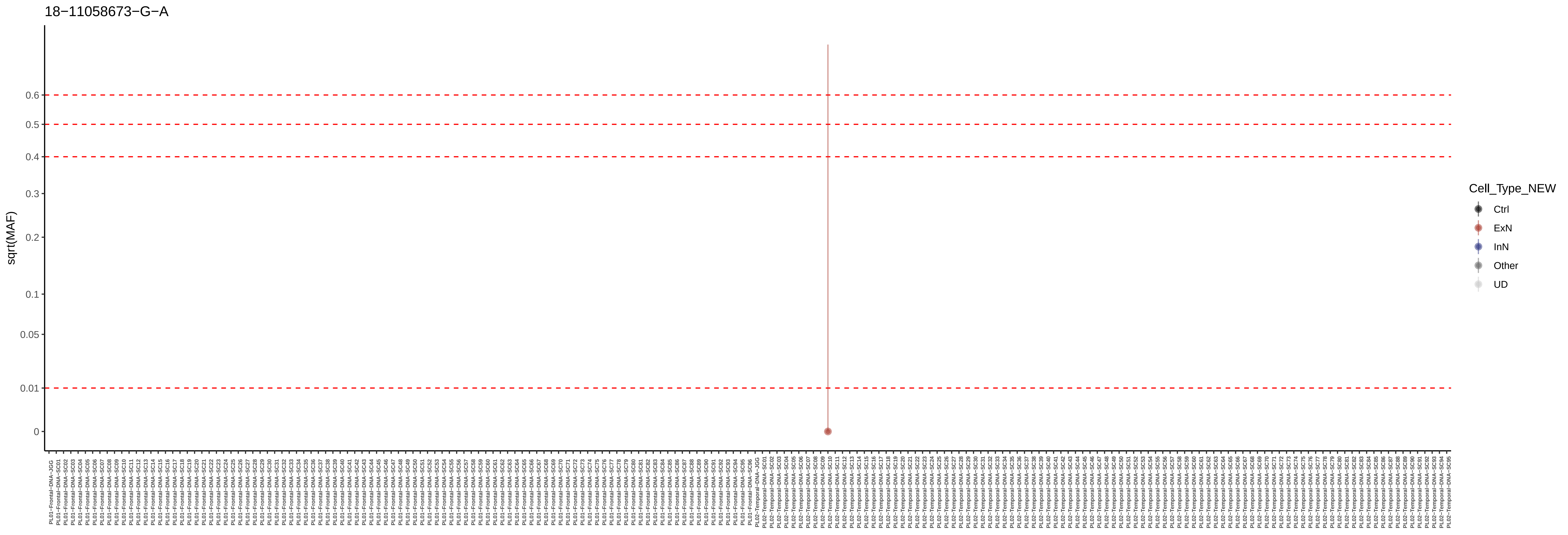

18−11058673−G−A
sqrt(MAF)
0.6
0.5
0.4
0.3
0.2
0.1
0.05
0.01
0
Cell_Type_NEW
Ctrl
ExN
InN
Other
UD

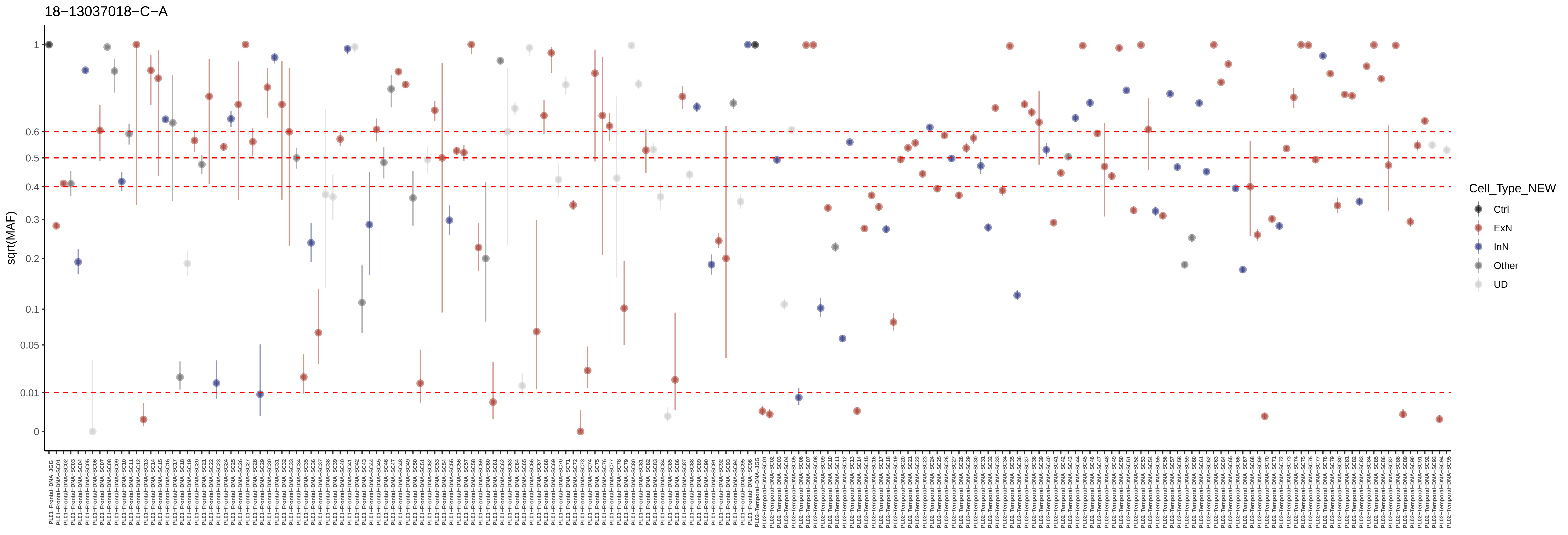
18−13037018−C−A
sqrt(MAF)
1
0.6
0.5
0.4
0.3
0.2
0.1
0.05
0.01
0
Cell_Type_NEW
Ctrl
ExN
InN
Other
UD

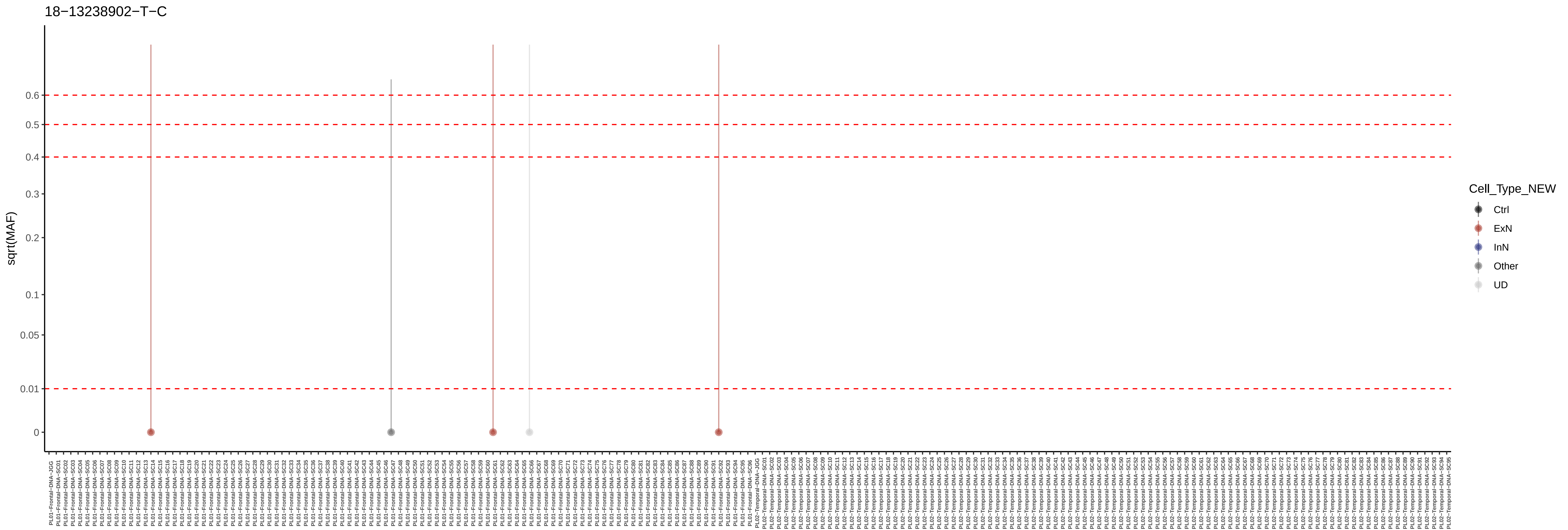

18−13238902−T−C
sqrt(MAF)
0.6
0.5
0.4
0.3
0.2
0.1
0.05
0.01
0
Cell_Type_NEW
Ctrl
ExN
InN
Other
UD

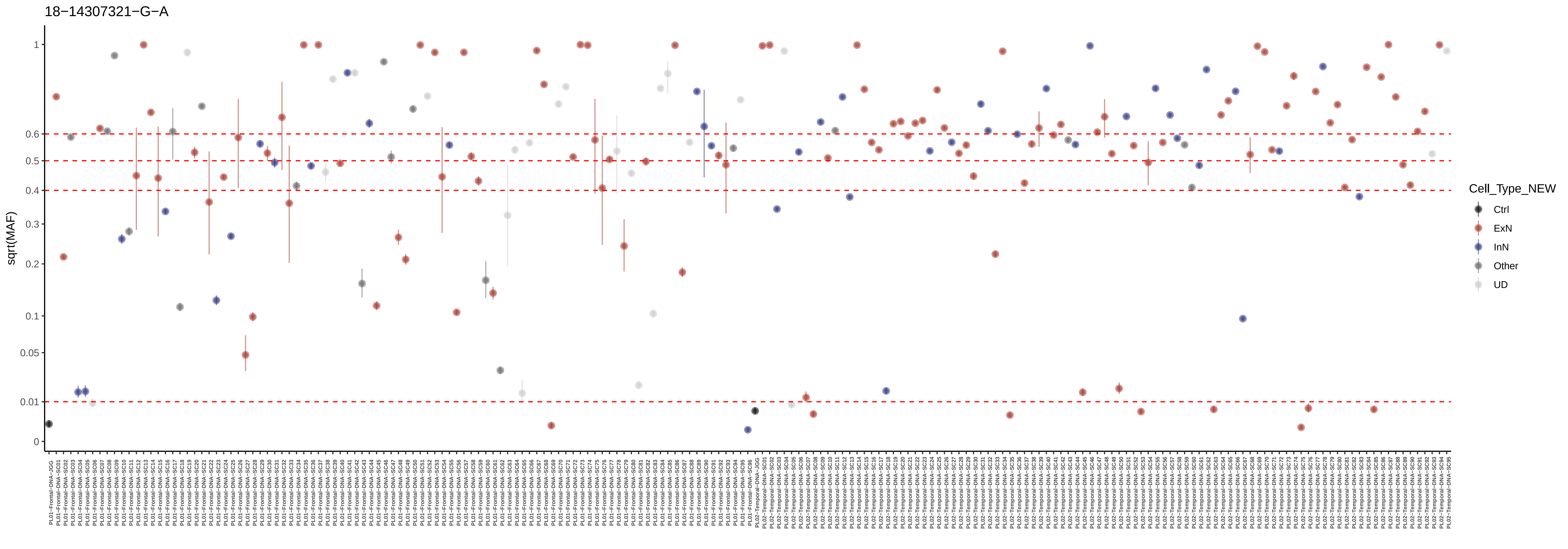
18−14307321−G−A
sqrt(MAF)
1
0.6
0.5
0.4
0.3
0.2
0.1
0.05
0.01
0
Cell_Type_NEW
Ctrl
ExN
InN
Other
UD

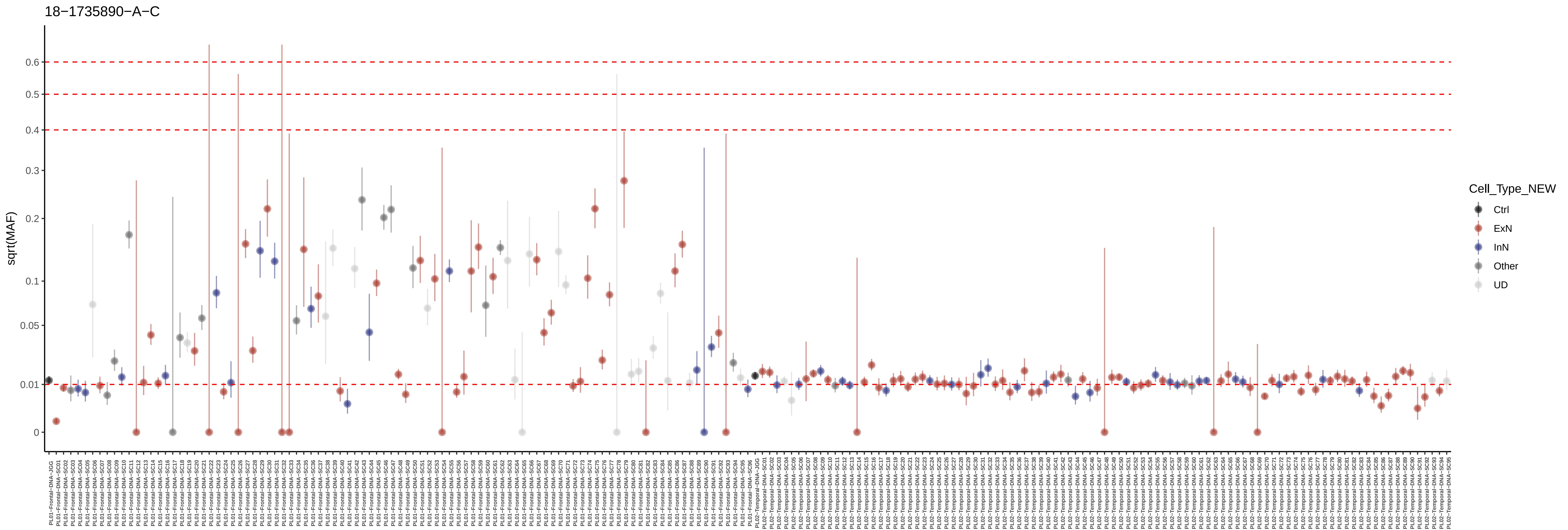
18−1735890−A−C
sqrt(MAF)
0.6
0.5
0.4
0.3
0.2
0.1
0.05
0.01
0
Cell_Type_NEW
Ctrl
ExN
InN
Other
UD

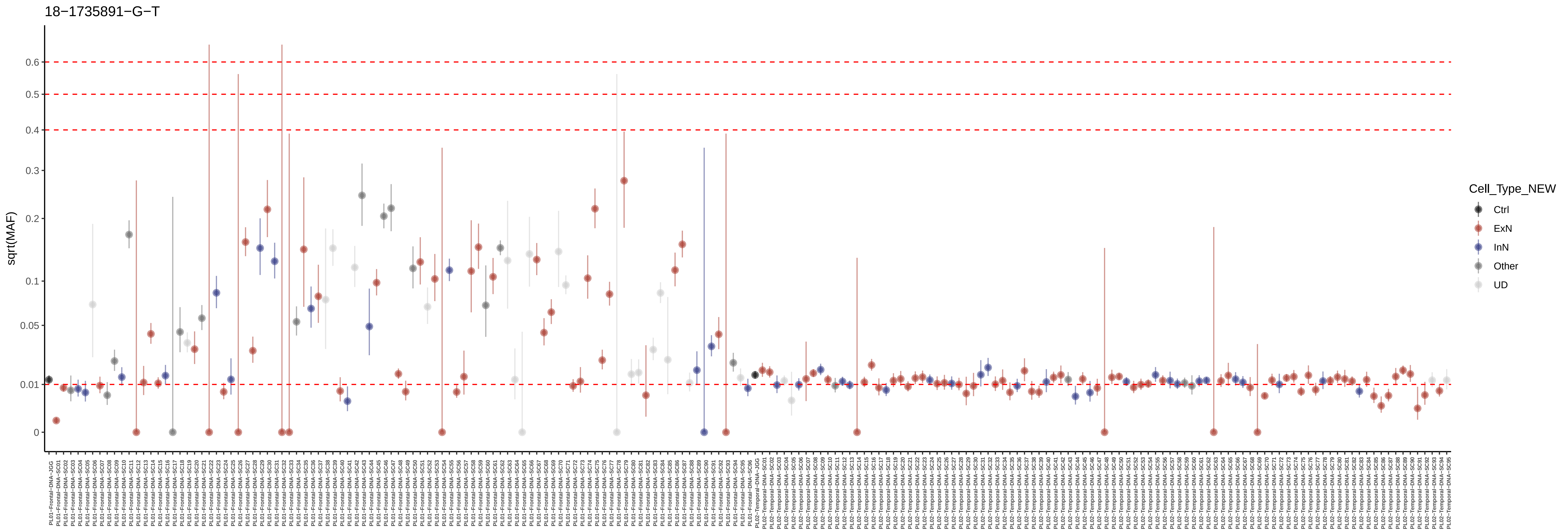

18−1735891−G−T
sqrt(MAF)
0.6
0.5
0.4
0.3
0.2
0.1
0.05
0.01
0
Cell_Type_NEW
Ctrl
ExN
InN
Other
UD

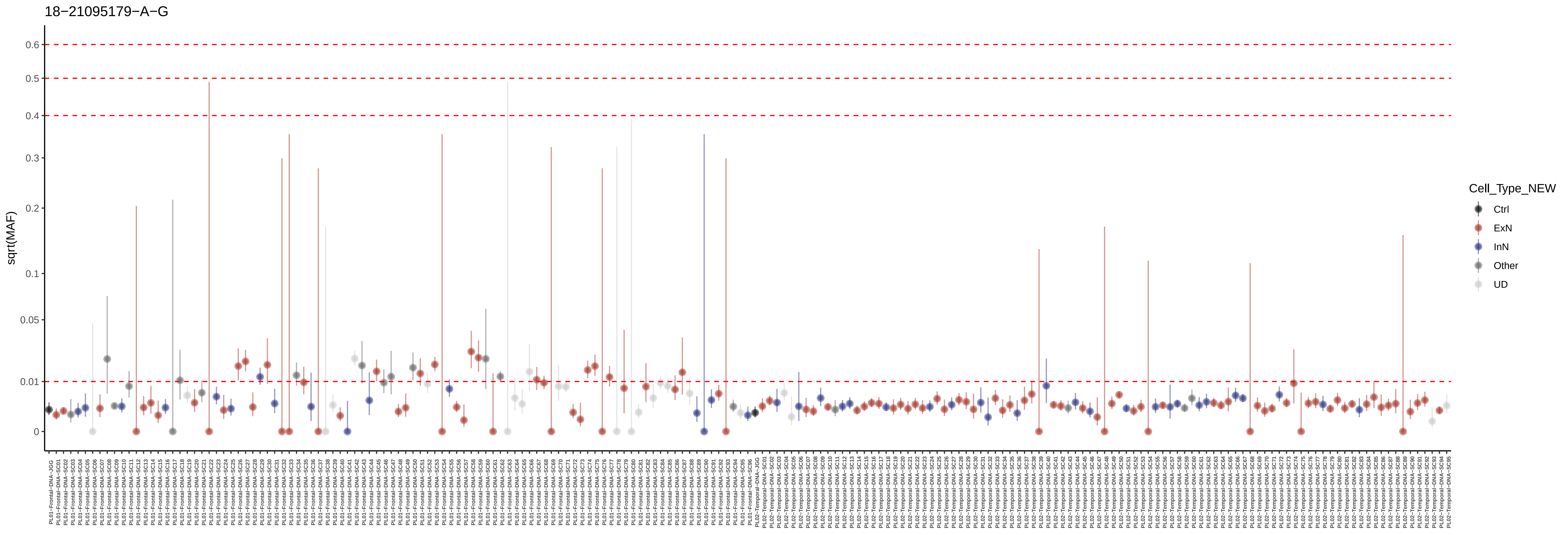

18−21095179−A−G
sqrt(MAF)
0.6
0.5
0.4
0.3
0.2
0.1
0.05
0.01
0
Cell_Type_NEW
Ctrl
ExN
InN
Other
UD

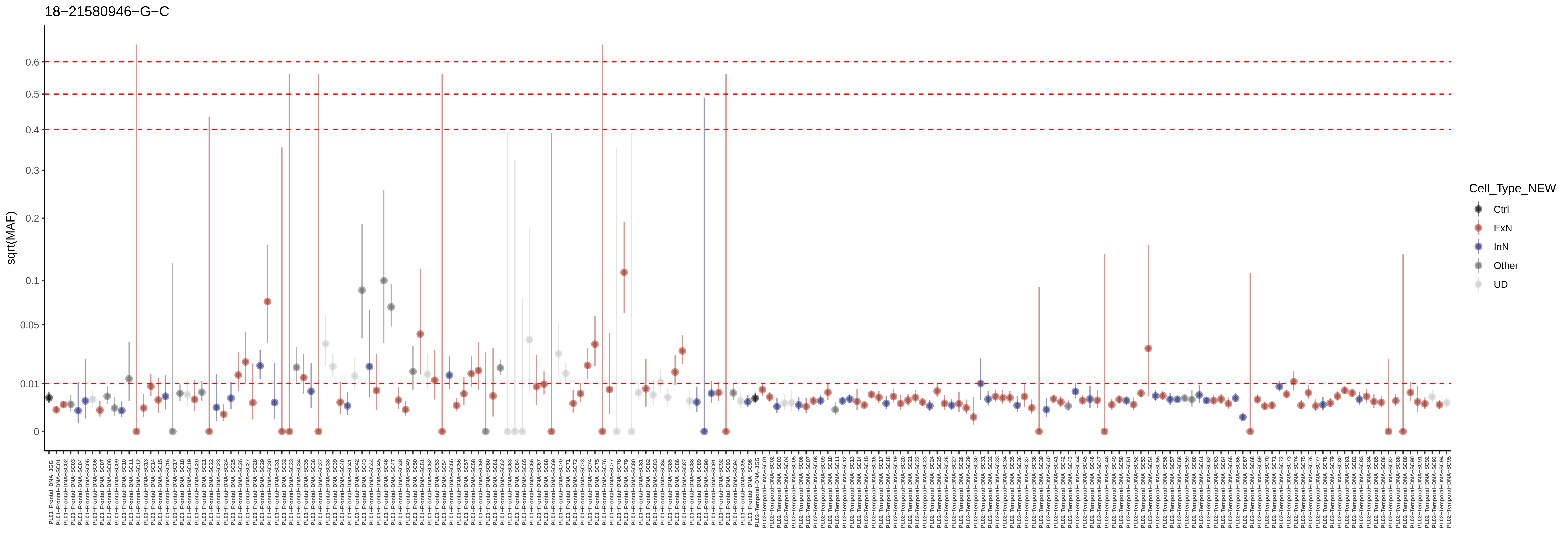

18−21580946−G−C
sqrt(MAF)
0.6
0.5
0.4
0.3
0.2
0.1
0.05
0.01
0
Cell_Type_NEW
Ctrl
ExN
InN
Other
UD

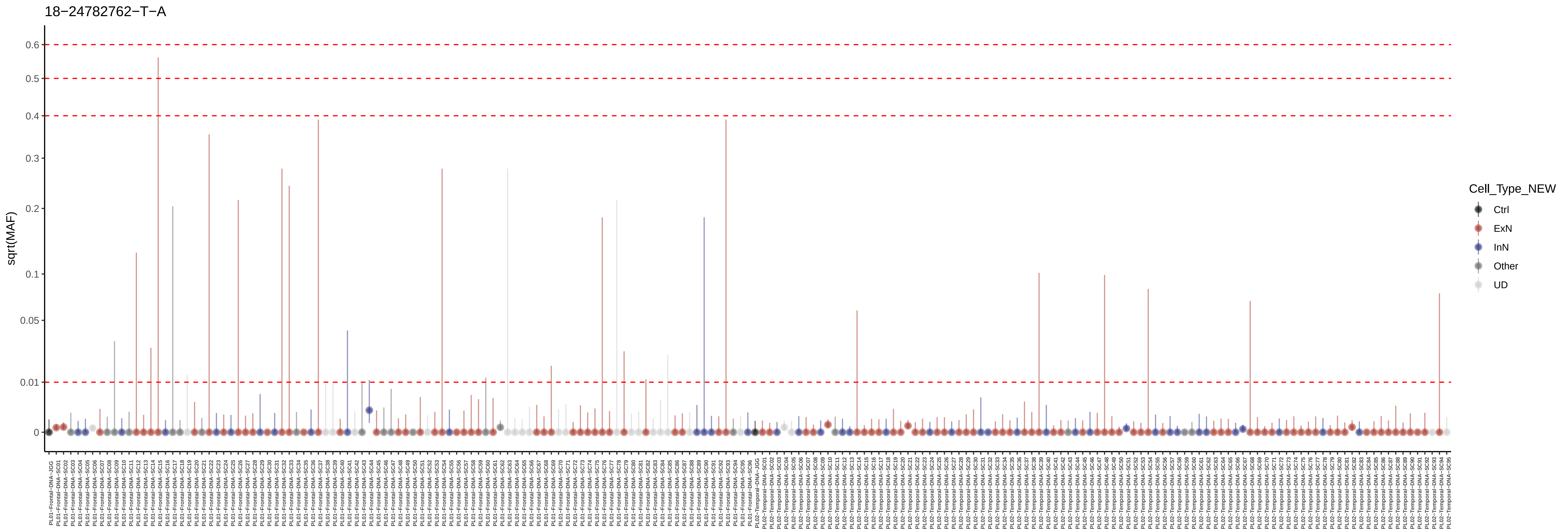
18−24782762−T−A
sqrt(MAF)
0.6
0.5
0.4
0.3
0.2
0.1
0.05
0.01
0
Cell_Type_NEW
Ctrl
ExN
InN
Other
UD

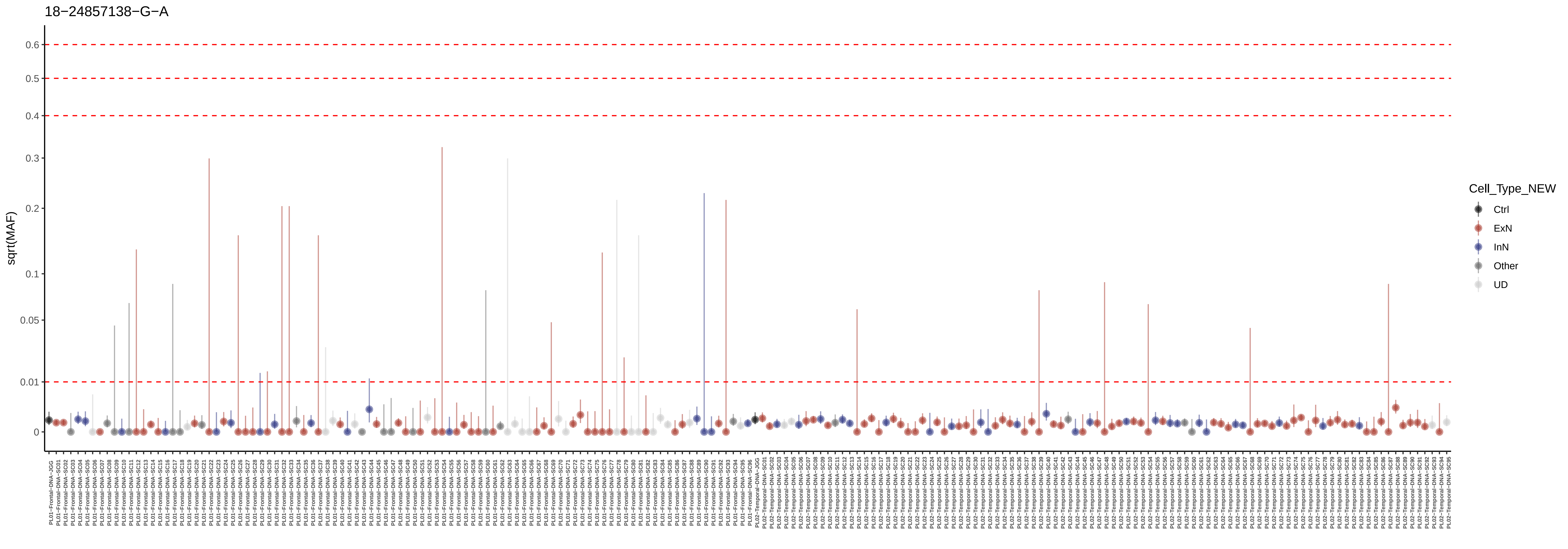
18−24857138−G−A
sqrt(MAF)
0.6
0.5
0.4
0.3
0.2
0.1
0.05
0.01
0
Cell_Type_NEW
Ctrl
ExN
InN
Other
UD

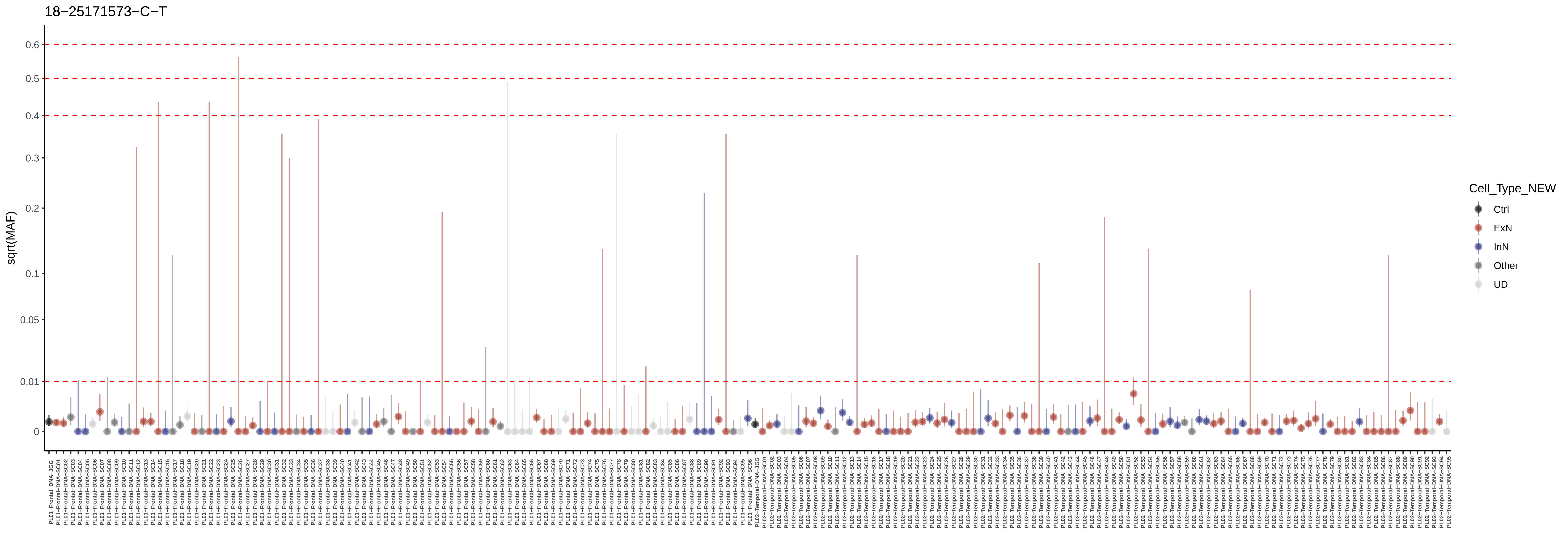

18−25171573−C−T
sqrt(MAF)
0.6
0.5
0.4
0.3
0.2
0.1
0.05
0.01
0
Cell_Type_NEW
Ctrl
ExN
InN
Other
UD

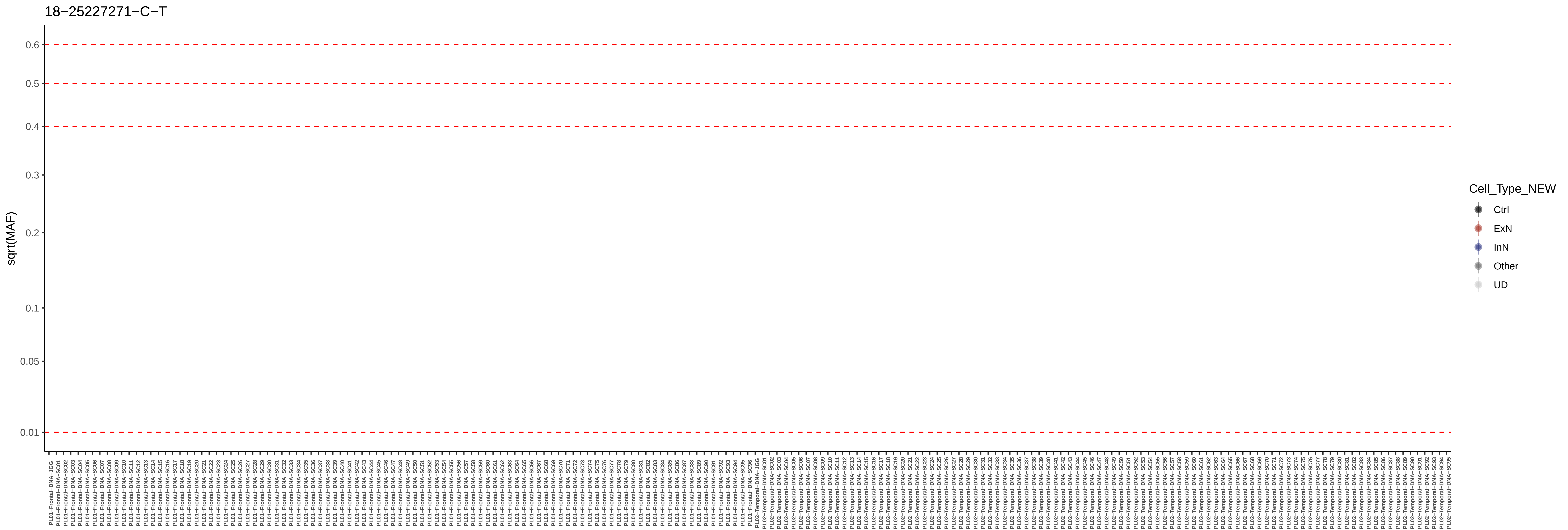

18−25227271−C−T
sqrt(MAF)
0.6
0.5
0.4
0.3
0.2
0.1
0.05
0.01
Cell_Type_NEW
Ctrl
ExN
InN
Other
UD

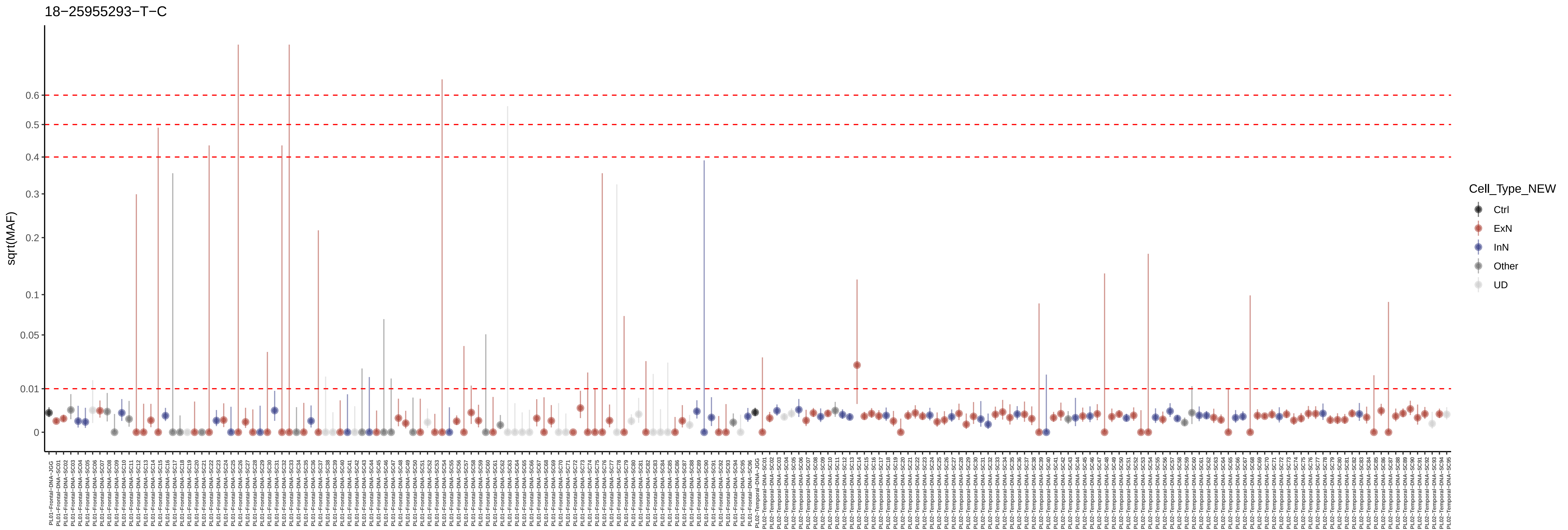
18−25955293−T−C
sqrt(MAF)
0.6
0.5
0.4
0.3
0.2
0.1
0.05
0.01
0
Cell_Type_NEW
Ctrl
ExN
InN
Other
UD

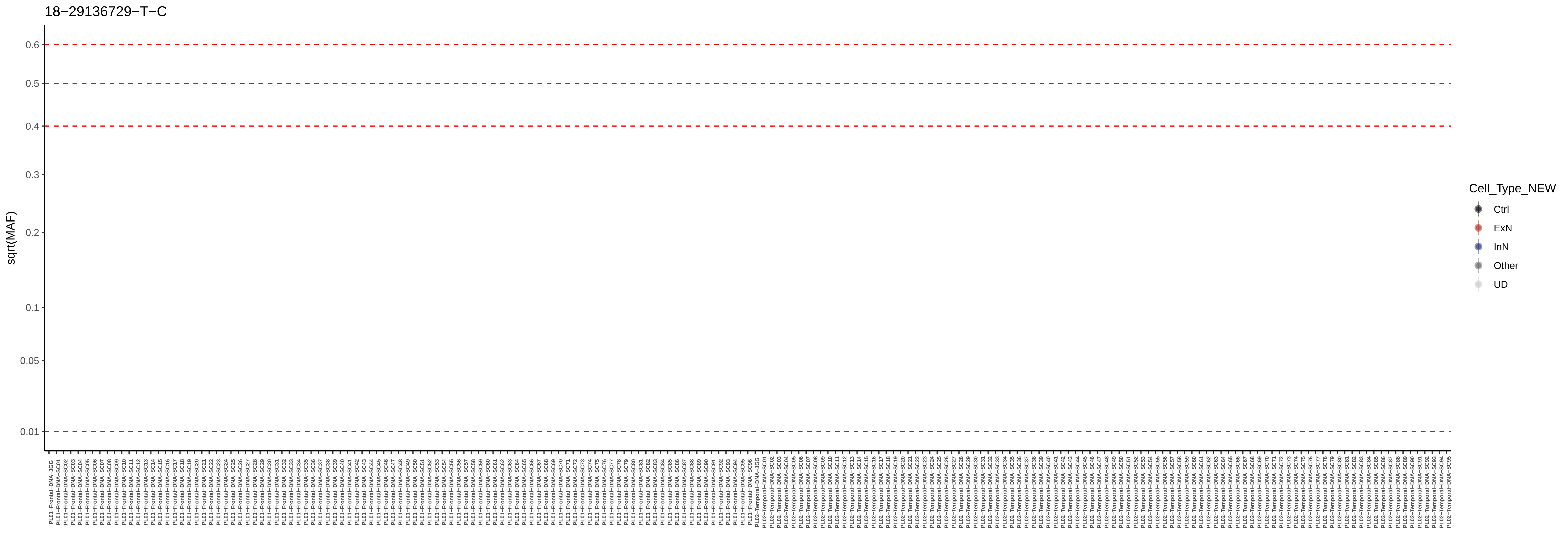
18−29136729−T−C
sqrt(MAF)
0.6
0.5
0.4
0.3
0.2
0.1
0.05
0.01
Cell_Type_NEW
Ctrl
ExN
InN
Other
UD

18−301430−T−C

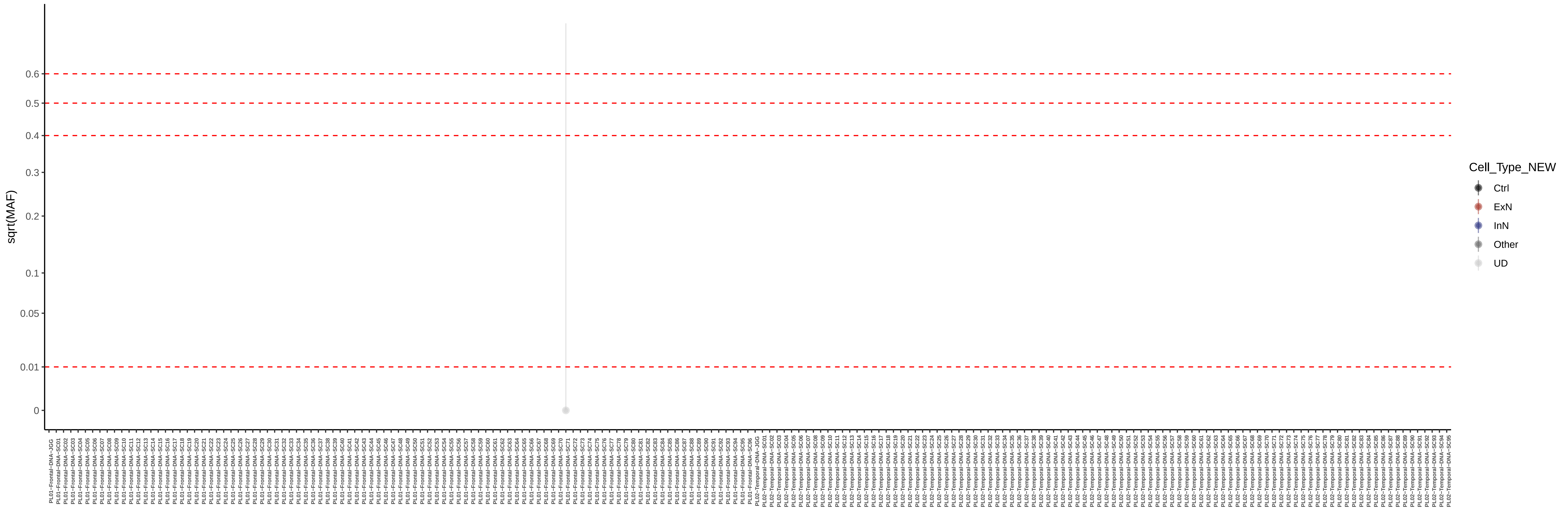

sqrt(MAF)
0.6
0.5
0.4
0.3
0.2
0.1
0.05
0.01
0
Cell_Type_NEW
Ctrl
ExN
InN
Other
UD

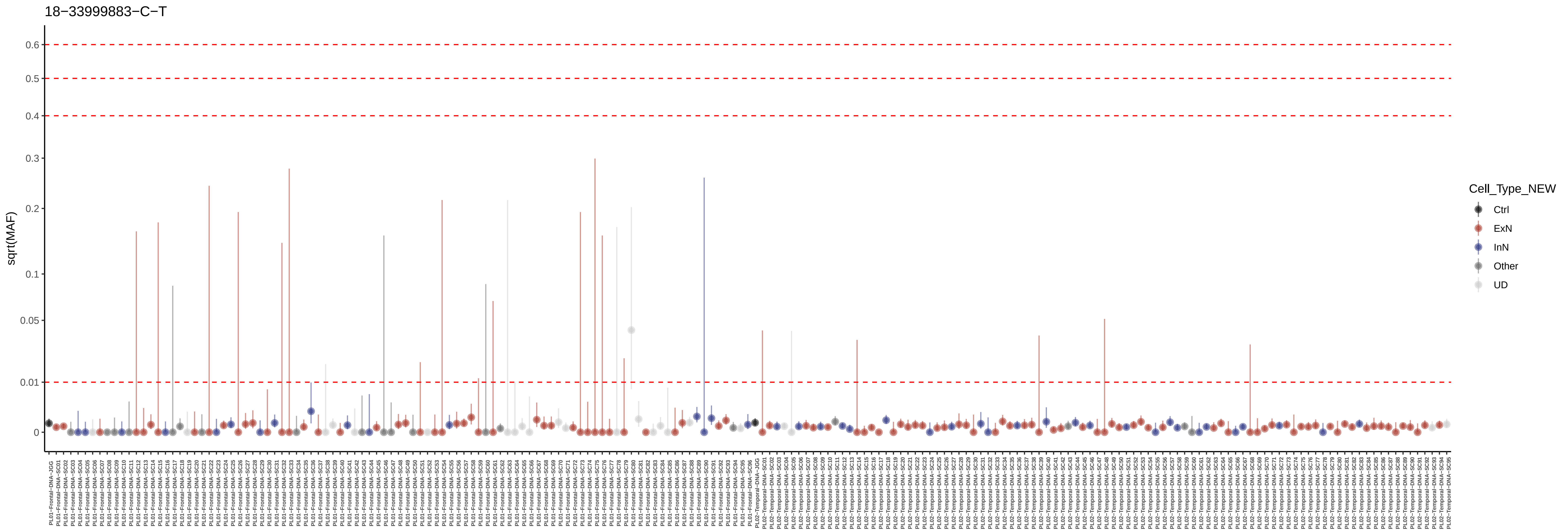
18−33999883−C−T
sqrt(MAF)
0.6
0.5
0.4
0.3
0.2
0.1
0.05
0.01
0
Cell_Type_NEW
Ctrl
ExN
InN
Other
UD

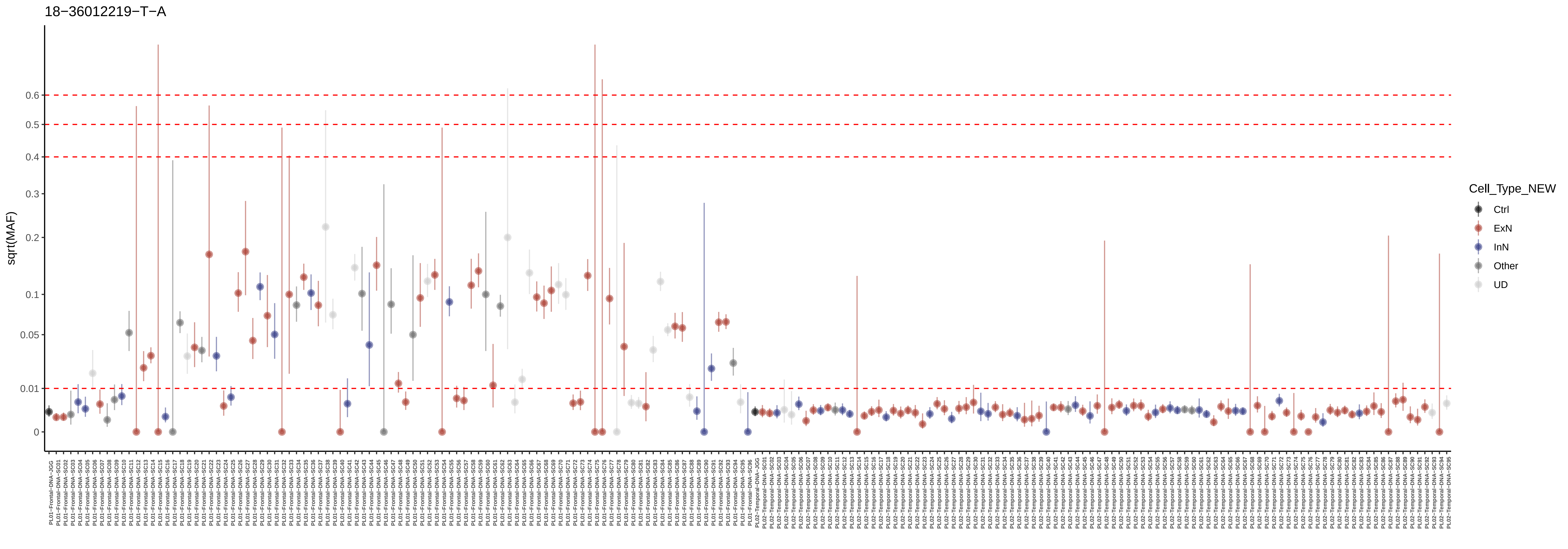
18−36012219−T−A
sqrt(MAF)
0.6
0.5
0.4
0.3
0.2
0.1
0.05
0.01
0
Cell_Type_NEW
Ctrl
ExN
InN
Other
UD

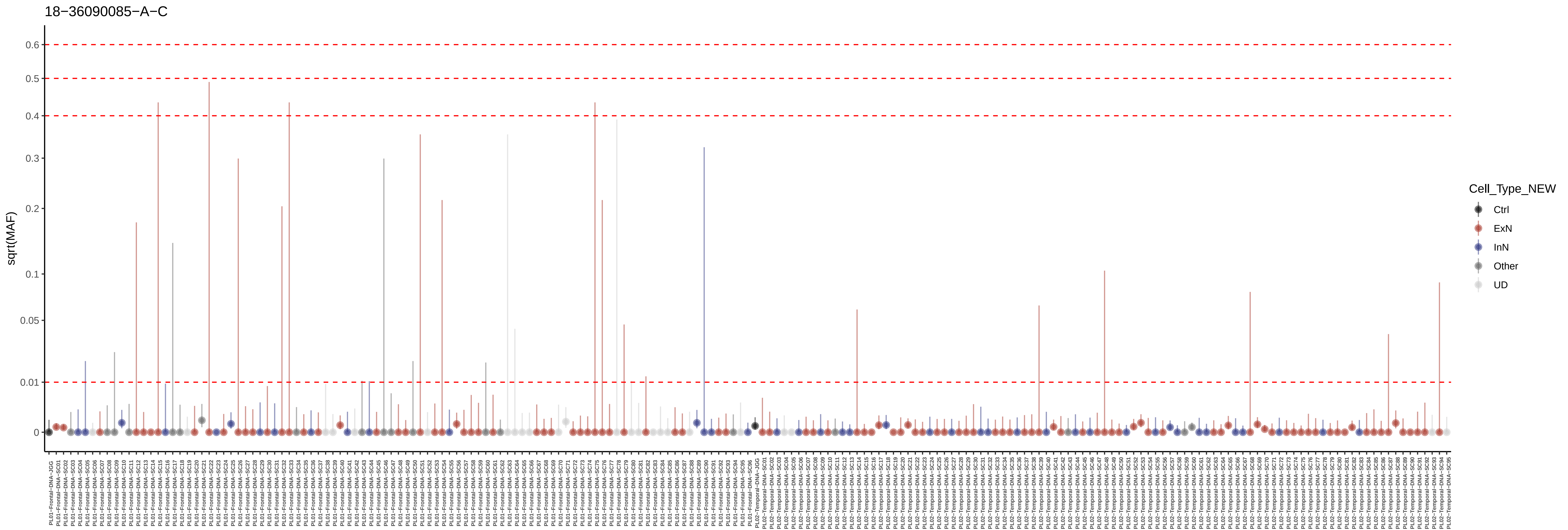
18−36090085−A−C
sqrt(MAF)
0.6
0.5
0.4
0.3
0.2
0.1
0.05
0.01
0
Cell_Type_NEW
Ctrl
ExN
InN
Other
UD

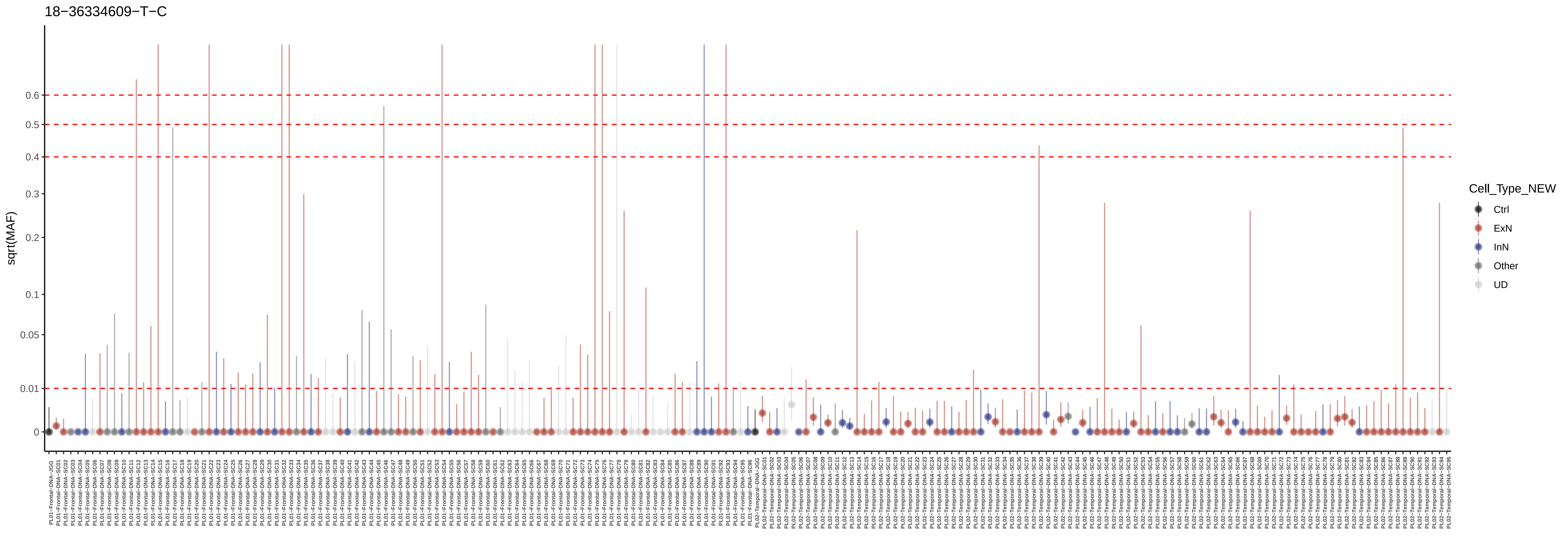
18−36334609−T−C
sqrt(MAF)
0.6
0.5
0.4
0.3
0.2
0.1
0.05
0.01
0
Cell_Type_NEW
Ctrl
ExN
InN
Other
UD

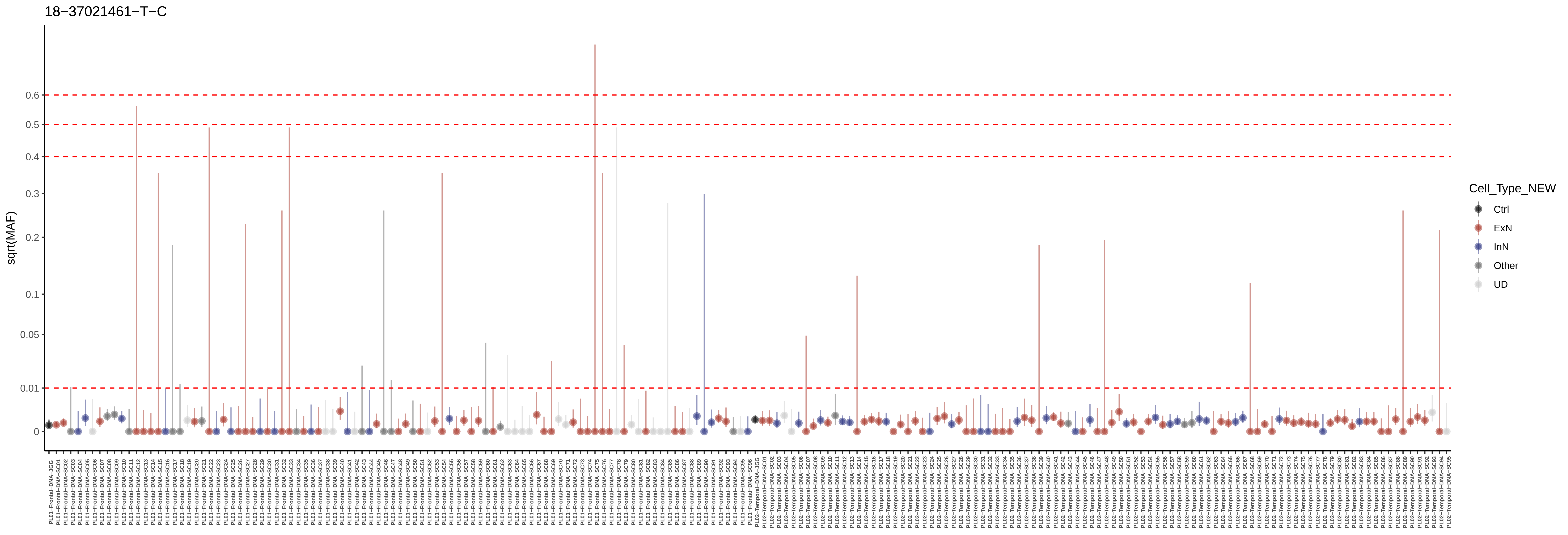
18−37021461−T−C
sqrt(MAF)
0.6
0.5
0.4
0.3
0.2
0.1
0.05
0.01
0
Cell_Type_NEW
Ctrl
ExN
InN
Other
UD

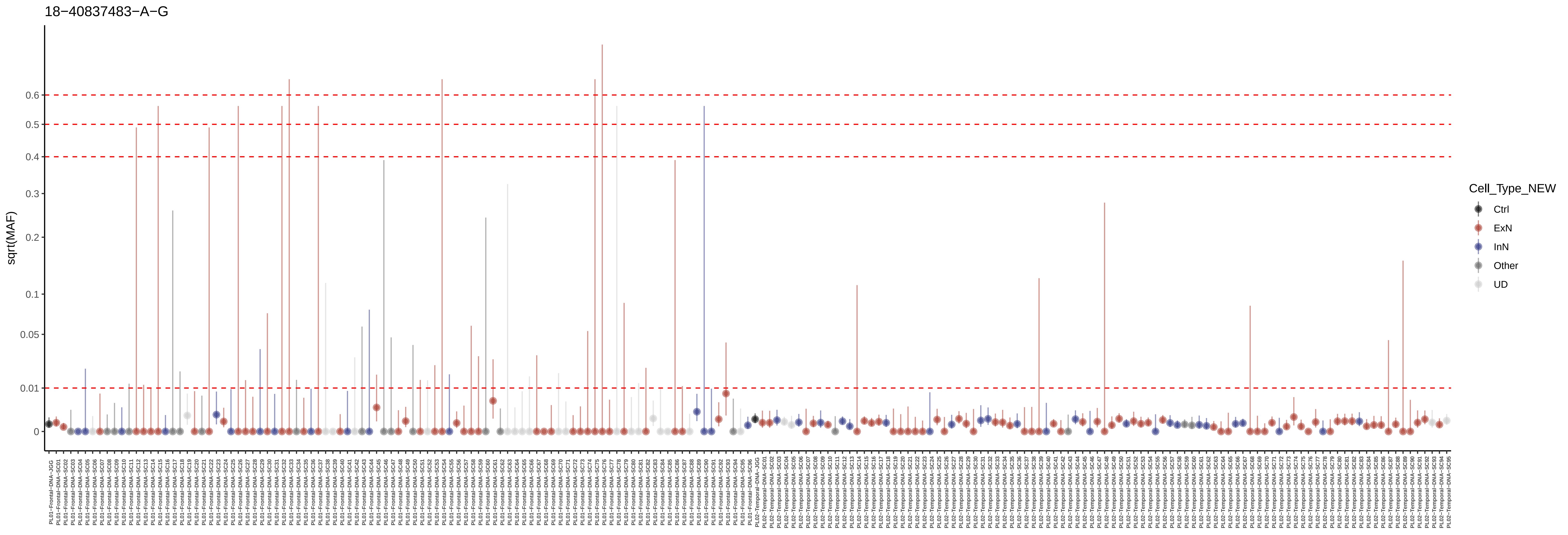
18−40837483−A−G
sqrt(MAF)
0.6
0.5
0.4
0.3
0.2
0.1
0.05
0.01
0
Cell_Type_NEW
Ctrl
ExN
InN
Other
UD

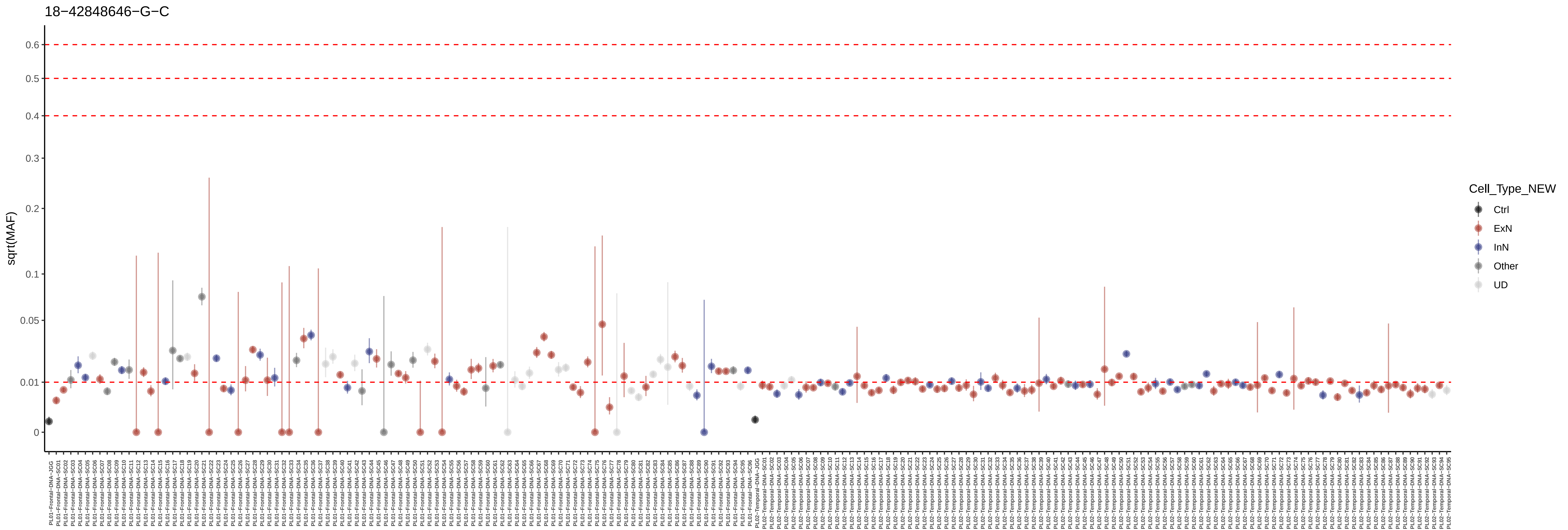
18−42848646−G−C
sqrt(MAF)
0.6
0.5
0.4
0.3
0.2
0.1
0.05
0.01
0
Cell_Type_NEW
Ctrl
ExN
InN
Other
UD

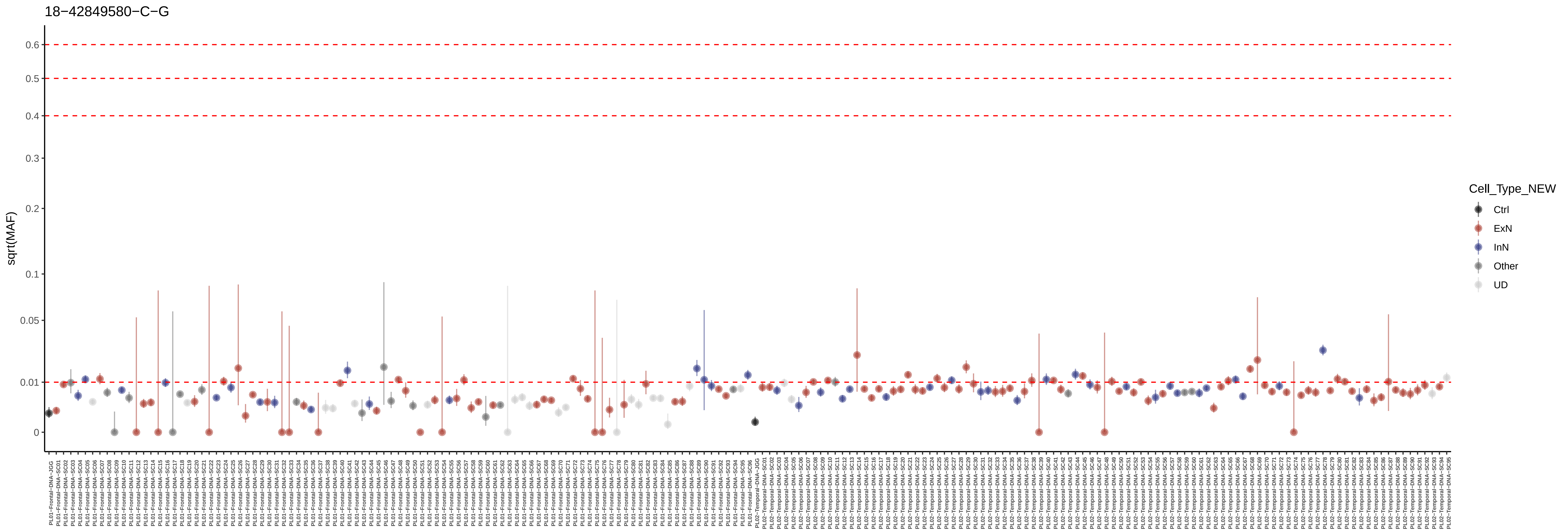

18−42849580−C−G
sqrt(MAF)
0.6
0.5
0.4
0.3
0.2
0.1
0.05
0.01
0
Cell_Type_NEW
Ctrl
ExN
InN
Other
UD

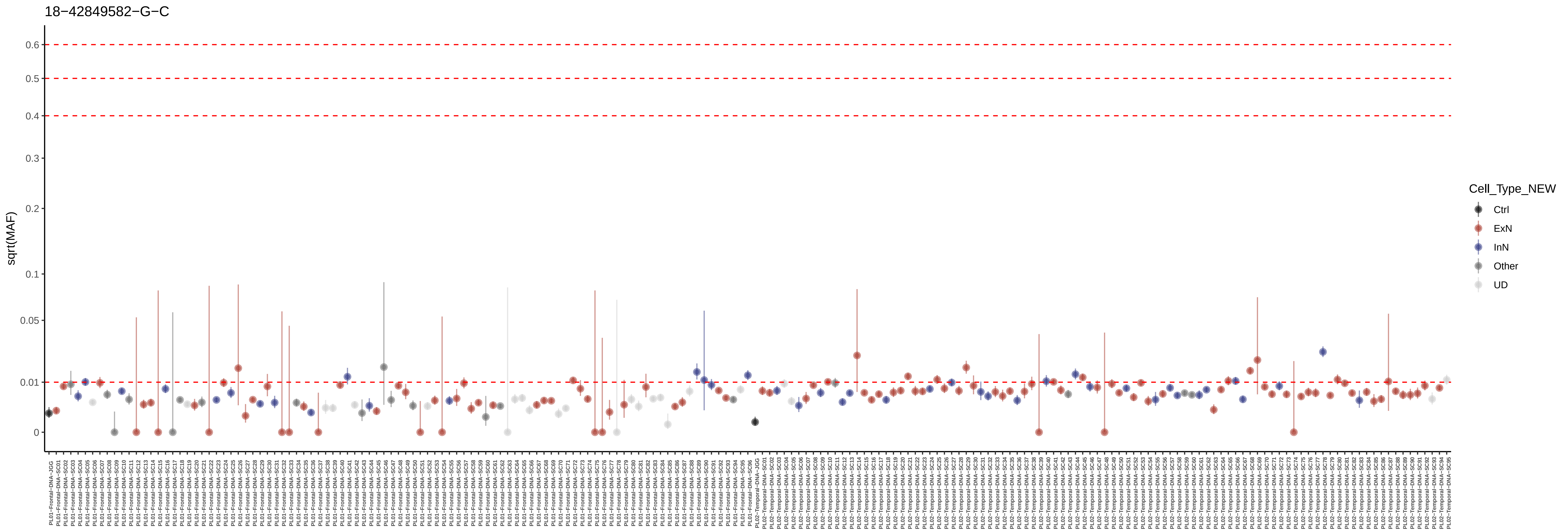

18−42849582−G−C
sqrt(MAF)
0.6
0.5
0.4
0.3
0.2
0.1
0.05
0.01
0
Cell_Type_NEW
Ctrl
ExN
InN
Other
UD

18−42932839−C−T

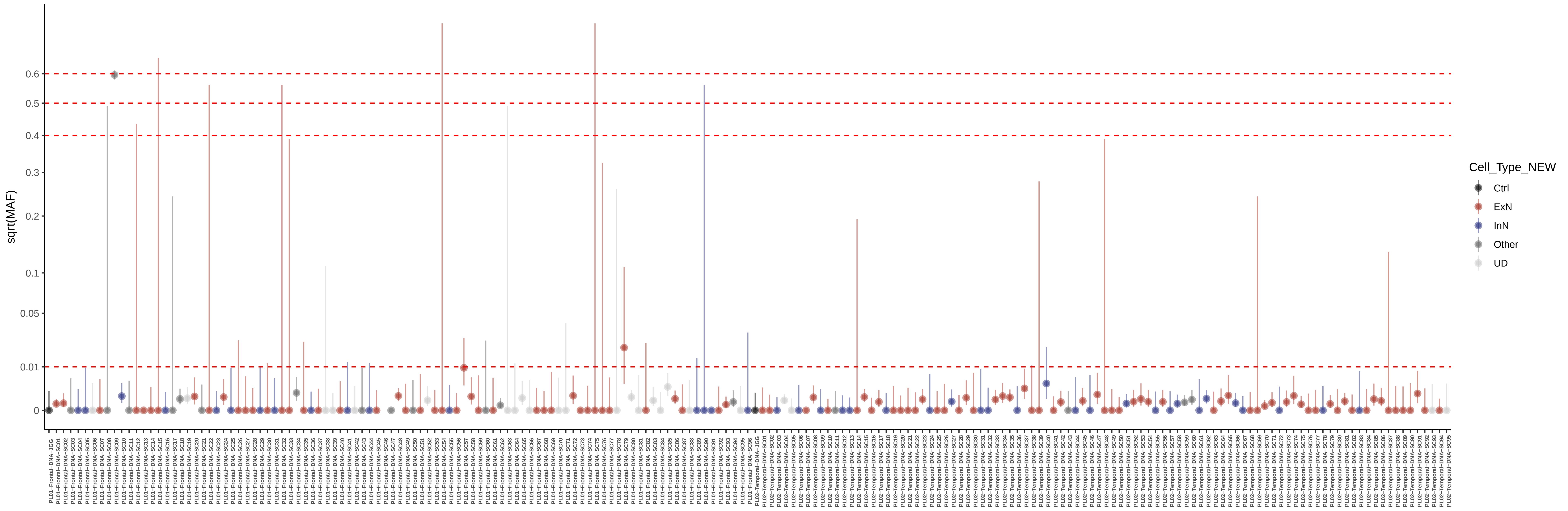

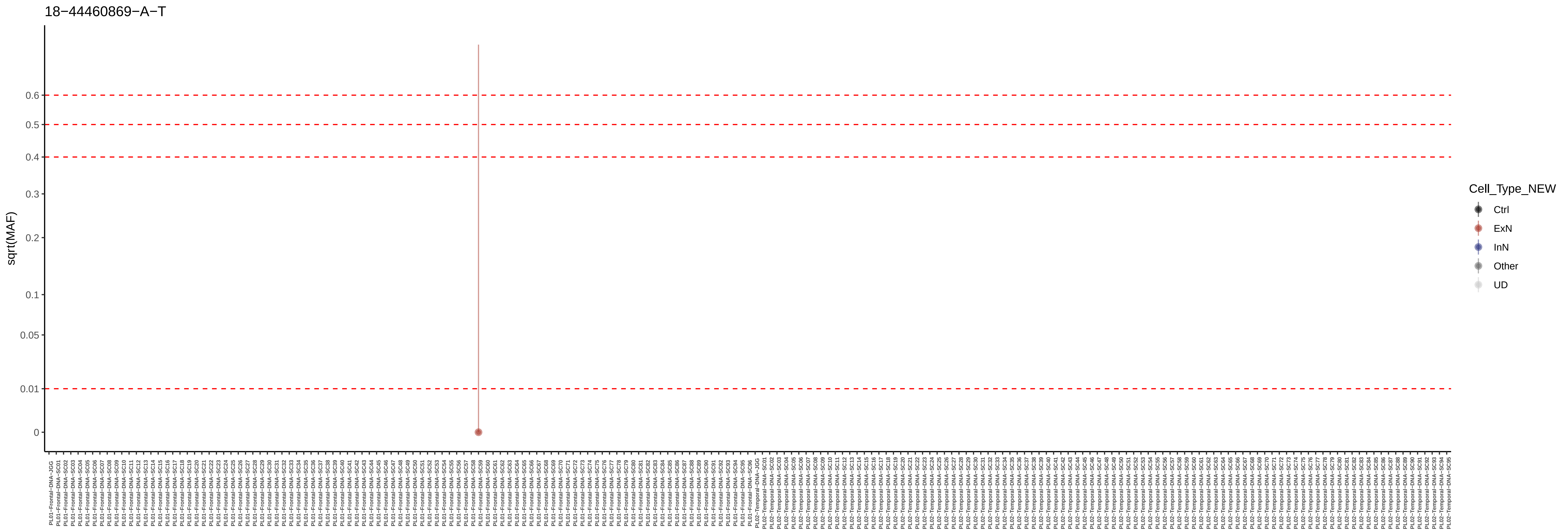
18−44460869−A−T
sqrt(MAF)
0.6
0.5
0.4
0.3
0.2
0.1
0.05
0.01
0
Cell_Type_NEW
Ctrl
ExN
InN
Other
UD

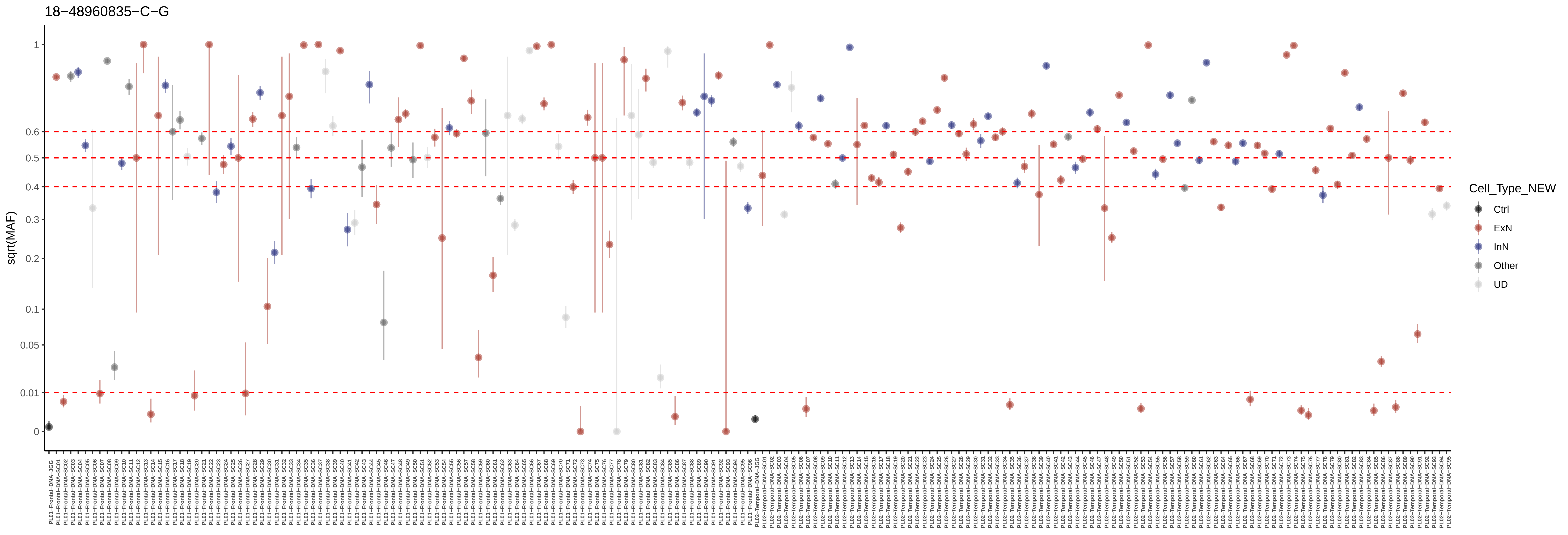
18−48960835−C−G
sqrt(MAF)
1
0.6
0.5
0.4
0.3
0.2
0.1
0.05
0.01
0
Cell_Type_NEW
Ctrl
ExN
InN
Other
UD

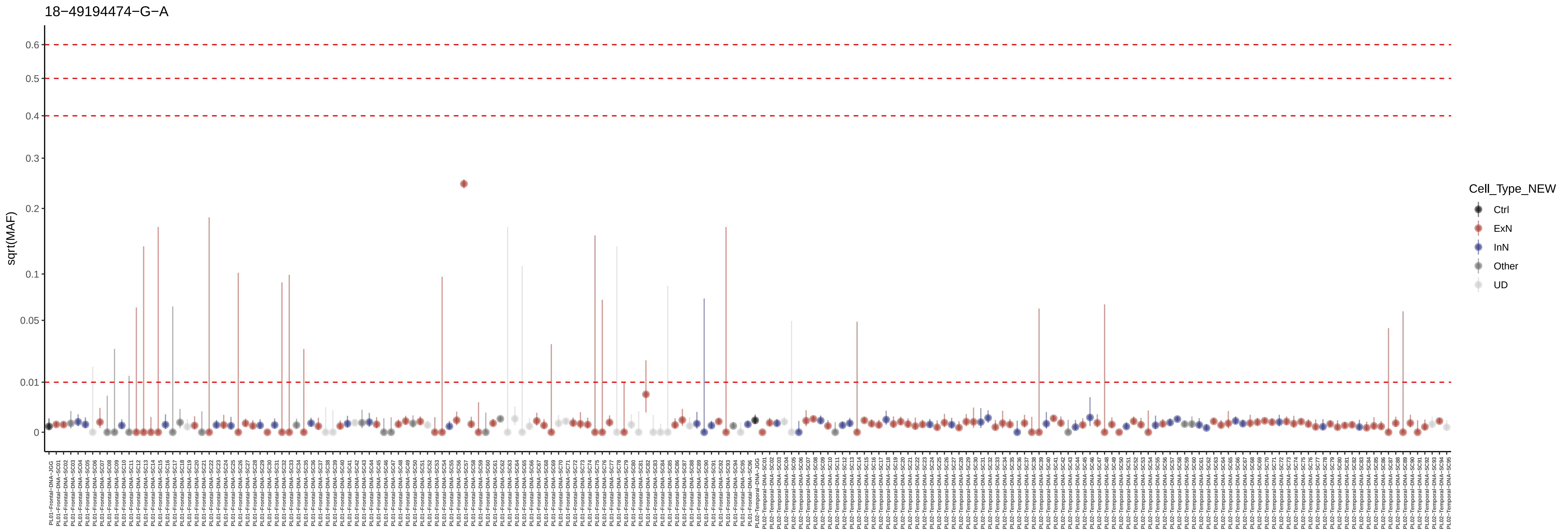

18−49194474−G−A
sqrt(MAF)
Cell_Type_NEW
Ctrl
ExN
InN
Other
UD

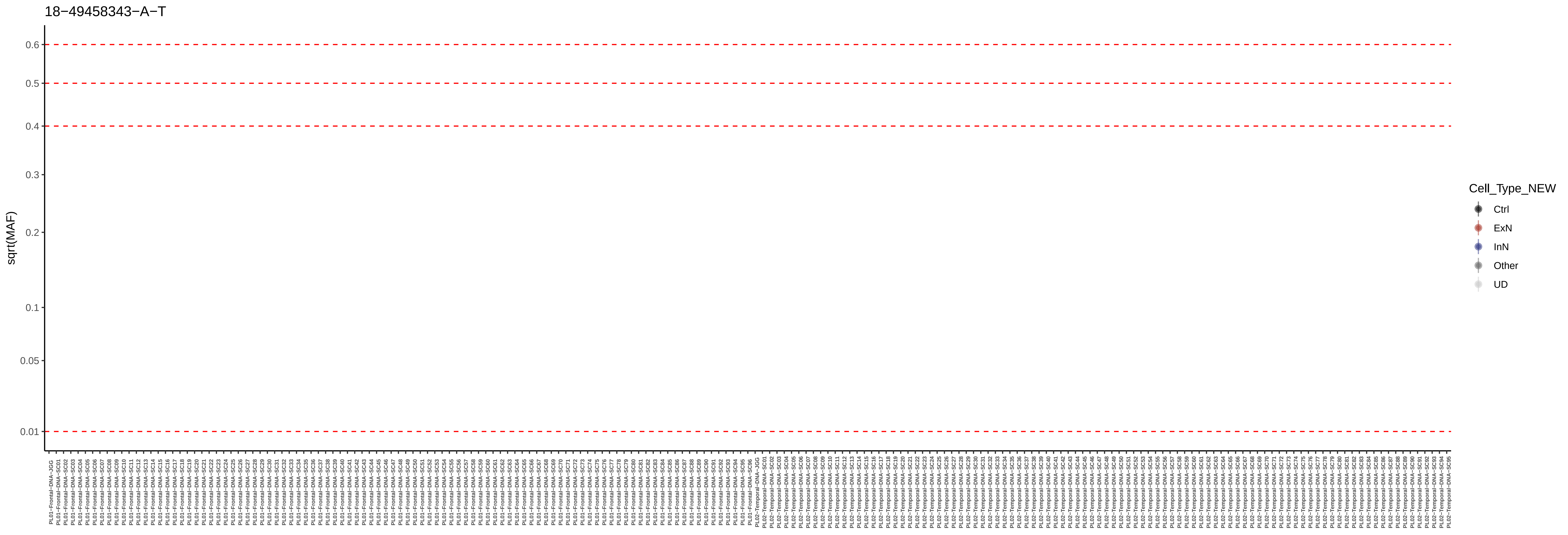
18−49458343−A−T
sqrt(MAF)
Cell_Type_NEW
Ctrl
ExN
InN
Other
UD

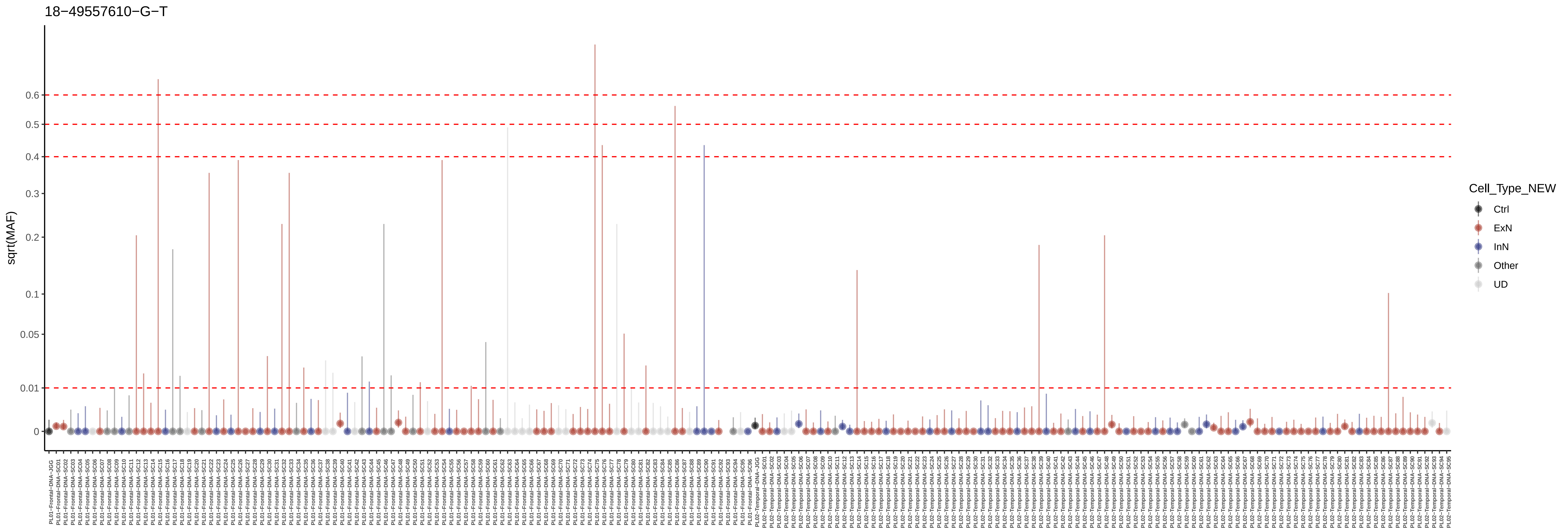
18−49557610−G−T
sqrt(MAF)
0.6
0.5
0.4
0.3
0.2
0.1
0.05
0.01
0
Cell_Type_NEW
Ctrl
ExN
InN
Other
UD

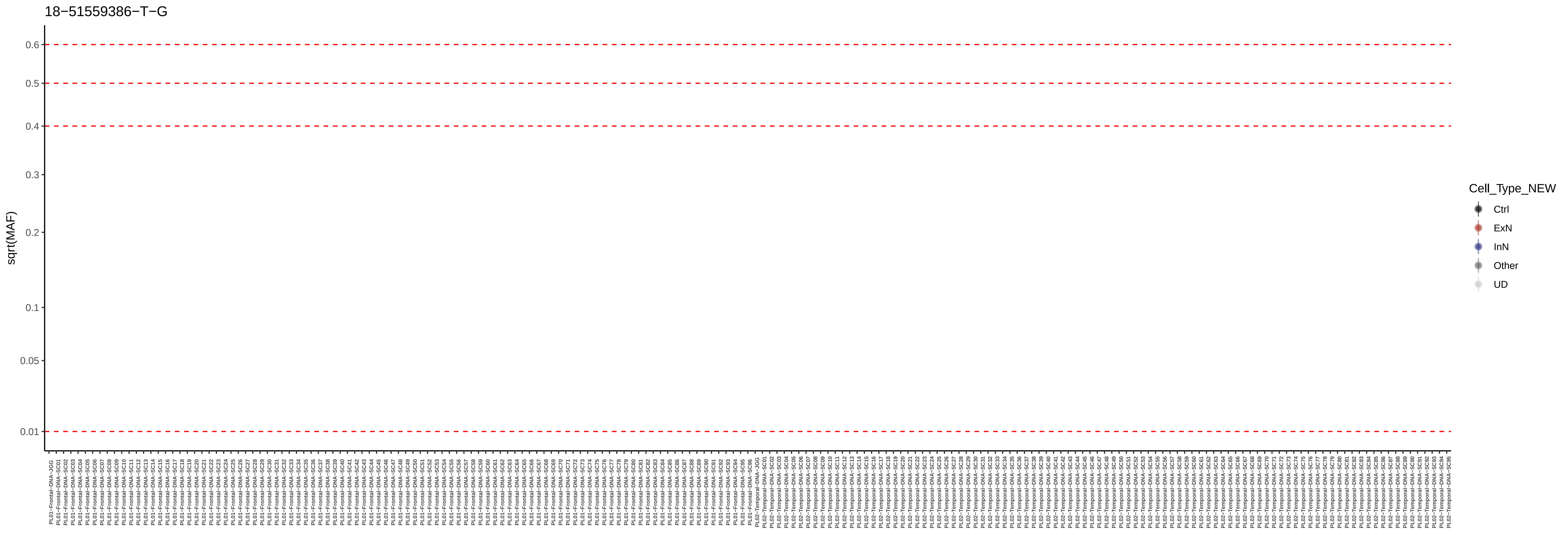
18−51559386−T−G
sqrt(MAF)
0.6
0.5
0.4
0.3
0.2
0.1
0.05
0.01
Cell_Type_NEW
Ctrl
ExN
InN
Other
UD

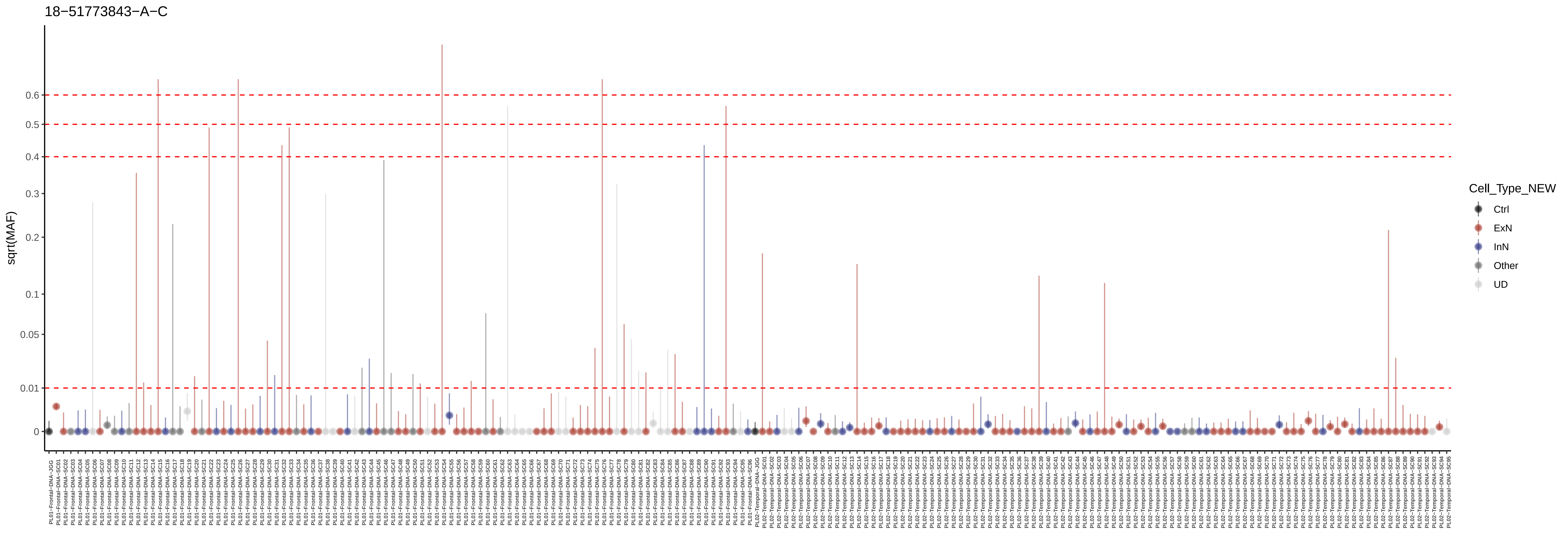
18−51773843−A−C
sqrt(MAF)
0.6
0.5
0.4
0.3
0.2
0.1
0.05
0.01
0
Cell_Type_NEW
Ctrl
ExN
InN
Other
UD

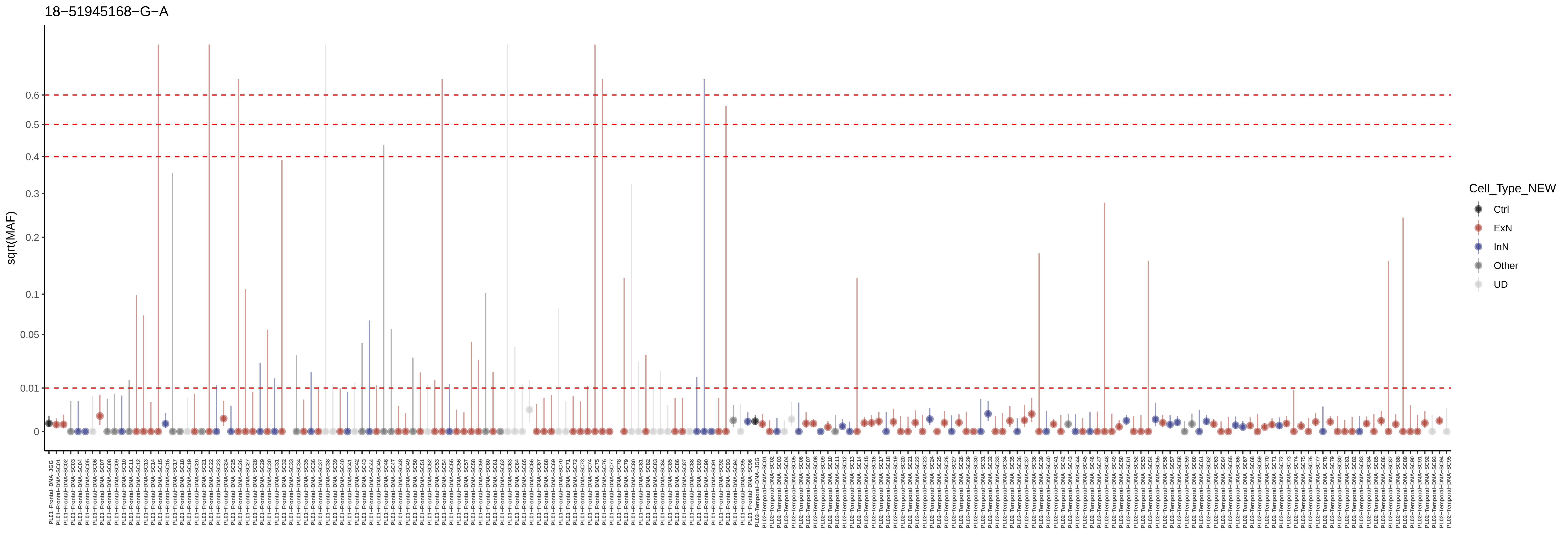
18−51945168−G−A
sqrt(MAF)
0
0.01
0.05
0.1
0.2
0.3
0.4
0.5
0.6
Cell_Type_NEW
Ctrl
ExN
InN
Other
UD

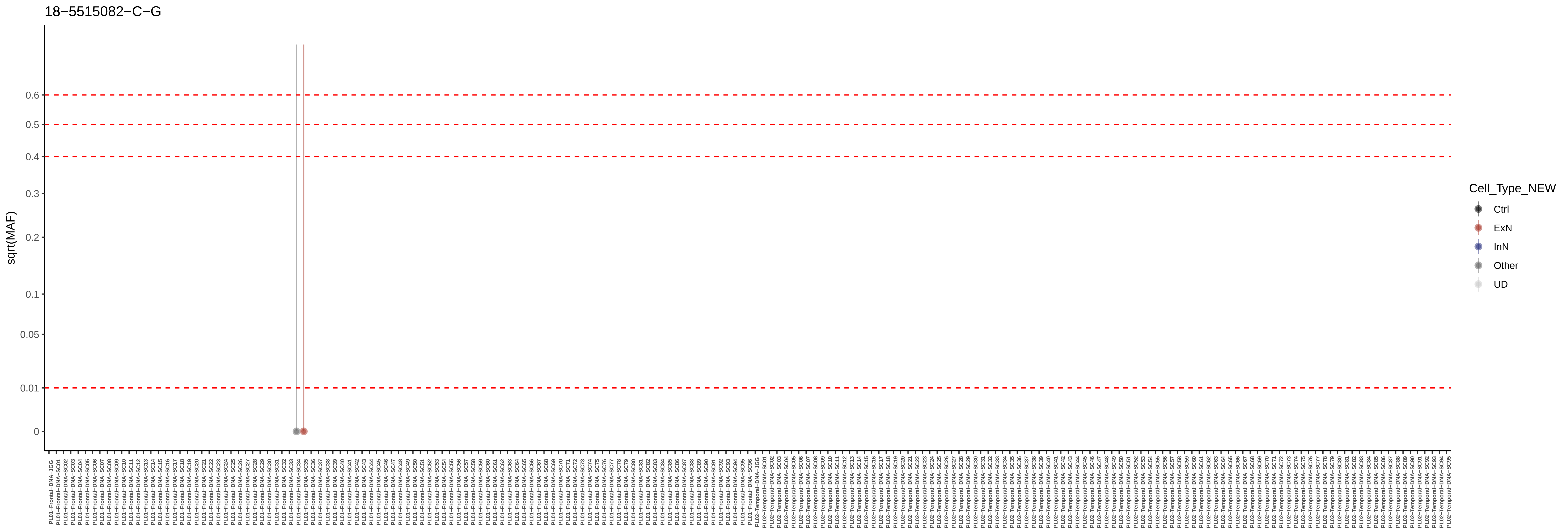
18−5515082−C−G
sqrt(MAF)
0.6
0.5
0.4
0.3
0.2
0.1
0.05
0.01
0
Cell_Type_NEW
Ctrl
ExN
InN
Other
UD

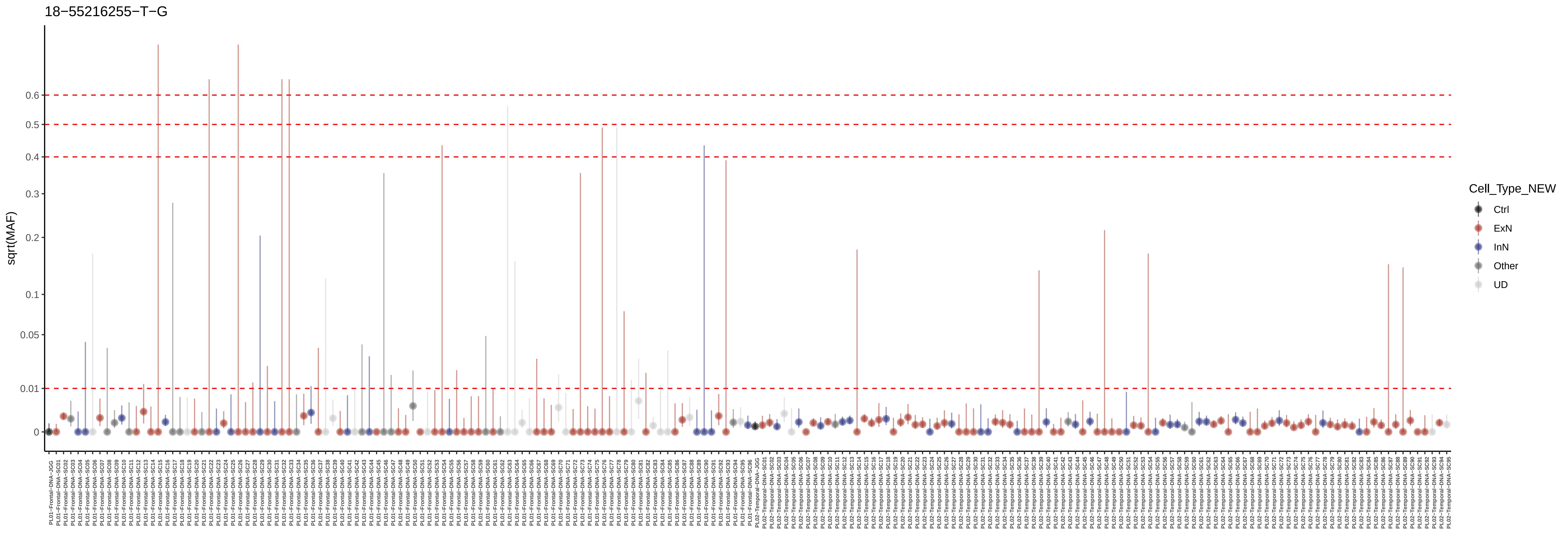
18−55216255−T−G
sqrt(MAF)
0.6
0.5
0.4
0.3
0.2
0.1
0.05
0.01
0
Cell_Type_NEW
Ctrl
ExN
InN
Other
UD

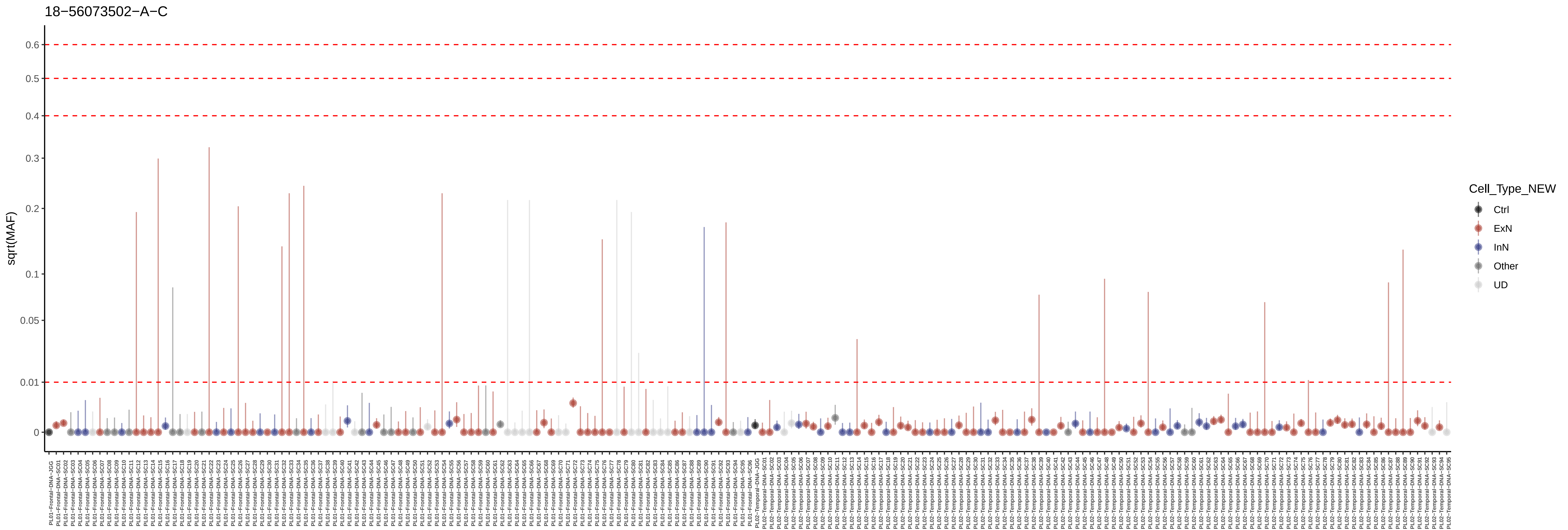
18−56073502−A−C
sqrt(MAF)
0.6
0.5
0.4
0.3
0.2
0.1
0.05
0.01
0
Cell_Type_NEW
Ctrl
ExN
InN
Other
UD

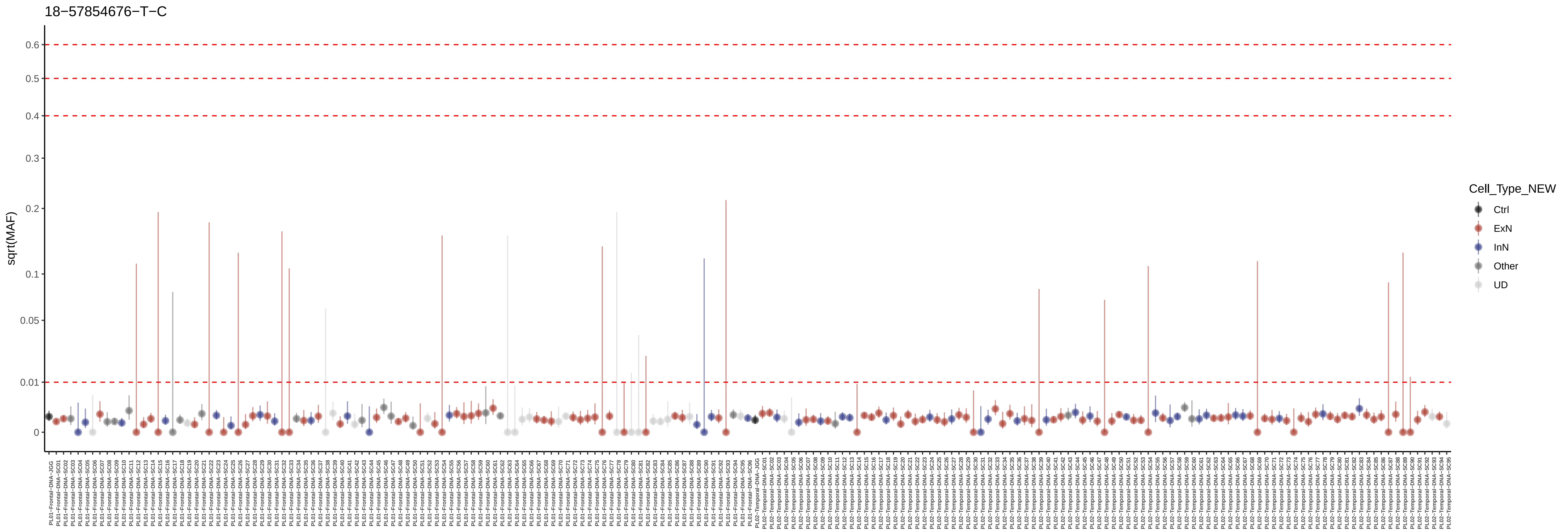
18−57854676−T−C
sqrt(MAF)
0.6
0.5
0.4
0.3
0.2
0.1
0.05
0.01
0
Cell_Type_NEW
Ctrl
ExN
InN
Other
UD

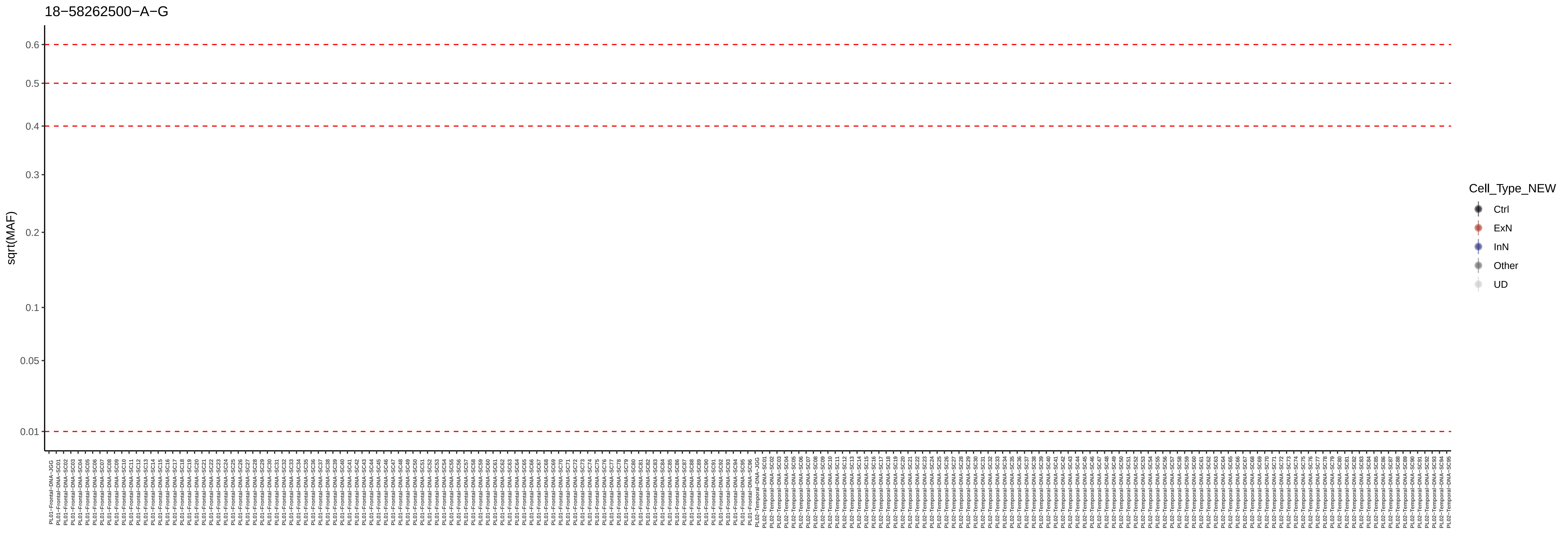
18−58262500−A−G
sqrt(MAF)
0.6
0.5
0.4
0.3
0.2
0.1
0.05
0.01
Cell_Type_NEW
Ctrl
ExN
InN
Other
UD

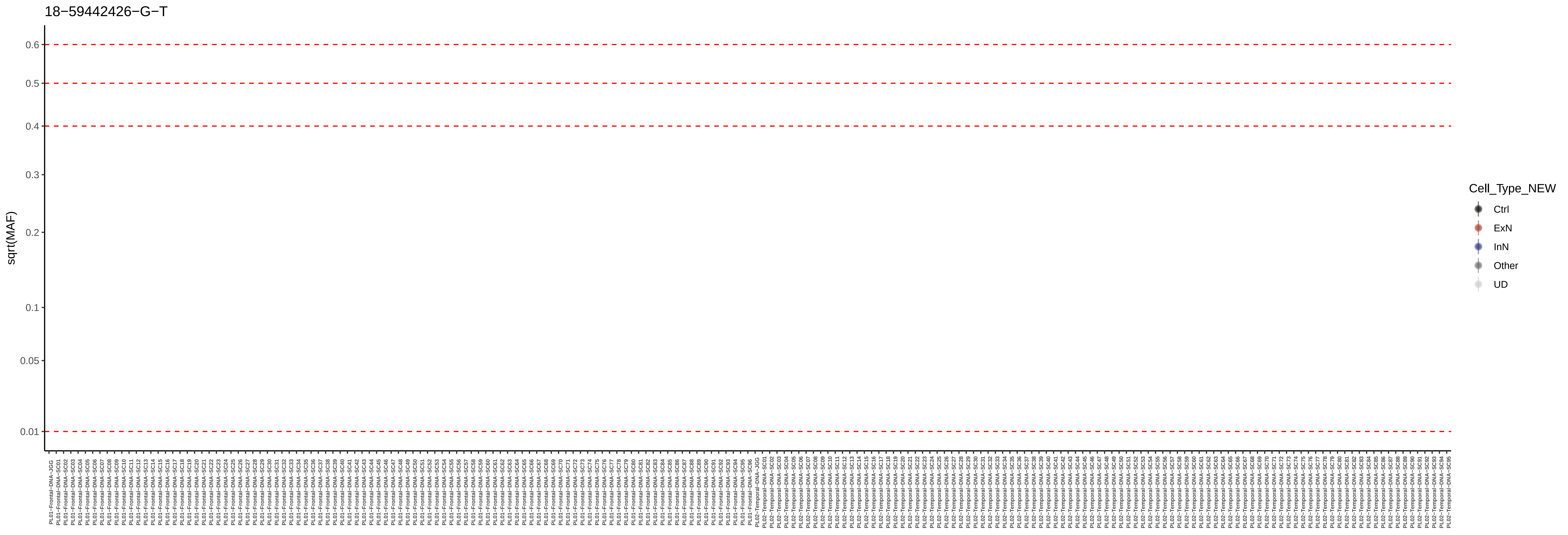
18−59442426−G−T
sqrt(MAF)
0.6
0.5
0.4
0.3
0.2
0.1
0.05
0.01
Cell_Type_NEW
Ctrl
ExN
InN
Other
UD

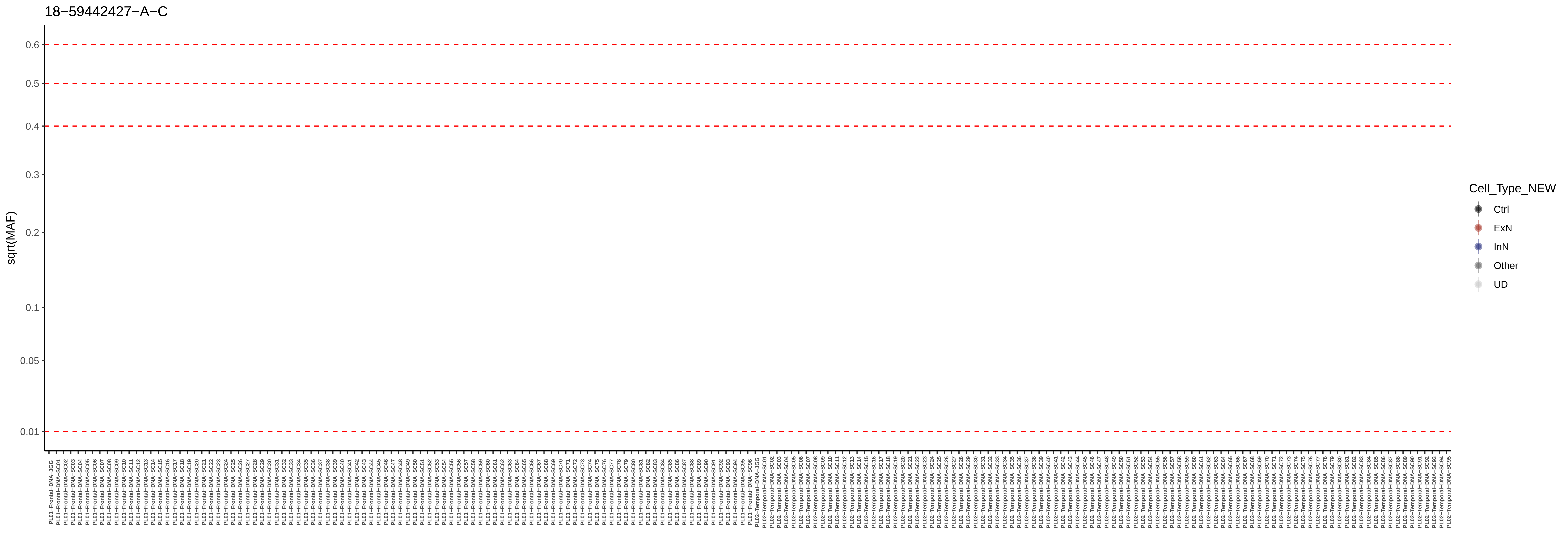

18−59442427−A−C
sqrt(MAF)
0.6
0.5
0.4
0.3
0.2
0.1
0.05
0.01
Cell_Type_NEW
Ctrl
ExN
InN
Other
UD

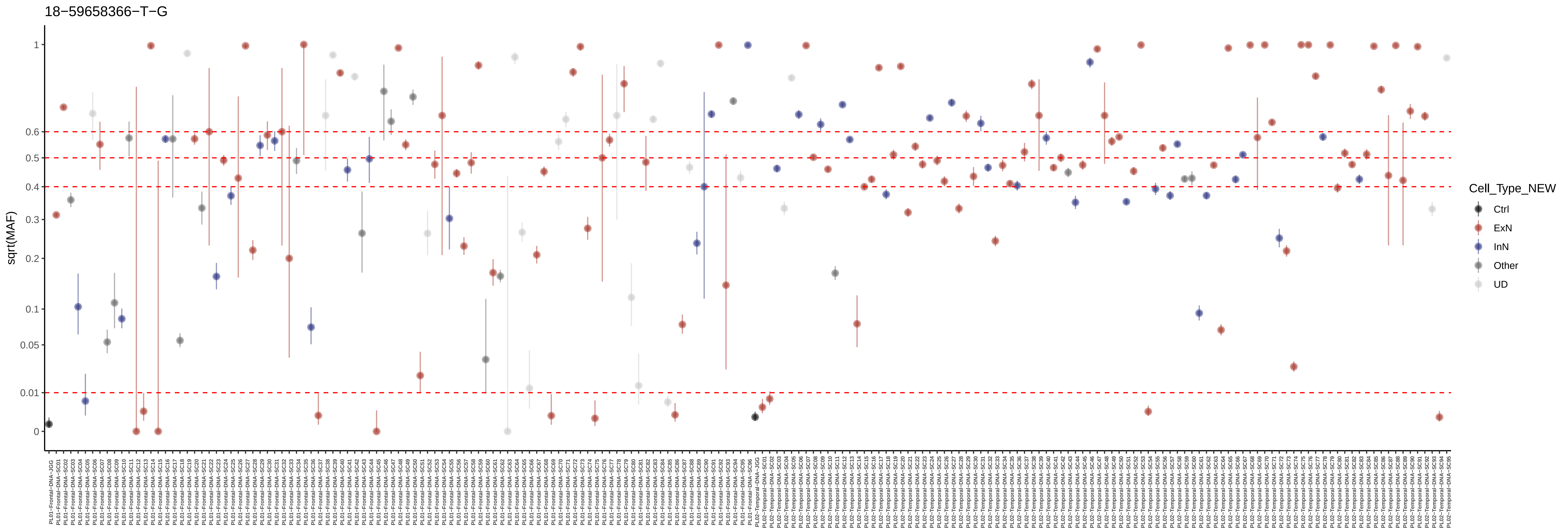
18−59658366−T−G
sqrt(MAF)
1
0.6
0.5
0.4
0.3
0.2
0.1
0.05
0.01
0
Cell_Type_NEW
Ctrl
ExN
InN
Other
UD

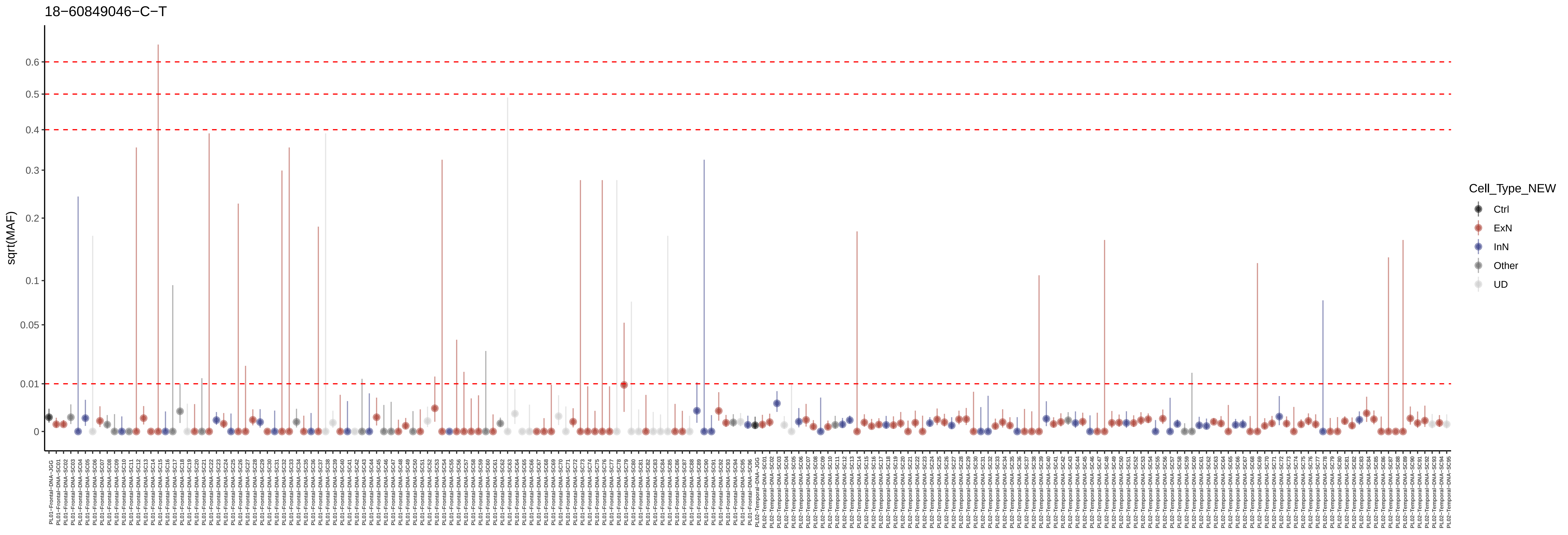

18−60849046−C−T
sqrt(MAF)
0.6
0.5
0.4
0.3
0.2
0.1
0.05
0.01
0
Cell_Type_NEW
Ctrl
ExN
InN
Other
UD

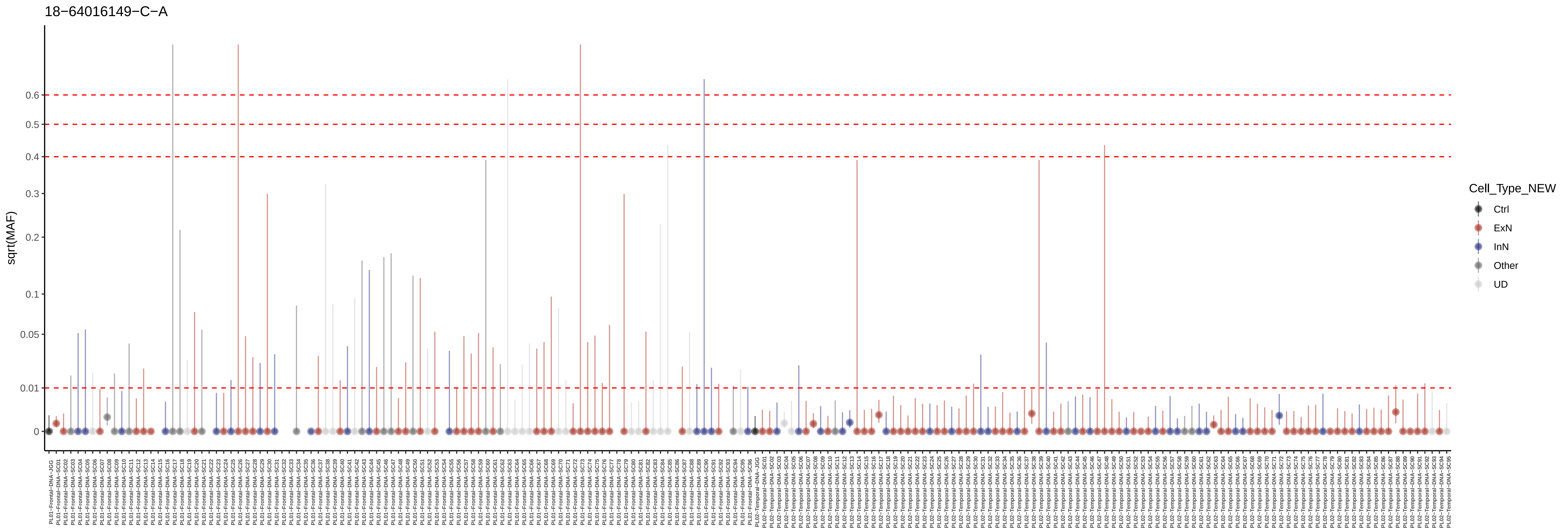
18−64016149−C−A
sqrt(MAF)
0.6
0.5
0.4
0.3
0.2
0.1
0.05
0.01
0
Cell_Type_NEW
Ctrl
ExN
InN
Other
UD

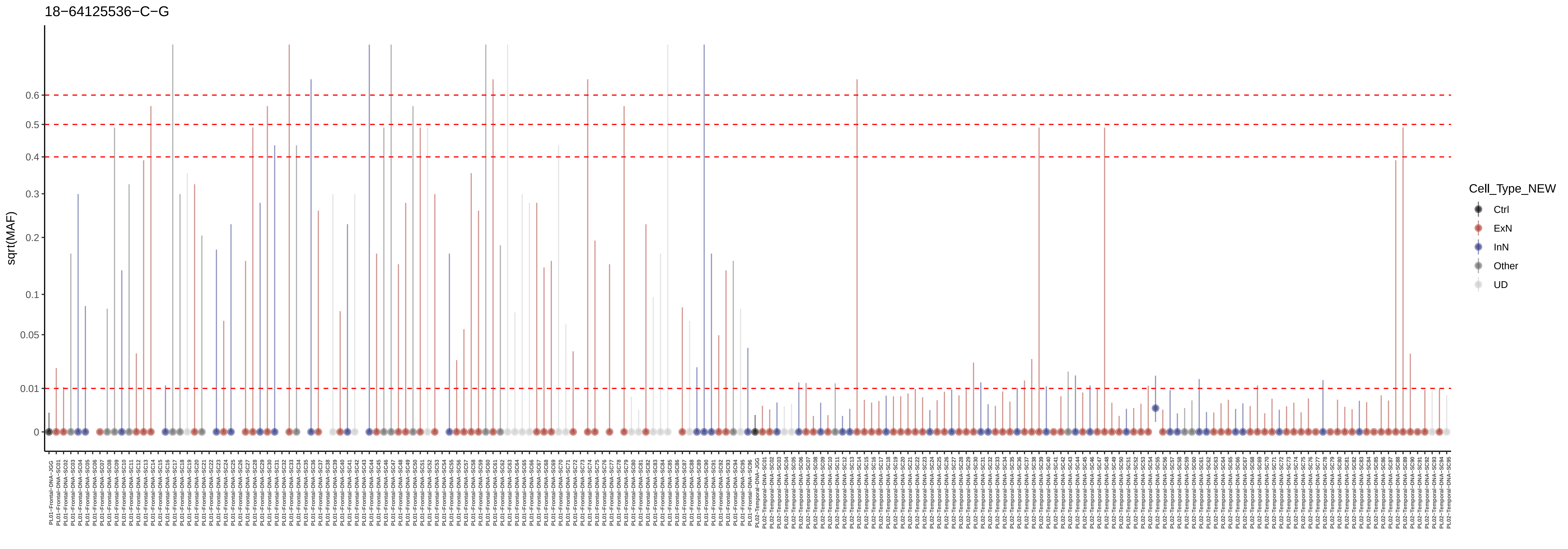
18−64125536−C−G
sqrt(MAF)
0.6
0.5
0.4
0.3
0.2
0.1
0.05
0.01
0
Cell_Type_NEW
Ctrl
ExN
InN
Other
UD

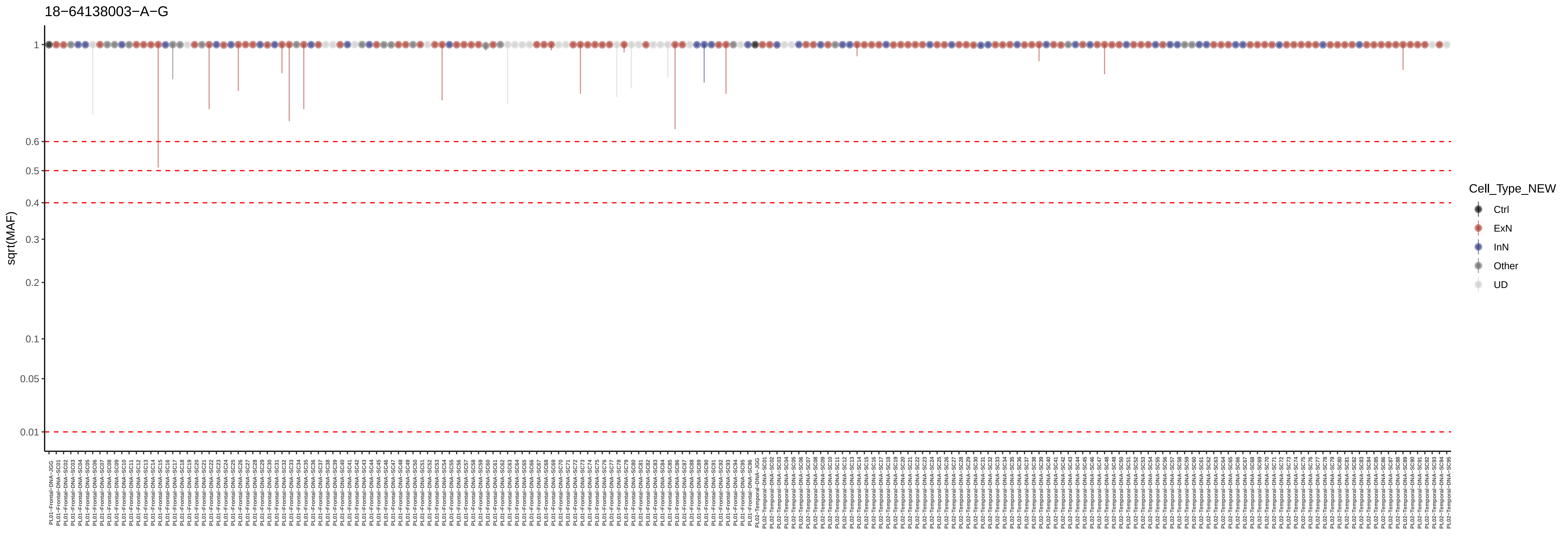

18−64138003−A−G
sqrt(MAF)
1
0.6
0.5
0.4
0.3
0.2
0.1
0.05
0.01
Cell_Type_NEW
Ctrl
ExN
InN
Other
UD

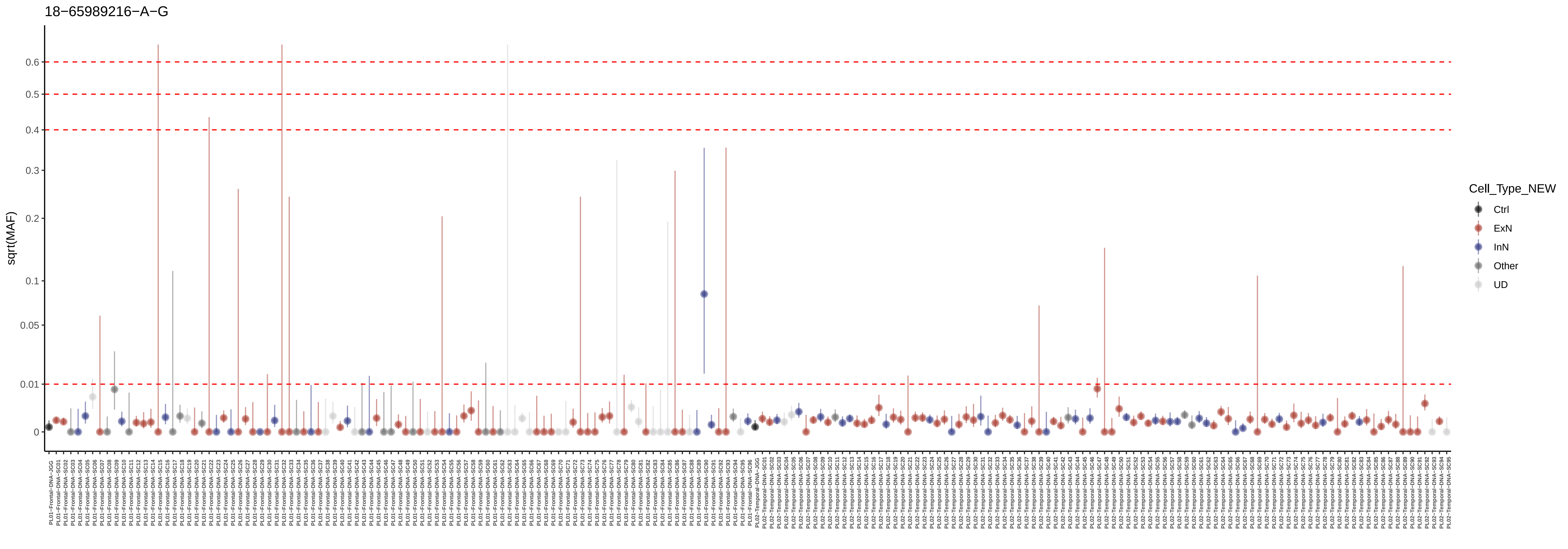

18−65989216−A−G
sqrt(MAF)
0.6
0.5
0.4
0.3
0.2
0.1
0.05
0.01
0
Cell_Type_NEW
Ctrl
ExN
InN
Other
UD

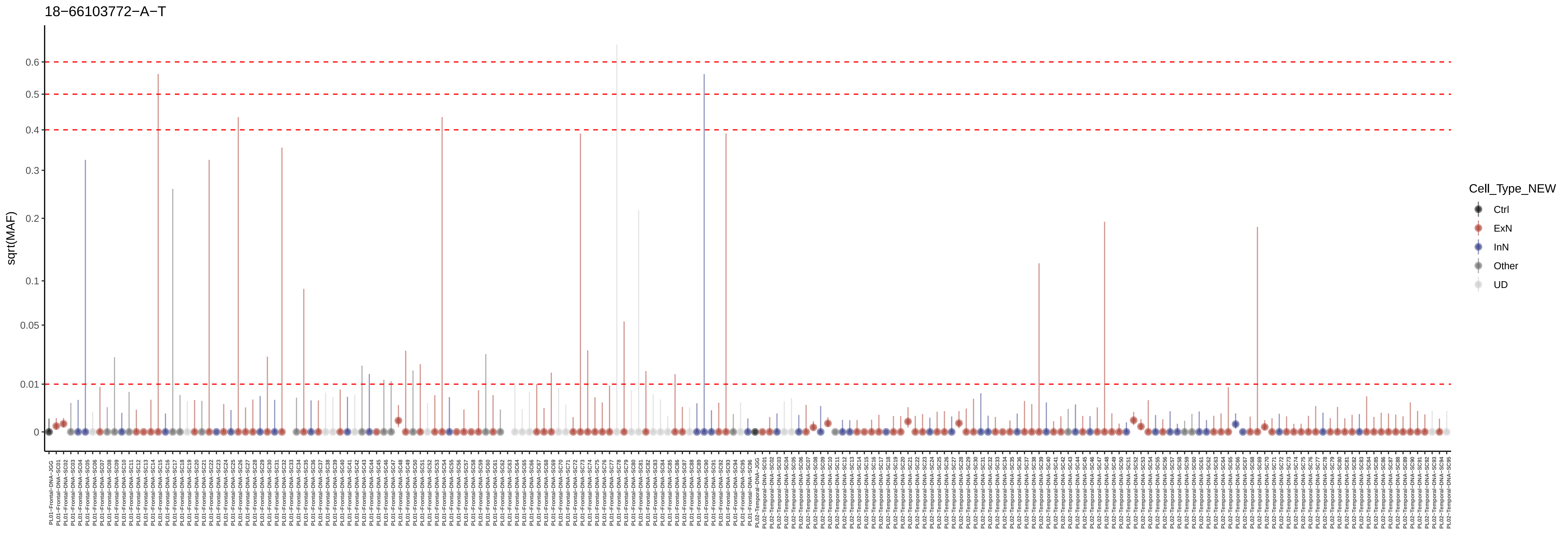
18−66103772−A−T
sqrt(MAF)
0.6
0.5
0.4
0.3
0.2
0.1
0.05
0.01
0
Cell_Type_NEW
Ctrl
ExN
InN
Other
UD

18−67366318−C−T

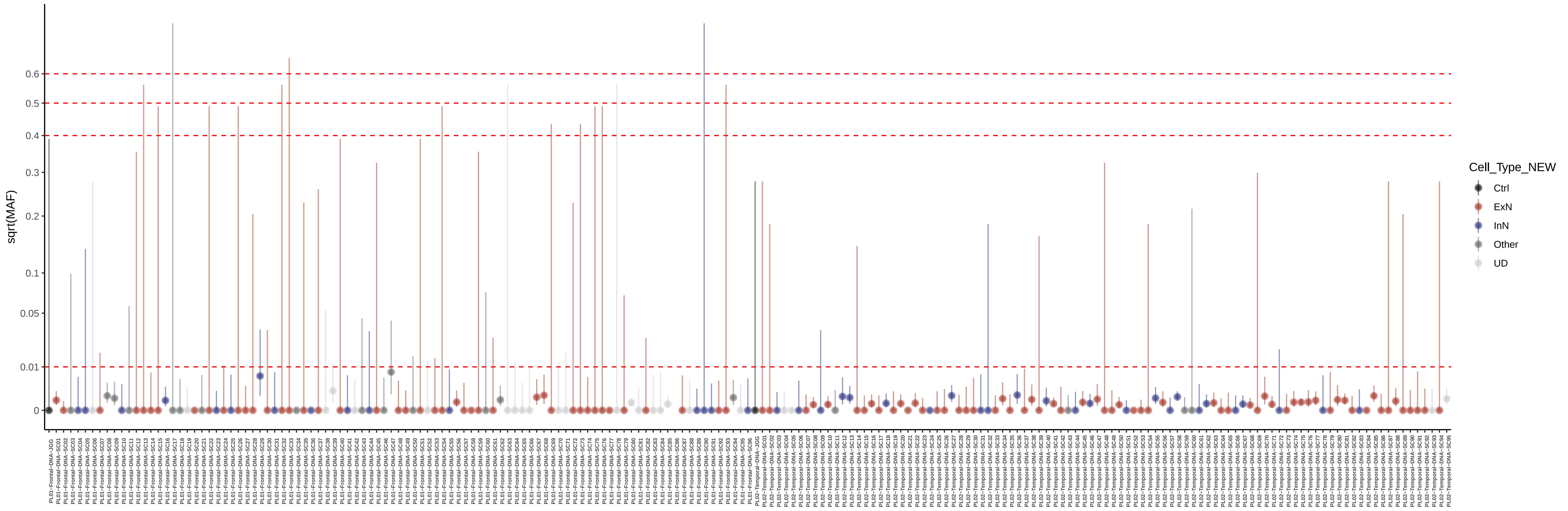

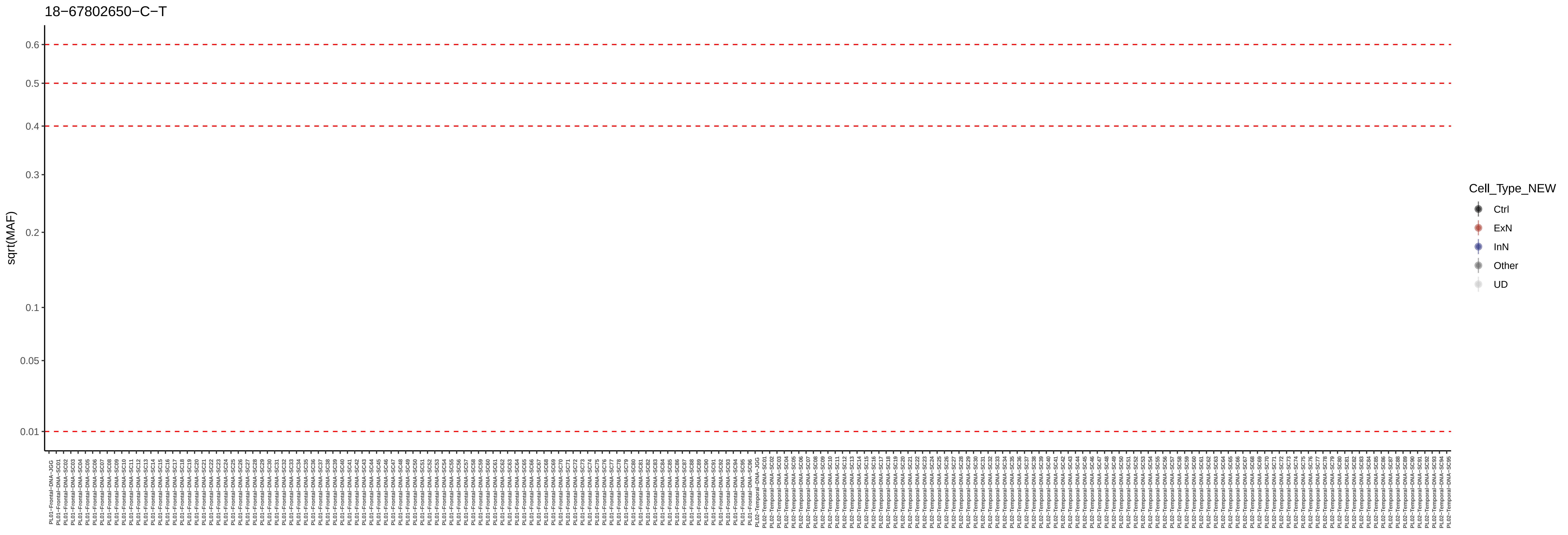
18−67802650−C−T
sqrt(MAF)
0.6
0.5
0.4
0.3
0.2
0.1
0.05
0.01
Cell_Type_NEW
Ctrl
ExN
InN
Other
UD

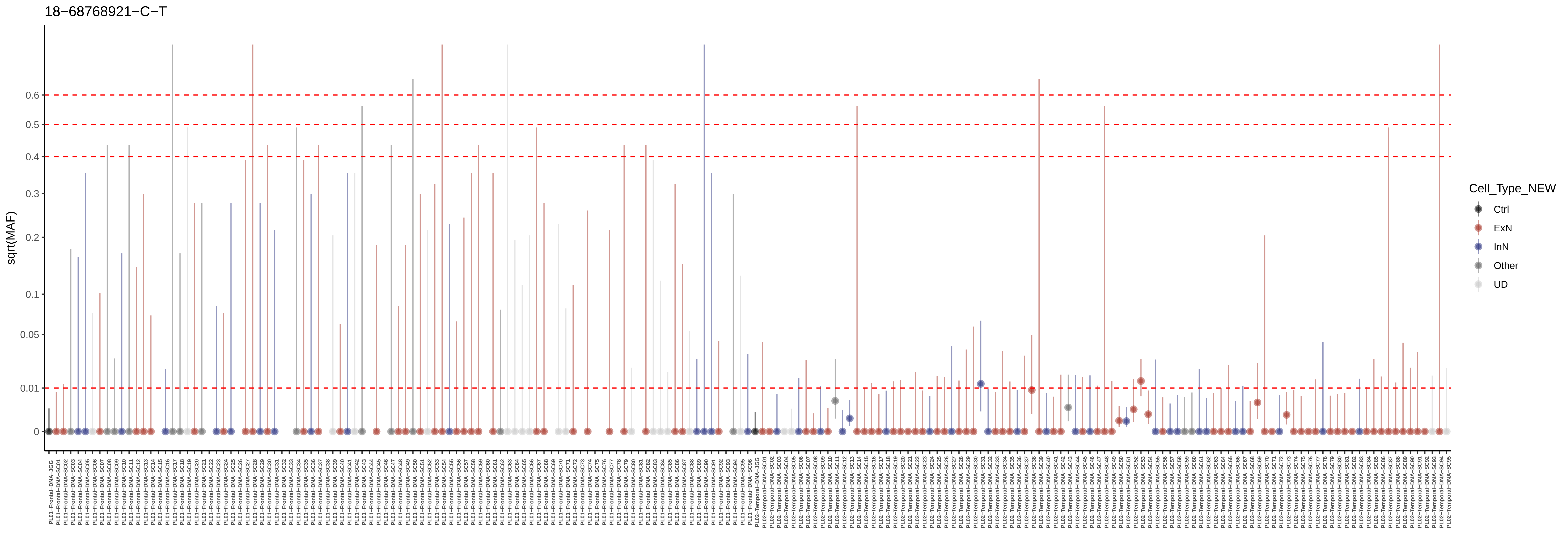
18−68768921−C−T
sqrt(MAF)
0.6
0.5
0.4
0.3
0.2
0.1
0.05
0.01
0
Cell_Type_NEW
Ctrl
ExN
InN
Other
UD

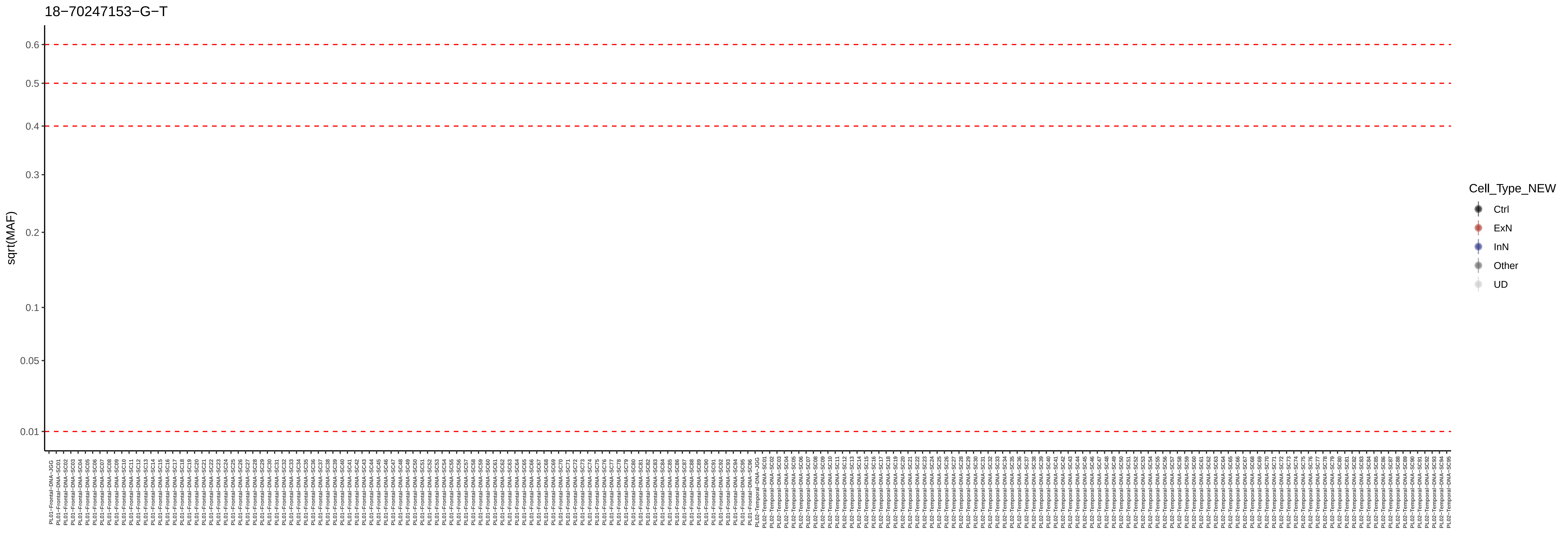
18−70247153−G−T
sqrt(MAF)
0.6
0.5
0.4
0.3
0.2
0.1
0.05
0.01
Cell_Type_NEW
Ctrl
ExN
InN
Other
UD

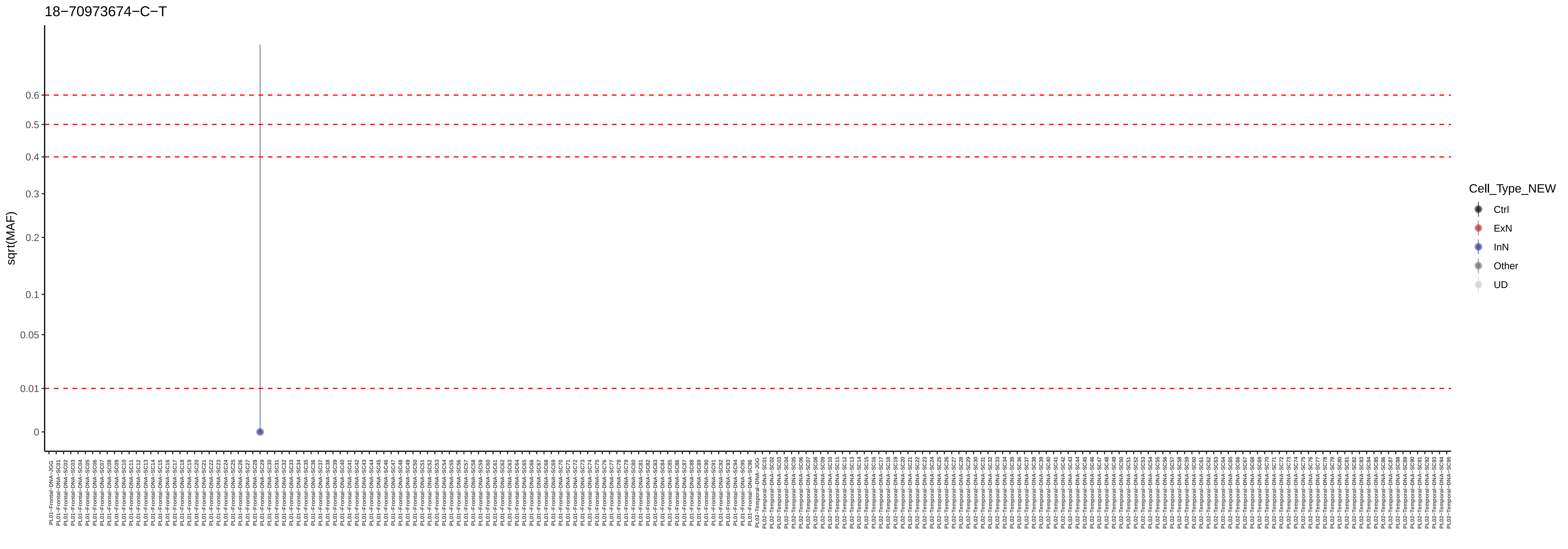
18−70973674−C−T
sqrt(MAF)
0.6
0.5
0.4
0.3
0.2
0.1
0.05
0.01
0
Cell_Type_NEW
Ctrl
ExN
InN
Other
UD

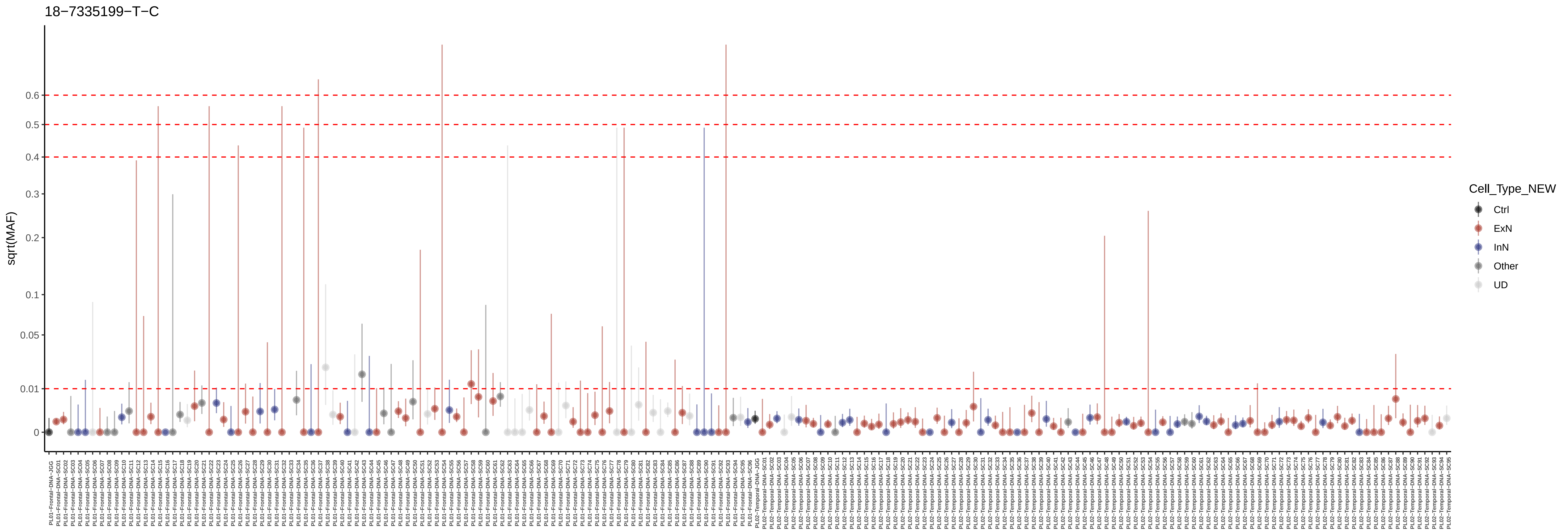
18−7335199−T−C
sqrt(MAF)
0.6
0.5
0.4
0.3
0.2
0.1
0.05
0.01
0
Cell_Type_NEW
Ctrl
ExN
InN
Other
UD

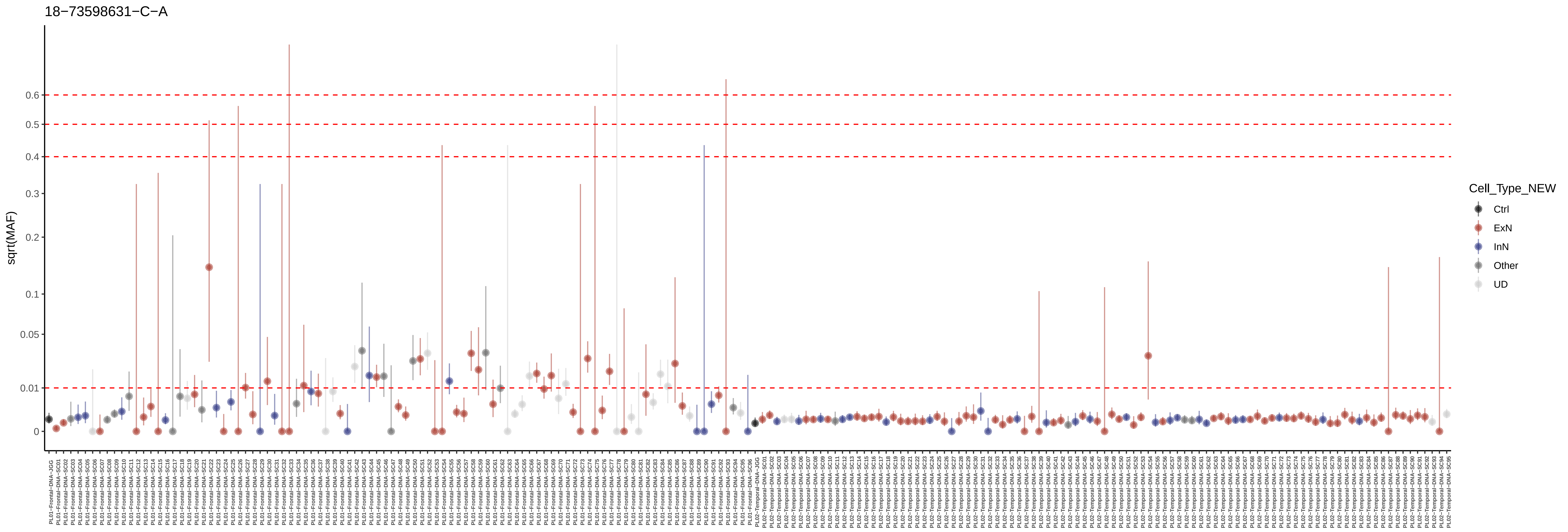
18−73598631−C−A
sqrt(MAF)
0.6
0.5
0.4
0.3
0.2
0.1
0.05
0.01
0
Cell_Type_NEW
Ctrl
ExN
InN
Other
UD

18−77870121−T−C

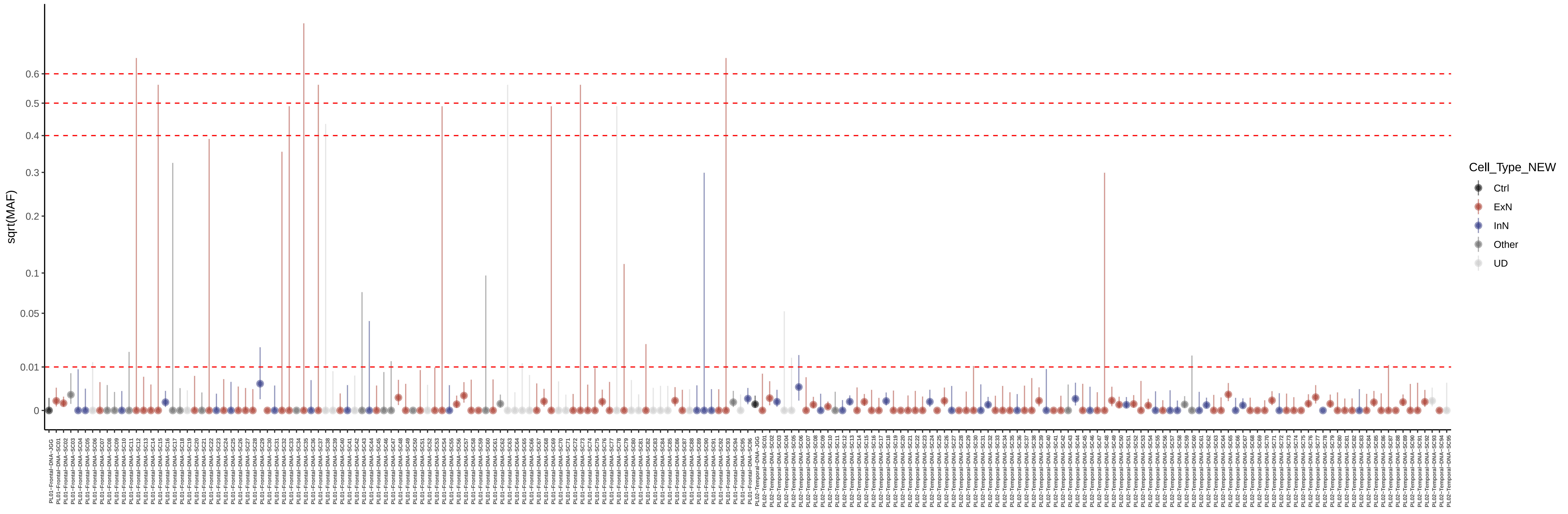

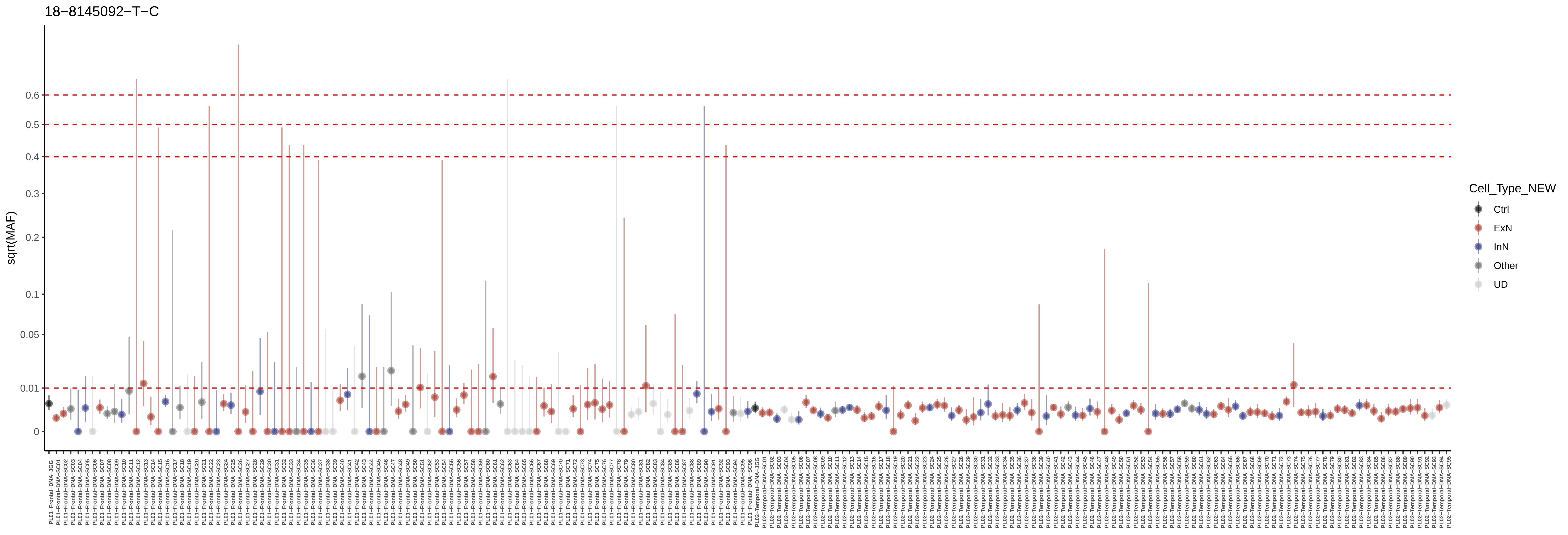

18−8145092−T−C
sqrt(MAF)
0.6
0.5
0.4
0.3
0.2
0.1
0.05
0.01
0
Cell_Type_NEW
Ctrl
ExN
InN
Other
UD

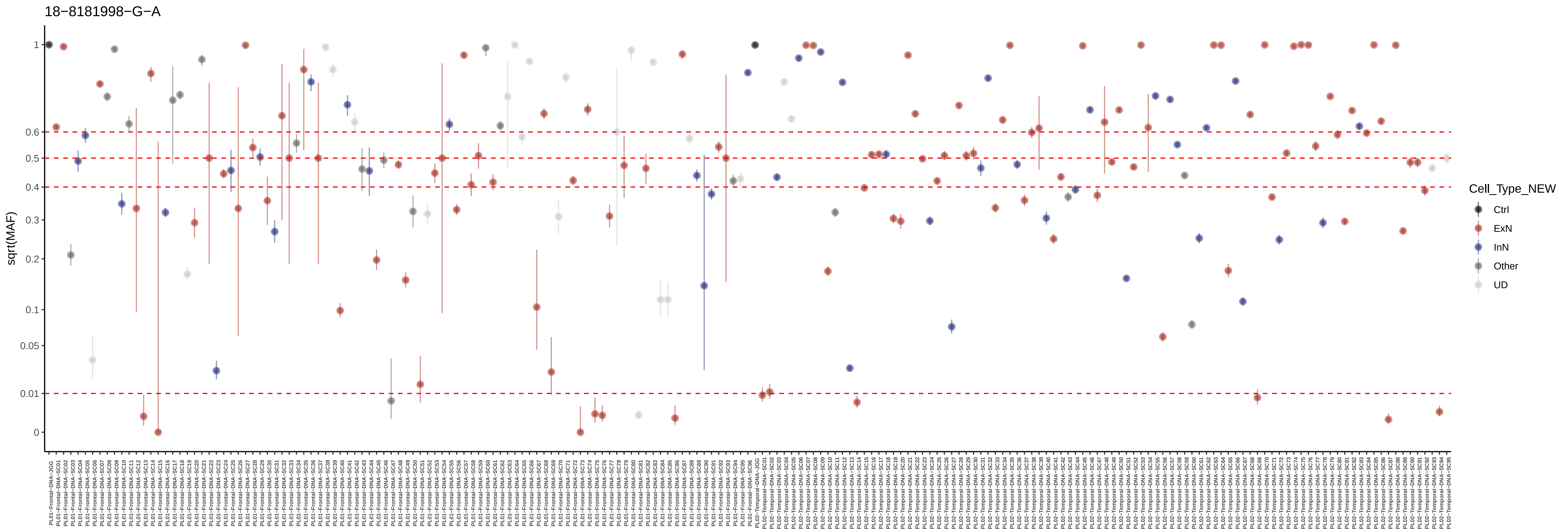
18−8181998−G−A
sqrt(MAF)
1
0.6
0.5
0.4
0.3
0.2
0.1
0.05
0.01
0
Cell_Type_NEW
Ctrl
ExN
InN
Other
UD

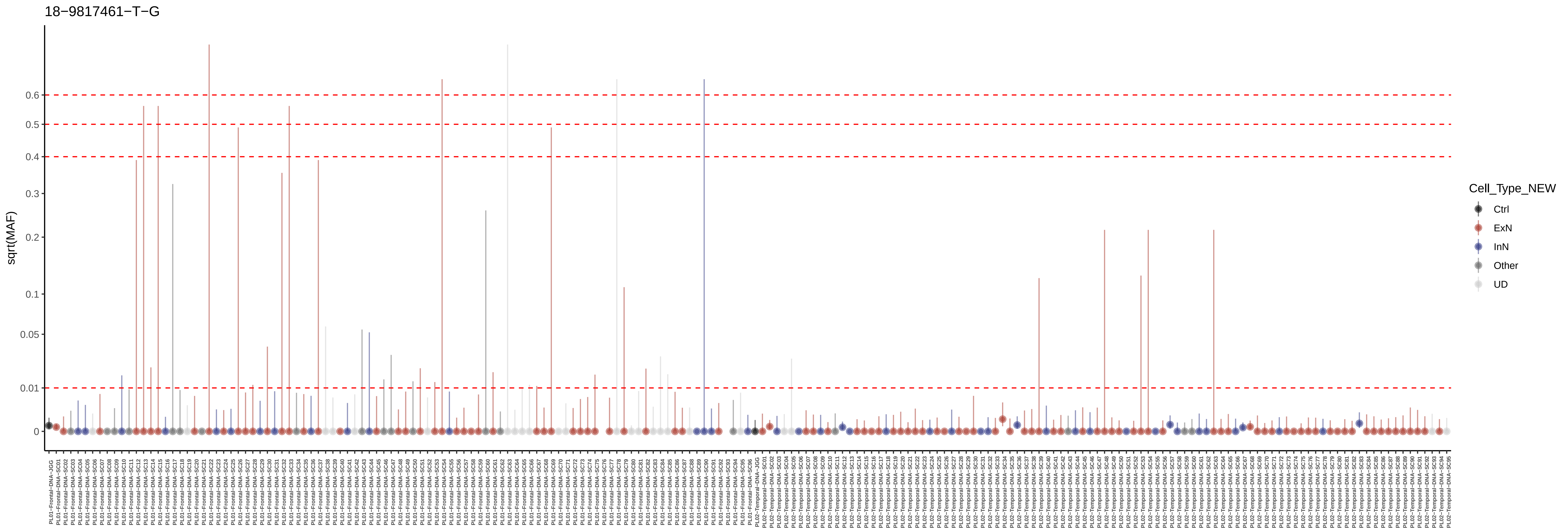
18−9817461−T−G
sqrt(MAF)
0.6
0.5
0.4
0.3
0.2
0.1
0.05
0.01
0
Cell_Type_NEW
Ctrl
ExN
InN
Other
UD

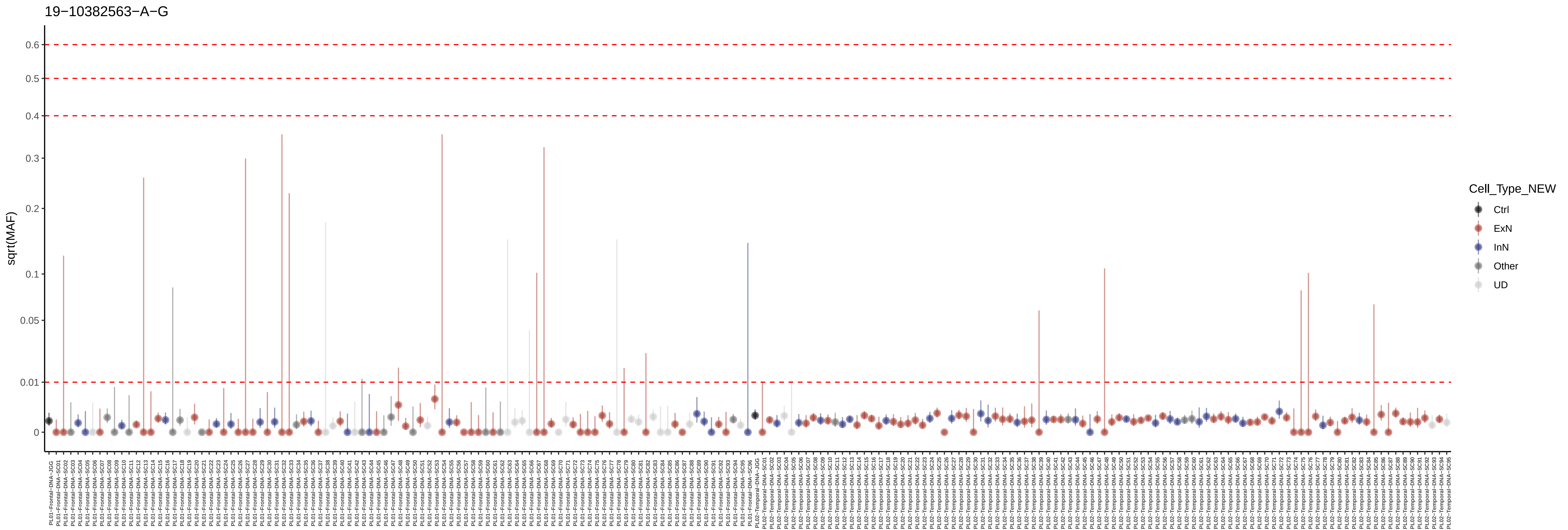

19−10382563−A−G
sqrt(MAF)
0.6
0.5
0.4
0.3
0.2
0.1
0.05
0.01
0
Cell_Type_NEW
Ctrl
ExN
InN
Other
UD

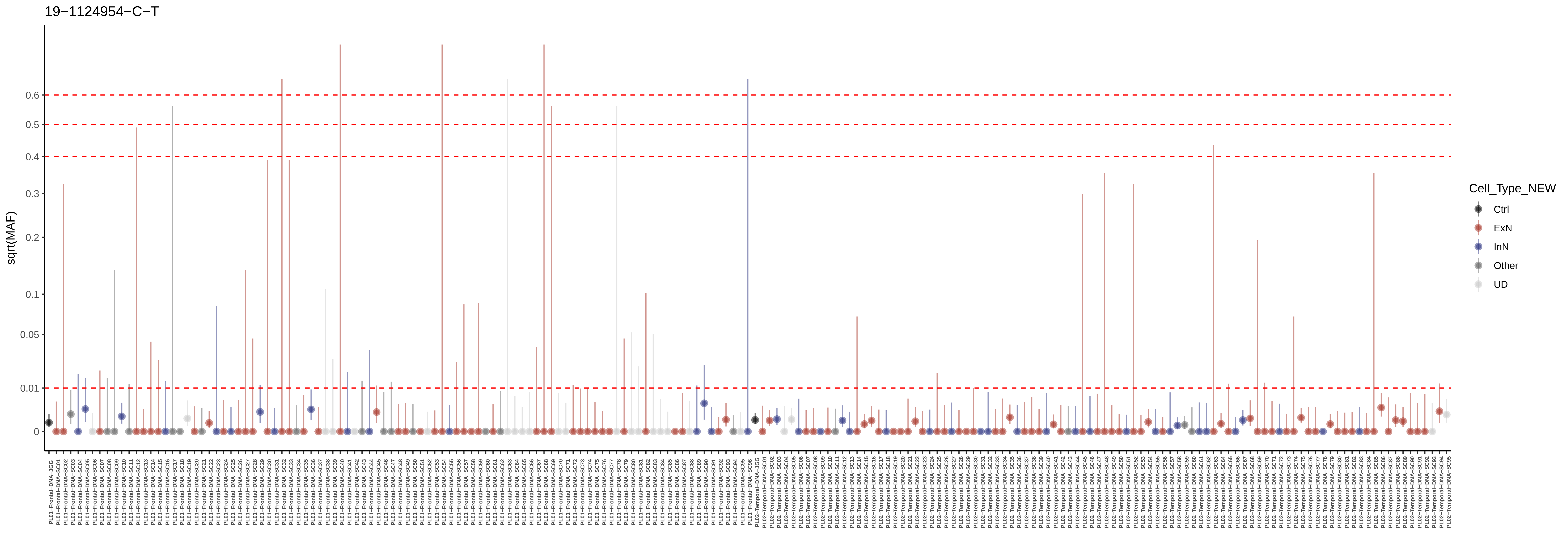
19−1124954−C−T
sqrt(MAF)
0.6
0.5
0.4
0.3
0.2
0.1
0.05
0.01
0
Cell_Type_NEW
Ctrl
ExN
InN
Other
UD

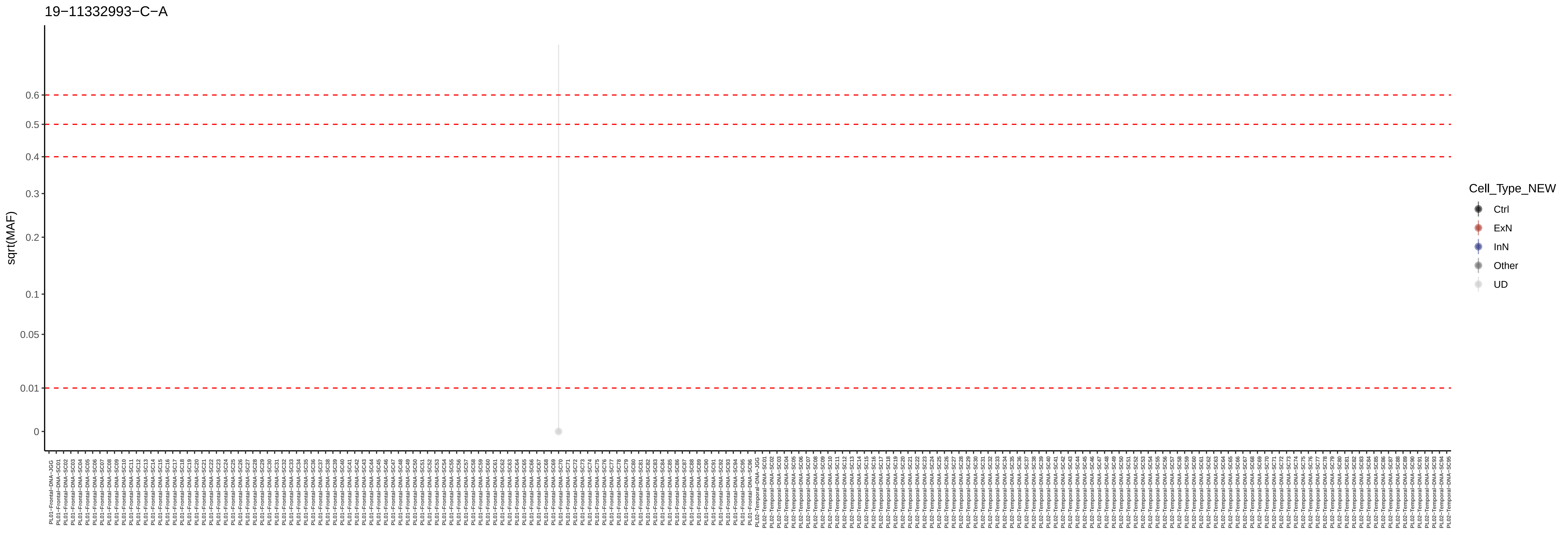

19−11332993−C−A
sqrt(MAF)
Cell_Type_NEW
Ctrl
ExN
InN
Other
UD

19−13941229−G−A

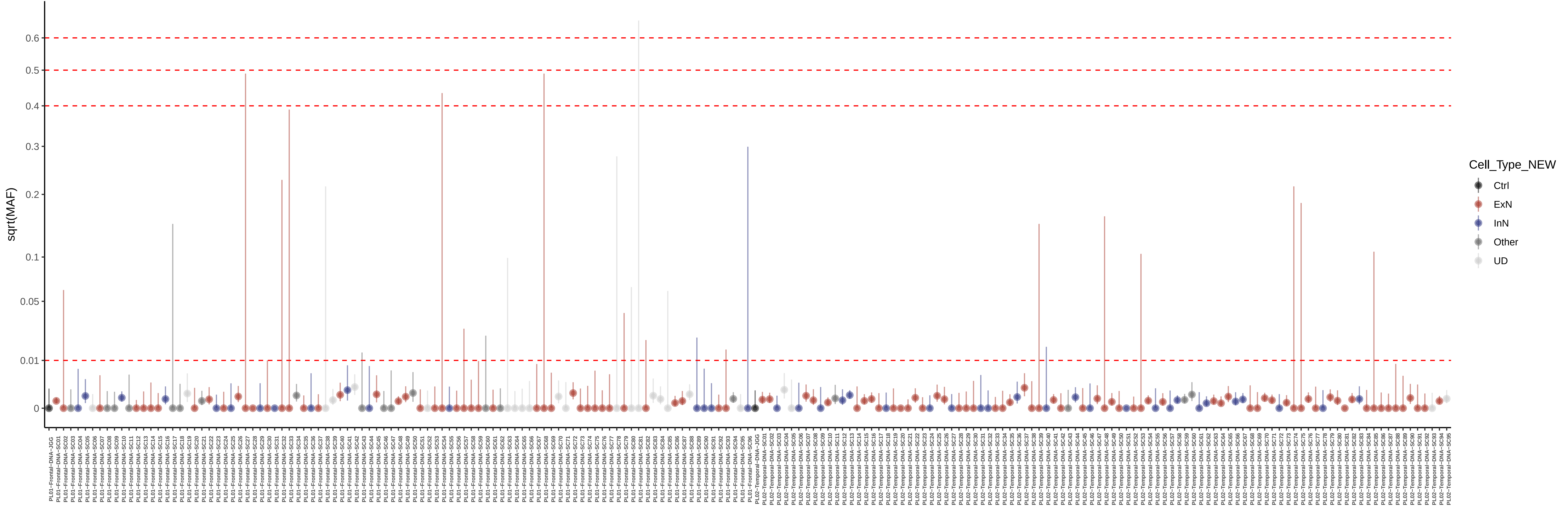

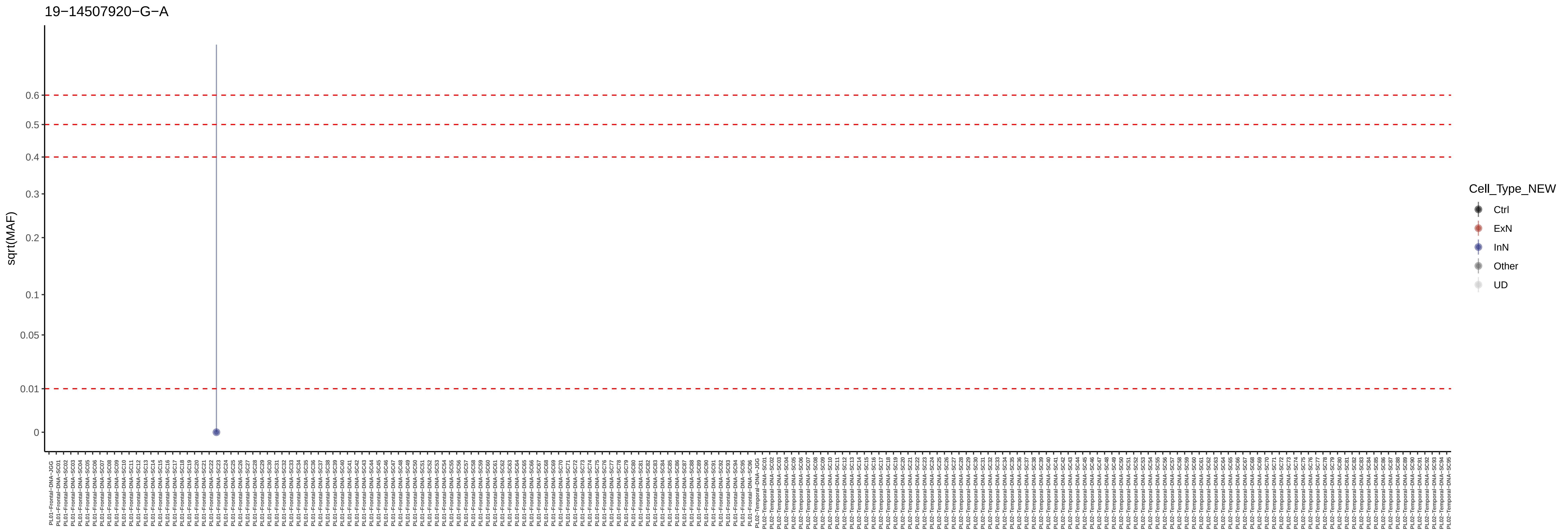

19−14507920−G−A
sqrt(MAF)
0.6
0.5
0.4
0.3
0.2
0.1
0.05
0.01
0
Cell_Type_NEW
Ctrl
ExN
InN
Other
UD

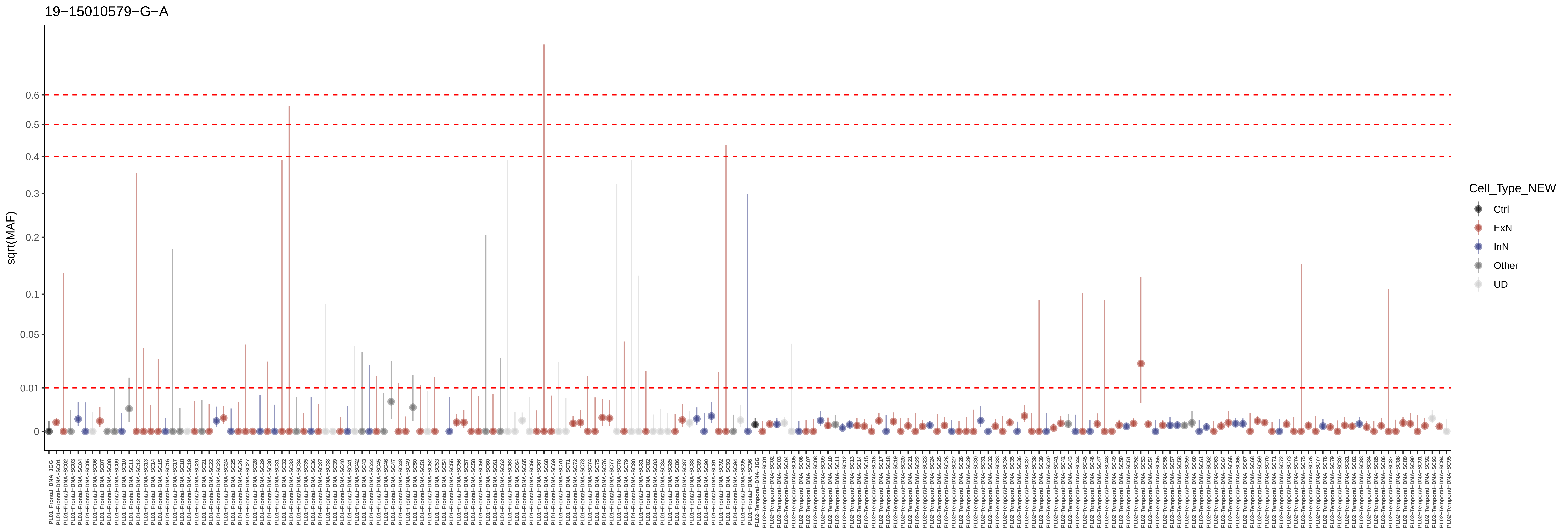
19−15010579−G−A
sqrt(MAF)
0.6
0.5
0.4
0.3
0.2
0.1
0.05
0.01
0
Cell_Type_NEW
Ctrl
ExN
InN
Other
UD

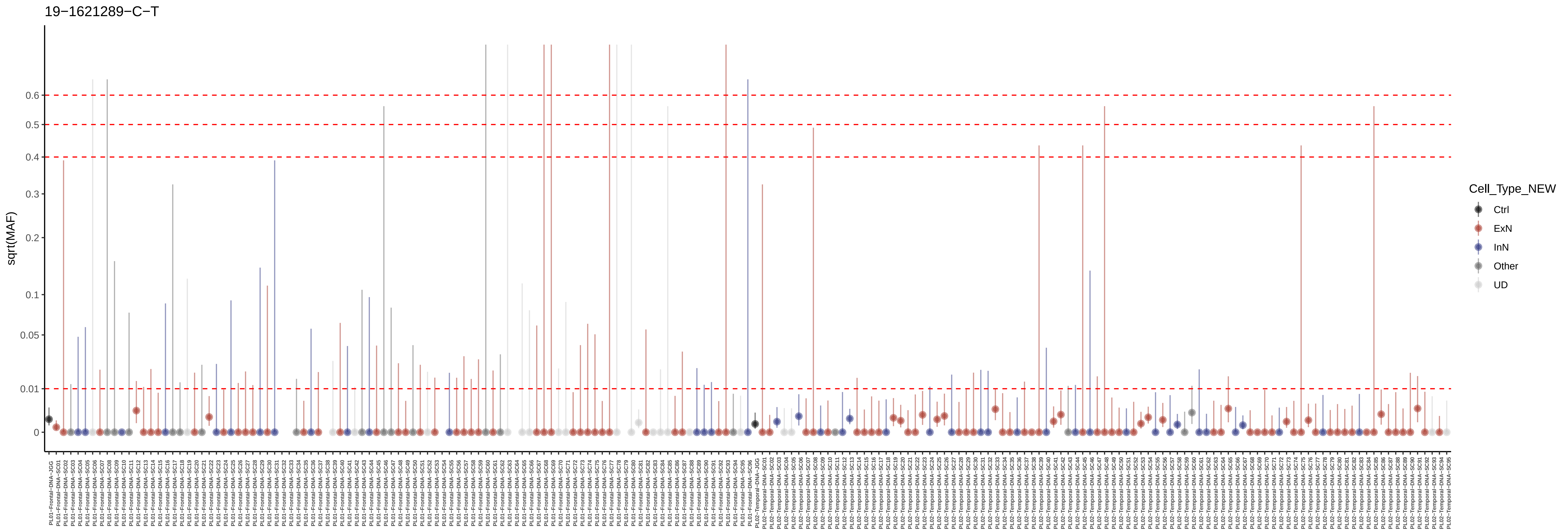
19−1621289−C−T
sqrt(MAF)
0.6
0.5
0.4
0.3
0.2
0.1
0.05
0.01
0
Cell_Type_NEW
Ctrl
ExN
InN
Other
UD

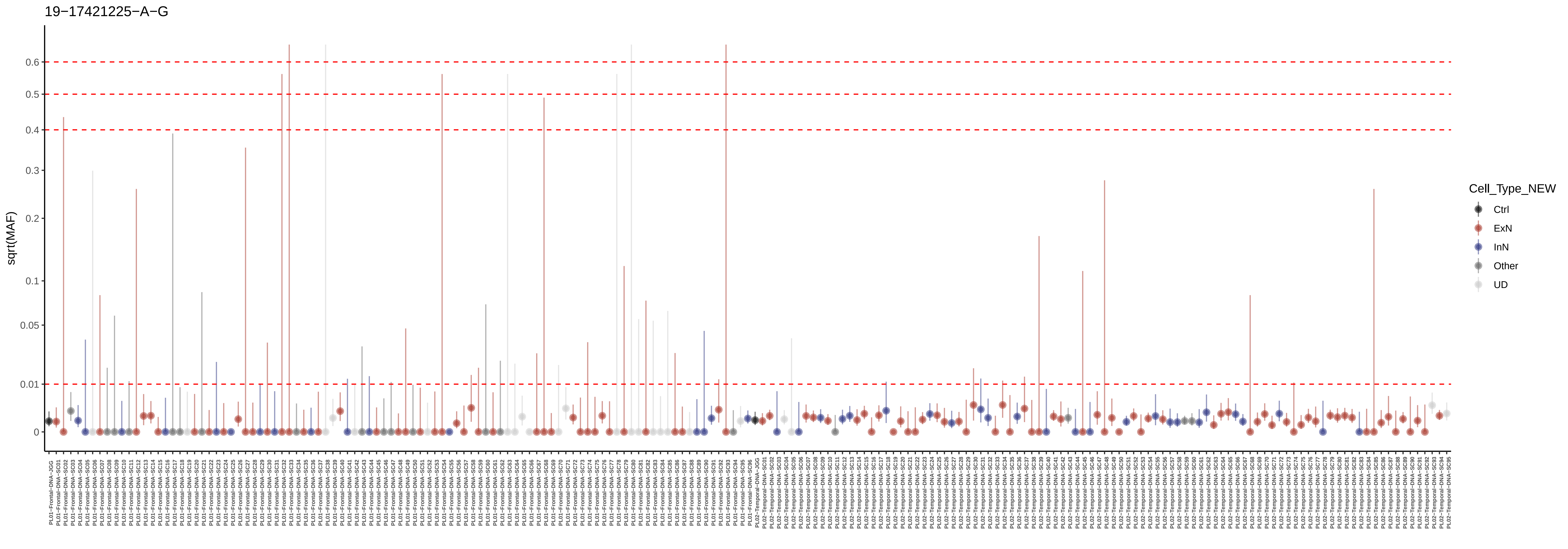

19−17421225−A−G
sqrt(MAF)
0.6
0.5
0.4
0.3
0.2
0.1
0.05
0.01
0
Cell_Type_NEW
Ctrl
ExN
InN
Other
UD

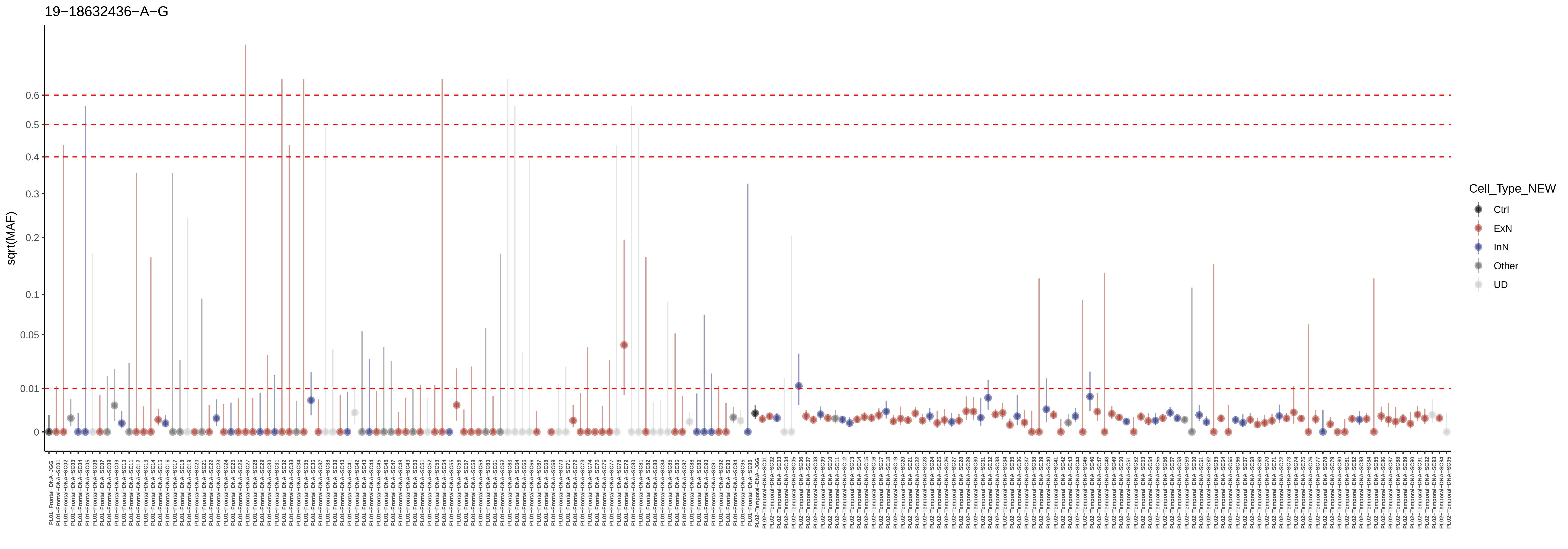
19−18632436−A−G
sqrt(MAF)
0.6
0.5
0.4
0.3
0.2
0.1
0.05
0.01
0
Cell_Type_NEW
Ctrl
ExN
InN
Other
UD

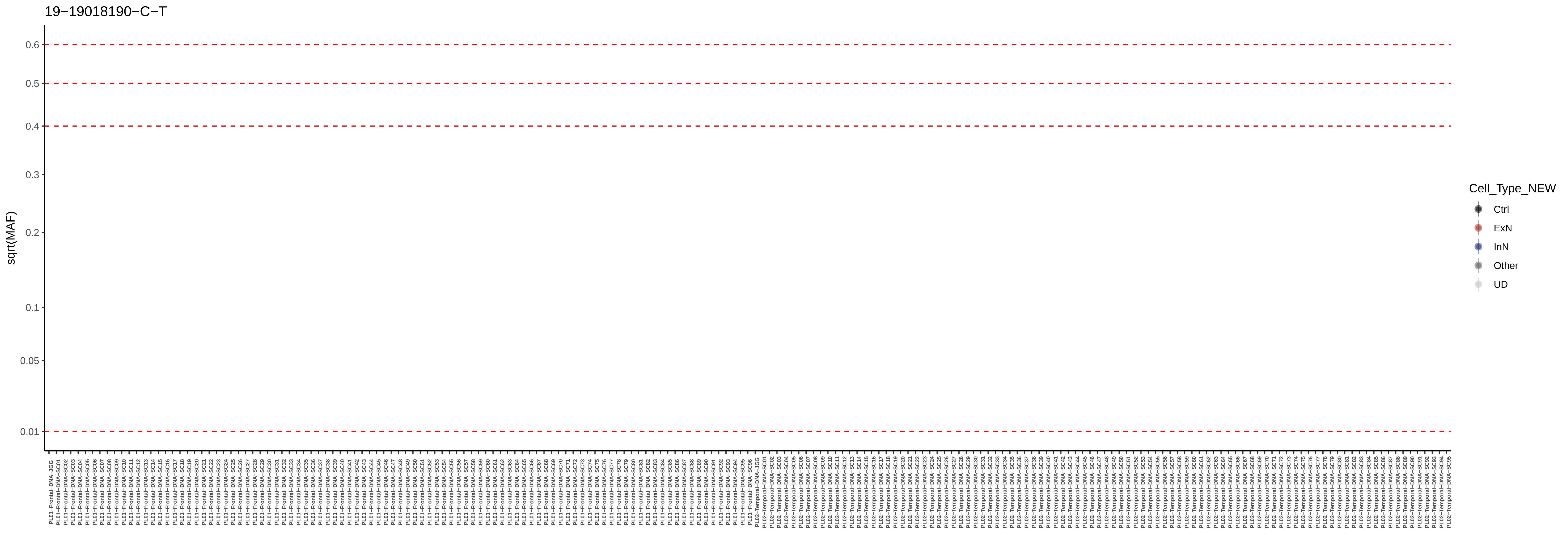

19−19018190−C−T
sqrt(MAF)
0.6
0.5
0.4
0.3
0.2
0.1
0.05
0.01
Cell_Type_NEW
Ctrl
ExN
InN
Other
UD

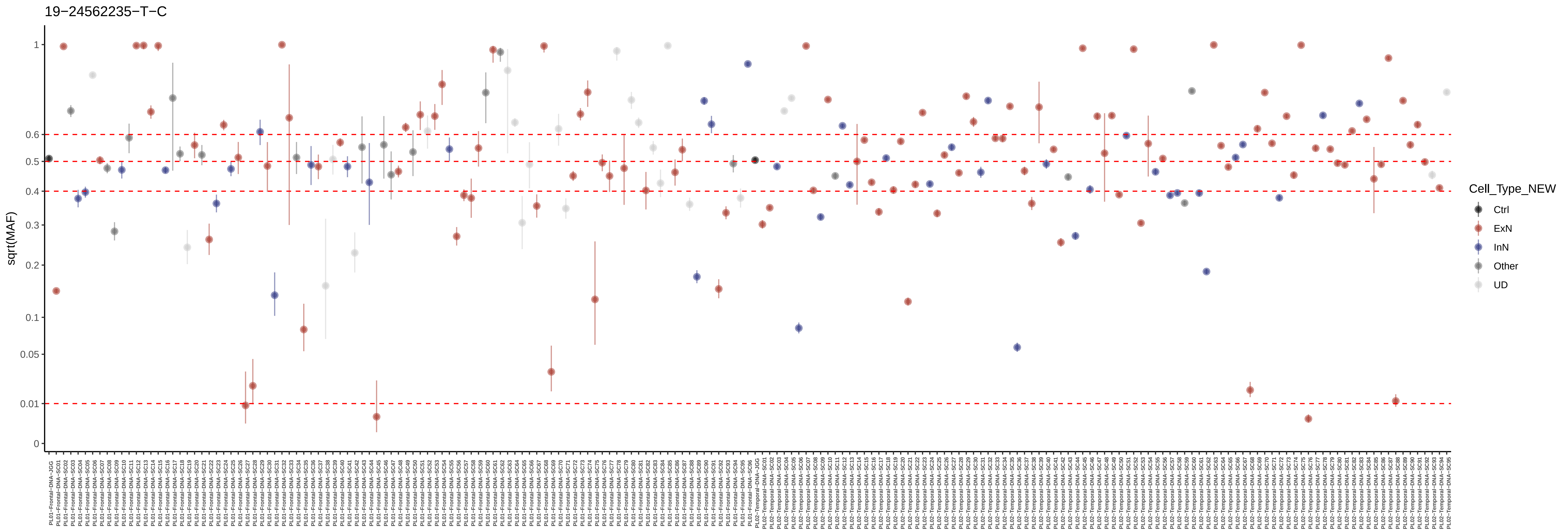

19−24562235−T−C
sqrt(MAF)
1
0.6
0.5
0.4
0.3
0.2
0.1
0.05
0.01
0
Cell_Type_NEW
Ctrl
ExN
InN
Other
UD

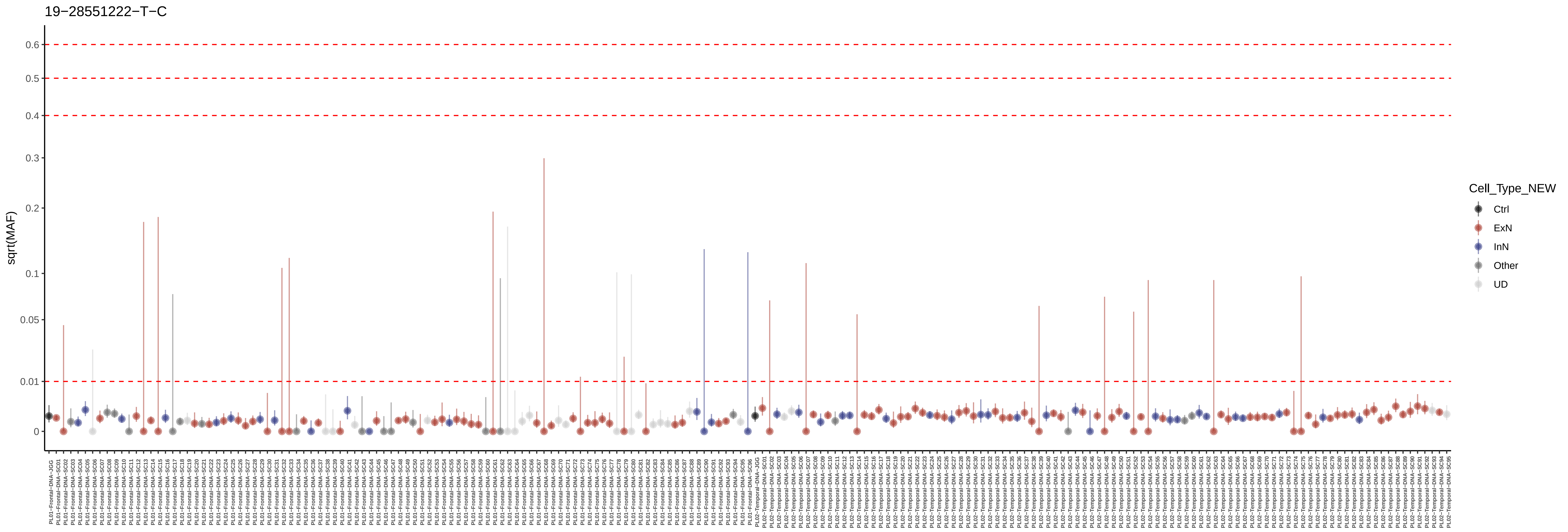
19−28551222−T−C
sqrt(MAF)
0.6
0.5
0.4
0.3
0.2
0.1
0.05
0.01
0
Cell_Type_NEW
Ctrl
ExN
InN
Other
UD

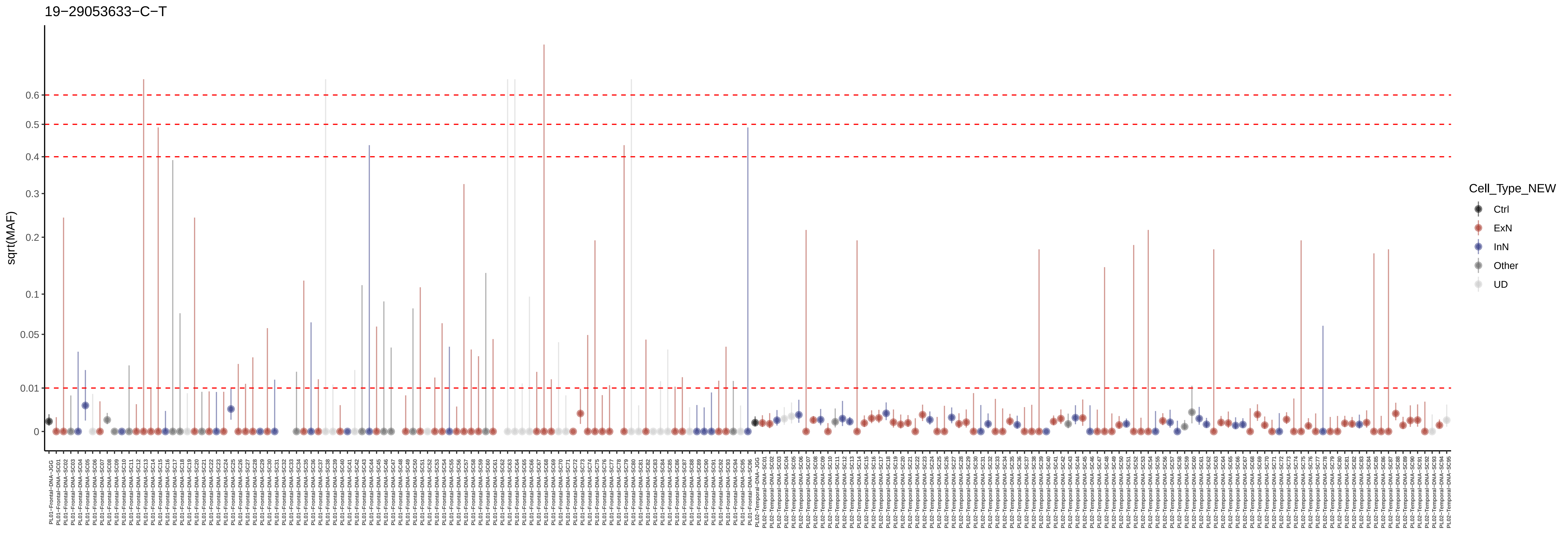
19−29053633−C−T
sqrt(MAF)
0.6
0.5
0.4
0.3
0.2
0.1
0.05
0.01
0
Cell_Type_NEW
Ctrl
ExN
InN
Other
UD

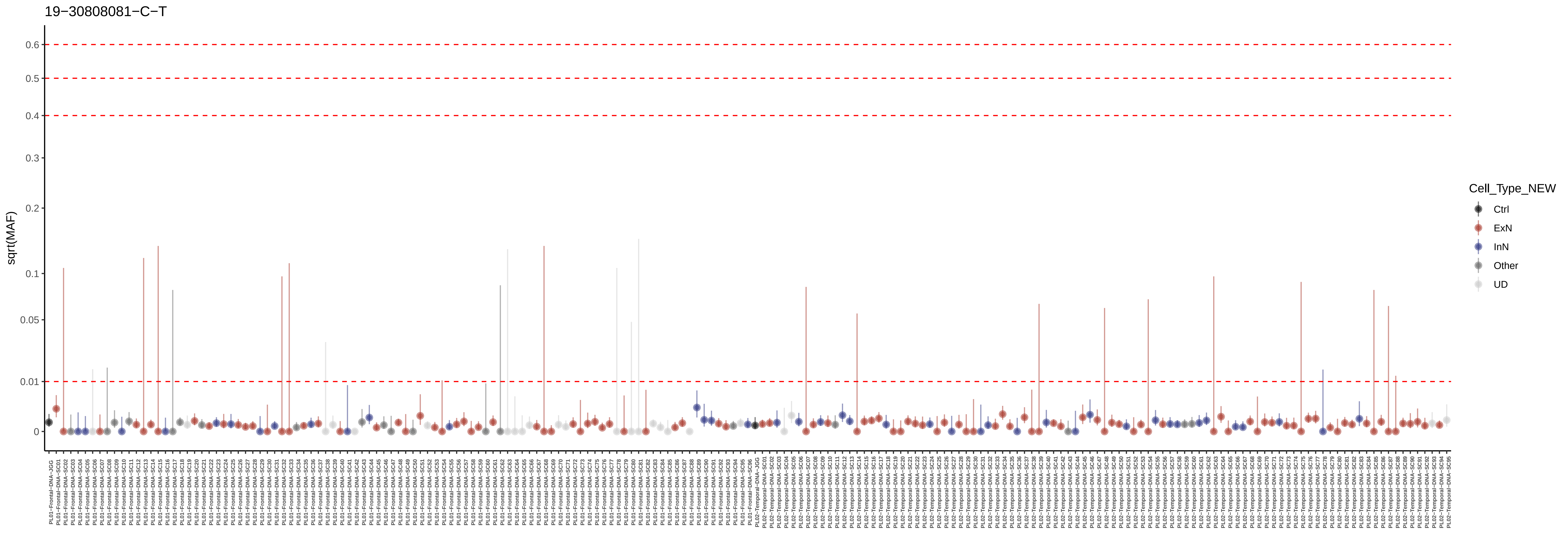

19−30808081−C−T
sqrt(MAF)
0.6
0.5
0.4
0.3
0.2
0.1
0.05
0.01
0
Cell_Type_NEW
Ctrl
ExN
InN
Other
UD

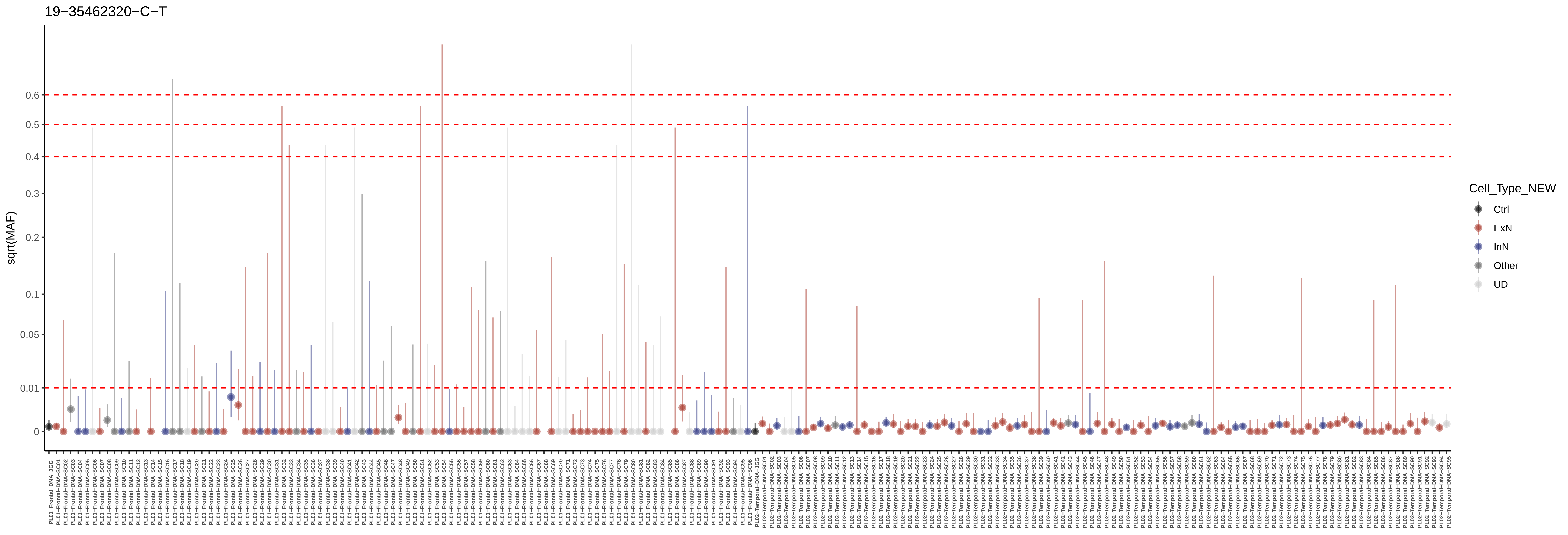
19−35462320−C−T
sqrt(MAF)
0.6
0.5
0.4
0.3
0.2
0.1
0.05
0.01
0
Cell_Type_NEW
Ctrl
ExN
InN
Other
UD

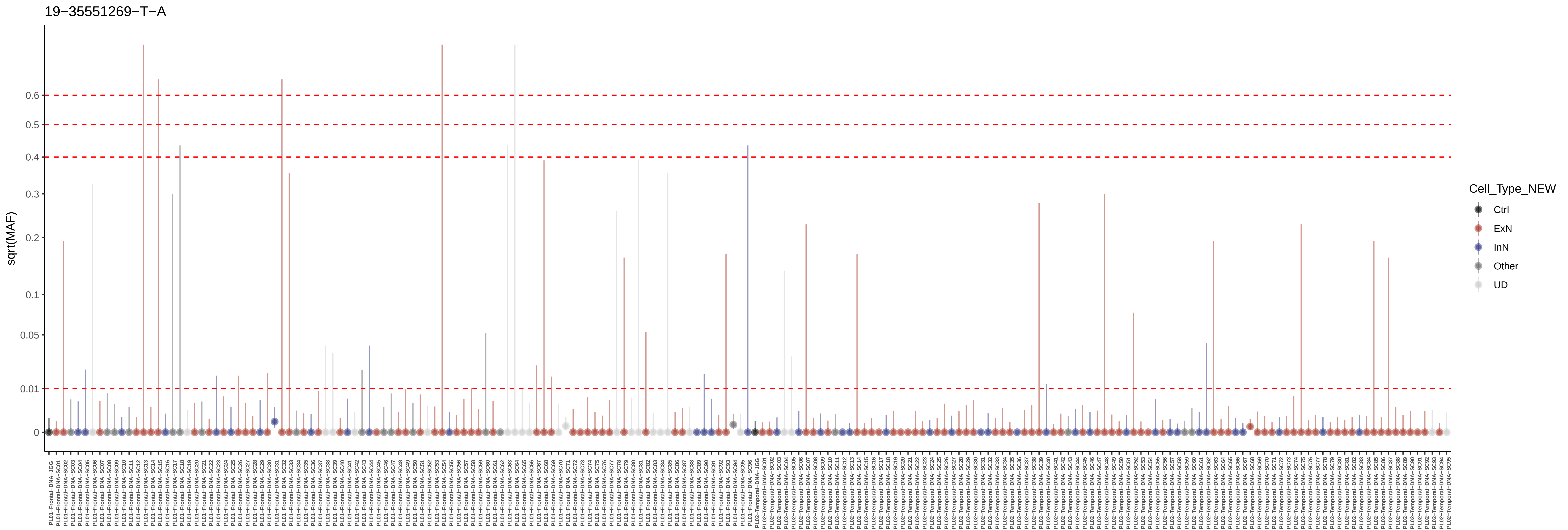
19−35551269−T−A
sqrt(MAF)
0.6
0.5
0.4
0.3
0.2
0.1
0.05
0.01
0
Cell_Type_NEW
Ctrl
ExN
InN
Other
UD

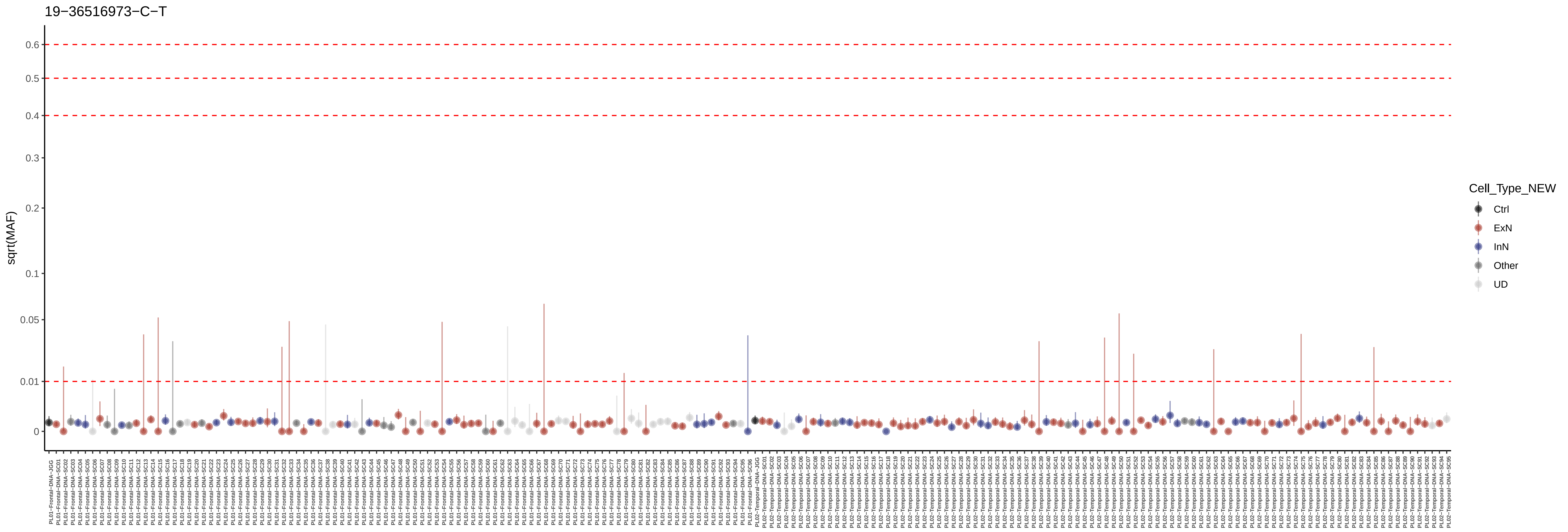
19−36516973−C−T
sqrt(MAF)
0.6
0.5
0.4
0.3
0.2
0.1
0.05
0.01
0
Cell_Type_NEW
Ctrl
ExN
InN
Other
UD

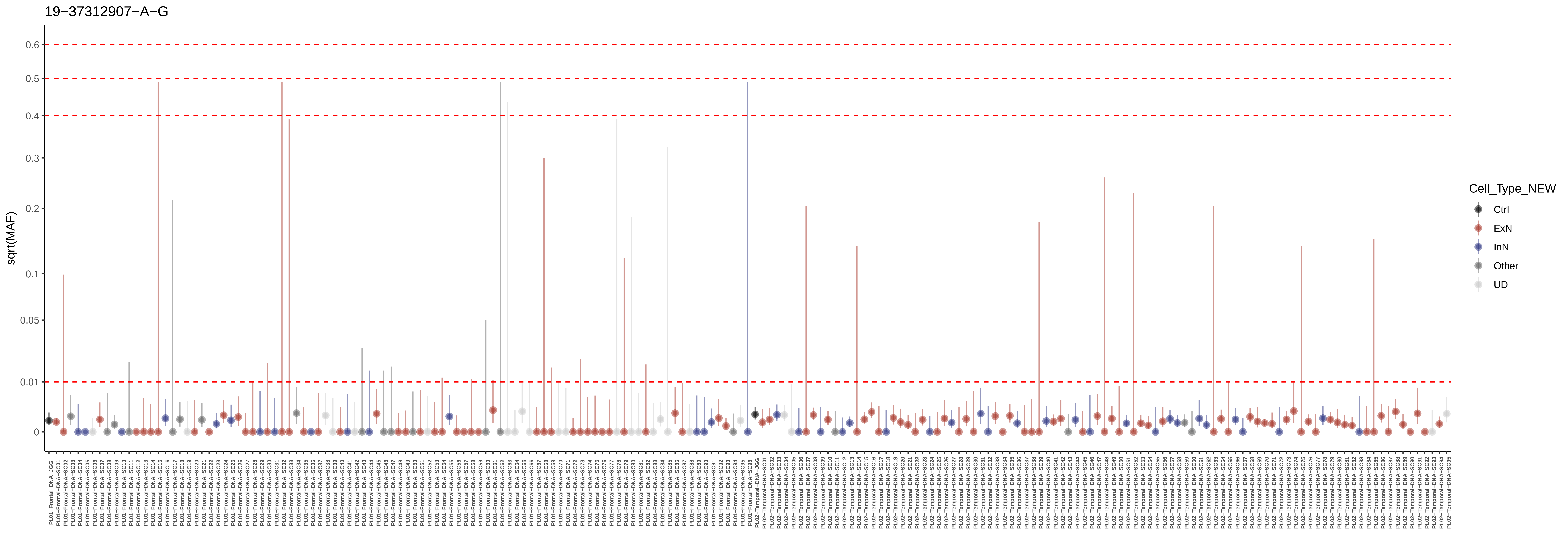
19−37312907−A−G
sqrt(MAF)
0.6
0.5
0.4
0.3
0.2
0.1
0.05
0.01
0
Cell_Type_NEW
Ctrl
ExN
InN
Other
UD

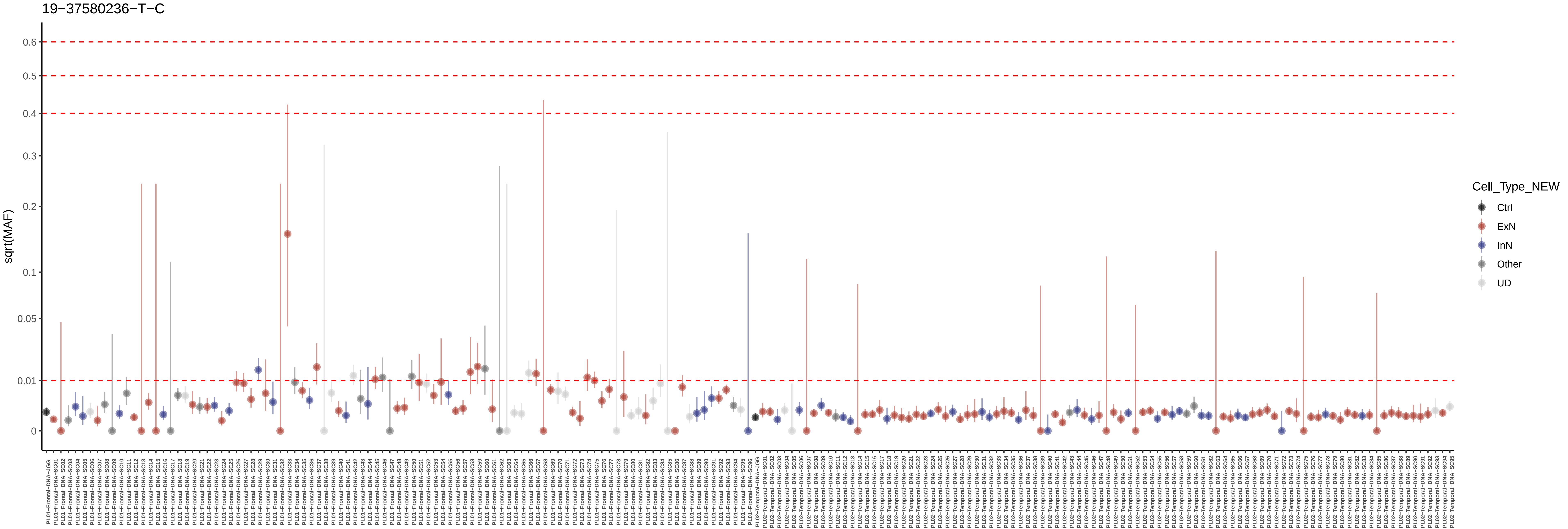
19−37580236−T−C
sqrt(MAF)
0.6
0.5
0.4
0.3
0.2
0.1
0.05
0.01
0
Cell_Type_NEW
Ctrl
ExN
InN
Other
UD

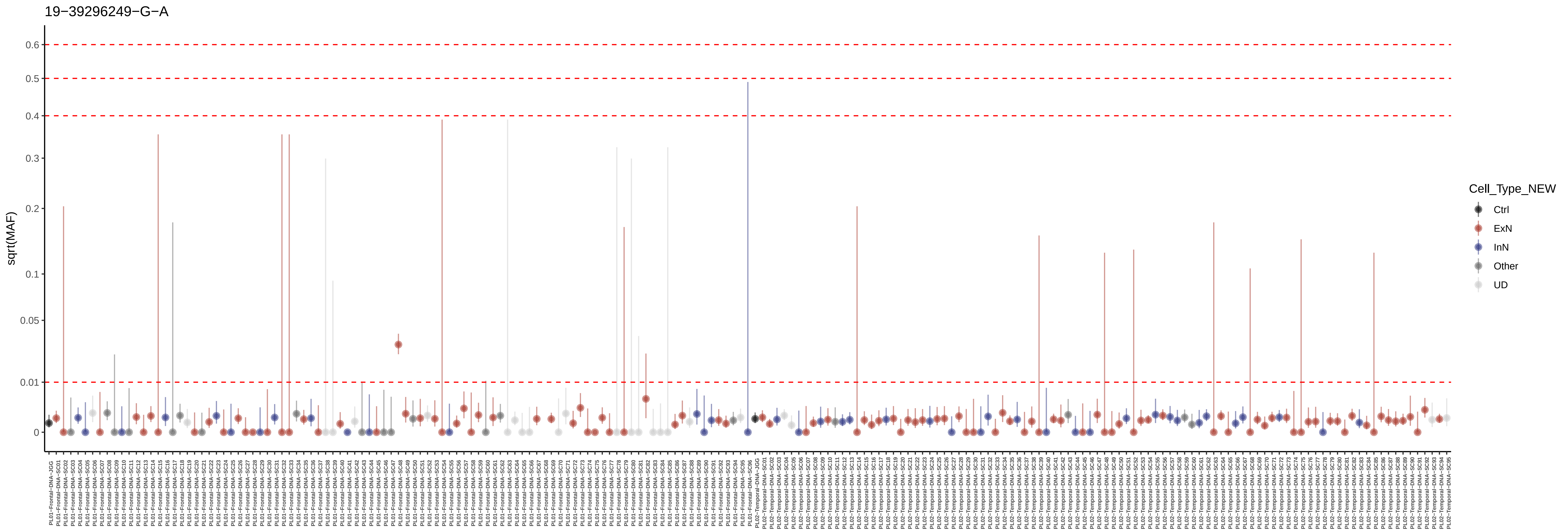
19−39296249−G−A
sqrt(MAF)
0.6
0.5
0.4
0.3
0.2
0.1
0.05
0.01
0
Cell_Type_NEW
Ctrl
ExN
InN
Other
UD

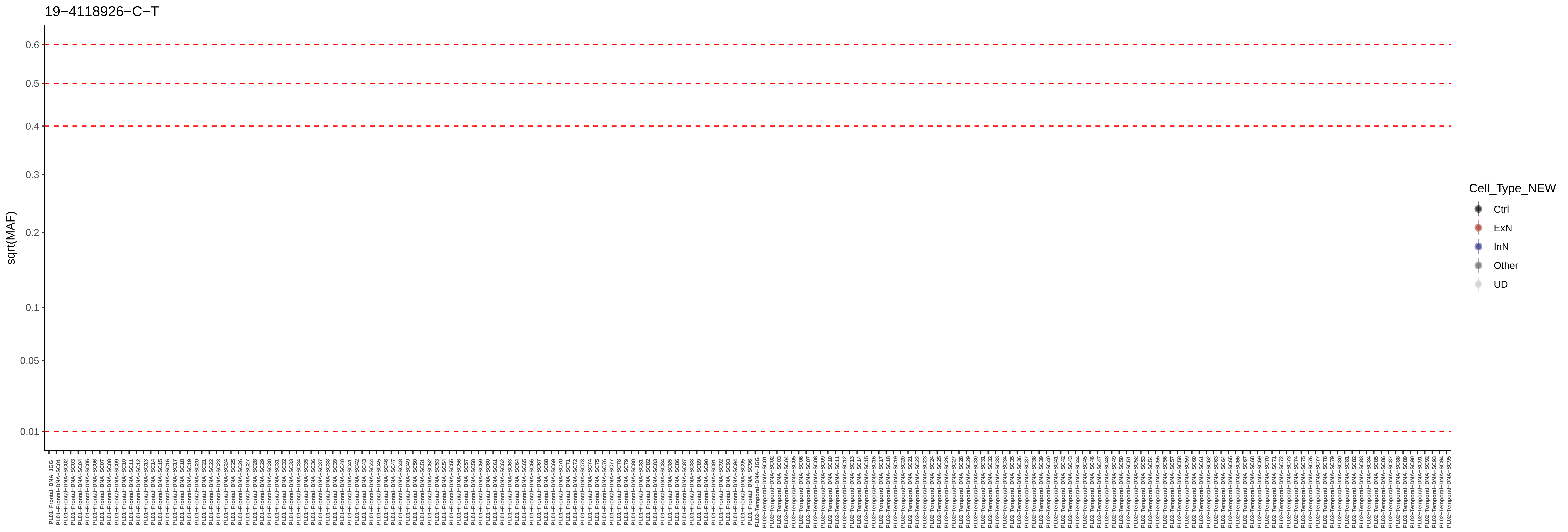
19−4118926−C−T
sqrt(MAF)
0.6
0.5
0.4
0.3
0.2
0.1
0.05
0.01
Cell_Type_NEW
Ctrl
ExN
InN
Other
UD

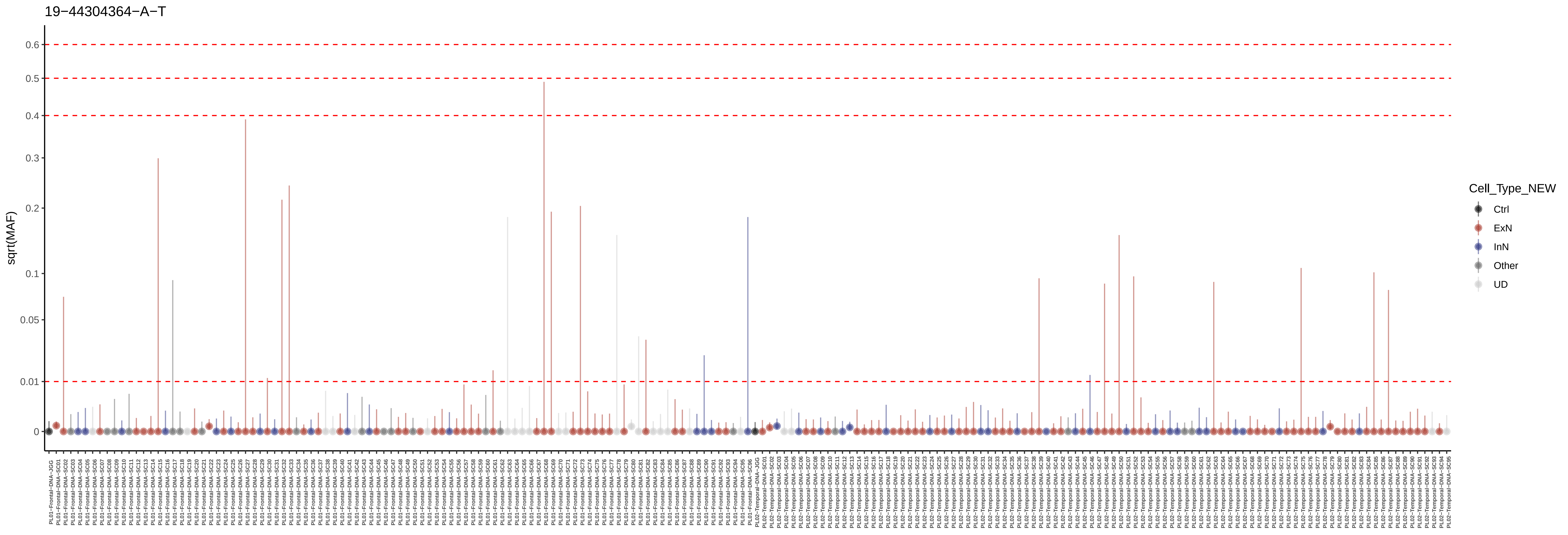
19−44304364−A−T
sqrt(MAF)
0.6
0.5
0.4
0.3
0.2
0.1
0.05
0.01
0
Cell_Type_NEW
Ctrl
ExN
InN
Other
UD

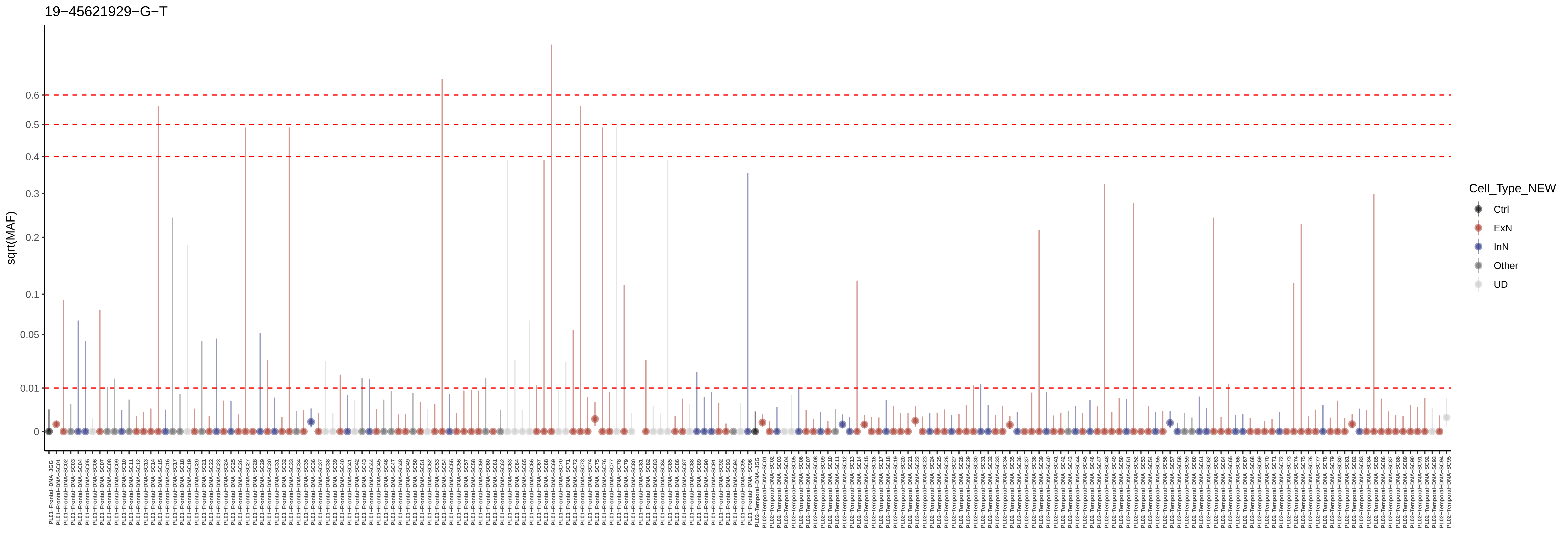
19−45621929−G−T
sqrt(MAF)
0.6
0.5
0.4
0.3
0.2
0.1
0.05
0.01
0
Cell_Type_NEW
Ctrl
ExN
InN
Other
UD

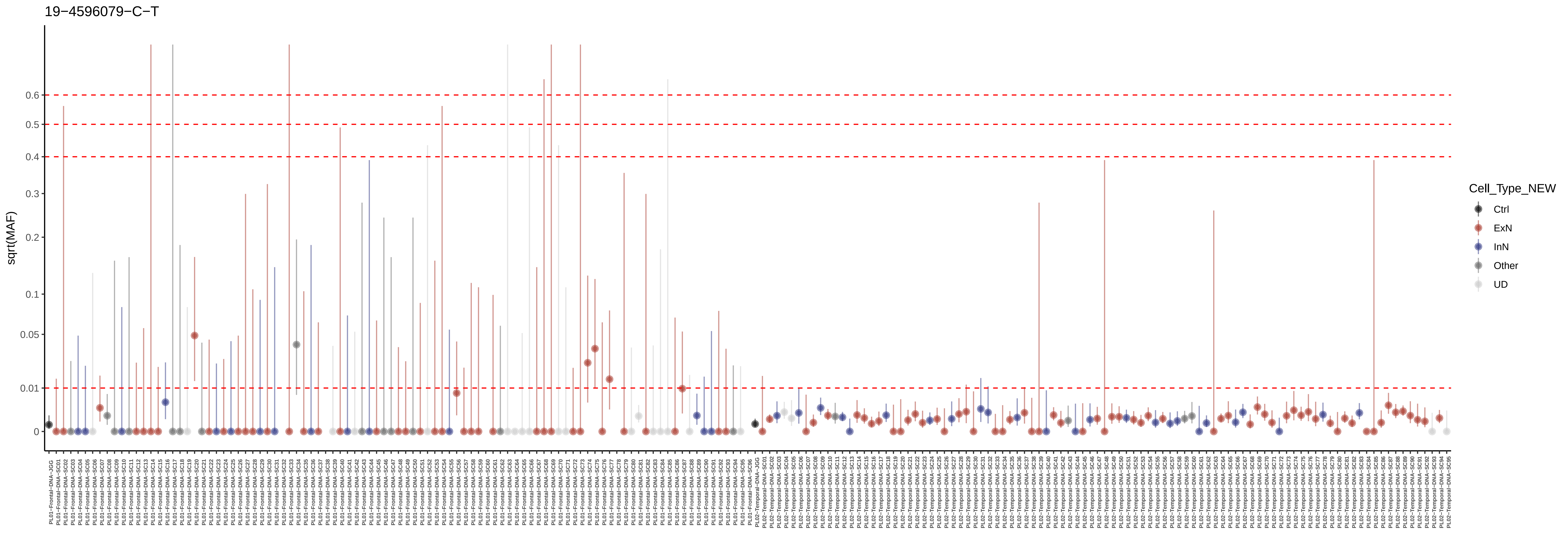
19−4596079−C−T
sqrt(MAF)
0.6
0.5
0.4
0.3
0.2
0.1
0.05
0.01
0
Cell_Type_NEW
Ctrl
ExN
InN
Other
UD

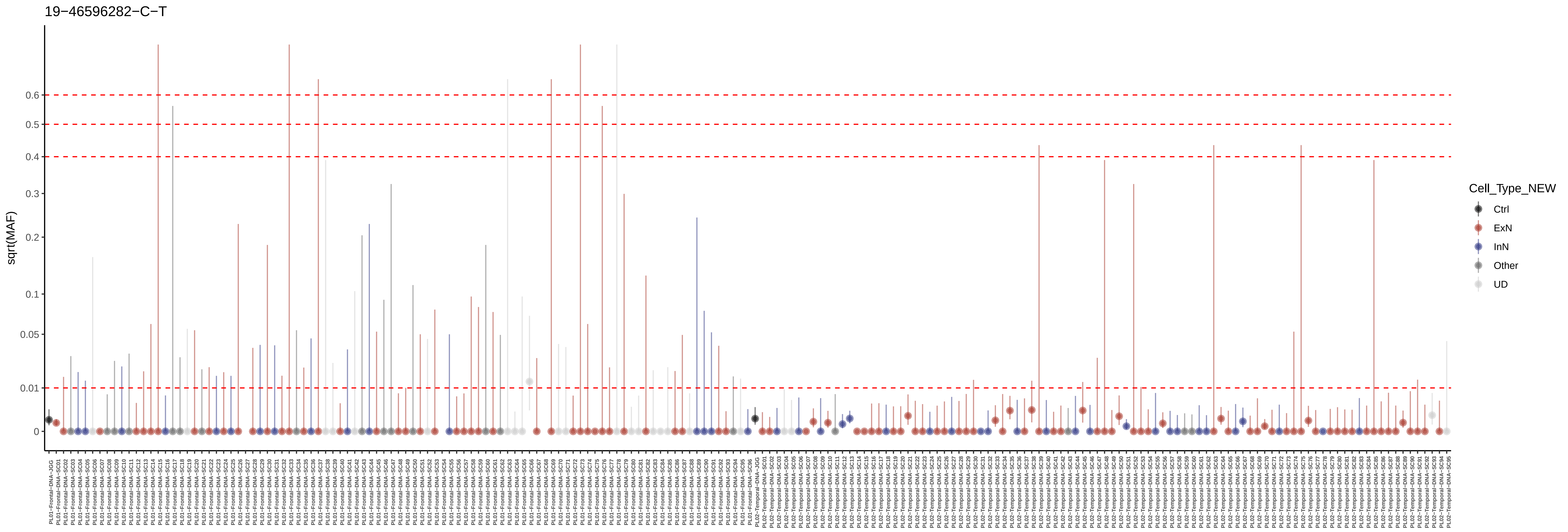

19−46596282−C−T
sqrt(MAF)
0.6
0.5
0.4
0.3
0.2
0.1
0.05
0.01
0
Cell_Type_NEW
Ctrl
ExN
InN
Other
UD

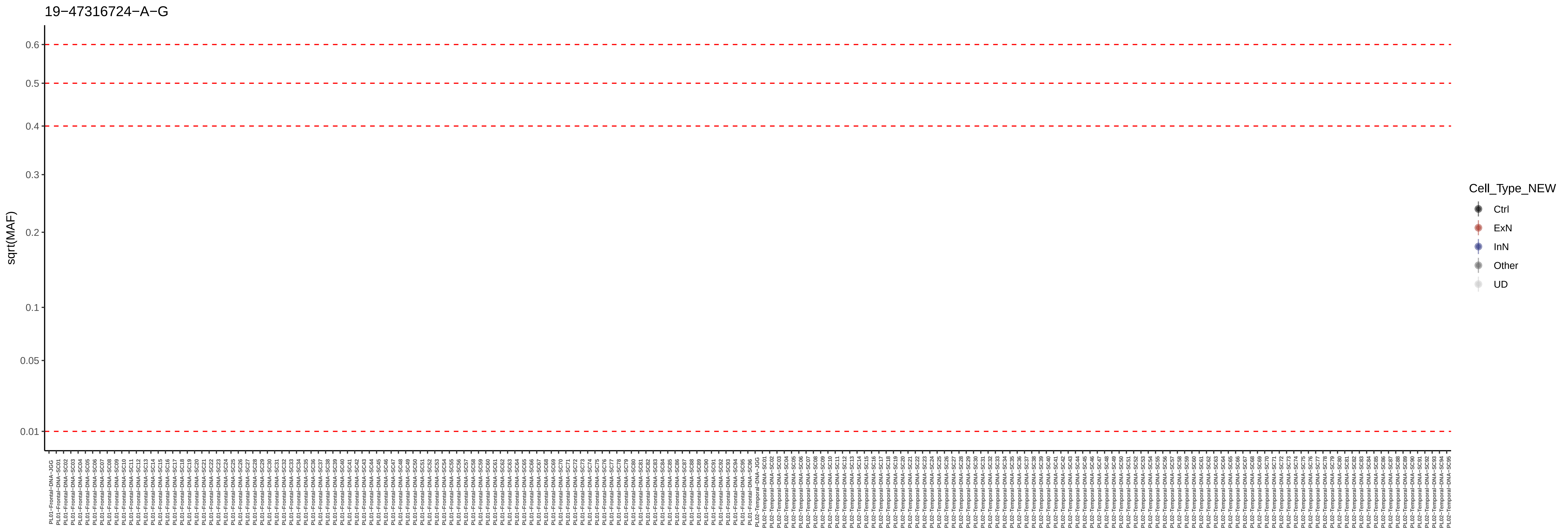

19−47316724−A−G
sqrt(MAF)
0.6
0.5
0.4
0.3
0.2
0.1
0.05
0.01
Cell_Type_NEW
Ctrl
ExN
InN
Other
UD

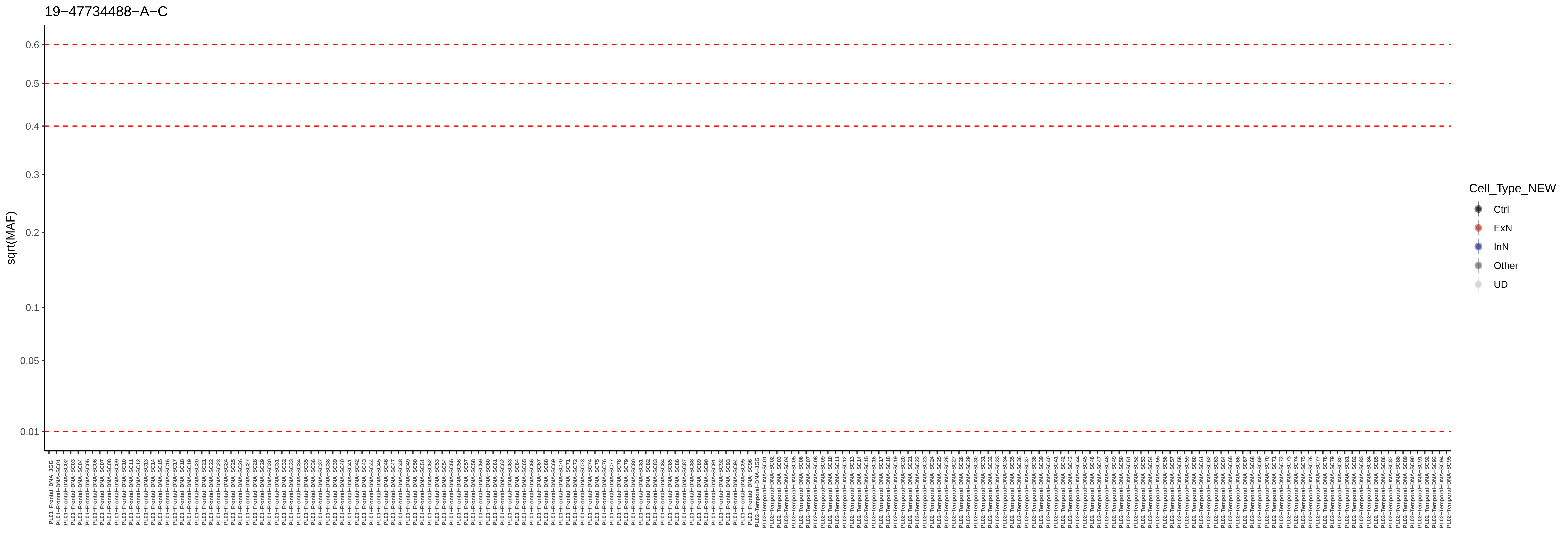
19−47734488−A−C
sqrt(MAF)
0.6
0.5
0.4
0.3
0.2
0.1
0.05
0.01
Cell_Type_NEW
Ctrl
ExN
InN
Other
UD

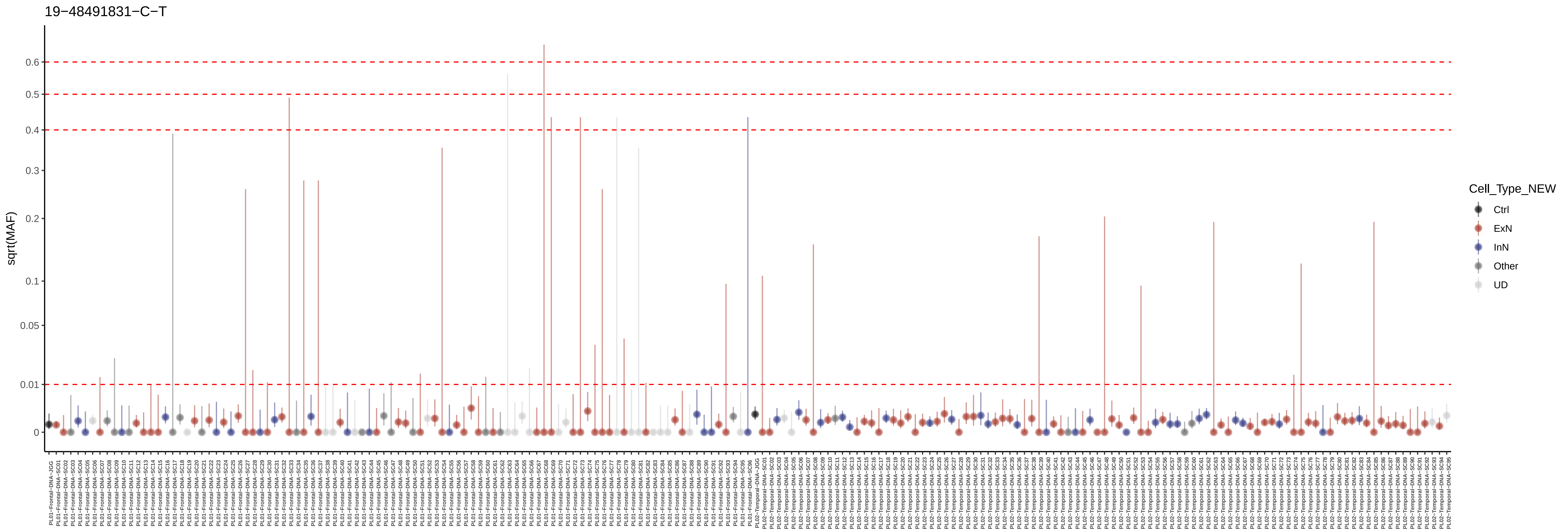
19−48491831−C−T
sqrt(MAF)
0.6
0.5
0.4
0.3
0.2
0.1
0.05
0.01
0
Cell_Type_NEW
Ctrl
ExN
InN
Other
UD

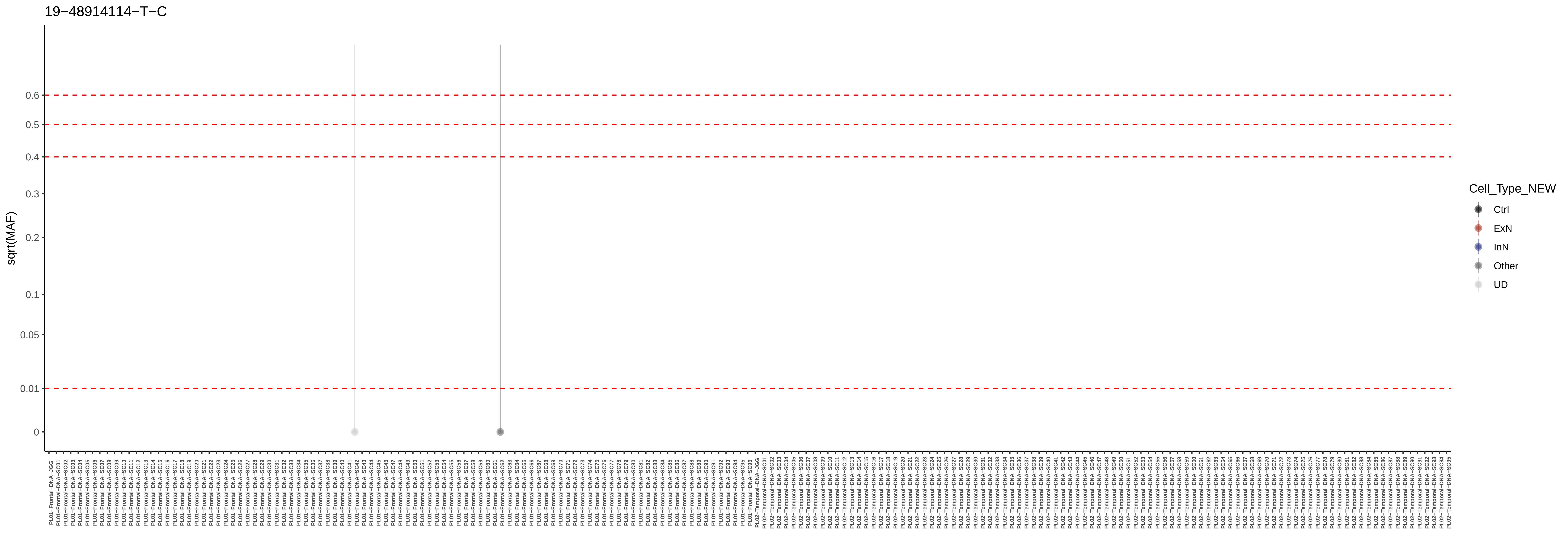
19−48914114−T−C
sqrt(MAF)
0.6
0.5
0.4
0.3
0.2
0.1
0.05
0.01
0
Cell_Type_NEW
Ctrl
ExN
InN
Other
UD

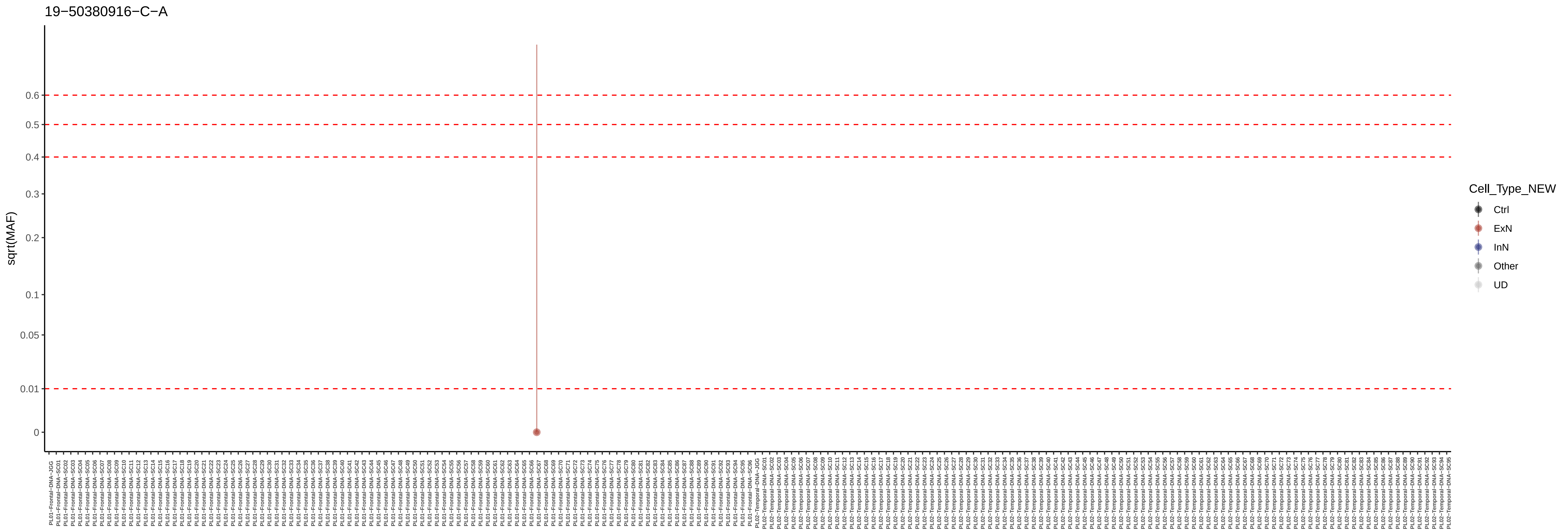

19−50380916−C−A
sqrt(MAF)
0.6
0.5
0.4
0.3
0.2
0.1
0.05
0.01
0
Cell_Type_NEW
Ctrl
ExN
InN
Other
UD

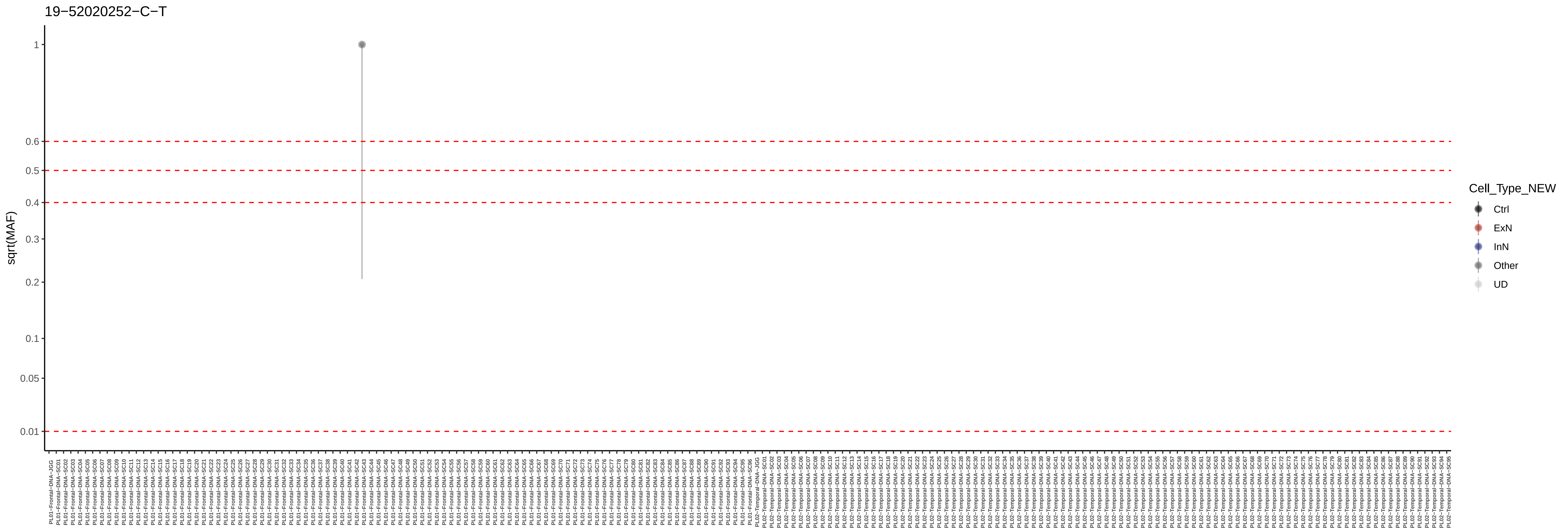
19−52020252−C−T
sqrt(MAF)
1
0.6
0.5
0.4
0.3
0.2
0.1
0.05
0.01
Cell_Type_NEW
Ctrl
ExN
InN
Other
UD

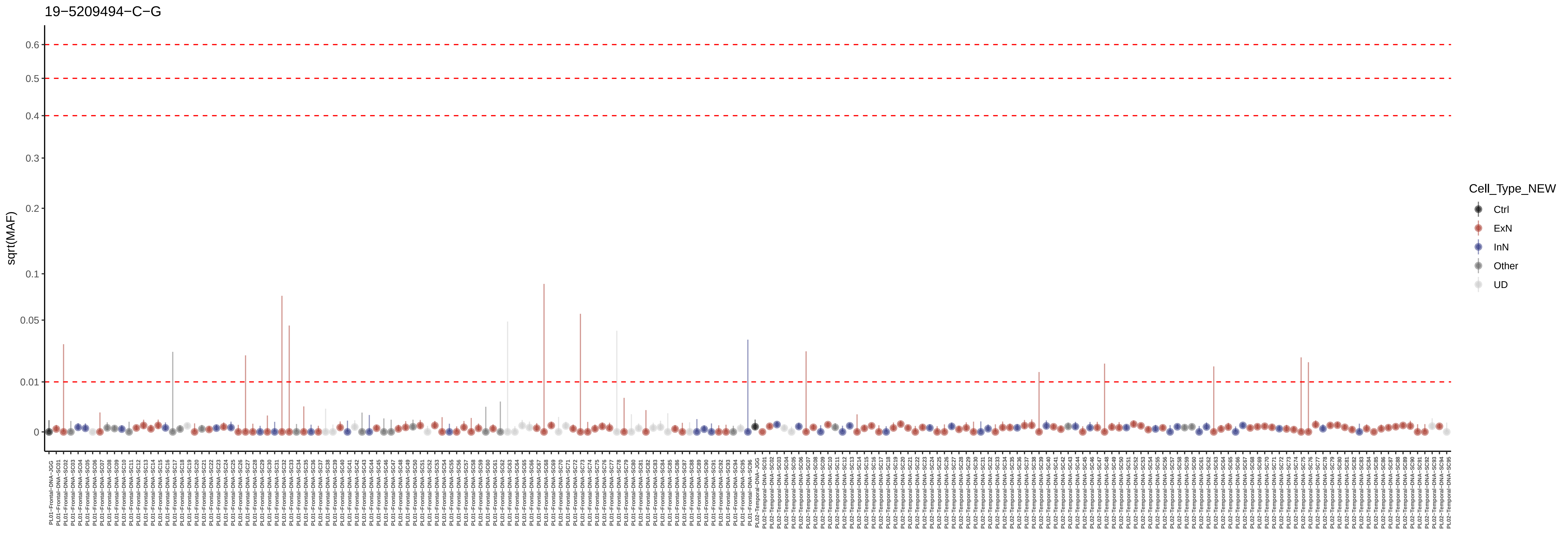
19−5209494−C−G
sqrt(MAF)
0.6
0.5
0.4
0.3
0.2
0.1
0.05
0.01
0
Cell_Type_NEW
Ctrl
ExN
InN
Other
UD

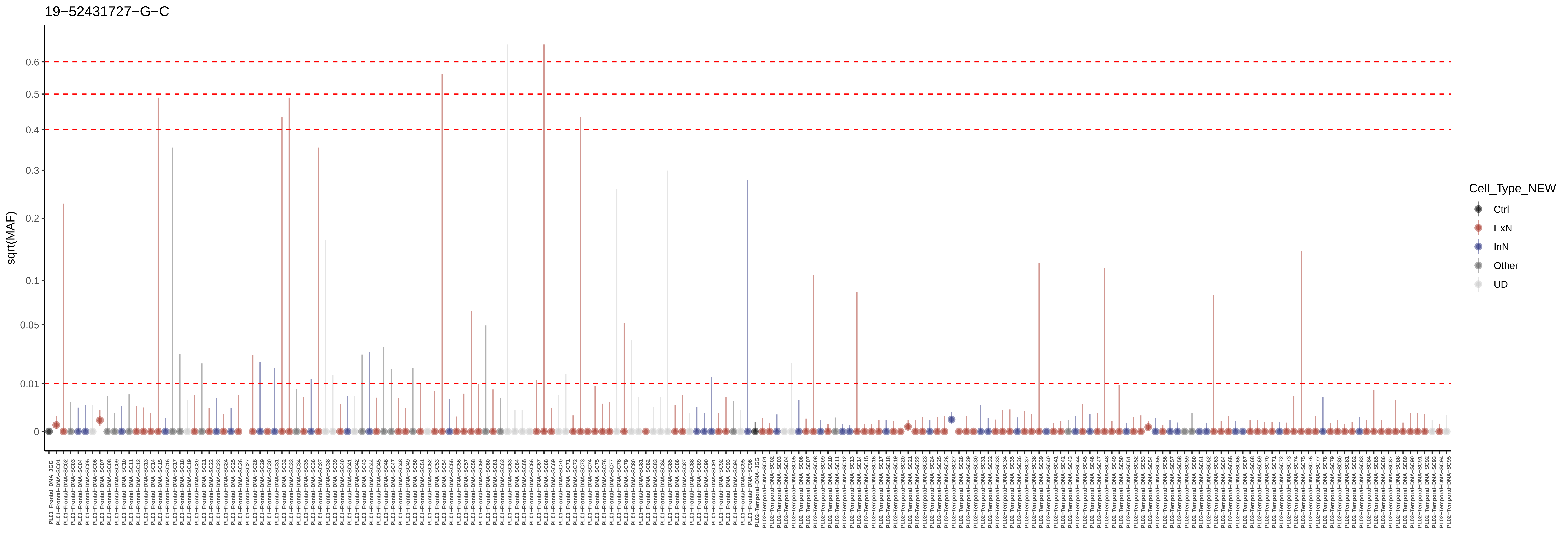
19−52431727−G−C
sqrt(MAF)
0.6
0.5
0.4
0.3
0.2
0.1
0.05
0.01
0
Cell_Type_NEW
Ctrl
ExN
InN
Other
UD

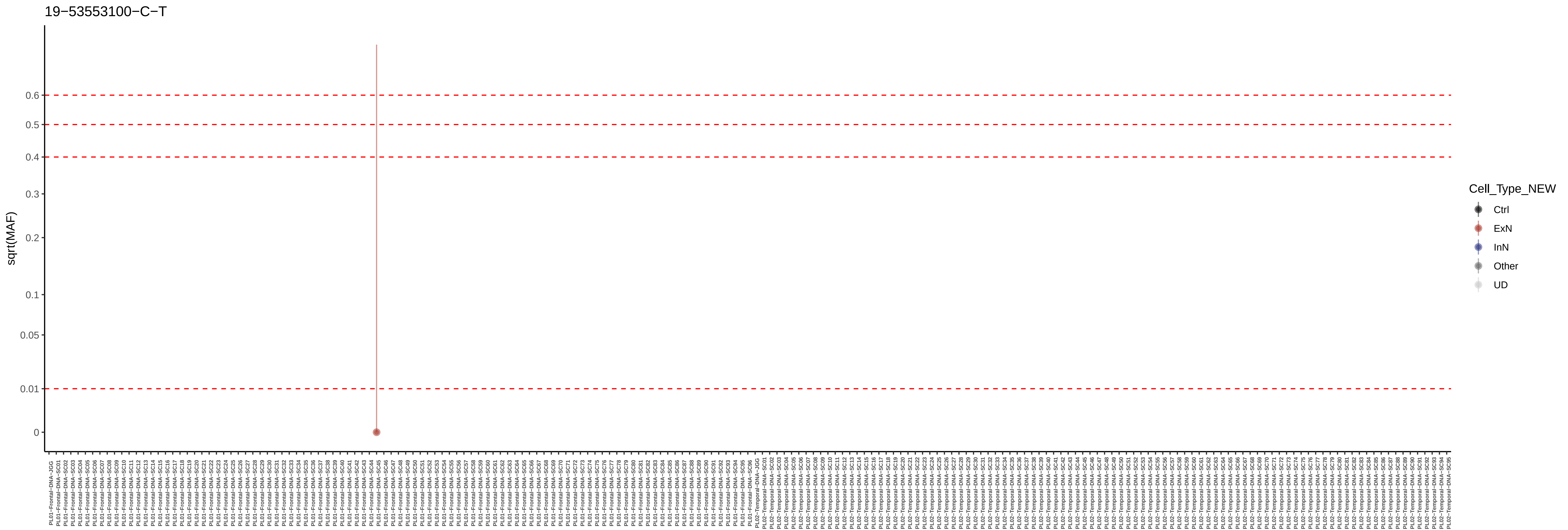

19−53553100−C−T
sqrt(MAF)
0.6
0.5
0.4
0.3
0.2
0.1
0.05
0.01
0
Cell_Type_NEW
Ctrl
ExN
InN
Other
UD

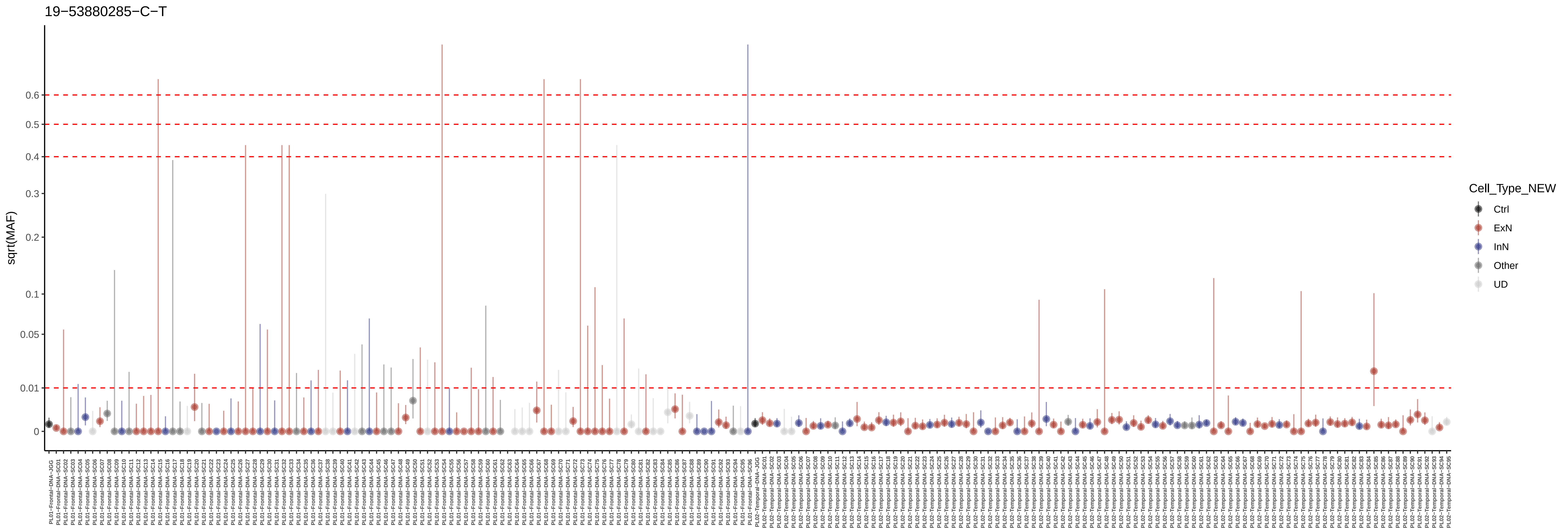

19−53880285−C−T
sqrt(MAF)
0.6
0.5
0.4
0.3
0.2
0.1
0.05
0.01
0
Cell_Type_NEW
Ctrl
ExN
InN
Other
UD

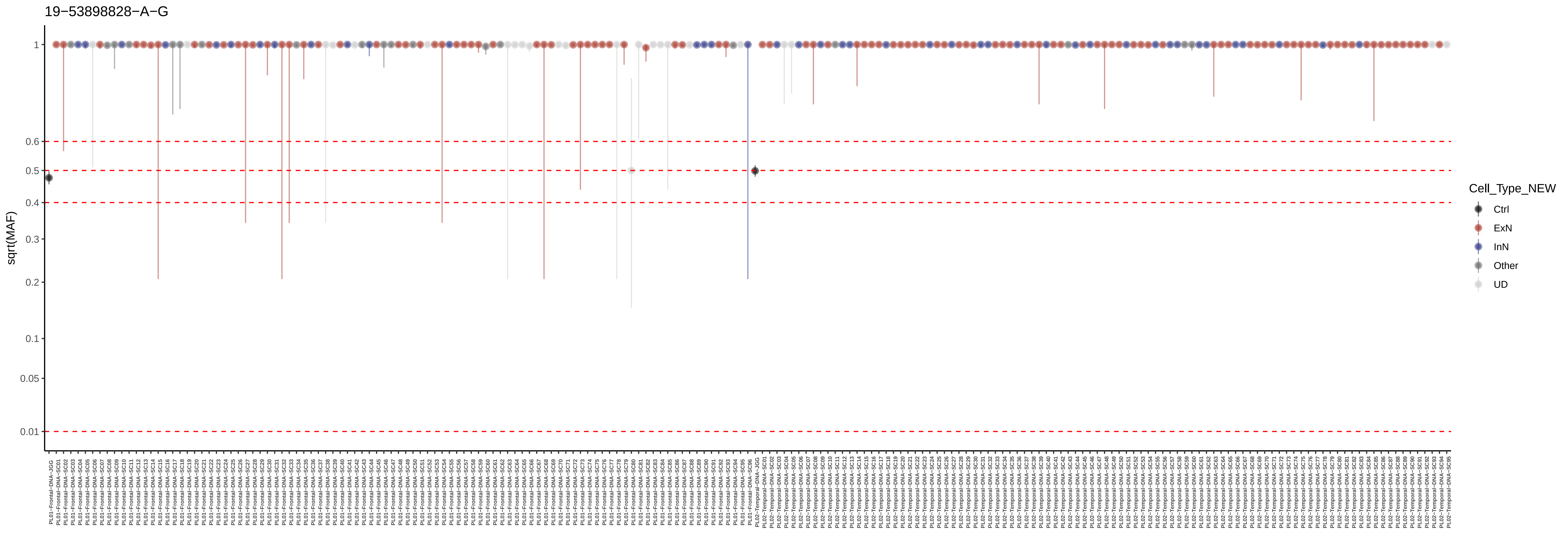
19−53898828−A−G
sqrt(MAF)
1
0.6
0.5
0.4
0.3
0.2
0.1
0.05
0.01
Cell_Type_NEW
Ctrl
ExN
InN
Other
UD

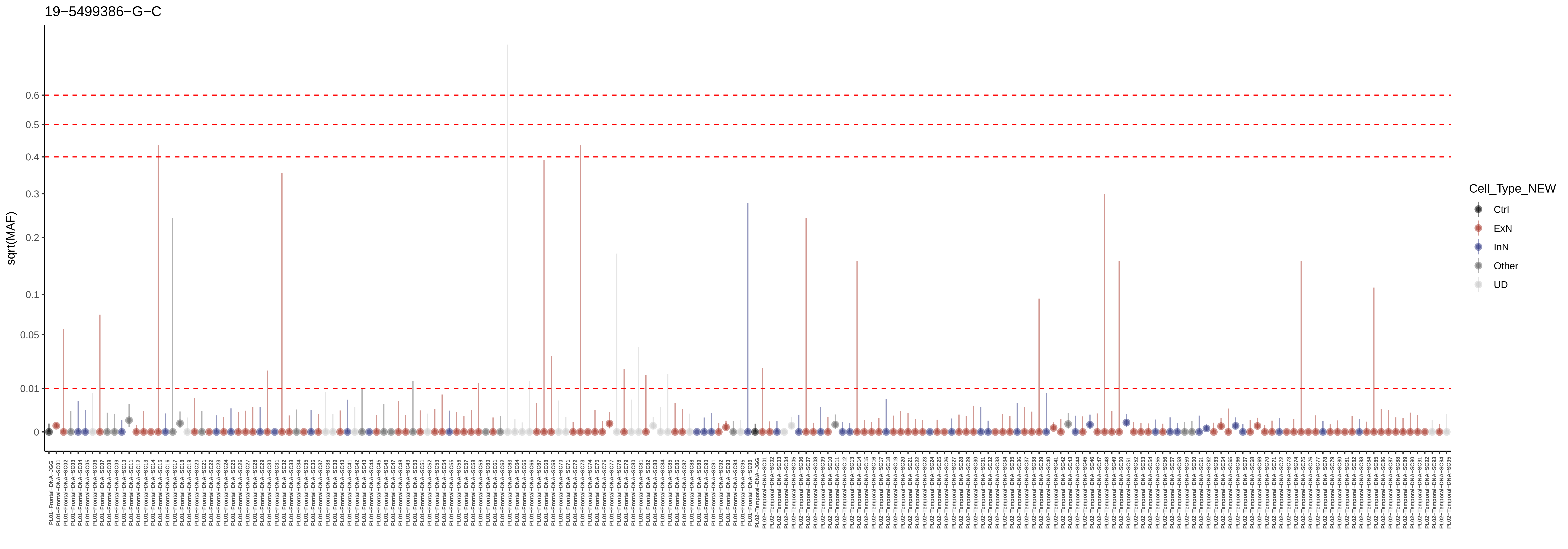

19−5499386−G−C
sqrt(MAF)
0.6
0.5
0.4
0.3
0.2
0.1
0.05
0.01
0
Cell_Type_NEW
Ctrl
ExN
InN
Other
UD

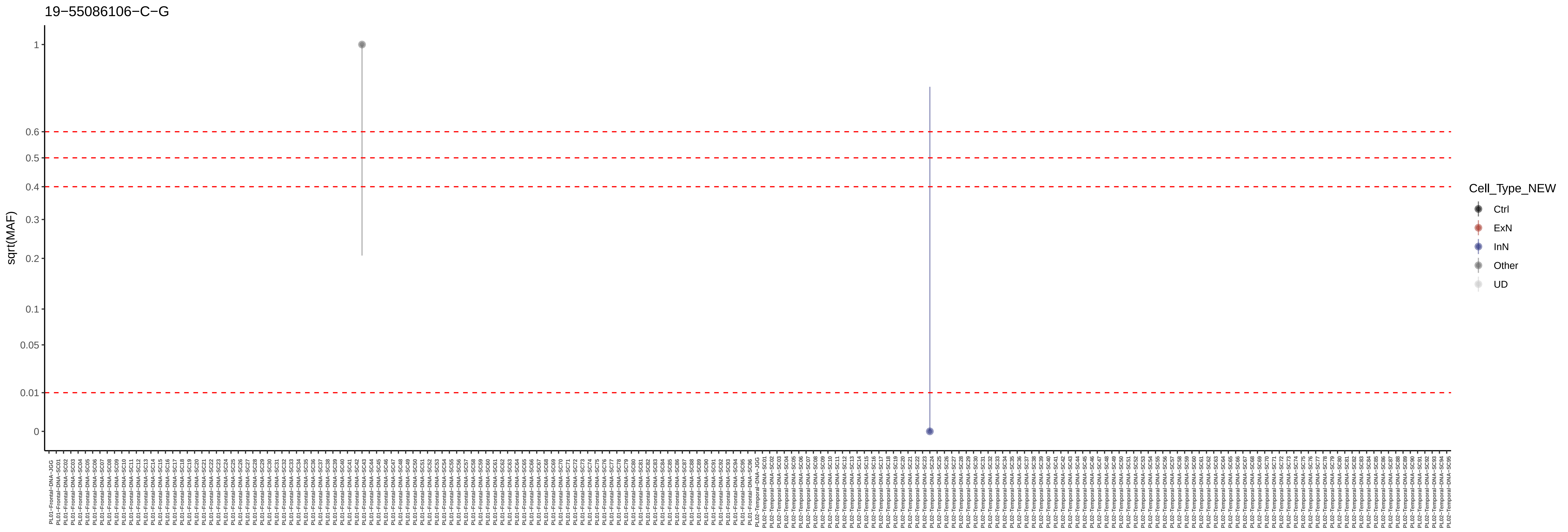

19−55086106−C−G
sqrt(MAF)
1
0.6
0.5
0.4
0.3
0.2
0.1
0.05
0.01
0
Cell_Type_NEW
Ctrl
ExN
InN
Other
UD

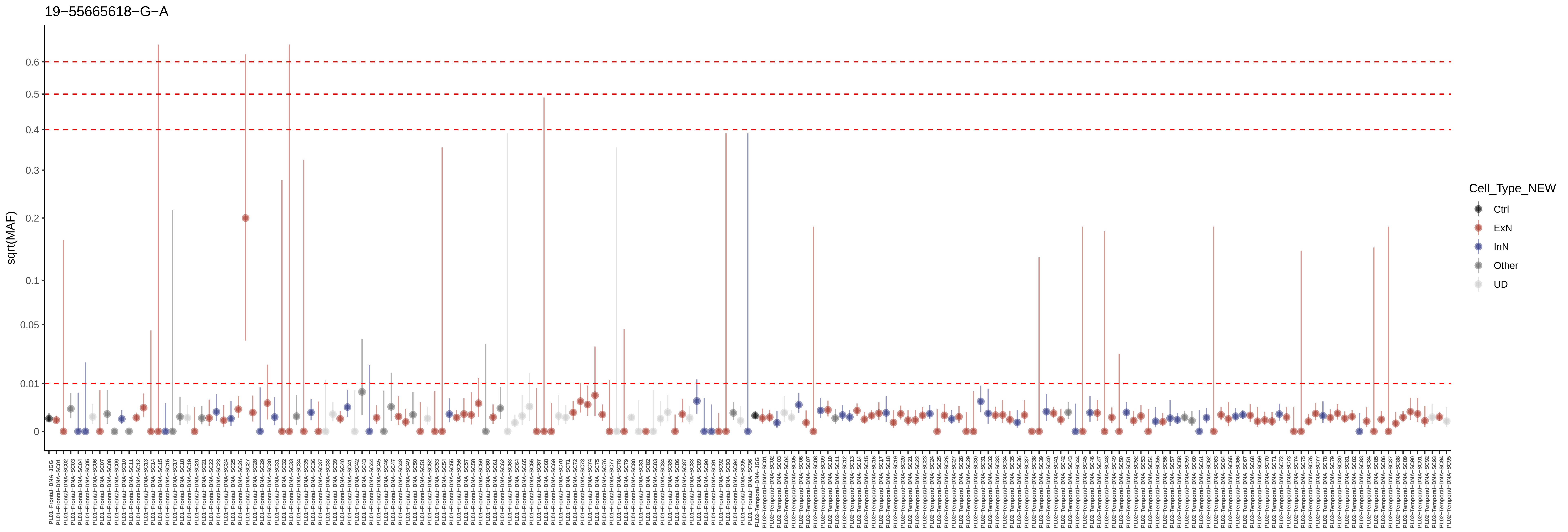
19−55665618−G−A
sqrt(MAF)
0.6
0.5
0.4
0.3
0.2
0.1
0.05
0.01
0
Cell_Type_NEW
Ctrl
ExN
InN
Other
UD

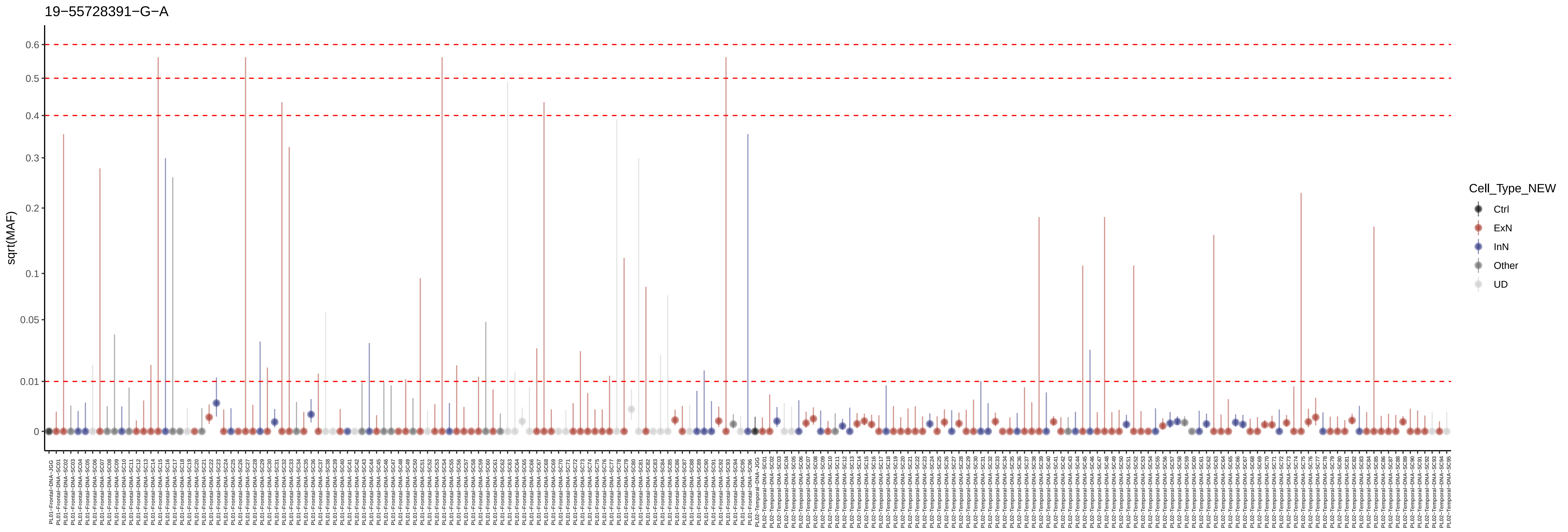
19−55728391−G−A
sqrt(MAF)
0.6
0.5
0.4
0.3
0.2
0.1
0.05
0.01
0
Cell_Type_NEW
Ctrl
ExN
InN
Other
UD

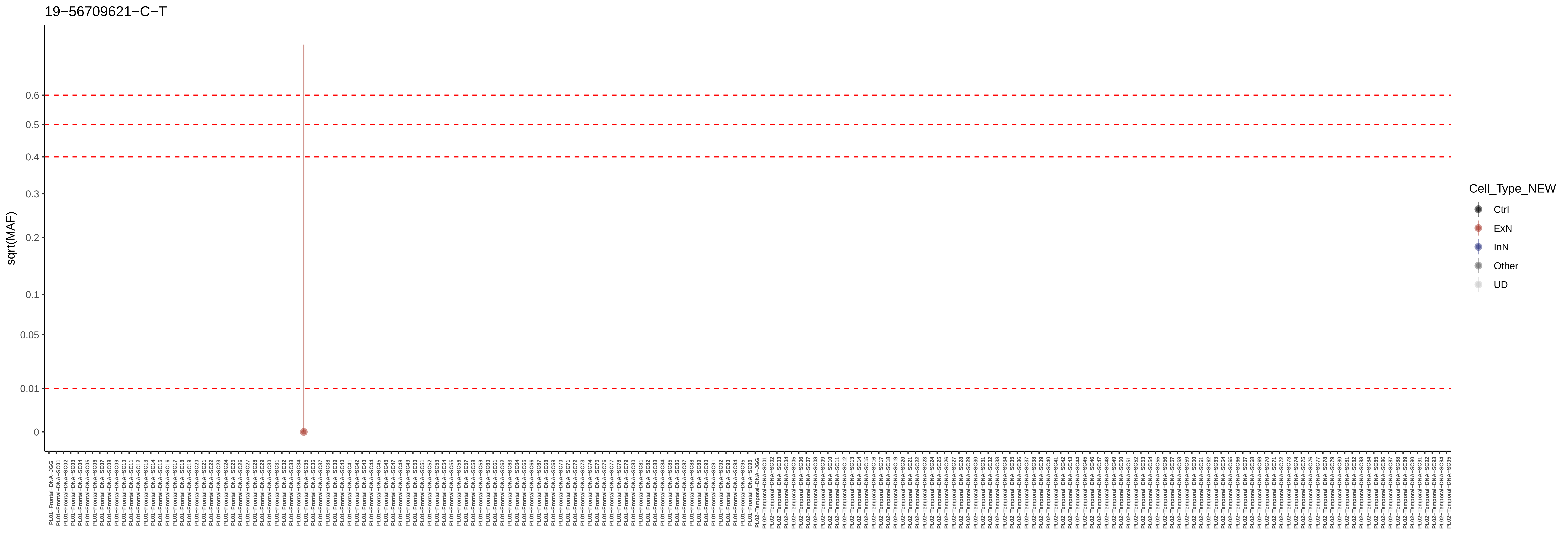
19−56709621−C−T
sqrt(MAF)
0.6
0.5
0.4
0.3
0.2
0.1
0.05
0.01
0
Cell_Type_NEW
Ctrl
ExN
InN
Other
UD

19−57342142−G−A

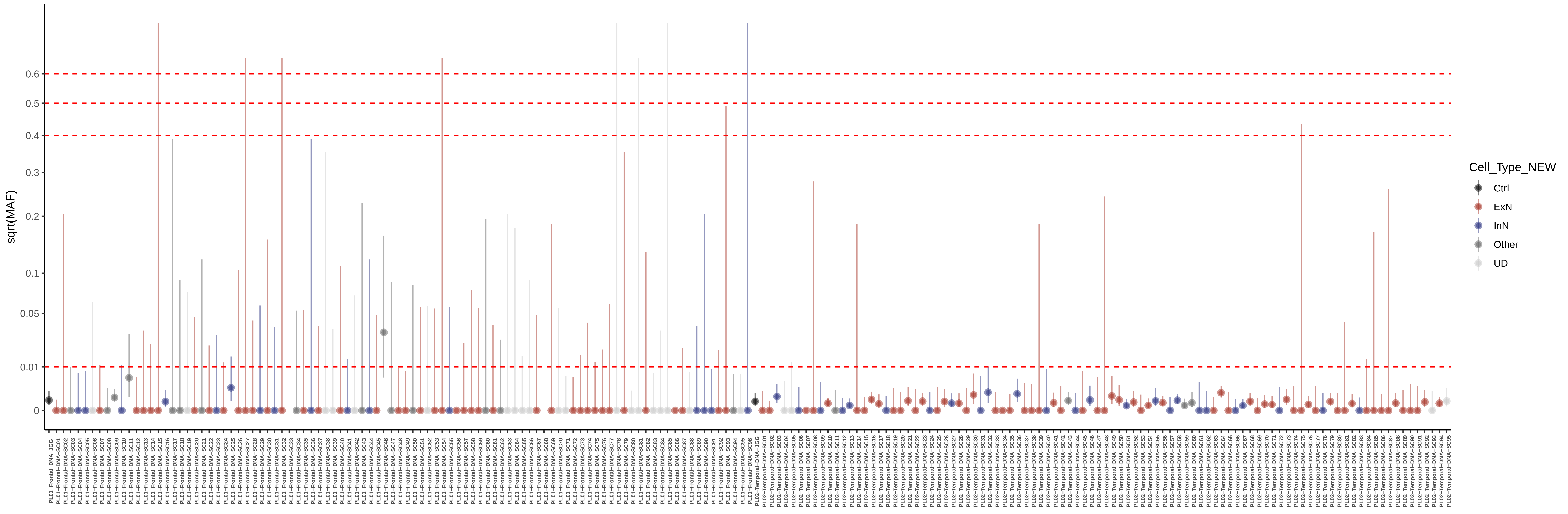

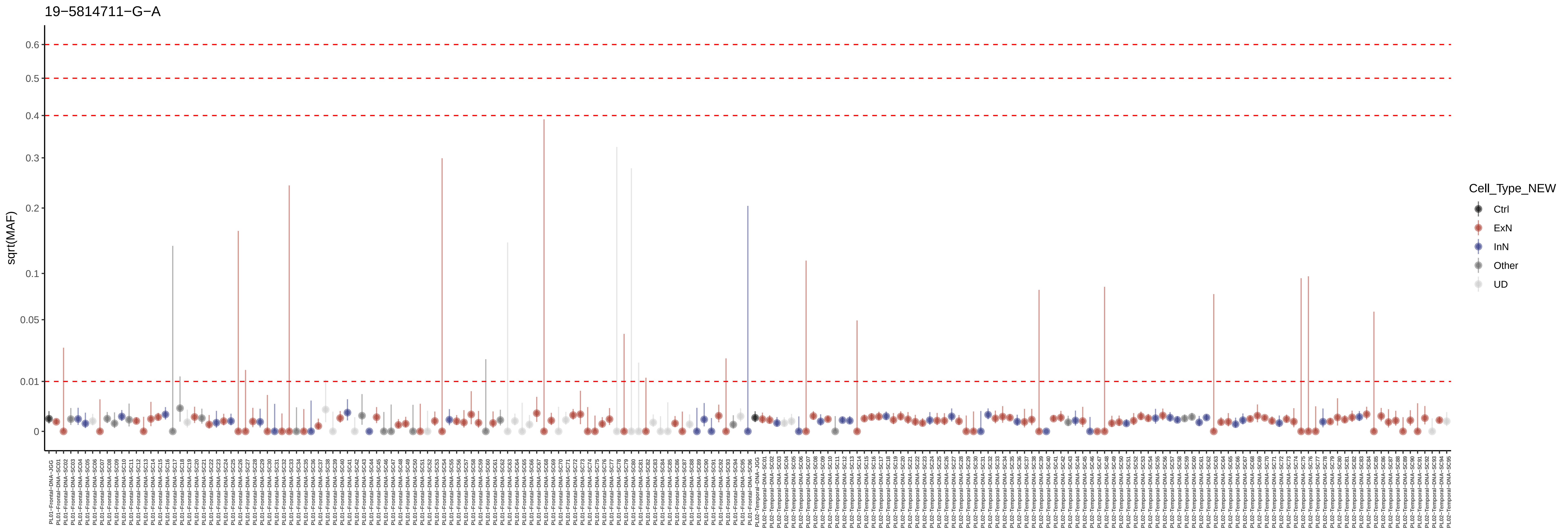
19−5814711−G−A
sqrt(MAF)
0.6
0.5
0.4
0.3
0.2
0.1
0.05
0.01
0
Cell_Type_NEW
Ctrl
ExN
InN
Other
UD

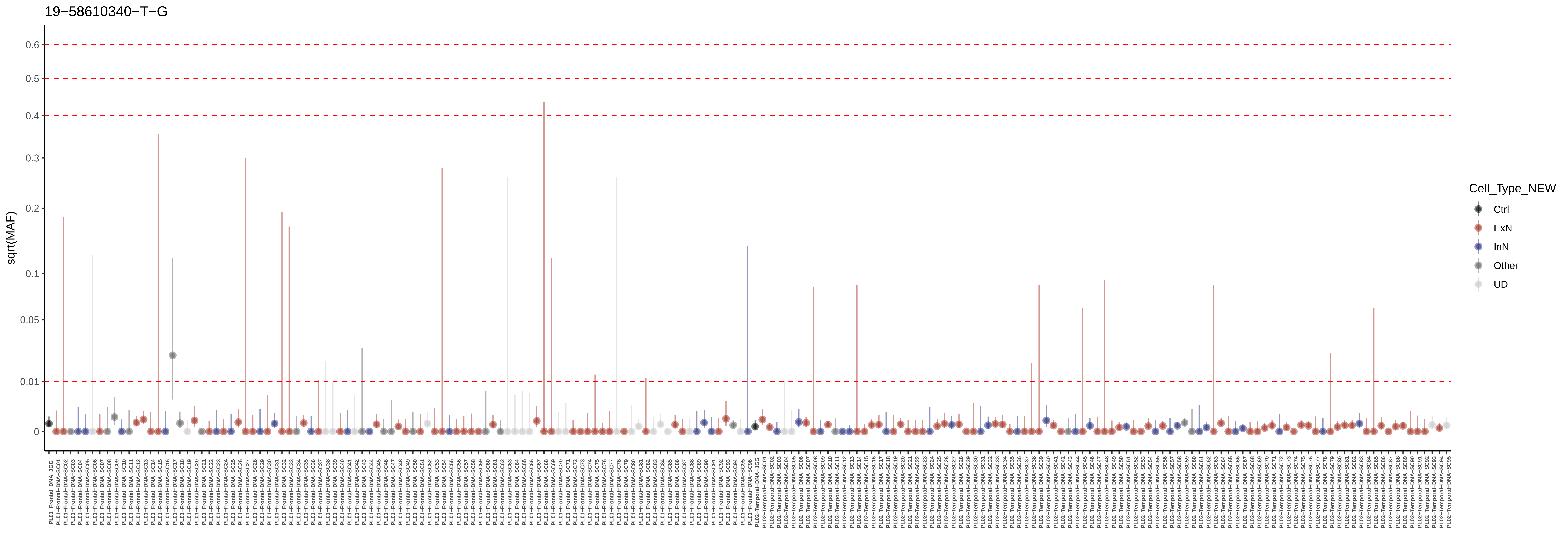

19−58610340−T−G
sqrt(MAF)
0.6
0.5
0.4
0.3
0.2
0.1
0.05
0.01
0
Cell_Type_NEW
Ctrl
ExN
InN
Other
UD

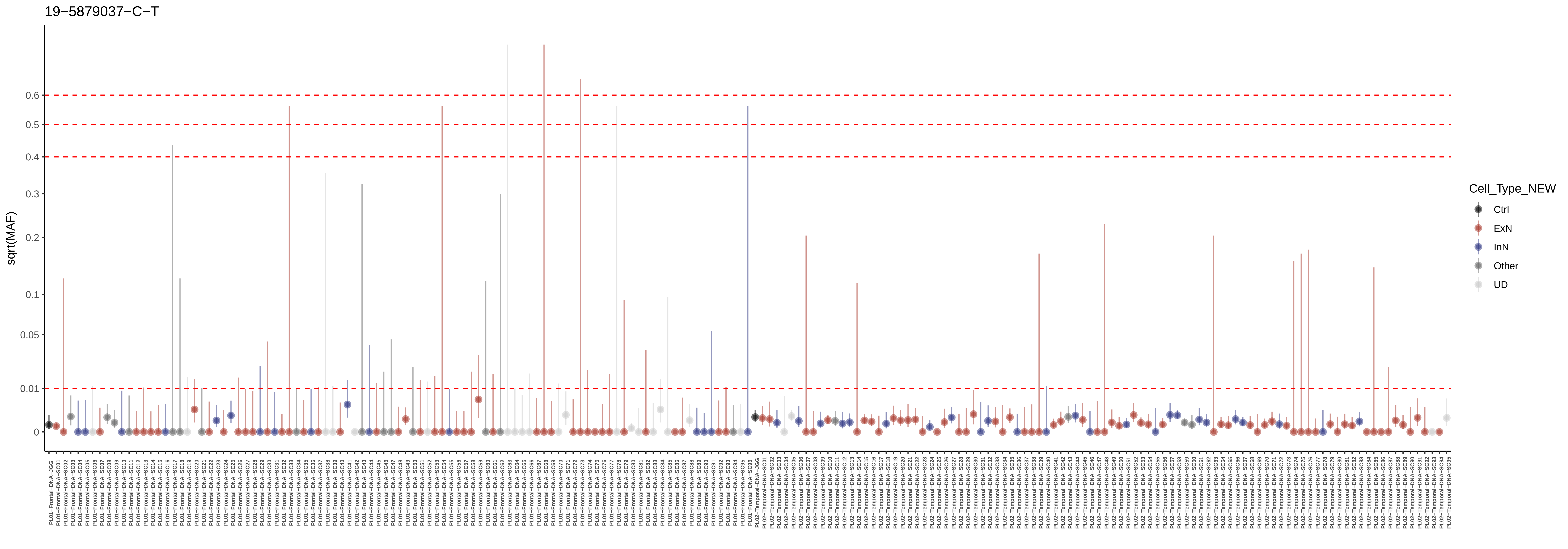
19−5879037−C−T
sqrt(MAF)
0.6
0.5
0.4
0.3
0.2
0.1
0.05
0.01
0
Cell_Type_NEW
Ctrl
ExN
InN
Other
UD

19−6263788−G−A

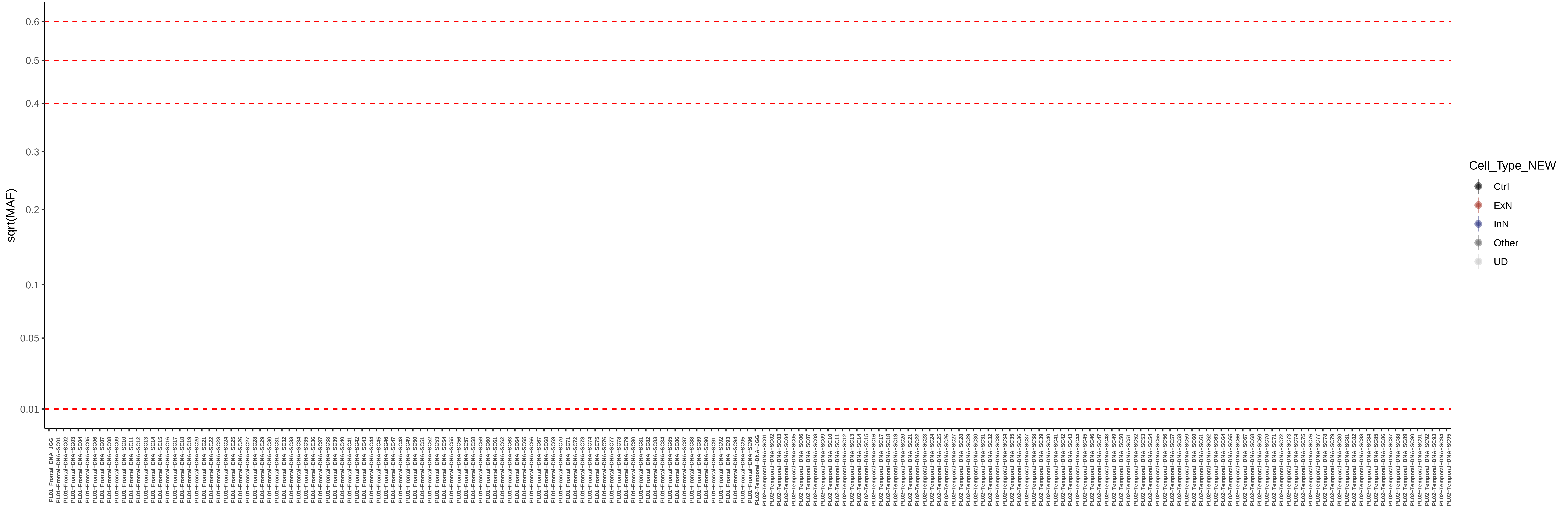

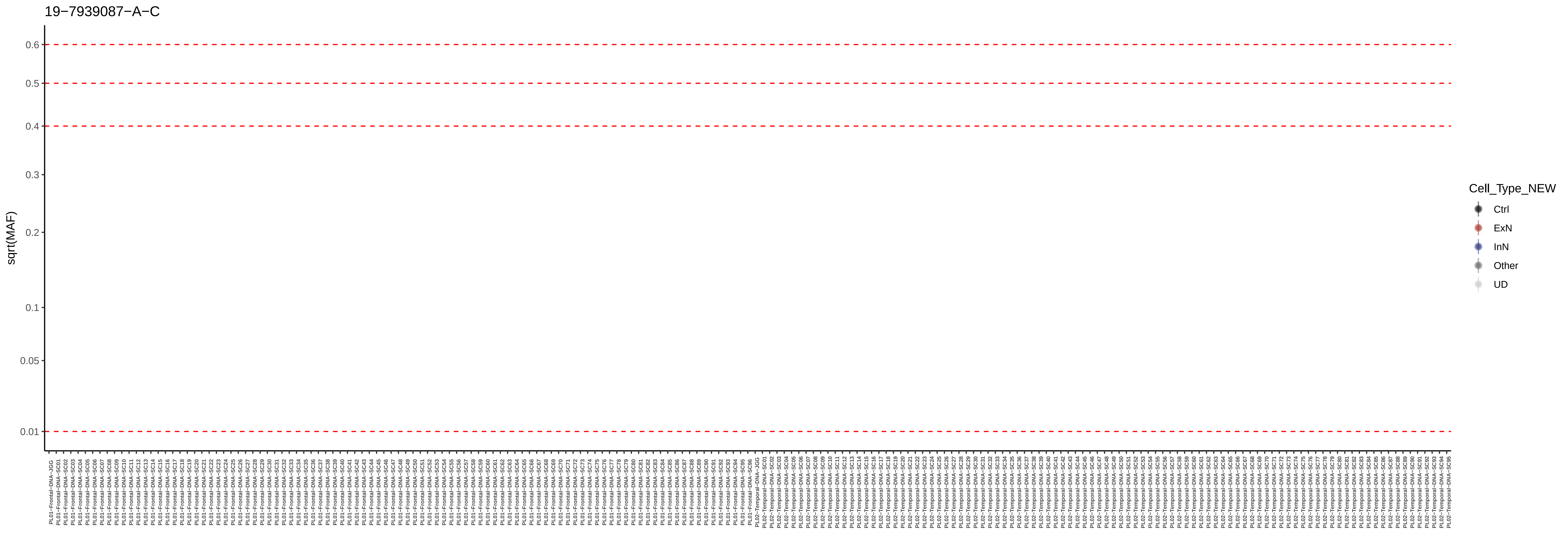

19−7939087−A−C
sqrt(MAF)
0.6
0.5
0.4
0.3
0.2
0.1
0.05
0.01
Cell_Type_NEW
Ctrl
ExN
InN
Other
UD

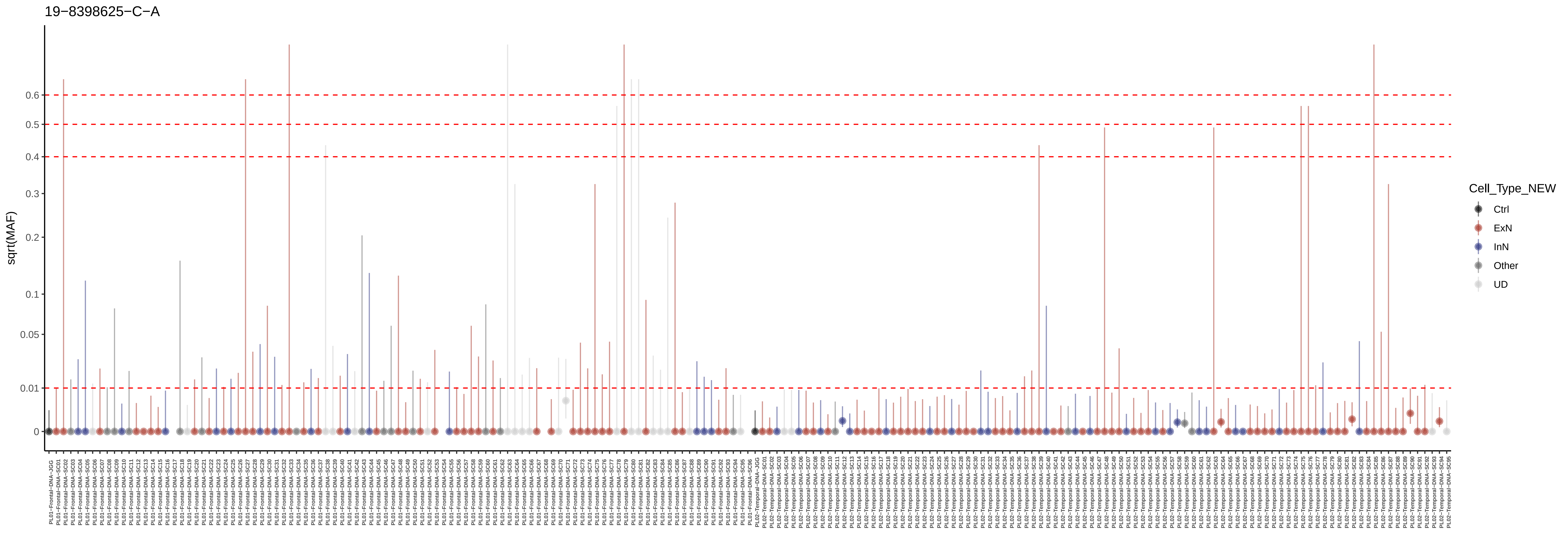
19−8398625−C−A
sqrt(MAF)
0.6
0.5
0.4
0.3
0.2
0.1
0.05
0.01
0
Cell_Type_NEW
Ctrl
ExN
InN
Other
UD

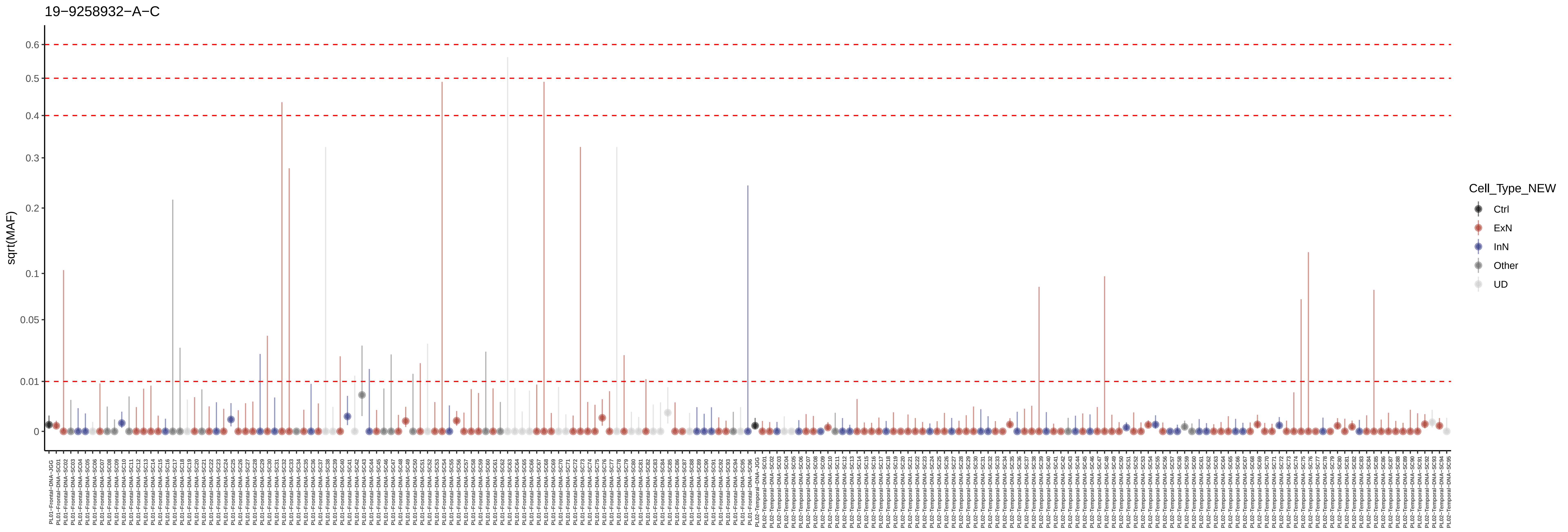
19−9258932−A−C
sqrt(MAF)
Cell_Type_NEW
Ctrl
ExN
InN
Other
UD

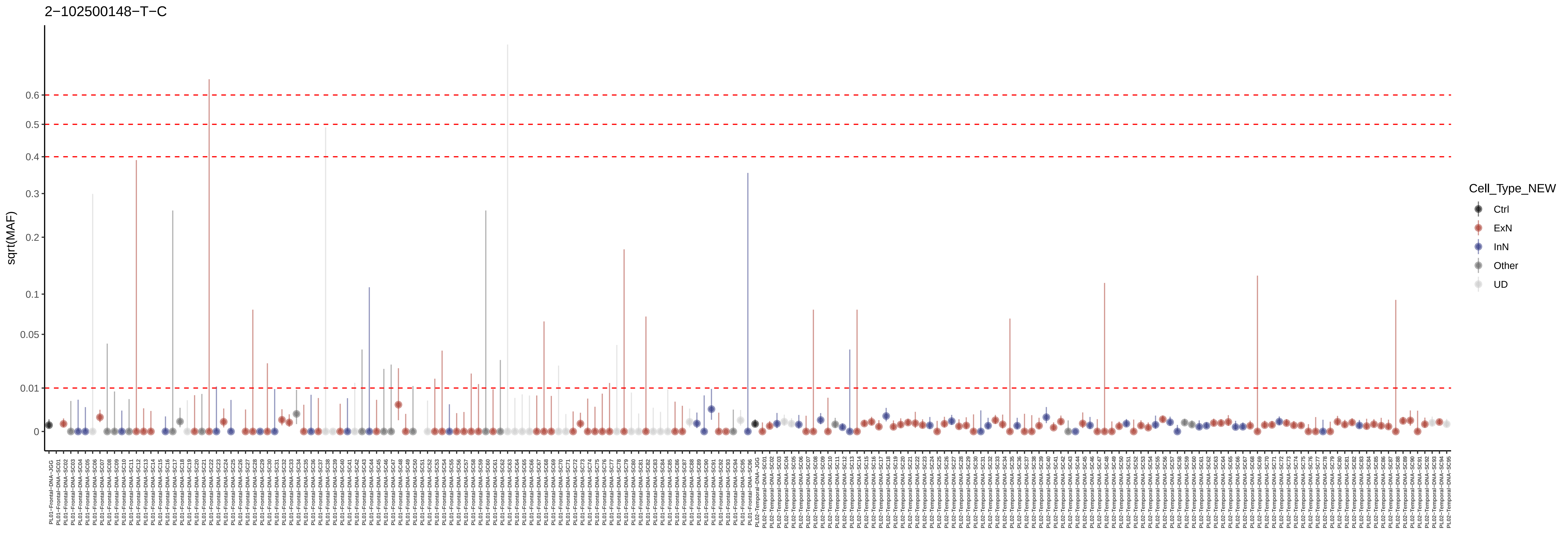
2−102500148−T−C
sqrt(MAF)
0.6
0.5
0.4
0.3
0.2
0.1
0.05
0.01
0
Cell_Type_NEW
Ctrl
ExN
InN
Other
UD

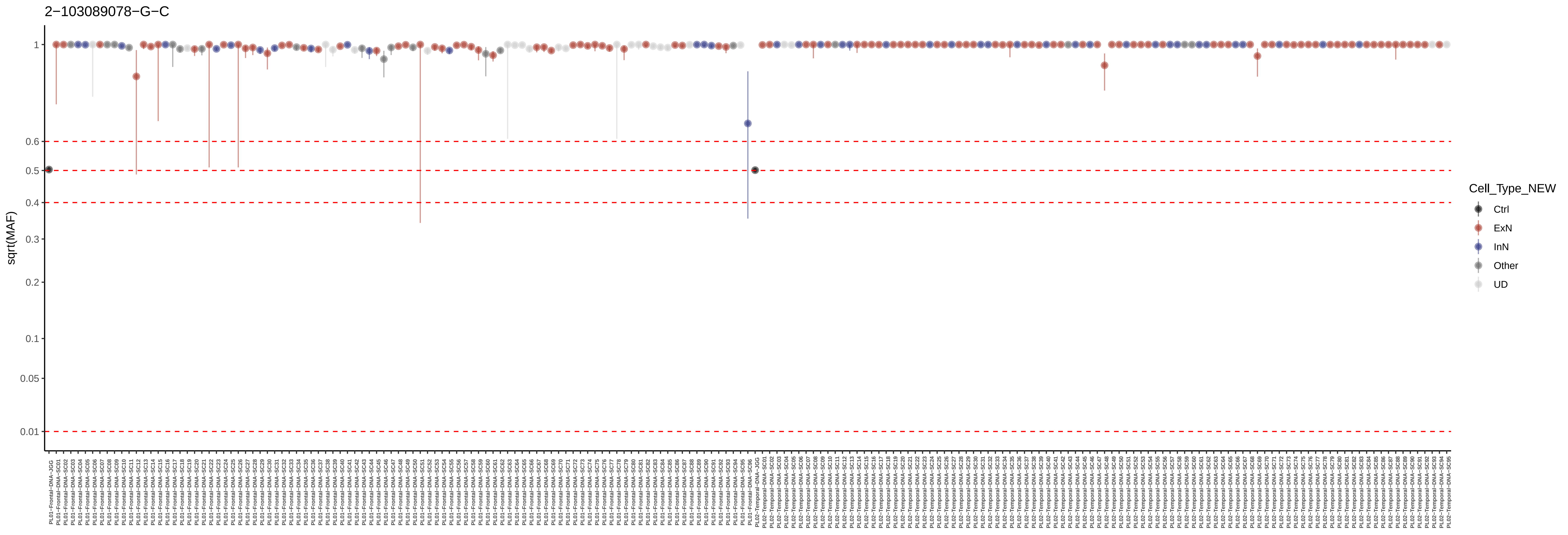
2−103089078−G−C
sqrt(MAF)
1
0.6
0.5
0.4
0.3
0.2
0.1
0.05
0.01
Cell_Type_NEW
Ctrl
ExN
InN
Other
UD

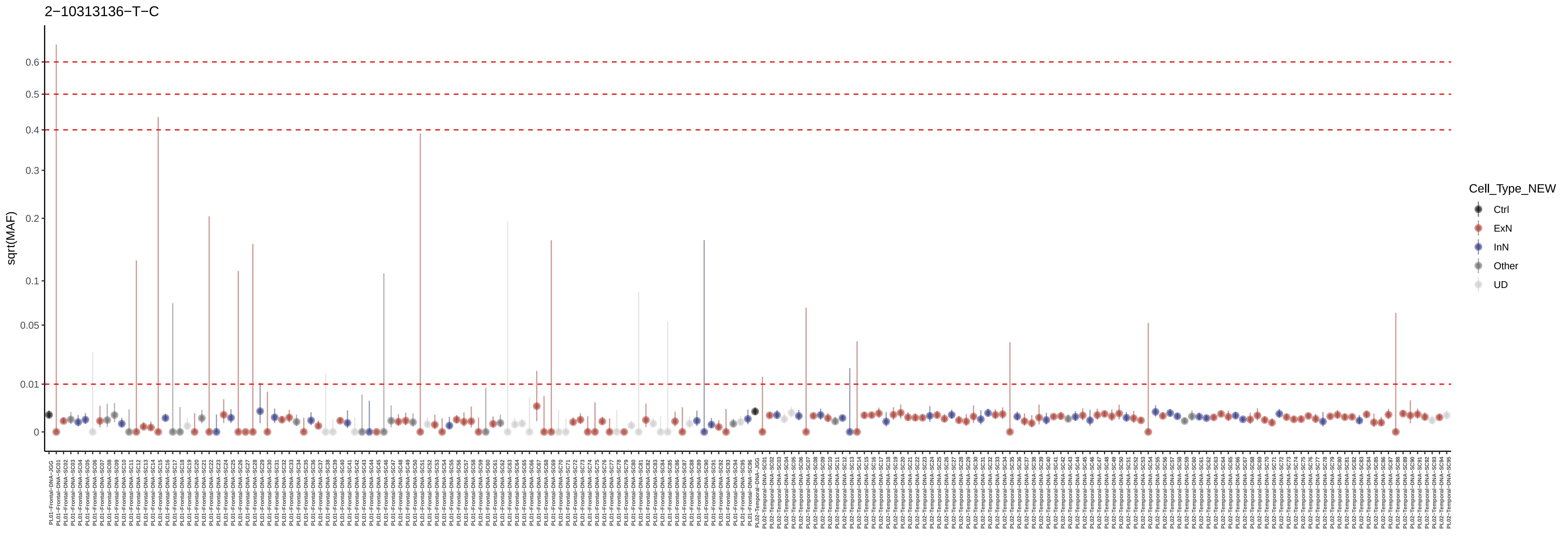

2−10313136−T−C
sqrt(MAF)
0.6
0.5
0.4
0.3
0.2
0.1
0.05
0.01
0
Cell_Type_NEW
Ctrl
ExN
InN
Other
UD

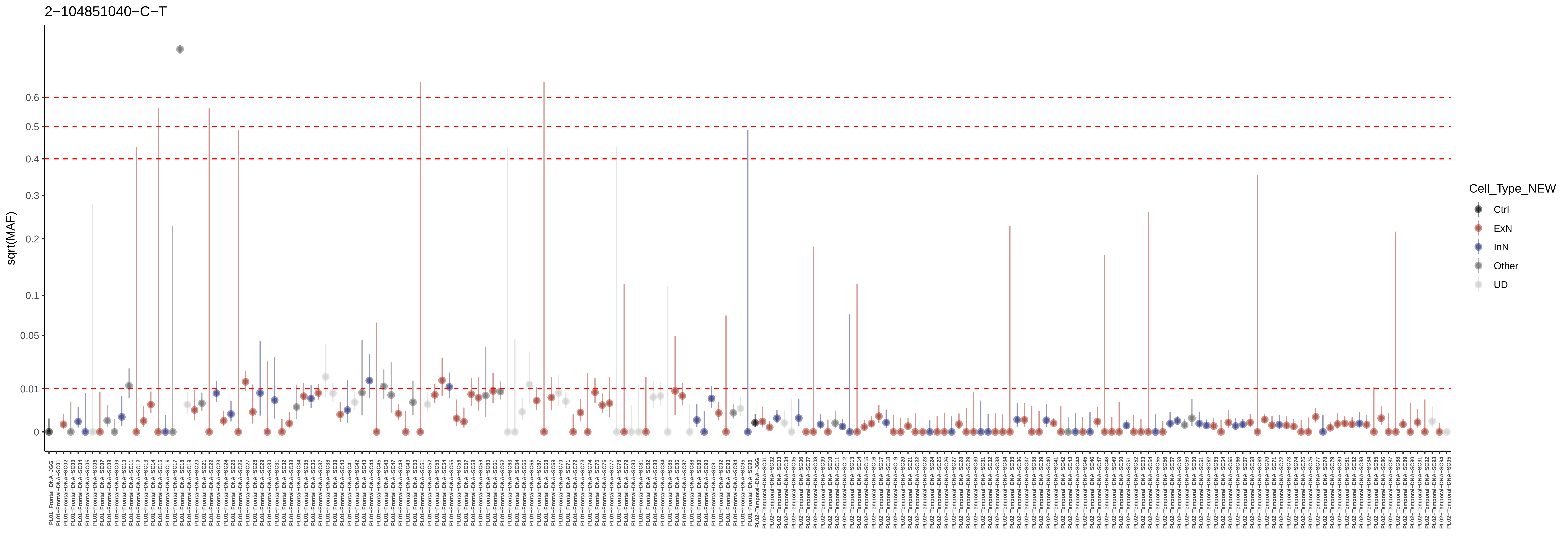

2−104851040−C−T
sqrt(MAF)
0.6
0.5
0.4
0.3
0.2
0.1
0.05
0.01
0
Cell_Type_NEW
Ctrl
ExN
InN
Other
UD

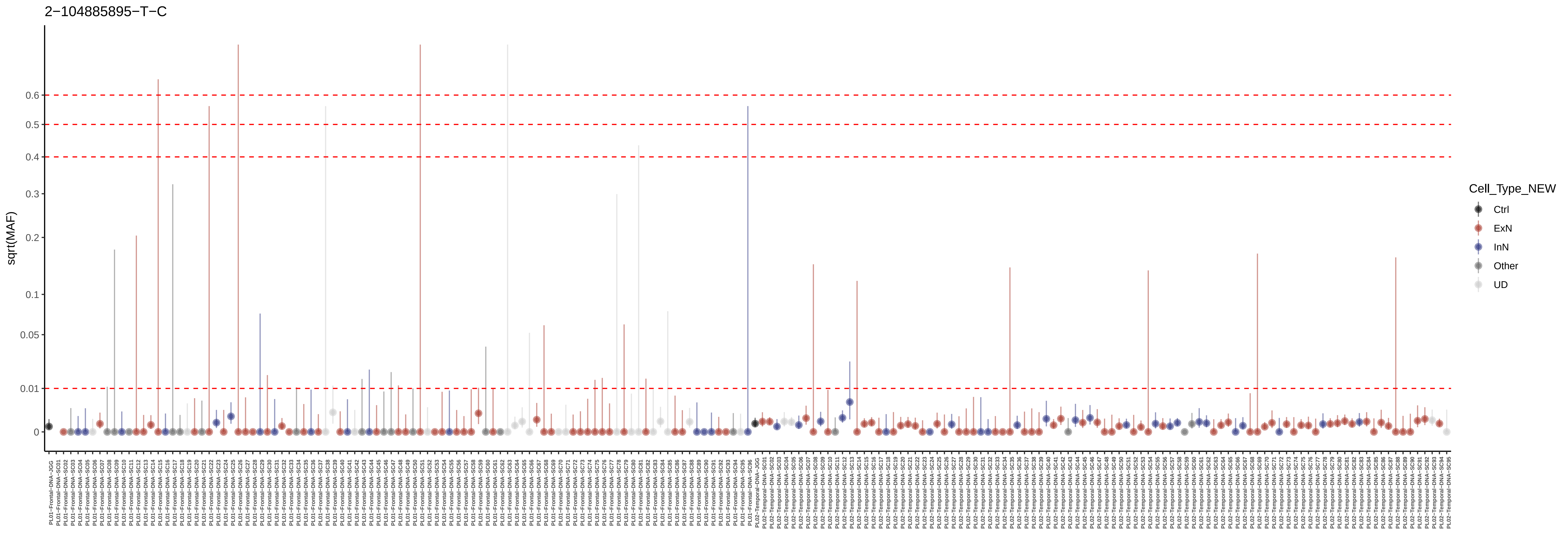

2−104885895−T−C
sqrt(MAF)
0.6
0.5
0.4
0.3
0.2
0.1
0.05
0.01
0
Cell_Type_NEW
Ctrl
ExN
InN
Other
UD

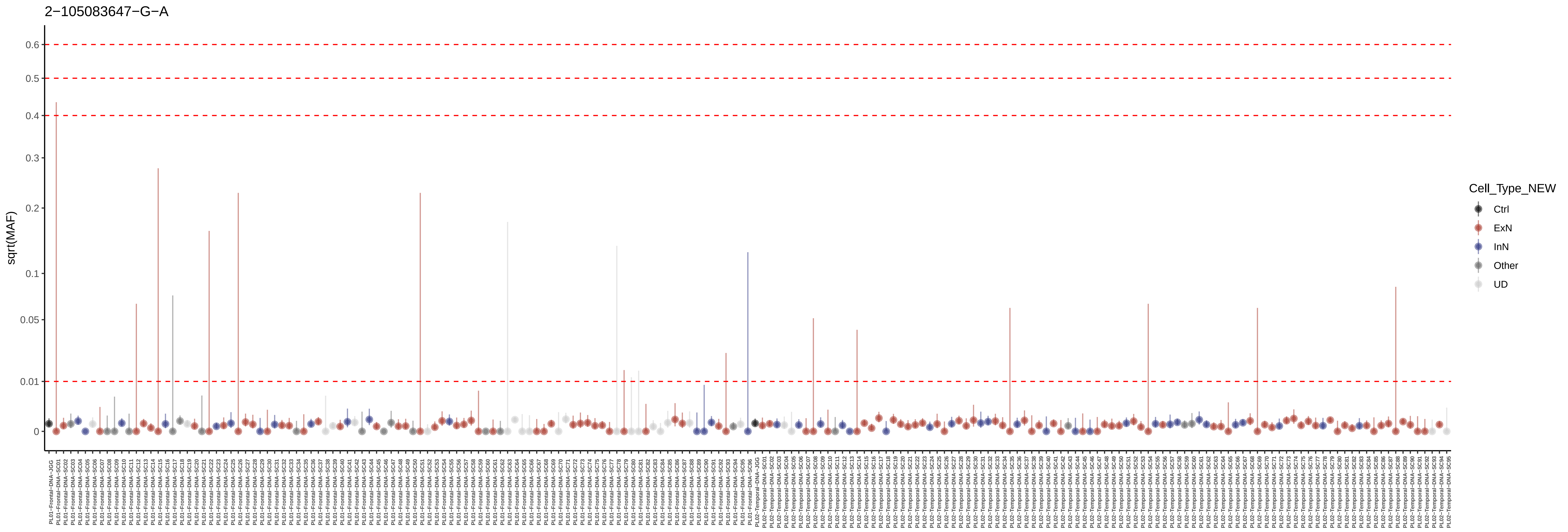
2−105083647−G−A
sqrt(MAF)
0.6
0.5
0.4
0.3
0.2
0.1
0.05
0.01
0
Cell_Type_NEW
Ctrl
ExN
InN
Other
UD

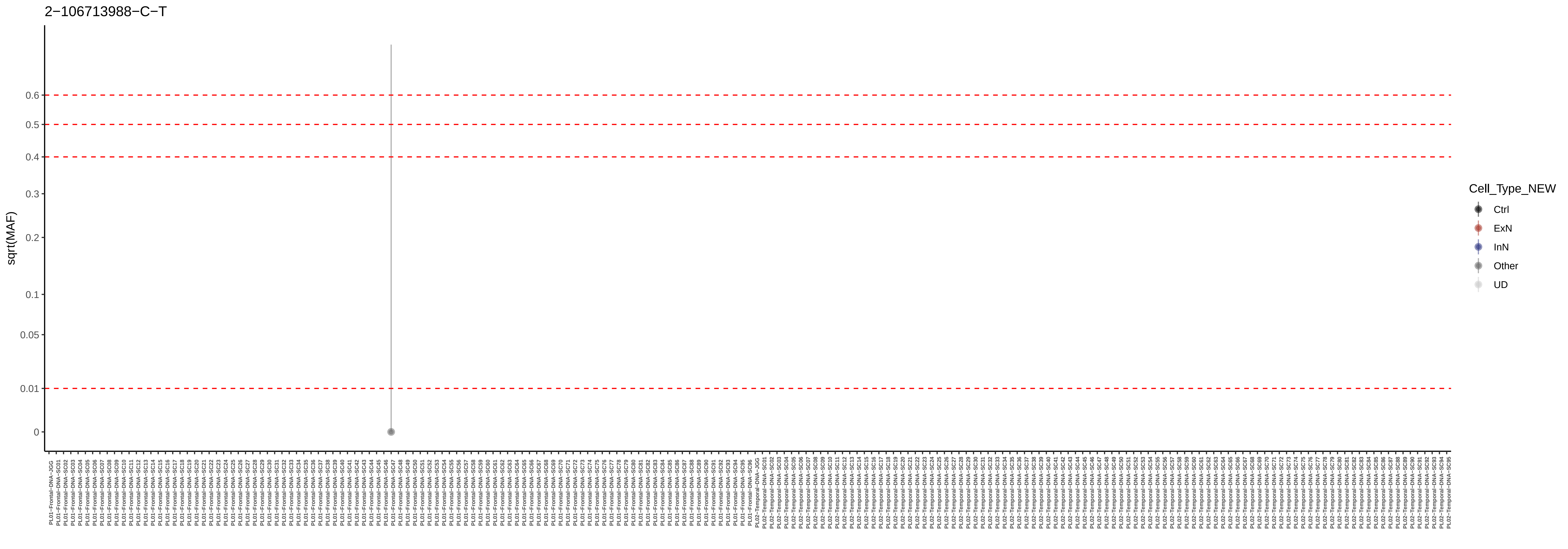
2−106713988−C−T
sqrt(MAF)
Cell_Type_NEW
Ctrl
ExN
InN
Other
UD

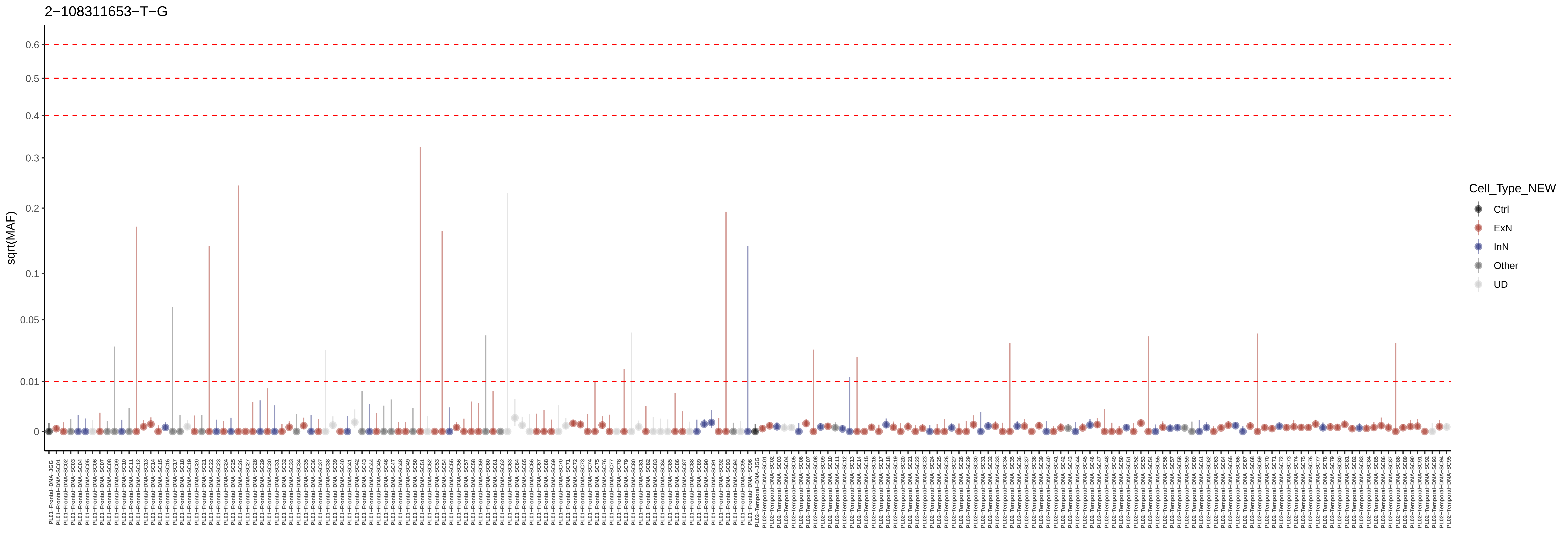
2−108311653−T−G
sqrt(MAF)
0.6
0.5
0.4
0.3
0.2
0.1
0.05
0.01
0
Cell_Type_NEW
Ctrl
ExN
InN
Other
UD

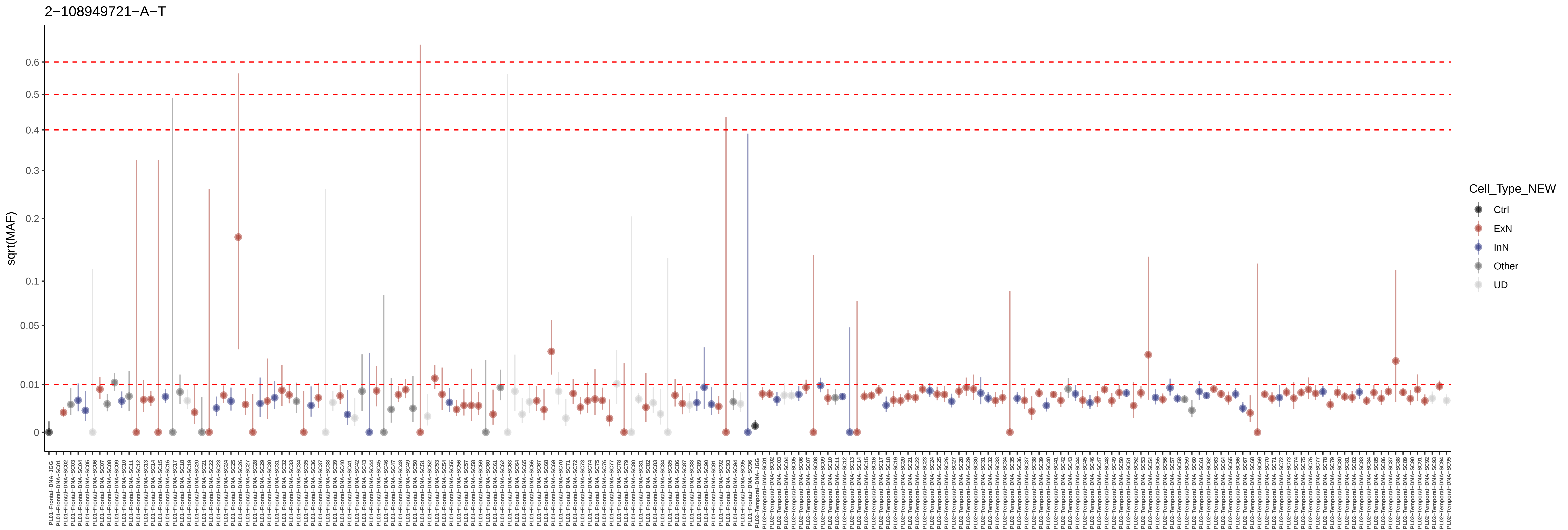

2−108949721−A−T
sqrt(MAF)
0.6
0.5
0.4
0.3
0.2
0.1
0.05
0.01
0
Cell_Type_NEW
Ctrl
ExN
InN
Other
UD

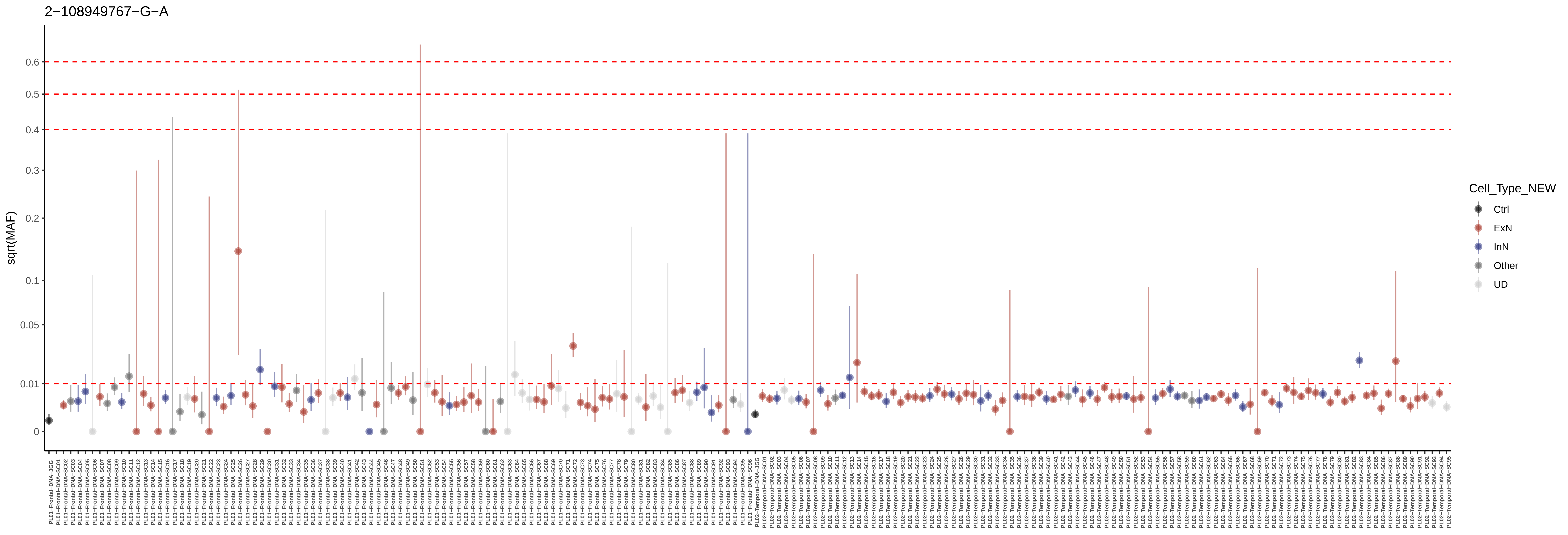
2−108949767−G−A
sqrt(MAF)
0.6
0.5
0.4
0.3
0.2
0.1
0.05
0.01
0
Cell_Type_NEW
Ctrl
ExN
InN
Other
UD

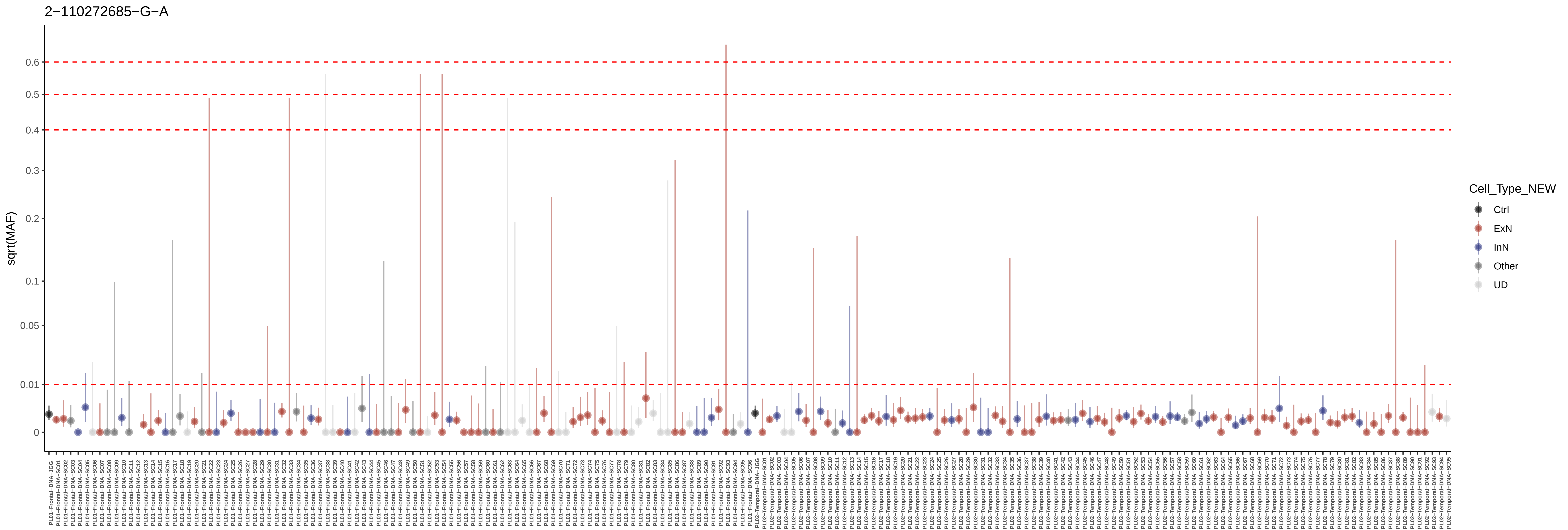
2−110272685−G−A
sqrt(MAF)
0.6
0.5
0.4
0.3
0.2
0.1
0.05
0.01
0
Cell_Type_NEW
Ctrl
ExN
InN
Other
UD

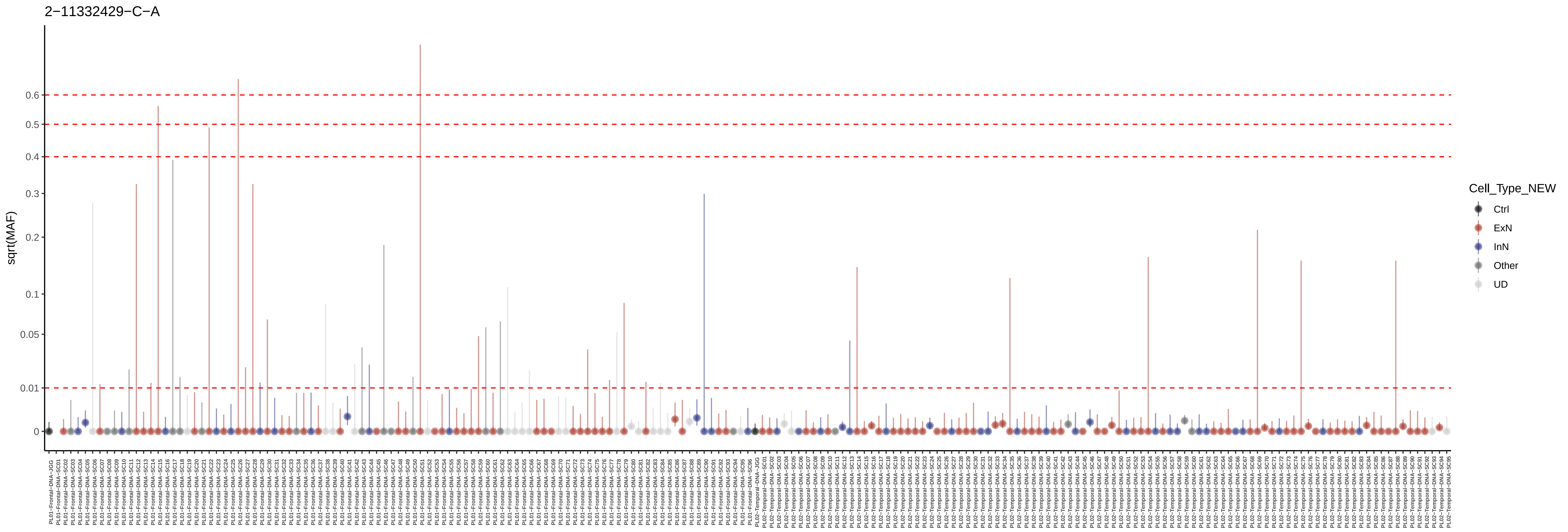
2−11332429−C−A
sqrt(MAF)
0.6
0.5
0.4
0.3
0.2
0.1
0.05
0.01
0
Cell_Type_NEW
Ctrl
ExN
InN
Other
UD

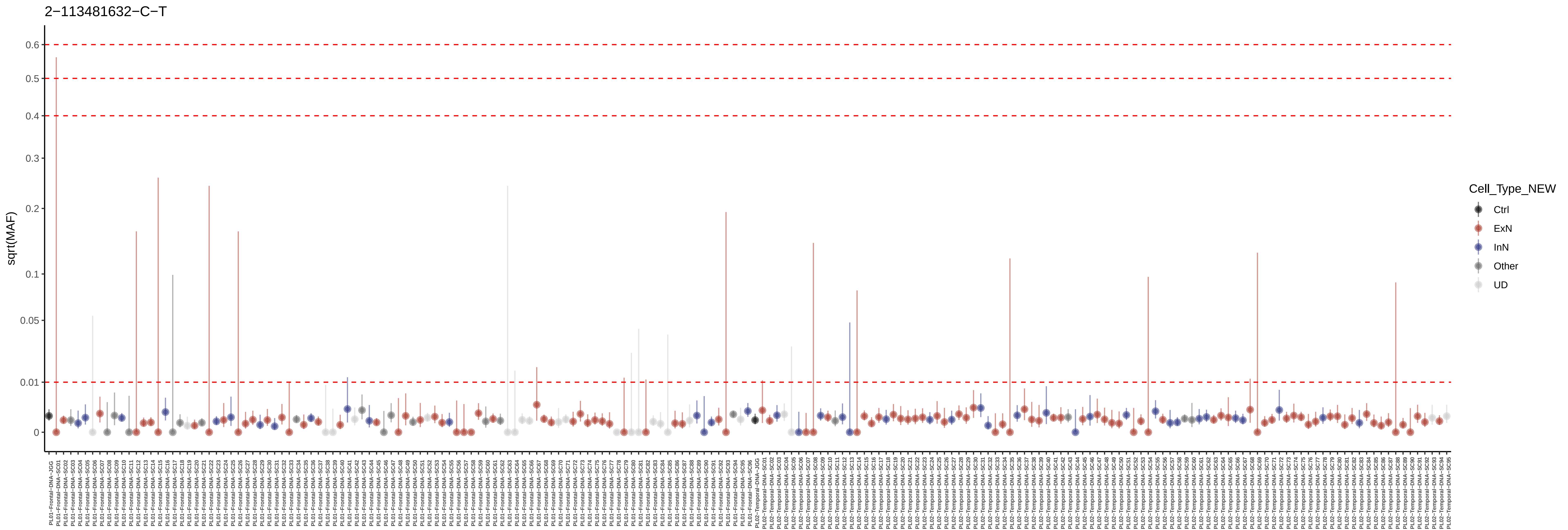

2−113481632−C−T
sqrt(MAF)
0.6
0.5
0.4
0.3
0.2
0.1
0.05
0.01
0
Cell_Type_NEW
Ctrl
ExN
InN
Other
UD

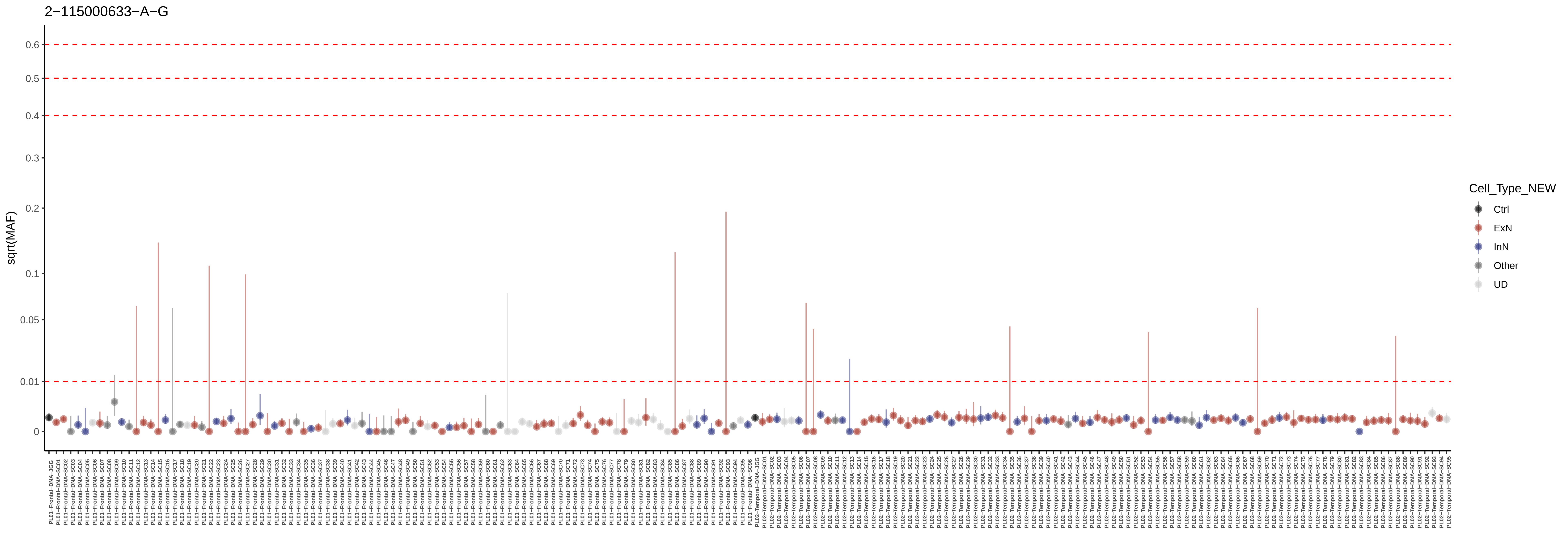
2−115000633−A−G
sqrt(MAF)
0.6
0.5
0.4
0.3
0.2
0.1
0.05
0.01
0
Cell_Type_NEW
Ctrl
ExN
InN
Other
UD

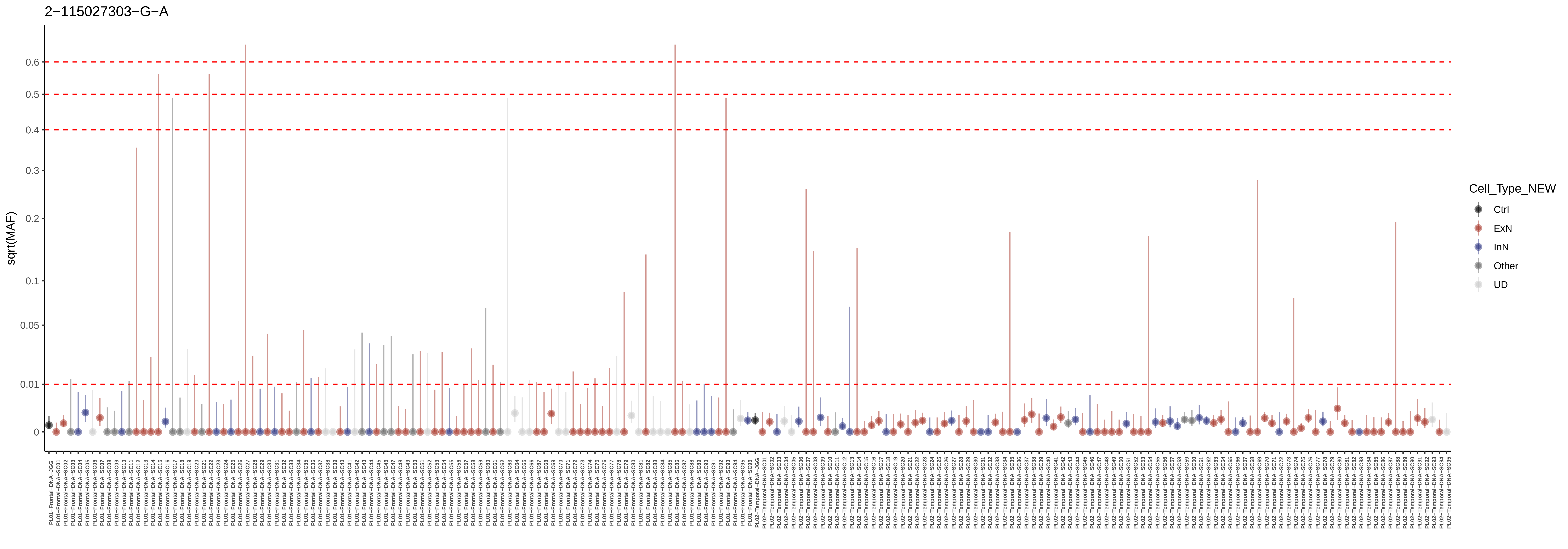
2−115027303−G−A
sqrt(MAF)
0.6
0.5
0.4
0.3
0.2
0.1
0.05
0.01
0
Cell_Type_NEW
Ctrl
ExN
InN
Other
UD

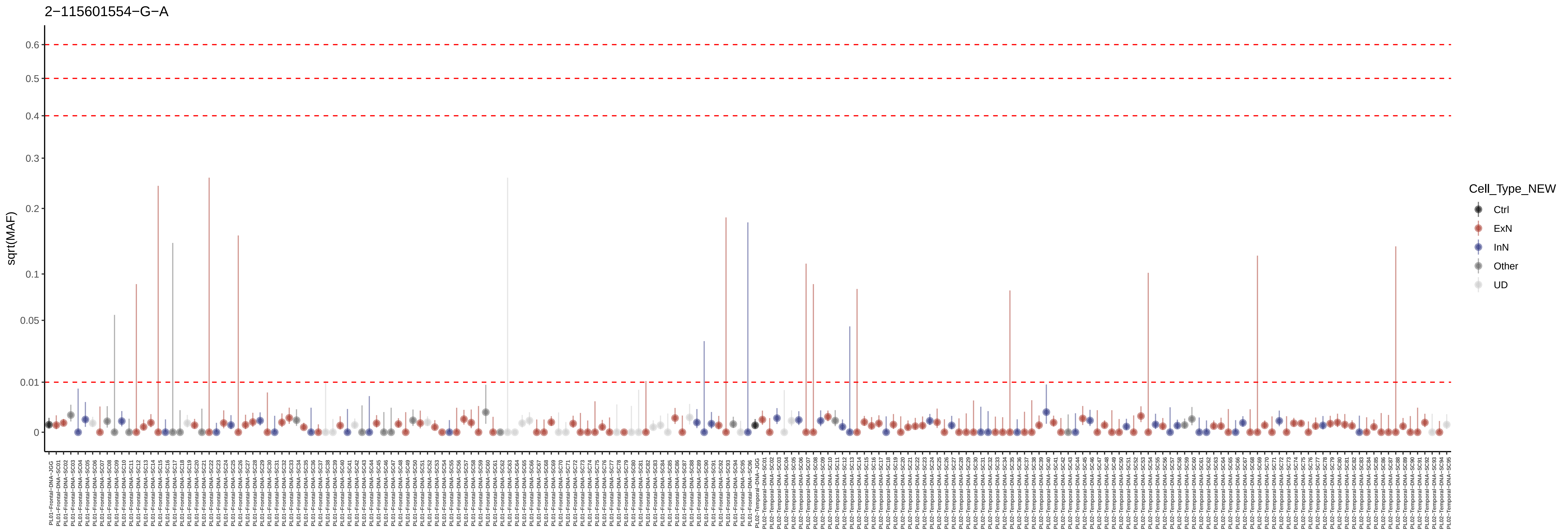
2−115601554−G−A
sqrt(MAF)
0.6
0.5
0.4
0.3
0.2
0.1
0.05
0.01
0
Cell_Type_NEW
Ctrl
ExN
InN
Other
UD

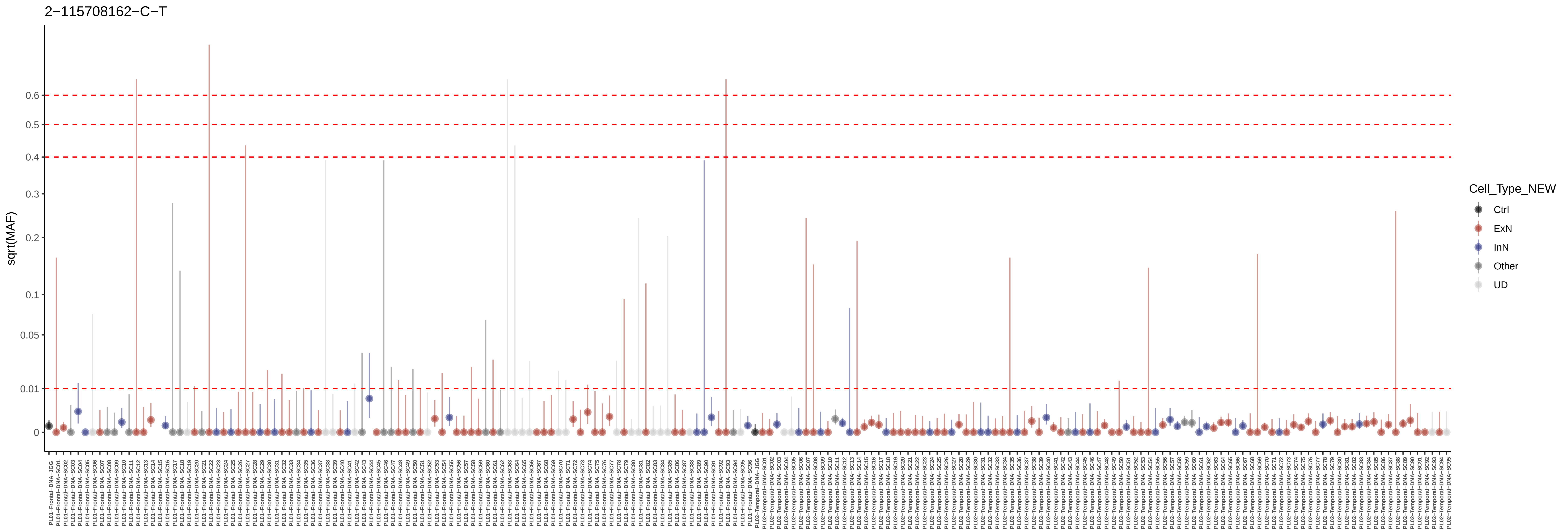

2−115708162−C−T
sqrt(MAF)
0.6
0.5
0.4
0.3
0.2
0.1
0.05
0.01
0
Cell_Type_NEW
Ctrl
ExN
InN
Other
UD

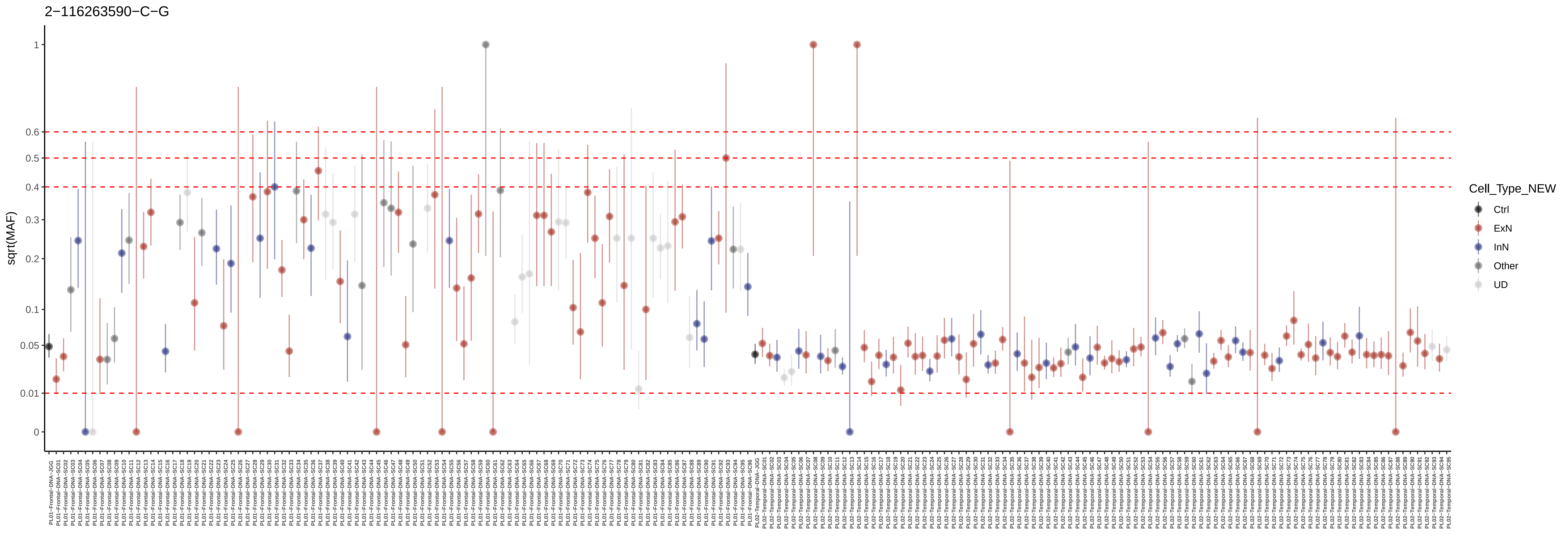
2−116263590−C−G
sqrt(MAF)
1
0.6
0.5
0.4
0.3
0.2
0.1
0.05
0.01
0
Cell_Type_NEW
Ctrl
ExN
InN
Other
UD

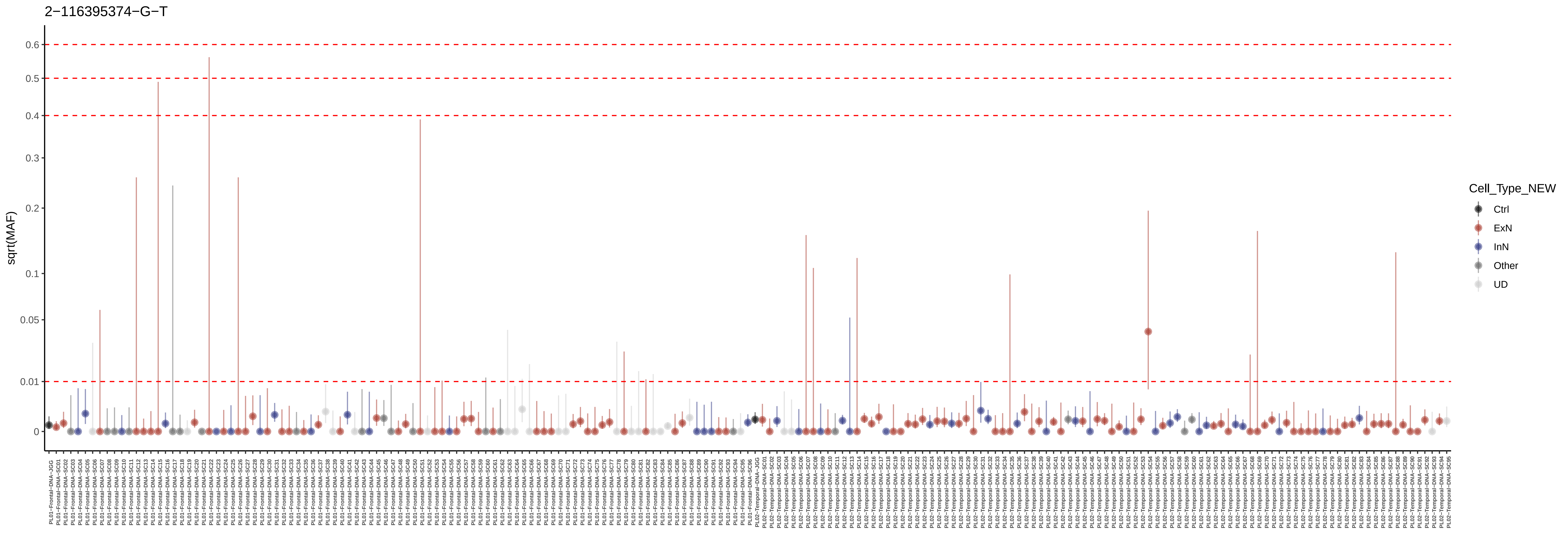
2−116395374−G−T
sqrt(MAF)
0.6
0.5
0.4
0.3
0.2
0.1
0.05
0.01
0
Cell_Type_NEW
Ctrl
ExN
InN
Other
UD

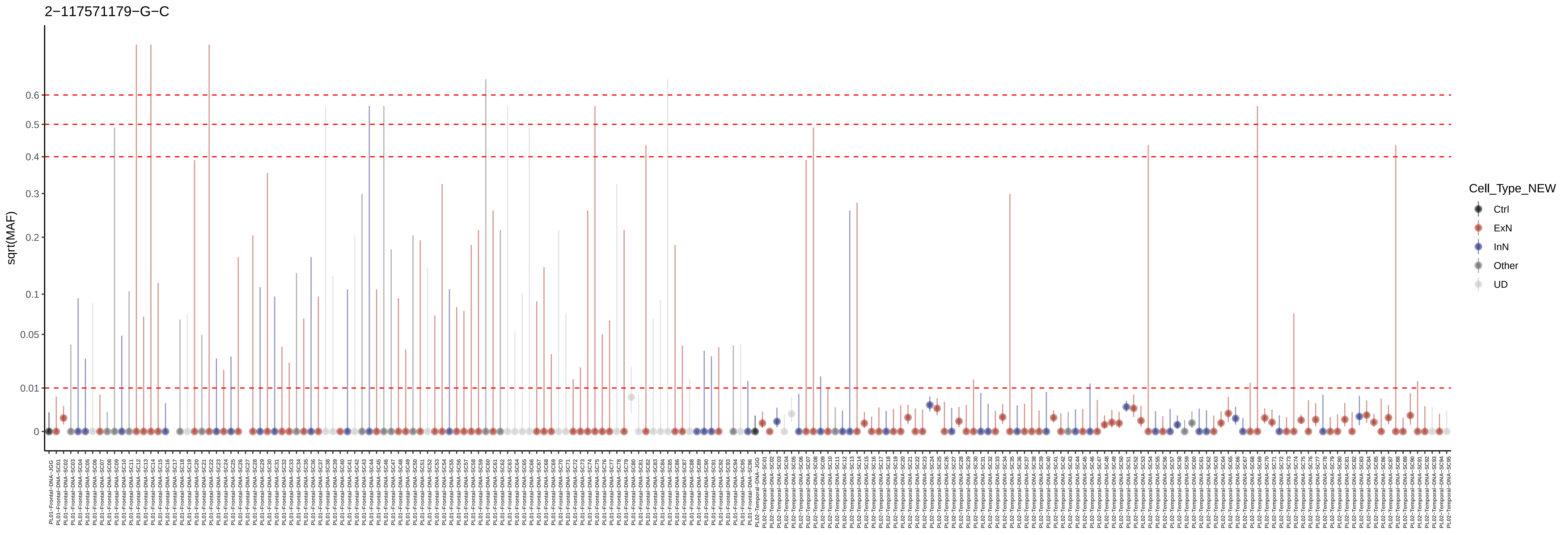
2−117571179−G−C
sqrt(MAF)
0.6
0.5
0.4
0.3
0.2
0.1
0.05
0.01
0
Cell_Type_NEW
Ctrl
ExN
InN
Other
UD

2−117571183−C−A

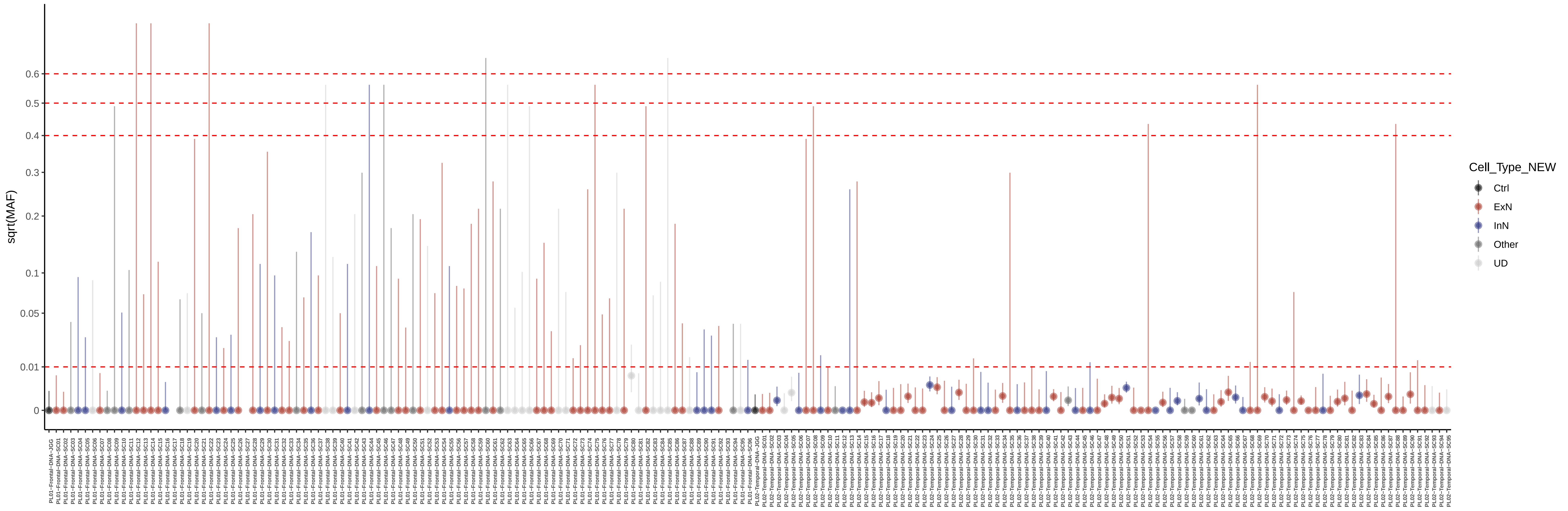

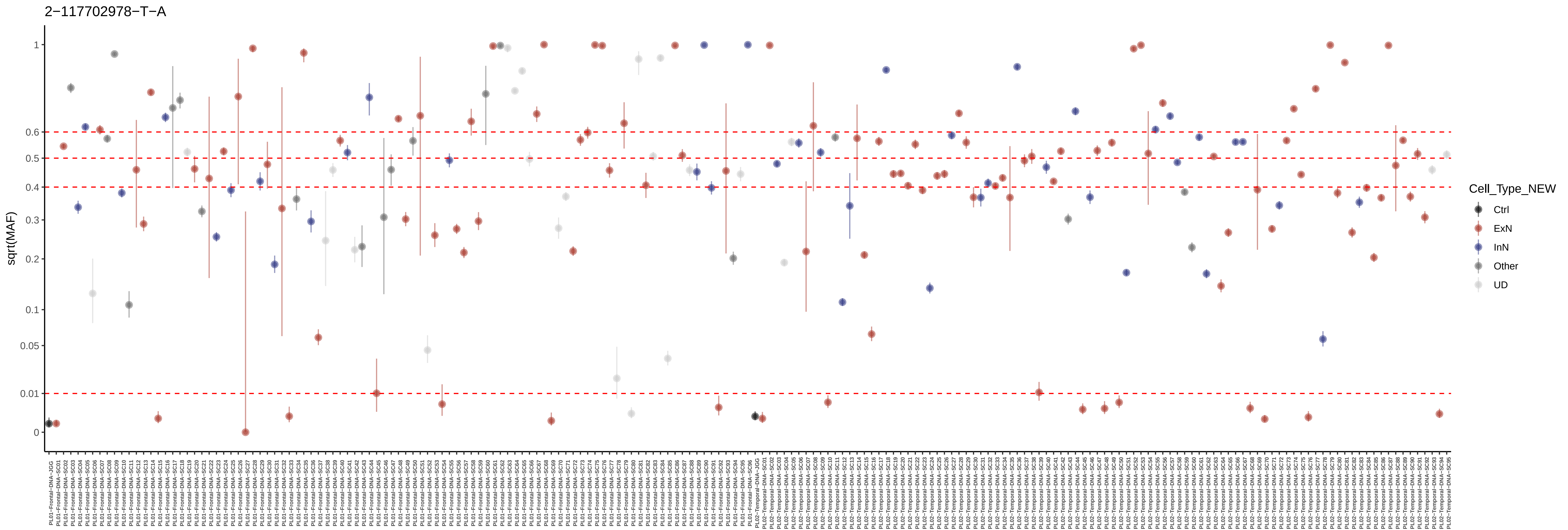
2−117702978−T−A
sqrt(MAF)
1
0.6
0.5
0.4
0.3
0.2
0.1
0.05
0.01
0
Cell_Type_NEW
Ctrl
ExN
InN
Other
UD

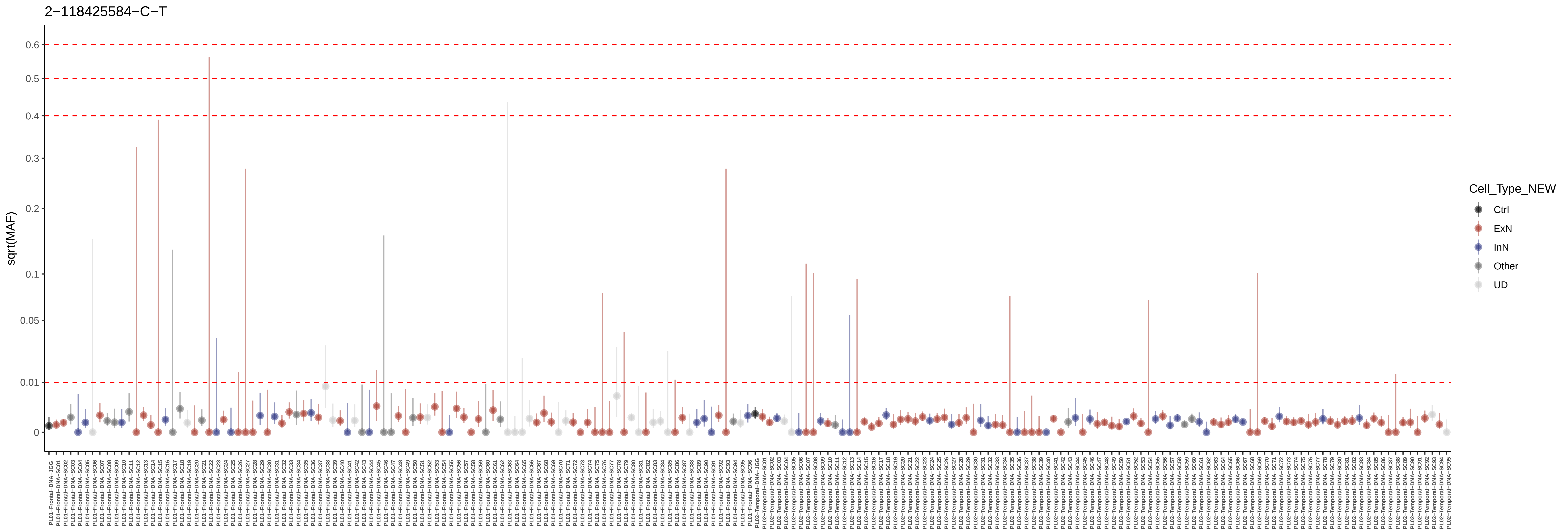
2−118425584−C−T
sqrt(MAF)
0.6
0.5
0.4
0.3
0.2
0.1
0.05
0.01
0
Cell_Type_NEW
Ctrl
ExN
InN
Other
UD

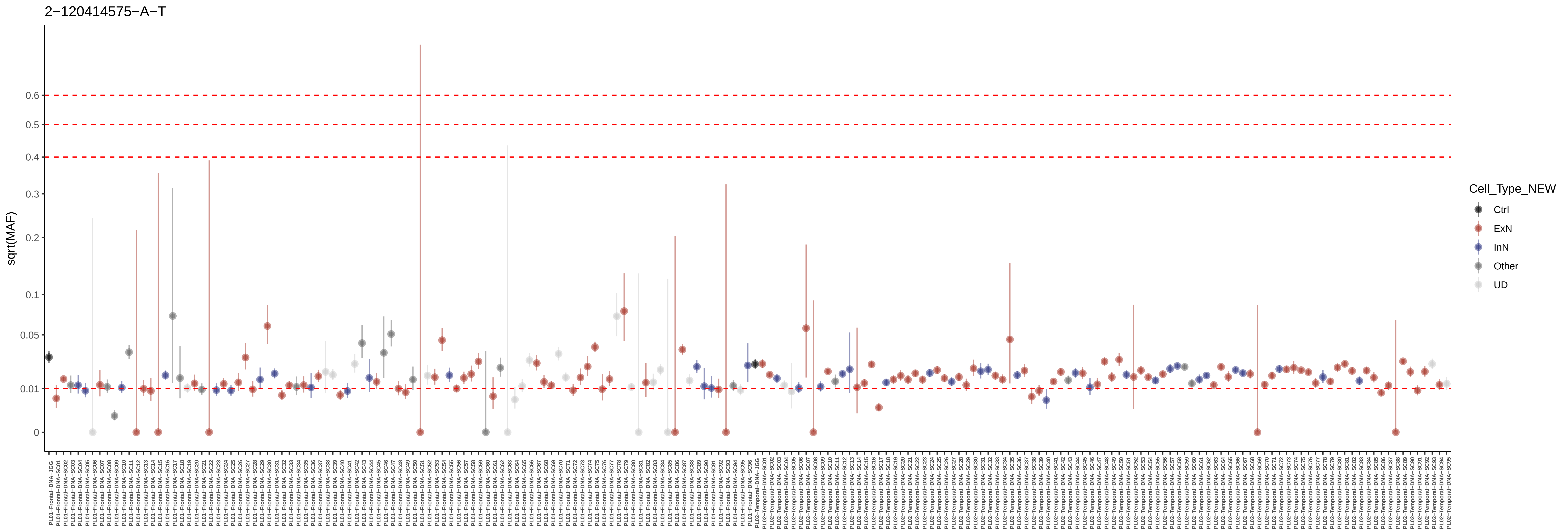
2−120414575−A−T
sqrt(MAF)
0.6
0.5
0.4
0.3
0.2
0.1
0.05
0.01
0
Cell_Type_NEW
Ctrl
ExN
InN
Other
UD

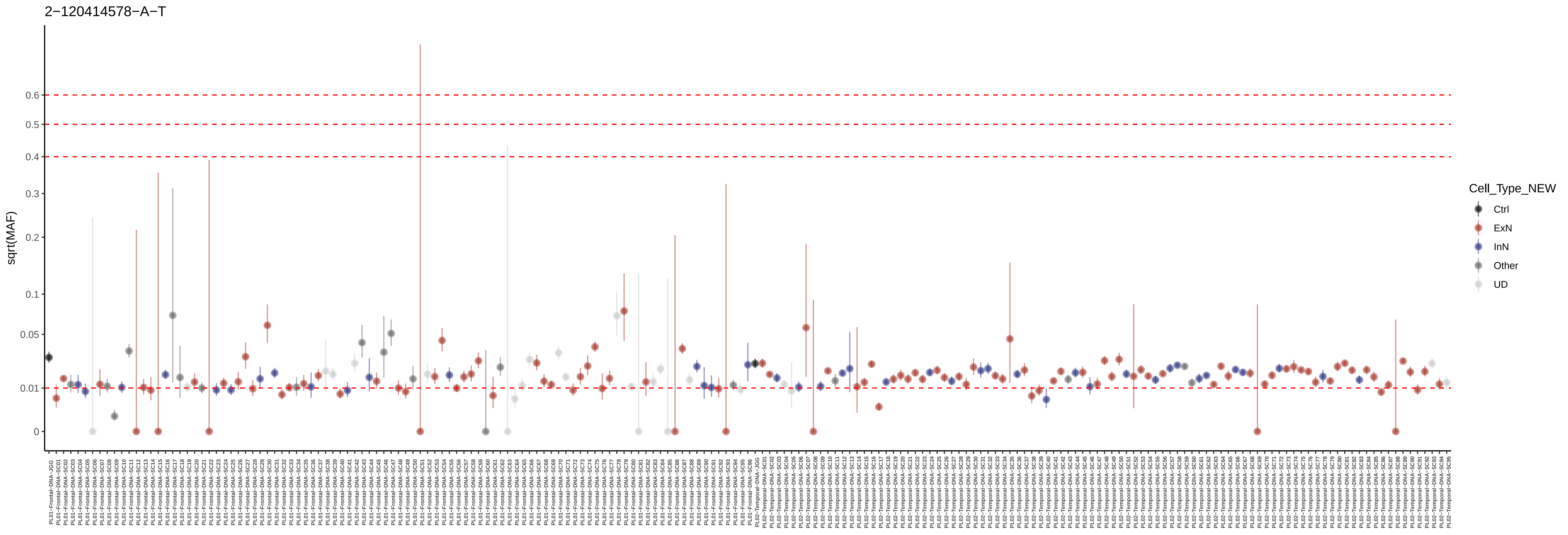
2−120414578−A−T
sqrt(MAF)
0.6
0.5
0.4
0.3
0.2
0.1
0.05
0.01
0
Cell_Type_NEW
Ctrl
ExN
InN
Other
UD

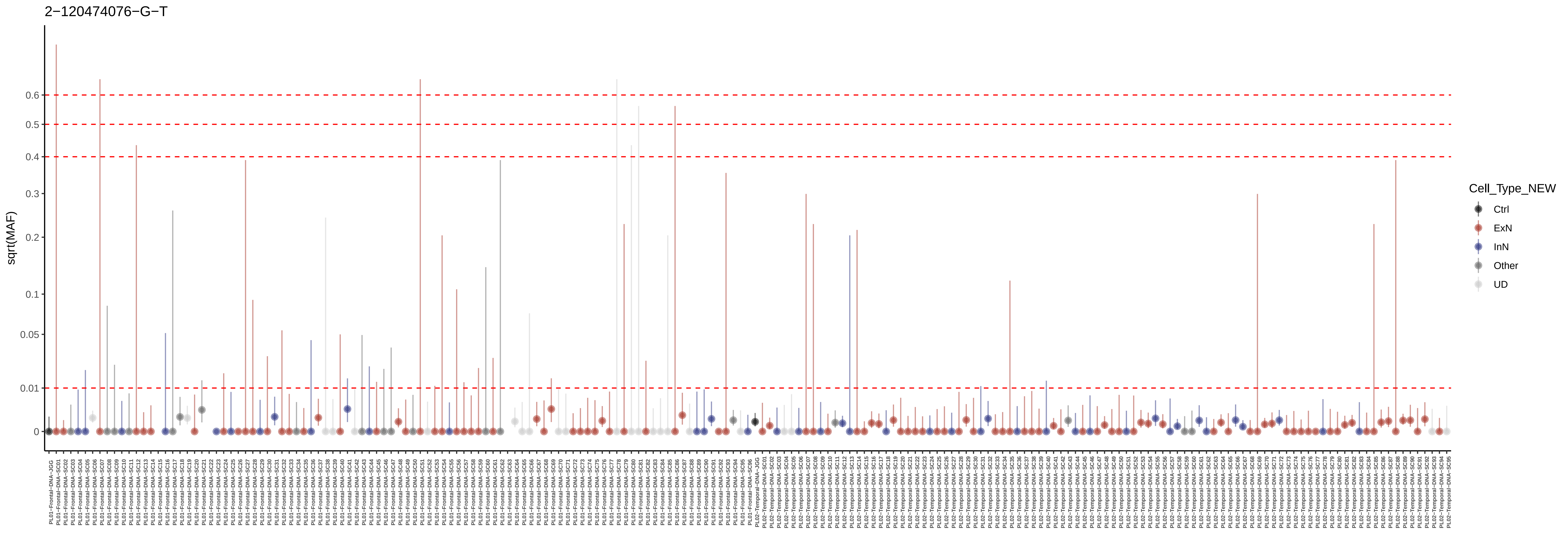
2−120474076−G−T
sqrt(MAF)
0.6
0.5
0.4
0.3
0.2
0.1
0.05
0.01
0
Cell_Type_NEW
Ctrl
ExN
InN
Other
UD

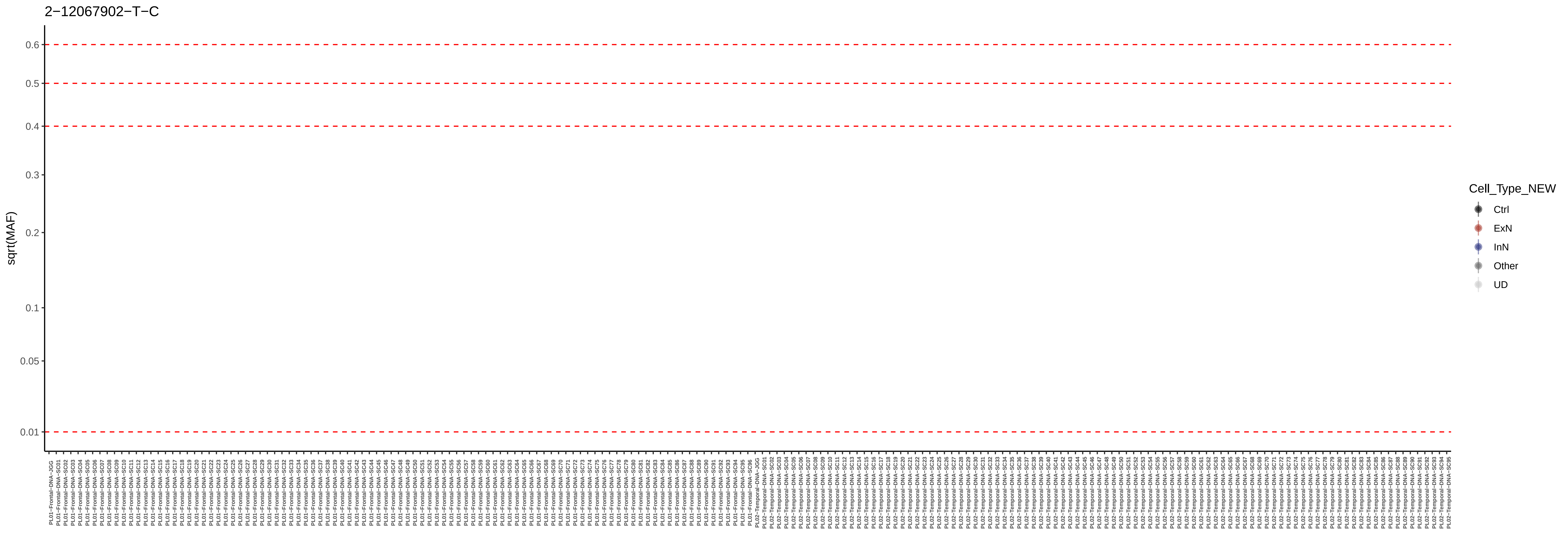
2−12067902−T−C
sqrt(MAF)
0.6
0.5
0.4
0.3
0.2
0.1
0.05
0.01
Cell_Type_NEW
Ctrl
ExN
InN
Other
UD

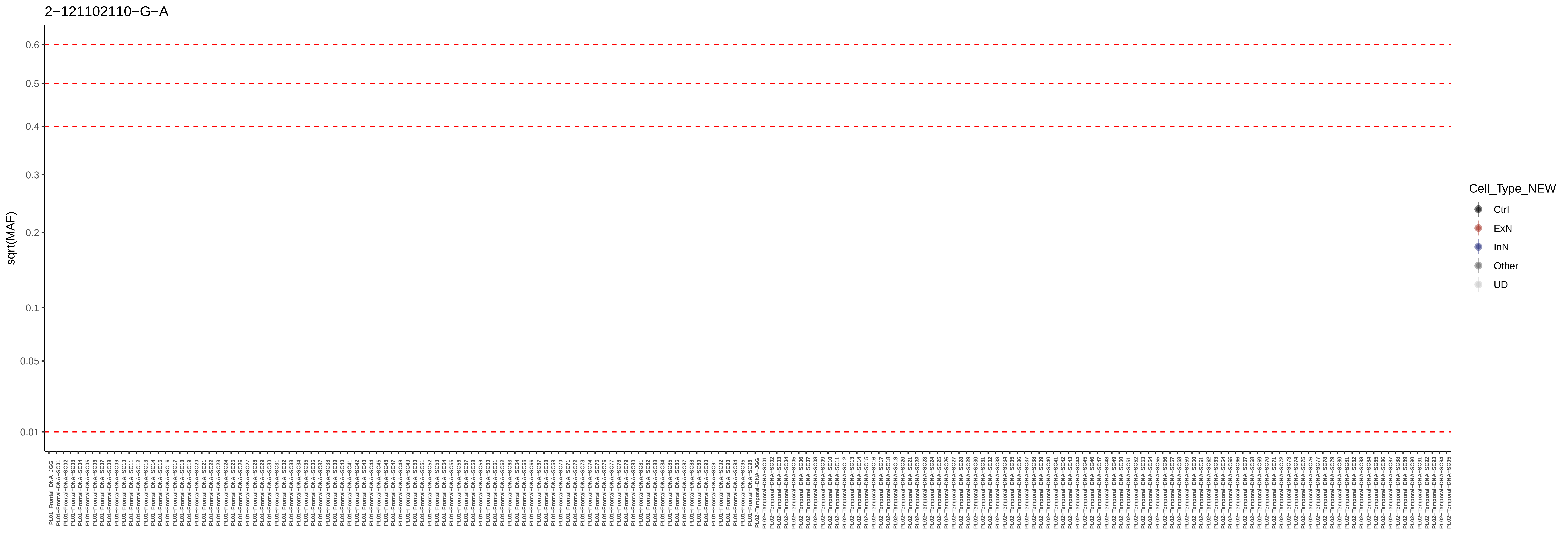
2−121102110−G−A
sqrt(MAF)
0.6
0.5
0.4
0.3
0.2
0.1
0.05
0.01
Cell_Type_NEW
Ctrl
ExN
InN
Other
UD

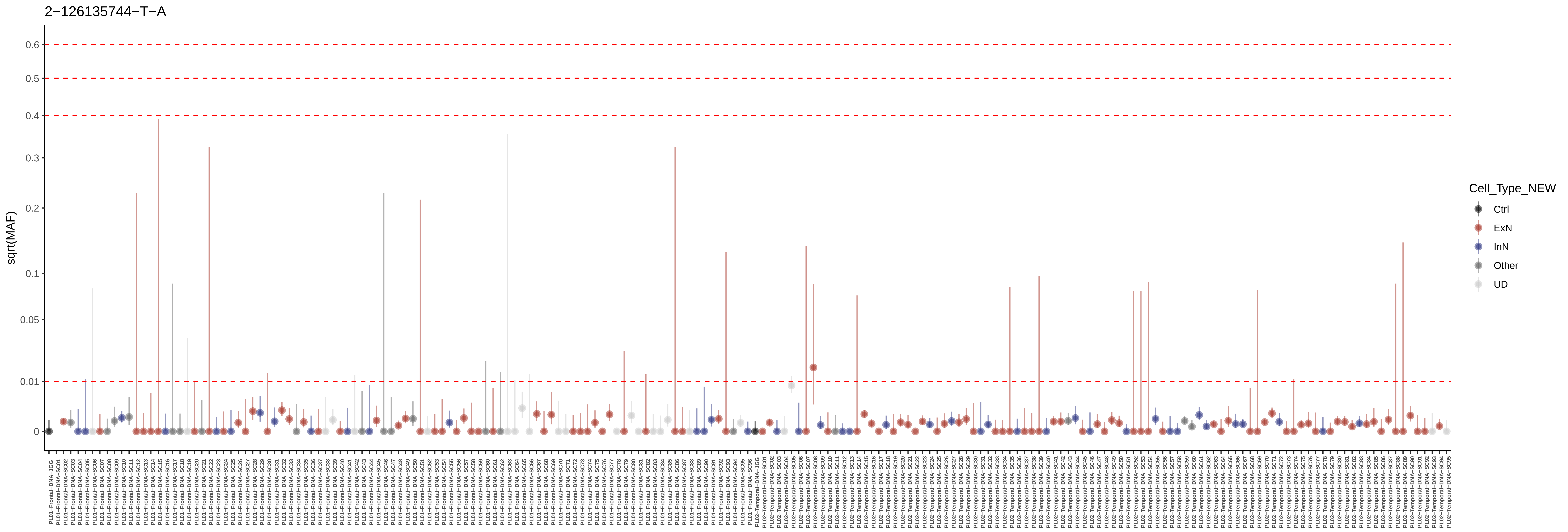

2−126135744−T−A
sqrt(MAF)
0.6
0.5
0.4
0.3
0.2
0.1
0.05
0.01
0
Cell_Type_NEW
Ctrl
ExN
InN
Other
UD

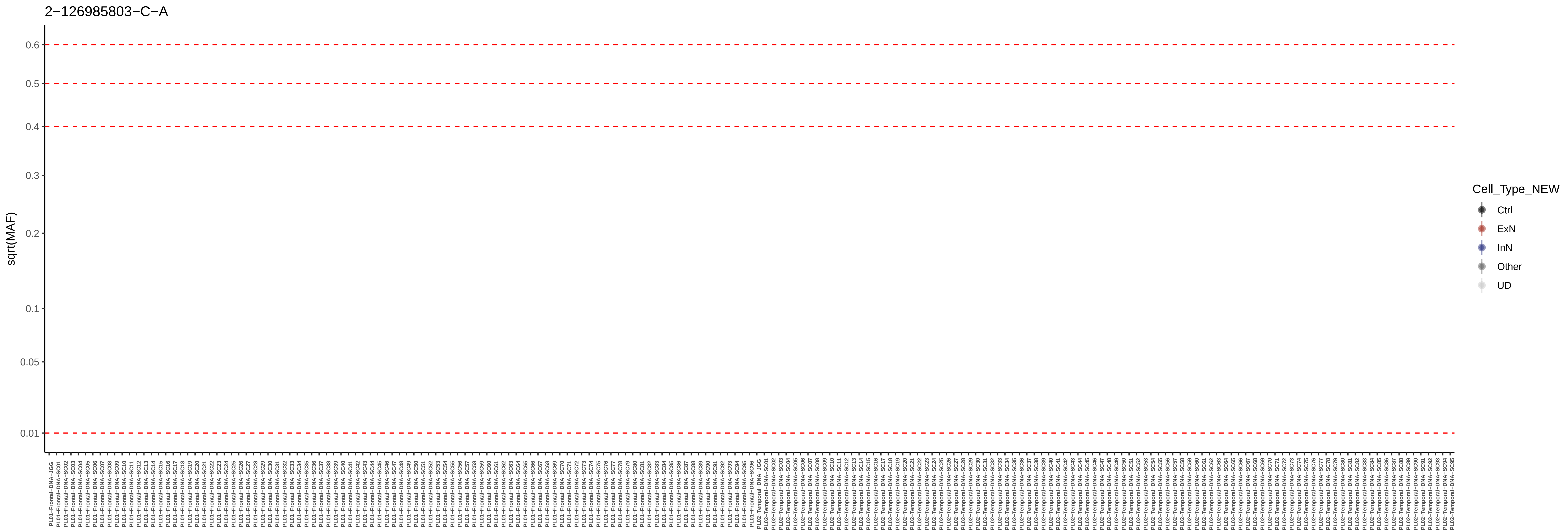

2−126985803−C−A
sqrt(MAF)
0.6
0.5
0.4
0.3
0.2
0.1
0.05
0.01
Cell_Type_NEW
Ctrl
ExN
InN
Other
UD

2−127028557−G−A

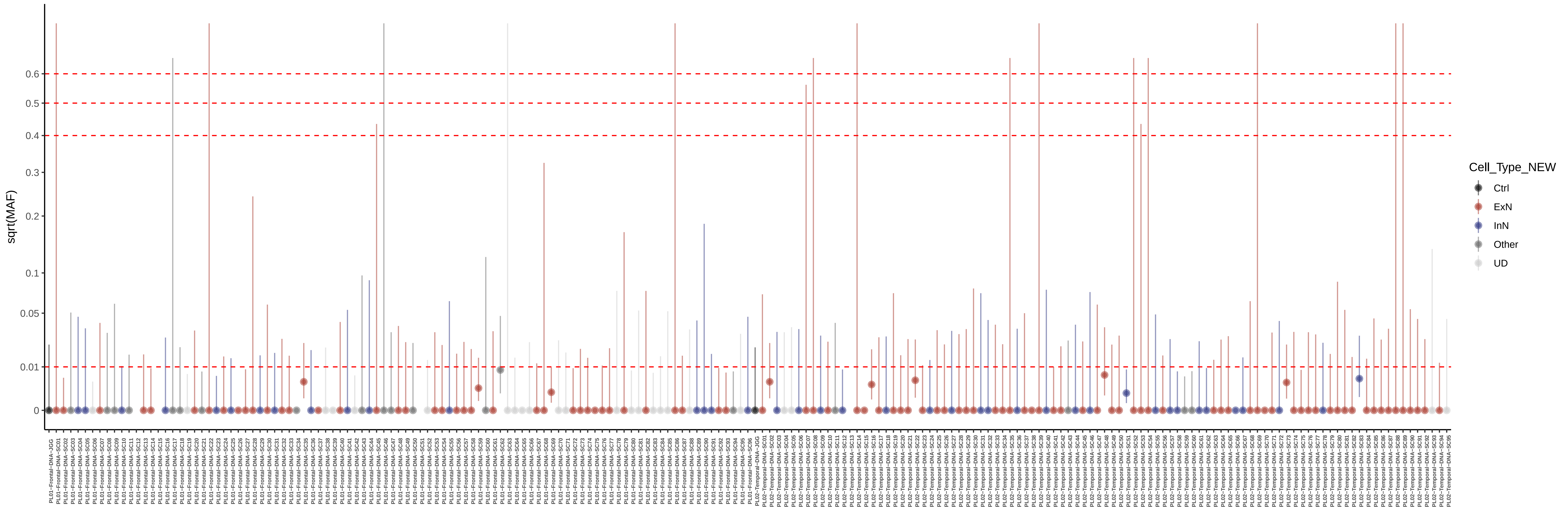

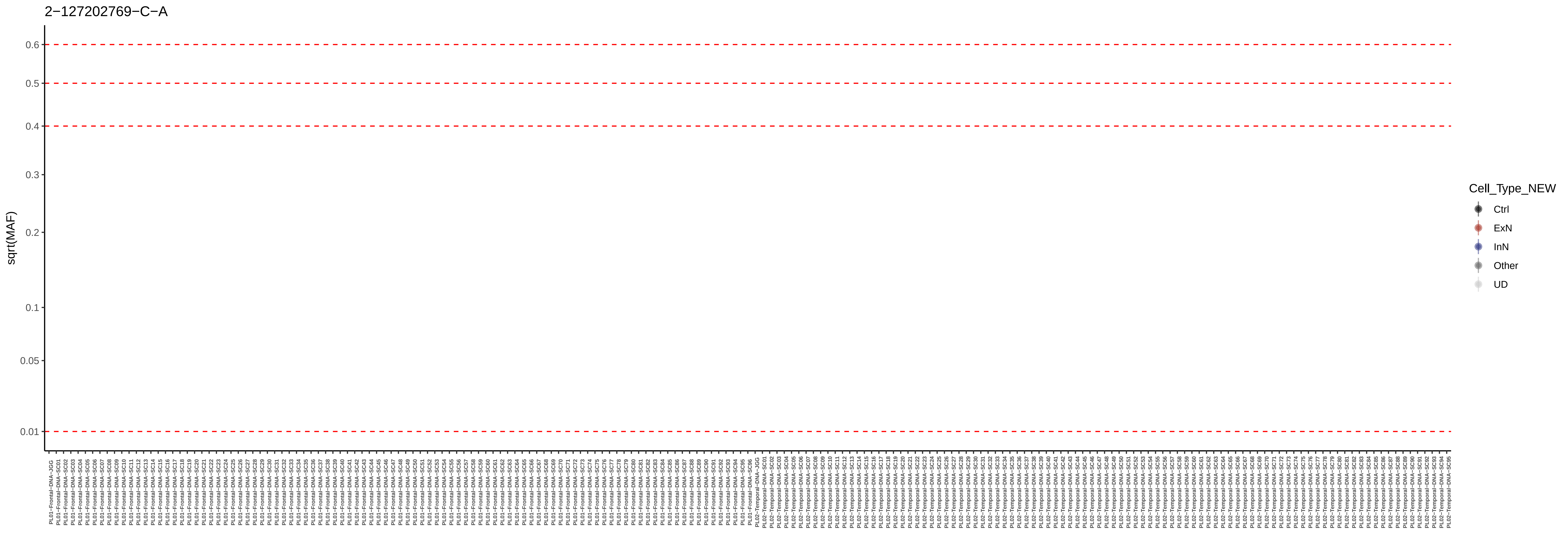
2−127202769−C−A
sqrt(MAF)
0.6
0.5
0.4
0.3
0.2
0.1
0.05
0.01
Cell_Type_NEW
Ctrl
ExN
InN
Other
UD

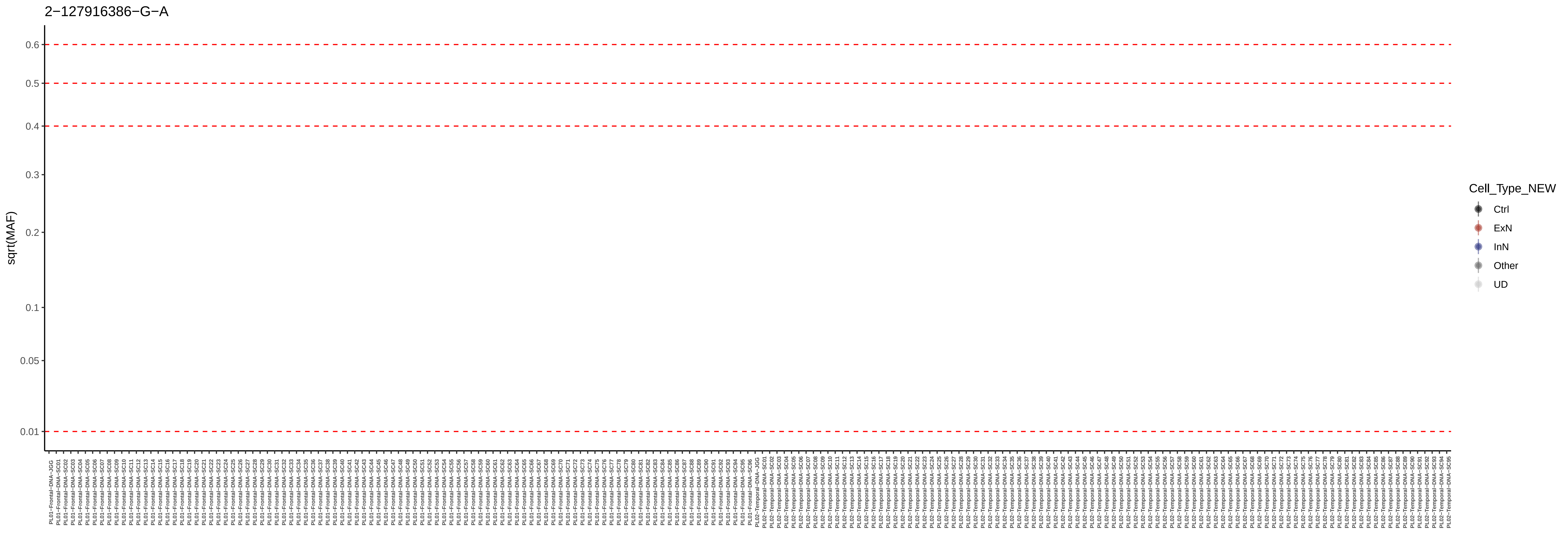
2−127916386−G−A
sqrt(MAF)
0.6
0.5
0.4
0.3
0.2
0.1
0.05
0.01
Cell_Type_NEW
Ctrl
ExN
InN
Other
UD

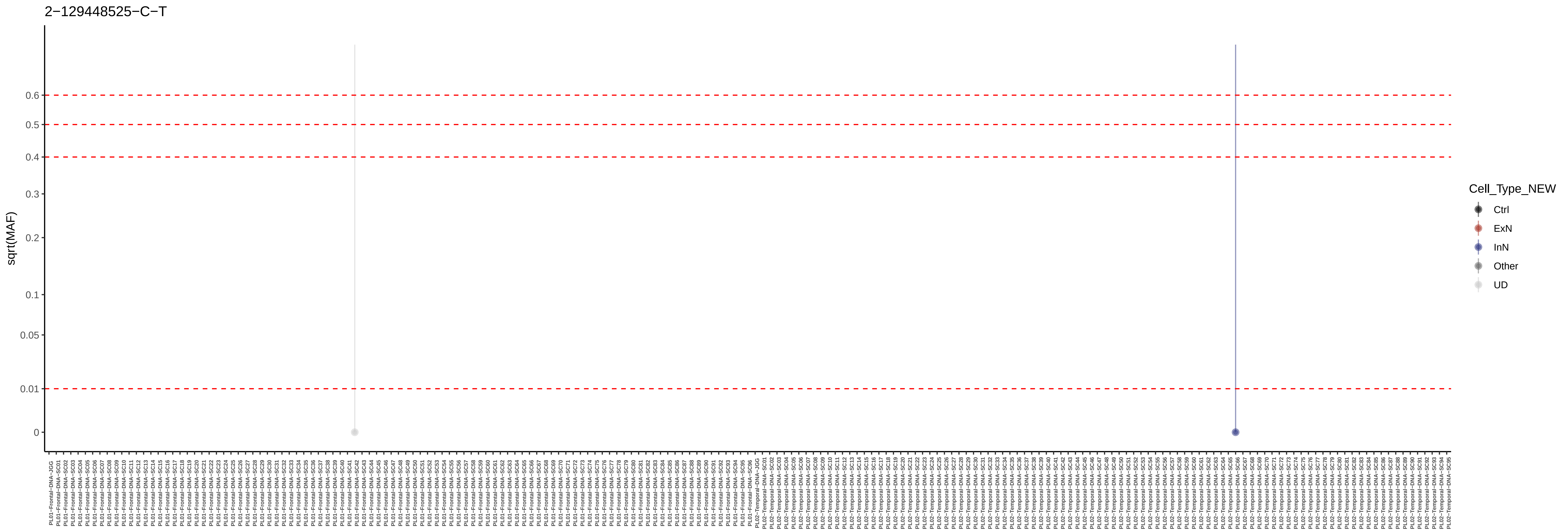
2−129448525−C−T
sqrt(MAF)
0.6
0.5
0.4
0.3
0.2
0.1
0.05
0.01
0
Cell_Type_NEW
Ctrl
ExN
InN
Other
UD

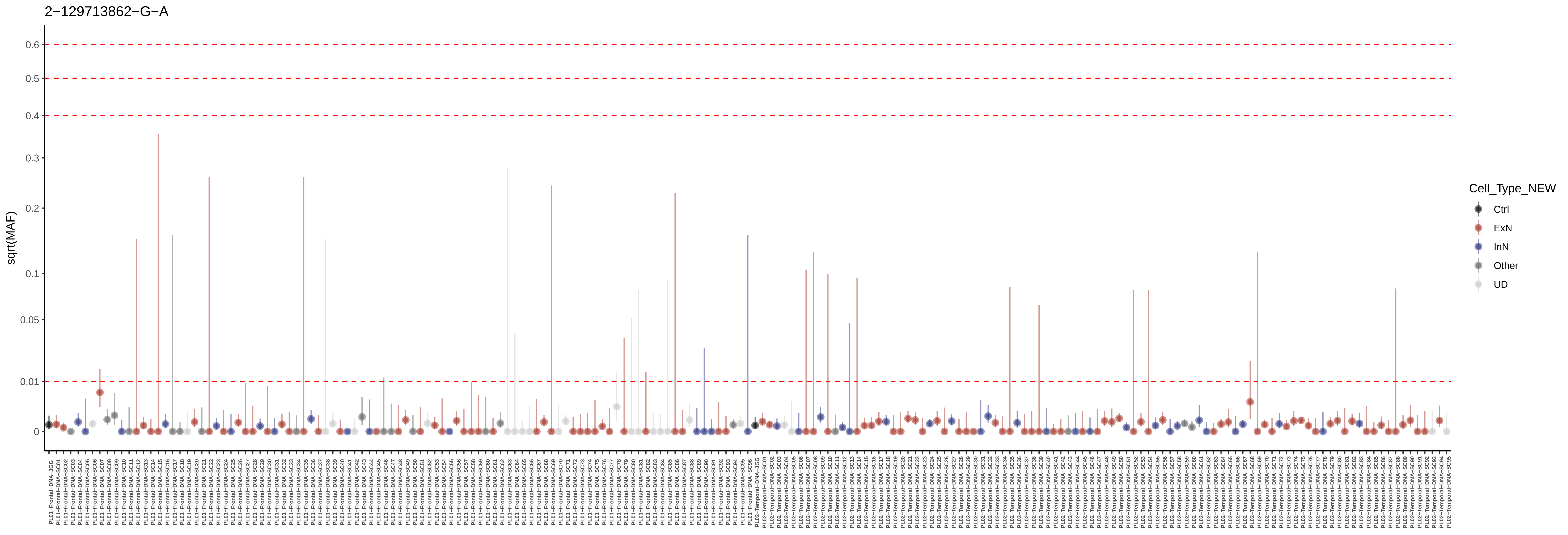

2−129713862−G−A
sqrt(MAF)
0.6
0.5
0.4
0.3
0.2
0.1
0.05
0.01
0
Cell_Type_NEW
Ctrl
ExN
InN
Other
UD

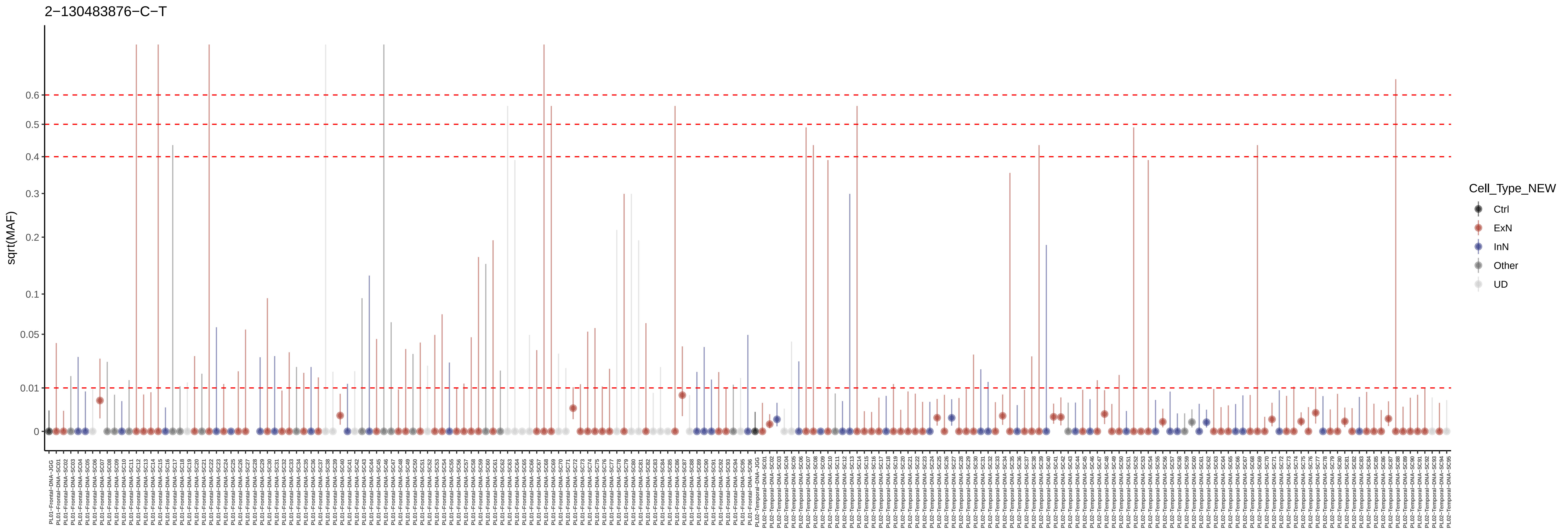
2−130483876−C−T
sqrt(MAF)
0.6
0.5
0.4
0.3
0.2
0.1
0.05
0.01
0
Cell_Type_NEW
Ctrl
ExN
InN
Other
UD

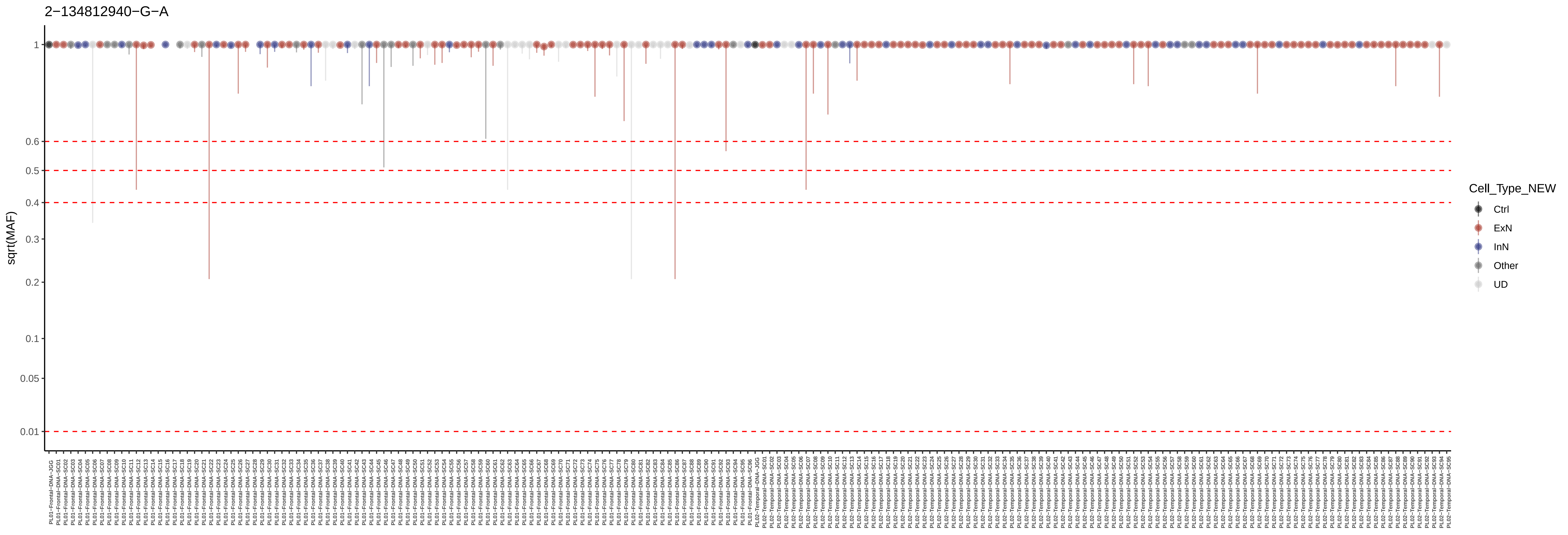

2−134812940−G−A
sqrt(MAF)
1
0.6
0.5
0.4
0.3
0.2
0.1
0.05
0.01
Cell_Type_NEW
Ctrl
ExN
InN
Other
UD

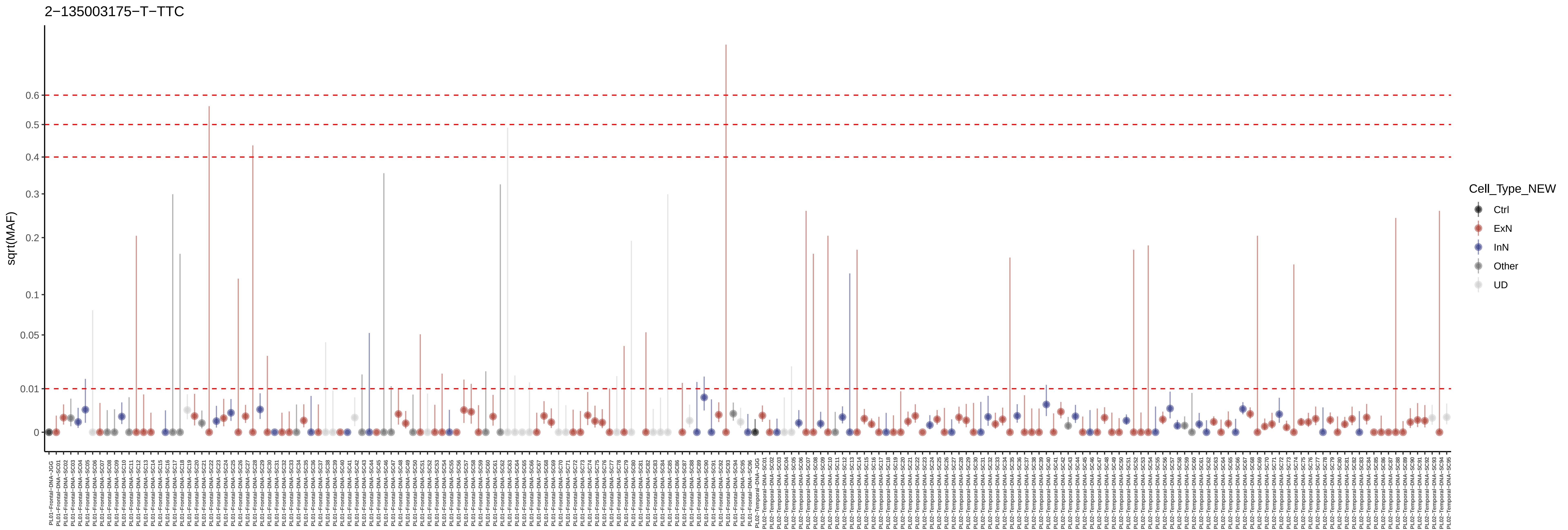
2−135003175−T−TTC
sqrt(MAF)
0.6
0.5
0.4
0.3
0.2
0.1
0.05
0.01
0
Cell_Type_NEW
Ctrl
ExN
InN
Other
UD

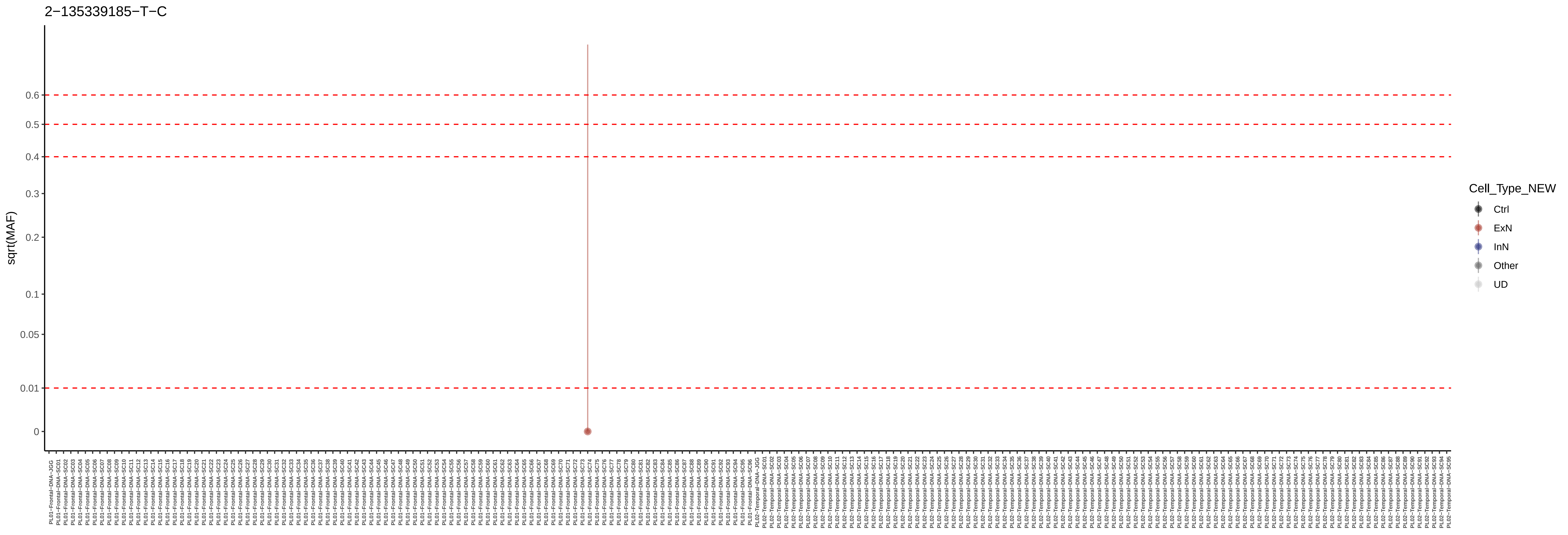
2−135339185−T−C
sqrt(MAF)
0.6
0.5
0.4
0.3
0.2
0.1
0.05
0.01
0
Cell_Type_NEW
Ctrl
ExN
InN
Other
UD

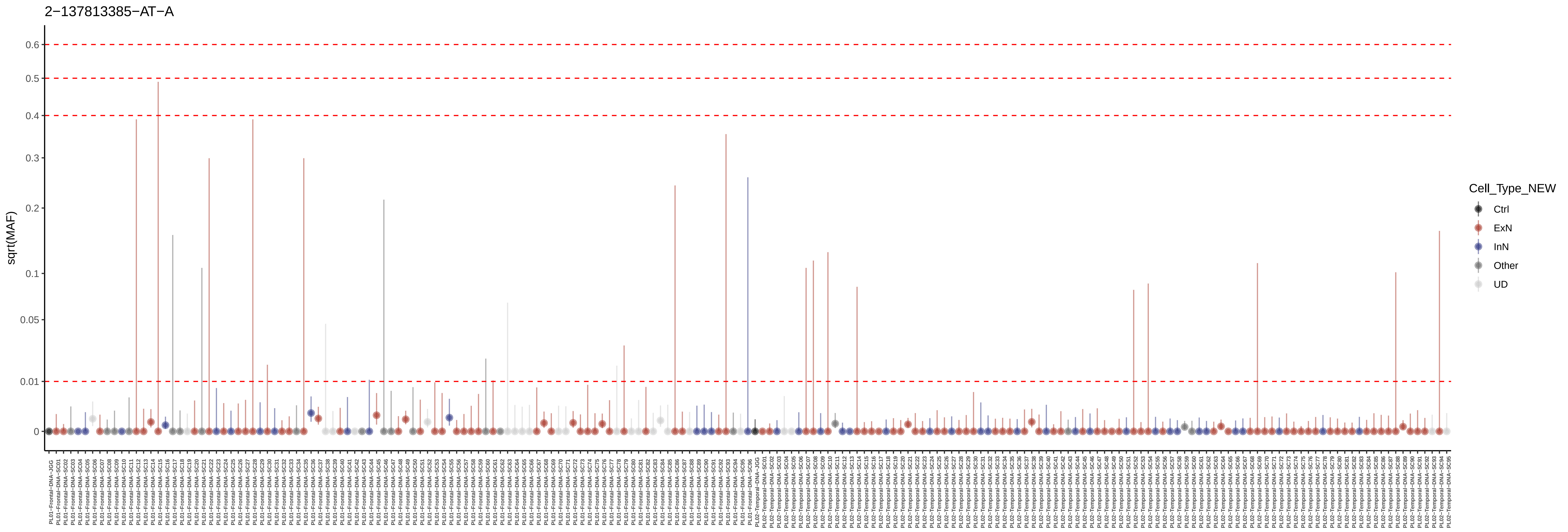
2−137813385−AT−A
sqrt(MAF)
0.6
0.5
0.4
0.3
0.2
0.1
0.05
0.01
0
Cell_Type_NEW
Ctrl
ExN
InN
Other
UD

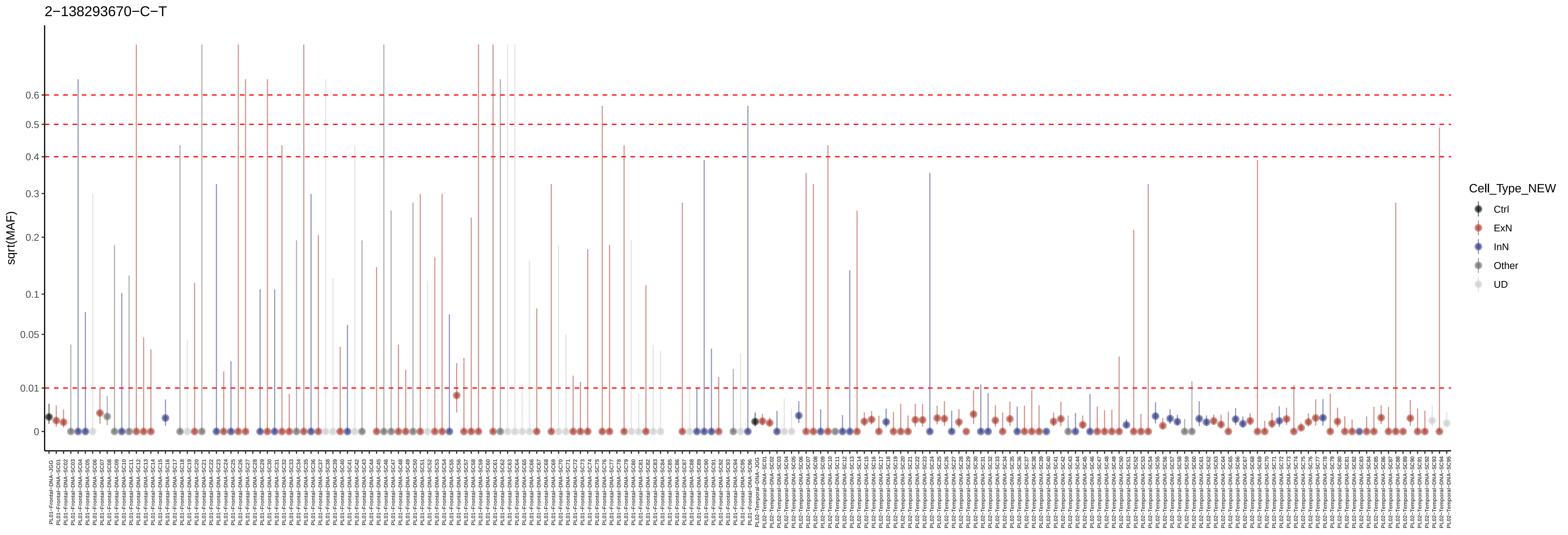
2−138293670−C−T
sqrt(MAF)
0.6
0.5
0.4
0.3
0.2
0.1
0.05
0.01
0
Cell_Type_NEW
Ctrl
ExN
InN
Other
UD

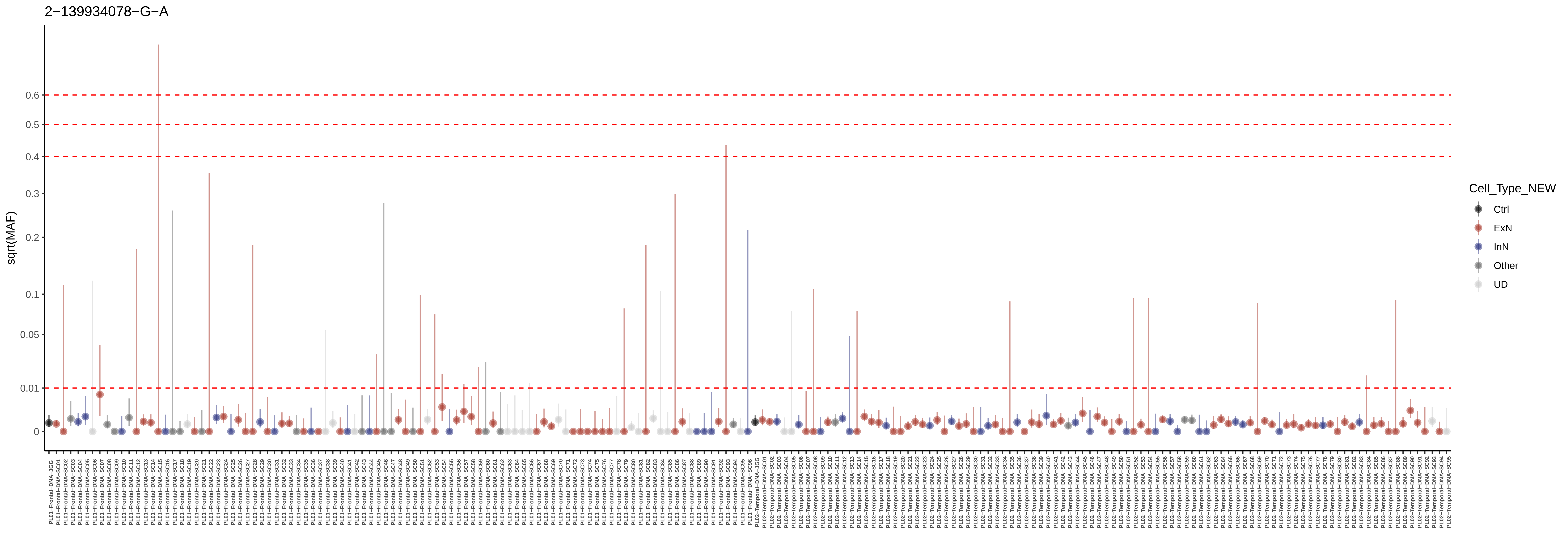

2−139934078−G−A
sqrt(MAF)
0.6
0.5
0.4
0.3
0.2
0.1
0.05
0.01
0
Cell_Type_NEW
Ctrl
ExN
InN
Other
UD

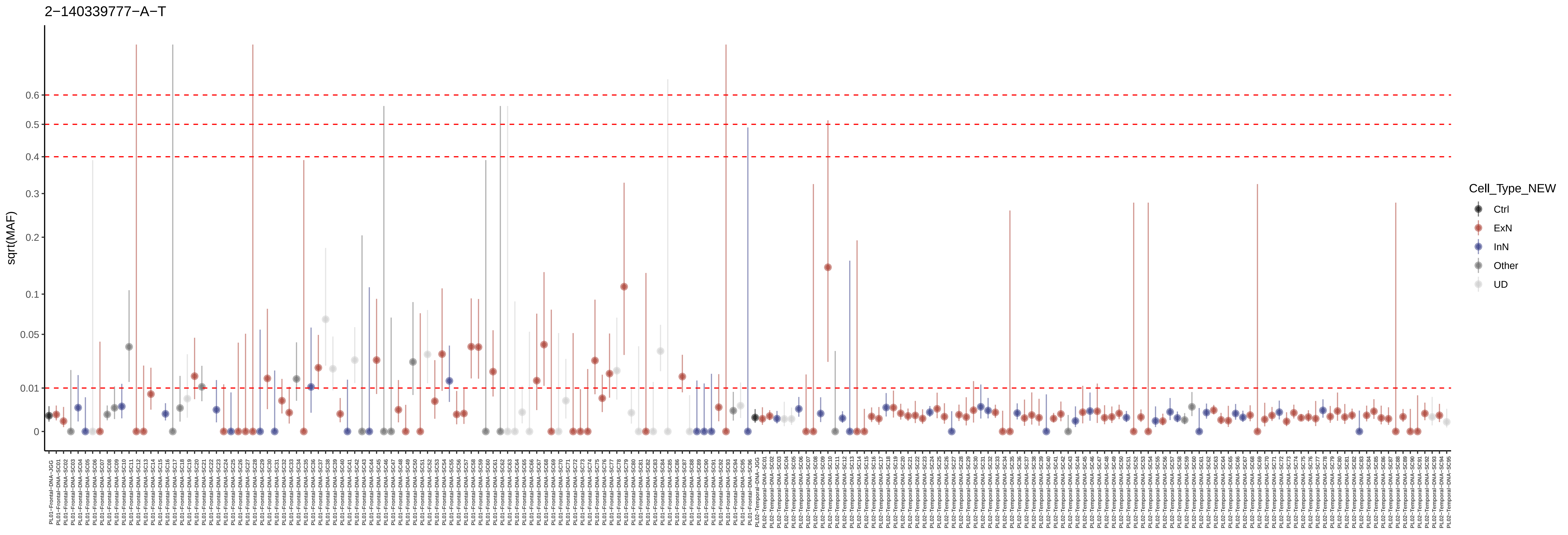

2−140339777−A−T
sqrt(MAF)
0.6
0.5
0.4
0.3
0.2
0.1
0.05
0.01
0
Cell_Type_NEW
Ctrl
ExN
InN
Other
UD

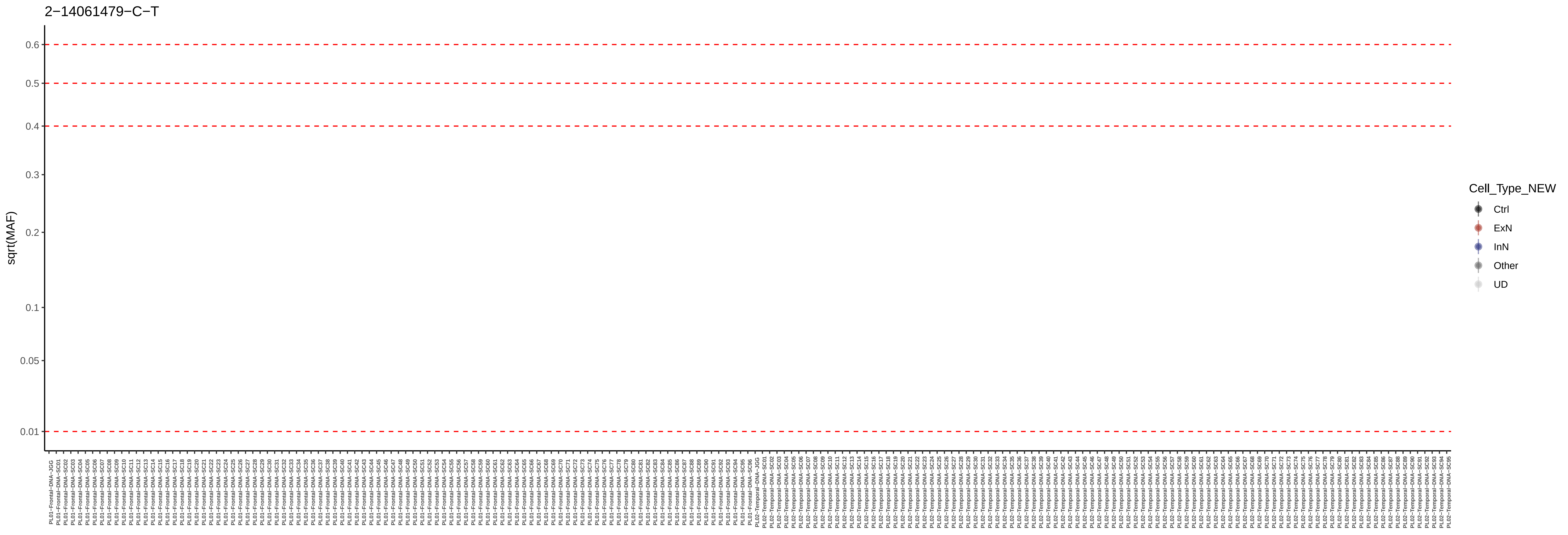
2−14061479−C−T
sqrt(MAF)
0.6
0.5
0.4
0.3
0.2
0.1
0.05
0.01
Cell_Type_NEW
Ctrl
ExN
InN
Other
UD

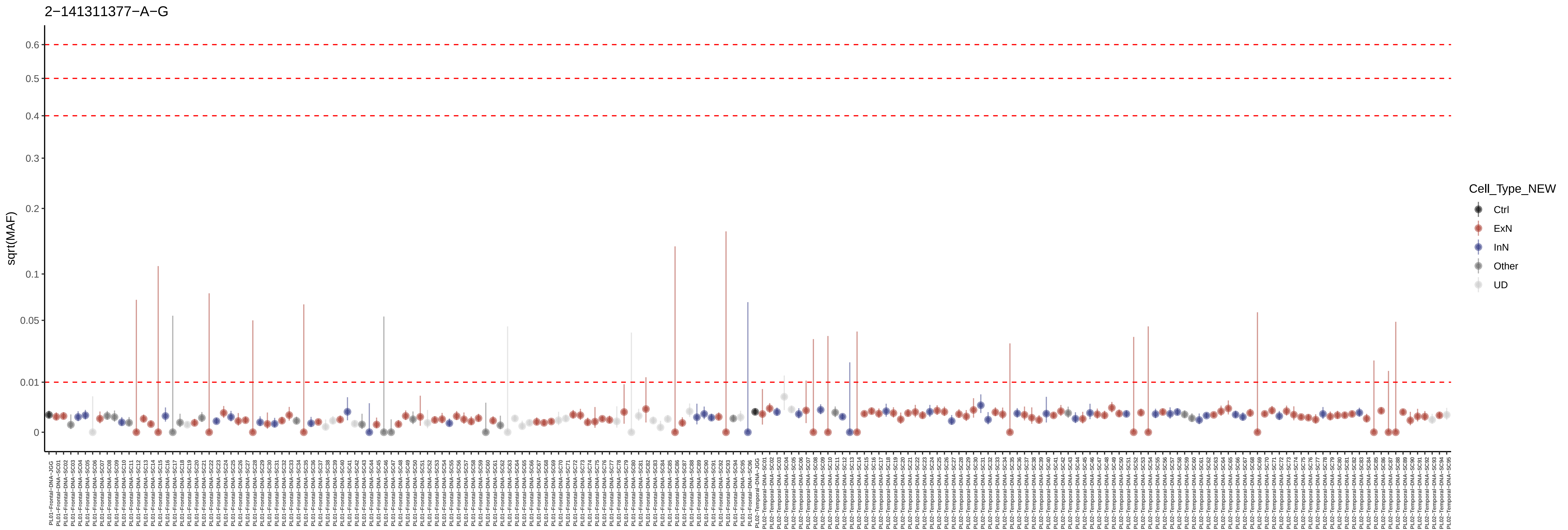

2−141311377−A−G
sqrt(MAF)
0.6
0.5
0.4
0.3
0.2
0.1
0.05
0.01
0
Cell_Type_NEW
Ctrl
ExN
InN
Other
UD

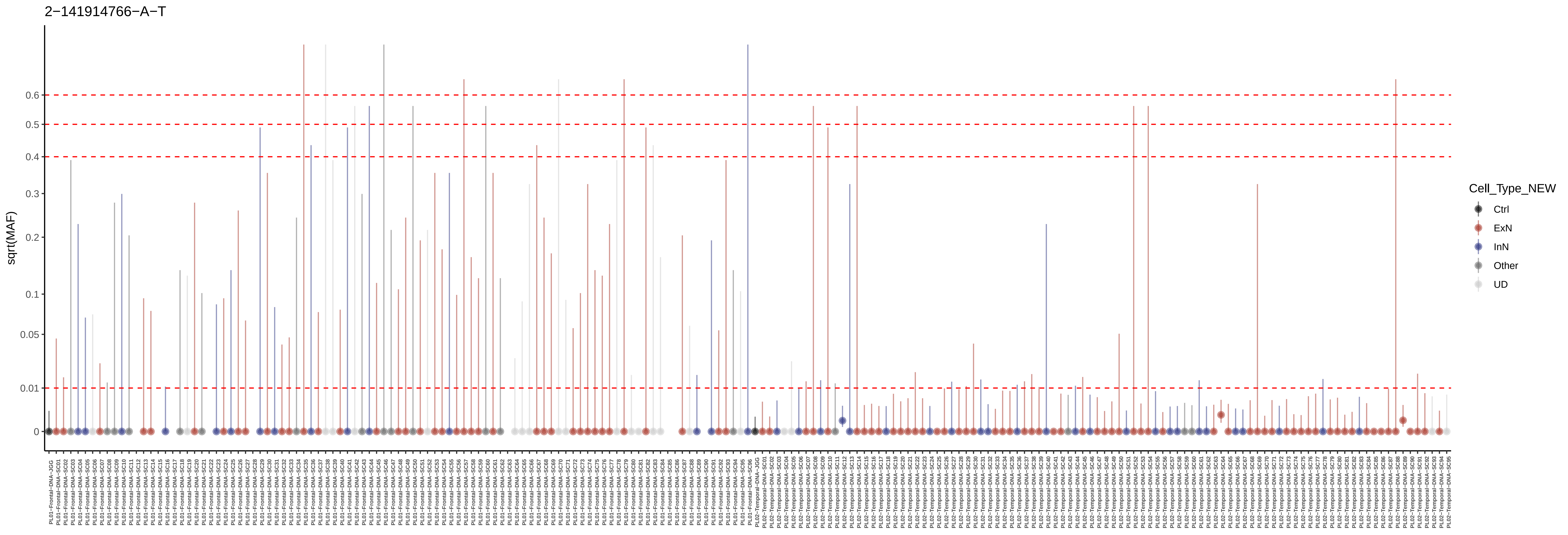

2−141914766−A−T
sqrt(MAF)
0.6
0.5
0.4
0.3
0.2
0.1
0.05
0.01
0
Cell_Type_NEW
Ctrl
ExN
InN
Other
UD

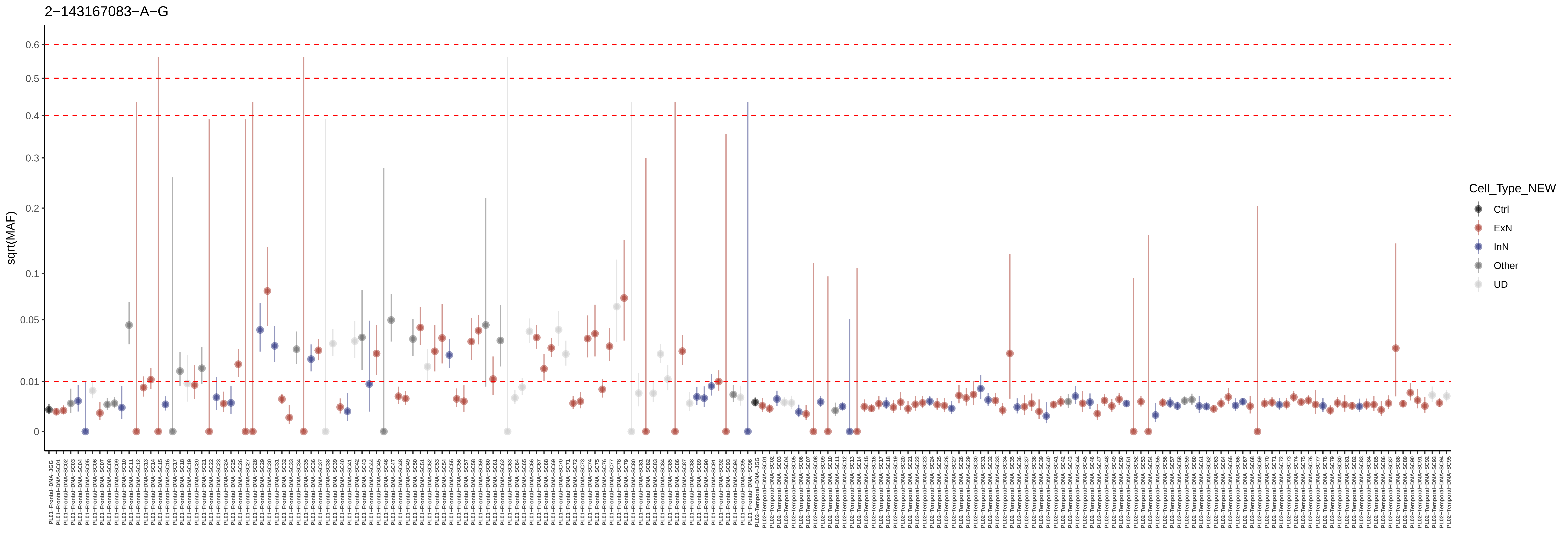
2−143167083−A−G
sqrt(MAF)
0.6
0.5
0.4
0.3
0.2
0.1
0.05
0.01
0
Cell_Type_NEW
Ctrl
ExN
InN
Other
UD

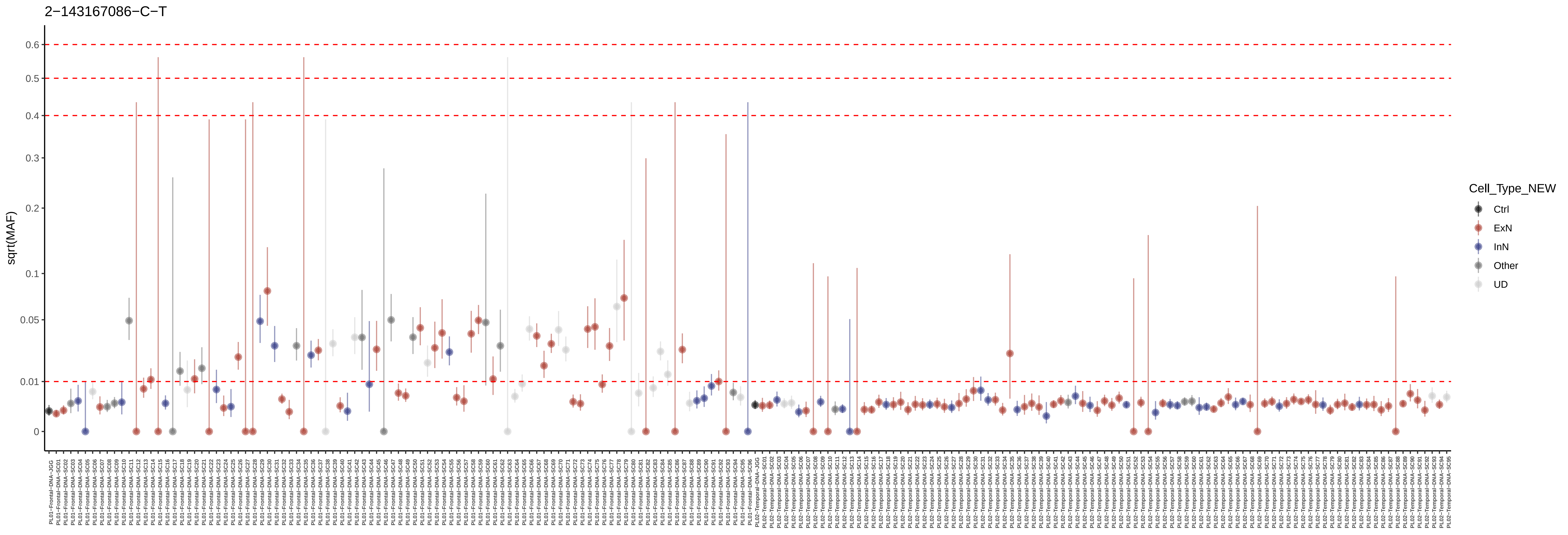

2−143167086−C−T
sqrt(MAF)
0.6
0.5
0.4
0.3
0.2
0.1
0.05
0.01
0
Cell_Type_NEW
Ctrl
ExN
InN
Other
UD

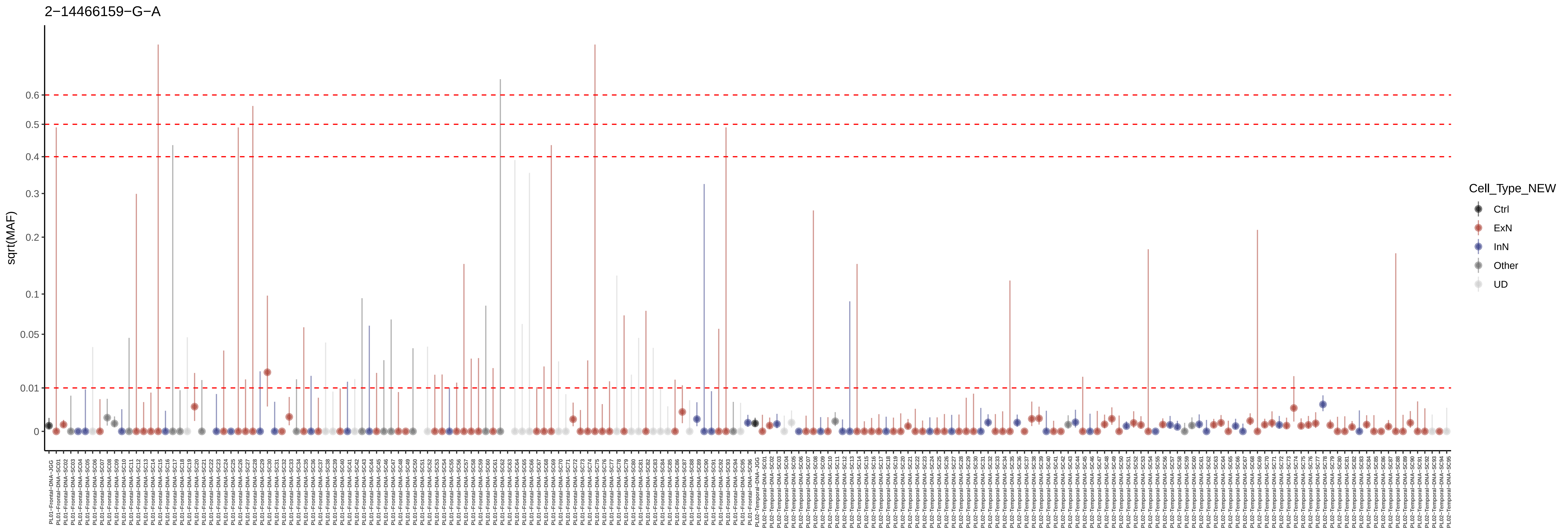
2−14466159−G−A
sqrt(MAF)
0.6
0.5
0.4
0.3
0.2
0.1
0.05
0.01
0
Cell_Type_NEW
Ctrl
ExN
InN
Other
UD

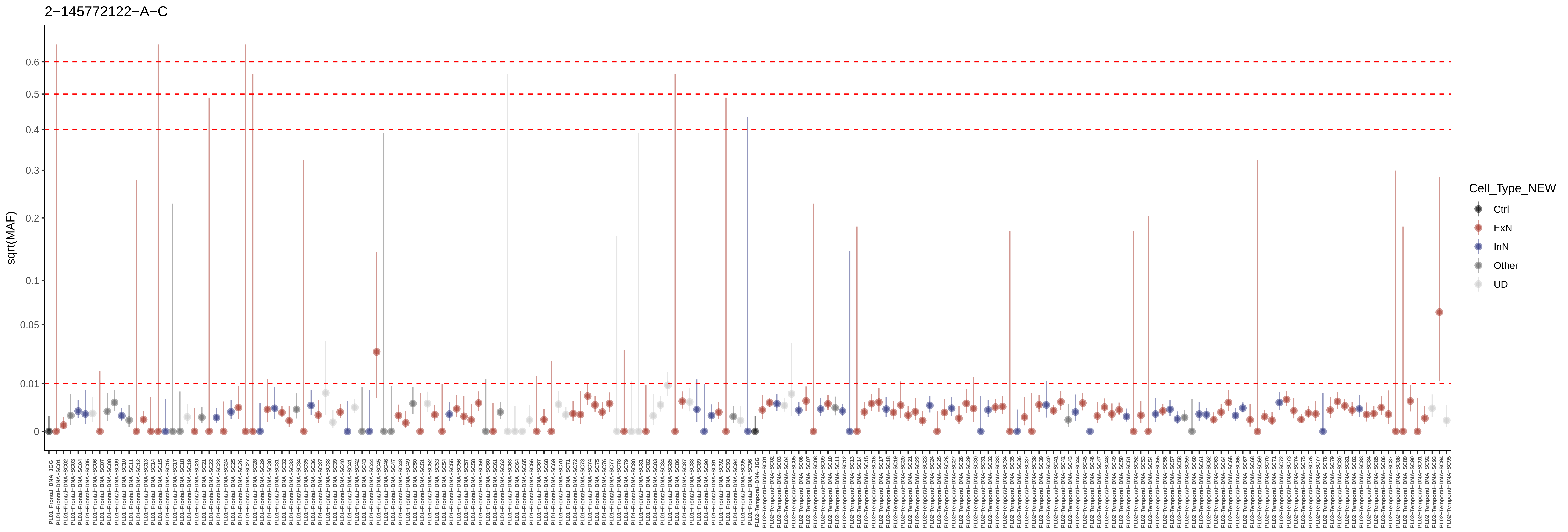

2−145772122−A−C
sqrt(MAF)
0.6
0.5
0.4
0.3
0.2
0.1
0.05
0.01
0
Cell_Type_NEW
Ctrl
ExN
InN
Other
UD

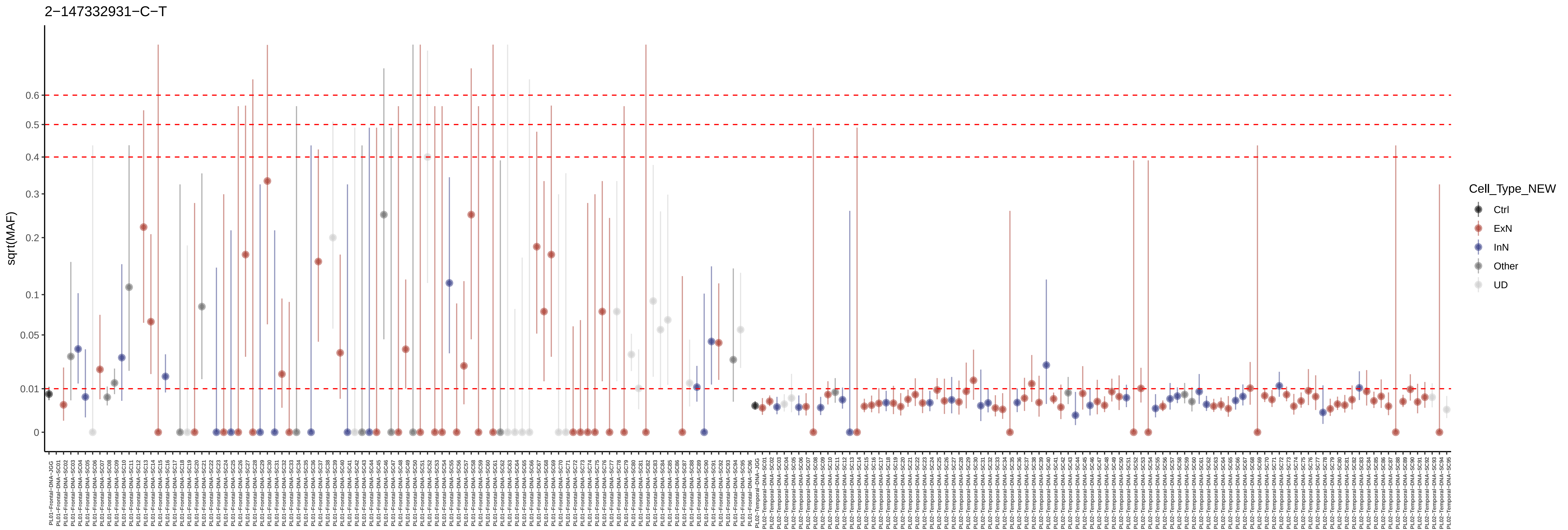
2−147332931−C−T
sqrt(MAF)
0.6
0.5
0.4
0.3
0.2
0.1
0.05
0.01
0
Cell_Type_NEW
Ctrl
ExN
InN
Other
UD

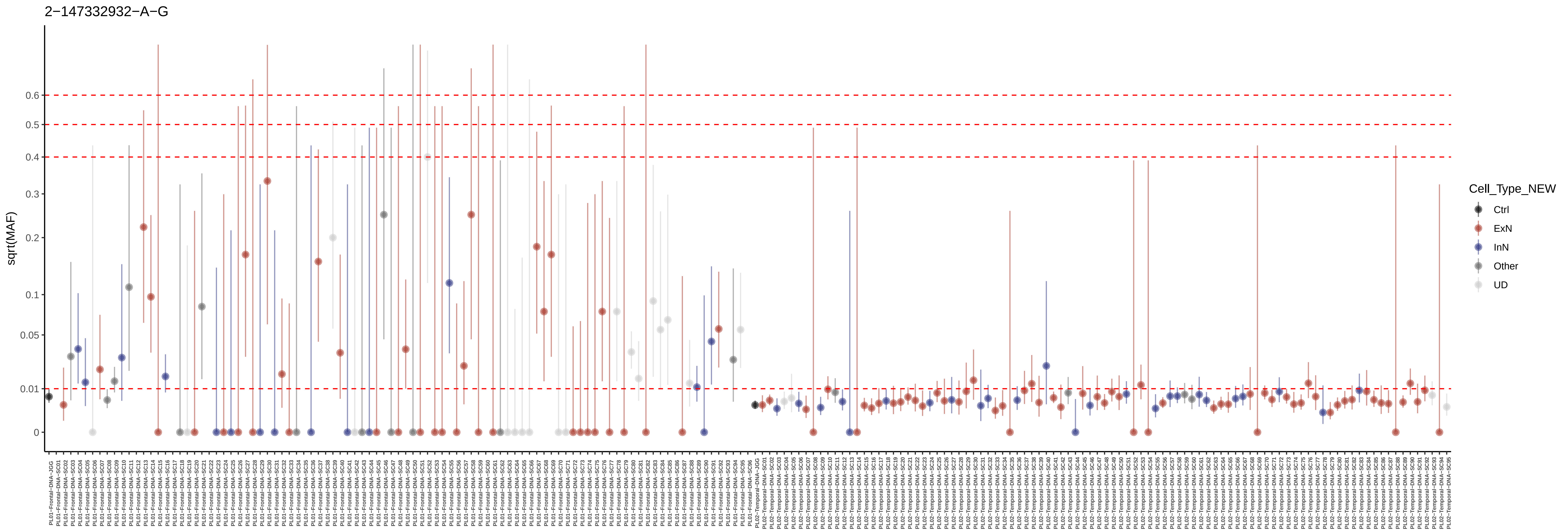
2−147332932−A−G
sqrt(MAF)
0.6
0.5
0.4
0.3
0.2
0.1
0.05
0.01
0
Cell_Type_NEW
Ctrl
ExN
InN
Other
UD

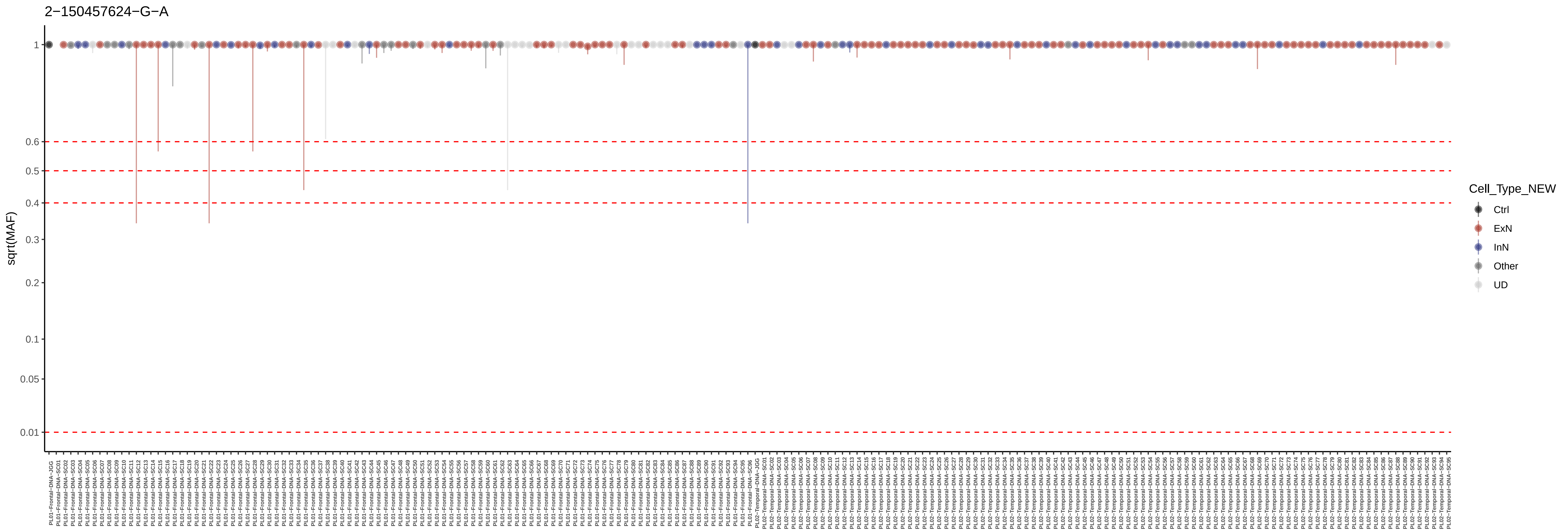

2−150457624−G−A
sqrt(MAF)
1
0.6
0.5
0.4
0.3
0.2
0.1
0.05
0.01
Cell_Type_NEW
Ctrl
ExN
InN
Other
UD

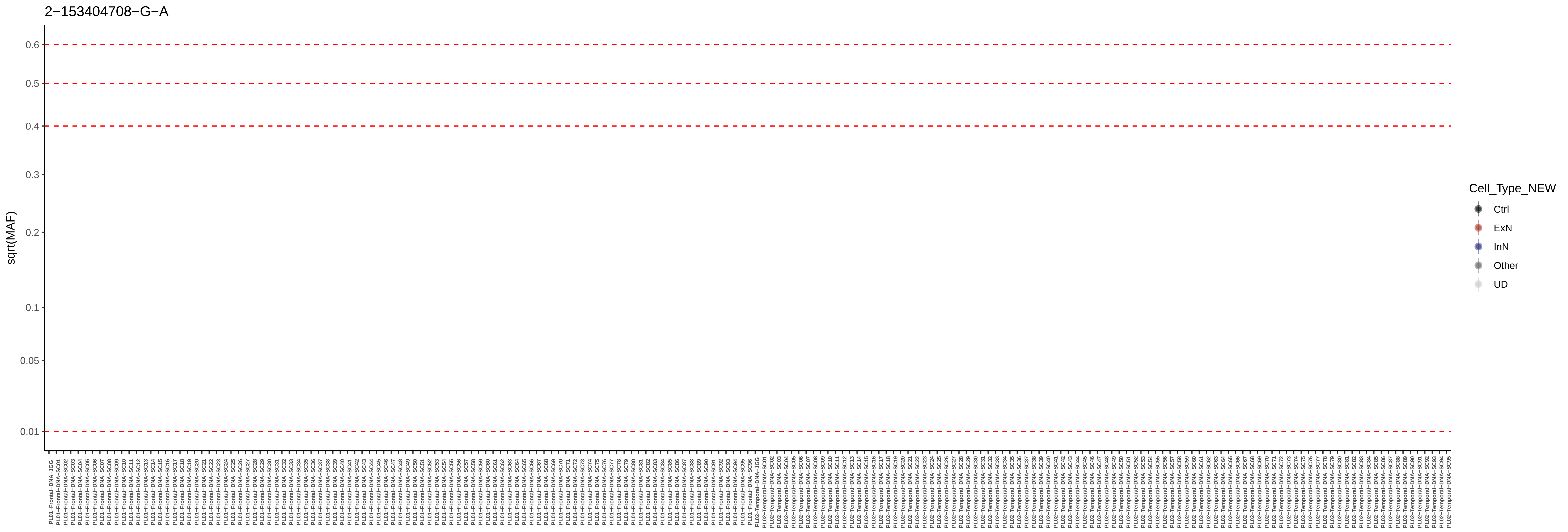
2−153404708−G−A
sqrt(MAF)
0.6
0.5
0.4
0.3
0.2
0.1
0.05
0.01
Cell_Type_NEW
Ctrl
ExN
InN
Other
UD

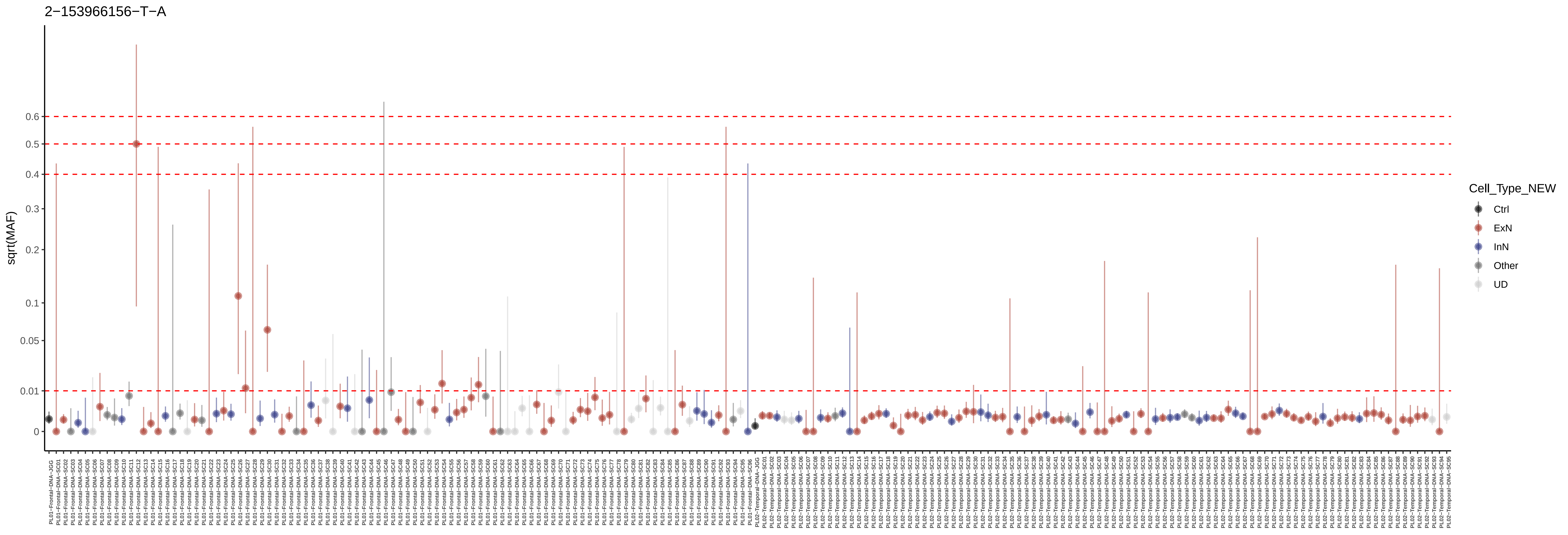
2−153966156−T−A
sqrt(MAF)
0.6
0.5
0.4
0.3
0.2
0.1
0.05
0.01
0
Cell_Type_NEW
Ctrl
ExN
InN
Other
UD

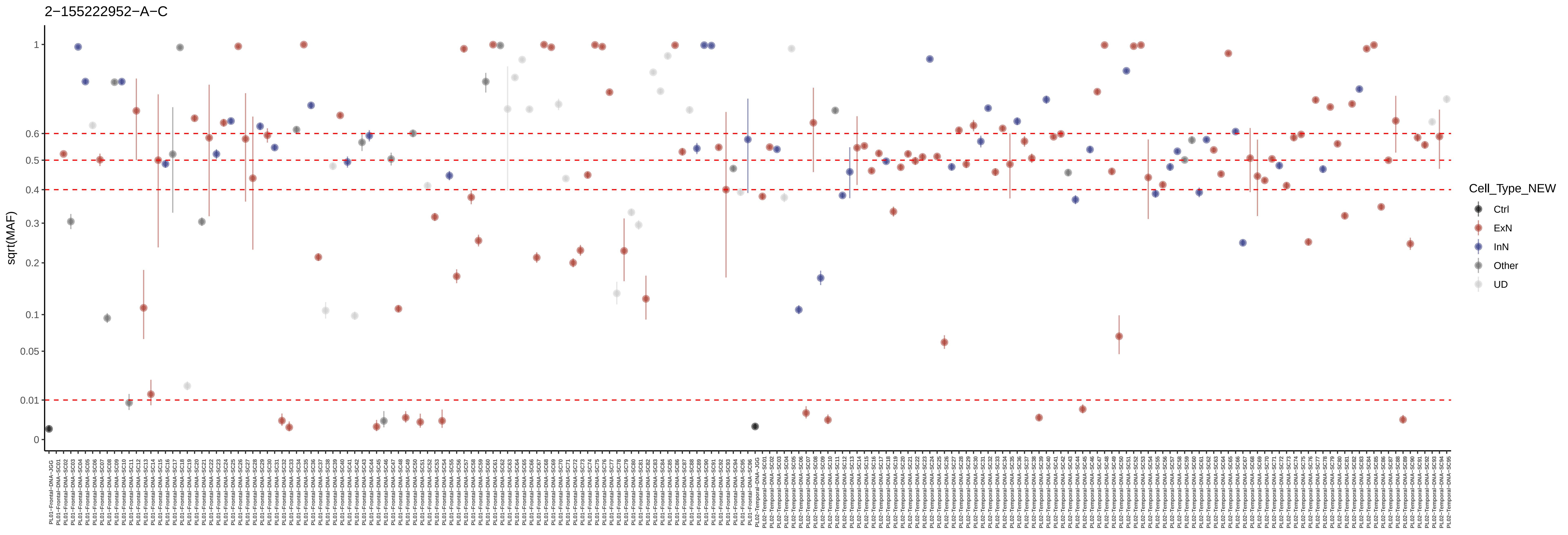
2−155222952−A−C
sqrt(MAF)
1
0.6
0.5
0.4
0.3
0.2
0.1
0.05
0.01
0
Cell_Type_NEW
Ctrl
ExN
InN
Other
UD

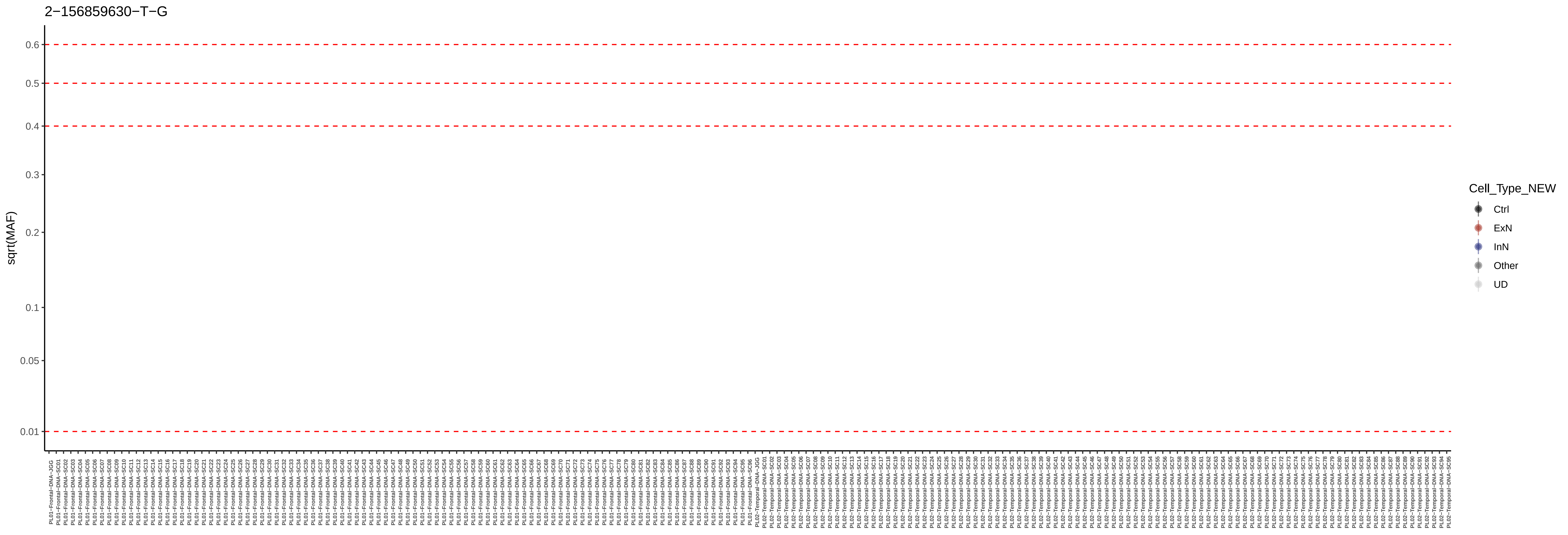
2−156859630−T−G
sqrt(MAF)
0.6
0.5
0.4
0.3
0.2
0.1
0.05
0.01
Cell_Type_NEW
Ctrl
ExN
InN
Other
UD

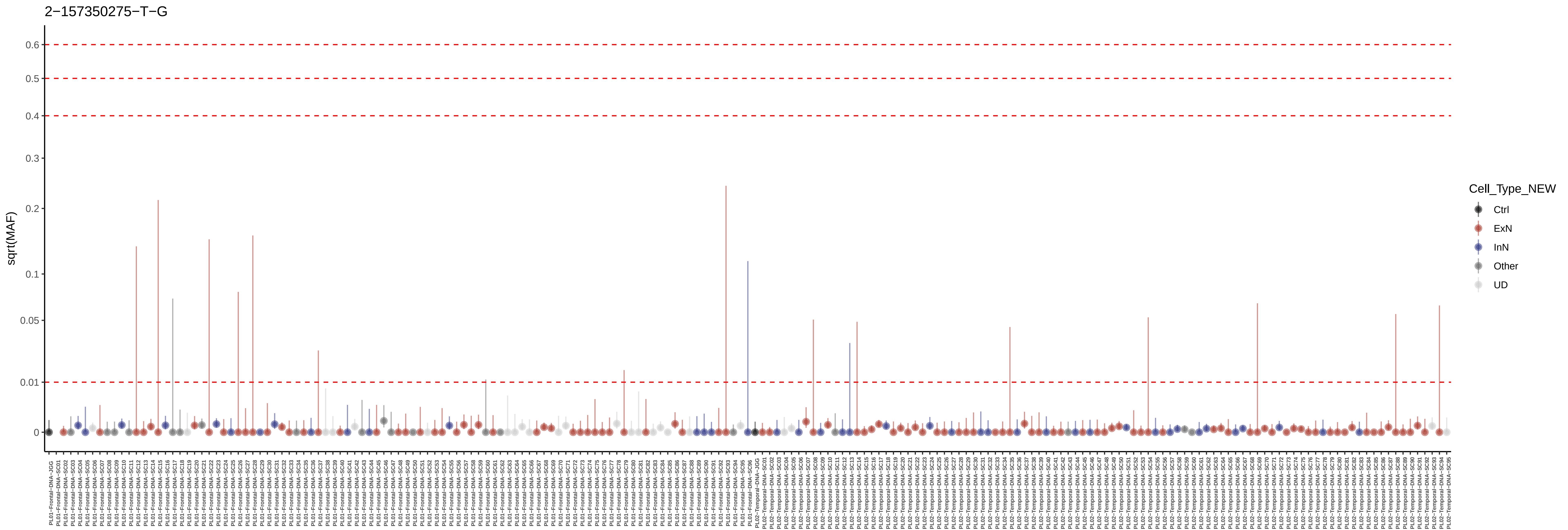

2−157350275−T−G
sqrt(MAF)
0.6
0.5
0.4
0.3
0.2
0.1
0.05
0.01
0
Cell_Type_NEW
Ctrl
ExN
InN
Other
UD

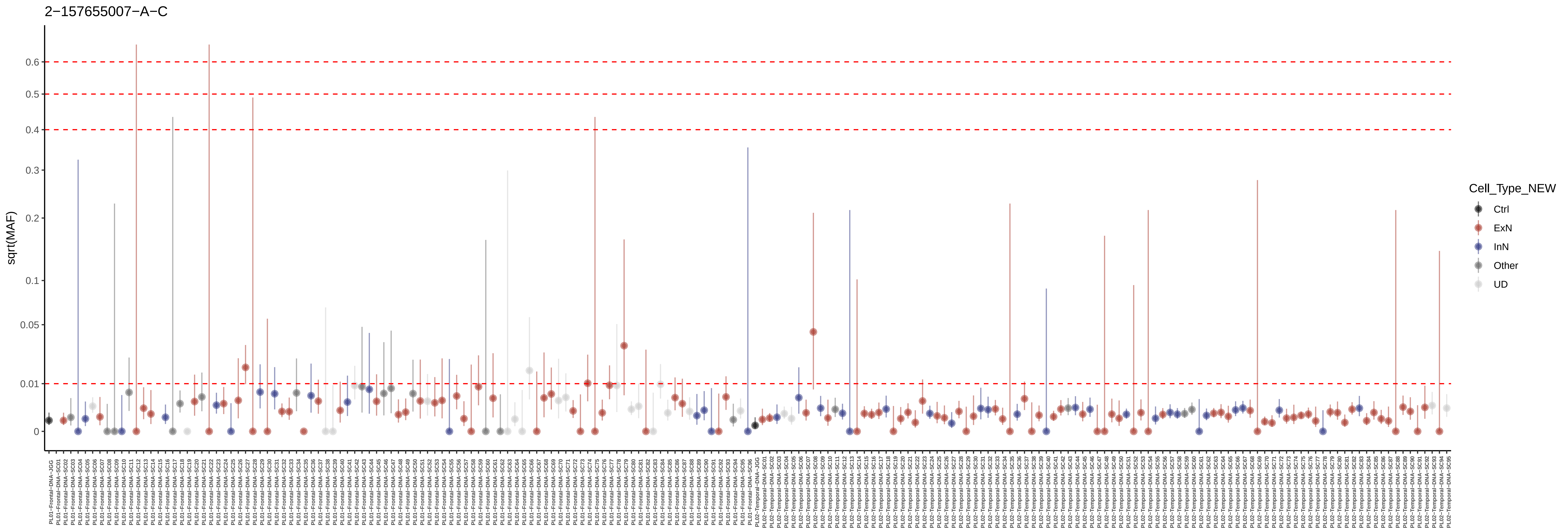
2−157655007−A−C
sqrt(MAF)
0.6
0.5
0.4
0.3
0.2
0.1
0.05
0.01
0
Cell_Type_NEW
Ctrl
ExN
InN
Other
UD

2−157655008−G−T

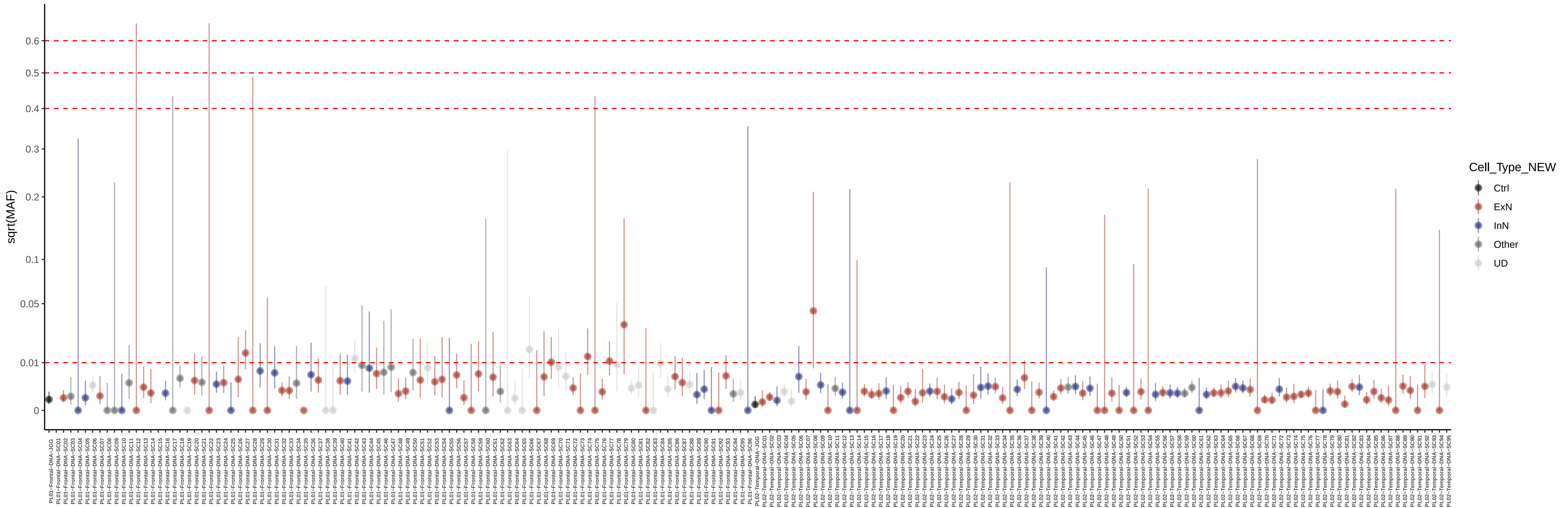

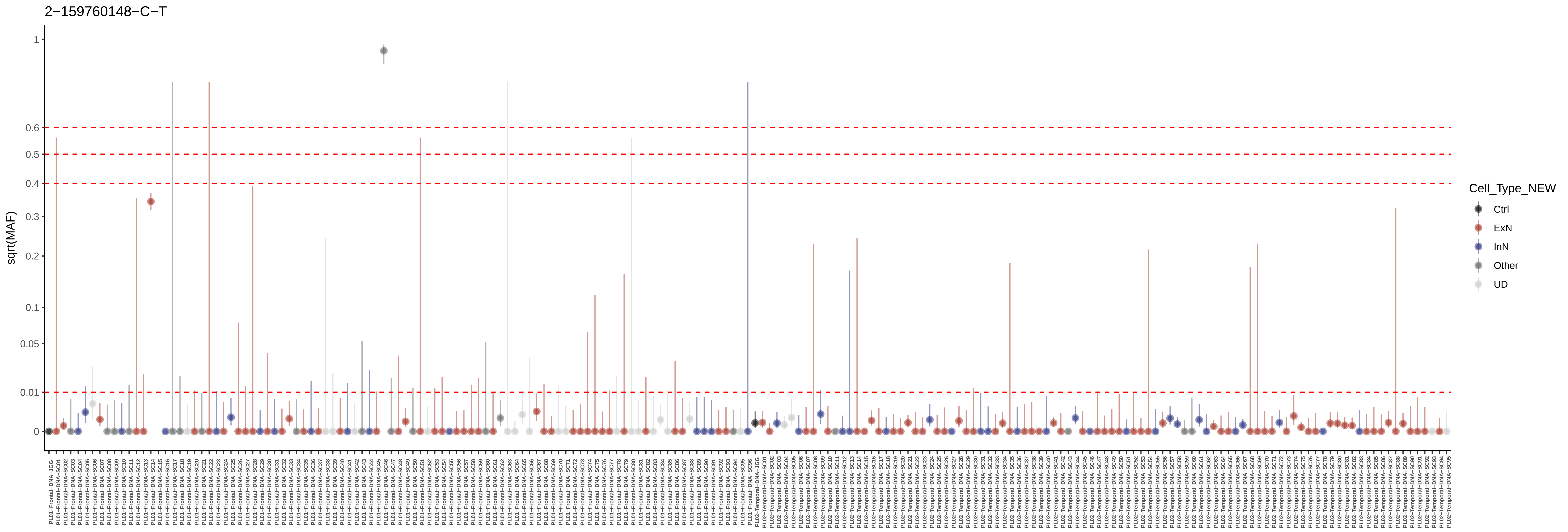
2−159760148−C−T
sqrt(MAF)
1
0.6
0.5
0.4
0.3
0.2
0.1
0.05
0.01
0
Cell_Type_NEW
Ctrl
ExN
InN
Other
UD

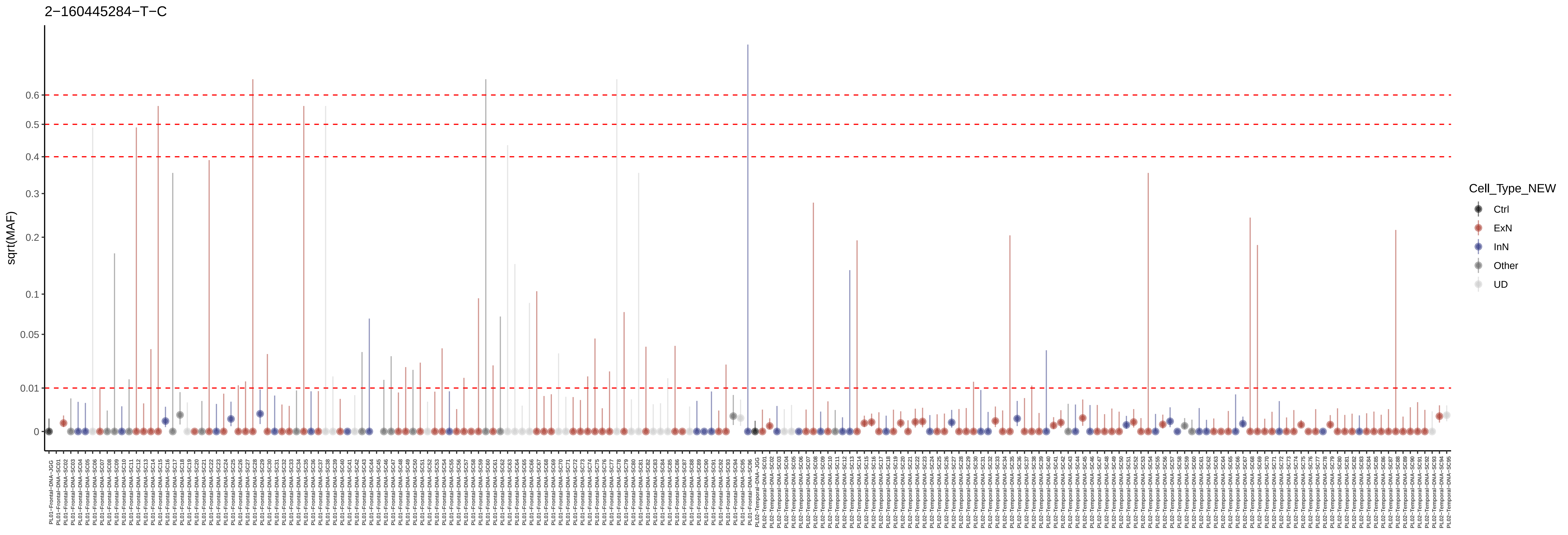

2−160445284−T−C
sqrt(MAF)
0.6
0.5
0.4
0.3
0.2
0.1
0.05
0.01
0
Cell_Type_NEW
Ctrl
ExN
InN
Other
UD

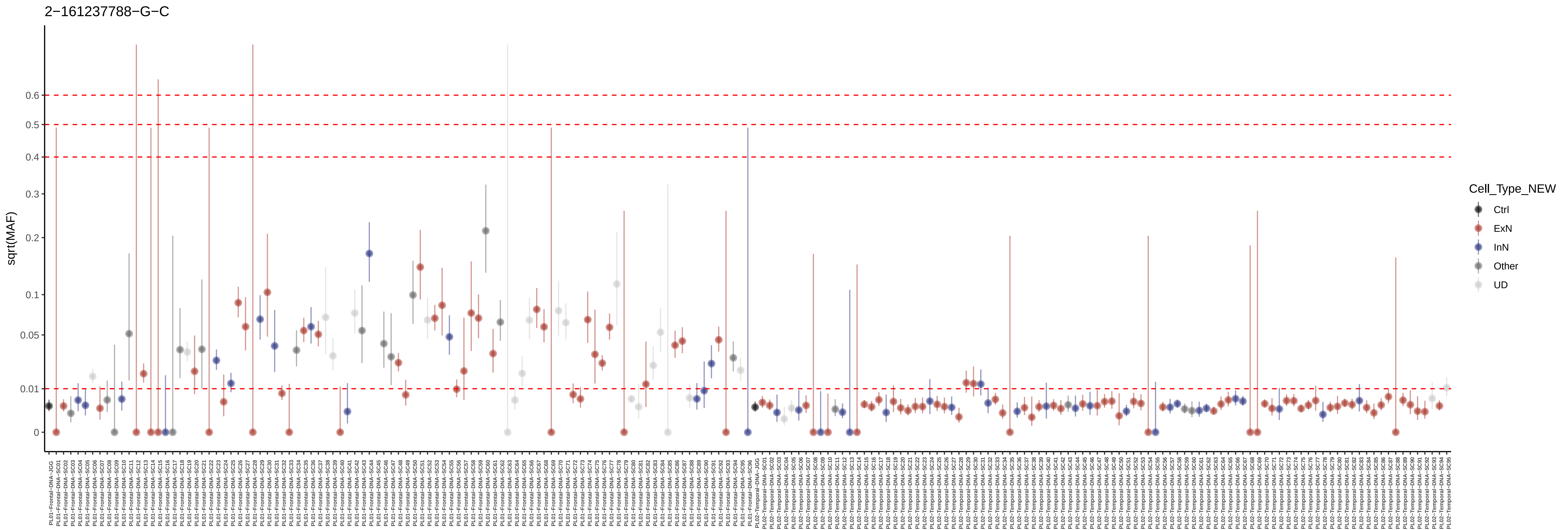

2−161237788−G−C
sqrt(MAF)
0
0.01
0.05
0.1
0.2
0.3
0.4
0.5
0.6
Cell_Type_NEW
Ctrl
ExN
InN
Other
UD

2−16269495−T−C

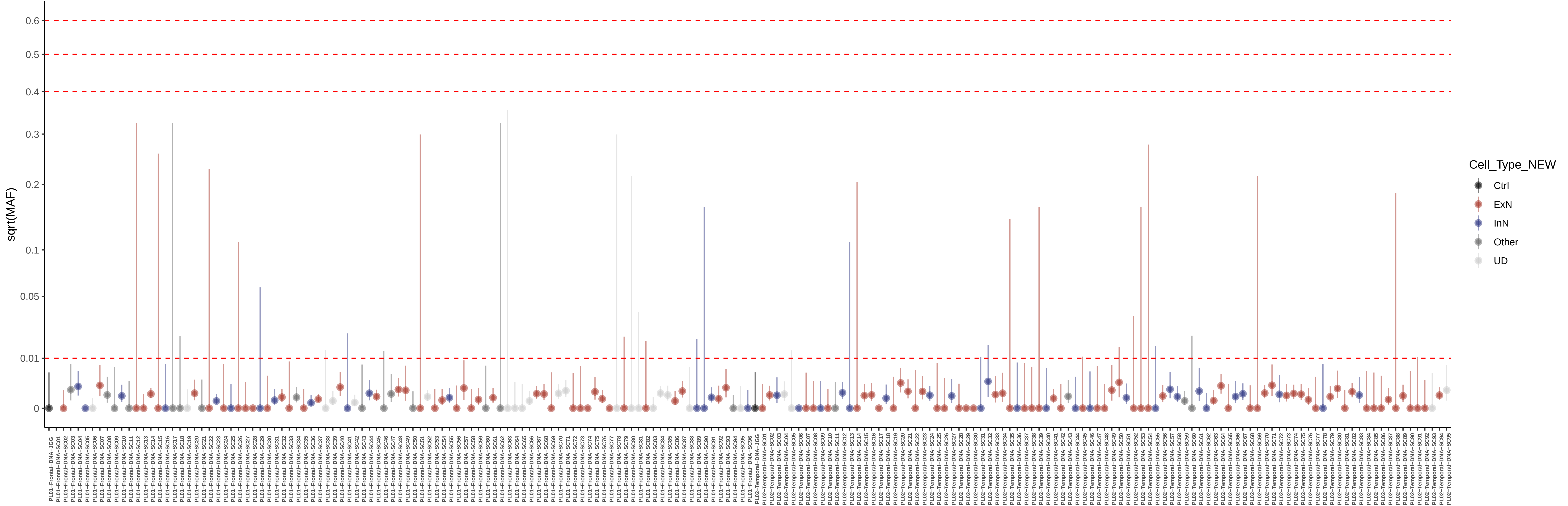

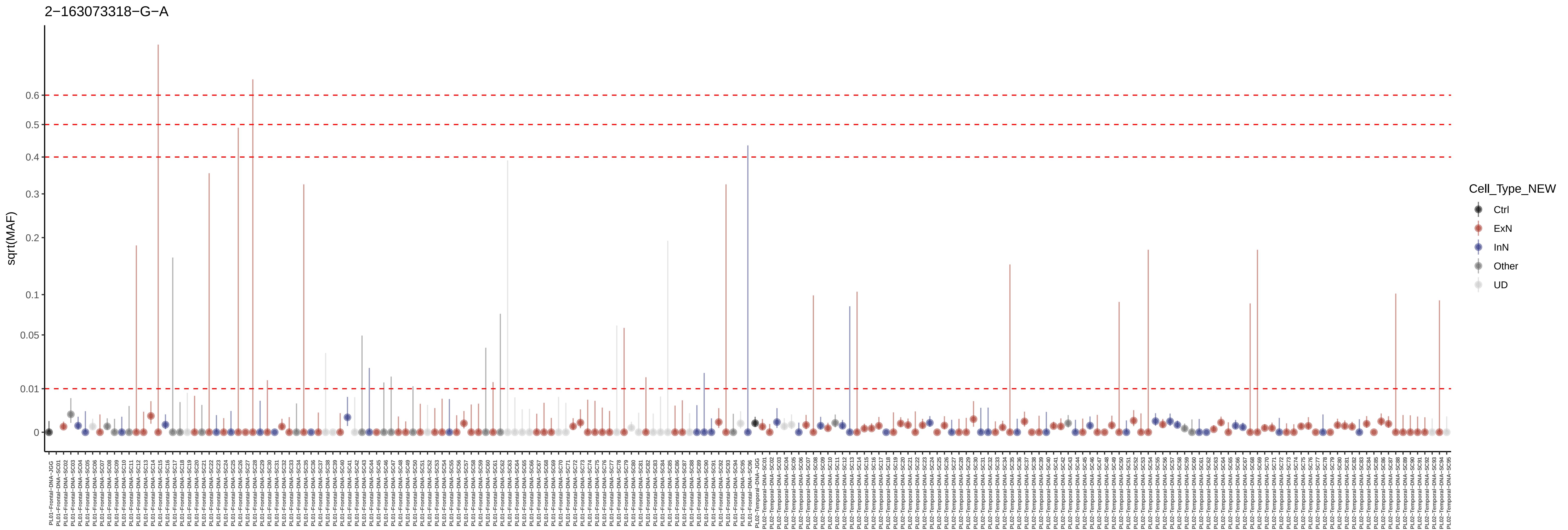

2−163073318−G−A
sqrt(MAF)
0.6
0.5
0.4
0.3
0.2
0.1
0.05
0.01
0
Cell_Type_NEW
Ctrl
ExN
InN
Other
UD

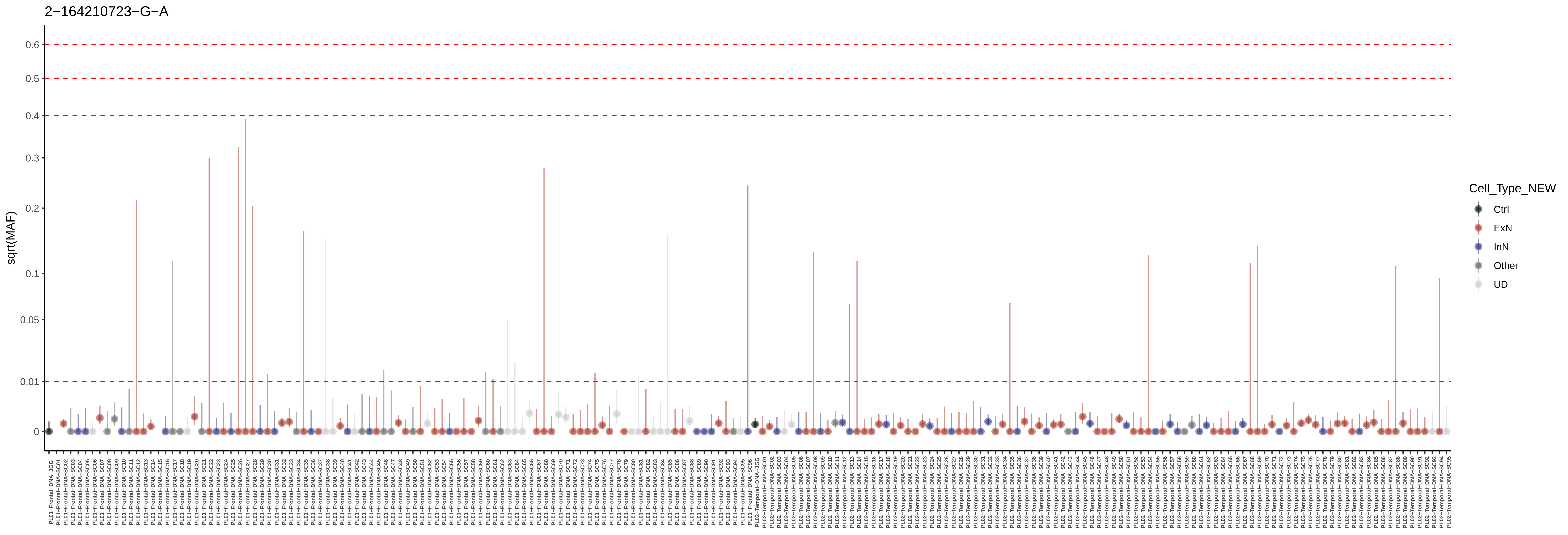

2−164210723−G−A
sqrt(MAF)
0.6
0.5
0.4
0.3
0.2
0.1
0.05
0.01
0
Cell_Type_NEW
Ctrl
ExN
InN
Other
UD

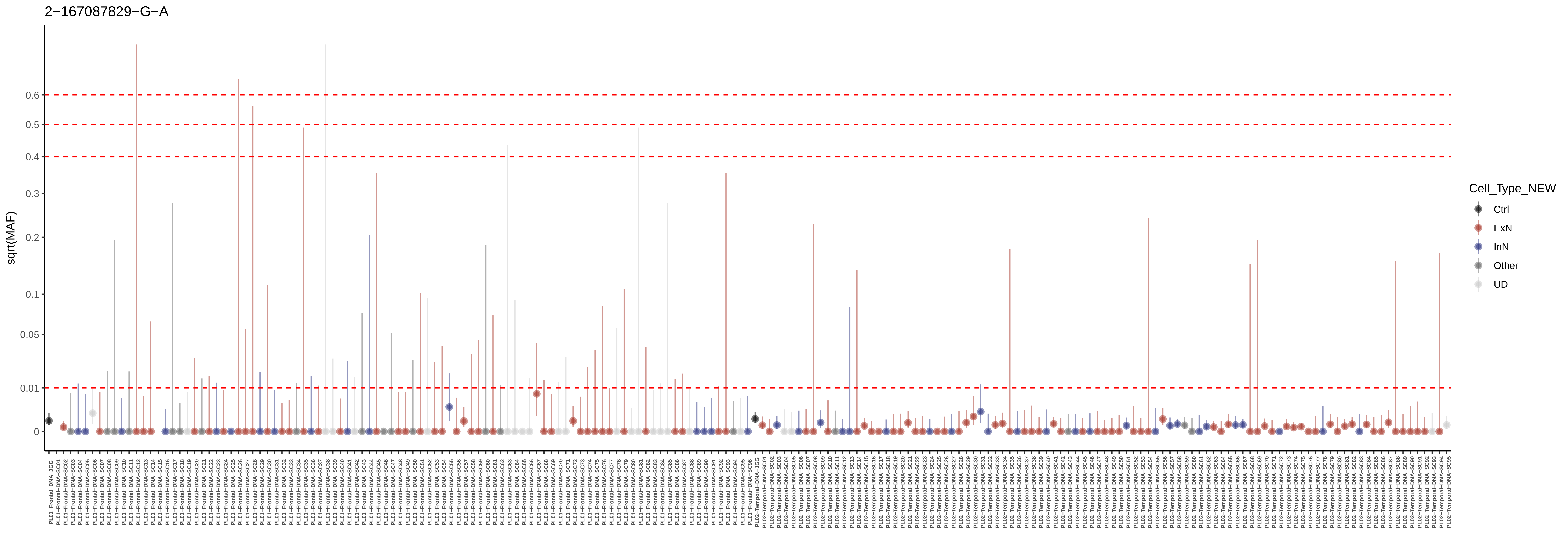

2−167087829−G−A
sqrt(MAF)
0.6
0.5
0.4
0.3
0.2
0.1
0.05
0.01
0
Cell_Type_NEW
Ctrl
ExN
InN
Other
UD

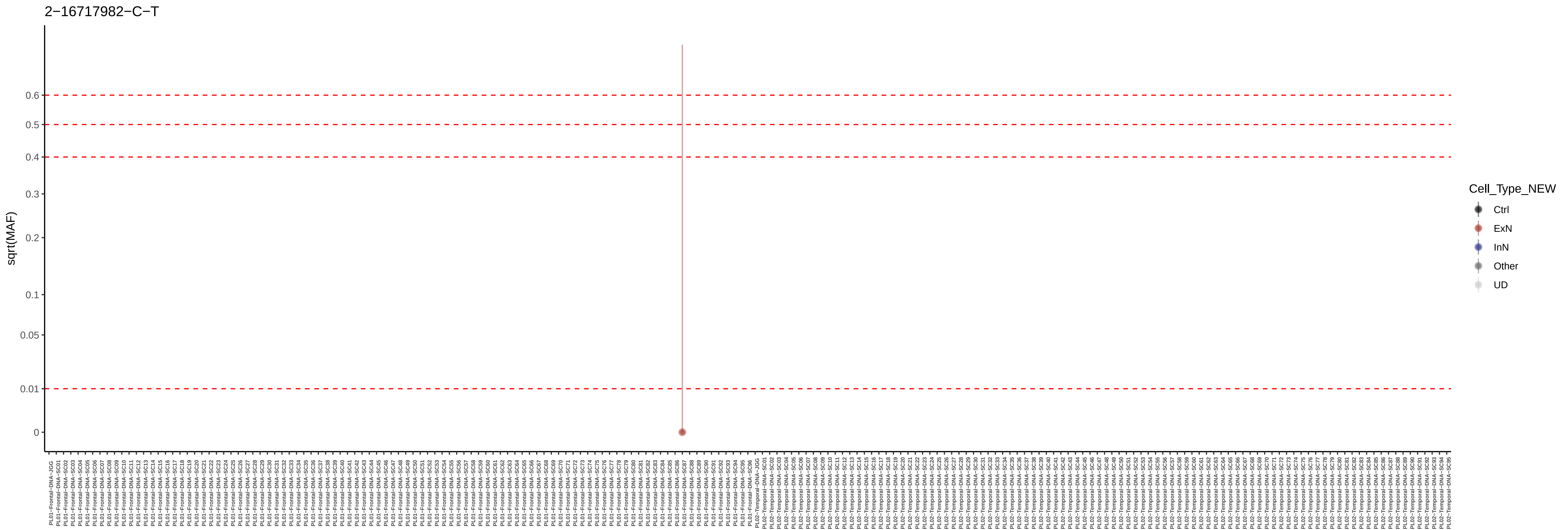

2−16717982−C−T
sqrt(MAF)
0.6
0.5
0.4
0.3
0.2
0.1
0.05
0.01
0
Cell_Type_NEW
Ctrl
ExN
InN
Other
UD

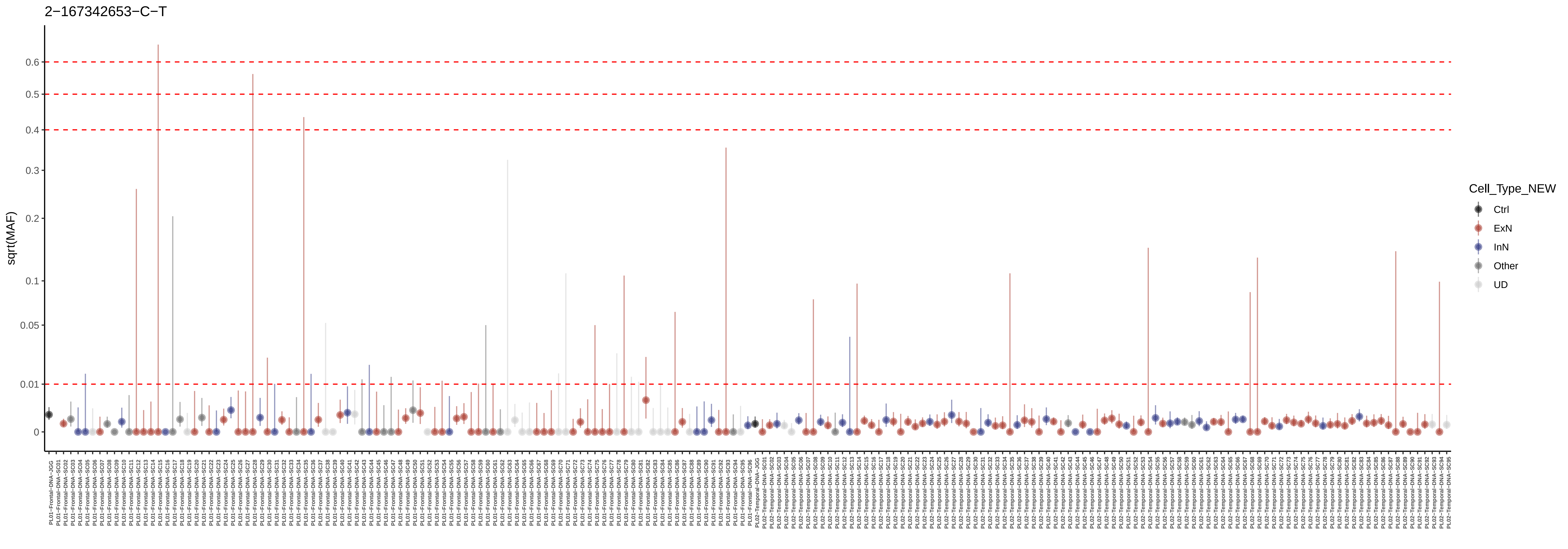

2−167342653−C−T
sqrt(MAF)
0.6
0.5
0.4
0.3
0.2
0.1
0.05
0.01
0
Cell_Type_NEW
Ctrl
ExN
InN
Other
UD

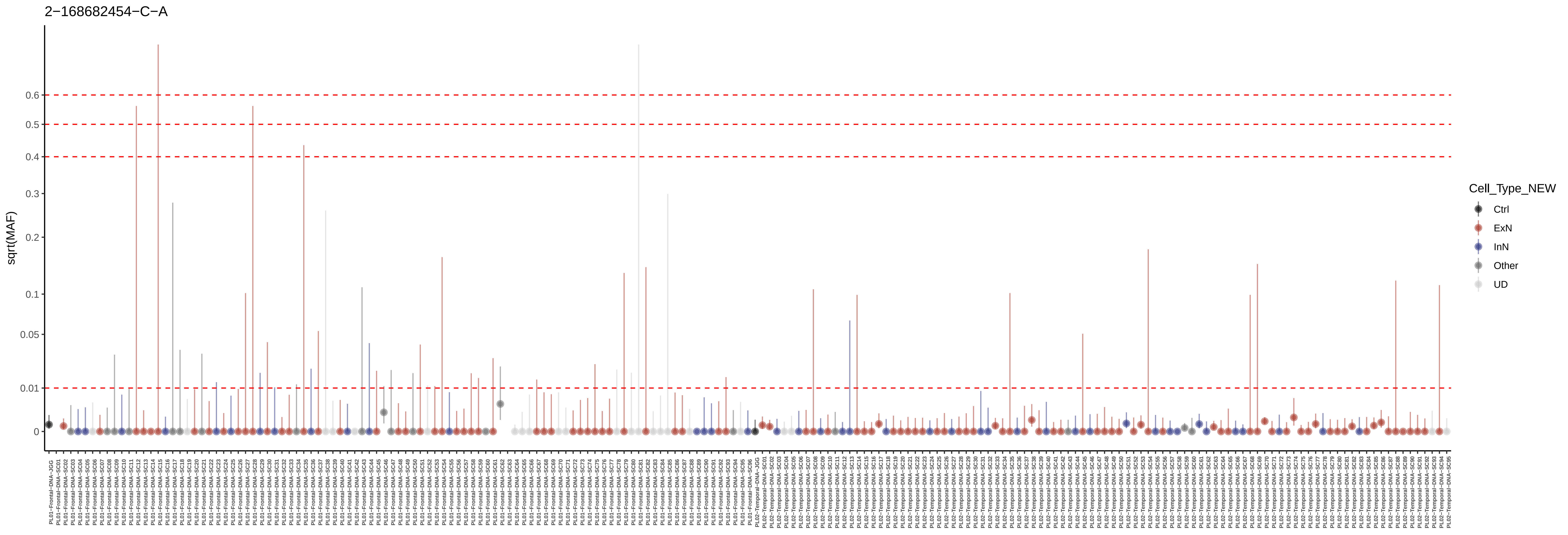

2−168682454−C−A
sqrt(MAF)
0.6
0.5
0.4
0.3
0.2
0.1
0.05
0.01
0
Cell_Type_NEW
Ctrl
ExN
InN
Other
UD

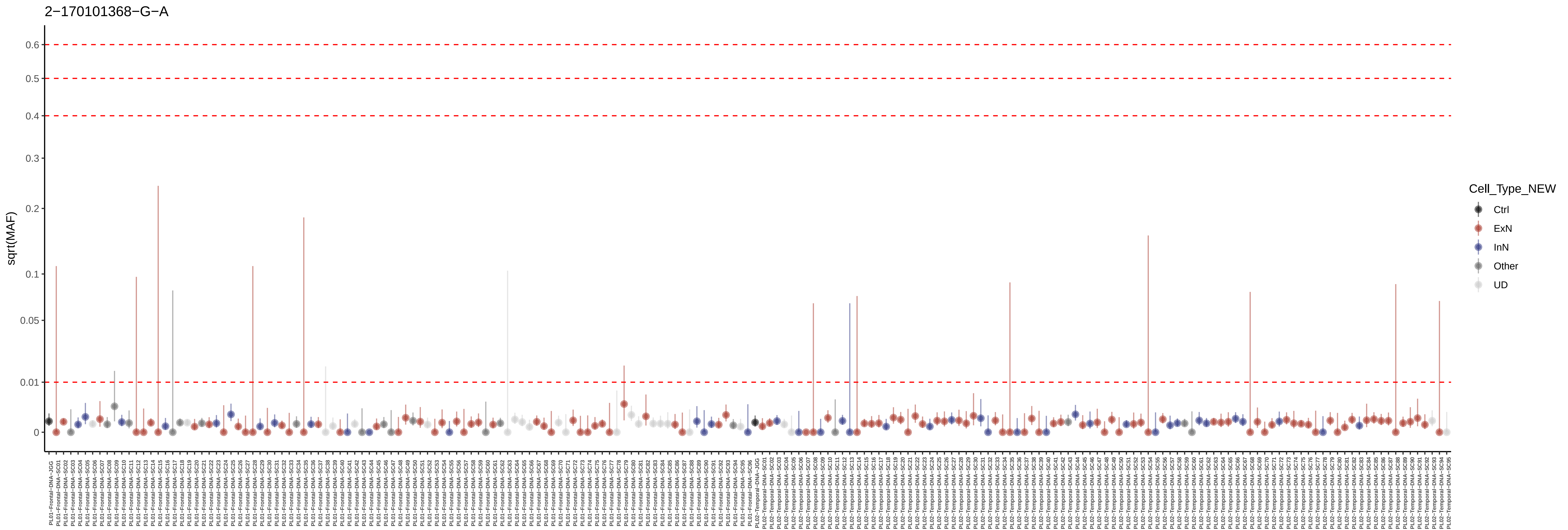

2−170101368−G−A
sqrt(MAF)
0.6
0.5
0.4
0.3
0.2
0.1
0.05
0.01
0
Cell_Type_NEW
Ctrl
ExN
InN
Other
UD

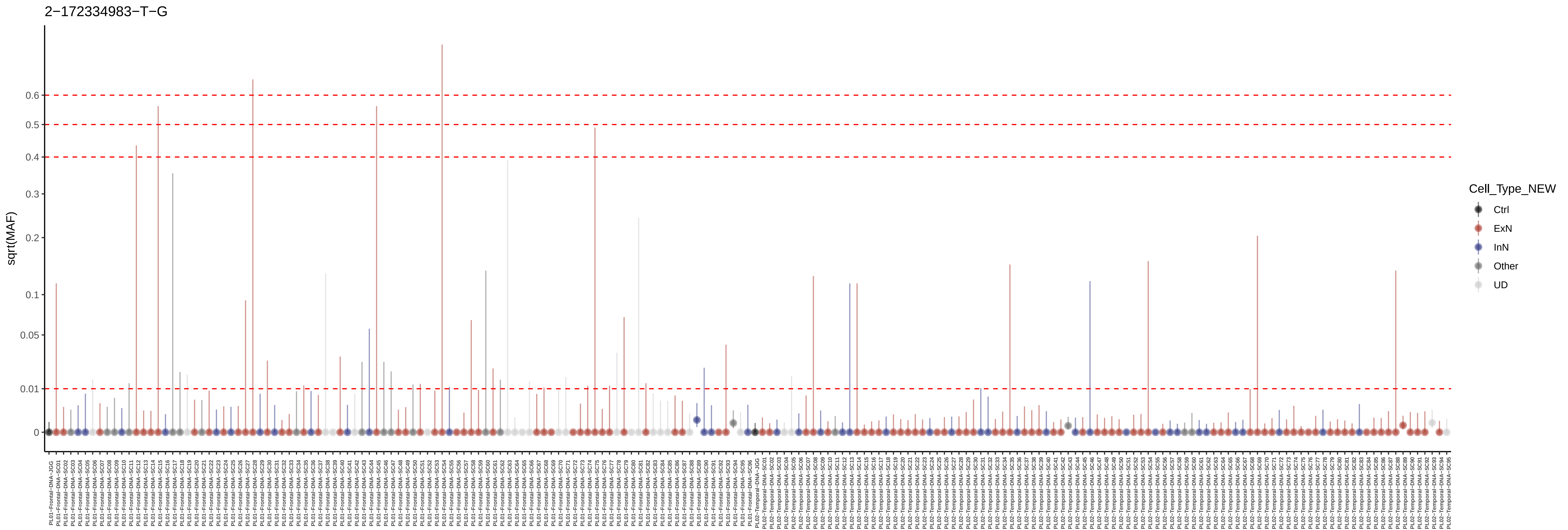
2−172334983−T−G
sqrt(MAF)
0
0.01
0.05
0.1
0.2
0.3
0.4
0.5
0.6
Cell_Type_NEW
Ctrl
ExN
InN
Other
UD

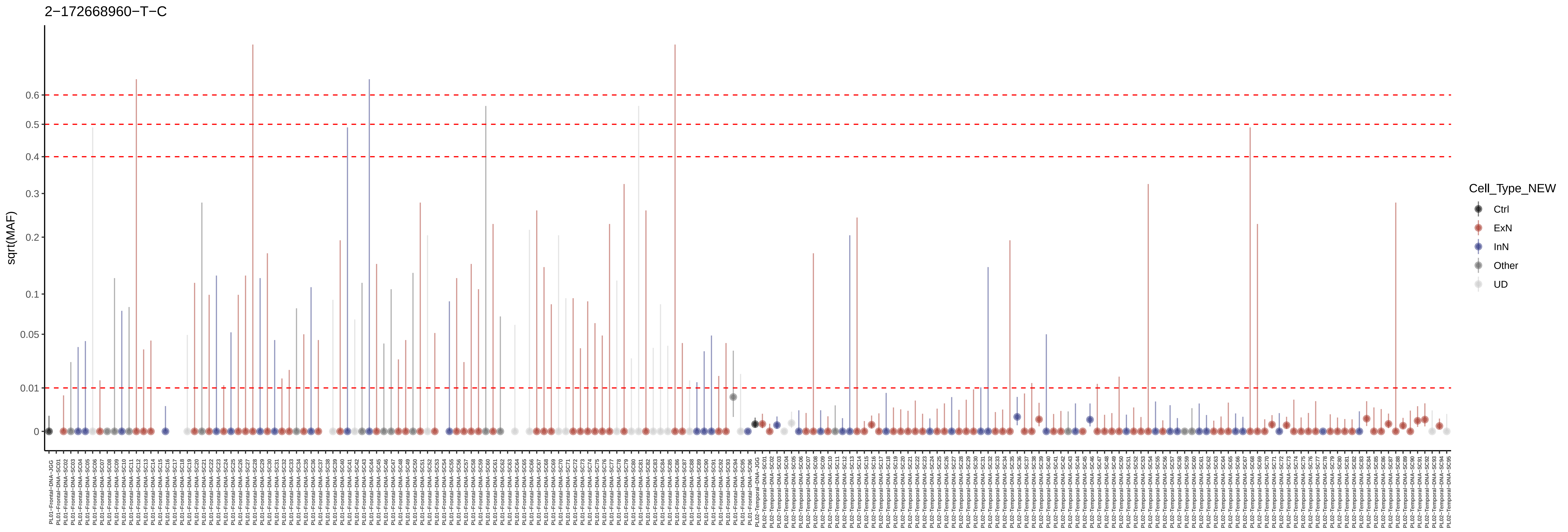
2−172668960−T−C
sqrt(MAF)
0.6
0.5
0.4
0.3
0.2
0.1
0.05
0.01
0
Cell_Type_NEW
Ctrl
ExN
InN
Other
UD

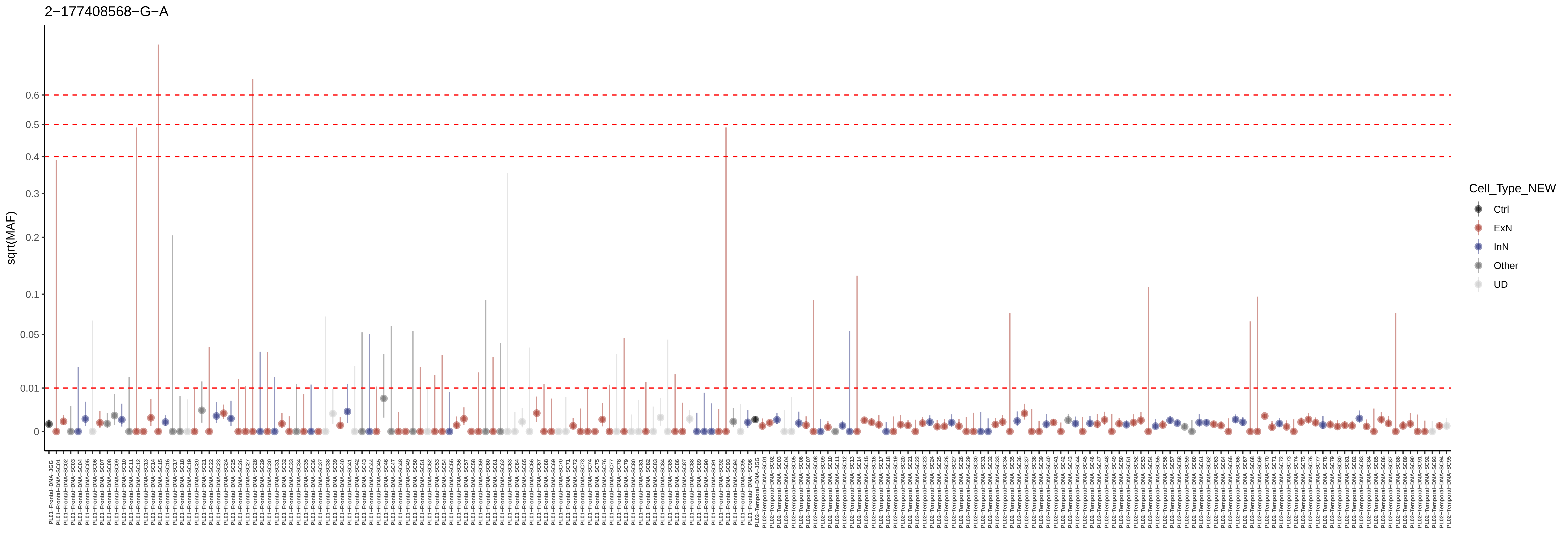
2−177408568−G−A
sqrt(MAF)
0.6
0.5
0.4
0.3
0.2
0.1
0.05
0.01
0
Cell_Type_NEW
Ctrl
ExN
InN
Other
UD

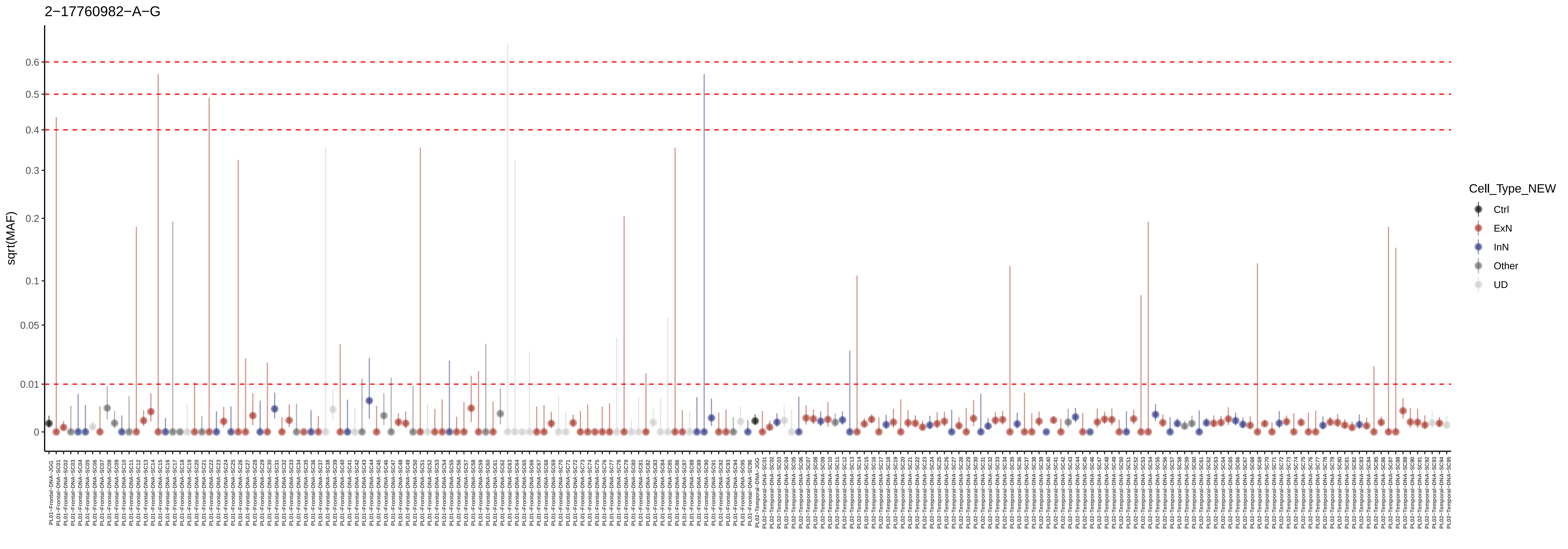

2−17760982−A−G
sqrt(MAF)
0.6
0.5
0.4
0.3
0.2
0.1
0.05
0.01
0
Cell_Type_NEW
Ctrl
ExN
InN
Other
UD

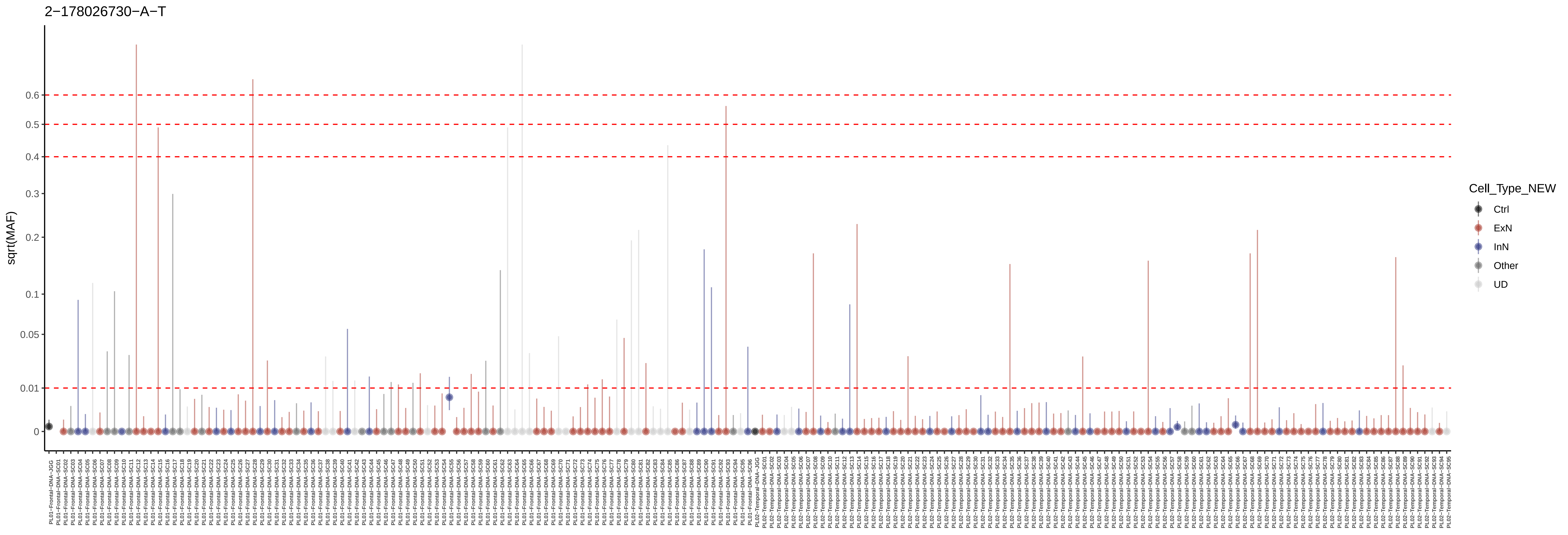
2−178026730−A−T
sqrt(MAF)
0.6
0.5
0.4
0.3
0.2
0.1
0.05
0.01
0
Cell_Type_NEW
Ctrl
ExN
InN
Other
UD

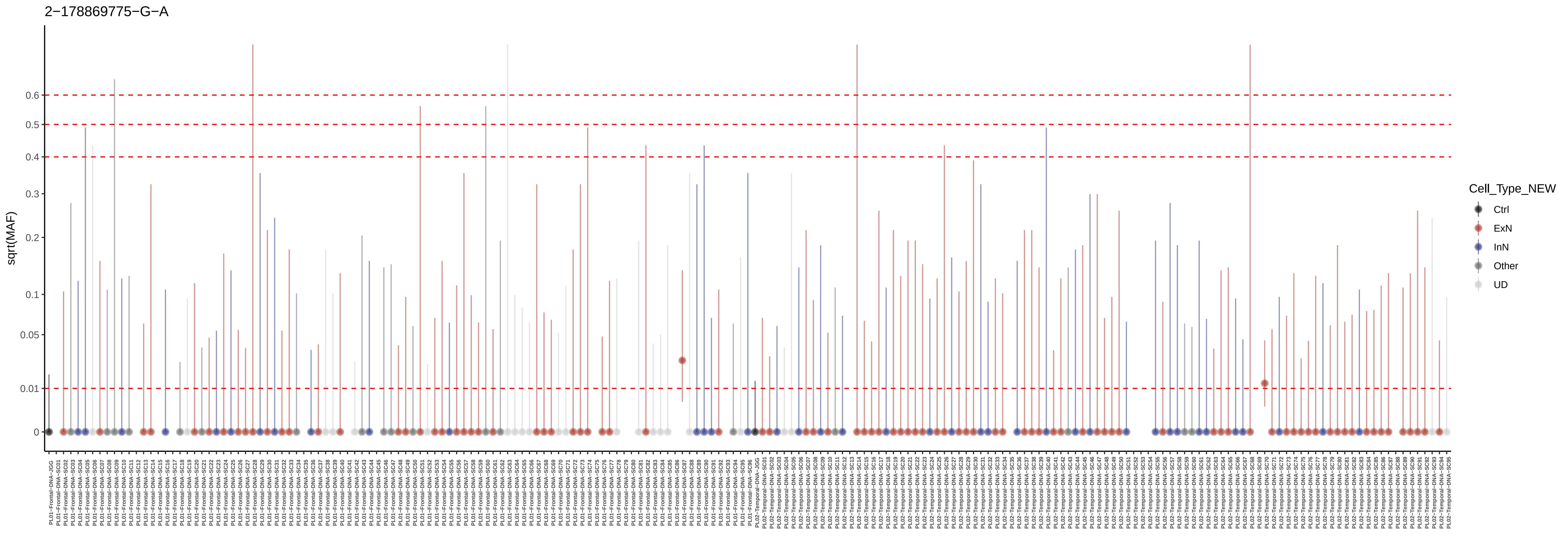

2−178869775−G−A
sqrt(MAF)
0
0.01
0.05
0.1
0.2
0.3
0.4
0.5
0.6
Cell_Type_NEW
Ctrl
ExN
InN
Other
UD

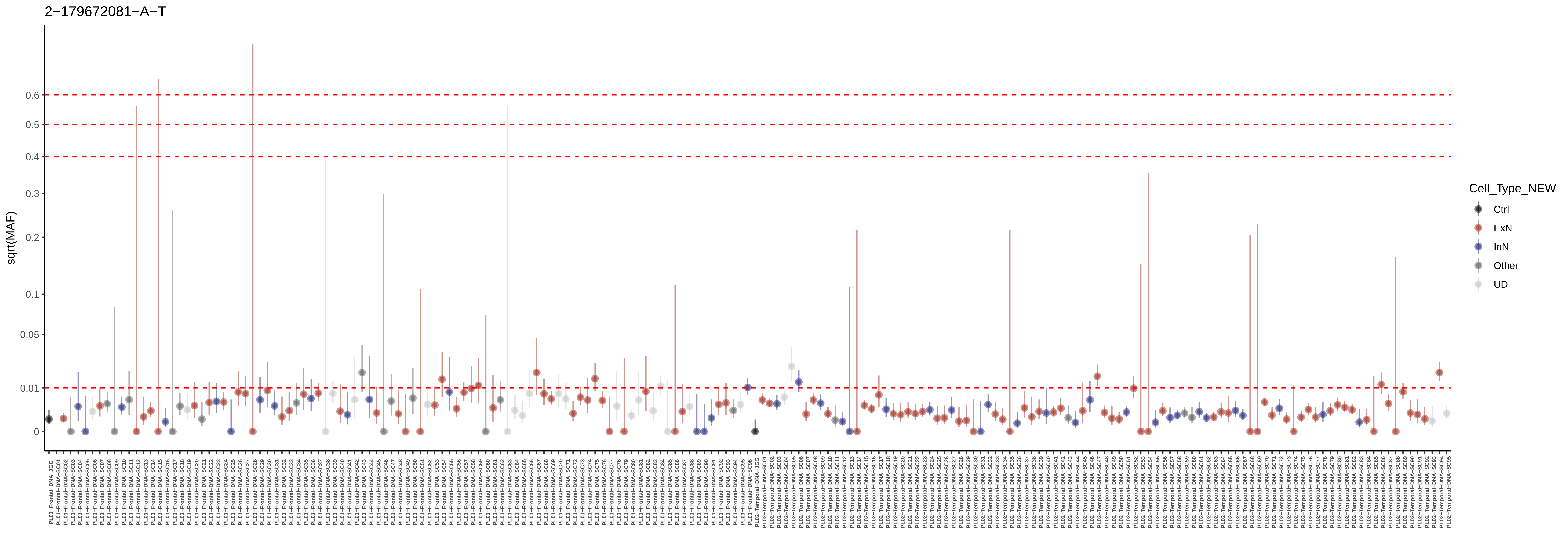
2−179672081−A−T
sqrt(MAF)
0.6
0.5
0.4
0.3
0.2
0.1
0.05
0.01
0
Cell_Type_NEW
Ctrl
ExN
InN
Other
UD

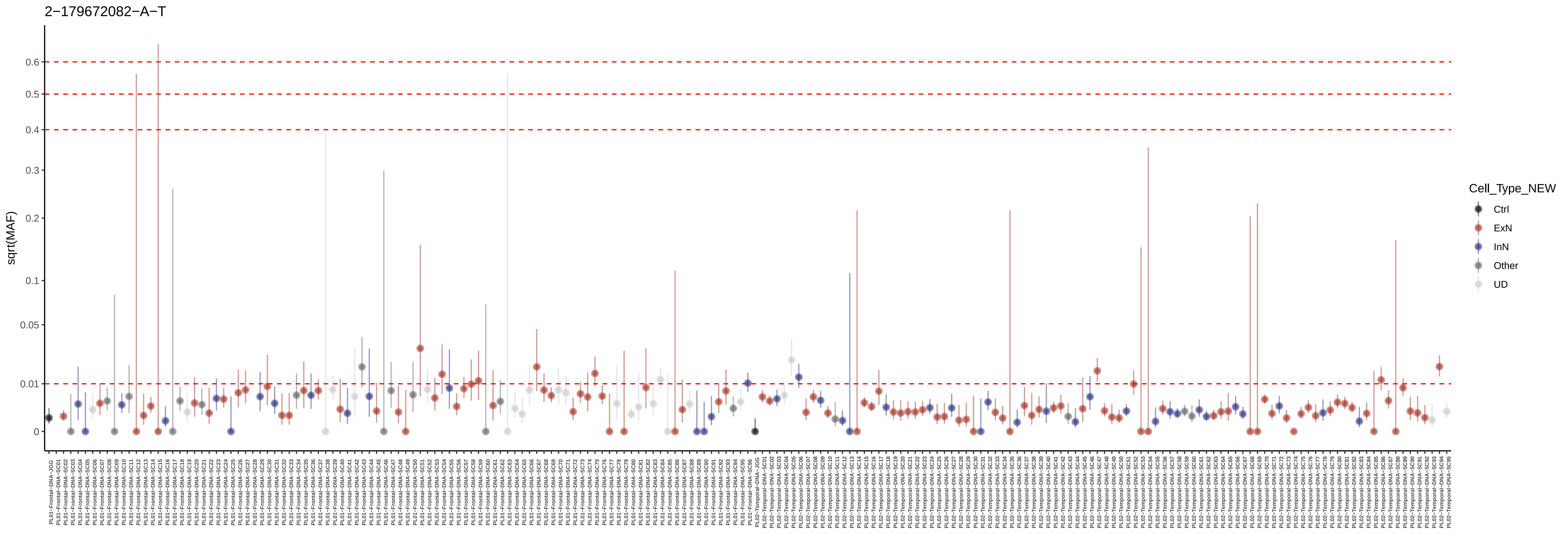
2−179672082−A−T
sqrt(MAF)
0.6
0.5
0.4
0.3
0.2
0.1
0.05
0.01
0
Cell_Type_NEW
Ctrl
ExN
InN
Other
UD

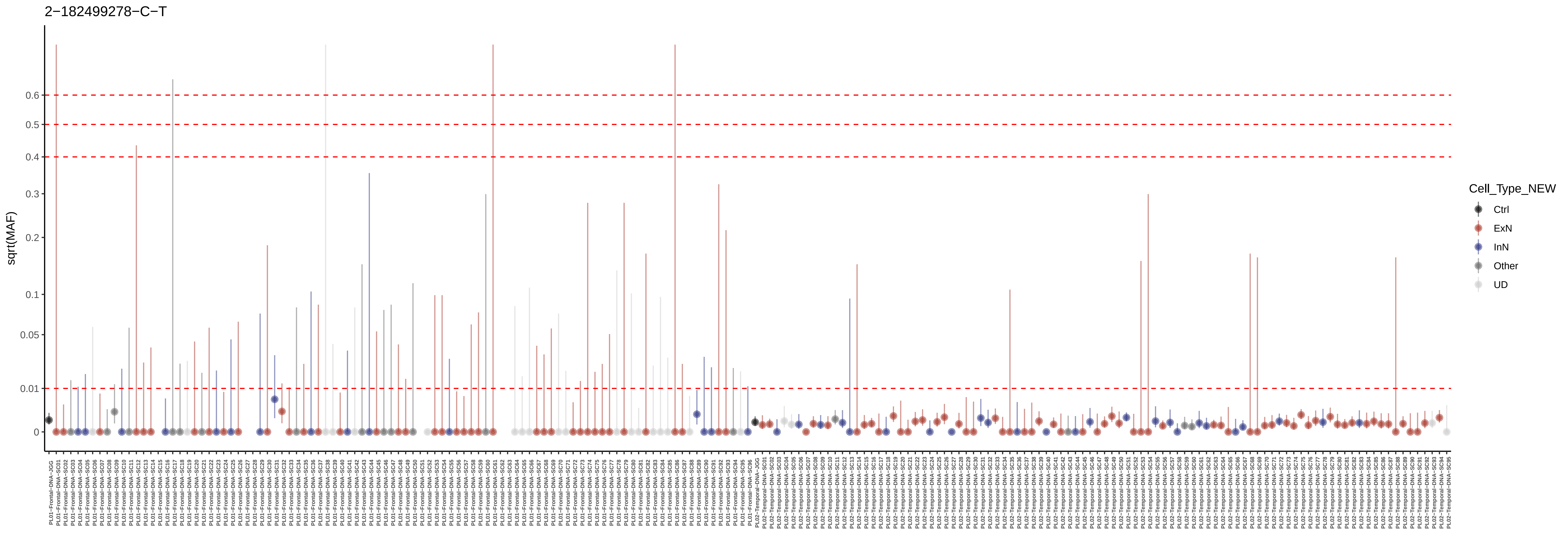

2−182499278−C−T
sqrt(MAF)
0.6
0.5
0.4
0.3
0.2
0.1
0.05
0.01
0
Cell_Type_NEW
Ctrl
ExN
InN
Other
UD

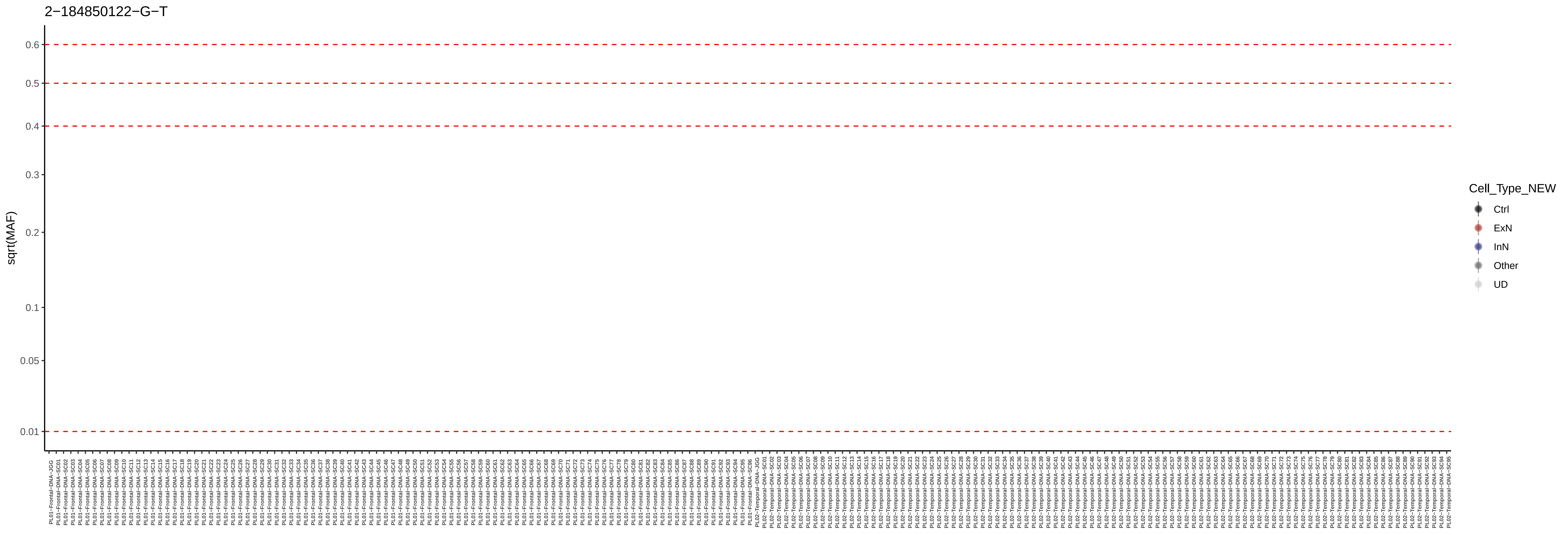
2−184850122−G−T
sqrt(MAF)
0.6
0.5
0.4
0.3
0.2
0.1
0.05
0.01
Cell_Type_NEW
Ctrl
ExN
InN
Other
UD

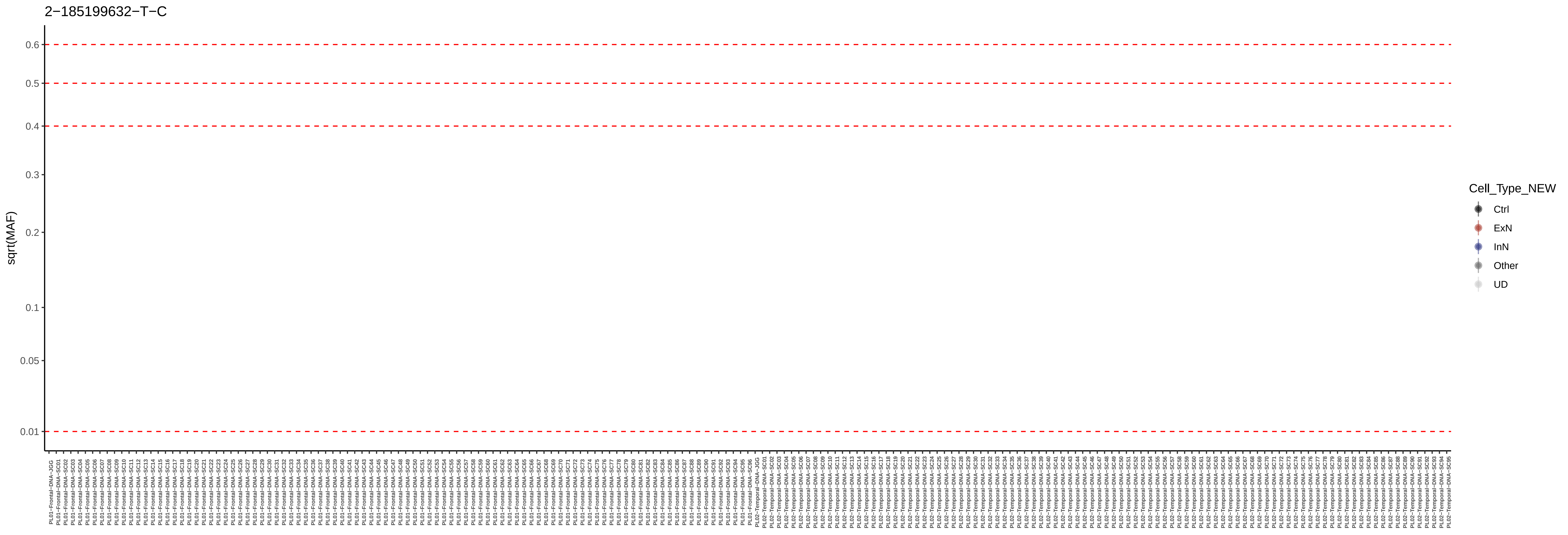
2−185199632−T−C
sqrt(MAF)
0.6
0.5
0.4
0.3
0.2
0.1
0.05
0.01
Cell_Type_NEW
Ctrl
ExN
InN
Other
UD

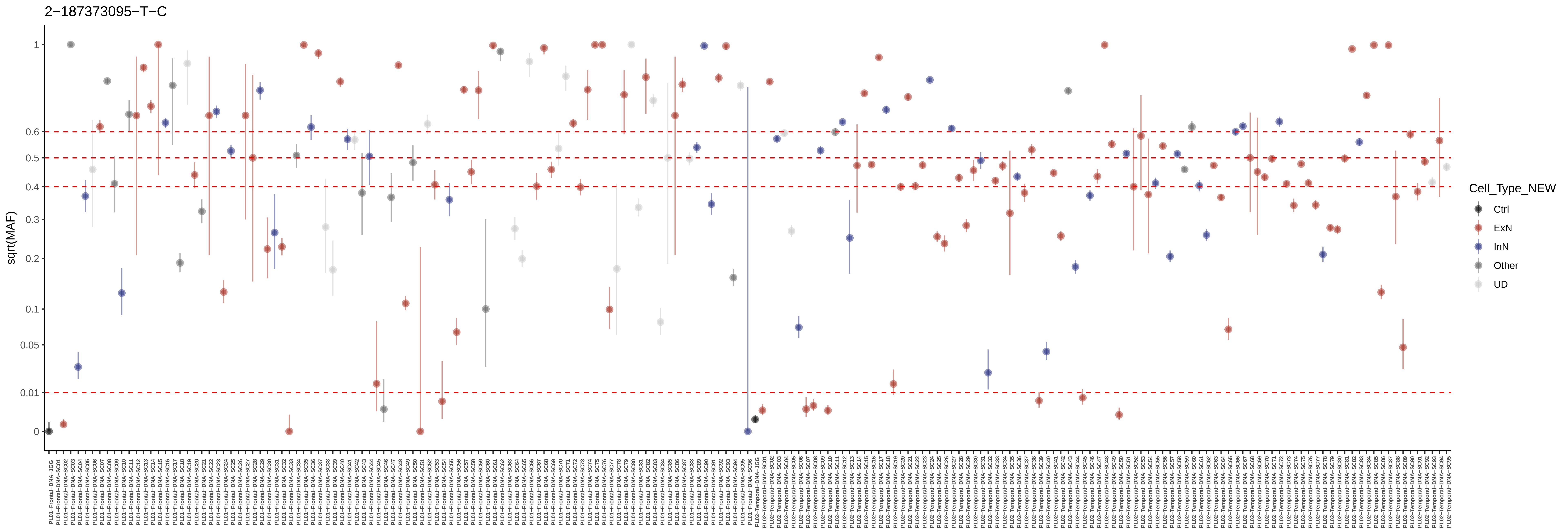
2−187373095−T−C
sqrt(MAF)
1
0.6
0.5
0.4
0.3
0.2
0.1
0.05
0.01
0
Cell_Type_NEW
Ctrl
ExN
InN
Other
UD

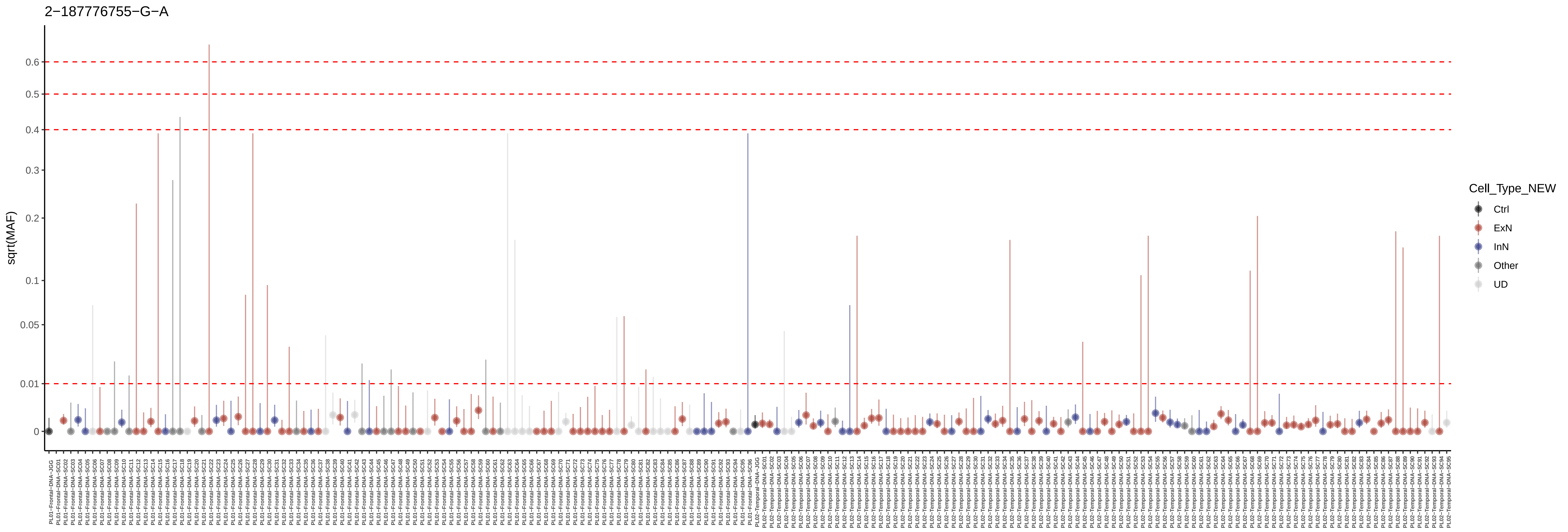
2−187776755−G−A
sqrt(MAF)
0.6
0.5
0.4
0.3
0.2
0.1
0.05
0.01
0
Cell_Type_NEW
Ctrl
ExN
InN
Other
UD

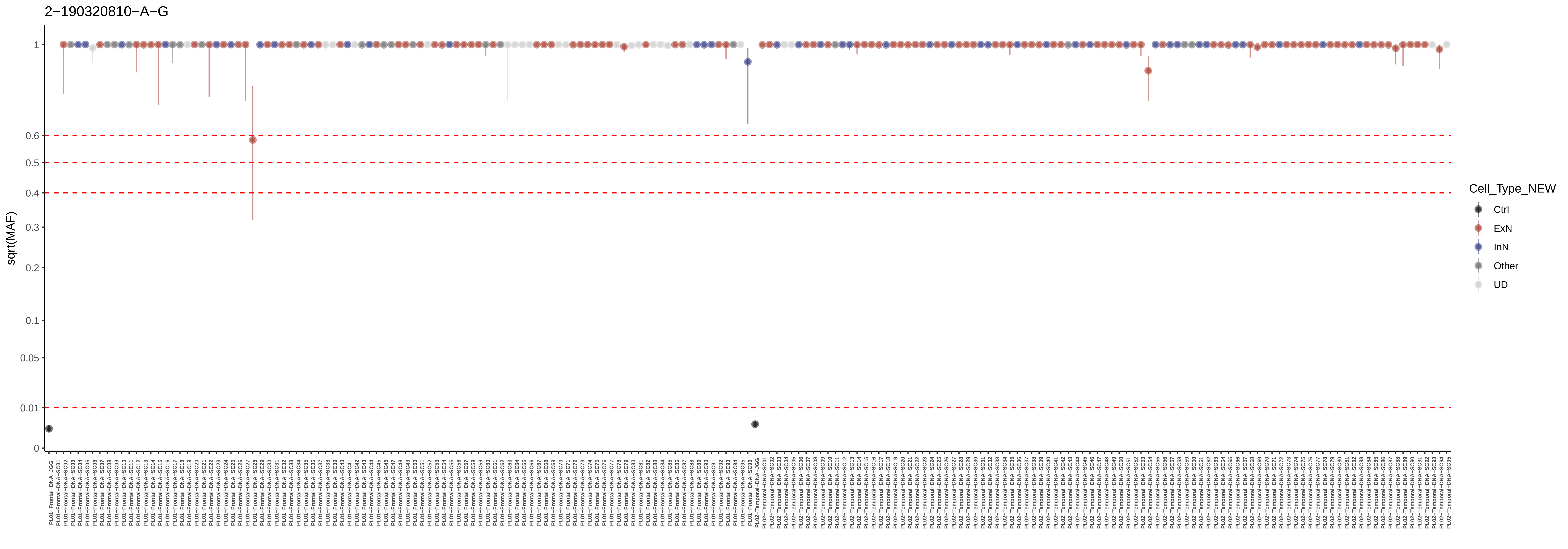
2−190320810−A−G
sqrt(MAF)
1
0.6
0.5
0.4
0.3
0.2
0.1
0.05
0.01
0
Cell_Type_NEW
Ctrl
ExN
InN
Other
UD

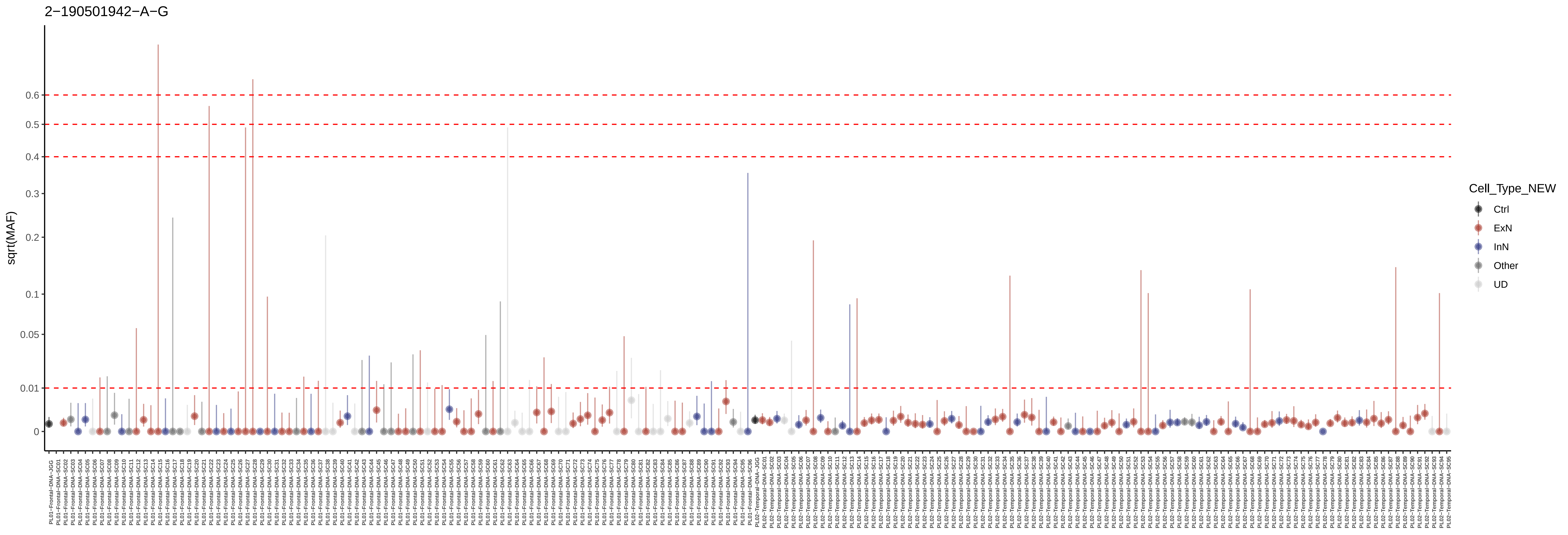
2−190501942−A−G
sqrt(MAF)
0.6
0.5
0.4
0.3
0.2
0.1
0.05
0.01
0
Cell_Type_NEW
Ctrl
ExN
InN
Other
UD

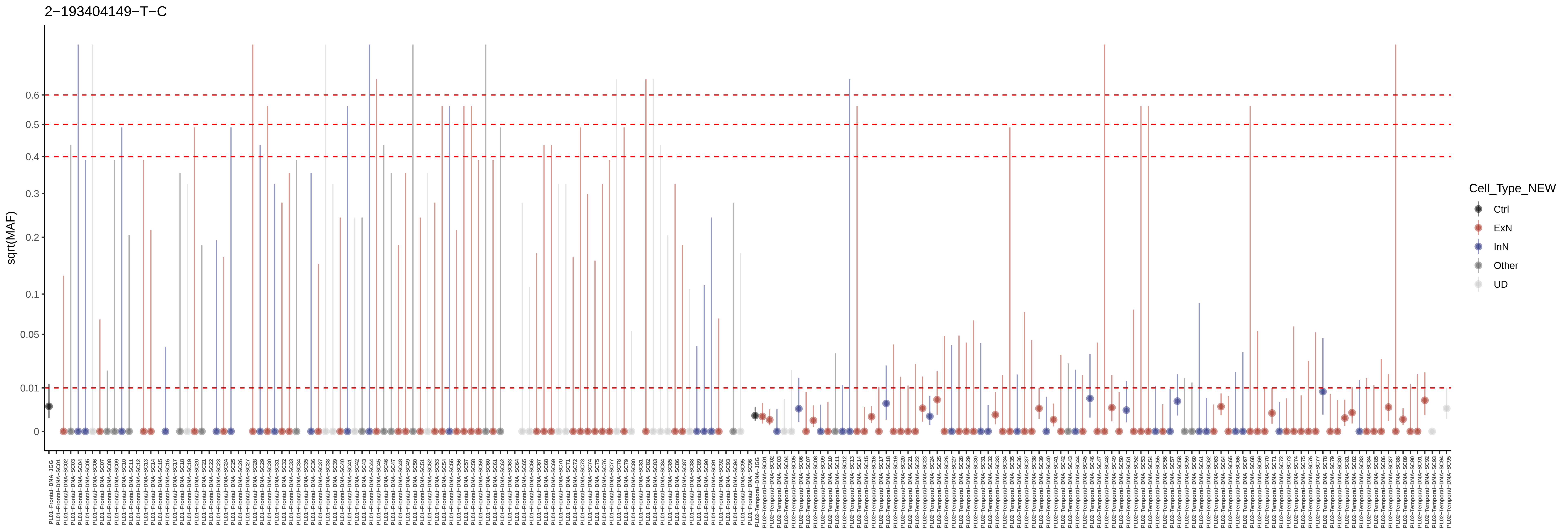
2−193404149−T−C
sqrt(MAF)
0.6
0.5
0.4
0.3
0.2
0.1
0.05
0.01
0
Cell_Type_NEW
Ctrl
ExN
InN
Other
UD

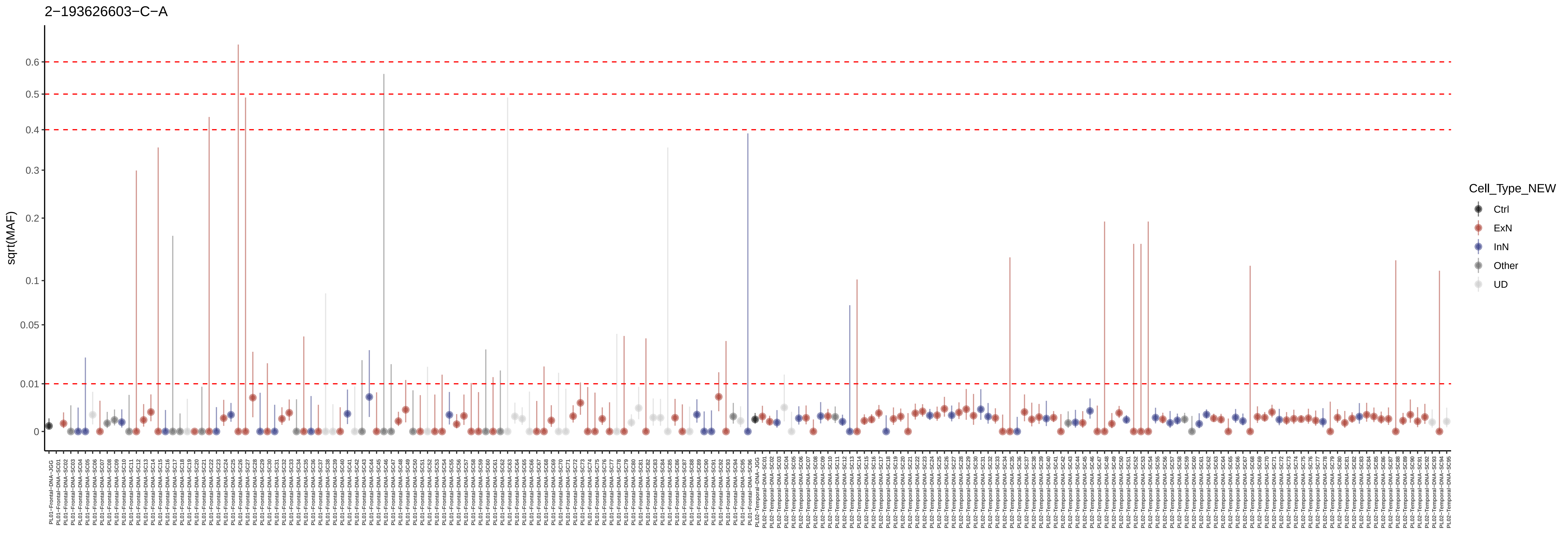

2−193626603−C−A
sqrt(MAF)
0.6
0.5
0.4
0.3
0.2
0.1
0.05
0.01
0
Cell_Type_NEW
Ctrl
ExN
InN
Other
UD

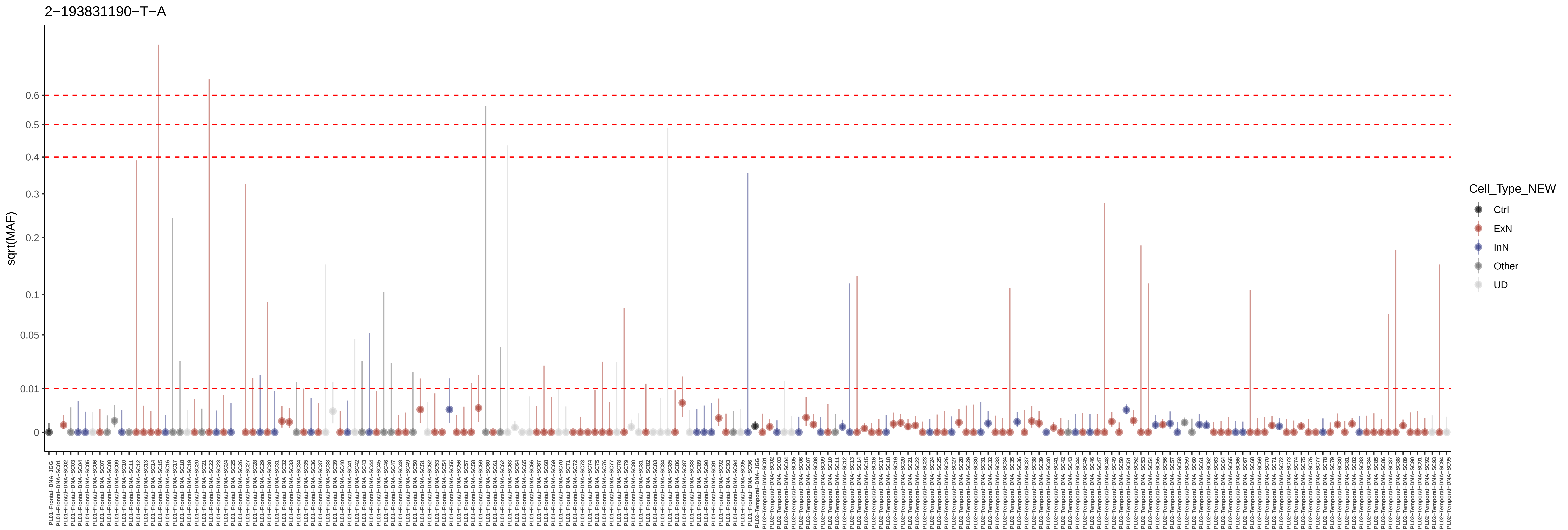
2−193831190−T−A
sqrt(MAF)
0.6
0.5
0.4
0.3
0.2
0.1
0.05
0.01
0
Cell_Type_NEW
Ctrl
ExN
InN
Other
UD

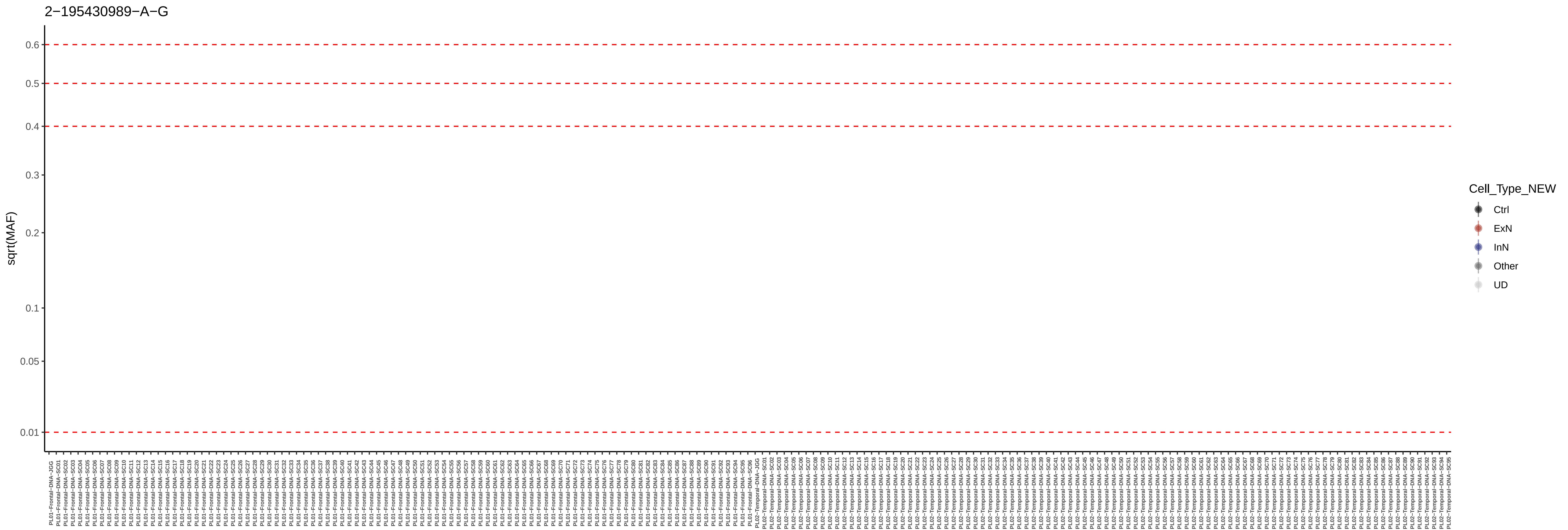

2−195430989−A−G
sqrt(MAF)
0.6
0.5
0.4
0.3
0.2
0.1
0.05
0.01
Cell_Type_NEW
Ctrl
ExN
InN
Other
UD

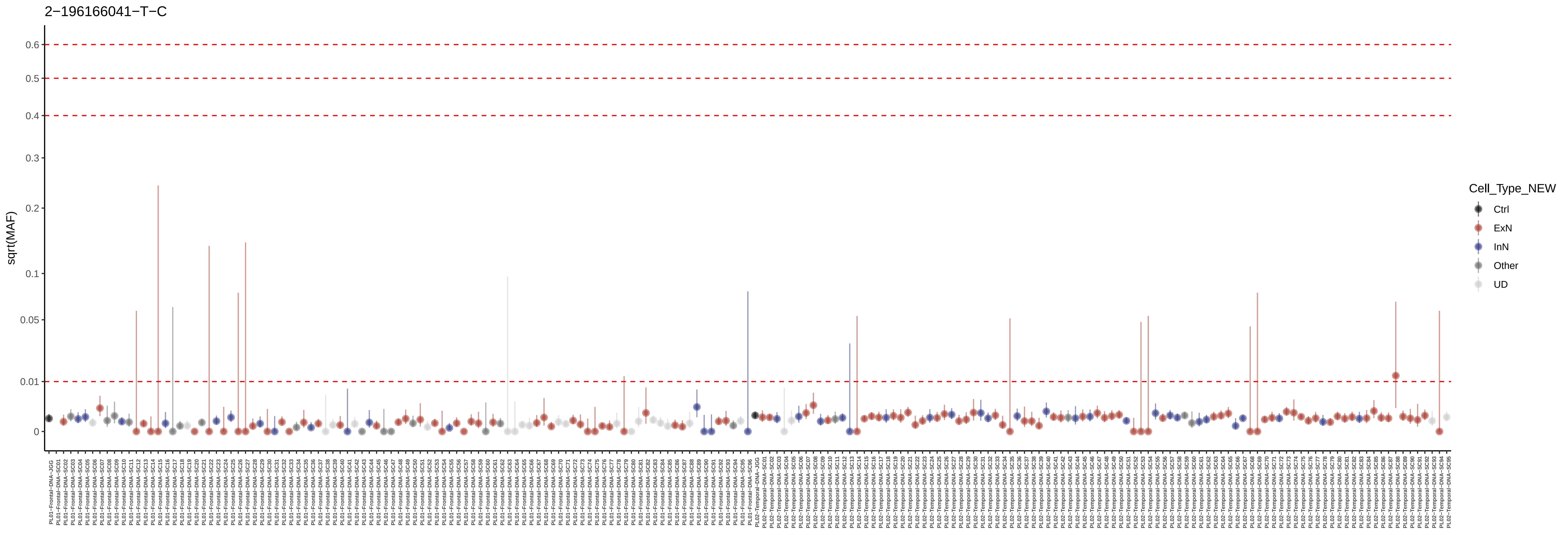

2−196166041−T−C
sqrt(MAF)
0.6
0.5
0.4
0.3
0.2
0.1
0.05
0.01
0
Cell_Type_NEW
Ctrl
ExN
InN
Other
UD

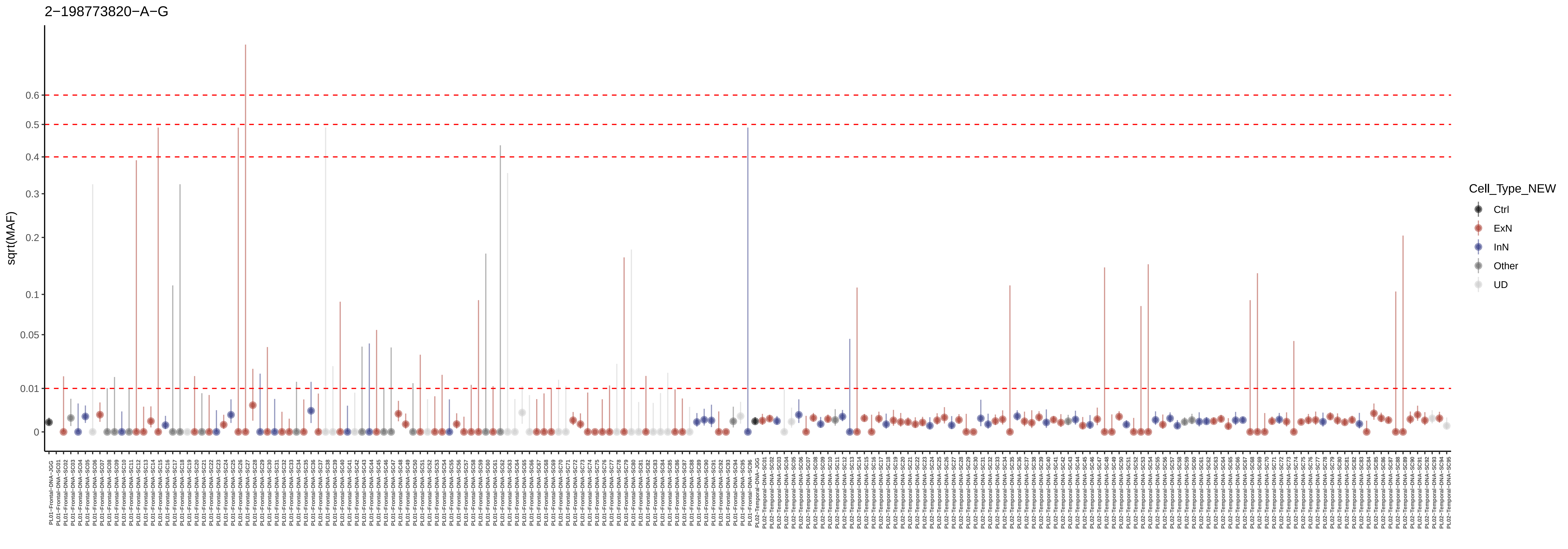
2−198773820−A−G
sqrt(MAF)
0.6
0.5
0.4
0.3
0.2
0.1
0.05
0.01
0
Cell_Type_NEW
Ctrl
ExN
InN
Other
UD

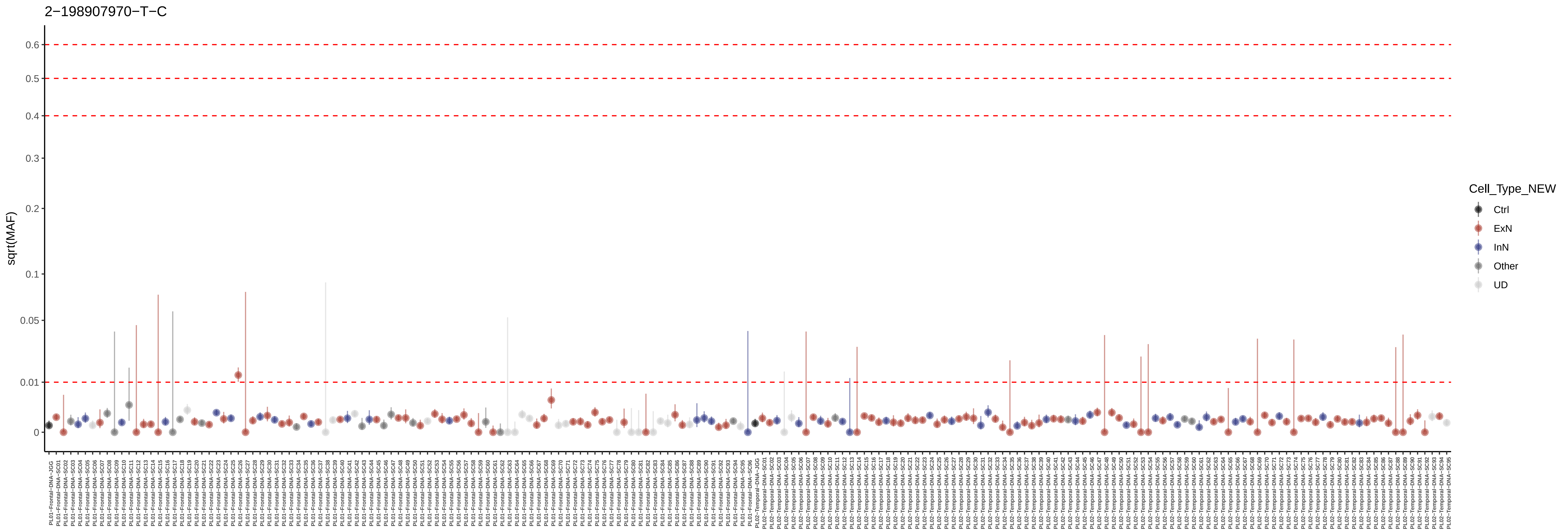
2−198907970−T−C
sqrt(MAF)
0.6
0.5
0.4
0.3
0.2
0.1
0.05
0.01
0
Cell_Type_NEW
Ctrl
ExN
InN
Other
UD

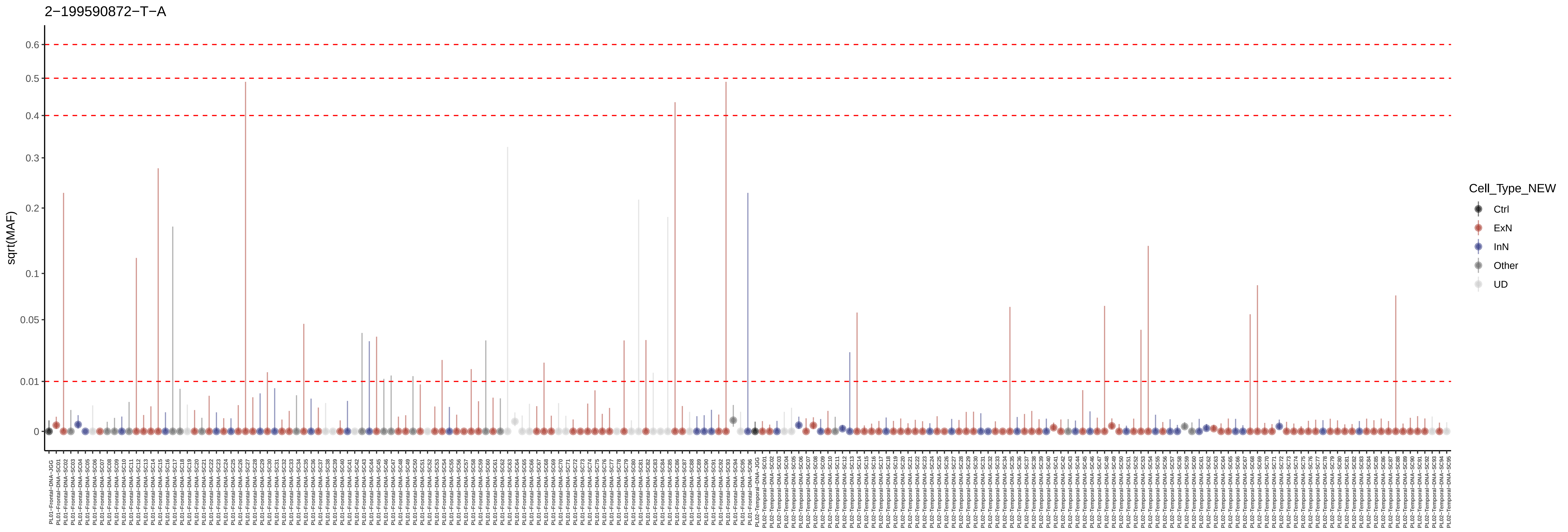

2−199590872−T−A
sqrt(MAF)
0.6
0.5
0.4
0.3
0.2
0.1
0.05
0.01
0
Cell_Type_NEW
Ctrl
ExN
InN
Other
UD

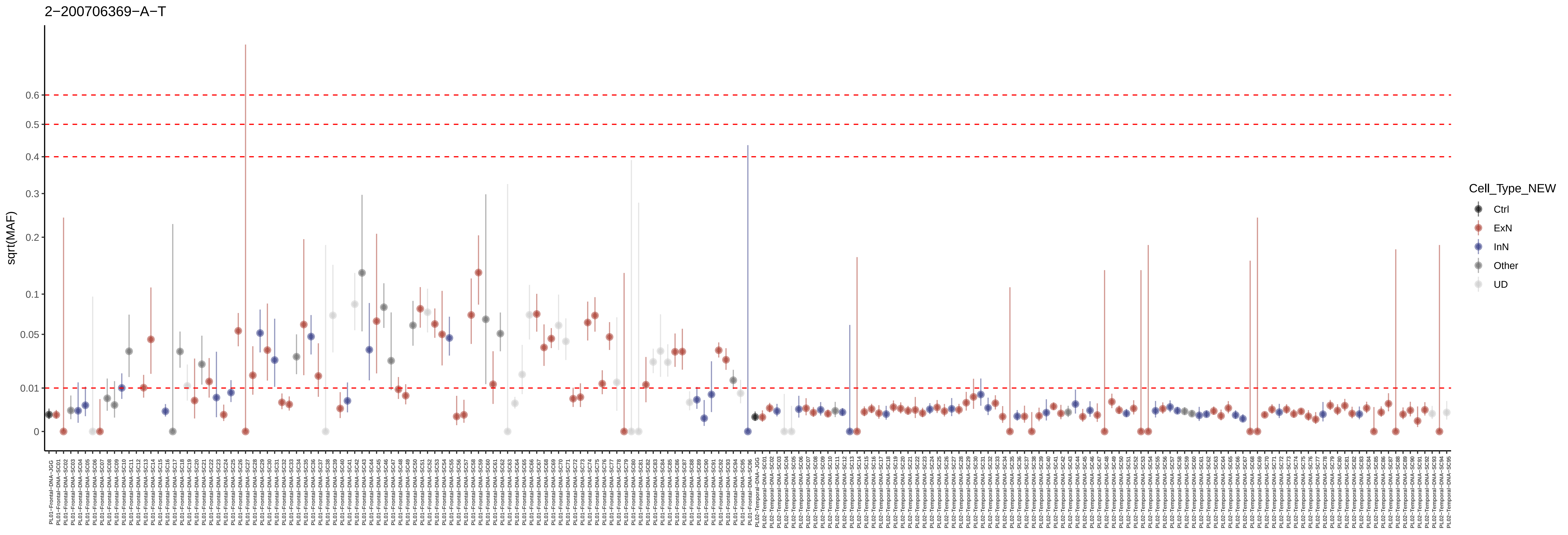
2−200706369−A−T
sqrt(MAF)
0.6
0.5
0.4
0.3
0.2
0.1
0.05
0.01
0
Cell_Type_NEW
Ctrl
ExN
InN
Other
UD

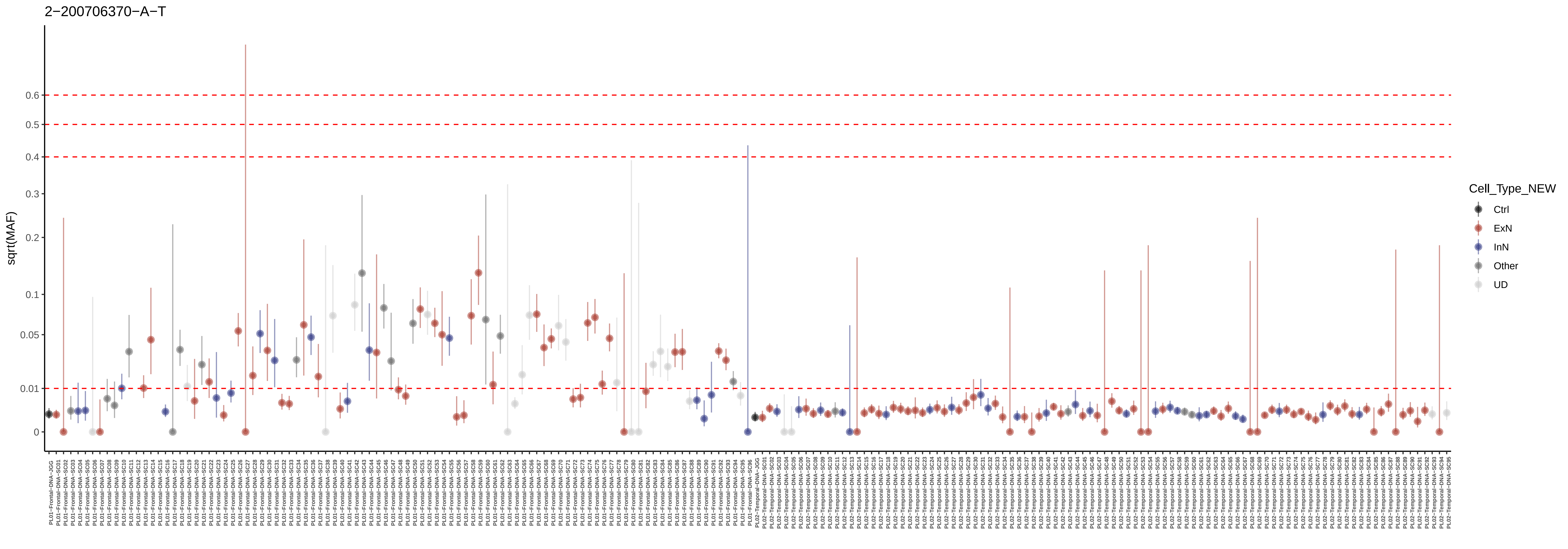
2−200706370−A−T
sqrt(MAF)
0.6
0.5
0.4
0.3
0.2
0.1
0.05
0.01
0
Cell_Type_NEW
Ctrl
ExN
InN
Other
UD

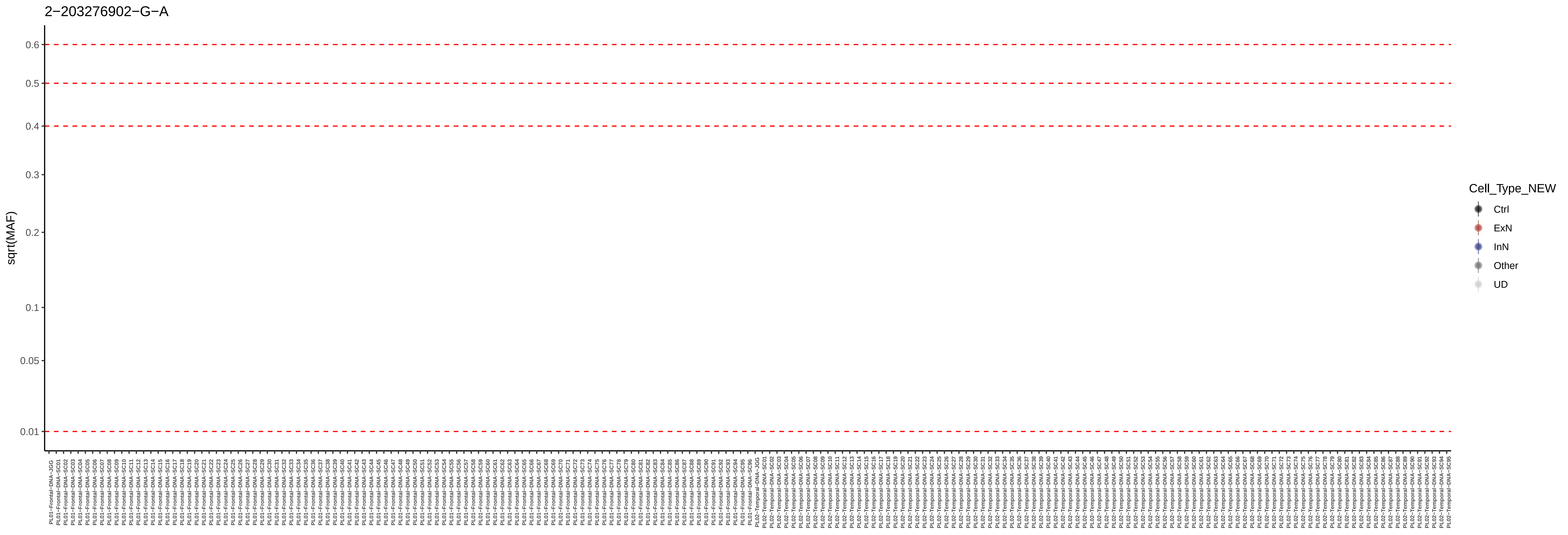

2−203276902−G−A
sqrt(MAF)
0.6
0.5
0.4
0.3
0.2
0.1
0.05
0.01
Cell_Type_NEW
Ctrl
ExN
InN
Other
UD

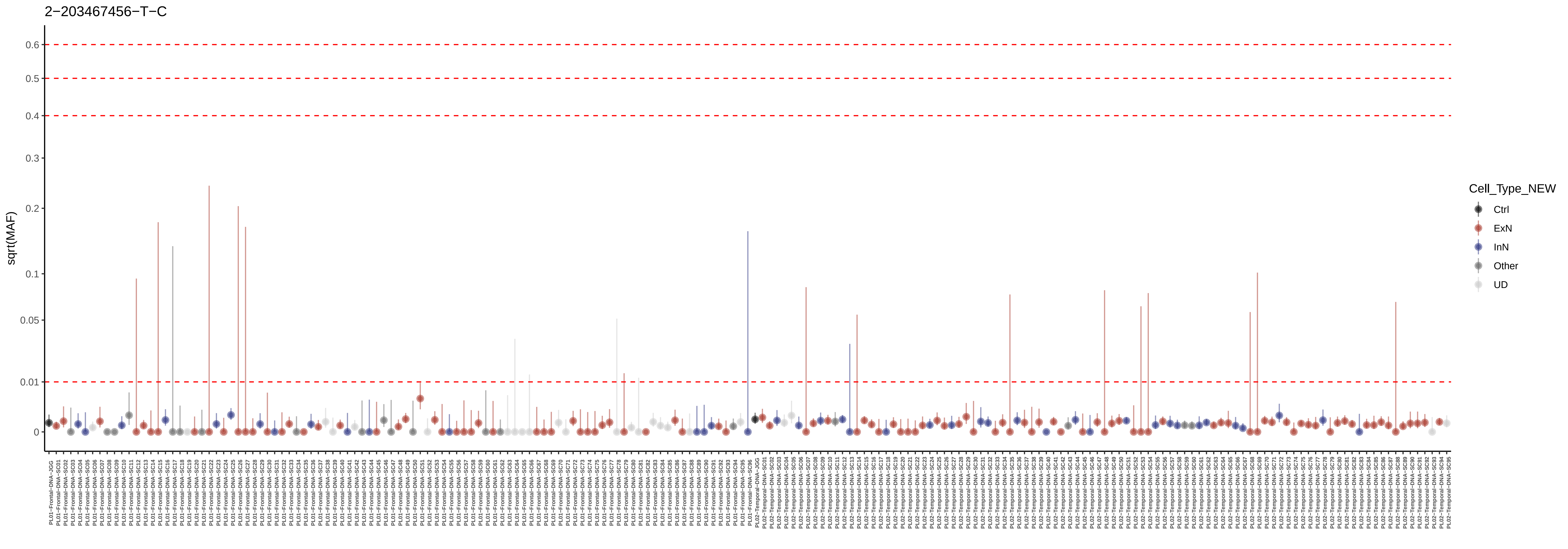
2−203467456−T−C
sqrt(MAF)
0.6
0.5
0.4
0.3
0.2
0.1
0.05
0.01
0
Cell_Type_NEW
Ctrl
ExN
InN
Other
UD

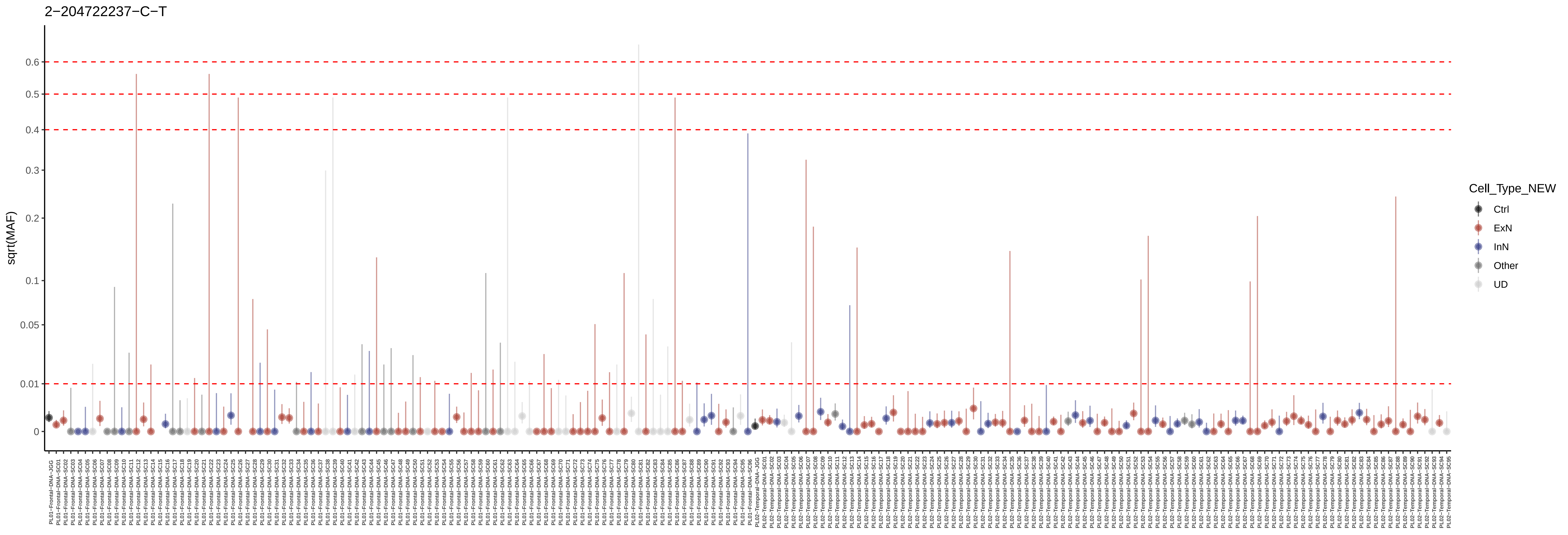
2−204722237−C−T
sqrt(MAF)
0.6
0.5
0.4
0.3
0.2
0.1
0.05
0.01
0
Cell_Type_NEW
Ctrl
ExN
InN
Other
UD

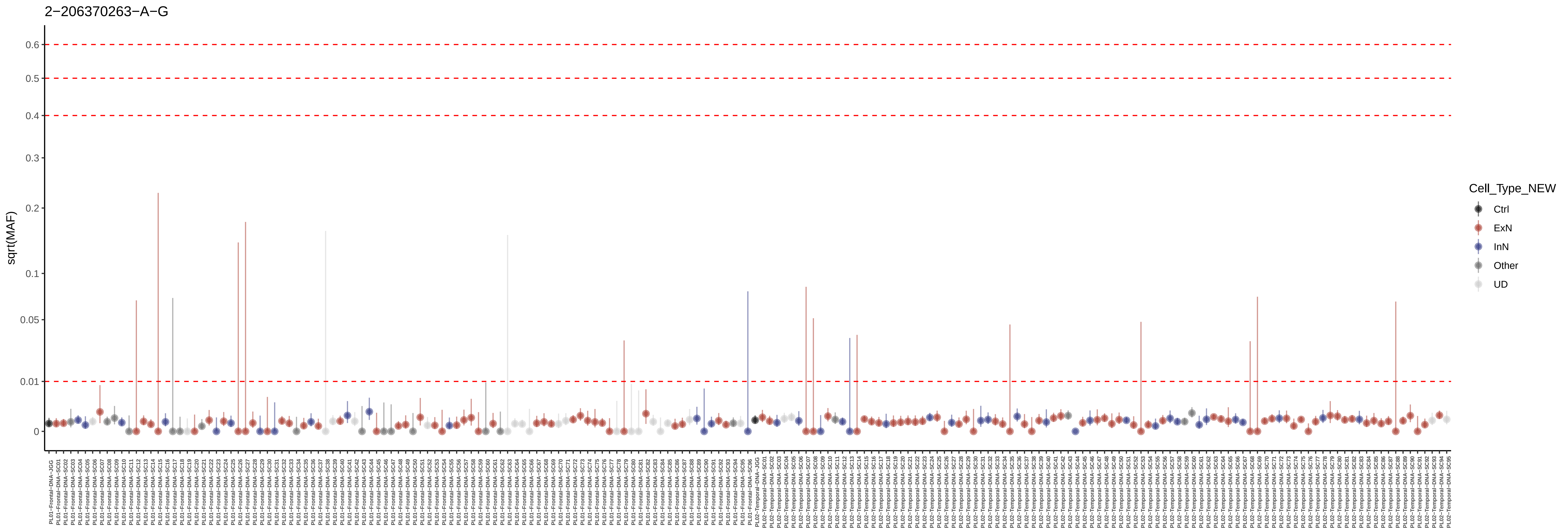
2−206370263−A−G
sqrt(MAF)
0.6
0.5
0.4
0.3
0.2
0.1
0.05
0.01
0
Cell_Type_NEW
Ctrl
ExN
InN
Other
UD

2−206998398−G−T

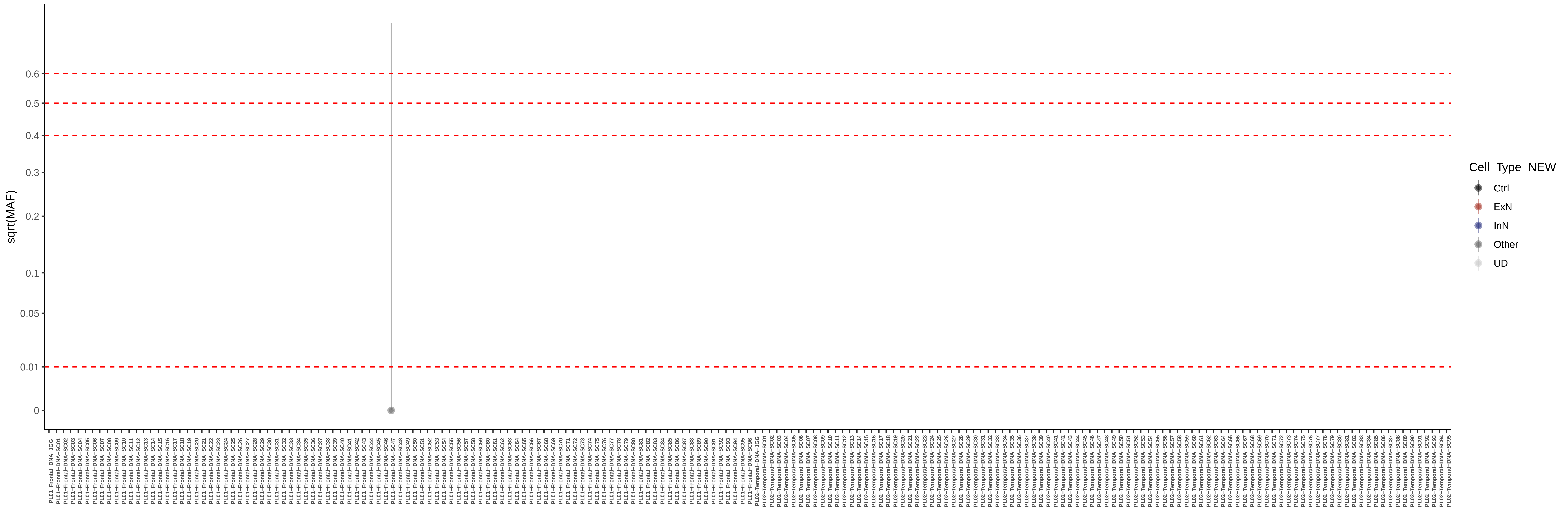

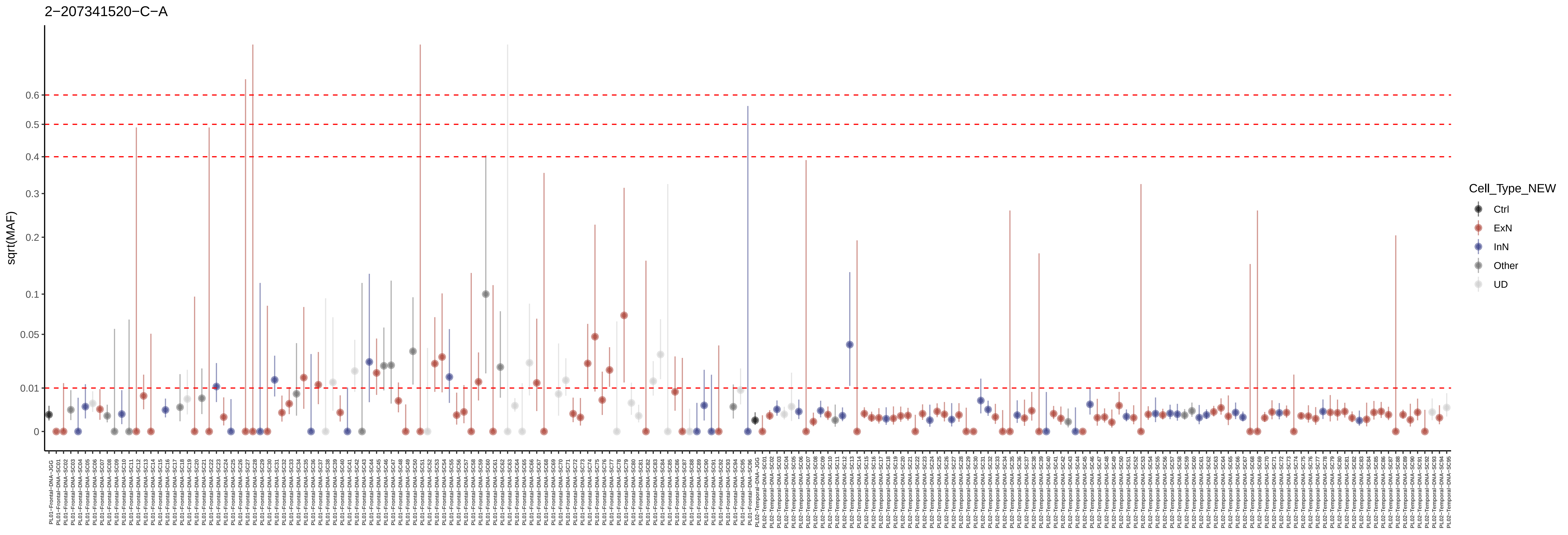
2−207341520−C−A
sqrt(MAF)
0.6
0.5
0.4
0.3
0.2
0.1
0.05
0.01
0
Cell_Type_NEW
Ctrl
ExN
InN
Other
UD

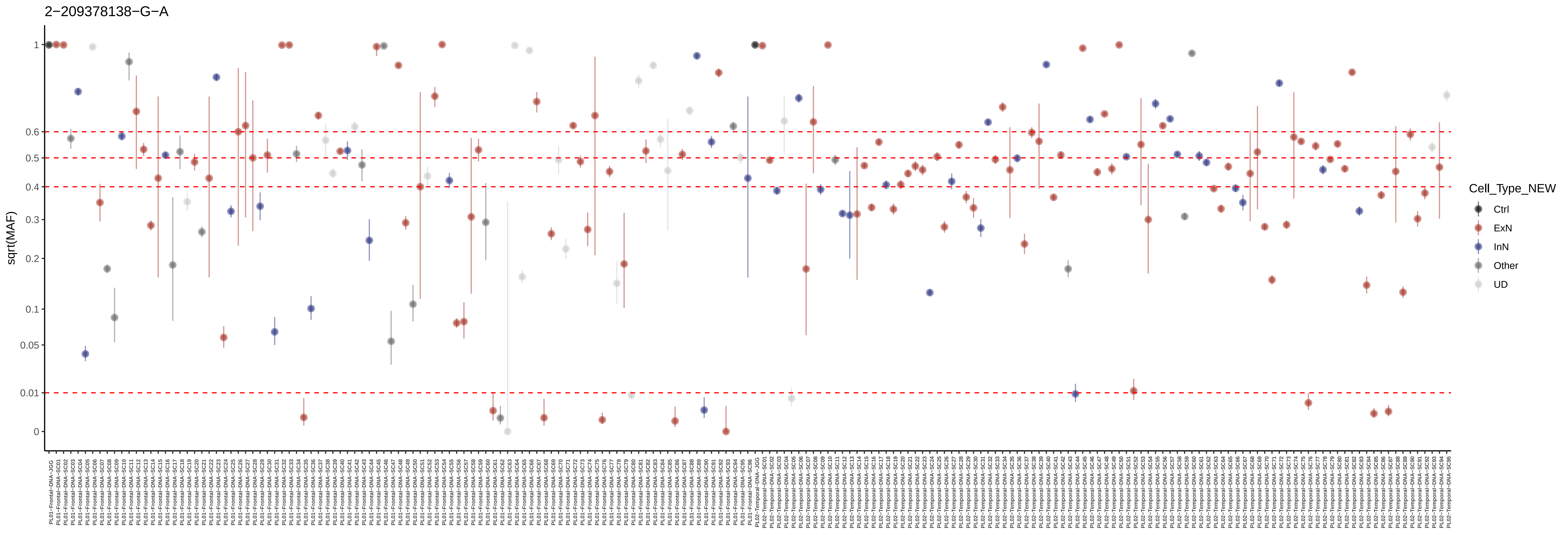
2−209378138−G−A
sqrt(MAF)
1
0.6
0.5
0.4
0.3
0.2
0.1
0.05
0.01
0
Cell_Type_NEW
Ctrl
ExN
InN
Other
UD

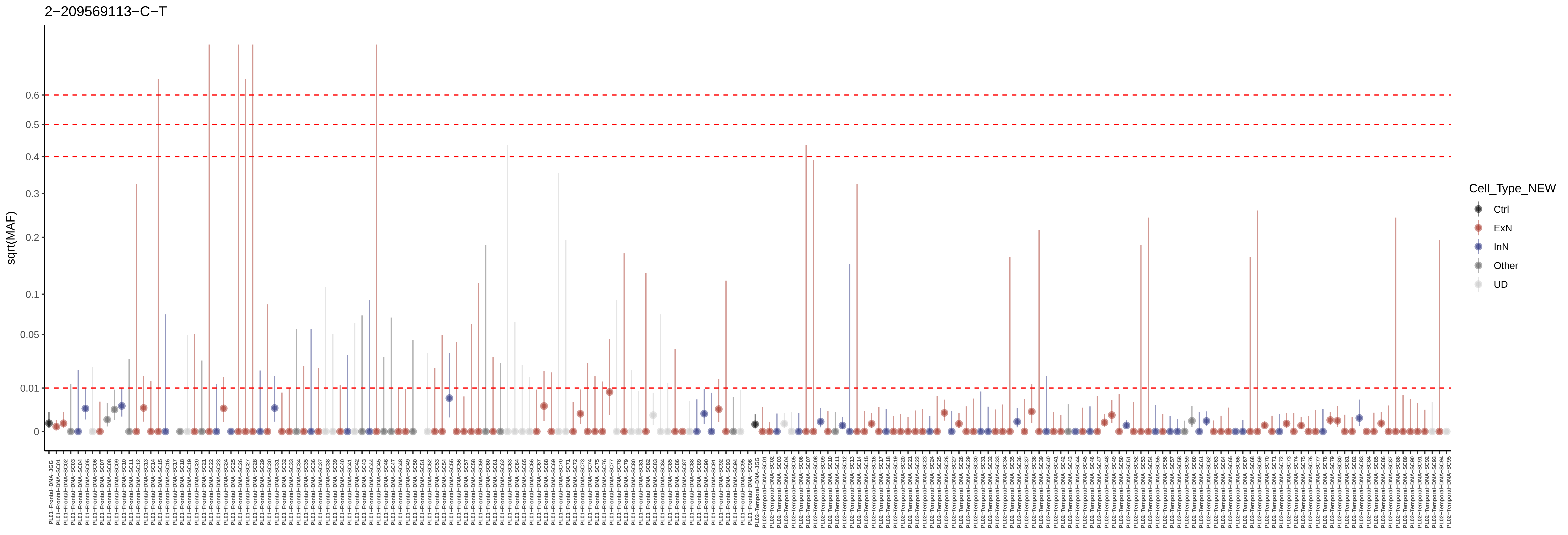
2−209569113−C−T
sqrt(MAF)
0.6
0.5
0.4
0.3
0.2
0.1
0.05
0.01
0
Cell_Type_NEW
Ctrl
ExN
InN
Other
UD

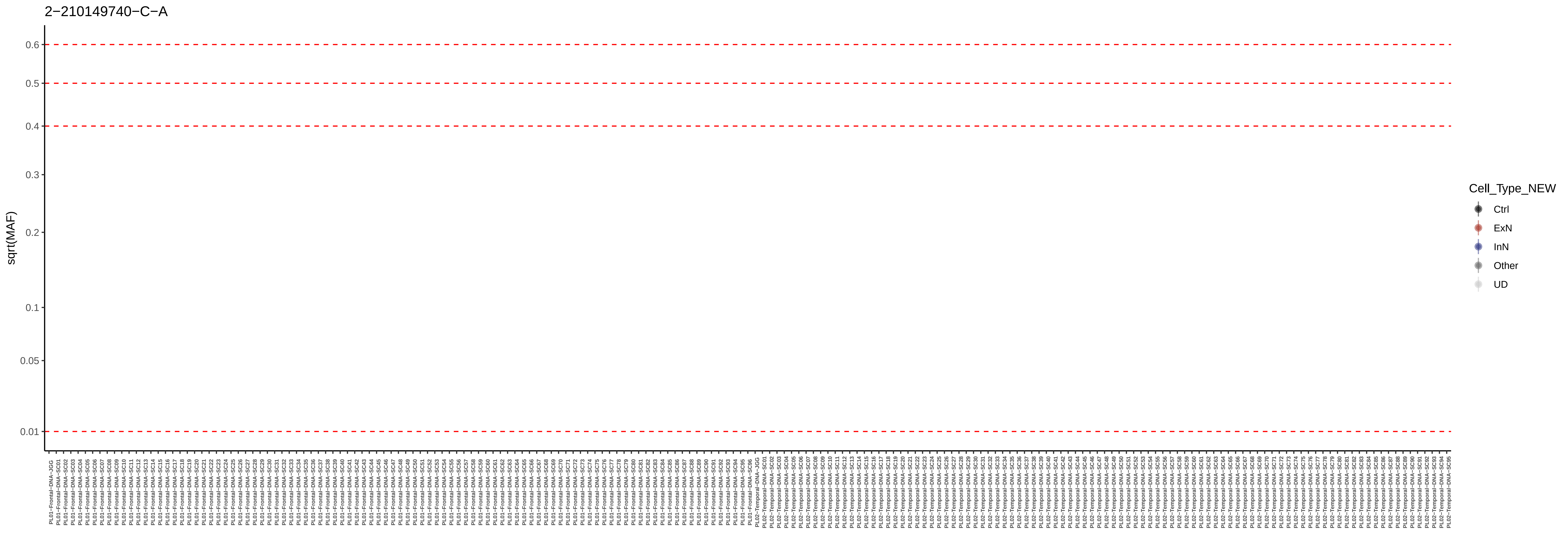
2−210149740−C−A
sqrt(MAF)
0.6
0.5
0.4
0.3
0.2
0.1
0.05
0.01
Cell_Type_NEW
Ctrl
ExN
InN
Other
UD

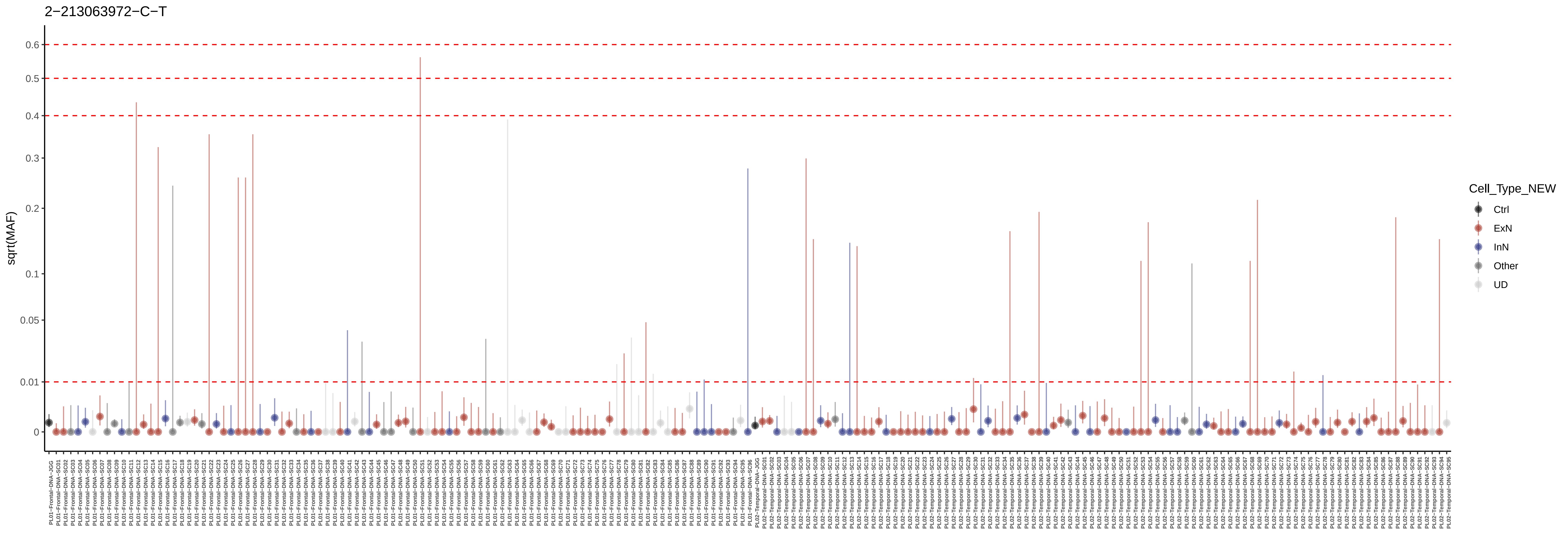
2−213063972−C−T
sqrt(MAF)
0.6
0.5
0.4
0.3
0.2
0.1
0.05
0.01
0
Cell_Type_NEW
Ctrl
ExN
InN
Other
UD

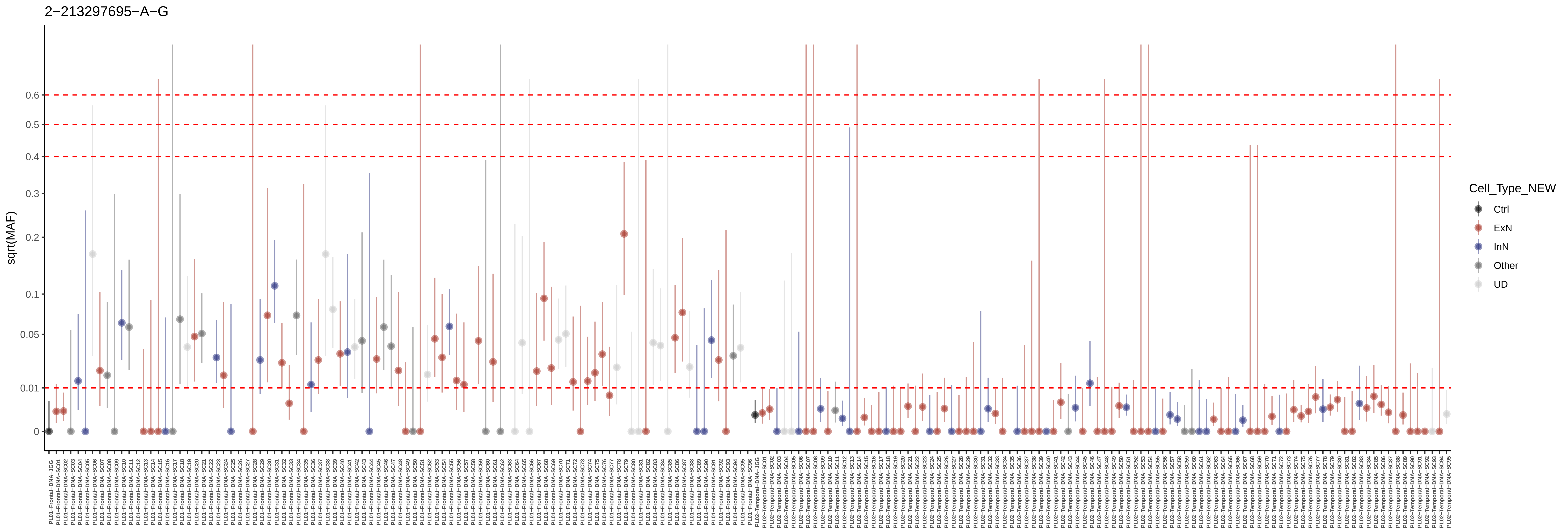
2−213297695−A−G
sqrt(MAF)
0.6
0.5
0.4
0.3
0.2
0.1
0.05
0.01
0
Cell_Type_NEW
Ctrl
ExN
InN
Other
UD

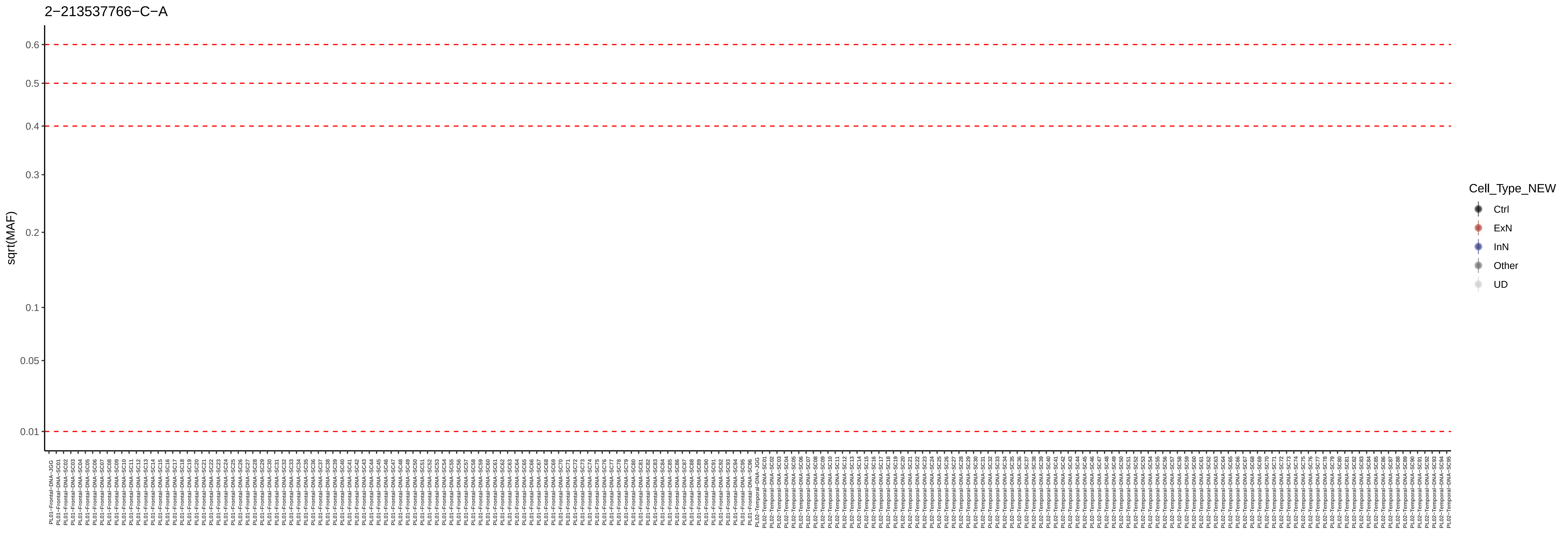
2−213537766−C−A
sqrt(MAF)
0.6
0.5
0.4
0.3
0.2
0.1
0.05
0.01
Cell_Type_NEW
Ctrl
ExN
InN
Other
UD

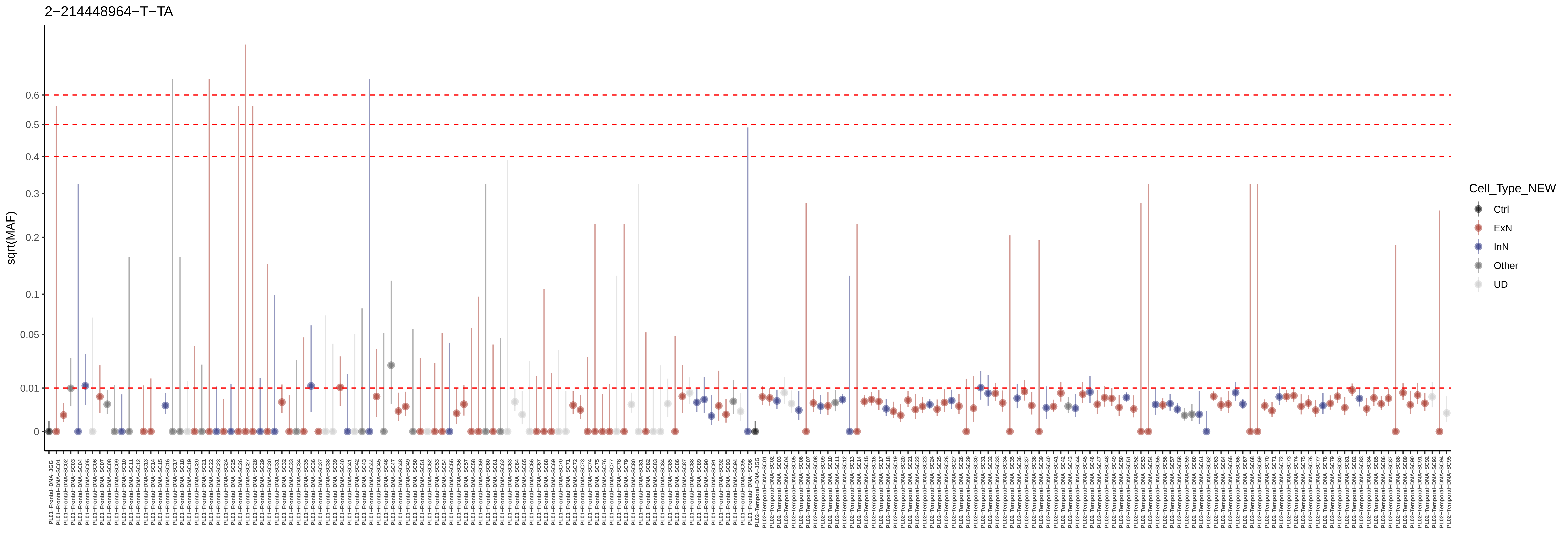

2−214448964−T−TA
sqrt(MAF)
0.6
0.5
0.4
0.3
0.2
0.1
0.05
0.01
0
Cell_Type_NEW
Ctrl
ExN
InN
Other
UD

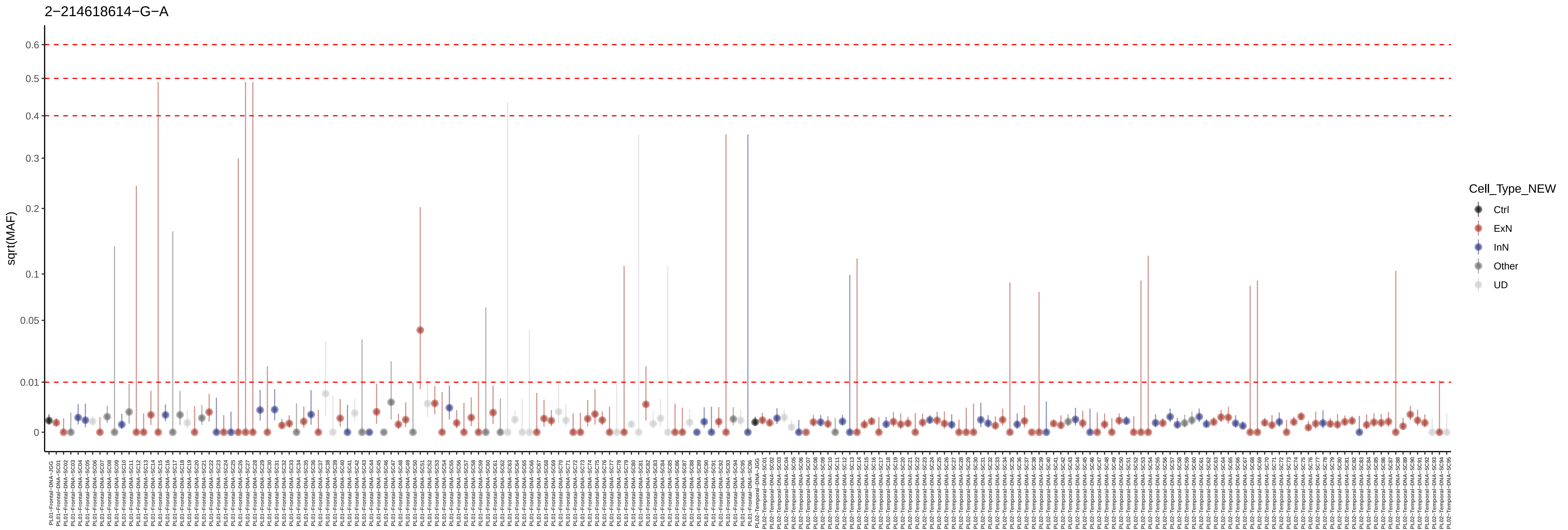

2−214618614−G−A
sqrt(MAF)
0.6
0.5
0.4
0.3
0.2
0.1
0.05
0.01
0
Cell_Type_NEW
Ctrl
ExN
InN
Other
UD

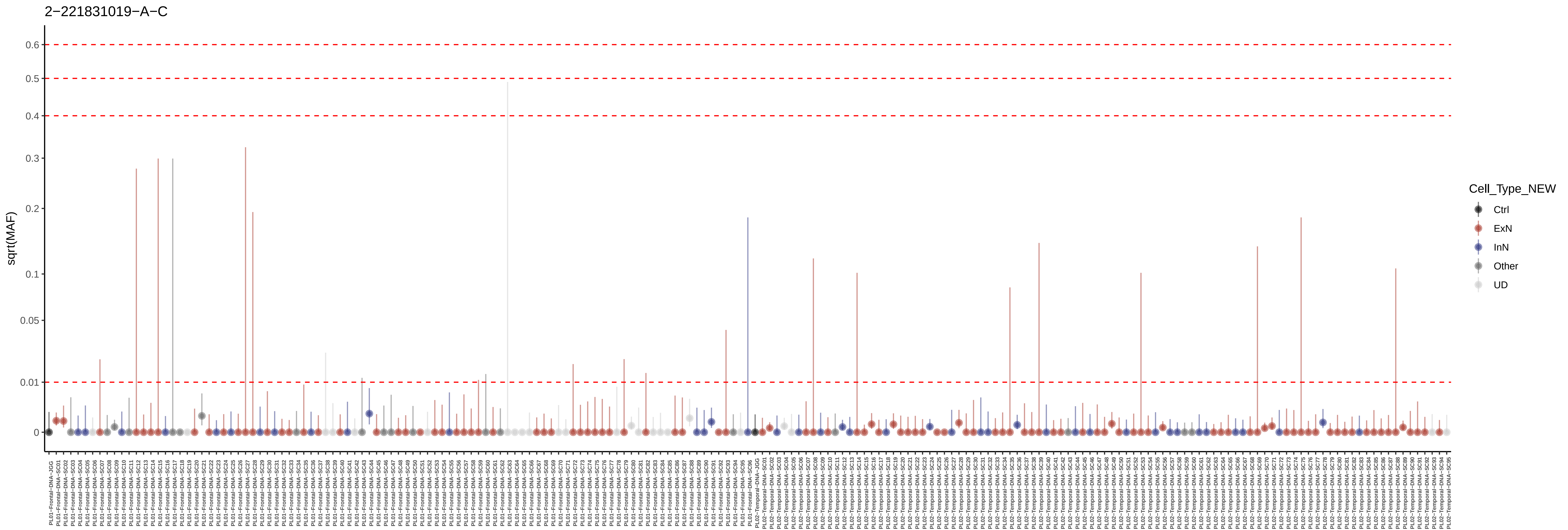
2−221831019−A−C
sqrt(MAF)
0.6
0.5
0.4
0.3
0.2
0.1
0.05
0.01
0
Cell_Type_NEW
Ctrl
ExN
InN
Other
UD

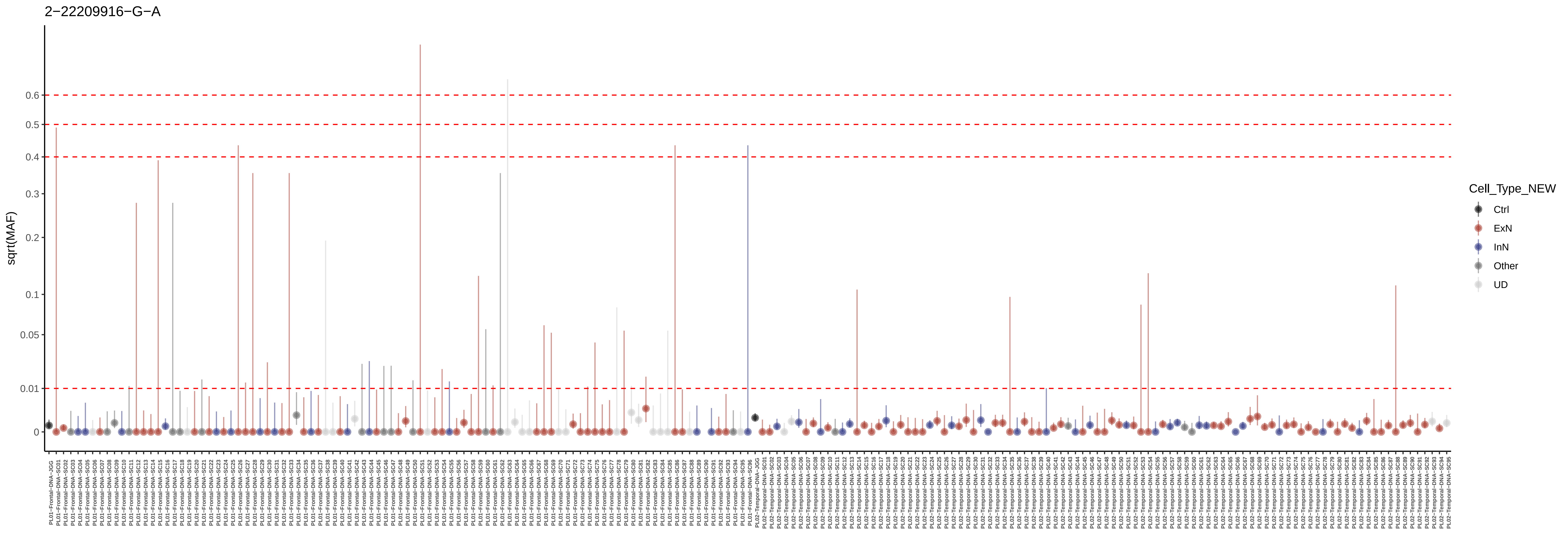

2−22209916−G−A
sqrt(MAF)
0.6
0.5
0.4
0.3
0.2
0.1
0.05
0.01
0
Cell_Type_NEW
Ctrl
ExN
InN
Other
UD

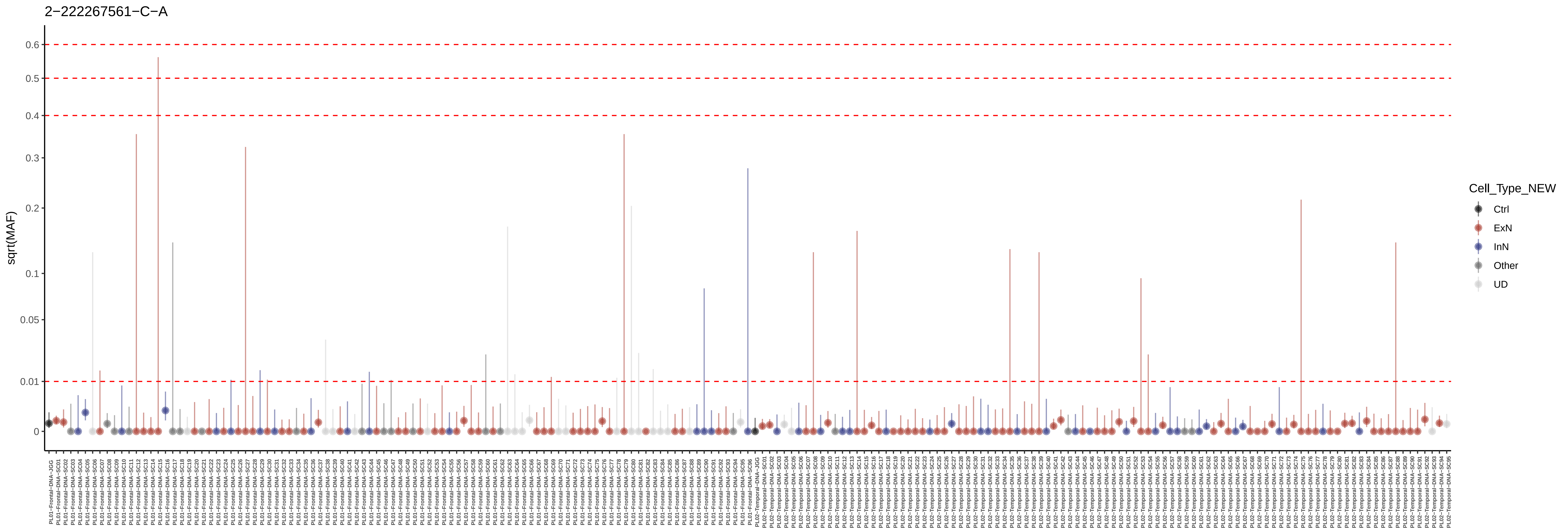

2−222267561−C−A
sqrt(MAF)
0.6
0.5
0.4
0.3
0.2
0.1
0.05
0.01
0
Cell_Type_NEW
Ctrl
ExN
InN
Other
UD

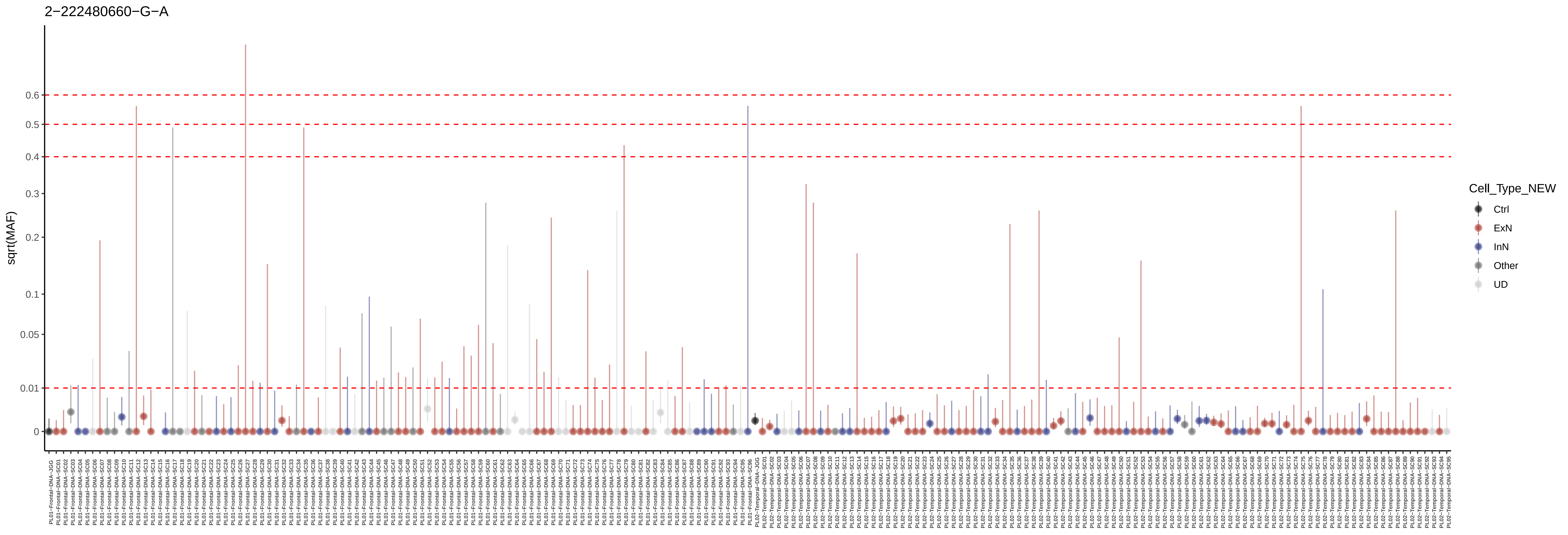
2−222480660−G−A
sqrt(MAF)
0.6
0.5
0.4
0.3
0.2
0.1
0.05
0.01
0
Cell_Type_NEW
Ctrl
ExN
InN
Other
UD

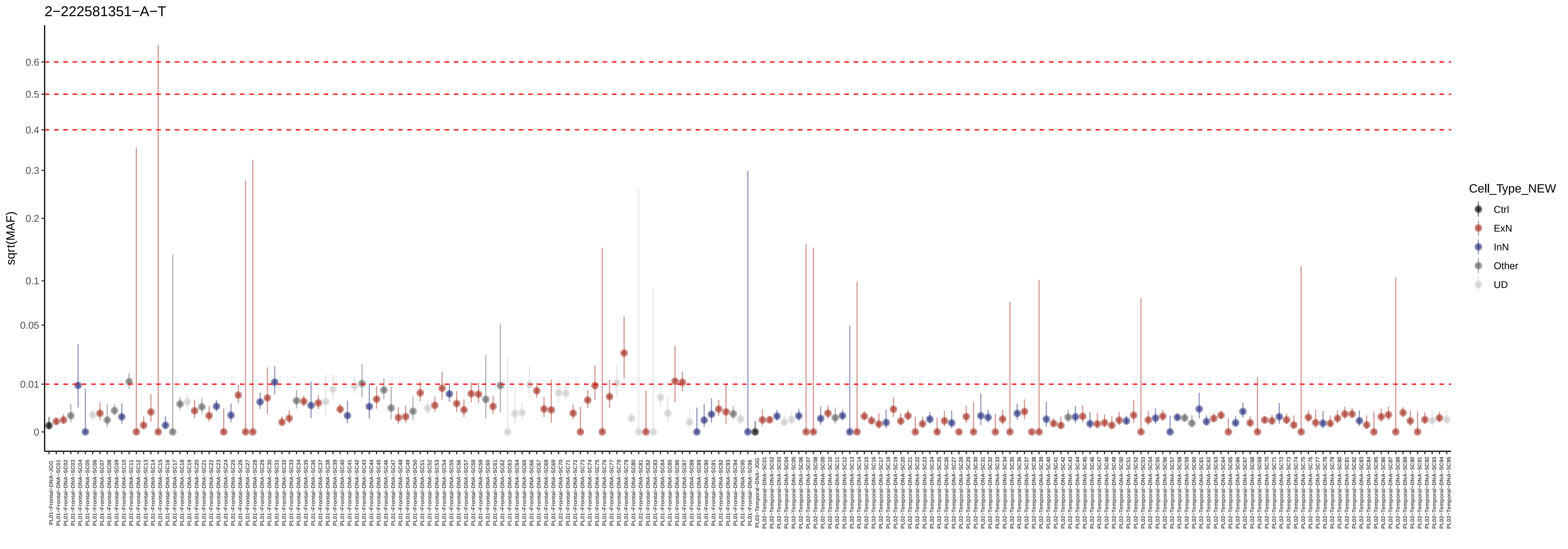
2−222581351−A−T
sqrt(MAF)
Cell_Type_NEW
Ctrl
ExN
InN
Other
UD

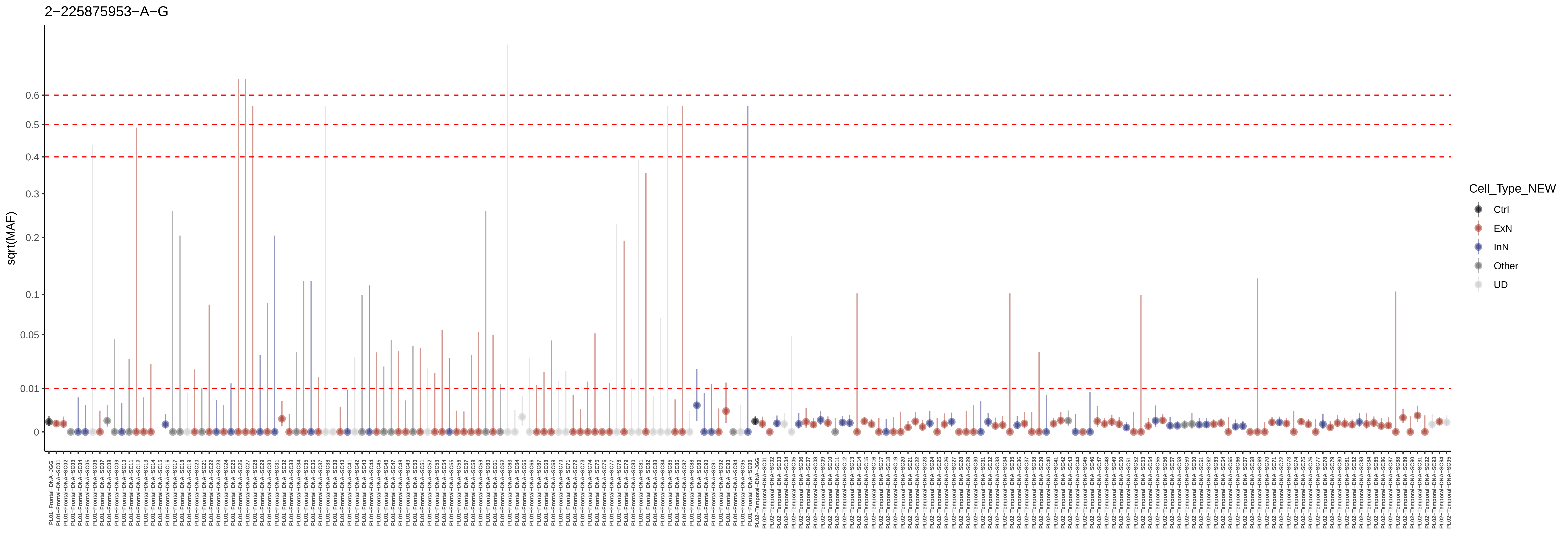

2−225875953−A−G
sqrt(MAF)
0.6
0.5
0.4
0.3
0.2
0.1
0.05
0.01
0
Cell_Type_NEW
Ctrl
ExN
InN
Other
UD

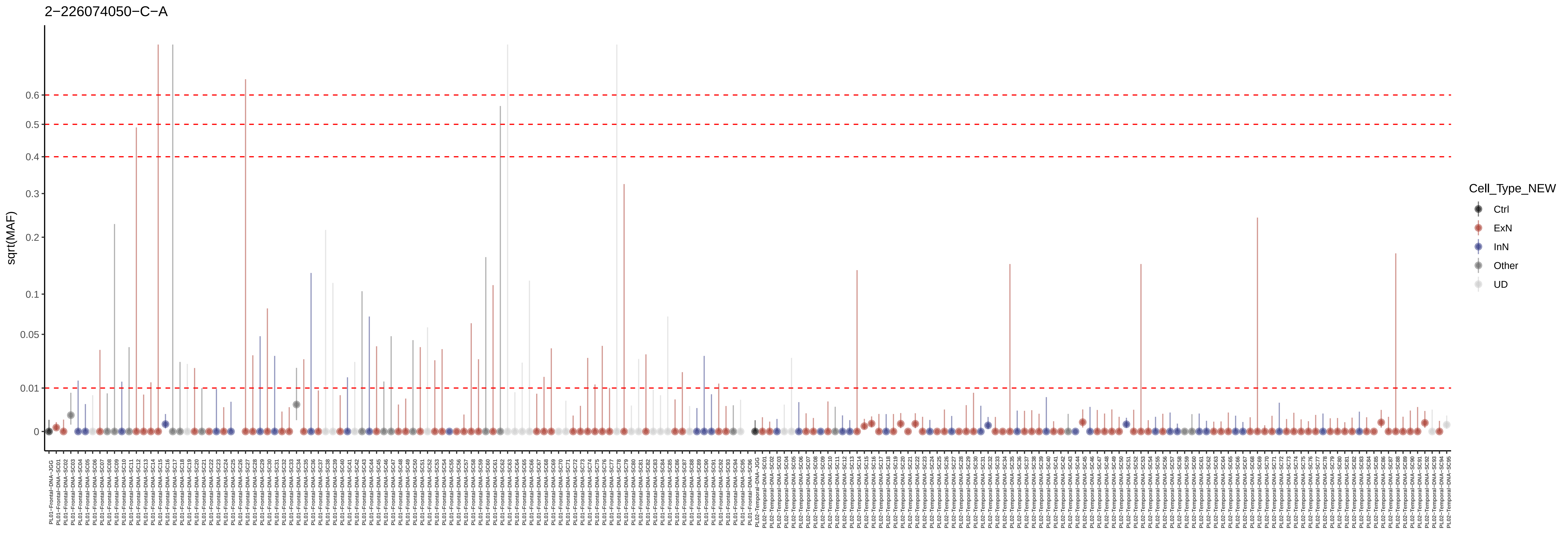

2−226074050−C−A
sqrt(MAF)
0.6
0.5
0.4
0.3
0.2
0.1
0.05
0.01
0
Cell_Type_NEW
Ctrl
ExN
InN
Other
UD

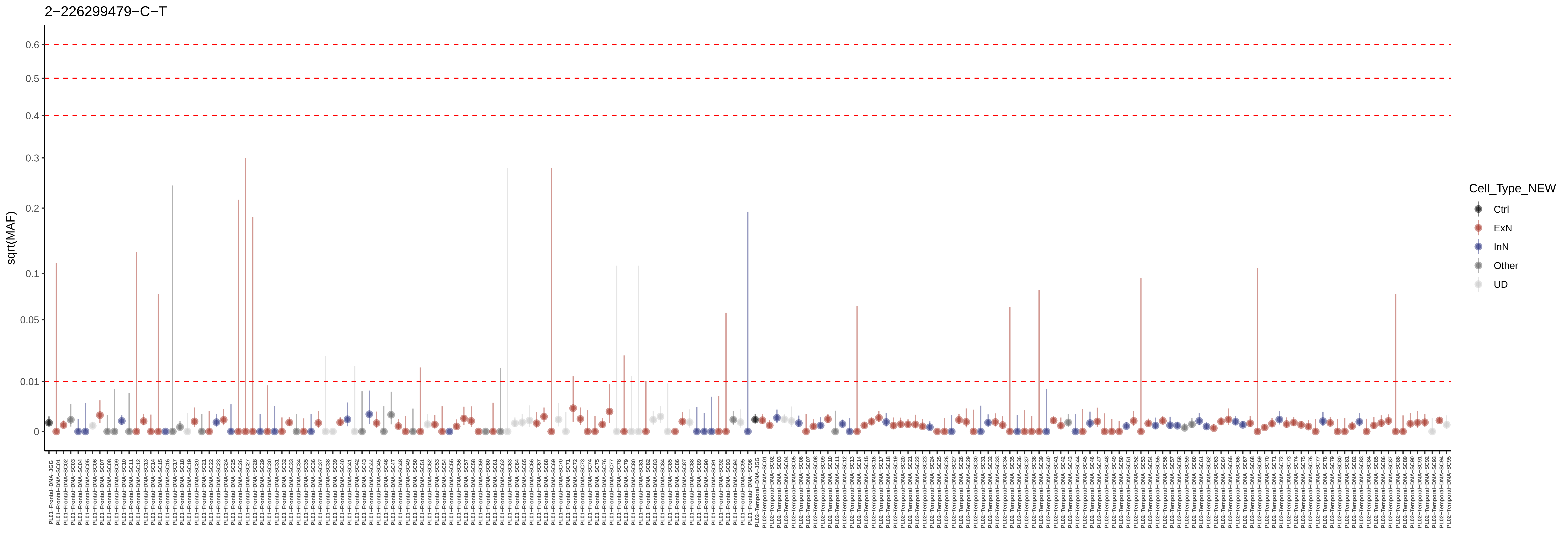

2−226299479−C−T
sqrt(MAF)
0.6
0.5
0.4
0.3
0.2
0.1
0.05
0.01
0
Cell_Type_NEW
Ctrl
ExN
InN
Other
UD

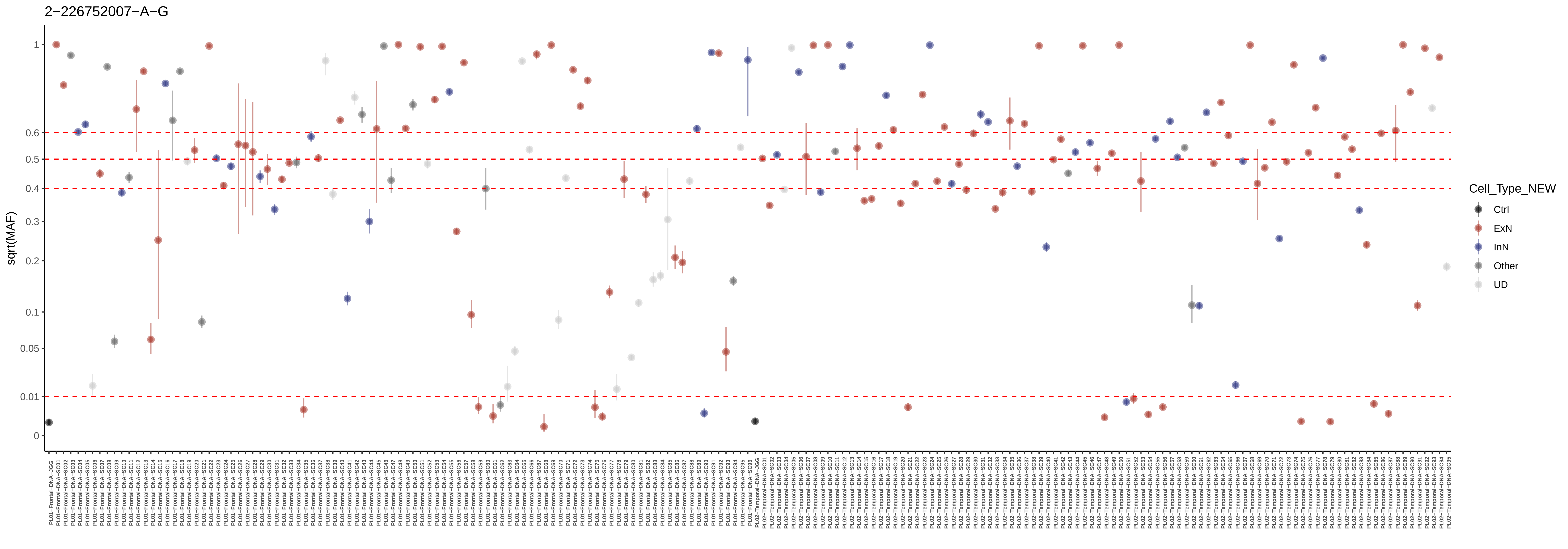
2−226752007−A−G
sqrt(MAF)
1
0.6
0.5
0.4
0.3
0.2
0.1
0.05
0.01
0
Cell_Type_NEW
Ctrl
ExN
InN
Other
UD

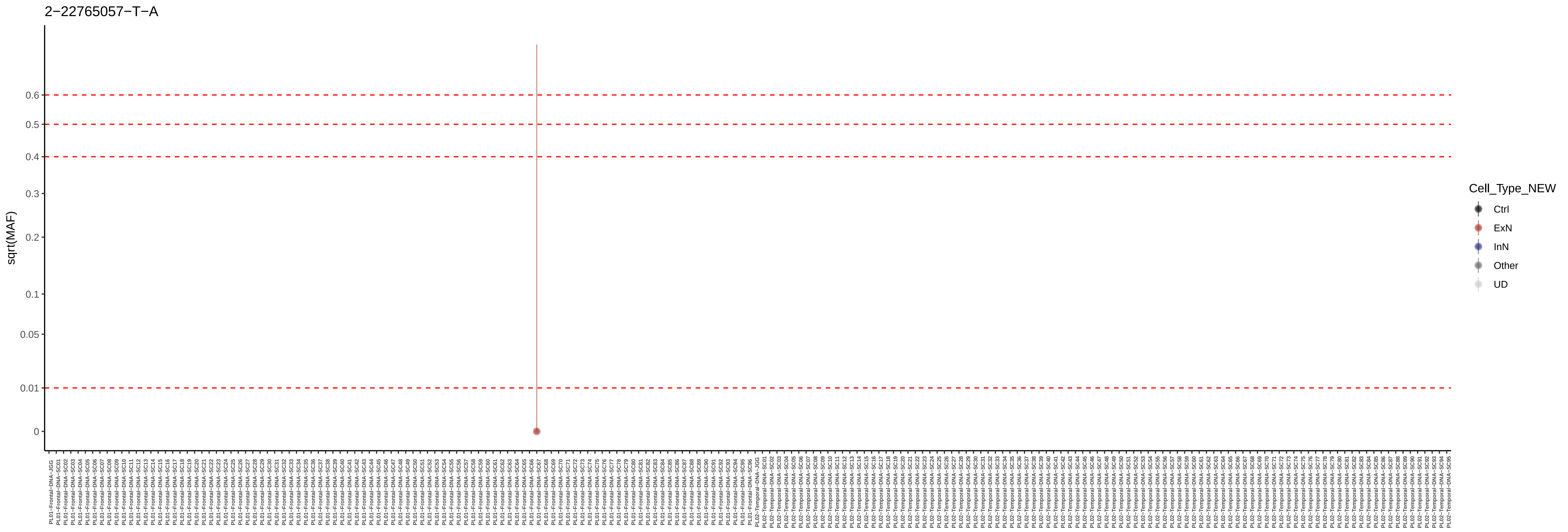

2−22765057−T−A
sqrt(MAF)
0.6
0.5
0.4
0.3
0.2
0.1
0.05
0.01
0
Cell_Type_NEW
Ctrl
ExN
InN
Other
UD

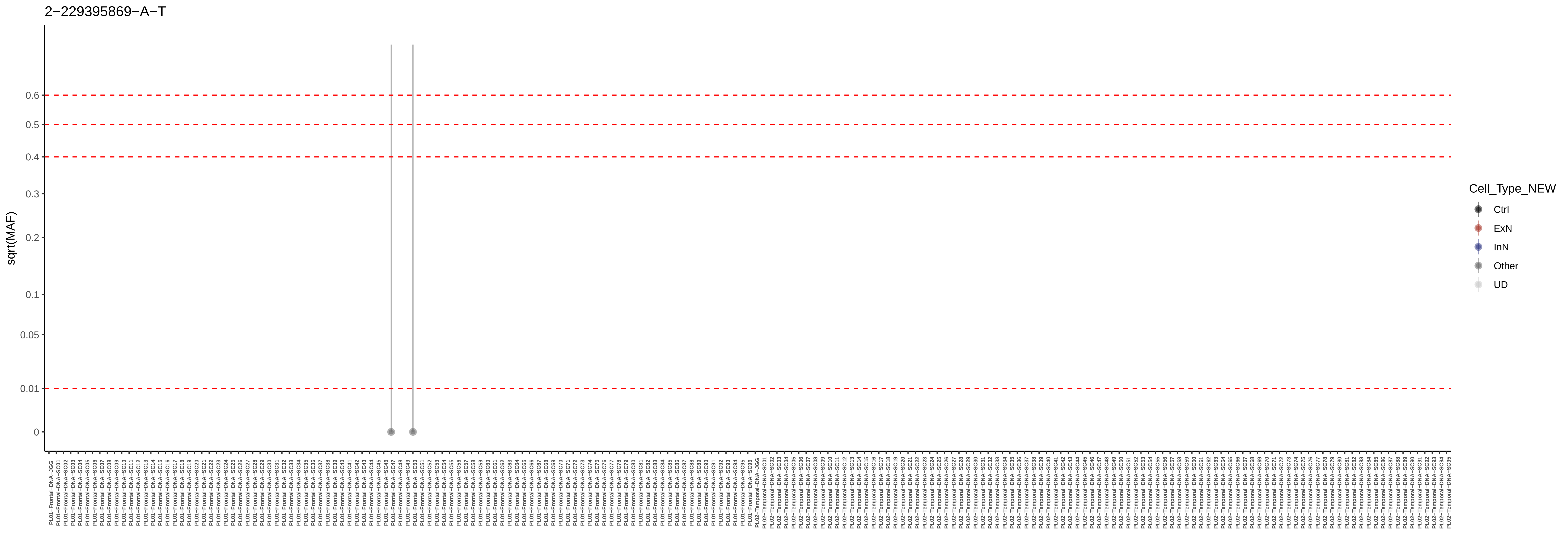

2−229395869−A−T
sqrt(MAF)
0.6
0.5
0.4
0.3
0.2
0.1
0.05
0.01
0
Cell_Type_NEW
Ctrl
ExN
InN
Other
UD

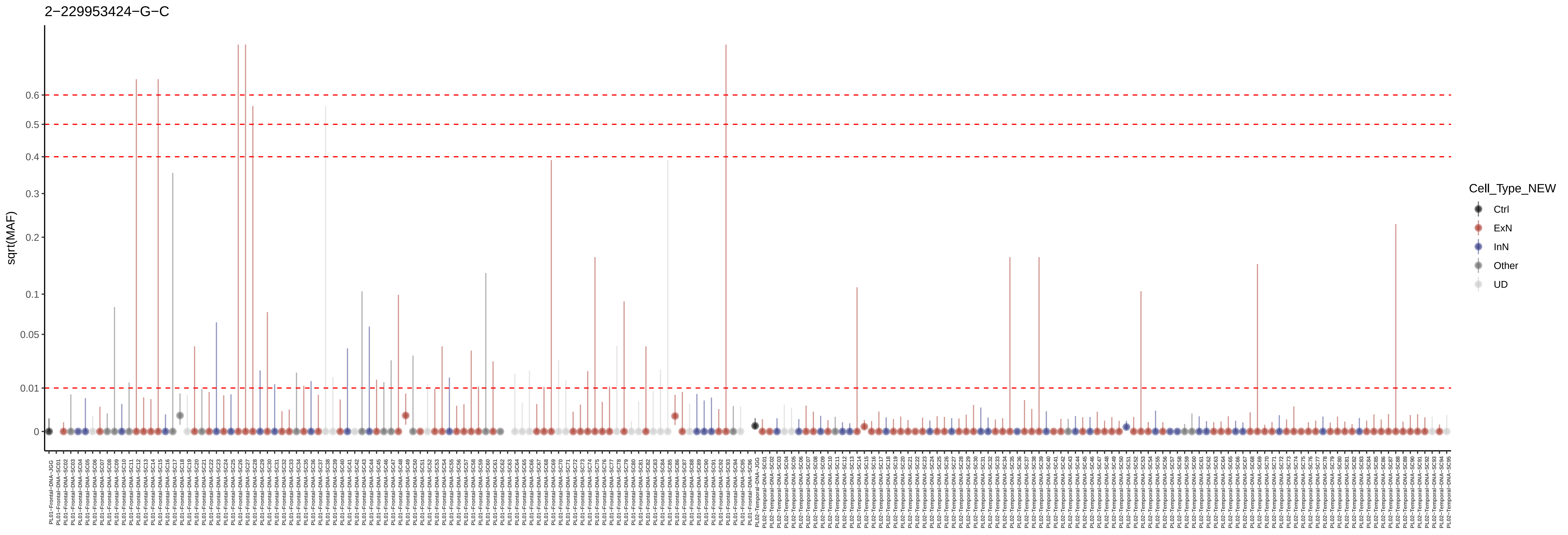
2−229953424−G−C
sqrt(MAF)
0.6
0.5
0.4
0.3
0.2
0.1
0.05
0.01
0
Cell_Type_NEW
Ctrl
ExN
InN
Other
UD

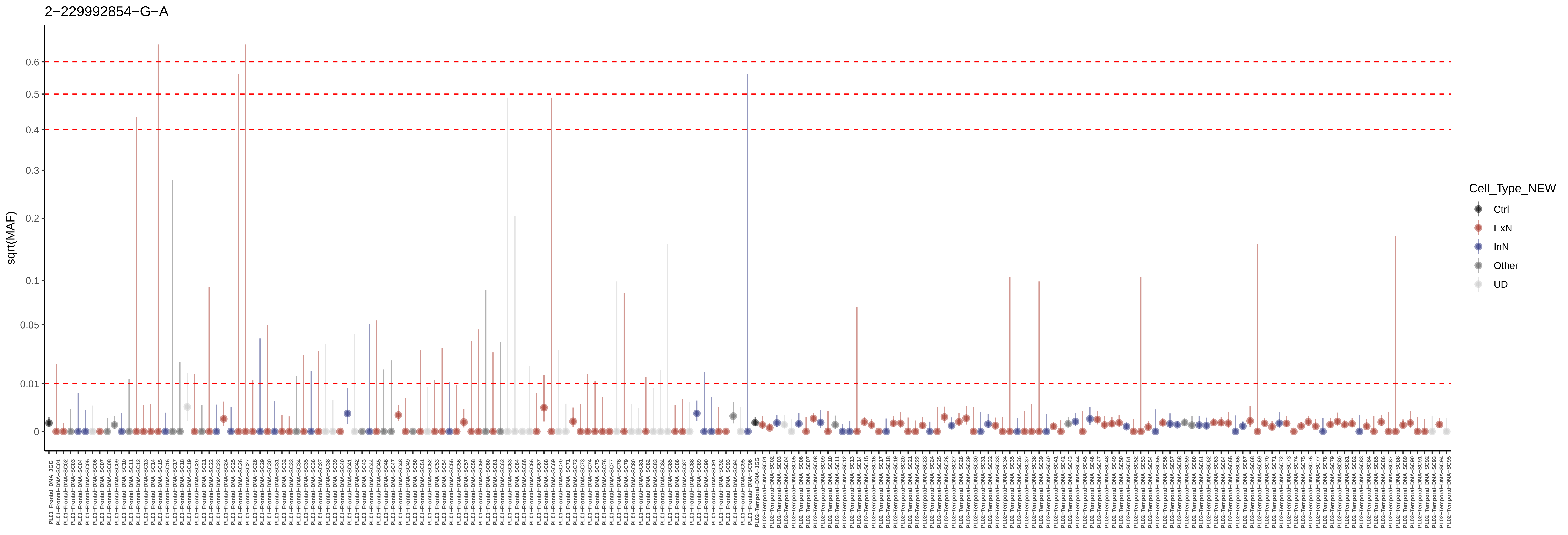

2−229992854−G−A
sqrt(MAF)
0.6
0.5
0.4
0.3
0.2
0.1
0.05
0.01
0
Cell_Type_NEW
Ctrl
ExN
InN
Other
UD

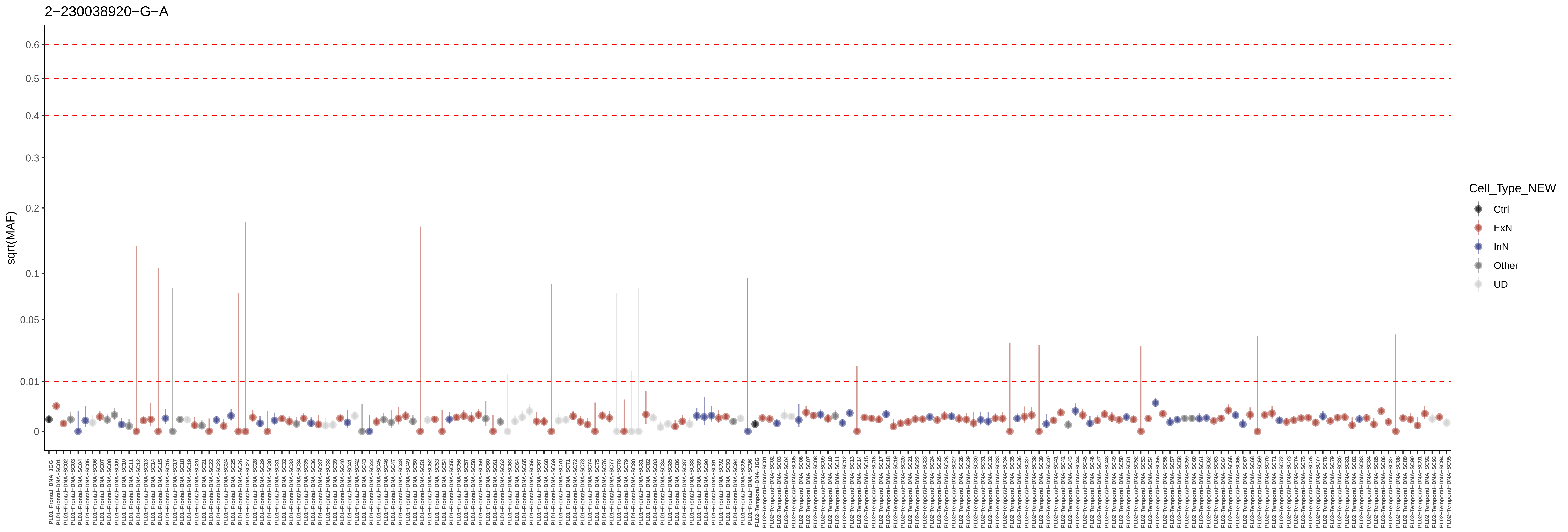

2−230038920−G−A
sqrt(MAF)
0.6
0.5
0.4
0.3
0.2
0.1
0.05
0.01
0
Cell_Type_NEW
Ctrl
ExN
InN
Other
UD

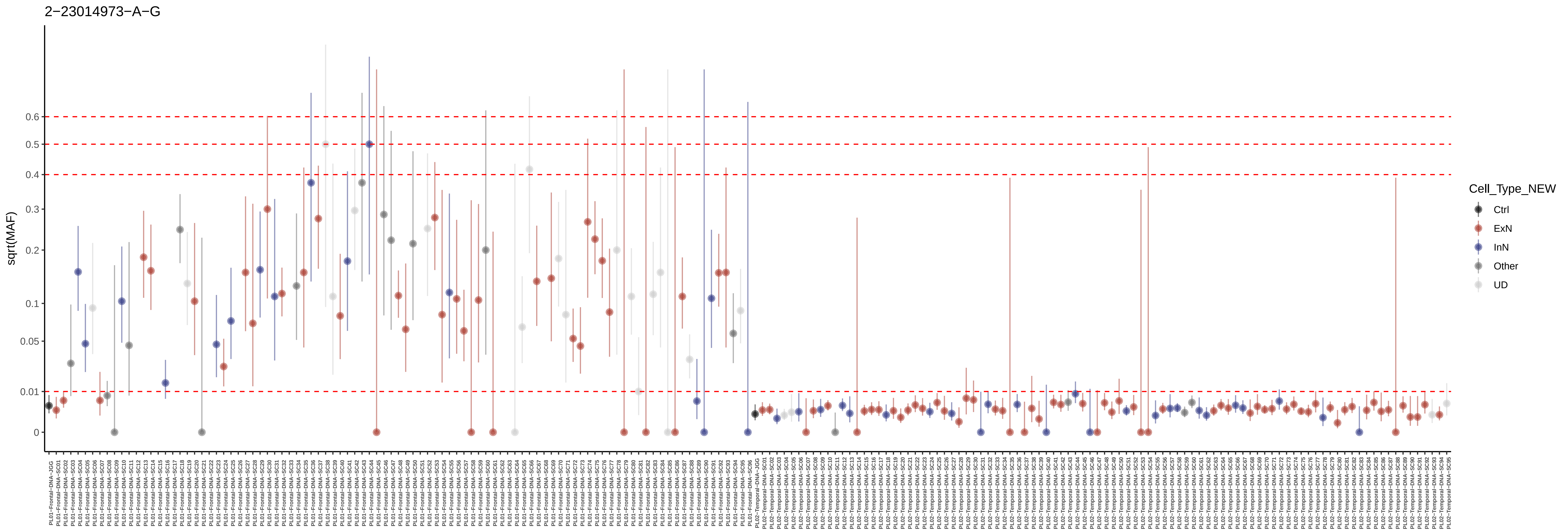

2−23014973−A−G
sqrt(MAF)
0.6
0.5
0.4
0.3
0.2
0.1
0.05
0.01
0
Cell_Type_NEW
Ctrl
ExN
InN
Other
UD

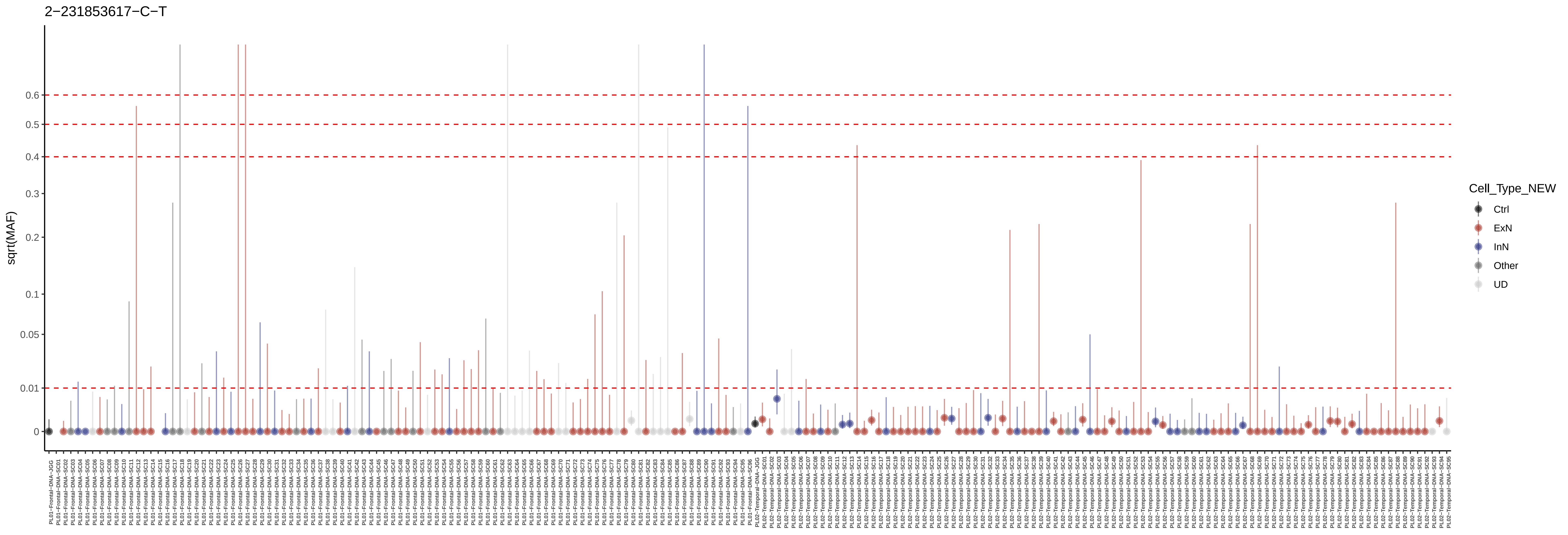
2−231853617−C−T
sqrt(MAF)
0.6
0.5
0.4
0.3
0.2
0.1
0.05
0.01
0
Cell_Type_NEW
Ctrl
ExN
InN
Other
UD

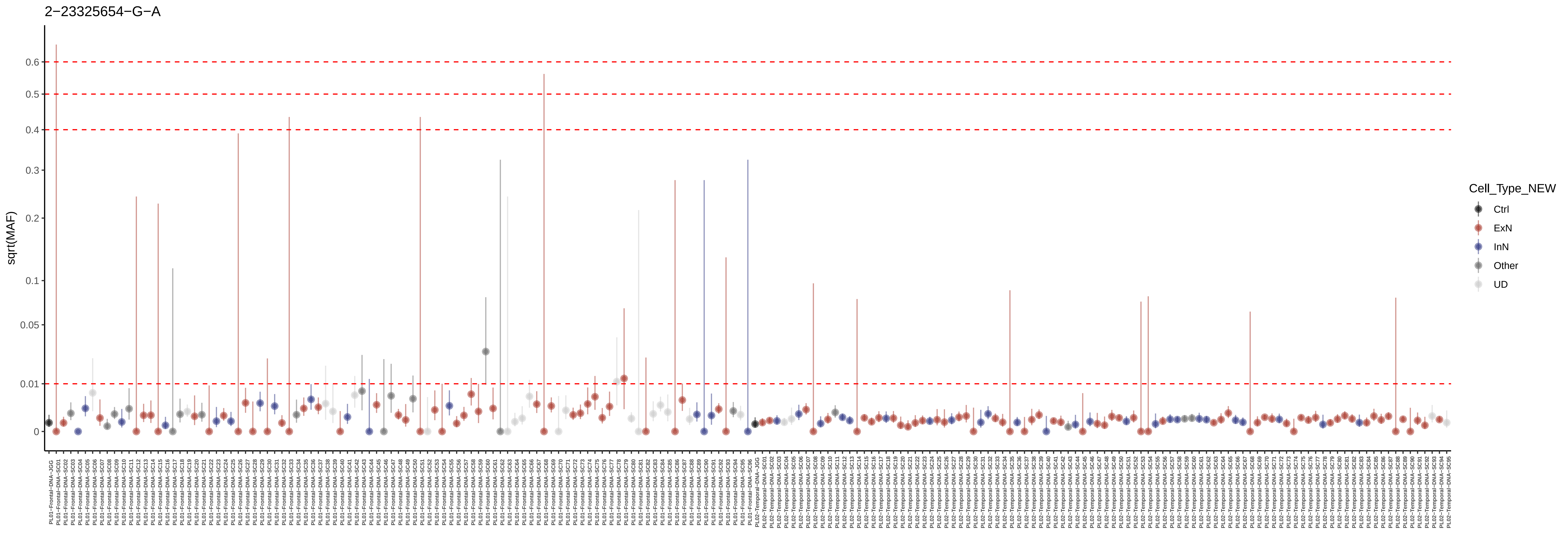
2−23325654−G−A
sqrt(MAF)
0.6
0.5
0.4
0.3
0.2
0.1
0.05
0.01
0
Cell_Type_NEW
Ctrl
ExN
InN
Other
UD

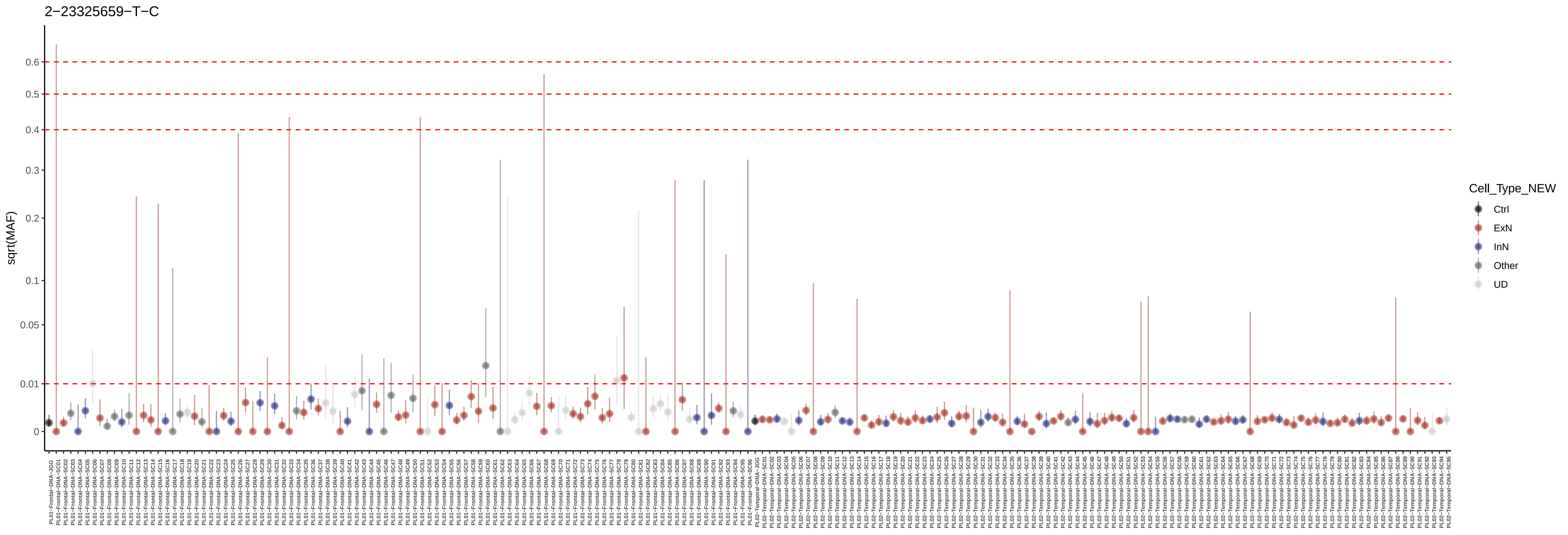
2−23325659−T−C
sqrt(MAF)
0.6
0.5
0.4
0.3
0.2
0.1
0.05
0.01
0
Cell_Type_NEW
Ctrl
ExN
InN
Other
UD

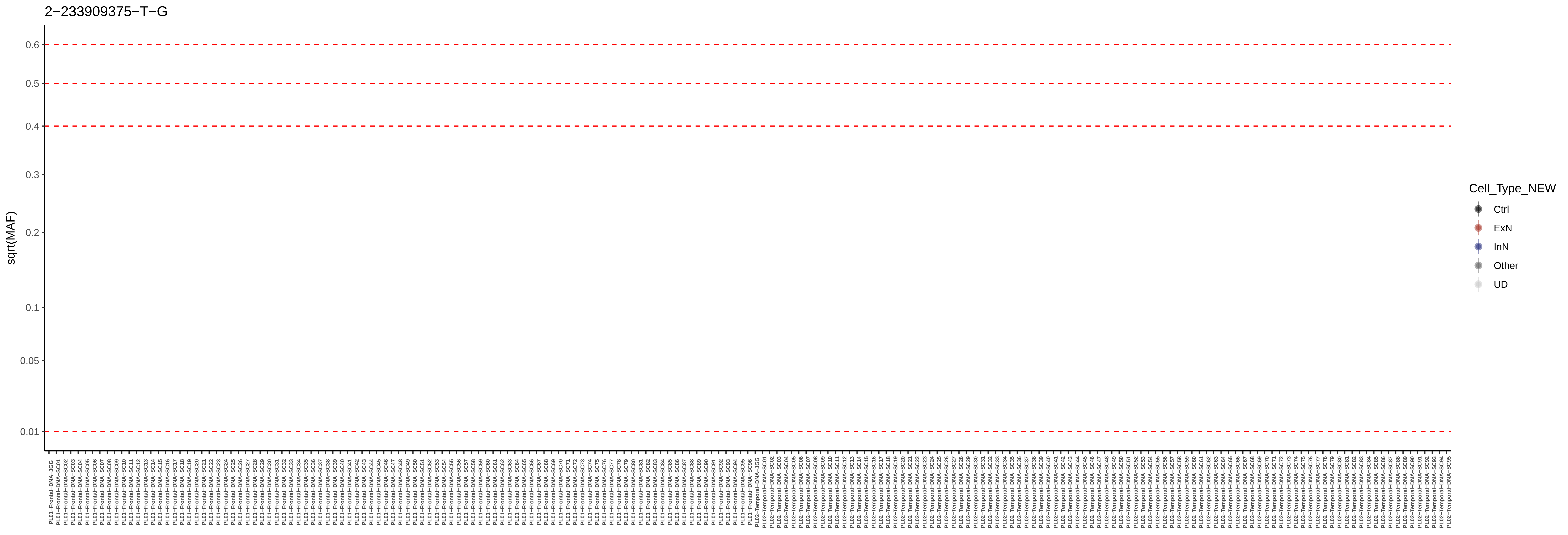

2−233909375−T−G
sqrt(MAF)
0.6
0.5
0.4
0.3
0.2
0.1
0.05
0.01
Cell_Type_NEW
Ctrl
ExN
InN
Other
UD

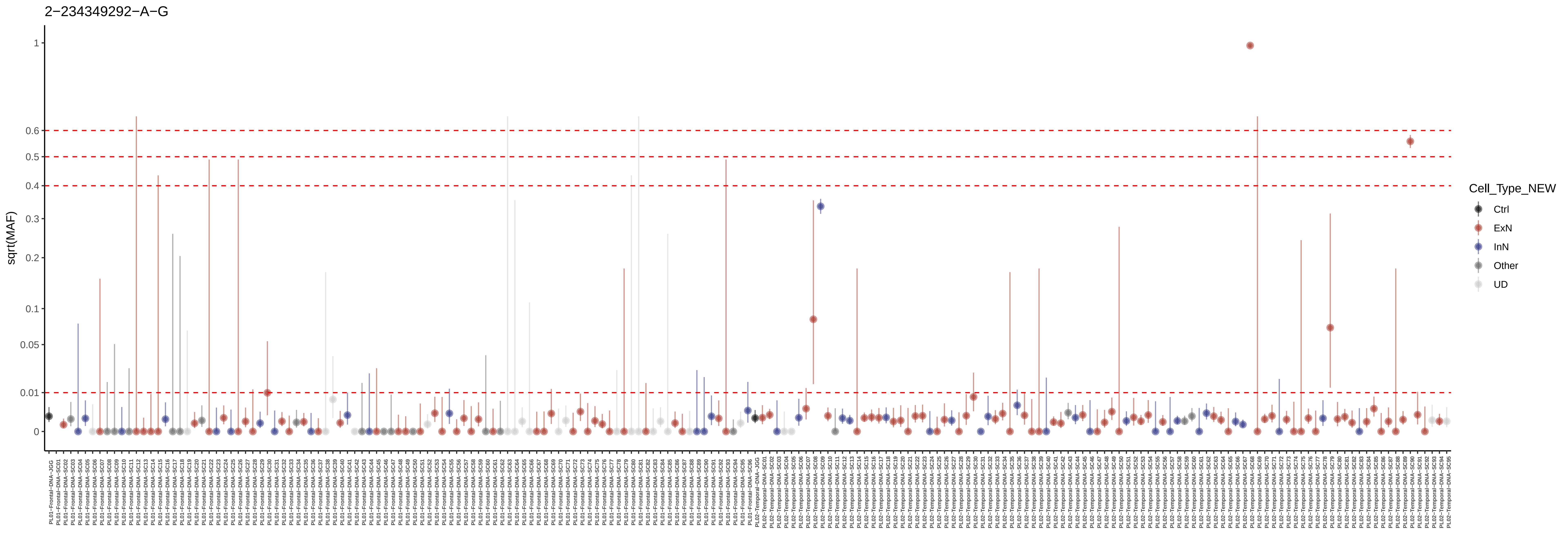

2−234349292−A−G
sqrt(MAF)
1
0.6
0.5
0.4
0.3
0.2
0.1
0.05
0.01
0
Cell_Type_NEW
Ctrl
ExN
InN
Other
UD

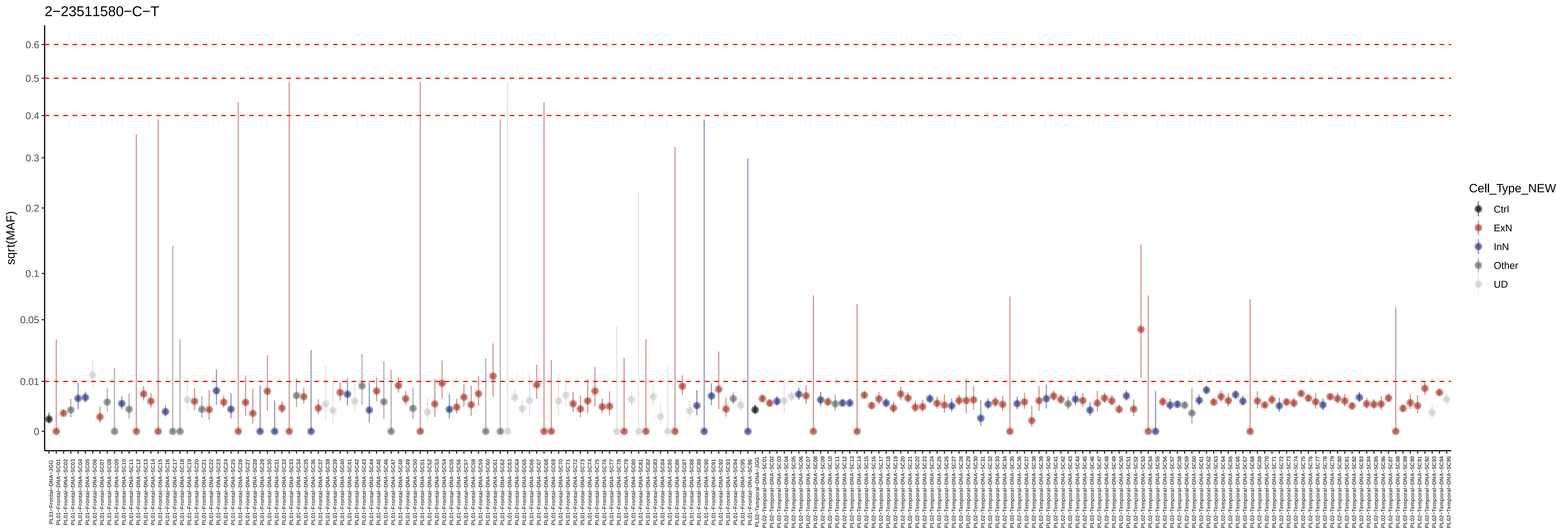

2−23511580−C−T
sqrt(MAF)
0.6
0.5
0.4
0.3
0.2
0.1
0.05
0.01
0
Cell_Type_NEW
Ctrl
ExN
InN
Other
UD

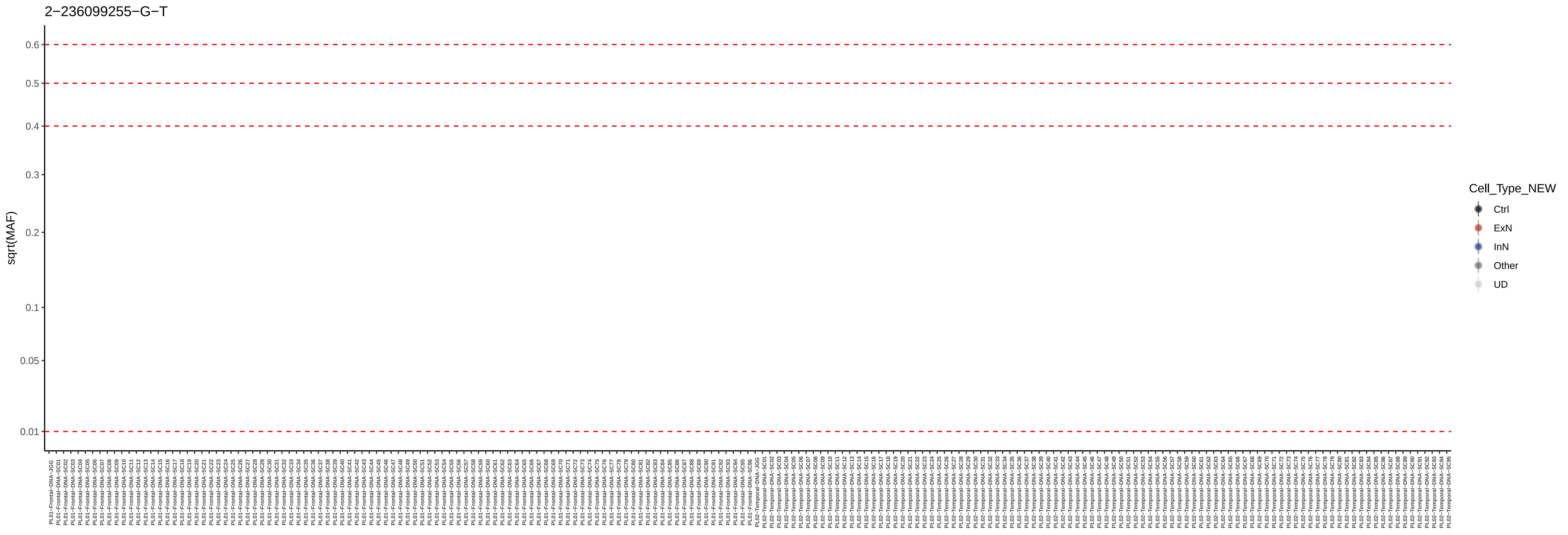
2−236099255−G−T
sqrt(MAF)
0.6
0.5
0.4
0.3
0.2
0.1
0.05
0.01
Cell_Type_NEW
Ctrl
ExN
InN
Other
UD

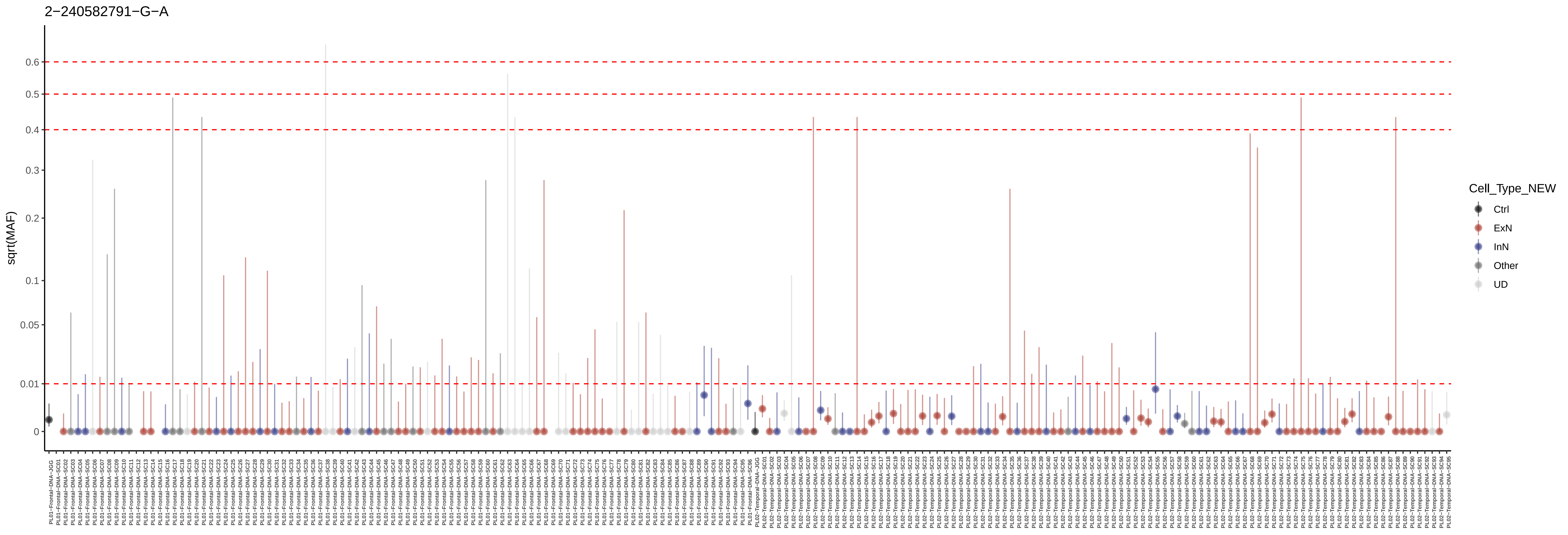
2−240582791−G−A
sqrt(MAF)
0.6
0.5
0.4
0.3
0.2
0.1
0.05
0.01
0
Cell_Type_NEW
Ctrl
ExN
InN
Other
UD

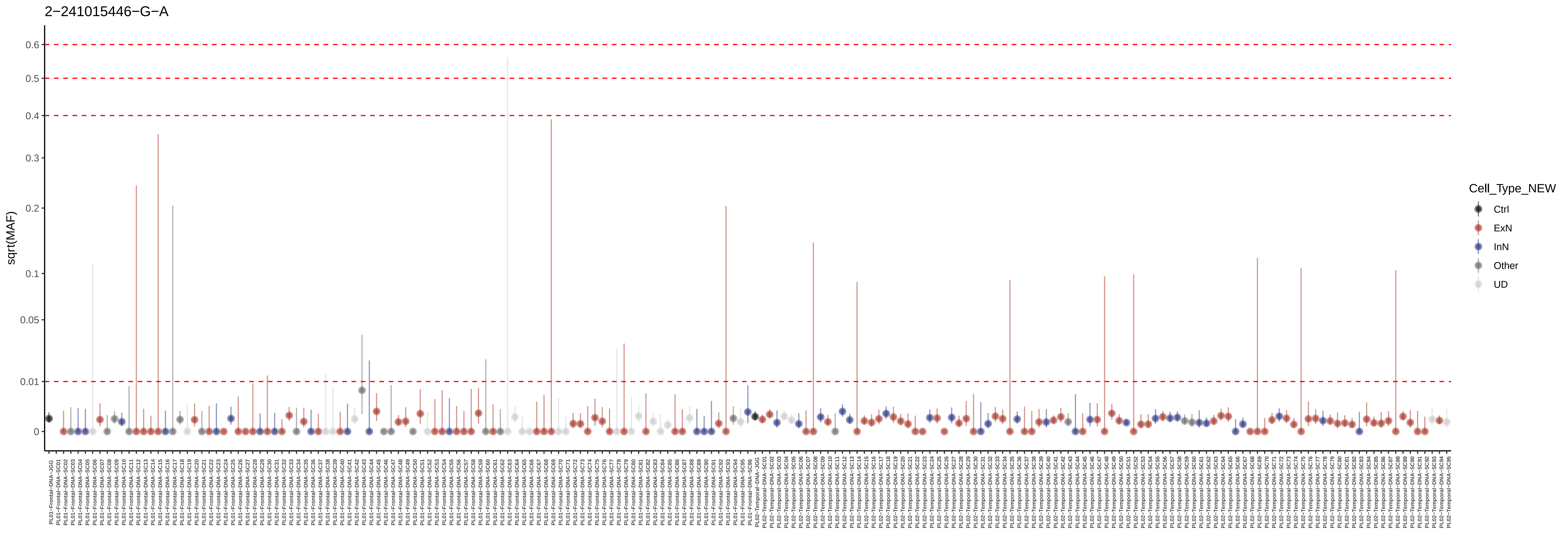
2−241015446−G−A
sqrt(MAF)
0.6
0.5
0.4
0.3
0.2
0.1
0.05
0.01
0
Cell_Type_NEW
Ctrl
ExN
InN
Other
UD

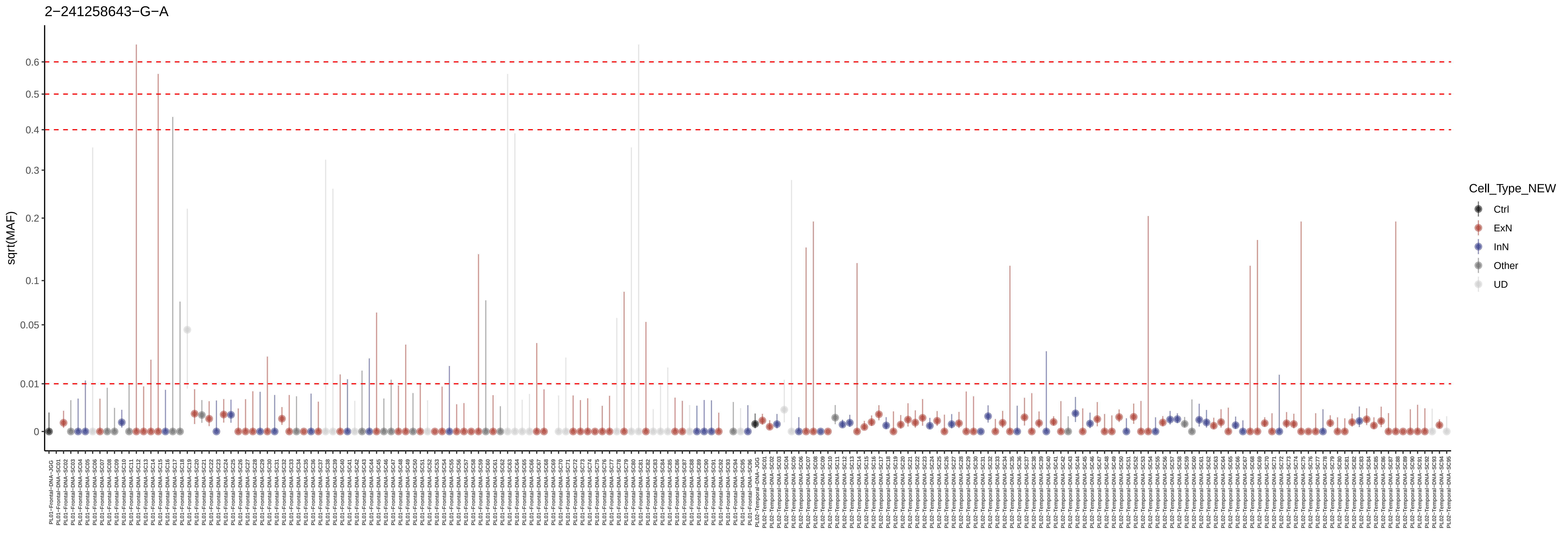
2−241258643−G−A
sqrt(MAF)
0.6
0.5
0.4
0.3
0.2
0.1
0.05
0.01
0
Cell_Type_NEW
Ctrl
ExN
InN
Other
UD

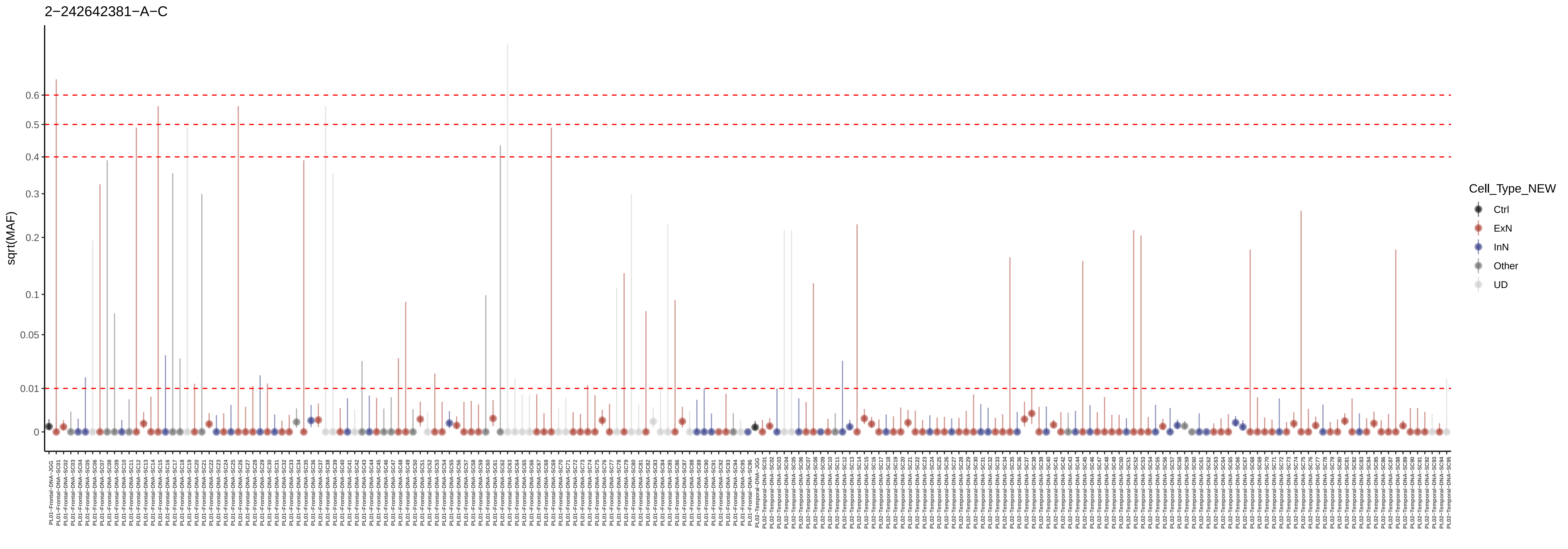
2−242642381−A−C
sqrt(MAF)
0.6
0.5
0.4
0.3
0.2
0.1
0.05
0.01
0
Cell_Type_NEW
Ctrl
ExN
InN
Other
UD

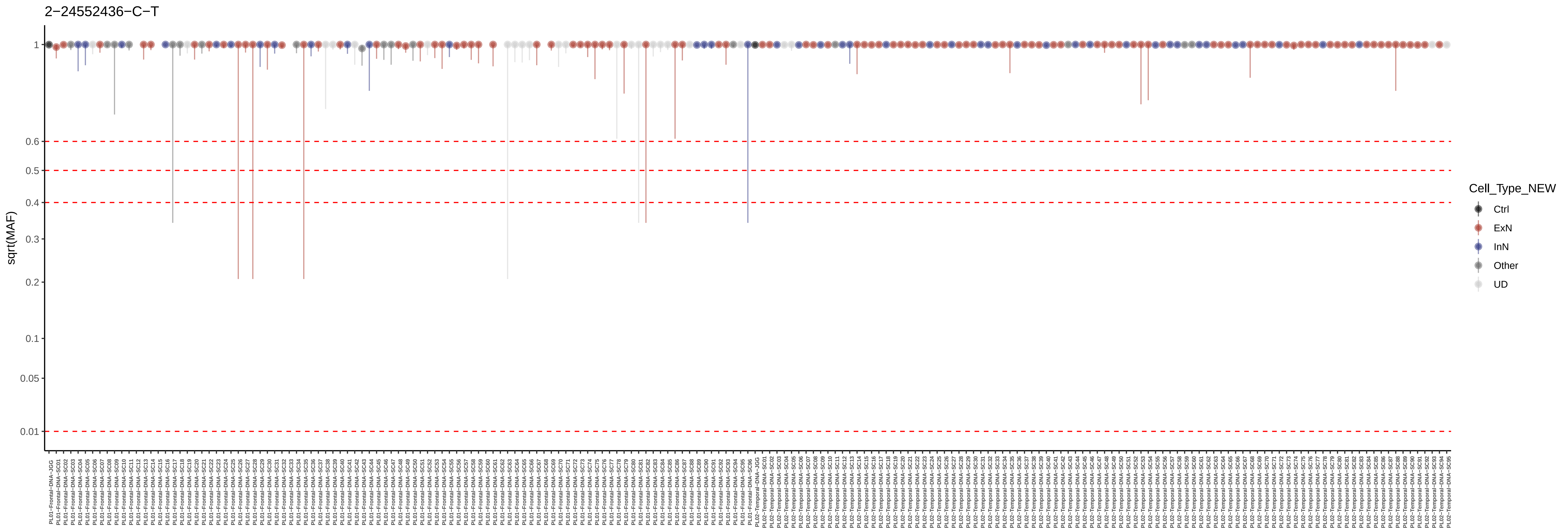

2−24552436−C−T
sqrt(MAF)
1
0.6
0.5
0.4
0.3
0.2
0.1
0.05
0.01
Cell_Type_NEW
Ctrl
ExN
InN
Other
UD

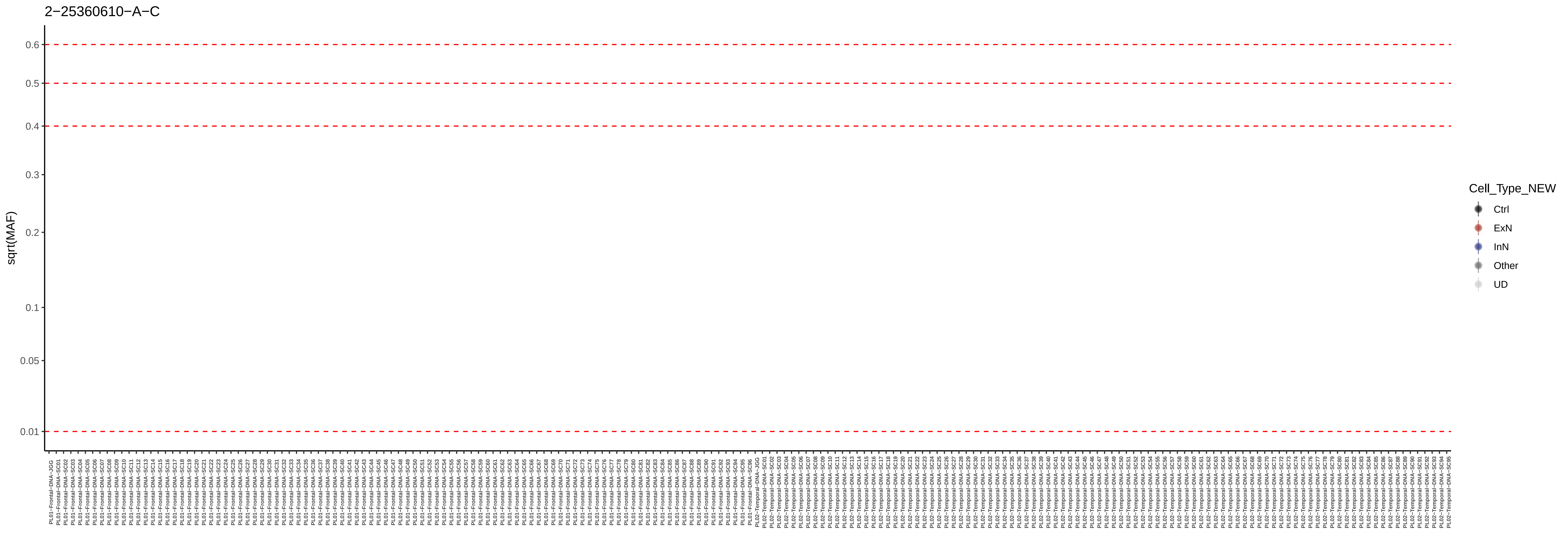
2−25360610−A−C
sqrt(MAF)
0.6
0.5
0.4
0.3
0.2
0.1
0.05
0.01
Cell_Type_NEW
Ctrl
ExN
InN
Other
UD

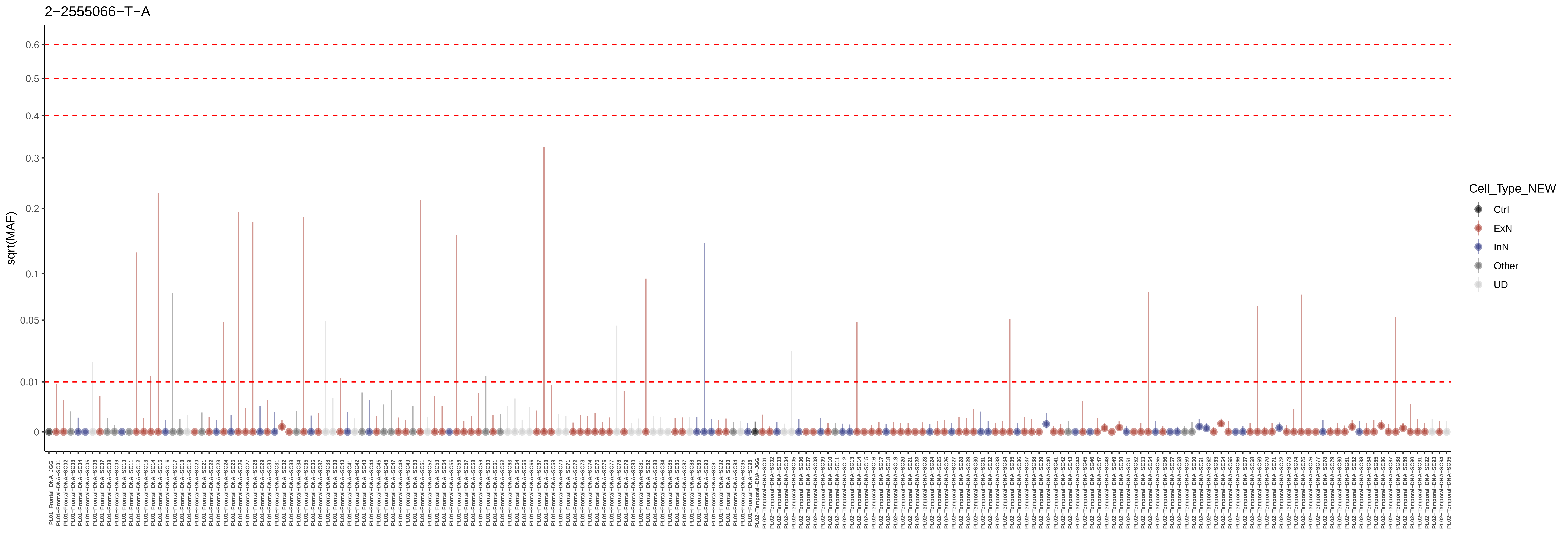

2−2555066−T−A
sqrt(MAF)
0.6
0.5
0.4
0.3
0.2
0.1
0.05
0.01
0
Cell_Type_NEW
Ctrl
ExN
InN
Other
UD

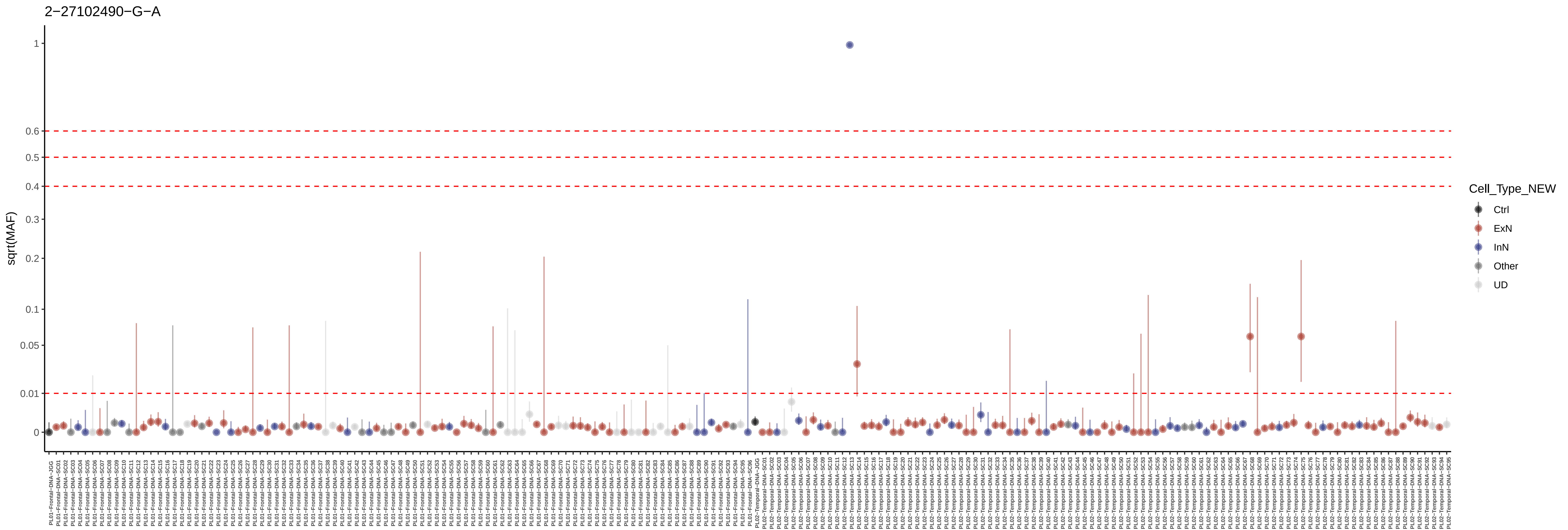

2−27102490−G−A
sqrt(MAF)
1
0.6
0.5
0.4
0.3
0.2
0.1
0.05
0.01
0
Cell_Type_NEW
Ctrl
ExN
InN
Other
UD

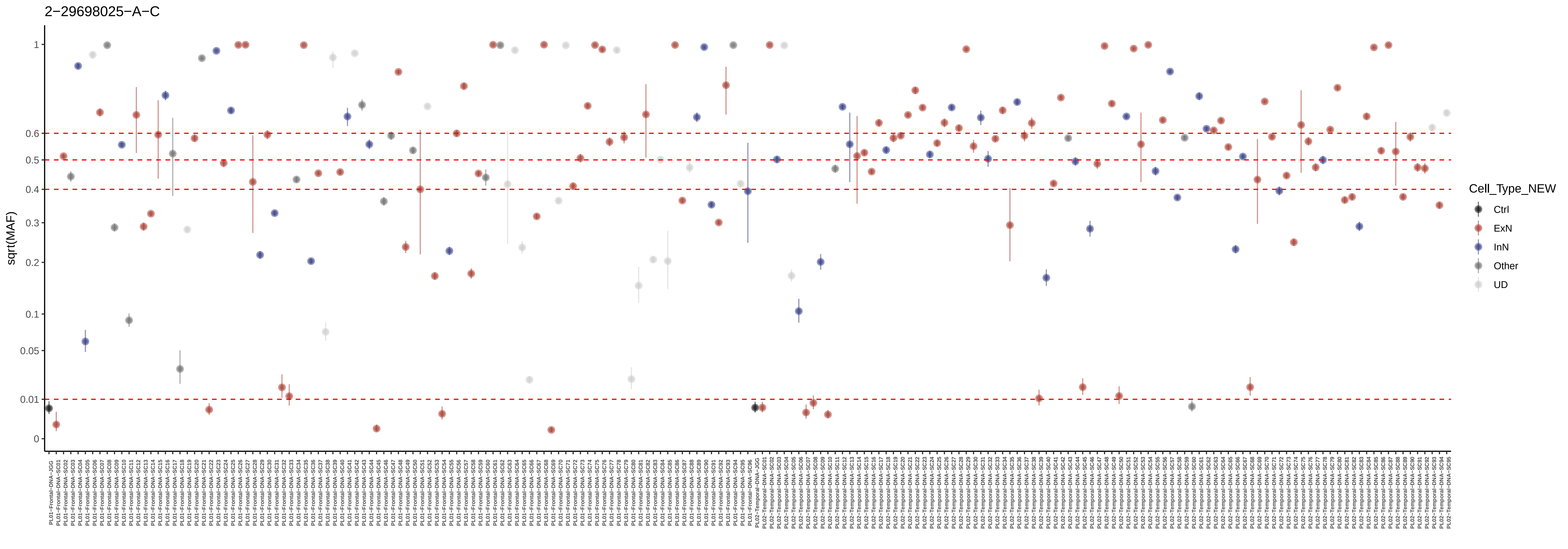

2−29698025−A−C
sqrt(MAF)
1
0.6
0.5
0.4
0.3
0.2
0.1
0.05
0.01
0
Cell_Type_NEW
Ctrl
ExN
InN
Other
UD

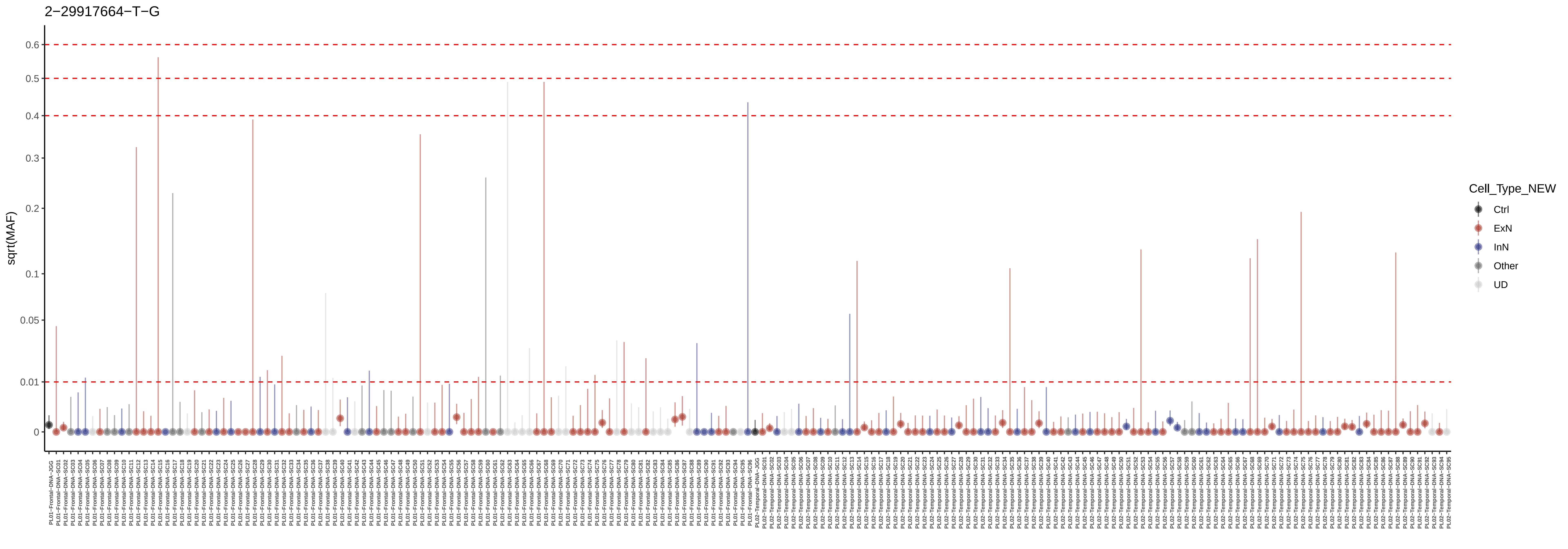
2−29917664−T−G
sqrt(MAF)
0.6
0.5
0.4
0.3
0.2
0.1
0.05
0.01
0
Cell_Type_NEW
Ctrl
ExN
InN
Other
UD

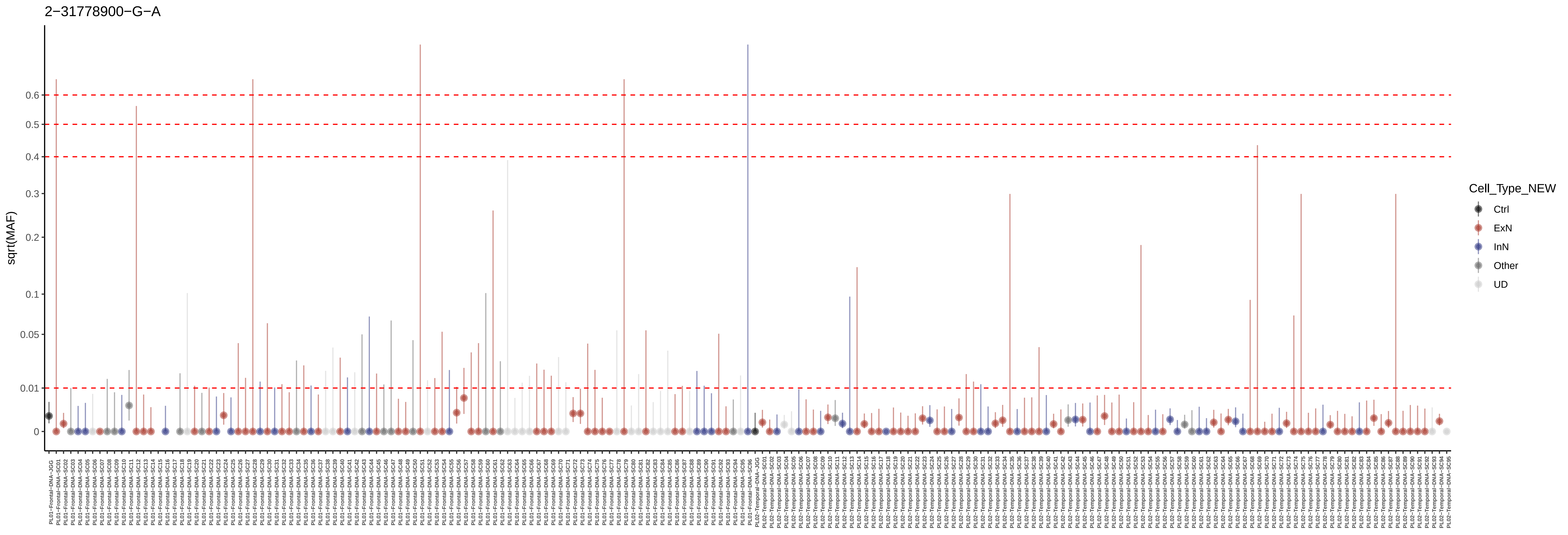

2−31778900−G−A
sqrt(MAF)
0.6
0.5
0.4
0.3
0.2
0.1
0.05
0.01
0
Cell_Type_NEW
Ctrl
ExN
InN
Other
UD

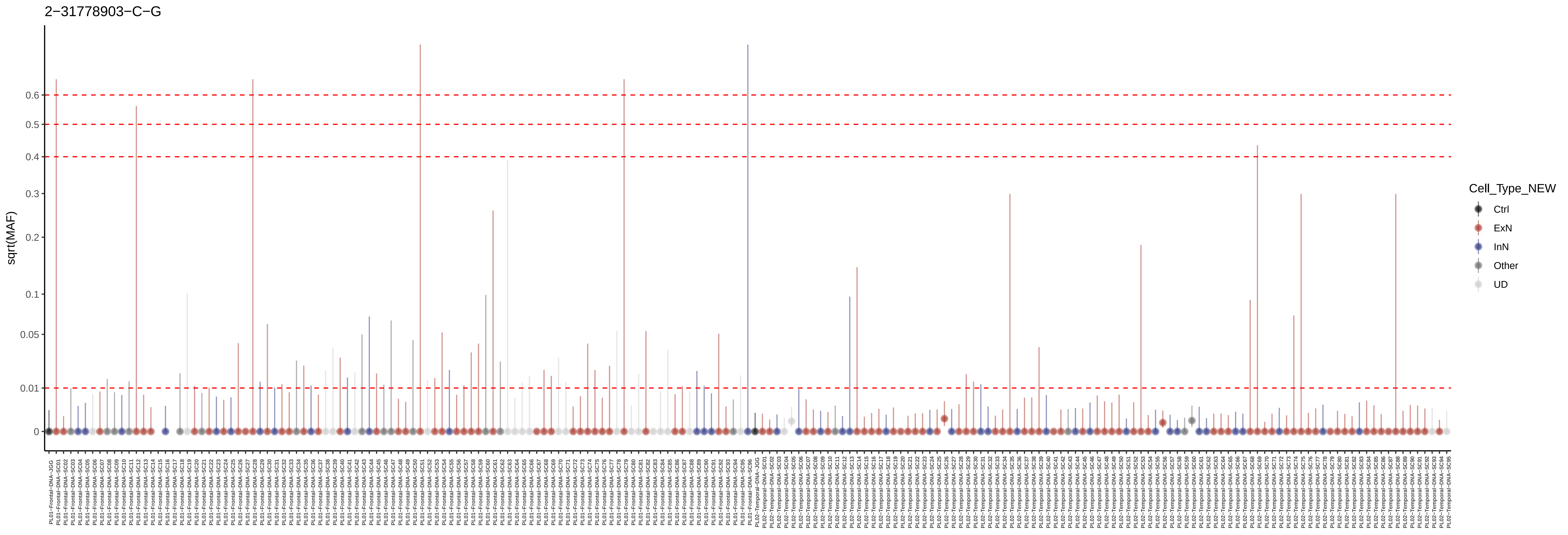
2−31778903−C−G
sqrt(MAF)
0.6
0.5
0.4
0.3
0.2
0.1
0.05
0.01
0
Cell_Type_NEW
Ctrl
ExN
InN
Other
UD

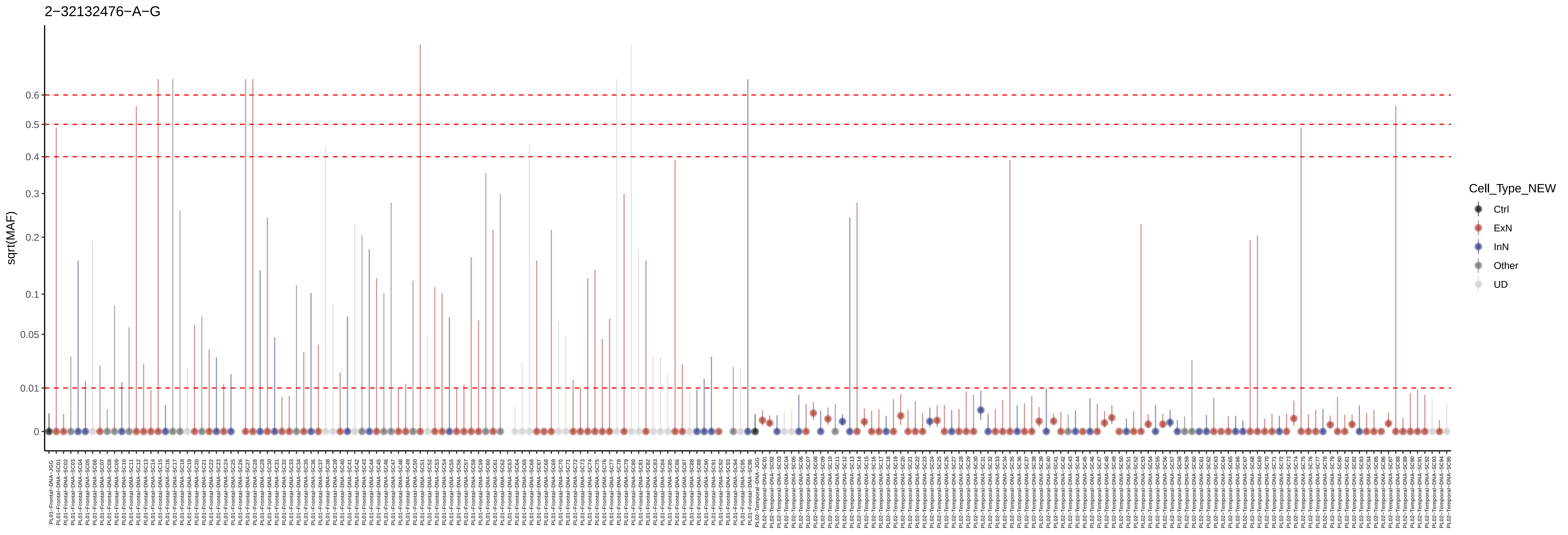

2−32132476−A−G
sqrt(MAF)
0.6
0.5
0.4
0.3
0.2
0.1
0.05
0.01
0
Cell_Type_NEW
Ctrl
ExN
InN
Other
UD

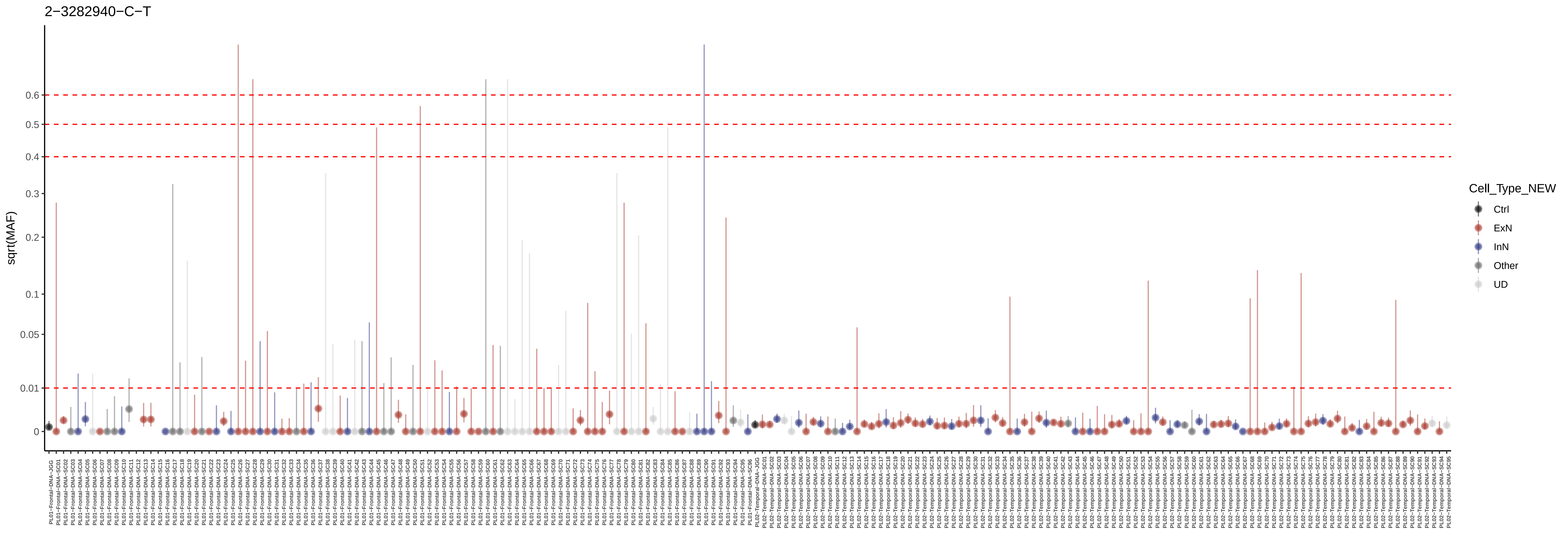

2−3282940−C−T
sqrt(MAF)
0.6
0.5
0.4
0.3
0.2
0.1
0.05
0.01
0
Cell_Type_NEW
Ctrl
ExN
InN
Other
UD

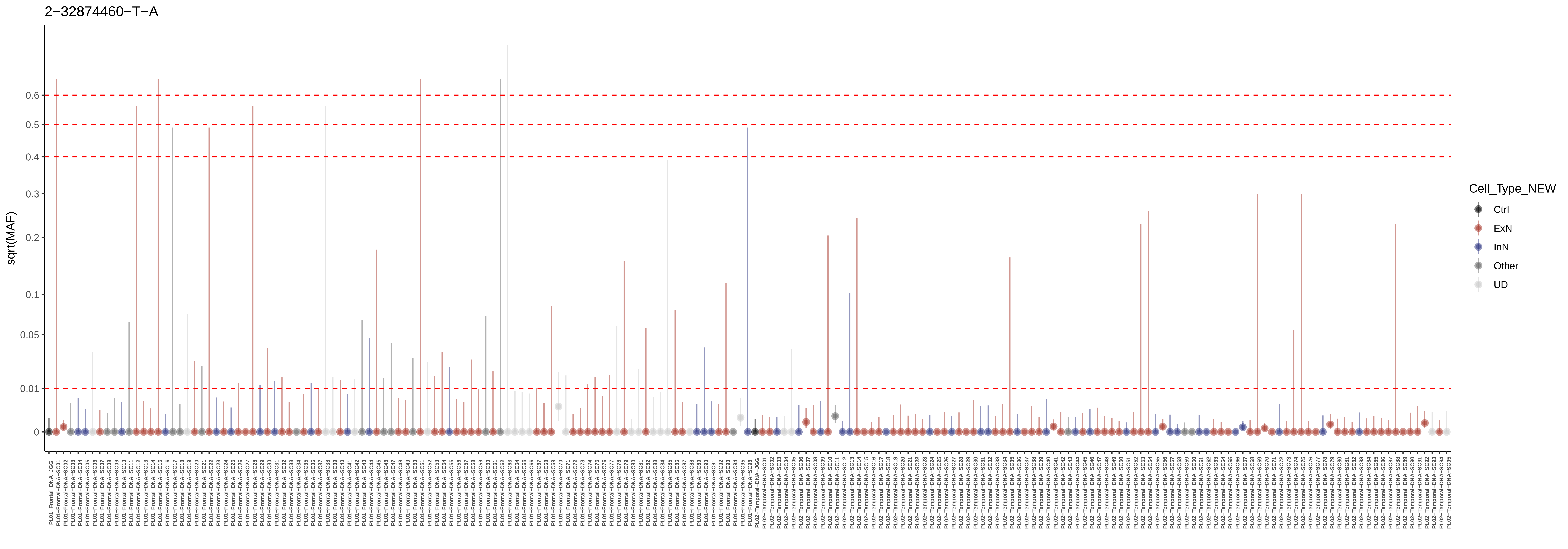
2−32874460−T−A
sqrt(MAF)
0.6
0.5
0.4
0.3
0.2
0.1
0.05
0.01
0
Cell_Type_NEW
Ctrl
ExN
InN
Other
UD

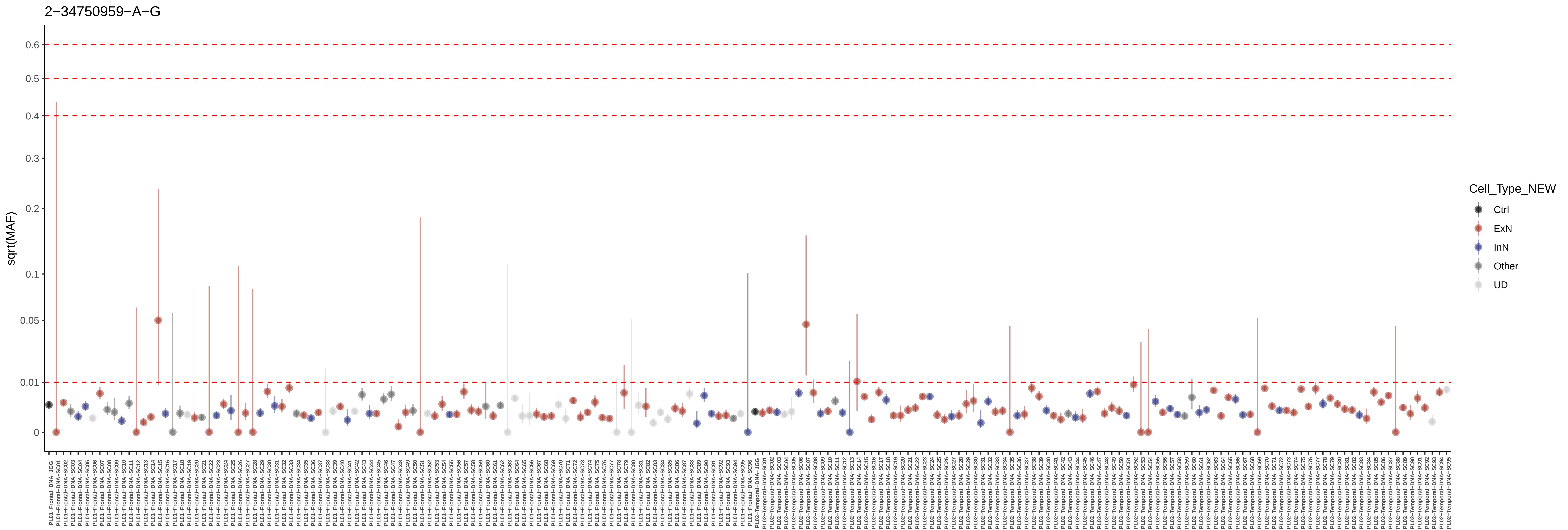
2−34750959−A−G
sqrt(MAF)
0.6
0.5
0.4
0.3
0.2
0.1
0.05
0.01
0
Cell_Type_NEW
Ctrl
ExN
InN
Other
UD

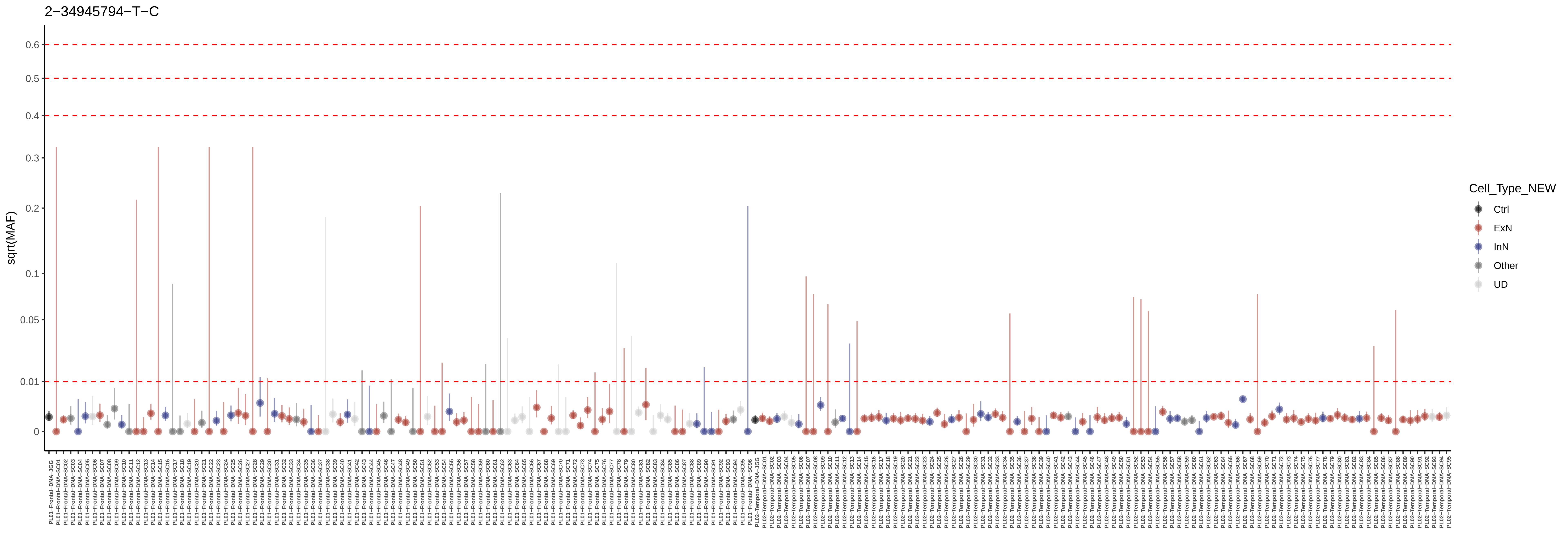
2−34945794−T−C
sqrt(MAF)
0.6
0.5
0.4
0.3
0.2
0.1
0.05
0.01
0
Cell_Type_NEW
Ctrl
ExN
InN
Other
UD

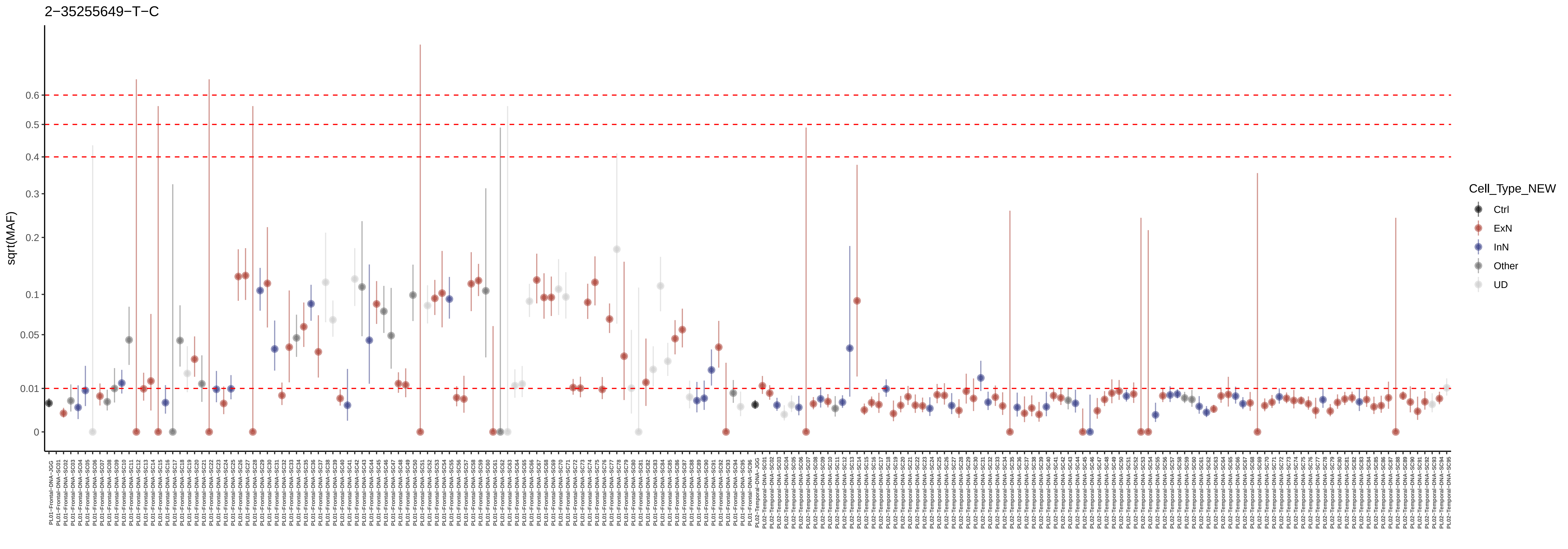

2−35255649−T−C
sqrt(MAF)
0
0.01
0.05
0.1
0.2
0.3
0.4
0.5
0.6
Cell_Type_NEW
Ctrl
ExN
InN
Other
UD

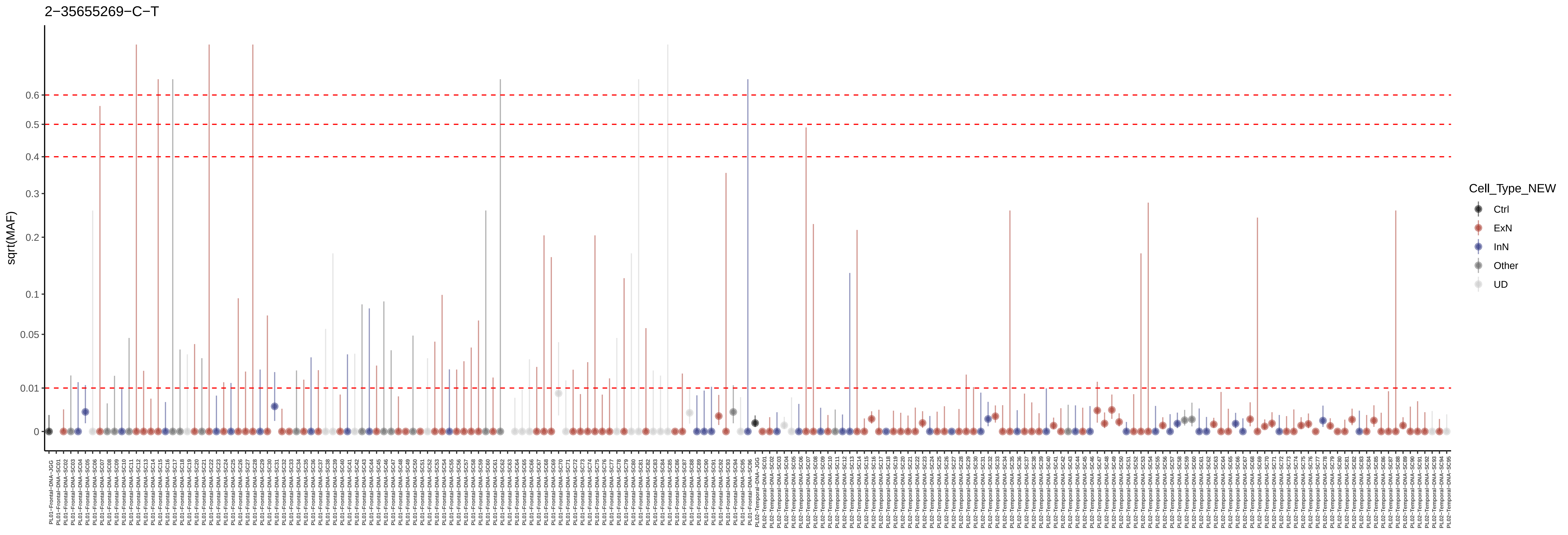
2−35655269−C−T
sqrt(MAF)
0.6
0.5
0.4
0.3
0.2
0.1
0.05
0.01
0
Cell_Type_NEW
Ctrl
ExN
InN
Other
UD

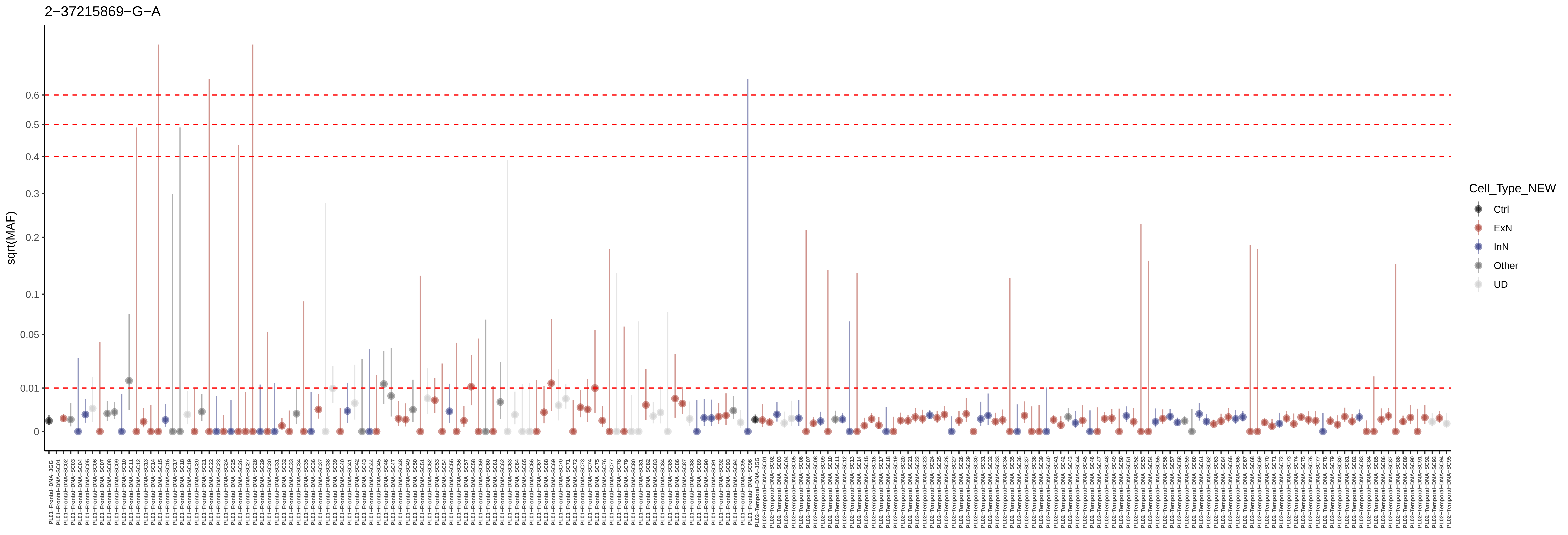
2−37215869−G−A
sqrt(MAF)
0.6
0.5
0.4
0.3
0.2
0.1
0.05
0.01
0
Cell_Type_NEW
Ctrl
ExN
InN
Other
UD

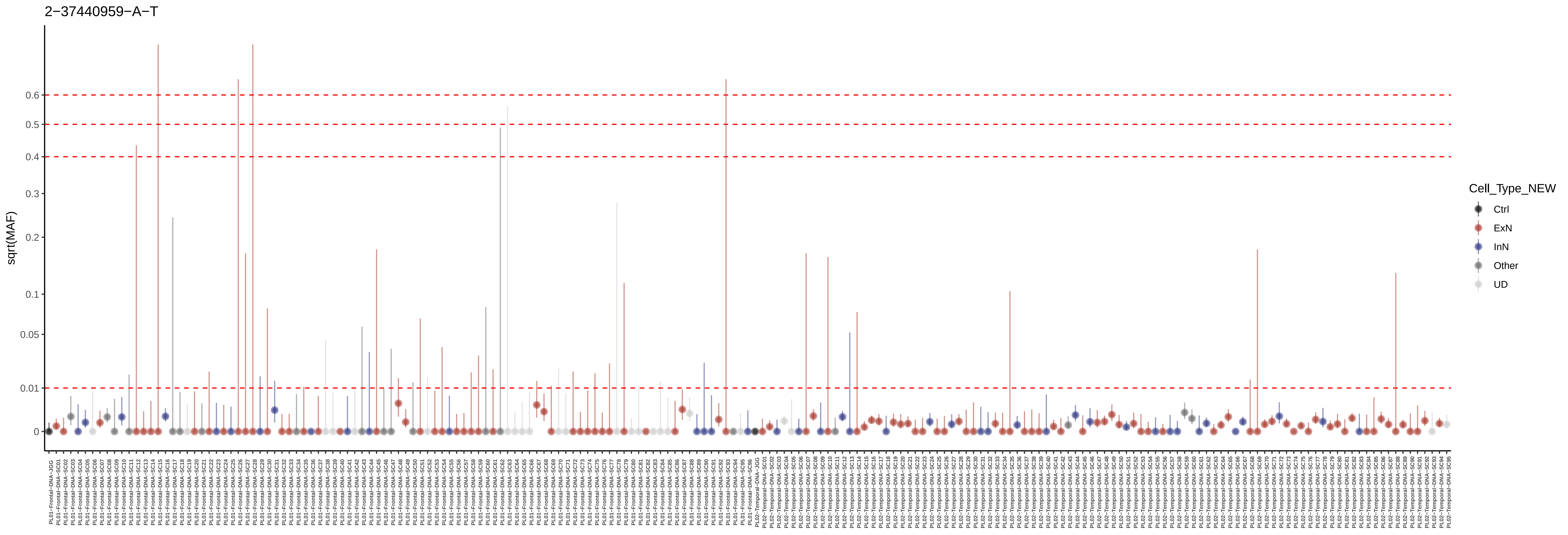
2-37440959-A-T
sqrt(MAF)
0.6
0.5
0.4
0.3
0.2
0.1
0.05
0.01
0
Cell_Type_NEW
Ctrl
ExN
InN
Other
UD

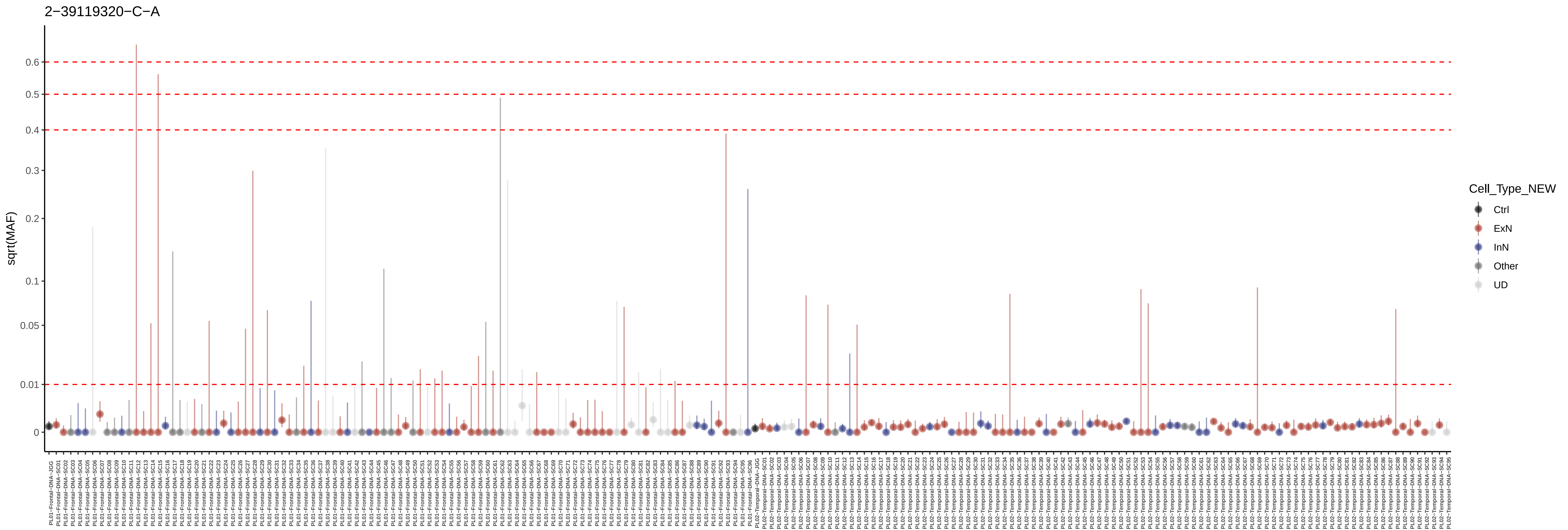

2−39119320−C−A
sqrt(MAF)
0.6
0.5
0.4
0.3
0.2
0.1
0.05
0.01
0
Cell_Type_NEW
Ctrl
ExN
InN
Other
UD

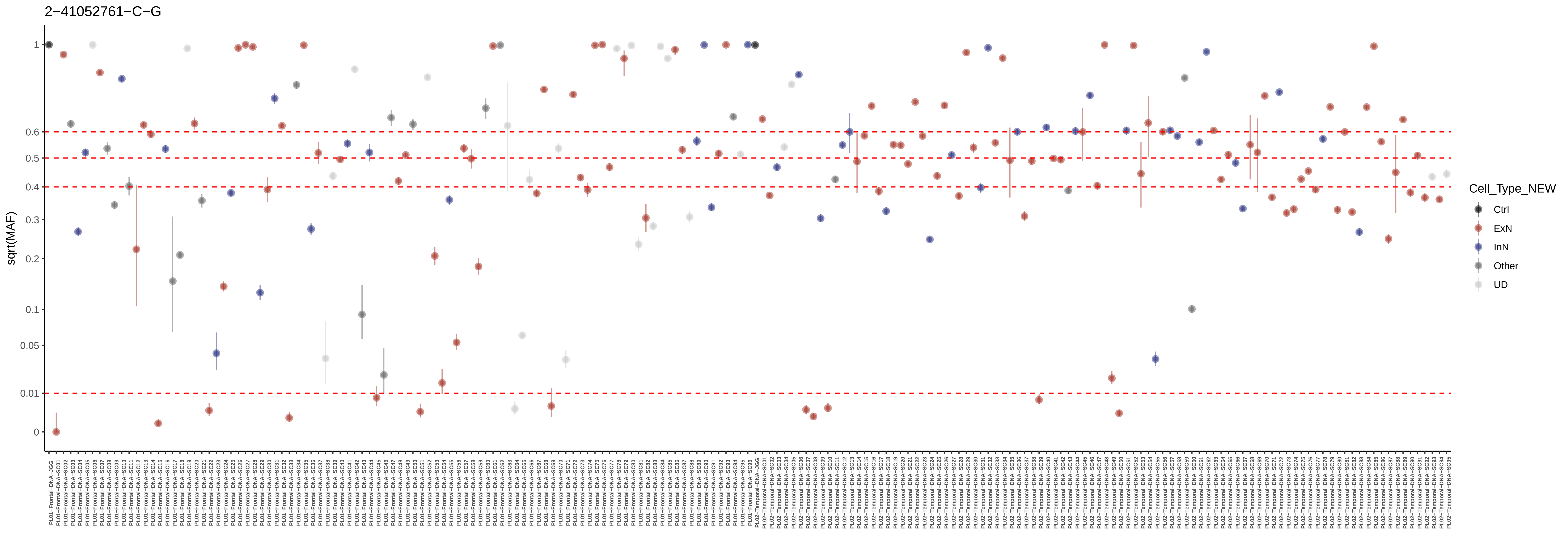
2−41052761−C−G
sqrt(MAF)
Cell_Type_NEW
Ctrl
ExN
InN
Other
UD

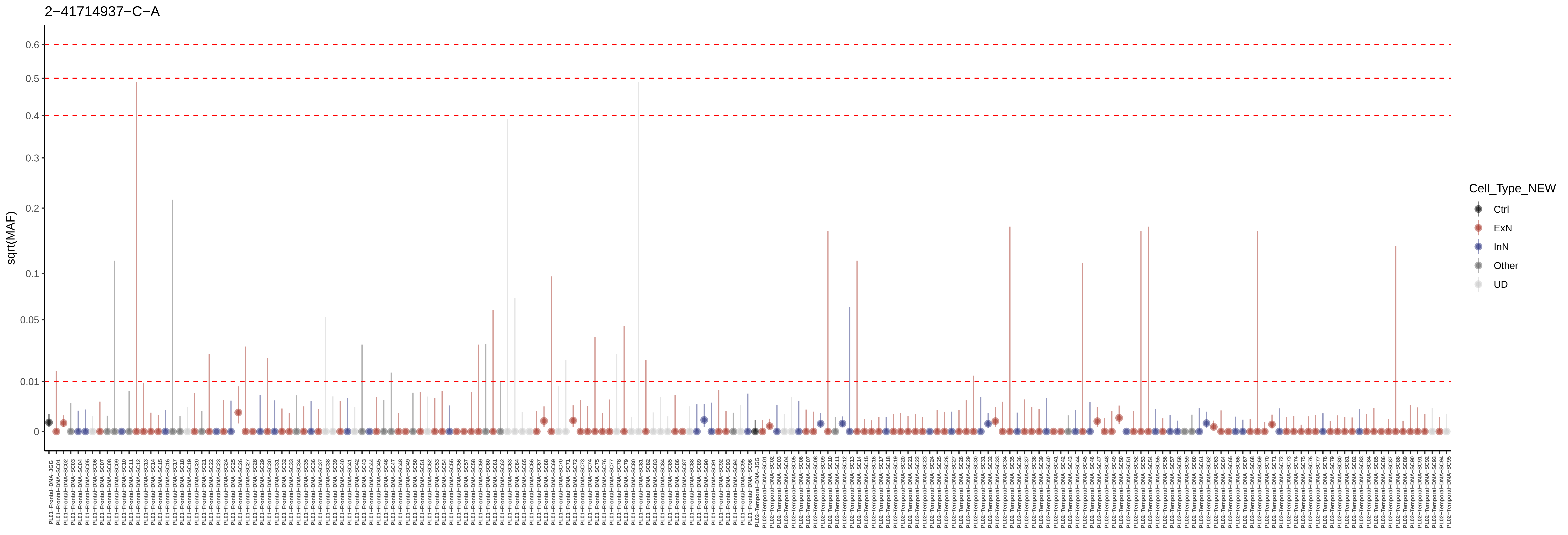

2−41714937−C−A
sqrt(MAF)
0.6
0.5
0.4
0.3
0.2
0.1
0.05
0.01
0
Cell_Type_NEW
Ctrl
ExN
InN
Other
UD

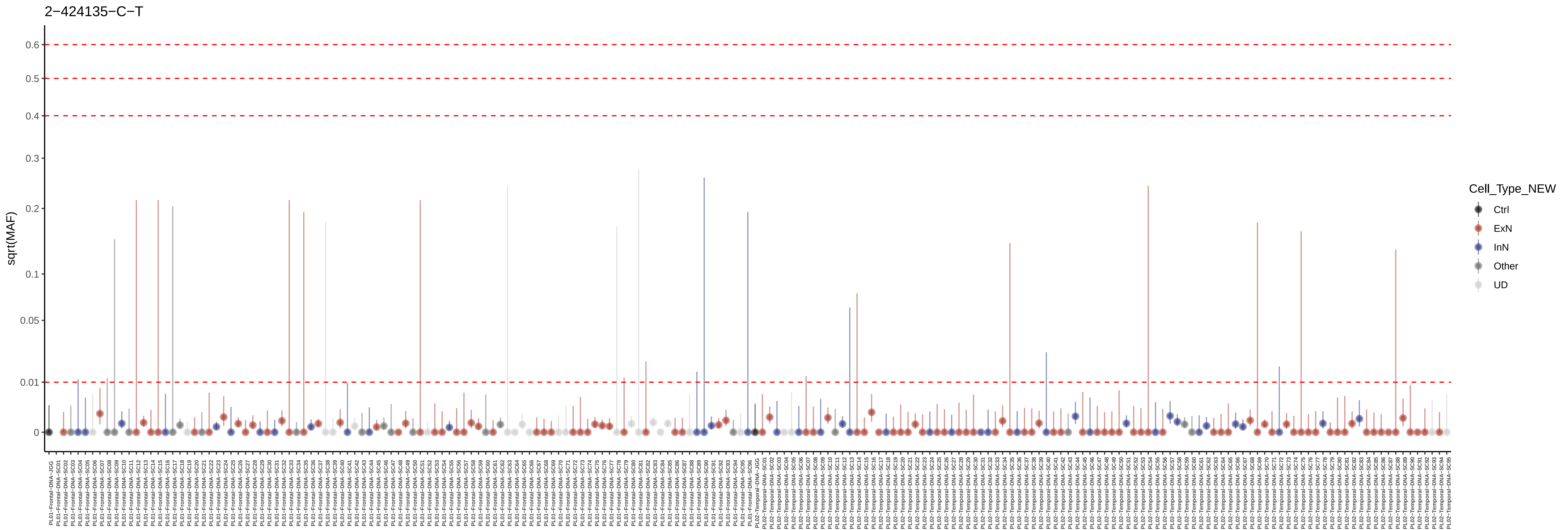
2−424135−C−T
sqrt(MAF)
0.6
0.5
0.4
0.3
0.2
0.1
0.05
0.01
0
Cell_Type_NEW
Ctrl
ExN
InN
Other
UD

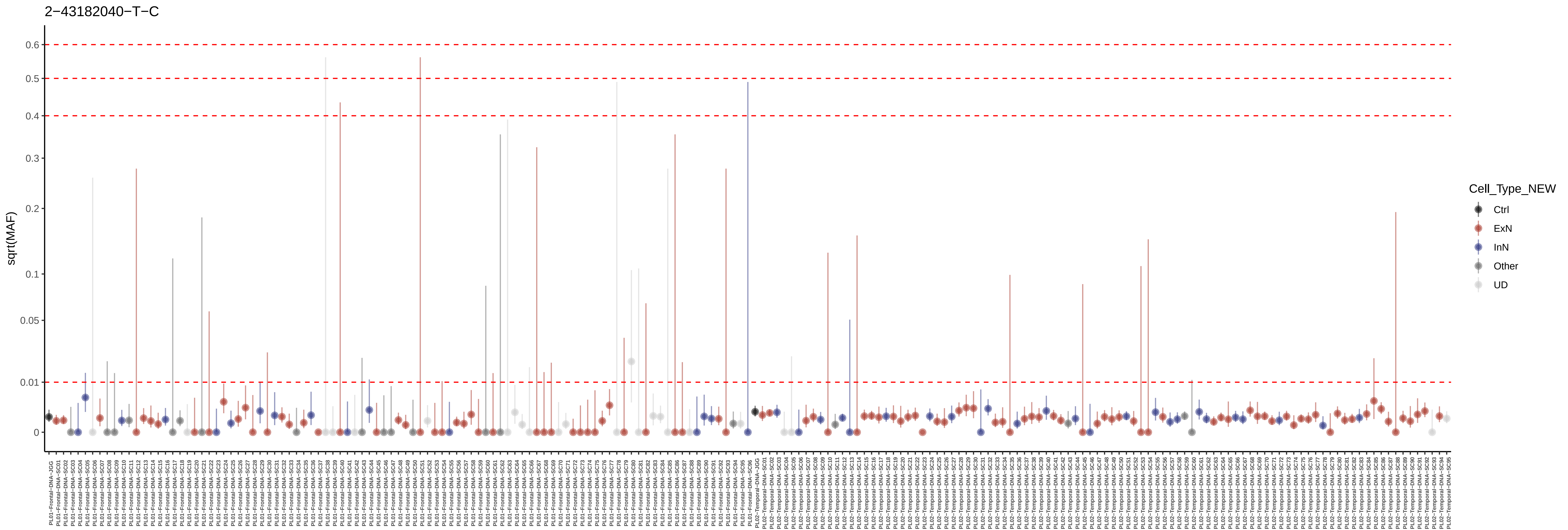

2−43182040−T−C
sqrt(MAF)
0.6
0.5
0.4
0.3
0.2
0.1
0.05
0.01
0
Cell_Type_NEW
Ctrl
ExN
InN
Other
UD

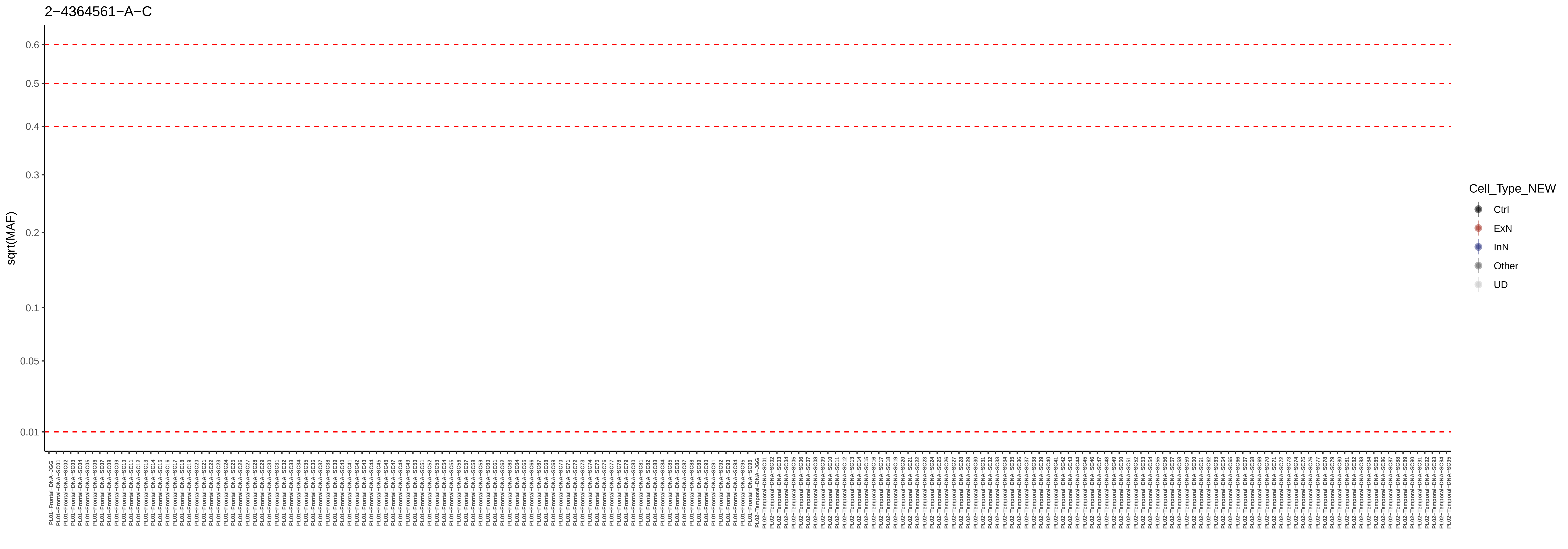

2−4364561−A−C
sqrt(MAF)
0.6
0.5
0.4
0.3
0.2
0.1
0.05
0.01
Cell_Type_NEW
Ctrl
ExN
InN
Other
UD

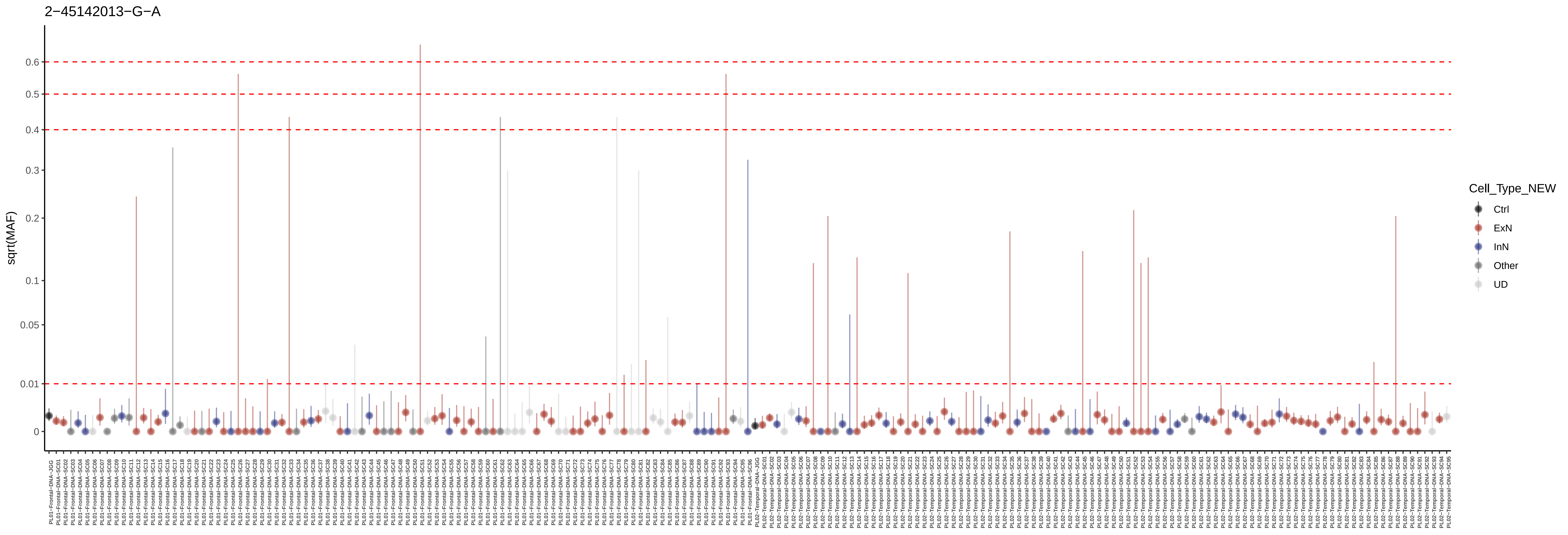

2−45142013−G−A
sqrt(MAF)
0.6
0.5
0.4
0.3
0.2
0.1
0.05
0.01
0
Cell_Type_NEW
Ctrl
ExN
InN
Other
UD

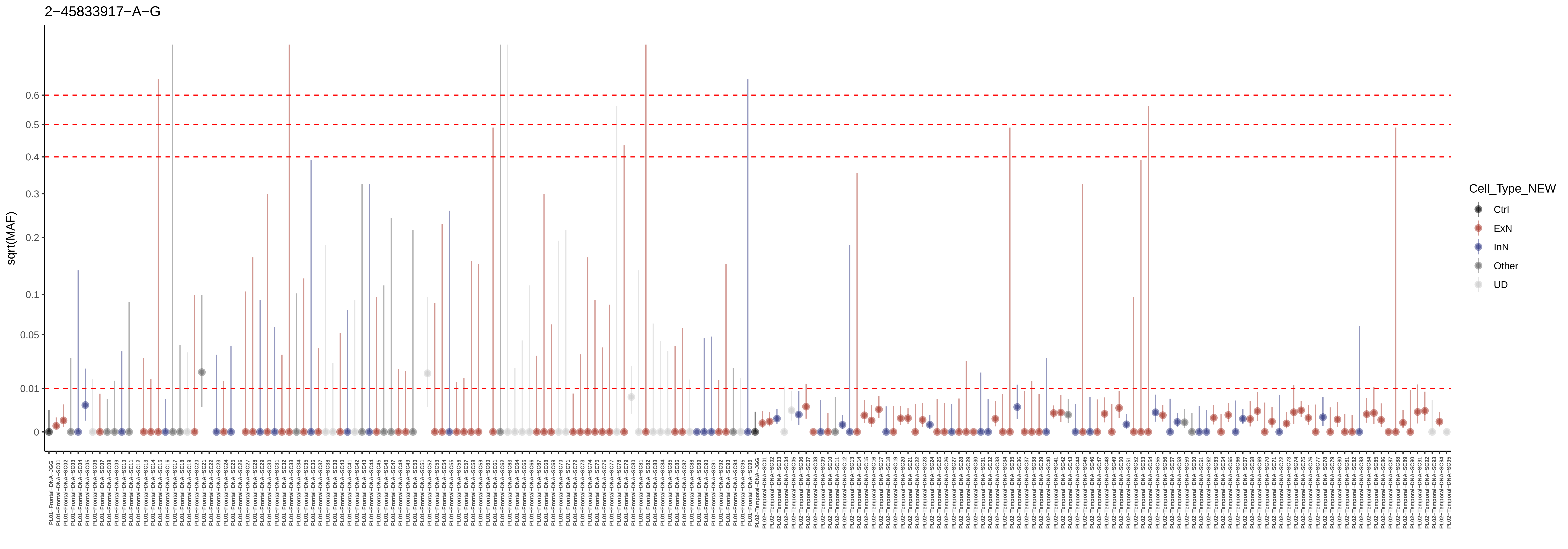
2−45833917−A−G
sqrt(MAF)
0.6
0.5
0.4
0.3
0.2
0.1
0.05
0.01
0
Cell_Type_NEW
Ctrl
ExN
InN
Other
UD

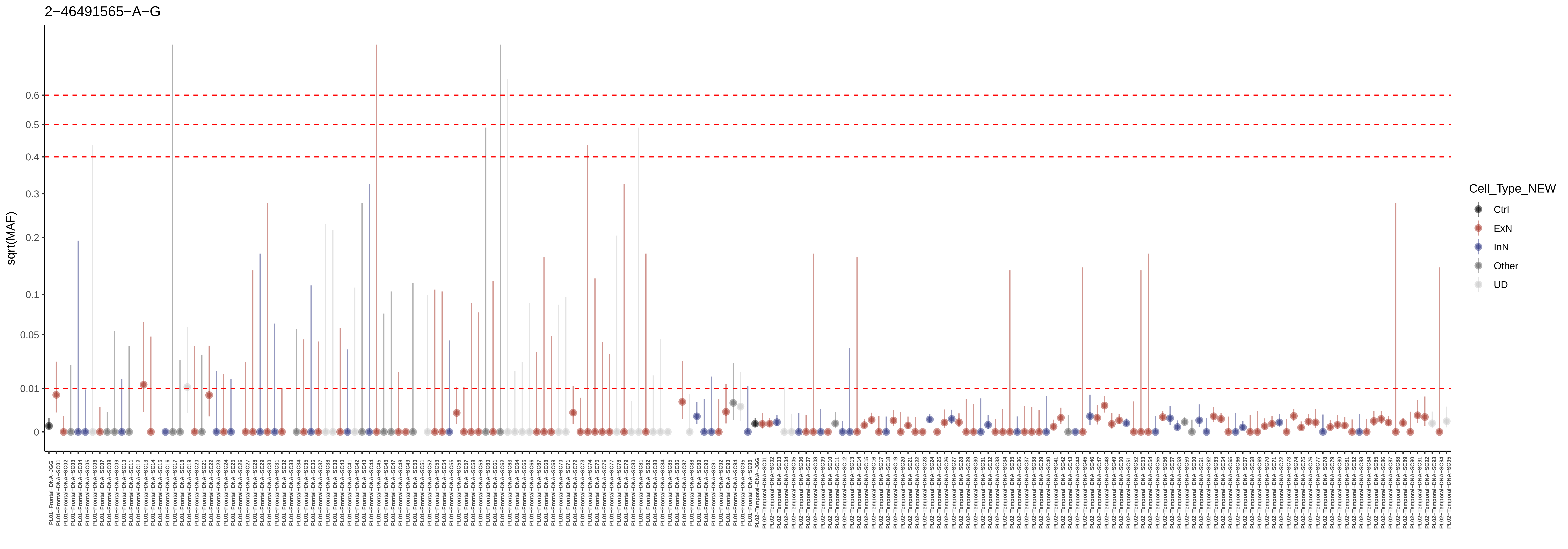
2−46491565−A−G
sqrt(MAF)
0.6
0.5
0.4
0.3
0.2
0.1
0.05
0.01
0
Cell_Type_NEW
Ctrl
ExN
InN
Other
UD

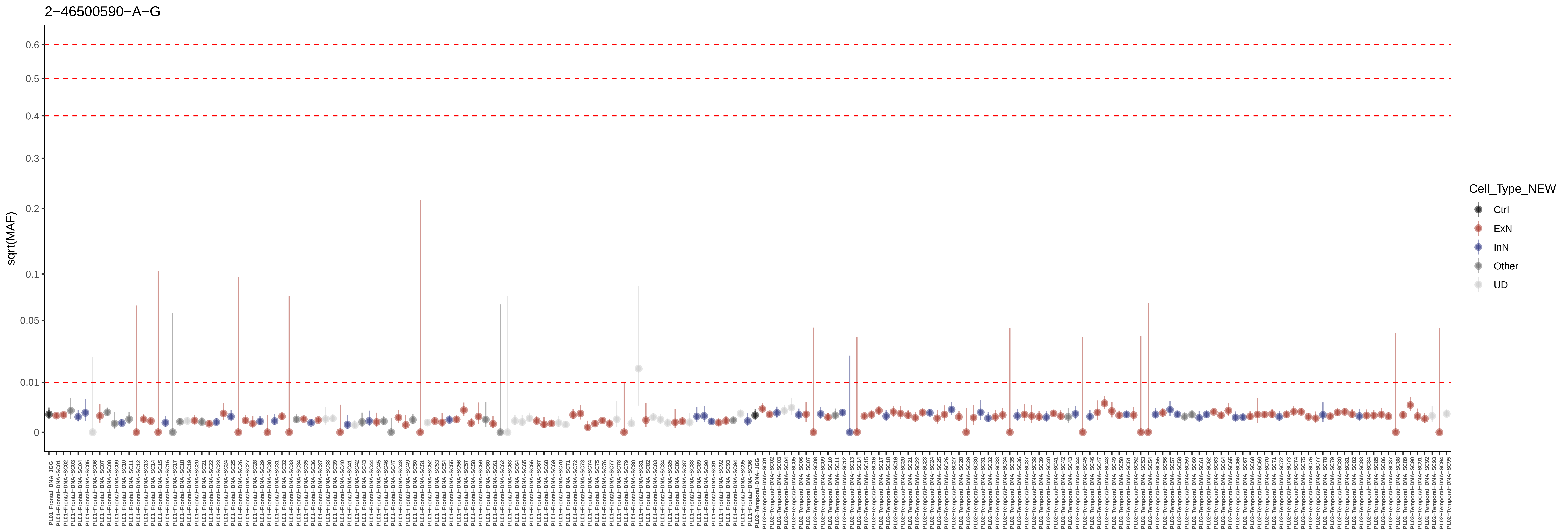
2−46500590−A−G
sqrt(MAF)
0.6
0.5
0.4
0.3
0.2
0.1
0.05
0.01
0
Cell_Type_NEW
Ctrl
ExN
InN
Other
UD

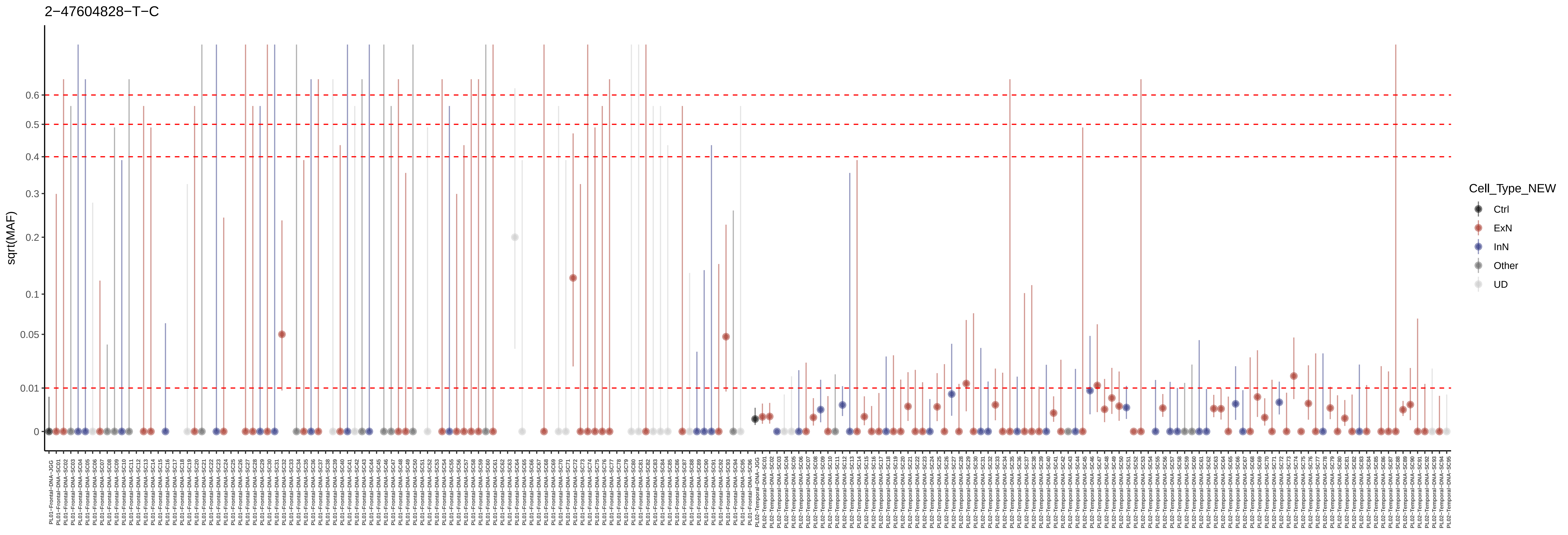
2−47604828−T−C
sqrt(MAF)
0.6
0.5
0.4
0.3
0.2
0.1
0.05
0.01
0
Cell_Type_NEW
Ctrl
ExN
InN
Other
UD

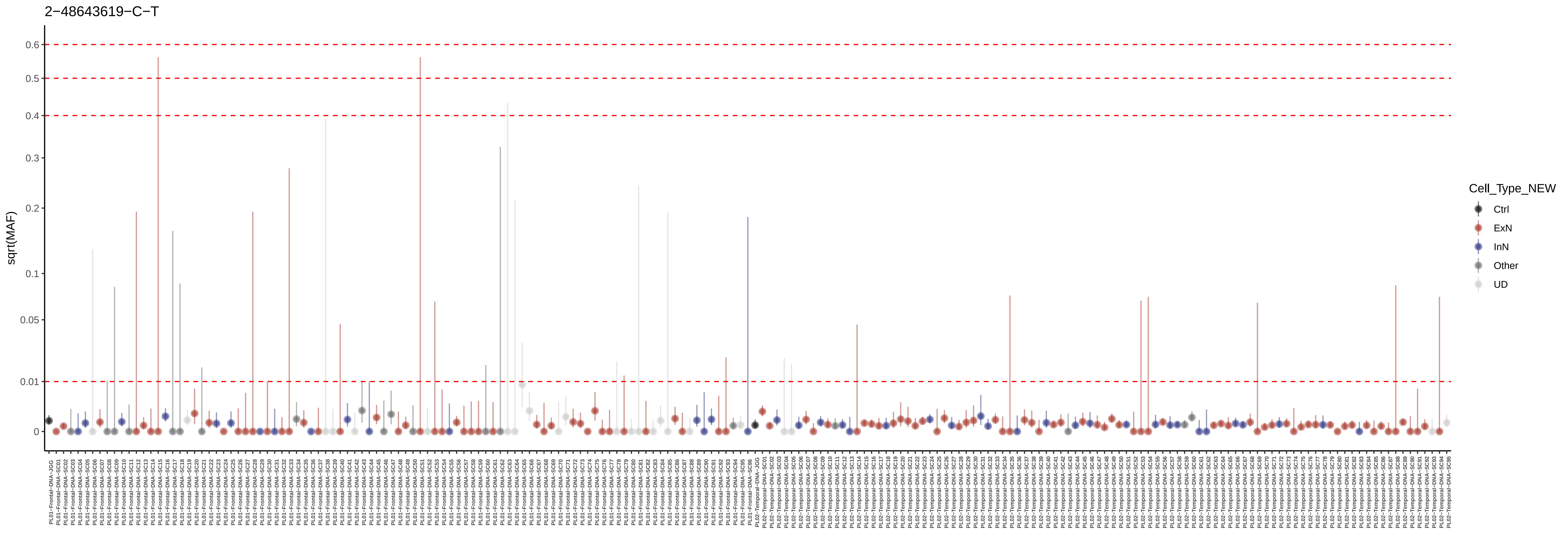

2−48643619−C−T
sqrt(MAF)
0.6
0.5
0.4
0.3
0.2
0.1
0.05
0.01
0
Cell_Type_NEW
Ctrl
ExN
InN
Other
UD

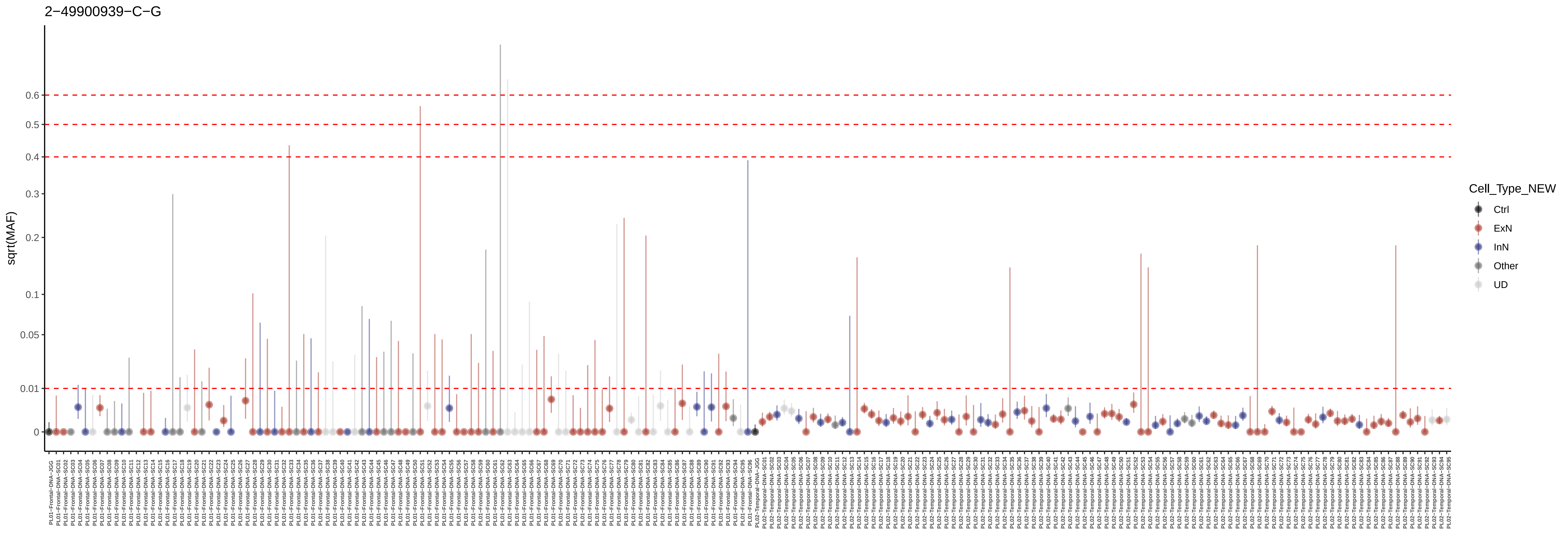

2−49900939−C−G
sqrt(MAF)
0
0.01
0.05
0.1
0.2
0.3
0.4
0.5
0.6
Cell_Type_NEW
Ctrl
ExN
InN
Other
UD

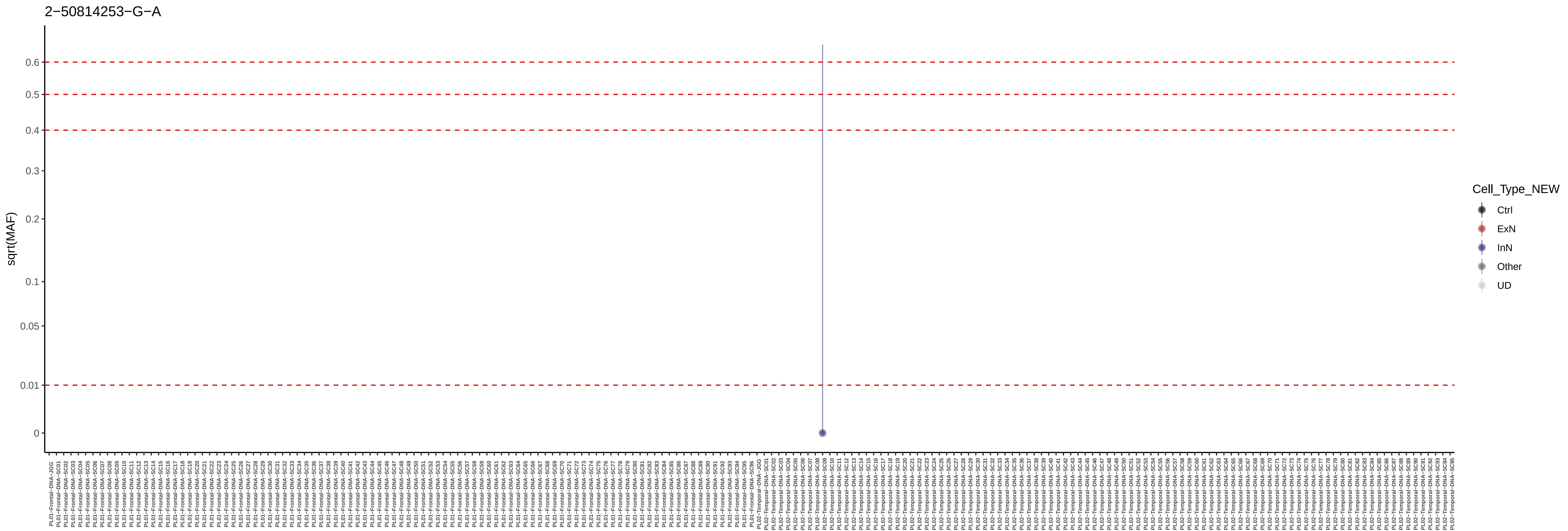

2−50814253−G−A
sqrt(MAF)
0.6
0.5
0.4
0.3
0.2
0.1
0.05
0.01
0
Cell_Type_NEW
Ctrl
ExN
InN
Other
UD

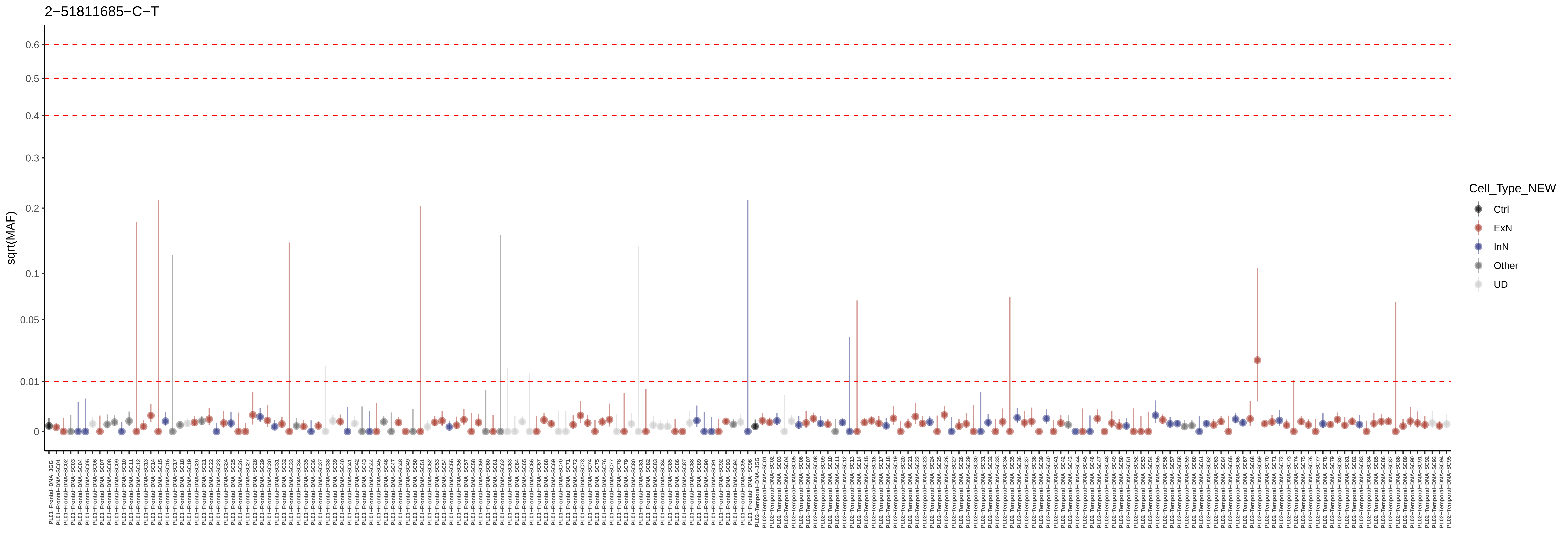

2−51811685−C−T
sqrt(MAF)
0.6
0.5
0.4
0.3
0.2
0.1
0.05
0.01
0
Cell_Type_NEW
Ctrl
ExN
InN
Other
UD

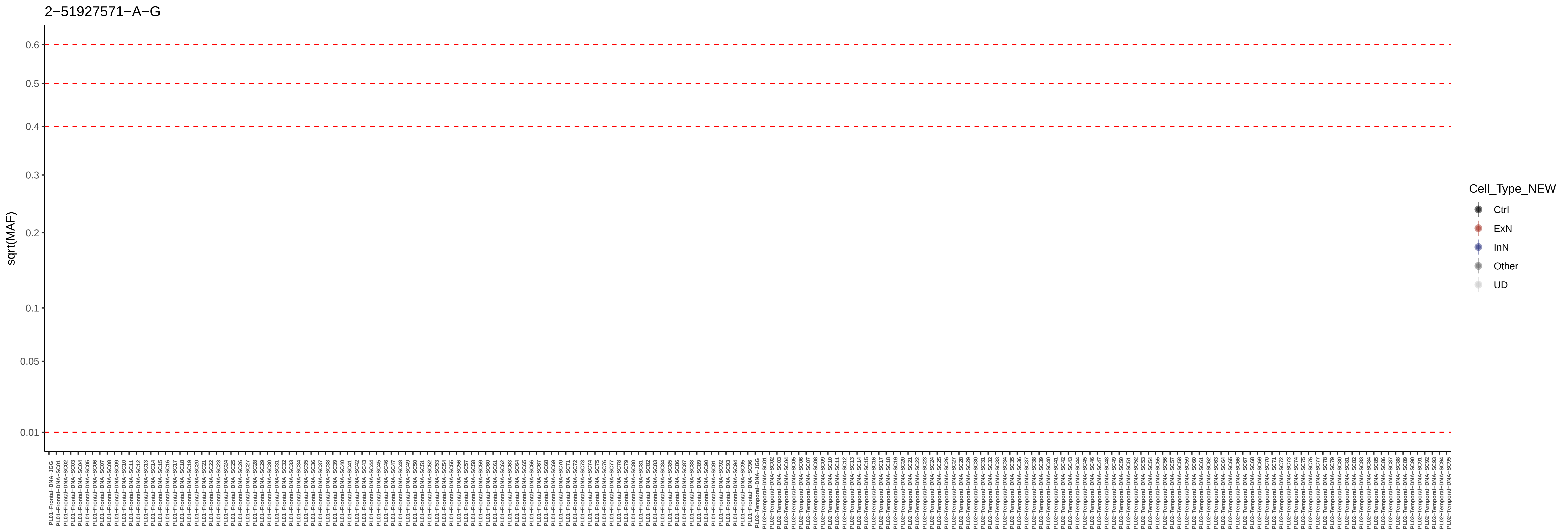
2−51927571−A−G
sqrt(MAF)
0.6
0.5
0.4
0.3
0.2
0.1
0.05
0.01
Cell_Type_NEW
Ctrl
ExN
InN
Other
UD

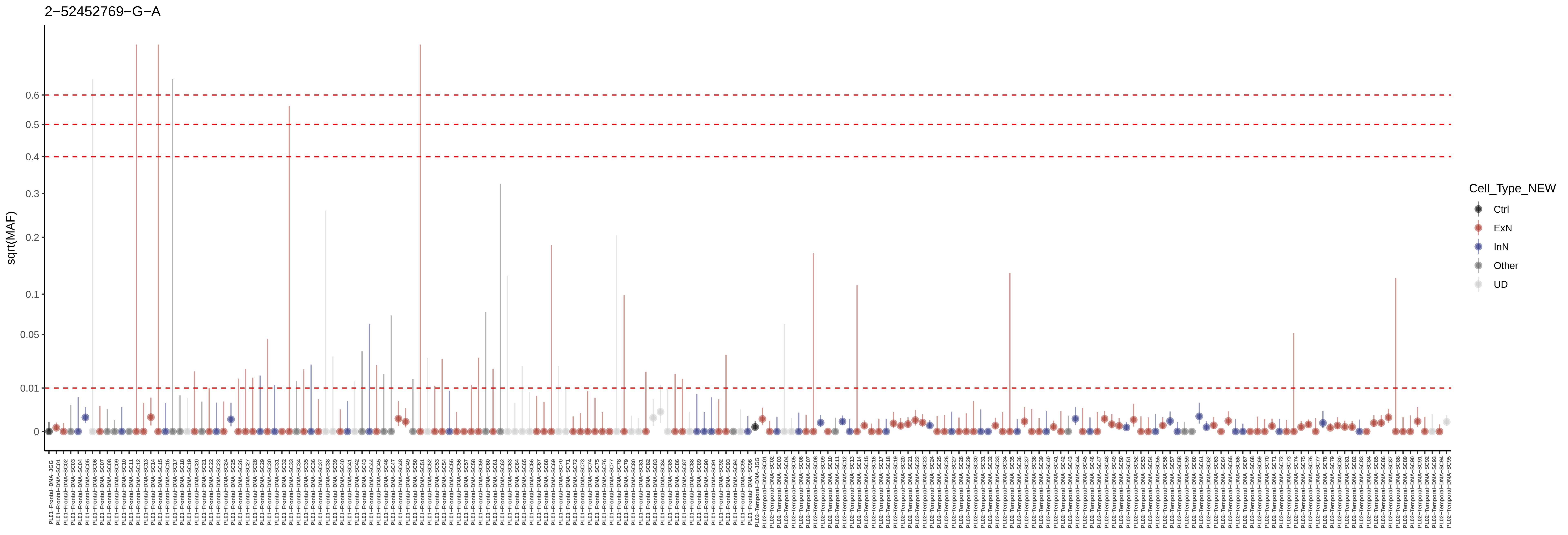

2−52452769−G−A
sqrt(MAF)
0.6
0.5
0.4
0.3
0.2
0.1
0.05
0.01
0
Cell_Type_NEW
Ctrl
ExN
InN
Other
UD

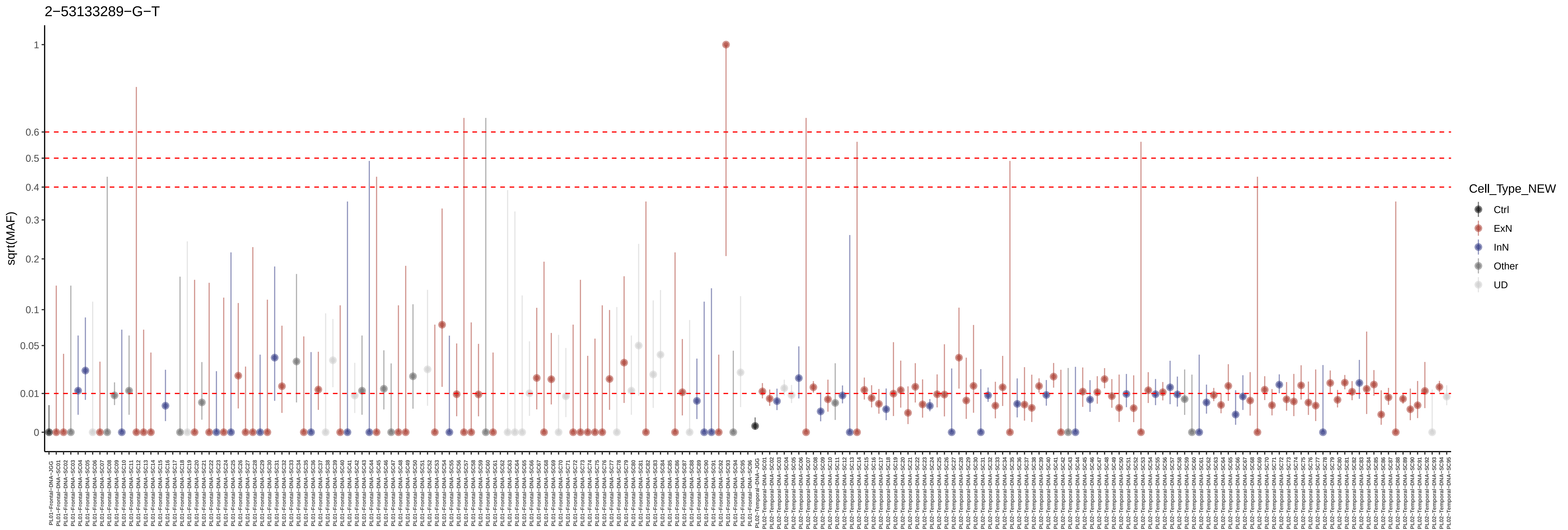

2−53133289−G−T
sqrt(MAF)
1
0.6
0.5
0.4
0.3
0.2
0.1
0.05
0.01
0
Cell_Type_NEW
Ctrl
ExN
InN
Other
UD

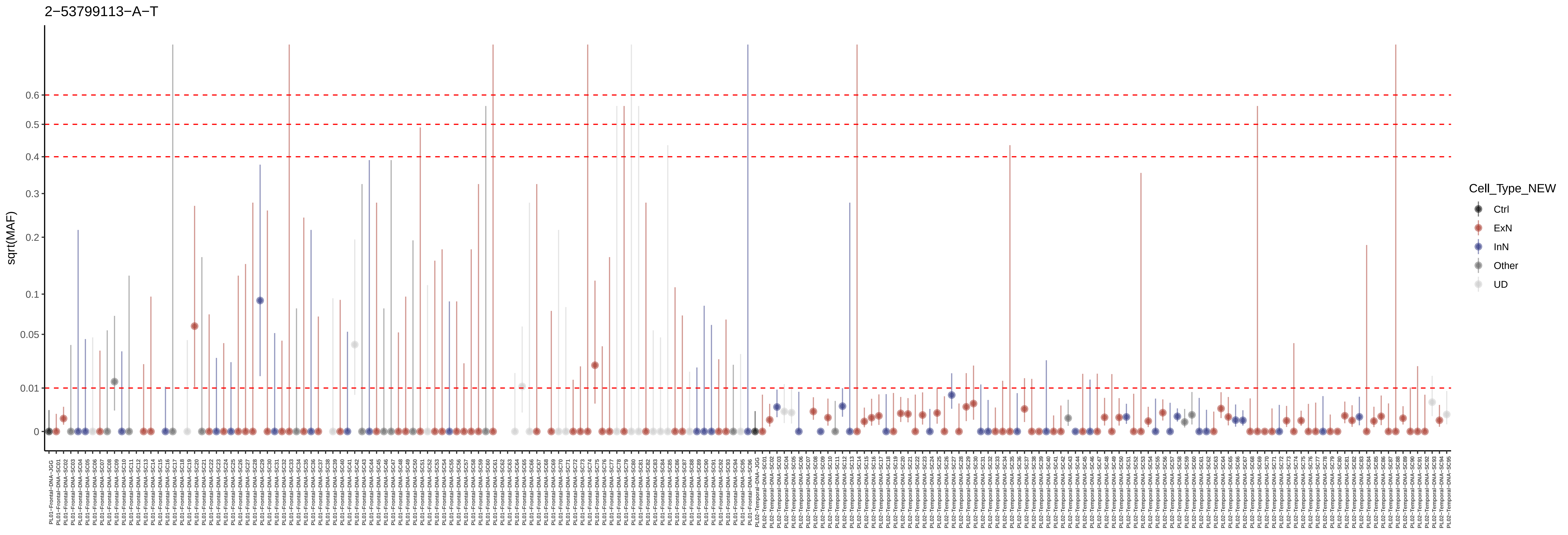
2−53799113−A−T
sqrt(MAF)
0.6
0.5
0.4
0.3
0.2
0.1
0.05
0.01
0
Cell_Type_NEW
Ctrl
ExN
InN
Other
UD

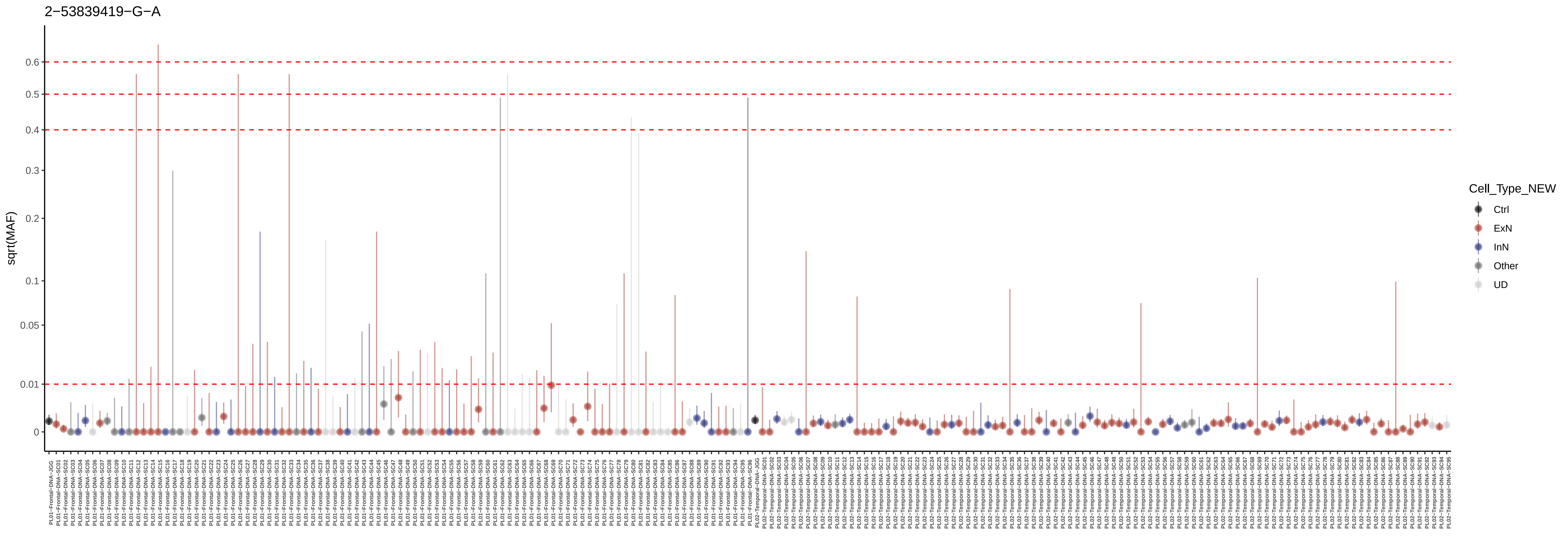

2−53839419−G−A
sqrt(MAF)
0.6
0.5
0.4
0.3
0.2
0.1
0.05
0.01
0
Cell_Type_NEW
Ctrl
ExN
InN
Other
UD

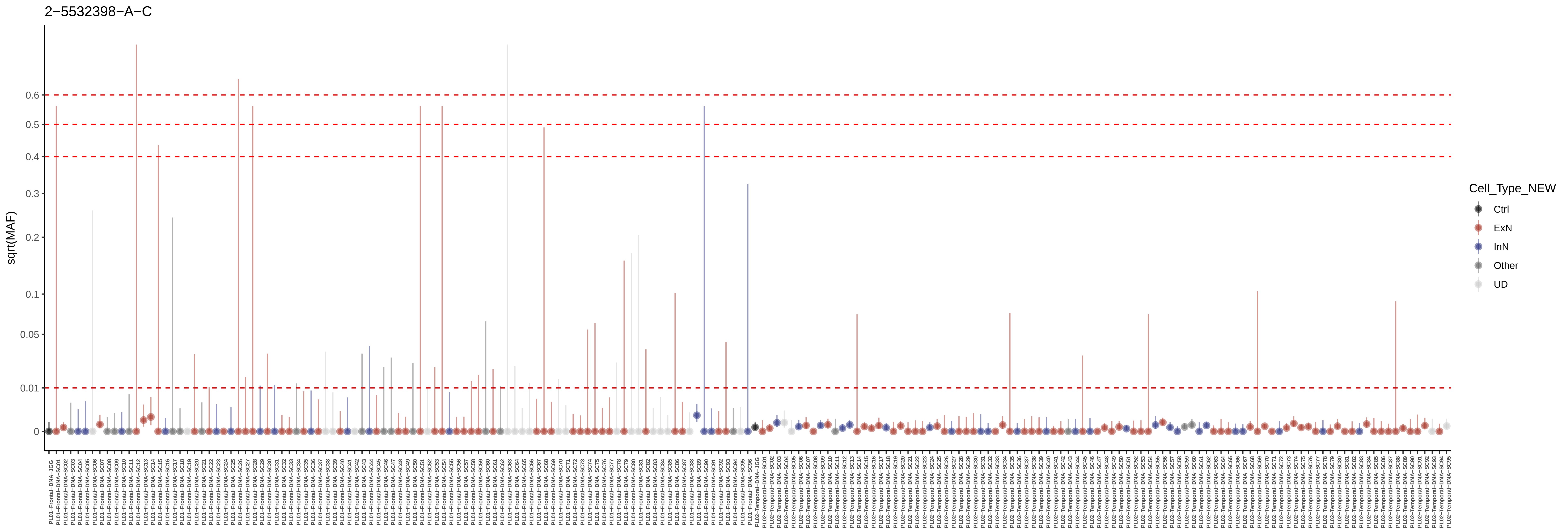

2−5532398−A−C
sqrt(MAF)
0.6
0.5
0.4
0.3
0.2
0.1
0.05
0.01
0
Cell_Type_NEW
Ctrl
ExN
InN
Other
UD

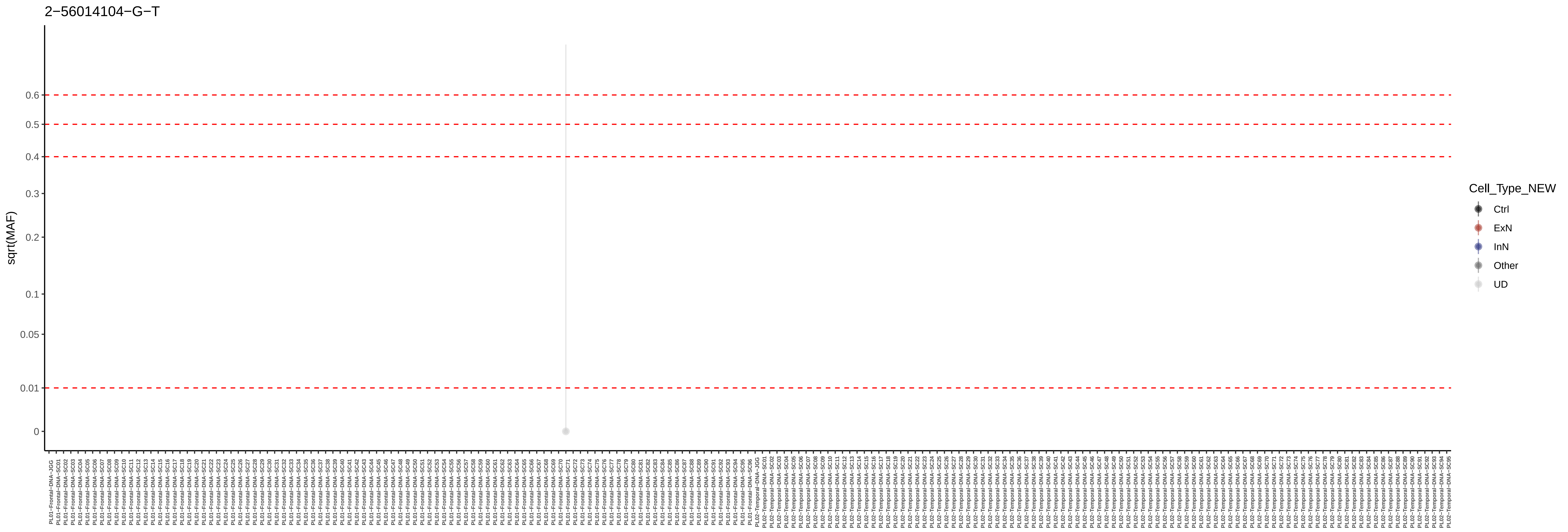
2−56014104−G−T
sqrt(MAF)
0.6
0.5
0.4
0.3
0.2
0.1
0.05
0.01
0
Cell_Type_NEW
Ctrl
ExN
InN
Other
UD

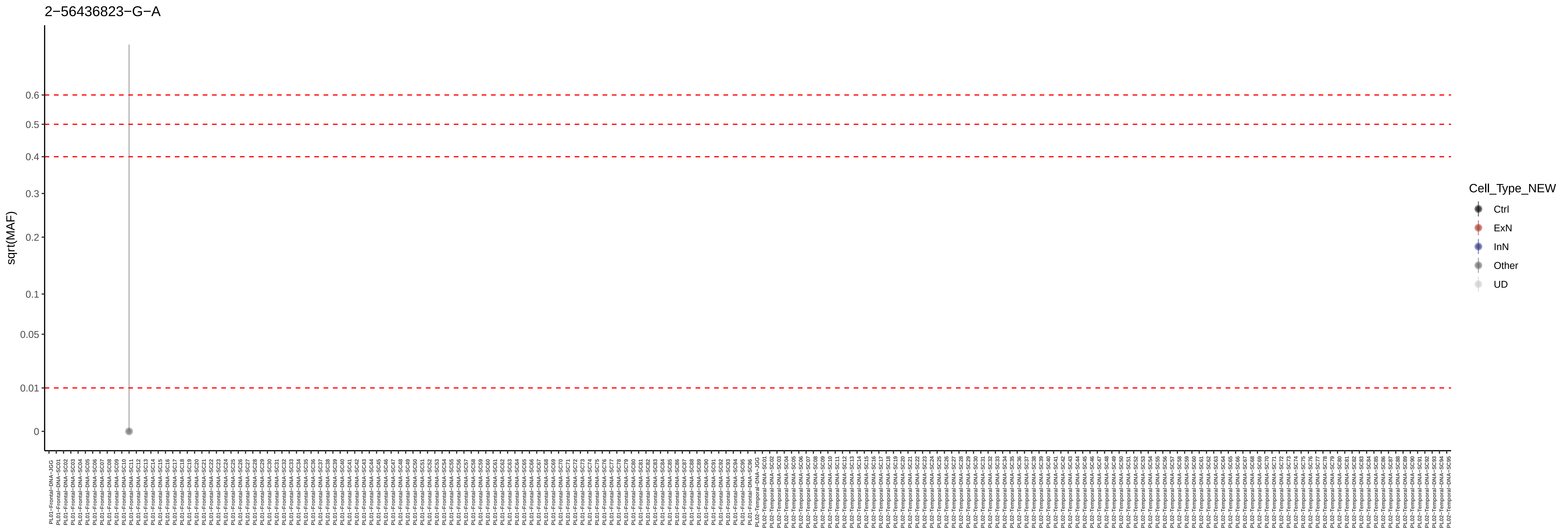
2−56436823−G−A
sqrt(MAF)
0.6
0.5
0.4
0.3
0.2
0.1
0.05
0.01
0
Cell_Type_NEW
Ctrl
ExN
InN
Other
UD

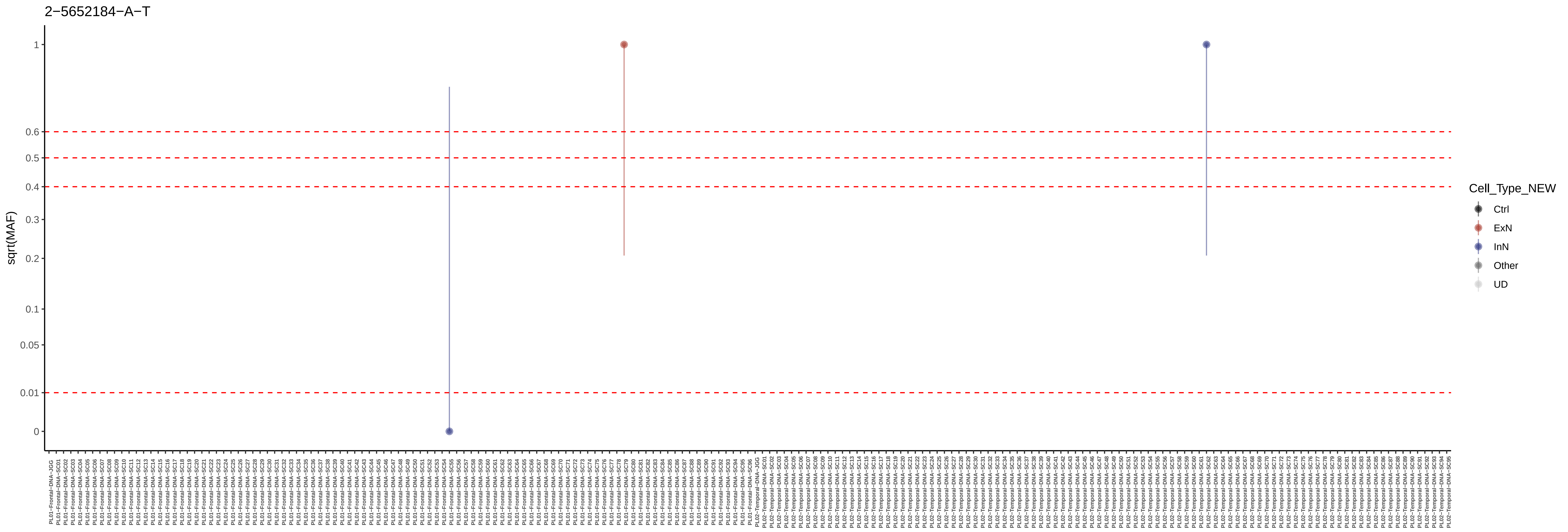
2−5652184−A−T
sqrt(MAF)
1
0.6
0.5
0.4
0.3
0.2
0.1
0.05
0.01
0
Cell_Type_NEW
Ctrl
ExN
InN
Other
UD

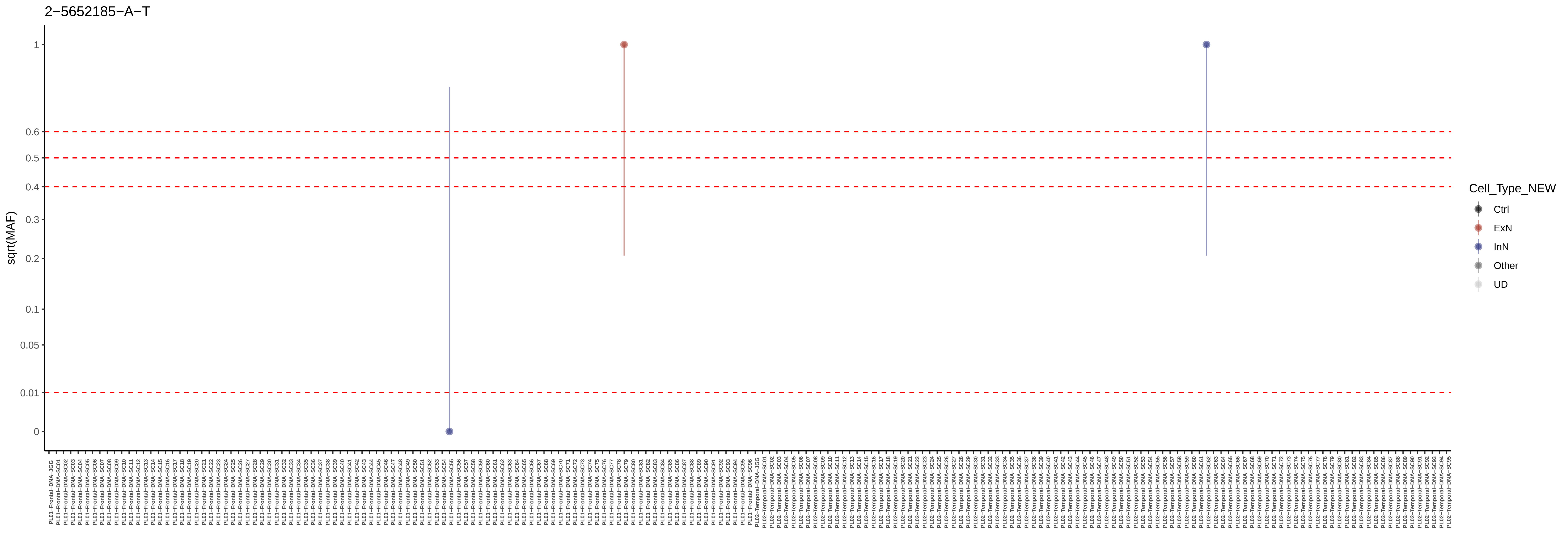
2−5652185−A−T
sqrt(MAF)
1
0.6
0.5
0.4
0.3
0.2
0.1
0.05
0.01
0
Cell_Type_NEW
Ctrl
ExN
InN
Other
UD

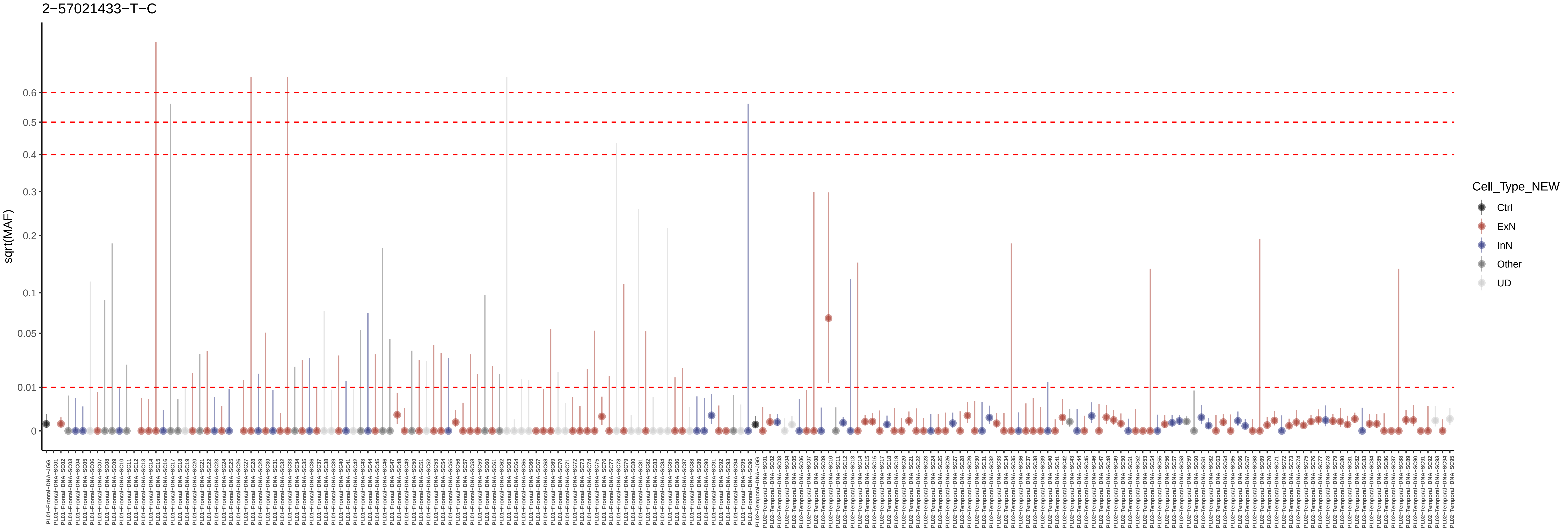
2−57021433−T−C
sqrt(MAF)
0.6
0.5
0.4
0.3
0.2
0.1
0.05
0.01
0
Cell_Type_NEW
Ctrl
ExN
InN
Other
UD

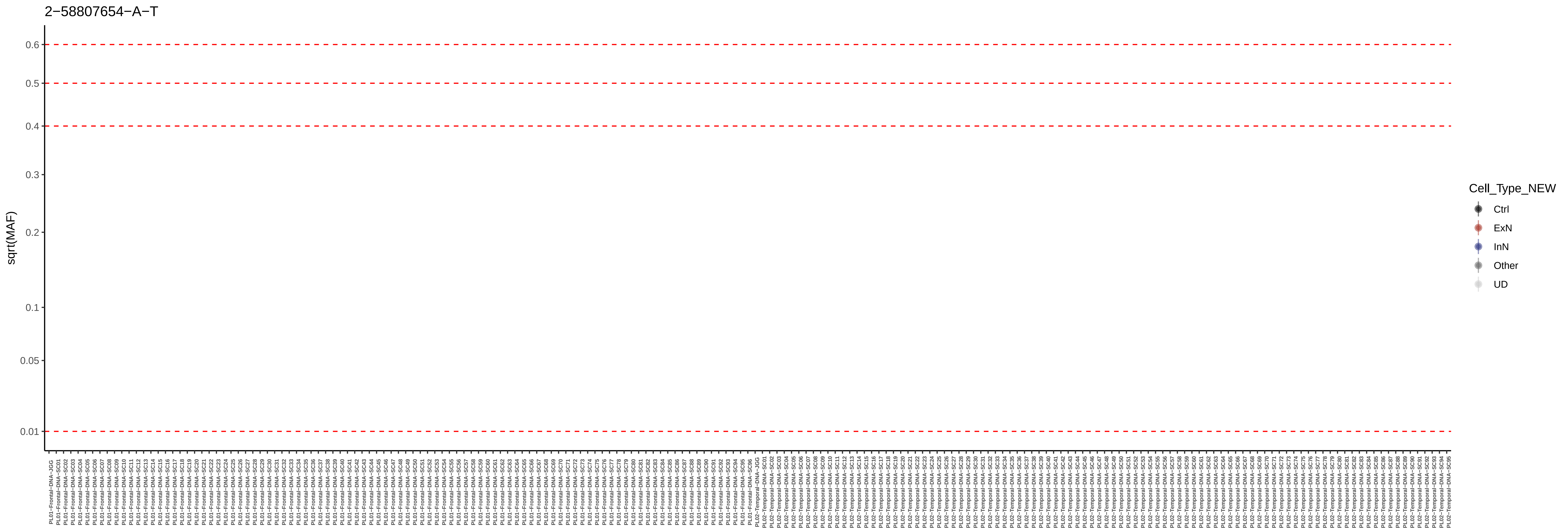
2−58807654−A−T
sqrt(MAF)
0.6
0.5
0.4
0.3
0.2
0.1
0.05
0.01
Cell_Type_NEW
Ctrl
ExN
InN
Other
UD

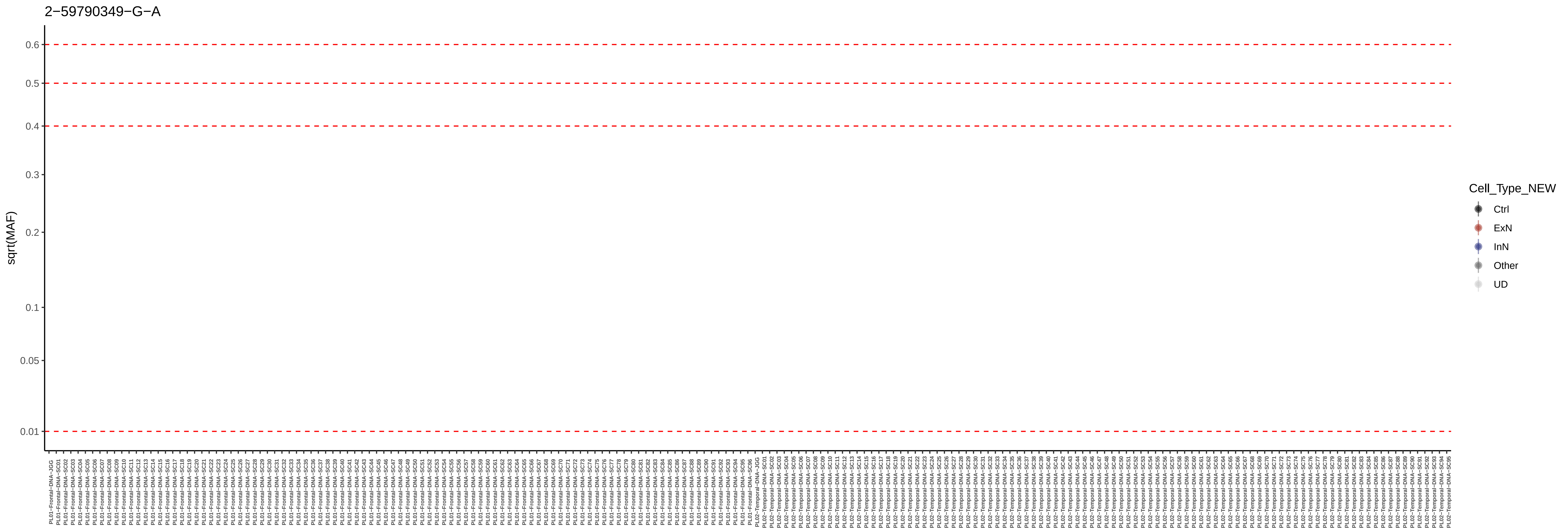
2−59790349−G−A
sqrt(MAF)
0.6
0.5
0.4
0.3
0.2
0.1
0.05
0.01
Cell_Type_NEW
Ctrl
ExN
InN
Other
UD

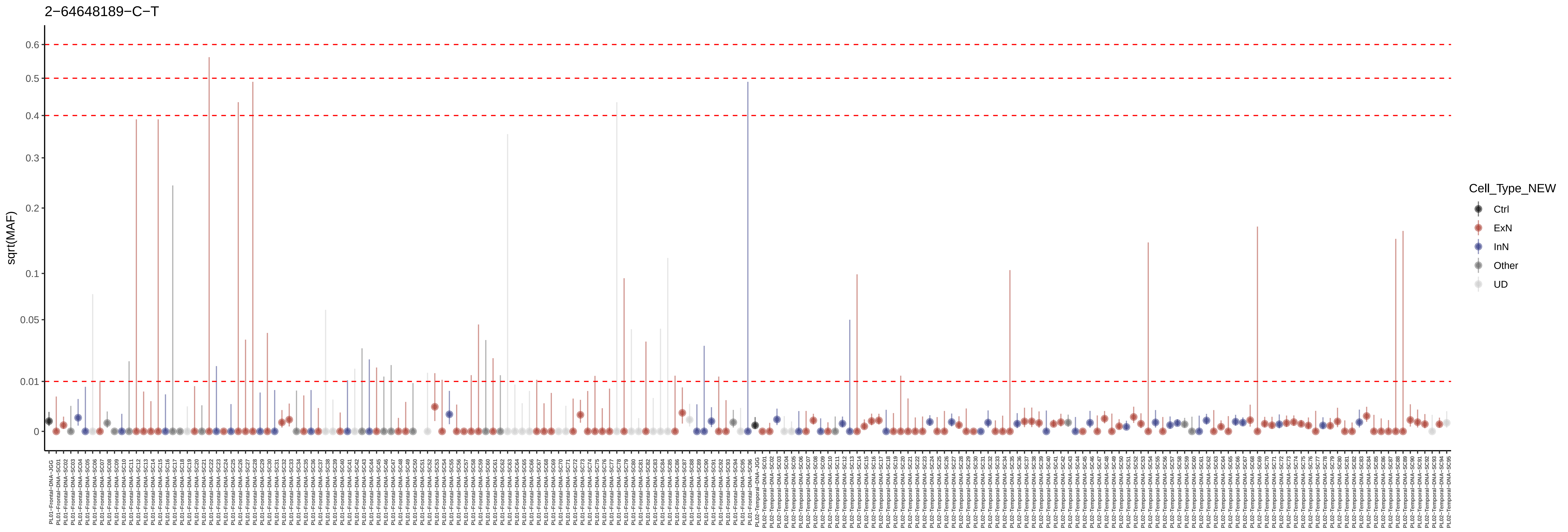
2−64648189−C−T
sqrt(MAF)
0.6
0.5
0.4
0.3
0.2
0.1
0.05
0.01
0
Cell_Type_NEW
Ctrl
ExN
InN
Other
UD

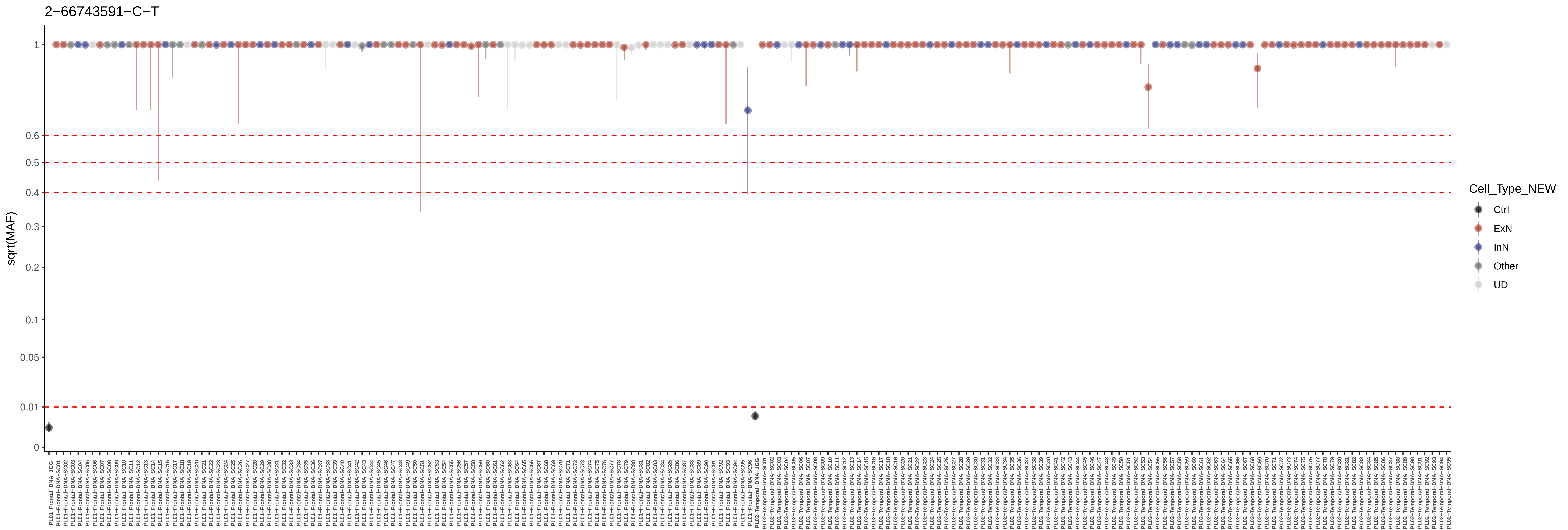

2−66743591−C−T
sqrt(MAF)
1
0.6
0.5
0.4
0.3
0.2
0.1
0.05
0.01
0
Cell_Type_NEW
Ctrl
ExN
InN
Other
UD

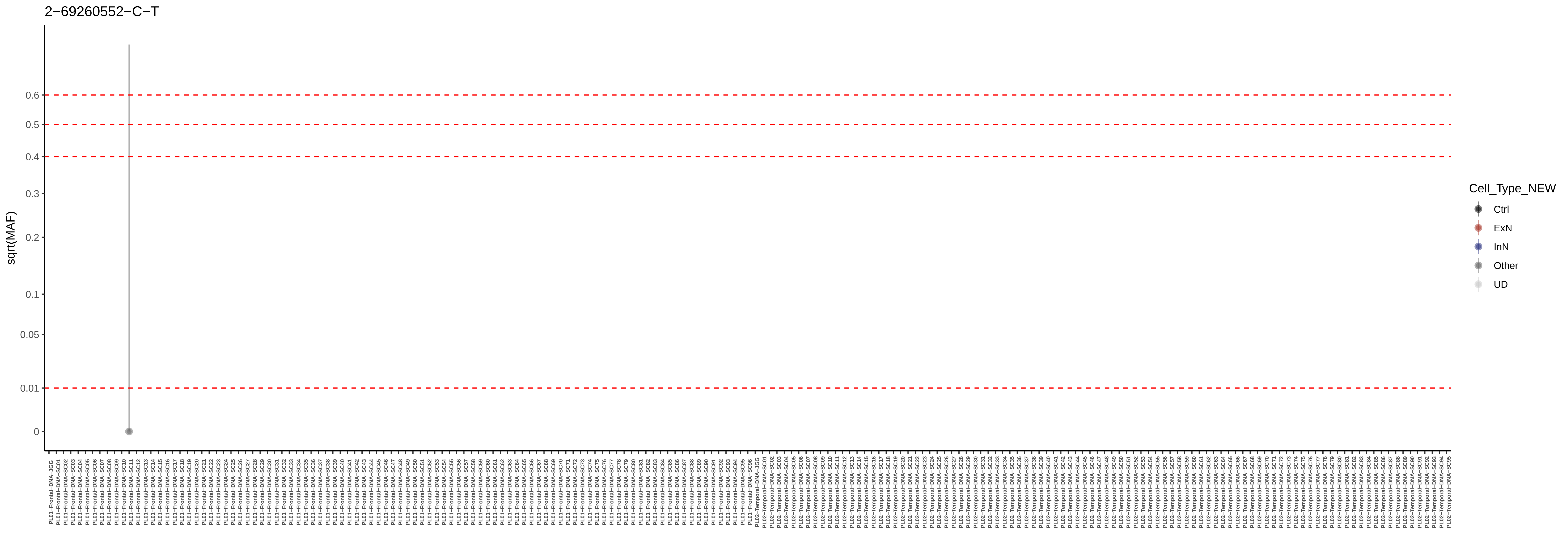
2−69260552−C−T
sqrt(MAF)
0.6
0.5
0.4
0.3
0.2
0.1
0.05
0.01
0
Cell_Type_NEW
Ctrl
ExN
InN
Other
UD

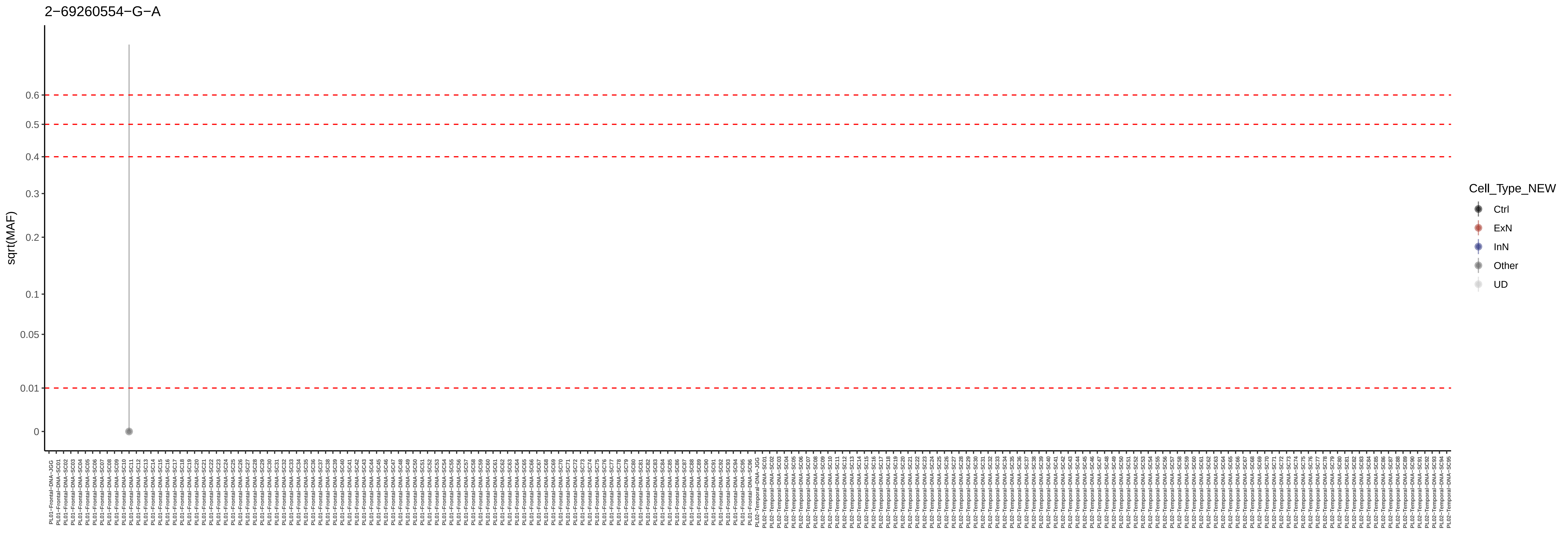
2−69260554−G−A
sqrt(MAF)
0.6
0.5
0.4
0.3
0.2
0.1
0.05
0.01
0
Cell_Type_NEW
Ctrl
ExN
InN
Other
UD

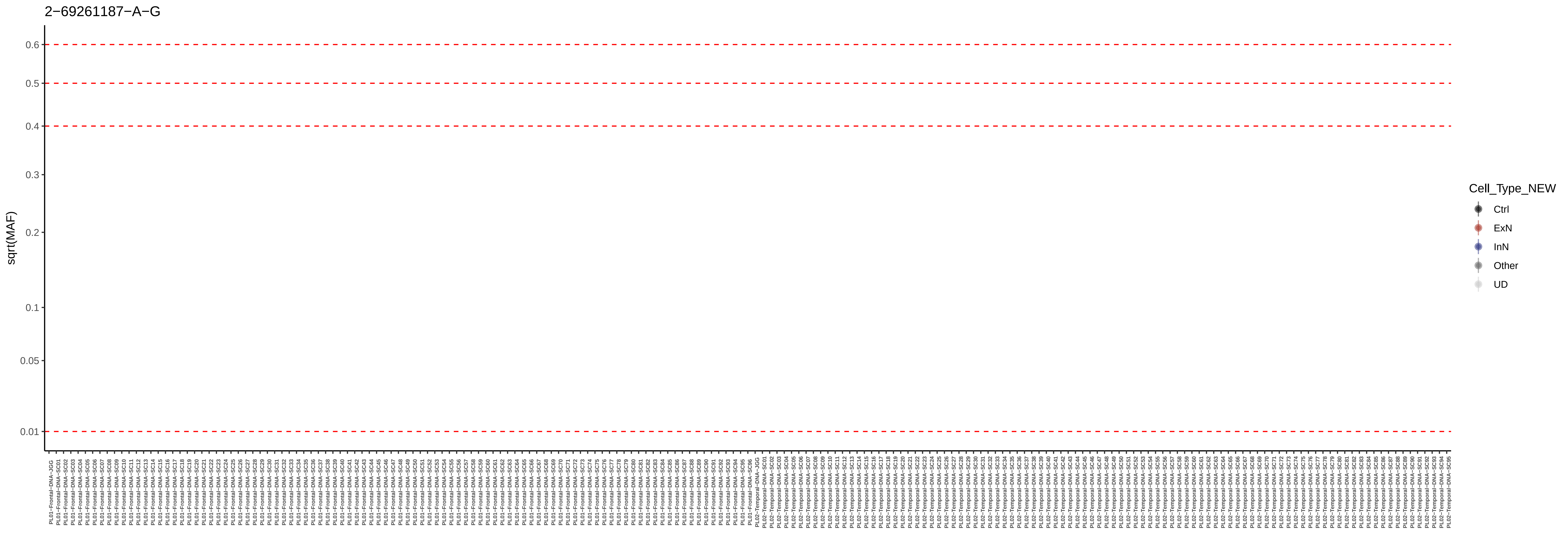
2−69261187−A−G
sqrt(MAF)
0.6
0.5
0.4
0.3
0.2
0.1
0.05
0.01
Cell_Type_NEW
Ctrl
ExN
InN
Other
UD

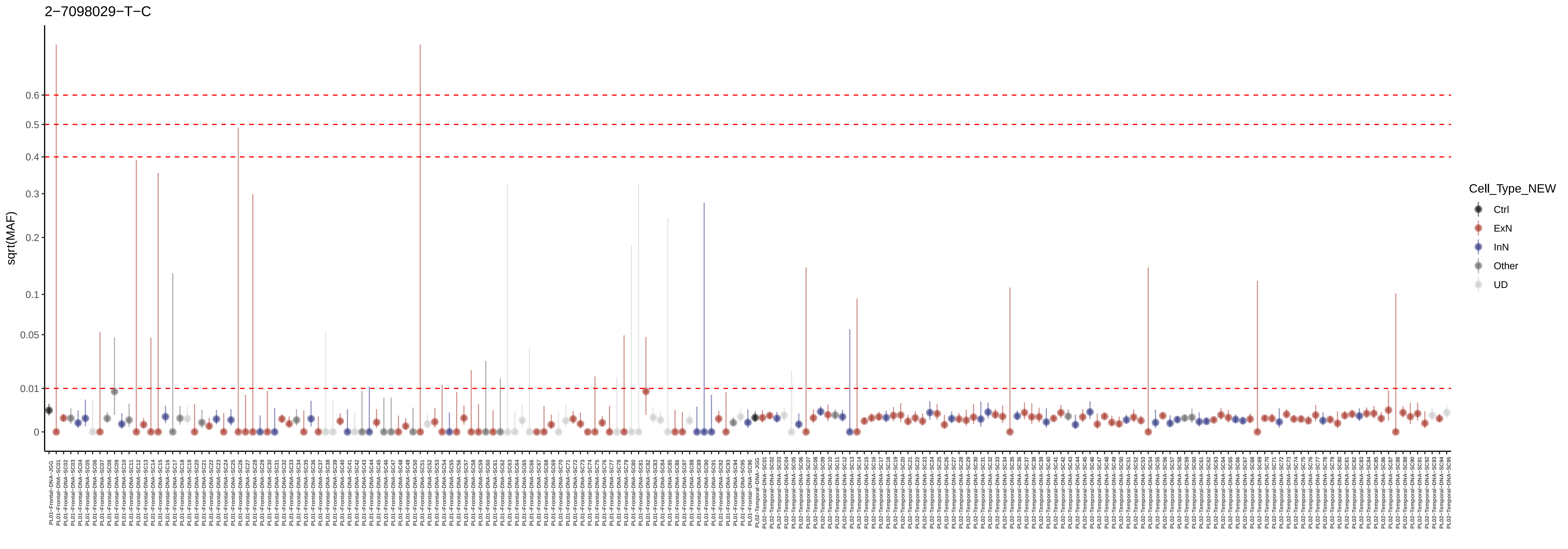

2−7098029−T−C
sqrt(MAF)
0.6
0.5
0.4
0.3
0.2
0.1
0.05
0.01
0
Cell_Type_NEW
Ctrl
ExN
InN
Other
UD

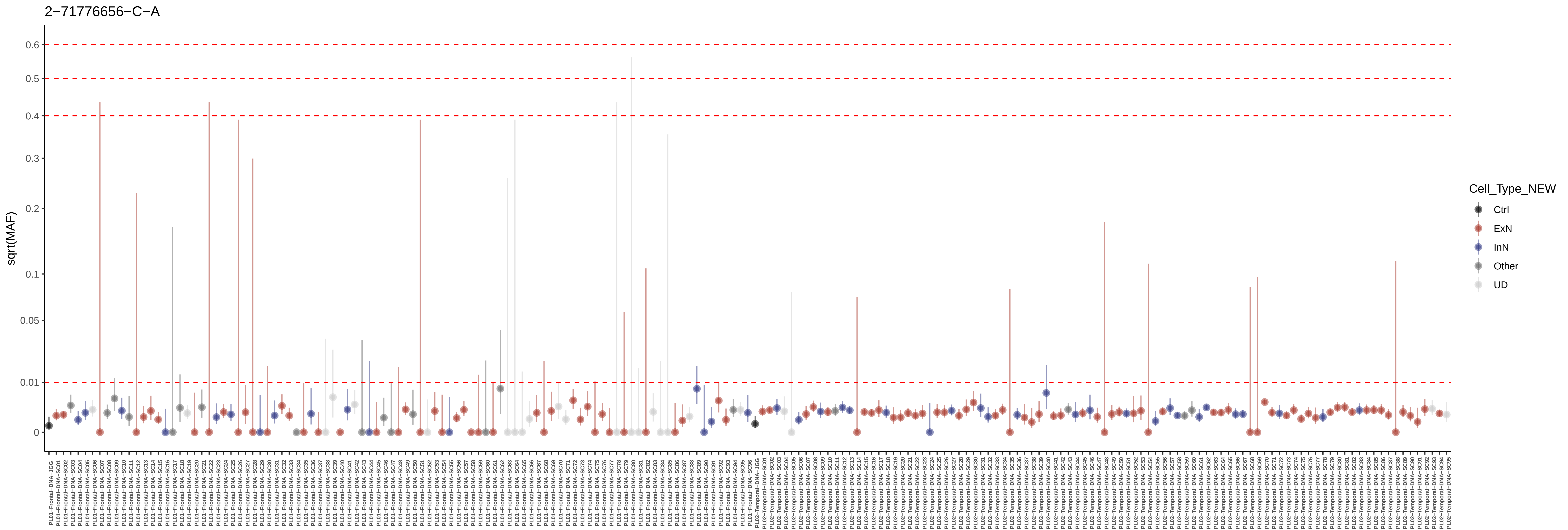

2−71776656−C−A
sqrt(MAF)
0.6
0.5
0.4
0.3
0.2
0.1
0.05
0.01
0
Cell_Type_NEW
Ctrl
ExN
InN
Other
UD

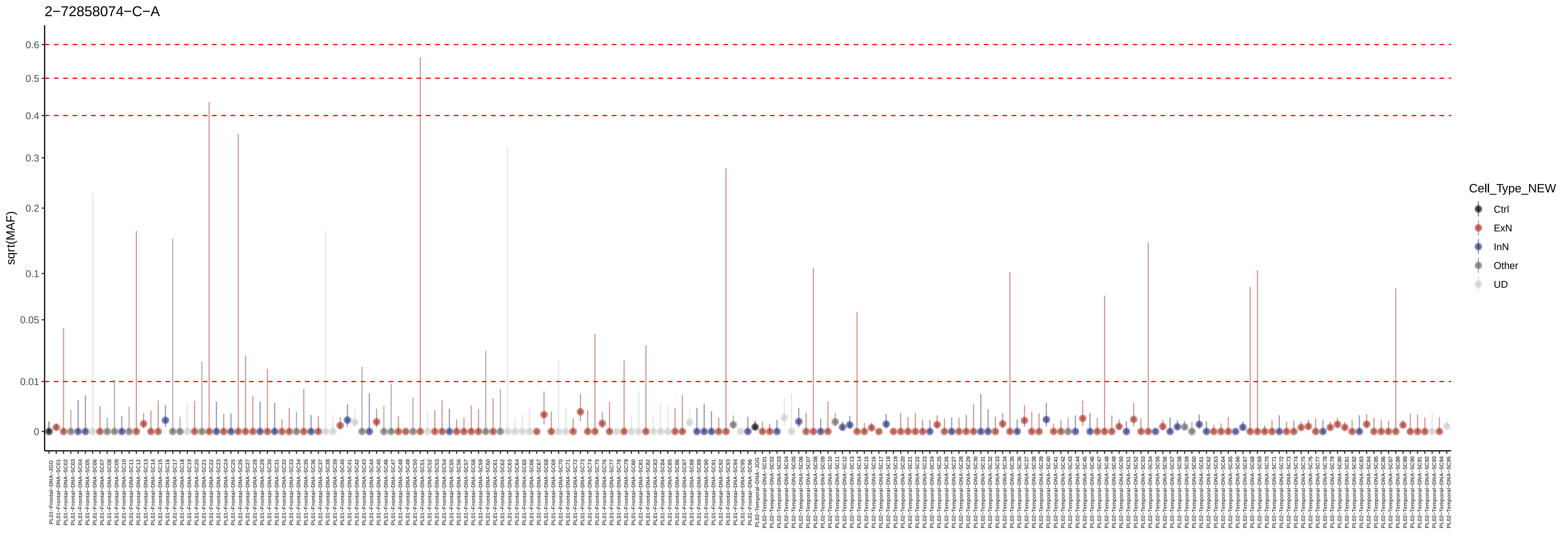

2−72858074−C−A
sqrt(MAF)
0.6
0.5
0.4
0.3
0.2
0.1
0.05
0.01
0
Cell_Type_NEW
Ctrl
ExN
InN
Other
UD

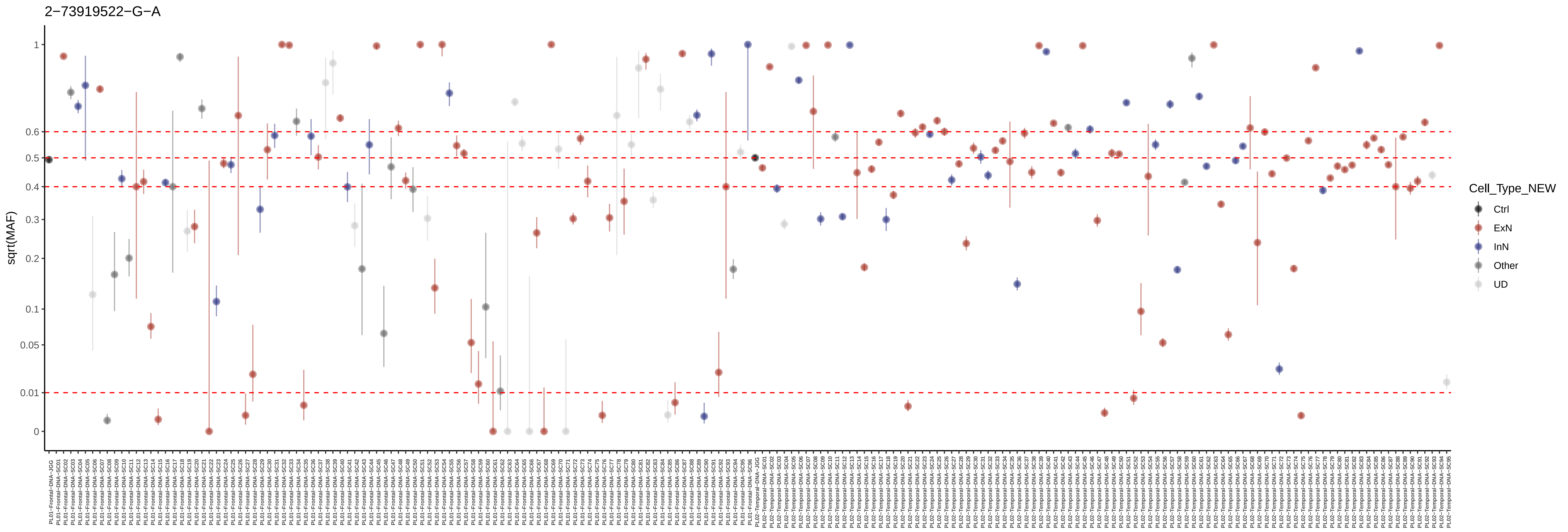
2−73919522−G−A
sqrt(MAF)
1
0.6
0.5
0.4
0.3
0.2
0.1
0.05
0.01
0
Cell_Type_NEW
Ctrl
ExN
InN
Other
UD

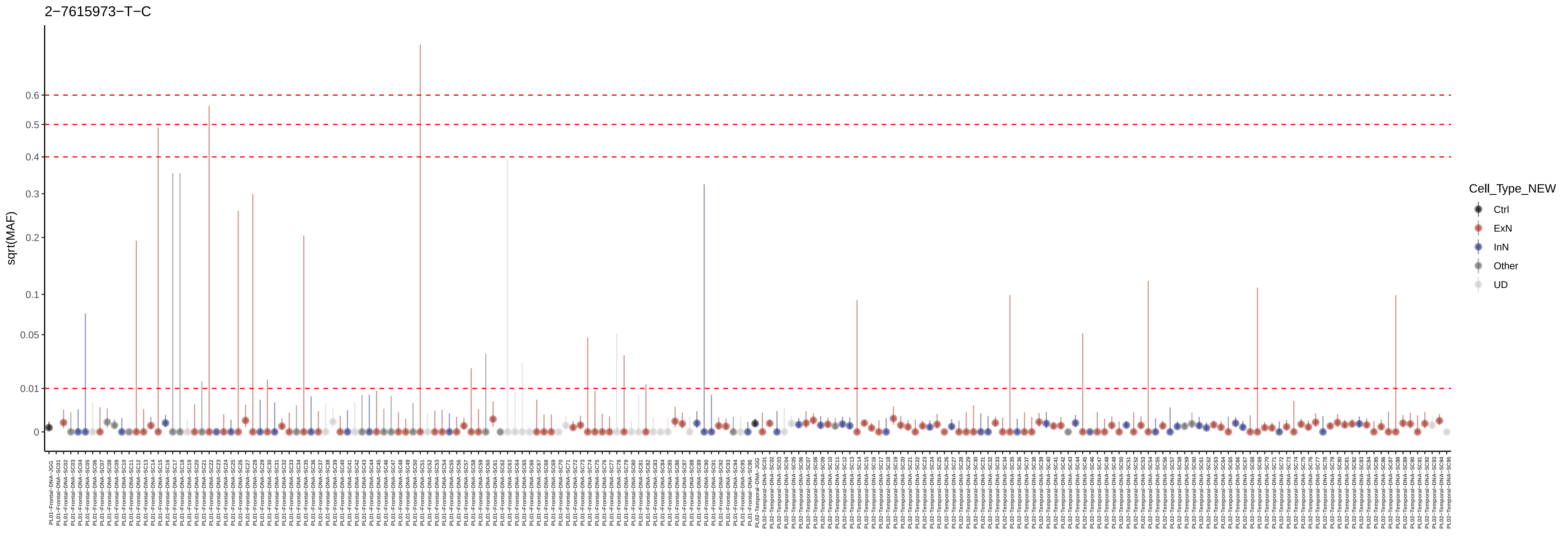
2−7615973−T−C
sqrt(MAF)
0.6
0.5
0.4
0.3
0.2
0.1
0.05
0.01
0
Cell_Type_NEW
Ctrl
ExN
InN
Other
UD

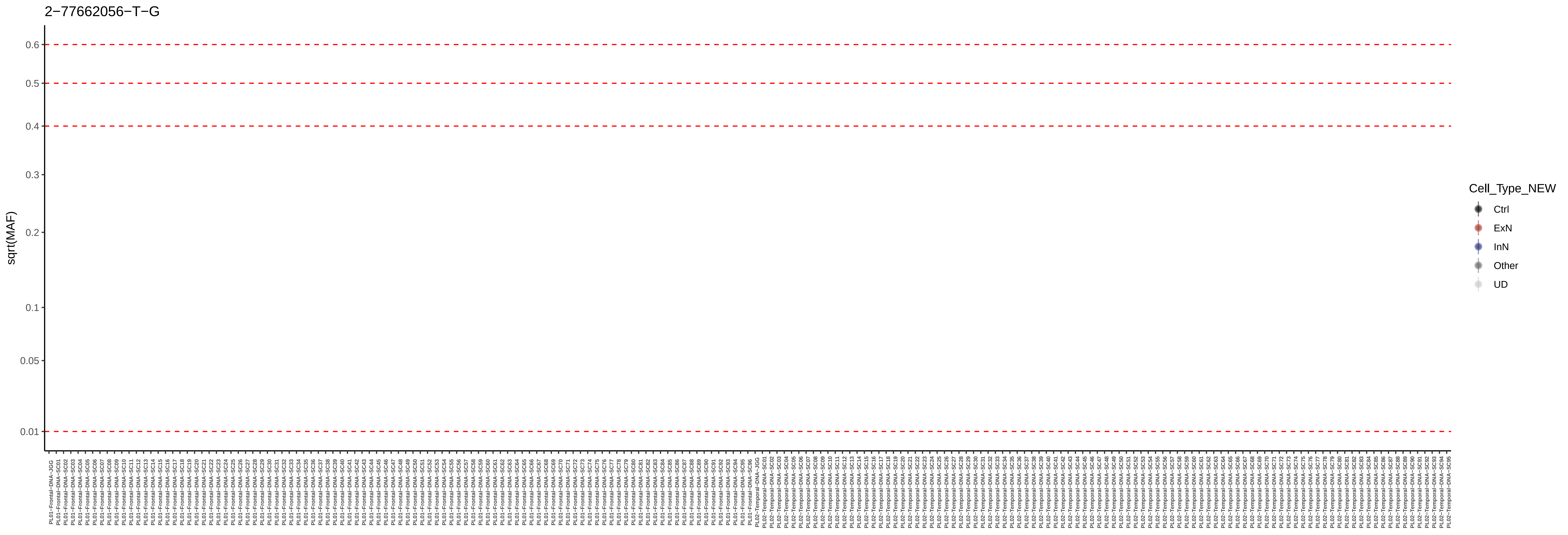

2−77662056−T−G
sqrt(MAF)
0.6
0.5
0.4
0.3
0.2
0.1
0.05
0.01
Cell_Type_NEW
Ctrl
ExN
InN
Other
UD

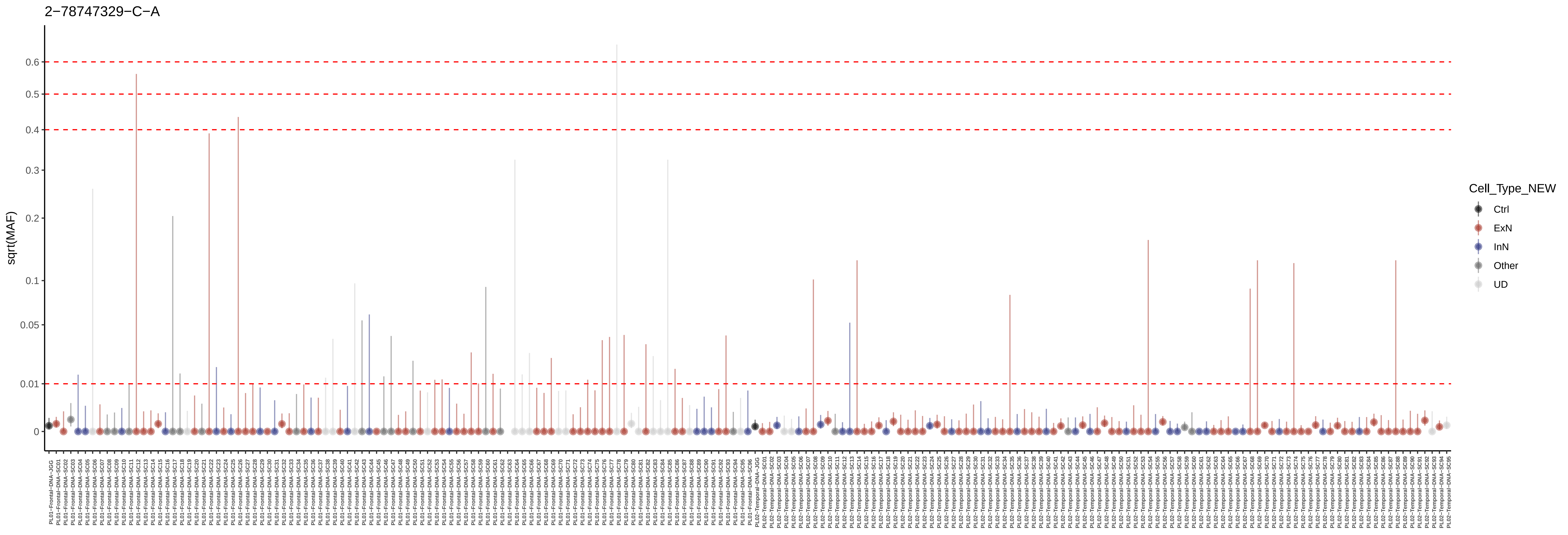
2−78747329−C−A
sqrt(MAF)
0.6
0.5
0.4
0.3
0.2
0.1
0.05
0.01
0
Cell_Type_NEW
Ctrl
ExN
InN
Other
UD

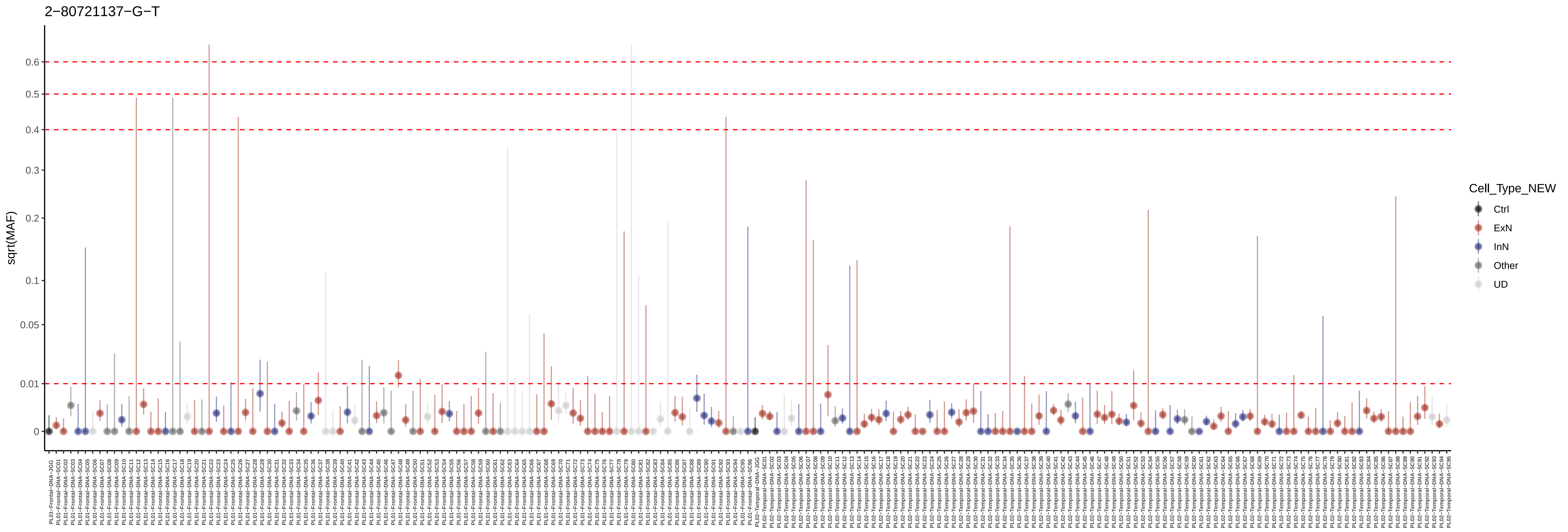
2−80721137−G−T
sqrt(MAF)
0.6
0.5
0.4
0.3
0.2
0.1
0.05
0.01
0
Cell_Type_NEW
Ctrl
ExN
InN
Other
UD

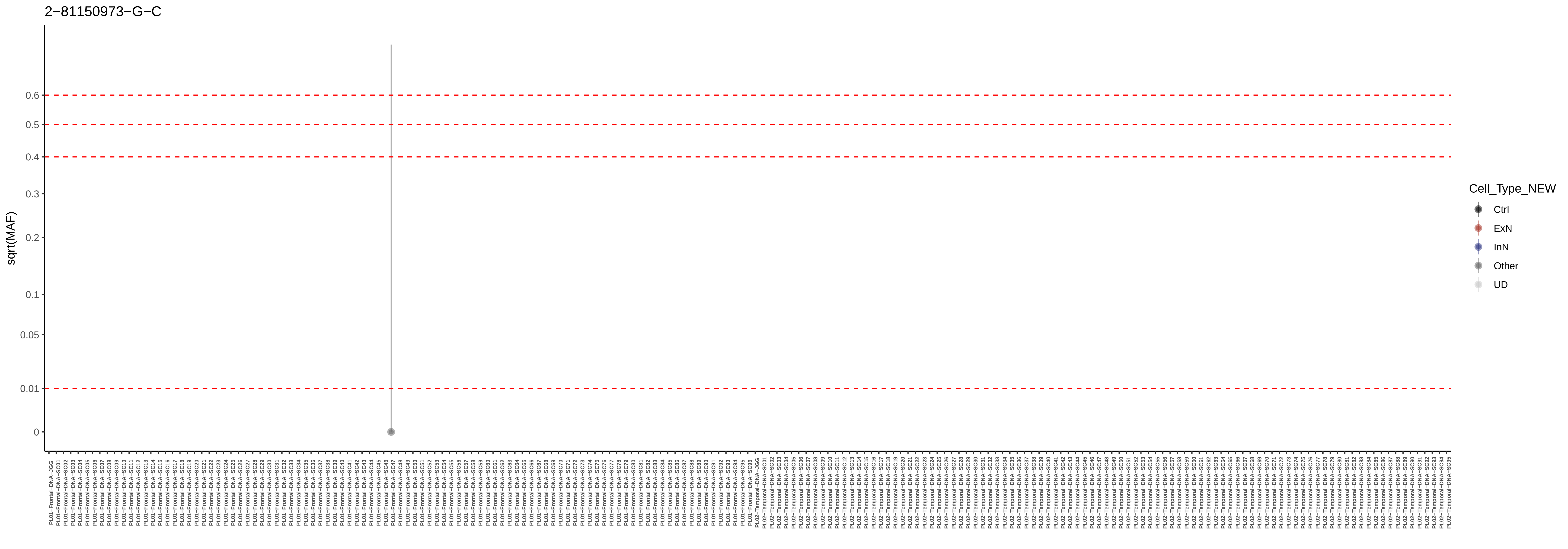
2−81150973−G−C
sqrt(MAF)
0.6
0.5
0.4
0.3
0.2
0.1
0.05
0.01
0
Cell_Type_NEW
Ctrl
ExN
InN
Other
UD

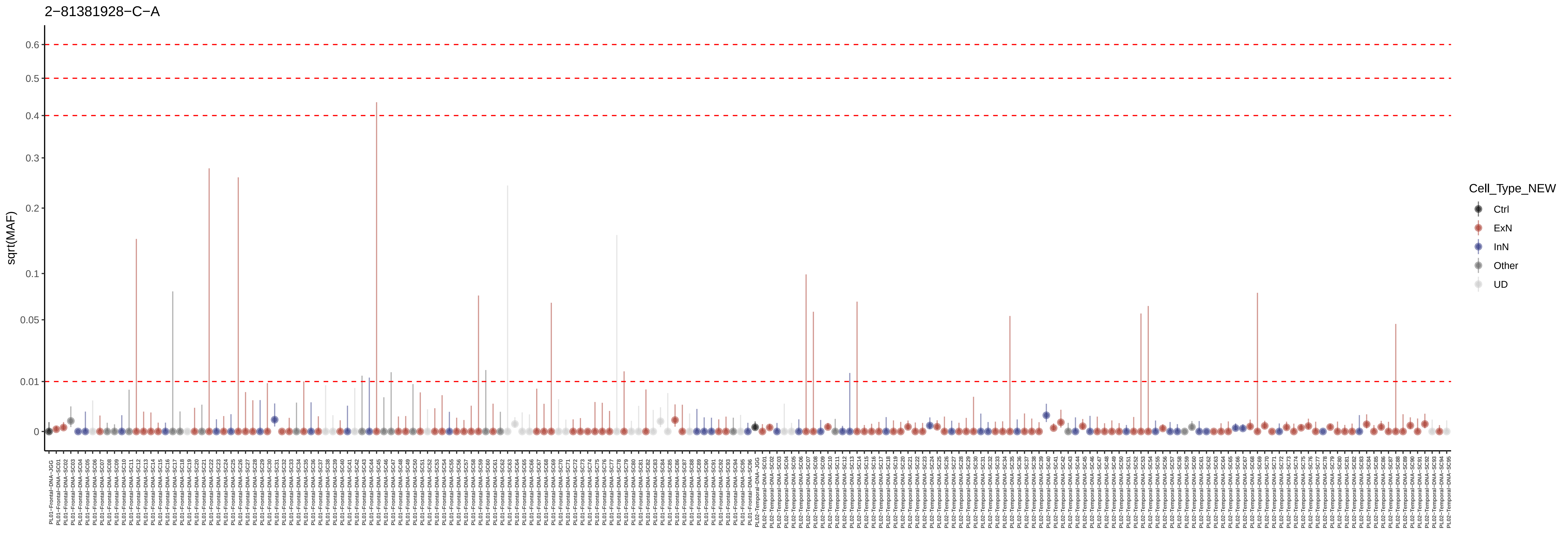

2−81381928−C−A
sqrt(MAF)
0.6
0.5
0.4
0.3
0.2
0.1
0.05
0.01
0
Cell_Type_NEW
Ctrl
ExN
InN
Other
UD

2−8175945−C−A

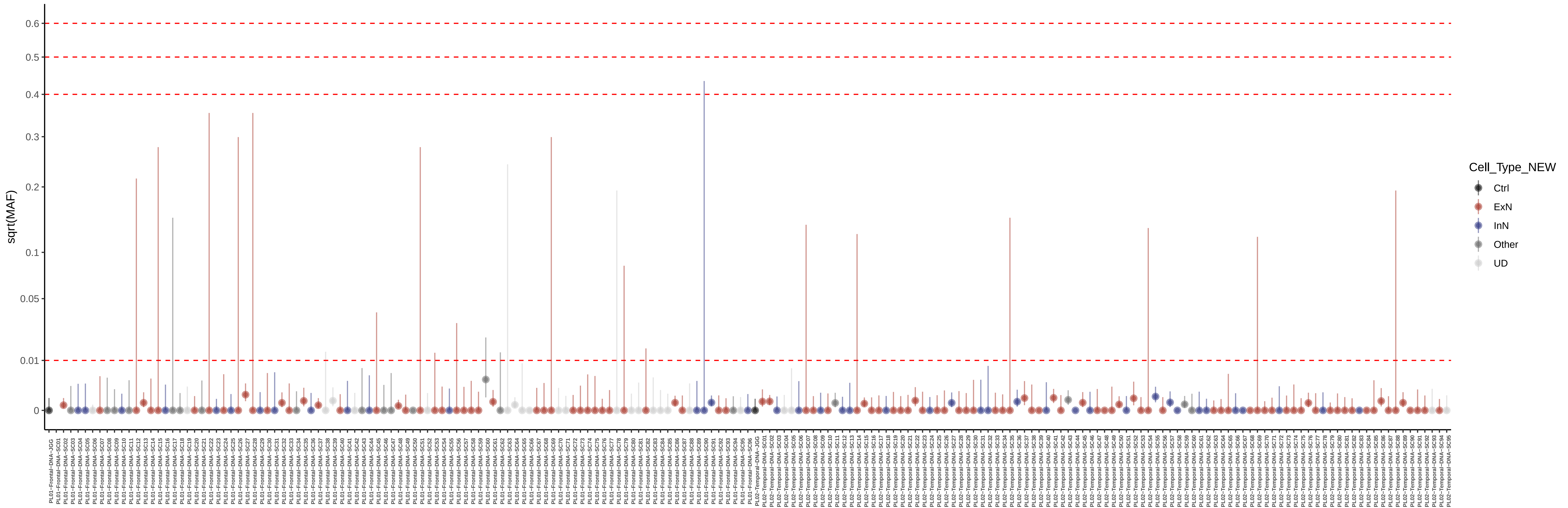
sqrt(MAF)
0.6
0.5
0.4
0.3
0.2
0.1
0.05
0.01
0
Cell_Type_NEW
Ctrl
ExN
InN
Other
UD

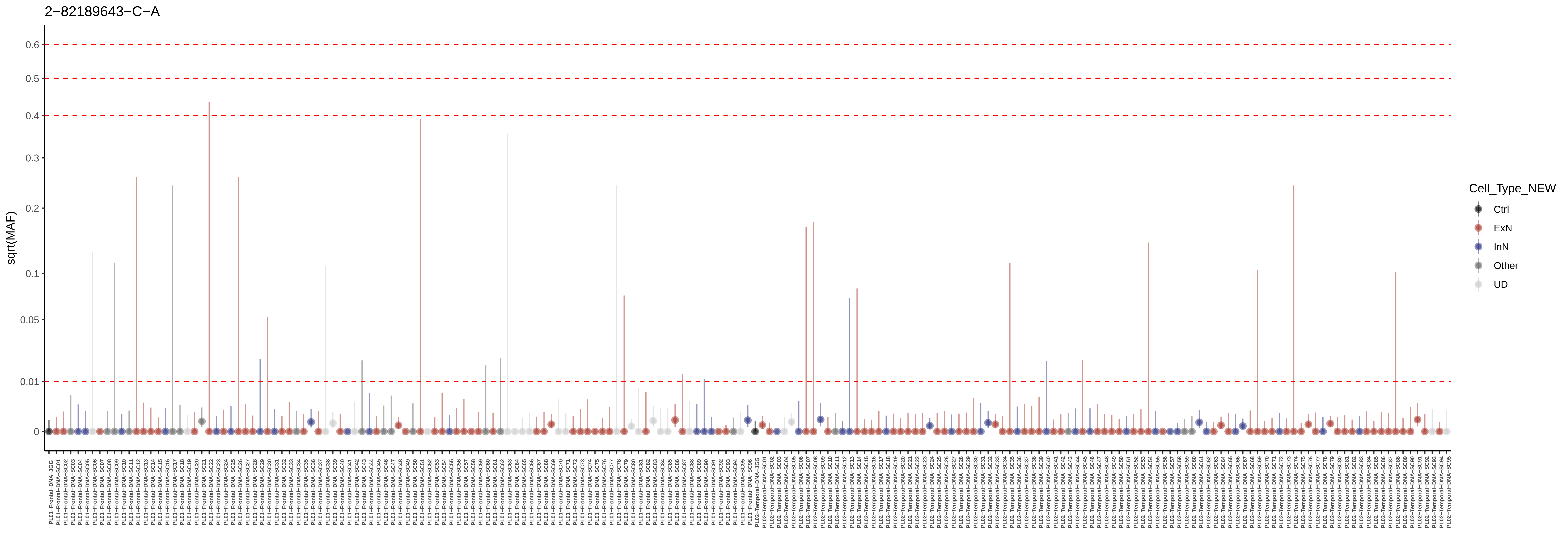
2−82189643−C−A
sqrt(MAF)
0.6
0.5
0.4
0.3
0.2
0.1
0.05
0.01
0
Cell_Type_NEW
Ctrl
ExN
InN
Other
UD

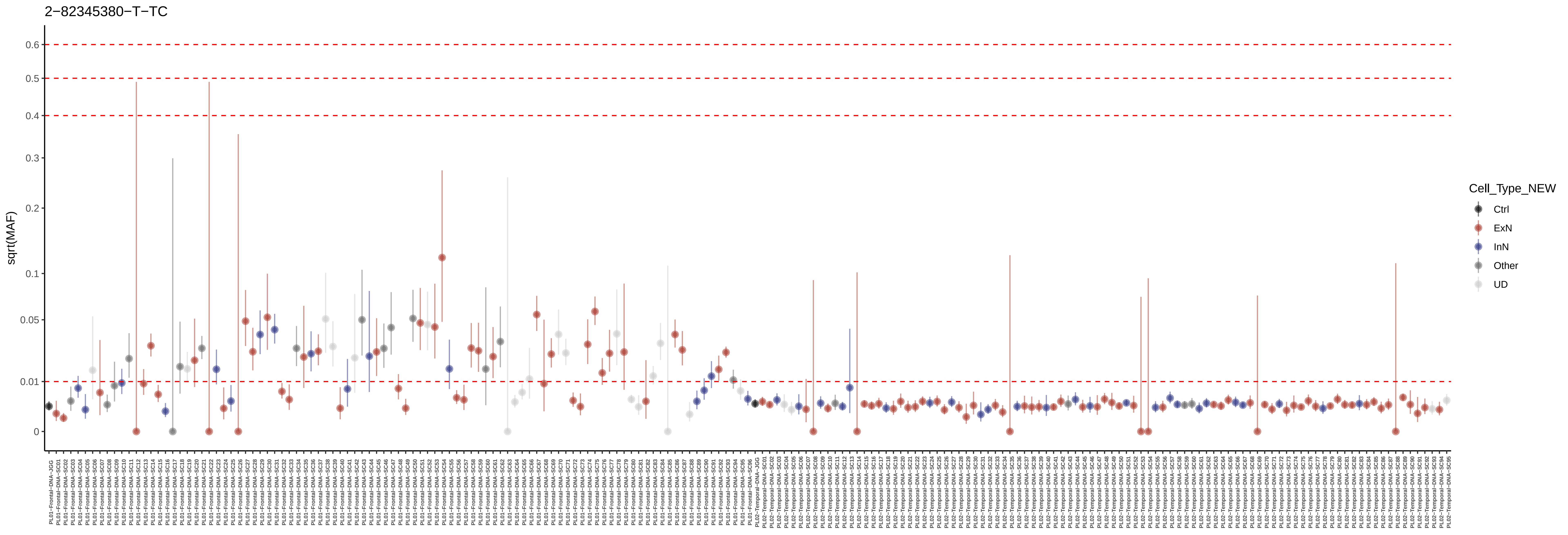
2−82345380−T−TC
sqrt(MAF)
0.6
0.5
0.4
0.3
0.2
0.1
0.05
0.01
0
Cell_Type_NEW
Ctrl
ExN
InN
Other
UD

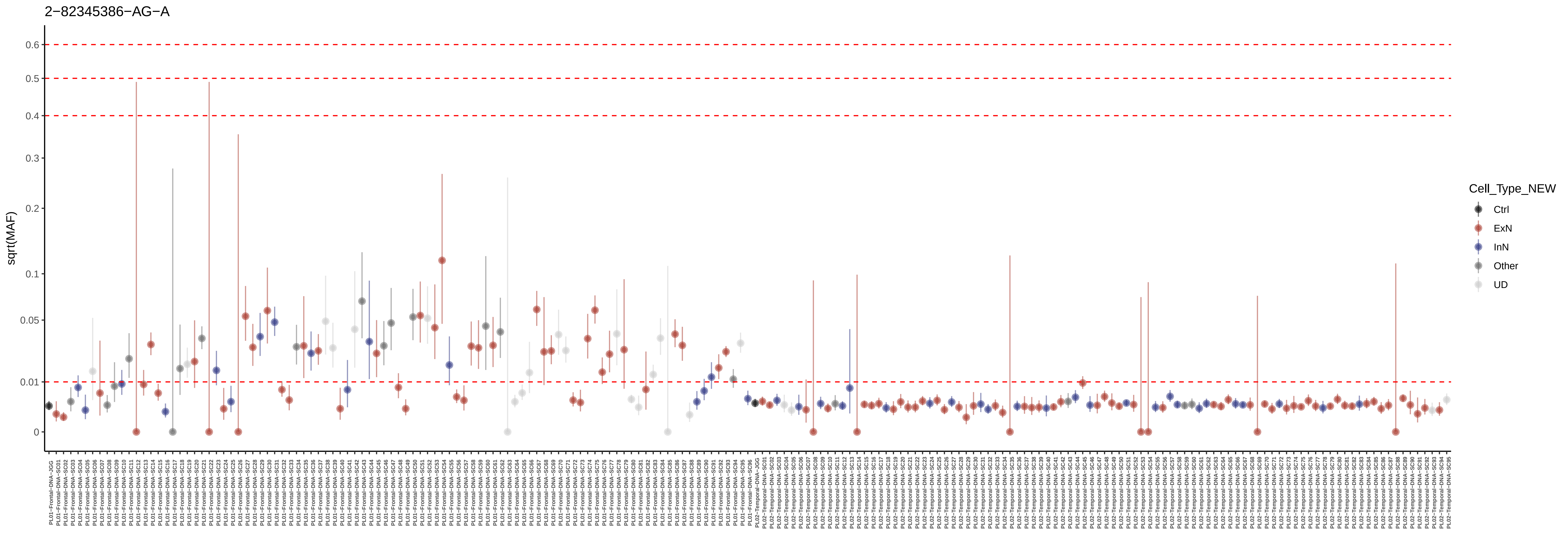

2−82345386−AG−A
sqrt(MAF)
0.6
0.5
0.4
0.3
0.2
0.1
0.05
0.01
0
Cell_Type_NEW
Ctrl
ExN
InN
Other
UD

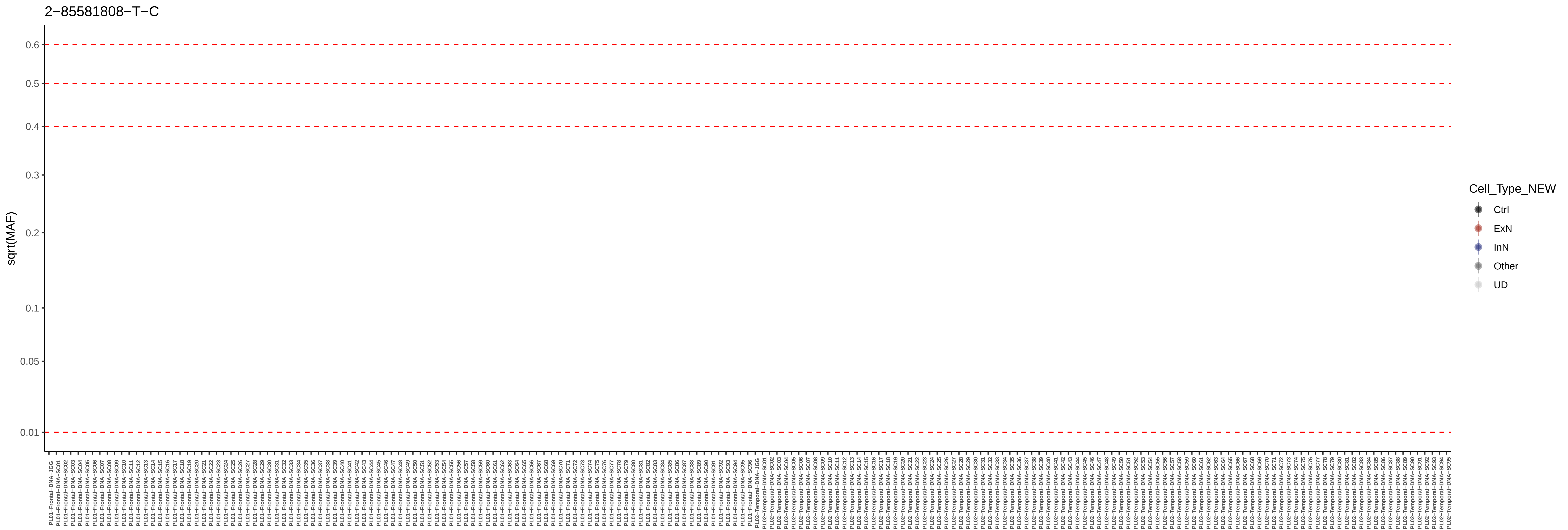

2−85581808−T−C
sqrt(MAF)
0.6
0.5
0.4
0.3
0.2
0.1
0.05
0.01
Cell_Type_NEW
Ctrl
ExN
InN
Other
UD

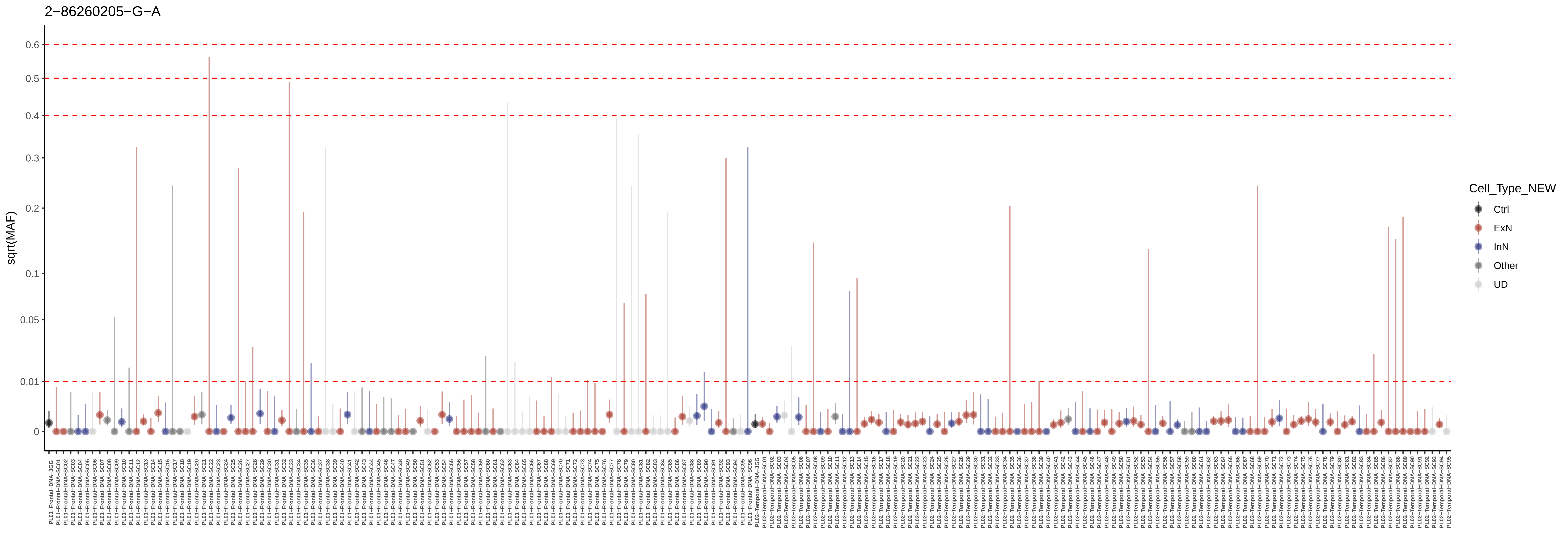

2−86260205−G−A
sqrt(MAF)
0.6
0.5
0.4
0.3
0.2
0.1
0.05
0.01
0
Cell_Type_NEW
Ctrl
ExN
InN
Other
UD

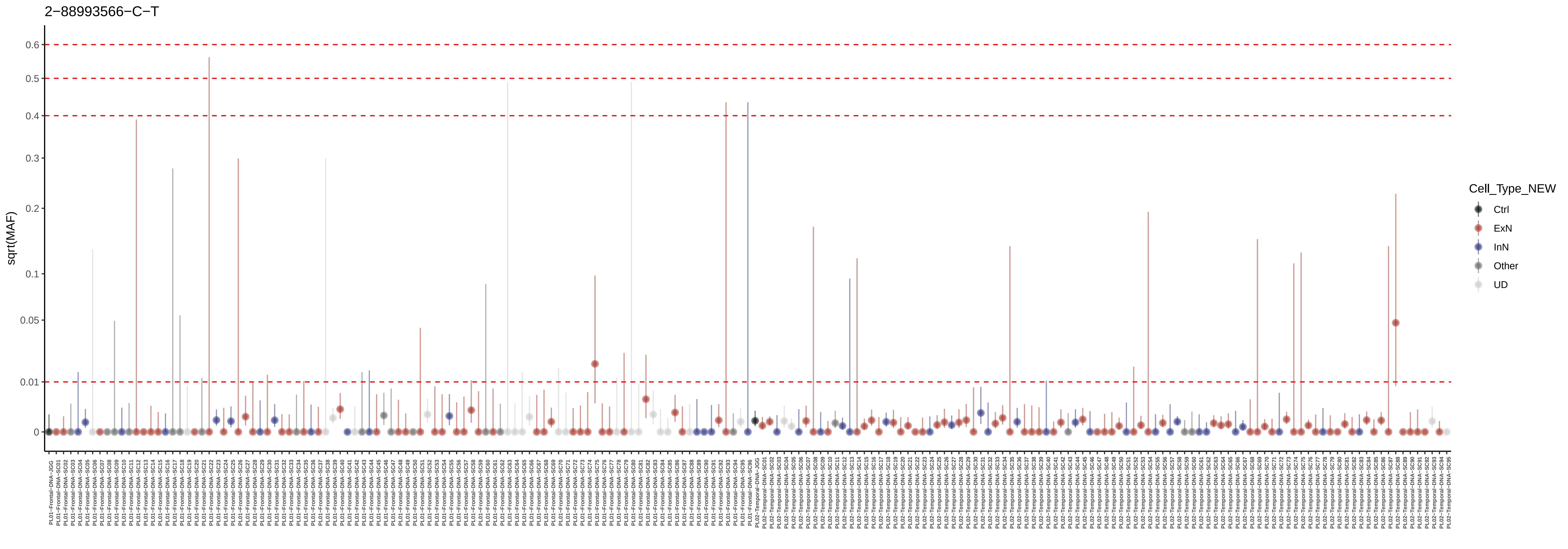
2−88993566−C−T
sqrt(MAF)
0.6
0.5
0.4
0.3
0.2
0.1
0.05
0.01
0
Cell_Type_NEW
Ctrl
ExN
InN
Other
UD

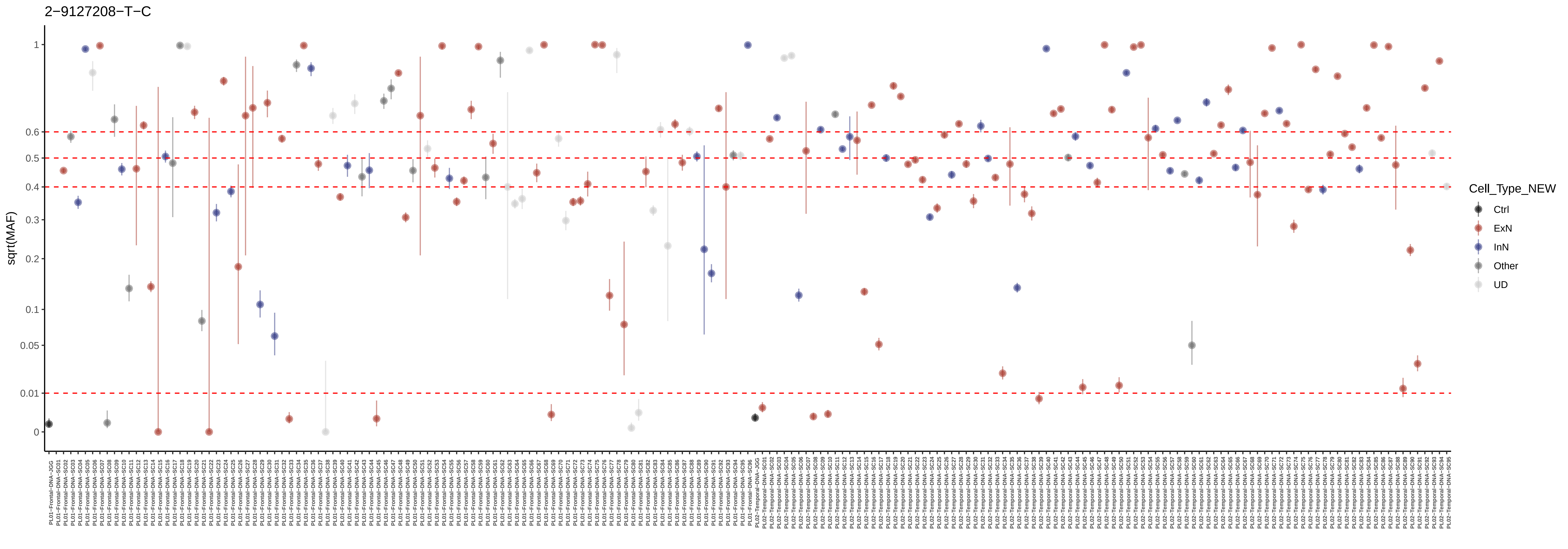

2−9127208−T−C
sqrt(MAF)
1
0.6
0.5
0.4
0.3
0.2
0.1
0.05
0.01
0
Cell_Type_NEW
Ctrl
ExN
InN
Other
UD

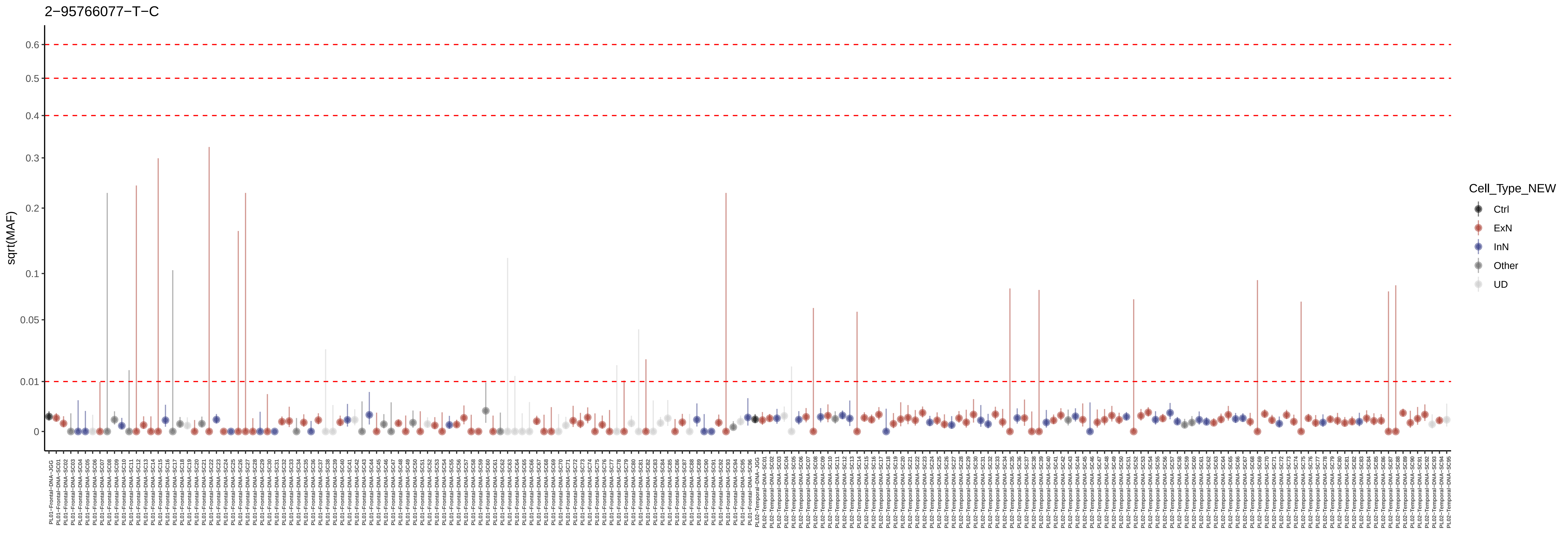

2−95766077−T−C
sqrt(MAF)
0.6
0.5
0.4
0.3
0.2
0.1
0.05
0.01
0
Cell_Type_NEW
Ctrl
ExN
InN
Other
UD

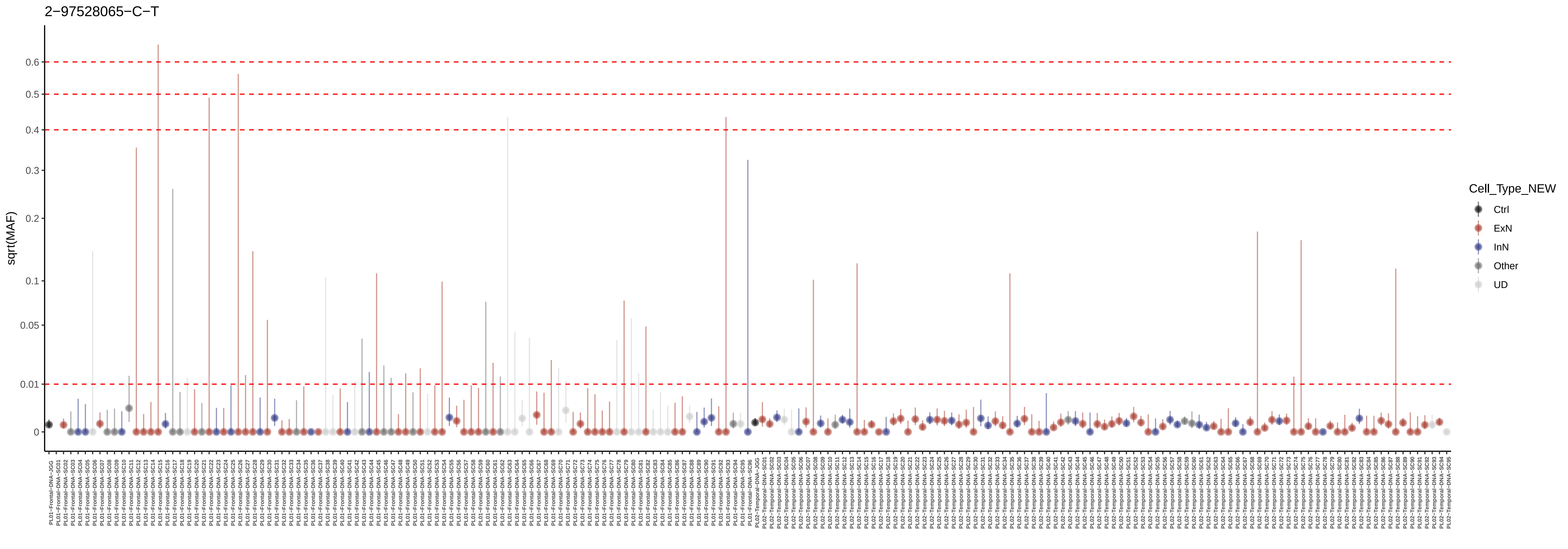

2−97528065−C−T
sqrt(MAF)
0.6
0.5
0.4
0.3
0.2
0.1
0.05
0.01
0
Cell_Type_NEW
Ctrl
ExN
InN
Other
UD

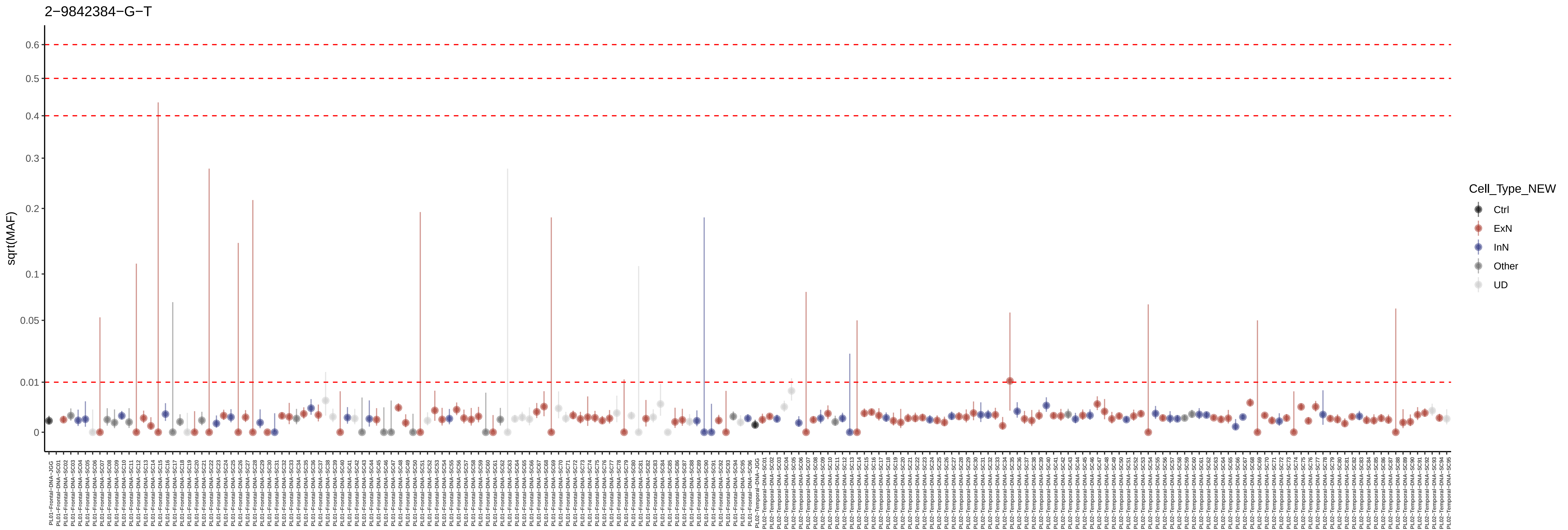
2−9842384−G−T
sqrt(MAF)
0.6
0.5
0.4
0.3
0.2
0.1
0.05
0.01
0
Cell_Type_NEW
Ctrl
ExN
InN
Other
UD

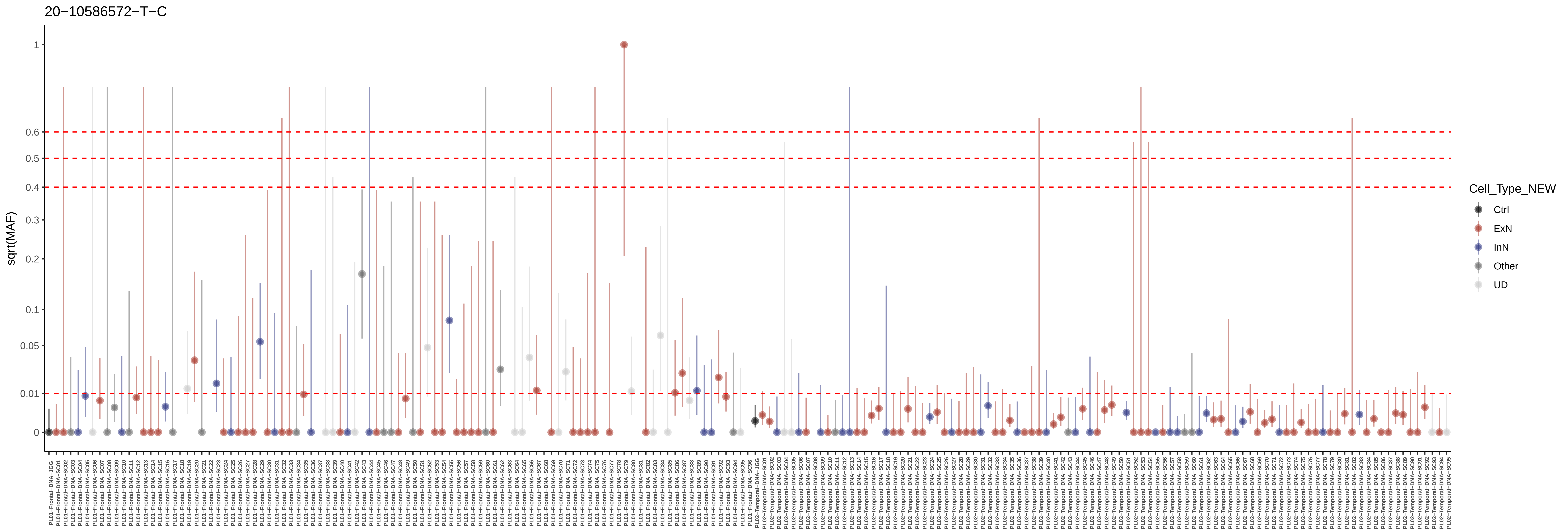
20−10586572−T−C
sqrt(MAF)
1
0.6
0.5
0.4
0.3
0.2
0.1
0.05
0.01
0
Cell_Type_NEW
Ctrl
ExN
InN
Other
UD

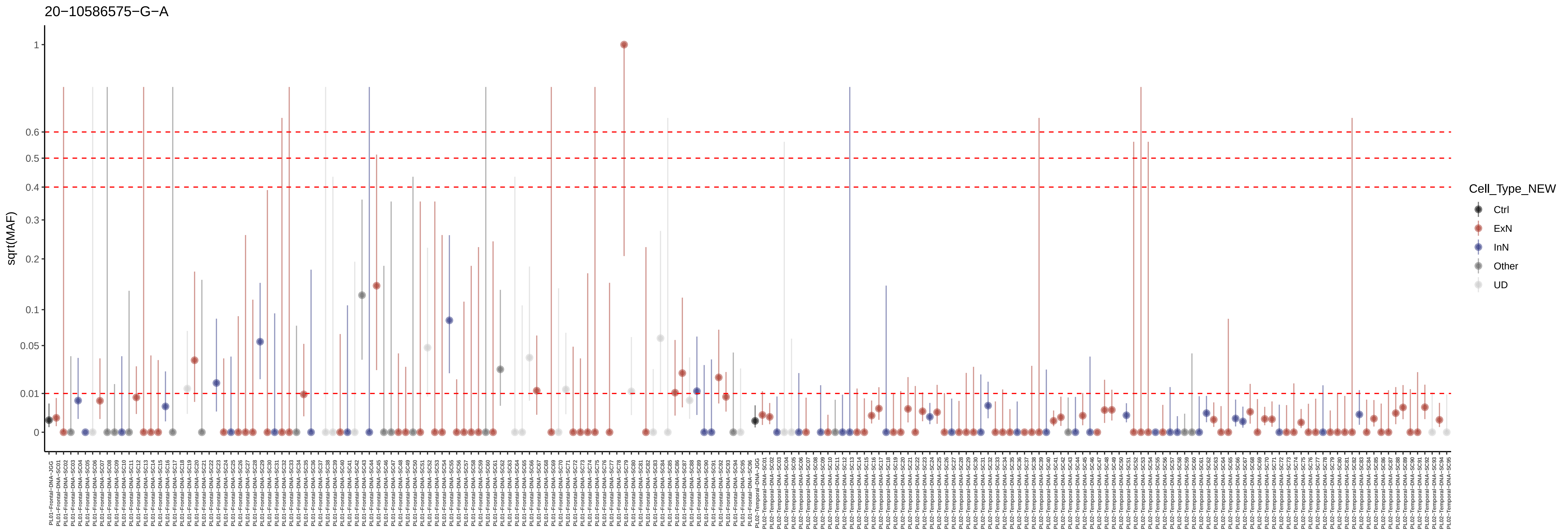
20−10586575−G−A
sqrt(MAF)
1
0.6
0.5
0.4
0.3
0.2
0.1
0.05
0.01
0
Cell_Type_NEW
Ctrl
ExN
InN
Other
UD

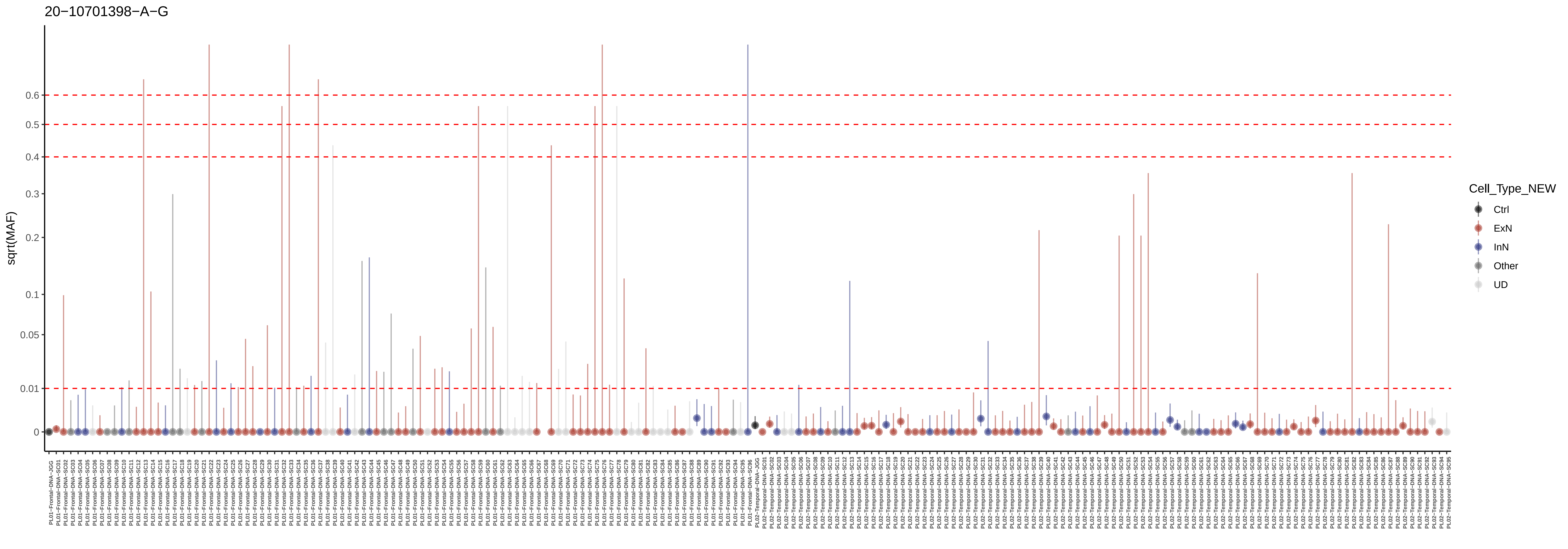
20−10701398−A−G
sqrt(MAF)
0.6
0.5
0.4
0.3
0.2
0.1
0.05
0.01
0
Cell_Type_NEW
Ctrl
ExN
InN
Other
UD

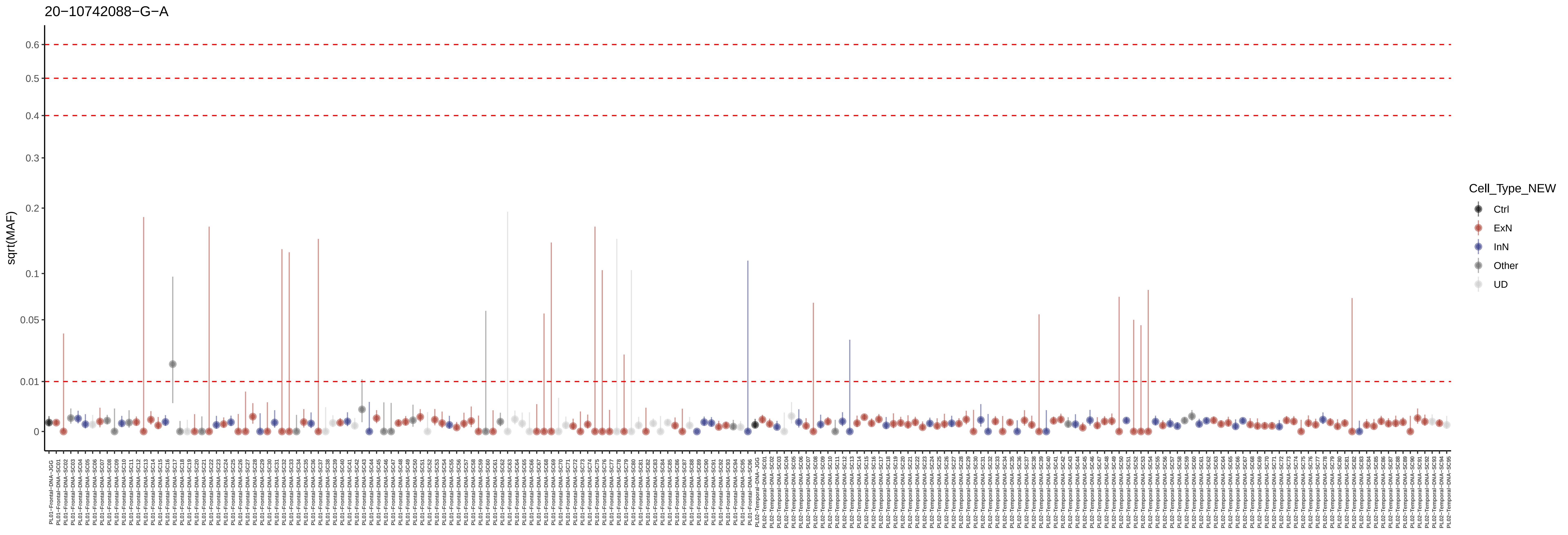
20−10742088−G−A
sqrt(MAF)
0.6
0.5
0.4
0.3
0.2
0.1
0.05
0.01
0
Cell_Type_NEW
Ctrl
ExN
InN
Other
UD

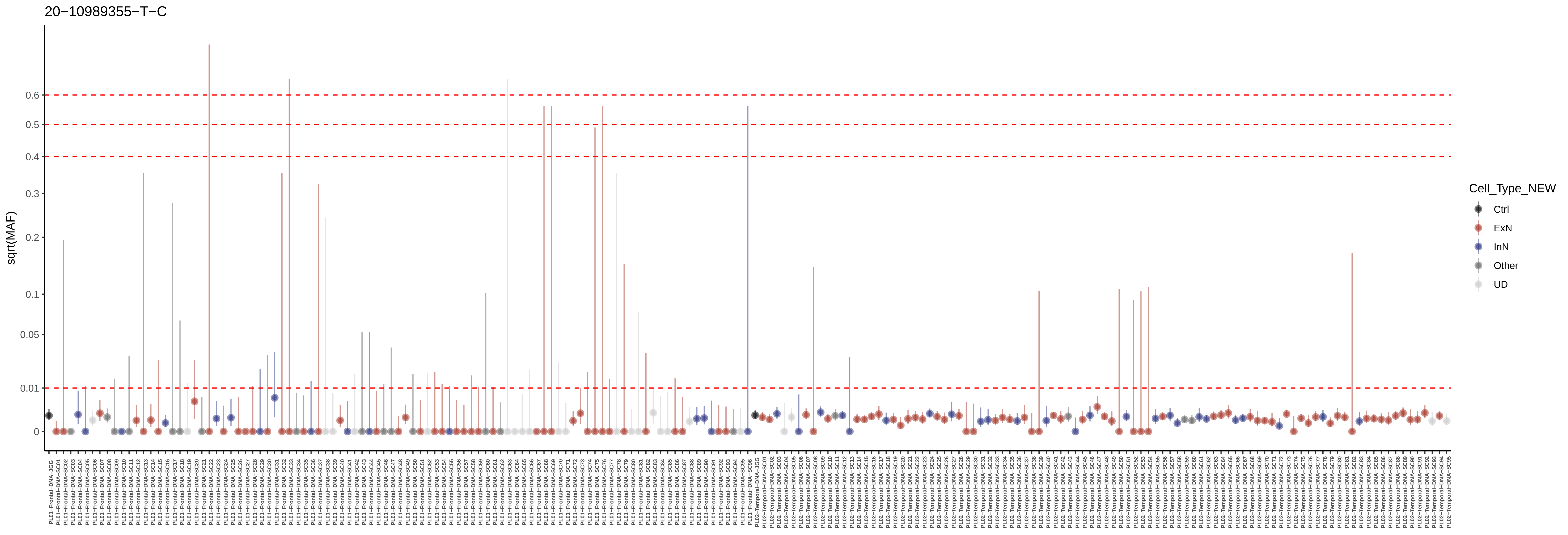

20−10989355−T−C
sqrt(MAF)
0.6
0.5
0.4
0.3
0.2
0.1
0.05
0.01
0
Cell_Type_NEW
Ctrl
ExN
InN
Other
UD

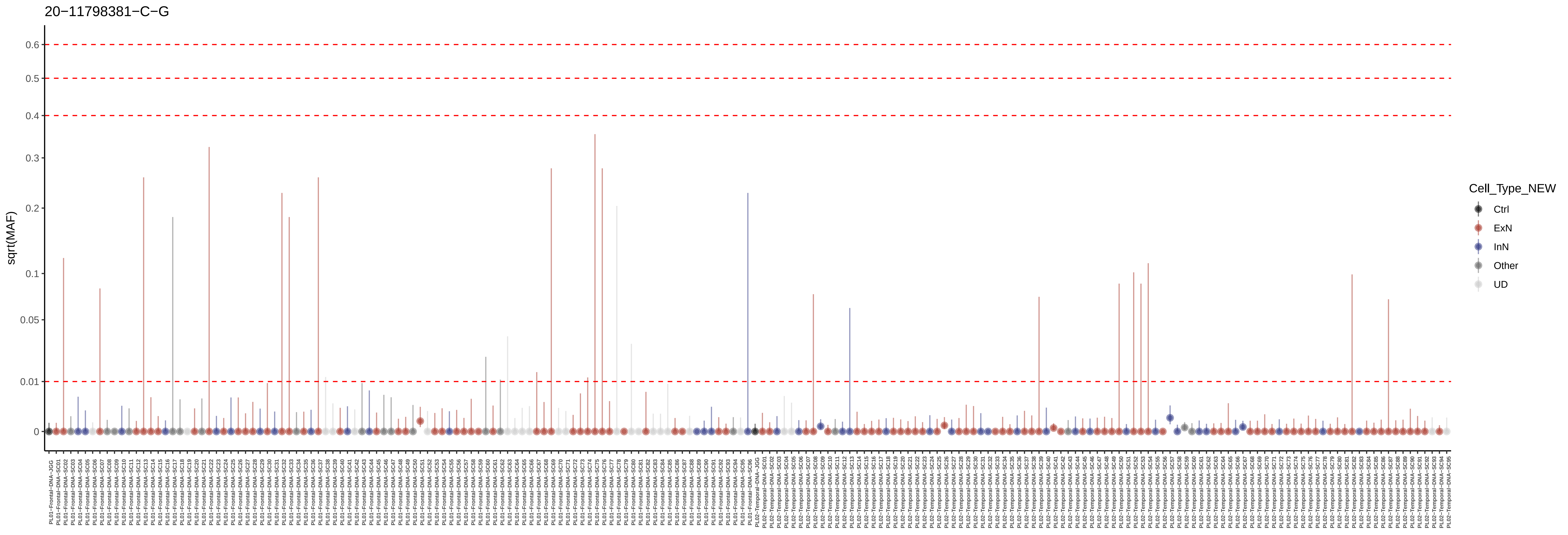

20−11798381−C−G
sqrt(MAF)
0.6
0.5
0.4
0.3
0.2
0.1
0.05
0.01
0
Cell_Type_NEW
Ctrl
ExN
InN
Other
UD

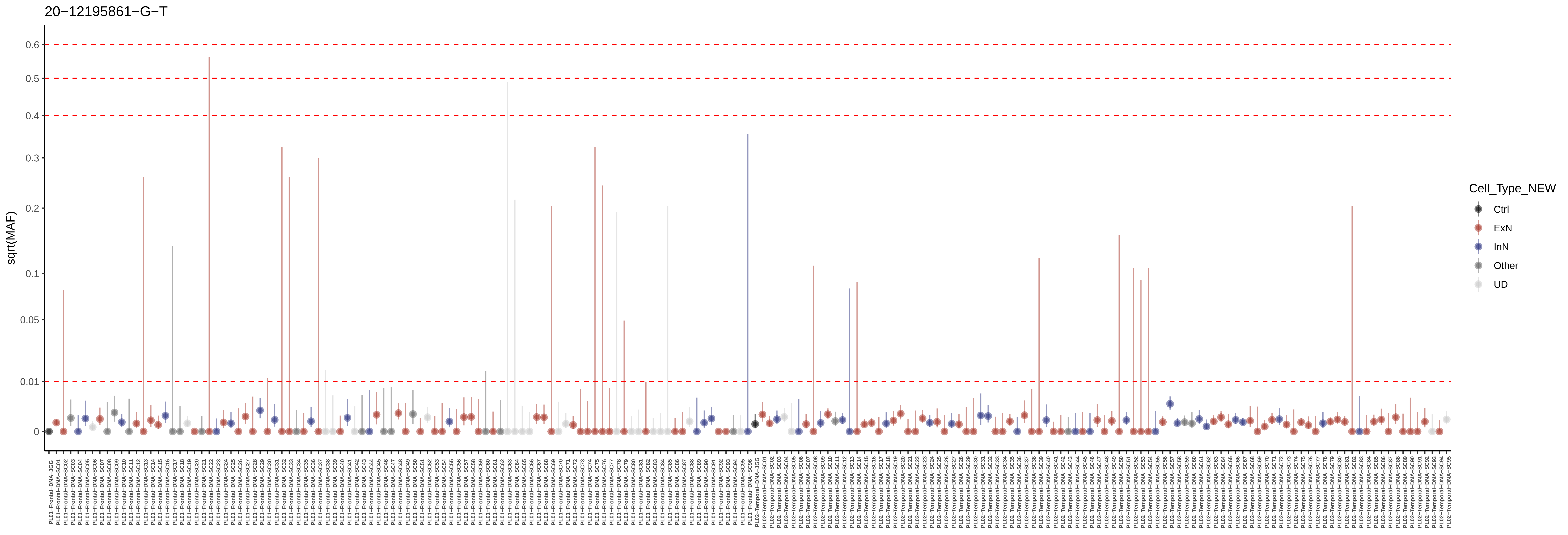
20−12195861−G−T
sqrt(MAF)
0.6
0.5
0.4
0.3
0.2
0.1
0.05
0.01
0
Cell_Type_NEW
Ctrl
ExN
InN
Other
UD

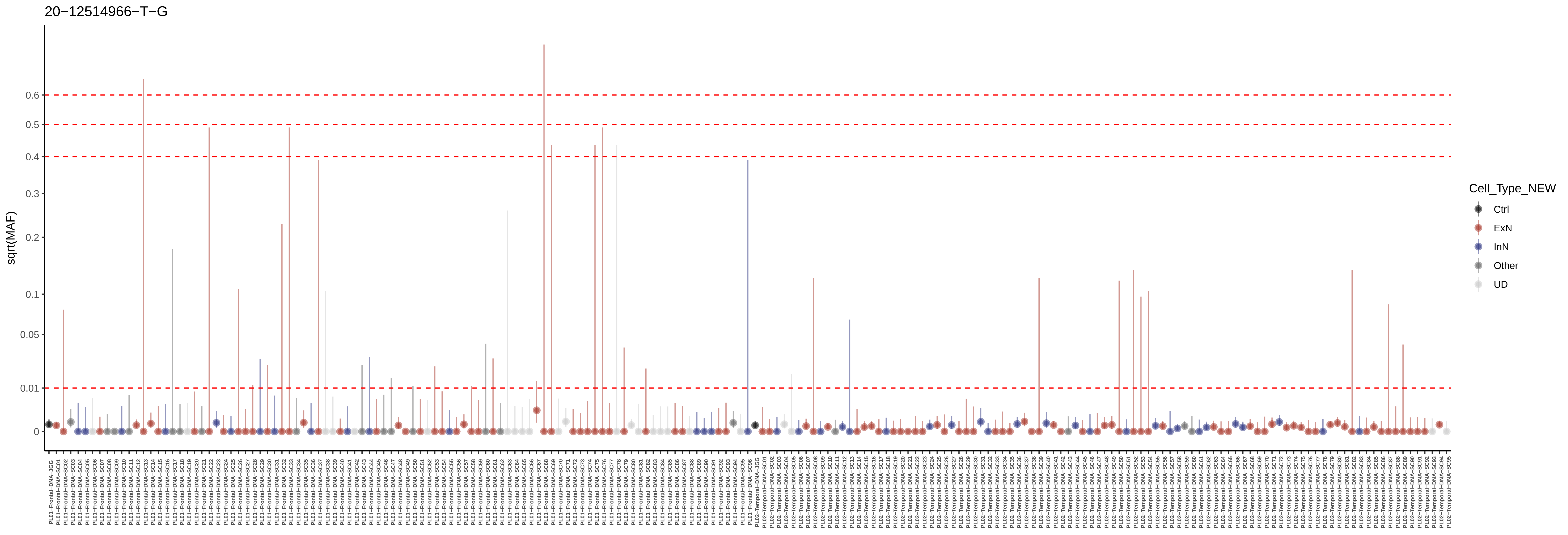

20−12514966−T−G
sqrt(MAF)
0.6
0.5
0.4
0.3
0.2
0.1
0.05
0.01
0
Cell_Type_NEW
Ctrl
ExN
InN
Other
UD

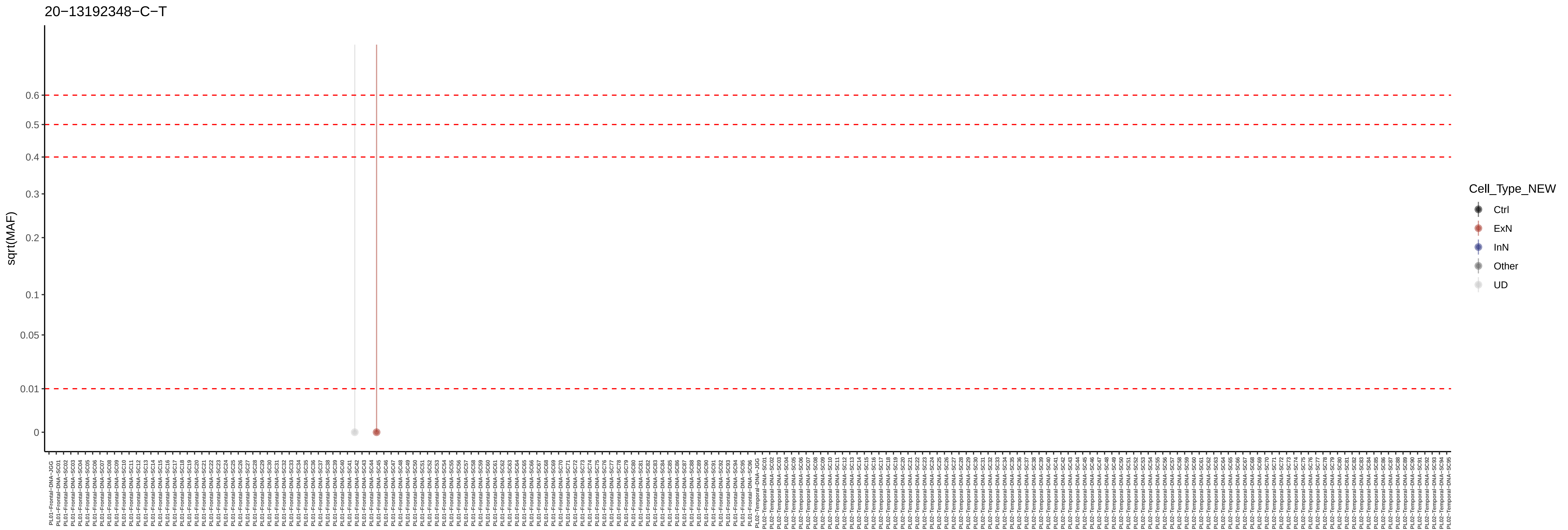
20−13192348−C−T
sqrt(MAF)
0.6
0.5
0.4
0.3
0.2
0.1
0.05
0.01
0
Cell_Type_NEW
Ctrl
ExN
InN
Other
UD

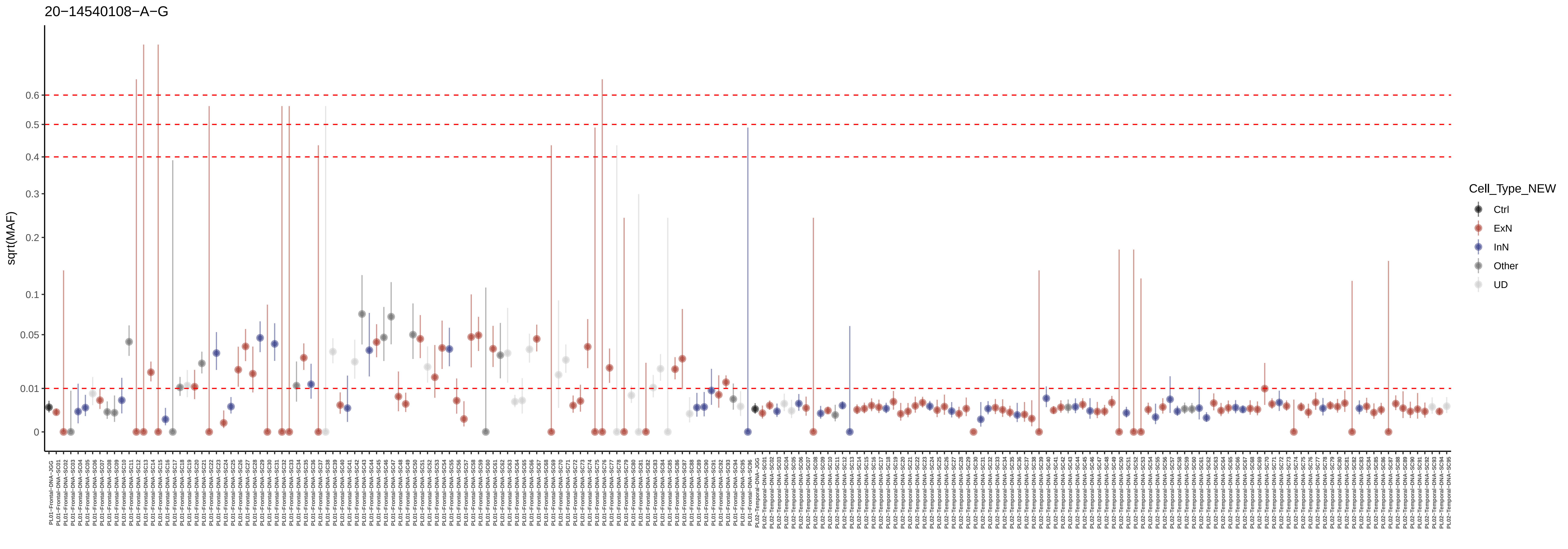
20−14540108−A−G
sqrt(MAF)
0.6
0.5
0.4
0.3
0.2
0.1
0.05
0.01
0
Cell_Type_NEW
Ctrl
ExN
InN
Other
UD

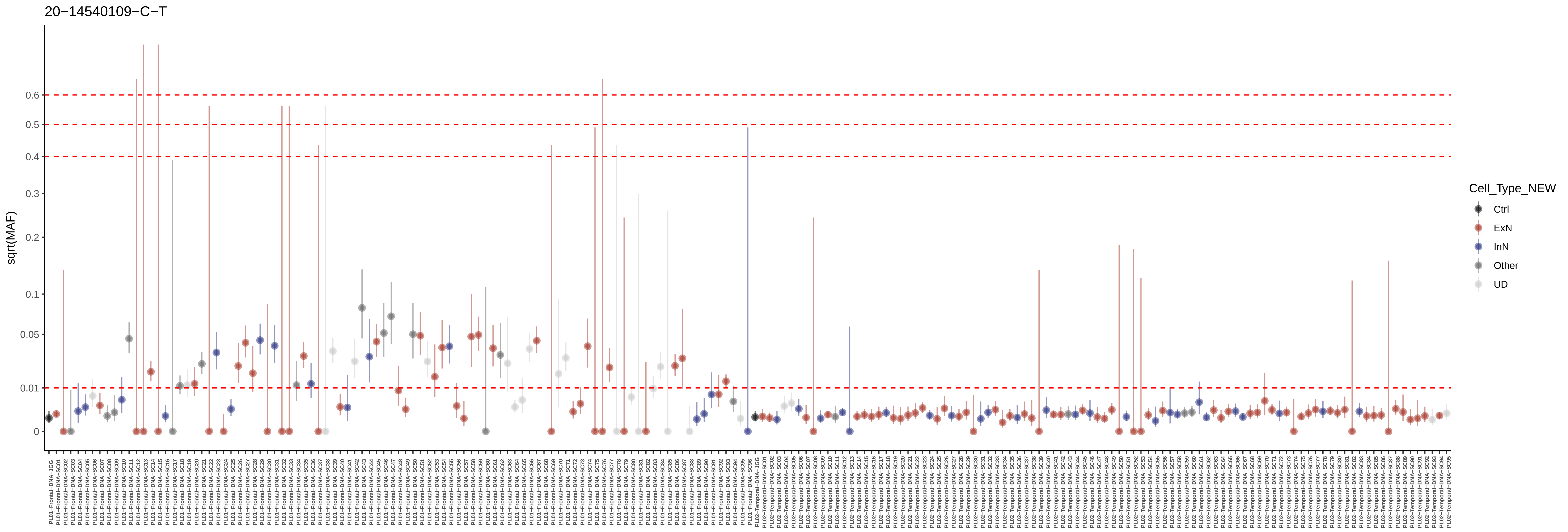
20−14540109−C−T
sqrt(MAF)
0.6
0.5
0.4
0.3
0.2
0.1
0.05
0.01
0
Cell_Type_NEW
Ctrl
ExN
InN
Other
UD

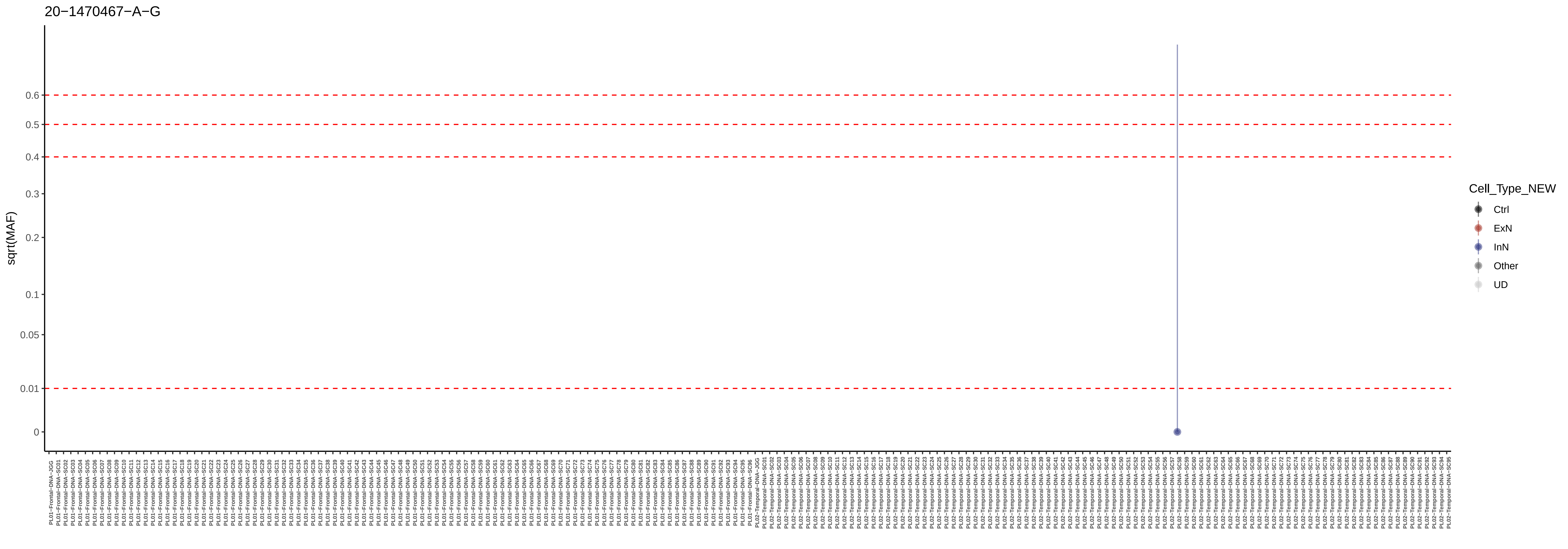
20−1470467−A−G
sqrt(MAF)
0.6
0.5
0.4
0.3
0.2
0.1
0.05
0.01
0
Cell_Type_NEW
Ctrl
ExN
InN
Other
UD

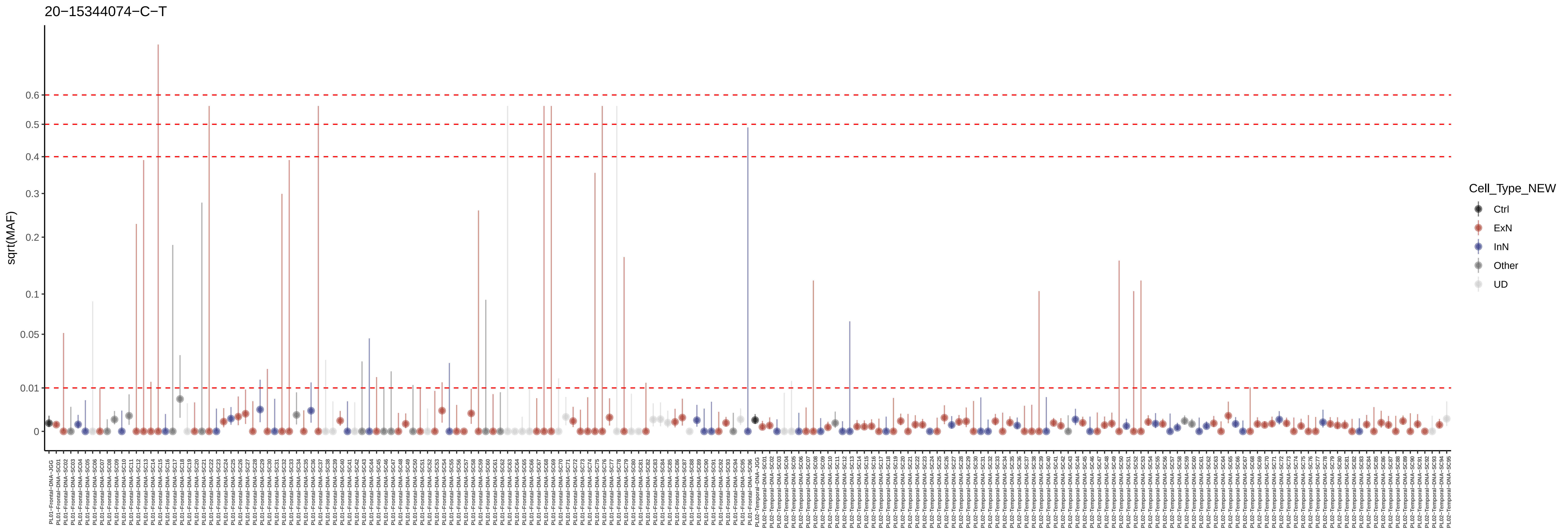

20−15344074−C−T
sqrt(MAF)
0.6
0.5
0.4
0.3
0.2
0.1
0.05
0.01
0
Cell_Type_NEW
Ctrl
ExN
InN
Other
UD

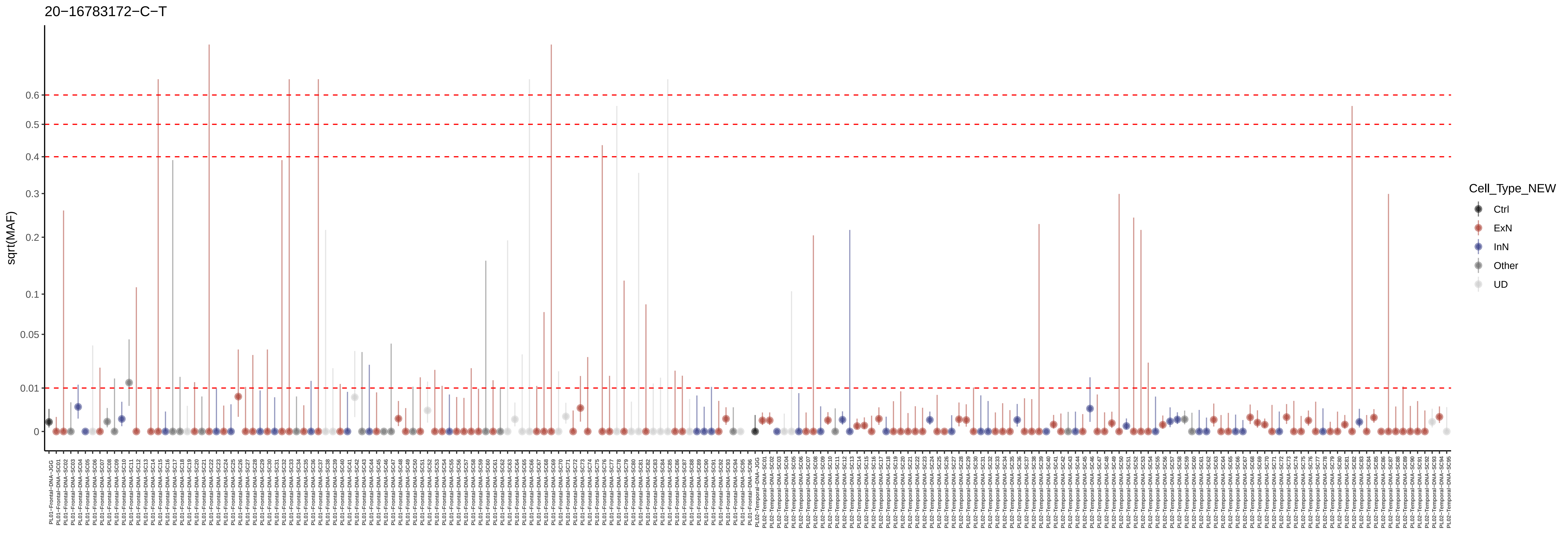
20−16783172−C−T
sqrt(MAF)
0.6
0.5
0.4
0.3
0.2
0.1
0.05
0.01
0
Cell_Type_NEW
Ctrl
ExN
InN
Other
UD

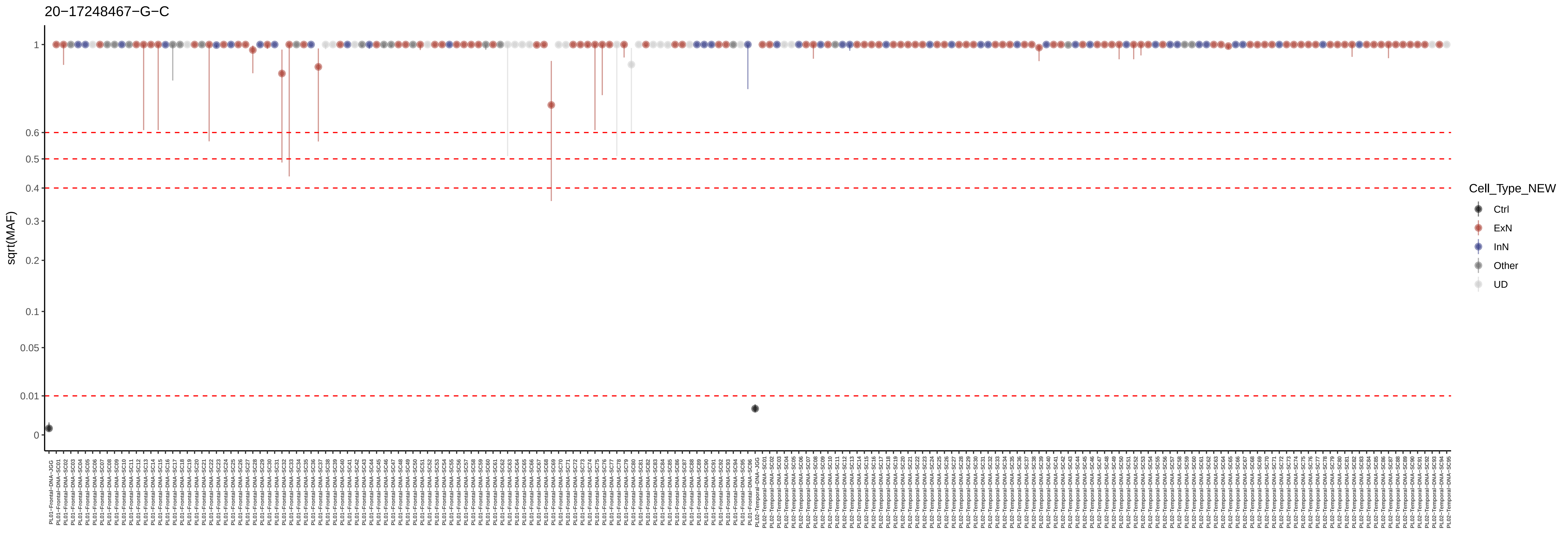
20−17248467−G−C
sqrt(MAF)
1
0.6
0.5
0.4
0.3
0.2
0.1
0.05
0.01
0
Cell_Type_NEW
Ctrl
ExN
InN
Other
UD

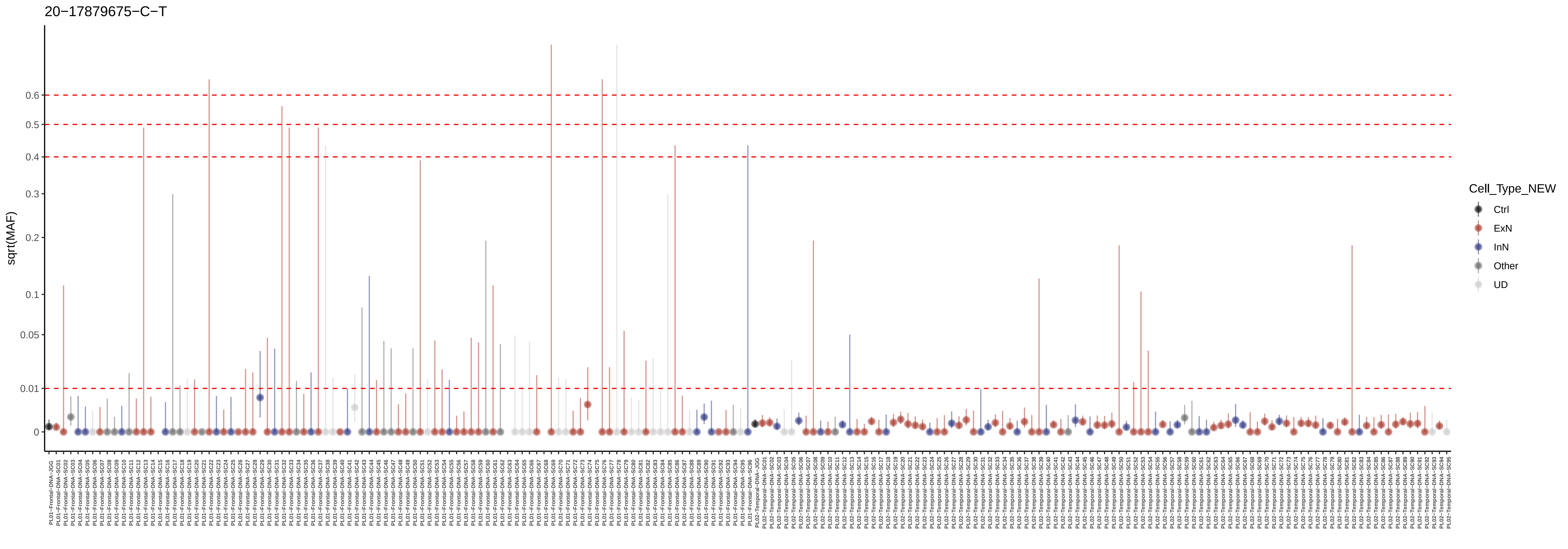

20−17879675−C−T
sqrt(MAF)
0
0.01
0.05
0.1
0.2
0.3
0.4
0.5
0.6
Cell_Type_NEW
Ctrl
ExN
InN
Other
UD

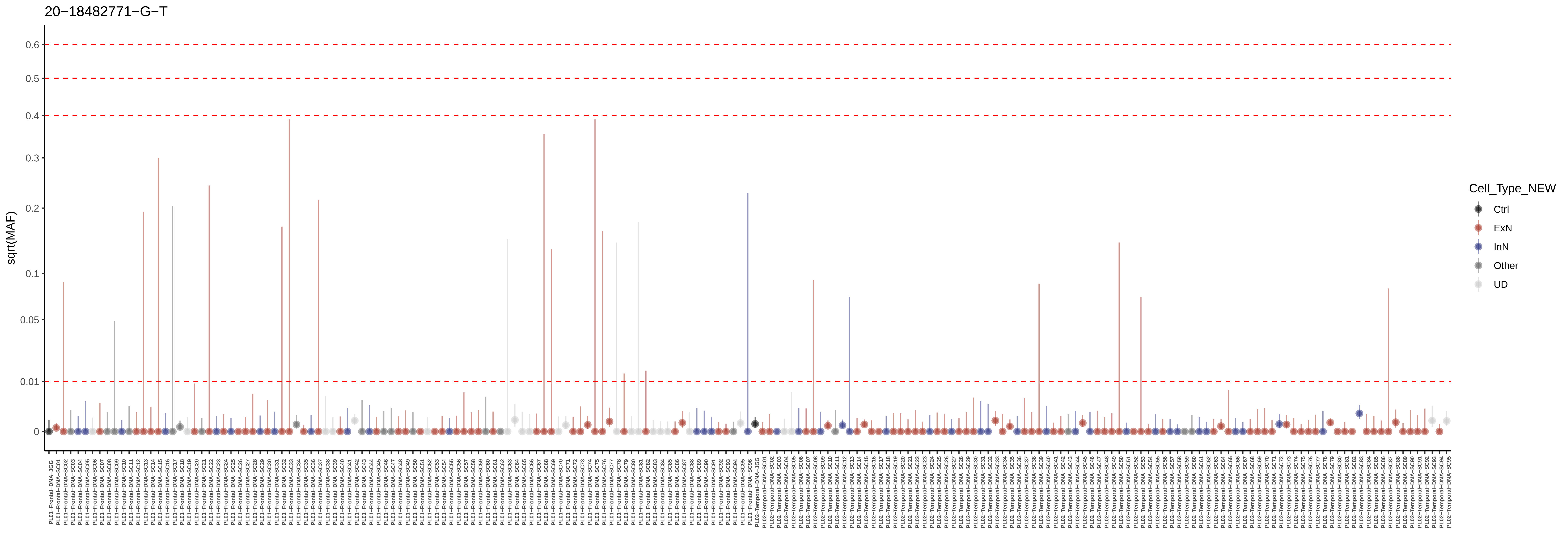
20−18482771−G−T
sqrt(MAF)
0.6
0.5
0.4
0.3
0.2
0.1
0.05
0.01
0
Cell_Type_NEW
Ctrl
ExN
InN
Other
UD

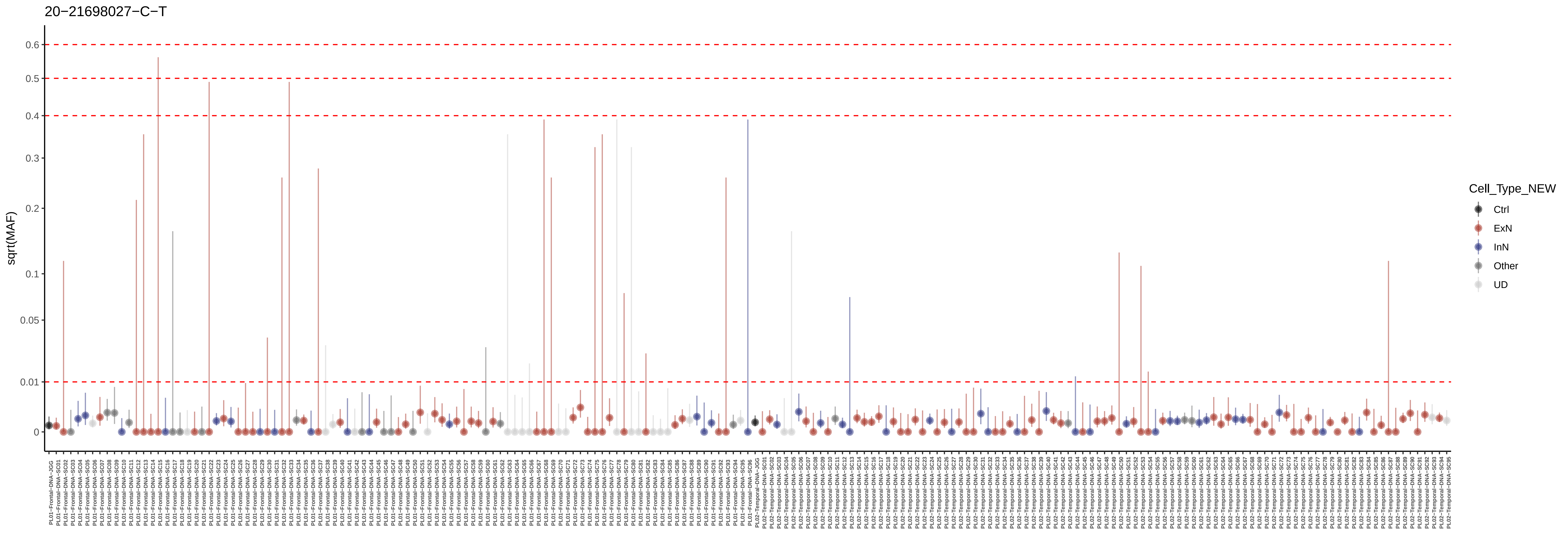
20−21698027−C−T
sqrt(MAF)
0.6
0.5
0.4
0.3
0.2
0.1
0.05
0.01
0
Cell_Type_NEW
Ctrl
ExN
InN
Other
UD

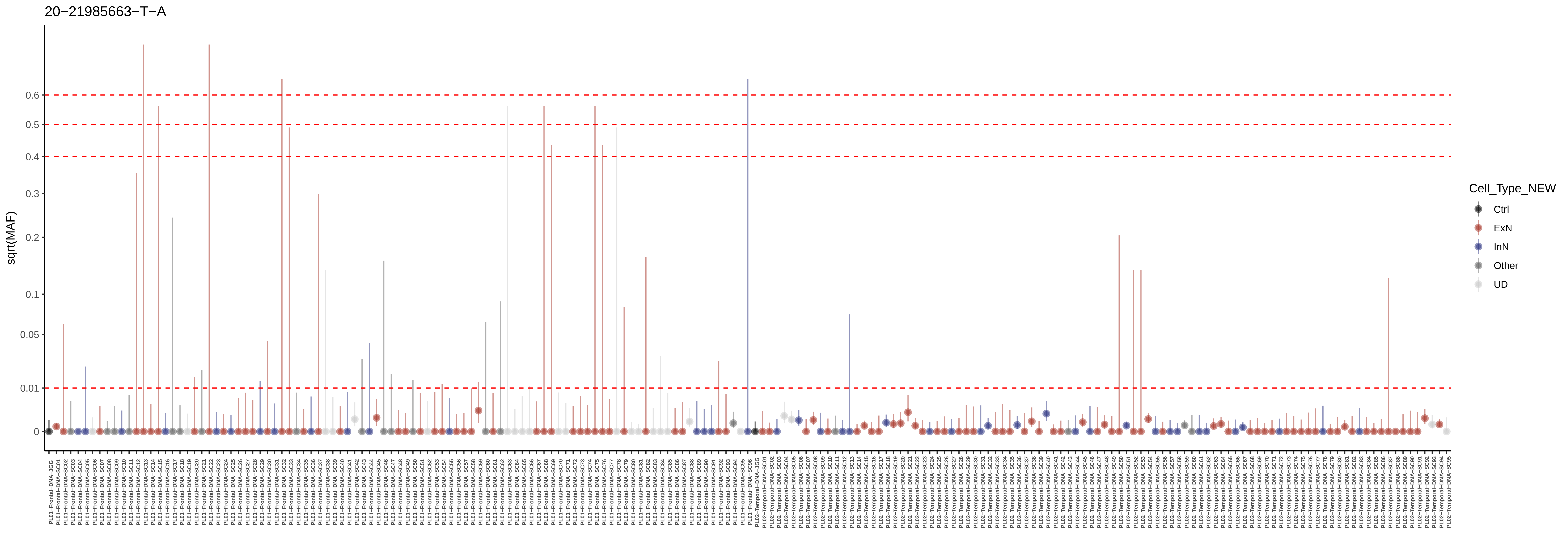

20−21985663−T−A
sqrt(MAF)
0.6
0.5
0.4
0.3
0.2
0.1
0.05
0.01
0
Cell_Type_NEW
Ctrl
ExN
InN
Other
UD

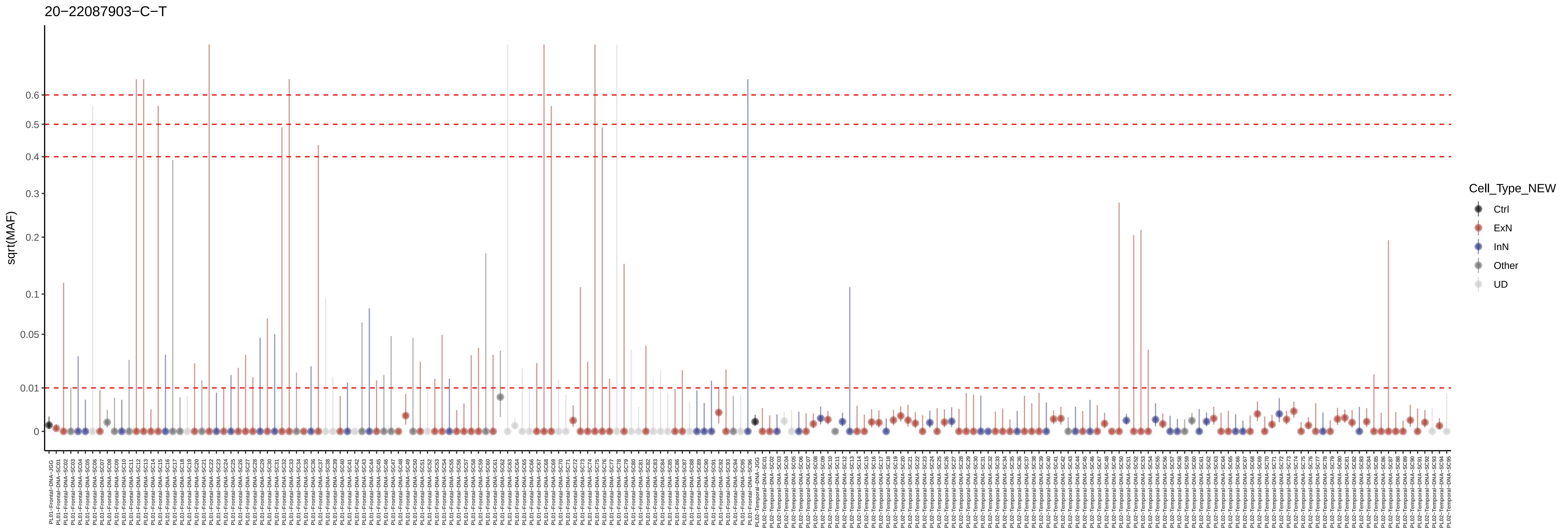
20−22087903−C−T
sqrt(MAF)
0.6
0.5
0.4
0.3
0.2
0.1
0.05
0.01
0
Cell_Type_NEW
Ctrl
ExN
InN
Other
UD

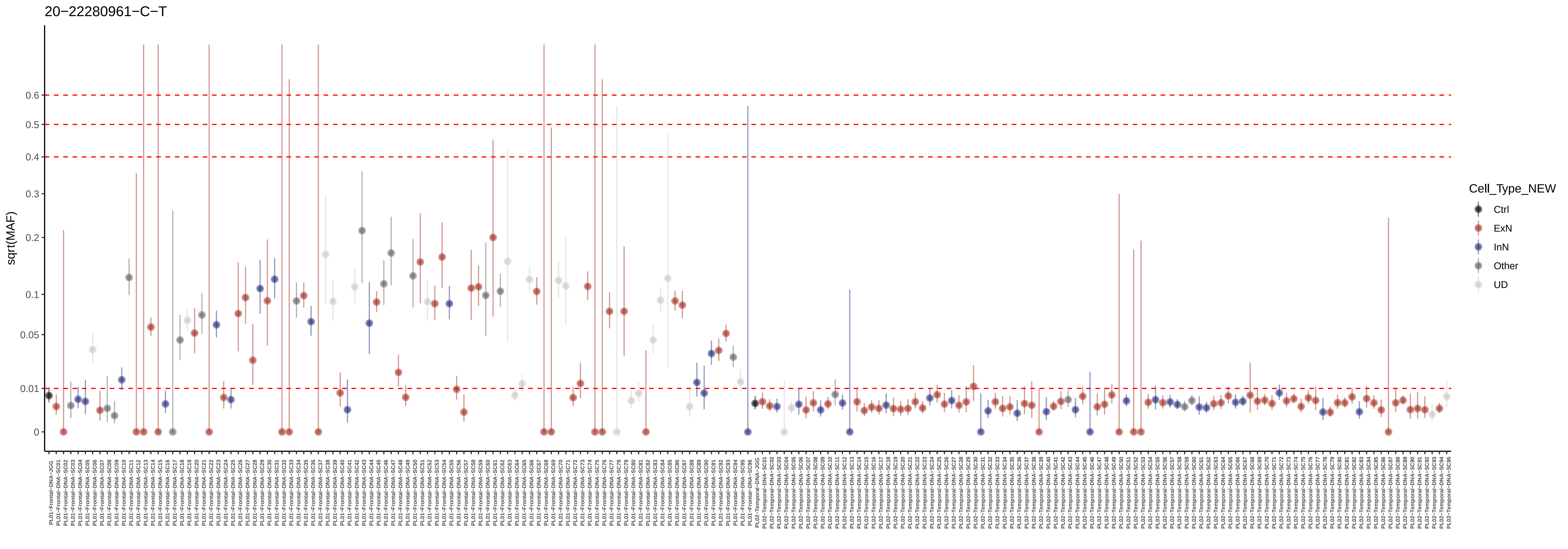

20−22280961−C−T
sqrt(MAF)
0.6
0.5
0.4
0.3
0.2
0.1
0.05
0.01
0
Cell_Type_NEW
Ctrl
ExN
InN
Other
UD

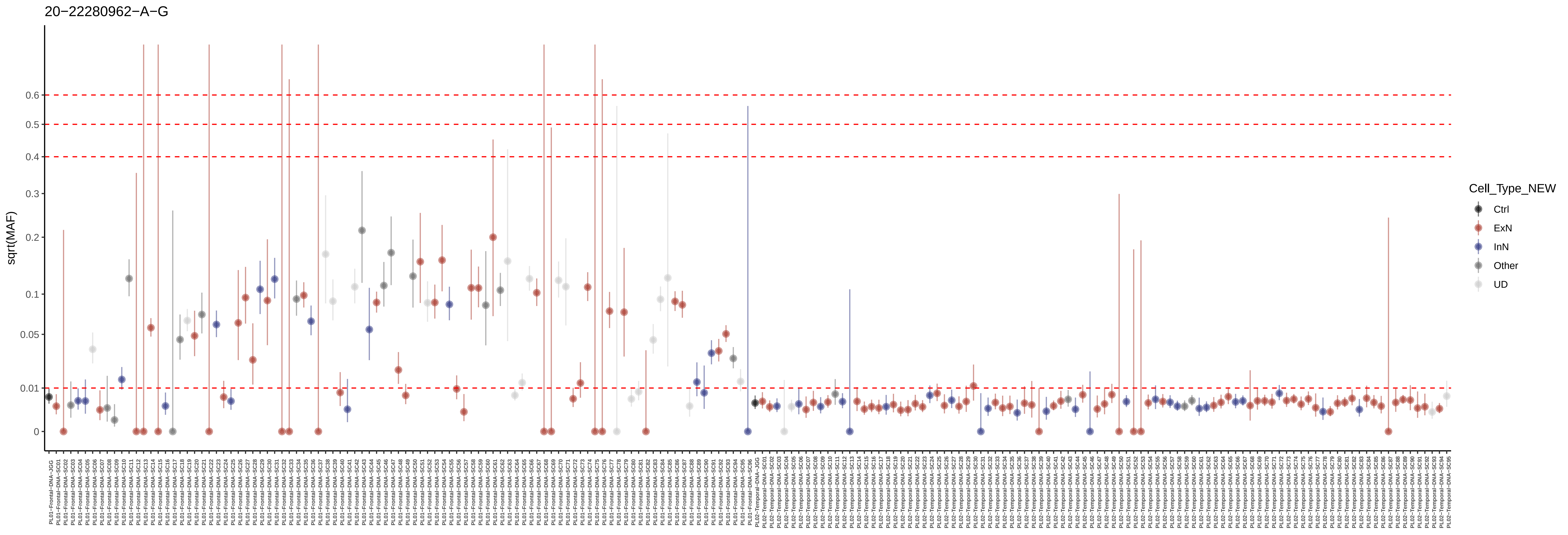

20−22280962−A−G
sqrt(MAF)
0
0.01
0.05
0.1
0.2
0.3
0.4
0.5
0.6
Cell_Type_NEW
Ctrl
ExN
InN
Other
UD

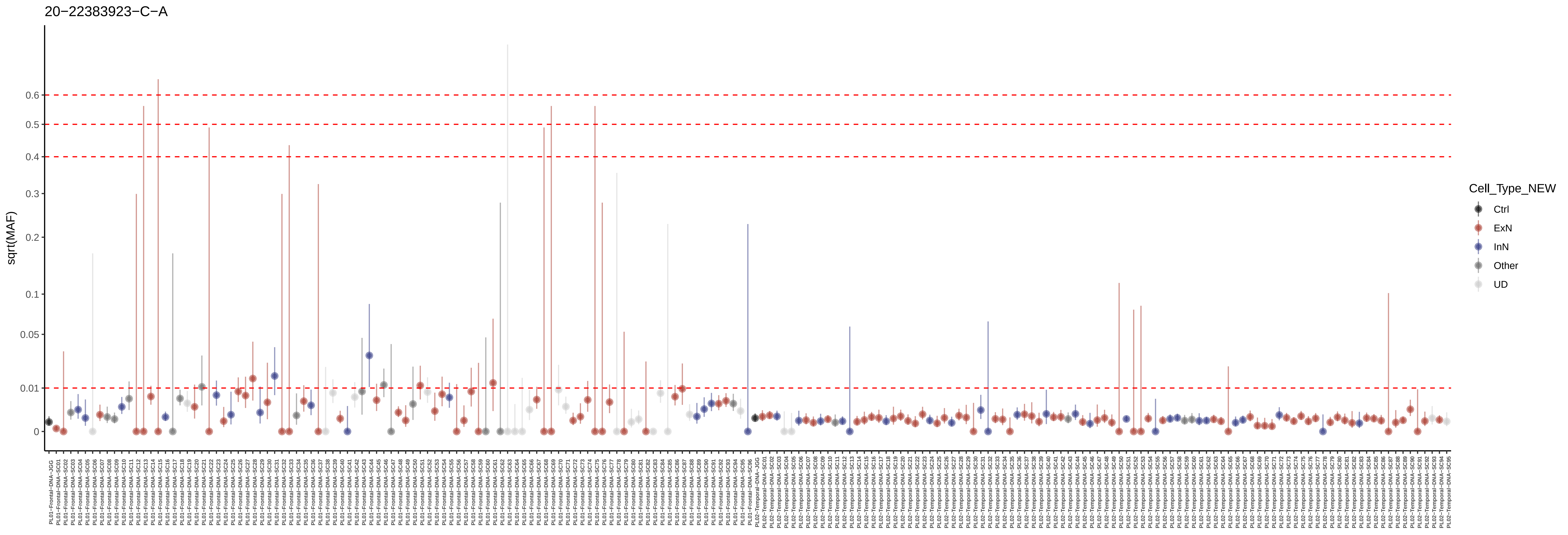
20−22383923−C−A
sqrt(MAF)
0.6
0.5
0.4
0.3
0.2
0.1
0.05
0.01
0
Cell_Type_NEW
Ctrl
ExN
InN
Other
UD

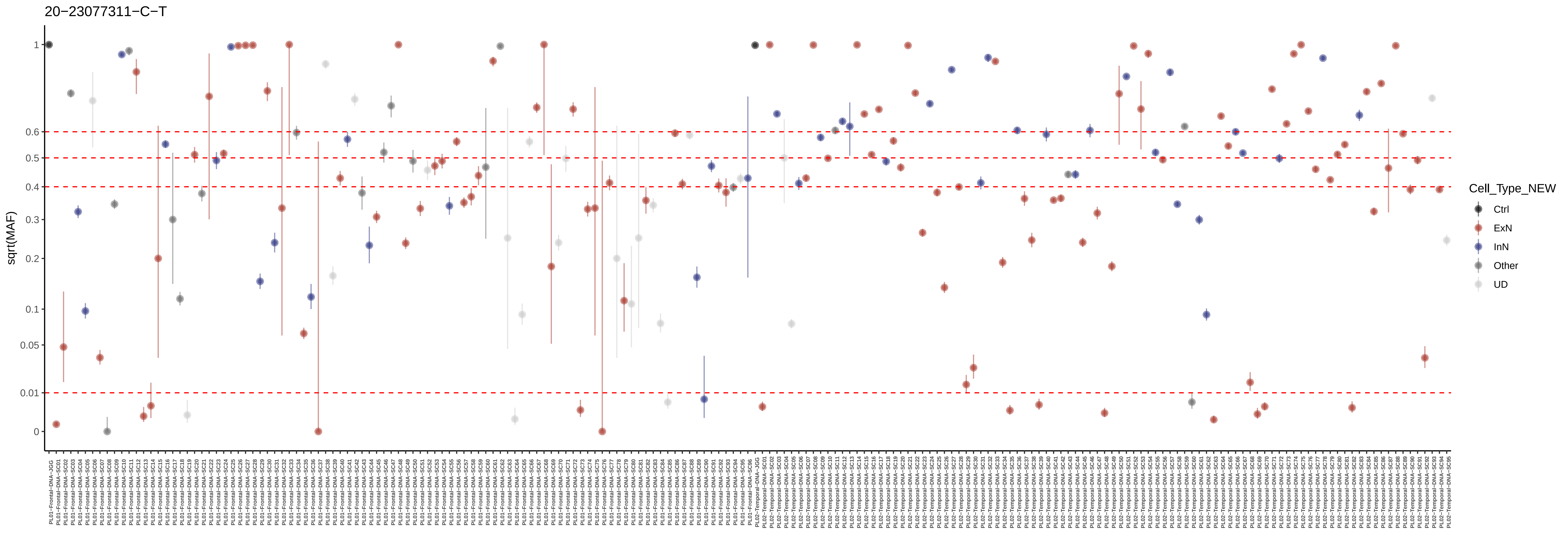
20−23077311−C−T
sqrt(MAF)
1
0.6
0.5
0.4
0.3
0.2
0.1
0.05
0.01
0
Cell_Type_NEW
Ctrl
ExN
InN
Other
UD

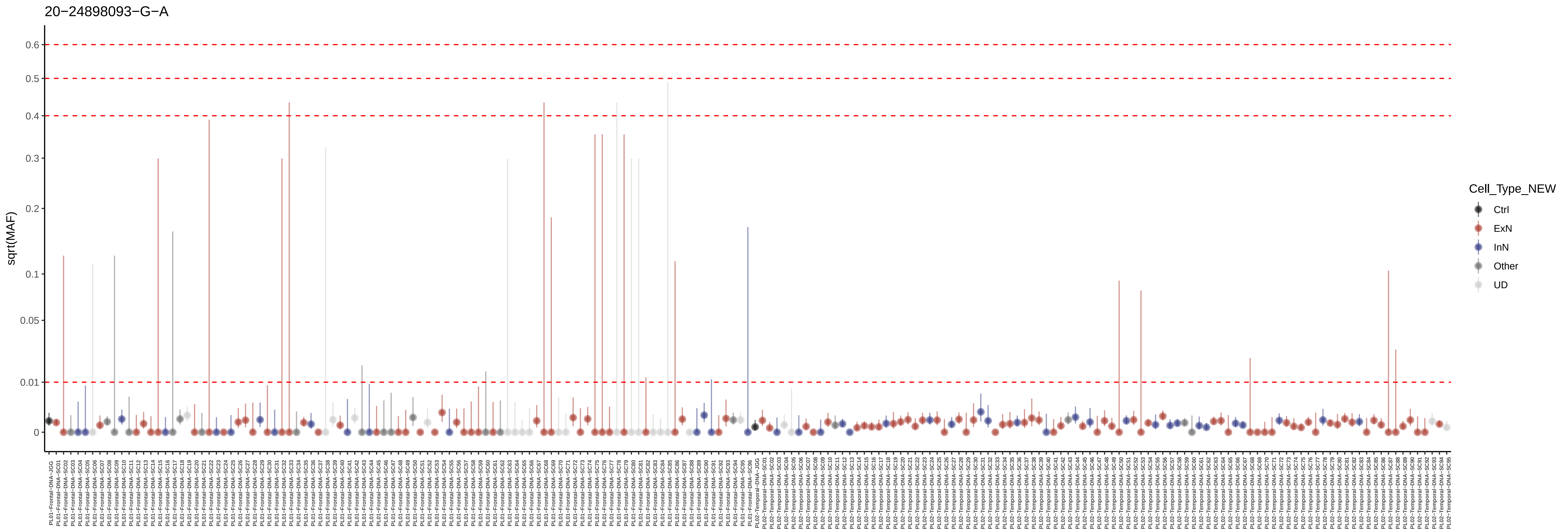

20−24898093−G−A
sqrt(MAF)
0.6
0.5
0.4
0.3
0.2
0.1
0.05
0.01
0
Cell_Type_NEW
Ctrl
ExN
InN
Other
UD

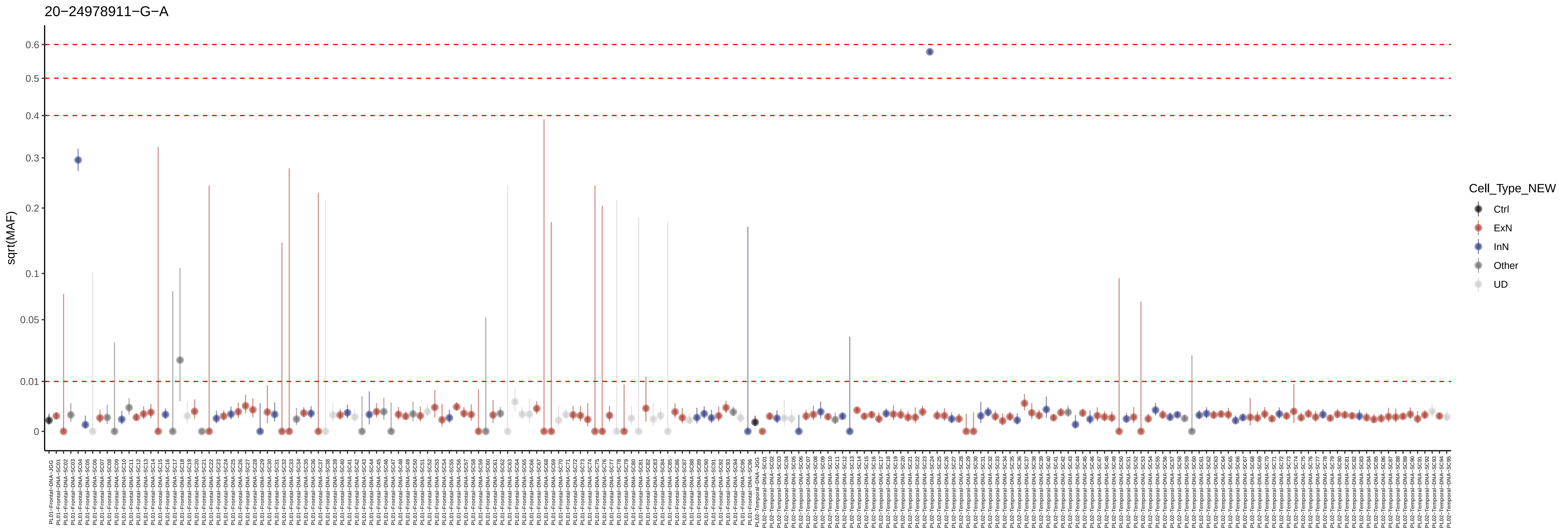
20−24978911−G−A
sqrt(MAF)
0.6
0.5
0.4
0.3
0.2
0.1
0.05
0.01
0
Cell_Type_NEW
Ctrl
ExN
InN
Other
UD

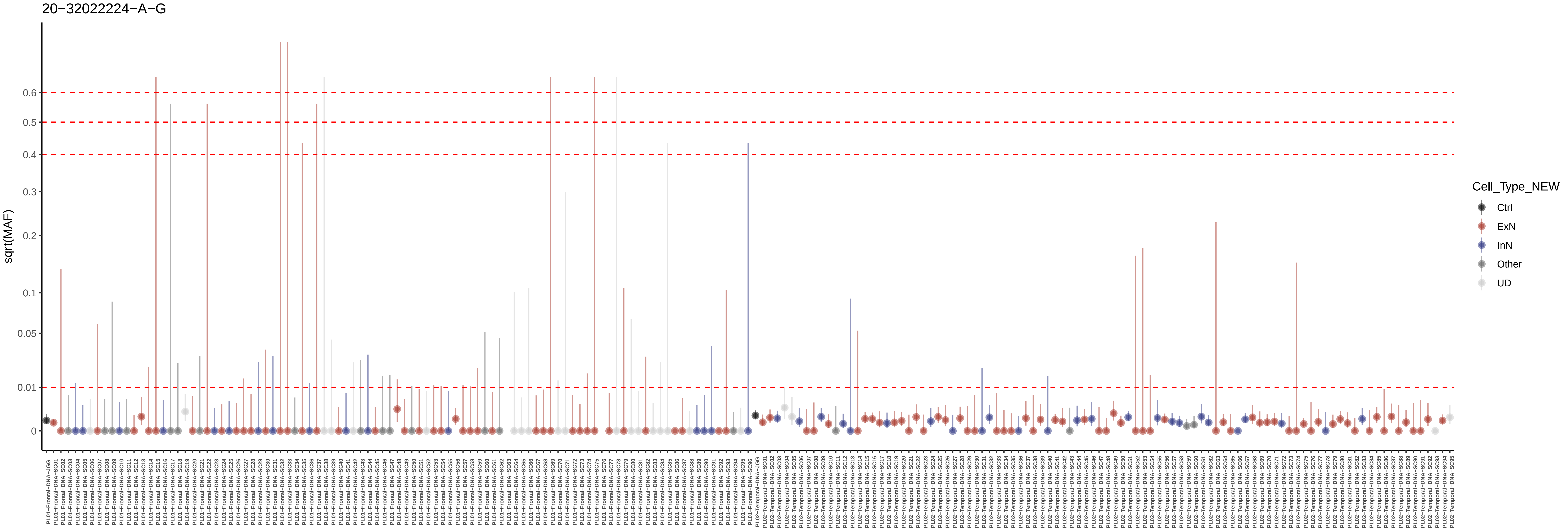
20−32022224−A−G
sqrt(MAF)
0.6
0.5
0.4
0.3
0.2
0.1
0.05
0.01
0
Cell_Type_NEW
Ctrl
ExN
InN
Other
UD

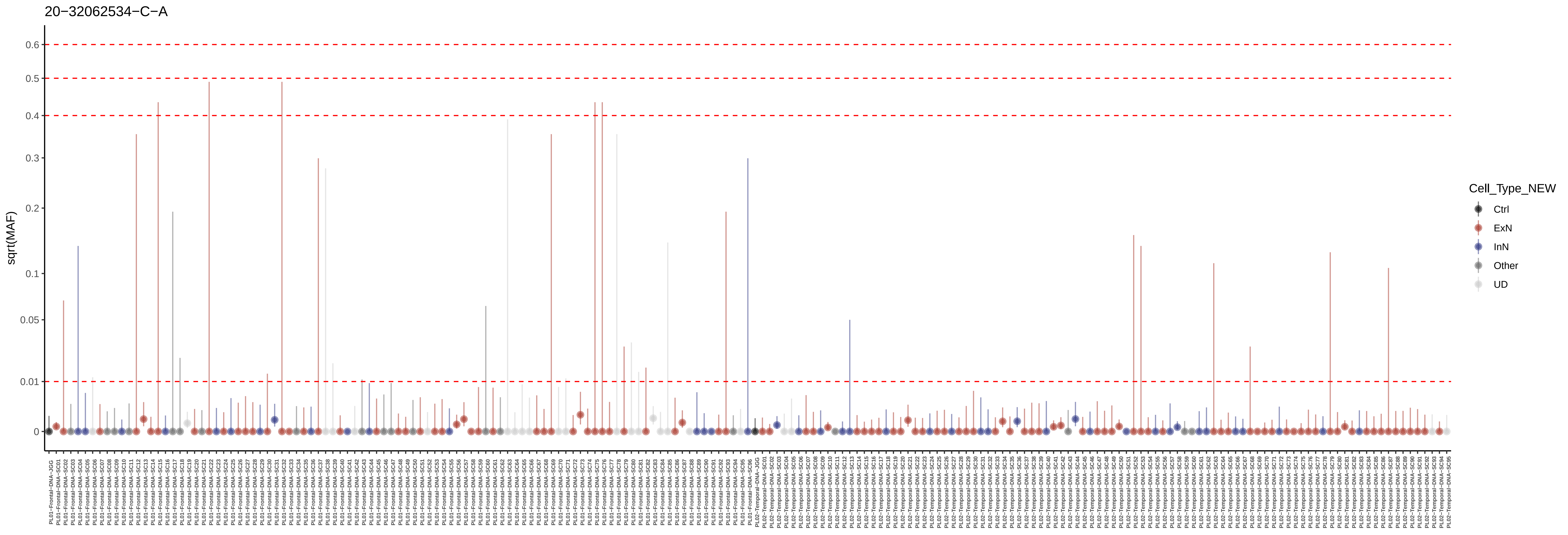
20−32062534−C−A
sqrt(MAF)
0.6
0.5
0.4
0.3
0.2
0.1
0.05
0.01
0
Cell_Type_NEW
Ctrl
ExN
InN
Other
UD

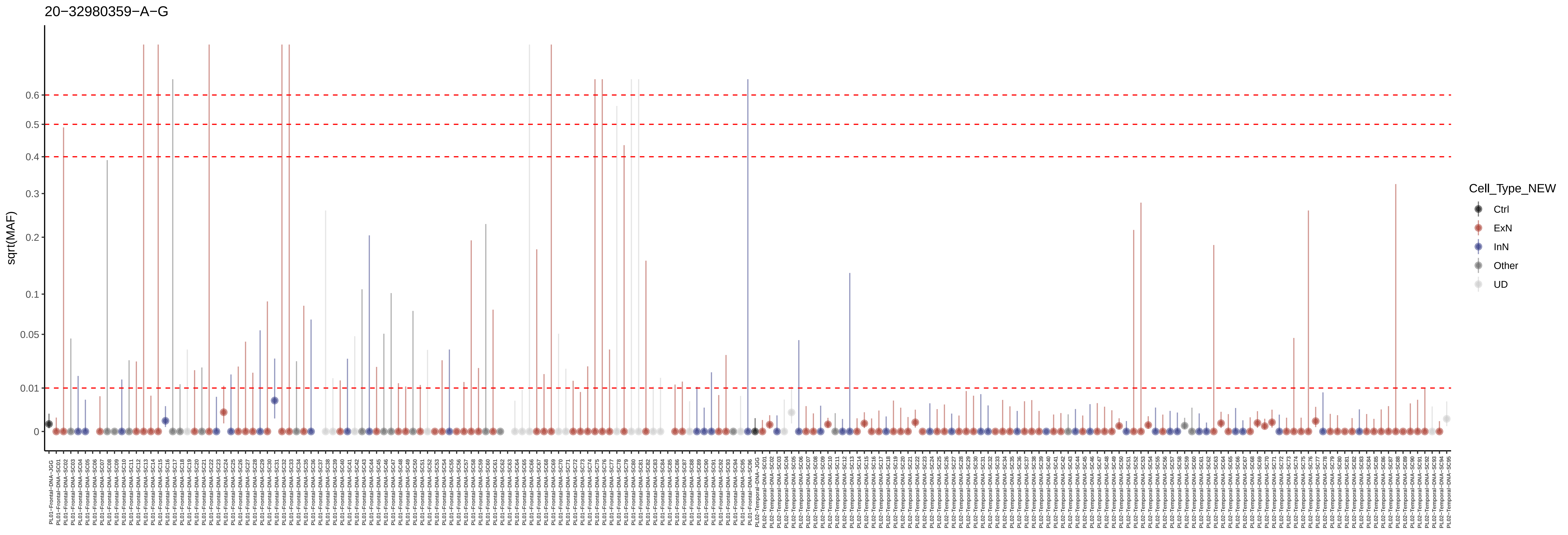

20−32980359−A−G
sqrt(MAF)
0.6
0.5
0.4
0.3
0.2
0.1
0.05
0.01
0
Cell_Type_NEW
Ctrl
ExN
InN
Other
UD

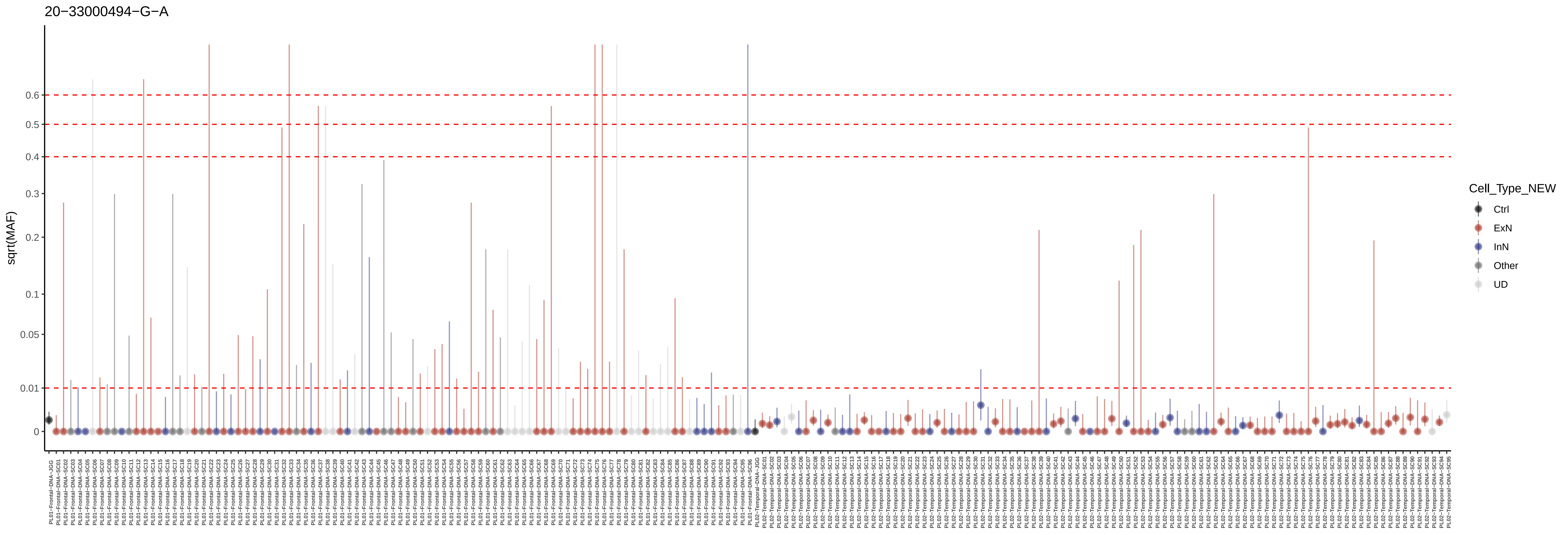

20−33000494−G−A
sqrt(MAF)
0.6
0.5
0.4
0.3
0.2
0.1
0.05
0.01
0
Cell_Type_NEW
Ctrl
ExN
InN
Other
UD

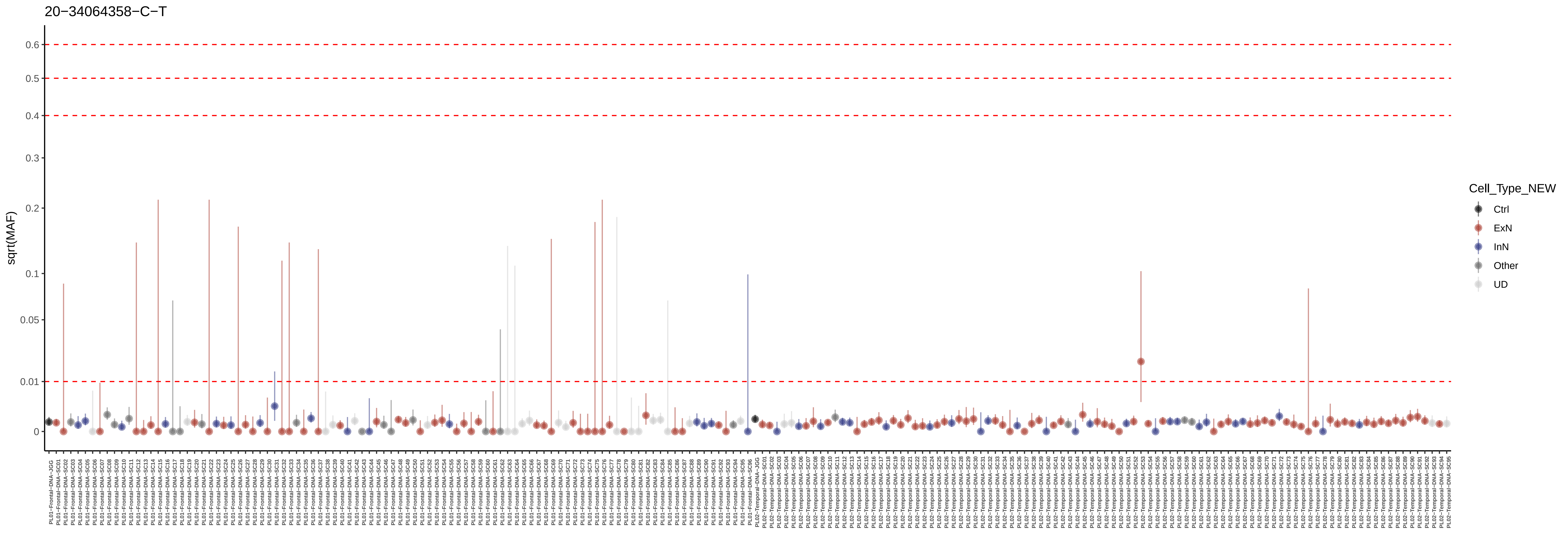

20−34064358−C−T
sqrt(MAF)
0.6
0.5
0.4
0.3
0.2
0.1
0.05
0.01
0
Cell_Type_NEW
Ctrl
ExN
InN
Other
UD

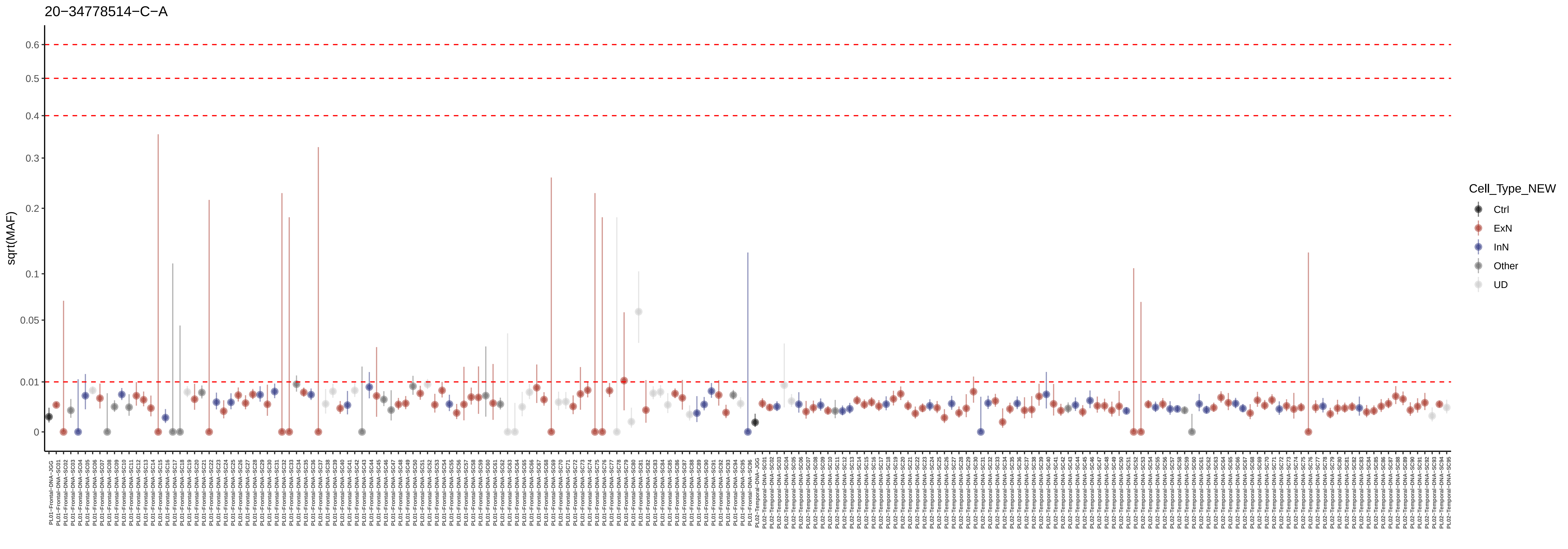

20−34778514−C−A
sqrt(MAF)
0.6
0.5
0.4
0.3
0.2
0.1
0.05
0.01
0
Cell_Type_NEW
Ctrl
ExN
InN
Other
UD

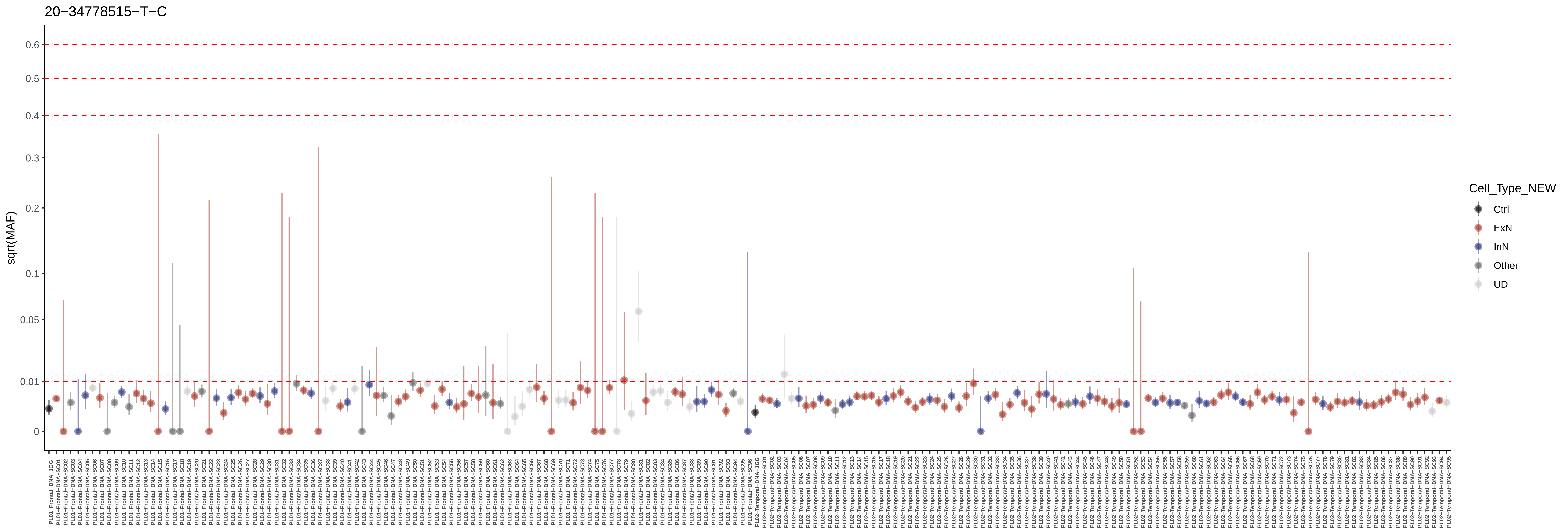
20−34778515−T−C
sqrt(MAF)
0.6
0.5
0.4
0.3
0.2
0.1
0.05
0.01
0
Cell_Type_NEW
Ctrl
ExN
InN
Other
UD

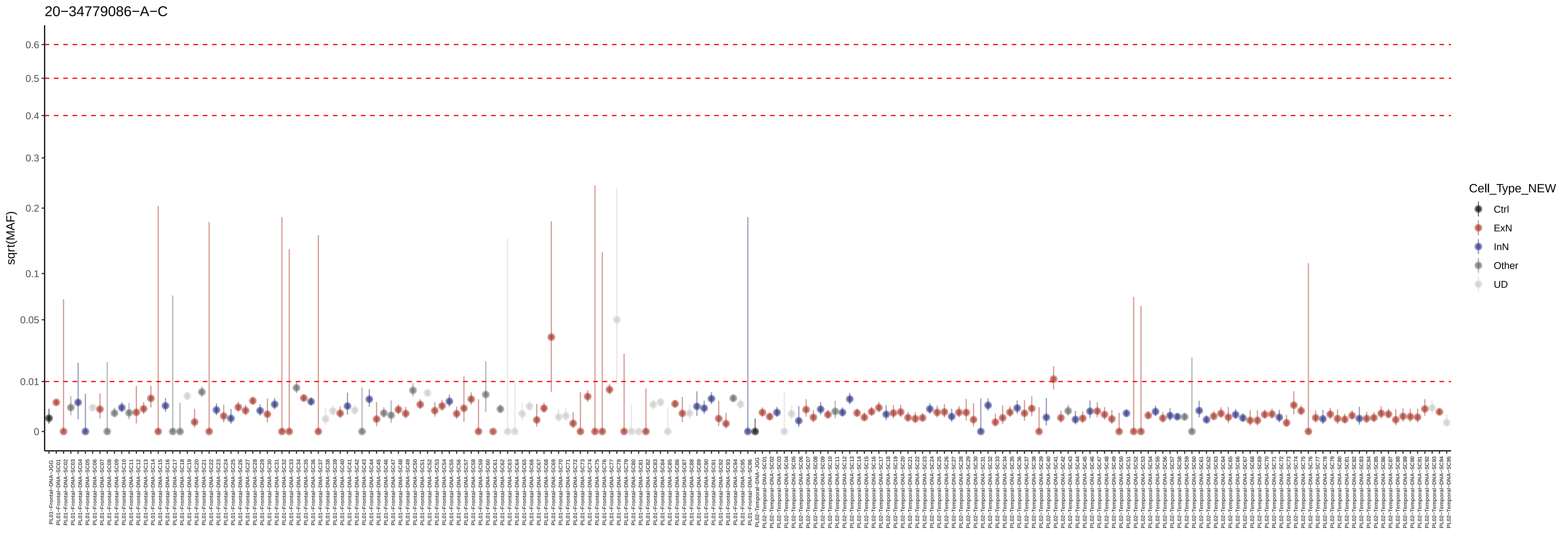

20−34779086−A−C
sqrt(MAF)
0.6
0.5
0.4
0.3
0.2
0.1
0.05
0.01
0
Cell_Type_NEW
Ctrl
ExN
InN
Other
UD

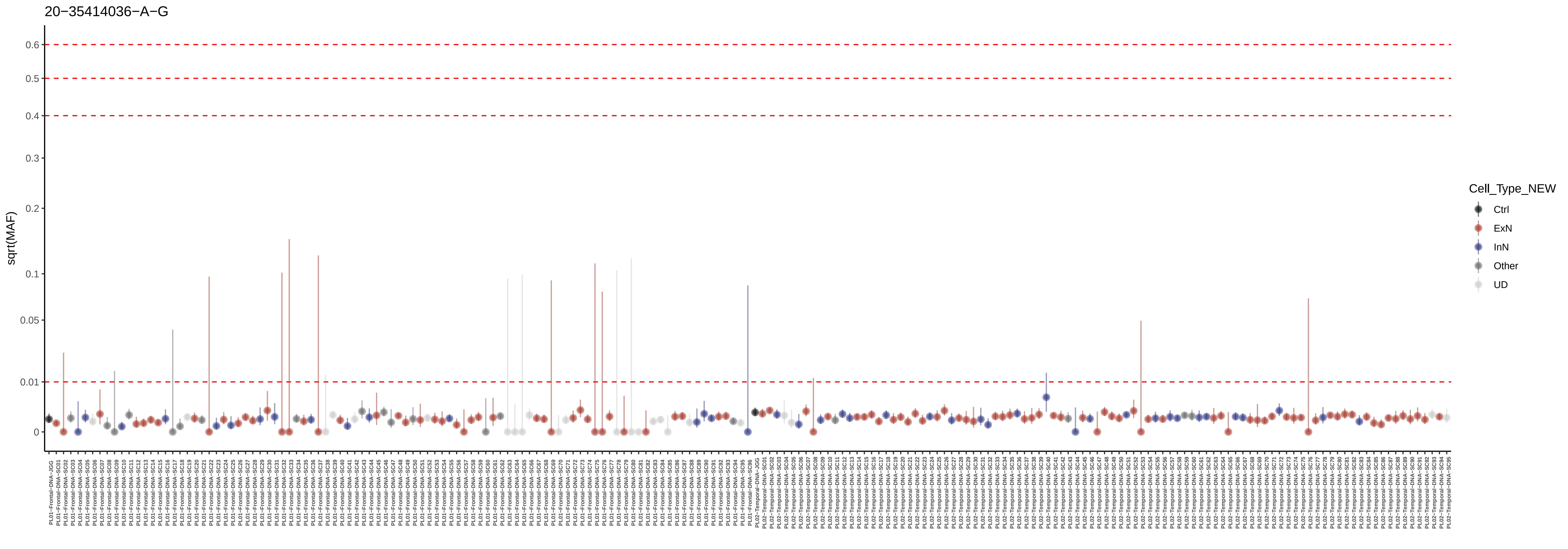

20−35414036−A−G
sqrt(MAF)
0.6
0.5
0.4
0.3
0.2
0.1
0.05
0.01
0
Cell_Type_NEW
Ctrl
ExN
InN
Other
UD

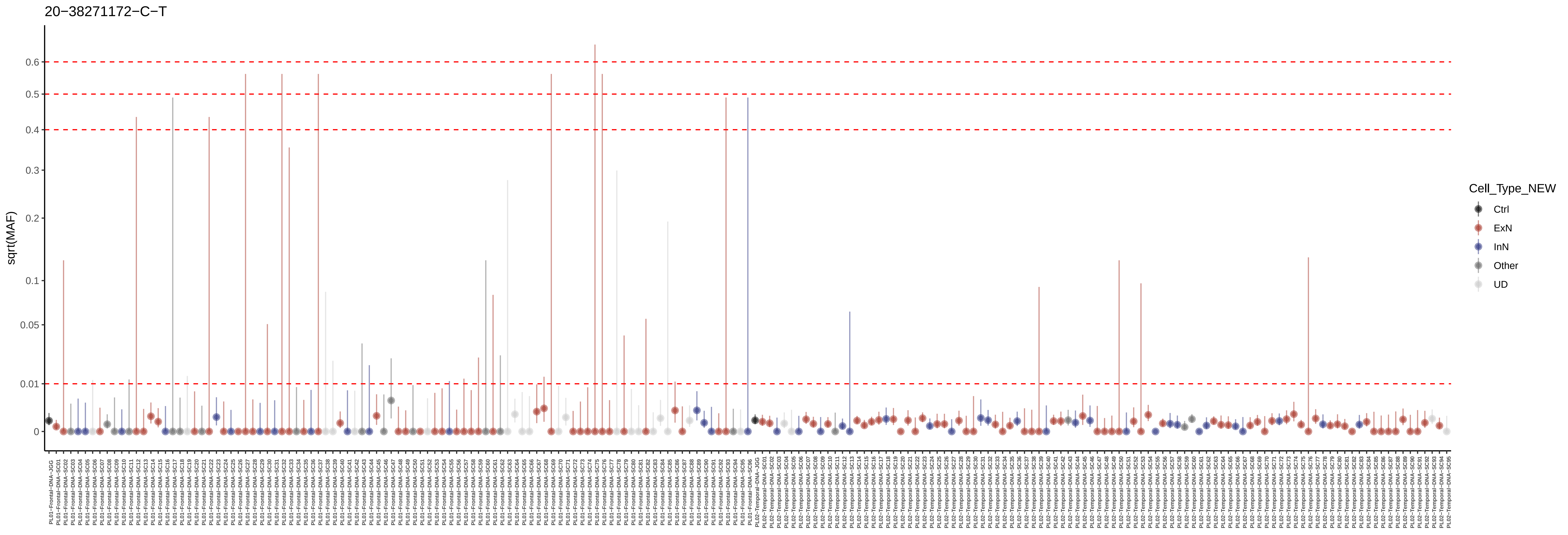
20−38271172−C−T
sqrt(MAF)
0.6
0.5
0.4
0.3
0.2
0.1
0.05
0.01
0
Cell_Type_NEW
Ctrl
ExN
InN
Other
UD

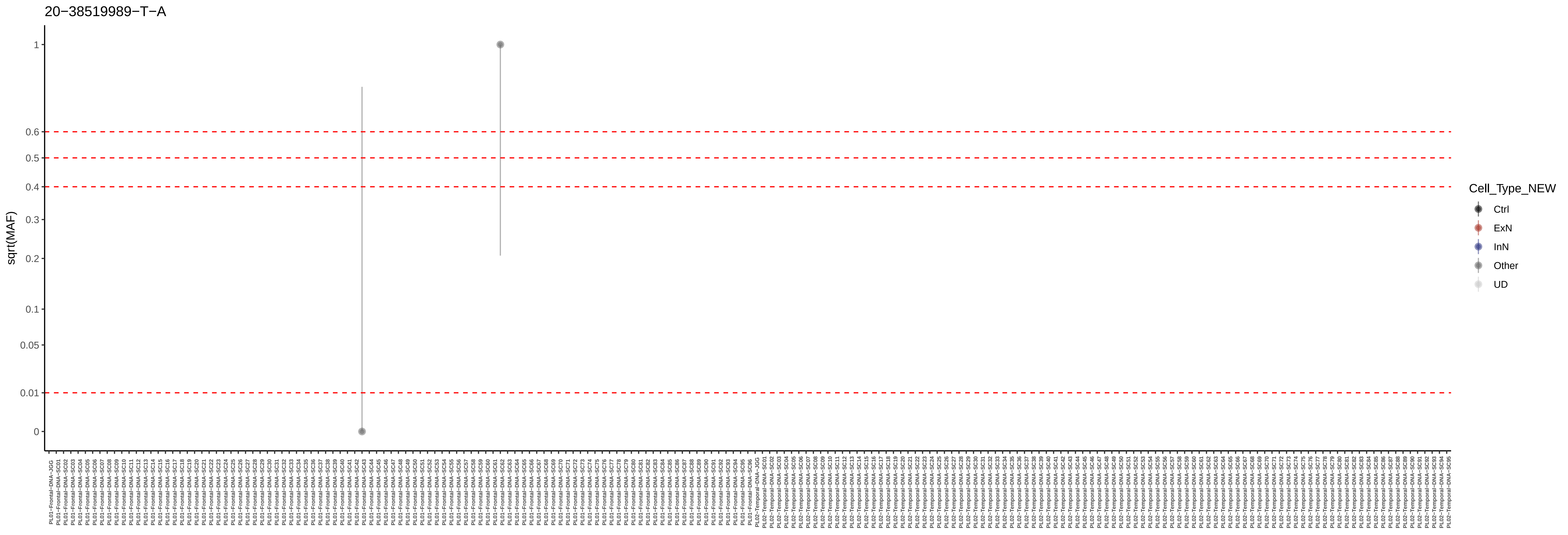
20−38519989−T−A
sqrt(MAF)
1
0.6
0.5
0.4
0.3
0.2
0.1
0.05
0.01
0
Cell_Type_NEW
Ctrl
ExN
InN
Other
UD

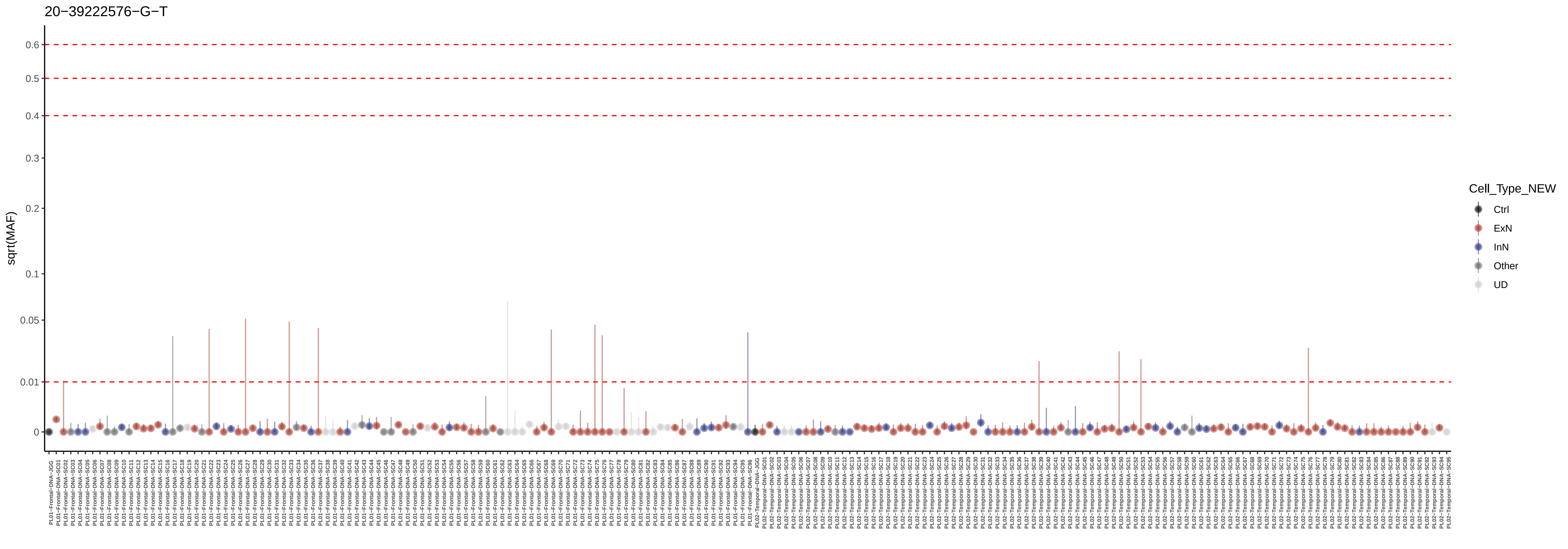
20−39222576−G−T
sqrt(MAF)
Cell_Type_NEW
Ctrl
ExN
InN
Other
UD

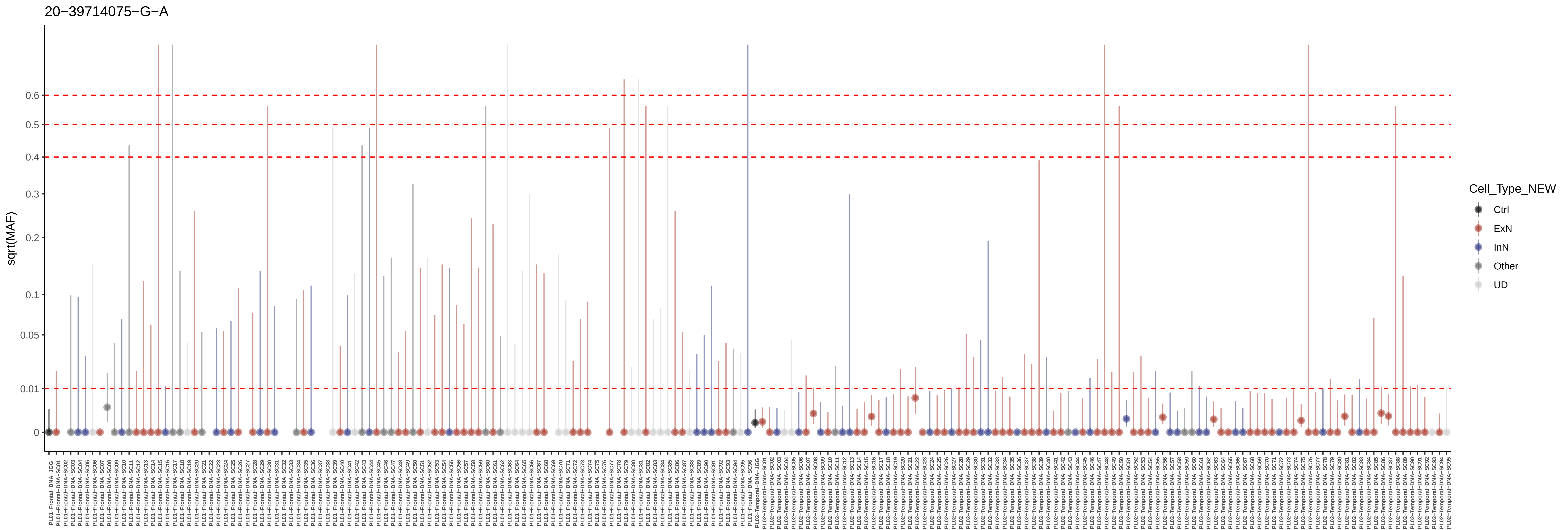
20−39714075−G−A
sqrt(MAF)
0.6
0.5
0.4
0.3
0.2
0.1
0.05
0.01
0
Cell_Type_NEW
Ctrl
ExN
InN
Other
UD

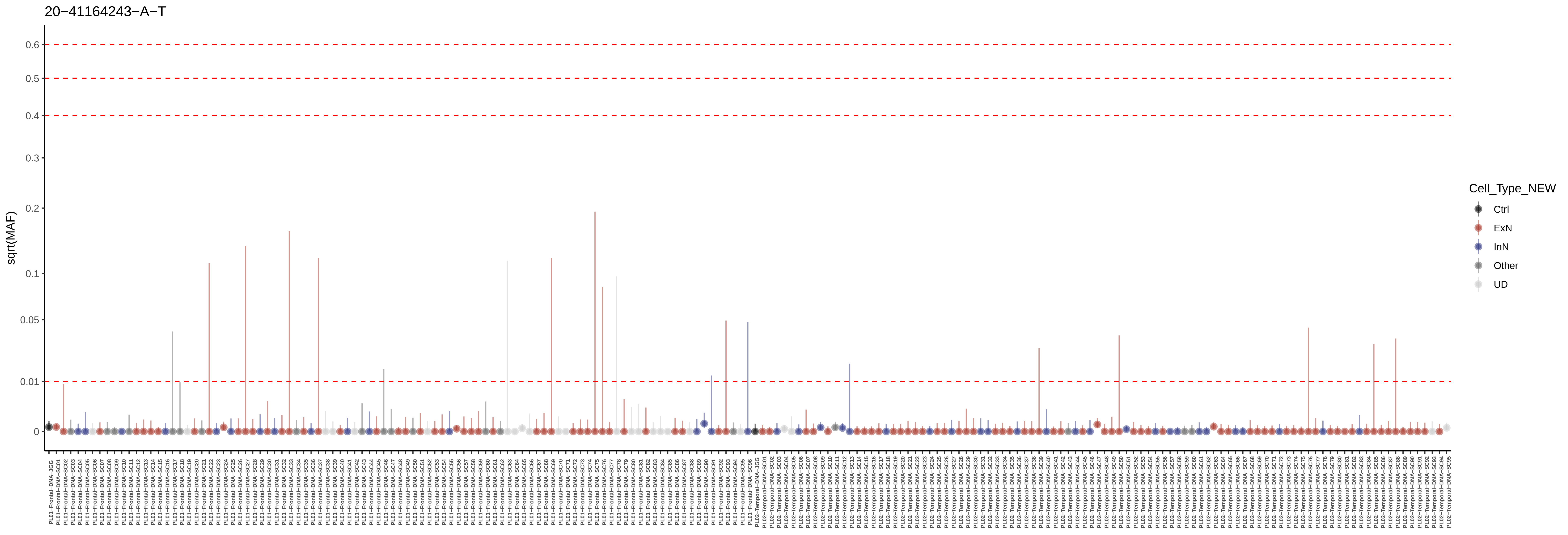

20−41164243−A−T
sqrt(MAF)
0.6
0.5
0.4
0.3
0.2
0.1
0.05
0.01
0
Cell_Type_NEW
Ctrl
ExN
InN
Other
UD

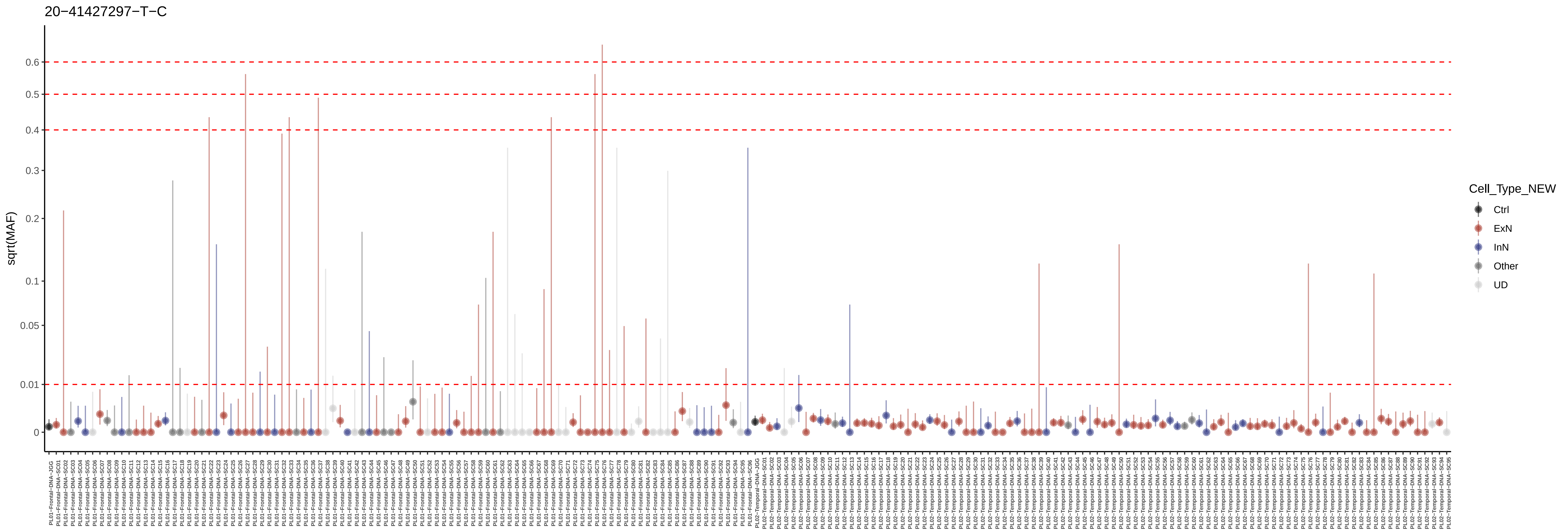

20−41427297−T−C
sqrt(MAF)
0.6
0.5
0.4
0.3
0.2
0.1
0.05
0.01
0
Cell_Type_NEW
Ctrl
ExN
InN
Other
UD

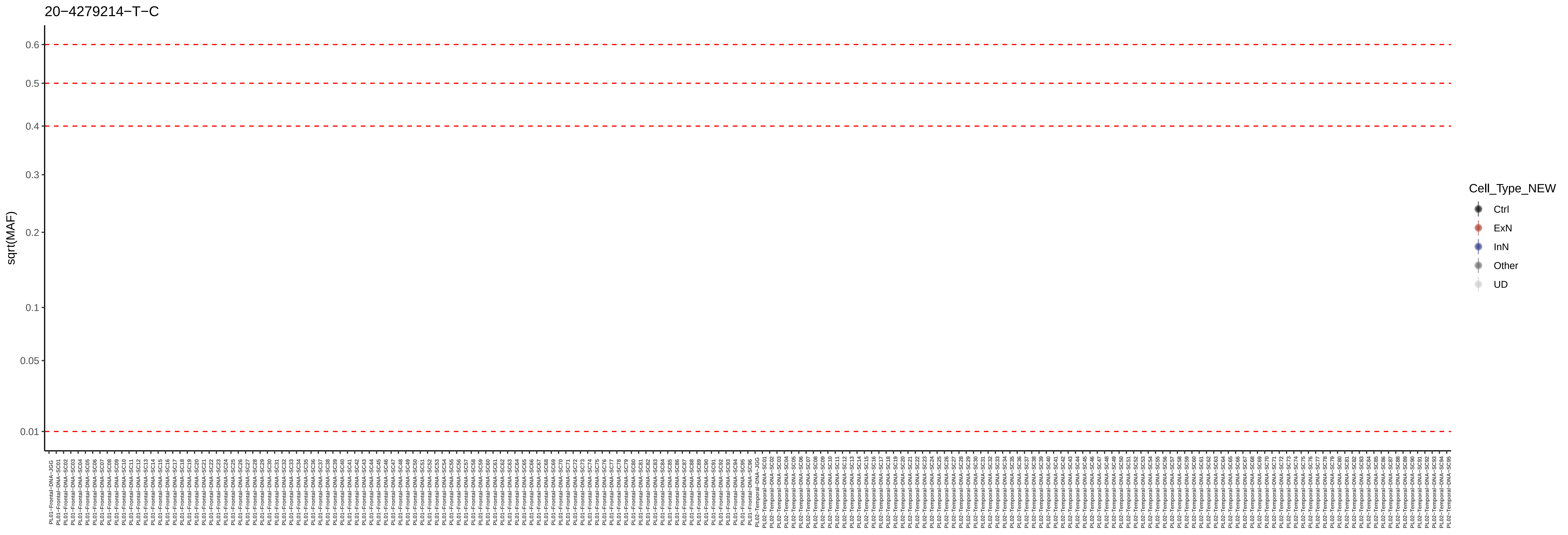

20−4279214−T−C
sqrt(MAF)
0.6
0.5
0.4
0.3
0.2
0.1
0.05
0.01
Cell_Type_NEW
Ctrl
ExN
InN
Other
UD

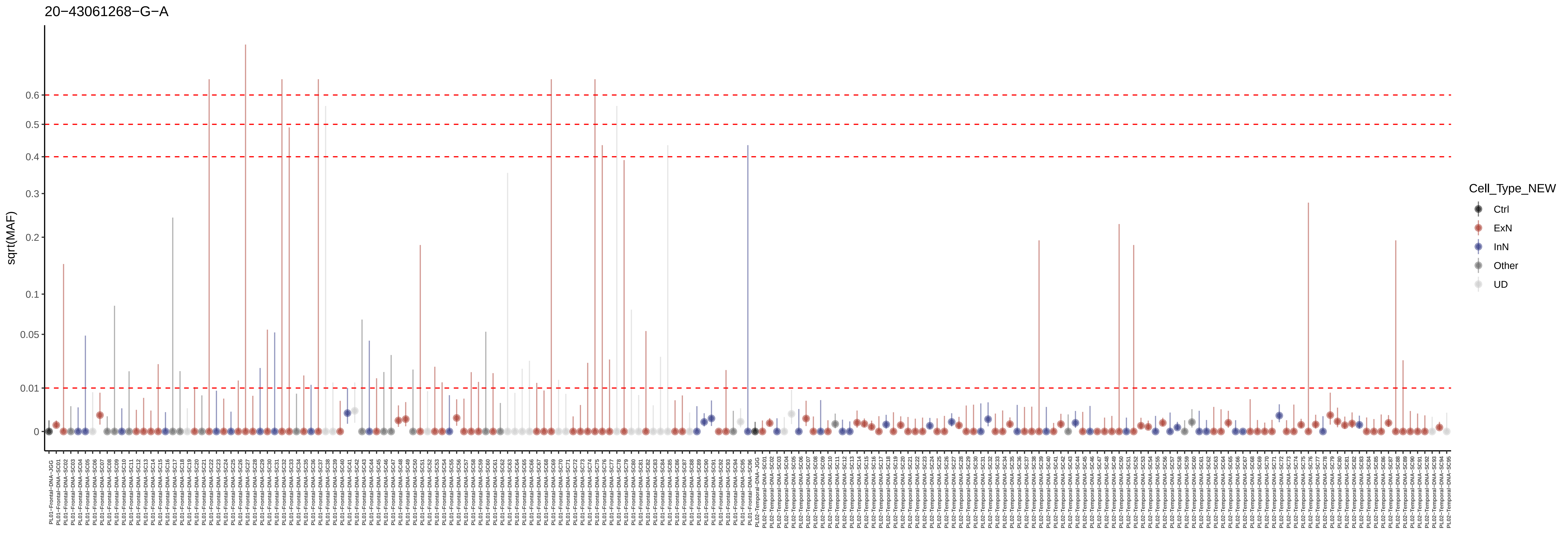
20−43061268−G−A
sqrt(MAF)
0.6
0.5
0.4
0.3
0.2
0.1
0.05
0.01
0
Cell_Type_NEW
Ctrl
ExN
InN
Other
UD

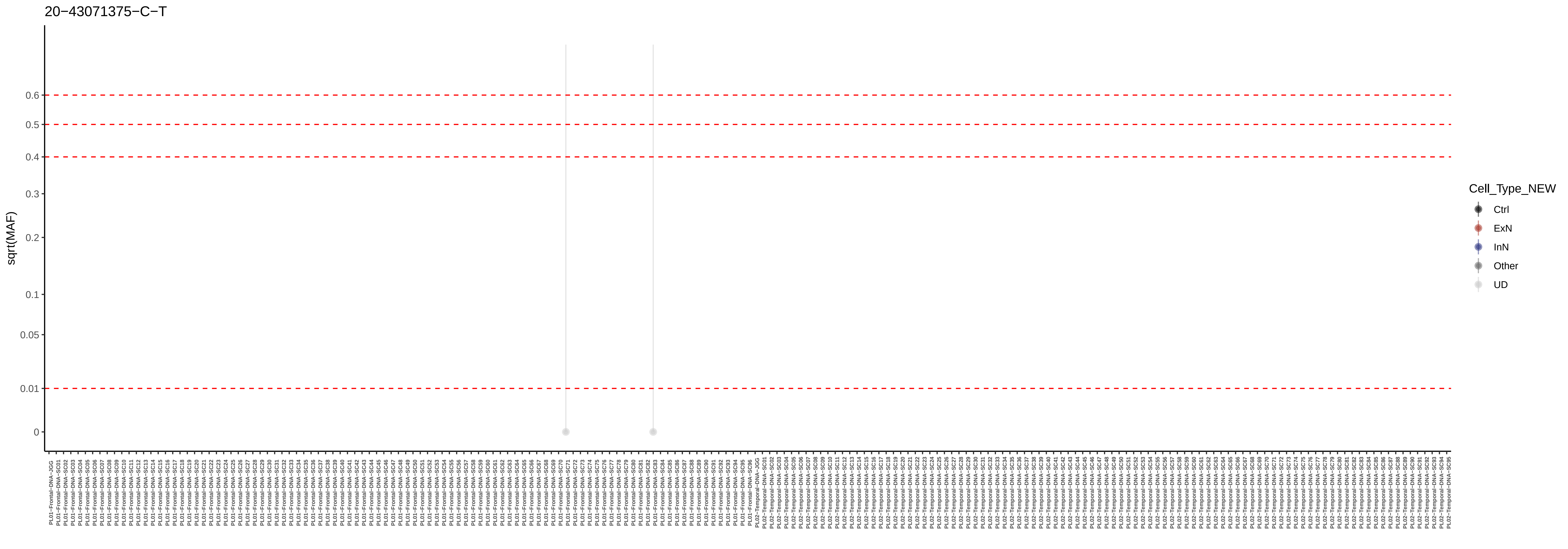

20−43071375−C−T
sqrt(MAF)
0.6
0.5
0.4
0.3
0.2
0.1
0.05
0.01
0
Cell_Type_NEW
Ctrl
ExN
InN
Other
UD

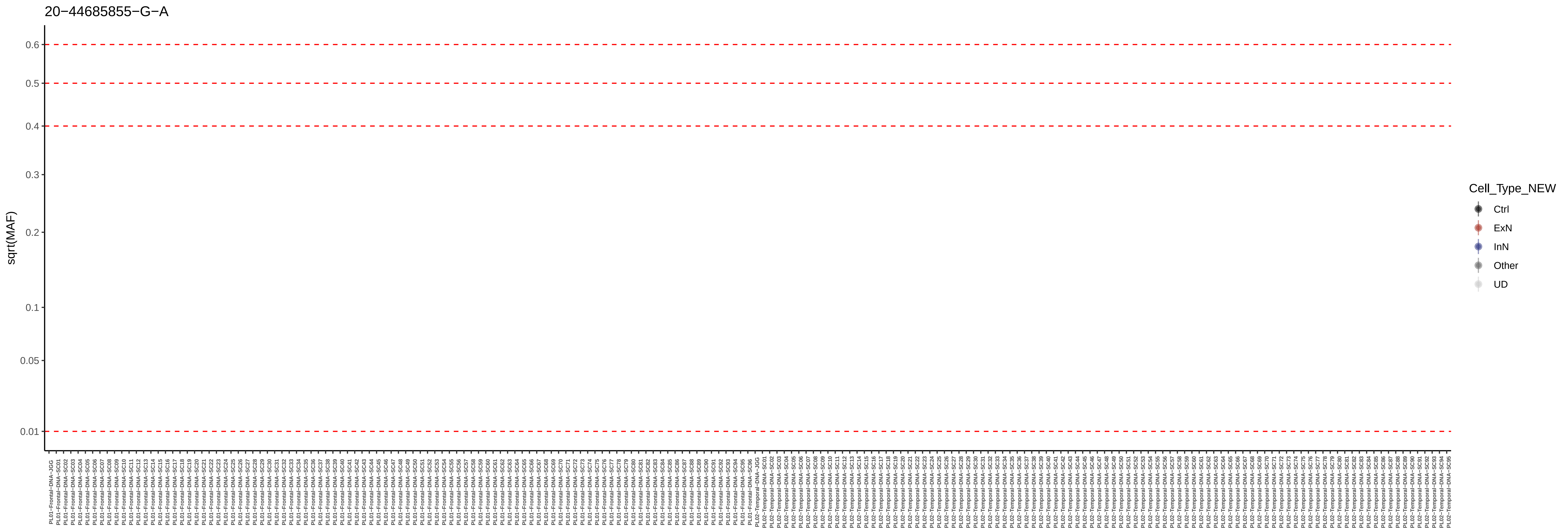
20−44685855−G−A
sqrt(MAF)
0.6
0.5
0.4
0.3
0.2
0.1
0.05
0.01
Cell_Type_NEW
Ctrl
ExN
InN
Other
UD

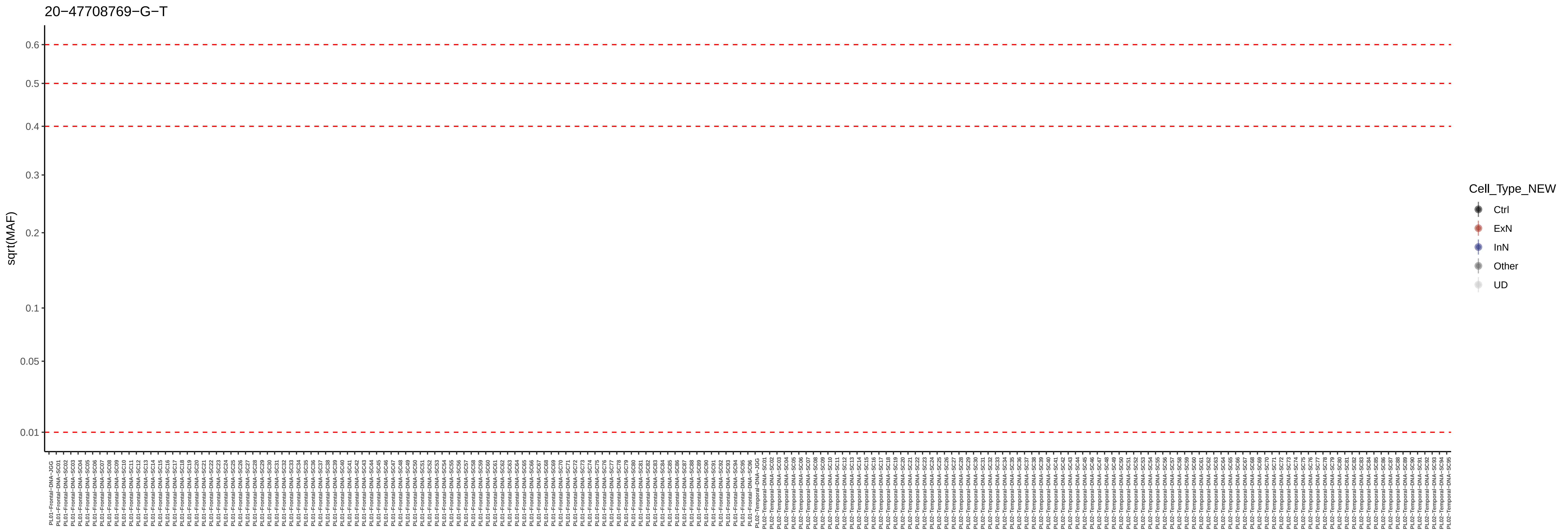
20−47708769−G−T
sqrt(MAF)
0.6
0.5
0.4
0.3
0.2
0.1
0.05
0.01
Cell_Type_NEW
Ctrl
ExN
InN
Other
UD

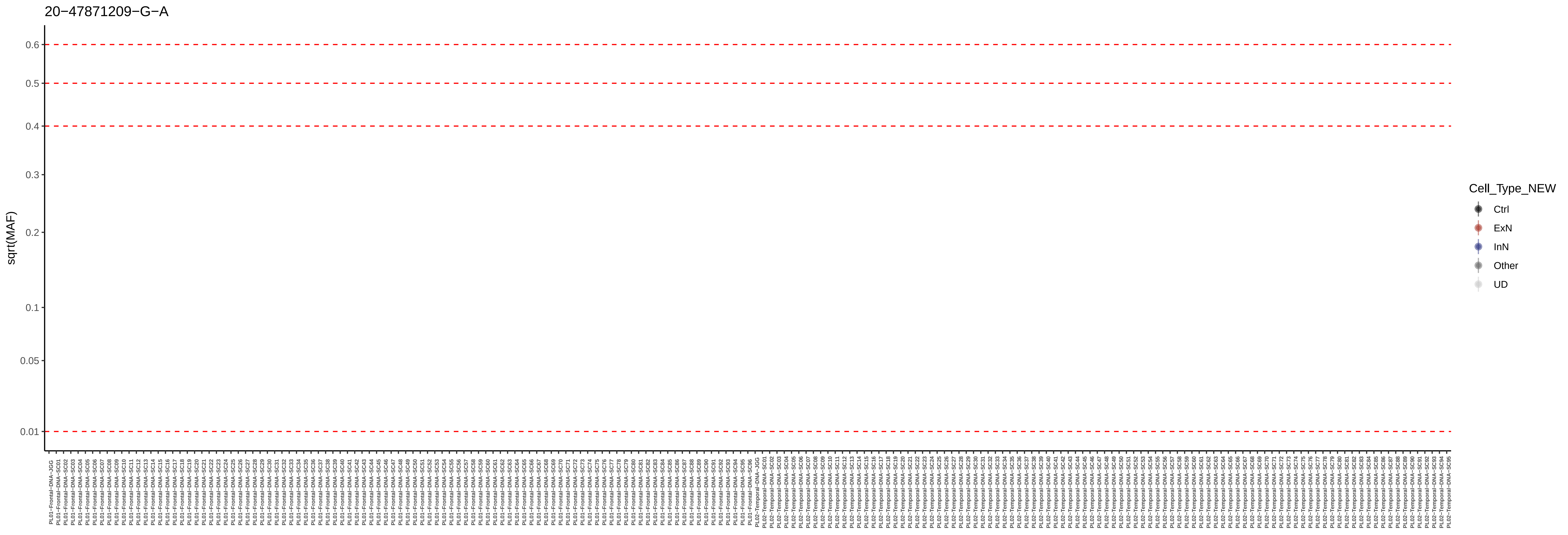
20−47871209−G−A
sqrt(MAF)
0.6
0.5
0.4
0.3
0.2
0.1
0.05
0.01
Cell_Type_NEW
Ctrl
ExN
InN
Other
UD

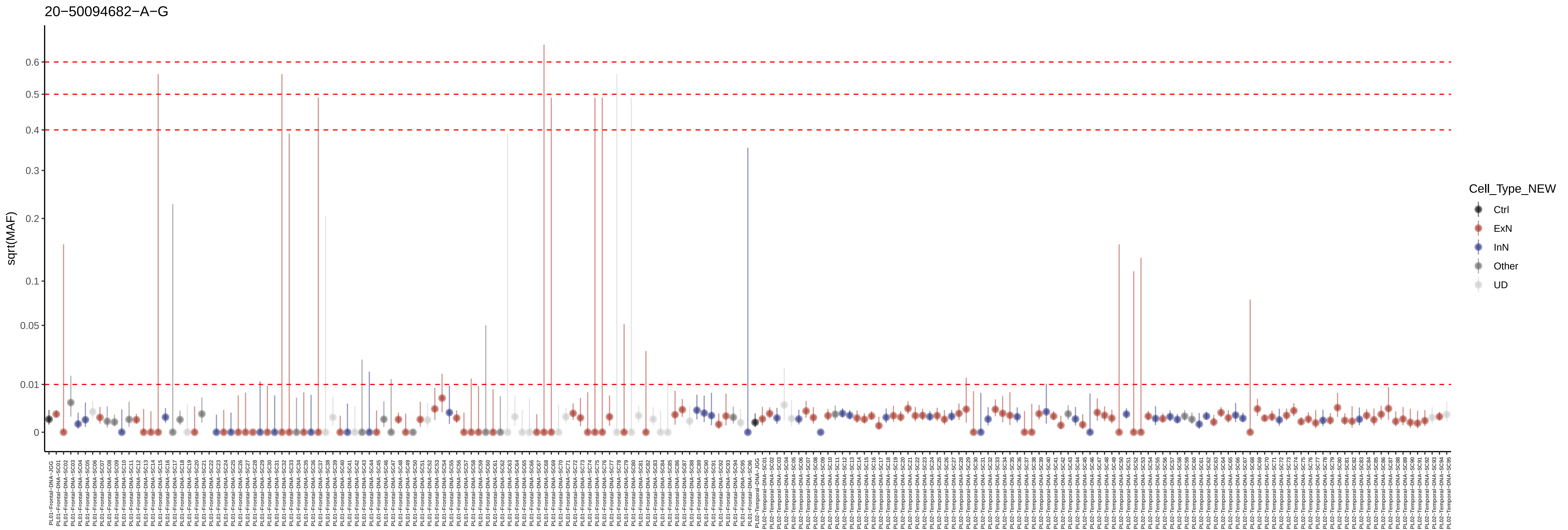

20−50094682−A−G
sqrt(MAF)
0.6
0.5
0.4
0.3
0.2
0.1
0.05
0.01
0
Cell_Type_NEW
Ctrl
ExN
InN
Other
UD

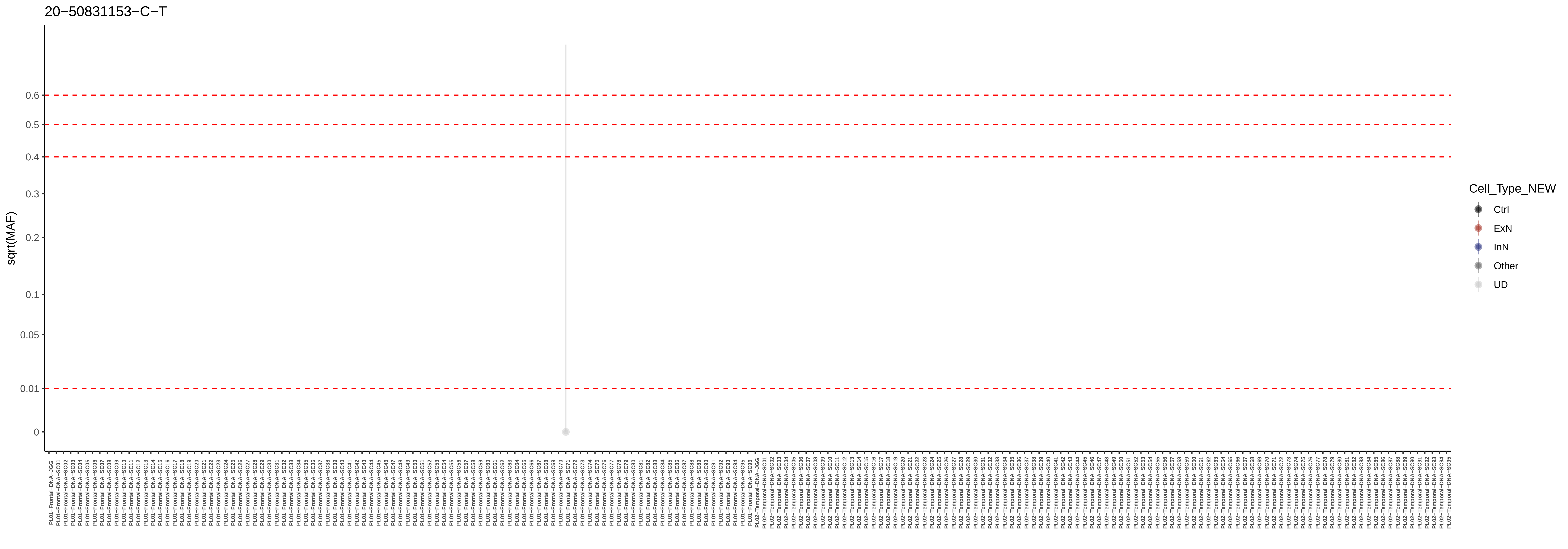
20−50831153−C−T
sqrt(MAF)
0.6
0.5
0.4
0.3
0.2
0.1
0.05
0.01
0
Cell_Type_NEW
Ctrl
ExN
InN
Other
UD

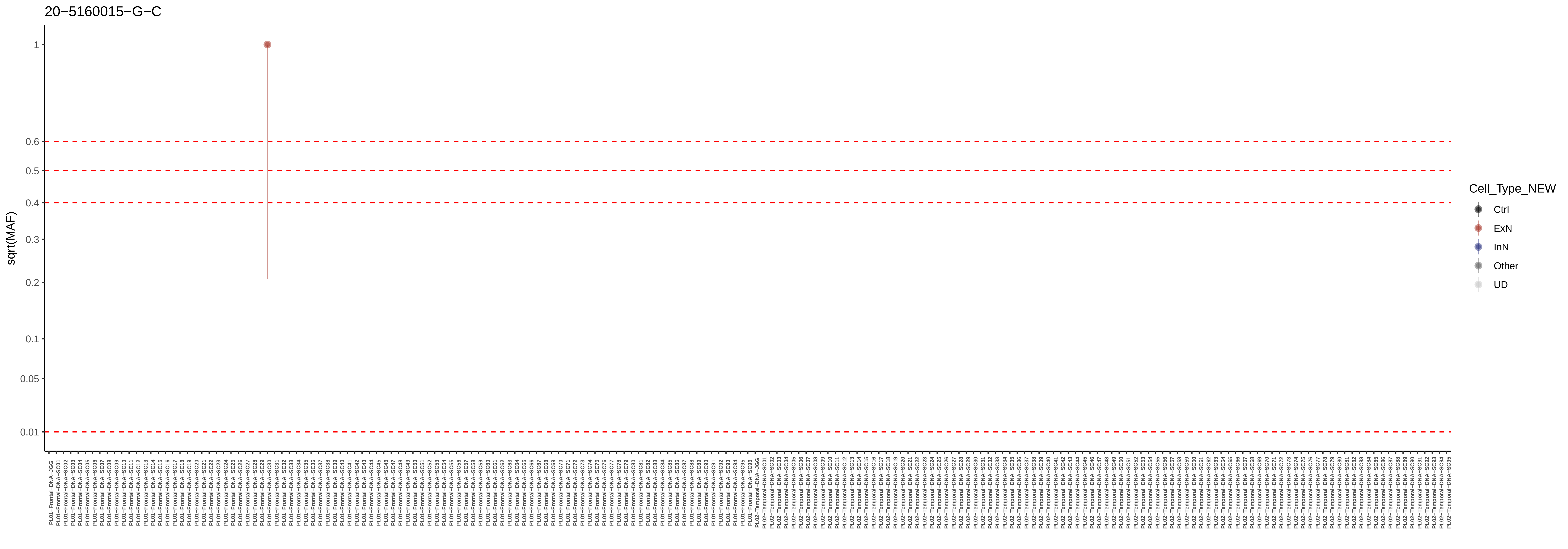

20−5160015−G−C
sqrt(MAF)
1
0.6
0.5
0.4
0.3
0.2
0.1
0.05
0.01
Cell_Type_NEW
Ctrl
ExN
InN
Other
UD

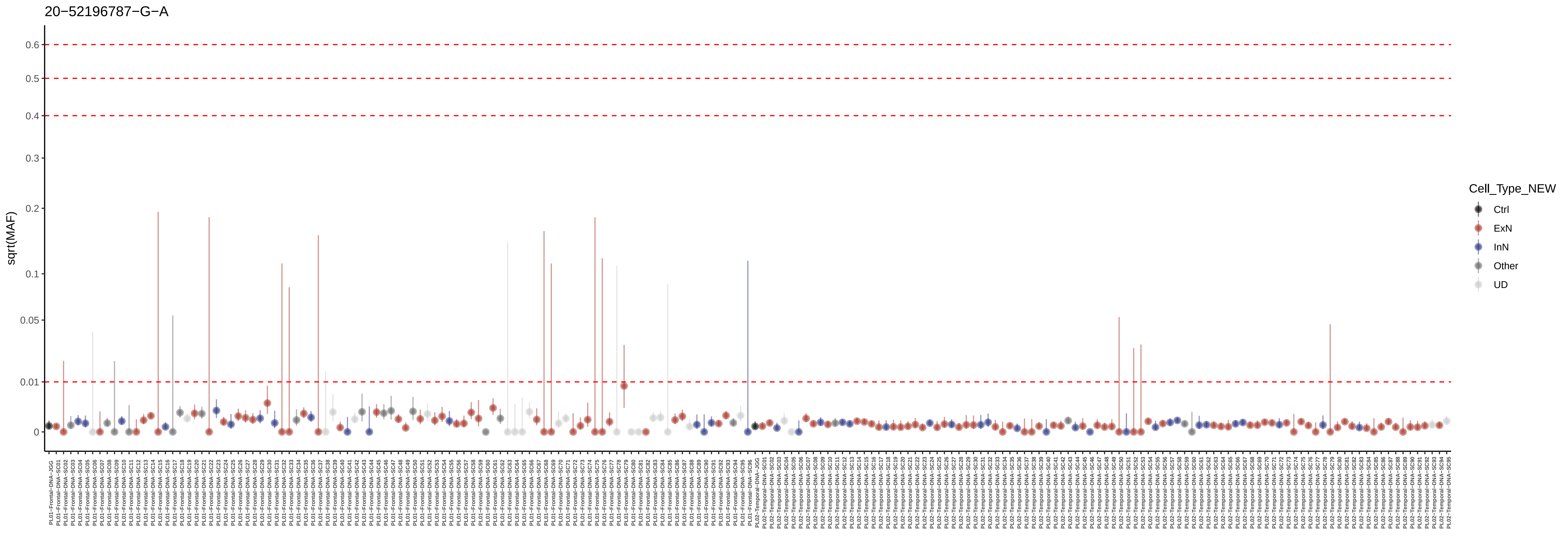
20−52196787−G−A
sqrt(MAF)
0.6
0.5
0.4
0.3
0.2
0.1
0.05
0.01
0
Cell_Type_NEW
Ctrl
ExN
InN
Other
UD

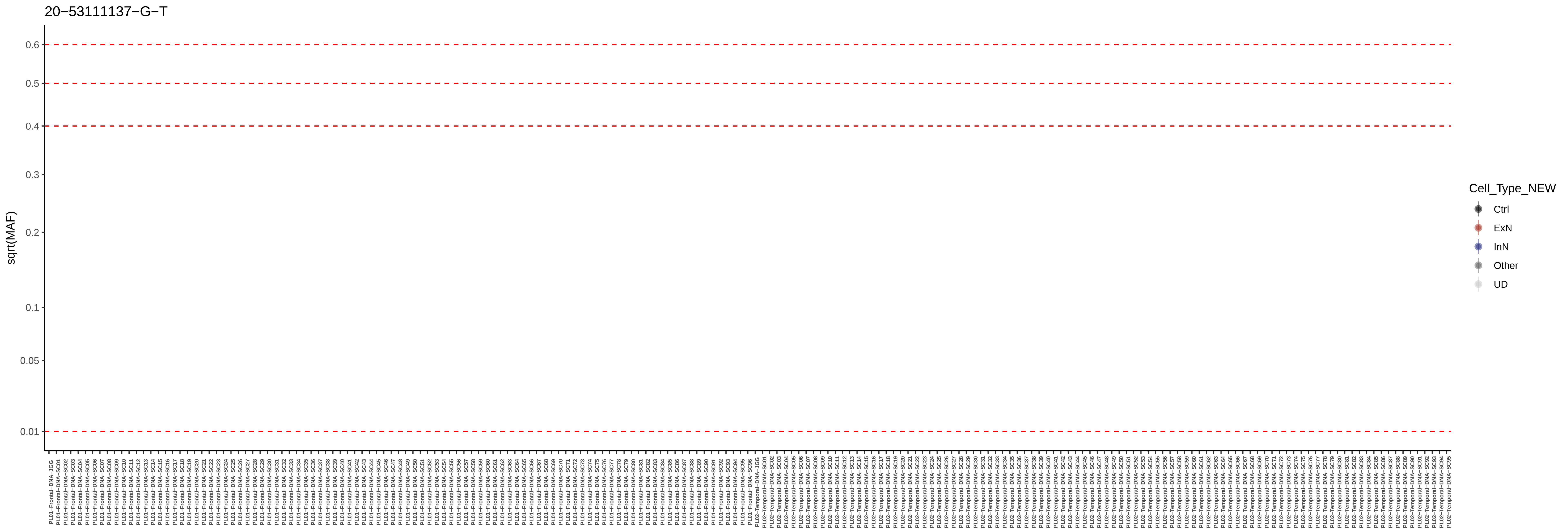
20−53111137−G−T
sqrt(MAF)
0.6
0.5
0.4
0.3
0.2
0.1
0.05
0.01
Cell_Type_NEW
Ctrl
ExN
InN
Other
UD

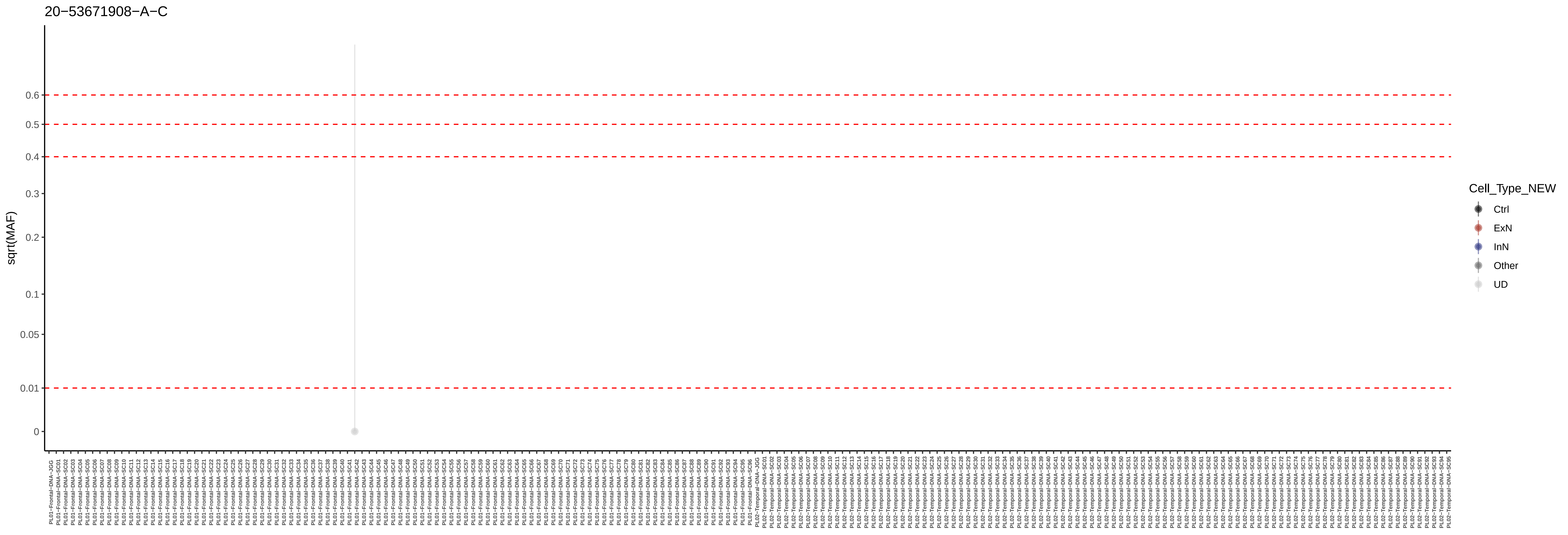

20−53671908−A−C
sqrt(MAF)
Cell_Type_NEW
Ctrl
ExN
InN
Other
UD

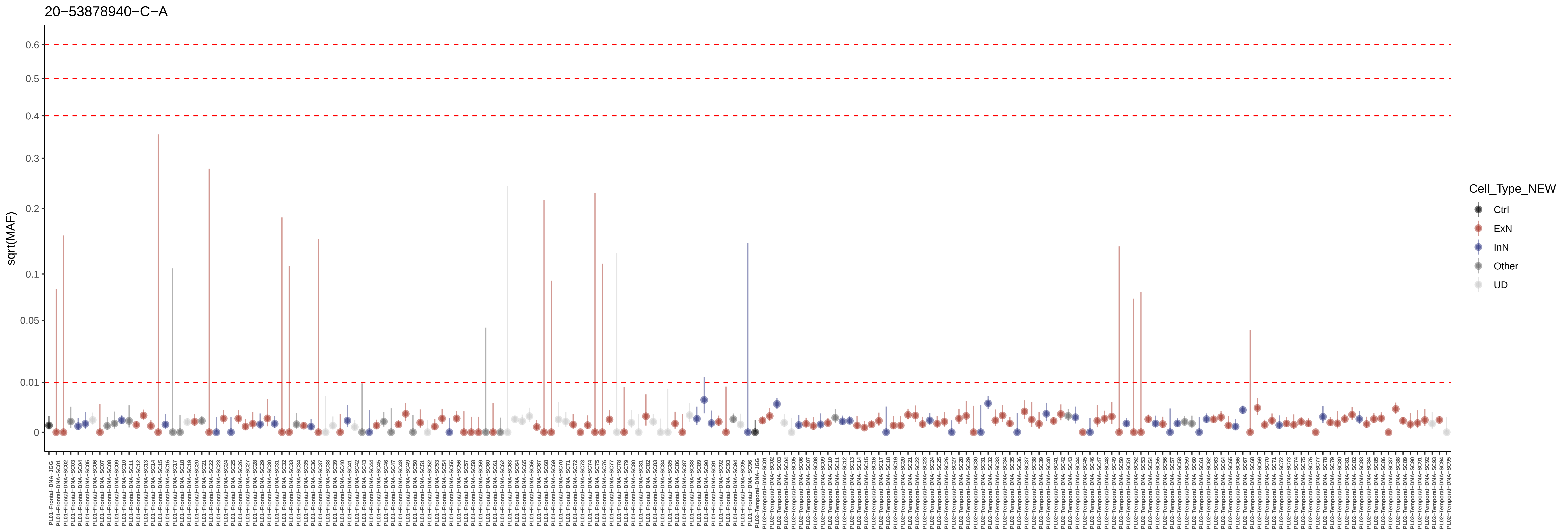

20−53878940−C−A
sqrt(MAF)
0.6
0.5
0.4
0.3
0.2
0.1
0.05
0.01
0
Cell_Type_NEW
Ctrl
ExN
InN
Other
UD

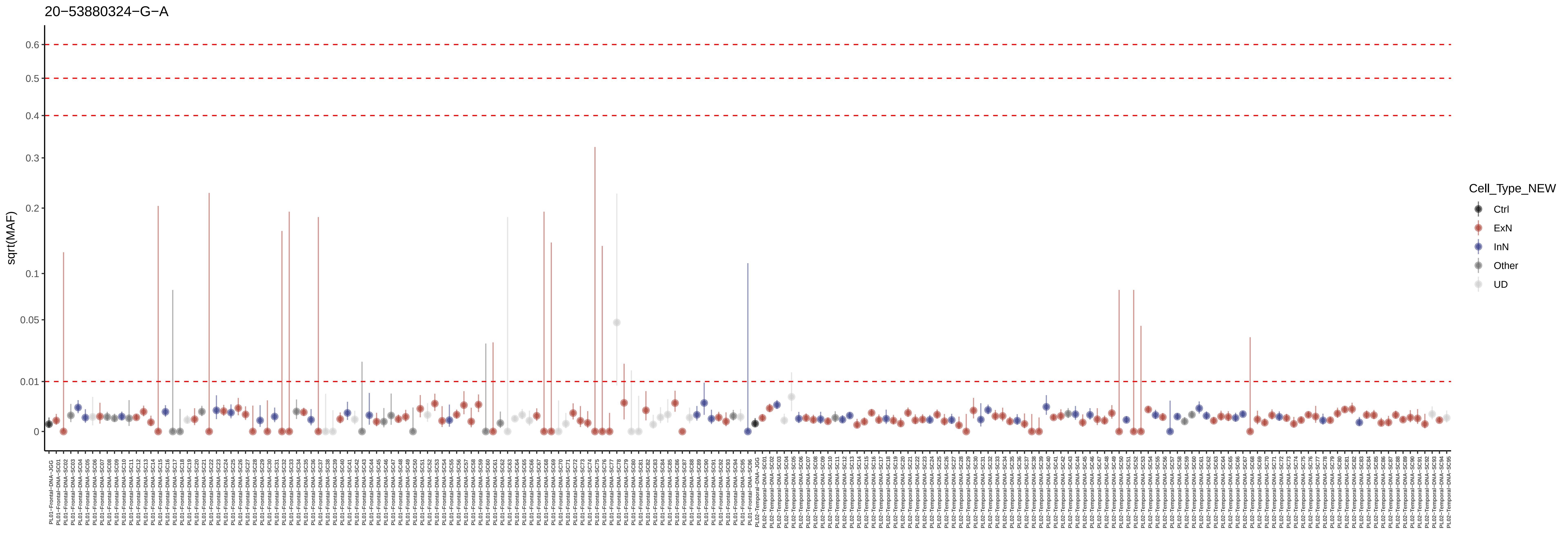

20−53880324−G−A
sqrt(MAF)
0.6
0.5
0.4
0.3
0.2
0.1
0.05
0.01
0
Cell_Type_NEW
Ctrl
ExN
InN
Other
UD

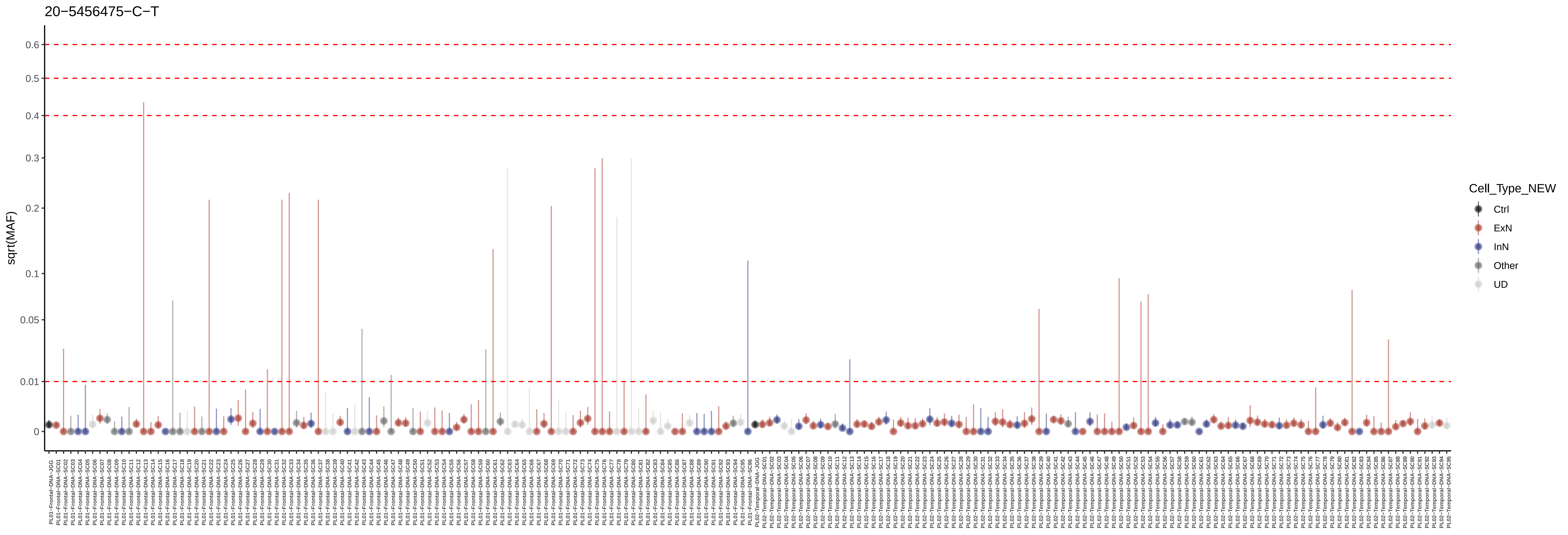

20−5456475−C−T
sqrt(MAF)
0.6
0.5
0.4
0.3
0.2
0.1
0.05
0.01
0
Cell_Type_NEW
Ctrl
ExN
InN
Other
UD

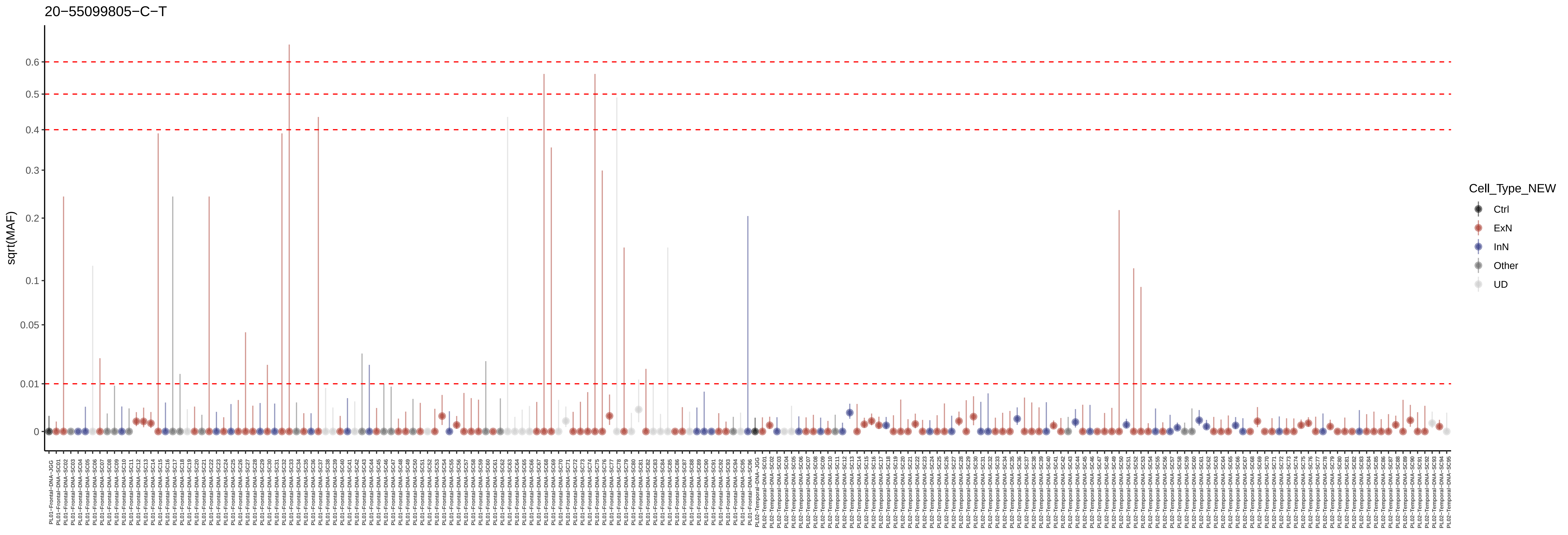
20−55099805−C−T
sqrt(MAF)
0.6
0.5
0.4
0.3
0.2
0.1
0.05
0.01
0
Cell_Type_NEW
Ctrl
ExN
InN
Other
UD

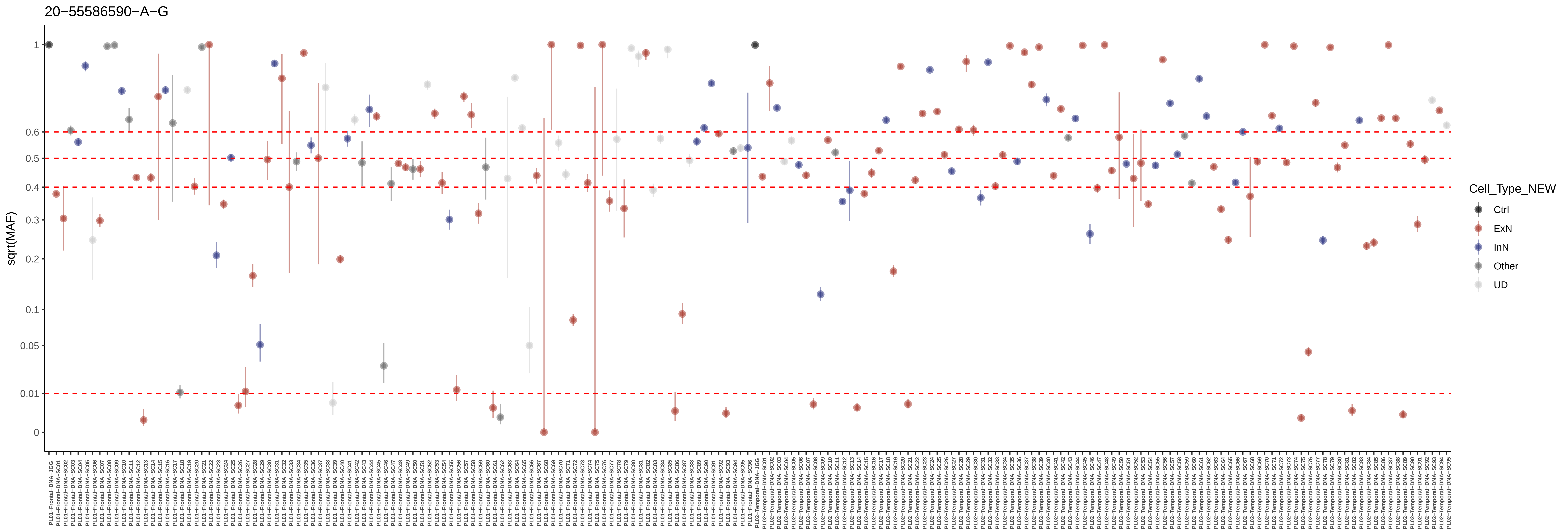
20−55586590−A−G
sqrt(MAF)
1
0.6
0.5
0.4
0.3
0.2
0.1
0.05
0.01
0
Cell_Type_NEW
Ctrl
ExN
InN
Other
UD

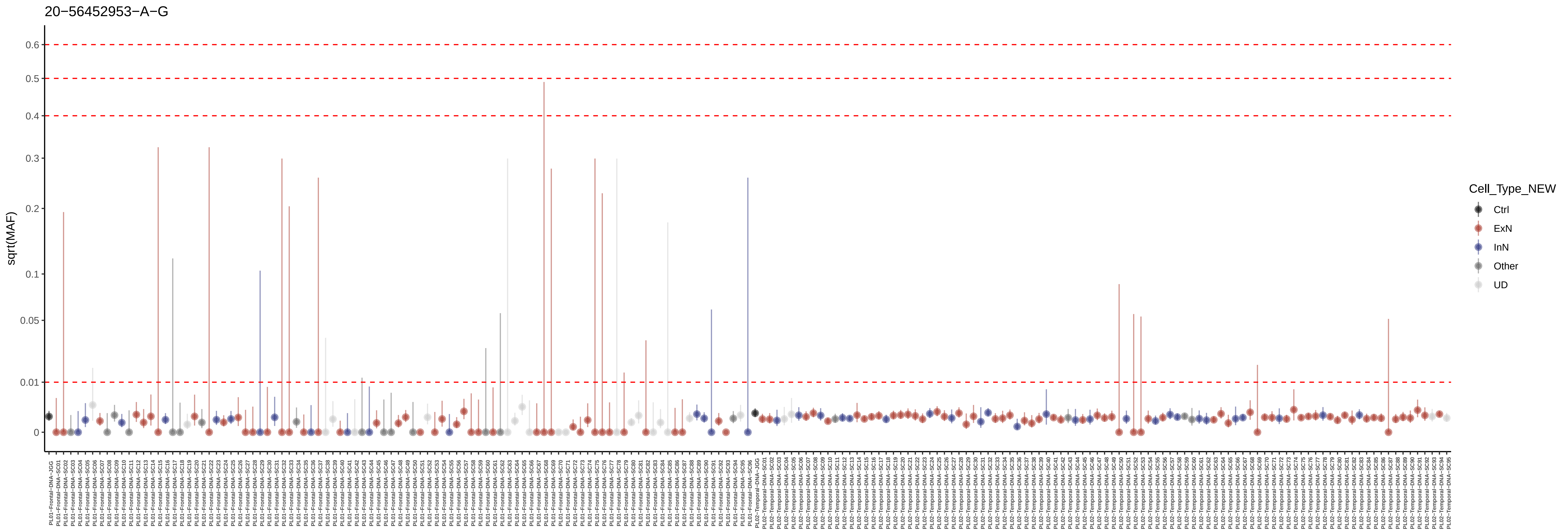
20−56452953−A−G
sqrt(MAF)
0.6
0.5
0.4
0.3
0.2
0.1
0.05
0.01
0
Cell_Type_NEW
Ctrl
ExN
InN
Other
UD

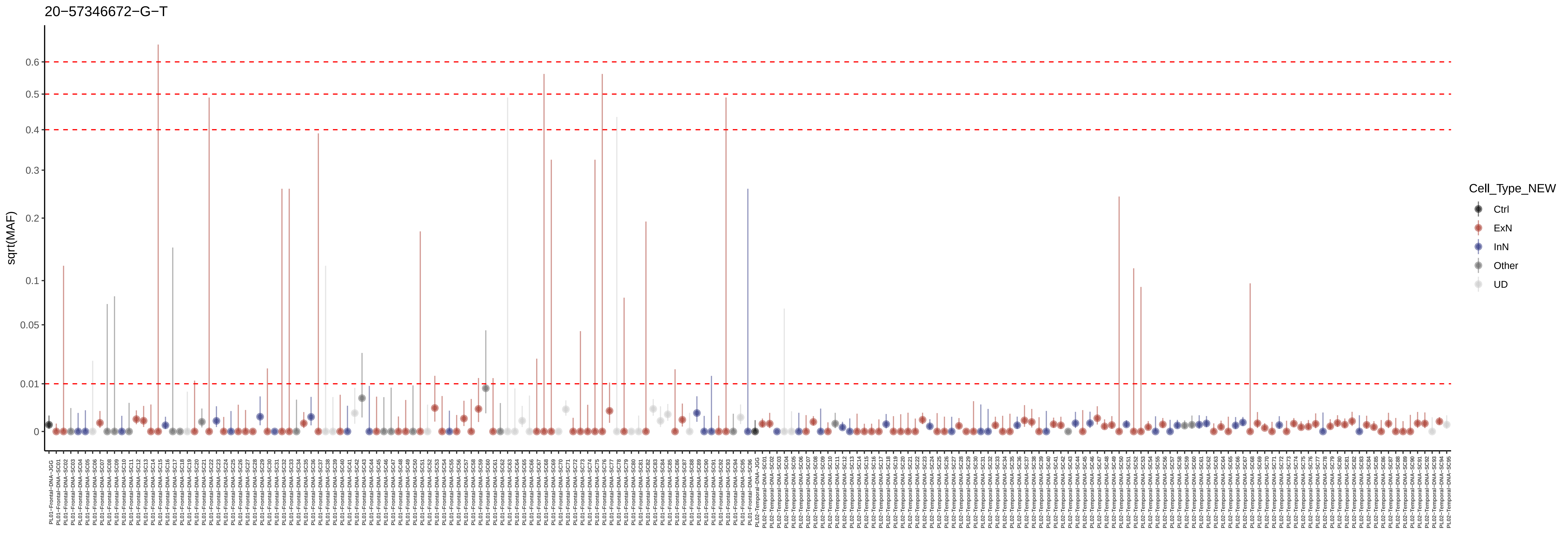

20−57346672−G−T
sqrt(MAF)
0.6
0.5
0.4
0.3
0.2
0.1
0.05
0.01
0
Cell_Type_NEW
Ctrl
ExN
InN
Other
UD

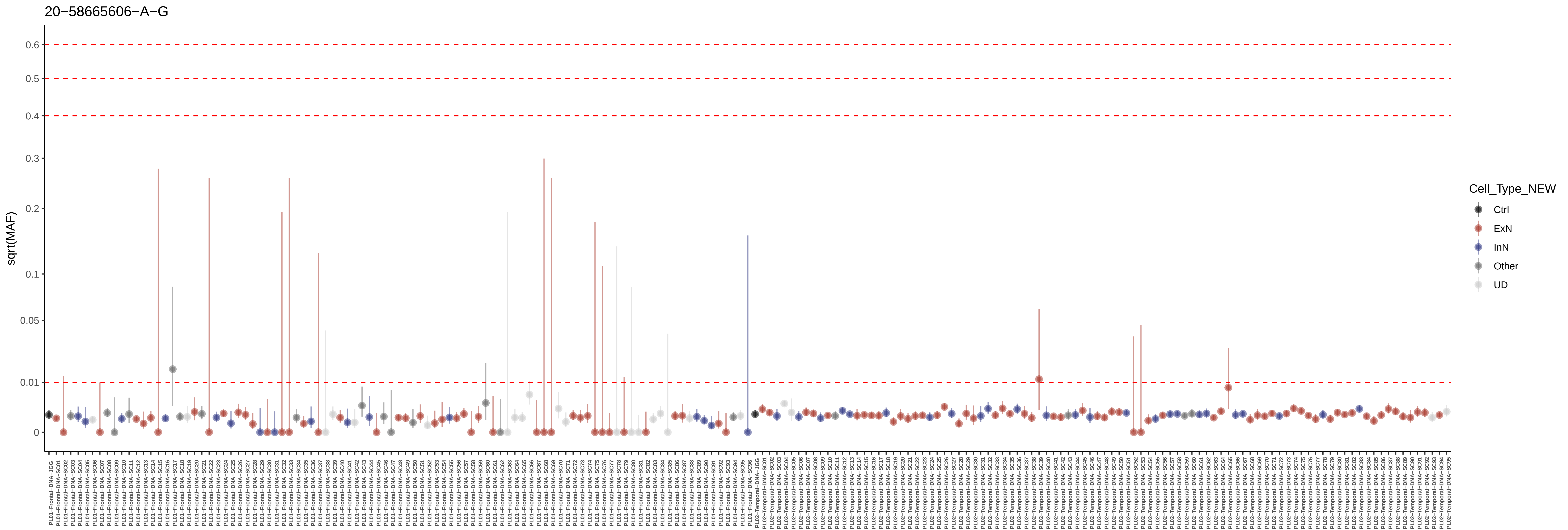
20−58665606−A−G
sqrt(MAF)
0.6
0.5
0.4
0.3
0.2
0.1
0.05
0.01
0
Cell_Type_NEW
Ctrl
ExN
InN
Other
UD

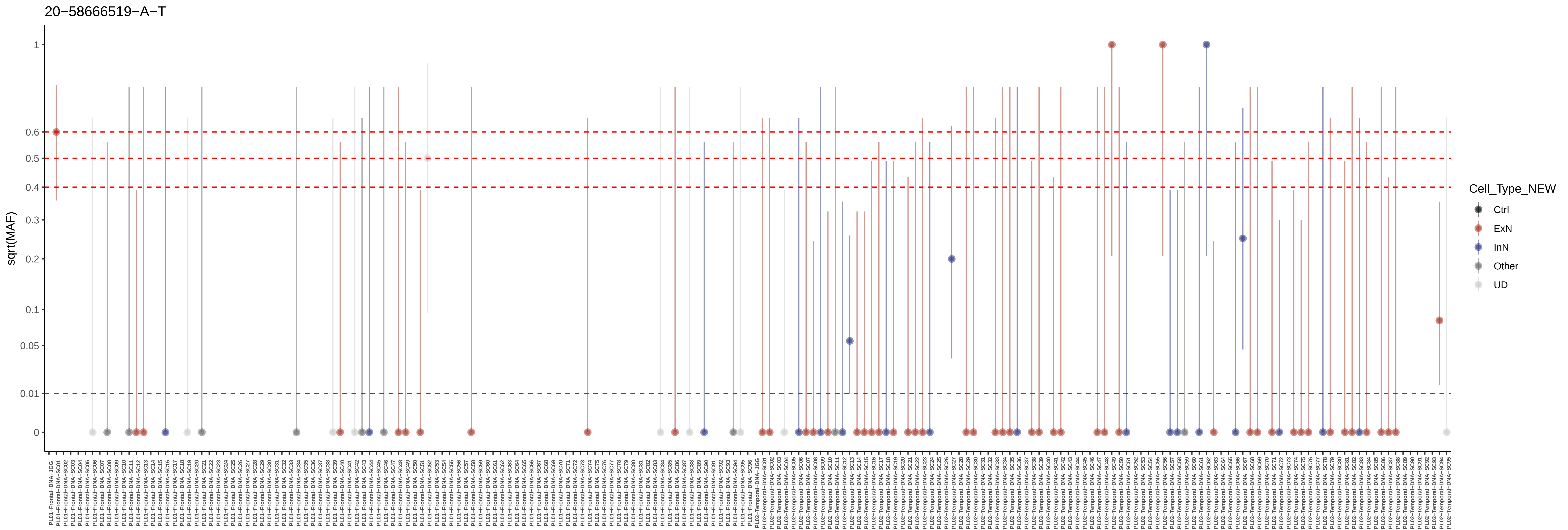
20−58666519−A−T
sqrt(MAF)
1
0.6
0.5
0.4
0.3
0.2
0.1
0.05
0.01
0
Cell_Type_NEW
Ctrl
ExN
InN
Other
UD

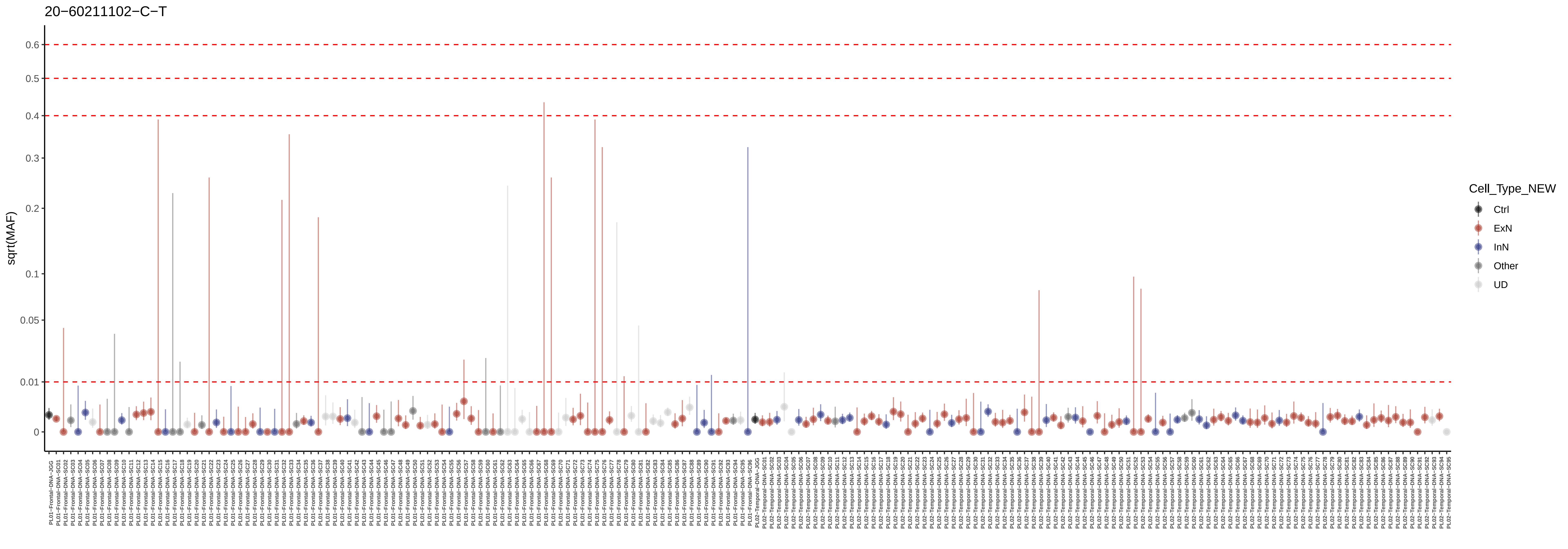
20−60211102−C−T
sqrt(MAF)
0.6
0.5
0.4
0.3
0.2
0.1
0.05
0.01
0
Cell_Type_NEW
Ctrl
ExN
InN
Other
UD

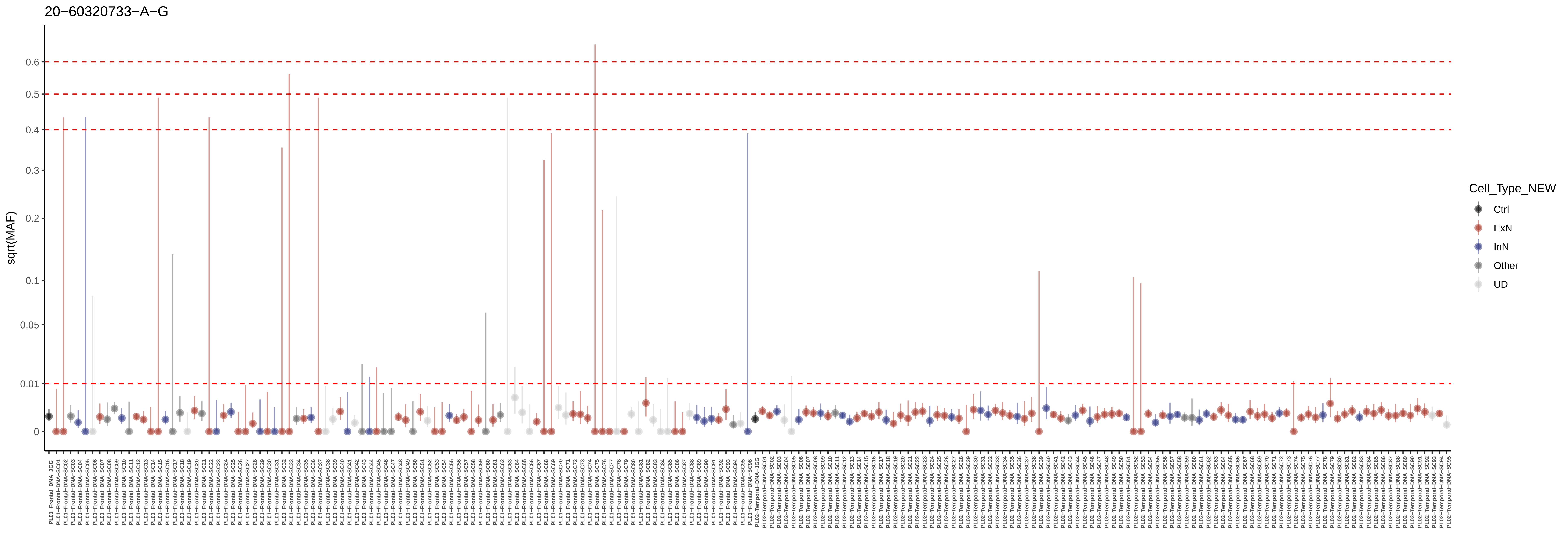
20−60320733−A−G
sqrt(MAF)
0.6
0.5
0.4
0.3
0.2
0.1
0.05
0.01
0
Cell_Type_NEW
Ctrl
ExN
InN
Other
UD

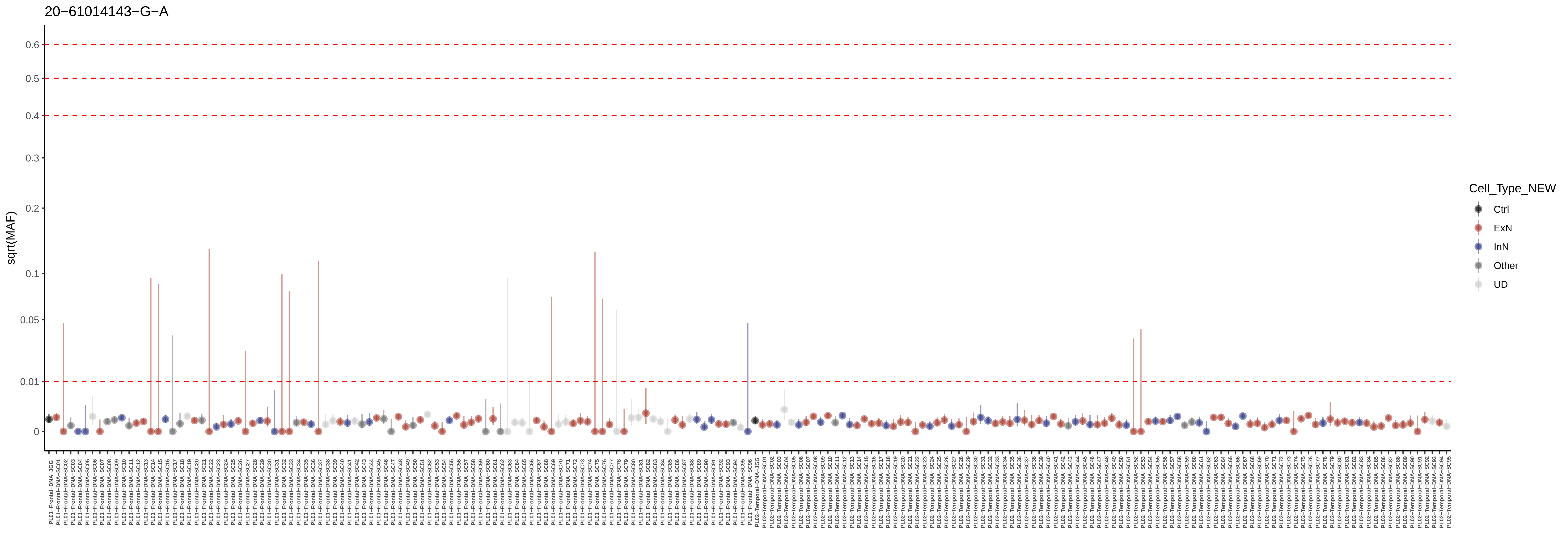
20−61014143−G−A
sqrt(MAF)
0.6
0.5
0.4
0.3
0.2
0.1
0.05
0.01
0
Cell_Type_NEW
Ctrl
ExN
InN
Other
UD

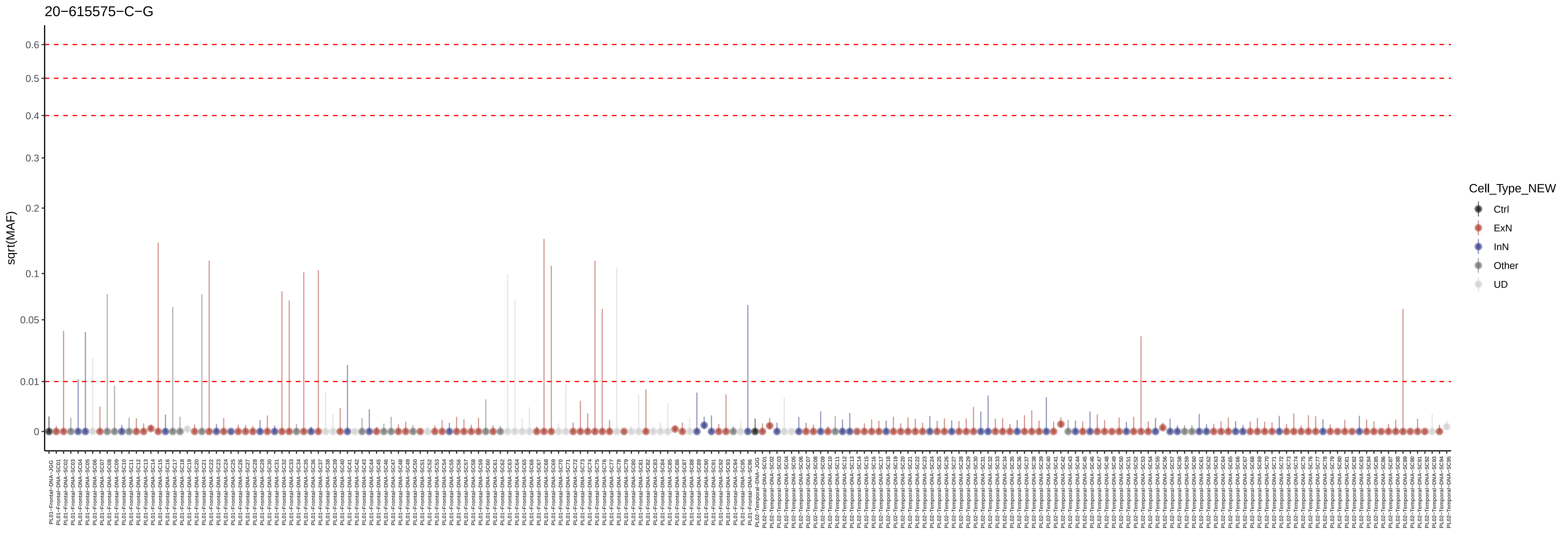
20−615575−C−G
sqrt(MAF)
0.6
0.5
0.4
0.3
0.2
0.1
0.05
0.01
0
Cell_Type_NEW
Ctrl
ExN
InN
Other
UD

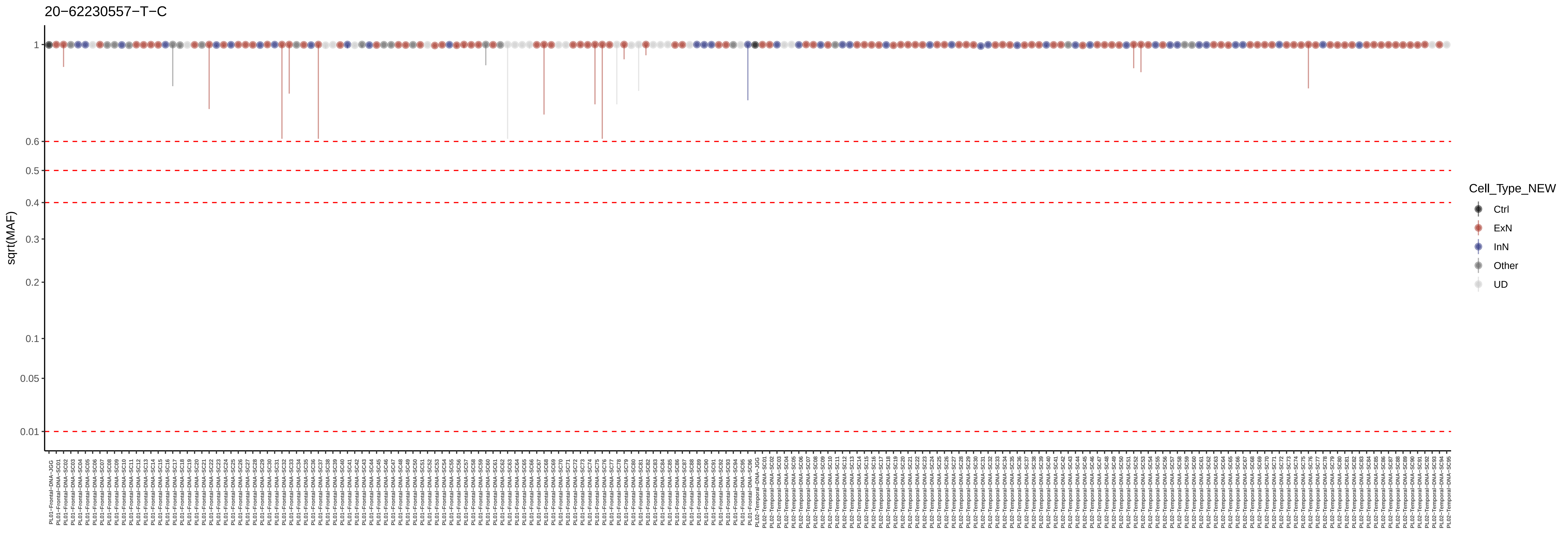
20−62230557−T−C
sqrt(MAF)
1
0.6
0.5
0.4
0.3
0.2
0.1
0.05
0.01
Cell_Type_NEW
Ctrl
ExN
InN
Other
UD

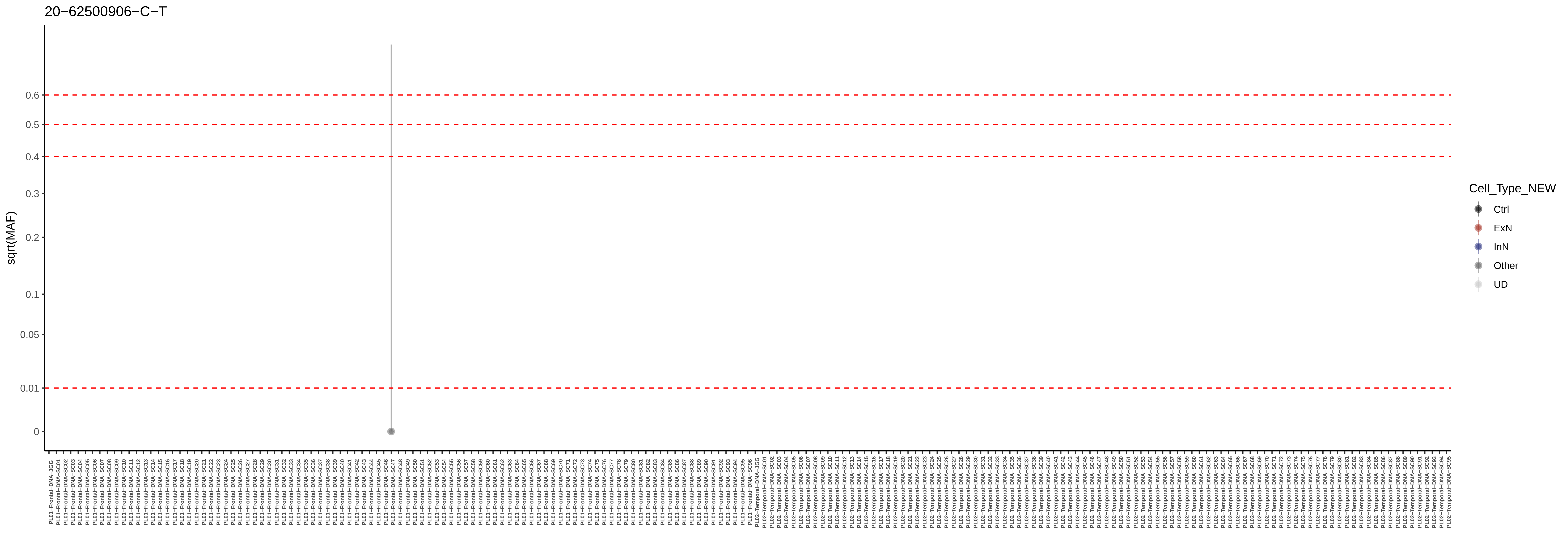
20−62500906−C−T
sqrt(MAF)
0.6
0.5
0.4
0.3
0.2
0.1
0.05
0.01
0
Cell_Type_NEW
Ctrl
ExN
InN
Other
UD

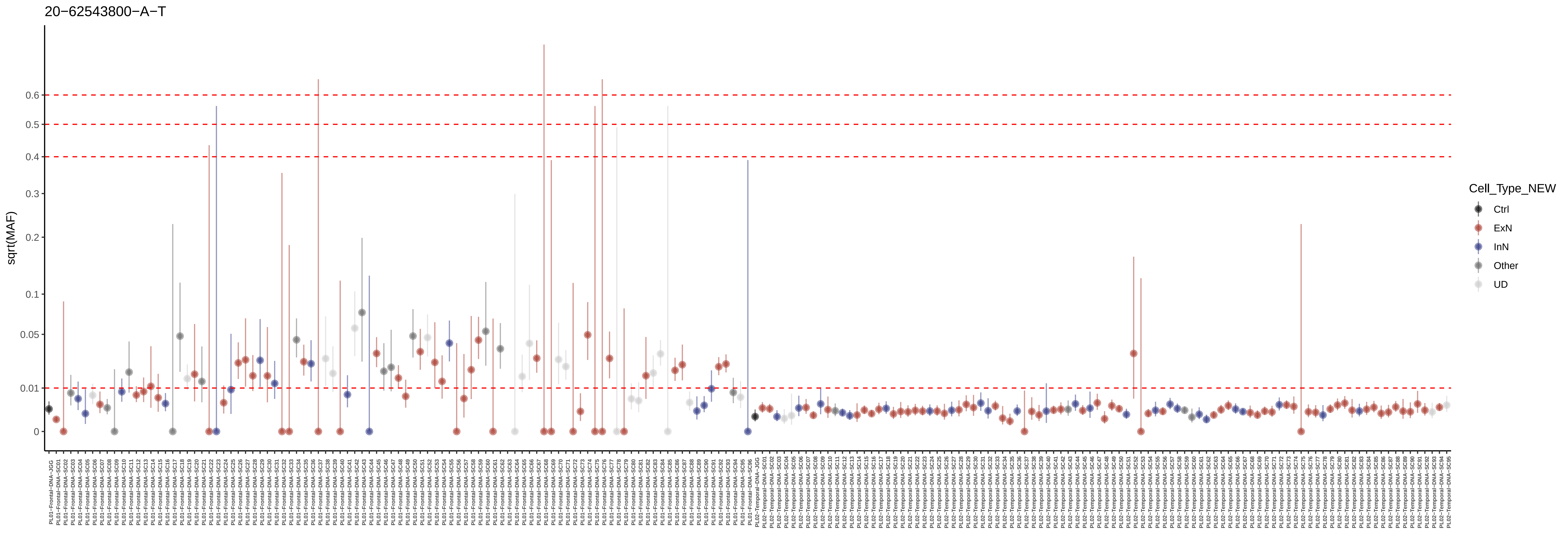

20−62543800−A−T
sqrt(MAF)
0.6
0.5
0.4
0.3
0.2
0.1
0.05
0.01
0
Cell_Type_NEW
Ctrl
ExN
InN
Other
UD

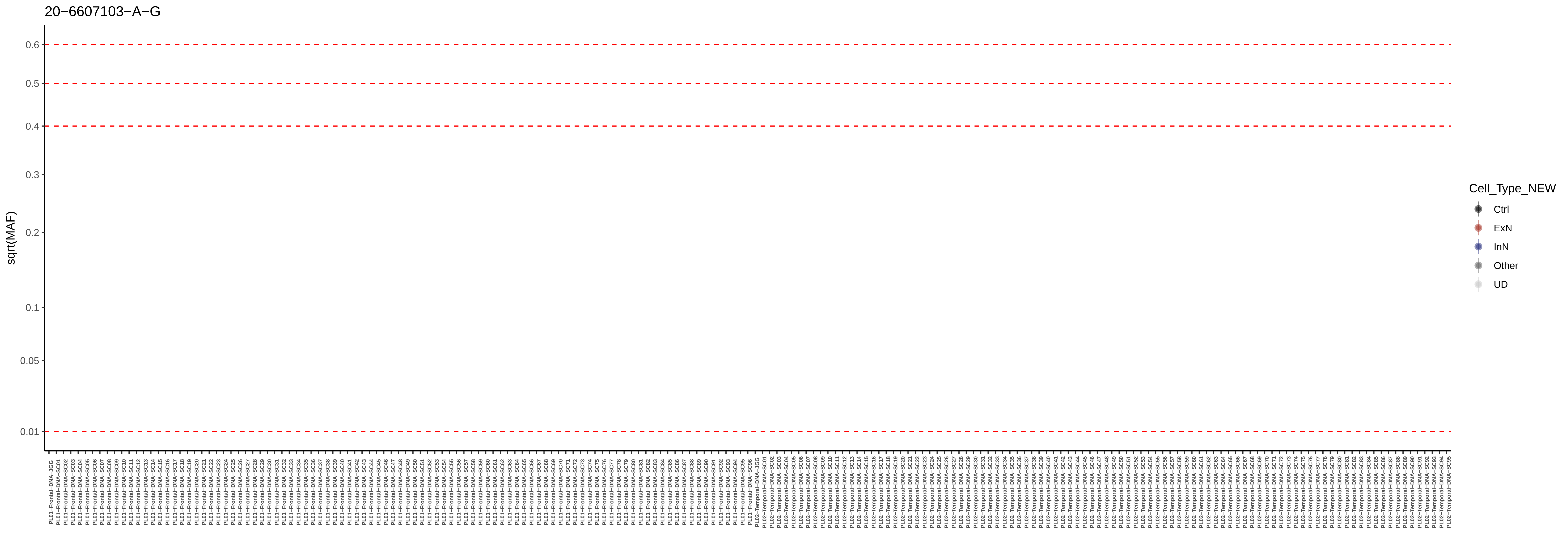

20−6607103−A−G
sqrt(MAF)
0.6
0.5
0.4
0.3
0.2
0.1
0.05
0.01
Cell_Type_NEW
Ctrl
ExN
InN
Other
UD

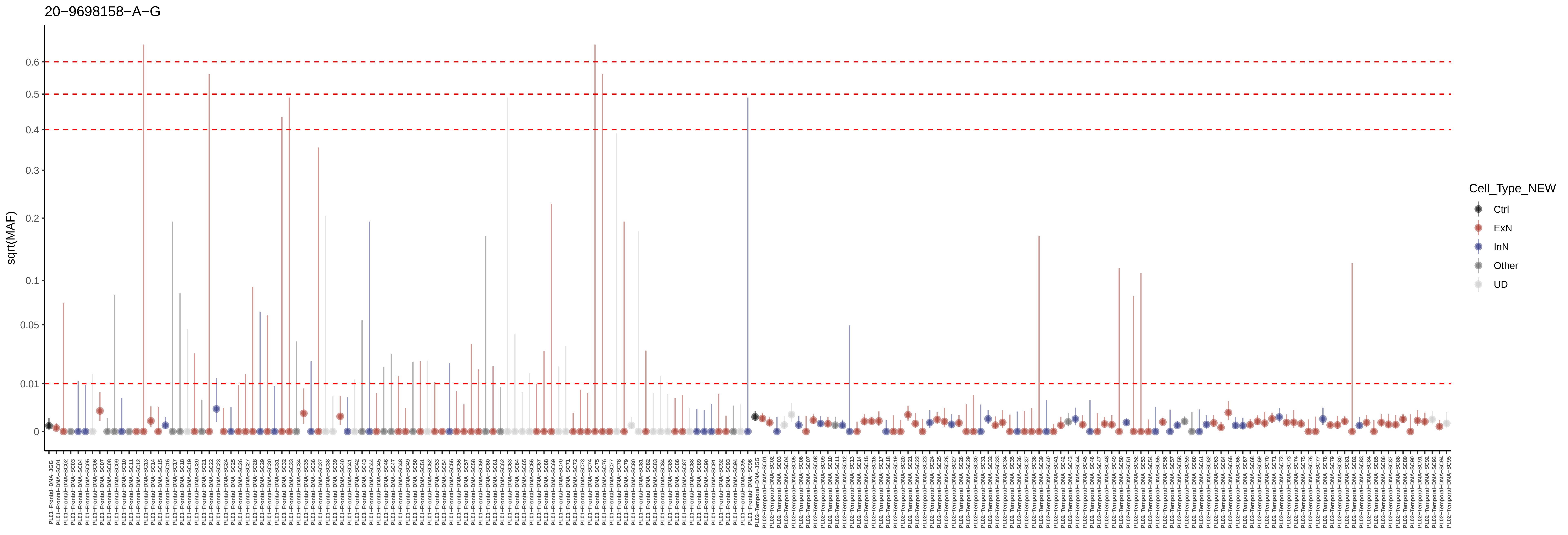
20−9698158−A−G
sqrt(MAF)
0.6
0.5
0.4
0.3
0.2
0.1
0.05
0.01
0
Cell_Type_NEW
Ctrl
ExN
InN
Other
UD

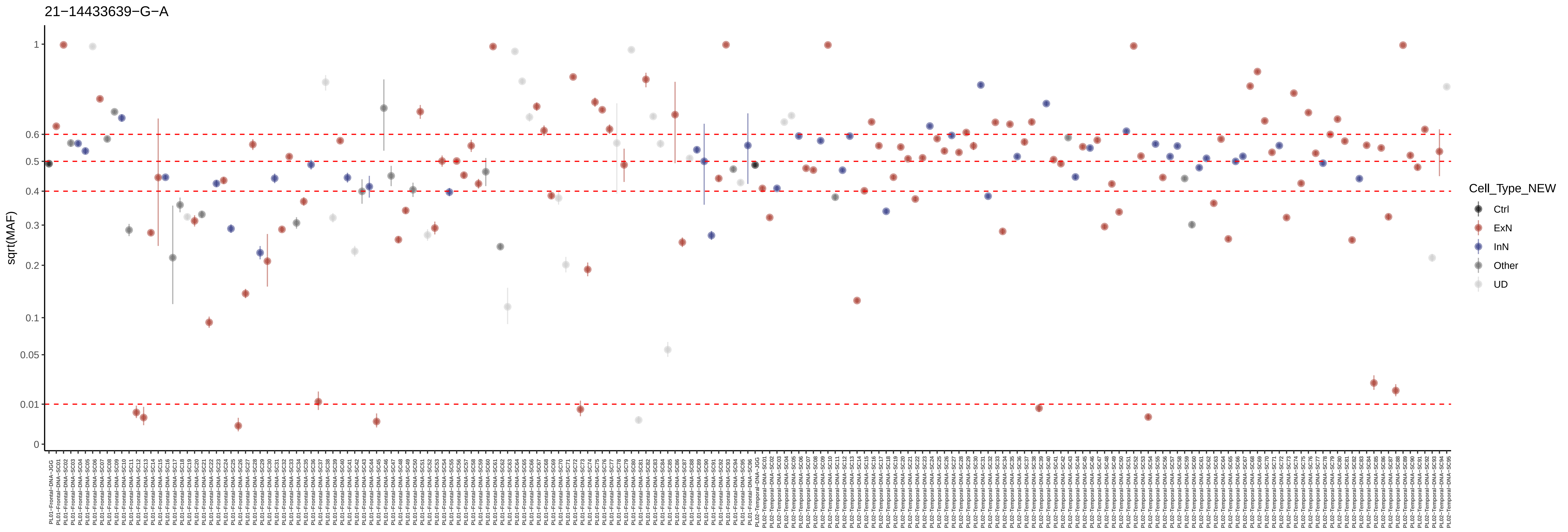
21−14433639−G−A
sqrt(MAF)
1
0.6
0.5
0.4
0.3
0.2
0.1
0.05
0.01
0
Cell_Type_NEW
Ctrl
ExN
InN
Other
UD

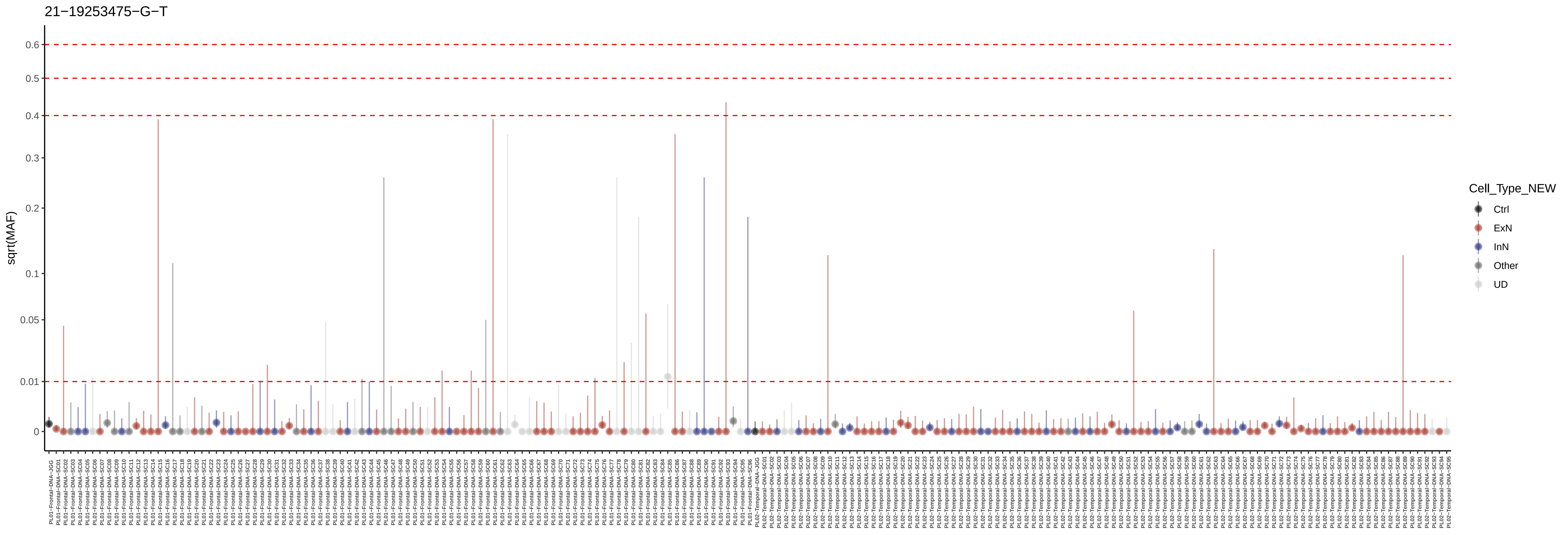
21−19253475−G−T
sqrt(MAF)
Cell_Type_NEW
Ctrl
ExN
InN
Other
UD

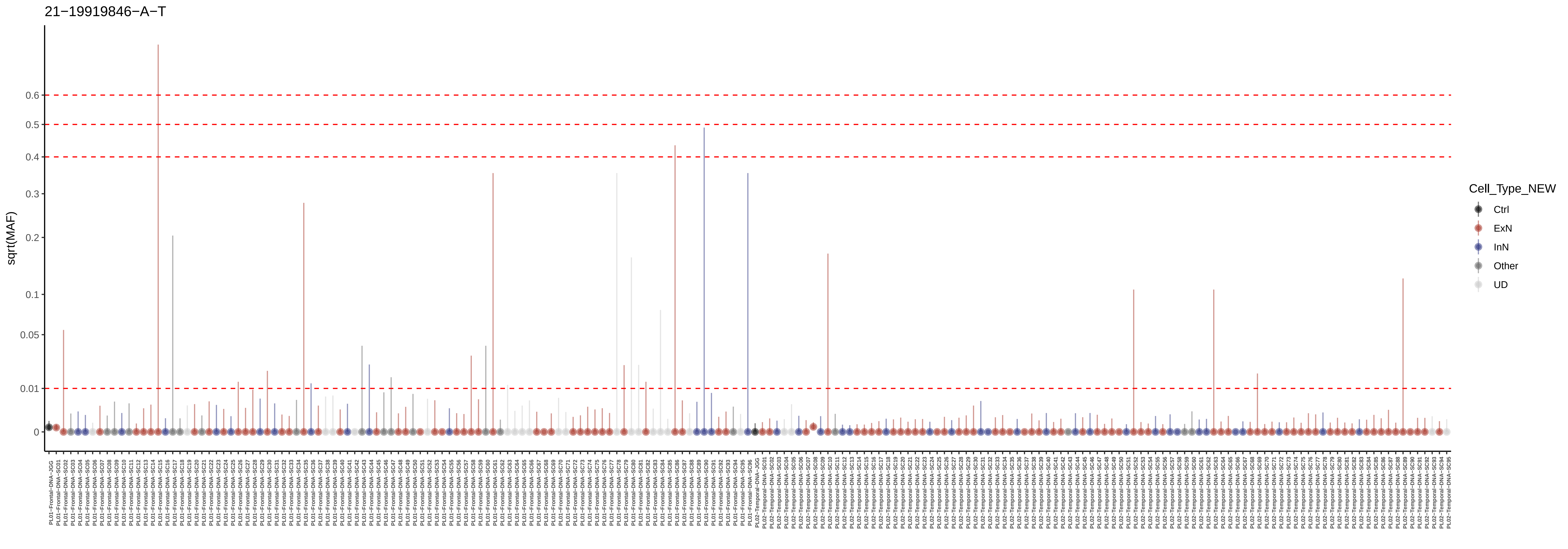
21−19919846−A−T
sqrt(MAF)
0.6
0.5
0.4
0.3
0.2
0.1
0.05
0.01
0
Cell_Type_NEW
Ctrl
ExN
InN
Other
UD

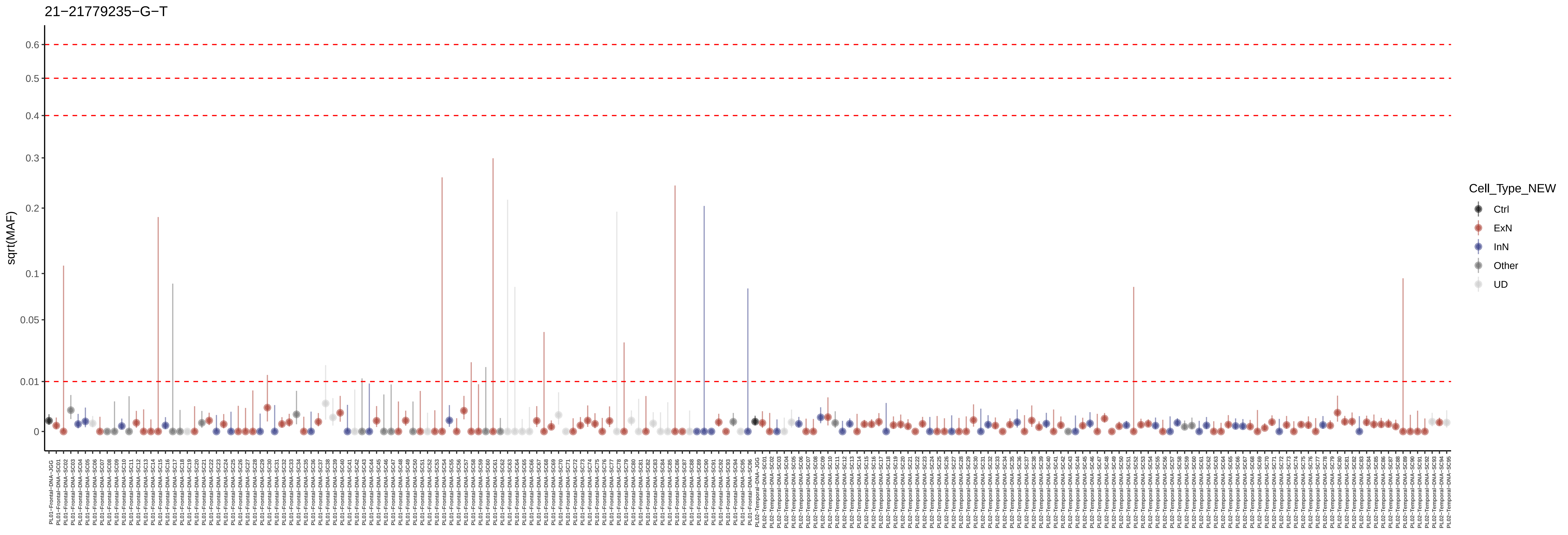
21−21779235−G−T
sqrt(MAF)
0.6
0.5
0.4
0.3
0.2
0.1
0.05
0.01
0
Cell_Type_NEW
Ctrl
ExN
InN
Other
UD

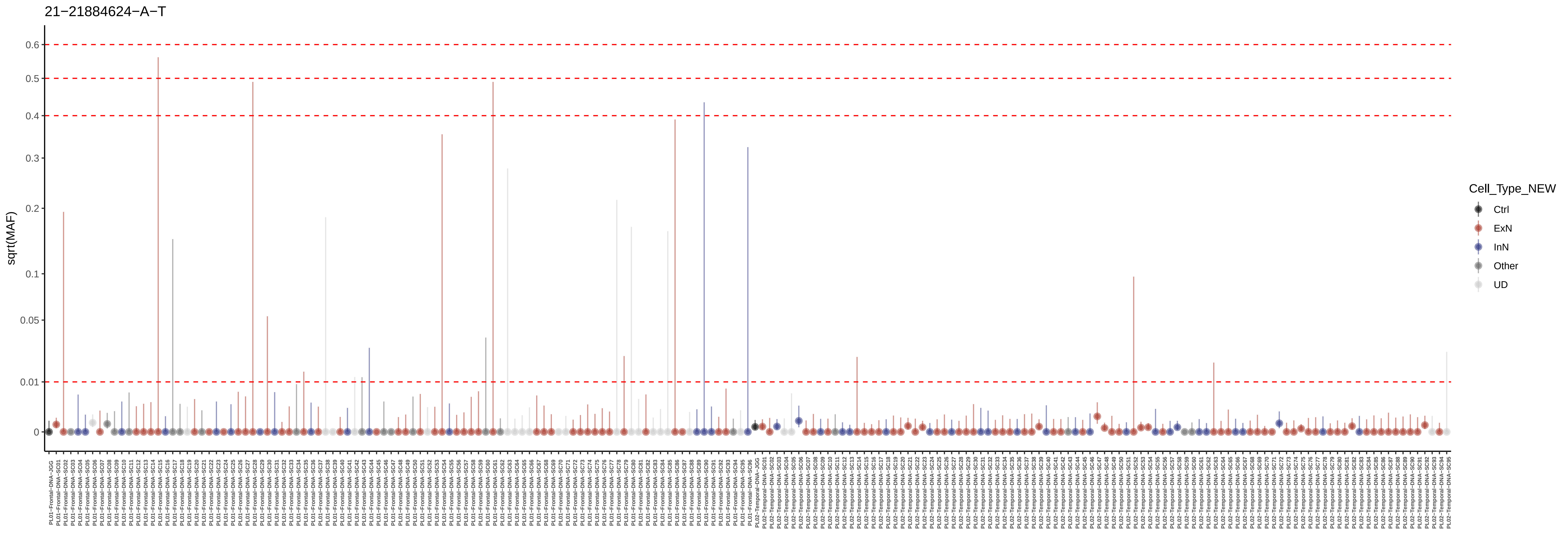
21−21884624−A−T
sqrt(MAF)
0.6
0.5
0.4
0.3
0.2
0.1
0.05
0.01
0
Cell_Type_NEW
Ctrl
ExN
InN
Other
UD

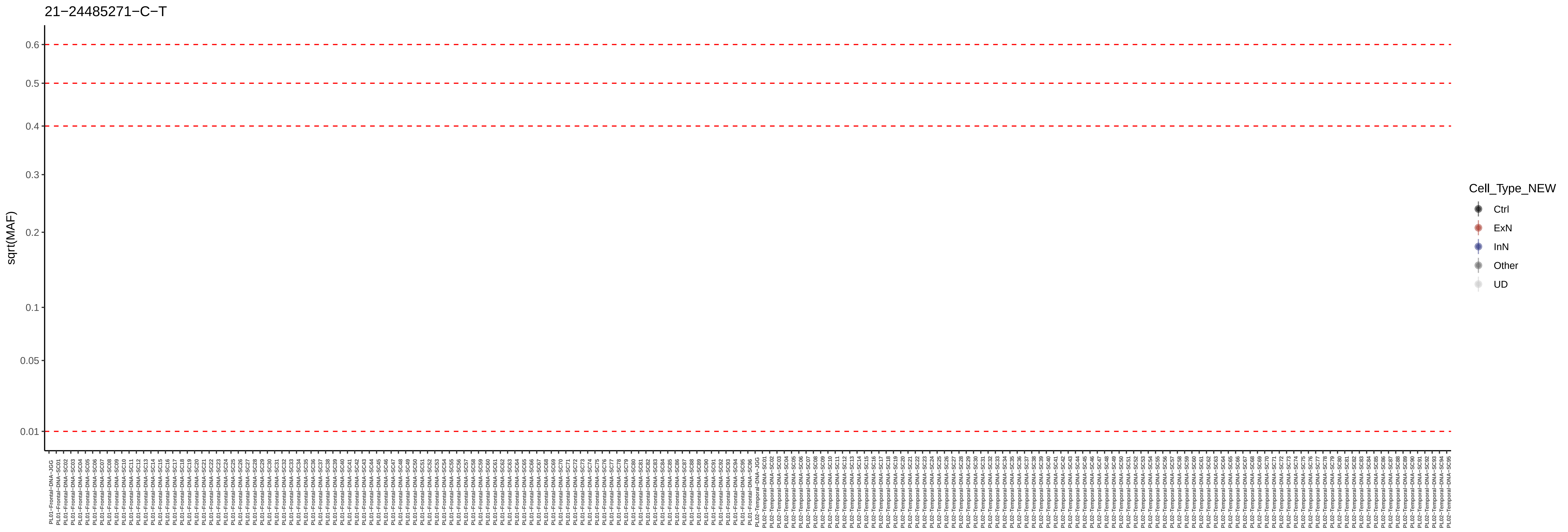
21−24485271−C−T
sqrt(MAF)
0.6
0.5
0.4
0.3
0.2
0.1
0.05
0.01
Cell_Type_NEW
Ctrl
ExN
InN
Other
UD

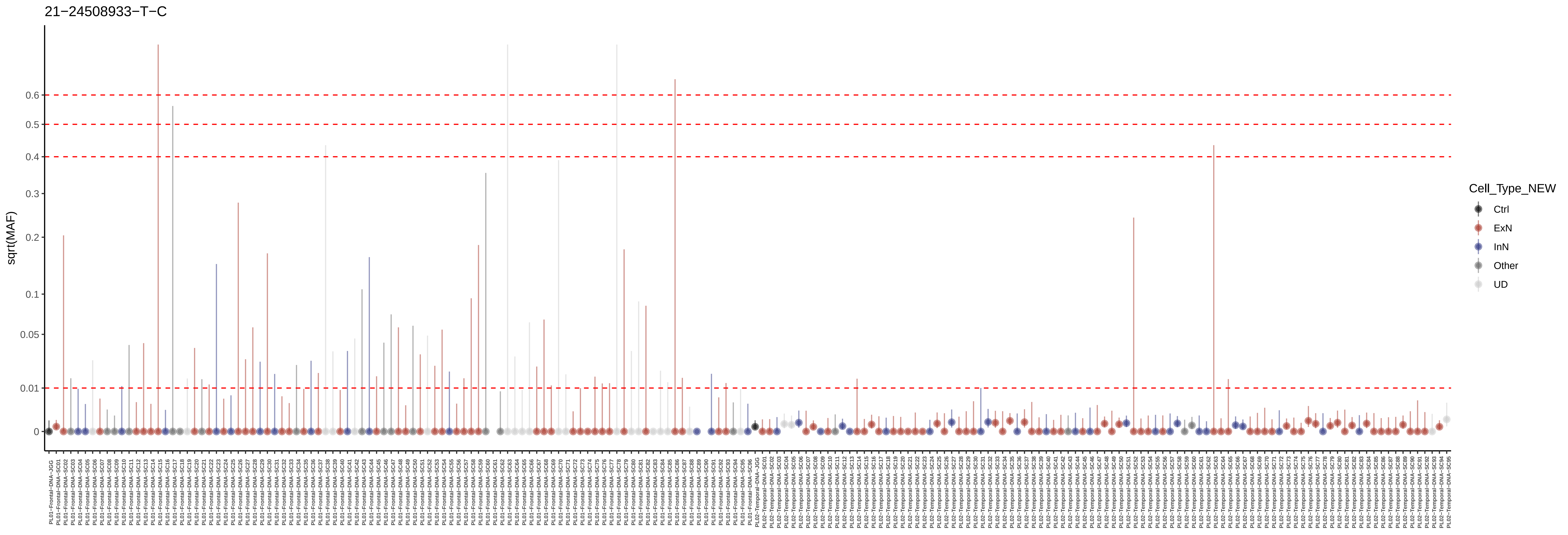

21−24508933−T−C
sqrt(MAF)
0.6
0.5
0.4
0.3
0.2
0.1
0.05
0.01
0
Cell_Type_NEW
Ctrl
ExN
InN
Other
UD

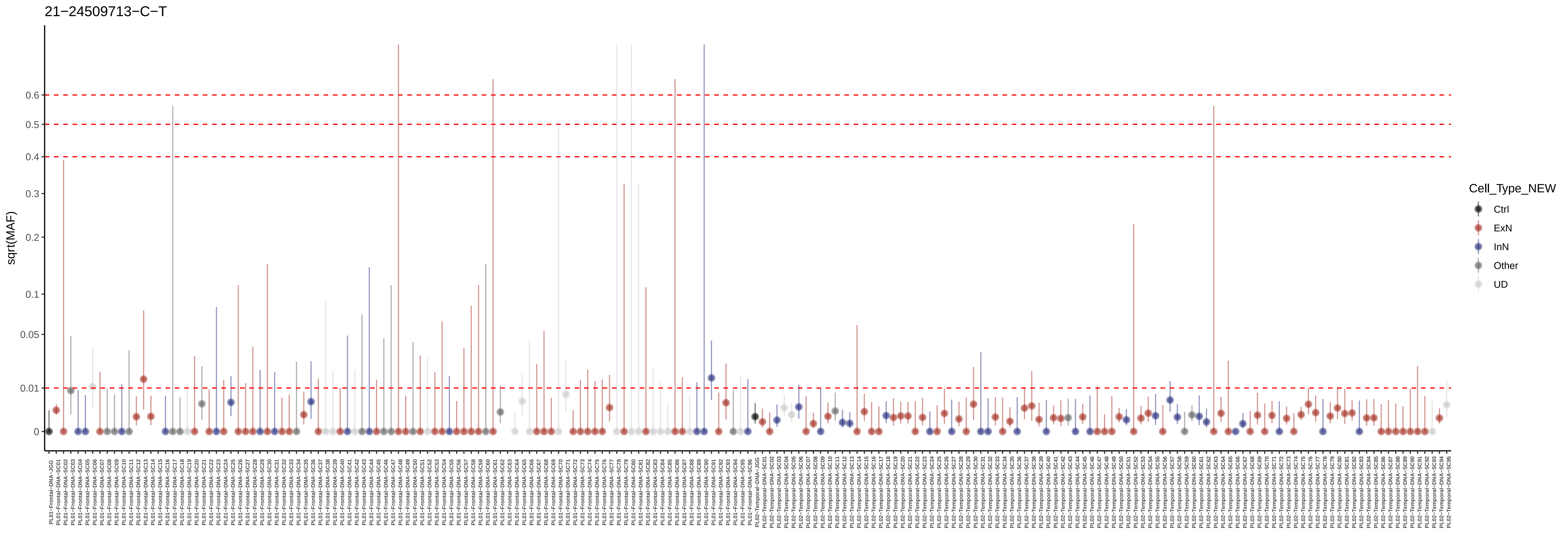
21−24509713−C−T
sqrt(MAF)
0.6
0.5
0.4
0.3
0.2
0.1
0.05
0.01
0
Cell_Type_NEW
Ctrl
ExN
InN
Other
UD

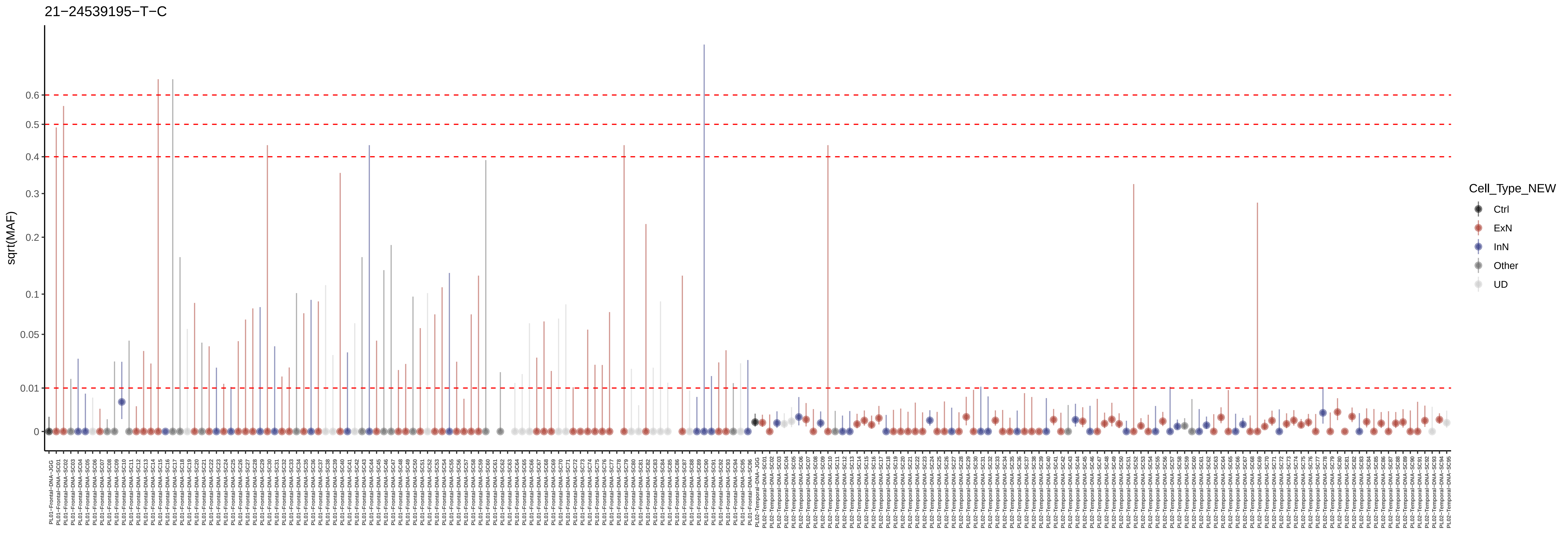

21−24539195−T−C
sqrt(MAF)
0.6
0.5
0.4
0.3
0.2
0.1
0.05
0.01
0
Cell_Type_NEW
Ctrl
ExN
InN
Other
UD

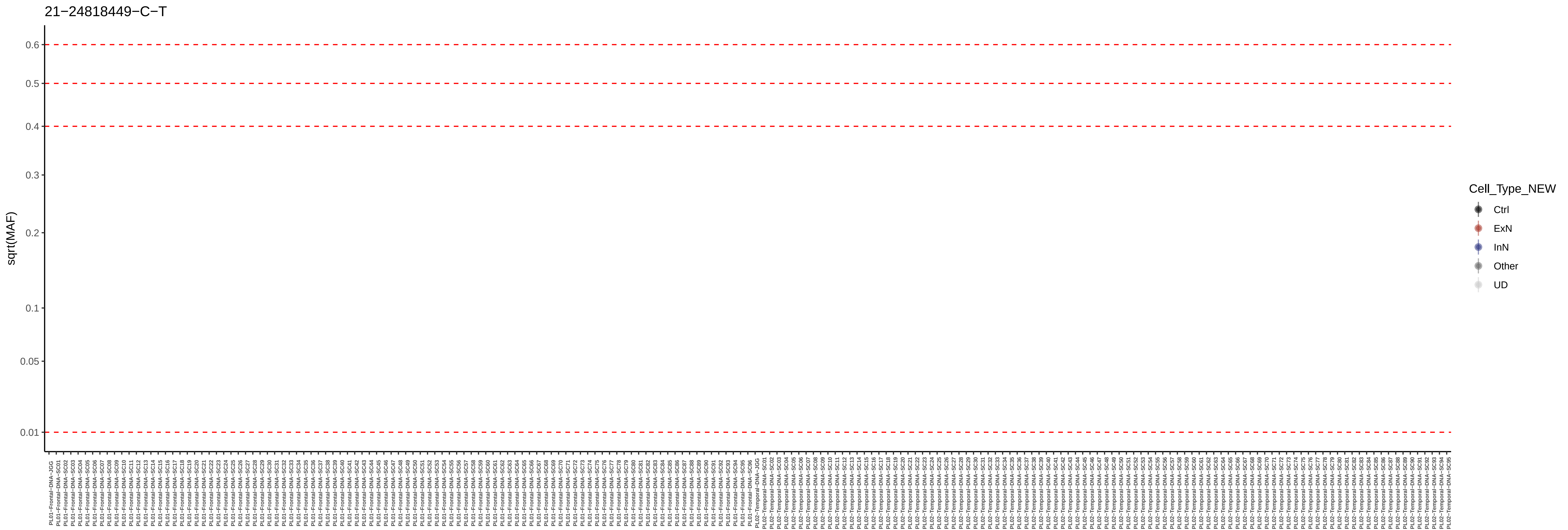

21−24818449−C−T
sqrt(MAF)
0.6
0.5
0.4
0.3
0.2
0.1
0.05
0.01
Cell_Type_NEW
Ctrl
ExN
InN
Other
UD

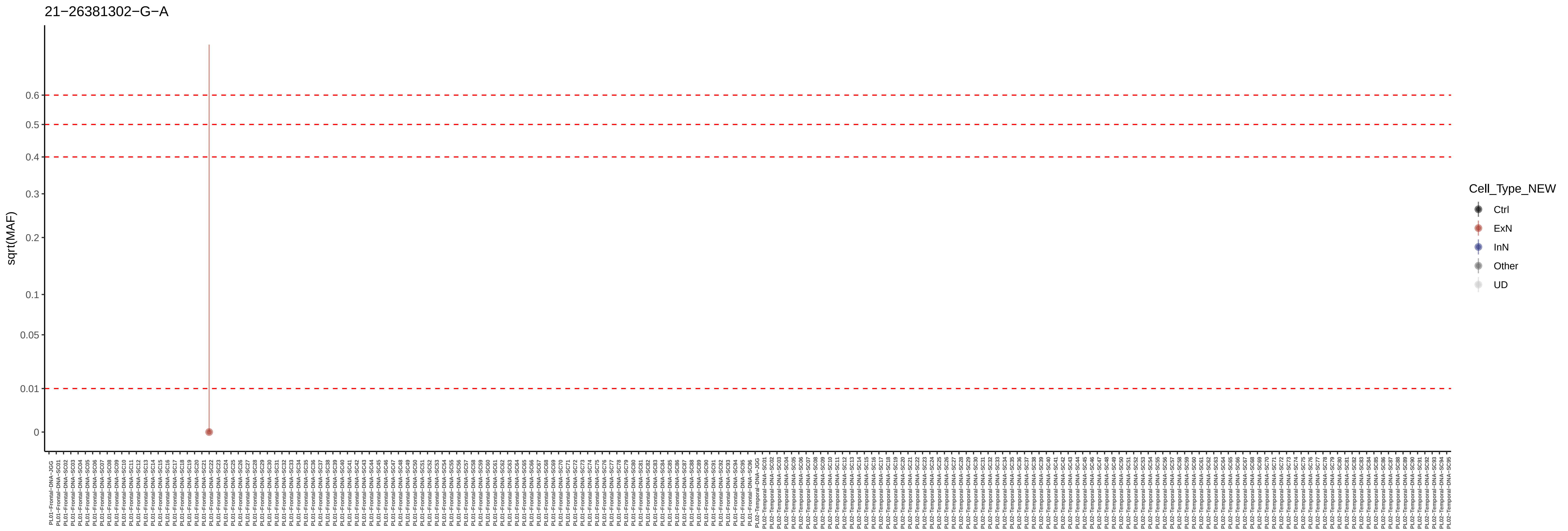
21−26381302−G−A
sqrt(MAF)
0.6
0.5
0.4
0.3
0.2
0.1
0.05
0.01
0
Cell_Type_NEW
Ctrl
ExN
InN
Other
UD

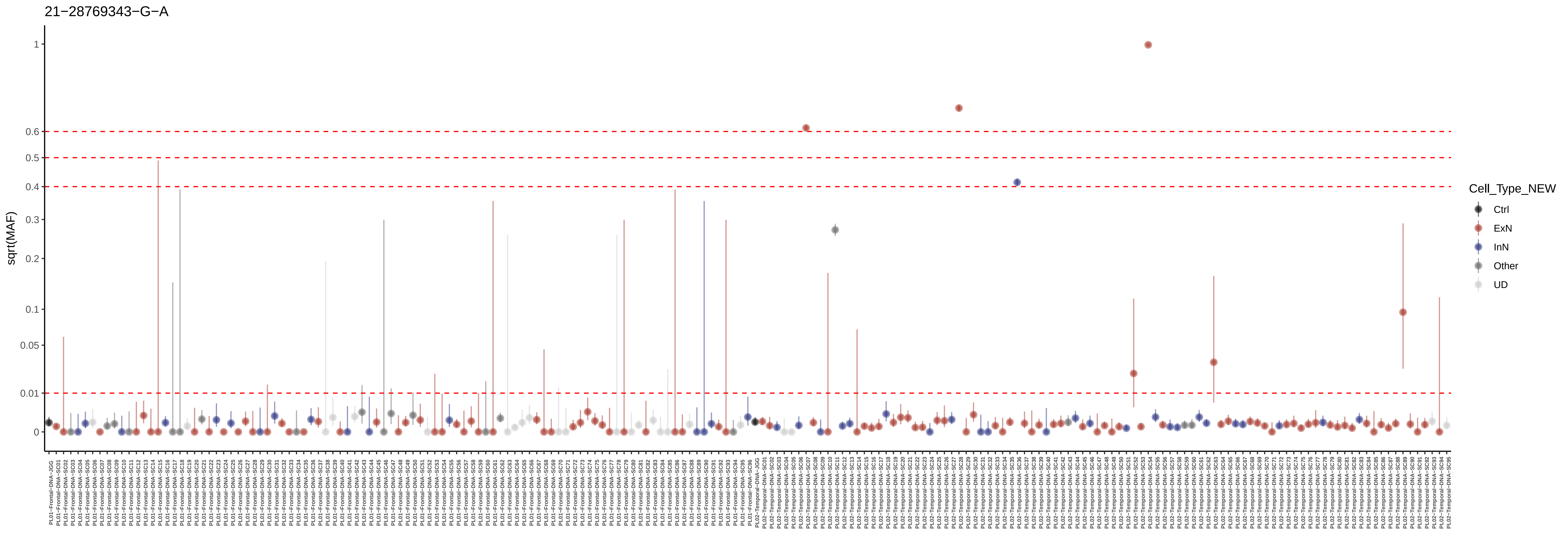
21−28769343−G−A
sqrt(MAF)
1
0.6
0.5
0.4
0.3
0.2
0.1
0.05
0.01
0
Cell_Type_NEW
Ctrl
ExN
InN
Other
UD

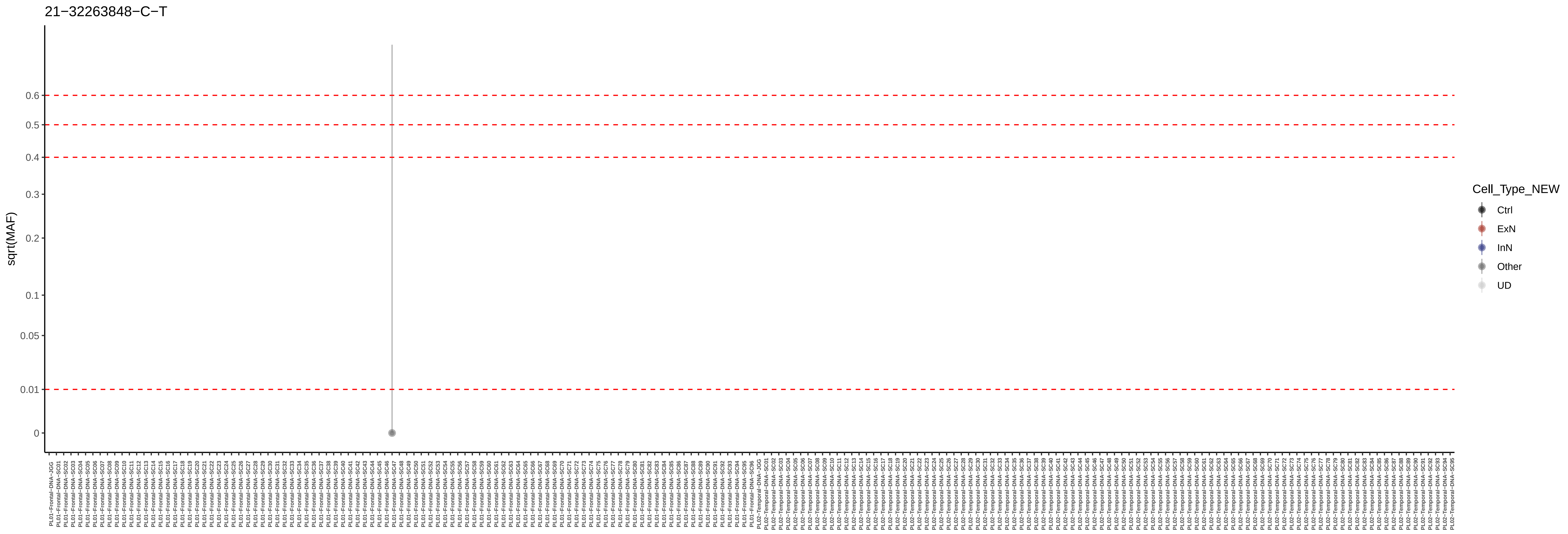
21−32263848−C−T
sqrt(MAF)
0.6
0.5
0.4
0.3
0.2
0.1
0.05
0.01
0
Cell_Type_NEW
Ctrl
ExN
InN
Other
UD

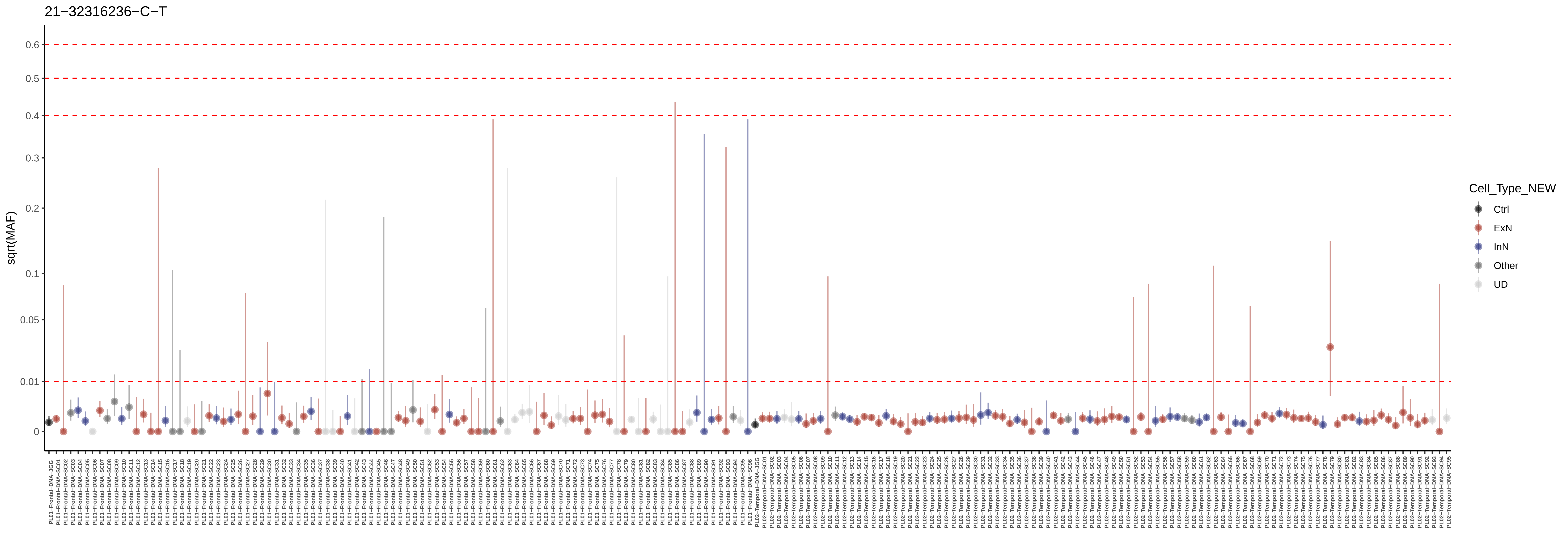

21−32316236−C−T
sqrt(MAF)
0.6
0.5
0.4
0.3
0.2
0.1
0.05
0.01
0
Cell_Type_NEW
Ctrl
ExN
InN
Other
UD

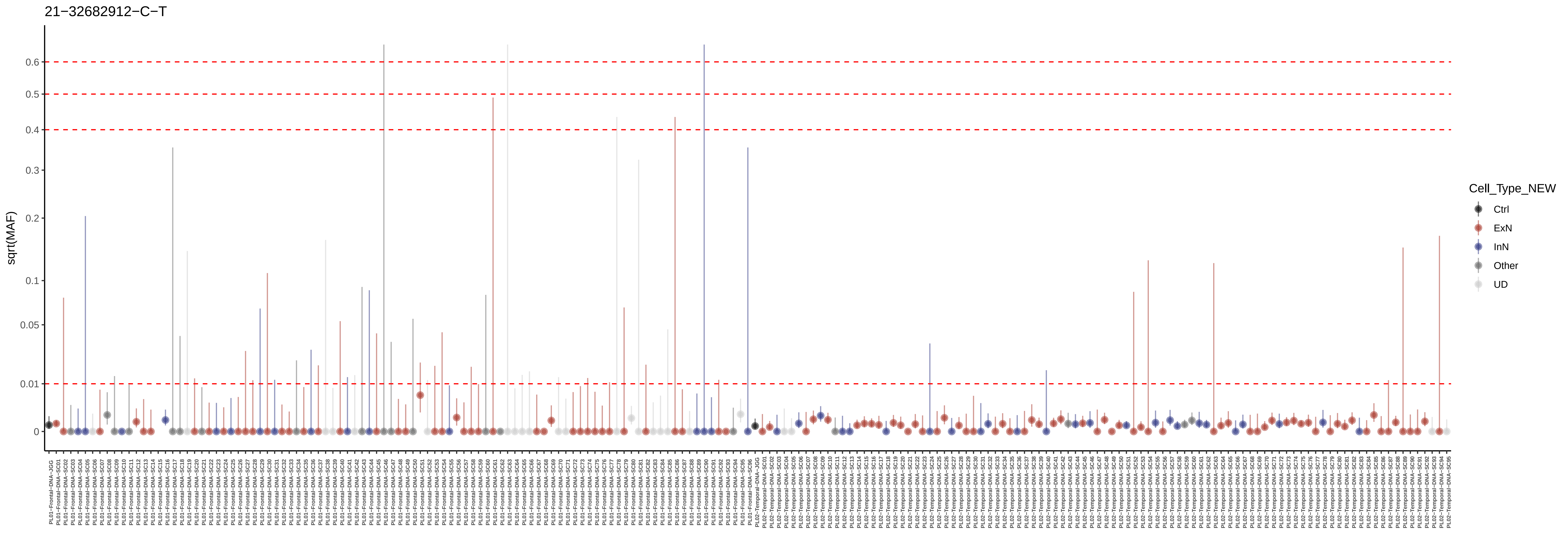

21−32682912−C−T
sqrt(MAF)
0.6
0.5
0.4
0.3
0.2
0.1
0.05
0.01
0
Cell_Type_NEW
Ctrl
ExN
InN
Other
UD

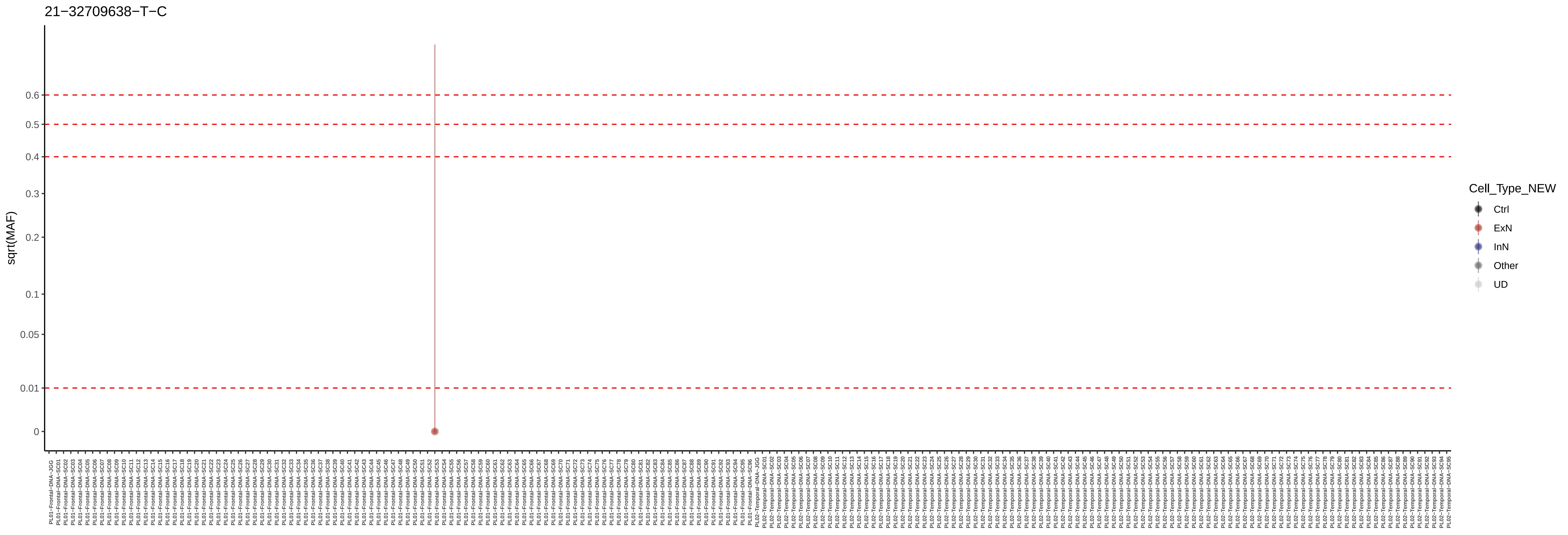

21−32709638−T−C
sqrt(MAF)
0.6
0.5
0.4
0.3
0.2
0.1
0.05
0.01
0
Cell_Type_NEW
Ctrl
ExN
InN
Other
UD

21−33248173−T−G

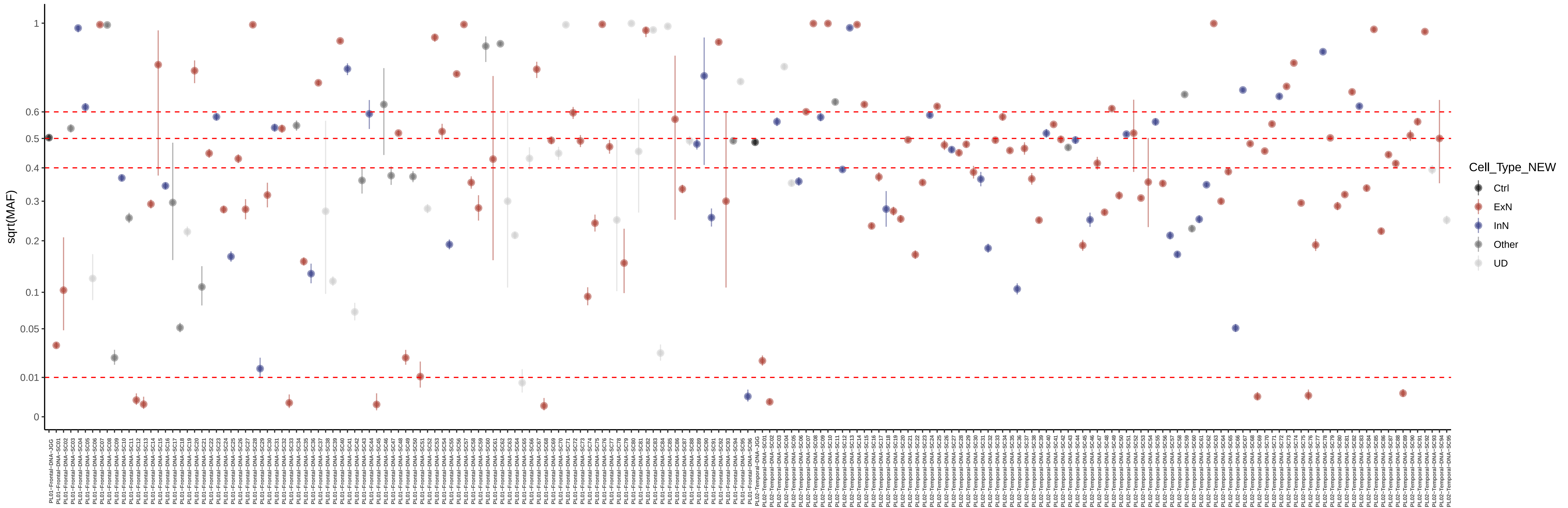

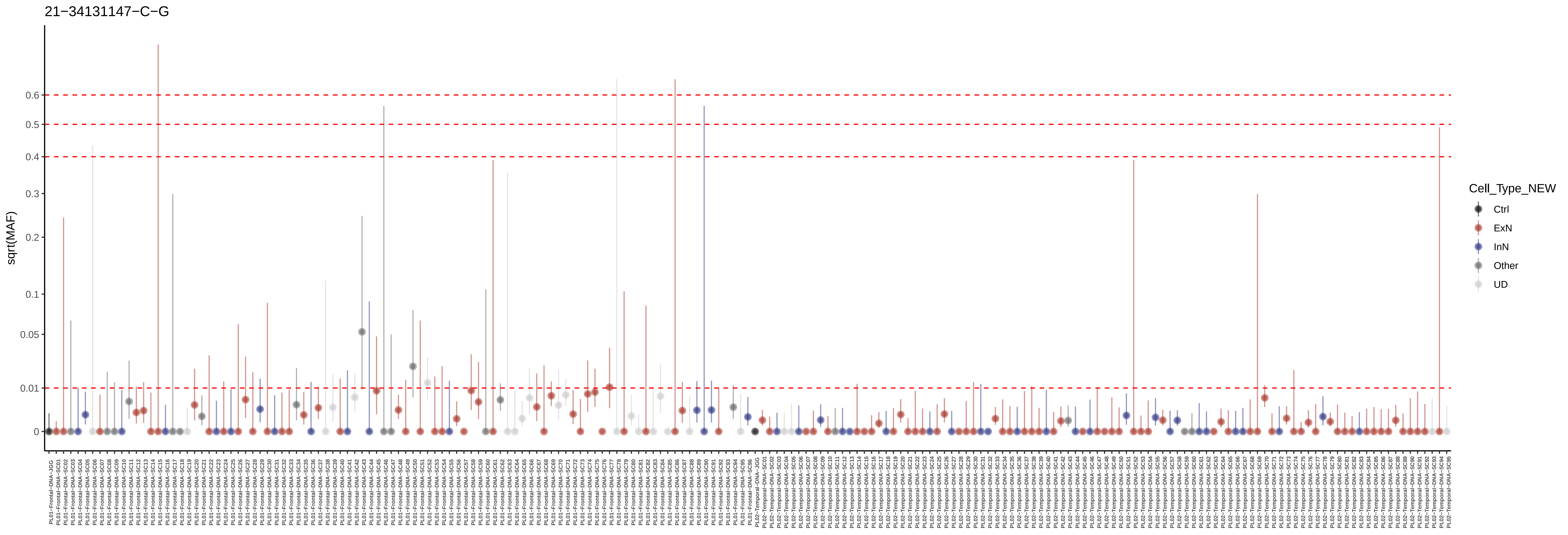

21−34131147−C−G
sqrt(MAF)
0.6
0.5
0.4
0.3
0.2
0.1
0.05
0.01
0
Cell_Type_NEW
Ctrl
ExN
InN
Other
UD

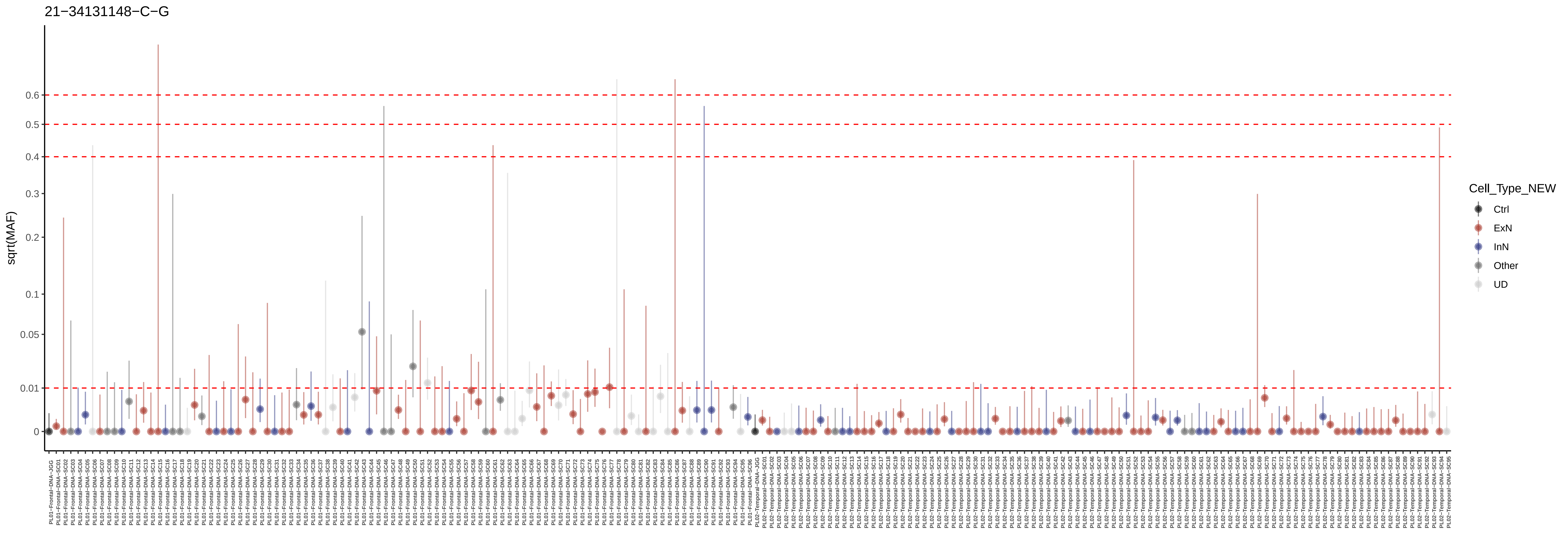
21−34131148−C−G
sqrt(MAF)
0.6
0.5
0.4
0.3
0.2
0.1
0.05
0.01
0
Cell_Type_NEW
Ctrl
ExN
InN
Other
UD

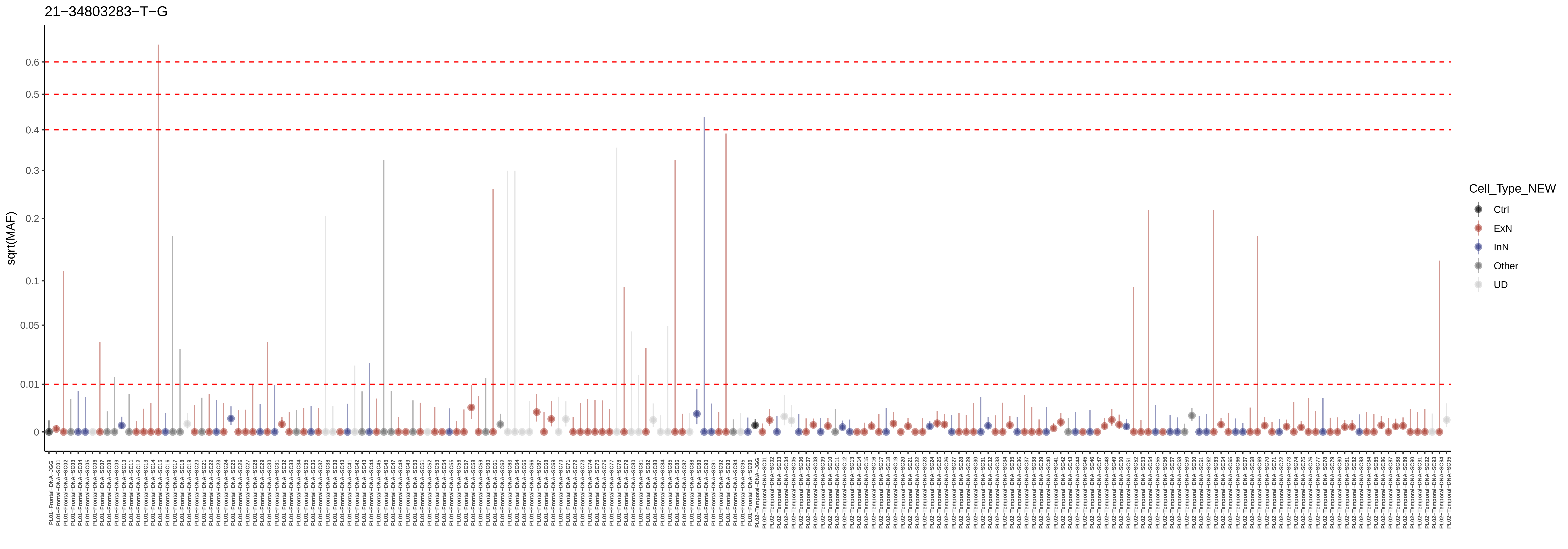
21−34803283−T−G
sqrt(MAF)
0.6
0.5
0.4
0.3
0.2
0.1
0.05
0.01
0
Cell_Type_NEW
Ctrl
ExN
InN
Other
UD

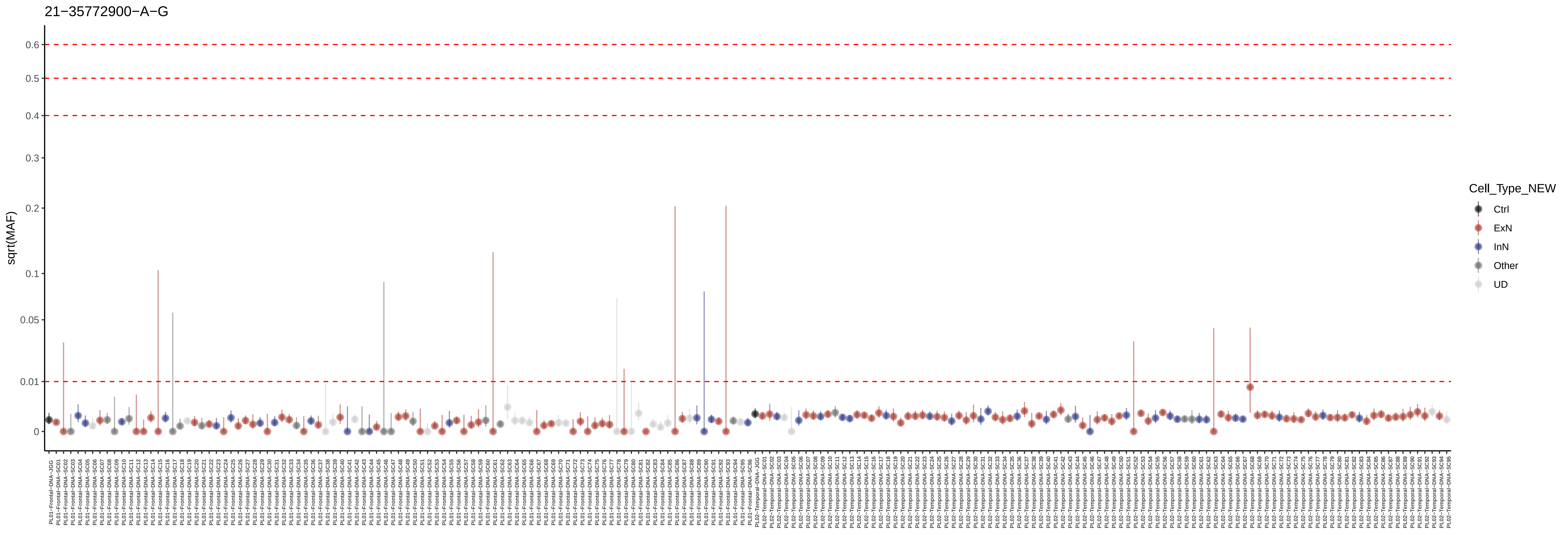
21−35772900−A−G
sqrt(MAF)
0.6
0.5
0.4
0.3
0.2
0.1
0.05
0.01
0
Cell_Type_NEW
Ctrl
ExN
InN
Other
UD

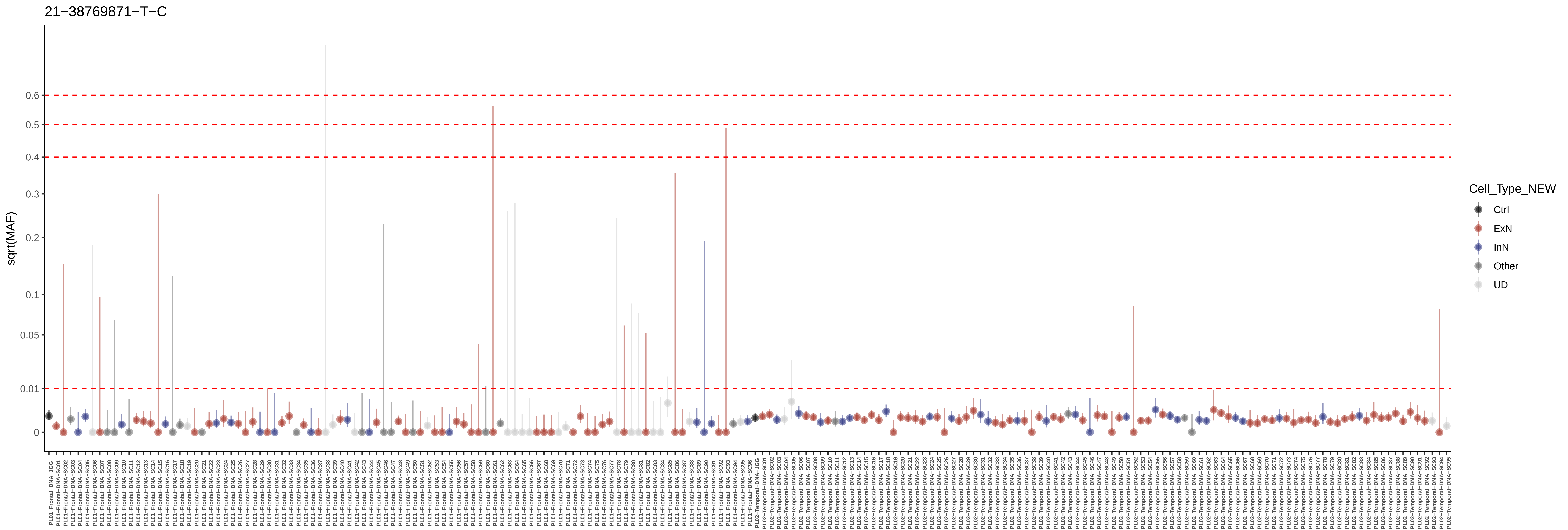

21−38769871−T−C
sqrt(MAF)
0.6
0.5
0.4
0.3
0.2
0.1
0.05
0.01
0
Cell_Type_NEW
Ctrl
ExN
InN
Other
UD

21−39917948−T−C

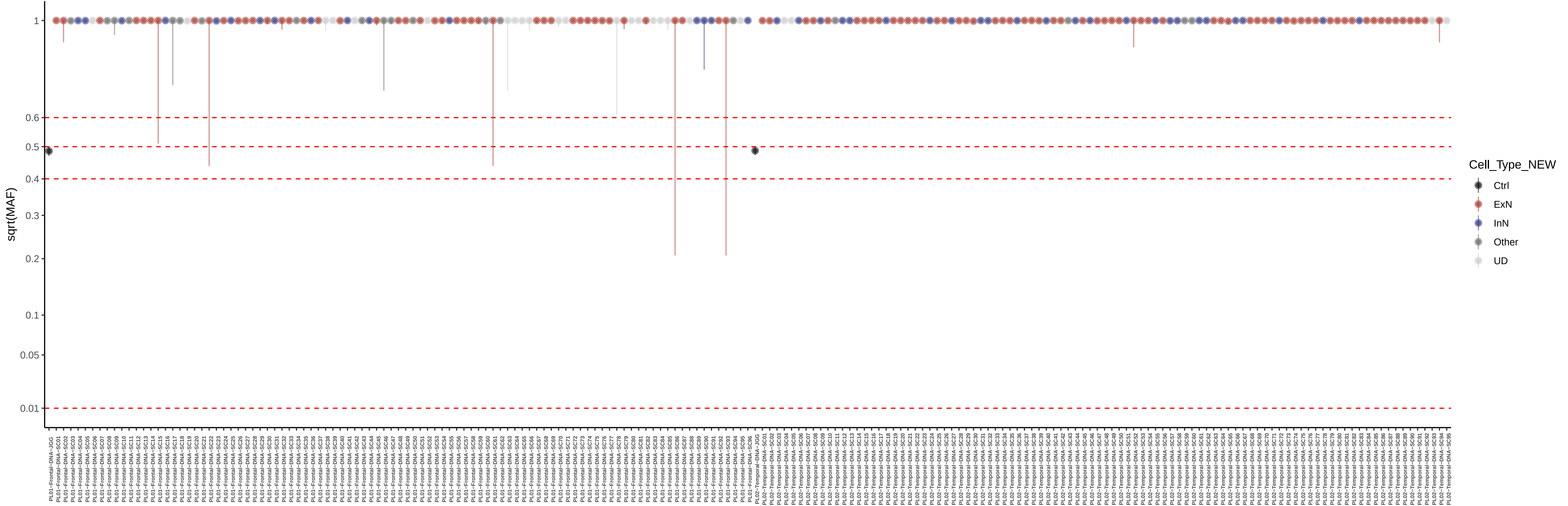

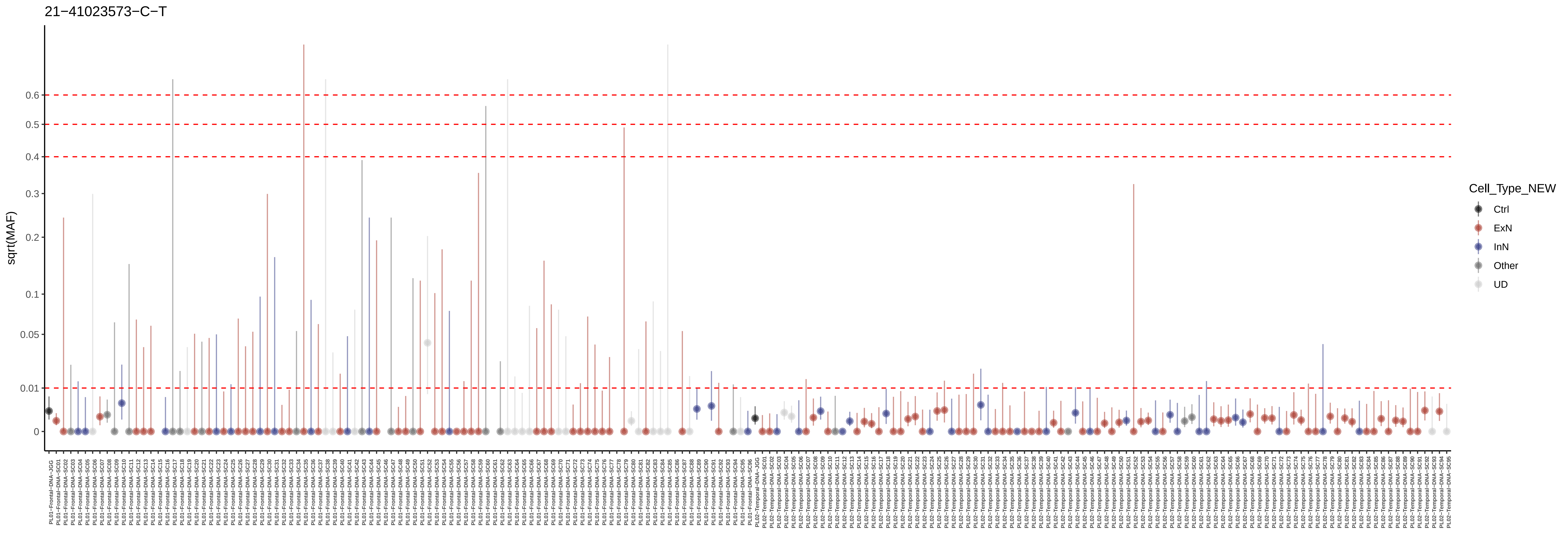
21−41023573−C−T
sqrt(MAF)
0.6
0.5
0.4
0.3
0.2
0.1
0.05
0.01
0
Cell_Type_NEW
Ctrl
ExN
InN
Other
UD

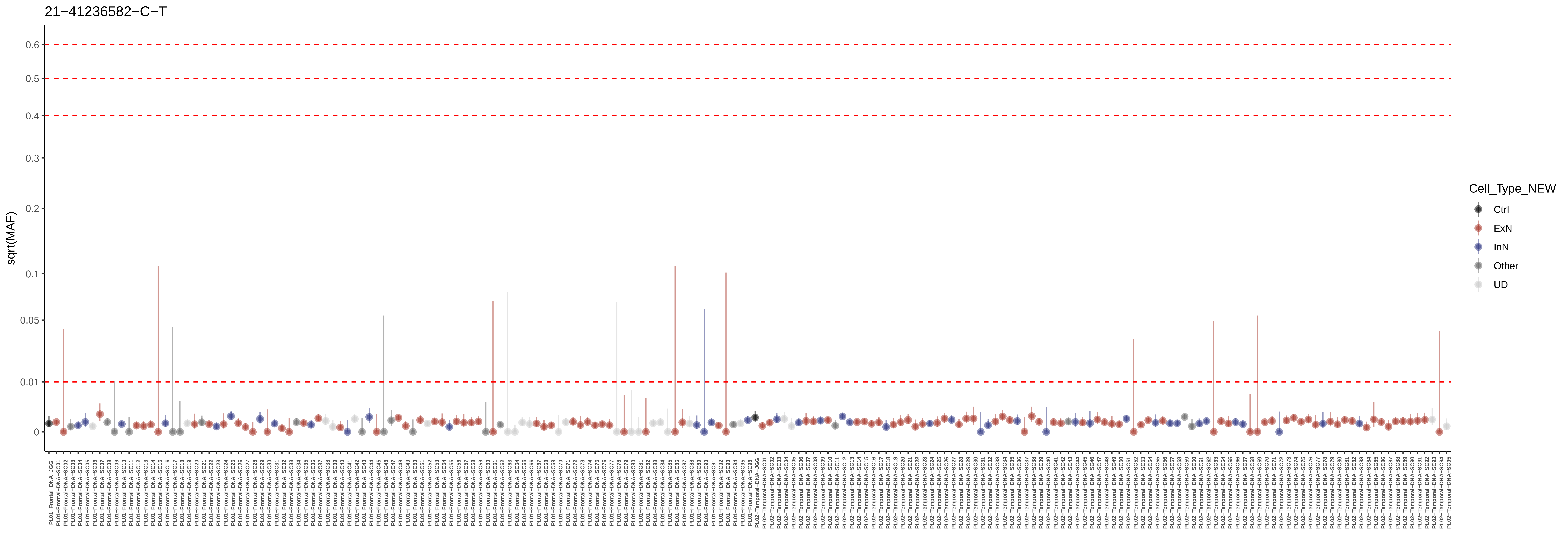
21−41236582−C−T
sqrt(MAF)
0.6
0.5
0.4
0.3
0.2
0.1
0.05
0.01
0
Cell_Type_NEW
Ctrl
ExN
InN
Other
UD

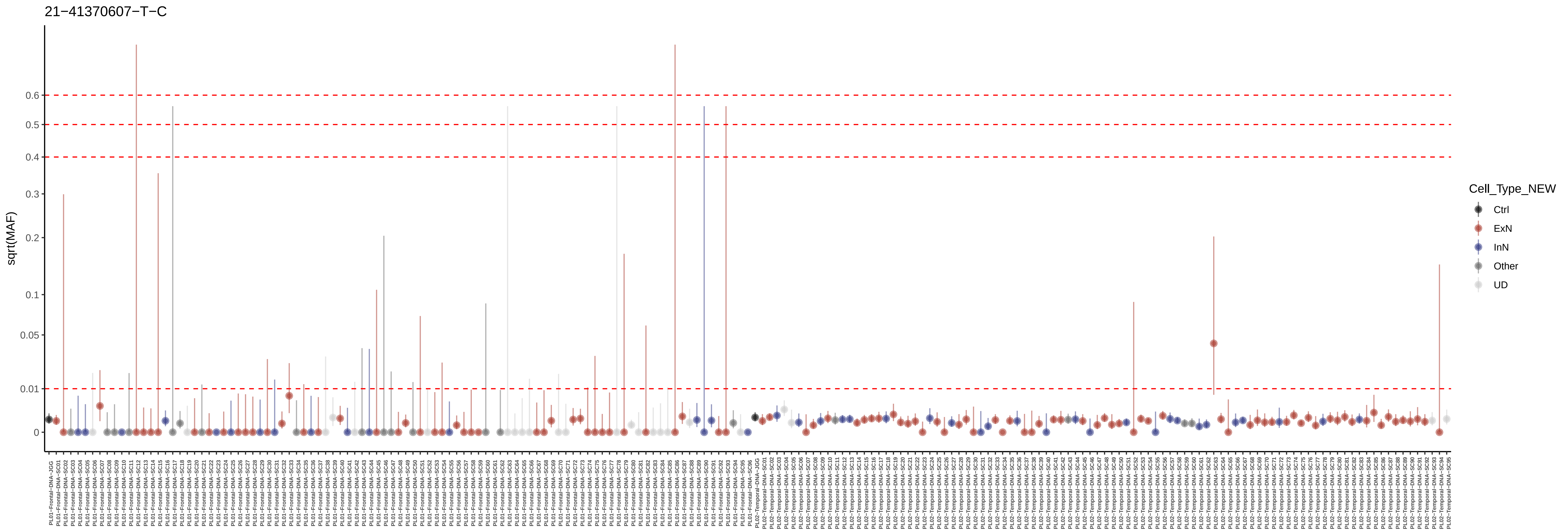
21−41370607−T−C
sqrt(MAF)
0.6
0.5
0.4
0.3
0.2
0.1
0.05
0.01
0
Cell_Type_NEW
Ctrl
ExN
InN
Other
UD

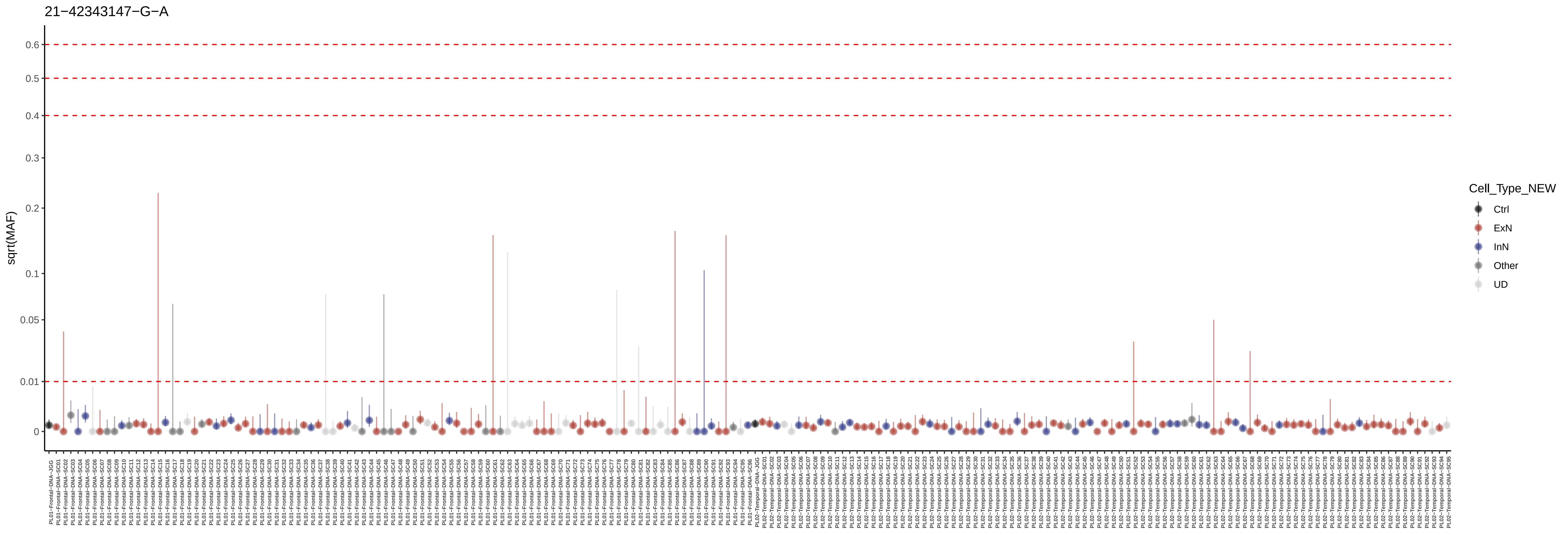

21−42343147−G−A
sqrt(MAF)
0.6
0.5
0.4
0.3
0.2
0.1
0.05
0.01
0
Cell_Type_NEW
Ctrl
ExN
InN
Other
UD

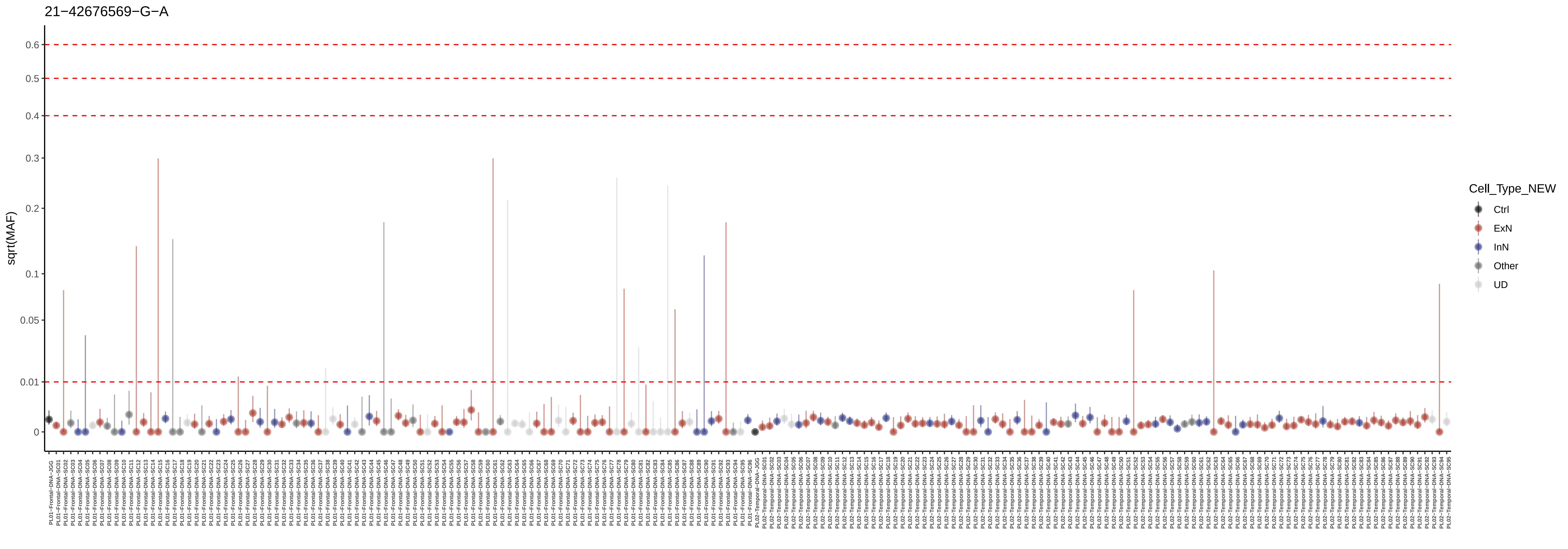

21−42676569−G−A
sqrt(MAF)
0.6
0.5
0.4
0.3
0.2
0.1
0.05
0.01
0
Cell_Type_NEW
Ctrl
ExN
InN
Other
UD

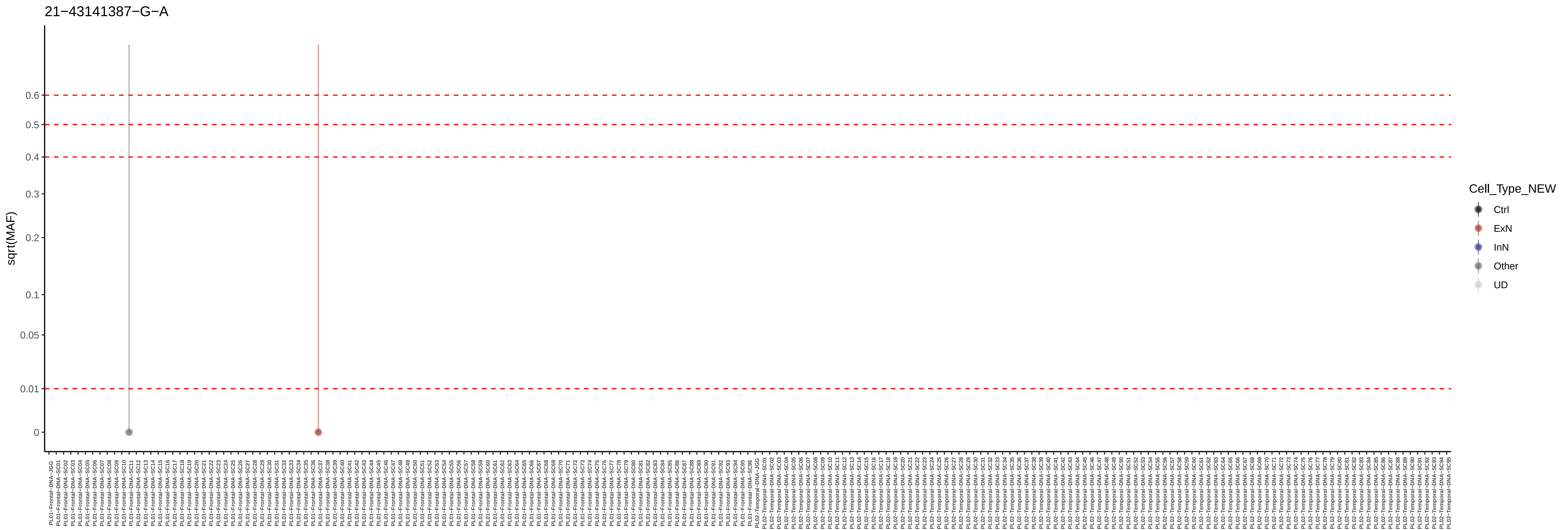

21−43141387−G−A
sqrt(MAF)
0.6
0.5
0.4
0.3
0.2
0.1
0.05
0.01
0
Cell_Type_NEW
Ctrl
ExN
InN
Other
UD

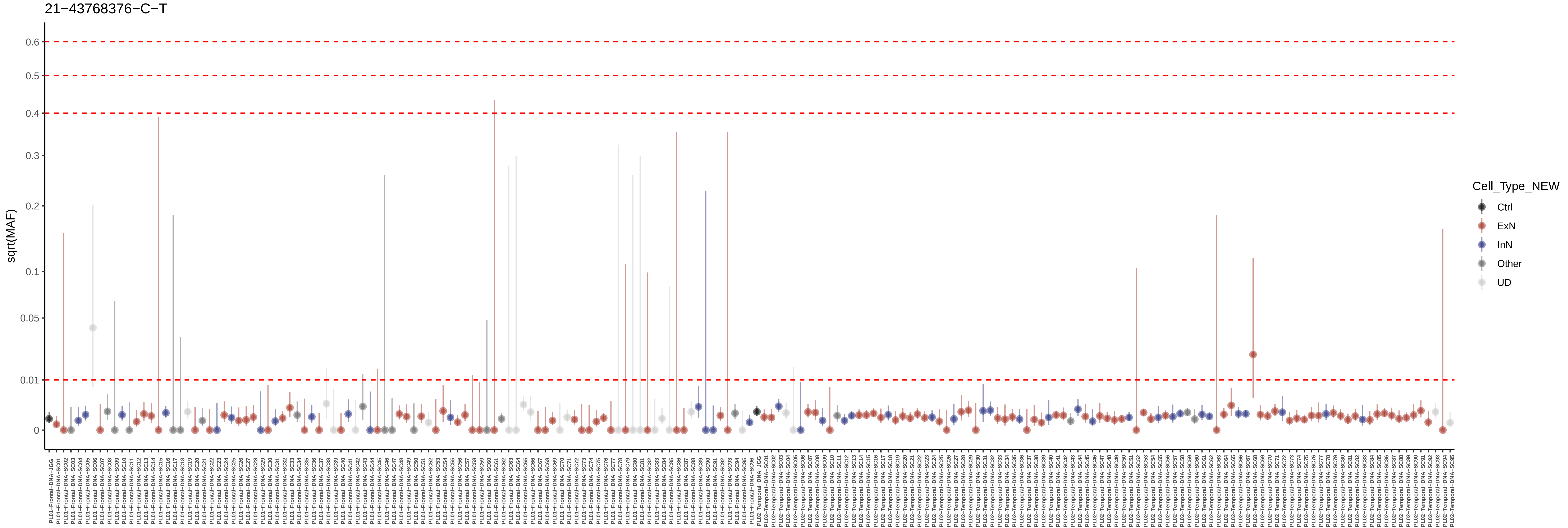

21−43768376−C−T
sqrt(MAF)
0.6
0.5
0.4
0.3
0.2
0.1
0.05
0.01
0
Cell_Type_NEW
Ctrl
ExN
InN
Other
UD

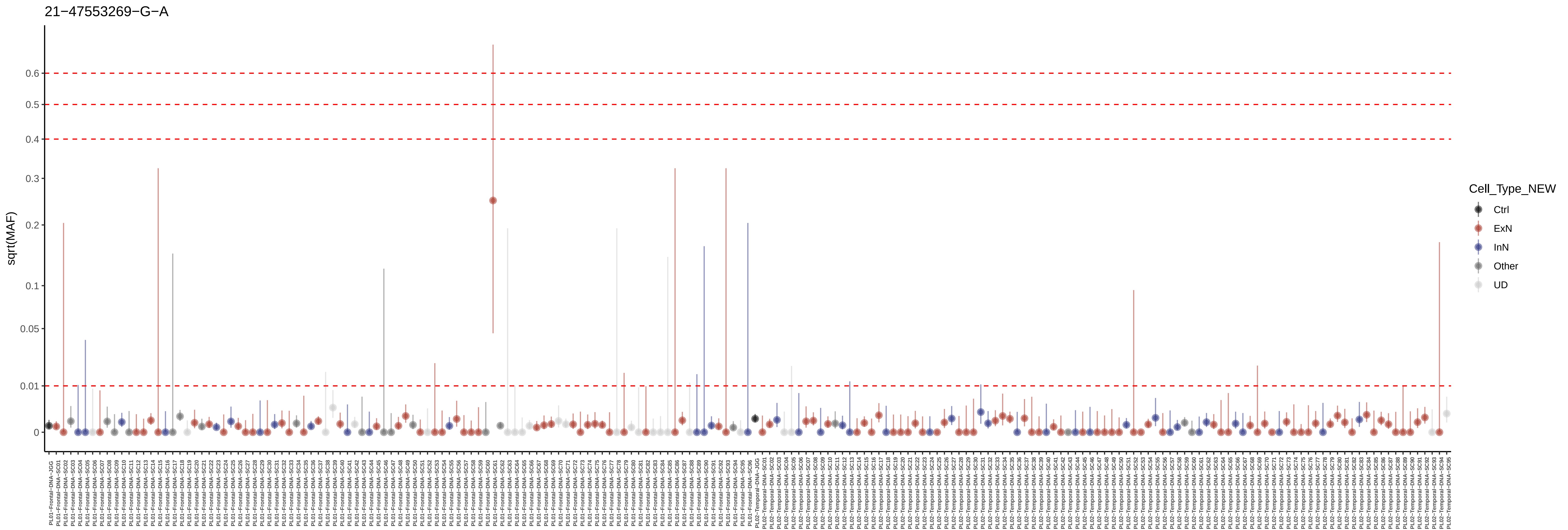
21−47553269−G−A
sqrt(MAF)
0.6
0.5
0.4
0.3
0.2
0.1
0.05
0.01
0
Cell_Type_NEW
Ctrl
ExN
InN
Other
UD

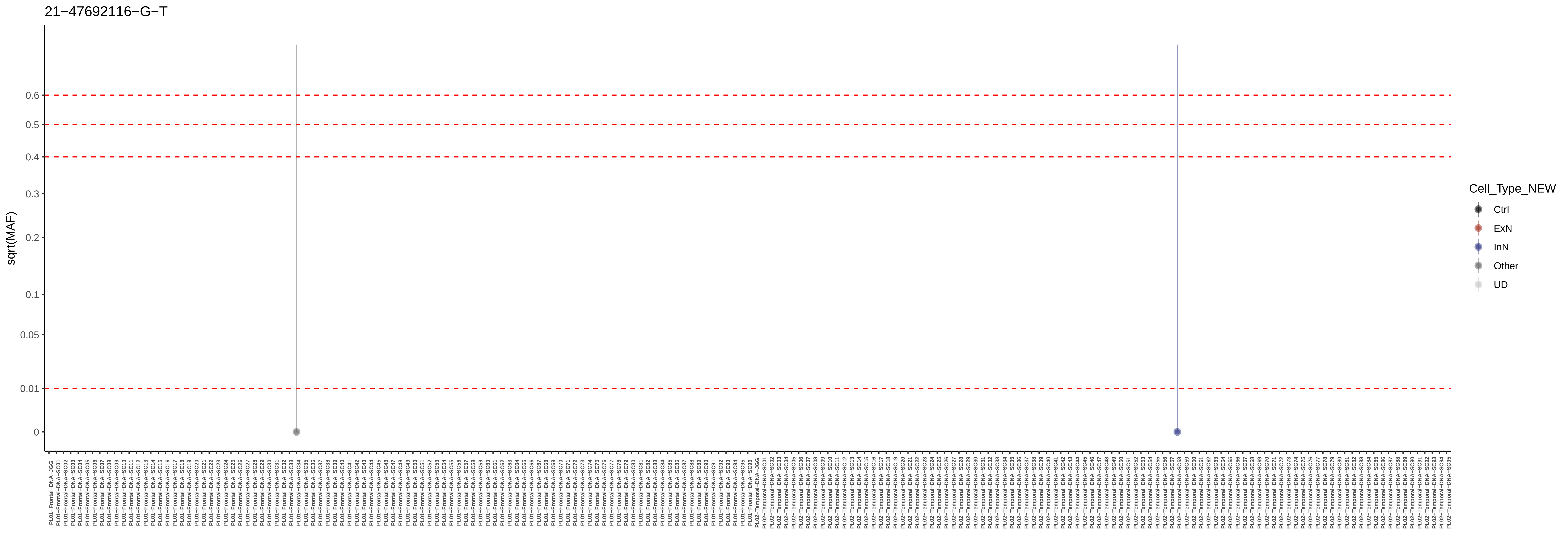
21−47692116−G−T
sqrt(MAF)
0.6
0.5
0.4
0.3
0.2
0.1
0.05
0.01
0
Cell_Type_NEW
Ctrl
ExN
InN
Other
UD

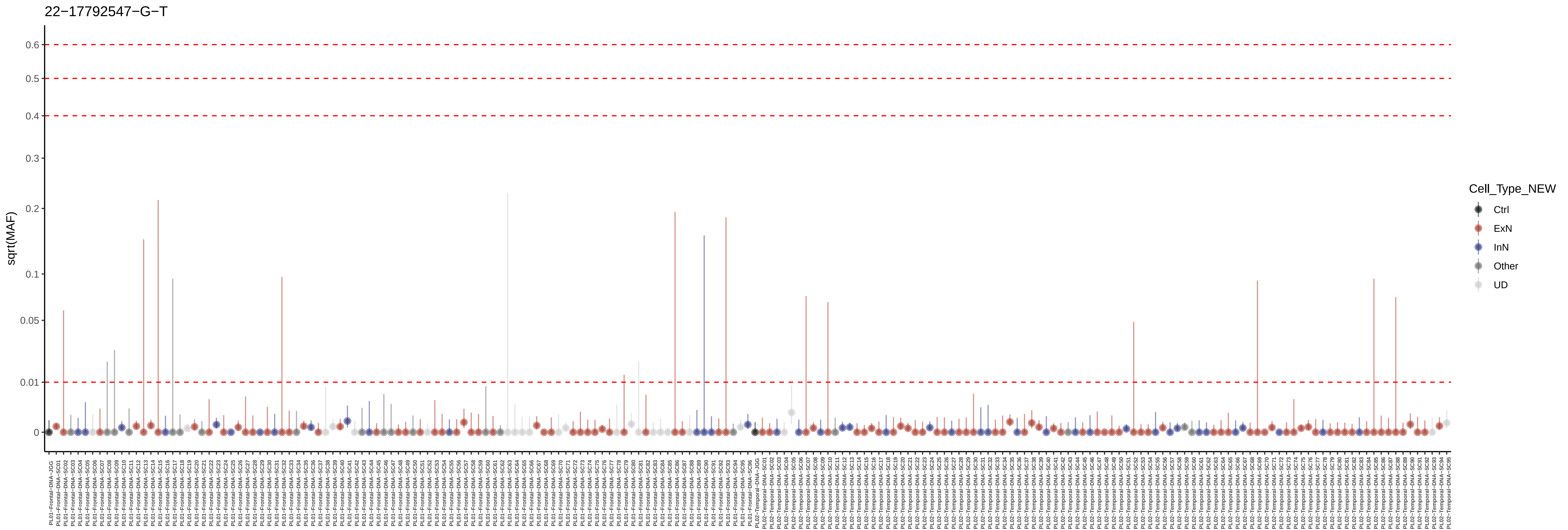

22−17792547−G−T
sqrt(MAF)
0.6
0.5
0.4
0.3
0.2
0.1
0.05
0.01
0
Cell_Type_NEW
Ctrl
ExN
InN
Other
UD

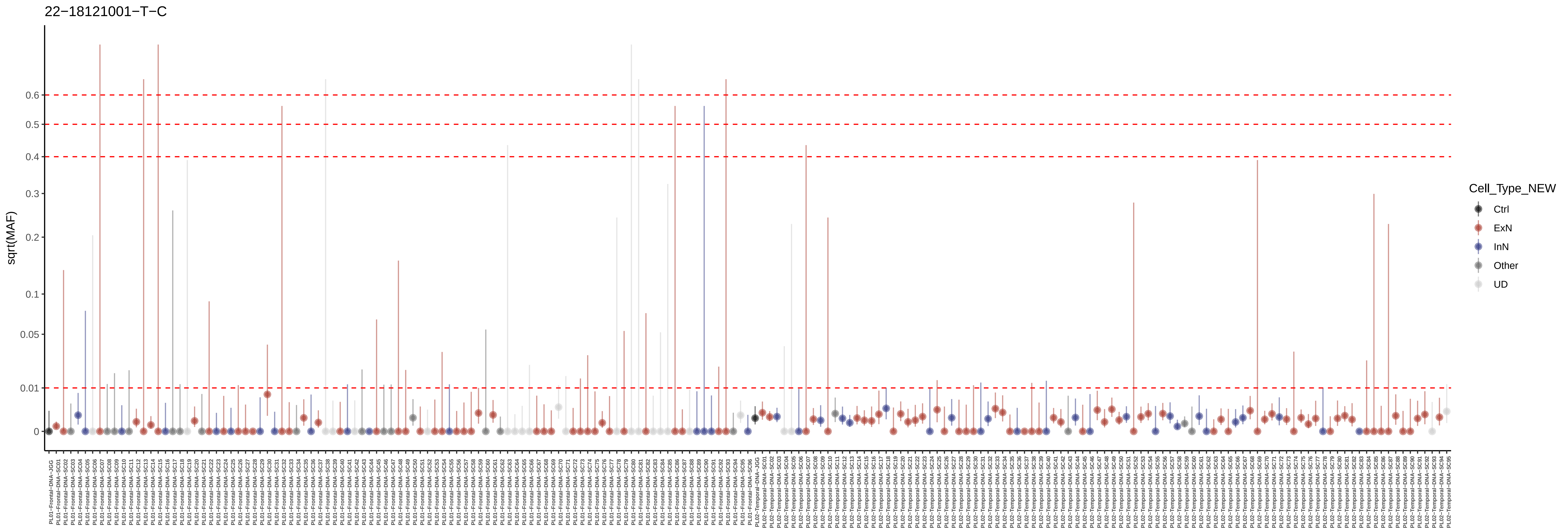
22−18121001−T−C
sqrt(MAF)
0.6
0.5
0.4
0.3
0.2
0.1
0.05
0.01
0
Cell_Type_NEW
Ctrl
ExN
InN
Other
UD

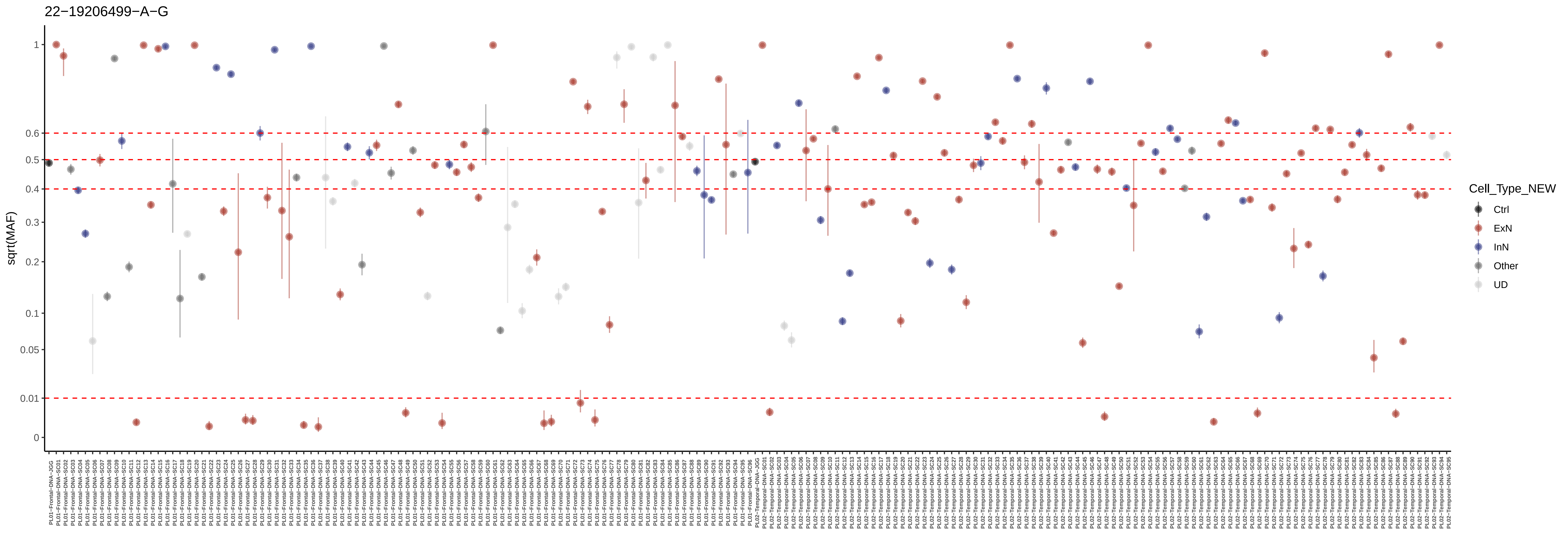

22−19206499−A−G
sqrt(MAF)
1
0.6
0.5
0.4
0.3
0.2
0.1
0.05
0.01
0
Cell_Type_NEW
Ctrl
ExN
InN
Other
UD

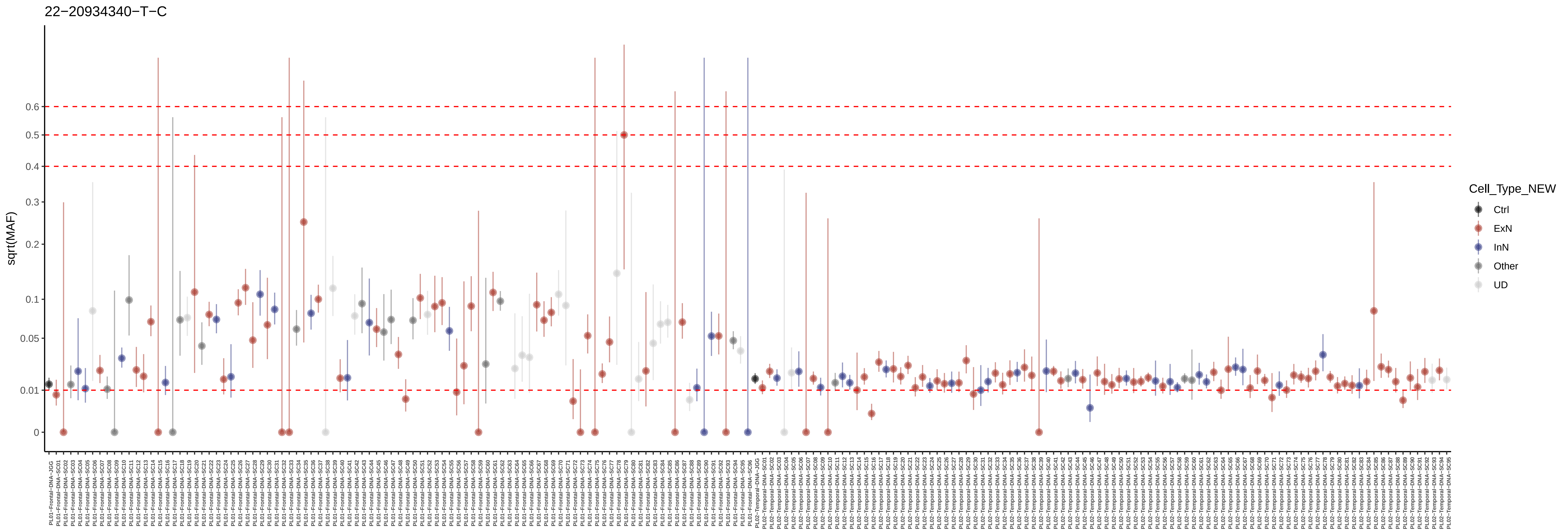
22−20934340−T−C
sqrt(MAF)
0.6
0.5
0.4
0.3
0.2
0.1
0.05
0.01
0
Cell_Type_NEW
Ctrl
ExN
InN
Other
UD

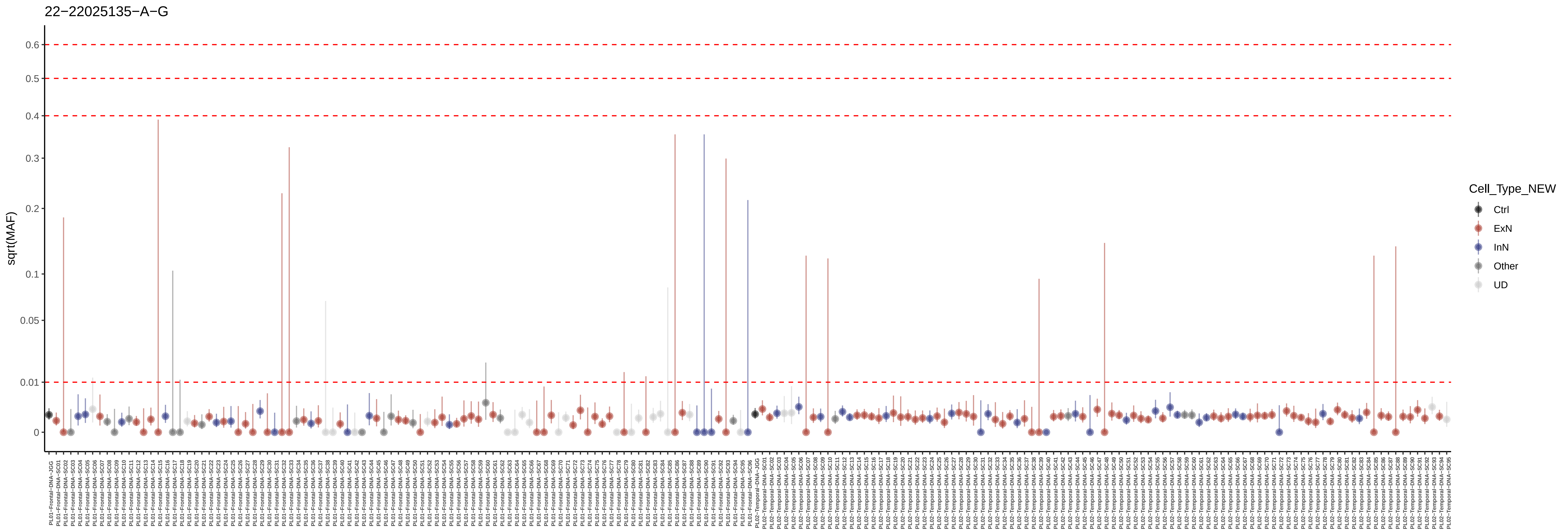

22−22025135−A−G
sqrt(MAF)
0.6
0.5
0.4
0.3
0.2
0.1
0.05
0.01
0
Cell_Type_NEW
Ctrl
ExN
InN
Other
UD

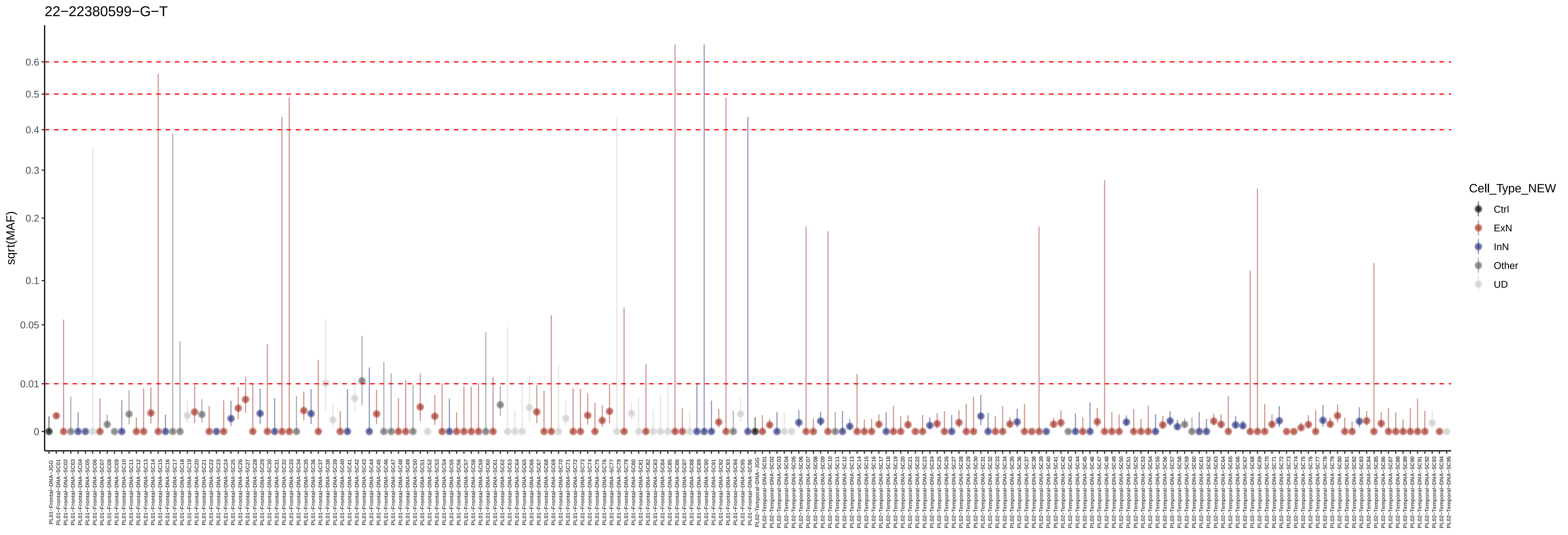

22−22380599−G−T
sqrt(MAF)
0.6
0.5
0.4
0.3
0.2
0.1
0.05
0.01
0
Cell_Type_NEW
Ctrl
ExN
InN
Other
UD

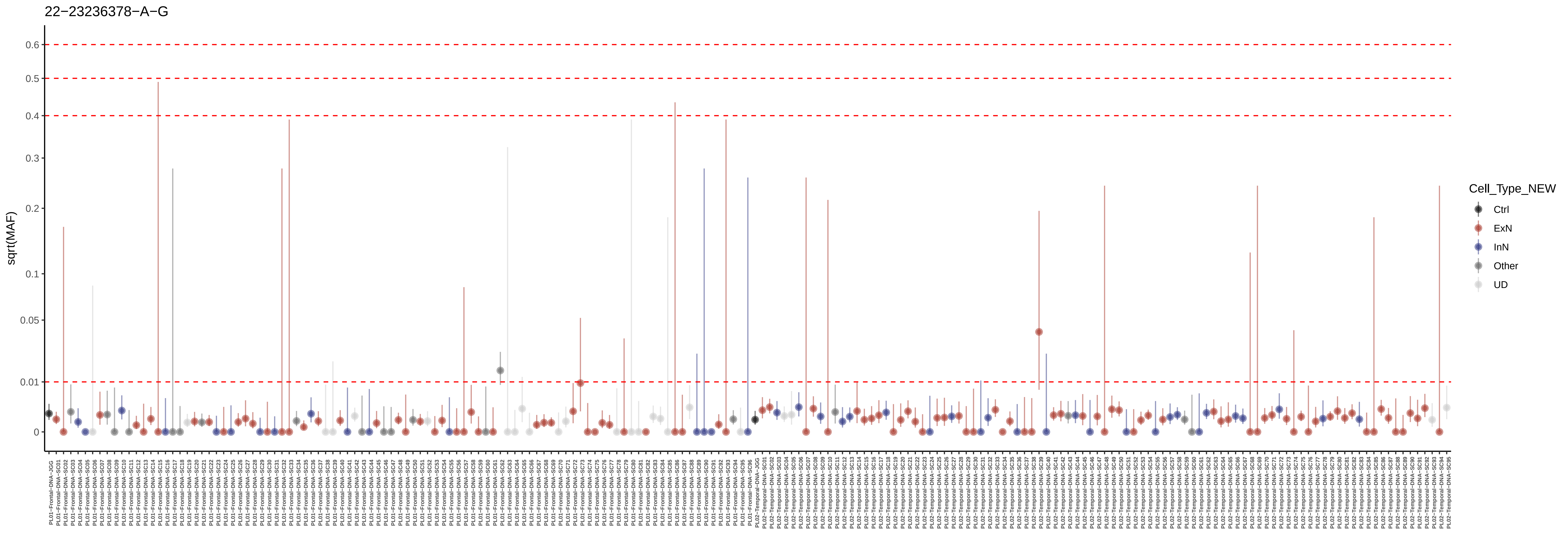
22−23236378−A−G
sqrt(MAF)
0.6
0.5
0.4
0.3
0.2
0.1
0.05
0.01
0
Cell_Type_NEW
Ctrl
ExN
InN
Other
UD

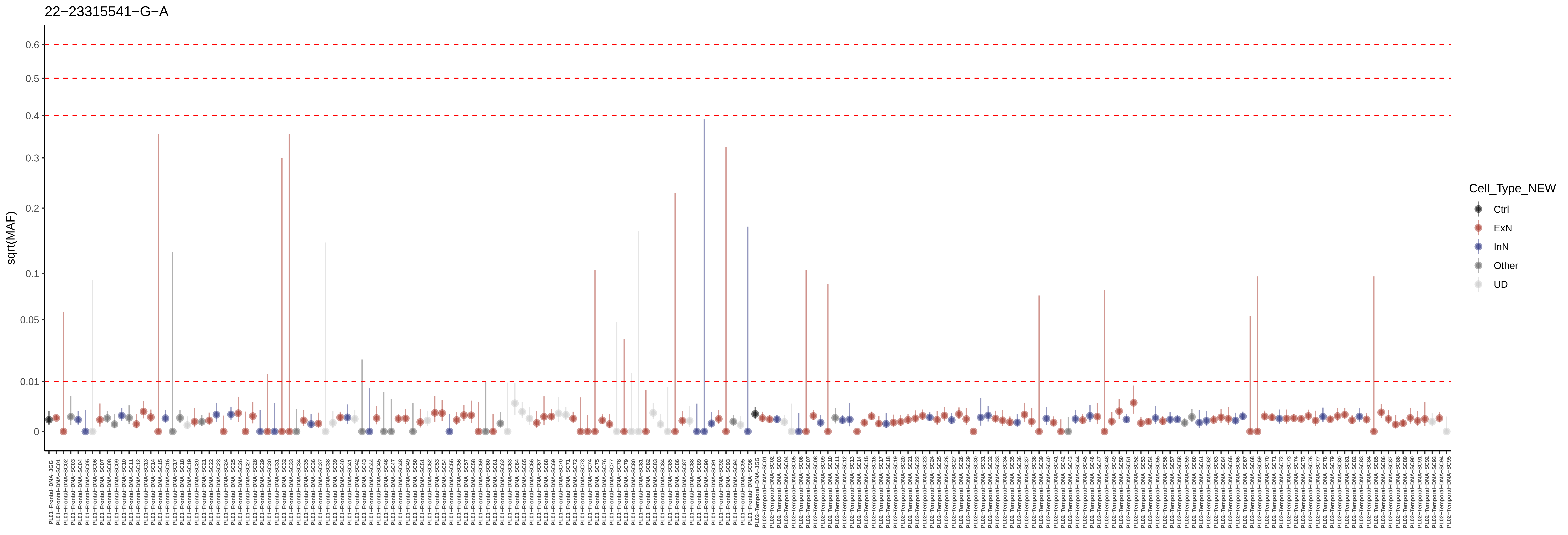
22−23315541−G−A
sqrt(MAF)
0.6
0.5
0.4
0.3
0.2
0.1
0.05
0.01
0
Cell_Type_NEW
Ctrl
ExN
InN
Other
UD

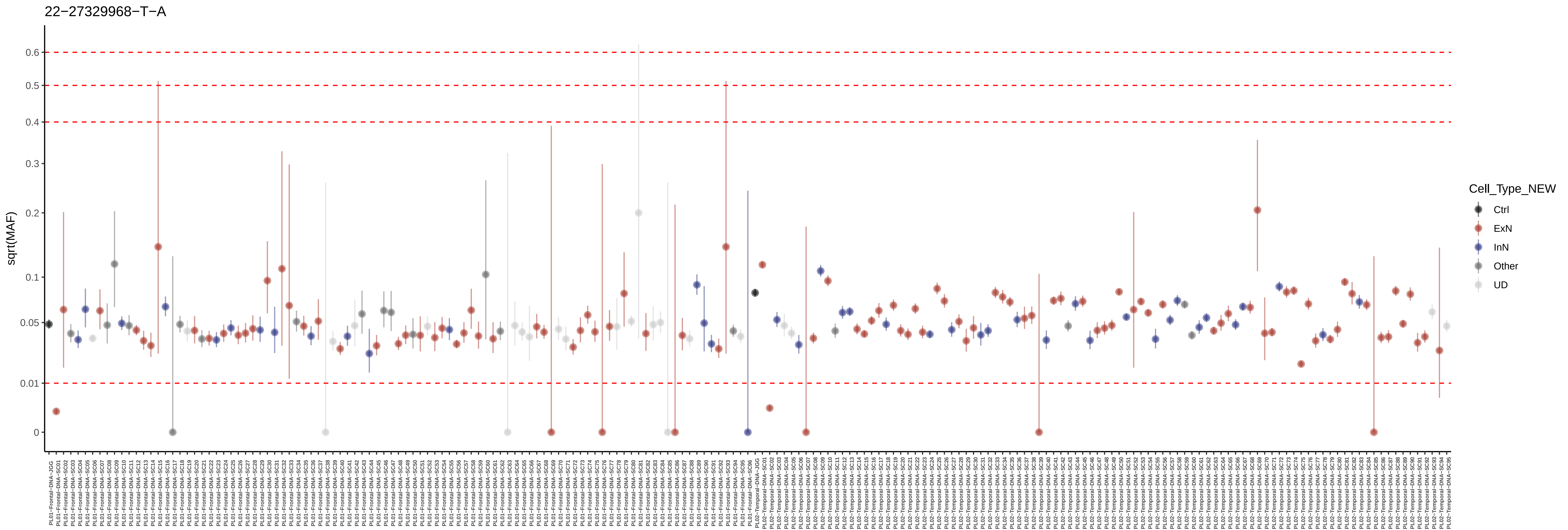

22−27329968−T−A
sqrt(MAF)
0.6
0.5
0.4
0.3
0.2
0.1
0.05
0.01
0
Cell_Type_NEW
Ctrl
ExN
InN
Other
UD

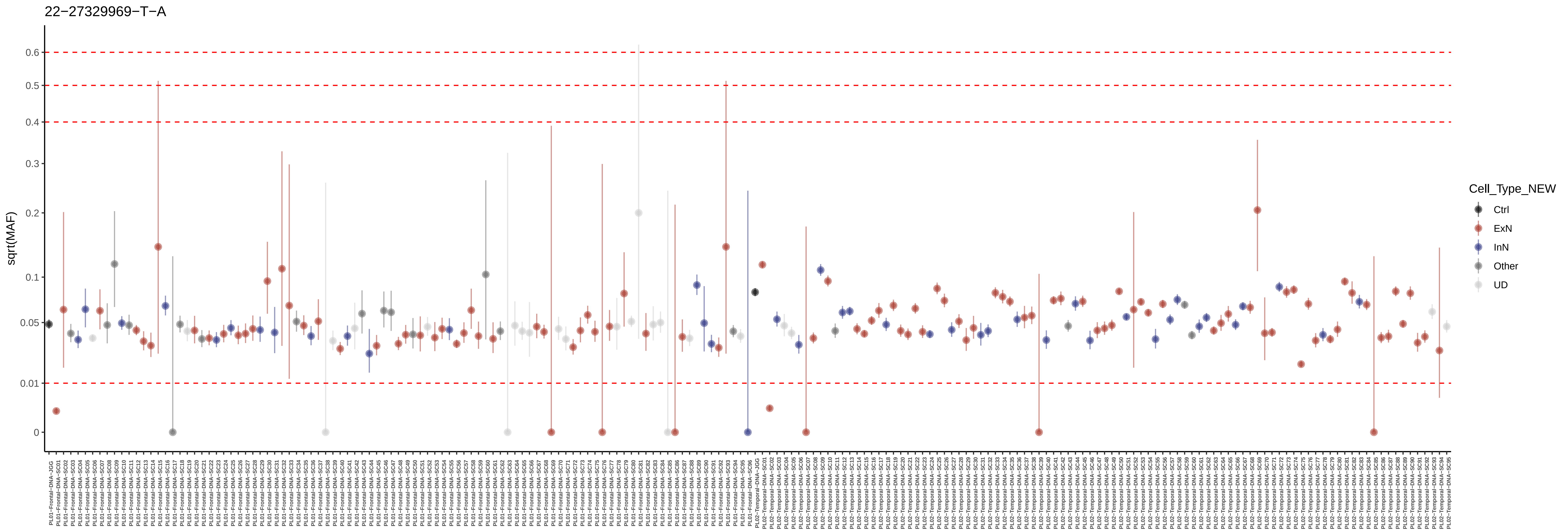
22−27329969−T−A
sqrt(MAF)
0.6
0.5
0.4
0.3
0.2
0.1
0.05
0.01
0
Cell_Type_NEW
Ctrl
ExN
InN
Other
UD

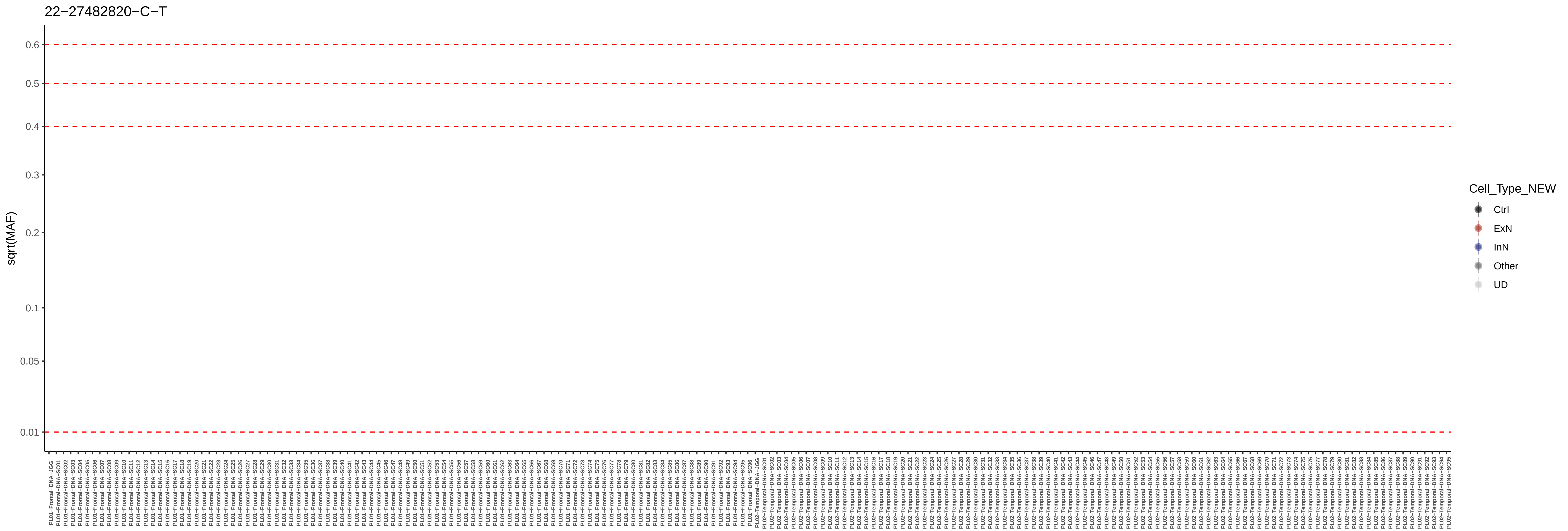

22−27482820−C−T
sqrt(MAF)
0.6
0.5
0.4
0.3
0.2
0.1
0.05
0.01
Cell_Type_NEW
Ctrl
ExN
InN
Other
UD

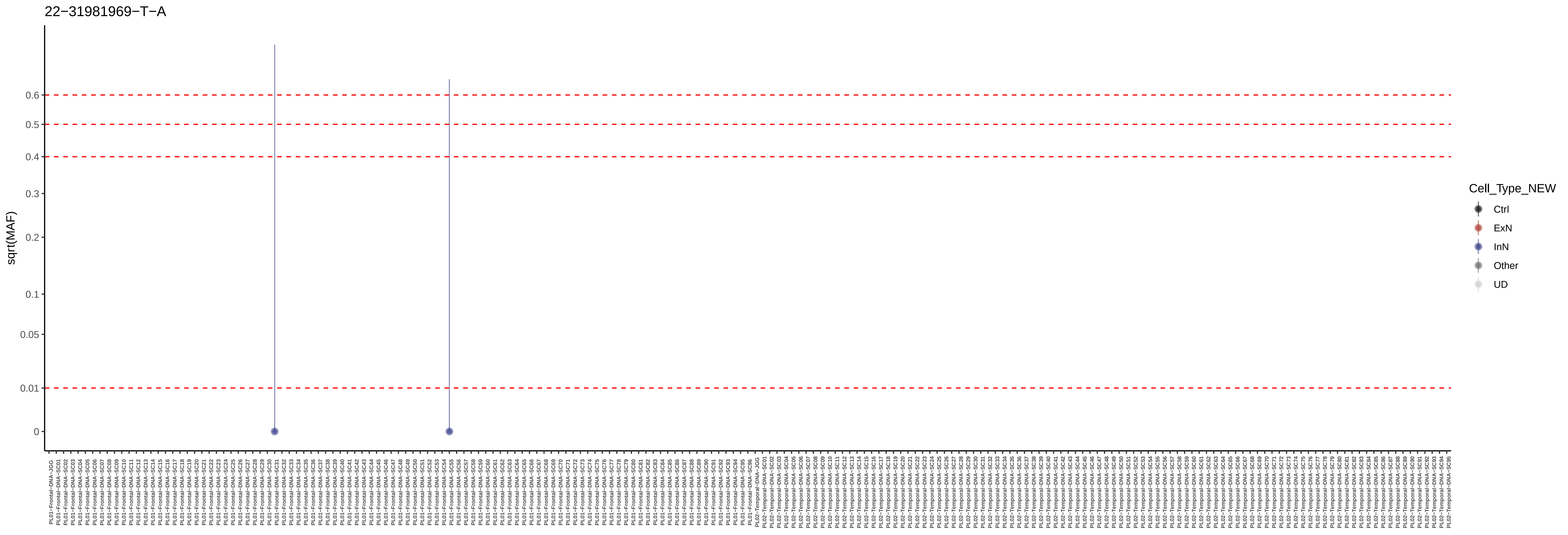
22−31981969−T−A
sqrt(MAF)
0.6
0.5
0.4
0.3
0.2
0.1
0.05
0.01
0
Cell_Type_NEW
Ctrl
ExN
InN
Other
UD

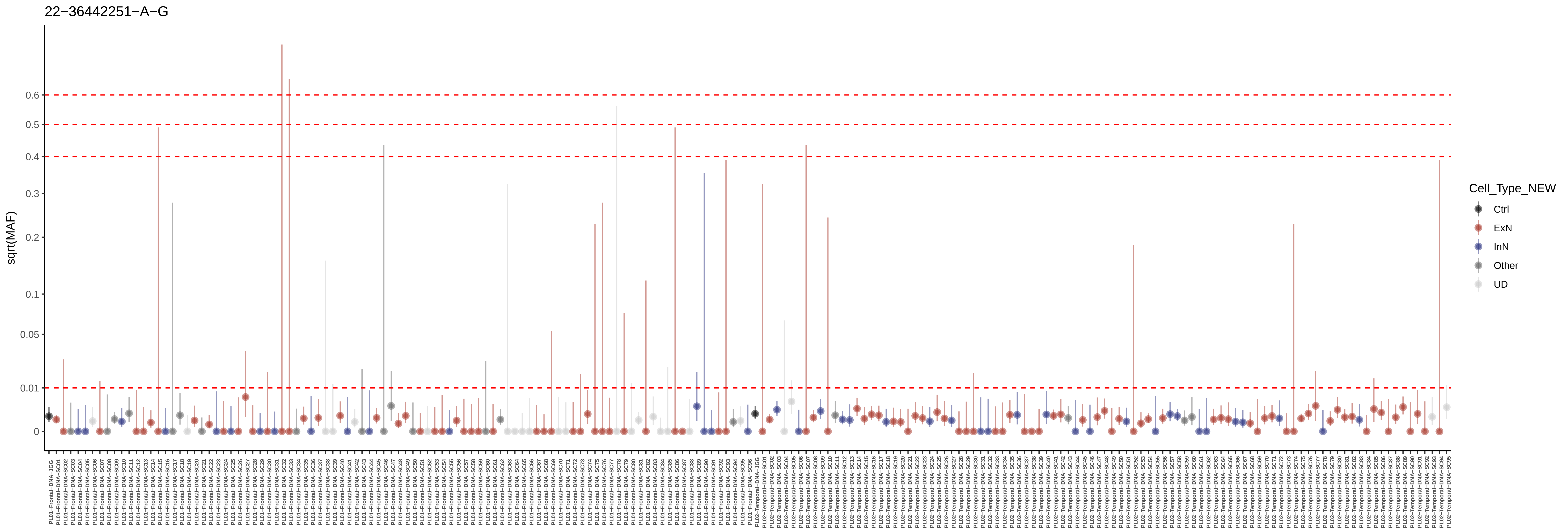

22−36442251−A−G
sqrt(MAF)
0.6
0.5
0.4
0.3
0.2
0.1
0.05
0.01
0
Cell_Type_NEW
Ctrl
ExN
InN
Other
UD

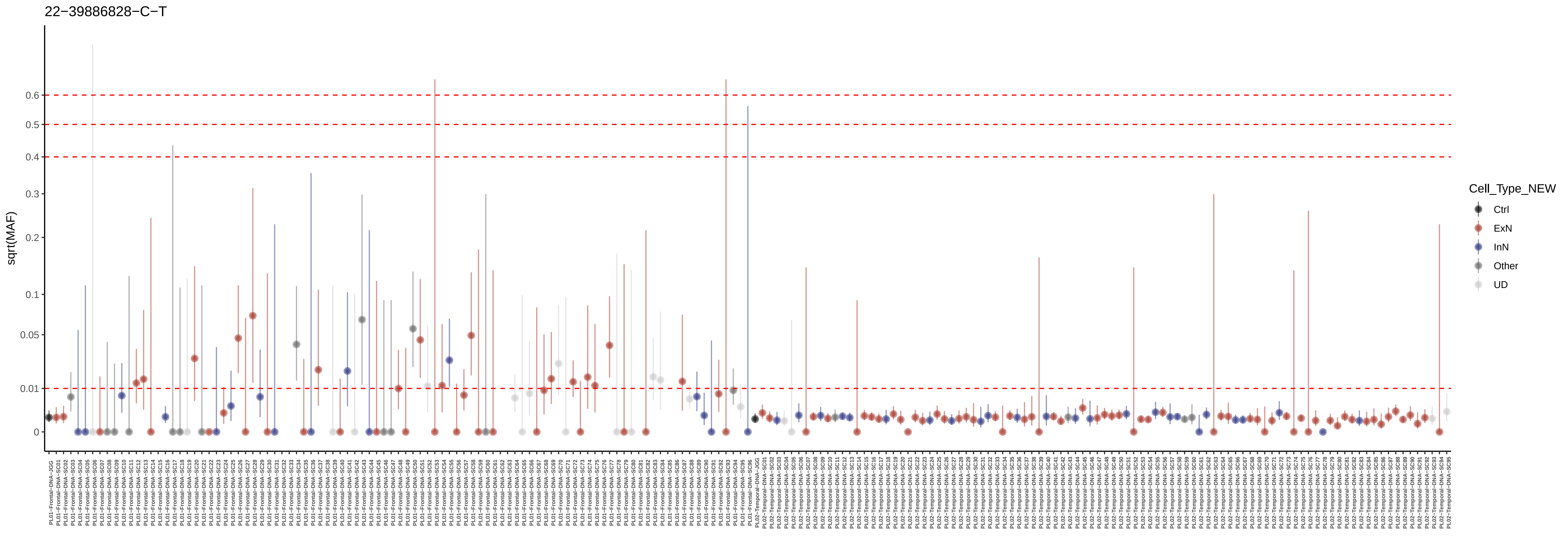
22−39886828−C−T
sqrt(MAF)
0.6
0.5
0.4
0.3
0.2
0.1
0.05
0.01
0
Cell_Type_NEW
Ctrl
ExN
InN
Other
UD

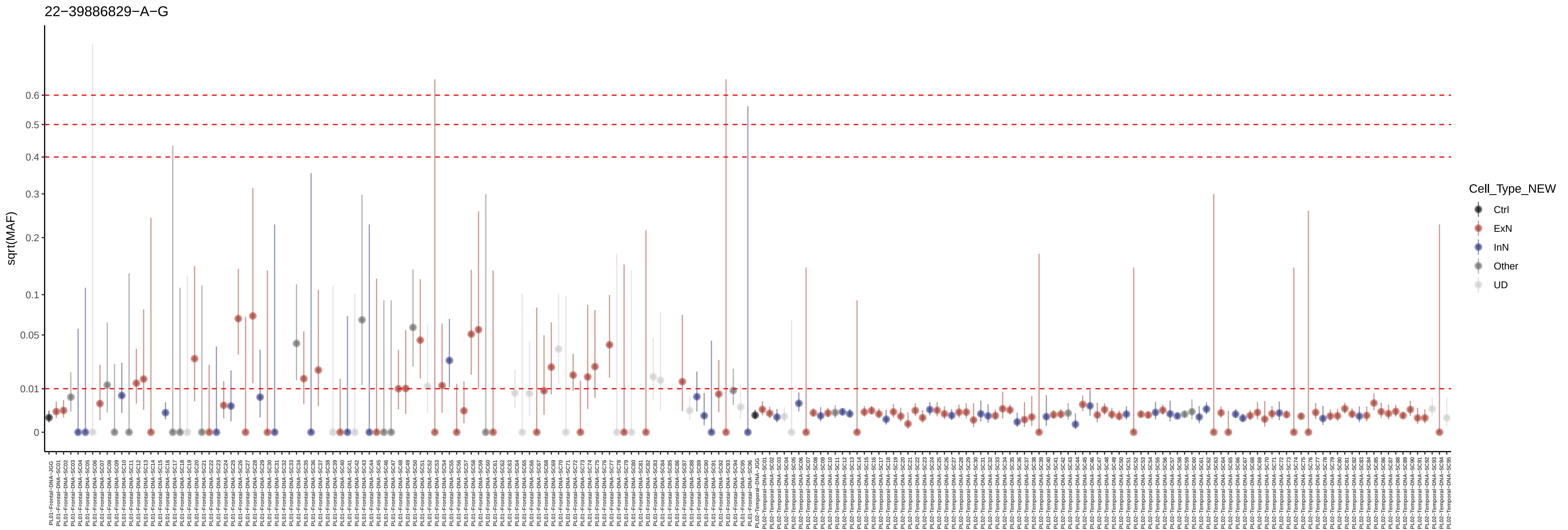

22−39886829−A−G
sqrt(MAF)
0.6
0.5
0.4
0.3
0.2
0.1
0.05
0.01
0
Cell_Type_NEW
Ctrl
ExN
InN
Other
UD

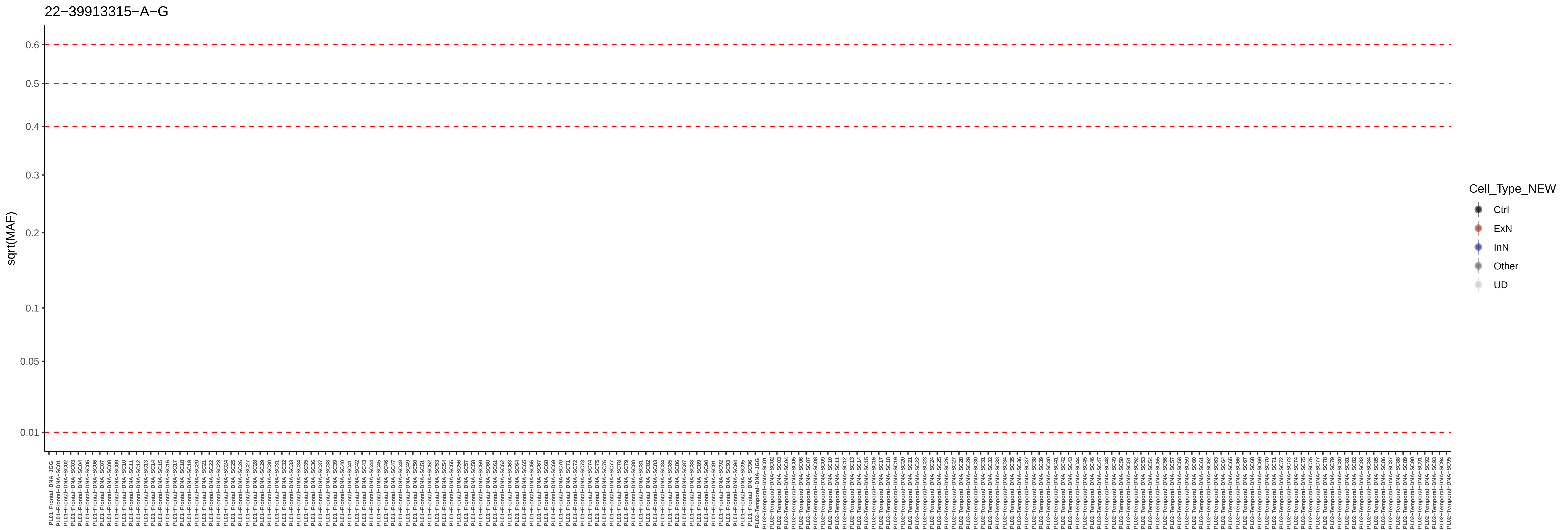

22−39913315−A−G
sqrt(MAF)
0.6
0.5
0.4
0.3
0.2
0.1
0.05
0.01
Cell_Type_NEW
Ctrl
ExN
InN
Other
UD

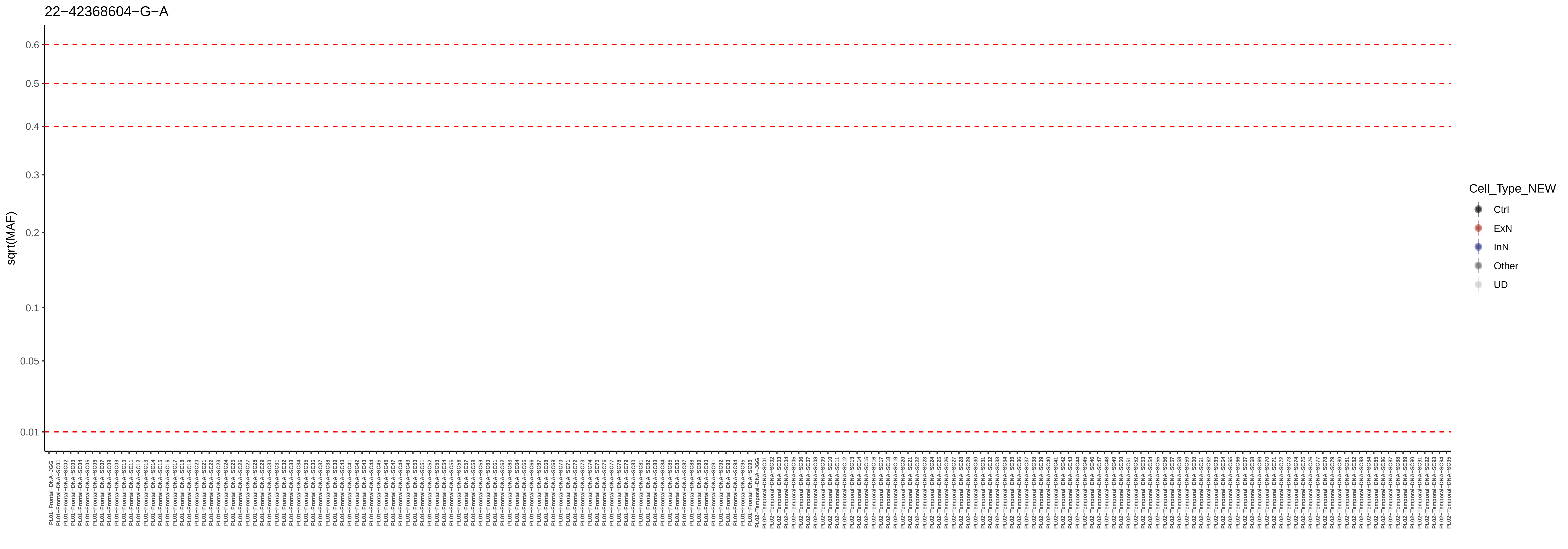
22−42368604−G−A
sqrt(MAF)
0.6
0.5
0.4
0.3
0.2
0.1
0.05
0.01
Cell_Type_NEW
Ctrl
ExN
InN
Other
UD

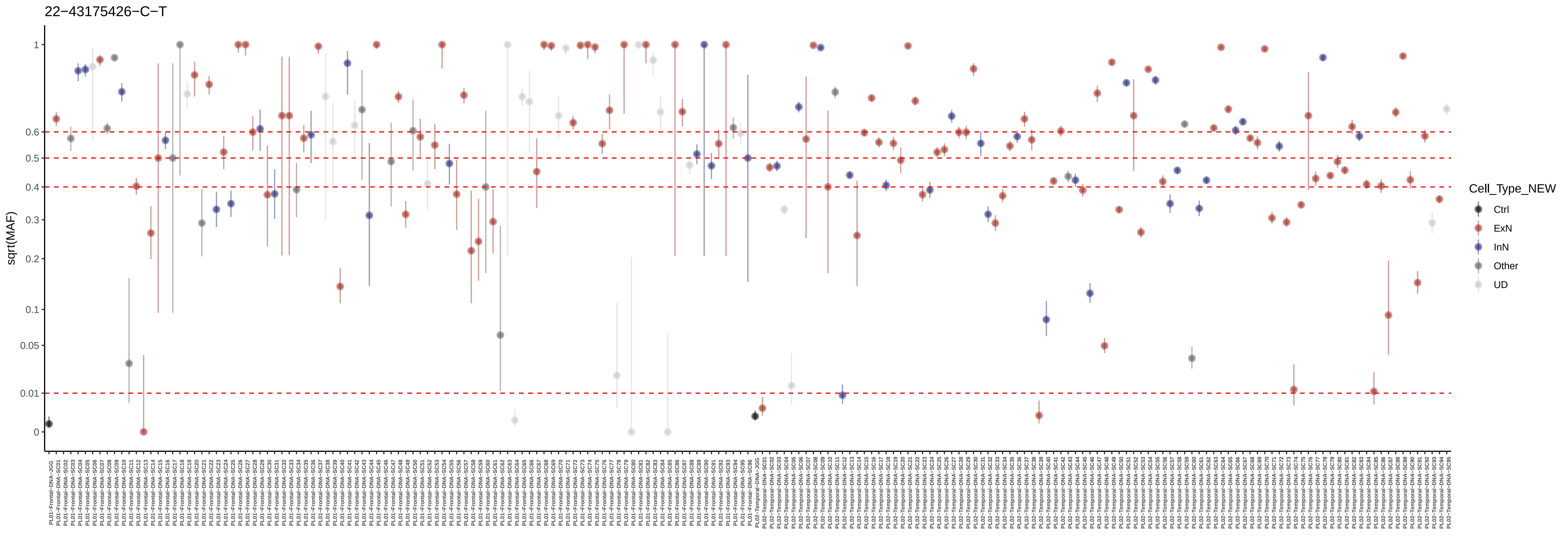

22−43175426−C−T
sqrt(MAF)
1
0.6
0.5
0.4
0.3
0.2
0.1
0.05
0.01
0
Cell_Type_NEW
Ctrl
ExN
InN
Other
UD

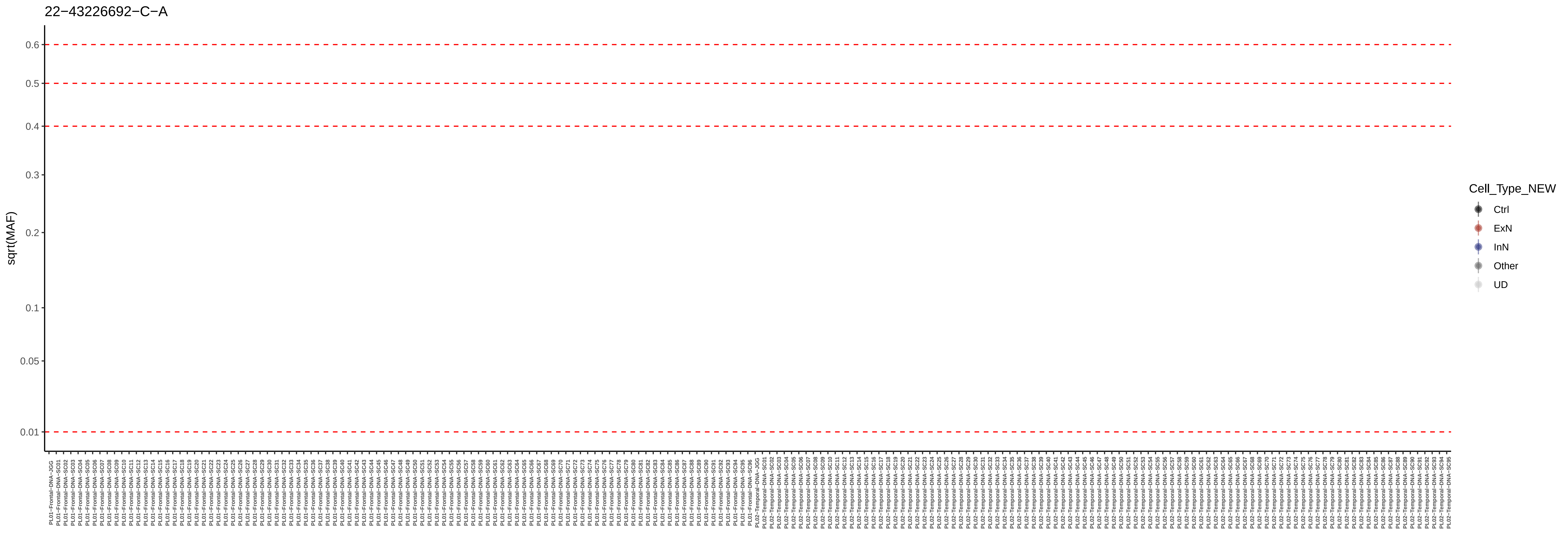
22−43226692−C−A
sqrt(MAF)
0.6
0.5
0.4
0.3
0.2
0.1
0.05
0.01
Cell_Type_NEW
Ctrl
ExN
InN
Other
UD

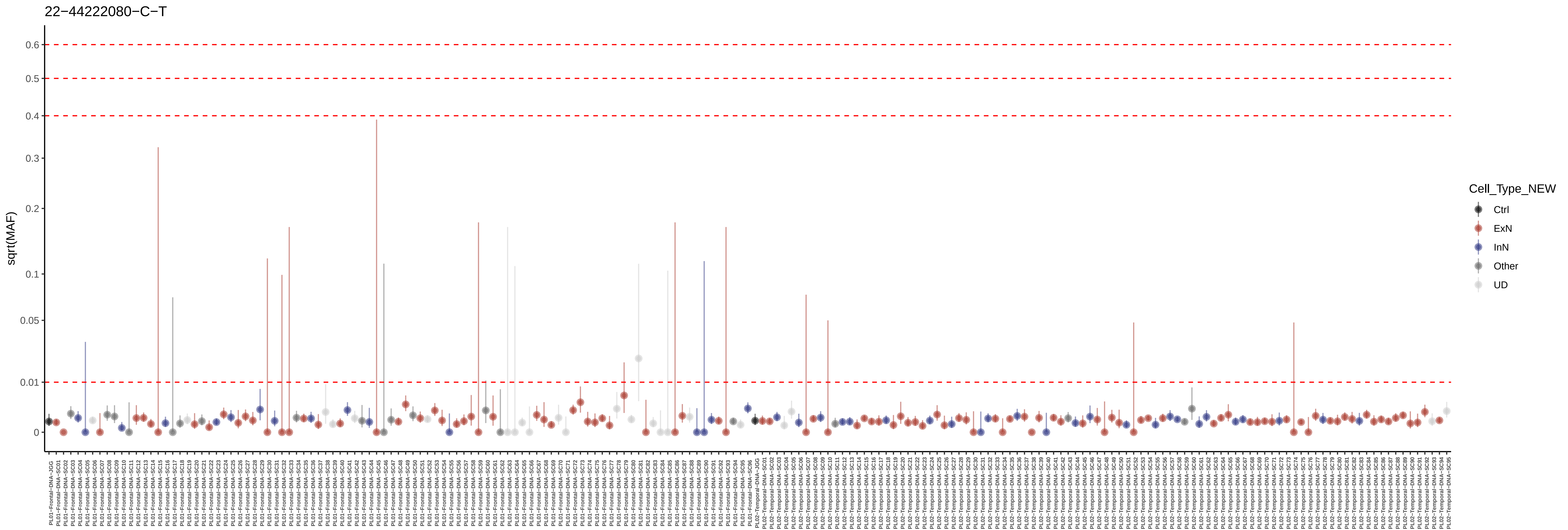
22−44222080−C−T
sqrt(MAF)
0.6
0.5
0.4
0.3
0.2
0.1
0.05
0.01
0
Cell_Type_NEW
Ctrl
ExN
InN
Other
UD

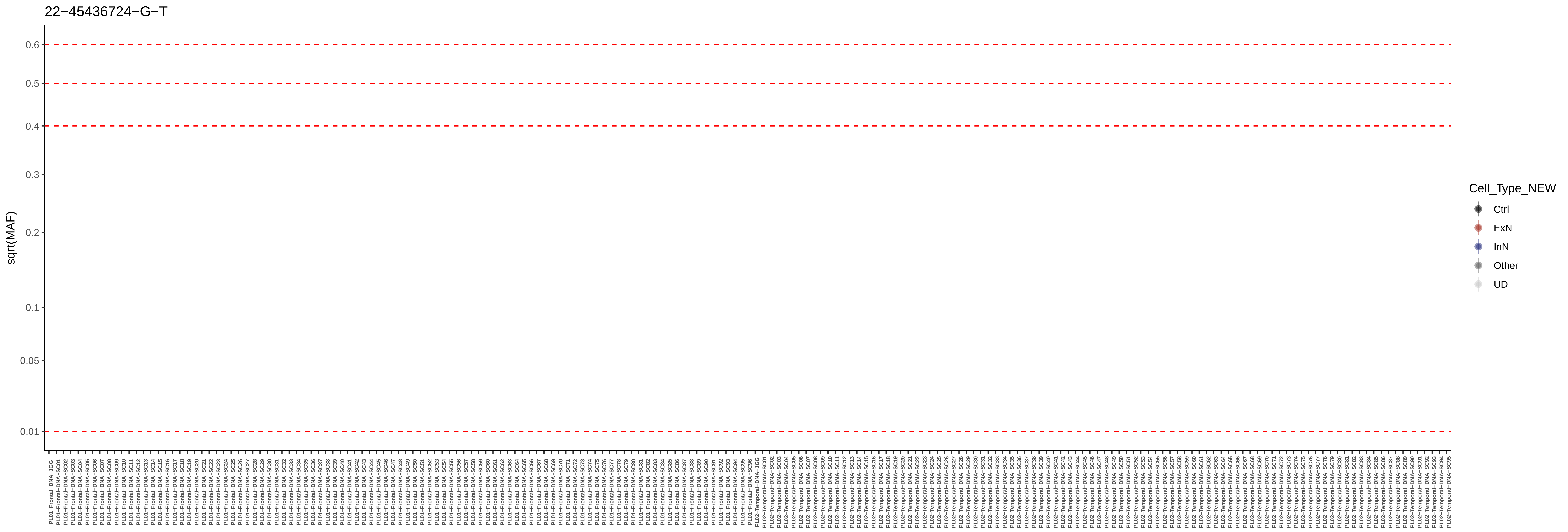
22−45436724−G−T
sqrt(MAF)
0.6
0.5
0.4
0.3
0.2
0.1
0.05
0.01
Cell_Type_NEW
Ctrl
ExN
InN
Other
UD

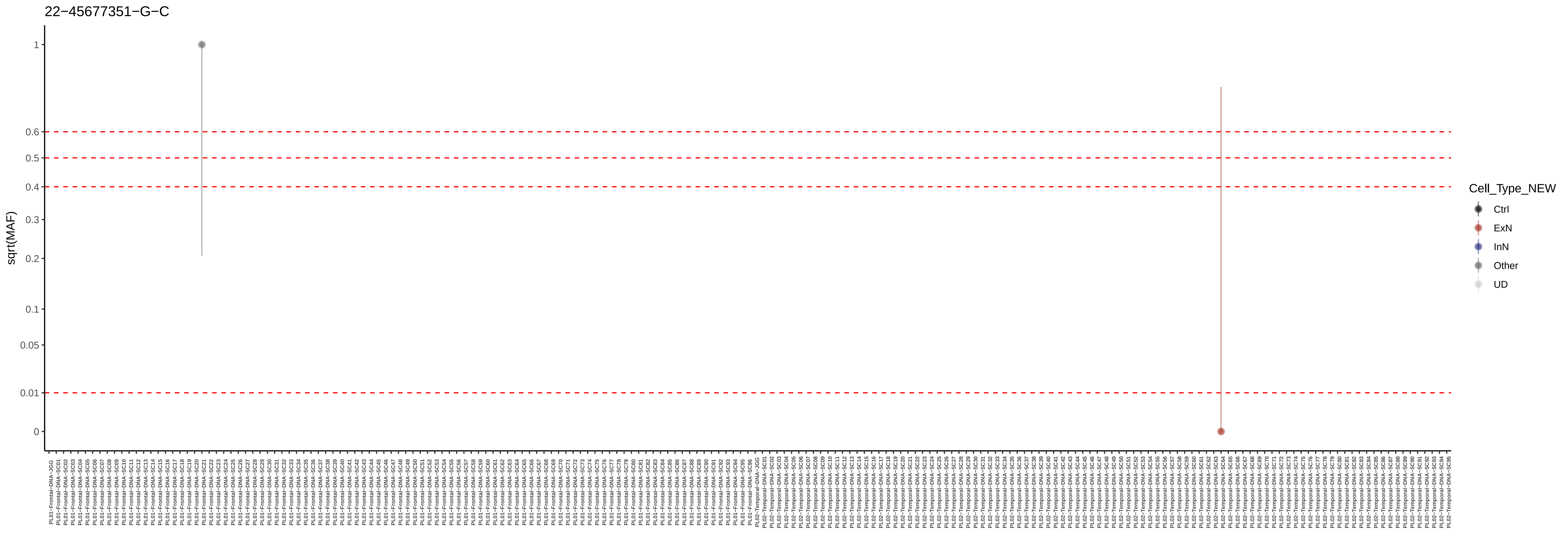
22−45677351−G−C
sqrt(MAF)
1
0.6
0.5
0.4
0.3
0.2
0.1
0.05
0.01
0
Cell_Type_NEW
Ctrl
ExN
InN
Other
UD

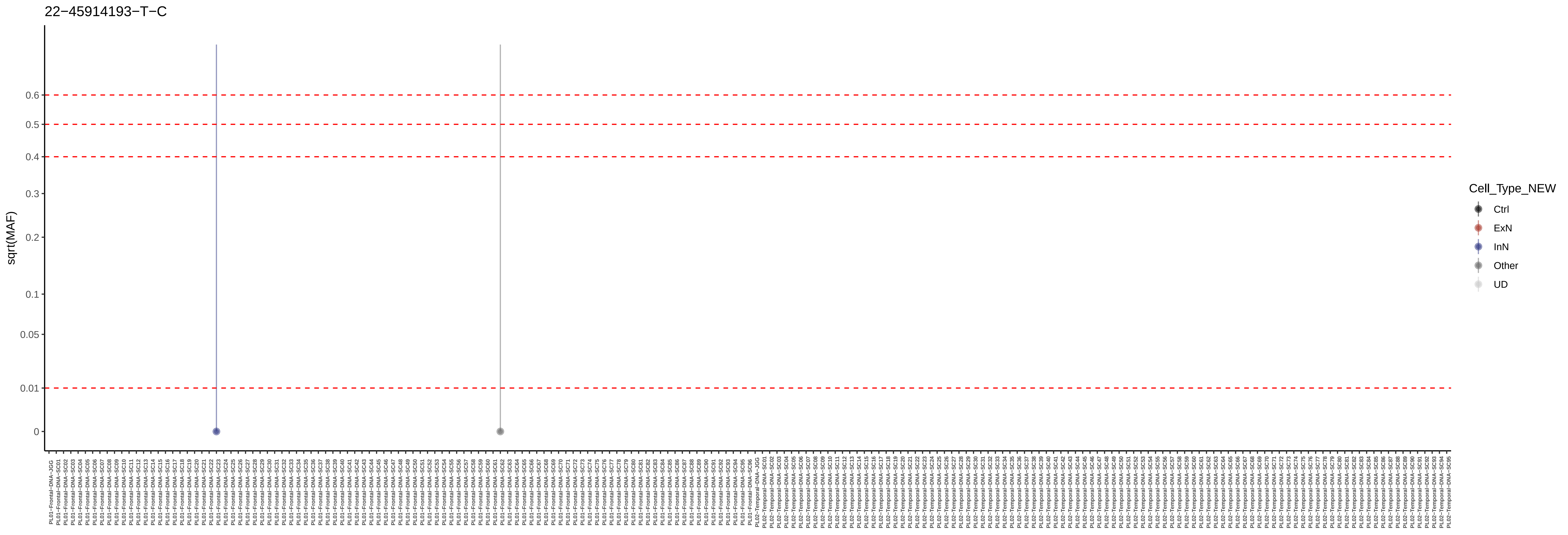
22−45914193−T−C
sqrt(MAF)
0.6
0.5
0.4
0.3
0.2
0.1
0.05
0.01
0
Cell_Type_NEW
Ctrl
ExN
InN
Other
UD

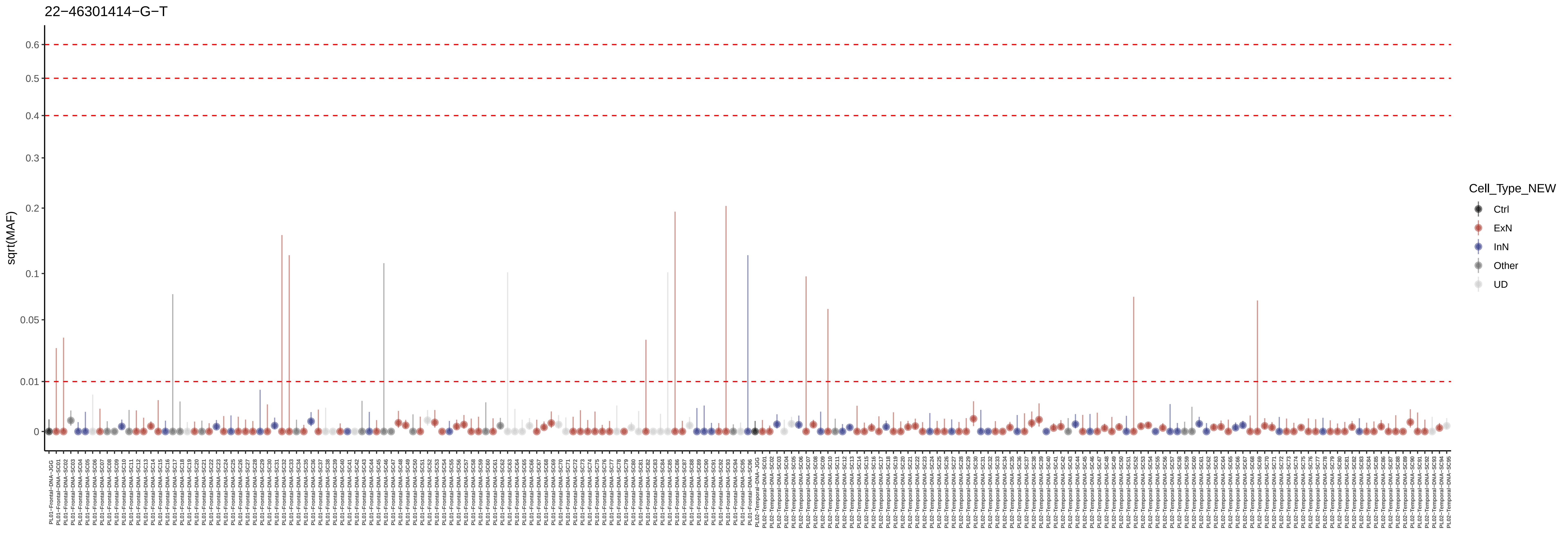

22−46301414−G−T
sqrt(MAF)
0.6
0.5
0.4
0.3
0.2
0.1
0.05
0.01
0
Cell_Type_NEW
Ctrl
ExN
InN
Other
UD

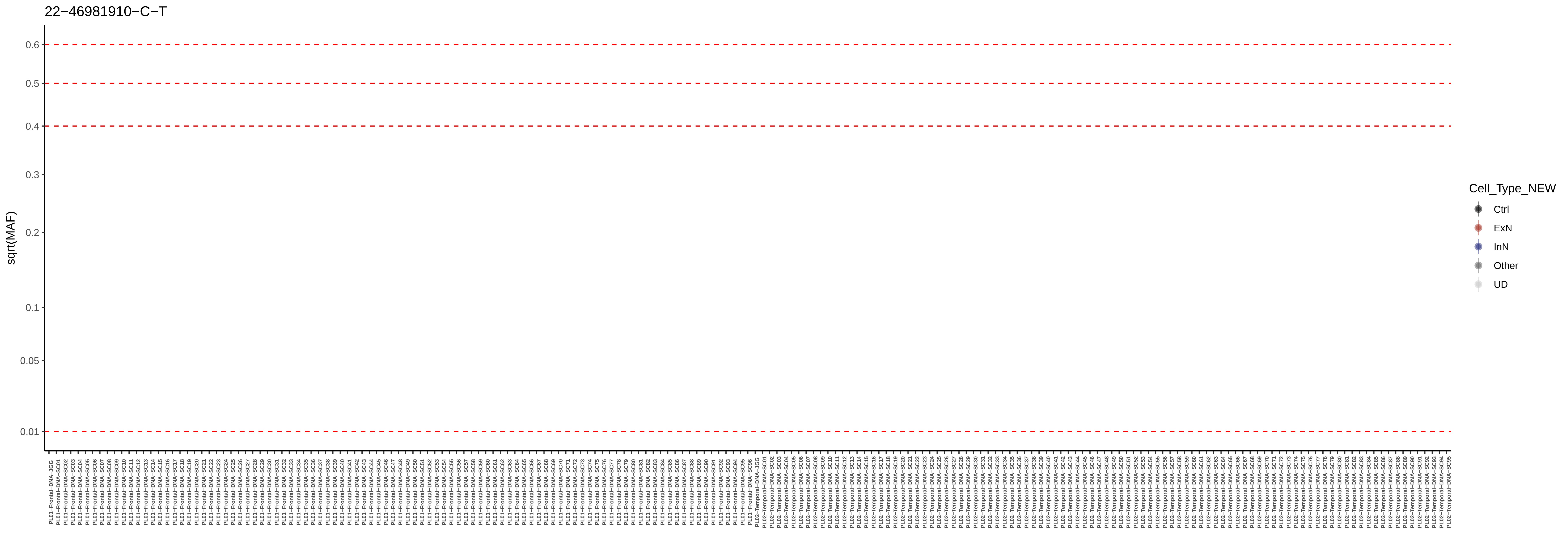

22−46981910−C−T
sqrt(MAF)
0.6
0.5
0.4
0.3
0.2
0.1
0.05
0.01
Cell_Type_NEW
Ctrl
ExN
InN
Other
UD

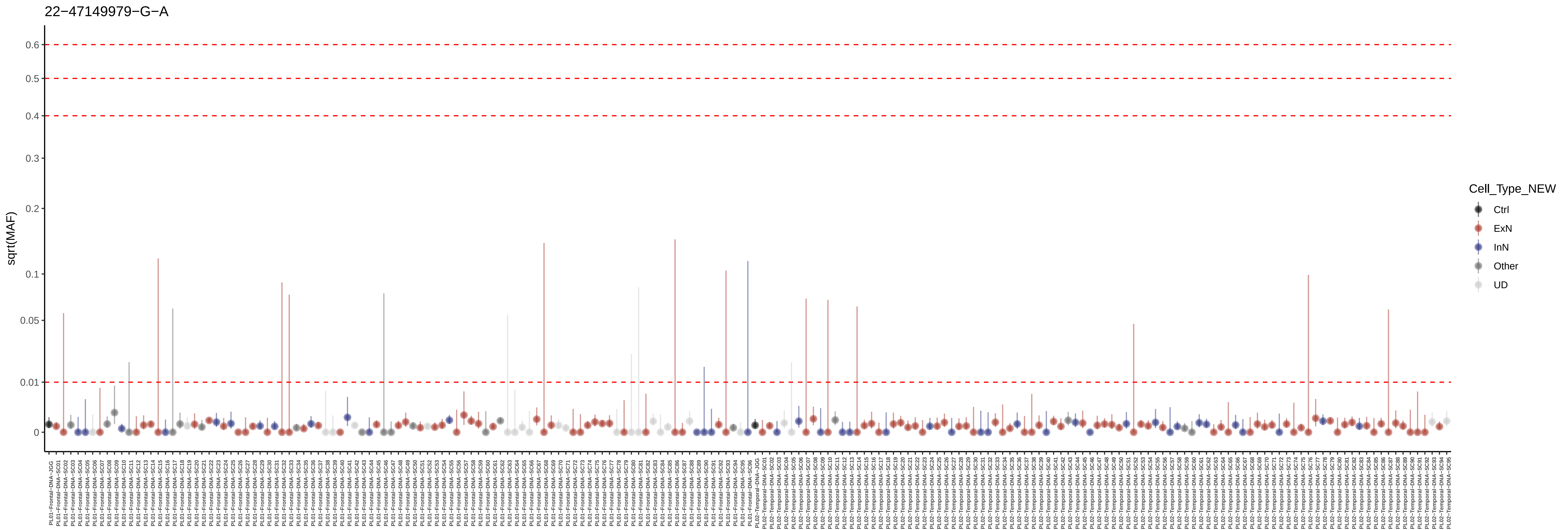
22−47149979−G−A
sqrt(MAF)
0.6
0.5
0.4
0.3
0.2
0.1
0.05
0.01
0
Cell_Type_NEW
Ctrl
ExN
InN
Other
UD

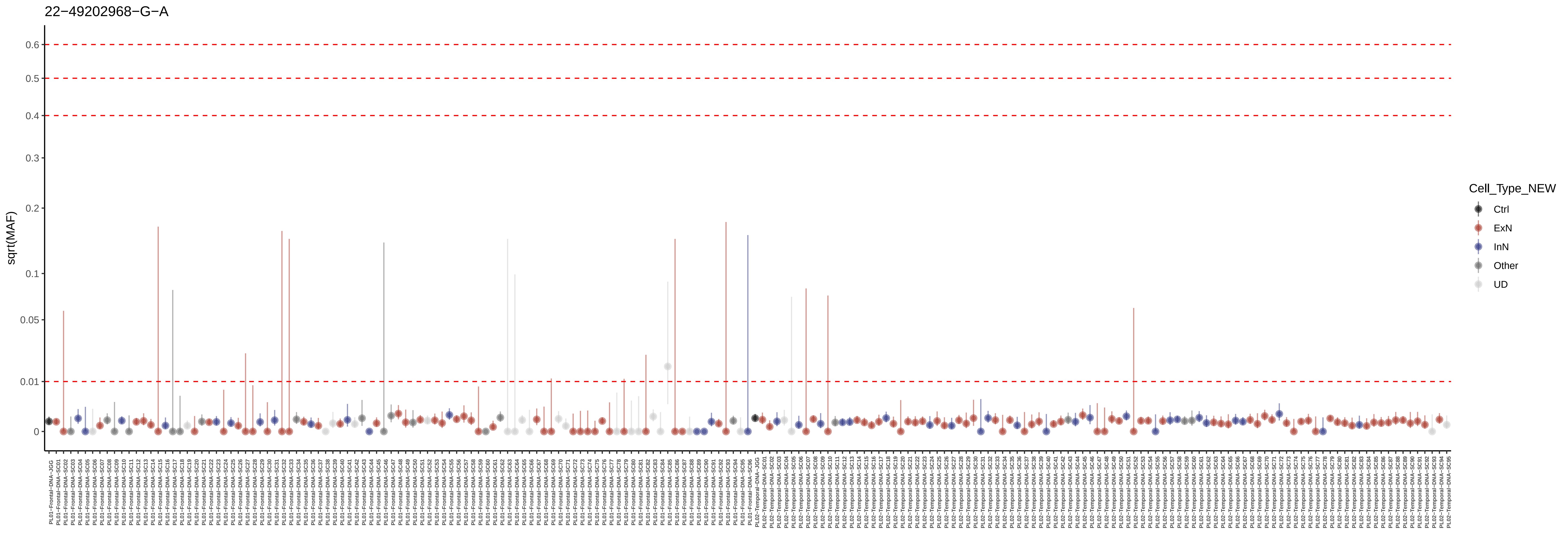

22−49202968−G−A
sqrt(MAF)
0.6
0.5
0.4
0.3
0.2
0.1
0.05
0.01
0
Cell_Type_NEW
Ctrl
ExN
InN
Other
UD

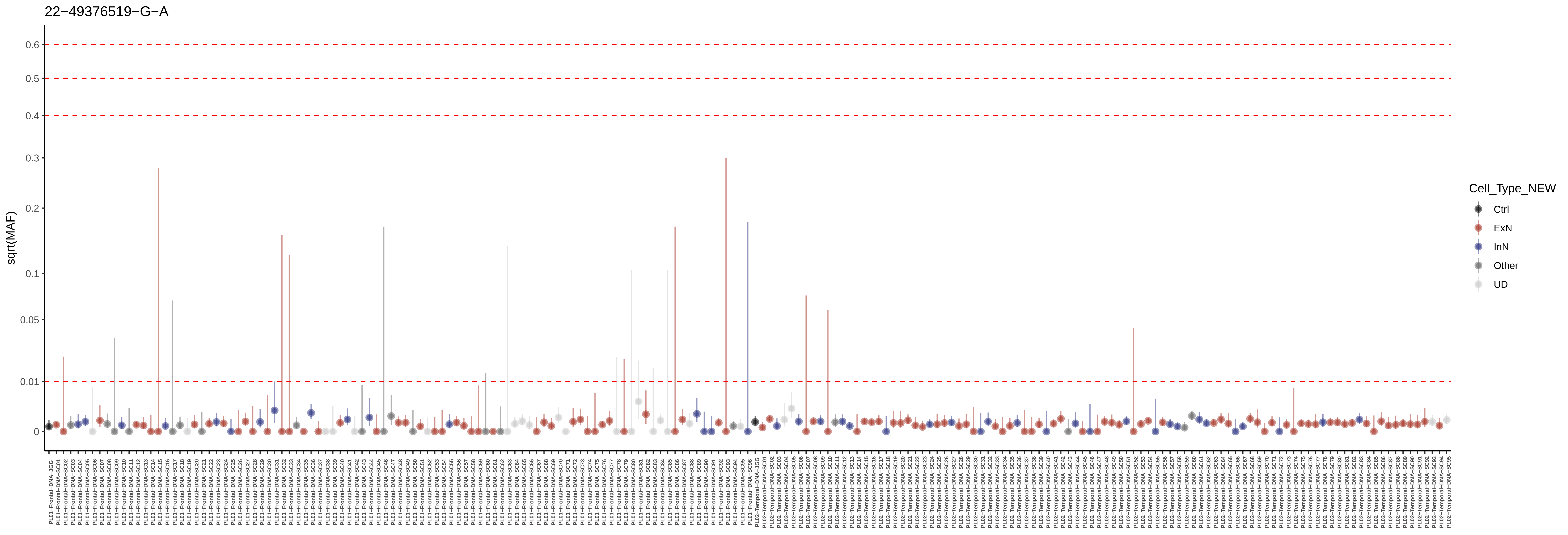
22−49376519−G−A
sqrt(MAF)
0.6
0.5
0.4
0.3
0.2
0.1
0.05
0.01
0
Cell_Type_NEW
Ctrl
ExN
InN
Other
UD

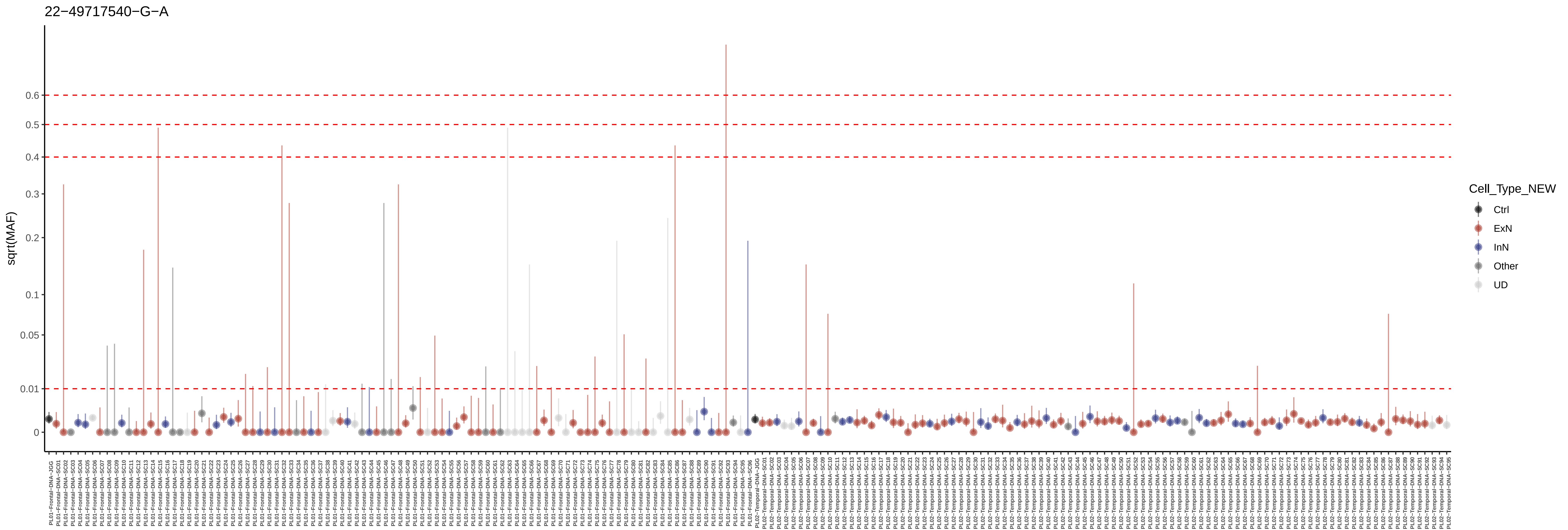
22−49717540−G−A
sqrt(MAF)
0.6
0.5
0.4
0.3
0.2
0.1
0.05
0.01
0
Cell_Type_NEW
Ctrl
ExN
InN
Other
UD

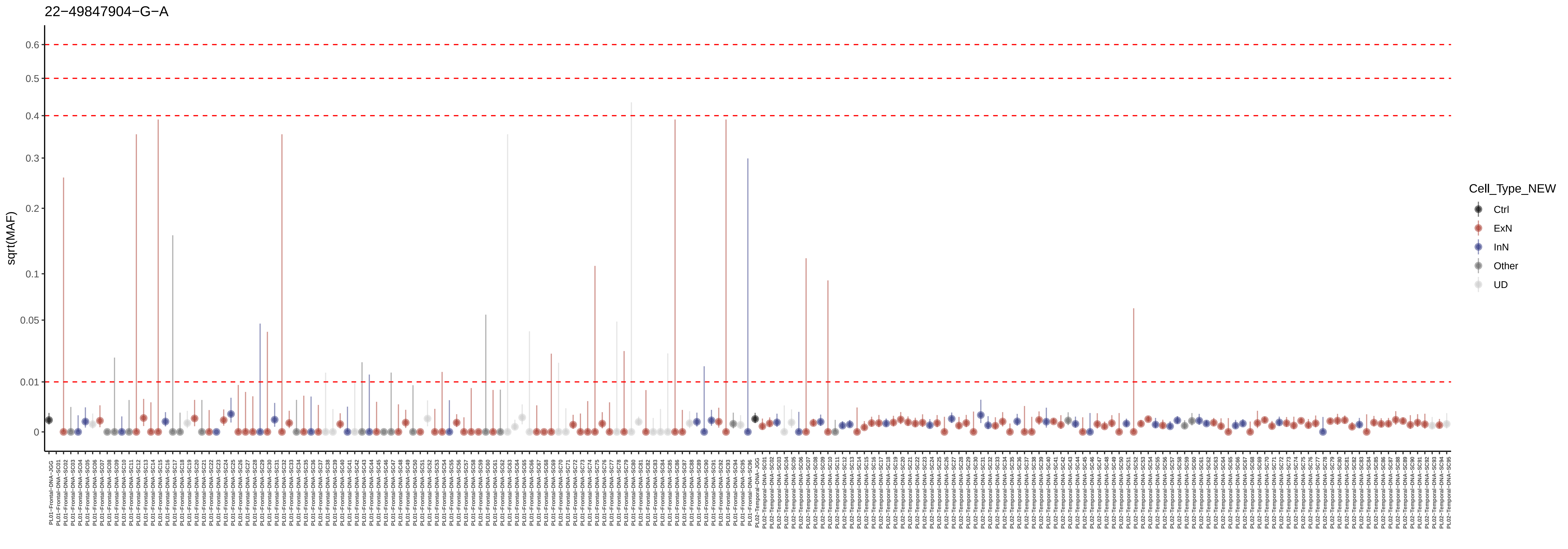

22−49847904−G−A
sqrt(MAF)
0.6
0.5
0.4
0.3
0.2
0.1
0.05
0.01
0
Cell_Type_NEW
Ctrl
ExN
InN
Other
UD

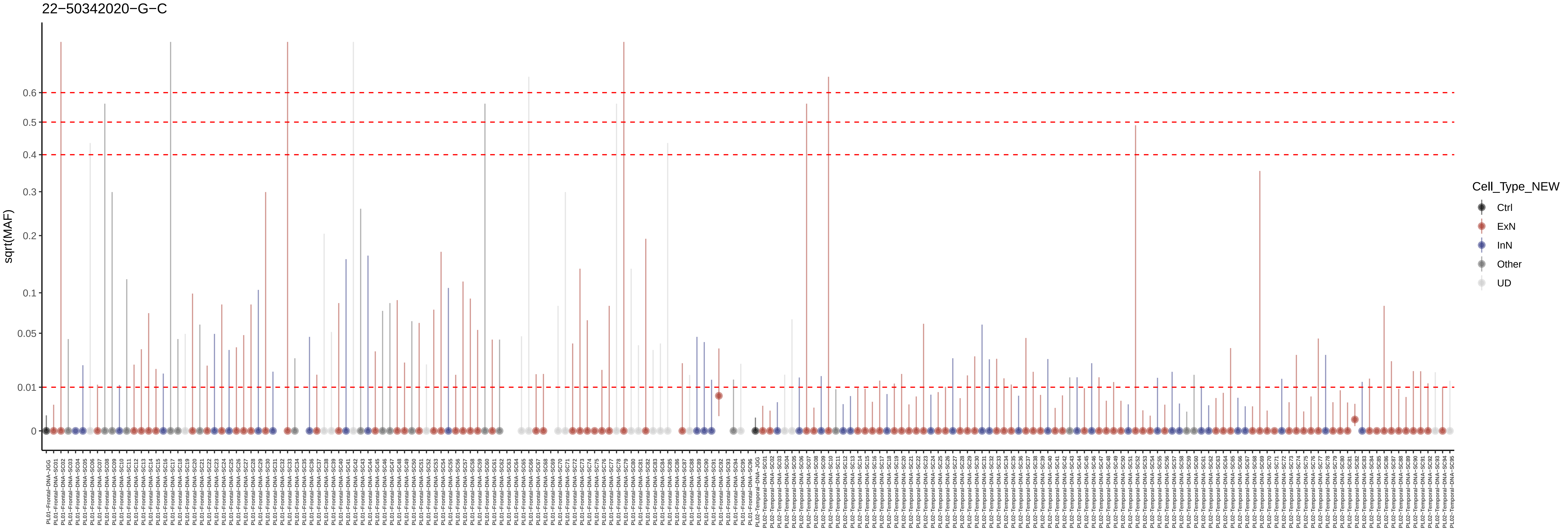
22−50342020−G−C
sqrt(MAF)
0.6
0.5
0.4
0.3
0.2
0.1
0.05
0.01
0
Cell_Type_NEW
Ctrl
ExN
InN
Other
UD

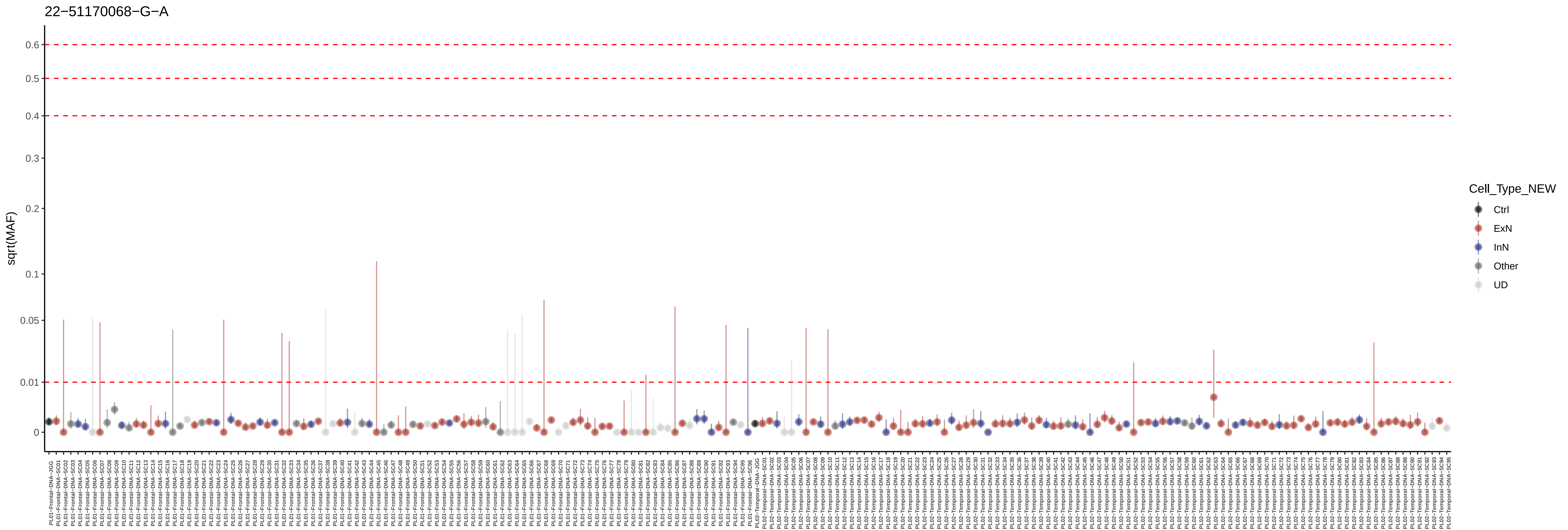

22−51170068−G−A
sqrt(MAF)
0.6
0.5
0.4
0.3
0.2
0.1
0.05
0.01
0
Cell_Type_NEW
Ctrl
ExN
InN
Other
UD

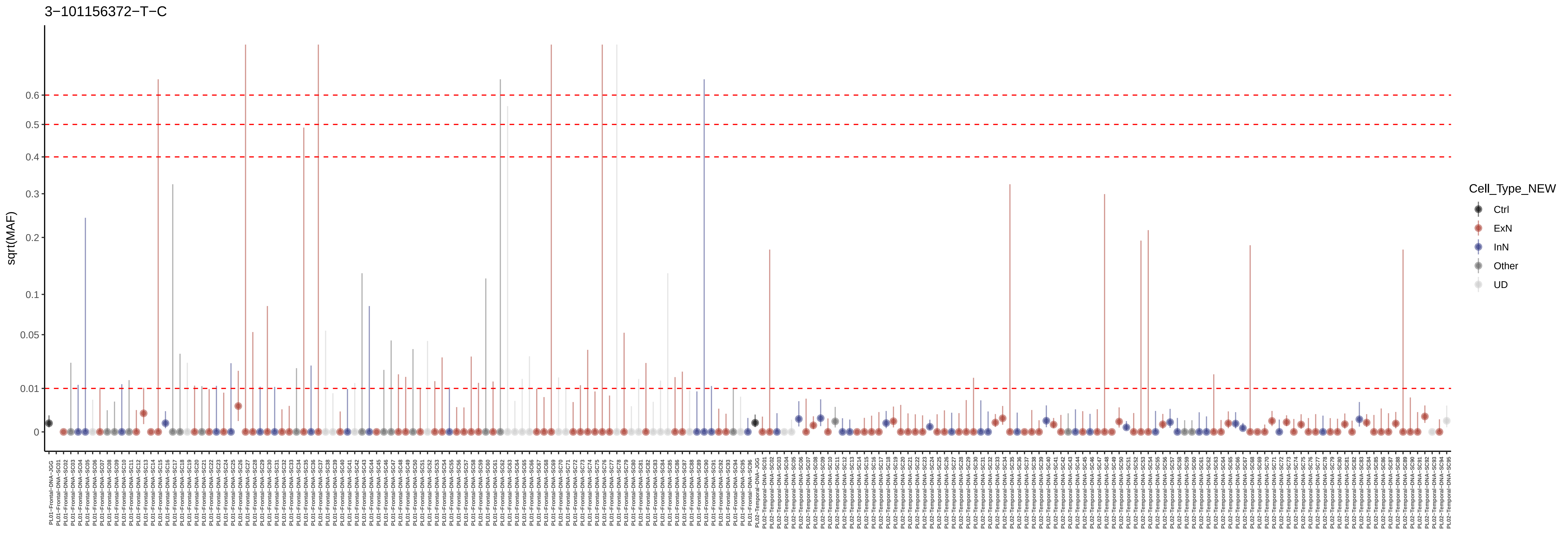

3−101156372−T−C
sqrt(MAF)
0.6
0.5
0.4
0.3
0.2
0.1
0.05
0.01
0
Cell_Type_NEW
Ctrl
ExN
InN
Other
UD

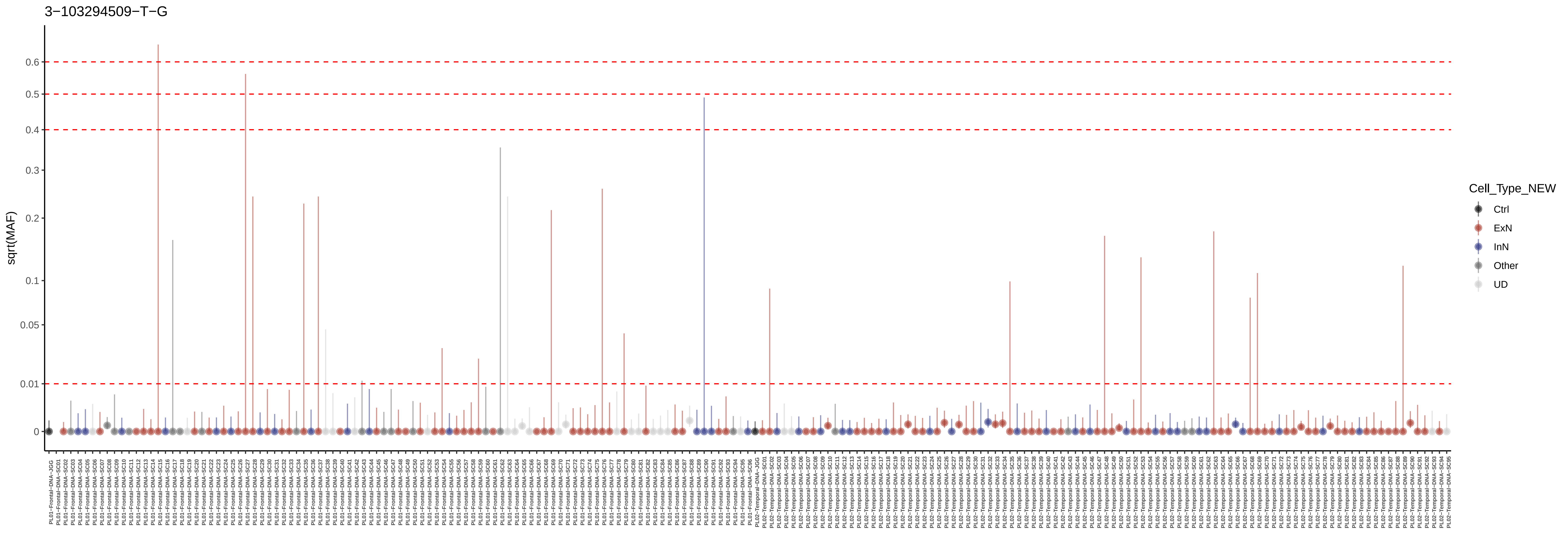

3−103294509−T−G
sqrt(MAF)
0.6
0.5
0.4
0.3
0.2
0.1
0.05
0.01
0
Cell_Type_NEW
Ctrl
ExN
InN
Other
UD

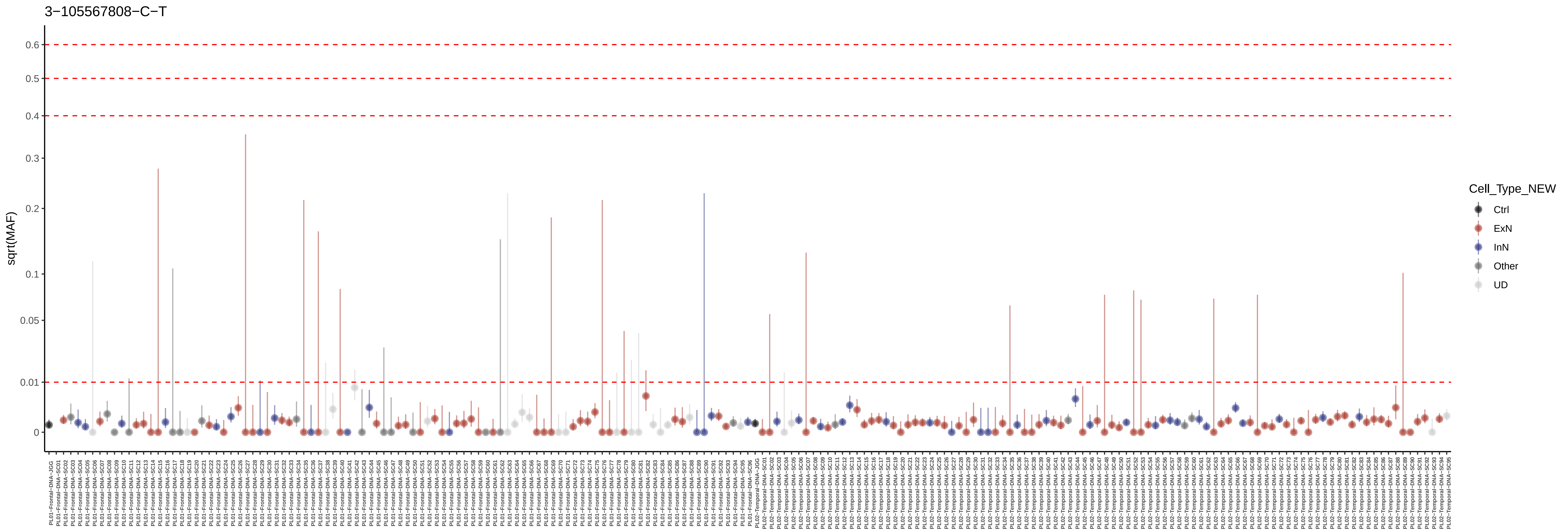

3−105567808−C−T
sqrt(MAF)
0.6
0.5
0.4
0.3
0.2
0.1
0.05
0.01
0
Cell_Type_NEW
Ctrl
ExN
InN
Other
UD

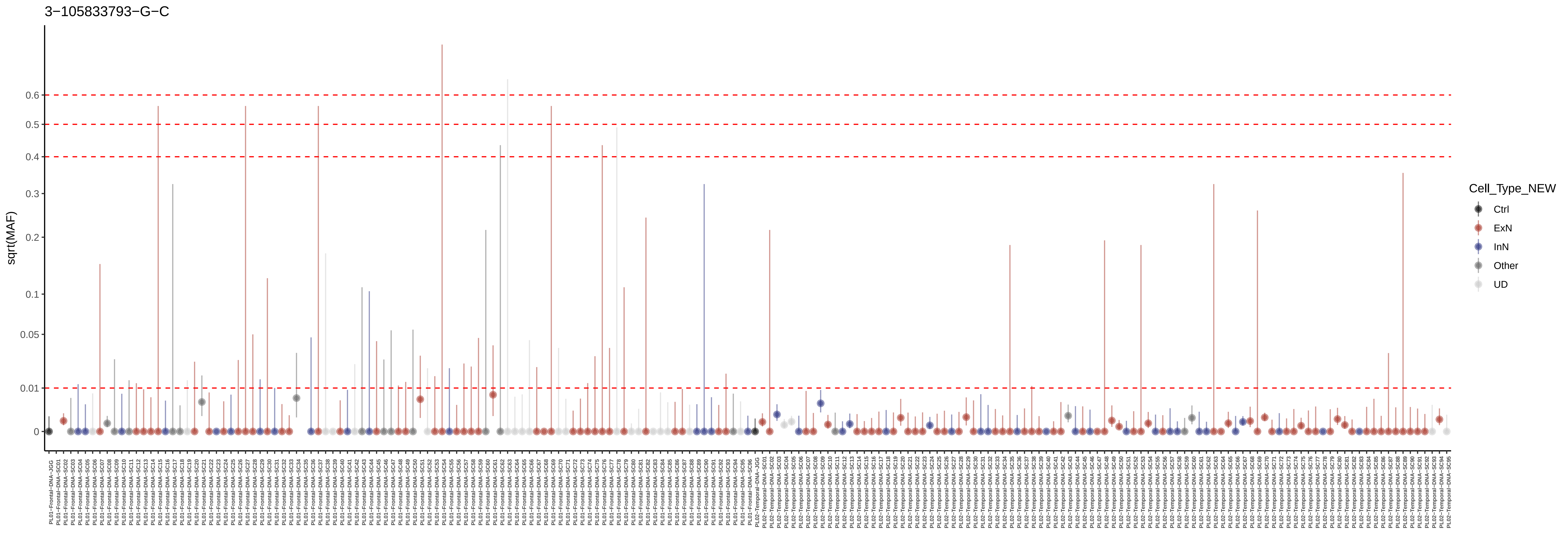

3−105833793−G−C
sqrt(MAF)
0.6
0.5
0.4
0.3
0.2
0.1
0.05
0.01
0
Cell_Type_NEW
Ctrl
ExN
InN
Other
UD

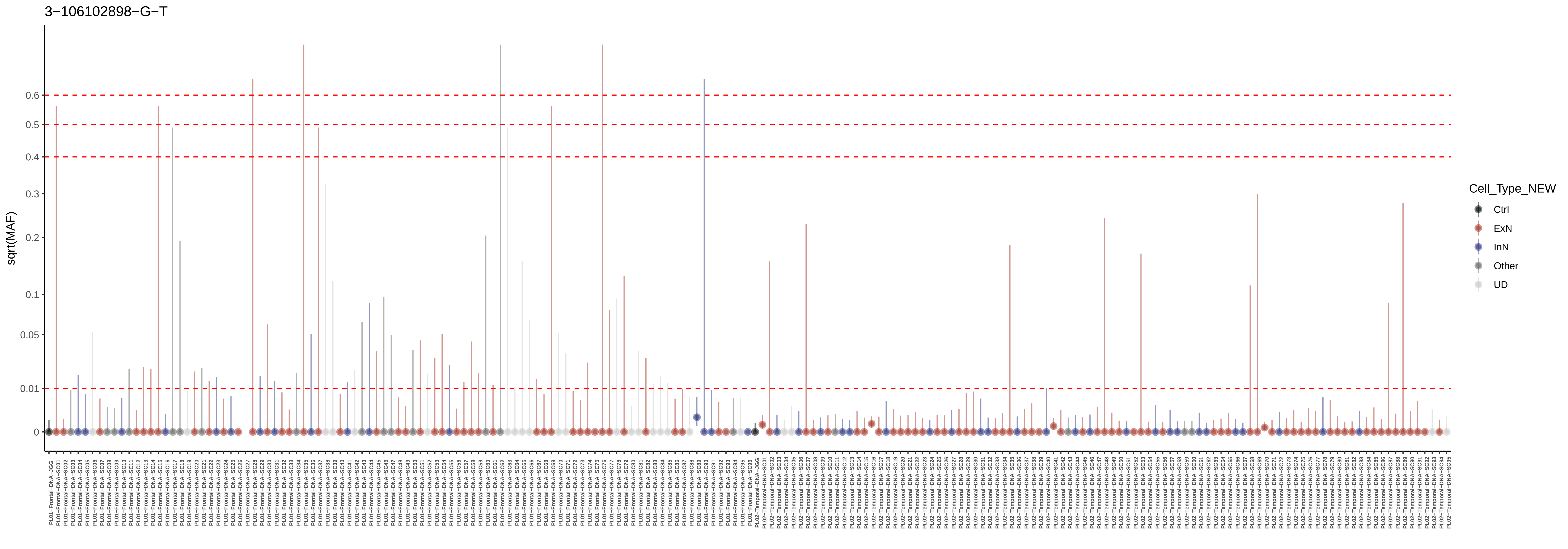
3−106102898−G−T
sqrt(MAF)
0.6
0.5
0.4
0.3
0.2
0.1
0.05
0.01
0
Cell_Type_NEW
Ctrl
ExN
InN
Other
UD

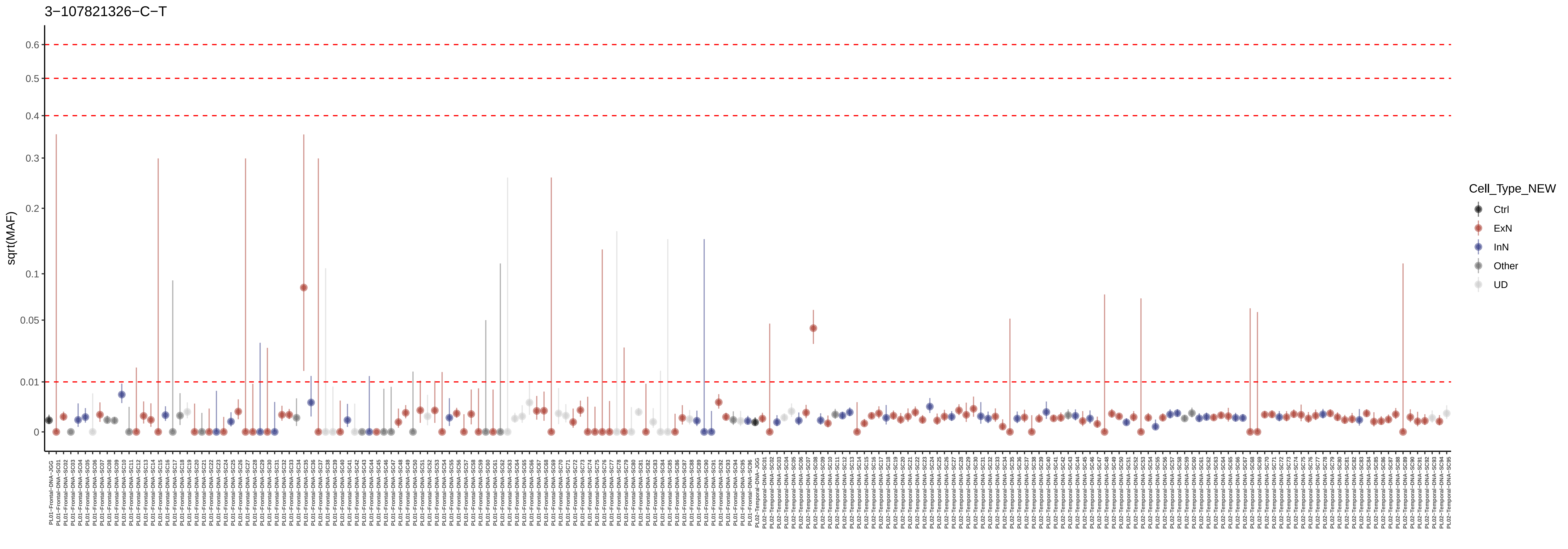

3−107821326−C−T
sqrt(MAF)
0.6
0.5
0.4
0.3
0.2
0.1
0.05
0.01
0
Cell_Type_NEW
Ctrl
ExN
InN
Other
UD

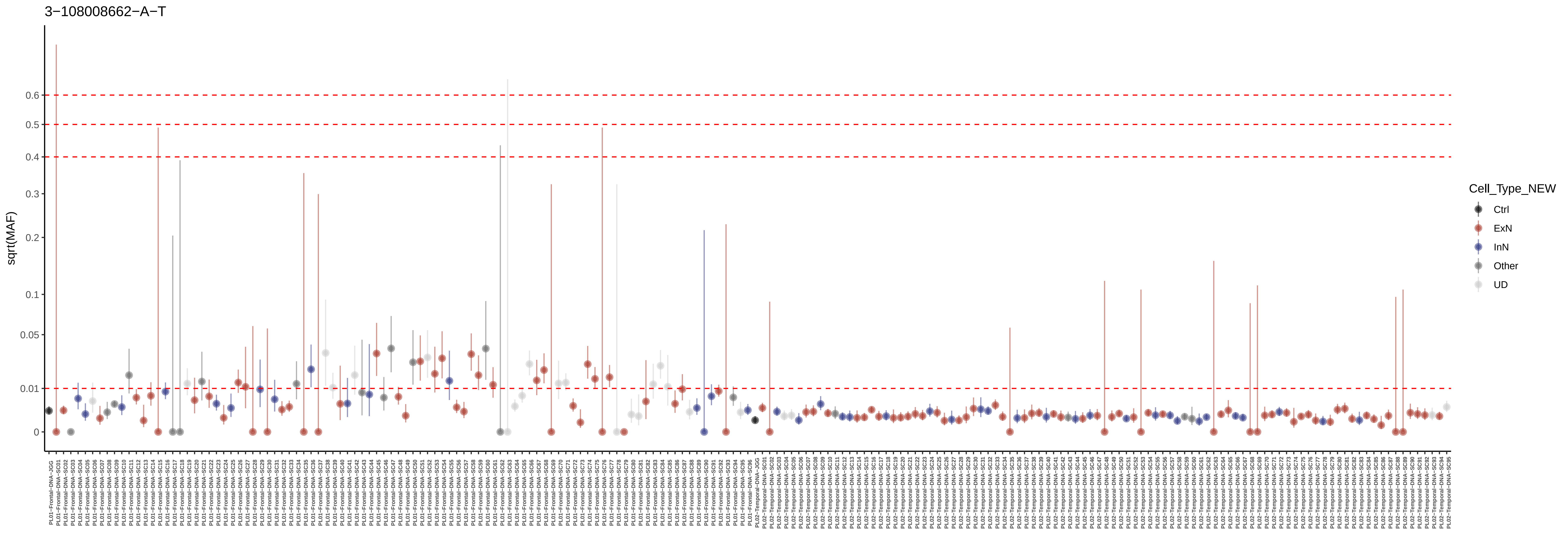

3−108008662−A−T
sqrt(MAF)
0.6
0.5
0.4
0.3
0.2
0.1
0.05
0.01
0
Cell_Type_NEW
Ctrl
ExN
InN
Other
UD

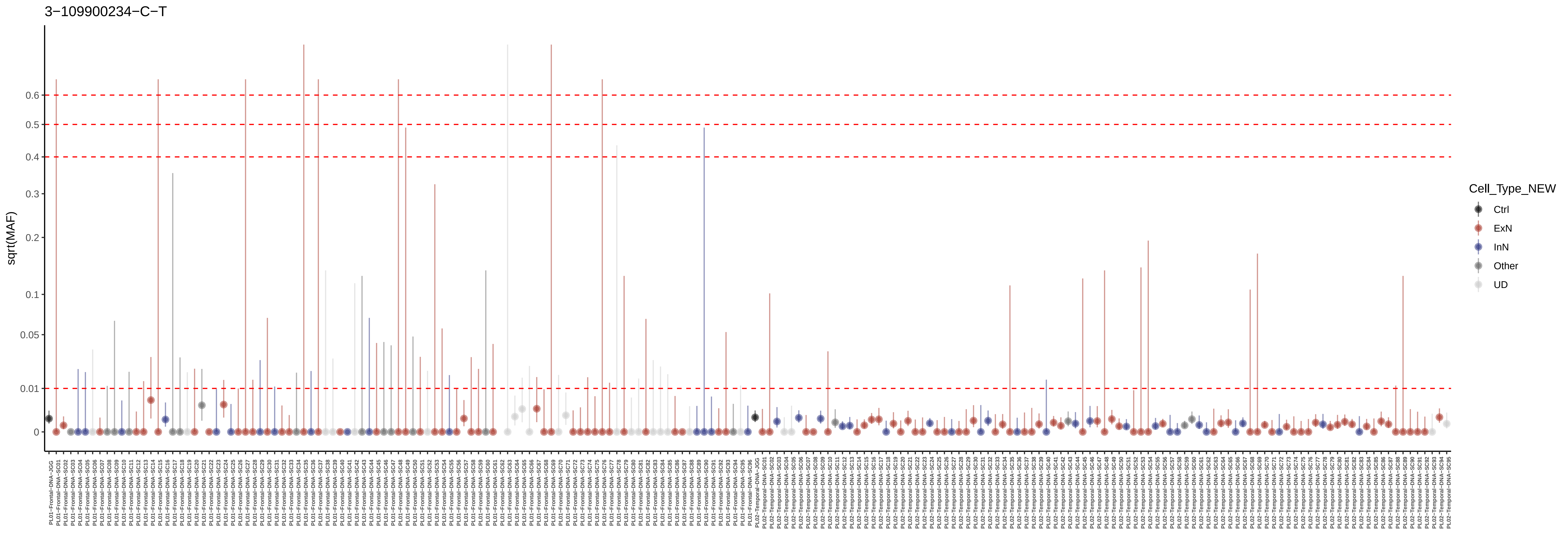

3−109900234−C−T
sqrt(MAF)
0.6
0.5
0.4
0.3
0.2
0.1
0.05
0.01
0
Cell_Type_NEW
Ctrl
ExN
InN
Other
UD

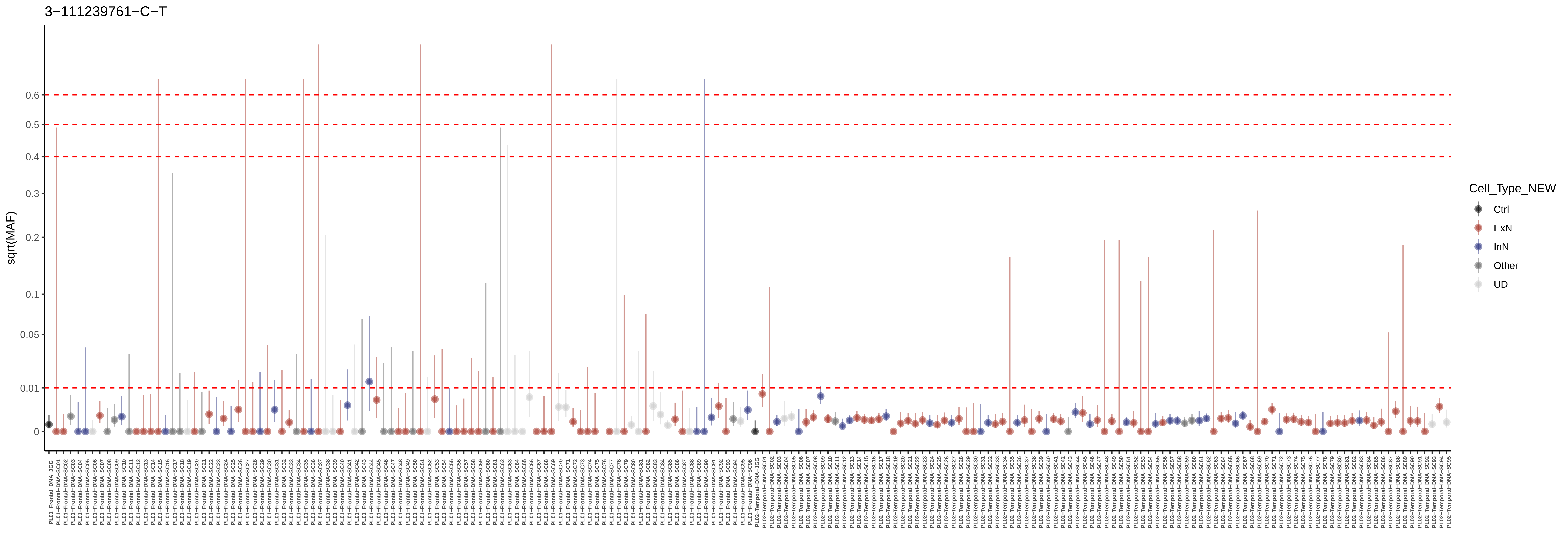
3−111239761−C−T
sqrt(MAF)
0.6
0.5
0.4
0.3
0.2
0.1
0.05
0.01
0
Cell_Type_NEW
Ctrl
ExN
InN
Other
UD

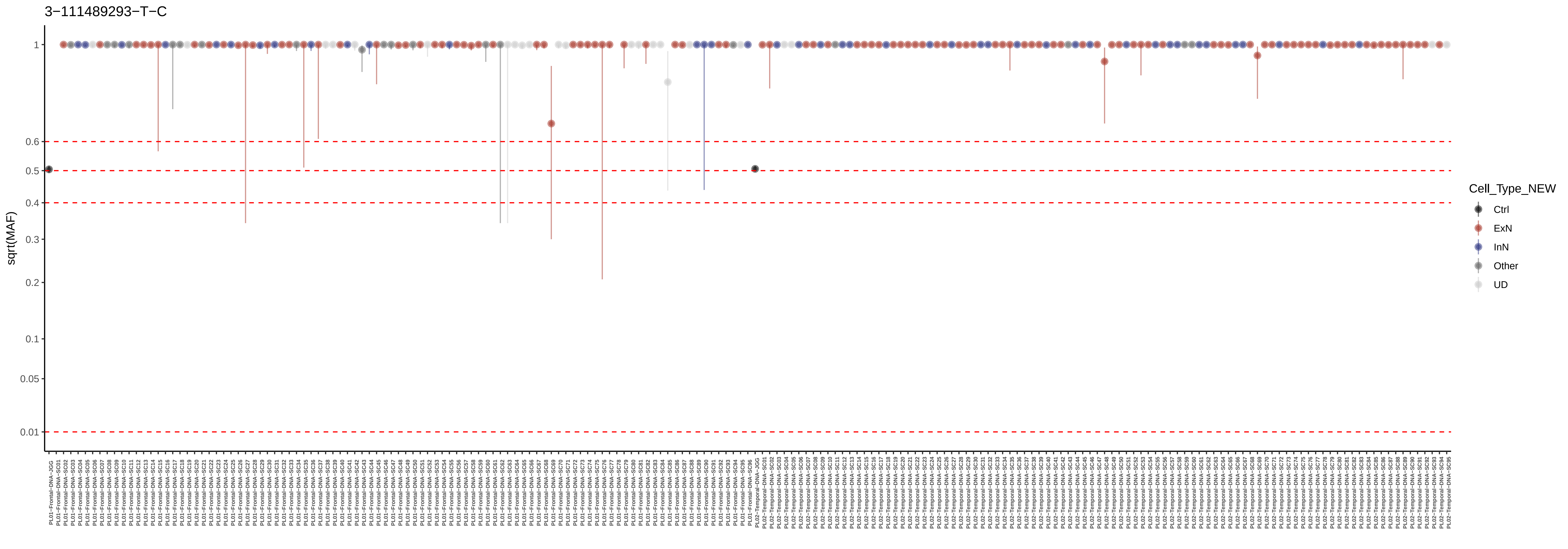

3−111489293−T−C
sqrt(MAF)
1
0.6
0.5
0.4
0.3
0.2
0.1
0.05
0.01
Cell_Type_NEW
Ctrl
ExN
InN
Other
UD

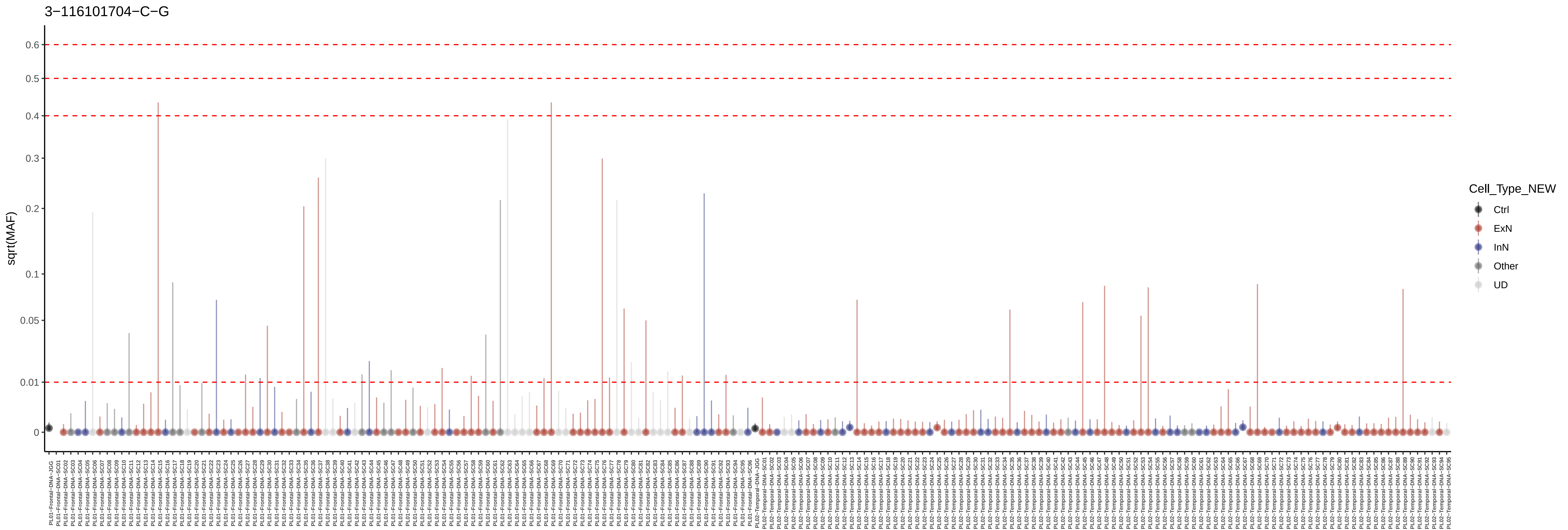
3−116101704−C−G
sqrt(MAF)
0.6
0.5
0.4
0.3
0.2
0.1
0.05
0.01
0
Cell_Type_NEW
Ctrl
ExN
InN
Other
UD

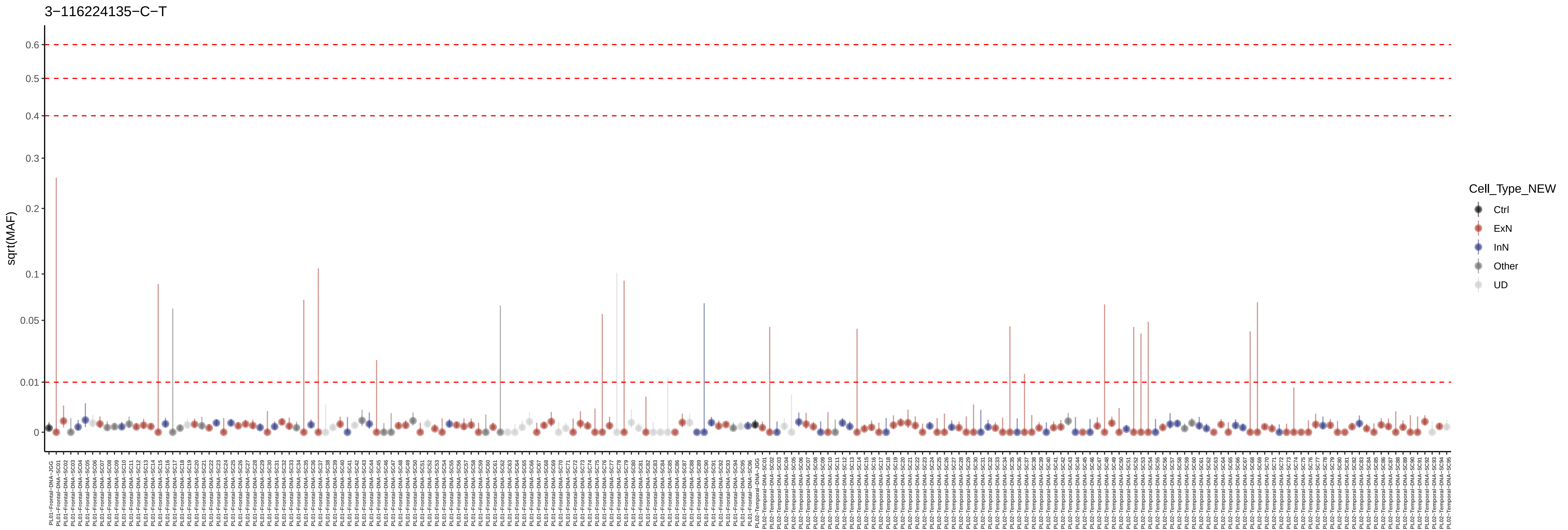

3−116224135−C−T
sqrt(MAF)
0.6
0.5
0.4
0.3
0.2
0.1
0.05
0.01
0
Cell_Type_NEW
Ctrl
ExN
InN
Other
UD

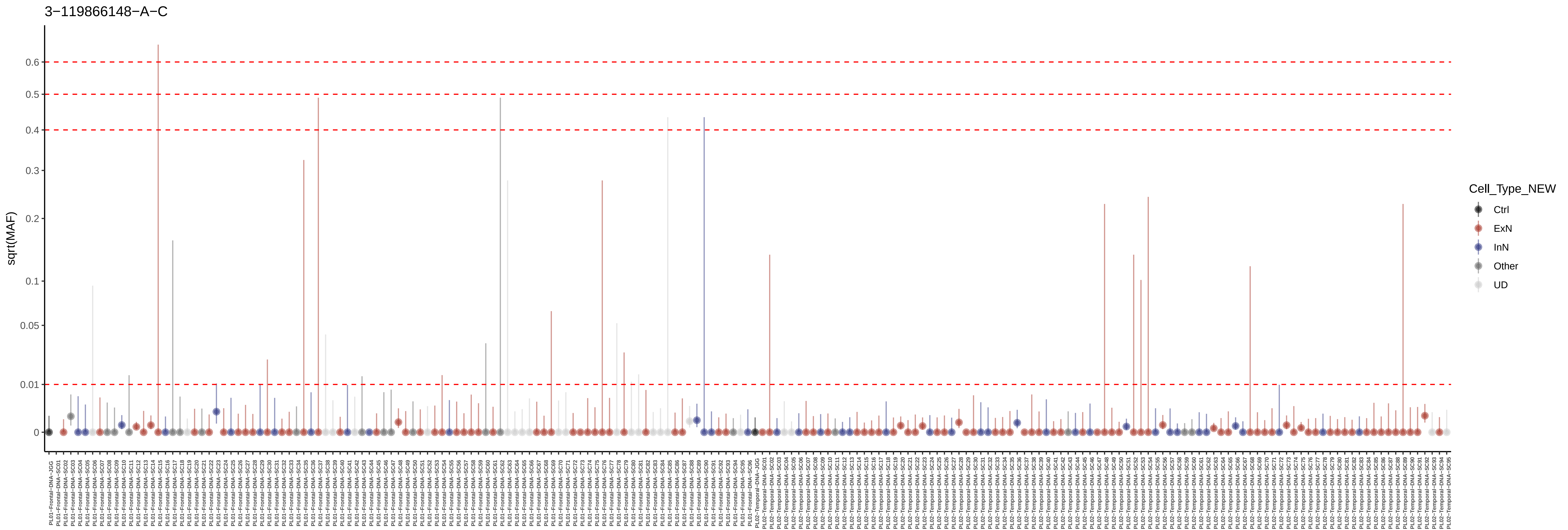
3−119866148−A−C
sqrt(MAF)
0.6
0.5
0.4
0.3
0.2
0.1
0.05
0.01
0
Cell_Type_NEW
Ctrl
ExN
InN
Other
UD

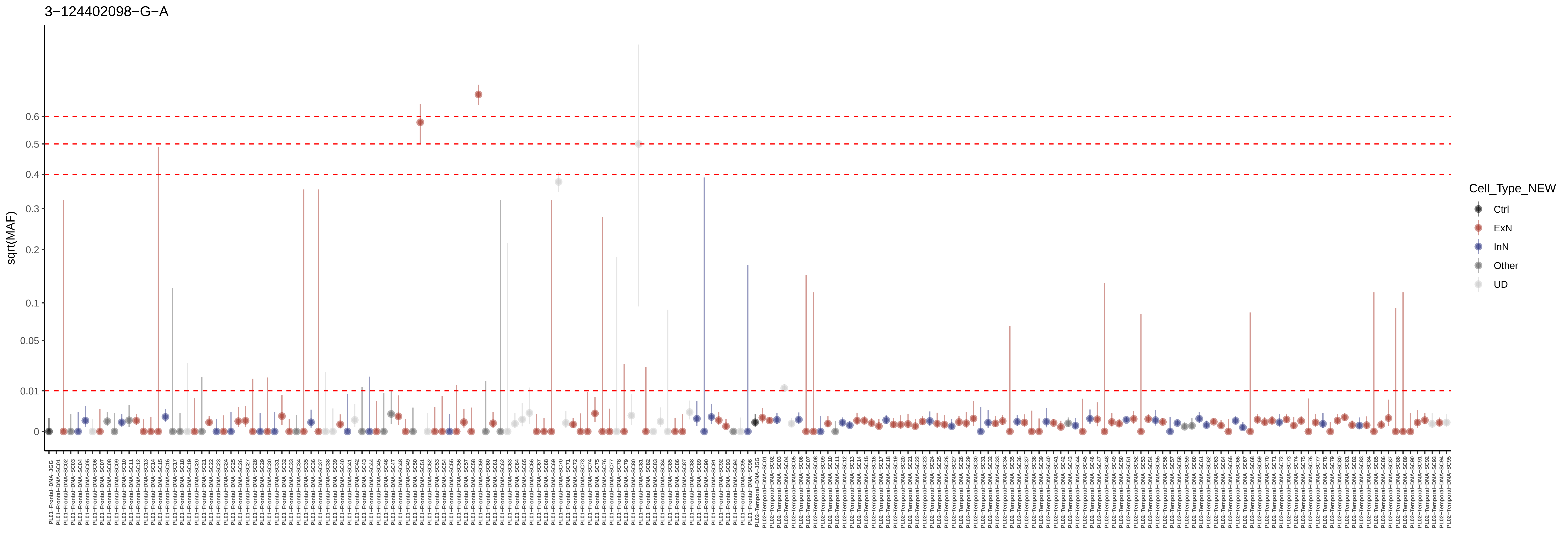
3−124402098−G−A
sqrt(MAF)
0.6
0.5
0.4
0.3
0.2
0.1
0.05
0.01
0
Cell_Type_NEW
Ctrl
ExN
InN
Other
UD

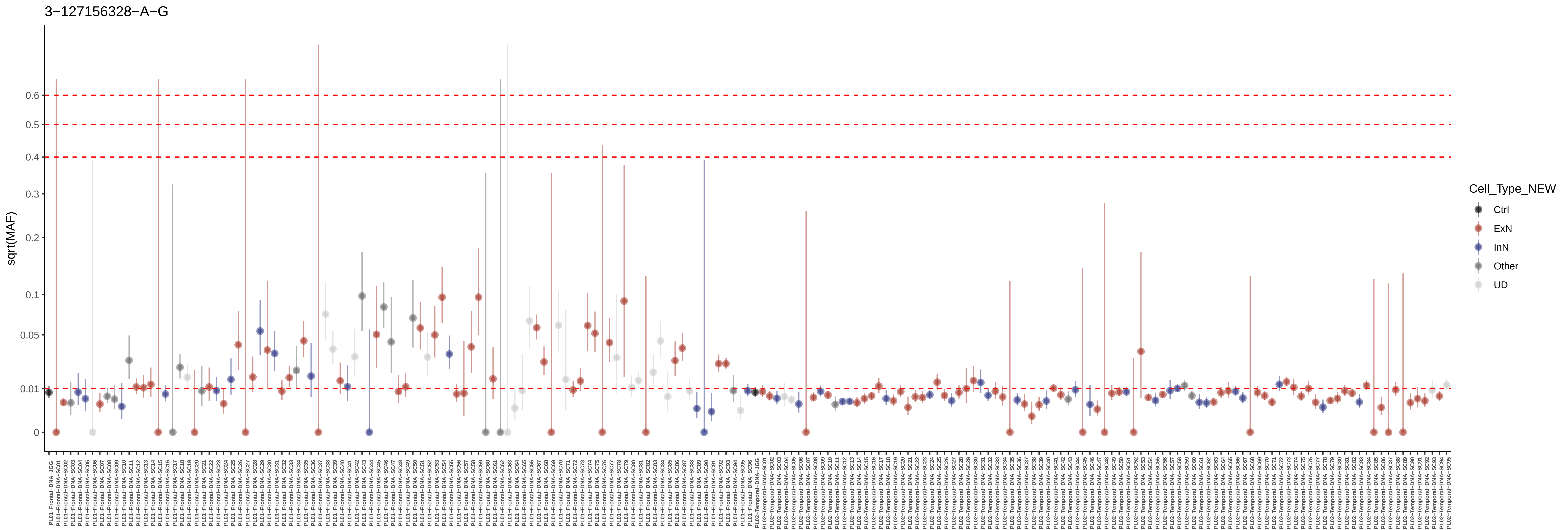
3−127156328−A−G
sqrt(MAF)
0.6
0.5
0.4
0.3
0.2
0.1
0.05
0.01
0
Cell_Type_NEW
Ctrl
ExN
InN
Other
UD

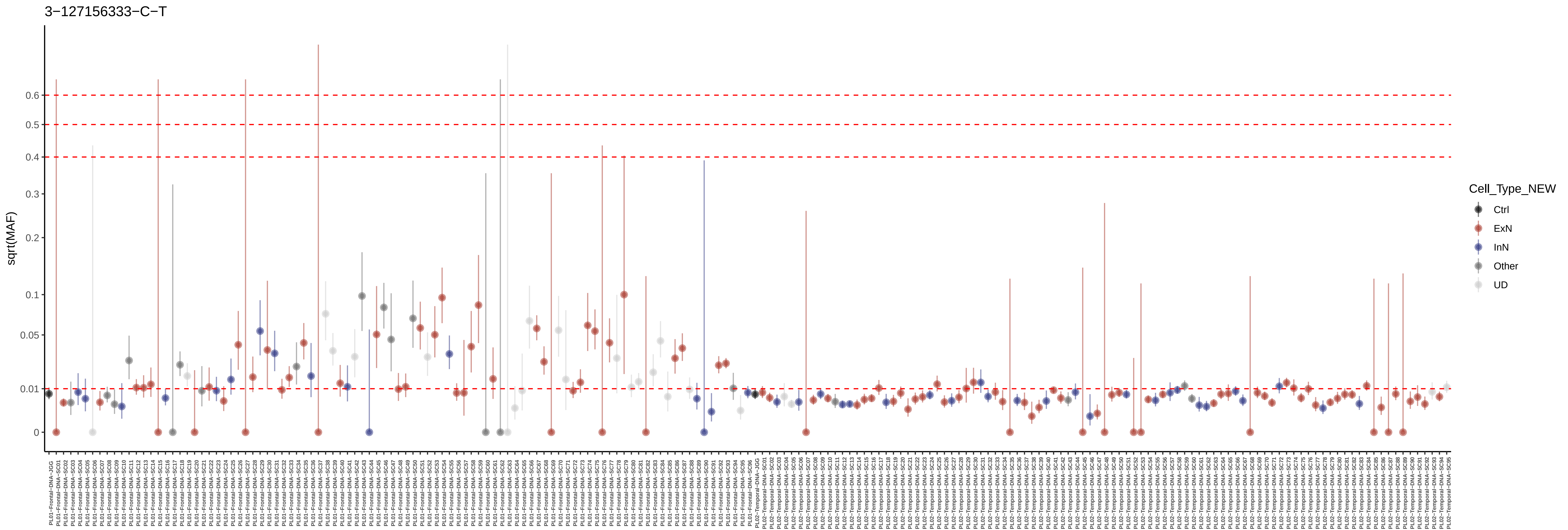
3−127156333−C−T
sqrt(MAF)
0.6
0.5
0.4
0.3
0.2
0.1
0.05
0.01
0
Cell_Type_NEW
Ctrl
ExN
InN
Other
UD

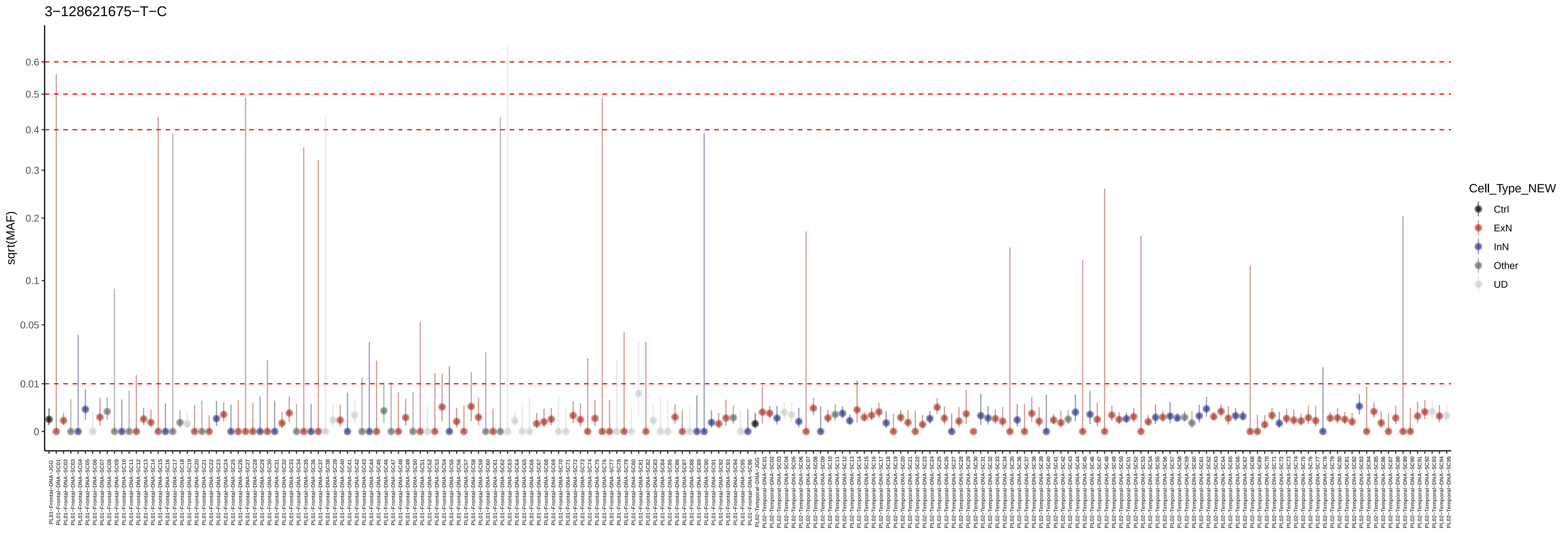
3−128621675−T−C
sqrt(MAF)
0.6
0.5
0.4
0.3
0.2
0.1
0.05
0.01
0
Cell_Type_NEW
Ctrl
ExN
InN
Other
UD

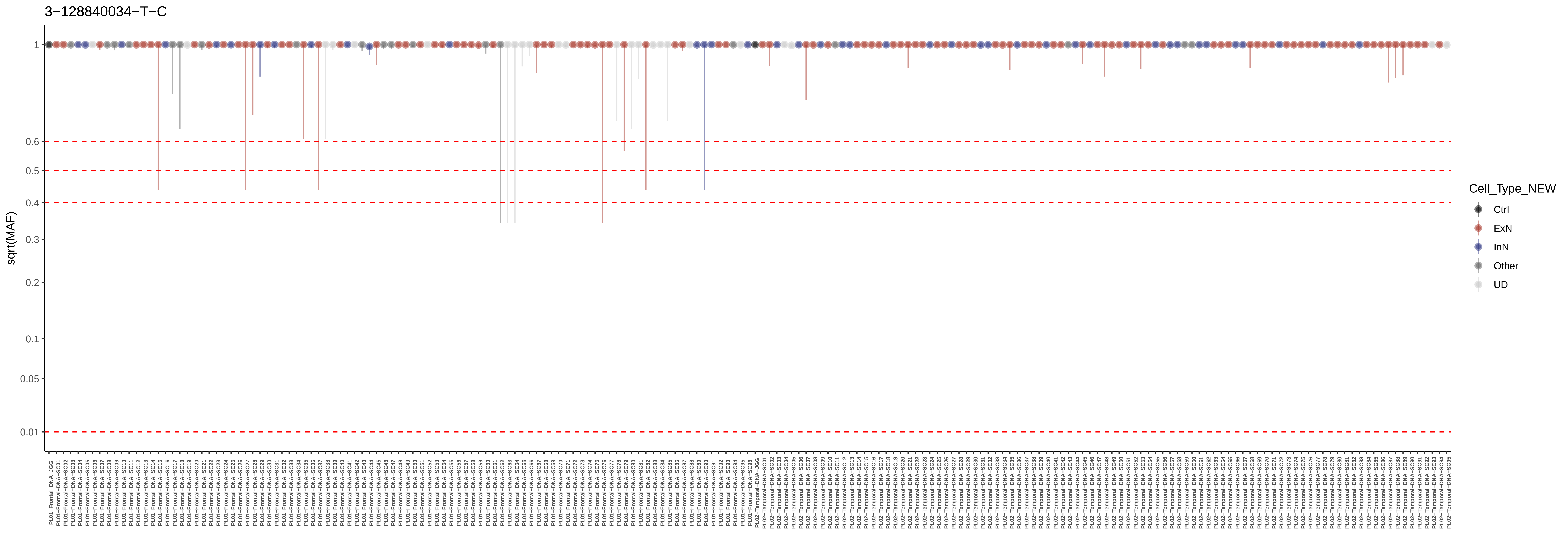

3−128840034−T−C
sqrt(MAF)
1
0.6
0.5
0.4
0.3
0.2
0.1
0.05
0.01
Cell_Type_NEW
Ctrl
ExN
InN
Other
UD

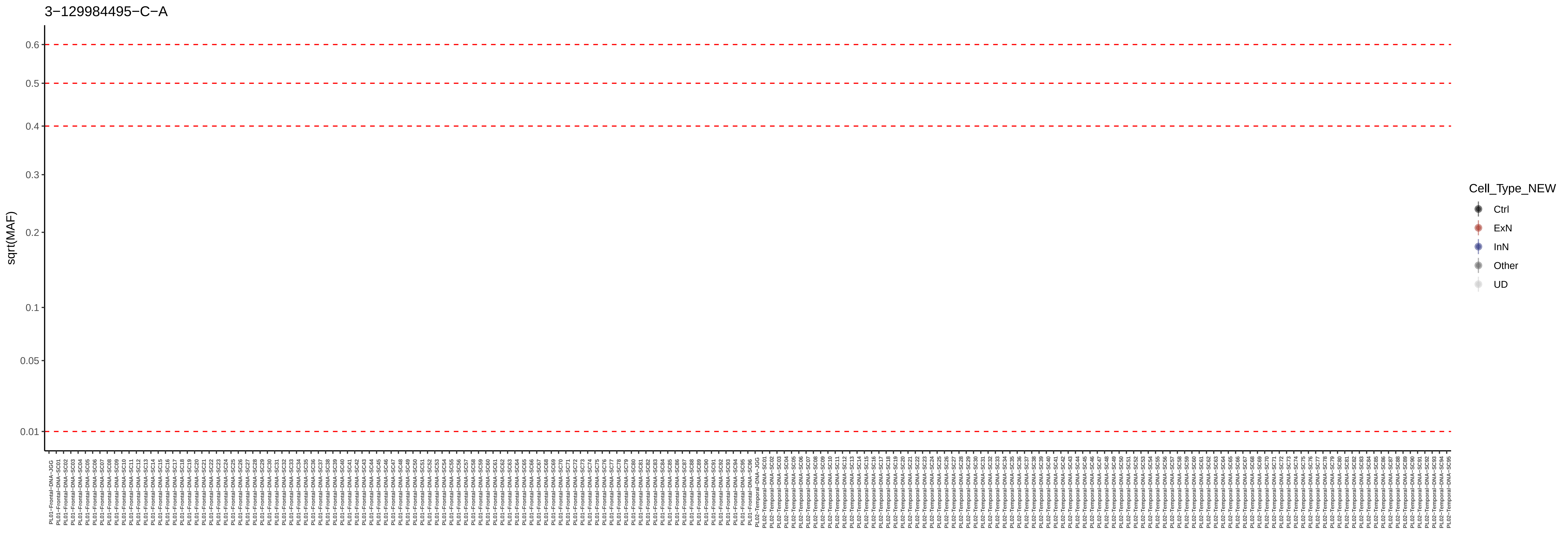
3−129984495−C−A
sqrt(MAF)
0.6
0.5
0.4
0.3
0.2
0.1
0.05
0.01
Cell_Type_NEW
Ctrl
ExN
InN
Other
UD

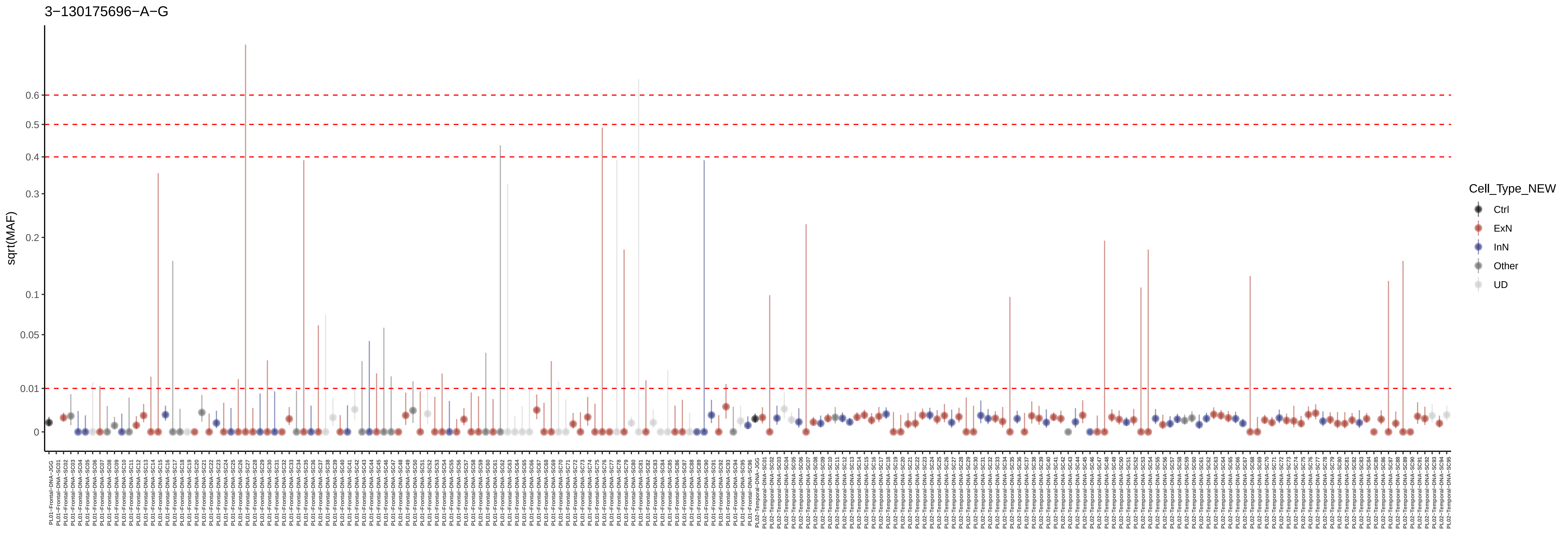

3−130175696−A−G
sqrt(MAF)
0.6
0.5
0.4
0.3
0.2
0.1
0.05
0.01
0
Cell_Type_NEW
Ctrl
ExN
InN
Other
UD

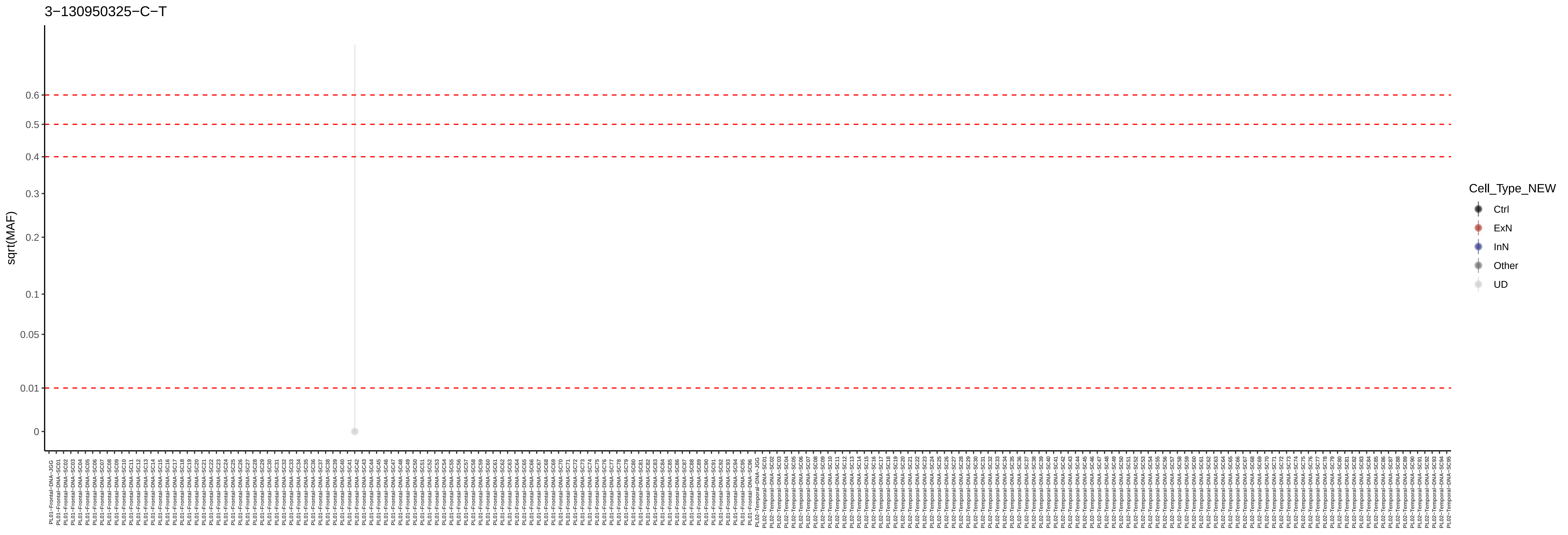
3−130950325−C−T
sqrt(MAF)
0.6
0.5
0.4
0.3
0.2
0.1
0.05
0.01
0
Cell_Type_NEW
Ctrl
ExN
InN
Other
UD

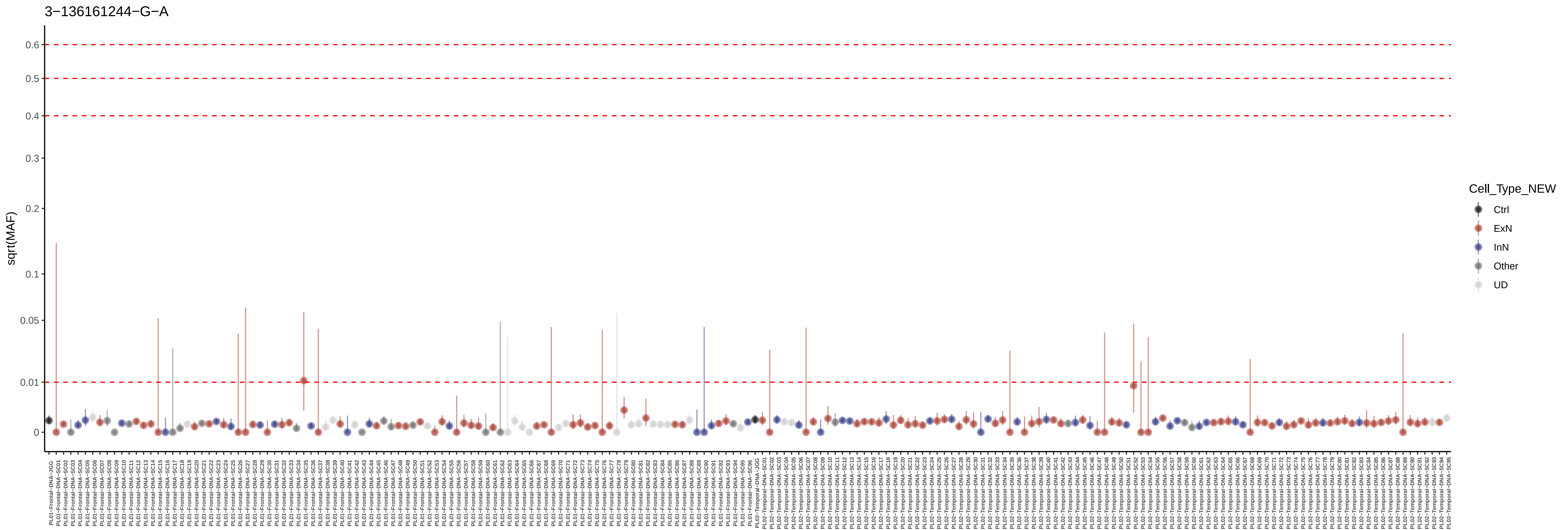

3−136161244−G−A
sqrt(MAF)
0.6
0.5
0.4
0.3
0.2
0.1
0.05
0.01
0
Cell_Type_NEW
Ctrl
ExN
InN
Other
UD

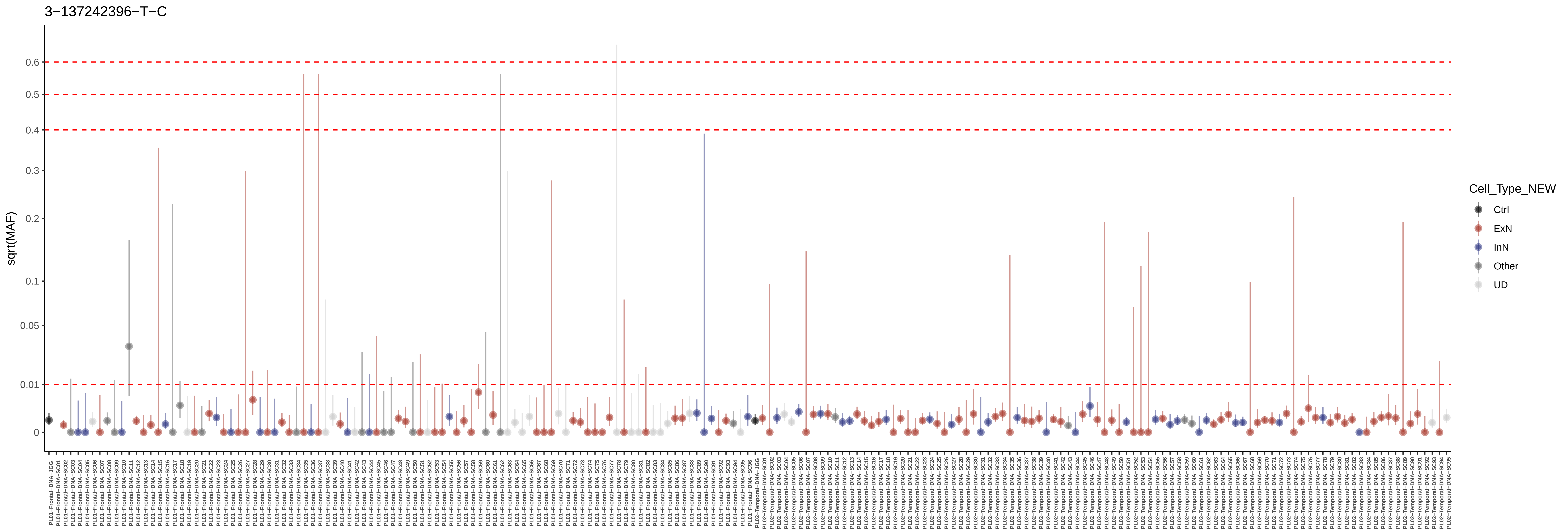
3−137242396−T−C
sqrt(MAF)
0.6
0.5
0.4
0.3
0.2
0.1
0.05
0.01
0
Cell_Type_NEW
Ctrl
ExN
InN
Other
UD

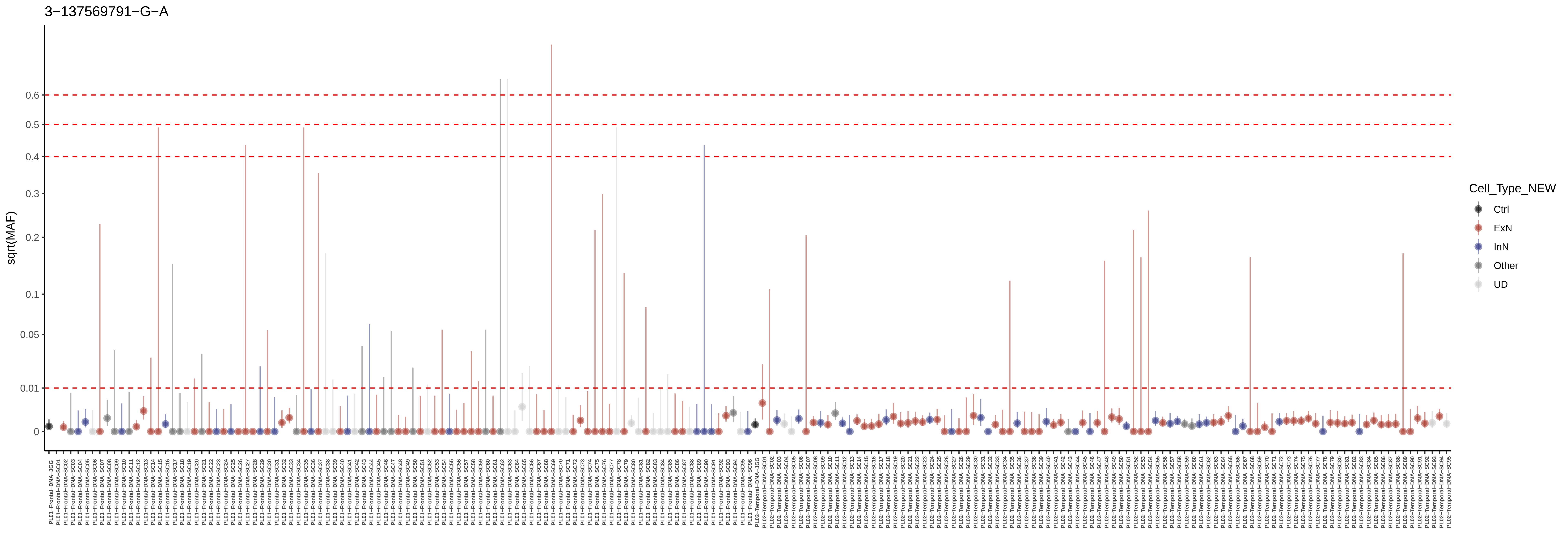
3−137569791−G−A
sqrt(MAF)
0.6
0.5
0.4
0.3
0.2
0.1
0.05
0.01
0
Cell_Type_NEW
Ctrl
ExN
InN
Other
UD

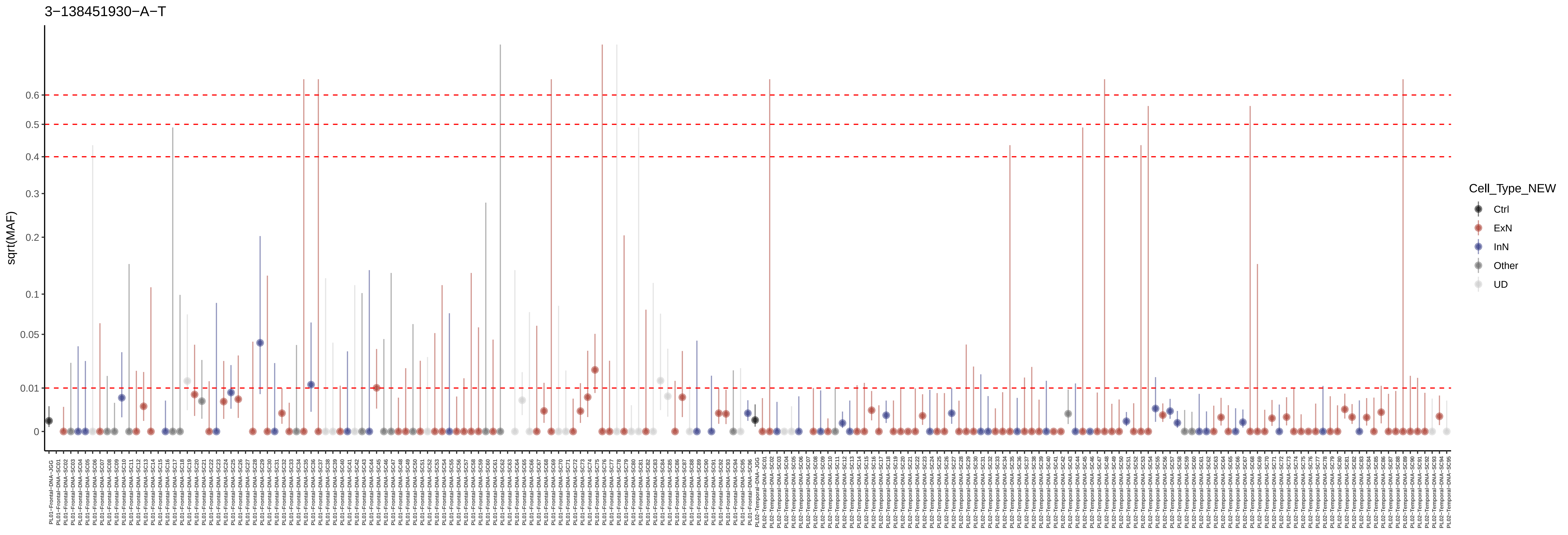
3−138451930−A−T
sqrt(MAF)
0.6
0.5
0.4
0.3
0.2
0.1
0.05
0.01
0
Cell_Type_NEW
Ctrl
ExN
InN
Other
UD

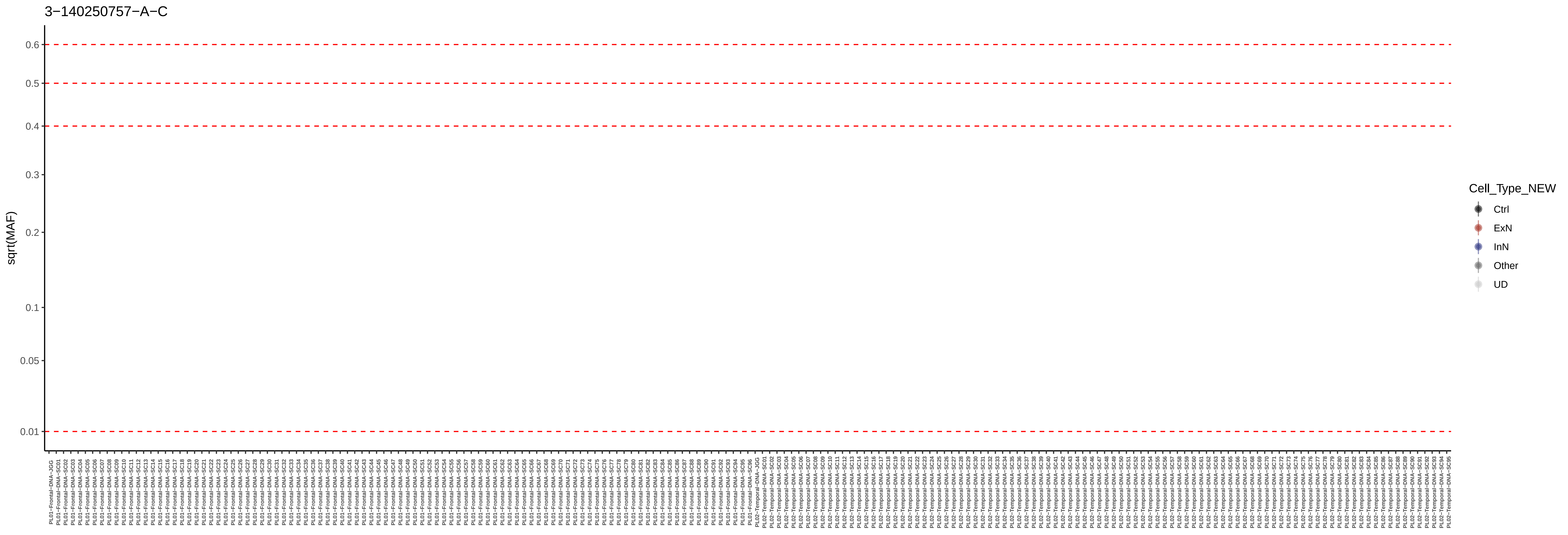
3−140250757−A−C
sqrt(MAF)
0.6
0.5
0.4
0.3
0.2
0.1
0.05
0.01
Cell_Type_NEW
Ctrl
ExN
InN
Other
UD

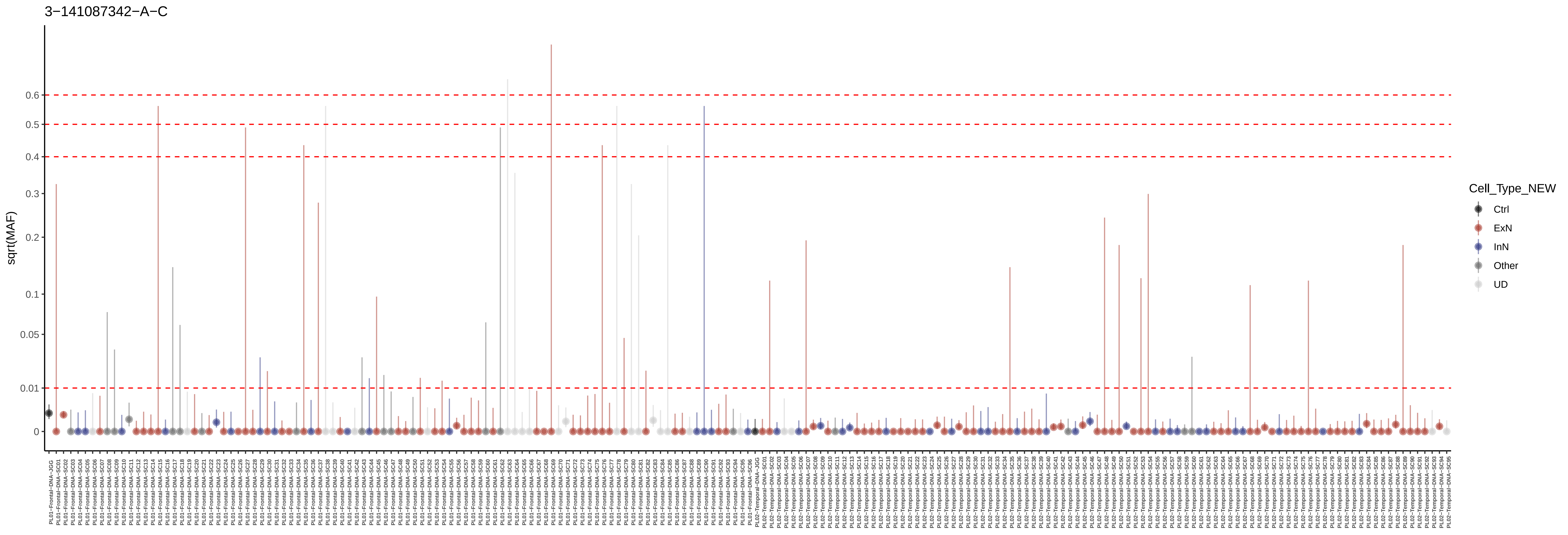
3−141087342−A−C
sqrt(MAF)
0.6
0.5
0.4
0.3
0.2
0.1
0.05
0.01
0
Cell_Type_NEW
Ctrl
ExN
InN
Other
UD

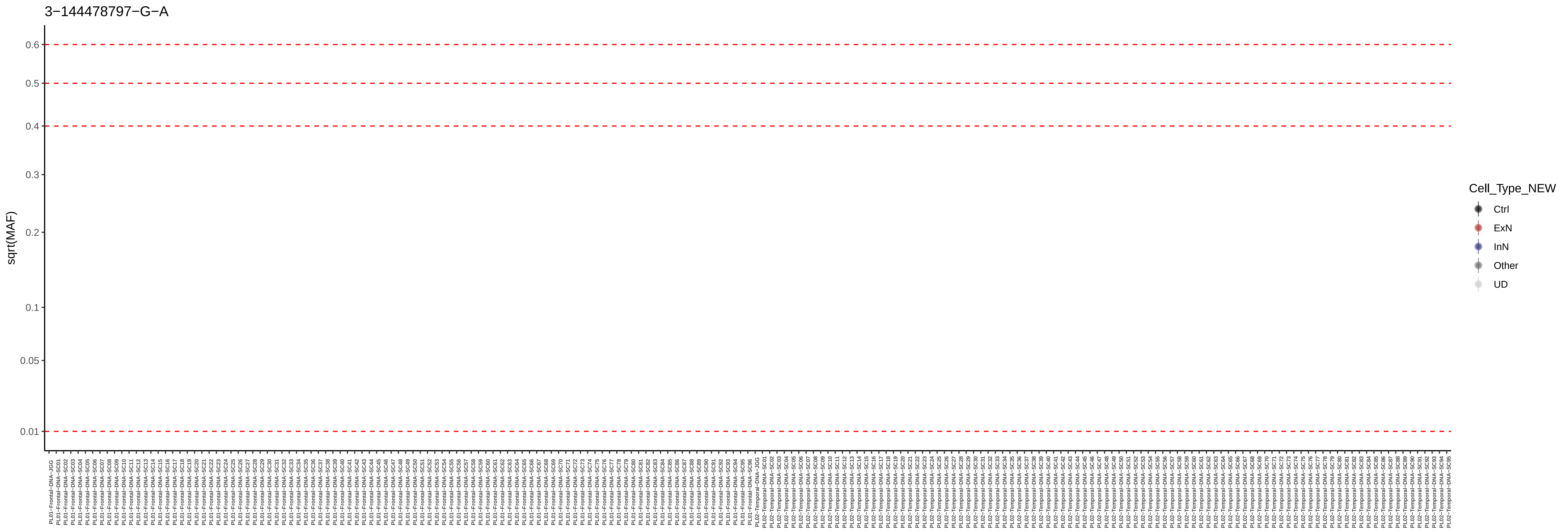
3−144478797−G−A
sqrt(MAF)
0.6
0.5
0.4
0.3
0.2
0.1
0.05
0.01
Cell_Type_NEW
Ctrl
ExN
InN
Other
UD

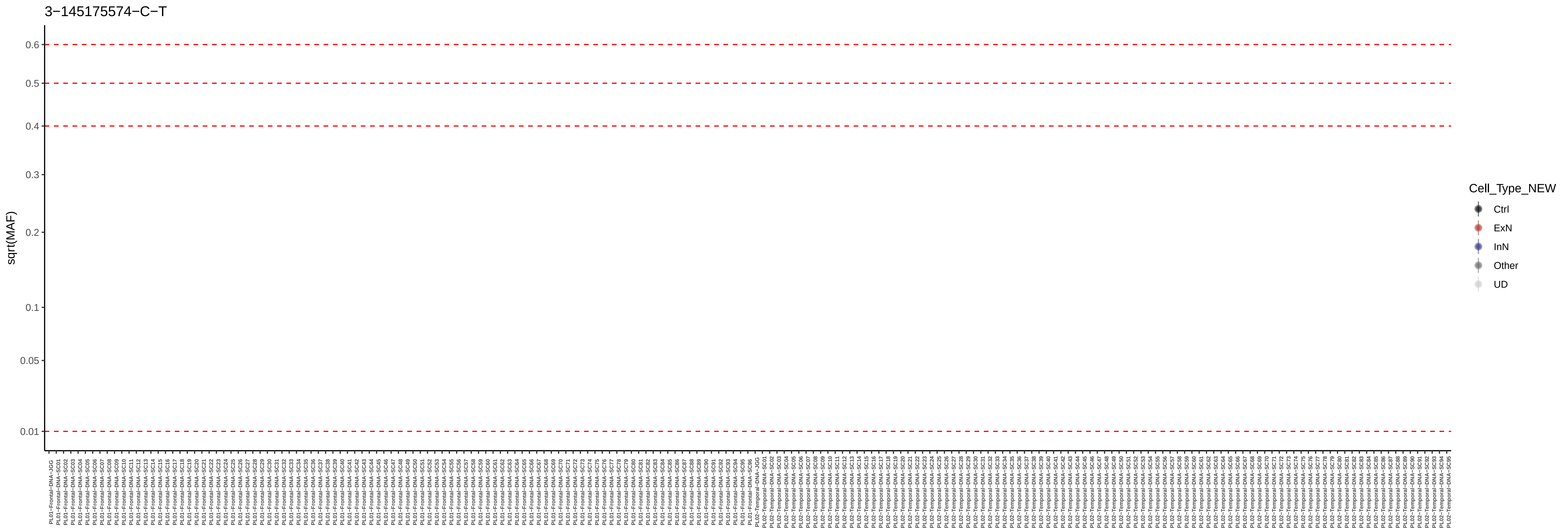
3−145175574−C−T
sqrt(MAF)
0.6
0.5
0.4
0.3
0.2
0.1
0.05
0.01
Cell_Type_NEW
Ctrl
ExN
InN
Other
UD

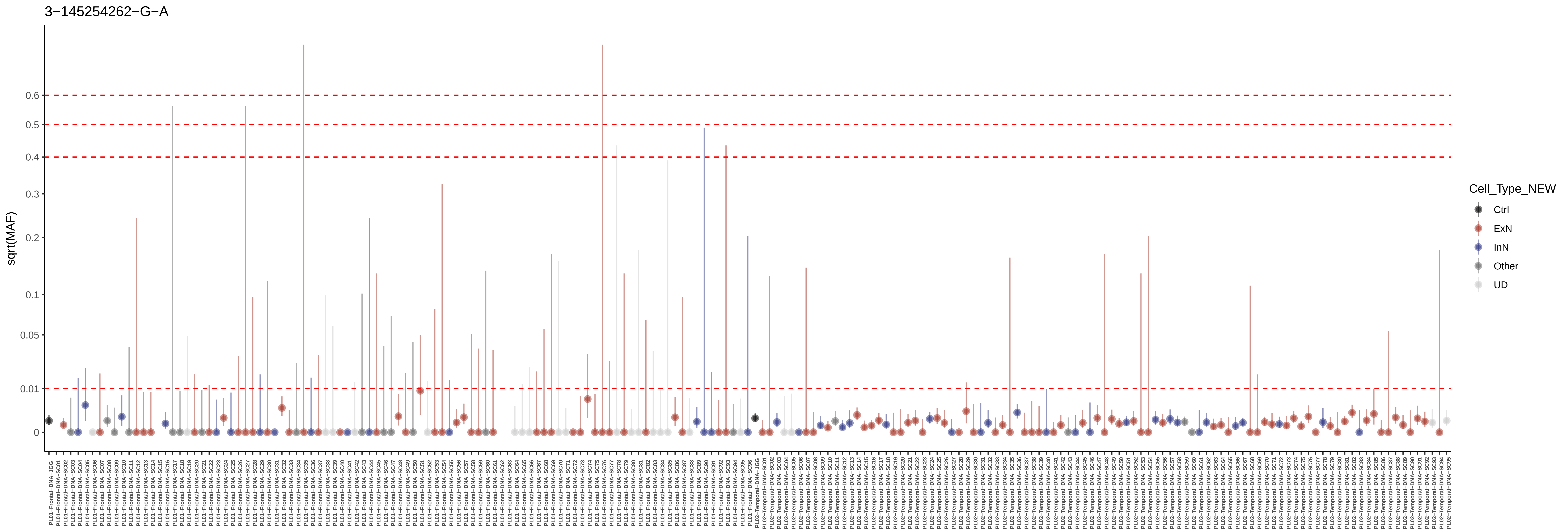

3−145254262−G−A
sqrt(MAF)
0.6
0.5
0.4
0.3
0.2
0.1
0.05
0.01
0
Cell_Type_NEW
Ctrl
ExN
InN
Other
UD

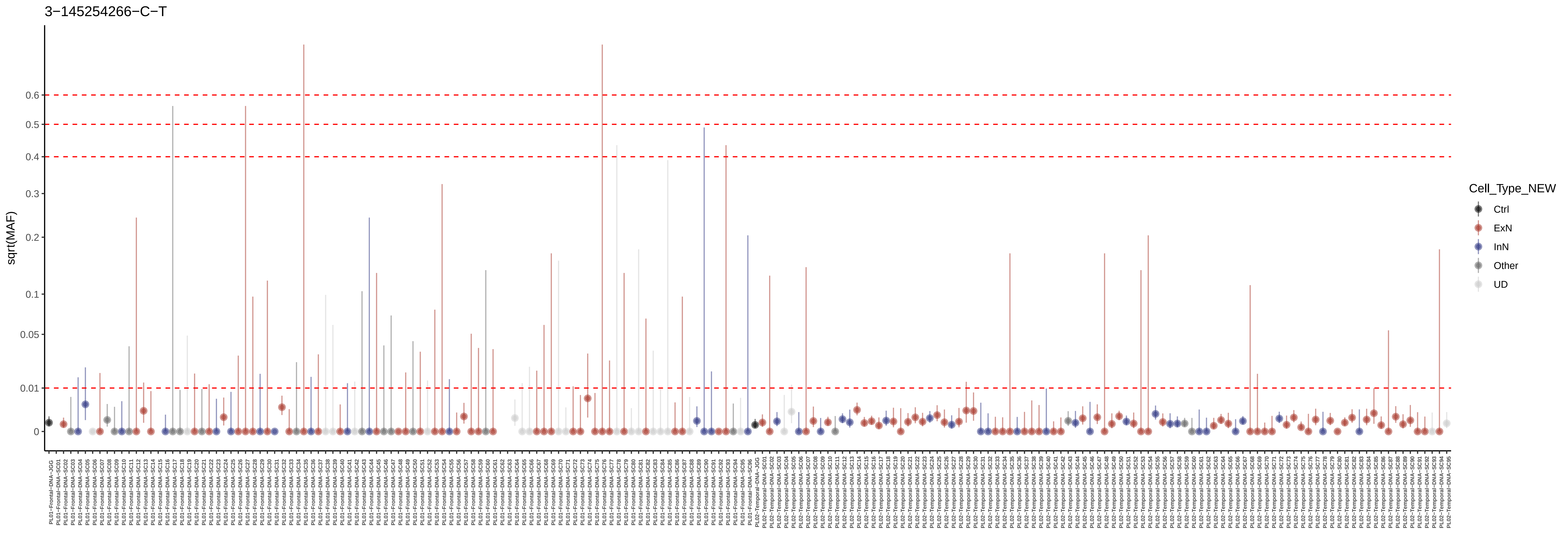
3−145254266−C−T
sqrt(MAF)
0.6
0.5
0.4
0.3
0.2
0.1
0.05
0.01
0
Cell_Type_NEW
Ctrl
ExN
InN
Other
UD

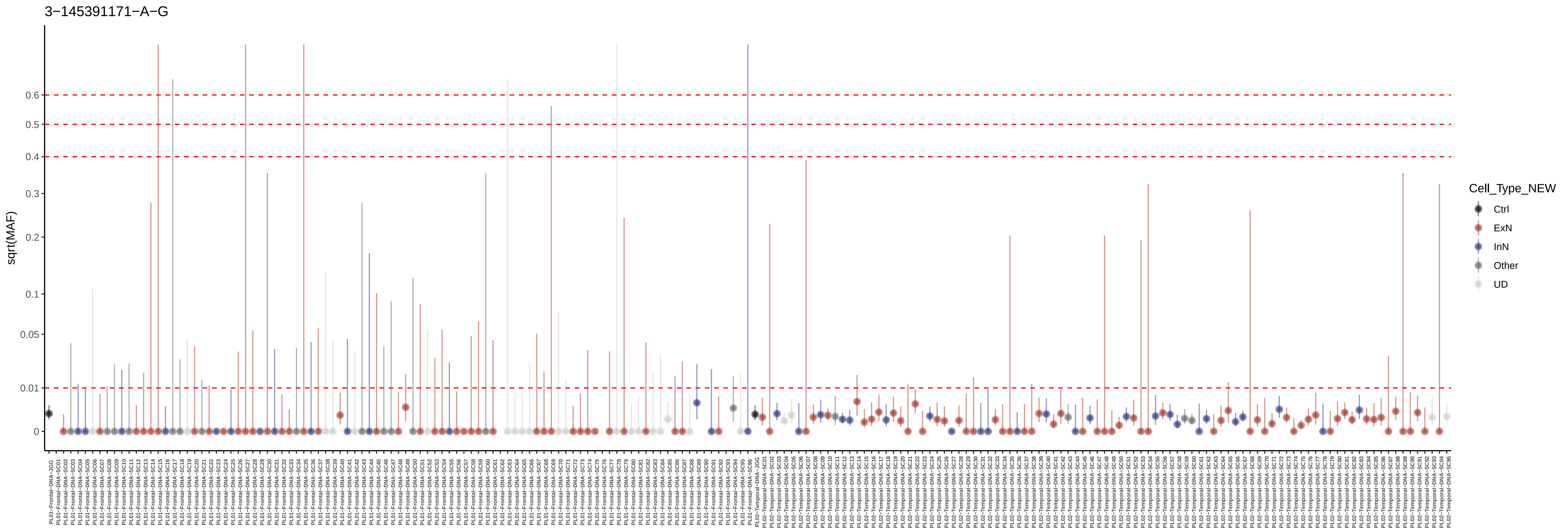
3−145391171−A−G
sqrt(MAF)
0.6
0.5
0.4
0.3
0.2
0.1
0.05
0.01
0
Cell_Type_NEW
Ctrl
ExN
InN
Other
UD

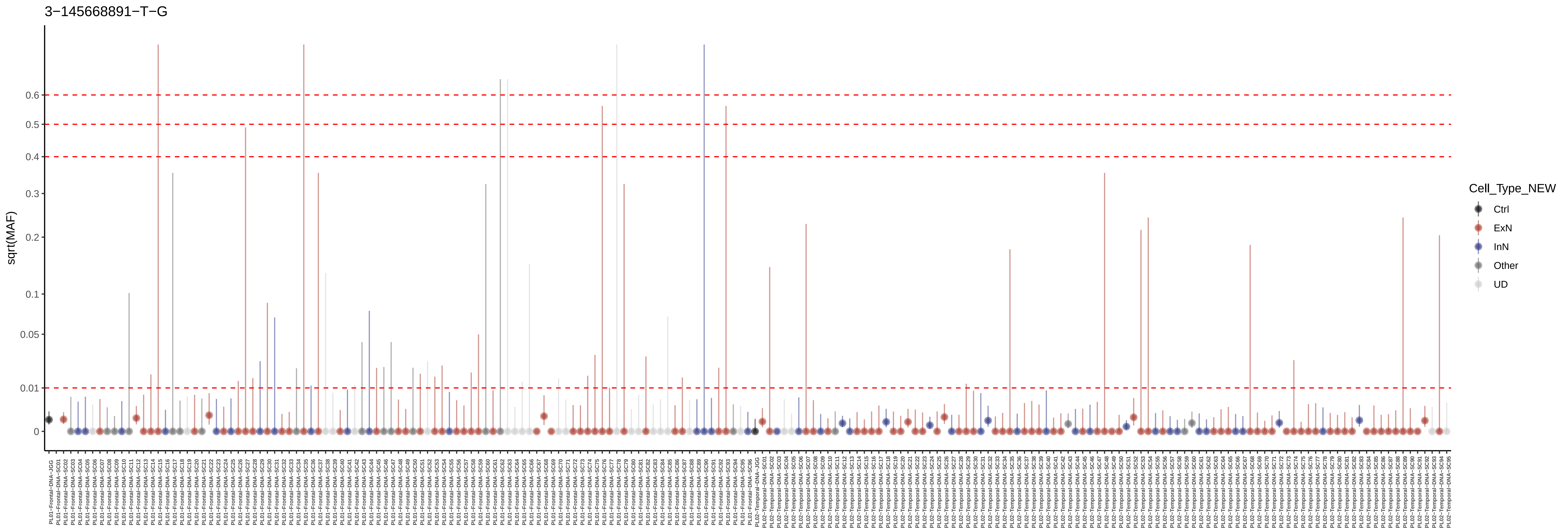
3−145668891−T−G
sqrt(MAF)
0.6
0.5
0.4
0.3
0.2
0.1
0.05
0.01
0
Cell_Type_NEW
Ctrl
ExN
InN
Other
UD

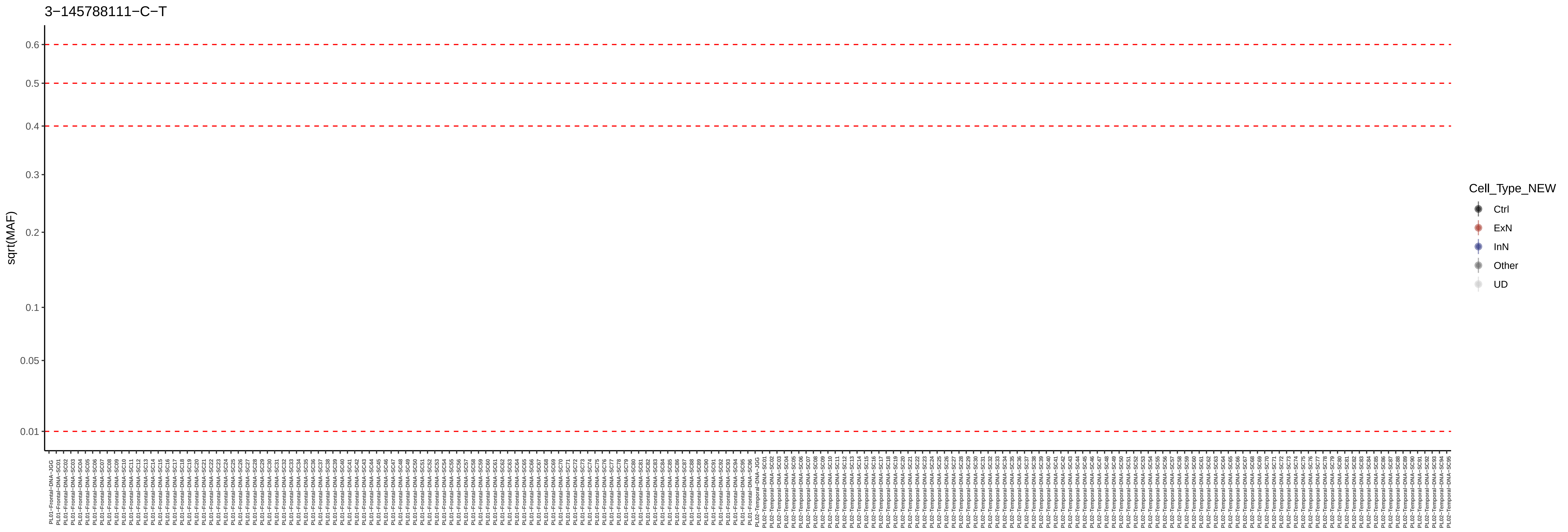
3−145788111−C−T
sqrt(MAF)
0.6
0.5
0.4
0.3
0.2
0.1
0.05
0.01
Cell_Type_NEW
Ctrl
ExN
InN
Other
UD

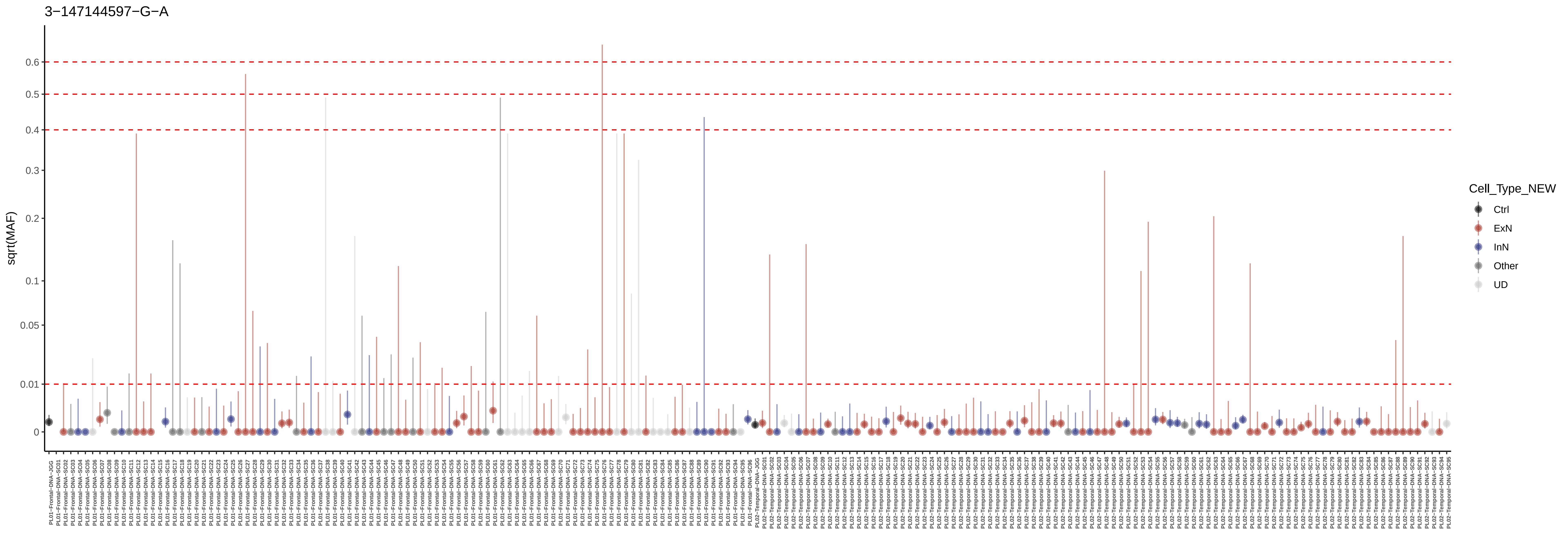

3−147144597−G−A
sqrt(MAF)
0.6
0.5
0.4
0.3
0.2
0.1
0.05
0.01
0
Cell_Type_NEW
Ctrl
ExN
InN
Other
UD

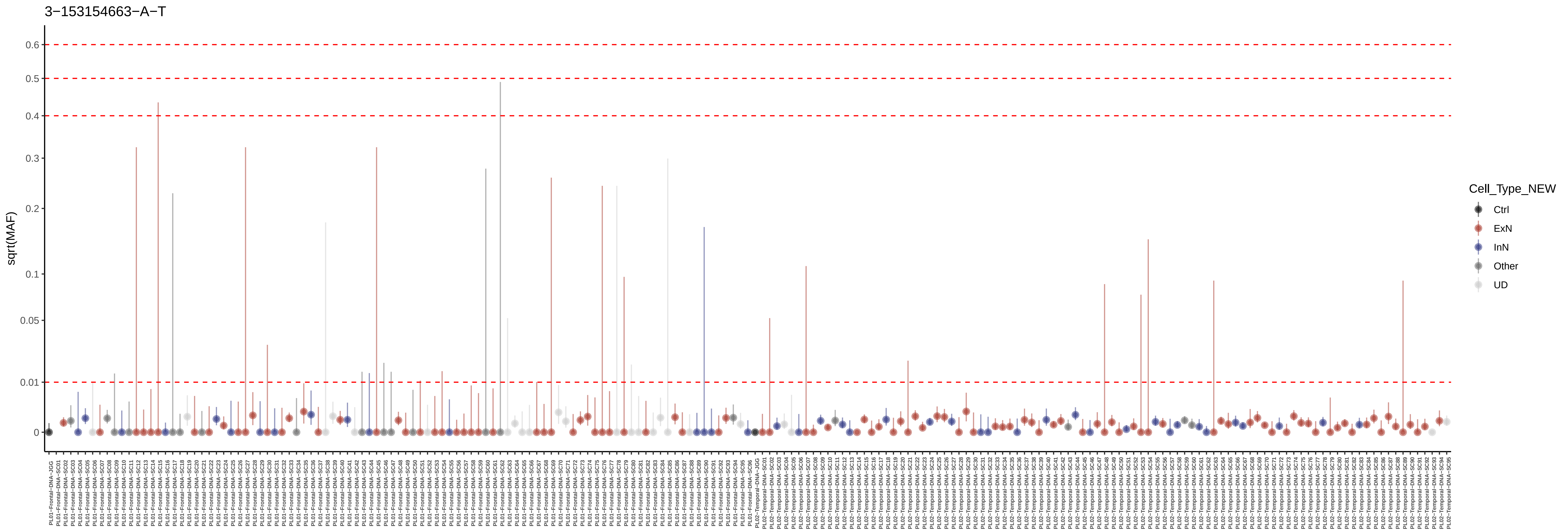
3−153154663−A−T
sqrt(MAF)
0.6
0.5
0.4
0.3
0.2
0.1
0.05
0.01
0
Cell_Type_NEW
Ctrl
ExN
InN
Other
UD

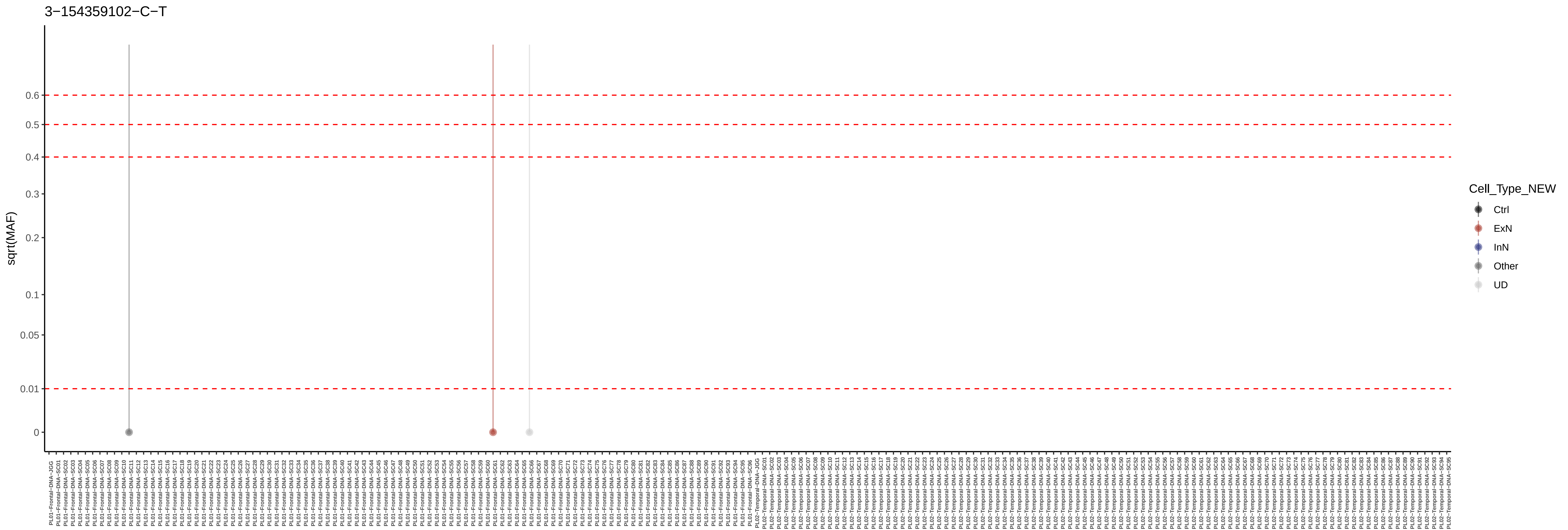
3−154359102−C−T
sqrt(MAF)
0.6
0.5
0.4
0.3
0.2
0.1
0.05
0.01
0
Cell_Type_NEW
Ctrl
ExN
InN
Other
UD

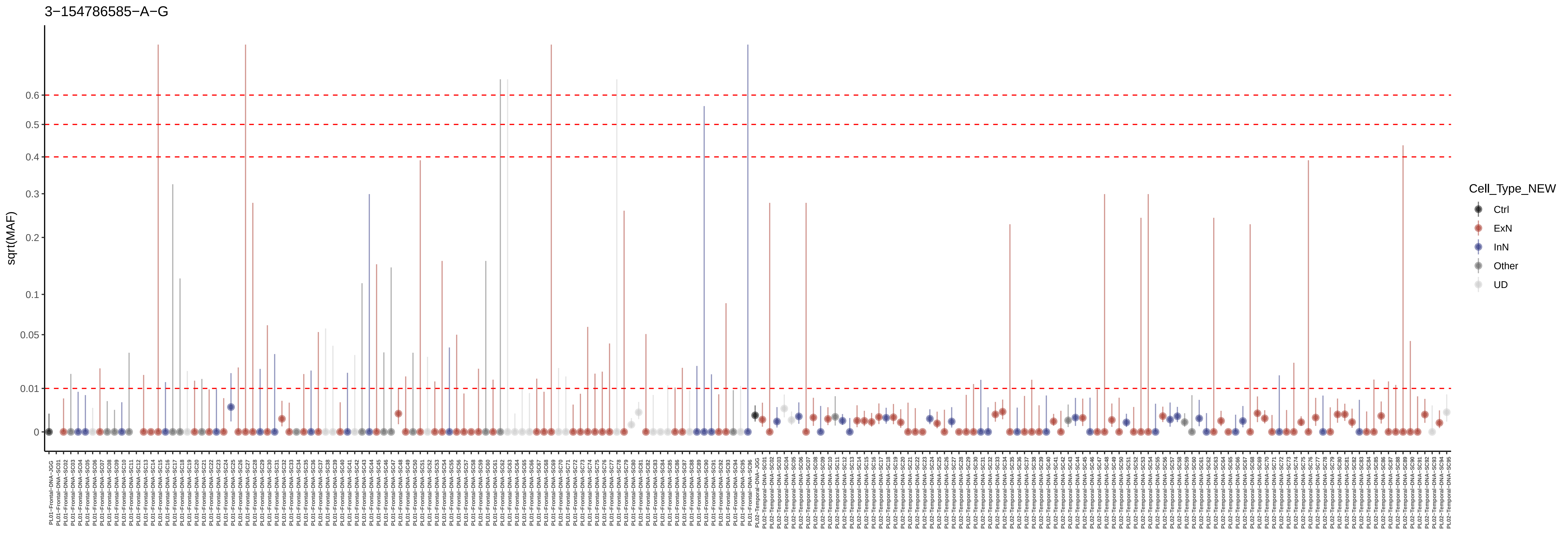
3−154786585−A−G
sqrt(MAF)
0.6
0.5
0.4
0.3
0.2
0.1
0.05
0.01
0
Cell_Type_NEW
Ctrl
ExN
InN
Other
UD

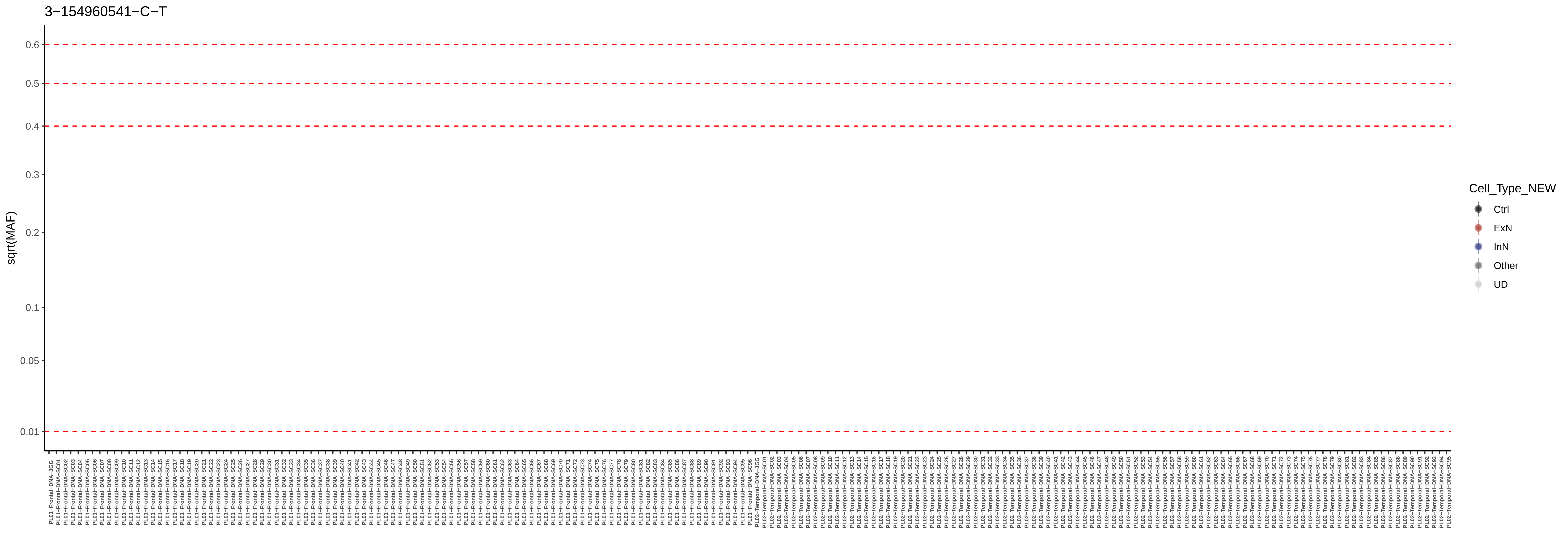
3−154960541−C−T
sqrt(MAF)
0.6
0.5
0.4
0.3
0.2
0.1
0.05
0.01
Cell_Type_NEW
Ctrl
ExN
InN
Other
UD

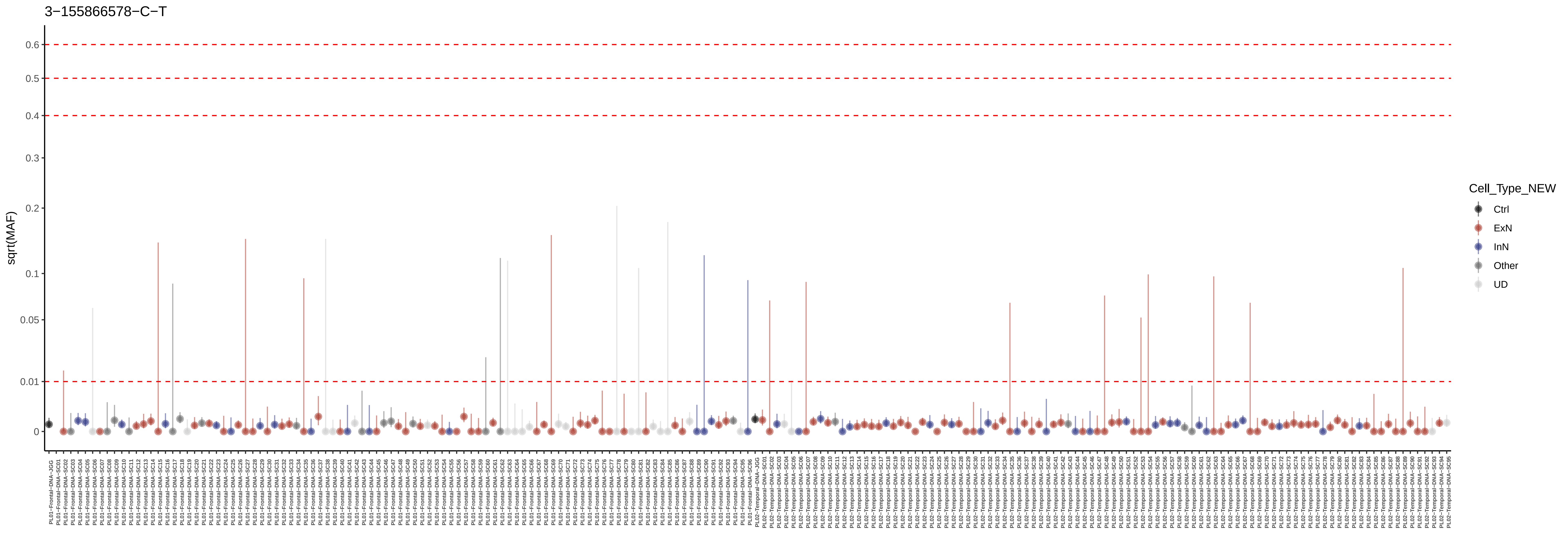

3−155866578−C−T
sqrt(MAF)
0.6
0.5
0.4
0.3
0.2
0.1
0.05
0.01
0
Cell_Type_NEW
Ctrl
ExN
InN
Other
UD

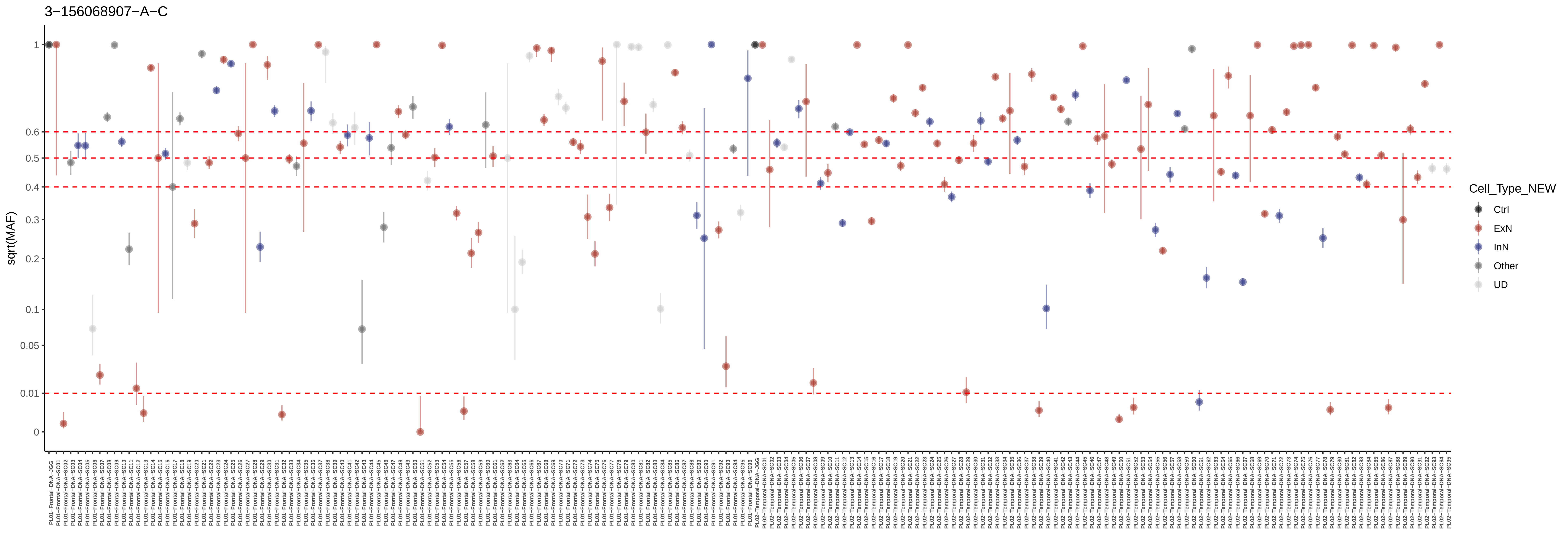
3−156068907−A−C
sqrt(MAF)
1
0.6
0.5
0.4
0.3
0.2
0.1
0.05
0.01
0
Cell_Type_NEW
Ctrl
ExN
InN
Other
UD

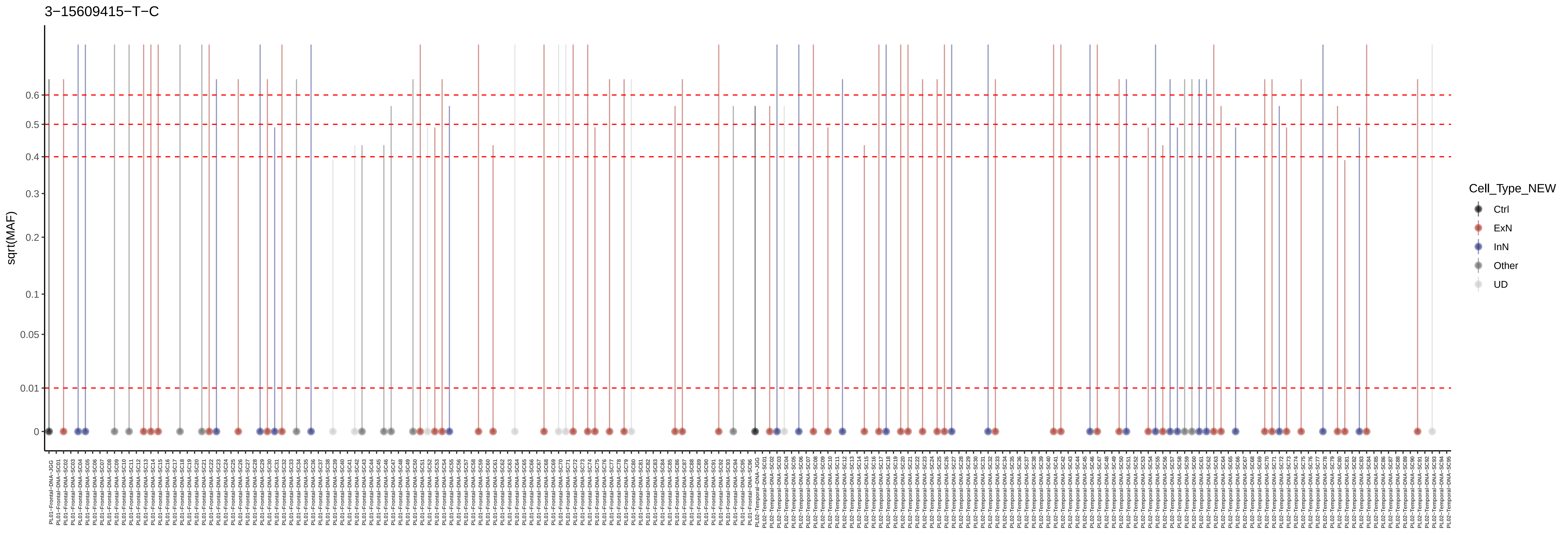
3−15609415−T−C
sqrt(MAF)
0.6
0.5
0.4
0.3
0.2
0.1
0.05
0.01
0
Cell_Type_NEW
Ctrl
ExN
InN
Other
UD

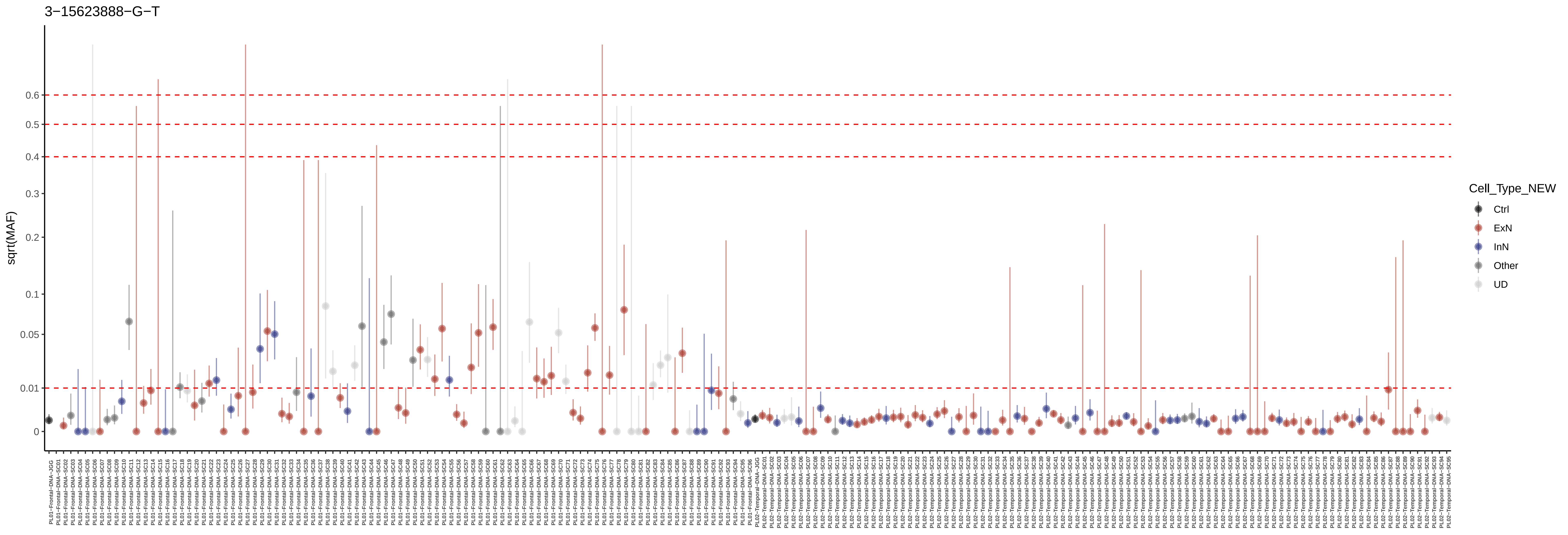
3−15623888−G−T
sqrt(MAF)
0.6
0.5
0.4
0.3
0.2
0.1
0.05
0.01
0
Cell_Type_NEW
Ctrl
ExN
InN
Other
UD

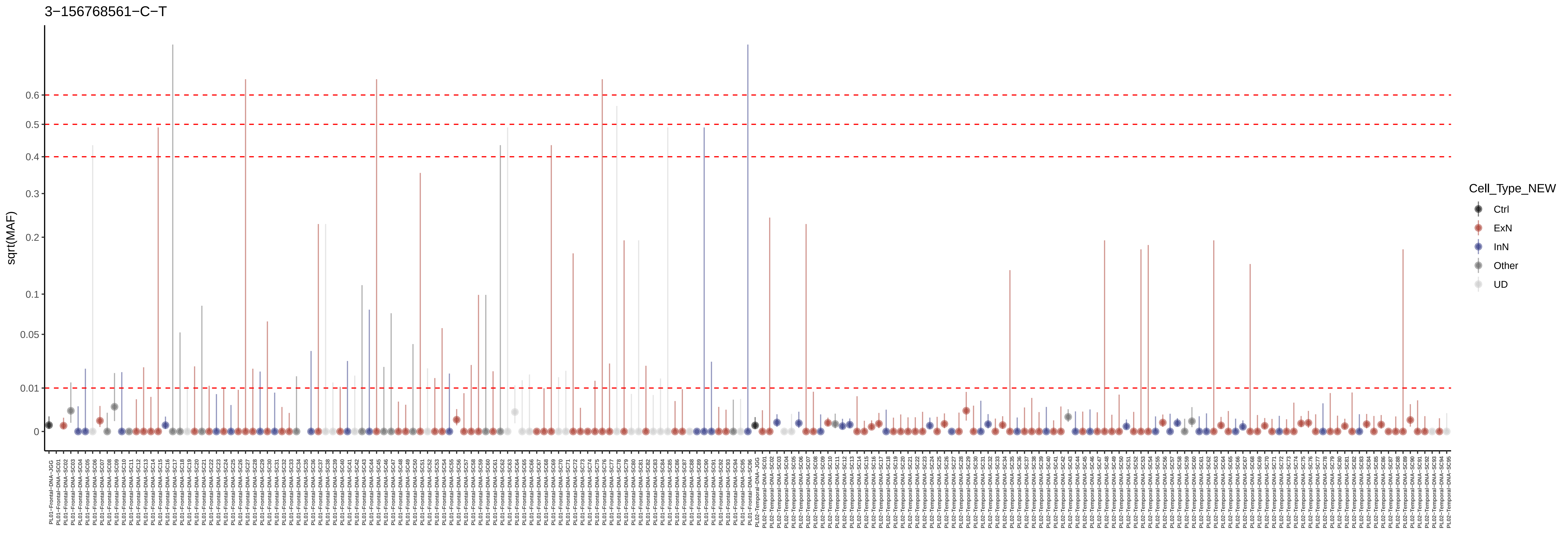
3−156768561−C−T
sqrt(MAF)
0
0.01
0.05
0.1
0.2
0.3
0.4
0.5
0.6
Cell_Type_NEW
Ctrl
ExN
InN
Other
UD

3−157131608−A−T

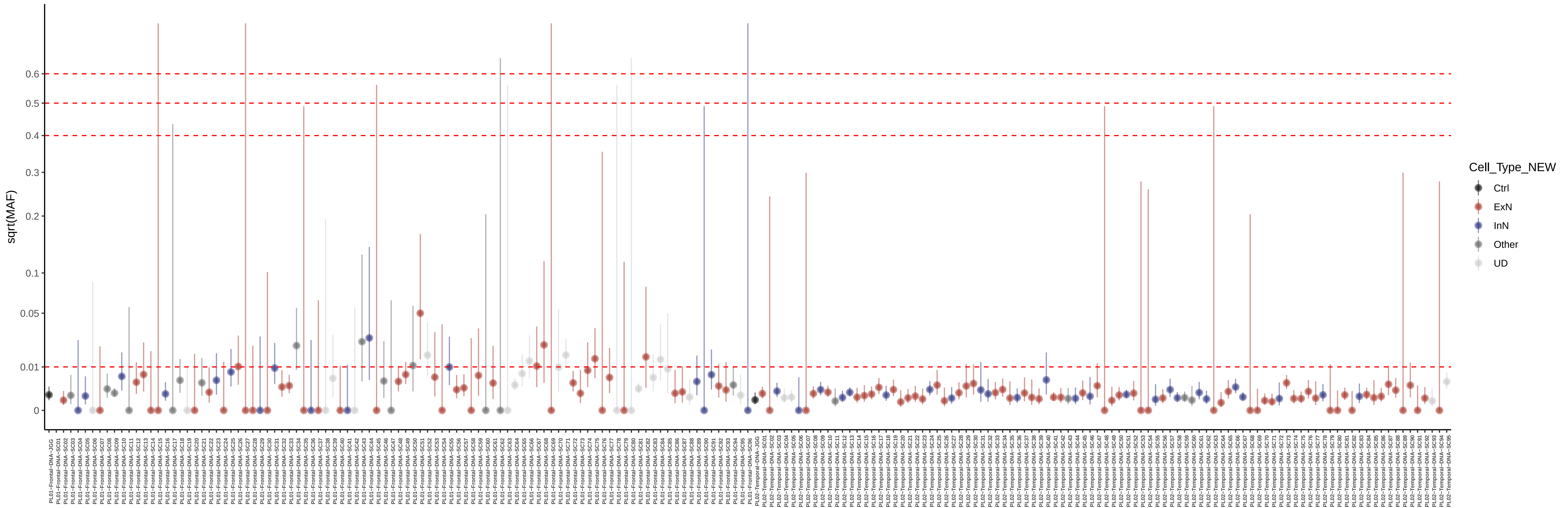

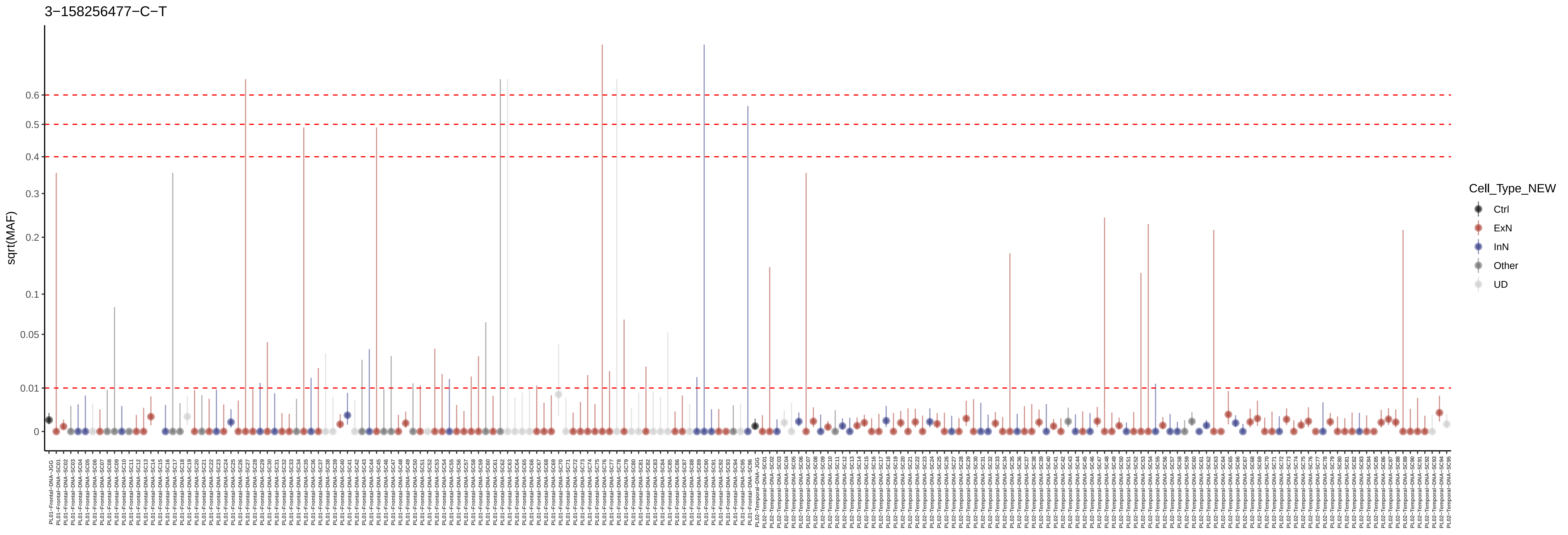

3−158256477−C−T
sqrt(MAF)
0.6
0.5
0.4
0.3
0.2
0.1
0.05
0.01
0
Cell_Type_NEW
Ctrl
ExN
InN
Other
UD

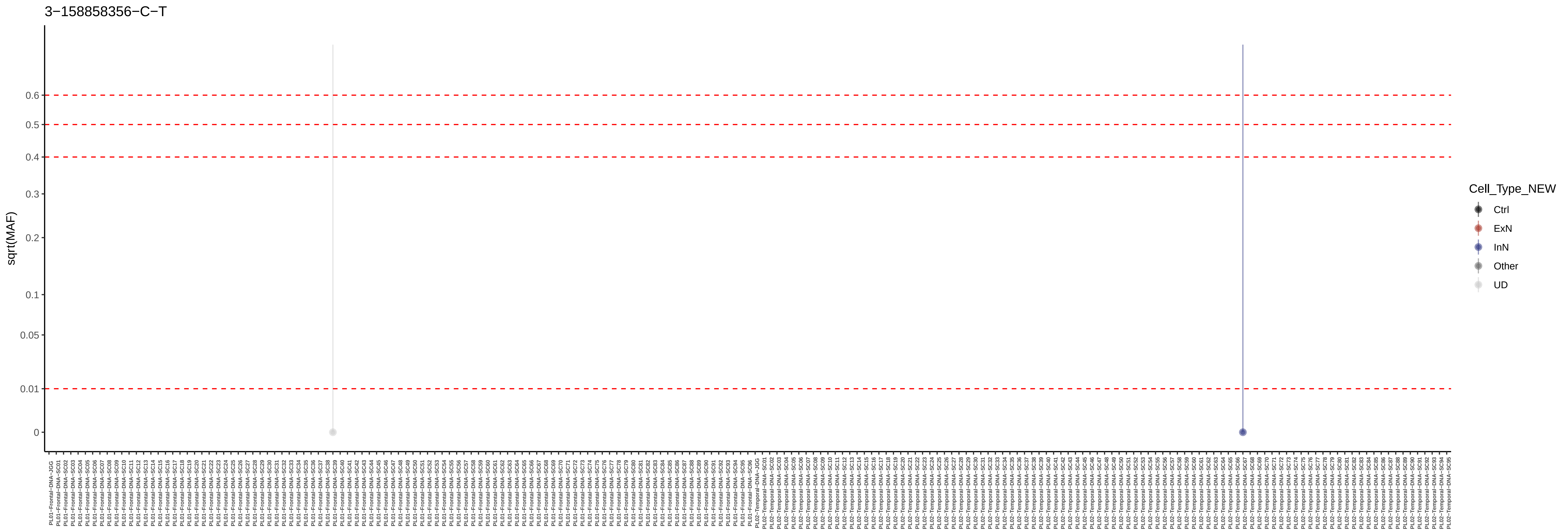
3−158858356−C−T
sqrt(MAF)
0.6
0.5
0.4
0.3
0.2
0.1
0.05
0.01
0
Cell_Type_NEW
Ctrl
ExN
InN
Other
UD

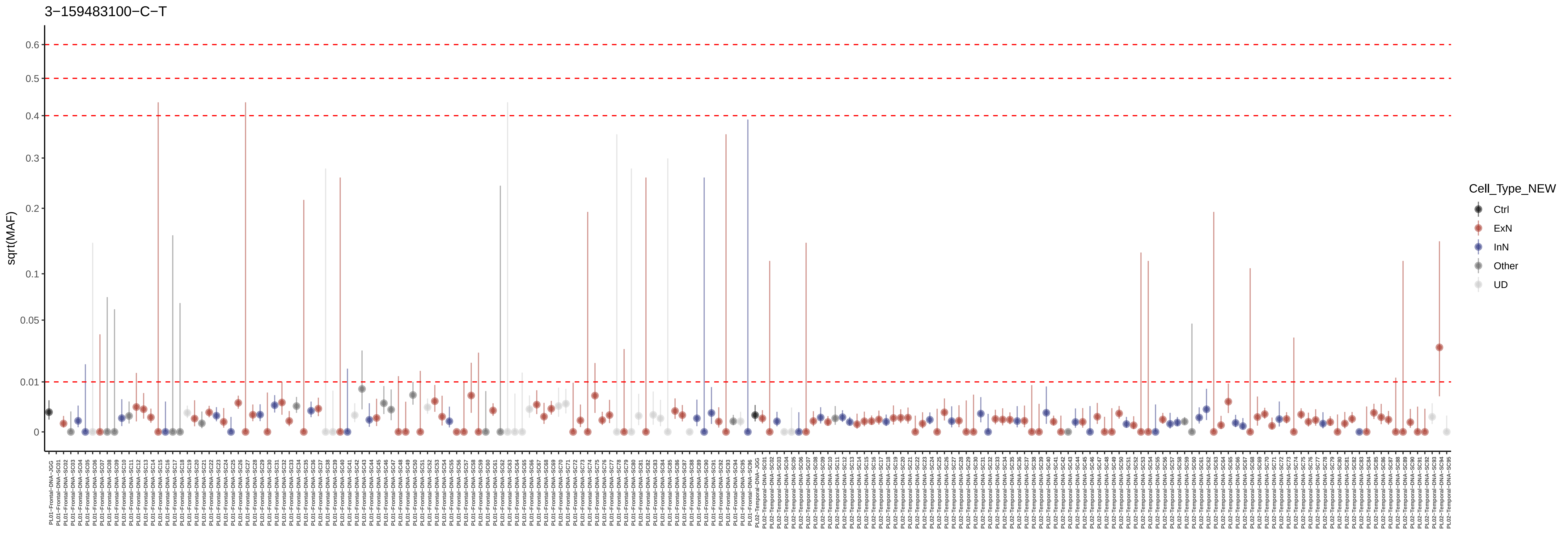

3−159483100−C−T
sqrt(MAF)
0.6
0.5
0.4
0.3
0.2
0.1
0.05
0.01
0
Cell_Type_NEW
Ctrl
ExN
InN
Other
UD

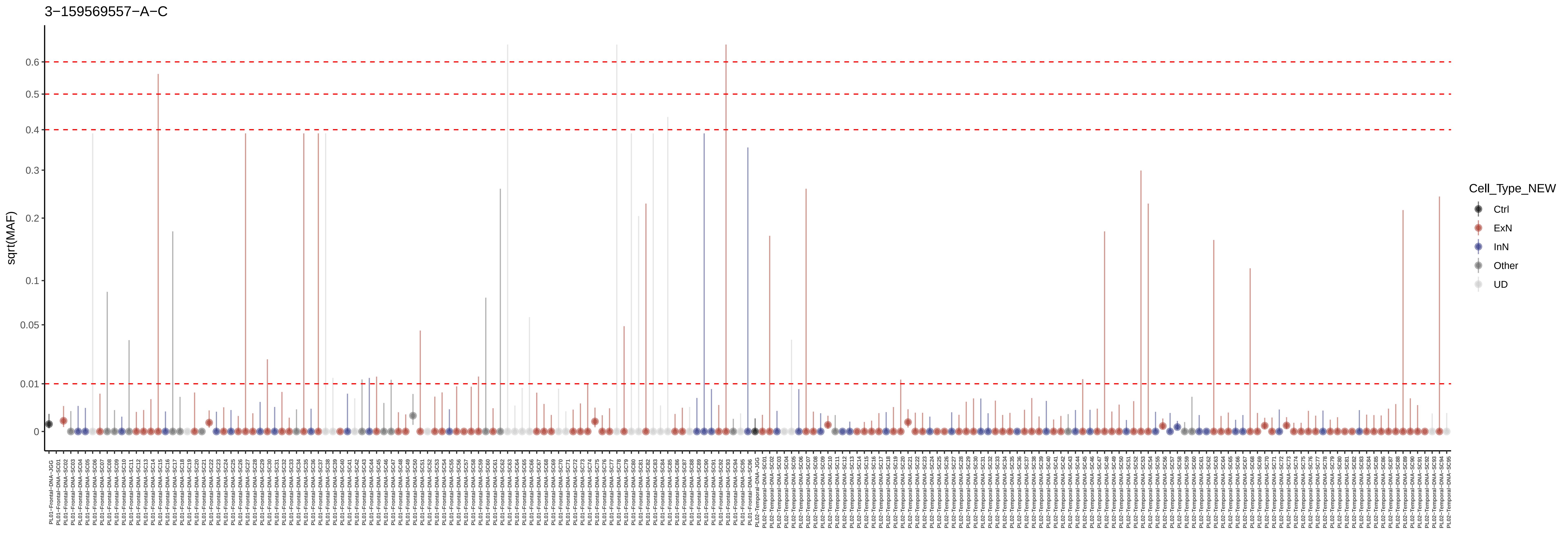
3−159569557−A−C
sqrt(MAF)
0.6
0.5
0.4
0.3
0.2
0.1
0.05
0.01
0
Cell_Type_NEW
Ctrl
ExN
InN
Other
UD

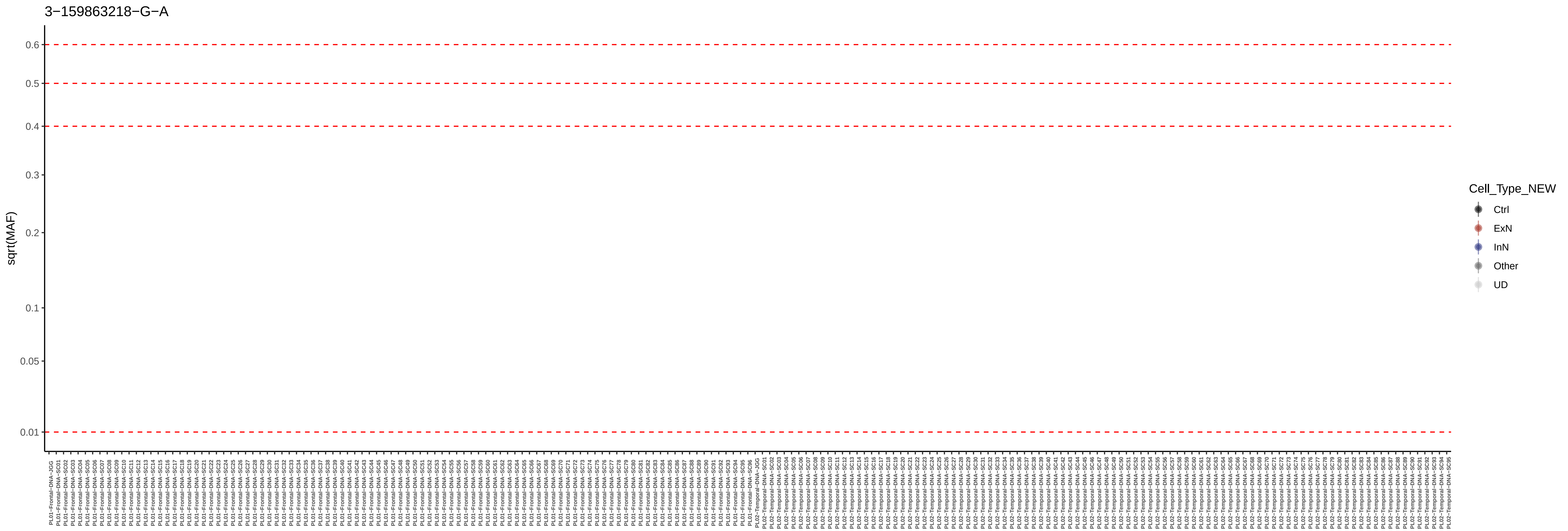
3−159863218−G−A
sqrt(MAF)
0.6
0.5
0.4
0.3
0.2
0.1
0.05
0.01
Cell_Type_NEW
Ctrl
ExN
InN
Other
UD

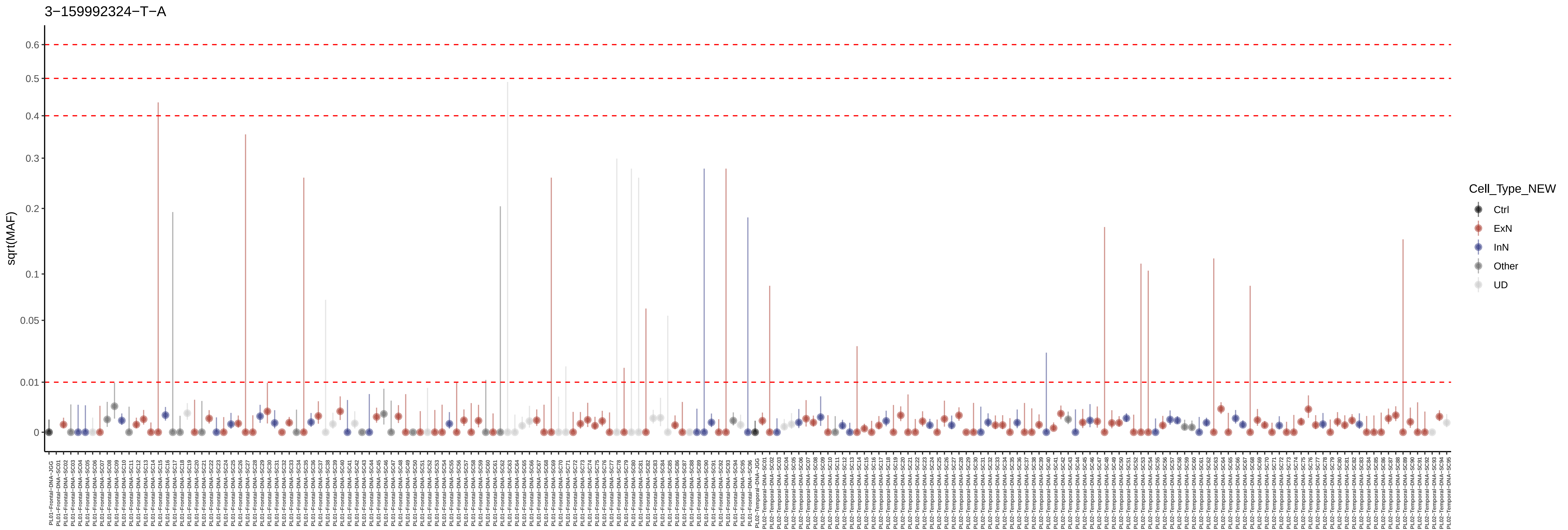
3−159992324−T−A
sqrt(MAF)
0.6
0.5
0.4
0.3
0.2
0.1
0.05
0.01
0
Cell_Type_NEW
Ctrl
ExN
InN
Other
UD

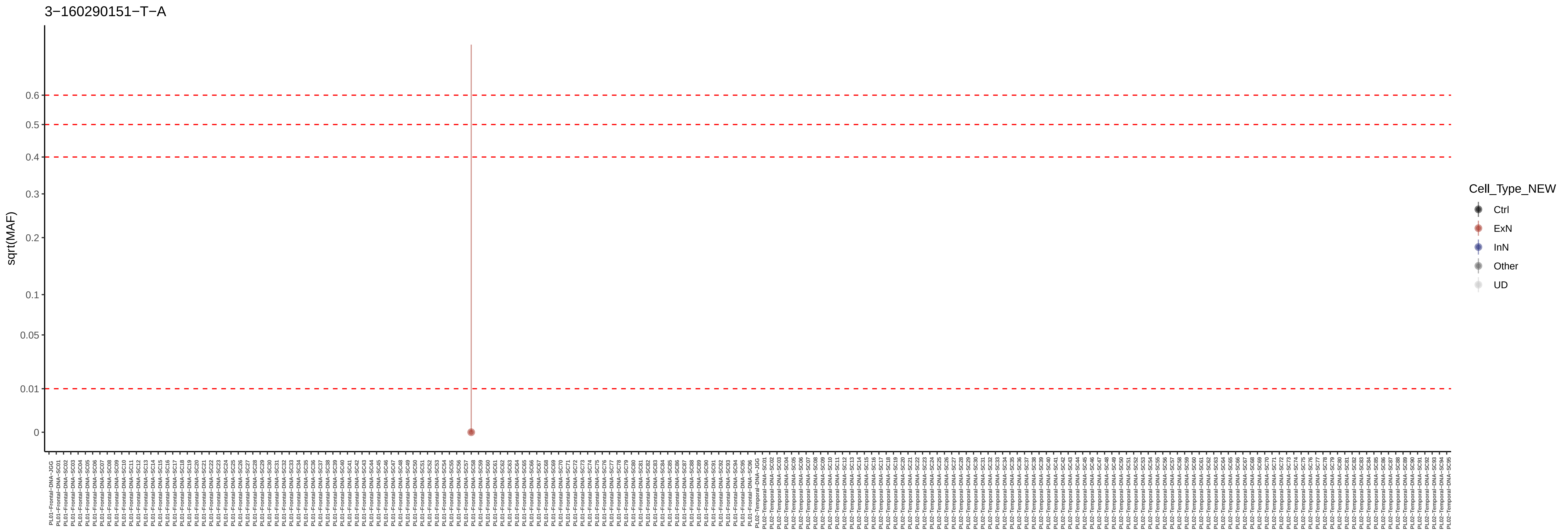

3−160290151−T−A
sqrt(MAF)
0.6
0.5
0.4
0.3
0.2
0.1
0.05
0.01
0
Cell_Type_NEW
Ctrl
ExN
InN
Other
UD

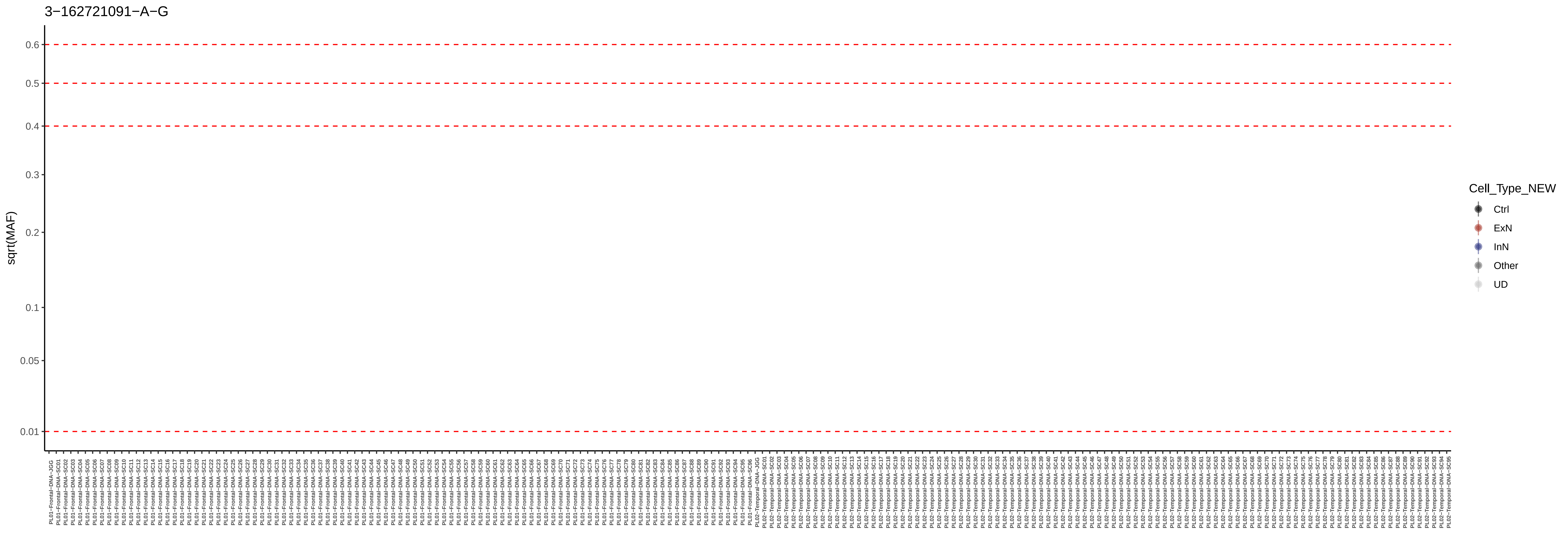
3−162721091−A−G
sqrt(MAF)
0.6
0.5
0.4
0.3
0.2
0.1
0.05
0.01
Cell_Type_NEW
Ctrl
ExN
InN
Other
UD

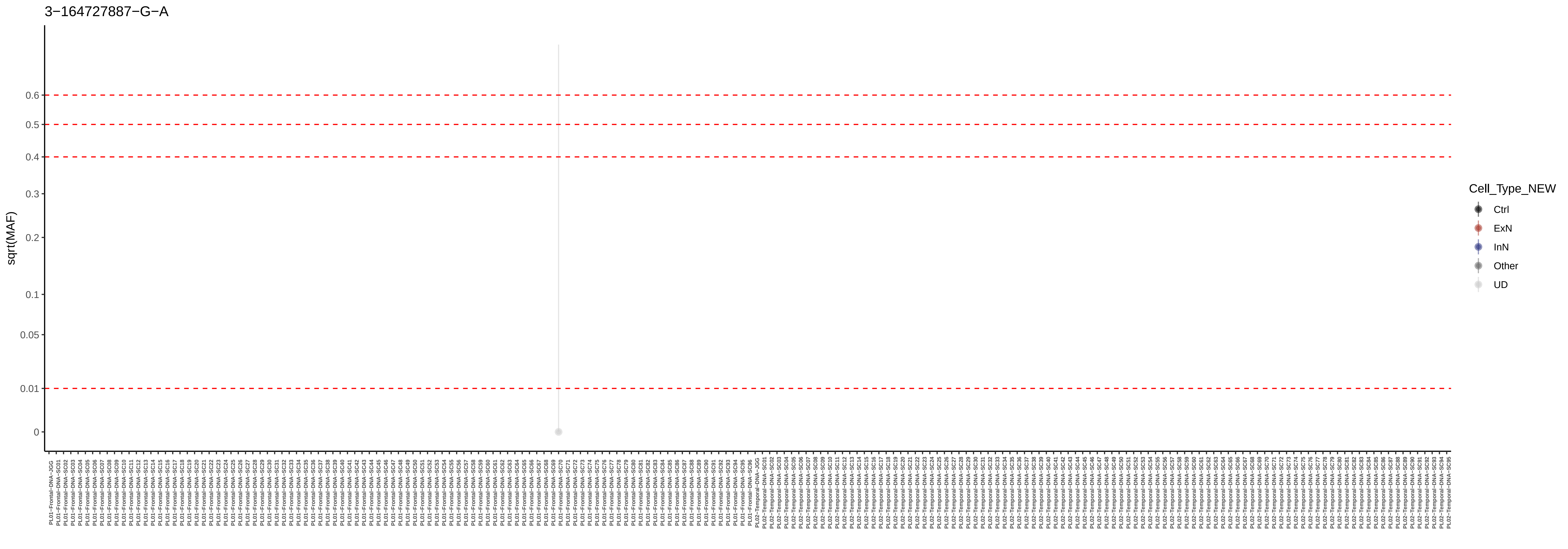

3−164727887−G−A
sqrt(MAF)
0.6
0.5
0.4
0.3
0.2
0.1
0.05
0.01
0
Cell_Type_NEW
Ctrl
ExN
InN
Other
UD

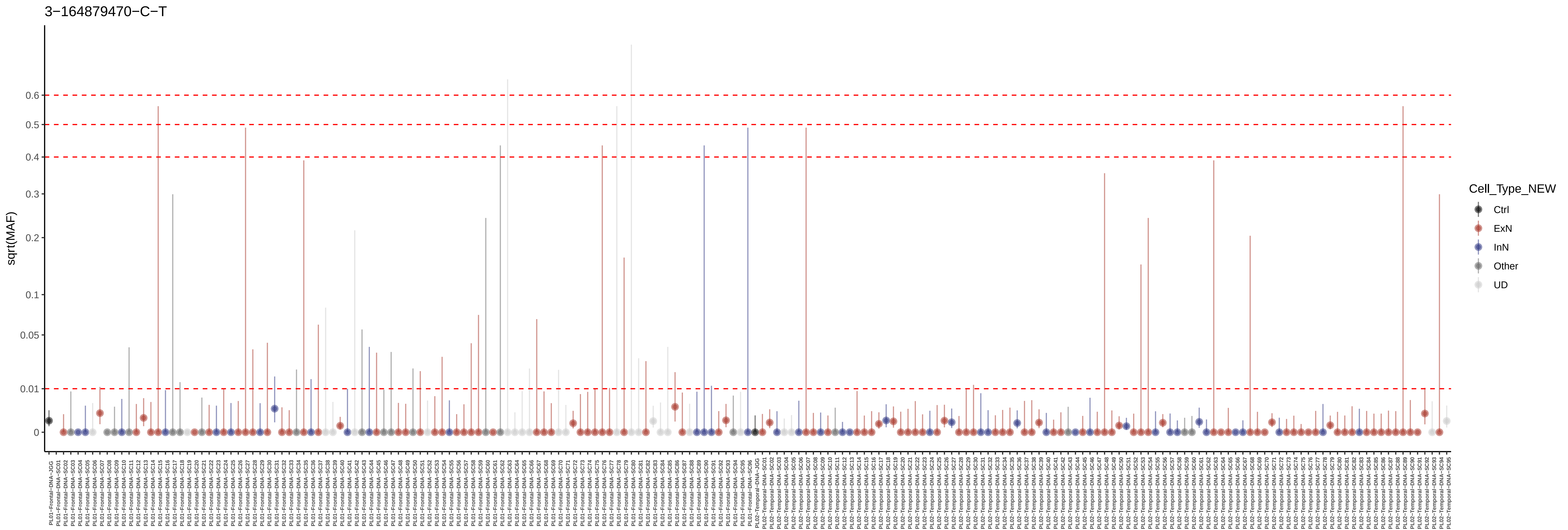

3−164879470−C−T
sqrt(MAF)
0.6
0.5
0.4
0.3
0.2
0.1
0.05
0.01
0
Cell_Type_NEW
Ctrl
ExN
InN
Other
UD

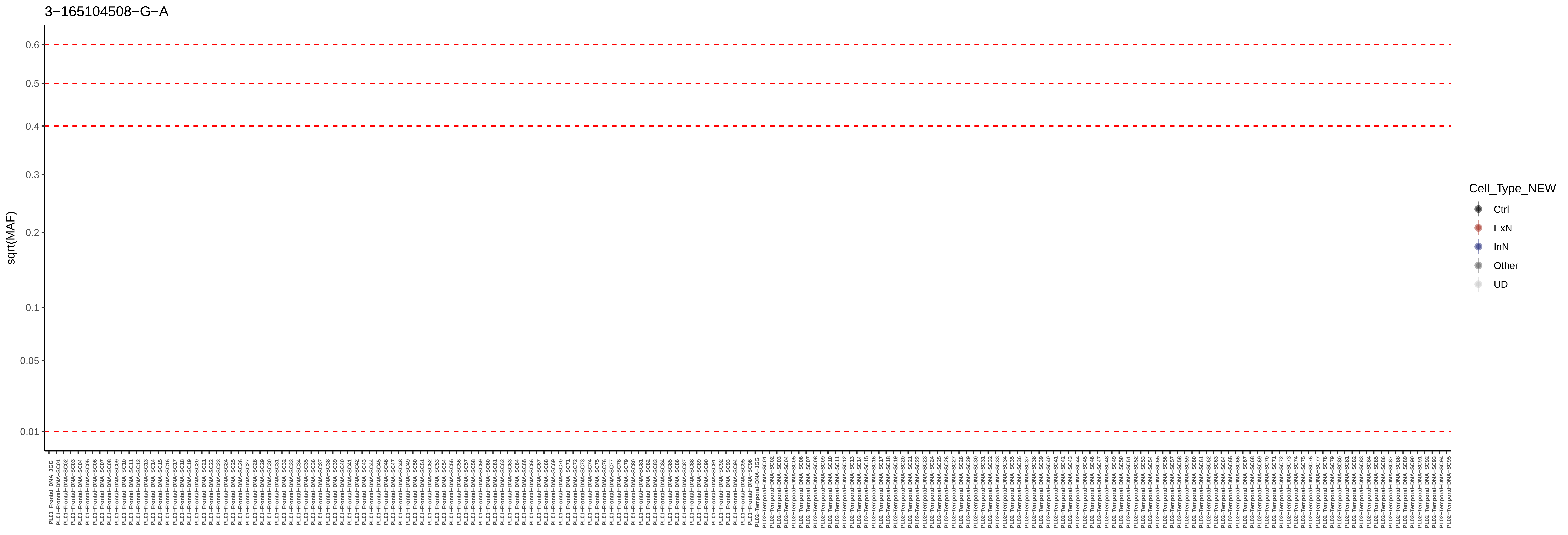

3−165104508−G−A
sqrt(MAF)
0.6
0.5
0.4
0.3
0.2
0.1
0.05
0.01
Cell_Type_NEW
Ctrl
ExN
InN
Other
UD

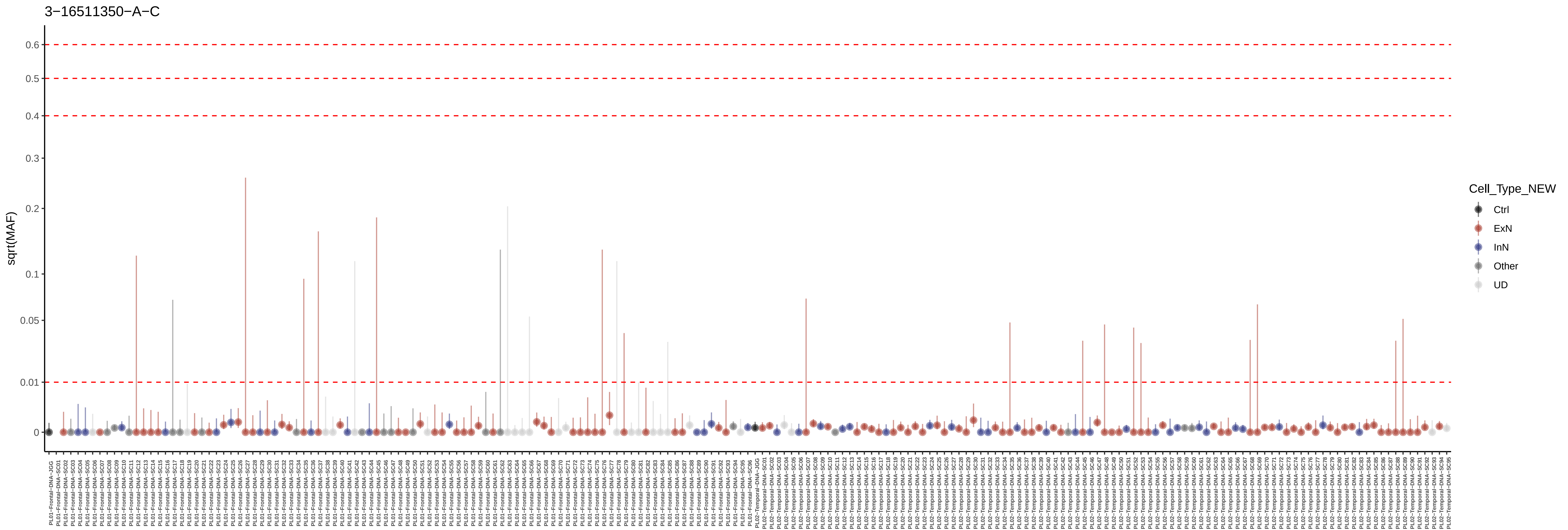
3−16511350−A−C
sqrt(MAF)
0.6
0.5
0.4
0.3
0.2
0.1
0.05
0.01
0
Cell_Type_NEW
Ctrl
ExN
InN
Other
UD

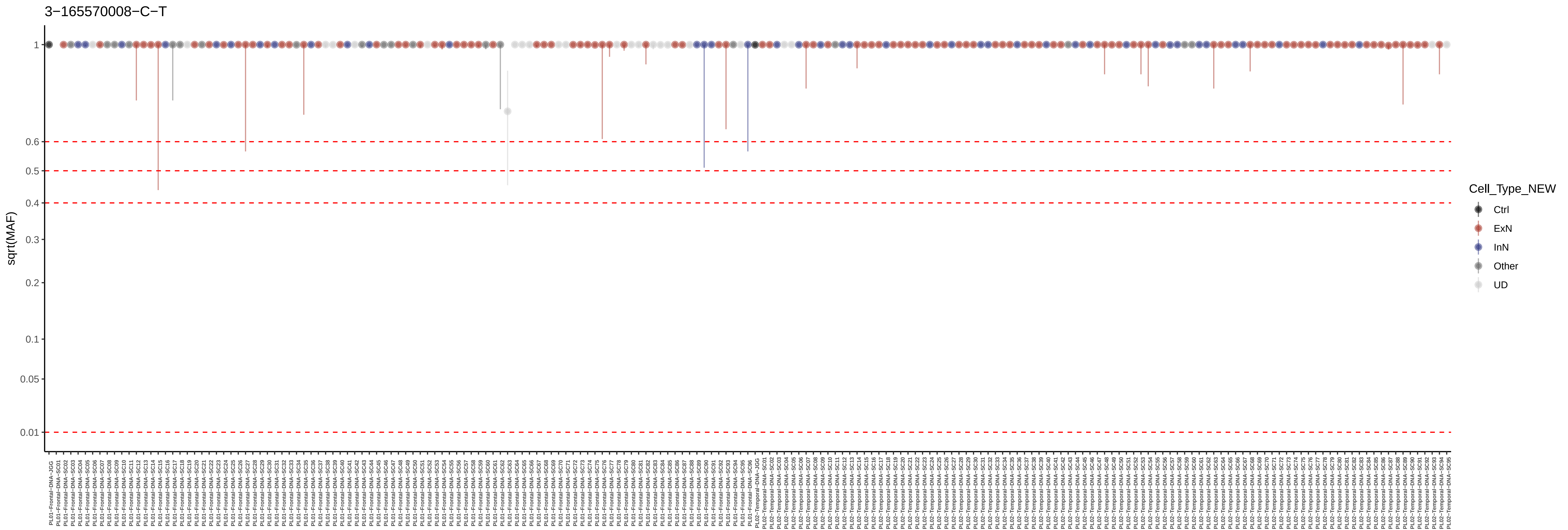

3−165570008−C−T
sqrt(MAF)
1
0.6
0.5
0.4
0.3
0.2
0.1
0.05
0.01
Cell_Type_NEW
Ctrl
ExN
InN
Other
UD

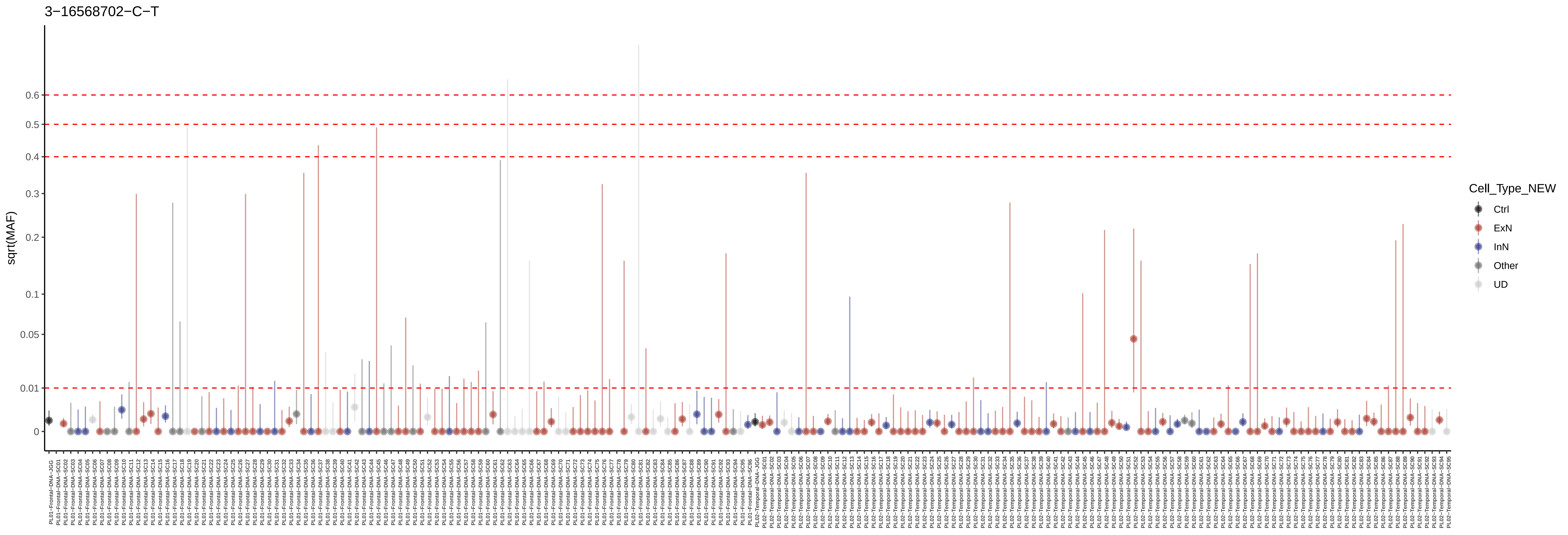
3−16568702−C−T
sqrt(MAF)
0.6
0.5
0.4
0.3
0.2
0.1
0.05
0.01
0
Cell_Type_NEW
Ctrl
ExN
InN
Other
UD

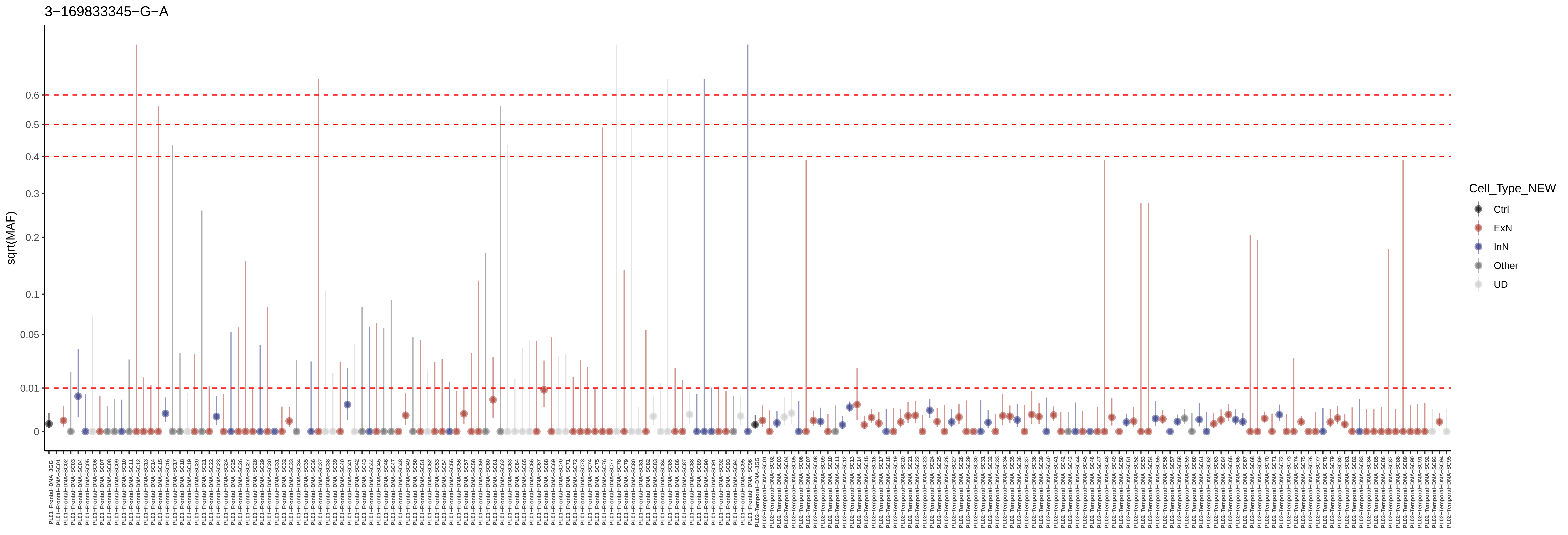
3−169833345−G−A
sqrt(MAF)
0.6
0.5
0.4
0.3
0.2
0.1
0.05
0.01
0
Cell_Type_NEW
Ctrl
ExN
InN
Other
UD

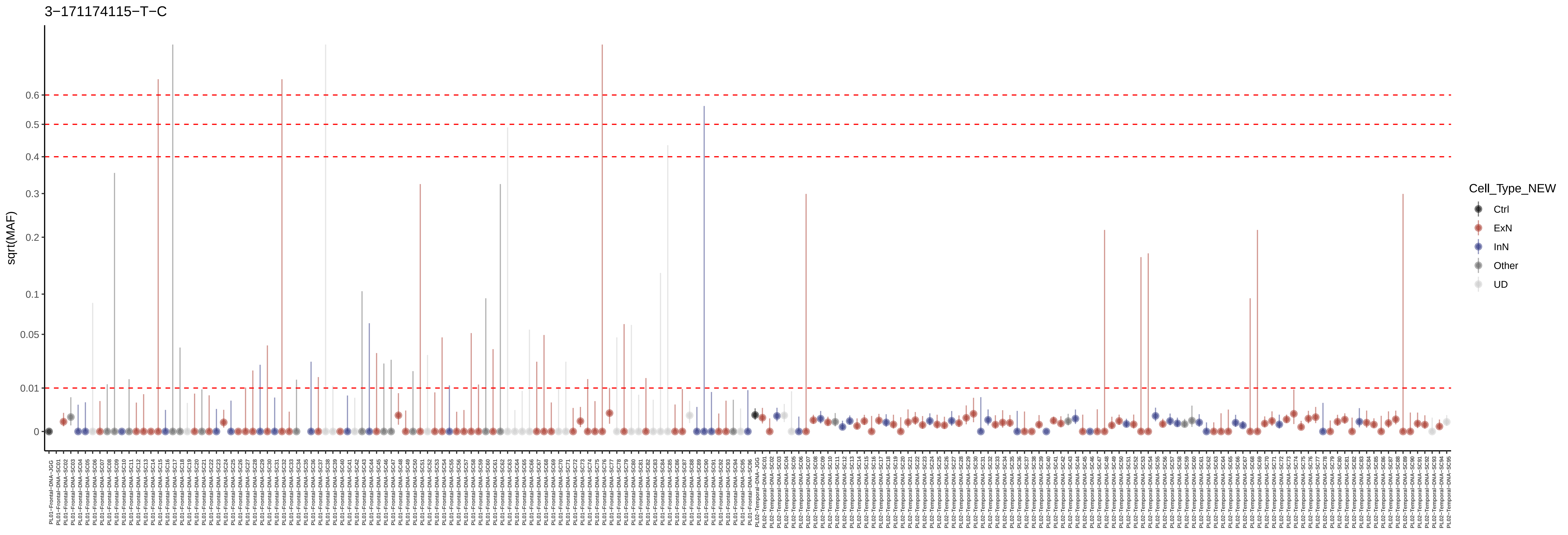

3−171174115−T−C
sqrt(MAF)
0
0.01
0.05
0.1
0.2
0.3
0.4
0.5
0.6
Cell_Type_NEW
Ctrl
ExN
InN
Other
UD

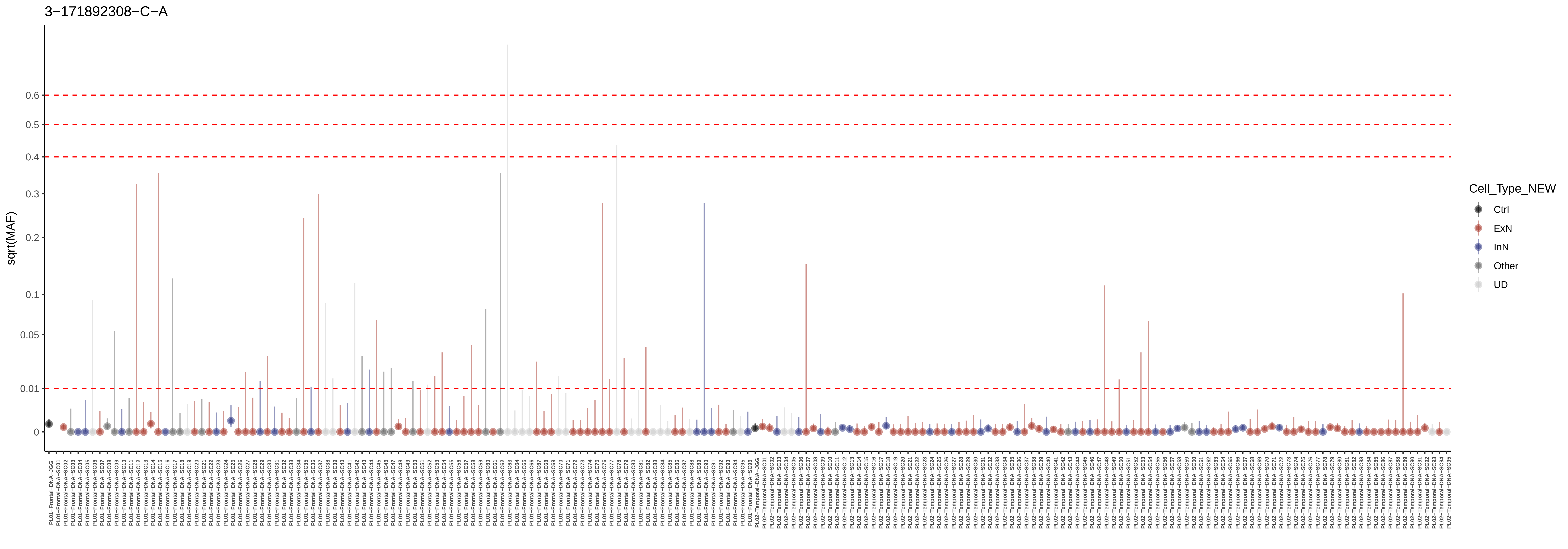

3−171892308−C−A
sqrt(MAF)
0.6
0.5
0.4
0.3
0.2
0.1
0.05
0.01
0
Cell_Type_NEW
Ctrl
ExN
InN
Other
UD

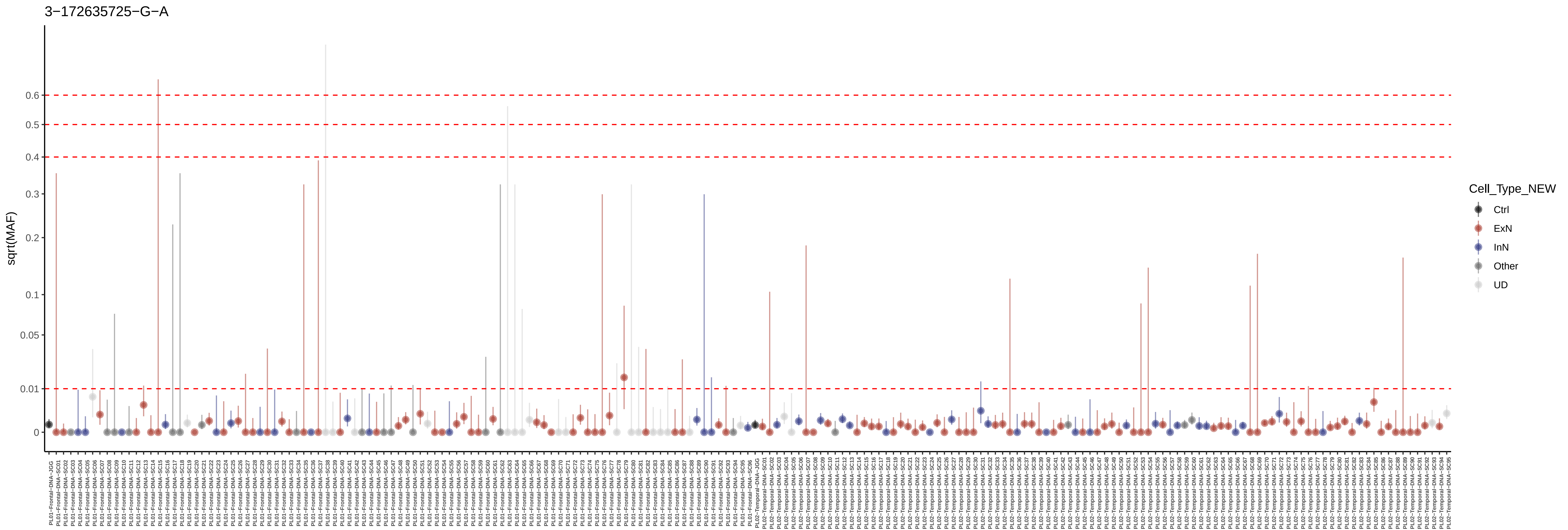
3−172635725−G−A
sqrt(MAF)
0.6
0.5
0.4
0.3
0.2
0.1
0.05
0.01
0
Cell_Type_NEW
Ctrl
ExN
InN
Other
UD

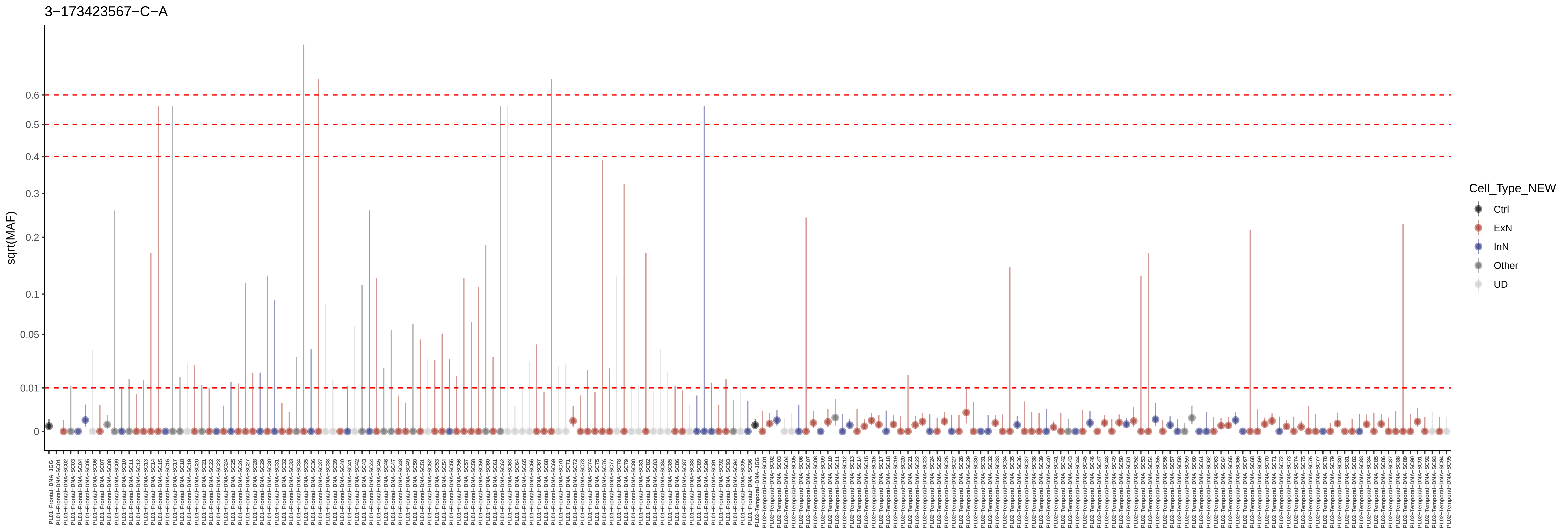
3−173423567−C−A
sqrt(MAF)
0.6
0.5
0.4
0.3
0.2
0.1
0.05
0.01
0
Cell_Type_NEW
Ctrl
ExN
InN
Other
UD

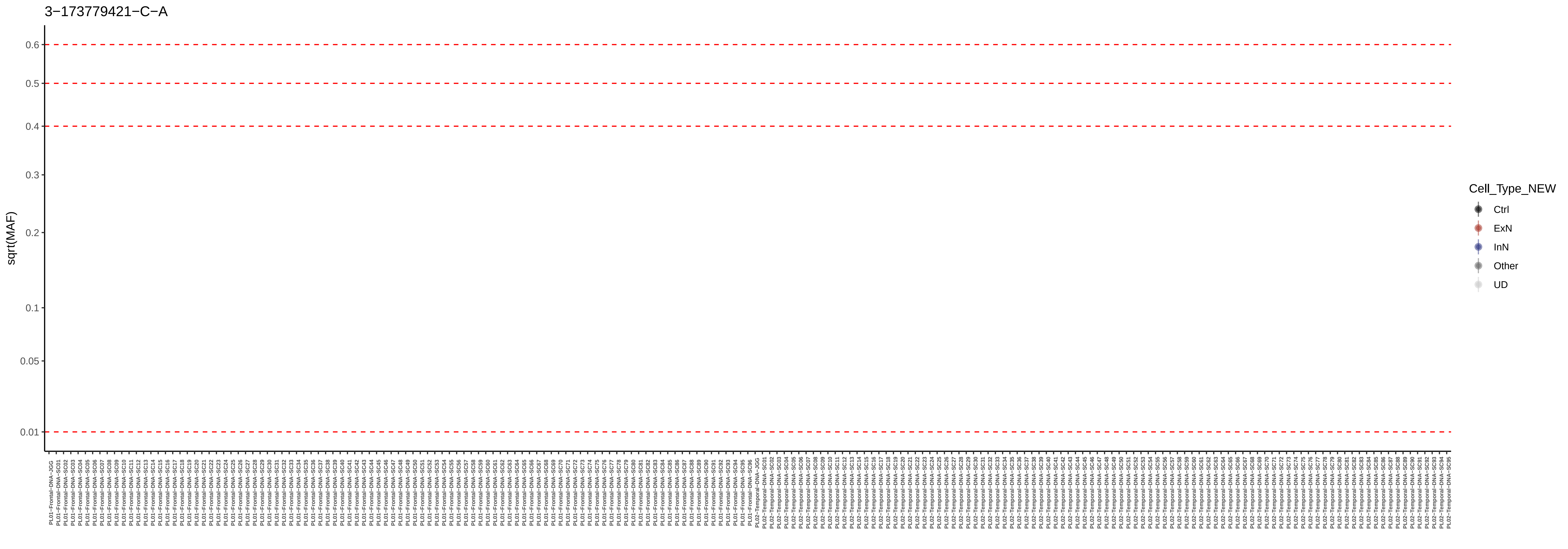
3−173779421−C−A
sqrt(MAF)
Cell_Type_NEW
Ctrl
ExN
InN
Other
UD

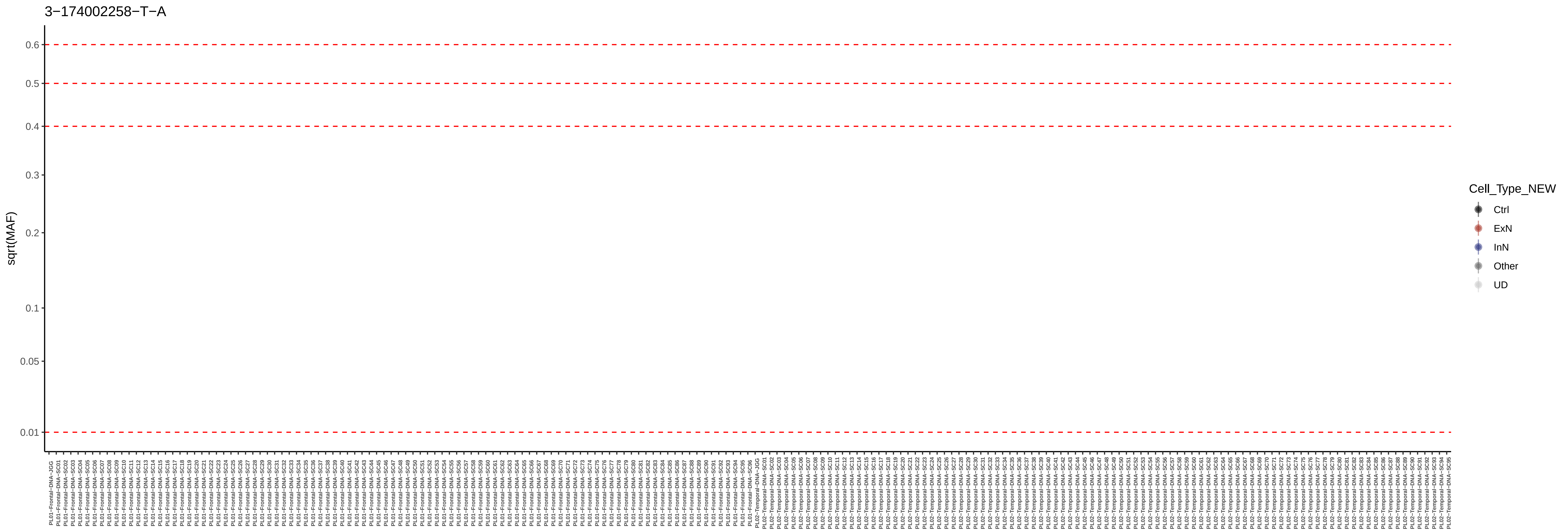
3−174002258−T−A
sqrt(MAF)
0.6
0.5
0.4
0.3
0.2
0.1
0.05
0.01
Cell_Type_NEW
Ctrl
ExN
InN
Other
UD

3−174523465−T−C

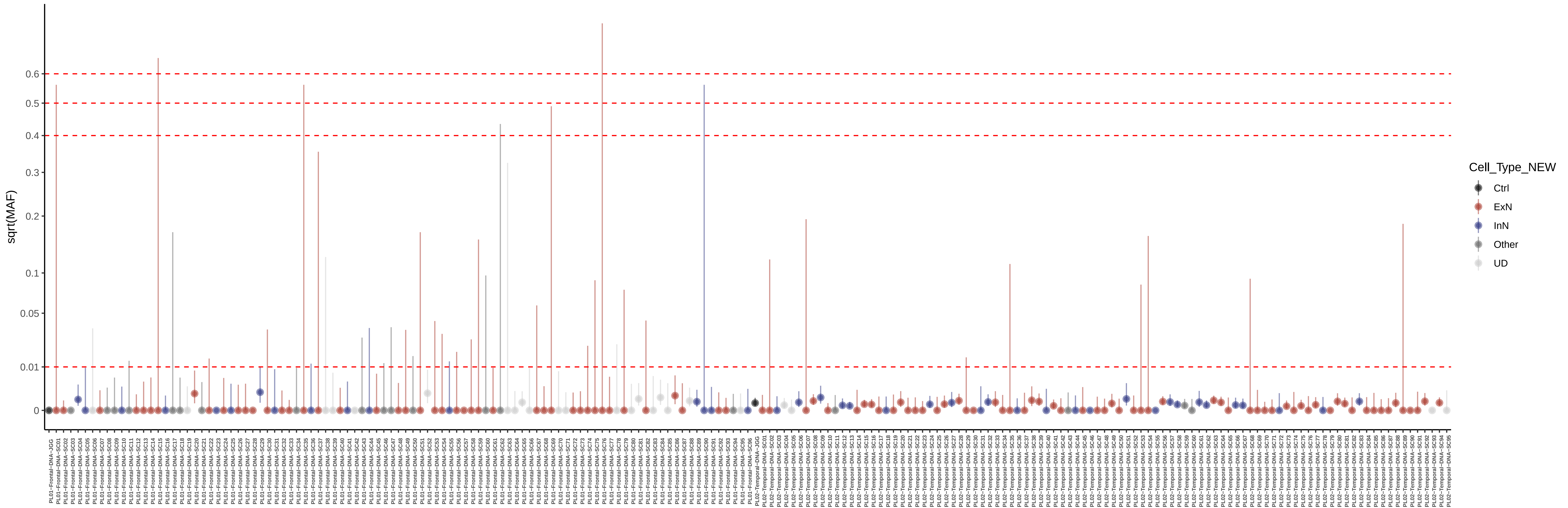

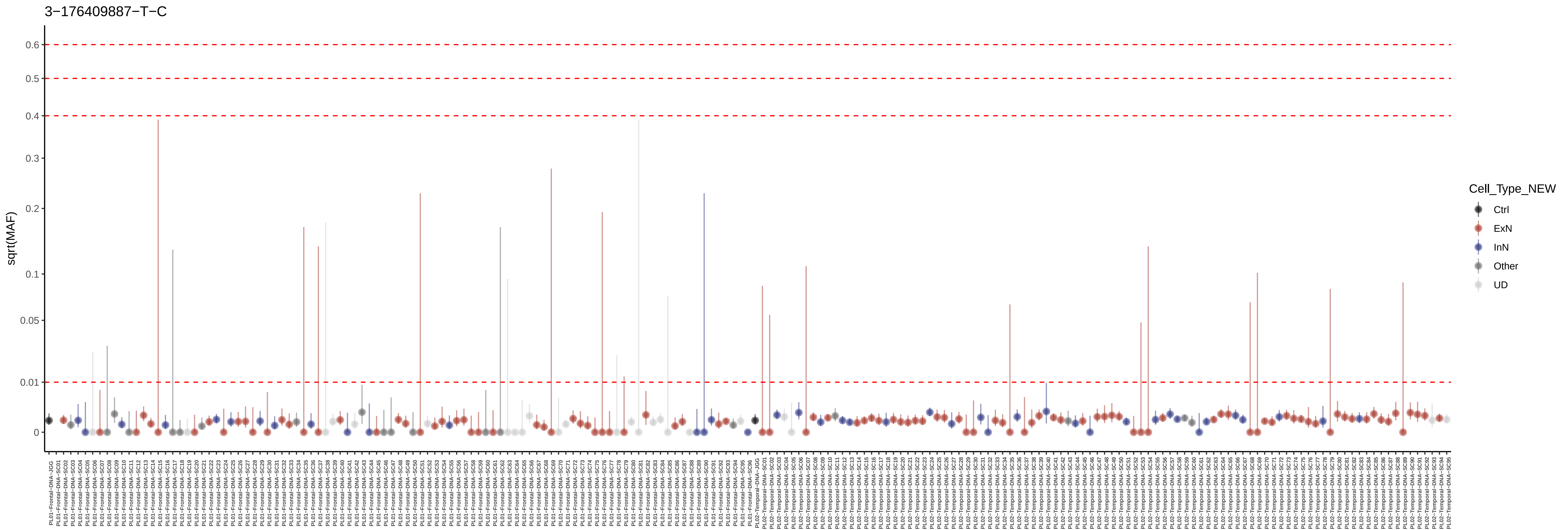
3−176409887−T−C
sqrt(MAF)
0.6
0.5
0.4
0.3
0.2
0.1
0.05
0.01
0
Cell_Type_NEW
Ctrl
ExN
InN
Other
UD

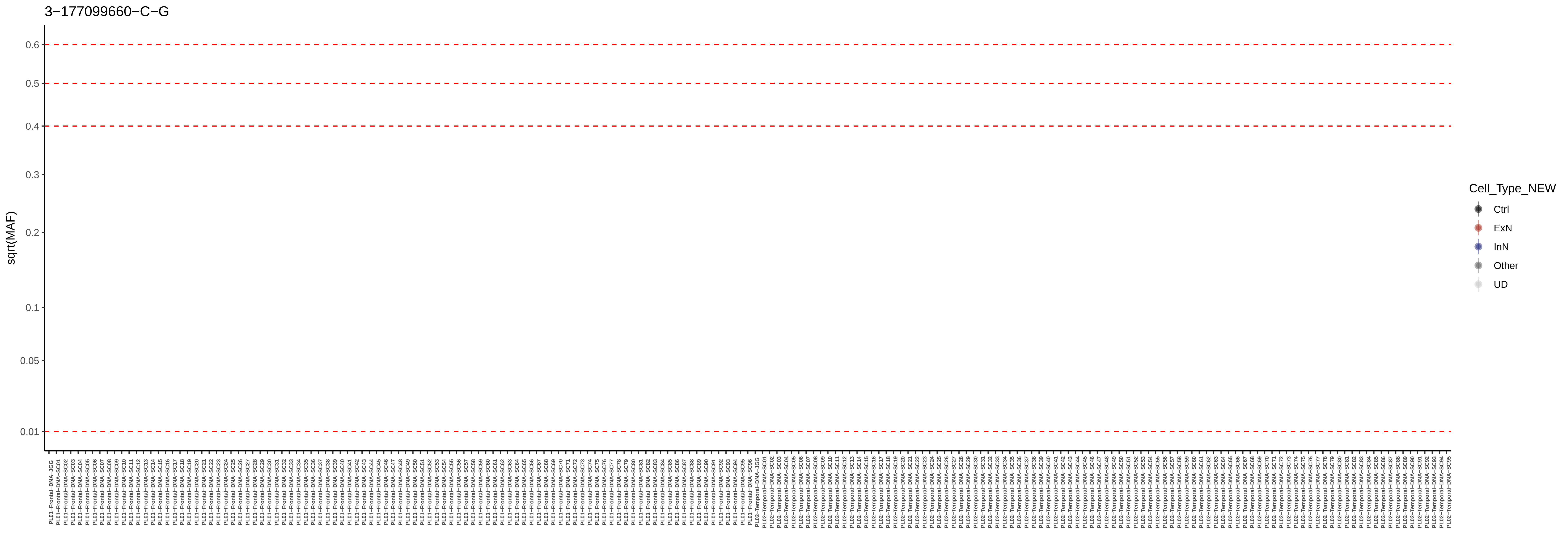
3−177099660−C−G
sqrt(MAF)
0.6
0.5
0.4
0.3
0.2
0.1
0.05
0.01
Cell_Type_NEW
Ctrl
ExN
InN
Other
UD
PL01−Frontal−DNA−JGG
PL01−Frontal−DNA−SC01
PL01−Frontal−DNA−SC02
PL01−Frontal−DNA−SC03
PL01−Frontal−DNA−SC04
PL01−Frontal−DNA−SC05
PL01−Frontal−DNA−SC06
PL01−Frontal−DNA−SC07
PL01−Frontal−DNA−SC08
PL01−Frontal−DNA−SC09
PL01−Frontal−DNA−SC10
PL01−Frontal−DNA−SC11
PL01−Frontal−DNA−SC12
PL01−Frontal−DNA−SC13
PL01−Frontal−DNA−SC14
PL01−Frontal−DNA−SC15
PL01−Frontal−DNA−SC16
PL01−Frontal−DNA−SC17
PL01−Frontal−DNA−SC18
PL01−Frontal−DNA−SC19
PL01−Frontal−DNA−SC20
PL01−Frontal−DNA−SC21
PL01−Frontal−DNA−SC22
PL01−Frontal−DNA−SC23
PL01−Frontal−DNA−SC24
PL01−Frontal−DNA−SC25
PL01−Frontal−DNA−SC26
PL01−Frontal−DNA−SC27
PL01−Frontal−DNA−SC28
PL01−Frontal−DNA−SC29
PL01−Frontal−DNA−SC30
PL01−Frontal−DNA−SC31
PL01−Frontal−DNA−SC32
PL01−Frontal−DNA−SC33
PL01−Frontal−DNA−SC34
PL01−Frontal−DNA−SC35
PL01−Frontal−DNA−SC36
PL01−Frontal−DNA−SC37
PL01−Frontal−DNA−SC38
PL01−Frontal−DNA−SC39
PL01−Frontal−DNA−SC40
PL01−Frontal−DNA−SC41
PL01−Frontal−DNA−SC42
PL01−Frontal−DNA−SC43
PL01−Frontal−DNA−SC44
PL01−Frontal−DNA−SC45
PL01−Frontal−DNA−SC46
PL01−Frontal−DNA−SC47
PL01−Frontal−DNA−SC48
PL01−Frontal−DNA−SC49
PL01−Frontal−DNA−SC50
PL01−Frontal−DNA−SC51
PL01−Frontal−DNA−SC52
PL01−Frontal−DNA−SC53
PL01−Frontal−DNA−SC54
PL01−Frontal−DNA−SC55
PL01−Frontal−DNA−SC56
PL01−Frontal−DNA−SC57
PL01−Frontal−DNA−SC58
PL01−Frontal−DNA−SC59
PL01−Frontal−DNA−SC60
PL01−Frontal−DNA−SC61
PL01−Frontal−DNA−SC62
PL01−Frontal−DNA−SC63
PL01−Frontal−DNA−SC64
PL01−Frontal−DNA−SC65
PL01−Frontal−DNA−SC66
PL01−Frontal−DNA−SC67
PL01−Frontal−DNA−SC68
PL01−Frontal−DNA−SC69
PL01−Frontal−DNA−SC70
PL01−Frontal−DNA−SC71
PL01−Frontal−DNA−SC72
PL01−Frontal−DNA−SC73
PL01−Frontal−DNA−SC74
PL01−Frontal−DNA−SC75
PL01−Frontal−DNA−SC76
PL01−Frontal−DNA−SC77
PL01−Frontal−DNA−SC78
PL01−Frontal−DNA−SC79
PL01−Frontal−DNA−SC80
PL01−Frontal−DNA−SC81
PL01−Frontal−DNA−SC82
PL01−Frontal−DNA−SC83
PL01−Frontal−DNA−SC84
PL01−Frontal−DNA−SC85
PL01−Frontal−DNA−SC86
PL01−Frontal−DNA−SC87
PL01−Frontal−DNA−SC88
PL01−Frontal−DNA−SC89
PL01−Frontal−DNA−SC90
PL01−Frontal−DNA−SC91
PL01−Frontal−DNA−SC92
PL01−Frontal−DNA−SC93
PL01−Frontal−DNA−SC94
PL01−Frontal−DNA−SC95
PL01−Frontal−DNA−SC96
PL02−Temporal−DNA−JGG
PL02−Temporal−DNA−SC01
PL02−Temporal−DNA−SC02
PL02−Temporal−DNA−SC03
PL02−Temporal−DNA−SC04
PL02−Temporal−DNA−SC05
PL02−Temporal−DNA−SC06
PL02−Temporal−DNA−SC07
PL02−Temporal−DNA−SC08
PL02−Temporal−DNA−SC09
PL02−Temporal−DNA−SC10
PL02−Temporal−DNA−SC11
PL02−Temporal−DNA−SC12
PL02−Temporal−DNA−SC13
PL02−Temporal−DNA−SC14
PL02−Temporal−DNA−SC15
PL02−Temporal−DNA−SC16
PL02−Temporal−DNA−SC17
PL02−Temporal−DNA−SC18
PL02−Temporal−DNA−SC19
PL02−Temporal−DNA−SC20
PL02−Temporal−DNA−SC21
PL02−Temporal−DNA−SC22
PL02−Temporal−DNA−SC23
PL02−Temporal−DNA−SC24
PL02−Temporal−DNA−SC25
PL02−Temporal−DNA−SC26
PL02−Temporal−DNA−SC27
PL02−Temporal−DNA−SC28
PL02−Temporal−DNA−SC29
PL02−Temporal−DNA−SC30
PL02−Temporal−DNA−SC31
PL02−Temporal−DNA−SC32
PL02−Temporal−DNA−SC33
PL02−Temporal−DNA−SC34
PL02−Temporal−DNA−SC35
PL02−Temporal−DNA−SC36
PL02−Temporal−DNA−SC37
PL02−Temporal−DNA−SC38
PL02−Temporal−DNA−SC39
PL02−Temporal−DNA−SC40
PL02−Temporal−DNA−SC41
PL02−Temporal−DNA−SC42
PL02−Temporal−DNA−SC43
PL02−Temporal−DNA−SC44
PL02−Temporal−DNA−SC45
PL02−Temporal−DNA−SC46
PL02−Temporal−DNA−SC47
PL02−Temporal−DNA−SC48
PL02−Temporal−DNA−SC49
PL02−Temporal−DNA−SC50
PL02−Temporal−DNA−SC51
PL02−Temporal−DNA−SC52
PL02−Temporal−DNA−SC53
PL02−Temporal−DNA−SC54
PL02−Temporal−DNA−SC55
PL02−Temporal−DNA−SC56
PL02−Temporal−DNA−SC57
PL02−Temporal−DNA−SC58
PL02−Temporal−DNA−SC59
PL02−Temporal−DNA−SC60
PL02−Temporal−DNA−SC61
PL02−Temporal−DNA−SC62
PL02−Temporal−DNA−SC63
PL02−Temporal−DNA−SC64
PL02−Temporal−DNA−SC65
PL02−Temporal−DNA−SC66
PL02−Temporal−DNA−SC67
PL02−Temporal−DNA−SC68
PL02−Temporal−DNA−SC69
PL02−Temporal−DNA−SC70
PL02−Temporal−DNA−SC71
PL02−Temporal−DNA−SC72
PL02−Temporal−DNA−SC73
PL02−Temporal−DNA−SC74
PL02−Temporal−DNA−SC75
PL02−Temporal−DNA−SC76
PL02−Temporal−DNA−SC77
PL02−Temporal−DNA−SC78
PL02−Temporal−DNA−SC79
PL02−Temporal−DNA−SC80
PL02−Temporal−DNA−SC81
PL02−Temporal−DNA−SC82
PL02−Temporal−DNA−SC83
PL02−Temporal−DNA−SC84
PL02−Temporal−DNA−SC85
PL02−Temporal−DNA−SC86
PL02−Temporal−DNA−SC87
PL02−Temporal−DNA−SC88
PL02−Temporal−DNA−SC89
PL02−Temporal−DNA−SC90
PL02−Temporal−DNA−SC91
PL02−Temporal−DNA−SC92
PL02−Temporal−DNA−SC93
PL02−Temporal−DNA−SC94
PL02−Temporal−DNA−SC95

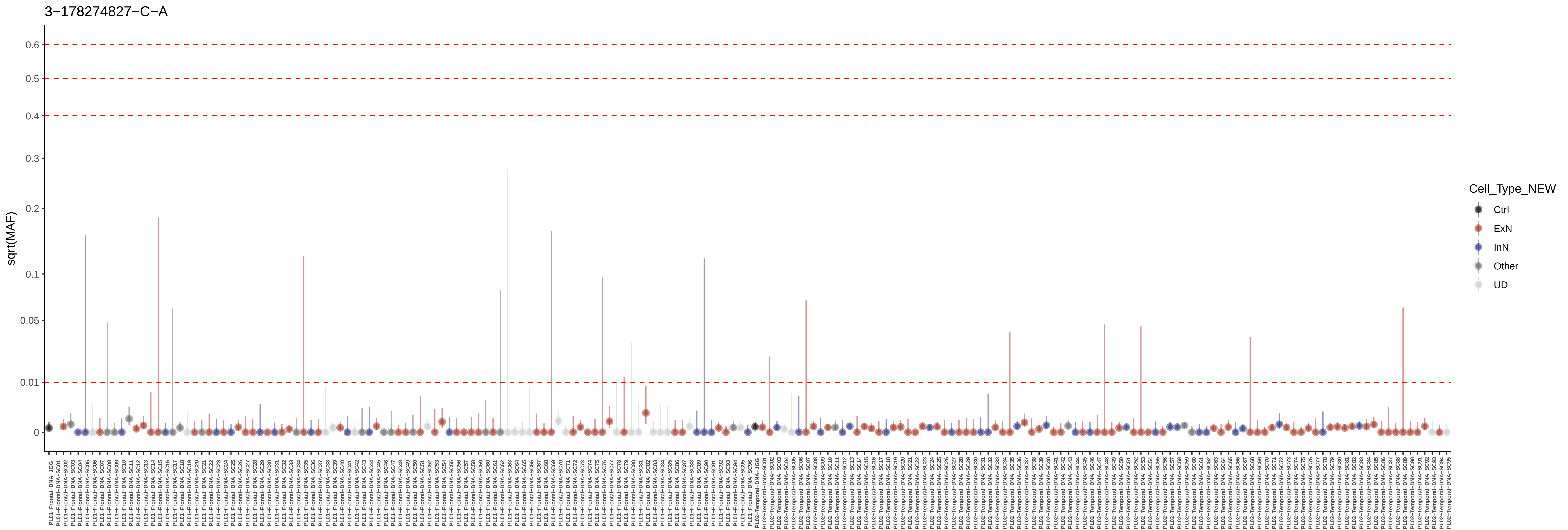

3−178274827−C−A
sqrt(MAF)
0.6
0.5
0.4
0.3
0.2
0.1
0.05
0.01
0
Cell_Type_NEW
Ctrl
ExN
InN
Other
UD

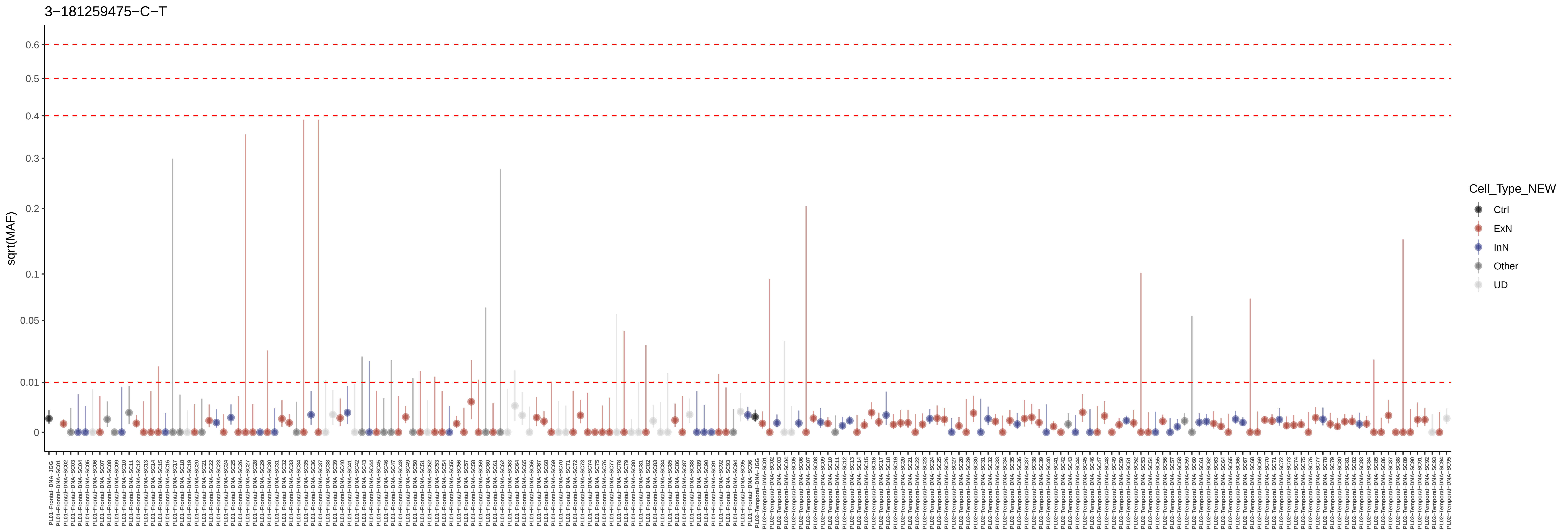
3−181259475−C−T
sqrt(MAF)
0.6
0.5
0.4
0.3
0.2
0.1
0.05
0.01
0
Cell_Type_NEW
Ctrl
ExN
InN
Other
UD

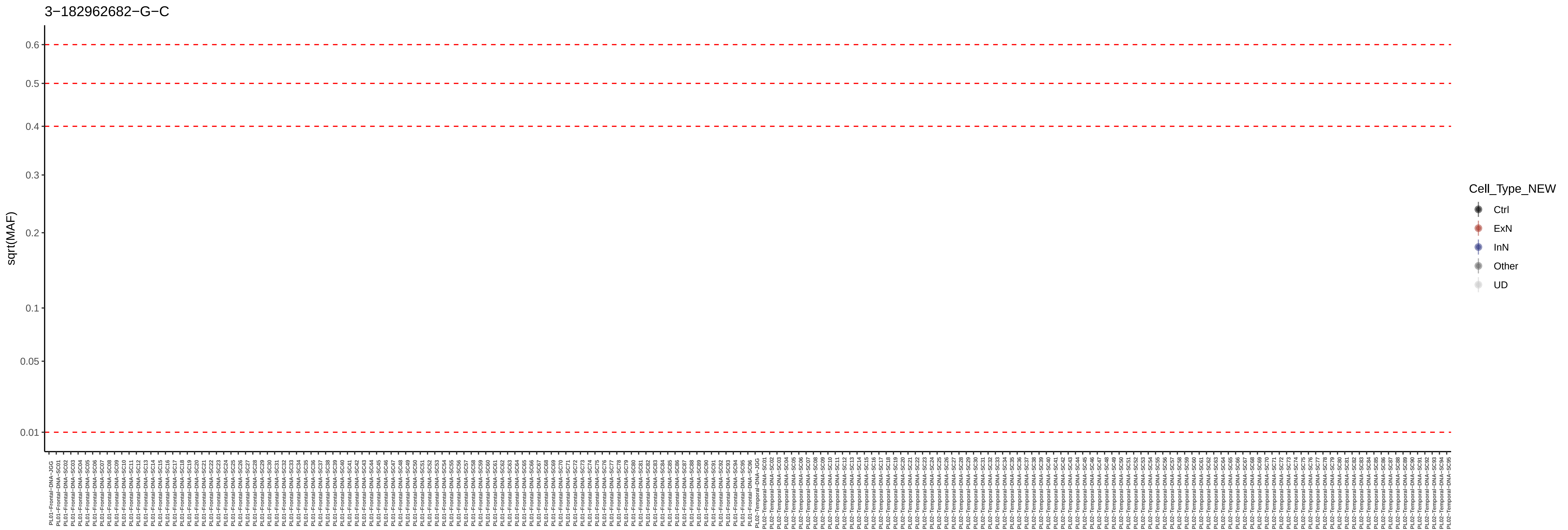

3−182962682−G−C
sqrt(MAF)
0.6
0.5
0.4
0.3
0.2
0.1
0.05
0.01
Cell_Type_NEW
Ctrl
ExN
InN
Other
UD

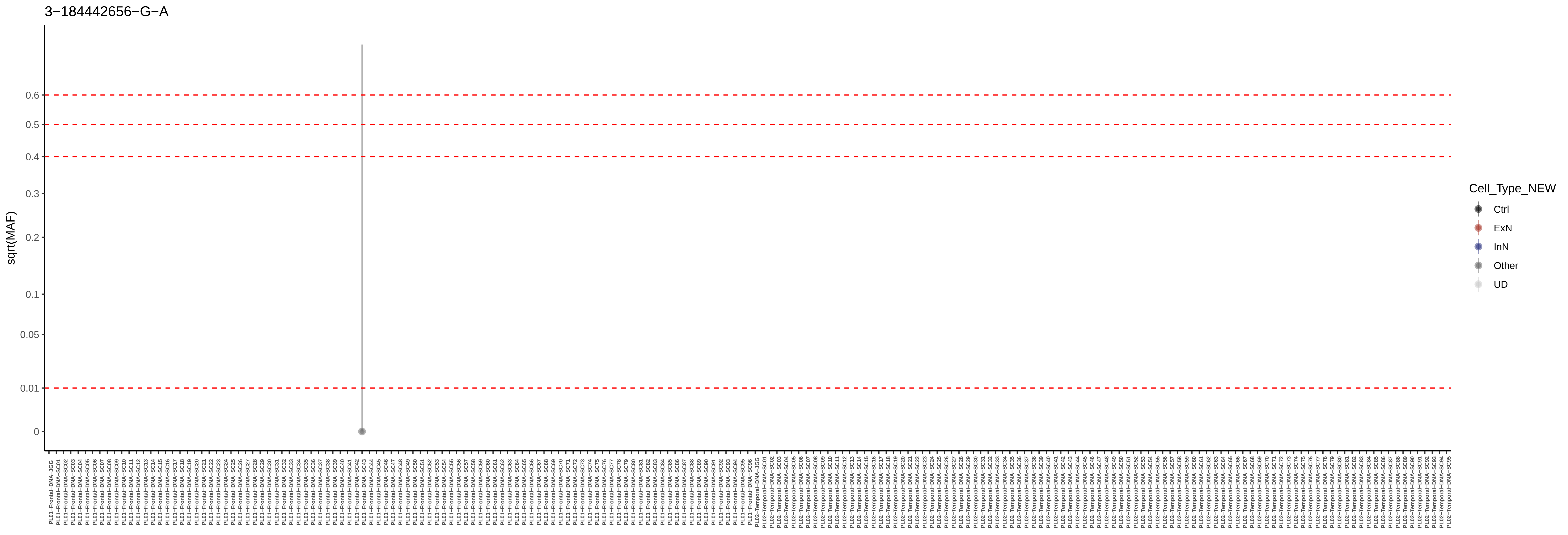
3−184442656−G−A
sqrt(MAF)
0.6
0.5
0.4
0.3
0.2
0.1
0.05
0.01
0
Cell_Type_NEW
Ctrl
ExN
InN
Other
UD

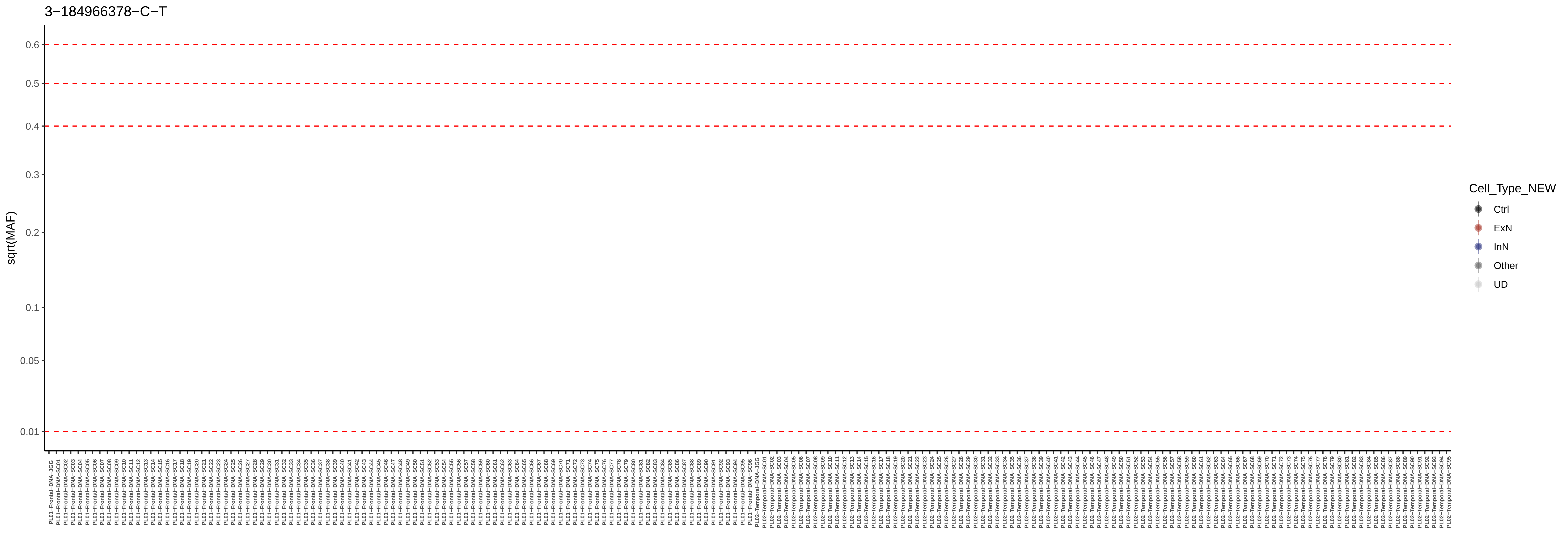

3−184966378−C−T
sqrt(MAF)
0.6
0.5
0.4
0.3
0.2
0.1
0.05
0.01
Cell_Type_NEW
Ctrl
ExN
InN
Other
UD

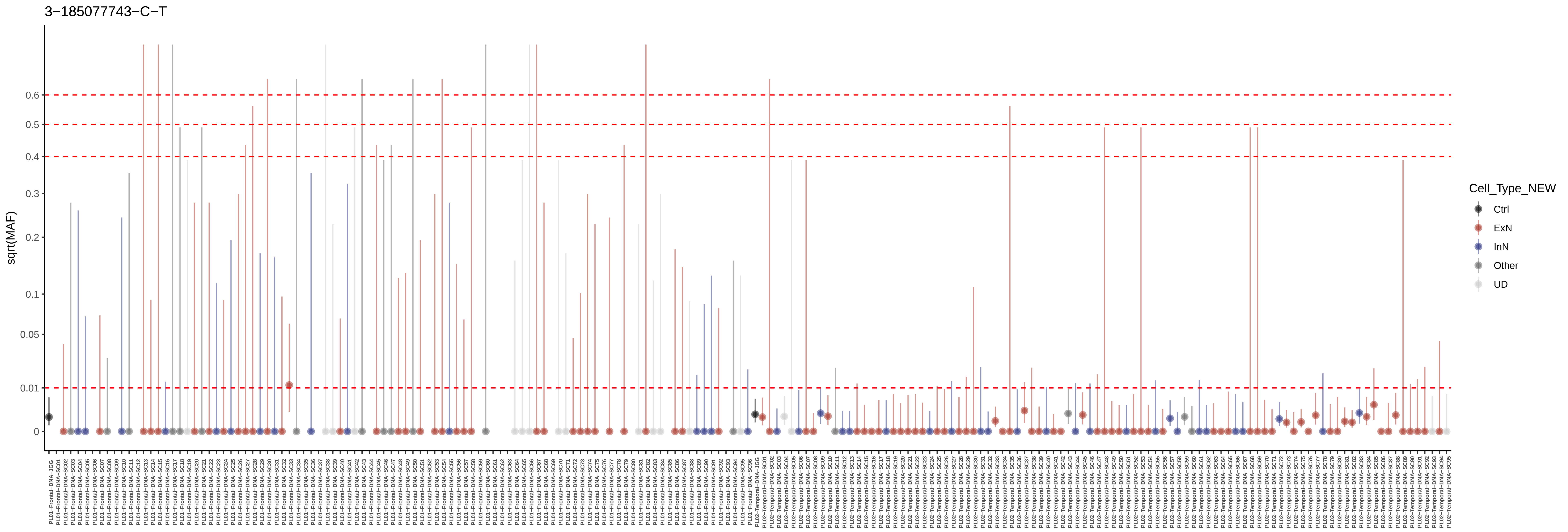
3−185077743−C−T
sqrt(MAF)
0.6
0.5
0.4
0.3
0.2
0.1
0.05
0.01
0
Cell_Type_NEW
Ctrl
ExN
InN
Other
UD

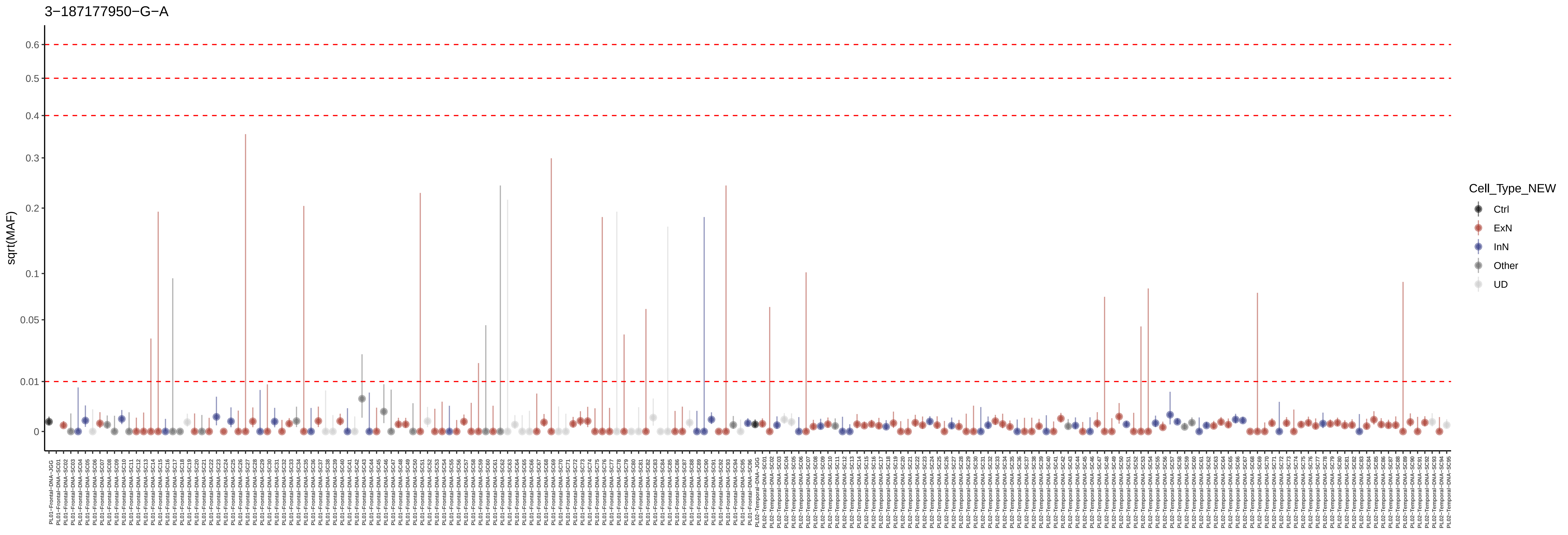

3−187177950−G−A
sqrt(MAF)
0.6
0.5
0.4
0.3
0.2
0.1
0.05
0.01
0
Cell_Type_NEW
Ctrl
ExN
InN
Other
UD

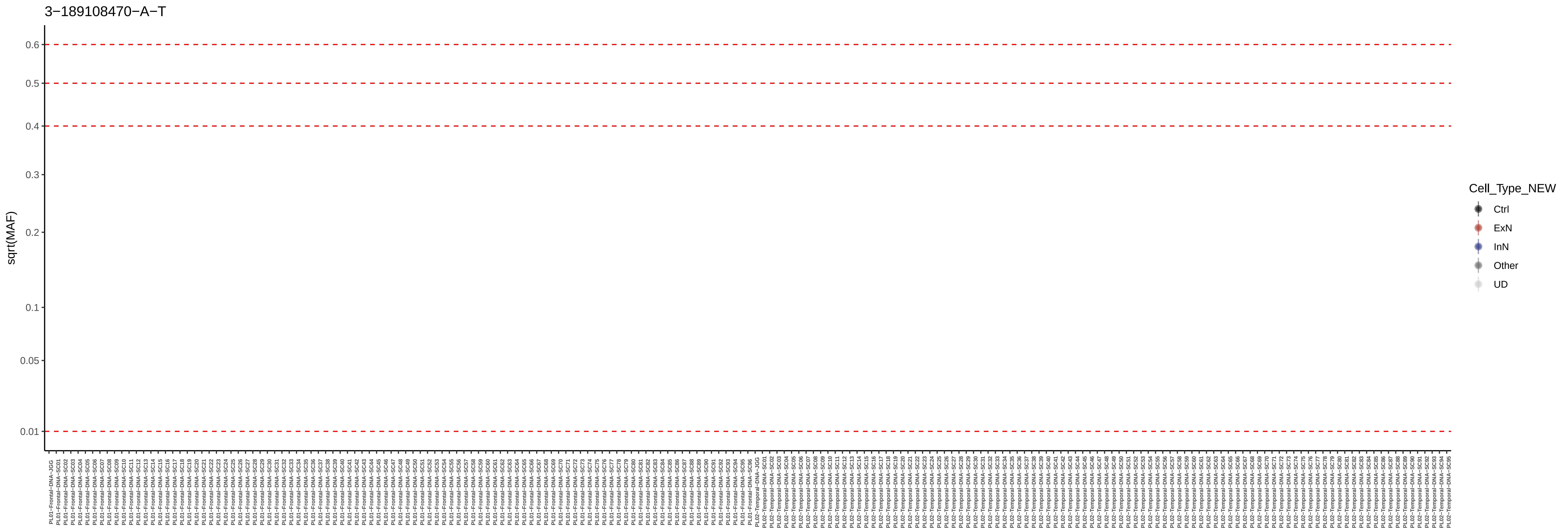

3−189108470−A−T
sqrt(MAF)
0.6
0.5
0.4
0.3
0.2
0.1
0.05
0.01
Cell_Type_NEW
Ctrl
ExN
InN
Other
UD

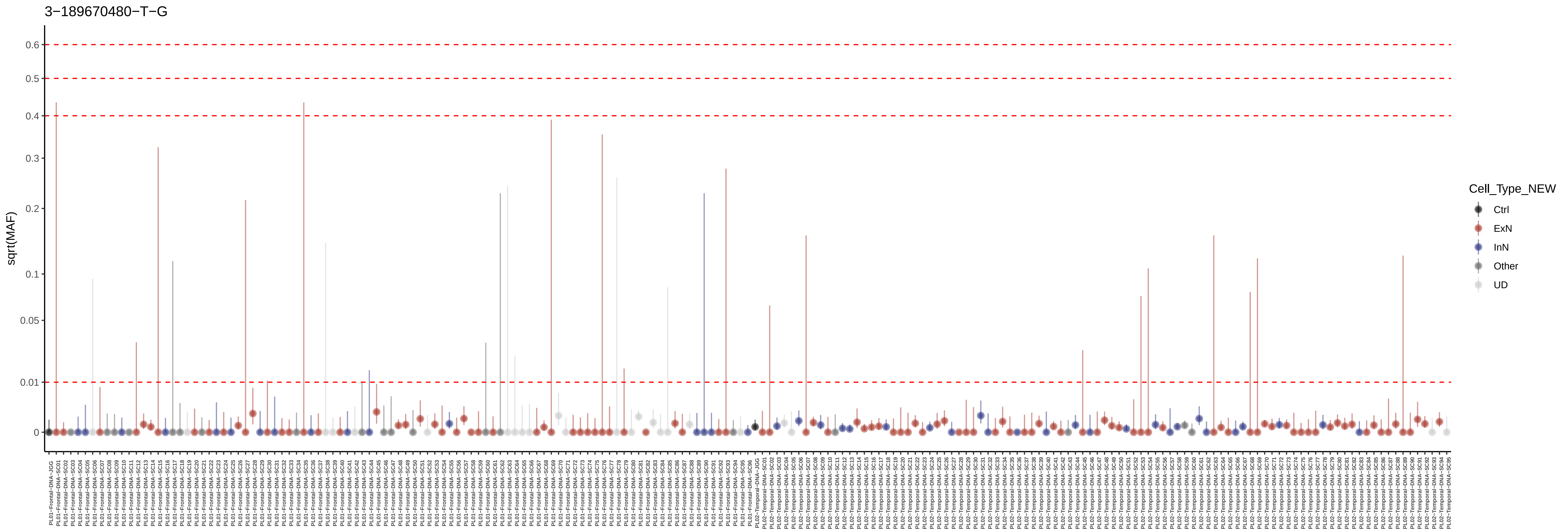

3−189670480−T−G
sqrt(MAF)
0.6
0.5
0.4
0.3
0.2
0.1
0.05
0.01
0
Cell_Type_NEW
Ctrl
ExN
InN
Other
UD

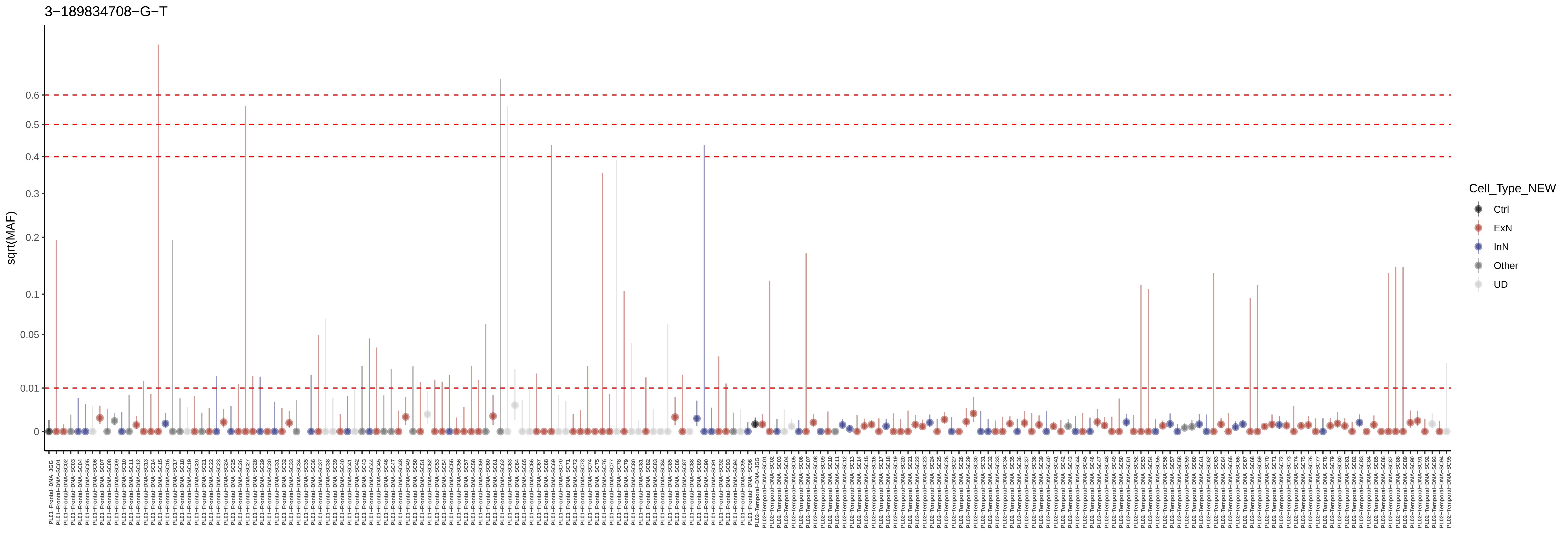

3−189834708−G−T
sqrt(MAF)
0.6
0.5
0.4
0.3
0.2
0.1
0.05
0.01
0
Cell_Type_NEW
Ctrl
ExN
InN
Other
UD

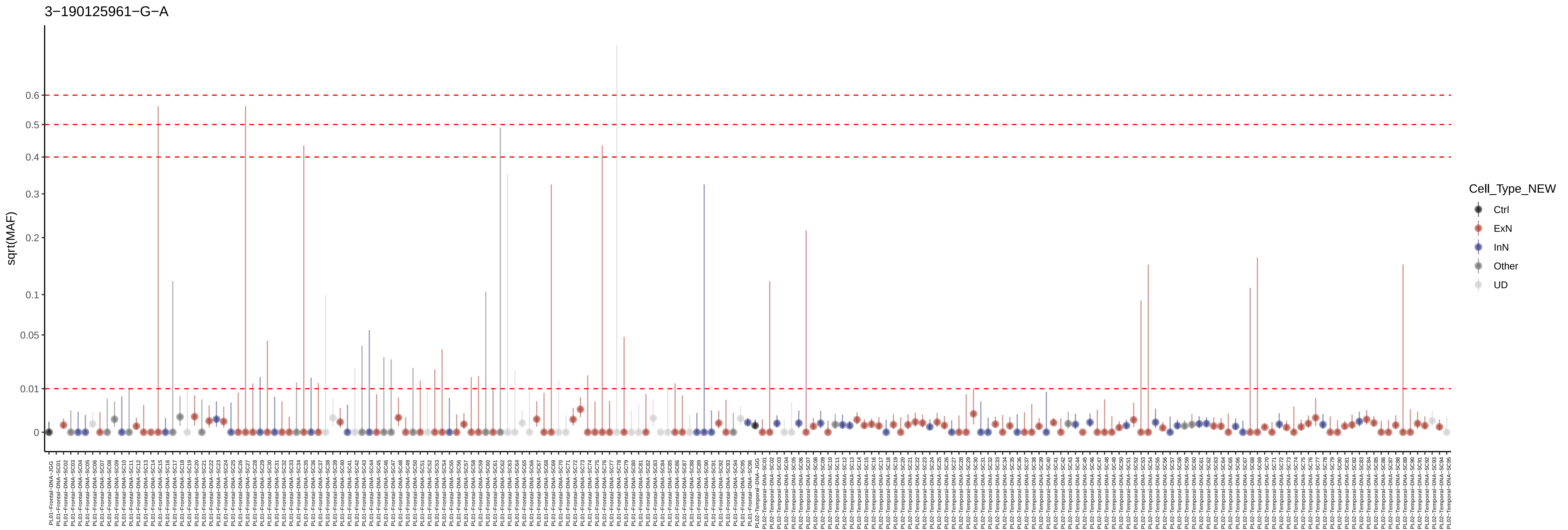

3−190125961−G−A
sqrt(MAF)
0.6
0.5
0.4
0.3
0.2
0.1
0.05
0.01
0
Cell_Type_NEW
Ctrl
ExN
InN
Other
UD

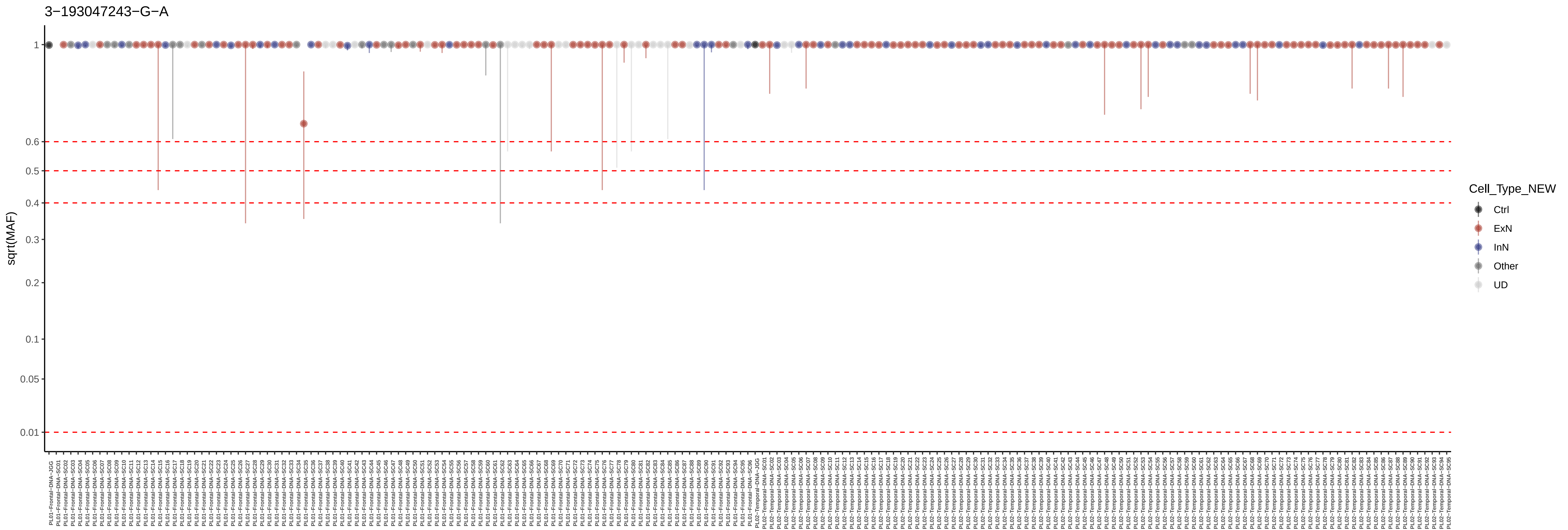
3−193047243−G−A
sqrt(MAF)
1
0.6
0.5
0.4
0.3
0.2
0.1
0.05
0.01
Cell_Type_NEW
Ctrl
ExN
InN
Other
UD

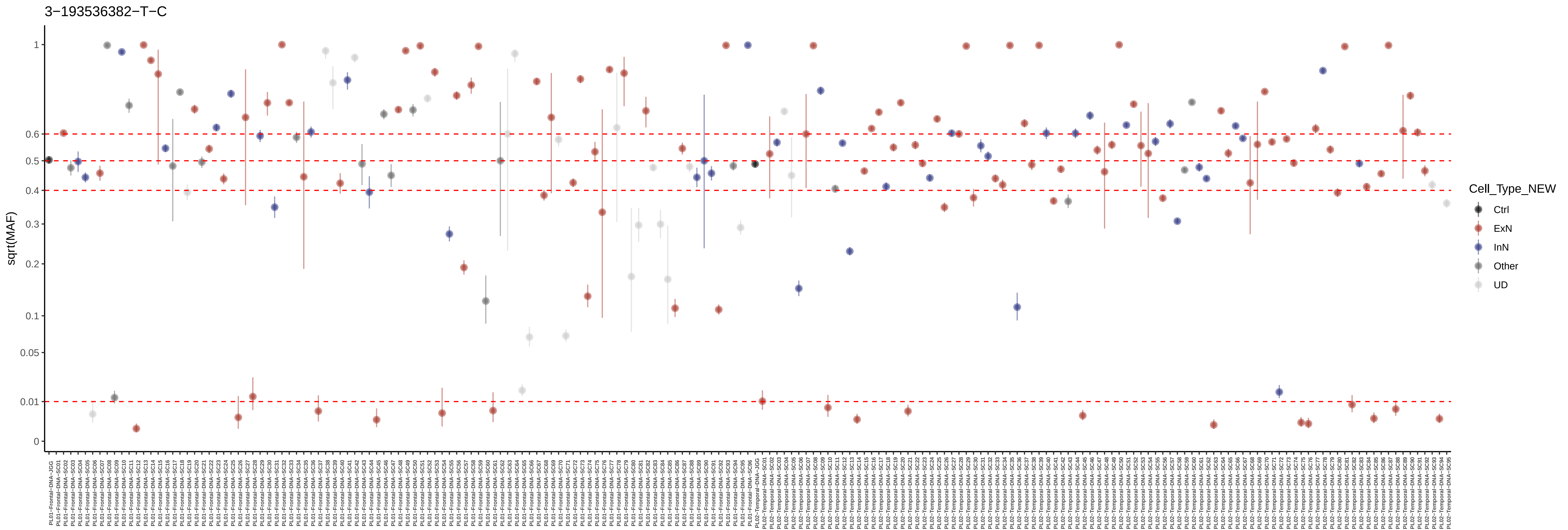
3−193536382−T−C
sqrt(MAF)
1
0.6
0.5
0.4
0.3
0.2
0.1
0.05
0.01
0
Cell_Type_NEW
Ctrl
ExN
InN
Other
UD

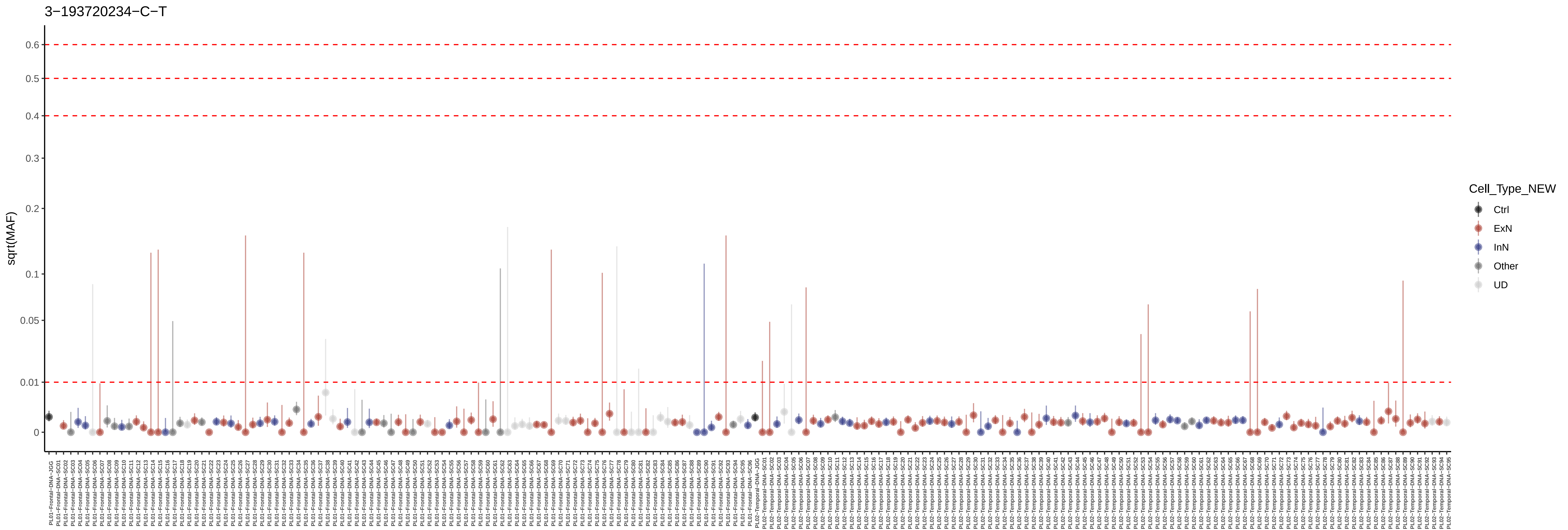
3−193720234−C−T
sqrt(MAF)
0.6
0.5
0.4
0.3
0.2
0.1
0.05
0.01
0
Cell_Type_NEW
Ctrl
ExN
InN
Other
UD

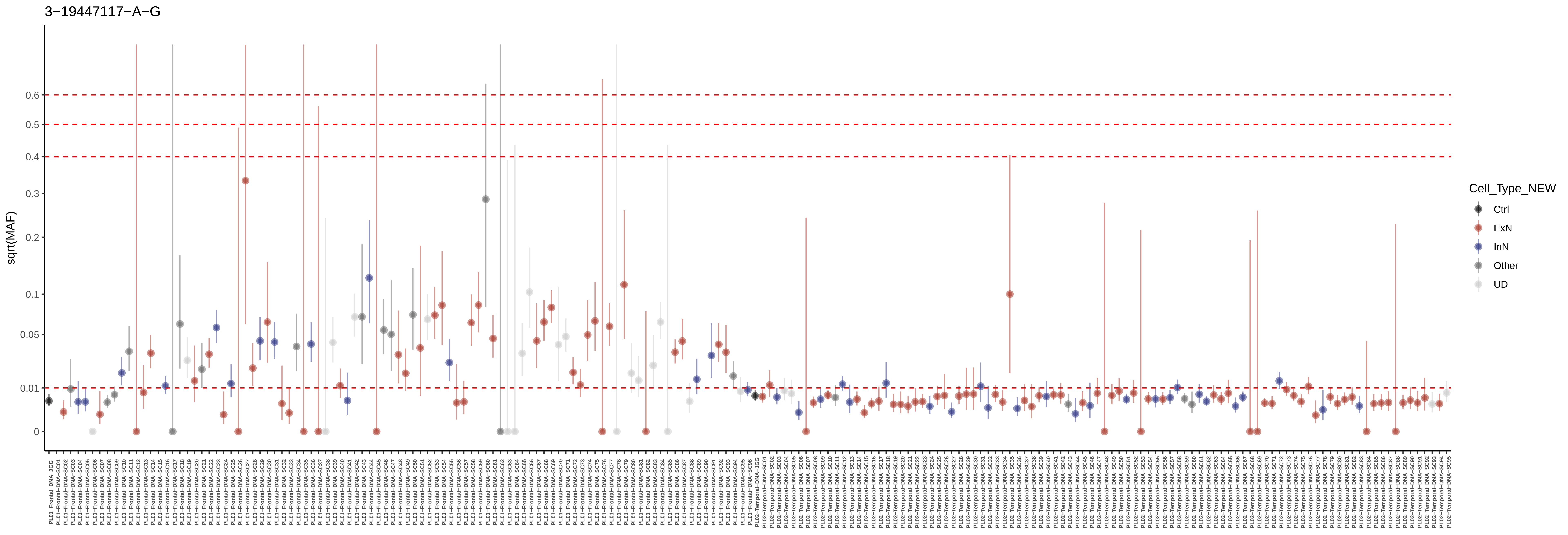

3−19447117−A−G
sqrt(MAF)
0.6
0.5
0.4
0.3
0.2
0.1
0.05
0.01
0
Cell_Type_NEW
Ctrl
ExN
InN
Other
UD

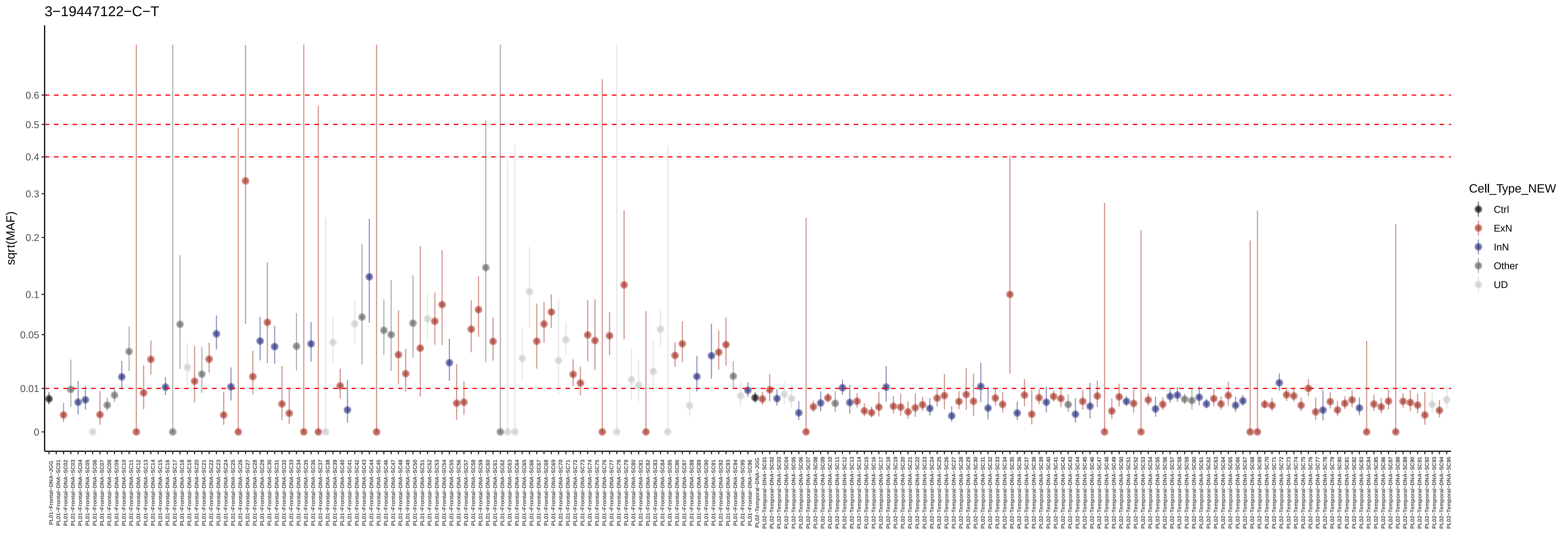

3−19447122−C−T
sqrt(MAF)
0.6
0.5
0.4
0.3
0.2
0.1
0.05
0.01
0
Cell_Type_NEW
Ctrl
ExN
InN
Other
UD

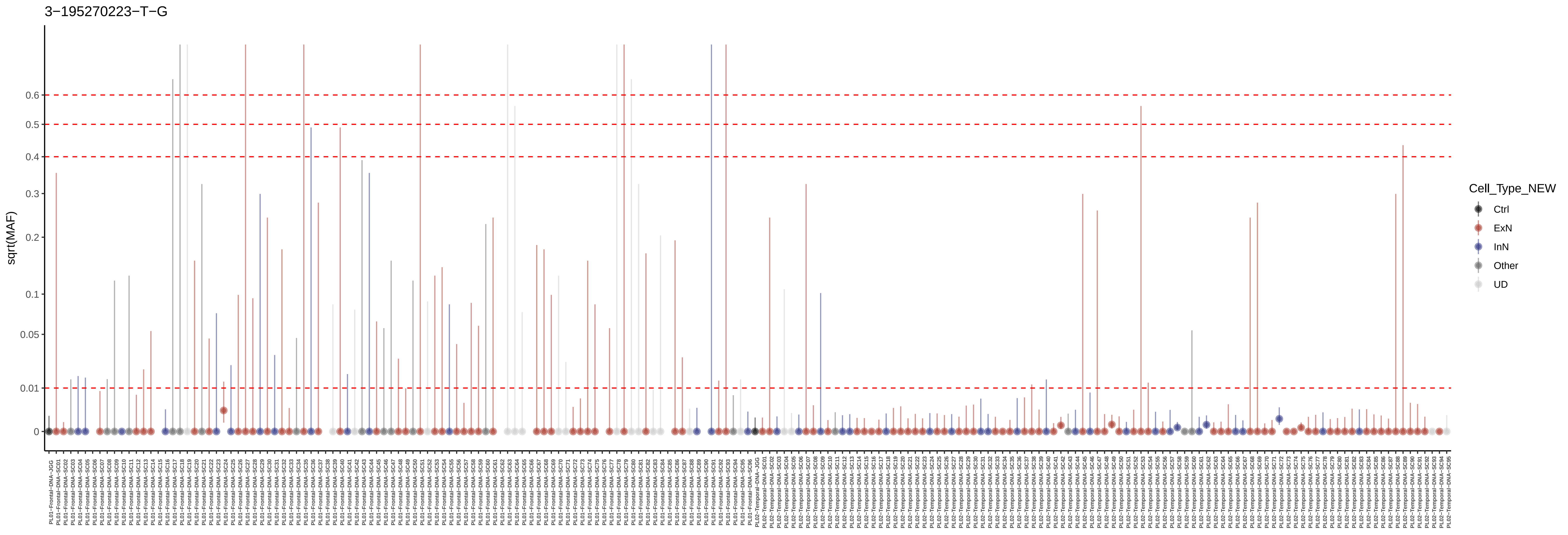
3−195270223−T−G
sqrt(MAF)
0.6
0.5
0.4
0.3
0.2
0.1
0.05
0.01
0
Cell_Type_NEW
Ctrl
ExN
InN
Other
UD

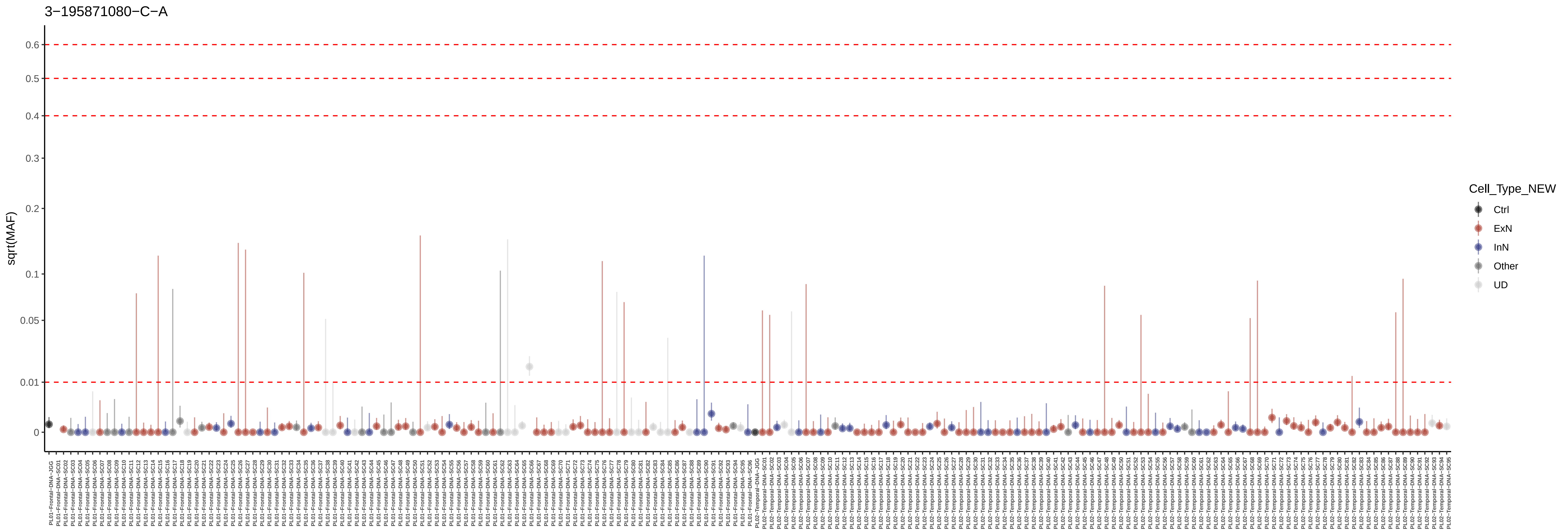

3−195871080−C−A
sqrt(MAF)
0.6
0.5
0.4
0.3
0.2
0.1
0.05
0.01
0
Cell_Type_NEW
Ctrl
ExN
InN
Other
UD

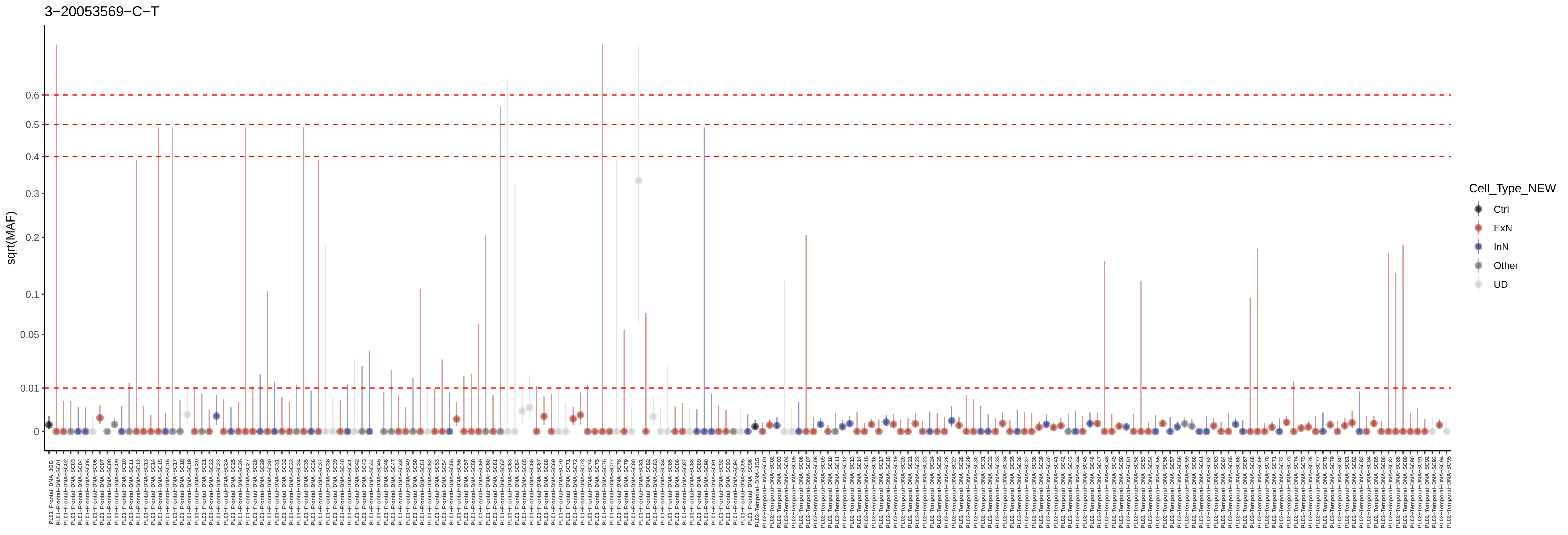
3−20053569−C−T
sqrt(MAF)
0.6
0.5
0.4
0.3
0.2
0.1
0.05
0.01
0
Cell_Type_NEW
Ctrl
ExN
InN
Other
UD

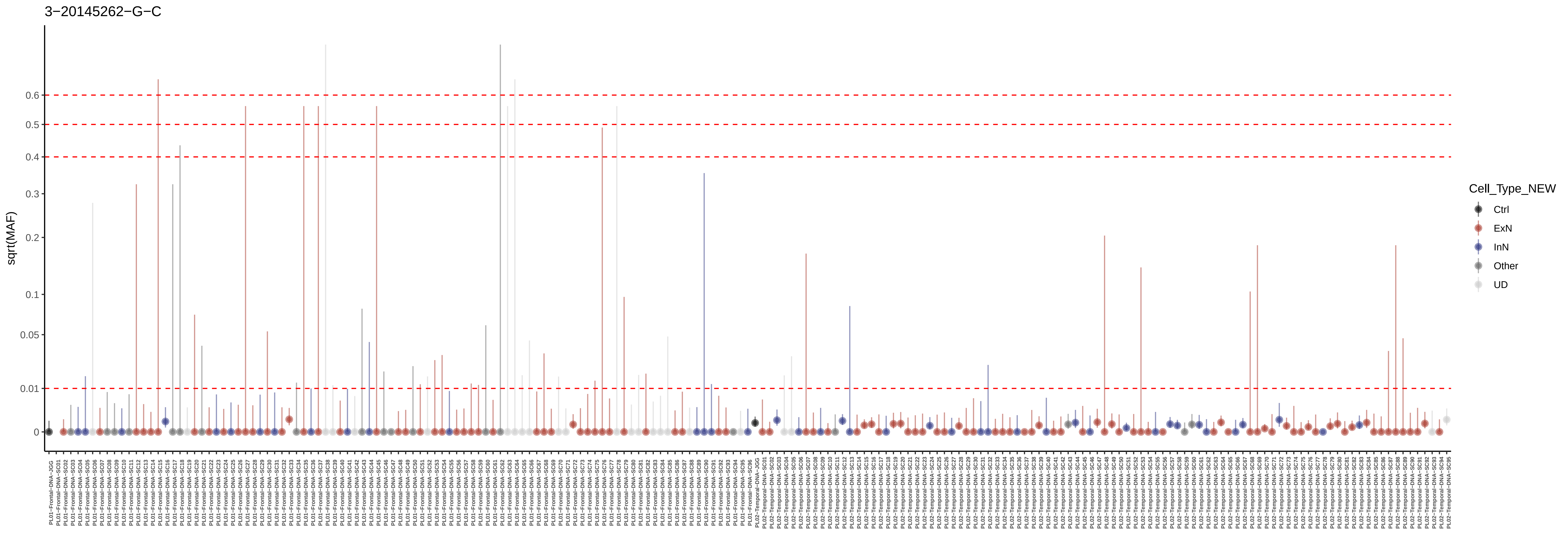
3−20145262−G−C
sqrt(MAF)
0.6
0.5
0.4
0.3
0.2
0.1
0.05
0.01
0
Cell_Type_NEW
Ctrl
ExN
InN
Other
UD

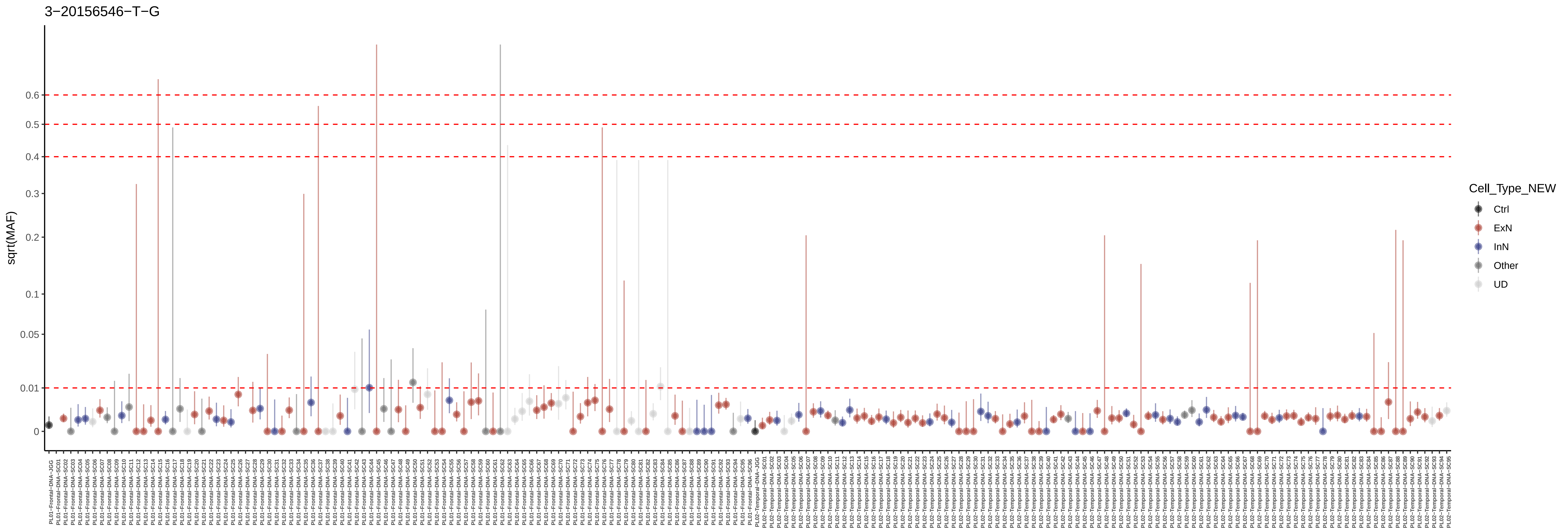
3−20156546−T−G
sqrt(MAF)
0.6
0.5
0.4
0.3
0.2
0.1
0.05
0.01
0
Cell_Type_NEW
Ctrl
ExN
InN
Other
UD

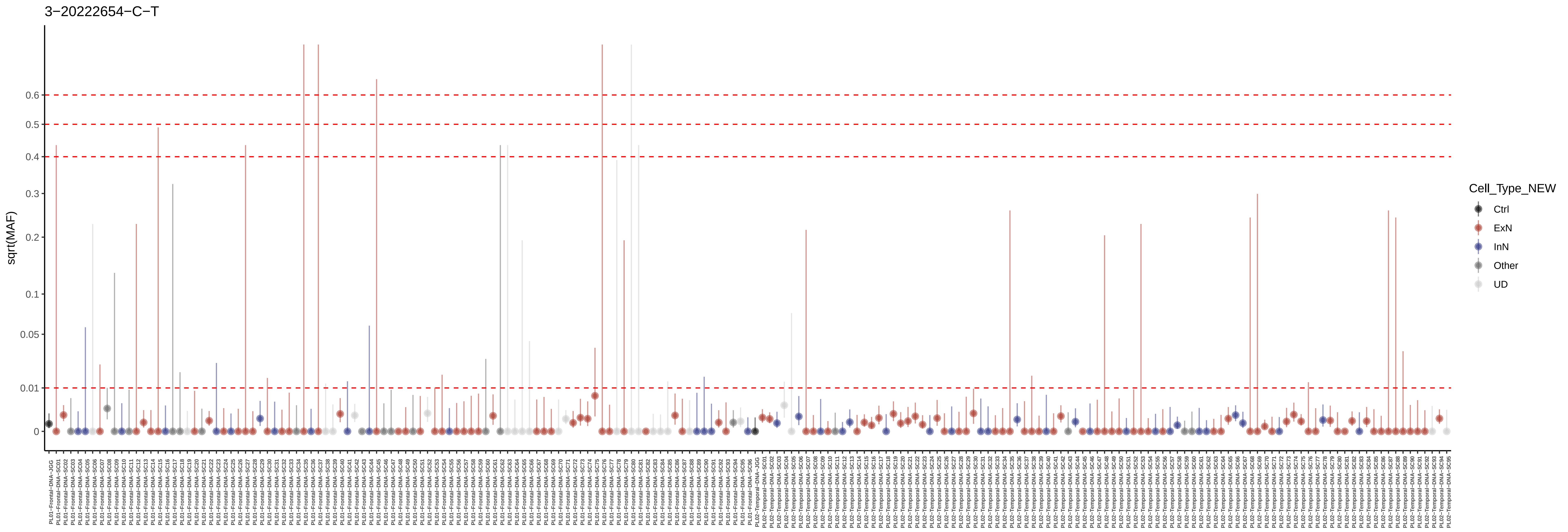
3−20222654−C−T
sqrt(MAF)
0.6
0.5
0.4
0.3
0.2
0.1
0.05
0.01
0
Cell_Type_NEW
Ctrl
ExN
InN
Other
UD

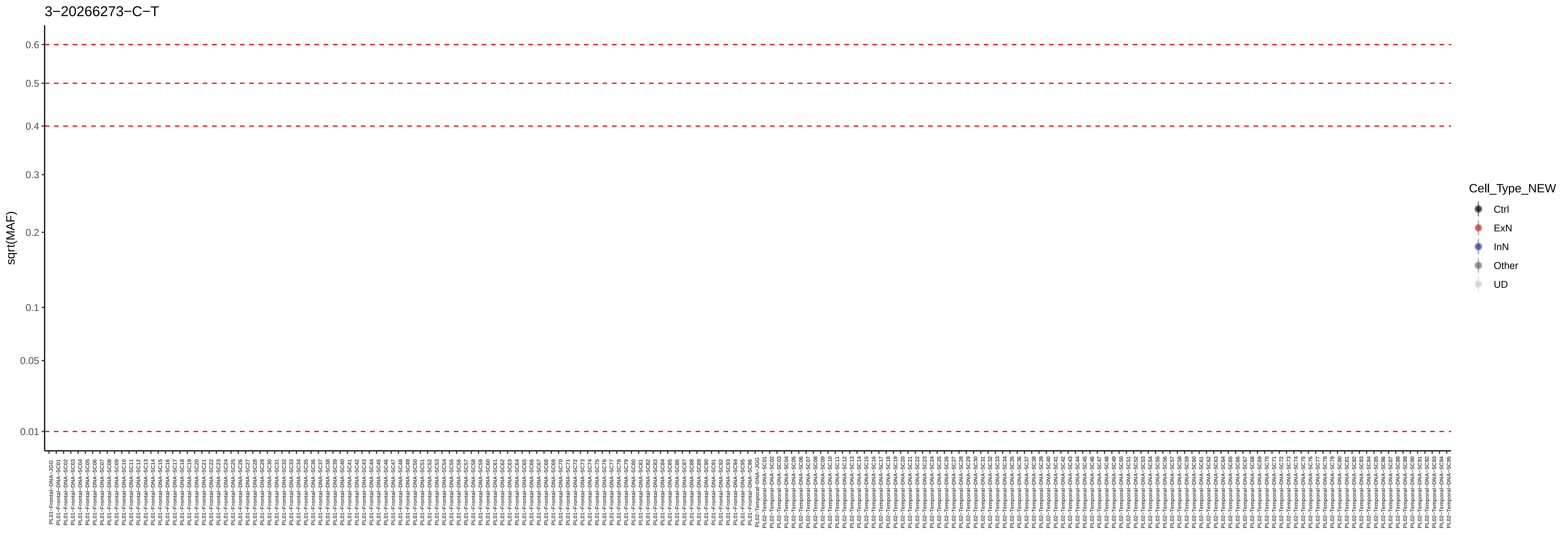
3−20266273−C−T
sqrt(MAF)
0.6
0.5
0.4
0.3
0.2
0.1
0.05
0.01
Cell_Type_NEW
Ctrl
ExN
InN
Other
UD

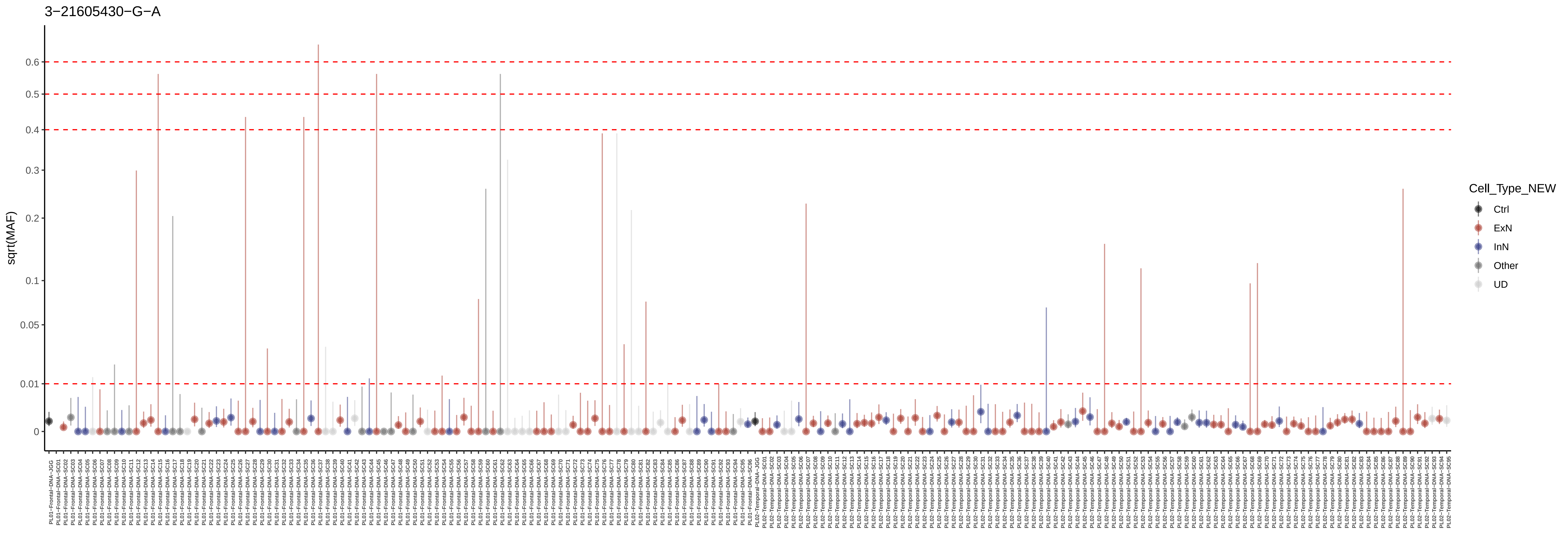
3−21605430−G−A
sqrt(MAF)
0.6
0.5
0.4
0.3
0.2
0.1
0.05
0.01
0
Cell_Type_NEW
Ctrl
ExN
InN
Other
UD

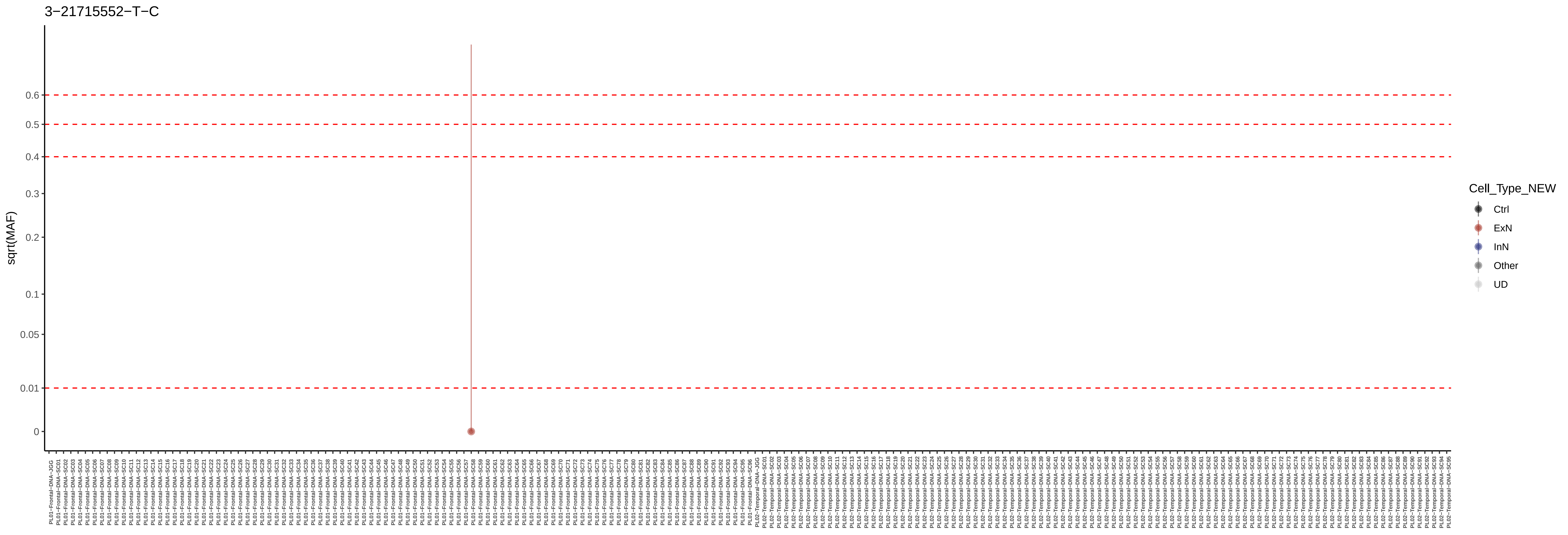
3−21715552−T−C
sqrt(MAF)
0.6
0.5
0.4
0.3
0.2
0.1
0.05
0.01
0
Cell_Type_NEW
Ctrl
ExN
InN
Other
UD

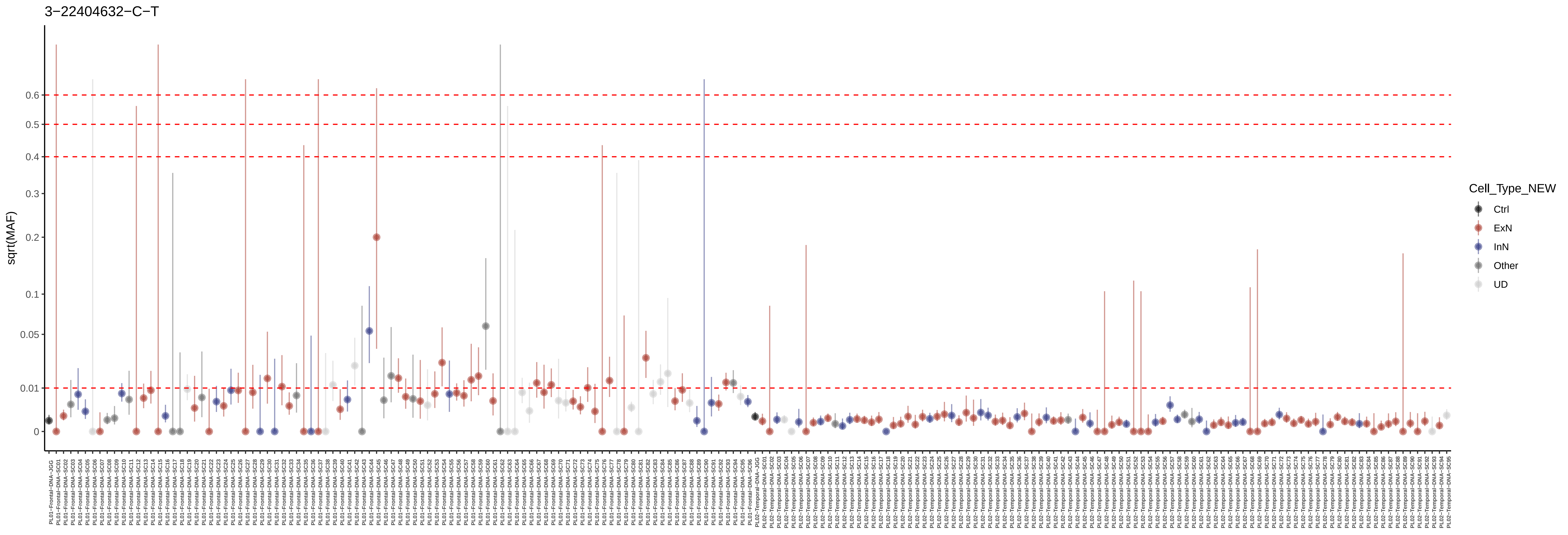
3−22404632−C−T
sqrt(MAF)
0.6
0.5
0.4
0.3
0.2
0.1
0.05
0.01
0
Cell_Type_NEW
Ctrl
ExN
InN
Other
UD

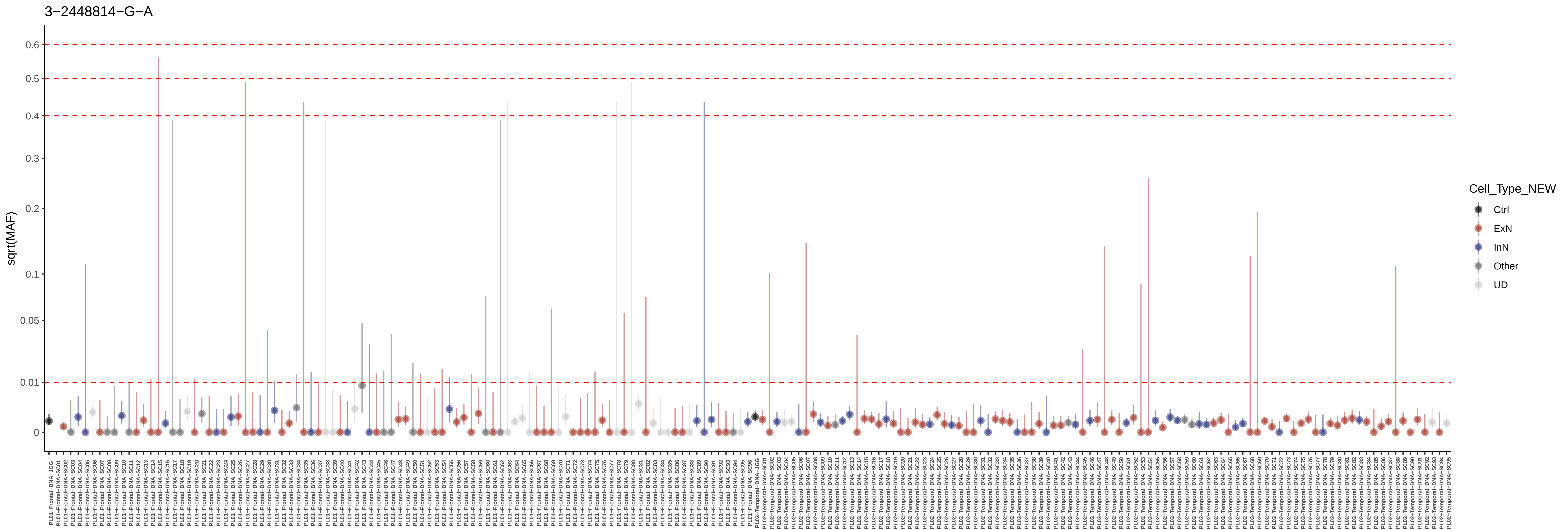

3−2448814−G−A
sqrt(MAF)
0.6
0.5
0.4
0.3
0.2
0.1
0.05
0.01
0
Cell_Type_NEW
Ctrl
ExN
InN
Other
UD

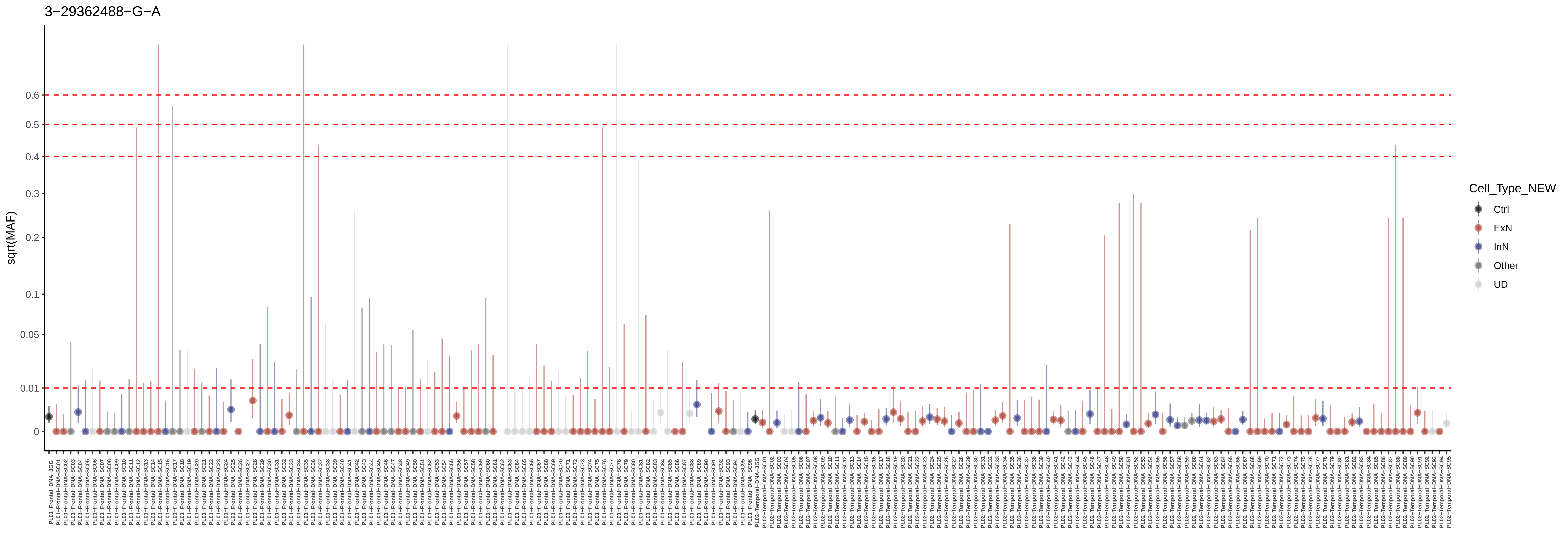

3−29362488−G−A
sqrt(MAF)
0.6
0.5
0.4
0.3
0.2
0.1
0.05
0.01
0
Cell_Type_NEW
Ctrl
ExN
InN
Other
UD

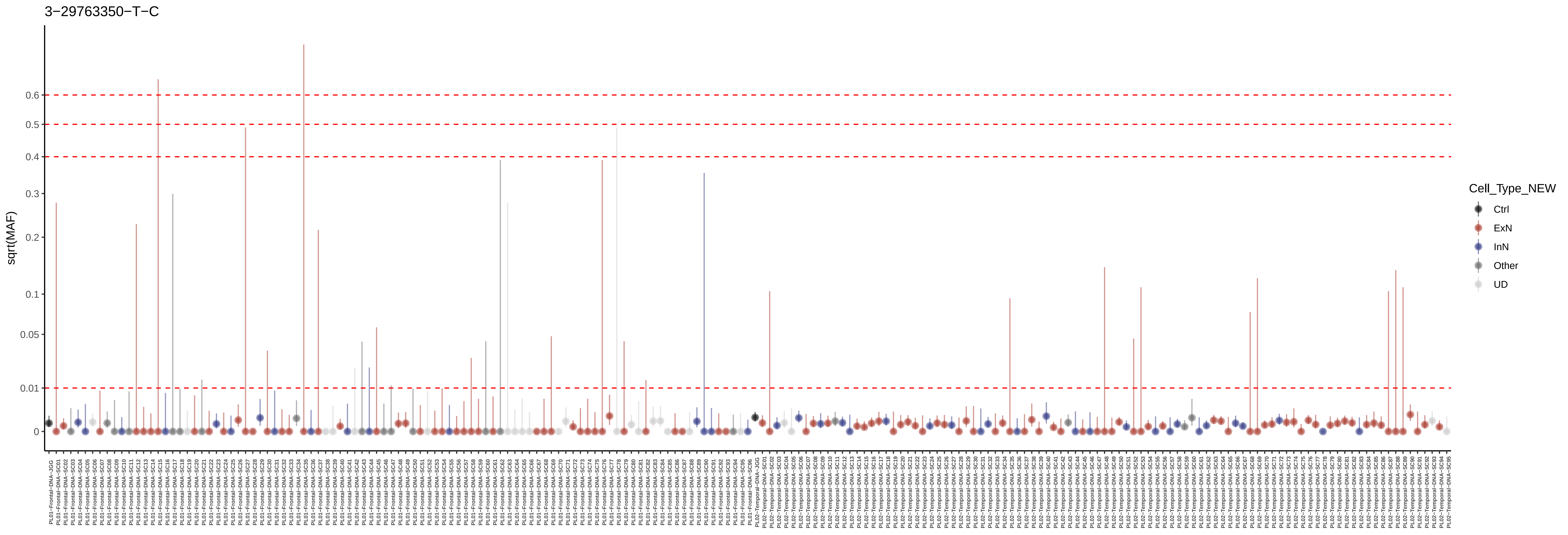

3−29763350−T−C
sqrt(MAF)
0.6
0.5
0.4
0.3
0.2
0.1
0.05
0.01
0
Cell_Type_NEW
Ctrl
ExN
InN
Other
UD

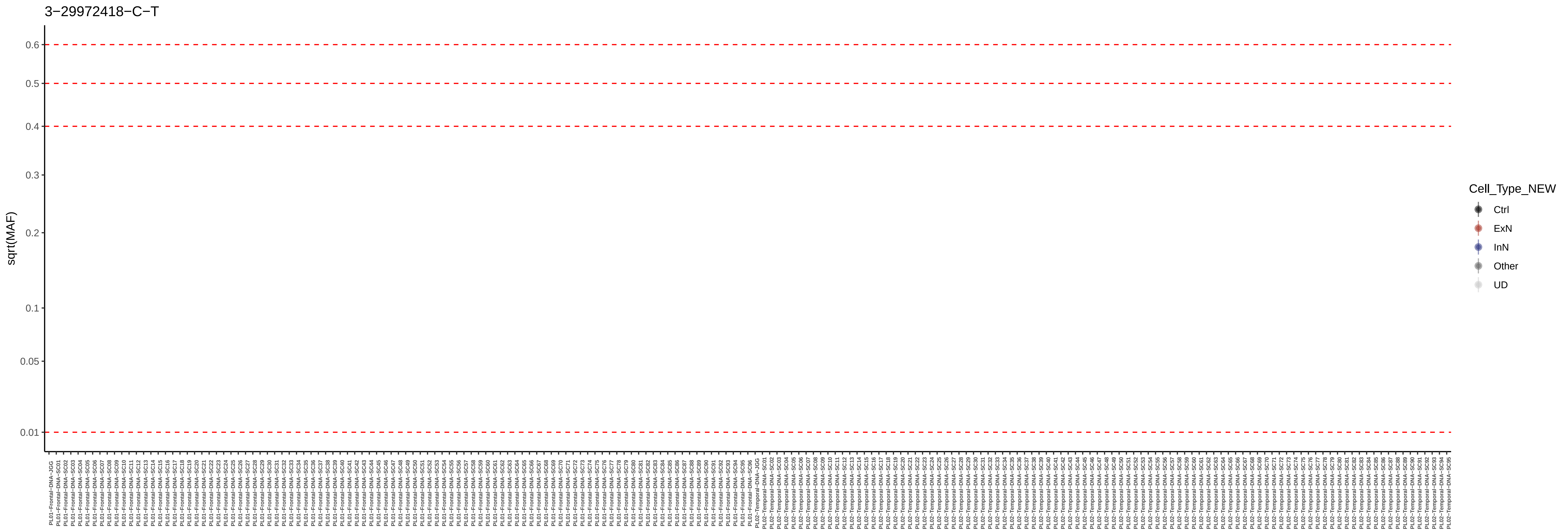
3−29972418−C−T
sqrt(MAF)
0.6
0.5
0.4
0.3
0.2
0.1
0.05
0.01
Cell_Type_NEW
Ctrl
ExN
InN
Other
UD

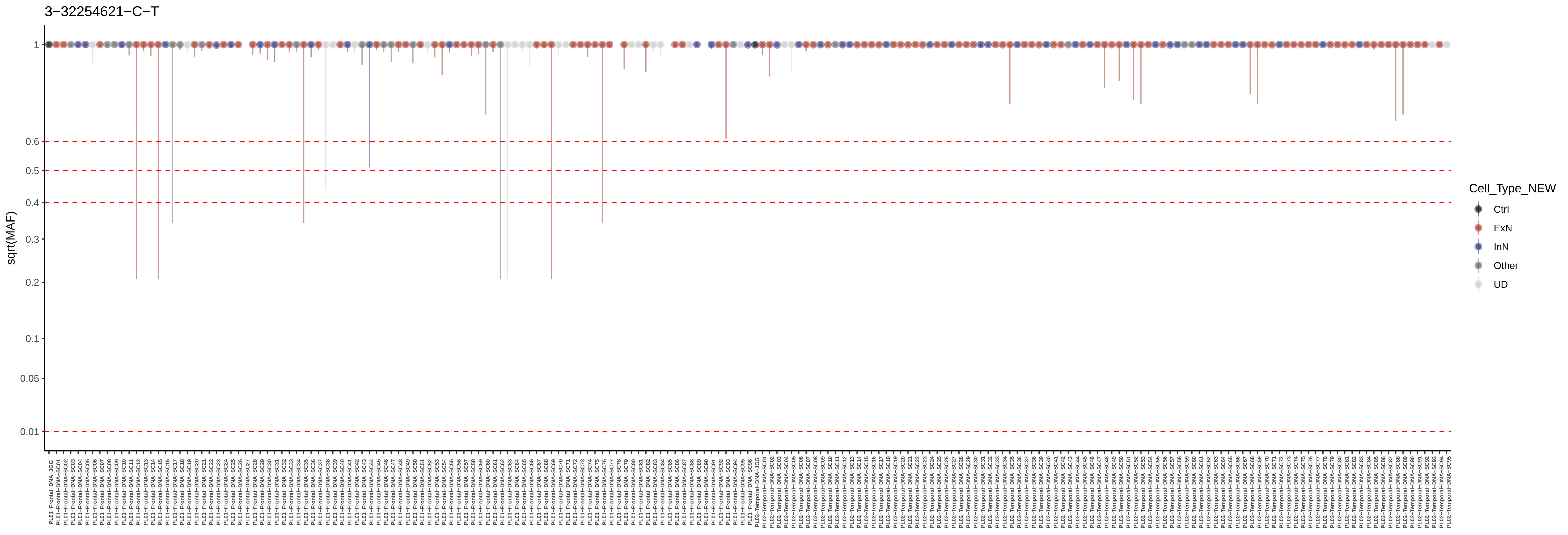

3−32254621−C−T
sqrt(MAF)
1
0.6
0.5
0.4
0.3
0.2
0.1
0.05
0.01
Cell_Type_NEW
Ctrl
ExN
InN
Other
UD

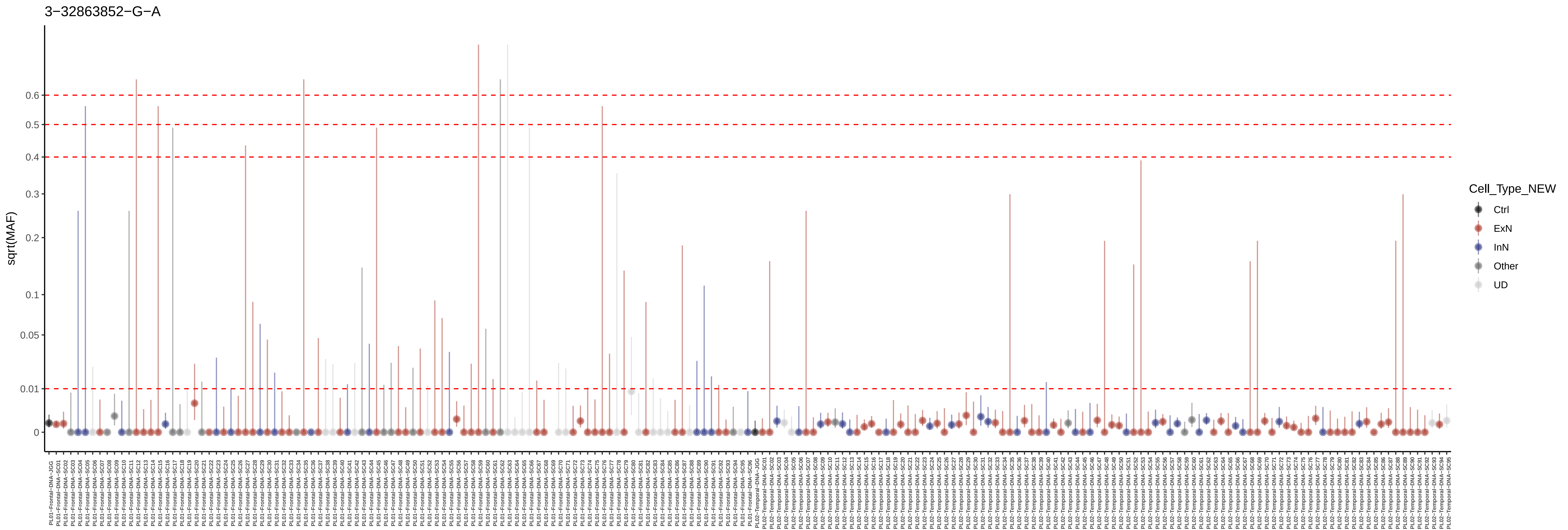
3−32863852−G−A
sqrt(MAF)
0.6
0.5
0.4
0.3
0.2
0.1
0.05
0.01
0
Cell_Type_NEW
Ctrl
ExN
InN
Other
UD

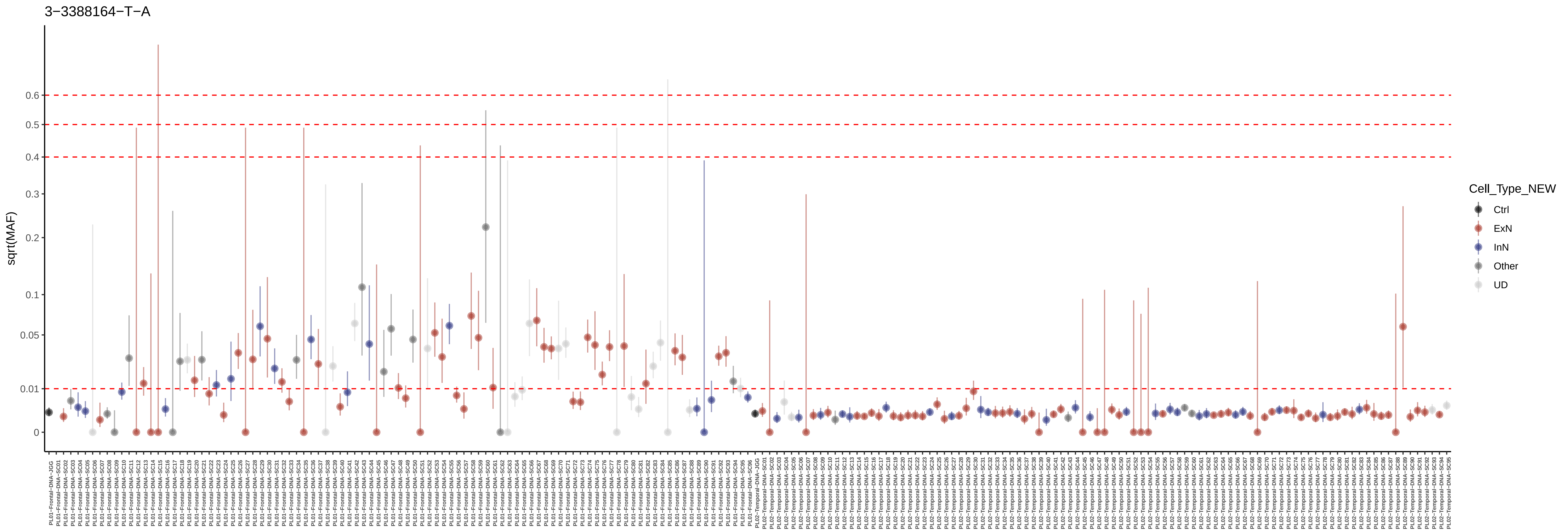
3−3388164−T−A
sqrt(MAF)
0.6
0.5
0.4
0.3
0.2
0.1
0.05
0.01
0
Cell_Type_NEW
Ctrl
ExN
InN
Other
UD

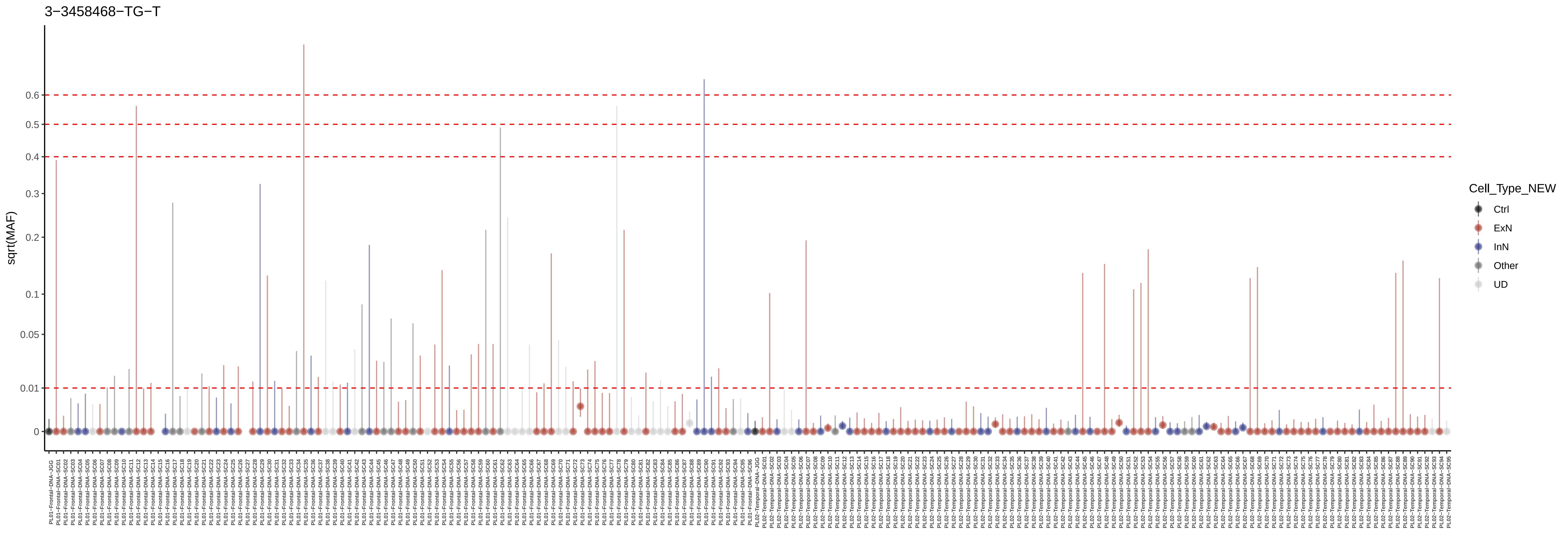
3−3458468−TG−T
sqrt(MAF)
0.6
0.5
0.4
0.3
0.2
0.1
0.05
0.01
0
Cell_Type_NEW
Ctrl
ExN
InN
Other
UD

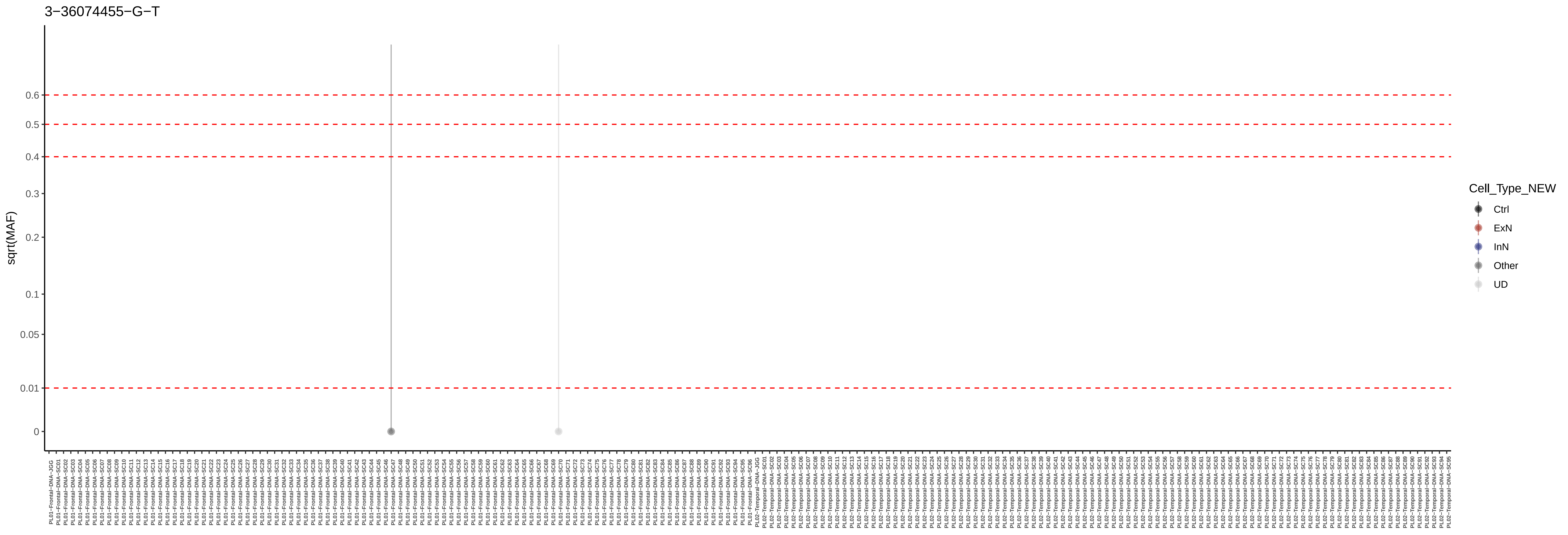
3−36074455−G−T
sqrt(MAF)
0.6
0.5
0.4
0.3
0.2
0.1
0.05
0.01
0
Cell_Type_NEW
Ctrl
ExN
InN
Other
UD

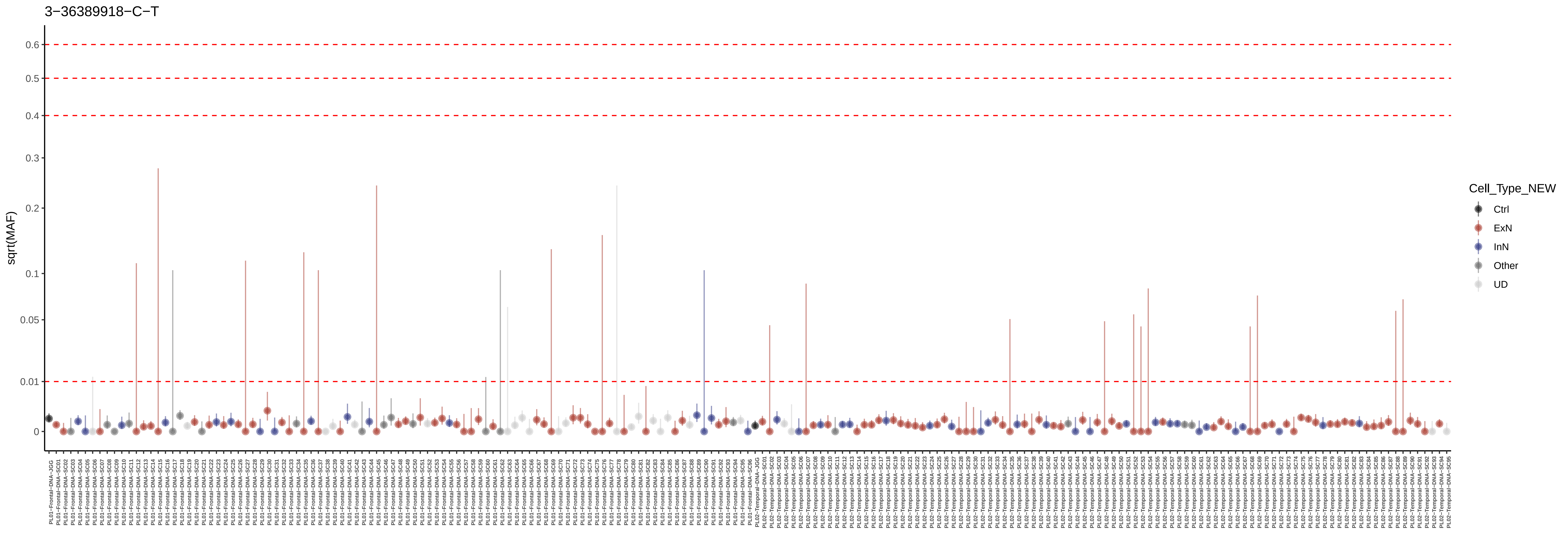

3−36389918−C−T
sqrt(MAF)
0.6
0.5
0.4
0.3
0.2
0.1
0.05
0.01
0
Cell_Type_NEW
Ctrl
ExN
InN
Other
UD

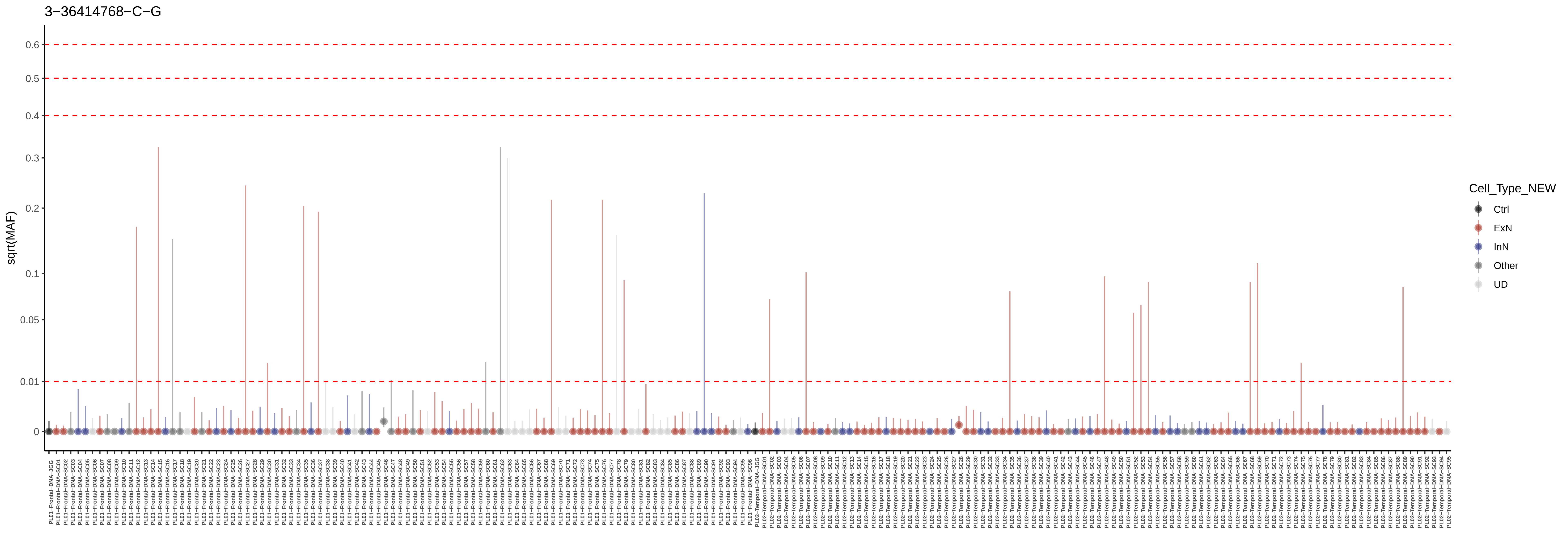
3−36414768−C−G
sqrt(MAF)
0.6
0.5
0.4
0.3
0.2
0.1
0.05
0.01
0
Cell_Type_NEW
Ctrl
ExN
InN
Other
UD

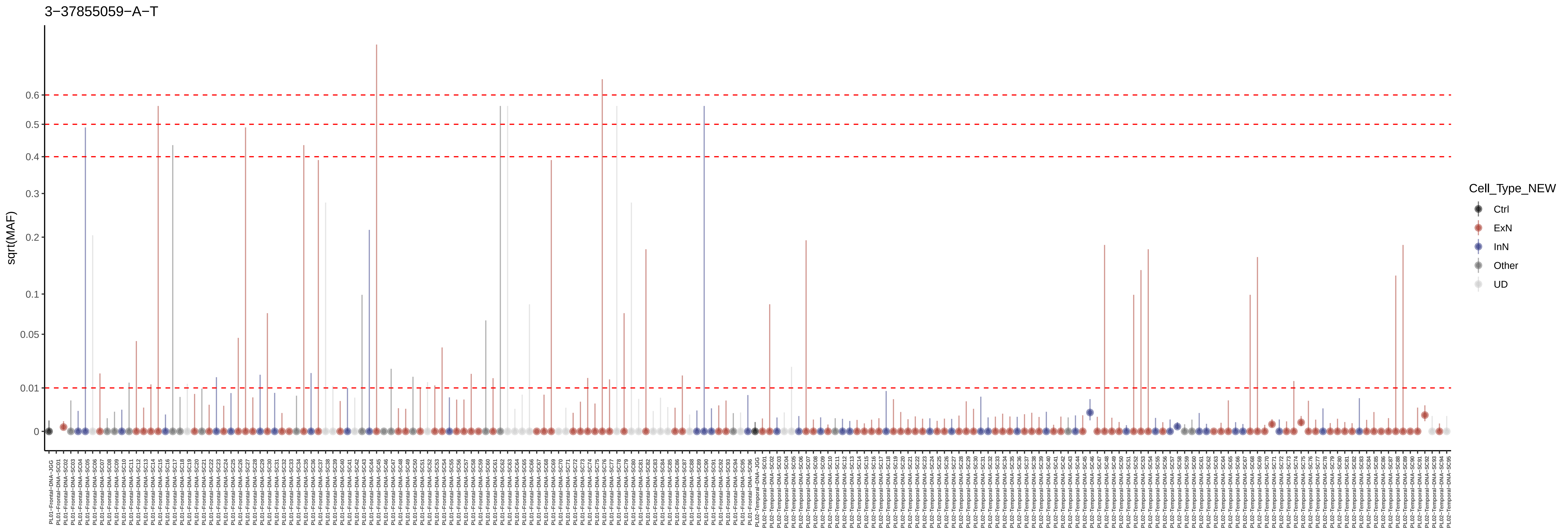
3−37855059−A−T
sqrt(MAF)
0.6
0.5
0.4
0.3
0.2
0.1
0.05
0.01
0
Cell_Type_NEW
Ctrl
ExN
InN
Other
UD

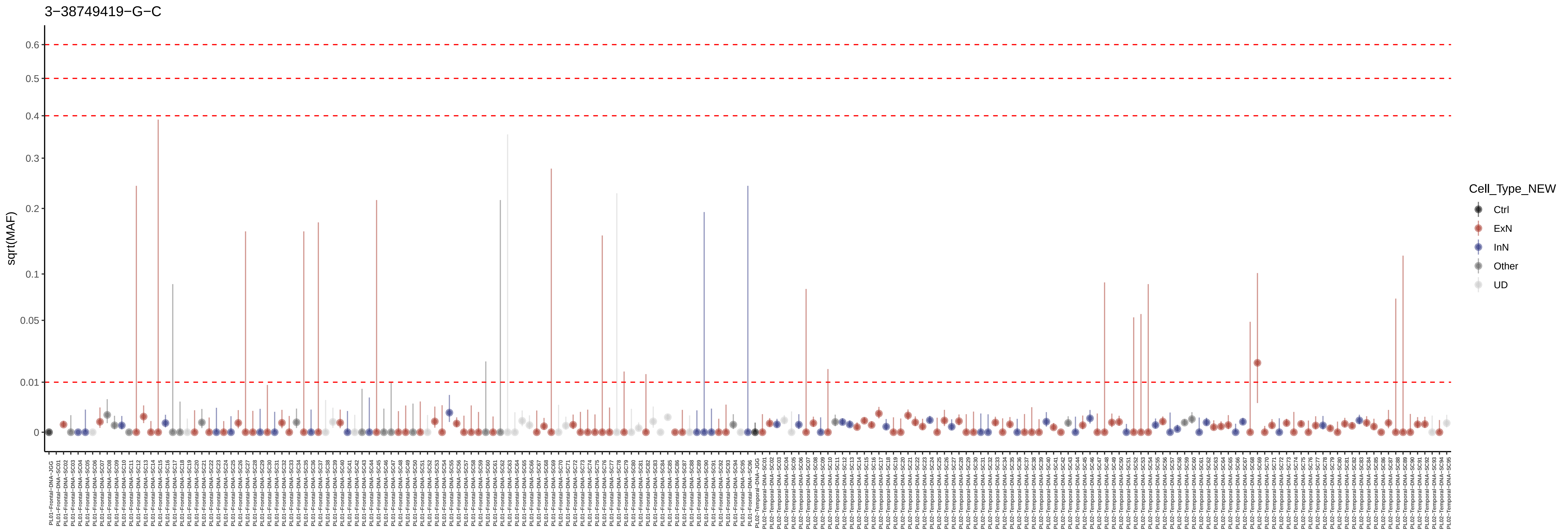

3−38749419−G−C
sqrt(MAF)
0.6
0.5
0.4
0.3
0.2
0.1
0.05
0.01
0
Cell_Type_NEW
Ctrl
ExN
InN
Other
UD

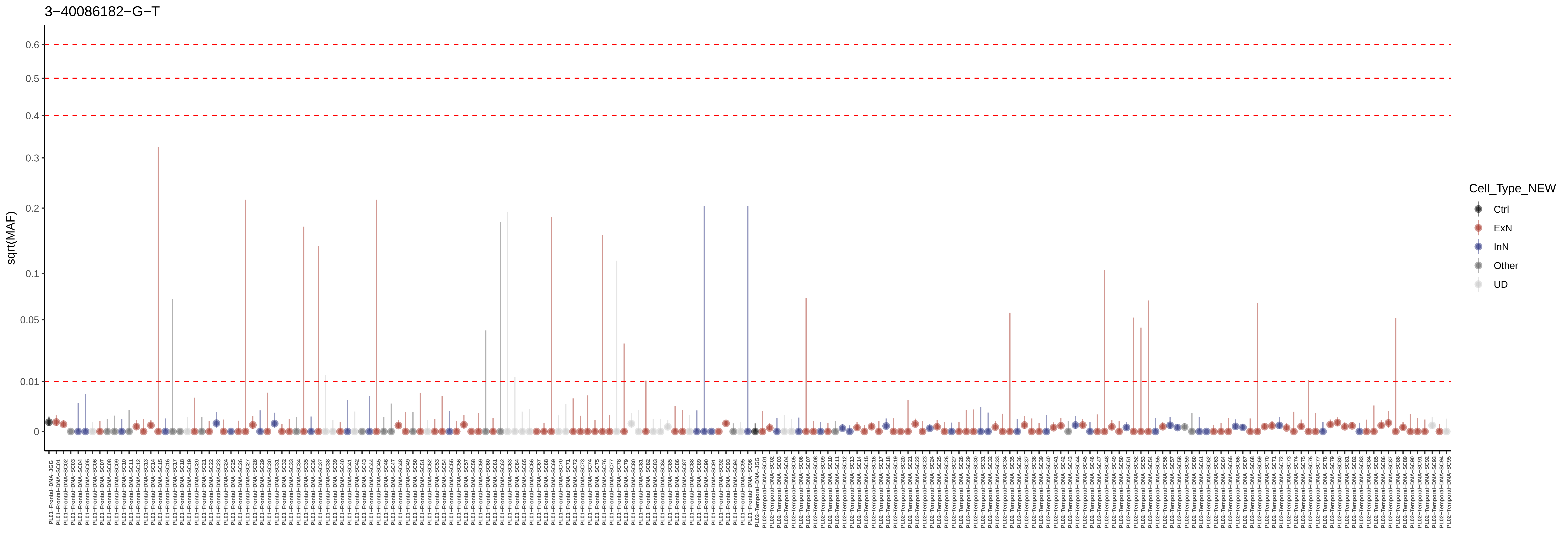

3−40086182−G−T
sqrt(MAF)
0.6
0.5
0.4
0.3
0.2
0.1
0.05
0.01
0
Cell_Type_NEW
Ctrl
ExN
InN
Other
UD

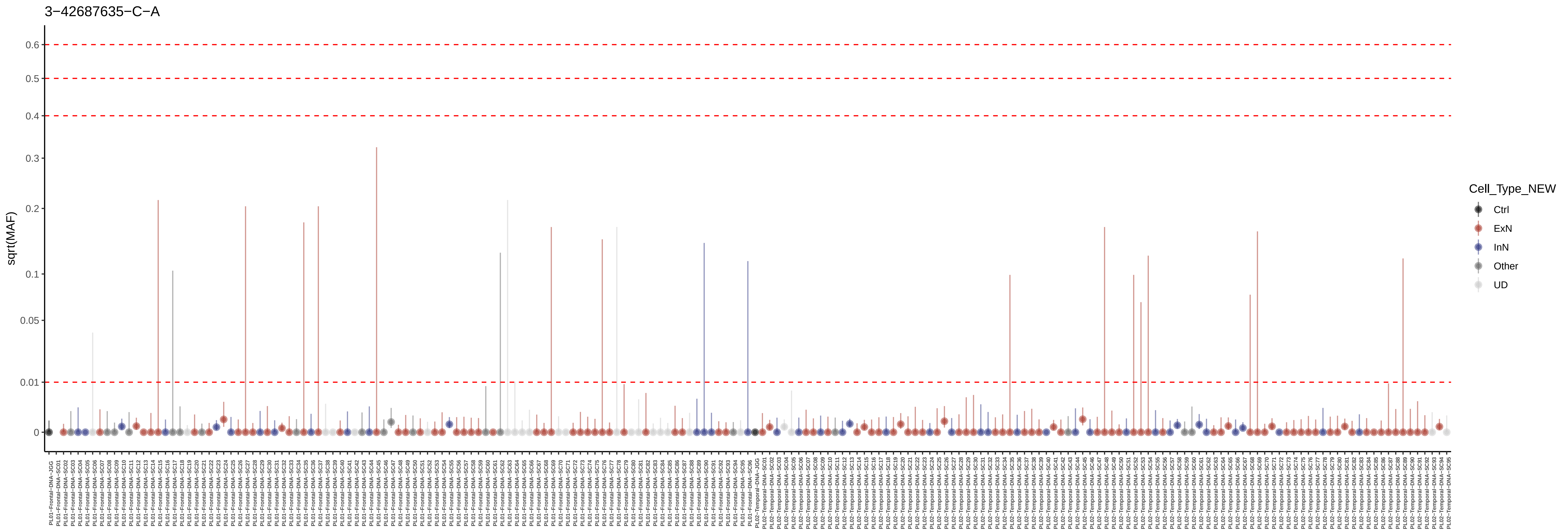

3−42687635−C−A
sqrt(MAF)
0.6
0.5
0.4
0.3
0.2
0.1
0.05
0.01
0
Cell_Type_NEW
Ctrl
ExN
InN
Other
UD

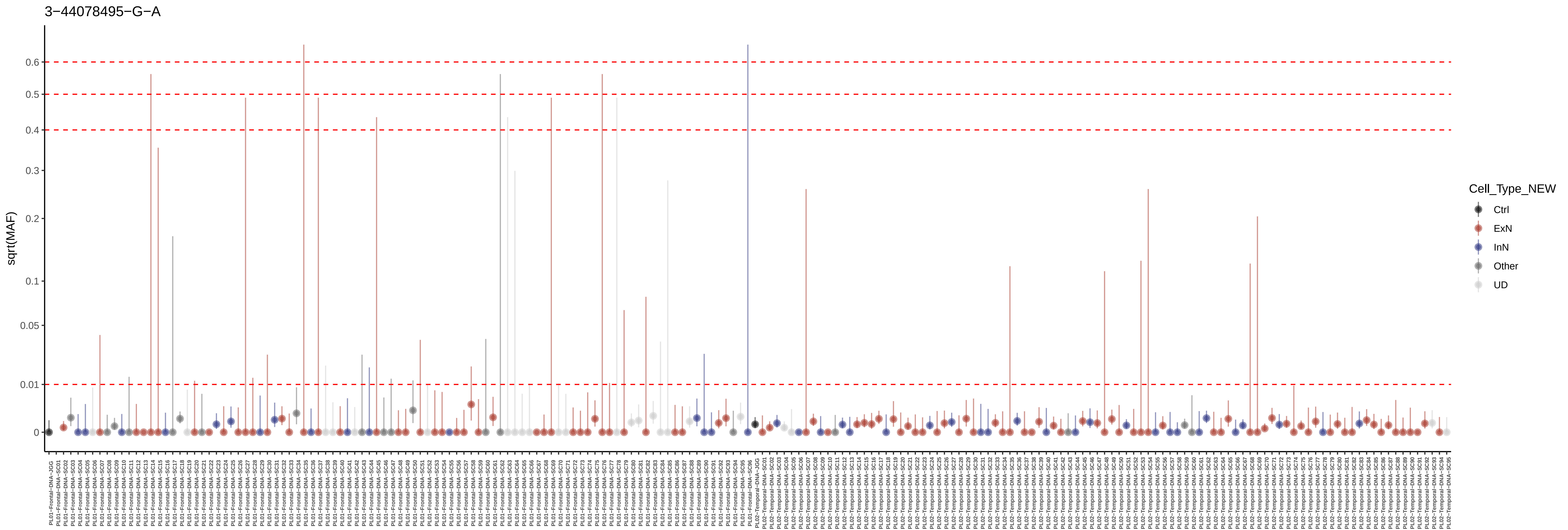
3−44078495−G−A
sqrt(MAF)
0.6
0.5
0.4
0.3
0.2
0.1
0.05
0.01
0
Cell_Type_NEW
Ctrl
ExN
InN
Other
UD

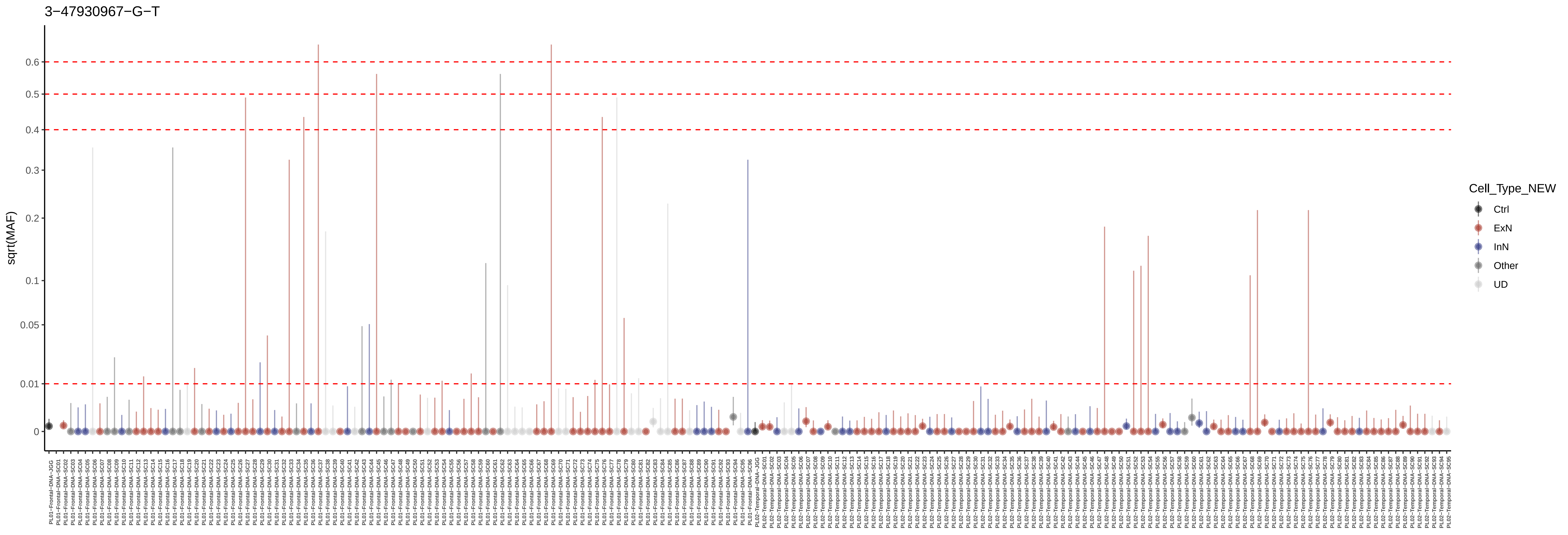
3−47930967−G−T
sqrt(MAF)
0.6
0.5
0.4
0.3
0.2
0.1
0.05
0.01
0
Cell_Type_NEW
Ctrl
ExN
InN
Other
UD

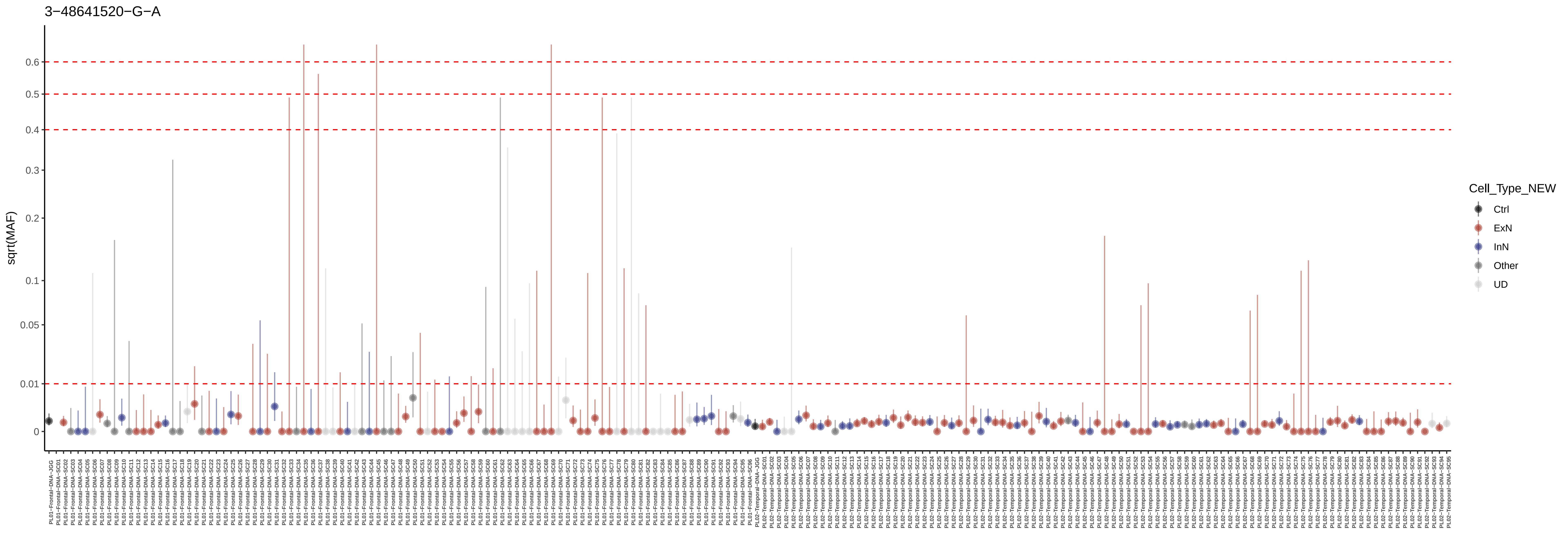
3−48641520−G−A
sqrt(MAF)
0.6
0.5
0.4
0.3
0.2
0.1
0.05
0.01
0
Cell_Type_NEW
Ctrl
ExN
InN
Other
UD

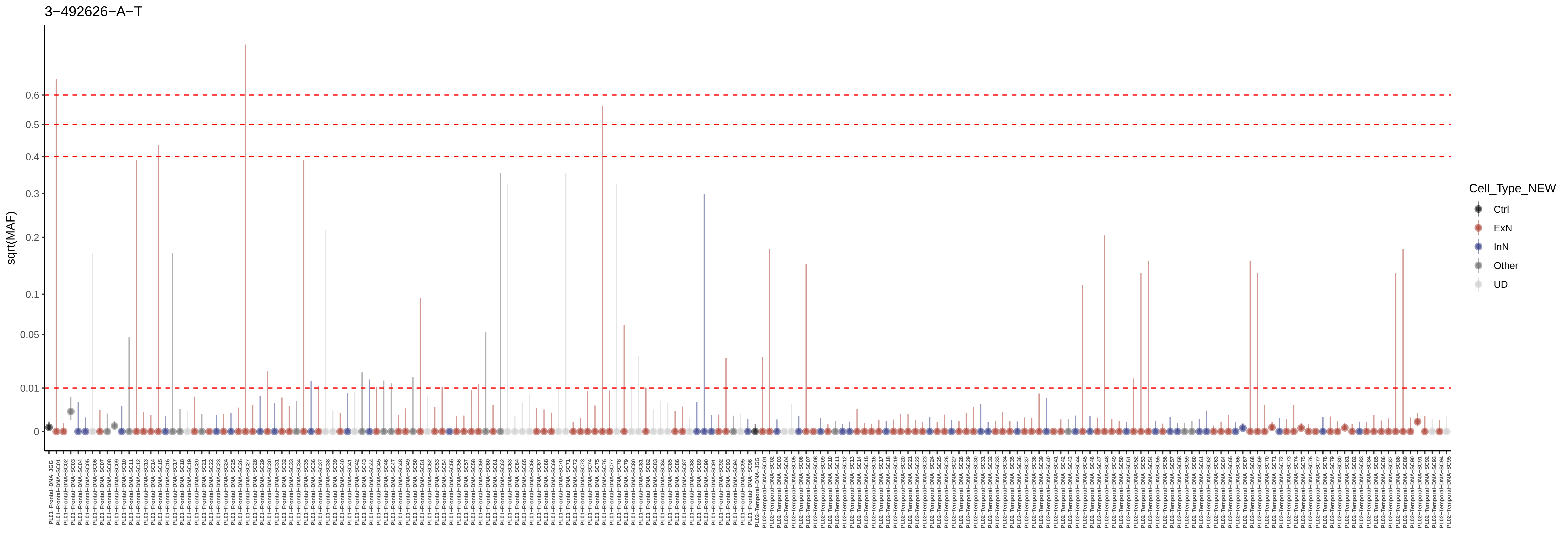
3−492626−A−T
sqrt(MAF)
0.6
0.5
0.4
0.3
0.2
0.1
0.05
0.01
0
Cell_Type_NEW
Ctrl
ExN
InN
Other
UD

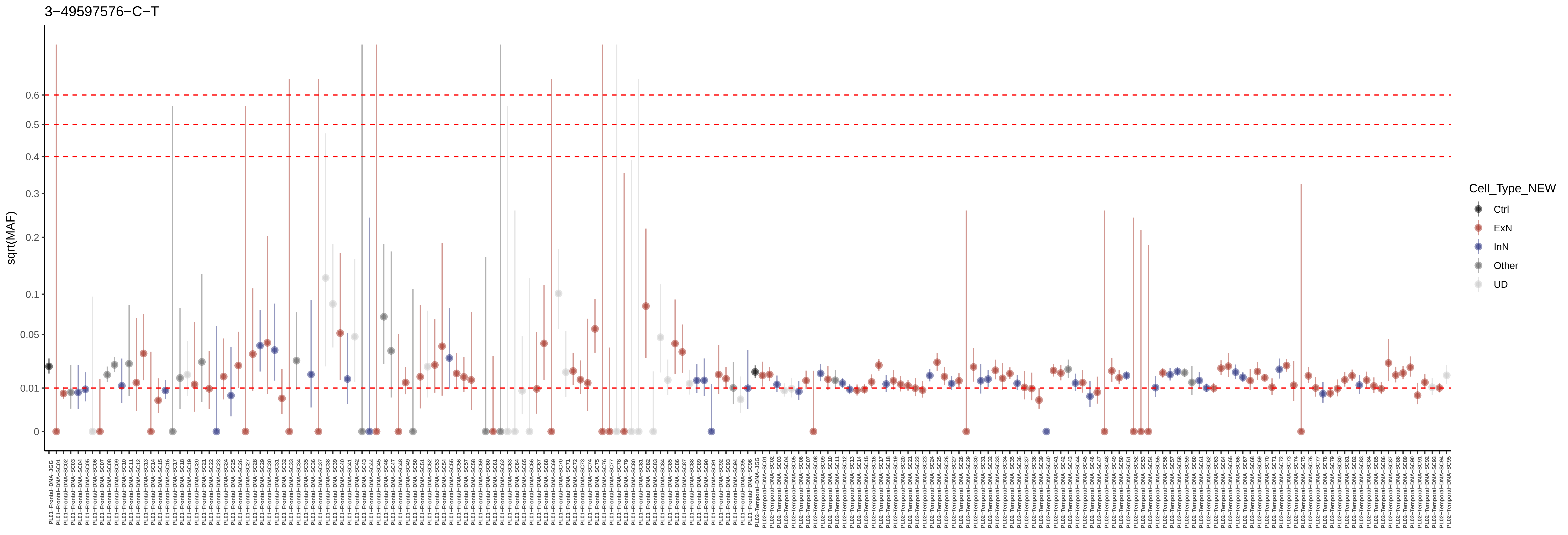

3−49597576−C−T
sqrt(MAF)
0.6
0.5
0.4
0.3
0.2
0.1
0.05
0.01
0
Cell_Type_NEW
Ctrl
ExN
InN
Other
UD

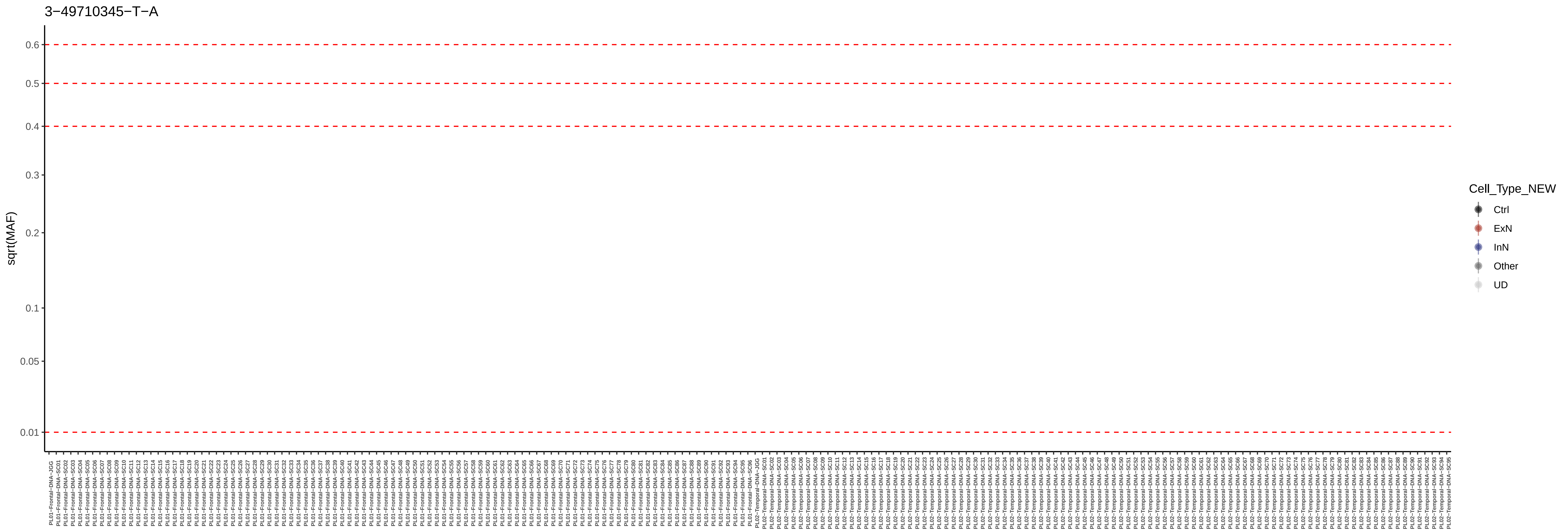

3−49710345−T−A
sqrt(MAF)
0.6
0.5
0.4
0.3
0.2
0.1
0.05
0.01
Cell_Type_NEW
Ctrl
ExN
InN
Other
UD

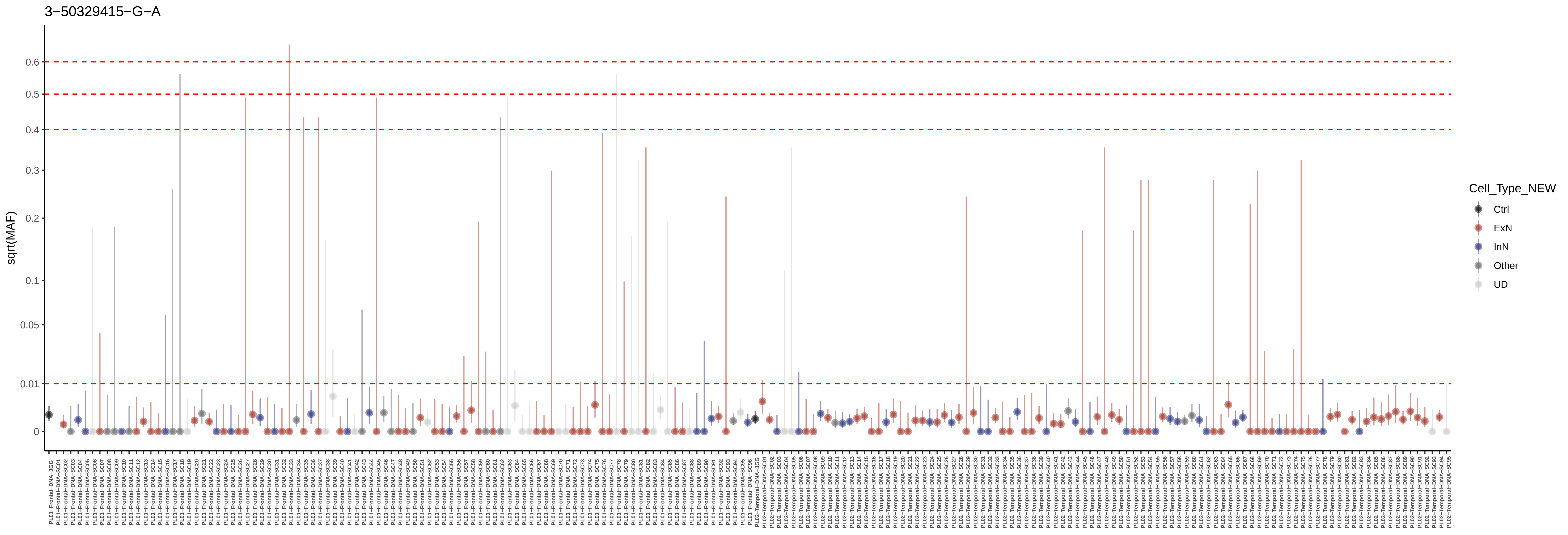
3−50329415−G−A
sqrt(MAF)
0.6
0.5
0.4
0.3
0.2
0.1
0.05
0.01
0
Cell_Type_NEW
Ctrl
ExN
InN
Other
UD

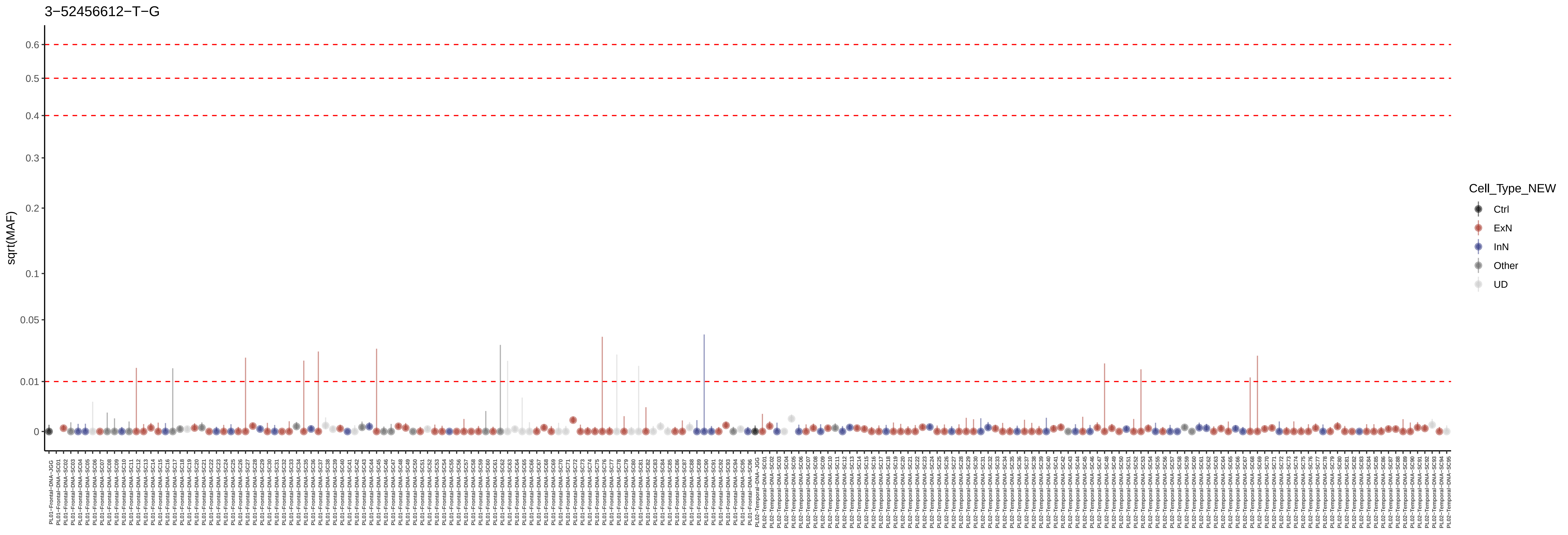
3−52456612−T−G
sqrt(MAF)
0.6
0.5
0.4
0.3
0.2
0.1
0.05
0.01
0
Cell_Type_NEW
Ctrl
ExN
InN
Other
UD

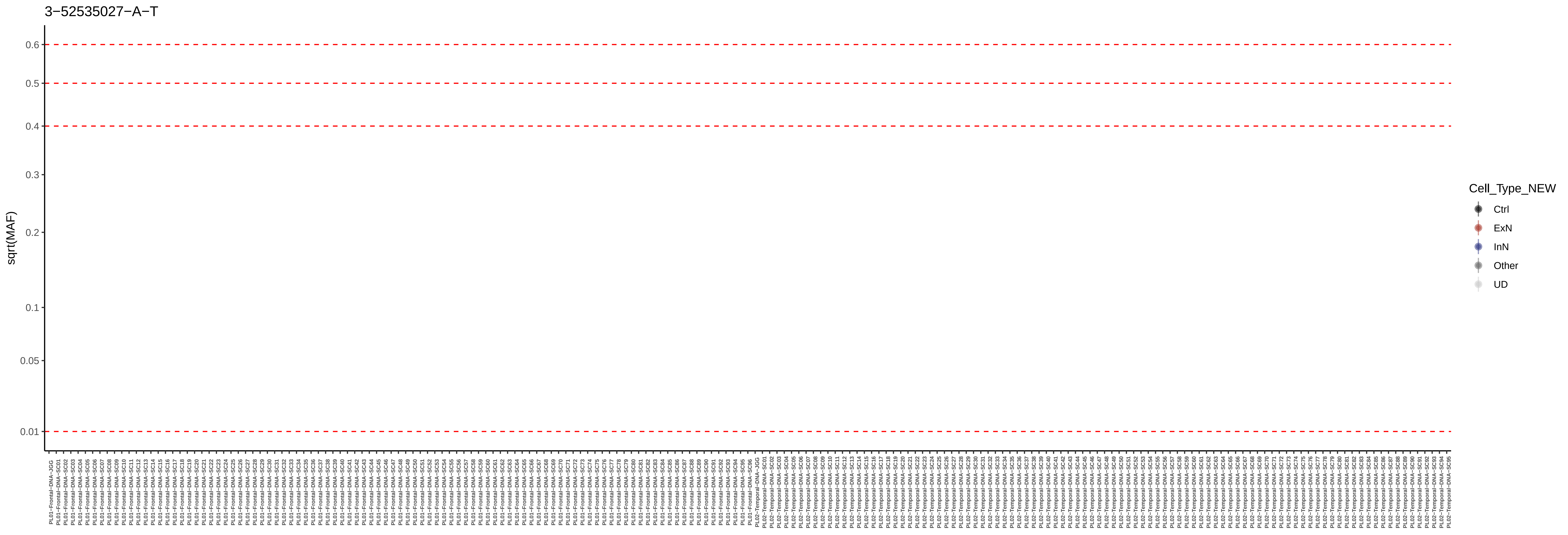

3−52535027−A−T
sqrt(MAF)
0.6
0.5
0.4
0.3
0.2
0.1
0.05
0.01
Cell_Type_NEW
Ctrl
ExN
InN
Other
UD

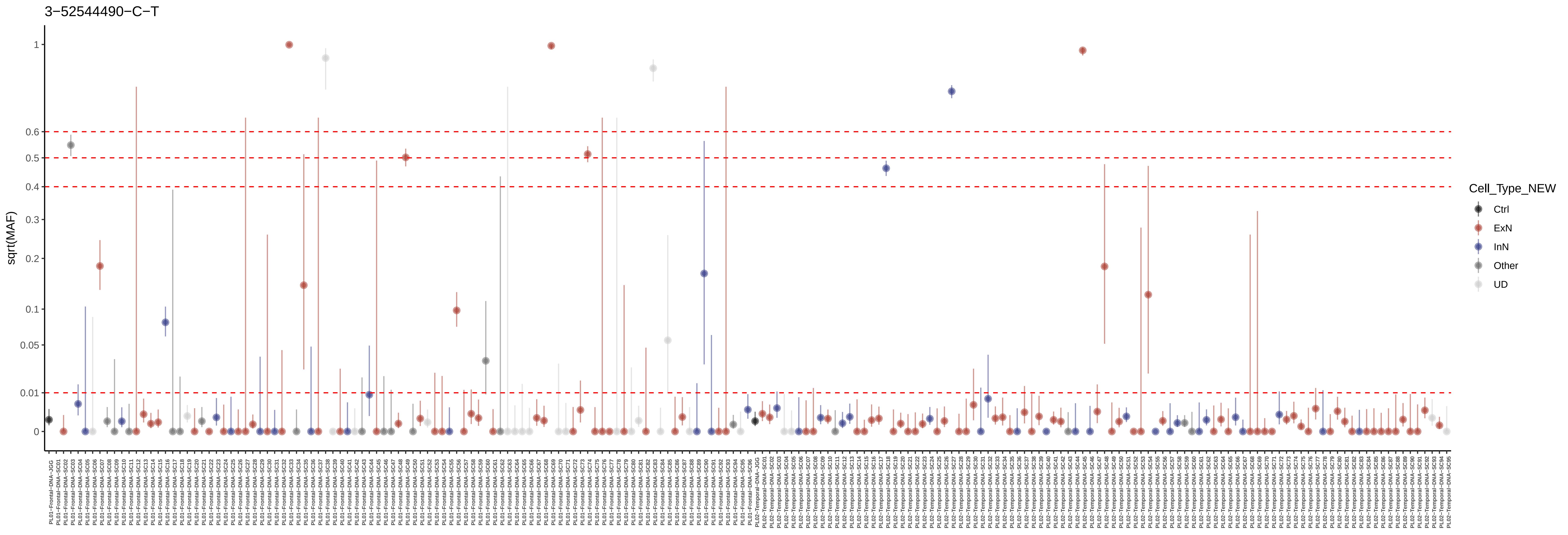
3−52544490−C−T
sqrt(MAF)
1
0.6
0.5
0.4
0.3
0.2
0.1
0.05
0.01
0
Cell_Type_NEW
Ctrl
ExN
InN
Other
UD

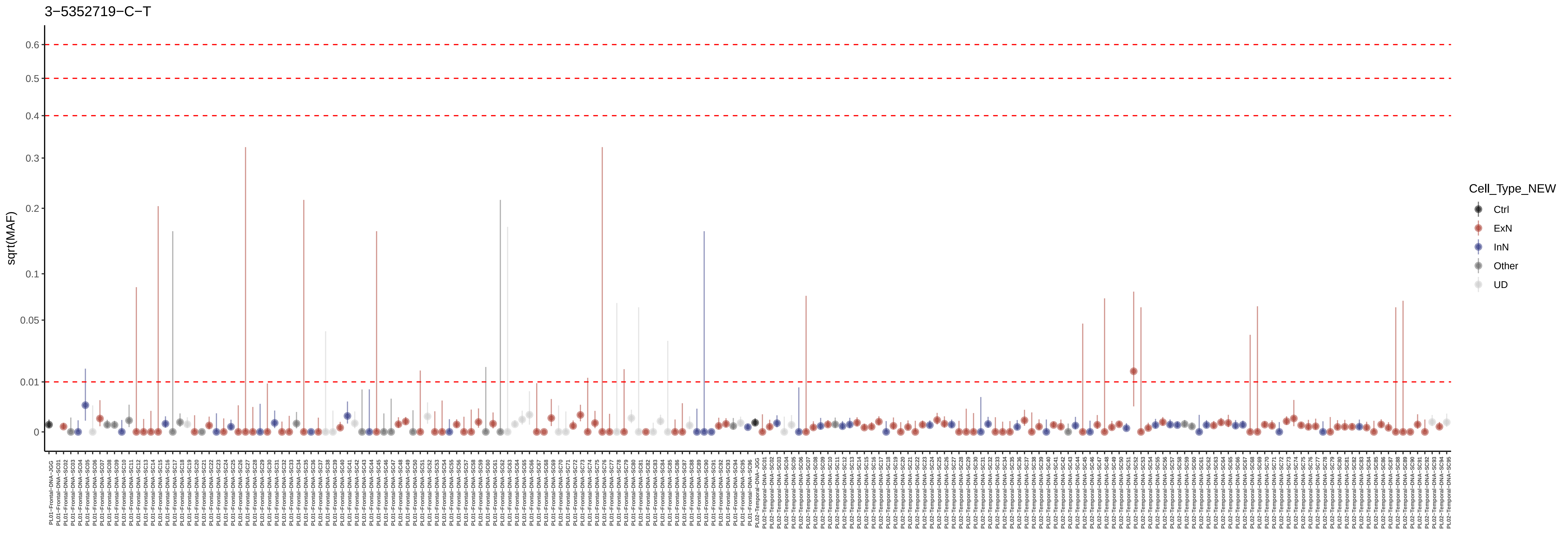

3−5352719−C−T
sqrt(MAF)
0.6
0.5
0.4
0.3
0.2
0.1
0.05
0.01
0
Cell_Type_NEW
Ctrl
ExN
InN
Other
UD

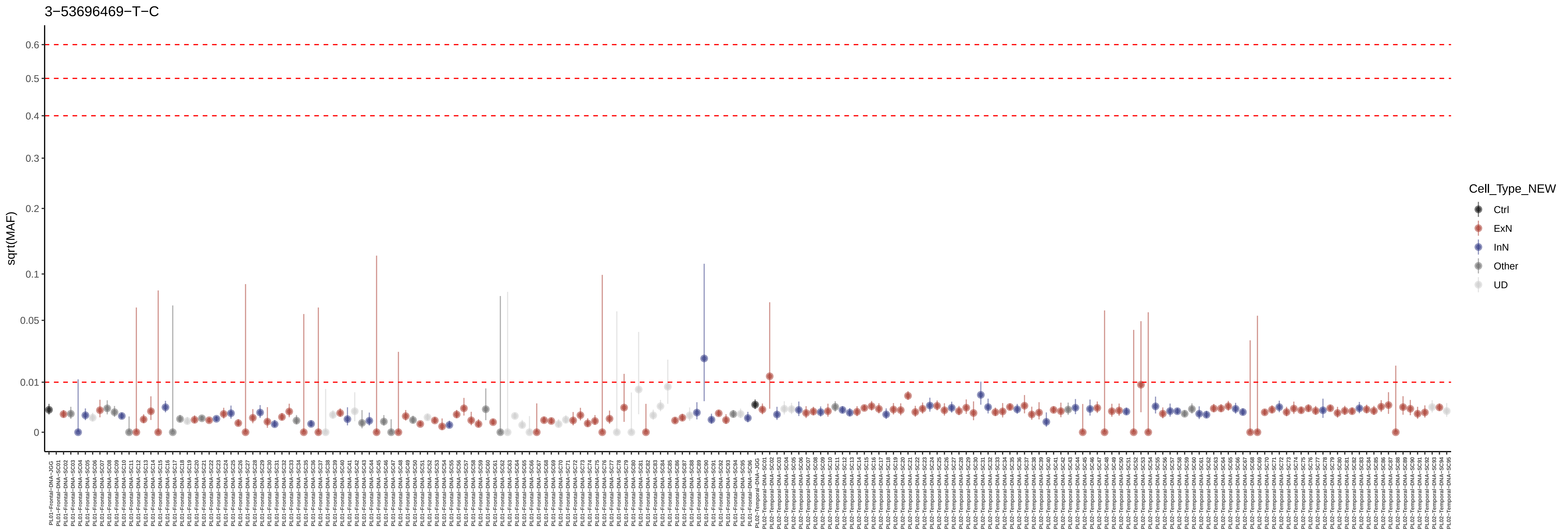
3−53696469−T−C
sqrt(MAF)
0.6
0.5
0.4
0.3
0.2
0.1
0.05
0.01
0
Cell_Type_NEW
Ctrl
ExN
InN
Other
UD

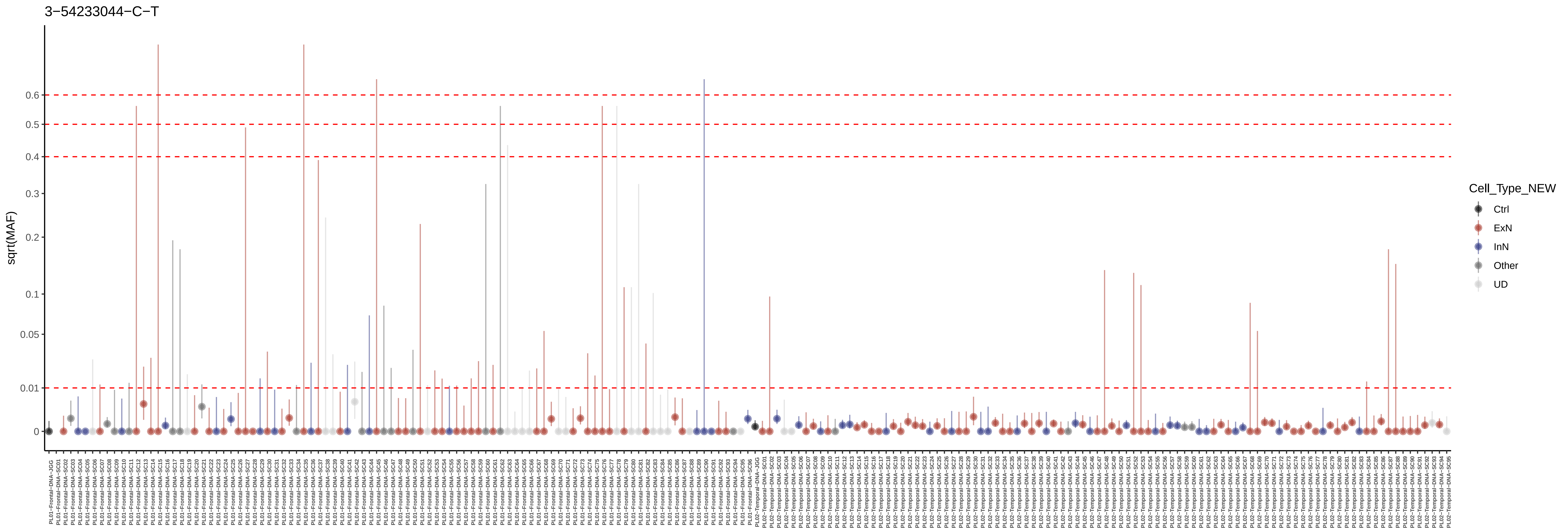

3−54233044−C−T
sqrt(MAF)
0.6
0.5
0.4
0.3
0.2
0.1
0.05
0.01
0
Cell_Type_NEW
Ctrl
ExN
InN
Other
UD

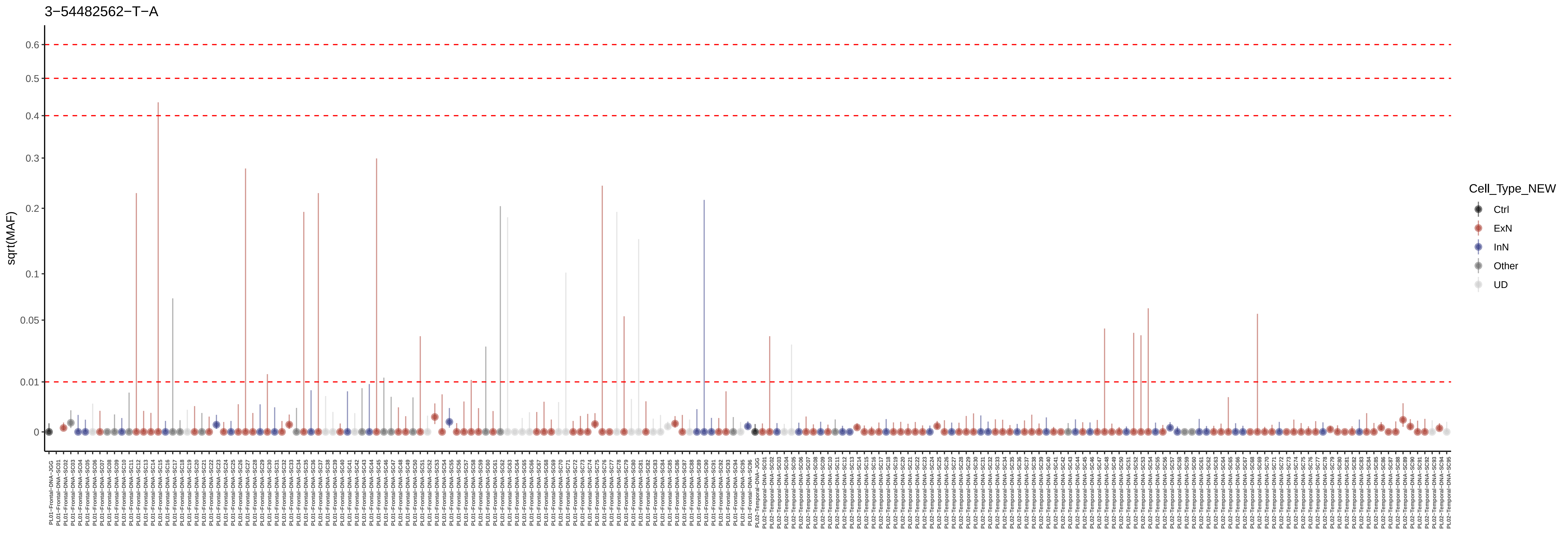
3−54482562−T−A
sqrt(MAF)
0.6
0.5
0.4
0.3
0.2
0.1
0.05
0.01
0
Cell_Type_NEW
Ctrl
ExN
InN
Other
UD

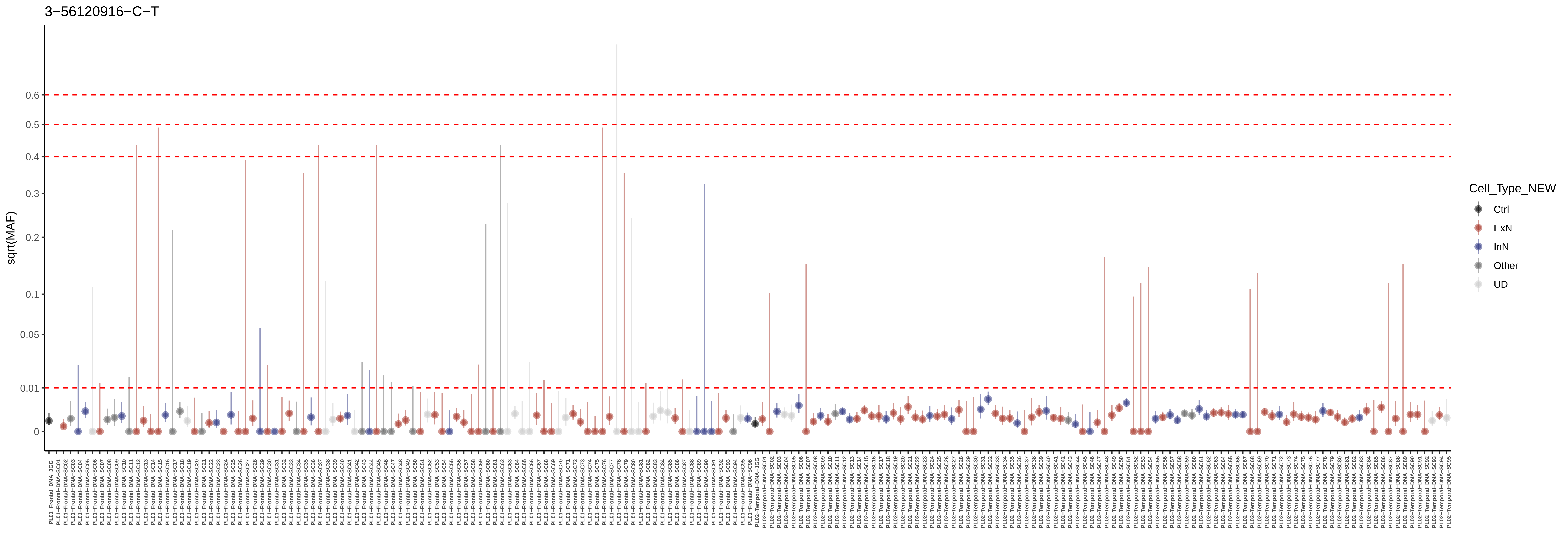
3−56120916−C−T
sqrt(MAF)
0.6
0.5
0.4
0.3
0.2
0.1
0.05
0.01
0
Cell_Type_NEW
Ctrl
ExN
InN
Other
UD

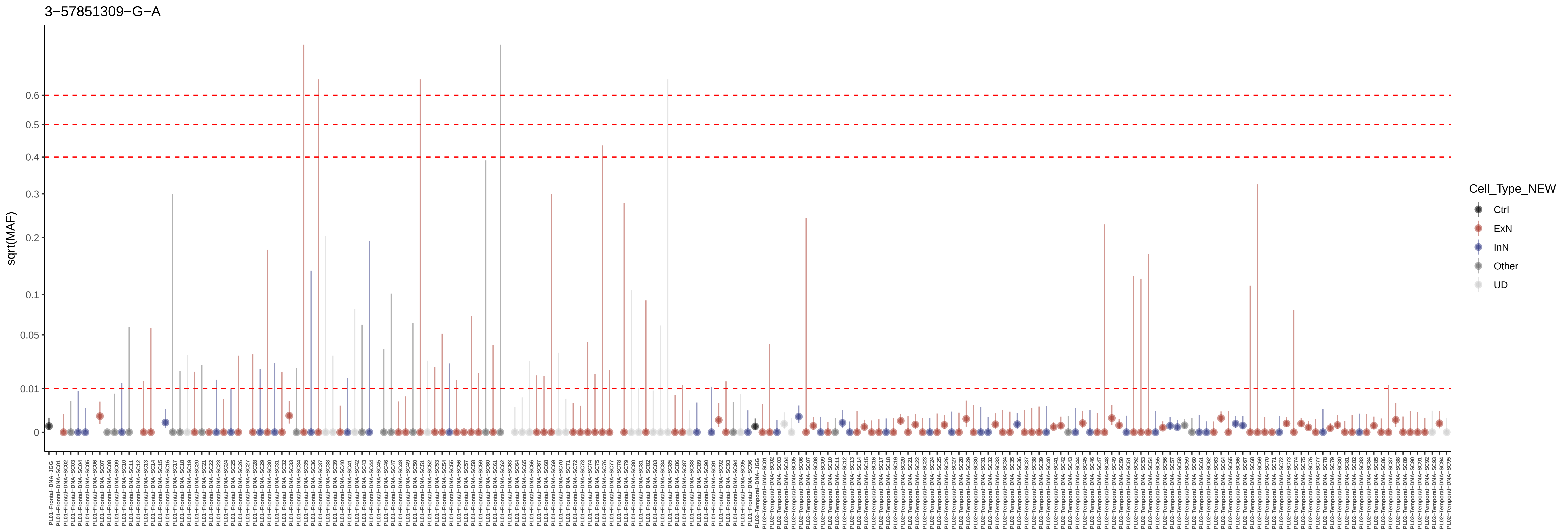
3−57851309−G−A
sqrt(MAF)
0.6
0.5
0.4
0.3
0.2
0.1
0.05
0.01
0
Cell_Type_NEW
Ctrl
ExN
InN
Other
UD

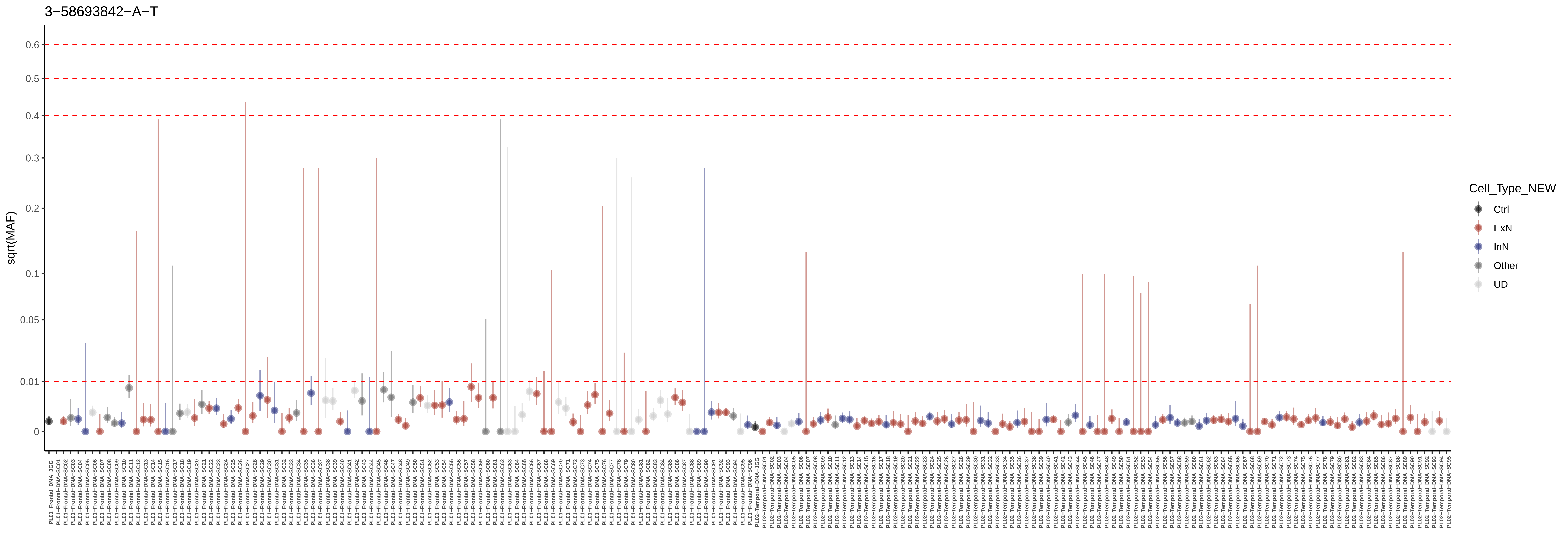
3−58693842−A−T
sqrt(MAF)
0.6
0.5
0.4
0.3
0.2
0.1
0.05
0.01
0
Cell_Type_NEW
Ctrl
ExN
InN
Other
UD

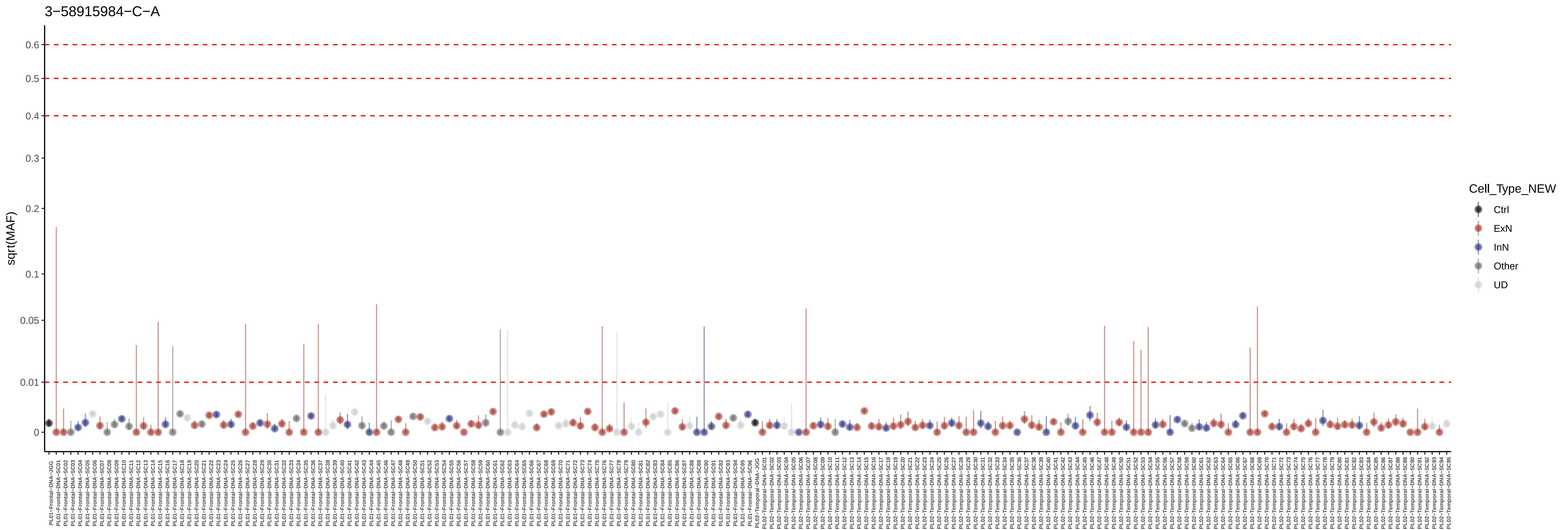
3−58915984−C−A
sqrt(MAF)
0.6
0.5
0.4
0.3
0.2
0.1
0.05
0.01
0
Cell_Type_NEW
Ctrl
ExN
InN
Other
UD

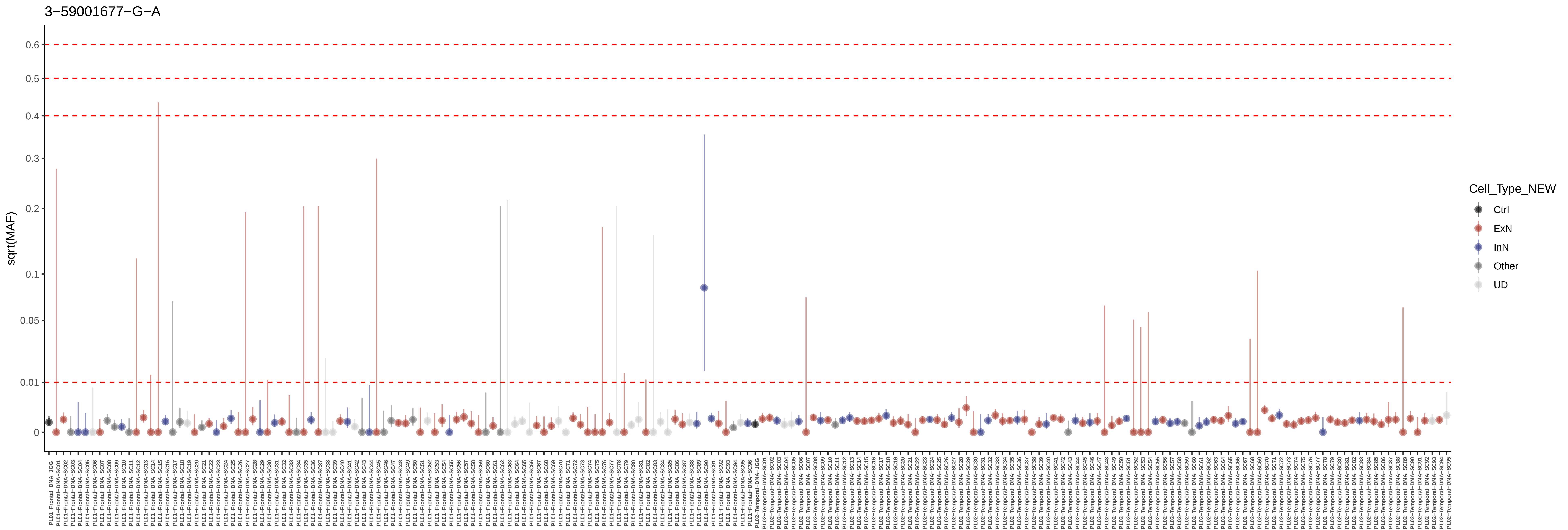
3−59001677−G−A
sqrt(MAF)
0
0.01
0.05
0.1
0.2
0.3
0.4
0.5
0.6
Cell_Type_NEW
Ctrl
ExN
InN
Other
UD

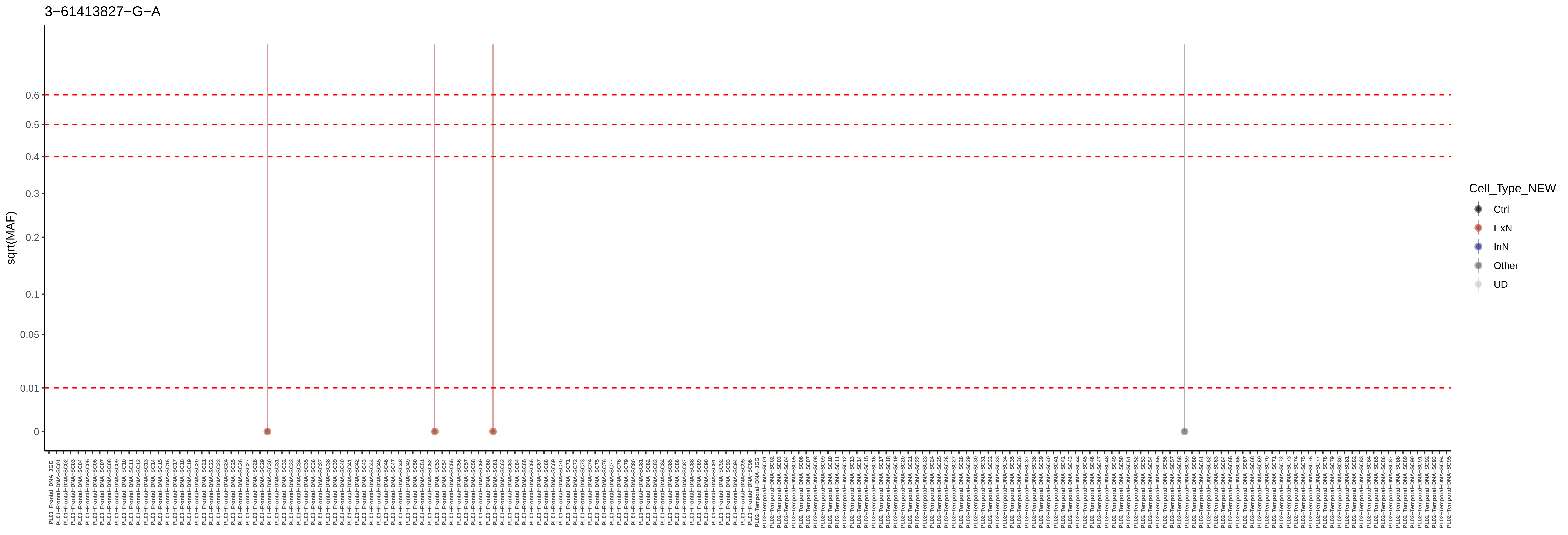

3−61413827−G−A
sqrt(MAF)
0.6
0.5
0.4
0.3
0.2
0.1
0.05
0.01
0
Cell_Type_NEW
Ctrl
ExN
InN
Other
UD

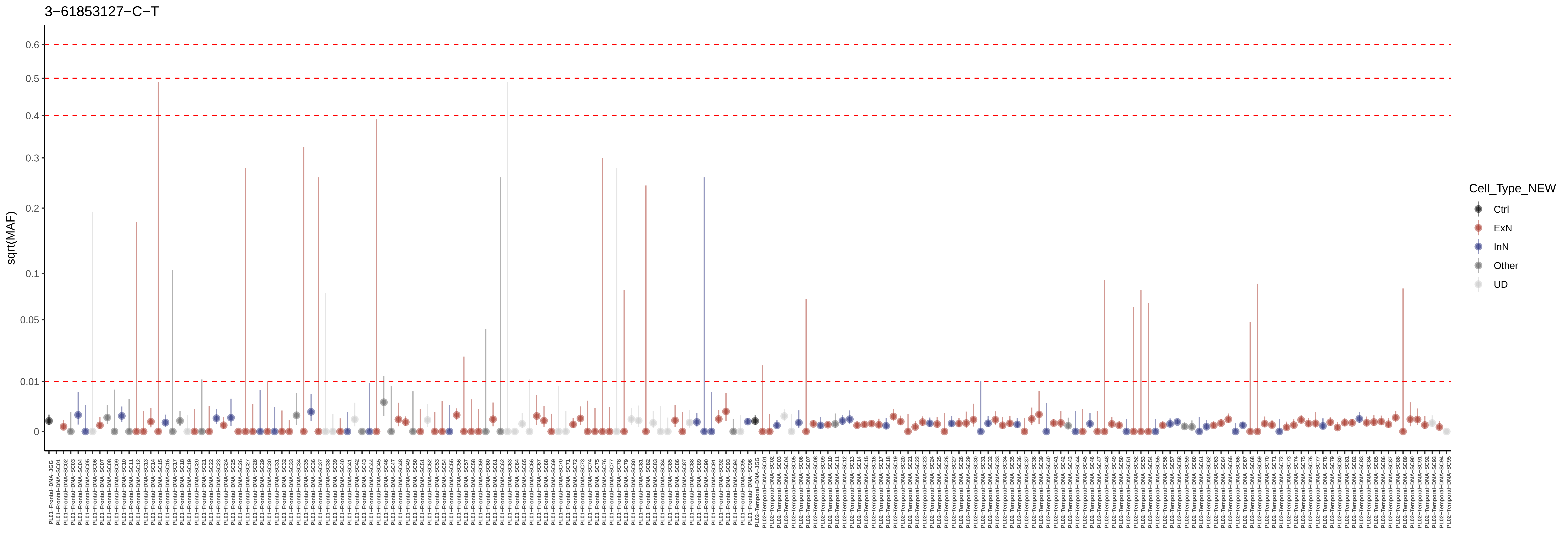
3−61853127−C−T
sqrt(MAF)
0.6
0.5
0.4
0.3
0.2
0.1
0.05
0.01
0
Cell_Type_NEW
Ctrl
ExN
InN
Other
UD

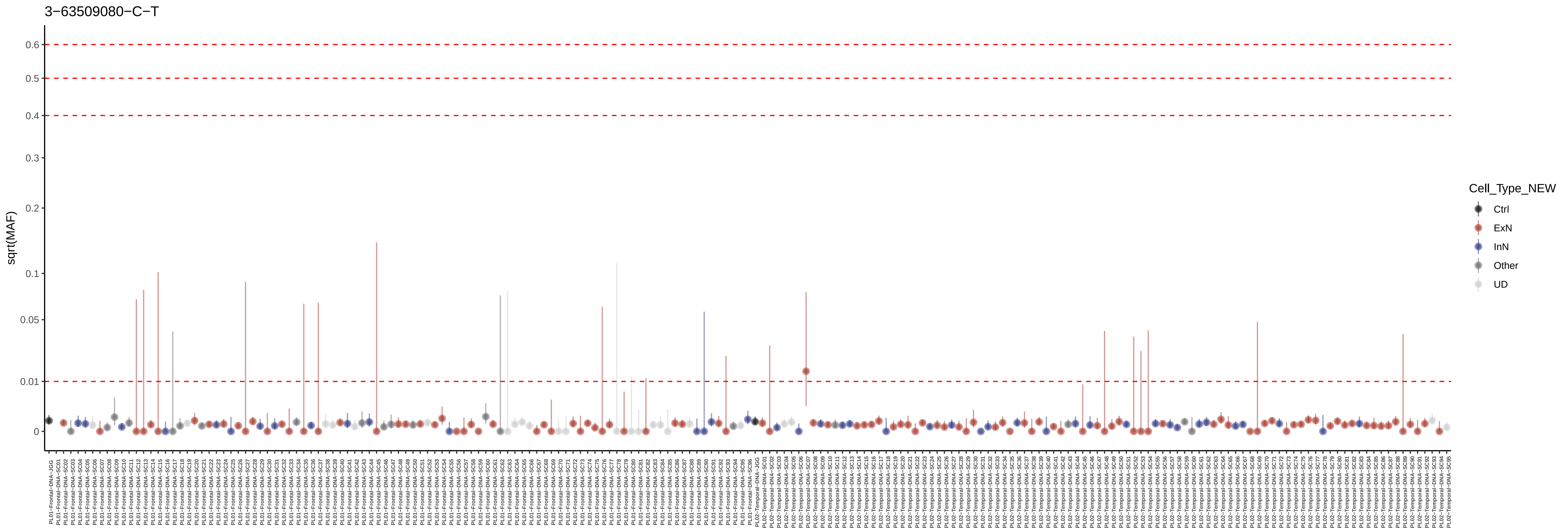
3−63509080−C−T
sqrt(MAF)
0.6
0.5
0.4
0.3
0.2
0.1
0.05
0.01
0
Cell_Type_NEW
Ctrl
ExN
InN
Other
UD

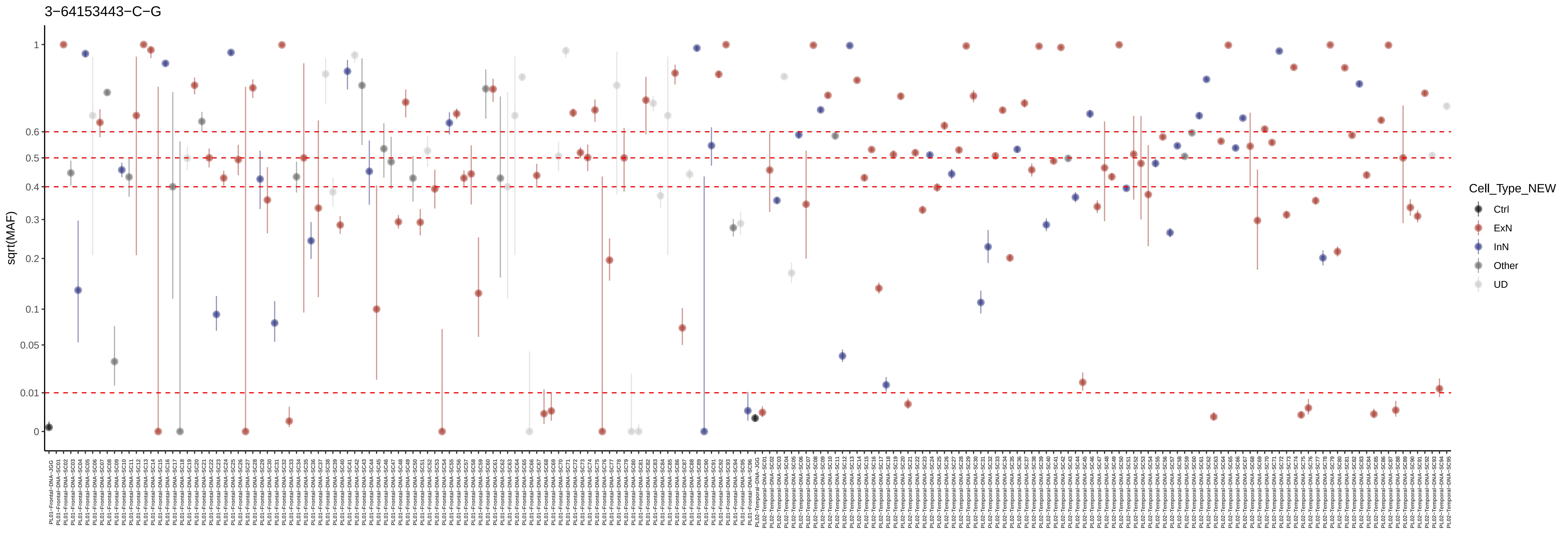
3−64153443−C−G
sqrt(MAF)
1
0.6
0.5
0.4
0.3
0.2
0.1
0.05
0.01
0
Cell_Type_NEW
Ctrl
ExN
InN
Other
UD

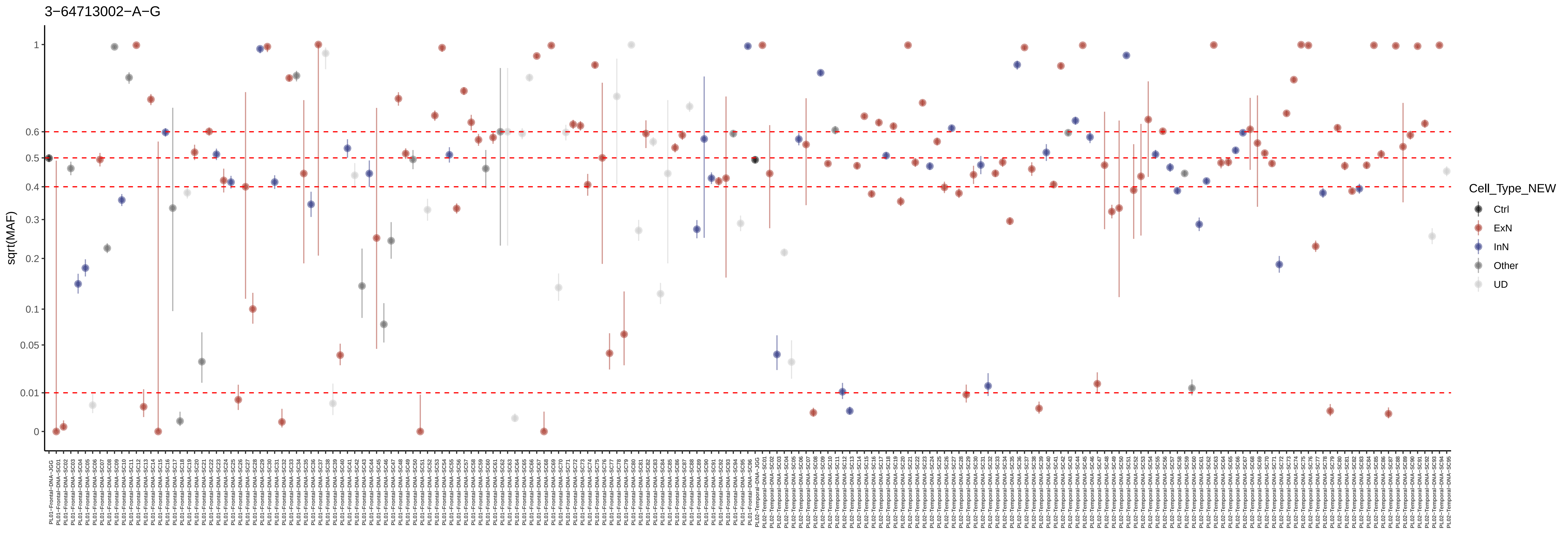
3−64713002−A−G
sqrt(MAF)
1
0.6
0.5
0.4
0.3
0.2
0.1
0.05
0.01
0
Cell_Type_NEW
Ctrl
ExN
InN
Other
UD

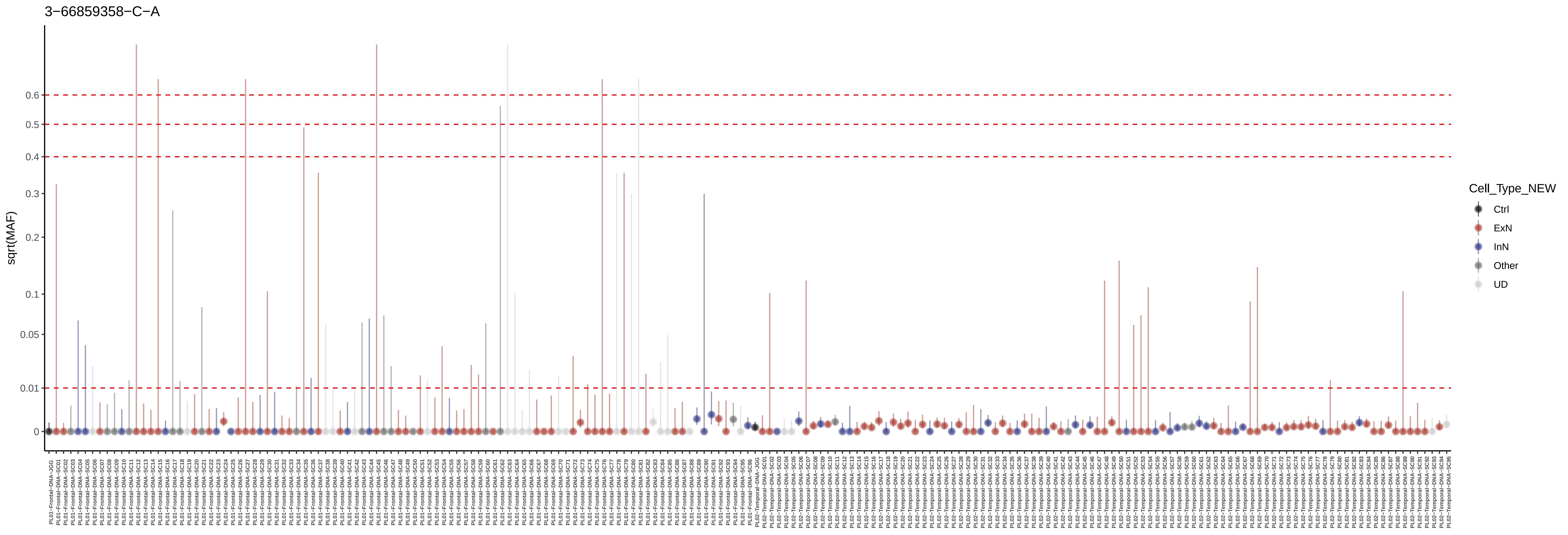
3−66859358−C−A
sqrt(MAF)
0.6
0.5
0.4
0.3
0.2
0.1
0.05
0.01
0
Cell_Type_NEW
Ctrl
ExN
InN
Other
UD

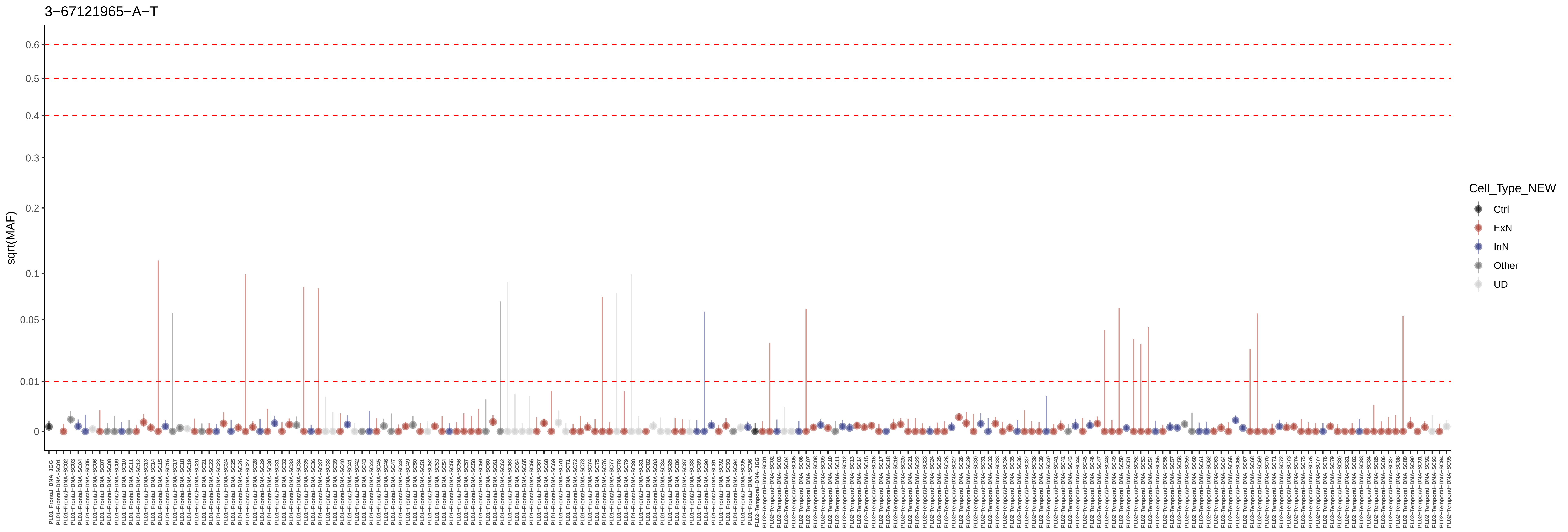
3−67121965−A−T
sqrt(MAF)
0.6
0.5
0.4
0.3
0.2
0.1
0.05
0.01
0
Cell_Type_NEW
Ctrl
ExN
InN
Other
UD

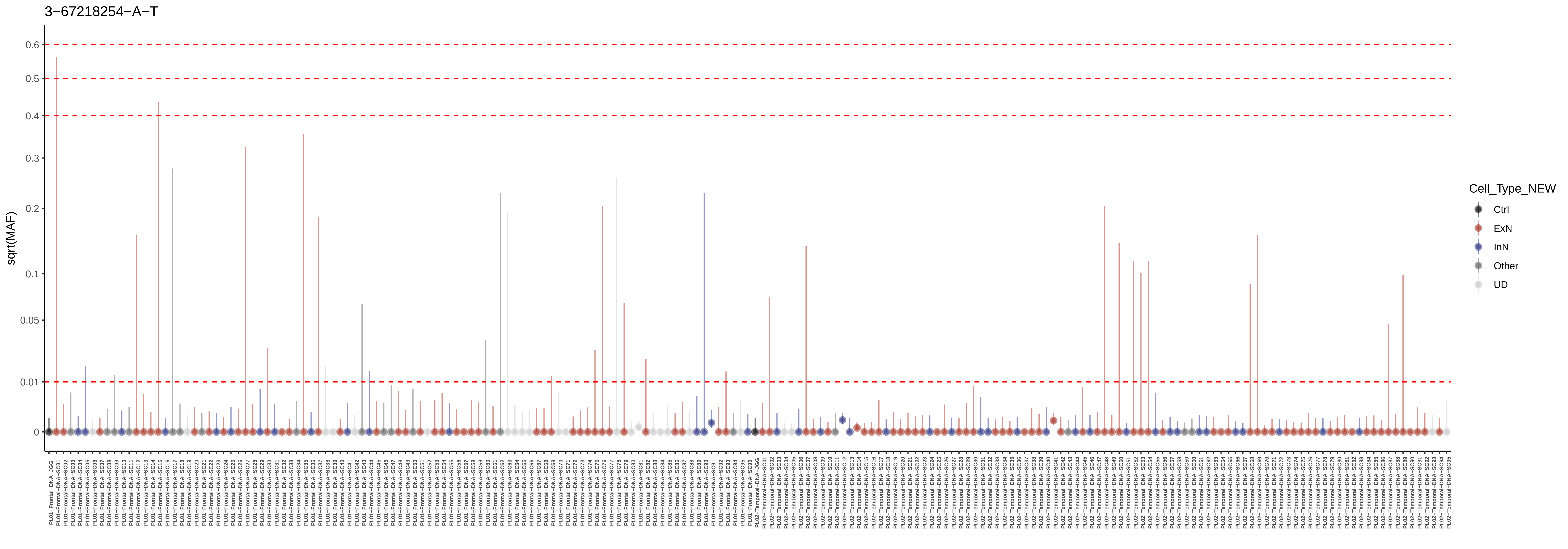

3−67218254−A−T
sqrt(MAF)
0.6
0.5
0.4
0.3
0.2
0.1
0.05
0.01
0
Cell_Type_NEW
Ctrl
ExN
InN
Other
UD

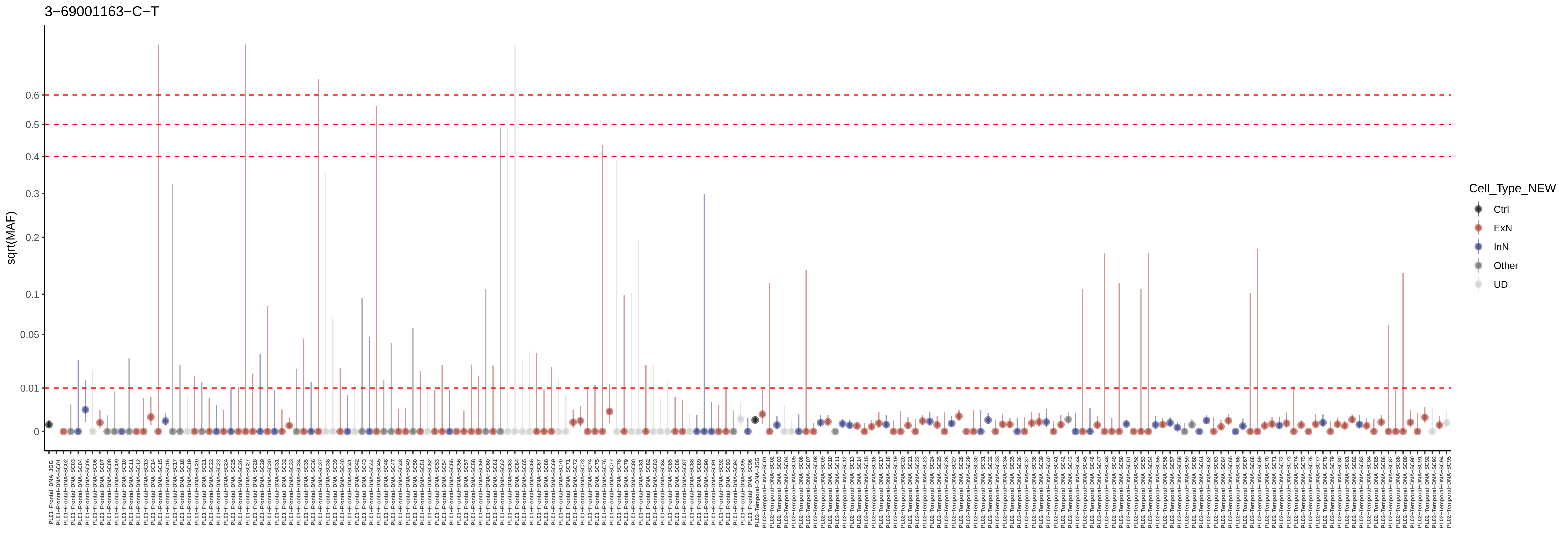
3−69001163−C−T
sqrt(MAF)
0.6
0.5
0.4
0.3
0.2
0.1
0.05
0.01
0
Cell_Type_NEW
Ctrl
ExN
InN
Other
UD

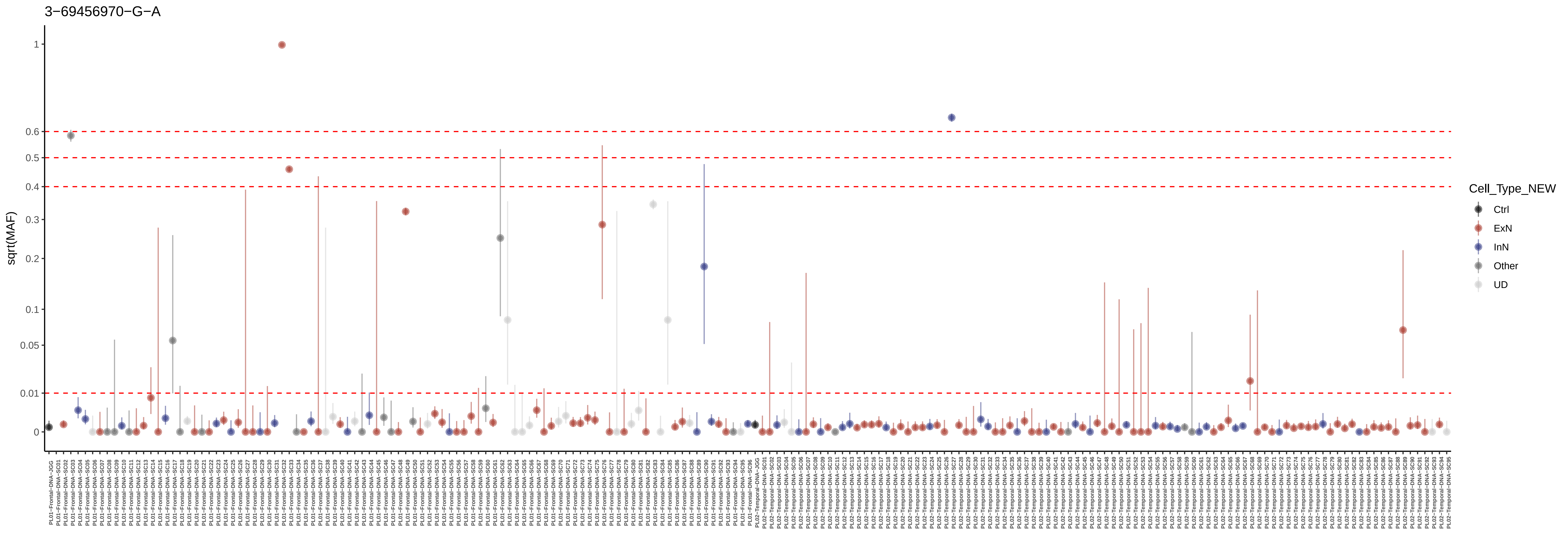

3−69456970−G−A
sqrt(MAF)
1
0.6
0.5
0.4
0.3
0.2
0.1
0.05
0.01
0
Cell_Type_NEW
Ctrl
ExN
InN
Other
UD

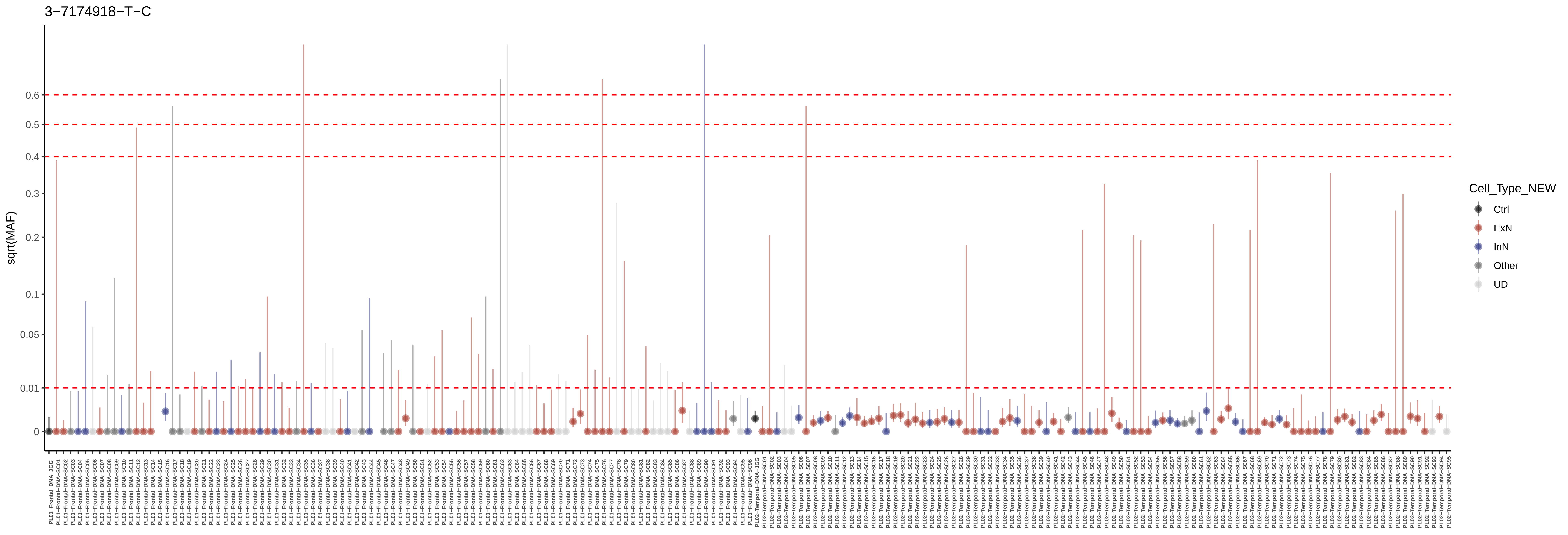
3−7174918−T−C
sqrt(MAF)
0.6
0.5
0.4
0.3
0.2
0.1
0.05
0.01
0
Cell_Type_NEW
Ctrl
ExN
InN
Other
UD

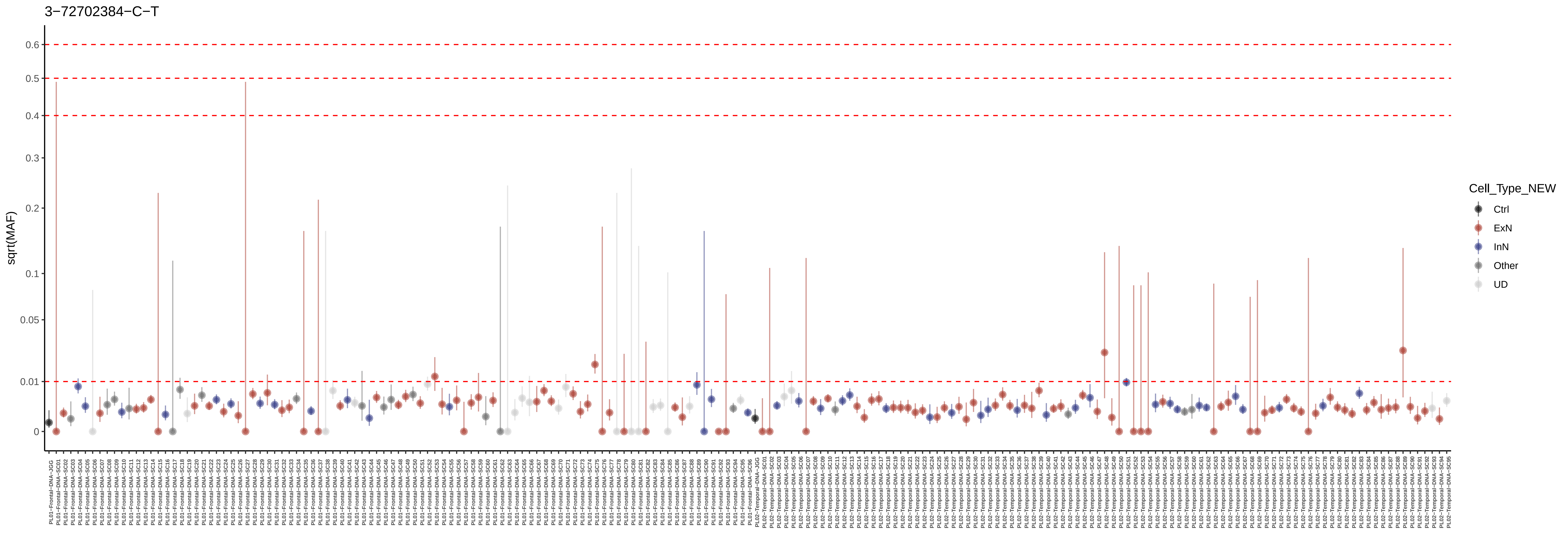
3−72702384−C−T
sqrt(MAF)
0.6
0.5
0.4
0.3
0.2
0.1
0.05
0.01
0
Cell_Type_NEW
Ctrl
ExN
InN
Other
UD

3−73073109−A−G

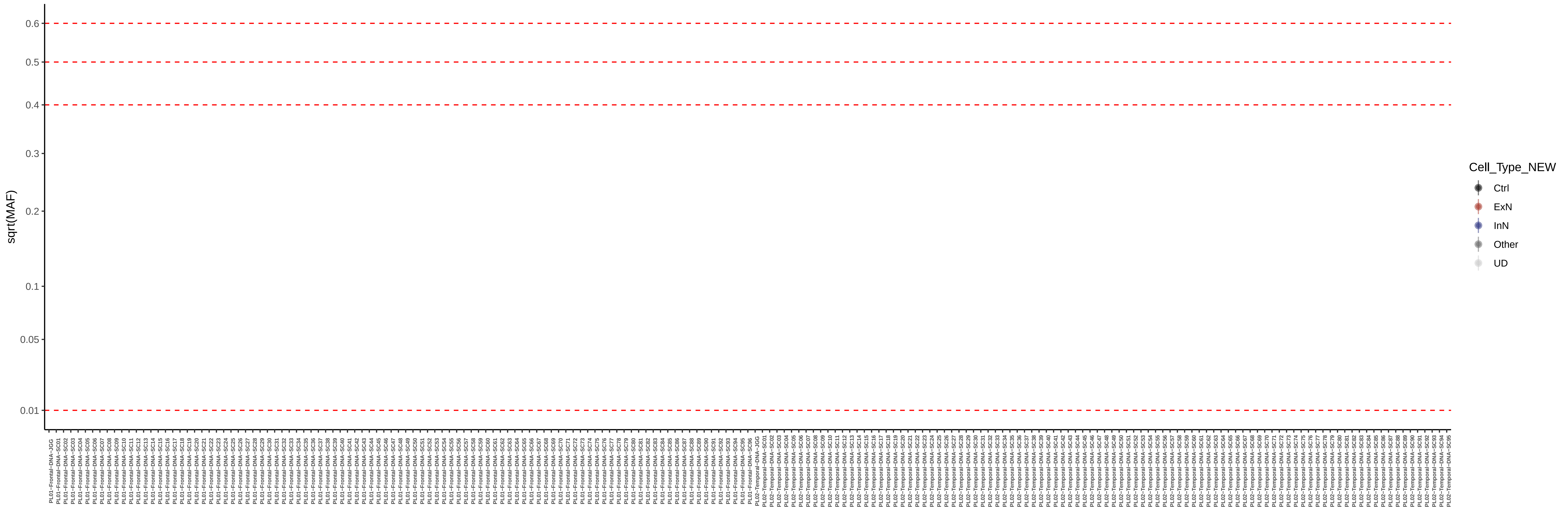

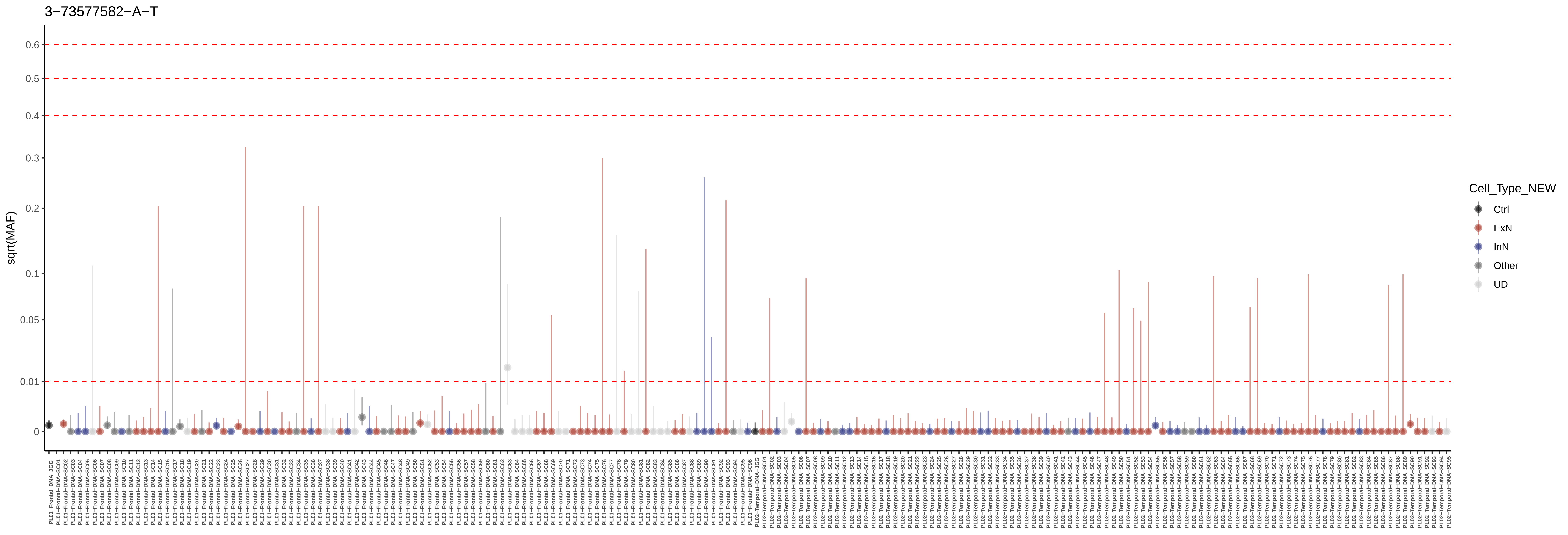

3−73577582−A−T
sqrt(MAF)
0.6
0.5
0.4
0.3
0.2
0.1
0.05
0.01
0
Cell_Type_NEW
Ctrl
ExN
InN
Other
UD

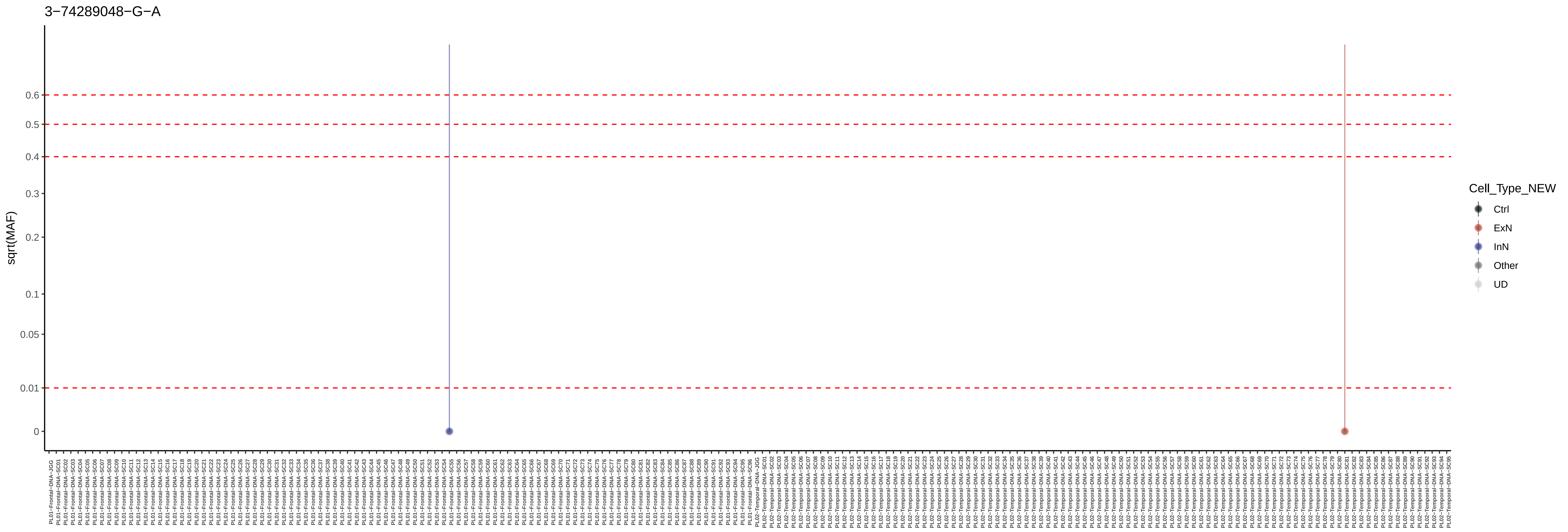
3−74289048−G−A
sqrt(MAF)
0.6
0.5
0.4
0.3
0.2
0.1
0.05
0.01
0
Cell_Type_NEW
Ctrl
ExN
InN
Other
UD

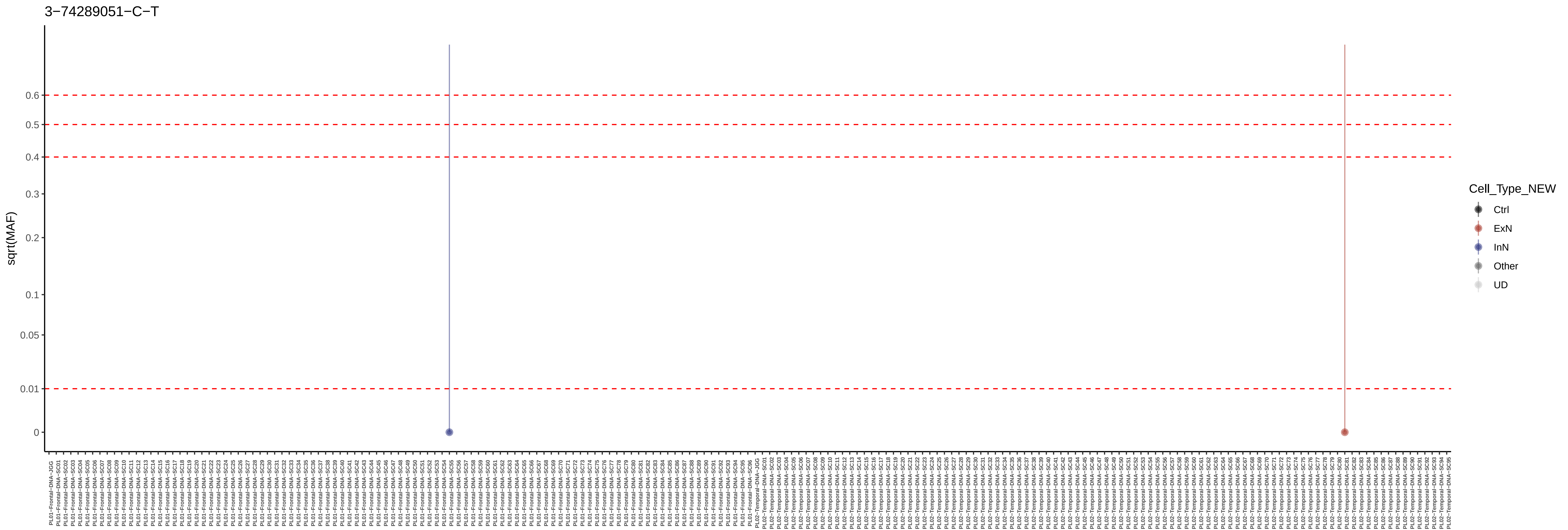
3−74289051−C−T
sqrt(MAF)
0.6
0.5
0.4
0.3
0.2
0.1
0.05
0.01
0
Cell_Type_NEW
Ctrl
ExN
InN
Other
UD

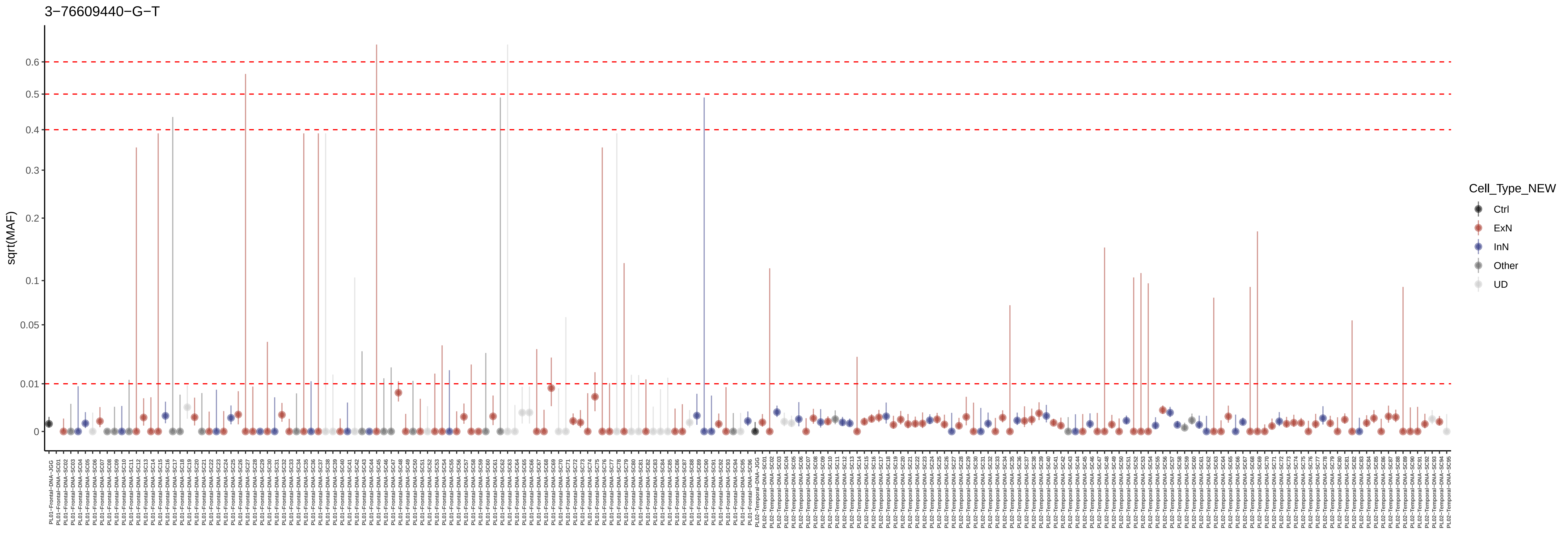

3−76609440−G−T
sqrt(MAF)
Cell_Type_NEW
Ctrl
ExN
InN
Other
UD

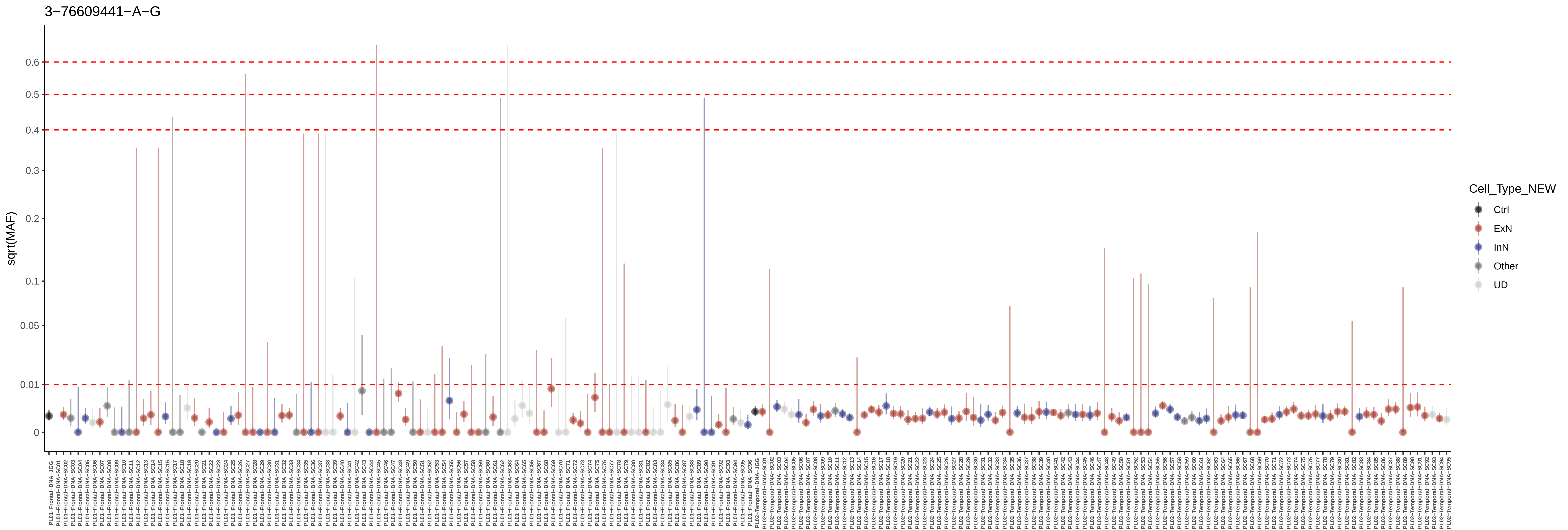
3−76609441−A−G
sqrt(MAF)
0.6
0.5
0.4
0.3
0.2
0.1
0.05
0.01
0
Cell_Type_NEW
Ctrl
ExN
InN
Other
UD

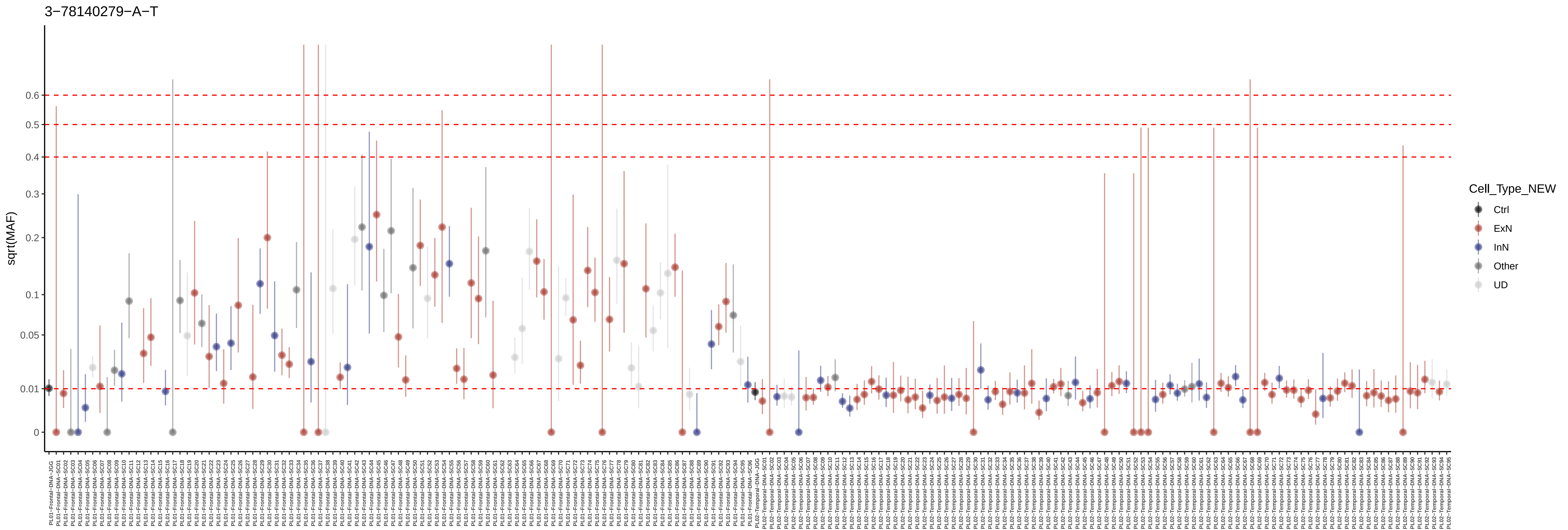
3−78140279−A−T
sqrt(MAF)
0.6
0.5
0.4
0.3
0.2
0.1
0.05
0.01
0
Cell_Type_NEW
Ctrl
ExN
InN
Other
UD

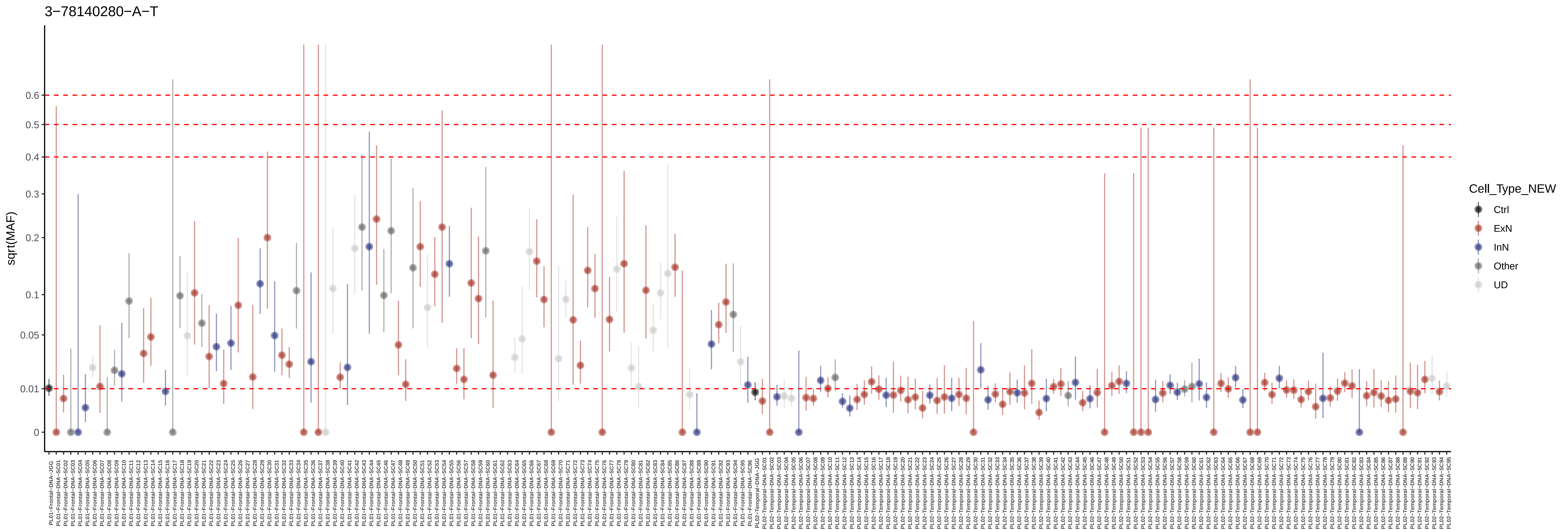
3−78140280−A−T
sqrt(MAF)
0.6
0.5
0.4
0.3
0.2
0.1
0.05
0.01
0
Cell_Type_NEW
Ctrl
ExN
InN
Other
UD

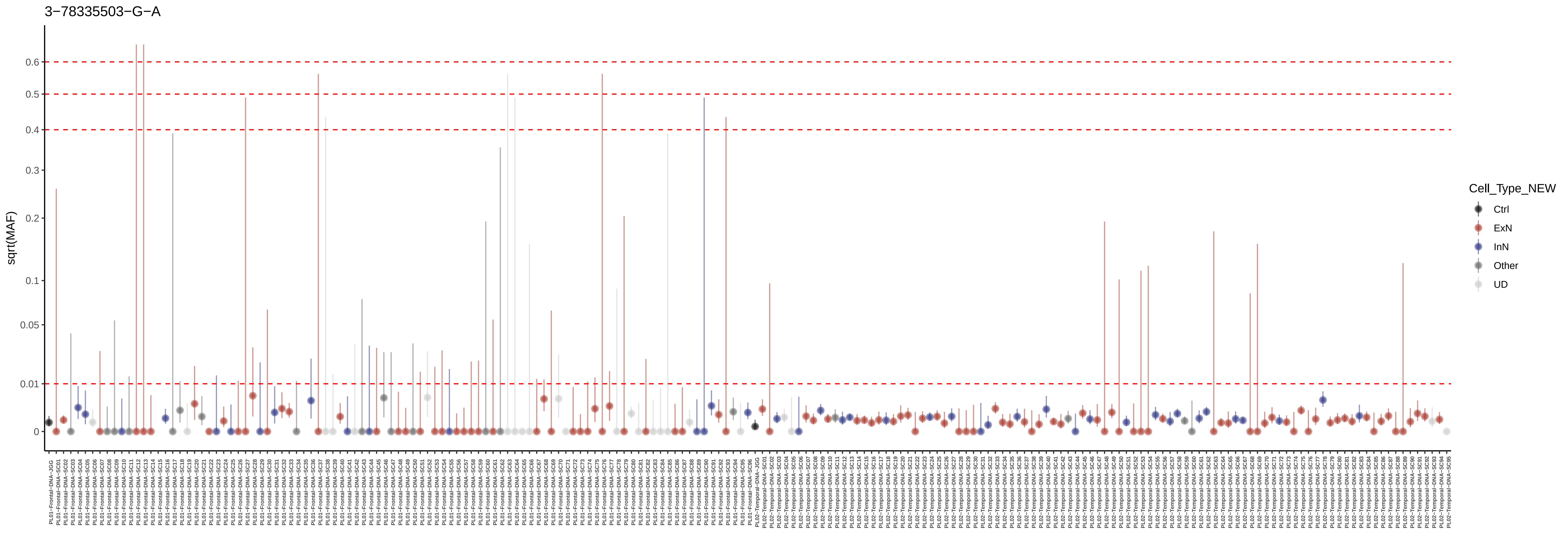

3−78335503−G−A
sqrt(MAF)
0.6
0.5
0.4
0.3
0.2
0.1
0.05
0.01
0
Cell_Type_NEW
Ctrl
ExN
InN
Other
UD

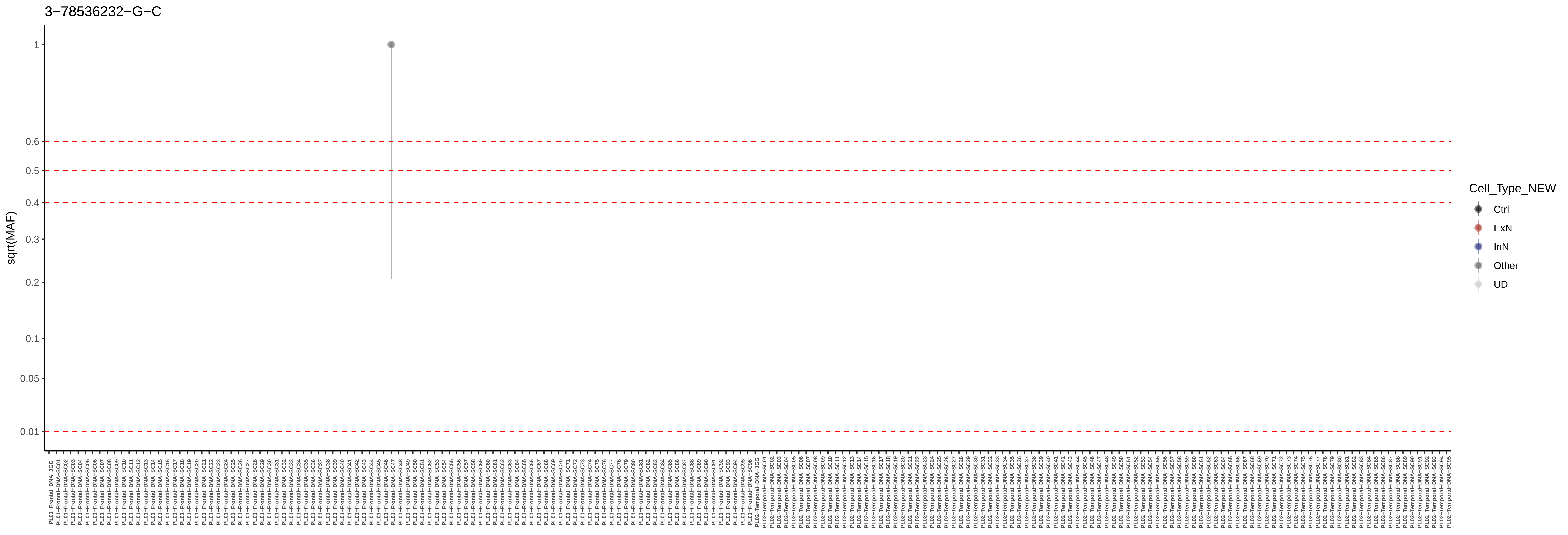
3−78536232−G−C
sqrt(MAF)
1
0.6
0.5
0.4
0.3
0.2
0.1
0.05
0.01
Cell_Type_NEW
Ctrl
ExN
InN
Other
UD

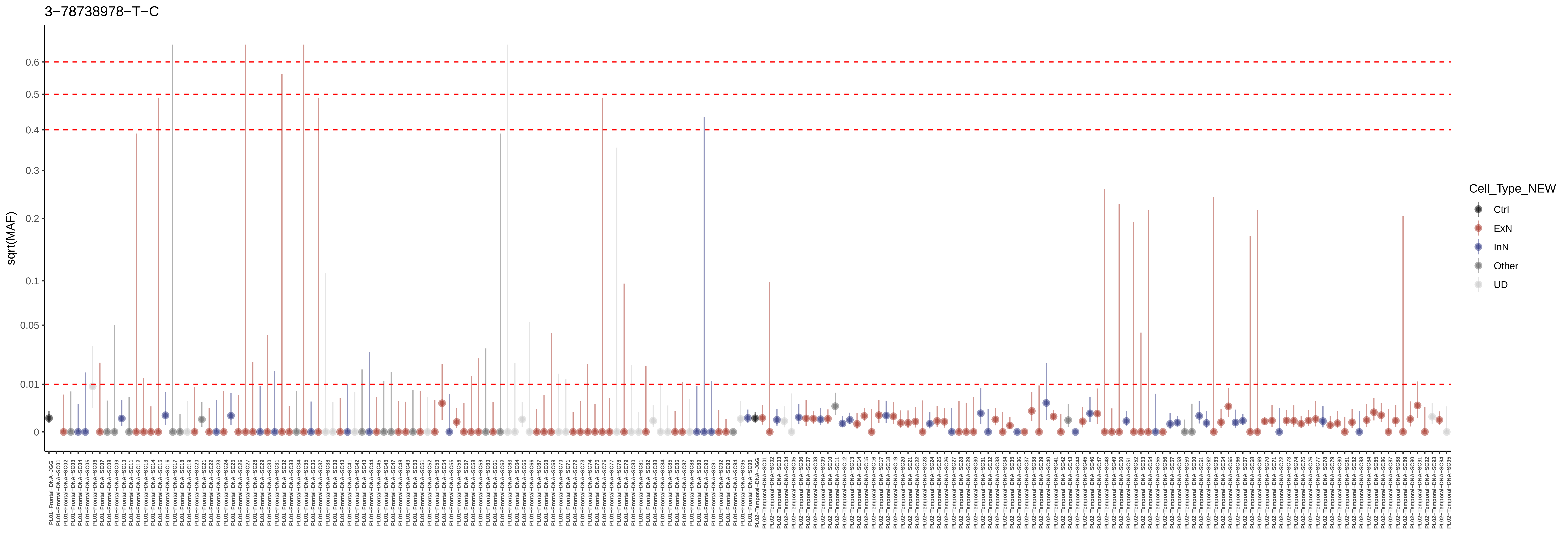
3−78738978−T−C
sqrt(MAF)
0.6
0.5
0.4
0.3
0.2
0.1
0.05
0.01
0
Cell_Type_NEW
Ctrl
ExN
InN
Other
UD

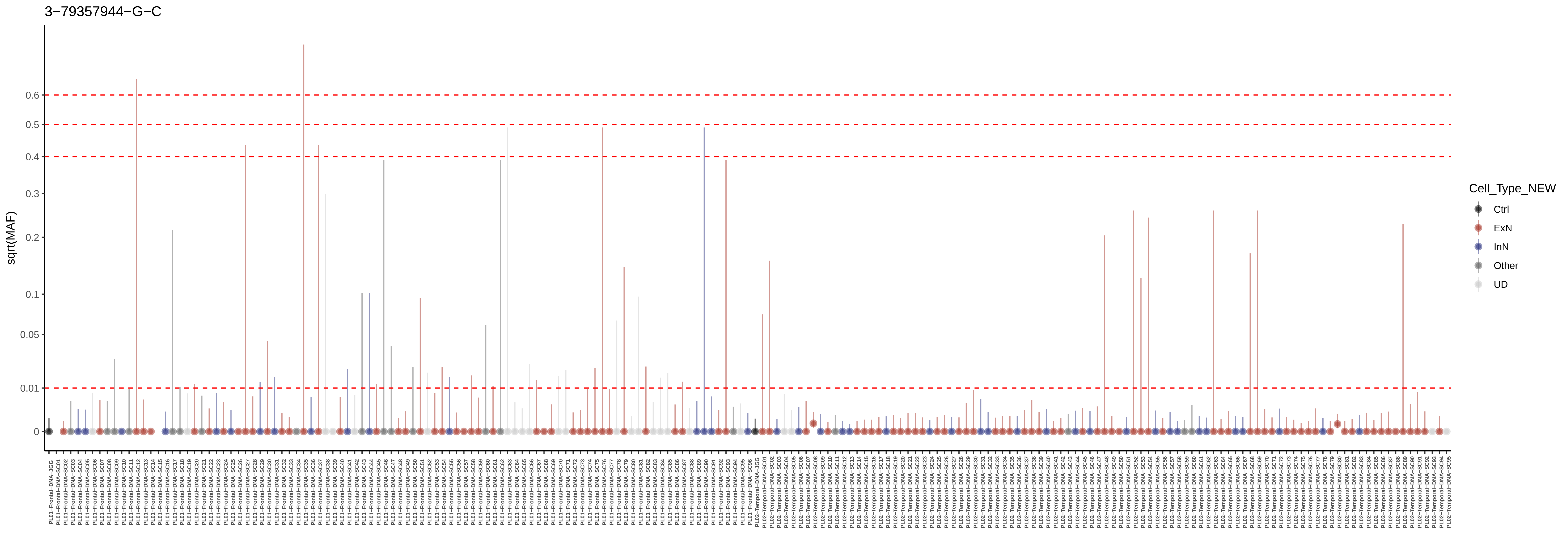
3−79357944−G−C
sqrt(MAF)
0.6
0.5
0.4
0.3
0.2
0.1
0.05
0.01
0
Cell_Type_NEW
Ctrl
ExN
InN
Other
UD

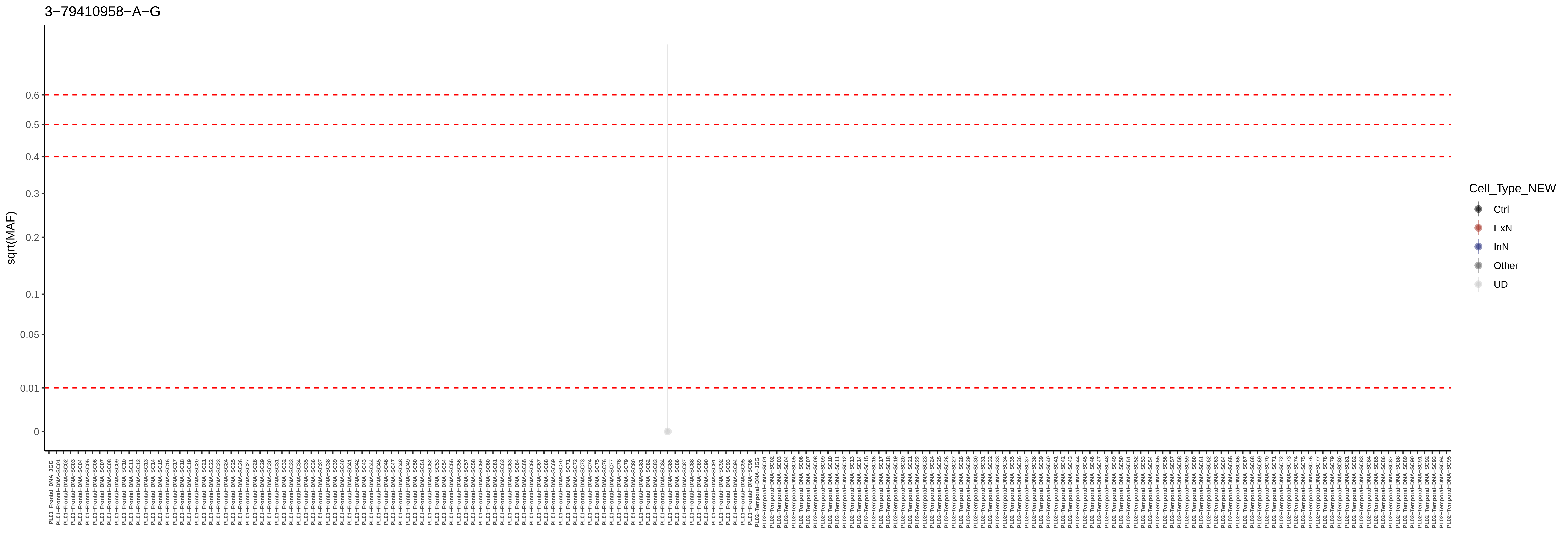

3−79410958−A−G
sqrt(MAF)
0.6
0.5
0.4
0.3
0.2
0.1
0.05
0.01
0
Cell_Type_NEW
Ctrl
ExN
InN
Other
UD

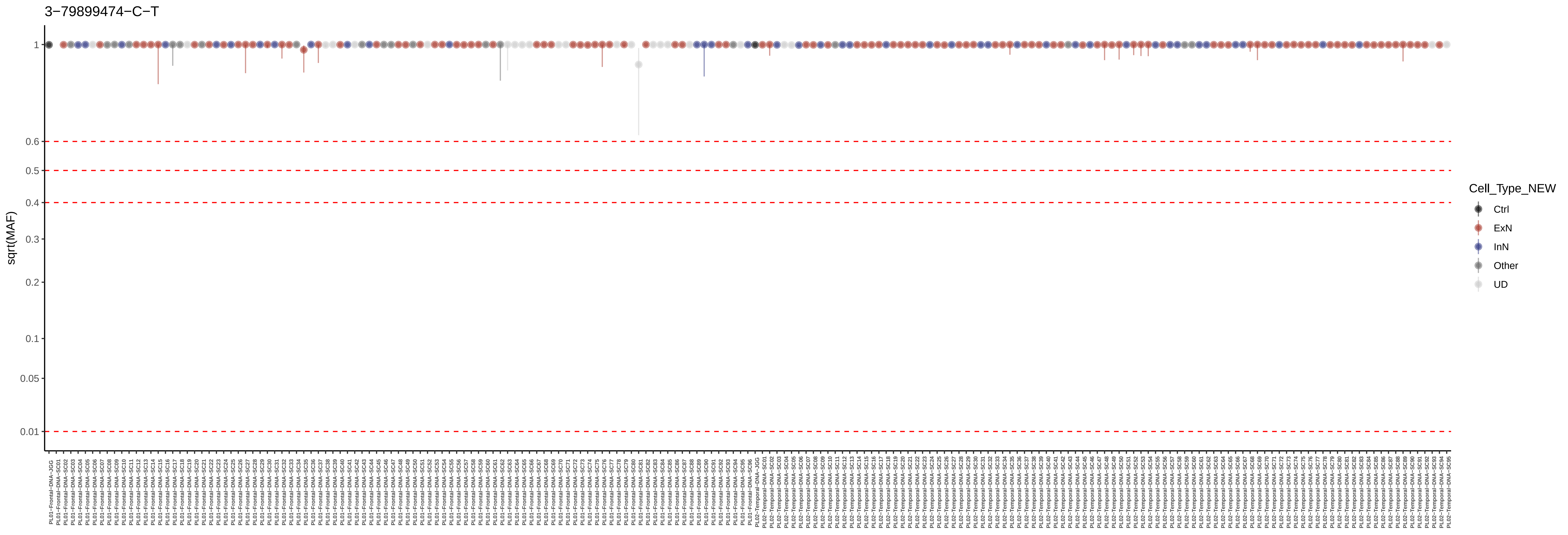

3−79899474−C−T
sqrt(MAF)
1
0.6
0.5
0.4
0.3
0.2
0.1
0.05
0.01
Cell_Type_NEW
Ctrl
ExN
InN
Other
UD

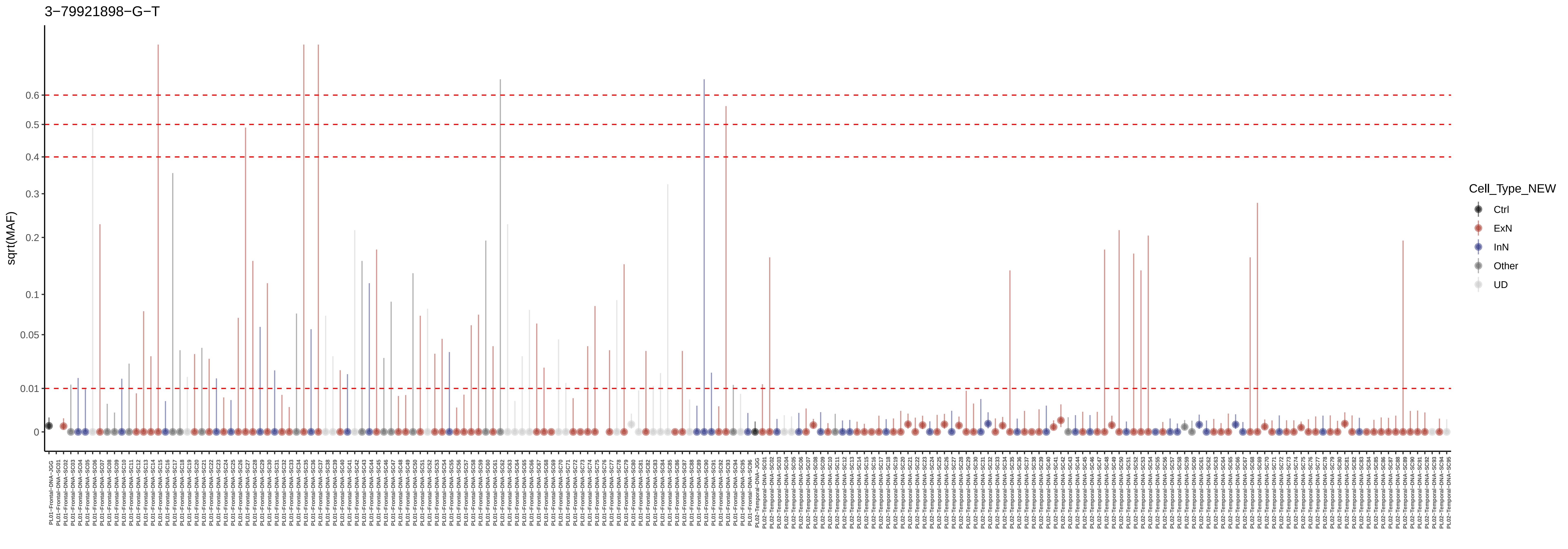
3−79921898−G−T
sqrt(MAF)
0.6
0.5
0.4
0.3
0.2
0.1
0.05
0.01
0
Cell_Type_NEW
Ctrl
ExN
InN
Other
UD

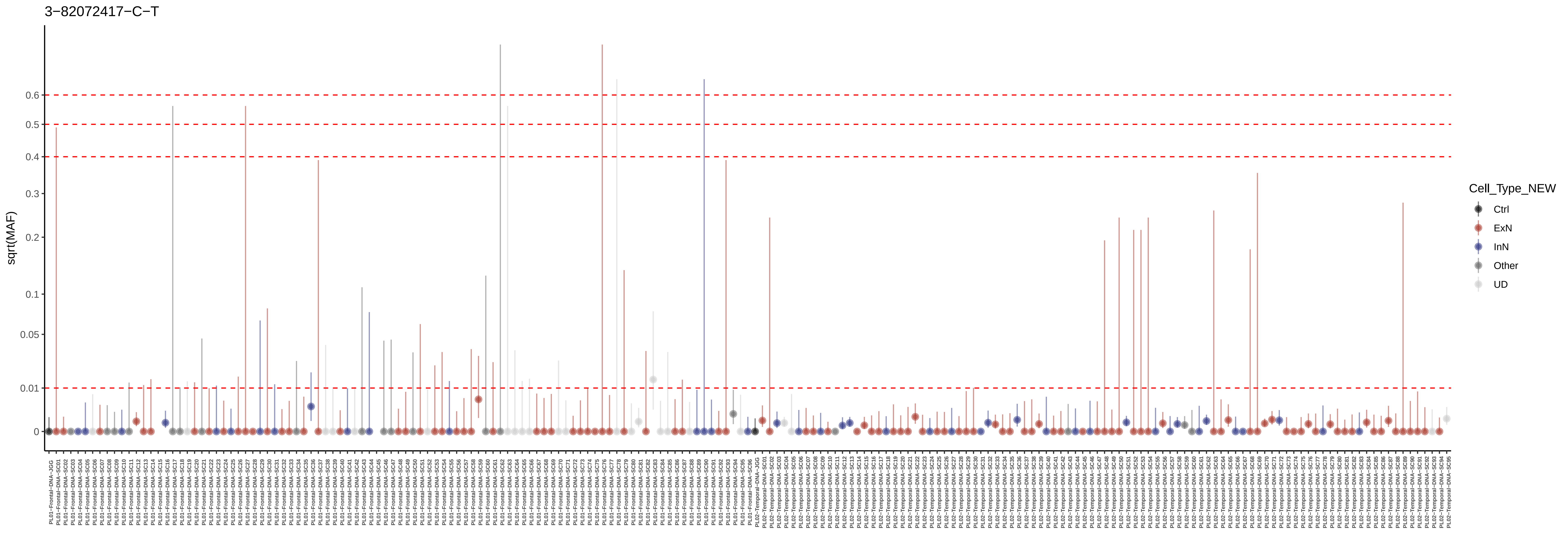
3−82072417−C−T
sqrt(MAF)
0.6
0.5
0.4
0.3
0.2
0.1
0.05
0.01
0
Cell_Type_NEW
Ctrl
ExN
InN
Other
UD

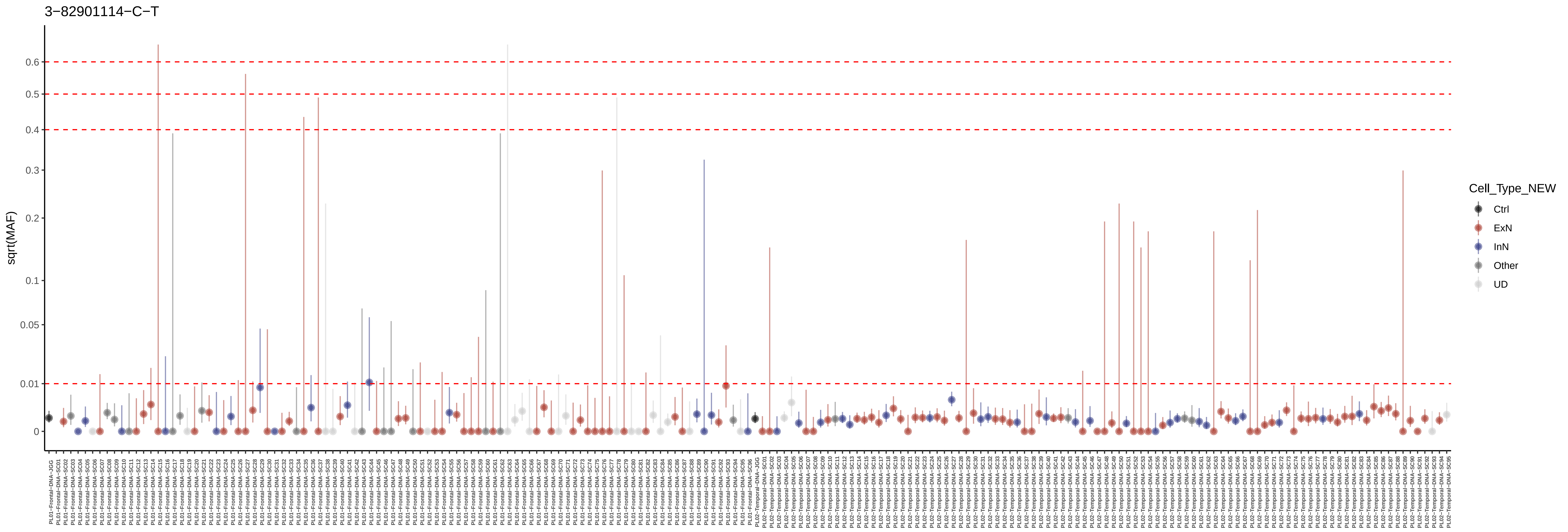
3−82901114−C−T
sqrt(MAF)
0.6
0.5
0.4
0.3
0.2
0.1
0.05
0.01
0
Cell_Type_NEW
Ctrl
ExN
InN
Other
UD

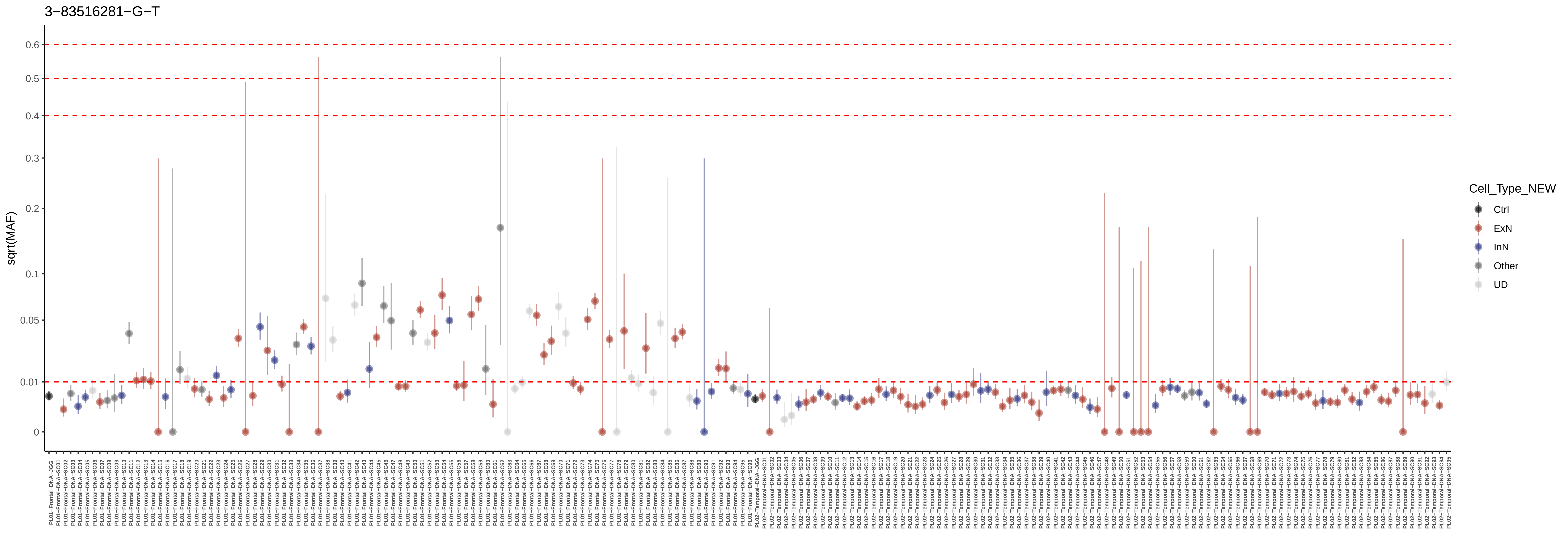

3−83516281−G−T
sqrt(MAF)
0.6
0.5
0.4
0.3
0.2
0.1
0.05
0.01
0
Cell_Type_NEW
Ctrl
ExN
InN
Other
UD

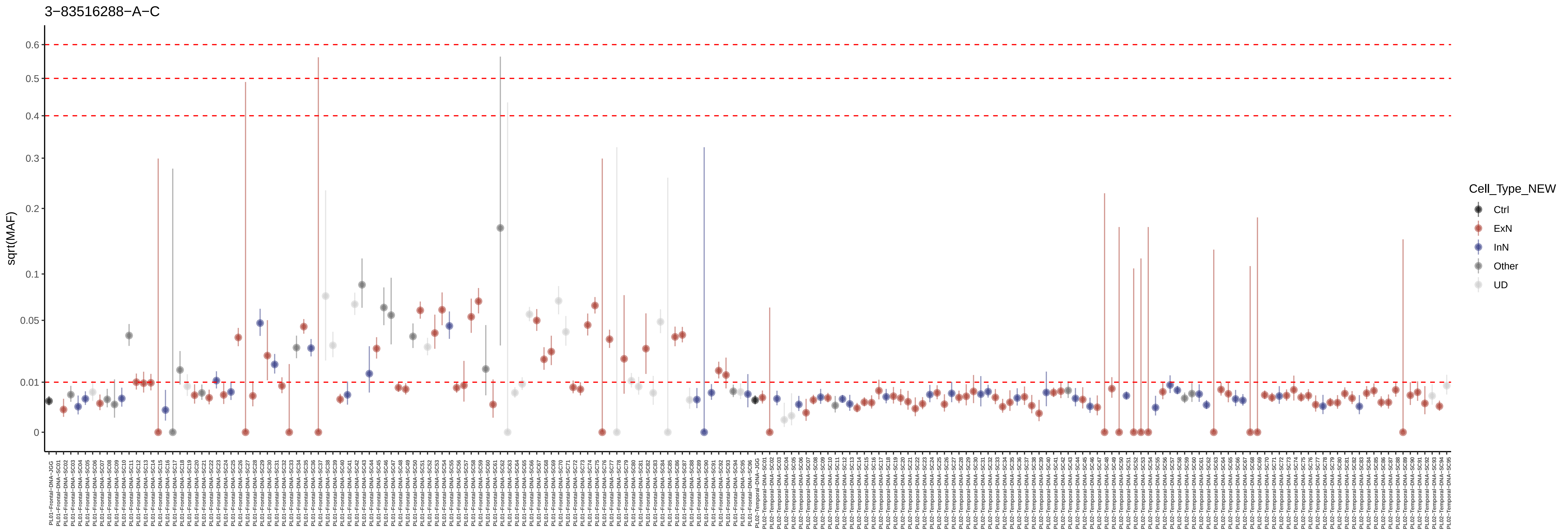
3−83516288−A−C
sqrt(MAF)
0.6
0.5
0.4
0.3
0.2
0.1
0.05
0.01
0
Cell_Type_NEW
Ctrl
ExN
InN
Other
UD

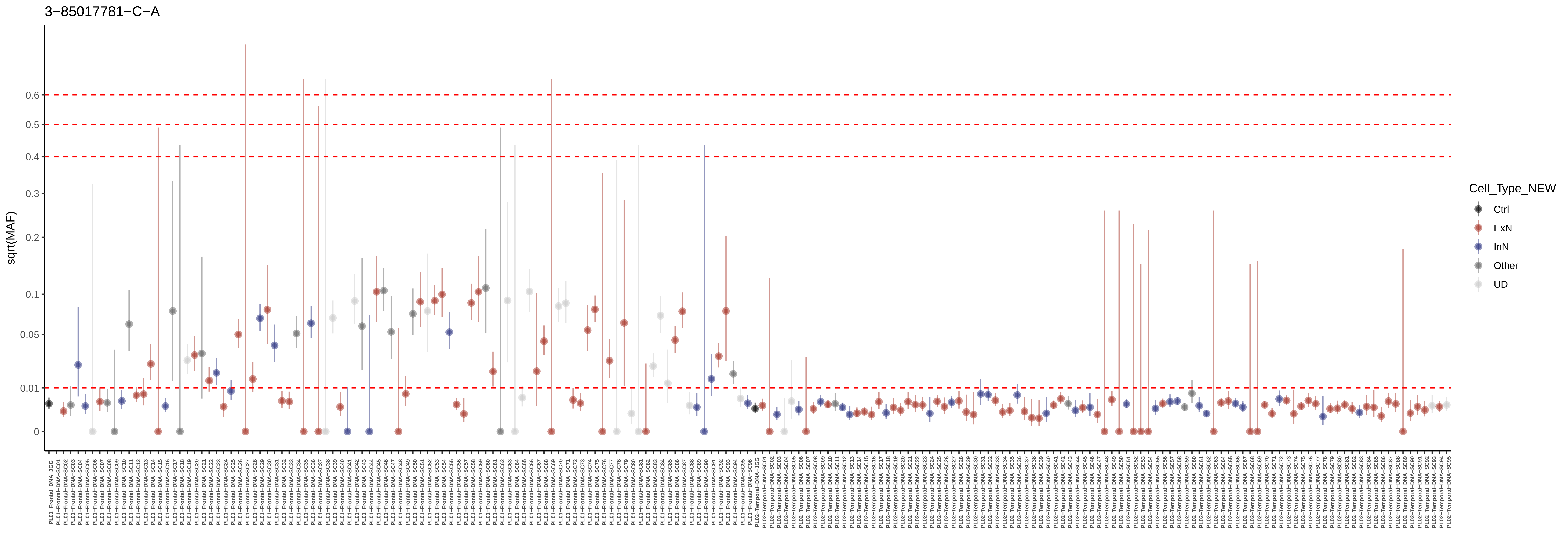

3−85017781−C−A
sqrt(MAF)
0
0.01
0.05
0.1
0.2
0.3
0.4
0.5
0.6
Cell_Type_NEW
Ctrl
ExN
InN
Other
UD

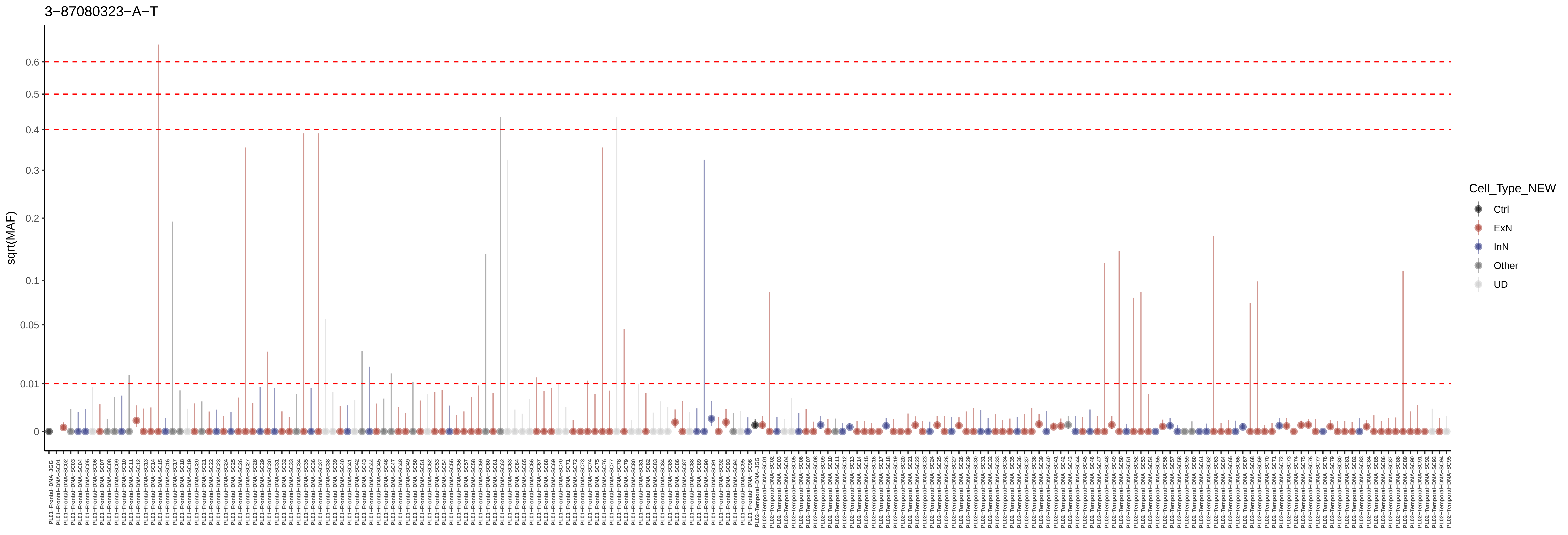

3−87080323−A−T
sqrt(MAF)
0.6
0.5
0.4
0.3
0.2
0.1
0.05
0.01
0
Cell_Type_NEW
Ctrl
ExN
InN
Other
UD

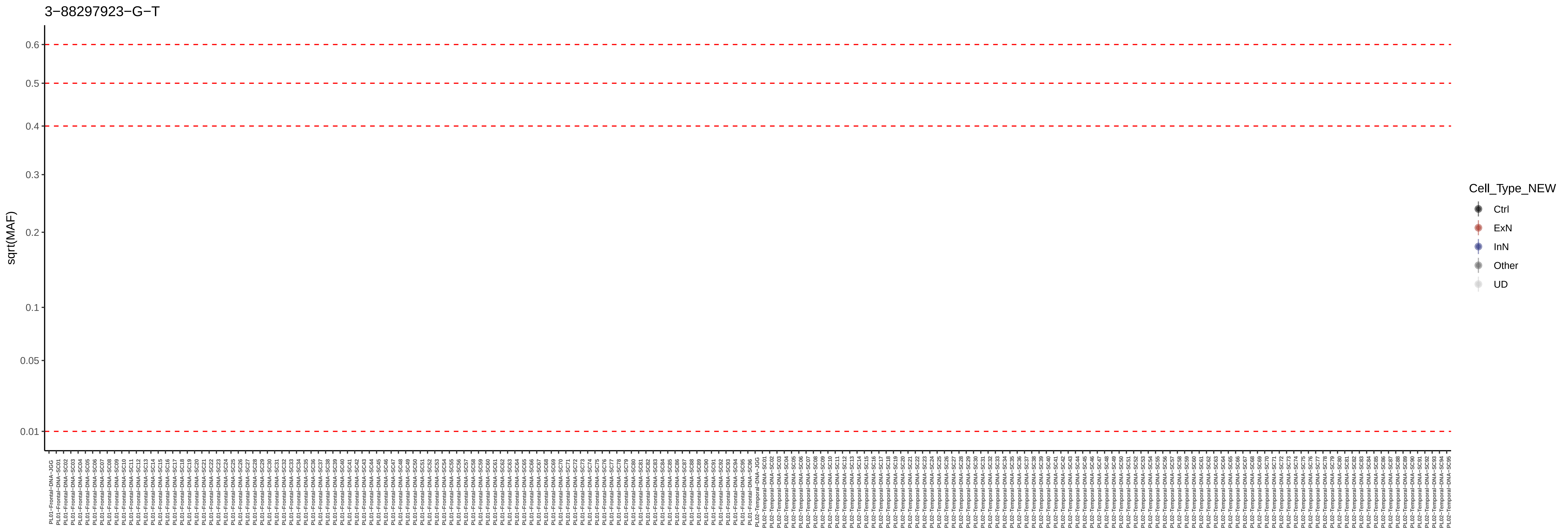
3−88297923−G−T
sqrt(MAF)
0.6
0.5
0.4
0.3
0.2
0.1
0.05
0.01
Cell_Type_NEW
Ctrl
ExN
InN
Other
UD

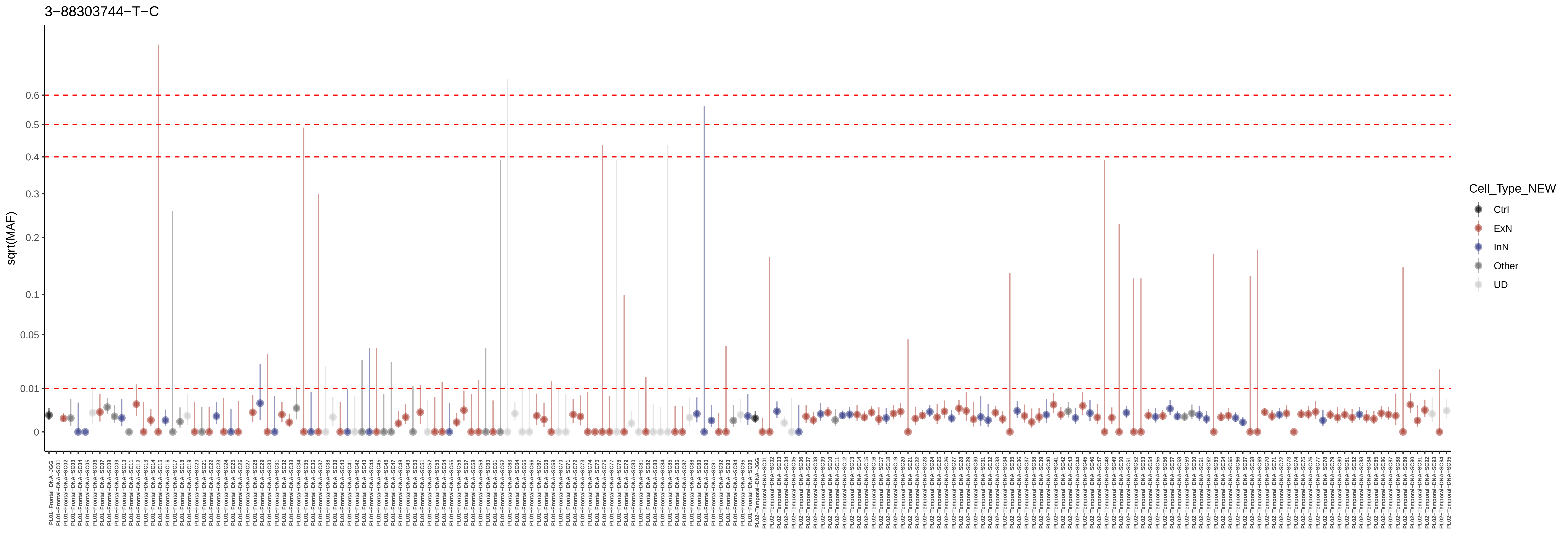

3−88303744−T−C
sqrt(MAF)
0.6
0.5
0.4
0.3
0.2
0.1
0.05
0.01
0
Cell_Type_NEW
Ctrl
ExN
InN
Other
UD

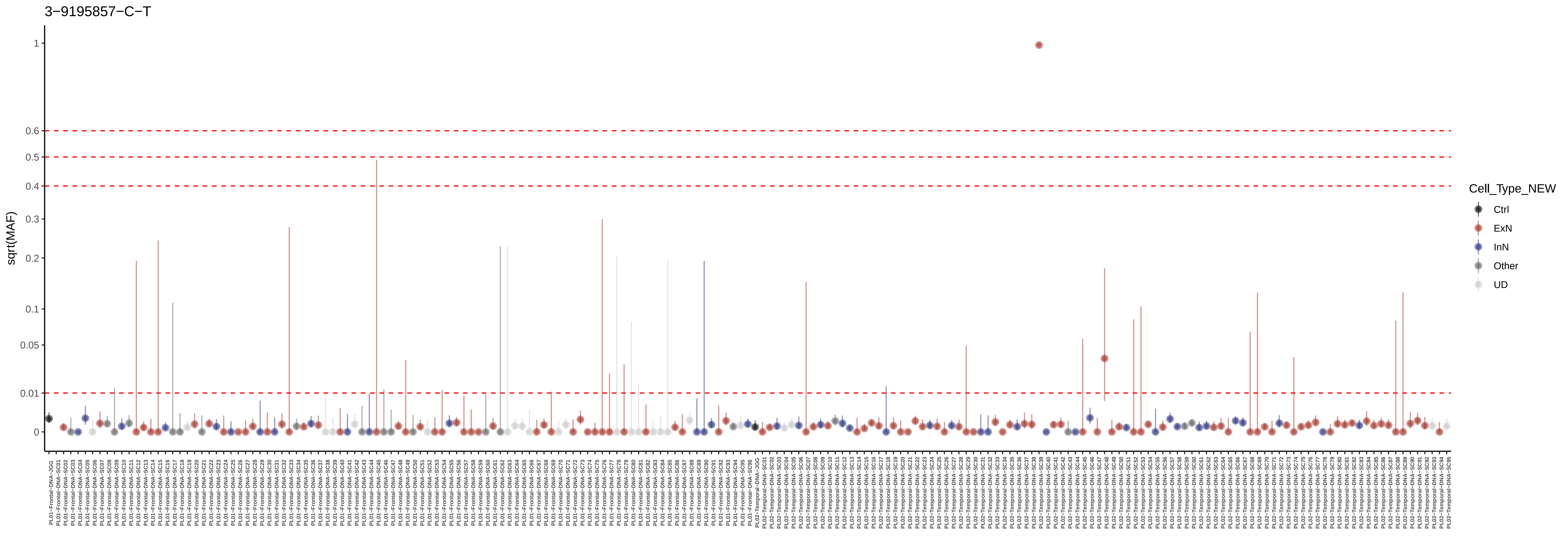
3−9195857−C−T
sqrt(MAF)
1
0.6
0.5
0.4
0.3
0.2
0.1
0.05
0.01
0
Cell_Type_NEW
Ctrl
ExN
InN
Other
UD

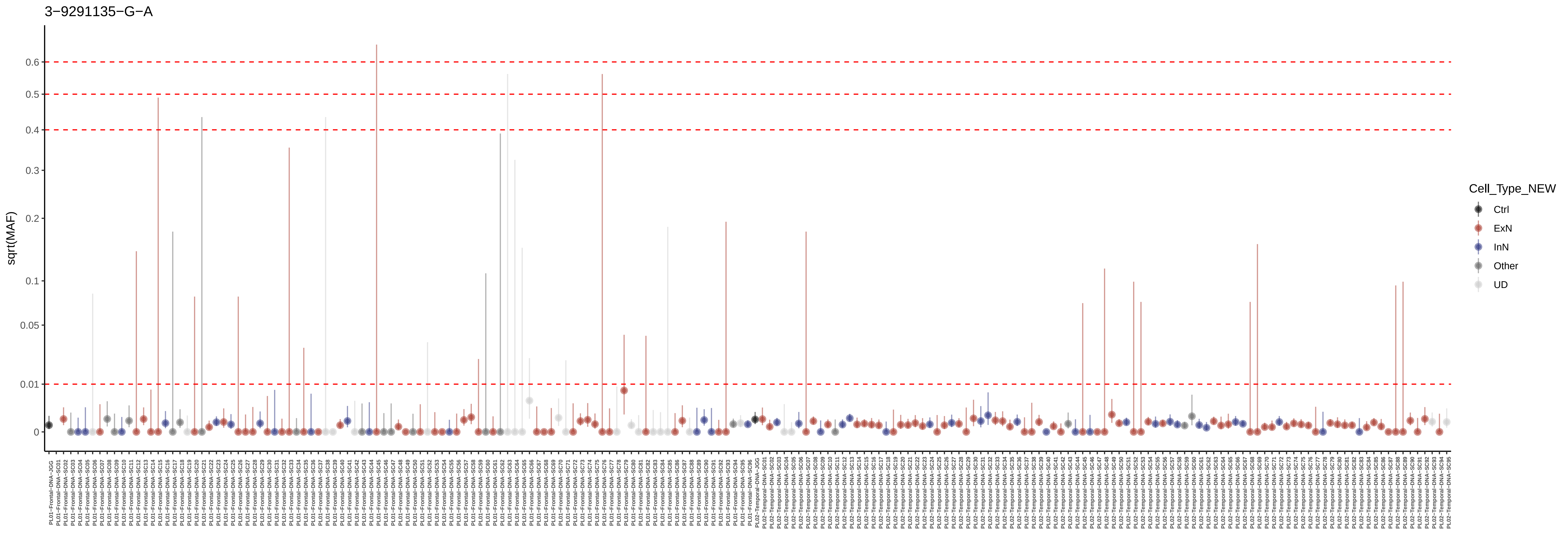
3−9291135−G−A
sqrt(MAF)
0.6
0.5
0.4
0.3
0.2
0.1
0.05
0.01
0
Cell_Type_NEW
Ctrl
ExN
InN
Other
UD

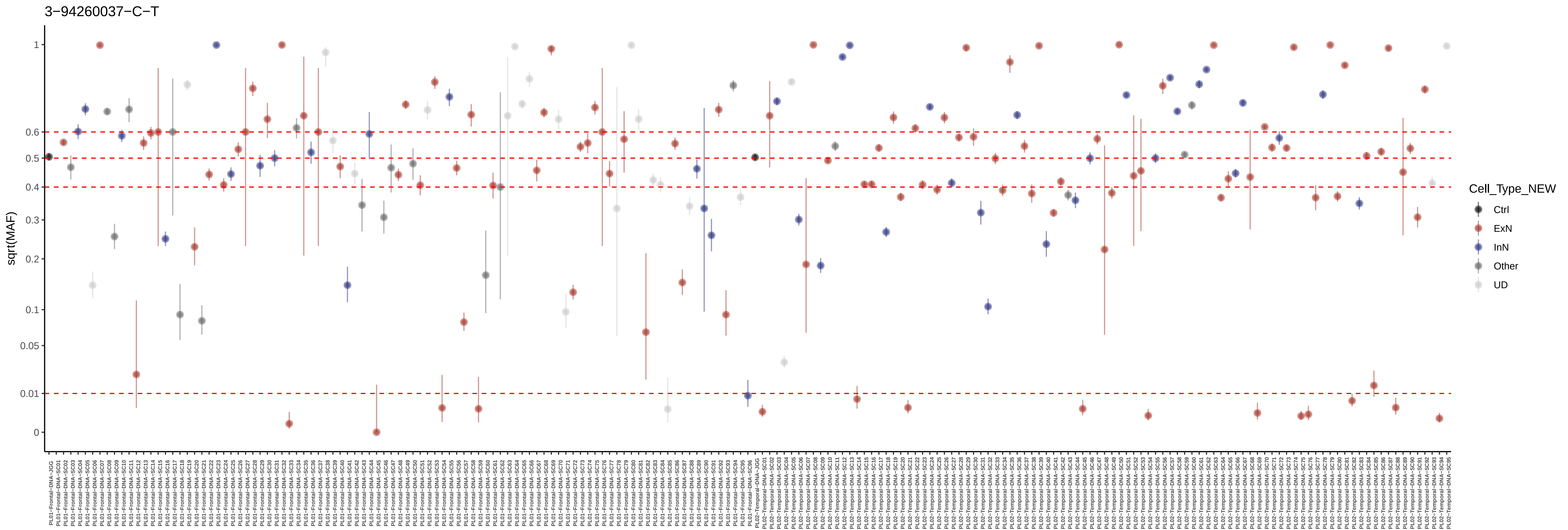
3−94260037−C−T
sqrt(MAF)
1
0.6
0.5
0.4
0.3
0.2
0.1
0.05
0.01
0
Cell_Type_NEW
Ctrl
ExN
InN
Other
UD

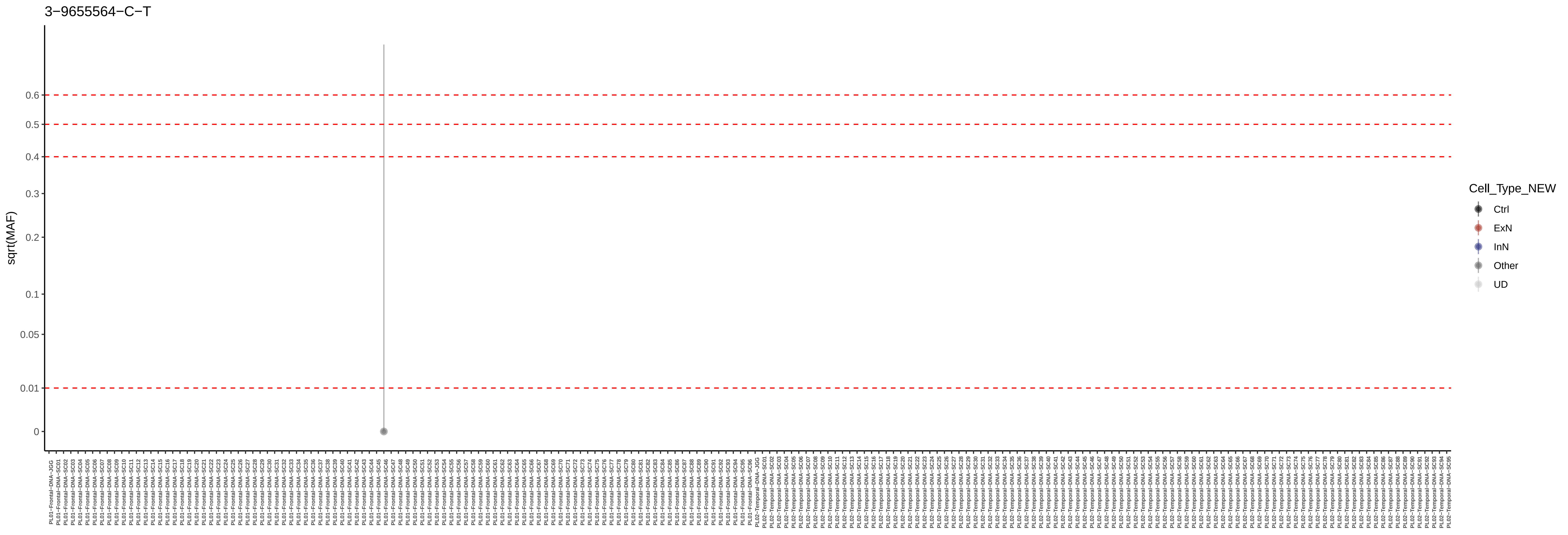

3−9655564−C−T
sqrt(MAF)
0.6
0.5
0.4
0.3
0.2
0.1
0.05
0.01
0
Cell_Type_NEW
Ctrl
ExN
InN
Other
UD

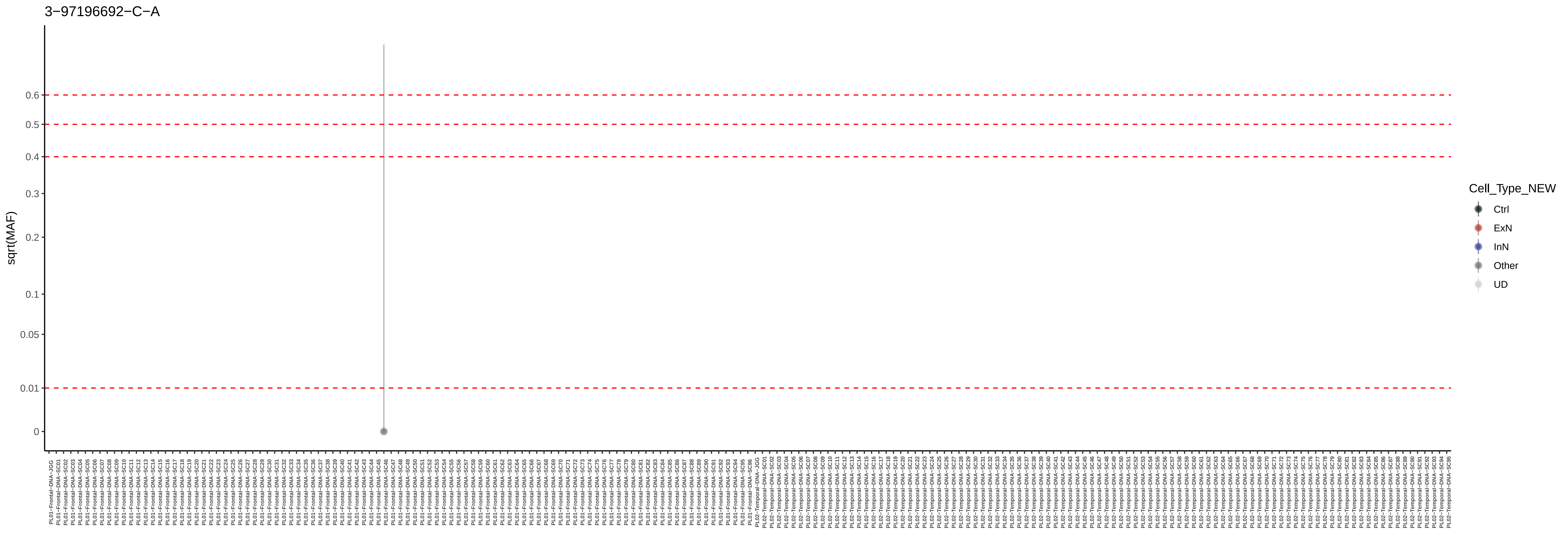
3−97196692−C−A
sqrt(MAF)
0.6
0.5
0.4
0.3
0.2
0.1
0.05
0.01
0
Cell_Type_NEW
Ctrl
ExN
InN
Other
UD

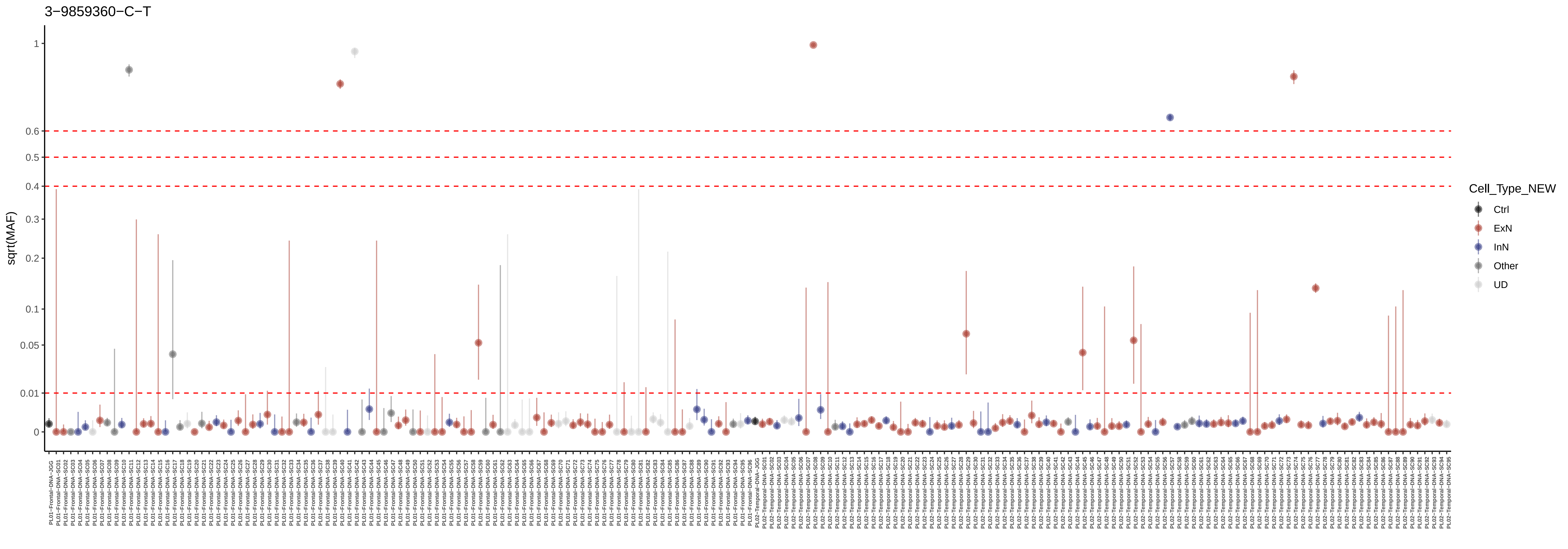
3−9859360−C−T
sqrt(MAF)
1
0.6
0.5
0.4
0.3
0.2
0.1
0.05
0.01
0
Cell_Type_NEW
Ctrl
ExN
InN
Other
UD

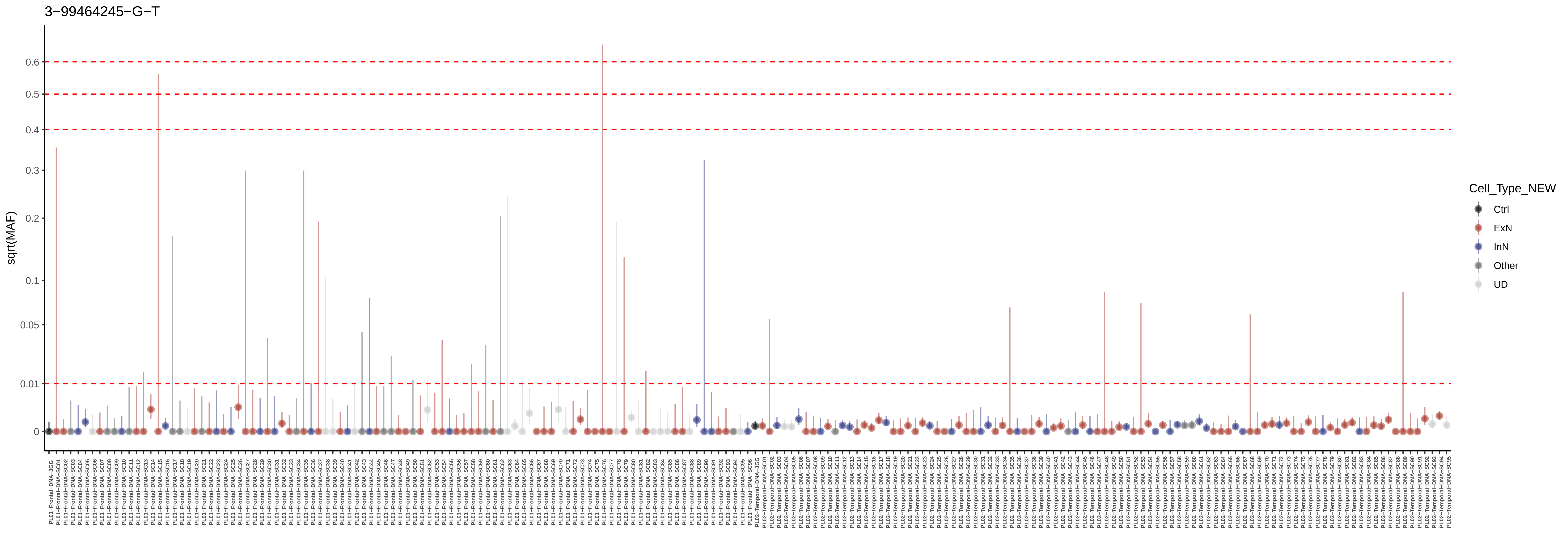

3−99464245−G−T
sqrt(MAF)
0.6
0.5
0.4
0.3
0.2
0.1
0.05
0.01
0
Cell_Type_NEW
Ctrl
ExN
InN
Other
UD

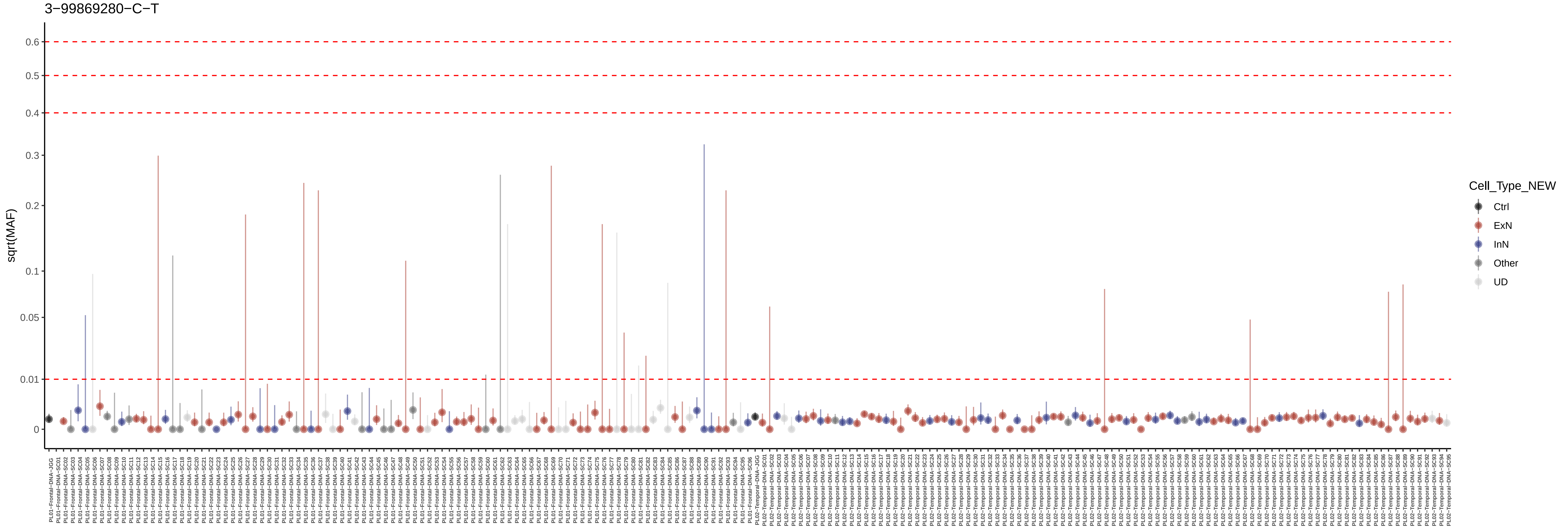

3−99869280−C−T
sqrt(MAF)
0.6
0.5
0.4
0.3
0.2
0.1
0.05
0.01
0
Cell_Type_NEW
Ctrl
ExN
InN
Other
UD

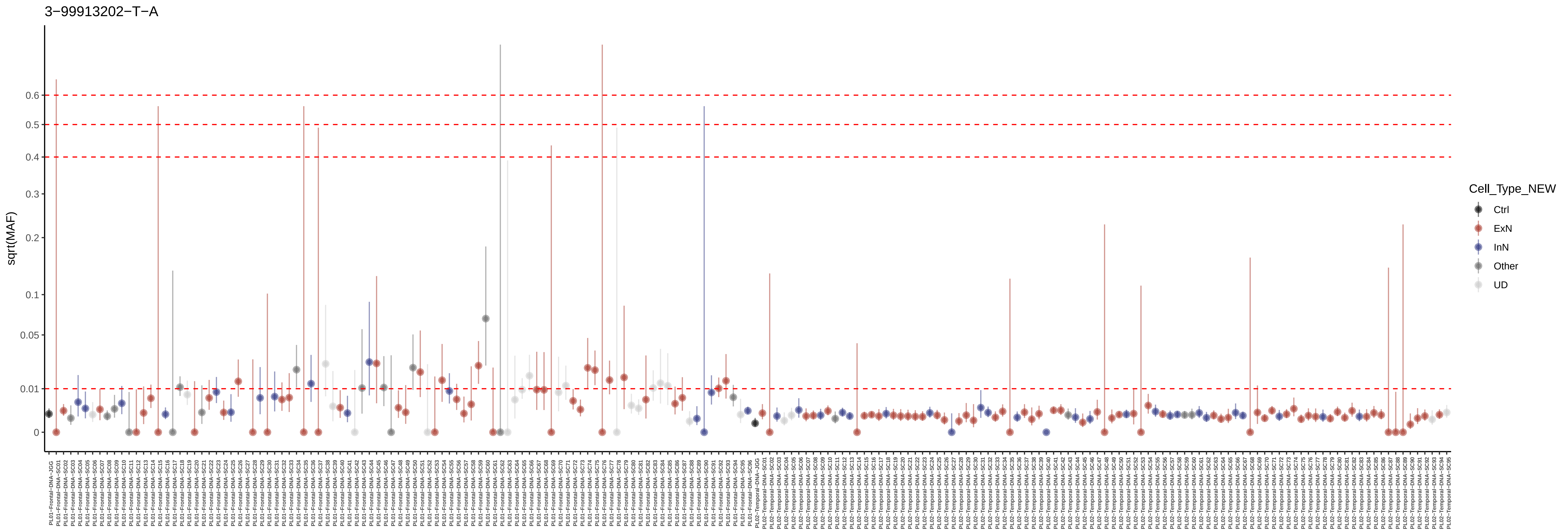

3−99913202−T−A
sqrt(MAF)
0.6
0.5
0.4
0.3
0.2
0.1
0.05
0.01
0
Cell_Type_NEW
Ctrl
ExN
InN
Other
UD

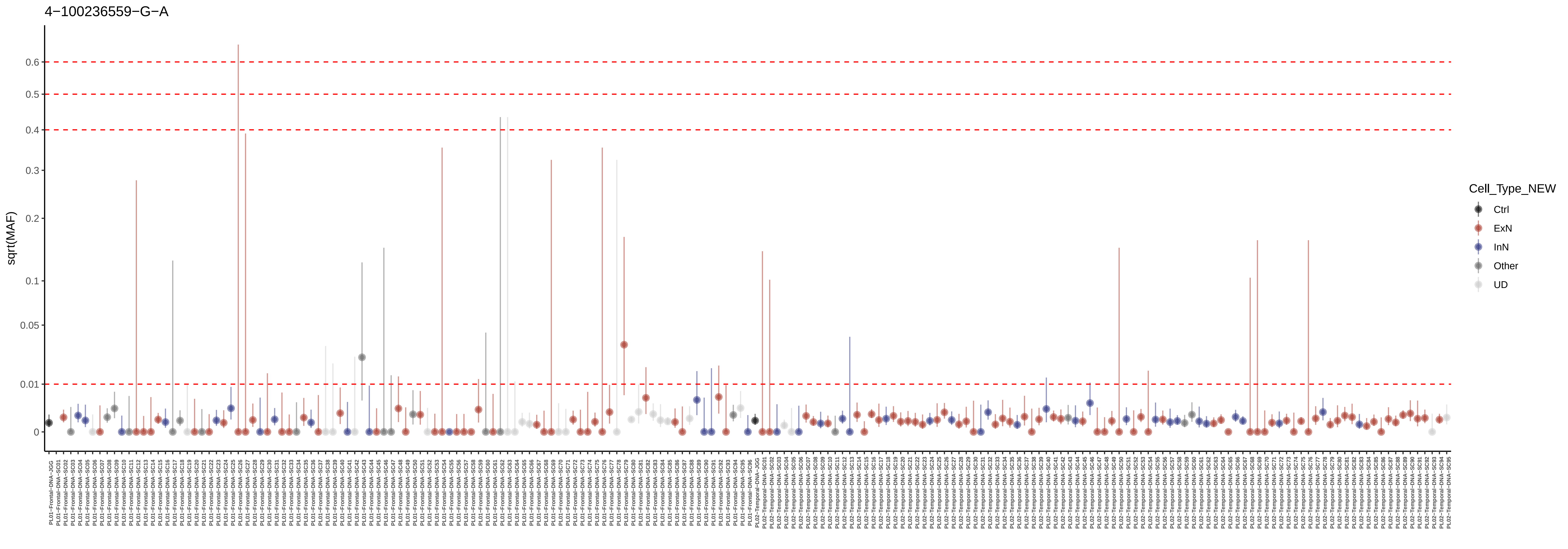
4−100236559−G−A
sqrt(MAF)
Cell_Type_NEW
Ctrl
ExN
InN
Other
UD

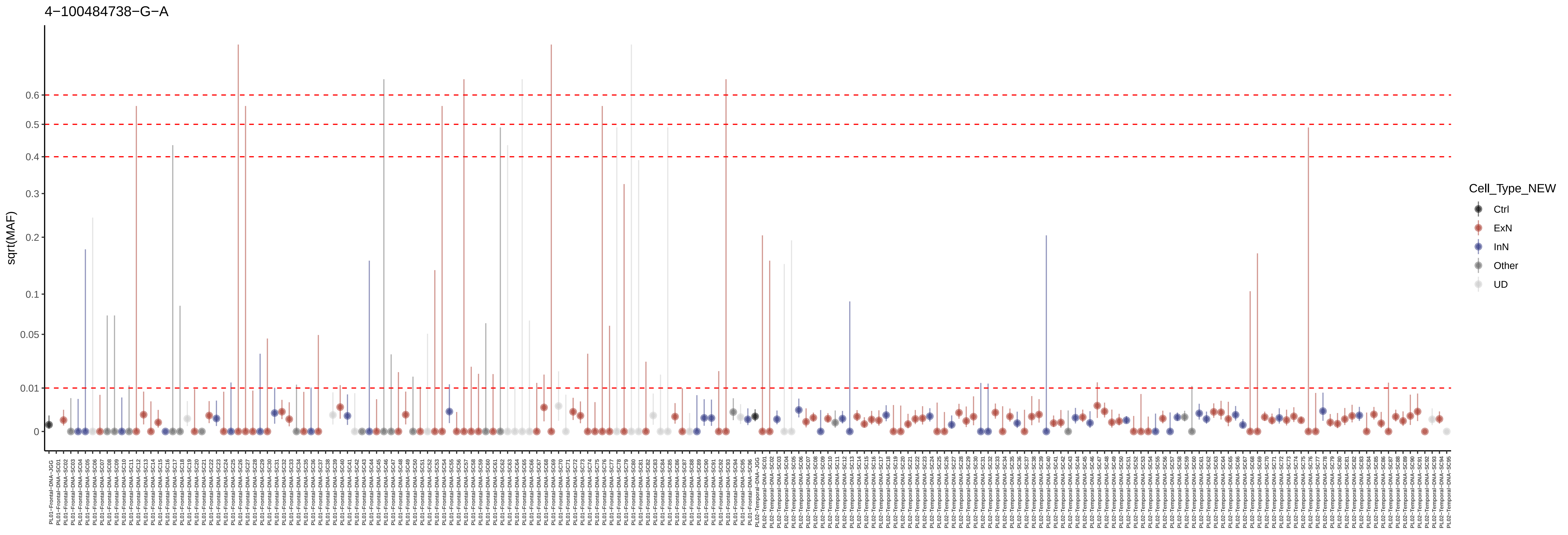
4−100484738−G−A
sqrt(MAF)
0.6
0.5
0.4
0.3
0.2
0.1
0.05
0.01
0
Cell_Type_NEW
Ctrl
ExN
InN
Other
UD

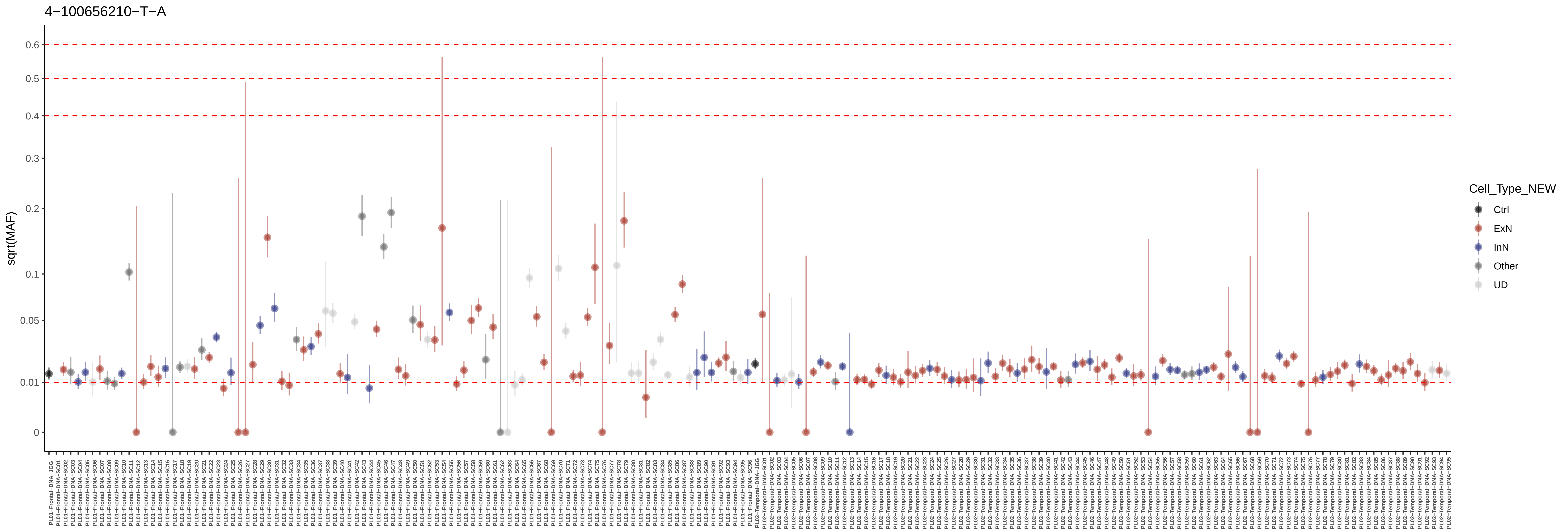

4−100656210−T−A
sqrt(MAF)
0.6
0.5
0.4
0.3
0.2
0.1
0.05
0.01
0
Cell_Type_NEW
Ctrl
ExN
InN
Other
UD

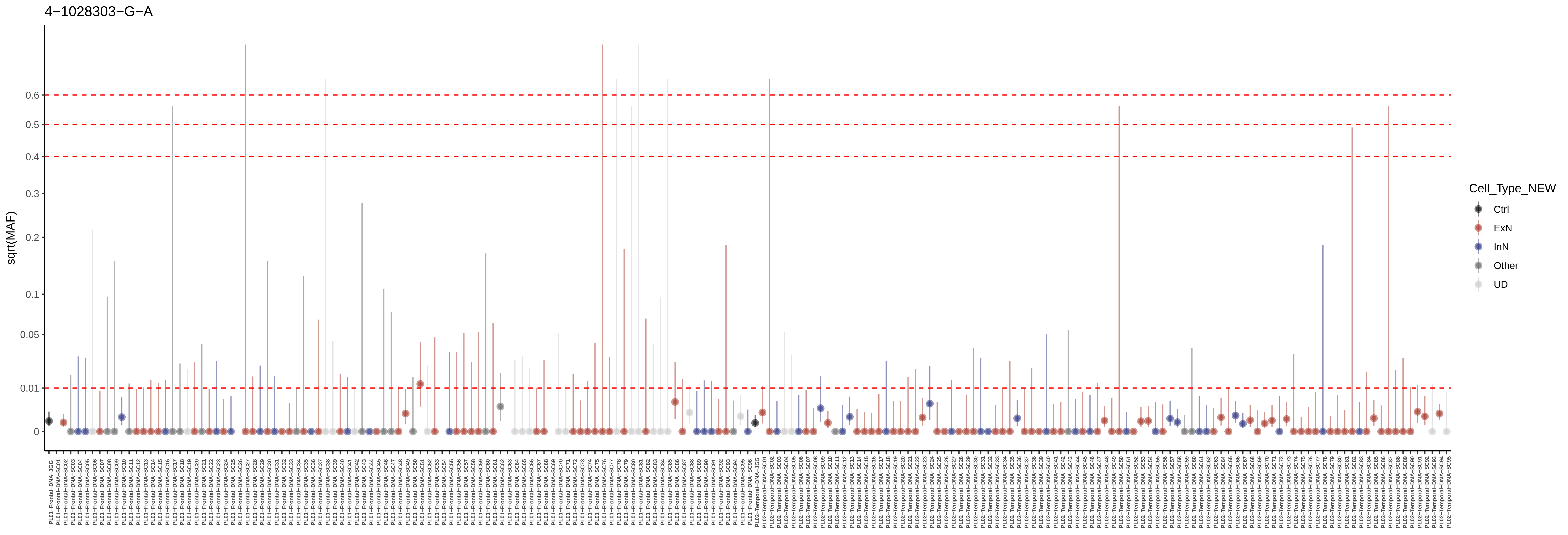

4−1028303−G−A
sqrt(MAF)
0.6
0.5
0.4
0.3
0.2
0.1
0.05
0.01
0
Cell_Type_NEW
Ctrl
ExN
InN
Other
UD

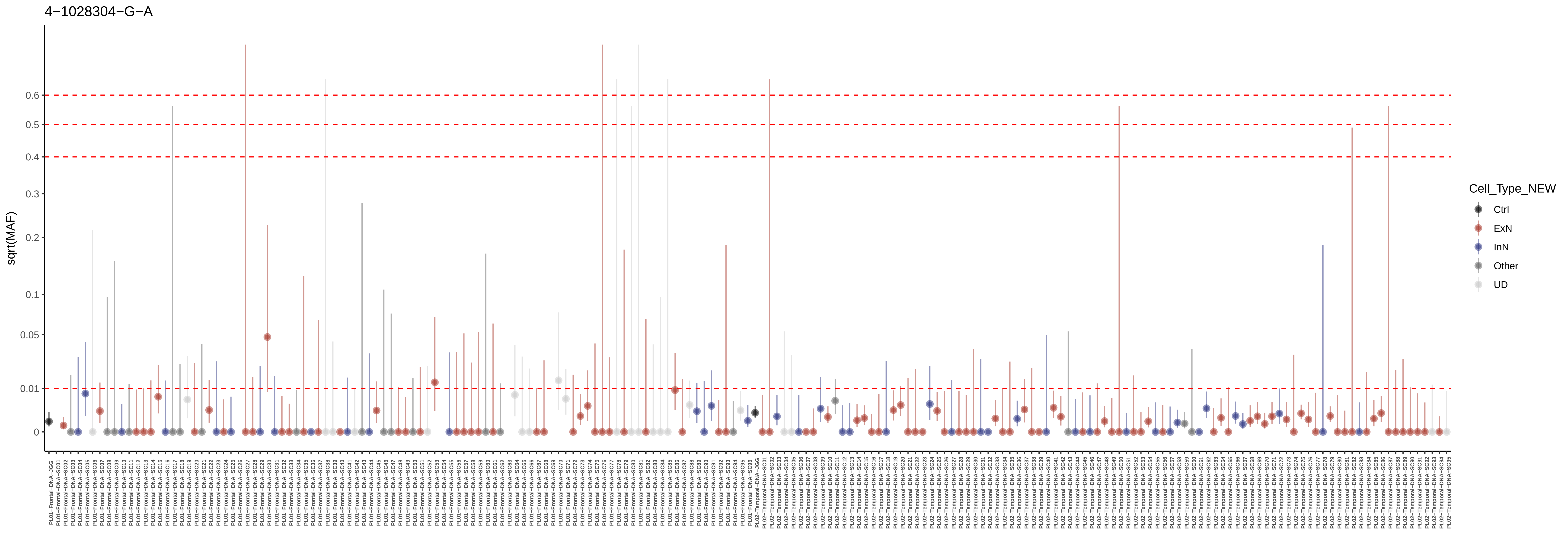
4−1028304−G−A
sqrt(MAF)
0.6
0.5
0.4
0.3
0.2
0.1
0.05
0.01
0
Cell_Type_NEW
Ctrl
ExN
InN
Other
UD

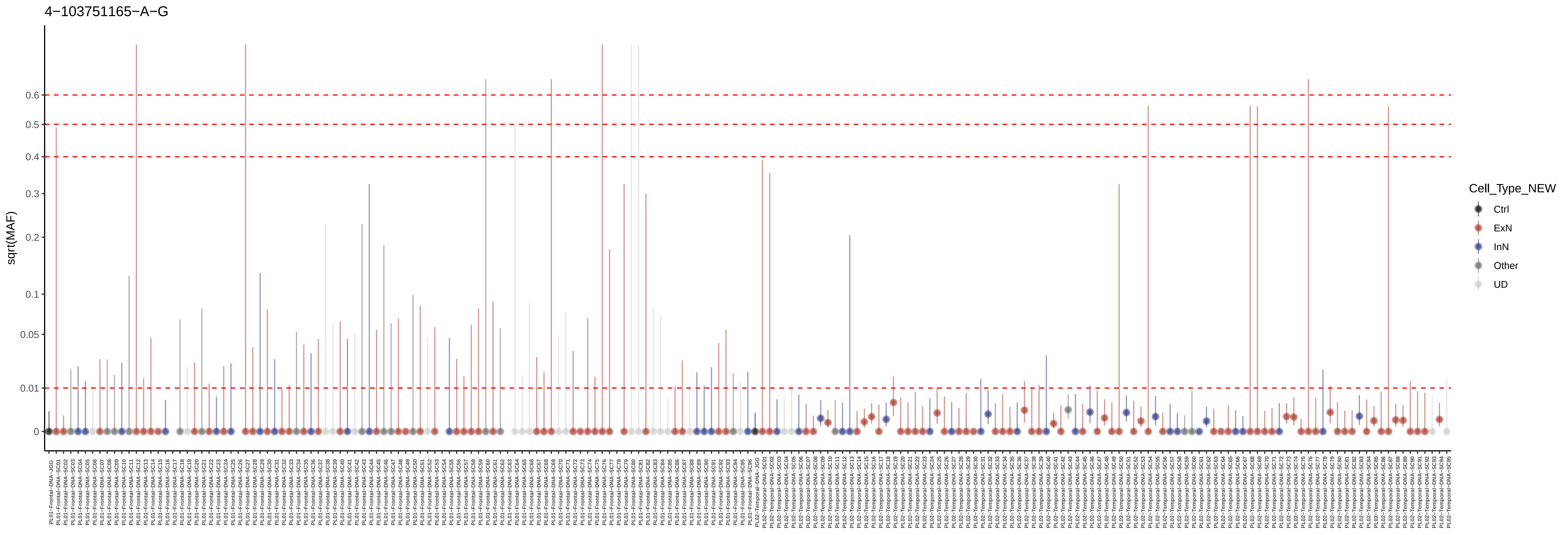

4−103751165−A−G
sqrt(MAF)
0.6
0.5
0.4
0.3
0.2
0.1
0.05
0.01
0
Cell_Type_NEW
Ctrl
ExN
InN
Other
UD

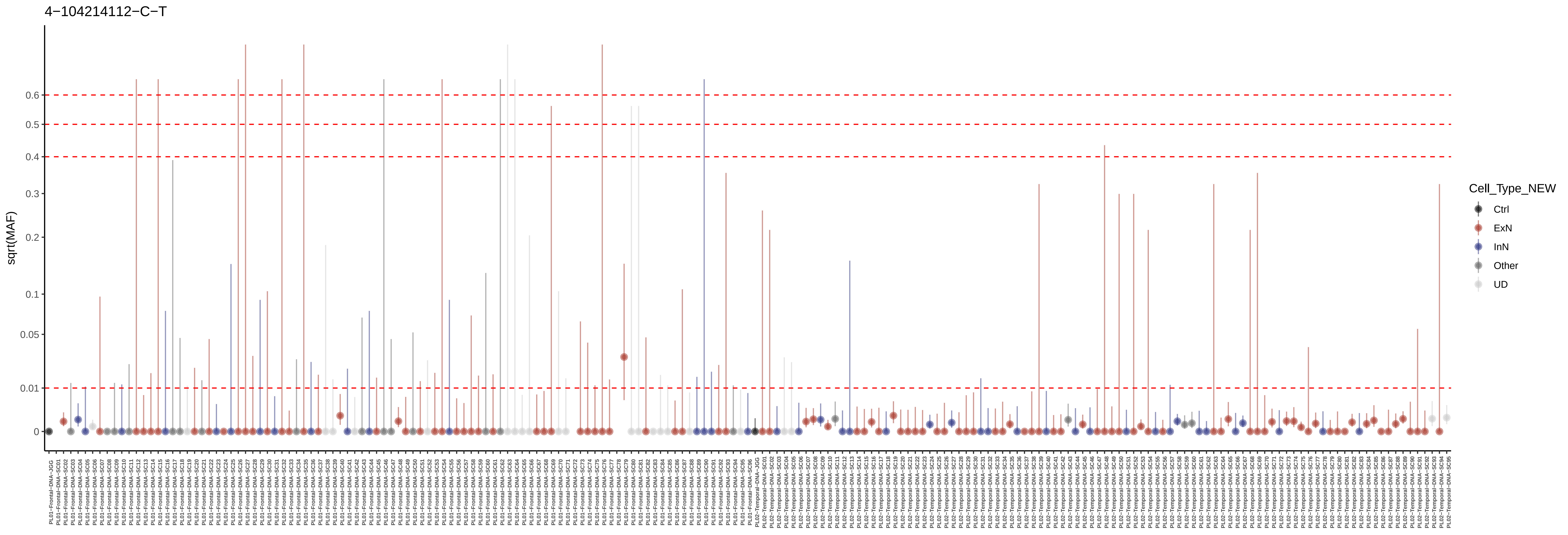
4−104214112−C−T
sqrt(MAF)
0.6
0.5
0.4
0.3
0.2
0.1
0.05
0.01
0
Cell_Type_NEW
Ctrl
ExN
InN
Other
UD

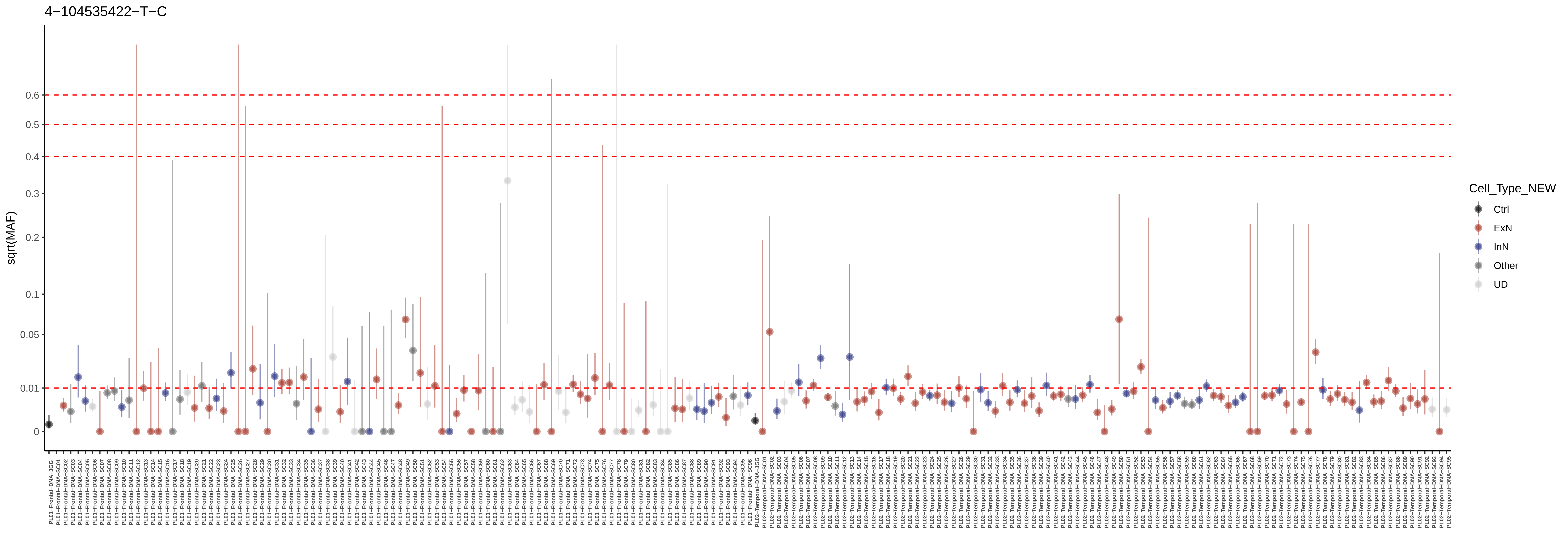

4−104535422−T−C
sqrt(MAF)
Cell_Type_NEW
Ctrl
ExN
InN
Other
UD

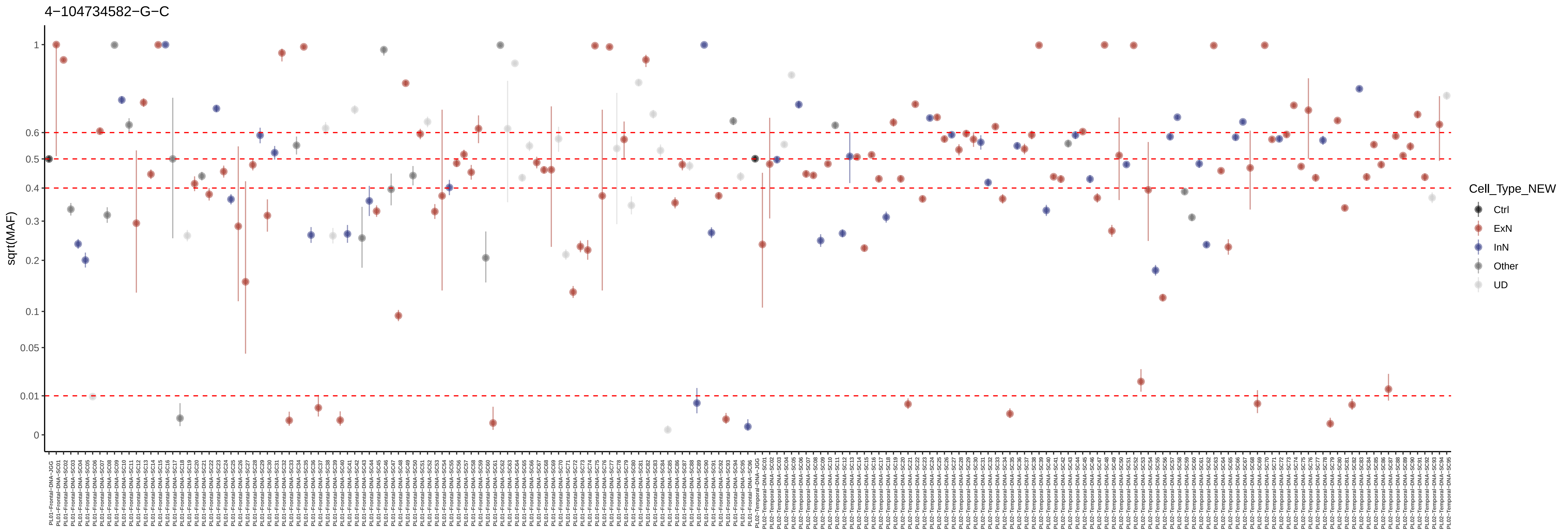

4−104734582−G−C
sqrt(MAF)
1
0.6
0.5
0.4
0.3
0.2
0.1
0.05
0.01
0
Cell_Type_NEW
Ctrl
ExN
InN
Other
UD

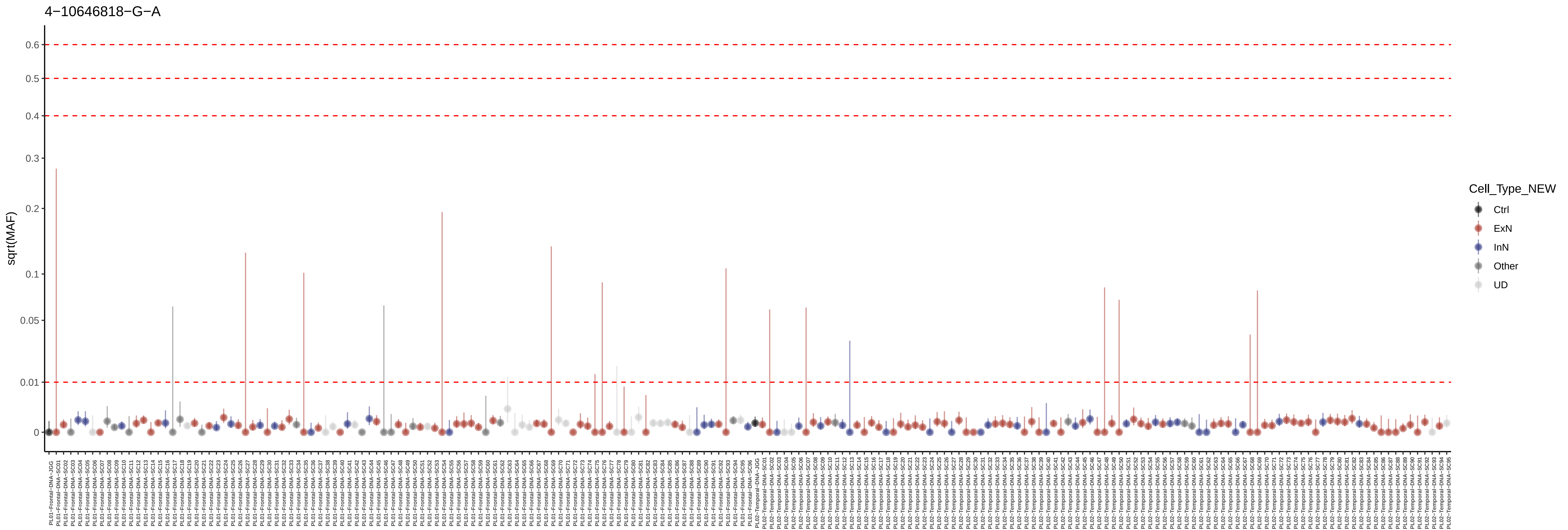

4−10646818−G−A
sqrt(MAF)
0.6
0.5
0.4
0.3
0.2
0.1
0.05
0.01
0
Cell_Type_NEW
Ctrl
ExN
InN
Other
UD

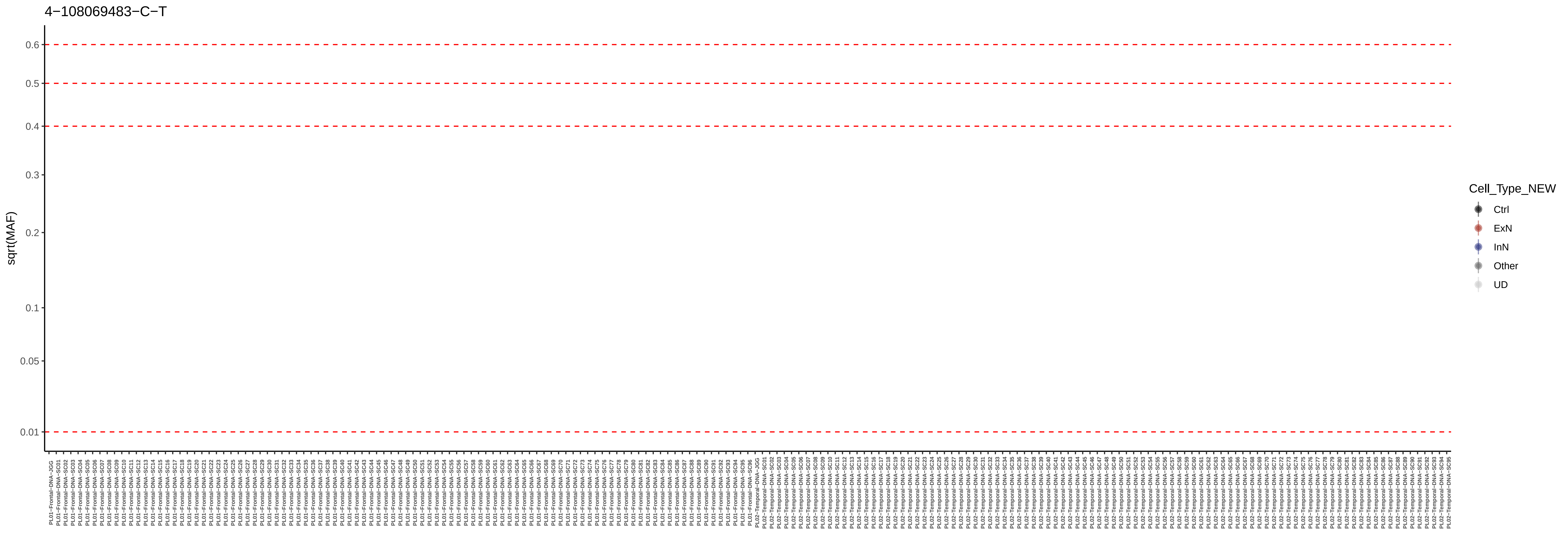

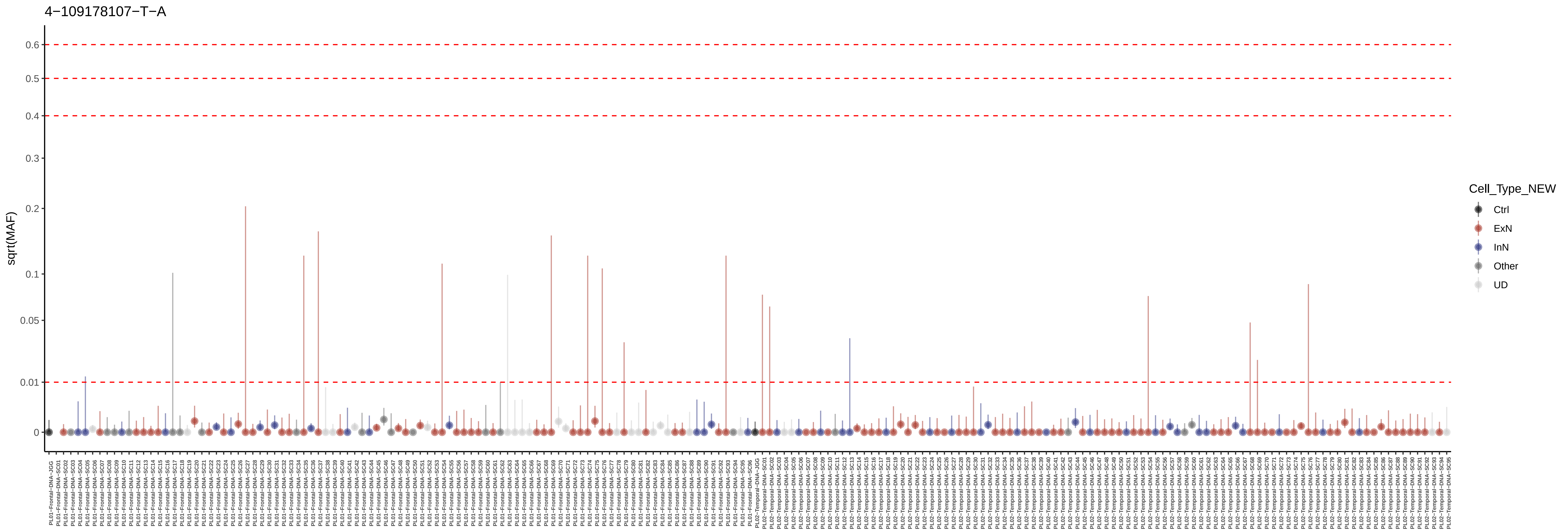
4−109178107−T−A
sqrt(MAF)
0.6
0.5
0.4
0.3
0.2
0.1
0.05
0.01
0
Cell_Type_NEW
Ctrl
ExN
InN
Other
UD

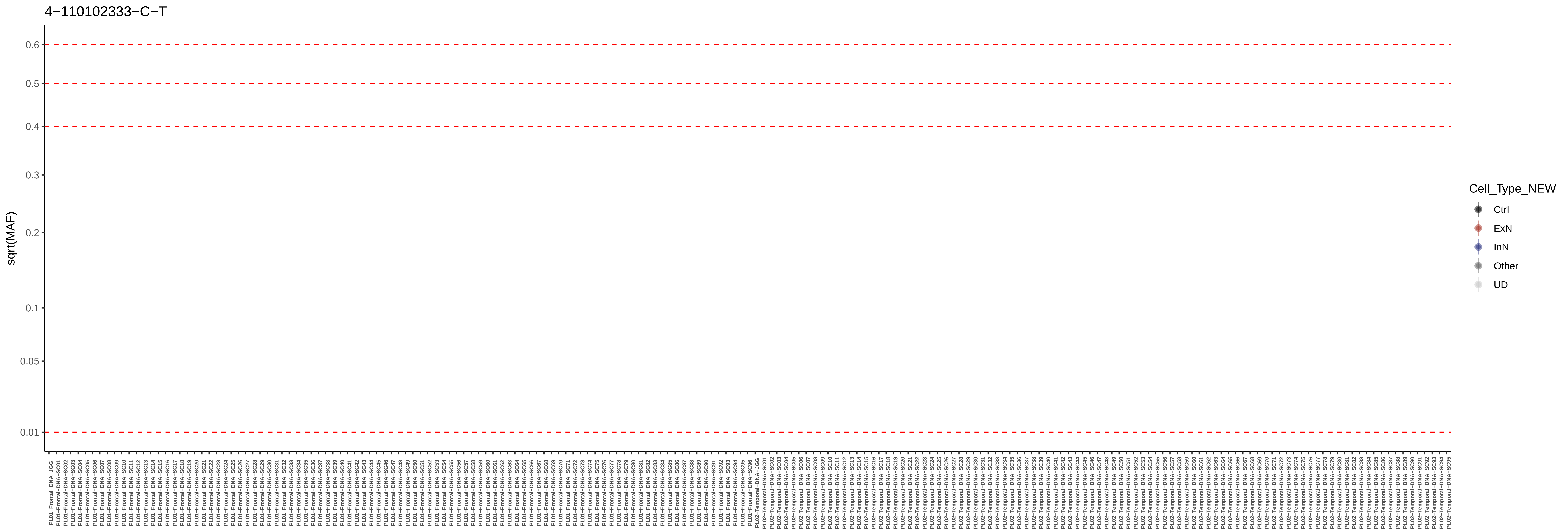

4−110102333−C−T
sqrt(MAF)
0.6
0.5
0.4
0.3
0.2
0.1
0.05
0.01
Cell_Type_NEW
Ctrl
ExN
InN
Other
UD

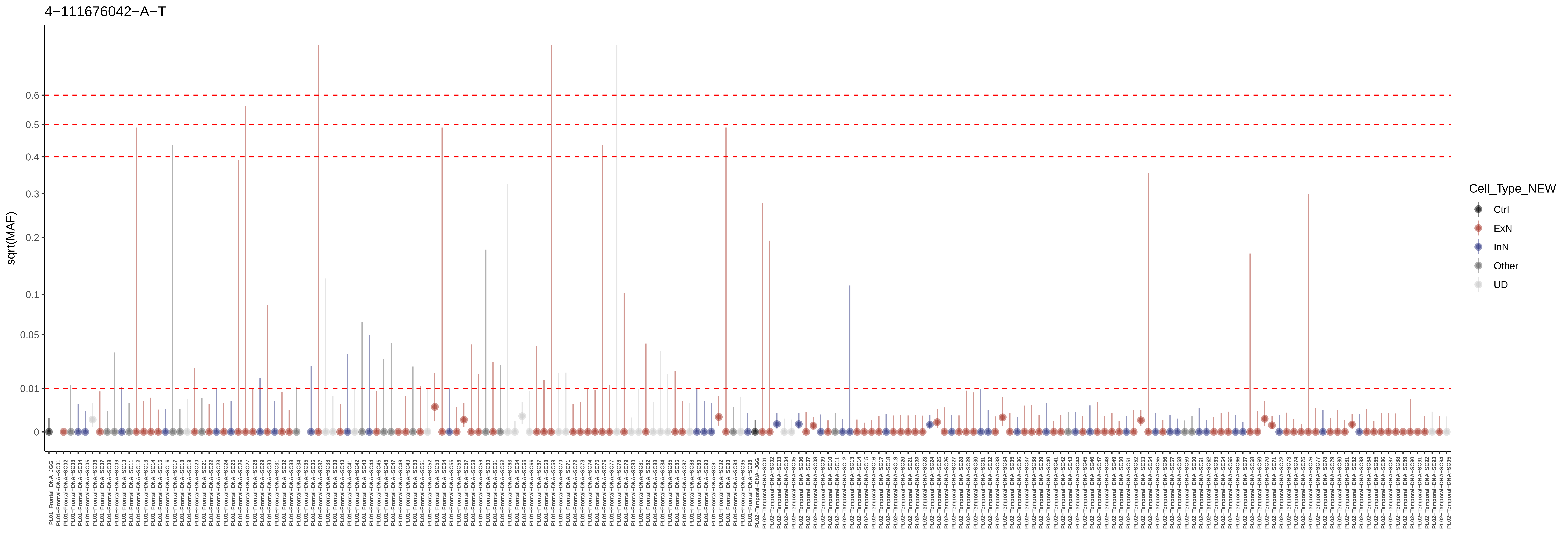
4−111676042−A−T
sqrt(MAF)
0.6
0.5
0.4
0.3
0.2
0.1
0.05
0.01
0
Cell_Type_NEW
Ctrl
ExN
InN
Other
UD

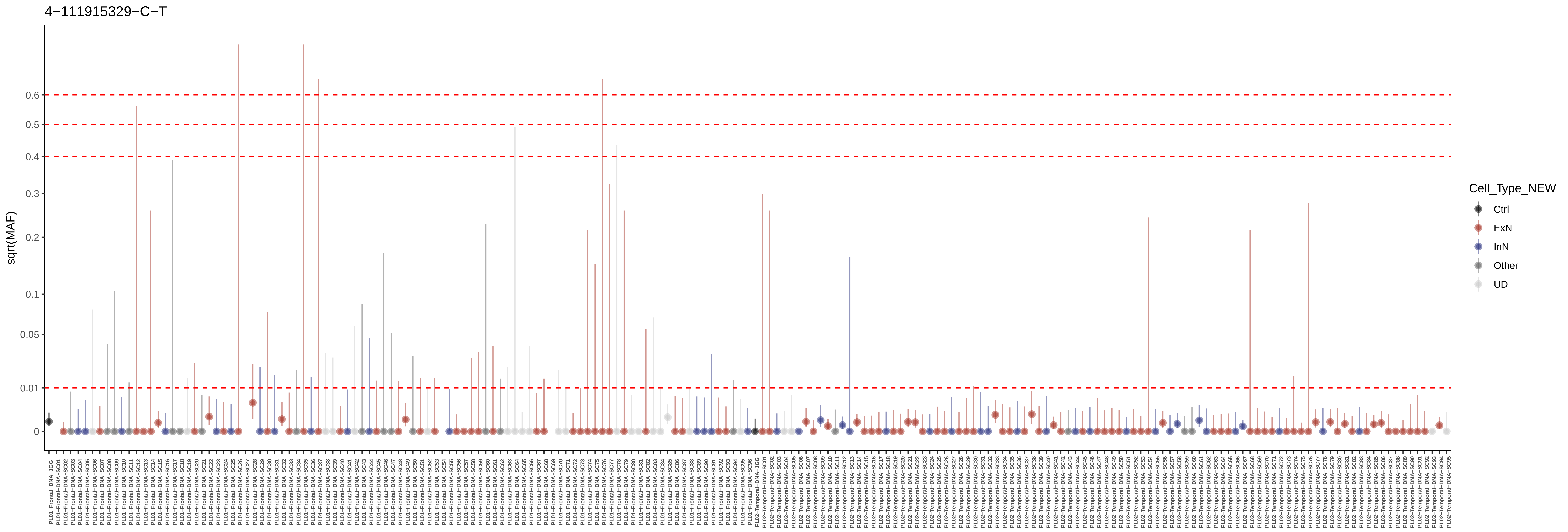
4−111915329−C−T
sqrt(MAF)
0.6
0.5
0.4
0.3
0.2
0.1
0.05
0.01
0
Cell_Type_NEW
Ctrl
ExN
InN
Other
UD

4−112266667−A−G

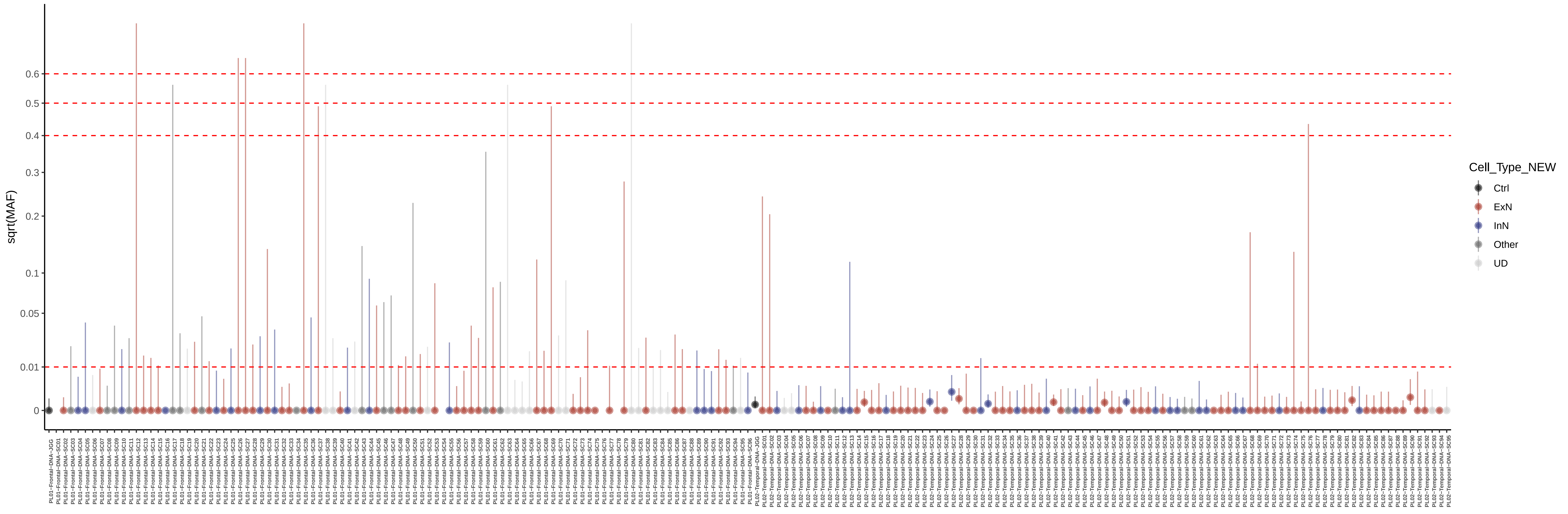

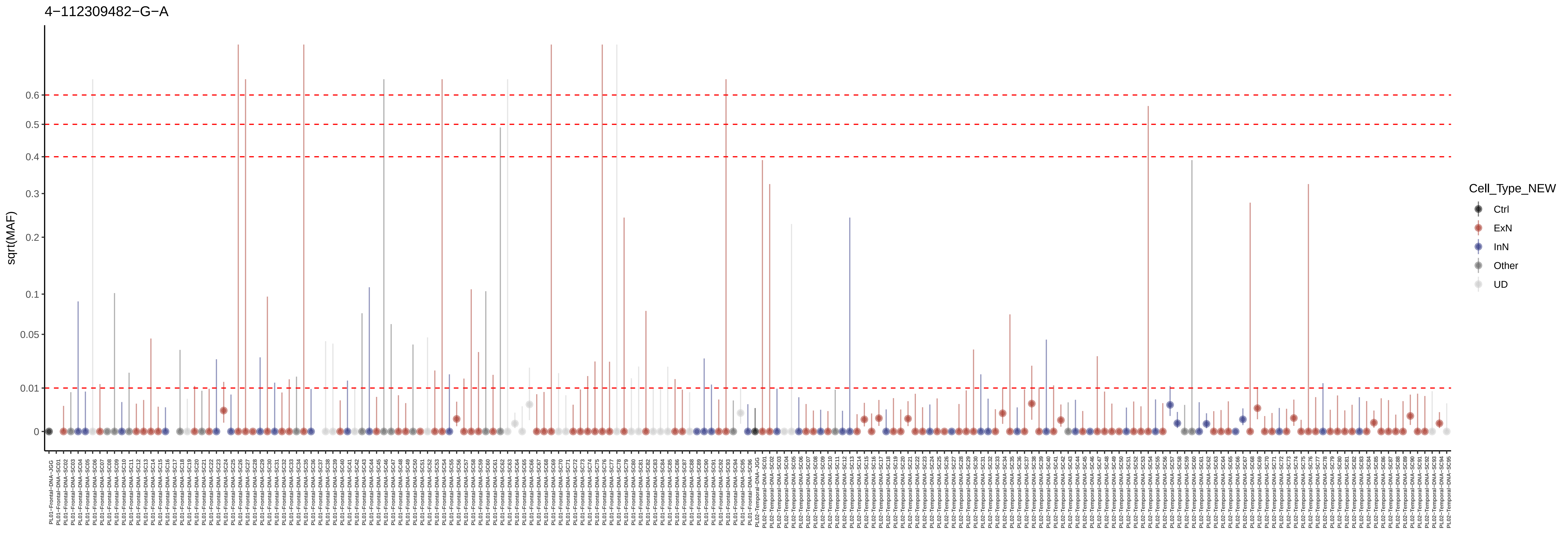

4−112309482−G−A
sqrt(MAF)
0.6
0.5
0.4
0.3
0.2
0.1
0.05
0.01
0
Cell_Type_NEW
Ctrl
ExN
InN
Other
UD

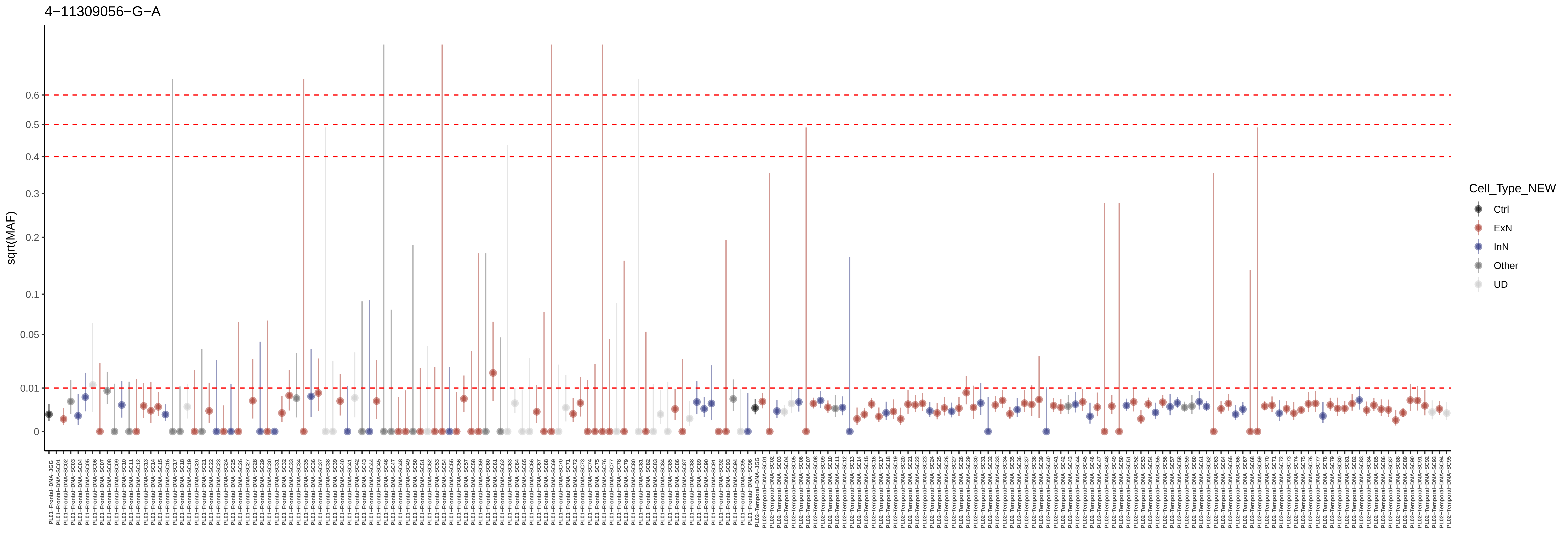

4−11309056−G−A
sqrt(MAF)
0.6
0.5
0.4
0.3
0.2
0.1
0.05
0.01
0
Cell_Type_NEW
Ctrl
ExN
InN
Other
UD

4−116228416−A−G

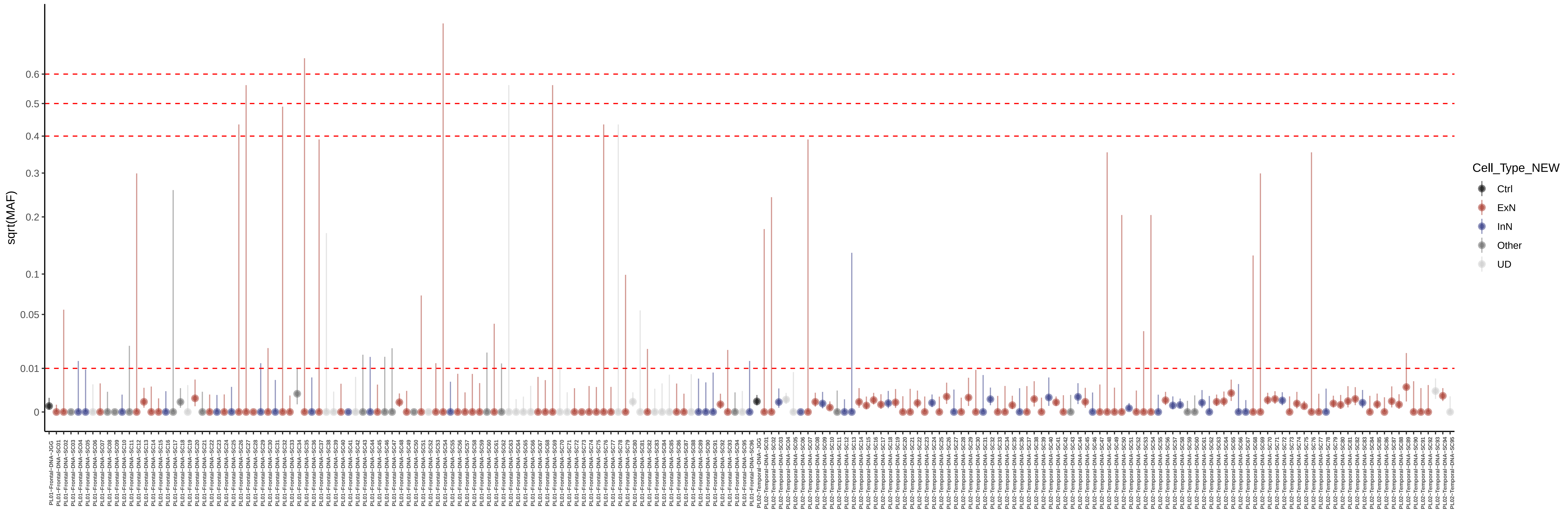

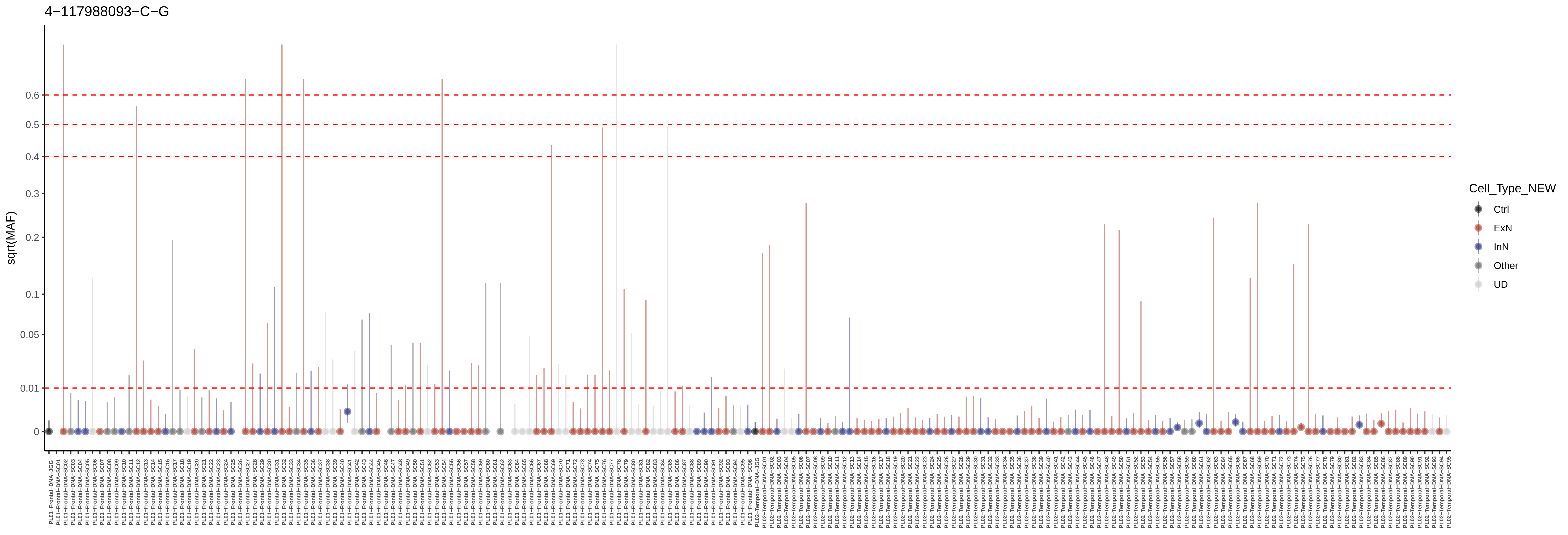
4−117988093−C−G
sqrt(MAF)
0.6
0.5
0.4
0.3
0.2
0.1
0.05
0.01
0
Cell_Type_NEW
Ctrl
ExN
InN
Other
UD

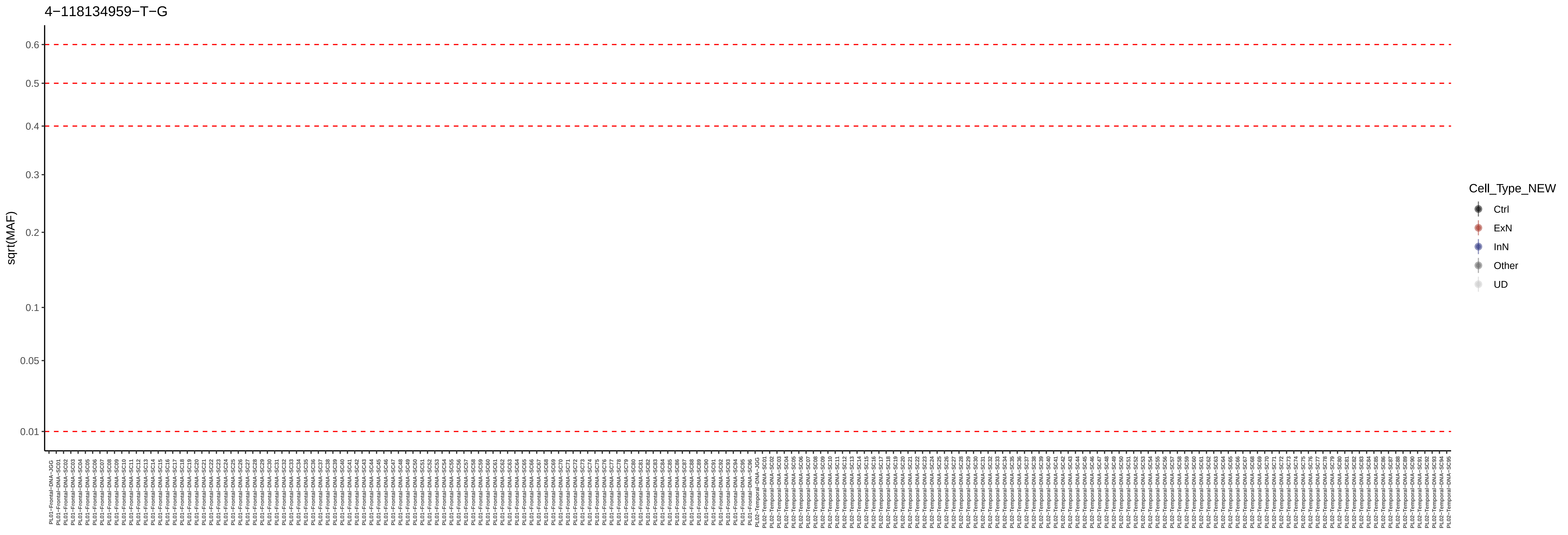
4−118134959−T−G
sqrt(MAF)
0.6
0.5
0.4
0.3
0.2
0.1
0.05
0.01
Cell_Type_NEW
Ctrl
ExN
InN
Other
UD

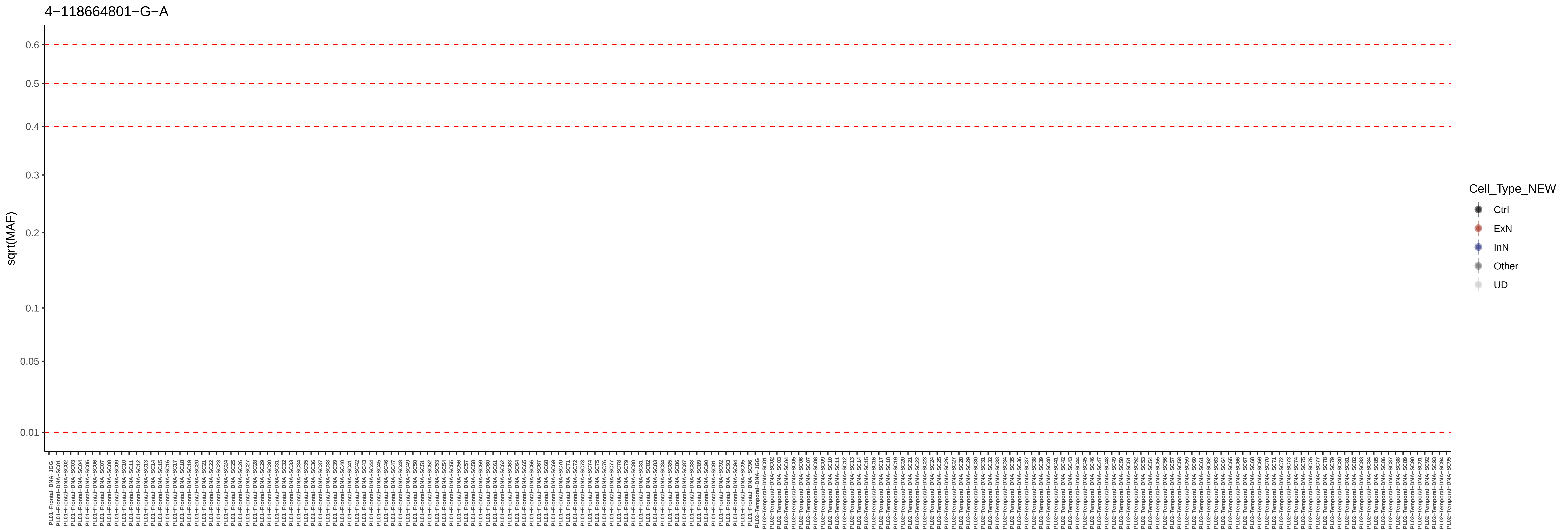

4−118664801−G−A
sqrt(MAF)
0.6
0.5
0.4
0.3
0.2
0.1
0.05
0.01
Cell_Type_NEW
Ctrl
ExN
InN
Other
UD

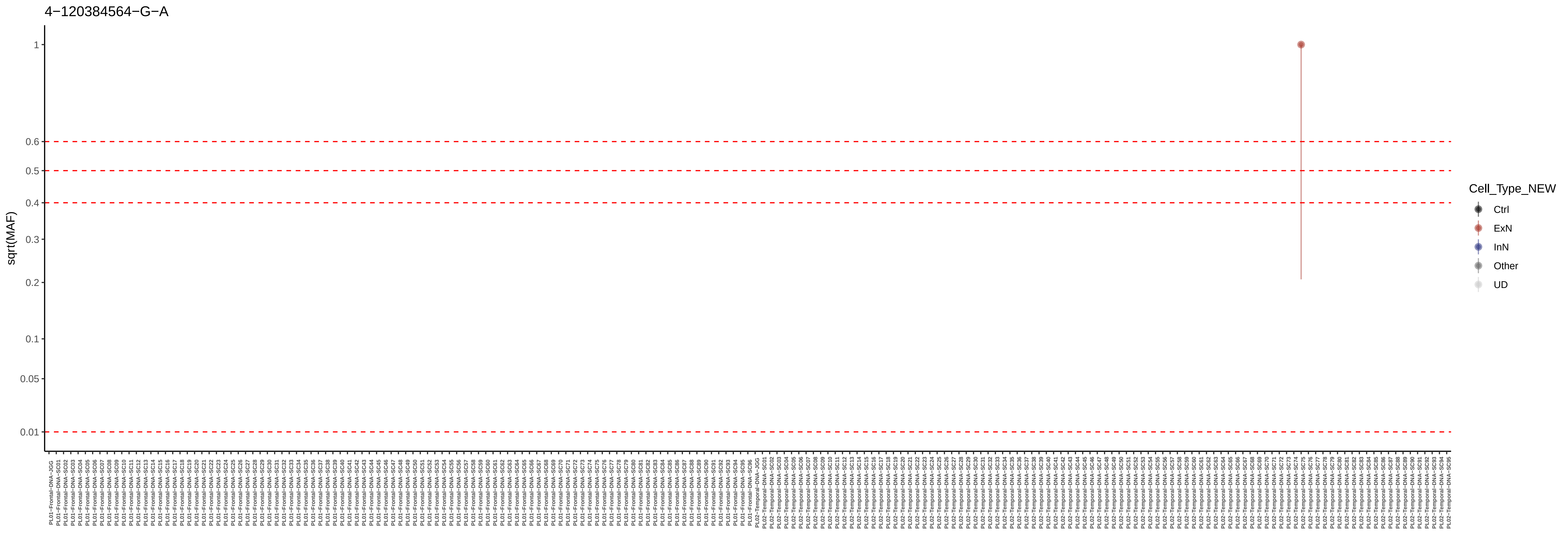
4−120384564−G−A
sqrt(MAF)
1
0.6
0.5
0.4
0.3
0.2
0.1
0.05
0.01
Cell_Type_NEW
Ctrl
ExN
InN
Other
UD

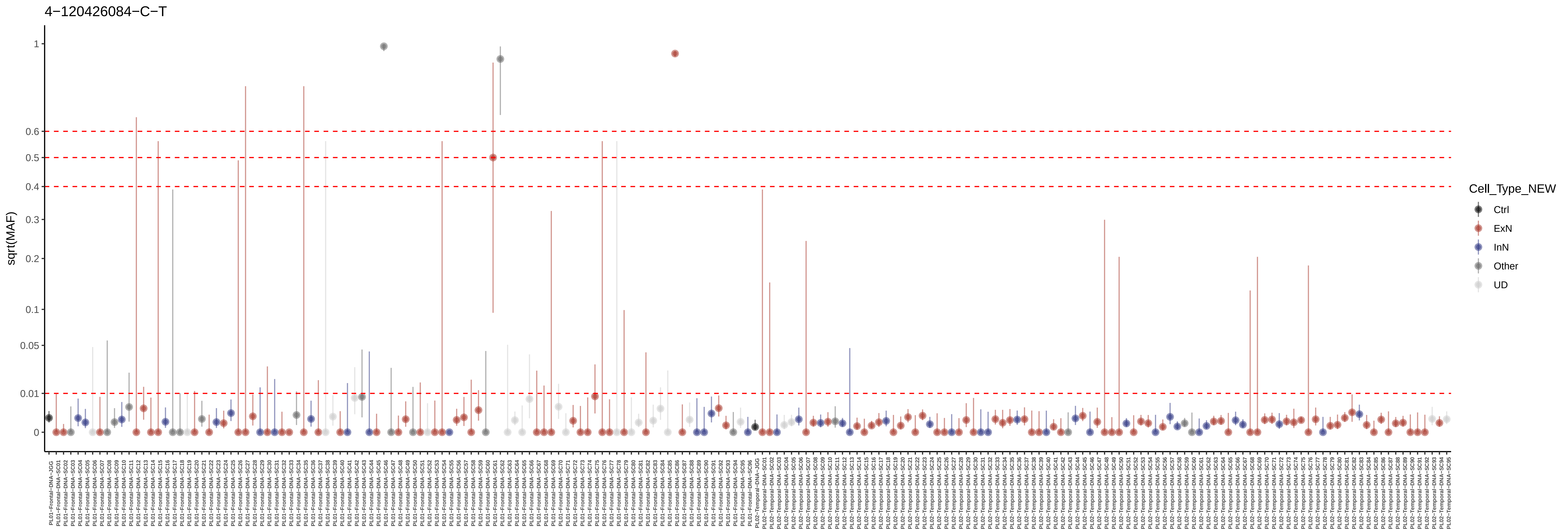
4−120426084−C−T
sqrt(MAF)
1
0.6
0.5
0.4
0.3
0.2
0.1
0.05
0.01
0
Cell_Type_NEW
Ctrl
ExN
InN
Other
UD

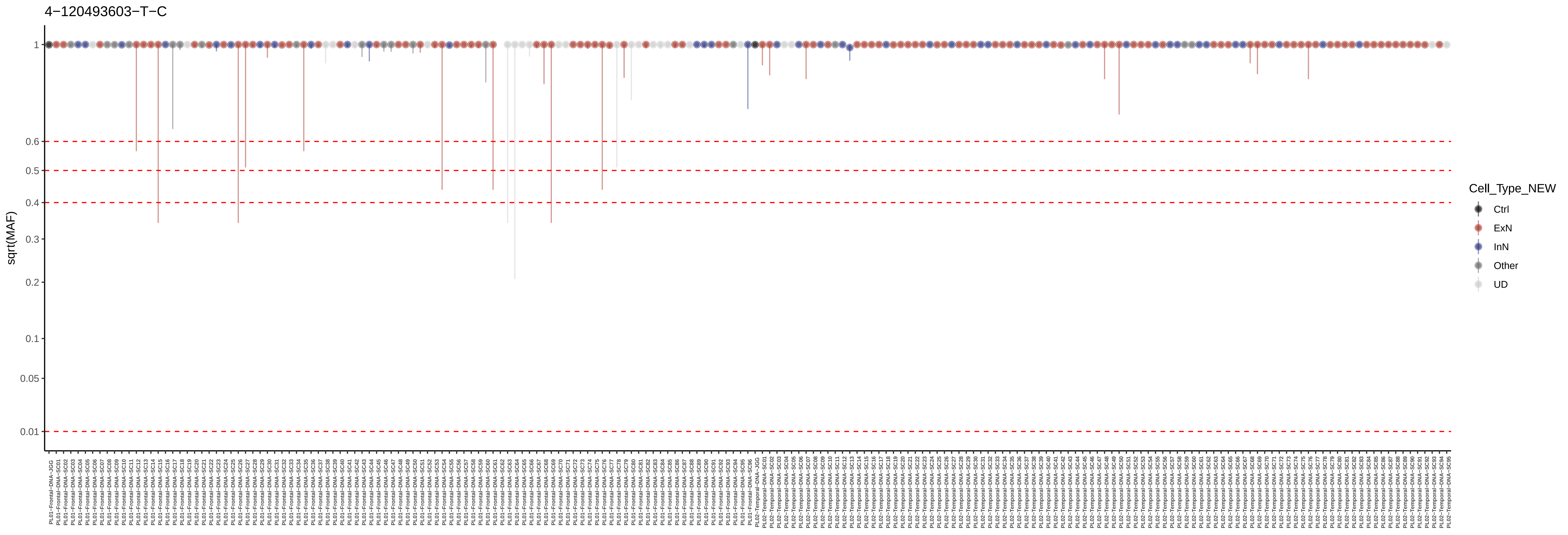
4−120493603−T−C
sqrt(MAF)
1
0.6
0.5
0.4
0.3
0.2
0.1
0.05
0.01
Cell_Type_NEW
Ctrl
ExN
InN
Other
UD

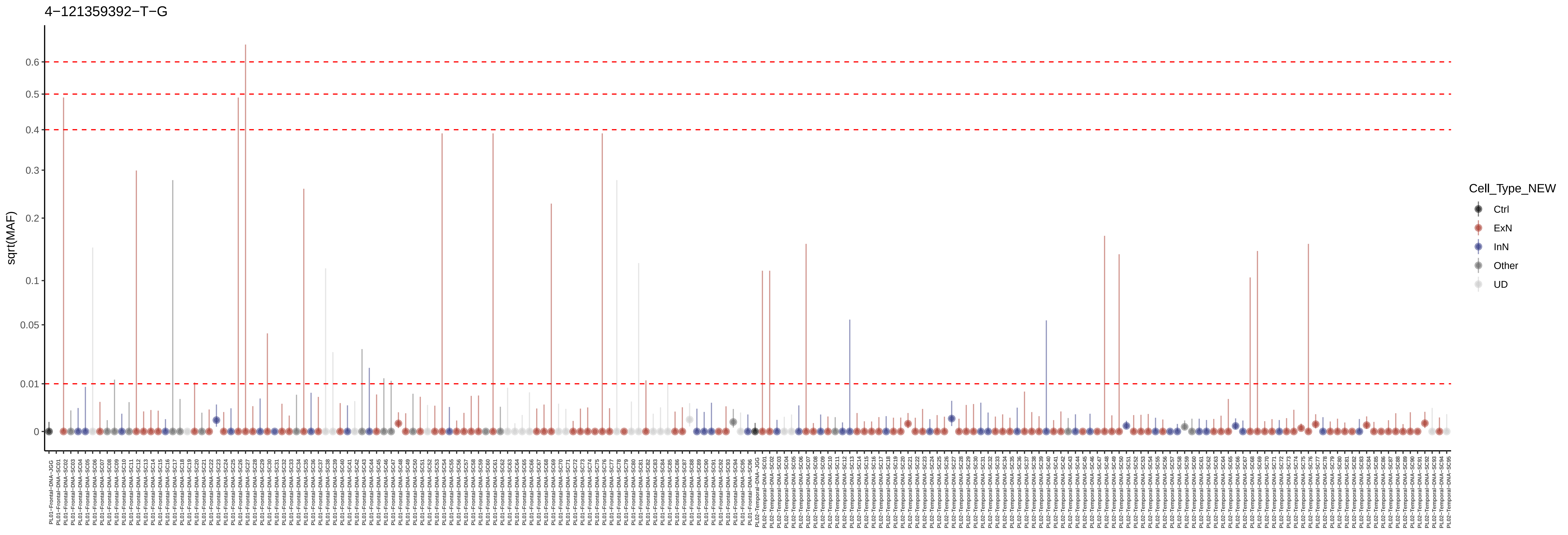

4−121359392−T−G
sqrt(MAF)
0.6
0.5
0.4
0.3
0.2
0.1
0.05
0.01
0
Cell_Type_NEW
Ctrl
ExN
InN
Other
UD

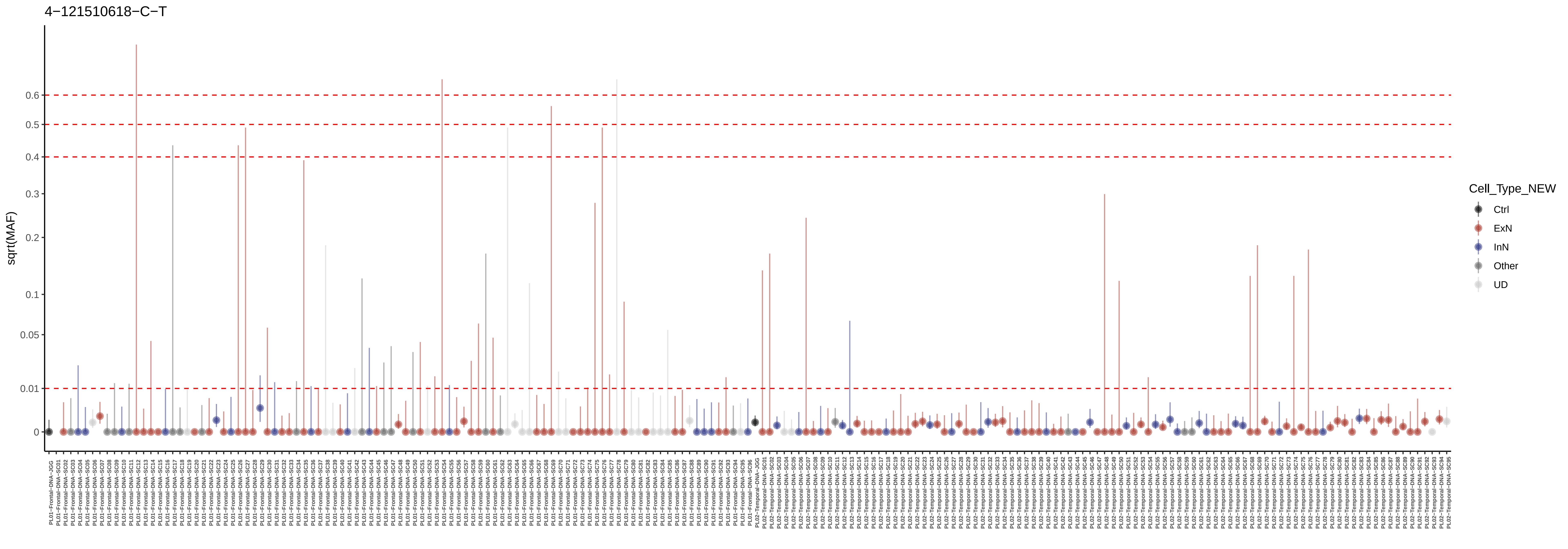
4−121510618−C−T
sqrt(MAF)
0.6
0.5
0.4
0.3
0.2
0.1
0.05
0.01
0
Cell_Type_NEW
Ctrl
ExN
InN
Other
UD

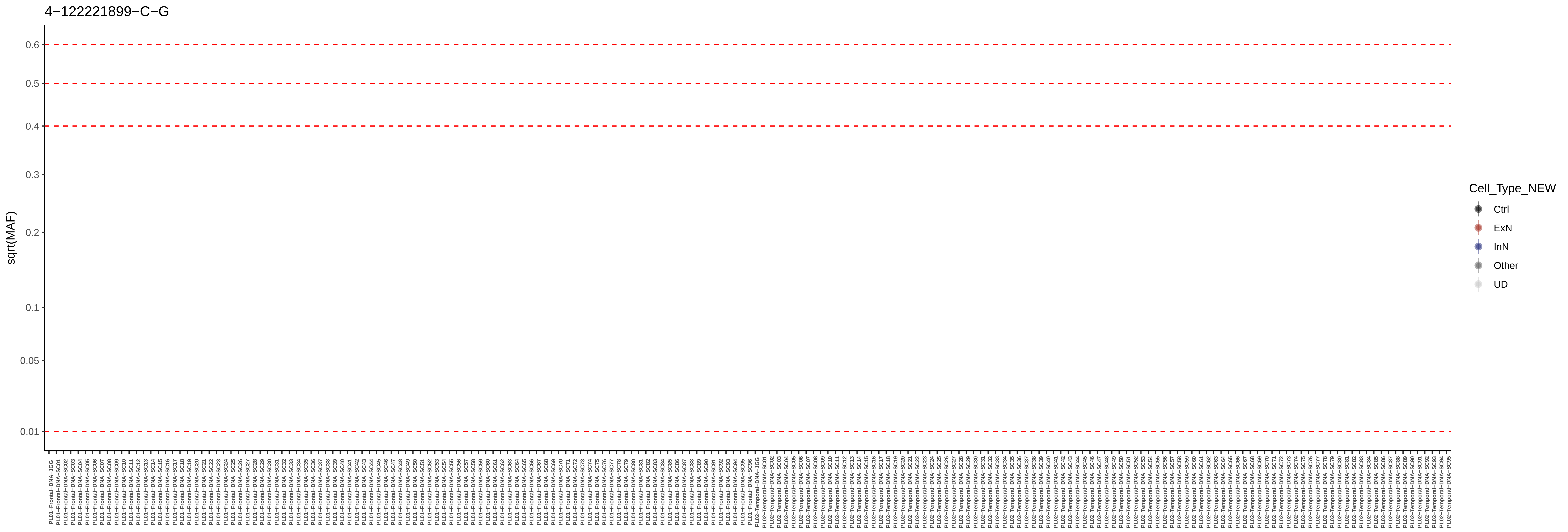
4−122221899−C−G
sqrt(MAF)
0.6
0.5
0.4
0.3
0.2
0.1
0.05
0.01
Cell_Type_NEW
Ctrl
ExN
InN
Other
UD

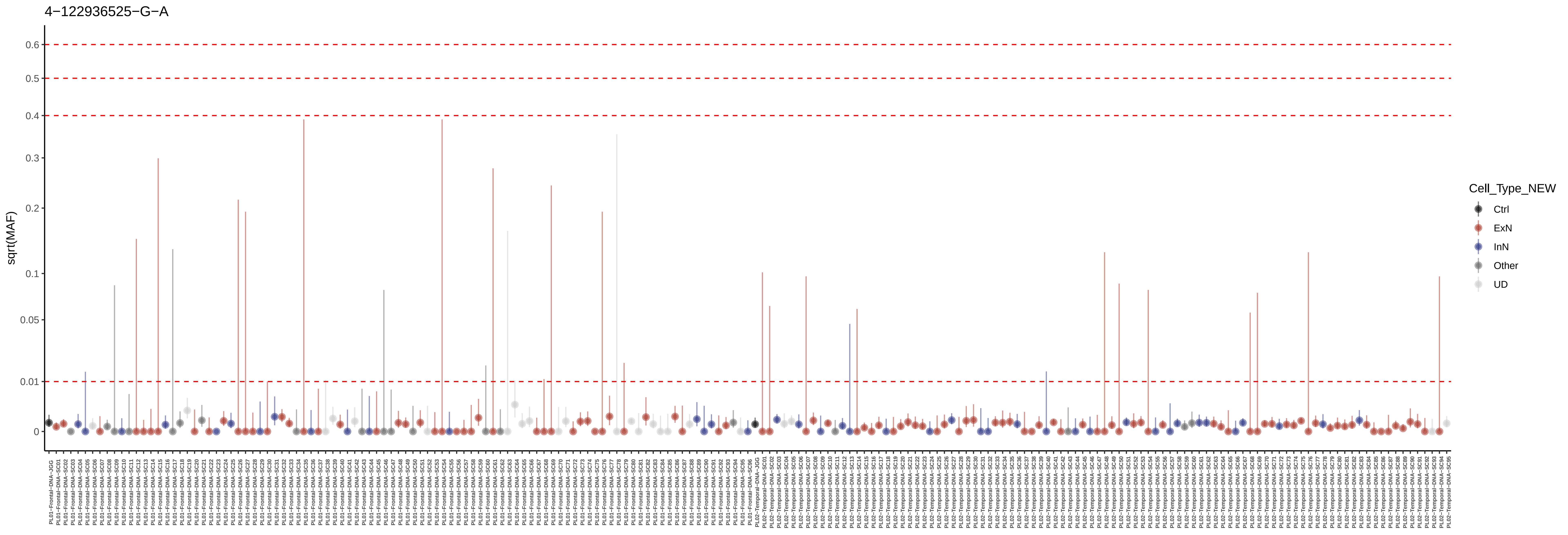

4−122936525−G−A
sqrt(MAF)
0.6
0.5
0.4
0.3
0.2
0.1
0.05
0.01
0
Cell_Type_NEW
Ctrl
ExN
InN
Other
UD

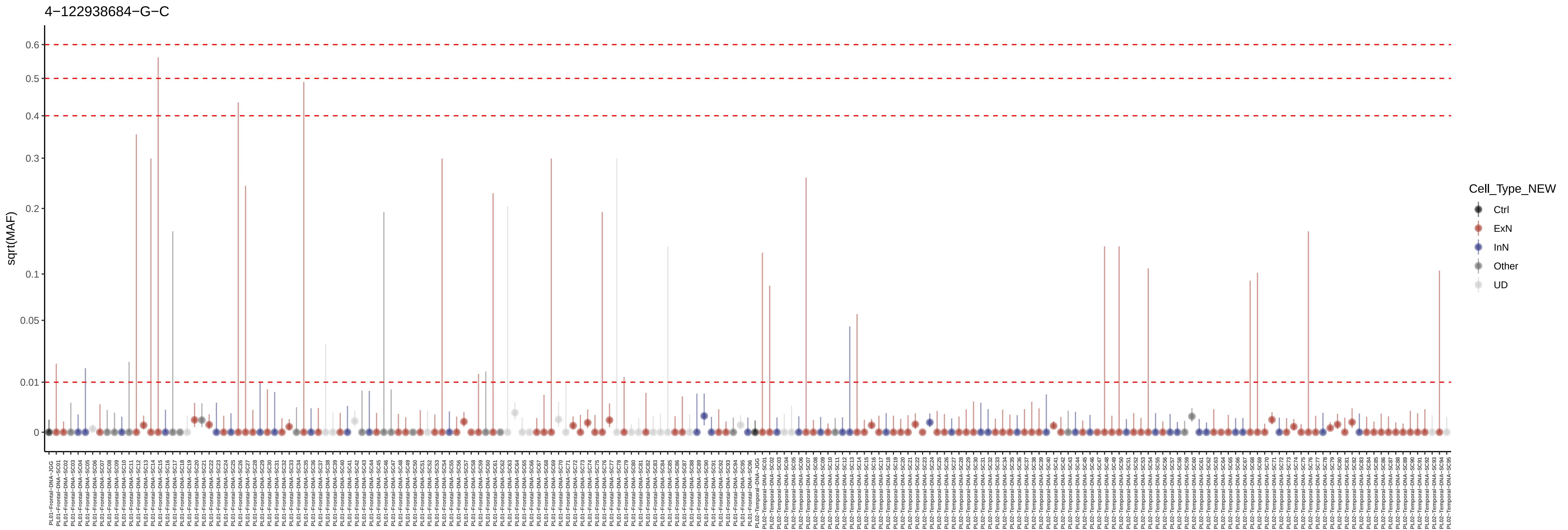
4−122938684−G−C
sqrt(MAF)
0.6
0.5
0.4
0.3
0.2
0.1
0.05
0.01
0
Cell_Type_NEW
Ctrl
ExN
InN
Other
UD

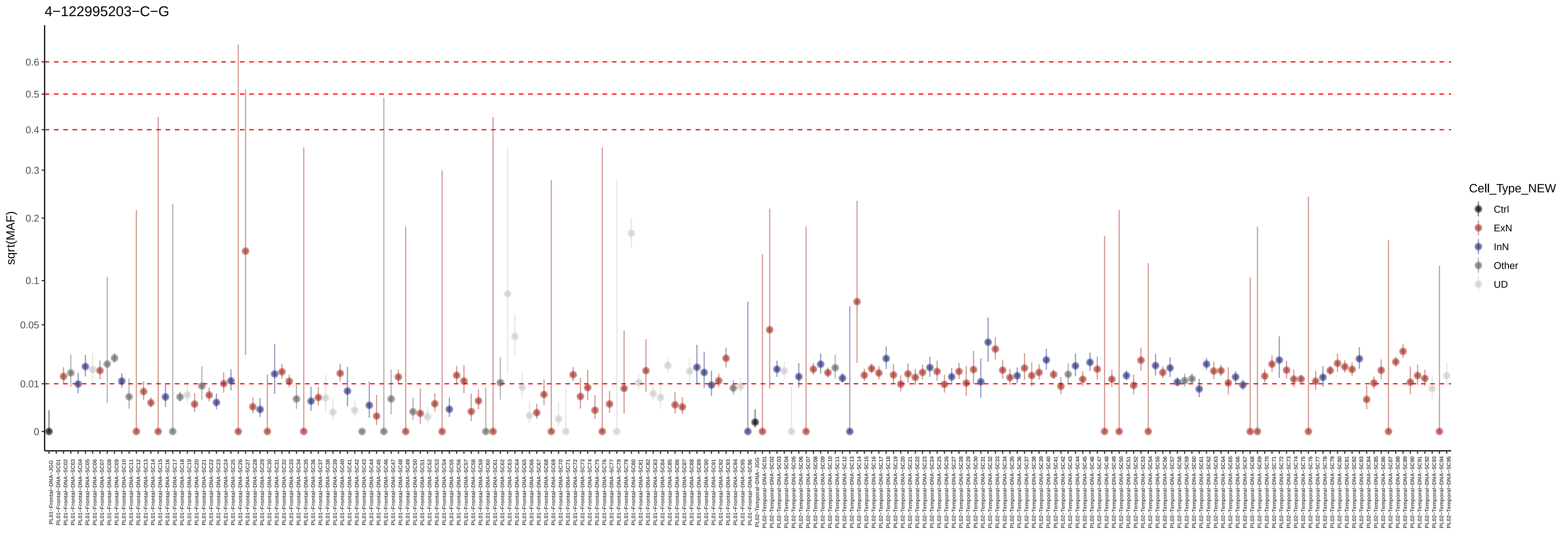

4−122995203−C−G
sqrt(MAF)
0.6
0.5
0.4
0.3
0.2
0.1
0.05
0.01
0
Cell_Type_NEW
Ctrl
ExN
InN
Other
UD

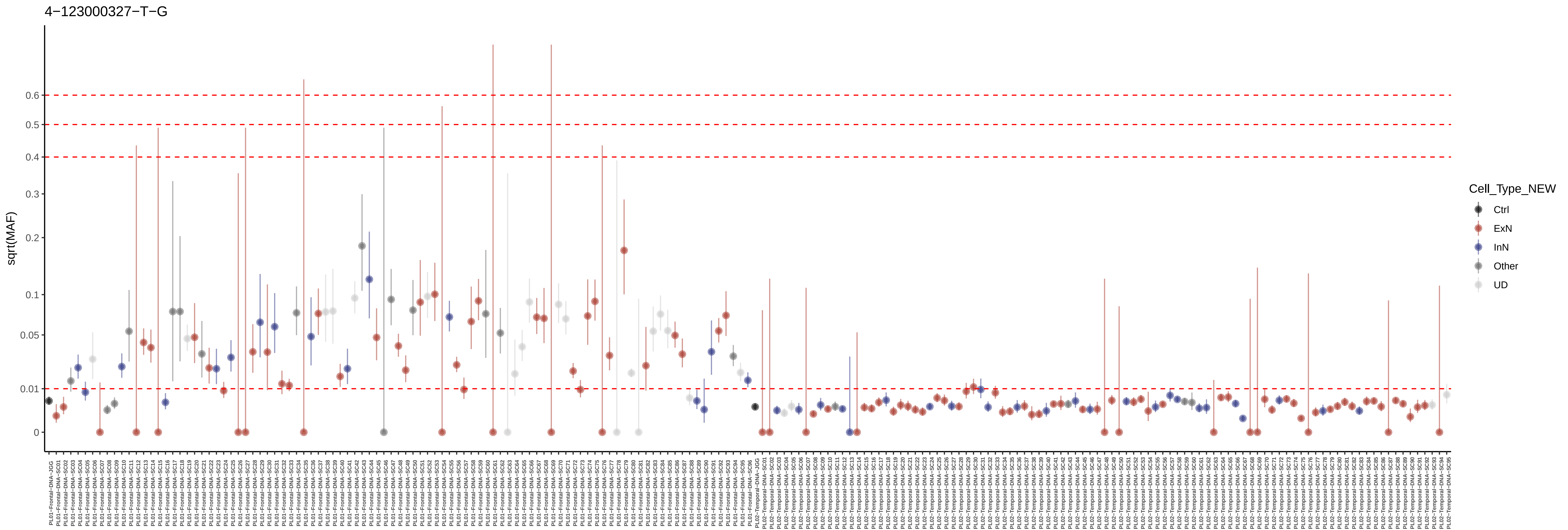
4−123000327−T−G
sqrt(MAF)
0.6
0.5
0.4
0.3
0.2
0.1
0.05
0.01
0
Cell_Type_NEW
Ctrl
ExN
InN
Other
UD

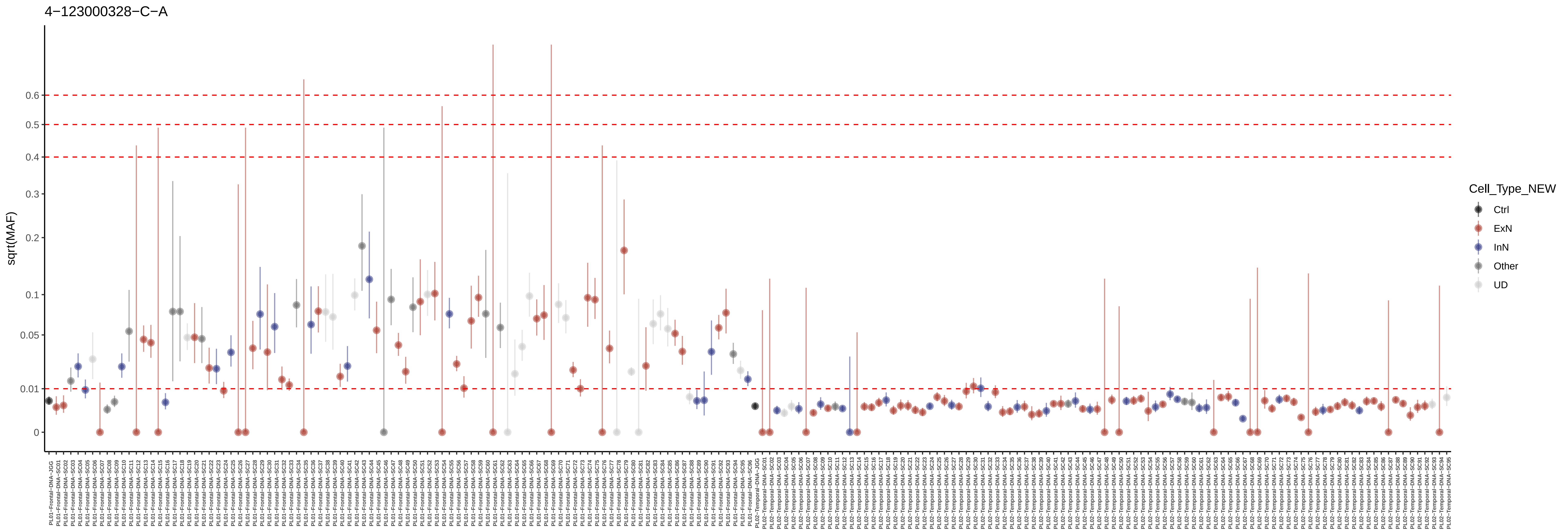

4−123000328−C−A
sqrt(MAF)
0.6
0.5
0.4
0.3
0.2
0.1
0.05
0.01
0
Cell_Type_NEW
Ctrl
ExN
InN
Other
UD

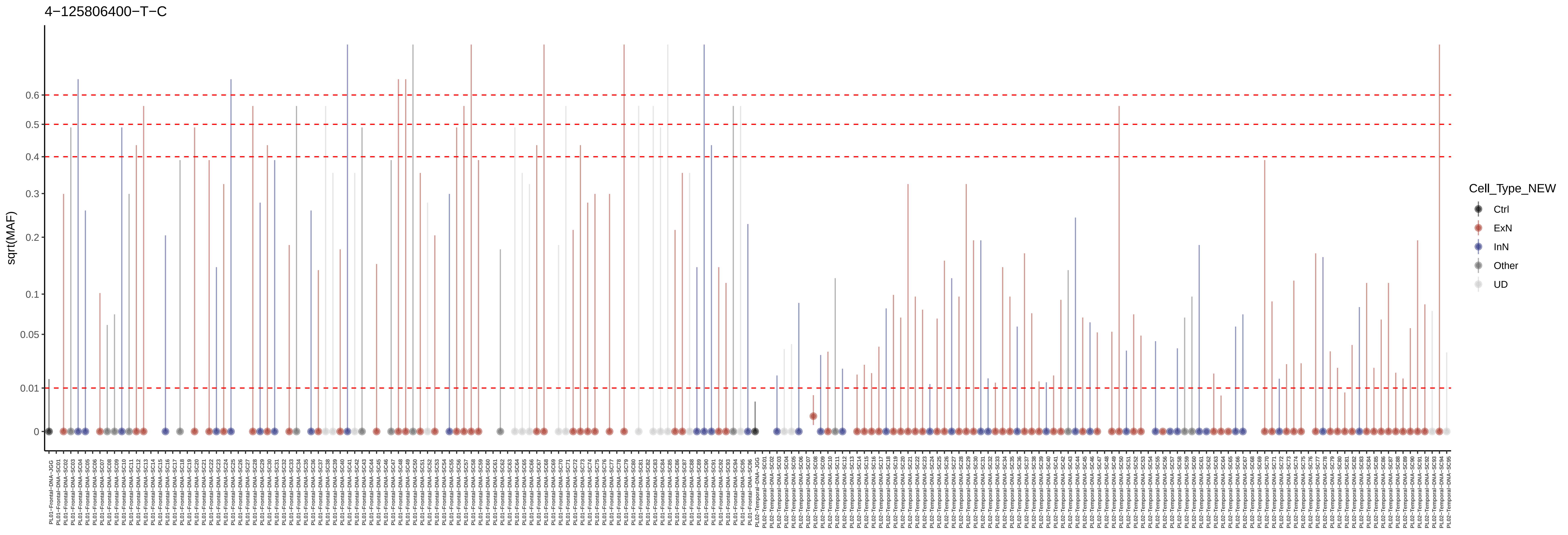

4−125806400−T−C
sqrt(MAF)
0
0.01
0.05
0.1
0.2
0.3
0.4
0.5
0.6
Cell_Type_NEW
Ctrl
ExN
InN
Other
UD

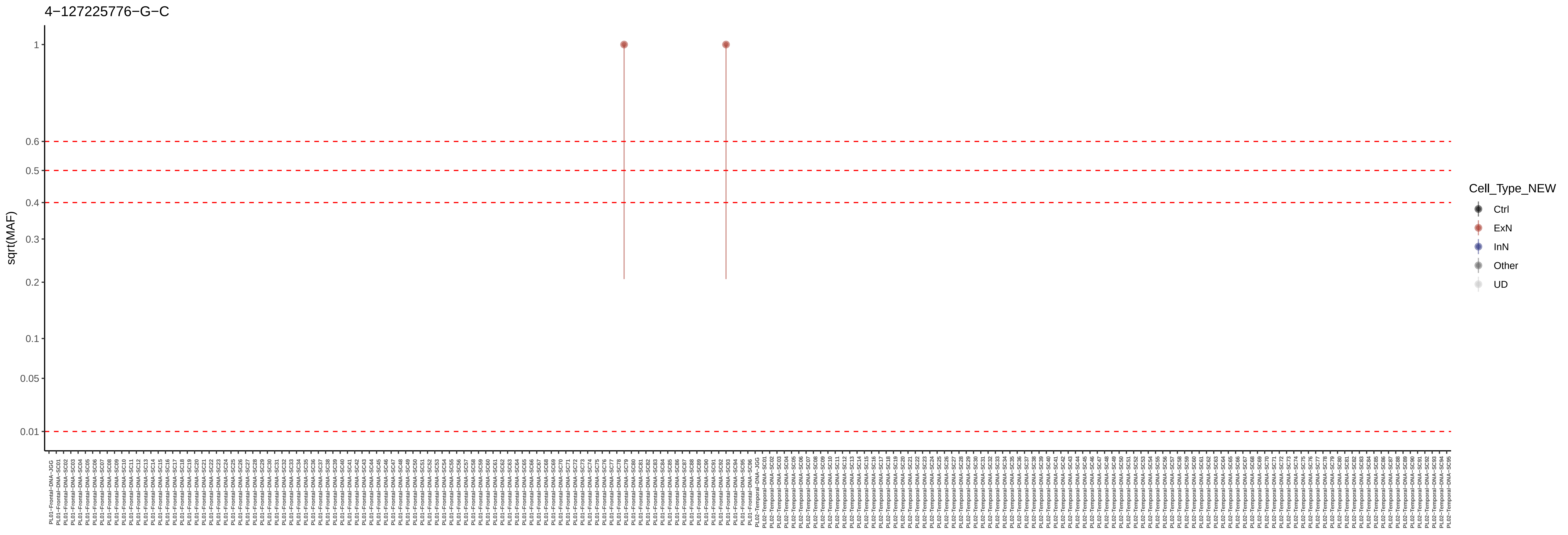

4−127225776−G−C
sqrt(MAF)
1
0.6
0.5
0.4
0.3
0.2
0.1
0.05
0.01
Cell_Type_NEW
Ctrl
ExN
InN
Other
UD

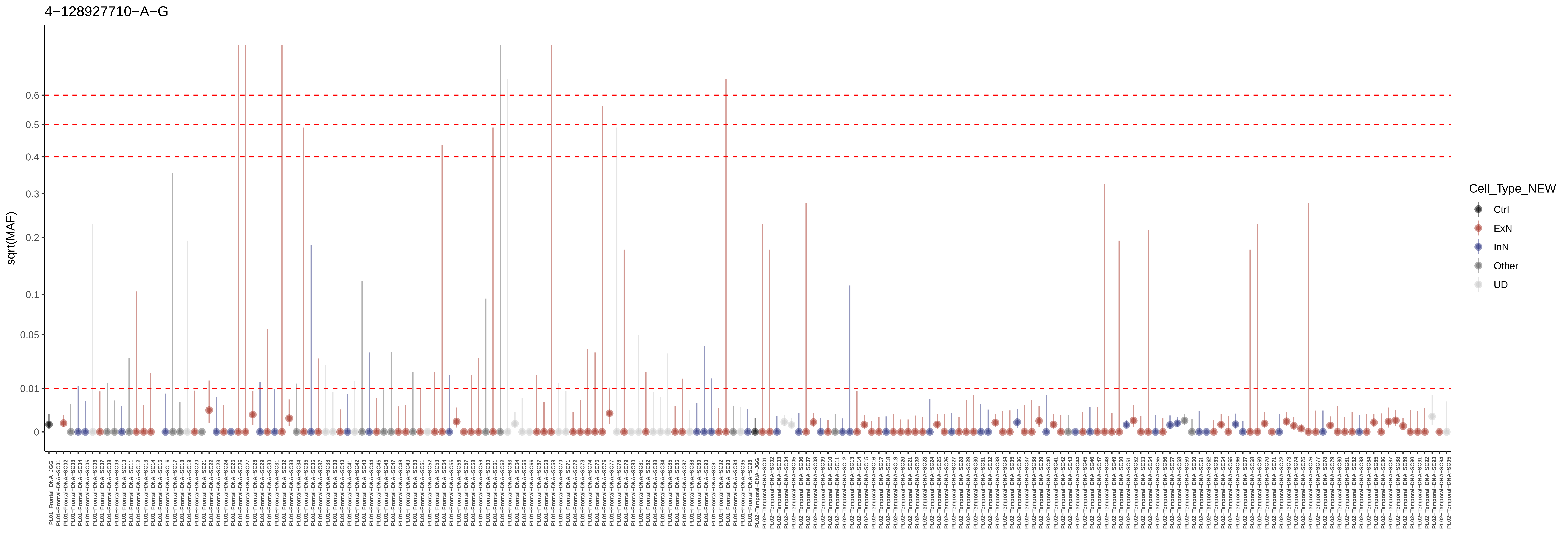
4−128927710−A−G
sqrt(MAF)
0.6
0.5
0.4
0.3
0.2
0.1
0.05
0.01
0
Cell_Type_NEW
Ctrl
ExN
InN
Other
UD

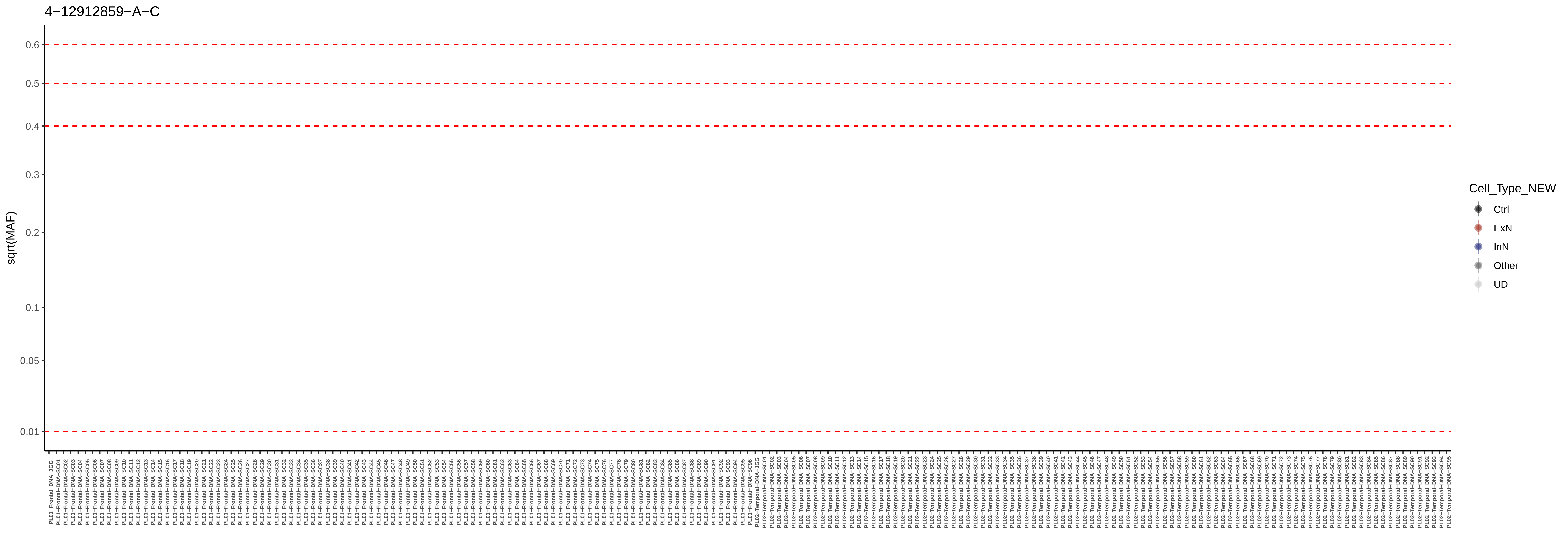
4−12912859−A−C
sqrt(MAF)
0.6
0.5
0.4
0.3
0.2
0.1
0.05
0.01
Cell_Type_NEW
Ctrl
ExN
InN
Other
UD

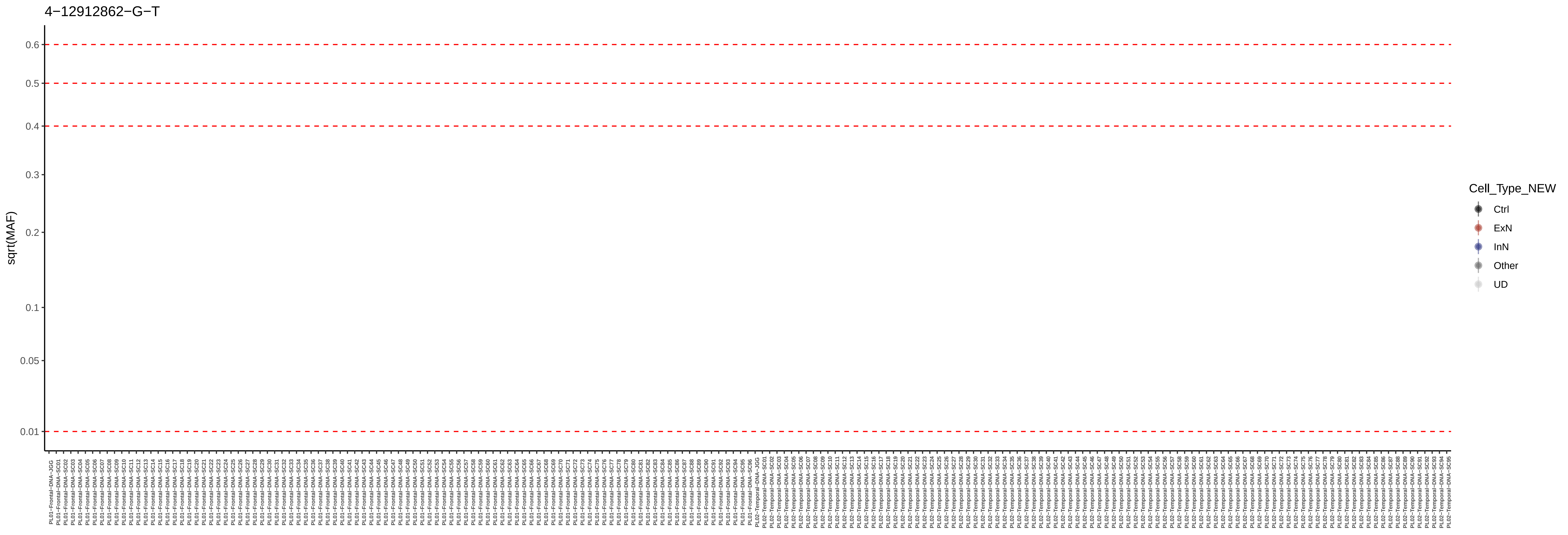
4−12912862−G−T
sqrt(MAF)
0.6
0.5
0.4
0.3
0.2
0.1
0.05
0.01
Cell_Type_NEW
Ctrl
ExN
InN
Other
UD

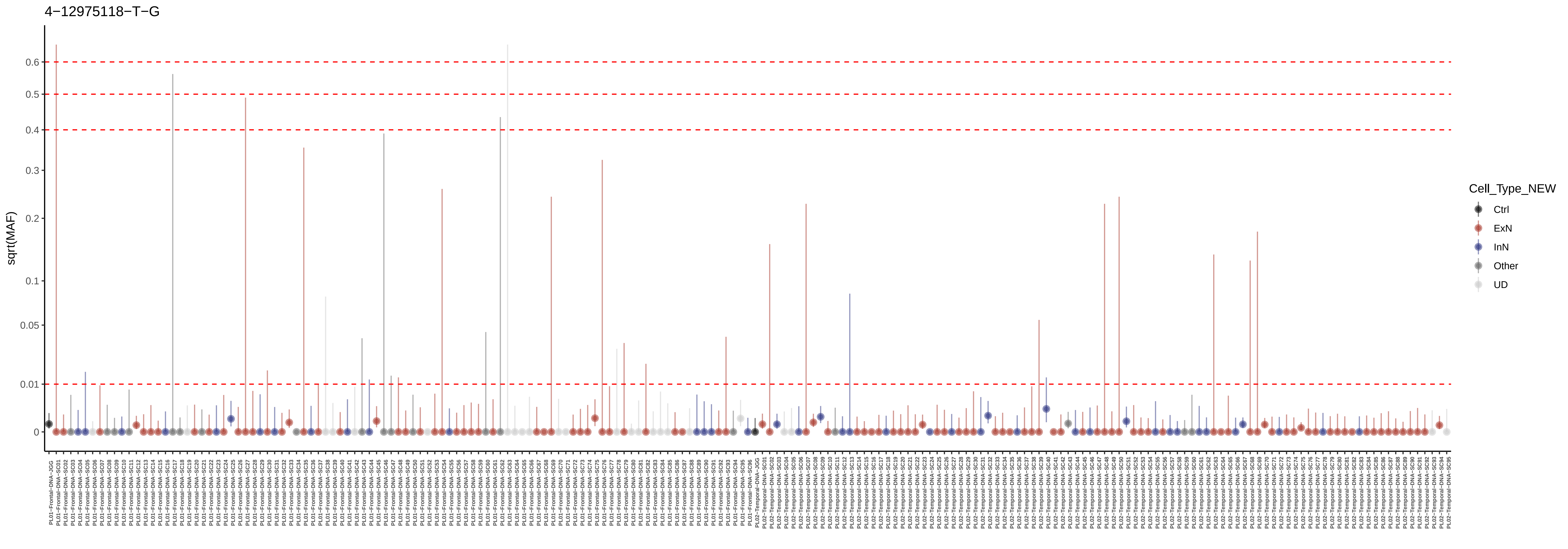

4−12975118−T−G
sqrt(MAF)
0.6
0.5
0.4
0.3
0.2
0.1
0.05
0.01
0
Cell_Type_NEW
Ctrl
ExN
InN
Other
UD

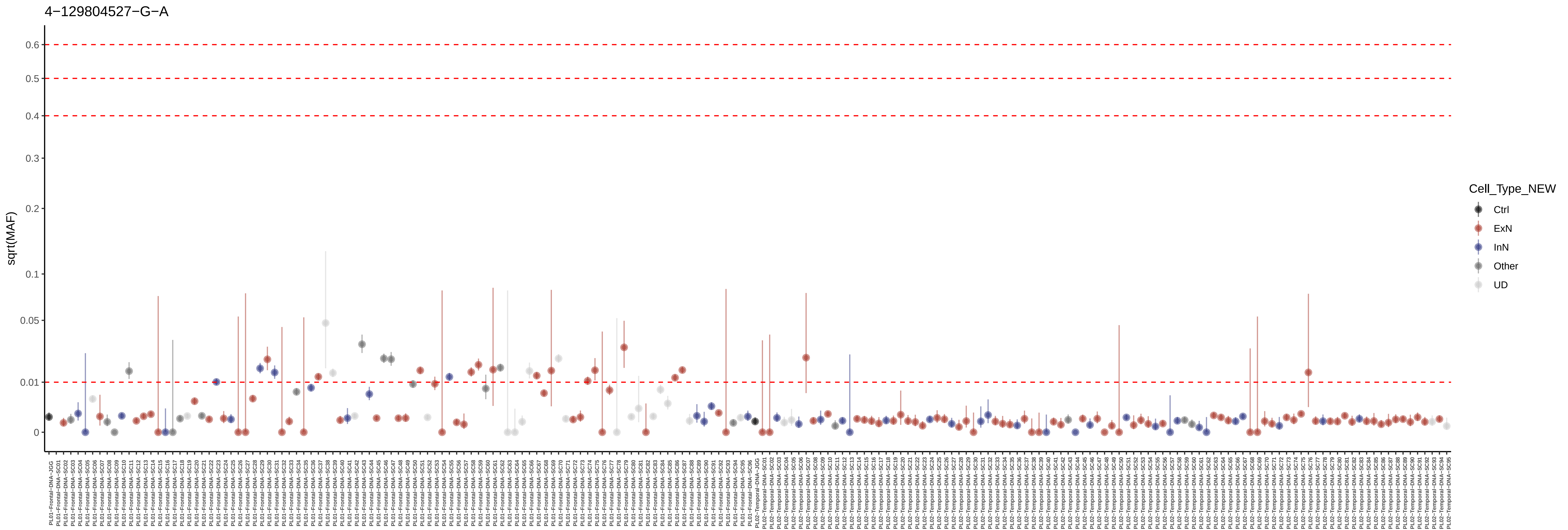

4−129804527−G−A
sqrt(MAF)
0.6
0.5
0.4
0.3
0.2
0.1
0.05
0.01
0
Cell_Type_NEW
Ctrl
ExN
InN
Other
UD

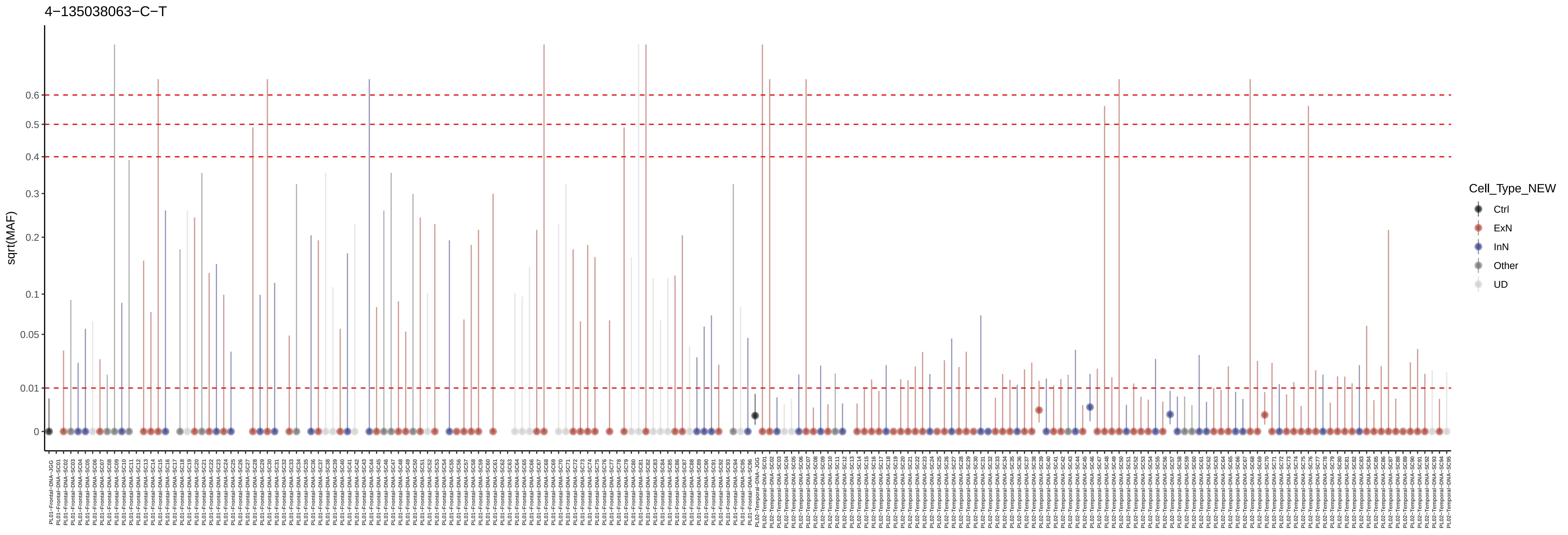
4−135038063−C−T
sqrt(MAF)
0.6
0.5
0.4
0.3
0.2
0.1
0.05
0.01
0
Cell_Type_NEW
Ctrl
ExN
InN
Other
UD

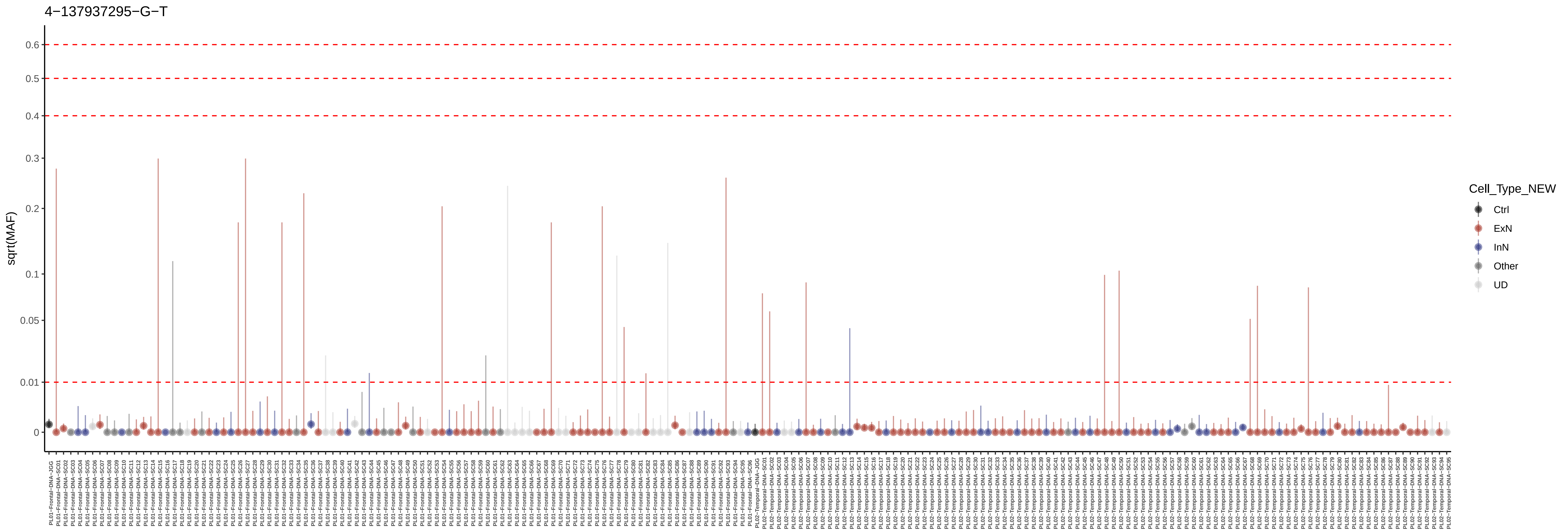
4−137937295−G−T
sqrt(MAF)
0.6
0.5
0.4
0.3
0.2
0.1
0.05
0.01
0
Cell_Type_NEW
Ctrl
ExN
InN
Other
UD

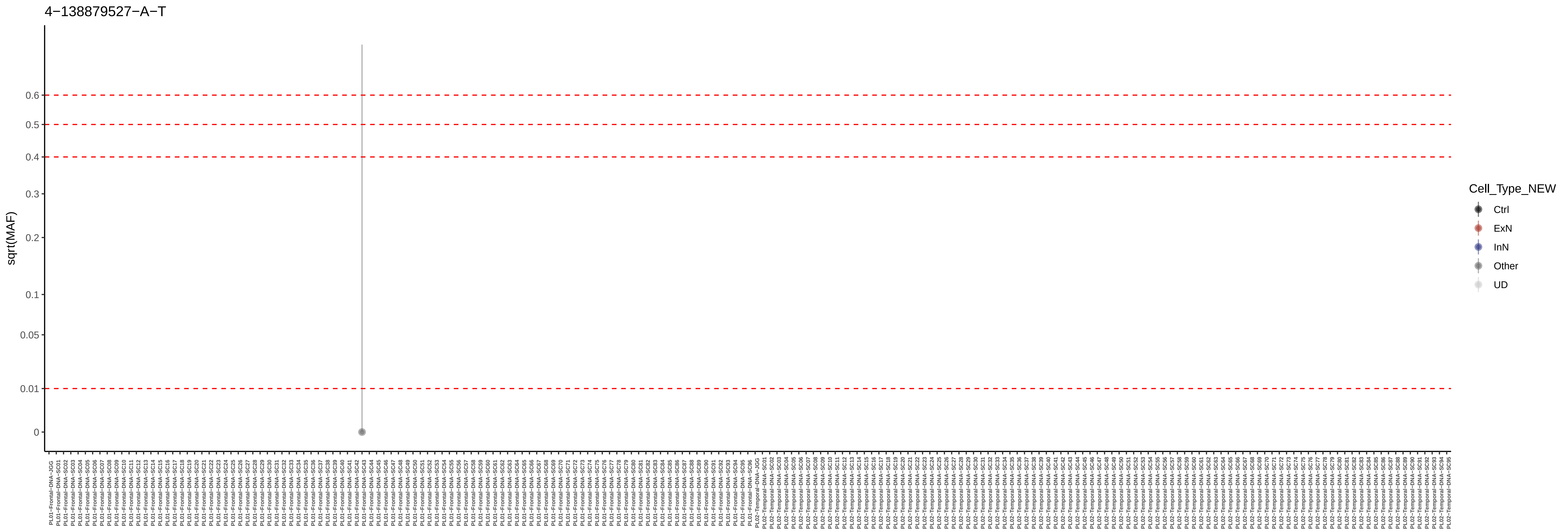
4−138879527−A−T
sqrt(MAF)
0.6
0.5
0.4
0.3
0.2
0.1
0.05
0.01
0
Cell_Type_NEW
Ctrl
ExN
InN
Other
UD

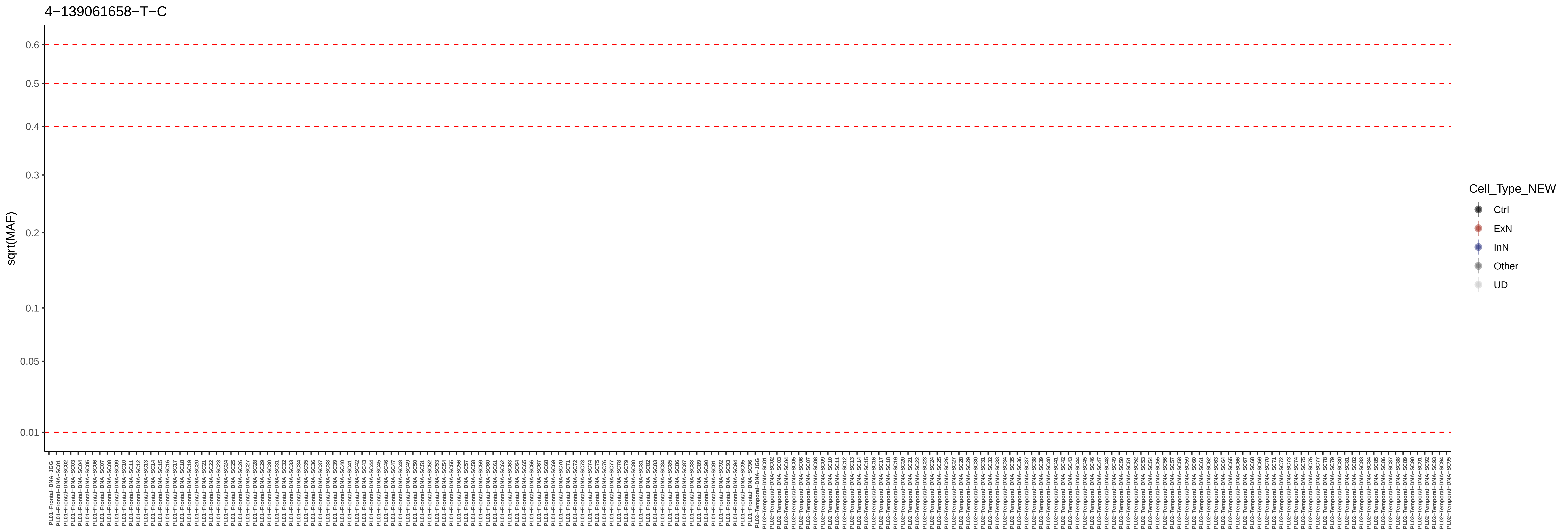
4−139061658−T−C
sqrt(MAF)
0.6
0.5
0.4
0.3
0.2
0.1
0.05
0.01
Cell_Type_NEW
Ctrl
ExN
InN
Other
UD

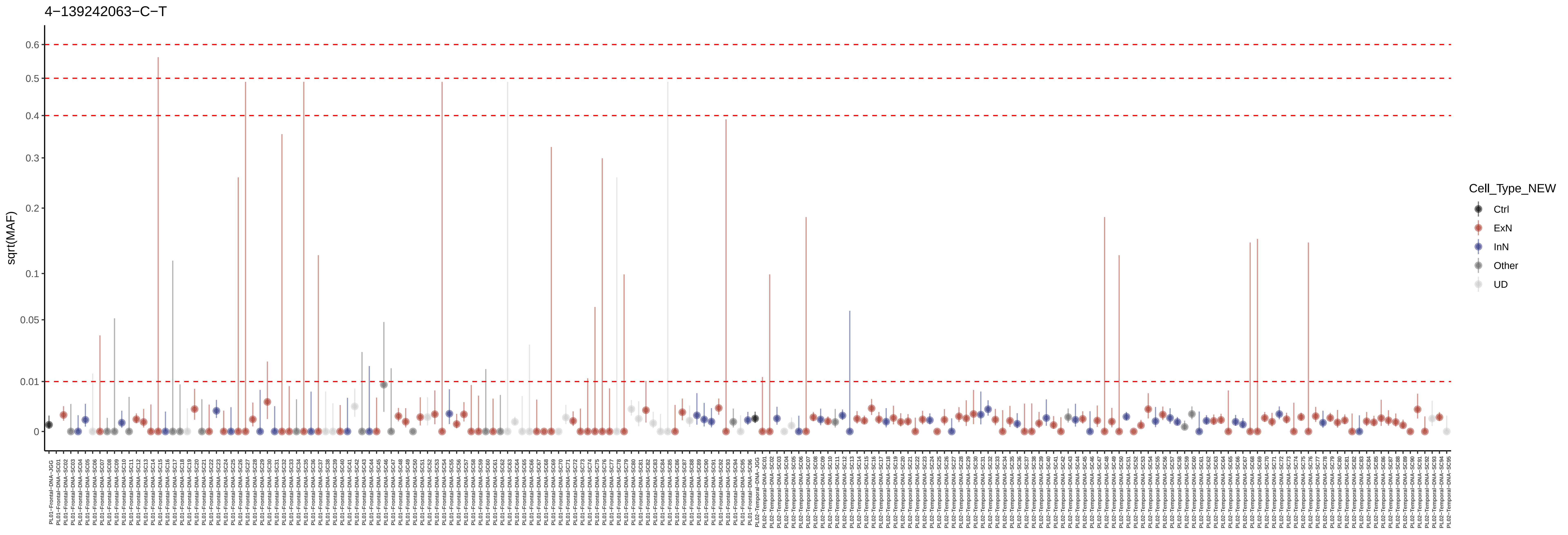
4−139242063−C−T
sqrt(MAF)
0.6
0.5
0.4
0.3
0.2
0.1
0.05
0.01
0
Cell_Type_NEW
Ctrl
ExN
InN
Other
UD

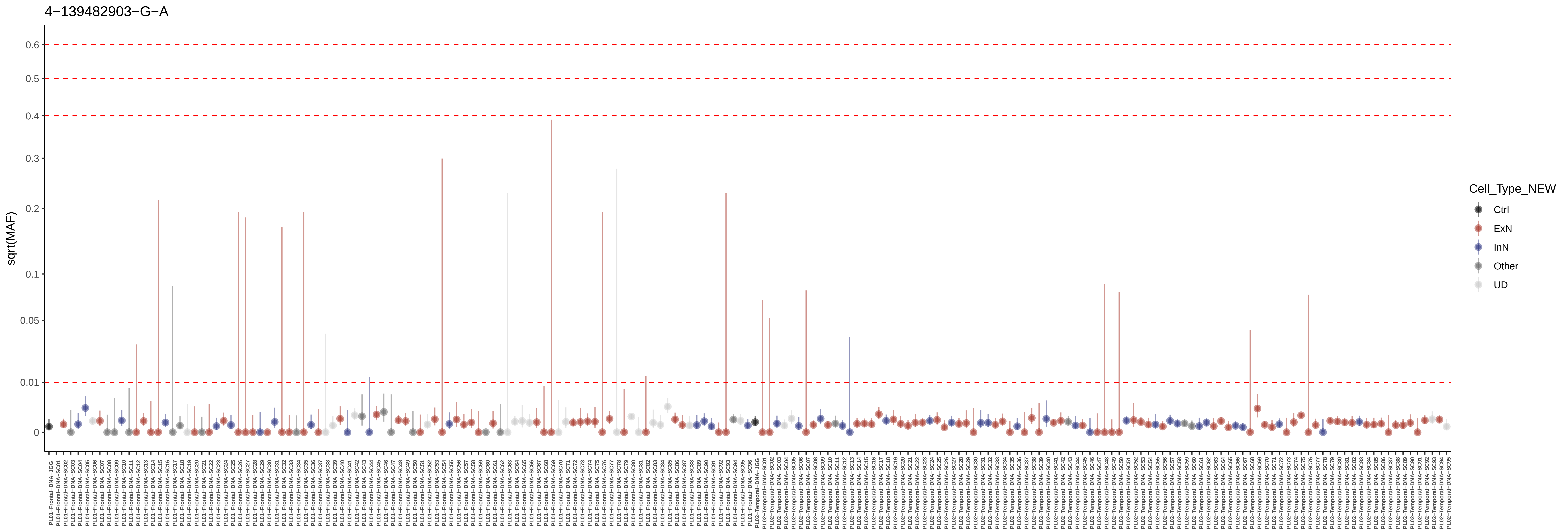

4−139482903−G−A
sqrt(MAF)
0.6
0.5
0.4
0.3
0.2
0.1
0.05
0.01
0
Cell_Type_NEW
Ctrl
ExN
InN
Other
UD

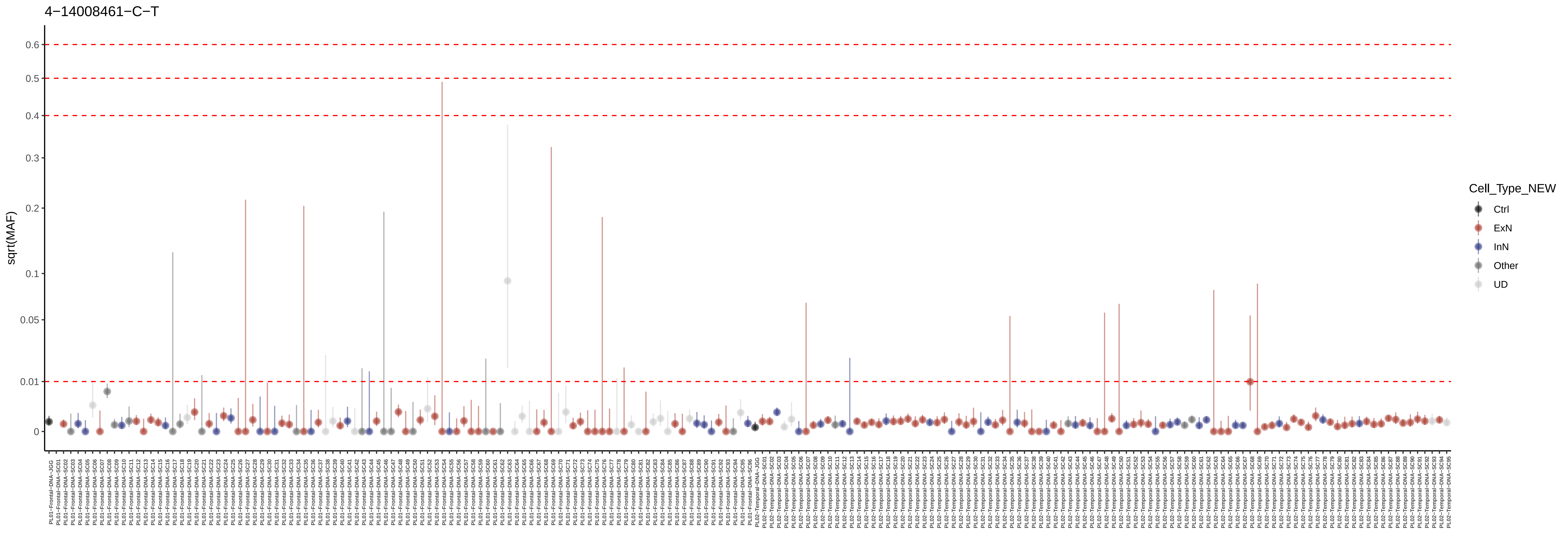
4−14008461−C−T
sqrt(MAF)
0.6
0.5
0.4
0.3
0.2
0.1
0.05
0.01
0
Cell_Type_NEW
Ctrl
ExN
InN
Other
UD

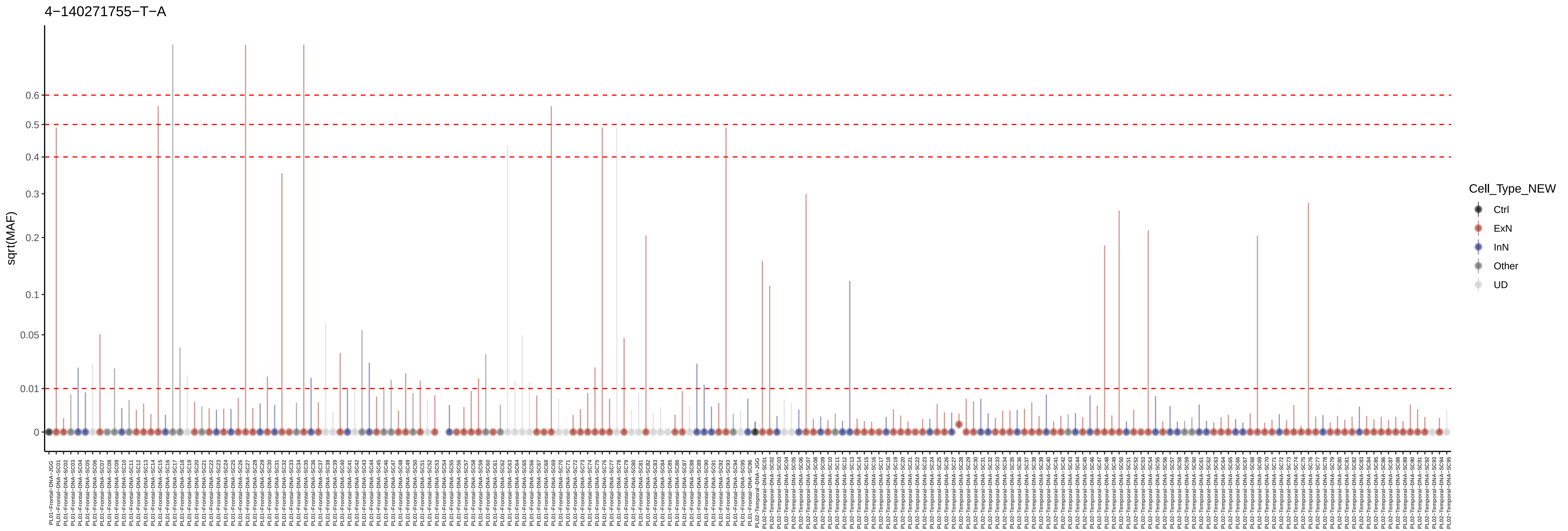
4−140271755−T−A
sqrt(MAF)
0.6
0.5
0.4
0.3
0.2
0.1
0.05
0.01
0
Cell_Type_NEW
Ctrl
ExN
InN
Other
UD

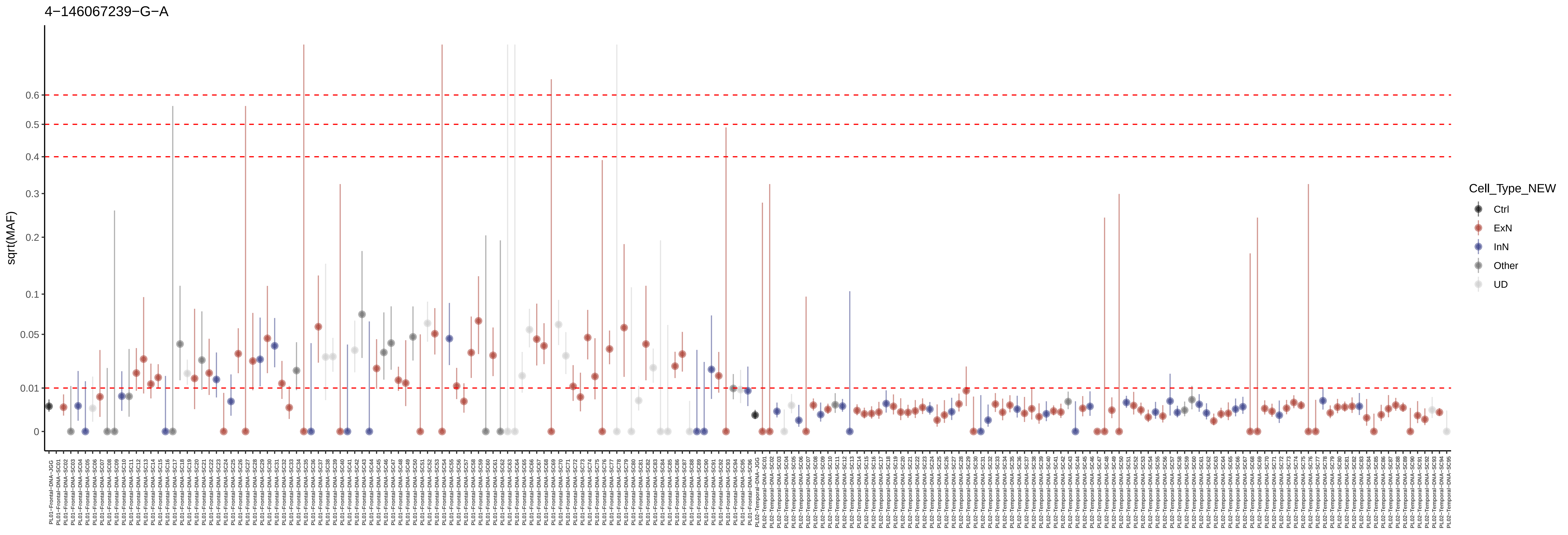

4−146067239−G−A
sqrt(MAF)
0.6
0.5
0.4
0.3
0.2
0.1
0.05
0.01
0
Cell_Type_NEW
Ctrl
ExN
InN
Other
UD

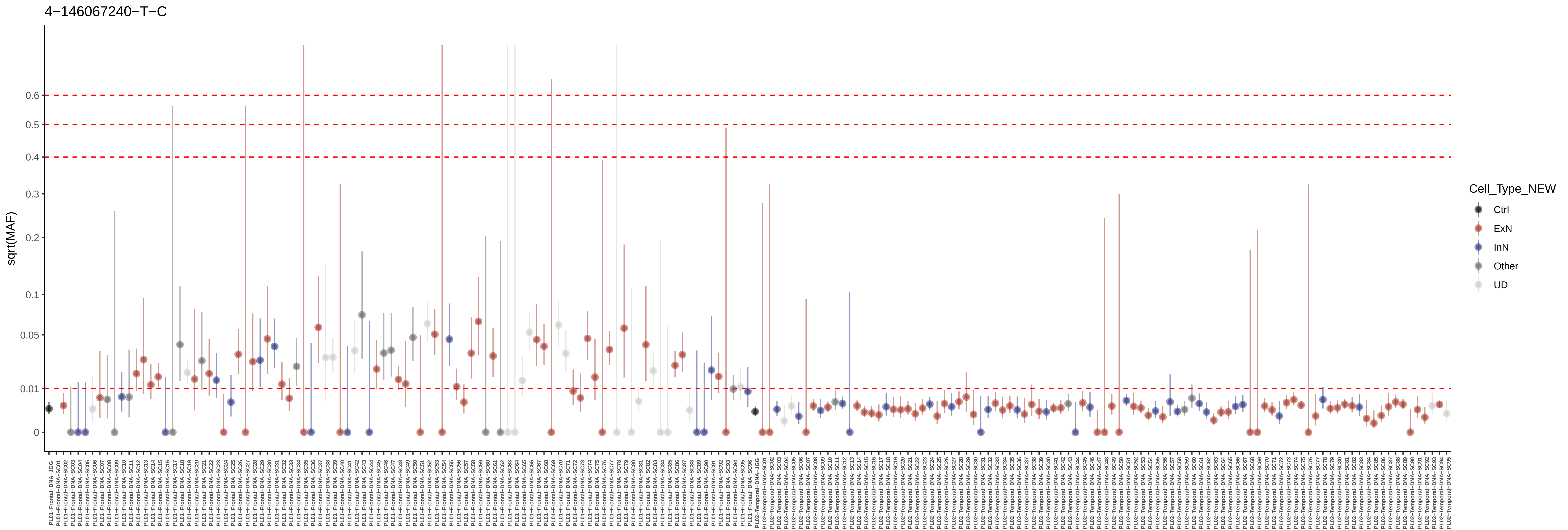

4−146067240−T−C
sqrt(MAF)
0.6
0.5
0.4
0.3
0.2
0.1
0.05
0.01
0
Cell_Type_NEW
Ctrl
ExN
InN
Other
UD

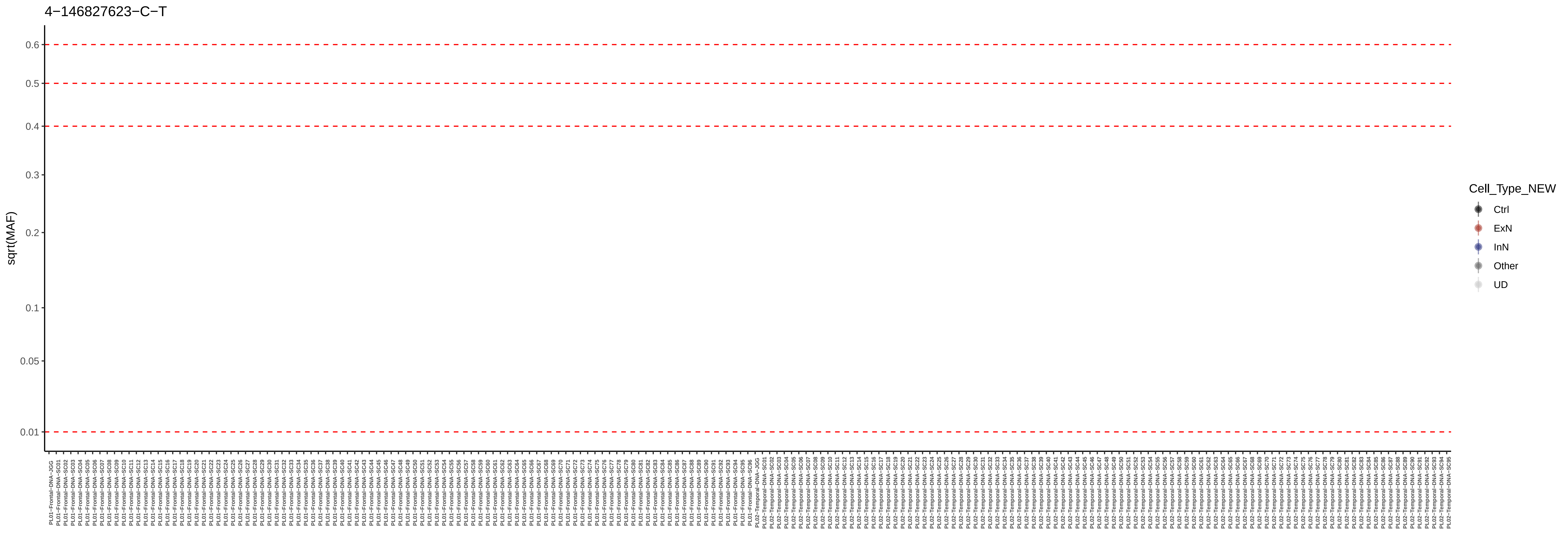

4−146827623−C−T
sqrt(MAF)
0.6
0.5
0.4
0.3
0.2
0.1
0.05
0.01
Cell_Type_NEW
Ctrl
ExN
InN
Other
UD

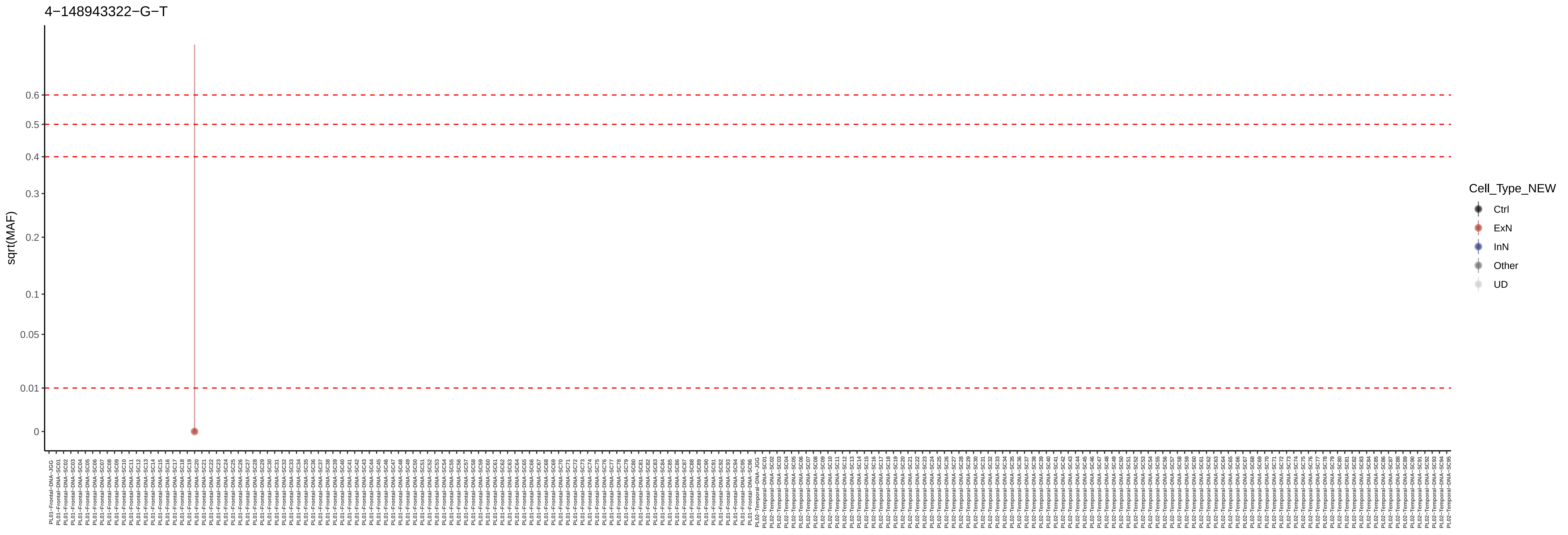

4−148943322−G−T
sqrt(MAF)
0.6
0.5
0.4
0.3
0.2
0.1
0.05
0.01
0
Cell_Type_NEW
Ctrl
ExN
InN
Other
UD

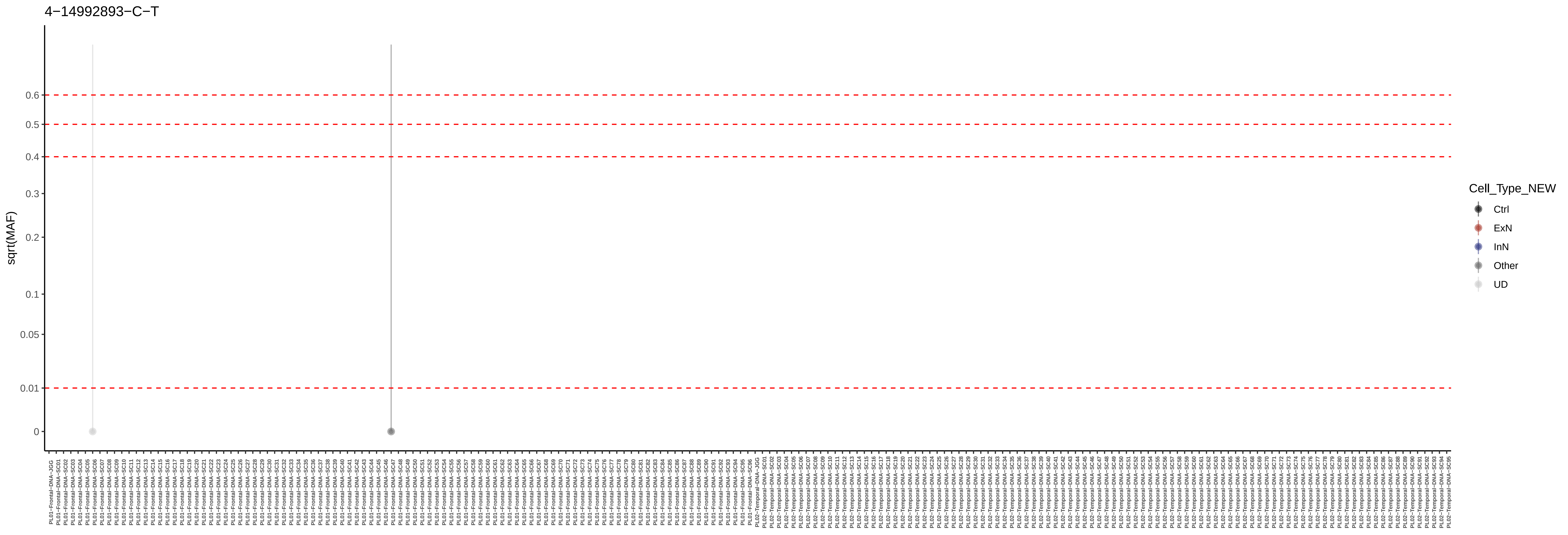

4−14992893−C−T
sqrt(MAF)
0.6
0.5
0.4
0.3
0.2
0.1
0.05
0.01
0
Cell_Type_NEW
Ctrl
ExN
InN
Other
UD

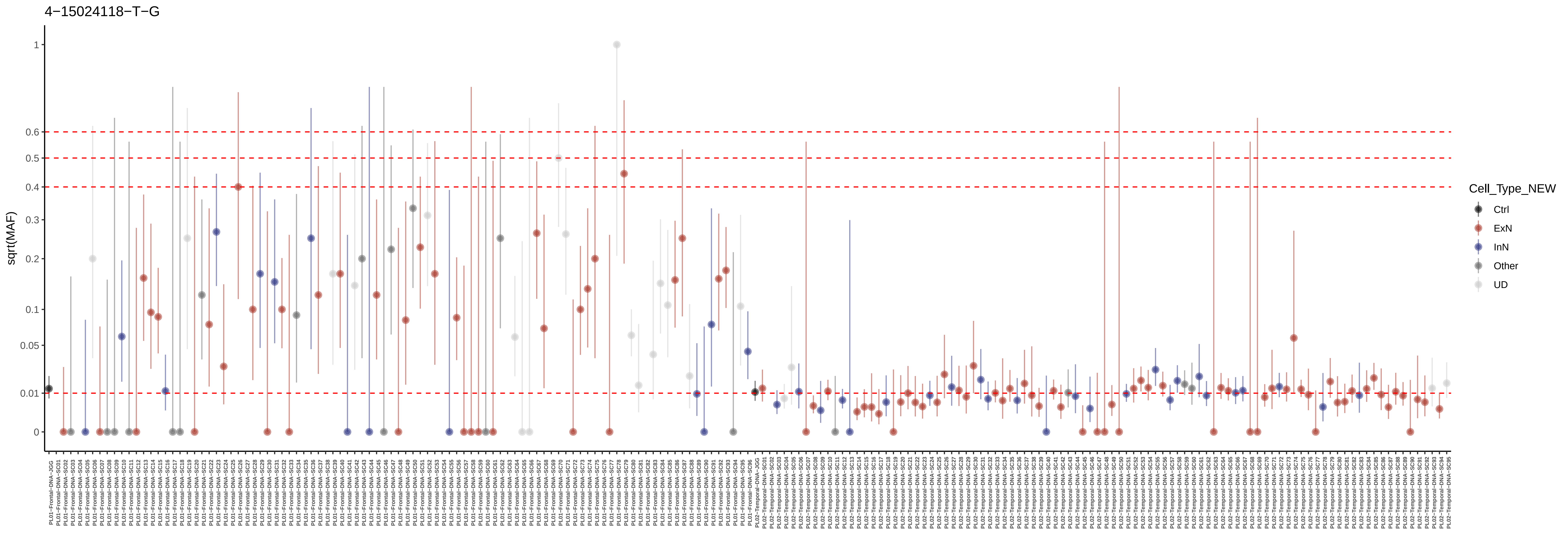

4−15024118−T−G
sqrt(MAF)
1
0.6
0.5
0.4
0.3
0.2
0.1
0.05
0.01
0
Cell_Type_NEW
Ctrl
ExN
InN
Other
UD

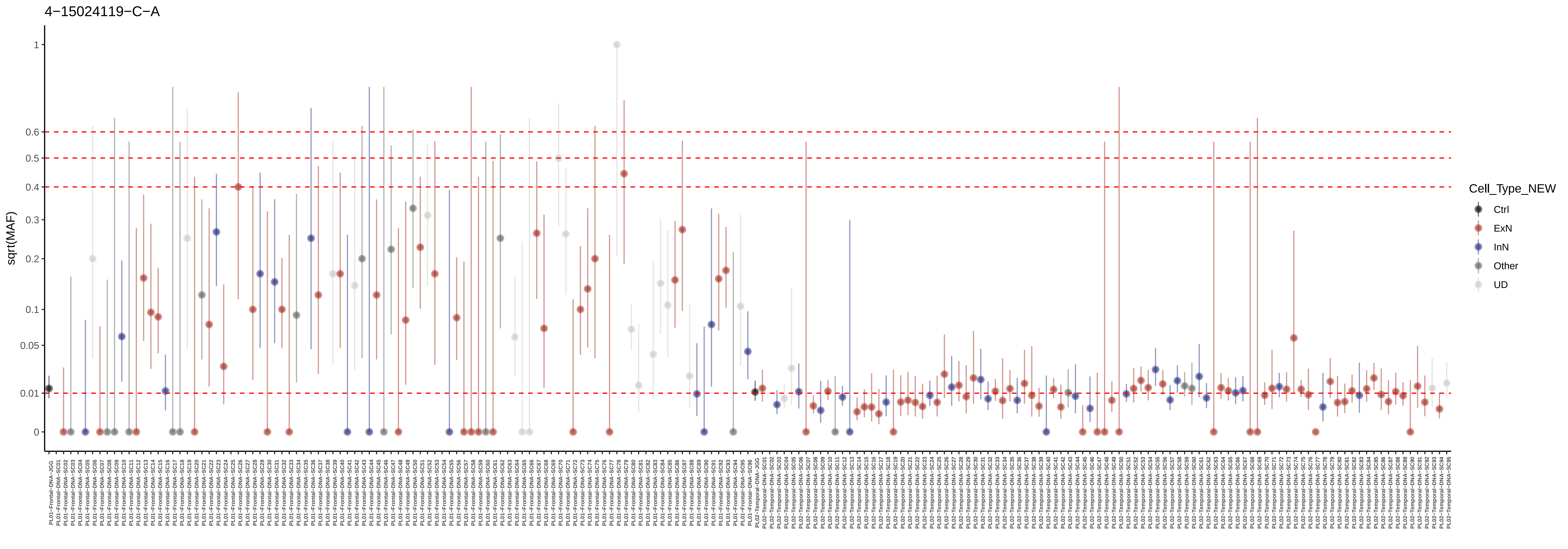

4−15024119−C−A
sqrt(MAF)
1
0.6
0.5
0.4
0.3
0.2
0.1
0.05
0.01
0
Cell_Type_NEW
Ctrl
ExN
InN
Other
UD

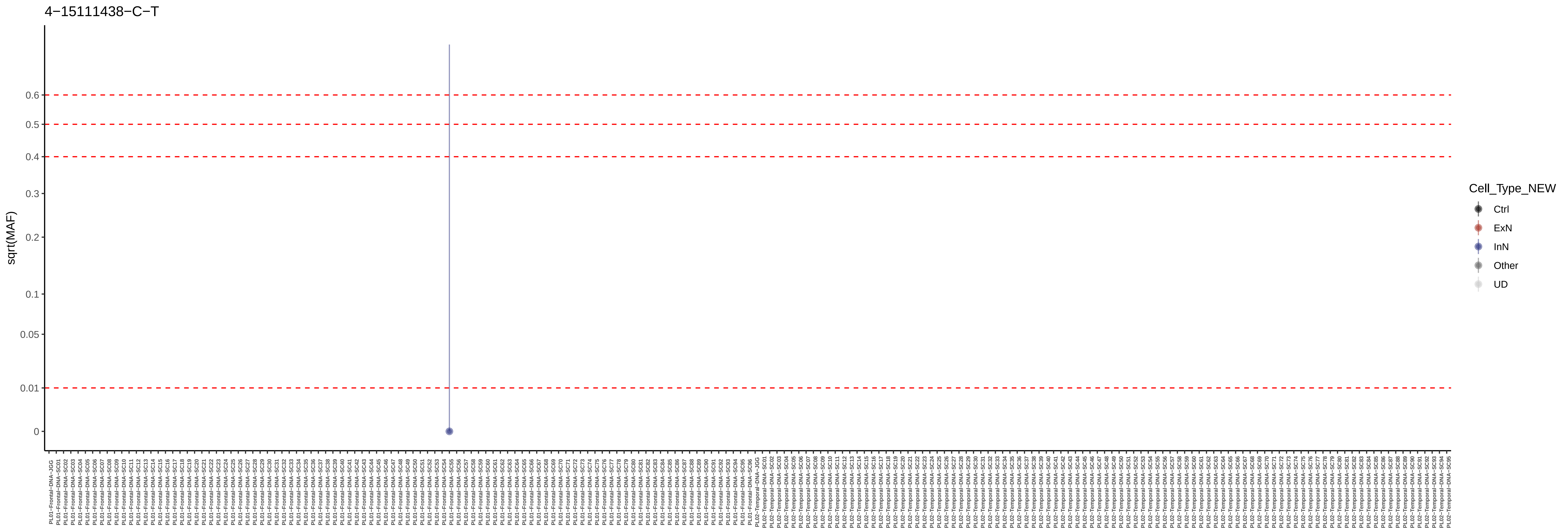
4−15111438−C−T
sqrt(MAF)
0.6
0.5
0.4
0.3
0.2
0.1
0.05
0.01
0
Cell_Type_NEW
Ctrl
ExN
InN
Other
UD

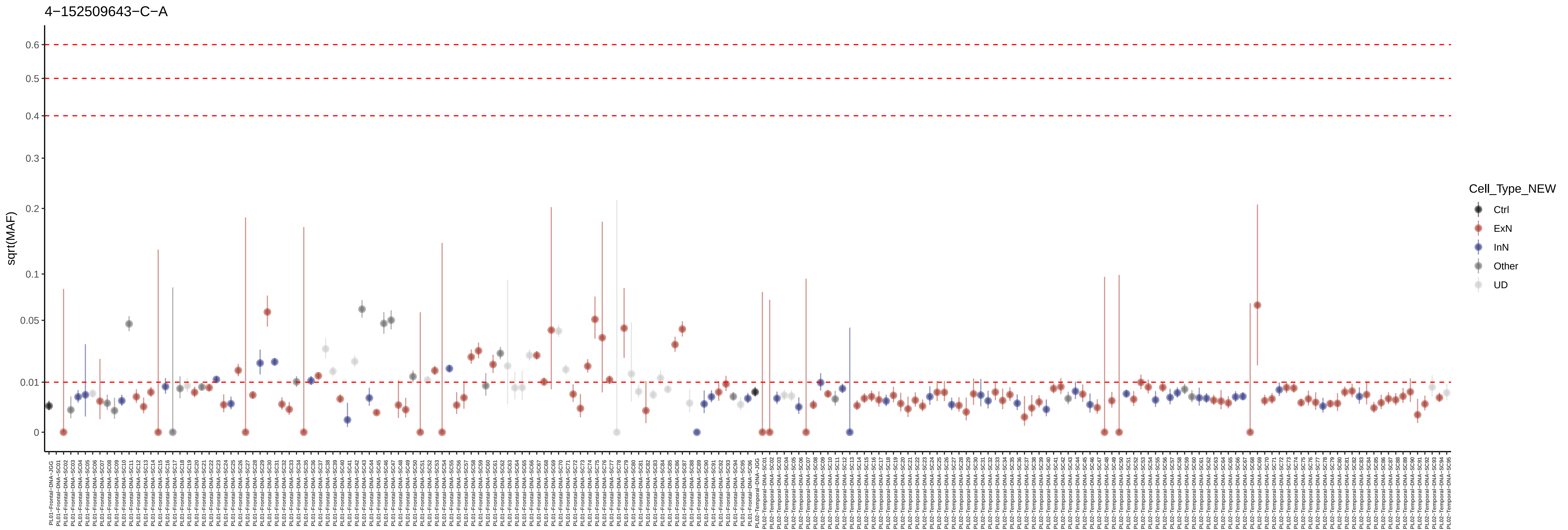

4−152509643−C−A
sqrt(MAF)
0.6
0.5
0.4
0.3
0.2
0.1
0.05
0.01
0
Cell_Type_NEW
Ctrl
ExN
InN
Other
UD

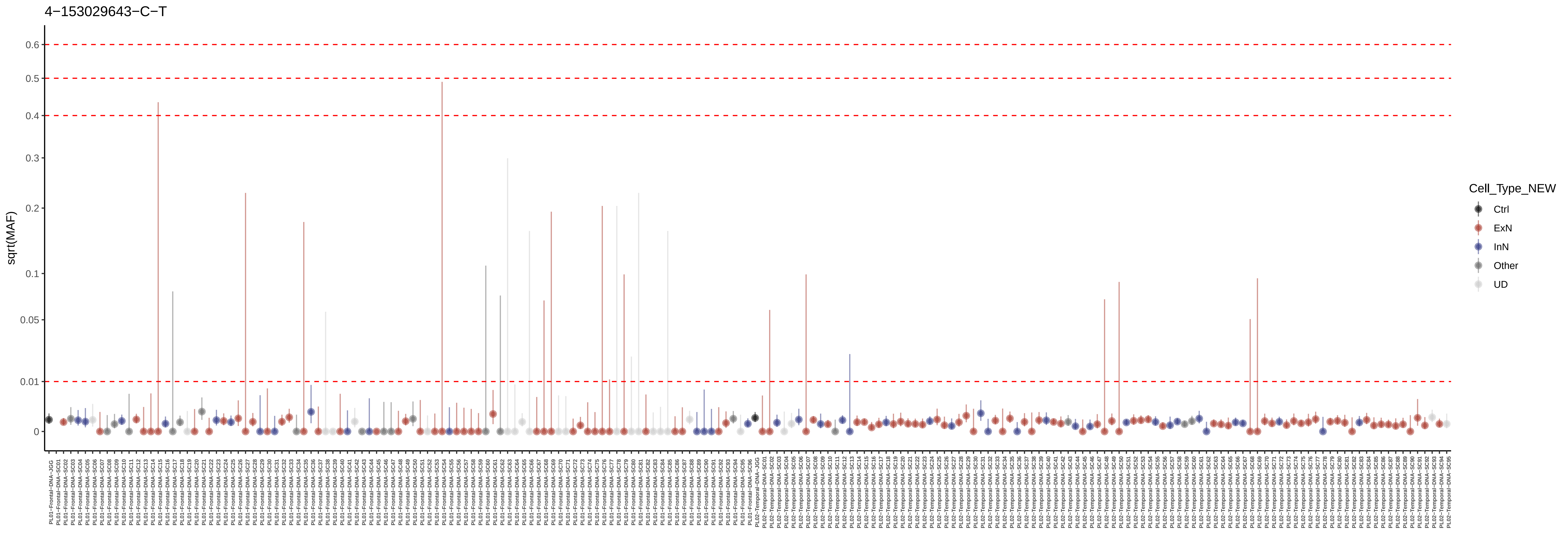
4−153029643−C−T
sqrt(MAF)
0.6
0.5
0.4
0.3
0.2
0.1
0.05
0.01
0
Cell_Type_NEW
Ctrl
ExN
InN
Other
UD

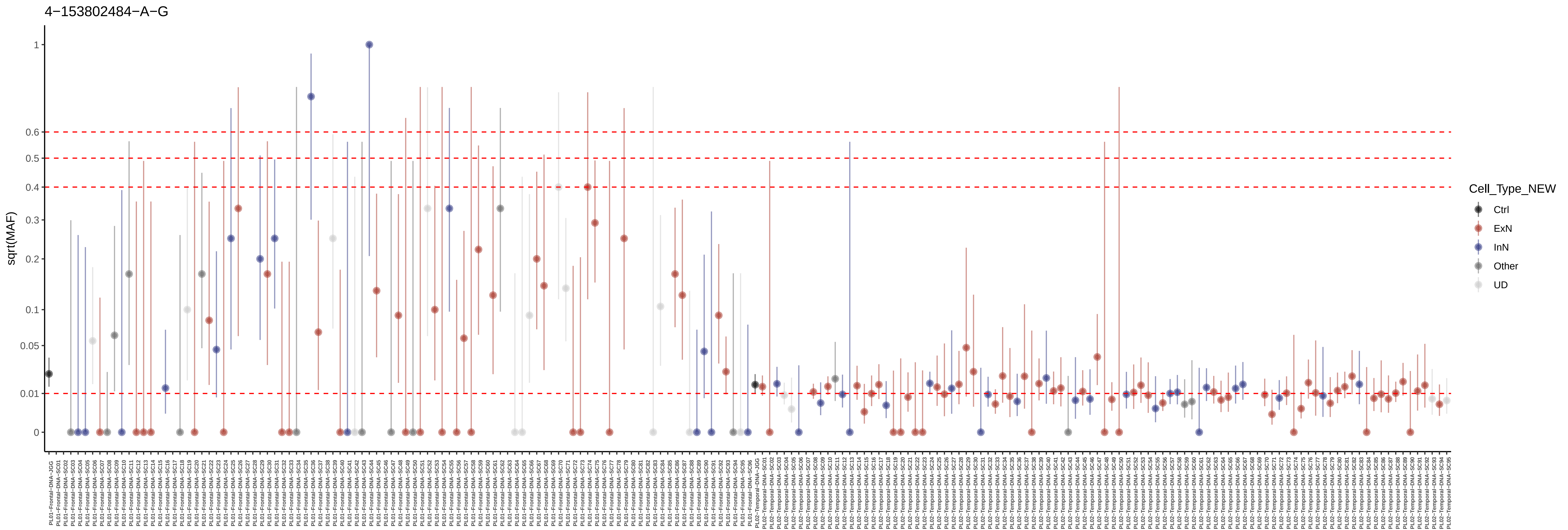

4−153802484−A−G
sqrt(MAF)
1
0.6
0.5
0.4
0.3
0.2
0.1
0.05
0.01
0
Cell_Type_NEW
Ctrl
ExN
InN
Other
UD

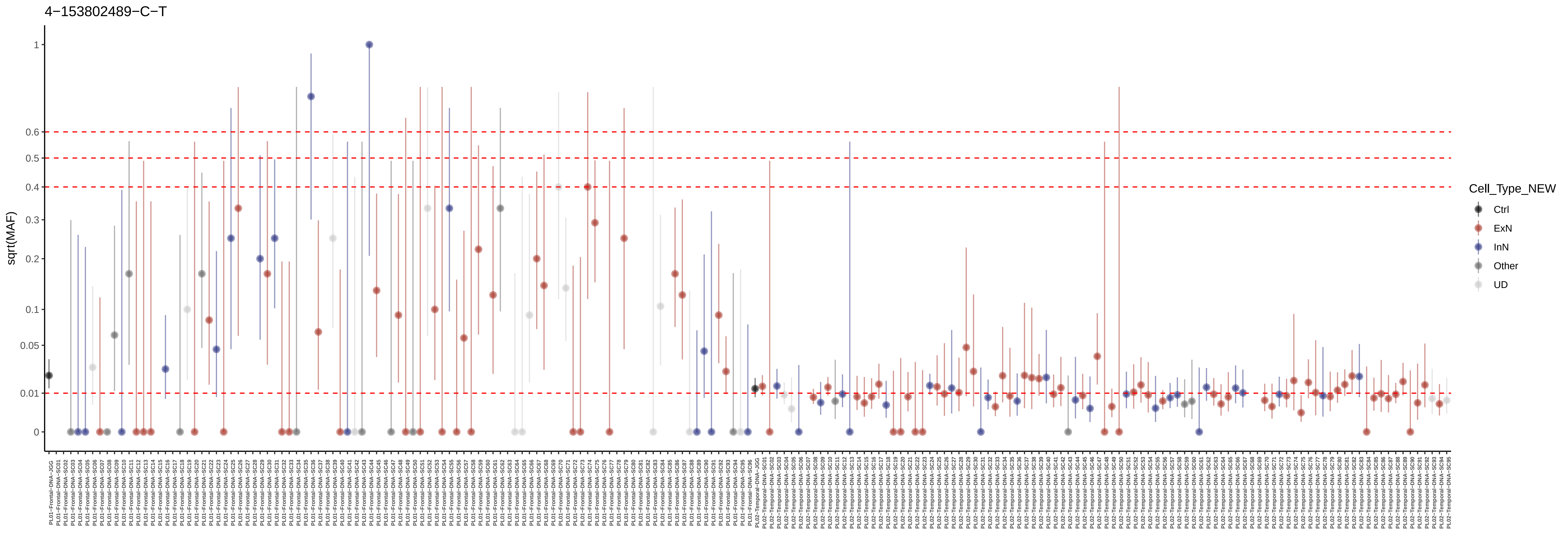

4−153802489−C−T
sqrt(MAF)
1
0.6
0.5
0.4
0.3
0.2
0.1
0.05
0.01
0
Cell_Type_NEW
Ctrl
ExN
InN
Other
UD

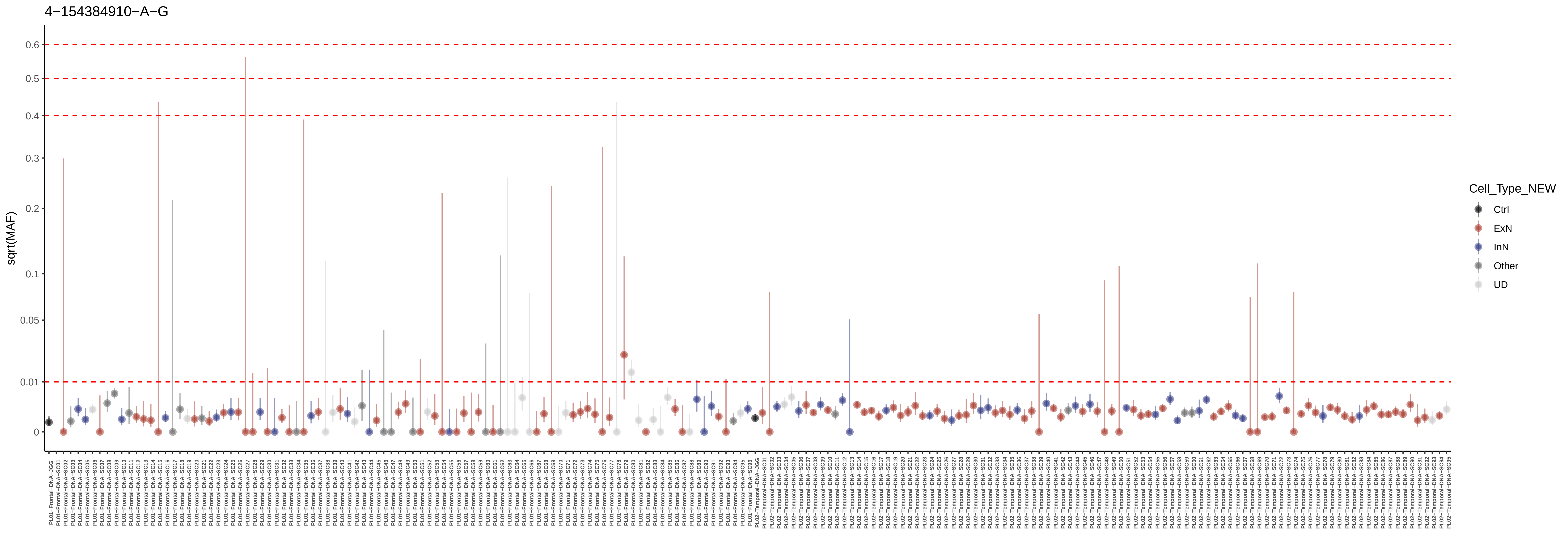

4−154384910−A−G
sqrt(MAF)
0.6
0.5
0.4
0.3
0.2
0.1
0.05
0.01
0
Cell_Type_NEW
Ctrl
ExN
InN
Other
UD

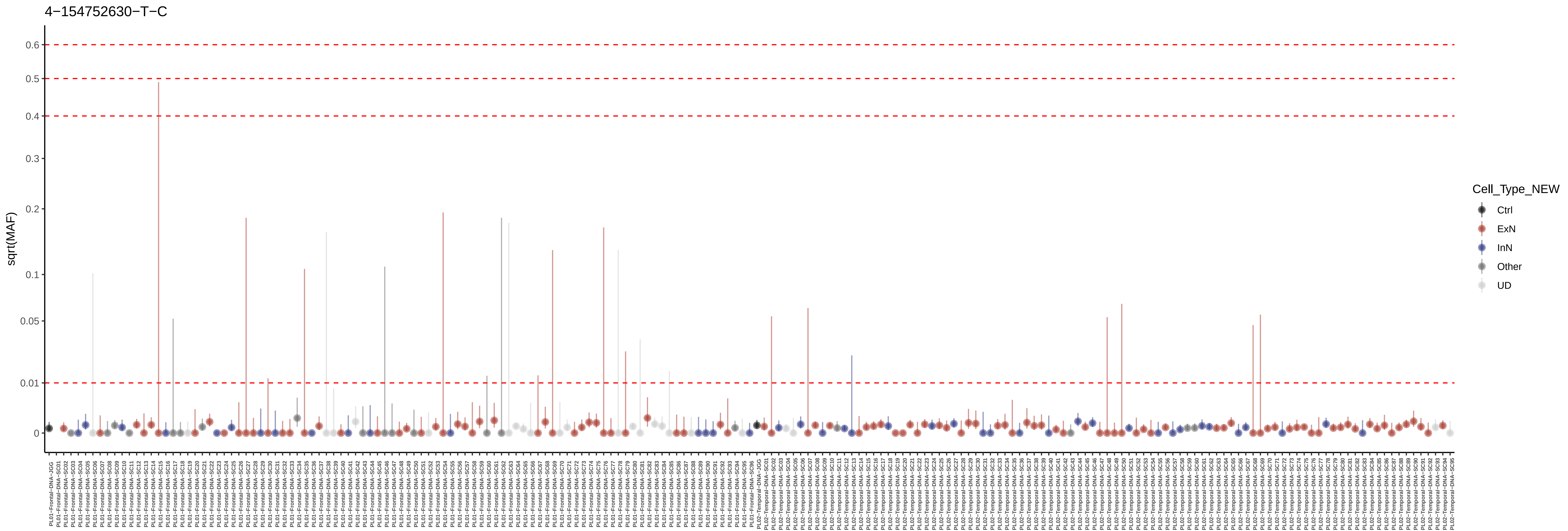

4−154752630−T−C
sqrt(MAF)
0.6
0.5
0.4
0.3
0.2
0.1
0.05
0.01
0
Cell_Type_NEW
Ctrl
ExN
InN
Other
UD

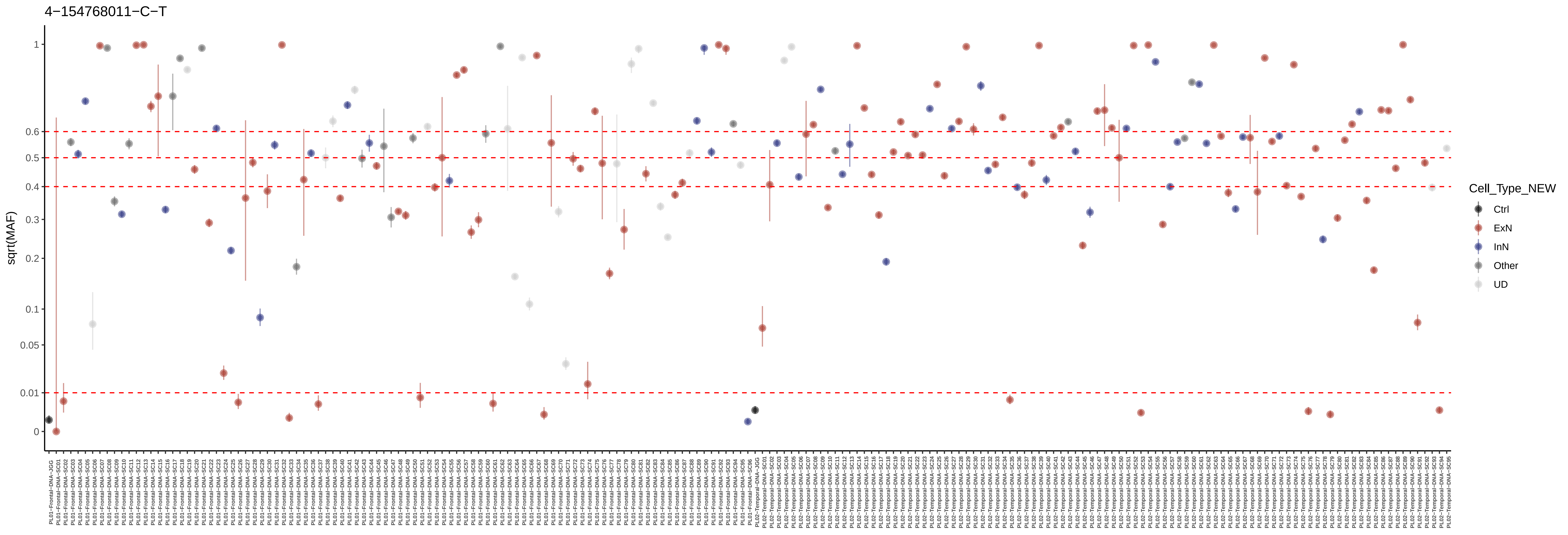

4−154768011−C−T
sqrt(MAF)
1
0.6
0.5
0.4
0.3
0.2
0.1
0.05
0.01
0
Cell_Type_NEW
Ctrl
ExN
InN
Other
UD

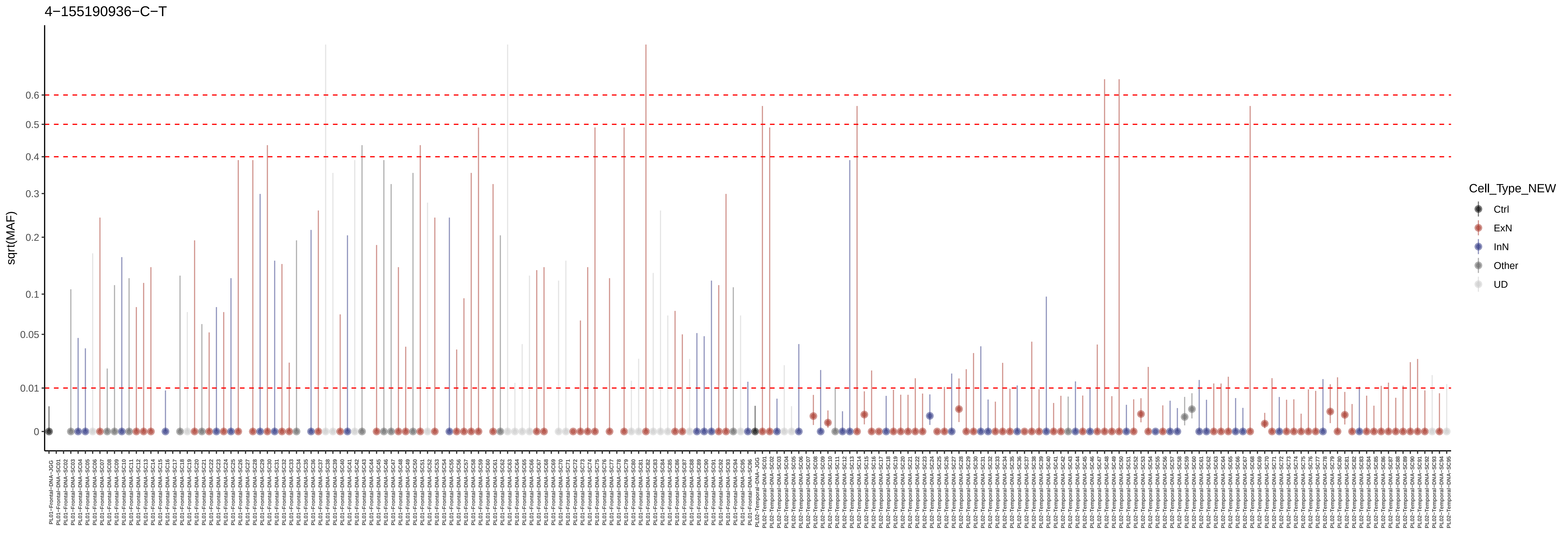
4−155190936−C−T
sqrt(MAF)
0.6
0.5
0.4
0.3
0.2
0.1
0.05
0.01
0
Cell_Type_NEW
Ctrl
ExN
InN
Other
UD

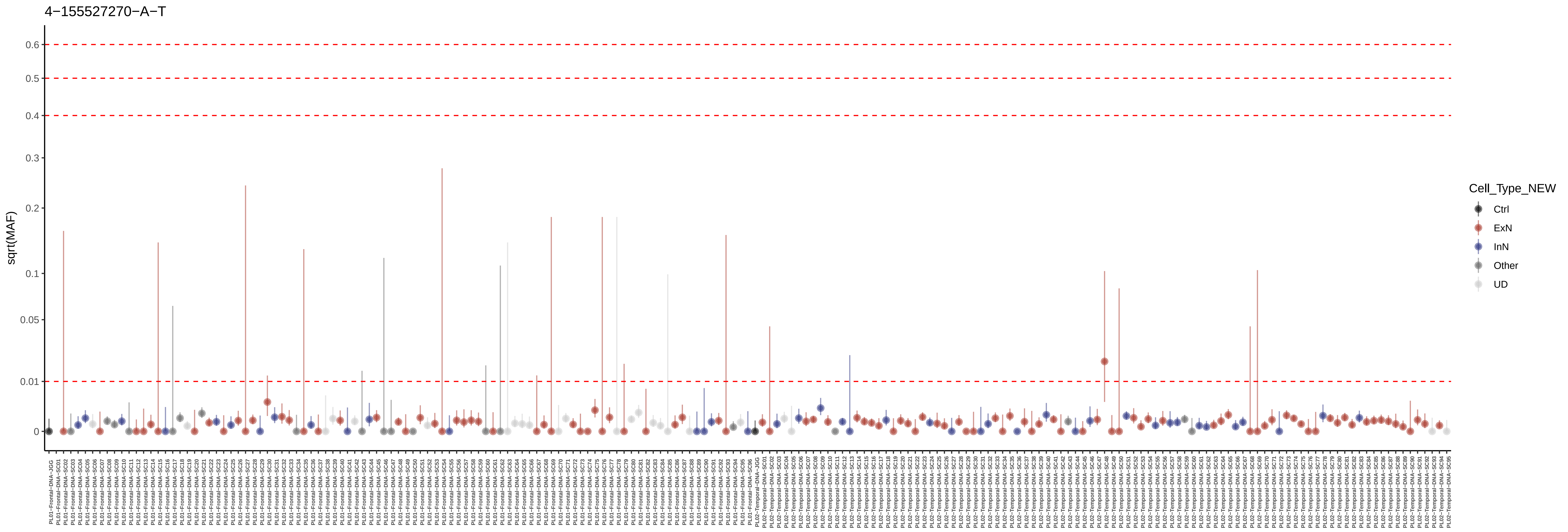
4−155527270−A−T
sqrt(MAF)
0.6
0.5
0.4
0.3
0.2
0.1
0.05
0.01
0
Cell_Type_NEW
Ctrl
ExN
InN
Other
UD

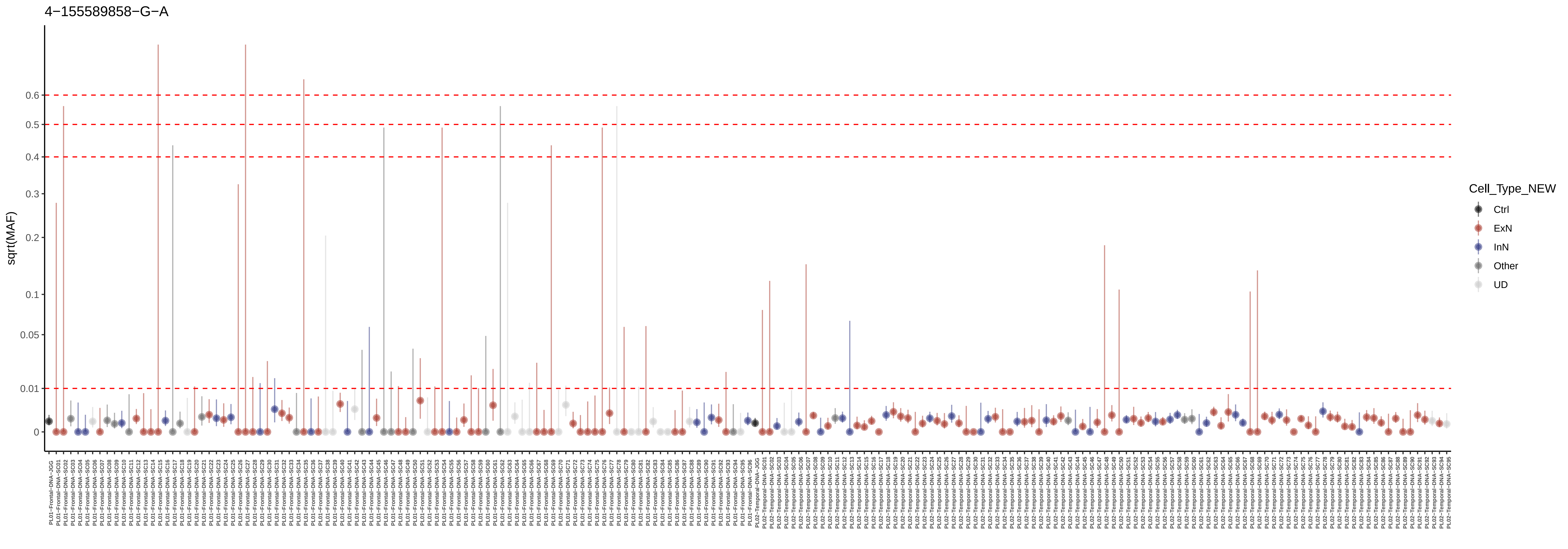

4−155589858−G−A
sqrt(MAF)
0.6
0.5
0.4
0.3
0.2
0.1
0.05
0.01
0
Cell_Type_NEW
Ctrl
ExN
InN
Other
UD

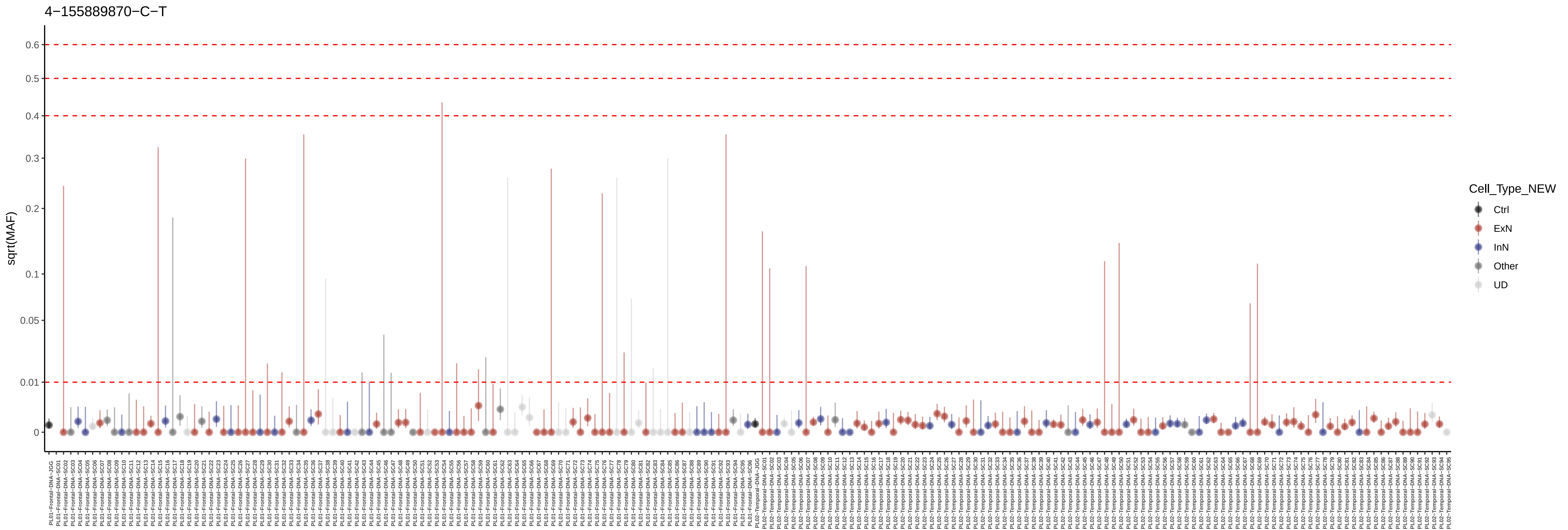

4−155889870−C−T
sqrt(MAF)
0.6
0.5
0.4
0.3
0.2
0.1
0.05
0.01
0
Cell_Type_NEW
Ctrl
ExN
InN
Other
UD

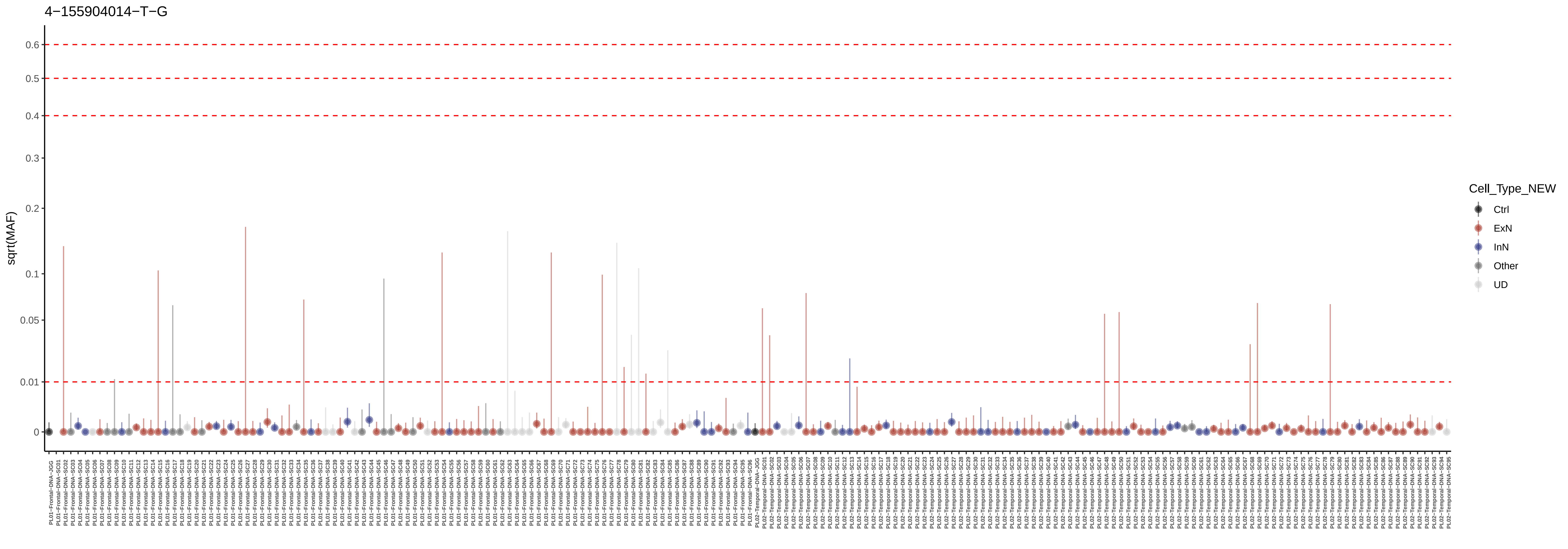
4−155904014−T−G
sqrt(MAF)
Cell_Type_NEW
Ctrl
ExN
InN
Other
UD

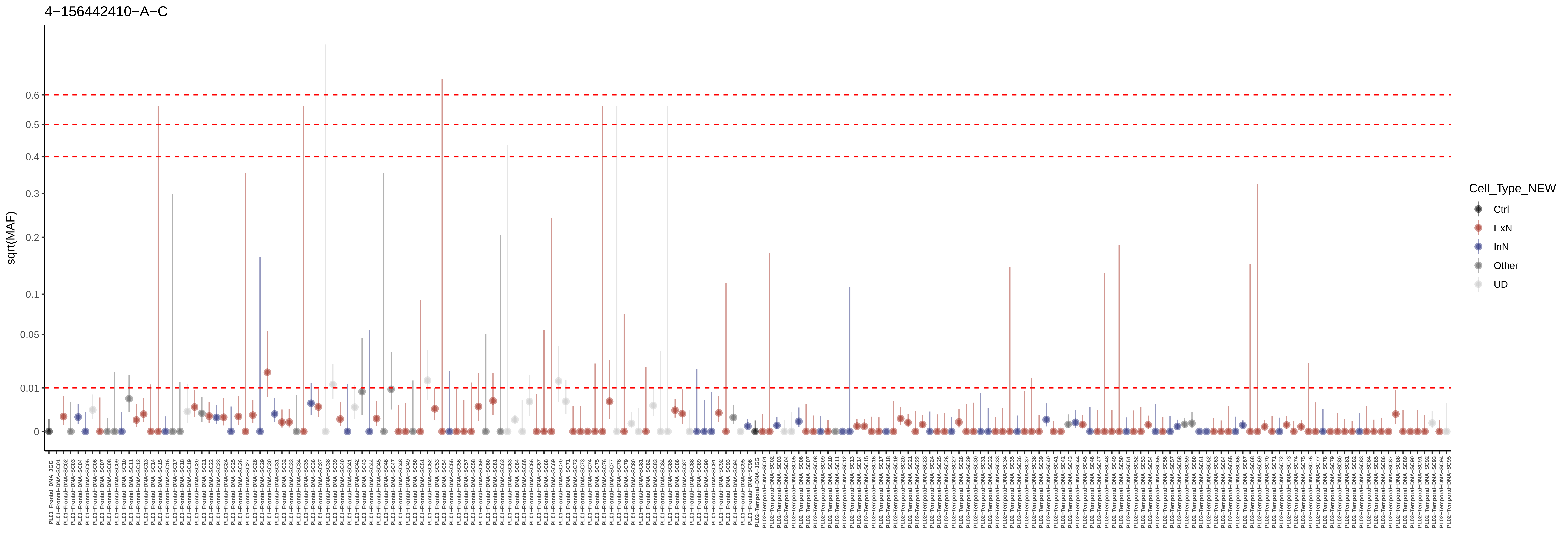
4−156442410−A−C
sqrt(MAF)
0.6
0.5
0.4
0.3
0.2
0.1
0.05
0.01
0
Cell_Type_NEW
Ctrl
ExN
InN
Other
UD

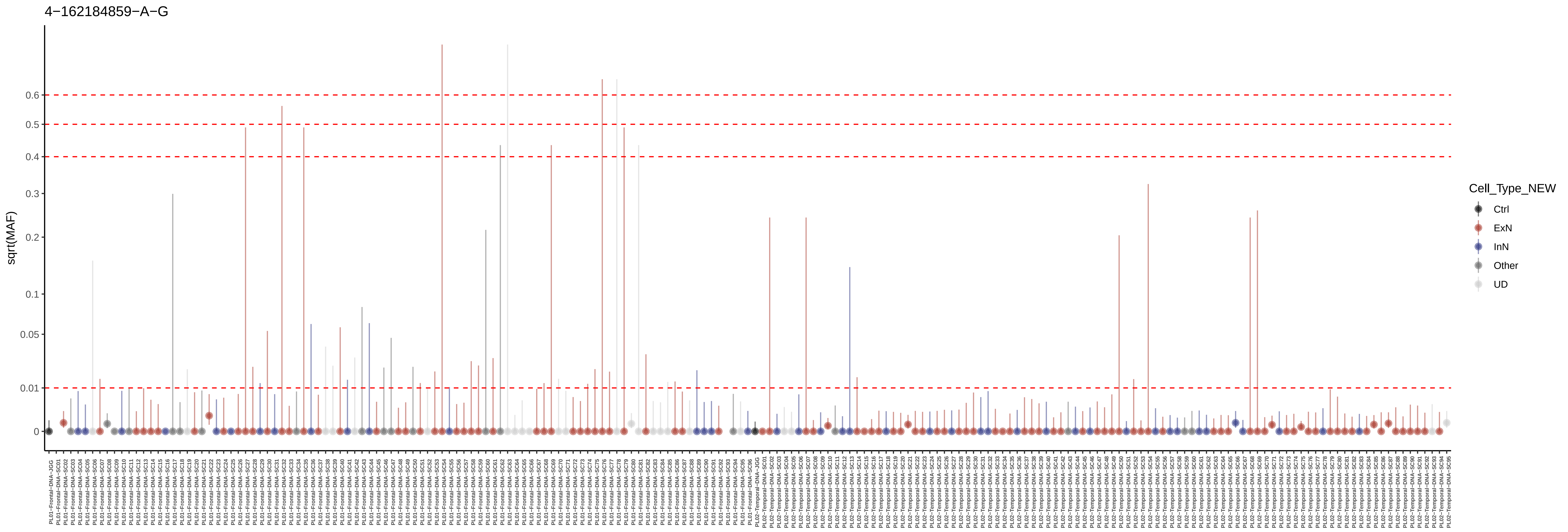

4−162184859−A−G
sqrt(MAF)
0.6
0.5
0.4
0.3
0.2
0.1
0.05
0.01
0
Cell_Type_NEW
Ctrl
ExN
InN
Other
UD

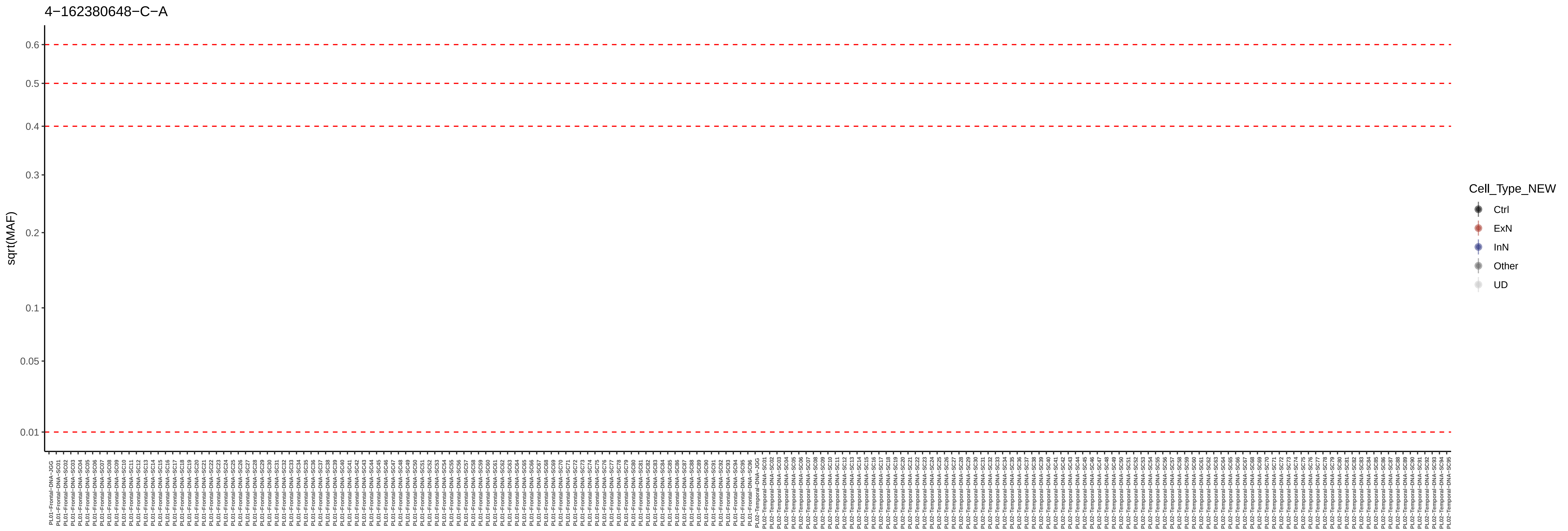
4−162380648−C−A
sqrt(MAF)
0.6
0.5
0.4
0.3
0.2
0.1
0.05
0.01
Cell_Type_NEW
Ctrl
ExN
InN
Other
UD

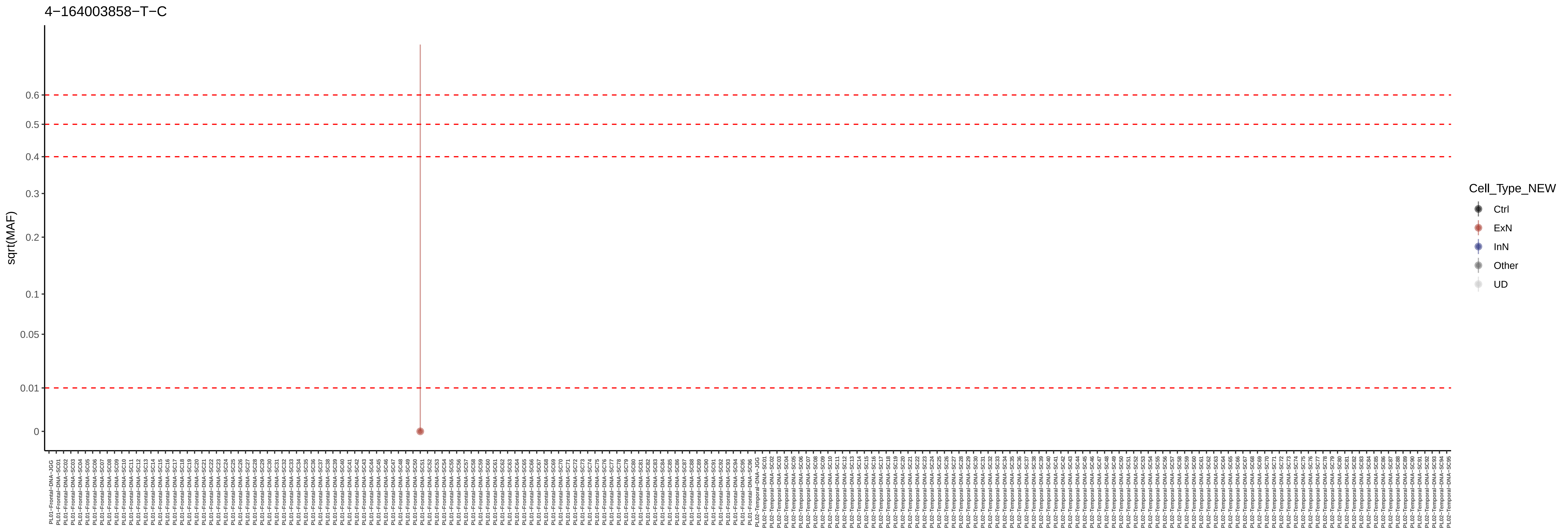
4−164003858−T−C
sqrt(MAF)
0.6
0.5
0.4
0.3
0.2
0.1
0.05
0.01
0
Cell_Type_NEW
Ctrl
ExN
InN
Other
UD

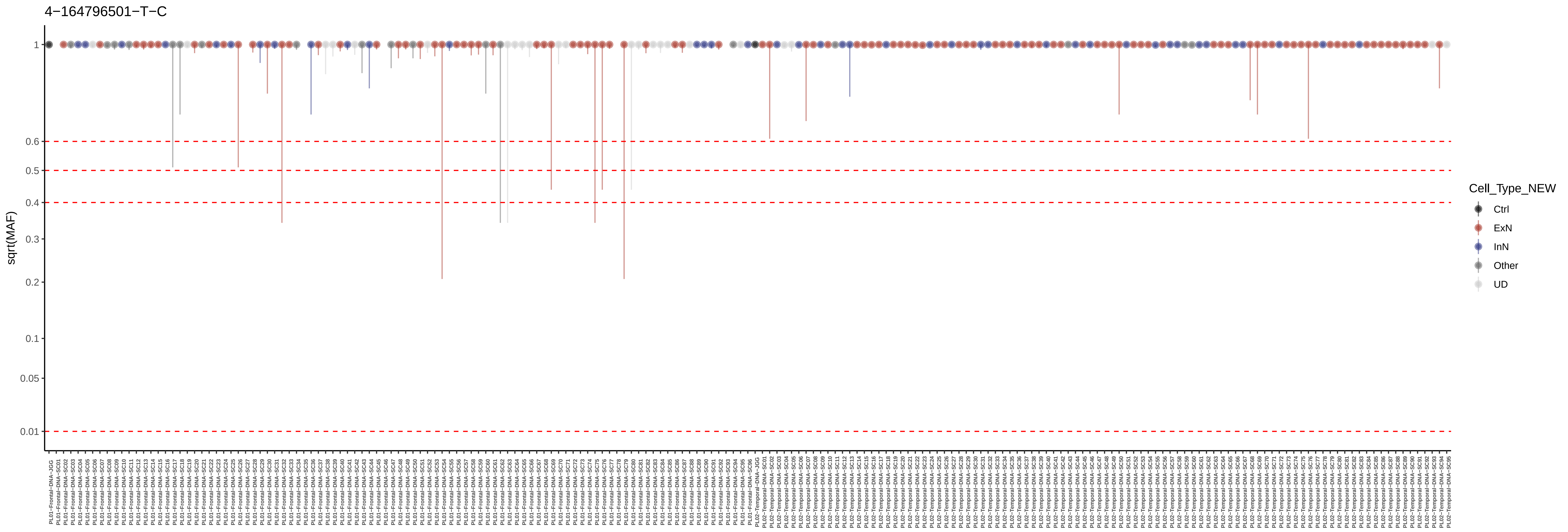
4−164796501−T−C
sqrt(MAF)
1
0.6
0.5
0.4
0.3
0.2
0.1
0.05
0.01
Cell_Type_NEW
Ctrl
ExN
InN
Other
UD

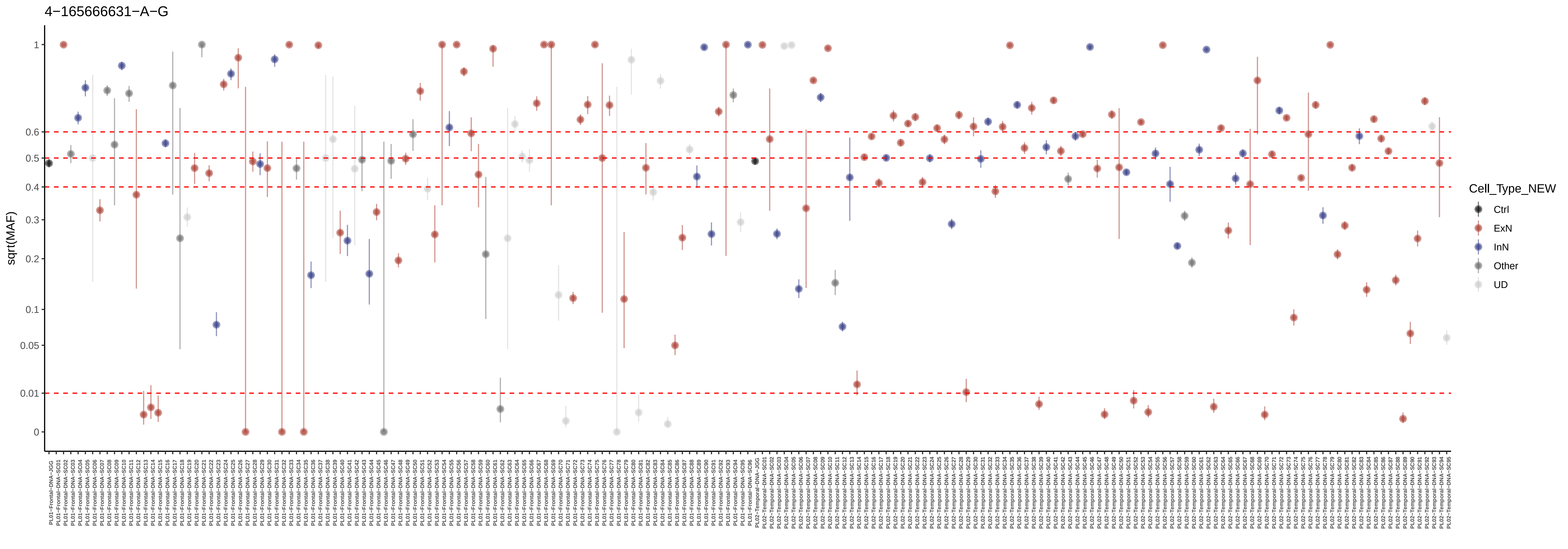
4−165666631−A−G
sqrt(MAF)
1
0.6
0.5
0.4
0.3
0.2
0.1
0.05
0.01
0
Cell_Type_NEW
Ctrl
ExN
InN
Other
UD

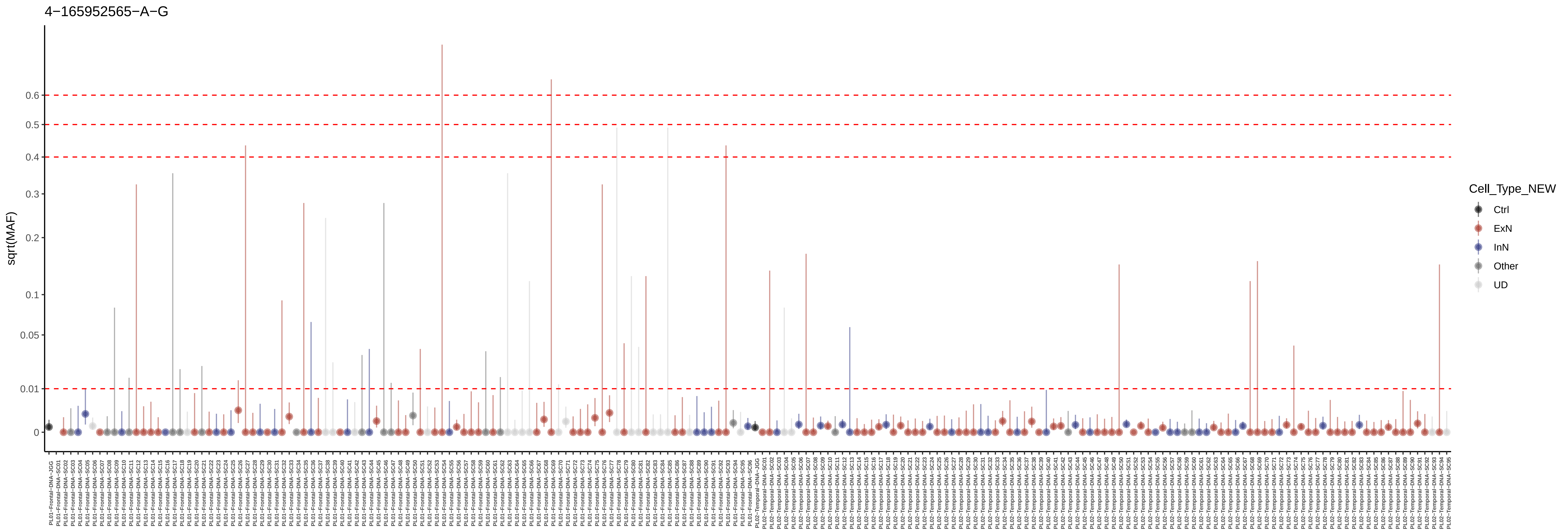

4−165952565−A−G
sqrt(MAF)
0.6
0.5
0.4
0.3
0.2
0.1
0.05
0.01
0
Cell_Type_NEW
Ctrl
ExN
InN
Other
UD

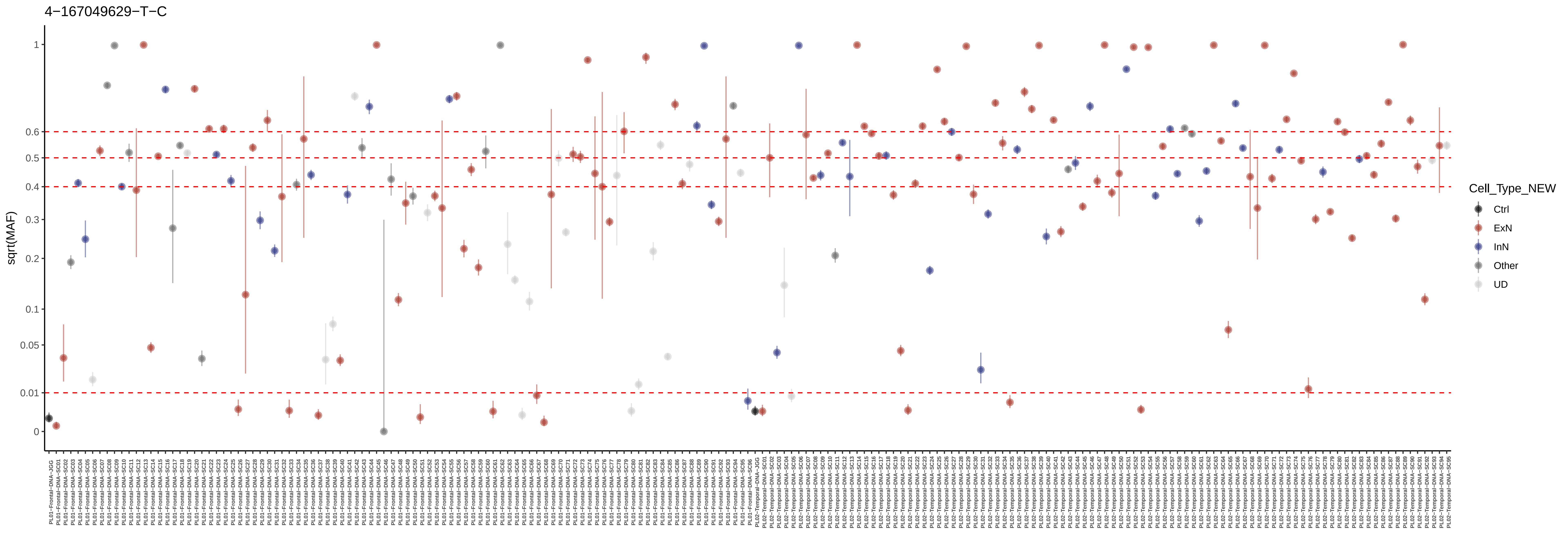

4−167049629−T−C
sqrt(MAF)
1
0.6
0.5
0.4
0.3
0.2
0.1
0.05
0.01
0
Cell_Type_NEW
Ctrl
ExN
InN
Other
UD

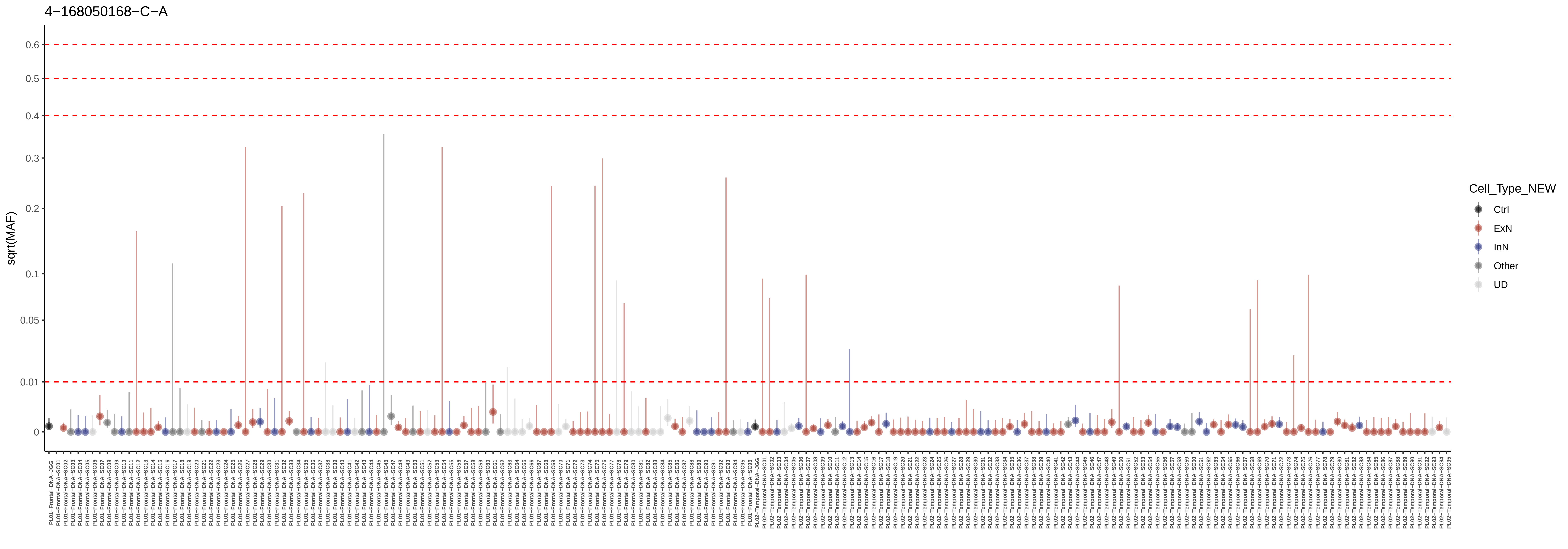
4−168050168−C−A
sqrt(MAF)
Cell_Type_NEW
Ctrl
ExN
InN
Other
UD

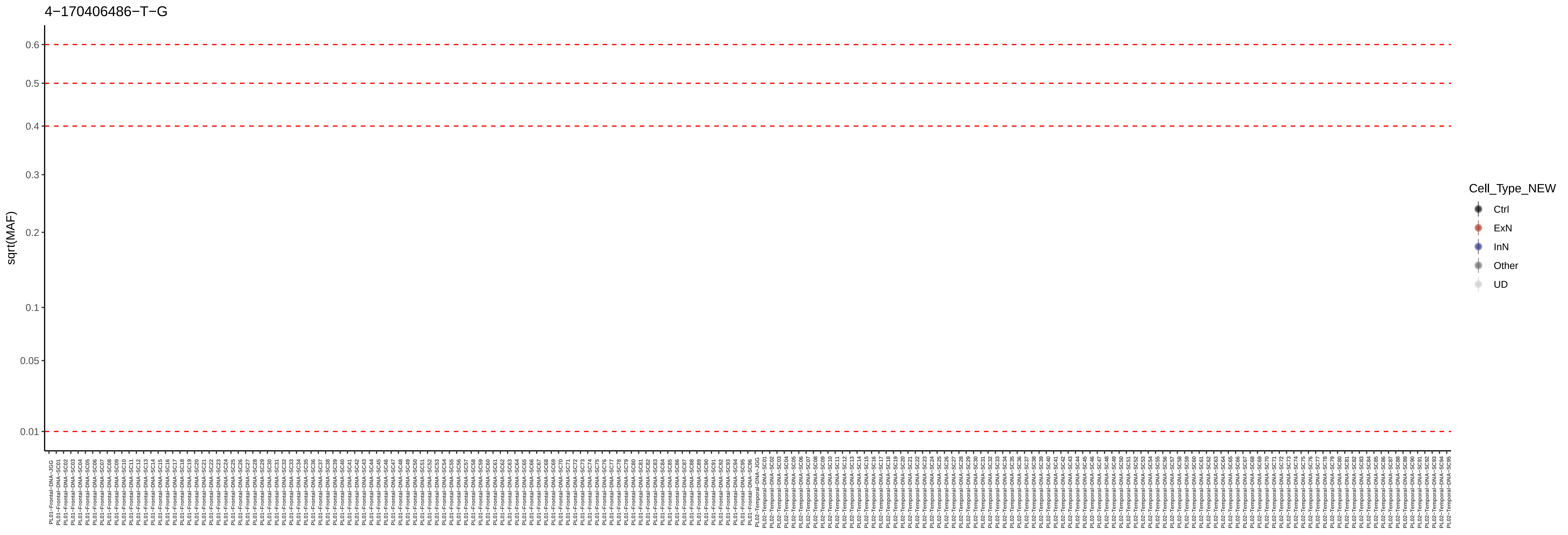
4−170406486−T−G
sqrt(MAF)
0.6
0.5
0.4
0.3
0.2
0.1
0.05
0.01
Cell_Type_NEW
Ctrl
ExN
InN
Other
UD

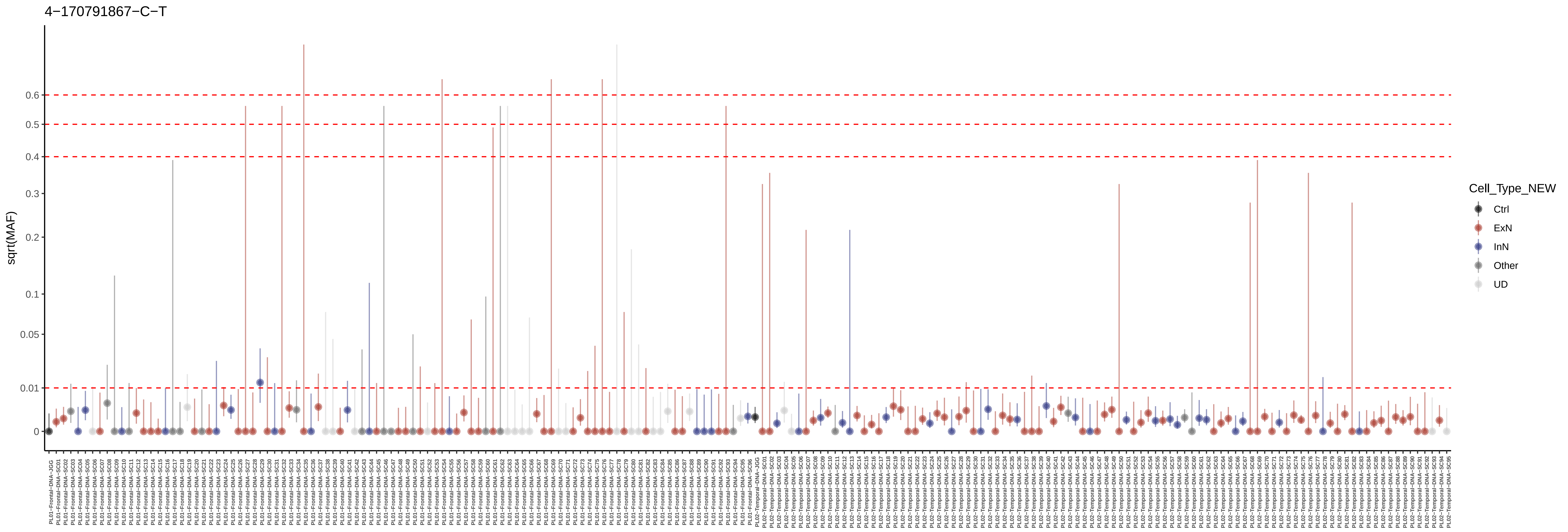

4−170791867−C−T
sqrt(MAF)
0.6
0.5
0.4
0.3
0.2
0.1
0.05
0.01
0
Cell_Type_NEW
Ctrl
ExN
InN
Other
UD

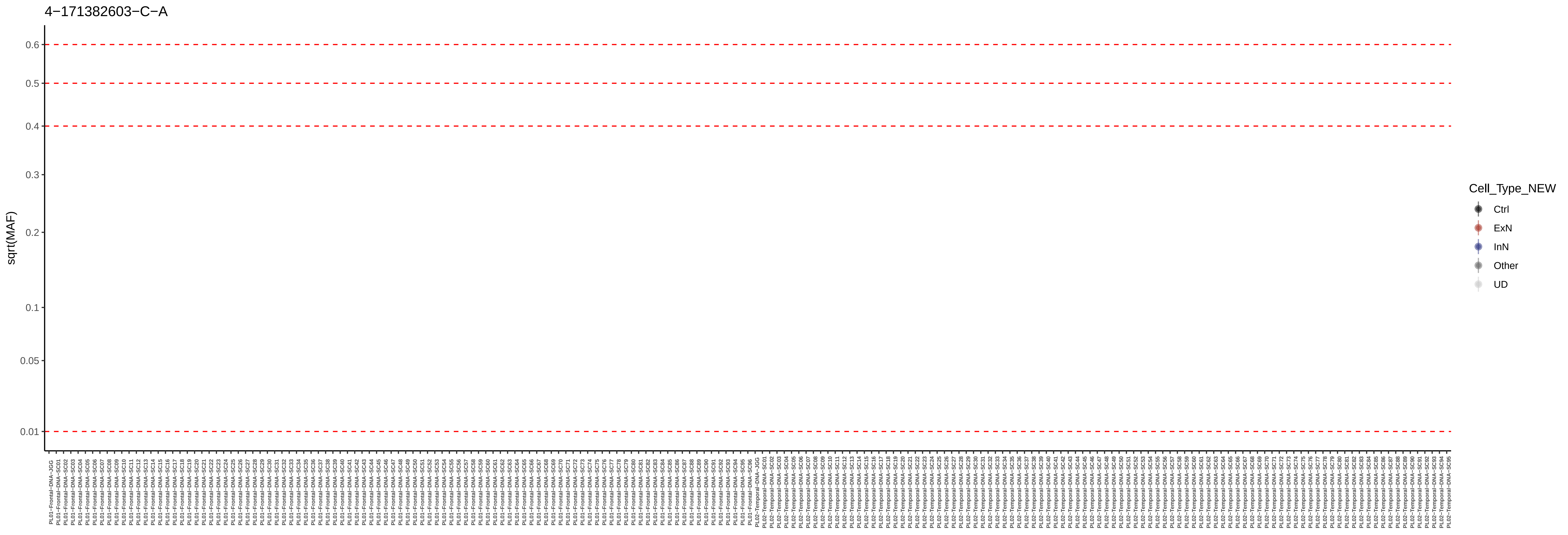
4−171382603−C−A
sqrt(MAF)
0.6
0.5
0.4
0.3
0.2
0.1
0.05
0.01
Cell_Type_NEW
Ctrl
ExN
InN
Other
UD

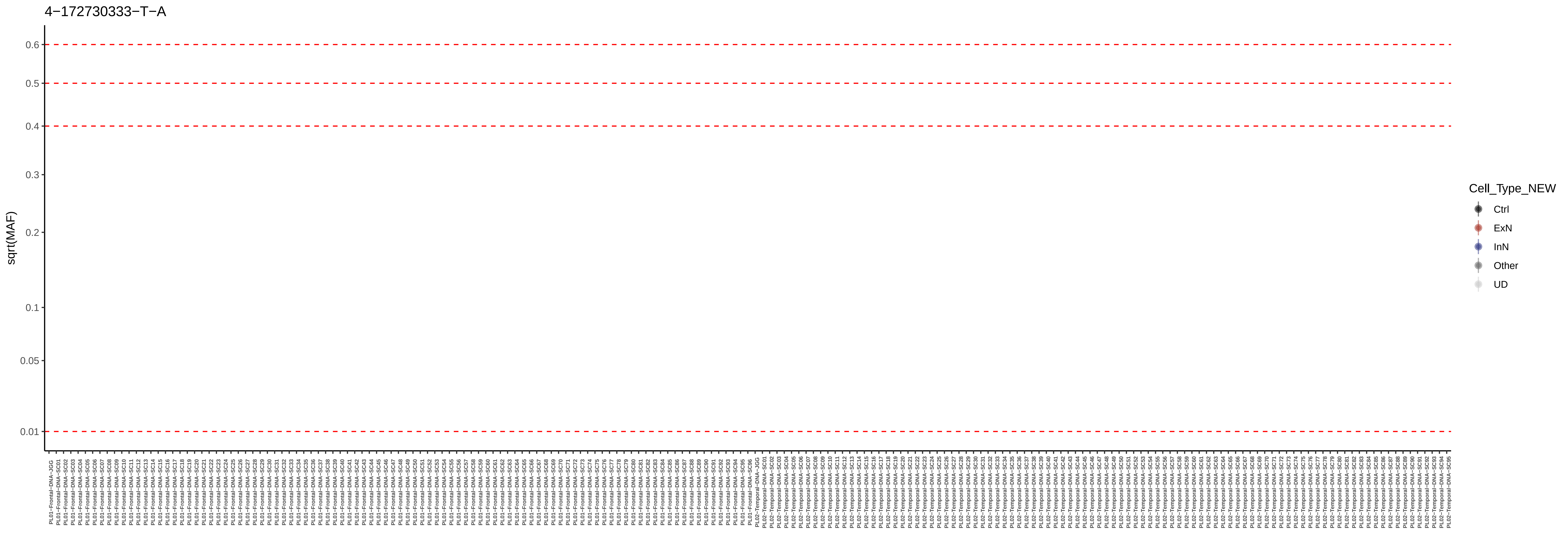
4−172730333−T−A
sqrt(MAF)
0.6
0.5
0.4
0.3
0.2
0.1
0.05
0.01
Cell_Type_NEW
Ctrl
ExN
InN
Other
UD

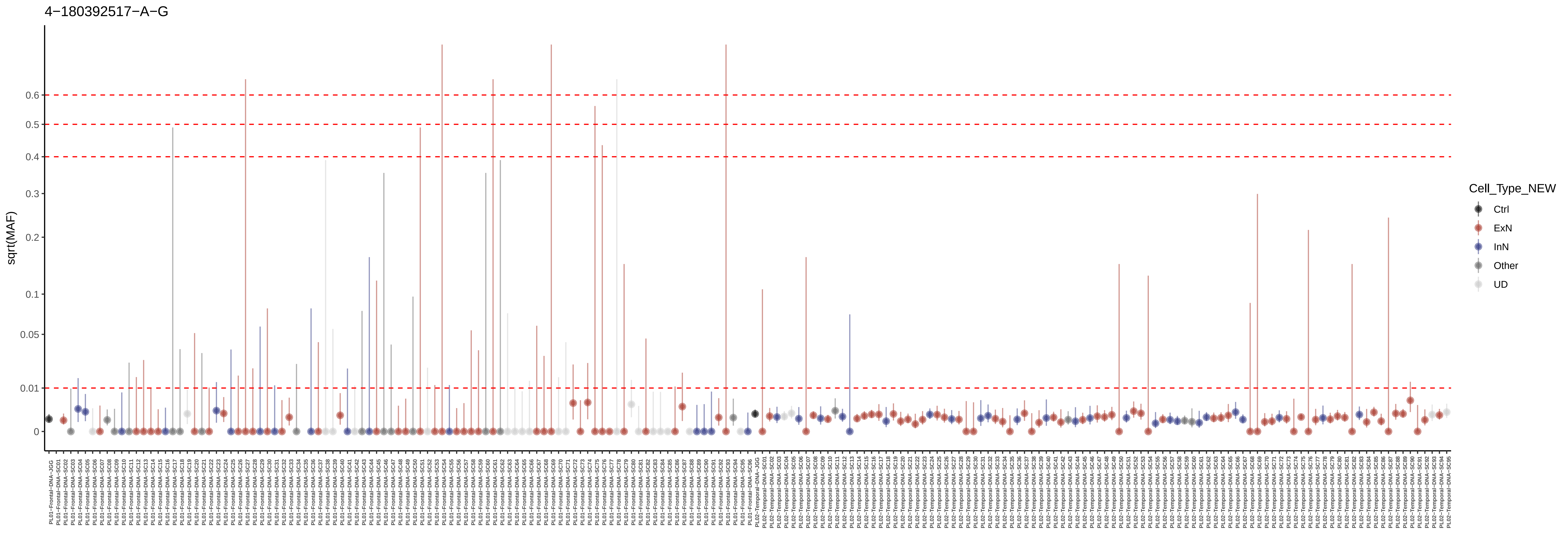

4−180392517−A−G
sqrt(MAF)
0.6
0.5
0.4
0.3
0.2
0.1
0.05
0.01
0
Cell_Type_NEW
Ctrl
ExN
InN
Other
UD

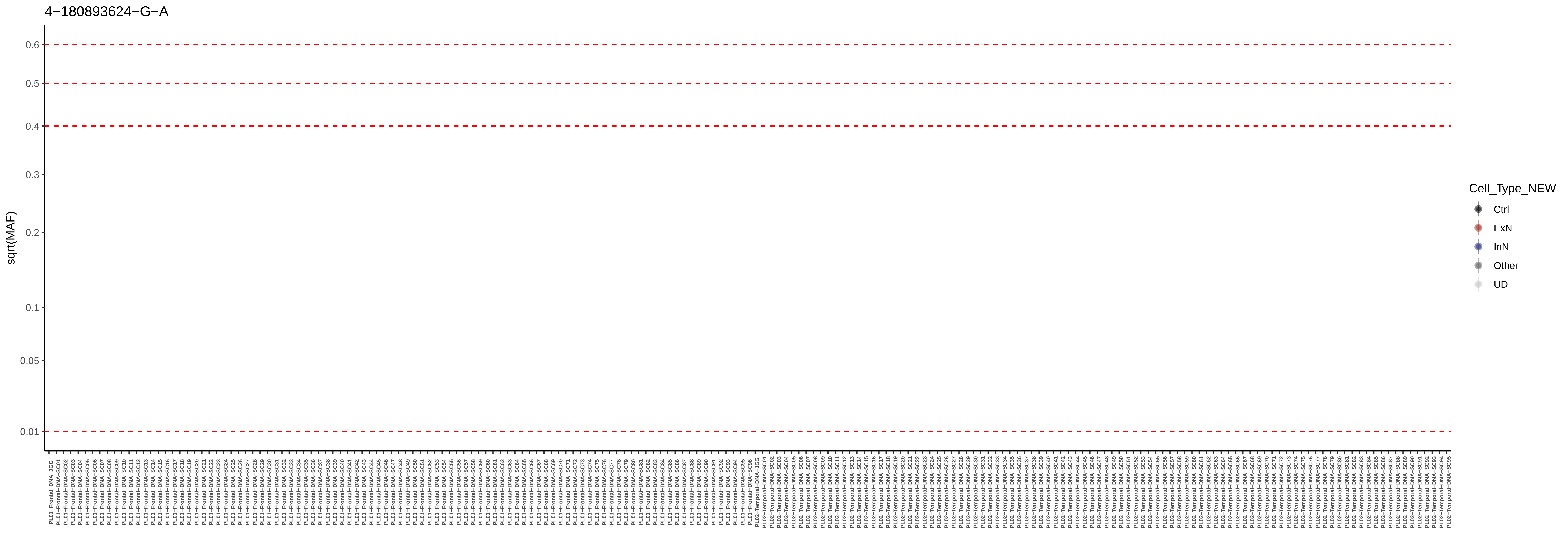

4−180893624−G−A
sqrt(MAF)
0.6
0.5
0.4
0.3
0.2
0.1
0.05
0.01
Cell_Type_NEW
Ctrl
ExN
InN
Other
UD

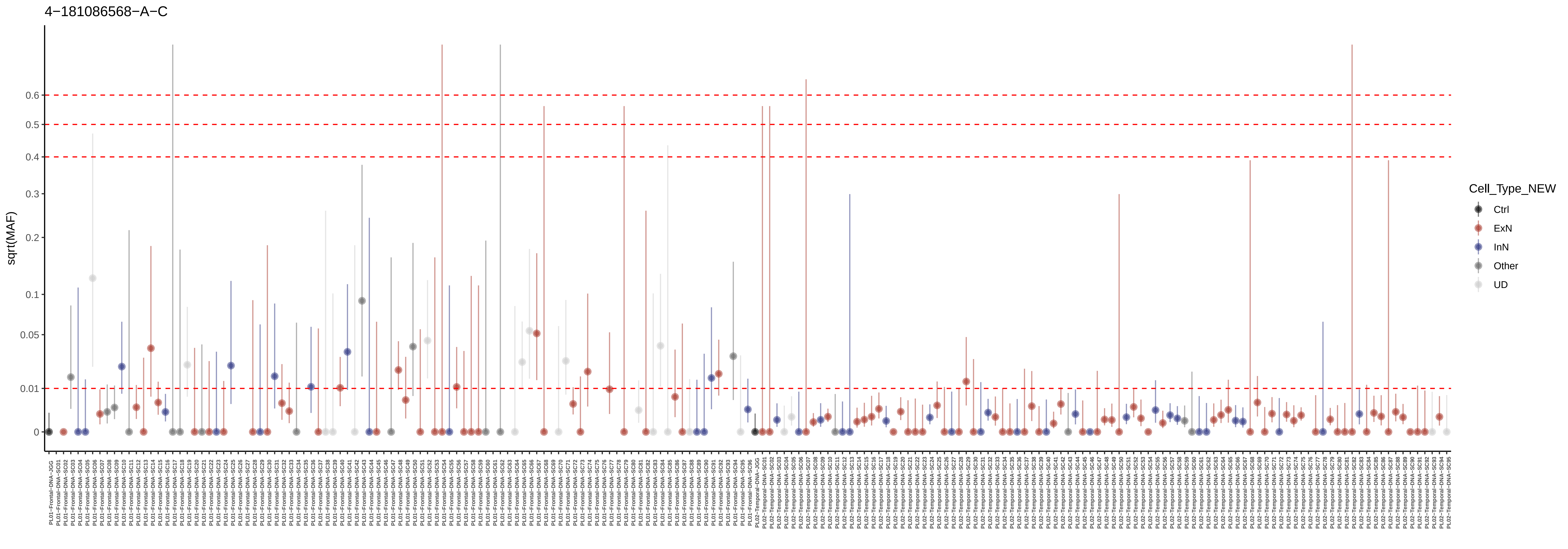

4−181086568−A−C
sqrt(MAF)
0.6
0.5
0.4
0.3
0.2
0.1
0.05
0.01
0
Cell_Type_NEW
Ctrl
ExN
InN
Other
UD

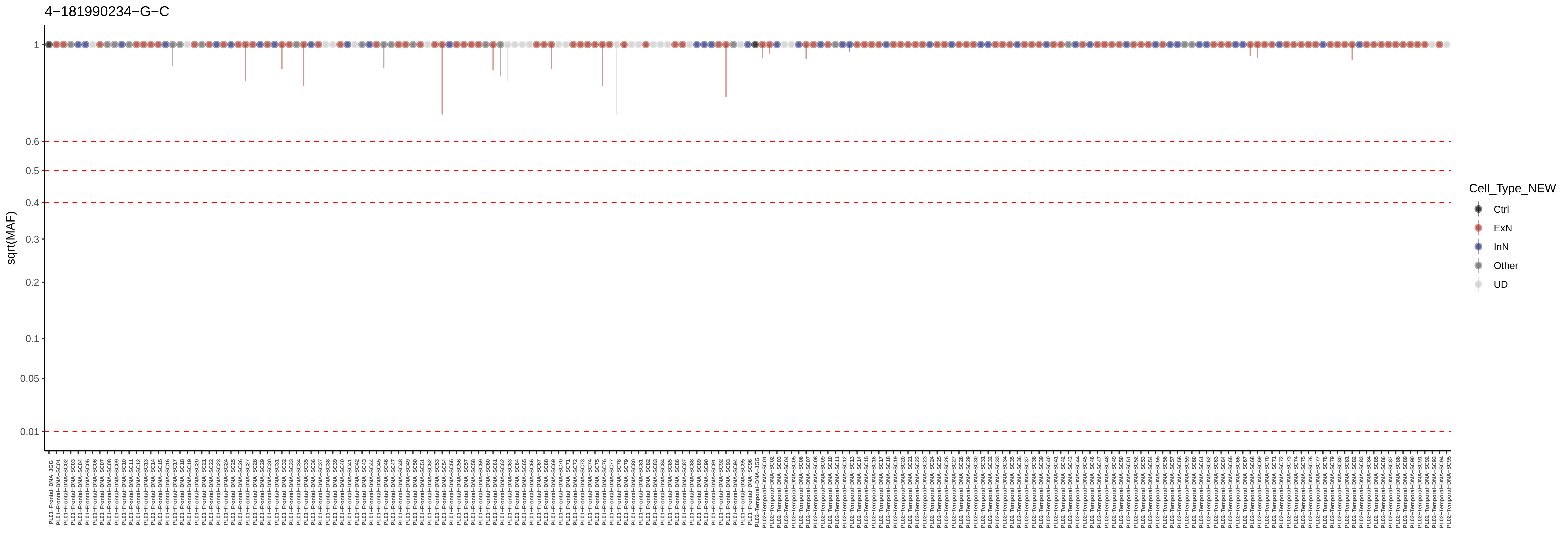

4−181990234−G−C
sqrt(MAF)
1
0.6
0.5
0.4
0.3
0.2
0.1
0.05
0.01
Cell_Type_NEW
Ctrl
ExN
InN
Other
UD

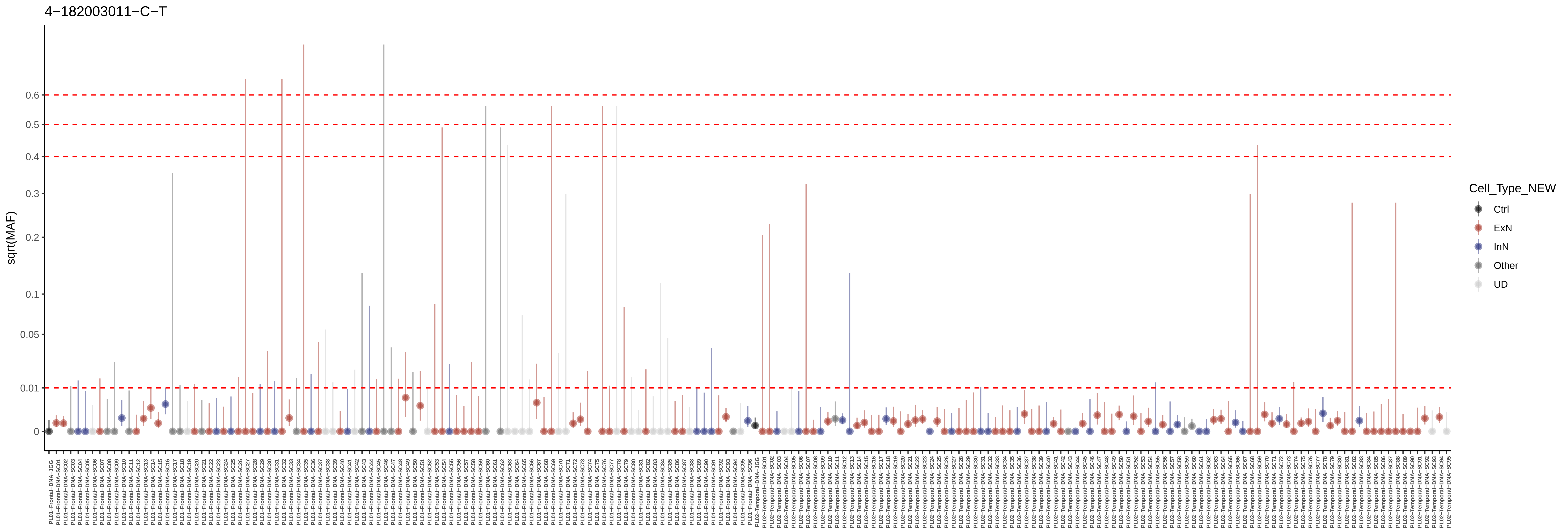
4−182003011−C−T
sqrt(MAF)
0.6
0.5
0.4
0.3
0.2
0.1
0.05
0.01
0
Cell_Type_NEW
Ctrl
ExN
InN
Other
UD

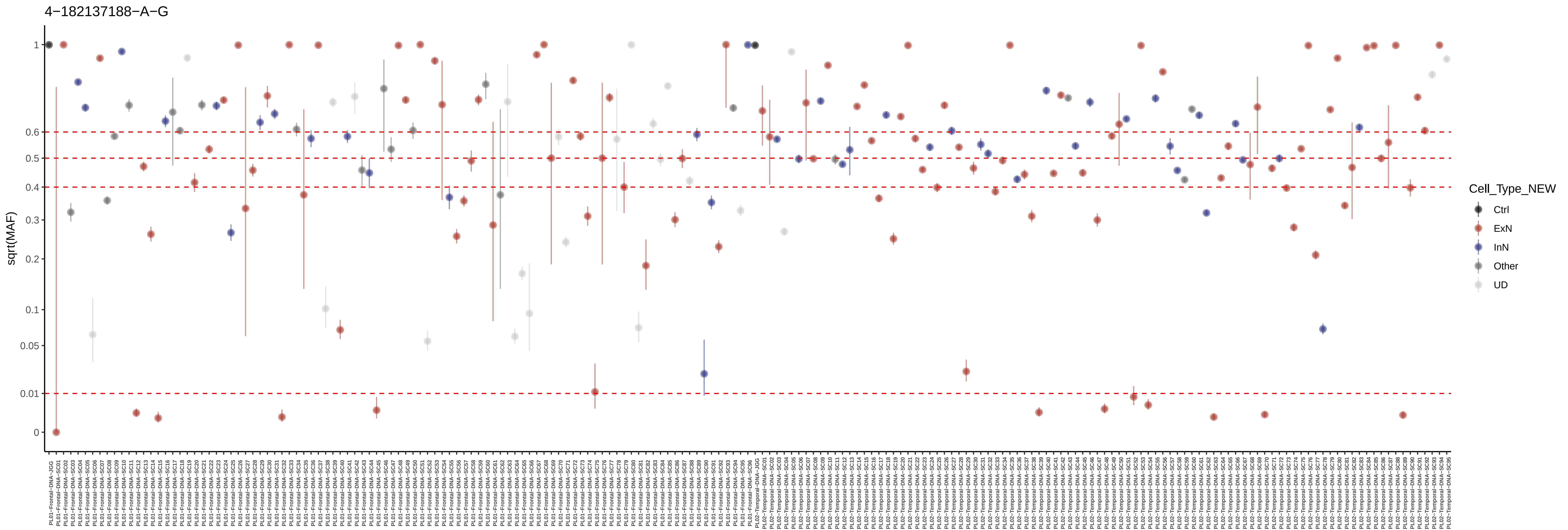

4−182137188−A−G
sqrt(MAF)
1
0.6
0.5
0.4
0.3
0.2
0.1
0.05
0.01
0
Cell_Type_NEW
Ctrl
ExN
InN
Other
UD

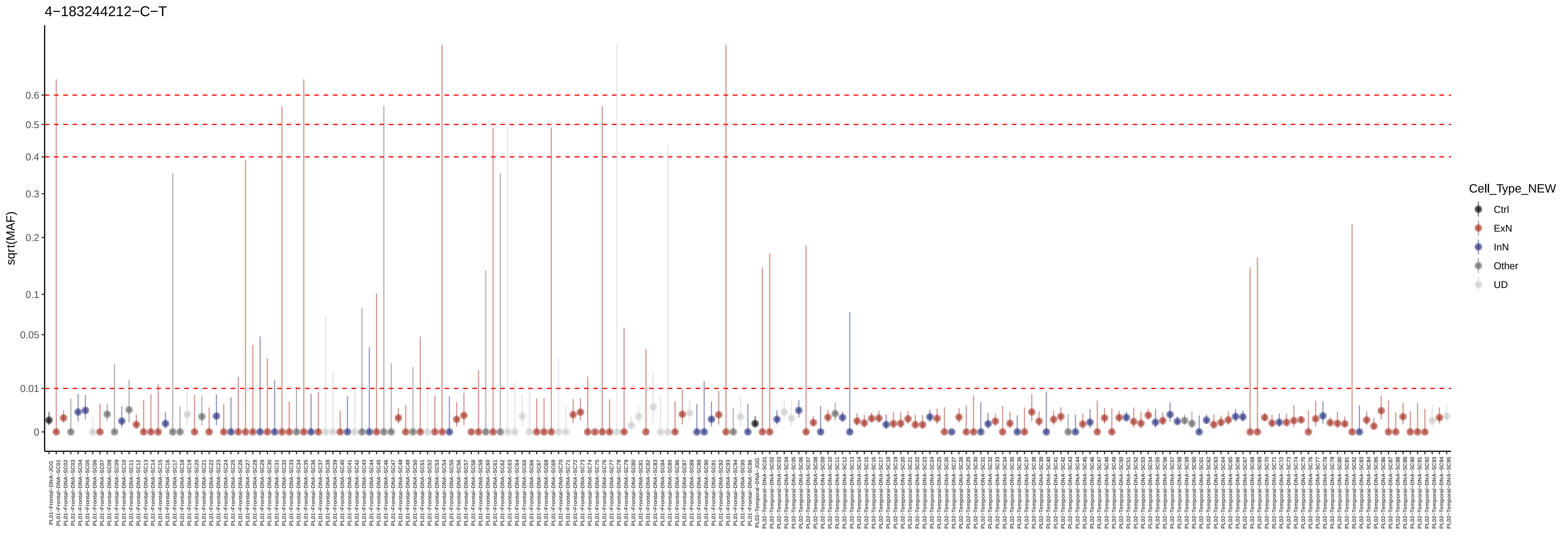
4−183244212−C−T
sqrt(MAF)
0.6
0.5
0.4
0.3
0.2
0.1
0.05
0.01
0
Cell_Type_NEW
Ctrl
ExN
InN
Other
UD

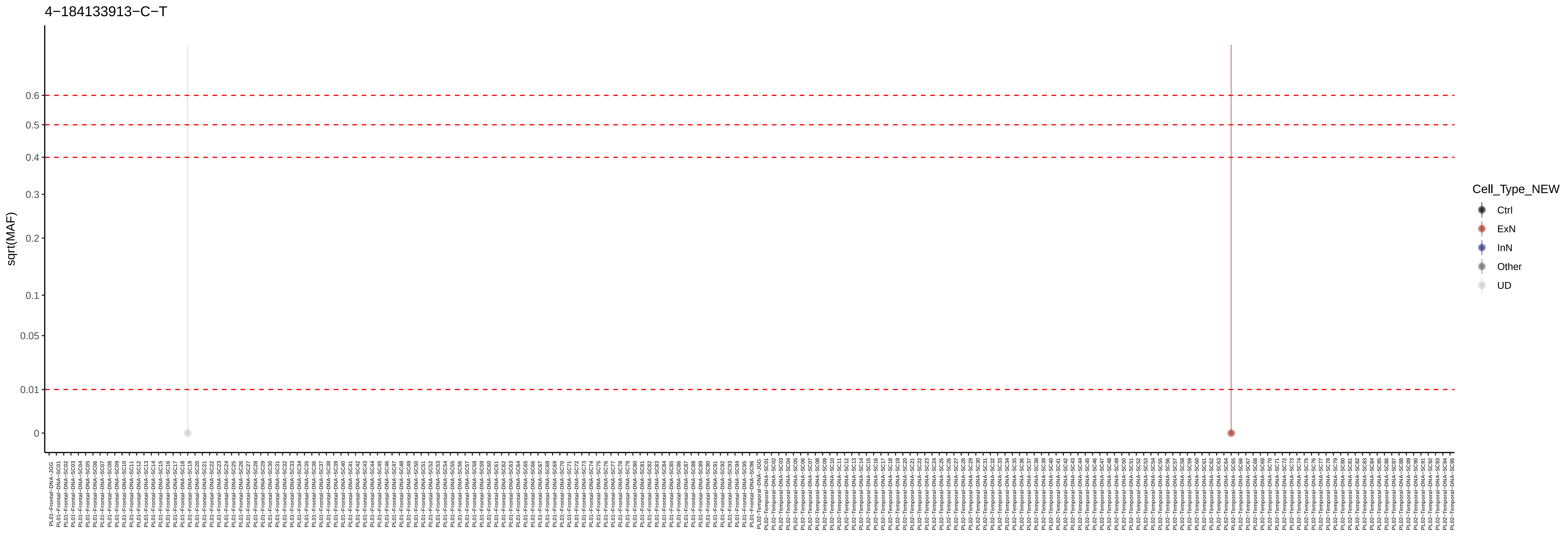
4−184133913−C−T
sqrt(MAF)
0.6
0.5
0.4
0.3
0.2
0.1
0.05
0.01
0
Cell_Type_NEW
Ctrl
ExN
InN
Other
UD

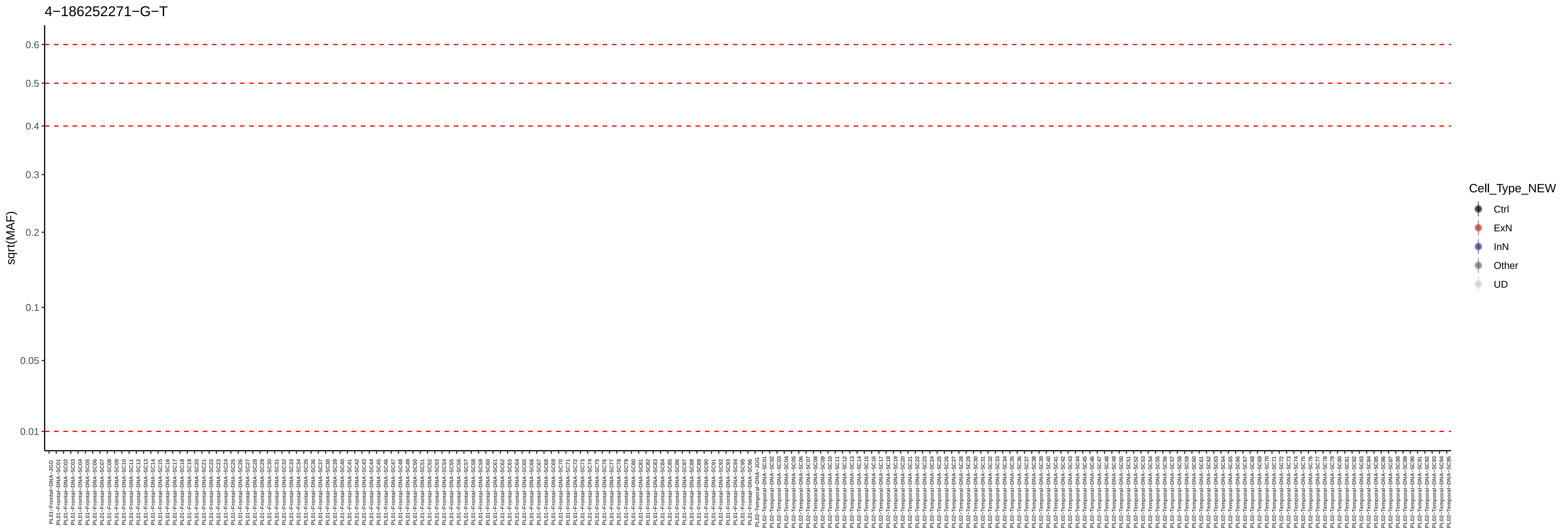
4−186252271−G−T
sqrt(MAF)
0.6
0.5
0.4
0.3
0.2
0.1
0.05
0.01
Cell_Type_NEW
Ctrl
ExN
InN
Other
UD

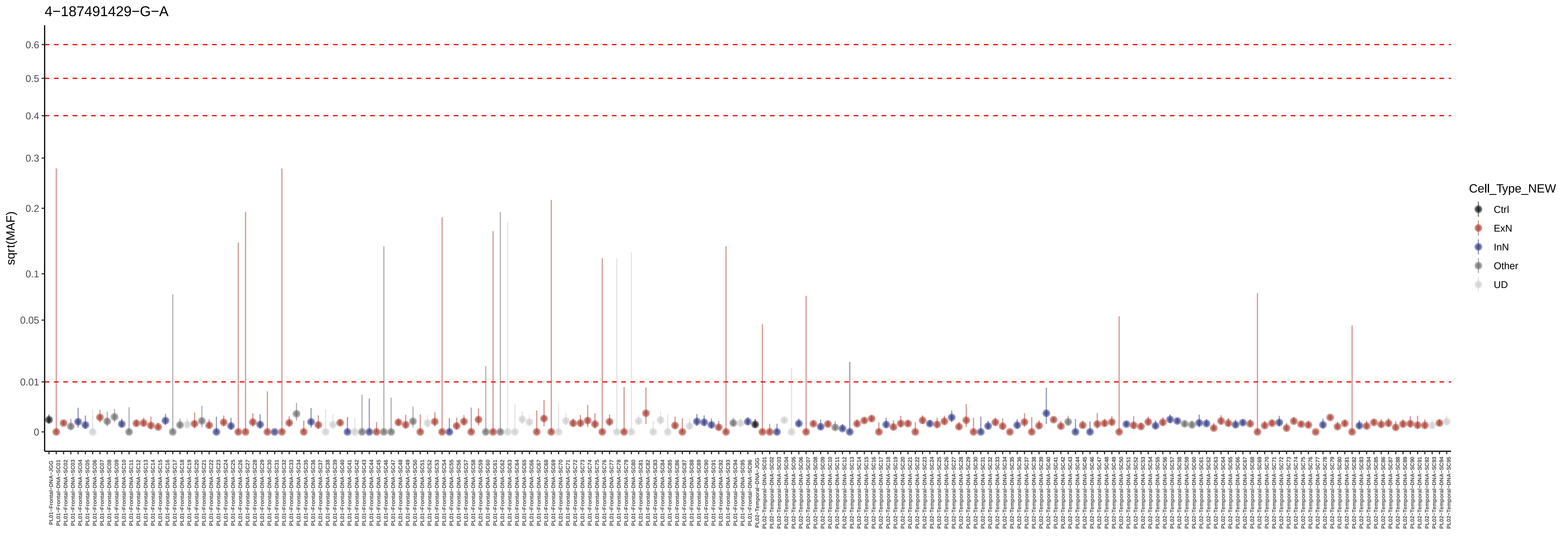
4−187491429−G−A
sqrt(MAF)
0.6
0.5
0.4
0.3
0.2
0.1
0.05
0.01
0
Cell_Type_NEW
Ctrl
ExN
InN
Other
UD

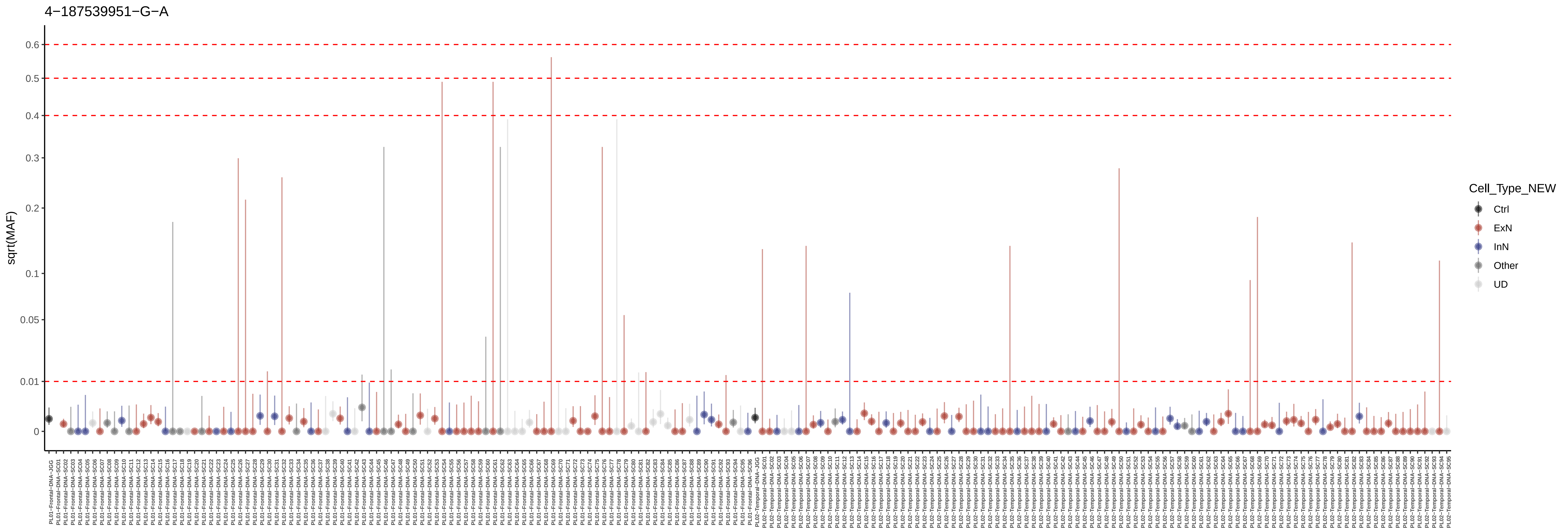
4−187539951−G−A
sqrt(MAF)
0.6
0.5
0.4
0.3
0.2
0.1
0.05
0.01
0
Cell_Type_NEW
Ctrl
ExN
InN
Other
UD

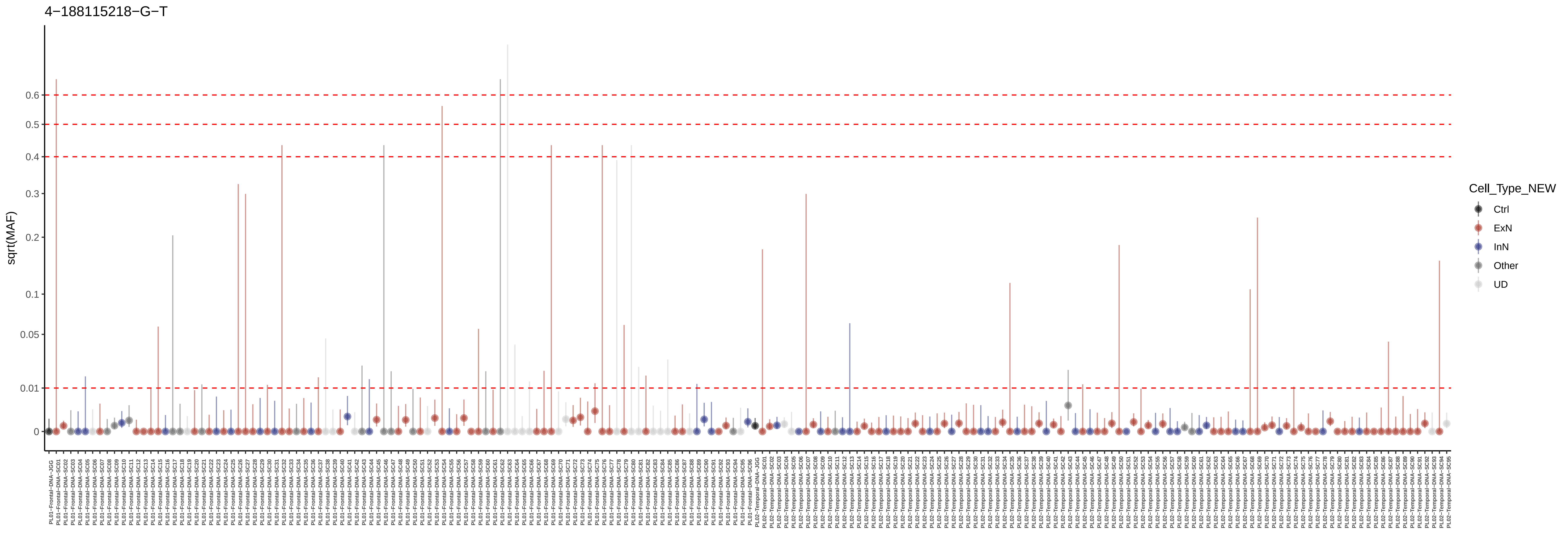
4−188115218−G−T
sqrt(MAF)
0.6
0.5
0.4
0.3
0.2
0.1
0.05
0.01
0
Cell_Type_NEW
Ctrl
ExN
InN
Other
UD

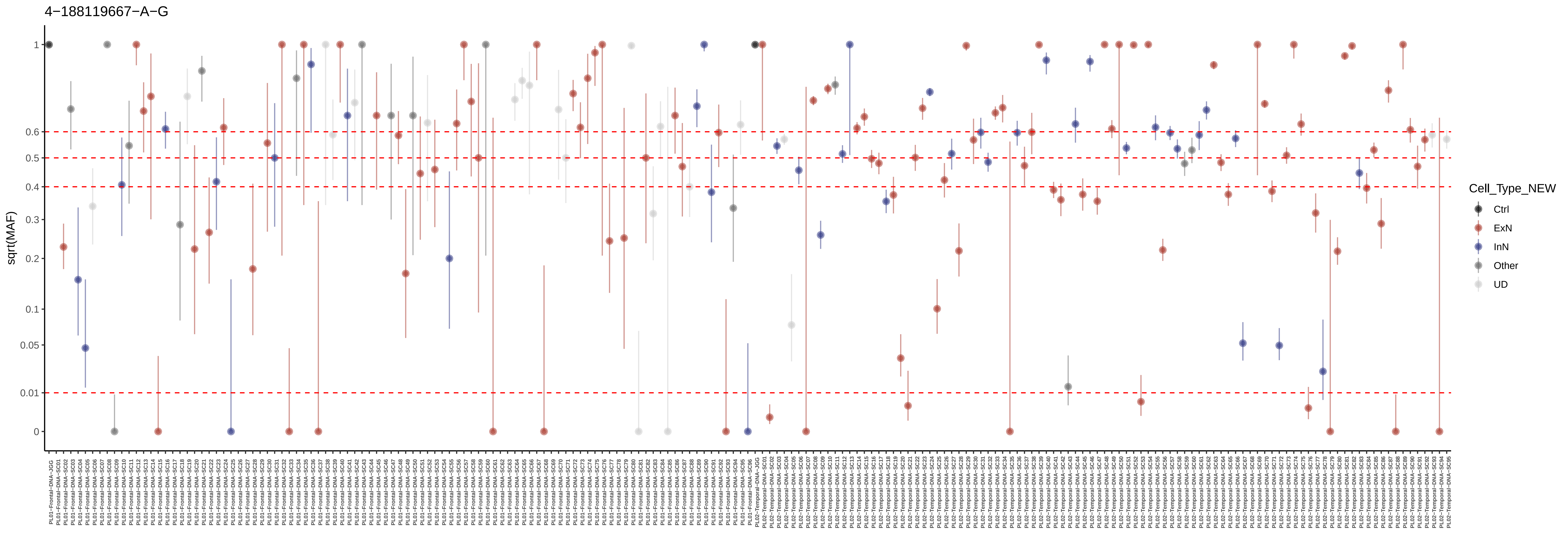

4−188119667−A−G
sqrt(MAF)
Cell_Type_NEW
Ctrl
ExN
InN
Other
UD

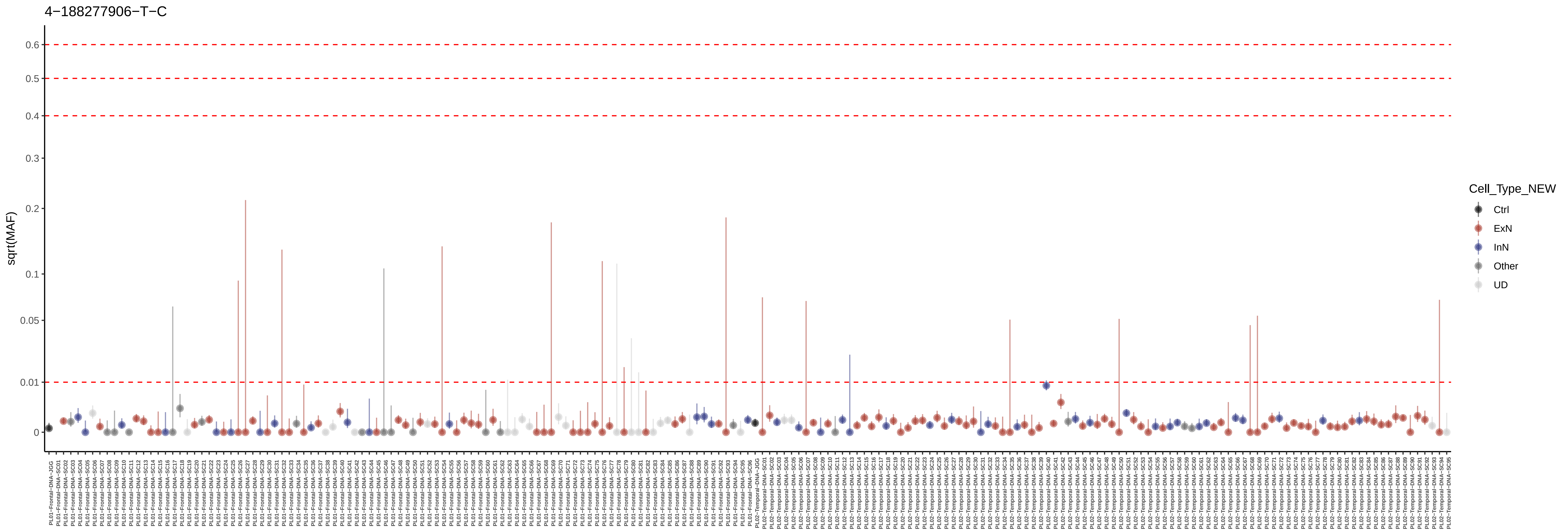
4−188277906−T−C
sqrt(MAF)
0.6
0.5
0.4
0.3
0.2
0.1
0.05
0.01
0
Cell_Type_NEW
Ctrl
ExN
InN
Other
UD

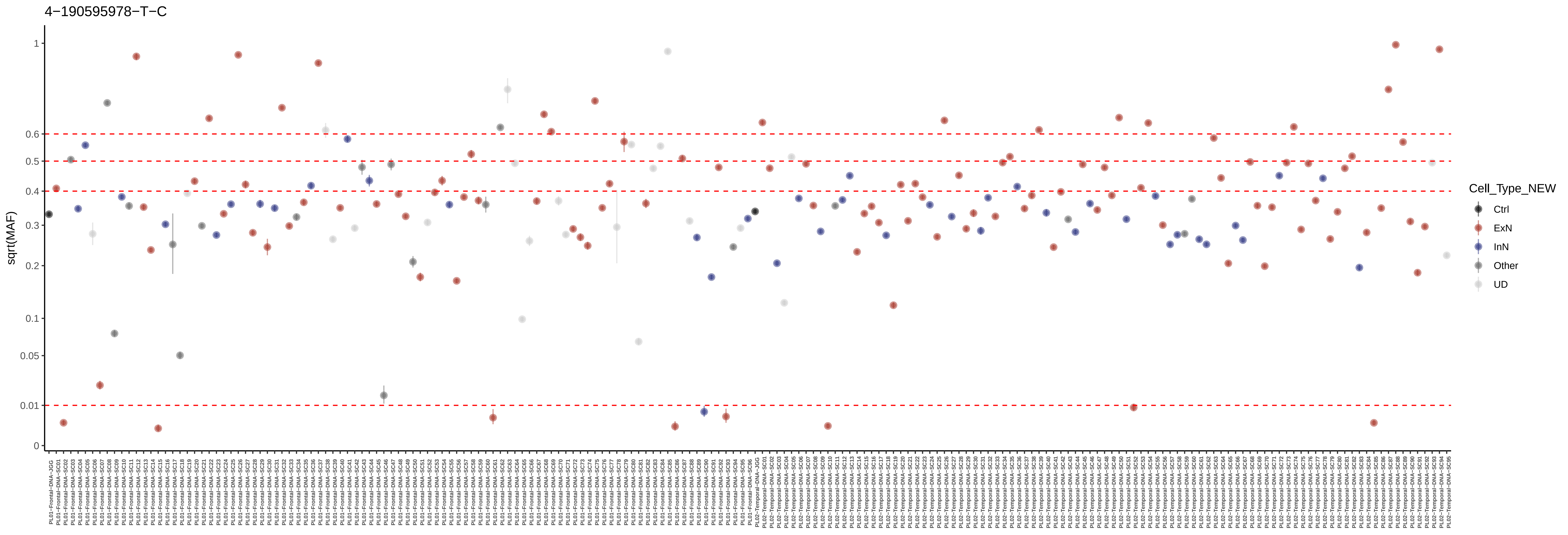
4−190595978−T−C
sqrt(MAF)
1
0.6
0.5
0.4
0.3
0.2
0.1
0.05
0.01
0
Cell_Type_NEW
Ctrl
ExN
InN
Other
UD

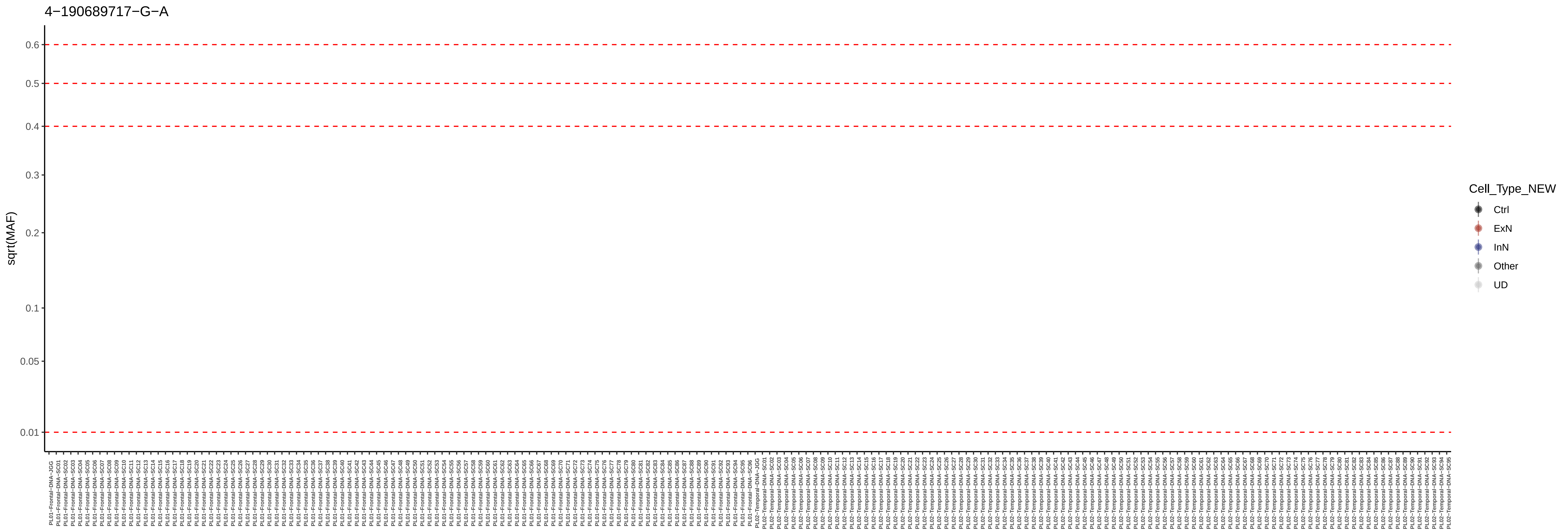
4−190689717−G−A
sqrt(MAF)
0.6
0.5
0.4
0.3
0.2
0.1
0.05
0.01
Cell_Type_NEW
Ctrl
ExN
InN
Other
UD

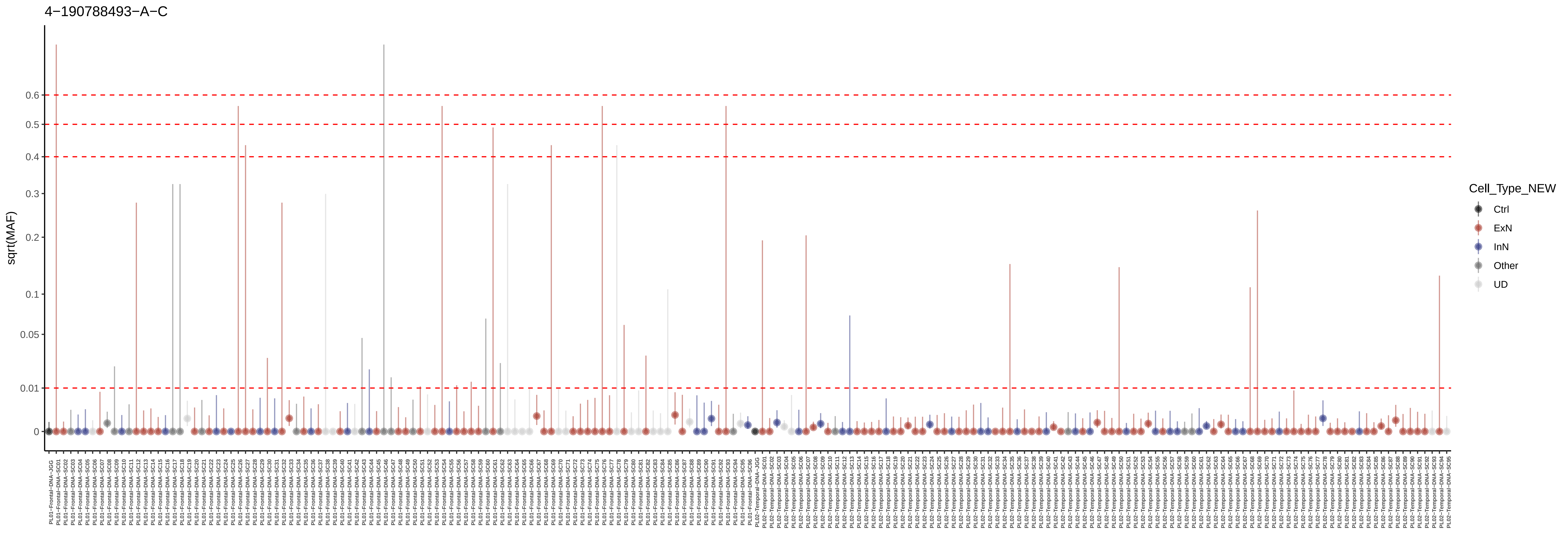

4−190788493−A−C
sqrt(MAF)
0.6
0.5
0.4
0.3
0.2
0.1
0.05
0.01
0
Cell_Type_NEW
Ctrl
ExN
InN
Other
UD

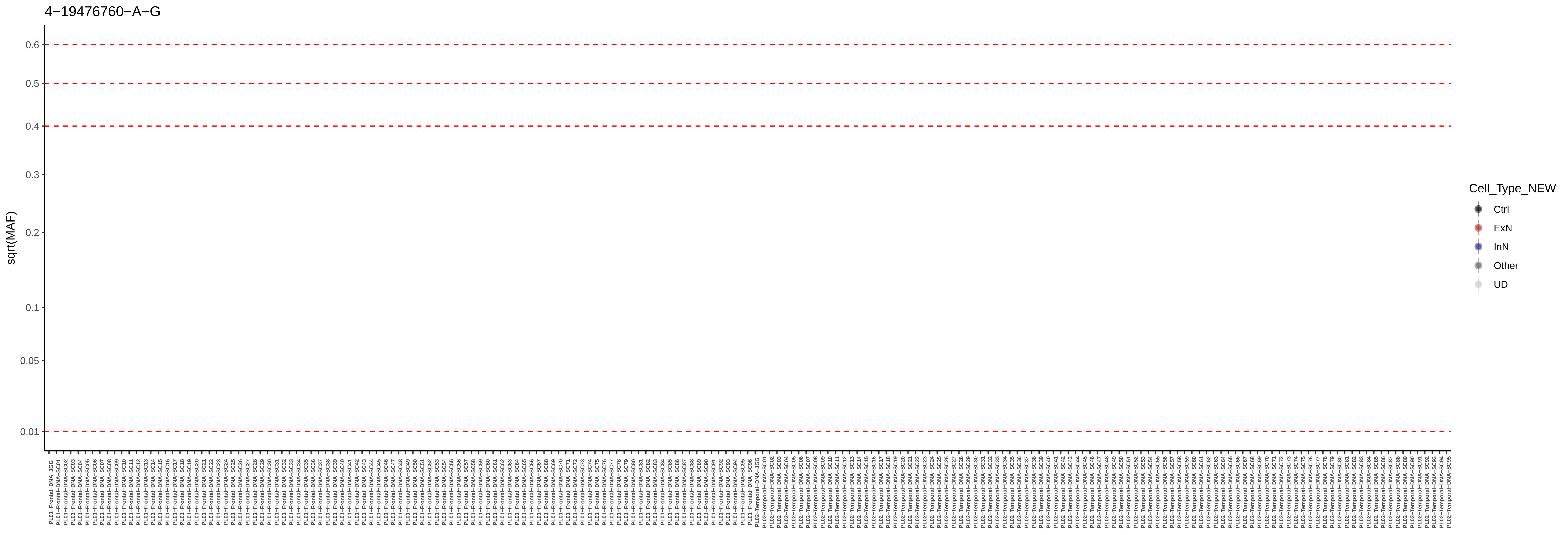
4−19476760−A−G
sqrt(MAF)
0.6
0.5
0.4
0.3
0.2
0.1
0.05
0.01
Cell_Type_NEW
Ctrl
ExN
InN
Other
UD

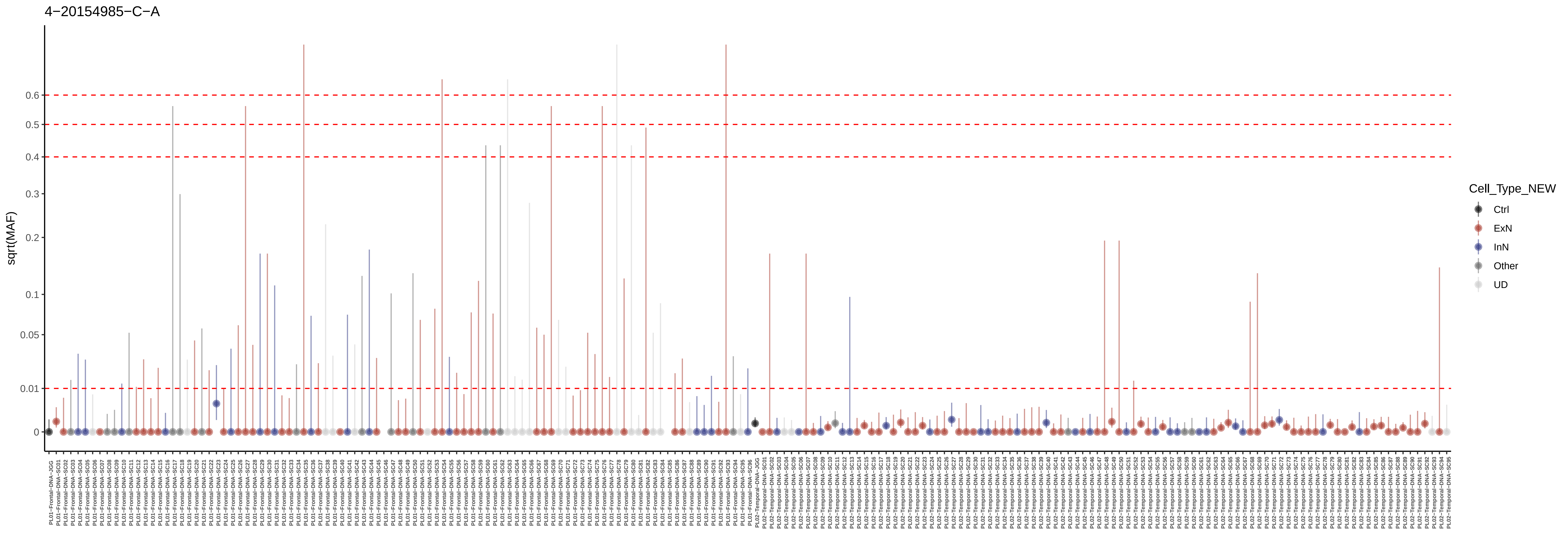
4−20154985−C−A
sqrt(MAF)
0.6
0.5
0.4
0.3
0.2
0.1
0.05
0.01
0
Cell_Type_NEW
Ctrl
ExN
InN
Other
UD

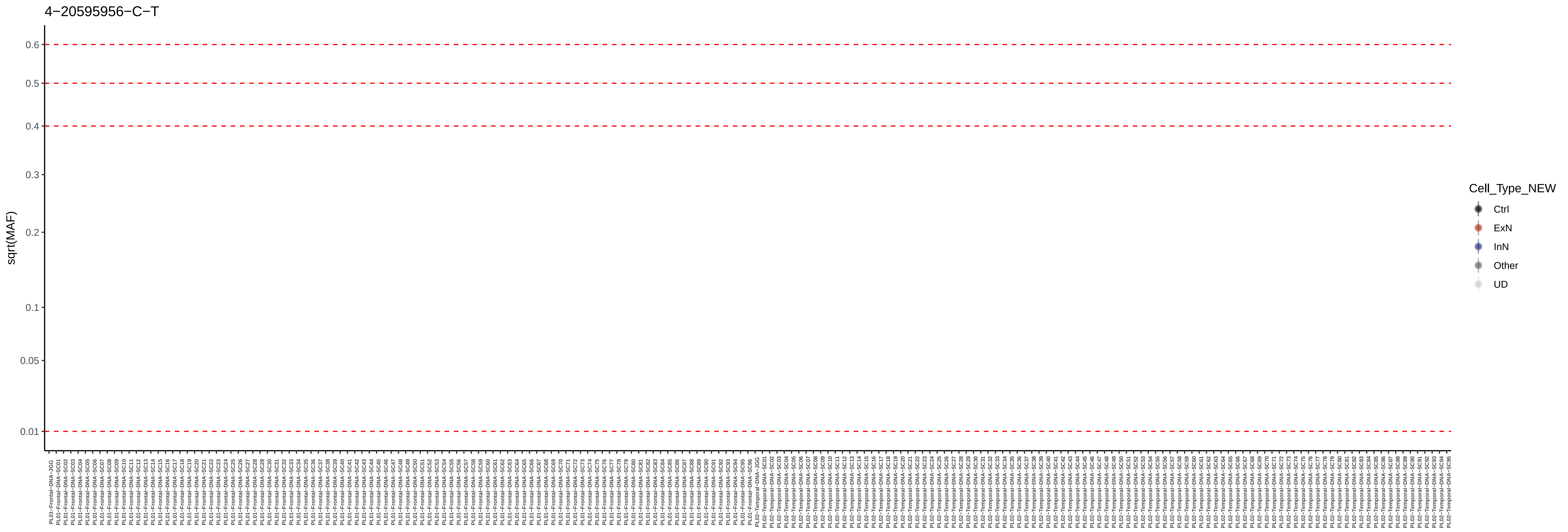
4−20595956−C−T
sqrt(MAF)
0.6
0.5
0.4
0.3
0.2
0.1
0.05
0.01
Cell_Type_NEW
Ctrl
ExN
InN
Other
UD

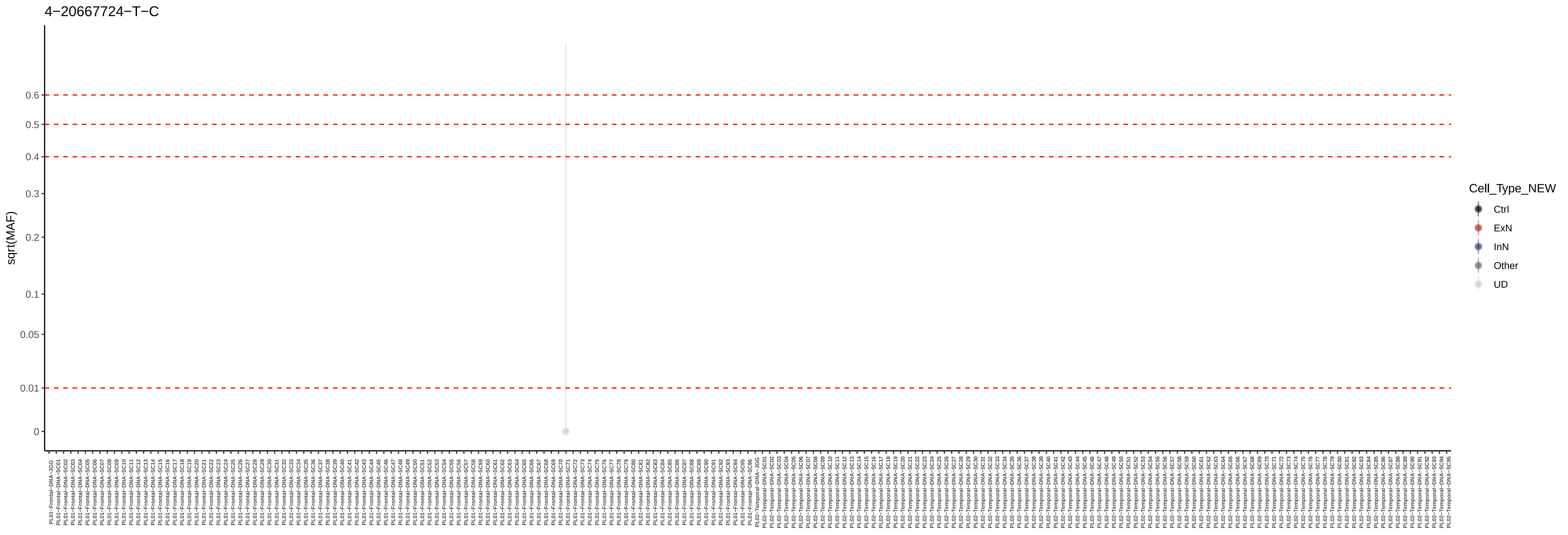
4−20667724−T−C
sqrt(MAF)
0.6
0.5
0.4
0.3
0.2
0.1
0.05
0.01
0
Cell_Type_NEW
Ctrl
ExN
InN
Other
UD

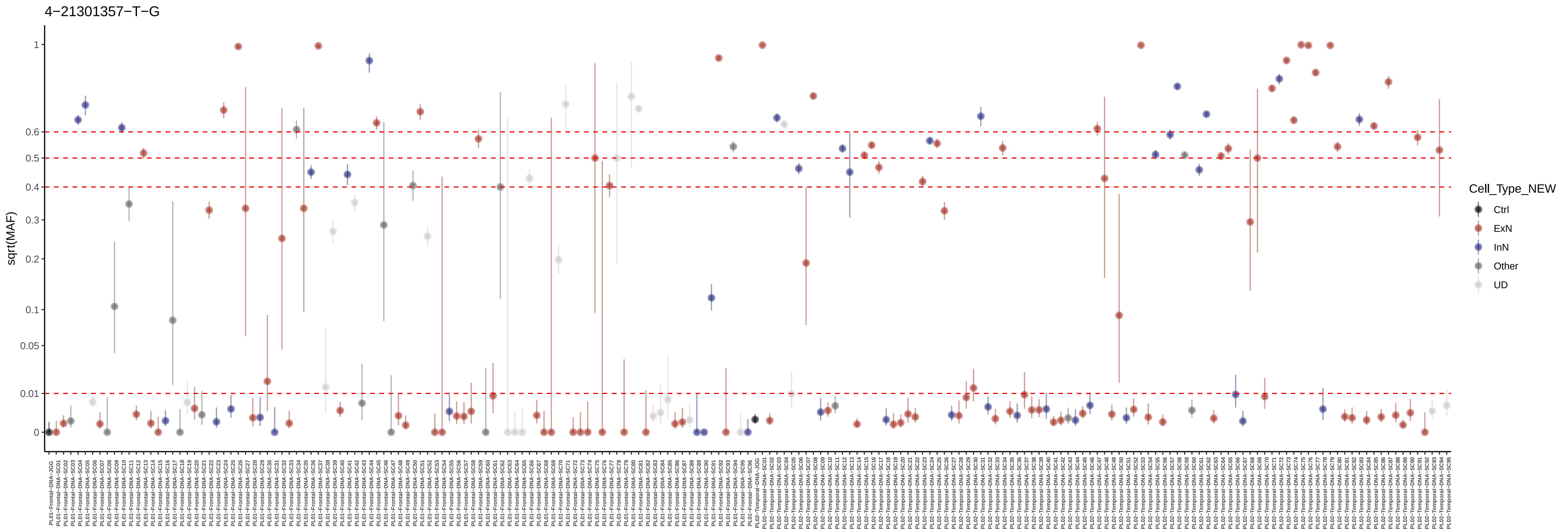
4−21301357−T−G
sqrt(MAF)
1
0.6
0.5
0.4
0.3
0.2
0.1
0.05
0.01
0
Cell_Type_NEW
Ctrl
ExN
InN
Other
UD

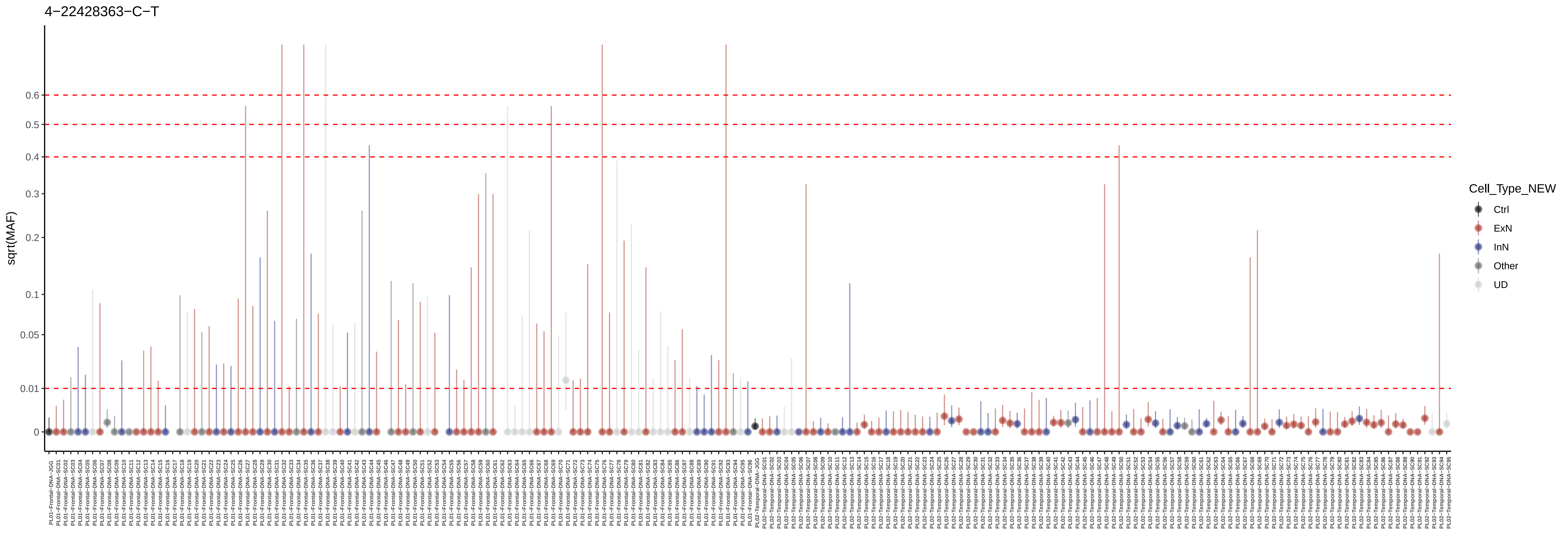
4−22428363−C−T
sqrt(MAF)
0.6
0.5
0.4
0.3
0.2
0.1
0.05
0.01
0
Cell_Type_NEW
Ctrl
ExN
InN
Other
UD

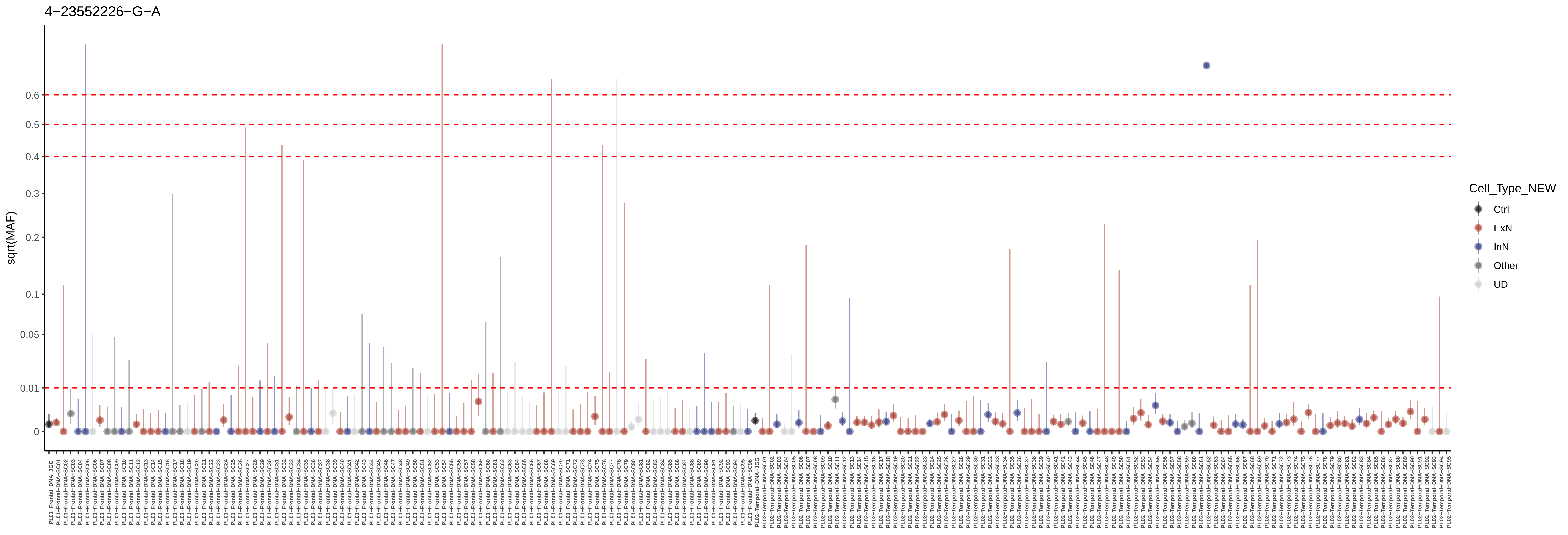

4−23552226−G−A
sqrt(MAF)
0
0.01
0.05
0.1
0.2
0.3
0.4
0.5
0.6
Cell_Type_NEW
Ctrl
ExN
InN
Other
UD

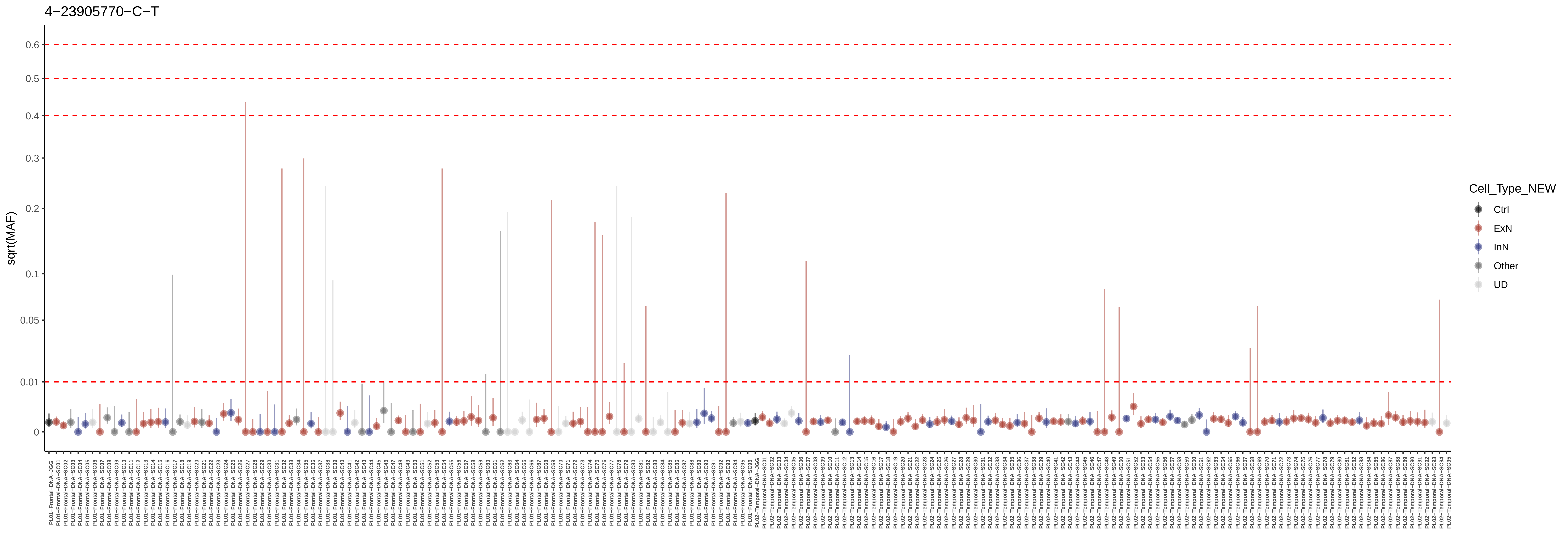
4−23905770−C−T
sqrt(MAF)
Cell_Type_NEW
Ctrl
ExN
InN
Other
UD

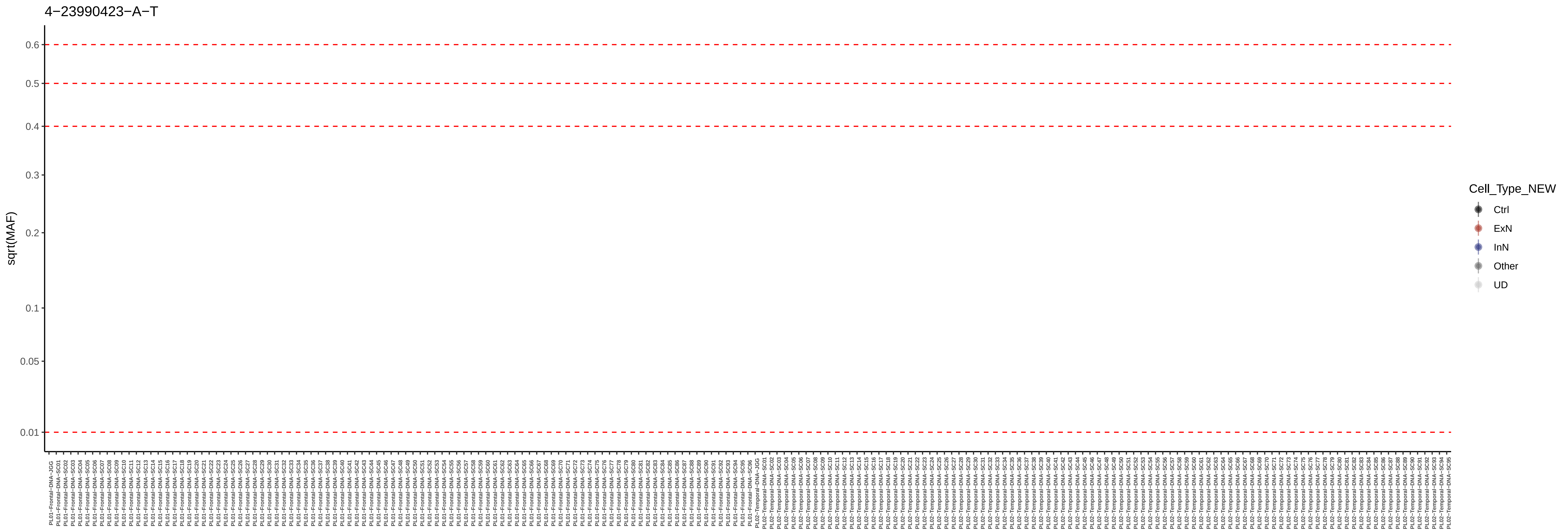
4−23990423−A−T
sqrt(MAF)
0.6
0.5
0.4
0.3
0.2
0.1
0.05
0.01
Cell_Type_NEW
Ctrl
ExN
InN
Other
UD
PL01−Frontal−DNA−JGG
PL01−Frontal−DNA−SC01
PL01−Frontal−DNA−SC02
PL01−Frontal−DNA−SC03
PL01−Frontal−DNA−SC04
PL01−Frontal−DNA−SC05
PL01−Frontal−DNA−SC06
PL01−Frontal−DNA−SC07
PL01−Frontal−DNA−SC08
PL01−Frontal−DNA−SC09
PL01−Frontal−DNA−SC10
PL01−Frontal−DNA−SC11
PL01−Frontal−DNA−SC12
PL01−Frontal−DNA−SC13
PL01−Frontal−DNA−SC14
PL01−Frontal−DNA−SC15
PL01−Frontal−DNA−SC16
PL01−Frontal−DNA−SC17
PL01−Frontal−DNA−SC18
PL01−Frontal−DNA−SC19
PL01−Frontal−DNA−SC20
PL01−Frontal−DNA−SC21
PL01−Frontal−DNA−SC22
PL01−Frontal−DNA−SC23
PL01−Frontal−DNA−SC24
PL01−Frontal−DNA−SC25
PL01−Frontal−DNA−SC26
PL01−Frontal−DNA−SC27
PL01−Frontal−DNA−SC28
PL01−Frontal−DNA−SC29
PL01−Frontal−DNA−SC30
PL01−Frontal−DNA−SC31
PL01−Frontal−DNA−SC32
PL01−Frontal−DNA−SC33
PL01−Frontal−DNA−SC34
PL01−Frontal−DNA−SC35
PL01−Frontal−DNA−SC36
PL01−Frontal−DNA−SC37
PL01−Frontal−DNA−SC38
PL01−Frontal−DNA−SC39
PL01−Frontal−DNA−SC40
PL01−Frontal−DNA−SC41
PL01−Frontal−DNA−SC42
PL01−Frontal−DNA−SC43
PL01−Frontal−DNA−SC44
PL01−Frontal−DNA−SC45
PL01−Frontal−DNA−SC46
PL01−Frontal−DNA−SC47
PL01−Frontal−DNA−SC48
PL01−Frontal−DNA−SC49
PL01−Frontal−DNA−SC50
PL01−Frontal−DNA−SC51
PL01−Frontal−DNA−SC52
PL01−Frontal−DNA−SC53
PL01−Frontal−DNA−SC54
PL01−Frontal−DNA−SC55
PL01−Frontal−DNA−SC56
PL01−Frontal−DNA−SC57
PL01−Frontal−DNA−SC58
PL01−Frontal−DNA−SC59
PL01−Frontal−DNA−SC60
PL01−Frontal−DNA−SC61
PL01−Frontal−DNA−SC62
PL01−Frontal−DNA−SC63
PL01−Frontal−DNA−SC64
PL01−Frontal−DNA−SC65
PL01−Frontal−DNA−SC66
PL01−Frontal−DNA−SC67
PL01−Frontal−DNA−SC68
PL01−Frontal−DNA−SC69
PL01−Frontal−DNA−SC70
PL01−Frontal−DNA−SC71
PL01−Frontal−DNA−SC72
PL01−Frontal−DNA−SC73
PL01−Frontal−DNA−SC74
PL01−Frontal−DNA−SC75
PL01−Frontal−DNA−SC76
PL01−Frontal−DNA−SC77
PL01−Frontal−DNA−SC78
PL01−Frontal−DNA−SC79
PL01−Frontal−DNA−SC80
PL01−Frontal−DNA−SC81
PL01−Frontal−DNA−SC82
PL01−Frontal−DNA−SC83
PL01−Frontal−DNA−SC84
PL01−Frontal−DNA−SC85
PL01−Frontal−DNA−SC86
PL01−Frontal−DNA−SC87
PL01−Frontal−DNA−SC88
PL01−Frontal−DNA−SC89
PL01−Frontal−DNA−SC90
PL01−Frontal−DNA−SC91
PL01−Frontal−DNA−SC92
PL01−Frontal−DNA−SC93
PL01−Frontal−DNA−SC94
PL01−Frontal−DNA−SC95
PL01−Frontal−DNA−SC96
PL02−Temporal−DNA−JGG
PL02−Temporal−DNA−SC01
PL02−Temporal−DNA−SC02
PL02−Temporal−DNA−SC03
PL02−Temporal−DNA−SC04
PL02−Temporal−DNA−SC05
PL02−Temporal−DNA−SC06
PL02−Temporal−DNA−SC07
PL02−Temporal−DNA−SC08
PL02−Temporal−DNA−SC09
PL02−Temporal−DNA−SC10
PL02−Temporal−DNA−SC11
PL02−Temporal−DNA−SC12
PL02−Temporal−DNA−SC13
PL02−Temporal−DNA−SC14
PL02−Temporal−DNA−SC15
PL02−Temporal−DNA−SC16
PL02−Temporal−DNA−SC17
PL02−Temporal−DNA−SC18
PL02−Temporal−DNA−SC19
PL02−Temporal−DNA−SC20
PL02−Temporal−DNA−SC21
PL02−Temporal−DNA−SC22
PL02−Temporal−DNA−SC23
PL02−Temporal−DNA−SC24
PL02−Temporal−DNA−SC25
PL02−Temporal−DNA−SC26
PL02−Temporal−DNA−SC27
PL02−Temporal−DNA−SC28
PL02−Temporal−DNA−SC29
PL02−Temporal−DNA−SC30
PL02−Temporal−DNA−SC31
PL02−Temporal−DNA−SC32
PL02−Temporal−DNA−SC33
PL02−Temporal−DNA−SC34
PL02−Temporal−DNA−SC35
PL02−Temporal−DNA−SC36
PL02−Temporal−DNA−SC37
PL02−Temporal−DNA−SC38
PL02−Temporal−DNA−SC39
PL02−Temporal−DNA−SC40
PL02−Temporal−DNA−SC41
PL02−Temporal−DNA−SC42
PL02−Temporal−DNA−SC43
PL02−Temporal−DNA−SC44
PL02−Temporal−DNA−SC45
PL02−Temporal−DNA−SC46
PL02−Temporal−DNA−SC47
PL02−Temporal−DNA−SC48
PL02−Temporal−DNA−SC49
PL02−Temporal−DNA−SC50
PL02−Temporal−DNA−SC51
PL02−Temporal−DNA−SC52
PL02−Temporal−DNA−SC53
PL02−Temporal−DNA−SC54
PL02−Temporal−DNA−SC55
PL02−Temporal−DNA−SC56
PL02−Temporal−DNA−SC57
PL02−Temporal−DNA−SC58
PL02−Temporal−DNA−SC59
PL02−Temporal−DNA−SC60
PL02−Temporal−DNA−SC61
PL02−Temporal−DNA−SC62
PL02−Temporal−DNA−SC63
PL02−Temporal−DNA−SC64
PL02−Temporal−DNA−SC65
PL02−Temporal−DNA−SC66
PL02−Temporal−DNA−SC67
PL02−Temporal−DNA−SC68
PL02−Temporal−DNA−SC69
PL02−Temporal−DNA−SC70
PL02−Temporal−DNA−SC71
PL02−Temporal−DNA−SC72
PL02−Temporal−DNA−SC73
PL02−Temporal−DNA−SC74
PL02−Temporal−DNA−SC75
PL02−Temporal−DNA−SC76
PL02−Temporal−DNA−SC77
PL02−Temporal−DNA−SC78
PL02−Temporal−DNA−SC79
PL02−Temporal−DNA−SC80
PL02−Temporal−DNA−SC81
PL02−Temporal−DNA−SC82
PL02−Temporal−DNA−SC83
PL02−Temporal−DNA−SC84
PL02−Temporal−DNA−SC85
PL02−Temporal−DNA−SC86
PL02−Temporal−DNA−SC87
PL02−Temporal−DNA−SC88
PL02−Temporal−DNA−SC89
PL02−Temporal−DNA−SC90
PL02−Temporal−DNA−SC91
PL02−Temporal−DNA−SC92
PL02−Temporal−DNA−SC93
PL02−Temporal−DNA−SC94
PL02−Temporal−DNA−SC95

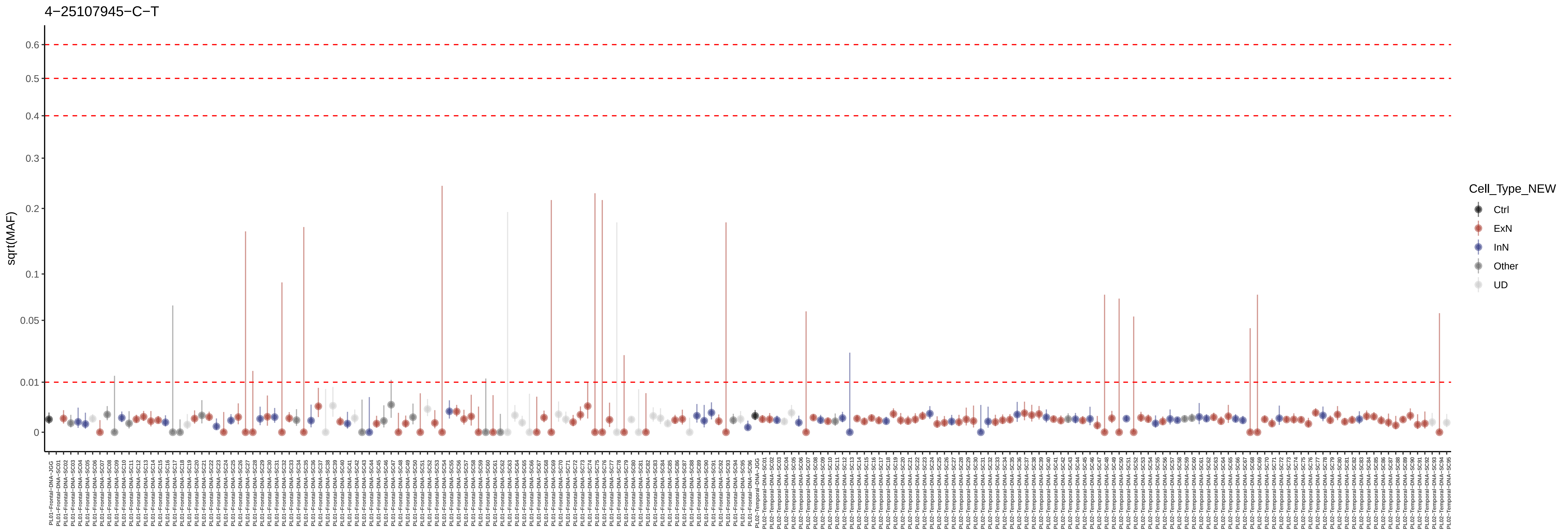

4−25107945−C−T
sqrt(MAF)
0.6
0.5
0.4
0.3
0.2
0.1
0.05
0.01
0
Cell_Type_NEW
Ctrl
ExN
InN
Other
UD

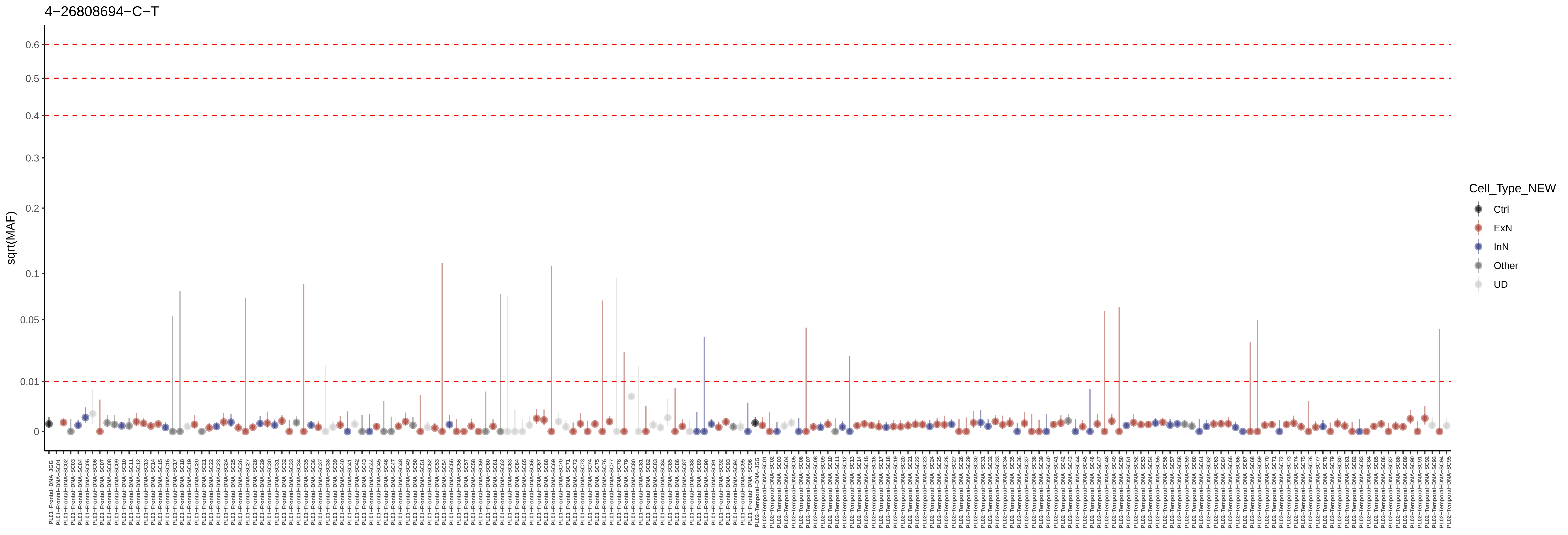

4−26808694−C−T
sqrt(MAF)
0.6
0.5
0.4
0.3
0.2
0.1
0.05
0.01
0
Cell_Type_NEW
Ctrl
ExN
InN
Other
UD

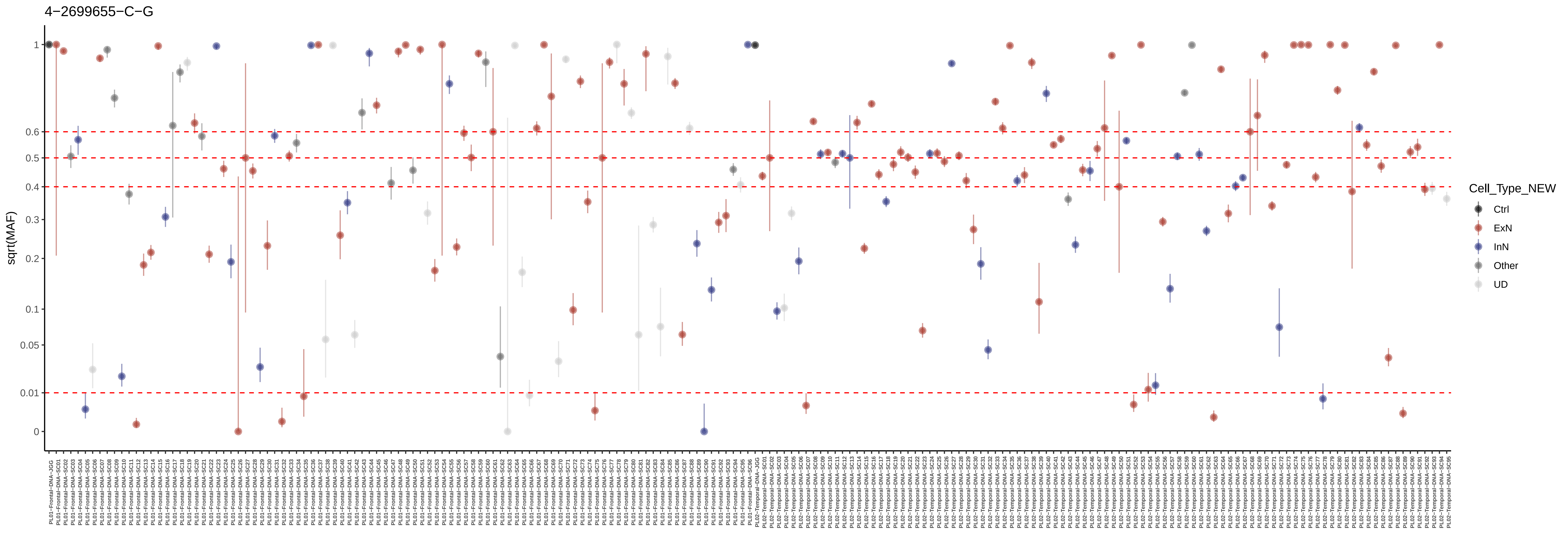

4−2699655−C−G
sqrt(MAF)
1
0.6
0.5
0.4
0.3
0.2
0.1
0.05
0.01
0
Cell_Type_NEW
Ctrl
ExN
InN
Other
UD

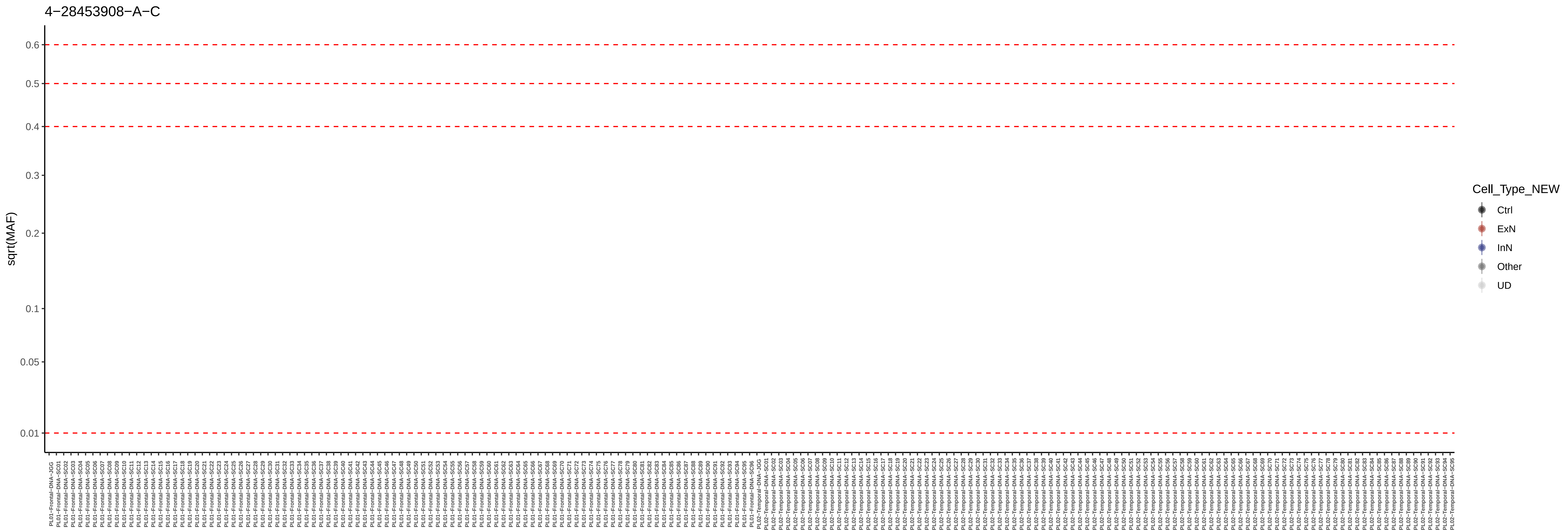
4−28453908−A−C
sqrt(MAF)
0.6
0.5
0.4
0.3
0.2
0.1
0.05
0.01
Cell_Type_NEW
Ctrl
ExN
InN
Other
UD

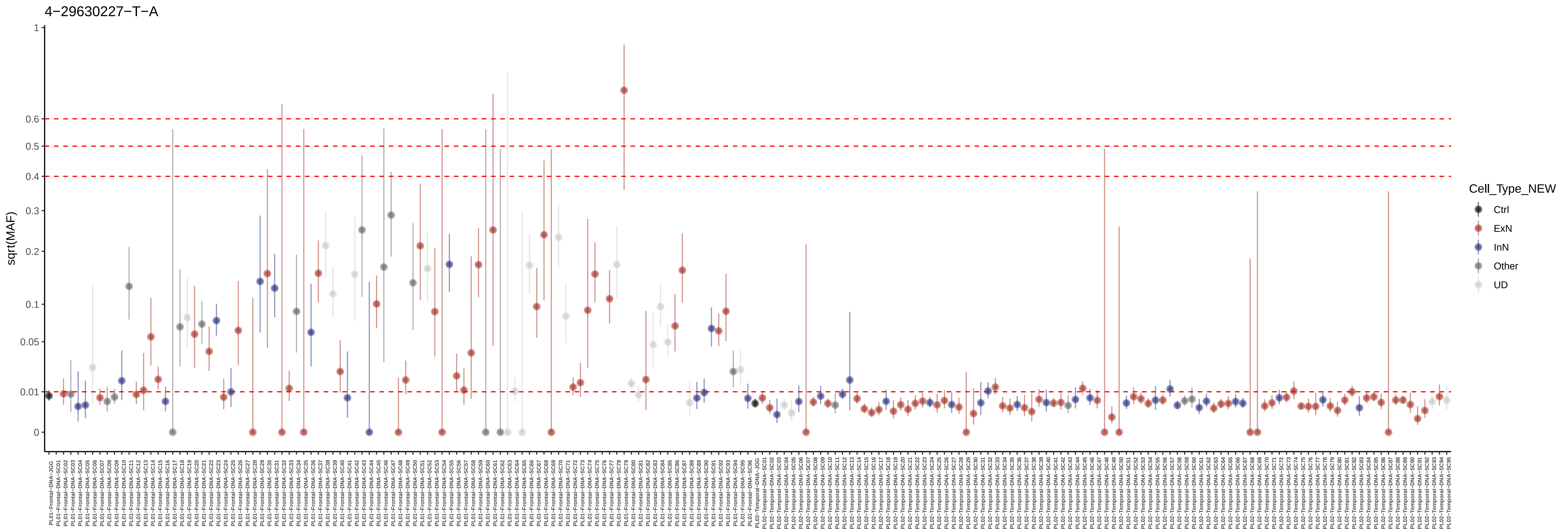

4−29630227−T−A
sqrt(MAF)
1
0.6
0.5
0.4
0.3
0.2
0.1
0.05
0.01
0
Cell_Type_NEW
Ctrl
ExN
InN
Other
UD

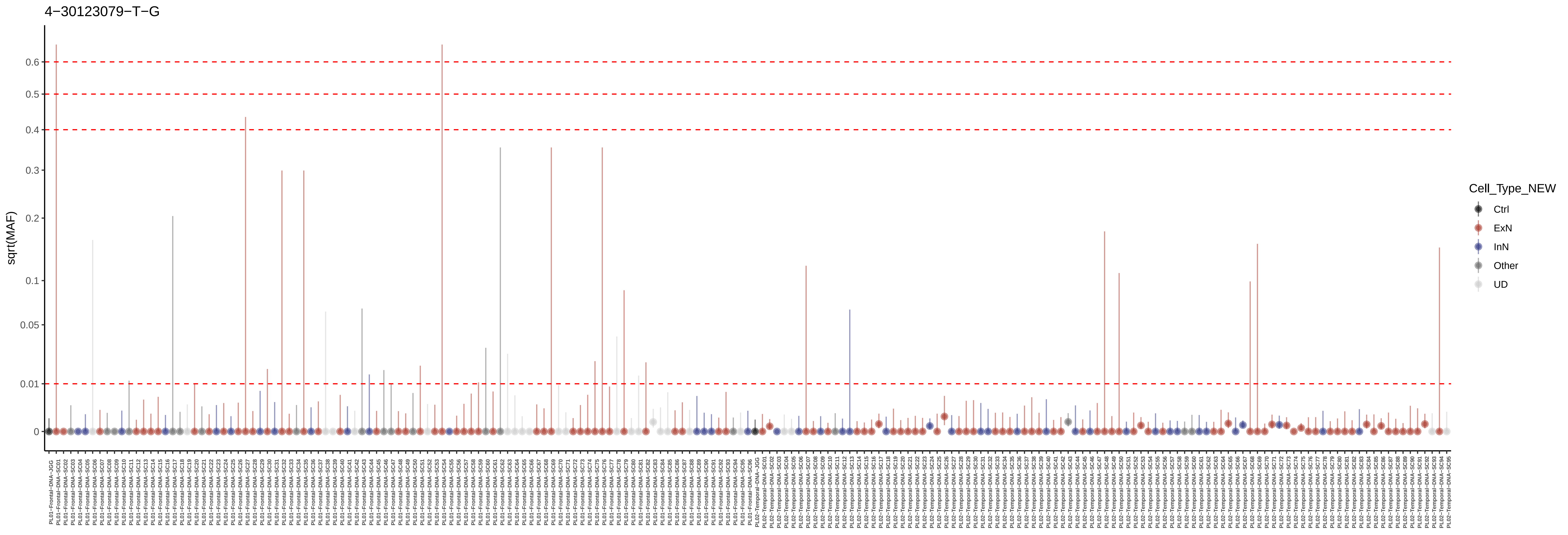
4−30123079−T−G
sqrt(MAF)
Cell_Type_NEW
Ctrl
ExN
InN
Other
UD

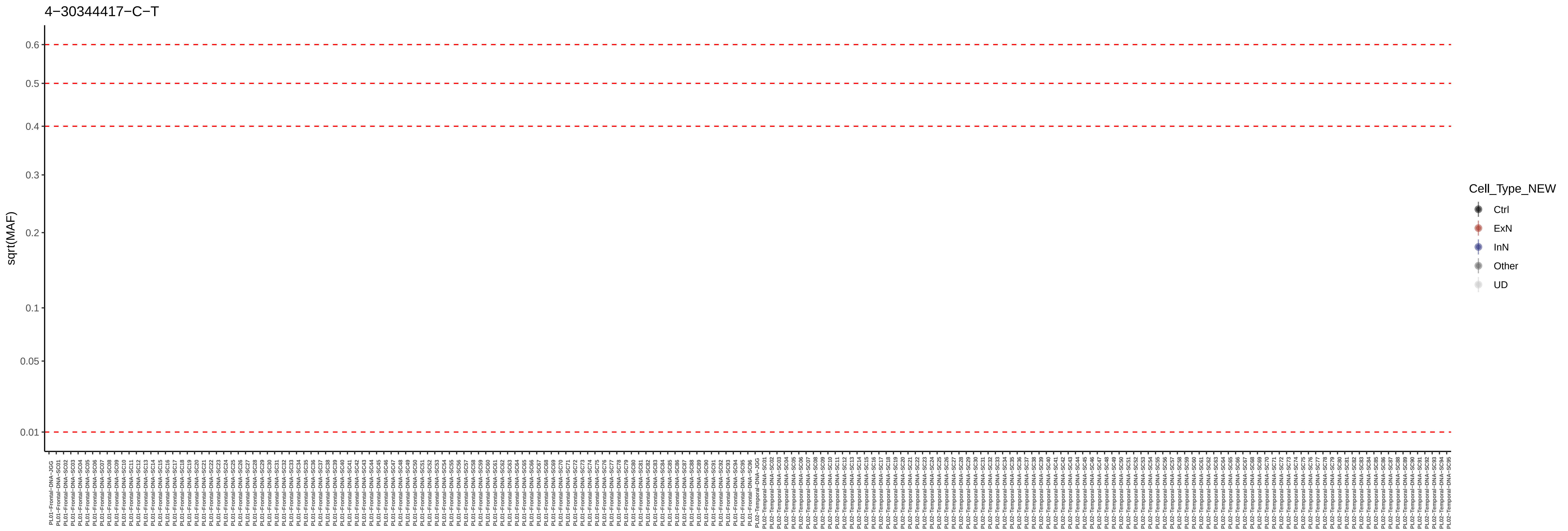
4−30344417−C−T
sqrt(MAF)
0.6
0.5
0.4
0.3
0.2
0.1
0.05
0.01
Cell_Type_NEW
Ctrl
ExN
InN
Other
UD

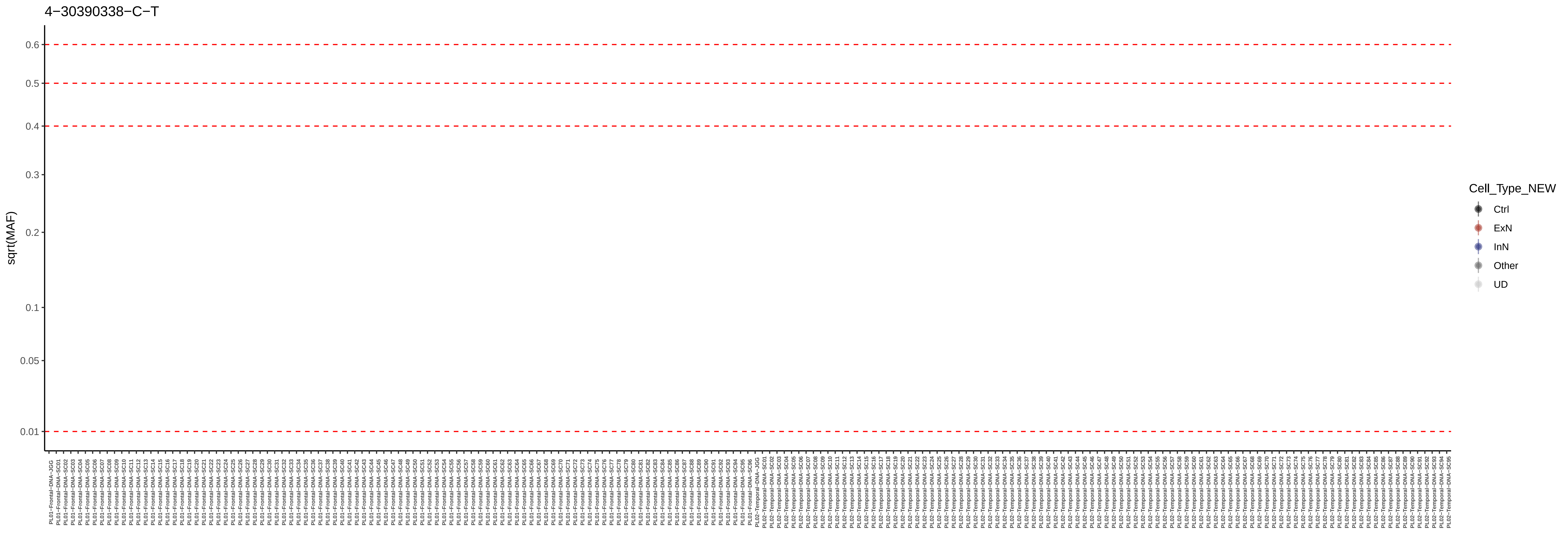
4−30390338−C−T
sqrt(MAF)
0.6
0.5
0.4
0.3
0.2
0.1
0.05
0.01
Cell_Type_NEW
Ctrl
ExN
InN
Other
UD

4−31555896−C−T

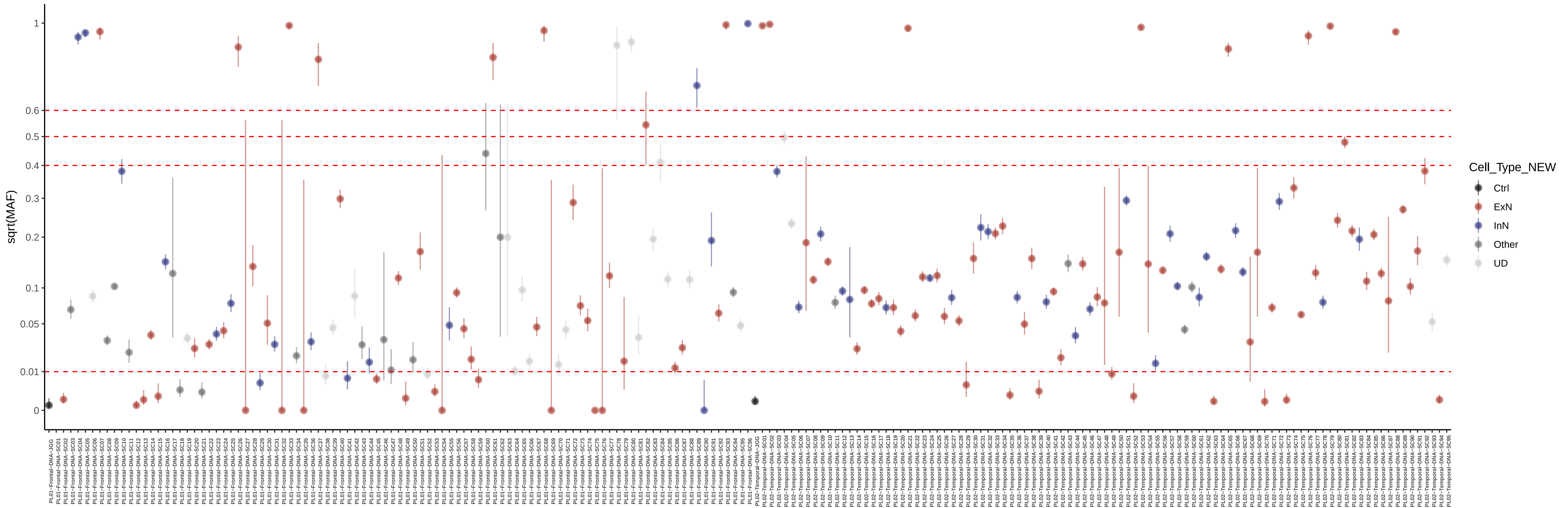

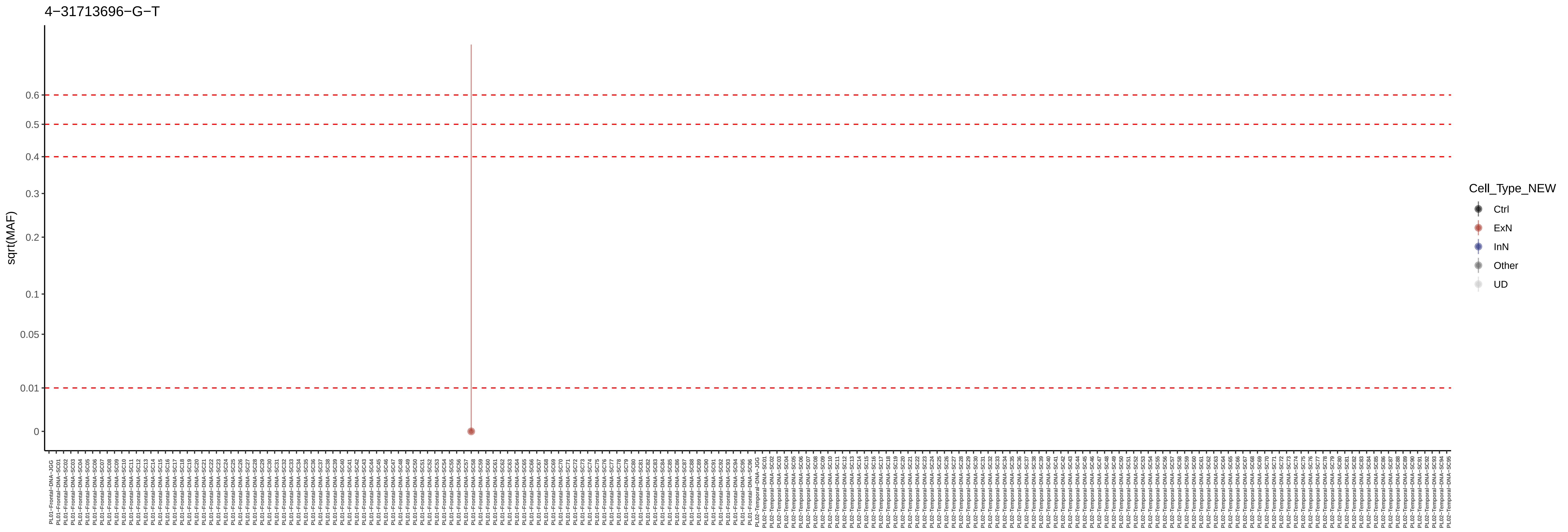
4−31713696−G−T
sqrt(MAF)
0.6
0.5
0.4
0.3
0.2
0.1
0.05
0.01
0
Cell_Type_NEW
Ctrl
ExN
InN
Other
UD

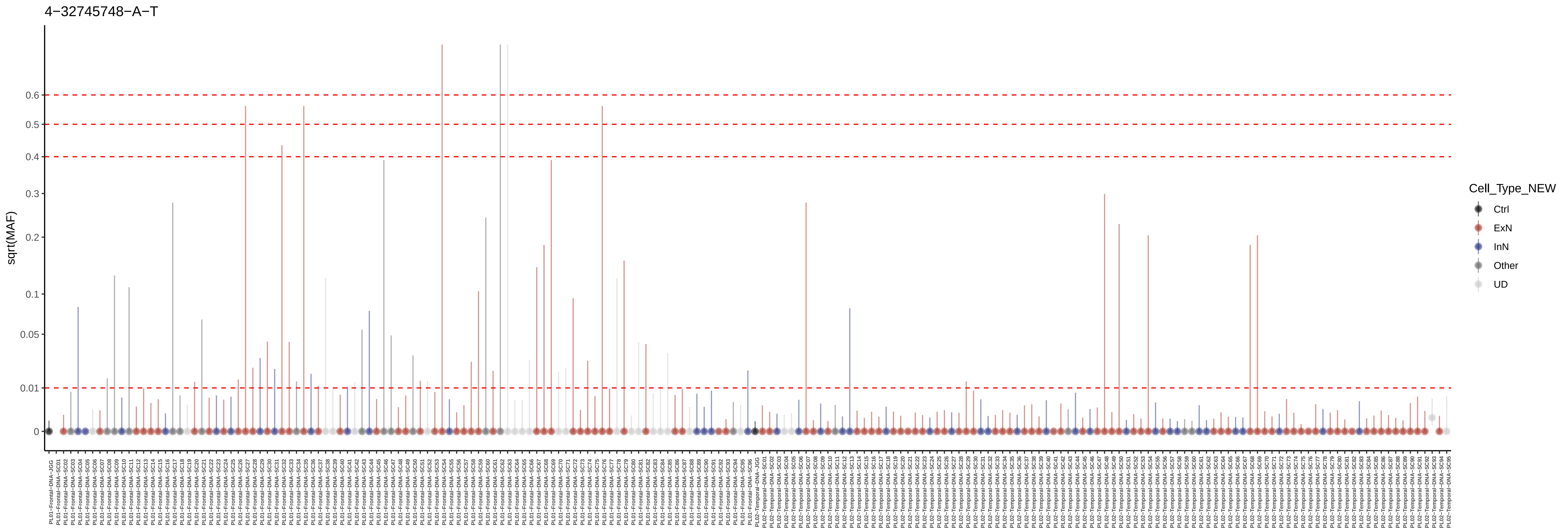

4−32745748−A−T
sqrt(MAF)
0.6
0.5
0.4
0.3
0.2
0.1
0.05
0.01
0
Cell_Type_NEW
Ctrl
ExN
InN
Other
UD

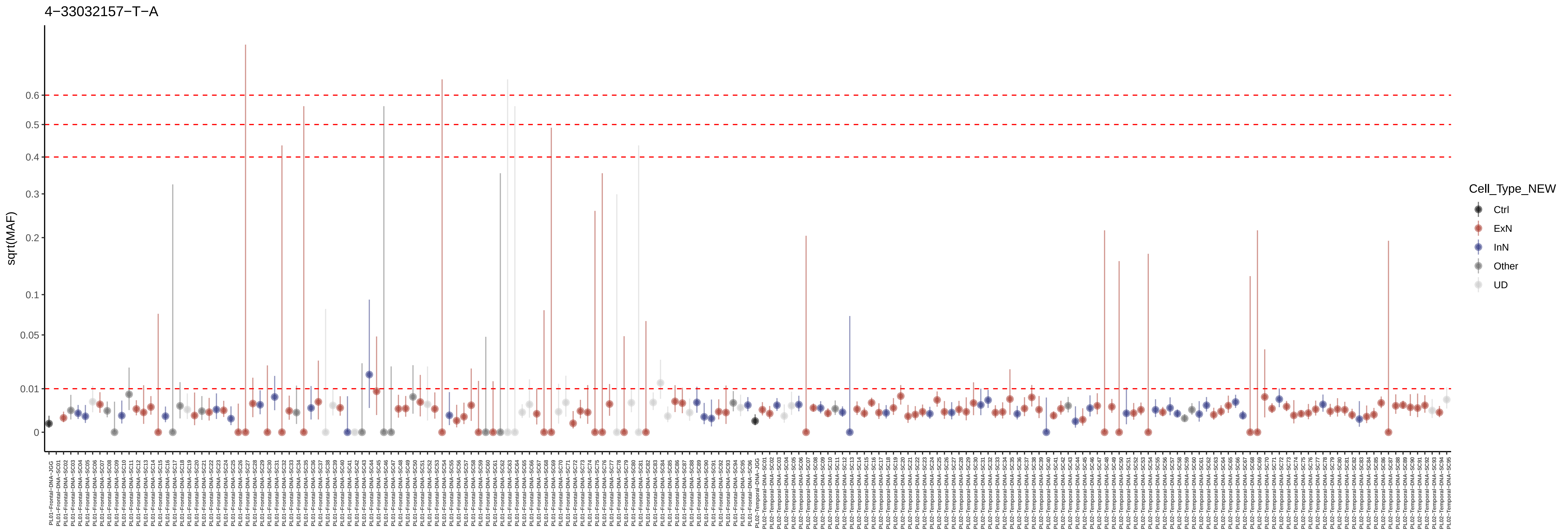

4−33032157−T−A
sqrt(MAF)
0.6
0.5
0.4
0.3
0.2
0.1
0.05
0.01
0
Cell_Type_NEW
Ctrl
ExN
InN
Other
UD

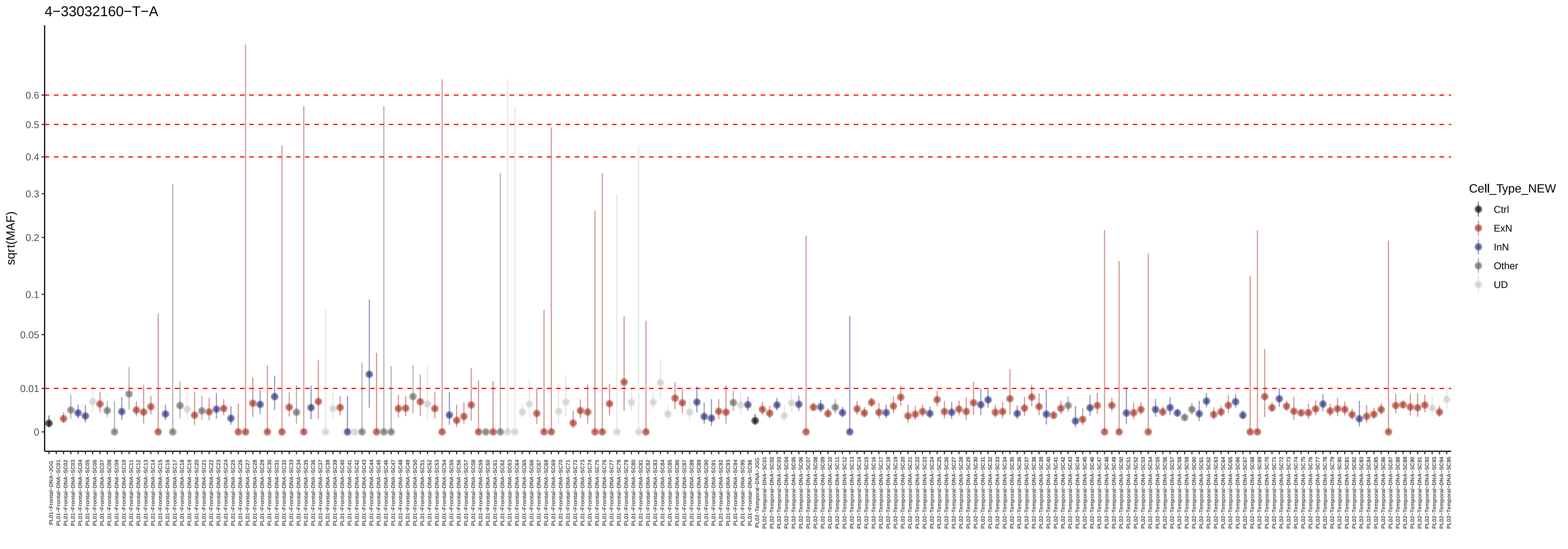

4−33032160−T−A
sqrt(MAF)
0.6
0.5
0.4
0.3
0.2
0.1
0.05
0.01
0
Cell_Type_NEW
Ctrl
ExN
InN
Other
UD

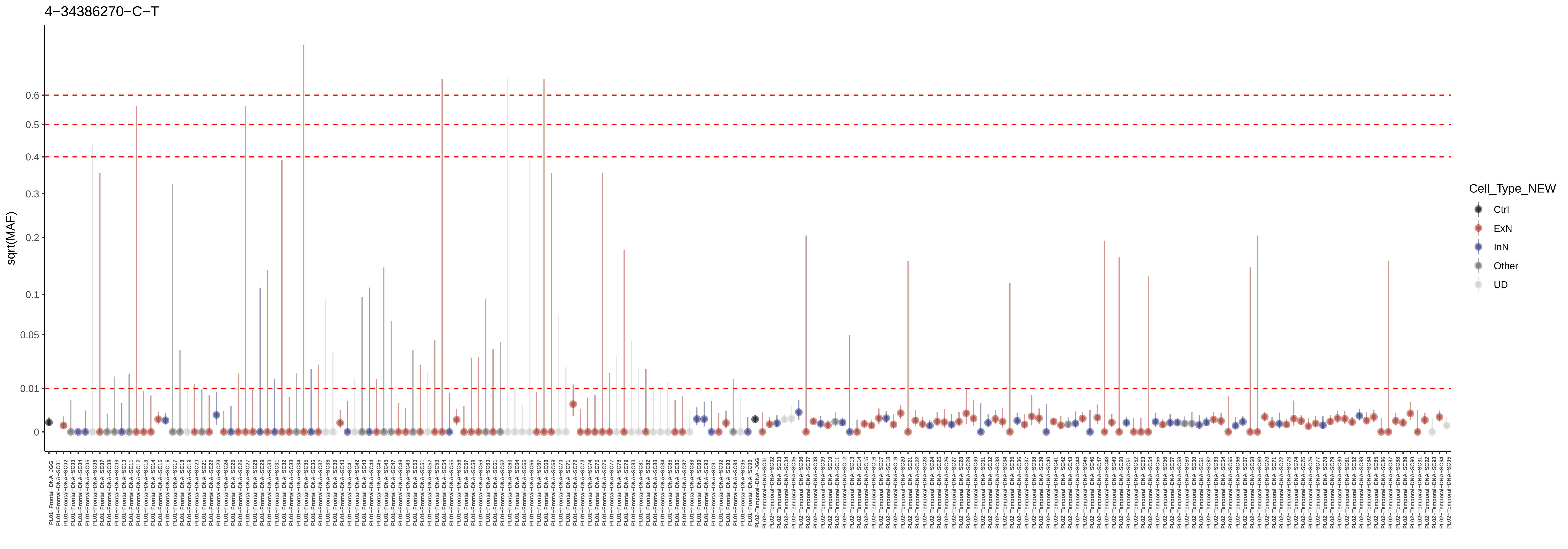

4−34386270−C−T
sqrt(MAF)
0.6
0.5
0.4
0.3
0.2
0.1
0.05
0.01
0
Cell_Type_NEW
Ctrl
ExN
InN
Other
UD

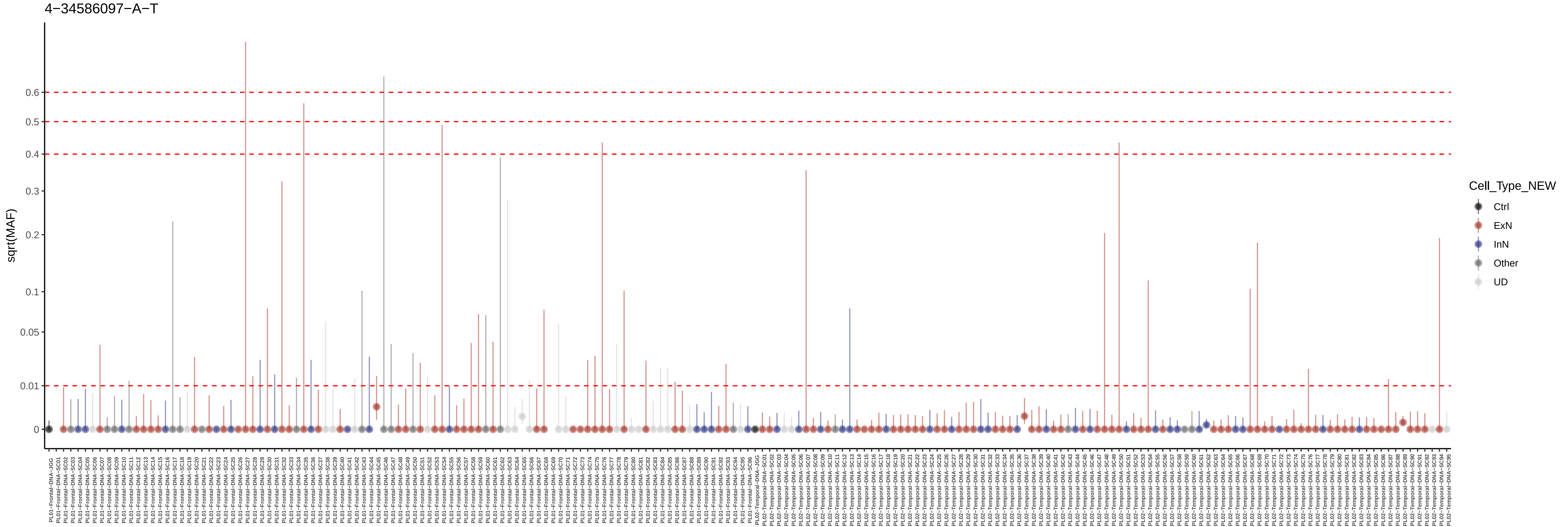
4−34586097−A−T
sqrt(MAF)
0.6
0.5
0.4
0.3
0.2
0.1
0.05
0.01
0
Cell_Type_NEW
Ctrl
ExN
InN
Other
UD

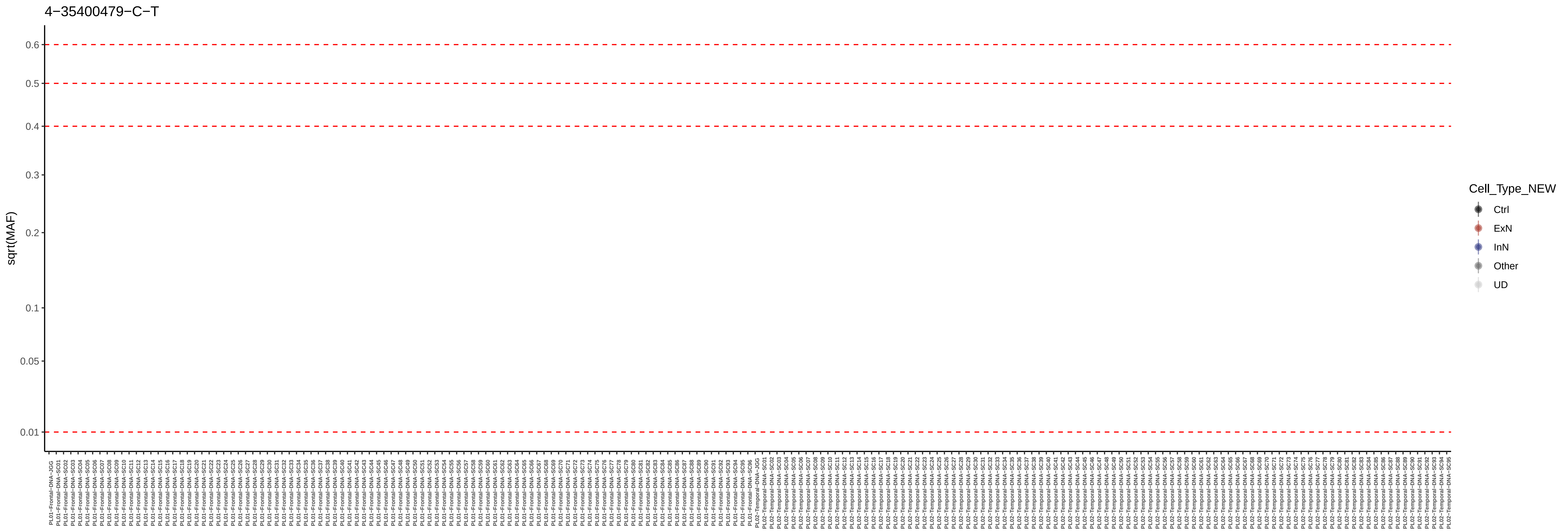
4−35400479−C−T
sqrt(MAF)
0.6
0.5
0.4
0.3
0.2
0.1
0.05
0.01
Cell_Type_NEW
Ctrl
ExN
InN
Other
UD

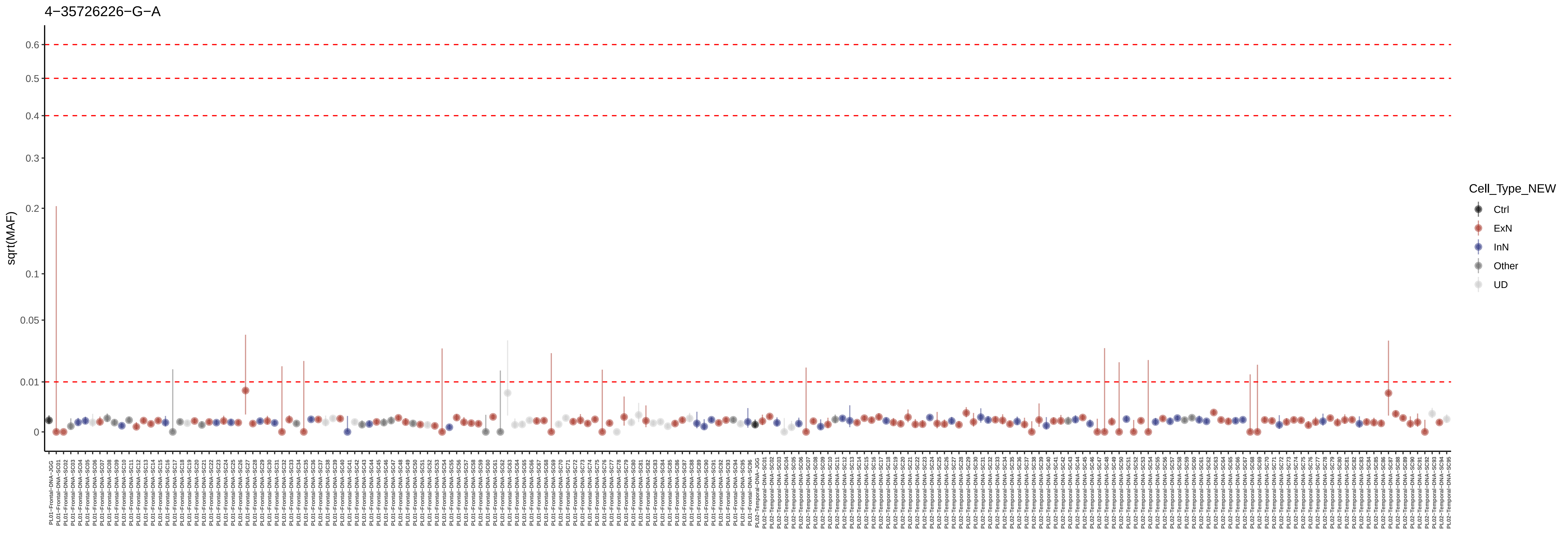

4−35726226−G−A
sqrt(MAF)
0.6
0.5
0.4
0.3
0.2
0.1
0.05
0.01
0
Cell_Type_NEW
Ctrl
ExN
InN
Other
UD

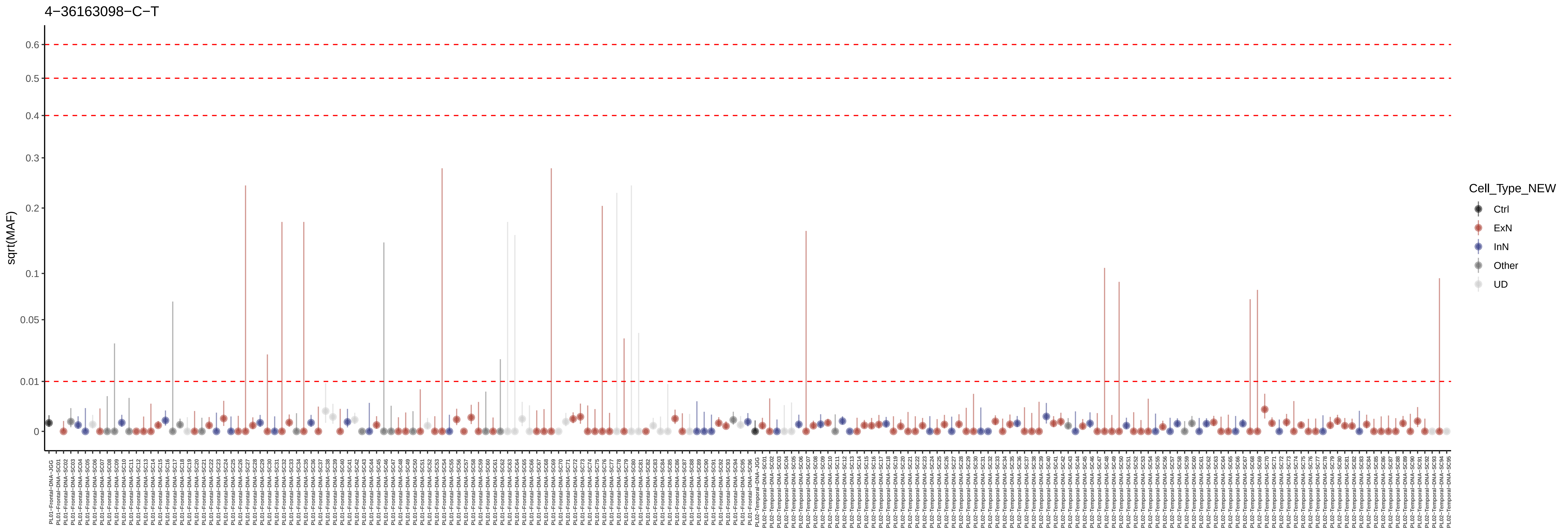
4−36163098−C−T
sqrt(MAF)
0.6
0.5
0.4
0.3
0.2
0.1
0.05
0.01
0
Cell_Type_NEW
Ctrl
ExN
InN
Other
UD

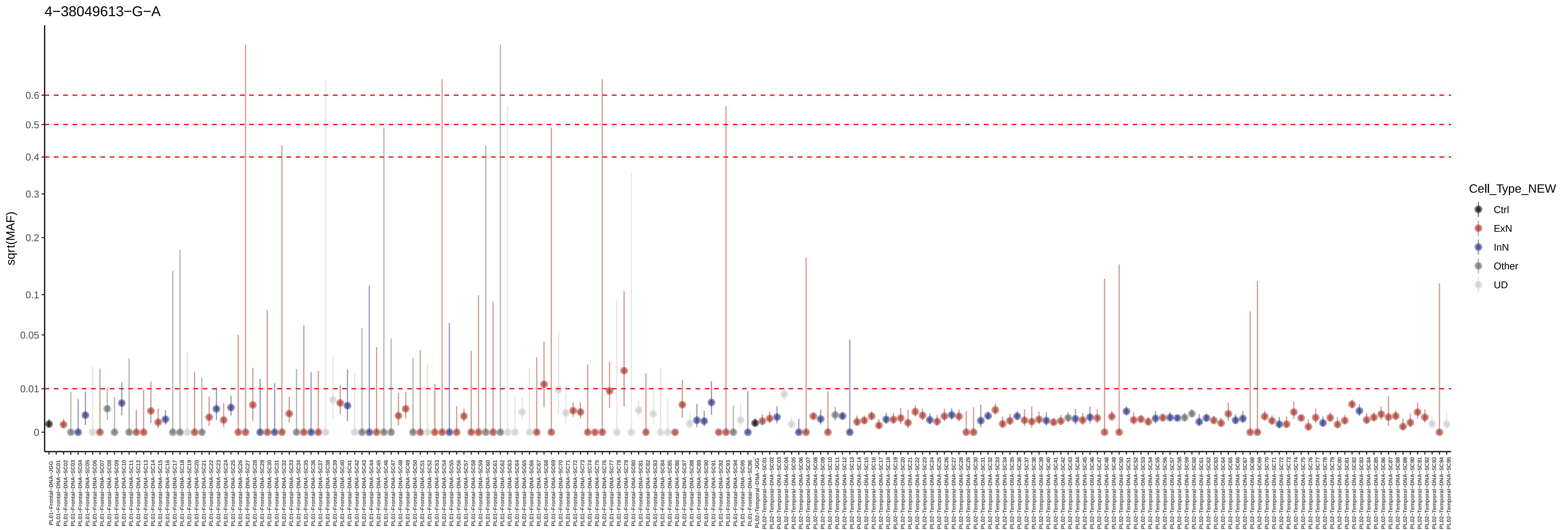

4−38049613−G−A
sqrt(MAF)
Cell_Type_NEW
Ctrl
ExN
InN
Other
UD

4−39424715−A−G

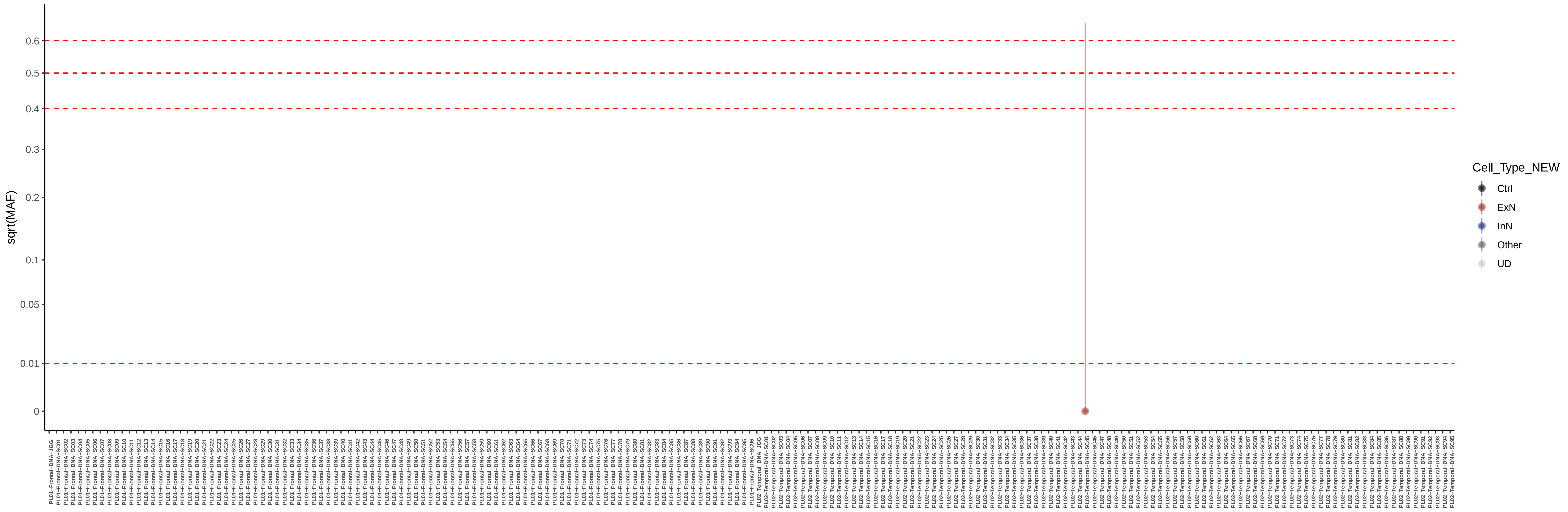

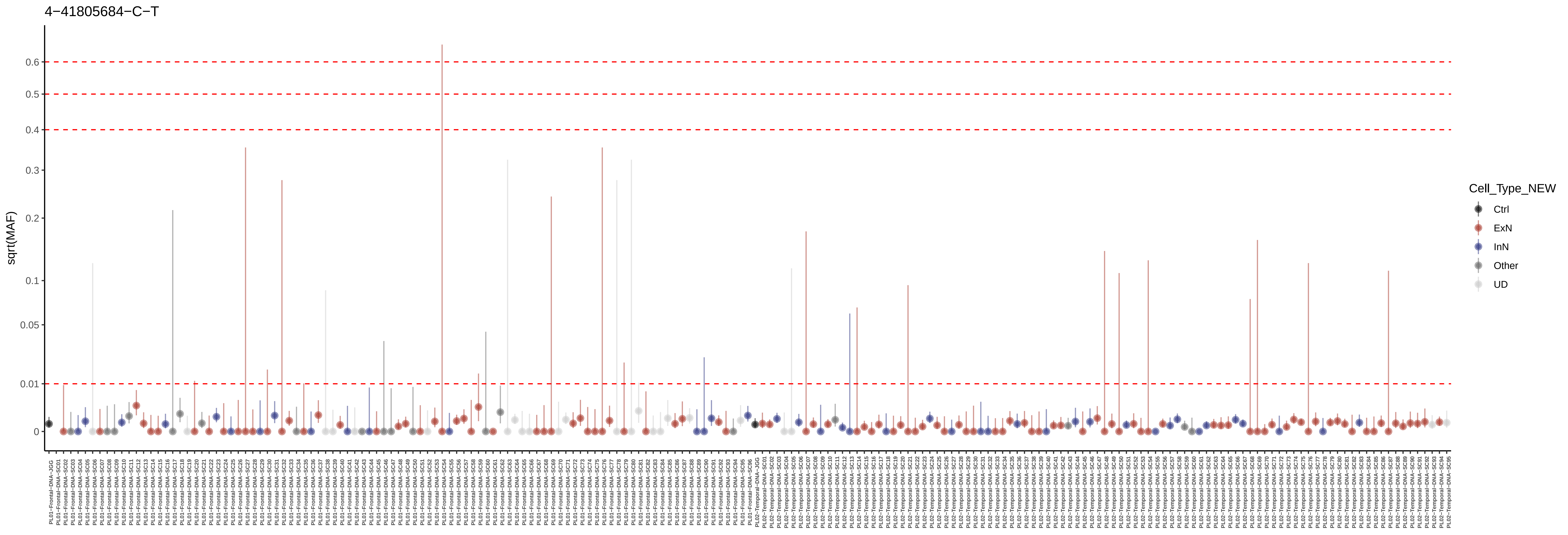
4−41805684−C−T
sqrt(MAF)
0.6
0.5
0.4
0.3
0.2
0.1
0.05
0.01
0
Cell_Type_NEW
Ctrl
ExN
InN
Other
UD

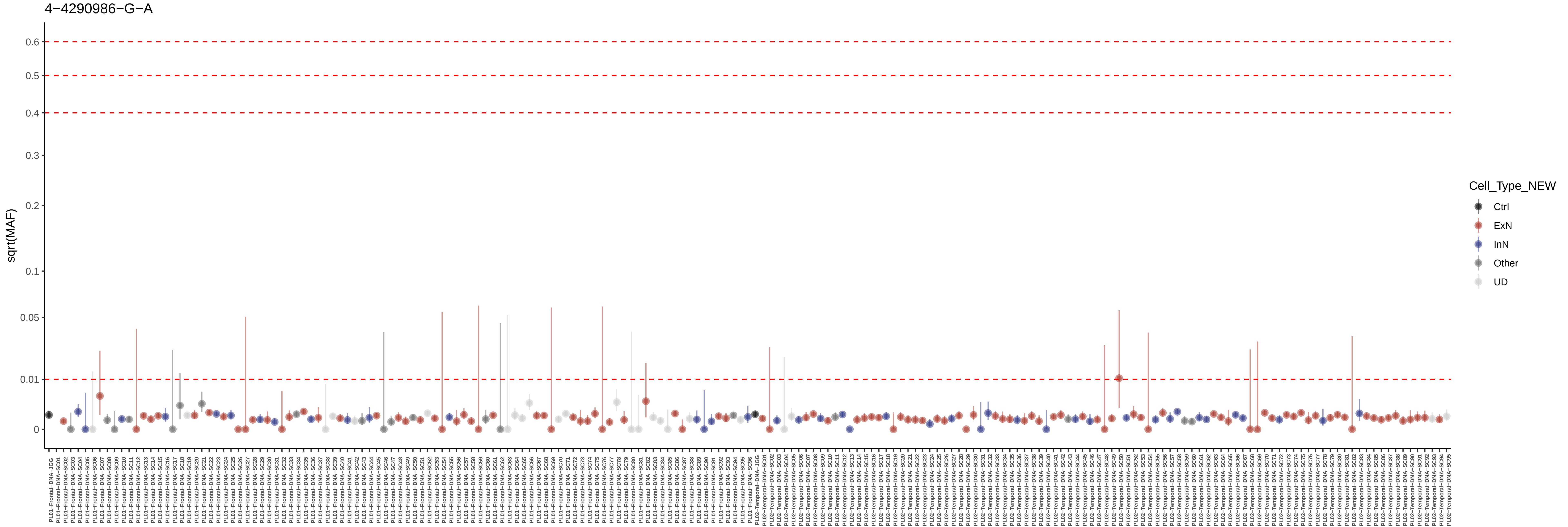

4−4290986−G−A
sqrt(MAF)
0.6
0.5
0.4
0.3
0.2
0.1
0.05
0.01
0
Cell_Type_NEW
Ctrl
ExN
InN
Other
UD

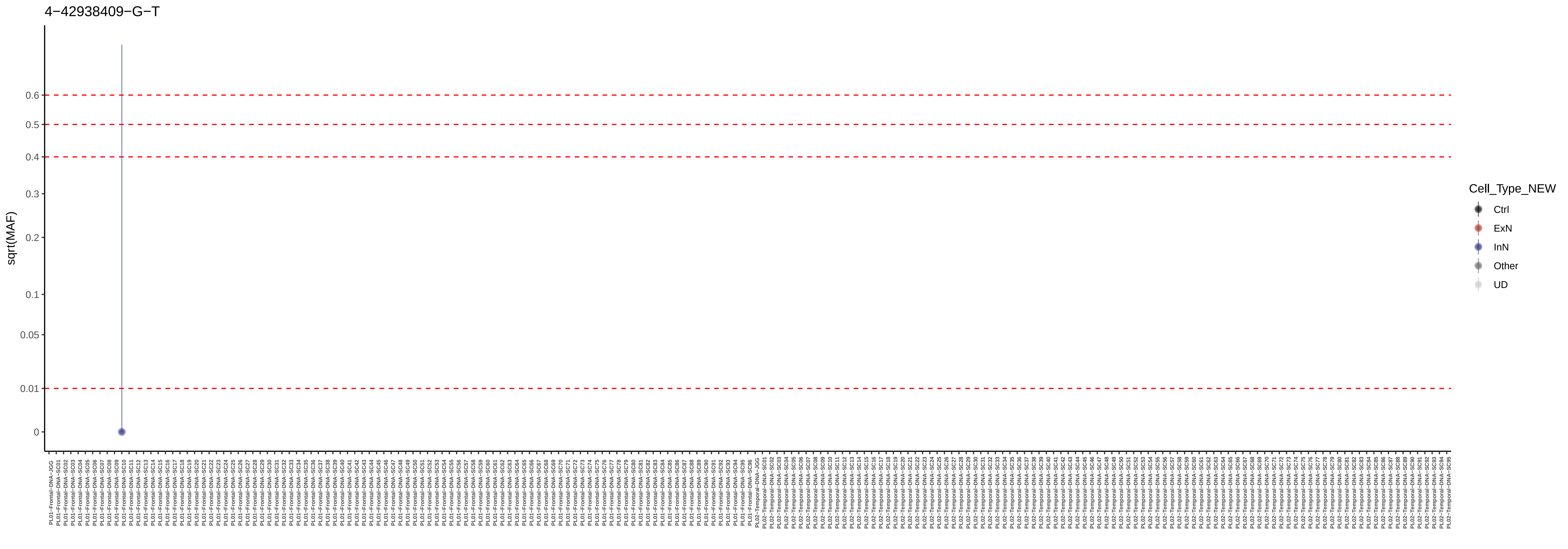

4−42938409−G−T
sqrt(MAF)
0.6
0.5
0.4
0.3
0.2
0.1
0.05
0.01
0
Cell_Type_NEW
Ctrl
ExN
InN
Other
UD

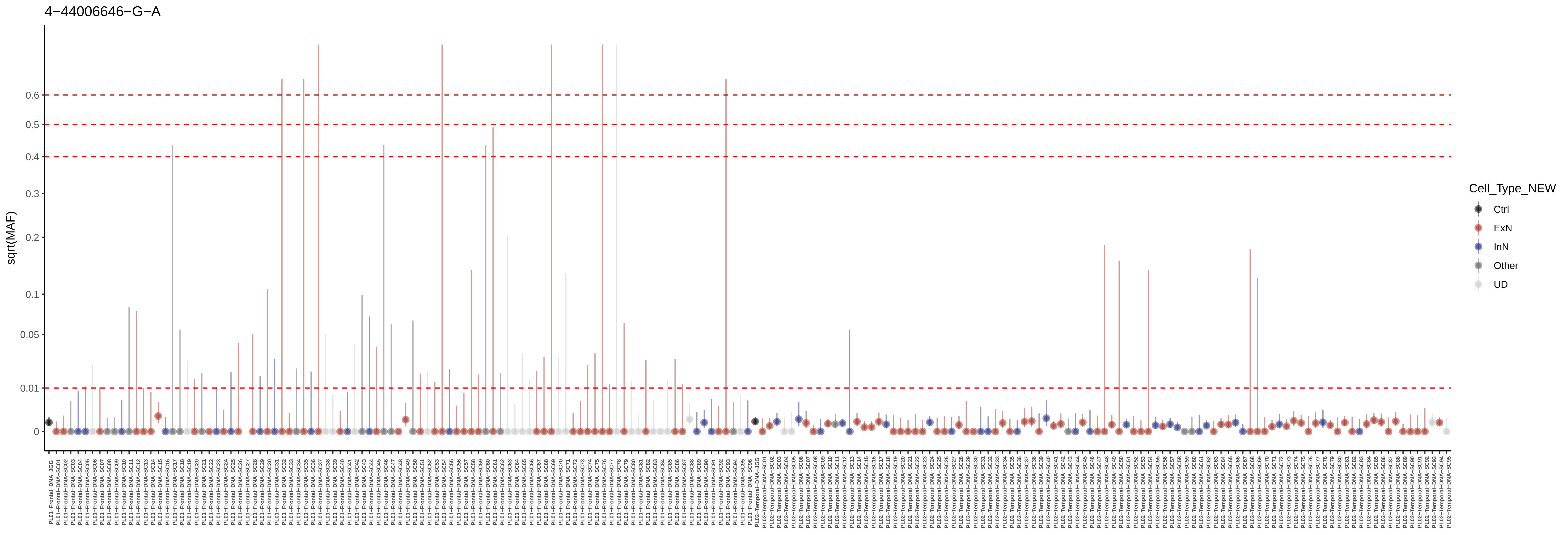
4−44006646−G−A
sqrt(MAF)
0.6
0.5
0.4
0.3
0.2
0.1
0.05
0.01
0
Cell_Type_NEW
Ctrl
ExN
InN
Other
UD

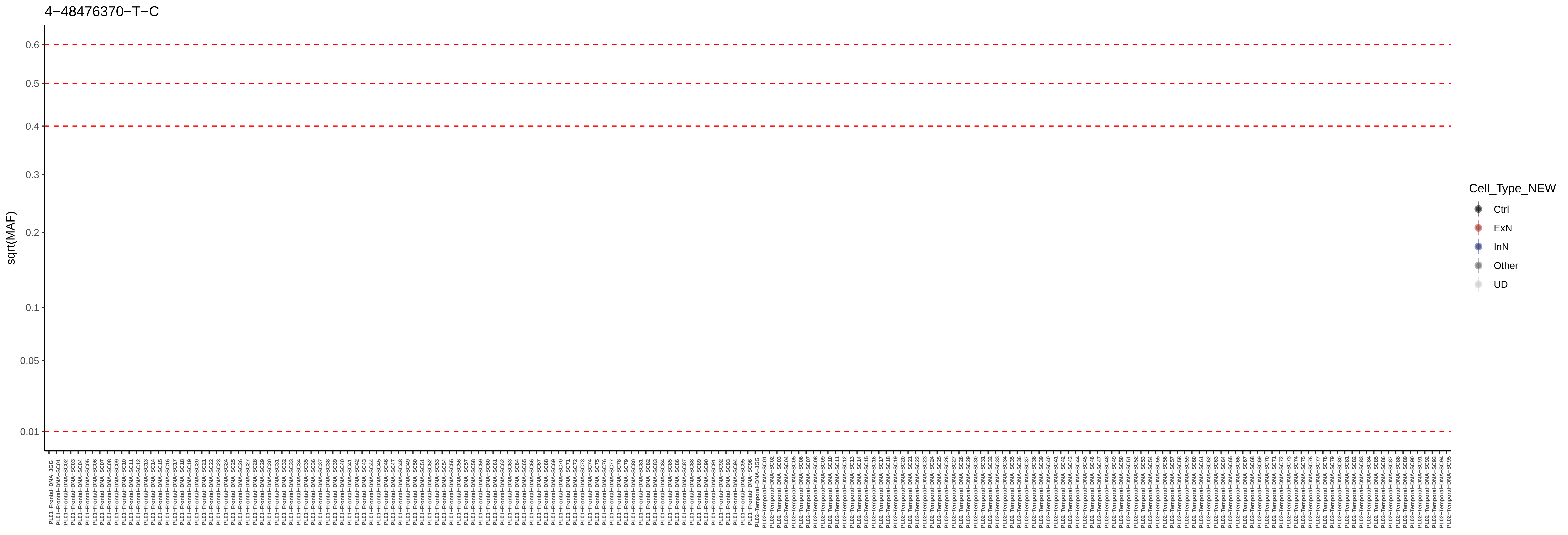
4−48476370−T−C
sqrt(MAF)
0.6
0.5
0.4
0.3
0.2
0.1
0.05
0.01
Cell_Type_NEW
Ctrl
ExN
InN
Other
UD

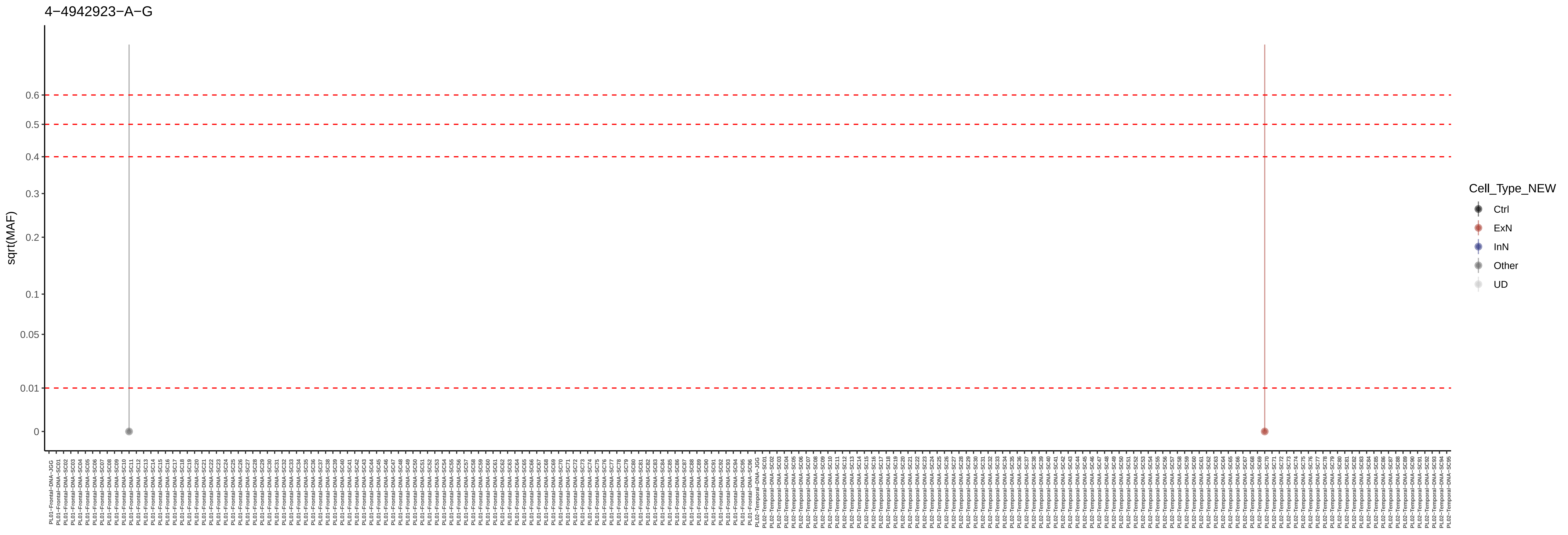
4−4942923−A−G
sqrt(MAF)
0.6
0.5
0.4
0.3
0.2
0.1
0.05
0.01
0
Cell_Type_NEW
Ctrl
ExN
InN
Other
UD

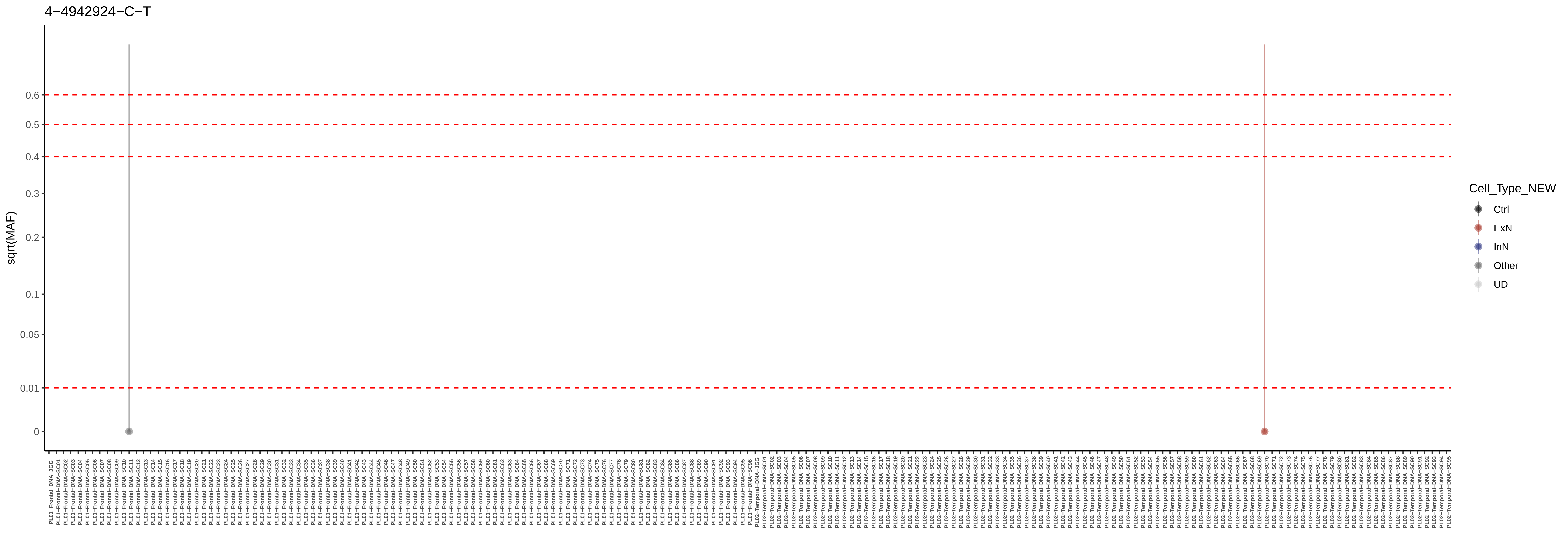
4−4942924−C−T
sqrt(MAF)
0.6
0.5
0.4
0.3
0.2
0.1
0.05
0.01
0
Cell_Type_NEW
Ctrl
ExN
InN
Other
UD

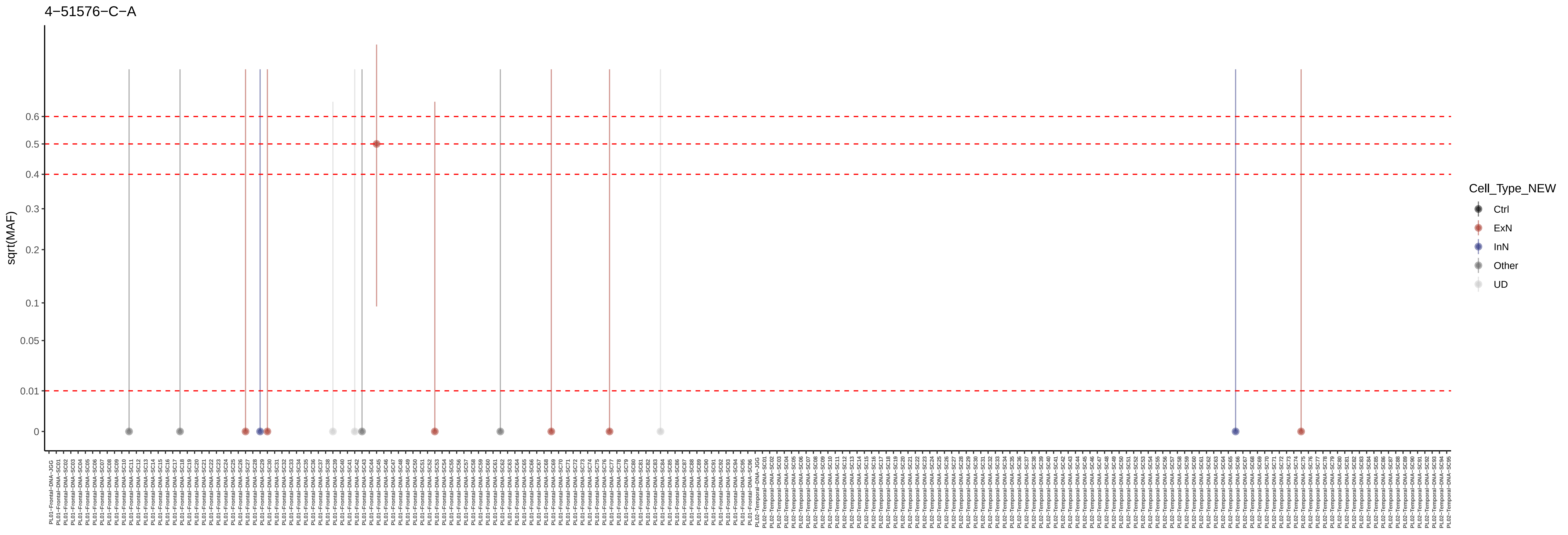
4−51576−C−A
sqrt(MAF)
0.6
0.5
0.4
0.3
0.2
0.1
0.05
0.01
0
Cell_Type_NEW
Ctrl
ExN
InN
Other
UD

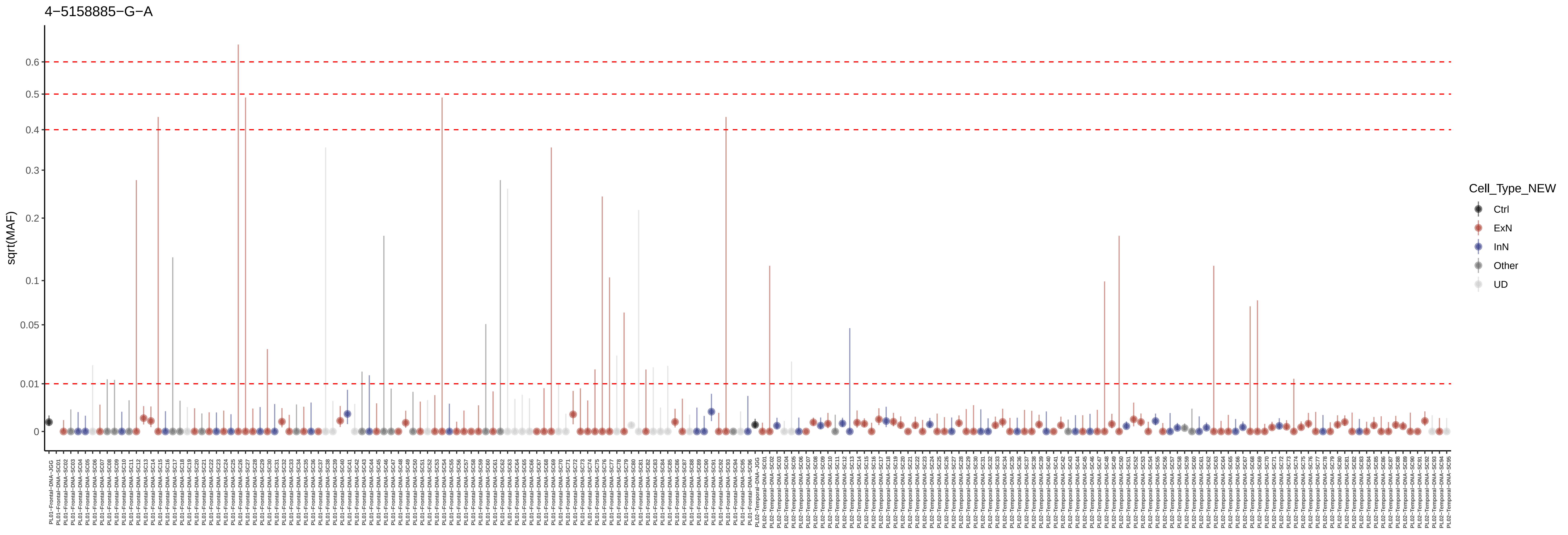

4−5158885−G−A
sqrt(MAF)
0.6
0.5
0.4
0.3
0.2
0.1
0.05
0.01
0
Cell_Type_NEW
Ctrl
ExN
InN
Other
UD

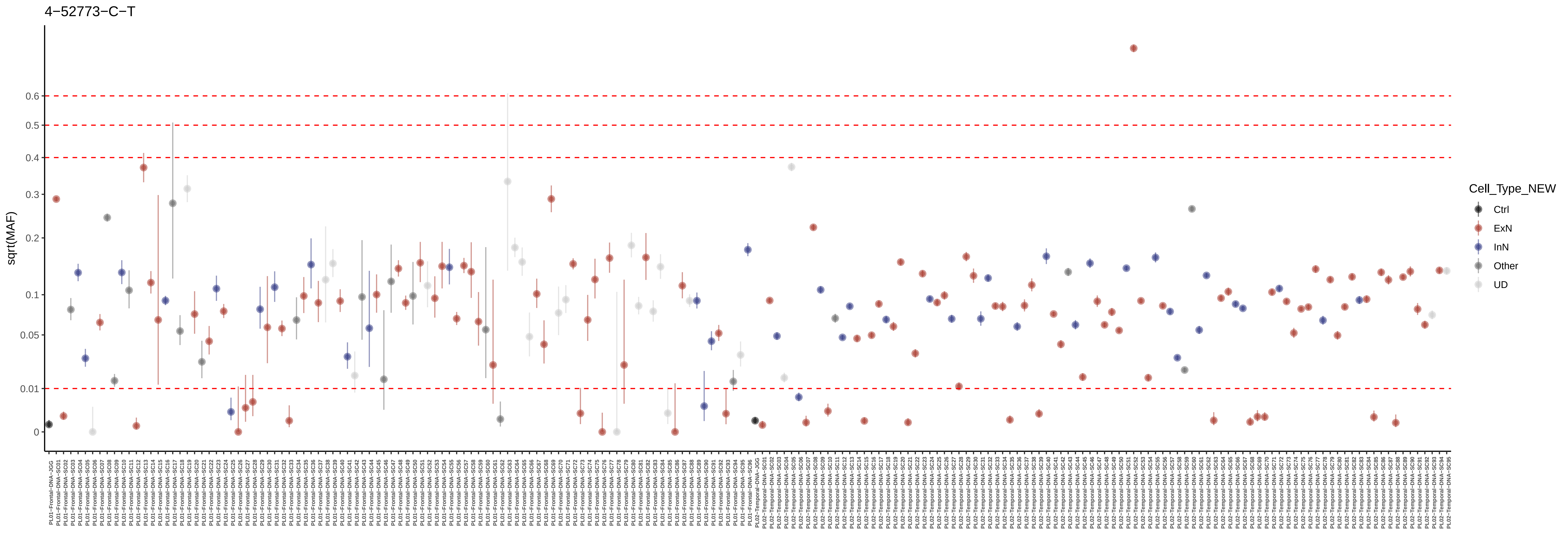

4−52773−C−T
sqrt(MAF)
0.6
0.5
0.4
0.3
0.2
0.1
0.05
0.01
0
Cell_Type_NEW
Ctrl
ExN
InN
Other
UD

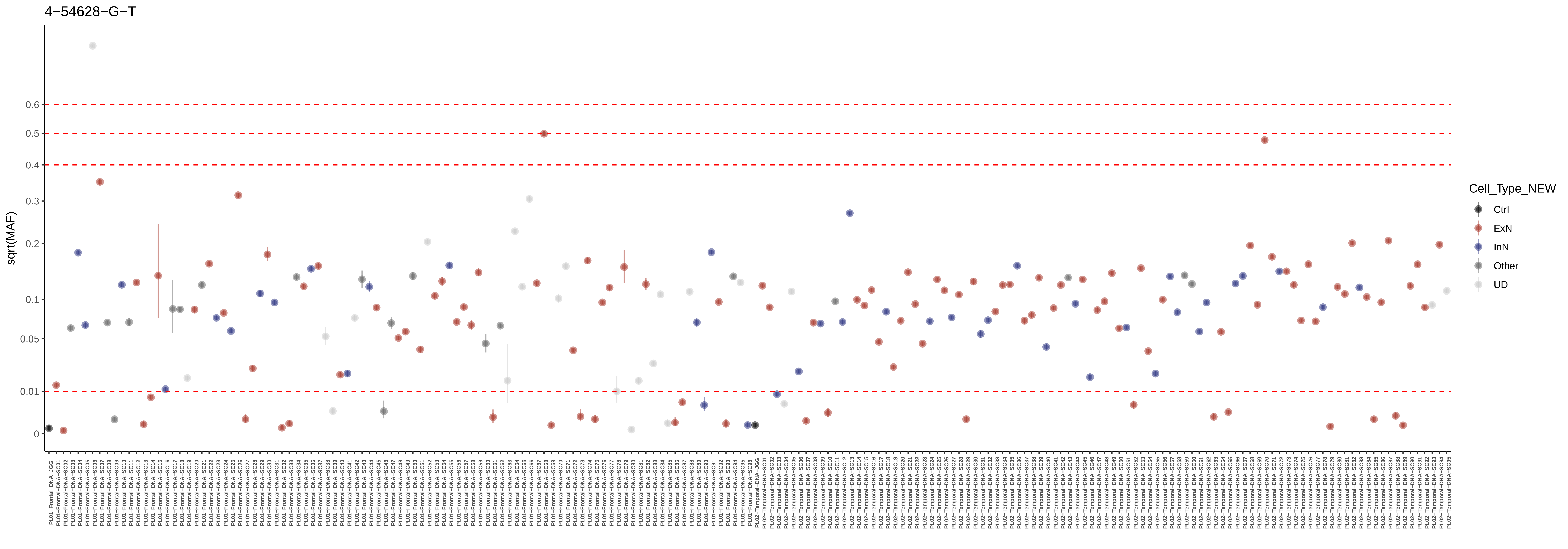

4−54628−G−T
sqrt(MAF)
0.6
0.5
0.4
0.3
0.2
0.1
0.05
0.01
0
Cell_Type_NEW
Ctrl
ExN
InN
Other
UD

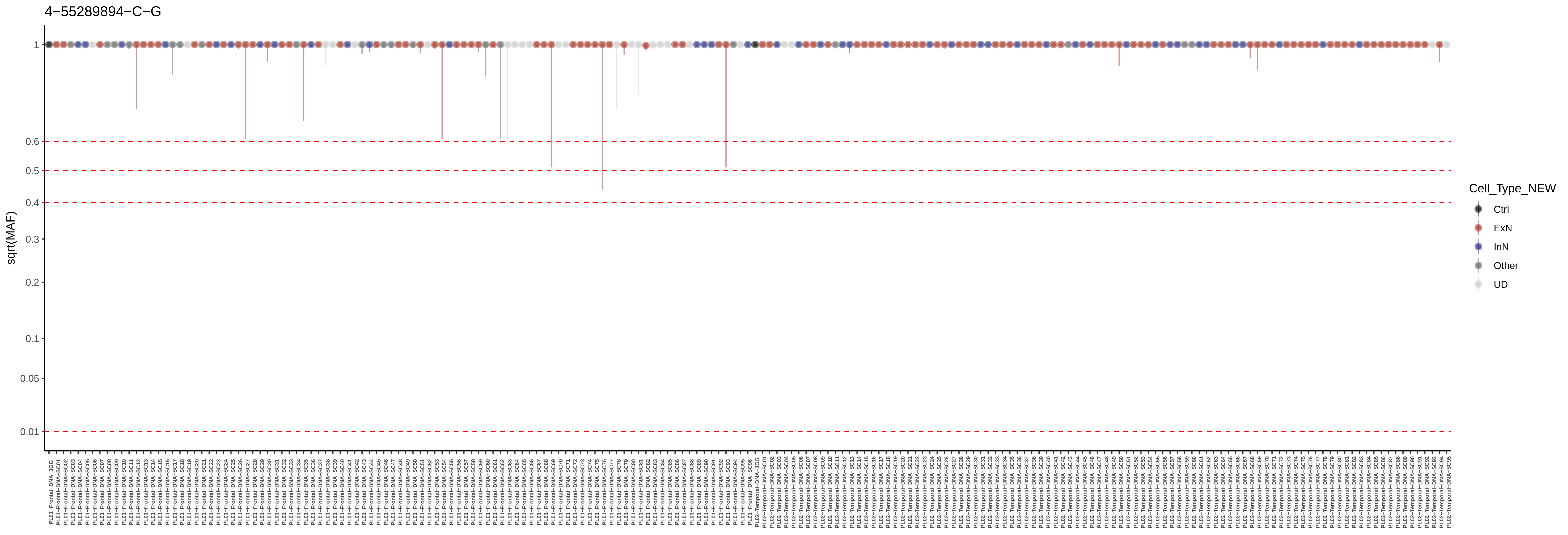

4−55289894−C−G
sqrt(MAF)
1
0.6
0.5
0.4
0.3
0.2
0.1
0.05
0.01
Cell_Type_NEW
Ctrl
ExN
InN
Other
UD

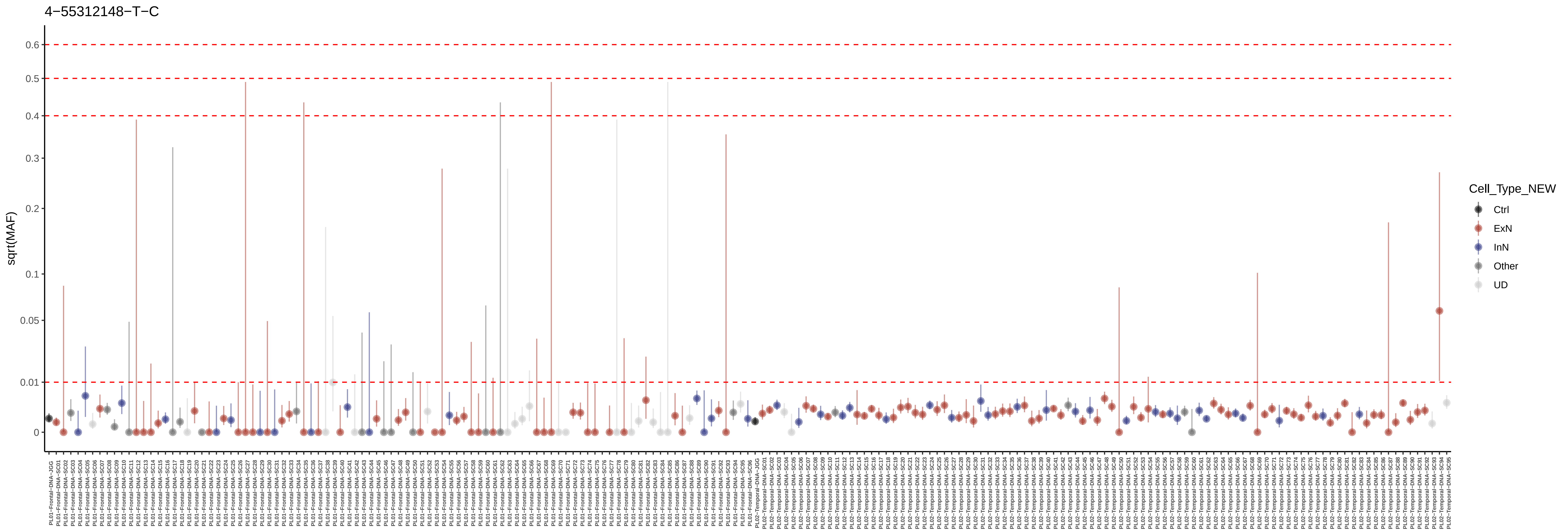

4−55312148−T−C
sqrt(MAF)
0.6
0.5
0.4
0.3
0.2
0.1
0.05
0.01
0
Cell_Type_NEW
Ctrl
ExN
InN
Other
UD

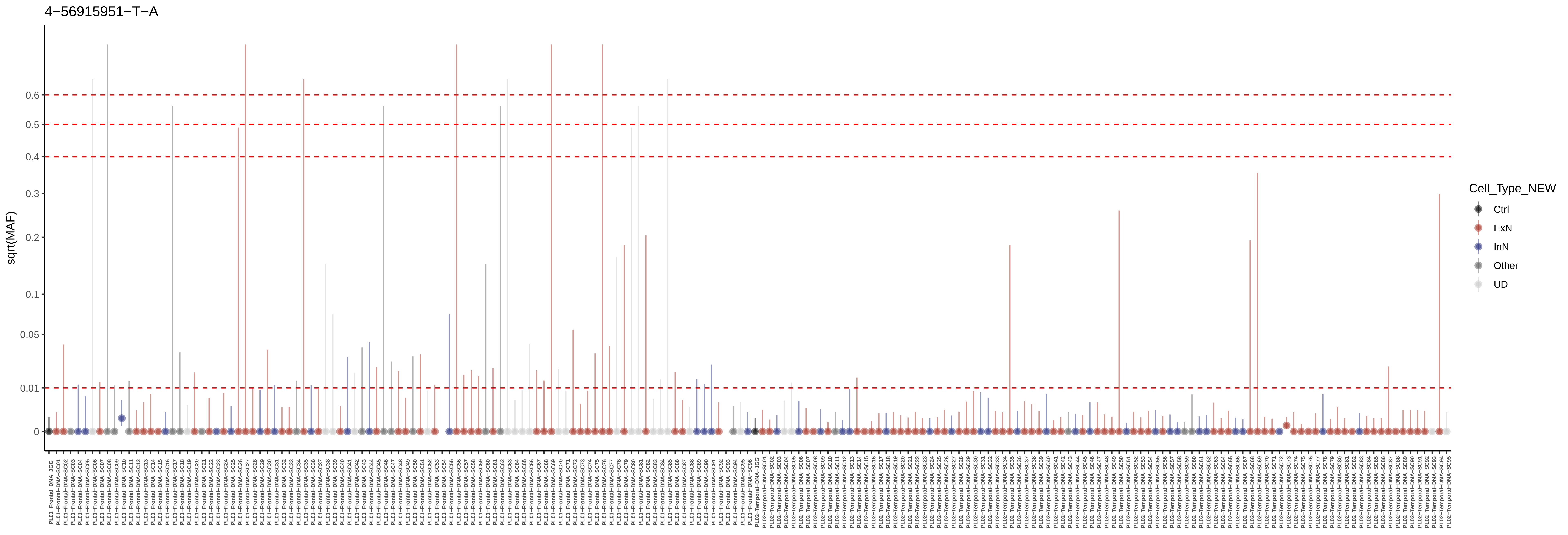

4−56915951−T−A
sqrt(MAF)
0.6
0.5
0.4
0.3
0.2
0.1
0.05
0.01
0
Cell_Type_NEW
Ctrl
ExN
InN
Other
UD

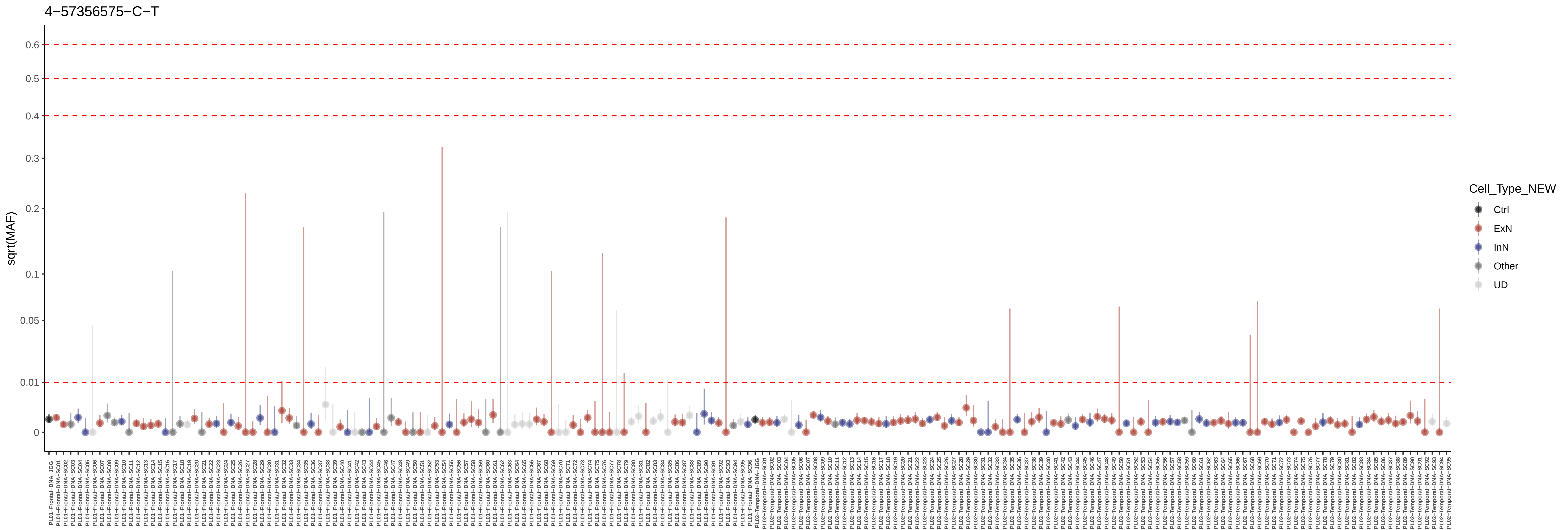

4−57356575−C−T
sqrt(MAF)
0.6
0.5
0.4
0.3
0.2
0.1
0.05
0.01
0
Cell_Type_NEW
Ctrl
ExN
InN
Other
UD

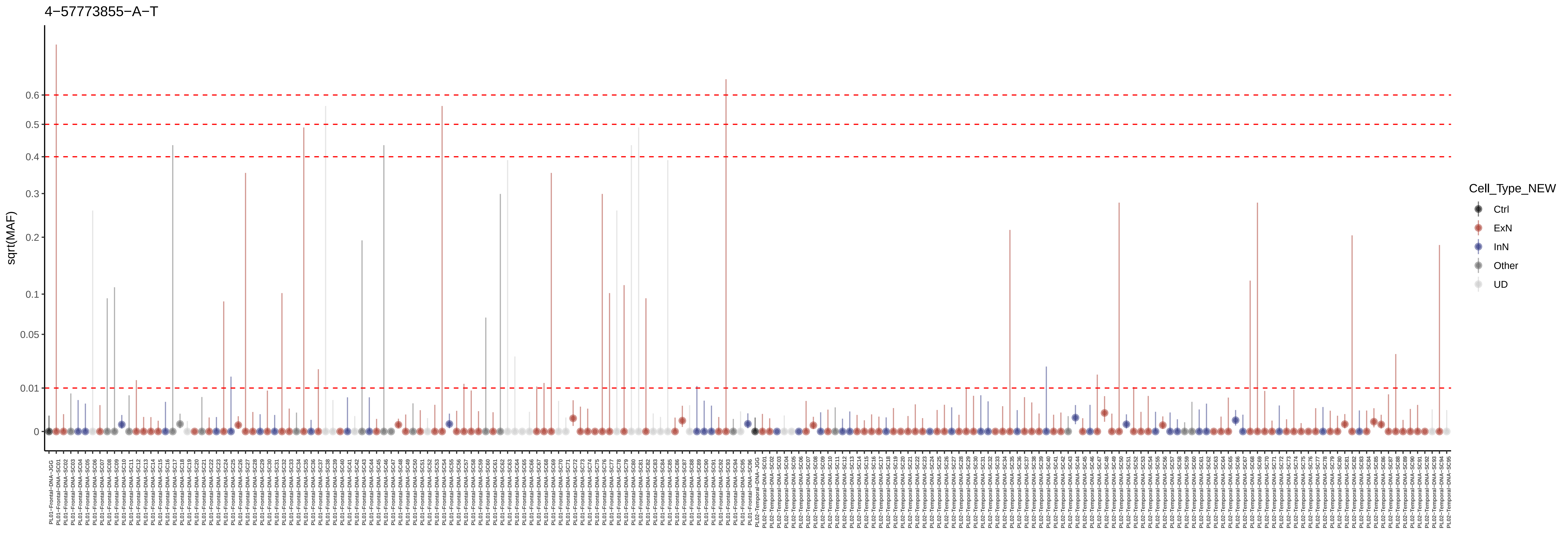
4−57773855−A−T
sqrt(MAF)
0.6
0.5
0.4
0.3
0.2
0.1
0.05
0.01
0
Cell_Type_NEW
Ctrl
ExN
InN
Other
UD

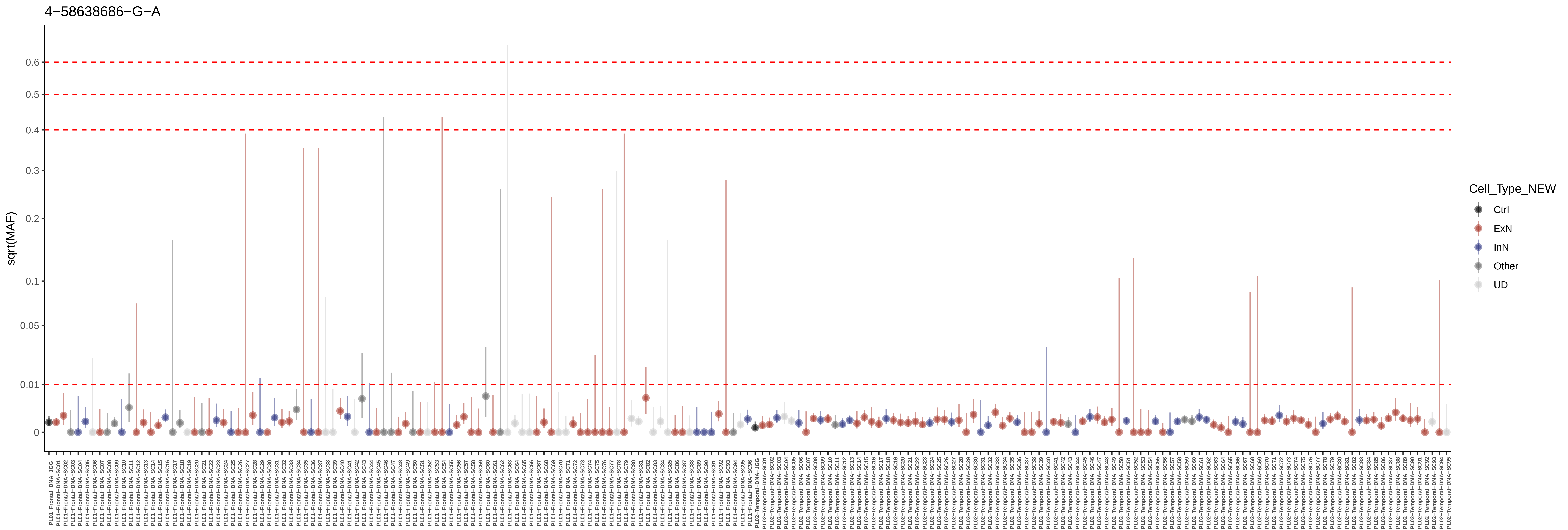
4−58638686−G−A
sqrt(MAF)
0.6
0.5
0.4
0.3
0.2
0.1
0.05
0.01
0
Cell_Type_NEW
Ctrl
ExN
InN
Other
UD

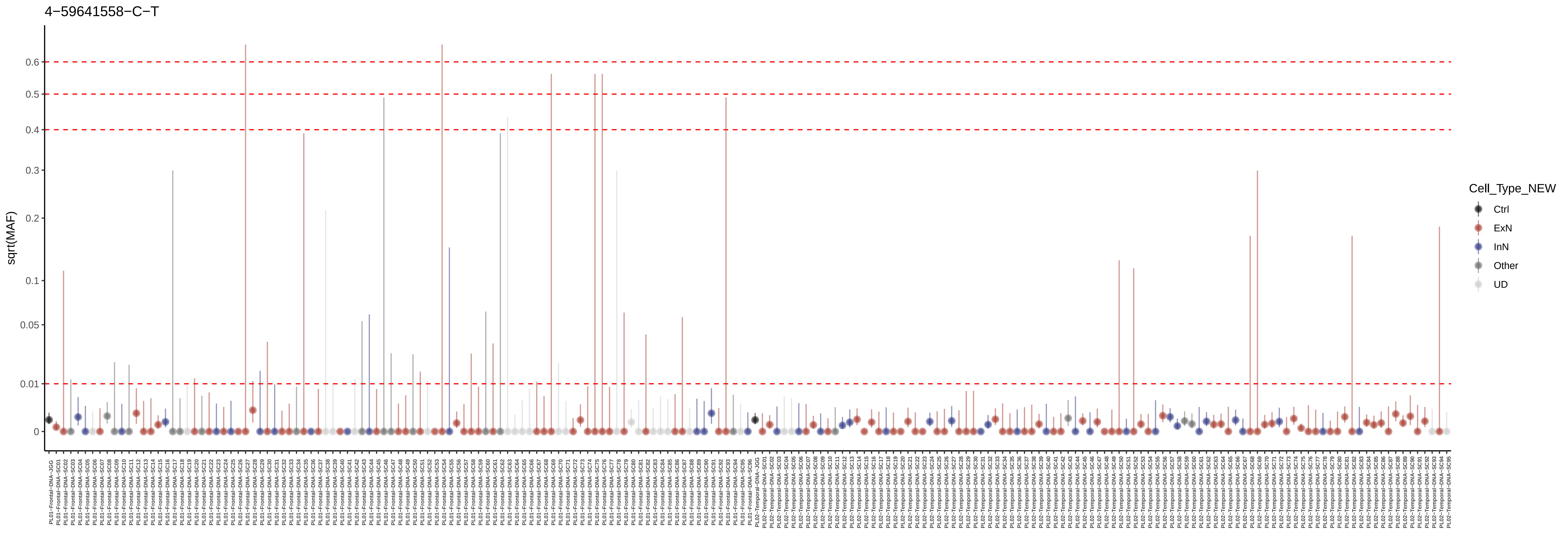
4−59641558−C−T
sqrt(MAF)
0.6
0.5
0.4
0.3
0.2
0.1
0.05
0.01
0
Cell_Type_NEW
Ctrl
ExN
InN
Other
UD

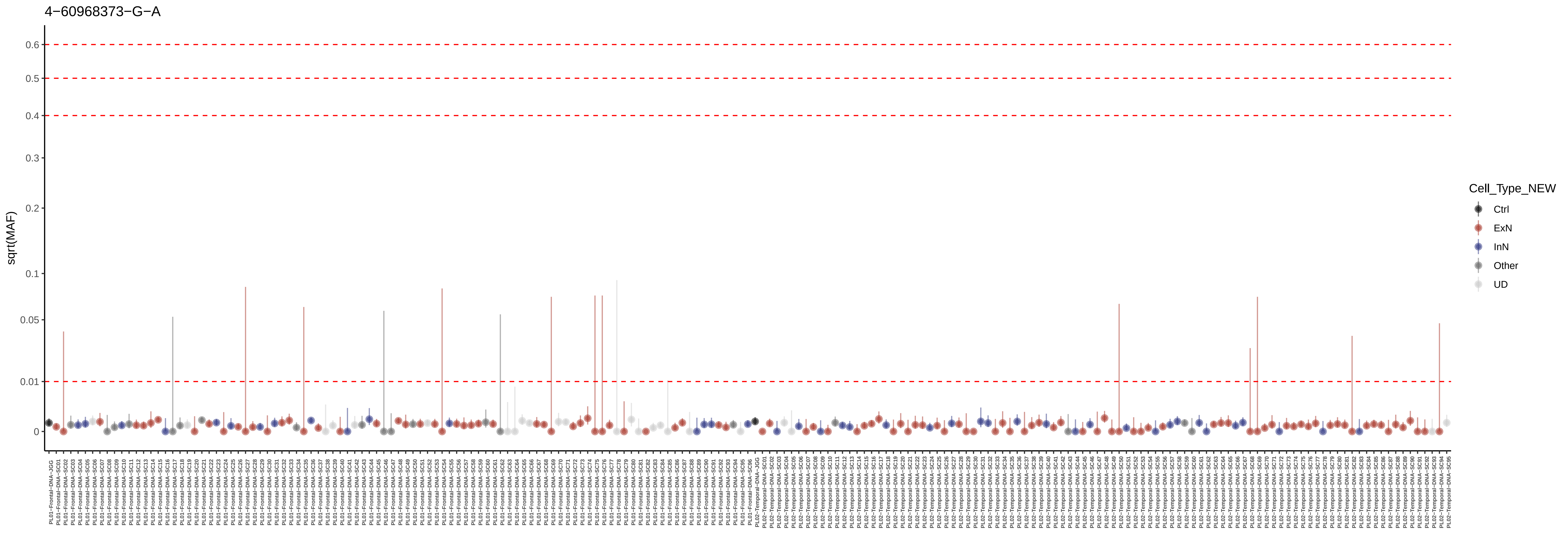
4−60968373−G−A
sqrt(MAF)
0.6
0.5
0.4
0.3
0.2
0.1
0.05
0.01
0
Cell_Type_NEW
Ctrl
ExN
InN
Other
UD

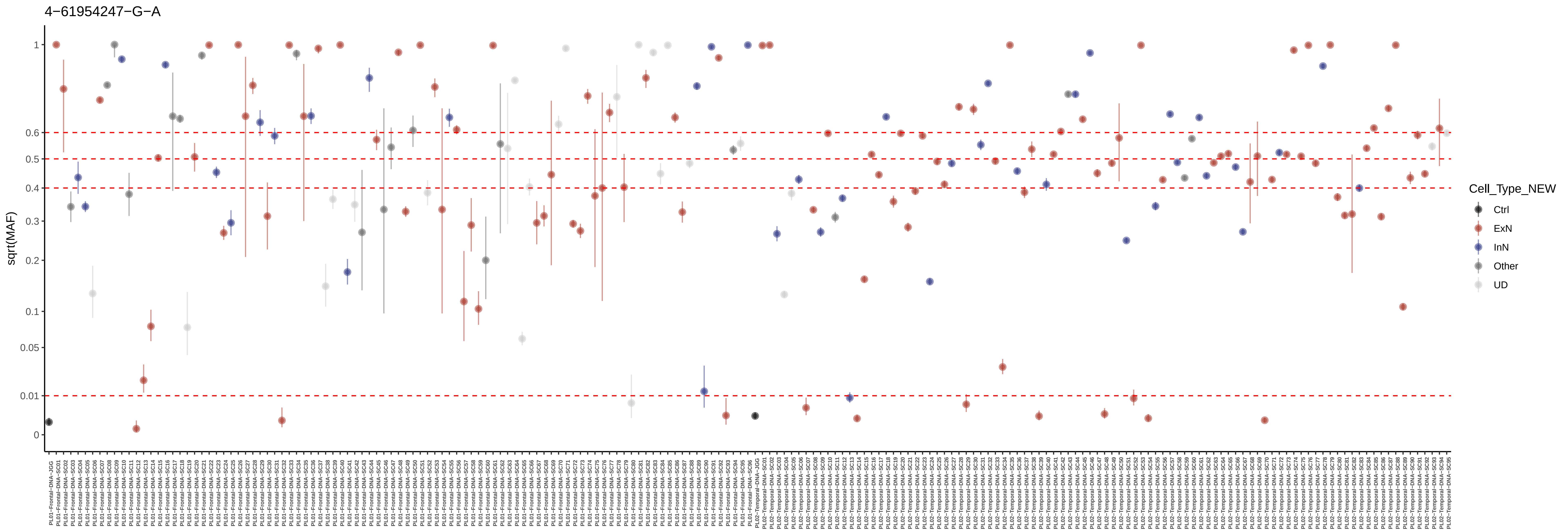
4−61954247−G−A
sqrt(MAF)
1
0.6
0.5
0.4
0.3
0.2
0.1
0.05
0.01
0
Cell_Type_NEW
Ctrl
ExN
InN
Other
UD

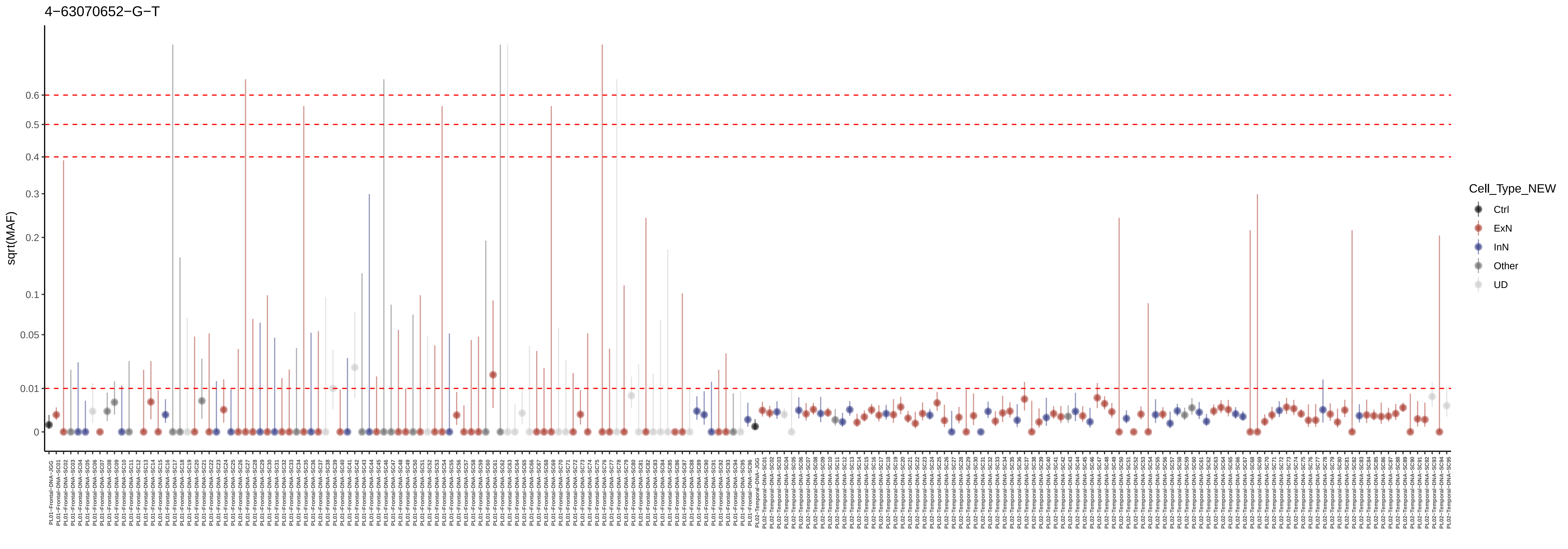

4−63070652−G−T
sqrt(MAF)
0
0.01
0.05
0.1
0.2
0.3
0.4
0.5
0.6
Cell_Type_NEW
Ctrl
ExN
InN
Other
UD

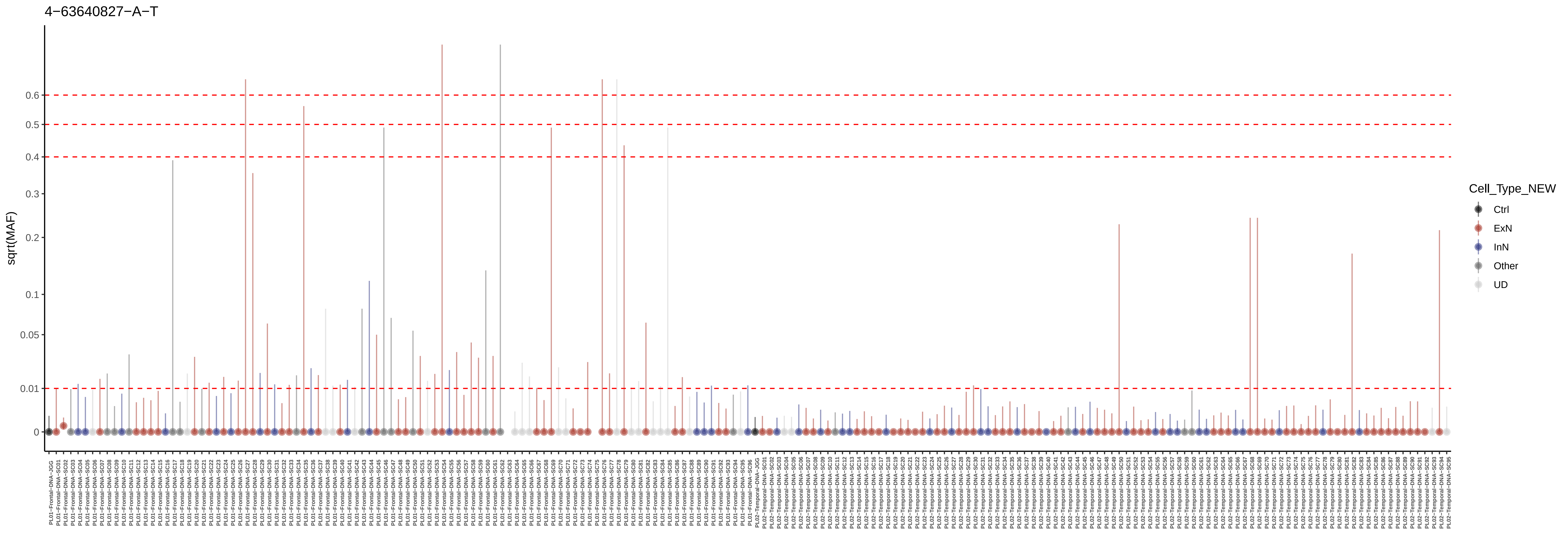

4−63640827−A−T
sqrt(MAF)
0.6
0.5
0.4
0.3
0.2
0.1
0.05
0.01
0
Cell_Type_NEW
Ctrl
ExN
InN
Other
UD

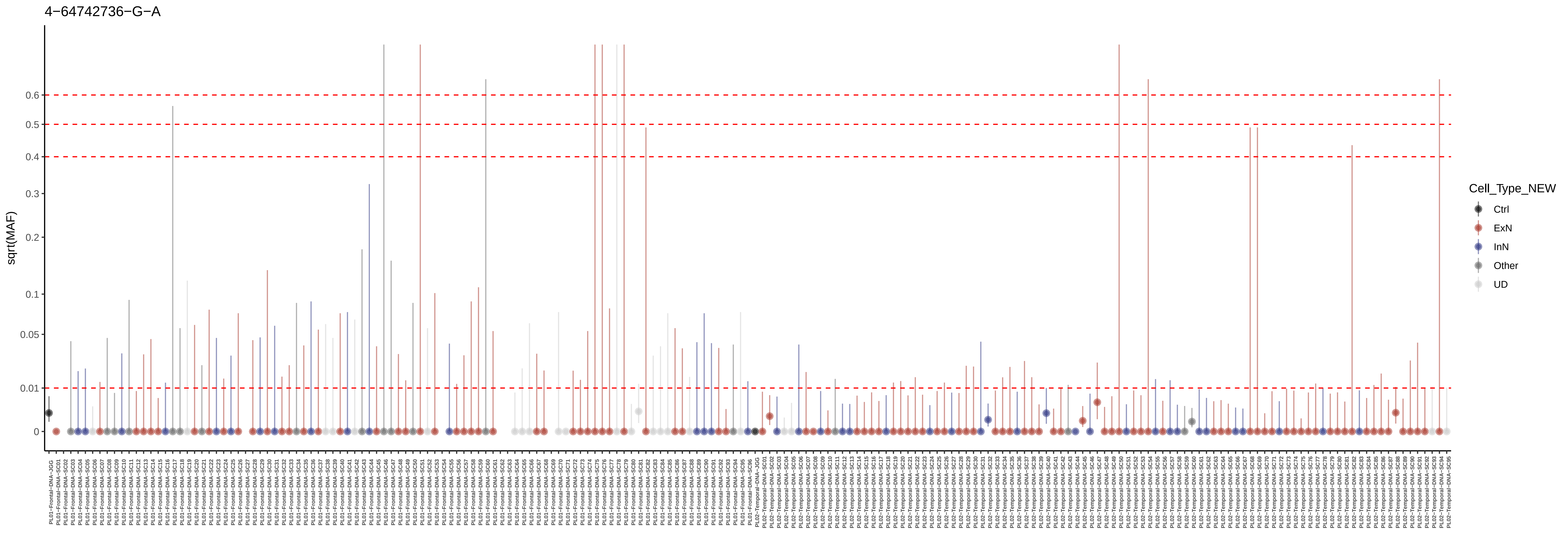
4−64742736−G−A
sqrt(MAF)
0.6
0.5
0.4
0.3
0.2
0.1
0.05
0.01
0
Cell_Type_NEW
Ctrl
ExN
InN
Other
UD

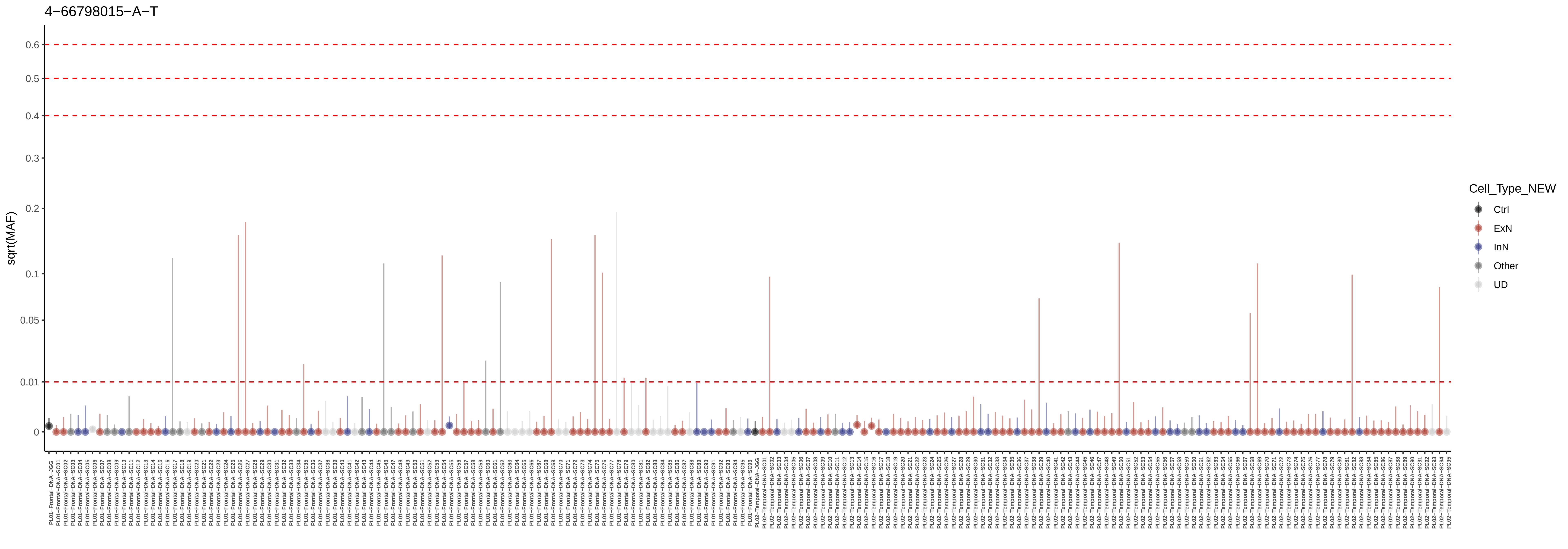
4−66798015−A−T
sqrt(MAF)
Cell_Type_NEW
Ctrl
ExN
InN
Other
UD

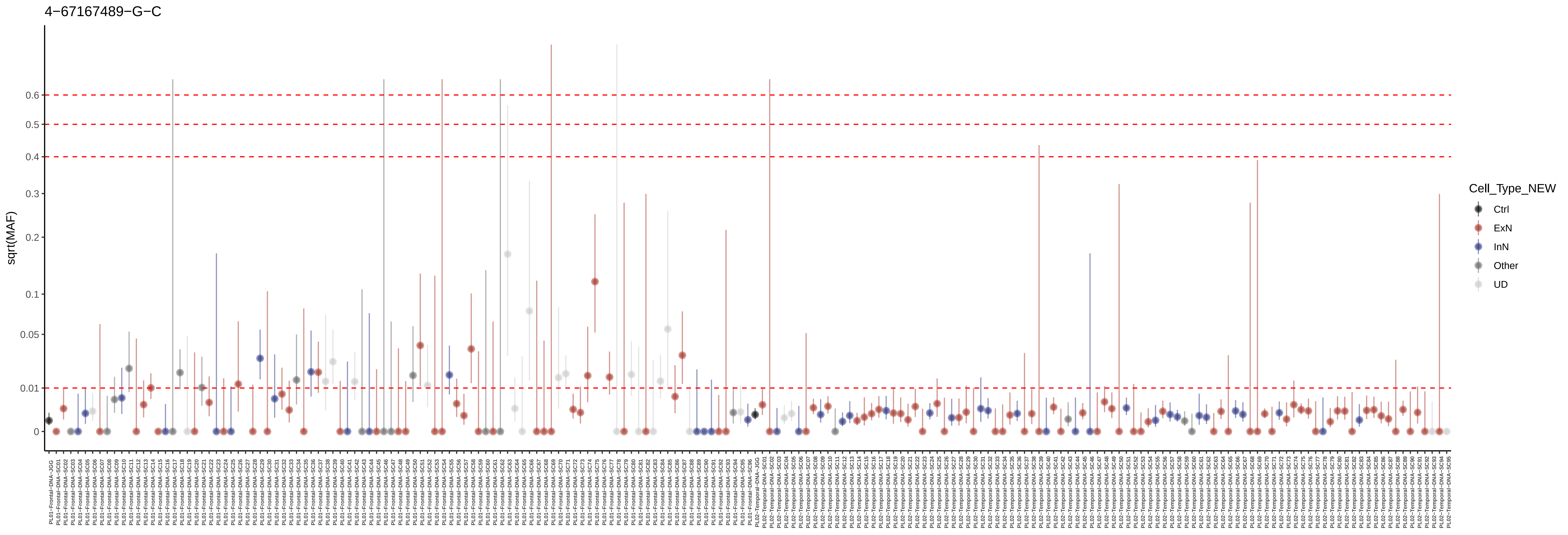

4−67167489−G−C
sqrt(MAF)
0.6
0.5
0.4
0.3
0.2
0.1
0.05
0.01
0
Cell_Type_NEW
Ctrl
ExN
InN
Other
UD

4−67729599−T−C

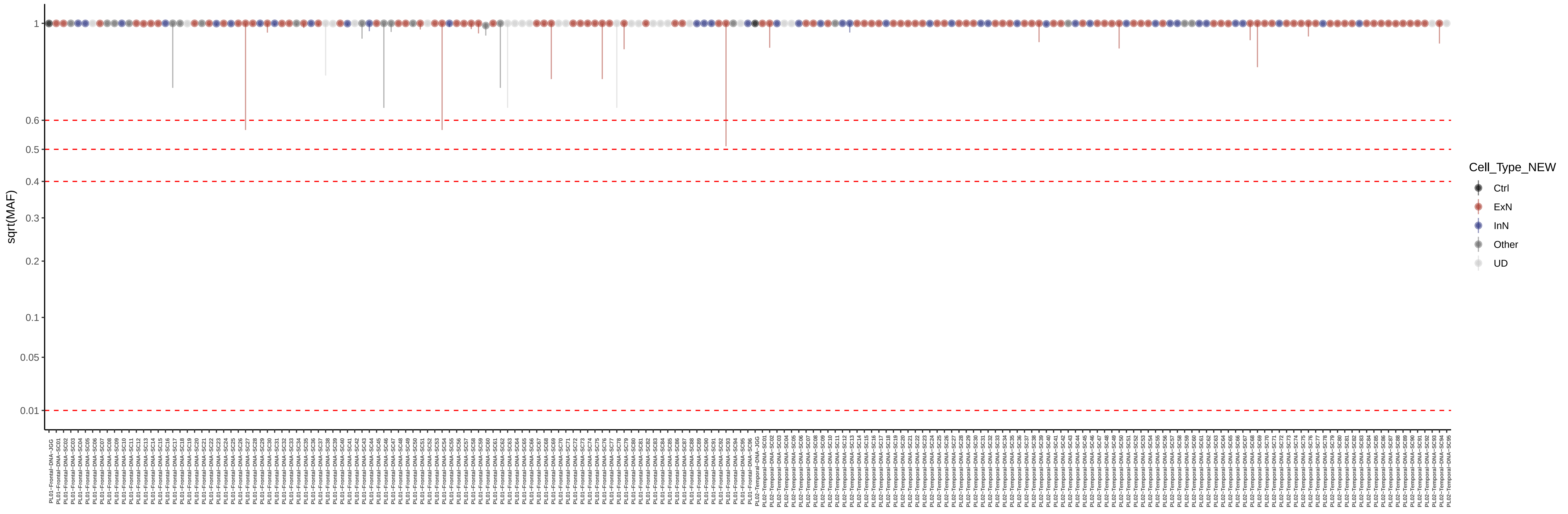

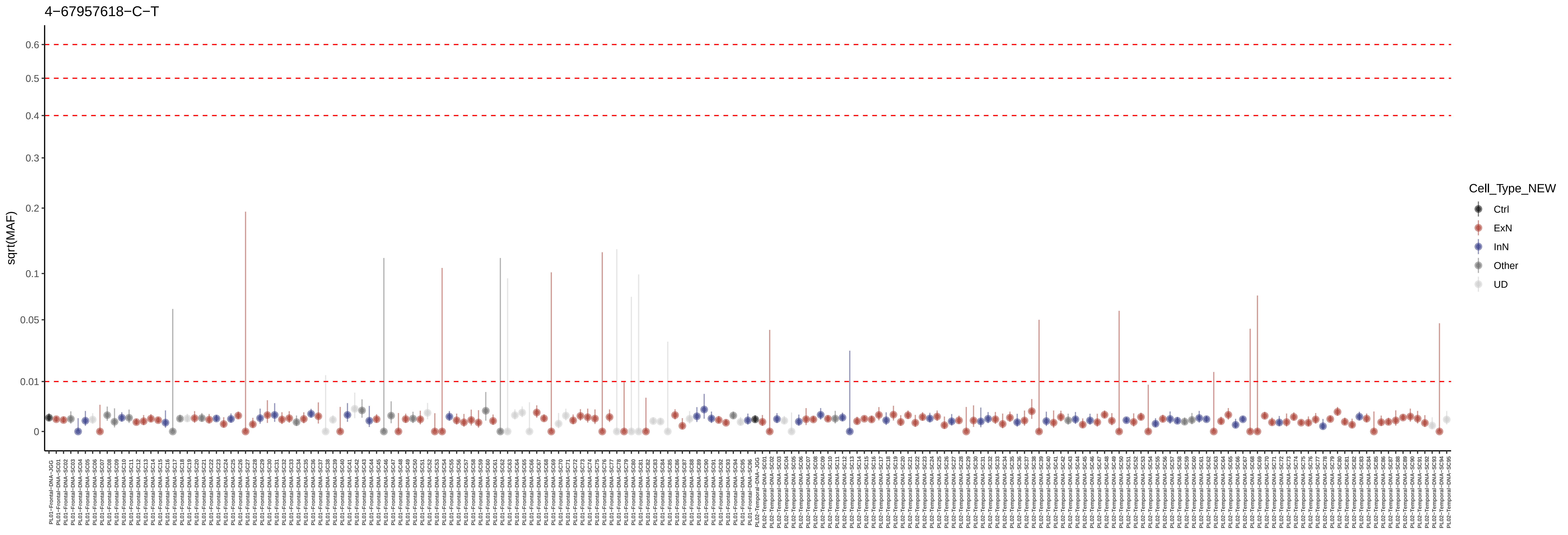
4−67957618−C−T
sqrt(MAF)
0.6
0.5
0.4
0.3
0.2
0.1
0.05
0.01
0
Cell_Type_NEW
Ctrl
ExN
InN
Other
UD

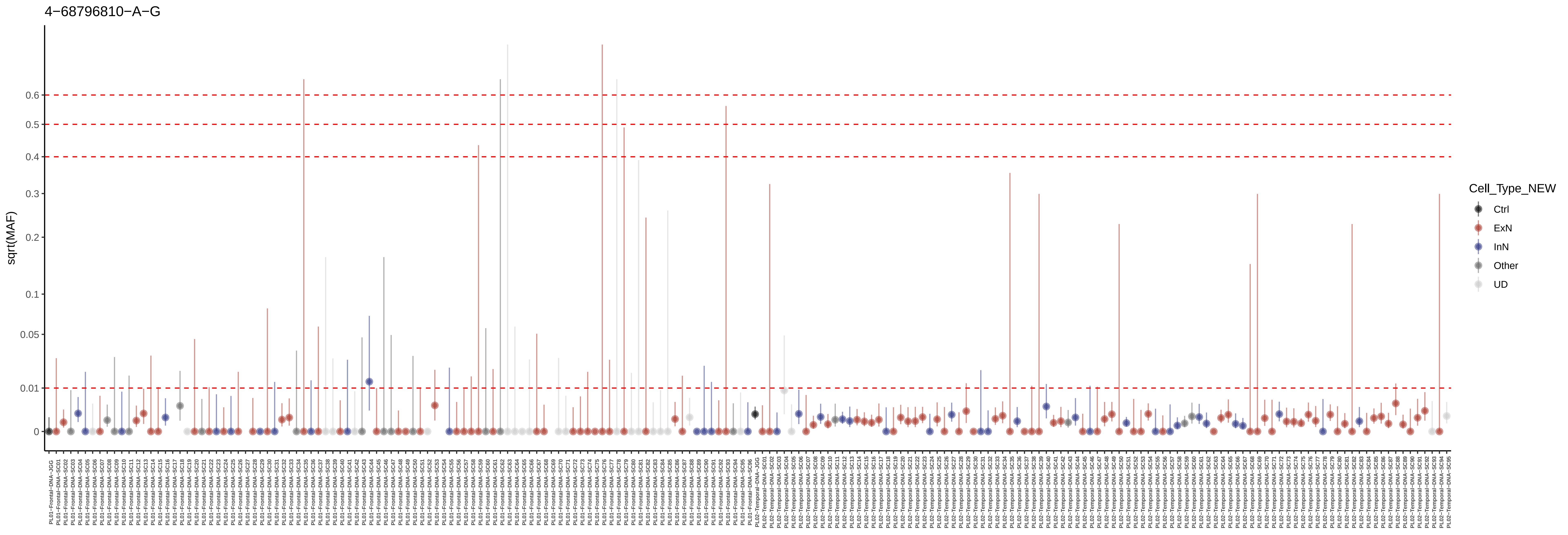
4−68796810−A−G
sqrt(MAF)
0.6
0.5
0.4
0.3
0.2
0.1
0.05
0.01
0
Cell_Type_NEW
Ctrl
ExN
InN
Other
UD

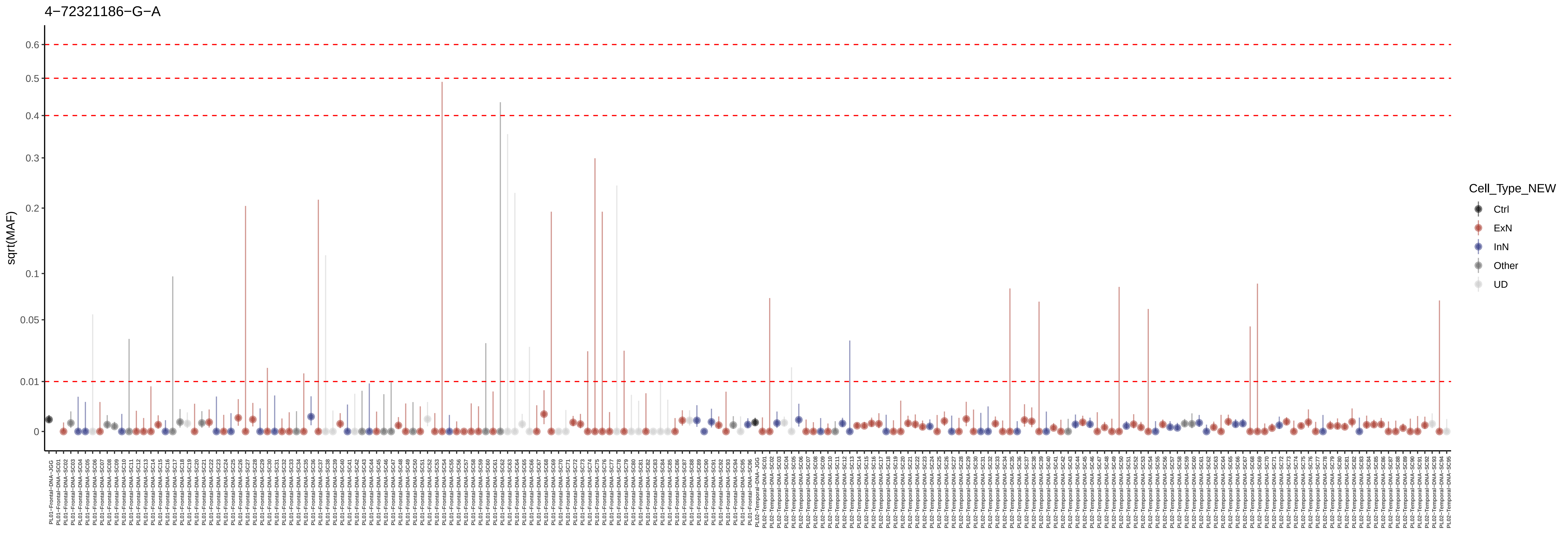
4−72321186−G−A
sqrt(MAF)
0.6
0.5
0.4
0.3
0.2
0.1
0.05
0.01
0
Cell_Type_NEW
Ctrl
ExN
InN
Other
UD

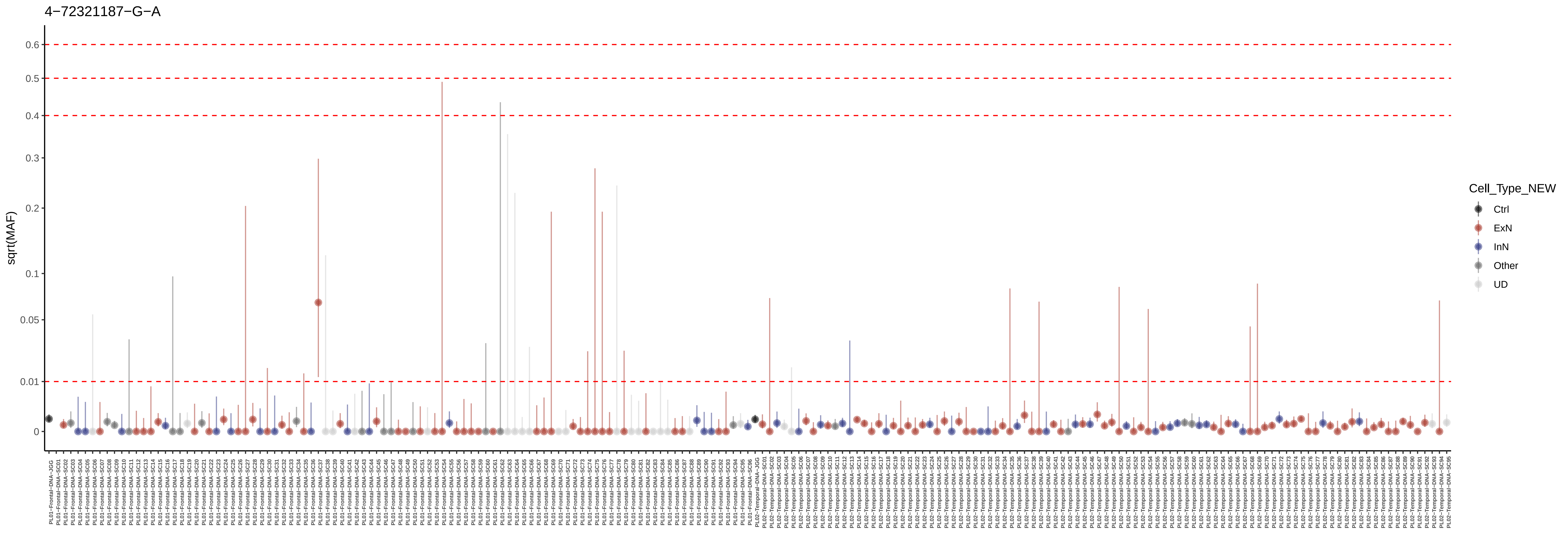
4−72321187−G−A
sqrt(MAF)
0.6
0.5
0.4
0.3
0.2
0.1
0.05
0.01
0
Cell_Type_NEW
Ctrl
ExN
InN
Other
UD

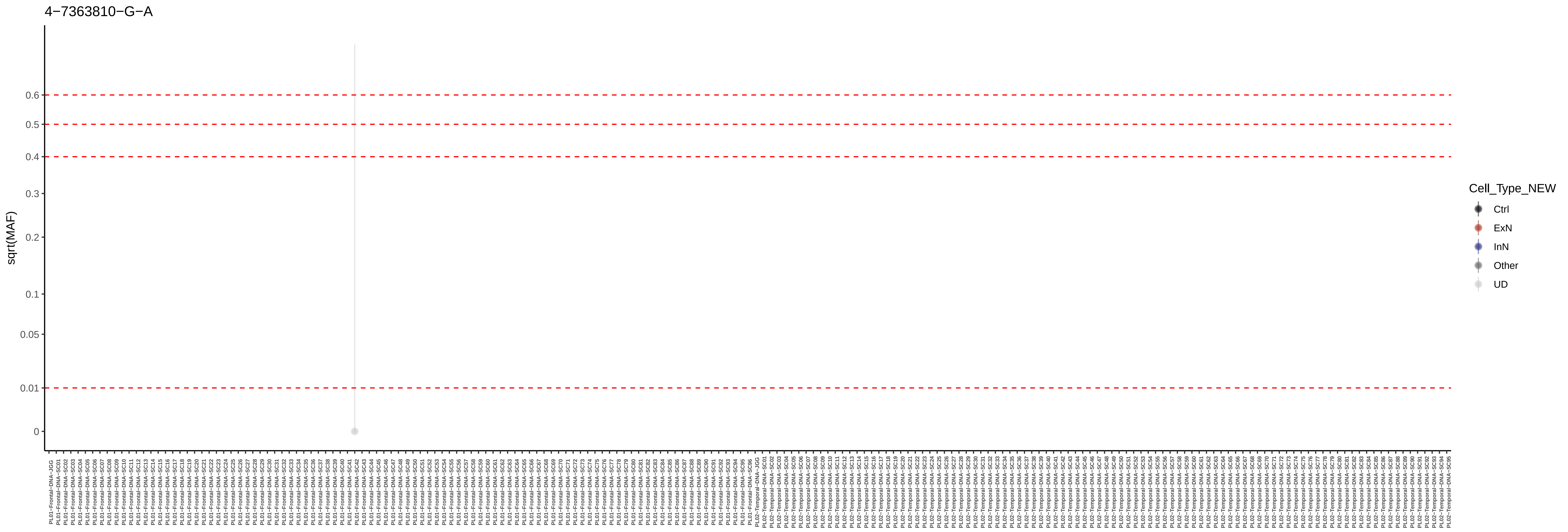
4−7363810−G−A
sqrt(MAF)
0.6
0.5
0.4
0.3
0.2
0.1
0.05
0.01
0
Cell_Type_NEW
Ctrl
ExN
InN
Other
UD

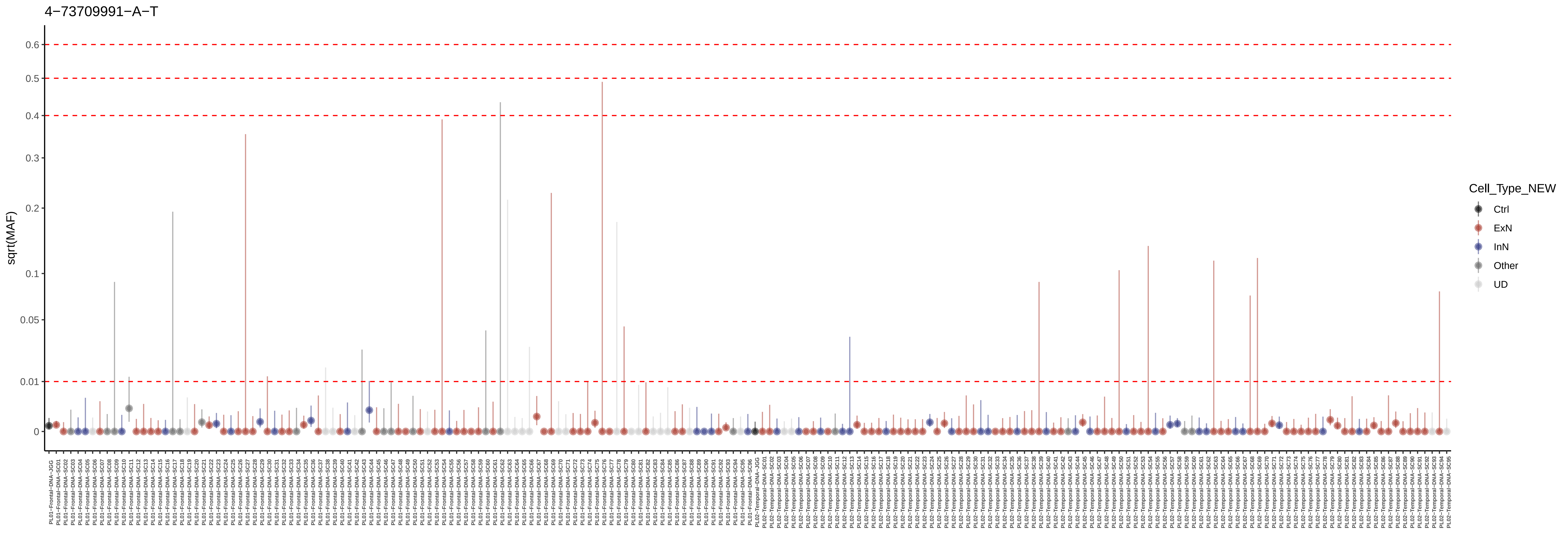

4−73709991−A−T
sqrt(MAF)
0.6
0.5
0.4
0.3
0.2
0.1
0.05
0.01
0
Cell_Type_NEW
Ctrl
ExN
InN
Other
UD

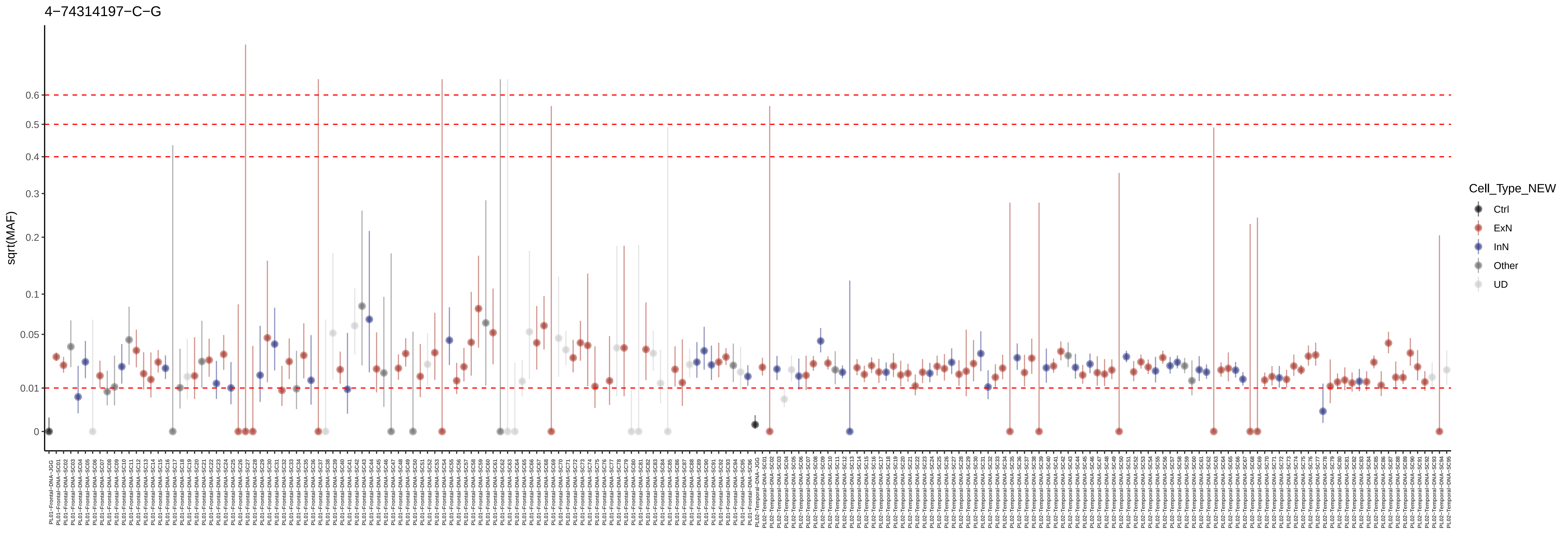

4−74314197−C−G
sqrt(MAF)
0.6
0.5
0.4
0.3
0.2
0.1
0.05
0.01
0
Cell_Type_NEW
Ctrl
ExN
InN
Other
UD

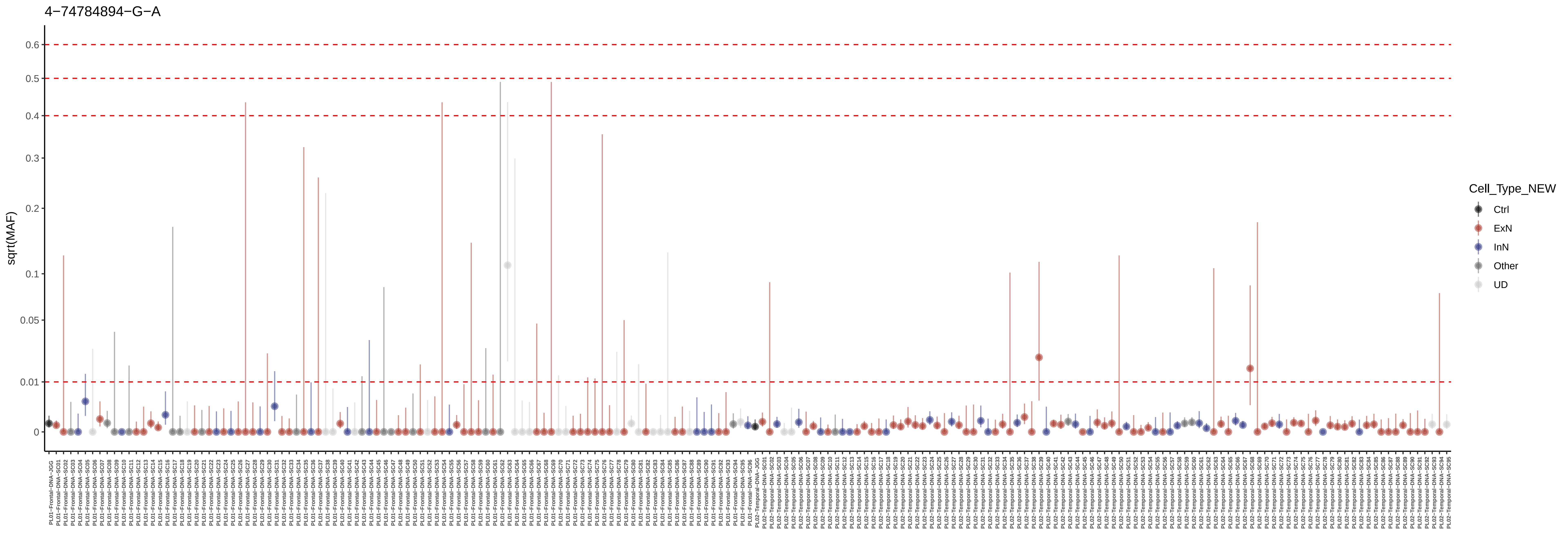

4−74784894−G−A
sqrt(MAF)
0.6
0.5
0.4
0.3
0.2
0.1
0.05
0.01
0
Cell_Type_NEW
Ctrl
ExN
InN
Other
UD

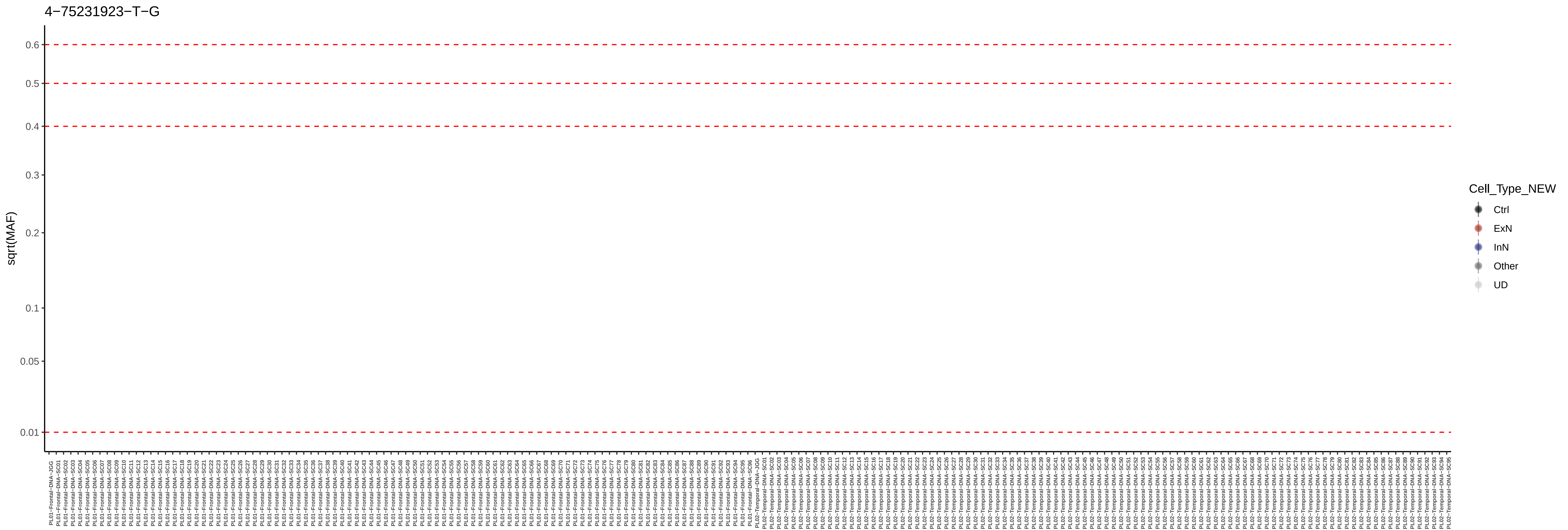
4−75231923−T−G
sqrt(MAF)
0.6
0.5
0.4
0.3
0.2
0.1
0.05
0.01
Cell_Type_NEW
Ctrl
ExN
InN
Other
UD

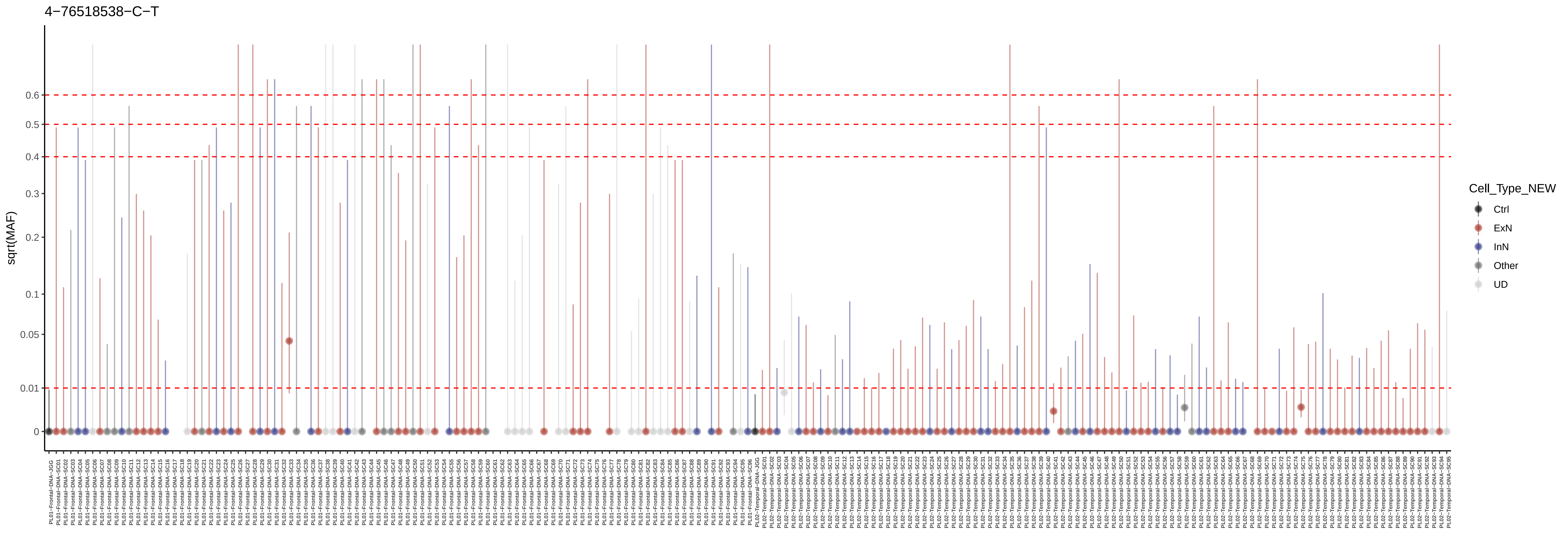
4−76518538−C−T
sqrt(MAF)
0.6
0.5
0.4
0.3
0.2
0.1
0.05
0.01
0
Cell_Type_NEW
Ctrl
ExN
InN
Other
UD

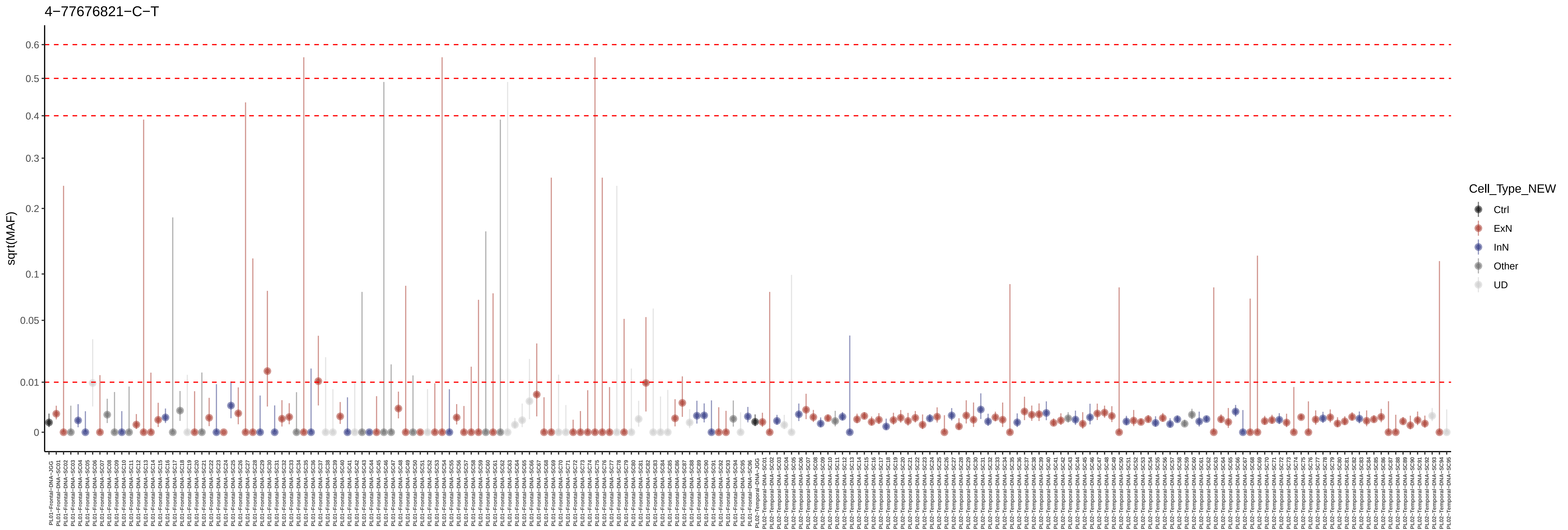

4−77676821−C−T
sqrt(MAF)
0.6
0.5
0.4
0.3
0.2
0.1
0.05
0.01
0
Cell_Type_NEW
Ctrl
ExN
InN
Other
UD

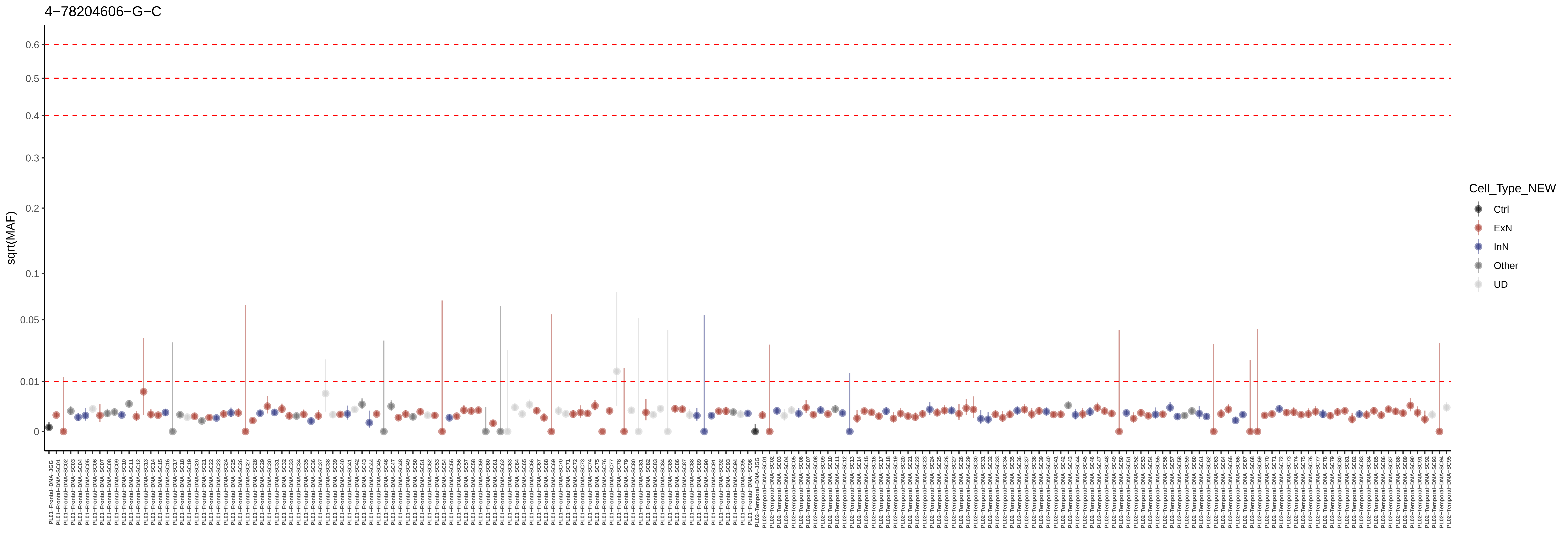

4−78204606−G−C
sqrt(MAF)
0.6
0.5
0.4
0.3
0.2
0.1
0.05
0.01
0
Cell_Type_NEW
Ctrl
ExN
InN
Other
UD

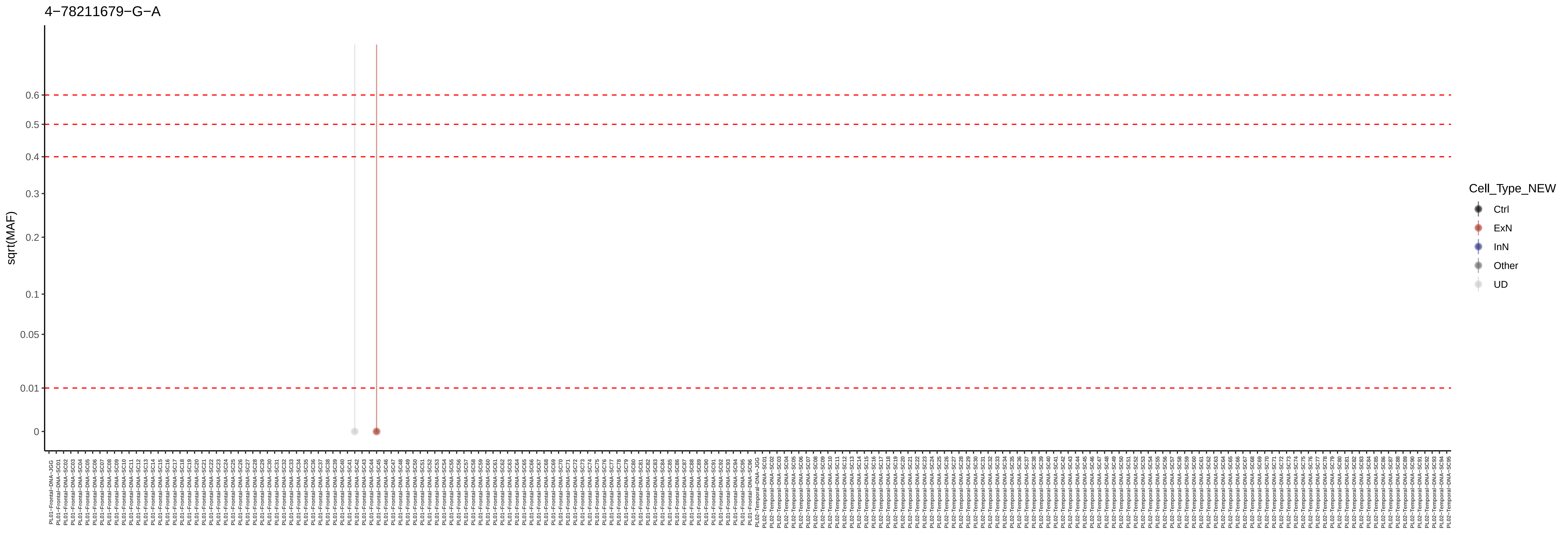
4−78211679−G−A
sqrt(MAF)
0.6
0.5
0.4
0.3
0.2
0.1
0.05
0.01
0
Cell_Type_NEW
Ctrl
ExN
InN
Other
UD

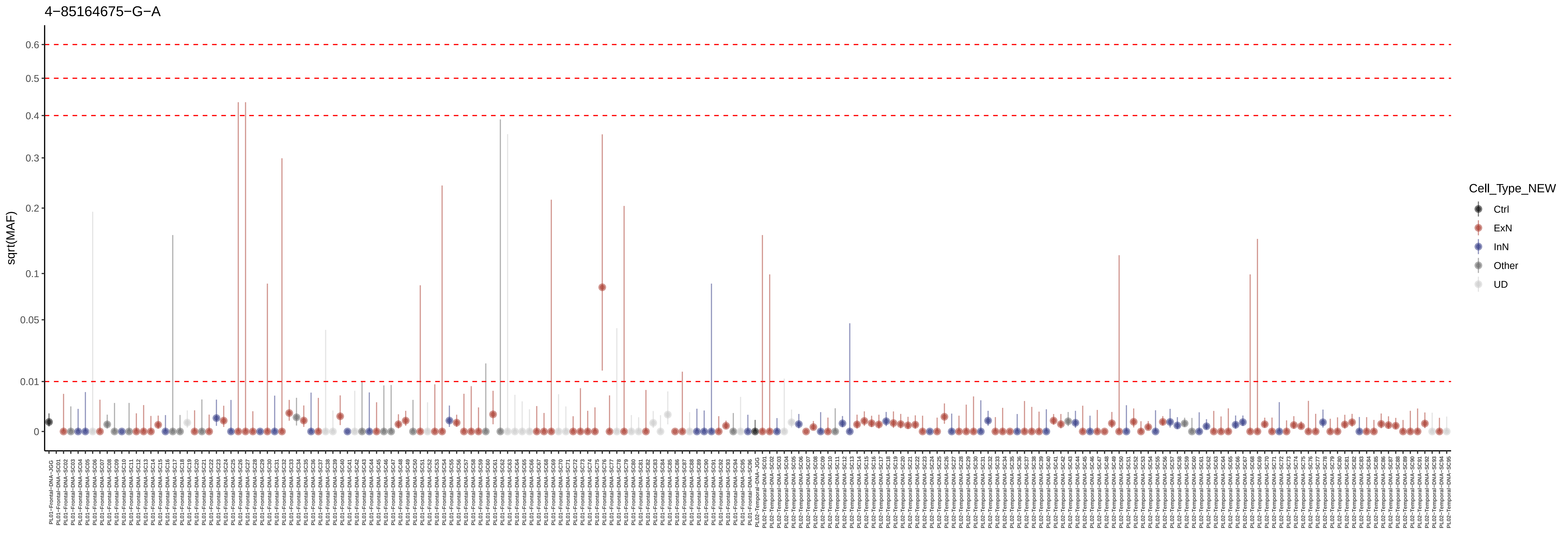
4−85164675−G−A
sqrt(MAF)
0.6
0.5
0.4
0.3
0.2
0.1
0.05
0.01
0
Cell_Type_NEW
Ctrl
ExN
InN
Other
UD

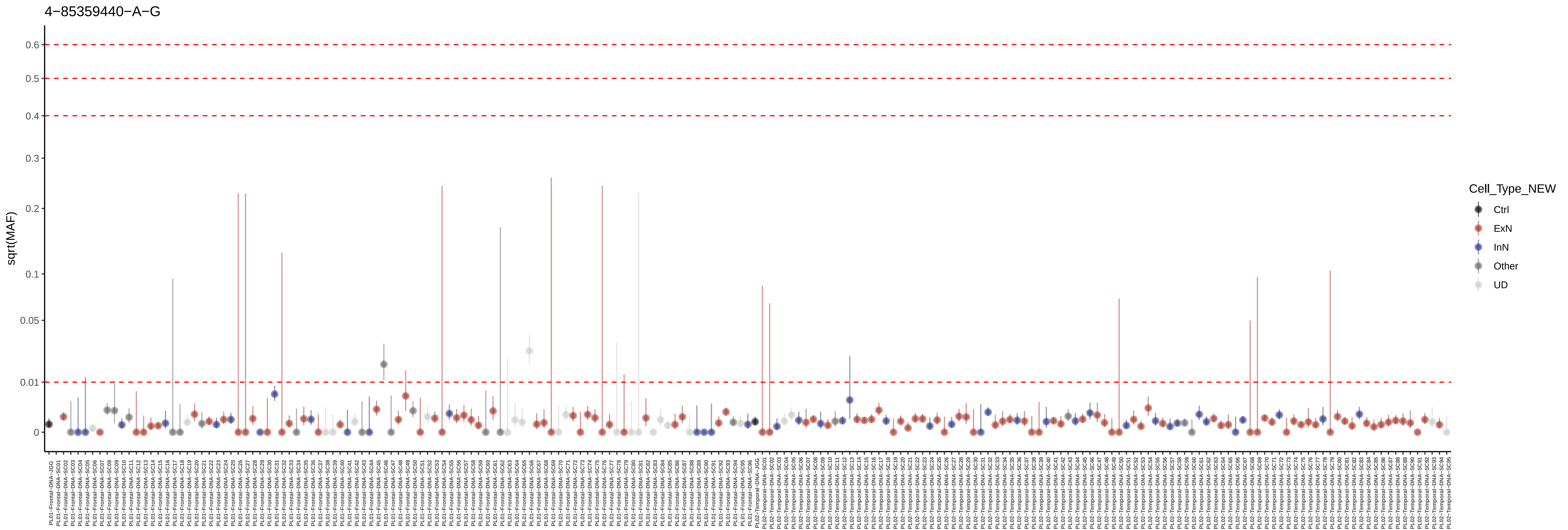
4−85359440−A−G
sqrt(MAF)
0.6
0.5
0.4
0.3
0.2
0.1
0.05
0.01
0
Cell_Type_NEW
Ctrl
ExN
InN
Other
UD

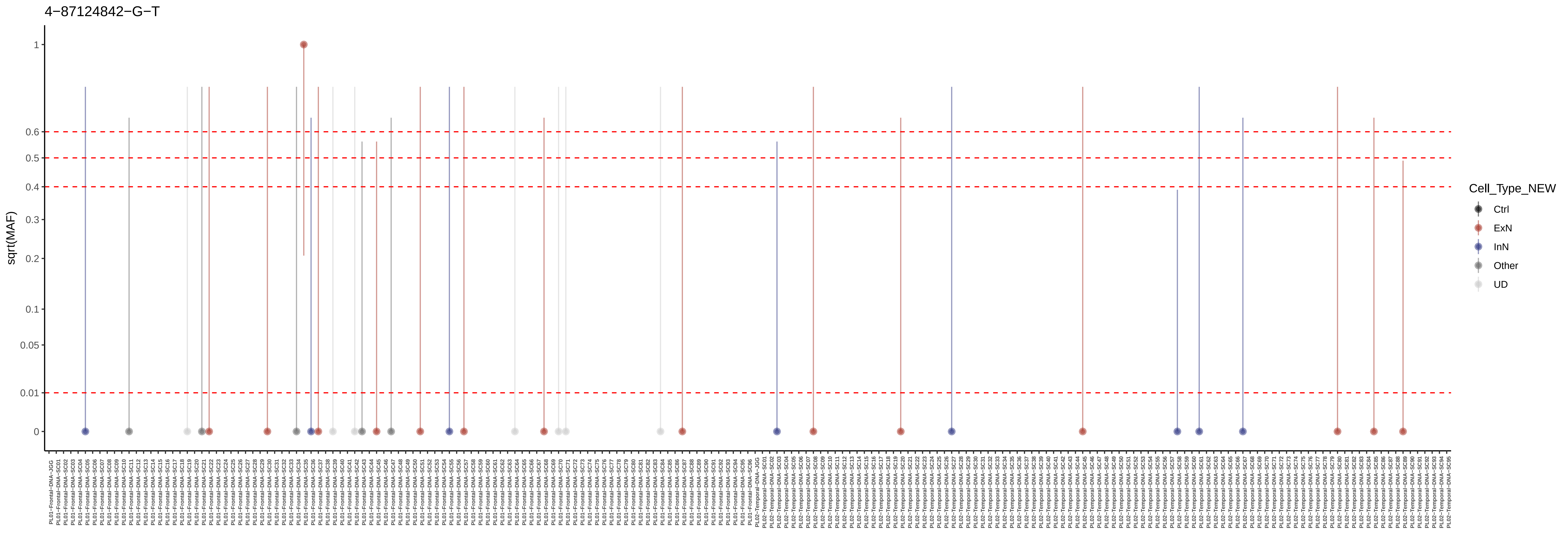
4−87124842−G−T
sqrt(MAF)
1
0.6
0.5
0.4
0.3
0.2
0.1
0.05
0.01
0
Cell_Type_NEW
Ctrl
ExN
InN
Other
UD

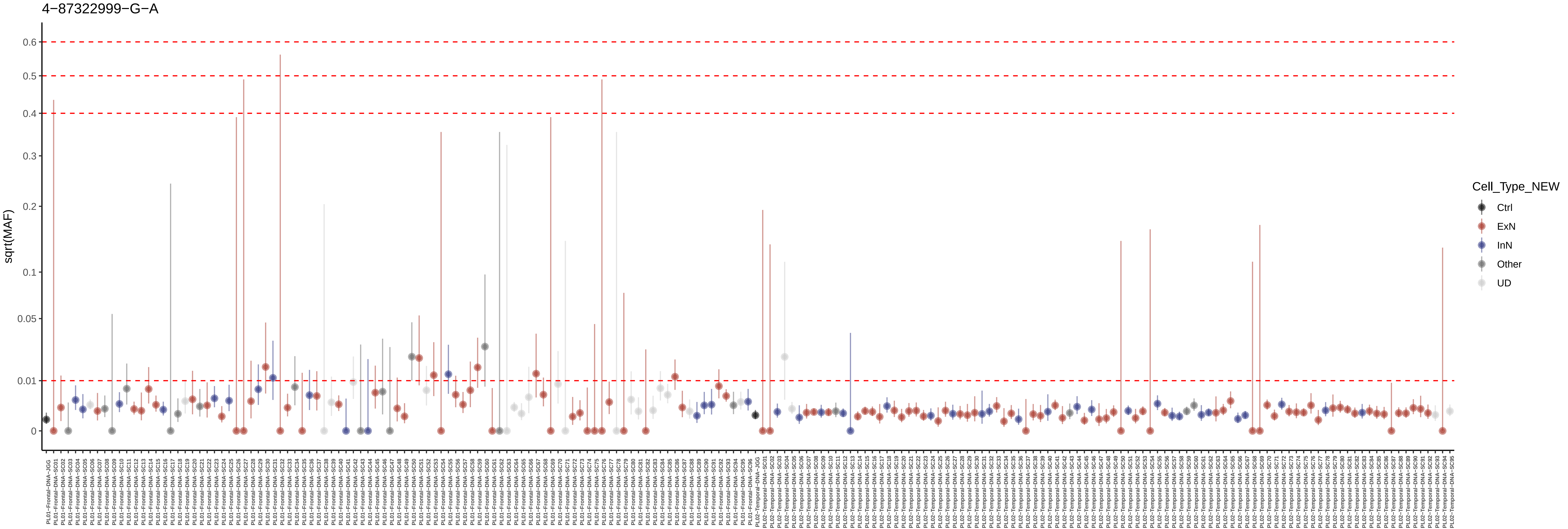
4−87322999−G−A
sqrt(MAF)
0.6
0.5
0.4
0.3
0.2
0.1
0.05
0.01
0
Cell_Type_NEW
Ctrl
ExN
InN
Other
UD

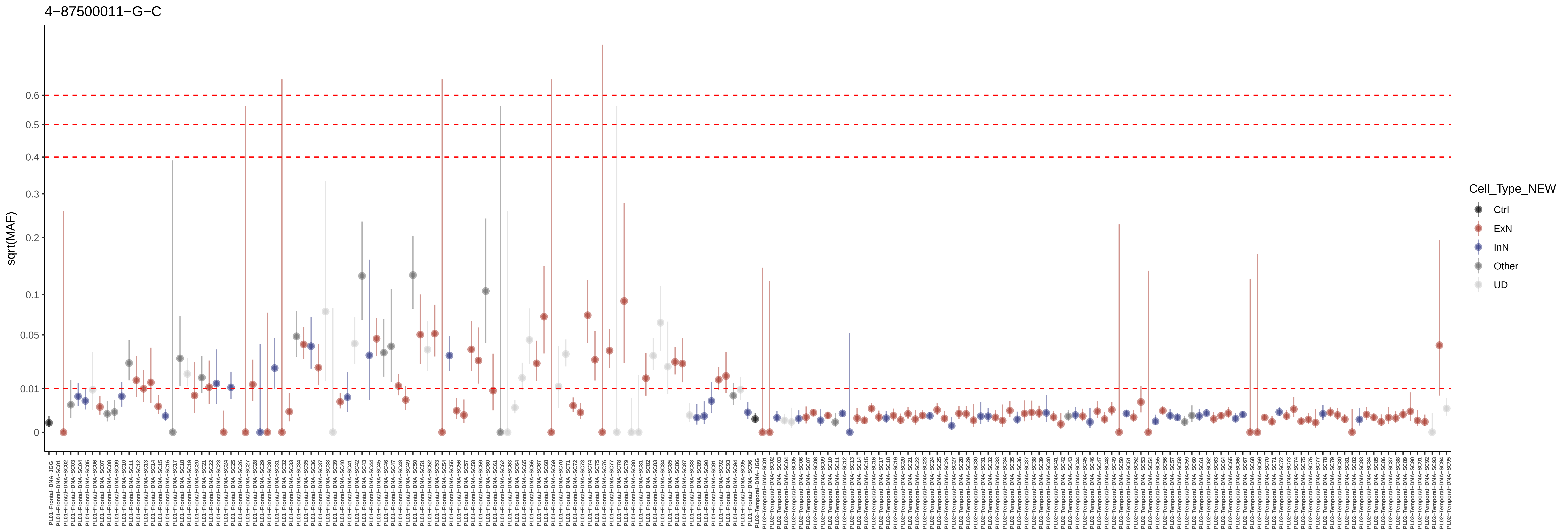
4−87500011−G−C
sqrt(MAF)
0.6
0.5
0.4
0.3
0.2
0.1
0.05
0.01
0
Cell_Type_NEW
Ctrl
ExN
InN
Other
UD

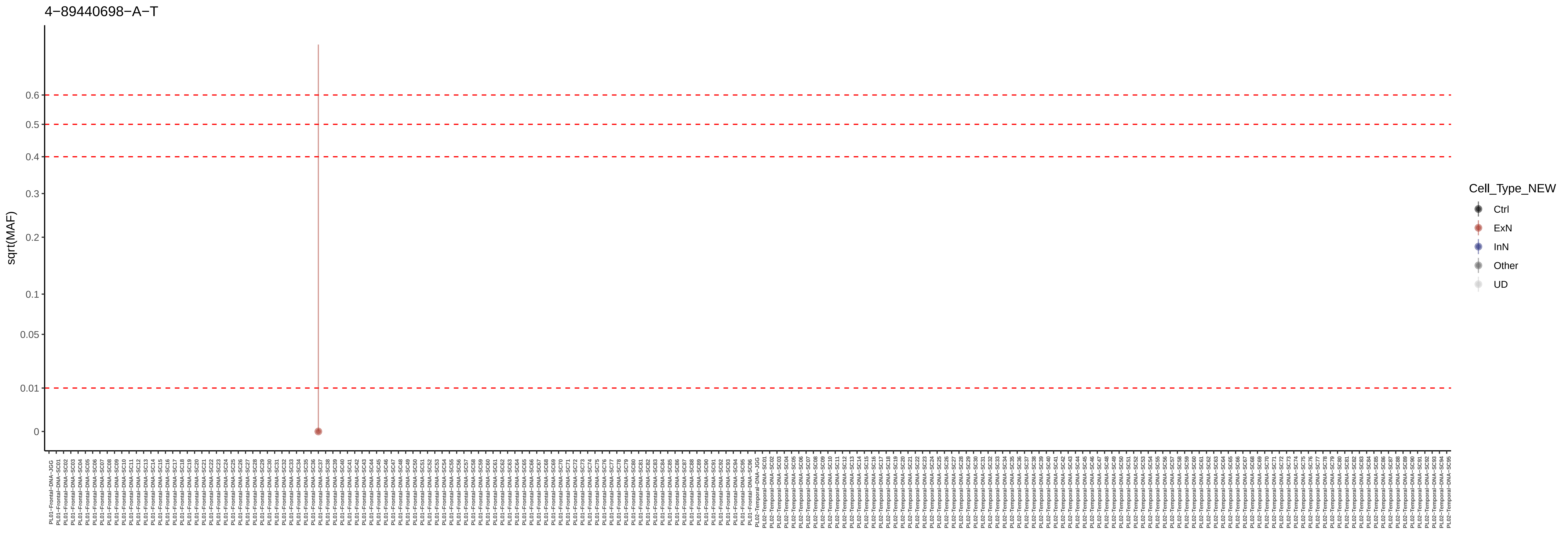
4−89440698−A−T
sqrt(MAF)
0.6
0.5
0.4
0.3
0.2
0.1
0.05
0.01
0
Cell_Type_NEW
Ctrl
ExN
InN
Other
UD

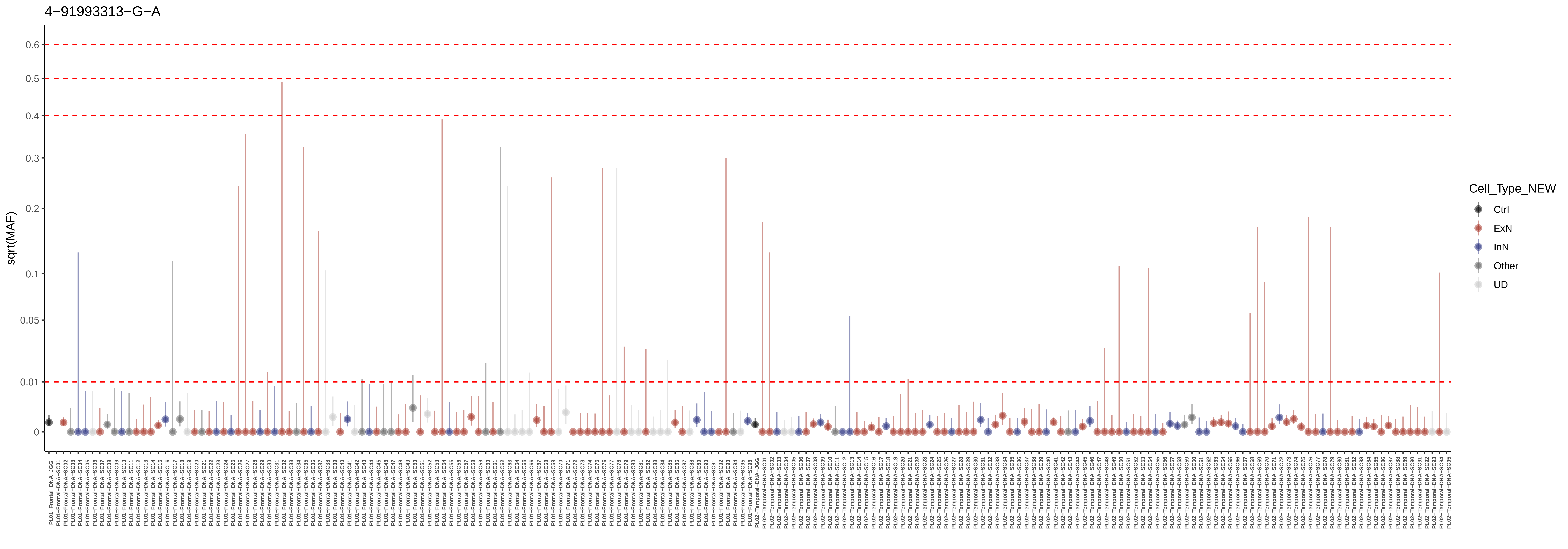

4−91993313−G−A
sqrt(MAF)
0.6
0.5
0.4
0.3
0.2
0.1
0.05
0.01
0
Cell_Type_NEW
Ctrl
ExN
InN
Other
UD

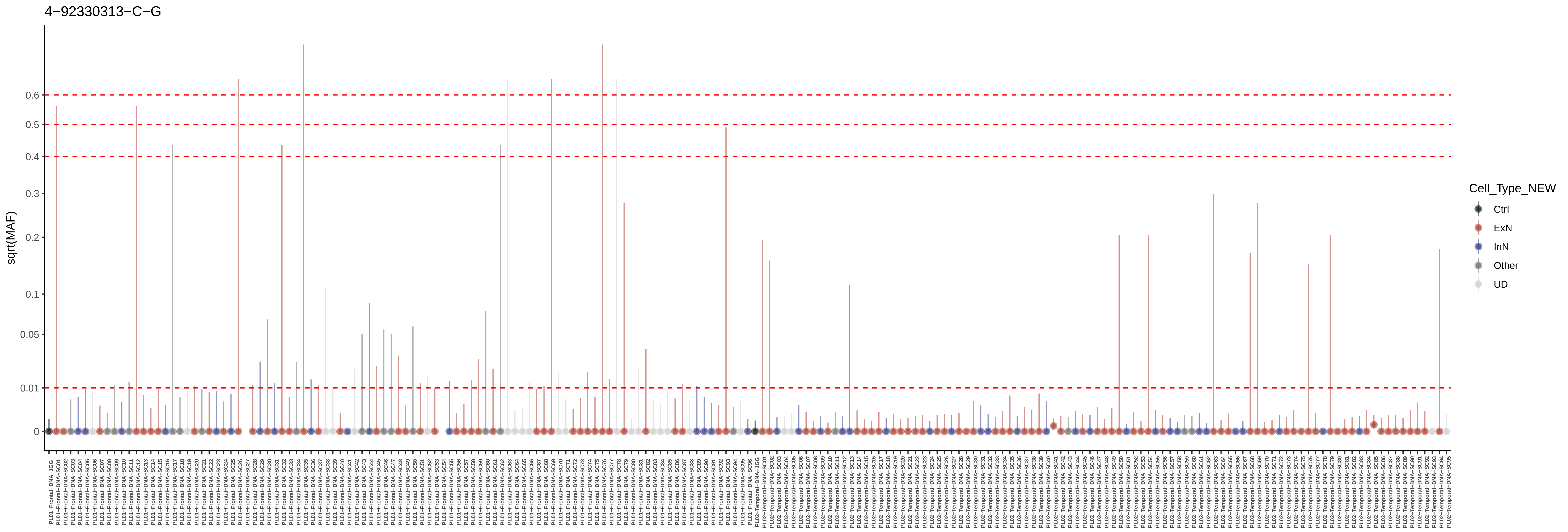
4−92330313−C−G
sqrt(MAF)
0.6
0.5
0.4
0.3
0.2
0.1
0.05
0.01
0
Cell_Type_NEW
Ctrl
ExN
InN
Other
UD

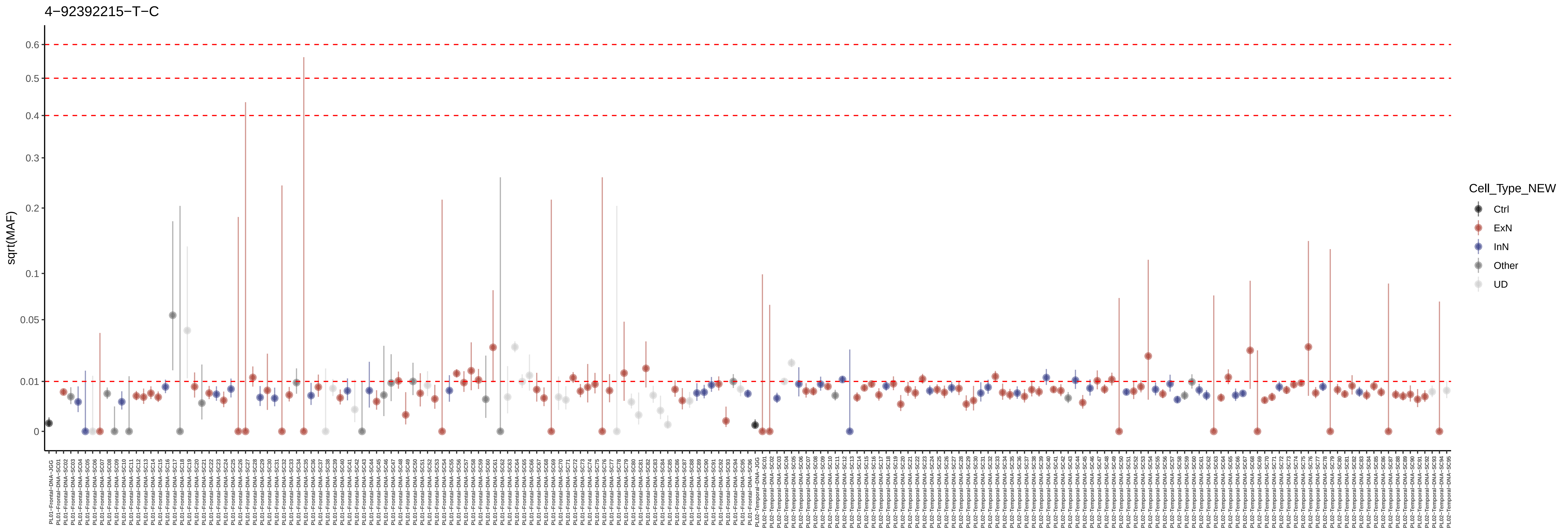
4−92392215−T−C
sqrt(MAF)
0.6
0.5
0.4
0.3
0.2
0.1
0.05
0.01
0
Cell_Type_NEW
Ctrl
ExN
InN
Other
UD

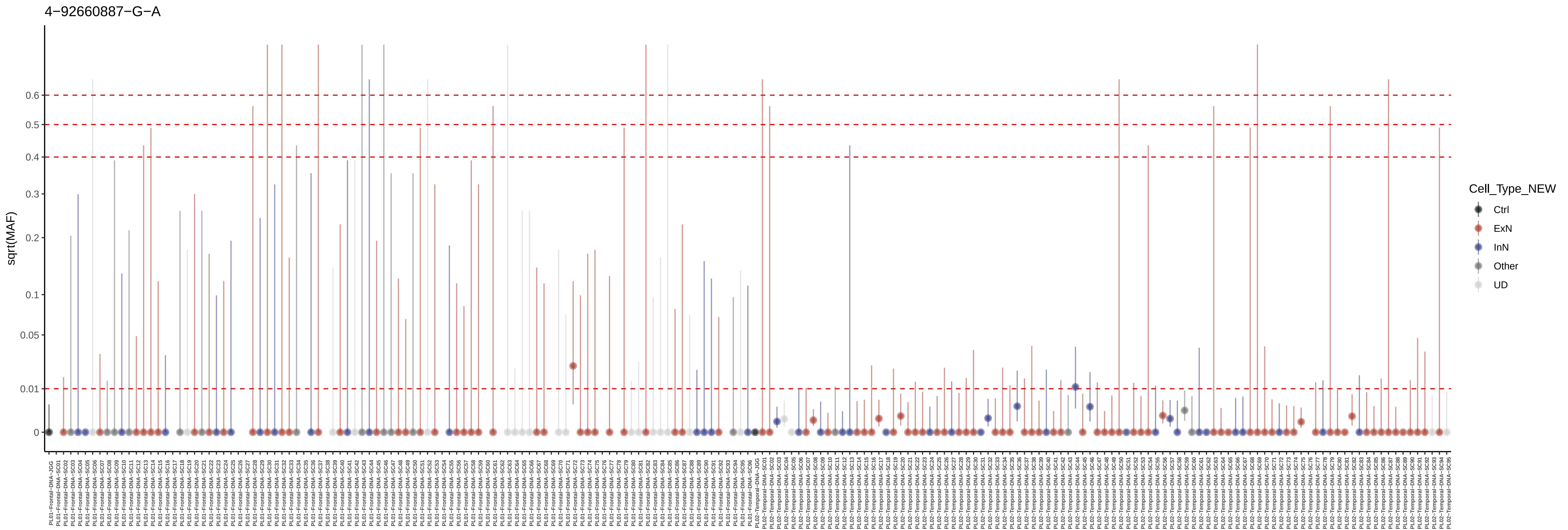

4−92660887−G−A
sqrt(MAF)
0.6
0.5
0.4
0.3
0.2
0.1
0.05
0.01
0
Cell_Type_NEW
Ctrl
ExN
InN
Other
UD

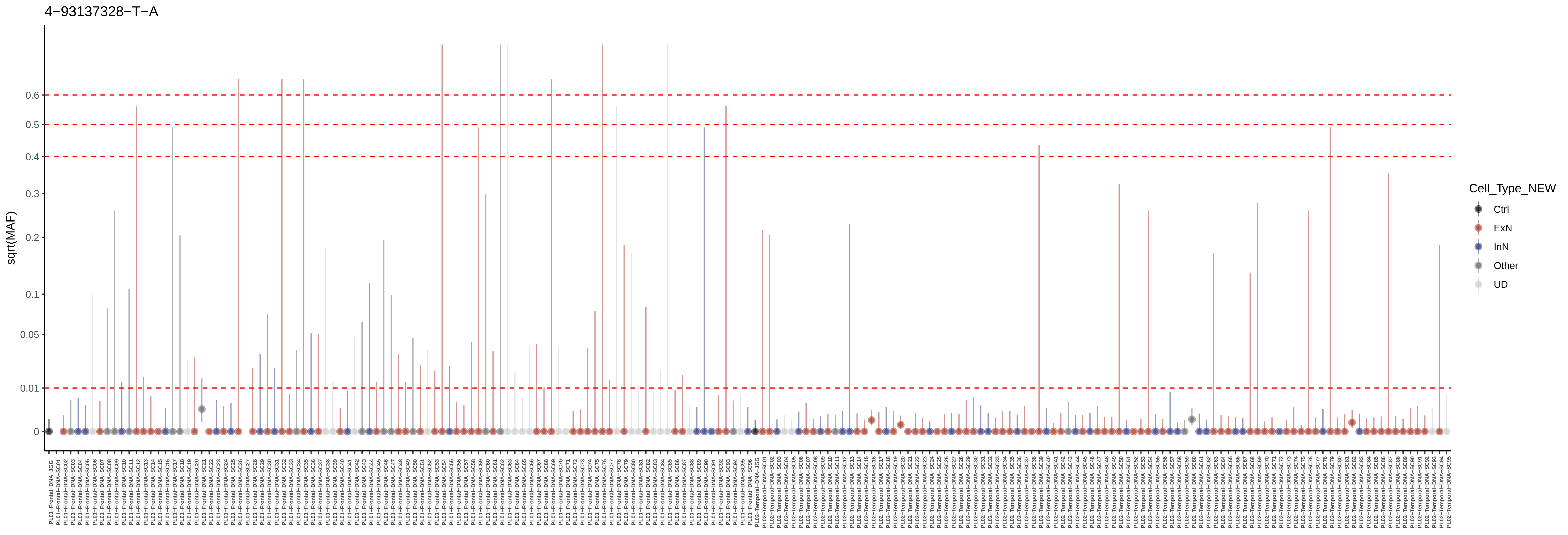
4−93137328−T−A
sqrt(MAF)
0.6
0.5
0.4
0.3
0.2
0.1
0.05
0.01
0
Cell_Type_NEW
Ctrl
ExN
InN
Other
UD

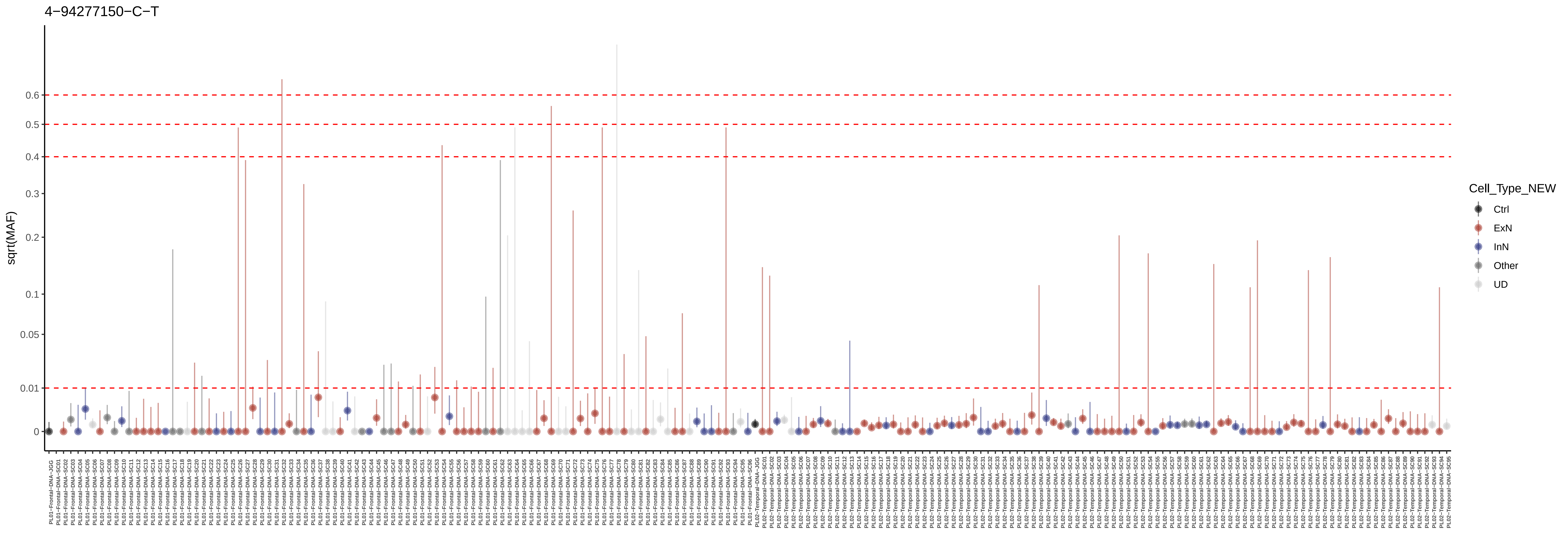
4−94277150−C−T
sqrt(MAF)
0.6
0.5
0.4
0.3
0.2
0.1
0.05
0.01
0
Cell_Type_NEW
Ctrl
ExN
InN
Other
UD

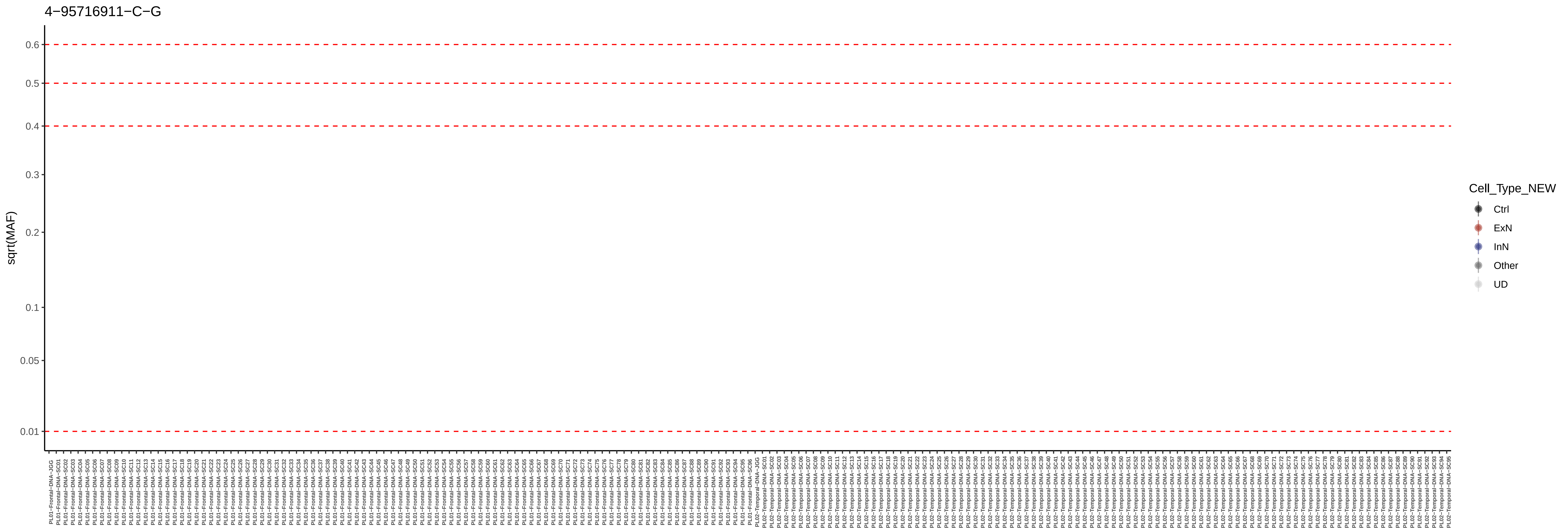
4−95716911−C−G
sqrt(MAF)
0.6
0.5
0.4
0.3
0.2
0.1
0.05
0.01
Cell_Type_NEW
Ctrl
ExN
InN
Other
UD

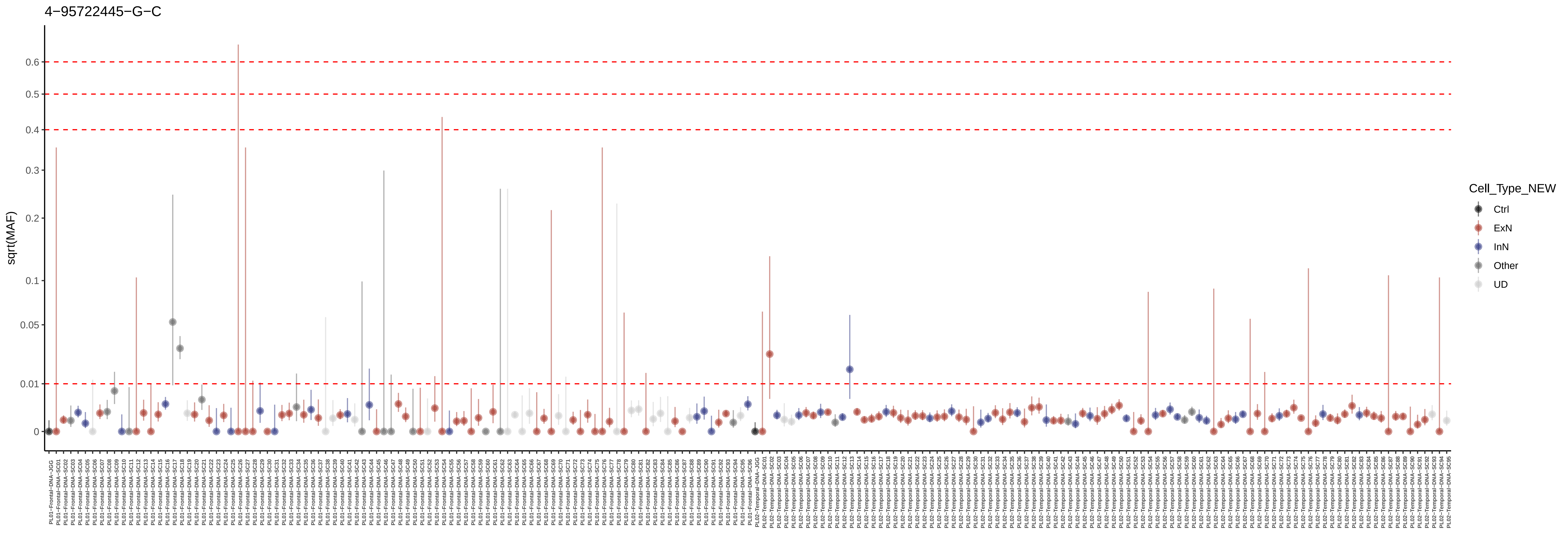

4−95722445−G−C
sqrt(MAF)
0.6
0.5
0.4
0.3
0.2
0.1
0.05
0.01
0
Cell_Type_NEW
Ctrl
ExN
InN
Other
UD

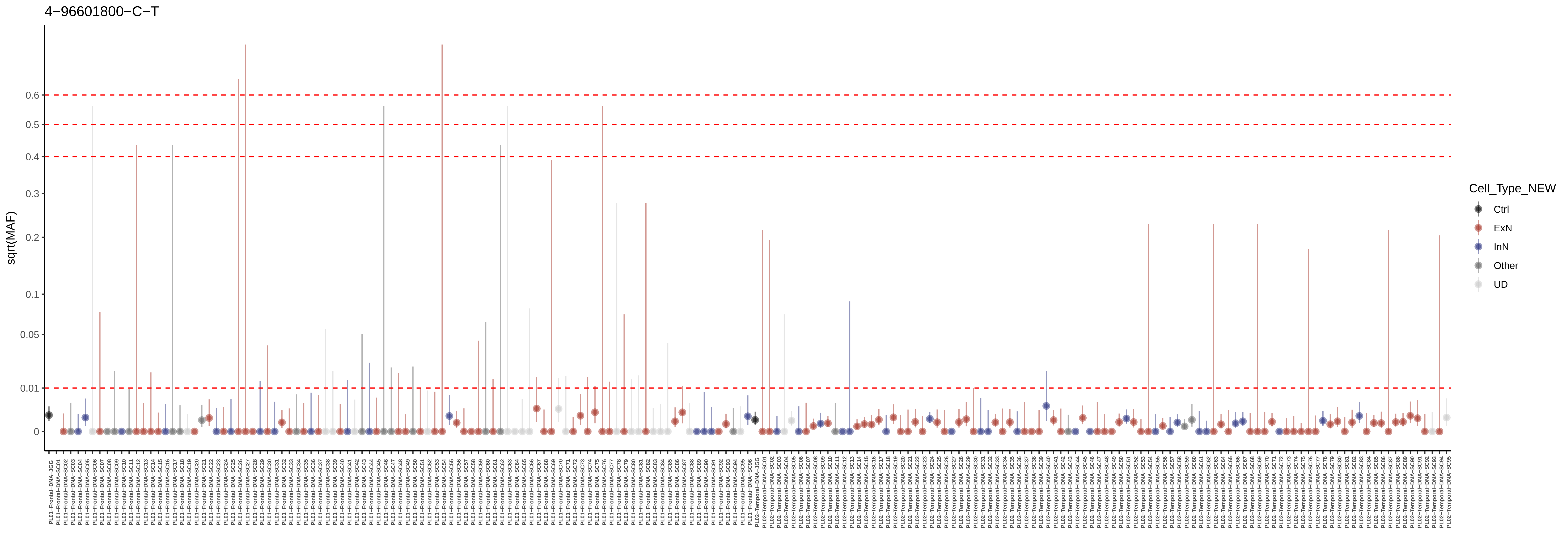
4−96601800−C−T
sqrt(MAF)
0.6
0.5
0.4
0.3
0.2
0.1
0.05
0.01
0
Cell_Type_NEW
Ctrl
ExN
InN
Other
UD

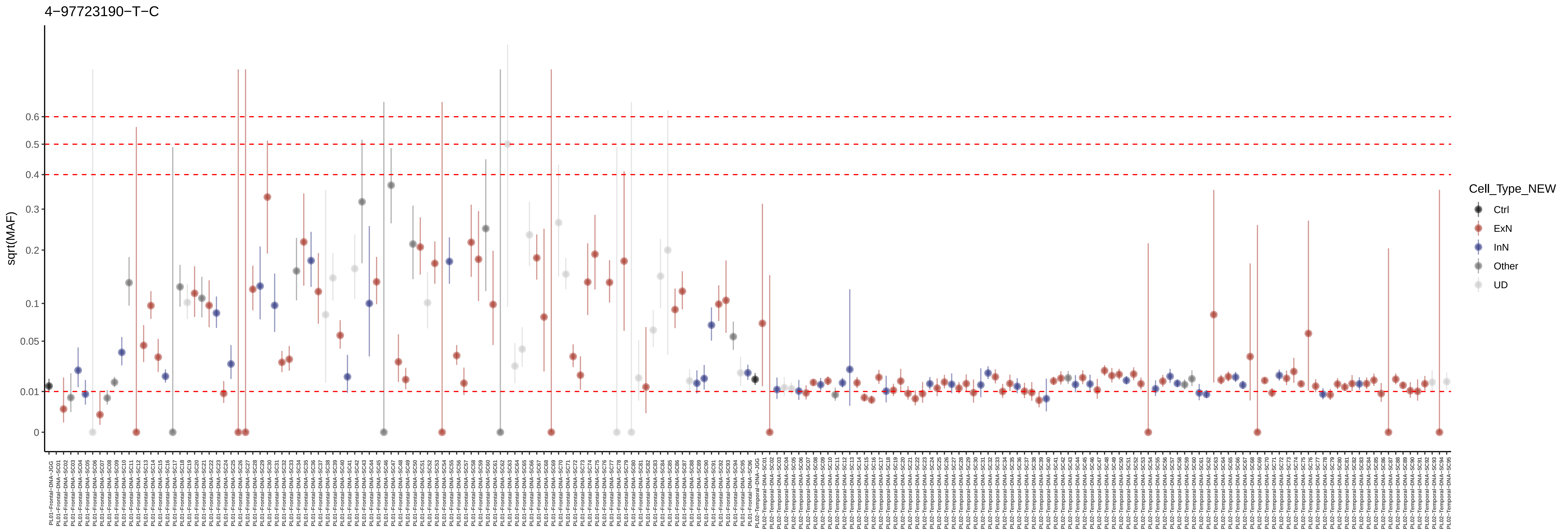
4−97723190−T−C
sqrt(MAF)
0.6
0.5
0.4
0.3
0.2
0.1
0.05
0.01
0
Cell_Type_NEW
Ctrl
ExN
InN
Other
UD

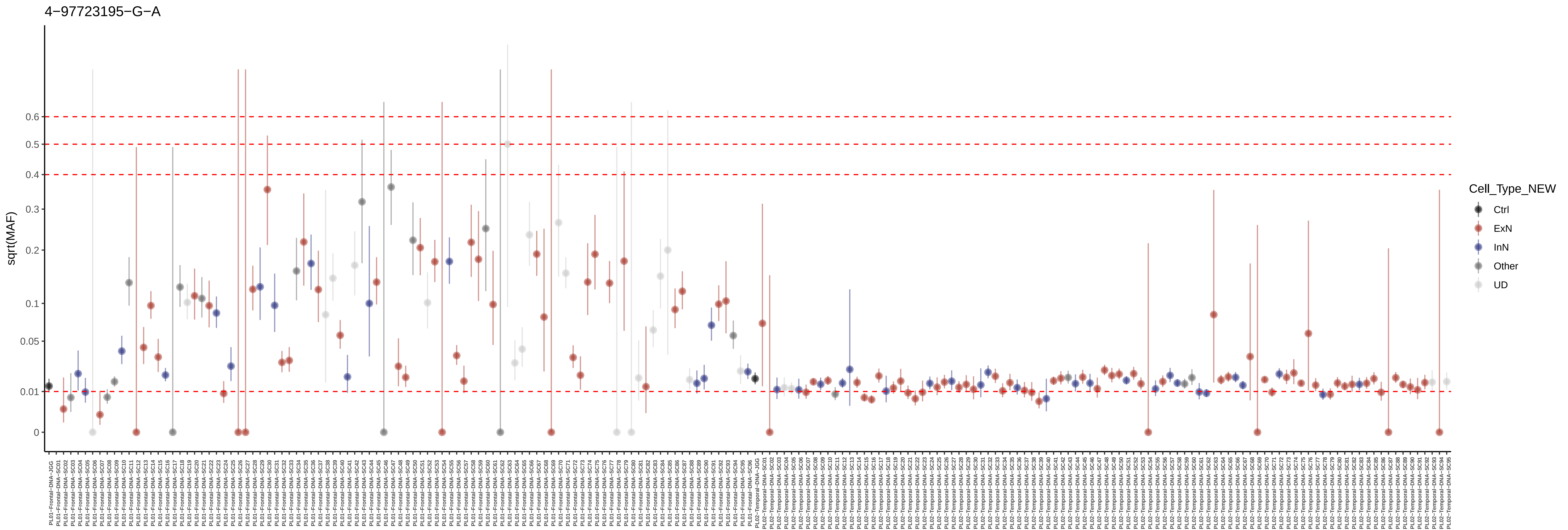
4−97723195−G−A
sqrt(MAF)
0.6
0.5
0.4
0.3
0.2
0.1
0.05
0.01
0
Cell_Type_NEW
Ctrl
ExN
InN
Other
UD

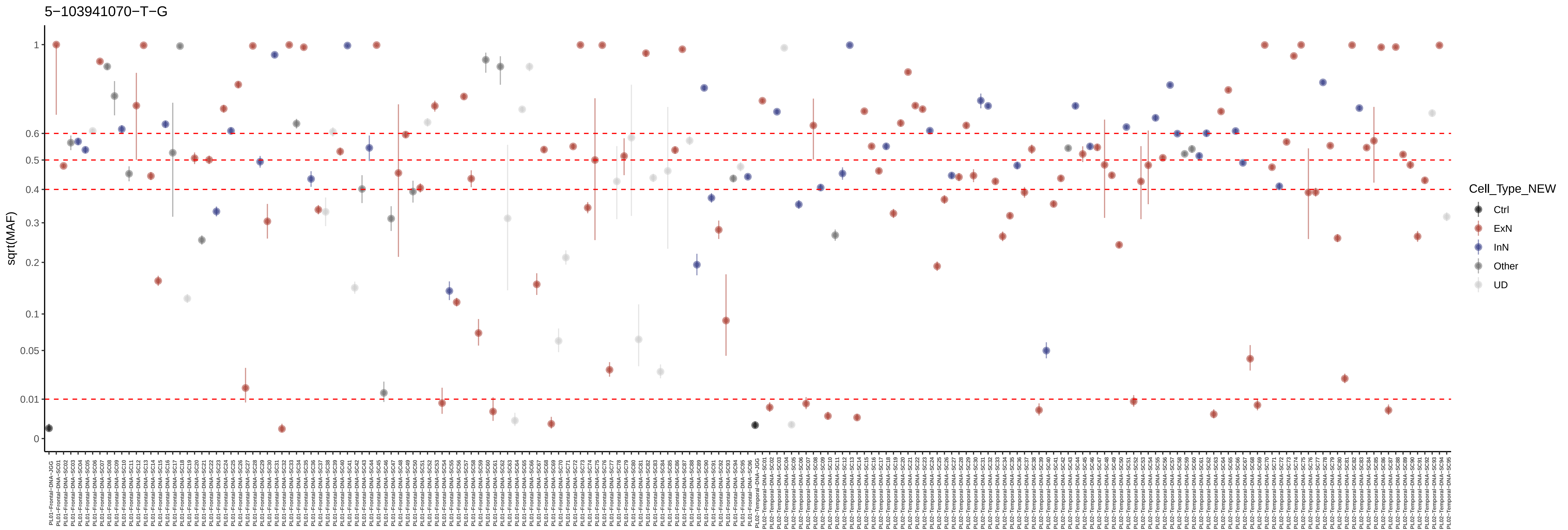
5−103941070−T−G
sqrt(MAF)
1
0.6
0.5
0.4
0.3
0.2
0.1
0.05
0.01
0
Cell_Type_NEW
Ctrl
ExN
InN
Other
UD

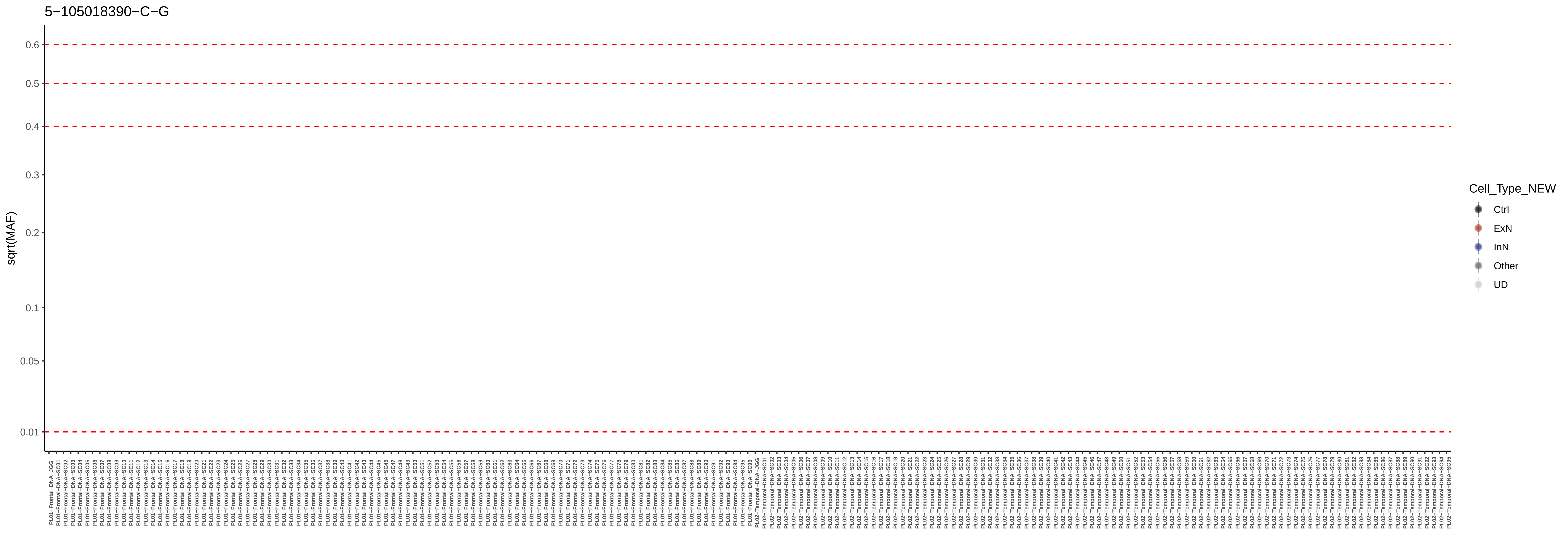
5−105018390−C−G
sqrt(MAF)
0.6
0.5
0.4
0.3
0.2
0.1
0.05
0.01
Cell_Type_NEW
Ctrl
ExN
InN
Other
UD

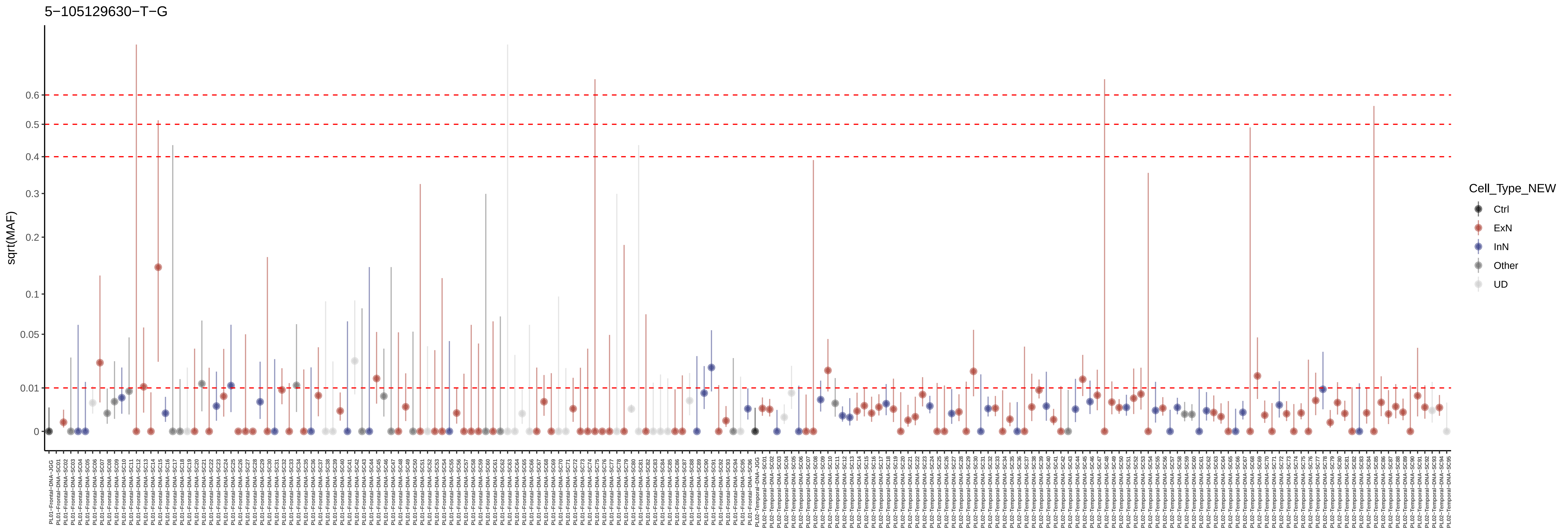
5−105129630−T−G
sqrt(MAF)
0.6
0.5
0.4
0.3
0.2
0.1
0.05
0.01
0
Cell_Type_NEW
Ctrl
ExN
InN
Other
UD

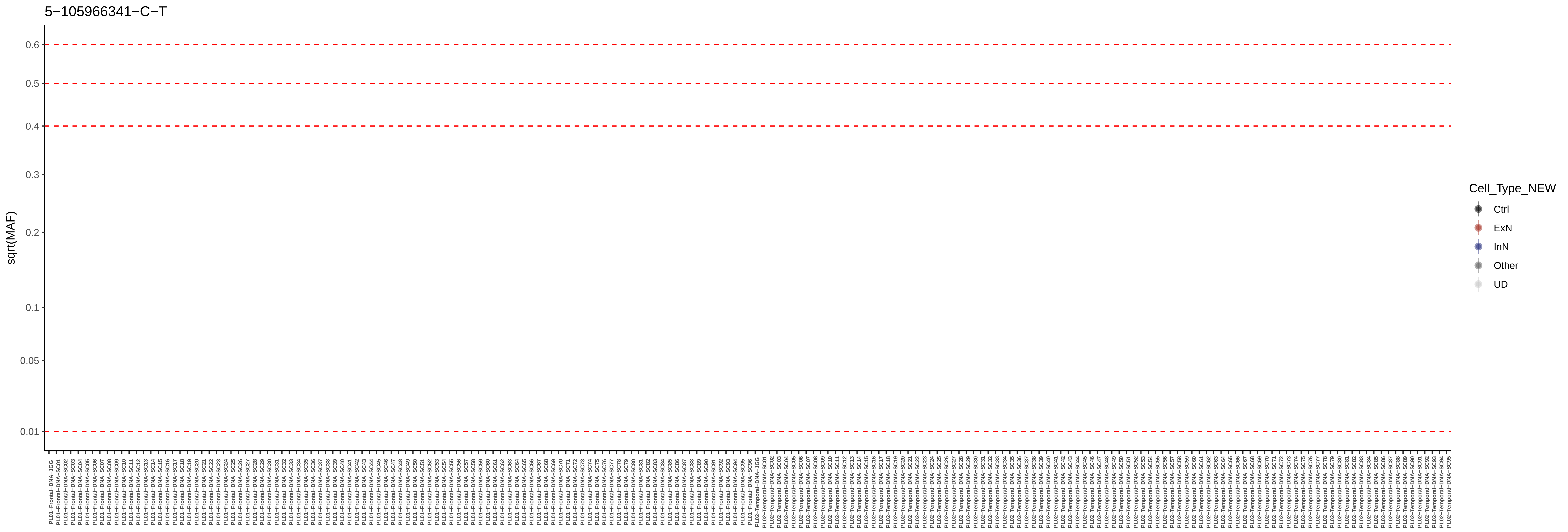
5−105966341−C−T
sqrt(MAF)
0.6
0.5
0.4
0.3
0.2
0.1
0.05
0.01
Cell_Type_NEW
Ctrl
ExN
InN
Other
UD

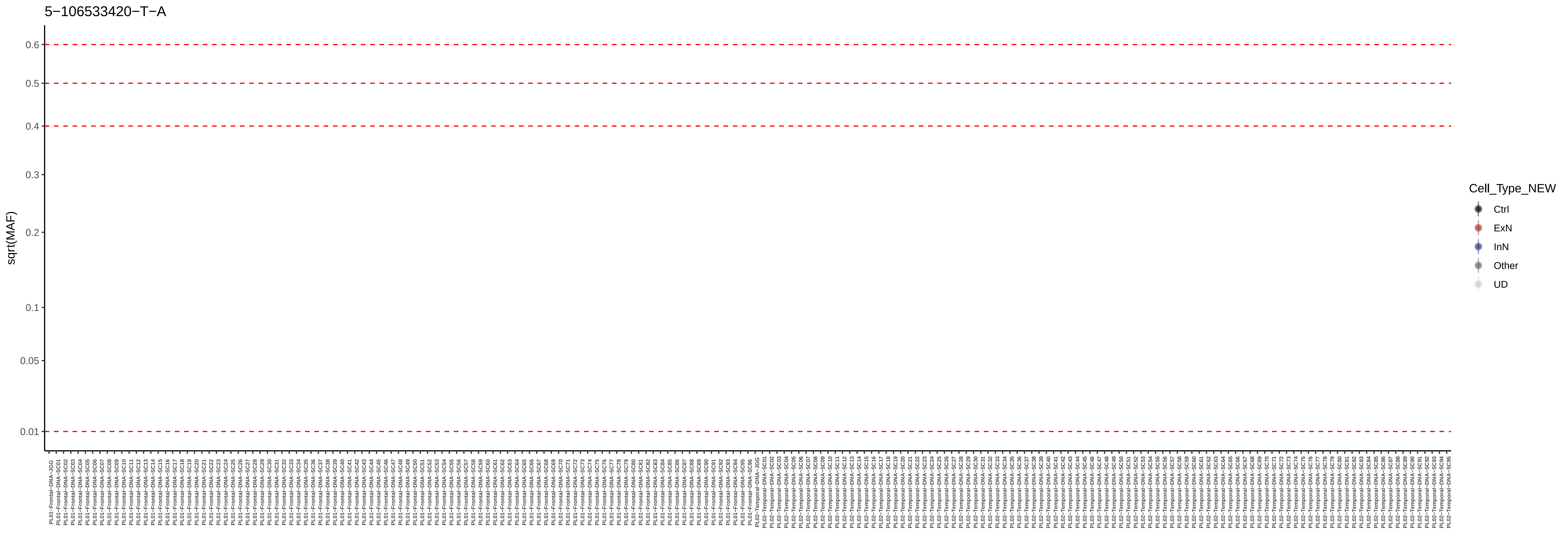

5−106533420−T−A
sqrt(MAF)
0.6
0.5
0.4
0.3
0.2
0.1
0.05
0.01
Cell_Type_NEW
Ctrl
ExN
InN
Other
UD

5−10808399−G−A

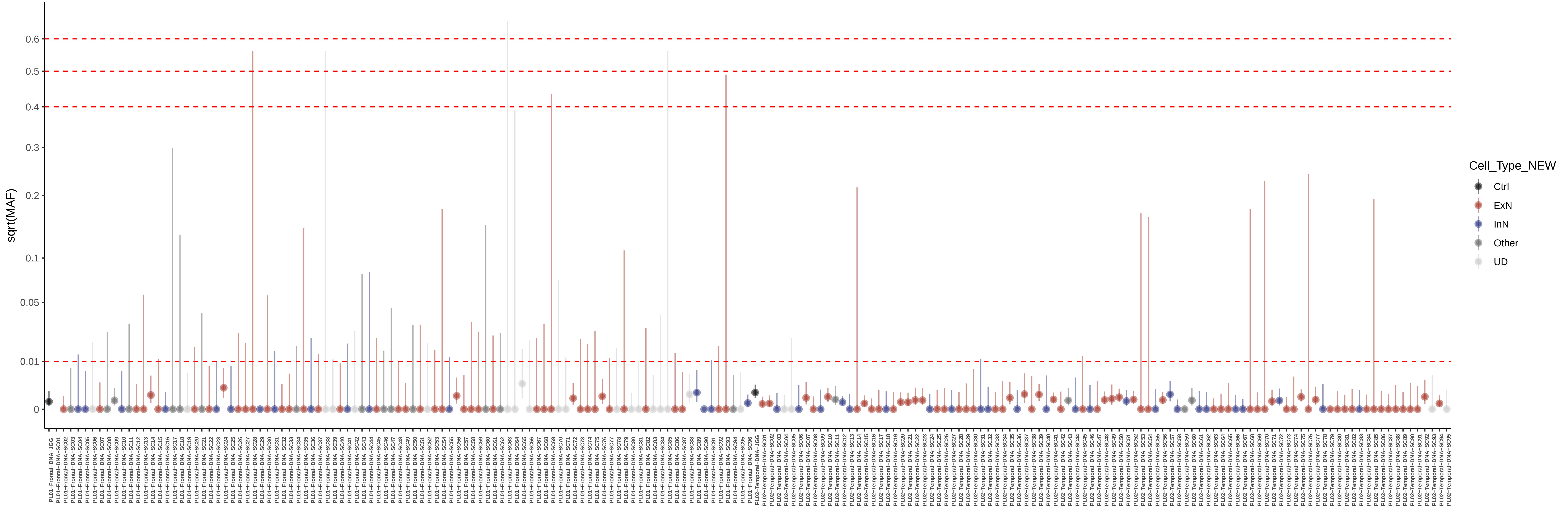

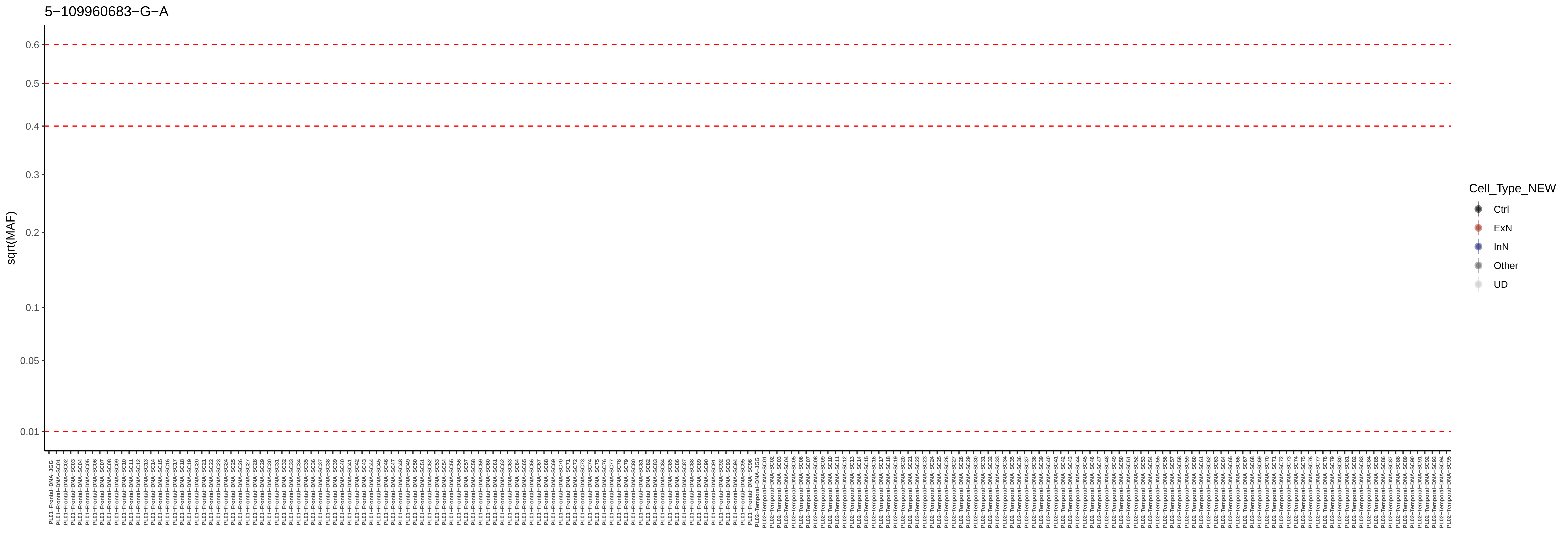

5−109960683−G−A
sqrt(MAF)
0.6
0.5
0.4
0.3
0.2
0.1
0.05
0.01
Cell_Type_NEW
Ctrl
ExN
InN
Other
UD

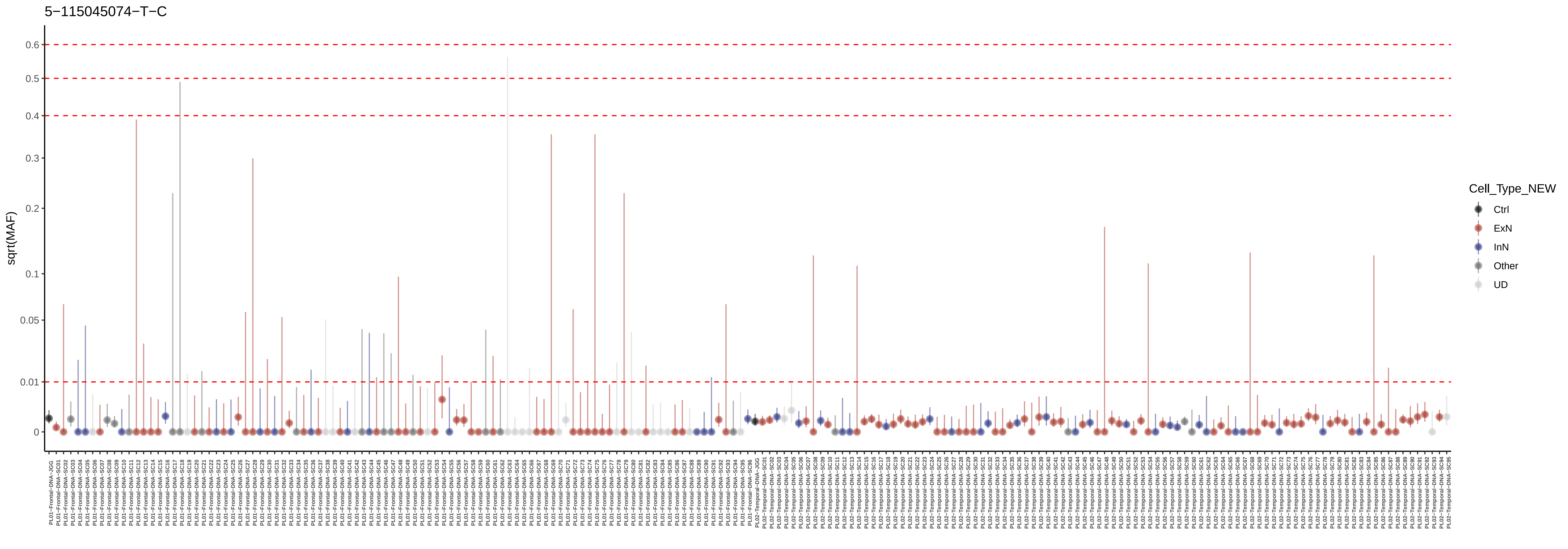

5−115045074−T−C
sqrt(MAF)
0.6
0.5
0.4
0.3
0.2
0.1
0.05
0.01
0
Cell_Type_NEW
Ctrl
ExN
InN
Other
UD

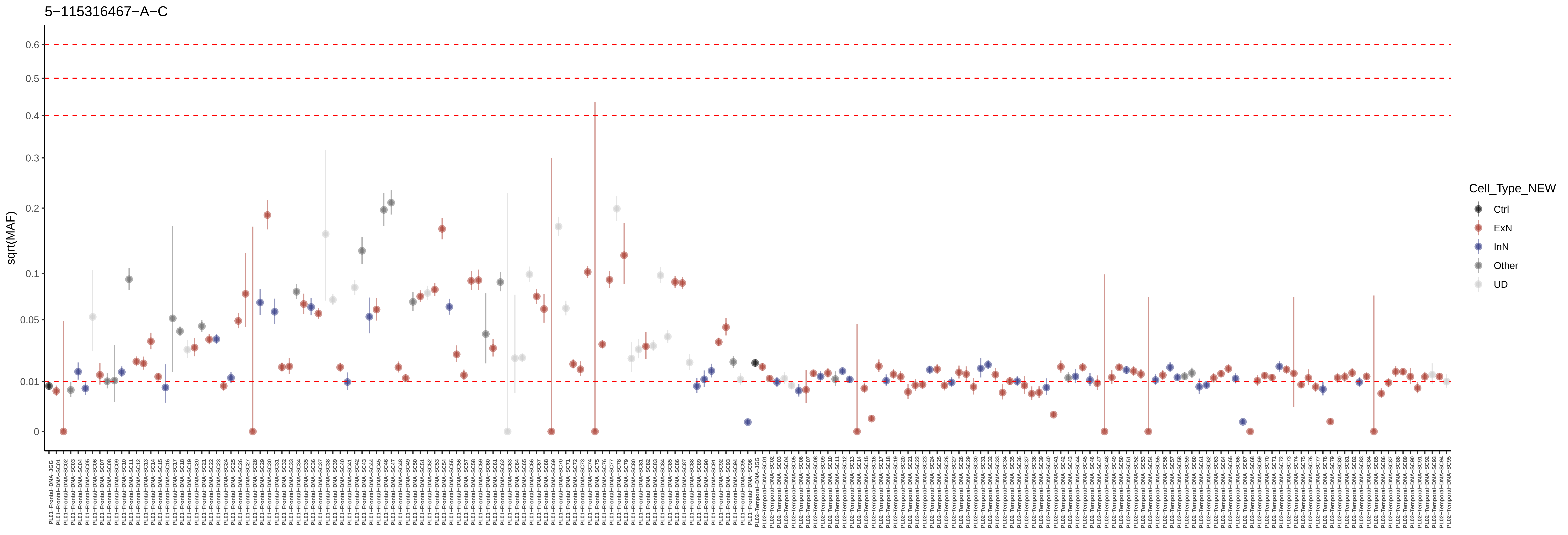
5−115316467−A−C
sqrt(MAF)
0.6
0.5
0.4
0.3
0.2
0.1
0.05
0.01
0
Cell_Type_NEW
Ctrl
ExN
InN
Other
UD

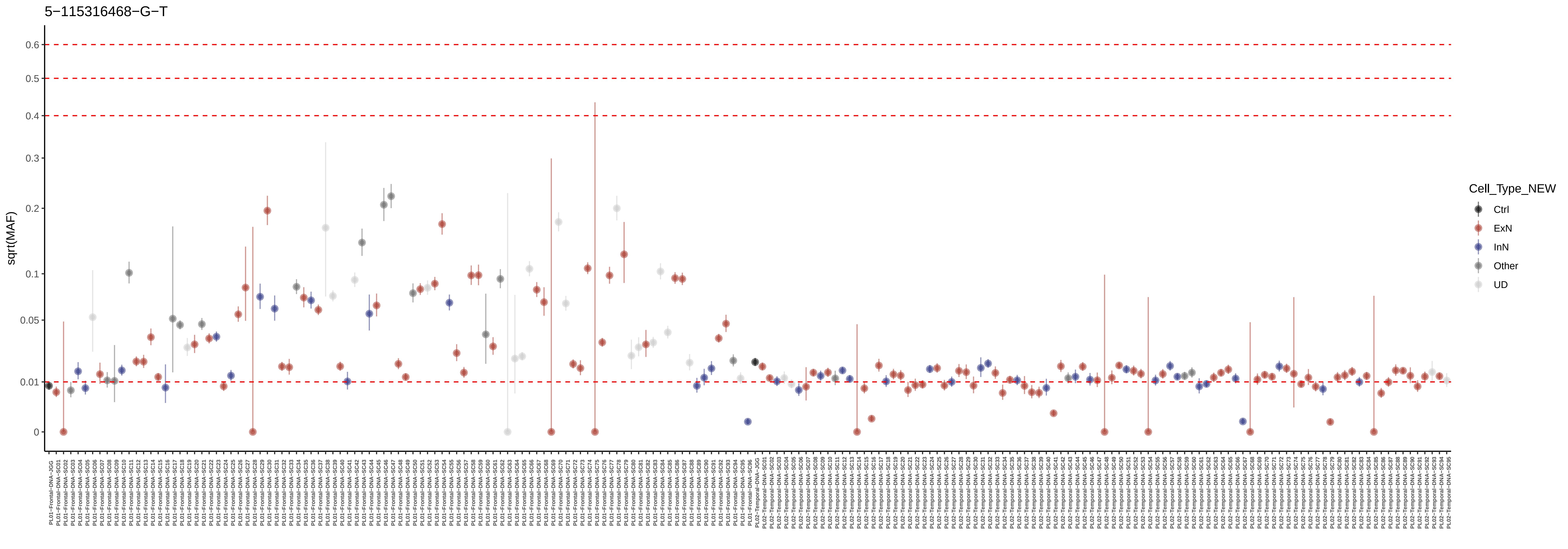
5−115316468−G−T
sqrt(MAF)
Cell_Type_NEW
Ctrl
ExN
InN
Other
UD

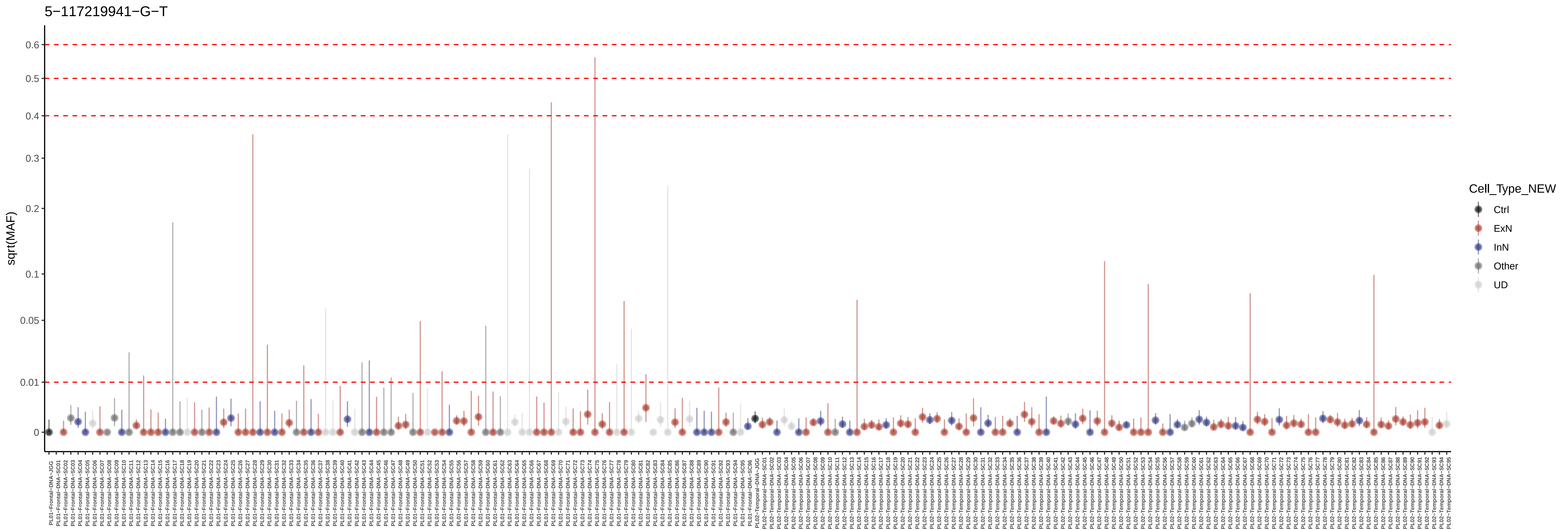
5−117219941−G−T
sqrt(MAF)
0.6
0.5
0.4
0.3
0.2
0.1
0.05
0.01
0
Cell_Type_NEW
Ctrl
ExN
InN
Other
UD

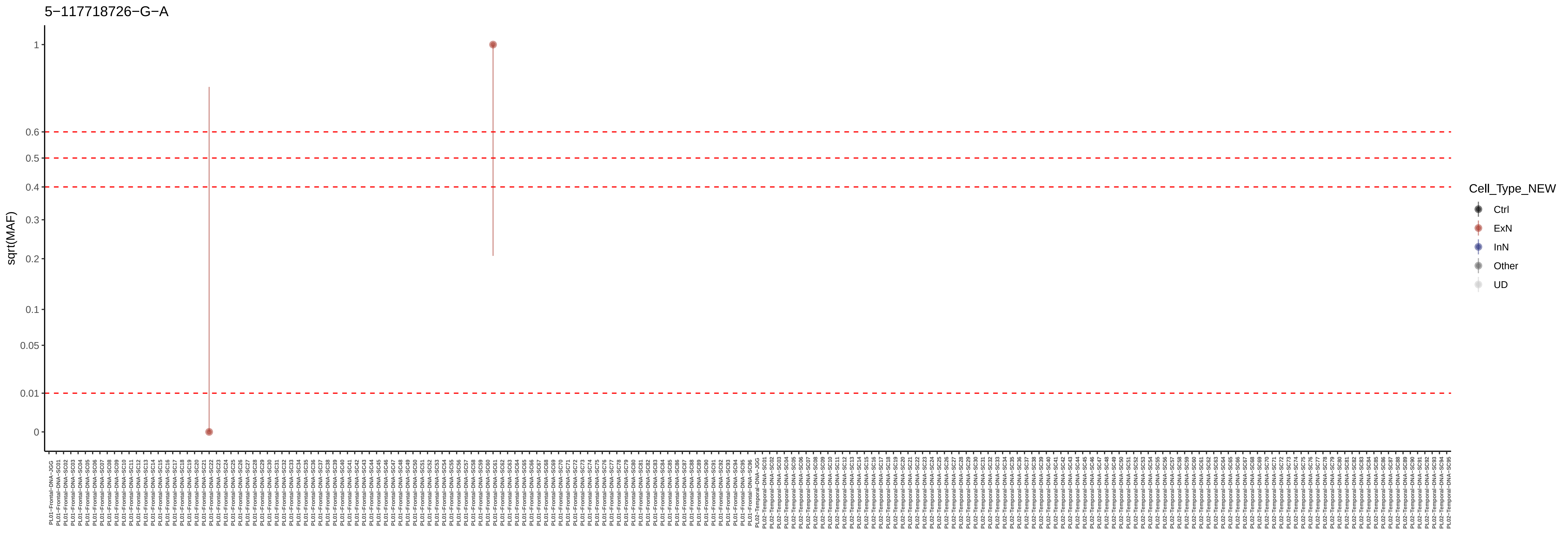
5−117718726−G−A
sqrt(MAF)
1
0.6
0.5
0.4
0.3
0.2
0.1
0.05
0.01
0
Cell_Type_NEW
Ctrl
ExN
InN
Other
UD

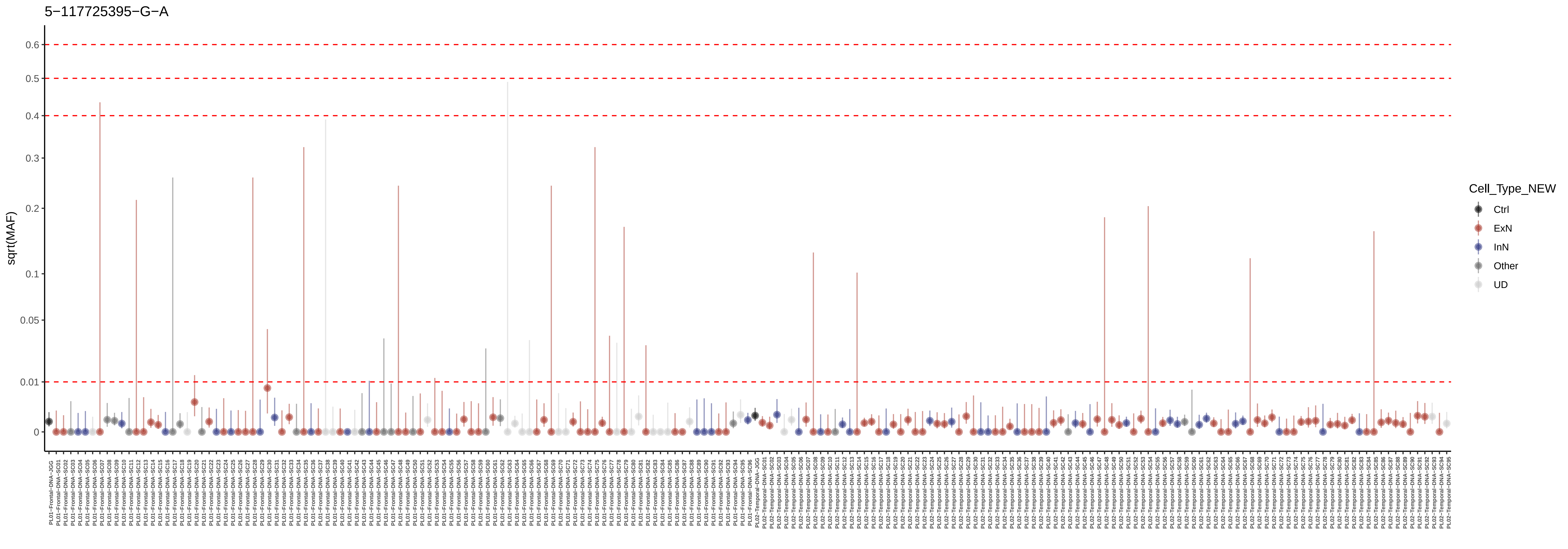

5−117725395−G−A
sqrt(MAF)
0.6
0.5
0.4
0.3
0.2
0.1
0.05
0.01
0
Cell_Type_NEW
Ctrl
ExN
InN
Other
UD

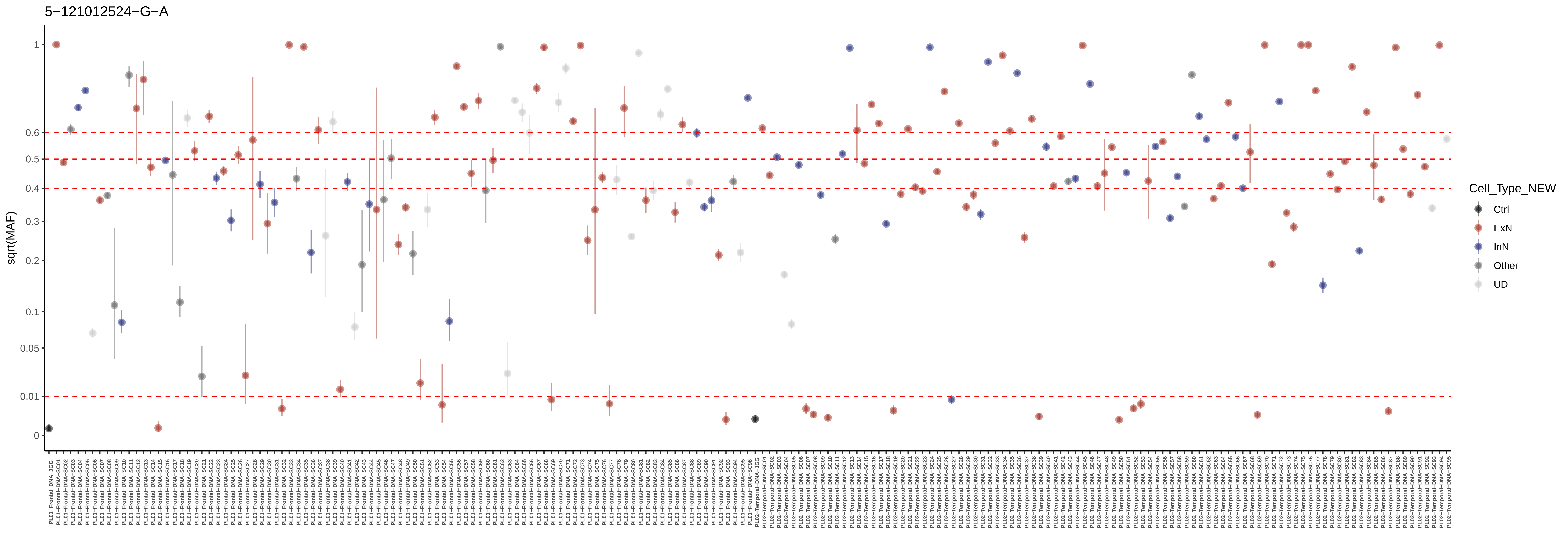
5−121012524−G−A
sqrt(MAF)
1
0.6
0.5
0.4
0.3
0.2
0.1
0.05
0.01
0
Cell_Type_NEW
Ctrl
ExN
InN
Other
UD

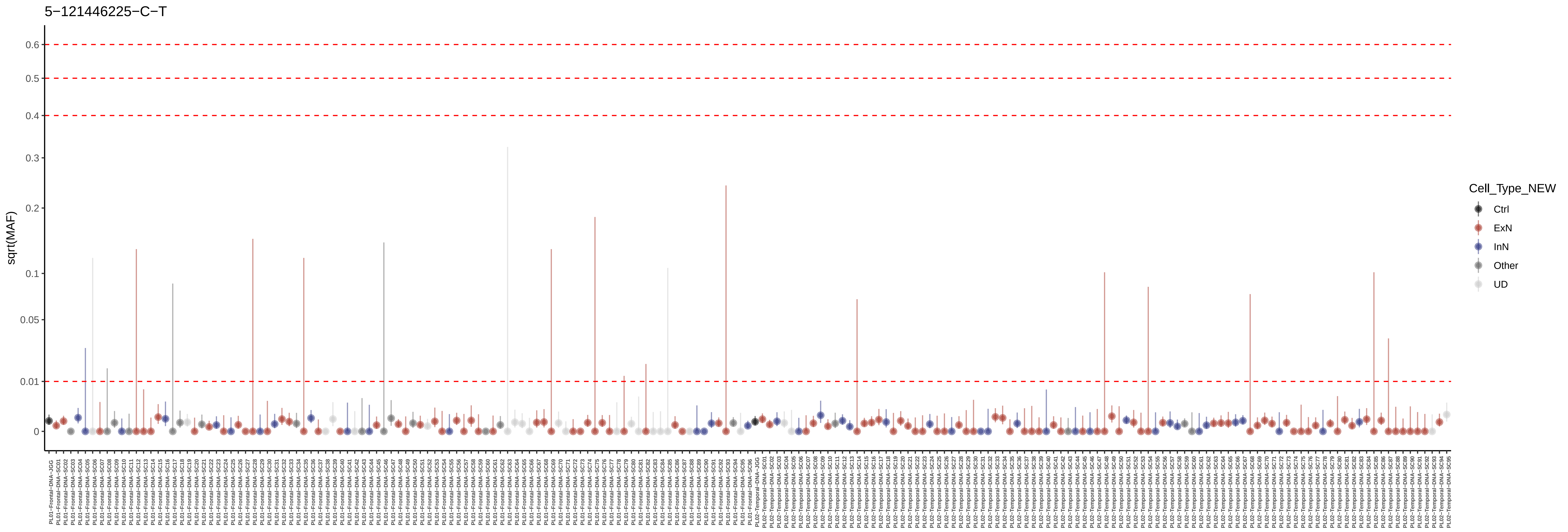
5−121446225−C−T
sqrt(MAF)
0.6
0.5
0.4
0.3
0.2
0.1
0.05
0.01
0
Cell_Type_NEW
Ctrl
ExN
InN
Other
UD

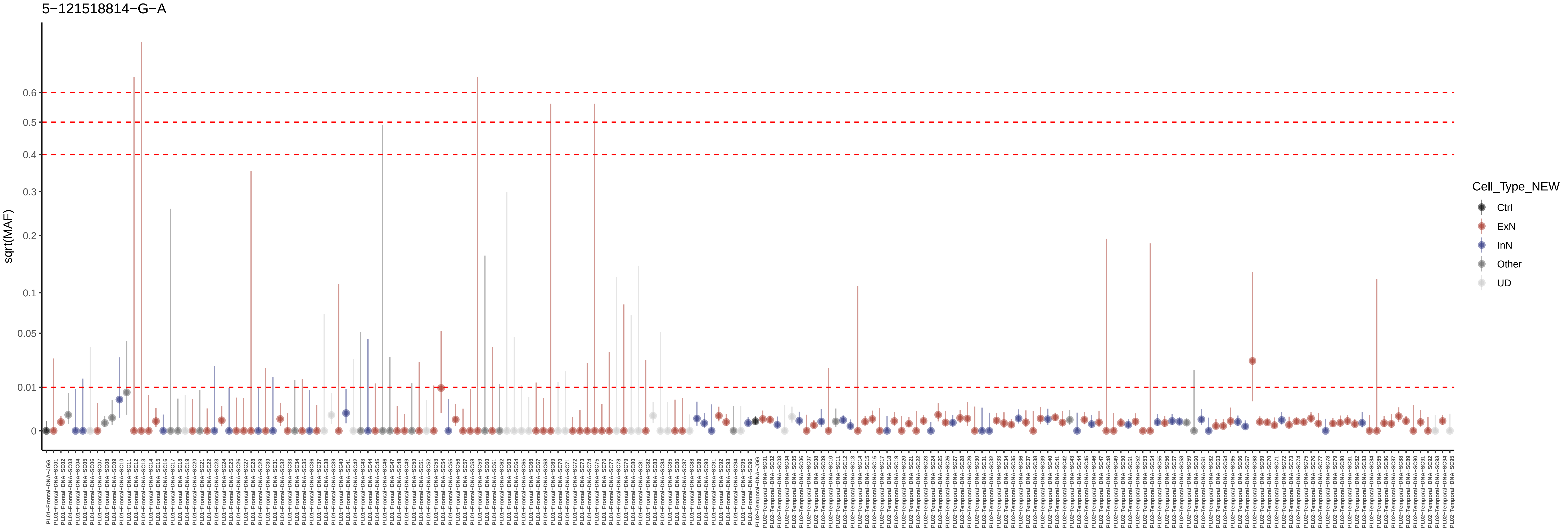
5−121518814−G−A
sqrt(MAF)
0.6
0.5
0.4
0.3
0.2
0.1
0.05
0.01
0
Cell_Type_NEW
Ctrl
ExN
InN
Other
UD

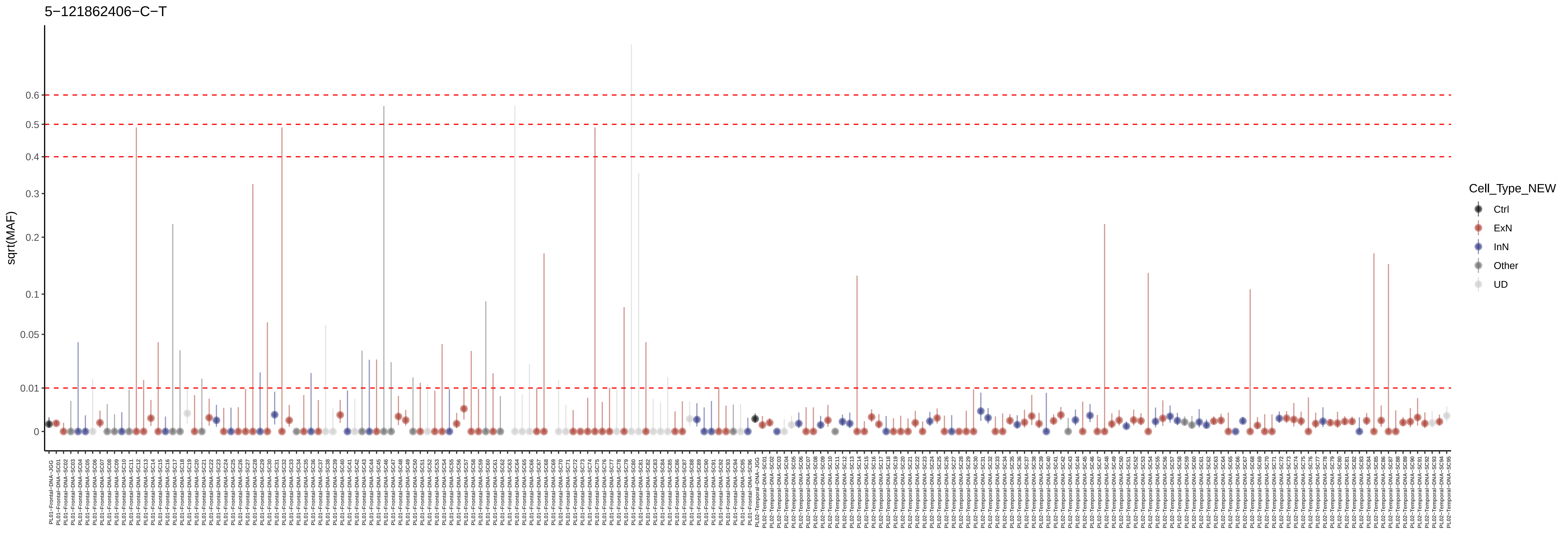

5−121862406−C−T
sqrt(MAF)
0.6
0.5
0.4
0.3
0.2
0.1
0.05
0.01
0
Cell_Type_NEW
Ctrl
ExN
InN
Other
UD

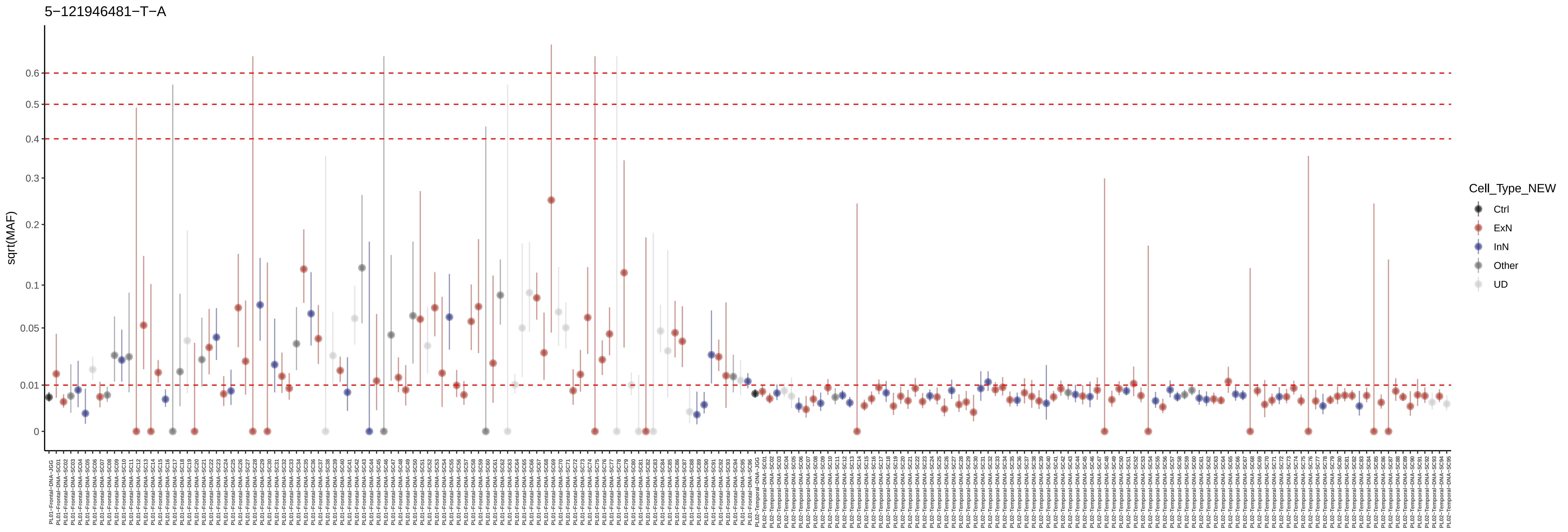
5−121946481−T−A
sqrt(MAF)
0.6
0.5
0.4
0.3
0.2
0.1
0.05
0.01
0
Cell_Type_NEW
Ctrl
ExN
InN
Other
UD

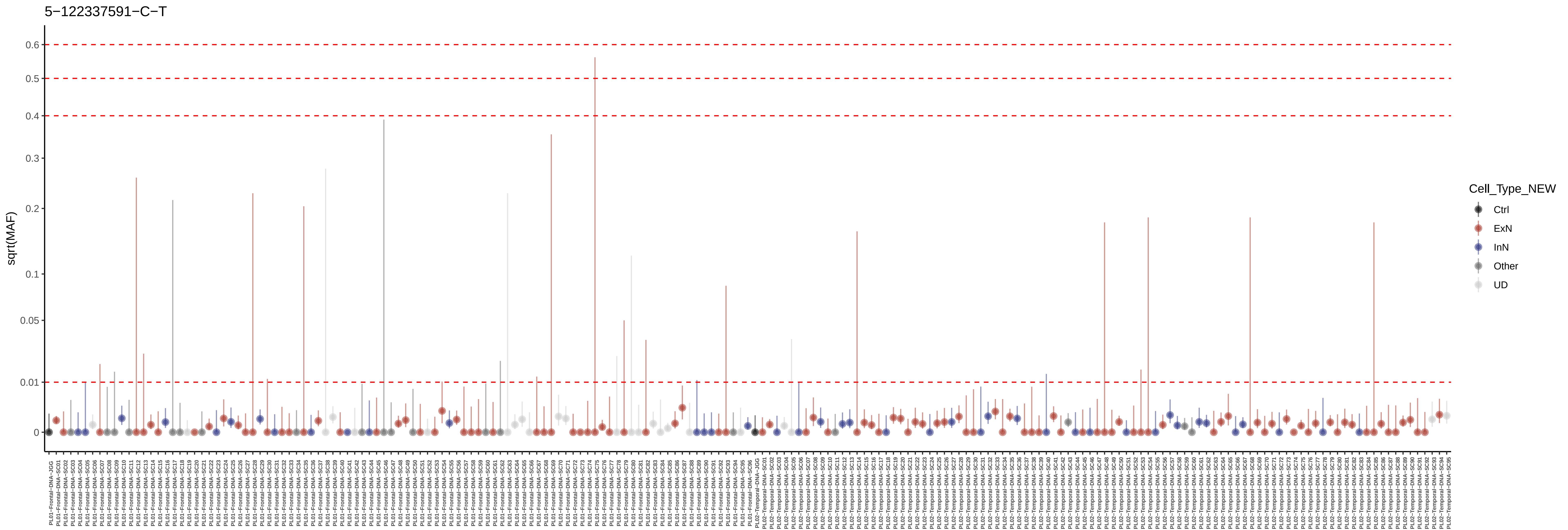

5−122337591−C−T
sqrt(MAF)
0.6
0.5
0.4
0.3
0.2
0.1
0.05
0.01
0
Cell_Type_NEW
Ctrl
ExN
InN
Other
UD

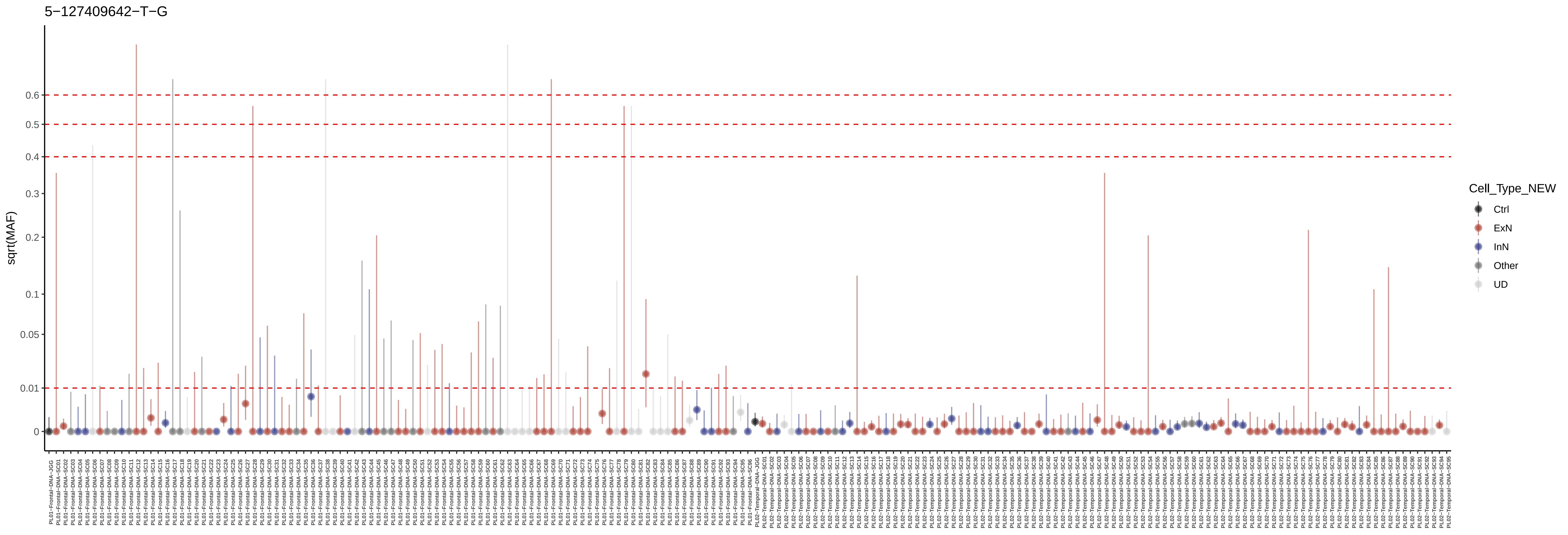
5−127409642−T−G
sqrt(MAF)
0.6
0.5
0.4
0.3
0.2
0.1
0.05
0.01
0
Cell_Type_NEW
Ctrl
ExN
InN
Other
UD

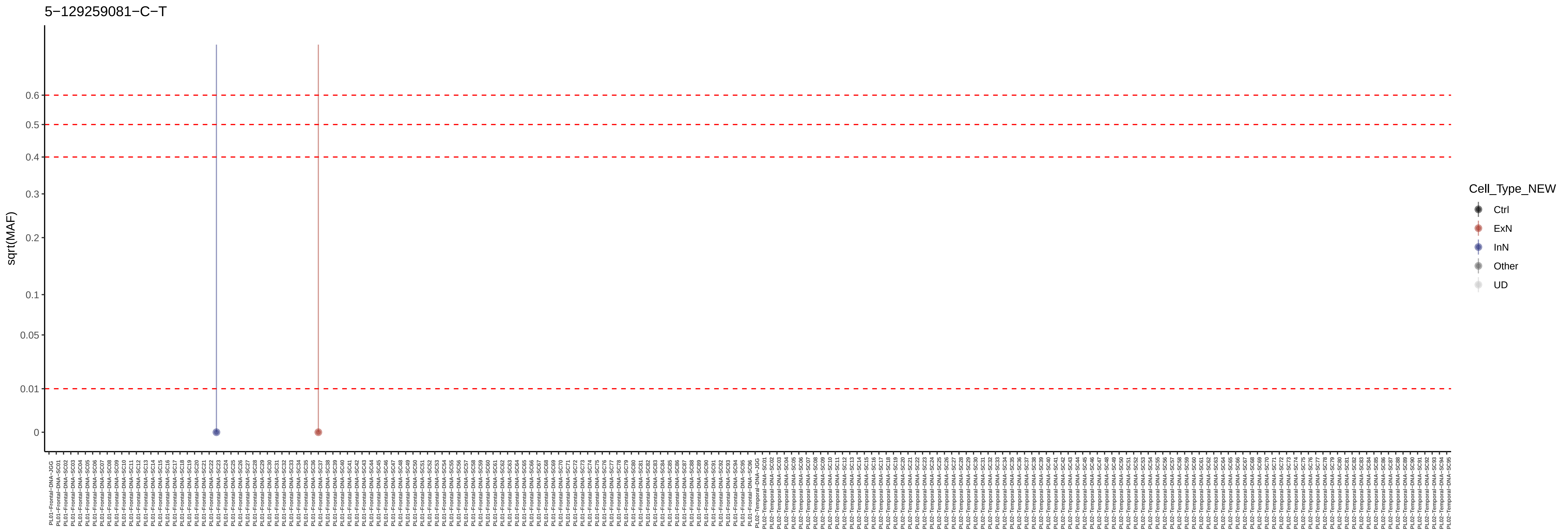

5−129259081−C−T
sqrt(MAF)
0.6
0.5
0.4
0.3
0.2
0.1
0.05
0.01
0
Cell_Type_NEW
Ctrl
ExN
InN
Other
UD

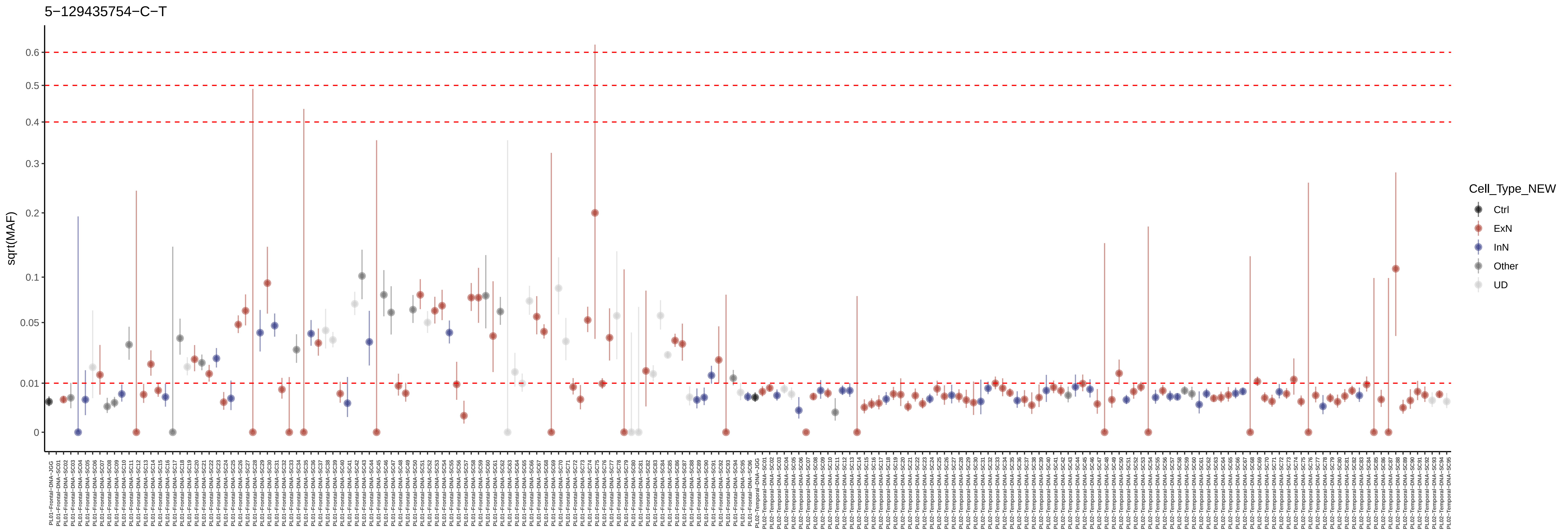

5−129435754−C−T
sqrt(MAF)
0.6
0.5
0.4
0.3
0.2
0.1
0.05
0.01
0
Cell_Type_NEW
Ctrl
ExN
InN
Other
UD

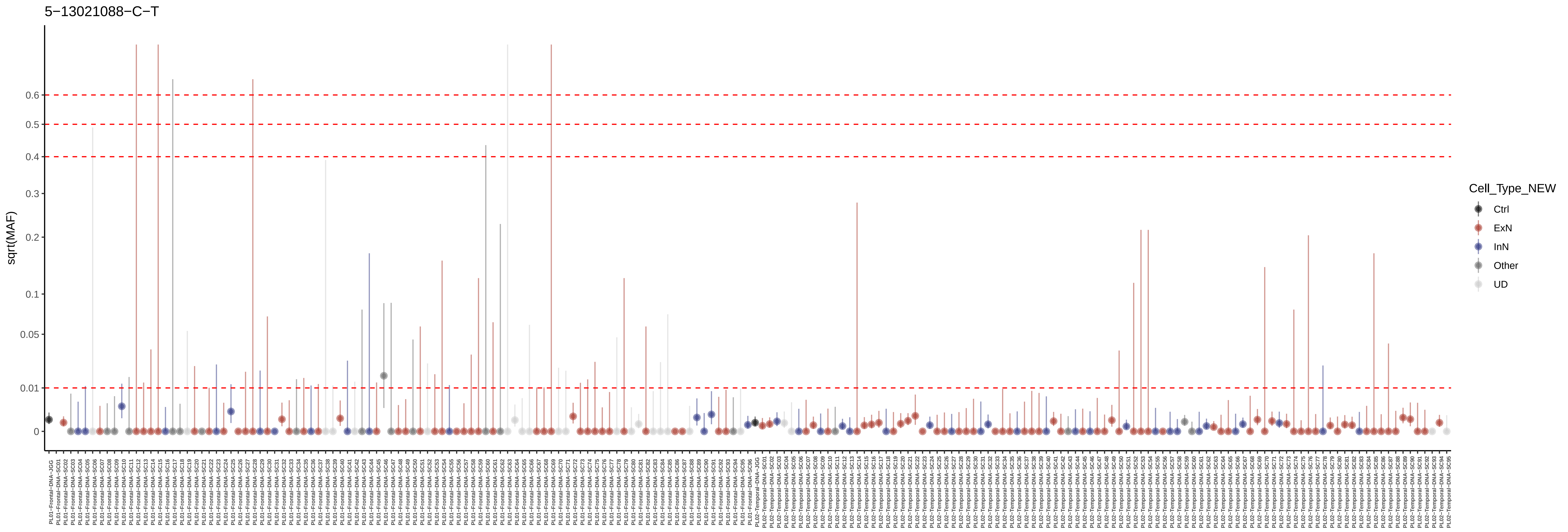
5−13021088−C−T
sqrt(MAF)
0.6
0.5
0.4
0.3
0.2
0.1
0.05
0.01
0
Cell_Type_NEW
Ctrl
ExN
InN
Other
UD

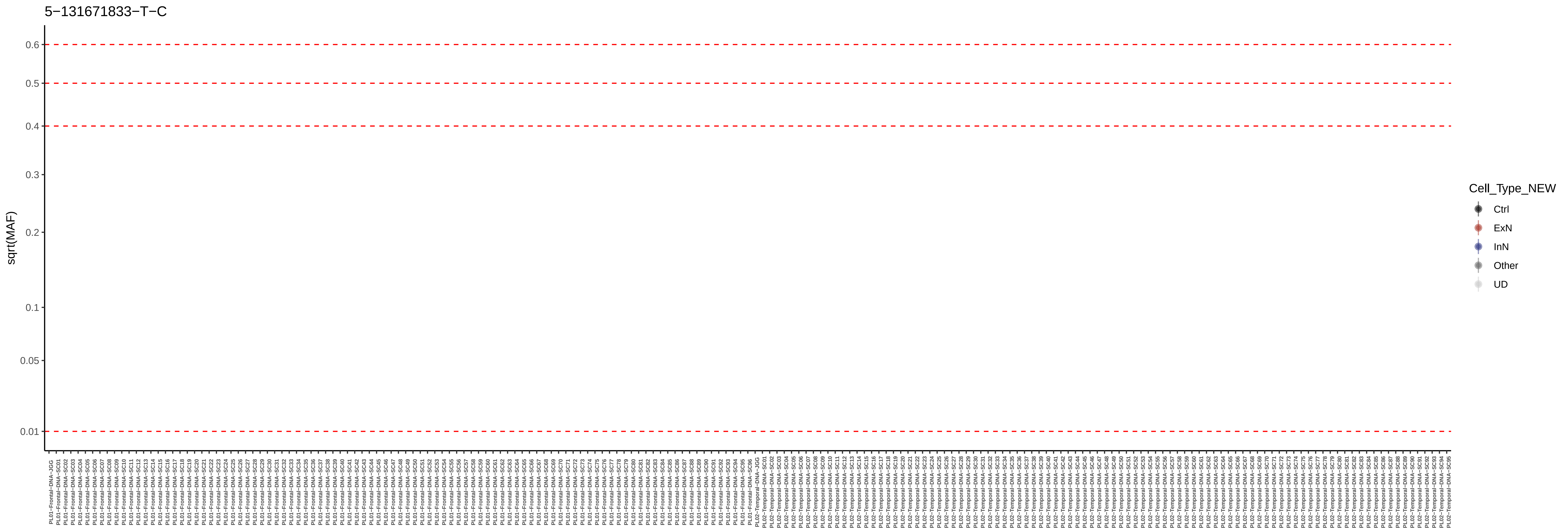
5−131671833−T−C
sqrt(MAF)
0.6
0.5
0.4
0.3
0.2
0.1
0.05
0.01
Cell_Type_NEW
Ctrl
ExN
InN
Other
UD

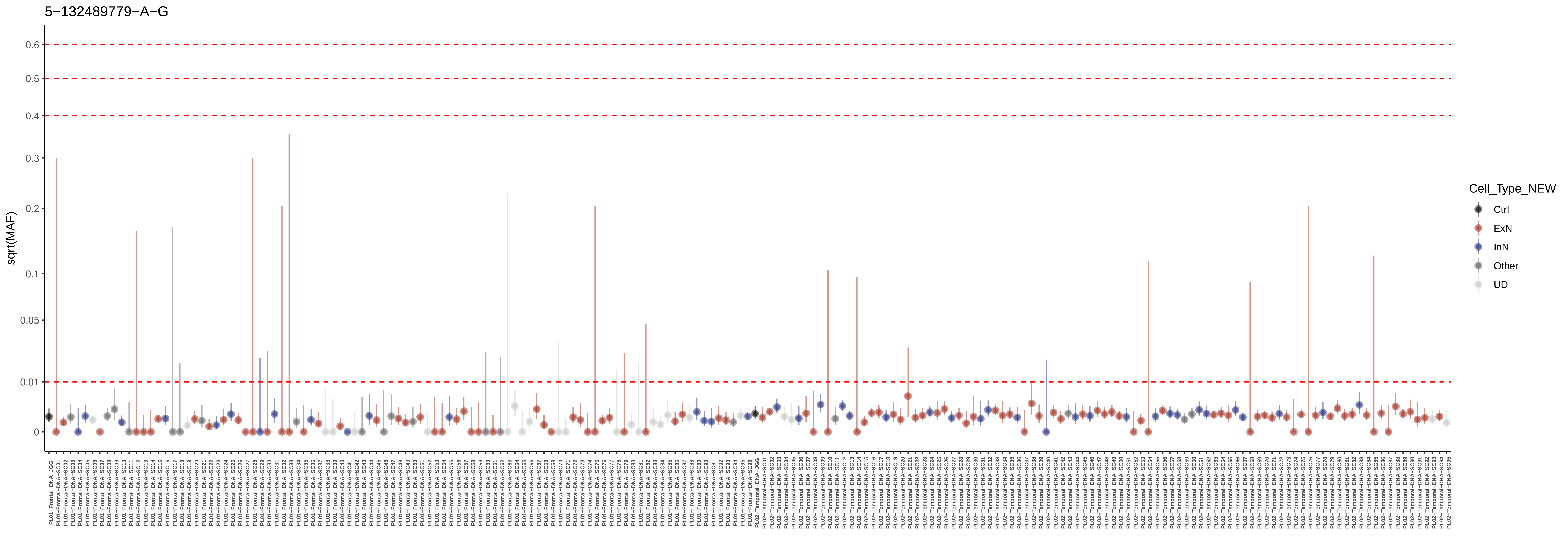

5−132489779−A−G
sqrt(MAF)
0.6
0.5
0.4
0.3
0.2
0.1
0.05
0.01
0
Cell_Type_NEW
Ctrl
ExN
InN
Other
UD

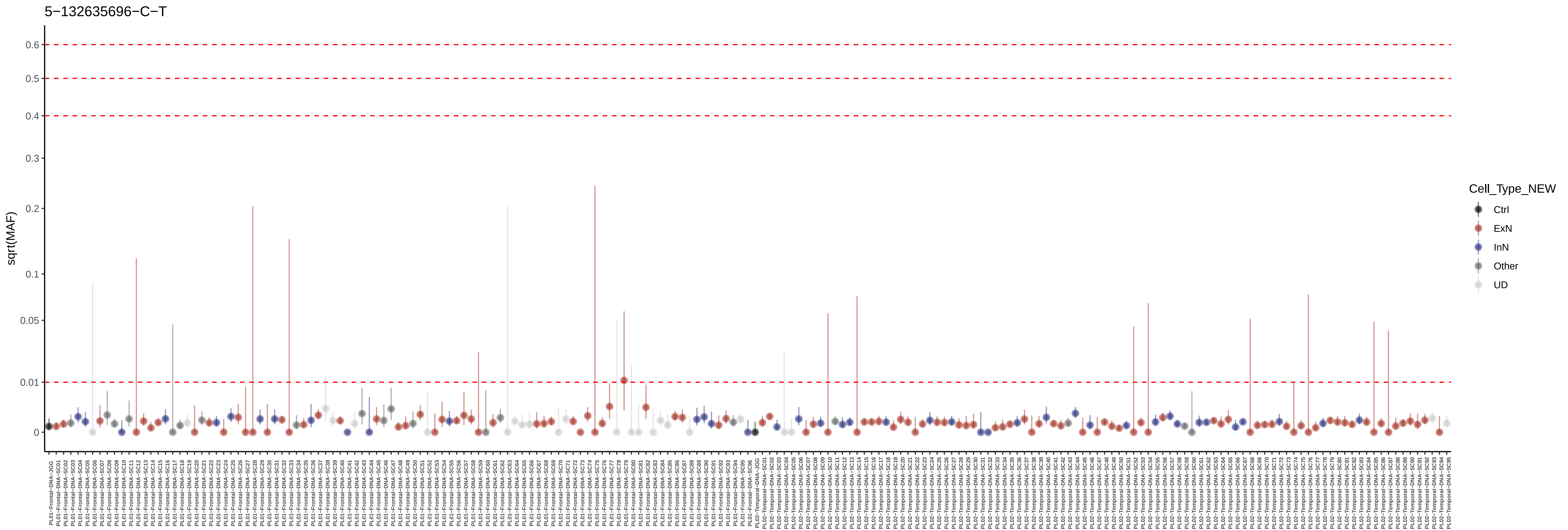
5−132635696−C−T
sqrt(MAF)
0.6
0.5
0.4
0.3
0.2
0.1
0.05
0.01
0
Cell_Type_NEW
Ctrl
ExN
InN
Other
UD

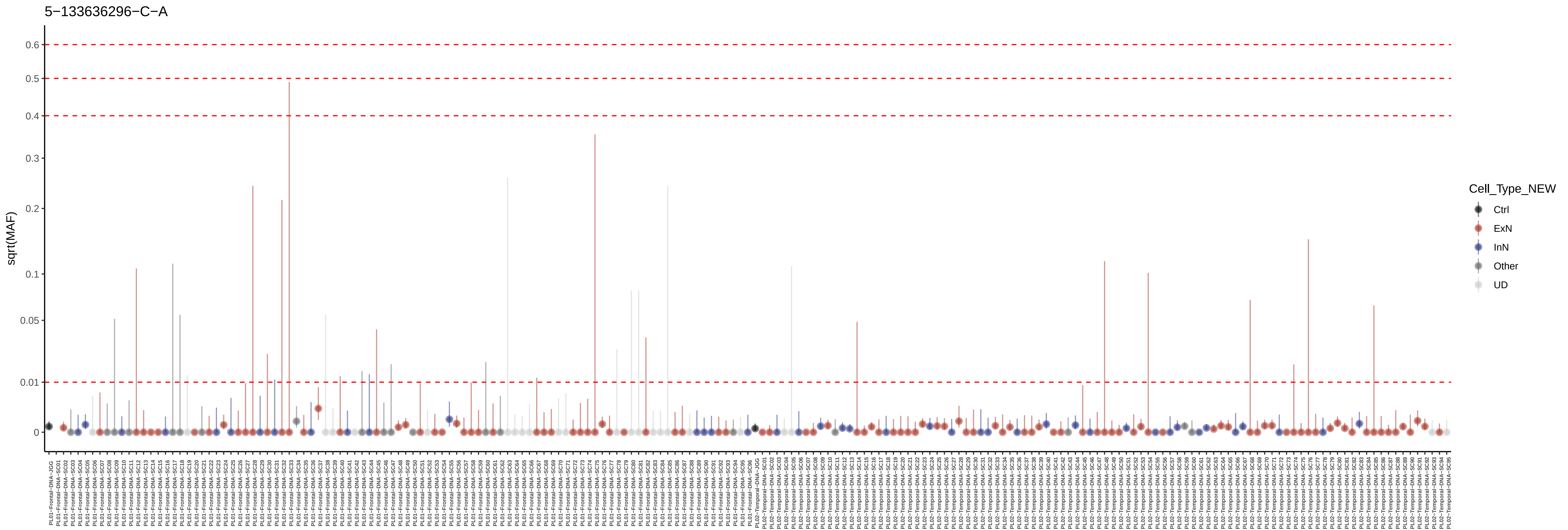
5−133636296−C−A
sqrt(MAF)
0.6
0.5
0.4
0.3
0.2
0.1
0.05
0.01
0
Cell_Type_NEW
Ctrl
ExN
InN
Other
UD

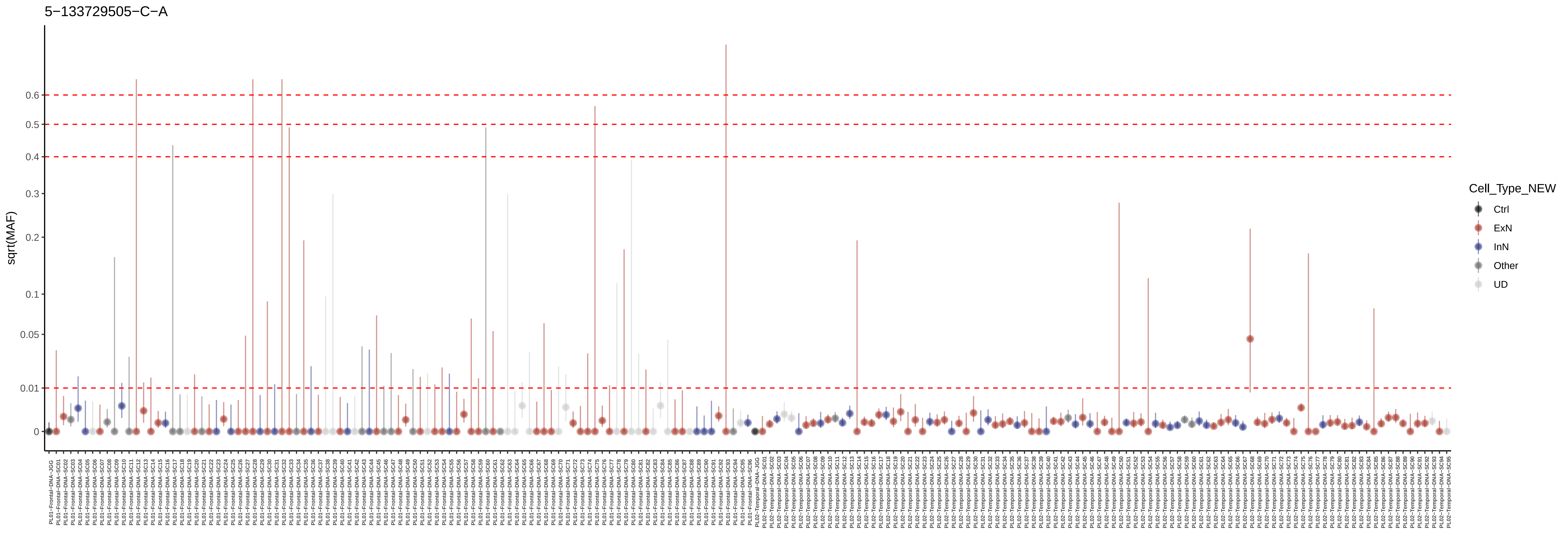
5−133729505−C−A
sqrt(MAF)
0
0.01
0.05
0.1
0.2
0.3
0.4
0.5
0.6
Cell_Type_NEW
Ctrl
ExN
InN
Other
UD

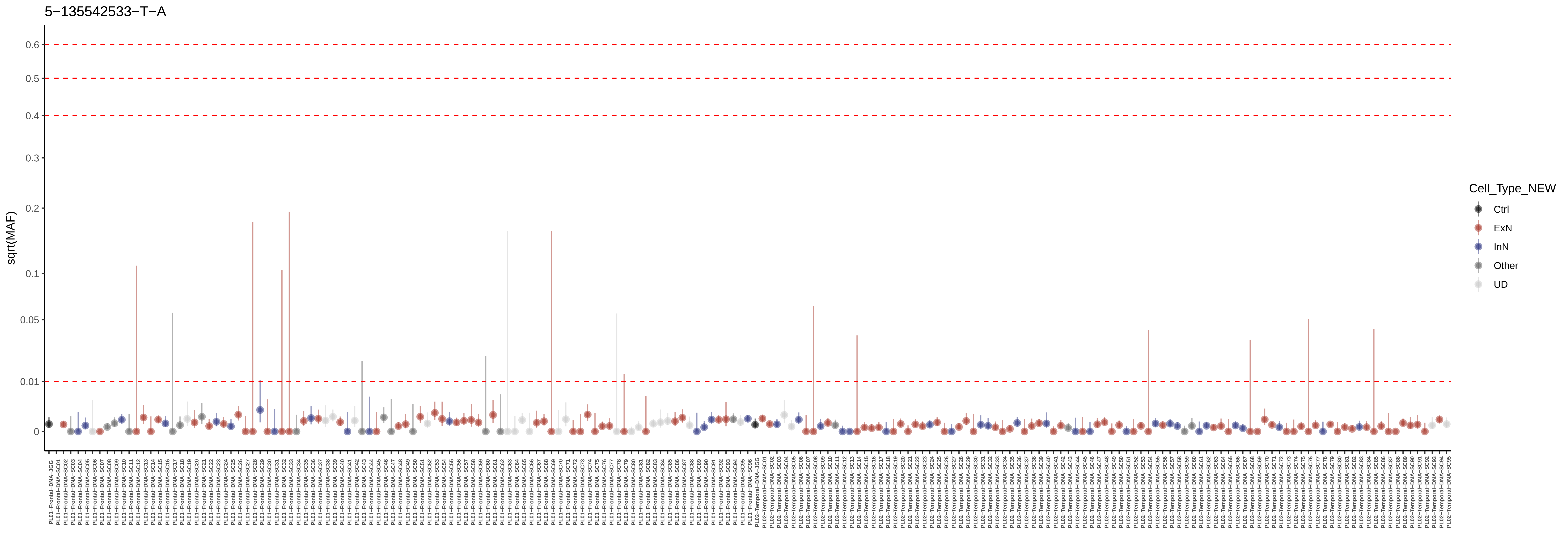
5−135542533−T−A
sqrt(MAF)
0.6
0.5
0.4
0.3
0.2
0.1
0.05
0.01
0
Cell_Type_NEW
Ctrl
ExN
InN
Other
UD

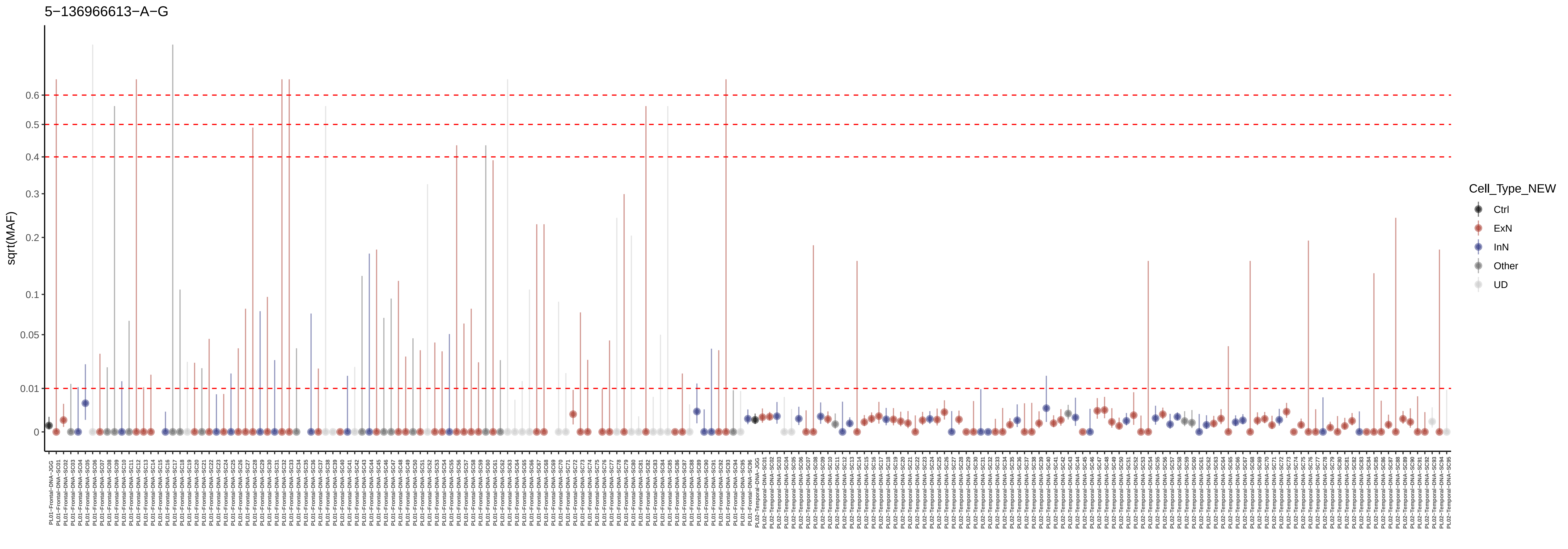
5−136966613−A−G
sqrt(MAF)
0.6
0.5
0.4
0.3
0.2
0.1
0.05
0.01
0
Cell_Type_NEW
Ctrl
ExN
InN
Other
UD

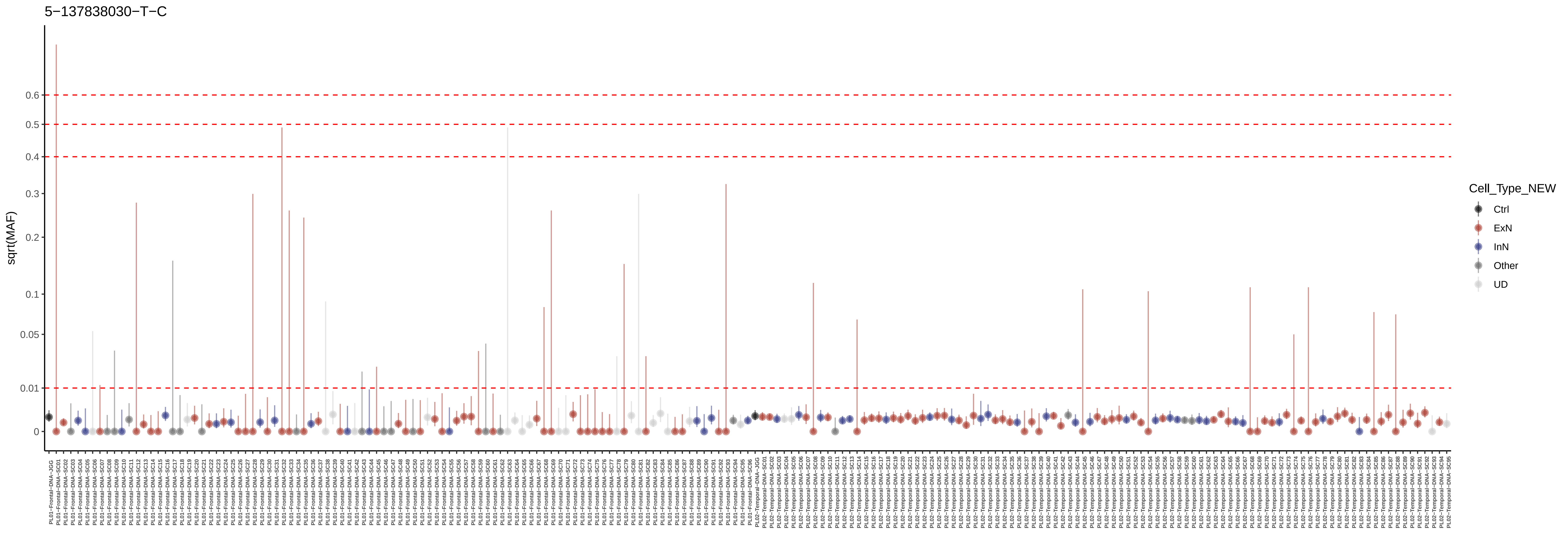

5−137838030−T−C
sqrt(MAF)
0.6
0.5
0.4
0.3
0.2
0.1
0.05
0.01
0
Cell_Type_NEW
Ctrl
ExN
InN
Other
UD

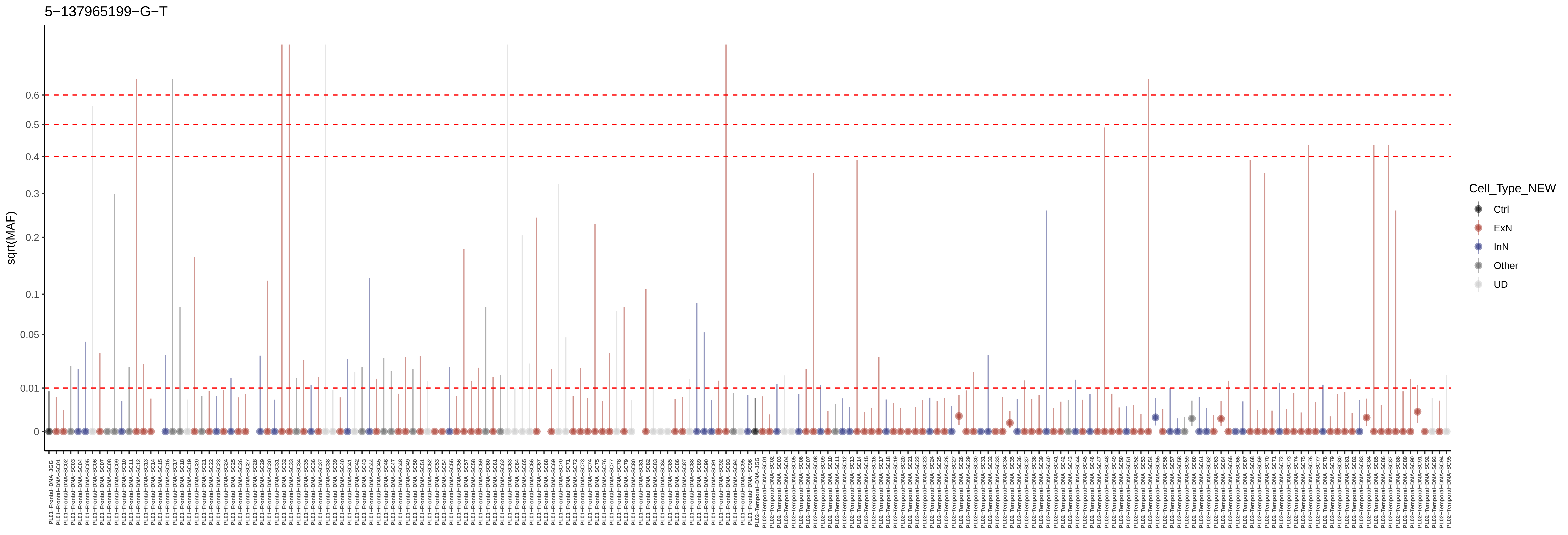
5−137965199−G−T
sqrt(MAF)
0.6
0.5
0.4
0.3
0.2
0.1
0.05
0.01
0
Cell_Type_NEW
Ctrl
ExN
InN
Other
UD

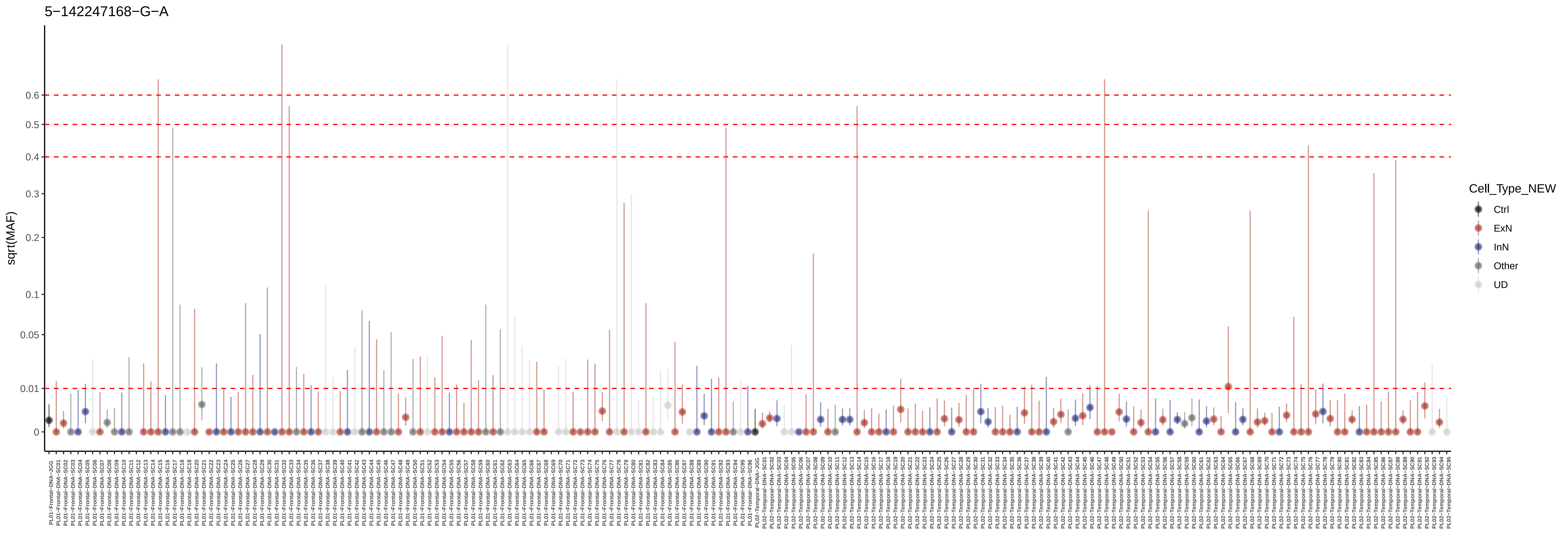

5−142247168−G−A
sqrt(MAF)
0.6
0.5
0.4
0.3
0.2
0.1
0.05
0.01
0
Cell_Type_NEW
Ctrl
ExN
InN
Other
UD

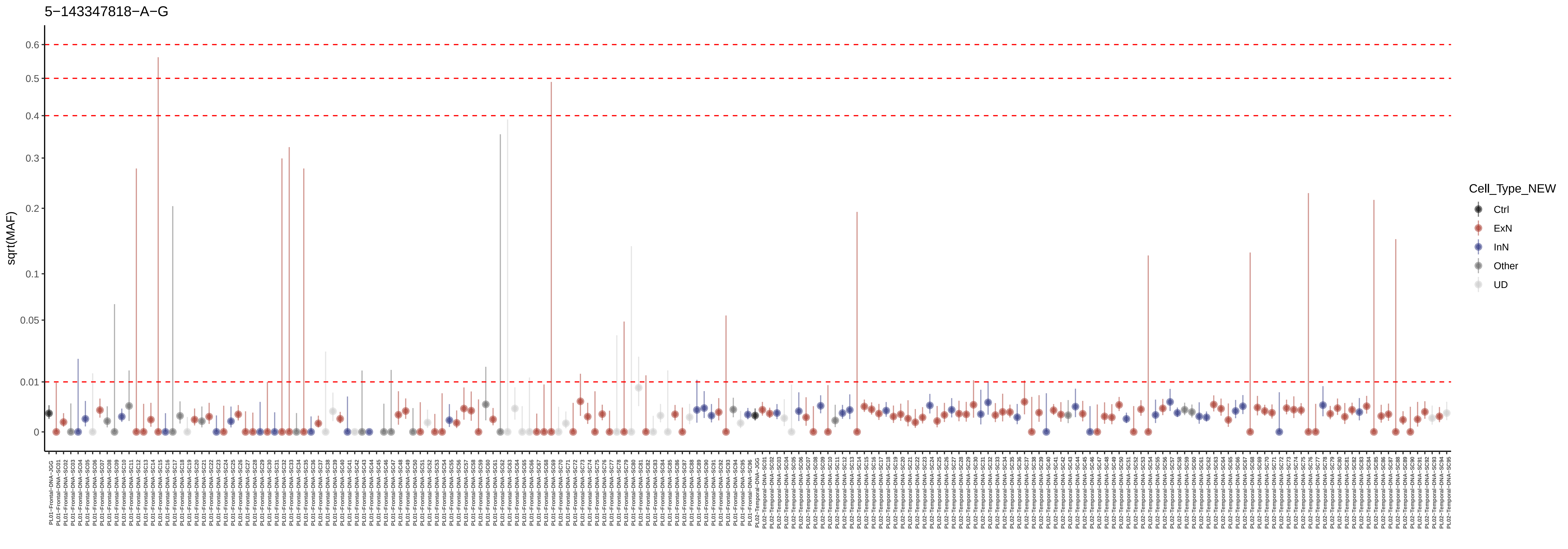
5−143347818−A−G
sqrt(MAF)
0.6
0.5
0.4
0.3
0.2
0.1
0.05
0.01
0
Cell_Type_NEW
Ctrl
ExN
InN
Other
UD

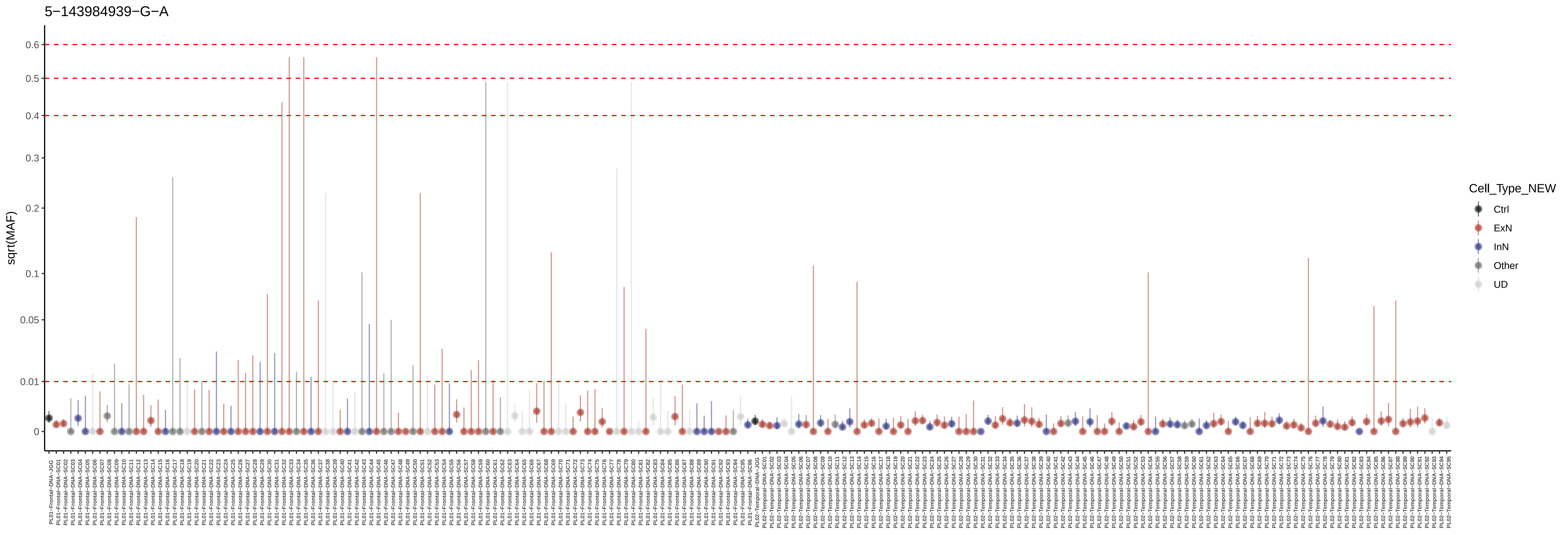
5−143984939−G−A
sqrt(MAF)
0.6
0.5
0.4
0.3
0.2
0.1
0.05
0.01
0
Cell_Type_NEW
Ctrl
ExN
InN
Other
UD

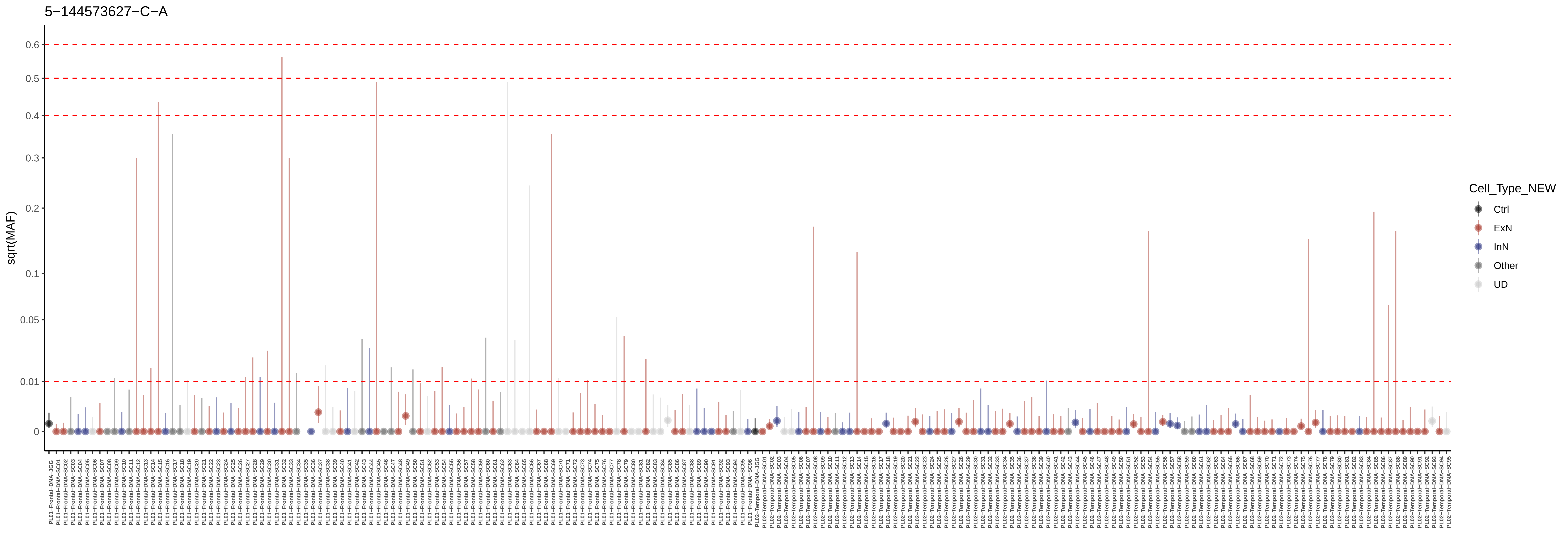
5−144573627−C−A
sqrt(MAF)
0.6
0.5
0.4
0.3
0.2
0.1
0.05
0.01
0
Cell_Type_NEW
Ctrl
ExN
InN
Other
UD

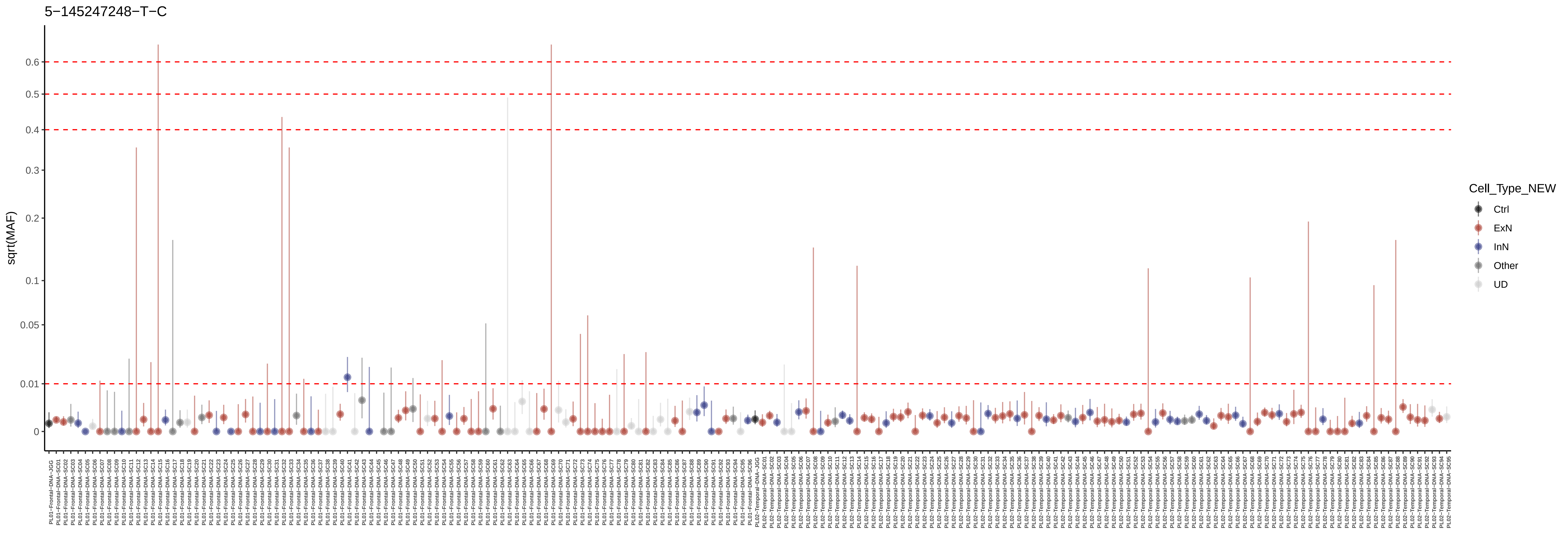

5−145247248−T−C
sqrt(MAF)
0.6
0.5
0.4
0.3
0.2
0.1
0.05
0.01
0
Cell_Type_NEW
Ctrl
ExN
InN
Other
UD

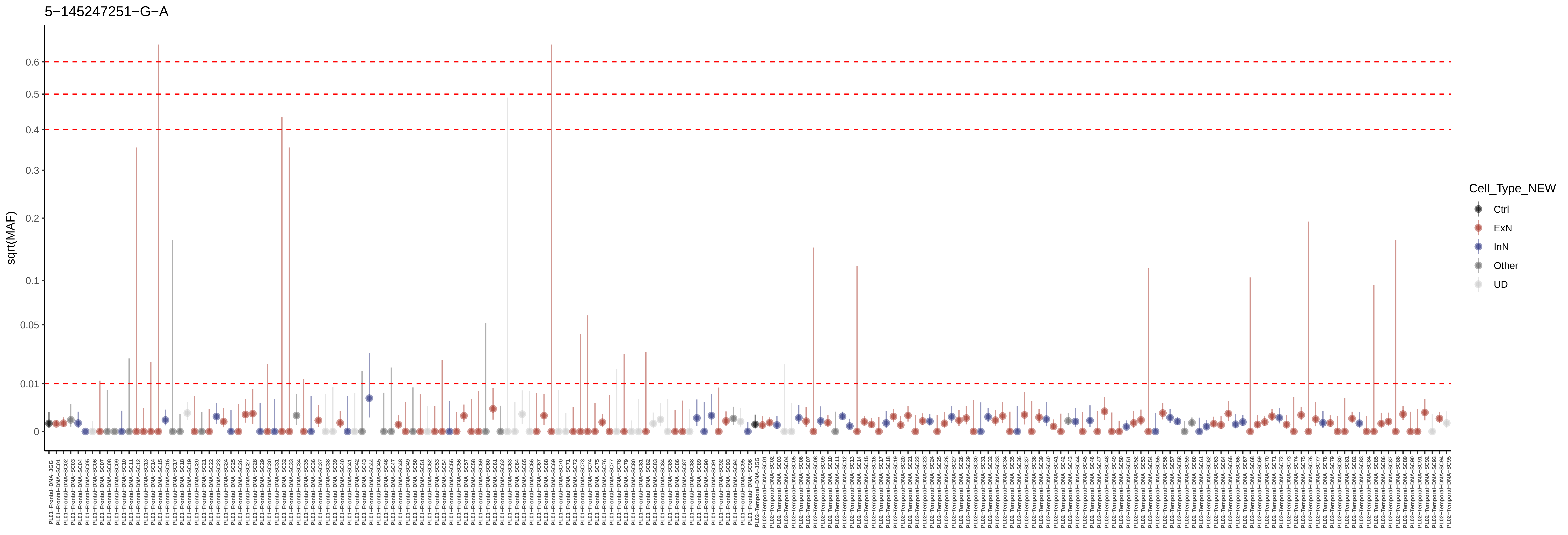
5−145247251−G−A
sqrt(MAF)
0.6
0.5
0.4
0.3
0.2
0.1
0.05
0.01
0
Cell_Type_NEW
Ctrl
ExN
InN
Other
UD

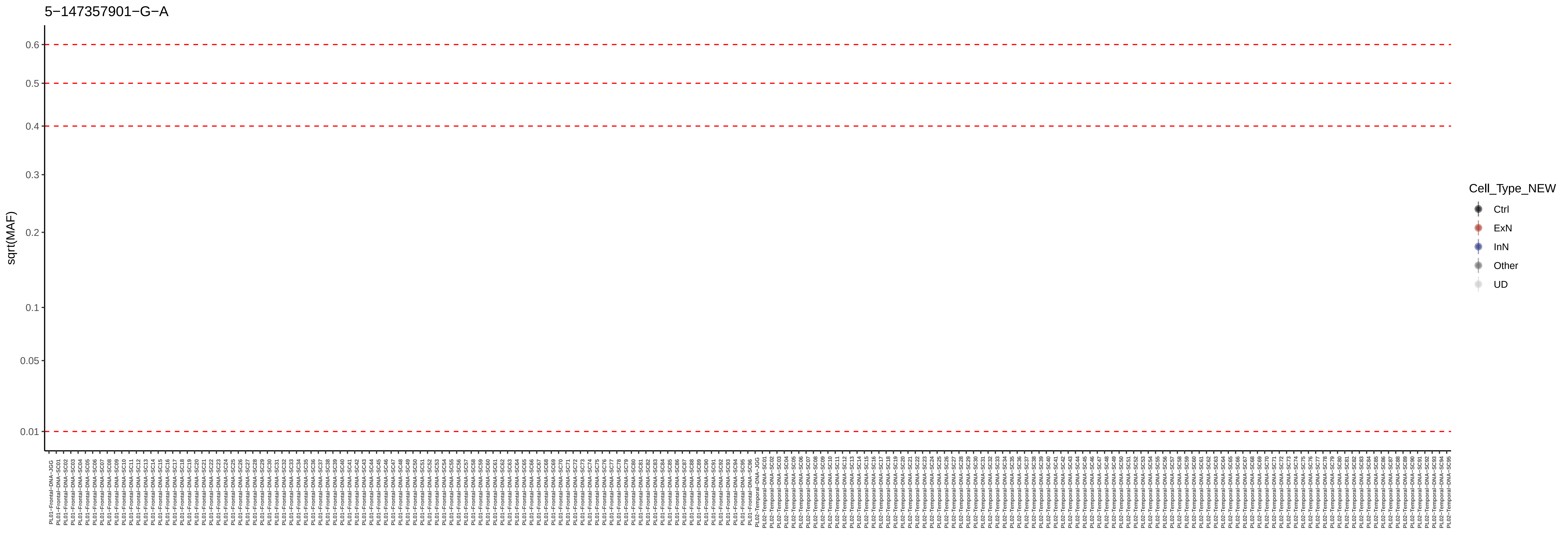
5−147357901−G−A
sqrt(MAF)
0.6
0.5
0.4
0.3
0.2
0.1
0.05
0.01
Cell_Type_NEW
Ctrl
ExN
InN
Other
UD

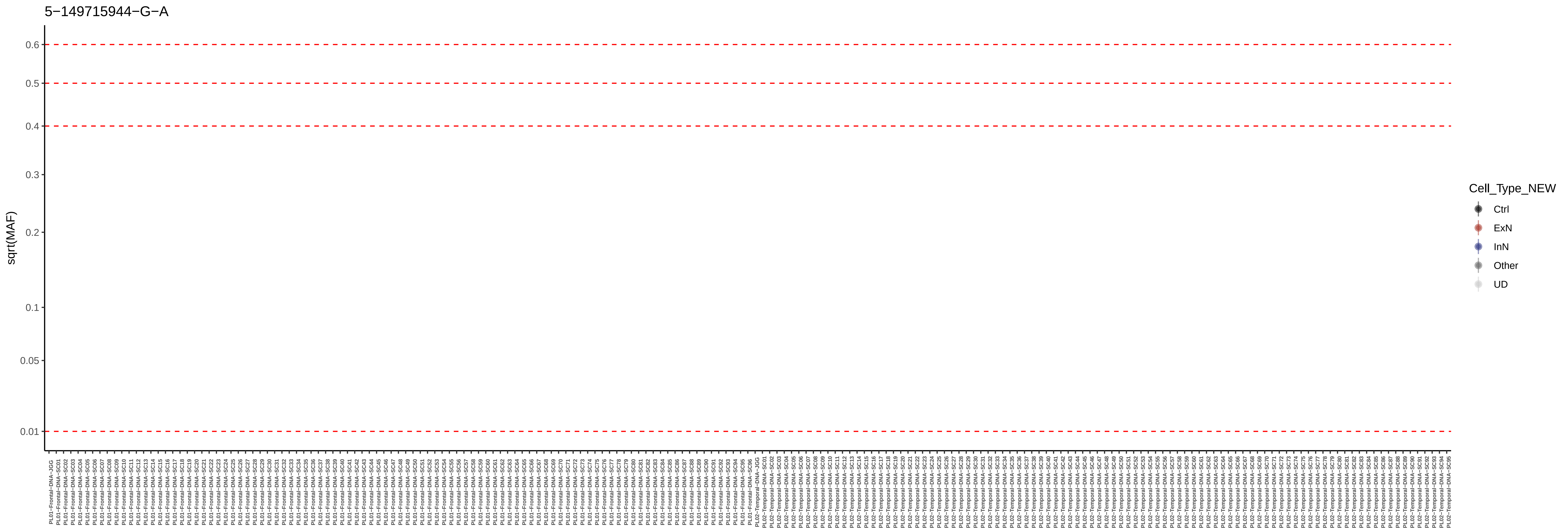
5−149715944−G−A
sqrt(MAF)
0.6
0.5
0.4
0.3
0.2
0.1
0.05
0.01
Cell_Type_NEW
Ctrl
ExN
InN
Other
UD

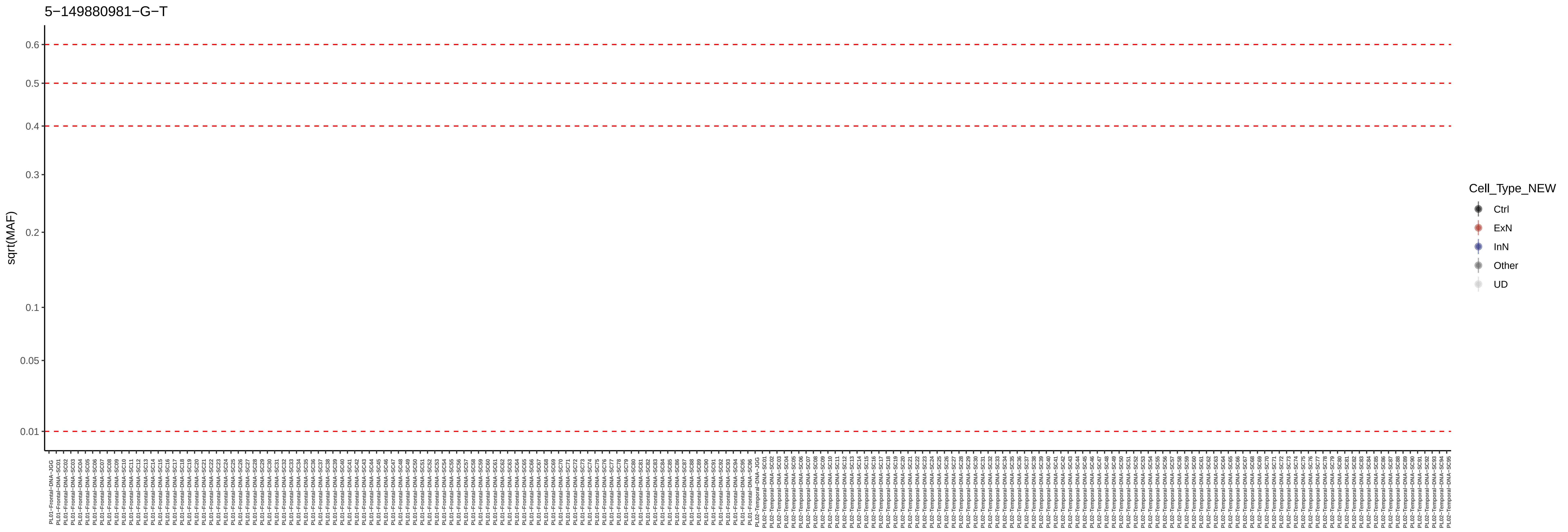

5−149880981−G−T
sqrt(MAF)
0.6
0.5
0.4
0.3
0.2
0.1
0.05
0.01
Cell_Type_NEW
Ctrl
ExN
InN
Other
UD

5−149967700−T−TAA

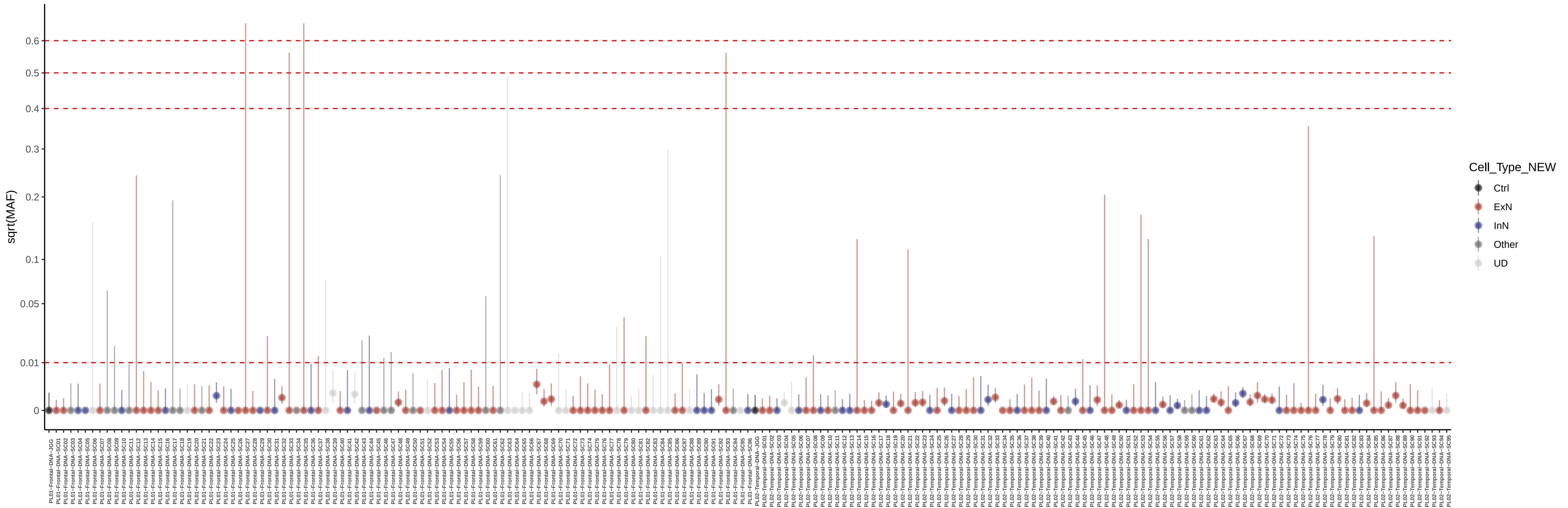

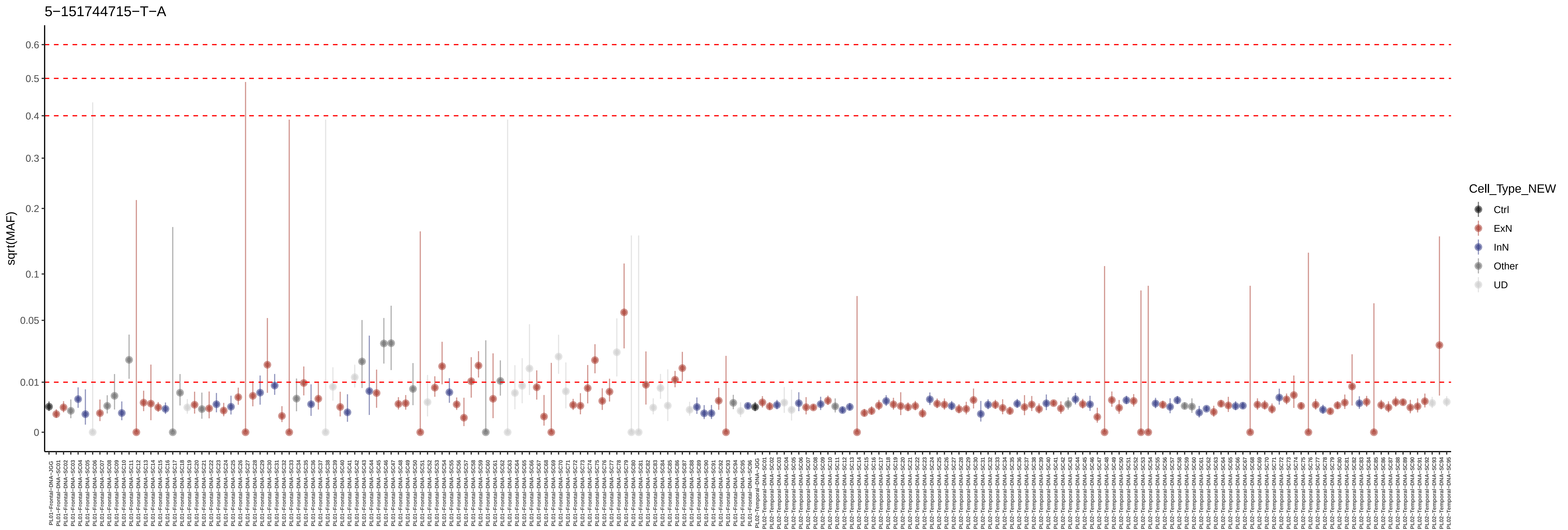
5−151744715−T−A
sqrt(MAF)
0.6
0.5
0.4
0.3
0.2
0.1
0.05
0.01
0
Cell_Type_NEW
Ctrl
ExN
InN
Other
UD
PL01−Frontal−DNA−JGG
PL01−Frontal−DNA−SC01 … PL01−Frontal−DNA−SC96
PL02−Temporal−DNA−JGG
PL02−Temporal−DNA−SC01 … PL02−Temporal−DNA−SC95

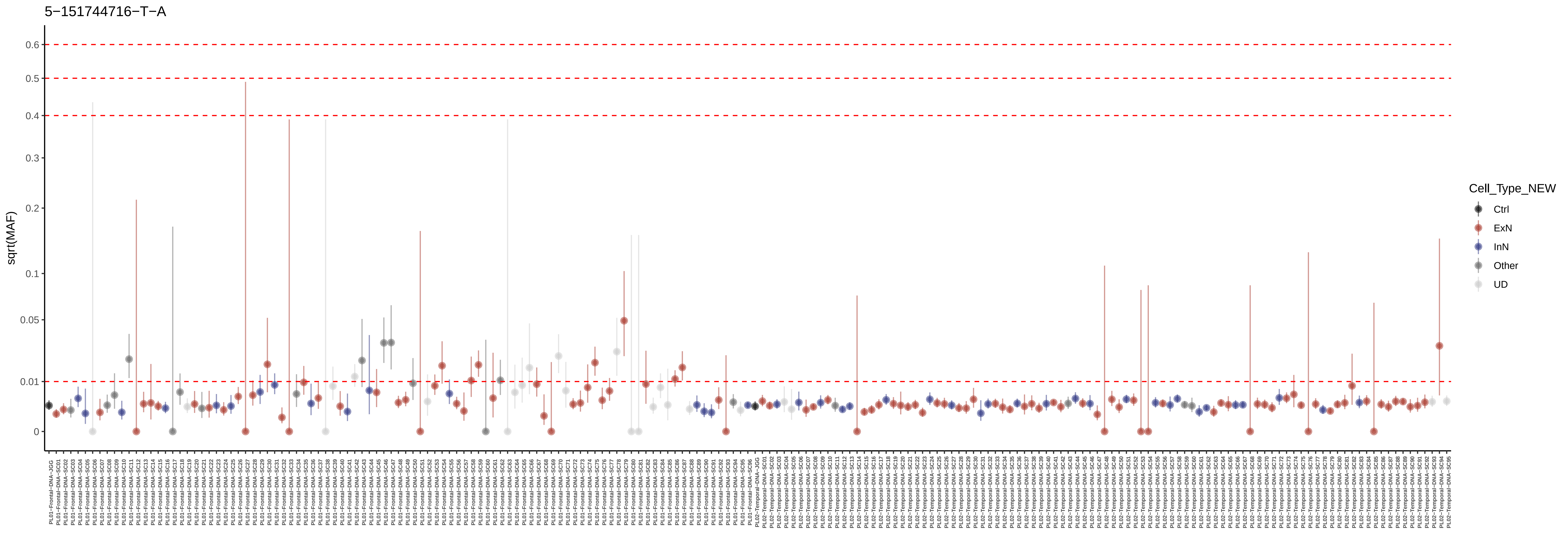
5−151744716−T−A
sqrt(MAF)
0.6
0.5
0.4
0.3
0.2
0.1
0.05
0.01
0
Cell_Type_NEW
Ctrl
ExN
InN
Other
UD

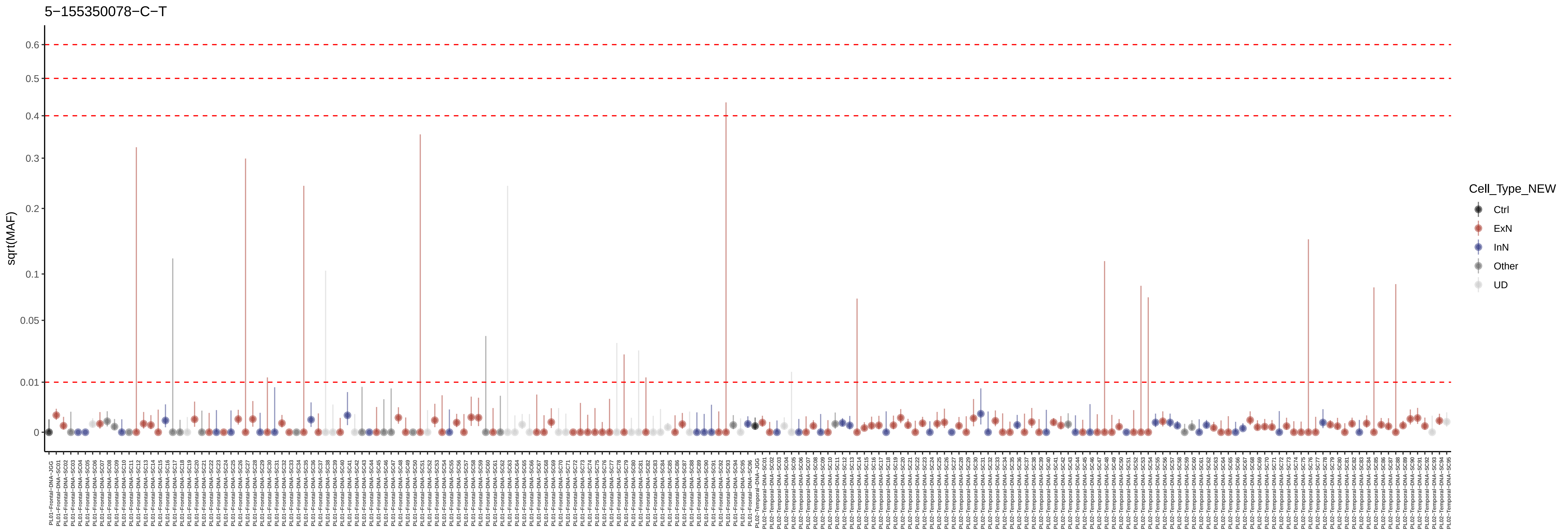
5−155350078−C−T
sqrt(MAF)
0.6
0.5
0.4
0.3
0.2
0.1
0.05
0.01
0
Cell_Type_NEW
Ctrl
ExN
InN
Other
UD

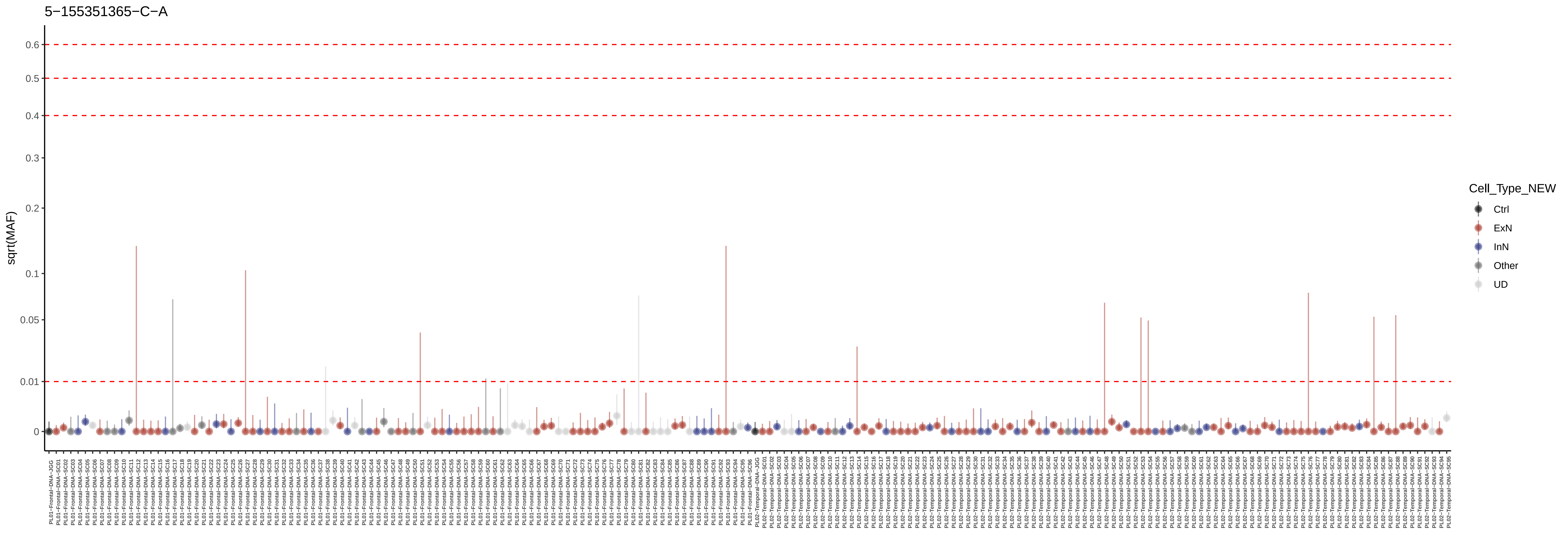
5−155351365−C−A
sqrt(MAF)
0.6
0.5
0.4
0.3
0.2
0.1
0.05
0.01
0
Cell_Type_NEW
Ctrl
ExN
InN
Other
UD

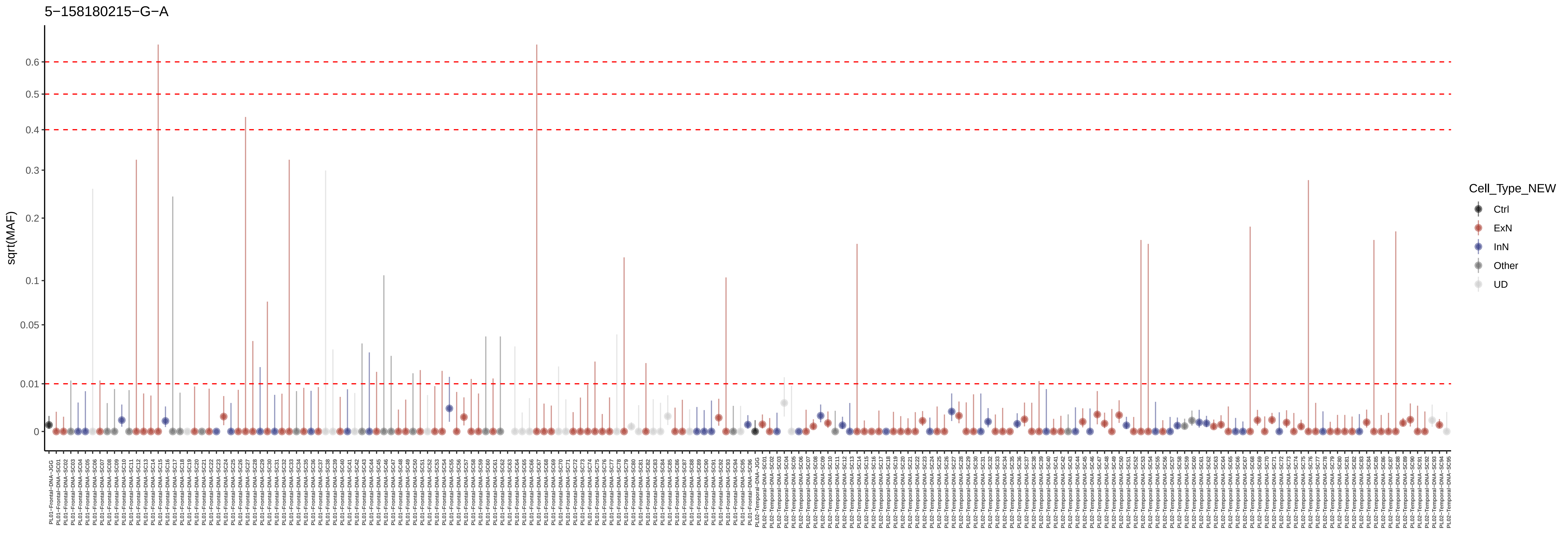
5−158180215−G−A
sqrt(MAF)
0.6
0.5
0.4
0.3
0.2
0.1
0.05
0.01
0
Cell_Type_NEW
Ctrl
ExN
InN
Other
UD

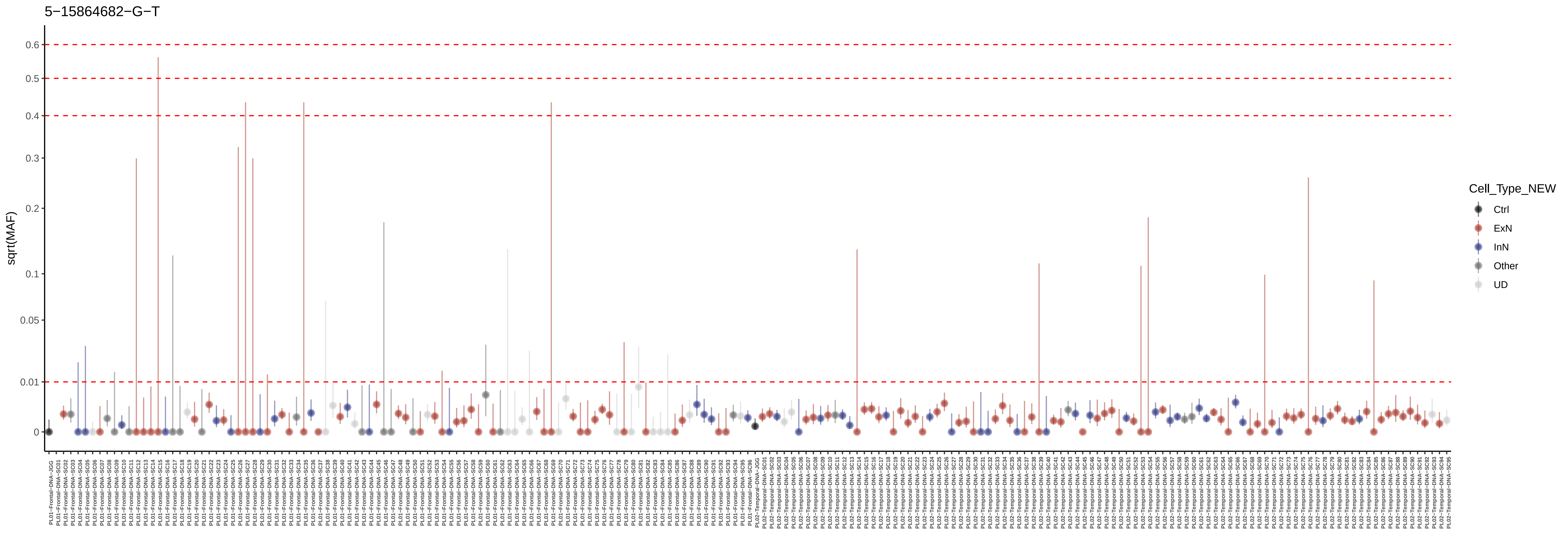

5−15864682−G−T
sqrt(MAF)
0.6
0.5
0.4
0.3
0.2
0.1
0.05
0.01
0
Cell_Type_NEW
Ctrl
ExN
InN
Other
UD

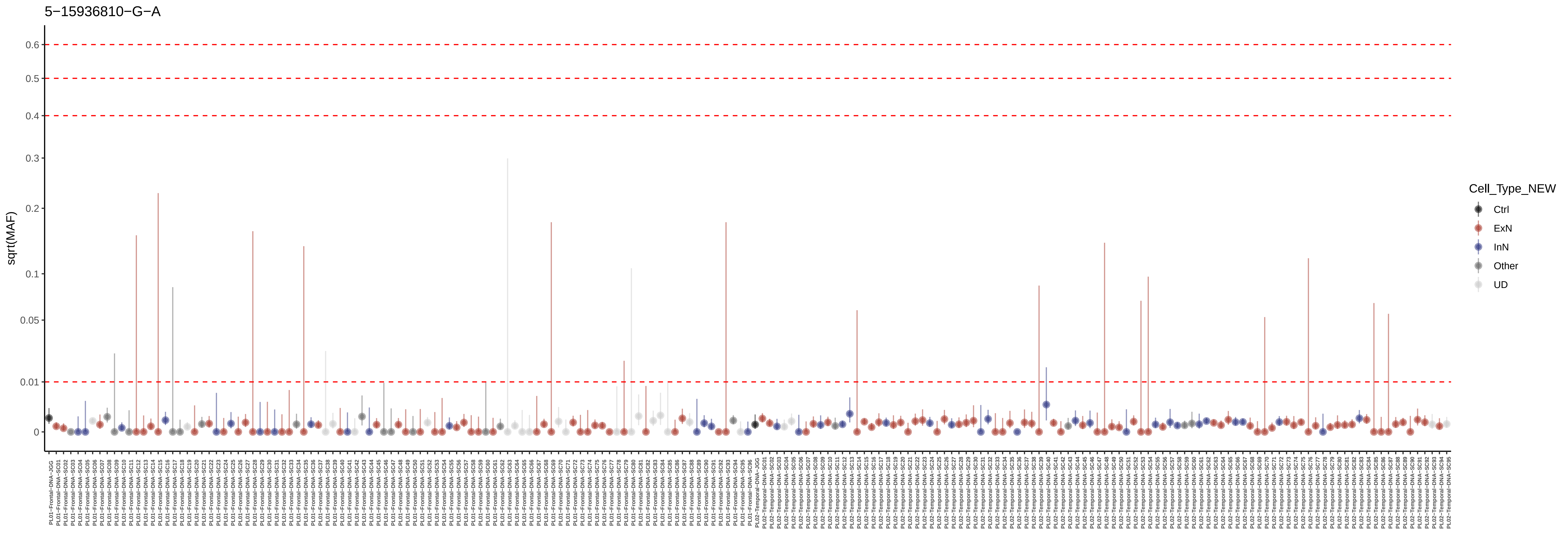
5−15936810−G−A
sqrt(MAF)
0.6
0.5
0.4
0.3
0.2
0.1
0.05
0.01
0
Cell_Type_NEW
Ctrl
ExN
InN
Other
UD

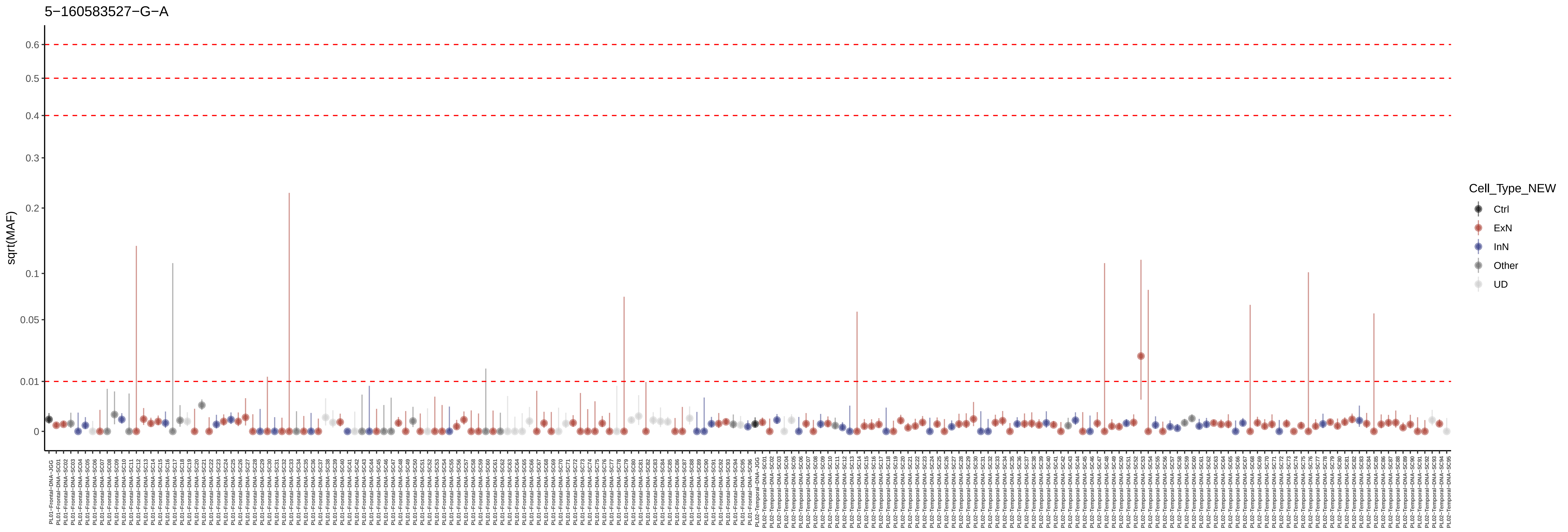

5−160583527−G−A
sqrt(MAF)
0.6
0.5
0.4
0.3
0.2
0.1
0.05
0.01
0
Cell_Type_NEW
Ctrl
ExN
InN
Other
UD

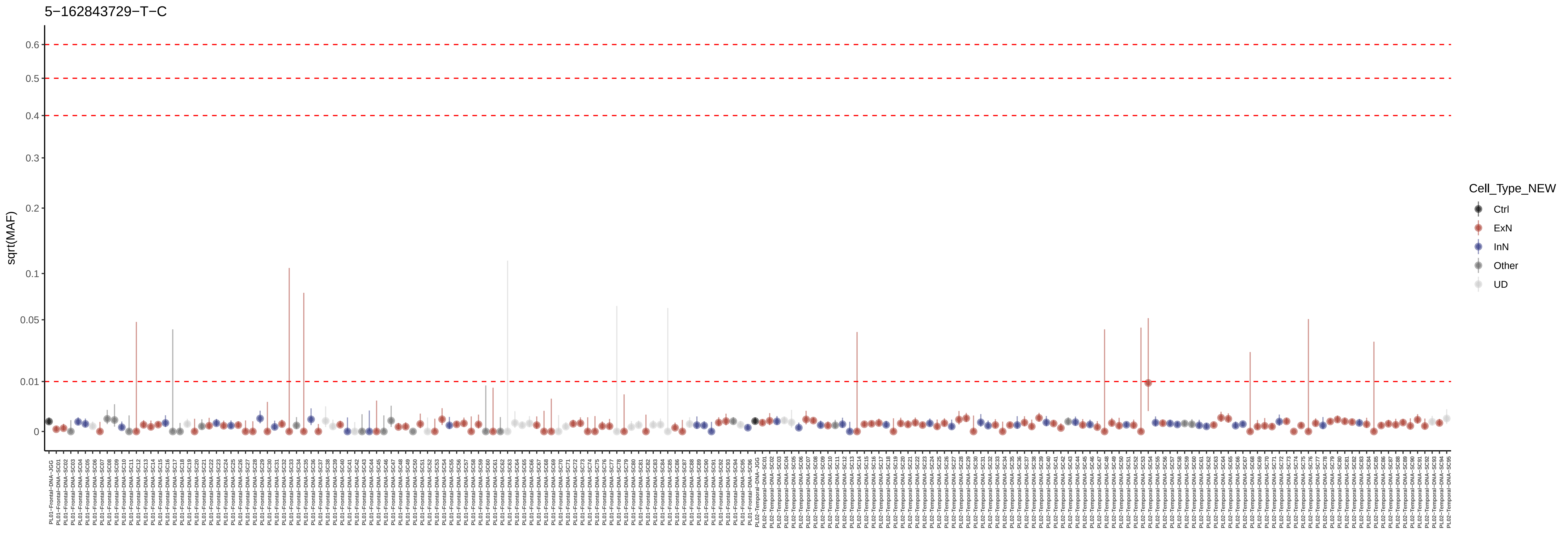

5−162843729−T−C
sqrt(MAF)
0.6
0.5
0.4
0.3
0.2
0.1
0.05
0.01
0
Cell_Type_NEW
Ctrl
ExN
InN
Other
UD

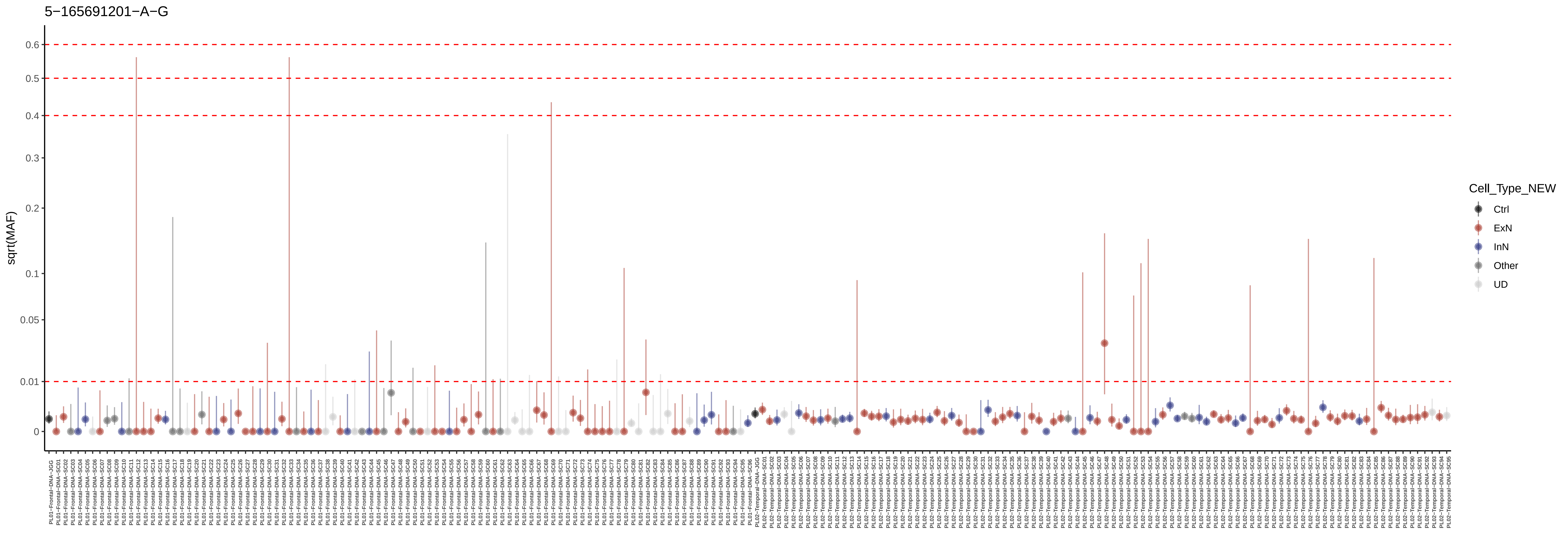
5−165691201−A−G
sqrt(MAF)
0.6
0.5
0.4
0.3
0.2
0.1
0.05
0.01
0
Cell_Type_NEW
Ctrl
ExN
InN
Other
UD

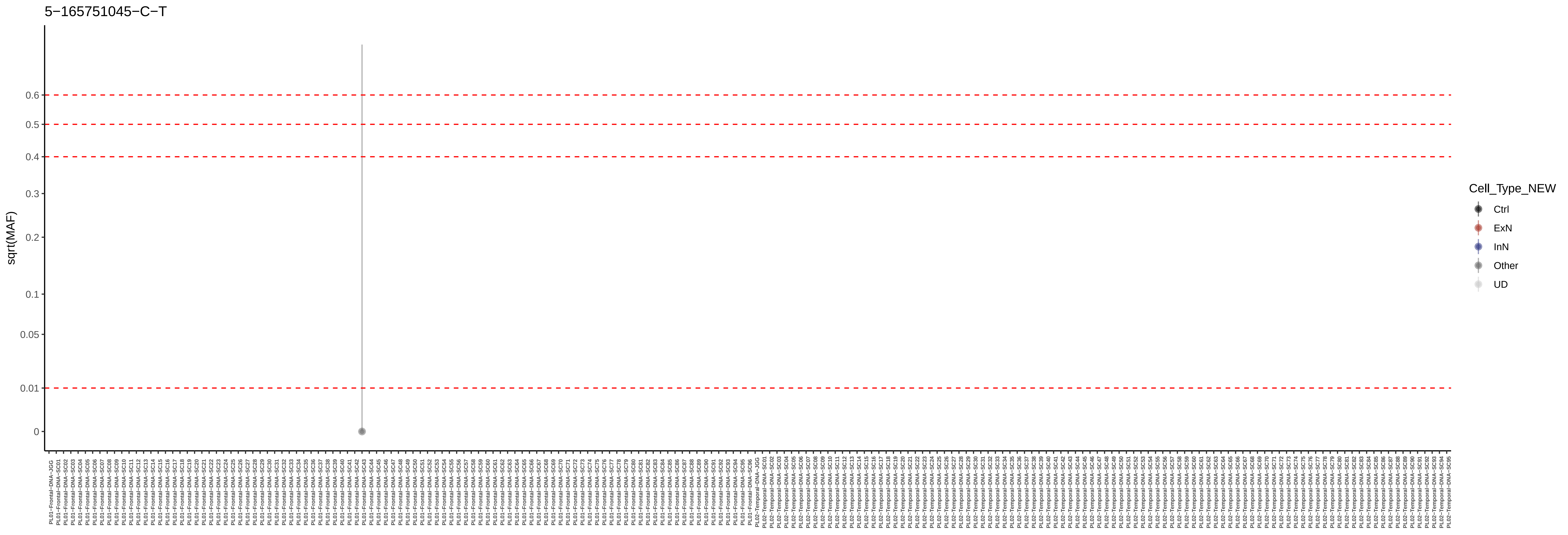

5−165751045−C−T
sqrt(MAF)
0.6
0.5
0.4
0.3
0.2
0.1
0.05
0.01
0
Cell_Type_NEW
Ctrl
ExN
InN
Other
UD

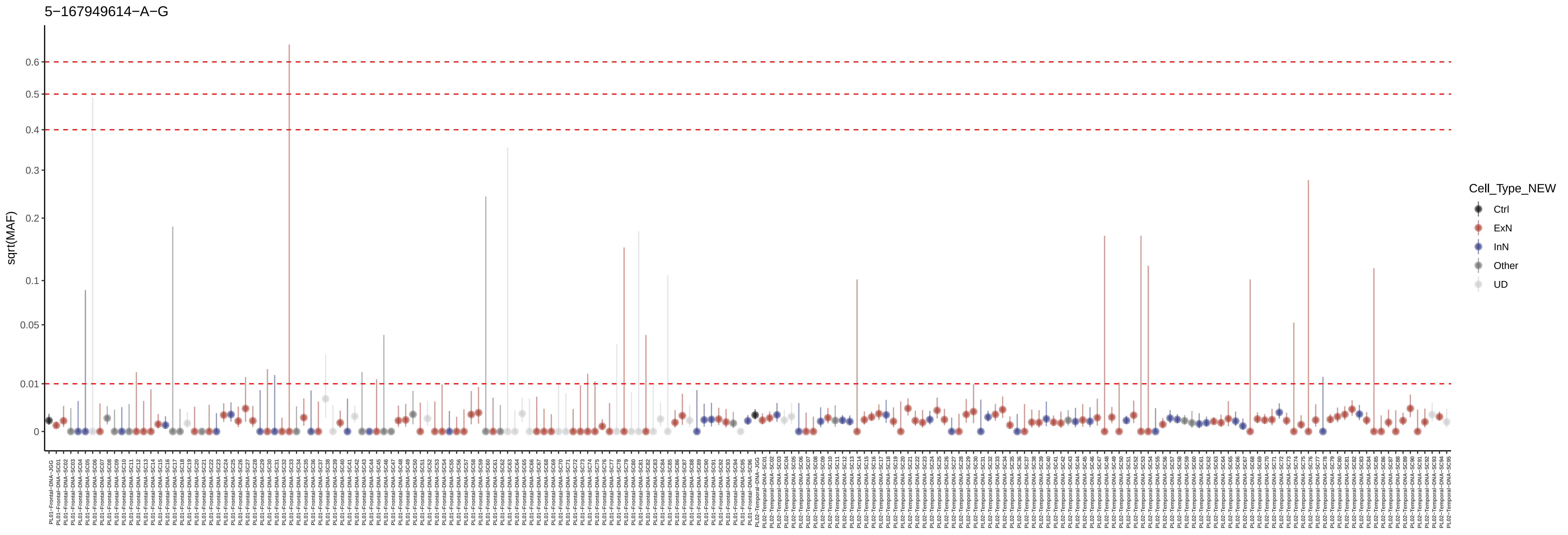
5−167949614−A−G
sqrt(MAF)
0.6
0.5
0.4
0.3
0.2
0.1
0.05
0.01
0
Cell_Type_NEW
Ctrl
ExN
InN
Other
UD

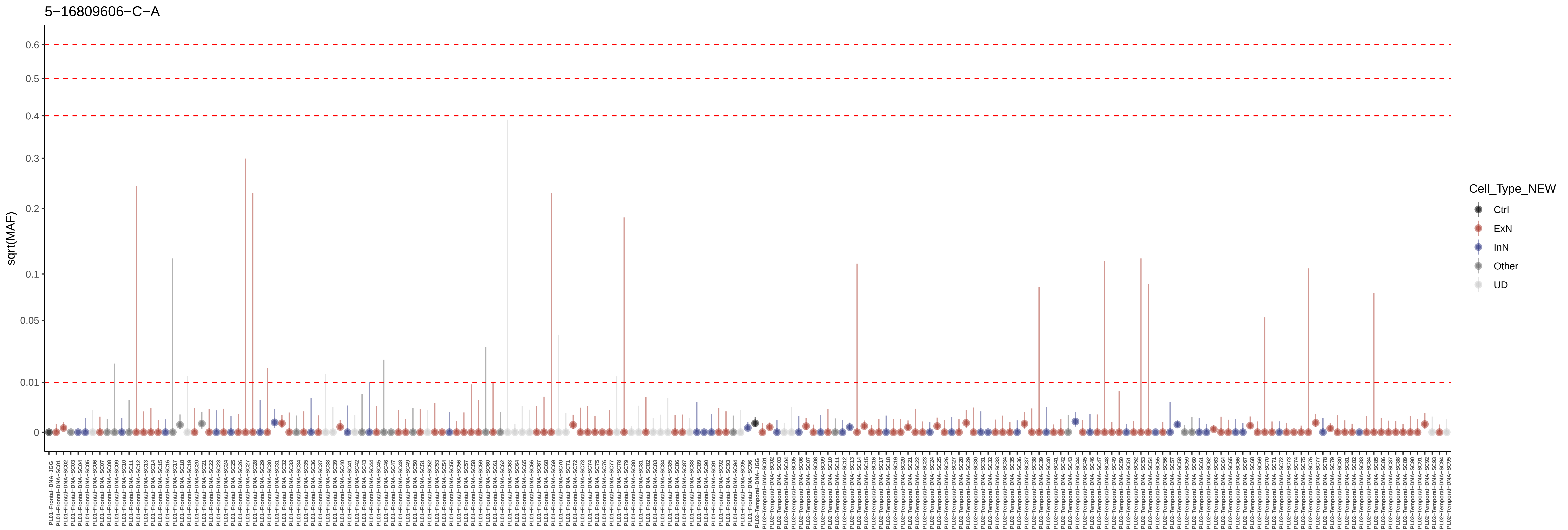

5−16809606−C−A
sqrt(MAF)
0.6
0.5
0.4
0.3
0.2
0.1
0.05
0.01
0
Cell_Type_NEW
Ctrl
ExN
InN
Other
UD

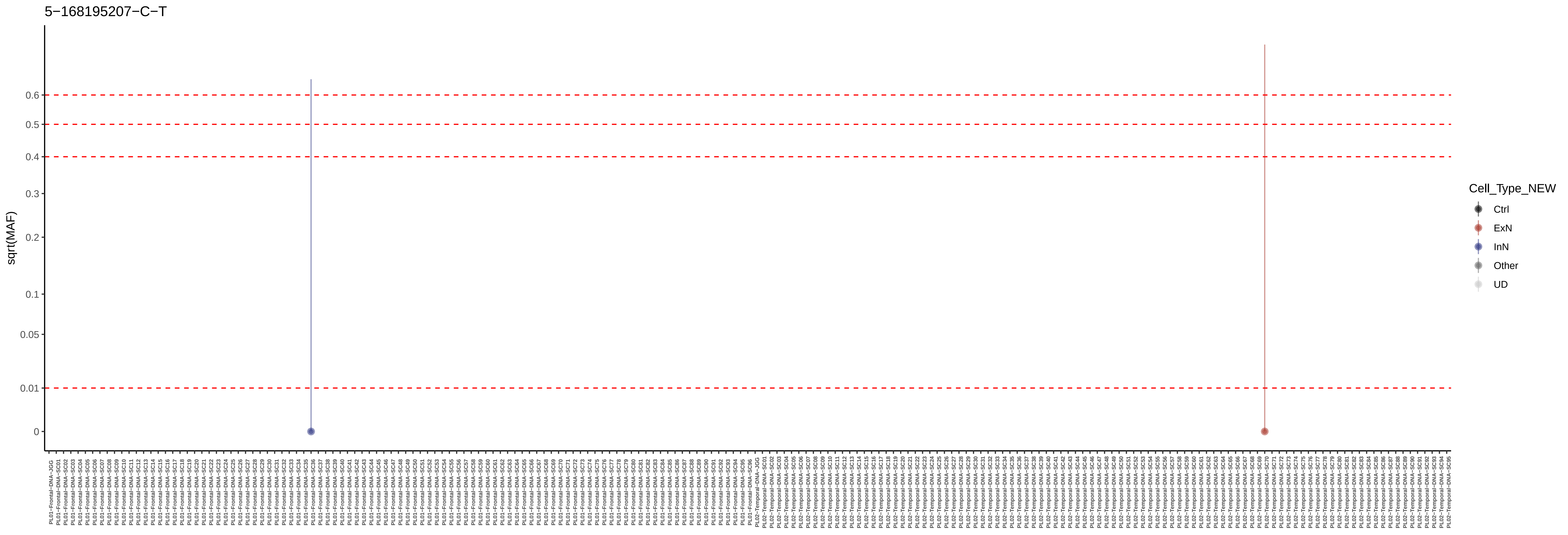
5−168195207−C−T
sqrt(MAF)
0.6
0.5
0.4
0.3
0.2
0.1
0.05
0.01
0
Cell_Type_NEW
Ctrl
ExN
InN
Other
UD

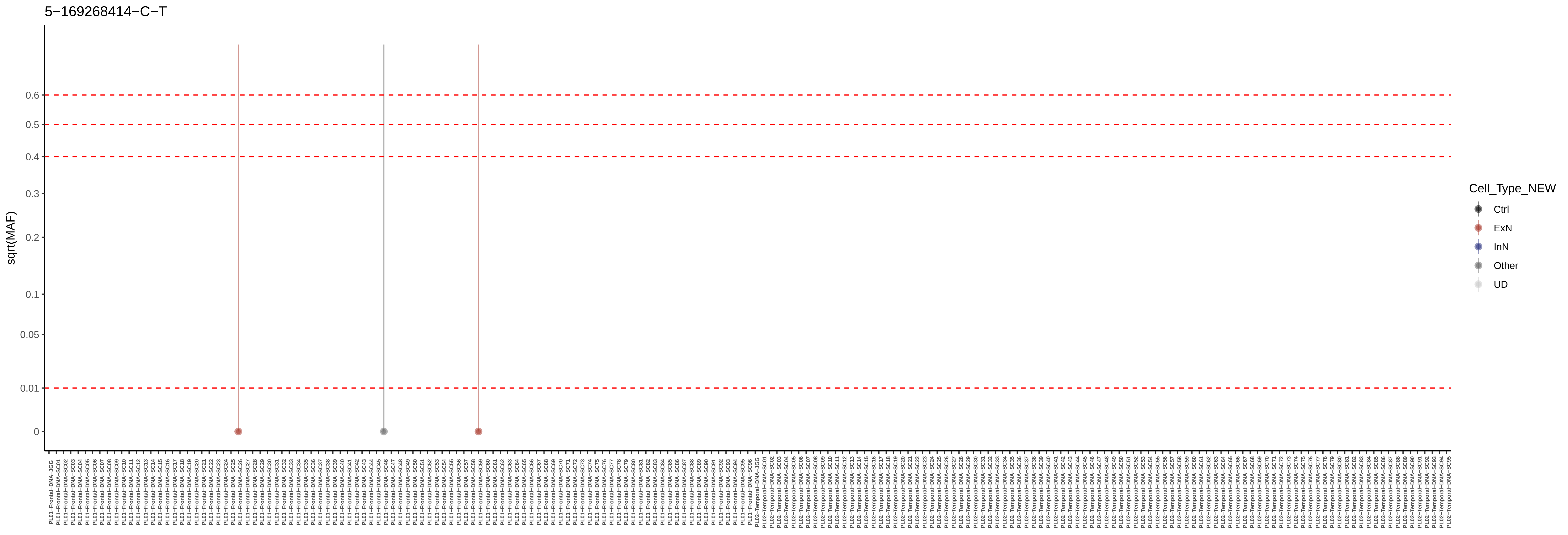
5−169268414−C−T
sqrt(MAF)
0.6
0.5
0.4
0.3
0.2
0.1
0.05
0.01
0
Cell_Type_NEW
Ctrl
ExN
InN
Other
UD

5−170372203−A−G

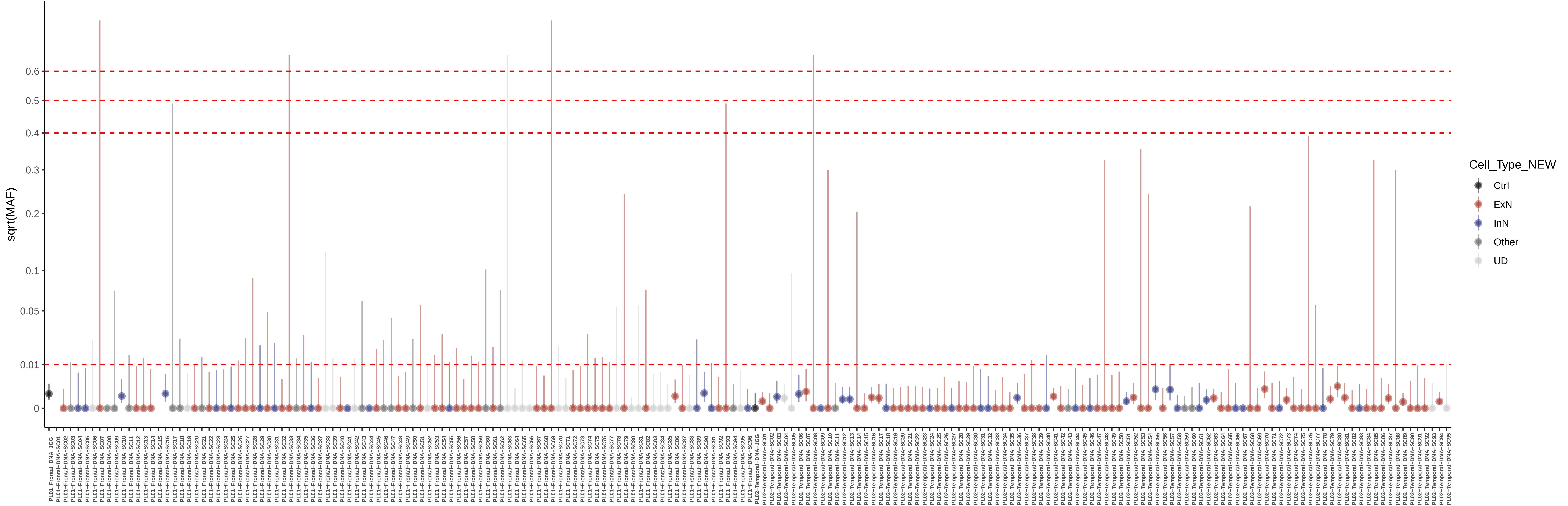

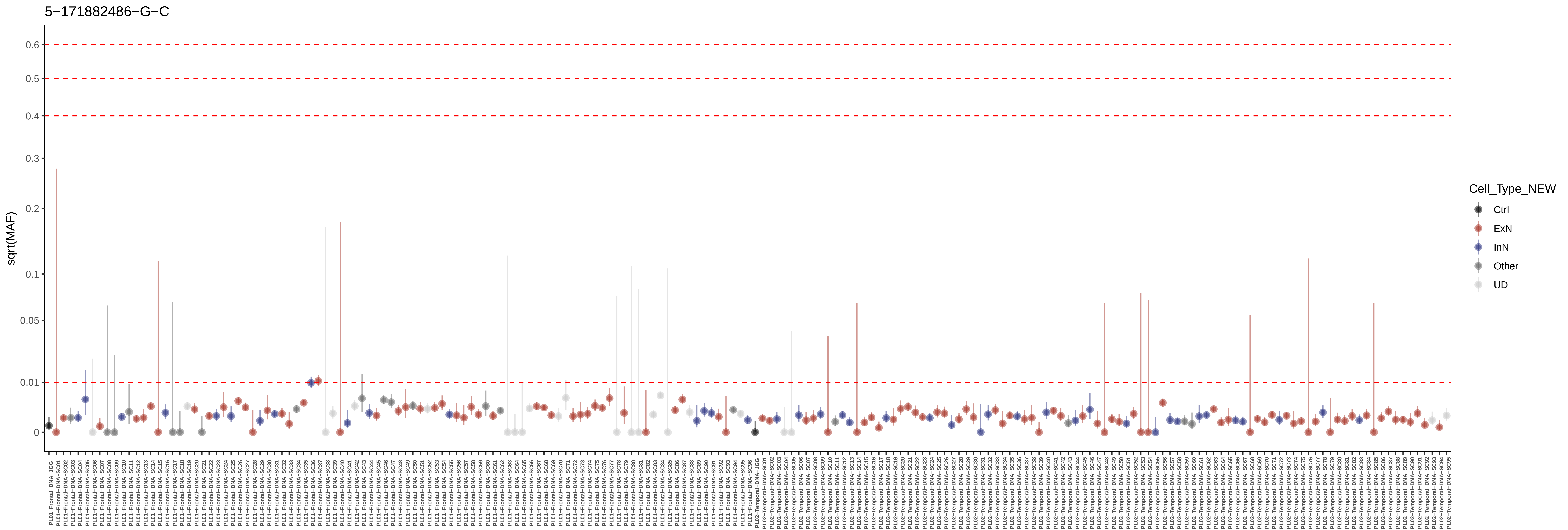

5−171882486−G−C
sqrt(MAF)
Cell_Type_NEW
Ctrl
ExN
InN
Other
UD

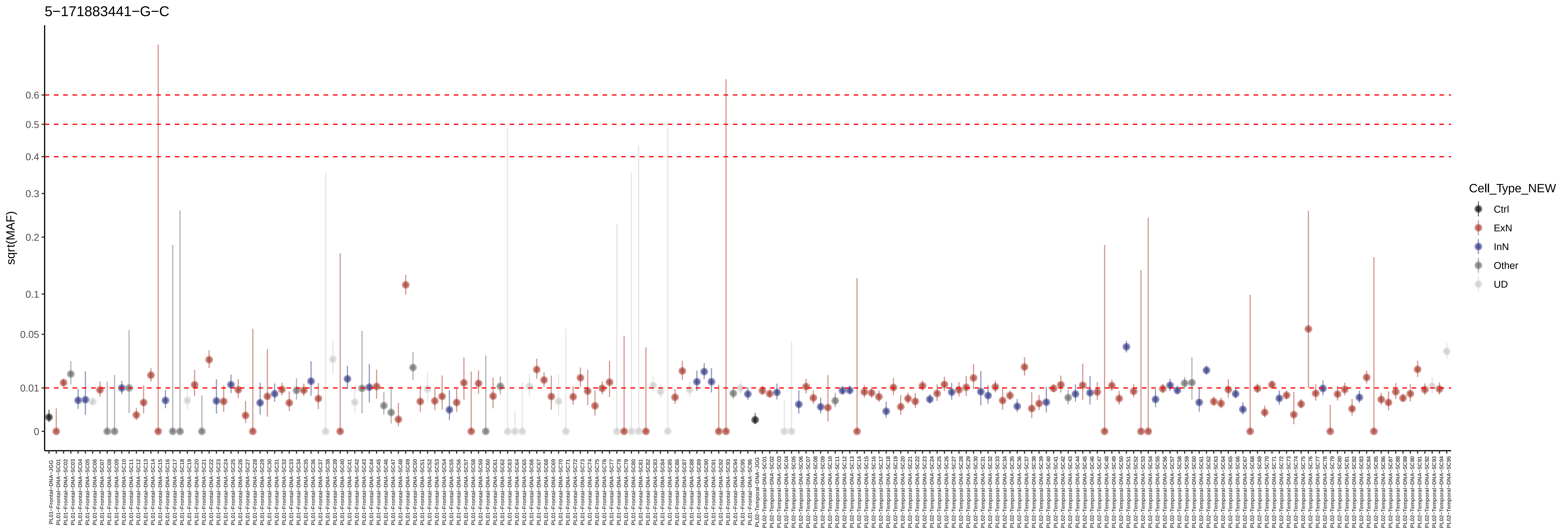

5−171883441−G−C
sqrt(MAF)
0.6
0.5
0.4
0.3
0.2
0.1
0.05
0.01
0
Cell_Type_NEW
Ctrl
ExN
InN
Other
UD

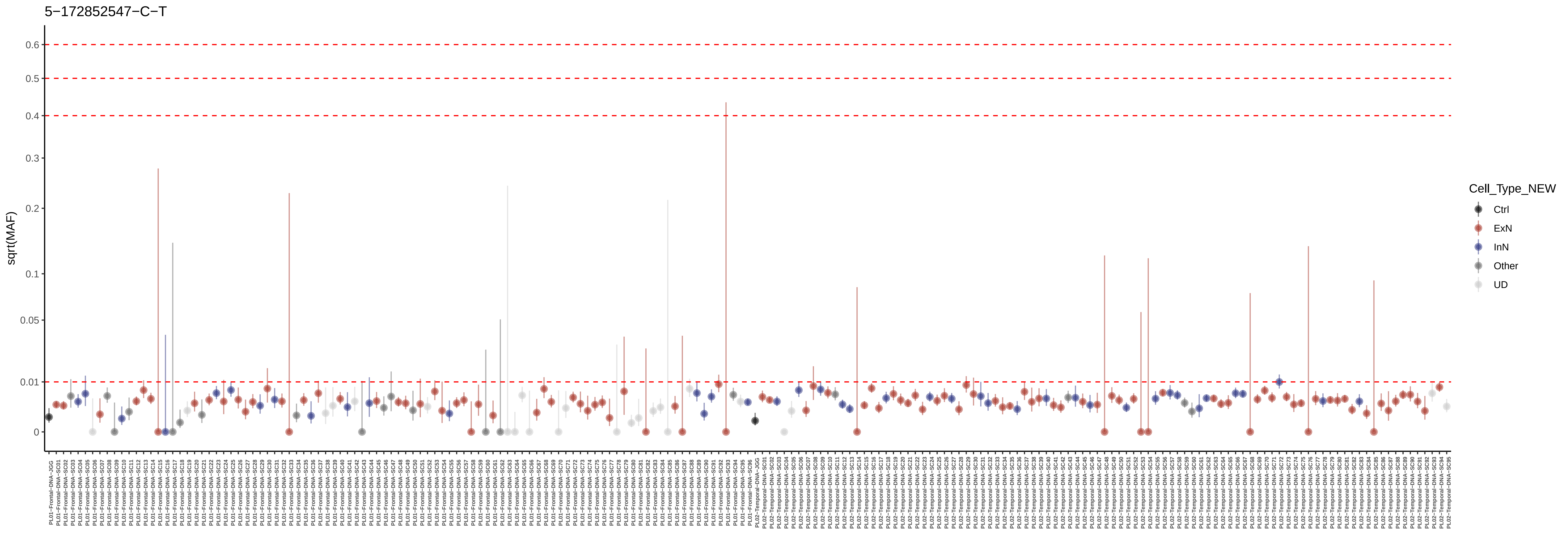
5−172852547−C−T
sqrt(MAF)
0.6
0.5
0.4
0.3
0.2
0.1
0.05
0.01
0
Cell_Type_NEW
Ctrl
ExN
InN
Other
UD

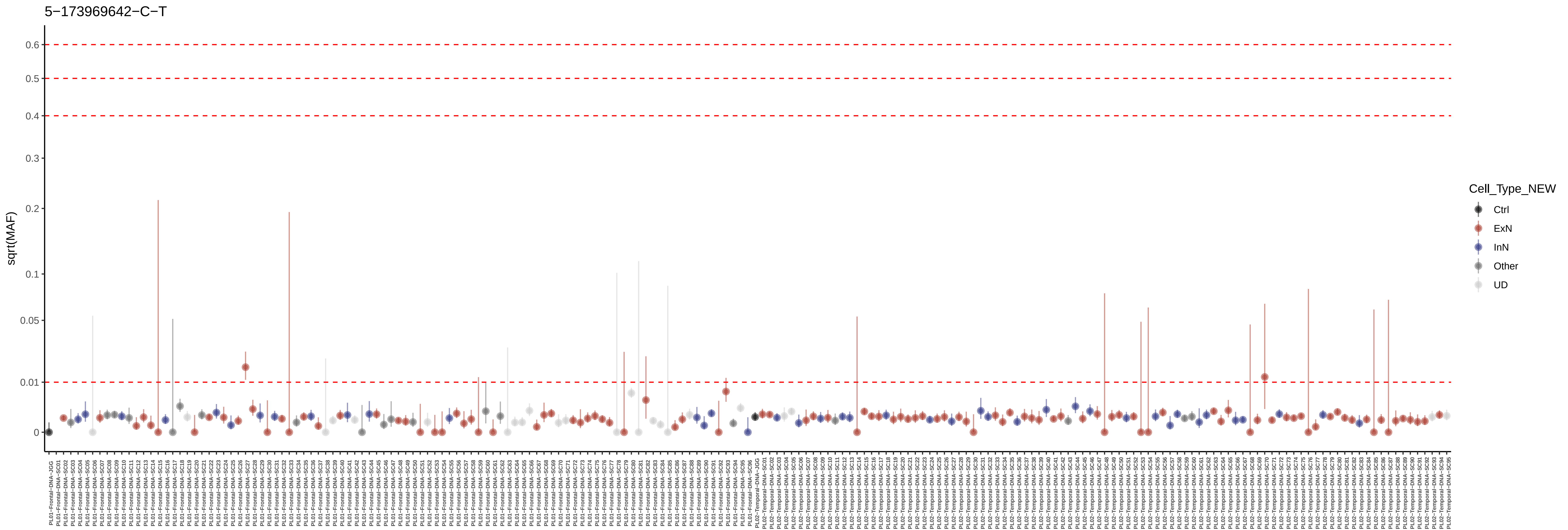

5−173969642−C−T
sqrt(MAF)
0.6
0.5
0.4
0.3
0.2
0.1
0.05
0.01
0
Cell_Type_NEW
Ctrl
ExN
InN
Other
UD

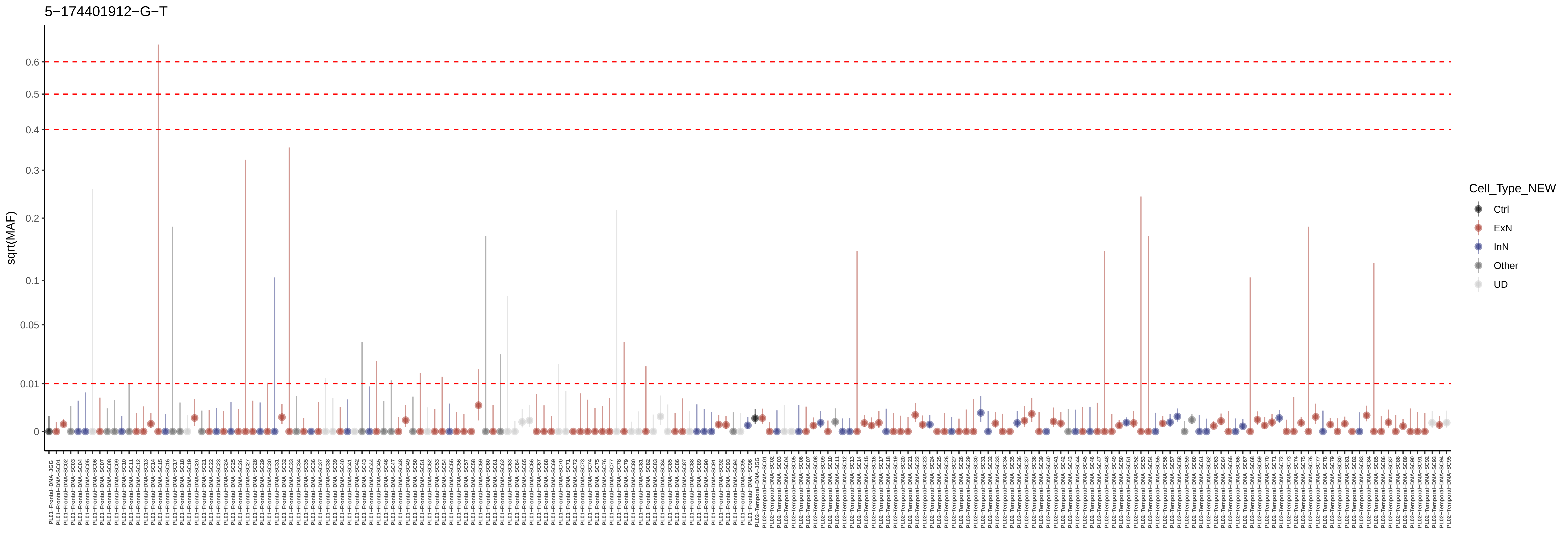
5−174401912−G−T
sqrt(MAF)
0.6
0.5
0.4
0.3
0.2
0.1
0.05
0.01
0
Cell_Type_NEW
Ctrl
ExN
InN
Other
UD

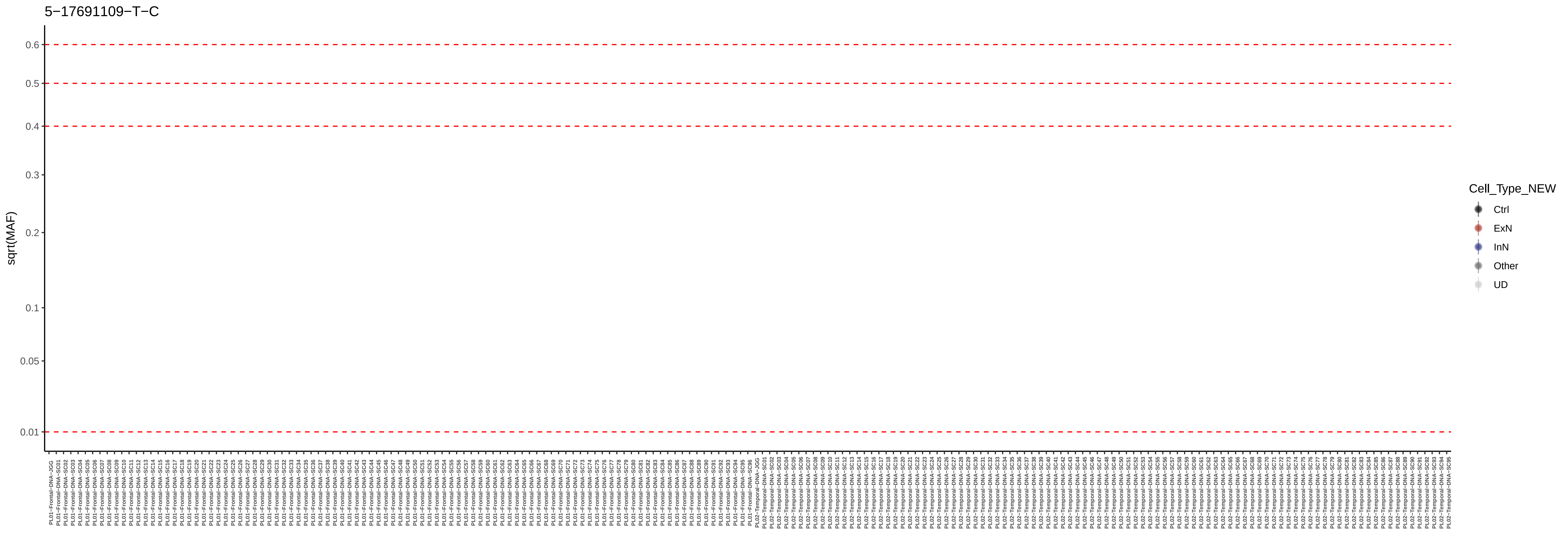
5−17691109−T−C
sqrt(MAF)
0.6
0.5
0.4
0.3
0.2
0.1
0.05
0.01
Cell_Type_NEW
Ctrl
ExN
InN
Other
UD

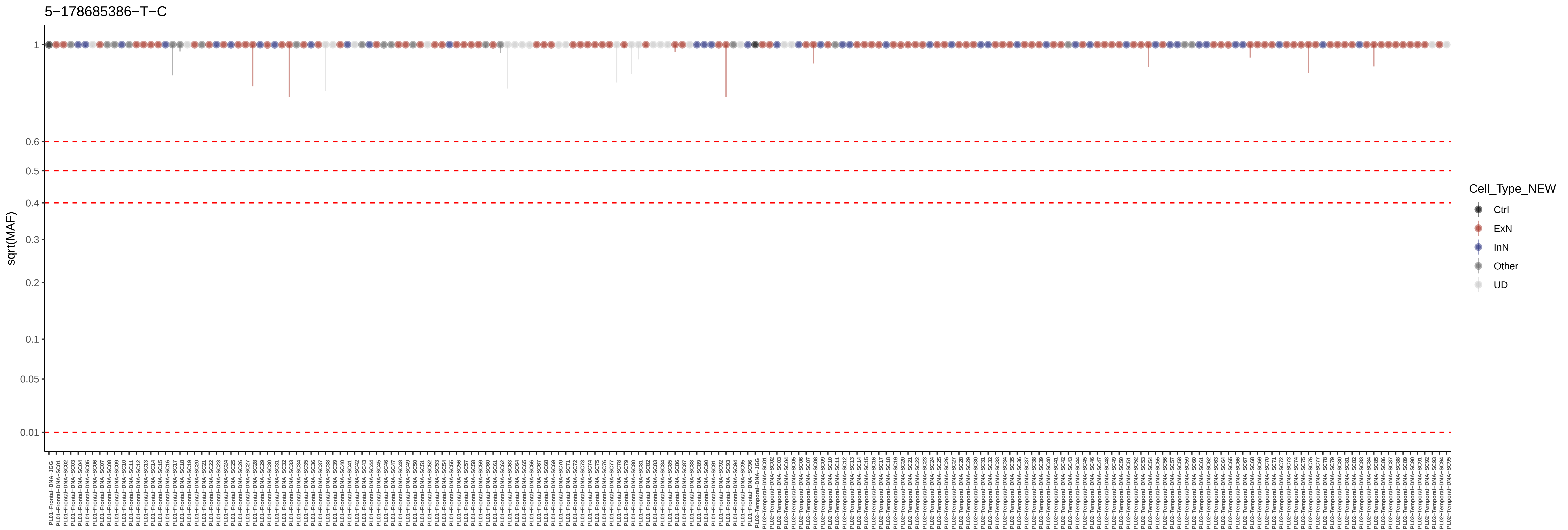

5−178685386−T−C
sqrt(MAF)
1
0.6
0.5
0.4
0.3
0.2
0.1
0.05
0.01
Cell_Type_NEW
Ctrl
ExN
InN
Other
UD

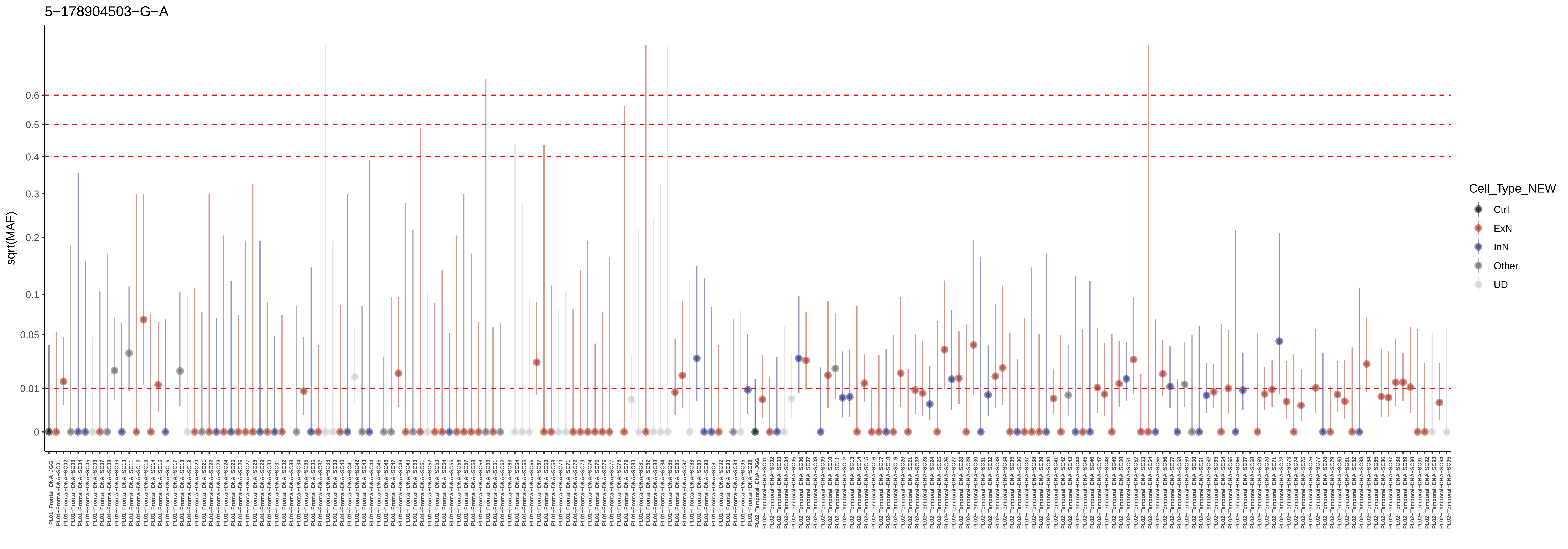
5−178904503−G−A
sqrt(MAF)
0.6
0.5
0.4
0.3
0.2
0.1
0.05
0.01
0
Cell_Type_NEW
Ctrl
ExN
InN
Other
UD

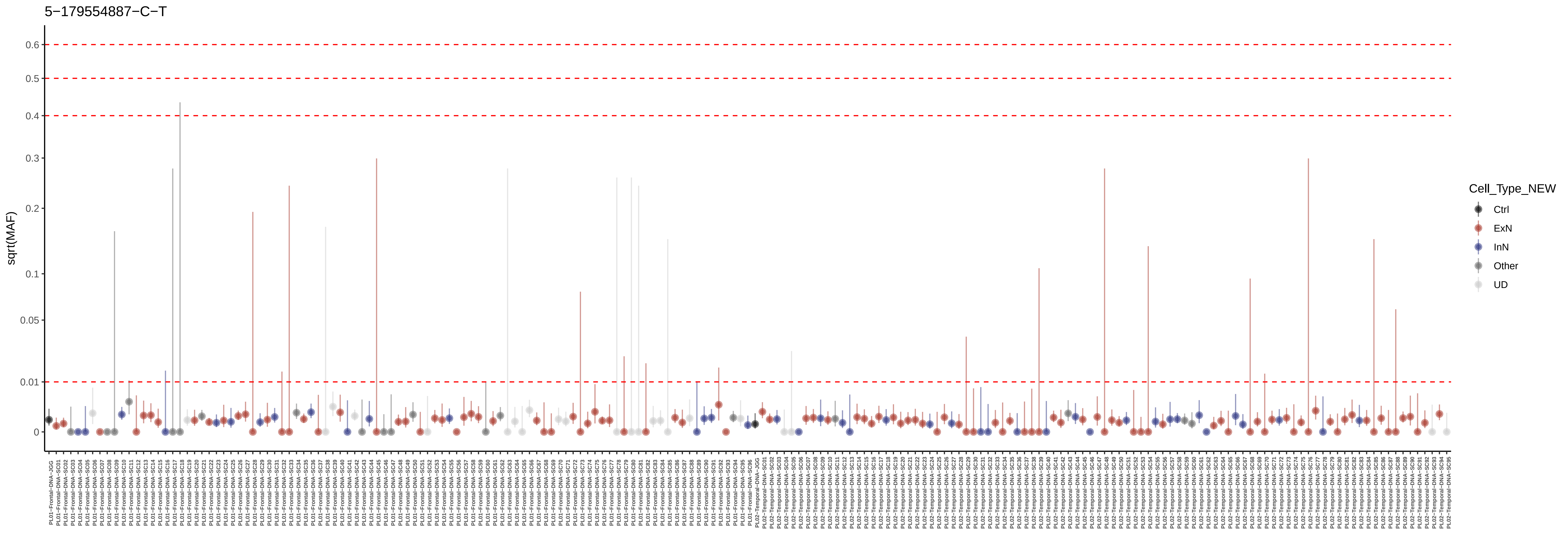

5−179554887−C−T
sqrt(MAF)
0.6
0.5
0.4
0.3
0.2
0.1
0.05
0.01
0
Cell_Type_NEW
Ctrl
ExN
InN
Other
UD

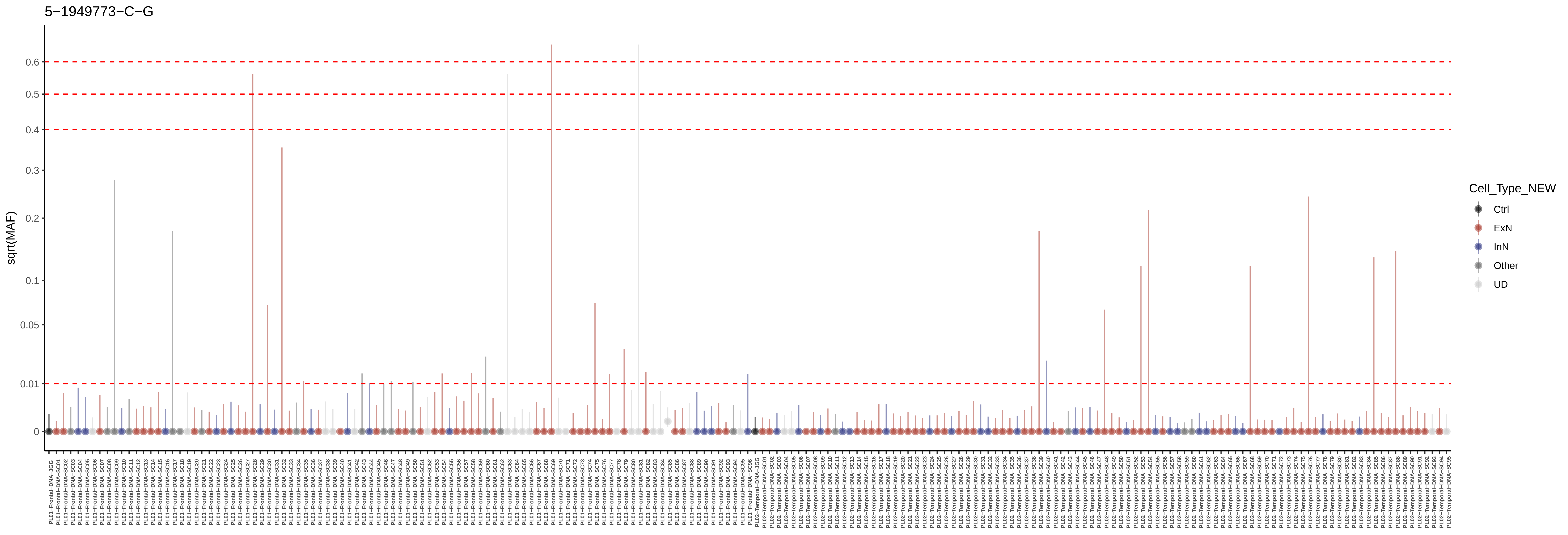

5−1949773−C−G
sqrt(MAF)
0.6
0.5
0.4
0.3
0.2
0.1
0.05
0.01
0
Cell_Type_NEW
Ctrl
ExN
InN
Other
UD

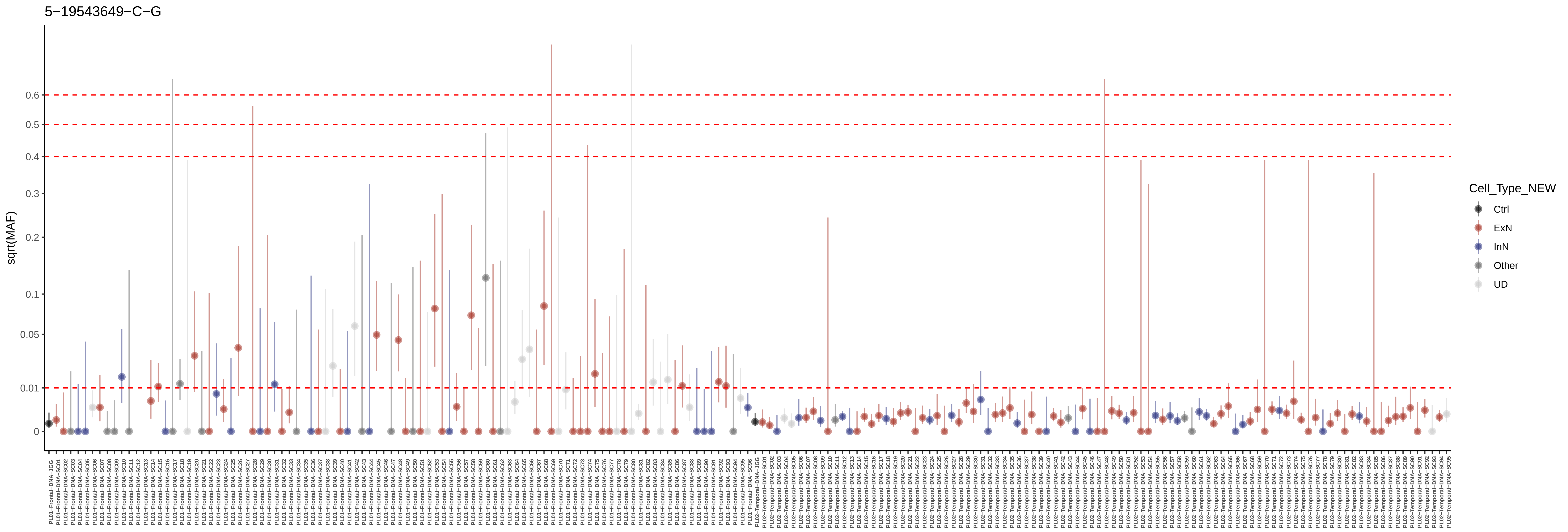
5−19543649−C−G
sqrt(MAF)
0.6
0.5
0.4
0.3
0.2
0.1
0.05
0.01
0
Cell_Type_NEW
Ctrl
ExN
InN
Other
UD

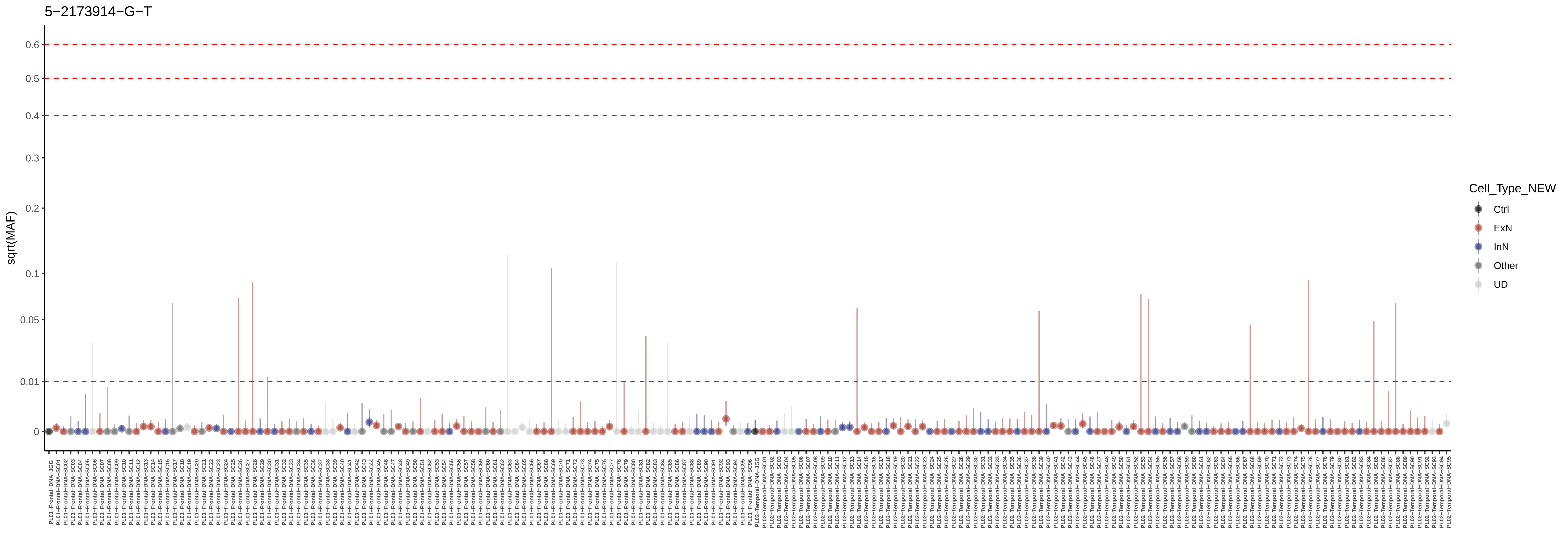
5−2173914−G−T
sqrt(MAF)
0.6
0.5
0.4
0.3
0.2
0.1
0.05
0.01
0
Cell_Type_NEW
Ctrl
ExN
InN
Other
UD

5−2376747−T−G

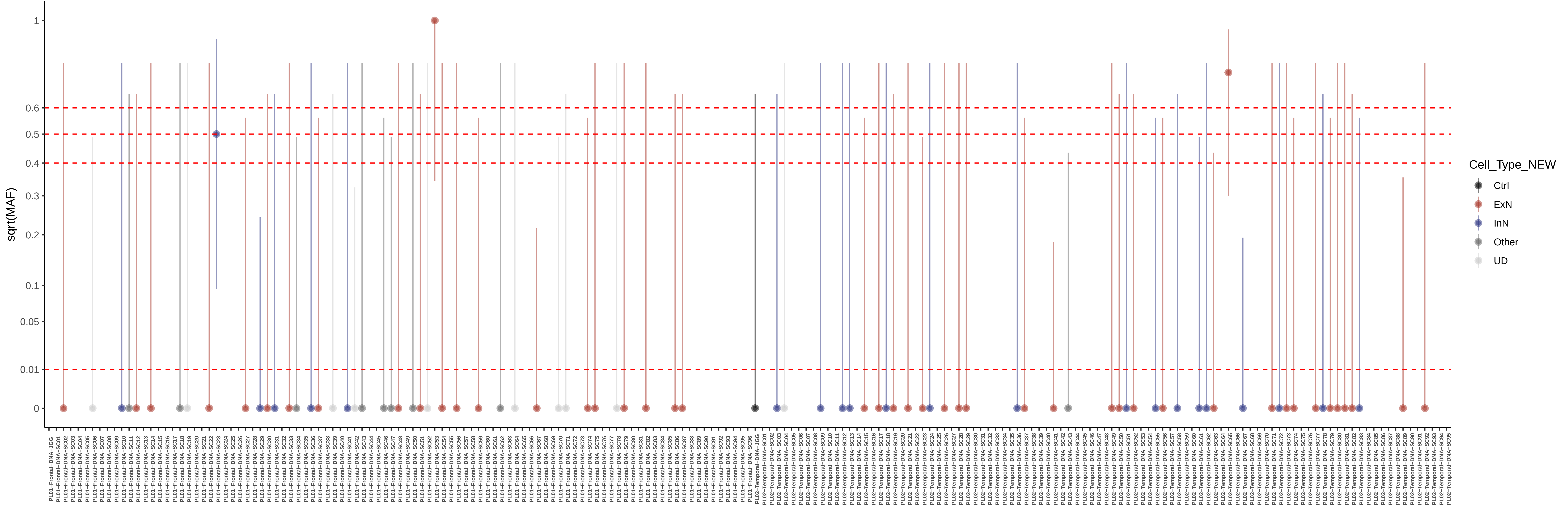
sqrt(MAF)
1
0.6
0.5
0.4
0.3
0.2
0.1
0.05
0.01
0
Cell_Type_NEW
Ctrl
ExN
InN
Other
UD

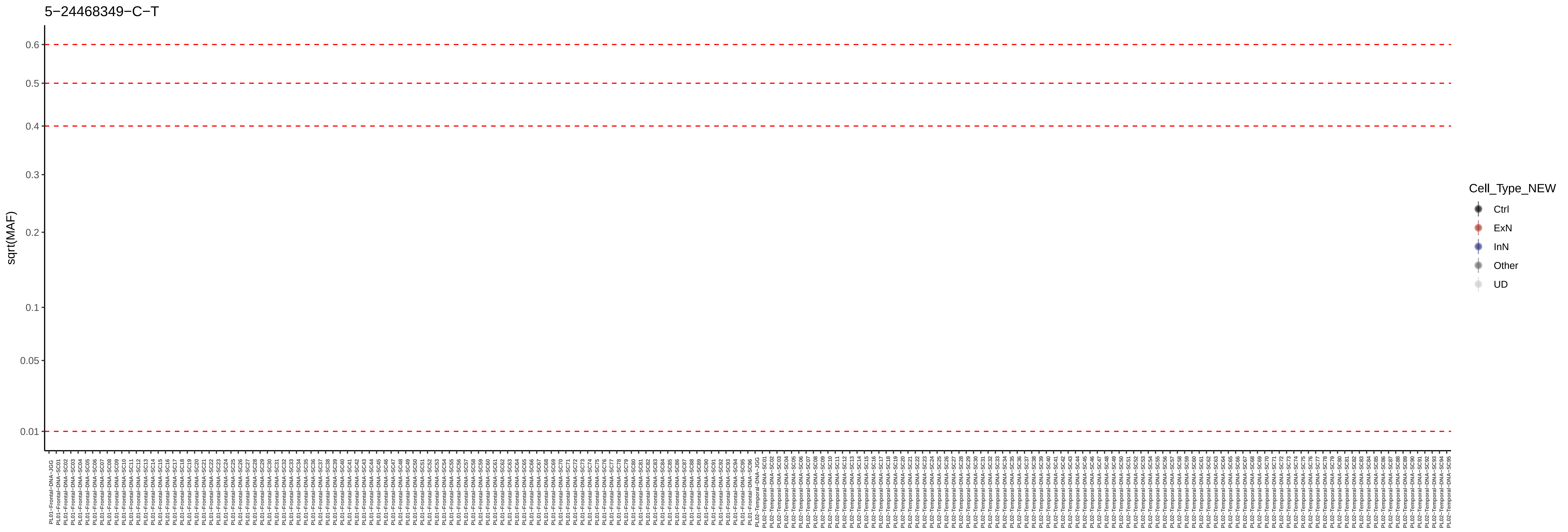
5−24468349−C−T
sqrt(MAF)
0.6
0.5
0.4
0.3
0.2
0.1
0.05
0.01
Cell_Type_NEW
Ctrl
ExN
InN
Other
UD

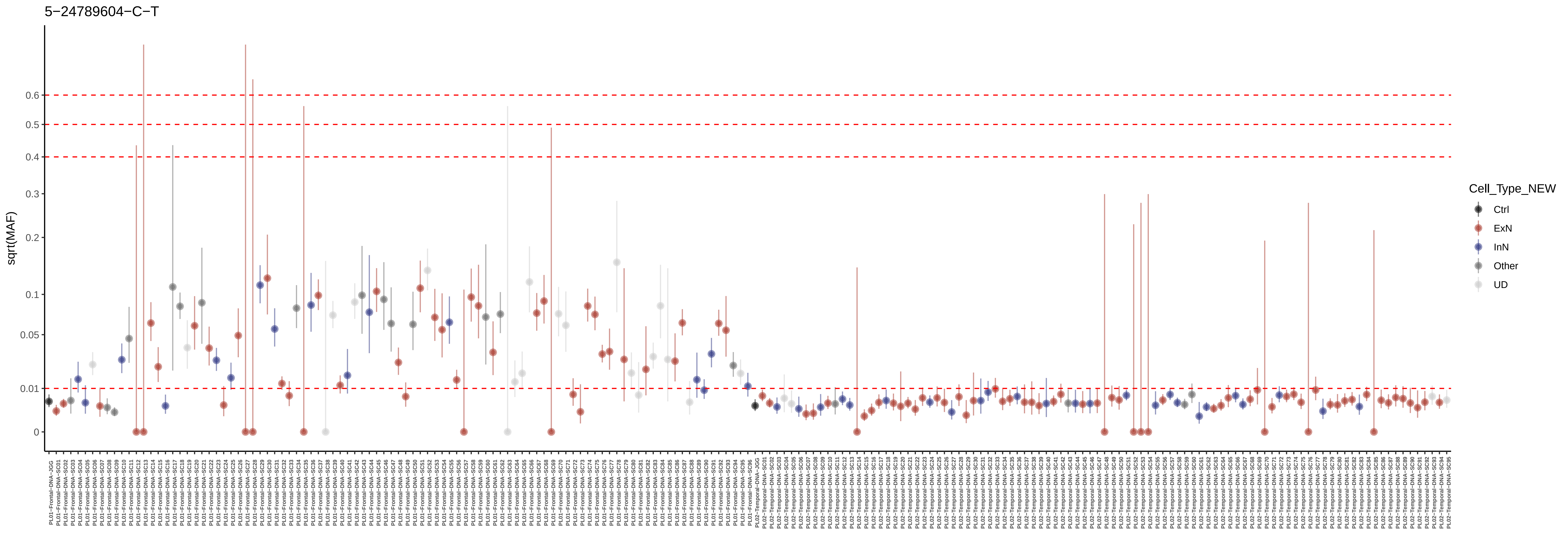
5−24789604−C−T
sqrt(MAF)
0.6
0.5
0.4
0.3
0.2
0.1
0.05
0.01
0
Cell_Type_NEW
Ctrl
ExN
InN
Other
UD

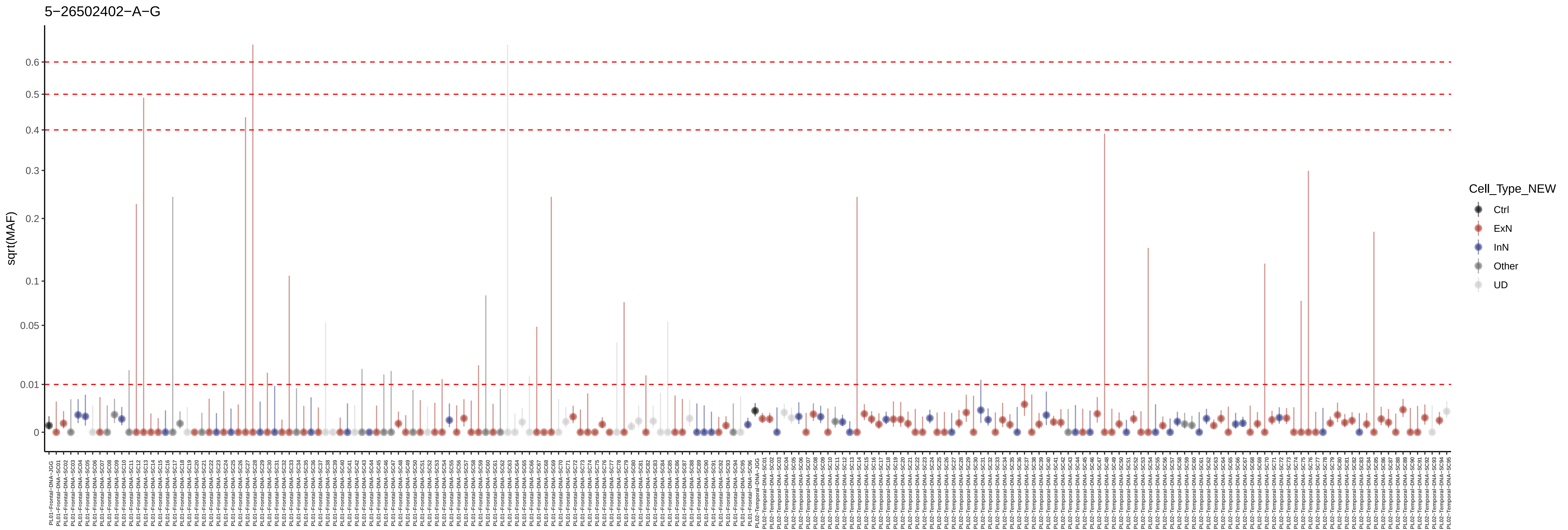

5−26502402−A−G
sqrt(MAF)
0.6
0.5
0.4
0.3
0.2
0.1
0.05
0.01
0
Cell_Type_NEW
Ctrl
ExN
InN
Other
UD

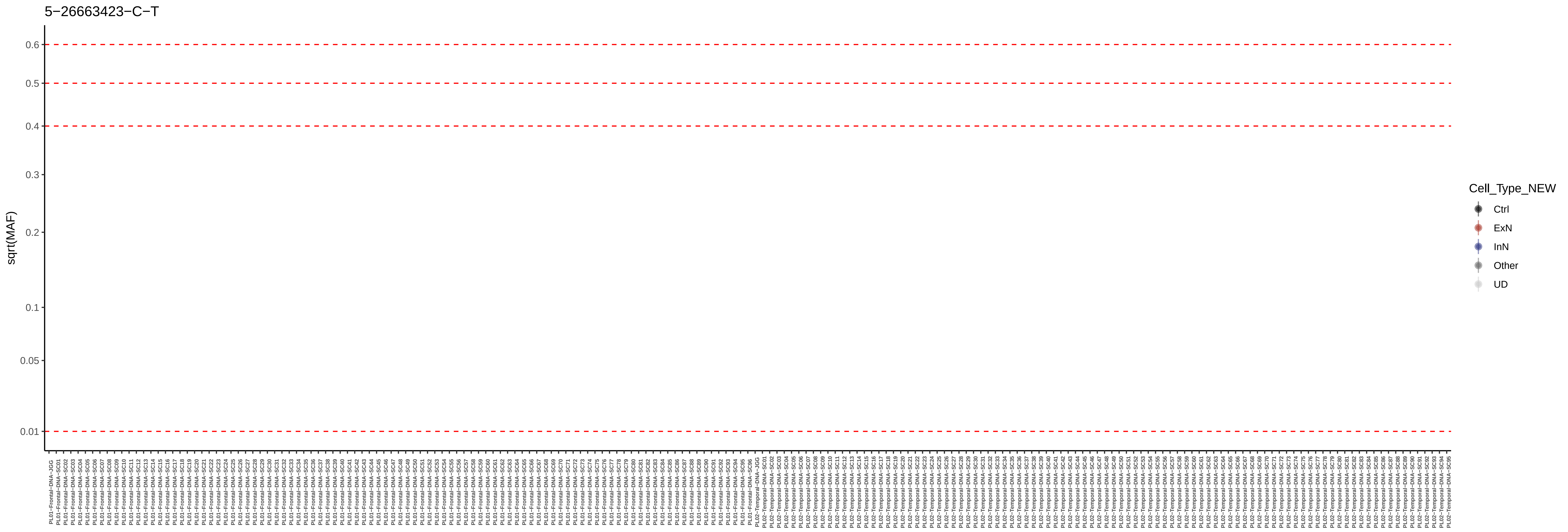
5−26663423−C−T
sqrt(MAF)
0.6
0.5
0.4
0.3
0.2
0.1
0.05
0.01
Cell_Type_NEW
Ctrl
ExN
InN
Other
UD

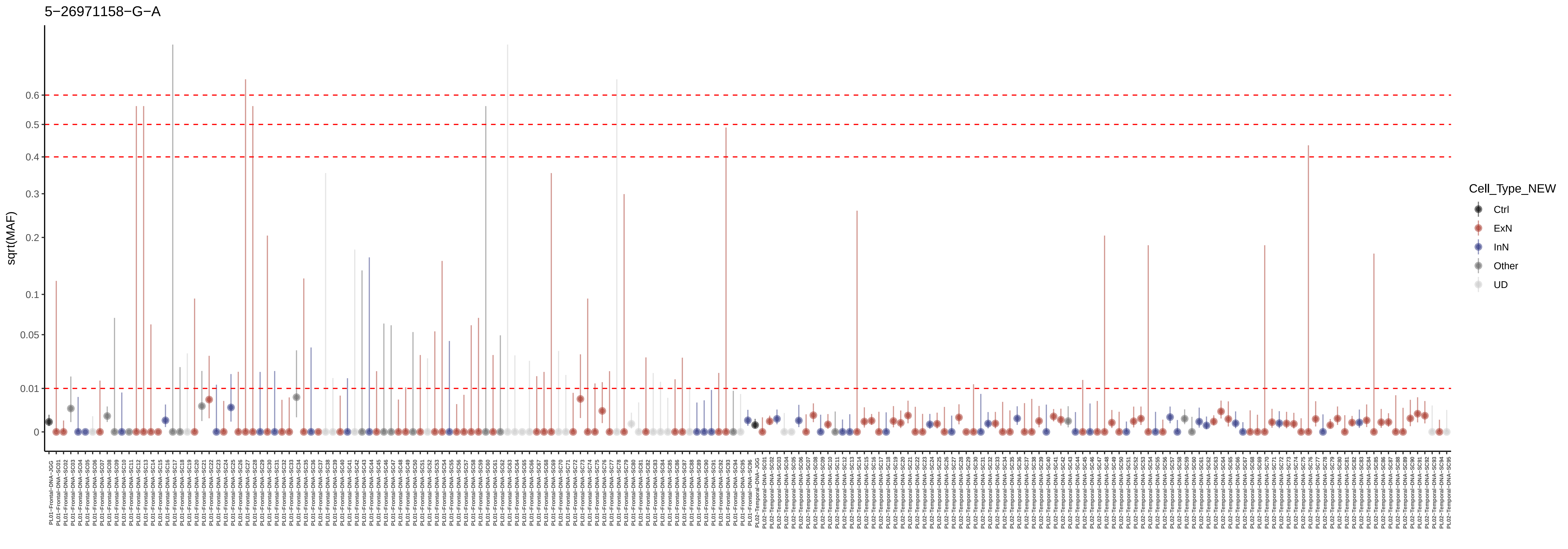
5−26971158−G−A
sqrt(MAF)
0.6
0.5
0.4
0.3
0.2
0.1
0.05
0.01
0
Cell_Type_NEW
Ctrl
ExN
InN
Other
UD

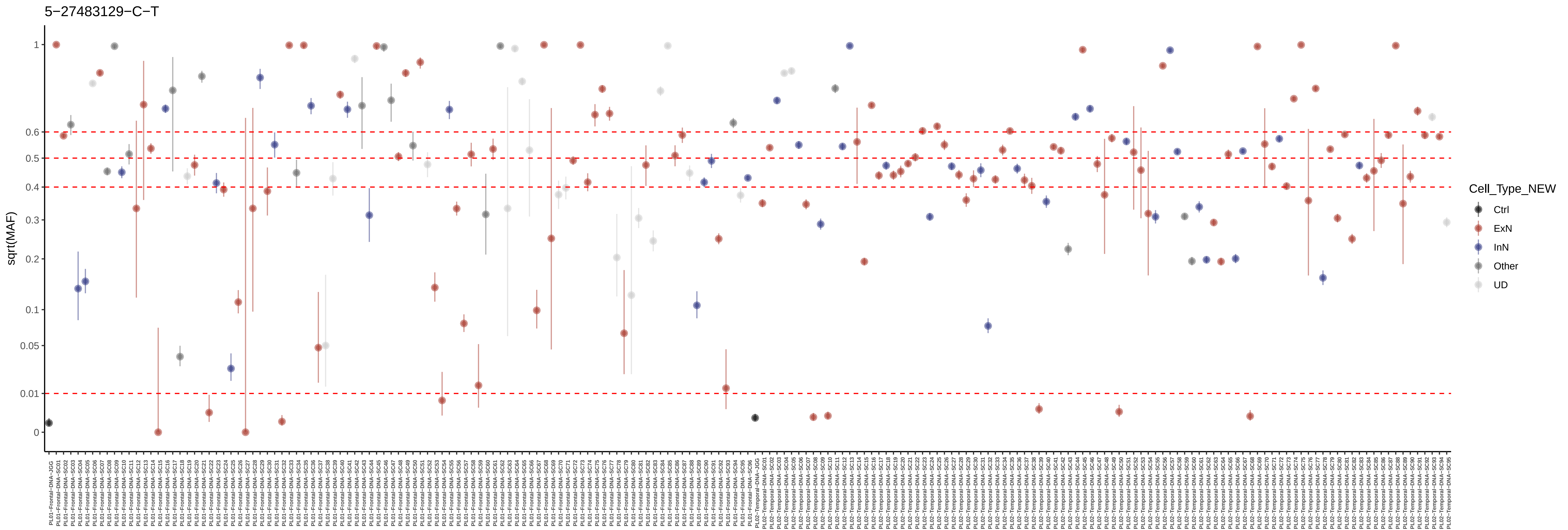

5−27483129−C−T
sqrt(MAF)
1
0.6
0.5
0.4
0.3
0.2
0.1
0.05
0.01
0
Cell_Type_NEW
Ctrl
ExN
InN
Other
UD

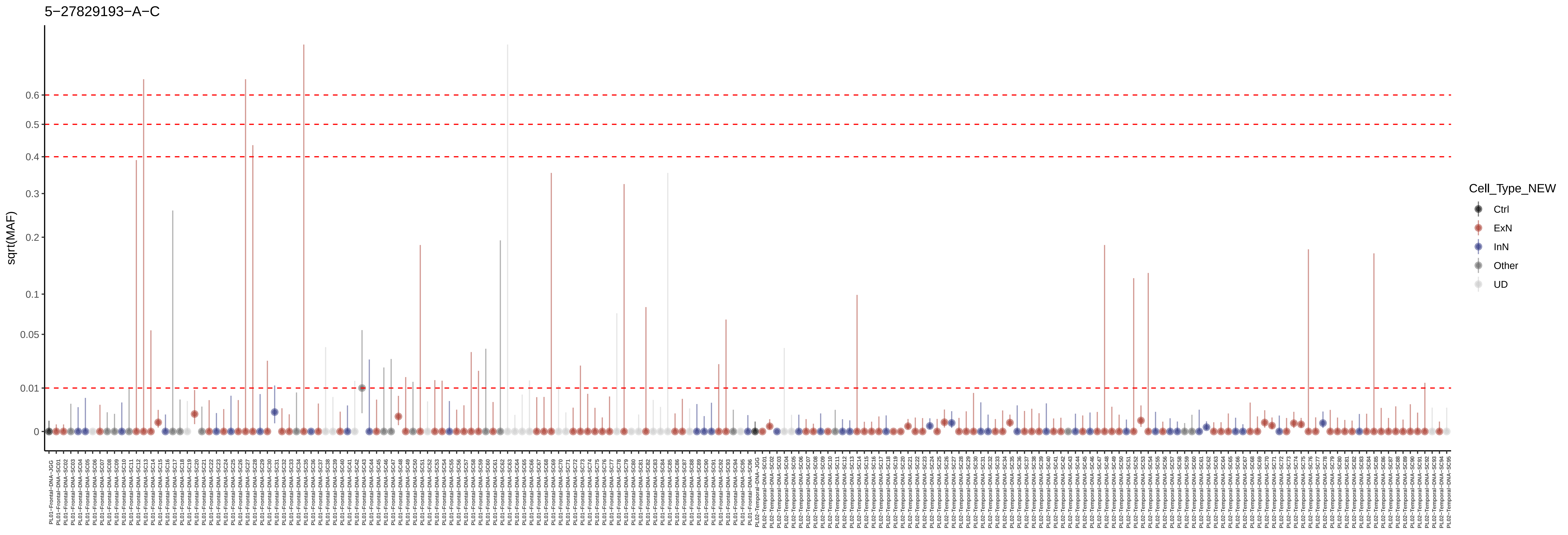
5−27829193−A−C
sqrt(MAF)
0.6
0.5
0.4
0.3
0.2
0.1
0.05
0.01
0
Cell_Type_NEW
Ctrl
ExN
InN
Other
UD

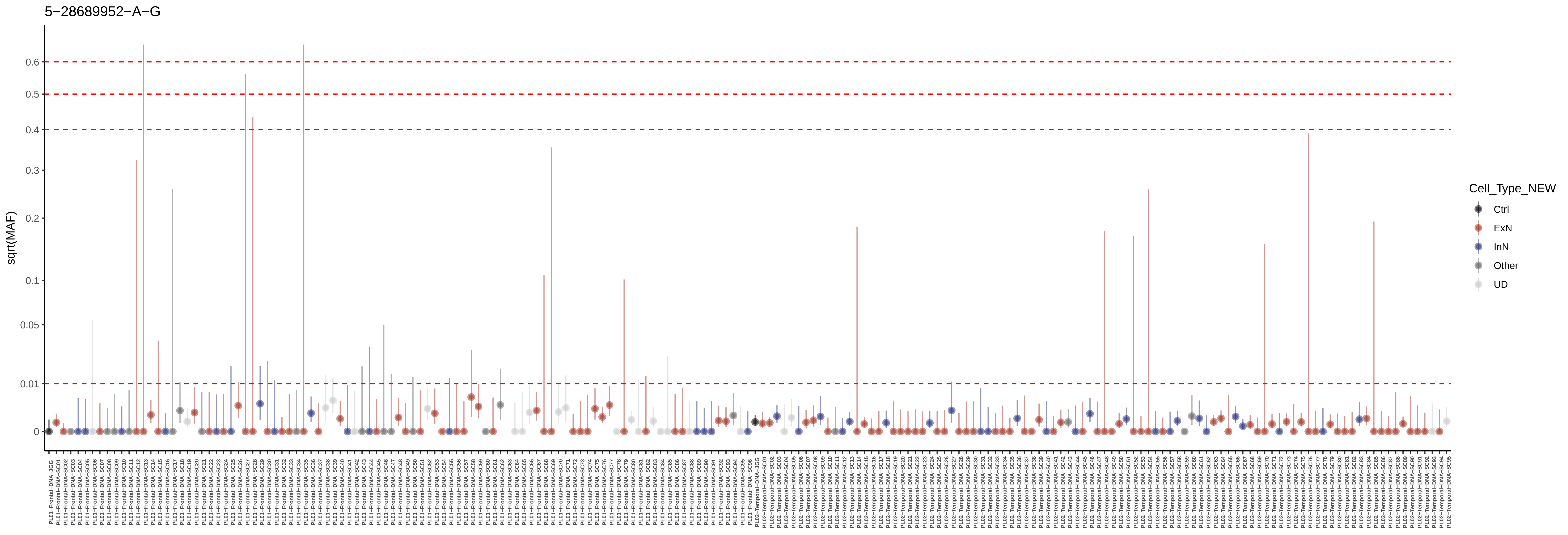
5−28689952−A−G
sqrt(MAF)
0.6
0.5
0.4
0.3
0.2
0.1
0.05
0.01
0
Cell_Type_NEW
Ctrl
ExN
InN
Other
UD

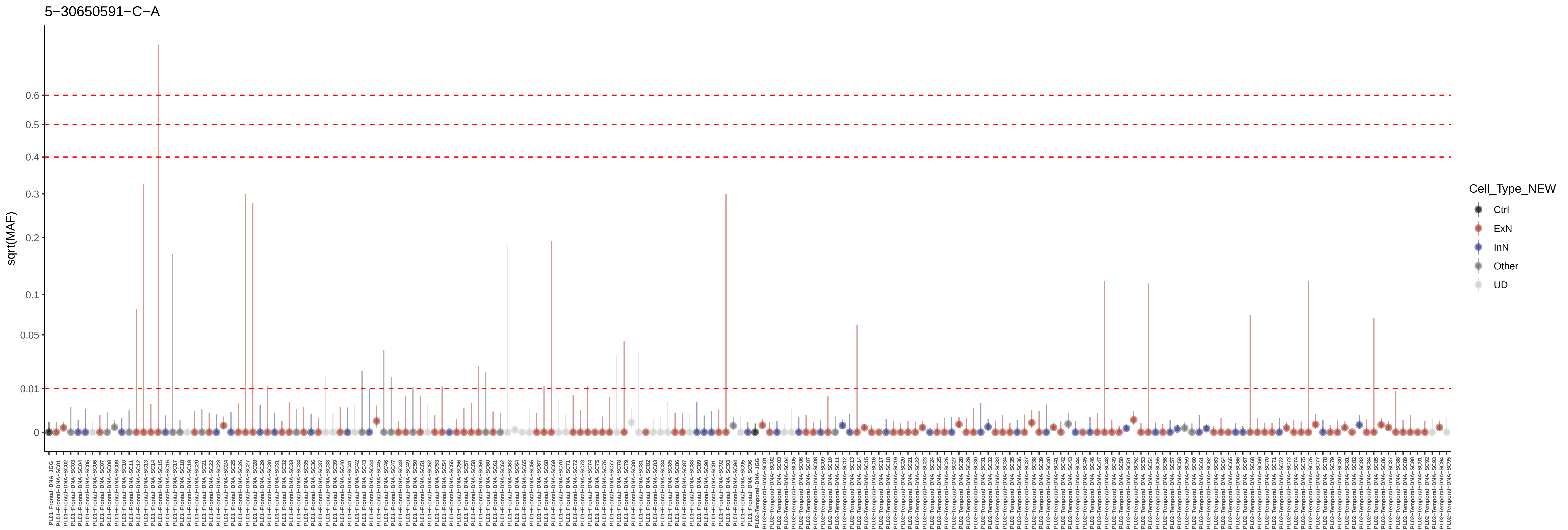
5−30650591−C−A
sqrt(MAF)
0.6
0.5
0.4
0.3
0.2
0.1
0.05
0.01
0
Cell_Type_NEW
Ctrl
ExN
InN
Other
UD

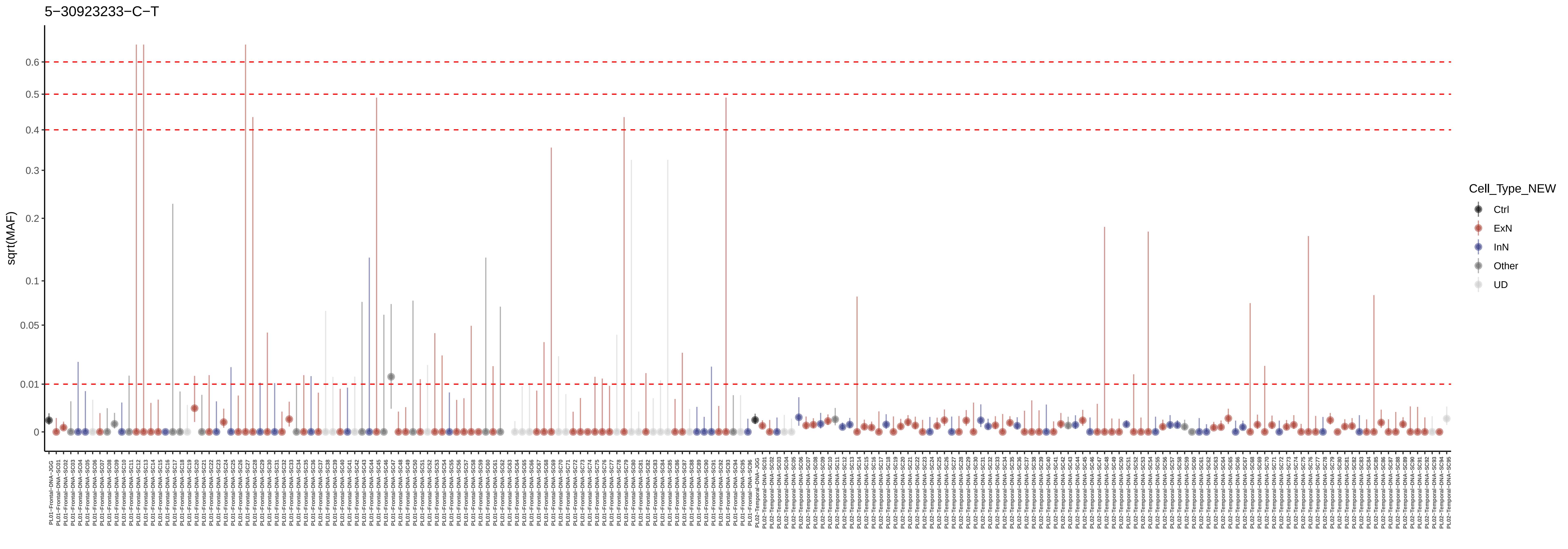
5−30923233−C−T
sqrt(MAF)
0.6
0.5
0.4
0.3
0.2
0.1
0.05
0.01
0
Cell_Type_NEW
Ctrl
ExN
InN
Other
UD

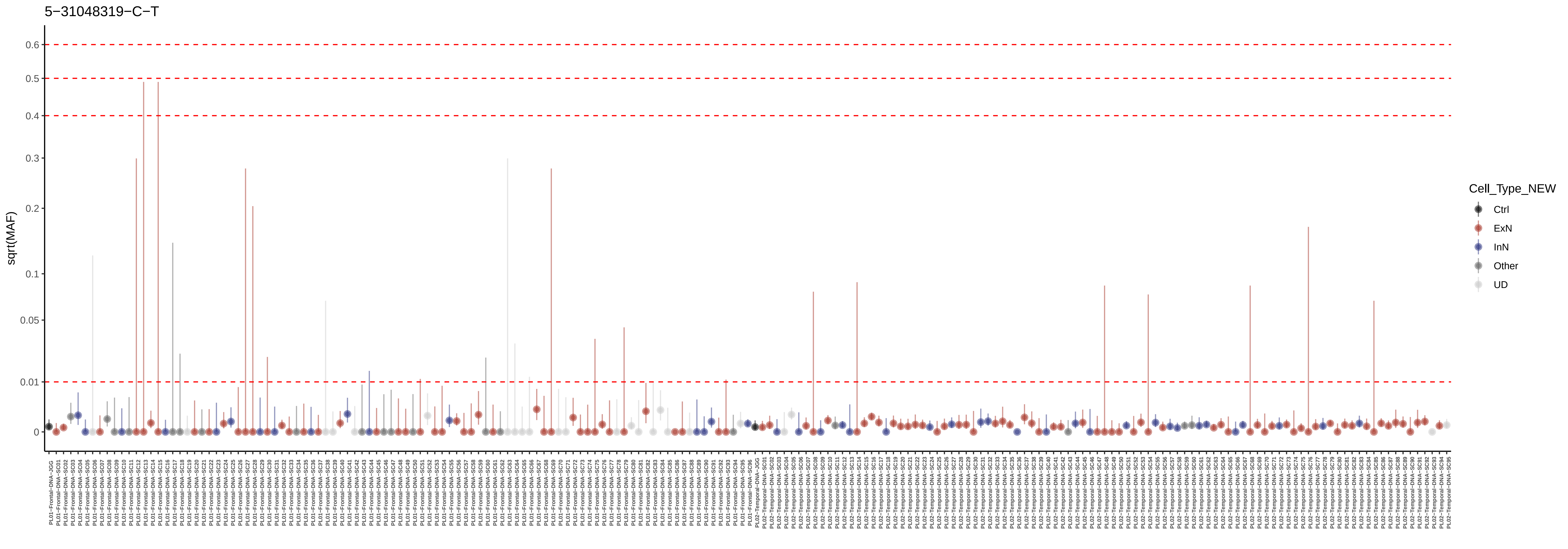
5−31048319−C−T
sqrt(MAF)
0.6
0.5
0.4
0.3
0.2
0.1
0.05
0.01
0
Cell_Type_NEW
Ctrl
ExN
InN
Other
UD

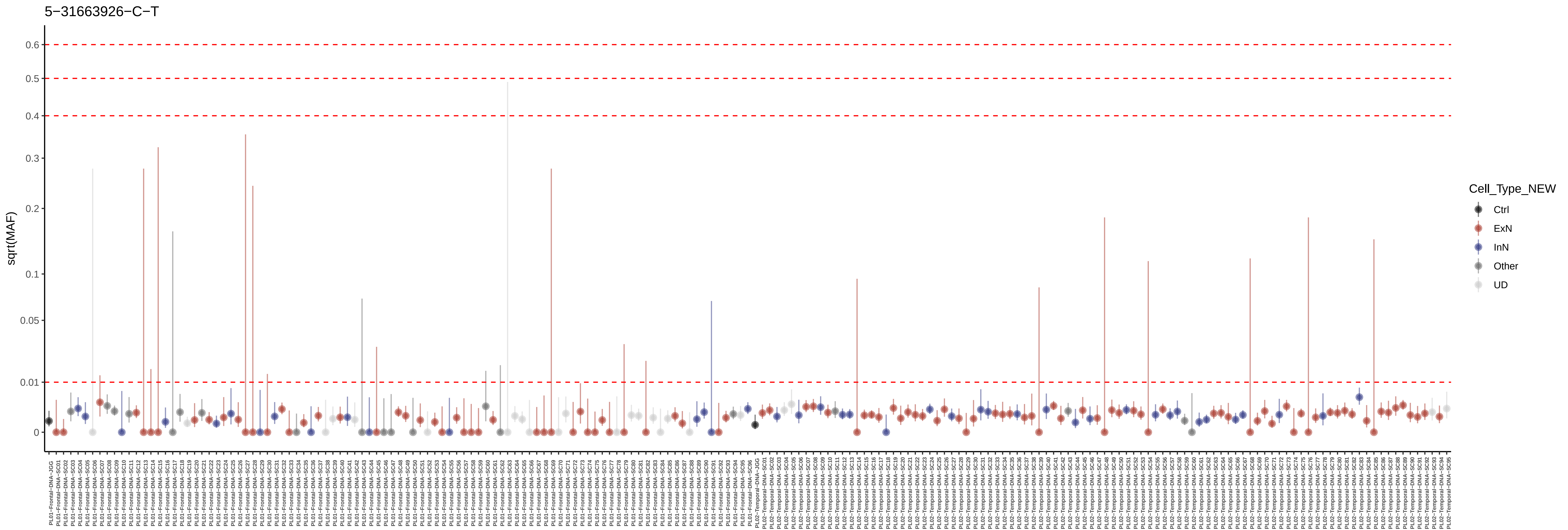

5−31663926−C−T
sqrt(MAF)
0.6
0.5
0.4
0.3
0.2
0.1
0.05
0.01
0
Cell_Type_NEW
Ctrl
ExN
InN
Other
UD

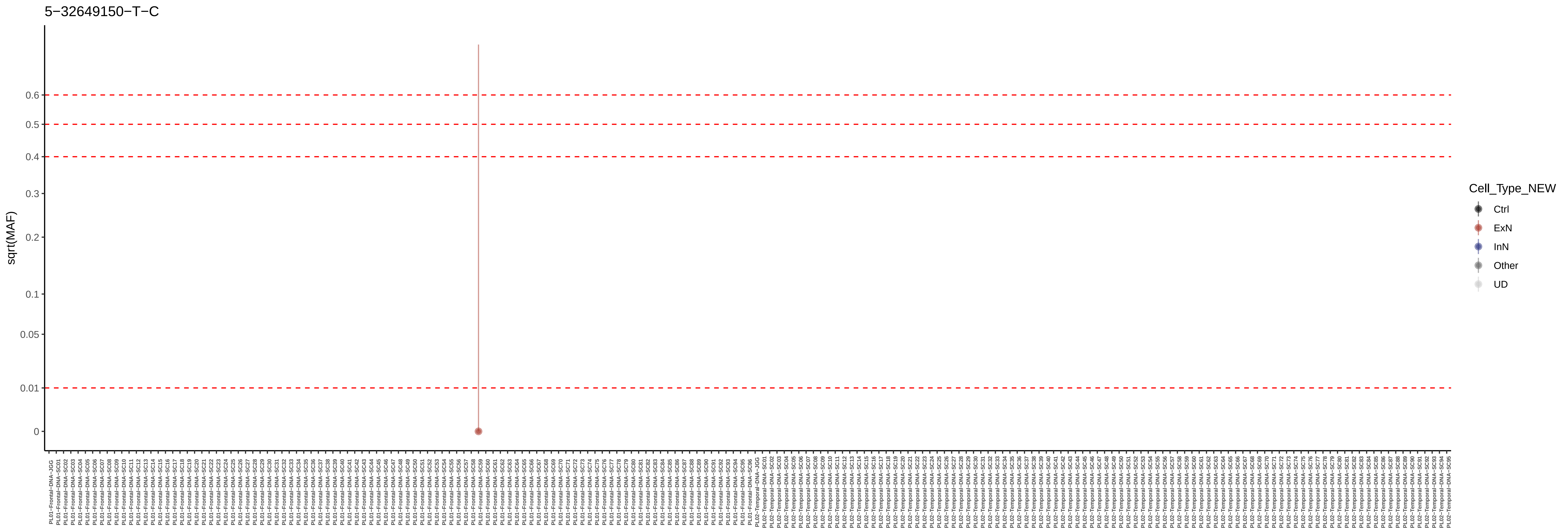
5−32649150−T−C
sqrt(MAF)
0.6
0.5
0.4
0.3
0.2
0.1
0.05
0.01
0
Cell_Type_NEW
Ctrl
ExN
InN
Other
UD

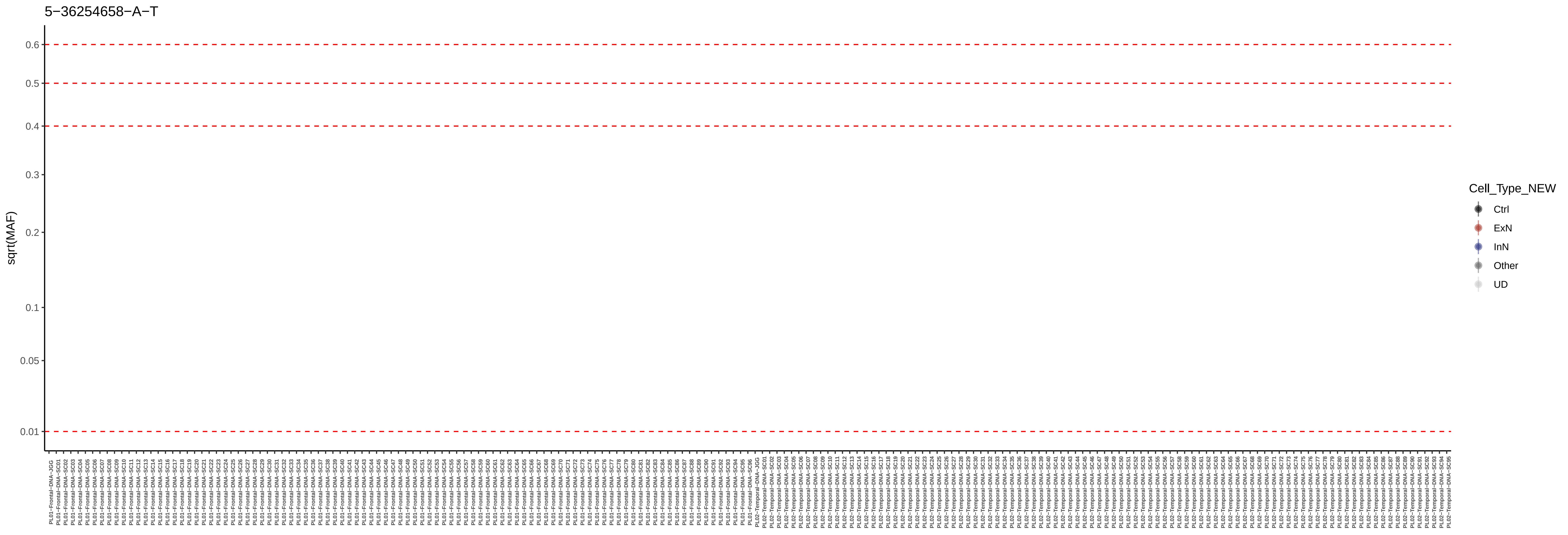
5−36254658−A−T
sqrt(MAF)
0.6
0.5
0.4
0.3
0.2
0.1
0.05
0.01
Cell_Type_NEW
Ctrl
ExN
InN
Other
UD

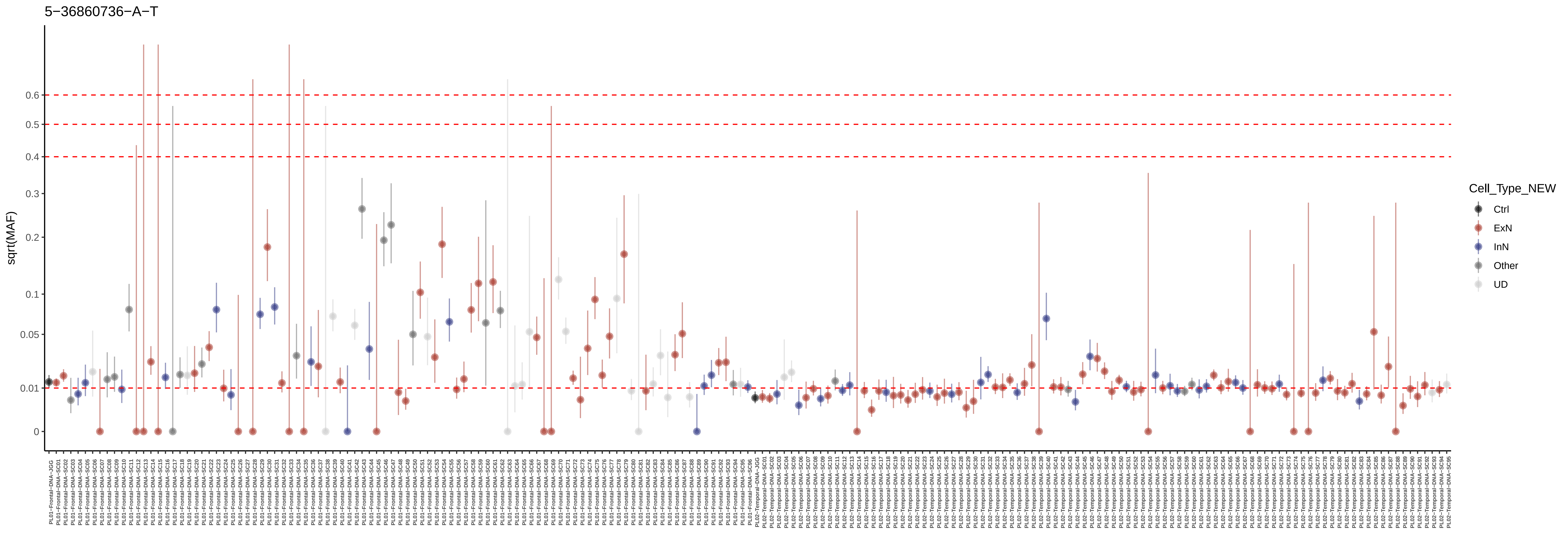
5−36860736−A−T
sqrt(MAF)
0
0.01
0.05
0.1
0.2
0.3
0.4
0.5
0.6
Cell_Type_NEW
Ctrl
ExN
InN
Other
UD

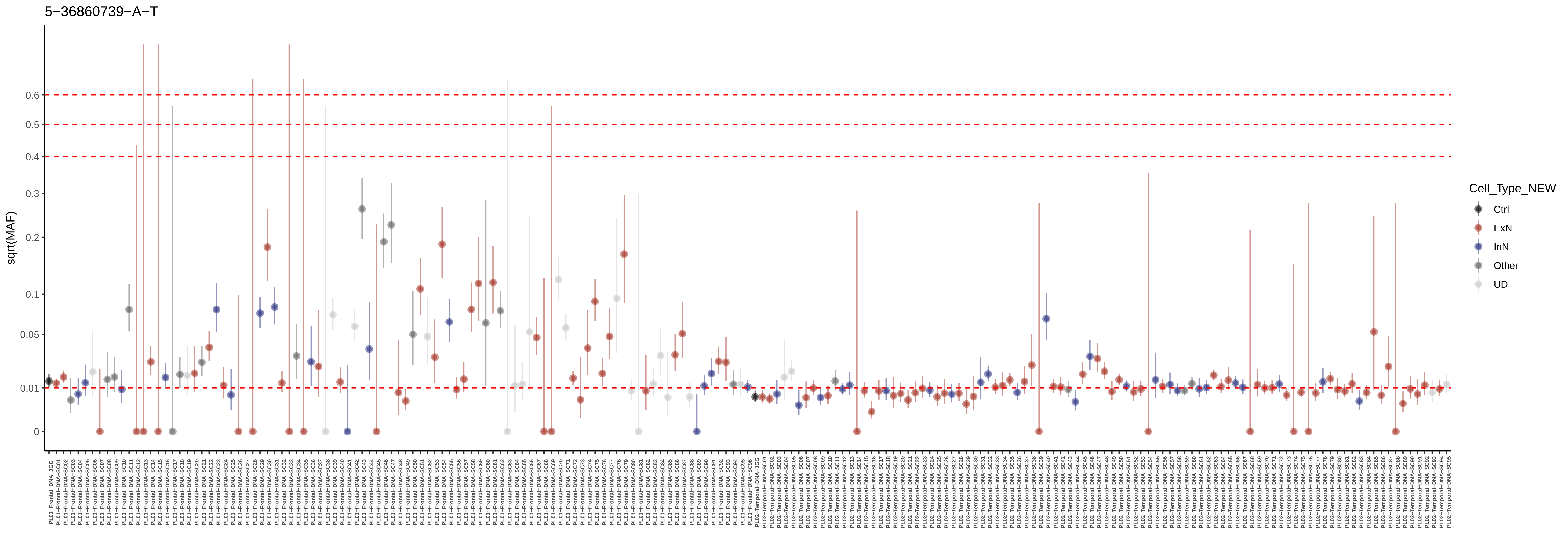

5−36860739−A−T
sqrt(MAF)
0.6
0.5
0.4
0.3
0.2
0.1
0.05
0.01
0
Cell_Type_NEW
Ctrl
ExN
InN
Other
UD

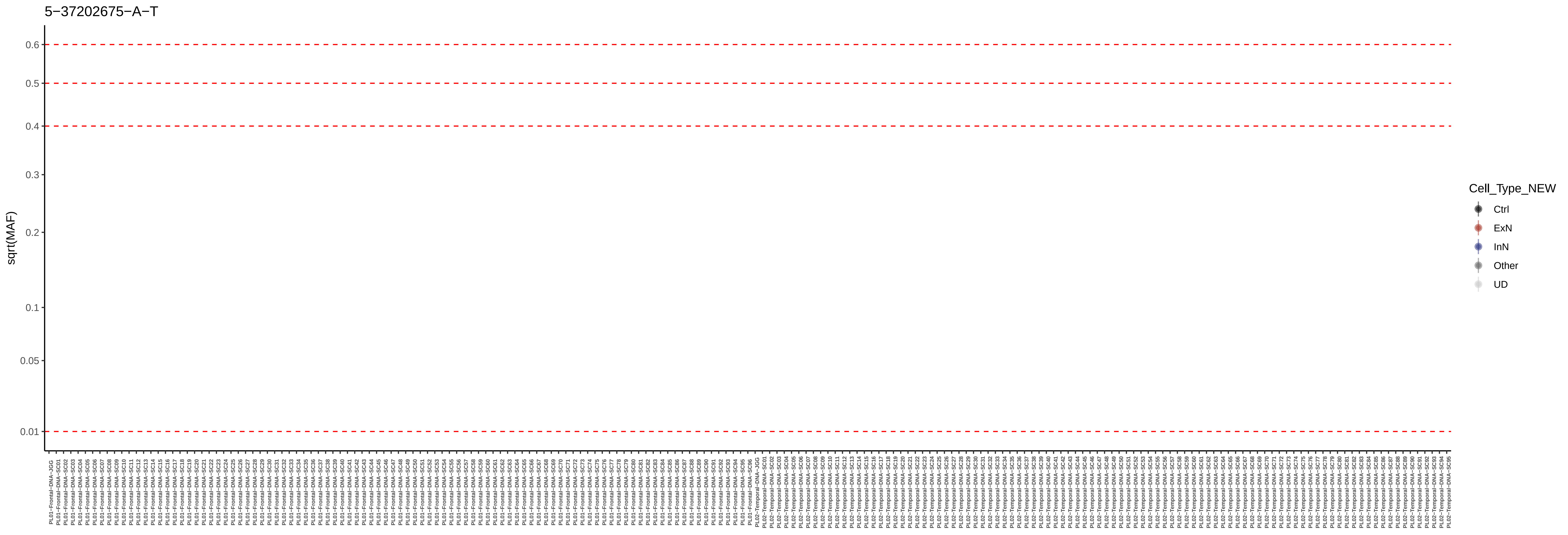
5−37202675−A−T
sqrt(MAF)
0.6
0.5
0.4
0.3
0.2
0.1
0.05
0.01
Cell_Type_NEW
Ctrl
ExN
InN
Other
UD

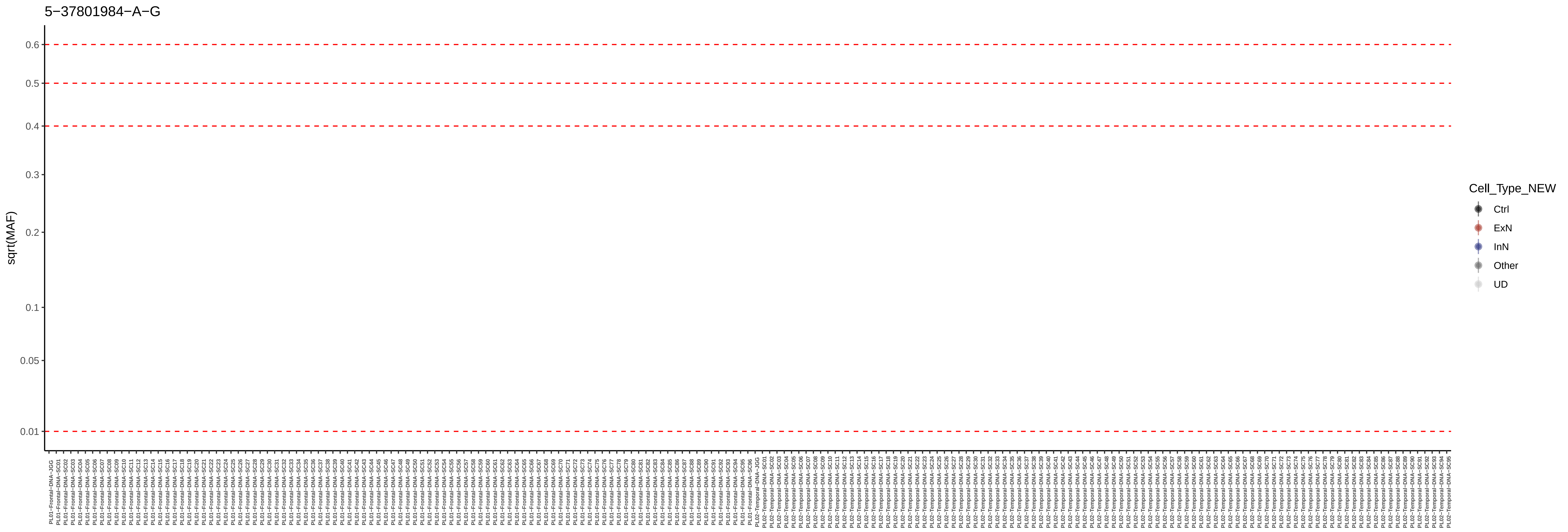
5−37801984−A−G
sqrt(MAF)
0.6
0.5
0.4
0.3
0.2
0.1
0.05
0.01
Cell_Type_NEW
Ctrl
ExN
InN
Other
UD

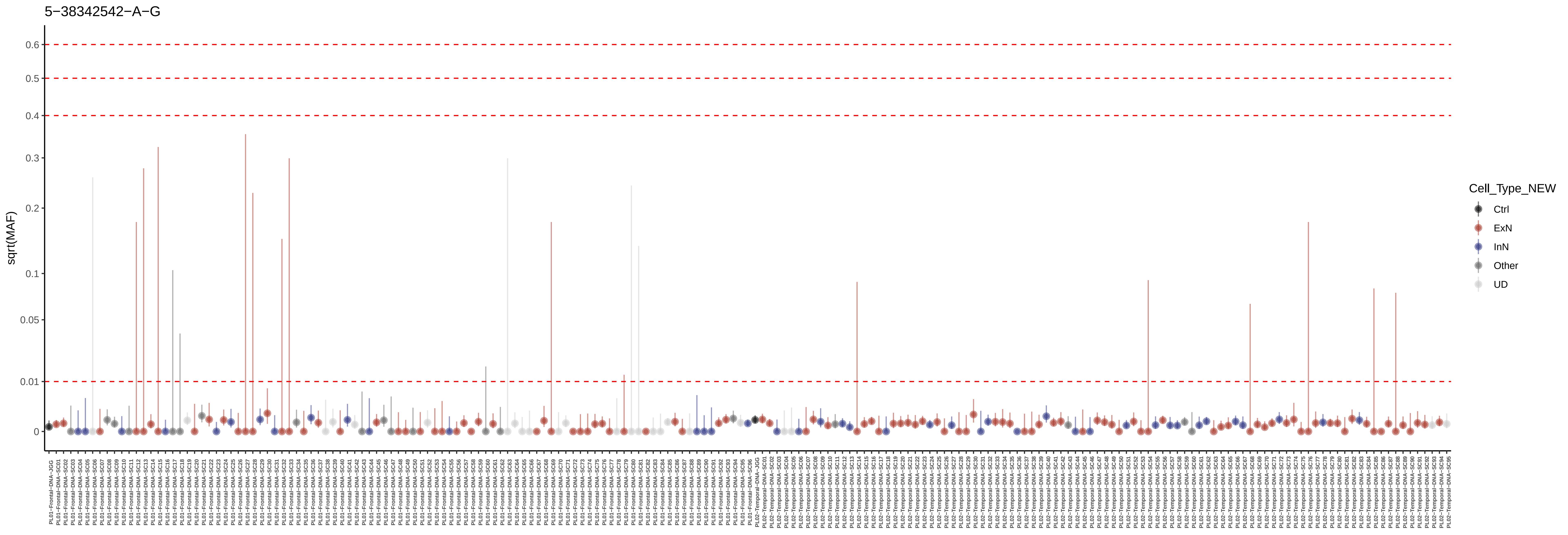

5−38342542−A−G
sqrt(MAF)
0.6
0.5
0.4
0.3
0.2
0.1
0.05
0.01
0
Cell_Type_NEW
Ctrl
ExN
InN
Other
UD

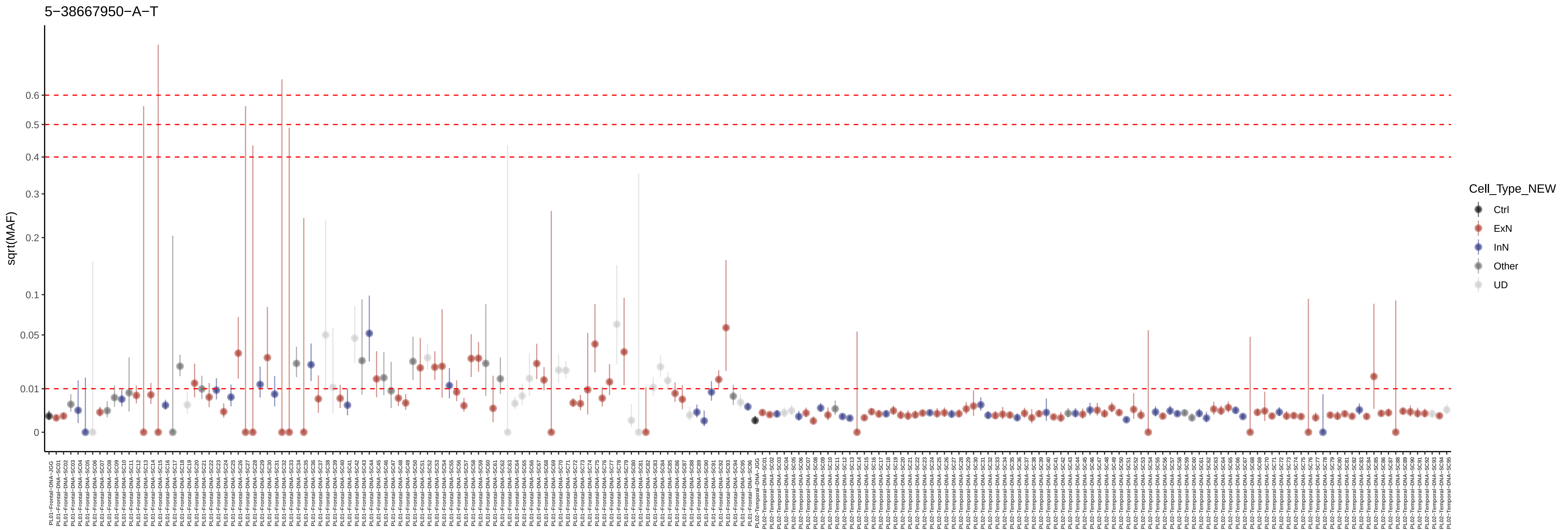

5−38667950−A−T
sqrt(MAF)
0.6
0.5
0.4
0.3
0.2
0.1
0.05
0.01
0
Cell_Type_NEW
Ctrl
ExN
InN
Other
UD

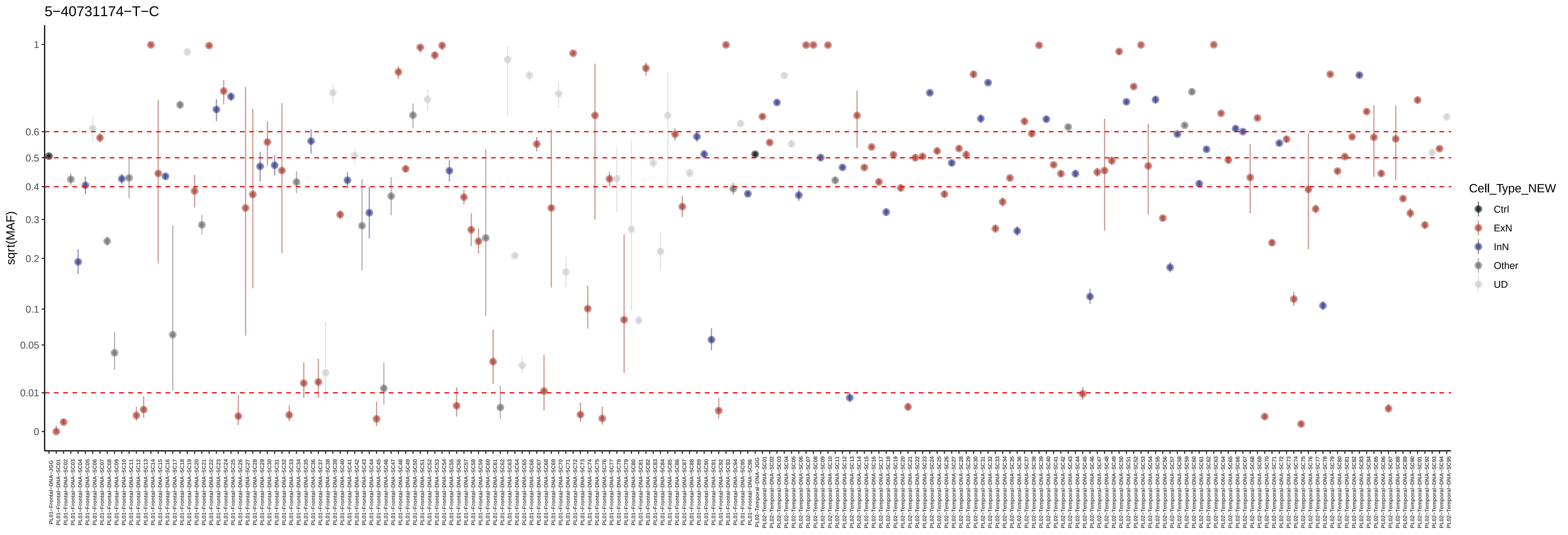

5−40731174−T−C
sqrt(MAF)
1
0.6
0.5
0.4
0.3
0.2
0.1
0.05
0.01
0
Cell_Type_NEW
Ctrl
ExN
InN
Other
UD

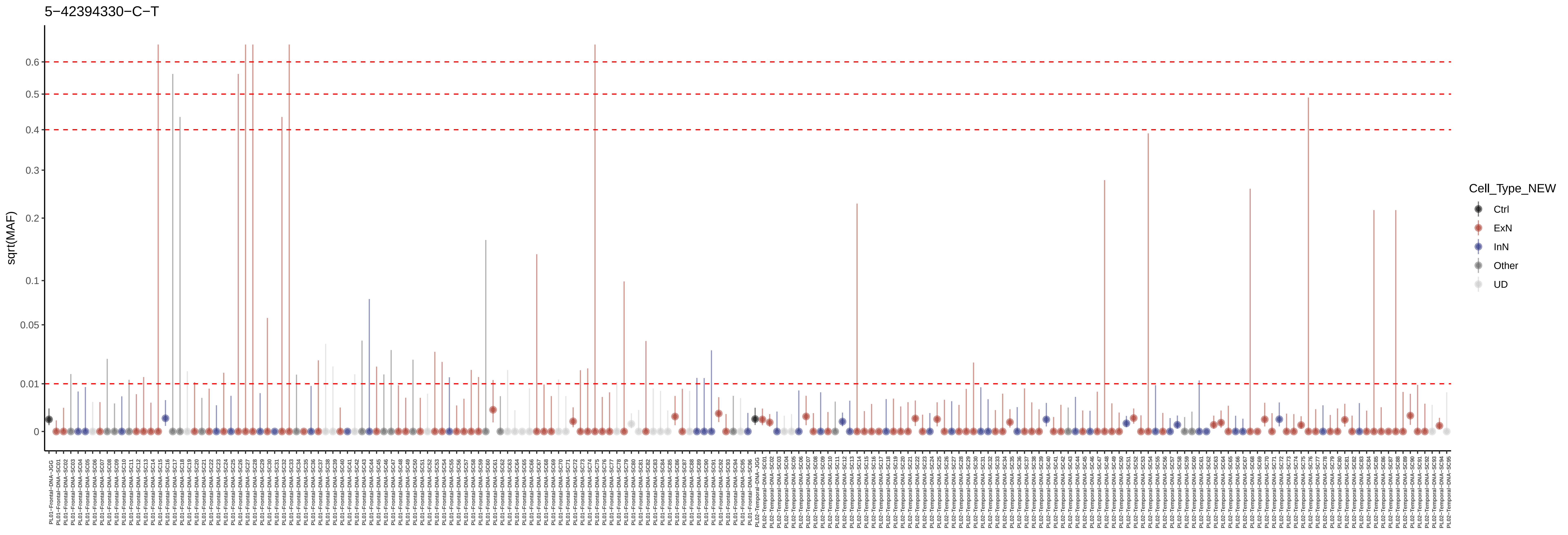

5−42394330−C−T
sqrt(MAF)
0.6
0.5
0.4
0.3
0.2
0.1
0.05
0.01
0
Cell_Type_NEW
Ctrl
ExN
InN
Other
UD

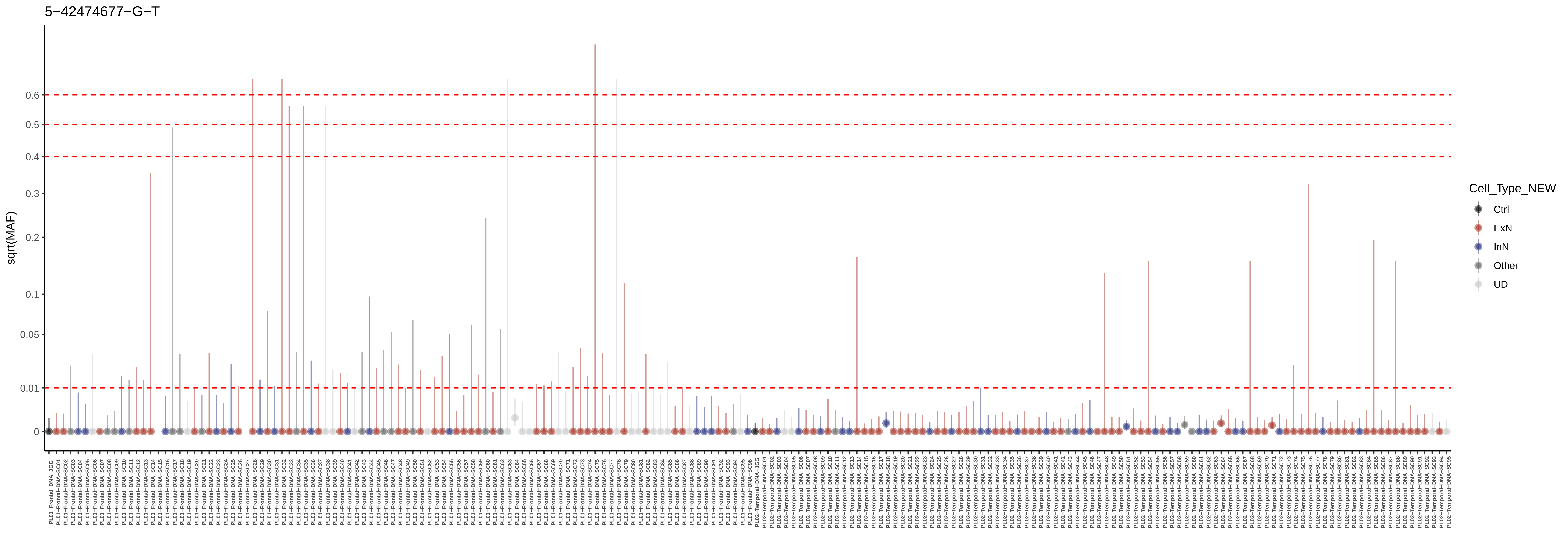

5−42474677−G−T
sqrt(MAF)
Cell_Type_NEW
Ctrl
ExN
InN
Other
UD

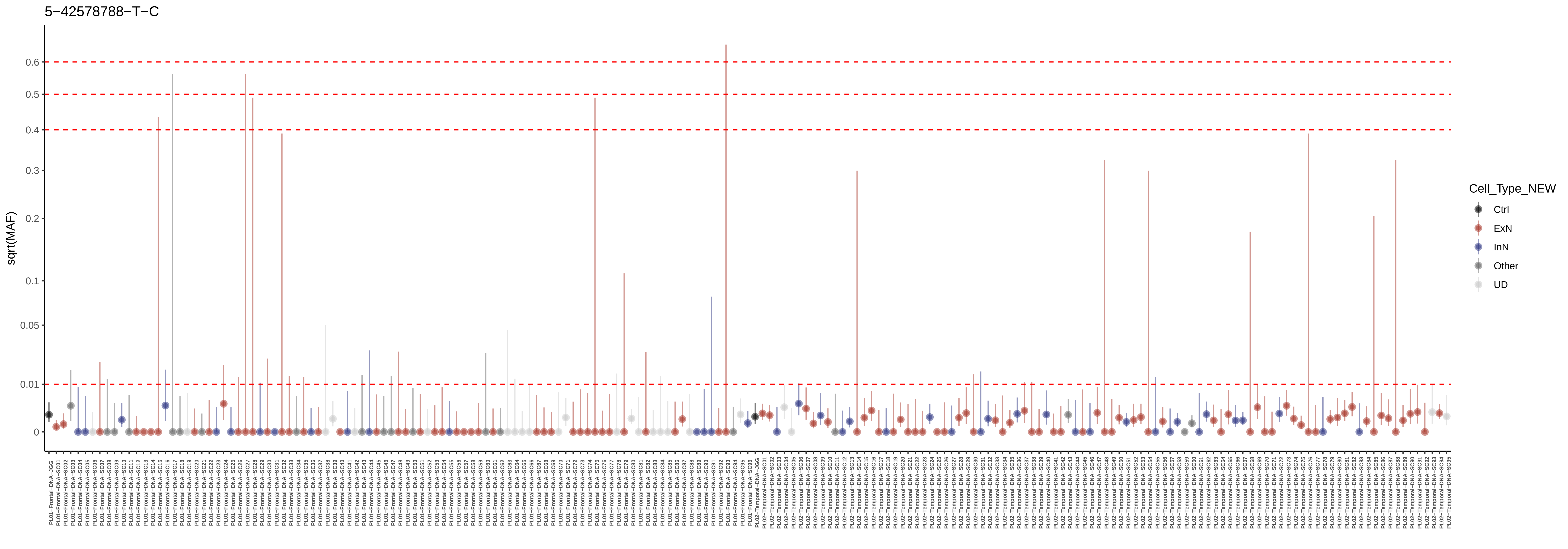
5−42578788−T−C
sqrt(MAF)
0.6
0.5
0.4
0.3
0.2
0.1
0.05
0.01
0
Cell_Type_NEW
Ctrl
ExN
InN
Other
UD

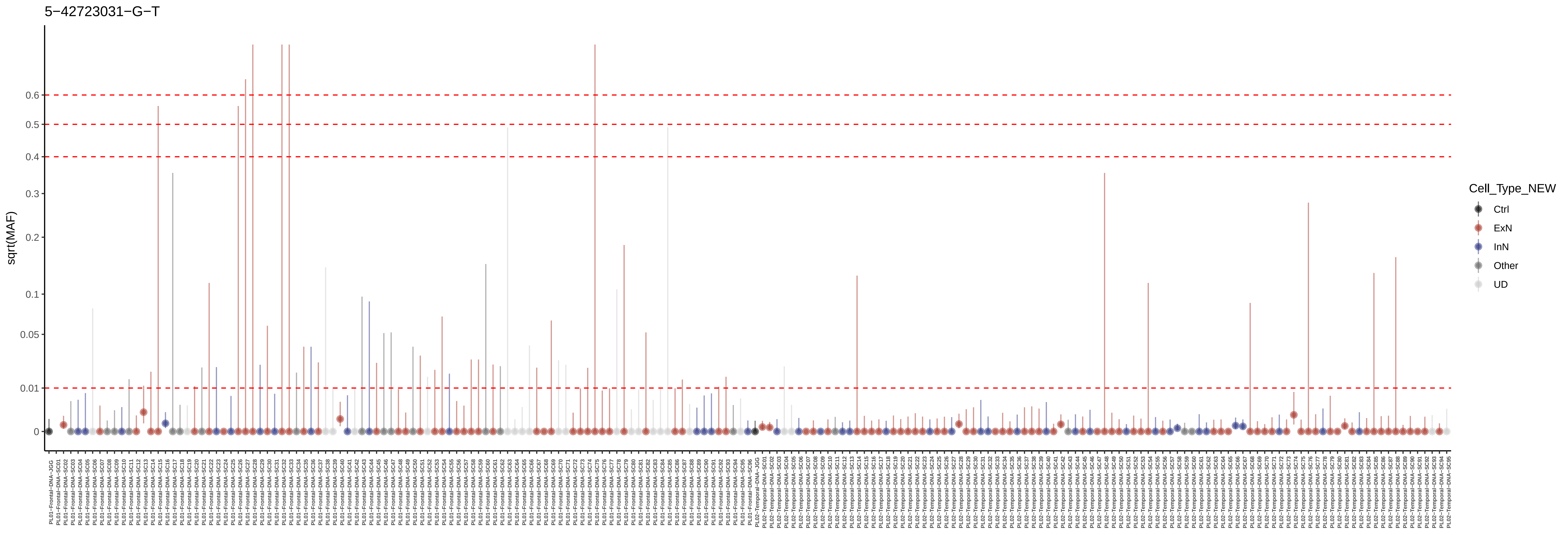

5−42723031−G−T
sqrt(MAF)
0.6
0.5
0.4
0.3
0.2
0.1
0.05
0.01
0
Cell_Type_NEW
Ctrl
ExN
InN
Other
UD

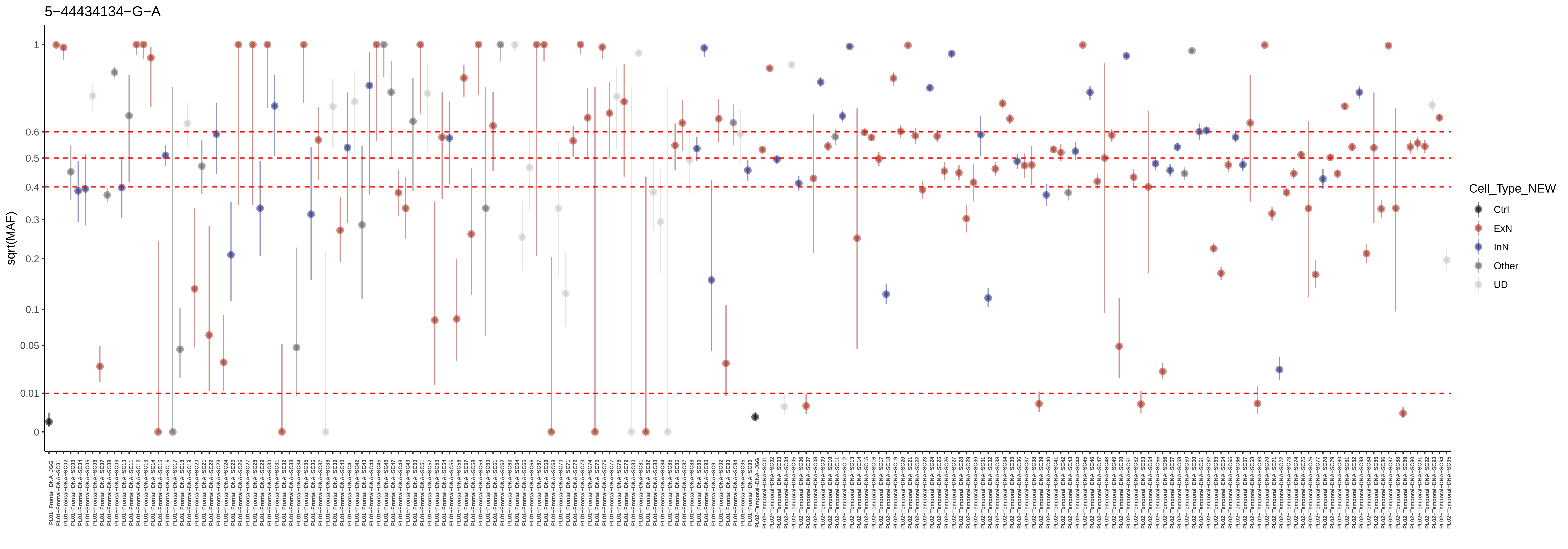

5−44434134−G−A
sqrt(MAF)
1
0.6
0.5
0.4
0.3
0.2
0.1
0.05
0.01
0
Cell_Type_NEW
Ctrl
ExN
InN
Other
UD

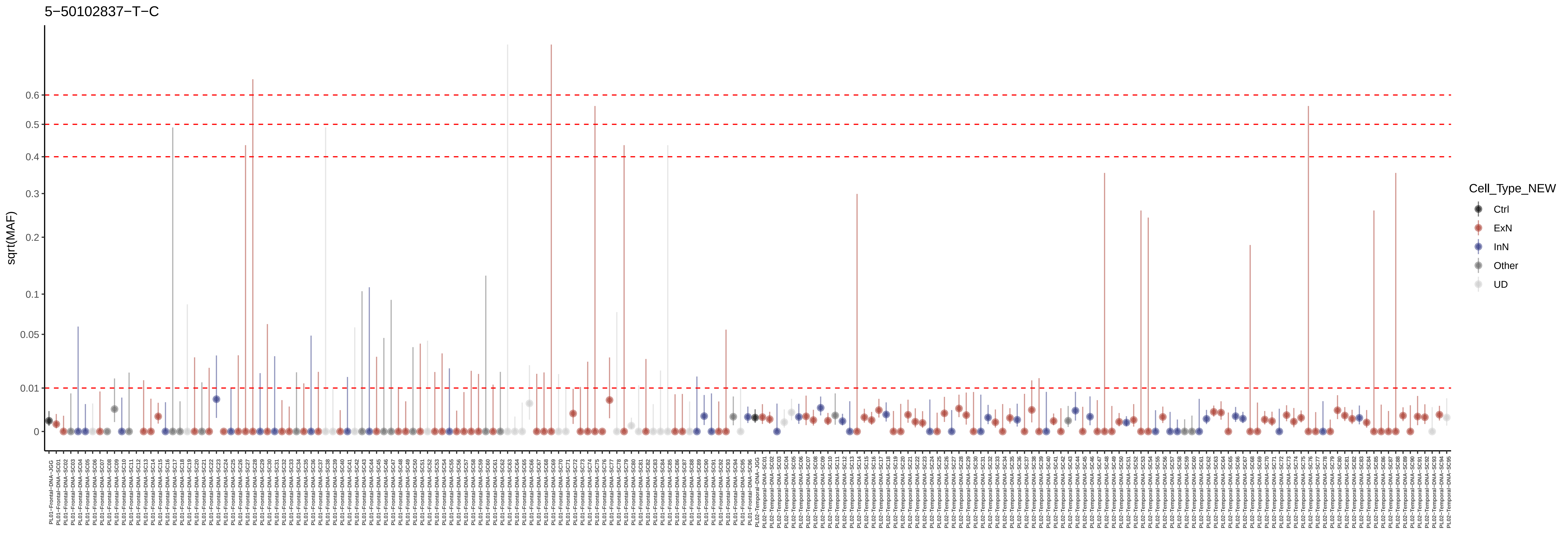

5−50102837−T−C
sqrt(MAF)
0.6
0.5
0.4
0.3
0.2
0.1
0.05
0.01
0
Cell_Type_NEW
Ctrl
ExN
InN
Other
UD

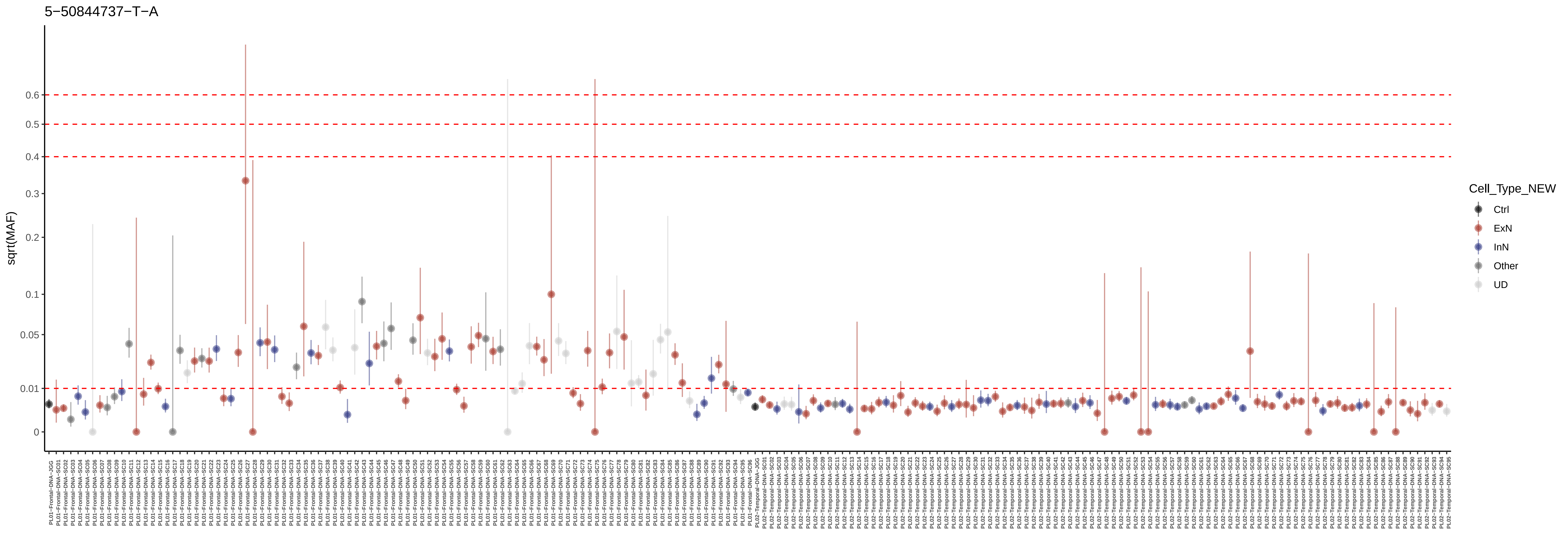
5−50844737−T−A
sqrt(MAF)
0.6
0.5
0.4
0.3
0.2
0.1
0.05
0.01
0
Cell_Type_NEW
Ctrl
ExN
InN
Other
UD

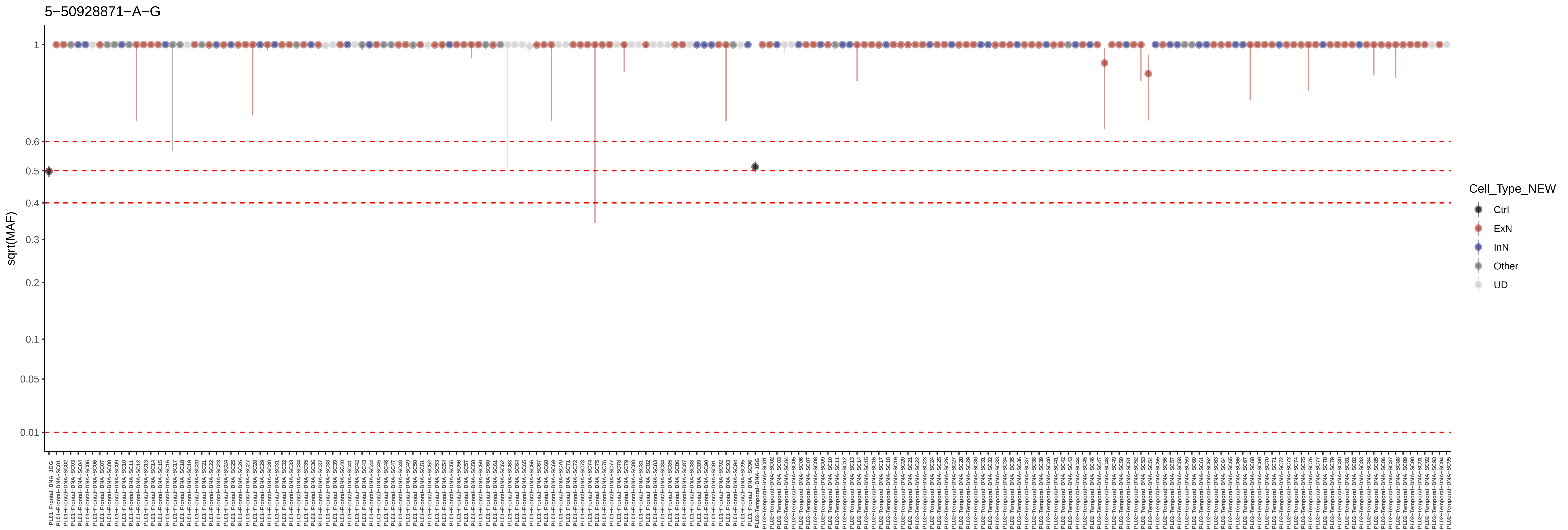
5−50928871−A−G
sqrt(MAF)
1
0.6
0.5
0.4
0.3
0.2
0.1
0.05
0.01
Cell_Type_NEW
Ctrl
ExN
InN
Other
UD

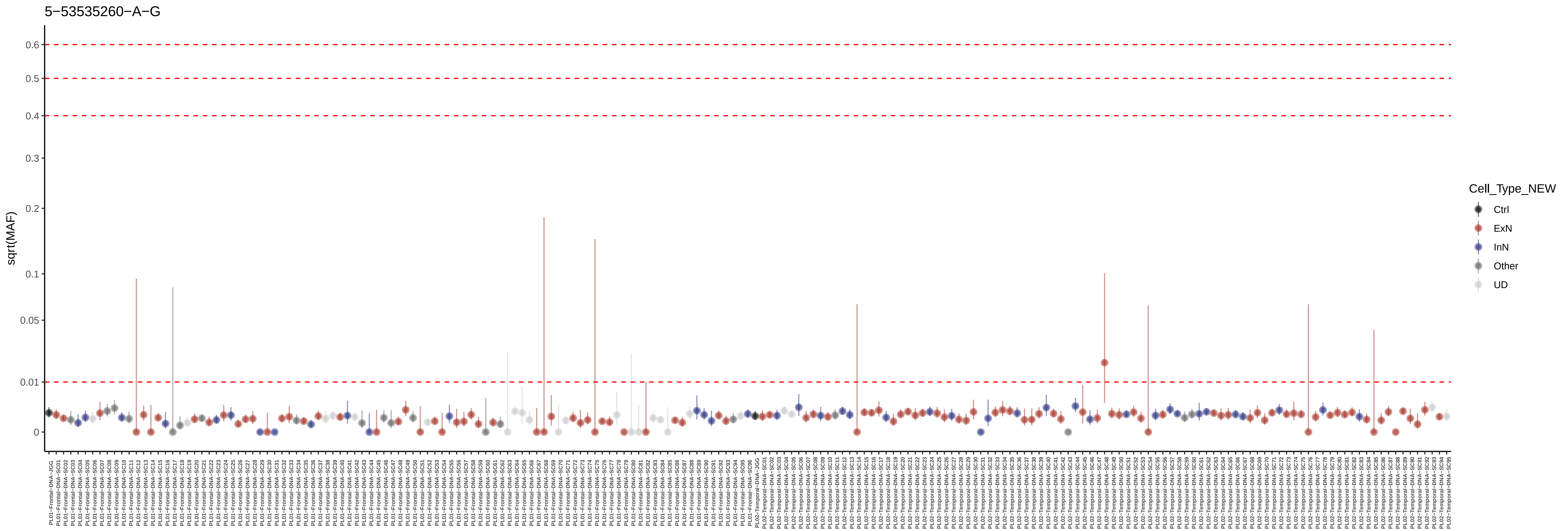
5−53535260−A−G
sqrt(MAF)
0.6
0.5
0.4
0.3
0.2
0.1
0.05
0.01
0
Cell_Type_NEW
Ctrl
ExN
InN
Other
UD

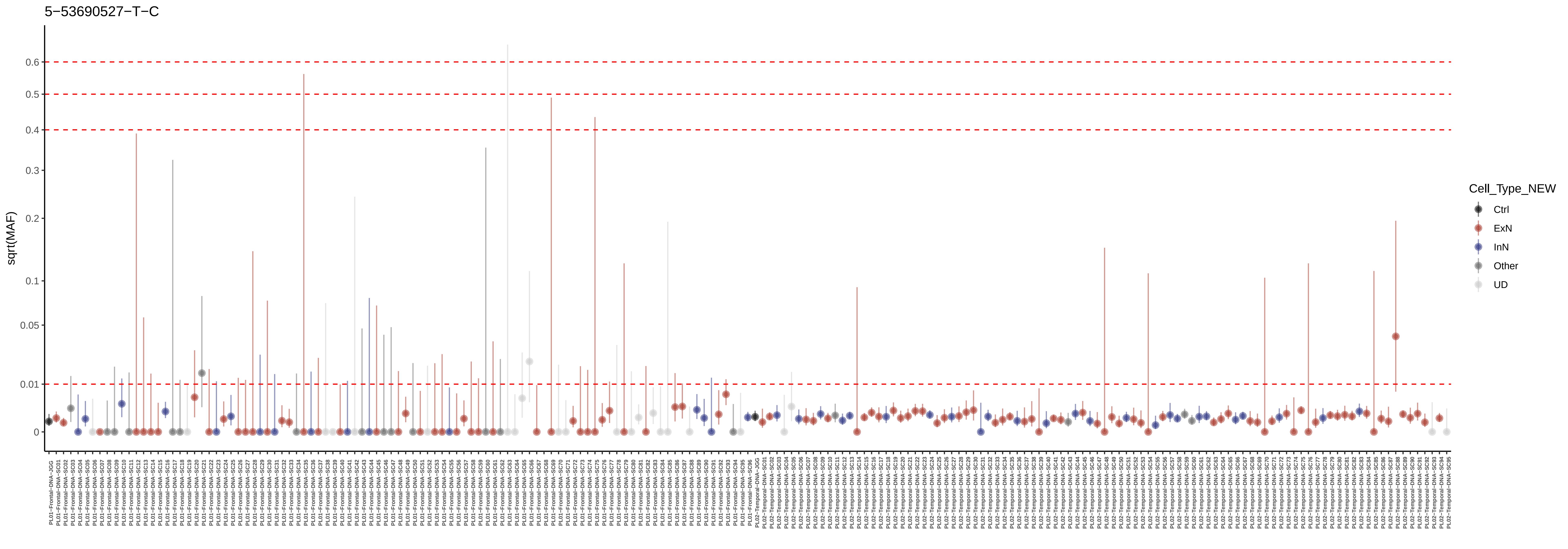
5−53690527−T−C
sqrt(MAF)
0.6
0.5
0.4
0.3
0.2
0.1
0.05
0.01
0
Cell_Type_NEW
Ctrl
ExN
InN
Other
UD

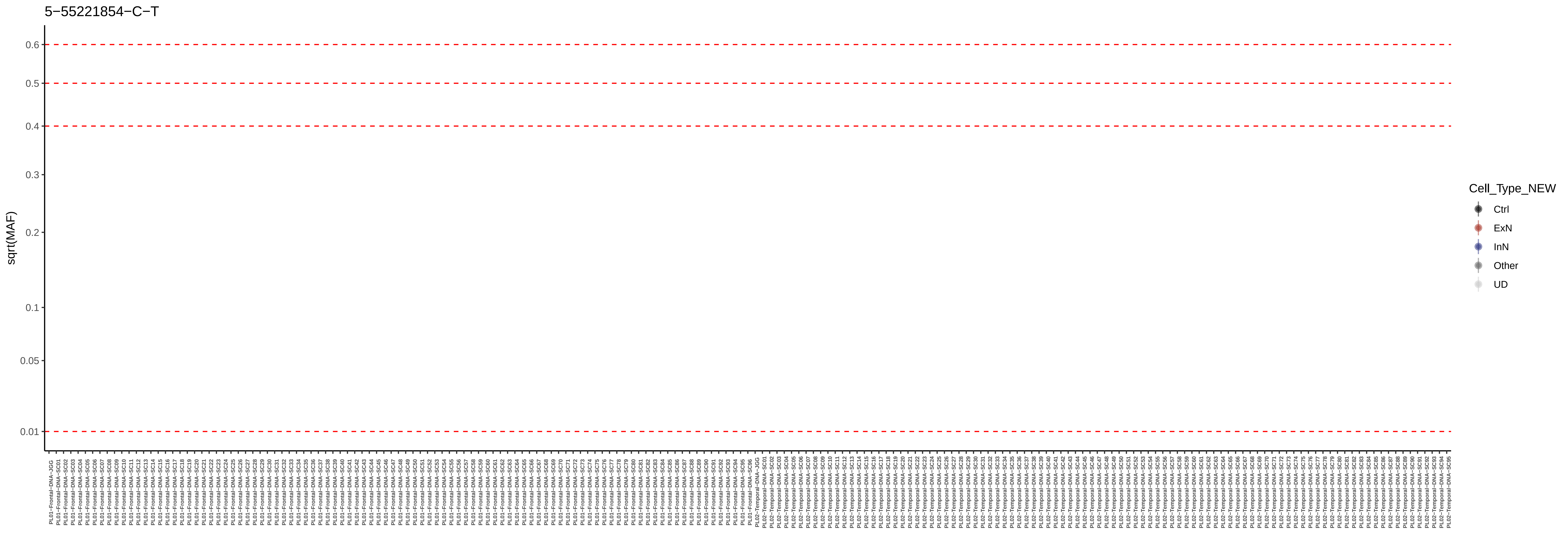

5−55221854−C−T
sqrt(MAF)
0.6
0.5
0.4
0.3
0.2
0.1
0.05
0.01
Cell_Type_NEW
Ctrl
ExN
InN
Other
UD

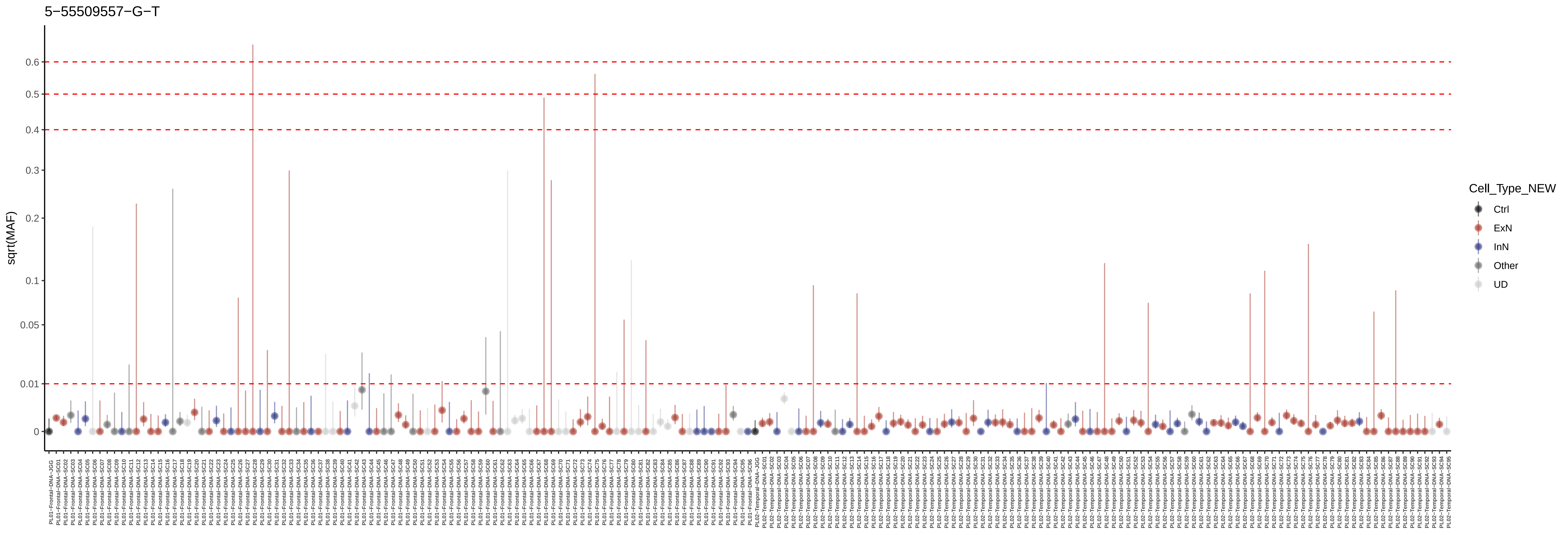
5−55509557−G−T
sqrt(MAF)
0.6
0.5
0.4
0.3
0.2
0.1
0.05
0.01
0
Cell_Type_NEW
Ctrl
ExN
InN
Other
UD

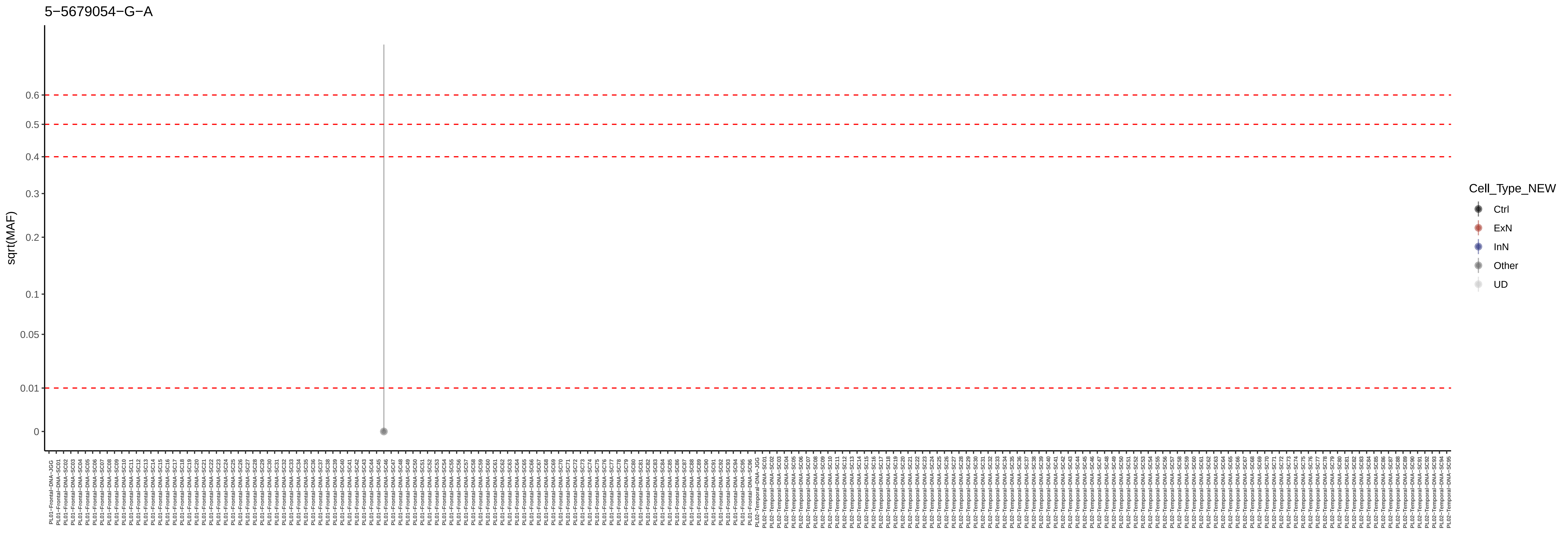

5−5679054−G−A
sqrt(MAF)
0.6
0.5
0.4
0.3
0.2
0.1
0.05
0.01
0
Cell_Type_NEW
Ctrl
ExN
InN
Other
UD

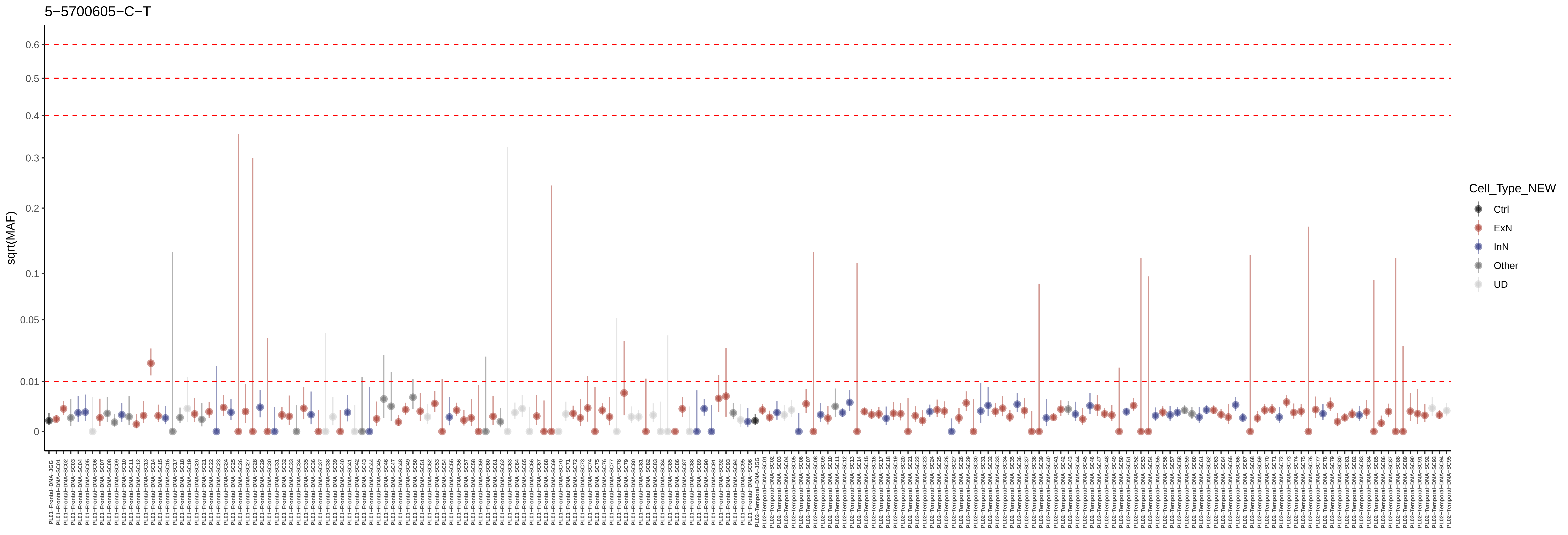

5−5700605−C−T
sqrt(MAF)
0.6
0.5
0.4
0.3
0.2
0.1
0.05
0.01
0
Cell_Type_NEW
Ctrl
ExN
InN
Other
UD

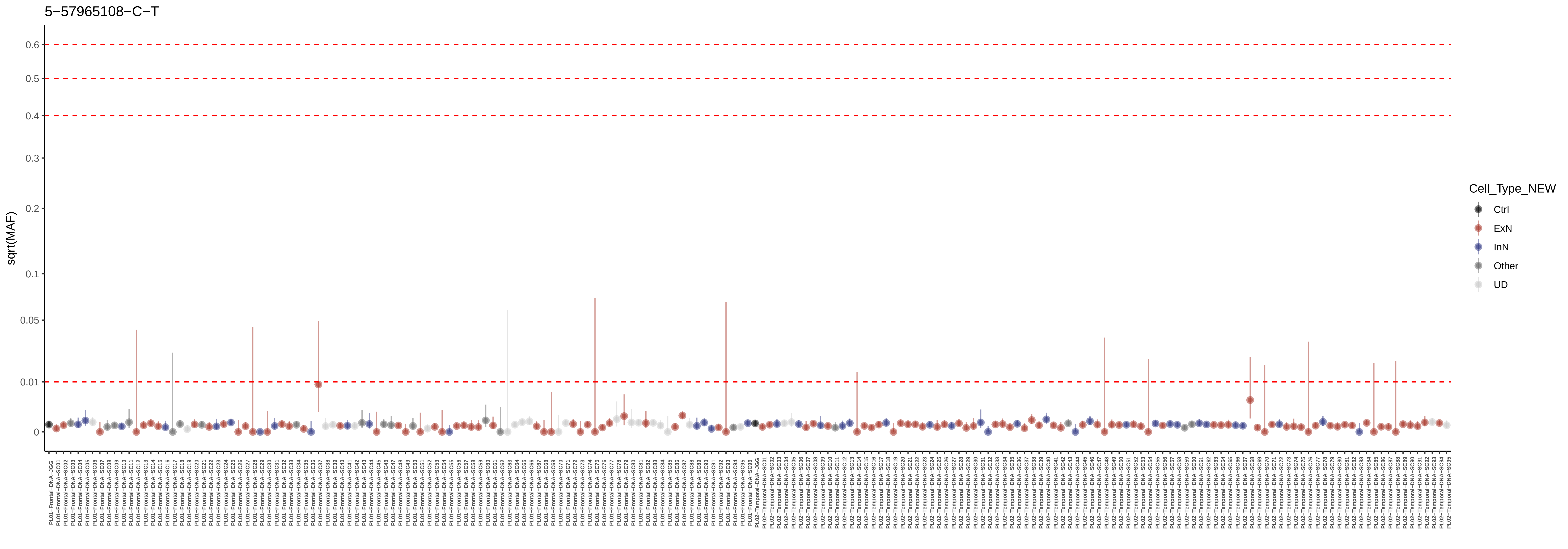

5−57965108−C−T
sqrt(MAF)
0.6
0.5
0.4
0.3
0.2
0.1
0.05
0.01
0
Cell_Type_NEW
Ctrl
ExN
InN
Other
UD

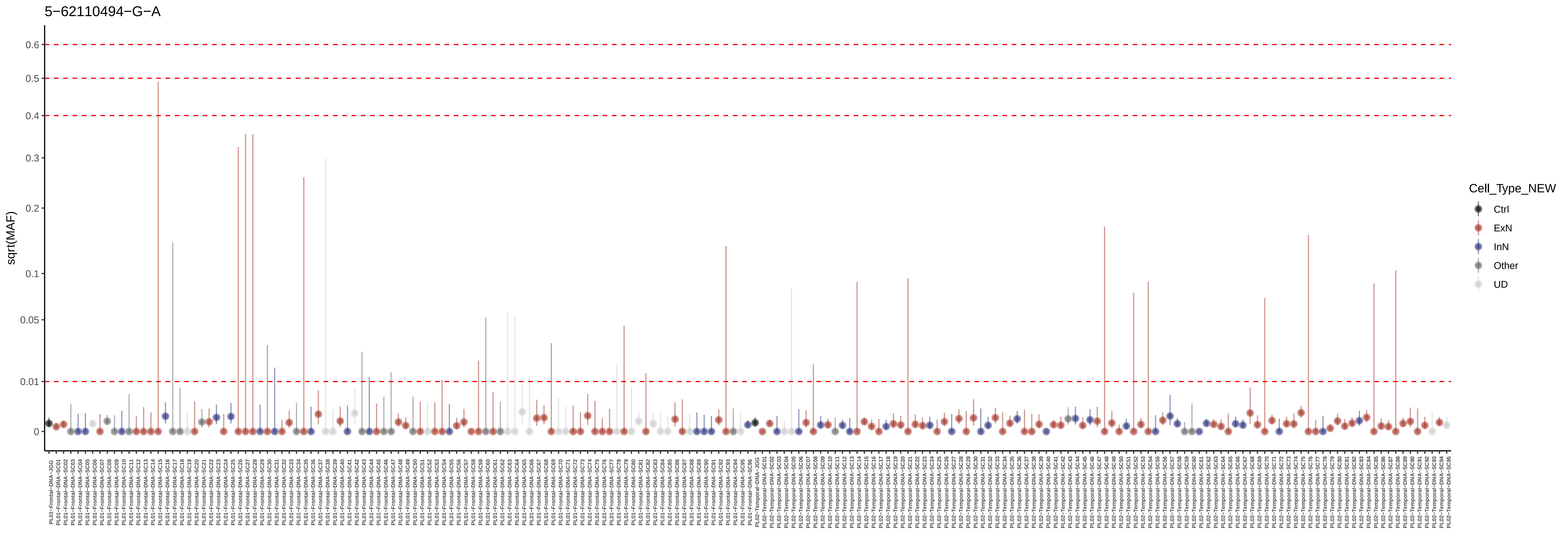

5−62110494−G−A
sqrt(MAF)
0.6
0.5
0.4
0.3
0.2
0.1
0.05
0.01
0
Cell_Type_NEW
Ctrl
ExN
InN
Other
UD

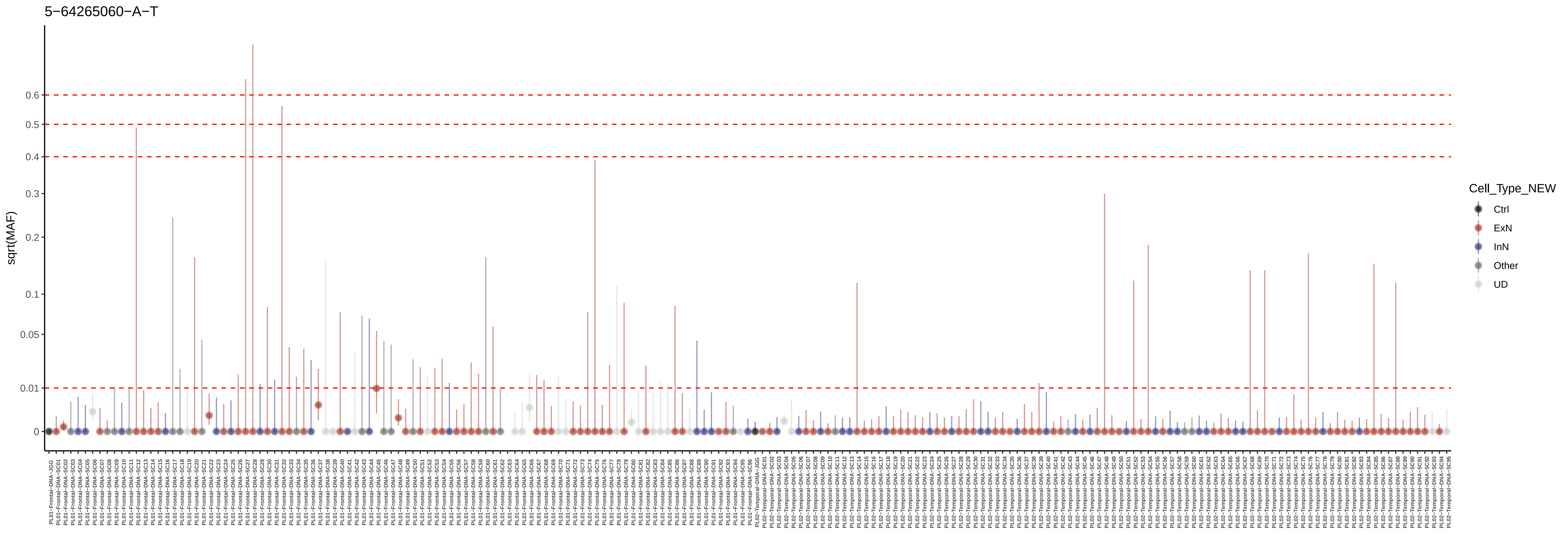

5−64265060−A−T
sqrt(MAF)
0.6
0.5
0.4
0.3
0.2
0.1
0.05
0.01
0
Cell_Type_NEW
Ctrl
ExN
InN
Other
UD

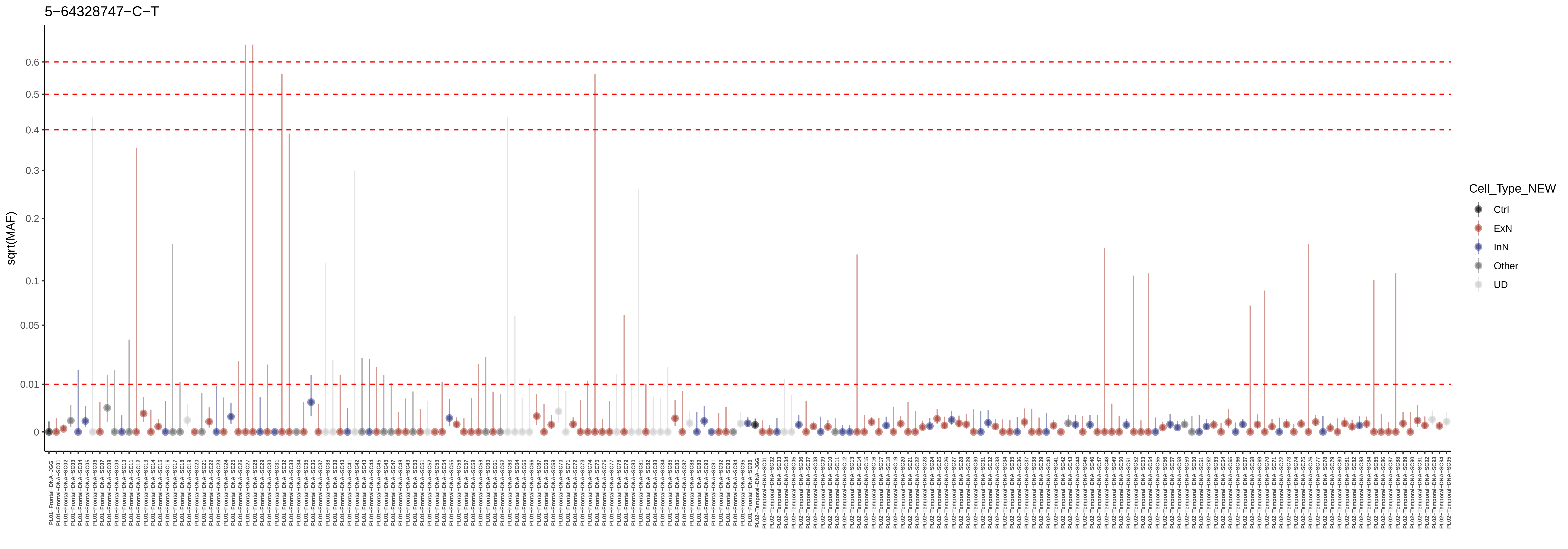
5−64328747−C−T
sqrt(MAF)
0.6
0.5
0.4
0.3
0.2
0.1
0.05
0.01
0
Cell_Type_NEW
Ctrl
ExN
InN
Other
UD

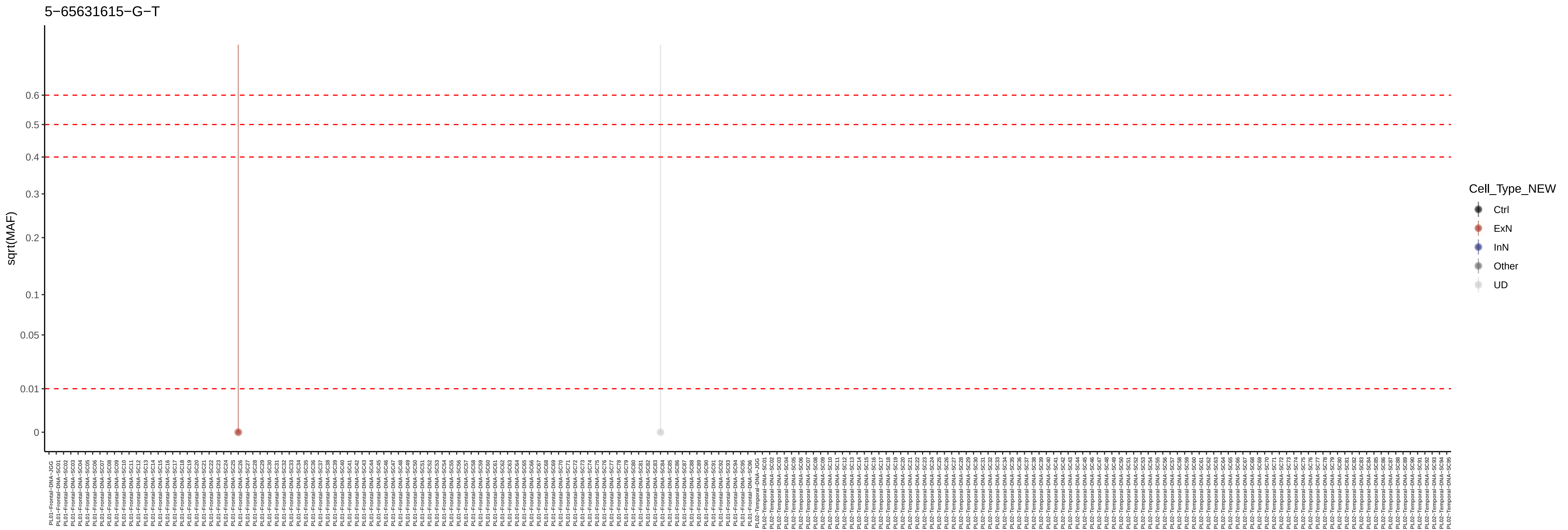
5−65631615−G−T
sqrt(MAF)
0.6
0.5
0.4
0.3
0.2
0.1
0.05
0.01
0
Cell_Type_NEW
Ctrl
ExN
InN
Other
UD

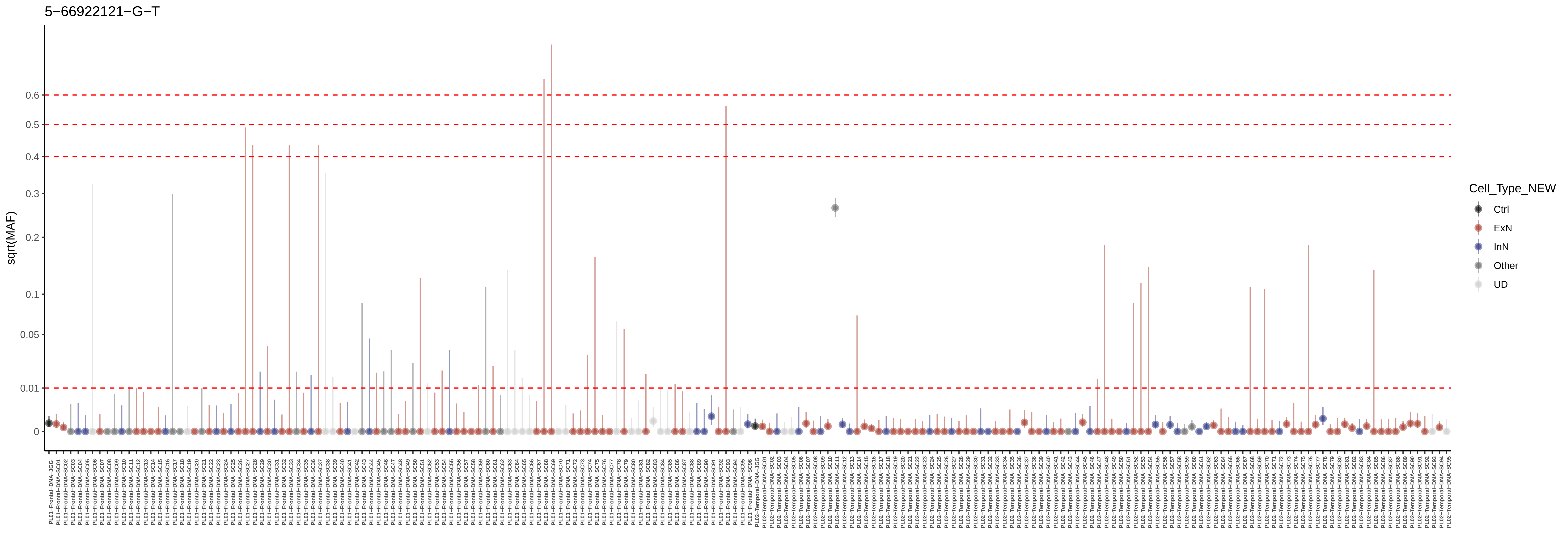
5−66922121−G−T
sqrt(MAF)
0.6
0.5
0.4
0.3
0.2
0.1
0.05
0.01
0
Cell_Type_NEW
Ctrl
ExN
InN
Other
UD

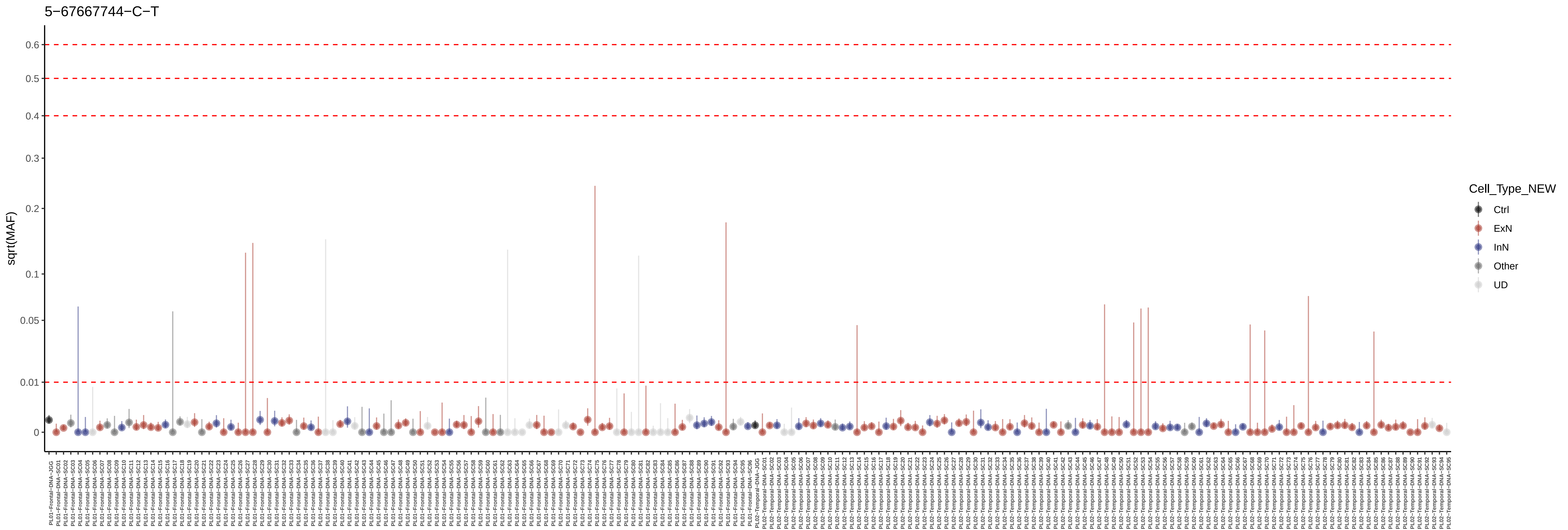
5−67667744−C−T
sqrt(MAF)
Cell_Type_NEW
Ctrl
ExN
InN
Other
UD

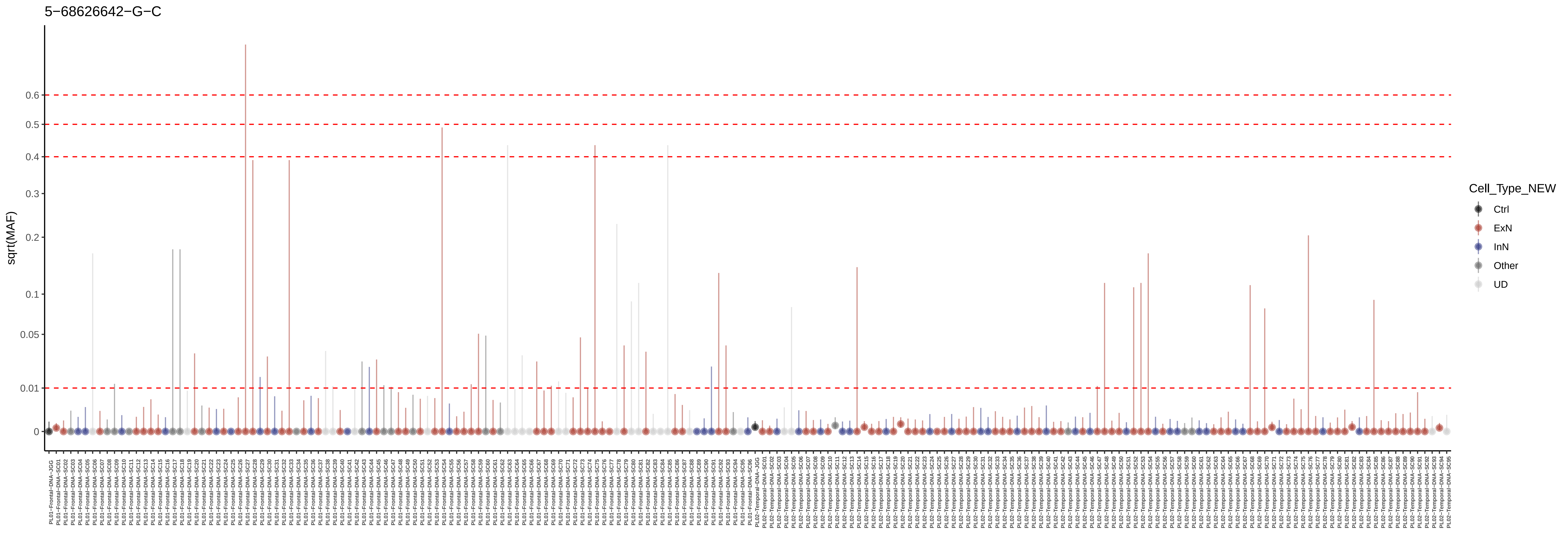
5−68626642−G−C
sqrt(MAF)
0.6
0.5
0.4
0.3
0.2
0.1
0.05
0.01
0
Cell_Type_NEW
Ctrl
ExN
InN
Other
UD

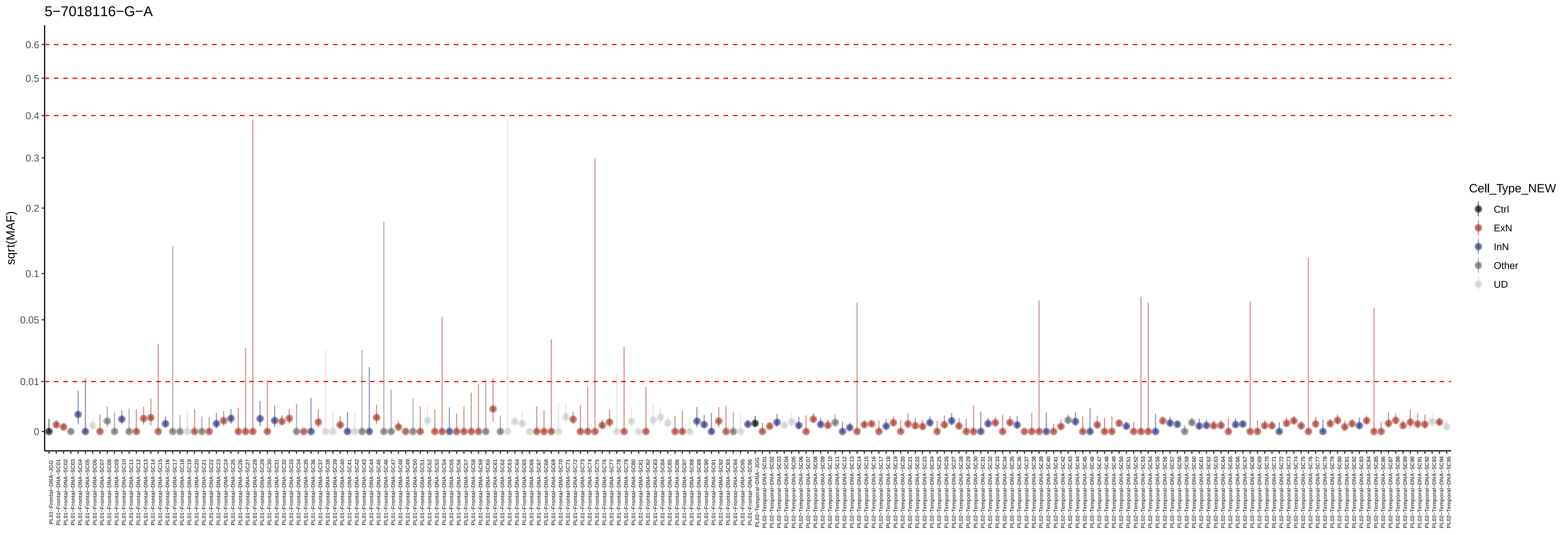
5−7018116−G−A
sqrt(MAF)
0.6
0.5
0.4
0.3
0.2
0.1
0.05
0.01
0
Cell_Type_NEW
Ctrl
ExN
InN
Other
UD

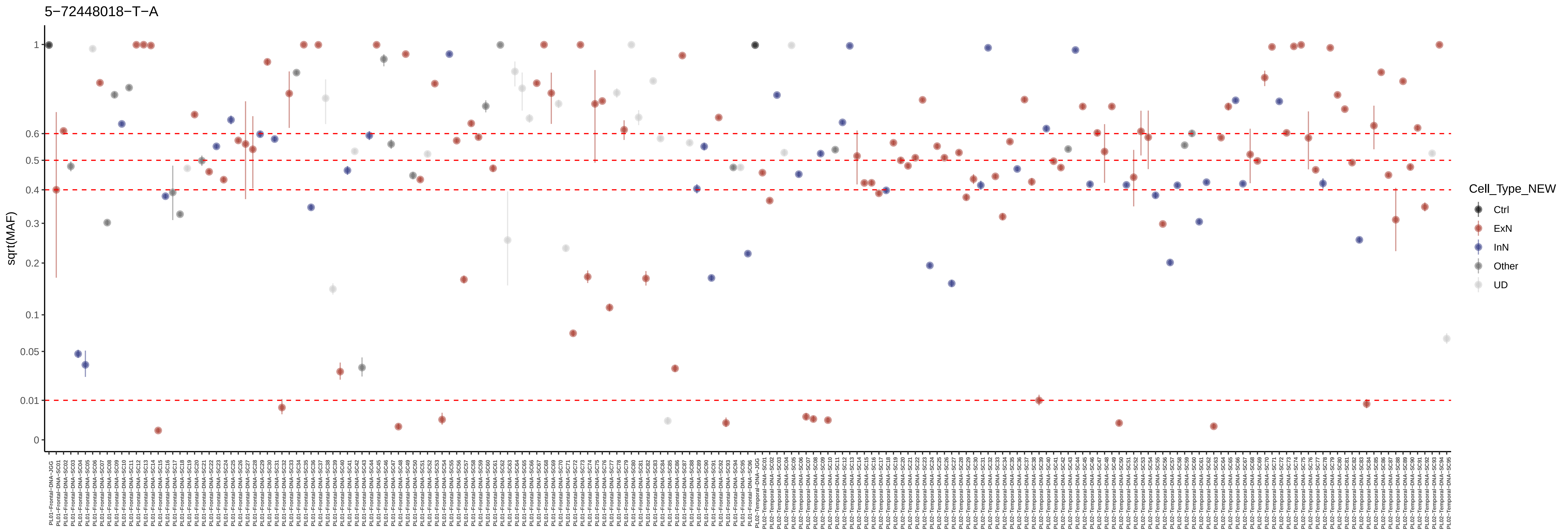

5−72448018−T−A
sqrt(MAF)
1
0.6
0.5
0.4
0.3
0.2
0.1
0.05
0.01
0
Cell_Type_NEW
Ctrl
ExN
InN
Other
UD

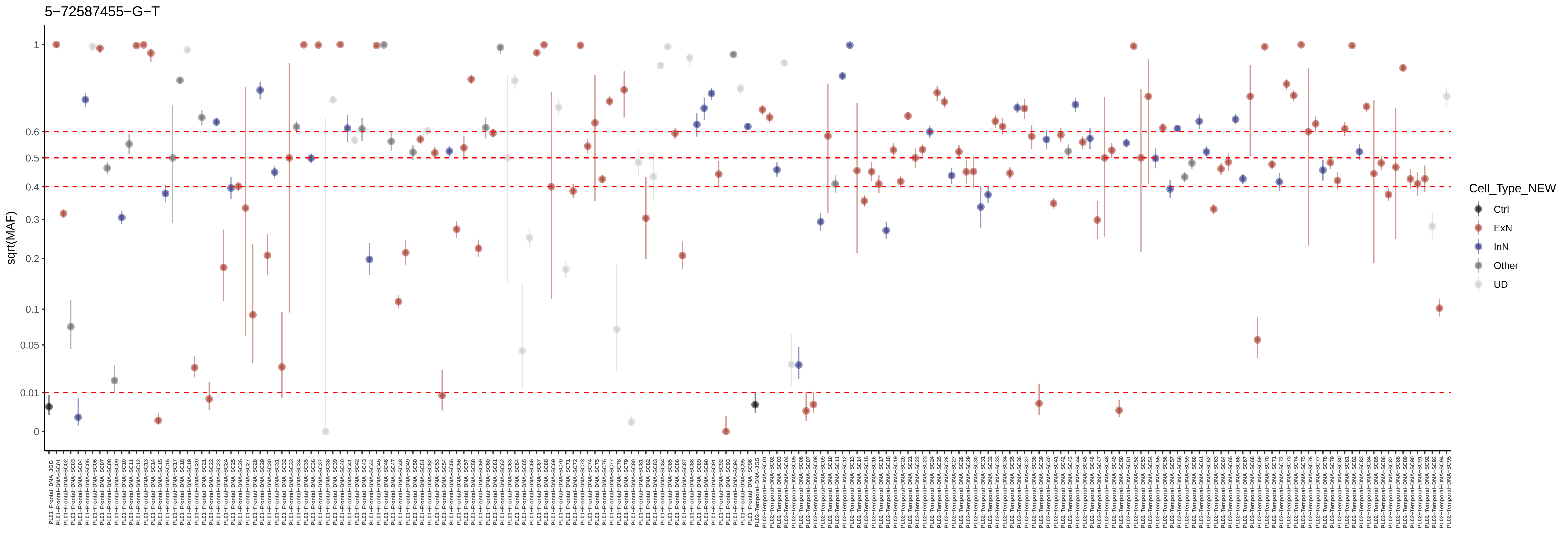
5−72587455−G−T
sqrt(MAF)
1
0.6
0.5
0.4
0.3
0.2
0.1
0.05
0.01
0
Cell_Type_NEW
Ctrl
ExN
InN
Other
UD

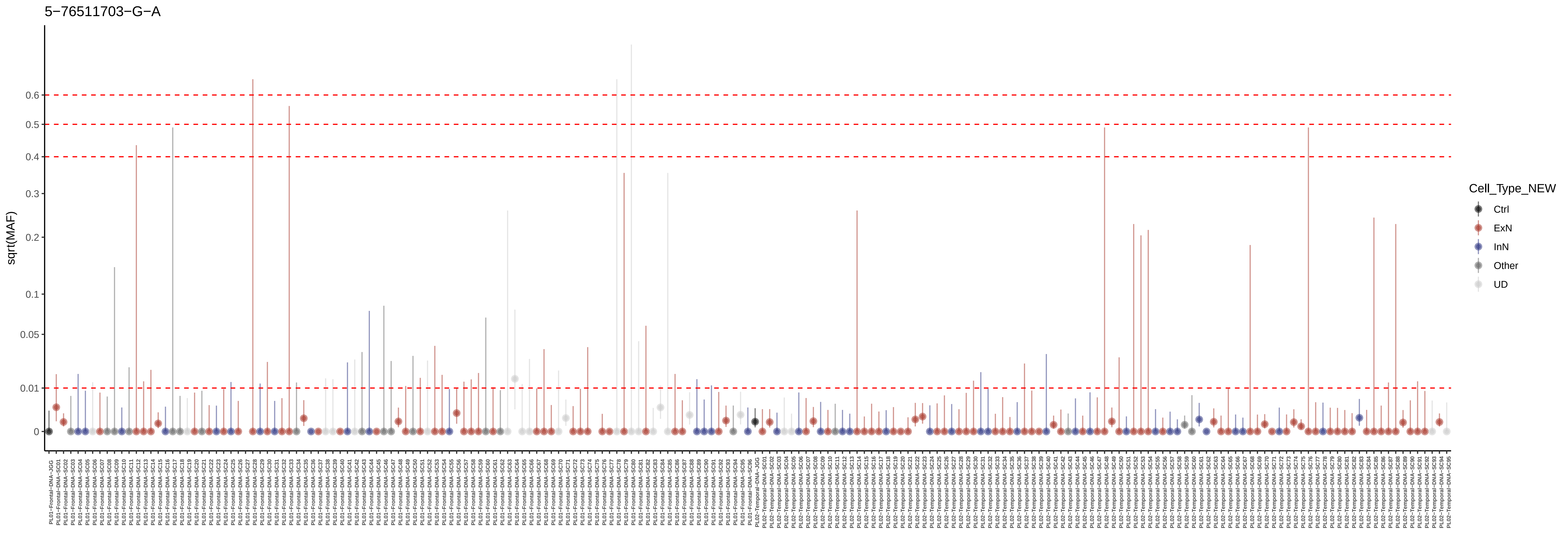

5−76511703−G−A
sqrt(MAF)
0.6
0.5
0.4
0.3
0.2
0.1
0.05
0.01
0
Cell_Type_NEW
Ctrl
ExN
InN
Other
UD

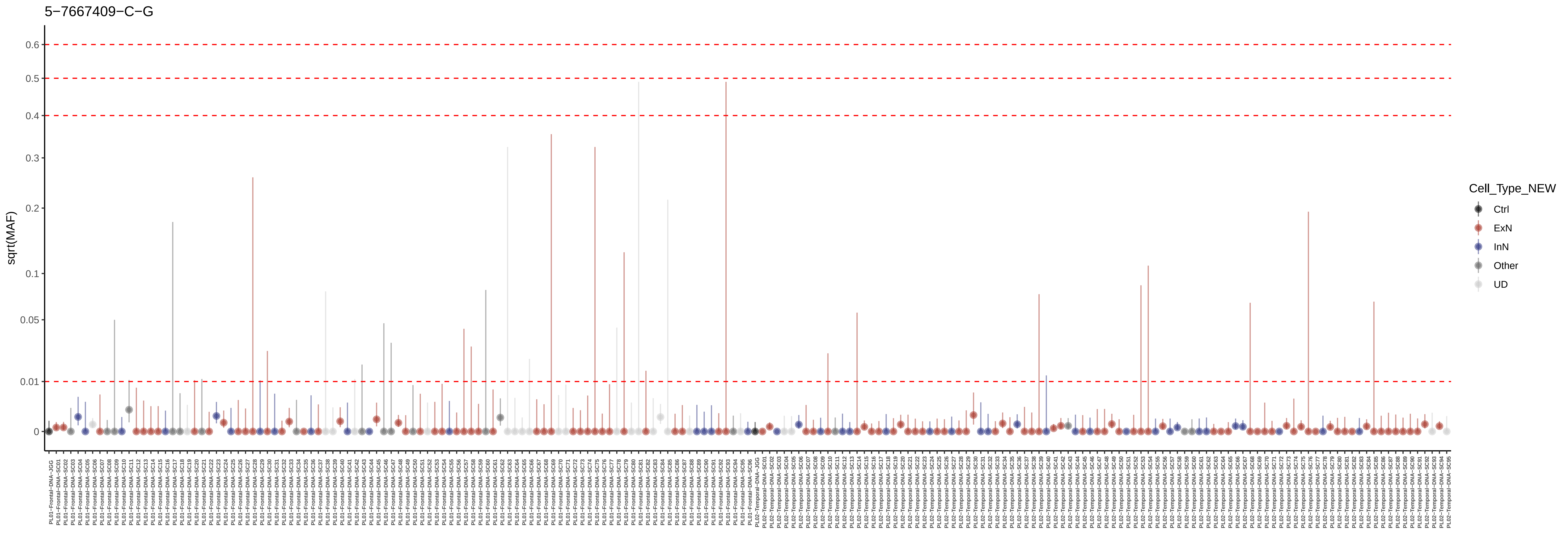

5−7667409−C−G
sqrt(MAF)
0.6
0.5
0.4
0.3
0.2
0.1
0.05
0.01
0
Cell_Type_NEW
Ctrl
ExN
InN
Other
UD

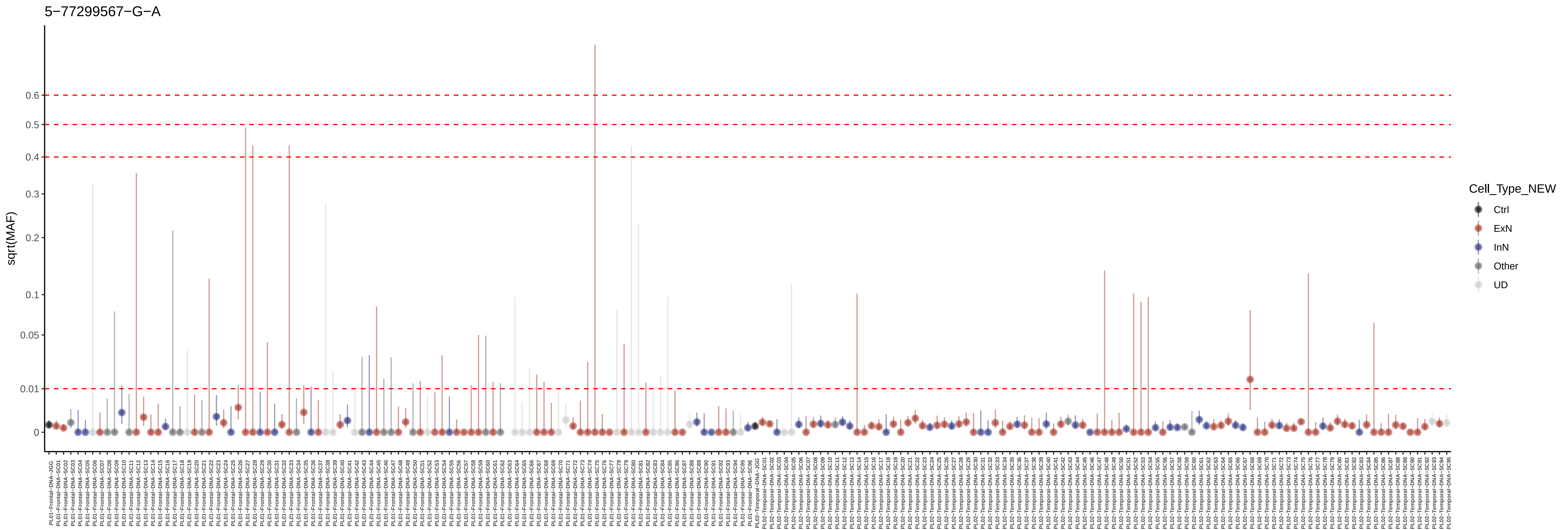
5−77299567−G−A
sqrt(MAF)
0.6
0.5
0.4
0.3
0.2
0.1
0.05
0.01
0
Cell_Type_NEW
Ctrl
ExN
InN
Other
UD

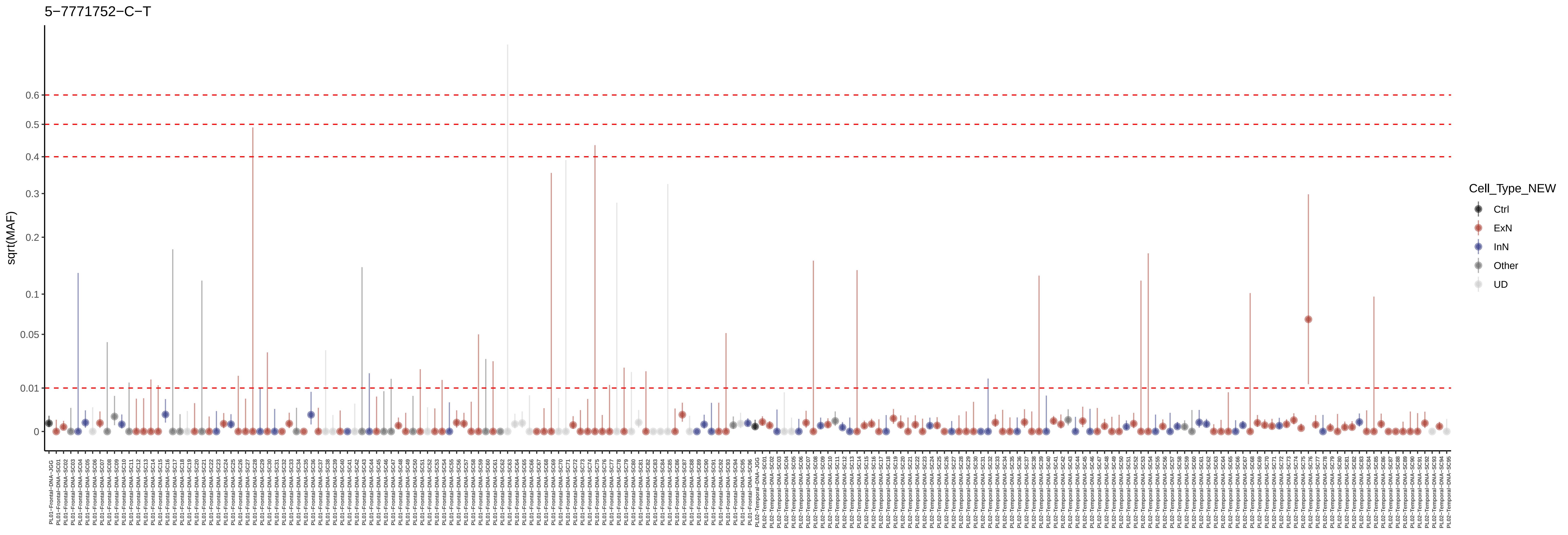
5−7771752−C−T
sqrt(MAF)
0.6
0.5
0.4
0.3
0.2
0.1
0.05
0.01
0
Cell_Type_NEW
Ctrl
ExN
InN
Other
UD

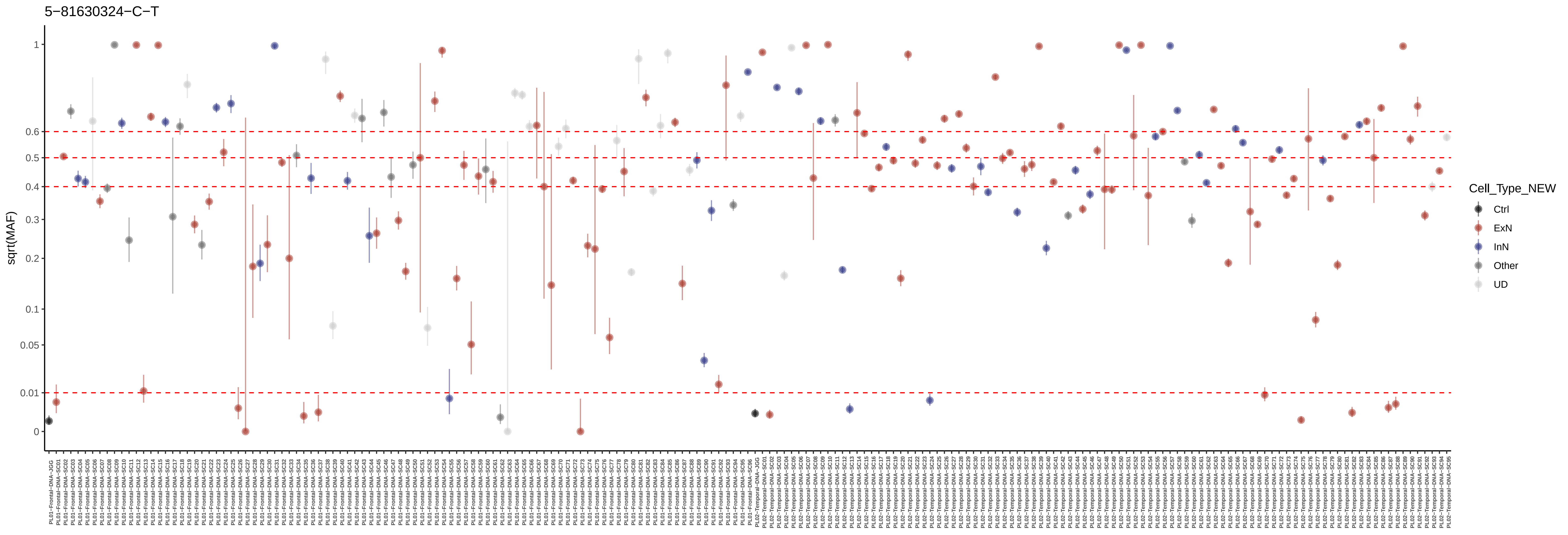

5−81630324−C−T
sqrt(MAF)
1
0.6
0.5
0.4
0.3
0.2
0.1
0.05
0.01
0
Cell_Type_NEW
Ctrl
ExN
InN
Other
UD

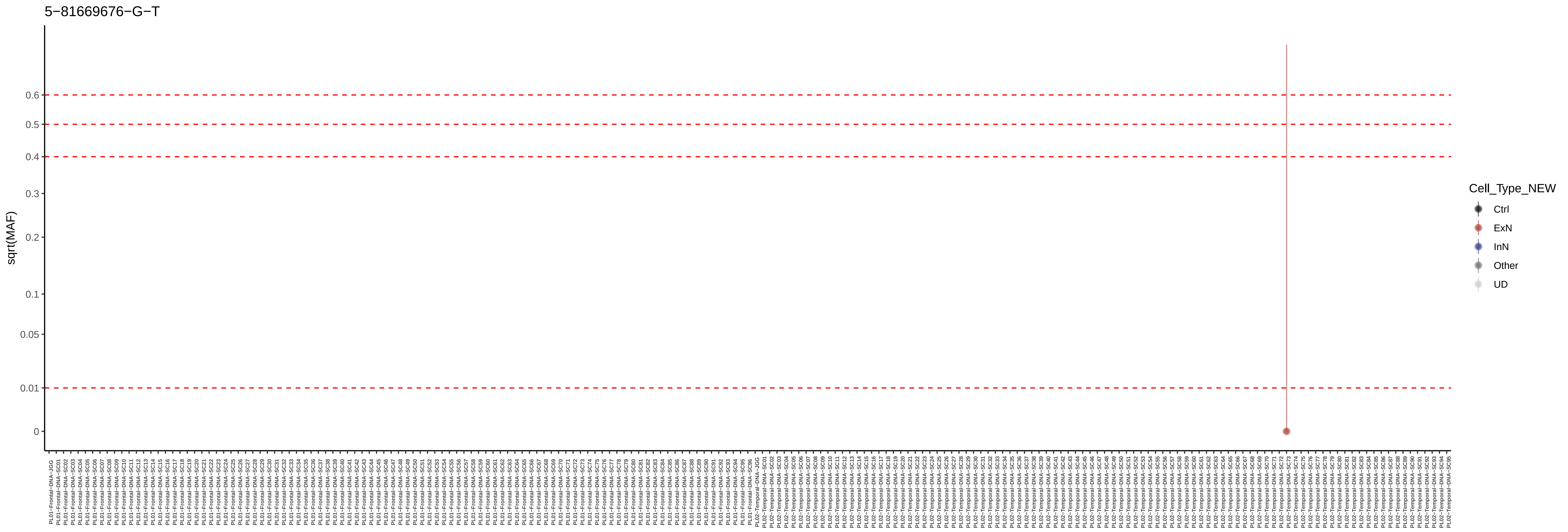

5−81669676−G−T
sqrt(MAF)
0.6
0.5
0.4
0.3
0.2
0.1
0.05
0.01
0
Cell_Type_NEW
Ctrl
ExN
InN
Other
UD

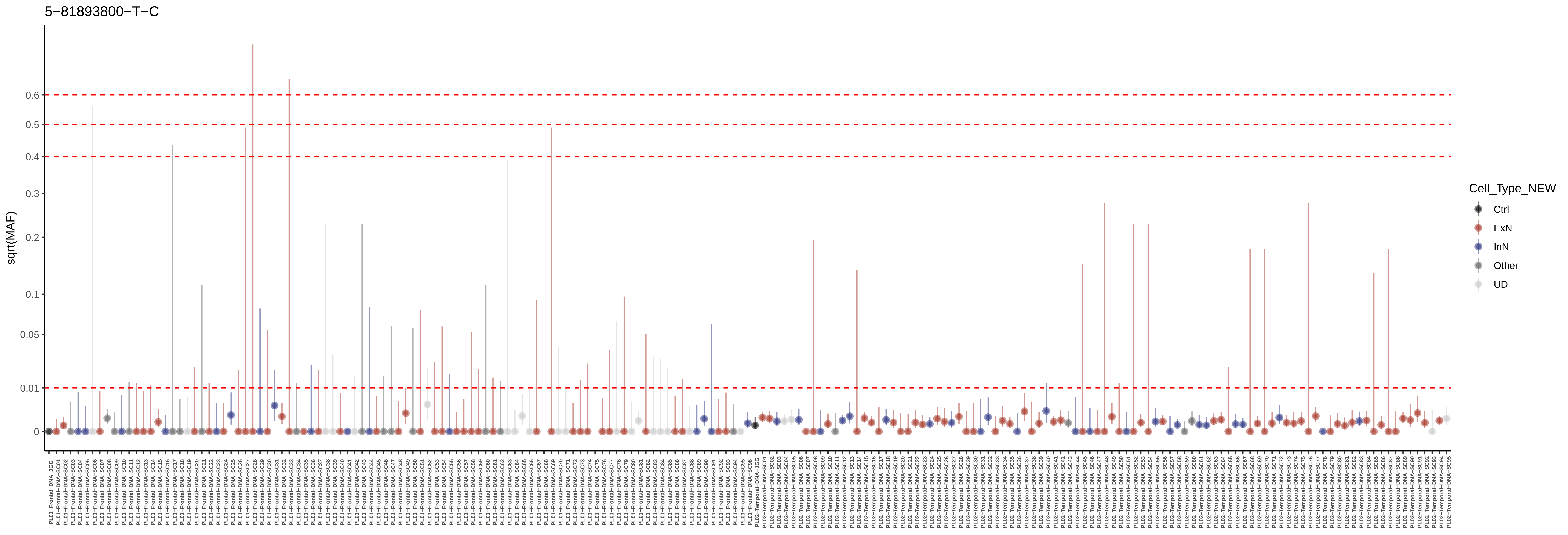
5−81893800−T−C
sqrt(MAF)
0.6
0.5
0.4
0.3
0.2
0.1
0.05
0.01
0
Cell_Type_NEW
Ctrl
ExN
InN
Other
UD

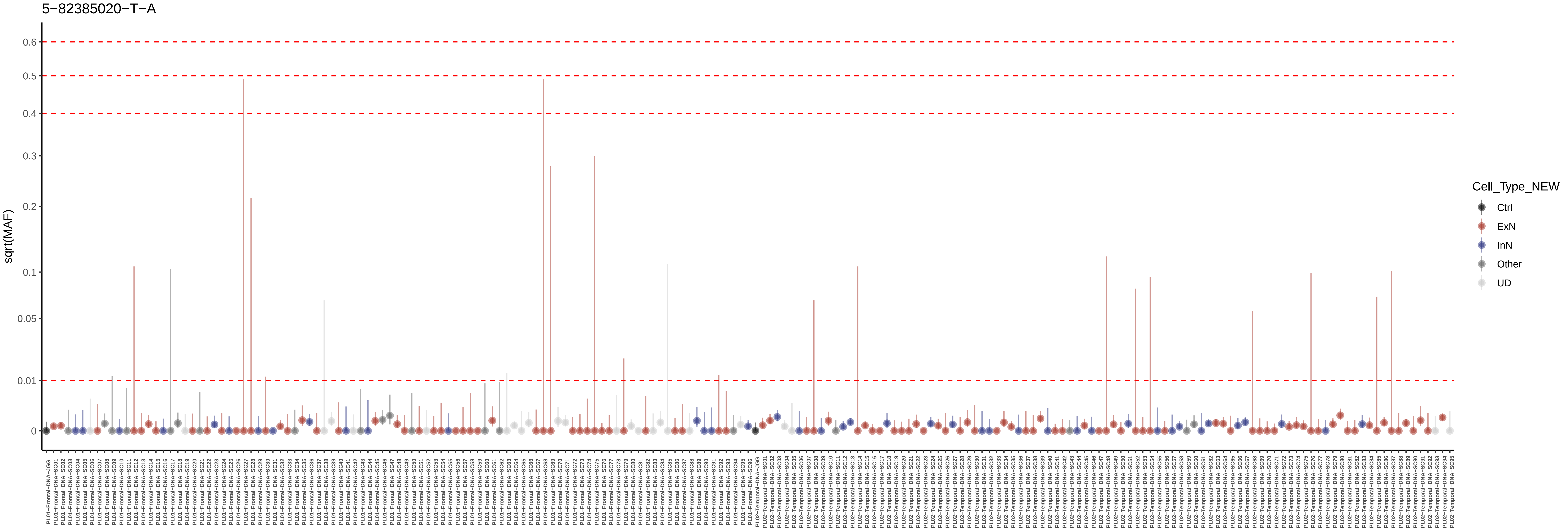

5−82385020−T−A
sqrt(MAF)
0.6
0.5
0.4
0.3
0.2
0.1
0.05
0.01
0
Cell_Type_NEW
Ctrl
ExN
InN
Other
UD

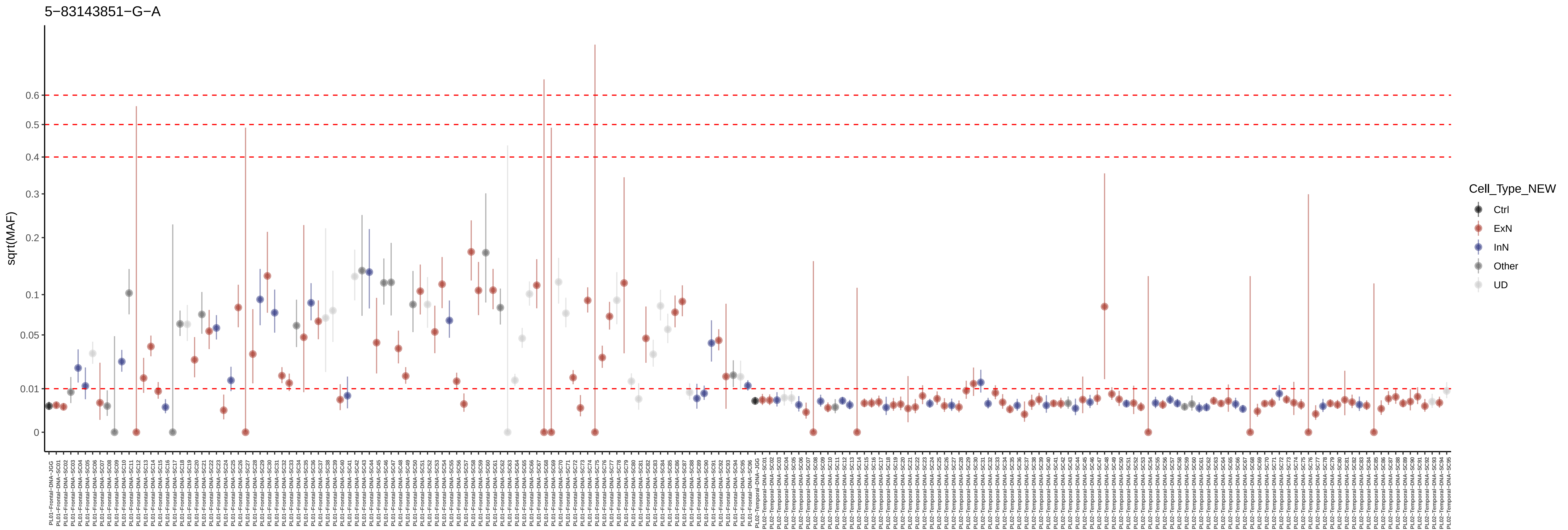
5−83143851−G−A
sqrt(MAF)
0.6
0.5
0.4
0.3
0.2
0.1
0.05
0.01
0
Cell_Type_NEW
Ctrl
ExN
InN
Other
UD

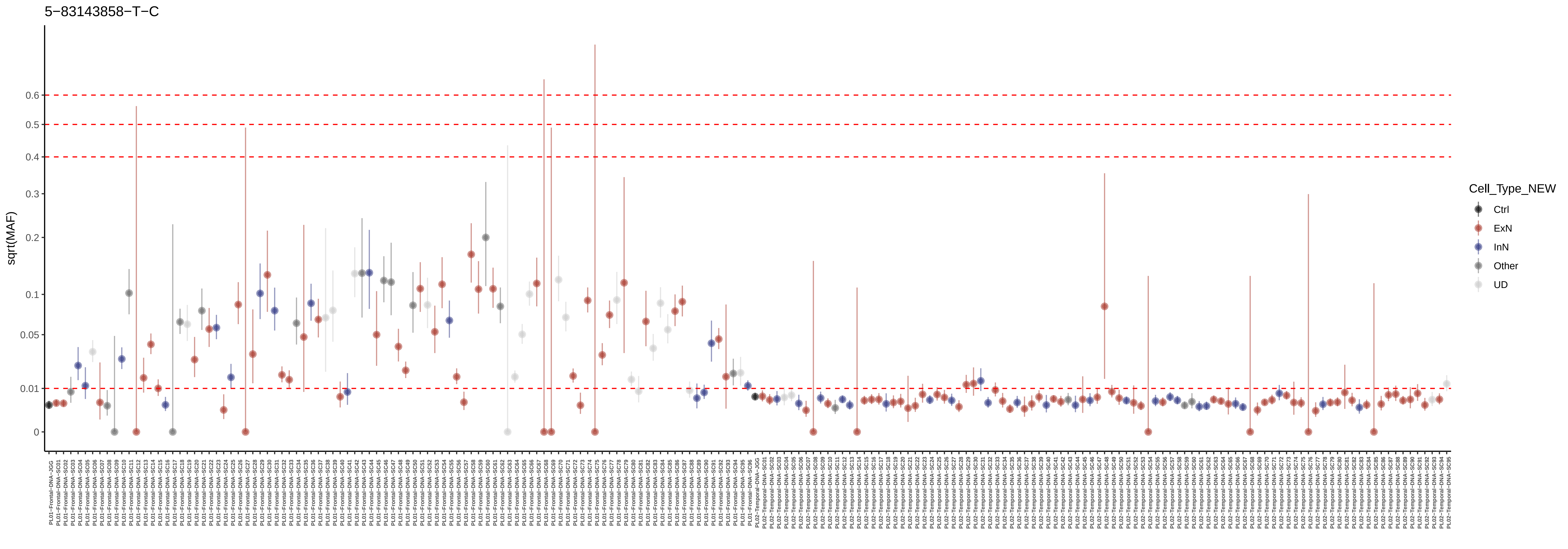

5−83143858−T−C
sqrt(MAF)
0
0.01
0.05
0.1
0.2
0.3
0.4
0.5
0.6
Cell_Type_NEW
Ctrl
ExN
InN
Other
UD

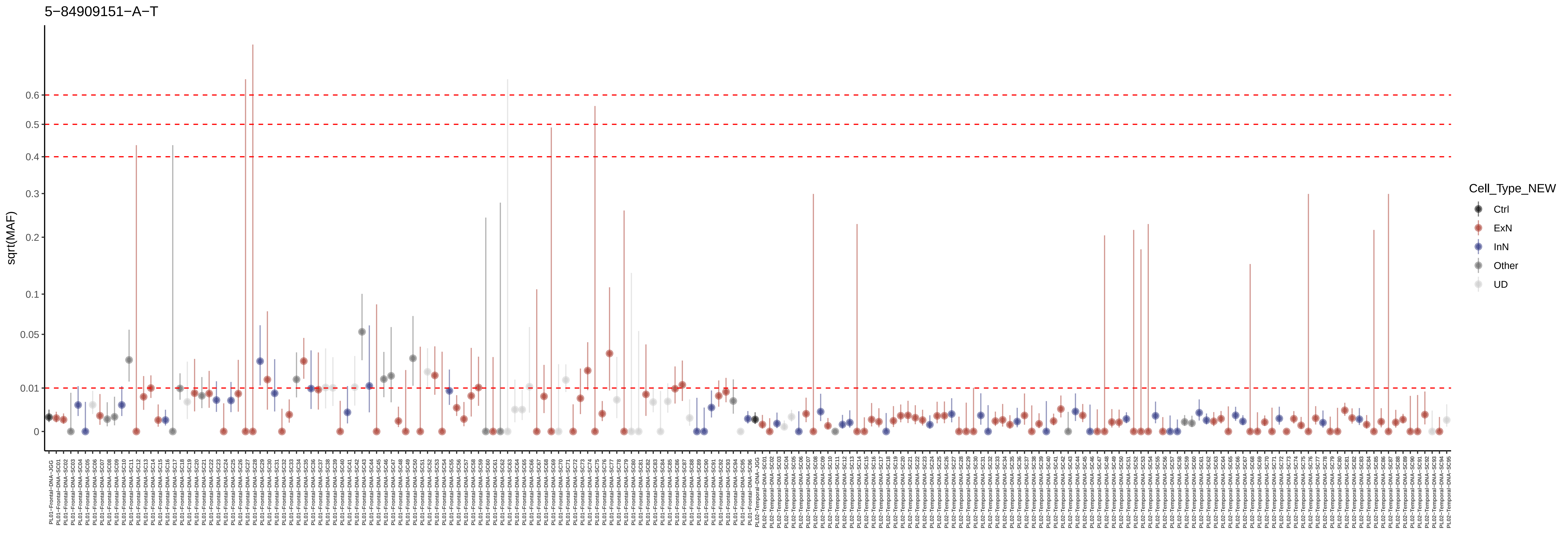

5−84909151−A−T
sqrt(MAF)
0.6
0.5
0.4
0.3
0.2
0.1
0.05
0.01
0
Cell_Type_NEW
Ctrl
ExN
InN
Other
UD

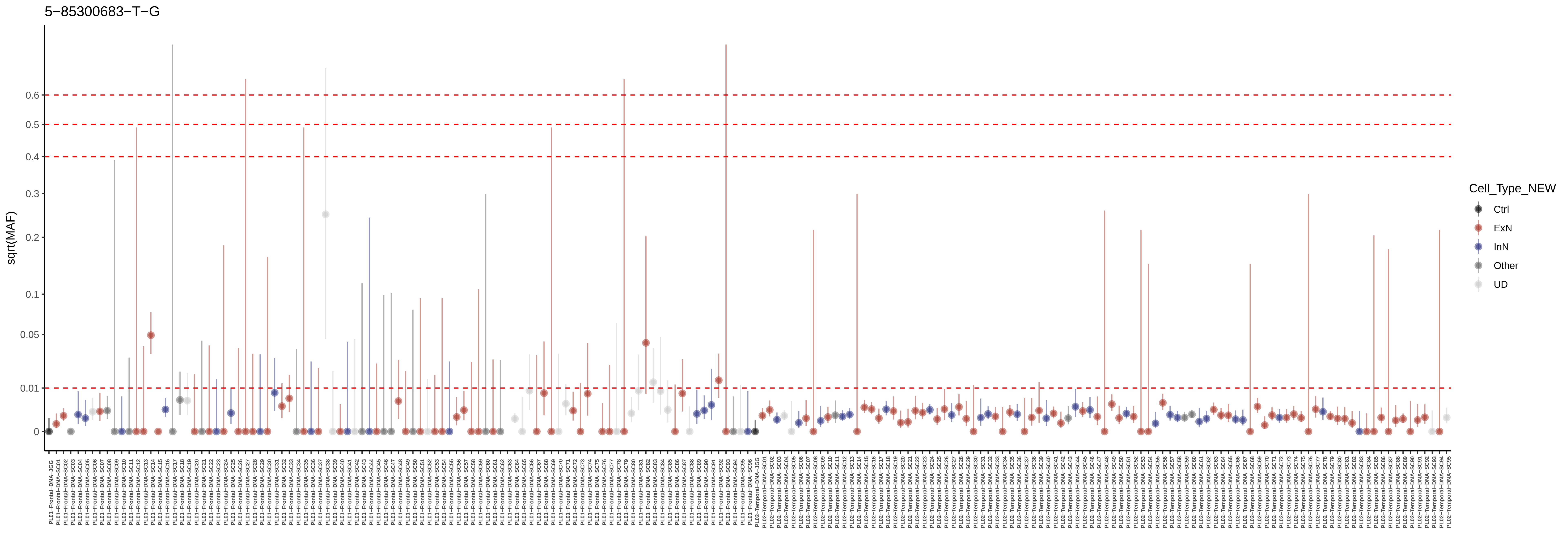
5−85300683−T−G
sqrt(MAF)
0.6
0.5
0.4
0.3
0.2
0.1
0.05
0.01
0
Cell_Type_NEW
Ctrl
ExN
InN
Other
UD

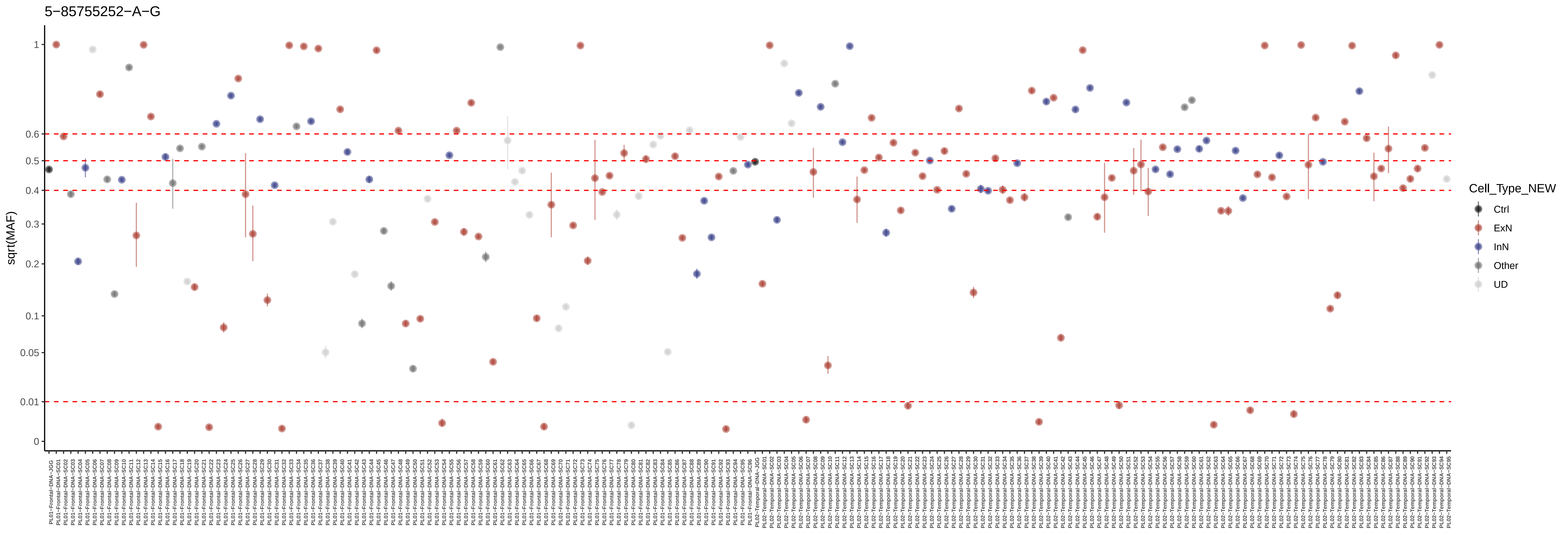
5−85755252−A−G
sqrt(MAF)
1
0.6
0.5
0.4
0.3
0.2
0.1
0.05
0.01
0
Cell_Type_NEW
Ctrl
ExN
InN
Other
UD

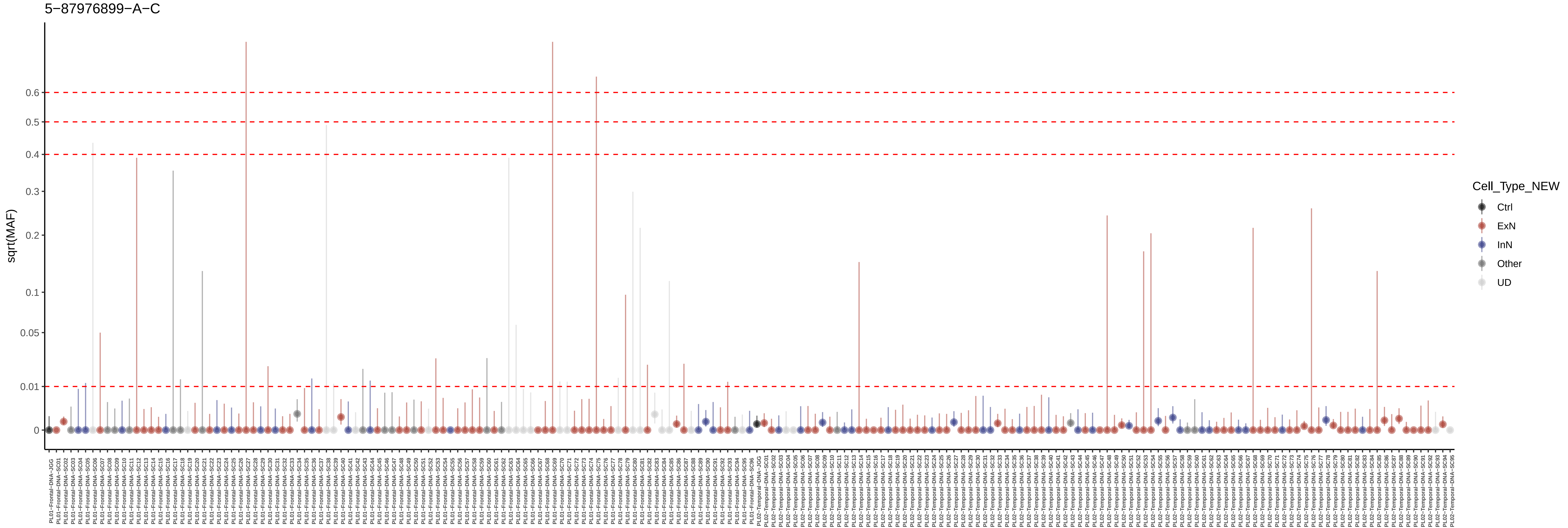
5−87976899−A−C
sqrt(MAF)
0.6
0.5
0.4
0.3
0.2
0.1
0.05
0.01
0
Cell_Type_NEW
Ctrl
ExN
InN
Other
UD

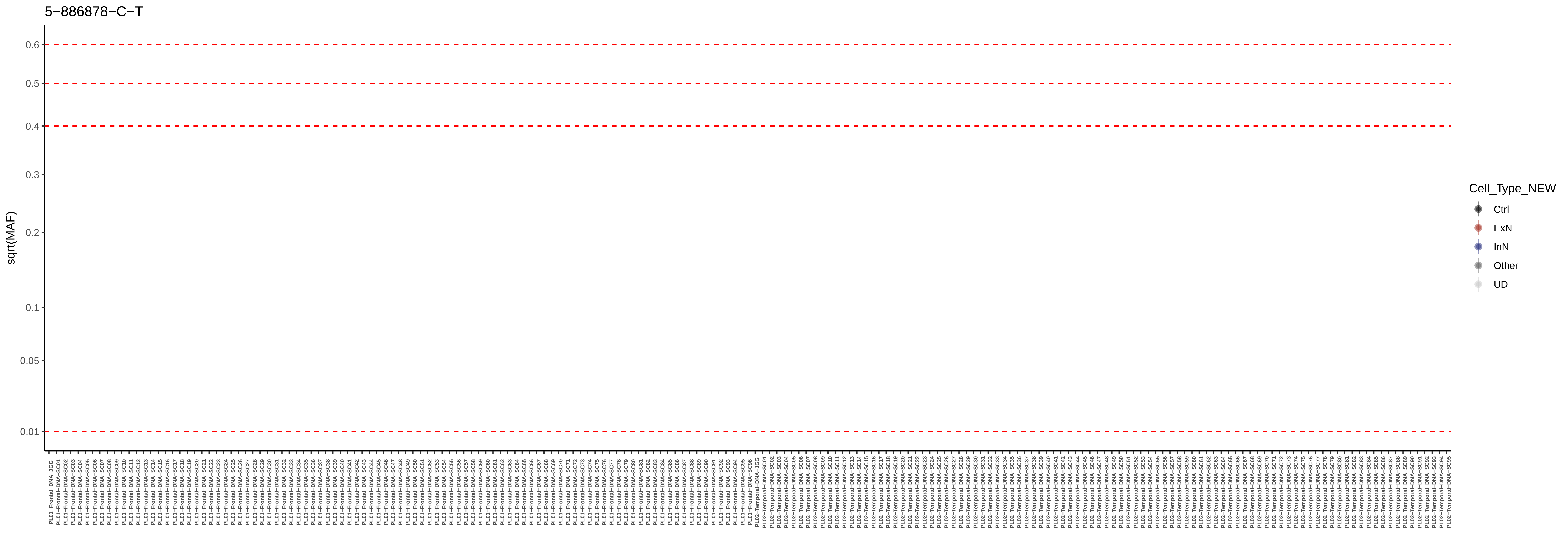
5−886878−C−T
sqrt(MAF)
0.6
0.5
0.4
0.3
0.2
0.1
0.05
0.01
Cell_Type_NEW
Ctrl
ExN
InN
Other
UD

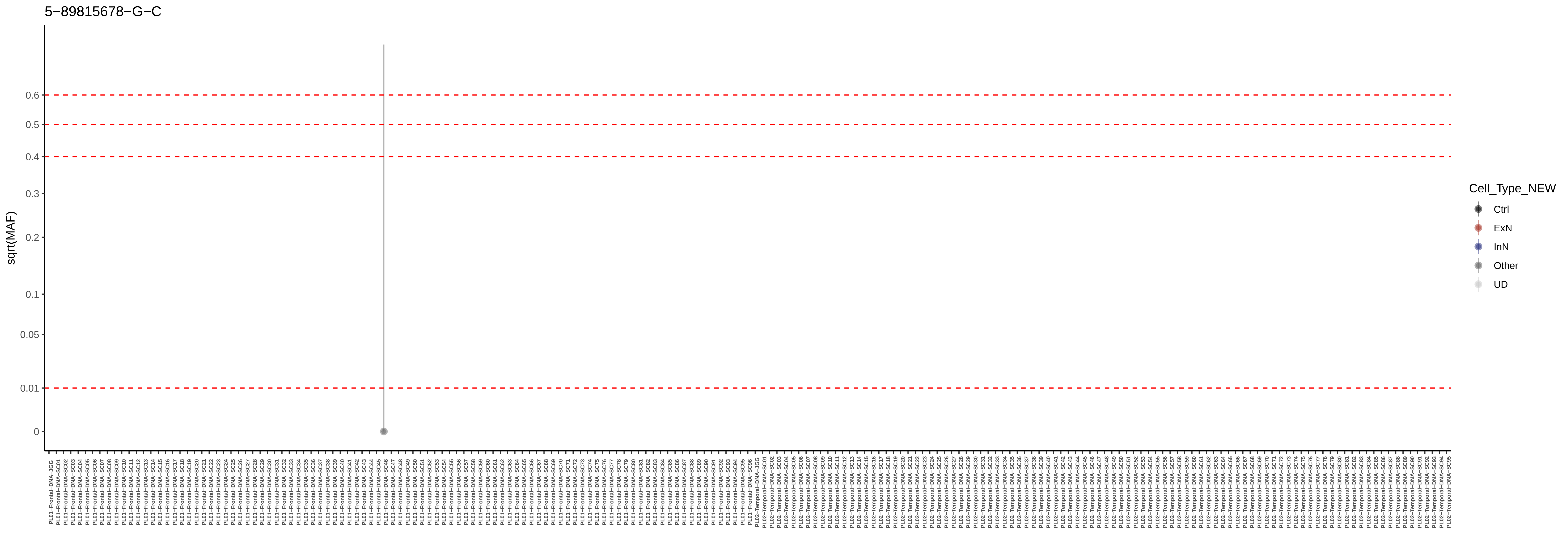
5−89815678−G−C
sqrt(MAF)
0.6
0.5
0.4
0.3
0.2
0.1
0.05
0.01
0
Cell_Type_NEW
Ctrl
ExN
InN
Other
UD

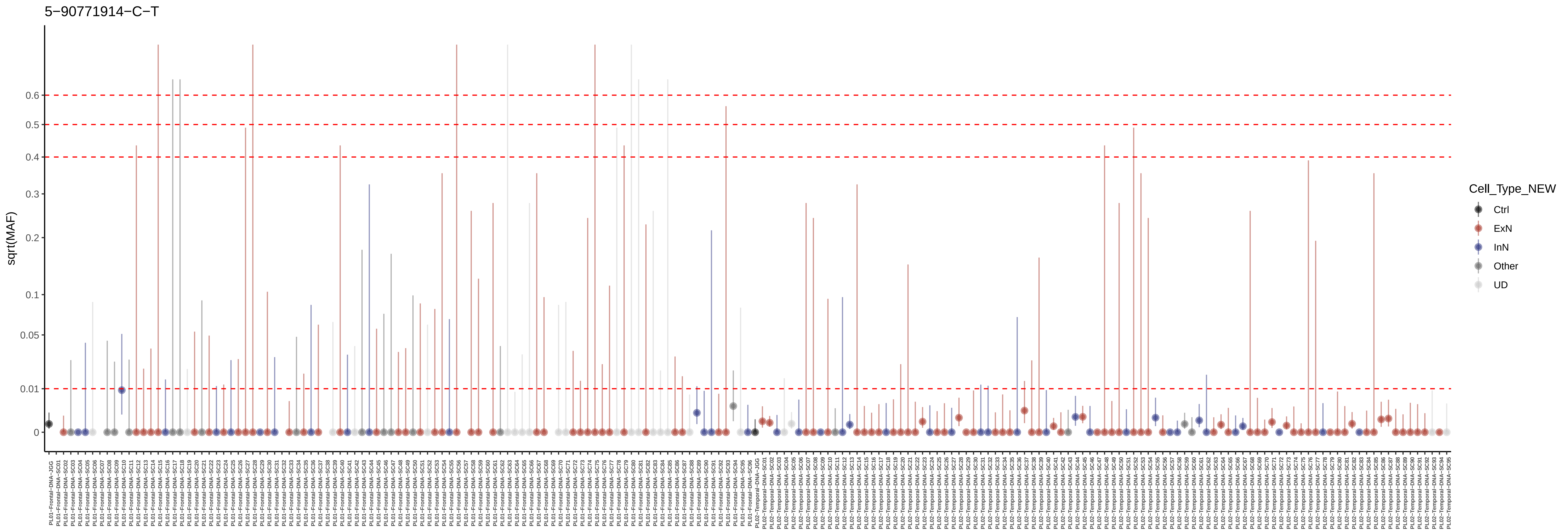

5−90771914−C−T
sqrt(MAF)
0.6
0.5
0.4
0.3
0.2
0.1
0.05
0.01
0
Cell_Type_NEW
Ctrl
ExN
InN
Other
UD

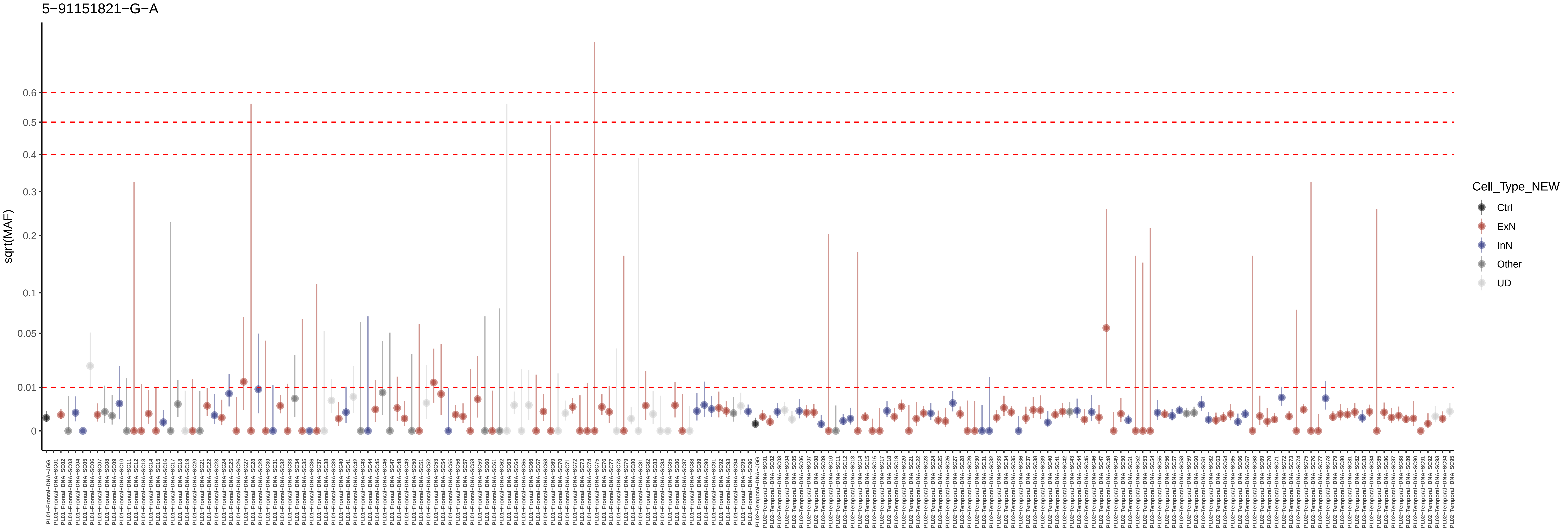
5−91151821−G−A
sqrt(MAF)
0.6
0.5
0.4
0.3
0.2
0.1
0.05
0.01
0
Cell_Type_NEW
Ctrl
ExN
InN
Other
UD

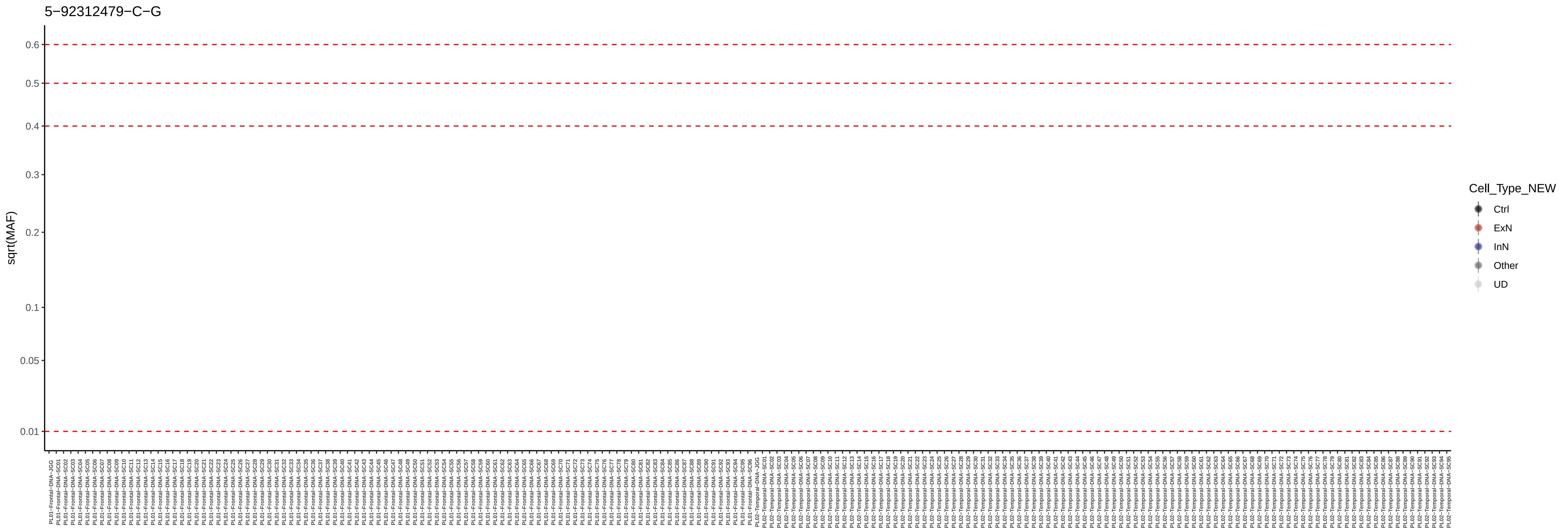
5−92312479−C−G
sqrt(MAF)
0.6
0.5
0.4
0.3
0.2
0.1
0.05
0.01
Cell_Type_NEW
Ctrl
ExN
InN
Other
UD

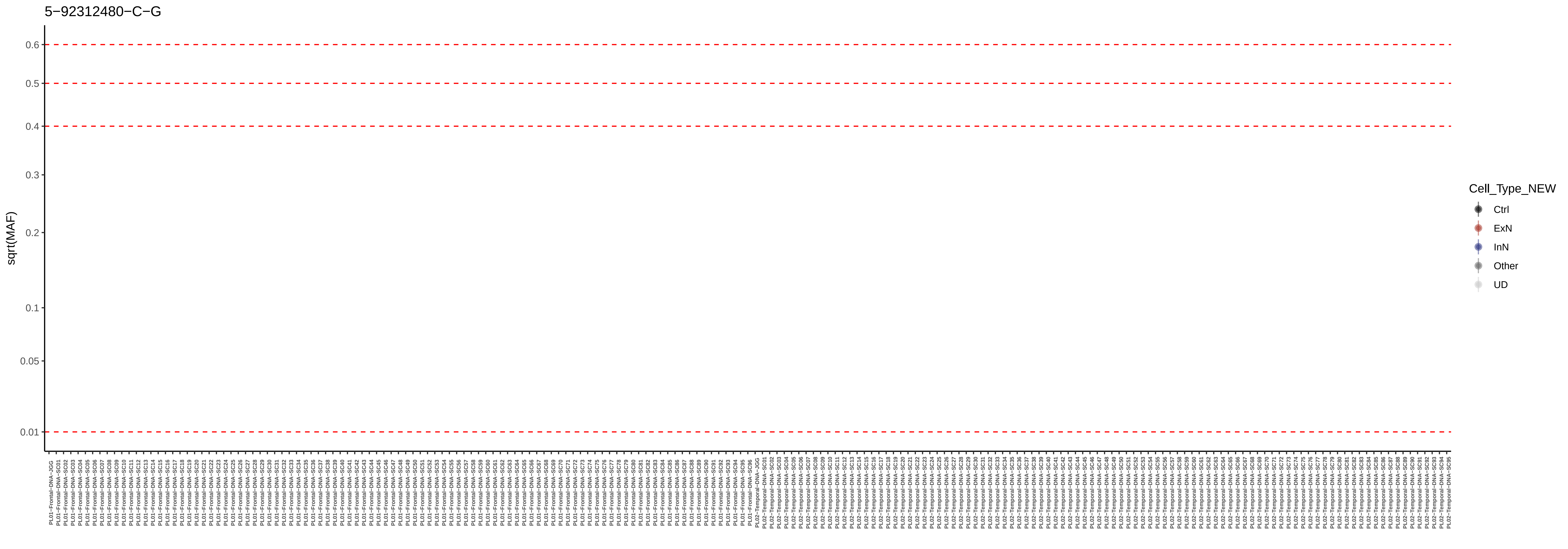
5−92312480−C−G
sqrt(MAF)
0.6
0.5
0.4
0.3
0.2
0.1
0.05
0.01
Cell_Type_NEW
Ctrl
ExN
InN
Other
UD

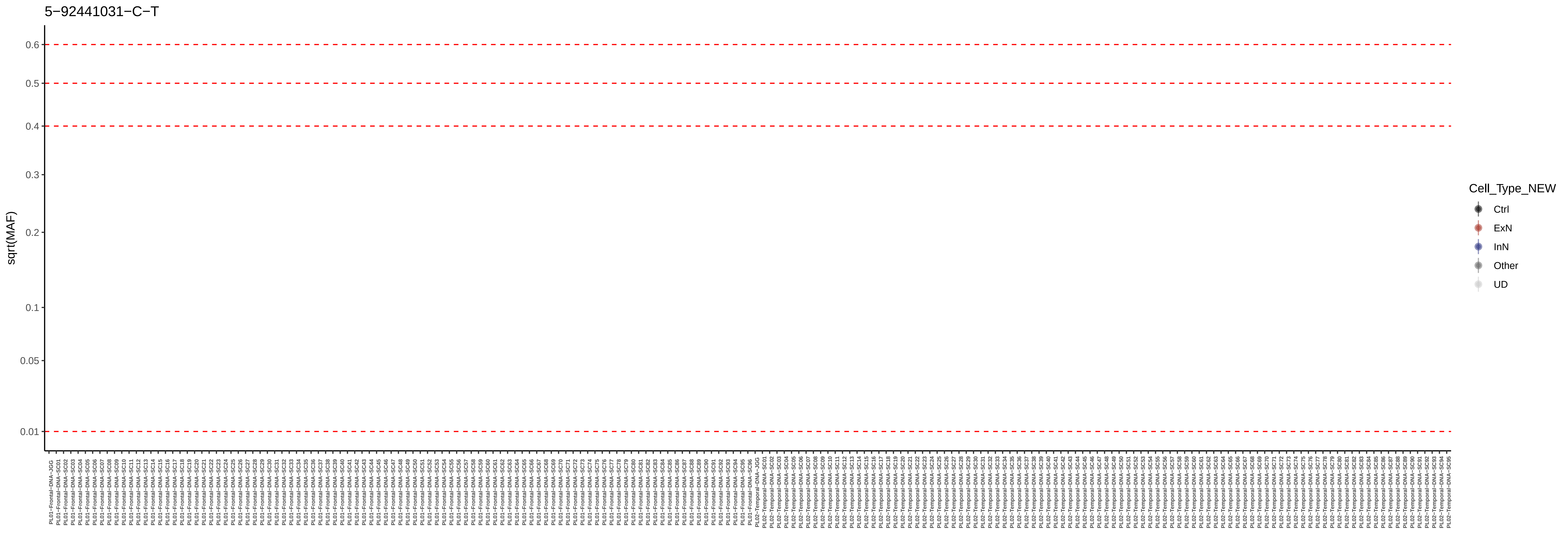

5−92441031−C−T
sqrt(MAF)
0.6
0.5
0.4
0.3
0.2
0.1
0.05
0.01
Cell_Type_NEW
Ctrl
ExN
InN
Other
UD

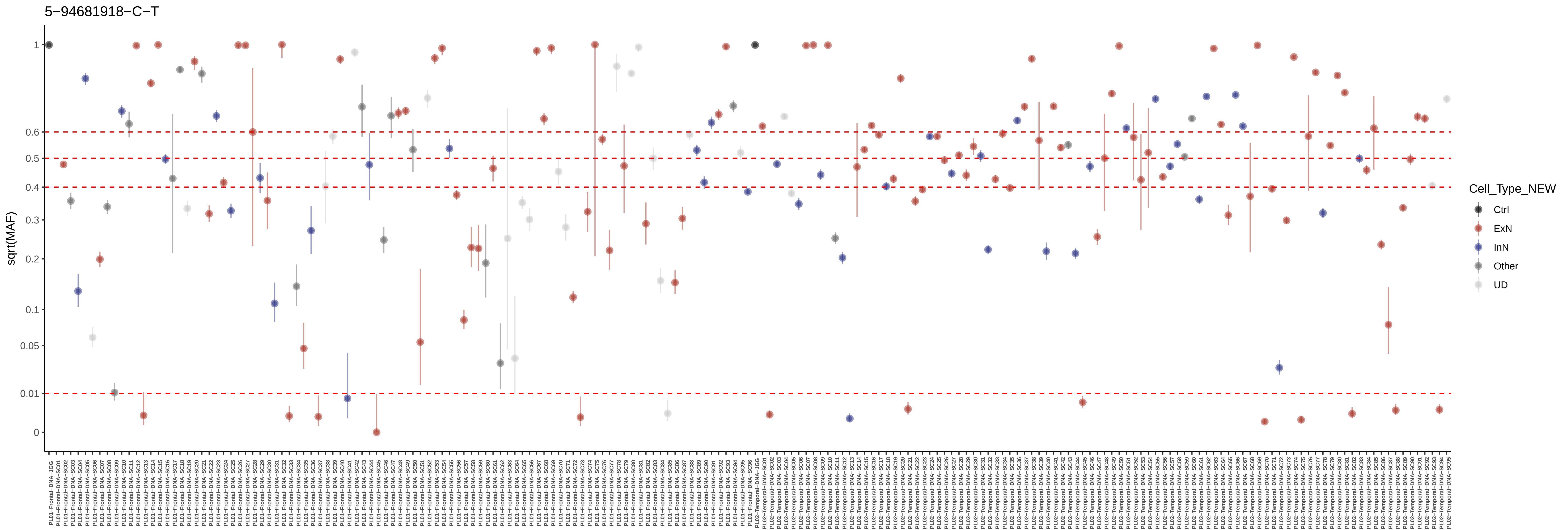
5−94681918−C−T
sqrt(MAF)
1
0.6
0.5
0.4
0.3
0.2
0.1
0.05
0.01
0
Cell_Type_NEW
Ctrl
ExN
InN
Other
UD

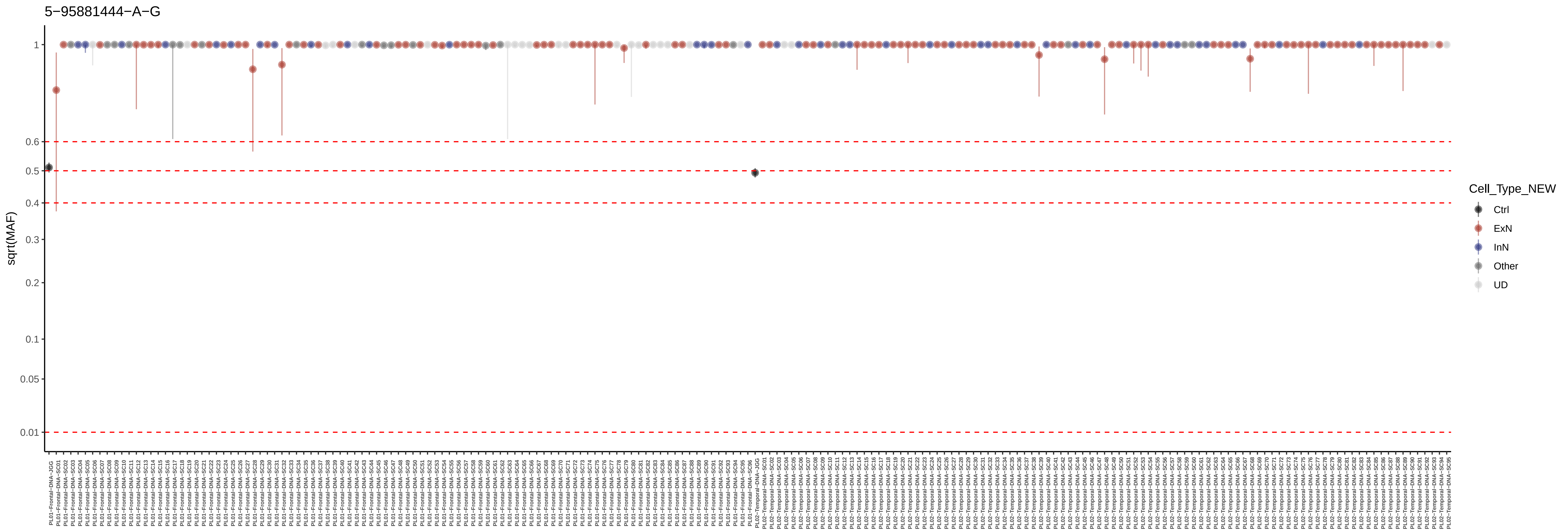

5−95881444−A−G
sqrt(MAF)
1
0.6
0.5
0.4
0.3
0.2
0.1
0.05
0.01
Cell_Type_NEW
Ctrl
ExN
InN
Other
UD

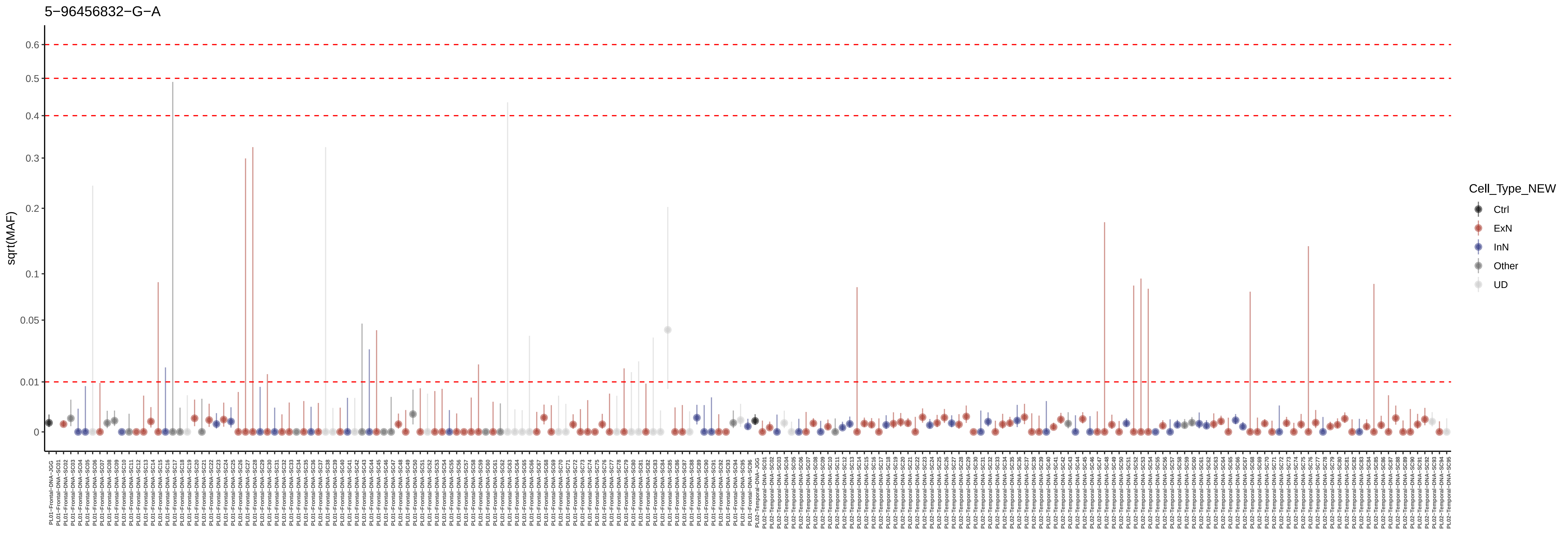
5−96456832−G−A
sqrt(MAF)
0.6
0.5
0.4
0.3
0.2
0.1
0.05
0.01
0
Cell_Type_NEW
Ctrl
ExN
InN
Other
UD

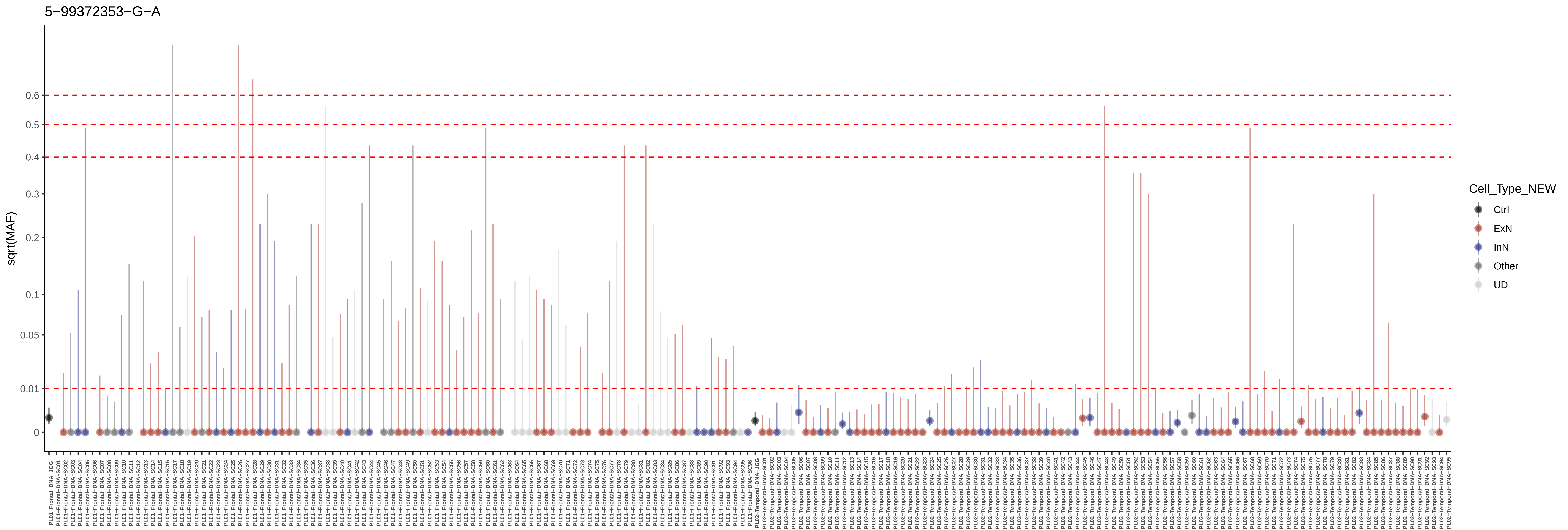

5−99372353−G−A
sqrt(MAF)
0.6
0.5
0.4
0.3
0.2
0.1
0.05
0.01
0
Cell_Type_NEW
Ctrl
ExN
InN
Other
UD

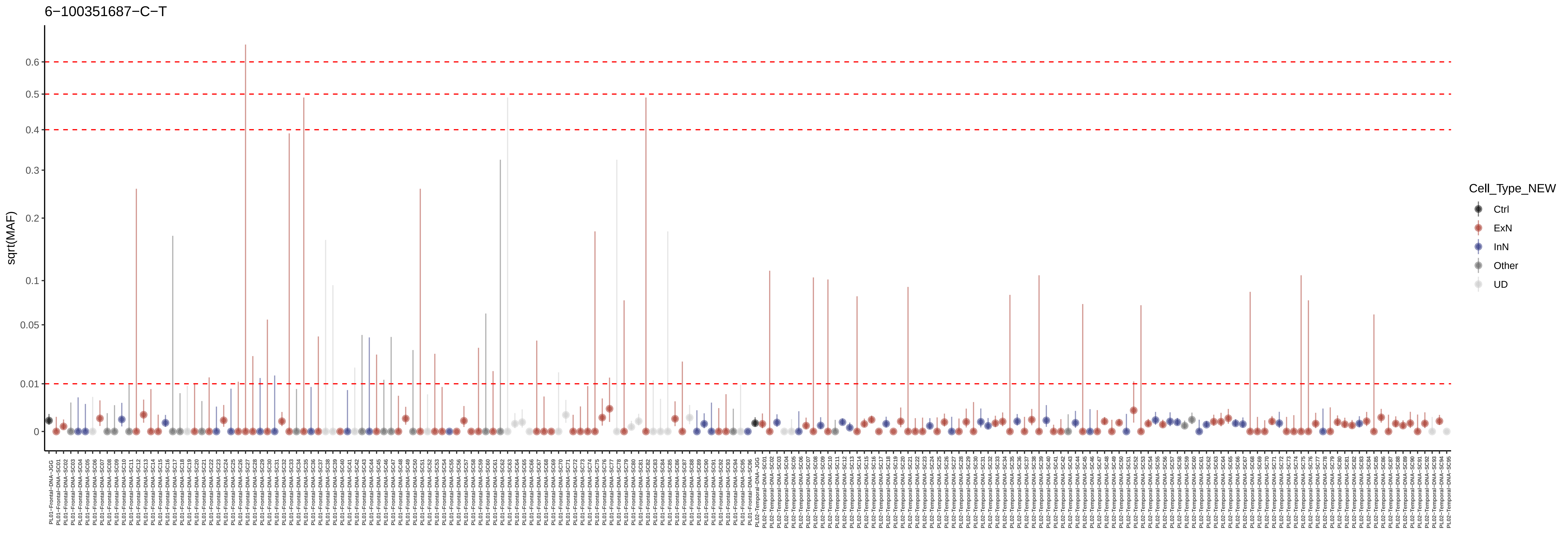

6−100351687−C−T
sqrt(MAF)
0.6
0.5
0.4
0.3
0.2
0.1
0.05
0.01
0
Cell_Type_NEW
Ctrl
ExN
InN
Other
UD

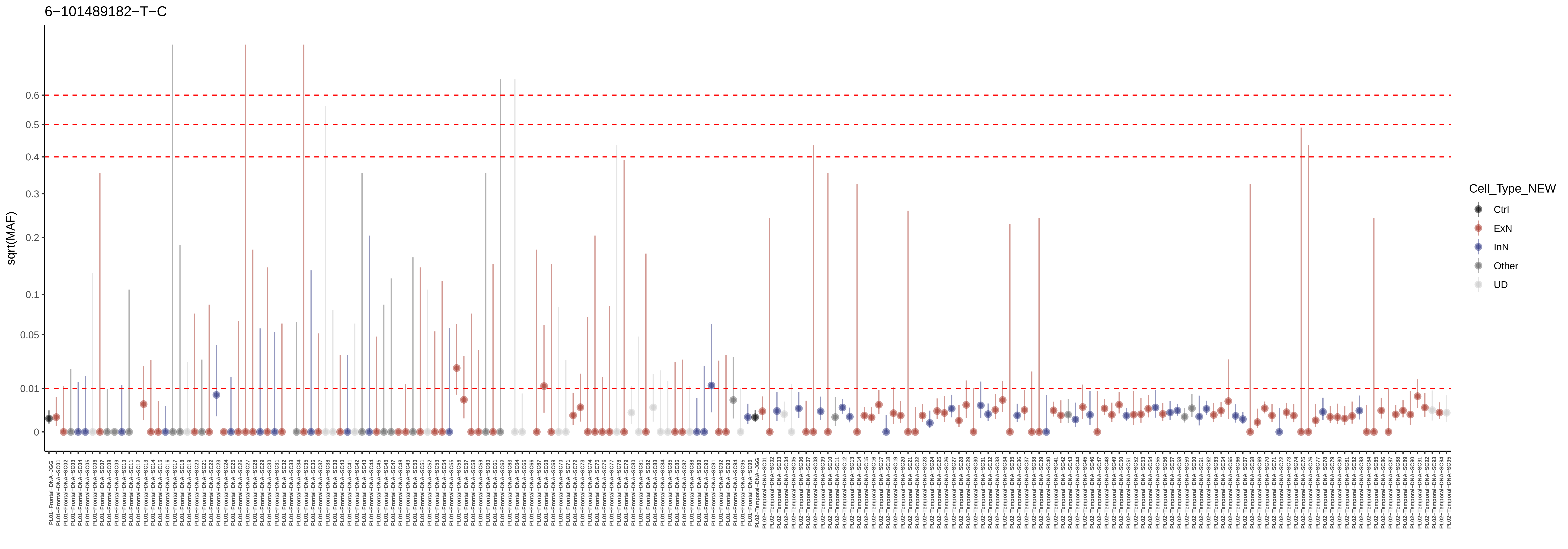

6−101489182−T−C
sqrt(MAF)
0.6
0.5
0.4
0.3
0.2
0.1
0.05
0.01
0
Cell_Type_NEW
Ctrl
ExN
InN
Other
UD

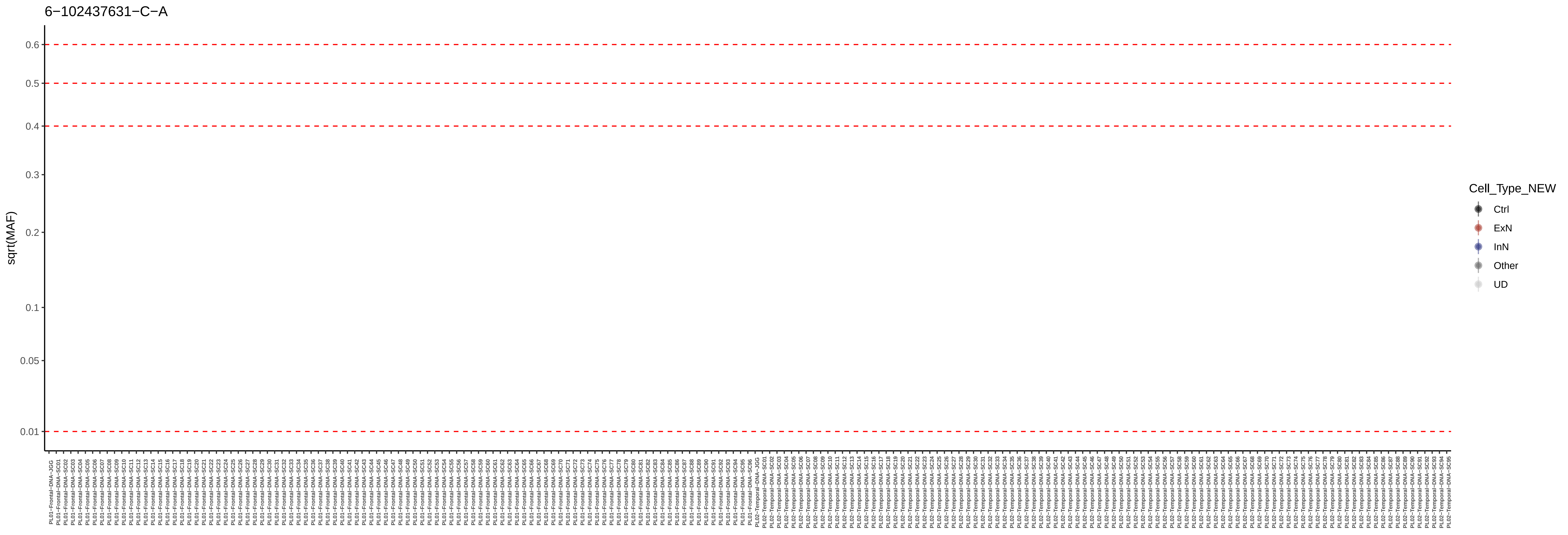

6−102437631−C−A
sqrt(MAF)
0.6
0.5
0.4
0.3
0.2
0.1
0.05
0.01
Cell_Type_NEW
Ctrl
ExN
InN
Other
UD

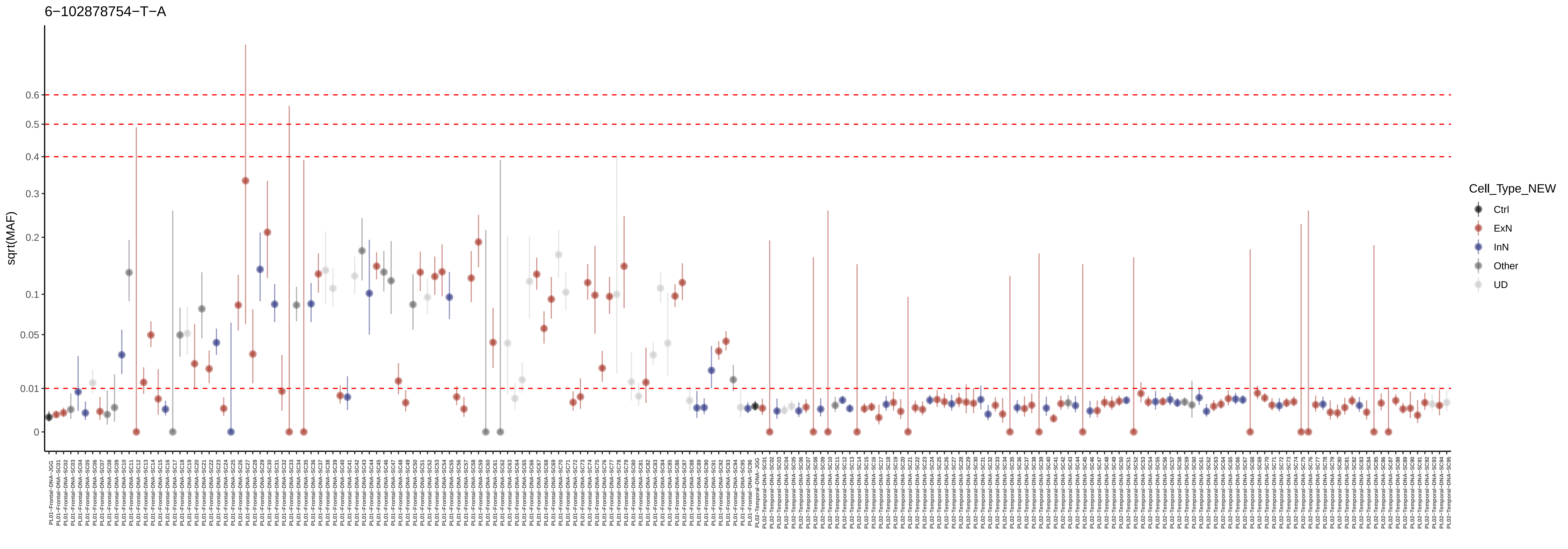
6−102878754−T−A
sqrt(MAF)
0.6
0.5
0.4
0.3
0.2
0.1
0.05
0.01
0
Cell_Type_NEW
Ctrl
ExN
InN
Other
UD

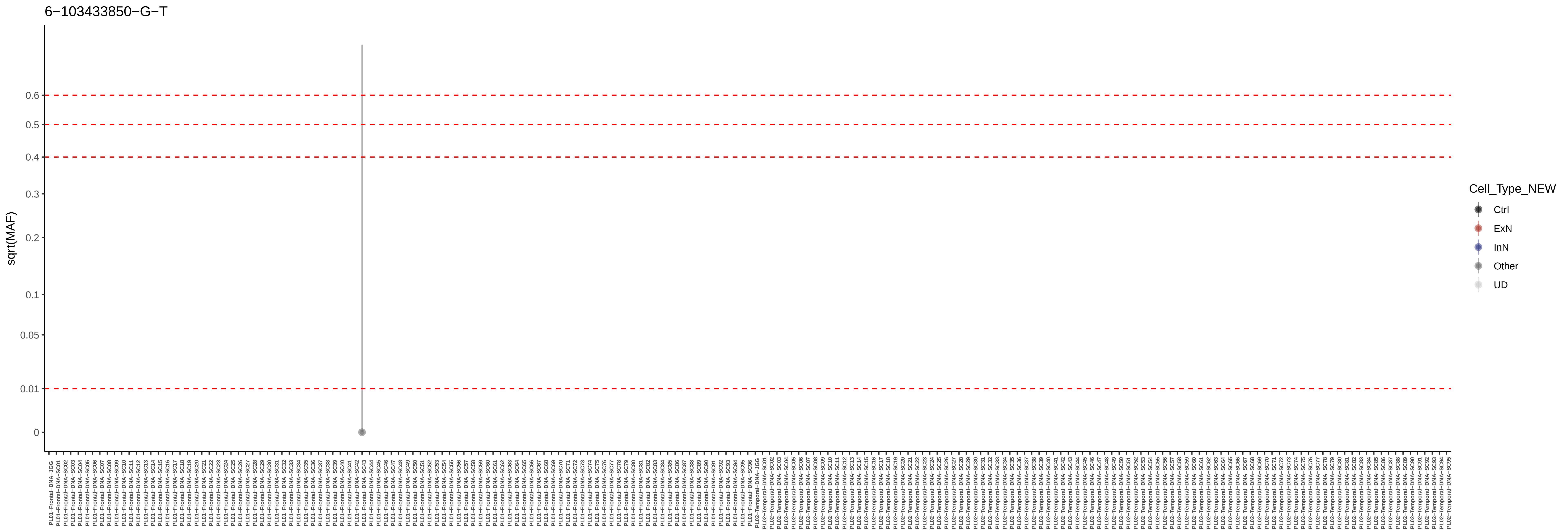

6−103433850−G−T
sqrt(MAF)
0.6
0.5
0.4
0.3
0.2
0.1
0.05
0.01
0
Cell_Type_NEW
Ctrl
ExN
InN
Other
UD

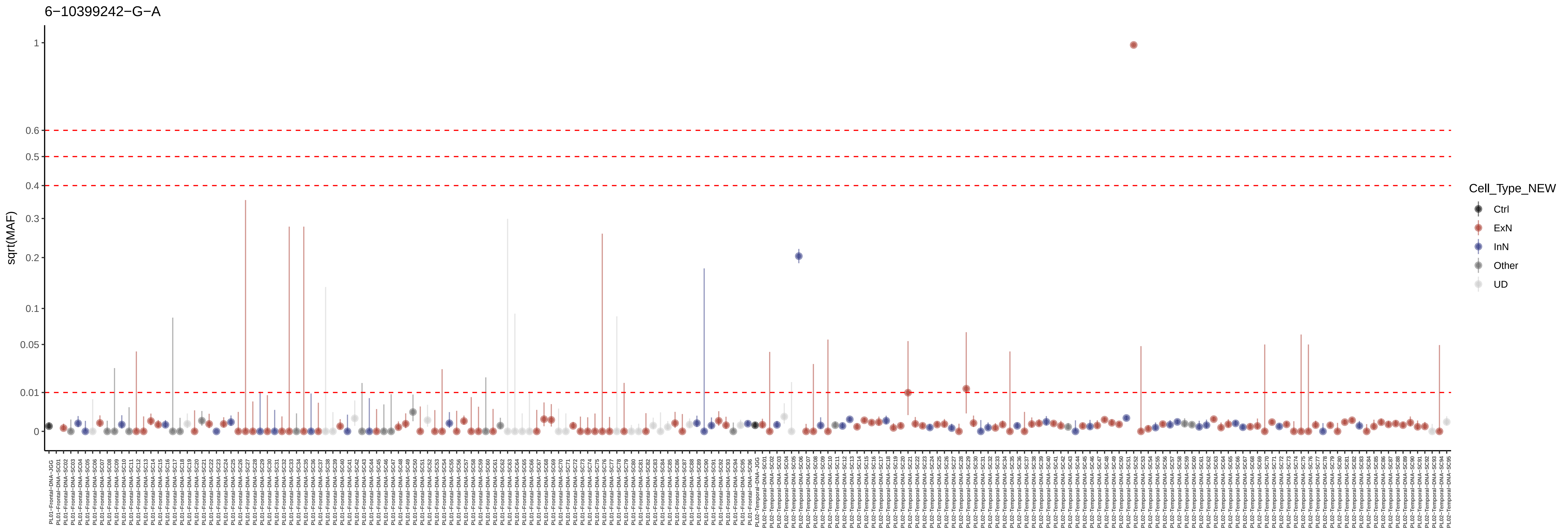
6−10399242−G−A
sqrt(MAF)
1
0.6
0.5
0.4
0.3
0.2
0.1
0.05
0.01
0
Cell_Type_NEW
Ctrl
ExN
InN
Other
UD

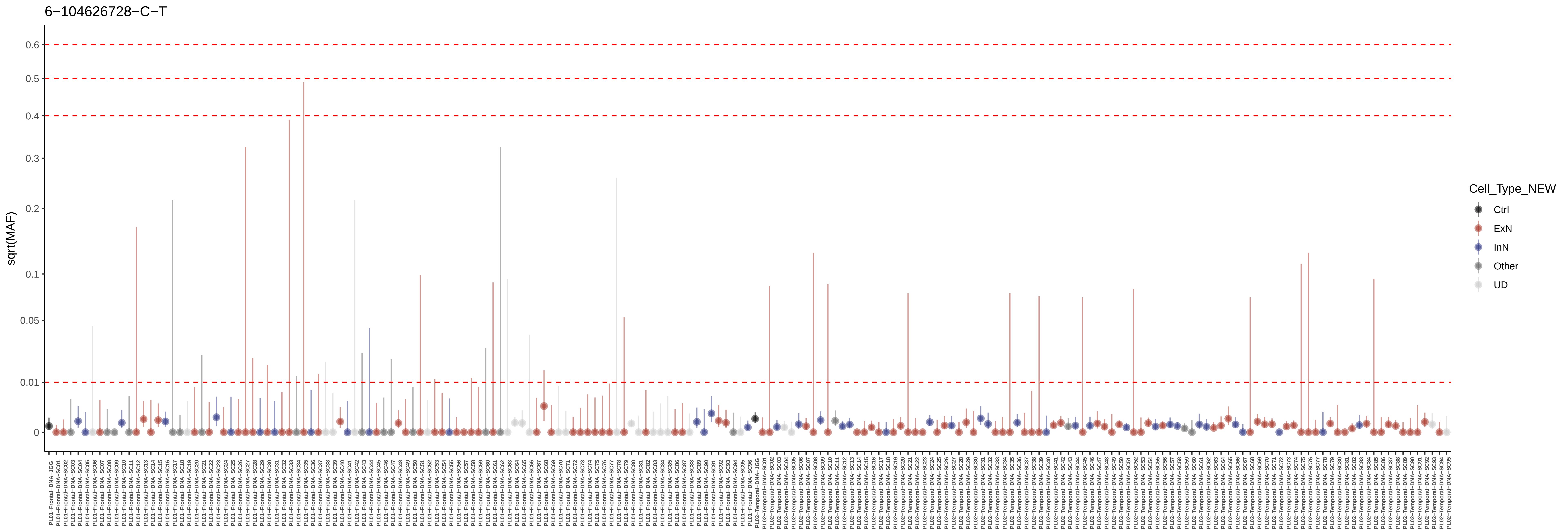

6−104626728−C−T
sqrt(MAF)
0.6
0.5
0.4
0.3
0.2
0.1
0.05
0.01
0
Cell_Type_NEW
Ctrl
ExN
InN
Other
UD

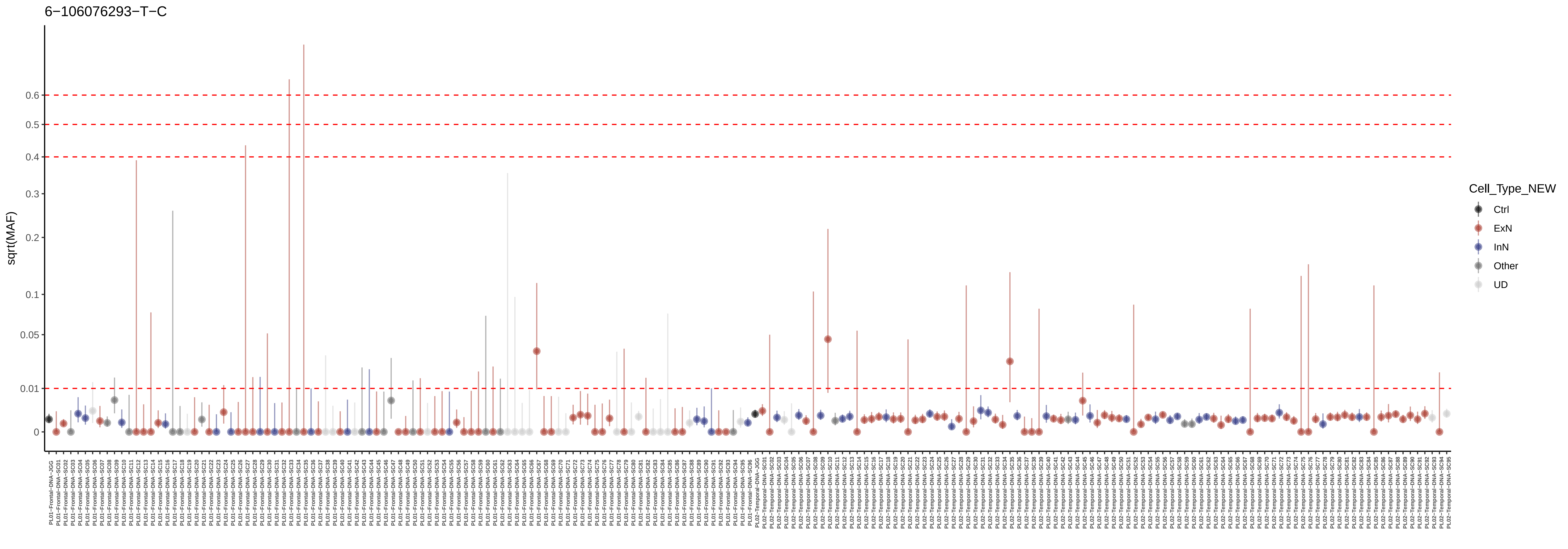

6−106076293−T−C
sqrt(MAF)
0.6
0.5
0.4
0.3
0.2
0.1
0.05
0.01
0
Cell_Type_NEW
Ctrl
ExN
InN
Other
UD

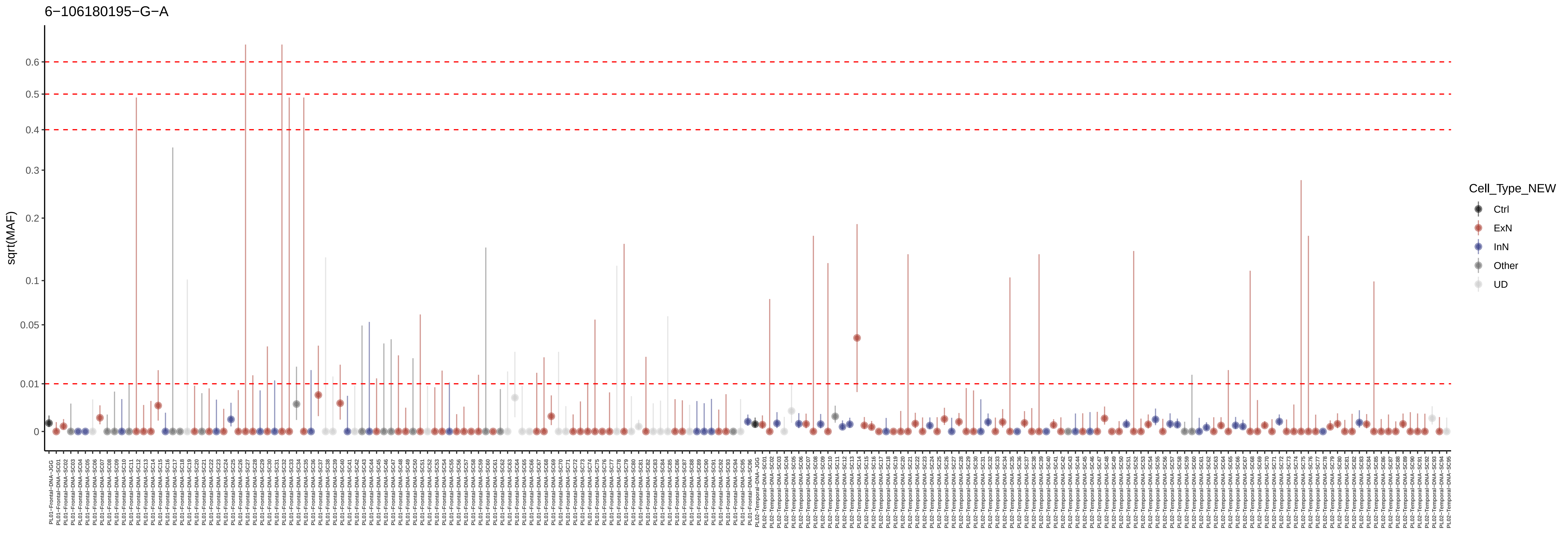

6−106180195−G−A
sqrt(MAF)
0.6
0.5
0.4
0.3
0.2
0.1
0.05
0.01
0
Cell_Type_NEW
Ctrl
ExN
InN
Other
UD

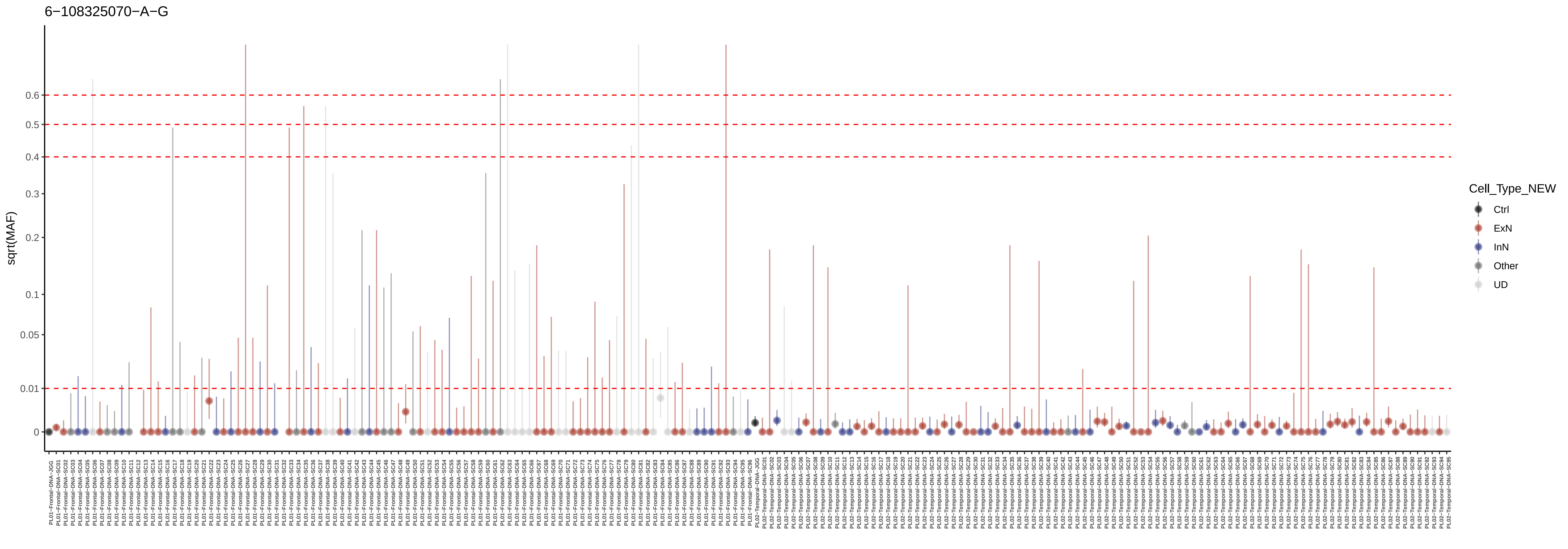
6−108325070−A−G
sqrt(MAF)
0
0.01
0.05
0.1
0.2
0.3
0.4
0.5
0.6
Cell_Type_NEW
Ctrl
ExN
InN
Other
UD

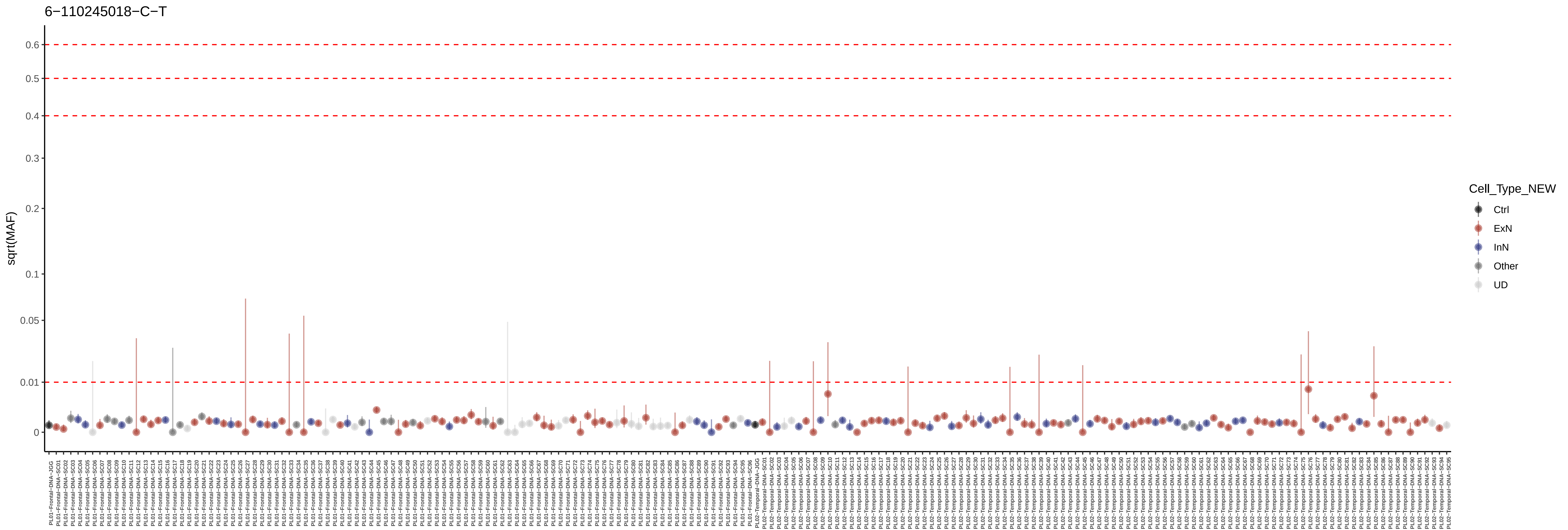
6−110245018−C−T
sqrt(MAF)
0.6
0.5
0.4
0.3
0.2
0.1
0.05
0.01
0
Cell_Type_NEW
Ctrl
ExN
InN
Other
UD

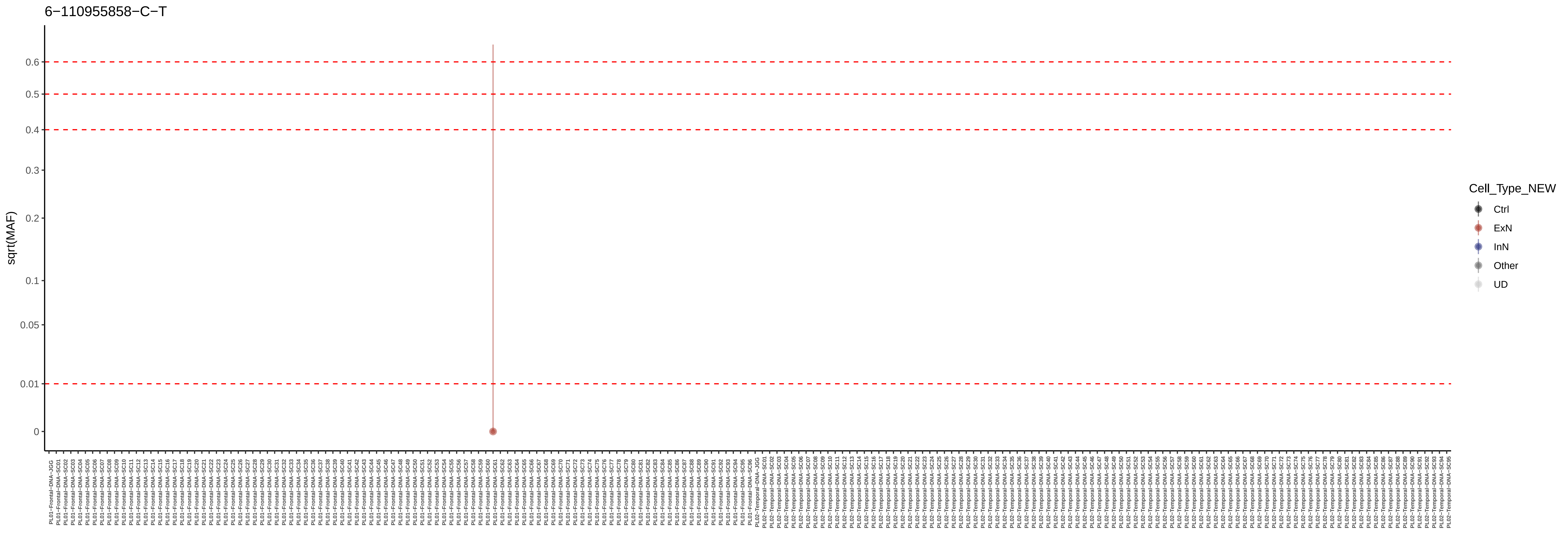
6−110955858−C−T
sqrt(MAF)
0.6
0.5
0.4
0.3
0.2
0.1
0.05
0.01
0
Cell_Type_NEW
Ctrl
ExN
InN
Other
UD

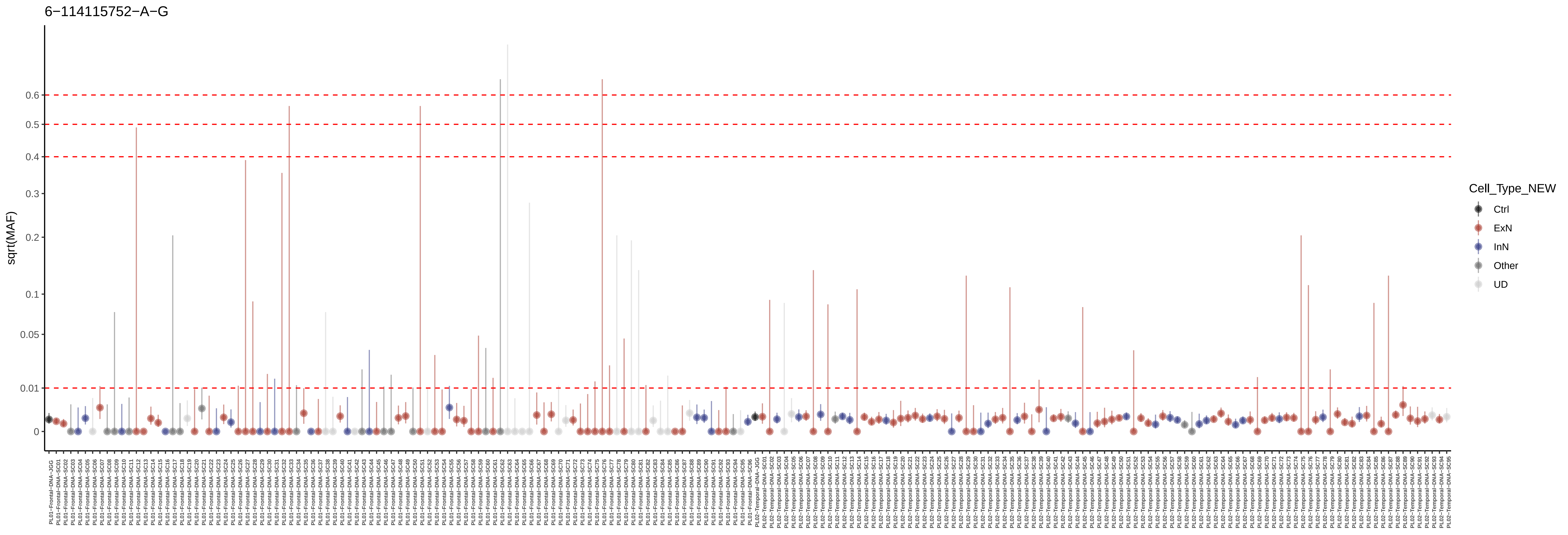

6−114115752−A−G
sqrt(MAF)
0.6
0.5
0.4
0.3
0.2
0.1
0.05
0.01
0
Cell_Type_NEW
Ctrl
ExN
InN
Other
UD

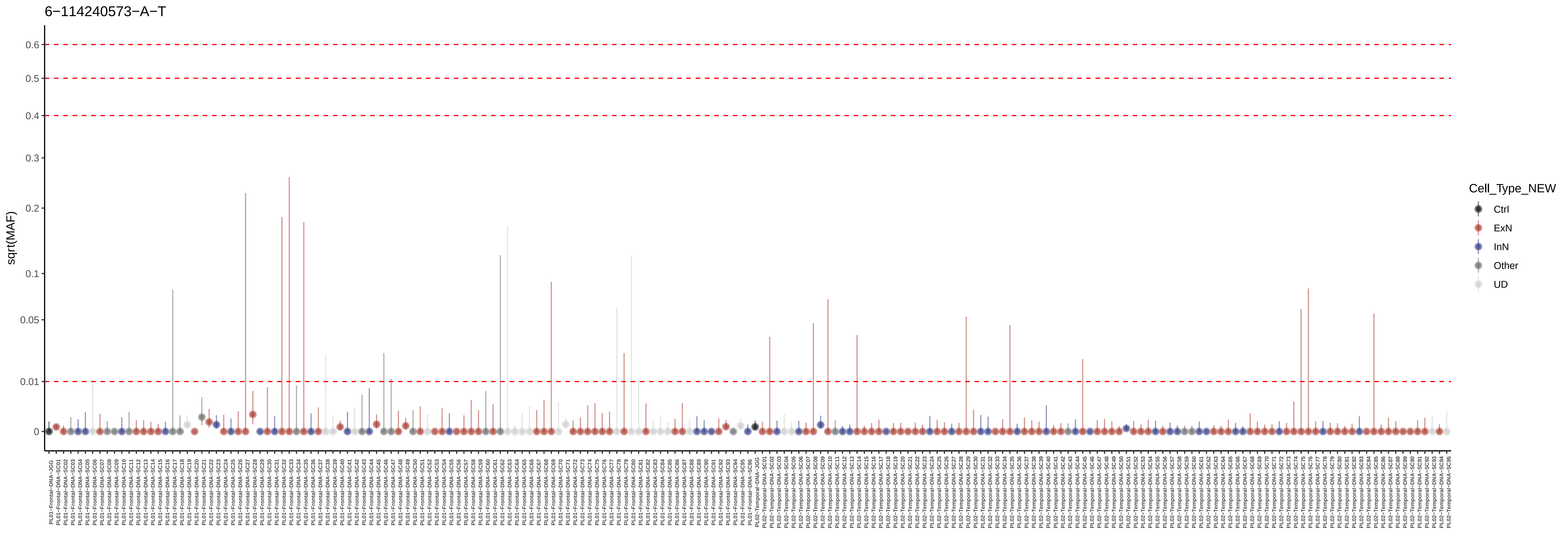

6−114240573−A−T
sqrt(MAF)
Cell_Type_NEW
Ctrl
ExN
InN
Other
UD

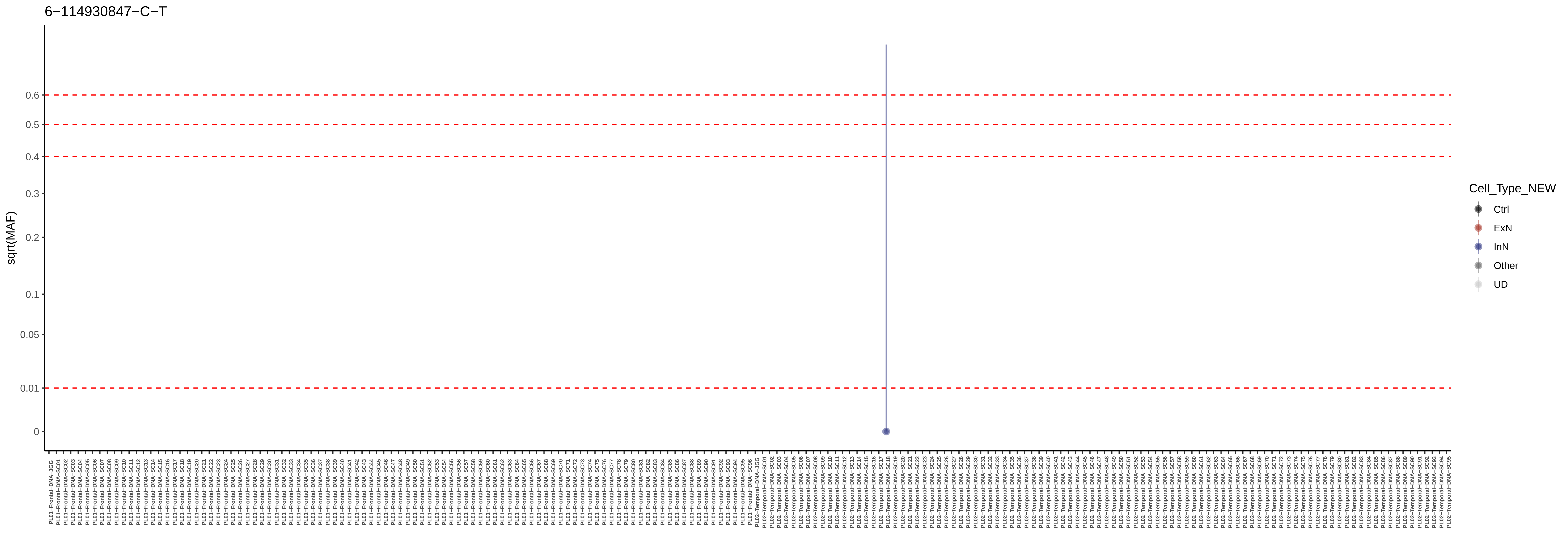
6−114930847−C−T
sqrt(MAF)
0.6
0.5
0.4
0.3
0.2
0.1
0.05
0.01
0
Cell_Type_NEW
Ctrl
ExN
InN
Other
UD

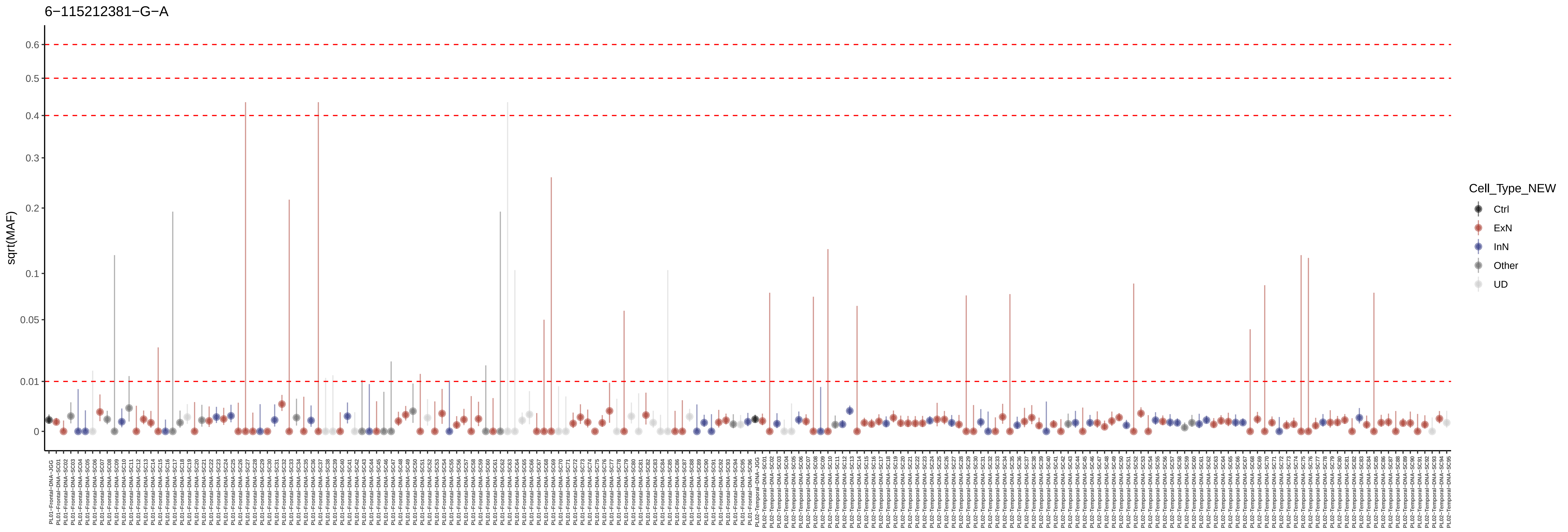
6−115212381−G−A
sqrt(MAF)
0.6
0.5
0.4
0.3
0.2
0.1
0.05
0.01
0
Cell_Type_NEW
Ctrl
ExN
InN
Other
UD

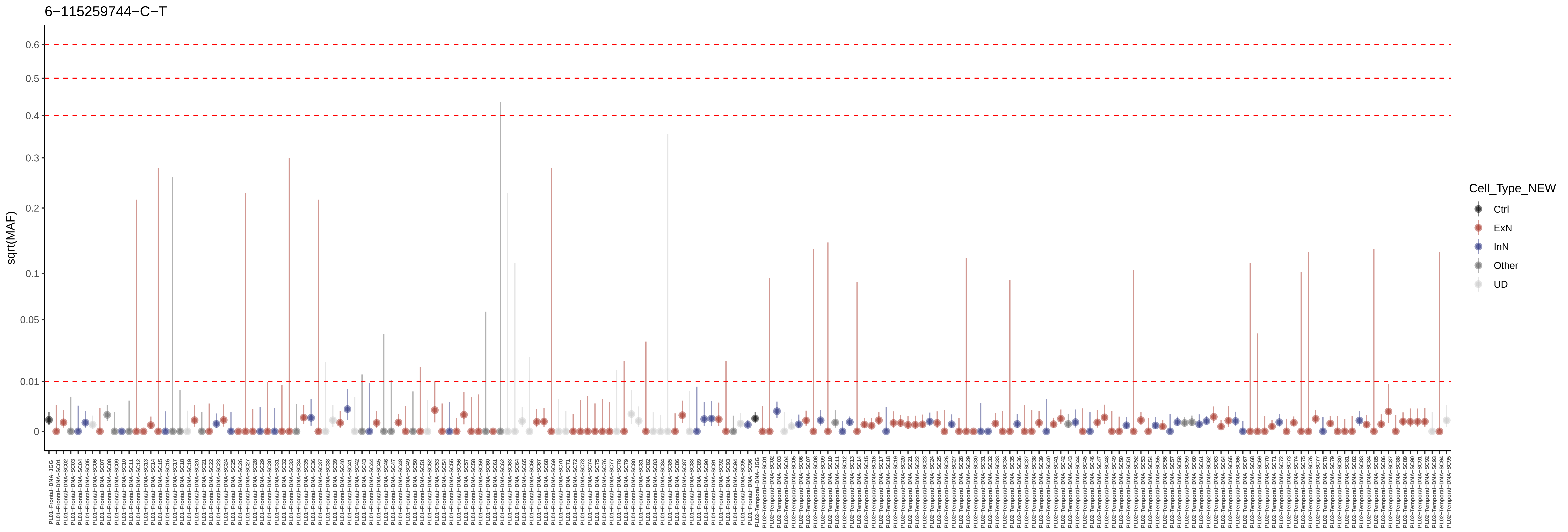

6−115259744−C−T
sqrt(MAF)
0.6
0.5
0.4
0.3
0.2
0.1
0.05
0.01
0
Cell_Type_NEW
Ctrl
ExN
InN
Other
UD

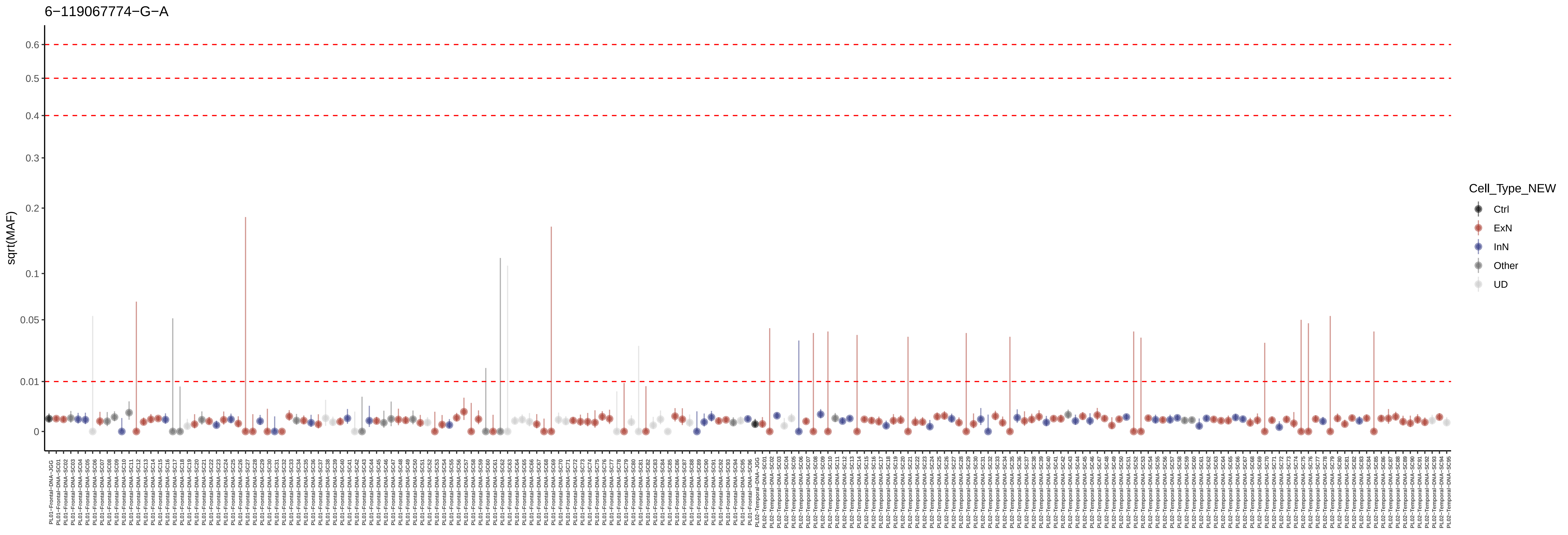
6−119067774−G−A
sqrt(MAF)
0.6
0.5
0.4
0.3
0.2
0.1
0.05
0.01
0
Cell_Type_NEW
Ctrl
ExN
InN
Other
UD

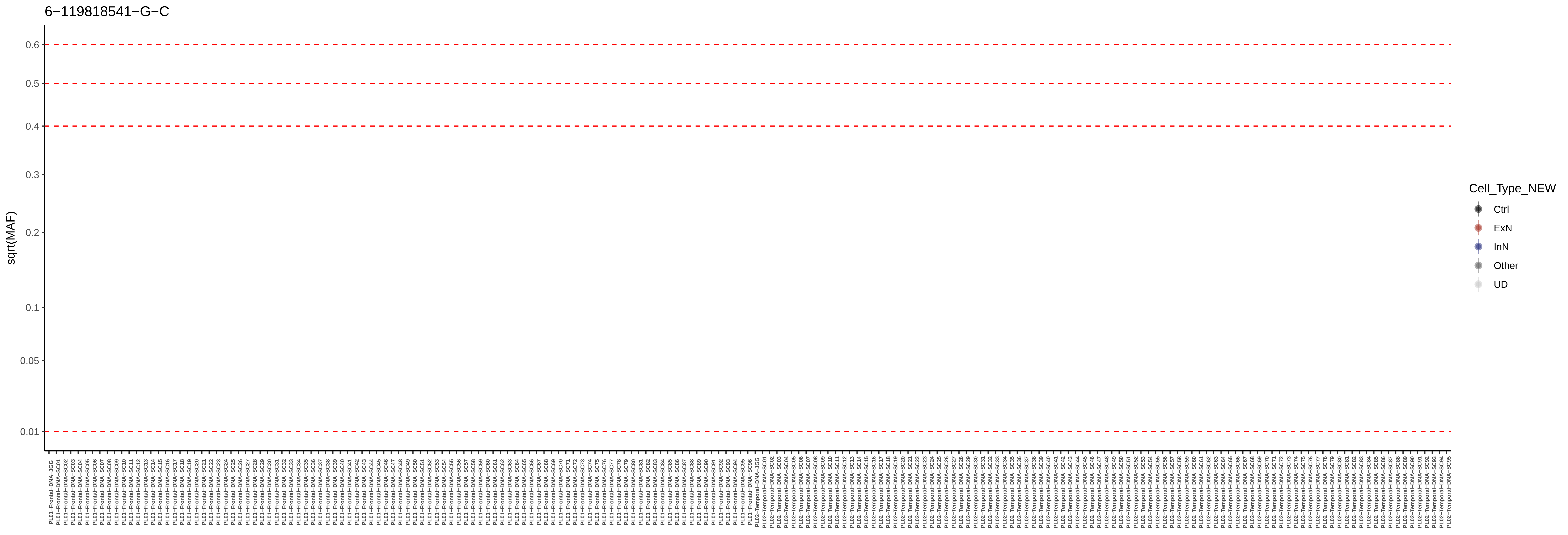
6−119818541−G−C
sqrt(MAF)
0.6
0.5
0.4
0.3
0.2
0.1
0.05
0.01
Cell_Type_NEW
Ctrl
ExN
InN
Other
UD

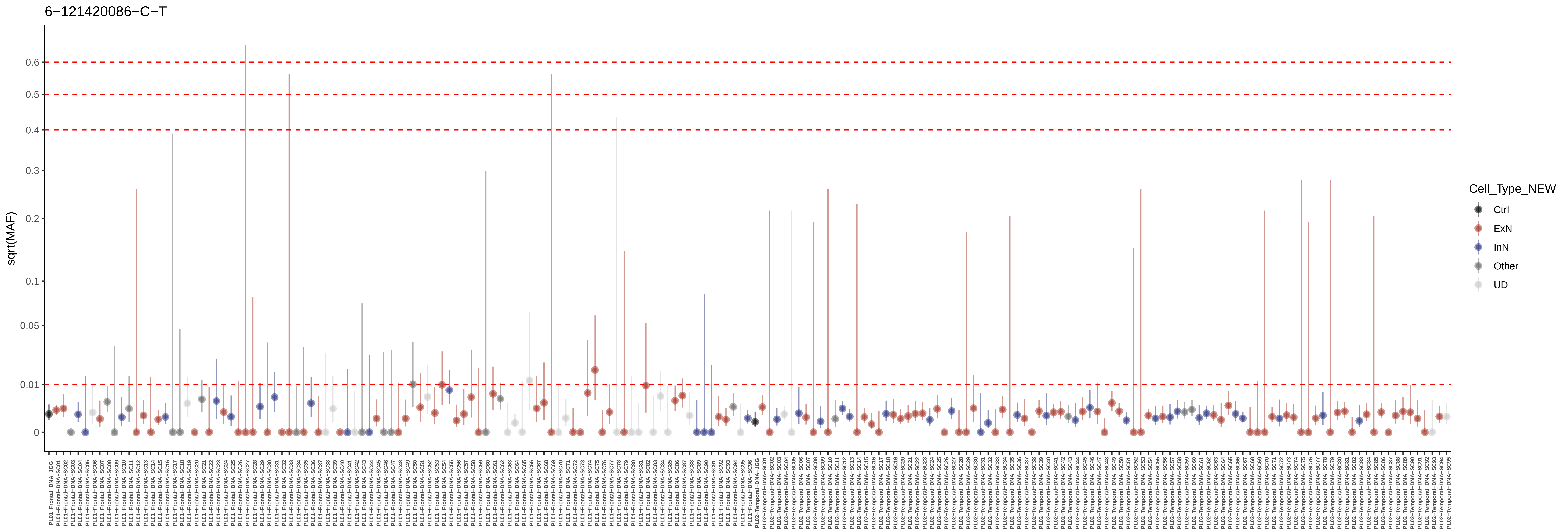

6−121420086−C−T
sqrt(MAF)
0
0.01
0.05
0.1
0.2
0.3
0.4
0.5
0.6
Cell_Type_NEW
Ctrl
ExN
InN
Other
UD

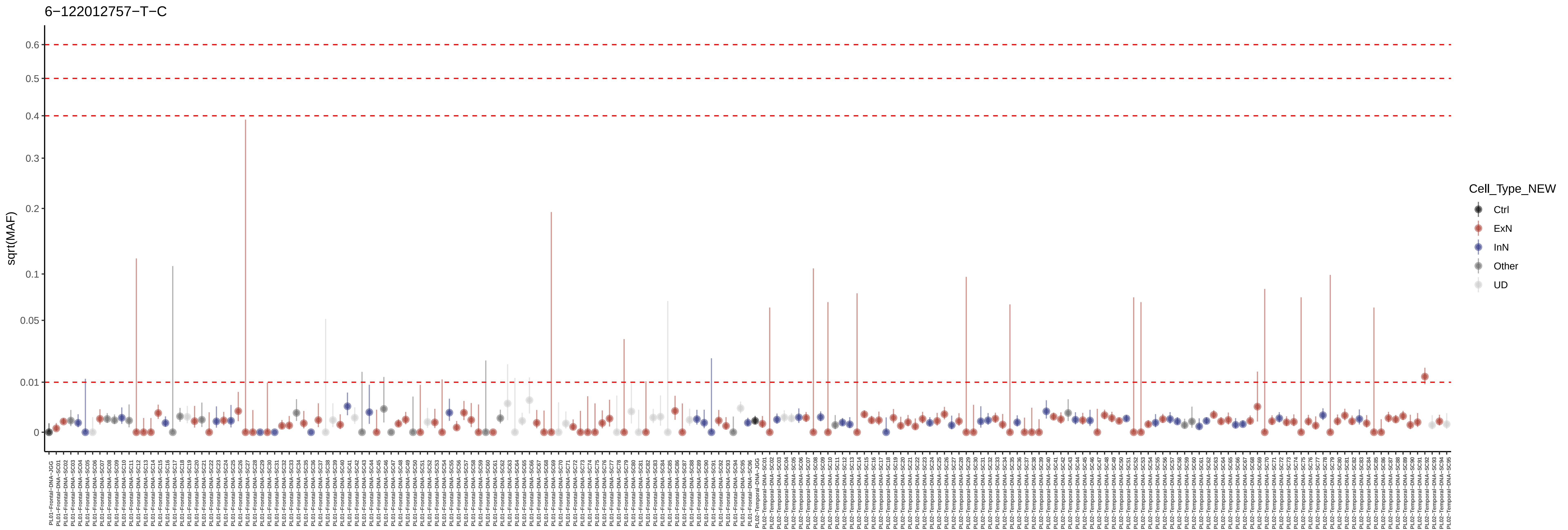
6−122012757−T−C
sqrt(MAF)
0.6
0.5
0.4
0.3
0.2
0.1
0.05
0.01
0
Cell_Type_NEW
Ctrl
ExN
InN
Other
UD

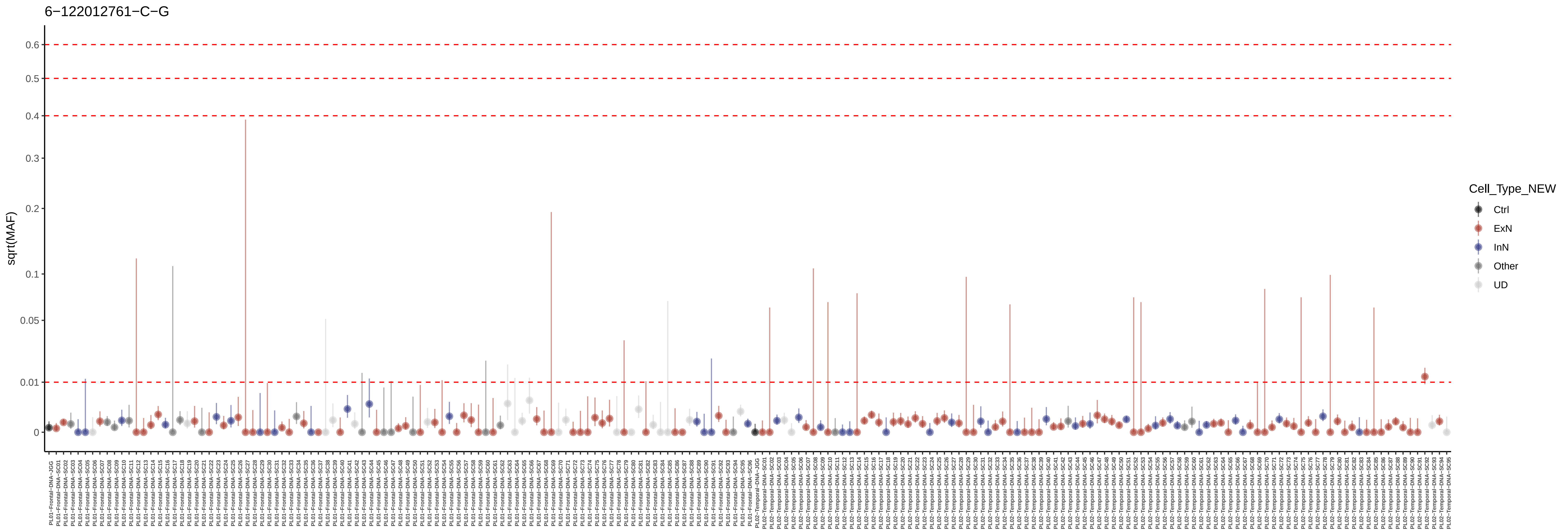

6−122012761−C−G
sqrt(MAF)
0.6
0.5
0.4
0.3
0.2
0.1
0.05
0.01
0
Cell_Type_NEW
Ctrl
ExN
InN
Other
UD

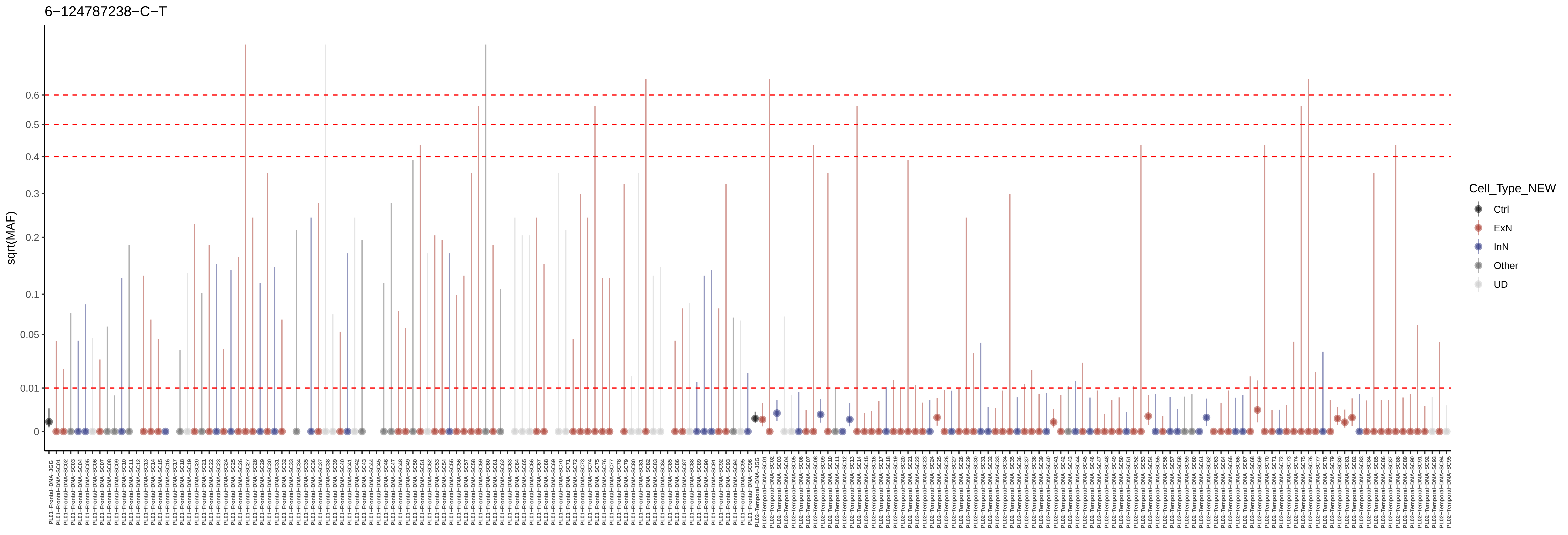
6−124787238−C−T
sqrt(MAF)
0.6
0.5
0.4
0.3
0.2
0.1
0.05
0.01
0
Cell_Type_NEW
Ctrl
ExN
InN
Other
UD

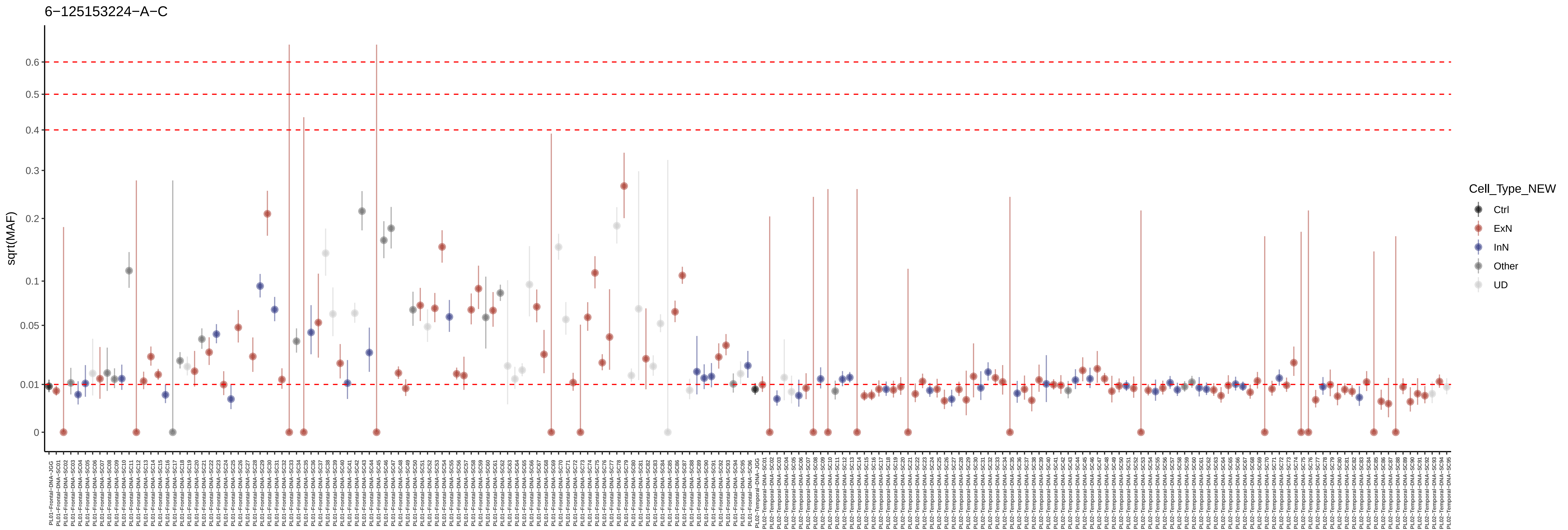
6−125153224−A−C
sqrt(MAF)
0.6
0.5
0.4
0.3
0.2
0.1
0.05
0.01
0
Cell_Type_NEW
Ctrl
ExN
InN
Other
UD

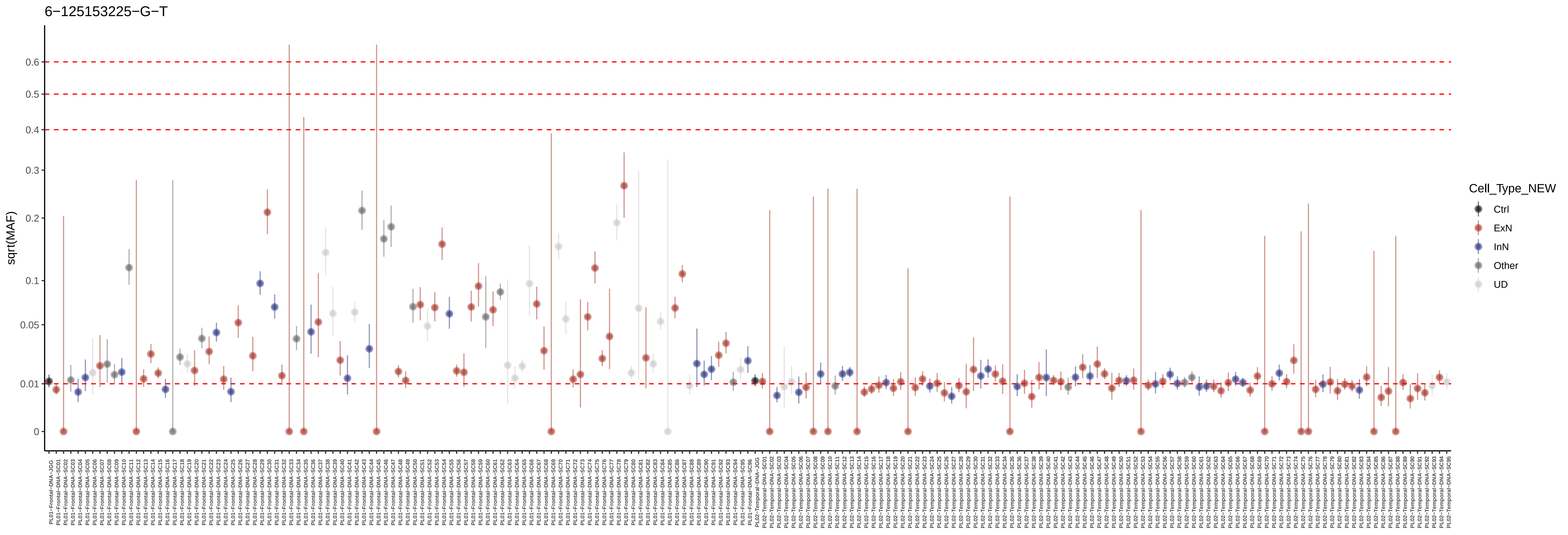
6−125153225−G−T
sqrt(MAF)
0.6
0.5
0.4
0.3
0.2
0.1
0.05
0.01
0
Cell_Type_NEW
Ctrl
ExN
InN
Other
UD

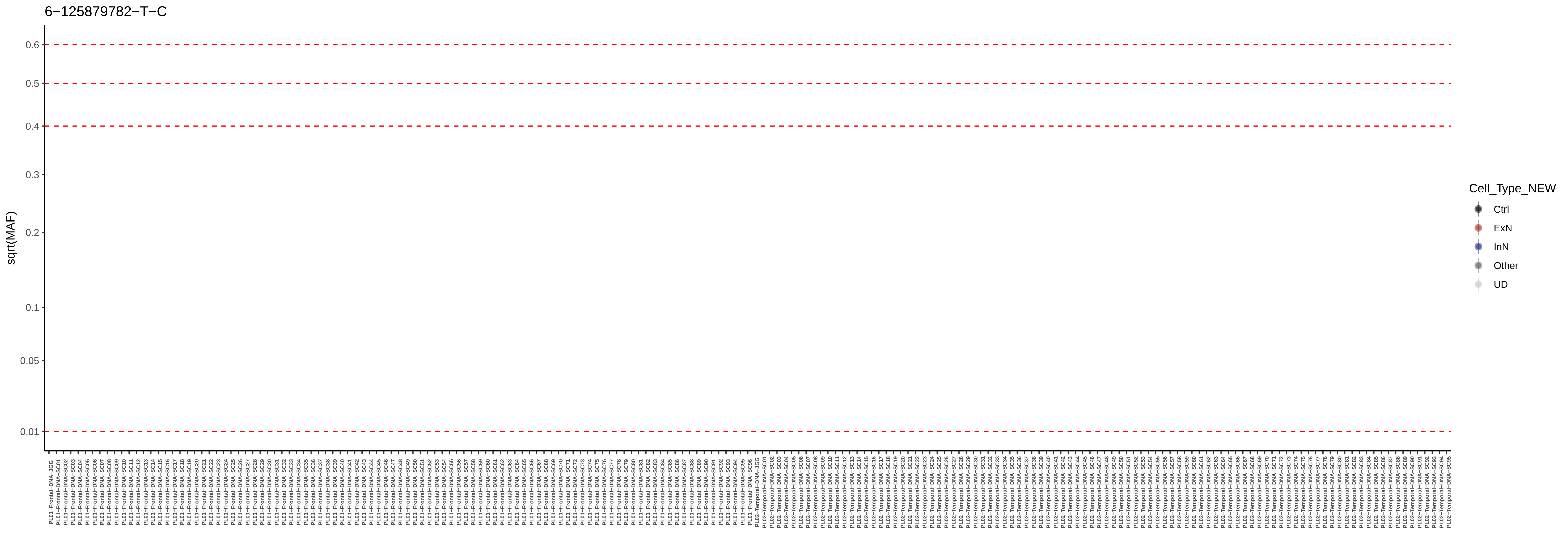
6−125879782−T−C
sqrt(MAF)
0.6
0.5
0.4
0.3
0.2
0.1
0.05
0.01
Cell_Type_NEW
Ctrl
ExN
InN
Other
UD

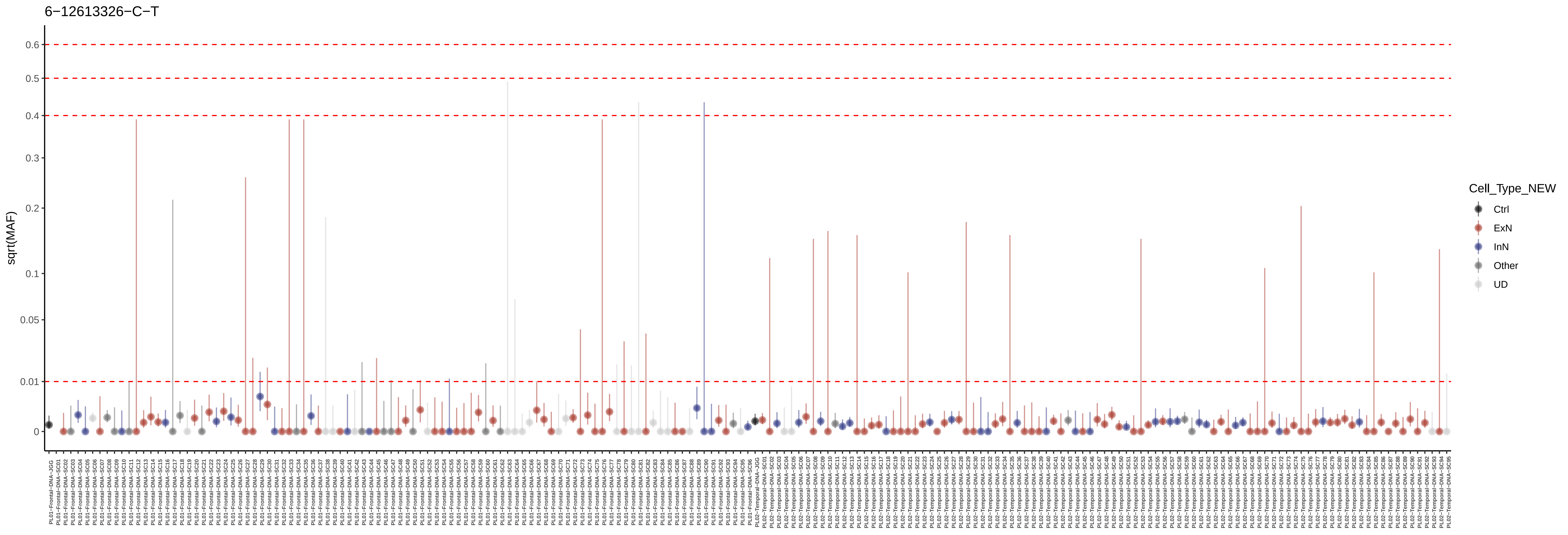
6−12613326−C−T
sqrt(MAF)
0.6
0.5
0.4
0.3
0.2
0.1
0.05
0.01
0
Cell_Type_NEW
Ctrl
ExN
InN
Other
UD

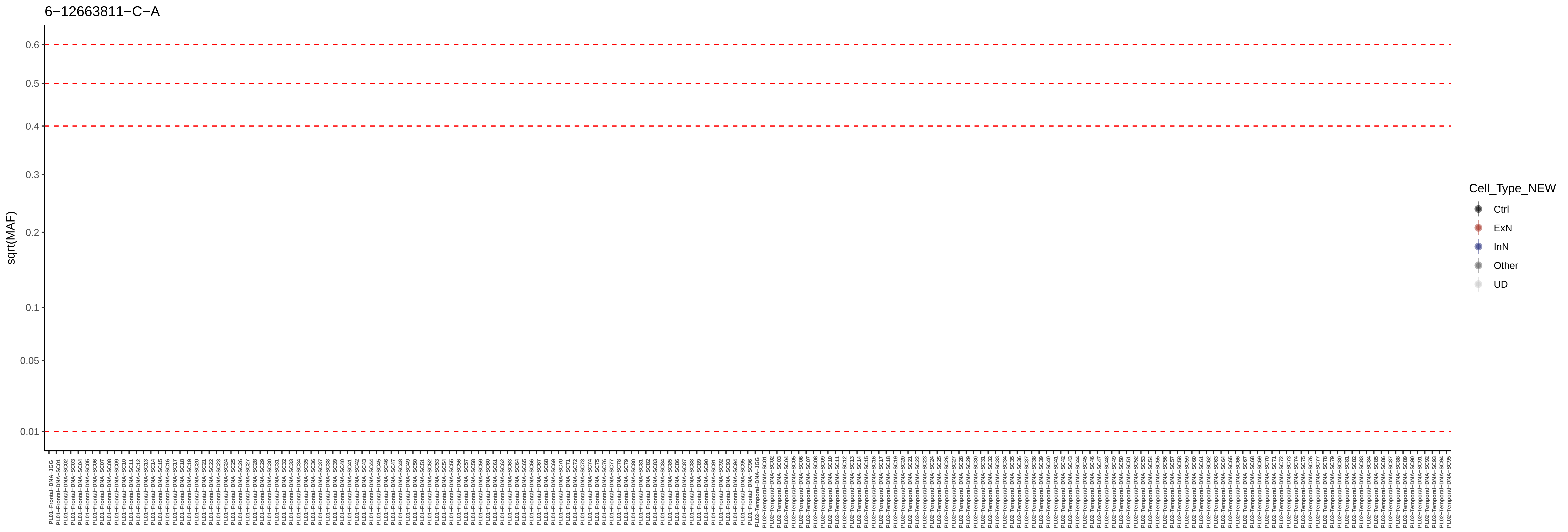
6−12663811−C−A
sqrt(MAF)
0.6
0.5
0.4
0.3
0.2
0.1
0.05
0.01
Cell_Type_NEW
Ctrl
ExN
InN
Other
UD

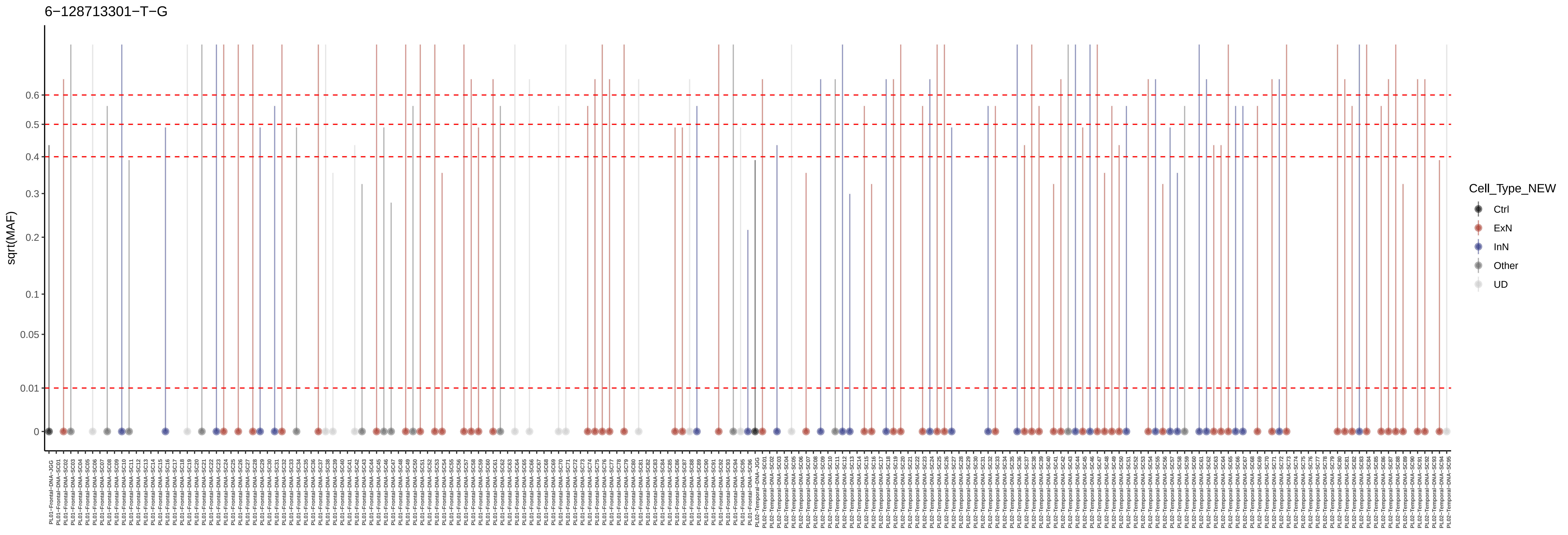

6−128713301−T−G
sqrt(MAF)
0.6
0.5
0.4
0.3
0.2
0.1
0.05
0.01
0
Cell_Type_NEW
Ctrl
ExN
InN
Other
UD

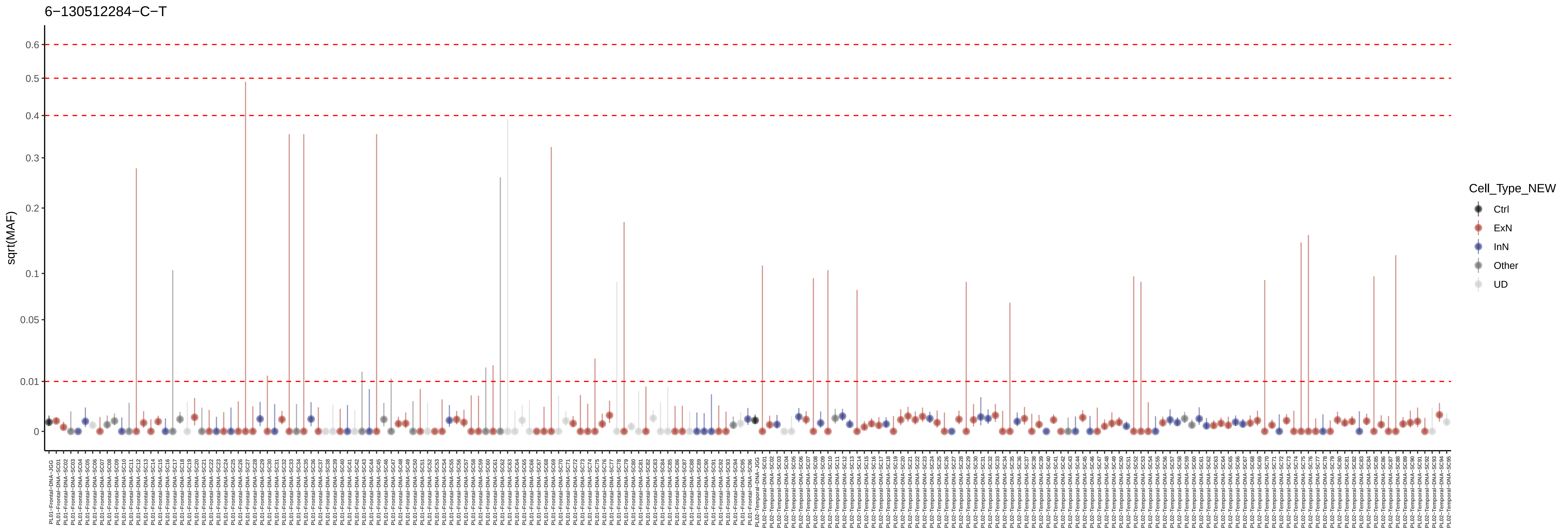
6−130512284−C−T
sqrt(MAF)
0.6
0.5
0.4
0.3
0.2
0.1
0.05
0.01
0
Cell_Type_NEW
Ctrl
ExN
InN
Other
UD

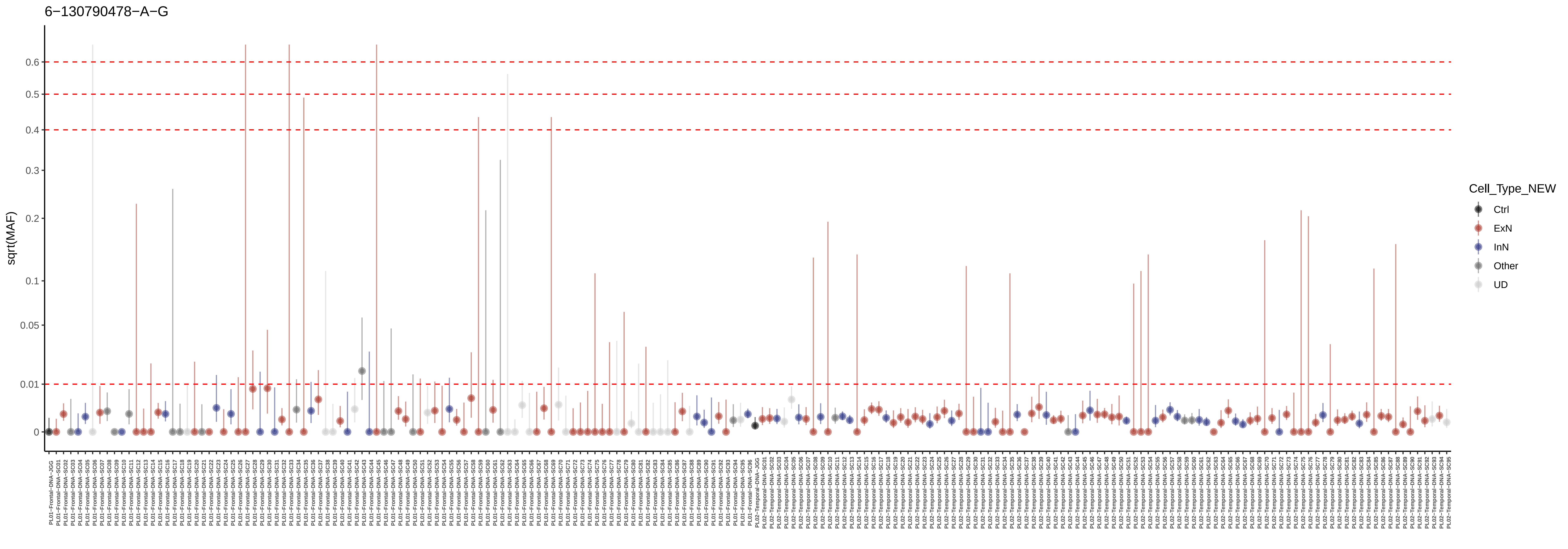
6−130790478−A−G
sqrt(MAF)
0.6
0.5
0.4
0.3
0.2
0.1
0.05
0.01
0
Cell_Type_NEW
Ctrl
ExN
InN
Other
UD

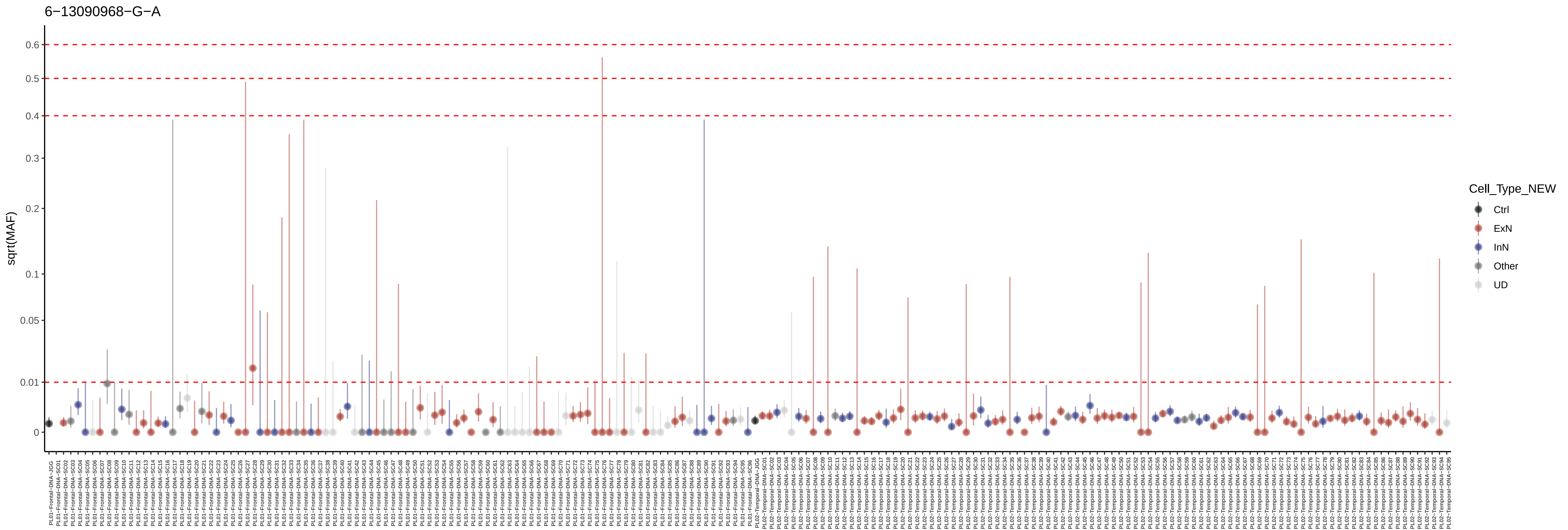
6−13090968−G−A
sqrt(MAF)
0.6
0.5
0.4
0.3
0.2
0.1
0.05
0.01
0
Cell_Type_NEW
Ctrl
ExN
InN
Other
UD

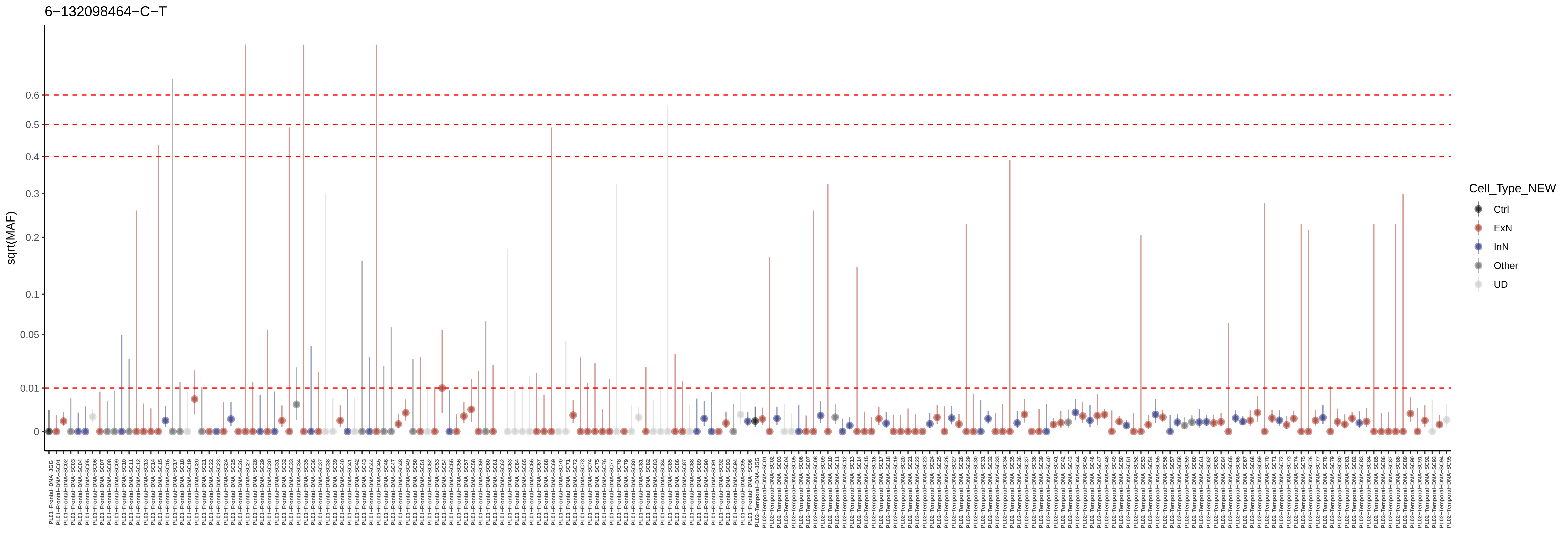

6−132098464−C−T
sqrt(MAF)
0.6
0.5
0.4
0.3
0.2
0.1
0.05
0.01
0
Cell_Type_NEW
Ctrl
ExN
InN
Other
UD

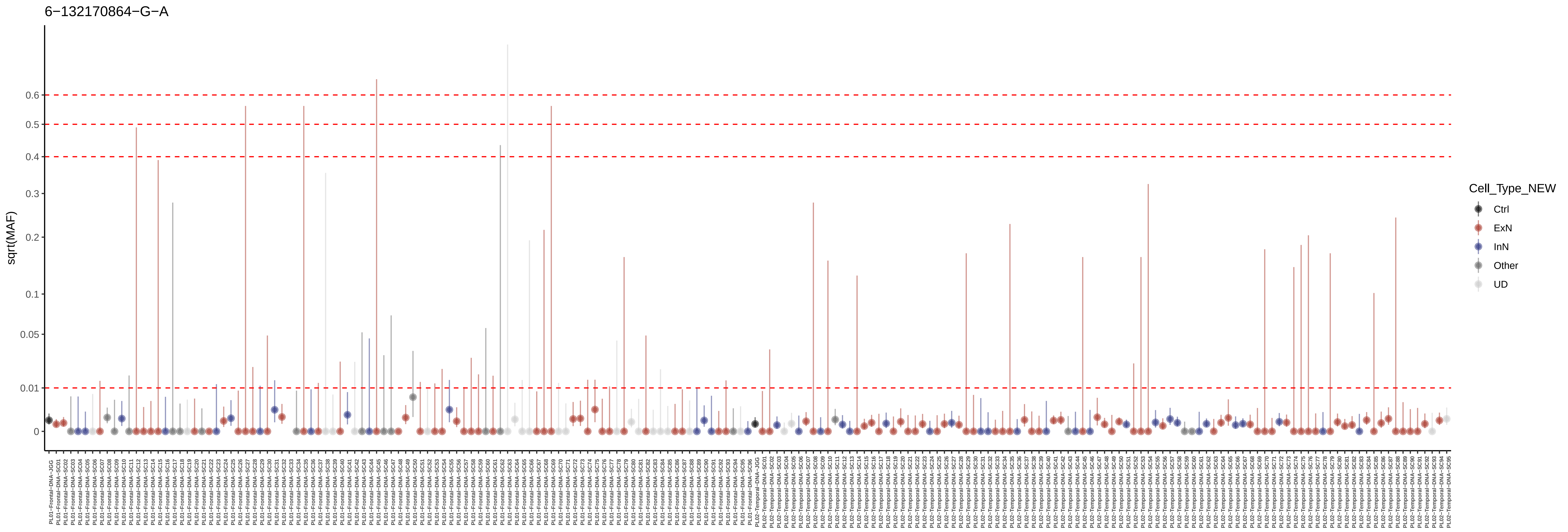
6−132170864−G−A
sqrt(MAF)
0.6
0.5
0.4
0.3
0.2
0.1
0.05
0.01
0
Cell_Type_NEW
Ctrl
ExN
InN
Other
UD

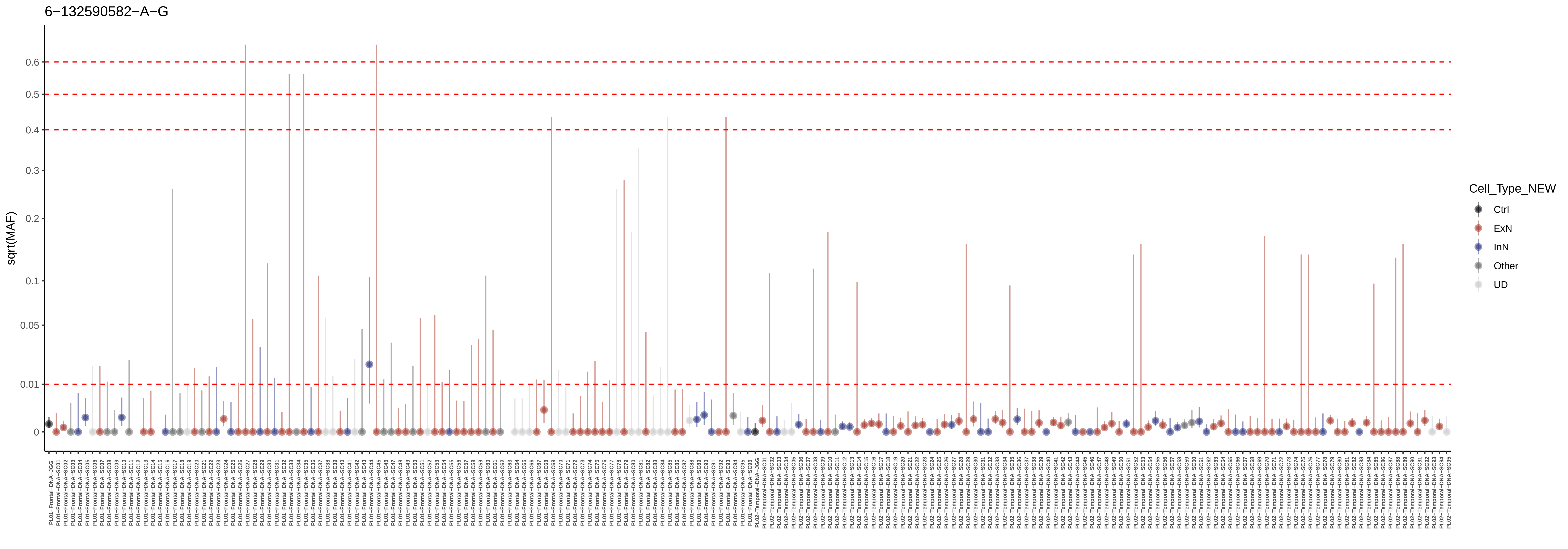

6−132590582−A−G
sqrt(MAF)
0.6
0.5
0.4
0.3
0.2
0.1
0.05
0.01
0
Cell_Type_NEW
Ctrl
ExN
InN
Other
UD

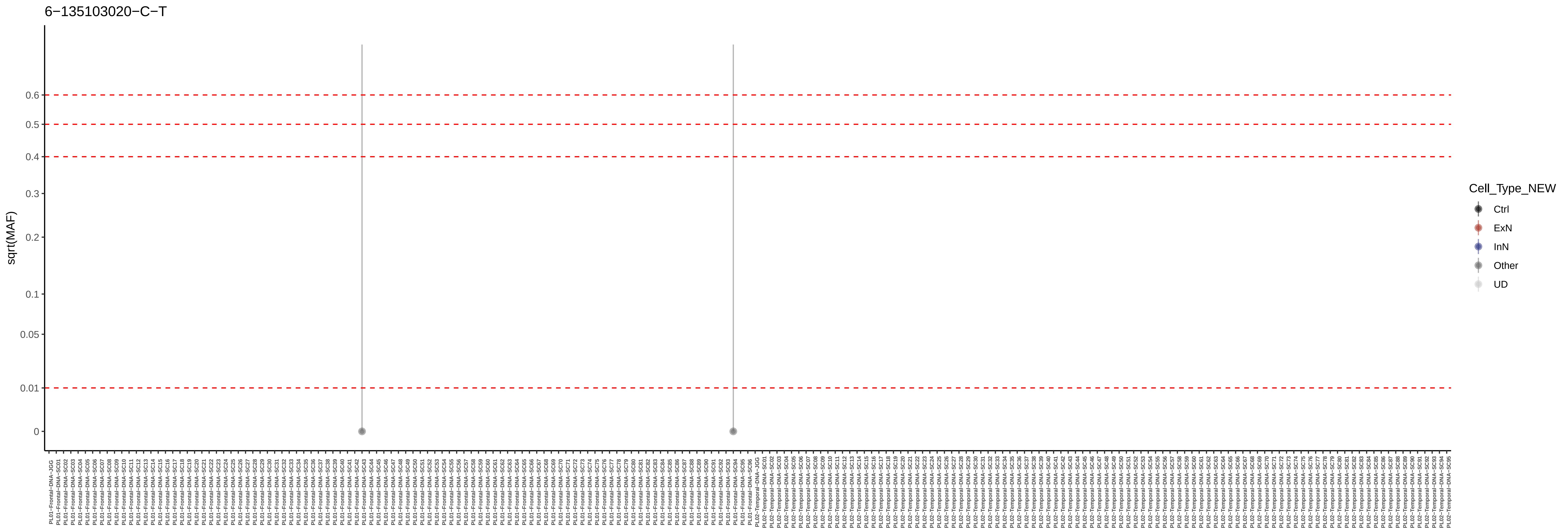
6−135103020−C−T
sqrt(MAF)
0.6
0.5
0.4
0.3
0.2
0.1
0.05
0.01
0
Cell_Type_NEW
Ctrl
ExN
InN
Other
UD

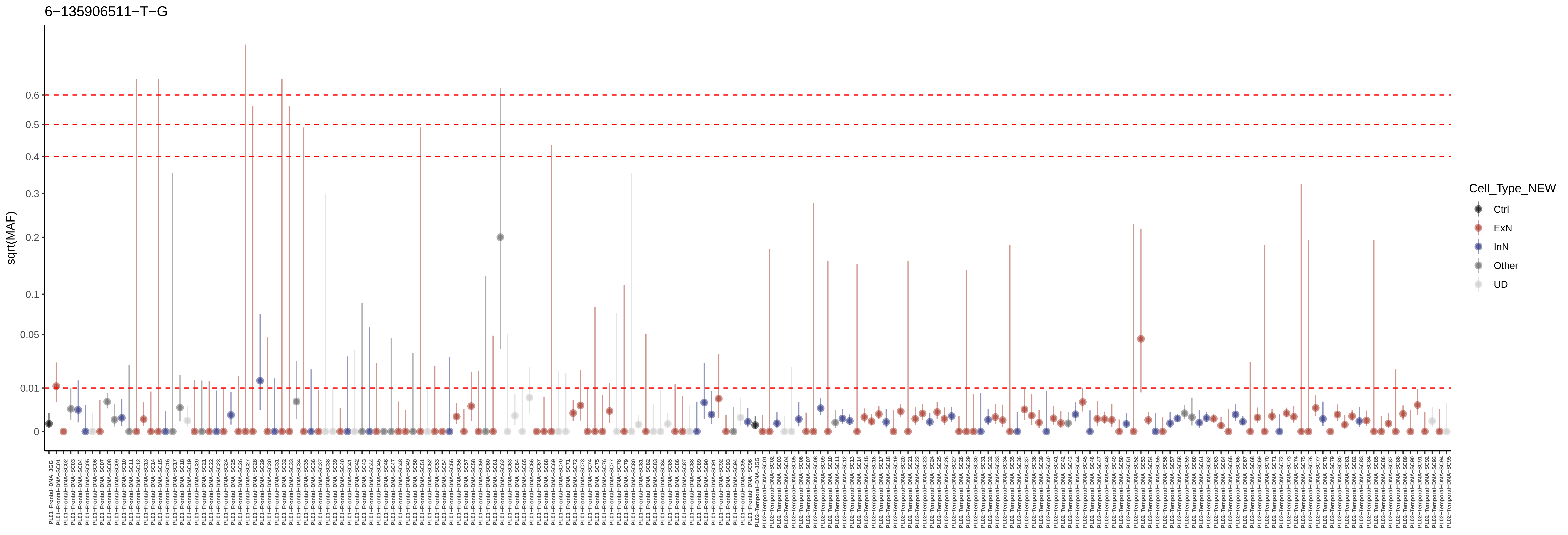
6−135906511−T−G
sqrt(MAF)
0.6
0.5
0.4
0.3
0.2
0.1
0.05
0.01
0
Cell_Type_NEW
Ctrl
ExN
InN
Other
UD

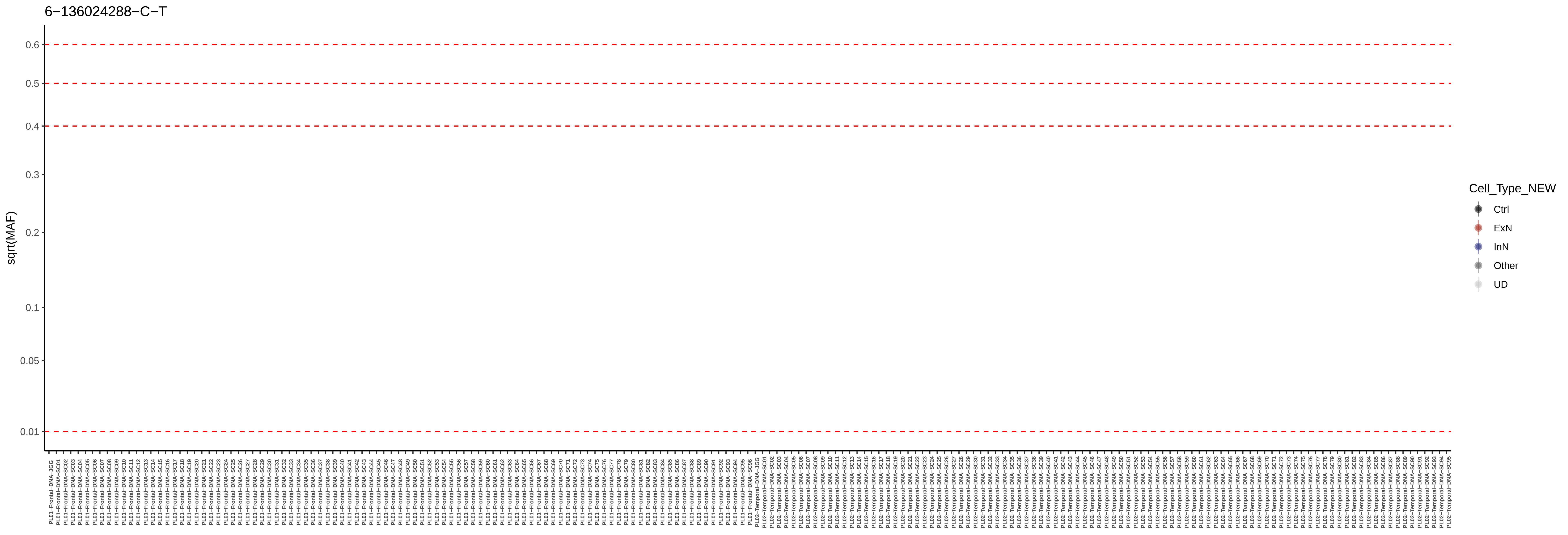
6−136024288−C−T
sqrt(MAF)
0.6
0.5
0.4
0.3
0.2
0.1
0.05
0.01
Cell_Type_NEW
Ctrl
ExN
InN
Other
UD

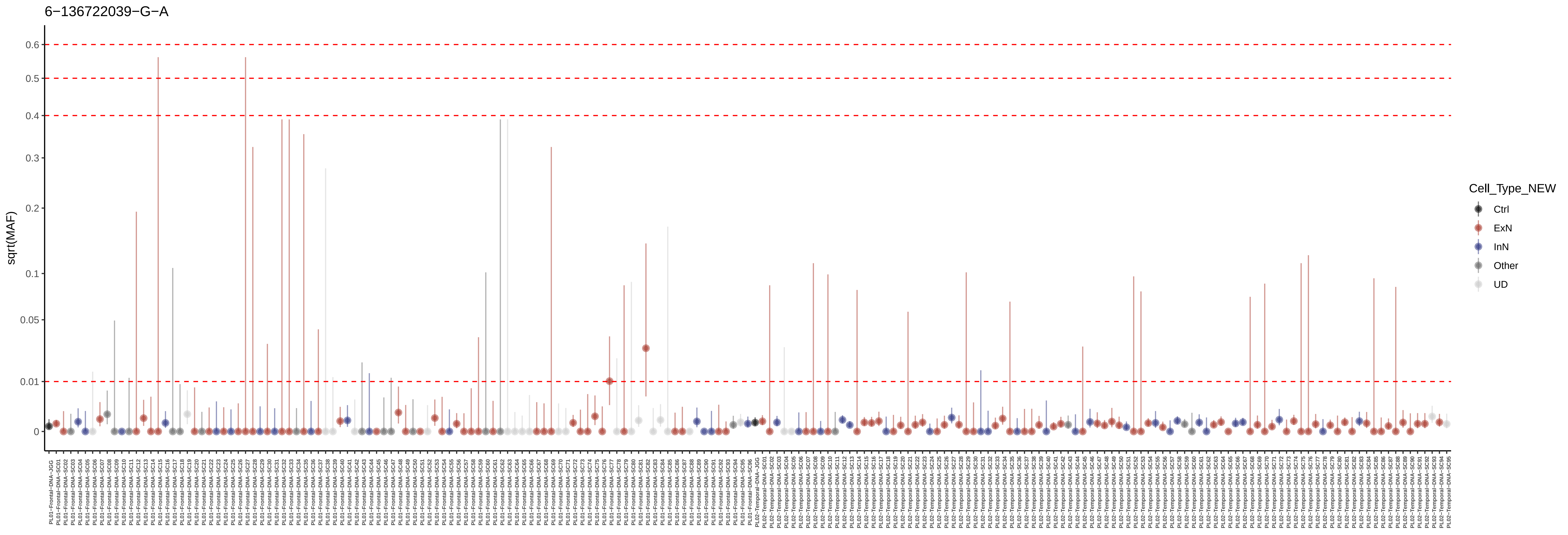
6−136722039−G−A
sqrt(MAF)
0.6
0.5
0.4
0.3
0.2
0.1
0.05
0.01
0
Cell_Type_NEW
Ctrl
ExN
InN
Other
UD

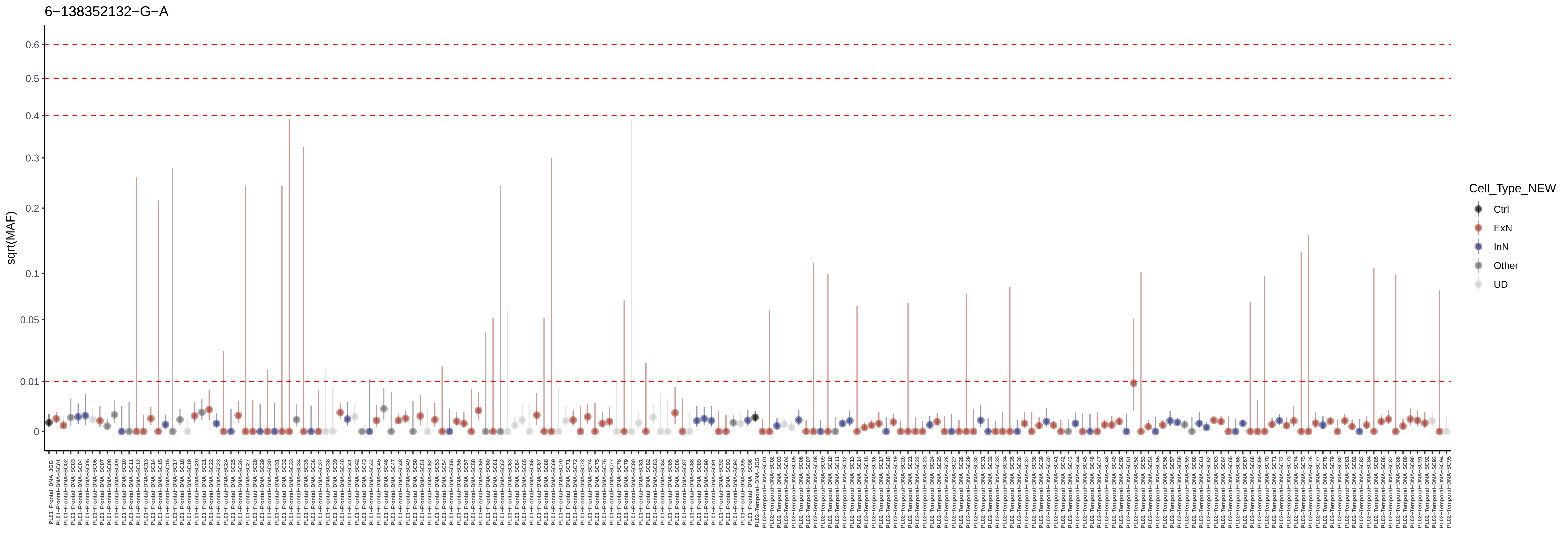
6−138352132−G−A
sqrt(MAF)
0.6
0.5
0.4
0.3
0.2
0.1
0.05
0.01
0
Cell_Type_NEW
Ctrl
ExN
InN
Other
UD

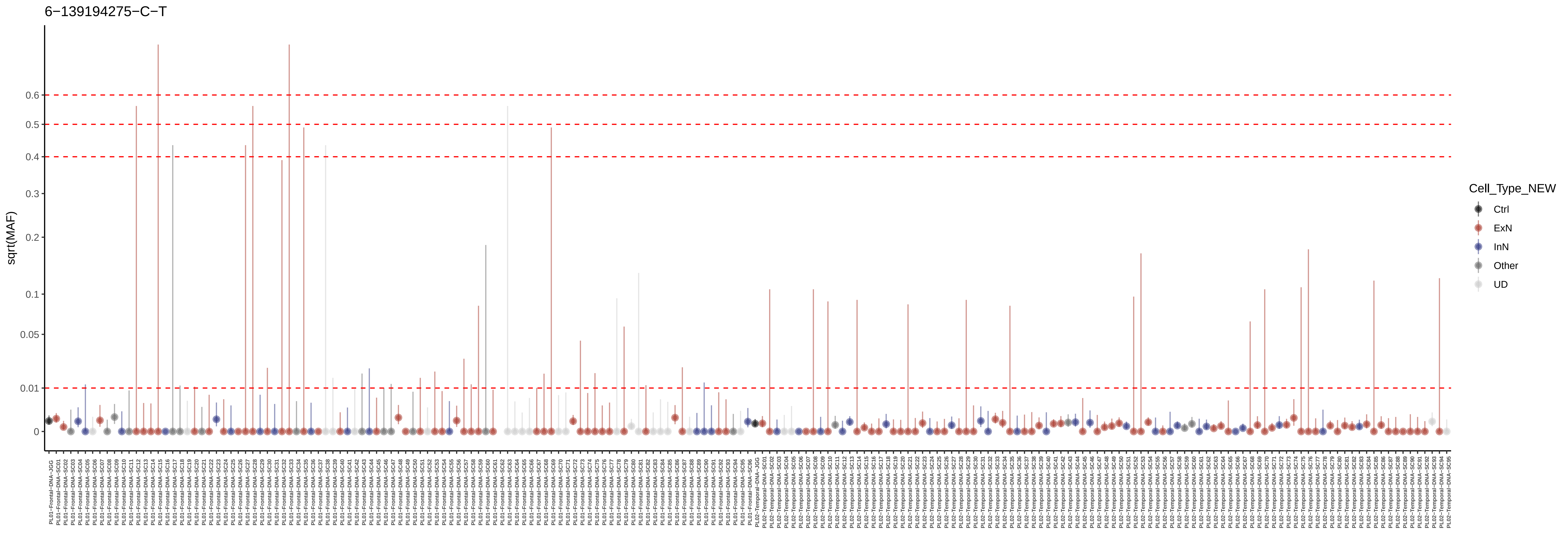
6−139194275−C−T
sqrt(MAF)
0.6
0.5
0.4
0.3
0.2
0.1
0.05
0.01
0
Cell_Type_NEW
Ctrl
ExN
InN
Other
UD

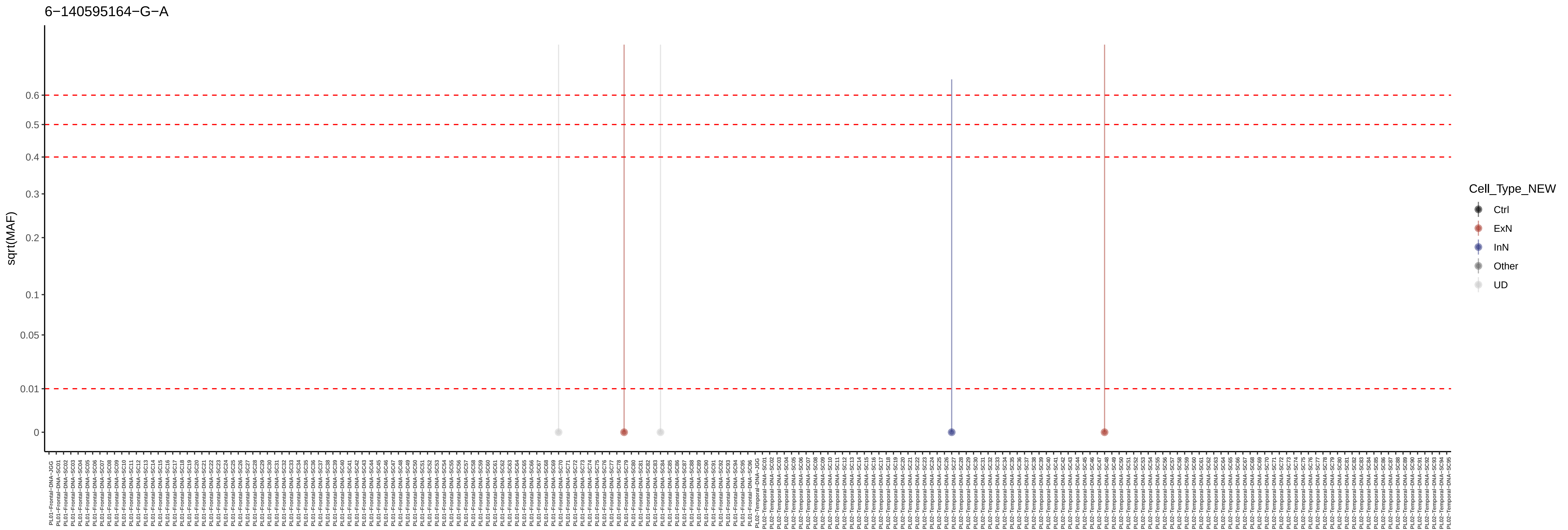
6−140595164−G−A
sqrt(MAF)
0
0.01
0.05
0.1
0.2
0.3
0.4
0.5
0.6
Cell_Type_NEW
Ctrl
ExN
InN
Other
UD

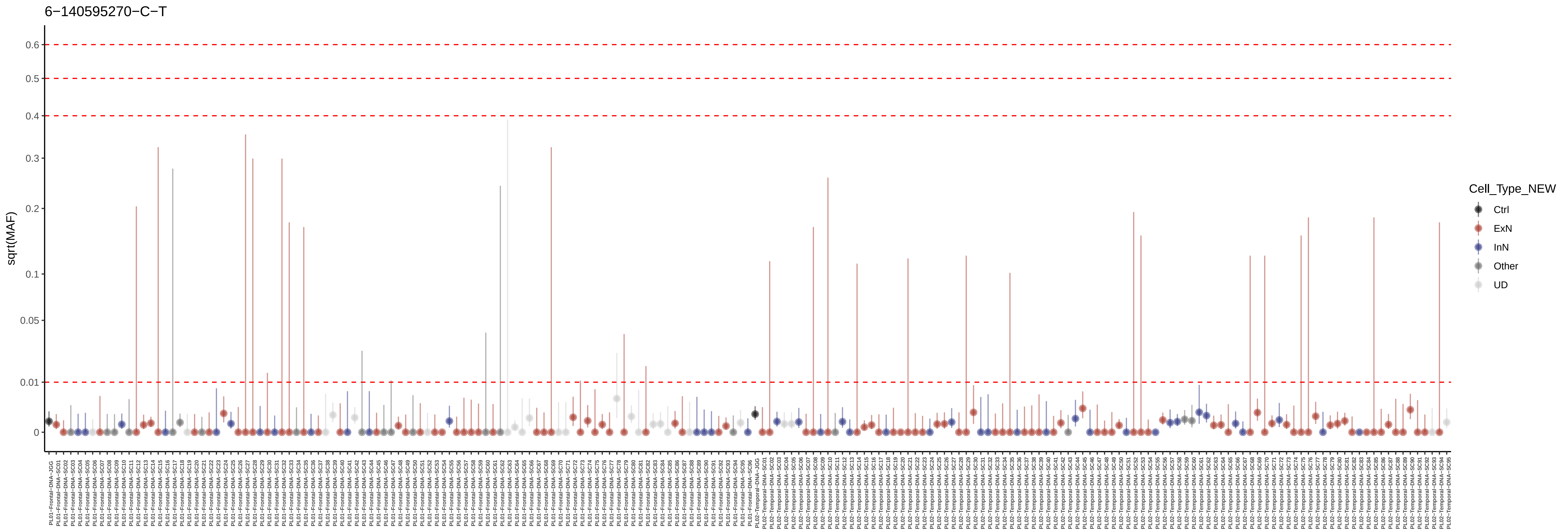
6−140595270−C−T
sqrt(MAF)
0.6
0.5
0.4
0.3
0.2
0.1
0.05
0.01
0
Cell_Type_NEW
Ctrl
ExN
InN
Other
UD

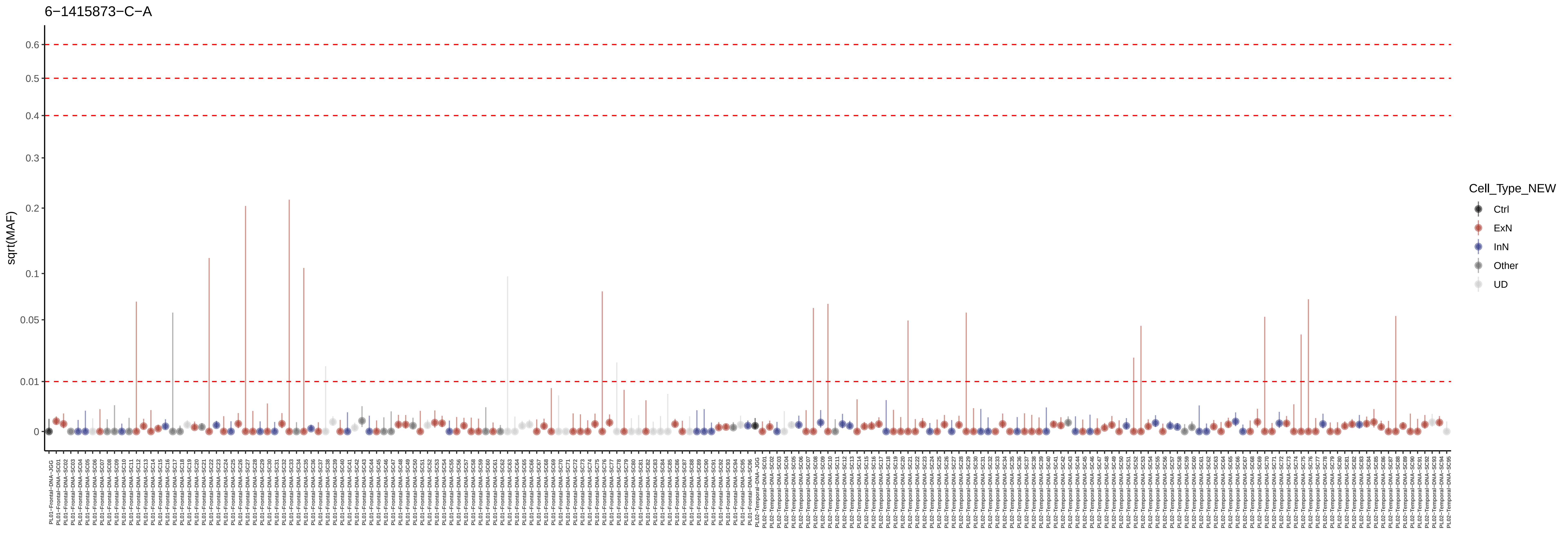

6−1415873−C−A
sqrt(MAF)
0.6
0.5
0.4
0.3
0.2
0.1
0.05
0.01
0
Cell_Type_NEW
Ctrl
ExN
InN
Other
UD

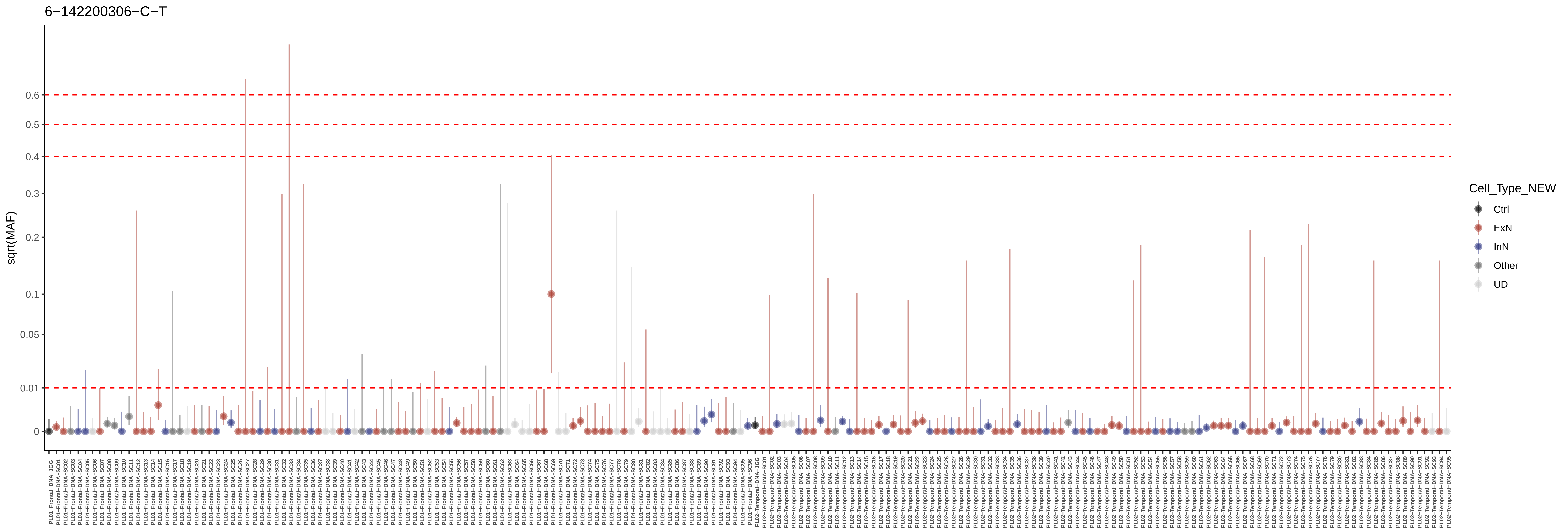
6−142200306−C−T
sqrt(MAF)
0.6
0.5
0.4
0.3
0.2
0.1
0.05
0.01
0
Cell_Type_NEW
Ctrl
ExN
InN
Other
UD

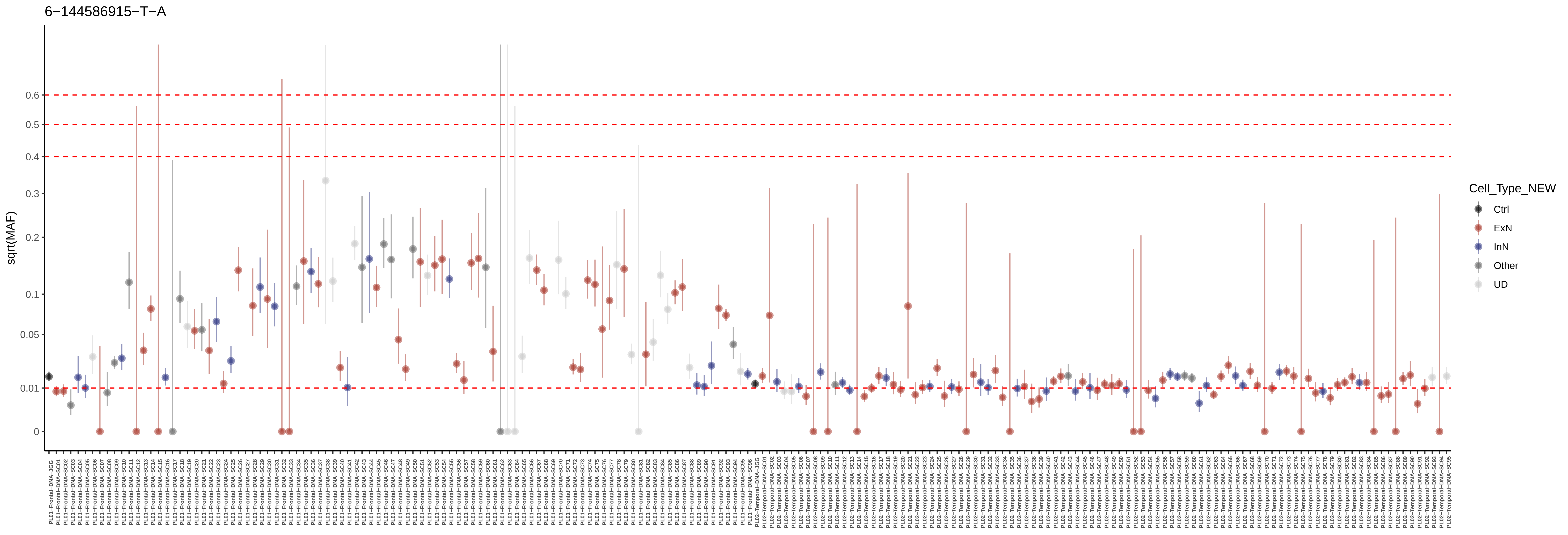
6−144586915−T−A
sqrt(MAF)
0.6
0.5
0.4
0.3
0.2
0.1
0.05
0.01
0
Cell_Type_NEW
Ctrl
ExN
InN
Other
UD

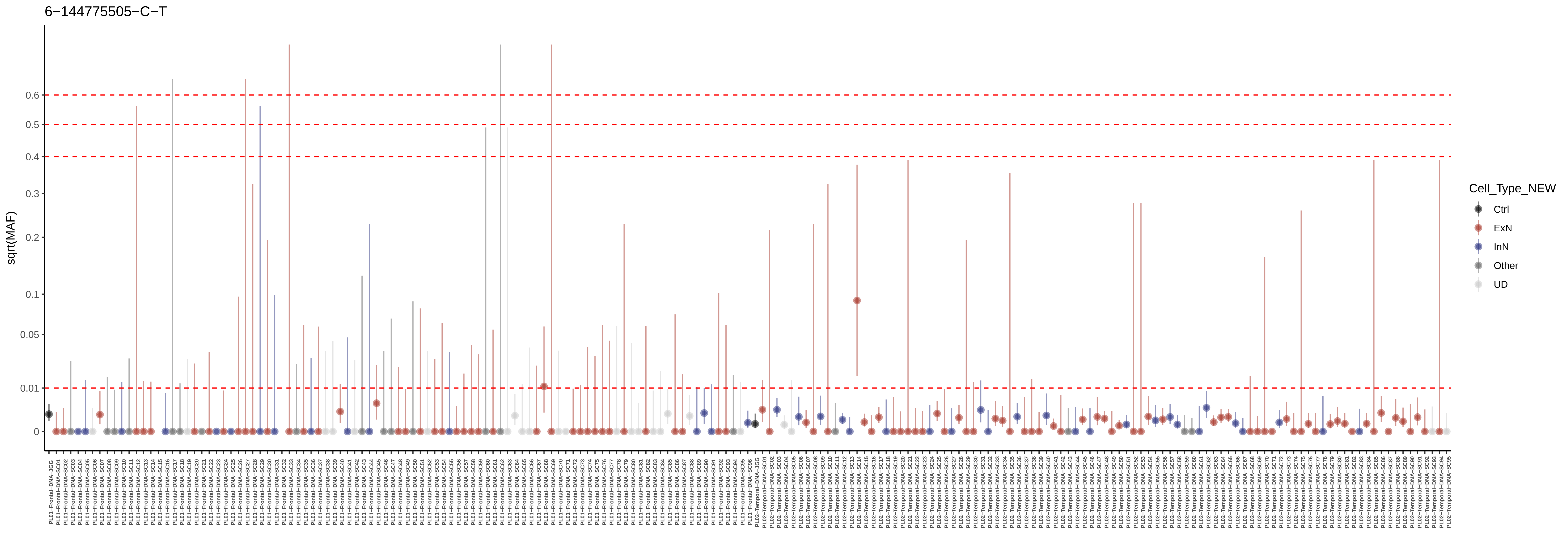
6−144775505−C−T
sqrt(MAF)
0.6
0.5
0.4
0.3
0.2
0.1
0.05
0.01
0
Cell_Type_NEW
Ctrl
ExN
InN
Other
UD

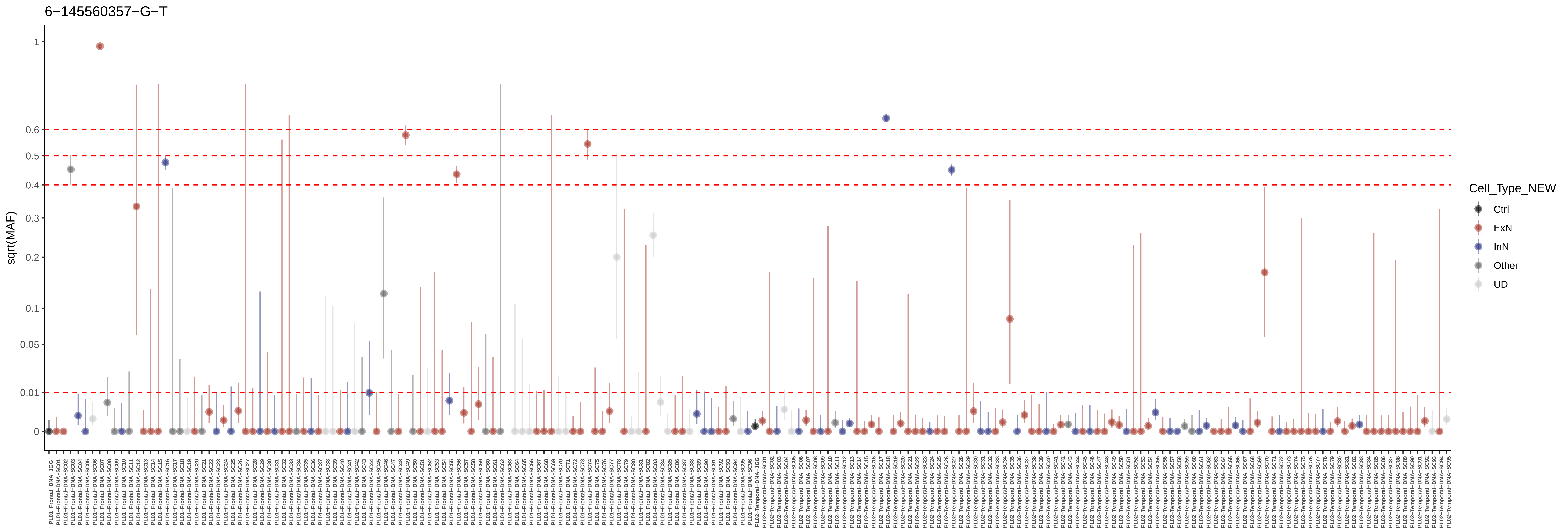
6−145560357−G−T
sqrt(MAF)
1
0.6
0.5
0.4
0.3
0.2
0.1
0.05
0.01
0
Cell_Type_NEW
Ctrl
ExN
InN
Other
UD

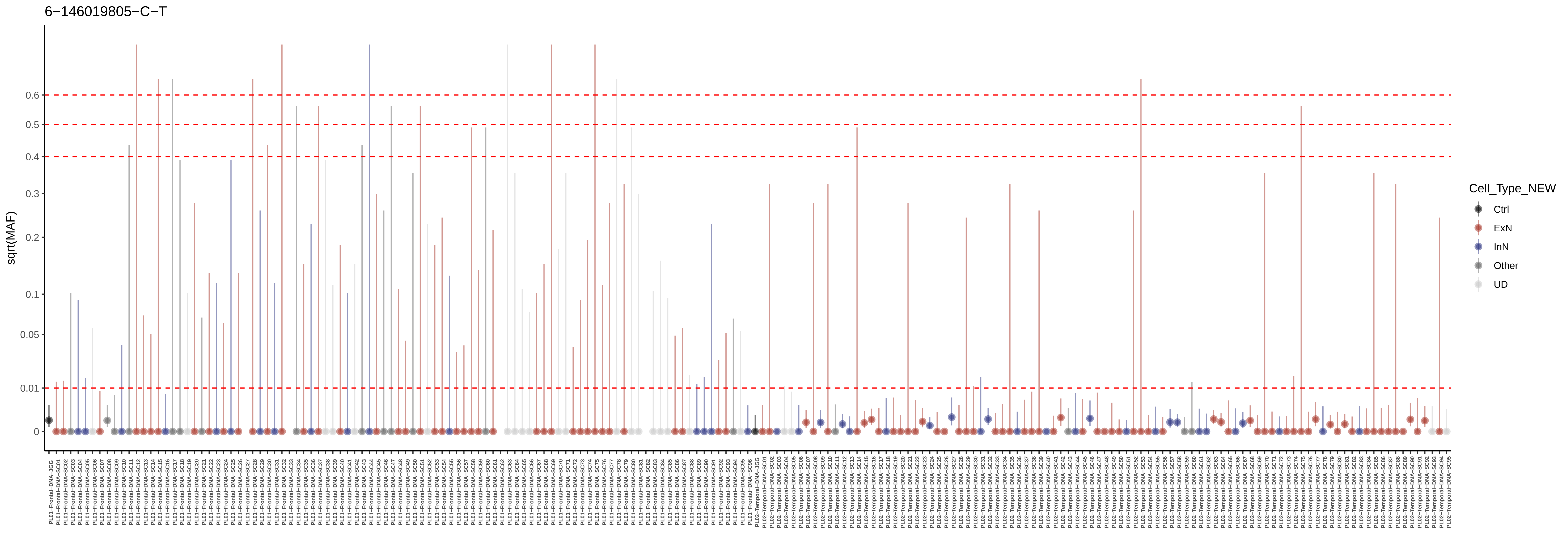

6−146019805−C−T
sqrt(MAF)
0
0.01
0.05
0.1
0.2
0.3
0.4
0.5
0.6
Cell_Type_NEW
Ctrl
ExN
InN
Other
UD

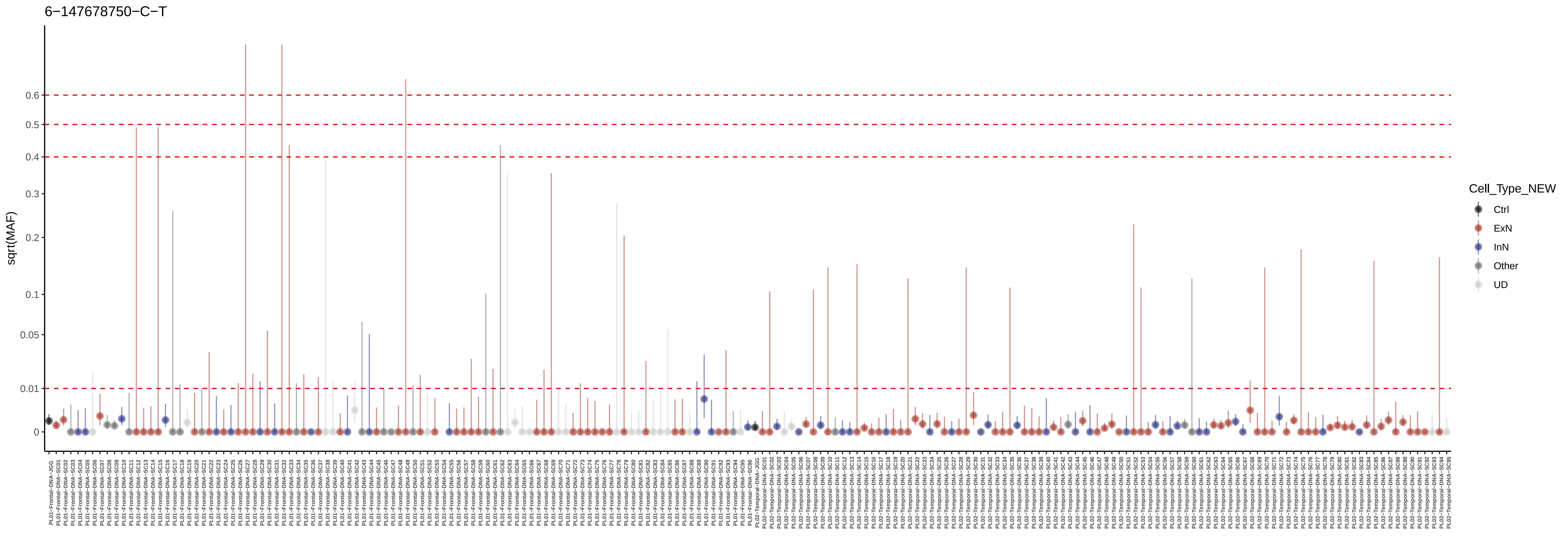

6−147678750−C−T
sqrt(MAF)
0.6
0.5
0.4
0.3
0.2
0.1
0.05
0.01
0
Cell_Type_NEW
Ctrl
ExN
InN
Other
UD

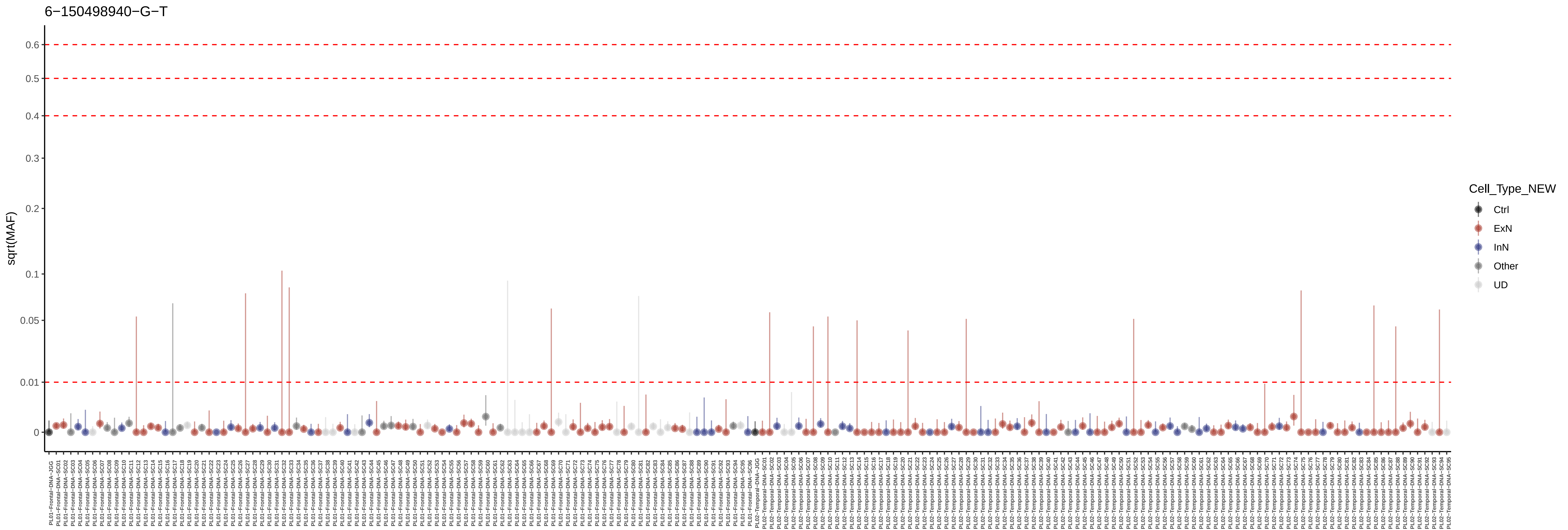
6−150498940−G−T
sqrt(MAF)
0.6
0.5
0.4
0.3
0.2
0.1
0.05
0.01
0
Cell_Type_NEW
Ctrl
ExN
InN
Other
UD

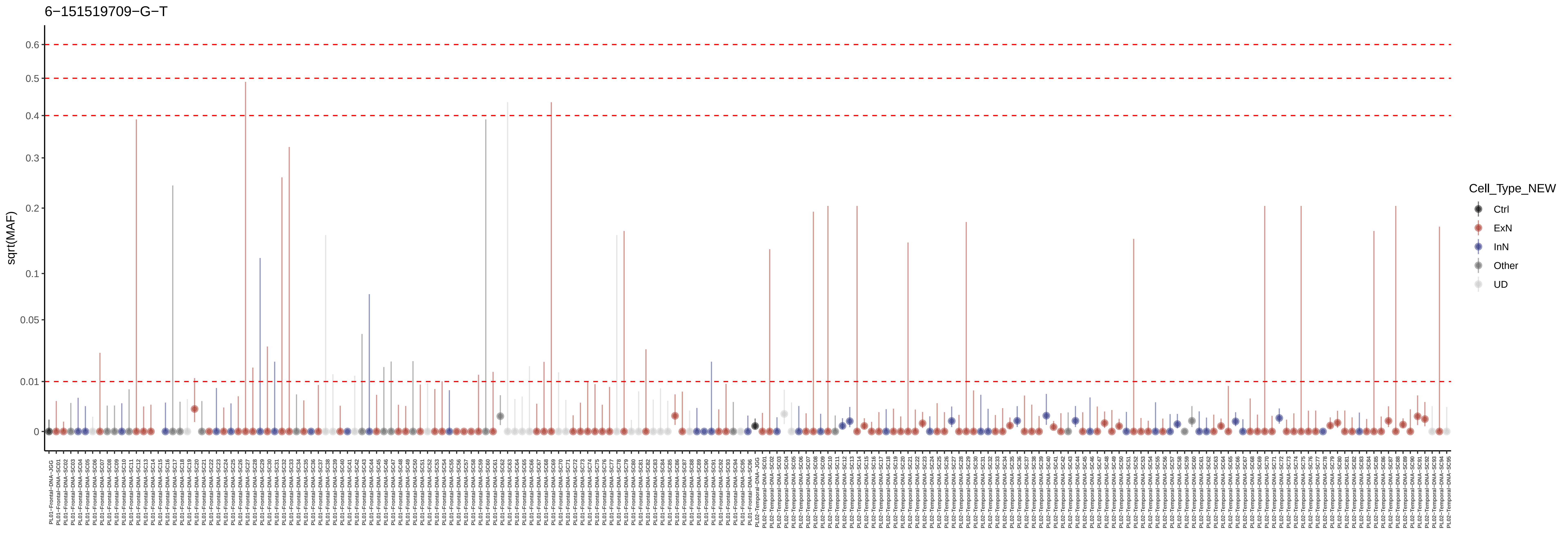

6−151519709−G−T
sqrt(MAF)
0.6
0.5
0.4
0.3
0.2
0.1
0.05
0.01
0
Cell_Type_NEW
Ctrl
ExN
InN
Other
UD

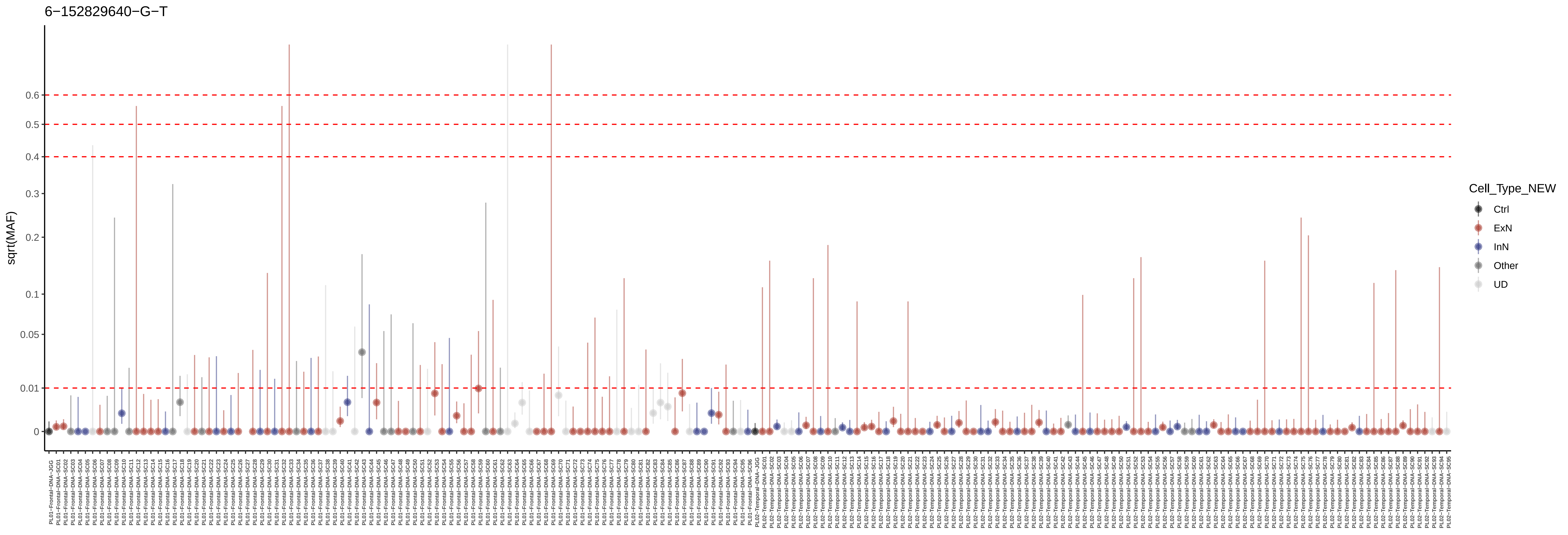
6−152829640−G−T
sqrt(MAF)
0.6
0.5
0.4
0.3
0.2
0.1
0.05
0.01
0
Cell_Type_NEW
Ctrl
ExN
InN
Other
UD

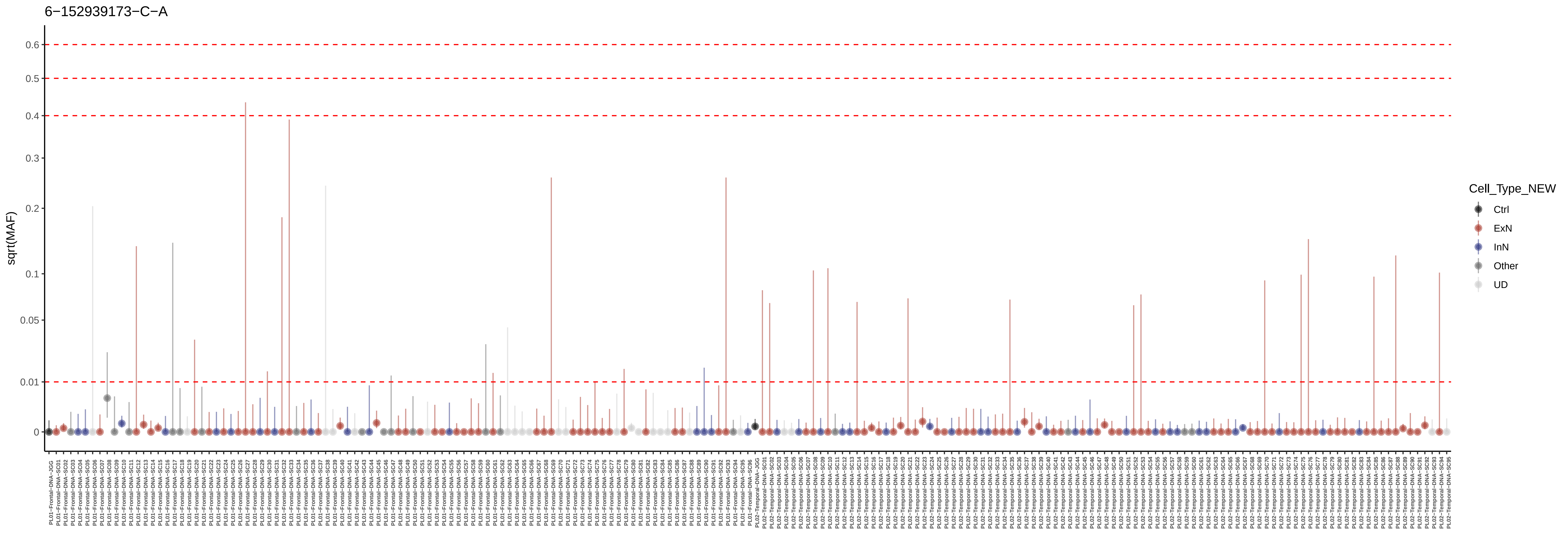

6−152939173−C−A
sqrt(MAF)
0.6
0.5
0.4
0.3
0.2
0.1
0.05
0.01
0
Cell_Type_NEW
Ctrl
ExN
InN
Other
UD

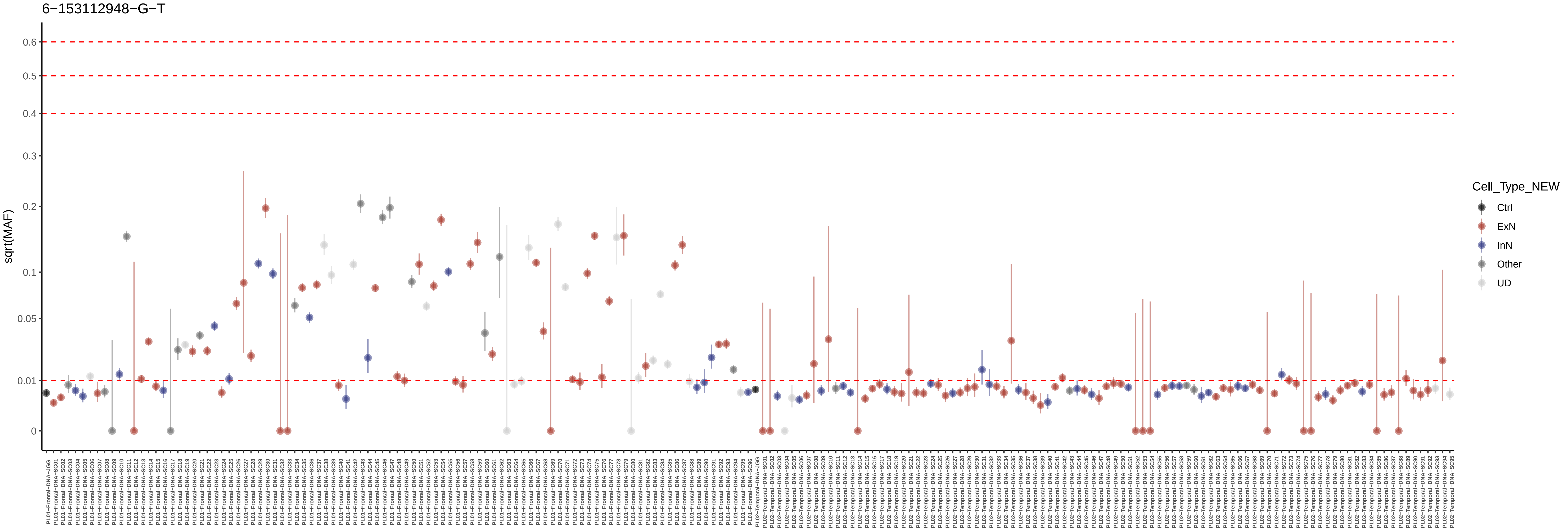
6−153112948−G−T
sqrt(MAF)
0.6
0.5
0.4
0.3
0.2
0.1
0.05
0.01
0
Cell_Type_NEW
Ctrl
ExN
InN
Other
UD

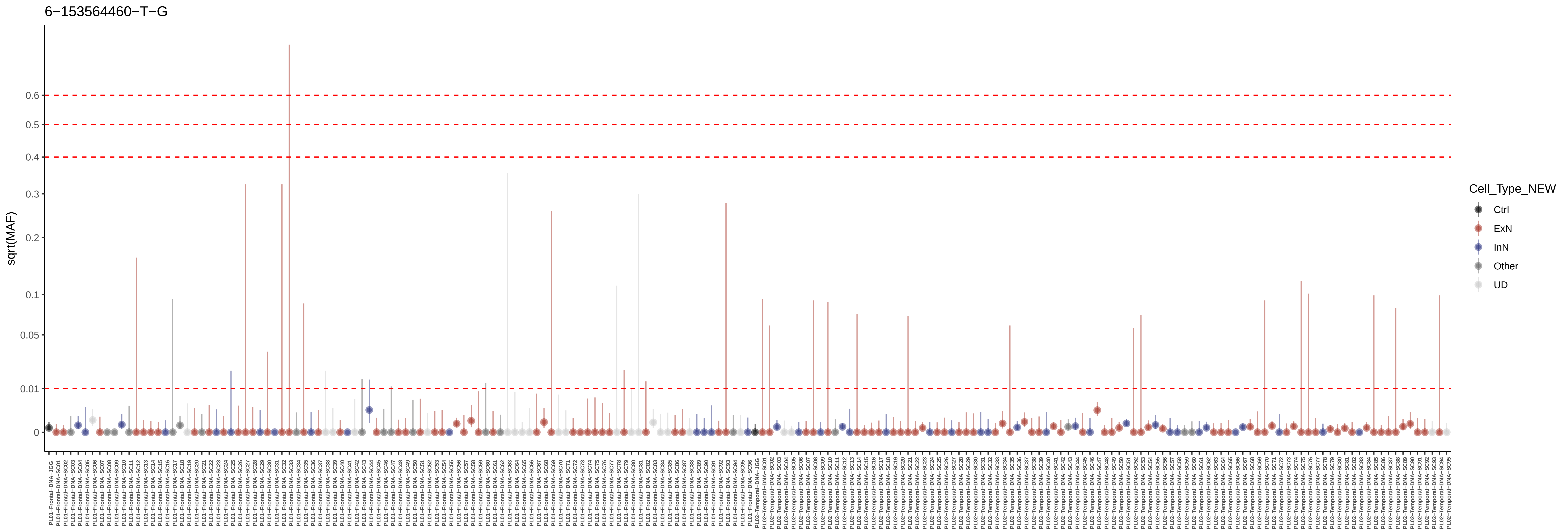
6−153564460−T−G
sqrt(MAF)
0.6
0.5
0.4
0.3
0.2
0.1
0.05
0.01
0
Cell_Type_NEW
Ctrl
ExN
InN
Other
UD

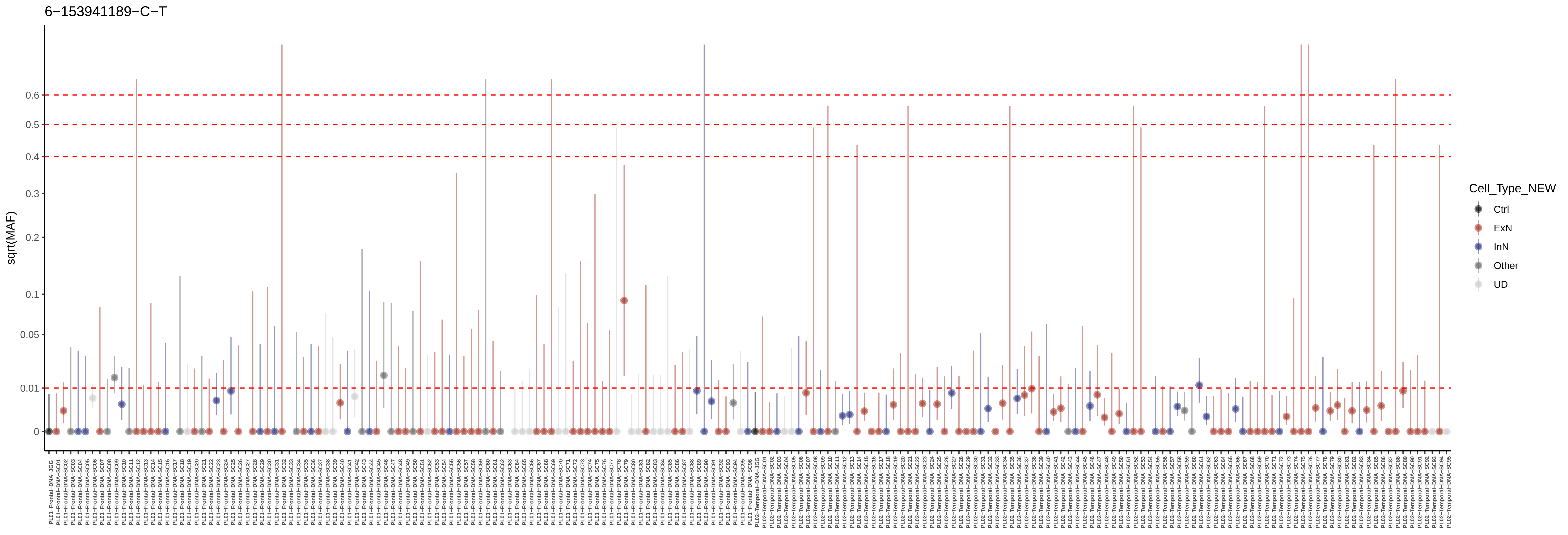

6−153941189−C−T
sqrt(MAF)
0.6
0.5
0.4
0.3
0.2
0.1
0.05
0.01
0
Cell_Type_NEW
Ctrl
ExN
InN
Other
UD

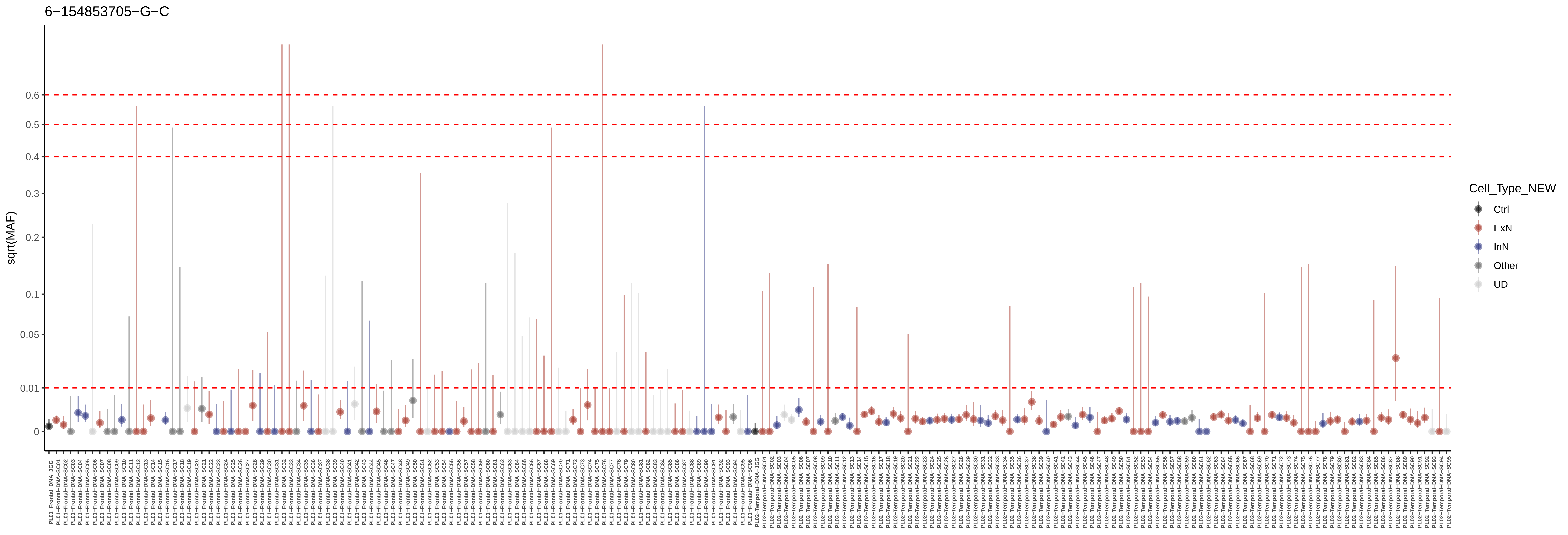
6−154853705−G−C
sqrt(MAF)
0.6
0.5
0.4
0.3
0.2
0.1
0.05
0.01
0
Cell_Type_NEW
Ctrl
ExN
InN
Other
UD

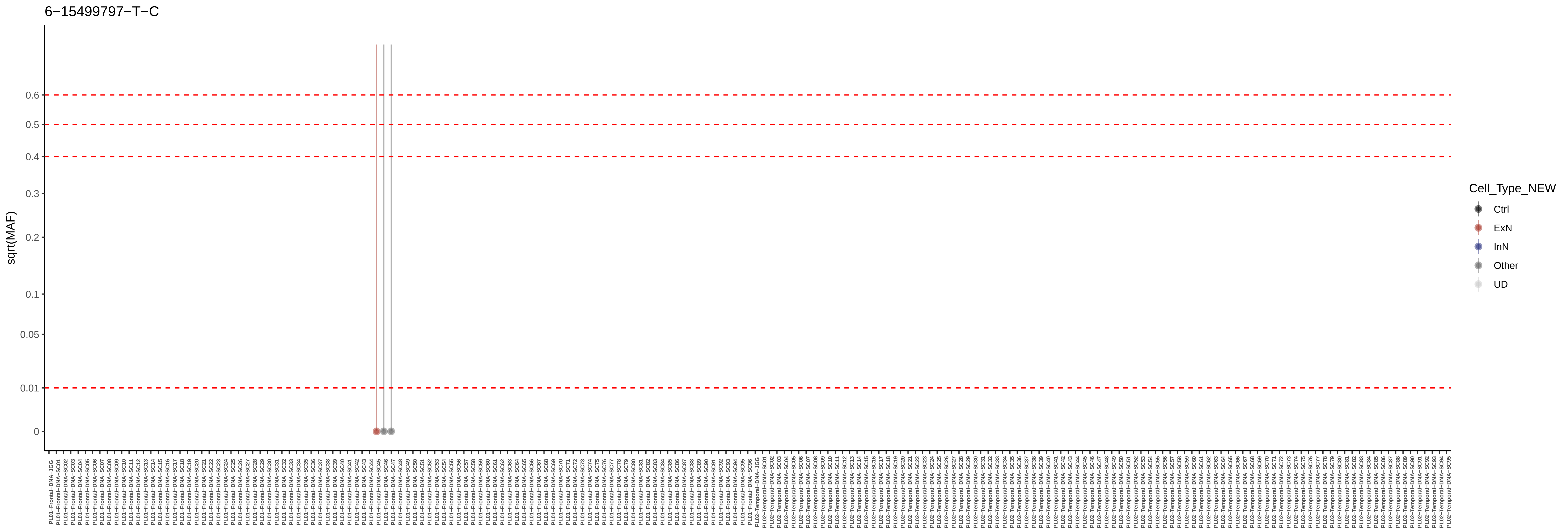
6−15499797−T−C
sqrt(MAF)
0.6
0.5
0.4
0.3
0.2
0.1
0.05
0.01
0
Cell_Type_NEW
Ctrl
ExN
InN
Other
UD

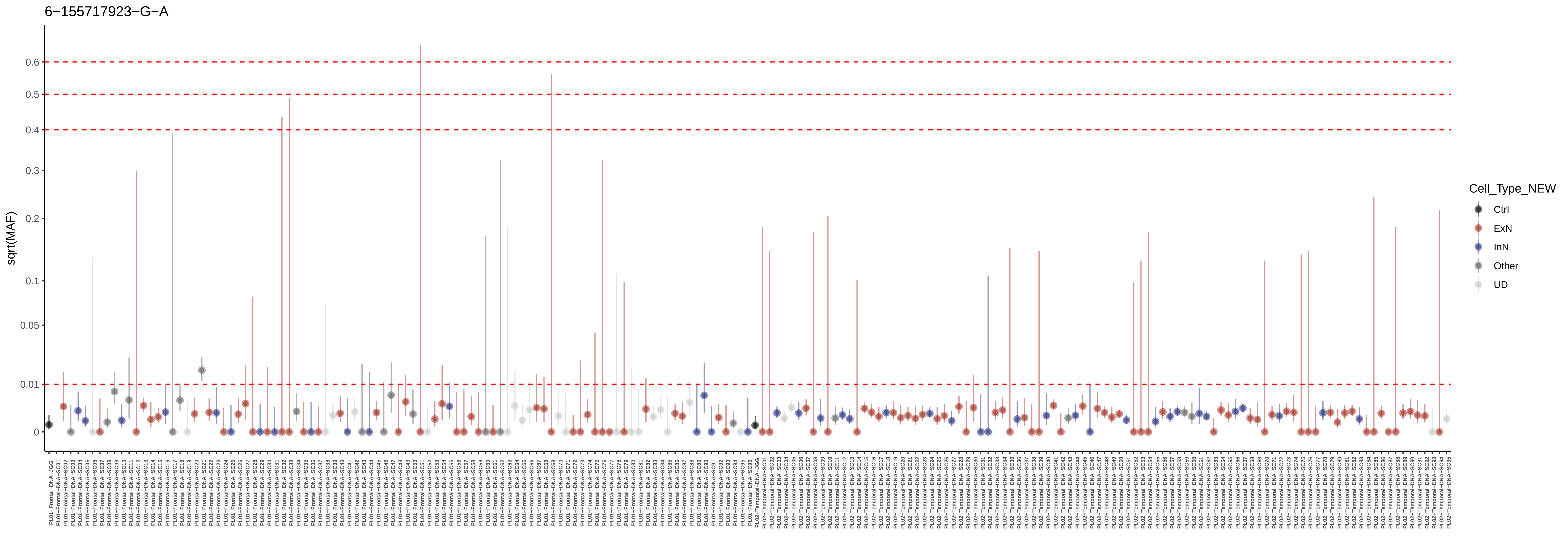
6−155717923−G−A
sqrt(MAF)
0.6
0.5
0.4
0.3
0.2
0.1
0.05
0.01
0
Cell_Type_NEW
Ctrl
ExN
InN
Other
UD

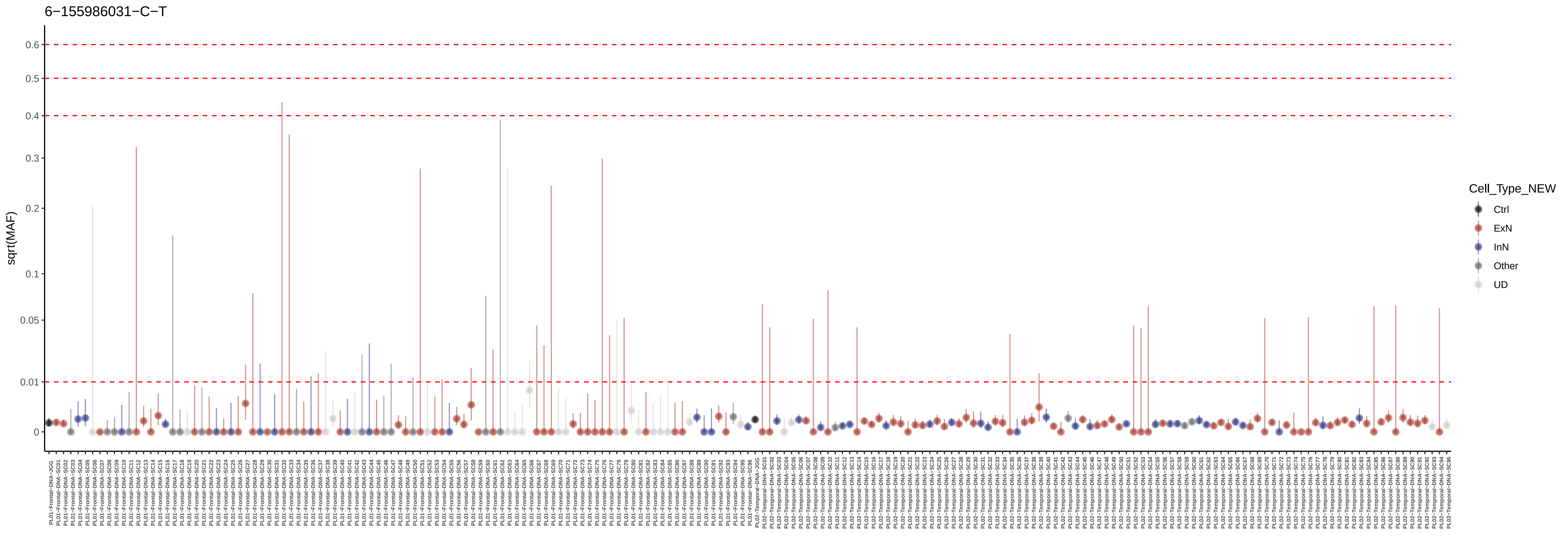
6−155986031−C−T
sqrt(MAF)
0.6
0.5
0.4
0.3
0.2
0.1
0.05
0.01
0
Cell_Type_NEW
Ctrl
ExN
InN
Other
UD

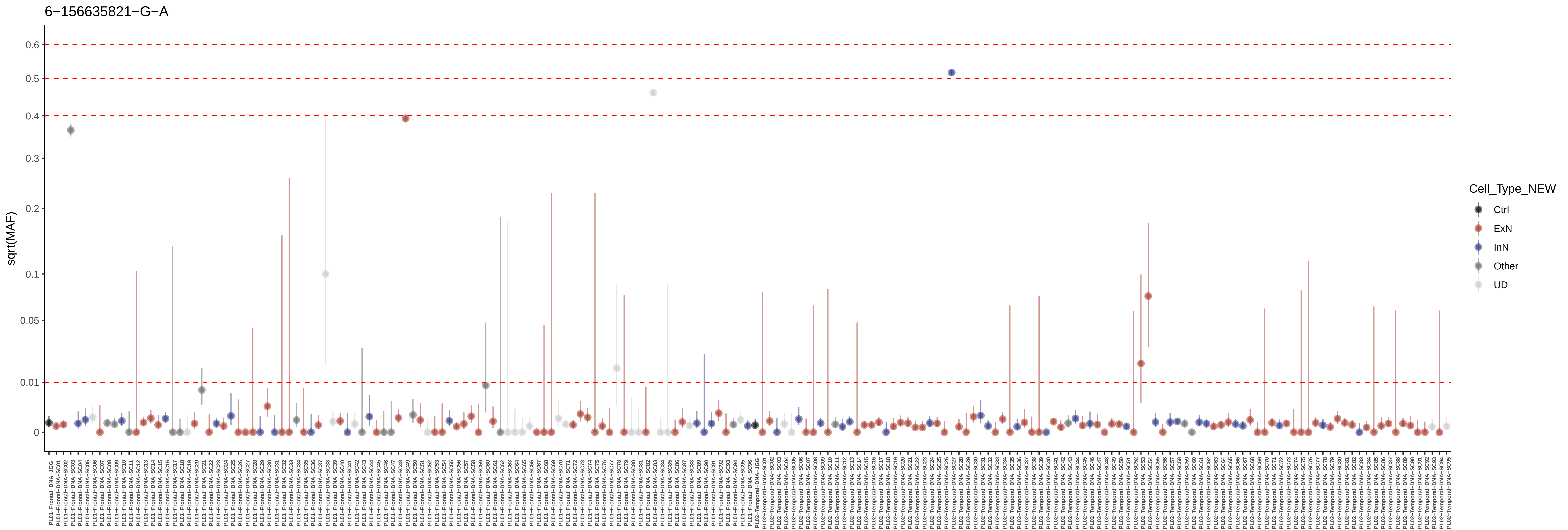
6−156635821−G−A
sqrt(MAF)
0.6
0.5
0.4
0.3
0.2
0.1
0.05
0.01
0
Cell_Type_NEW
Ctrl
ExN
InN
Other
UD

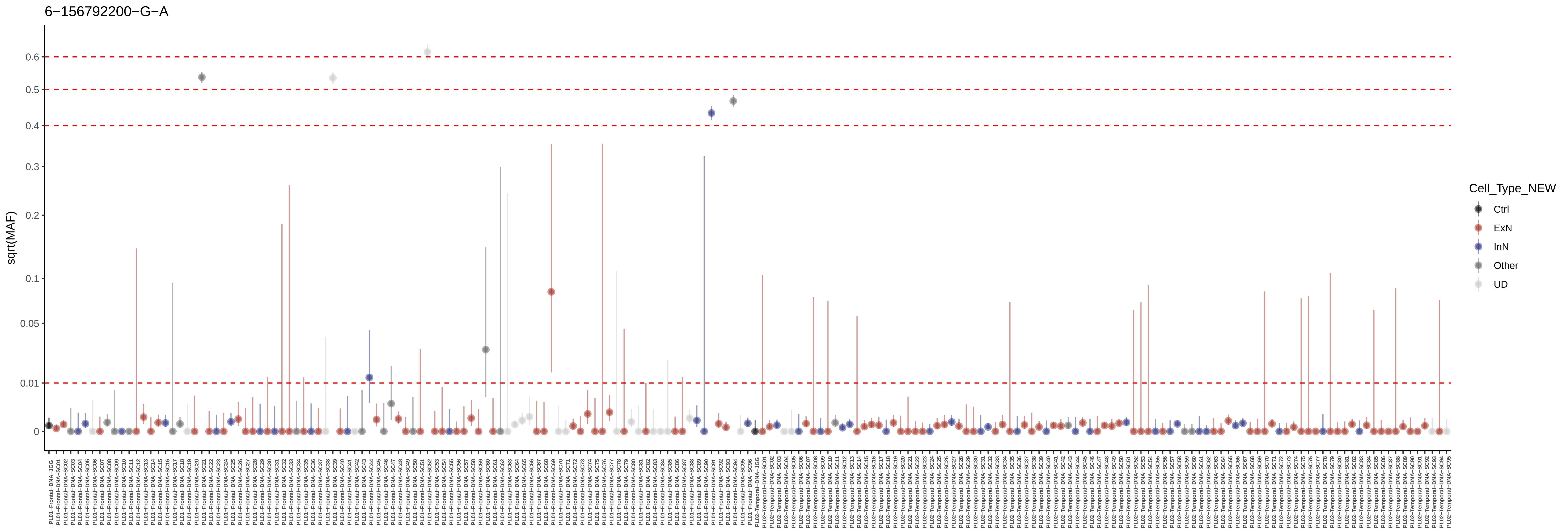
6−156792200−G−A
sqrt(MAF)
0.6
0.5
0.4
0.3
0.2
0.1
0.05
0.01
0
Cell_Type_NEW
Ctrl
ExN
InN
Other
UD

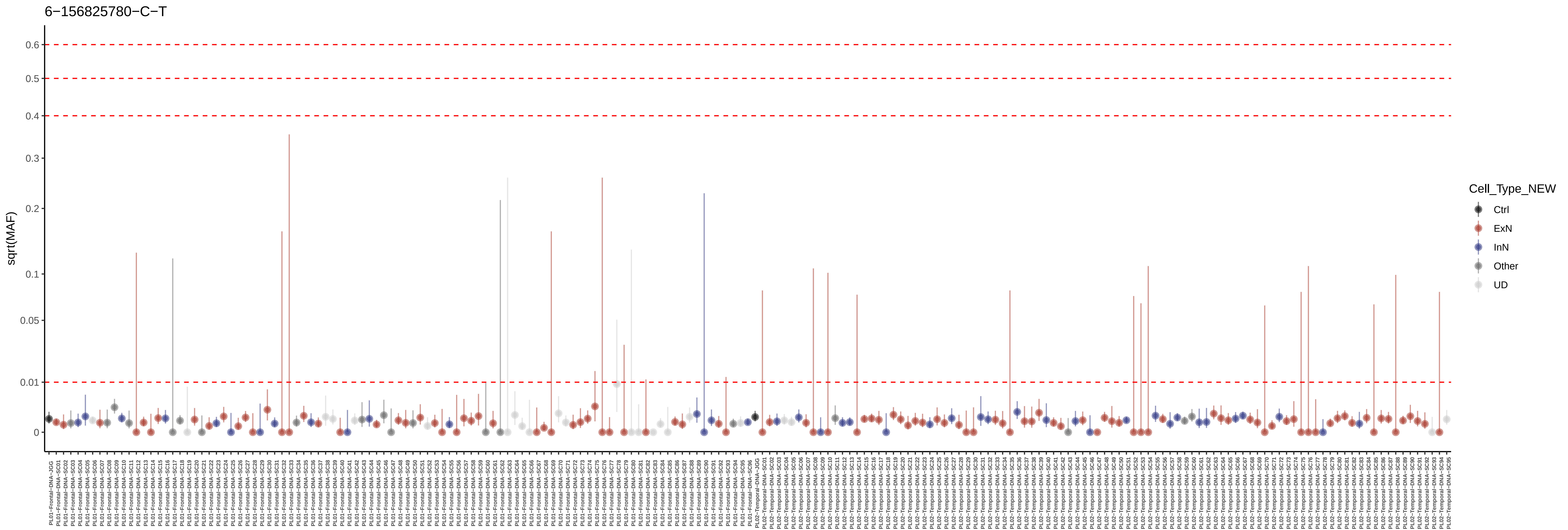
6−156825780−C−T
sqrt(MAF)
0.6
0.5
0.4
0.3
0.2
0.1
0.05
0.01
0
Cell_Type_NEW
Ctrl
ExN
InN
Other
UD

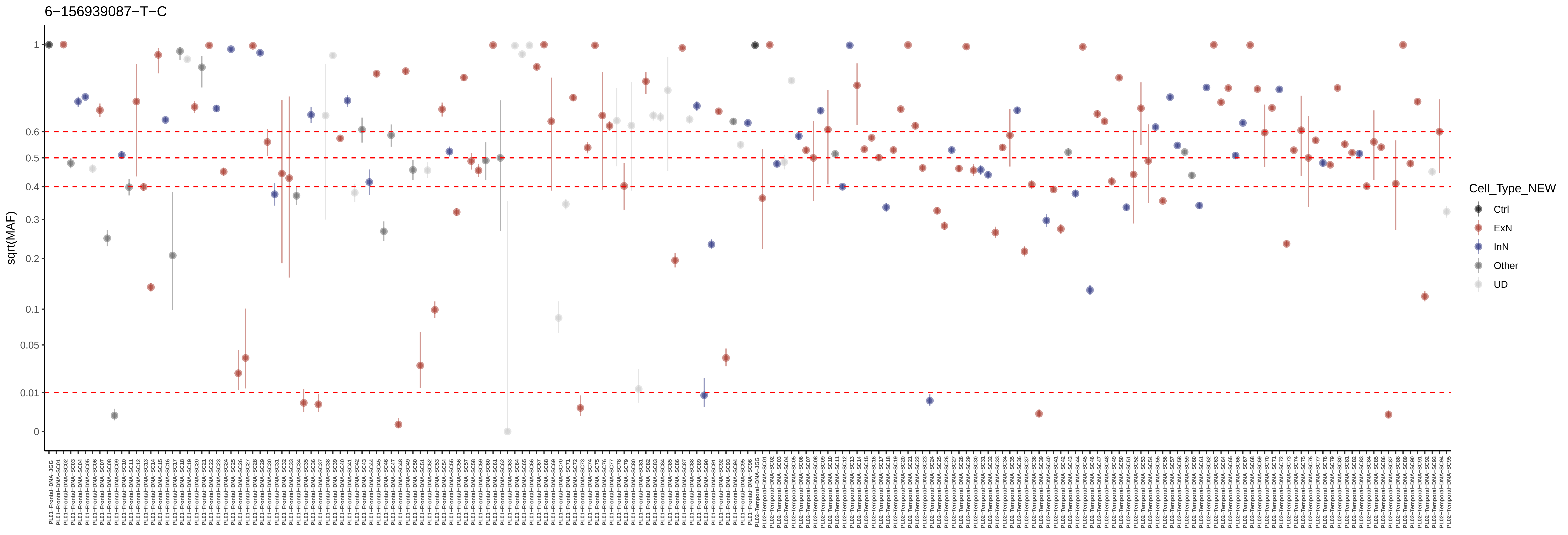
6−156939087−T−C
sqrt(MAF)
1
0.6
0.5
0.4
0.3
0.2
0.1
0.05
0.01
0
Cell_Type_NEW
Ctrl
ExN
InN
Other
UD

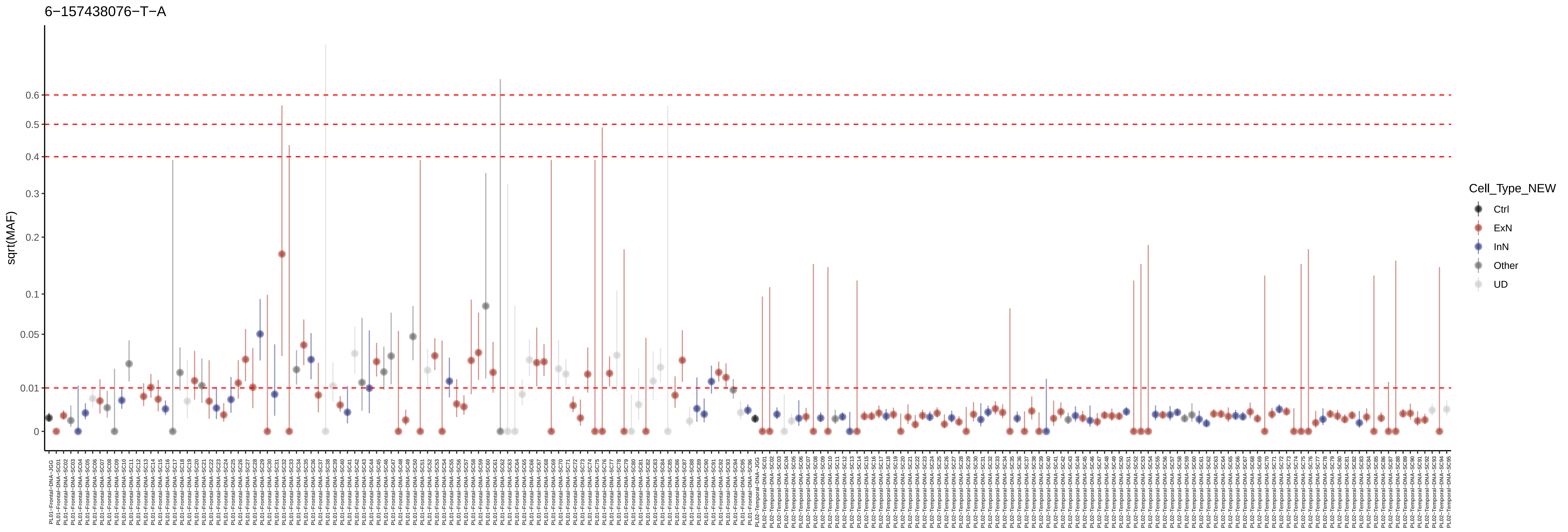
6−157438076−T−A
sqrt(MAF)
0.6
0.5
0.4
0.3
0.2
0.1
0.05
0.01
0
Cell_Type_NEW
Ctrl
ExN
InN
Other
UD

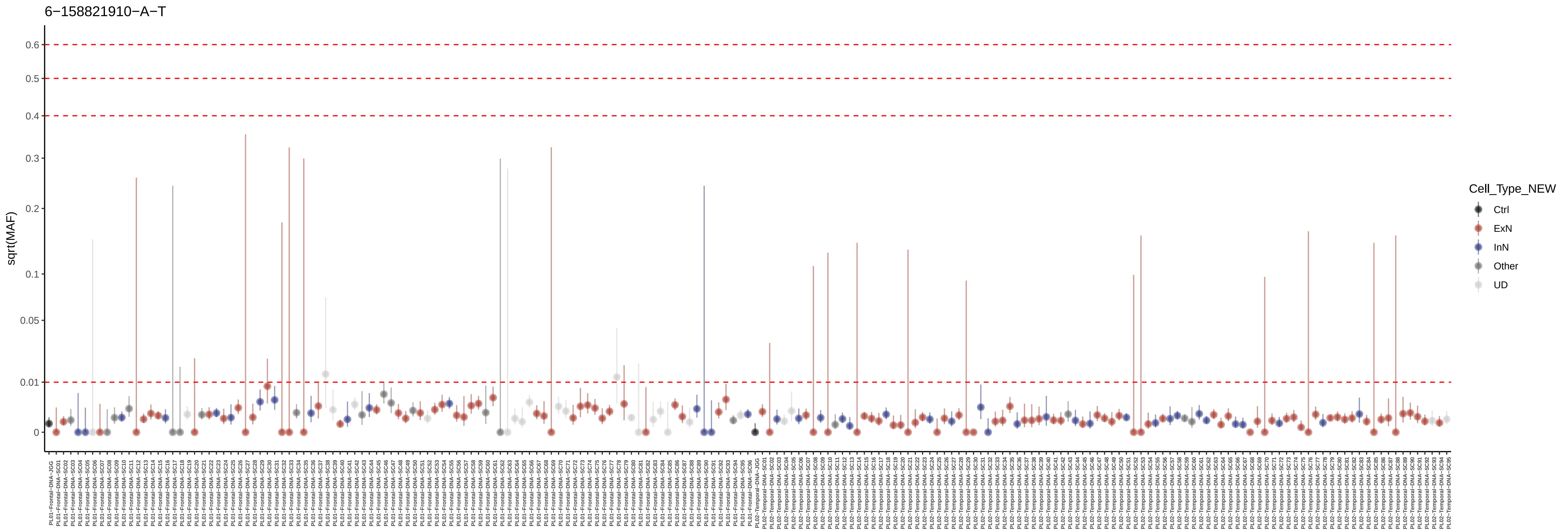
6−158821910−A−T
sqrt(MAF)
0.6
0.5
0.4
0.3
0.2
0.1
0.05
0.01
0
Cell_Type_NEW
Ctrl
ExN
InN
Other
UD

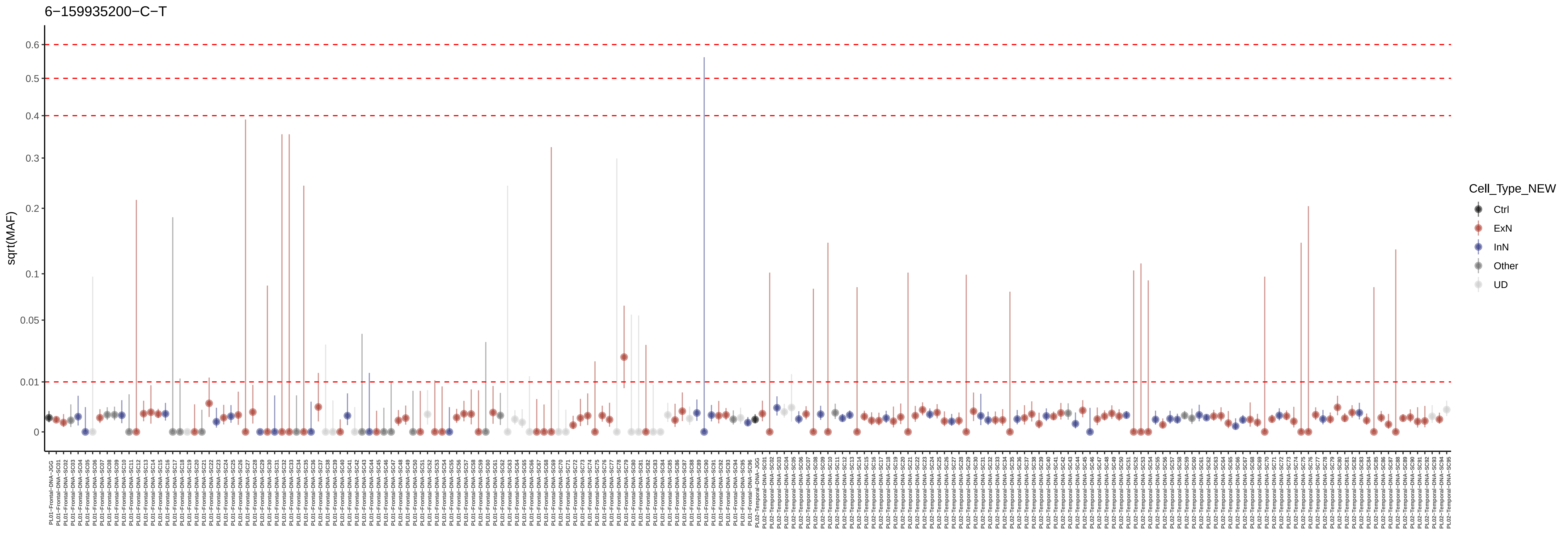
6−159935200−C−T
sqrt(MAF)
Cell_Type_NEW
Ctrl
ExN
InN
Other
UD

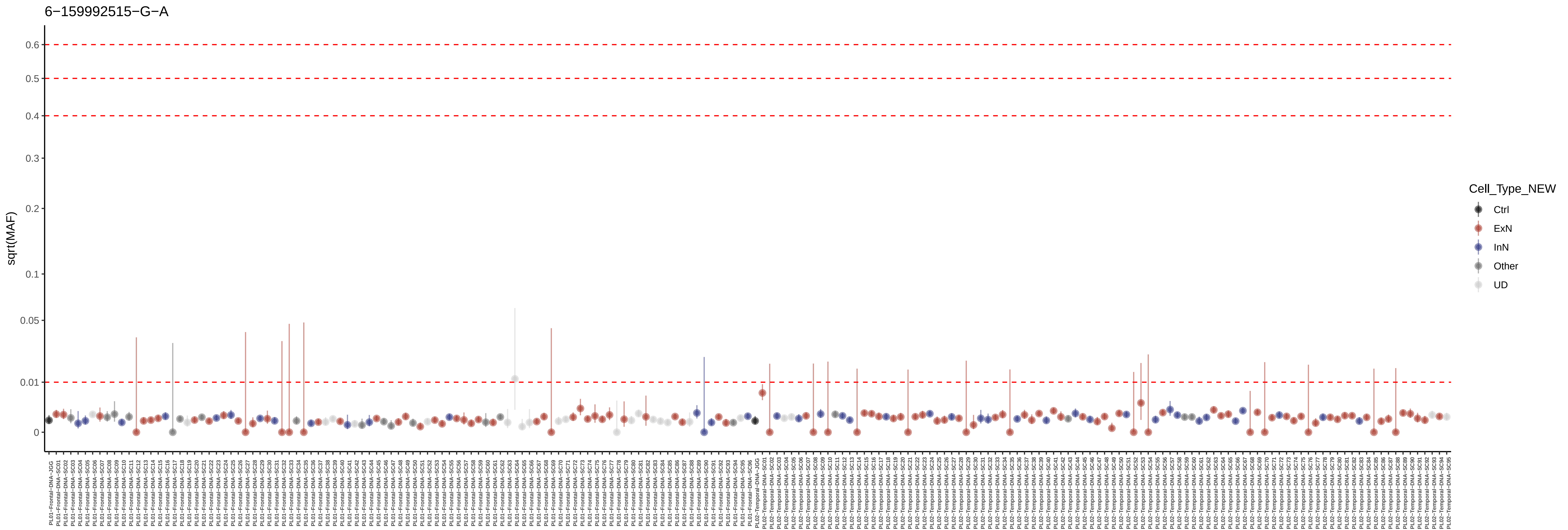
6−159992515−G−A
sqrt(MAF)
0.6
0.5
0.4
0.3
0.2
0.1
0.05
0.01
0
Cell_Type_NEW
Ctrl
ExN
InN
Other
UD

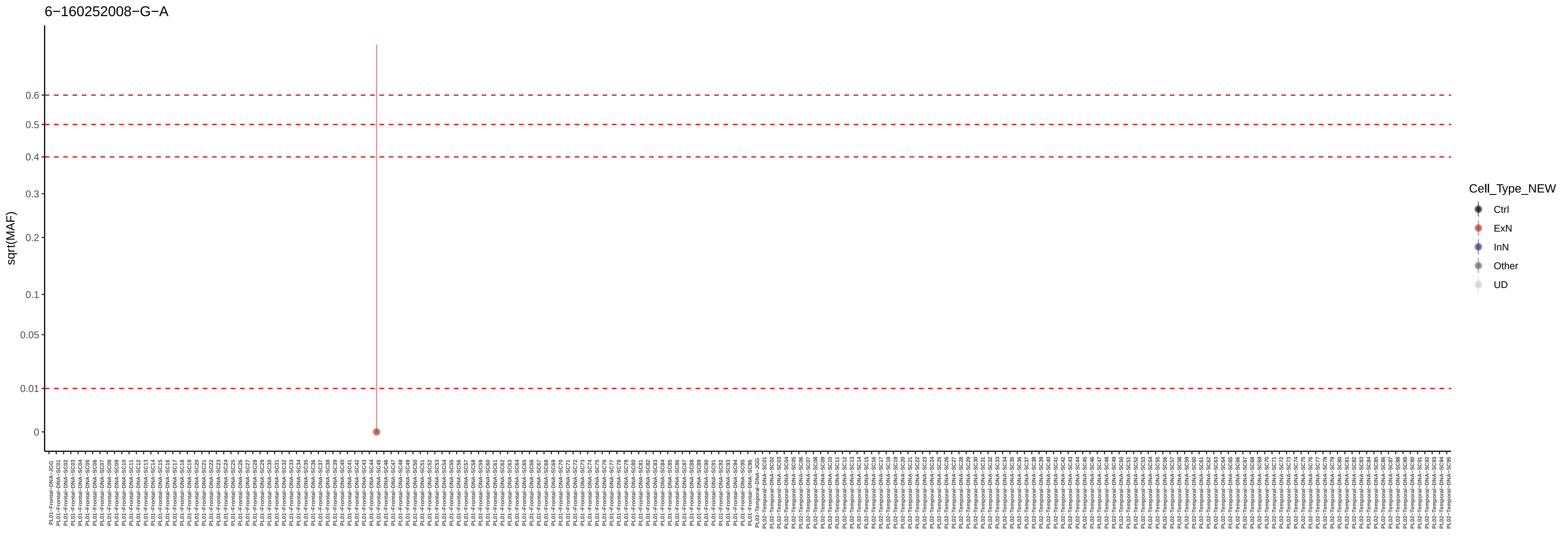

6−160252008−G−A
sqrt(MAF)
0.6
0.5
0.4
0.3
0.2
0.1
0.05
0.01
0
Cell_Type_NEW
Ctrl
ExN
InN
Other
UD

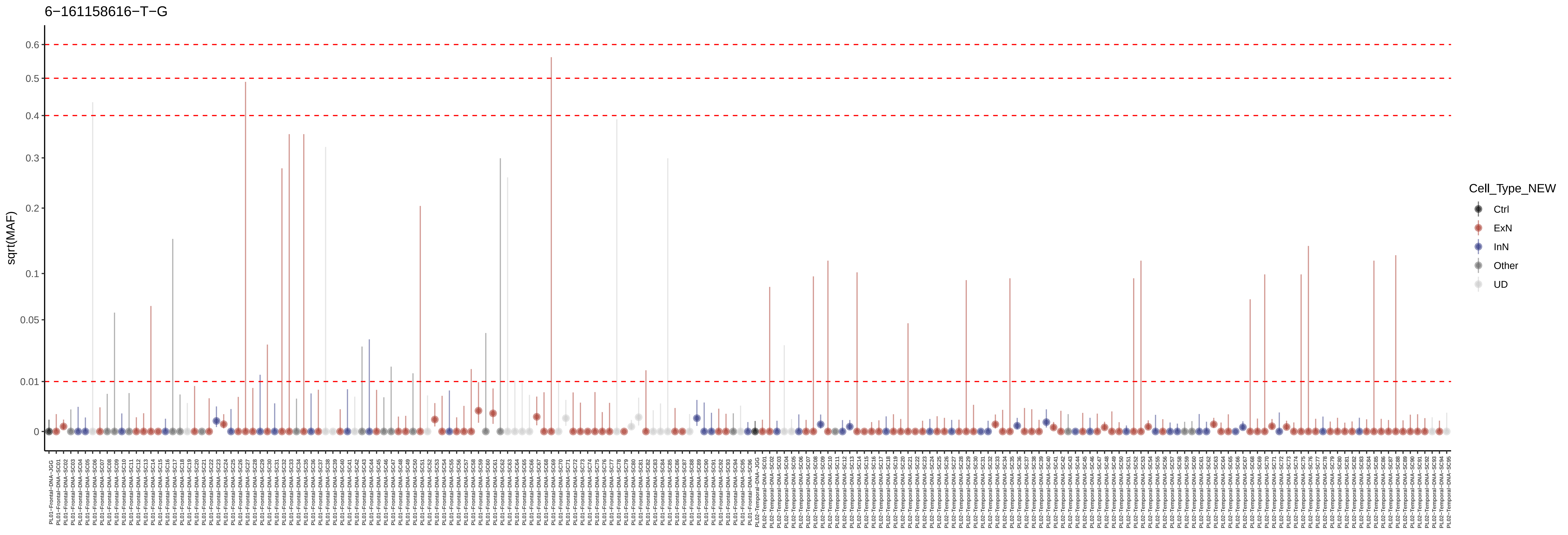

6−161158616−T−G
sqrt(MAF)
0.6
0.5
0.4
0.3
0.2
0.1
0.05
0.01
0
Cell_Type_NEW
Ctrl
ExN
InN
Other
UD

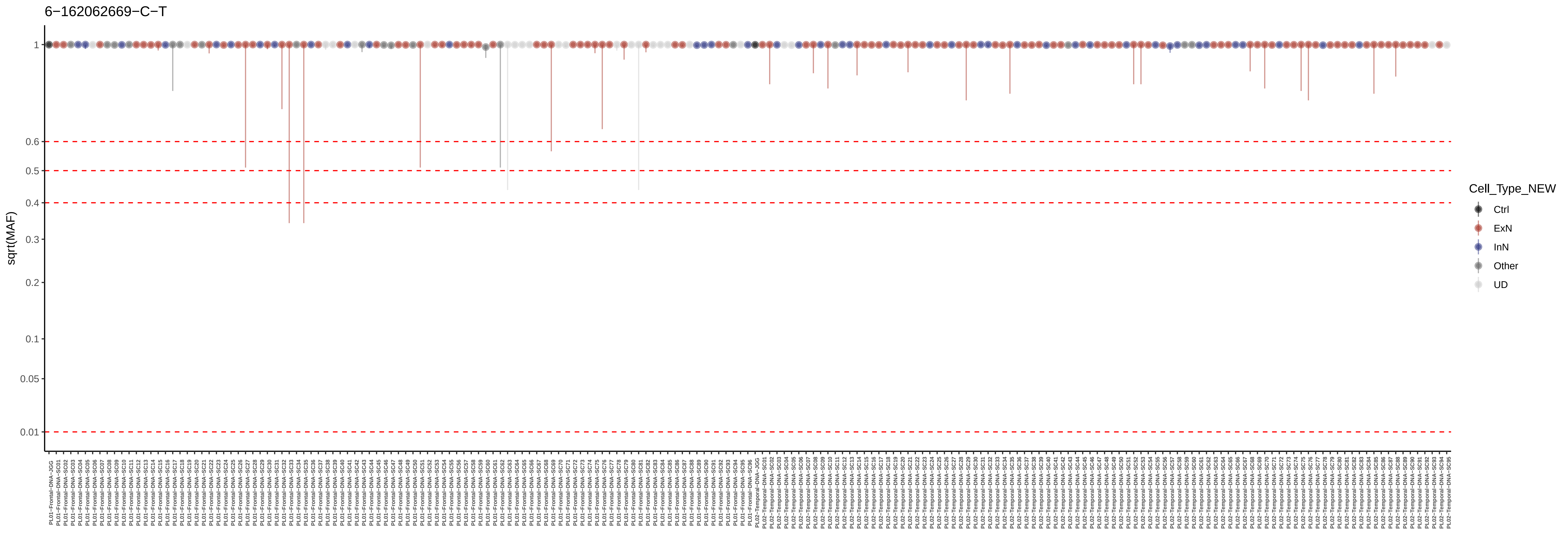
6−162062669−C−T
sqrt(MAF)
1
0.6
0.5
0.4
0.3
0.2
0.1
0.05
0.01
Cell_Type_NEW
Ctrl
ExN
InN
Other
UD

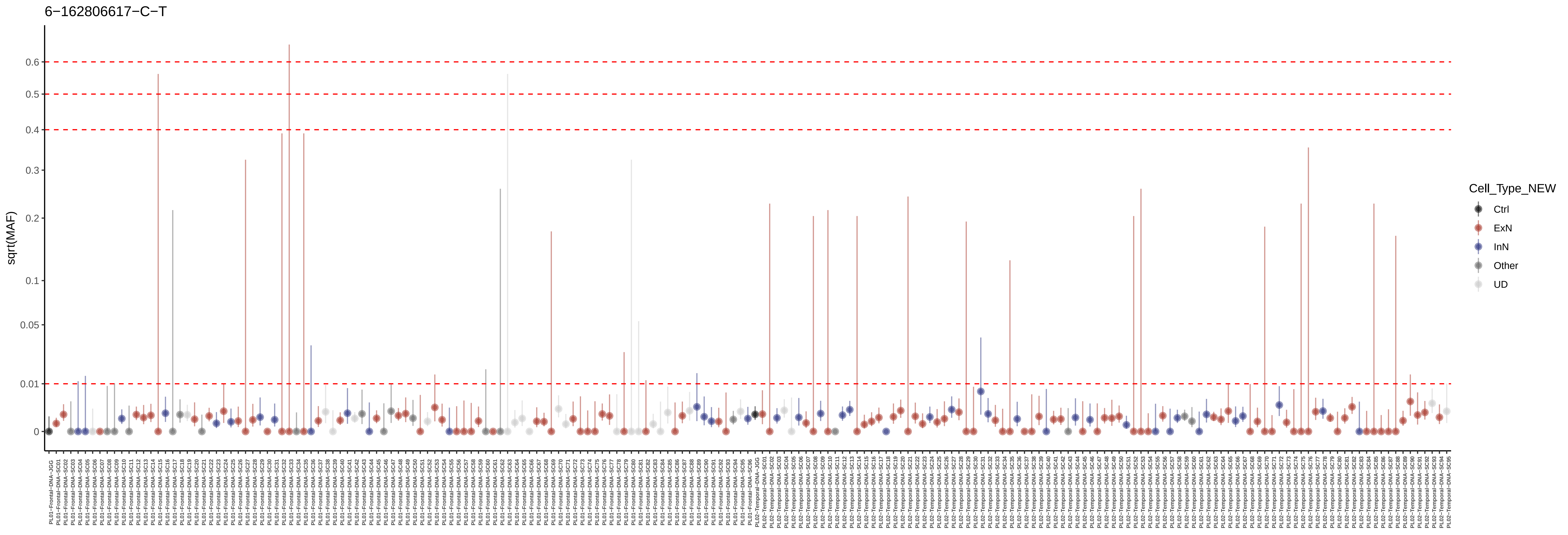
6−162806617−C−T
sqrt(MAF)
0.6
0.5
0.4
0.3
0.2
0.1
0.05
0.01
0
Cell_Type_NEW
Ctrl
ExN
InN
Other
UD

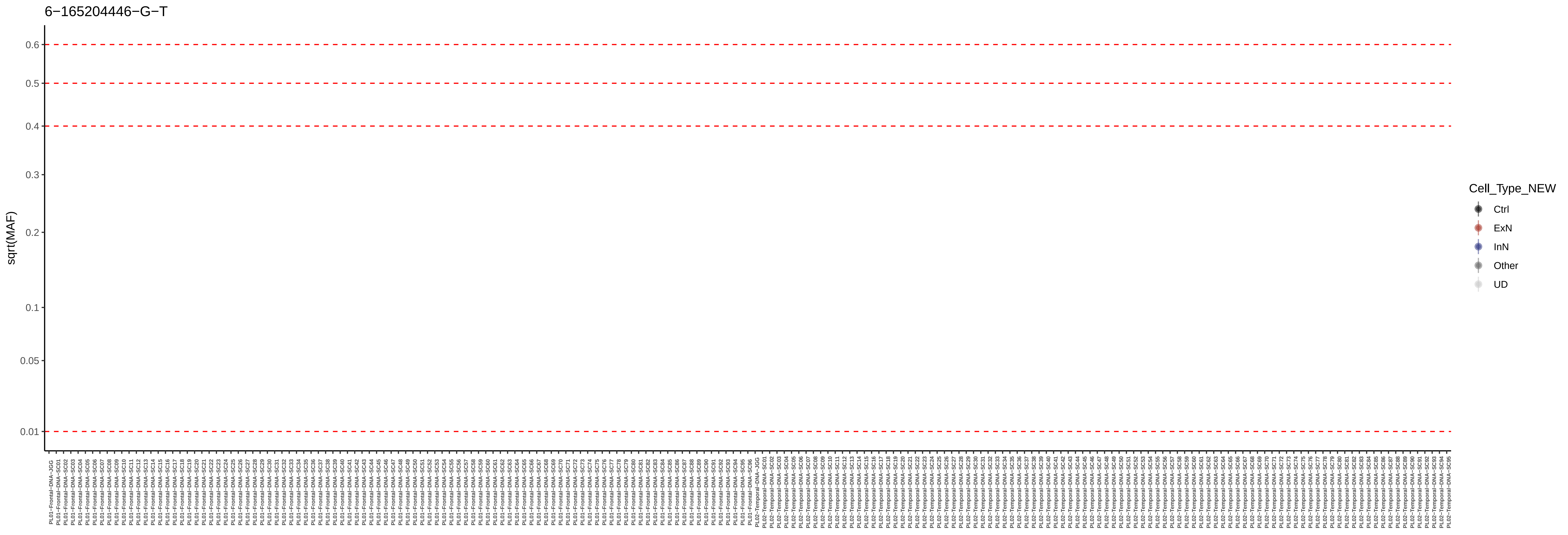
6−165204446−G−T
sqrt(MAF)
0.6
0.5
0.4
0.3
0.2
0.1
0.05
0.01
Cell_Type_NEW
Ctrl
ExN
InN
Other
UD

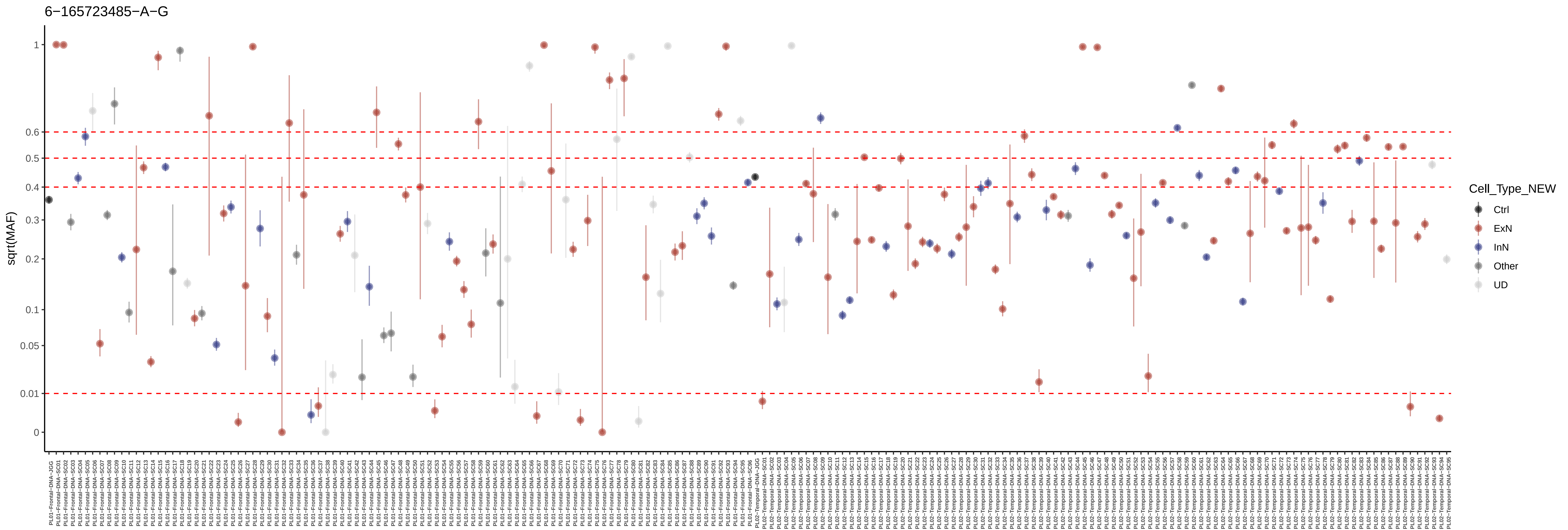
6−165723485−A−G
sqrt(MAF)
1
0.6
0.5
0.4
0.3
0.2
0.1
0.05
0.01
0
Cell_Type_NEW
Ctrl
ExN
InN
Other
UD

6−1661827−C−T

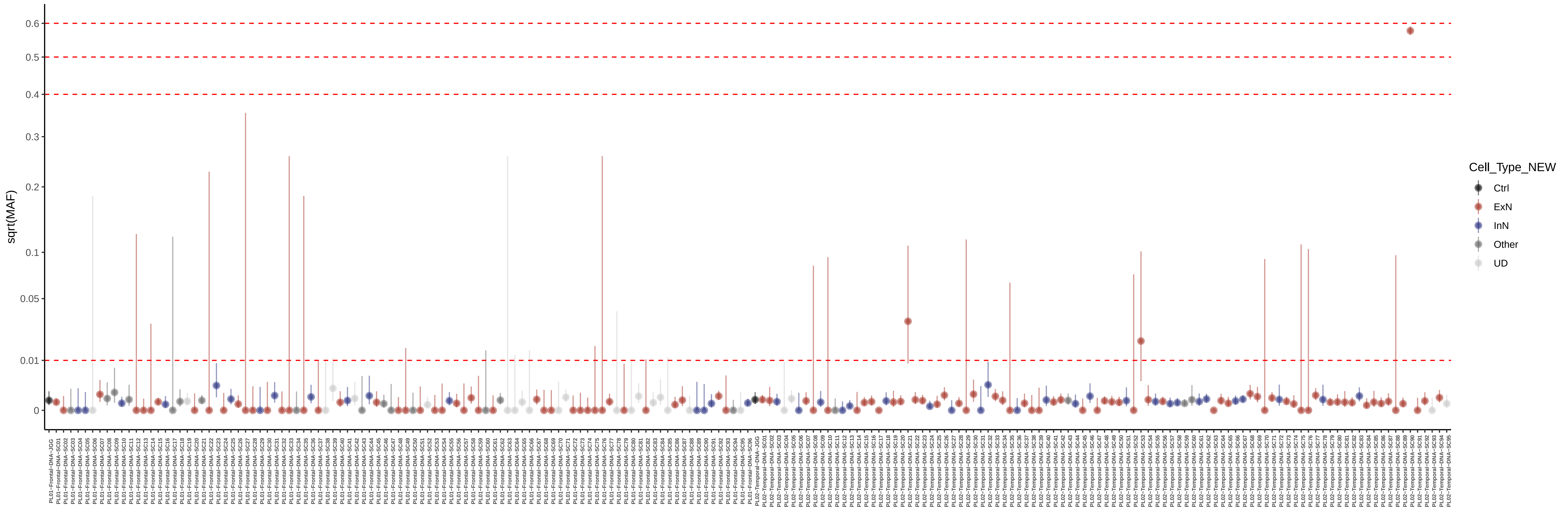
sqrt(MAF)
0.6
0.5
0.4
0.3
0.2
0.1
0.05
0.01
0
Cell_Type_NEW
Ctrl
ExN
InN
Other
UD

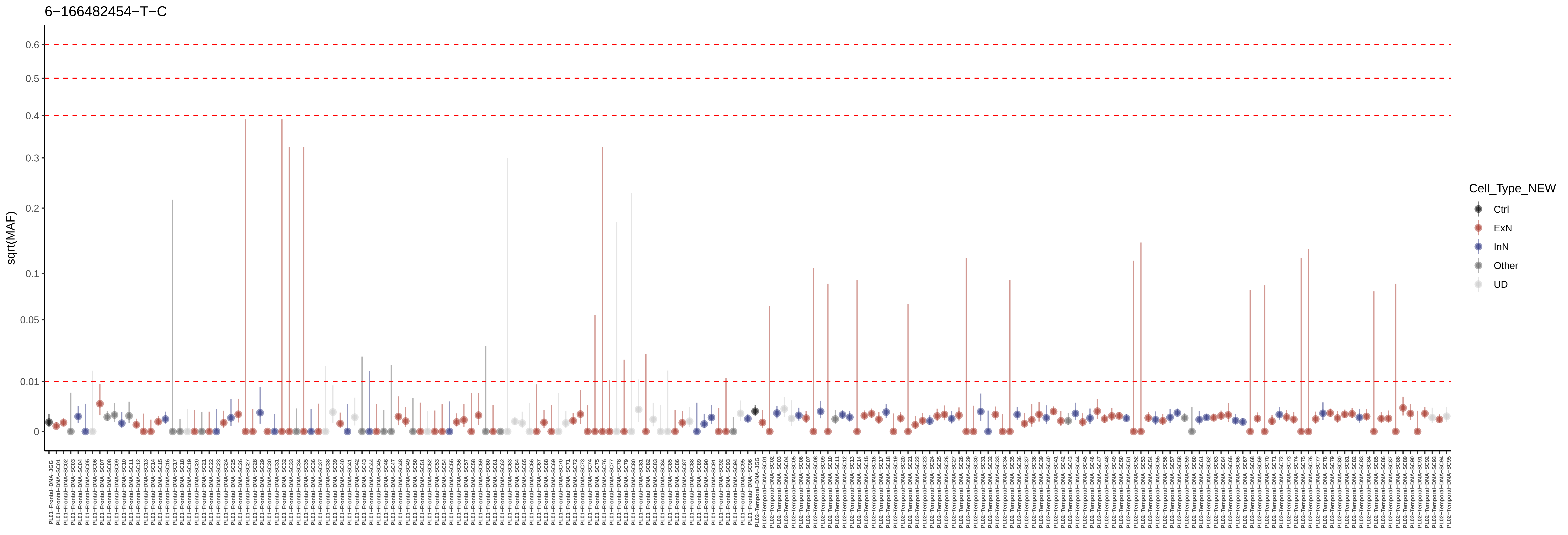
6−166482454−T−C
sqrt(MAF)
0.6
0.5
0.4
0.3
0.2
0.1
0.05
0.01
0
Cell_Type_NEW
Ctrl
ExN
InN
Other
UD

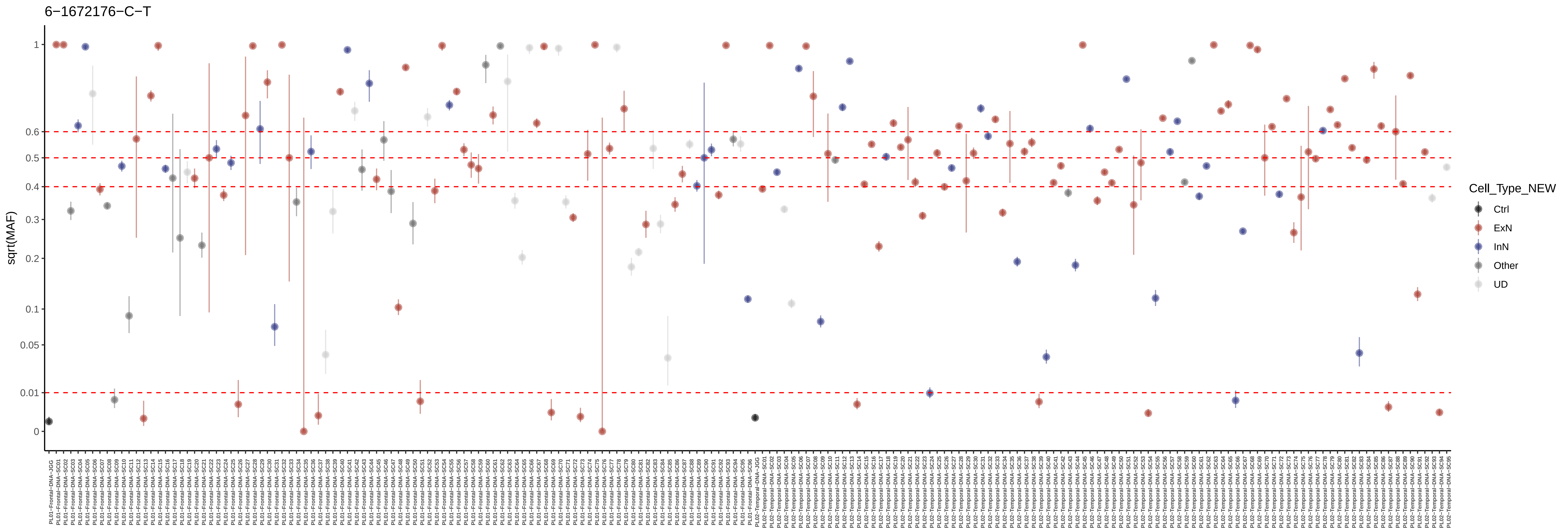
6−1672176−C−T
sqrt(MAF)
1
0.6
0.5
0.4
0.3
0.2
0.1
0.05
0.01
0
Cell_Type_NEW
Ctrl
ExN
InN
Other
UD

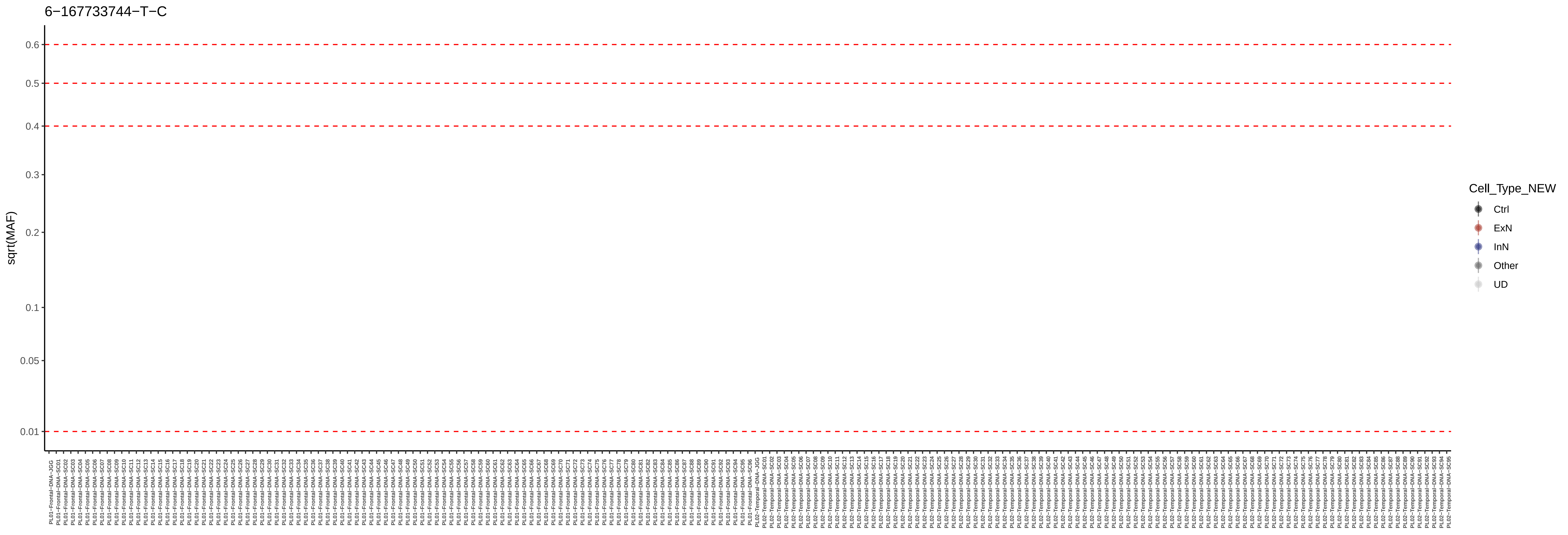
6−167733744−T−C
sqrt(MAF)
0.6
0.5
0.4
0.3
0.2
0.1
0.05
0.01
Cell_Type_NEW
Ctrl
ExN
InN
Other
UD

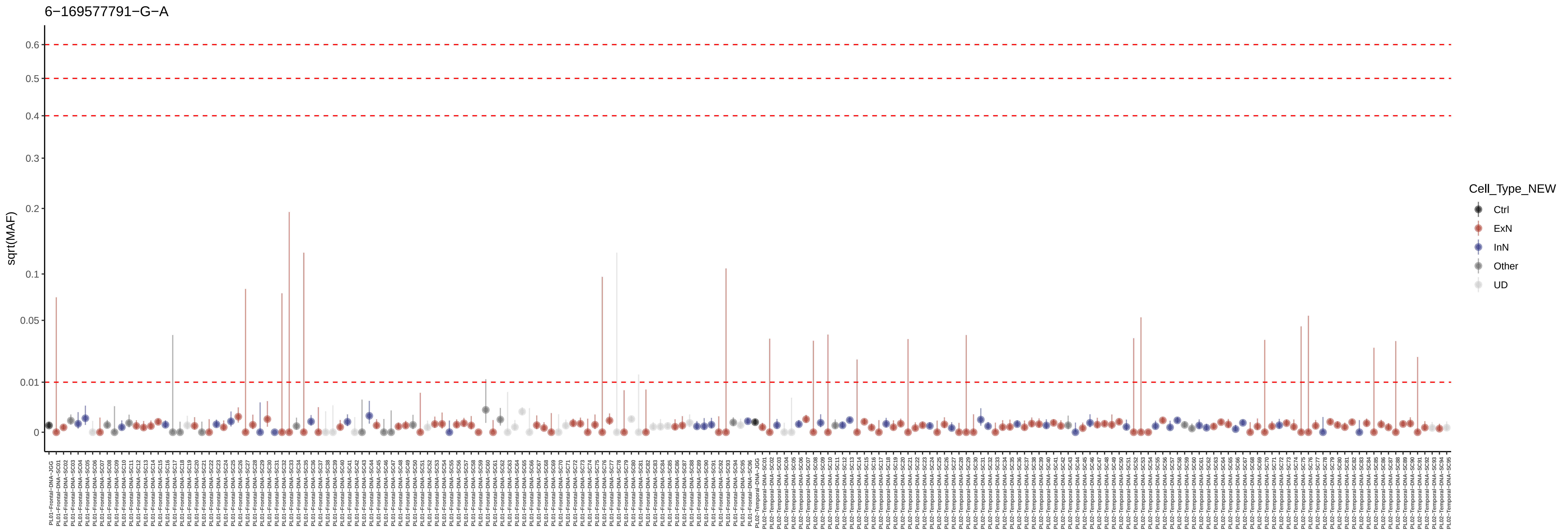

6−169577791−G−A
sqrt(MAF)
0.6
0.5
0.4
0.3
0.2
0.1
0.05
0.01
0
Cell_Type_NEW
Ctrl
ExN
InN
Other
UD

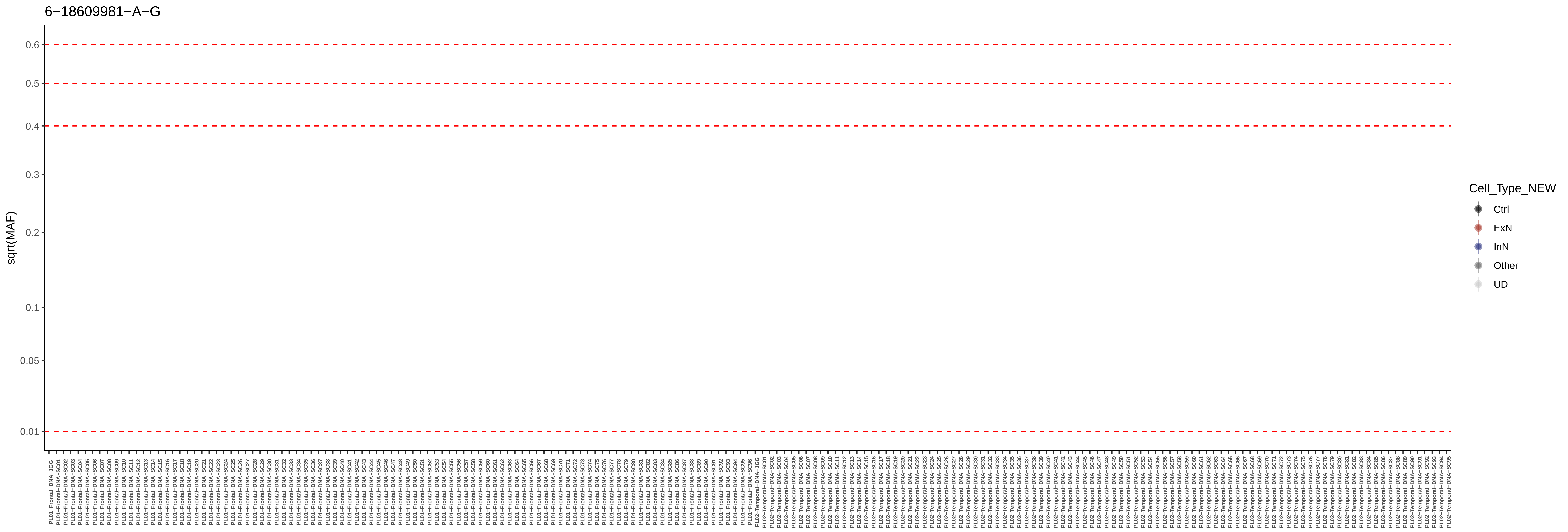
6−18609981−A−G
sqrt(MAF)
0.6
0.5
0.4
0.3
0.2
0.1
0.05
0.01
Cell_Type_NEW
Ctrl
ExN
InN
Other
UD

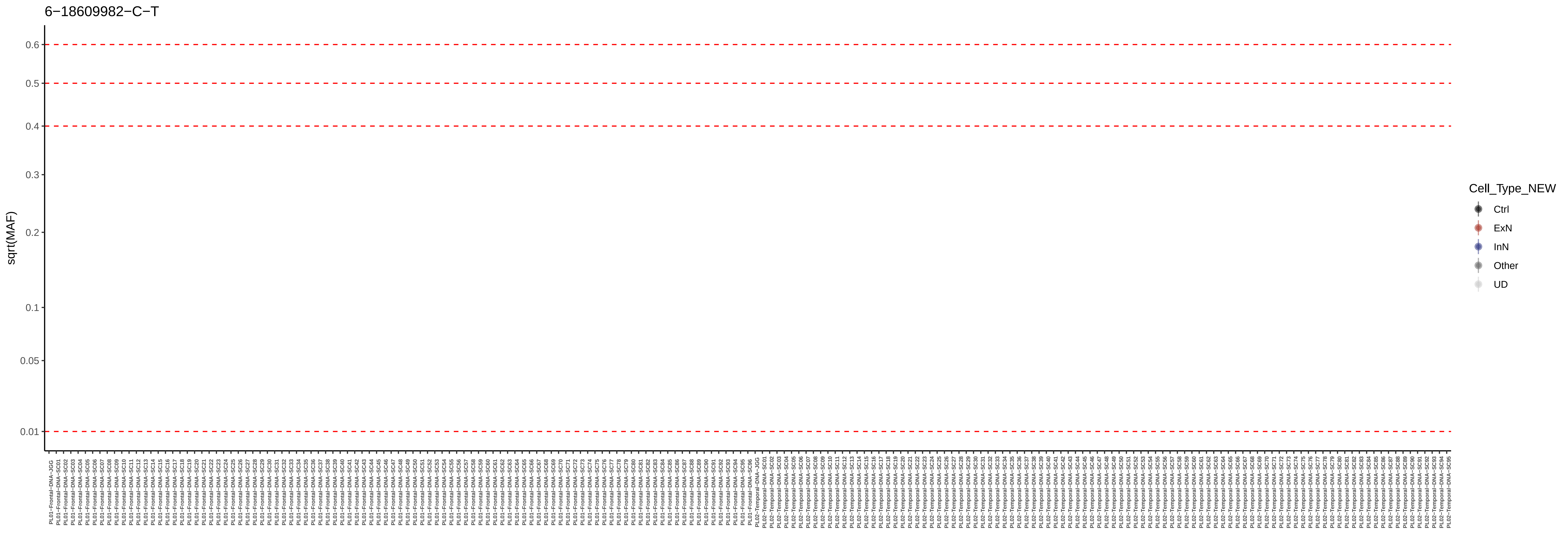
6−18609982−C−T
sqrt(MAF)
Cell_Type_NEW
Ctrl
ExN
InN
Other
UD

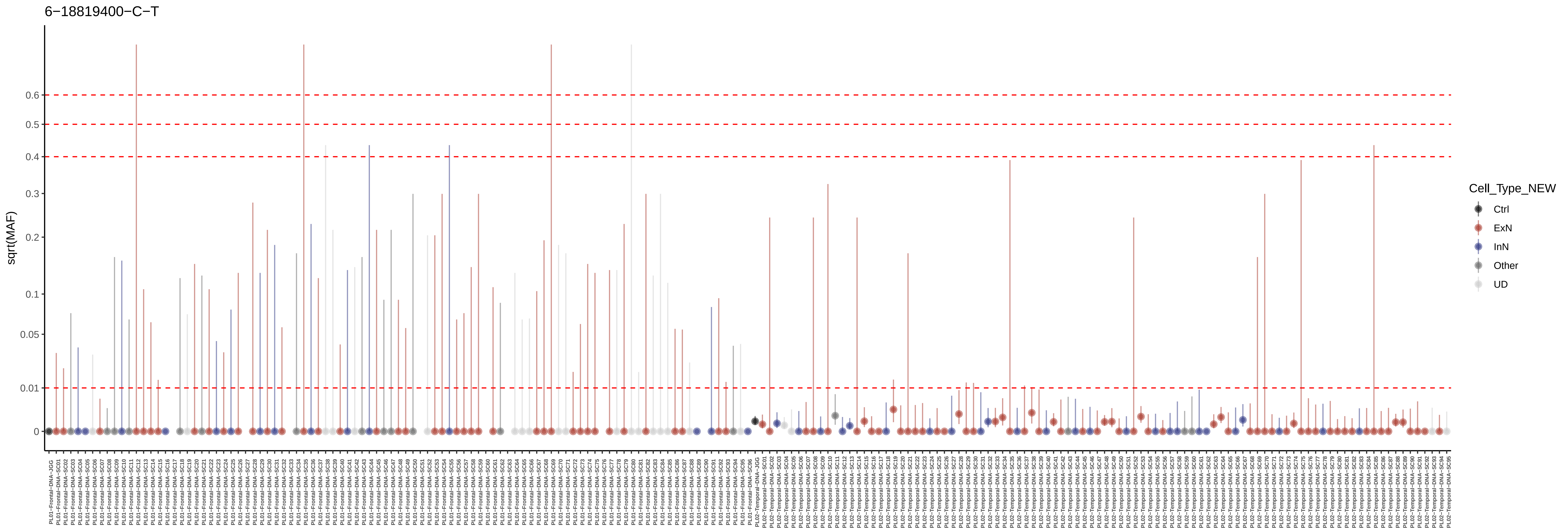
6−18819400−C−T
sqrt(MAF)
0.6
0.5
0.4
0.3
0.2
0.1
0.05
0.01
0
Cell_Type_NEW
Ctrl
ExN
InN
Other
UD

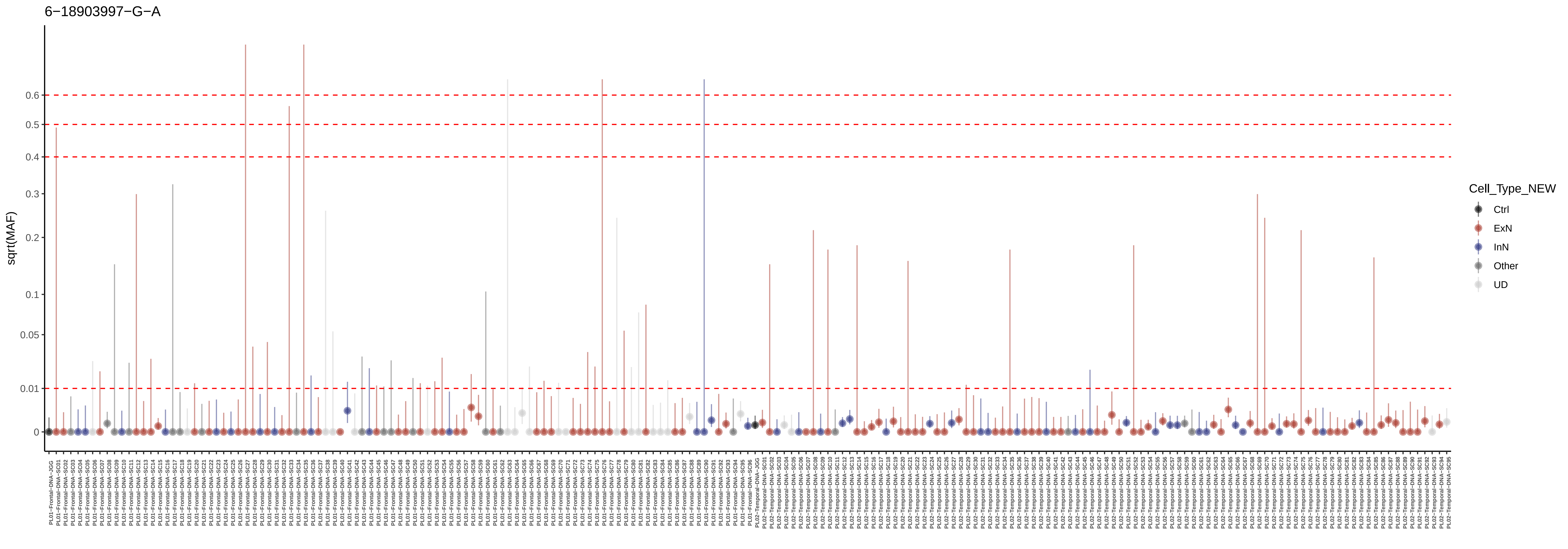

6−18903997−G−A
sqrt(MAF)
0.6
0.5
0.4
0.3
0.2
0.1
0.05
0.01
0
Cell_Type_NEW
Ctrl
ExN
InN
Other
UD

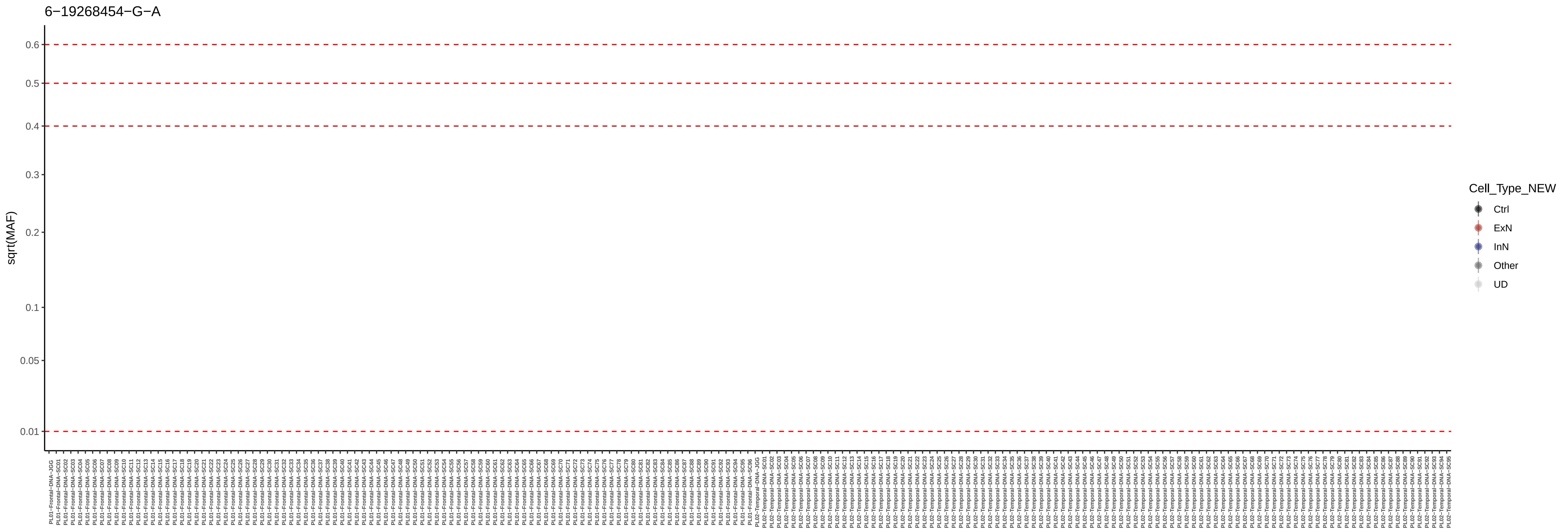
6−19268454−G−A
sqrt(MAF)
0.6
0.5
0.4
0.3
0.2
0.1
0.05
0.01
Cell_Type_NEW
Ctrl
ExN
InN
Other
UD

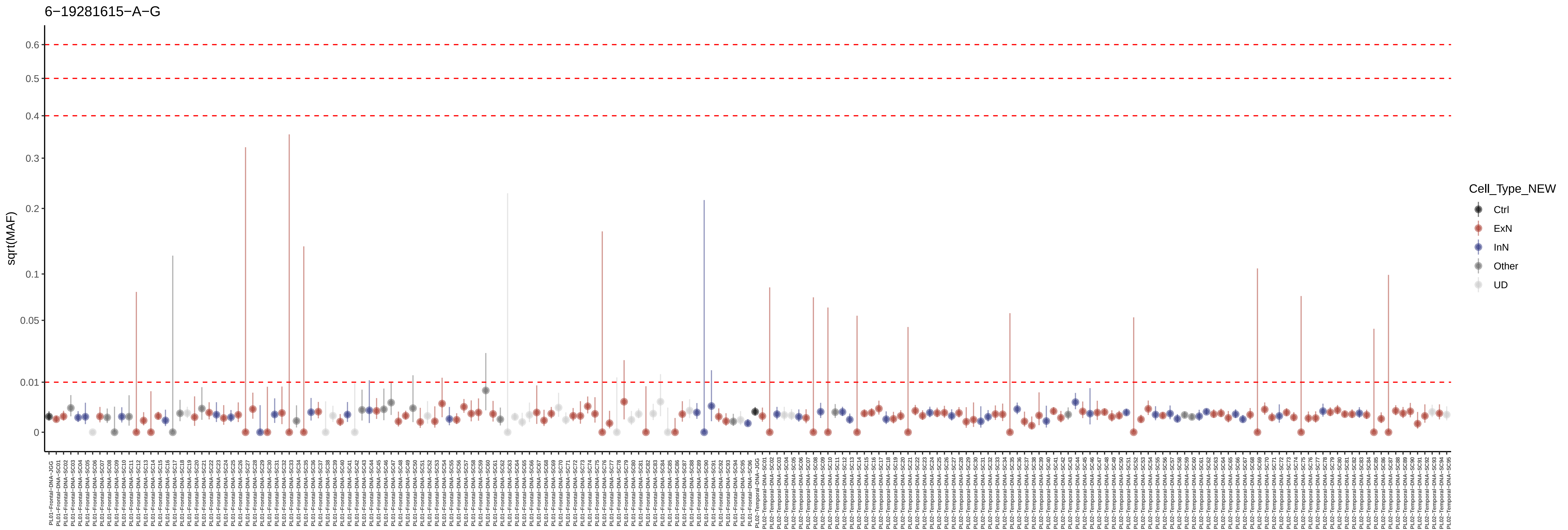

6−19281615−A−G
sqrt(MAF)
0.6
0.5
0.4
0.3
0.2
0.1
0.05
0.01
0
Cell_Type_NEW
Ctrl
ExN
InN
Other
UD

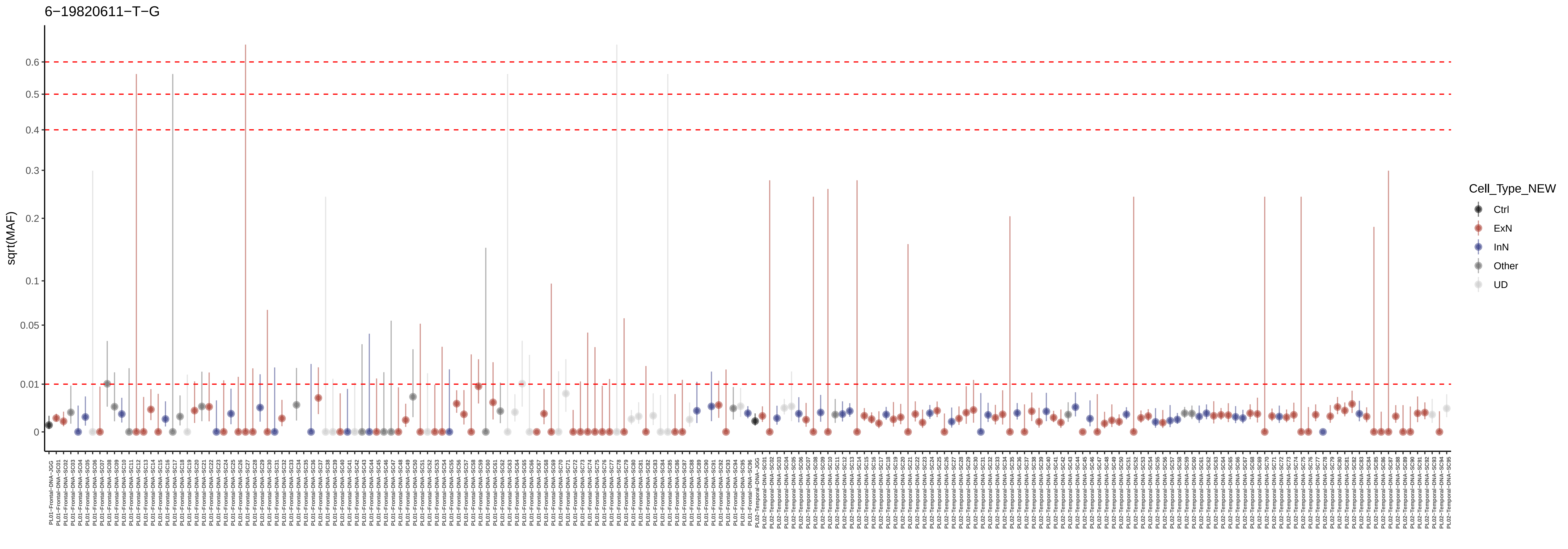

6−19820611−T−G
sqrt(MAF)
0
0.01
0.05
0.1
0.2
0.3
0.4
0.5
0.6
Cell_Type_NEW
Ctrl
ExN
InN
Other
UD

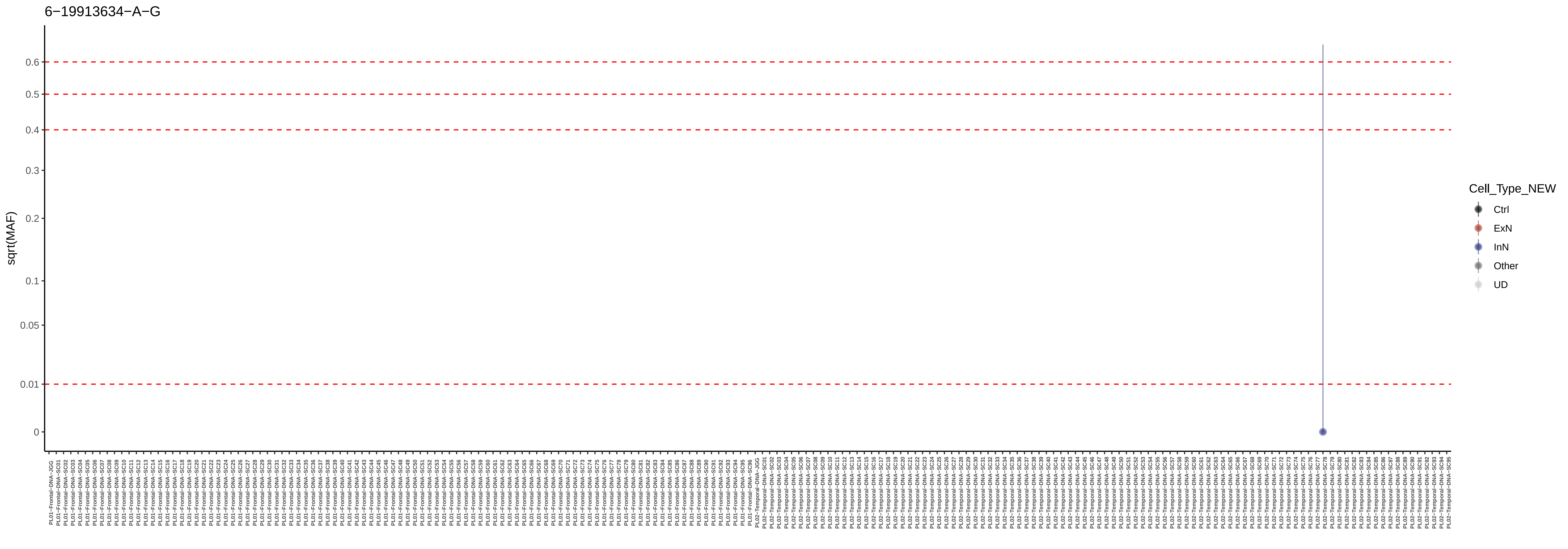

6−19913634−A−G
sqrt(MAF)
0.6
0.5
0.4
0.3
0.2
0.1
0.05
0.01
0
Cell_Type_NEW
Ctrl
ExN
InN
Other
UD

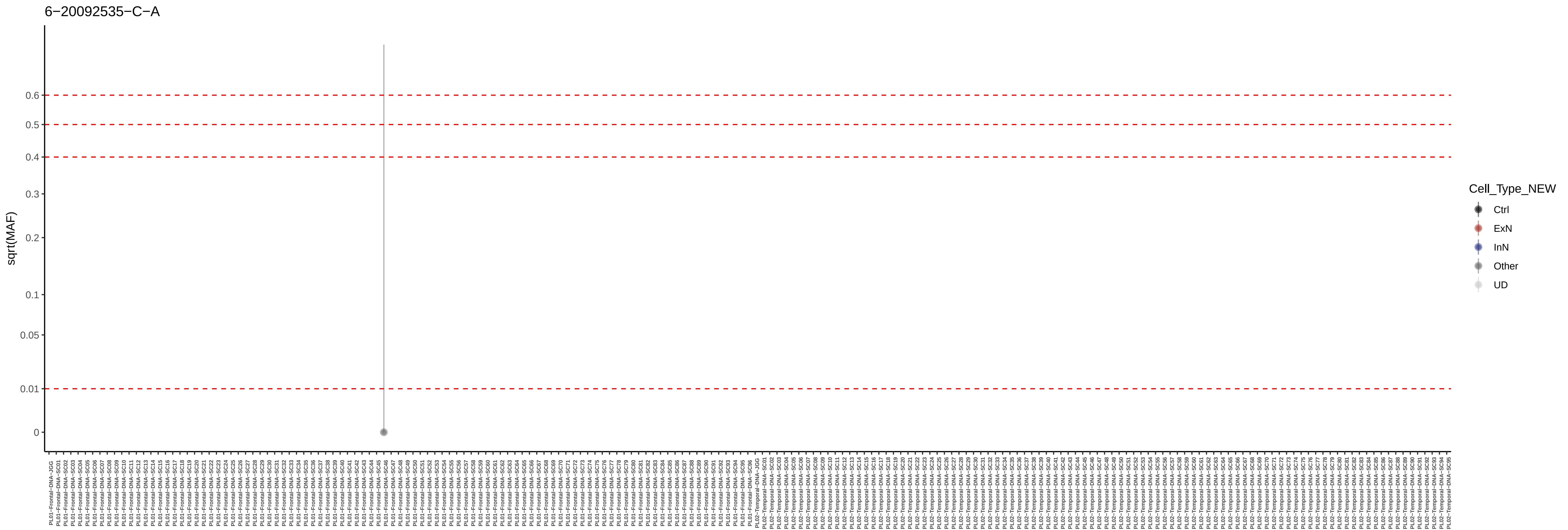

6−20092535−C−A
sqrt(MAF)
0.6
0.5
0.4
0.3
0.2
0.1
0.05
0.01
0
Cell_Type_NEW
Ctrl
ExN
InN
Other
UD

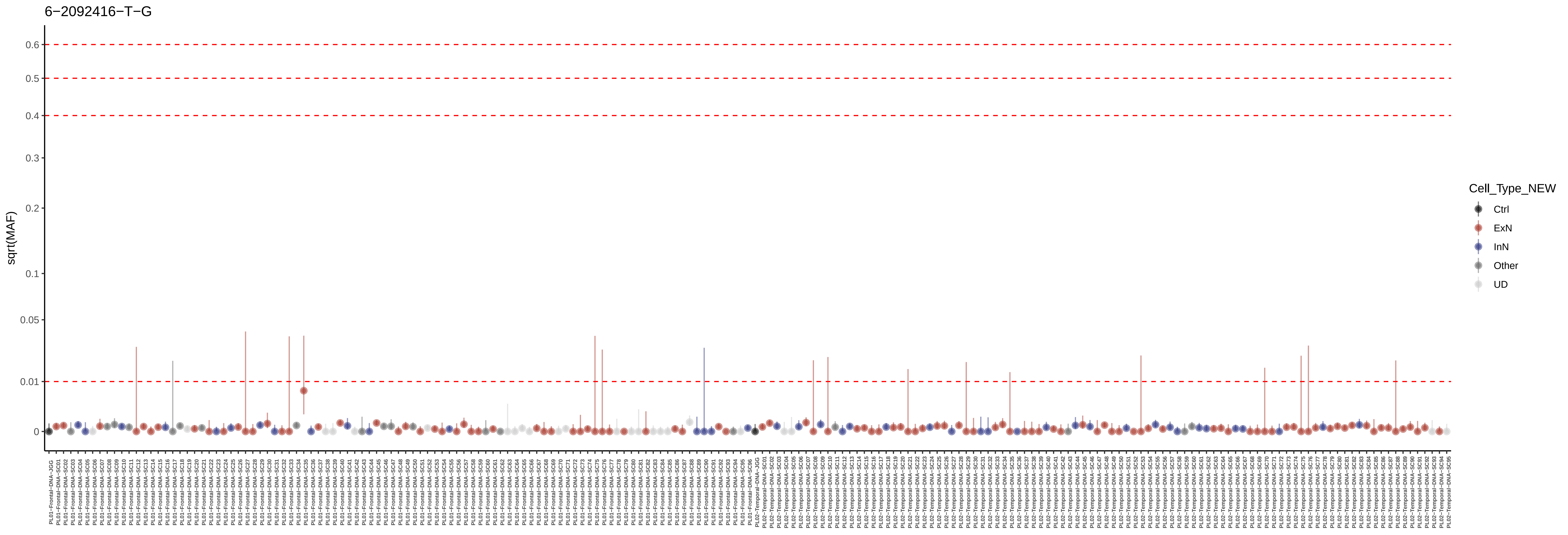

6−2092416−T−G
sqrt(MAF)
0.6
0.5
0.4
0.3
0.2
0.1
0.05
0.01
0
Cell_Type_NEW
Ctrl
ExN
InN
Other
UD

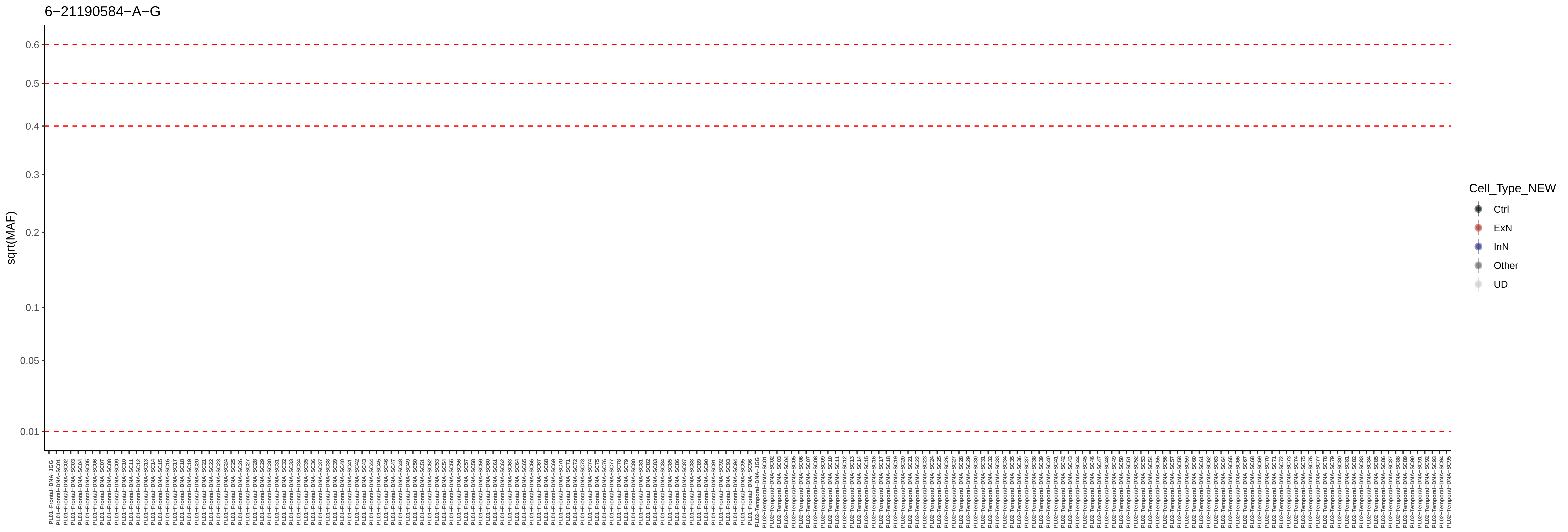
6−21190584−A−G
sqrt(MAF)
0.6
0.5
0.4
0.3
0.2
0.1
0.05
0.01
Cell_Type_NEW
Ctrl
ExN
InN
Other
UD

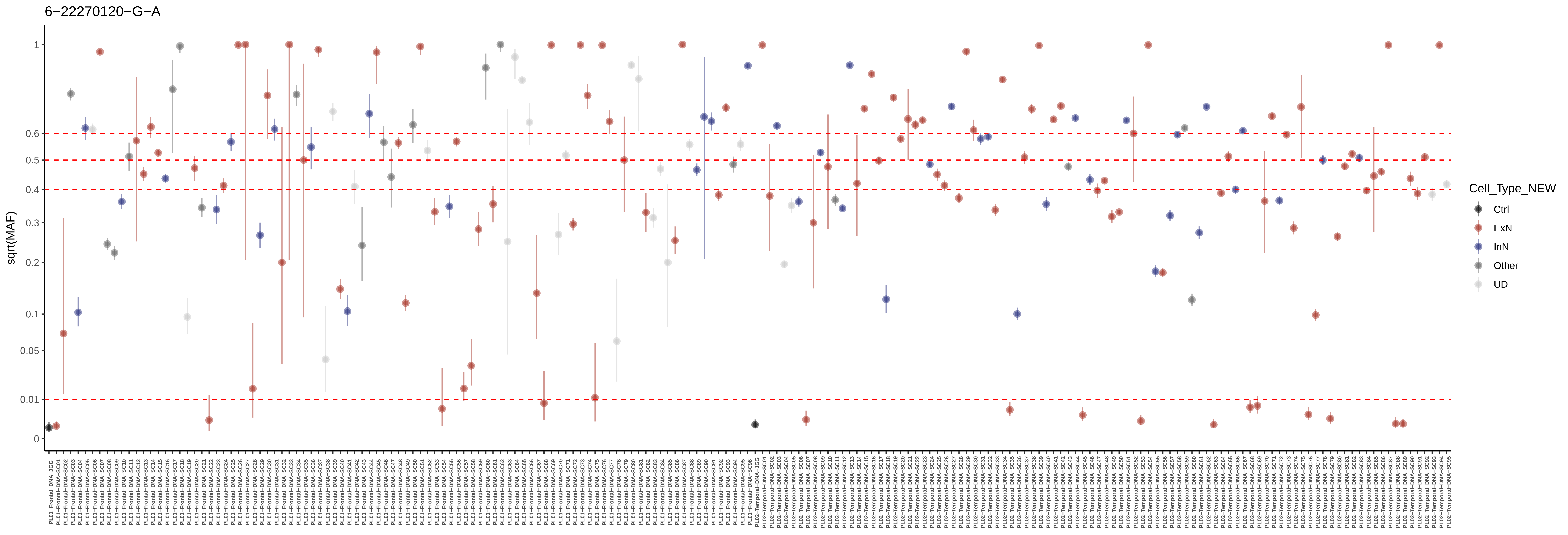

6−22270120−G−A
sqrt(MAF)
1
0.6
0.5
0.4
0.3
0.2
0.1
0.05
0.01
0
Cell_Type_NEW
Ctrl
ExN
InN
Other
UD

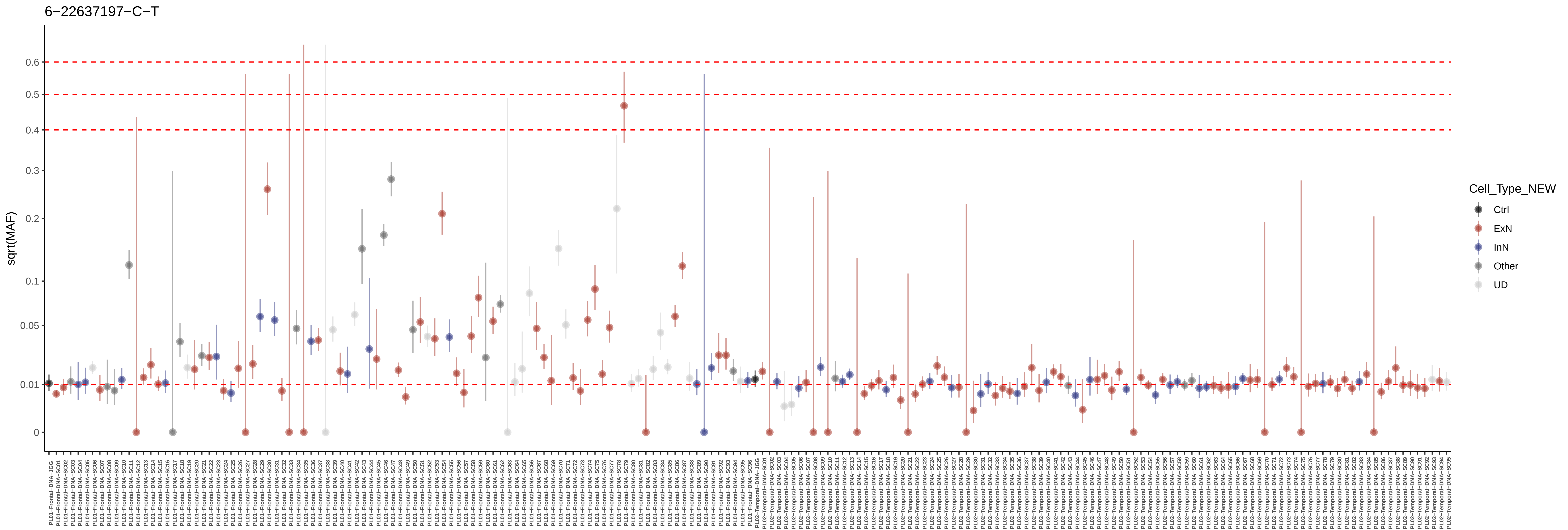
6−22637197−C−T
sqrt(MAF)
0.6
0.5
0.4
0.3
0.2
0.1
0.05
0.01
0
Cell_Type_NEW
Ctrl
ExN
InN
Other
UD

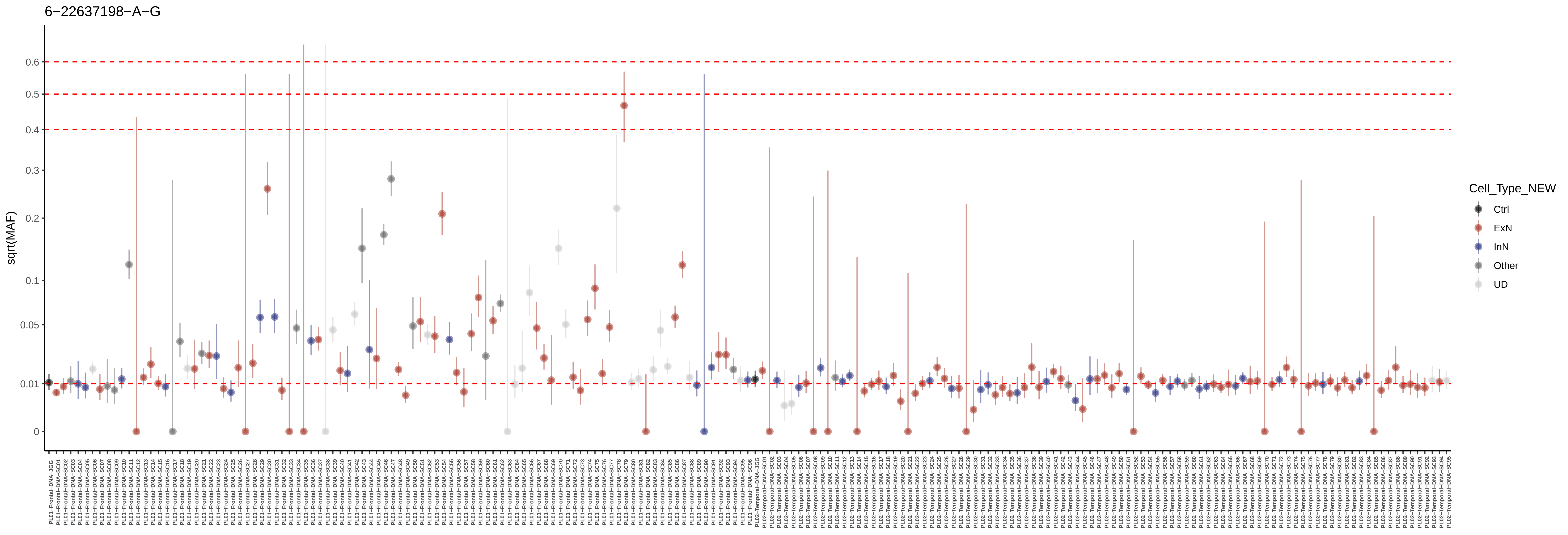

6−22637198−A−G
sqrt(MAF)
0.6
0.5
0.4
0.3
0.2
0.1
0.05
0.01
0
Cell_Type_NEW
Ctrl
ExN
InN
Other
UD

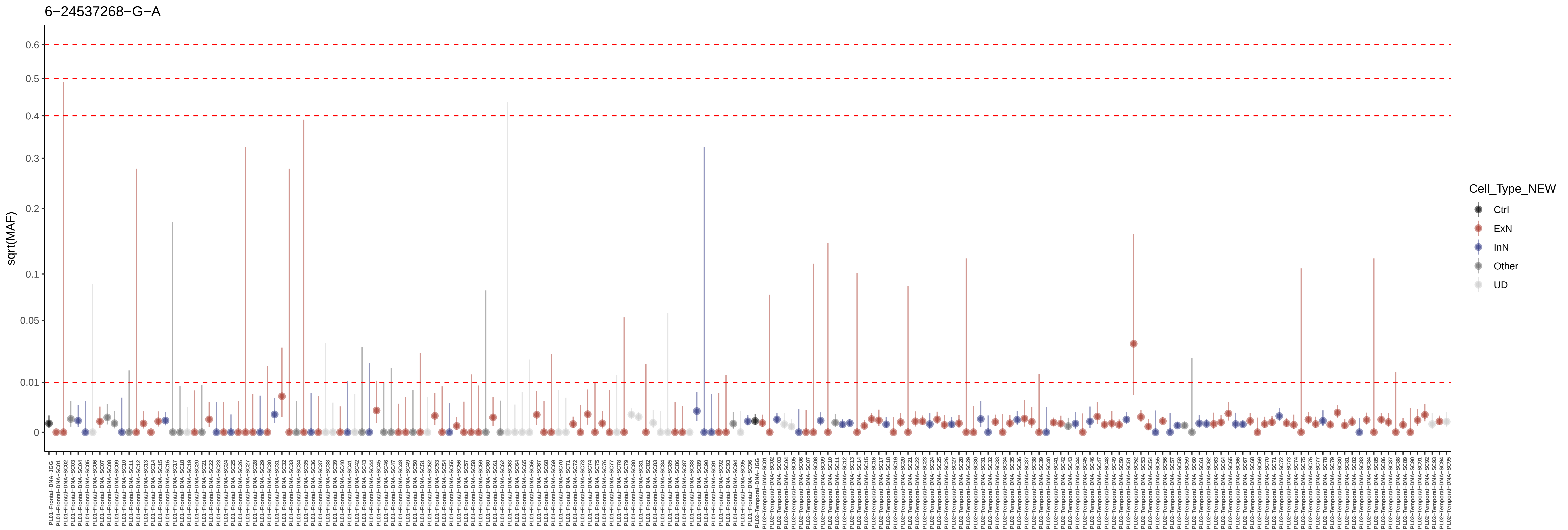
6−24537268−G−A
sqrt(MAF)
0.6
0.5
0.4
0.3
0.2
0.1
0.05
0.01
0
Cell_Type_NEW
Ctrl
ExN
InN
Other
UD

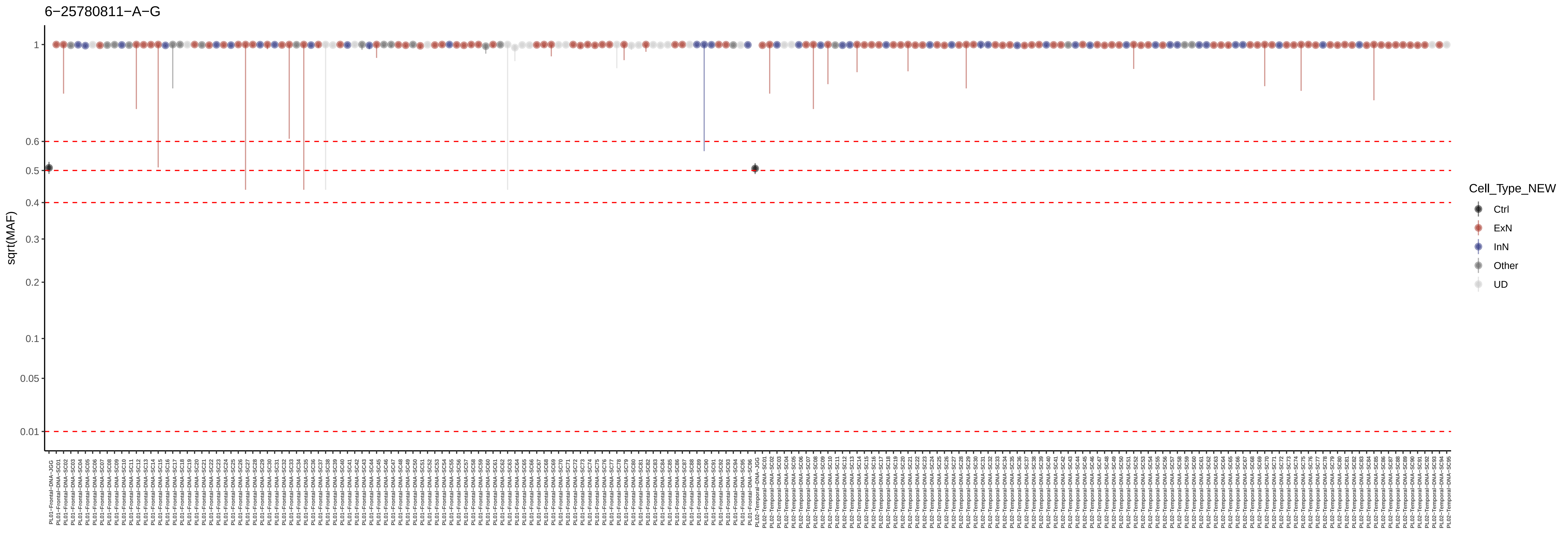

6−25780811−A−G
sqrt(MAF)
1
0.6
0.5
0.4
0.3
0.2
0.1
0.05
0.01
Cell_Type_NEW
Ctrl
ExN
InN
Other
UD

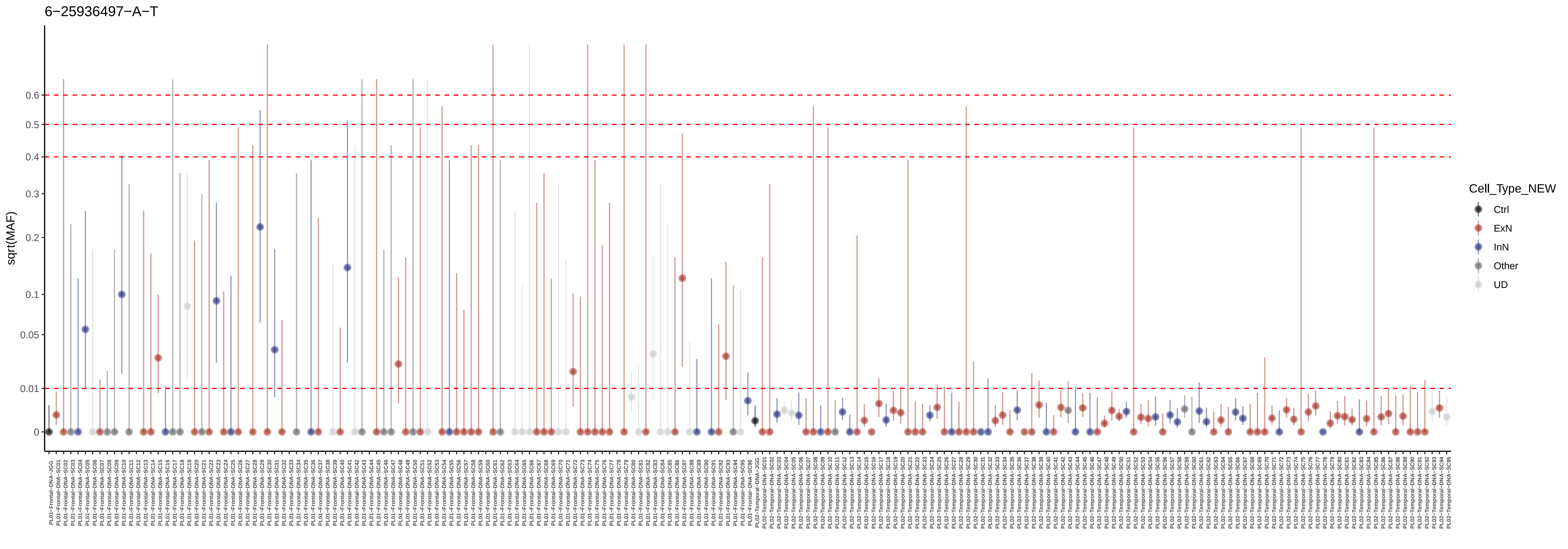

6−25936497−A−T
sqrt(MAF)
0.6
0.5
0.4
0.3
0.2
0.1
0.05
0.01
0
Cell_Type_NEW
Ctrl
ExN
InN
Other
UD

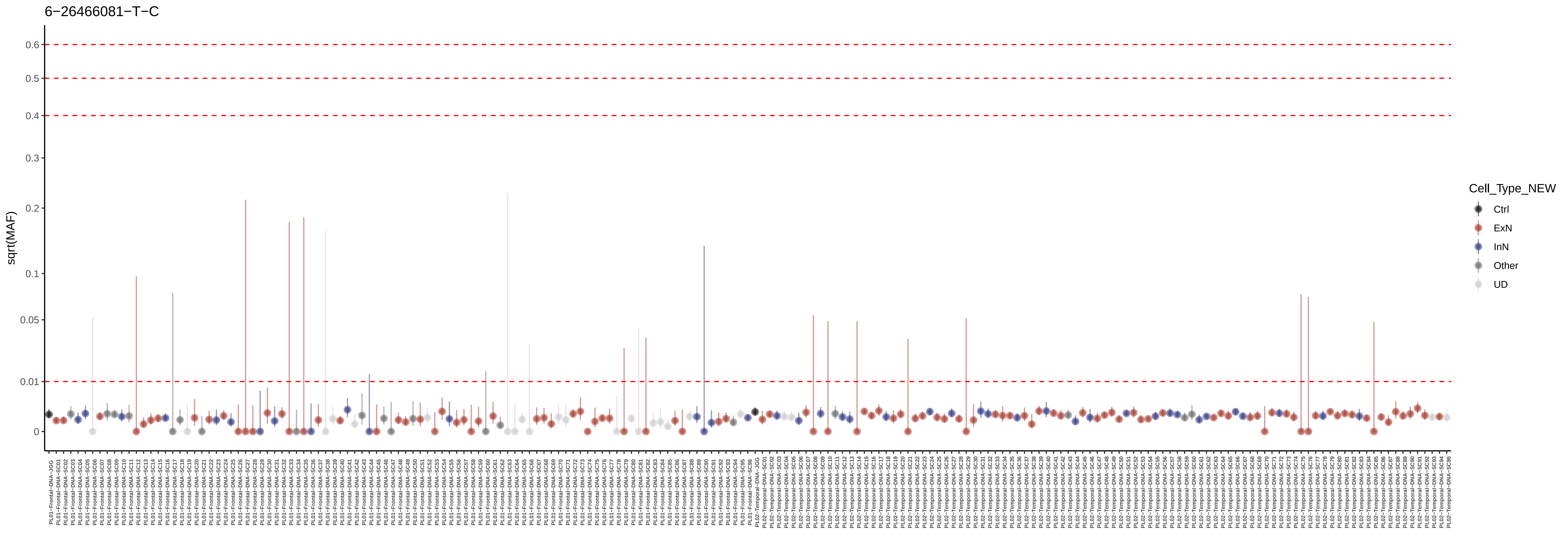

6−26466081−T−C
sqrt(MAF)
0.6
0.5
0.4
0.3
0.2
0.1
0.05
0.01
0
Cell_Type_NEW
Ctrl
ExN
InN
Other
UD

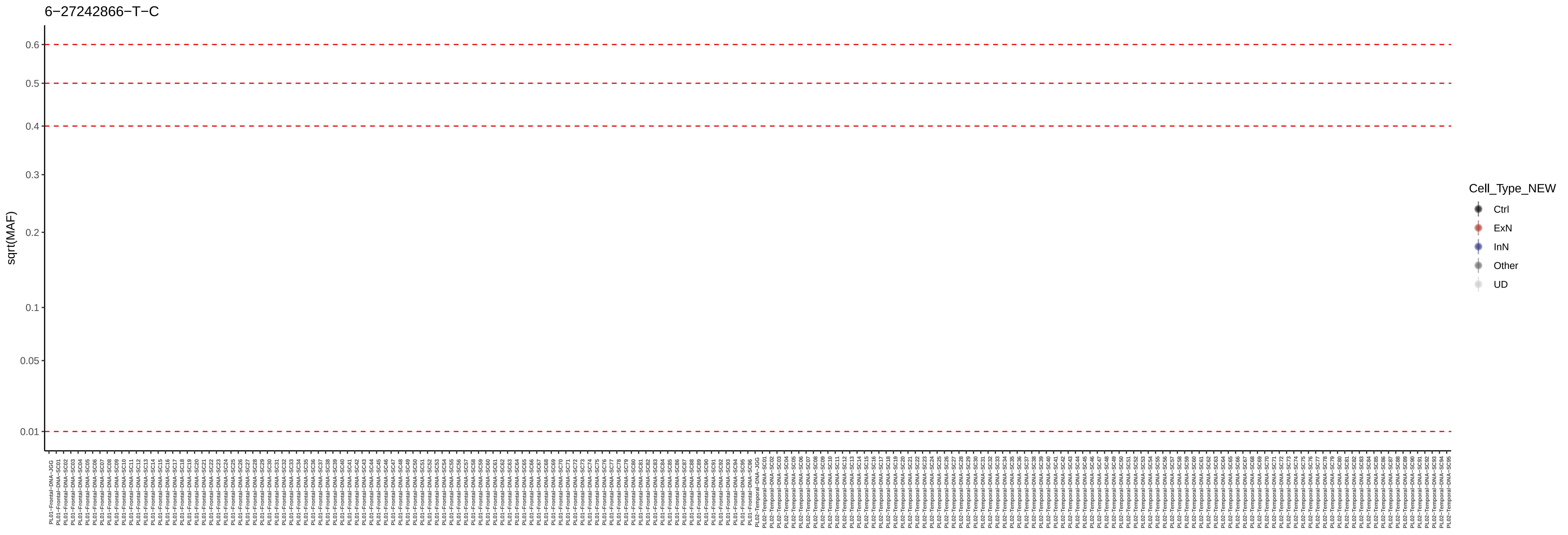
6−27242866−T−C
sqrt(MAF)
0.6
0.5
0.4
0.3
0.2
0.1
0.05
0.01
Cell_Type_NEW
Ctrl
ExN
InN
Other
UD

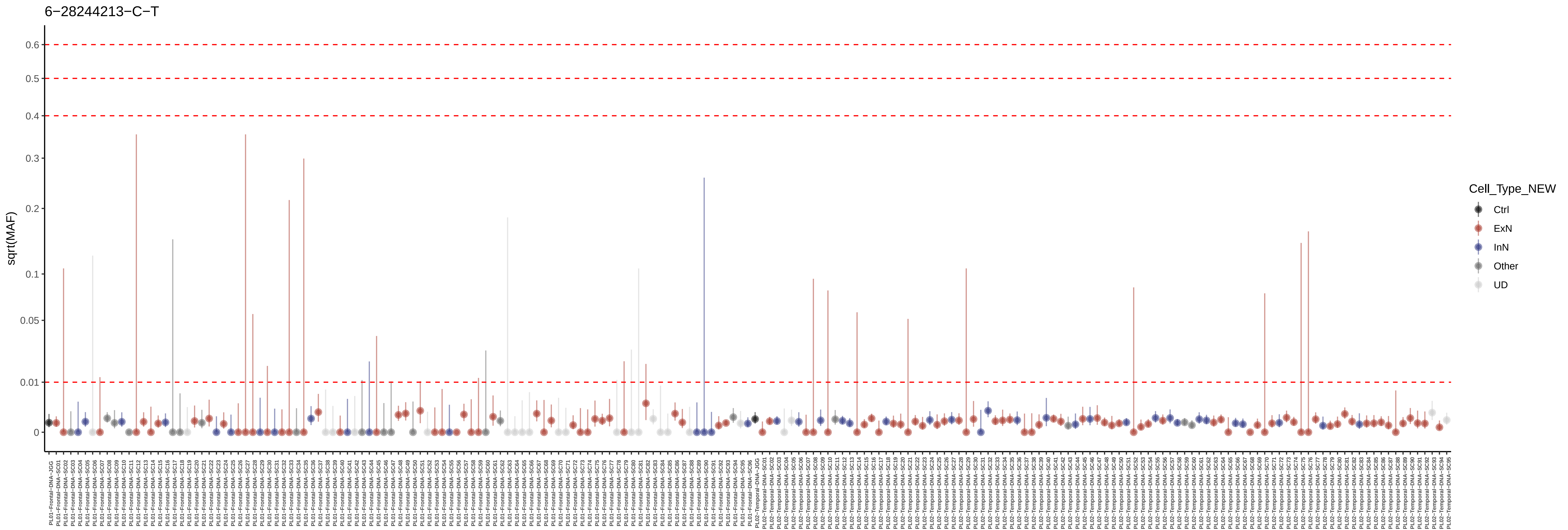
6−28244213−C−T
sqrt(MAF)
0.6
0.5
0.4
0.3
0.2
0.1
0.05
0.01
0
Cell_Type_NEW
Ctrl
ExN
InN
Other
UD

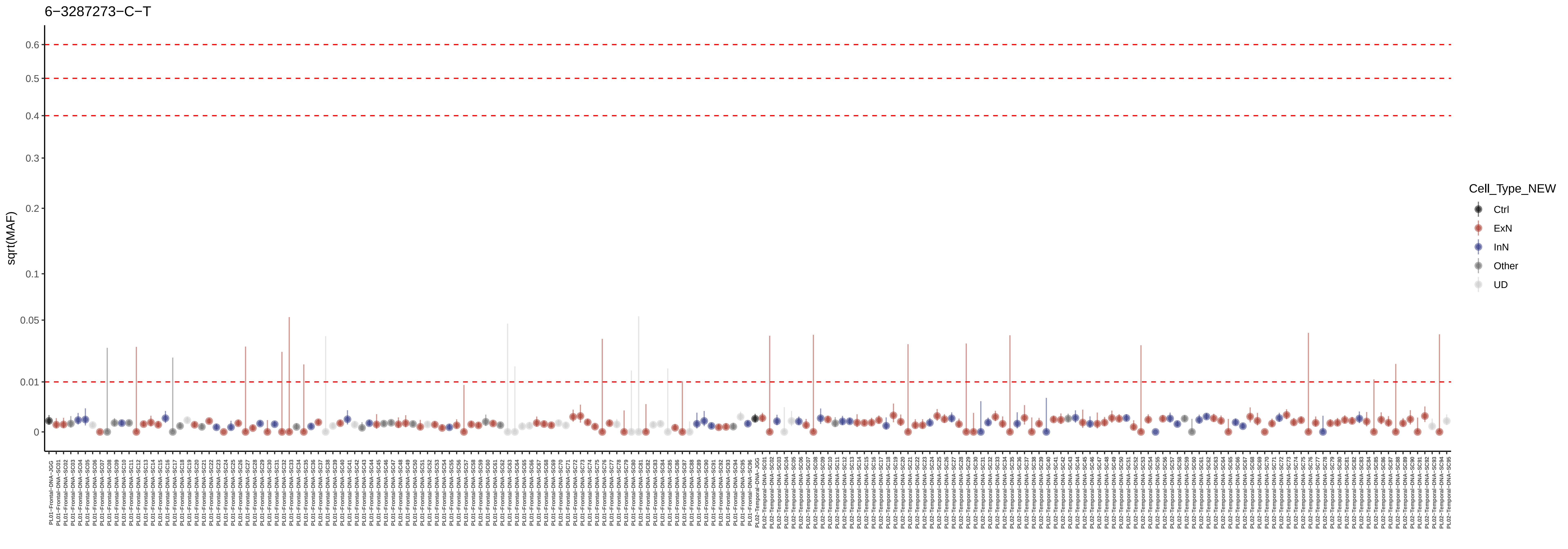
6−3287273−C−T
sqrt(MAF)
0.6
0.5
0.4
0.3
0.2
0.1
0.05
0.01
0
Cell_Type_NEW
Ctrl
ExN
InN
Other
UD

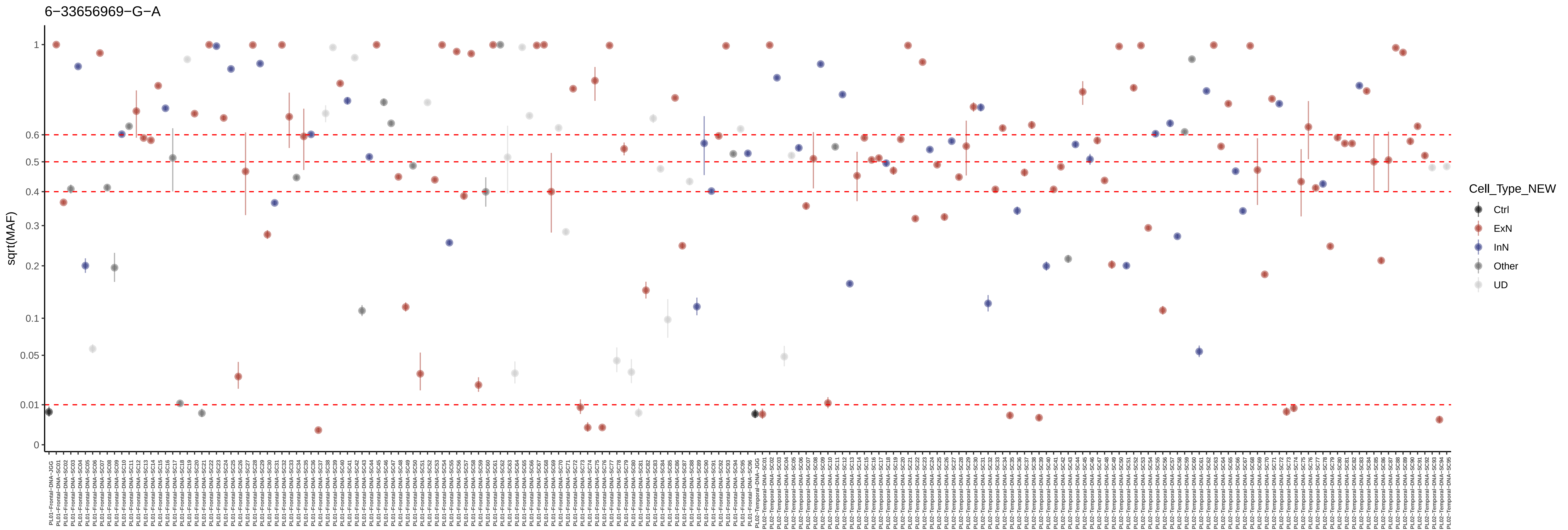

6−33656969−G−A
sqrt(MAF)
1
0.6
0.5
0.4
0.3
0.2
0.1
0.05
0.01
0
Cell_Type_NEW
Ctrl
ExN
InN
Other
UD

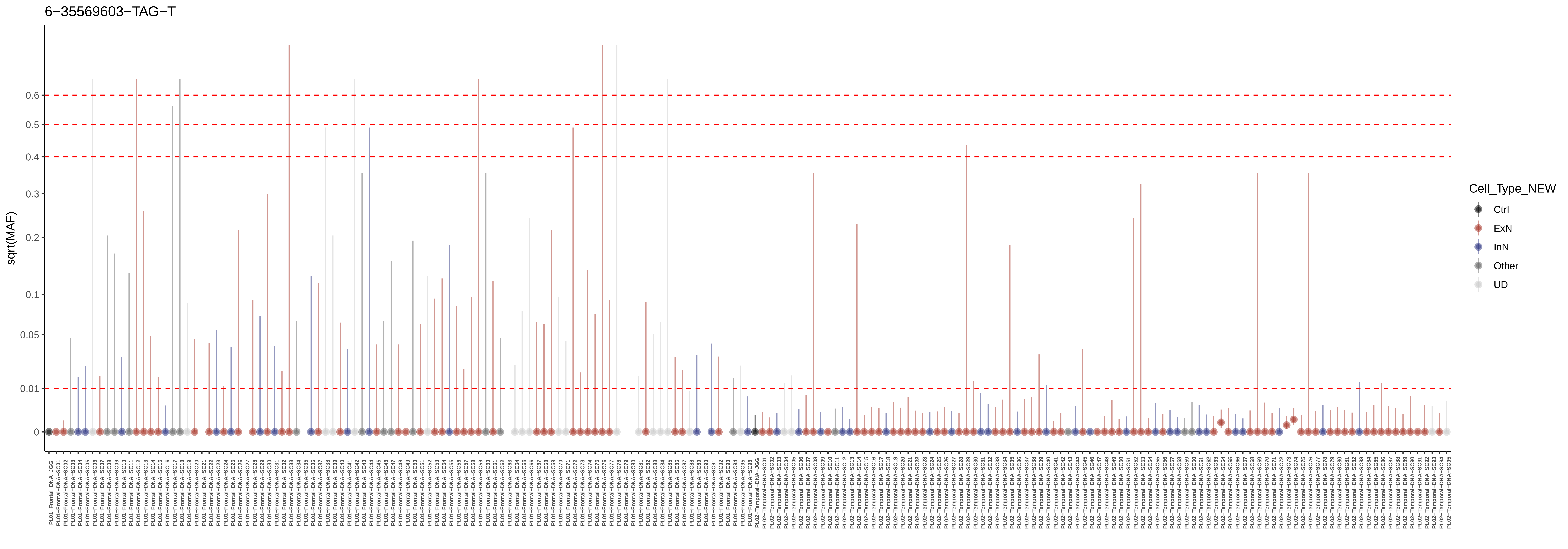
6−35569603−TAG−T
sqrt(MAF)
0.6
0.5
0.4
0.3
0.2
0.1
0.05
0.01
0
Cell_Type_NEW
Ctrl
ExN
InN
Other
UD

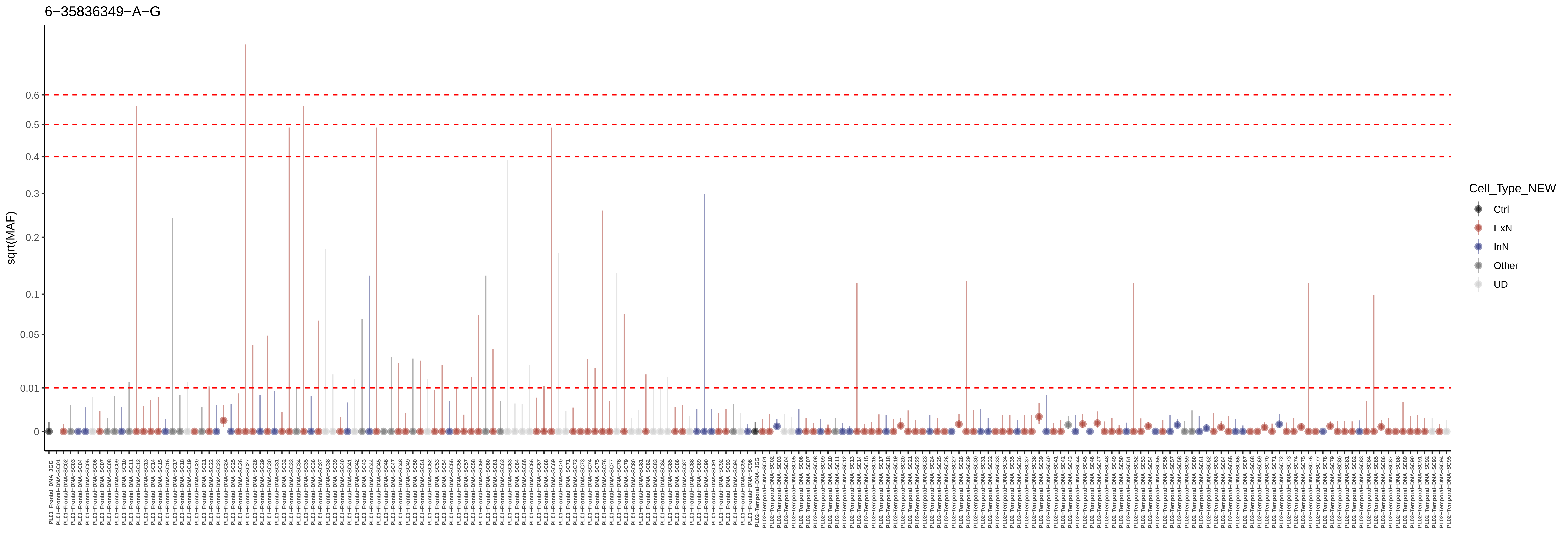
6−35836349−A−G
sqrt(MAF)
0.6
0.5
0.4
0.3
0.2
0.1
0.05
0.01
0
Cell_Type_NEW
Ctrl
ExN
InN
Other
UD

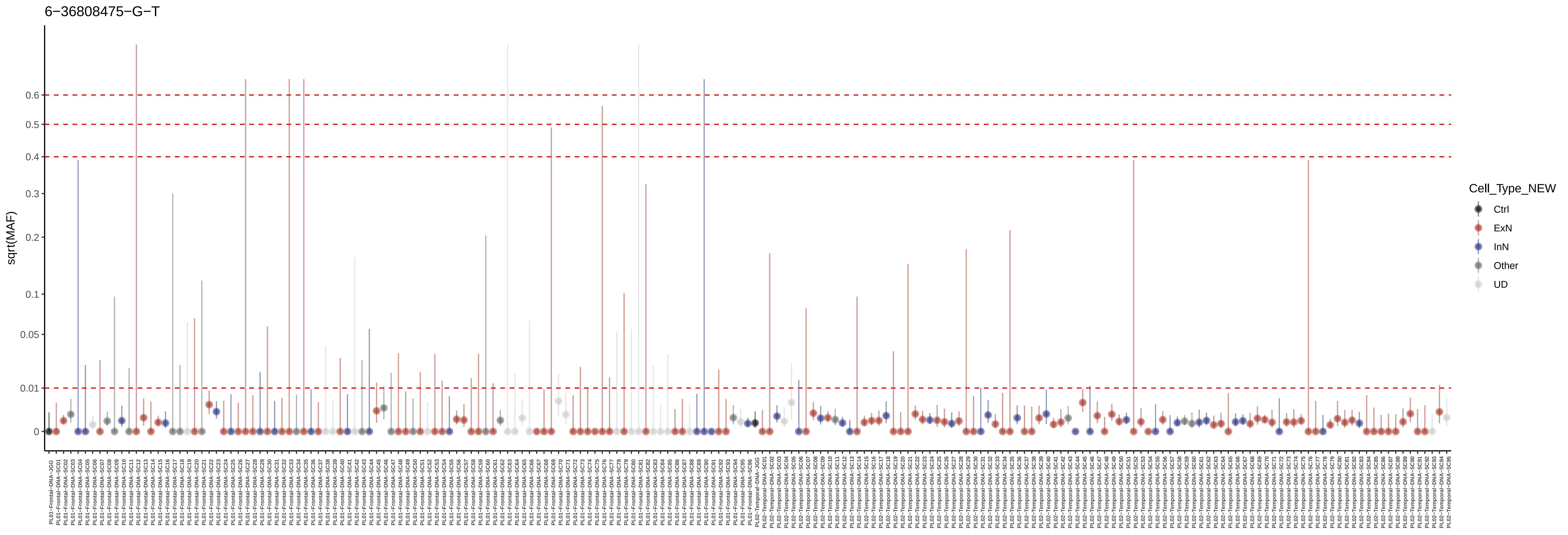
6−36808475−G−T
sqrt(MAF)
0.6
0.5
0.4
0.3
0.2
0.1
0.05
0.01
0
Cell_Type_NEW
Ctrl
ExN
InN
Other
UD

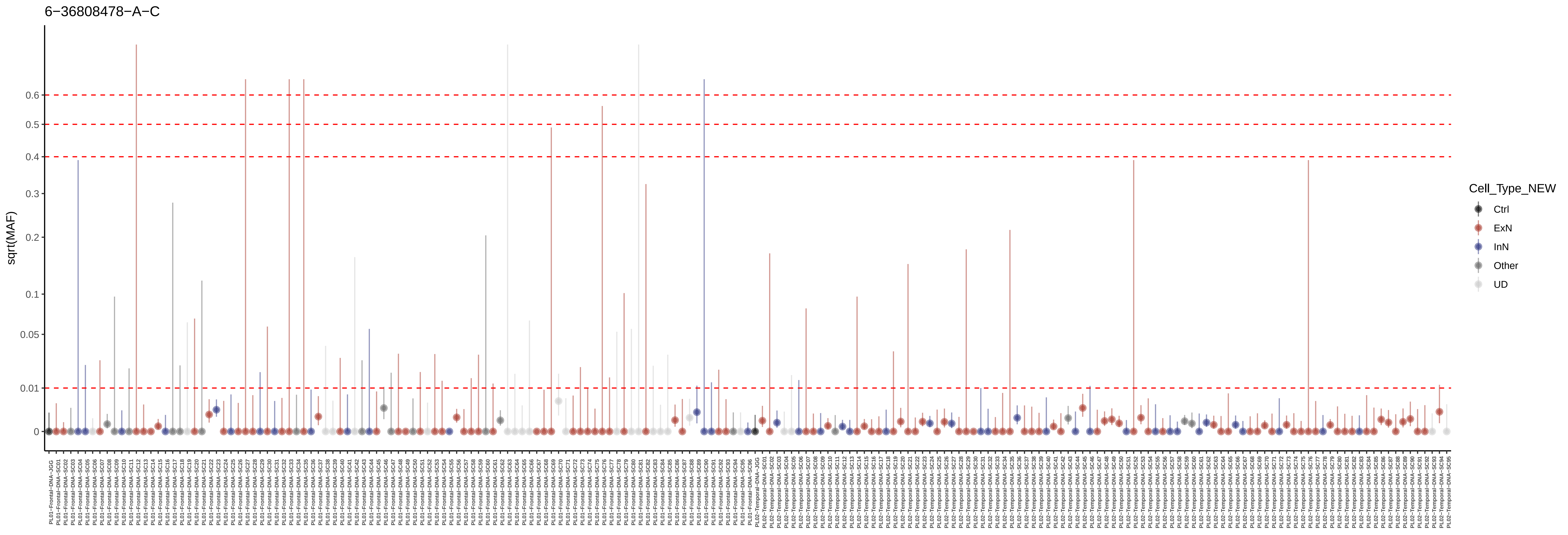

6−36808478−A−C
sqrt(MAF)
0.6
0.5
0.4
0.3
0.2
0.1
0.05
0.01
0
Cell_Type_NEW
Ctrl
ExN
InN
Other
UD

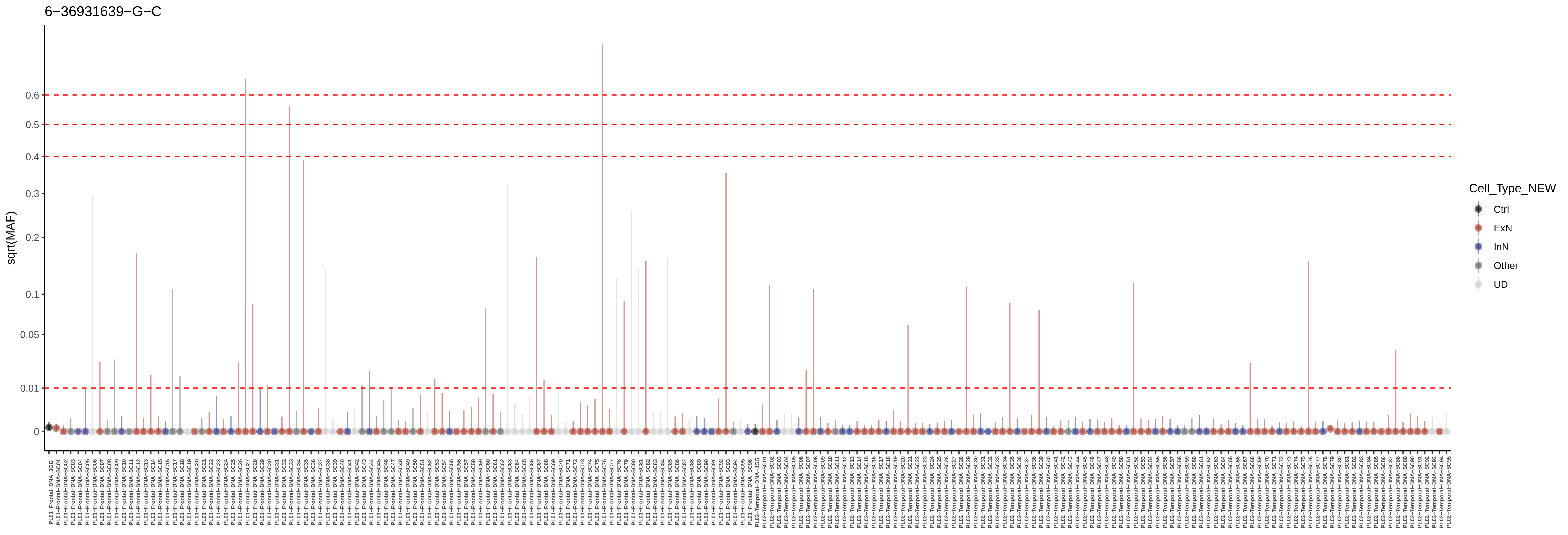
6−36931639−G−C
sqrt(MAF)
0.6
0.5
0.4
0.3
0.2
0.1
0.05
0.01
0
Cell_Type_NEW
Ctrl
ExN
InN
Other
UD

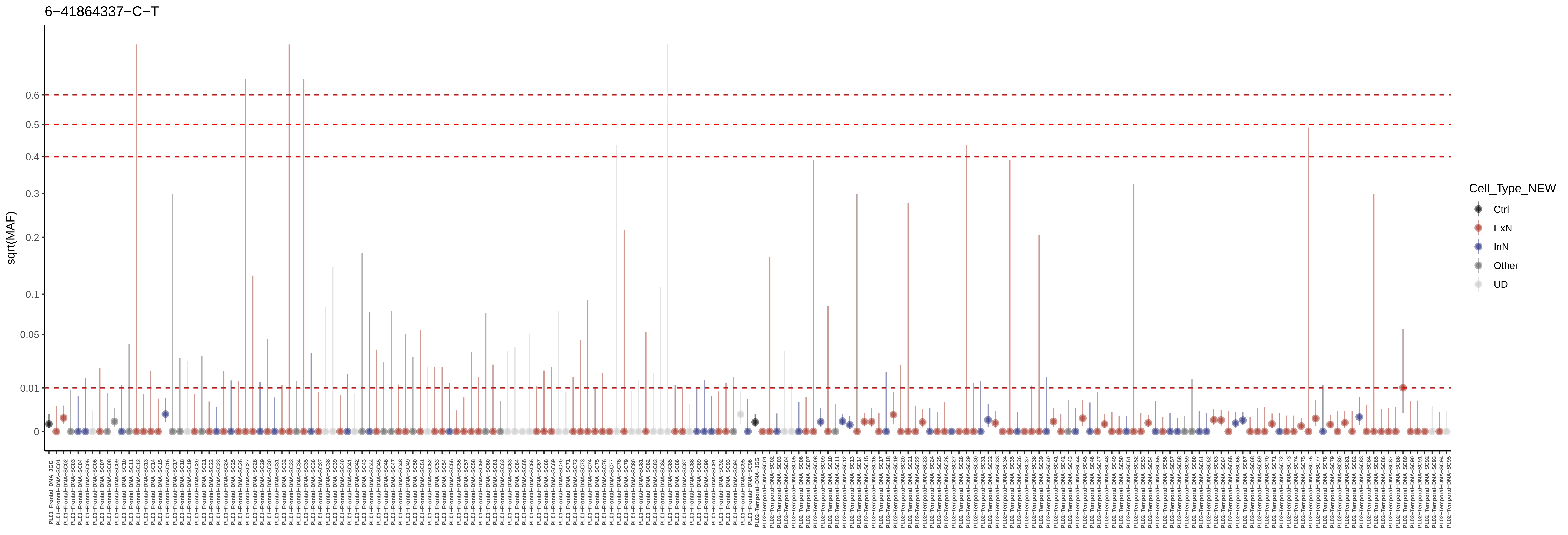
6−41864337−C−T
sqrt(MAF)
0.6
0.5
0.4
0.3
0.2
0.1
0.05
0.01
0
Cell_Type_NEW
Ctrl
ExN
InN
Other
UD

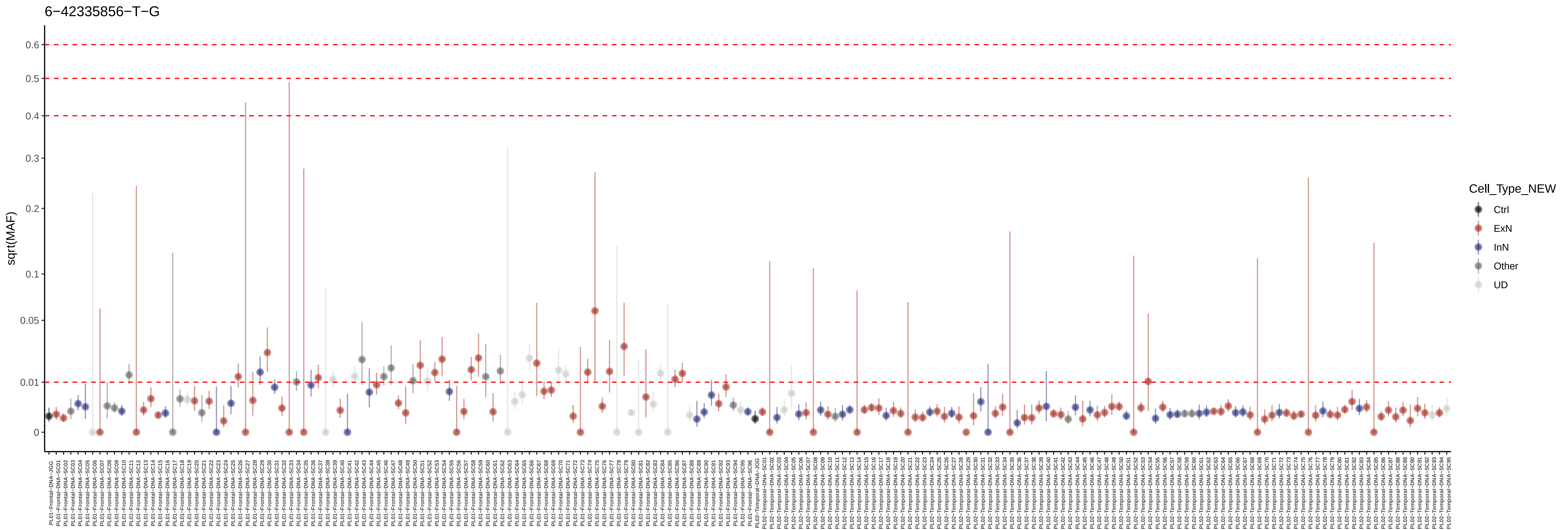
6−42335856−T−G
sqrt(MAF)
0.6
0.5
0.4
0.3
0.2
0.1
0.05
0.01
0
Cell_Type_NEW
Ctrl
ExN
InN
Other
UD

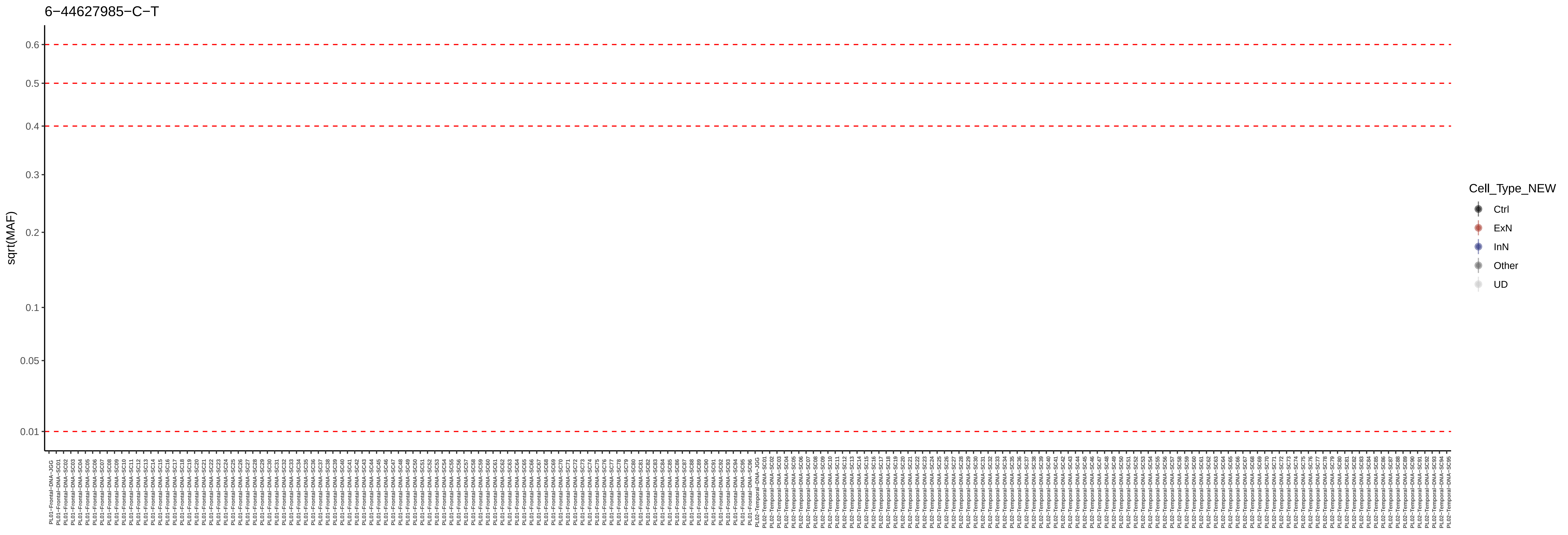
6−44627985−C−T
sqrt(MAF)
0.6
0.5
0.4
0.3
0.2
0.1
0.05
0.01
Cell_Type_NEW
Ctrl
ExN
InN
Other
UD

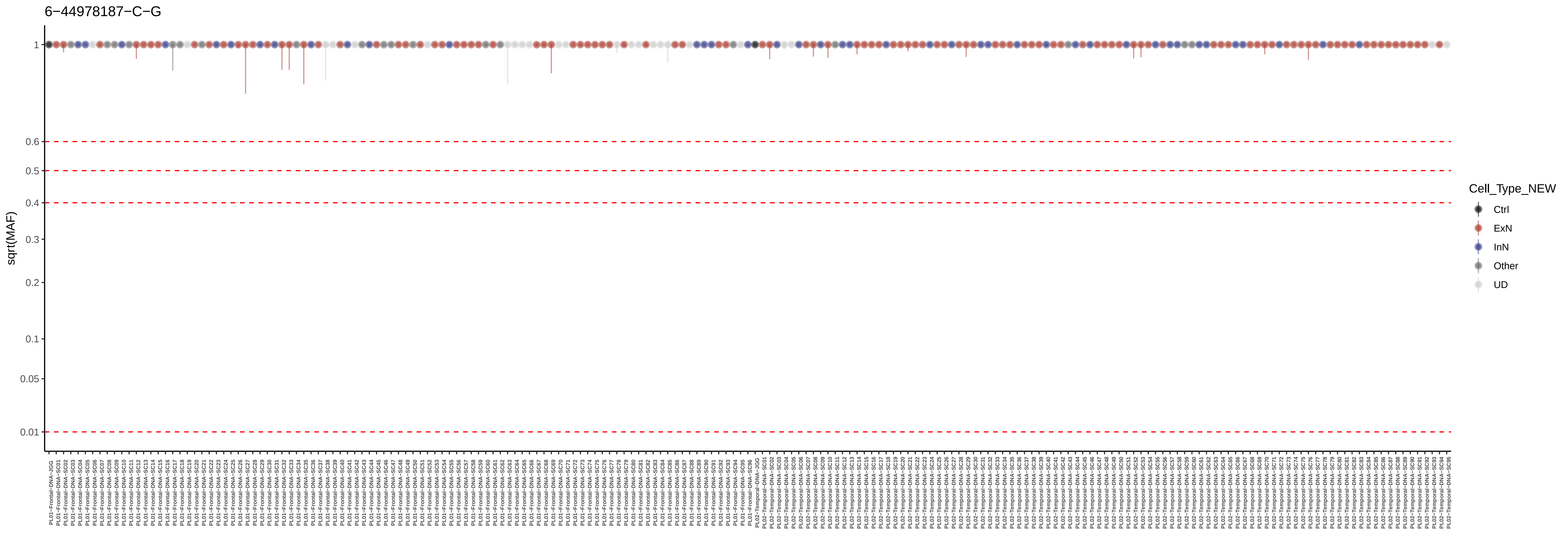
6−44978187−C−G
sqrt(MAF)
Cell_Type_NEW
Ctrl
ExN
InN
Other
UD

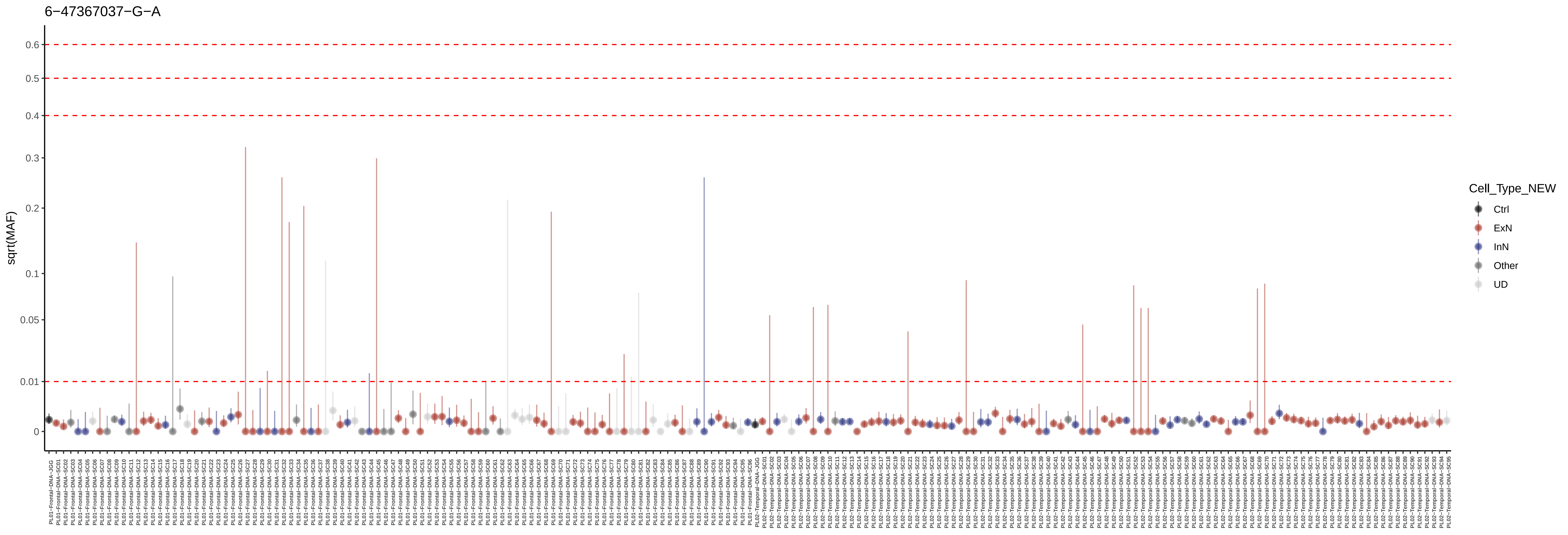
6−47367037−G−A
sqrt(MAF)
0.6
0.5
0.4
0.3
0.2
0.1
0.05
0.01
0
Cell_Type_NEW
Ctrl
ExN
InN
Other
UD

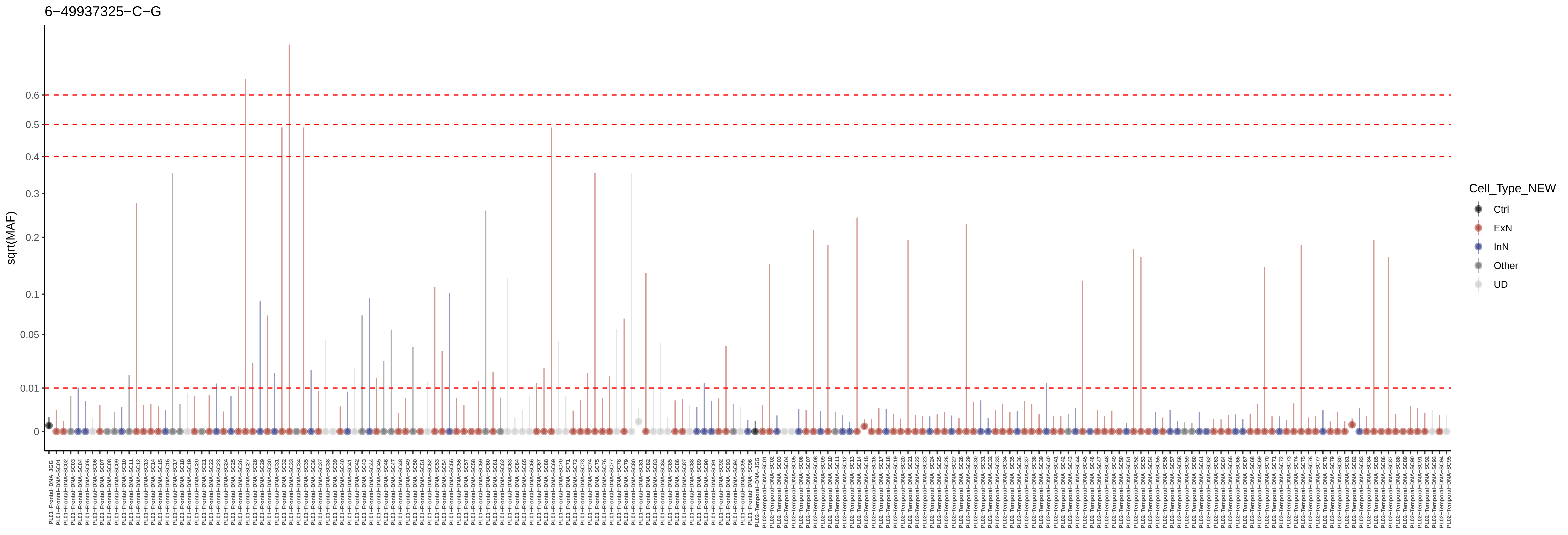

6−49937325−C−G
sqrt(MAF)
0.6
0.5
0.4
0.3
0.2
0.1
0.05
0.01
0
Cell_Type_NEW
Ctrl
ExN
InN
Other
UD

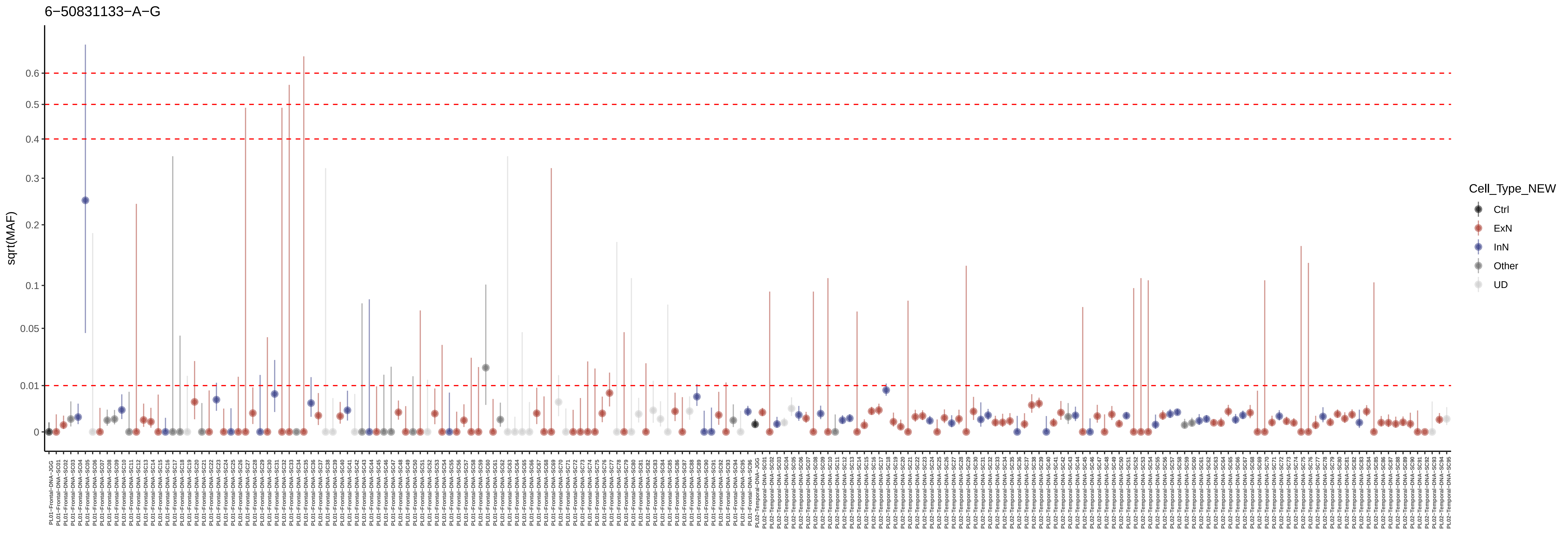

6−50831133−A−G
sqrt(MAF)
0.6
0.5
0.4
0.3
0.2
0.1
0.05
0.01
0
Cell_Type_NEW
Ctrl
ExN
InN
Other
UD

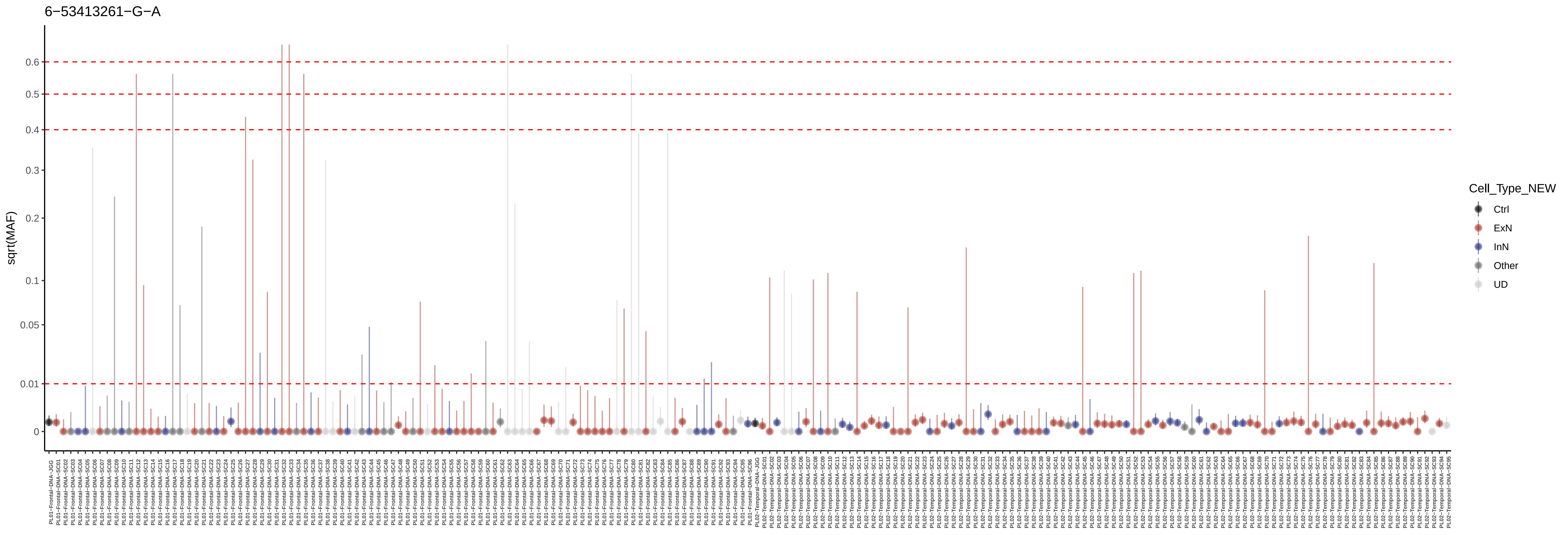
6−53413261−G−A
sqrt(MAF)
0.6
0.5
0.4
0.3
0.2
0.1
0.05
0.01
0
Cell_Type_NEW
Ctrl
ExN
InN
Other
UD

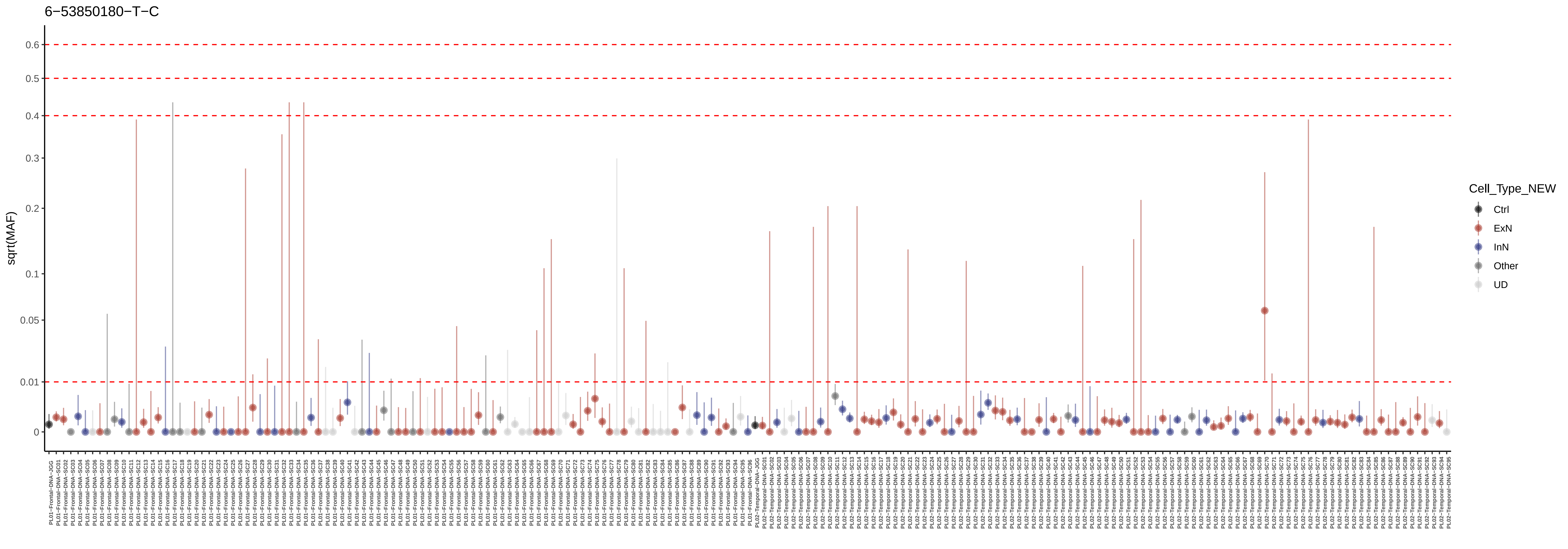
6−53850180−T−C
sqrt(MAF)
0.6
0.5
0.4
0.3
0.2
0.1
0.05
0.01
0
Cell_Type_NEW
Ctrl
ExN
InN
Other
UD

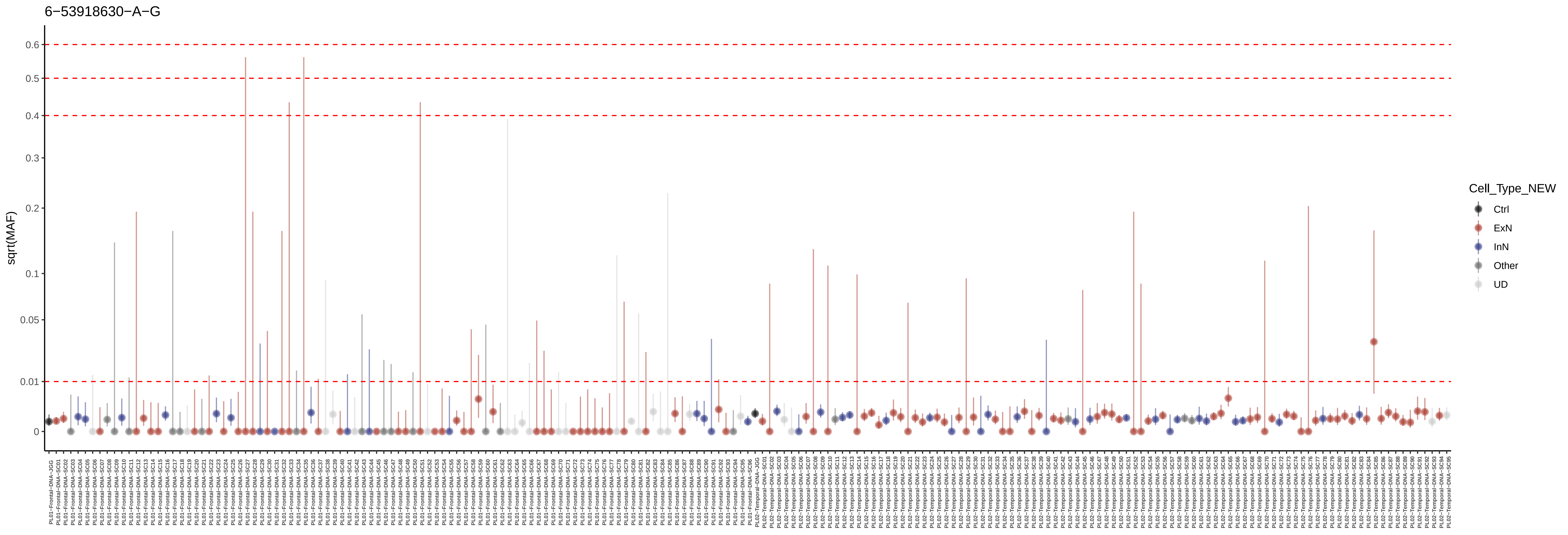

6−53918630−A−G
sqrt(MAF)
0.6
0.5
0.4
0.3
0.2
0.1
0.05
0.01
0
Cell_Type_NEW
Ctrl
ExN
InN
Other
UD

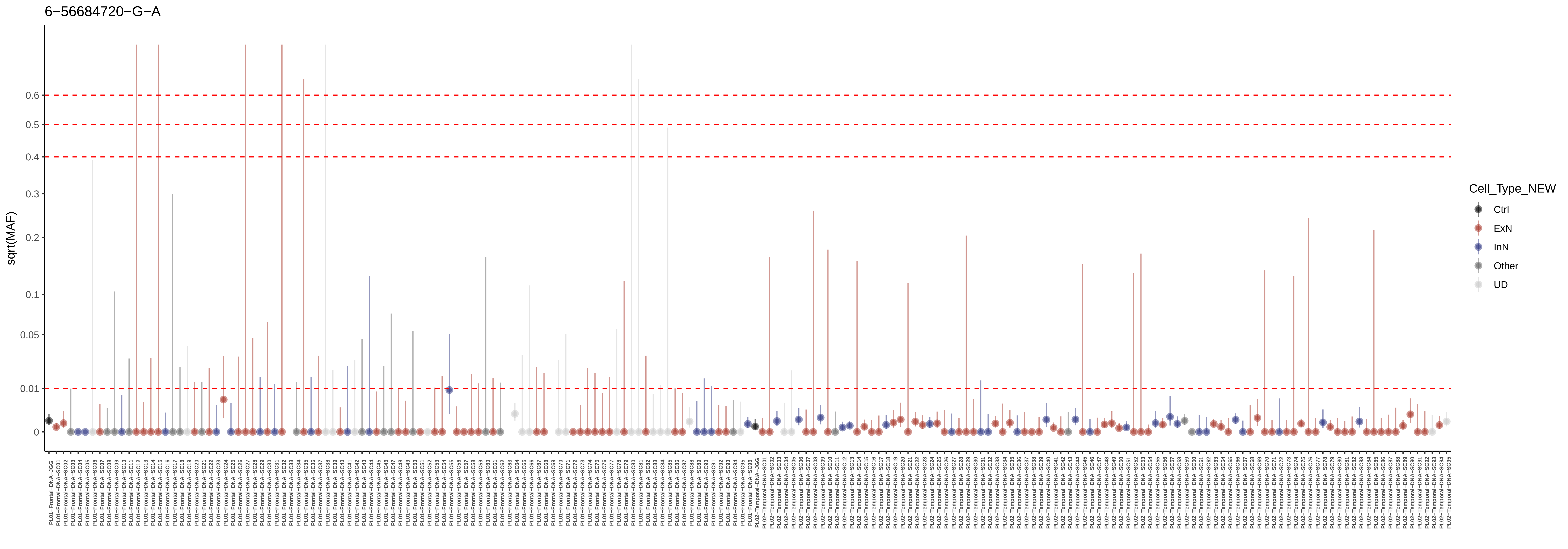
6−56684720−G−A
sqrt(MAF)
0.6
0.5
0.4
0.3
0.2
0.1
0.05
0.01
0
Cell_Type_NEW
Ctrl
ExN
InN
Other
UD

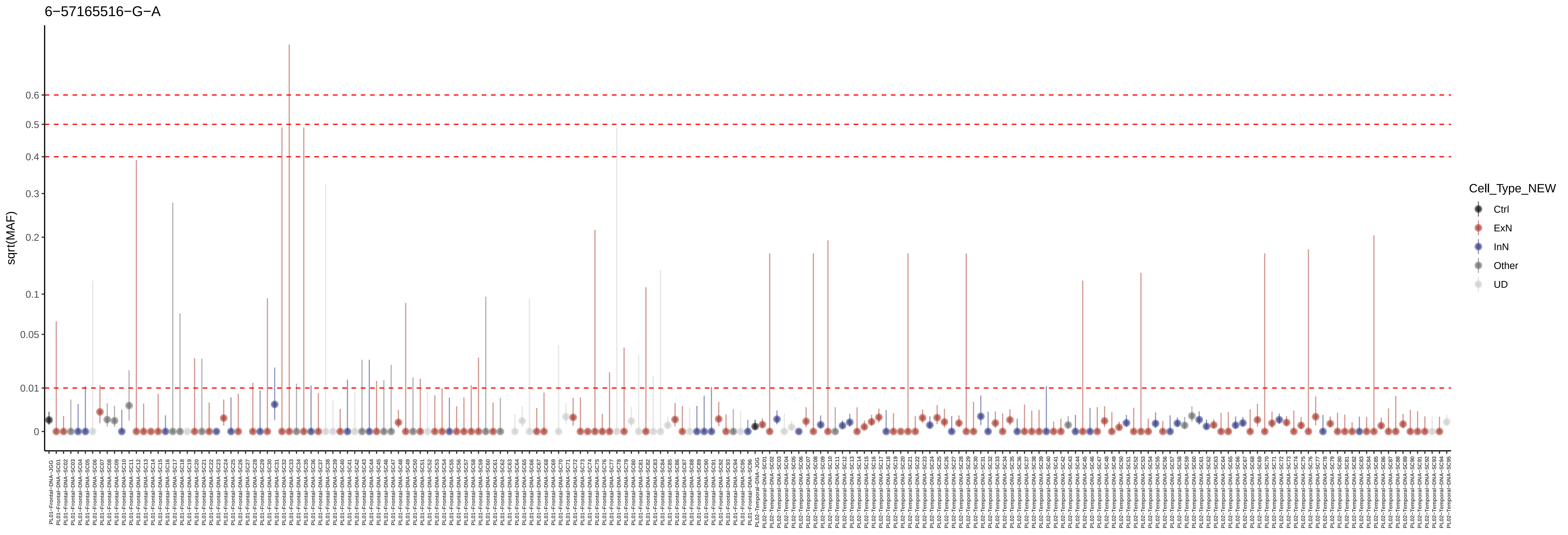
6−57165516−G−A
sqrt(MAF)
0.6
0.5
0.4
0.3
0.2
0.1
0.05
0.01
0
Cell_Type_NEW
Ctrl
ExN
InN
Other
UD

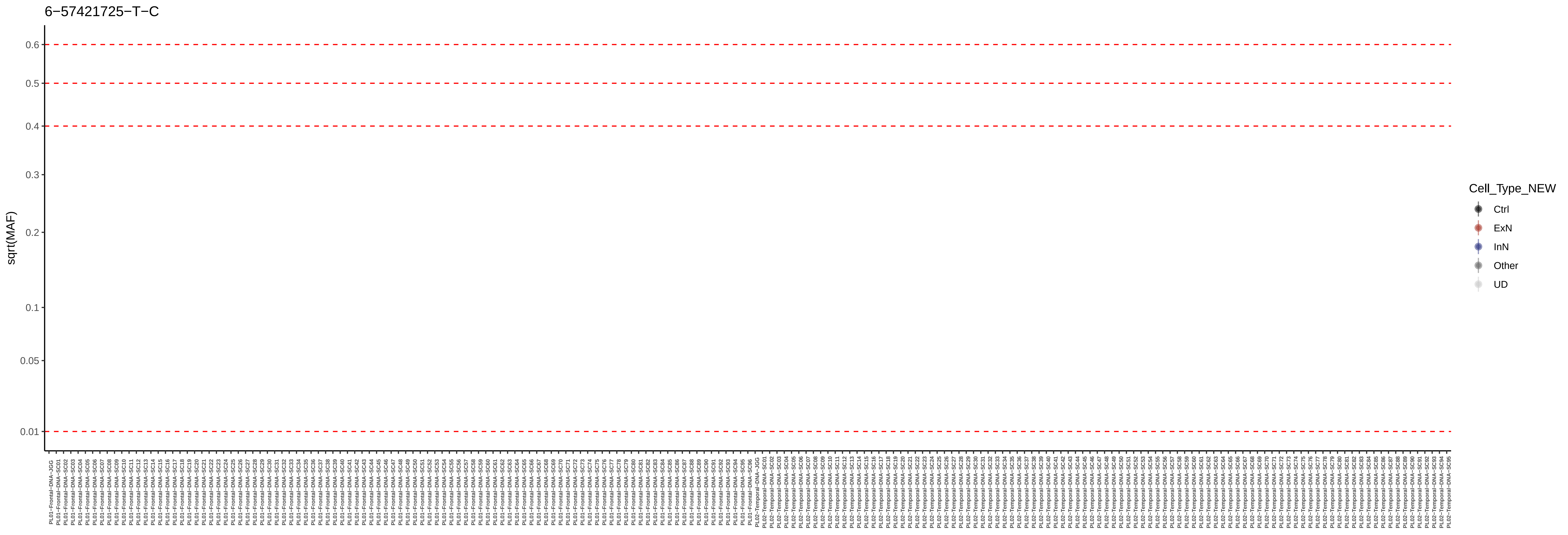
6−57421725−T−C
sqrt(MAF)
0.6
0.5
0.4
0.3
0.2
0.1
0.05
0.01
Cell_Type_NEW
Ctrl
ExN
InN
Other
UD

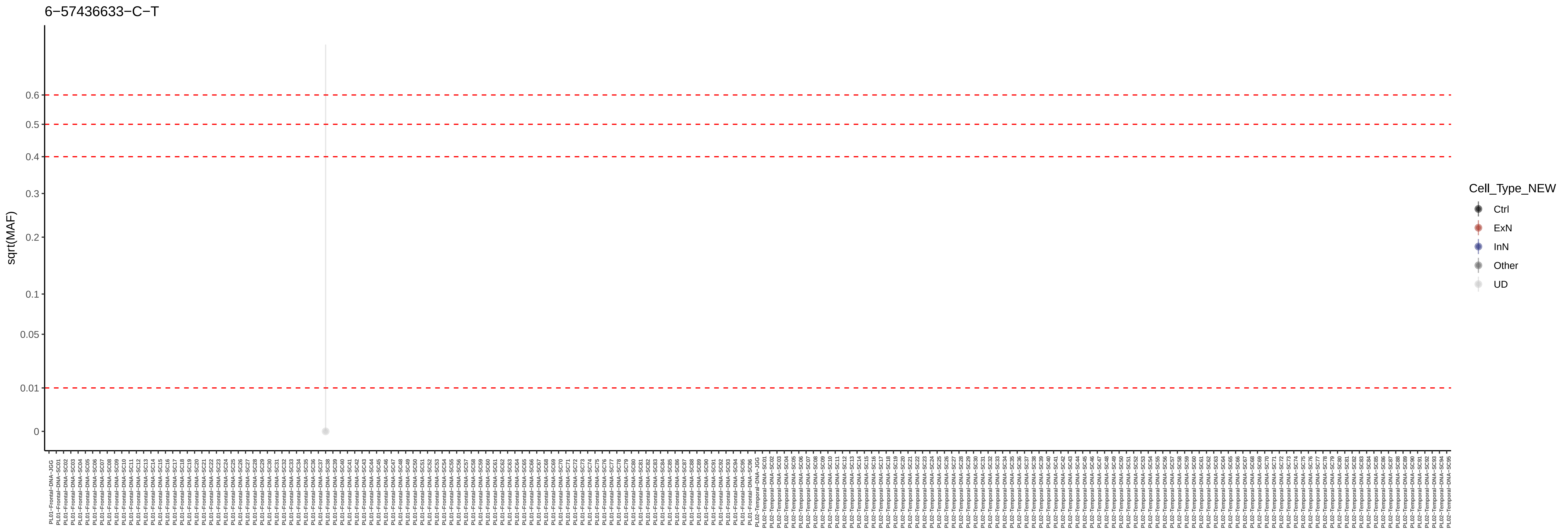

6−57436633−C−T
sqrt(MAF)
0.6
0.5
0.4
0.3
0.2
0.1
0.05
0.01
0
Cell_Type_NEW
Ctrl
ExN
InN
Other
UD

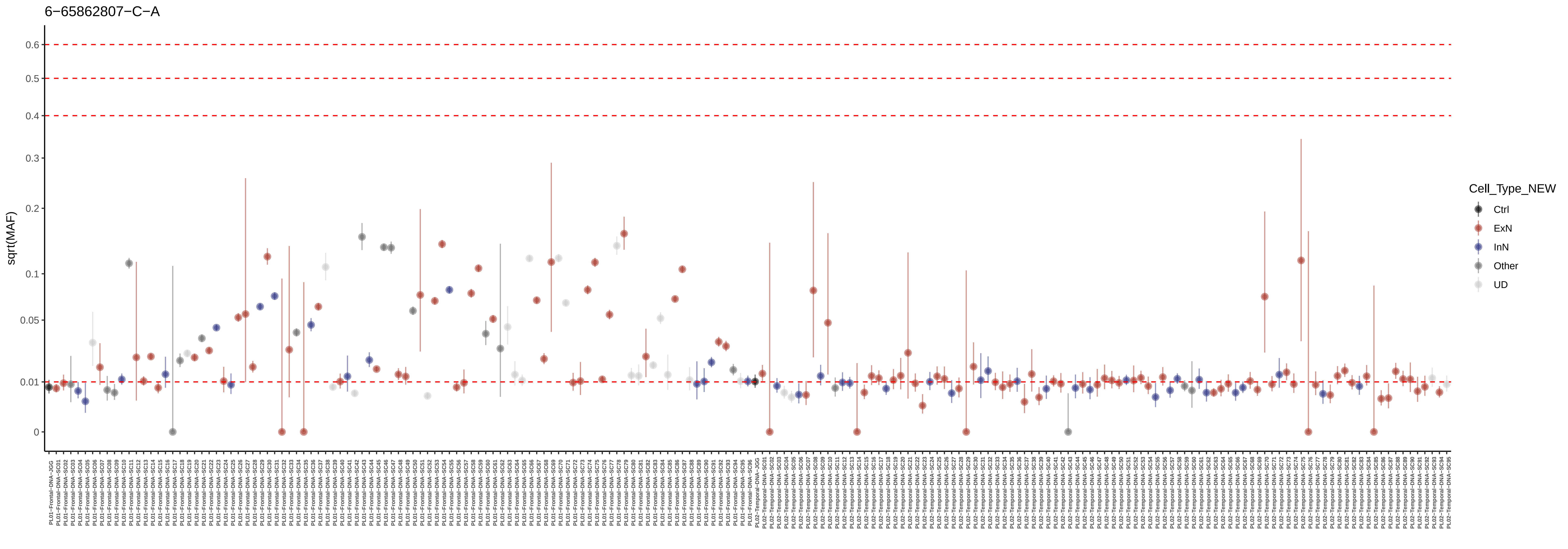
6−65862807−C−A
sqrt(MAF)
0.6
0.5
0.4
0.3
0.2
0.1
0.05
0.01
0
Cell_Type_NEW
Ctrl
ExN
InN
Other
UD

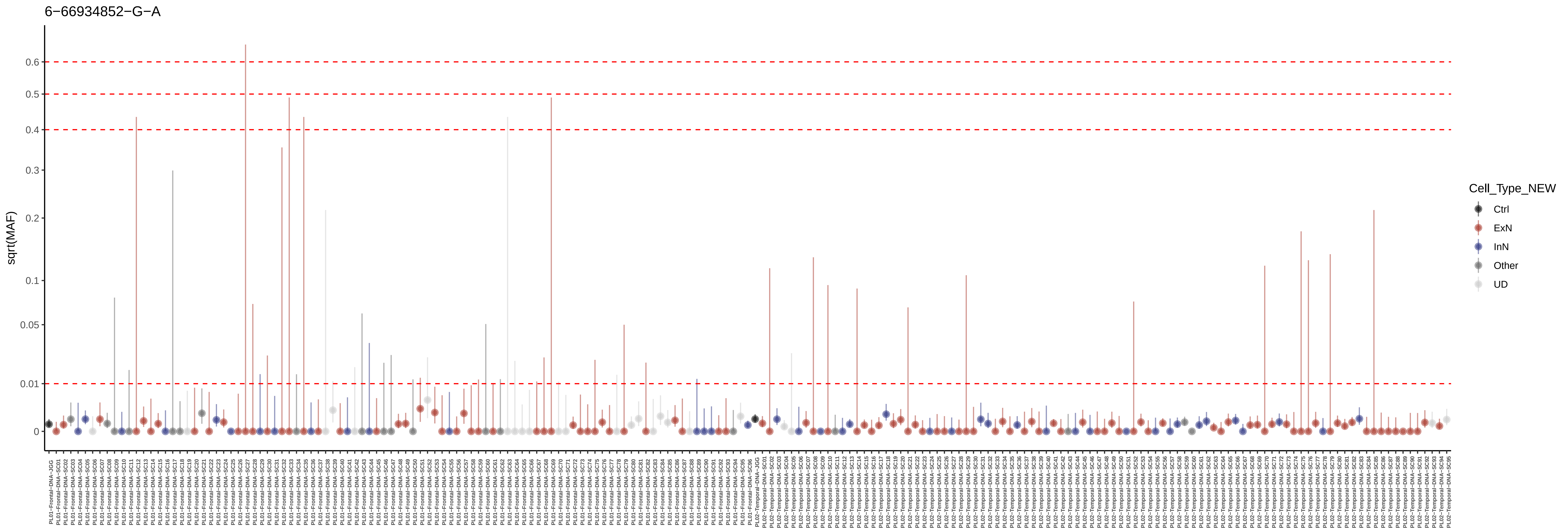
6−66934852−G−A
sqrt(MAF)
0.6
0.5
0.4
0.3
0.2
0.1
0.05
0.01
0
Cell_Type_NEW
Ctrl
ExN
InN
Other
UD

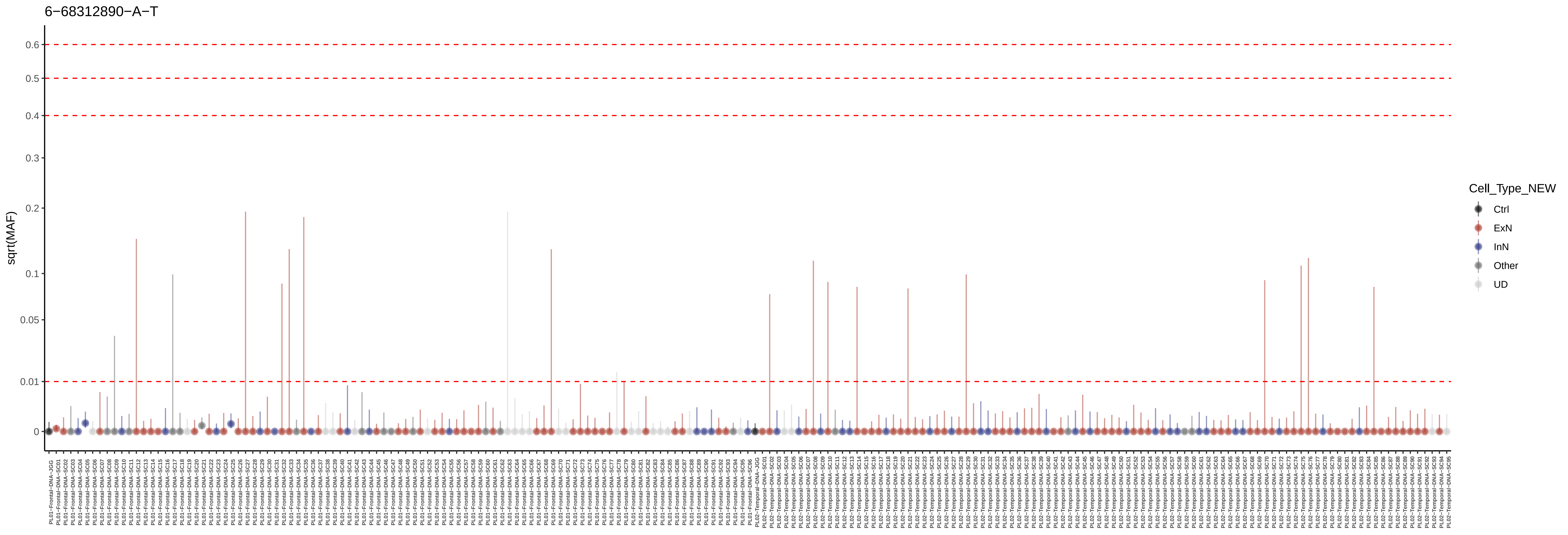
6−68312890−A−T
sqrt(MAF)
0.6
0.5
0.4
0.3
0.2
0.1
0.05
0.01
0
Cell_Type_NEW
Ctrl
ExN
InN
Other
UD

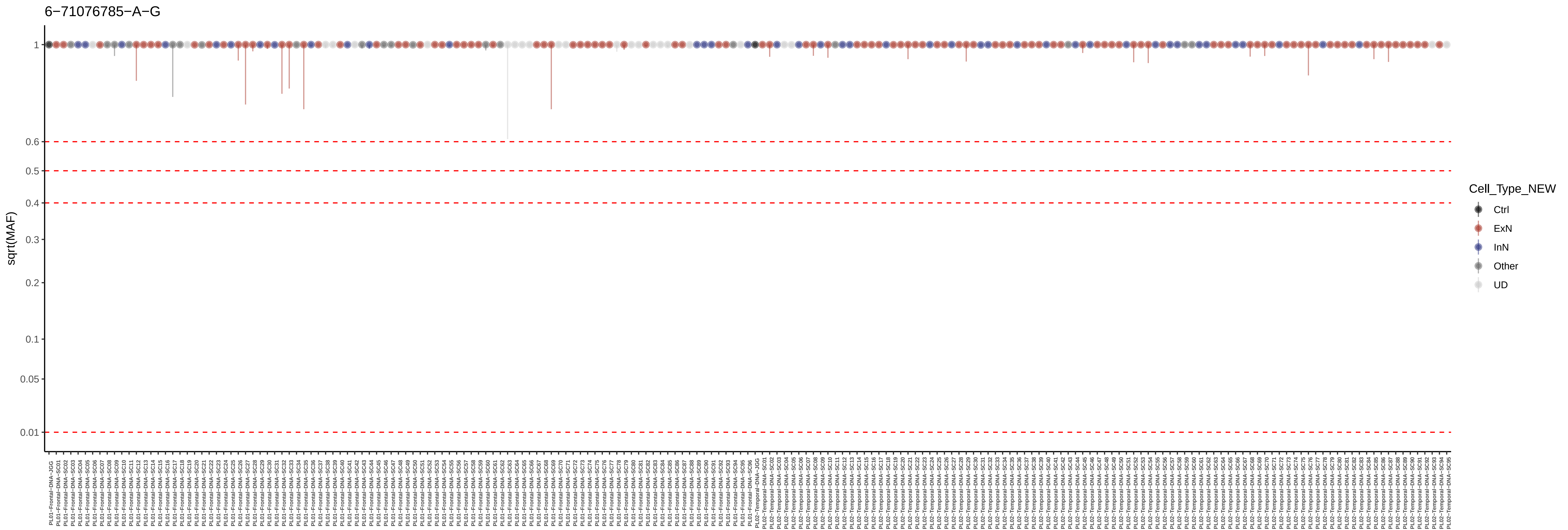
6−71076785−A−G
sqrt(MAF)
1
0.6
0.5
0.4
0.3
0.2
0.1
0.05
0.01
Cell_Type_NEW
Ctrl
ExN
InN
Other
UD

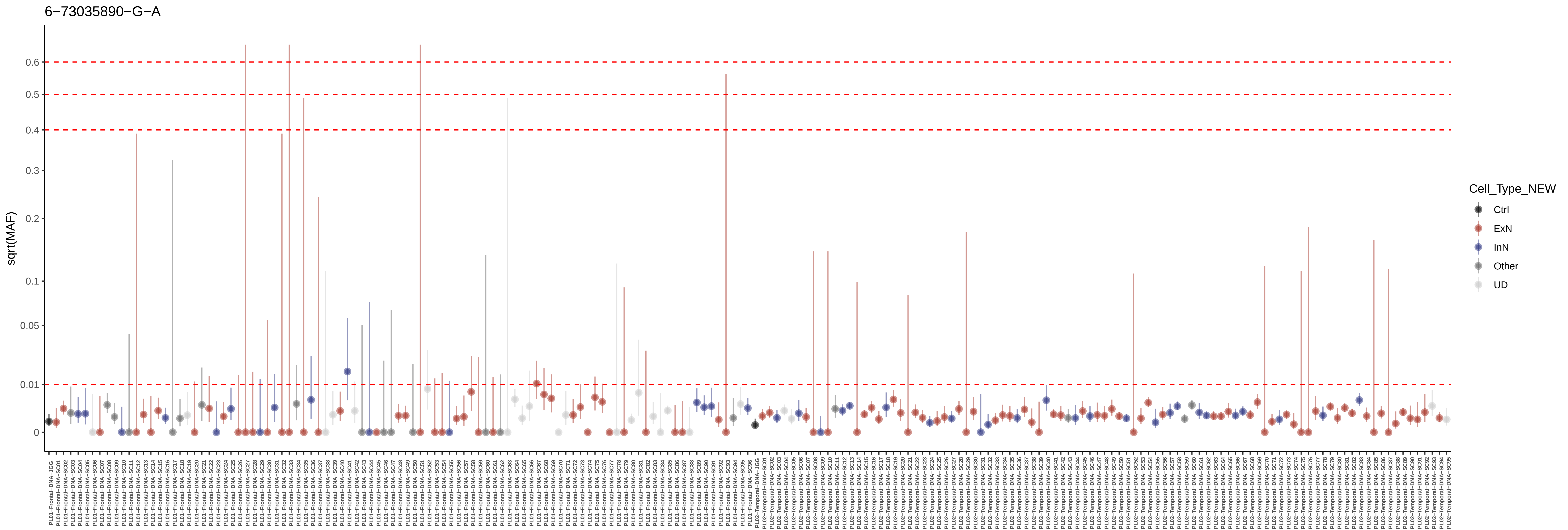
6−73035890−G−A
sqrt(MAF)
0.6
0.5
0.4
0.3
0.2
0.1
0.05
0.01
0
Cell_Type_NEW
Ctrl
ExN
InN
Other
UD

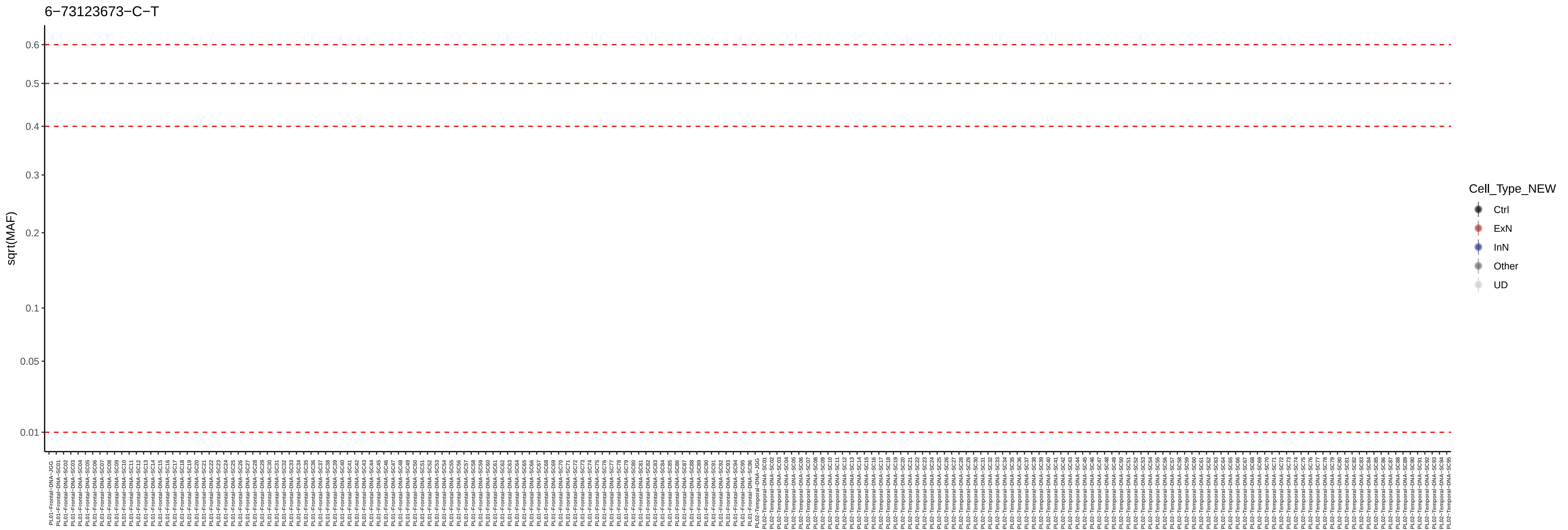

6−73123673−C−T
sqrt(MAF)
0.6
0.5
0.4
0.3
0.2
0.1
0.05
0.01
Cell_Type_NEW
Ctrl
ExN
InN
Other
UD

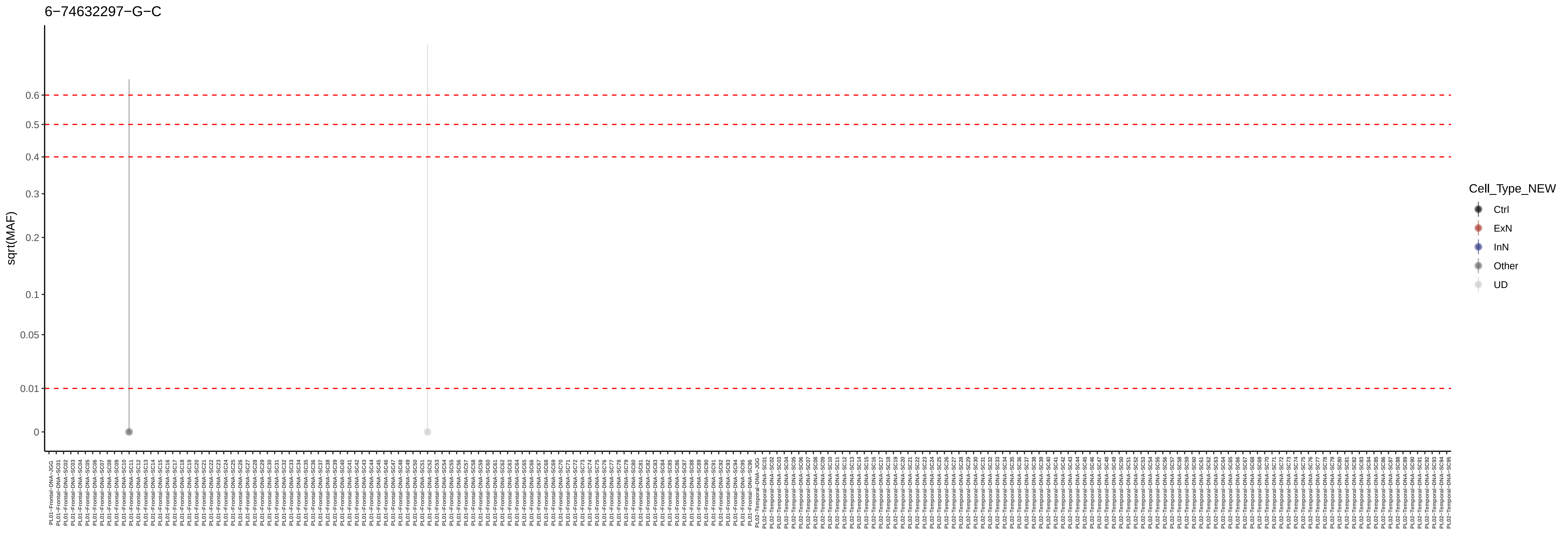
6−74632297−G−C
sqrt(MAF)
0.6
0.5
0.4
0.3
0.2
0.1
0.05
0.01
0
Cell_Type_NEW
Ctrl
ExN
InN
Other
UD

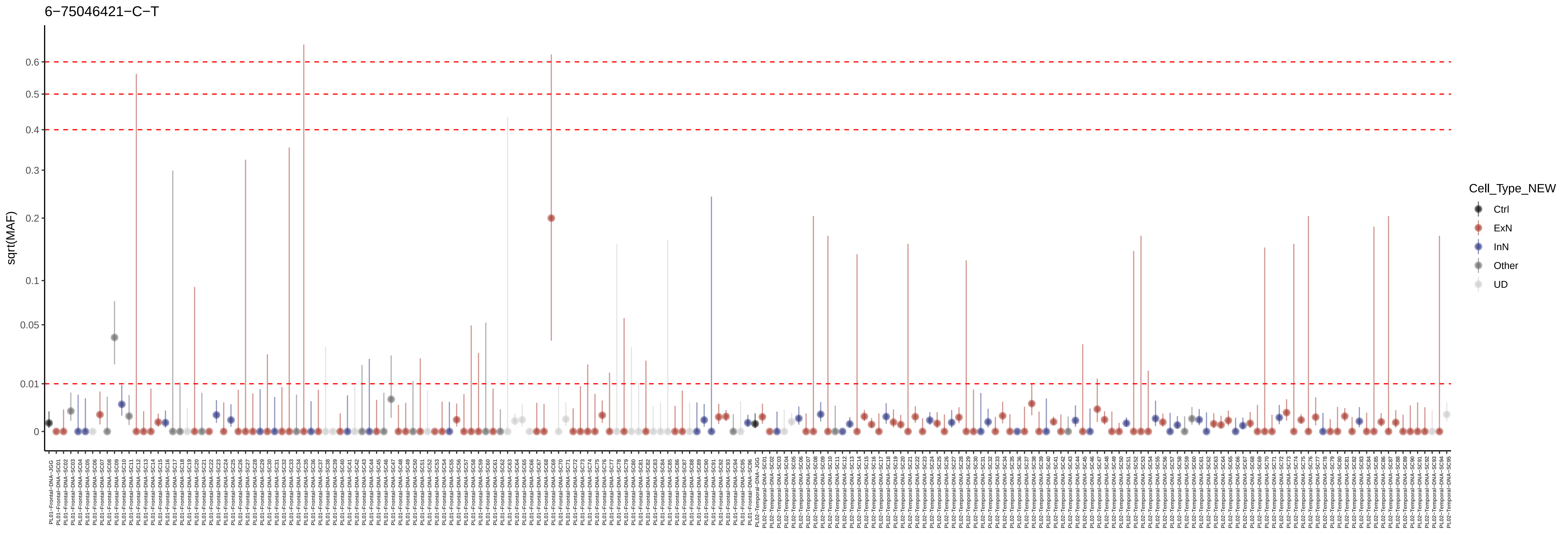

6−75046421−C−T
sqrt(MAF)
0.6
0.5
0.4
0.3
0.2
0.1
0.05
0.01
0
Cell_Type_NEW
Ctrl
ExN
InN
Other
UD

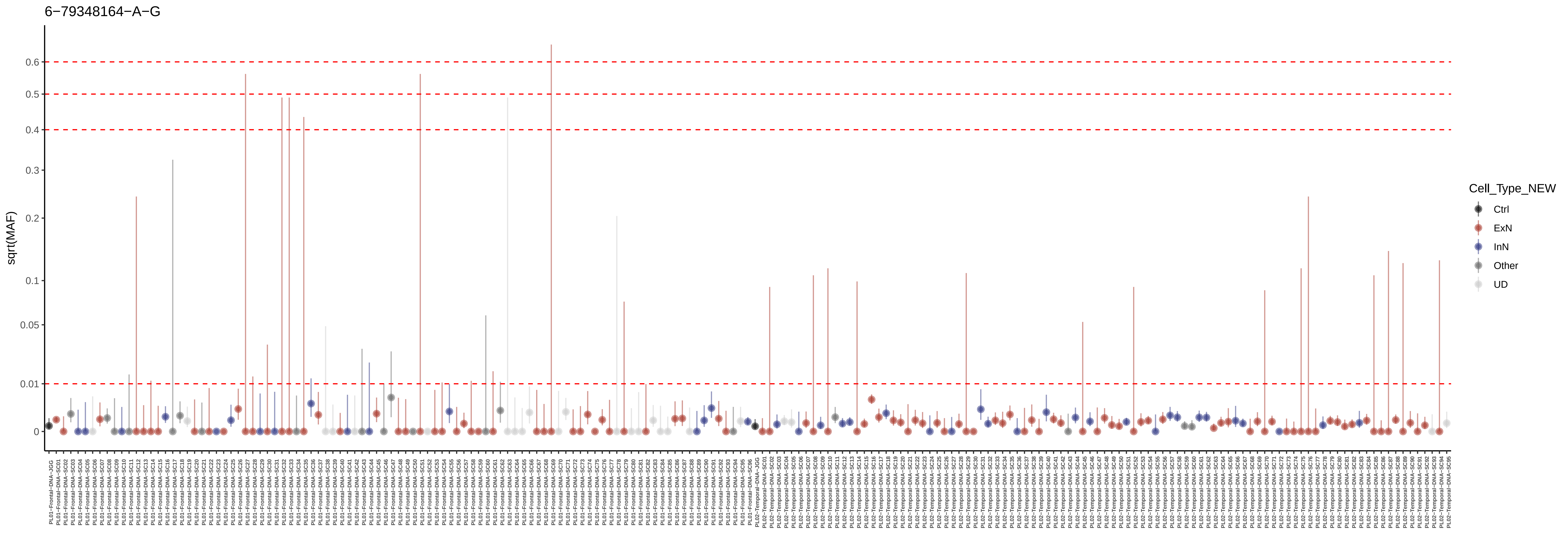
6−79348164−A−G
sqrt(MAF)
0.6
0.5
0.4
0.3
0.2
0.1
0.05
0.01
0
Cell_Type_NEW
Ctrl
ExN
InN
Other
UD

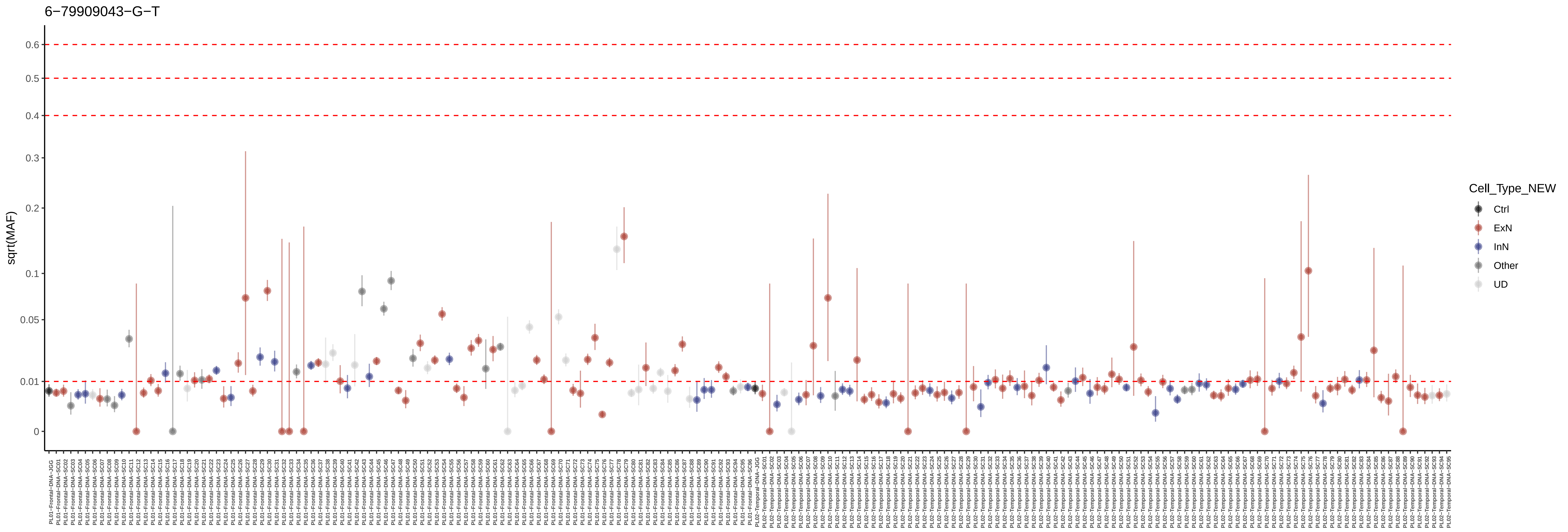
6−79909043−G−T
sqrt(MAF)
0.6
0.5
0.4
0.3
0.2
0.1
0.05
0.01
0
Cell_Type_NEW
Ctrl
ExN
InN
Other
UD

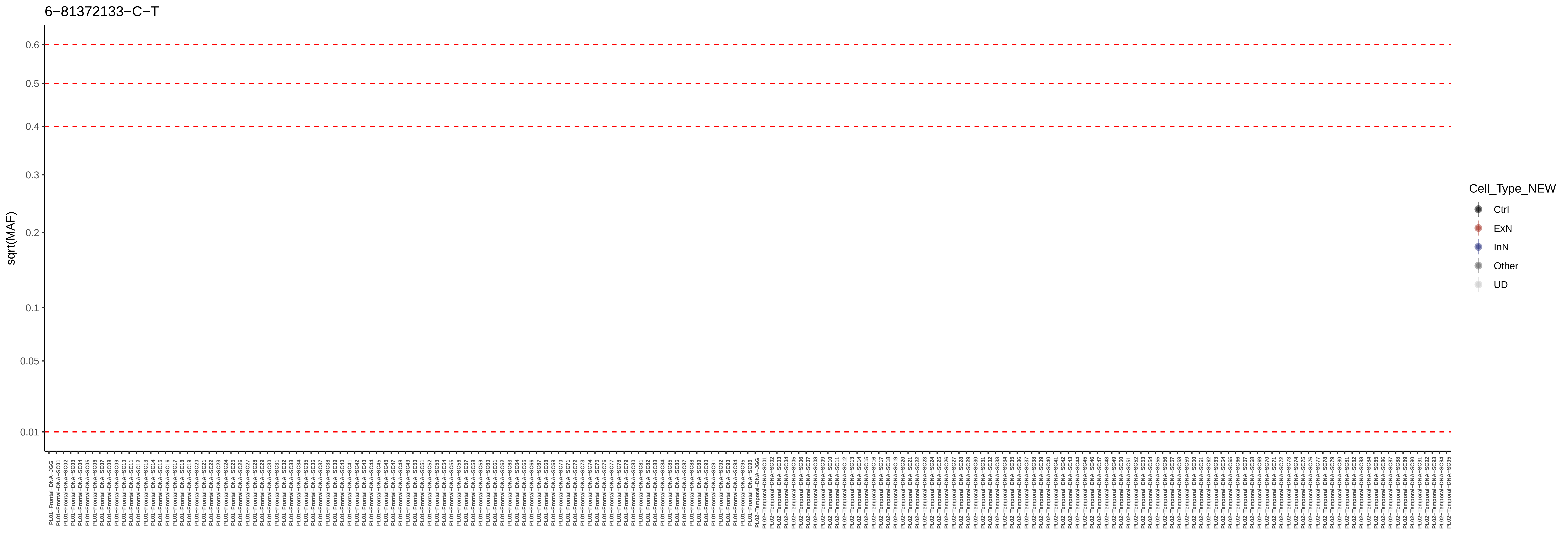

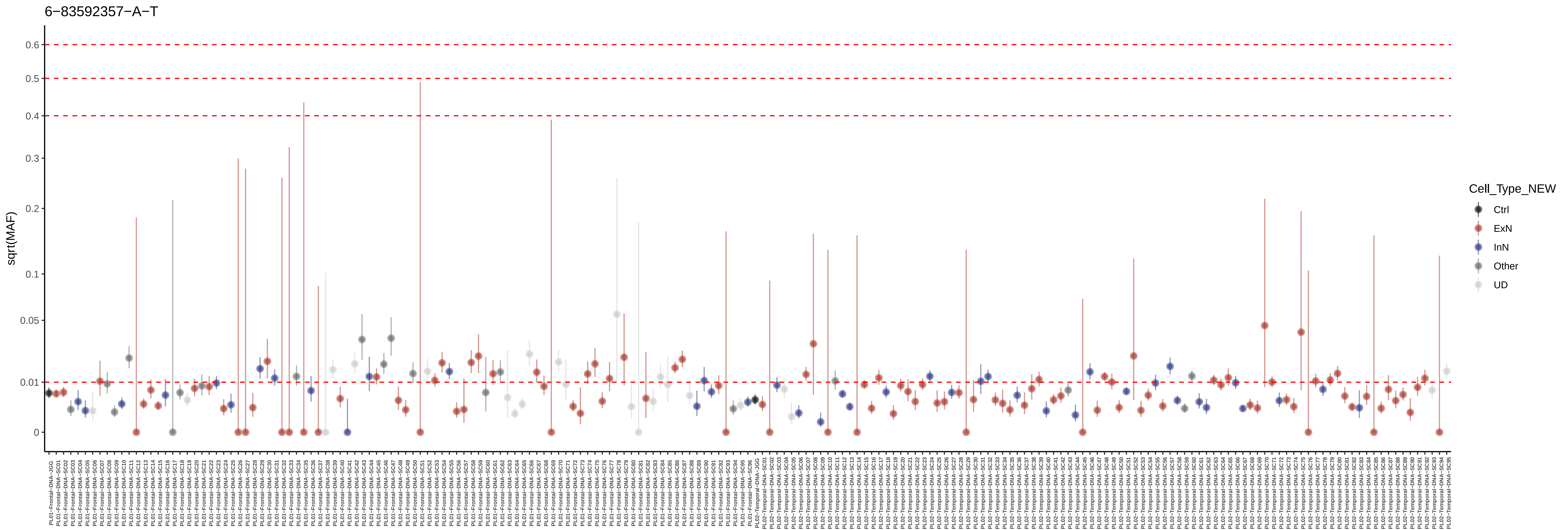

6−83592357−A−T
sqrt(MAF)
0.6
0.5
0.4
0.3
0.2
0.1
0.05
0.01
0
Cell_Type_NEW
Ctrl
ExN
InN
Other
UD

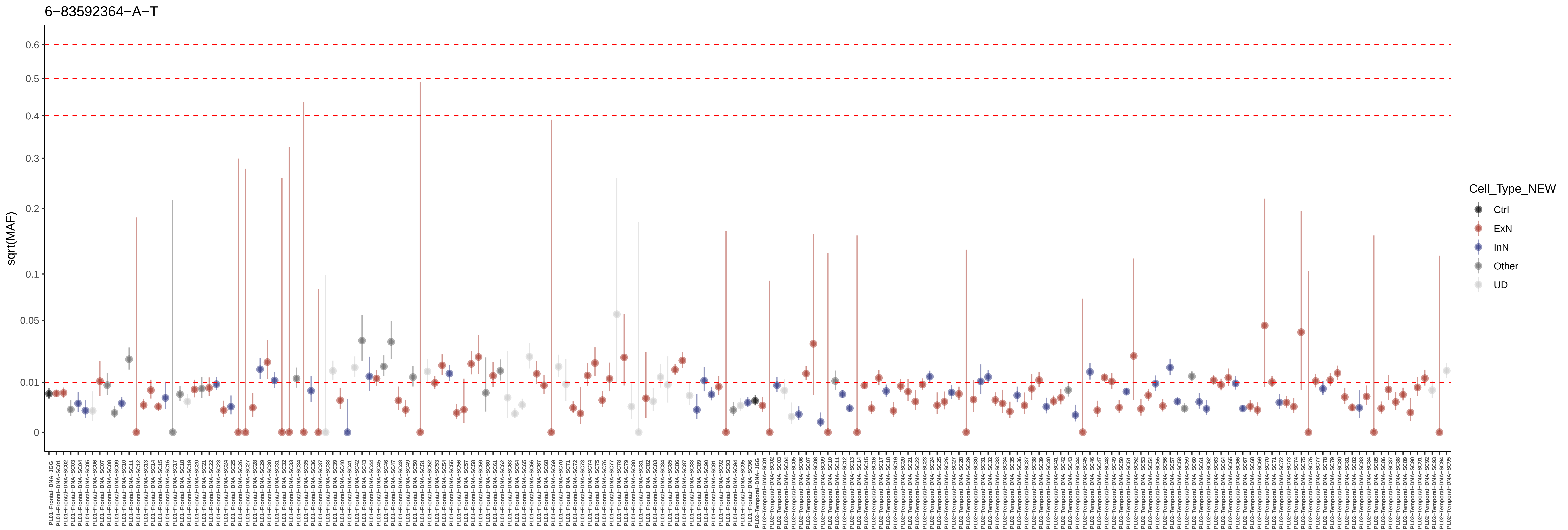

6−83592364−A−T
sqrt(MAF)
0.6
0.5
0.4
0.3
0.2
0.1
0.05
0.01
0
Cell_Type_NEW
Ctrl
ExN
InN
Other
UD

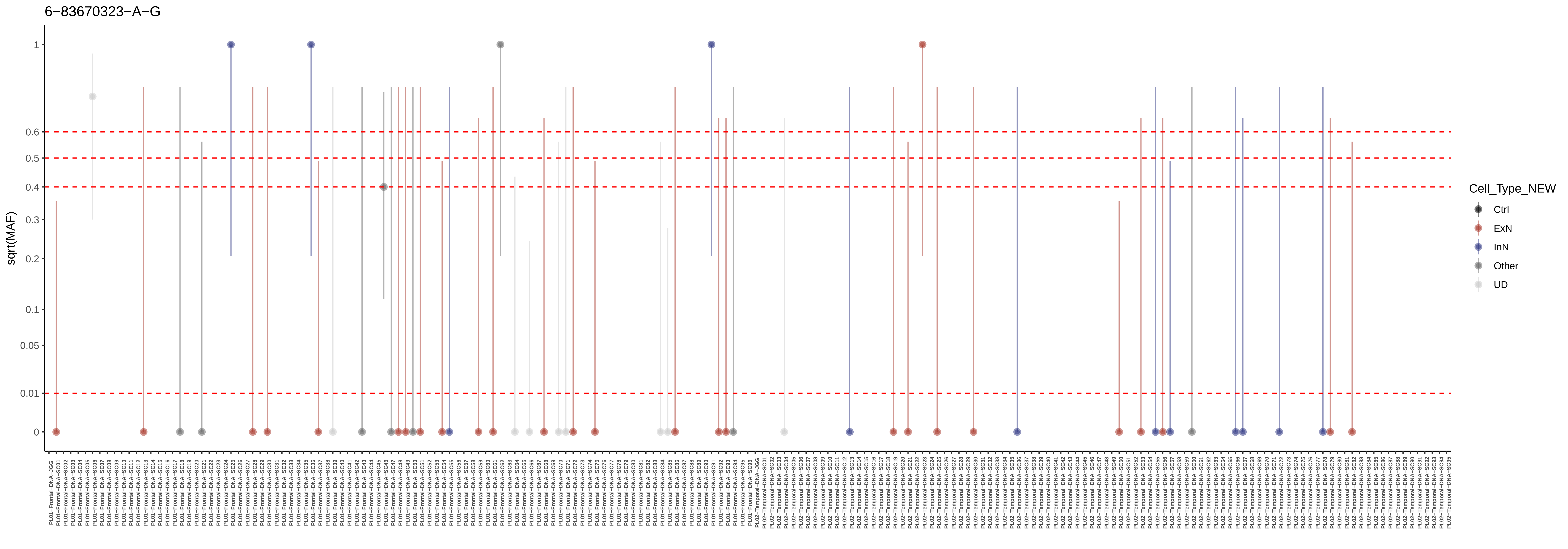
6−83670323−A−G
sqrt(MAF)
1
0.6
0.5
0.4
0.3
0.2
0.1
0.05
0.01
0
Cell_Type_NEW
Ctrl
ExN
InN
Other
UD

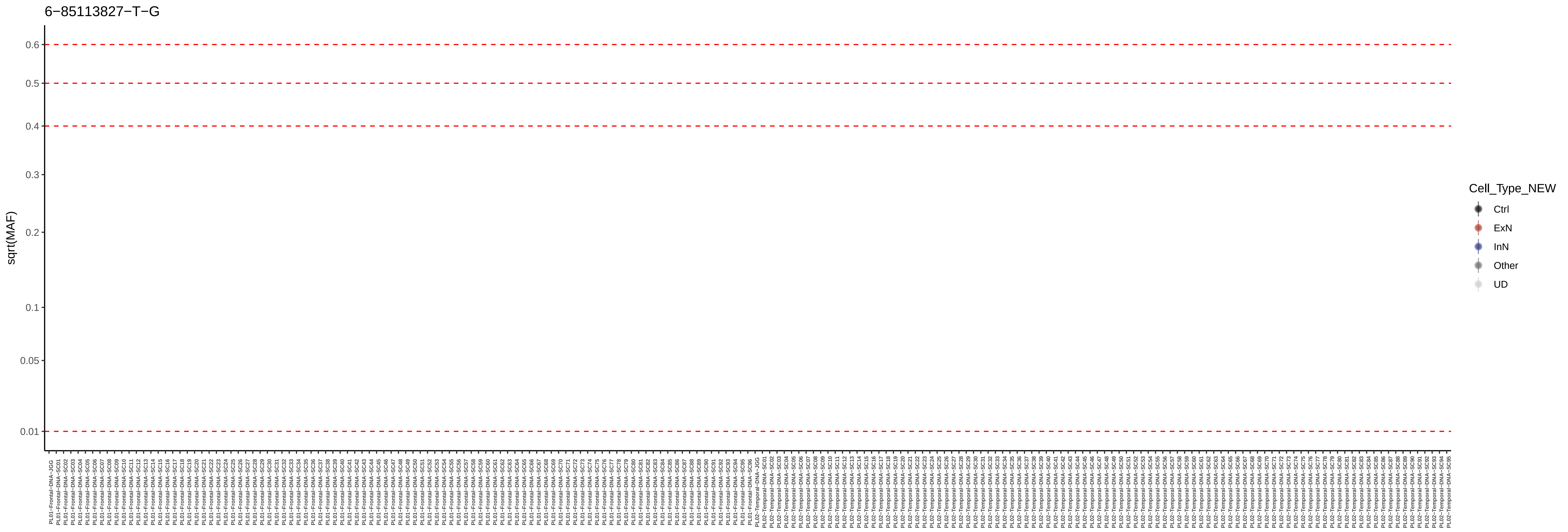

6−85113827−T−G
sqrt(MAF)
0.6
0.5
0.4
0.3
0.2
0.1
0.05
0.01
Cell_Type_NEW
Ctrl
ExN
InN
Other
UD

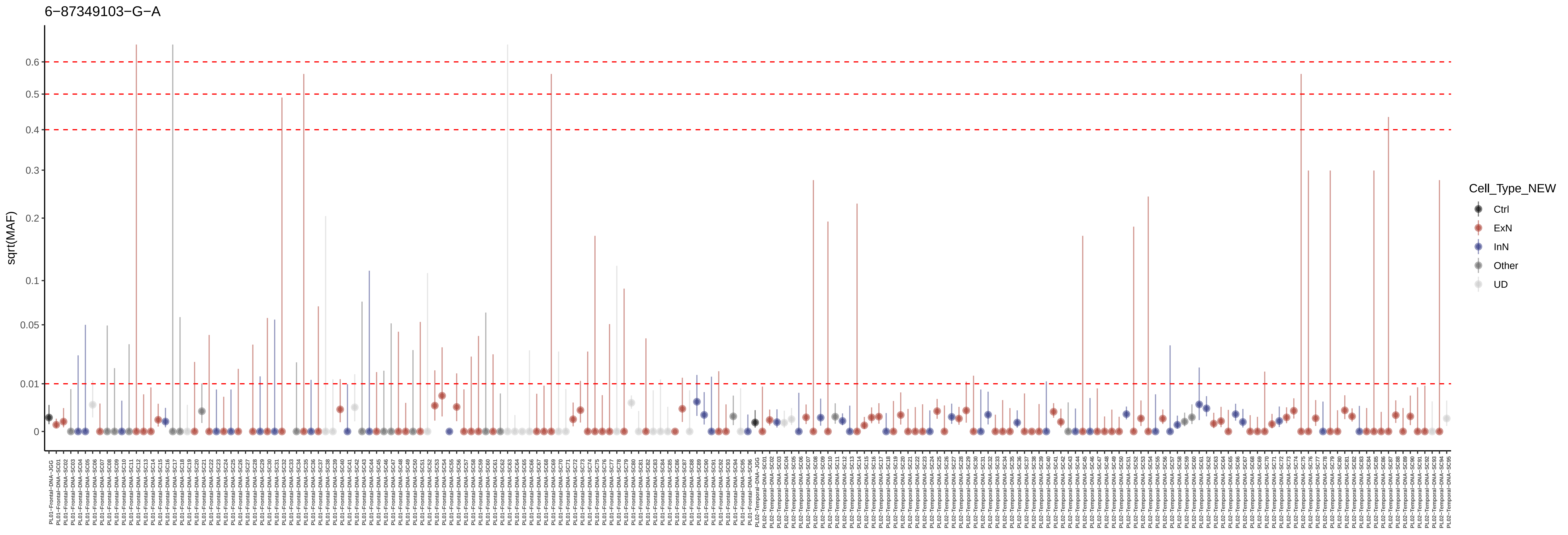
6−87349103−G−A
sqrt(MAF)
0.6
0.5
0.4
0.3
0.2
0.1
0.05
0.01
0
Cell_Type_NEW
Ctrl
ExN
InN
Other
UD

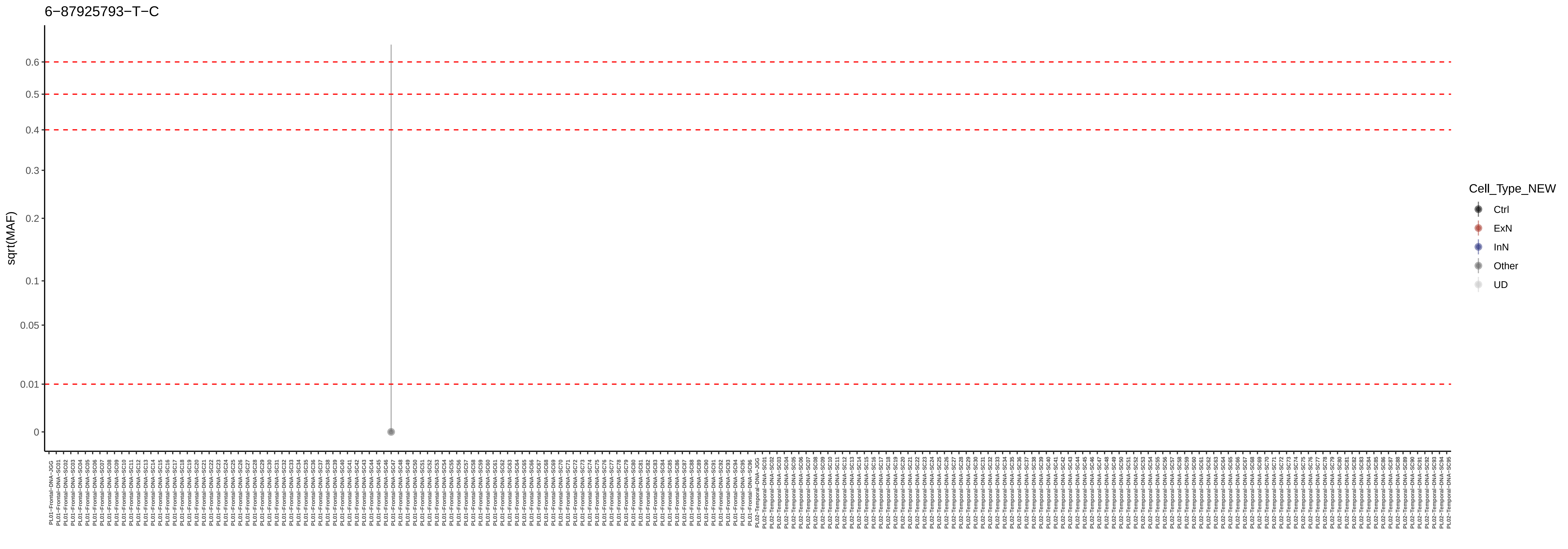
6−87925793−T−C
sqrt(MAF)
0.6
0.5
0.4
0.3
0.2
0.1
0.05
0.01
0
Cell_Type_NEW
Ctrl
ExN
InN
Other
UD

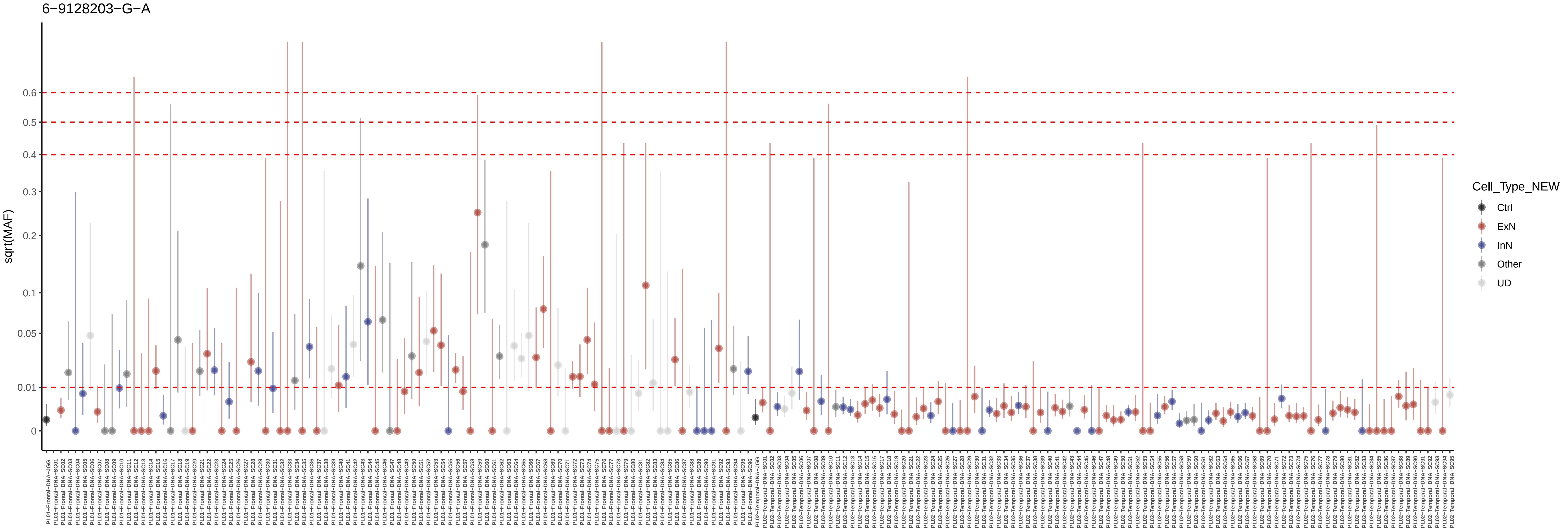
6−9128203−G−A
sqrt(MAF)
0.6
0.5
0.4
0.3
0.2
0.1
0.05
0.01
0
Cell_Type_NEW
Ctrl
ExN
InN
Other
UD

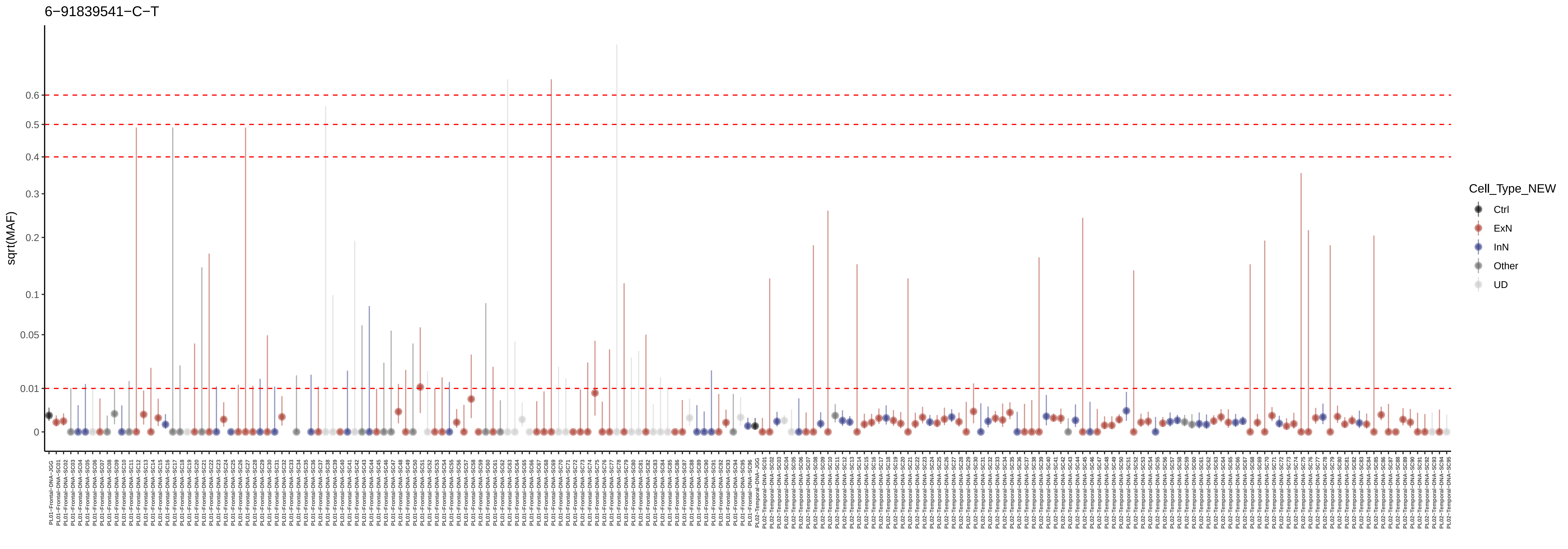
6−91839541−C−T
sqrt(MAF)
0.6
0.5
0.4
0.3
0.2
0.1
0.05
0.01
0
Cell_Type_NEW
Ctrl
ExN
InN
Other
UD

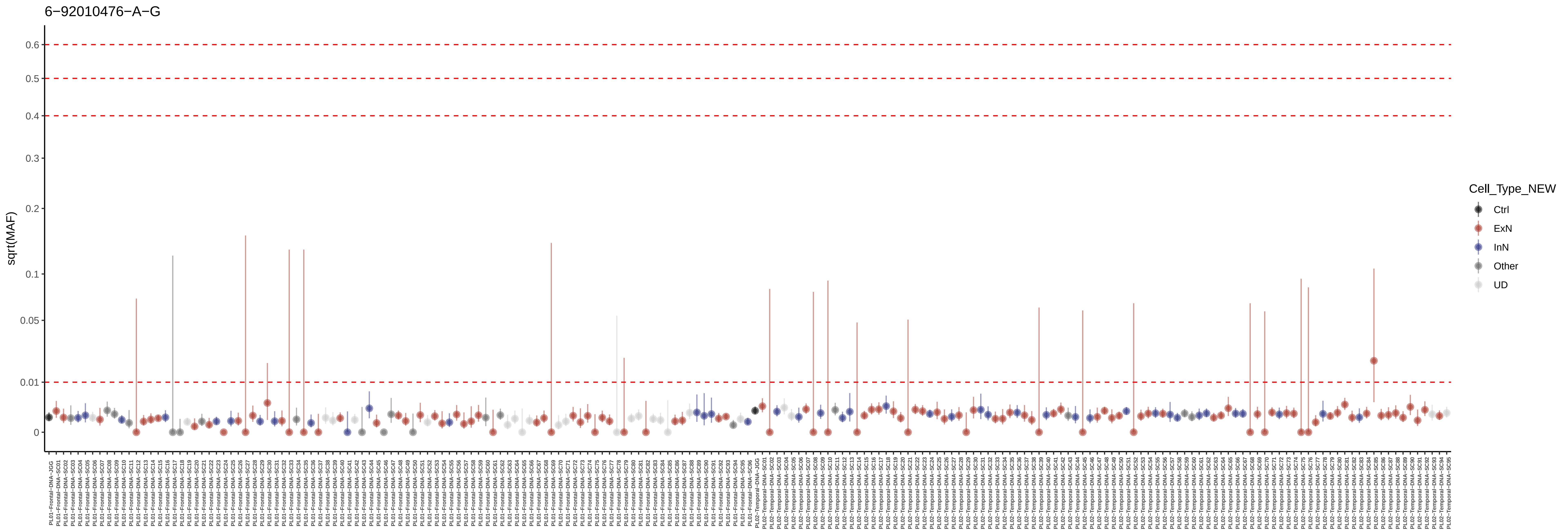
6−92010476−A−G
sqrt(MAF)
0.6
0.5
0.4
0.3
0.2
0.1
0.05
0.01
0
Cell_Type_NEW
Ctrl
ExN
InN
Other
UD

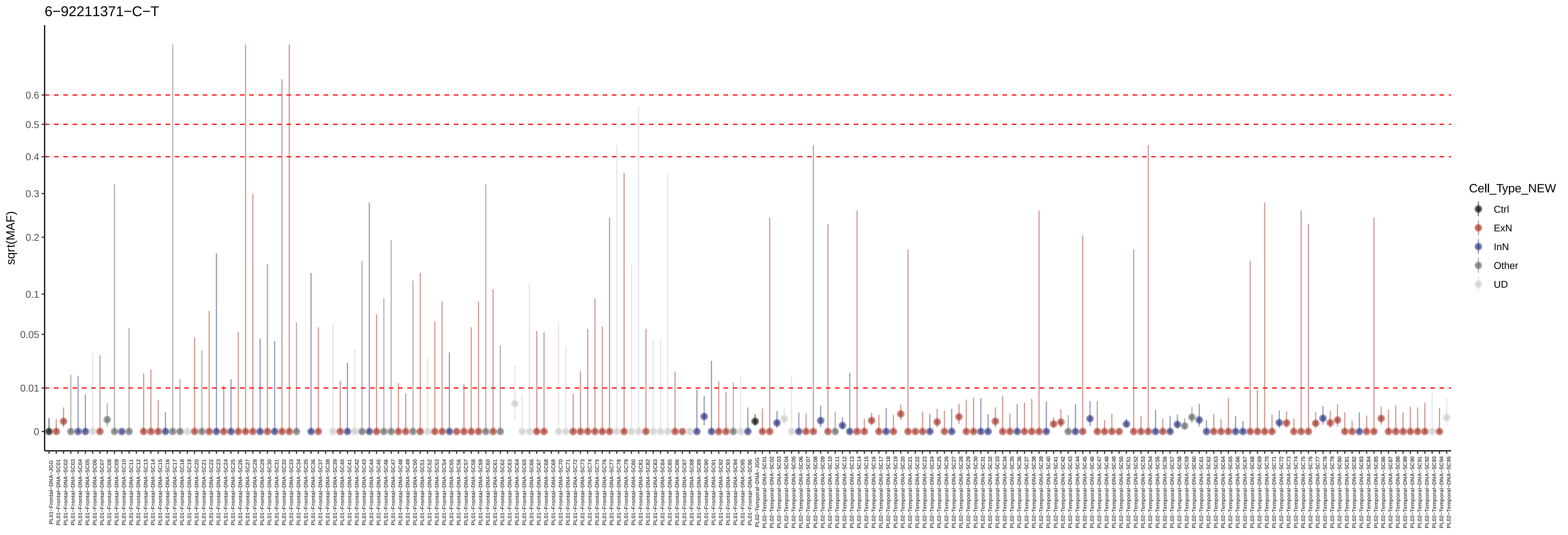
6−92211371−C−T
sqrt(MAF)
0.6
0.5
0.4
0.3
0.2
0.1
0.05
0.01
0
Cell_Type_NEW
Ctrl
ExN
InN
Other
UD

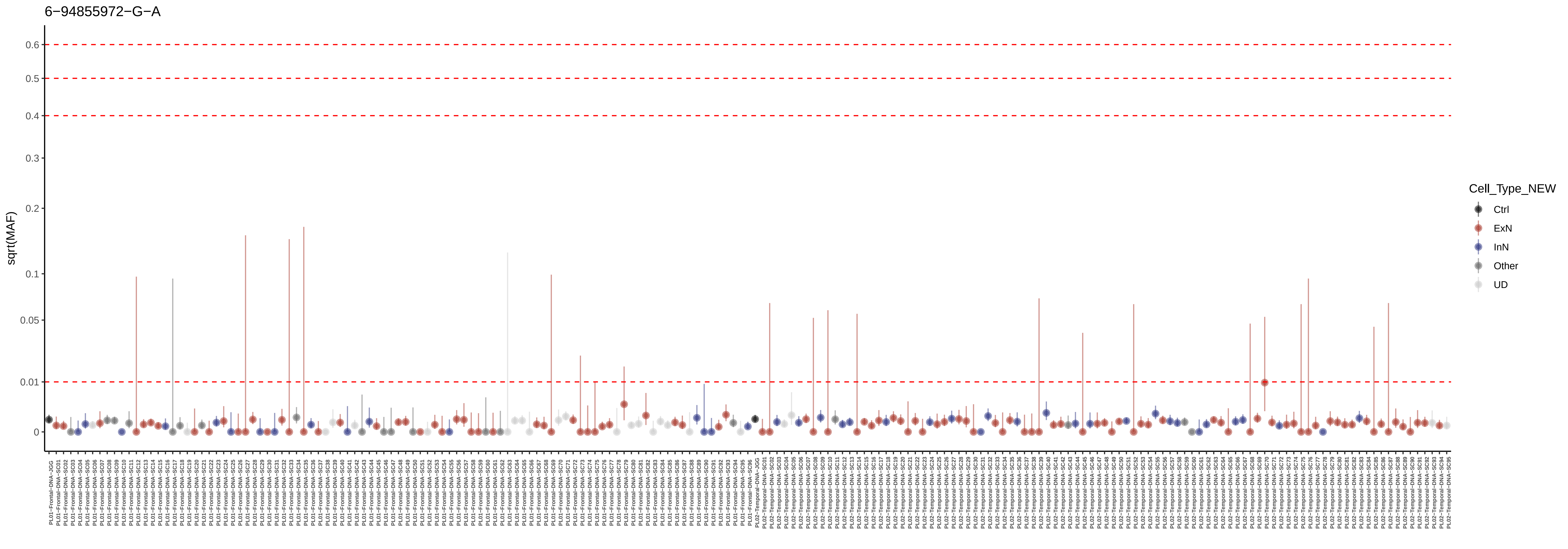

6−94855972−G−A
sqrt(MAF)
Cell_Type_NEW
Ctrl
ExN
InN
Other
UD

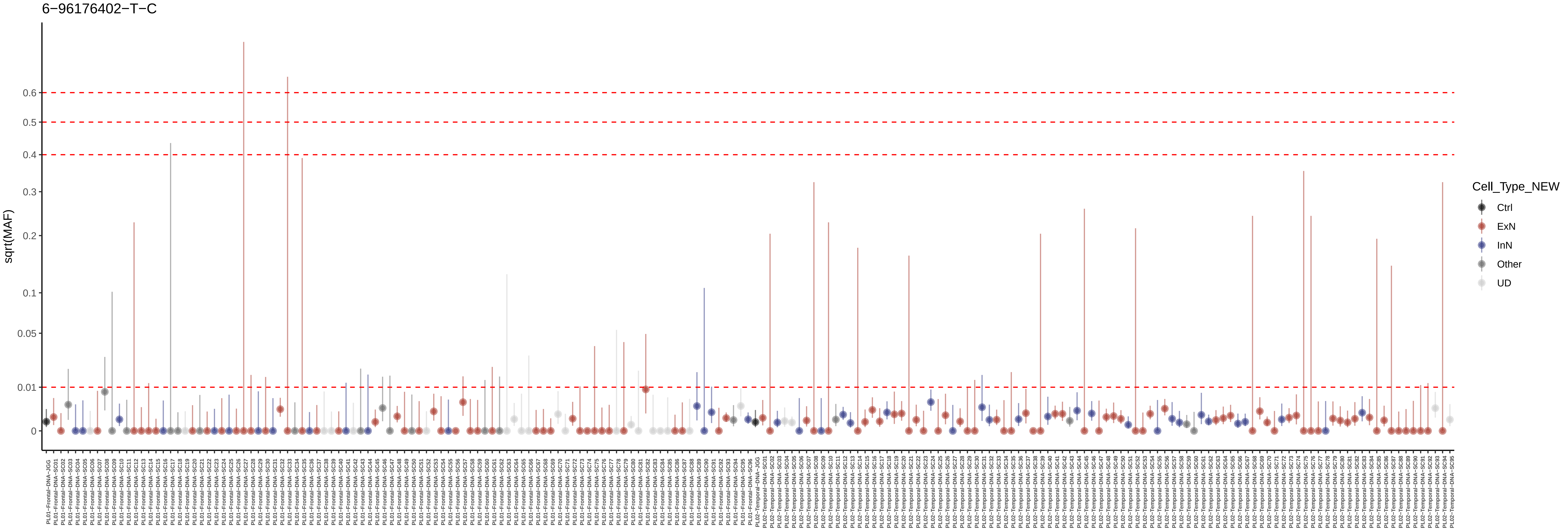
6−96176402−T−C
sqrt(MAF)
0.6
0.5
0.4
0.3
0.2
0.1
0.05
0.01
0
Cell_Type_NEW
Ctrl
ExN
InN
Other
UD

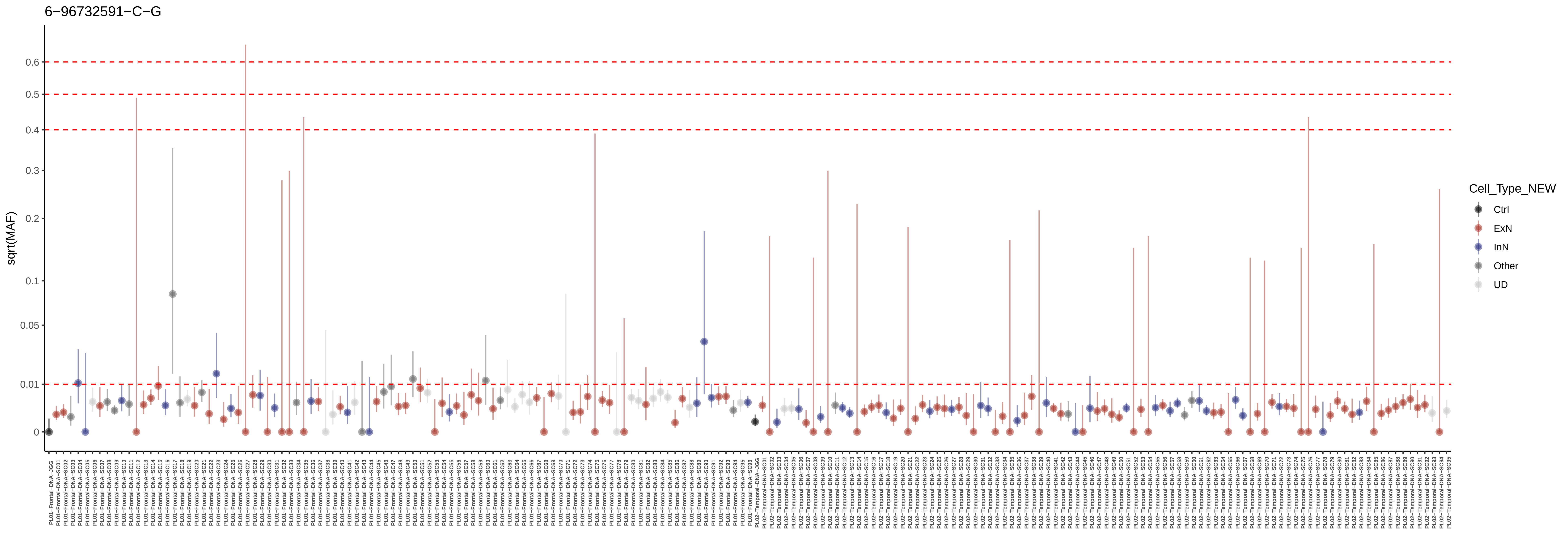
6−96732591−C−G
sqrt(MAF)
0.6
0.5
0.4
0.3
0.2
0.1
0.05
0.01
0
Cell_Type_NEW
Ctrl
ExN
InN
Other
UD

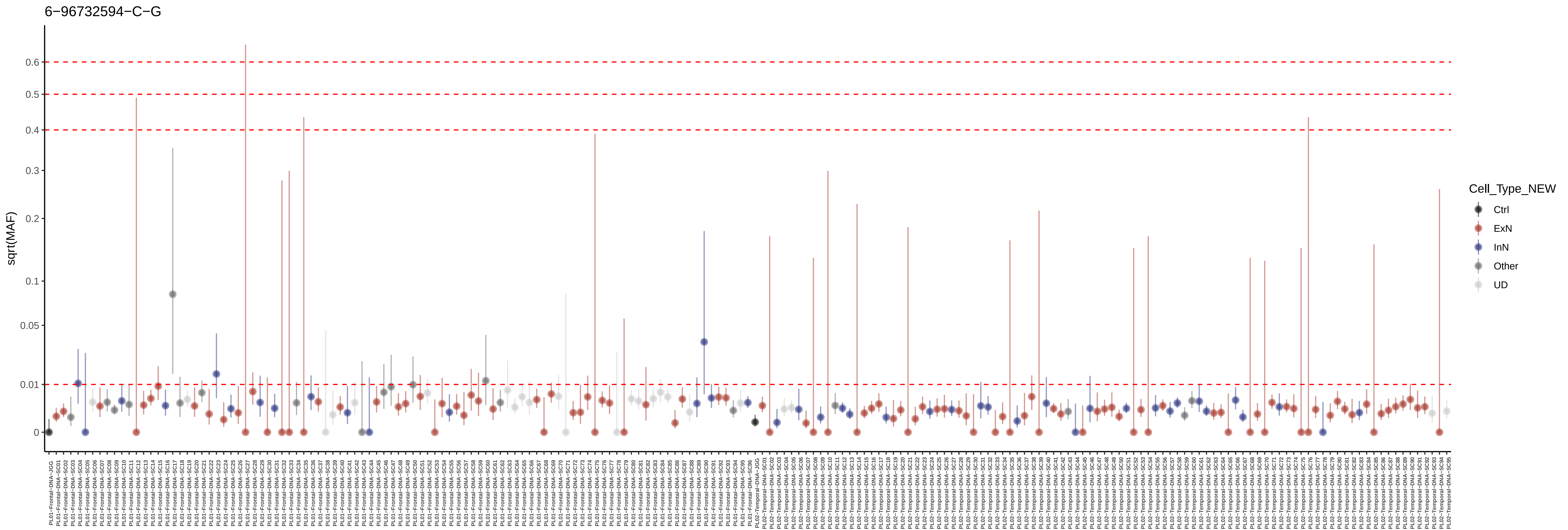
6−96732594−C−G
sqrt(MAF)
0.6
0.5
0.4
0.3
0.2
0.1
0.05
0.01
0
Cell_Type_NEW
Ctrl
ExN
InN
Other
UD

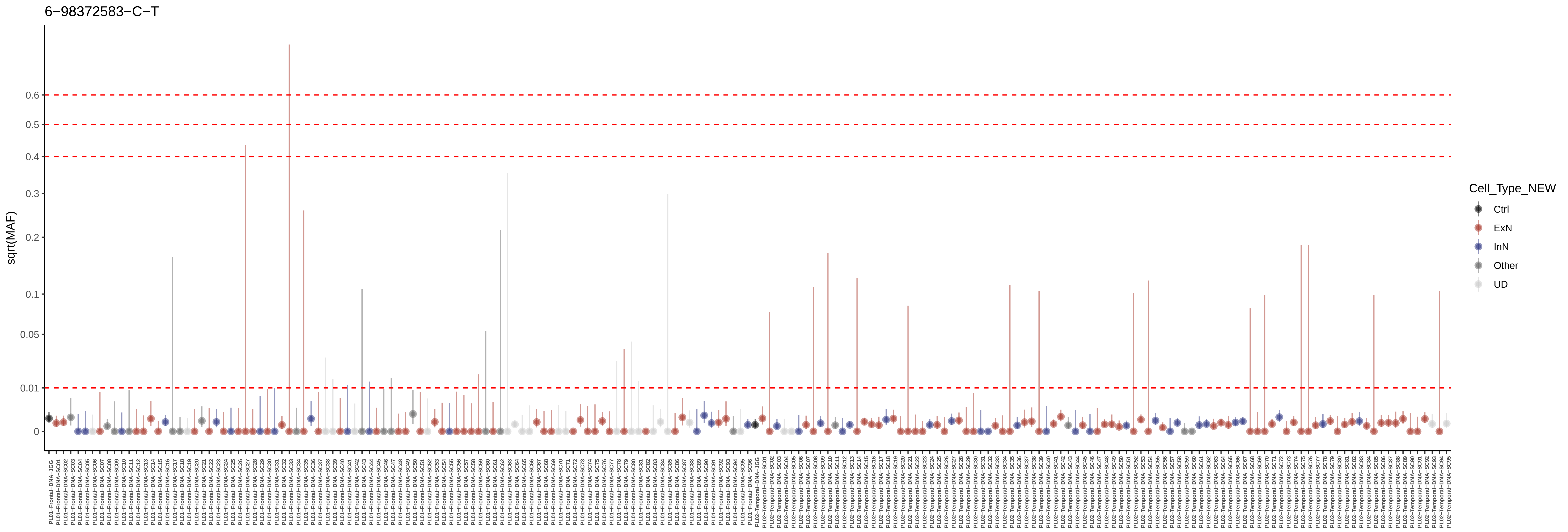
6−98372583−C−T
sqrt(MAF)
0.6
0.5
0.4
0.3
0.2
0.1
0.05
0.01
0
Cell_Type_NEW
Ctrl
ExN
InN
Other
UD

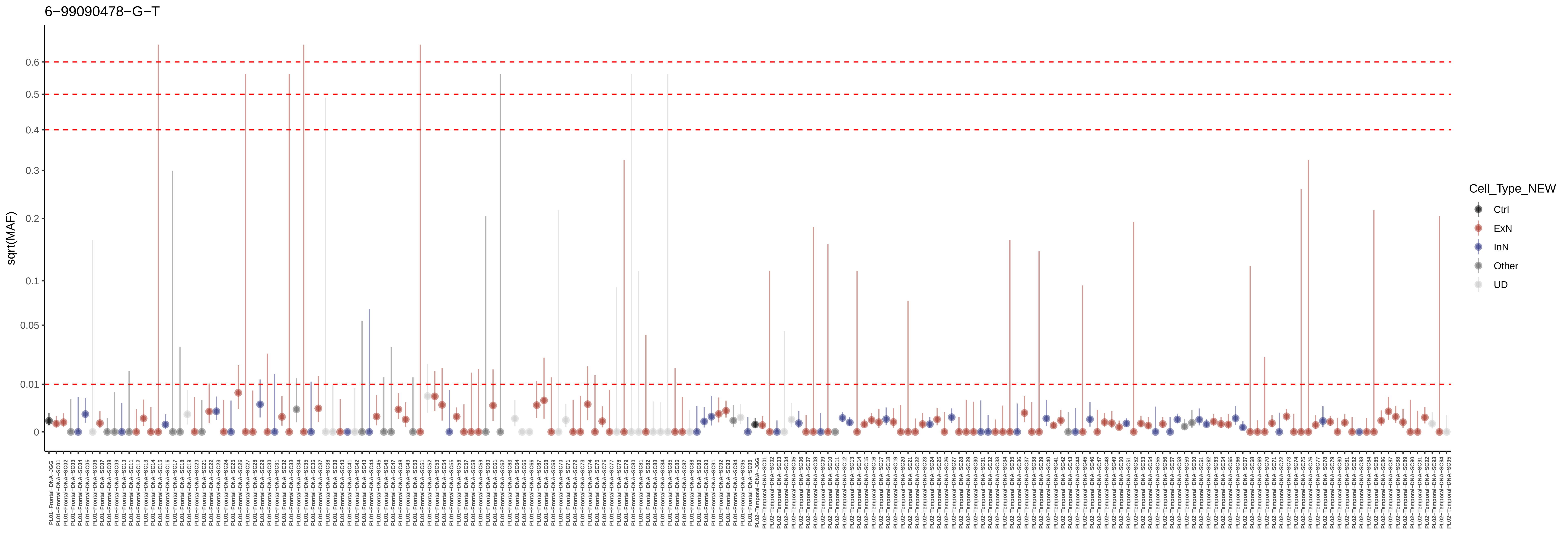
6−99090478−G−T
sqrt(MAF)
0.6
0.5
0.4
0.3
0.2
0.1
0.05
0.01
0
Cell_Type_NEW
Ctrl
ExN
InN
Other
UD

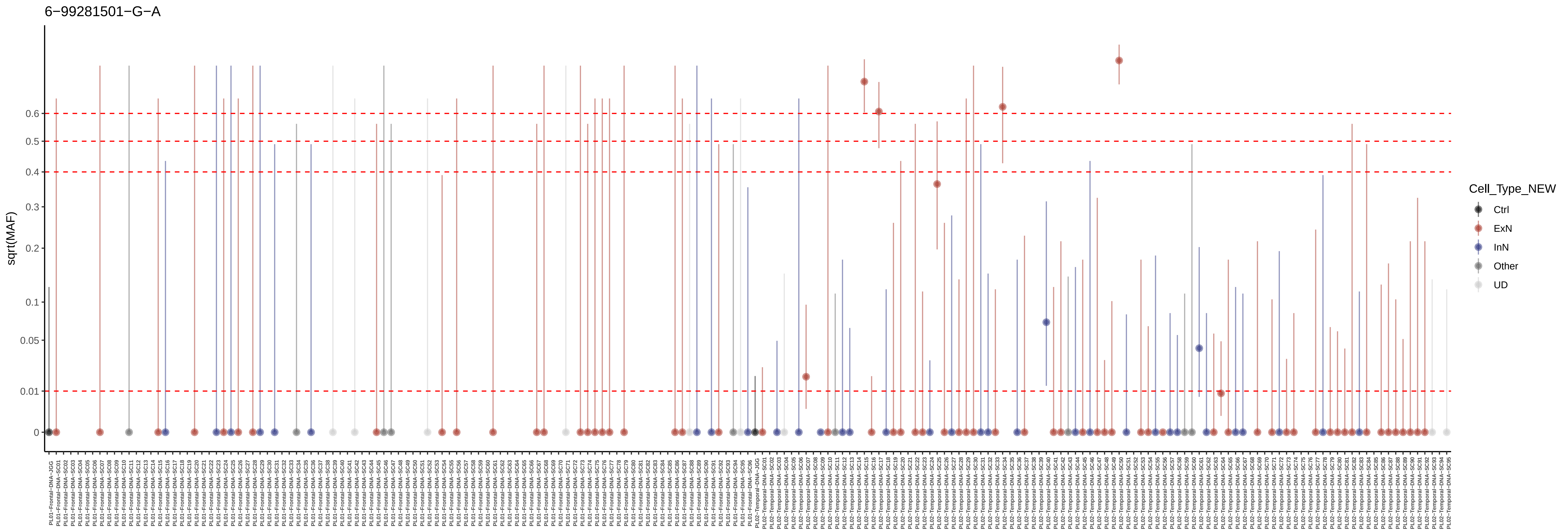
6−99281501−G−A
sqrt(MAF)
0
0.01
0.05
0.1
0.2
0.3
0.4
0.5
0.6
Cell_Type_NEW
Ctrl
ExN
InN
Other
UD

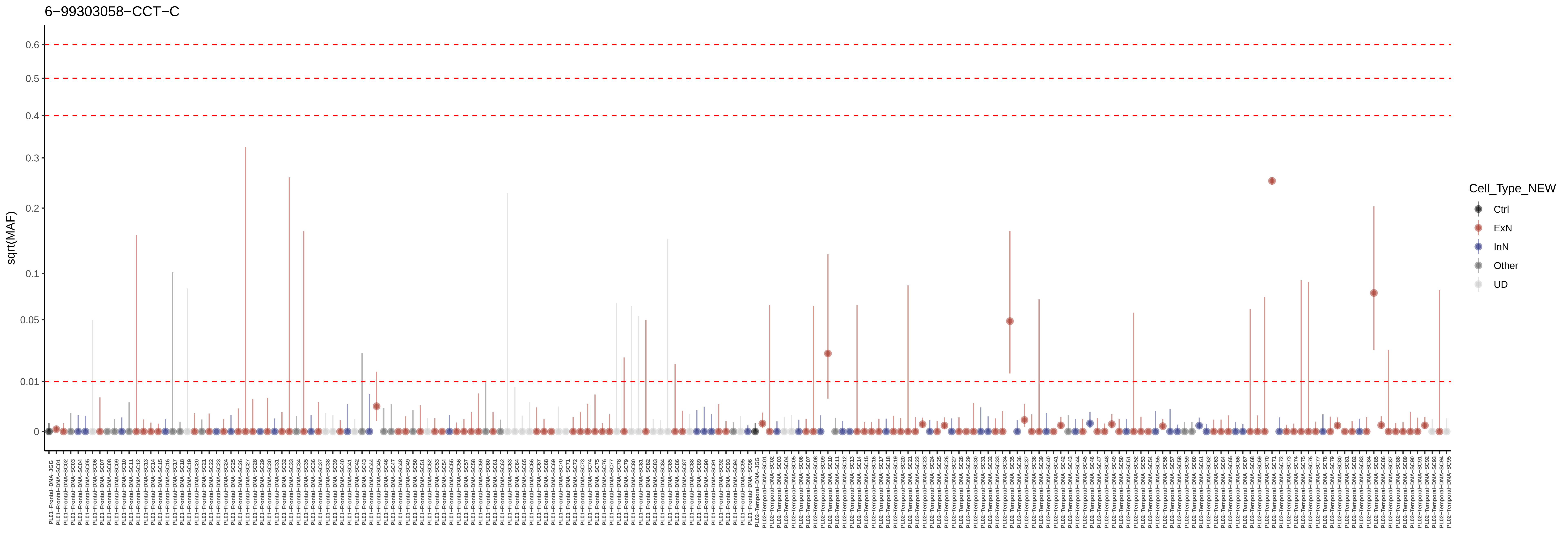
6−99303058−CCT−C
sqrt(MAF)
0.6
0.5
0.4
0.3
0.2
0.1
0.05
0.01
0
Cell_Type_NEW
Ctrl
ExN
InN
Other
UD

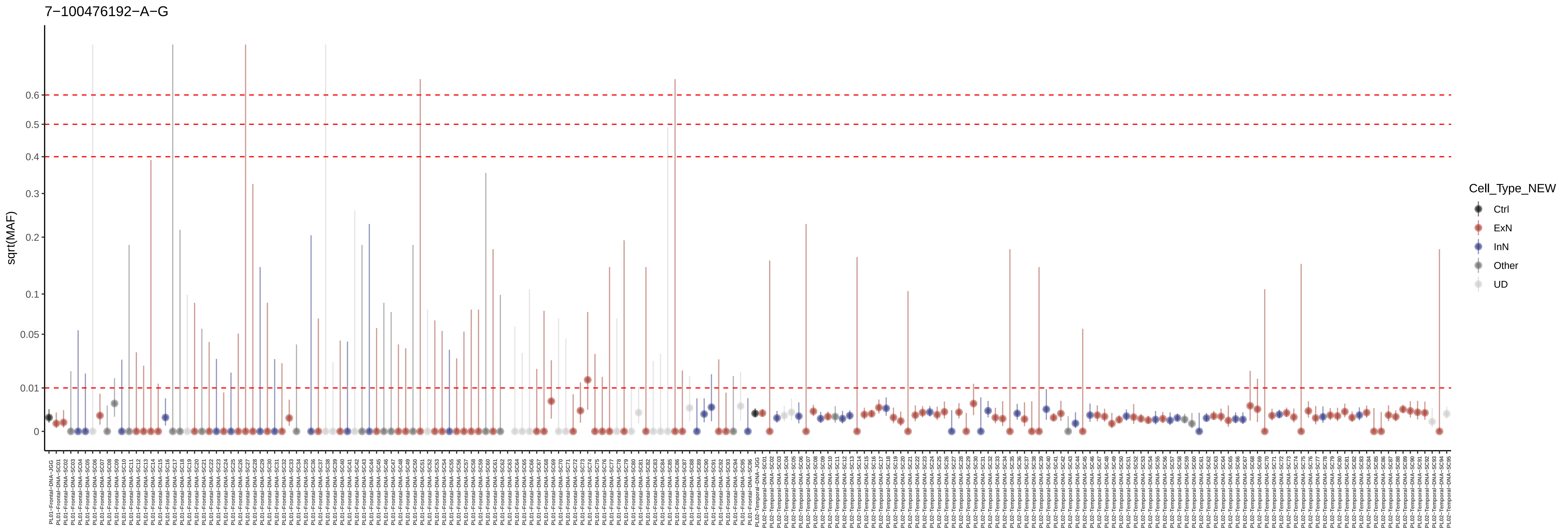
7−100476192−A−G
sqrt(MAF)
0.6
0.5
0.4
0.3
0.2
0.1
0.05
0.01
0
Cell_Type_NEW
Ctrl
ExN
InN
Other
UD

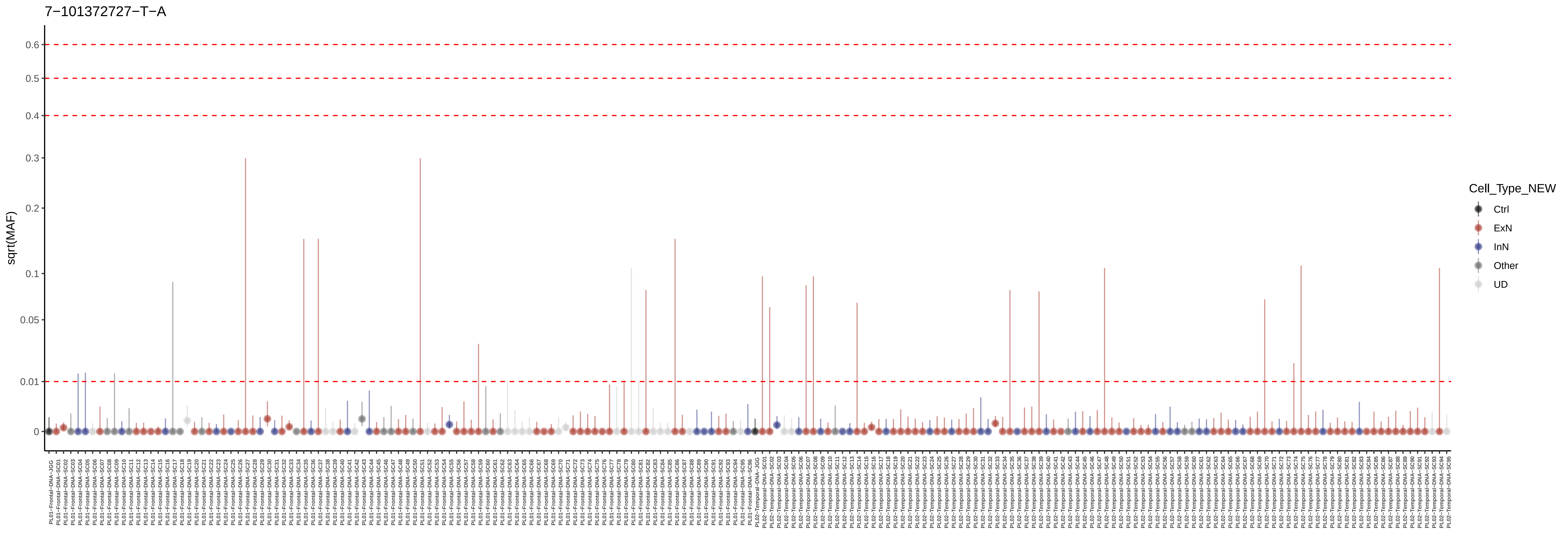
7−101372727−T−A
sqrt(MAF)
0.6
0.5
0.4
0.3
0.2
0.1
0.05
0.01
0
Cell_Type_NEW
Ctrl
ExN
InN
Other
UD

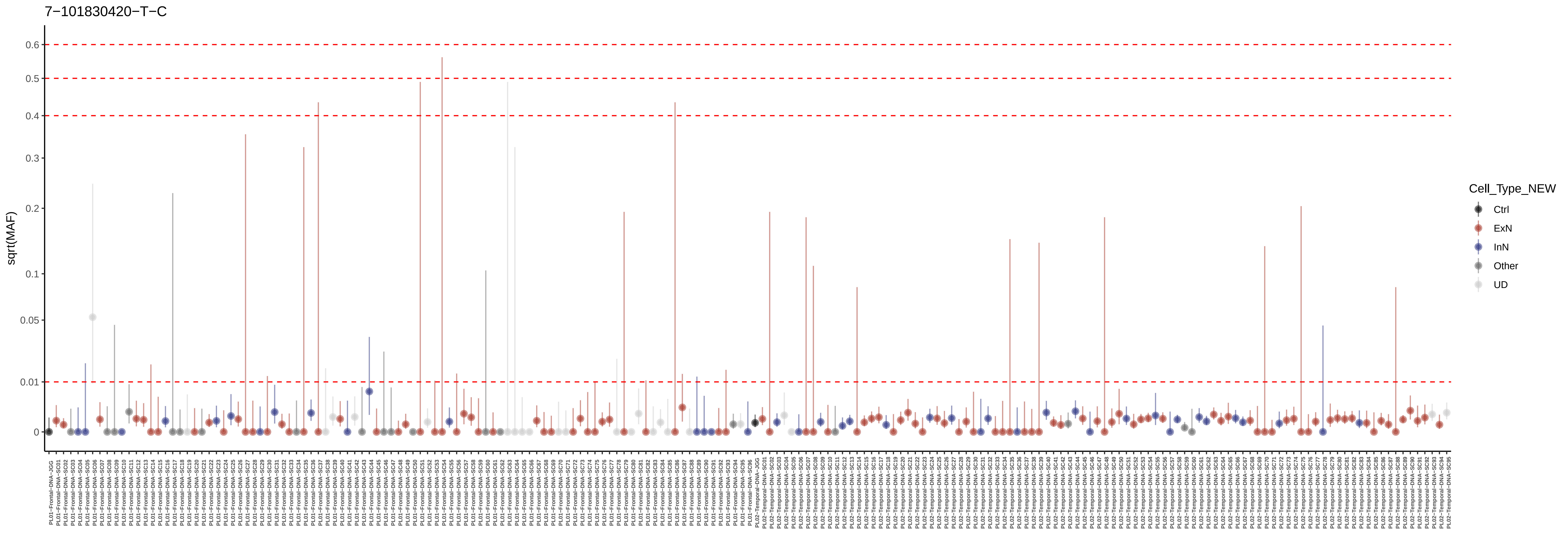

7−101830420−T−C
sqrt(MAF)
0.6
0.5
0.4
0.3
0.2
0.1
0.05
0.01
0
Cell_Type_NEW
Ctrl
ExN
InN
Other
UD

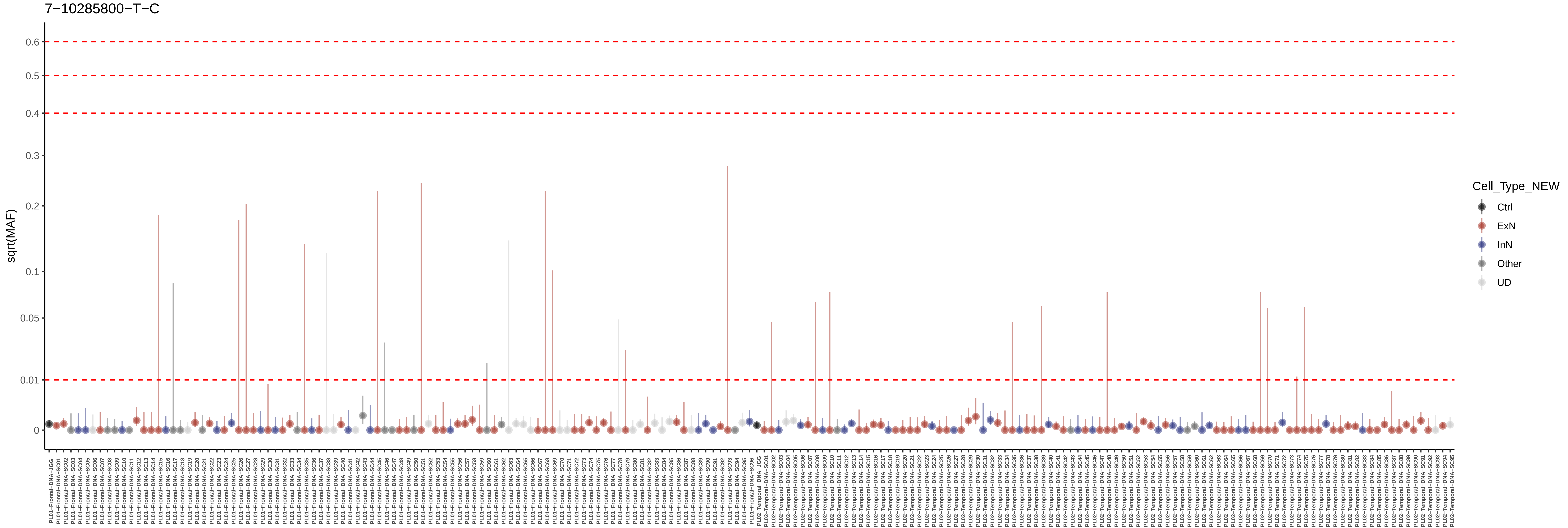
7−10285800−T−C
sqrt(MAF)
0.6
0.5
0.4
0.3
0.2
0.1
0.05
0.01
0
Cell_Type_NEW
Ctrl
ExN
InN
Other
UD

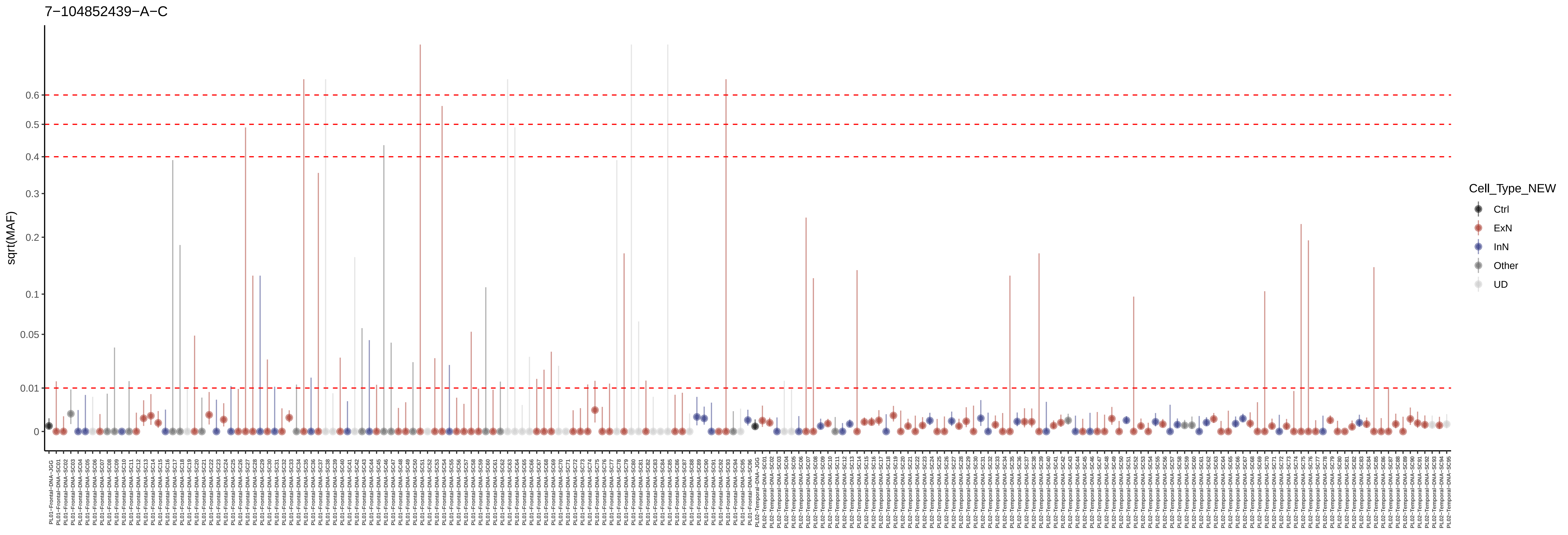
7−104852439−A−C
sqrt(MAF)
0.6
0.5
0.4
0.3
0.2
0.1
0.05
0.01
0
Cell_Type_NEW
Ctrl
ExN
InN
Other
UD

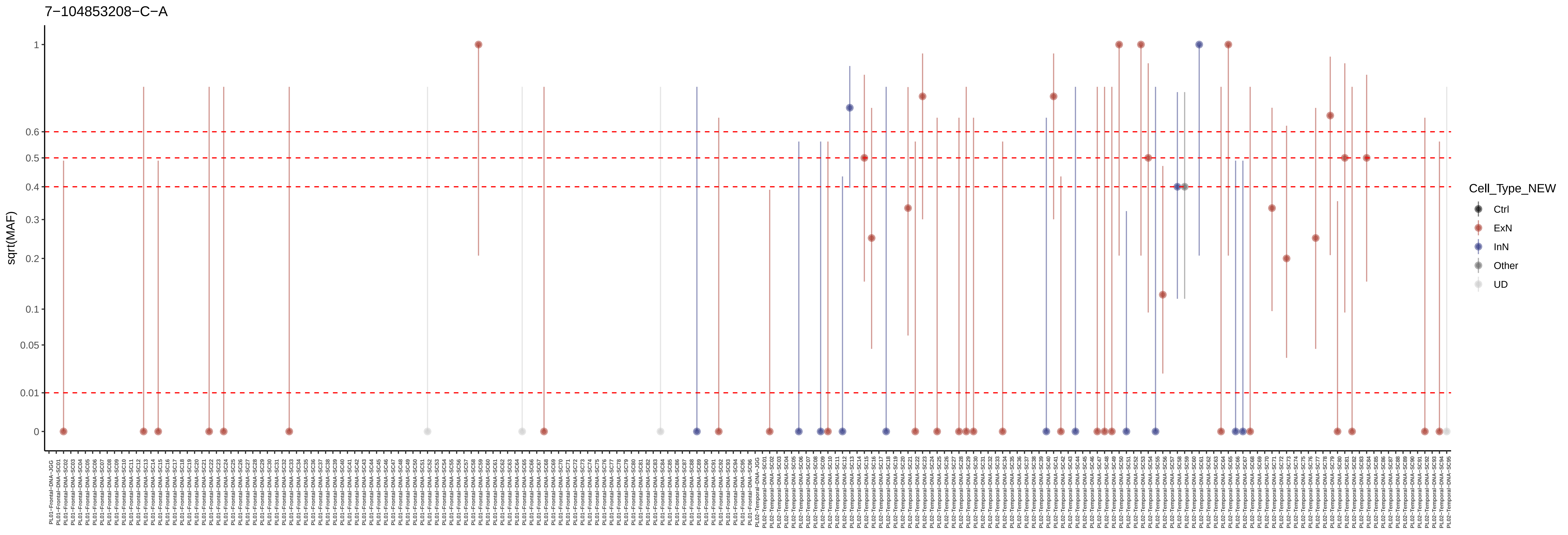

7−104853208−C−A
sqrt(MAF)
1
0.6
0.5
0.4
0.3
0.2
0.1
0.05
0.01
0
Cell_Type_NEW
Ctrl
ExN
InN
Other
UD

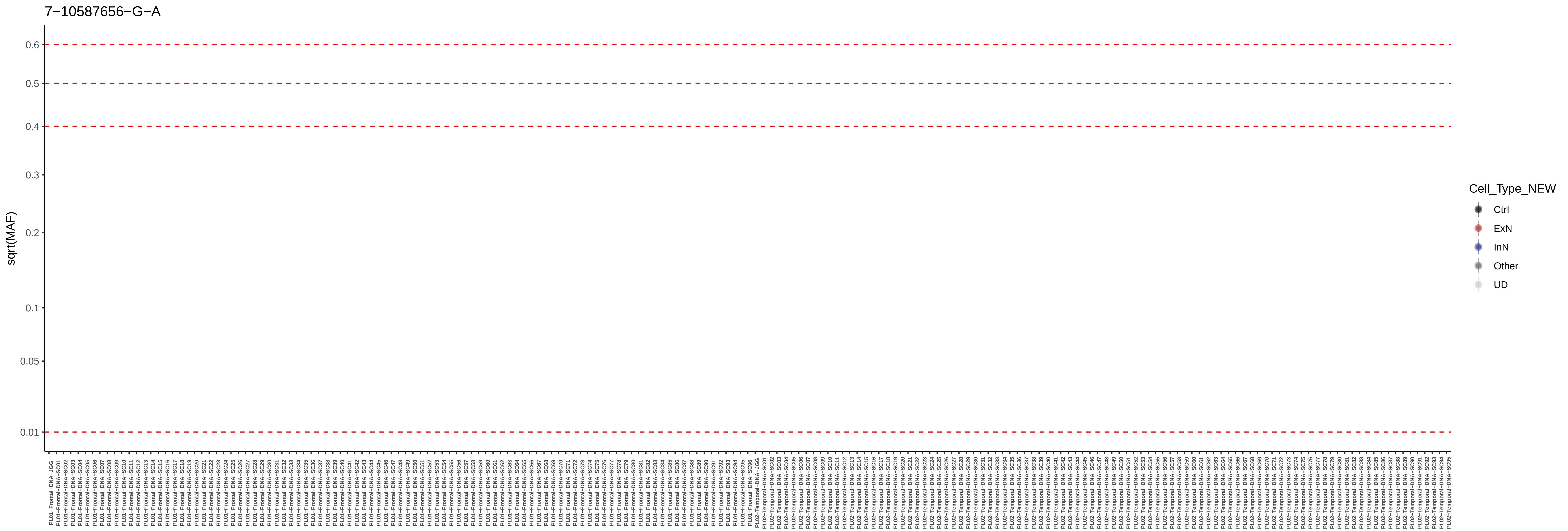
7−10587656−G−A
sqrt(MAF)
0.6
0.5
0.4
0.3
0.2
0.1
0.05
0.01
Cell_Type_NEW
Ctrl
ExN
InN
Other
UD

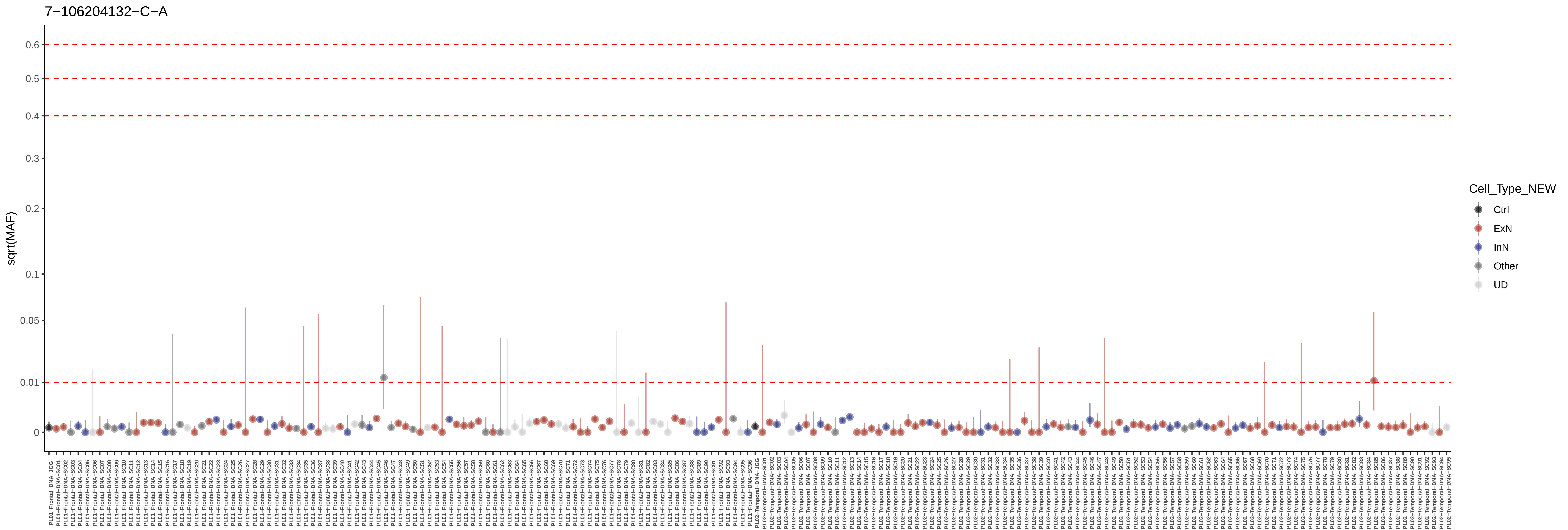

7−106204132−C−A
sqrt(MAF)
0.6
0.5
0.4
0.3
0.2
0.1
0.05
0.01
0
Cell_Type_NEW
Ctrl
ExN
InN
Other
UD

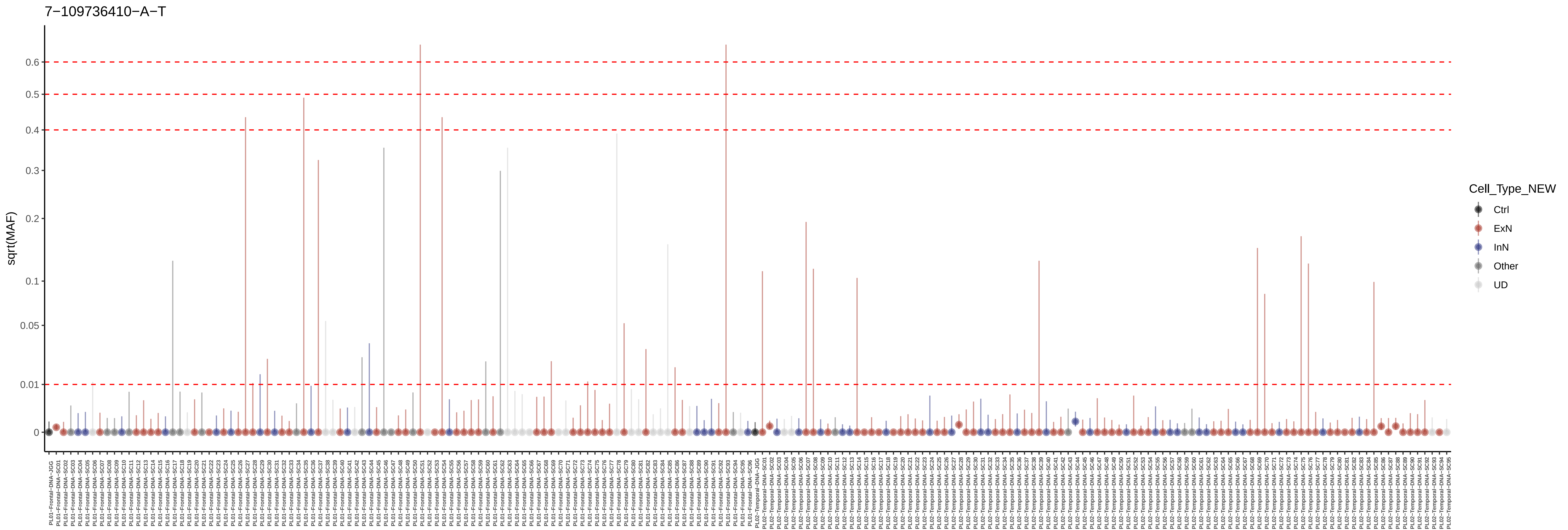

7−109736410−A−T
sqrt(MAF)
0.6
0.5
0.4
0.3
0.2
0.1
0.05
0.01
0
Cell_Type_NEW
Ctrl
ExN
InN
Other
UD

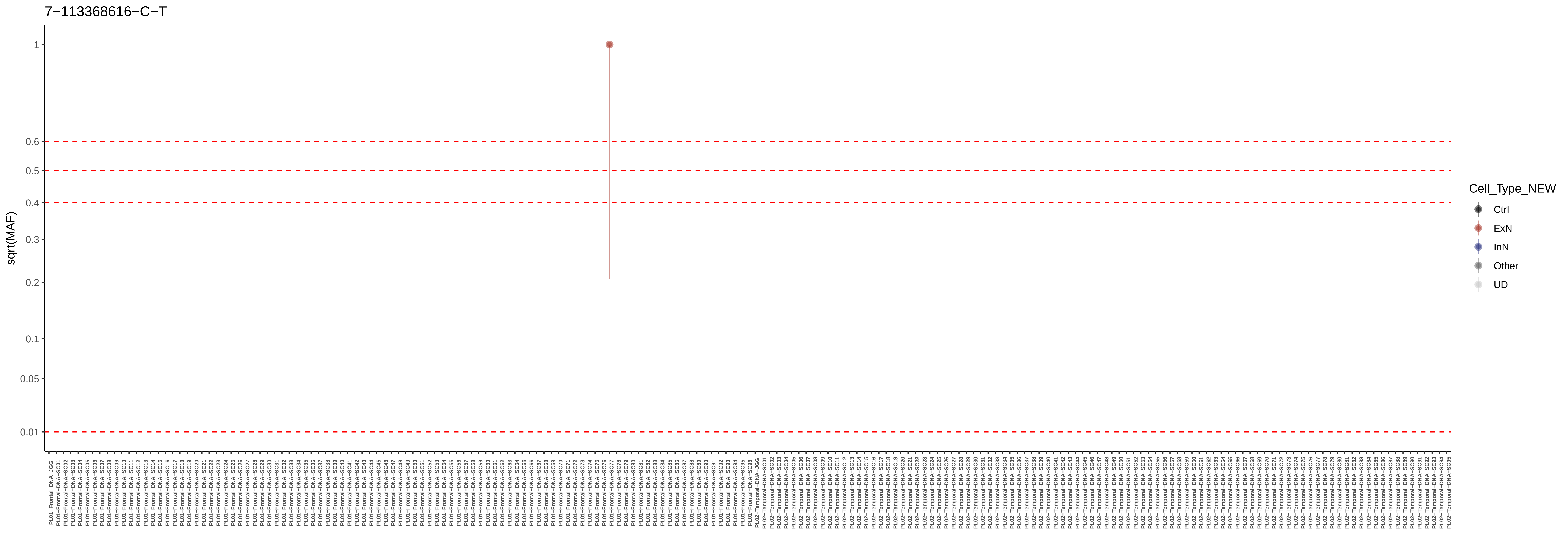
7−113368616−C−T
sqrt(MAF)
1
0.6
0.5
0.4
0.3
0.2
0.1
0.05
0.01
Cell_Type_NEW
Ctrl
ExN
InN
Other
UD

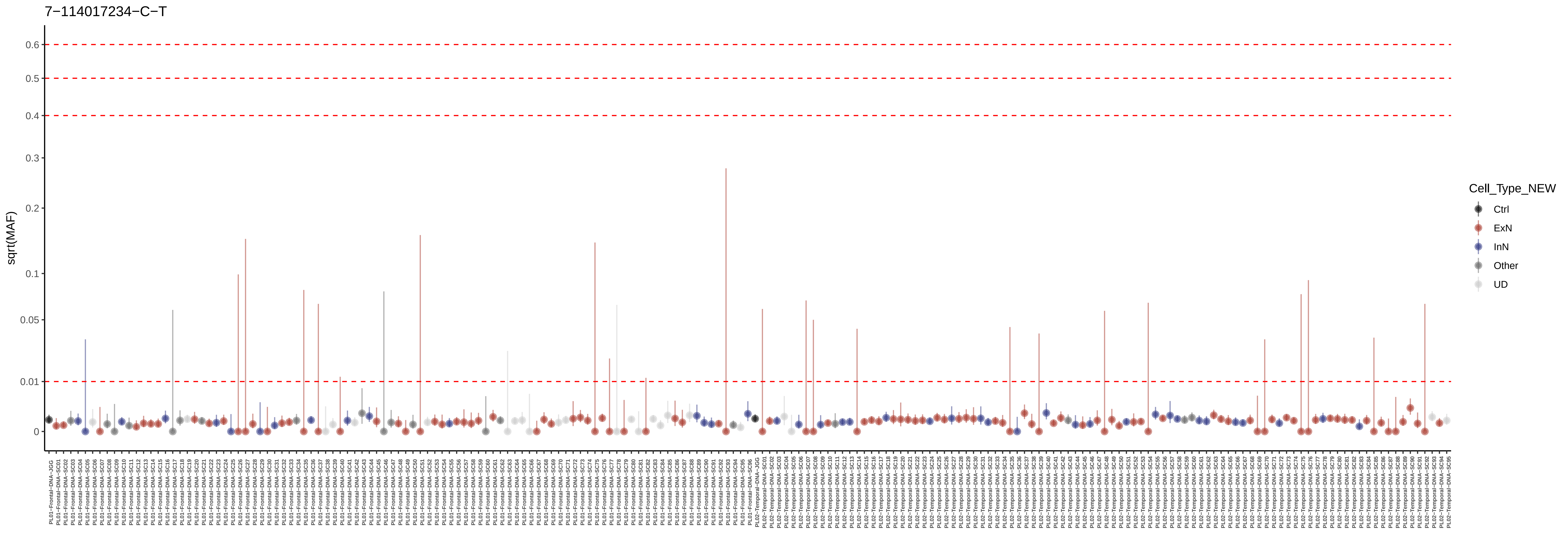

7−114017234−C−T
sqrt(MAF)
0.6
0.5
0.4
0.3
0.2
0.1
0.05
0.01
0
Cell_Type_NEW
Ctrl
ExN
InN
Other
UD

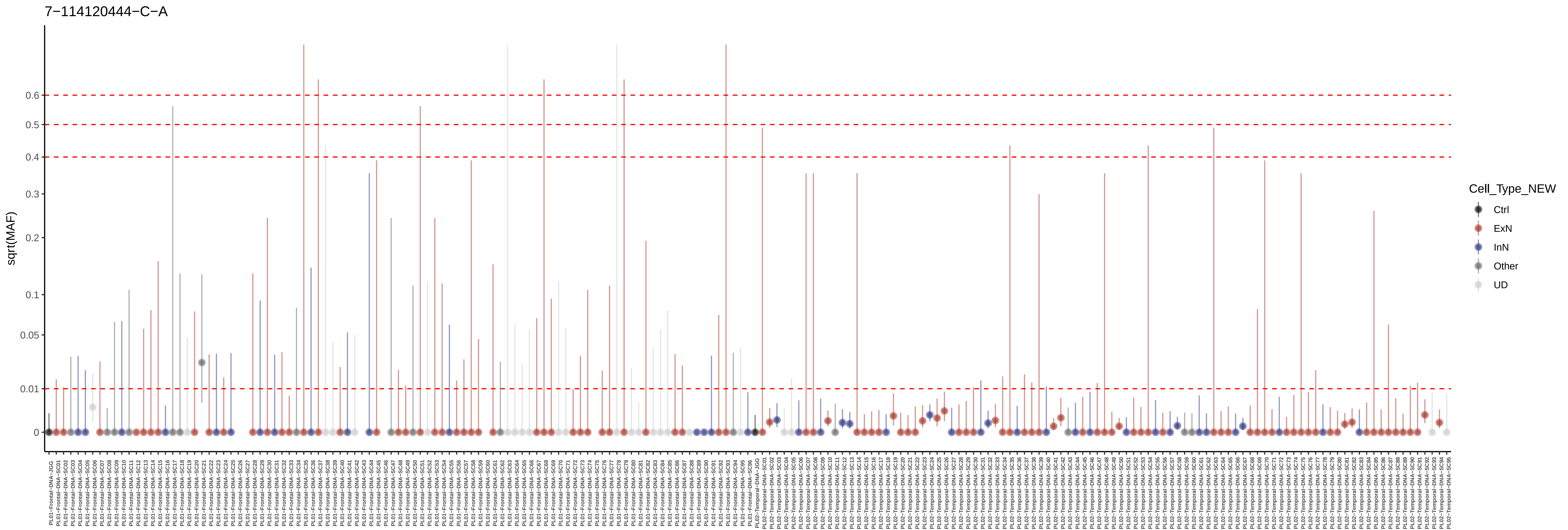
7−114120444−C−A
sqrt(MAF)
0.6
0.5
0.4
0.3
0.2
0.1
0.05
0.01
0
Cell_Type_NEW
Ctrl
ExN
InN
Other
UD

7−114658860−A−G

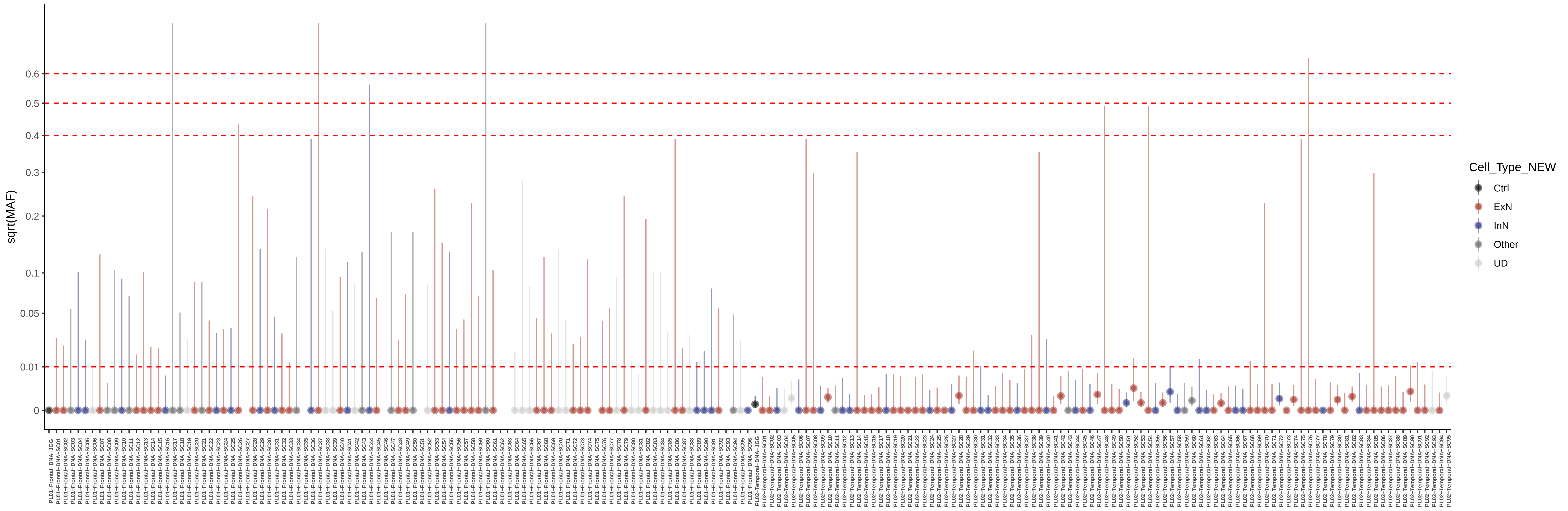

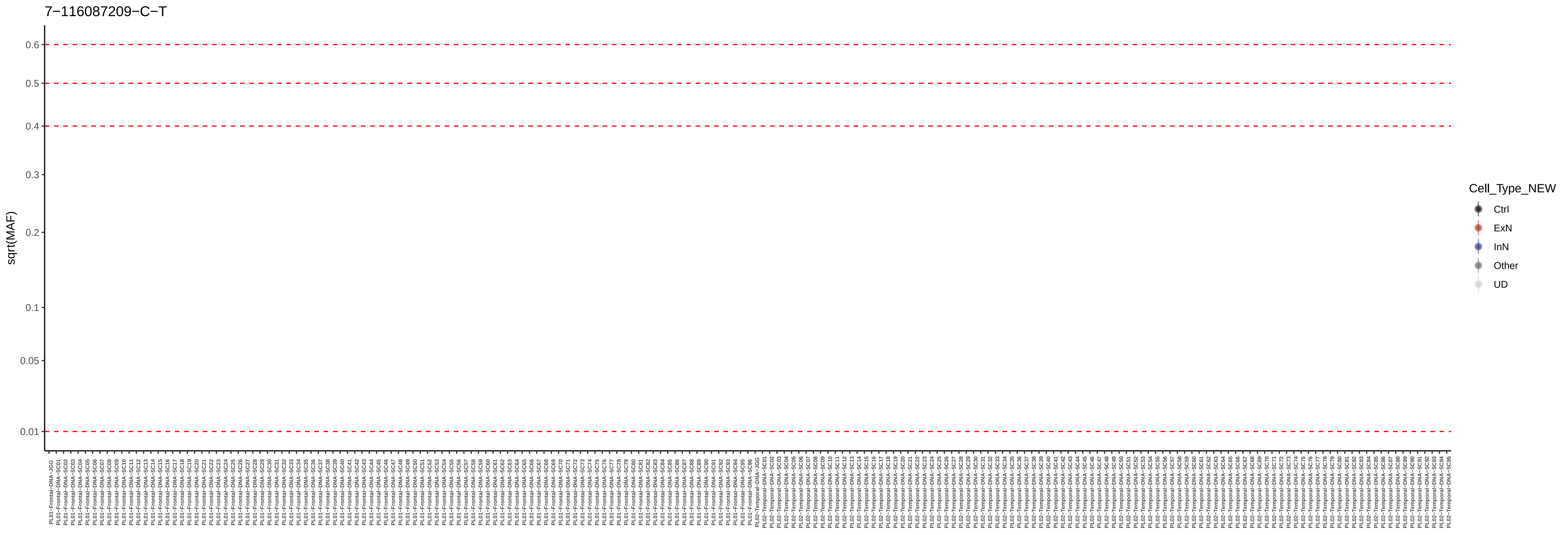
7−116087209−C−T
sqrt(MAF)
0.6
0.5
0.4
0.3
0.2
0.1
0.05
0.01
Cell_Type_NEW
Ctrl
ExN
InN
Other
UD

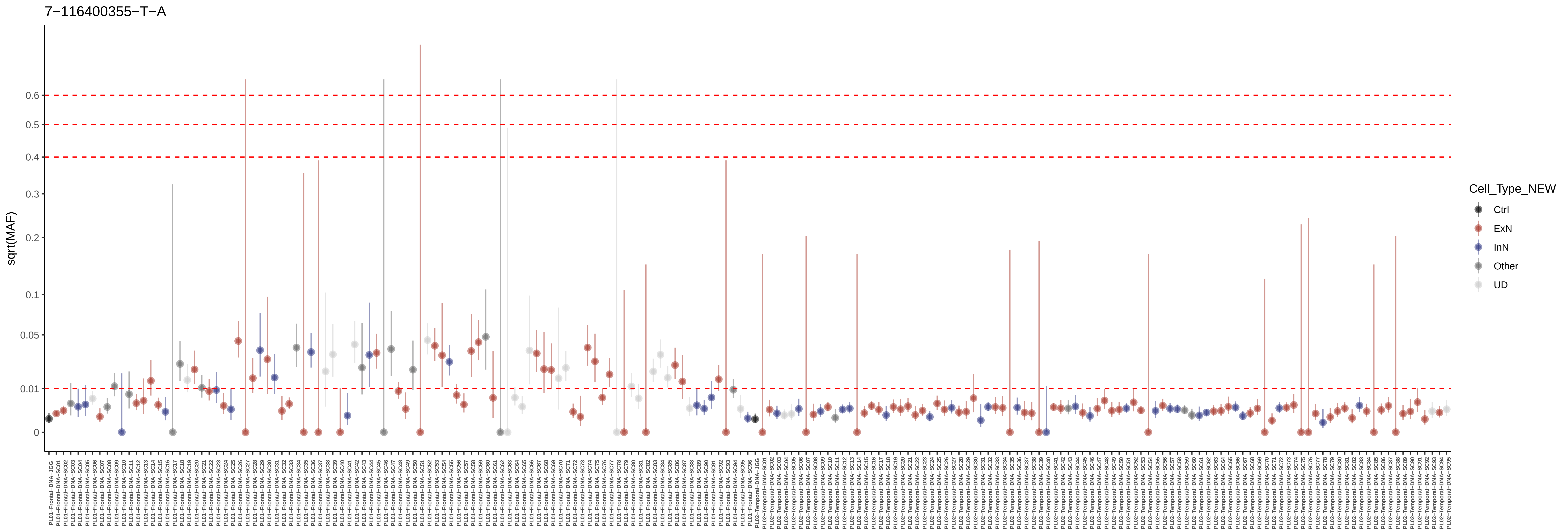
7−116400355−T−A
sqrt(MAF)
0.6
0.5
0.4
0.3
0.2
0.1
0.05
0.01
0
Cell_Type_NEW
Ctrl
ExN
InN
Other
UD

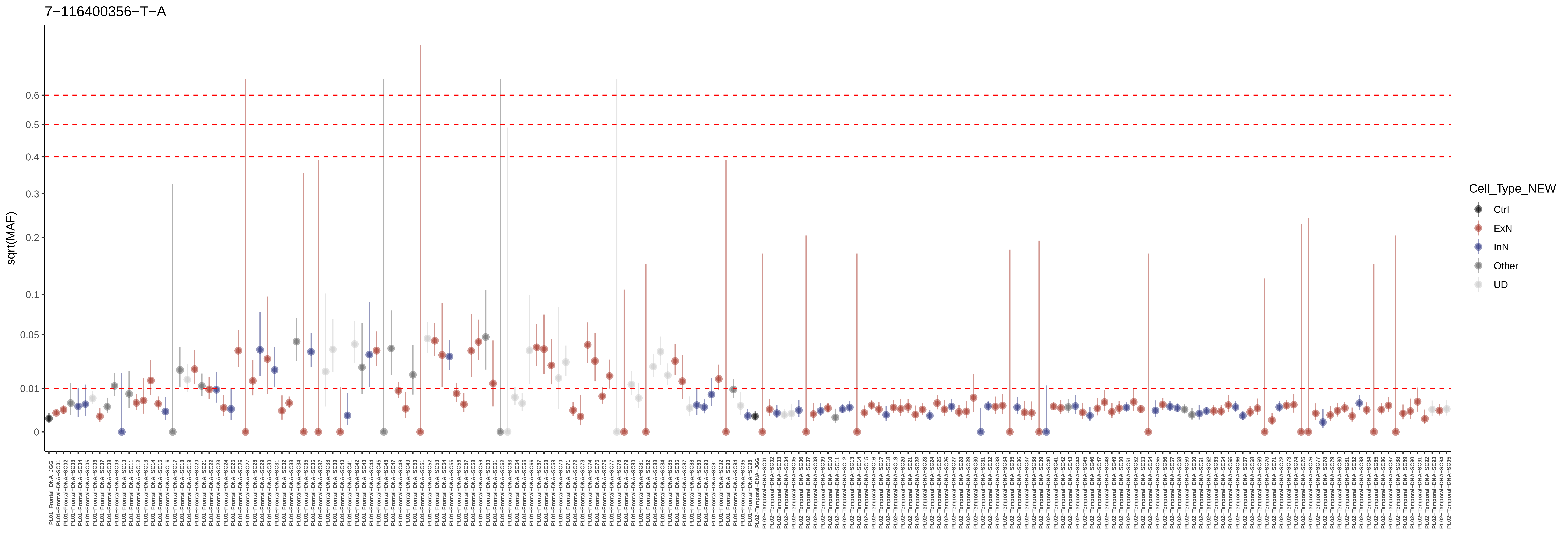
7−116400356−T−A
sqrt(MAF)
0.6
0.5
0.4
0.3
0.2
0.1
0.05
0.01
0
Cell_Type_NEW
Ctrl
ExN
InN
Other
UD

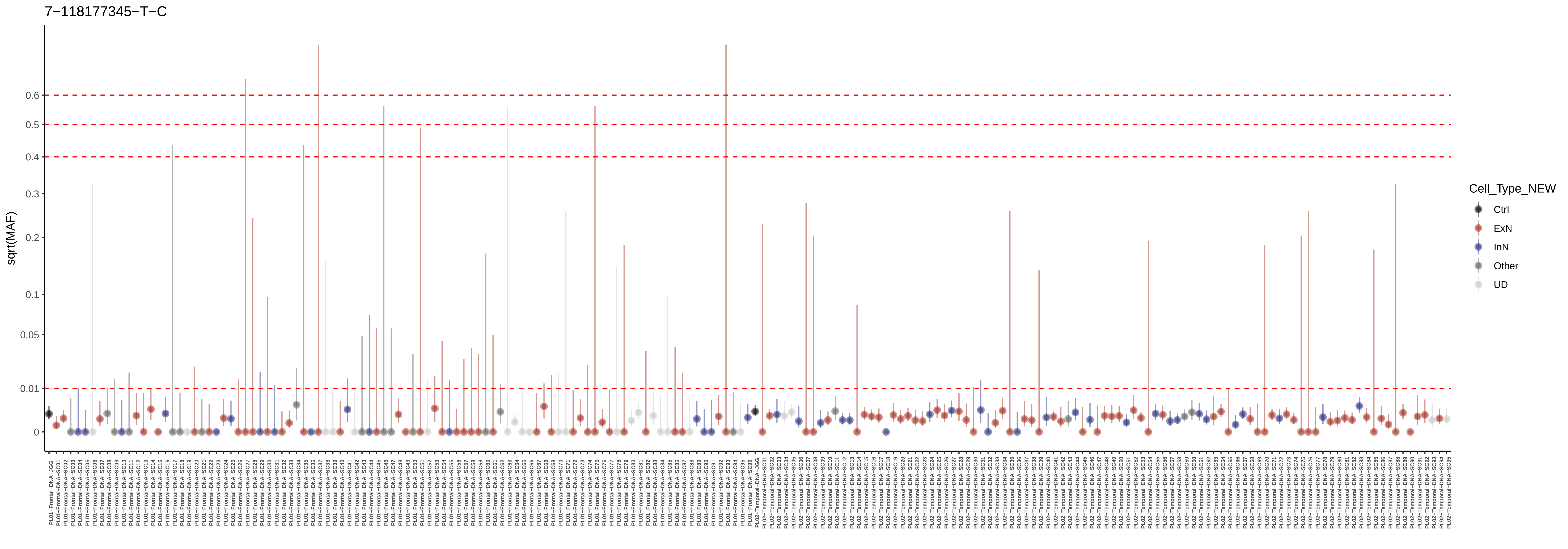
7−118177345−T−C
sqrt(MAF)
0.6
0.5
0.4
0.3
0.2
0.1
0.05
0.01
0
Cell_Type_NEW
Ctrl
ExN
InN
Other
UD

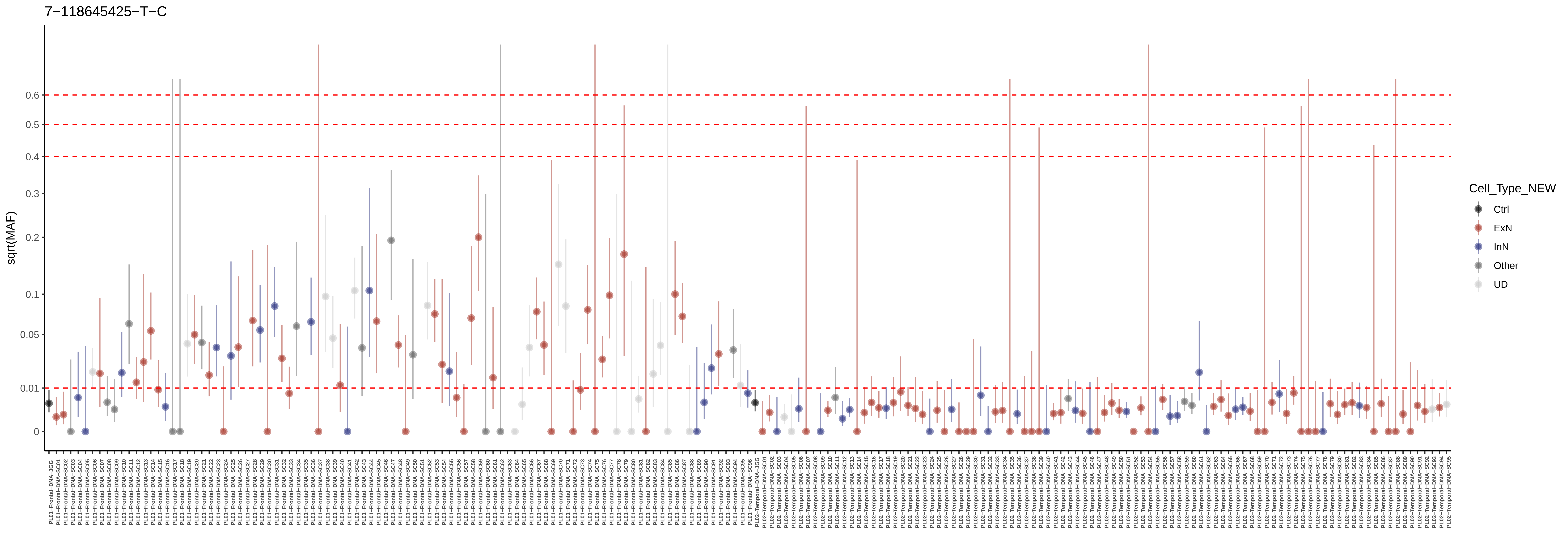
7−118645425−T−C
sqrt(MAF)
0.6
0.5
0.4
0.3
0.2
0.1
0.05
0.01
0
Cell_Type_NEW
Ctrl
ExN
InN
Other
UD

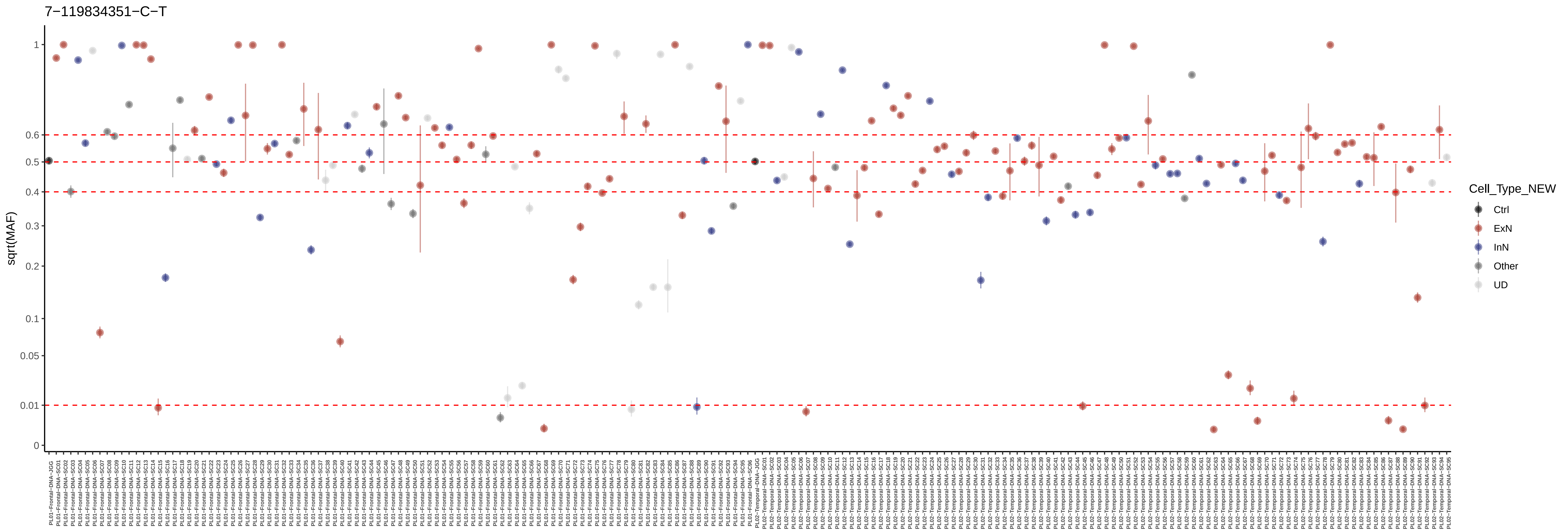

7−119834351−C−T
sqrt(MAF)
1
0.6
0.5
0.4
0.3
0.2
0.1
0.05
0.01
0
Cell_Type_NEW
Ctrl
ExN
InN
Other
UD

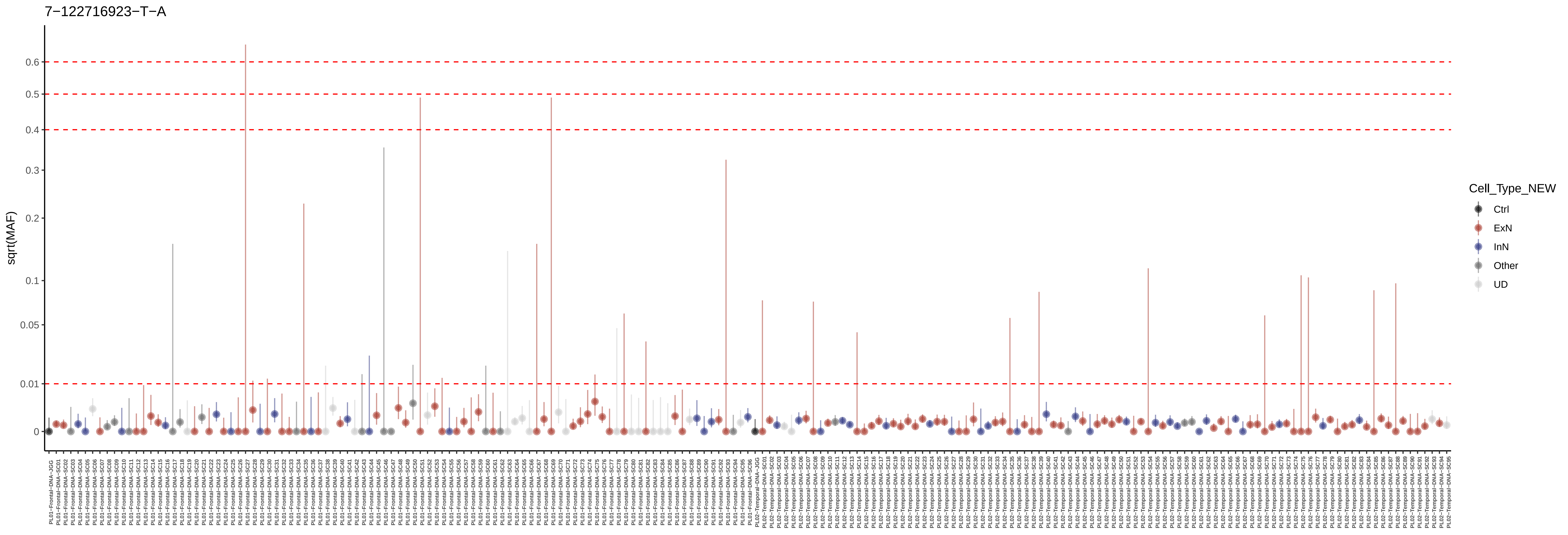

7−122716923−T−A
sqrt(MAF)
0.6
0.5
0.4
0.3
0.2
0.1
0.05
0.01
0
Cell_Type_NEW
Ctrl
ExN
InN
Other
UD

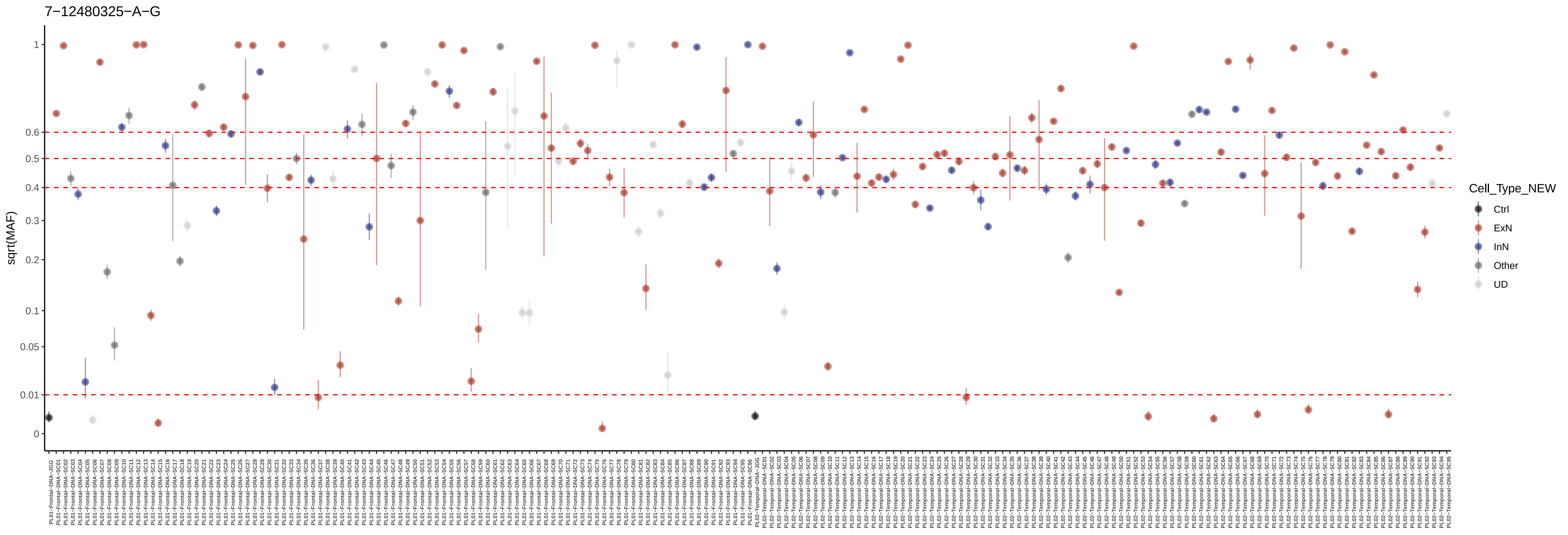
7−12480325−A−G
sqrt(MAF)
1
0.6
0.5
0.4
0.3
0.2
0.1
0.05
0.01
0
Cell_Type_NEW
Ctrl
ExN
InN
Other
UD

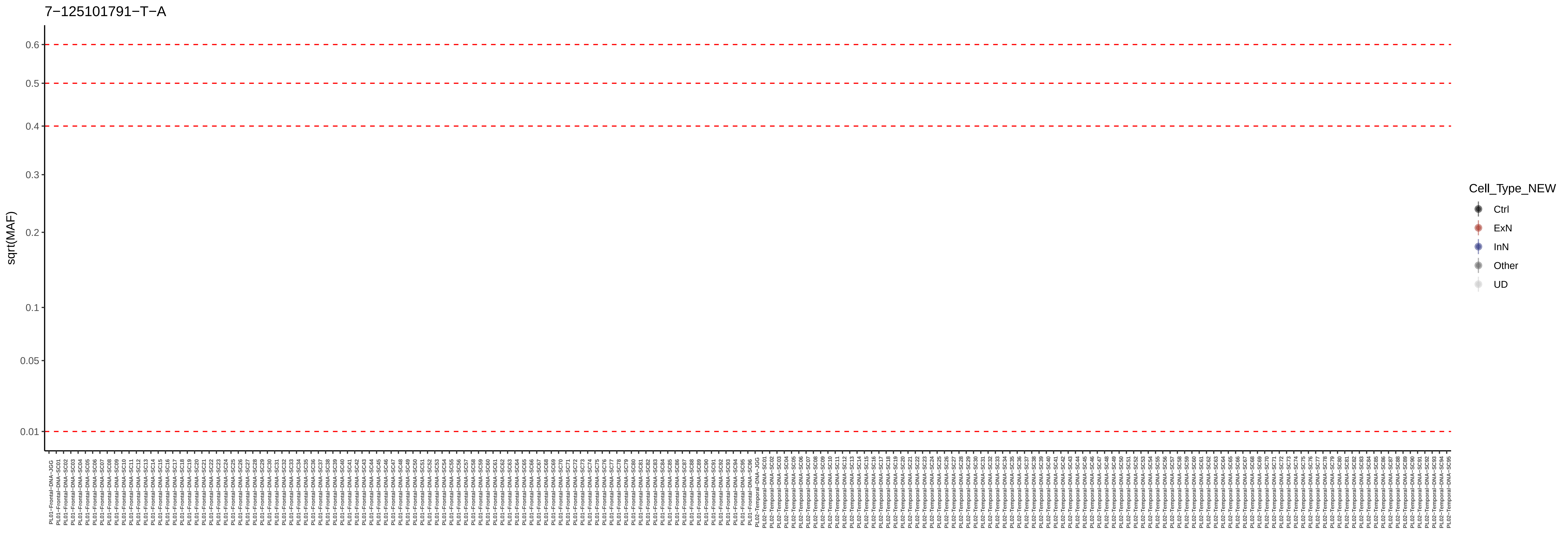

7−125101791−T−A
sqrt(MAF)
0.6
0.5
0.4
0.3
0.2
0.1
0.05
0.01
Cell_Type_NEW
Ctrl
ExN
InN
Other
UD

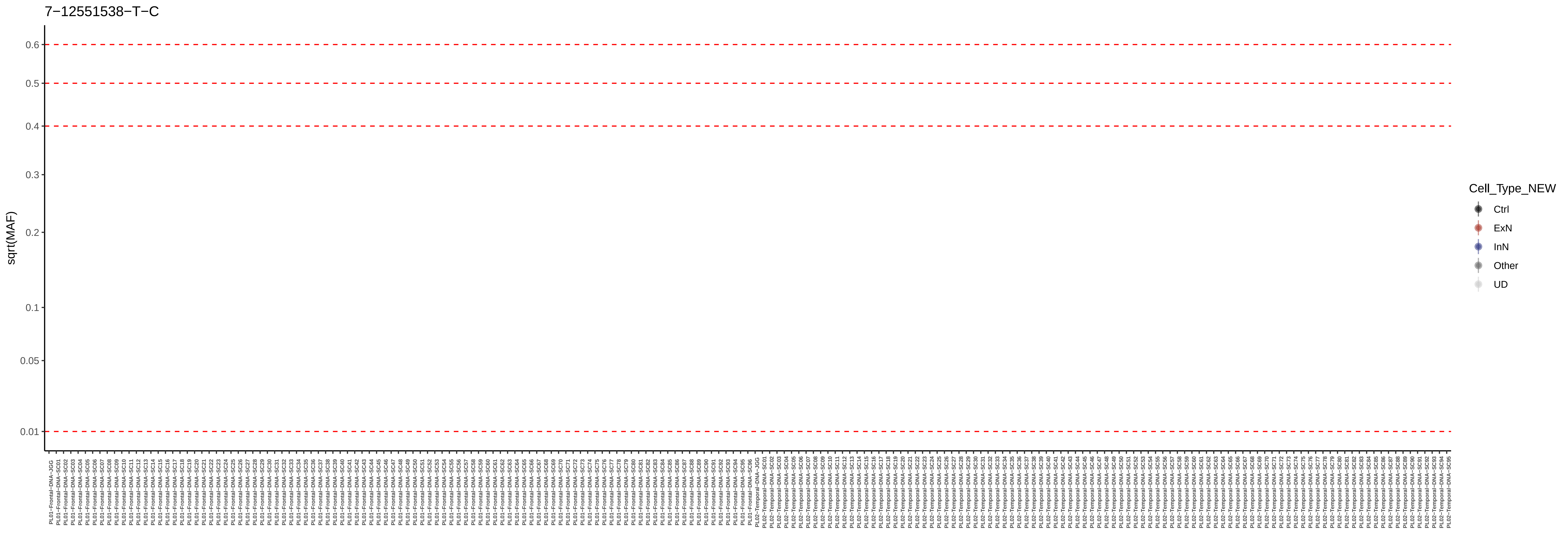

7−12551538−T−C
sqrt(MAF)
0.6
0.5
0.4
0.3
0.2
0.1
0.05
0.01
Cell_Type_NEW
Ctrl
ExN
InN
Other
UD

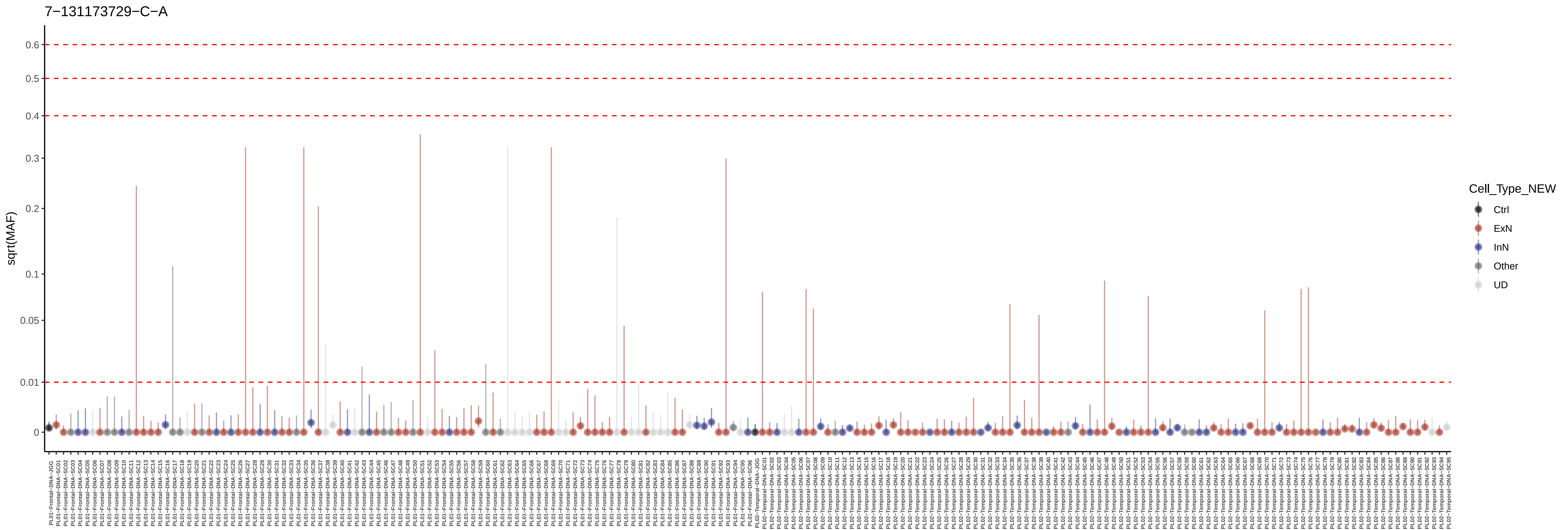
7−131173729−C−A
sqrt(MAF)
0
0.01
0.05
0.1
0.2
0.3
0.4
0.5
0.6
Cell_Type_NEW
Ctrl
ExN
InN
Other
UD

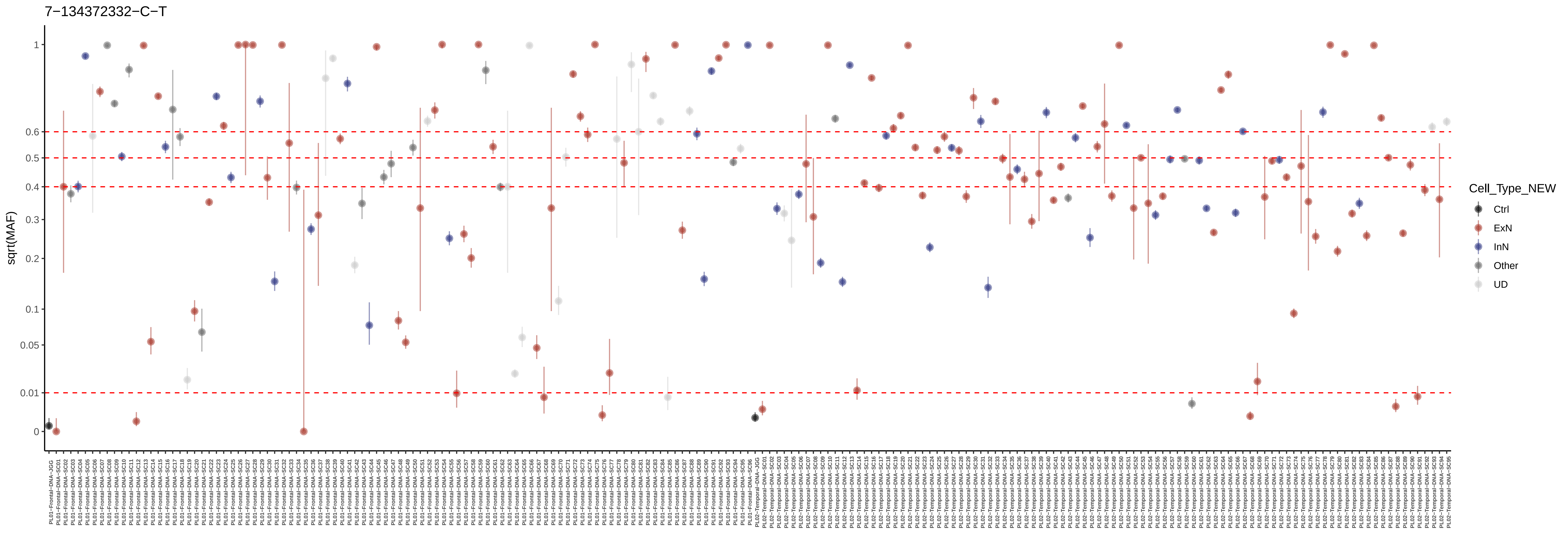

7−134372332−C−T
sqrt(MAF)
1
0.6
0.5
0.4
0.3
0.2
0.1
0.05
0.01
0
Cell_Type_NEW
Ctrl
ExN
InN
Other
UD

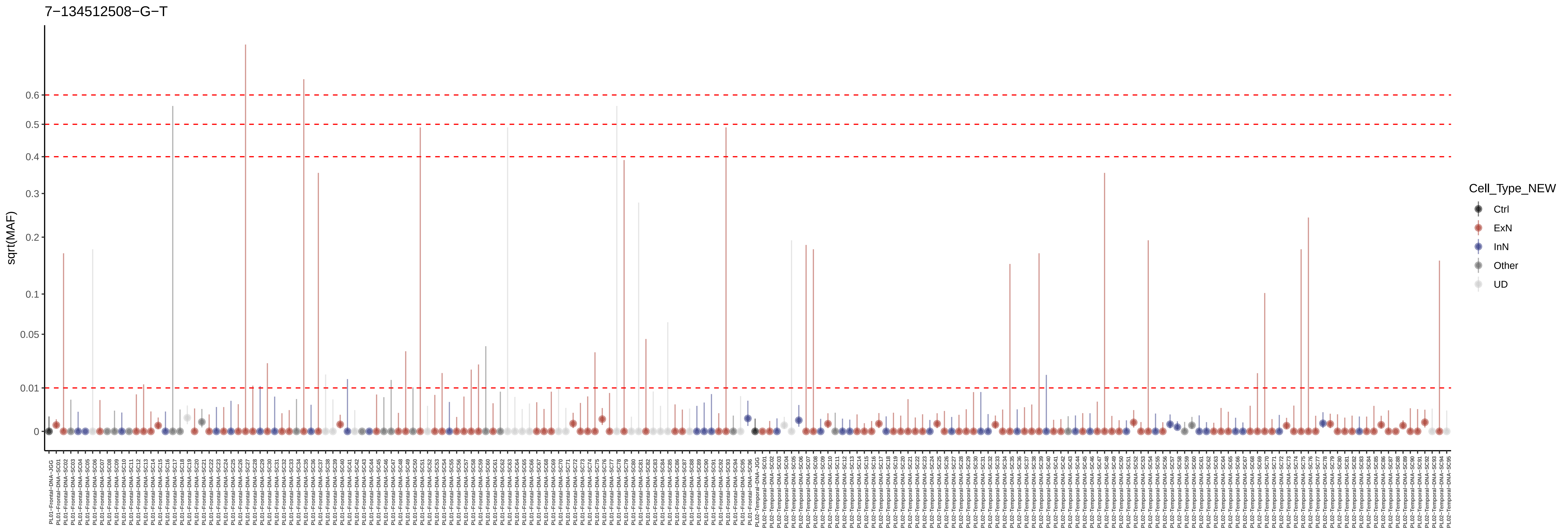
7−134512508−G−T
sqrt(MAF)
0.6
0.5
0.4
0.3
0.2
0.1
0.05
0.01
0
Cell_Type_NEW
Ctrl
ExN
InN
Other
UD

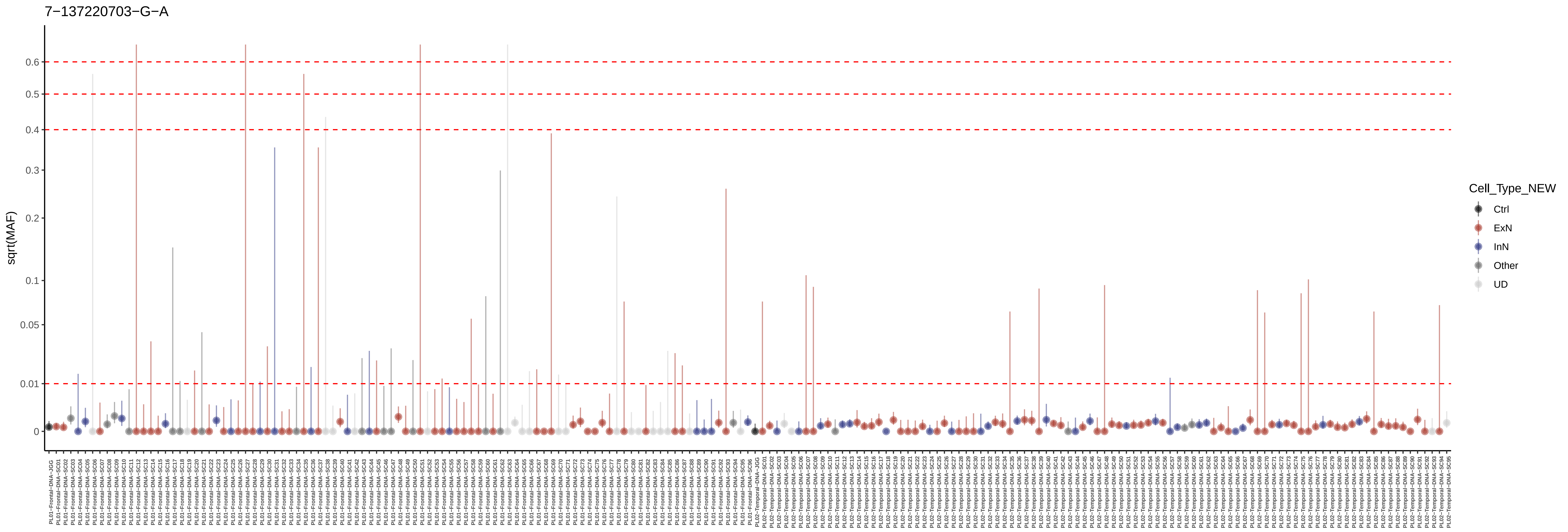
7−137220703−G−A
sqrt(MAF)
0.6
0.5
0.4
0.3
0.2
0.1
0.05
0.01
0
Cell_Type_NEW
Ctrl
ExN
InN
Other
UD

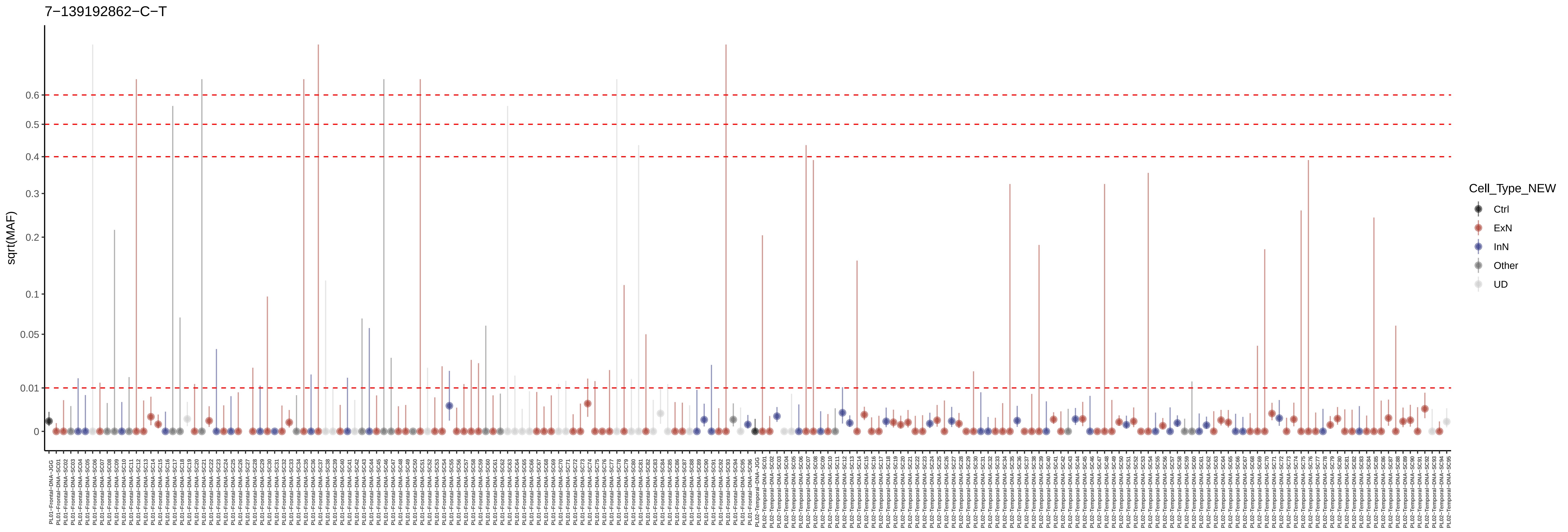
7−139192862−C−T
sqrt(MAF)
0.6
0.5
0.4
0.3
0.2
0.1
0.05
0.01
0
Cell_Type_NEW
Ctrl
ExN
InN
Other
UD

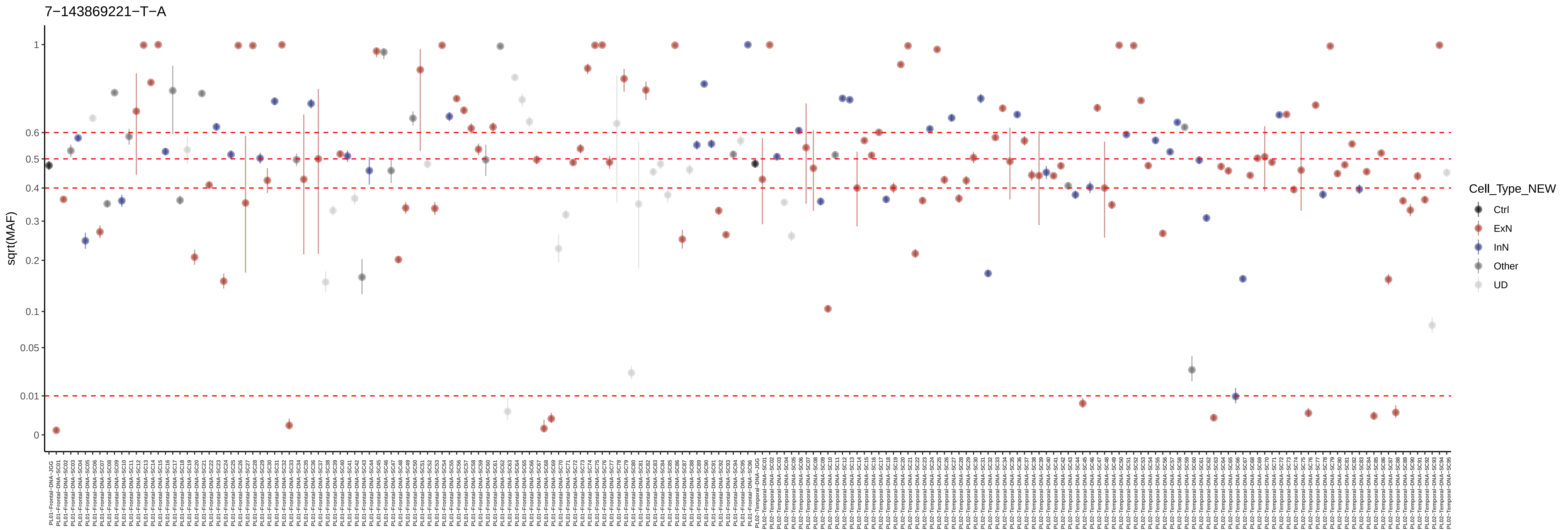
7−143869221−T−A
sqrt(MAF)
1
0.6
0.5
0.4
0.3
0.2
0.1
0.05
0.01
0
Cell_Type_NEW
Ctrl
ExN
InN
Other
UD

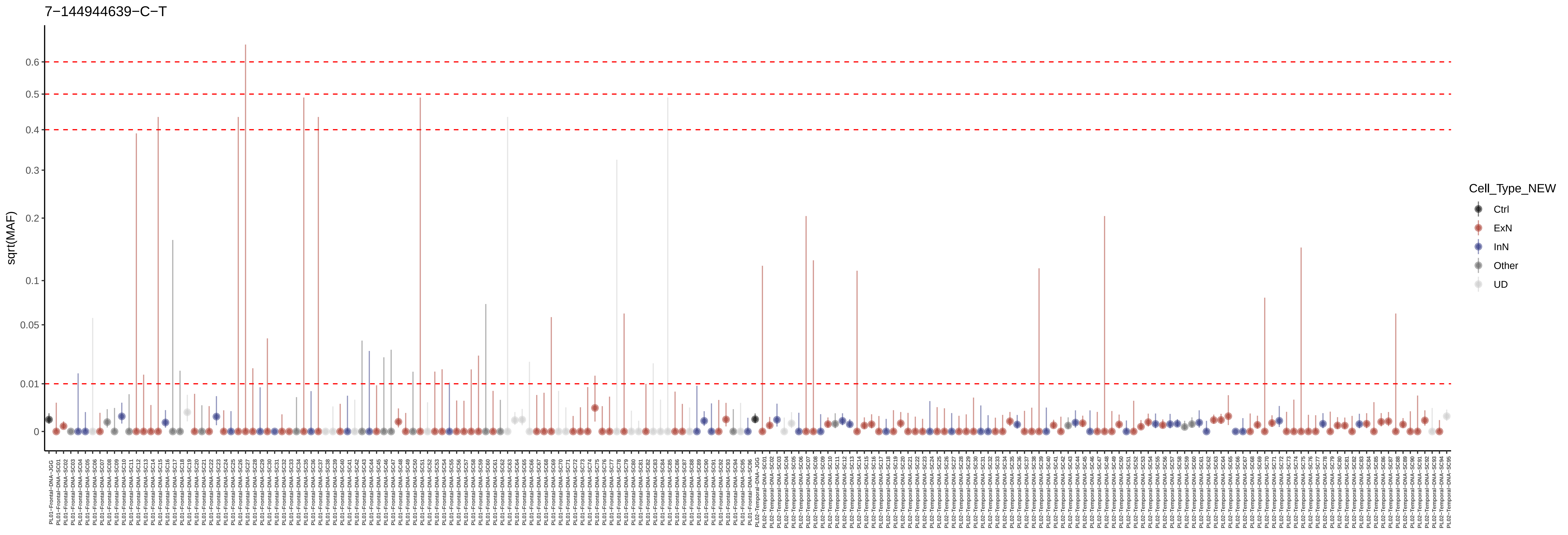
7−144944639−C−T
sqrt(MAF)
0.6
0.5
0.4
0.3
0.2
0.1
0.05
0.01
0
Cell_Type_NEW
Ctrl
ExN
InN
Other
UD

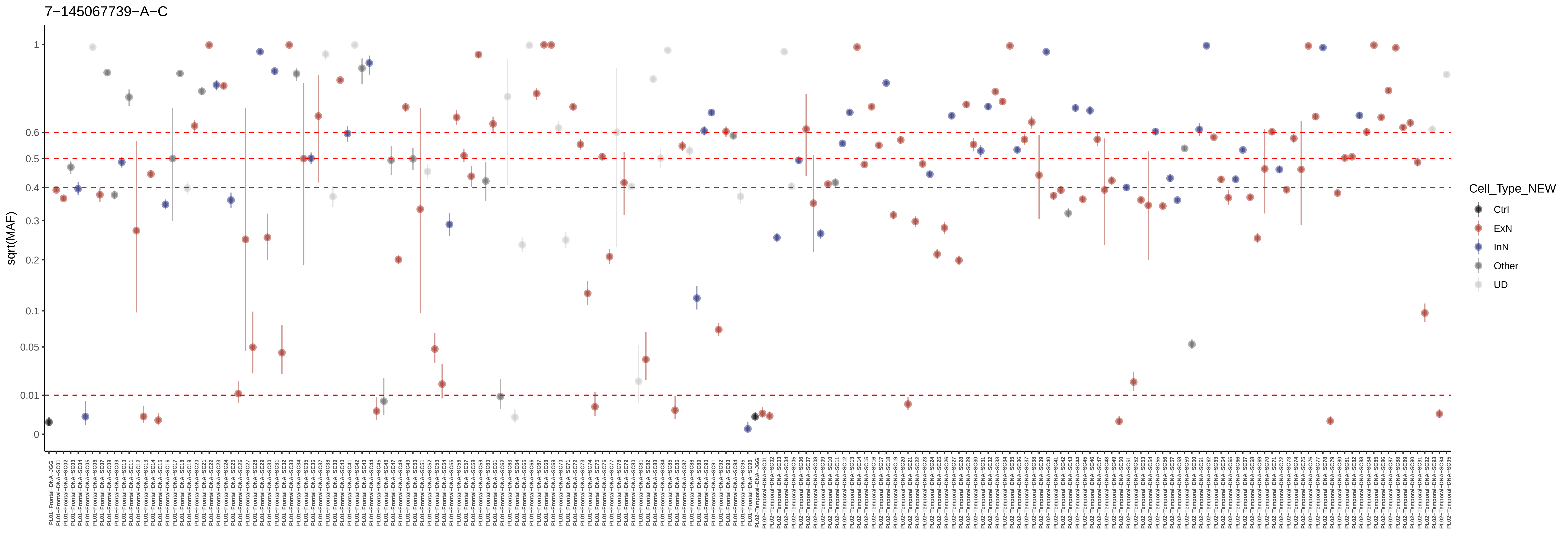
7−145067739−A−C
sqrt(MAF)
1
0.6
0.5
0.4
0.3
0.2
0.1
0.05
0.01
0
Cell_Type_NEW
Ctrl
ExN
InN
Other
UD

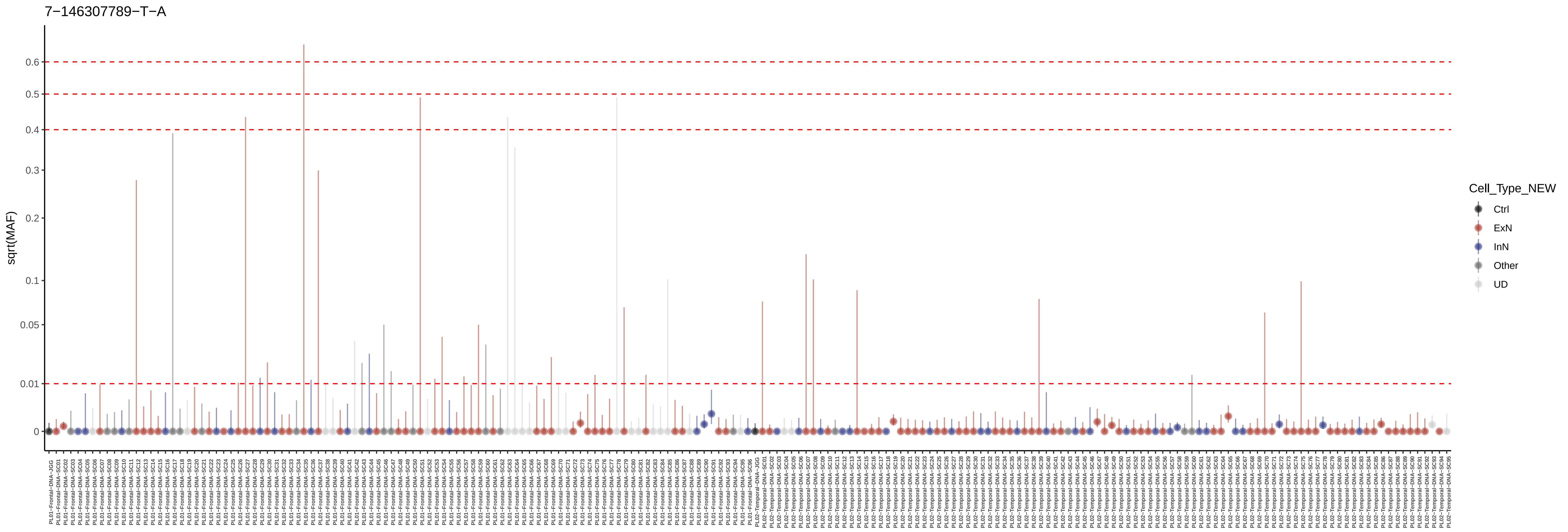
7−146307789−T−A
sqrt(MAF)
0.6
0.5
0.4
0.3
0.2
0.1
0.05
0.01
0
Cell_Type_NEW
Ctrl
ExN
InN
Other
UD

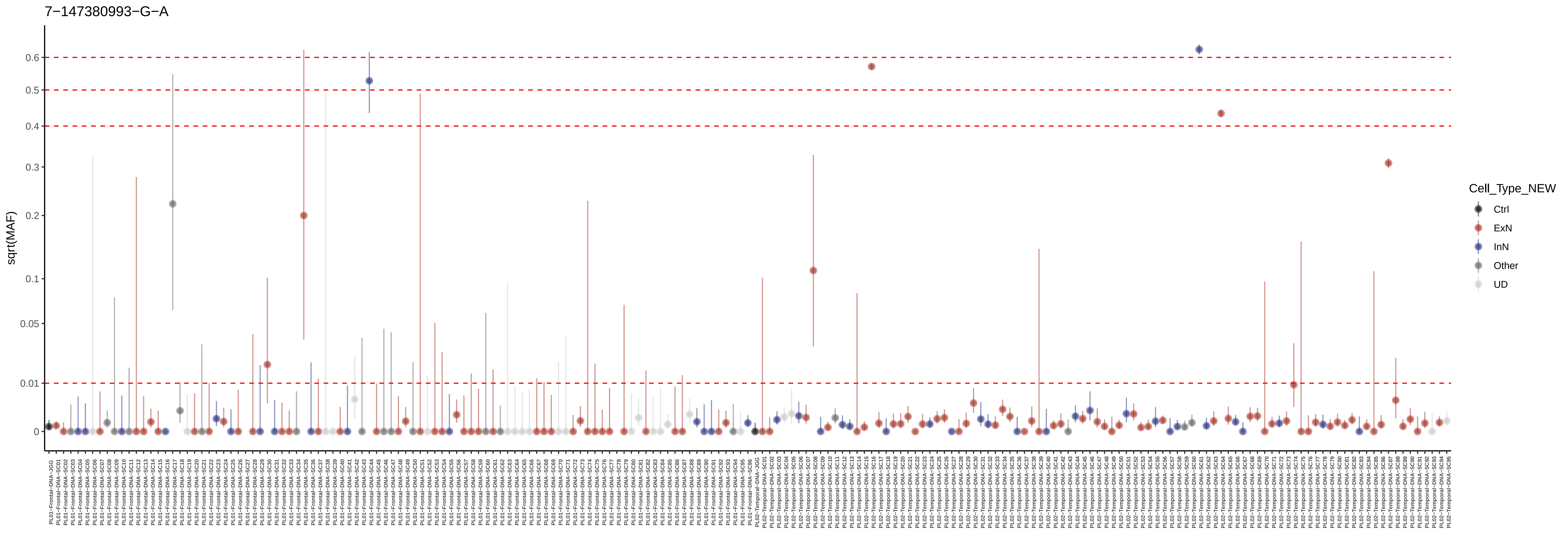
7−147380993−G−A
sqrt(MAF)
0.6
0.5
0.4
0.3
0.2
0.1
0.05
0.01
0
Cell_Type_NEW
Ctrl
ExN
InN
Other
UD

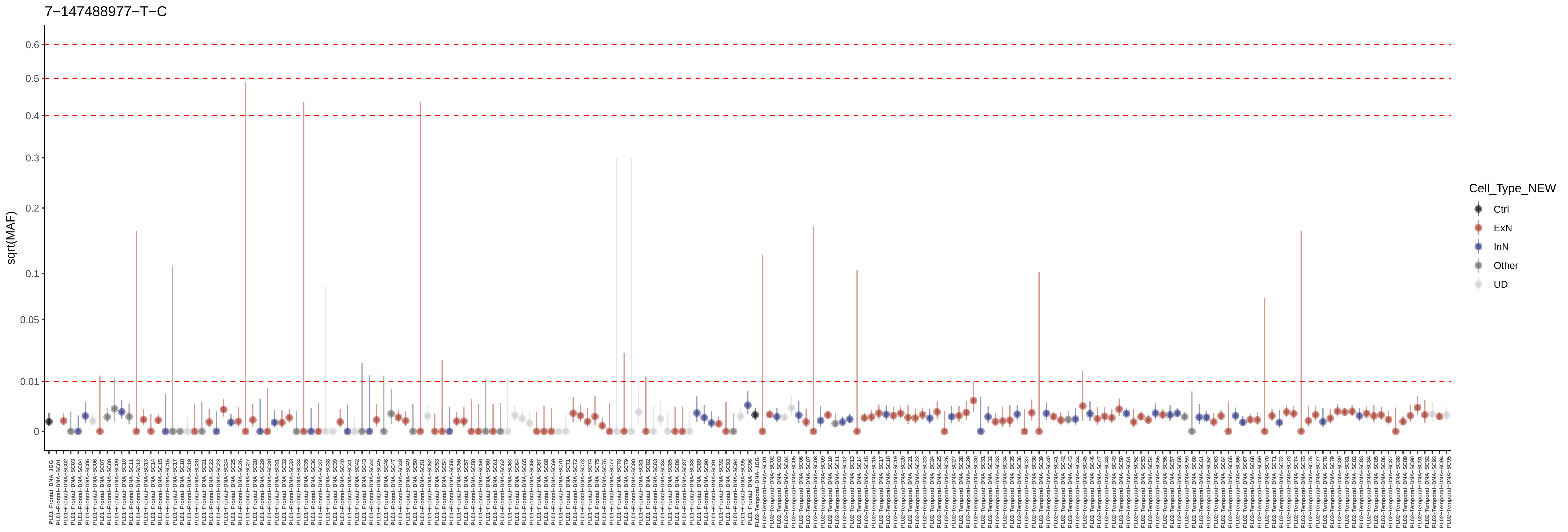
7−147488977−T−C
sqrt(MAF)
0.6
0.5
0.4
0.3
0.2
0.1
0.05
0.01
0
Cell_Type_NEW
Ctrl
ExN
InN
Other
UD

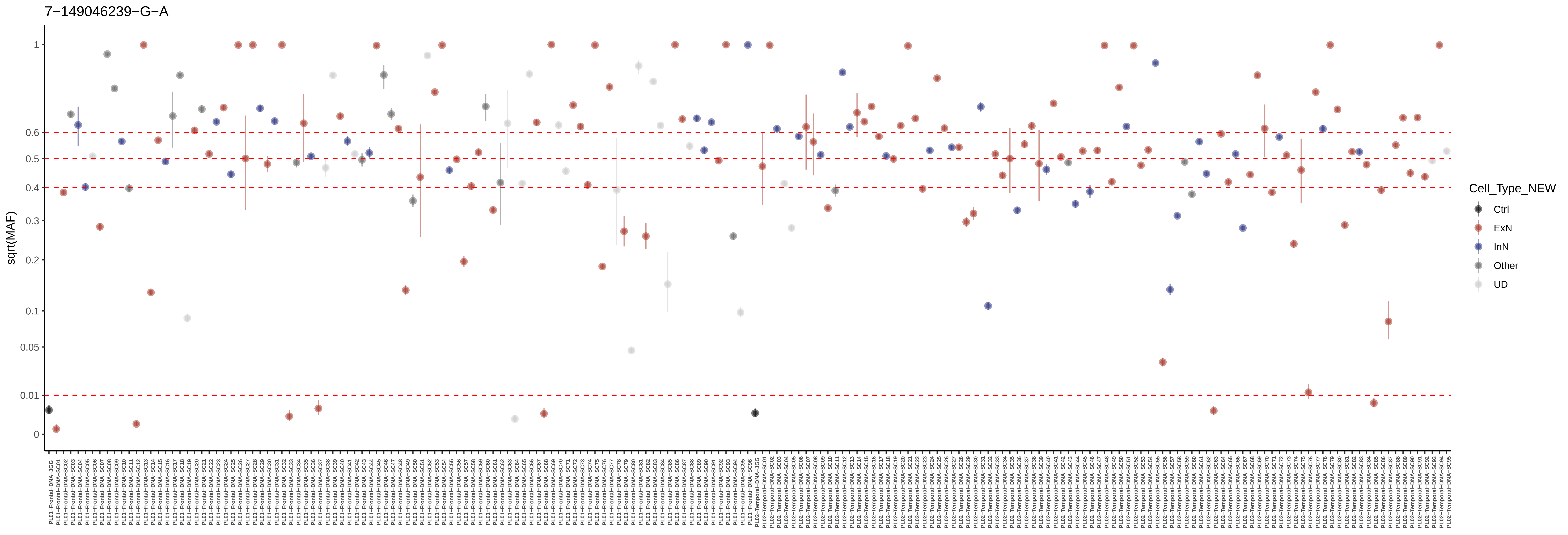
7−149046239−G−A
sqrt(MAF)
1
0.6
0.5
0.4
0.3
0.2
0.1
0.05
0.01
0
Cell_Type_NEW
Ctrl
ExN
InN
Other
UD

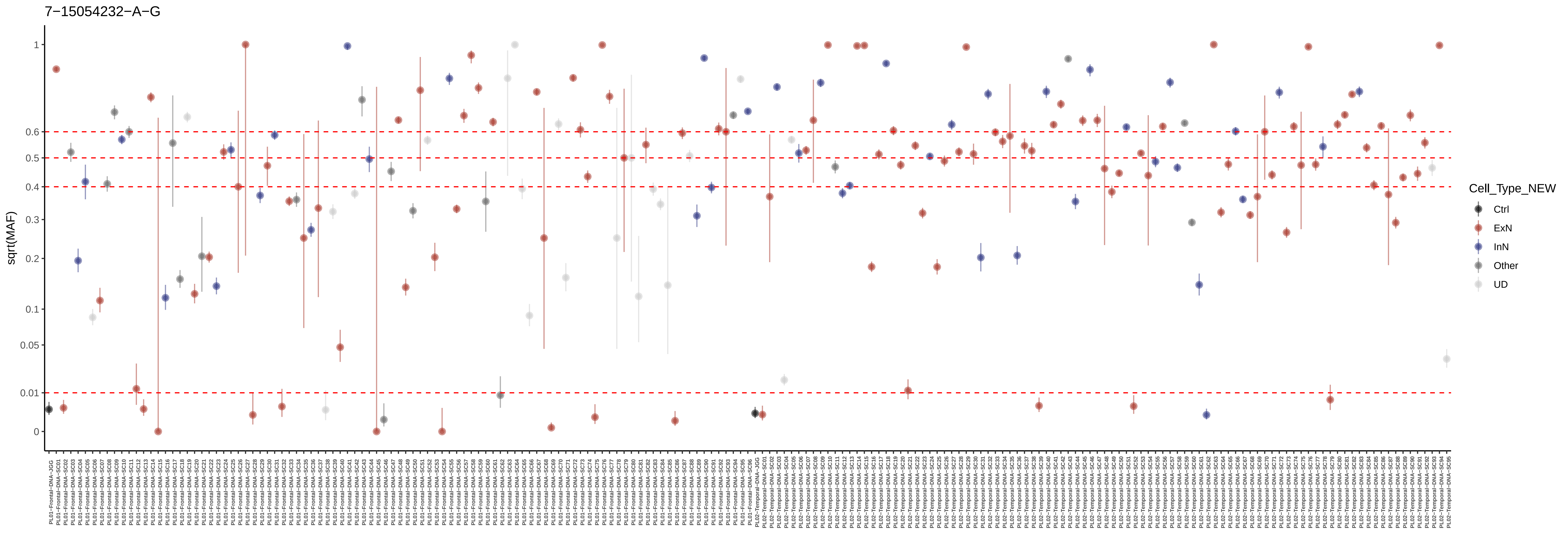

7−15054232−A−G
sqrt(MAF)
1
0.6
0.5
0.4
0.3
0.2
0.1
0.05
0.01
0
Cell_Type_NEW
Ctrl
ExN
InN
Other
UD

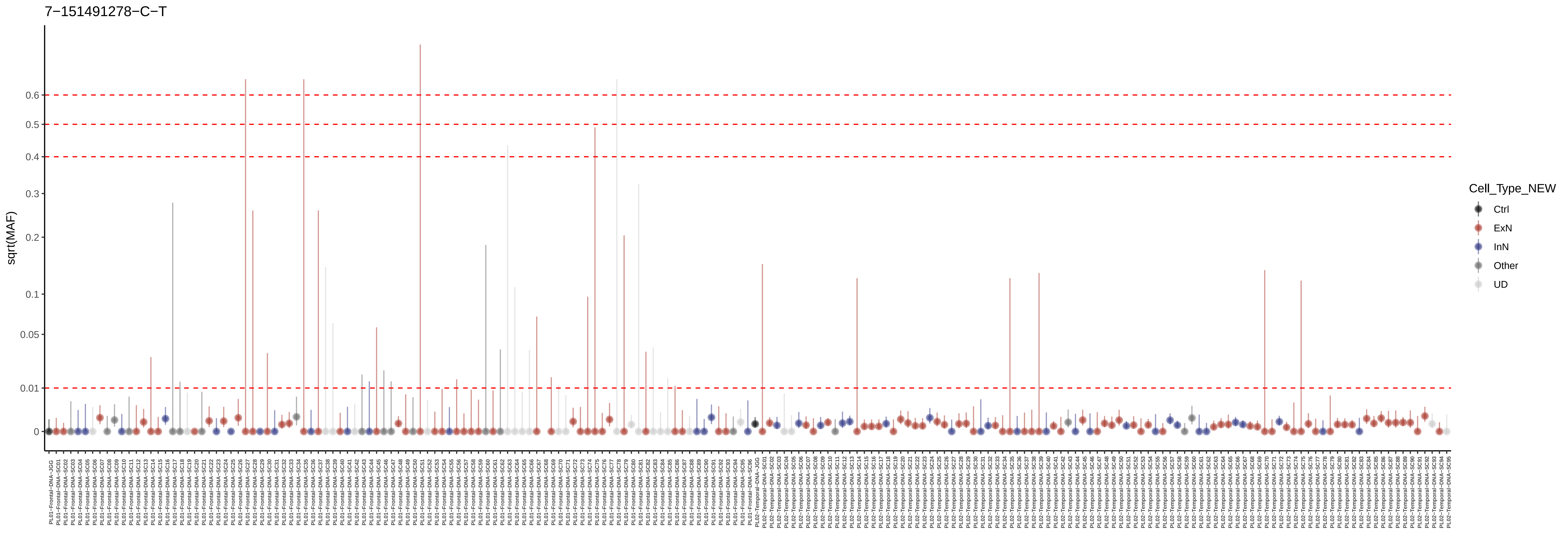

7−151491278−C−T
sqrt(MAF)
0.6
0.5
0.4
0.3
0.2
0.1
0.05
0.01
0
Cell_Type_NEW
Ctrl
ExN
InN
Other
UD

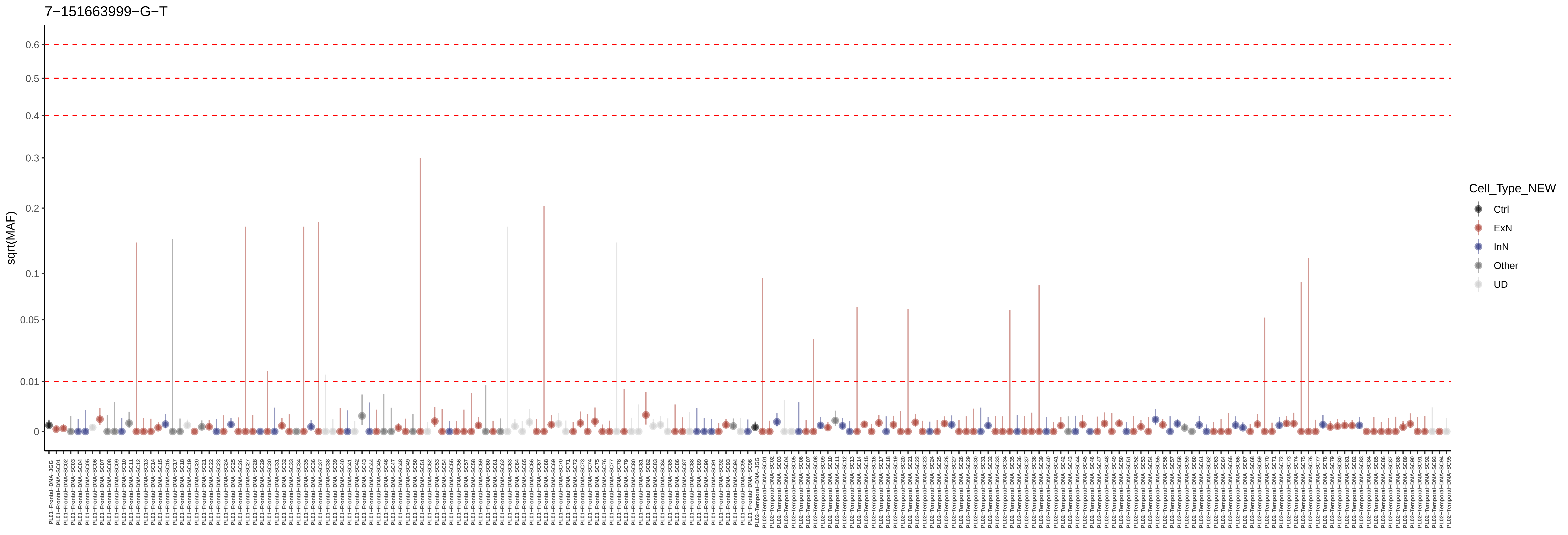
7−151663999−G−T
sqrt(MAF)
0.6
0.5
0.4
0.3
0.2
0.1
0.05
0.01
0
Cell_Type_NEW
Ctrl
ExN
InN
Other
UD

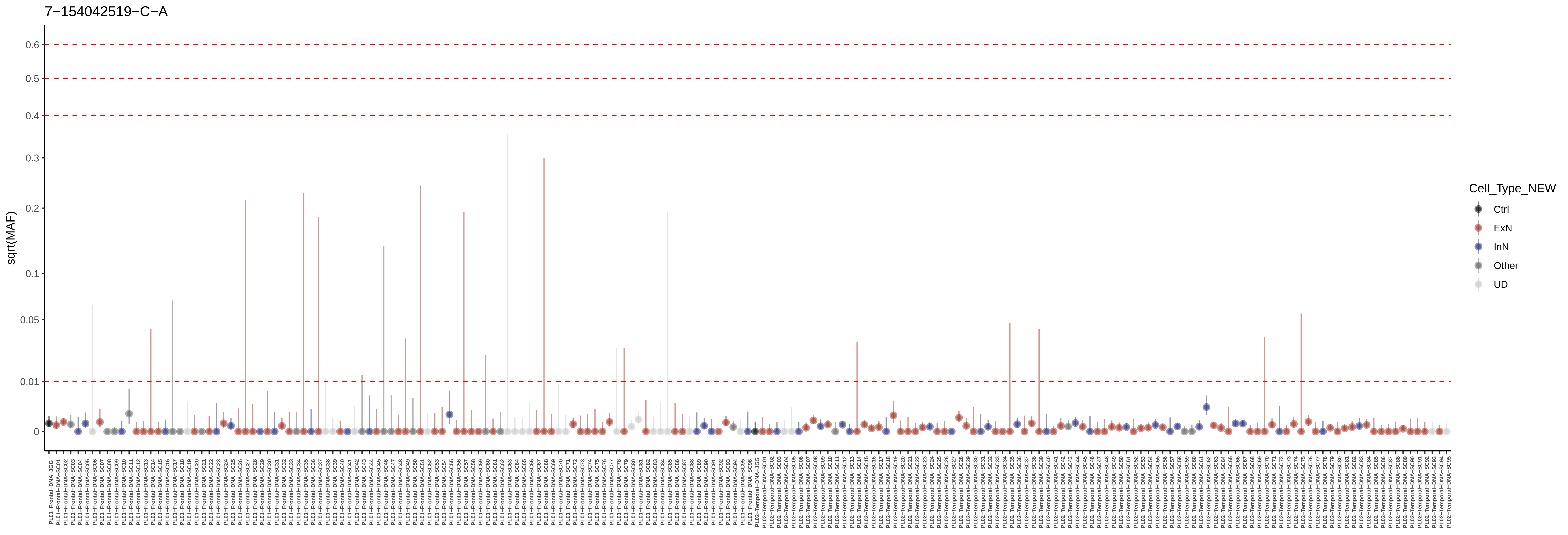
7−154042519−C−A
sqrt(MAF)
0.6
0.5
0.4
0.3
0.2
0.1
0.05
0.01
0
Cell_Type_NEW
Ctrl
ExN
InN
Other
UD

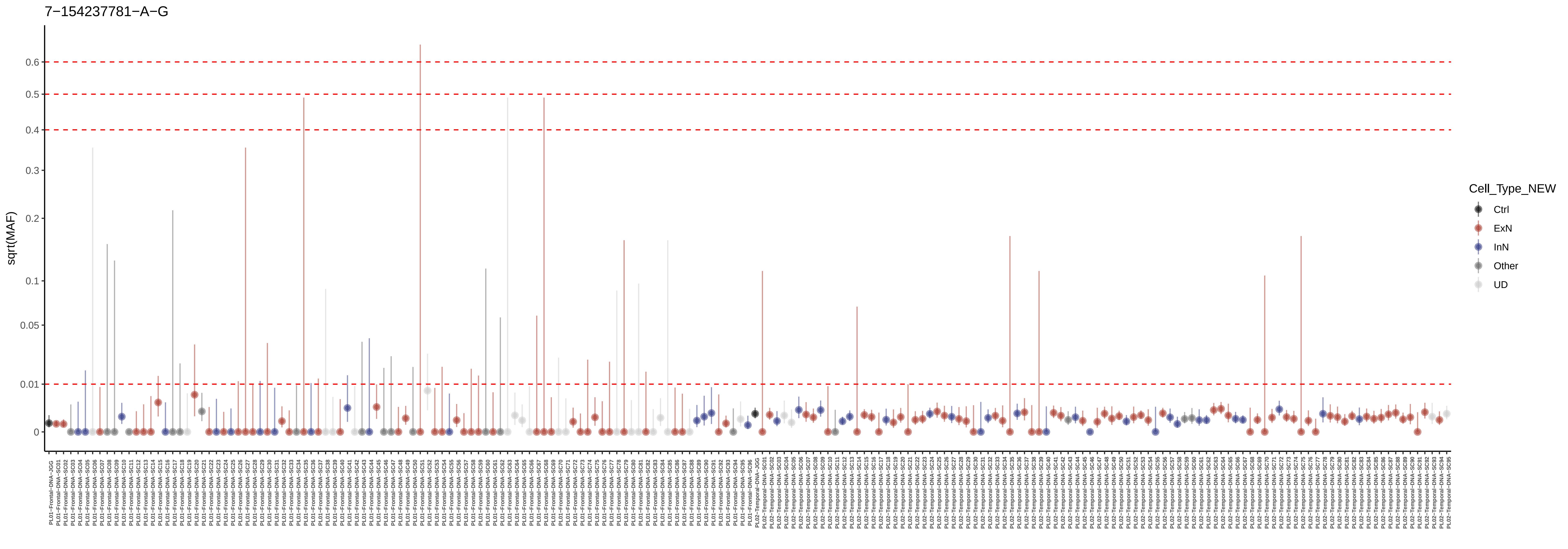

7−154237781−A−G
sqrt(MAF)
0.6
0.5
0.4
0.3
0.2
0.1
0.05
0.01
0
Cell_Type_NEW
Ctrl
ExN
InN
Other
UD

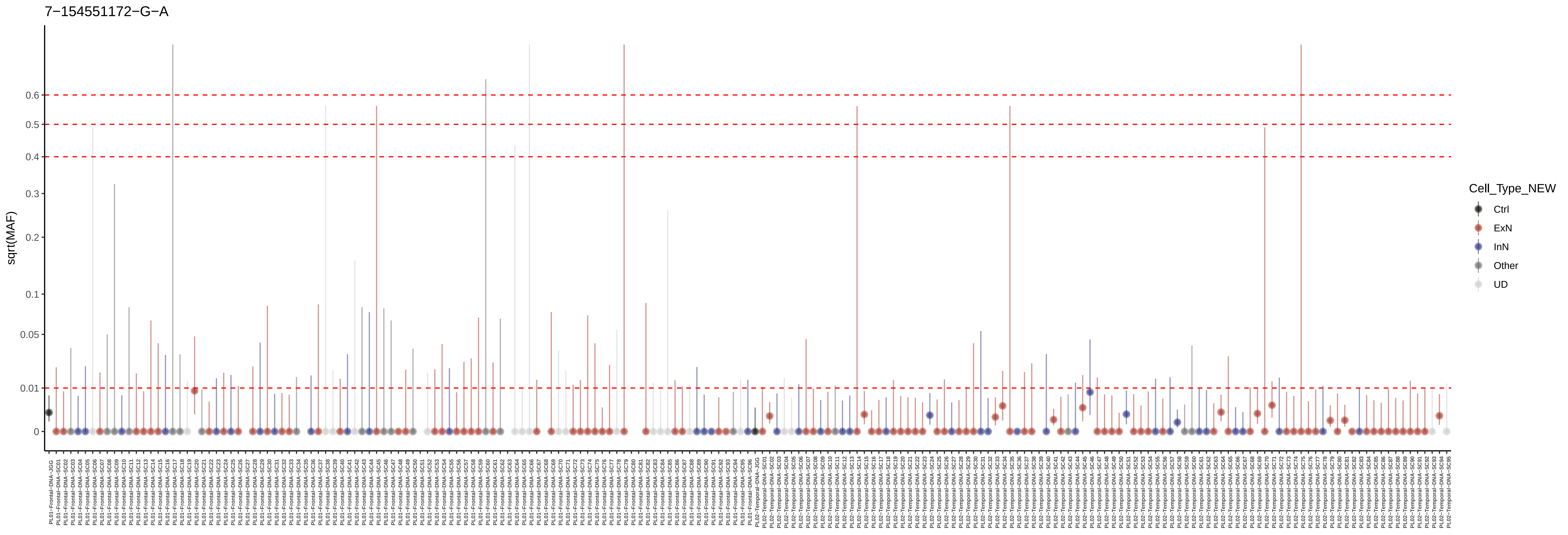

7−154551172−G−A
sqrt(MAF)
0.6
0.5
0.4
0.3
0.2
0.1
0.05
0.01
0
Cell_Type_NEW
Ctrl
ExN
InN
Other
UD

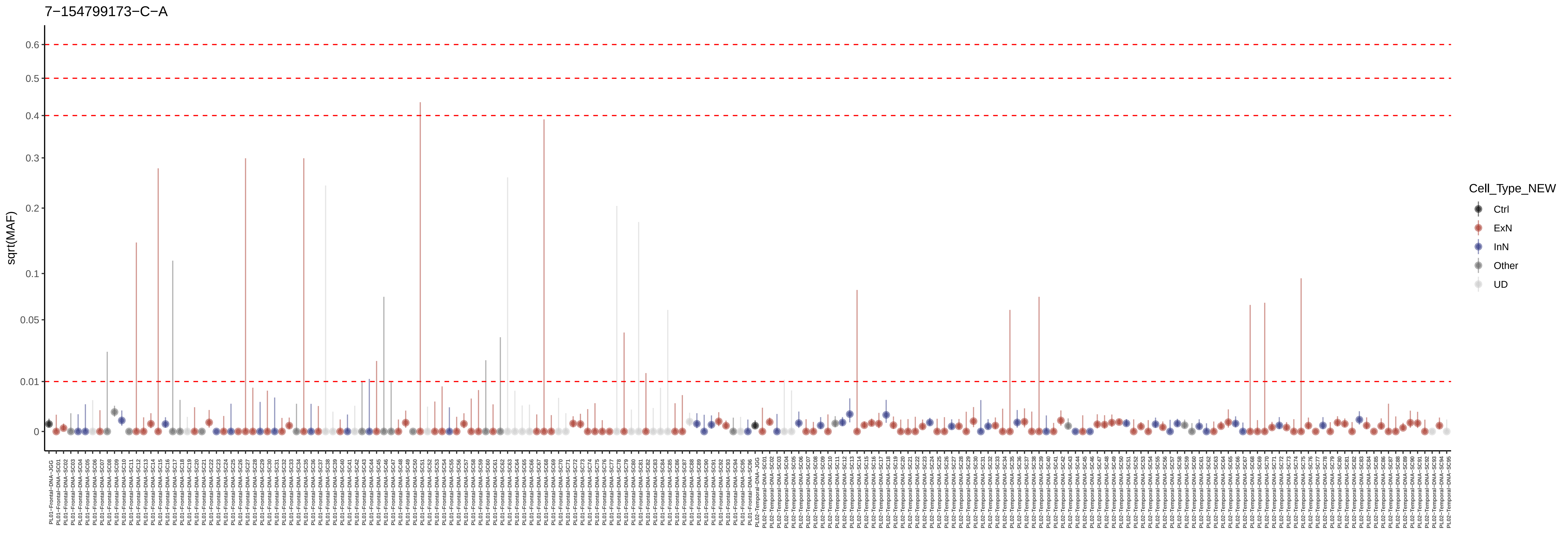
7−154799173−C−A
sqrt(MAF)
0.6
0.5
0.4
0.3
0.2
0.1
0.05
0.01
0
Cell_Type_NEW
Ctrl
ExN
InN
Other
UD

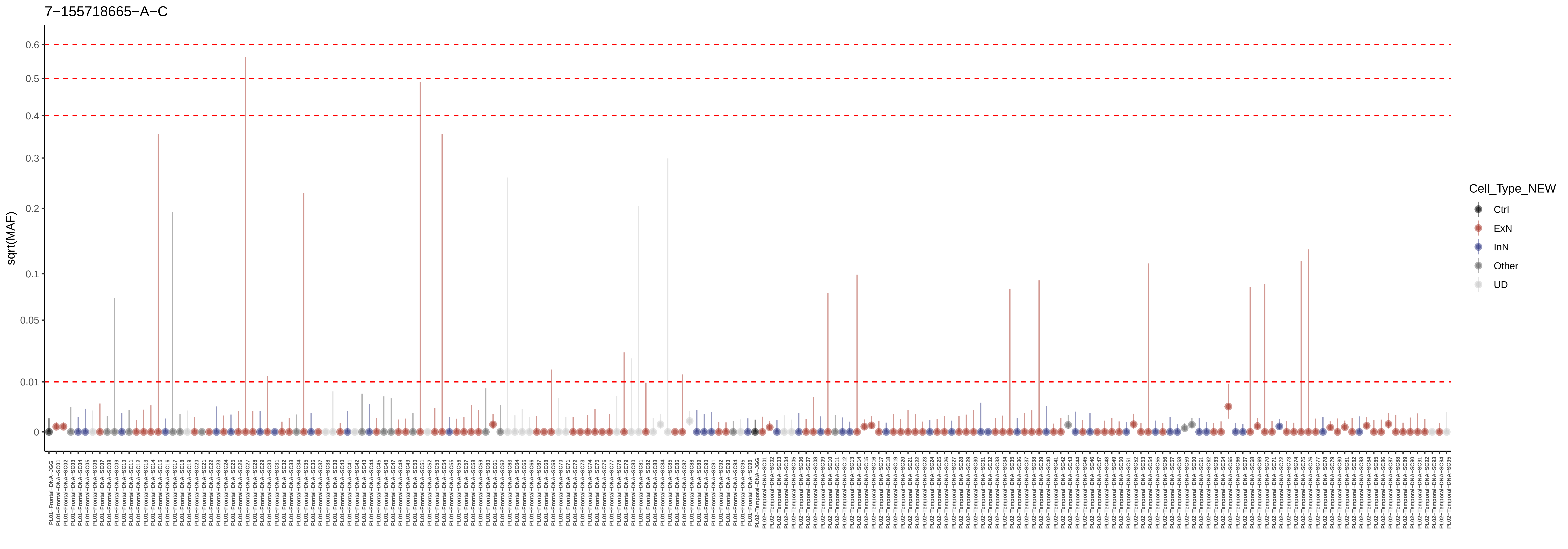
7−155718665−A−C
sqrt(MAF)
0.6
0.5
0.4
0.3
0.2
0.1
0.05
0.01
0
Cell_Type_NEW
Ctrl
ExN
InN
Other
UD

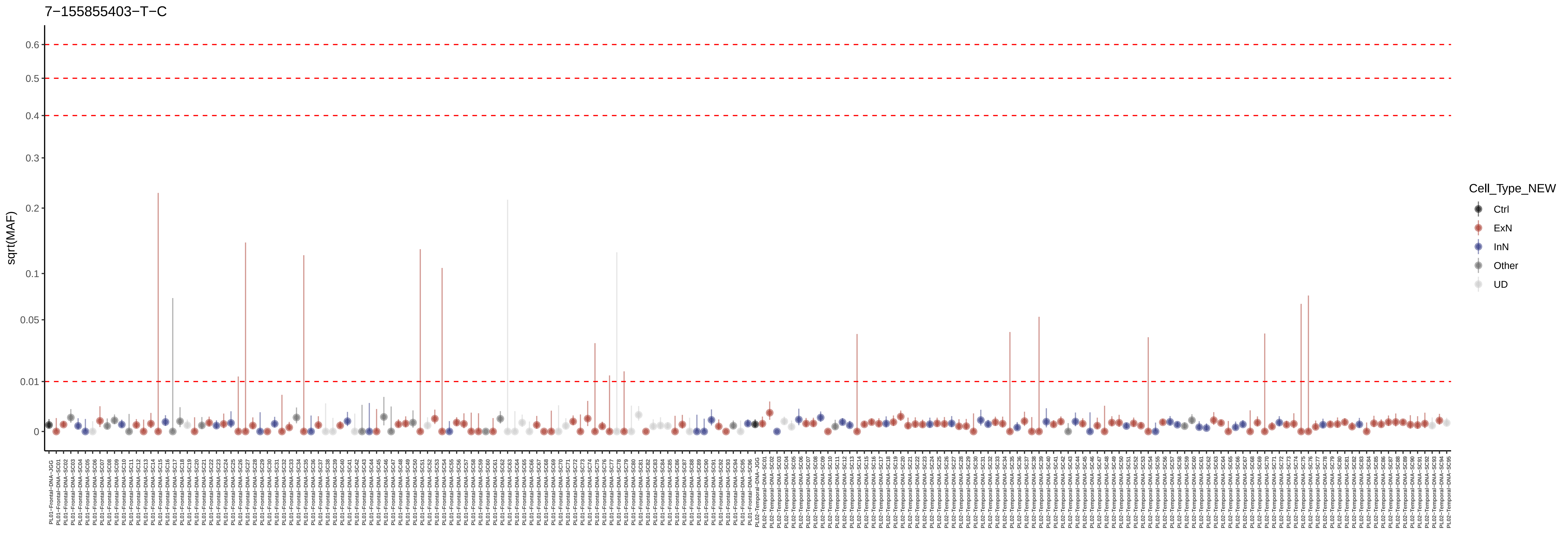

7−155855403−T−C
sqrt(MAF)
0.6
0.5
0.4
0.3
0.2
0.1
0.05
0.01
0
Cell_Type_NEW
Ctrl
ExN
InN
Other
UD

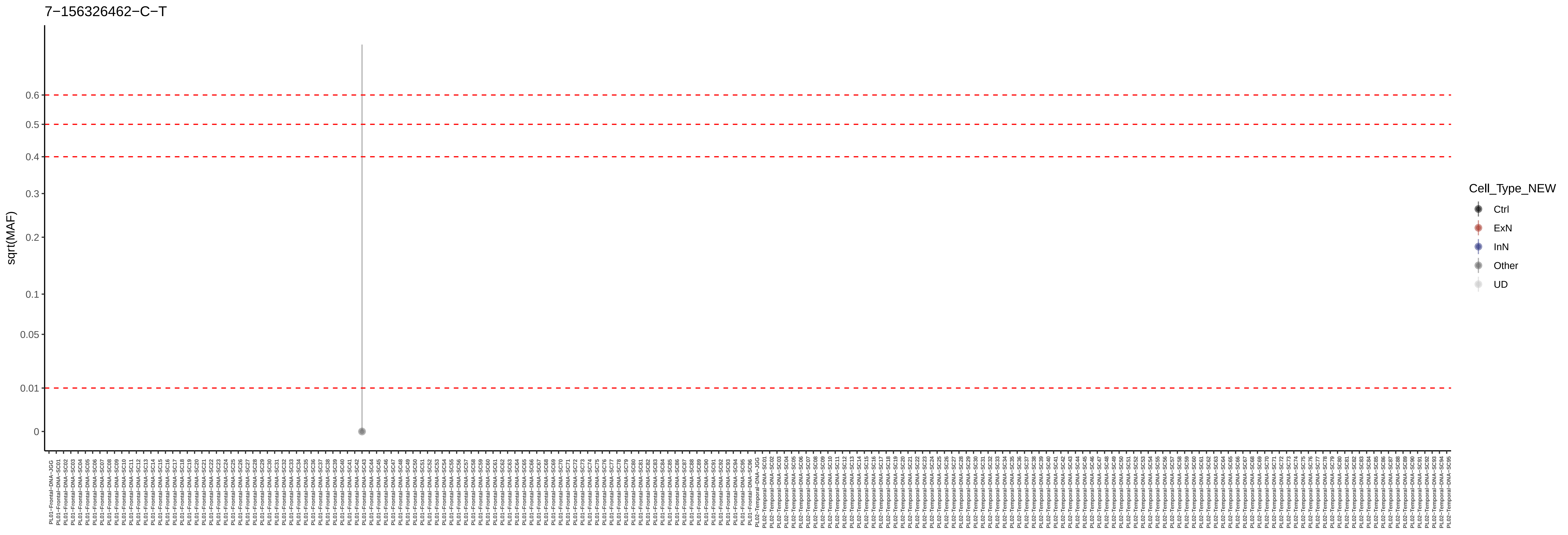
7−156326462−C−T
sqrt(MAF)
0.6
0.5
0.4
0.3
0.2
0.1
0.05
0.01
0
Cell_Type_NEW
Ctrl
ExN
InN
Other
UD

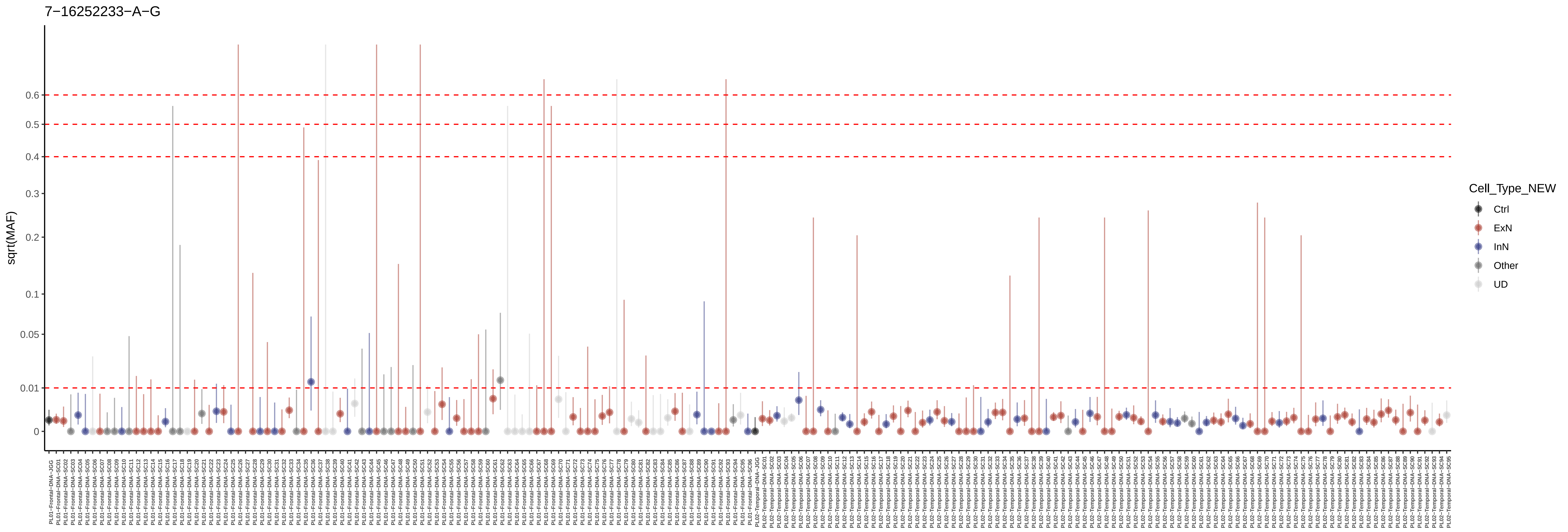
7−16252233−A−G
sqrt(MAF)
0.6
0.5
0.4
0.3
0.2
0.1
0.05
0.01
0
Cell_Type_NEW
Ctrl
ExN
InN
Other
UD

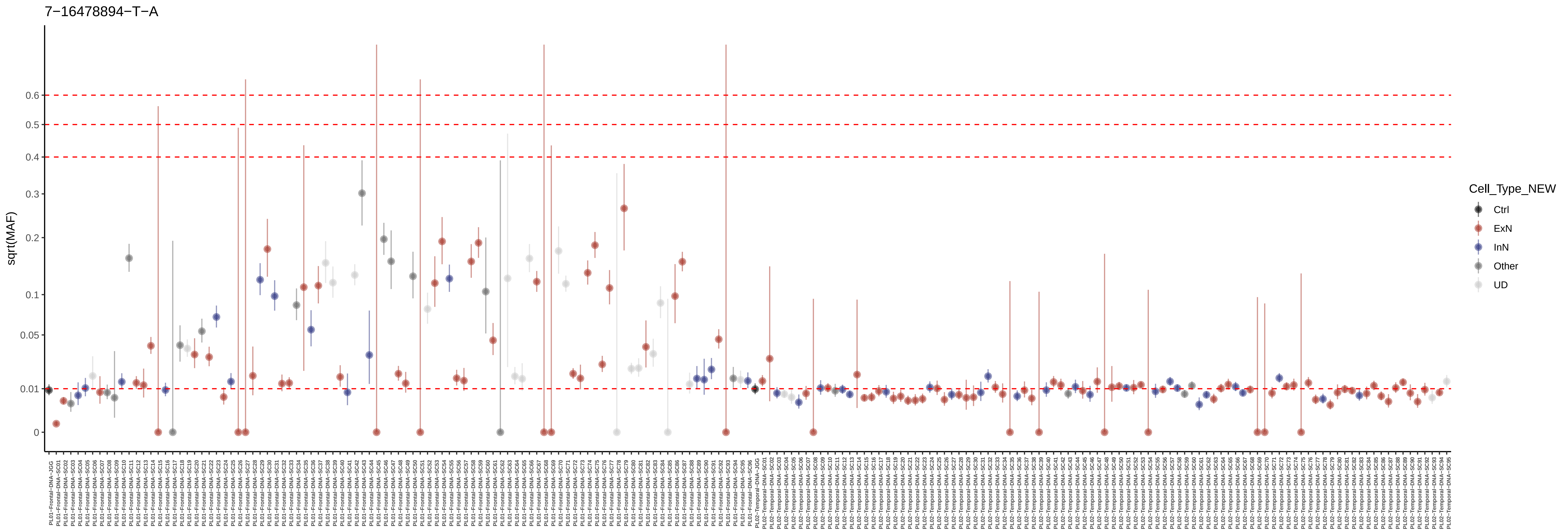

7−16478894−T−A
sqrt(MAF)
0.6
0.5
0.4
0.3
0.2
0.1
0.05
0.01
0
Cell_Type_NEW
Ctrl
ExN
InN
Other
UD

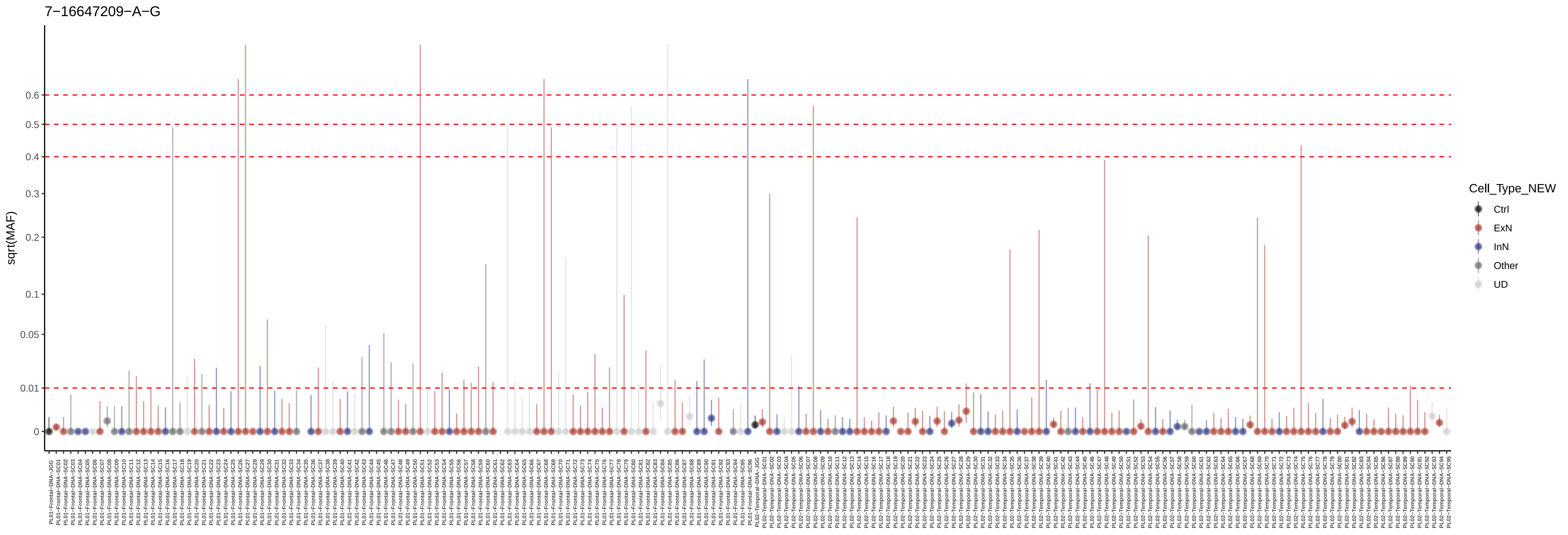

7−16647209−A−G
sqrt(MAF)
0.6
0.5
0.4
0.3
0.2
0.1
0.05
0.01
0
Cell_Type_NEW
Ctrl
ExN
InN
Other
UD

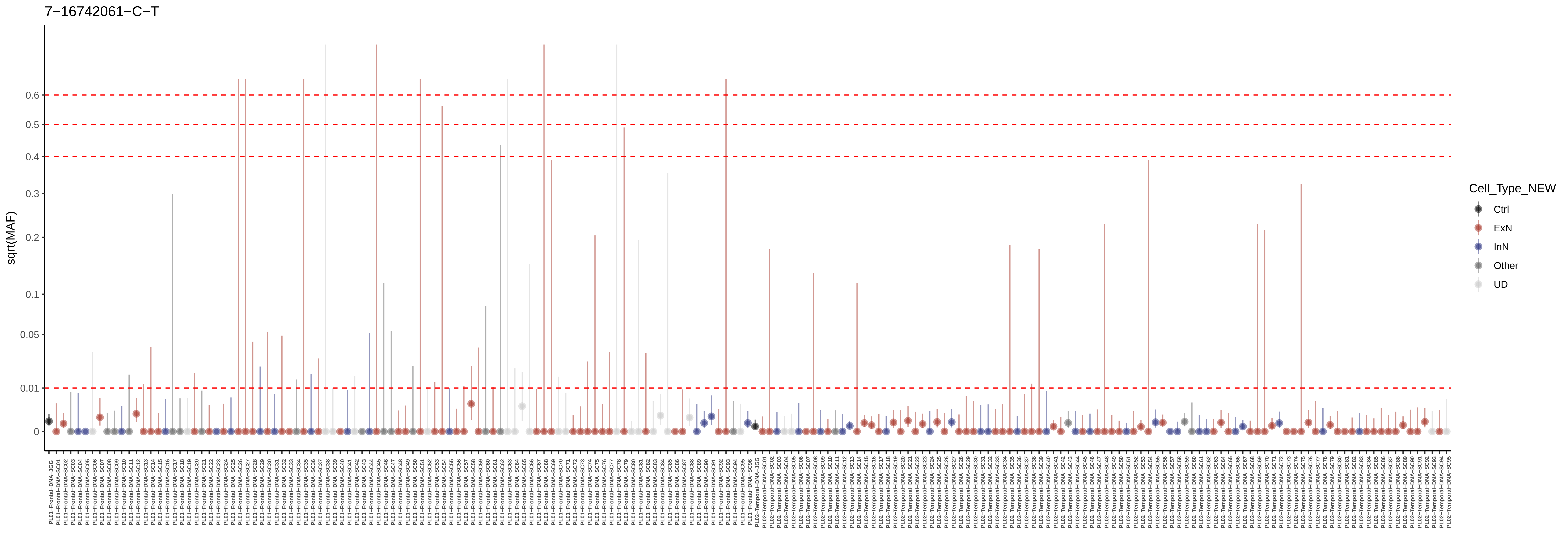
7−16742061−C−T
sqrt(MAF)
0.6
0.5
0.4
0.3
0.2
0.1
0.05
0.01
0
Cell_Type_NEW
Ctrl
ExN
InN
Other
UD

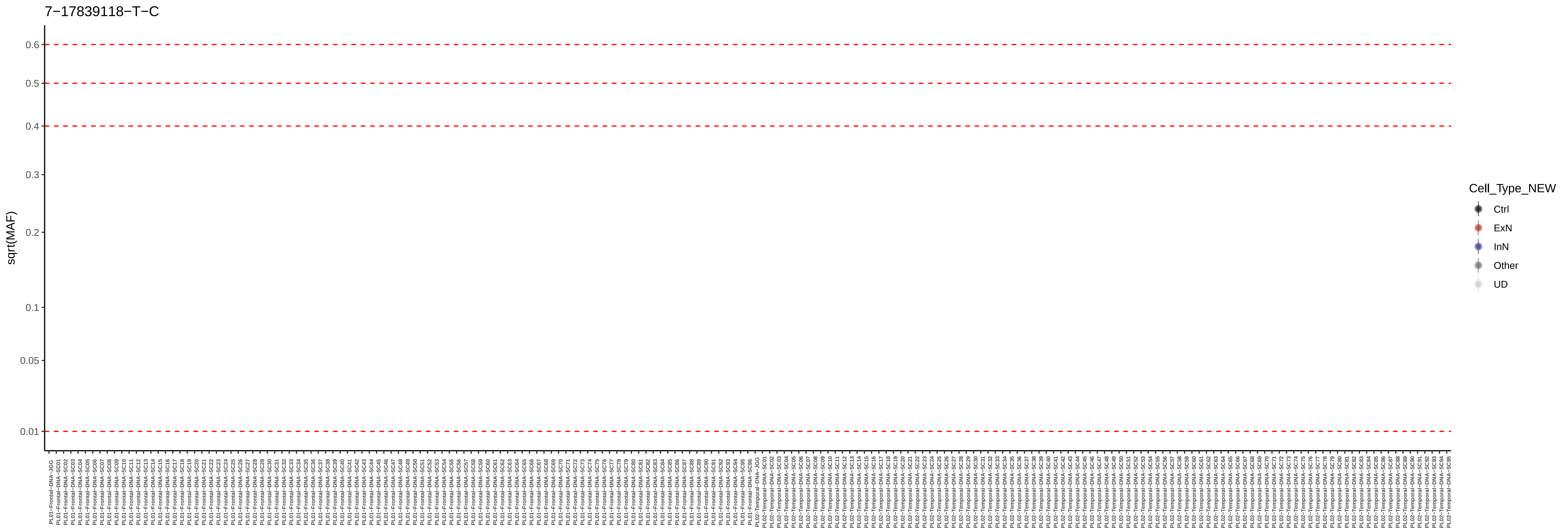
7−17839118−T−C
sqrt(MAF)
0.6
0.5
0.4
0.3
0.2
0.1
0.05
0.01
Cell_Type_NEW
Ctrl
ExN
InN
Other
UD

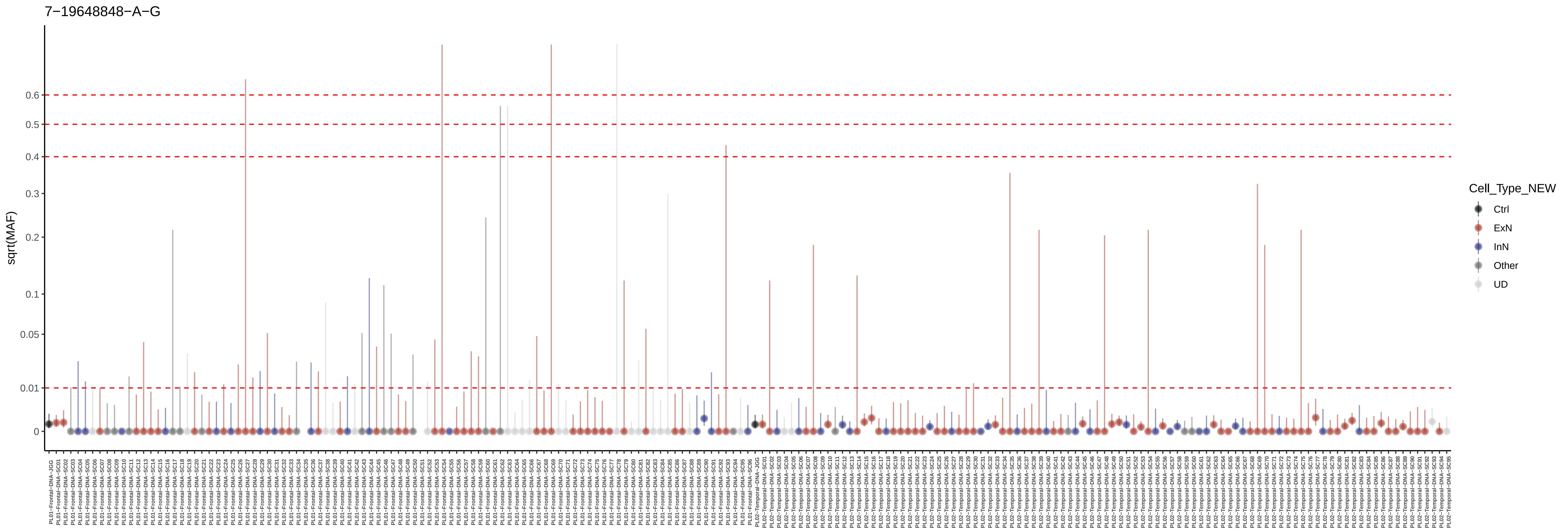
7−19648848−A−G
sqrt(MAF)
0
0.01
0.05
0.1
0.2
0.3
0.4
0.5
0.6
Cell_Type_NEW
Ctrl
ExN
InN
Other
UD

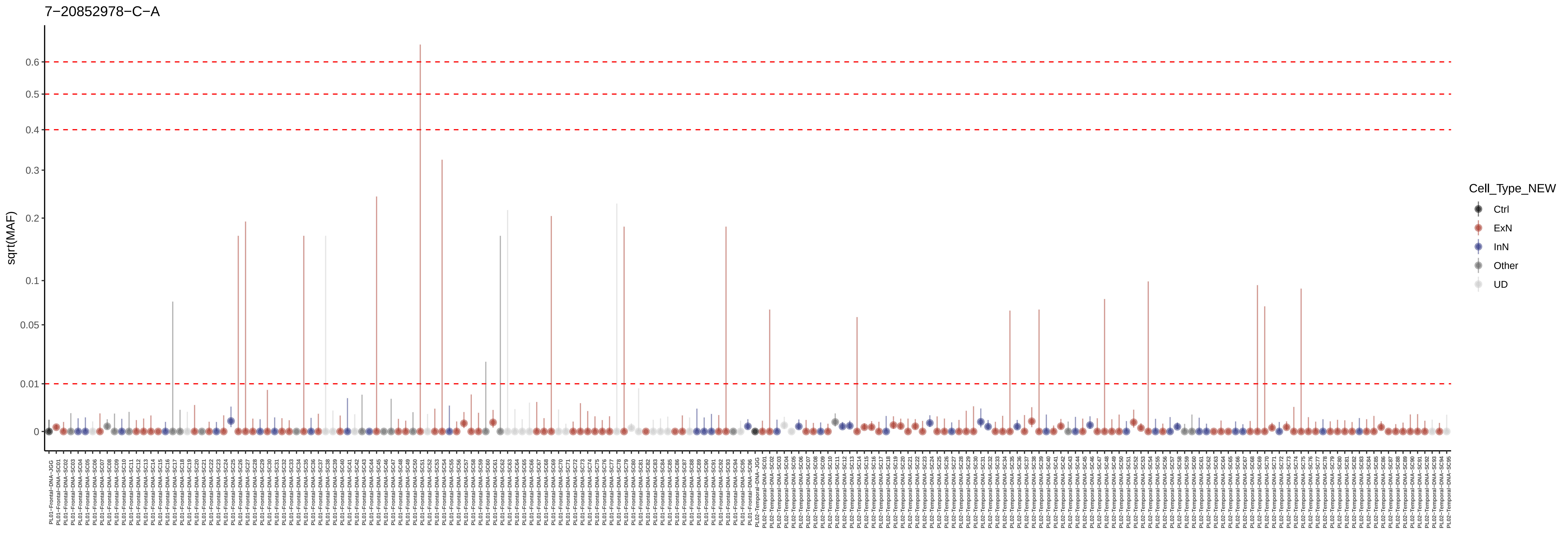
7−20852978−C−A
sqrt(MAF)
0.6
0.5
0.4
0.3
0.2
0.1
0.05
0.01
0
Cell_Type_NEW
Ctrl
ExN
InN
Other
UD

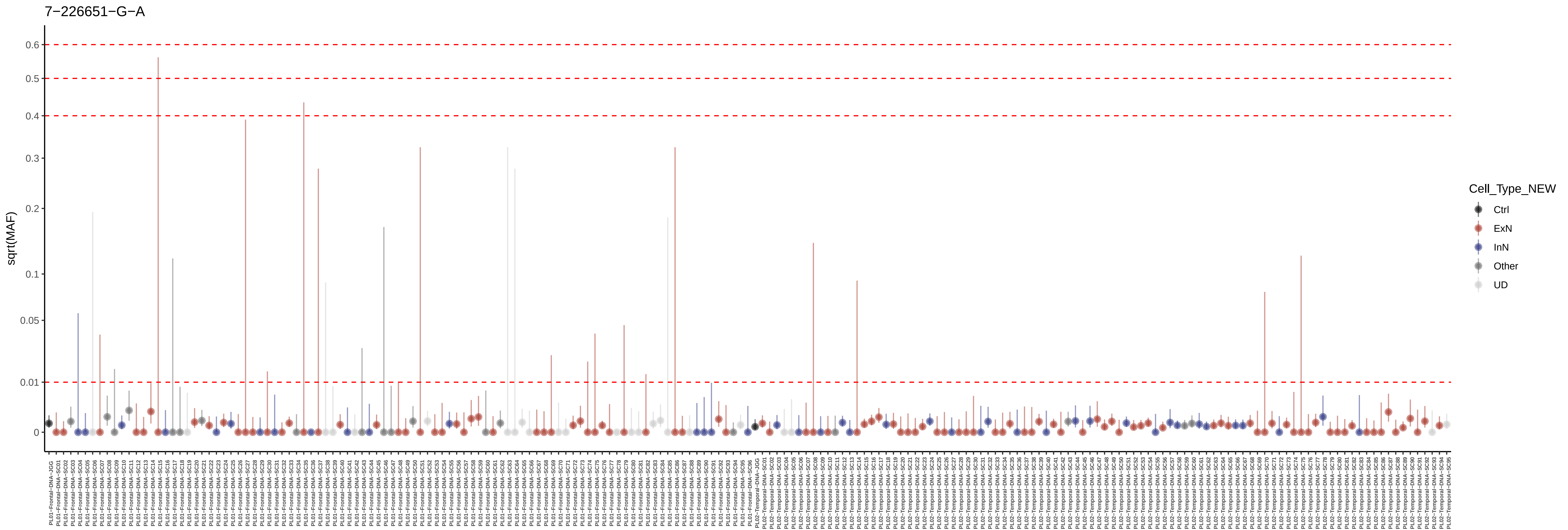
7−226651−G−A
sqrt(MAF)
0.6
0.5
0.4
0.3
0.2
0.1
0.05
0.01
0
Cell_Type_NEW
Ctrl
ExN
InN
Other
UD

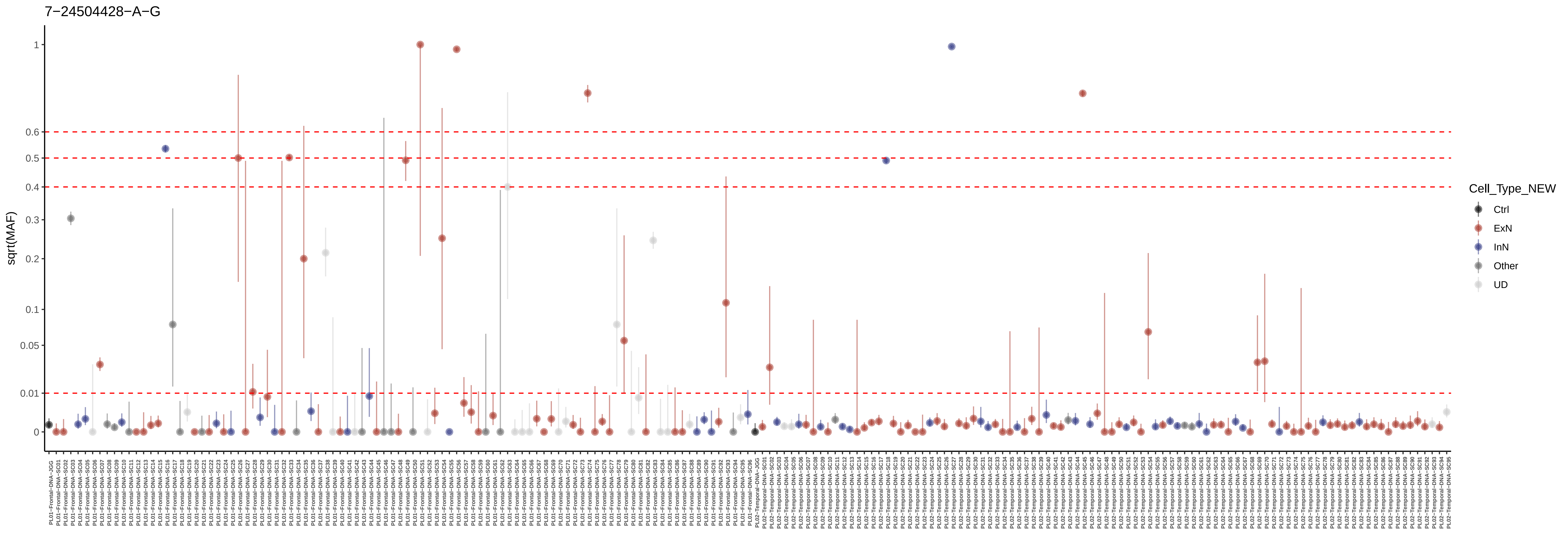
7−24504428−A−G
sqrt(MAF)
1
0.6
0.5
0.4
0.3
0.2
0.1
0.05
0.01
0
Cell_Type_NEW
Ctrl
ExN
InN
Other
UD

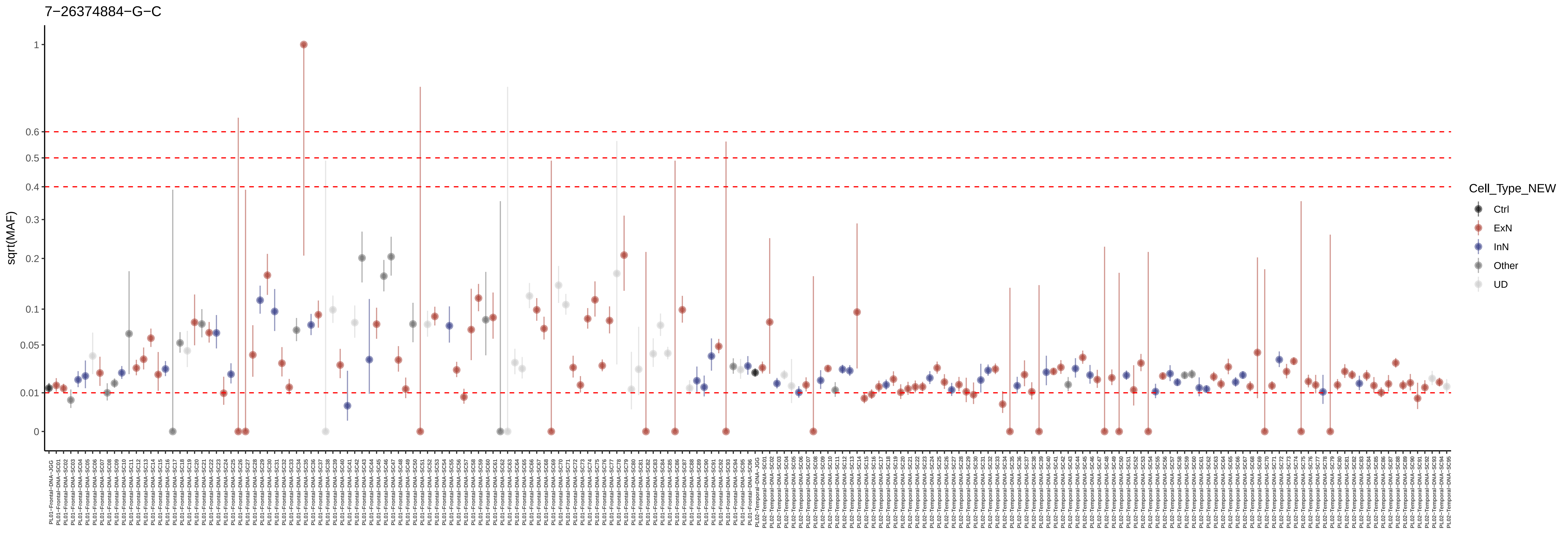

7−26374884−G−C
sqrt(MAF)
1
0.6
0.5
0.4
0.3
0.2
0.1
0.05
0.01
0
Cell_Type_NEW
Ctrl
ExN
InN
Other
UD

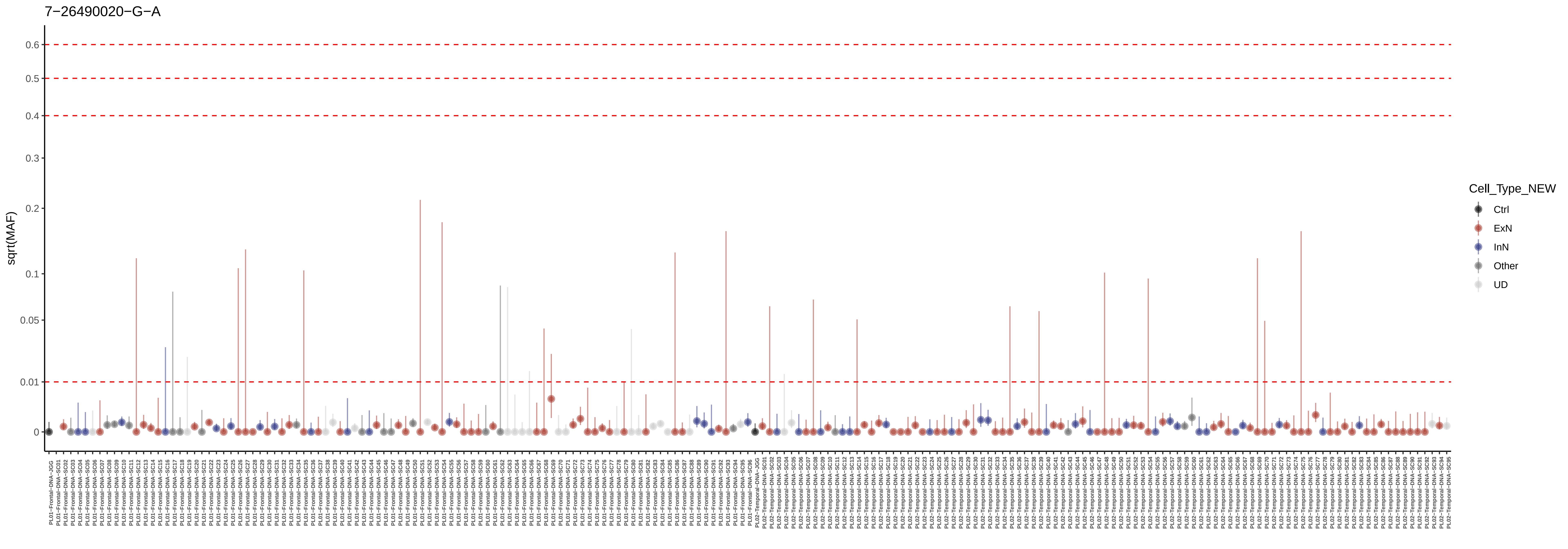
7−26490020−G−A
sqrt(MAF)
0.6
0.5
0.4
0.3
0.2
0.1
0.05
0.01
0
Cell_Type_NEW
Ctrl
ExN
InN
Other
UD

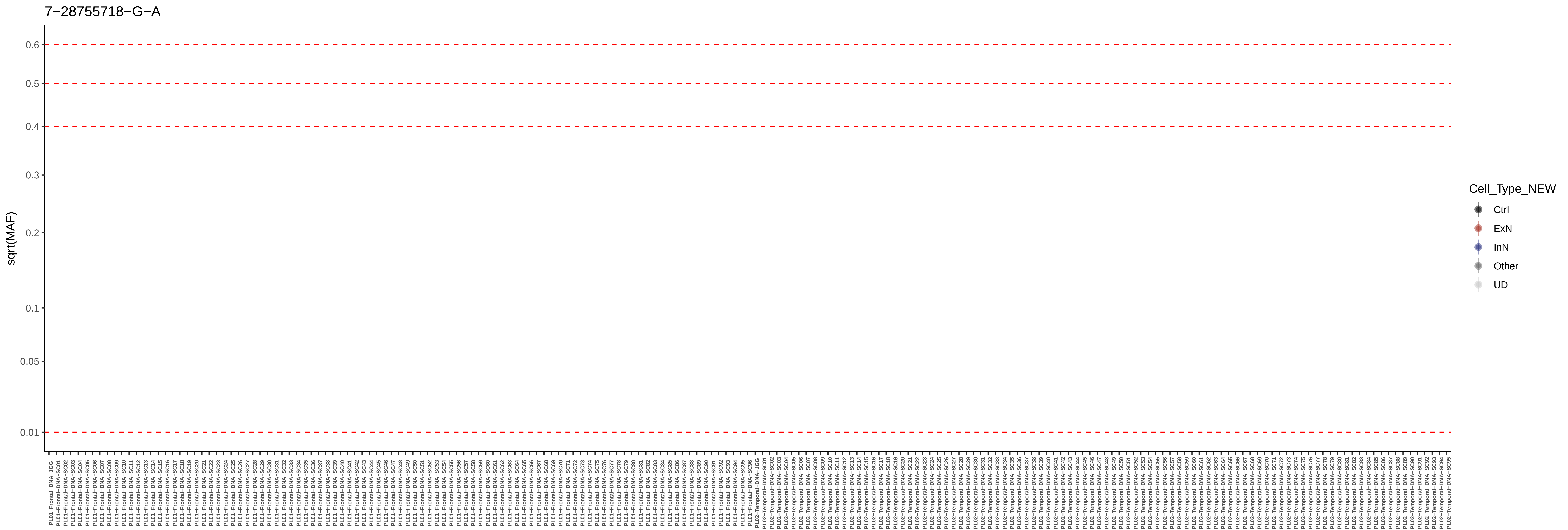
7−28755718−G−A
sqrt(MAF)
0.6
0.5
0.4
0.3
0.2
0.1
0.05
0.01
Cell_Type_NEW
Ctrl
ExN
InN
Other
UD

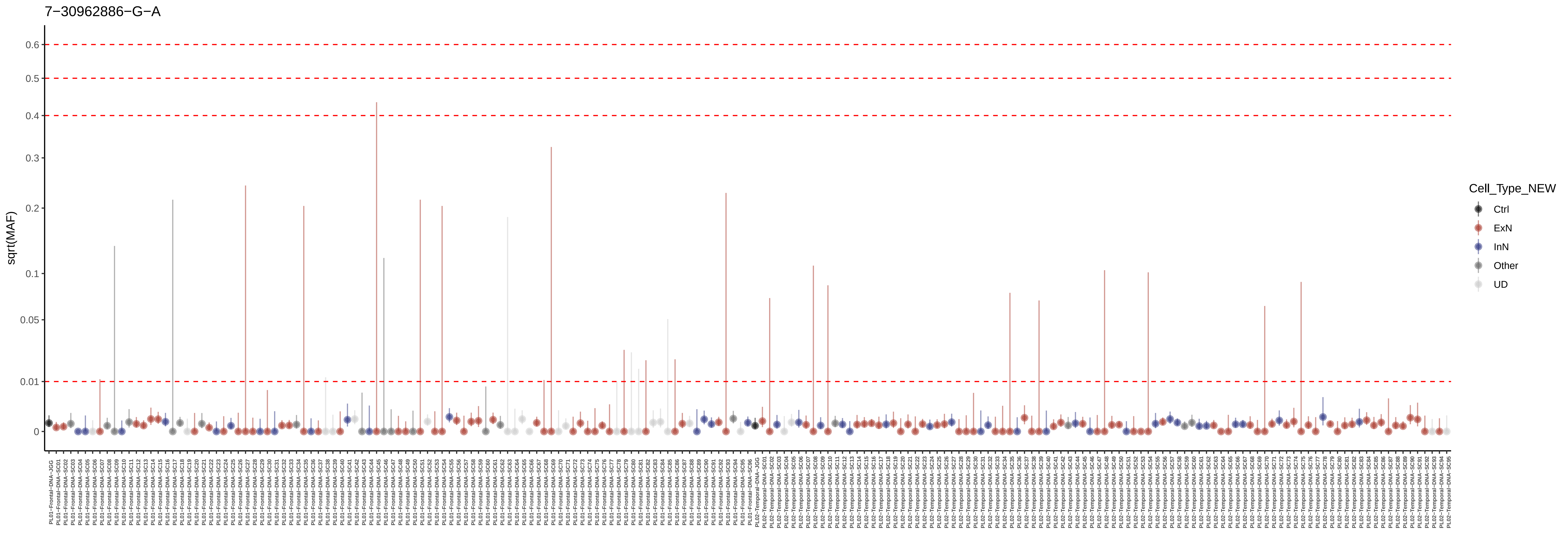
7−30962886−G−A
sqrt(MAF)
0.6
0.5
0.4
0.3
0.2
0.1
0.05
0.01
0
Cell_Type_NEW
Ctrl
ExN
InN
Other
UD

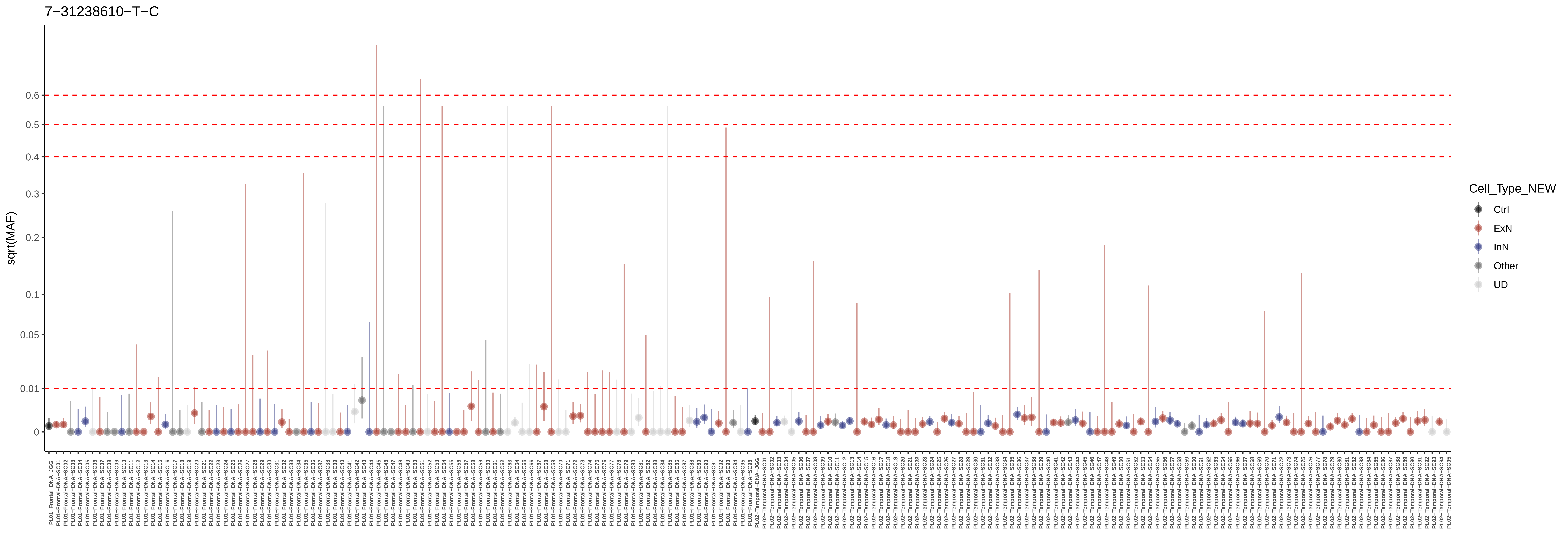
7−31238610−T−C
sqrt(MAF)
0.6
0.5
0.4
0.3
0.2
0.1
0.05
0.01
0
Cell_Type_NEW
Ctrl
ExN
InN
Other
UD

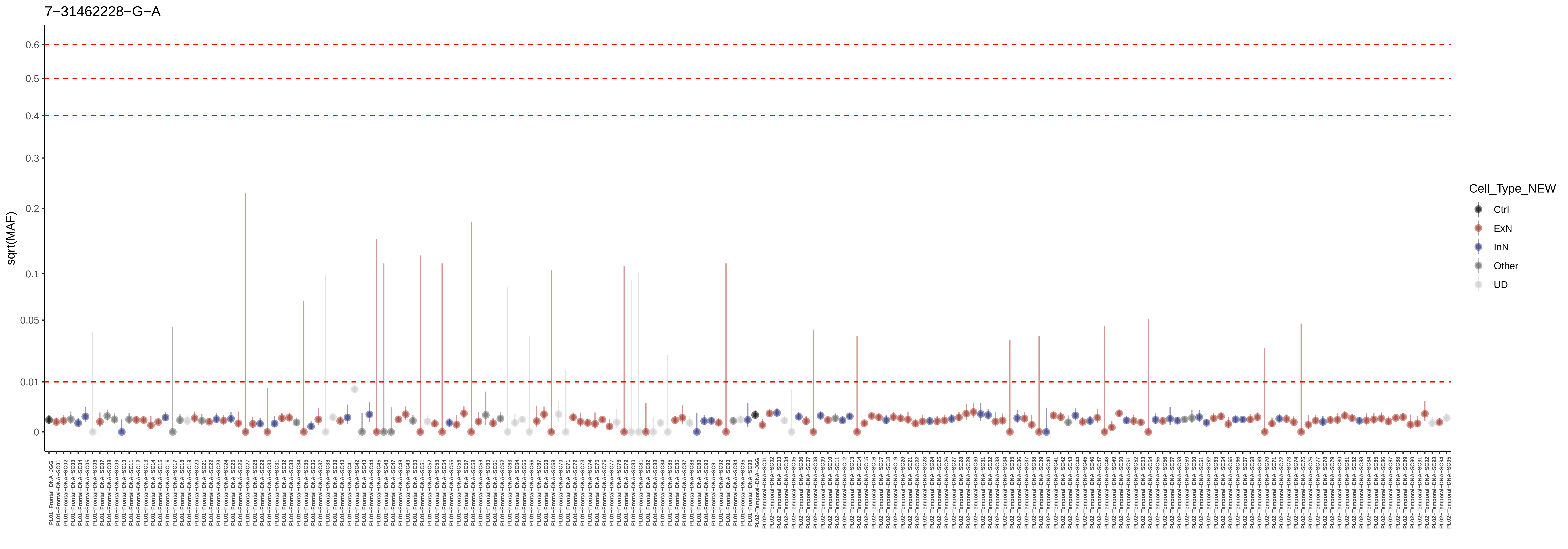
7−31462228−G−A
sqrt(MAF)
0.6
0.5
0.4
0.3
0.2
0.1
0.05
0.01
0
Cell_Type_NEW
Ctrl
ExN
InN
Other
UD

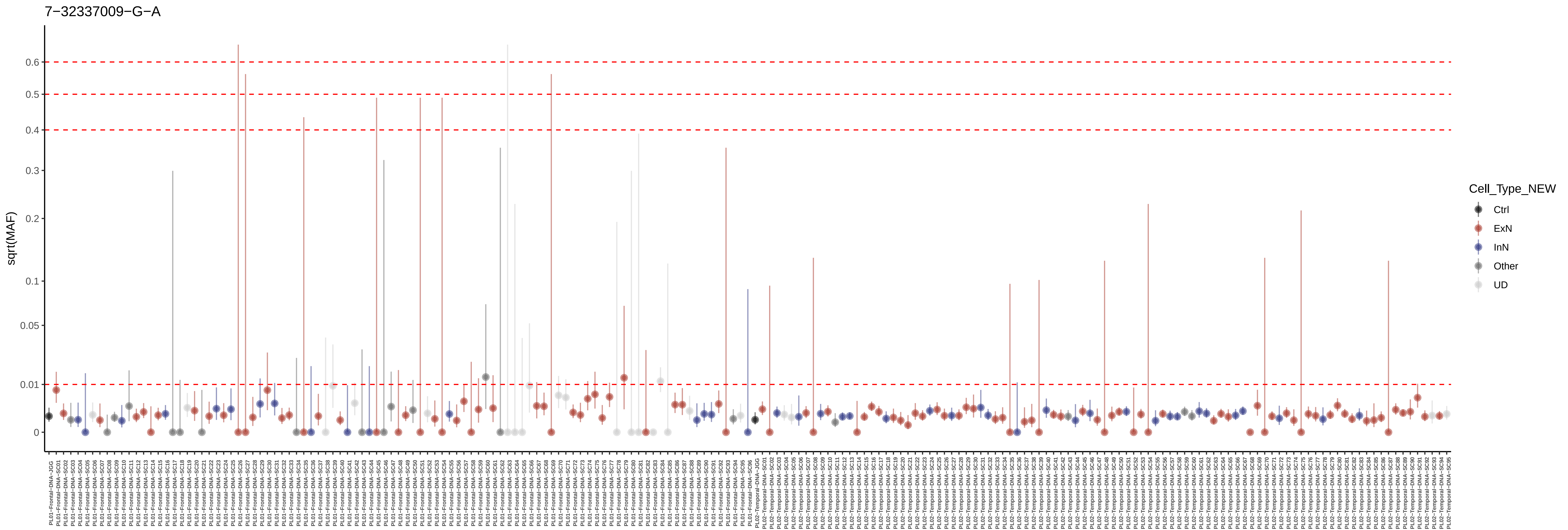
7−32337009−G−A
sqrt(MAF)
0.6
0.5
0.4
0.3
0.2
0.1
0.05
0.01
0
Cell_Type_NEW
Ctrl
ExN
InN
Other
UD

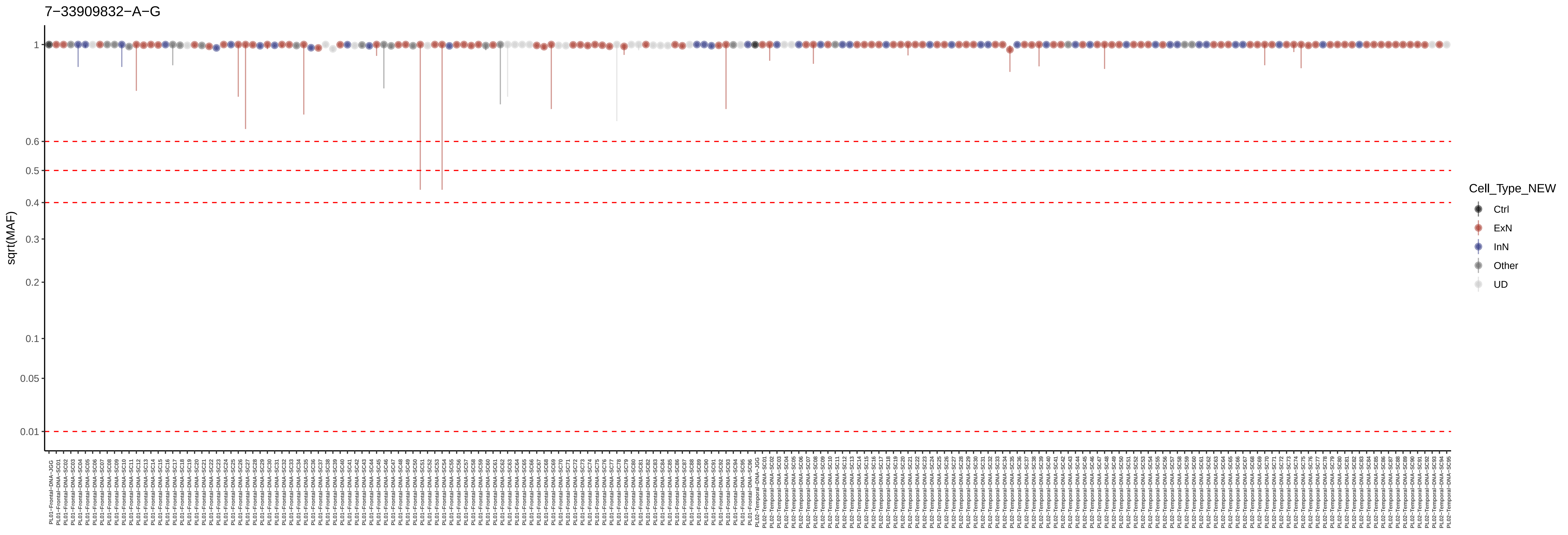
7−33909832−A−G
sqrt(MAF)
1
0.6
0.5
0.4
0.3
0.2
0.1
0.05
0.01
Cell_Type_NEW
Ctrl
ExN
InN
Other
UD

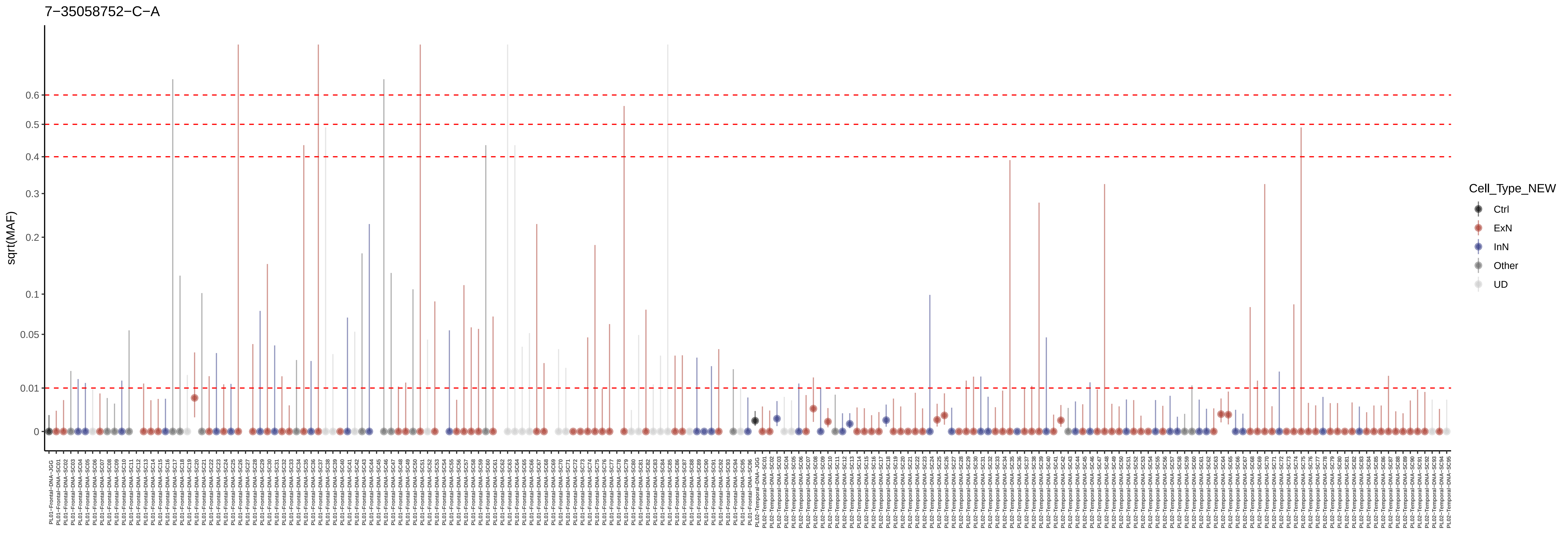

7−35058752−C−A
sqrt(MAF)
0.6
0.5
0.4
0.3
0.2
0.1
0.05
0.01
0
Cell_Type_NEW
Ctrl
ExN
InN
Other
UD

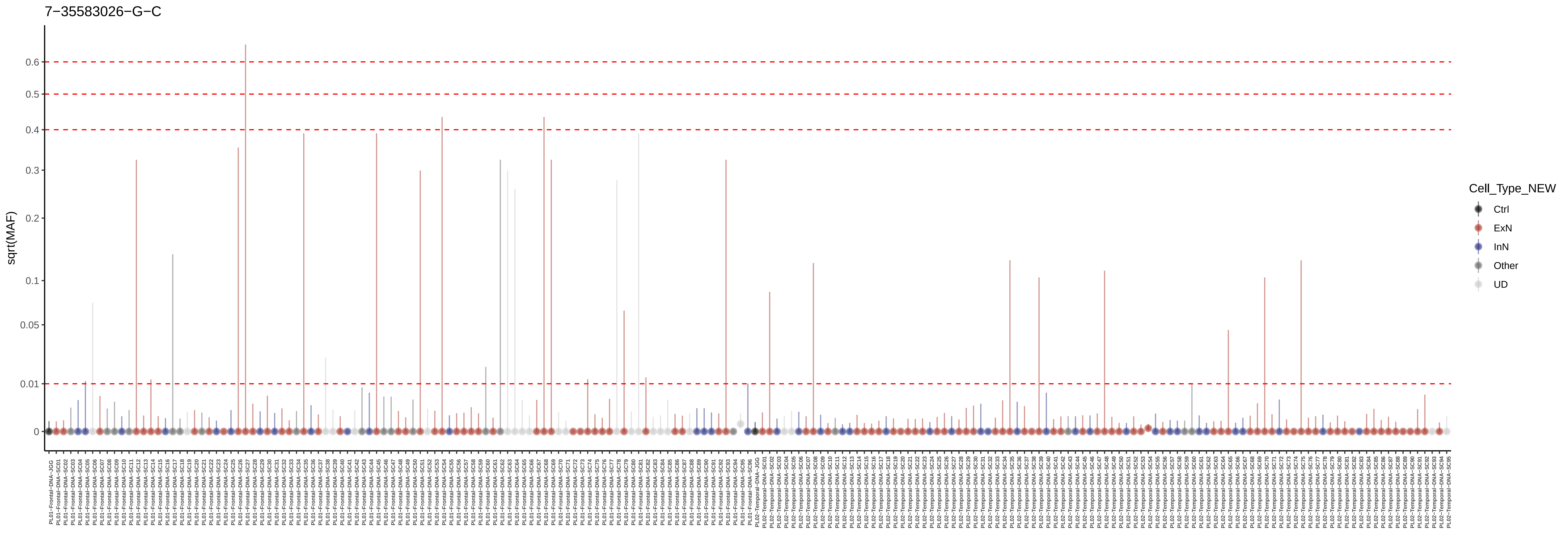

7−35583026−G−C
sqrt(MAF)
0.6
0.5
0.4
0.3
0.2
0.1
0.05
0.01
0
Cell_Type_NEW
Ctrl
ExN
InN
Other
UD

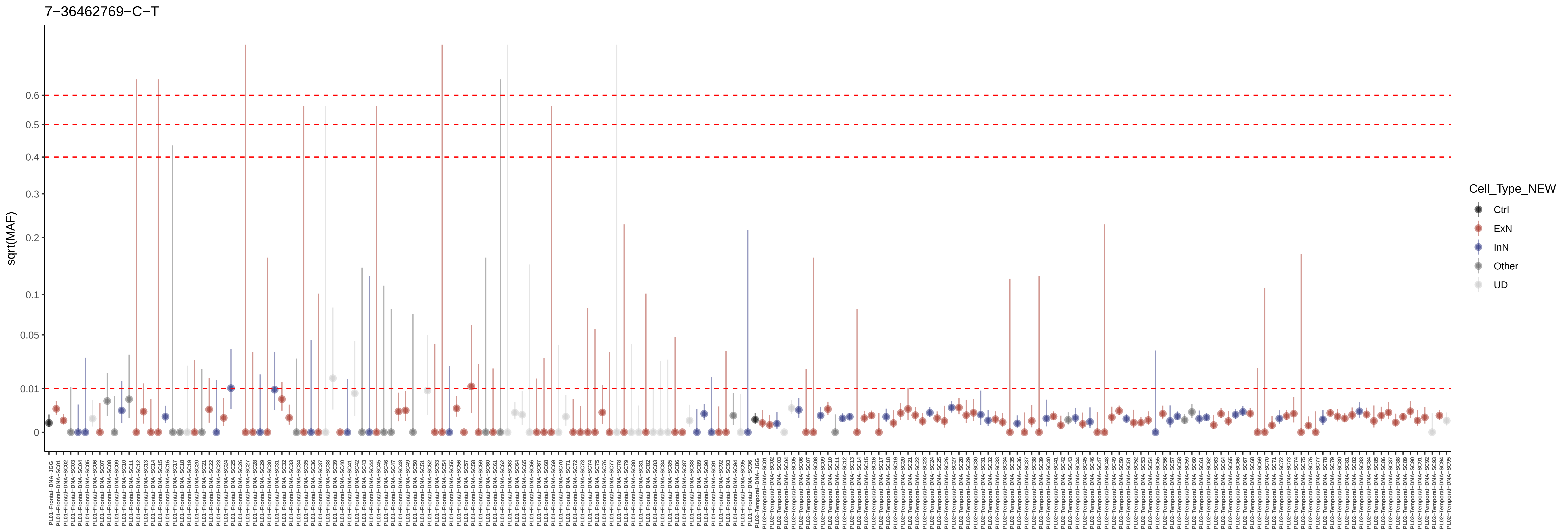
7−36462769−C−T
sqrt(MAF)
0.6
0.5
0.4
0.3
0.2
0.1
0.05
0.01
0
Cell_Type_NEW
Ctrl
ExN
InN
Other
UD

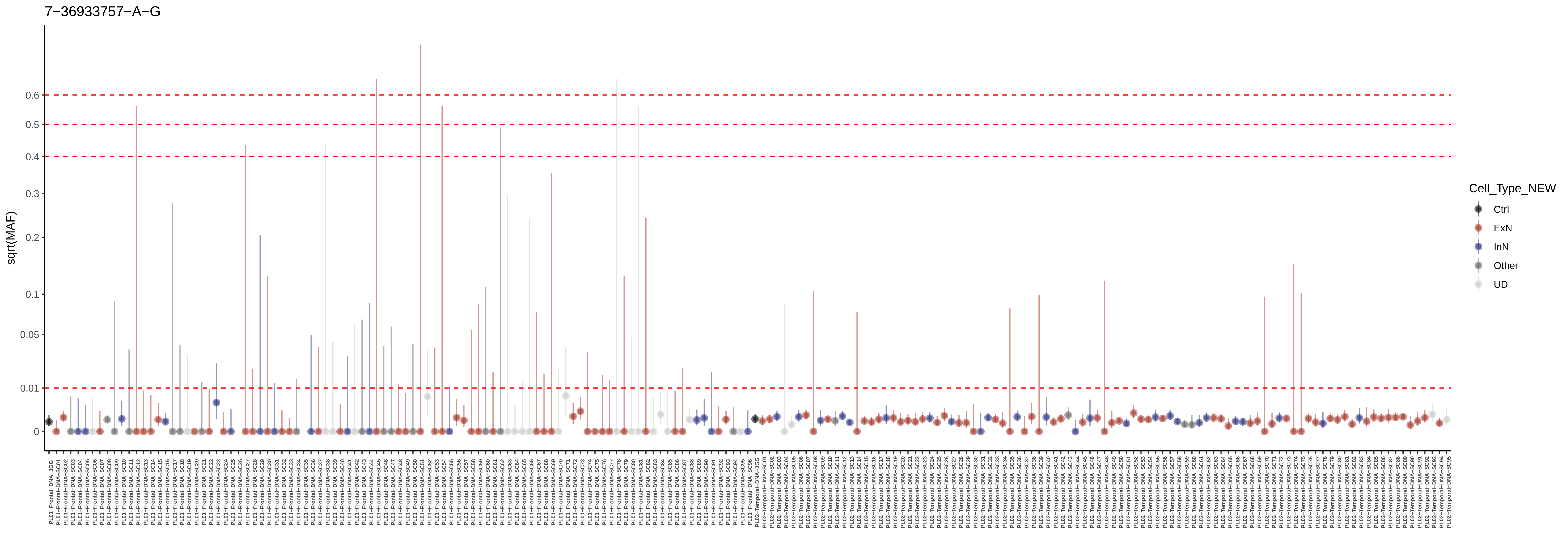

7−36933757−A−G
sqrt(MAF)
0.6
0.5
0.4
0.3
0.2
0.1
0.05
0.01
0
Cell_Type_NEW
Ctrl
ExN
InN
Other
UD

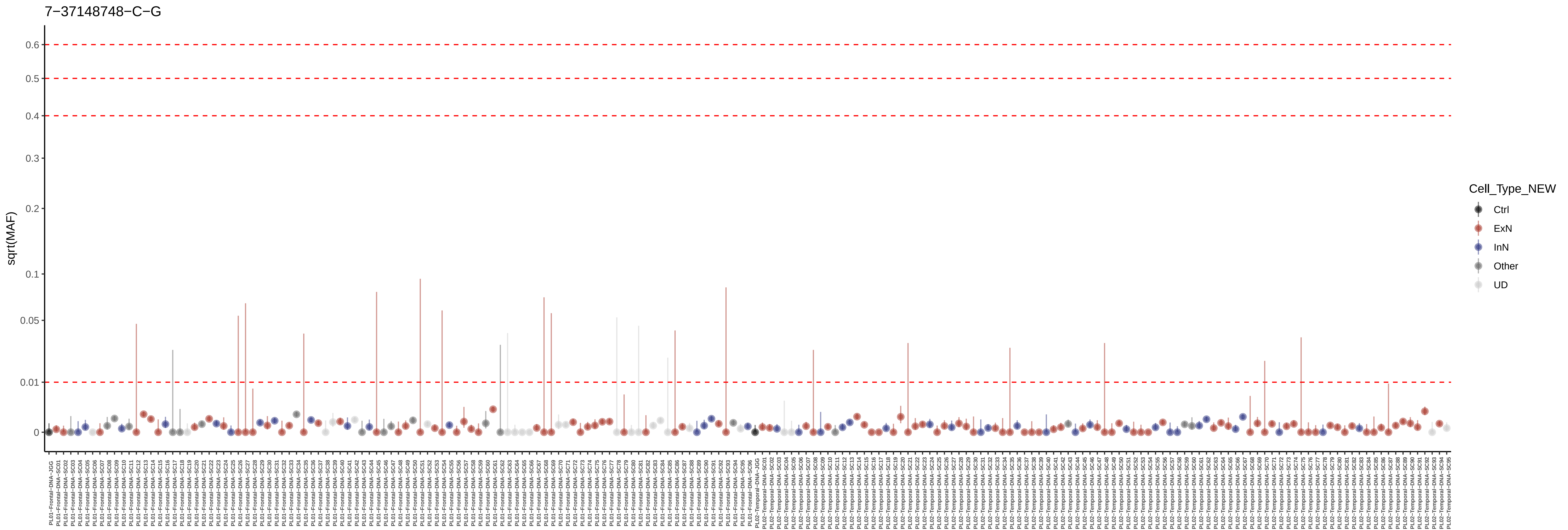
7−37148748−C−G
sqrt(MAF)
0.6
0.5
0.4
0.3
0.2
0.1
0.05
0.01
0
Cell_Type_NEW
Ctrl
ExN
InN
Other
UD

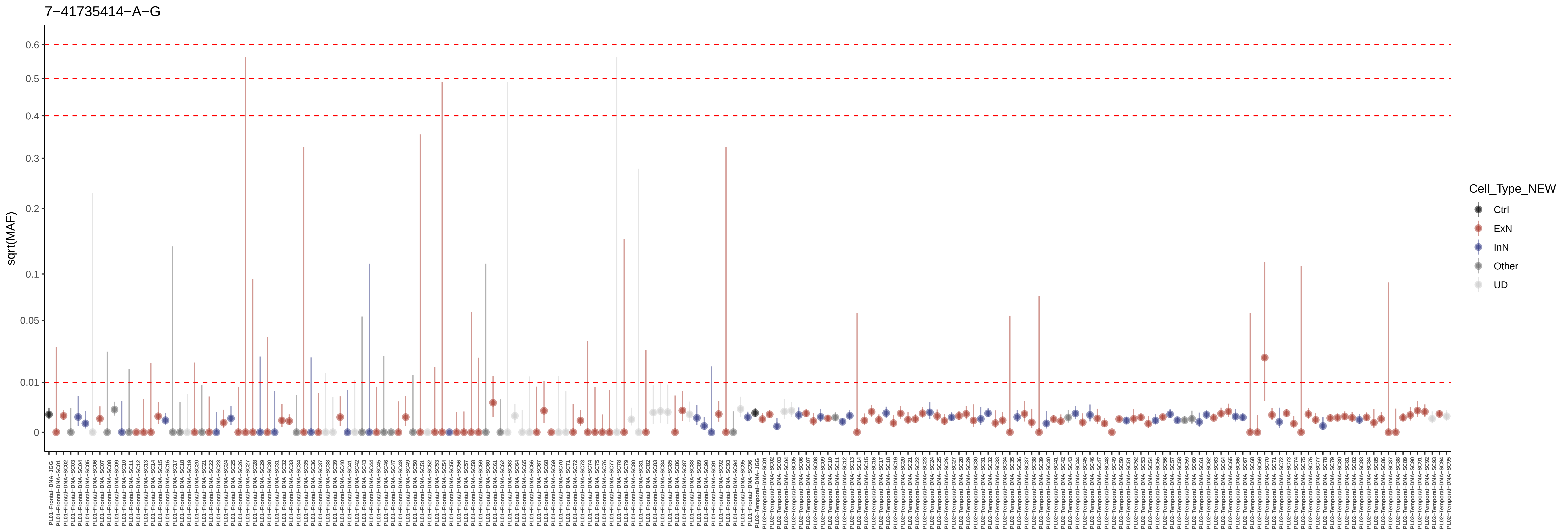
7−41735414−A−G
sqrt(MAF)
0.6
0.5
0.4
0.3
0.2
0.1
0.05
0.01
0
Cell_Type_NEW
Ctrl
ExN
InN
Other
UD

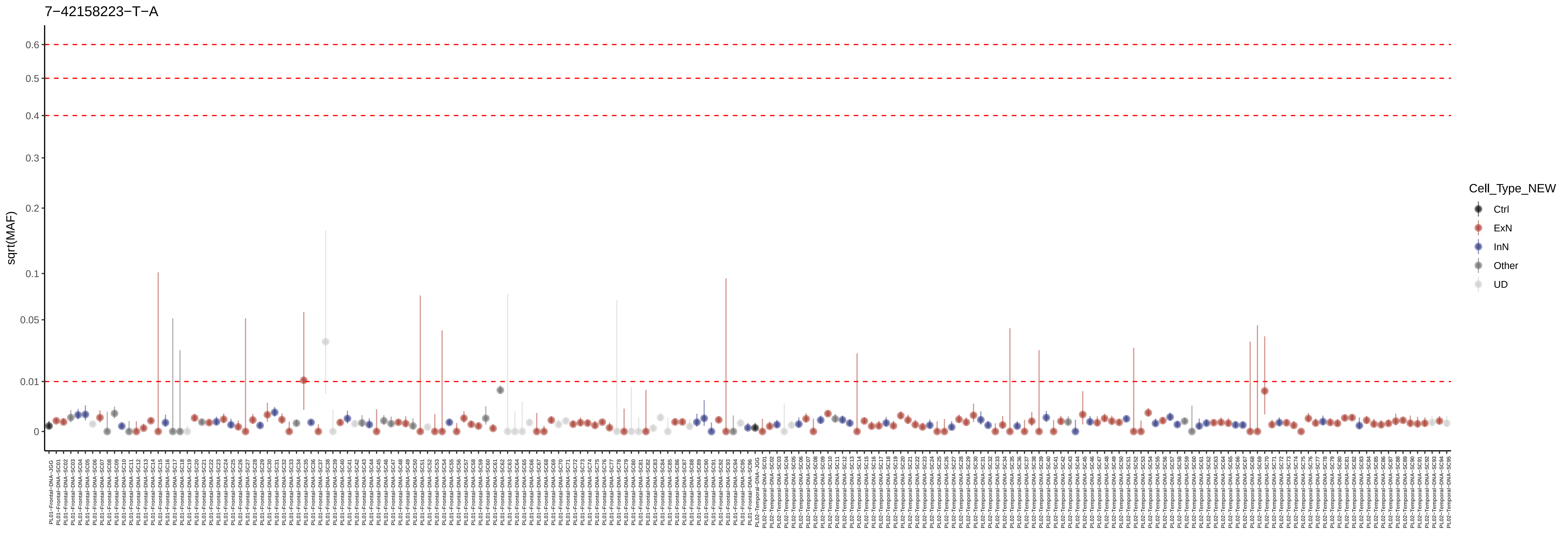
7−42158223−T−A
sqrt(MAF)
0.6
0.5
0.4
0.3
0.2
0.1
0.05
0.01
0
Cell_Type_NEW
Ctrl
ExN
InN
Other
UD

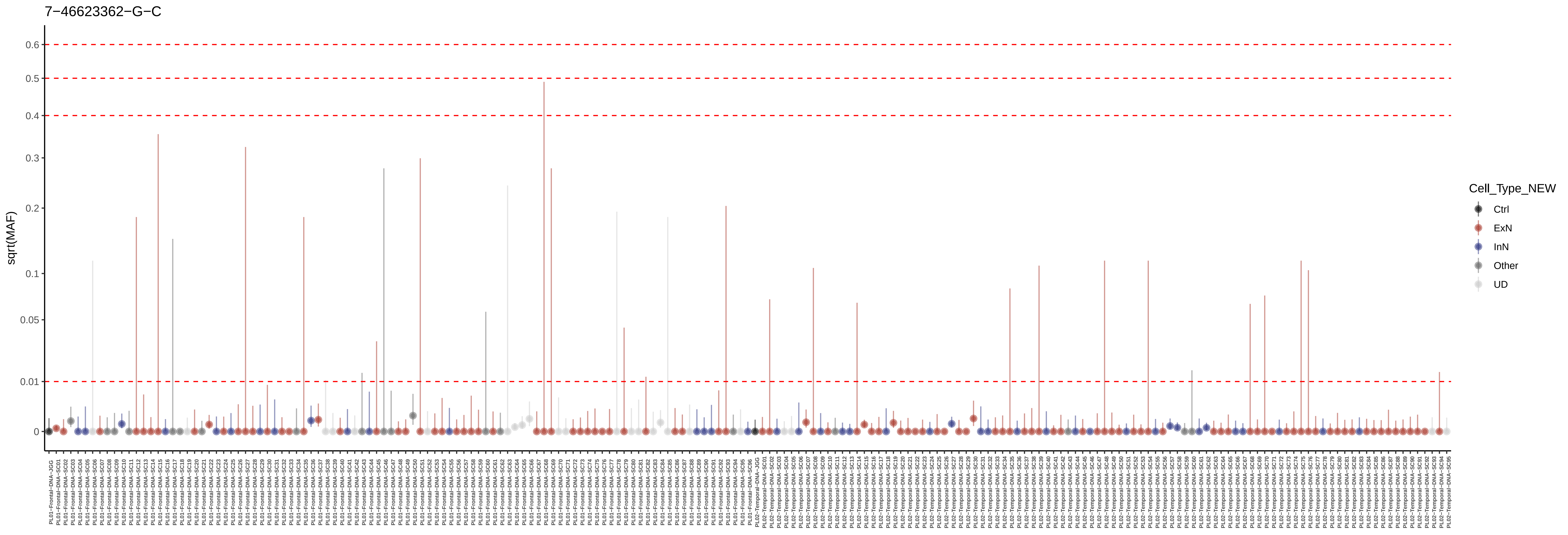

7−46623362−G−C
sqrt(MAF)
0.6
0.5
0.4
0.3
0.2
0.1
0.05
0.01
0
Cell_Type_NEW
Ctrl
ExN
InN
Other
UD

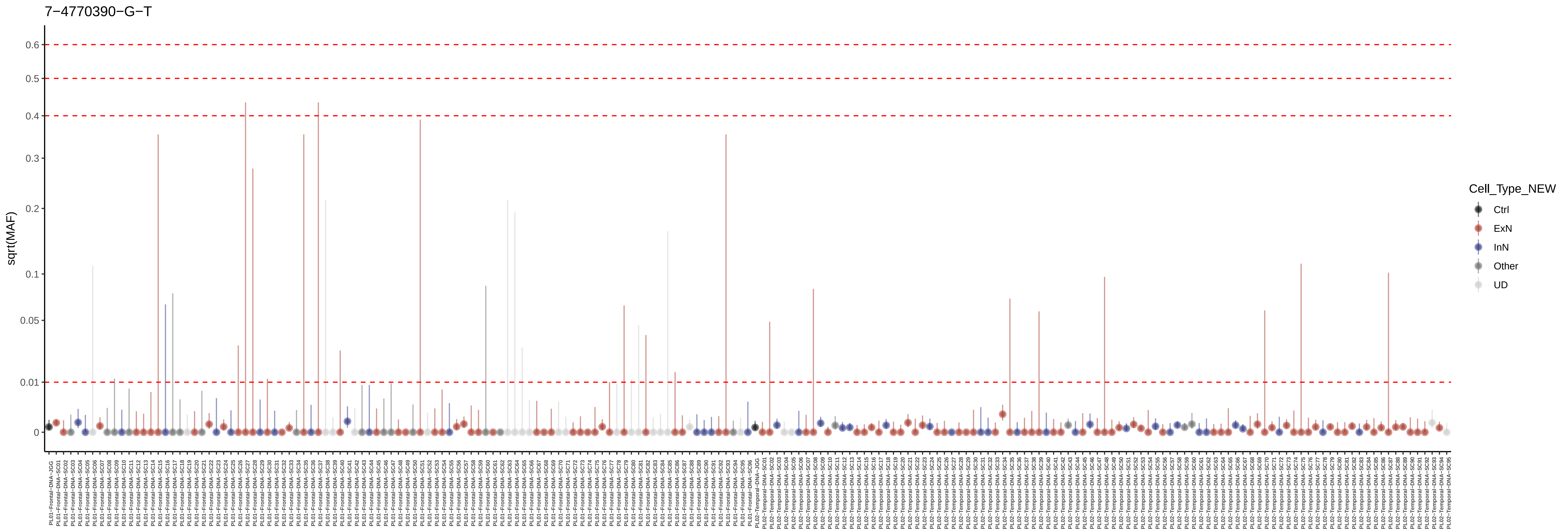

7−4770390−G−T
sqrt(MAF)
0.6
0.5
0.4
0.3
0.2
0.1
0.05
0.01
0
Cell_Type_NEW
Ctrl
ExN
InN
Other
UD

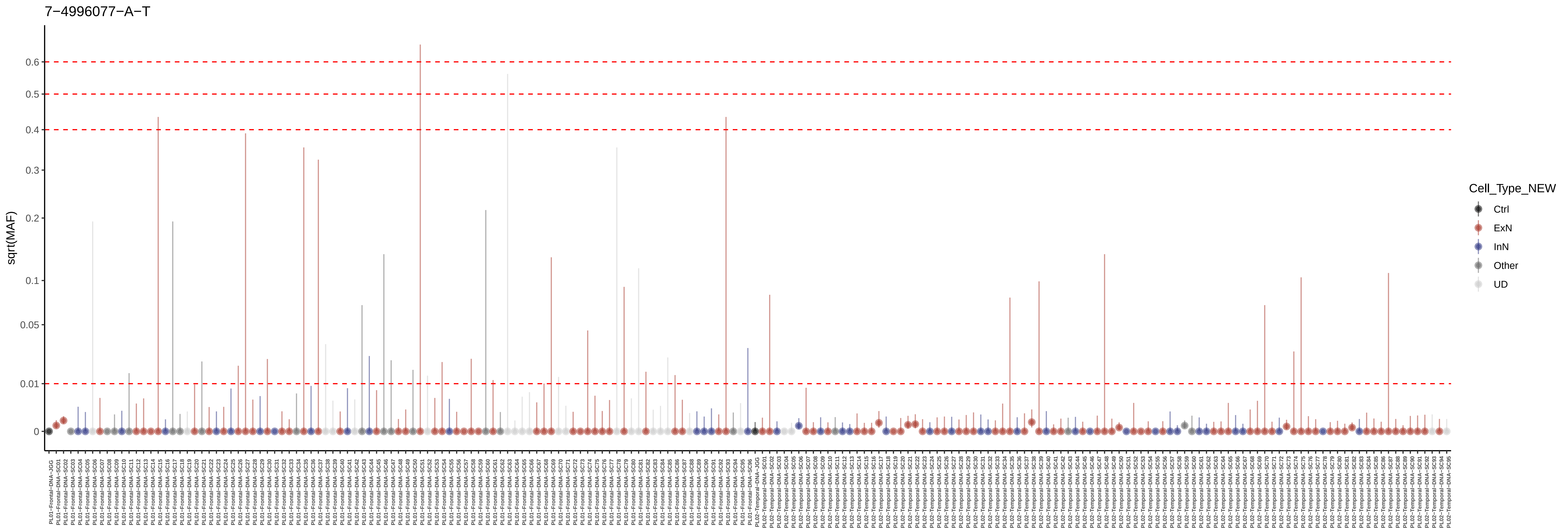
7−4996077−A−T
sqrt(MAF)
0.6
0.5
0.4
0.3
0.2
0.1
0.05
0.01
0
Cell_Type_NEW
Ctrl
ExN
InN
Other
UD

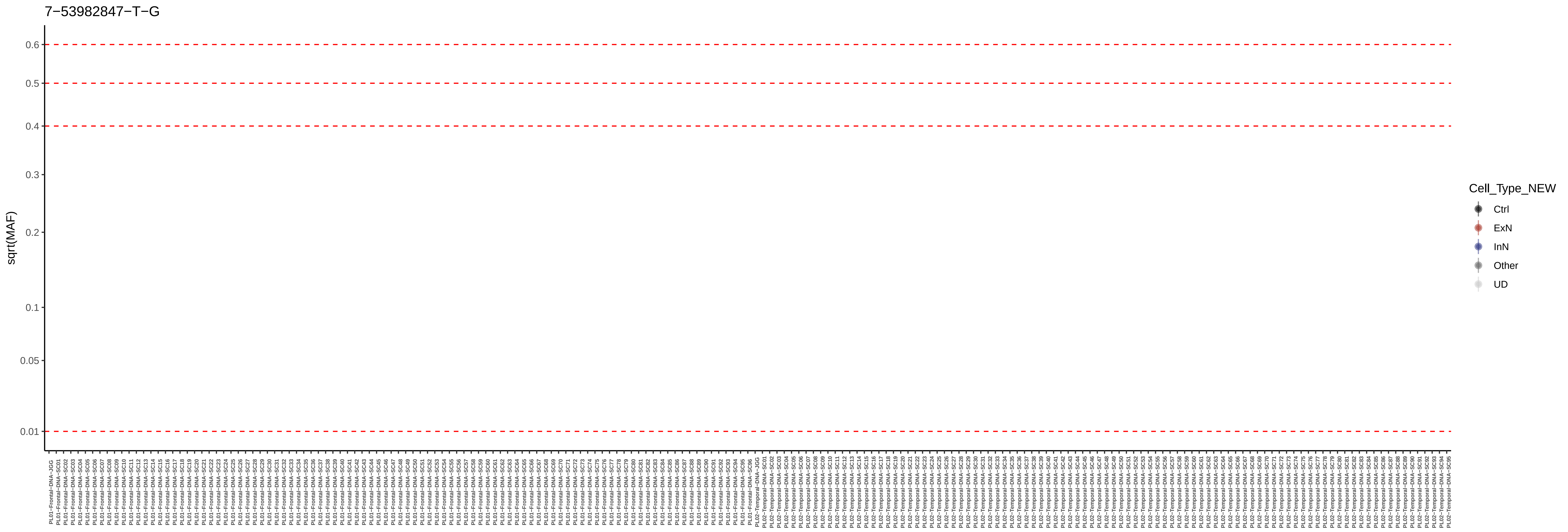
7−53982847−T−G
sqrt(MAF)
0.6
0.5
0.4
0.3
0.2
0.1
0.05
0.01
Cell_Type_NEW
Ctrl
ExN
InN
Other
UD

# 7−54130903−T−G

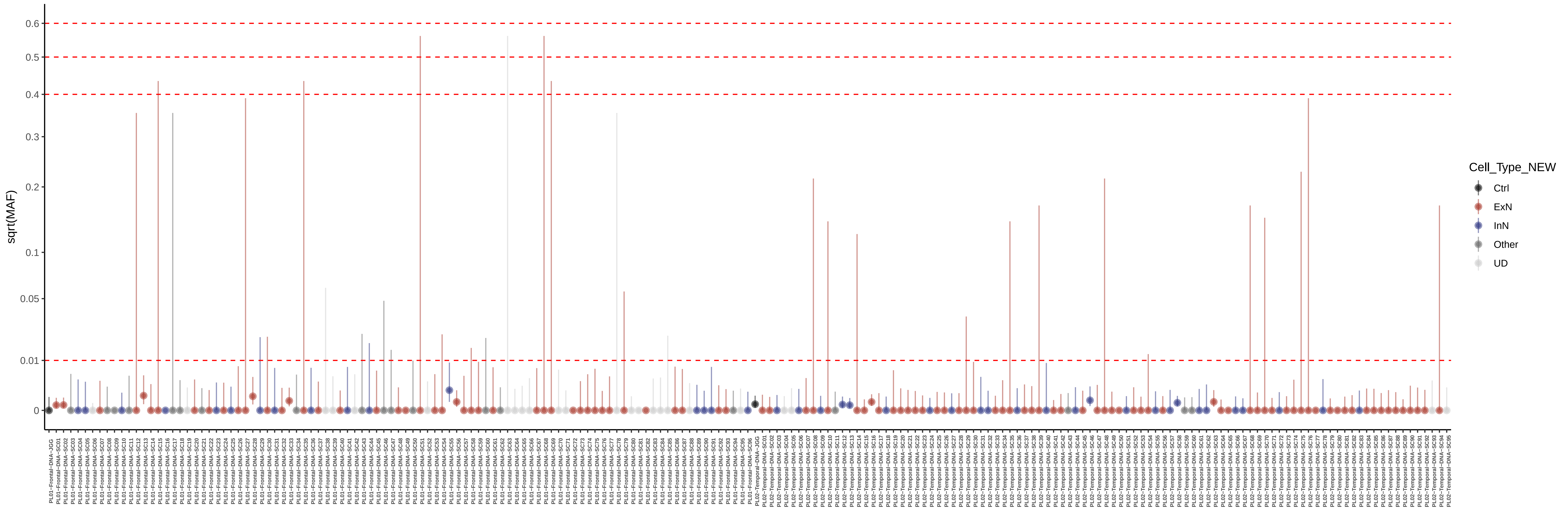

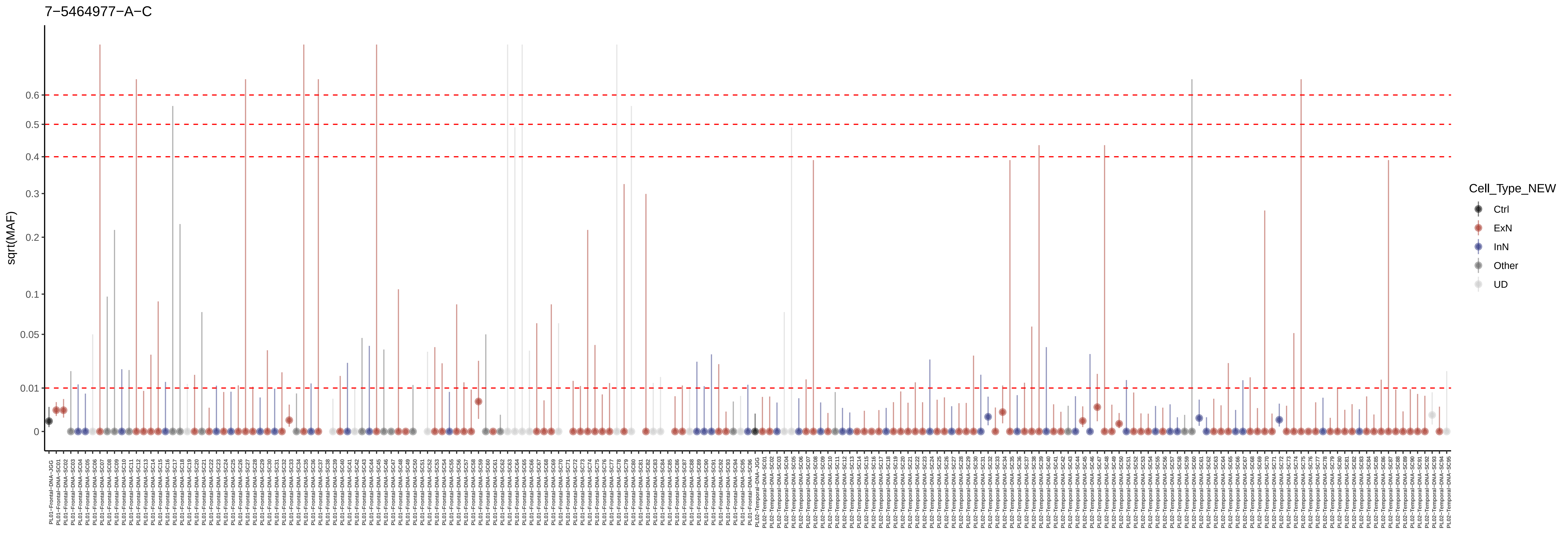
7−5464977−A−C
sqrt(MAF)
0.6
0.5
0.4
0.3
0.2
0.1
0.05
0.01
0
Cell_Type_NEW
Ctrl
ExN
InN
Other
UD

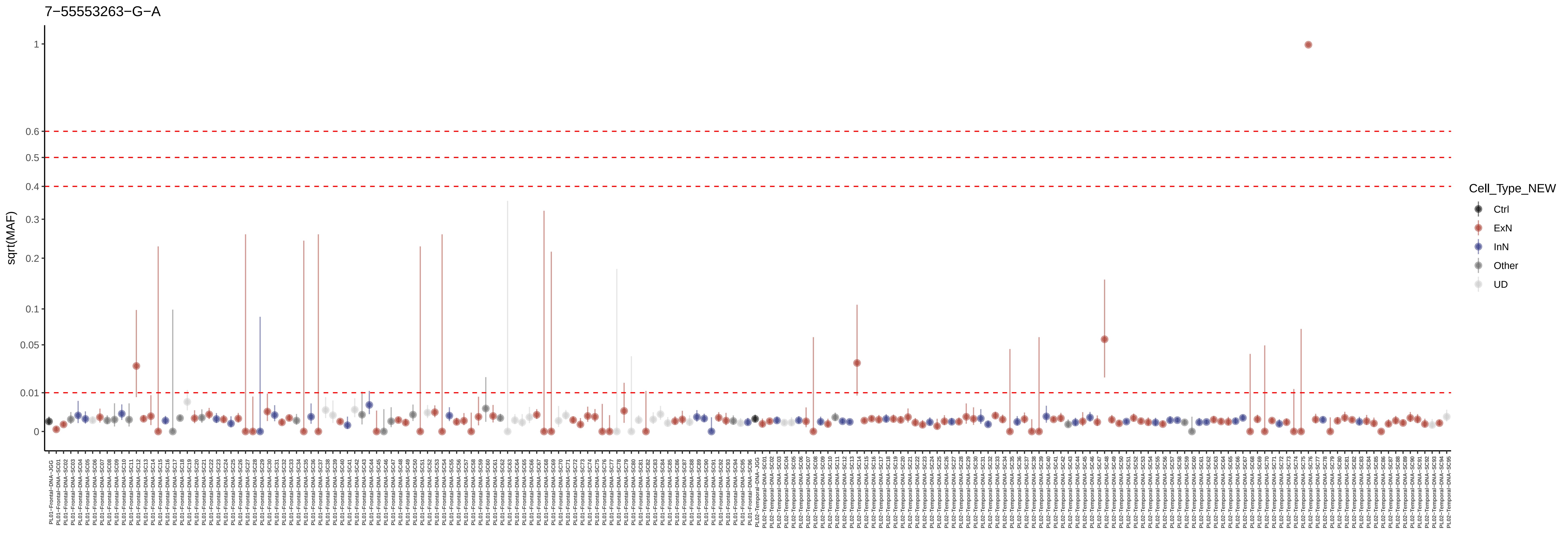
7−55553263−G−A
sqrt(MAF)
1
0.6
0.5
0.4
0.3
0.2
0.1
0.05
0.01
0
Cell_Type_NEW
Ctrl
ExN
InN
Other
UD

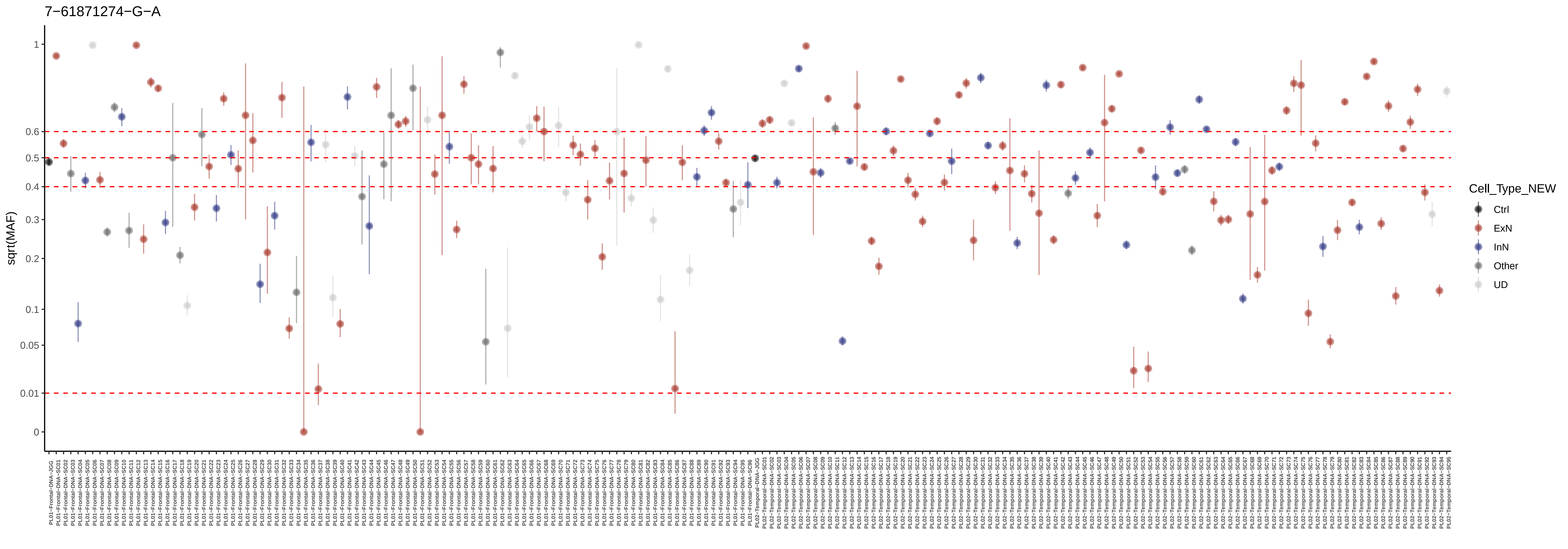
7−61871274−G−A
sqrt(MAF)
1
0.6
0.5
0.4
0.3
0.2
0.1
0.05
0.01
0
Cell_Type_NEW
Ctrl
ExN
InN
Other
UD

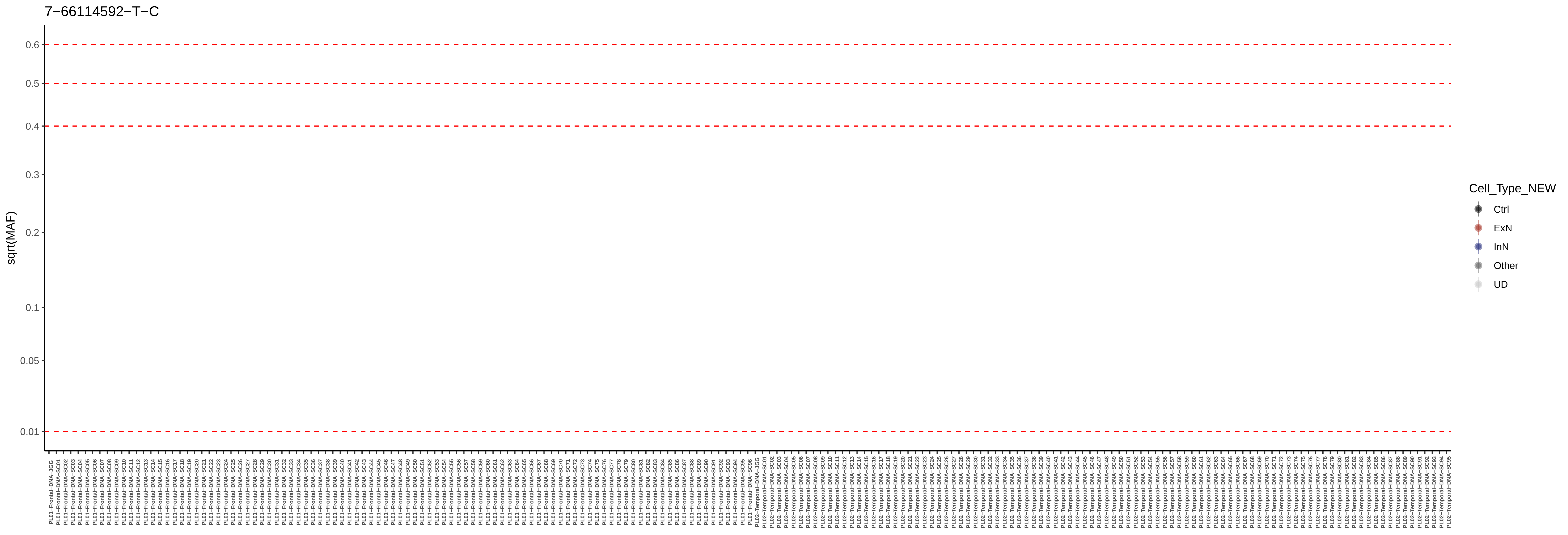
7−66114592−T−C
sqrt(MAF)
0.6
0.5
0.4
0.3
0.2
0.1
0.05
0.01
Cell_Type_NEW
Ctrl
ExN
InN
Other
UD

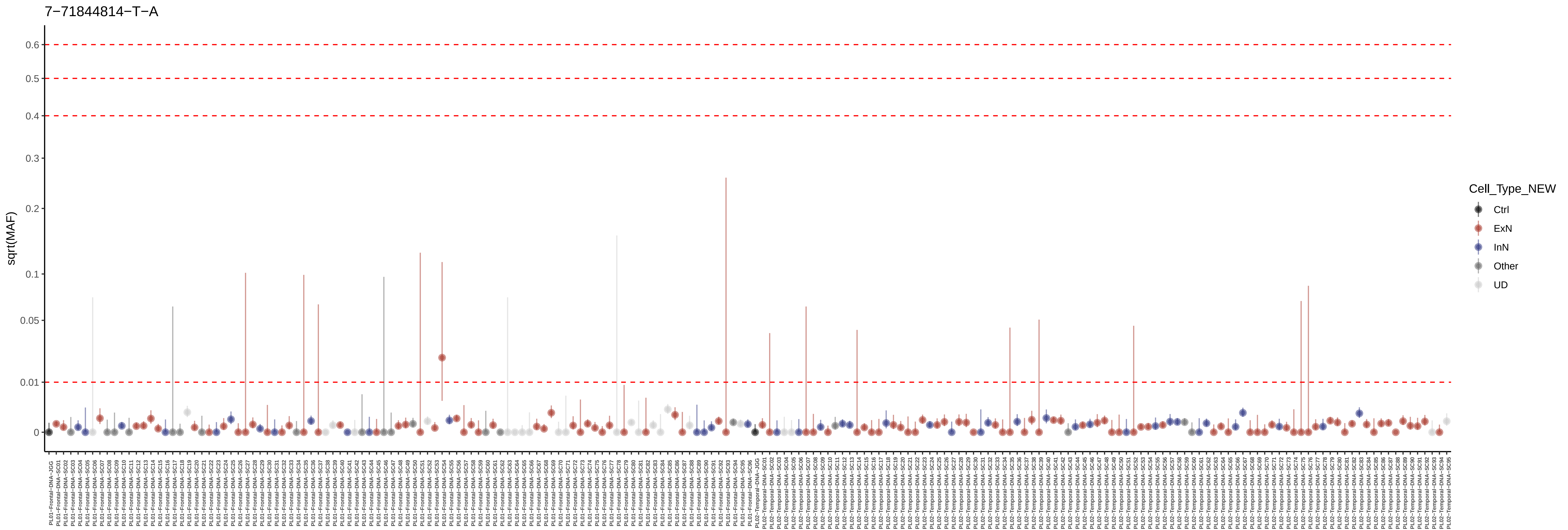
7−71844814−T−A
sqrt(MAF)
0.6
0.5
0.4
0.3
0.2
0.1
0.05
0.01
0
Cell_Type_NEW
Ctrl
ExN
InN
Other
UD

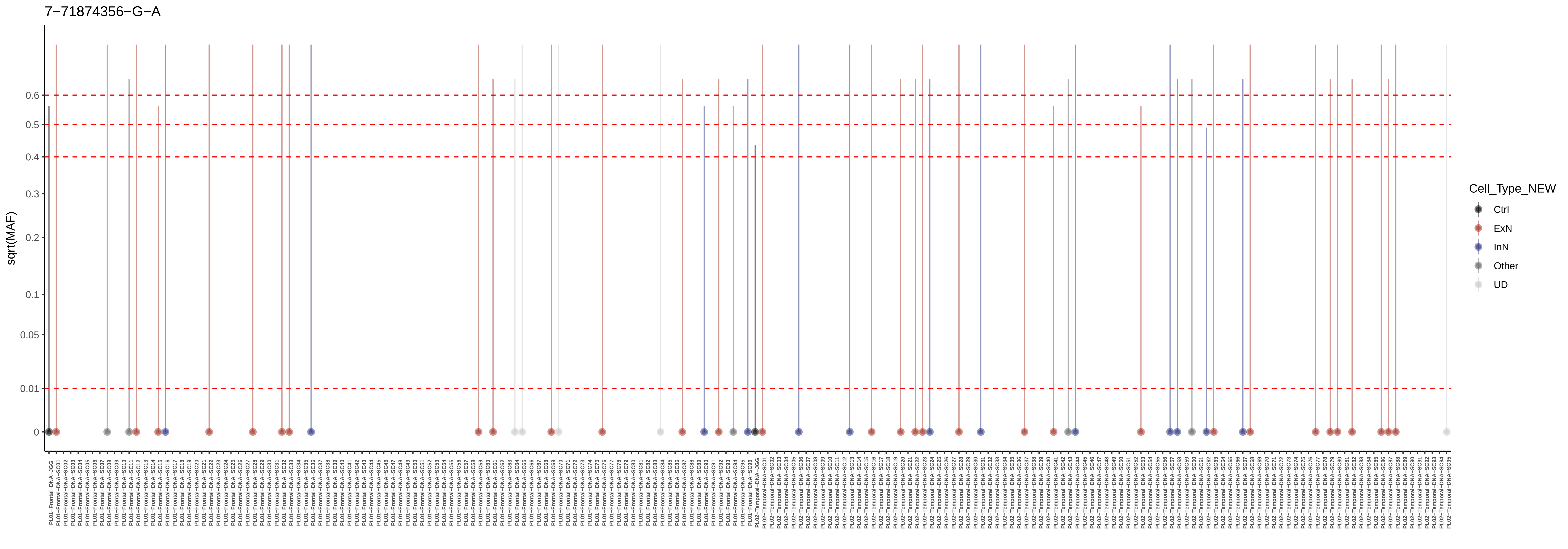

7−71874356−G−A
sqrt(MAF)
0.6
0.5
0.4
0.3
0.2
0.1
0.05
0.01
0
Cell_Type_NEW
Ctrl
ExN
InN
Other
UD

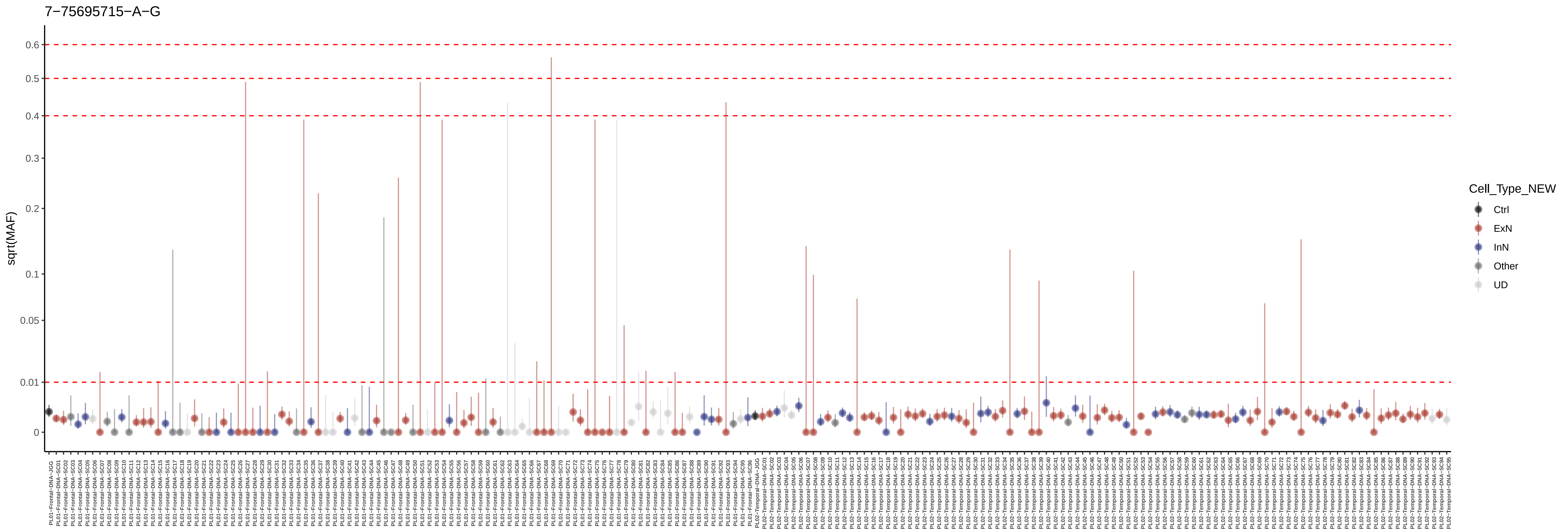

7−75695715−A−G
sqrt(MAF)
0.6
0.5
0.4
0.3
0.2
0.1
0.05
0.01
0
Cell_Type_NEW
Ctrl
ExN
InN
Other
UD

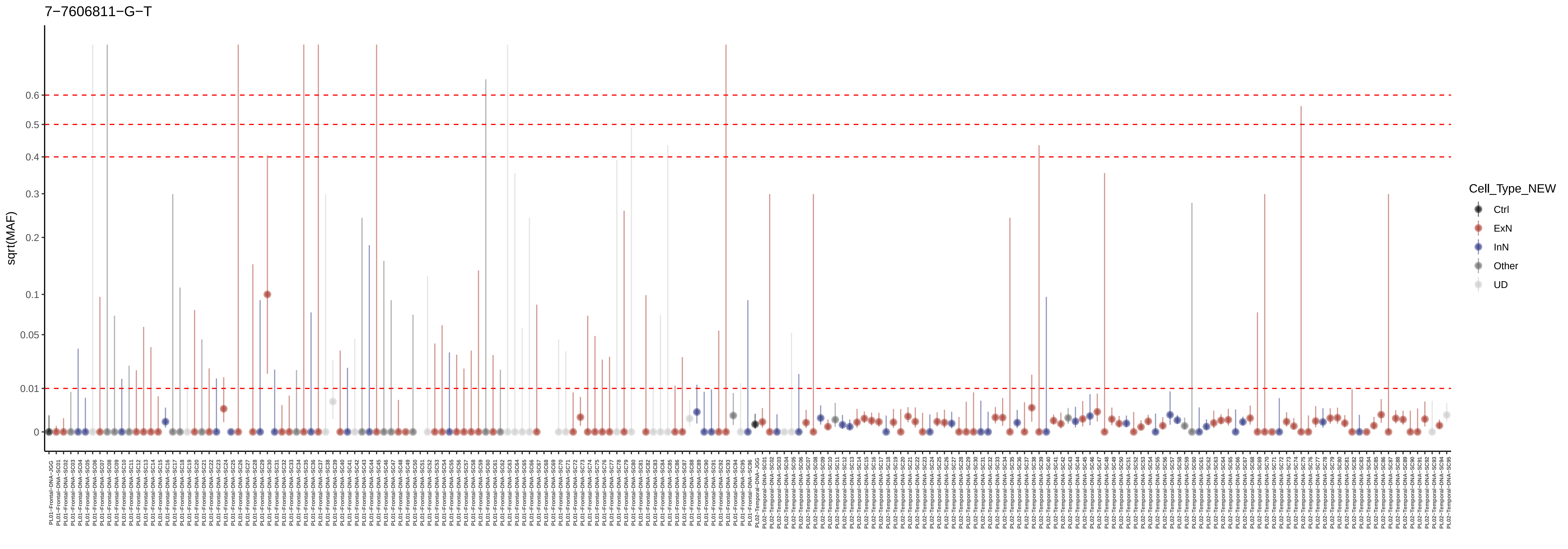
7−7606811−G−T
sqrt(MAF)
0.6
0.5
0.4
0.3
0.2
0.1
0.05
0.01
0
Cell_Type_NEW
Ctrl
ExN
InN
Other
UD

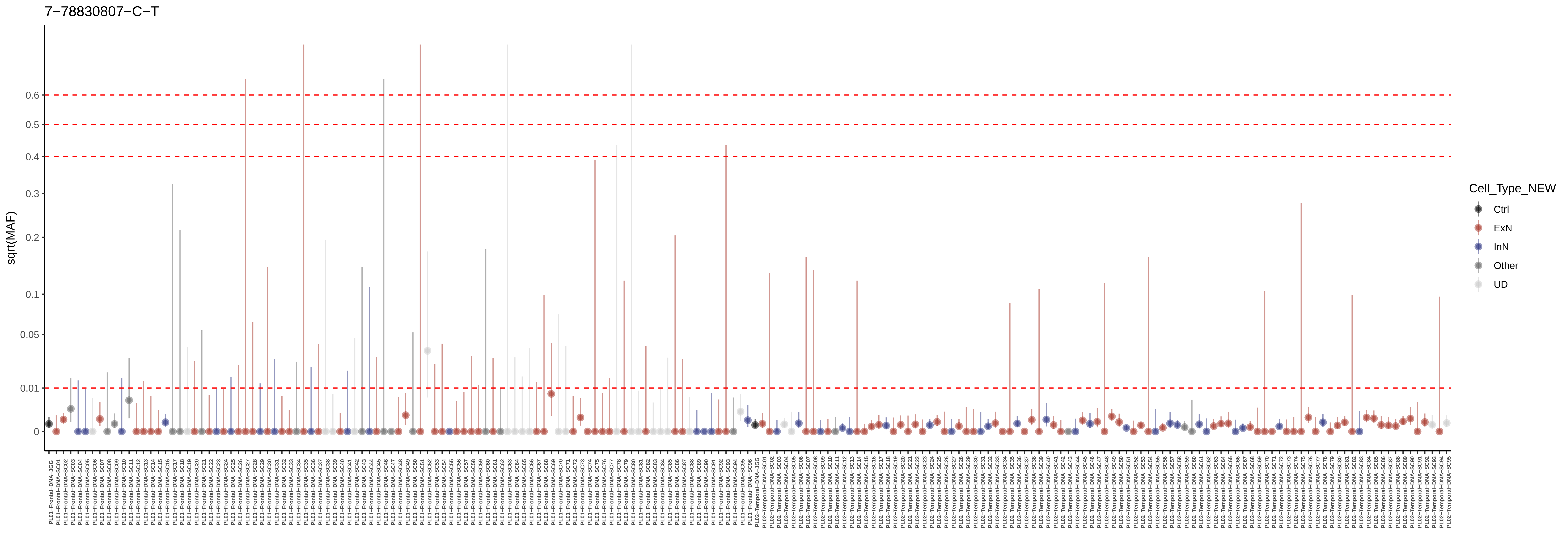
7−78830807−C−T
sqrt(MAF)
0.6
0.5
0.4
0.3
0.2
0.1
0.05
0.01
0
Cell_Type_NEW
Ctrl
ExN
InN
Other
UD

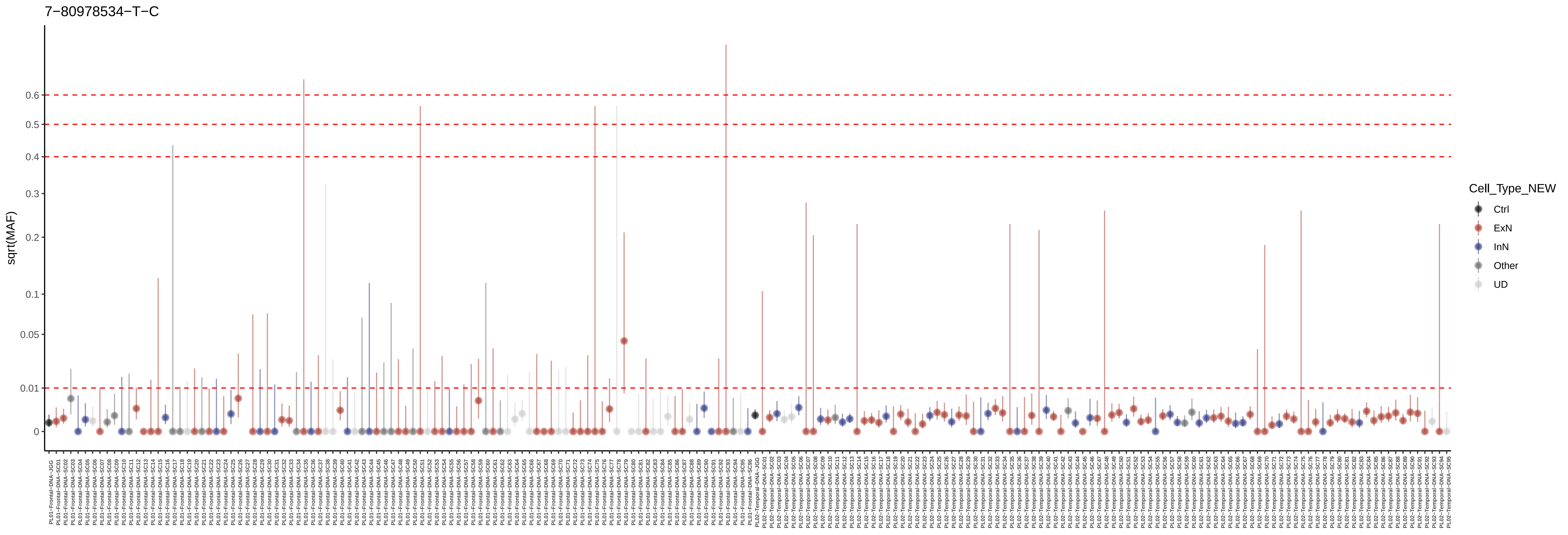
7−80978534−T−C
sqrt(MAF)
0.6
0.5
0.4
0.3
0.2
0.1
0.05
0.01
0
Cell_Type_NEW
Ctrl
ExN
InN
Other
UD

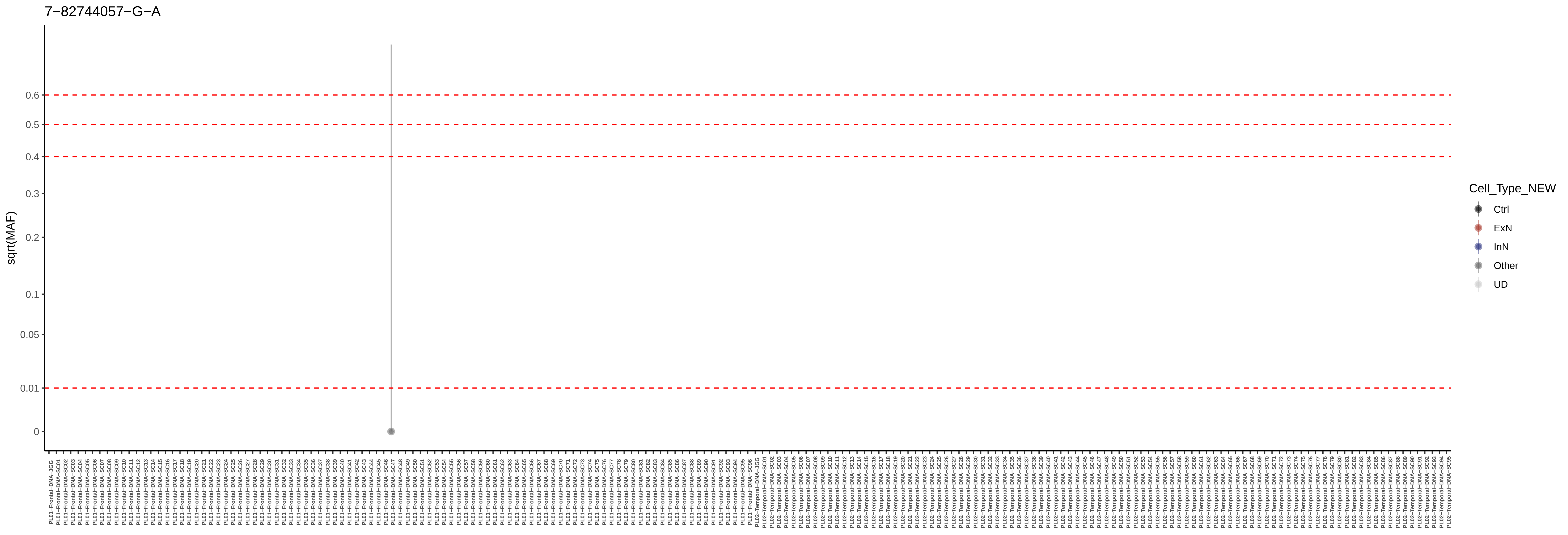
7−82744057−G−A
sqrt(MAF)
0
0.01
0.05
0.1
0.2
0.3
0.4
0.5
0.6
Cell_Type_NEW
Ctrl
ExN
InN
Other
UD

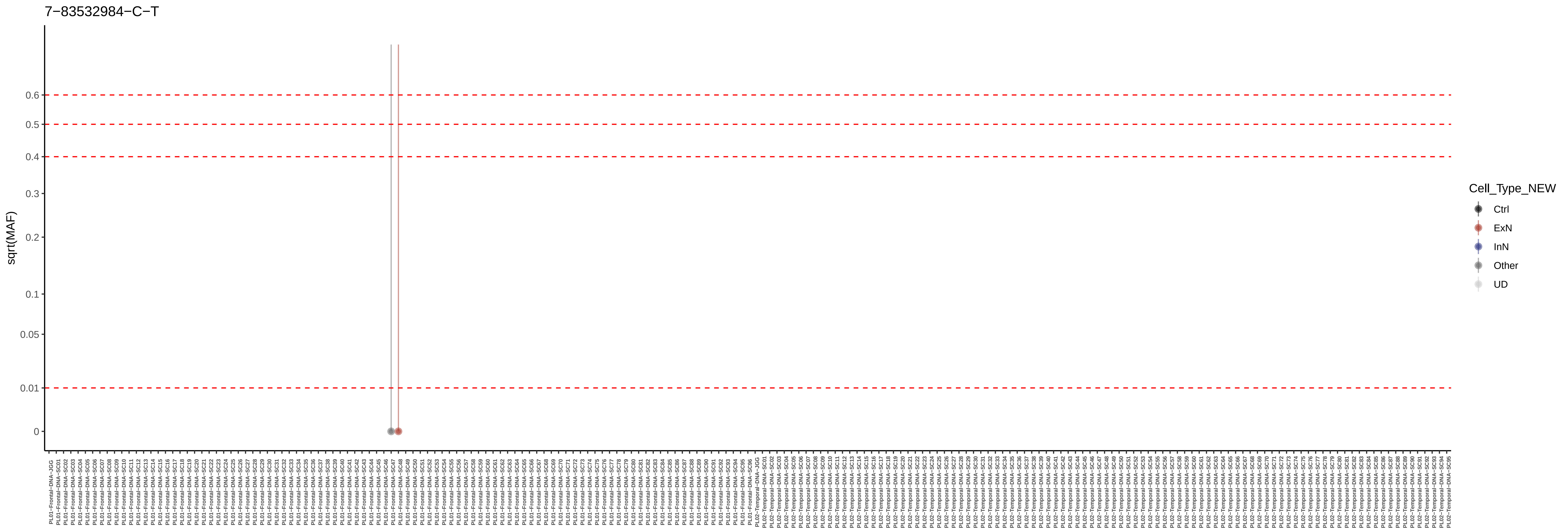
7−83532984−C−T
sqrt(MAF)
0.6
0.5
0.4
0.3
0.2
0.1
0.05
0.01
0
Cell_Type_NEW
Ctrl
ExN
InN
Other
UD

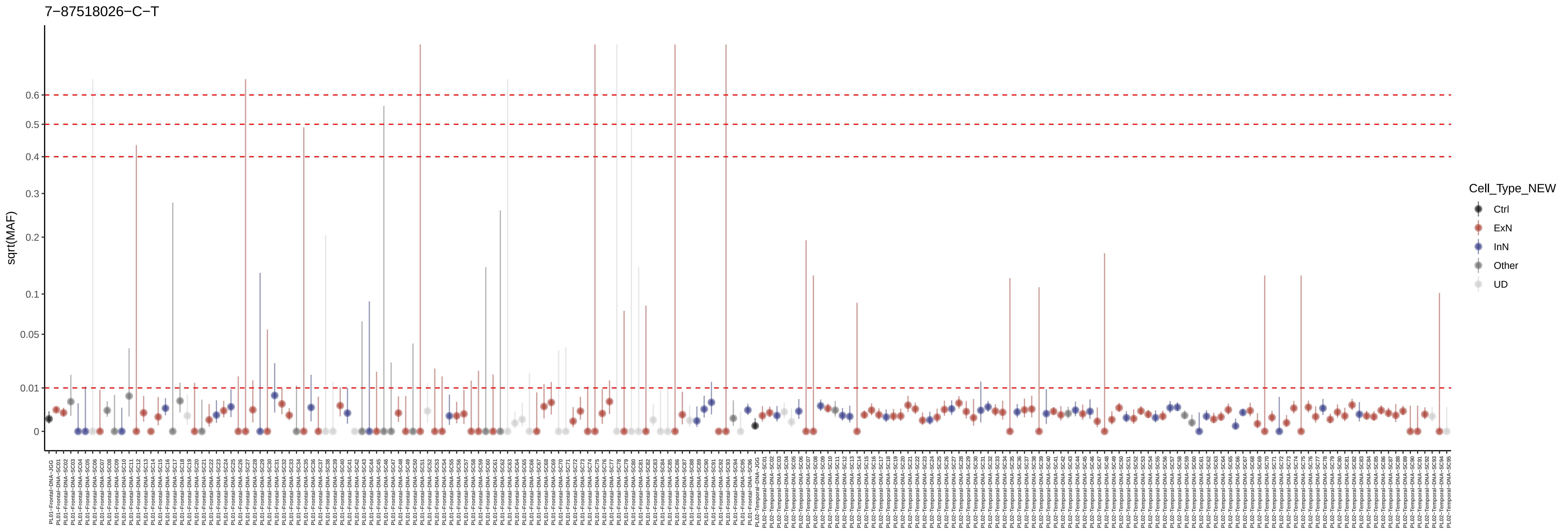
7−87518026−C−T
sqrt(MAF)
0.6
0.5
0.4
0.3
0.2
0.1
0.05
0.01
0
Cell_Type_NEW
Ctrl
ExN
InN
Other
UD

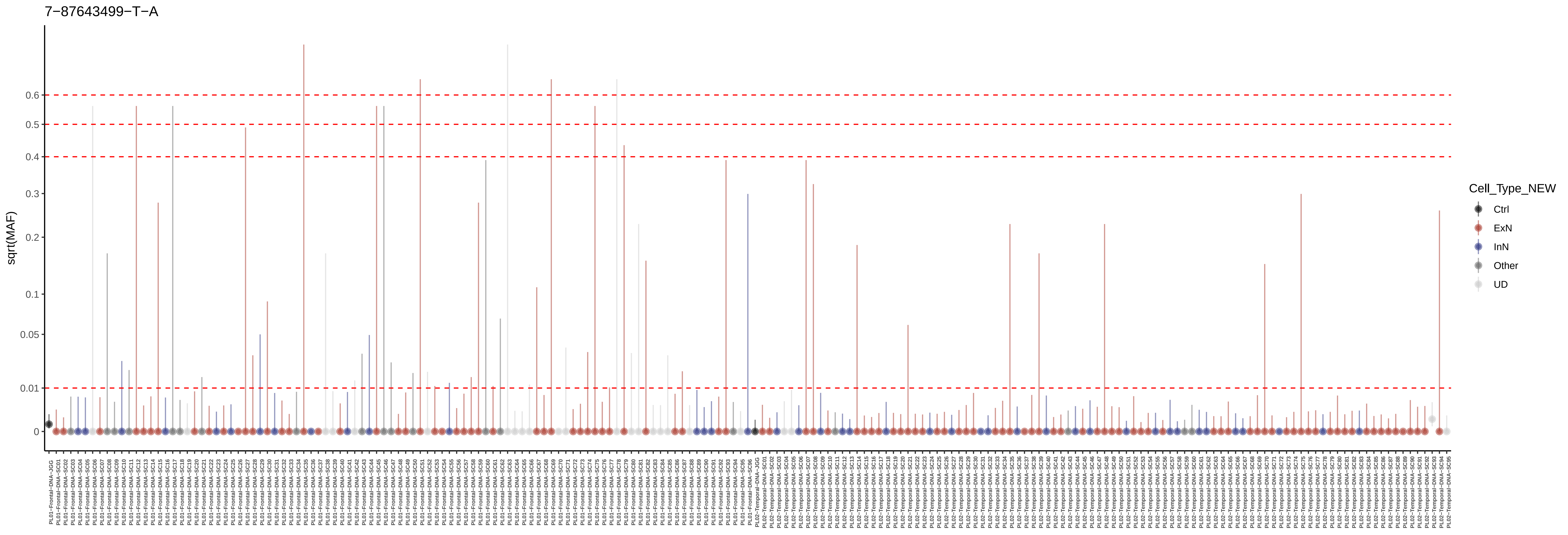
7−87643499−T−A
sqrt(MAF)
0.6
0.5
0.4
0.3
0.2
0.1
0.05
0.01
0
Cell_Type_NEW
Ctrl
ExN
InN
Other
UD

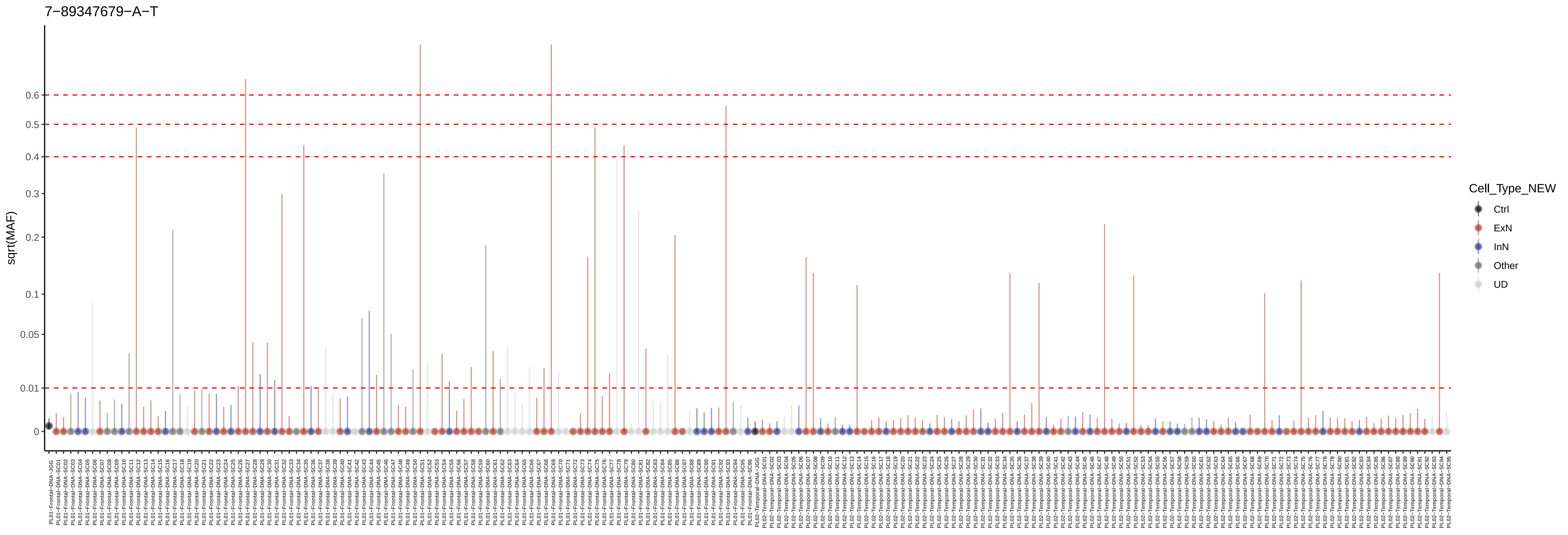

7−89347679−A−T
sqrt(MAF)
0.6
0.5
0.4
0.3
0.2
0.1
0.05
0.01
0
Cell_Type_NEW
Ctrl
ExN
InN
Other
UD

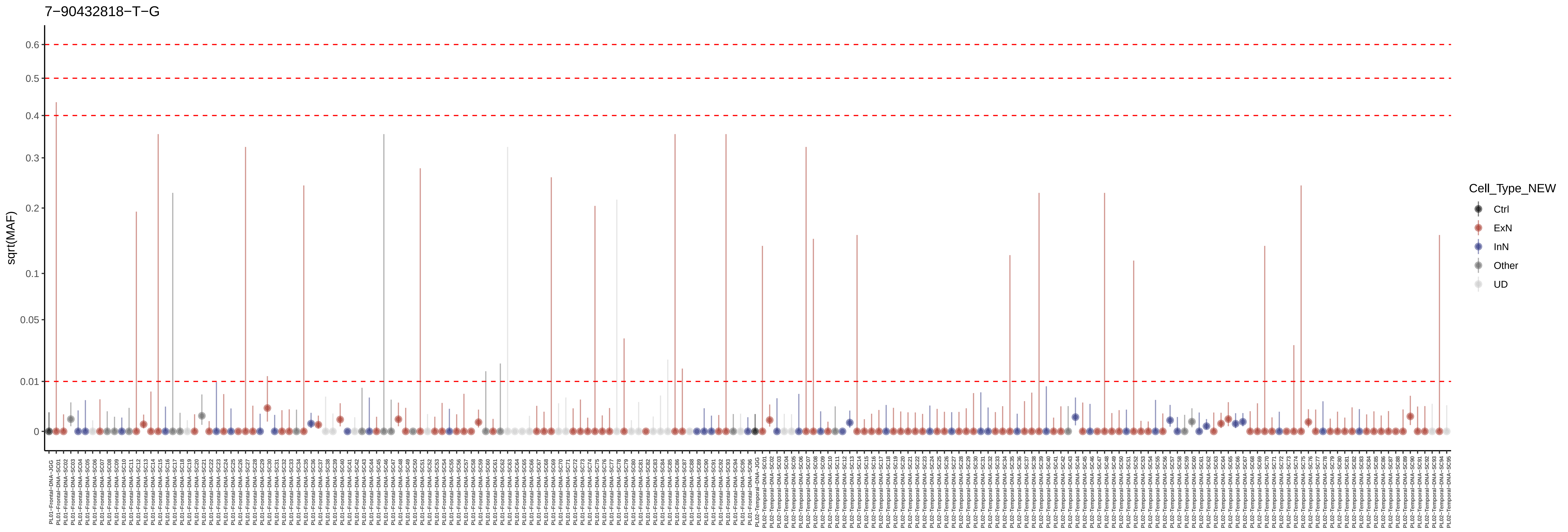
7−90432818−T−G
sqrt(MAF)
0.6
0.5
0.4
0.3
0.2
0.1
0.05
0.01
0
Cell_Type_NEW
Ctrl
ExN
InN
Other
UD

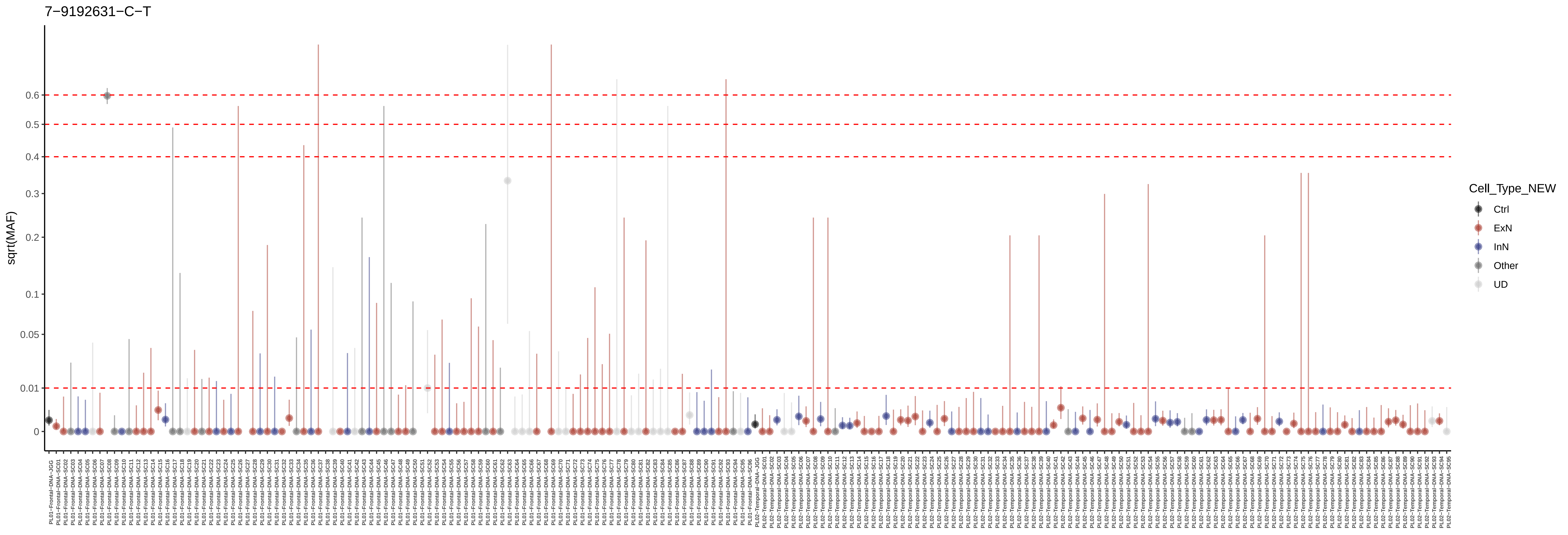

7−9192631−C−T
sqrt(MAF)
0.6
0.5
0.4
0.3
0.2
0.1
0.05
0.01
0
Cell_Type_NEW
Ctrl
ExN
InN
Other
UD

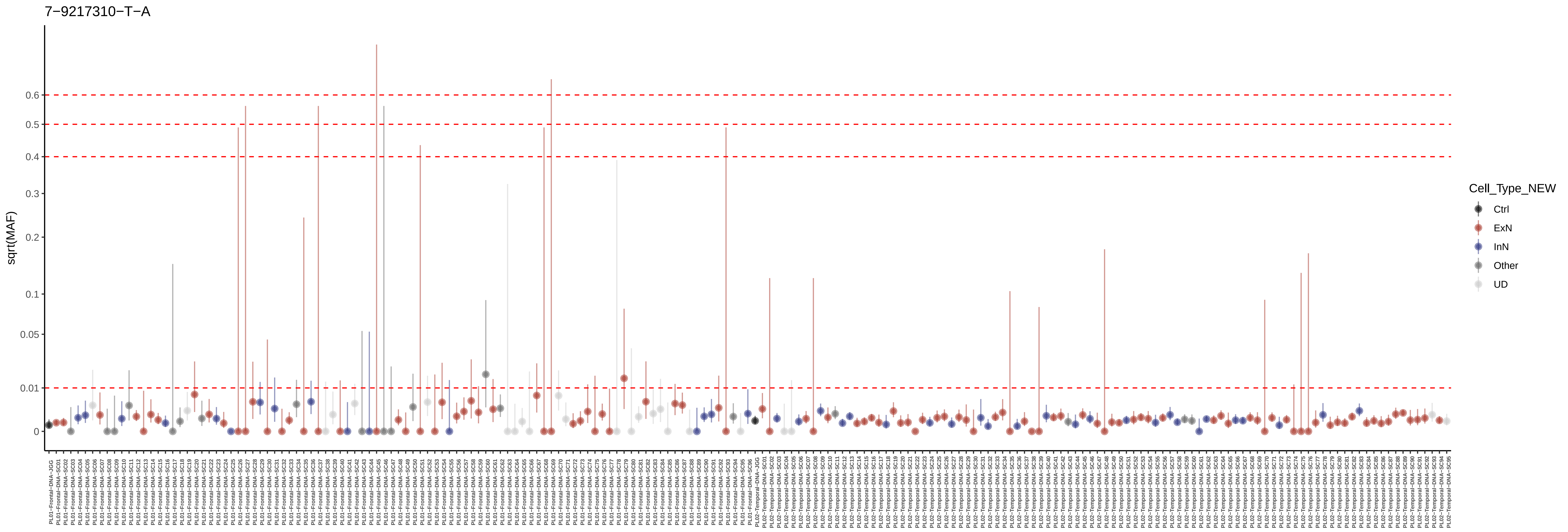
7−9217310−T−A
sqrt(MAF)
0.6
0.5
0.4
0.3
0.2
0.1
0.05
0.01
0
Cell_Type_NEW
Ctrl
ExN
InN
Other
UD

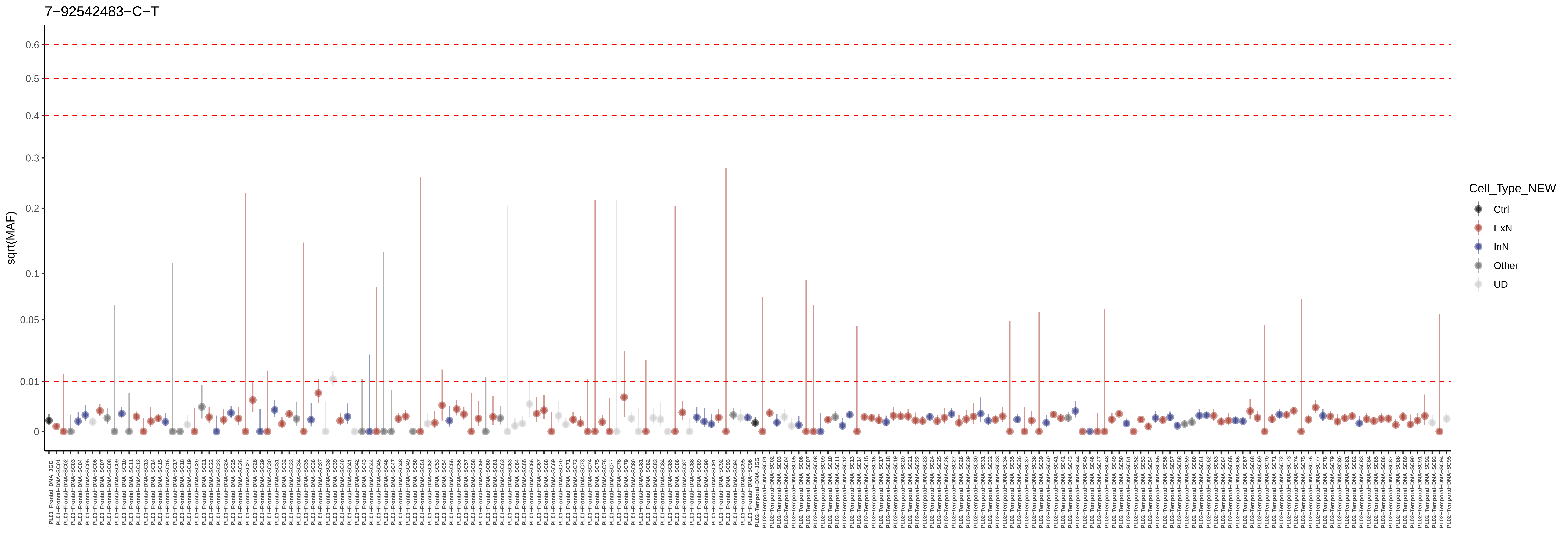

7−92542483−C−T
sqrt(MAF)
0.6
0.5
0.4
0.3
0.2
0.1
0.05
0.01
0
Cell_Type_NEW
Ctrl
ExN
InN
Other
UD

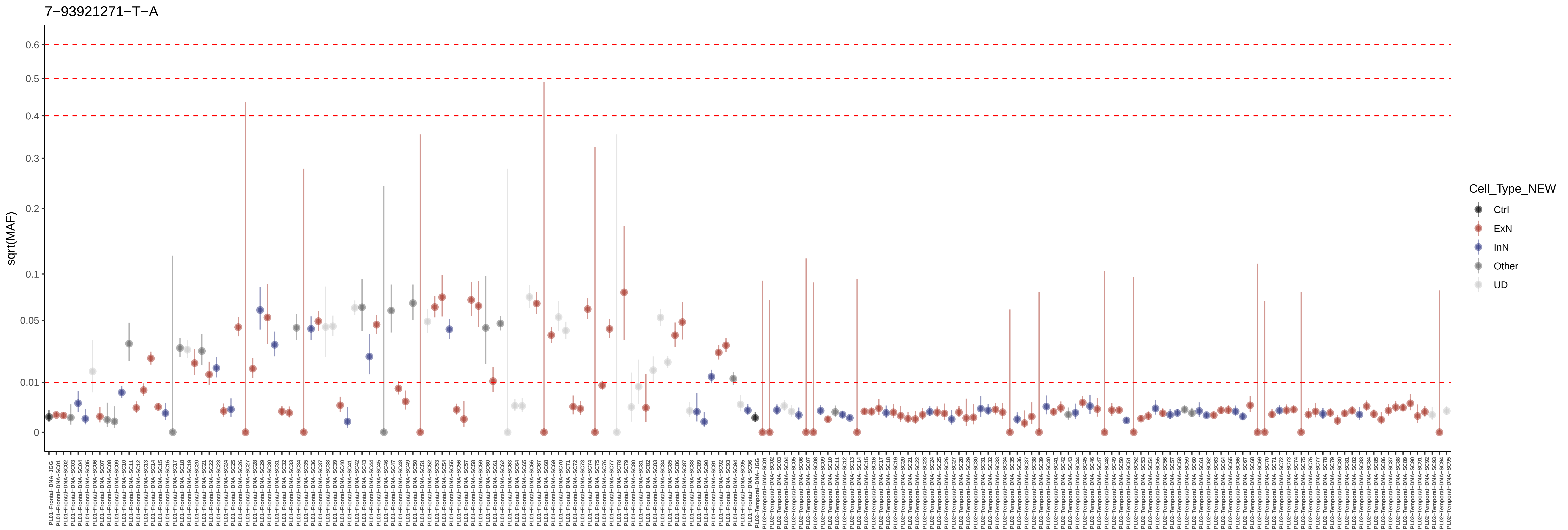

7−93921271−T−A
sqrt(MAF)
0.6
0.5
0.4
0.3
0.2
0.1
0.05
0.01
0
Cell_Type_NEW
Ctrl
ExN
InN
Other
UD

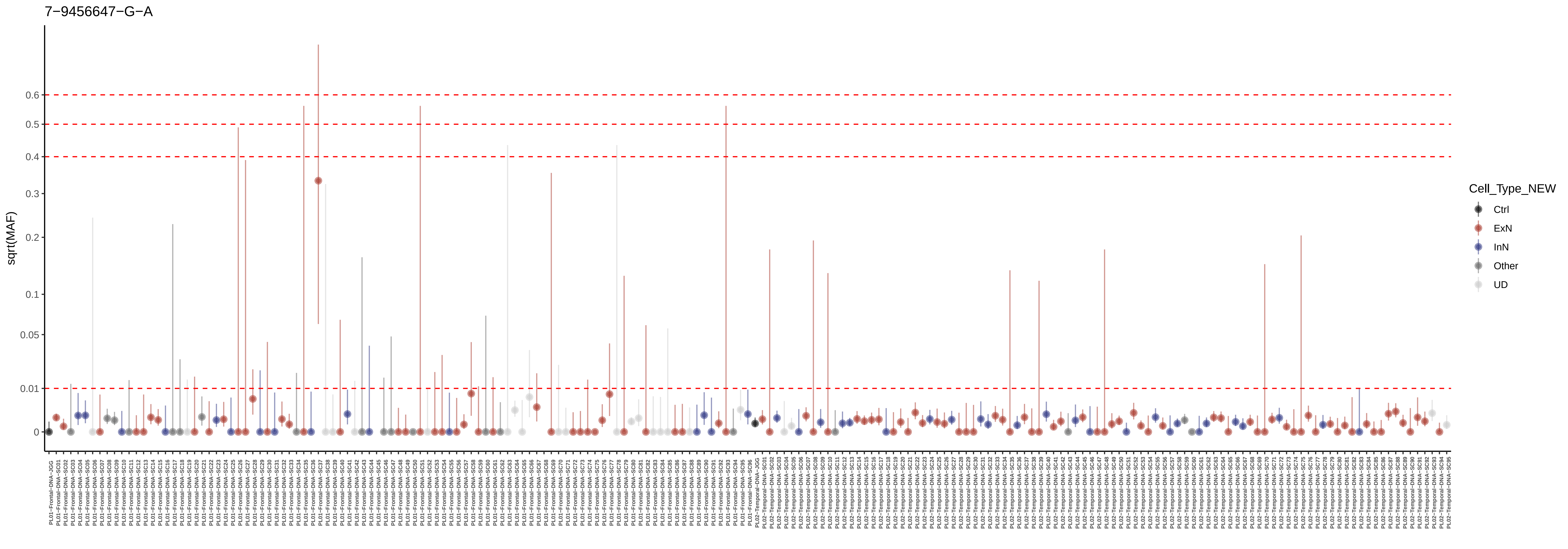
7−9456647−G−A
sqrt(MAF)
0.6
0.5
0.4
0.3
0.2
0.1
0.05
0.01
0
Cell_Type_NEW
Ctrl
ExN
InN
Other
UD

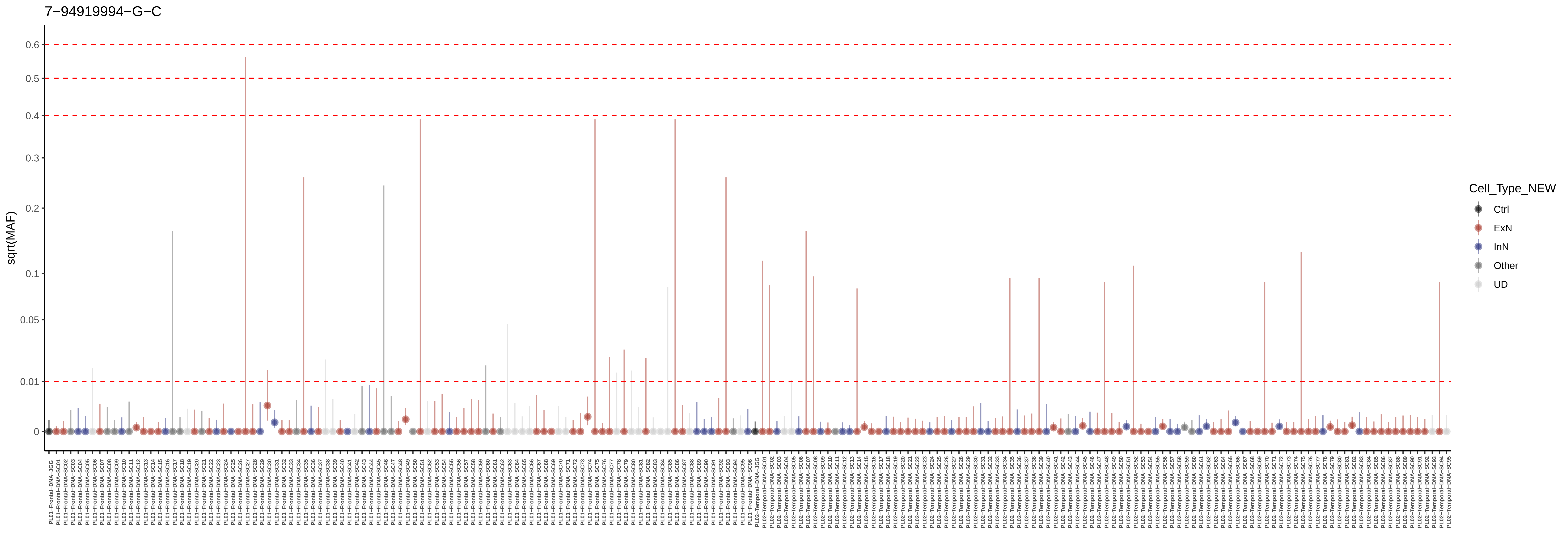

7−94919994−G−C
sqrt(MAF)
0.6
0.5
0.4
0.3
0.2
0.1
0.05
0.01
0
Cell_Type_NEW
Ctrl
ExN
InN
Other
UD

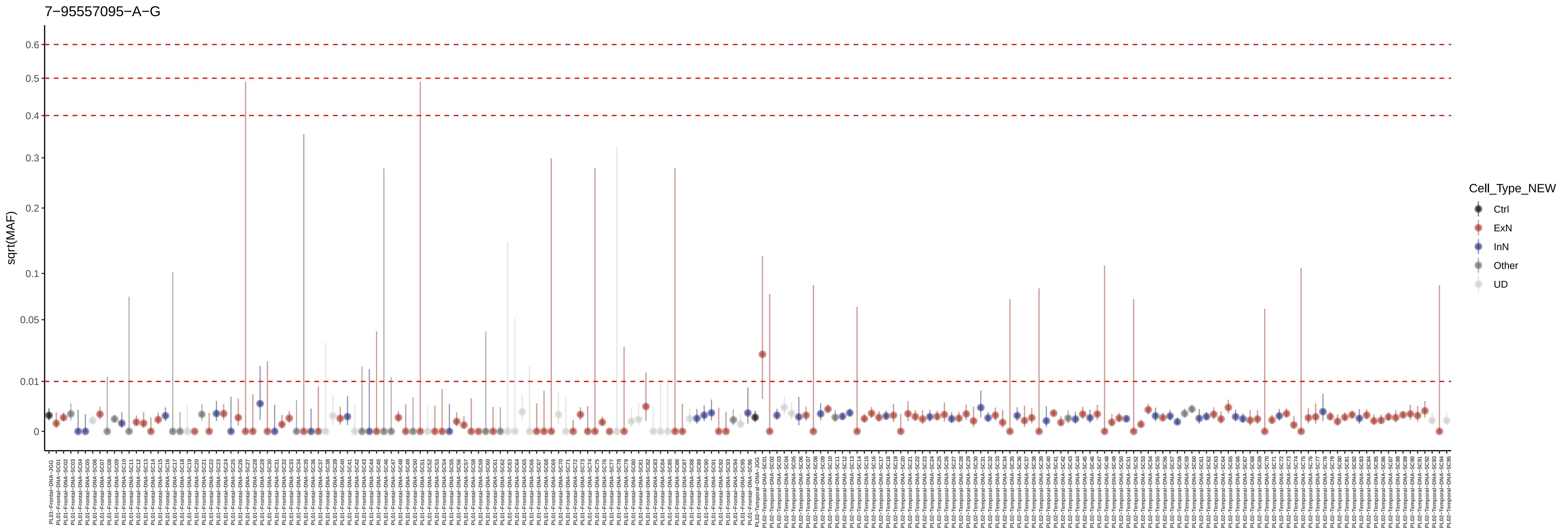
7−95557095−A−G
sqrt(MAF)
0.6
0.5
0.4
0.3
0.2
0.1
0.05
0.01
0
Cell_Type_NEW
Ctrl
ExN
InN
Other
UD

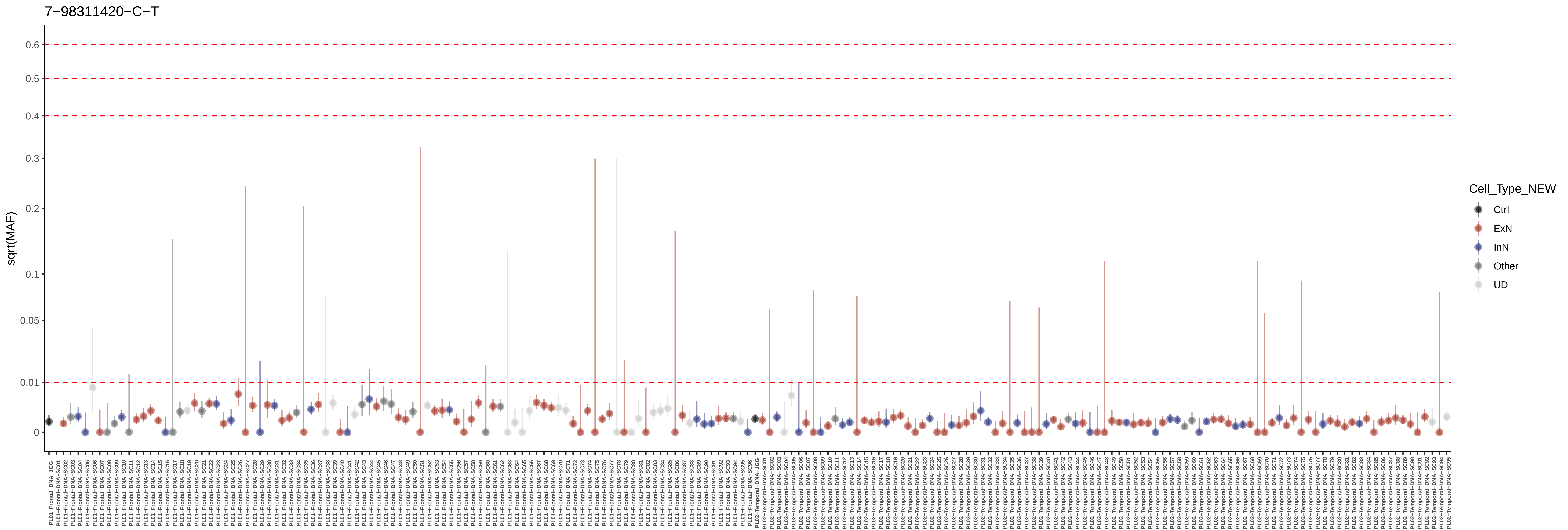

7−98311420−C−T
sqrt(MAF)
0.6
0.5
0.4
0.3
0.2
0.1
0.05
0.01
0
Cell_Type_NEW
Ctrl
ExN
InN
Other
UD

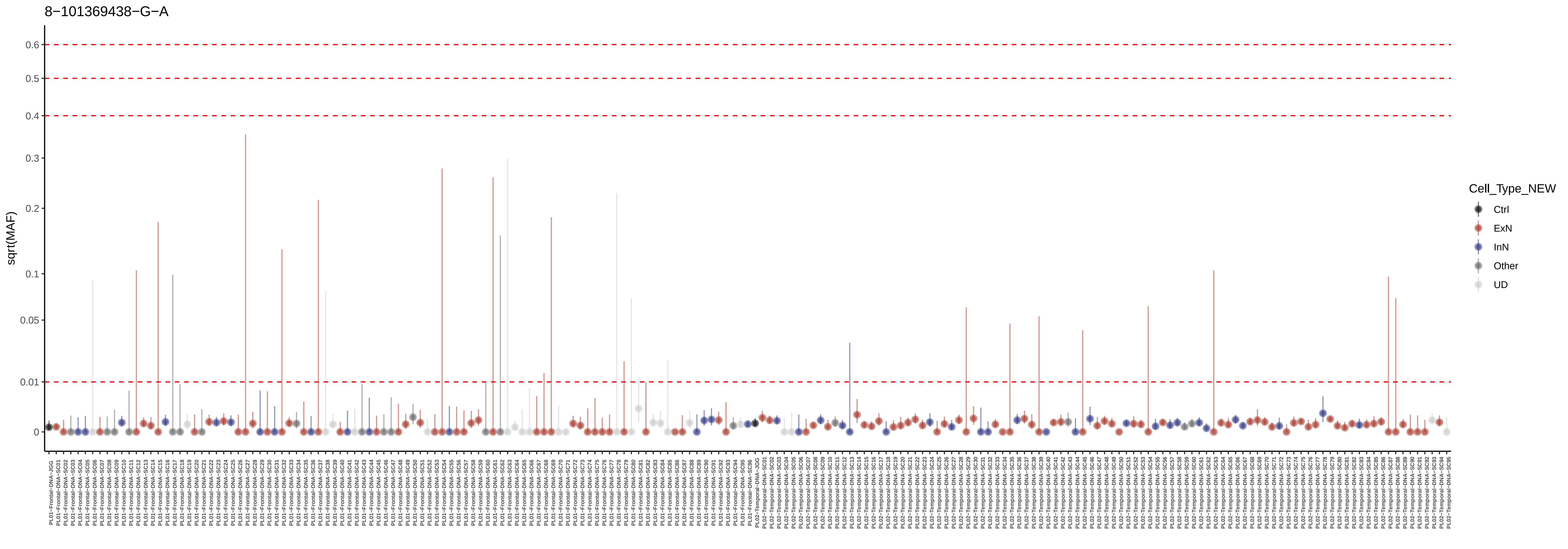

8−101369438−G−A
sqrt(MAF)
0.6
0.5
0.4
0.3
0.2
0.1
0.05
0.01
0
Cell_Type_NEW
Ctrl
ExN
InN
Other
UD

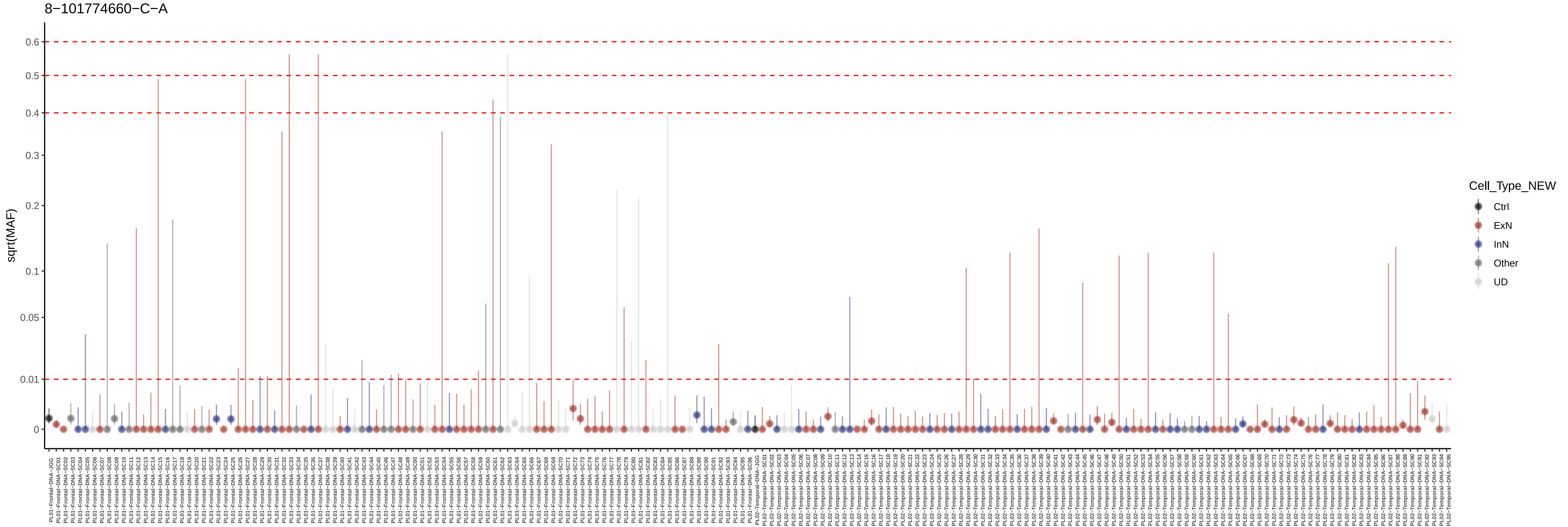
8−101774660−C−A
sqrt(MAF)
0.6
0.5
0.4
0.3
0.2
0.1
0.05
0.01
0
Cell_Type_NEW
Ctrl
ExN
InN
Other
UD

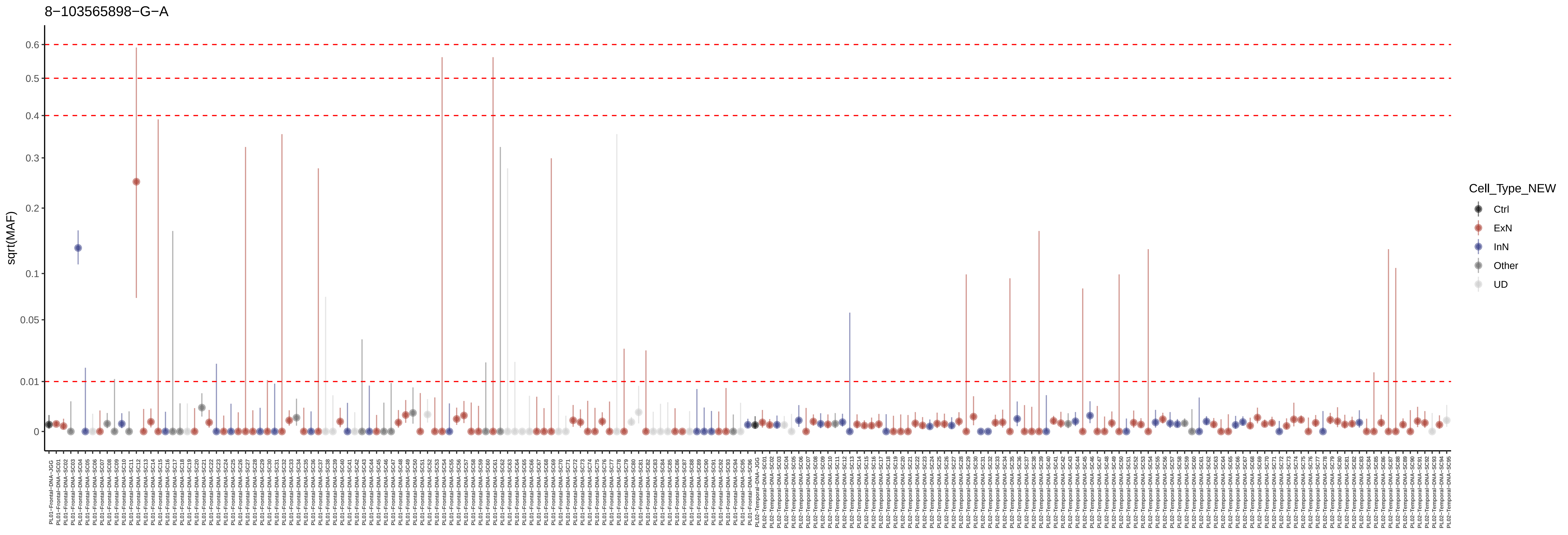
8−103565898−G−A
sqrt(MAF)
0.6
0.5
0.4
0.3
0.2
0.1
0.05
0.01
0
Cell_Type_NEW
Ctrl
ExN
InN
Other
UD

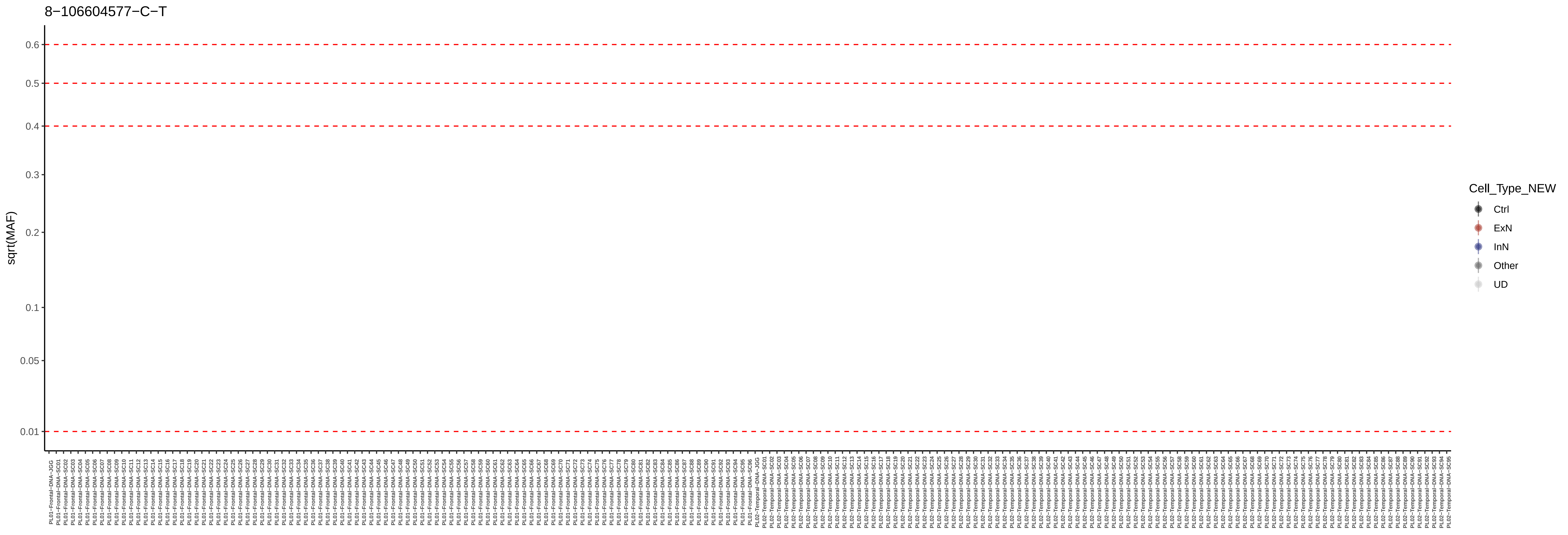
8−106604577−C−T
sqrt(MAF)
0.6
0.5
0.4
0.3
0.2
0.1
0.05
0.01
Cell_Type_NEW
Ctrl
ExN
InN
Other
UD

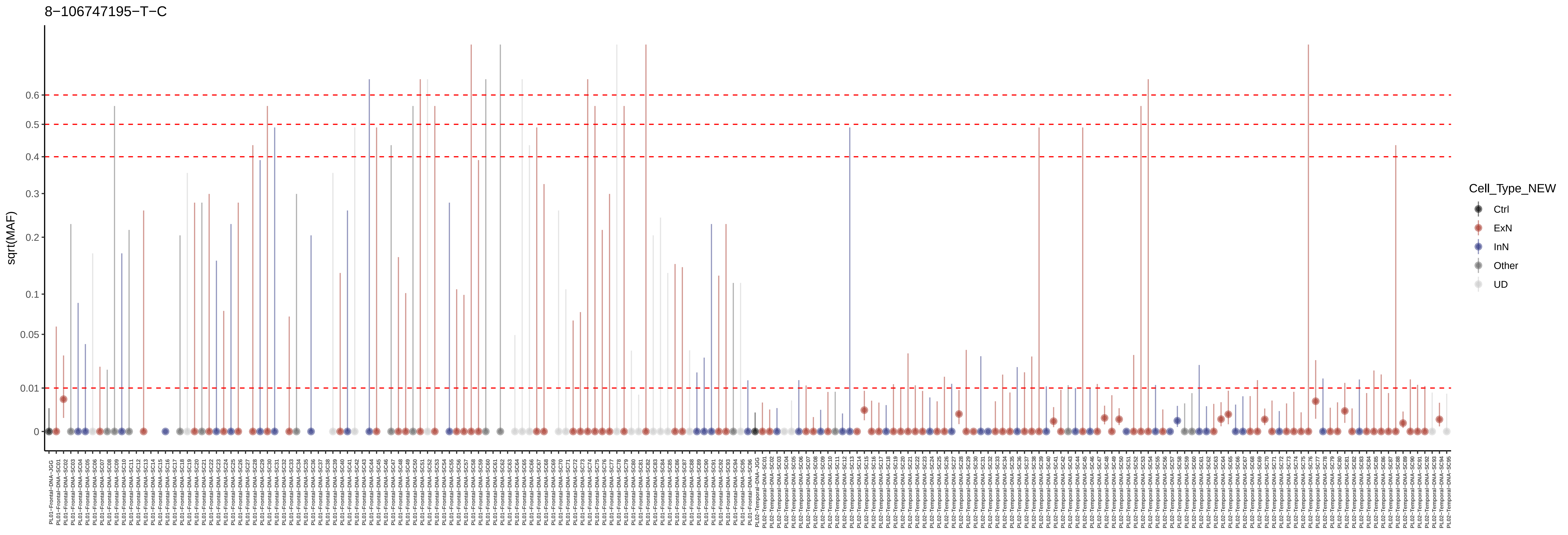
8−106747195−T−C
sqrt(MAF)
0.6
0.5
0.4
0.3
0.2
0.1
0.05
0.01
0
Cell_Type_NEW
Ctrl
ExN
InN
Other
UD

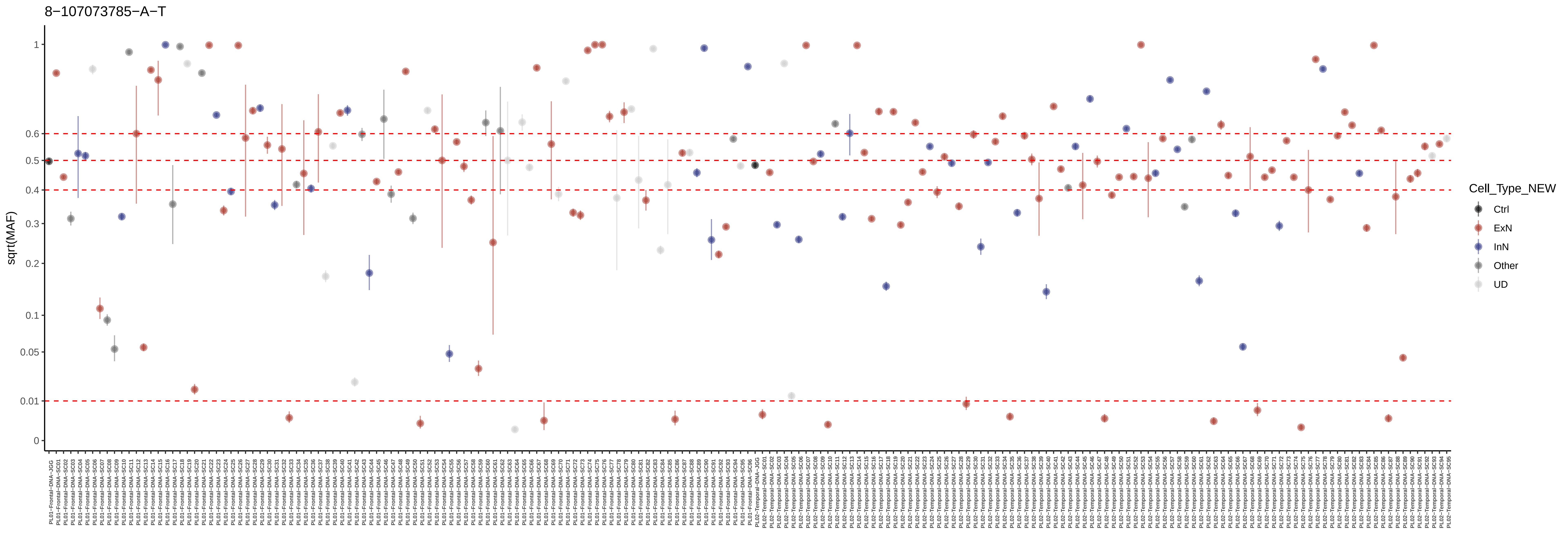

8−107073785−A−T
sqrt(MAF)
1
0.6
0.5
0.4
0.3
0.2
0.1
0.05
0.01
0
Cell_Type_NEW
Ctrl
ExN
InN
Other
UD

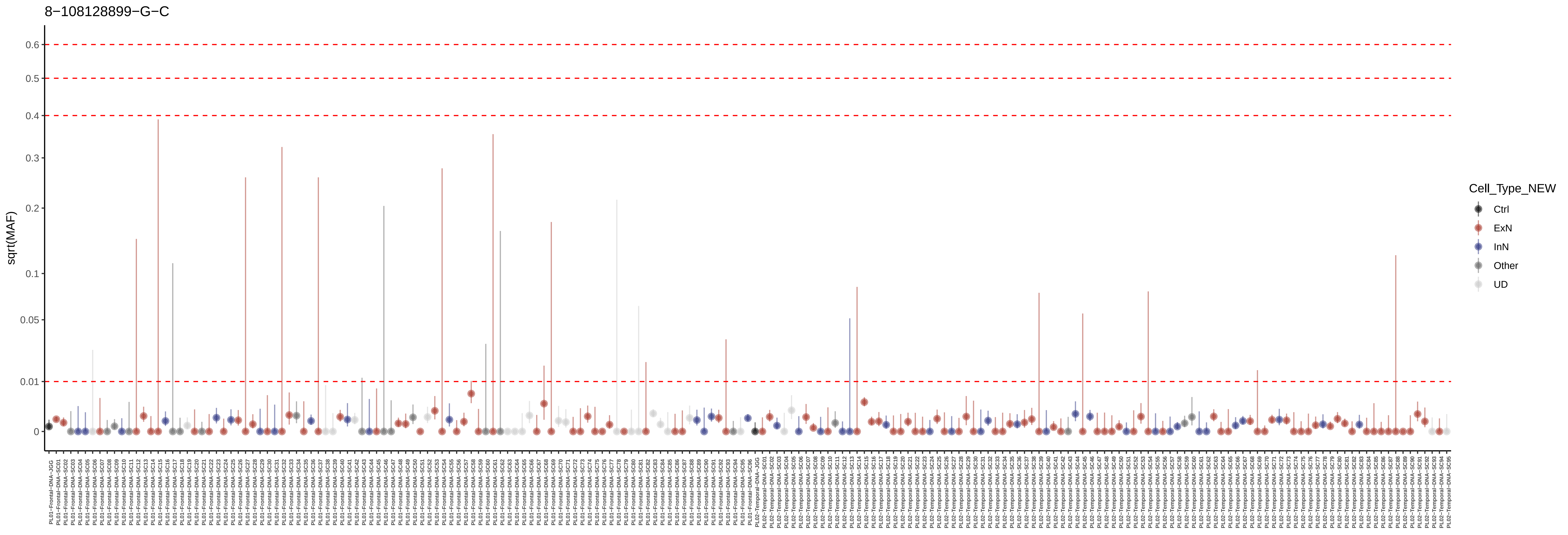
8−108128899−G−C
sqrt(MAF)
0.6
0.5
0.4
0.3
0.2
0.1
0.05
0.01
0
Cell_Type_NEW
Ctrl
ExN
InN
Other
UD

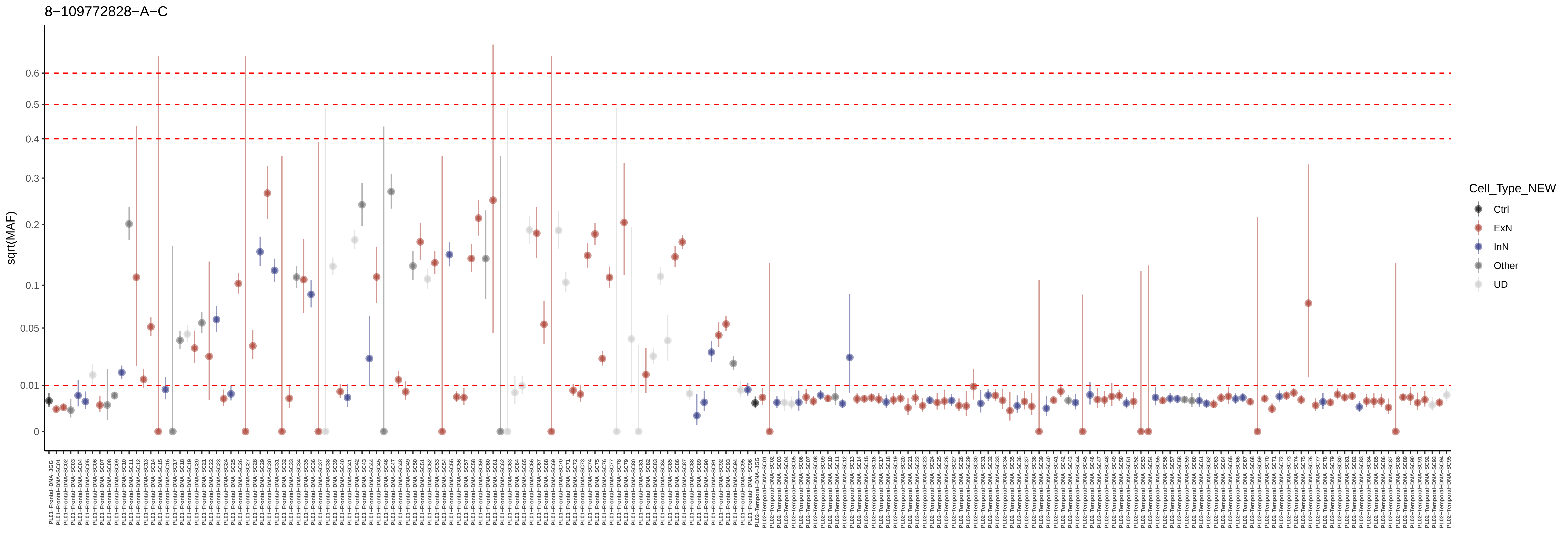
8−109772828−A−C
sqrt(MAF)
0.6
0.5
0.4
0.3
0.2
0.1
0.05
0.01
0
Cell_Type_NEW
Ctrl
ExN
InN
Other
UD

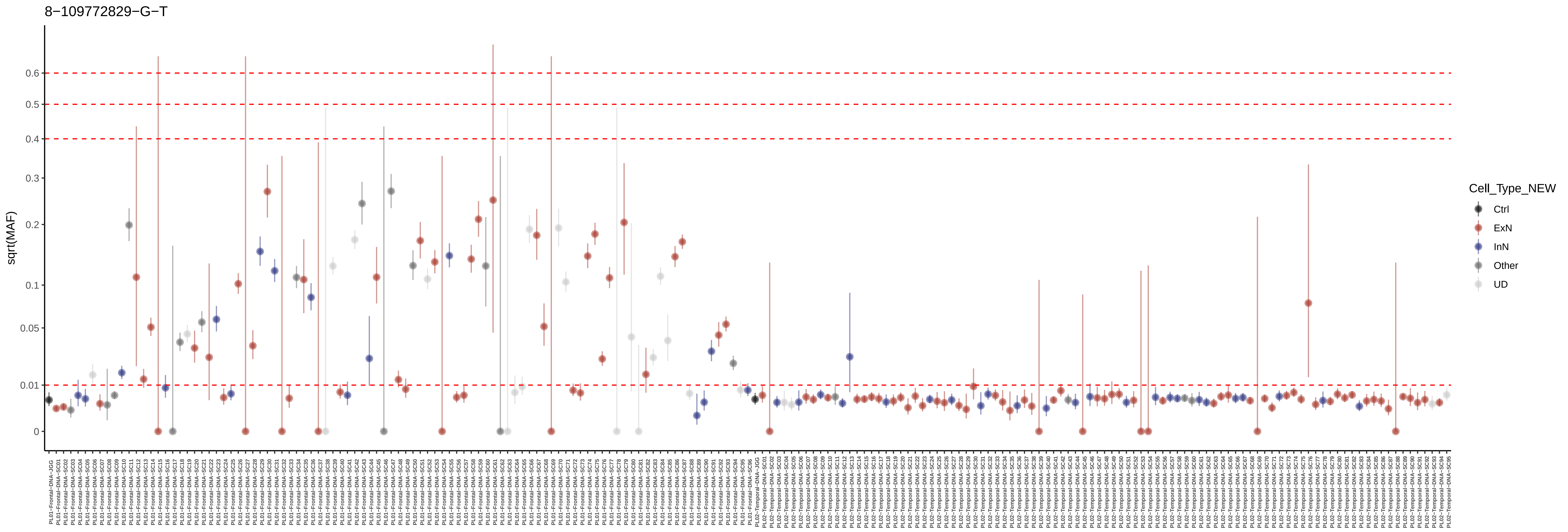
8−109772829−G−T
sqrt(MAF)
0.6
0.5
0.4
0.3
0.2
0.1
0.05
0.01
0
Cell_Type_NEW
Ctrl
ExN
InN
Other
UD

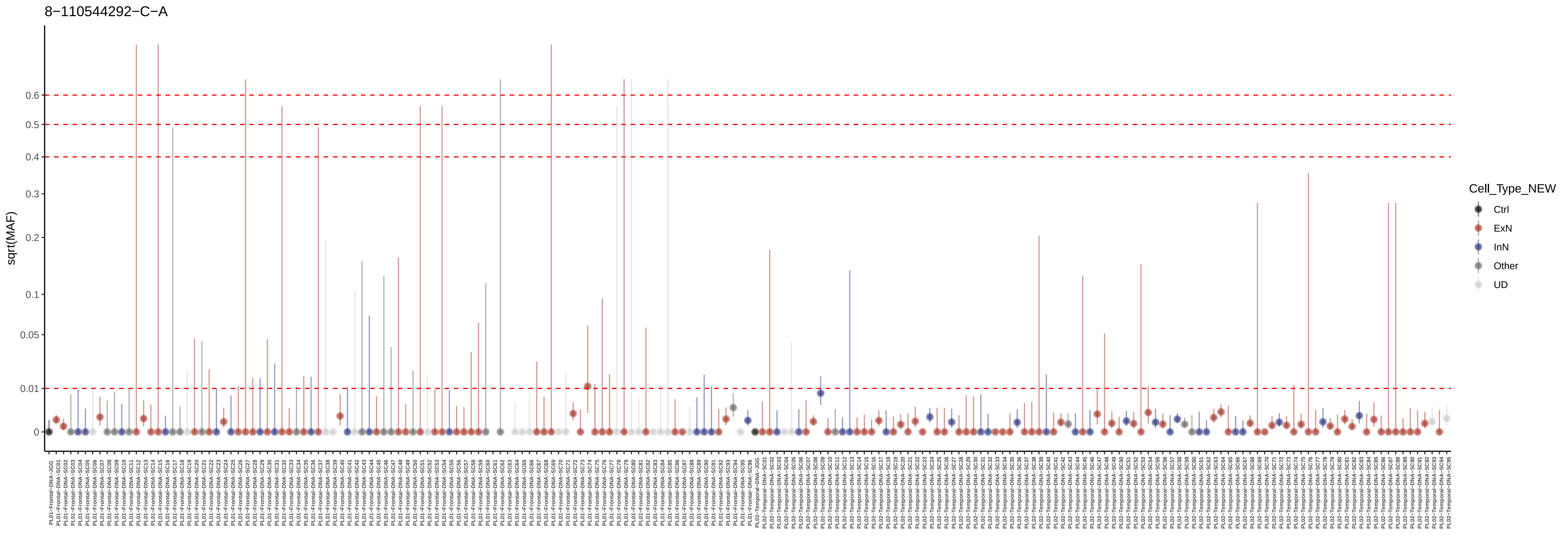
8−110544292−C−A
sqrt(MAF)
0.6
0.5
0.4
0.3
0.2
0.1
0.05
0.01
0
Cell_Type_NEW
Ctrl
ExN
InN
Other
UD

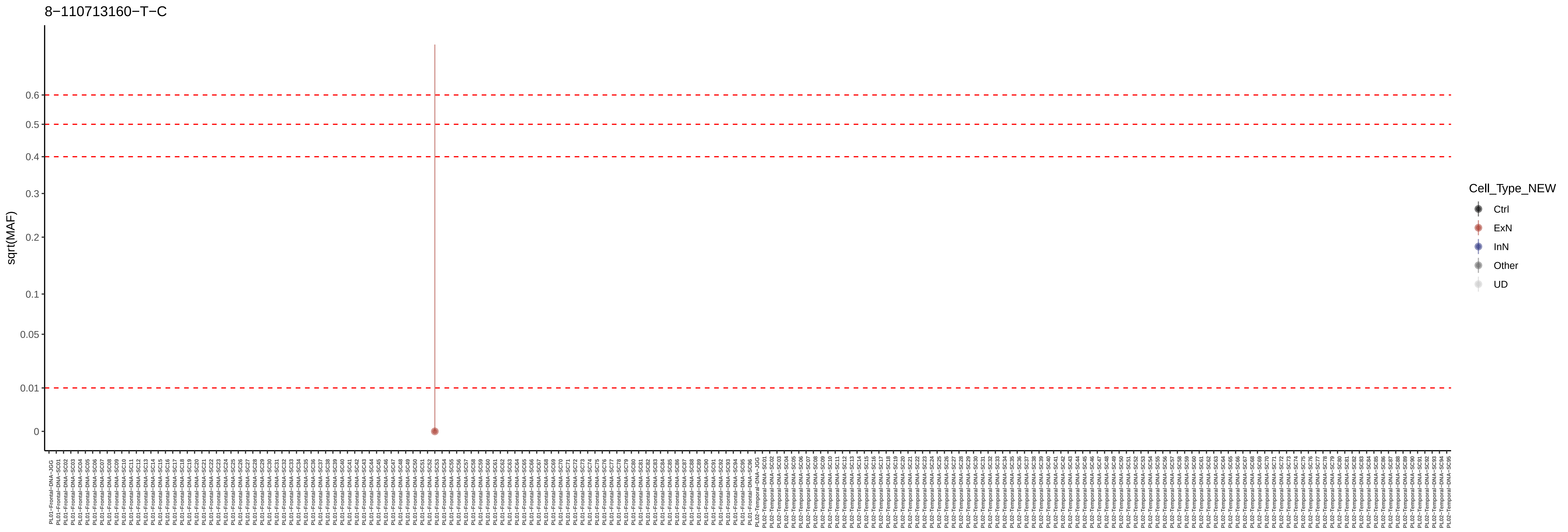

8−110713160−T−C
sqrt(MAF)
0.6
0.5
0.4
0.3
0.2
0.1
0.05
0.01
0
Cell_Type_NEW
Ctrl
ExN
InN
Other
UD

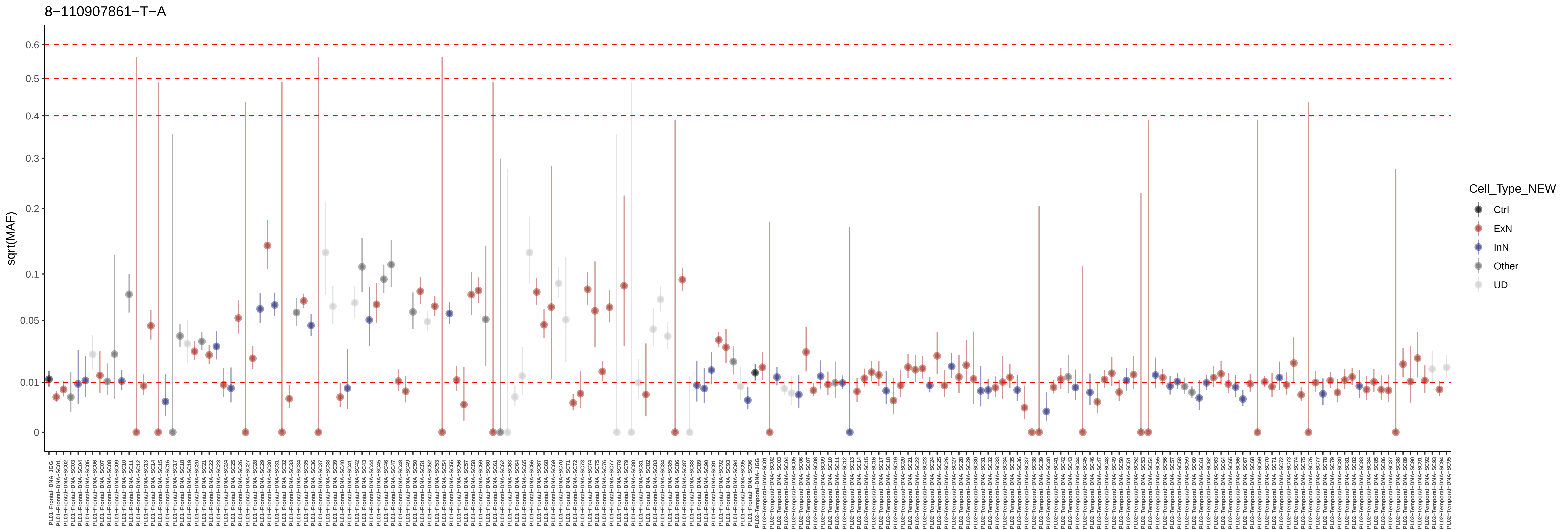

8−110907861−T−A
sqrt(MAF)
Cell_Type_NEW
Ctrl
ExN
InN
Other
UD

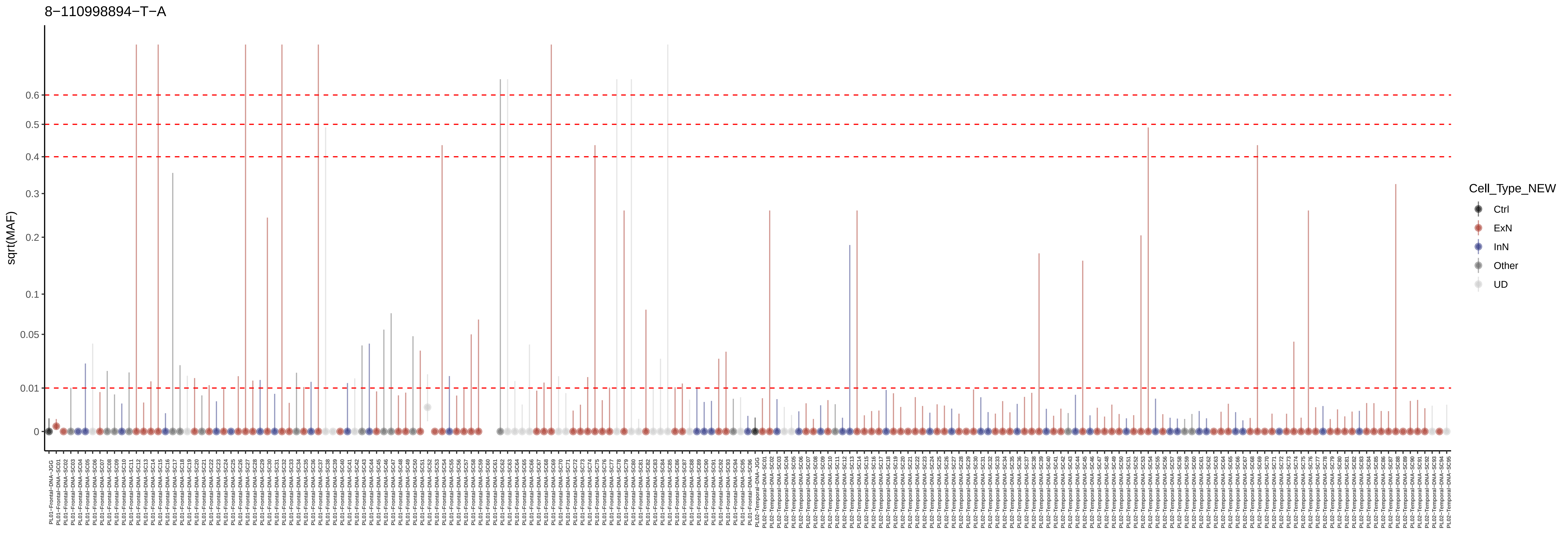

8−110998894−T−A
sqrt(MAF)
0.6
0.5
0.4
0.3
0.2
0.1
0.05
0.01
0
Cell_Type_NEW
Ctrl
ExN
InN
Other
UD

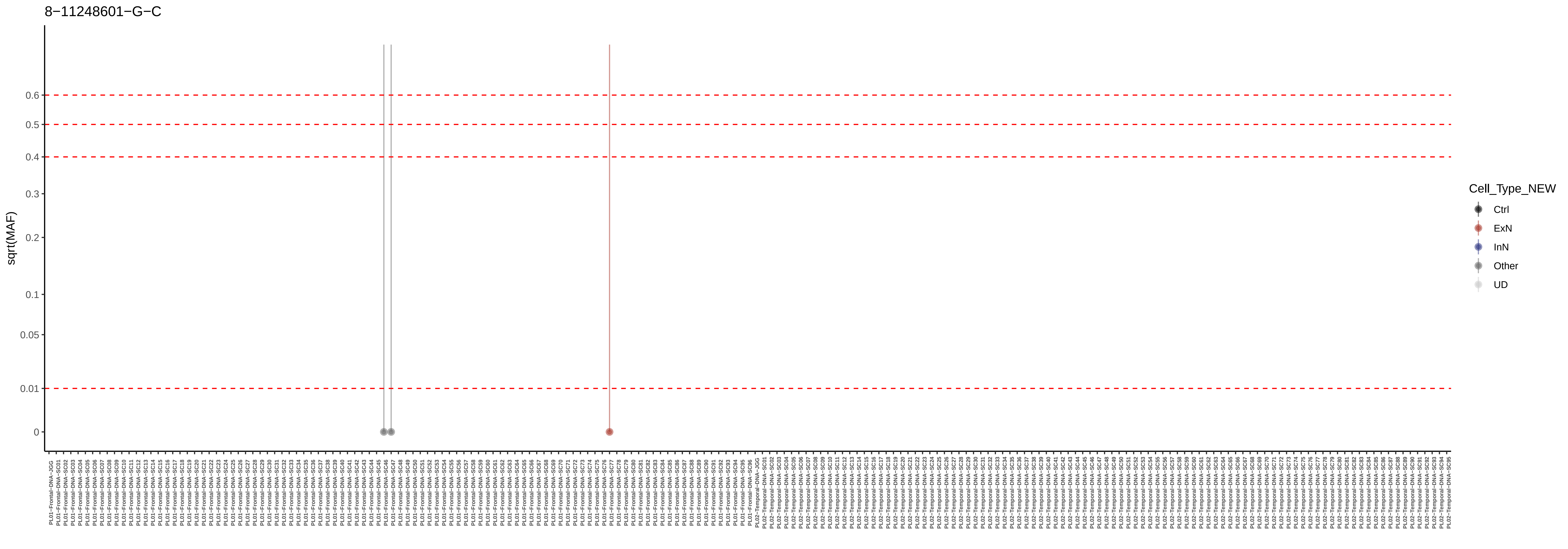

8−11248601−G−C
sqrt(MAF)
0.6
0.5
0.4
0.3
0.2
0.1
0.05
0.01
0
Cell_Type_NEW
Ctrl
ExN
InN
Other
UD

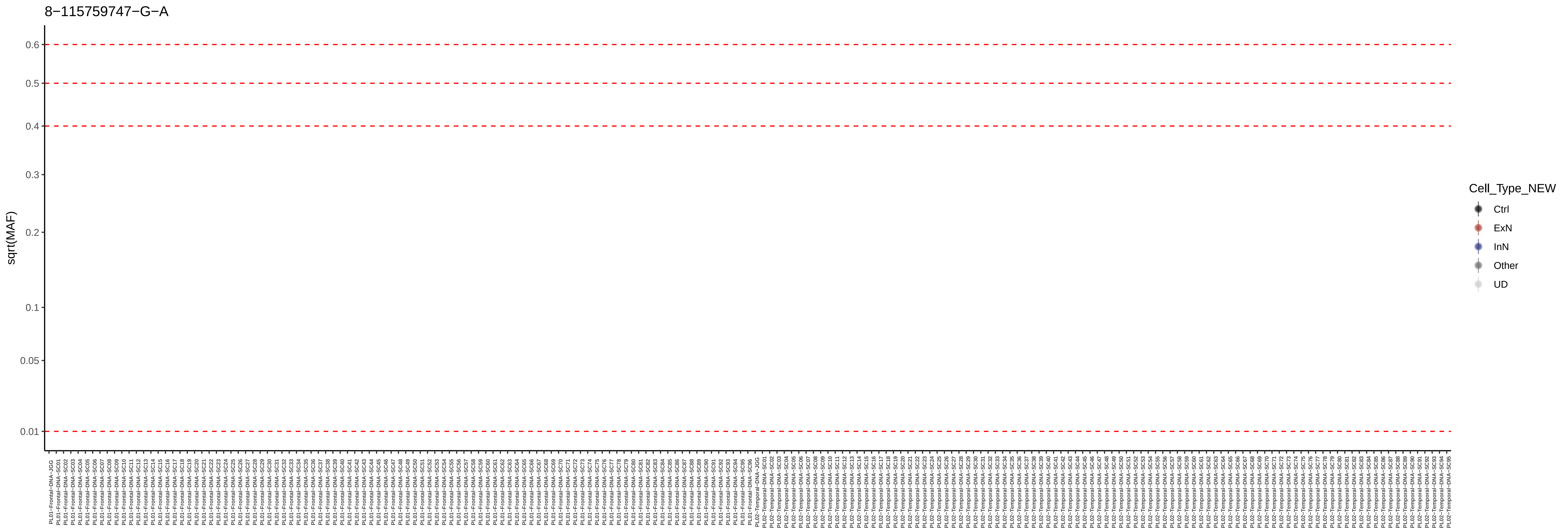
8−115759747−G−A
sqrt(MAF)
0.6
0.5
0.4
0.3
0.2
0.1
0.05
0.01
Cell_Type_NEW
Ctrl
ExN
InN
Other
UD

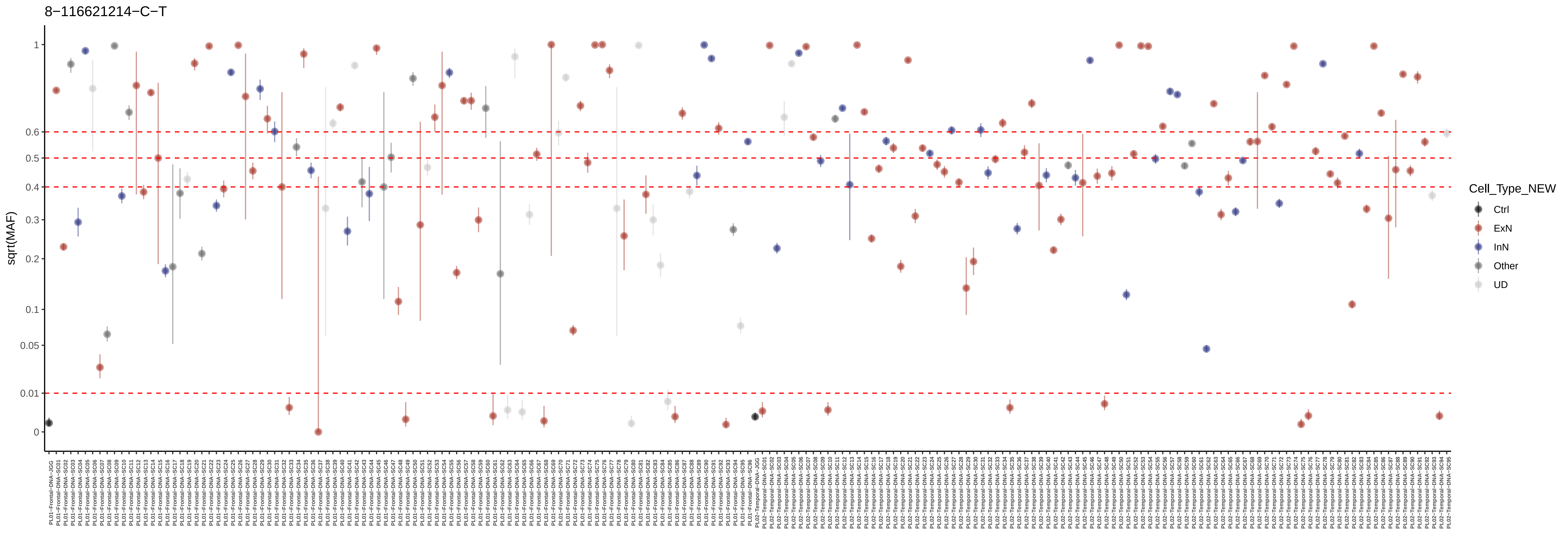
8−116621214−C−T
sqrt(MAF)
1
0.6
0.5
0.4
0.3
0.2
0.1
0.05
0.01
0
Cell_Type_NEW
Ctrl
ExN
InN
Other
UD

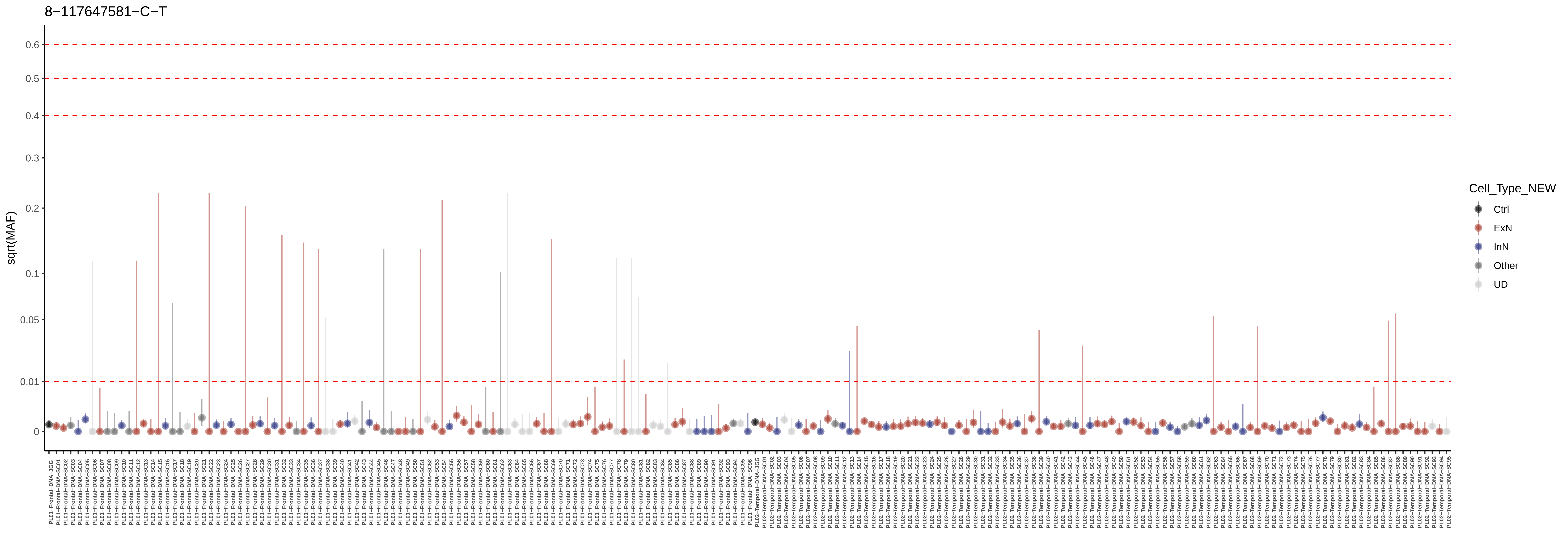
8−117647581−C−T
sqrt(MAF)
0.6
0.5
0.4
0.3
0.2
0.1
0.05
0.01
0
Cell_Type_NEW
Ctrl
ExN
InN
Other
UD

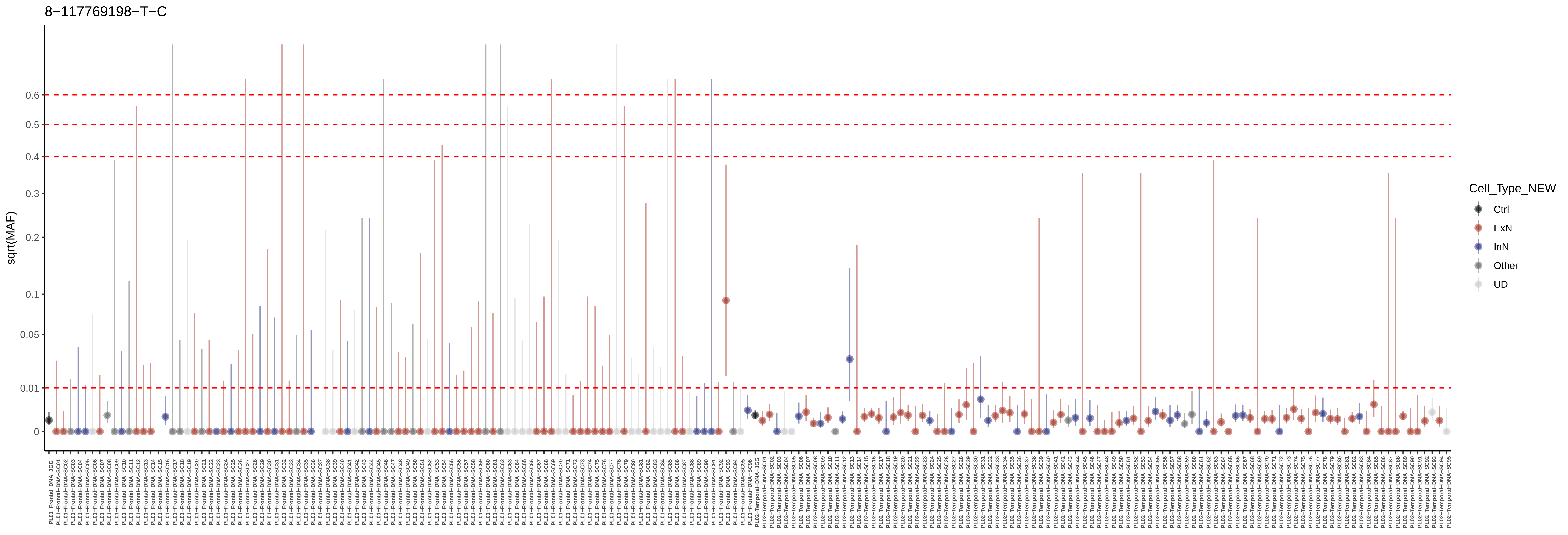
8−117769198−T−C
sqrt(MAF)
0.6
0.5
0.4
0.3
0.2
0.1
0.05
0.01
0
Cell_Type_NEW
Ctrl
ExN
InN
Other
UD

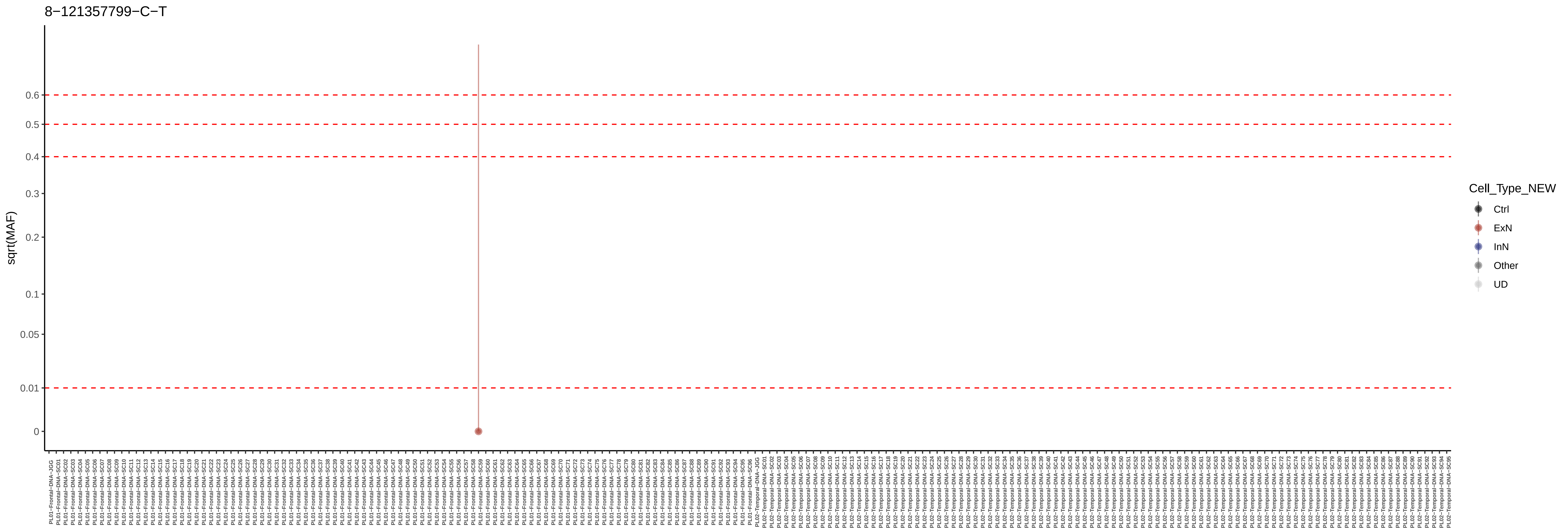

8−121357799−C−T
sqrt(MAF)
0.6
0.5
0.4
0.3
0.2
0.1
0.05
0.01
0
Cell_Type_NEW
Ctrl
ExN
InN
Other
UD

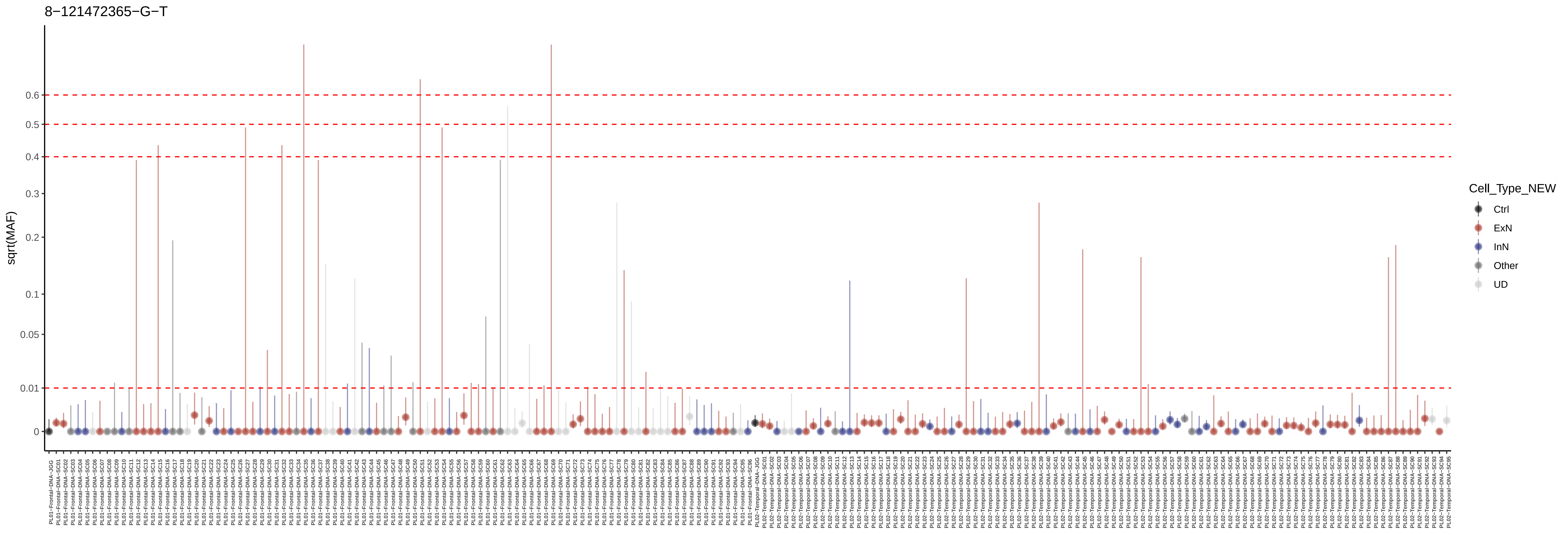
8−121472365−G−T
sqrt(MAF)
0.6
0.5
0.4
0.3
0.2
0.1
0.05
0.01
0
Cell_Type_NEW
Ctrl
ExN
InN
Other
UD

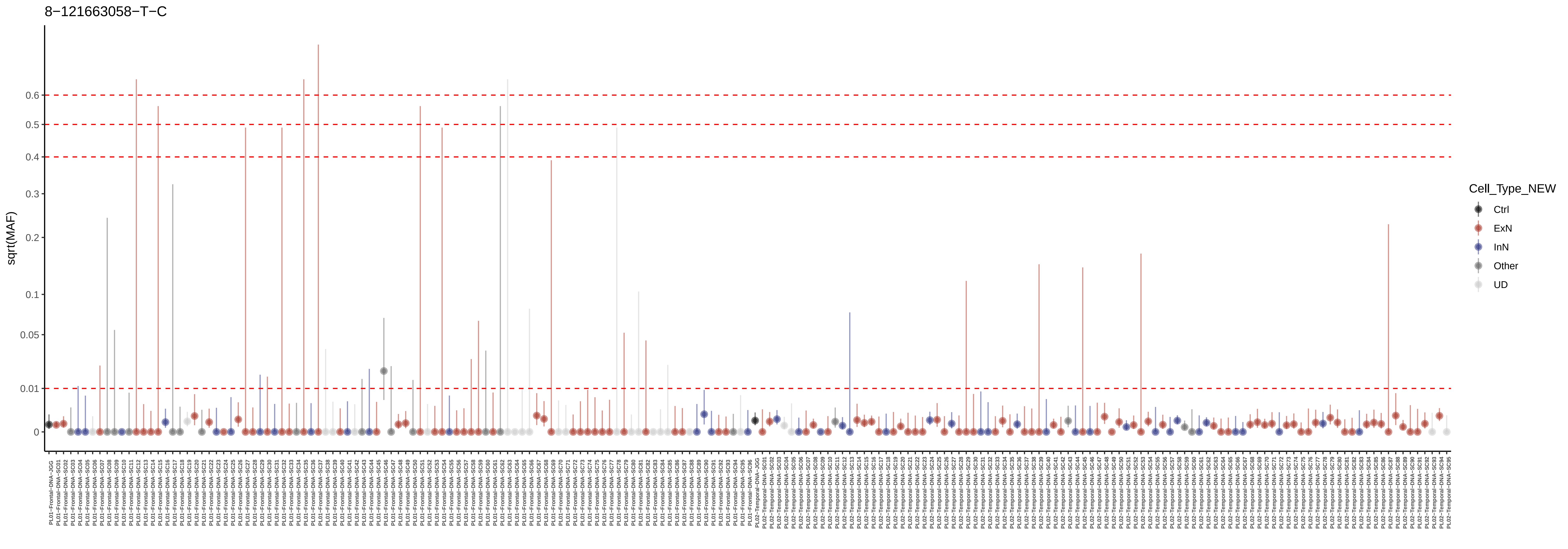
8−121663058−T−C
sqrt(MAF)
0.6
0.5
0.4
0.3
0.2
0.1
0.05
0.01
0
Cell_Type_NEW
Ctrl
ExN
InN
Other
UD

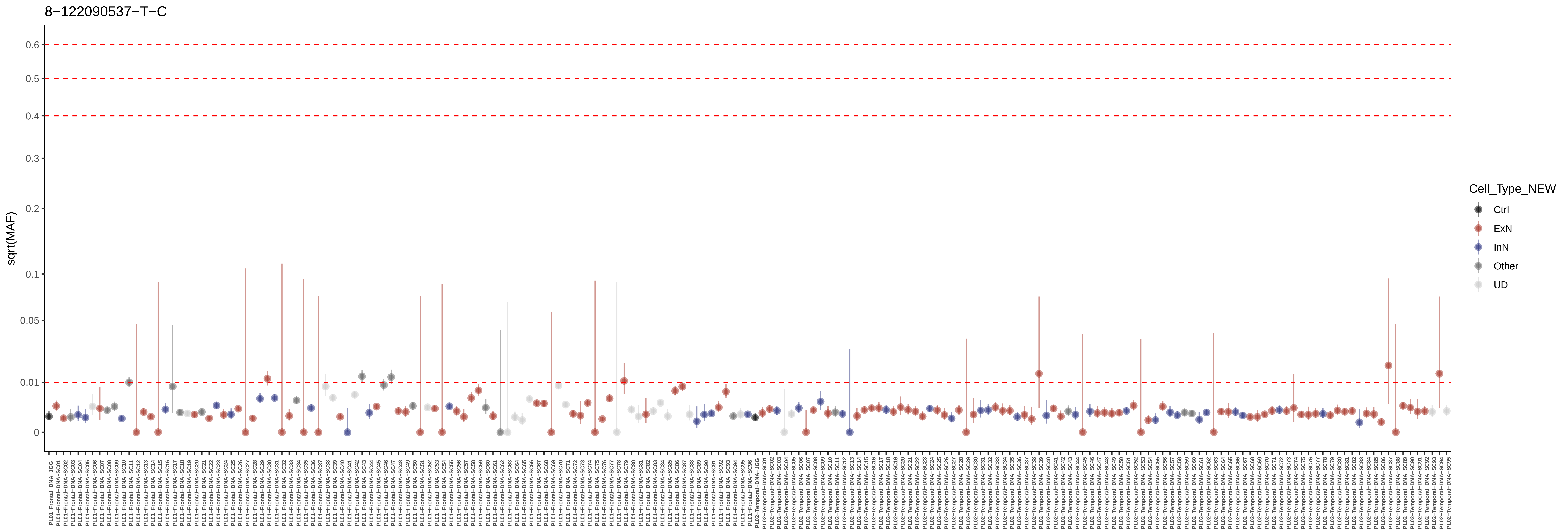

8−122090537−T−C
sqrt(MAF)
0.6
0.5
0.4
0.3
0.2
0.1
0.05
0.01
0
Cell_Type_NEW
Ctrl
ExN
InN
Other
UD

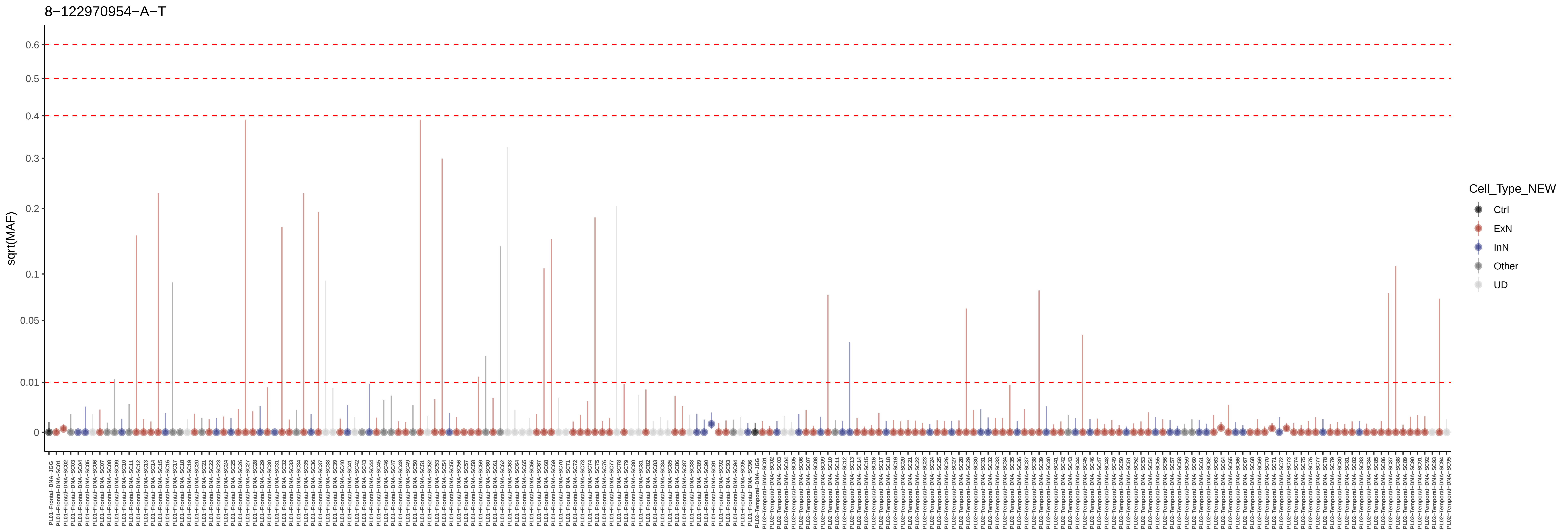
8−122970954−A−T
sqrt(MAF)
0.6
0.5
0.4
0.3
0.2
0.1
0.05
0.01
0
Cell_Type_NEW
Ctrl
ExN
InN
Other
UD

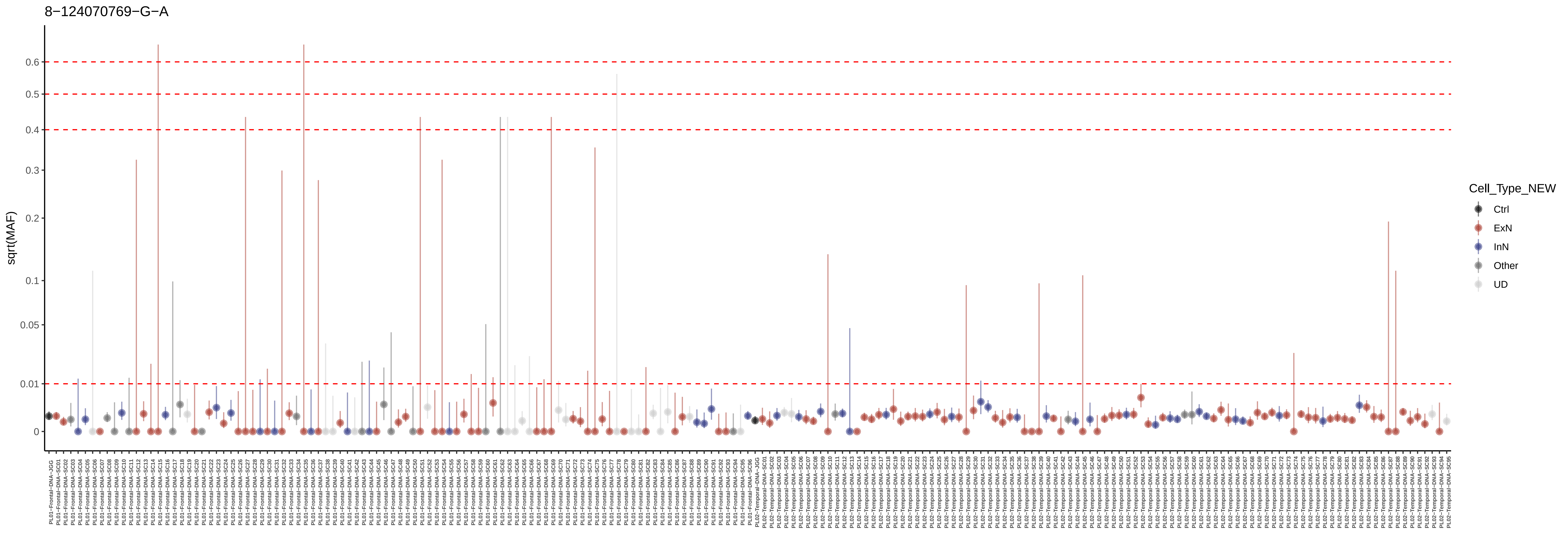
8−124070769−G−A
sqrt(MAF)
0.6
0.5
0.4
0.3
0.2
0.1
0.05
0.01
0
Cell_Type_NEW
Ctrl
ExN
InN
Other
UD

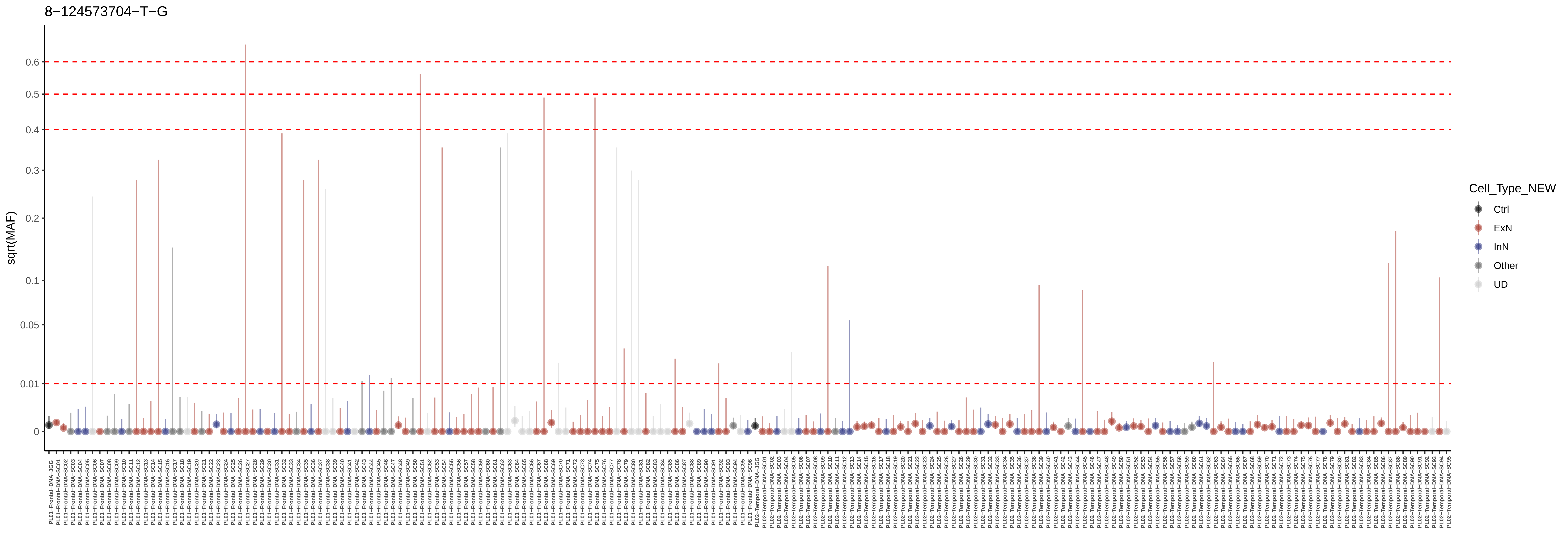

8−124573704−T−G
sqrt(MAF)
0.6
0.5
0.4
0.3
0.2
0.1
0.05
0.01
0
Cell_Type_NEW
Ctrl
ExN
InN
Other
UD

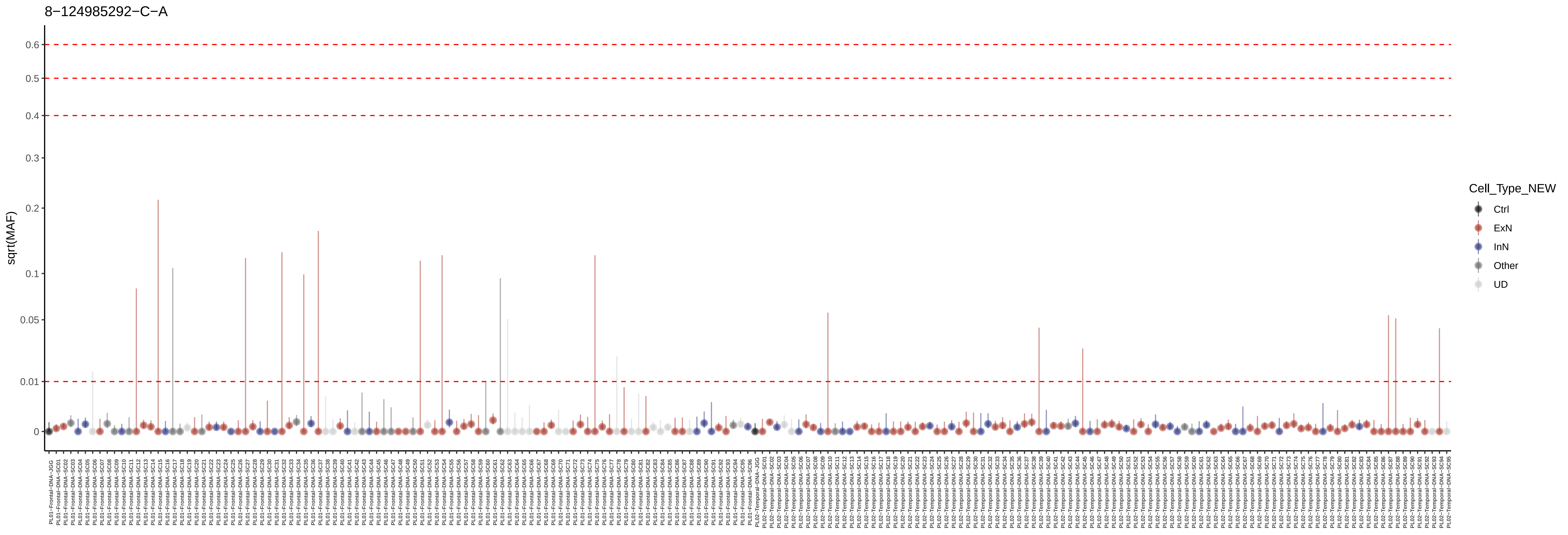

8−124985292−C−A
sqrt(MAF)
0.6
0.5
0.4
0.3
0.2
0.1
0.05
0.01
0
Cell_Type_NEW
Ctrl
ExN
InN
Other
UD

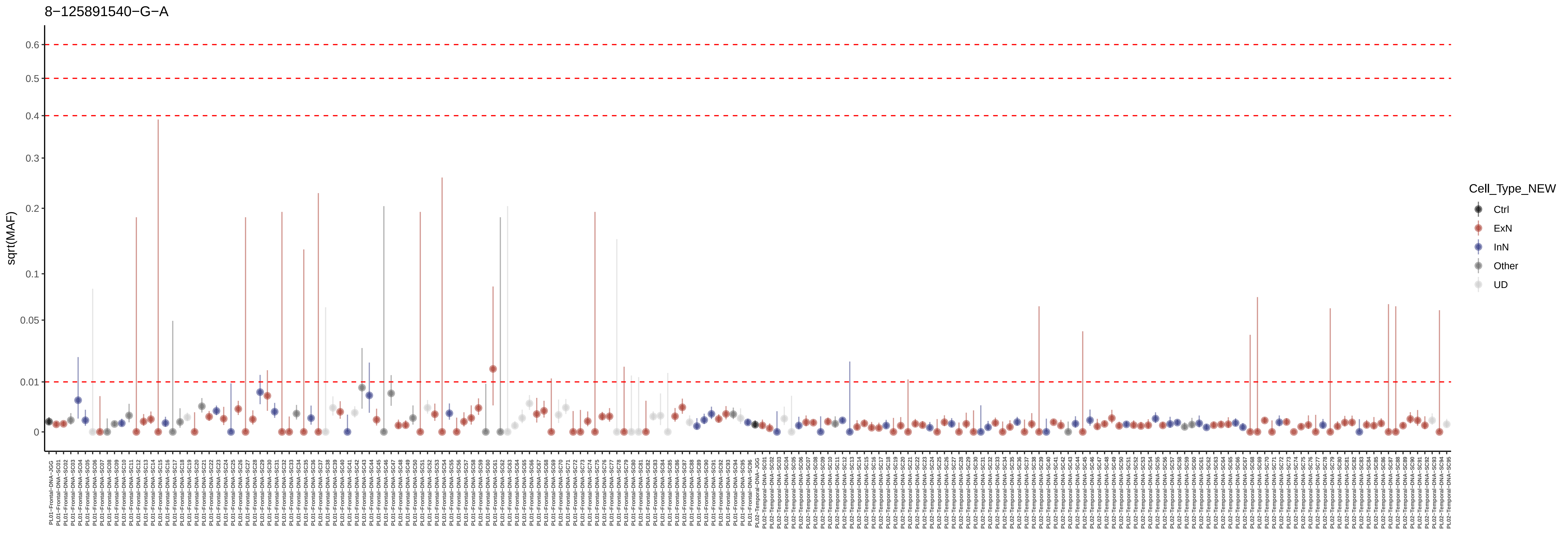
8−125891540−G−A
sqrt(MAF)
0.6
0.5
0.4
0.3
0.2
0.1
0.05
0.01
0
Cell_Type_NEW
Ctrl
ExN
InN
Other
UD

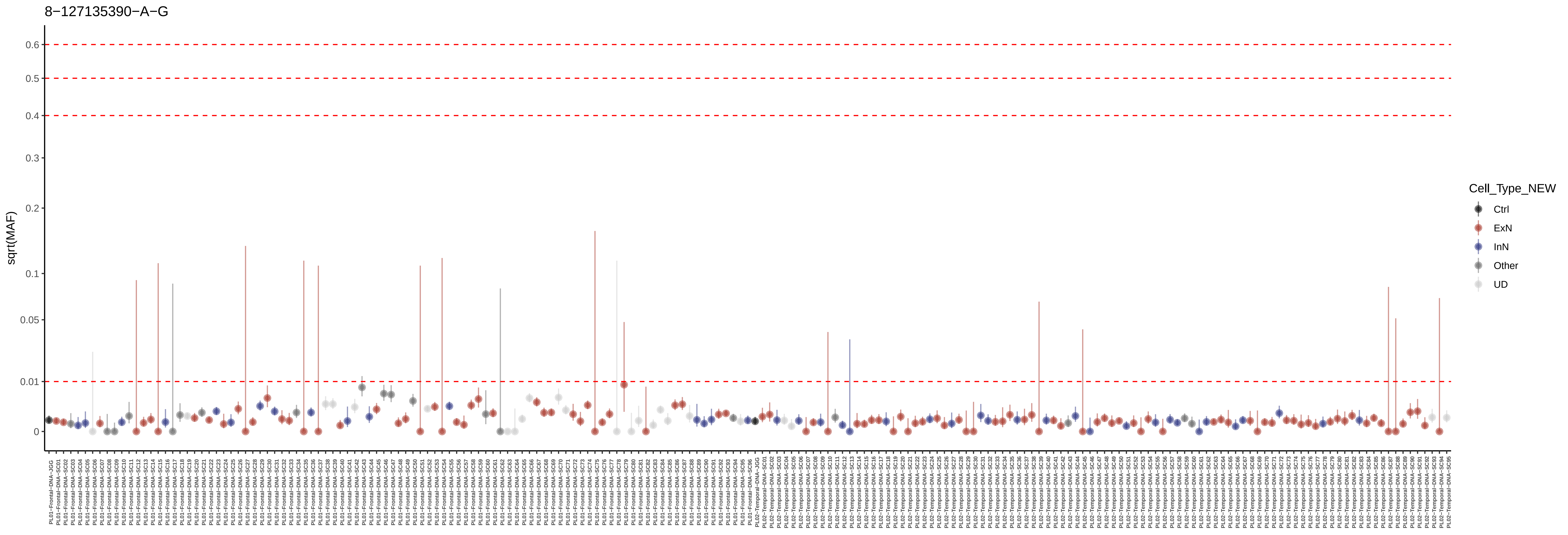
8−127135390−A−G
sqrt(MAF)
0.6
0.5
0.4
0.3
0.2
0.1
0.05
0.01
0
Cell_Type_NEW
Ctrl
ExN
InN
Other
UD

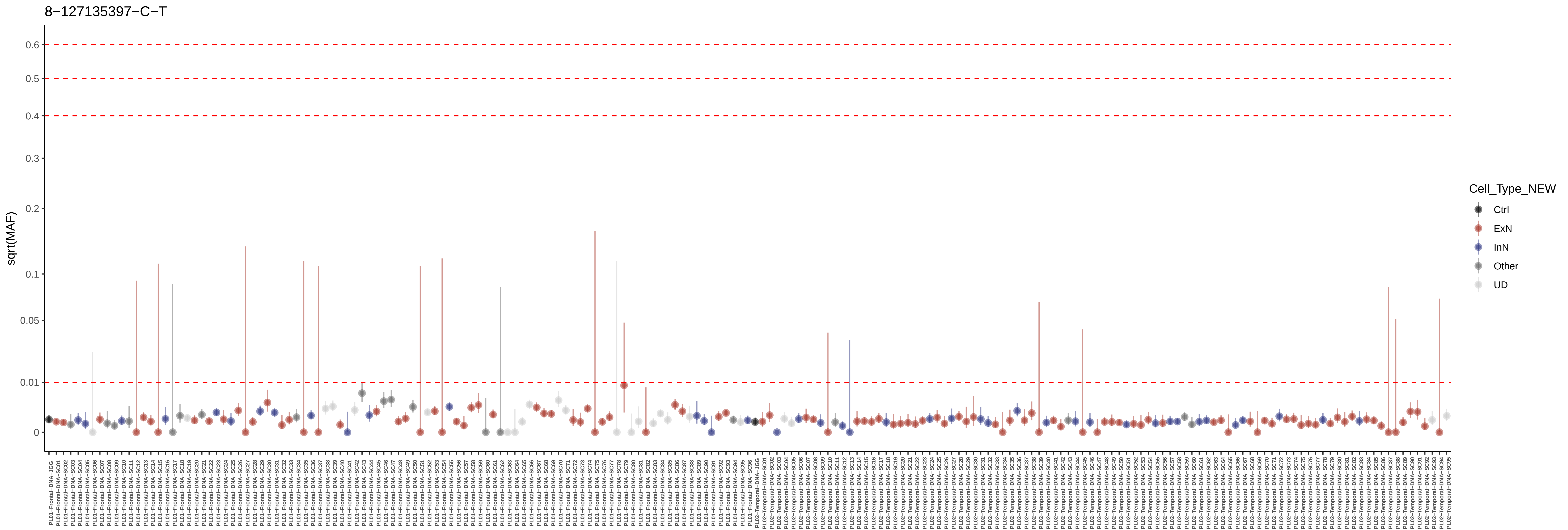
8−127135397−C−T
sqrt(MAF)
0.6
0.5
0.4
0.3
0.2
0.1
0.05
0.01
0
Cell_Type_NEW
Ctrl
ExN
InN
Other
UD

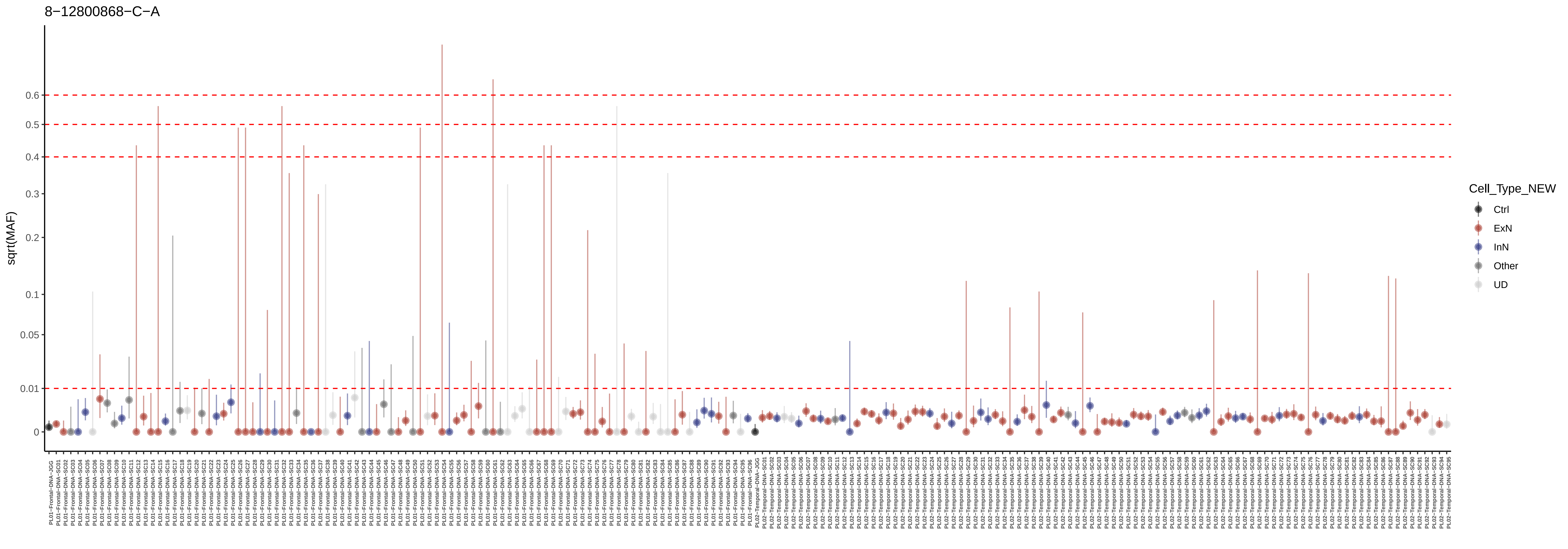

8−12800868−C−A
sqrt(MAF)
0.6
0.5
0.4
0.3
0.2
0.1
0.05
0.01
0
Cell_Type_NEW
Ctrl
ExN
InN
Other
UD

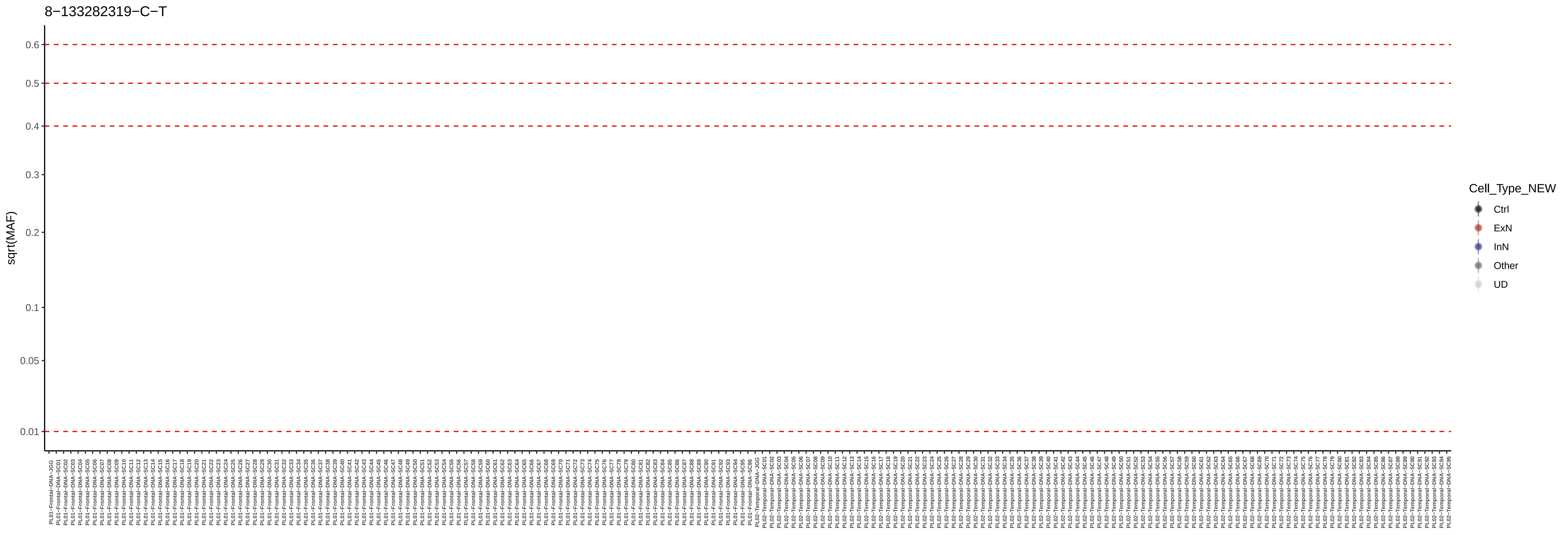
8−133282319−C−T
sqrt(MAF)
0.6
0.5
0.4
0.3
0.2
0.1
0.05
0.01
Cell_Type_NEW
Ctrl
ExN
InN
Other
UD

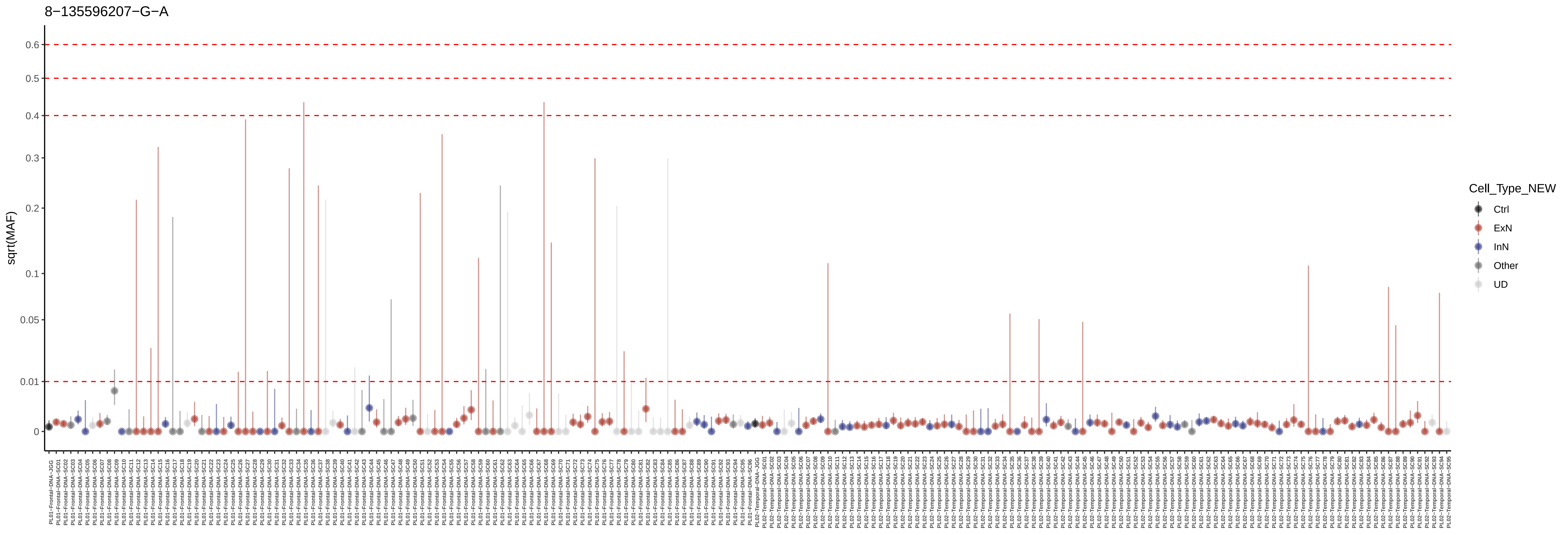

8−135596207−G−A
sqrt(MAF)
0.6
0.5
0.4
0.3
0.2
0.1
0.05
0.01
0
Cell_Type_NEW
Ctrl
ExN
InN
Other
UD

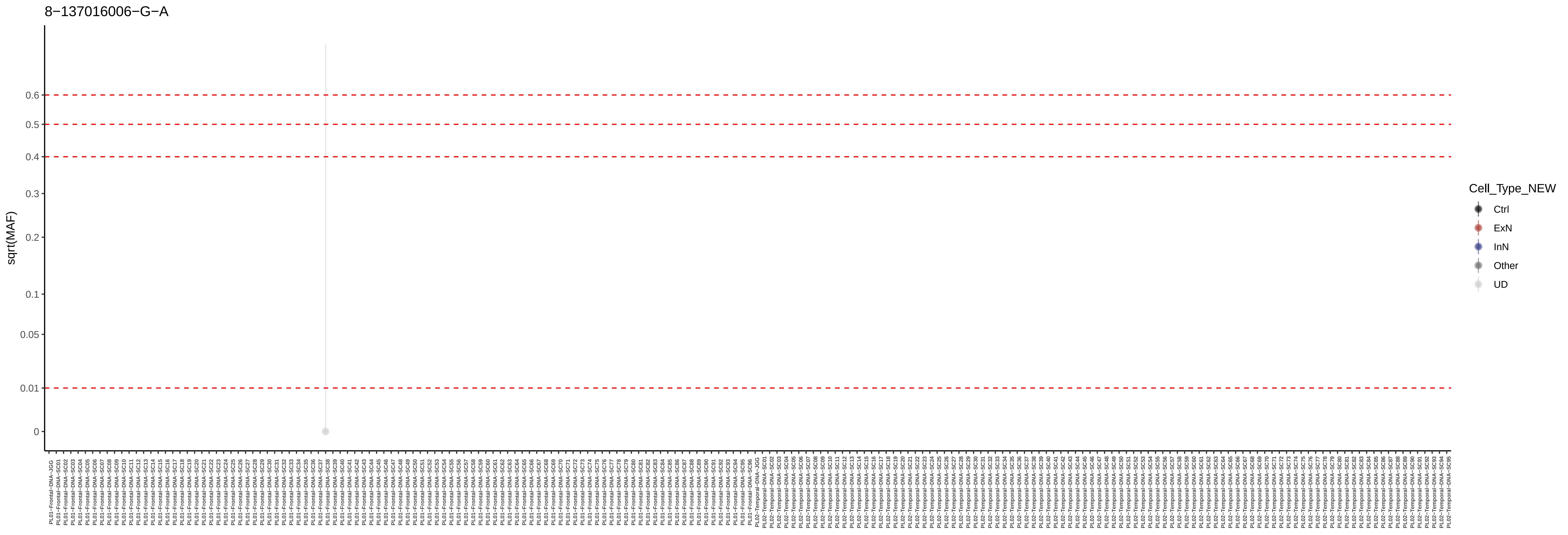

8−137016006−G−A
sqrt(MAF)
0.6
0.5
0.4
0.3
0.2
0.1
0.05
0.01
0
Cell_Type_NEW
Ctrl
ExN
InN
Other
UD

8−137016007−T−C

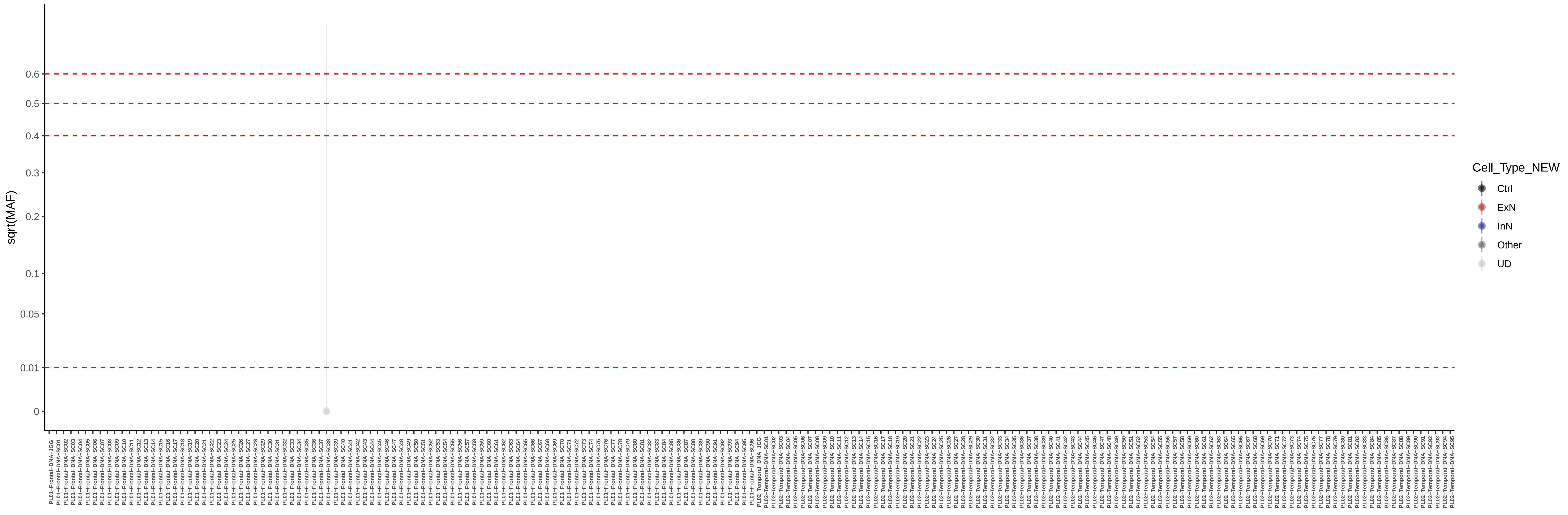

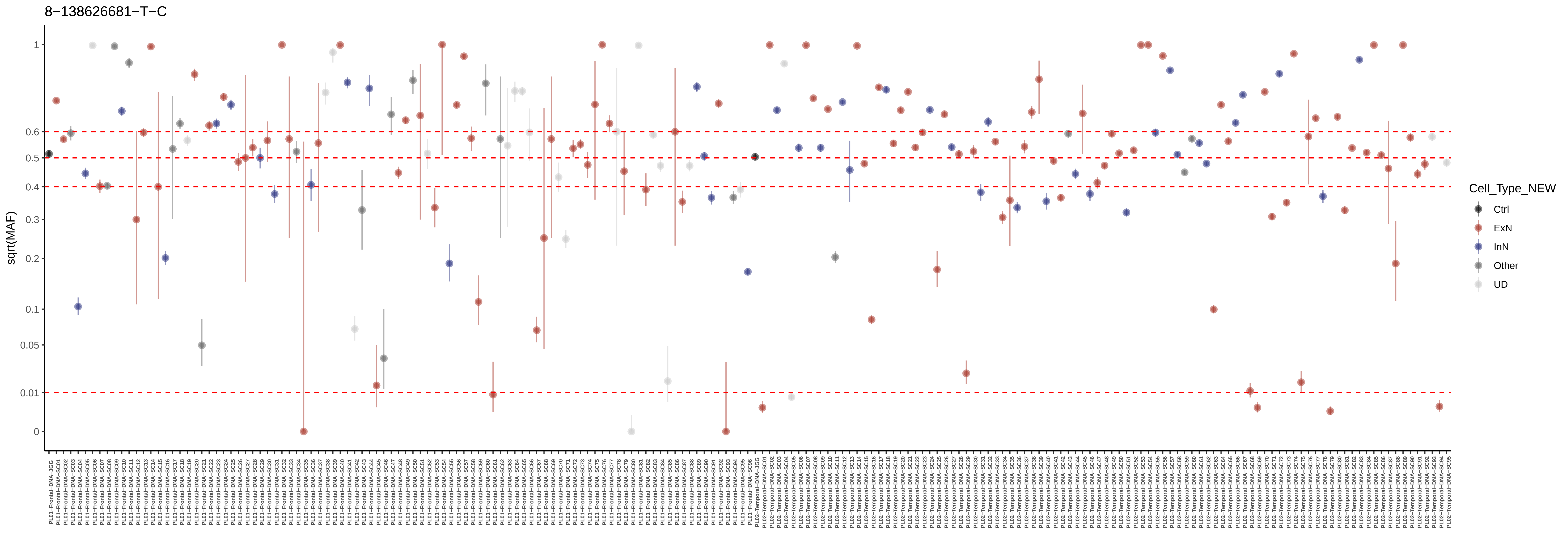
8−138626681−T−C
sqrt(MAF)
1
0.6
0.5
0.4
0.3
0.2
0.1
0.05
0.01
0
Cell_Type_NEW
Ctrl
ExN
InN
Other
UD

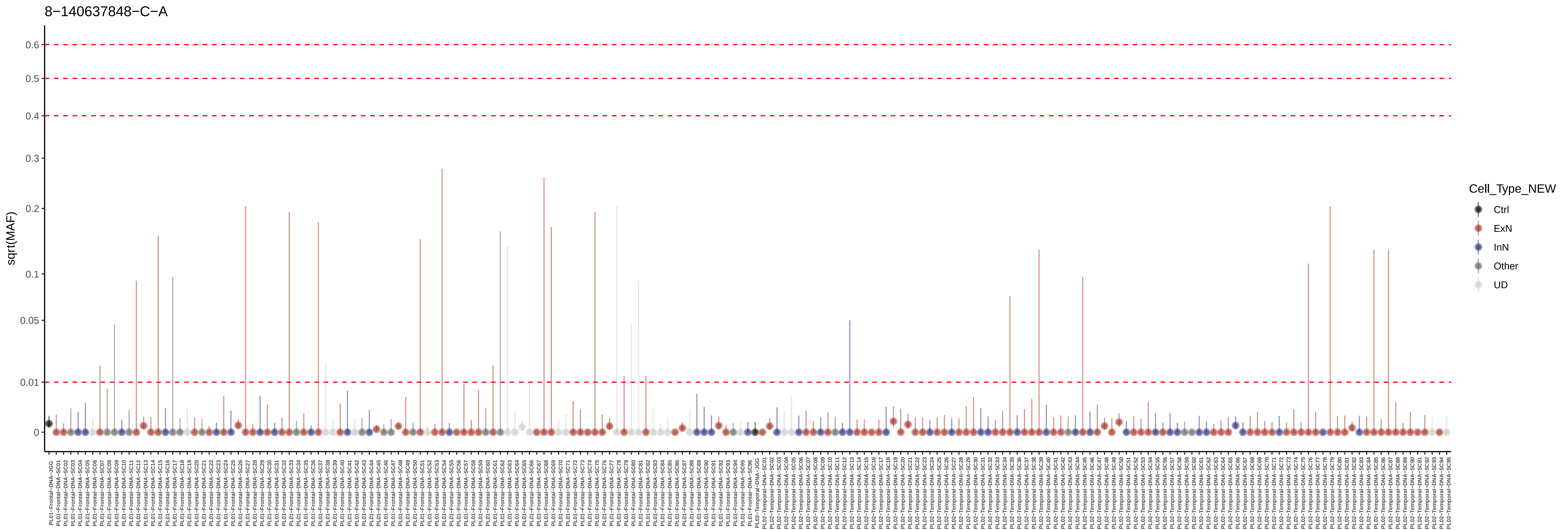
8−140637848−C−A
sqrt(MAF)
Cell_Type_NEW
Ctrl
ExN
InN
Other
UD

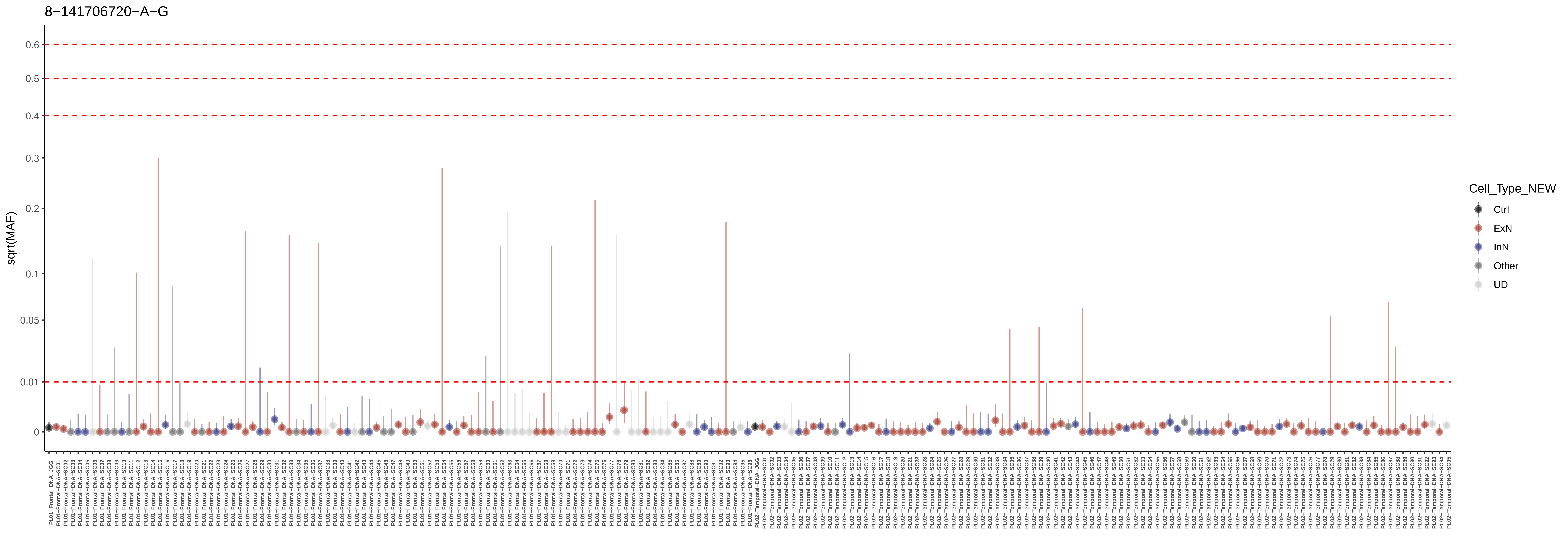

8−141706720−A−G
sqrt(MAF)
0.6
0.5
0.4
0.3
0.2
0.1
0.05
0.01
0
Cell_Type_NEW
Ctrl
ExN
InN
Other
UD

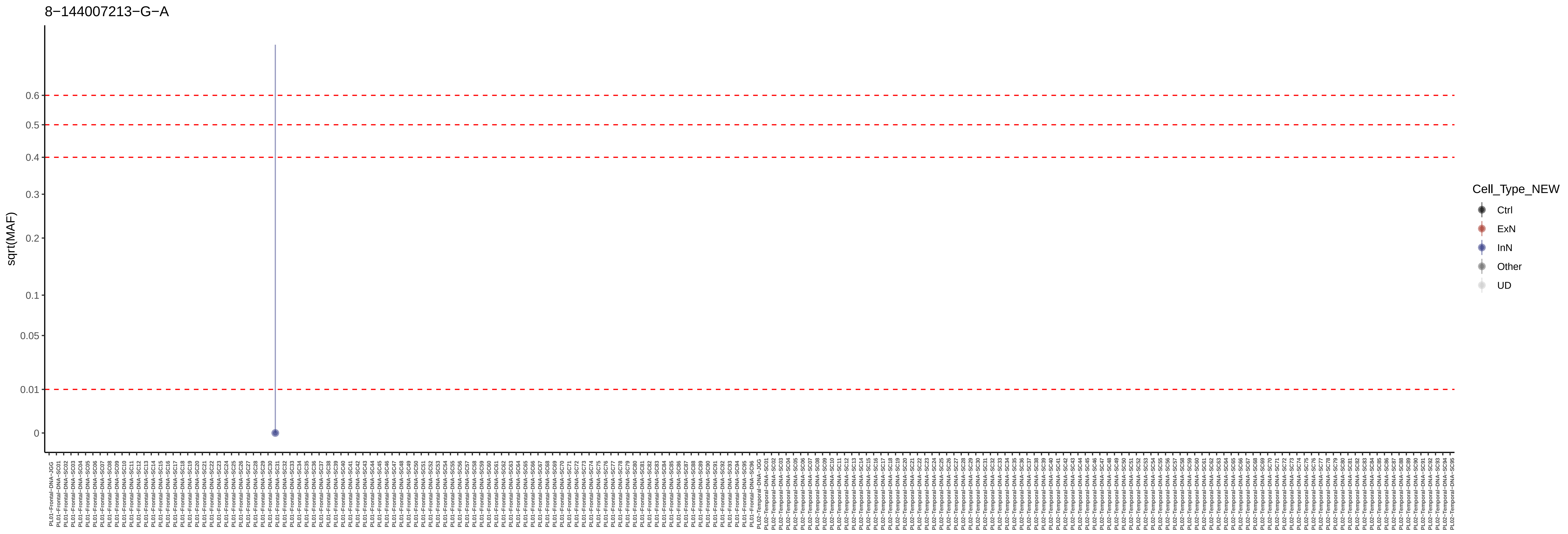

8−144007213−G−A
sqrt(MAF)
0.6
0.5
0.4
0.3
0.2
0.1
0.05
0.01
0
Cell_Type_NEW
Ctrl
ExN
InN
Other
UD

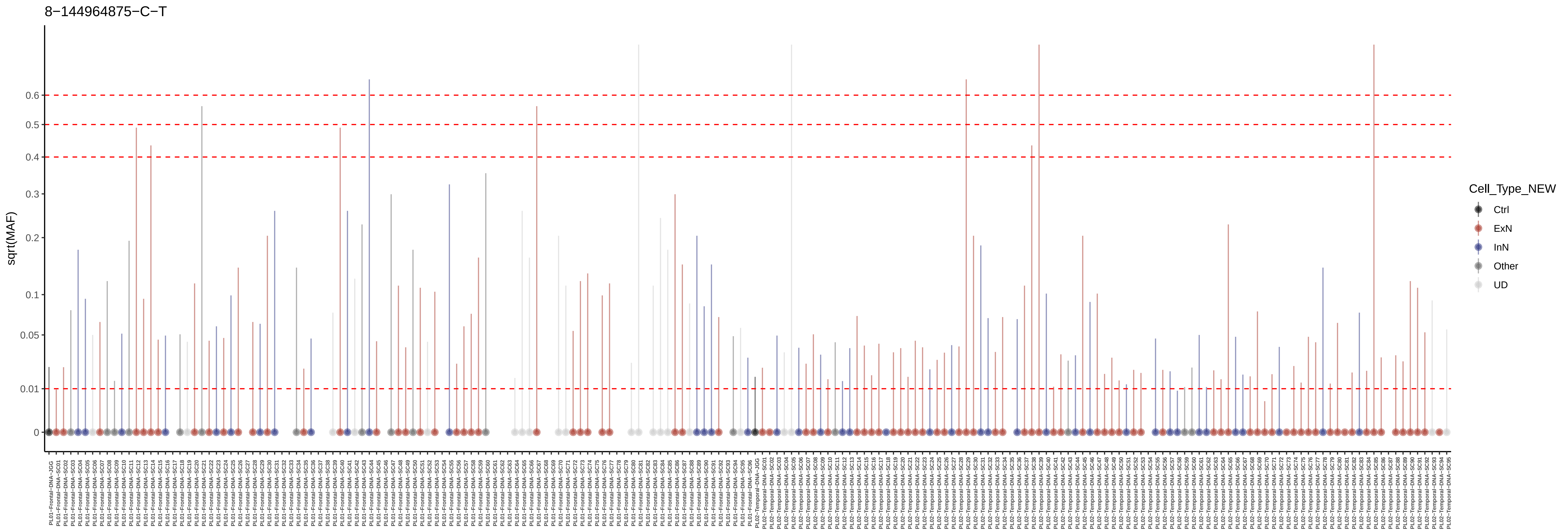

8−144964875−C−T
sqrt(MAF)
0.6
0.5
0.4
0.3
0.2
0.1
0.05
0.01
0
Cell_Type_NEW
Ctrl
ExN
InN
Other
UD

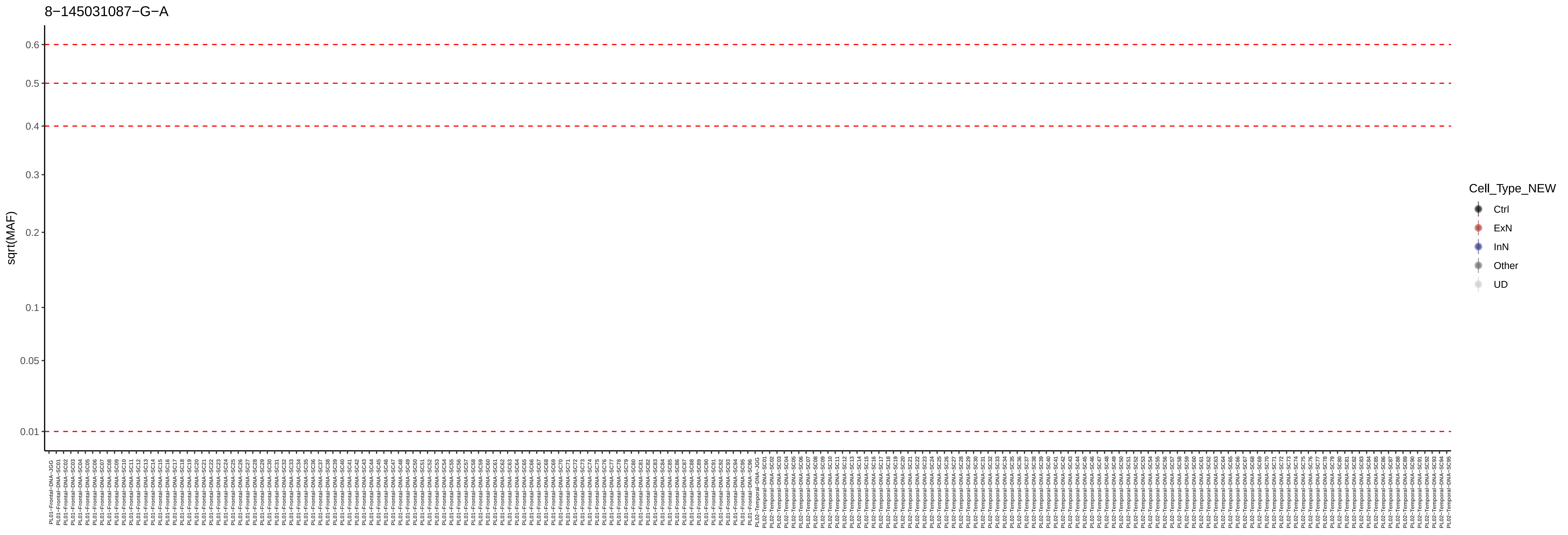

8−145031087−G−A
sqrt(MAF)
0.6
0.5
0.4
0.3
0.2
0.1
0.05
0.01
Cell_Type_NEW
Ctrl
ExN
InN
Other
UD

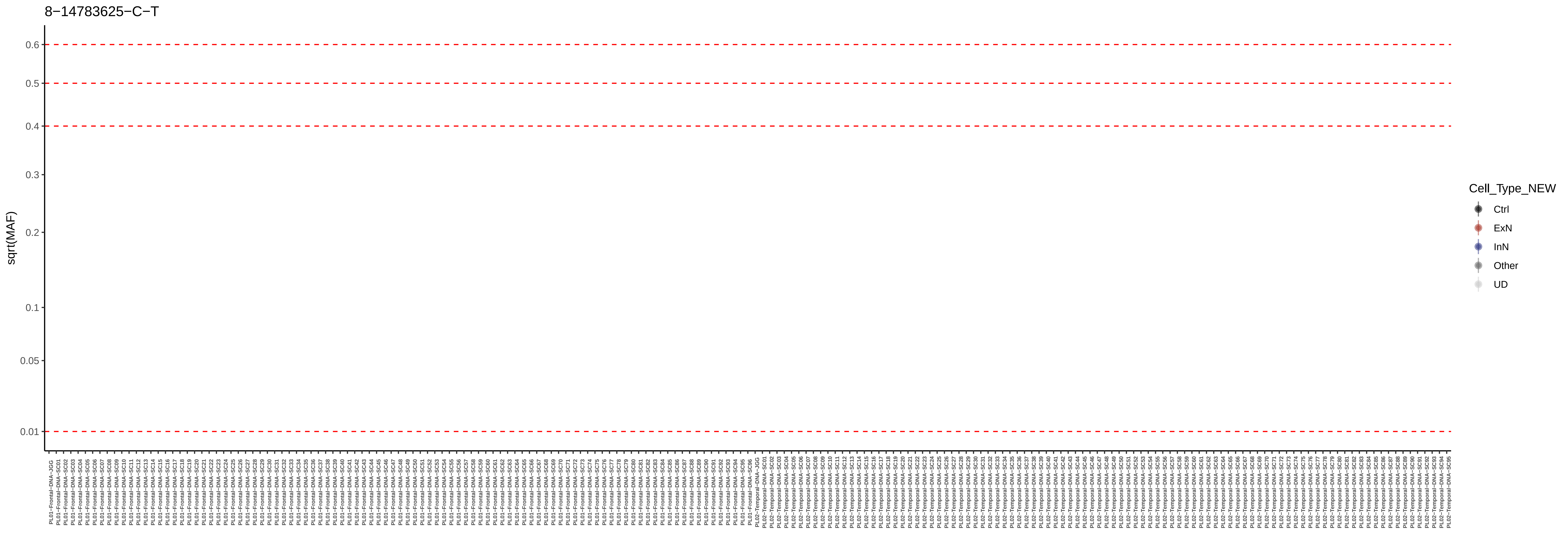
8−14783625−C−T
sqrt(MAF)
0.6
0.5
0.4
0.3
0.2
0.1
0.05
0.01
Cell_Type_NEW
Ctrl
ExN
InN
Other
UD

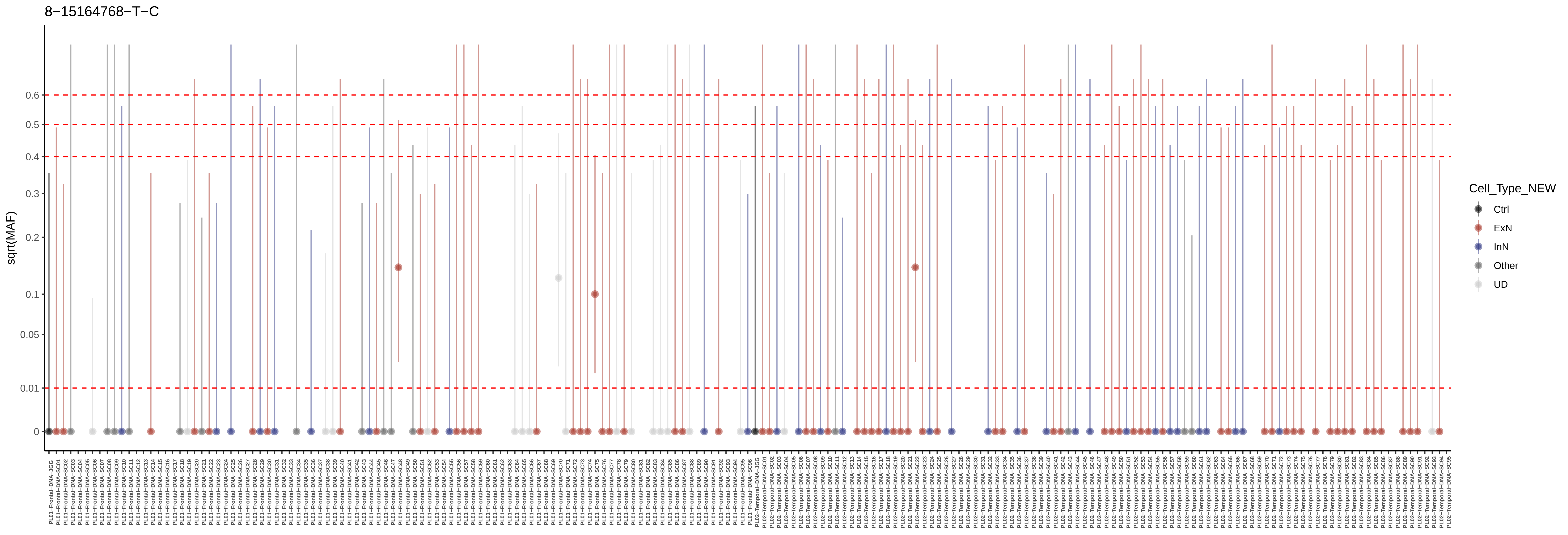

8−15164768−T−C
sqrt(MAF)
0.6
0.5
0.4
0.3
0.2
0.1
0.05
0.01
0
Cell_Type_NEW
Ctrl
ExN
InN
Other
UD

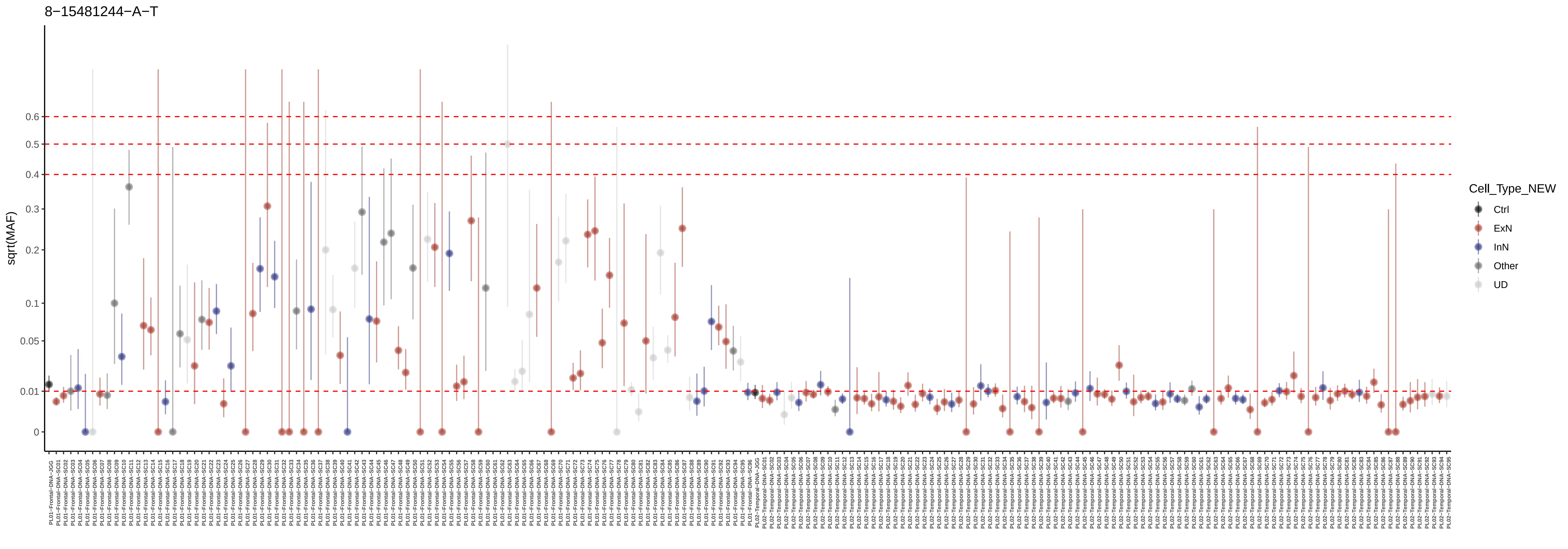

8−15481244−A−T
sqrt(MAF)
0.6
0.5
0.4
0.3
0.2
0.1
0.05
0.01
0
Cell_Type_NEW
Ctrl
ExN
InN
Other
UD

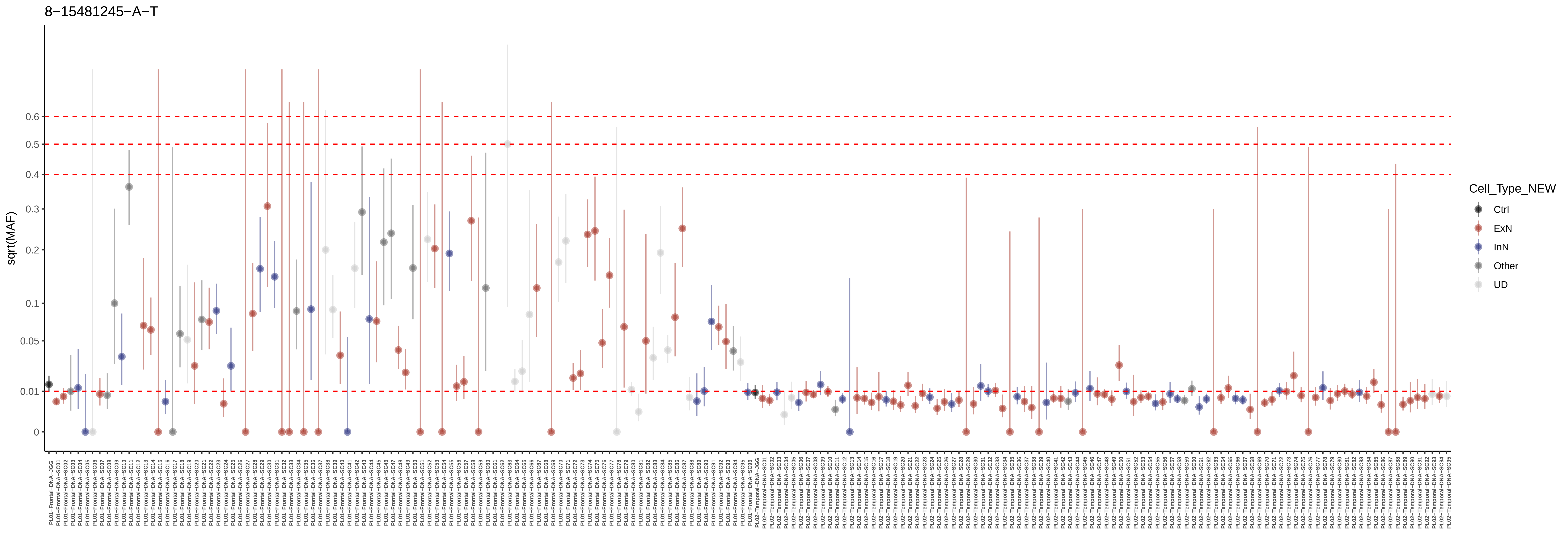
8−15481245−A−T
sqrt(MAF)
0.6
0.5
0.4
0.3
0.2
0.1
0.05
0.01
0
Cell_Type_NEW
Ctrl
ExN
InN
Other
UD

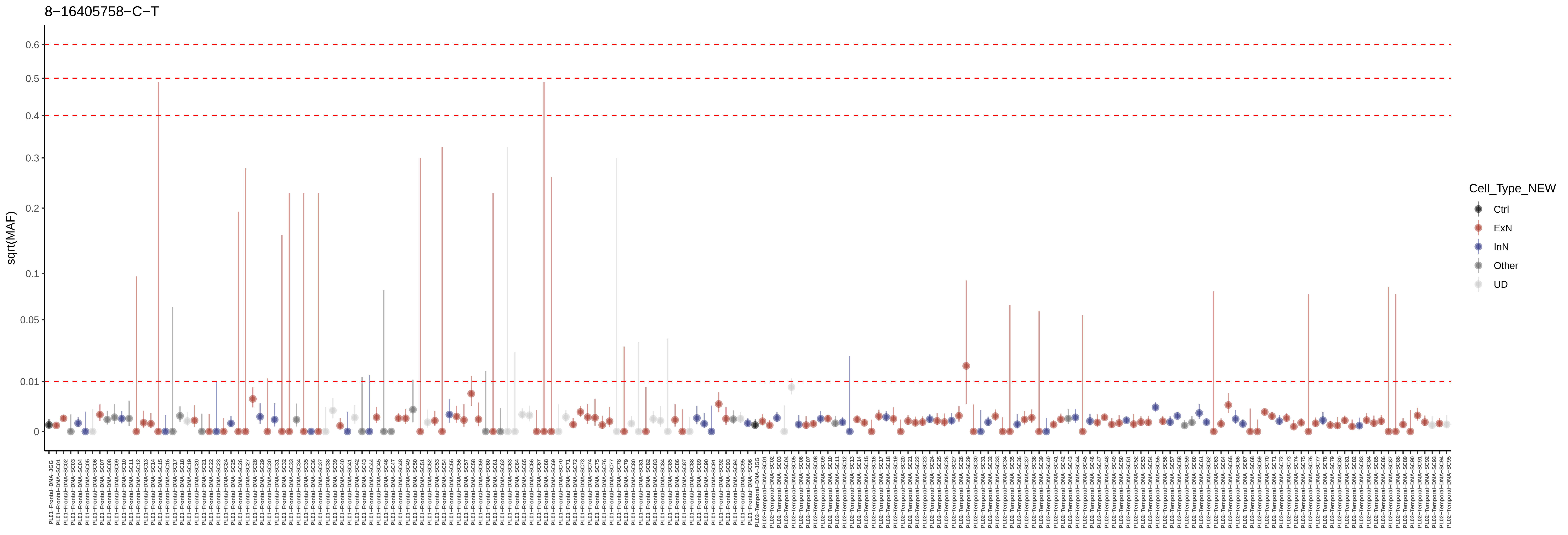

8−16405758−C−T
sqrt(MAF)
0.6
0.5
0.4
0.3
0.2
0.1
0.05
0.01
0
Cell_Type_NEW
Ctrl
ExN
InN
Other
UD

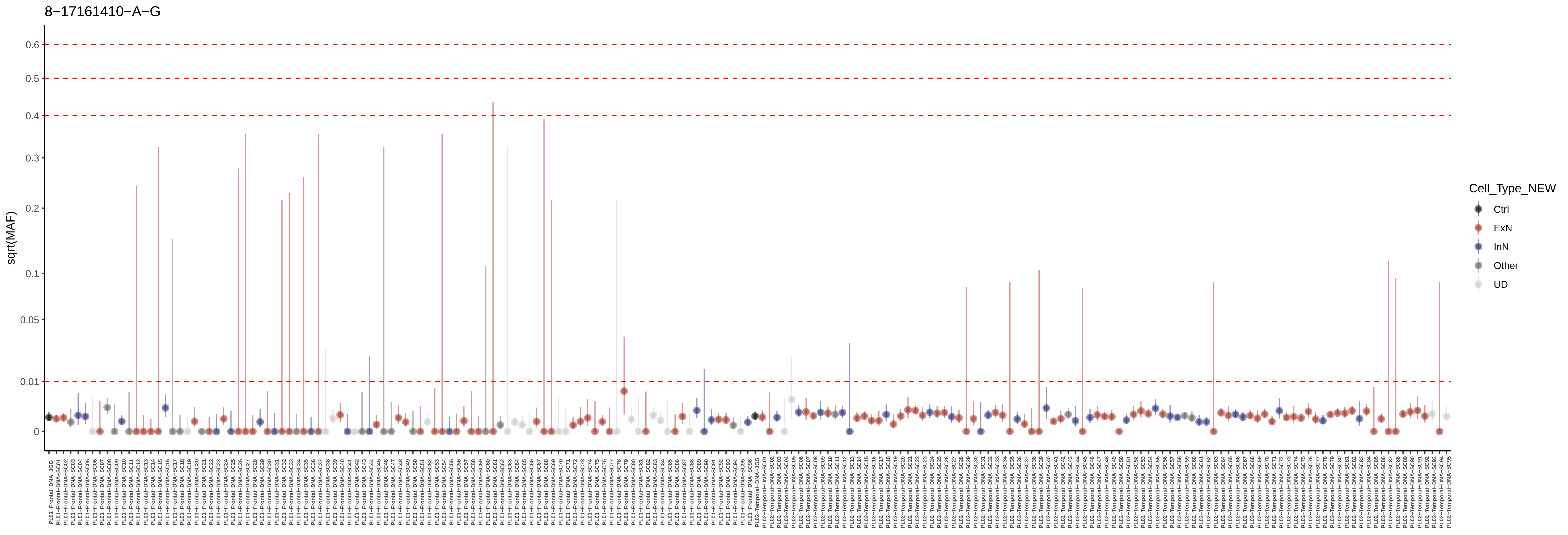
8−17161410−A−G
sqrt(MAF)
0.6
0.5
0.4
0.3
0.2
0.1
0.05
0.01
0
Cell_Type_NEW
Ctrl
ExN
InN
Other
UD

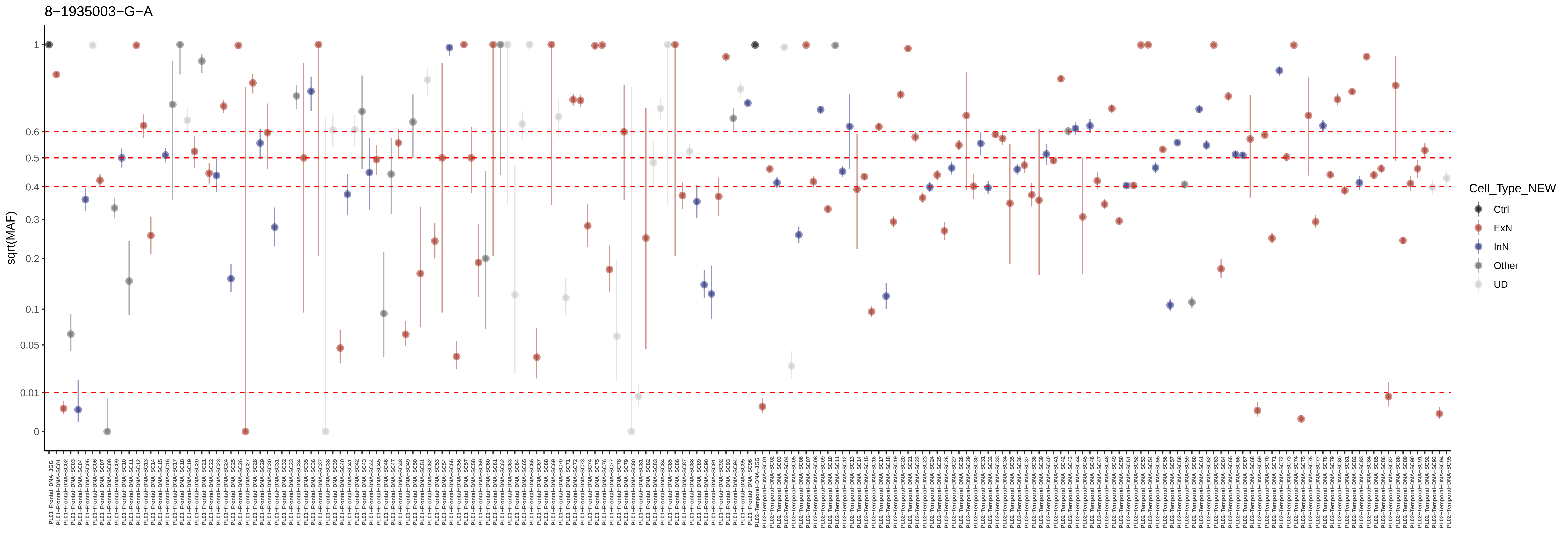

8−1935003−G−A
sqrt(MAF)
1
0.6
0.5
0.4
0.3
0.2
0.1
0.05
0.01
0
Cell_Type_NEW
Ctrl
ExN
InN
Other
UD

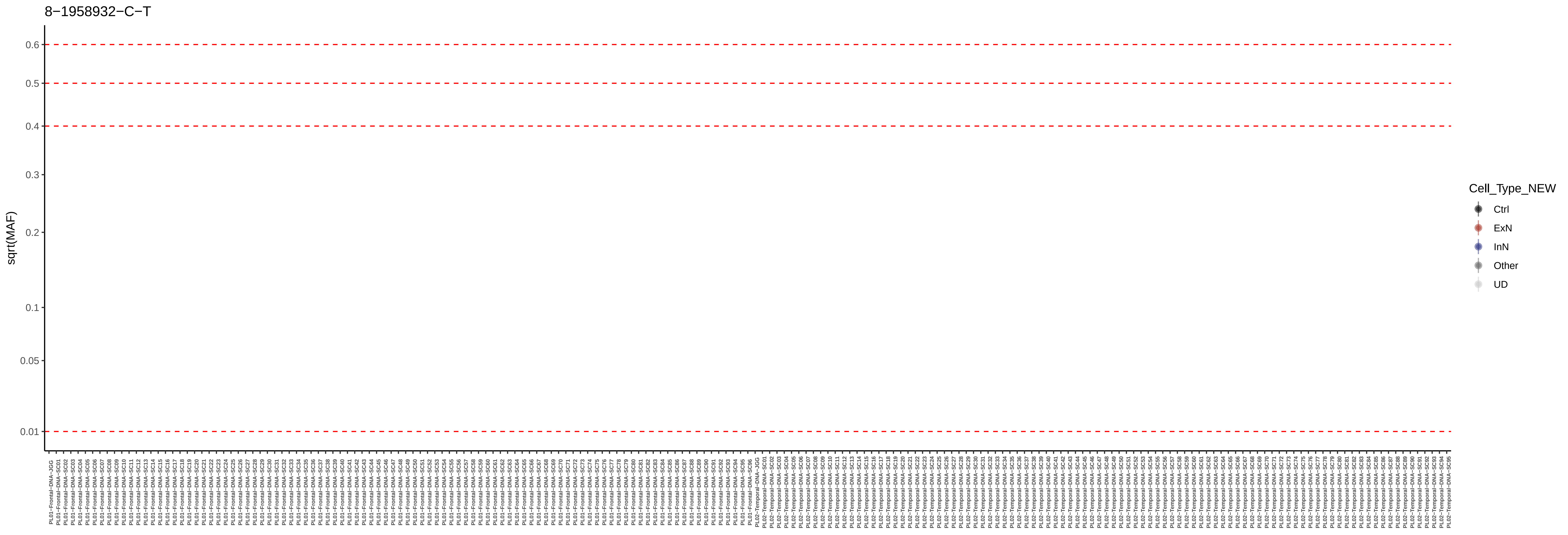
8−1958932−C−T
sqrt(MAF)
0.6
0.5
0.4
0.3
0.2
0.1
0.05
0.01
Cell_Type_NEW
Ctrl
ExN
InN
Other
UD

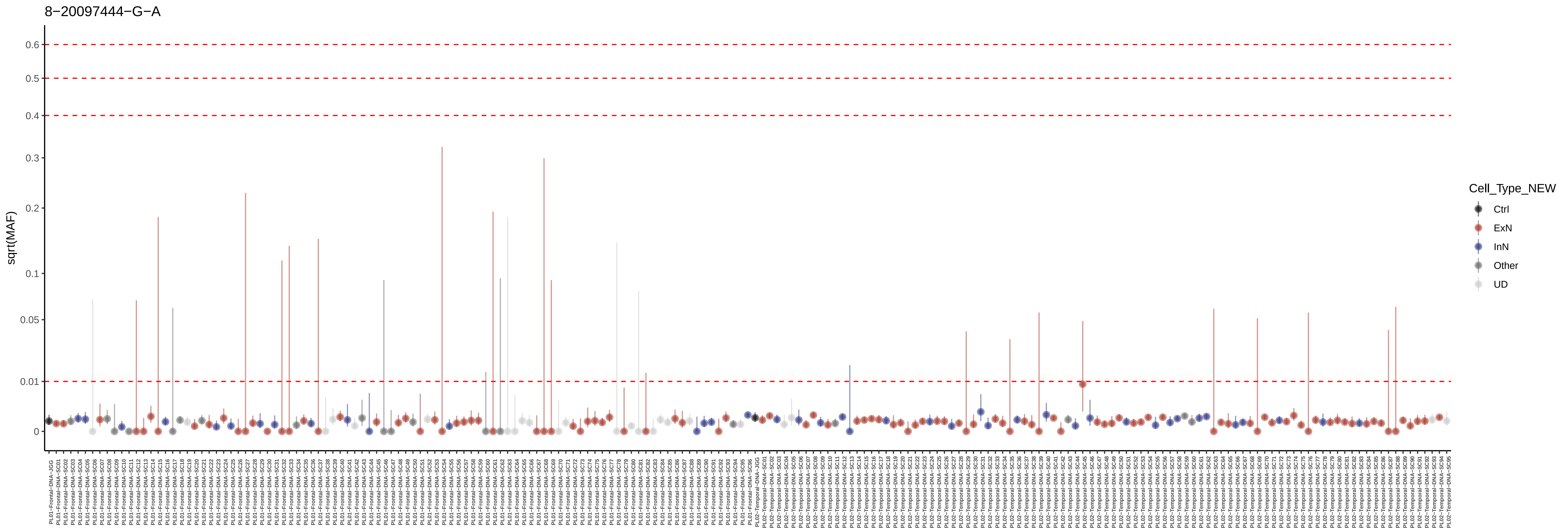
8−20097444−G−A
sqrt(MAF)
0.6
0.5
0.4
0.3
0.2
0.1
0.05
0.01
0
Cell_Type_NEW
Ctrl
ExN
InN
Other
UD

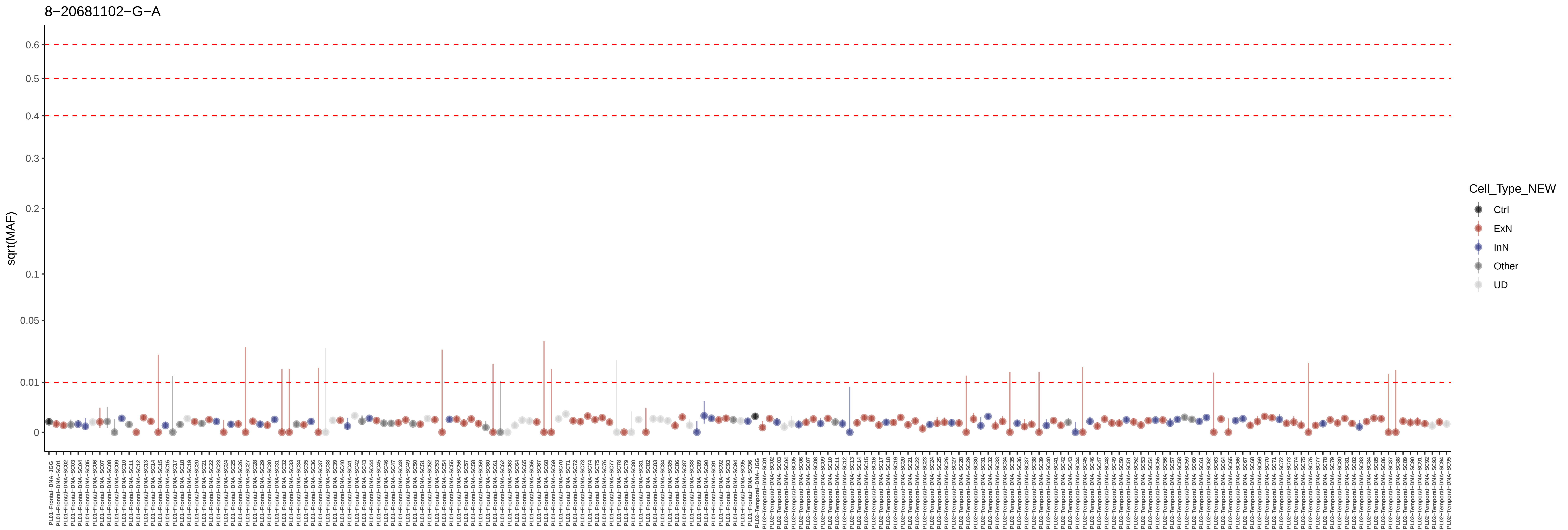

8−20681102−G−A
sqrt(MAF)
0.6
0.5
0.4
0.3
0.2
0.1
0.05
0.01
0
Cell_Type_NEW
Ctrl
ExN
InN
Other
UD

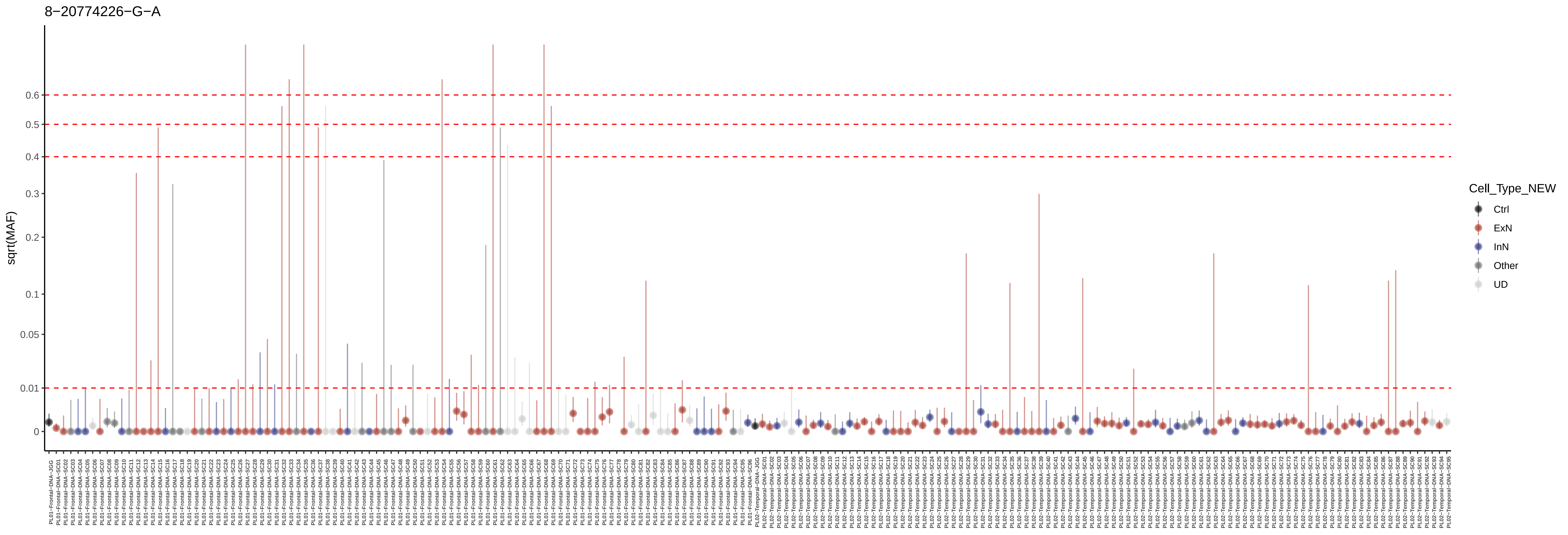

8−20774226−G−A
sqrt(MAF)
0.6
0.5
0.4
0.3
0.2
0.1
0.05
0.01
0
Cell_Type_NEW
Ctrl
ExN
InN
Other
UD

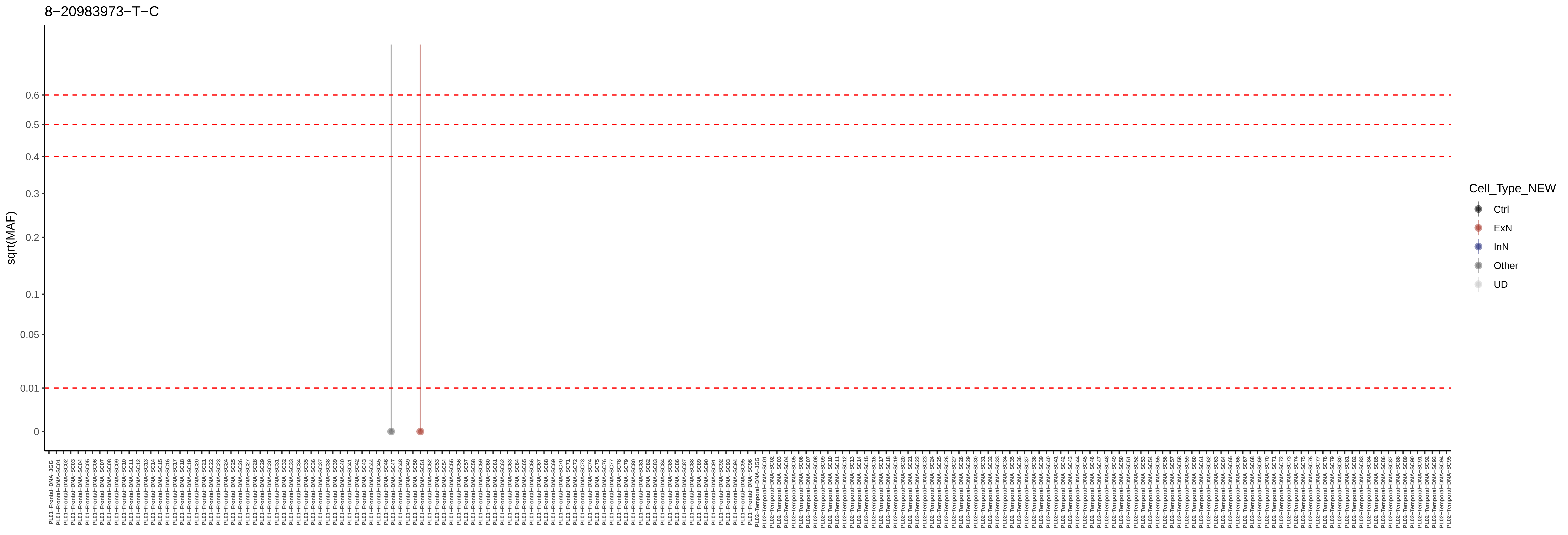
8−20983973−T−C
sqrt(MAF)
0.6
0.5
0.4
0.3
0.2
0.1
0.05
0.01
0
Cell_Type_NEW
Ctrl
ExN
InN
Other
UD

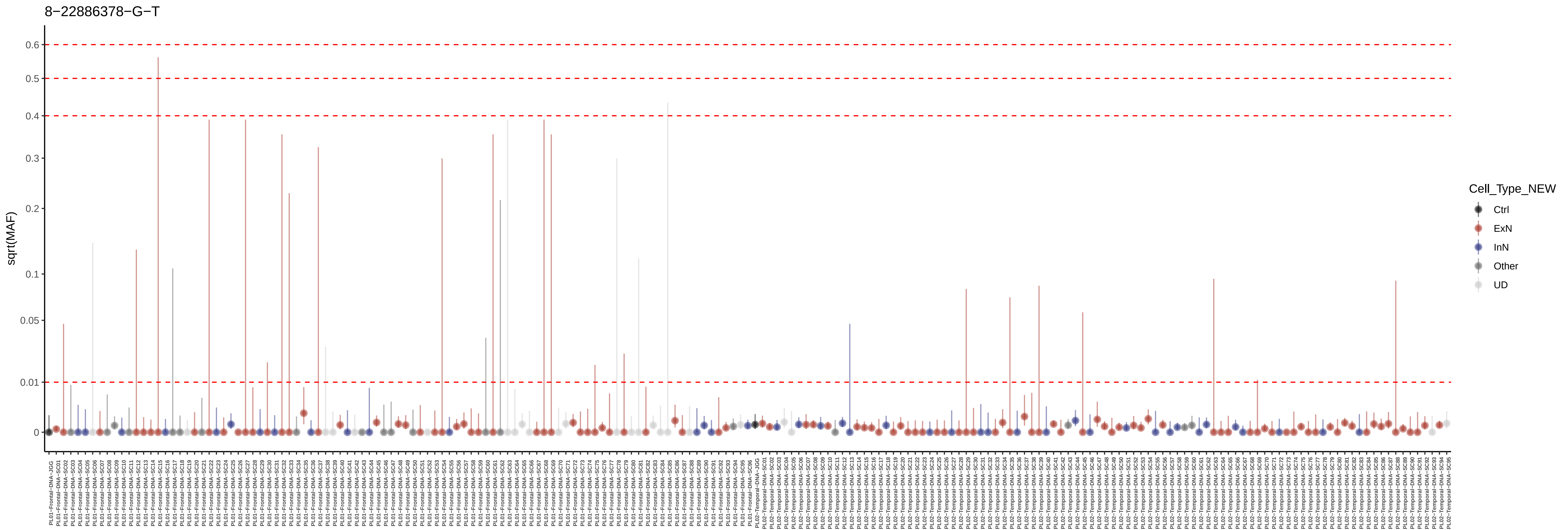

8−22886378−G−T
sqrt(MAF)
0.6
0.5
0.4
0.3
0.2
0.1
0.05
0.01
0
Cell_Type_NEW
Ctrl
ExN
InN
Other
UD

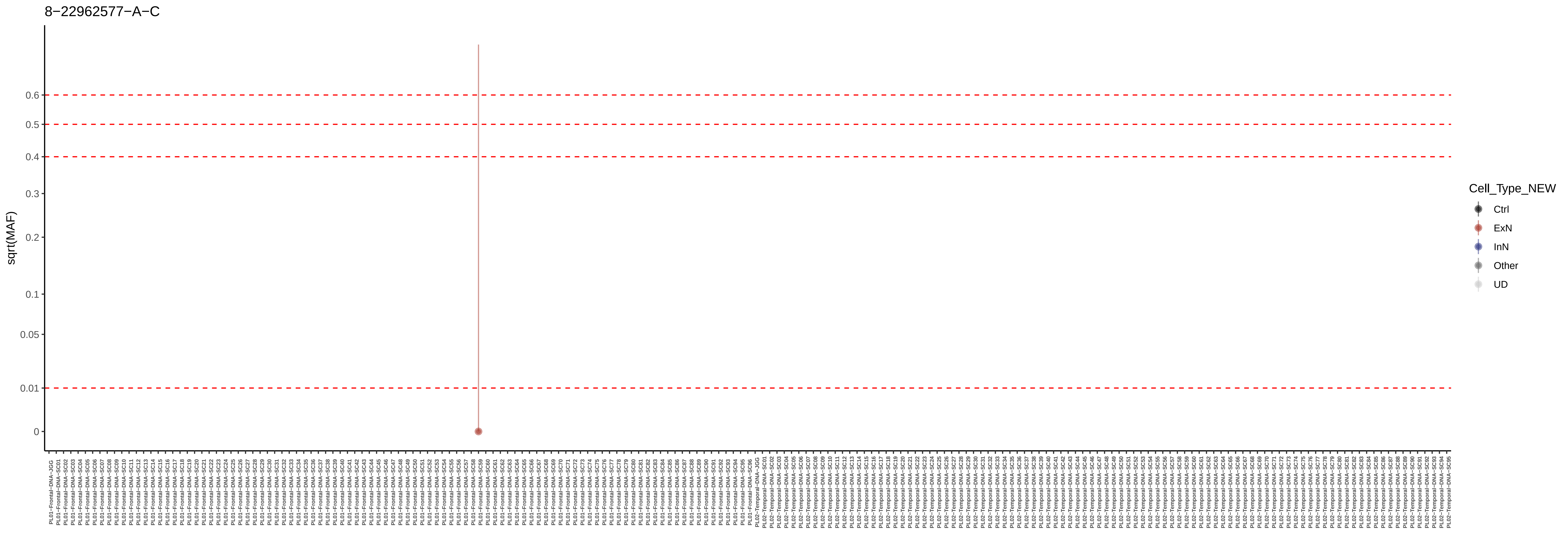

8−22962577−A−C
sqrt(MAF)
0.6
0.5
0.4
0.3
0.2
0.1
0.05
0.01
0
Cell_Type_NEW
Ctrl
ExN
InN
Other
UD

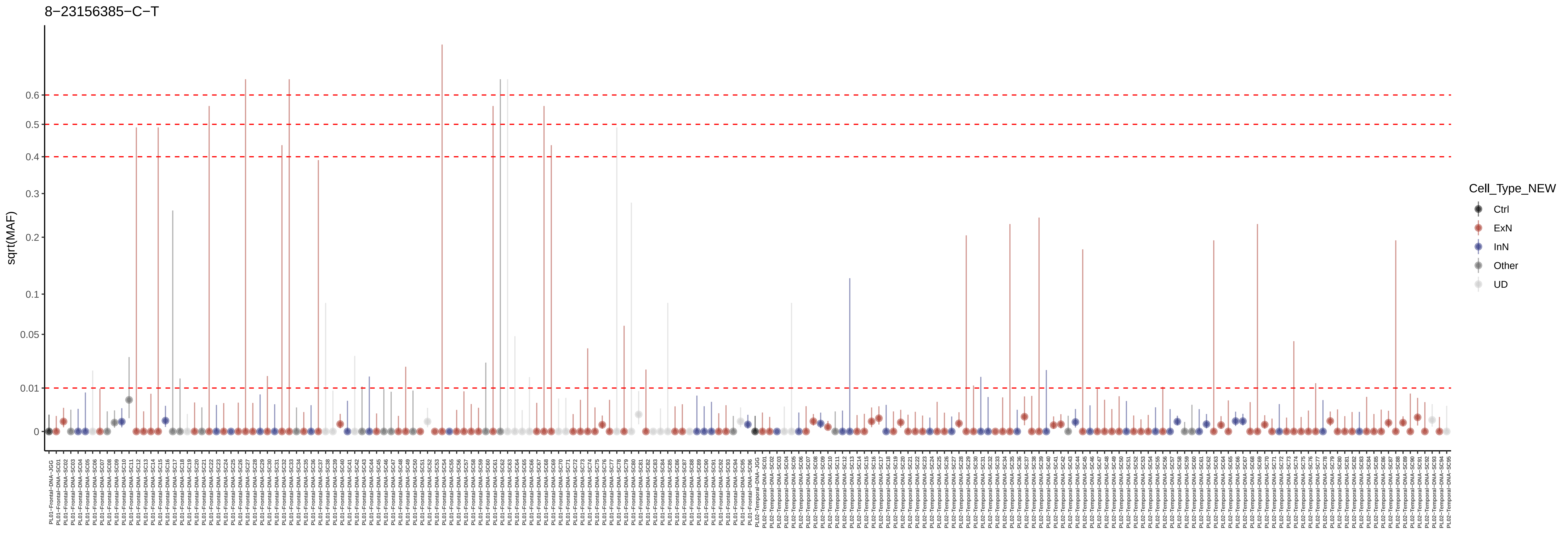
8−23156385−C−T
sqrt(MAF)
0.6
0.5
0.4
0.3
0.2
0.1
0.05
0.01
0
Cell_Type_NEW
Ctrl
ExN
InN
Other
UD

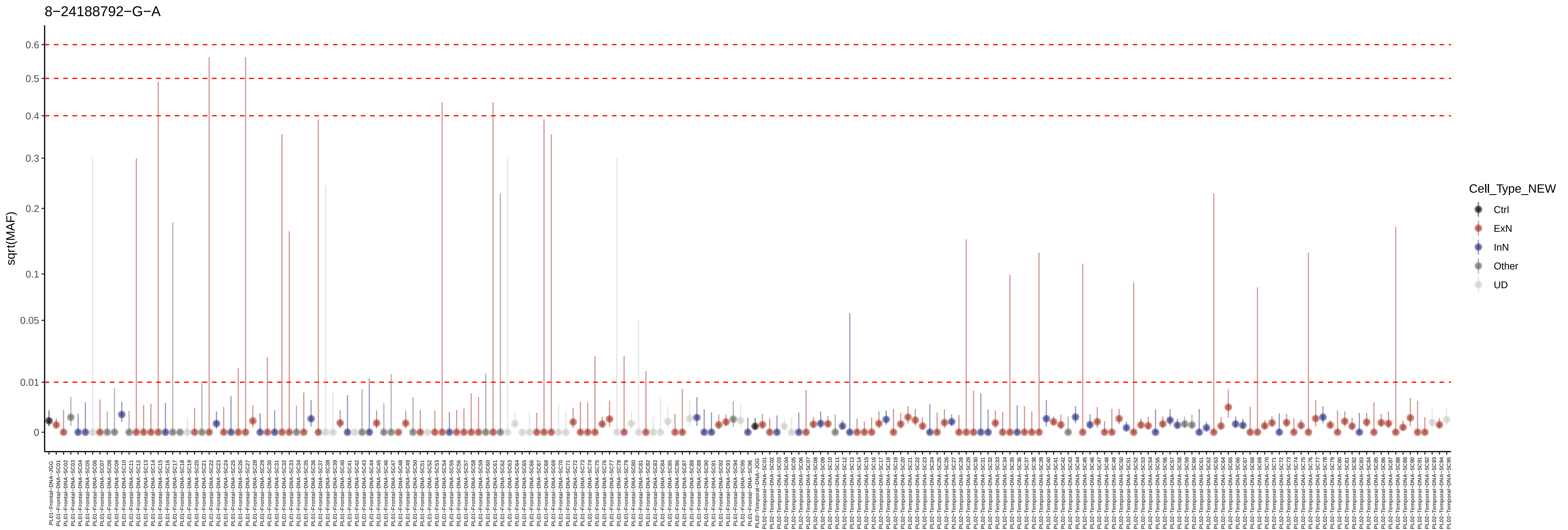

8−24188792−G−A
sqrt(MAF)
0.6
0.5
0.4
0.3
0.2
0.1
0.05
0.01
0
Cell_Type_NEW
Ctrl
ExN
InN
Other
UD

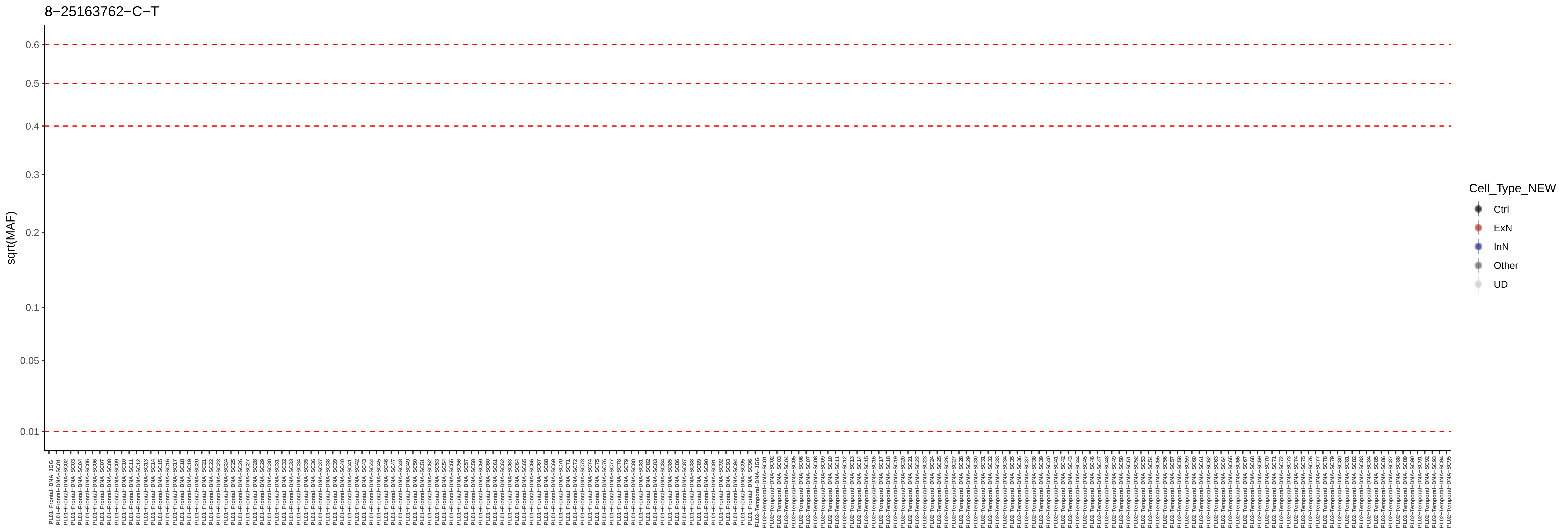
8−25163762−C−T
sqrt(MAF)
0.6
0.5
0.4
0.3
0.2
0.1
0.05
0.01
Cell_Type_NEW
Ctrl
ExN
InN
Other
UD
PL01−Frontal−DNA−JGG
PL01−Frontal−DNA−SC01
PL01−Frontal−DNA−SC02
PL01−Frontal−DNA−SC03
PL01−Frontal−DNA−SC04
PL01−Frontal−DNA−SC05
PL01−Frontal−DNA−SC06
PL01−Frontal−DNA−SC07
PL01−Frontal−DNA−SC08
PL01−Frontal−DNA−SC09
PL01−Frontal−DNA−SC10
PL01−Frontal−DNA−SC11
PL01−Frontal−DNA−SC12
PL01−Frontal−DNA−SC13
PL01−Frontal−DNA−SC14
PL01−Frontal−DNA−SC15
PL01−Frontal−DNA−SC16
PL01−Frontal−DNA−SC17
PL01−Frontal−DNA−SC18
PL01−Frontal−DNA−SC19
PL01−Frontal−DNA−SC20
PL01−Frontal−DNA−SC21
PL01−Frontal−DNA−SC22
PL01−Frontal−DNA−SC23
PL01−Frontal−DNA−SC24
PL01−Frontal−DNA−SC25
PL01−Frontal−DNA−SC26
PL01−Frontal−DNA−SC27
PL01−Frontal−DNA−SC28
PL01−Frontal−DNA−SC29
PL01−Frontal−DNA−SC30
PL01−Frontal−DNA−SC31
PL01−Frontal−DNA−SC32
PL01−Frontal−DNA−SC33
PL01−Frontal−DNA−SC34
PL01−Frontal−DNA−SC35
PL01−Frontal−DNA−SC36
PL01−Frontal−DNA−SC37
PL01−Frontal−DNA−SC38
PL01−Frontal−DNA−SC39
PL01−Frontal−DNA−SC40
PL01−Frontal−DNA−SC41
PL01−Frontal−DNA−SC42
PL01−Frontal−DNA−SC43
PL01−Frontal−DNA−SC44
PL01−Frontal−DNA−SC45
PL01−Frontal−DNA−SC46
PL01−Frontal−DNA−SC47
PL01−Frontal−DNA−SC48
PL01−Frontal−DNA−SC49
PL01−Frontal−DNA−SC50
PL01−Frontal−DNA−SC51
PL01−Frontal−DNA−SC52
PL01−Frontal−DNA−SC53
PL01−Frontal−DNA−SC54
PL01−Frontal−DNA−SC55
PL01−Frontal−DNA−SC56
PL01−Frontal−DNA−SC57
PL01−Frontal−DNA−SC58
PL01−Frontal−DNA−SC59
PL01−Frontal−DNA−SC60
PL01−Frontal−DNA−SC61
PL01−Frontal−DNA−SC62
PL01−Frontal−DNA−SC63
PL01−Frontal−DNA−SC64
PL01−Frontal−DNA−SC65
PL01−Frontal−DNA−SC66
PL01−Frontal−DNA−SC67
PL01−Frontal−DNA−SC68
PL01−Frontal−DNA−SC69
PL01−Frontal−DNA−SC70
PL01−Frontal−DNA−SC71
PL01−Frontal−DNA−SC72
PL01−Frontal−DNA−SC73
PL01−Frontal−DNA−SC74
PL01−Frontal−DNA−SC75
PL01−Frontal−DNA−SC76
PL01−Frontal−DNA−SC77
PL01−Frontal−DNA−SC78
PL01−Frontal−DNA−SC79
PL01−Frontal−DNA−SC80
PL01−Frontal−DNA−SC81
PL01−Frontal−DNA−SC82
PL01−Frontal−DNA−SC83
PL01−Frontal−DNA−SC84
PL01−Frontal−DNA−SC85
PL01−Frontal−DNA−SC86
PL01−Frontal−DNA−SC87
PL01−Frontal−DNA−SC88
PL01−Frontal−DNA−SC89
PL01−Frontal−DNA−SC90
PL01−Frontal−DNA−SC91
PL01−Frontal−DNA−SC92
PL01−Frontal−DNA−SC93
PL01−Frontal−DNA−SC94
PL01−Frontal−DNA−SC95
PL01−Frontal−DNA−SC96
PL02−Temporal−DNA−JGG
PL02−Temporal−DNA−SC01
PL02−Temporal−DNA−SC02
PL02−Temporal−DNA−SC03
PL02−Temporal−DNA−SC04
PL02−Temporal−DNA−SC05
PL02−Temporal−DNA−SC06
PL02−Temporal−DNA−SC07
PL02−Temporal−DNA−SC08
PL02−Temporal−DNA−SC09
PL02−Temporal−DNA−SC10
PL02−Temporal−DNA−SC11
PL02−Temporal−DNA−SC12
PL02−Temporal−DNA−SC13
PL02−Temporal−DNA−SC14
PL02−Temporal−DNA−SC15
PL02−Temporal−DNA−SC16
PL02−Temporal−DNA−SC17
PL02−Temporal−DNA−SC18
PL02−Temporal−DNA−SC19
PL02−Temporal−DNA−SC20
PL02−Temporal−DNA−SC21
PL02−Temporal−DNA−SC22
PL02−Temporal−DNA−SC23
PL02−Temporal−DNA−SC24
PL02−Temporal−DNA−SC25
PL02−Temporal−DNA−SC26
PL02−Temporal−DNA−SC27
PL02−Temporal−DNA−SC28
PL02−Temporal−DNA−SC29
PL02−Temporal−DNA−SC30
PL02−Temporal−DNA−SC31
PL02−Temporal−DNA−SC32
PL02−Temporal−DNA−SC33
PL02−Temporal−DNA−SC34
PL02−Temporal−DNA−SC35
PL02−Temporal−DNA−SC36
PL02−Temporal−DNA−SC37
PL02−Temporal−DNA−SC38
PL02−Temporal−DNA−SC39
PL02−Temporal−DNA−SC40
PL02−Temporal−DNA−SC41
PL02−Temporal−DNA−SC42
PL02−Temporal−DNA−SC43
PL02−Temporal−DNA−SC44
PL02−Temporal−DNA−SC45
PL02−Temporal−DNA−SC46
PL02−Temporal−DNA−SC47
PL02−Temporal−DNA−SC48
PL02−Temporal−DNA−SC49
PL02−Temporal−DNA−SC50
PL02−Temporal−DNA−SC51
PL02−Temporal−DNA−SC52
PL02−Temporal−DNA−SC53
PL02−Temporal−DNA−SC54
PL02−Temporal−DNA−SC55
PL02−Temporal−DNA−SC56
PL02−Temporal−DNA−SC57
PL02−Temporal−DNA−SC58
PL02−Temporal−DNA−SC59
PL02−Temporal−DNA−SC60
PL02−Temporal−DNA−SC61
PL02−Temporal−DNA−SC62
PL02−Temporal−DNA−SC63
PL02−Temporal−DNA−SC64
PL02−Temporal−DNA−SC65
PL02−Temporal−DNA−SC66
PL02−Temporal−DNA−SC67
PL02−Temporal−DNA−SC68
PL02−Temporal−DNA−SC69
PL02−Temporal−DNA−SC70
PL02−Temporal−DNA−SC71
PL02−Temporal−DNA−SC72
PL02−Temporal−DNA−SC73
PL02−Temporal−DNA−SC74
PL02−Temporal−DNA−SC75
PL02−Temporal−DNA−SC76
PL02−Temporal−DNA−SC77
PL02−Temporal−DNA−SC78
PL02−Temporal−DNA−SC79
PL02−Temporal−DNA−SC80
PL02−Temporal−DNA−SC81
PL02−Temporal−DNA−SC82
PL02−Temporal−DNA−SC83
PL02−Temporal−DNA−SC84
PL02−Temporal−DNA−SC85
PL02−Temporal−DNA−SC86
PL02−Temporal−DNA−SC87
PL02−Temporal−DNA−SC88
PL02−Temporal−DNA−SC89
PL02−Temporal−DNA−SC90
PL02−Temporal−DNA−SC91
PL02−Temporal−DNA−SC92
PL02−Temporal−DNA−SC93
PL02−Temporal−DNA−SC94
PL02−Temporal−DNA−SC95

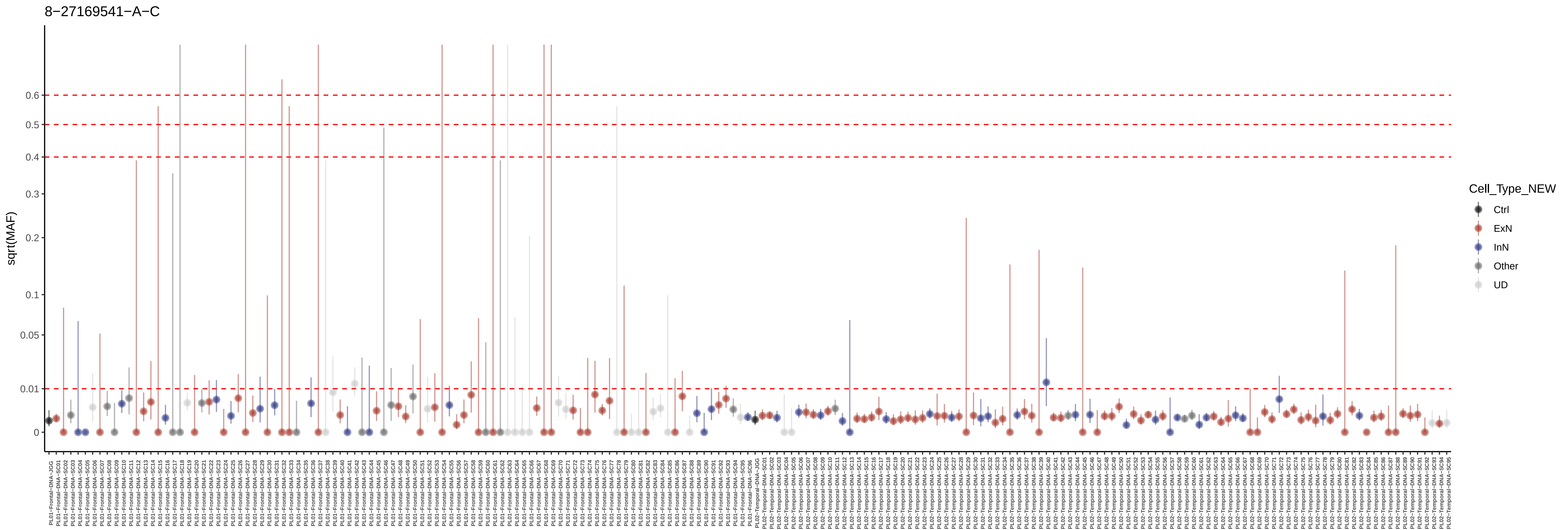

8−27169541−A−C
sqrt(MAF)
0.6
0.5
0.4
0.3
0.2
0.1
0.05
0.01
0
Cell_Type_NEW
Ctrl
ExN
InN
Other
UD

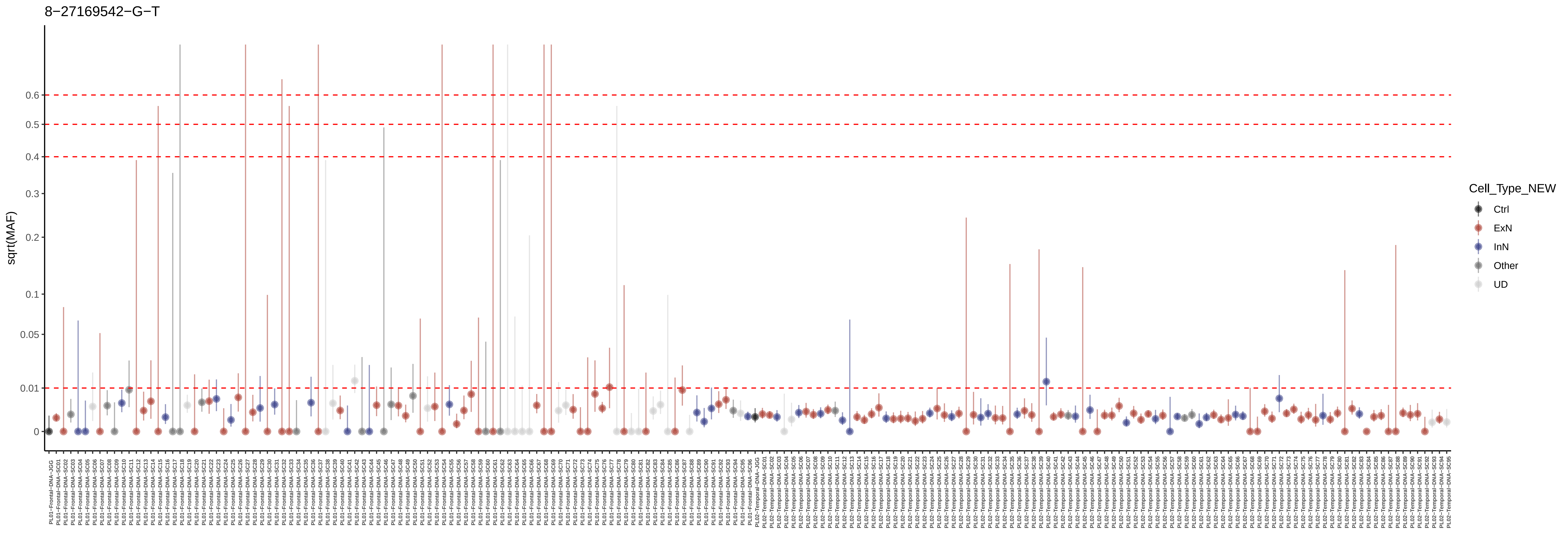
8−27169542−G−T
sqrt(MAF)
0.6
0.5
0.4
0.3
0.2
0.1
0.05
0.01
0
Cell_Type_NEW
Ctrl
ExN
InN
Other
UD

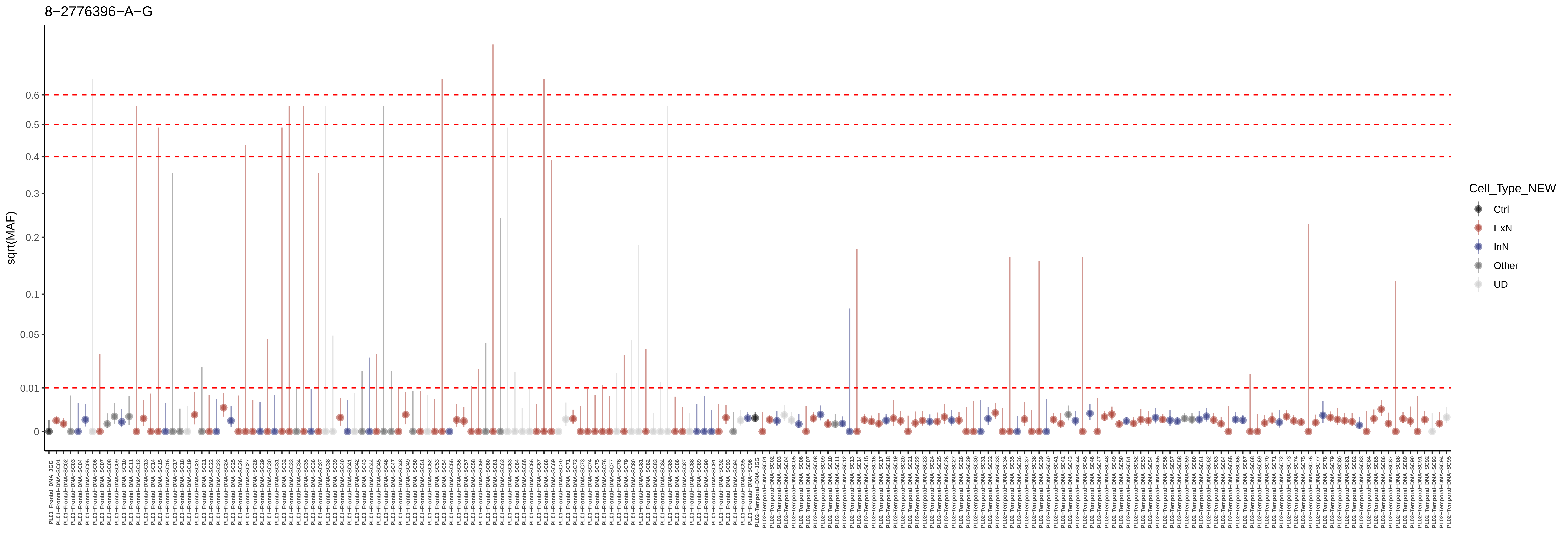

8-2776396-A-G
sqrt(MAF)
Cell_Type_NEW
Ctrl
ExN
InN
Other
UD

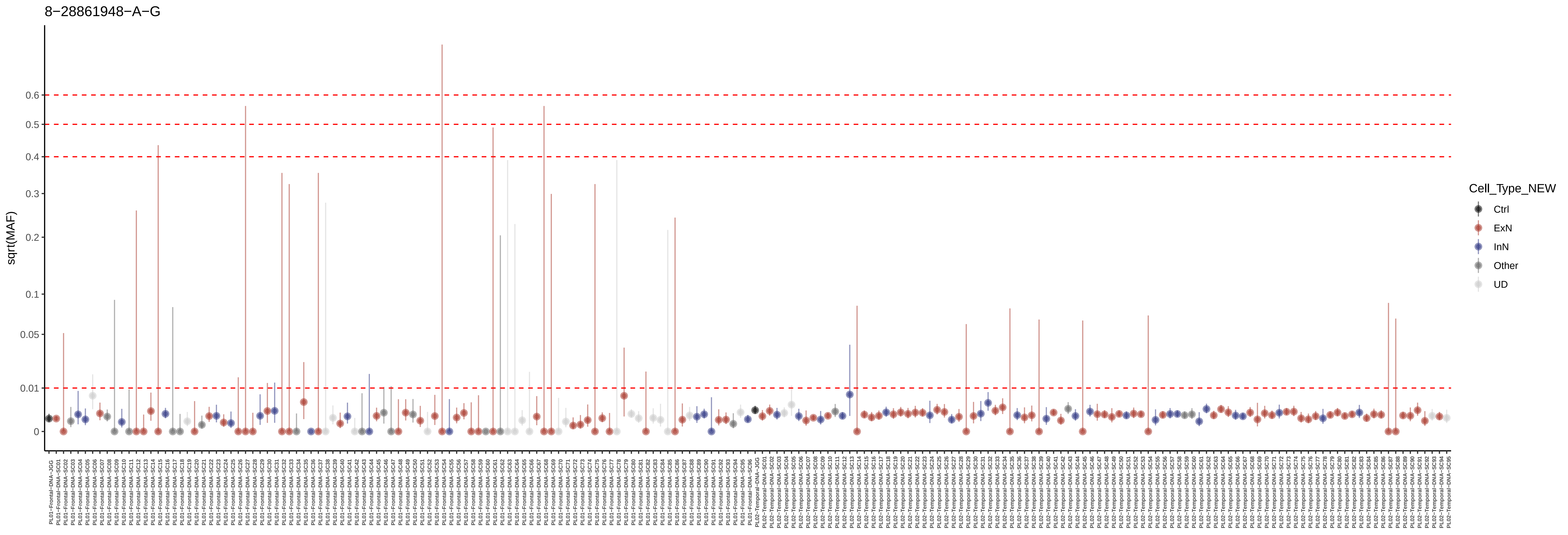
8−28861948−A−G
sqrt(MAF)
0.6
0.5
0.4
0.3
0.2
0.1
0.05
0.01
0
Cell_Type_NEW
Ctrl
ExN
InN
Other
UD

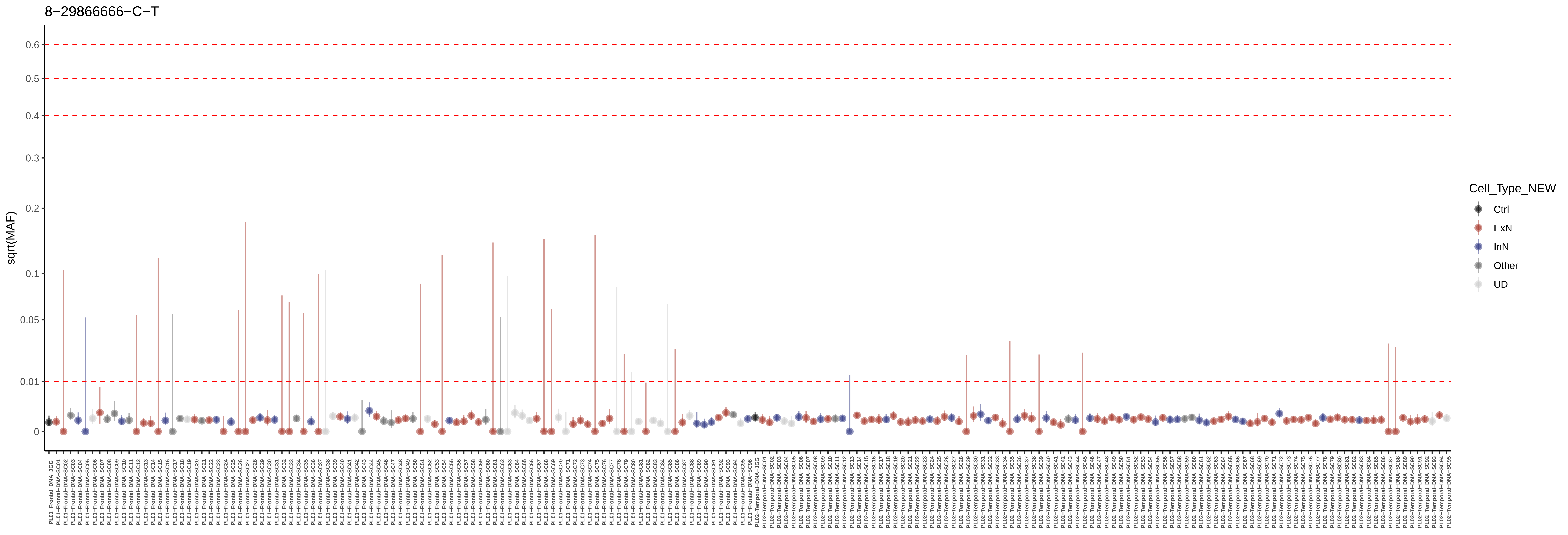

8−29866666−C−T
sqrt(MAF)
0.6
0.5
0.4
0.3
0.2
0.1
0.05
0.01
0
Cell_Type_NEW
Ctrl
ExN
InN
Other
UD

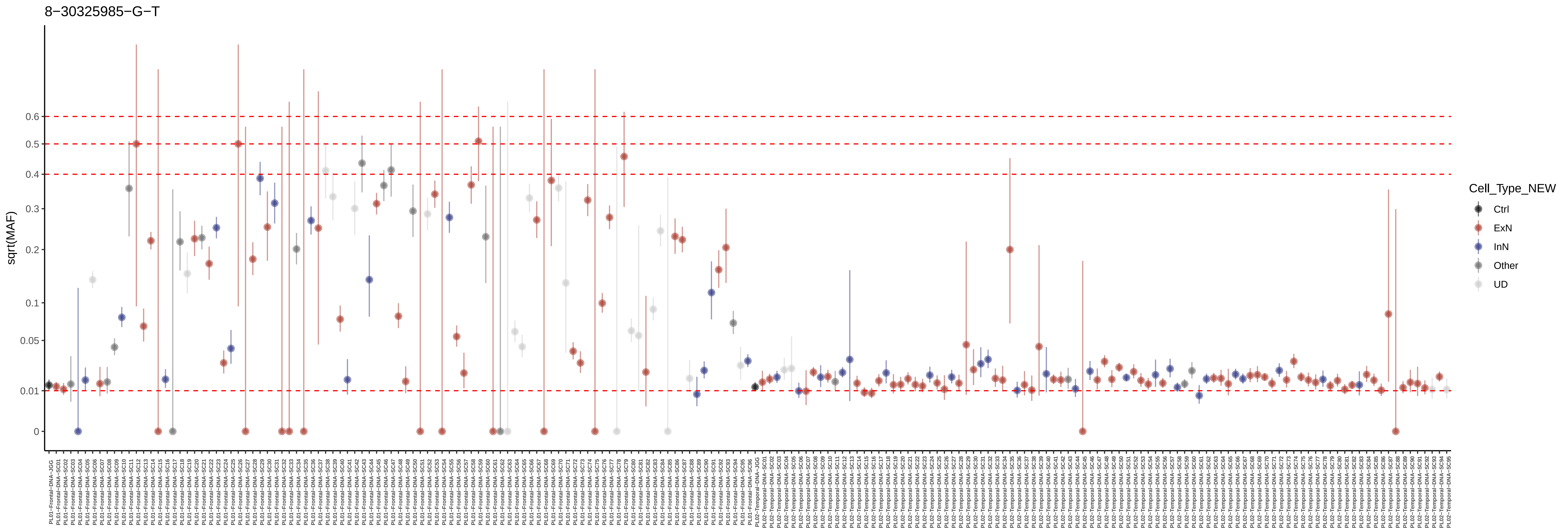
8−30325985−G−T
sqrt(MAF)
0.6
0.5
0.4
0.3
0.2
0.1
0.05
0.01
0
Cell_Type_NEW
Ctrl
ExN
InN
Other
UD

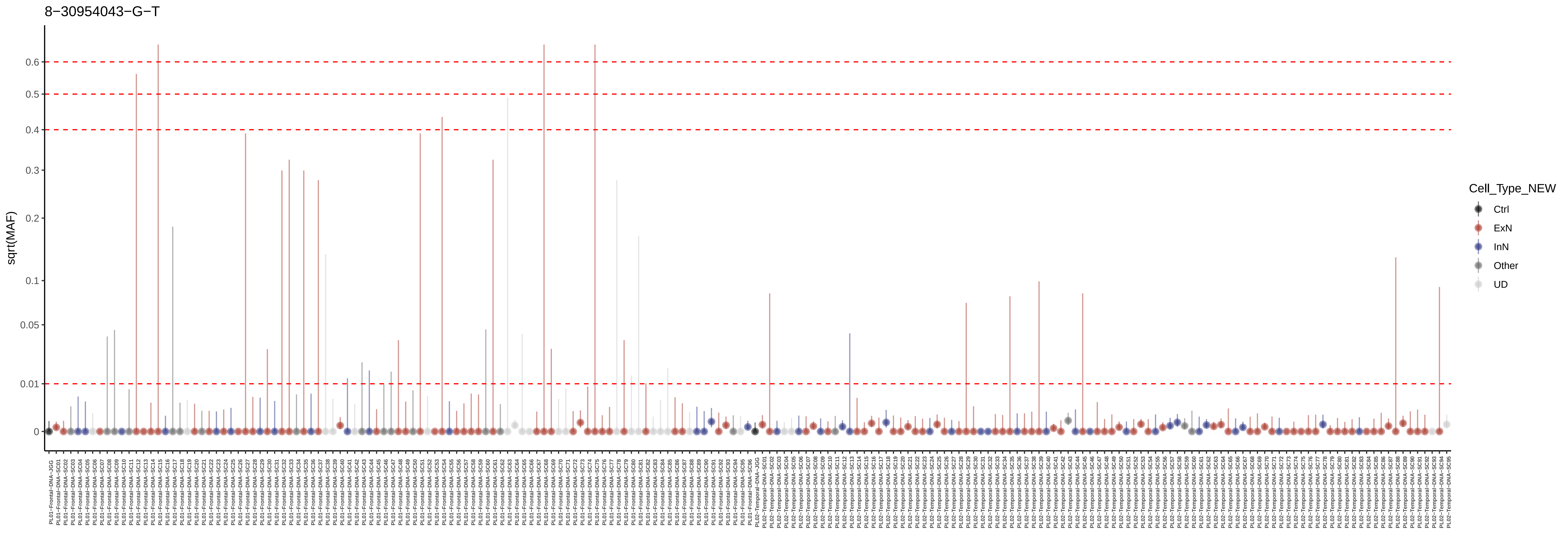
8−30954043−G−T
sqrt(MAF)
0.6
0.5
0.4
0.3
0.2
0.1
0.05
0.01
0
Cell_Type_NEW
Ctrl
ExN
InN
Other
UD

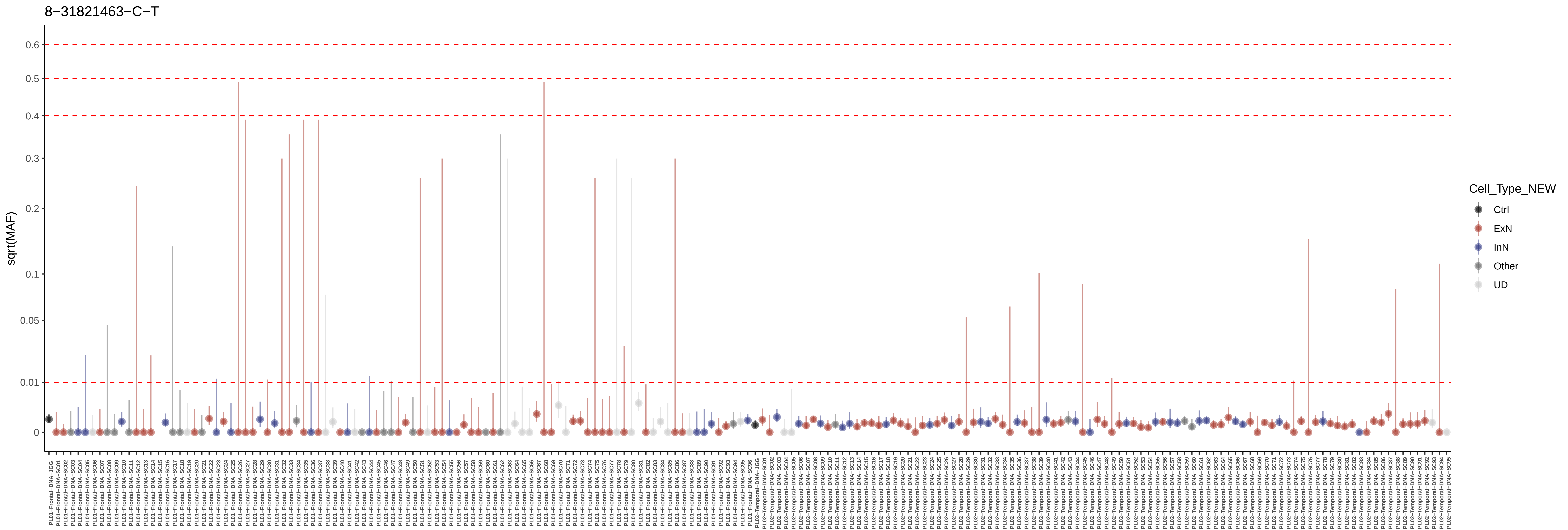

8−31821463−C−T
sqrt(MAF)
0.6
0.5
0.4
0.3
0.2
0.1
0.05
0.01
0
Cell_Type_NEW
Ctrl
ExN
InN
Other
UD

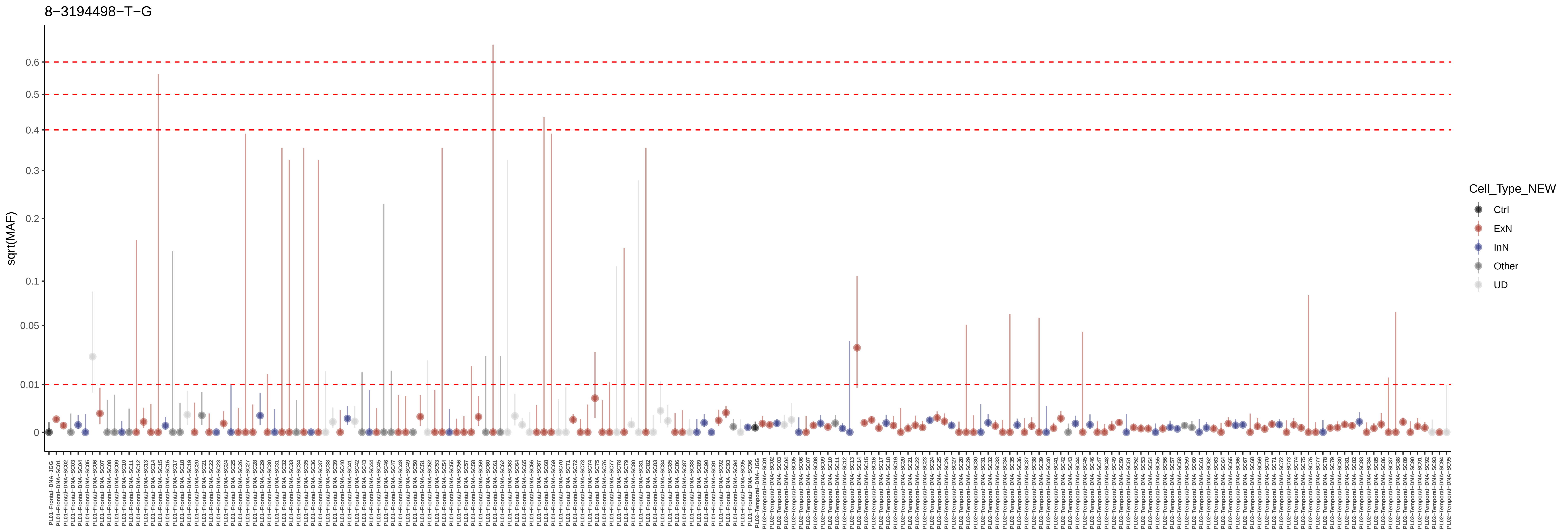
8−3194498−T−G
sqrt(MAF)
0.6
0.5
0.4
0.3
0.2
0.1
0.05
0.01
0
Cell_Type_NEW
Ctrl
ExN
InN
Other
UD

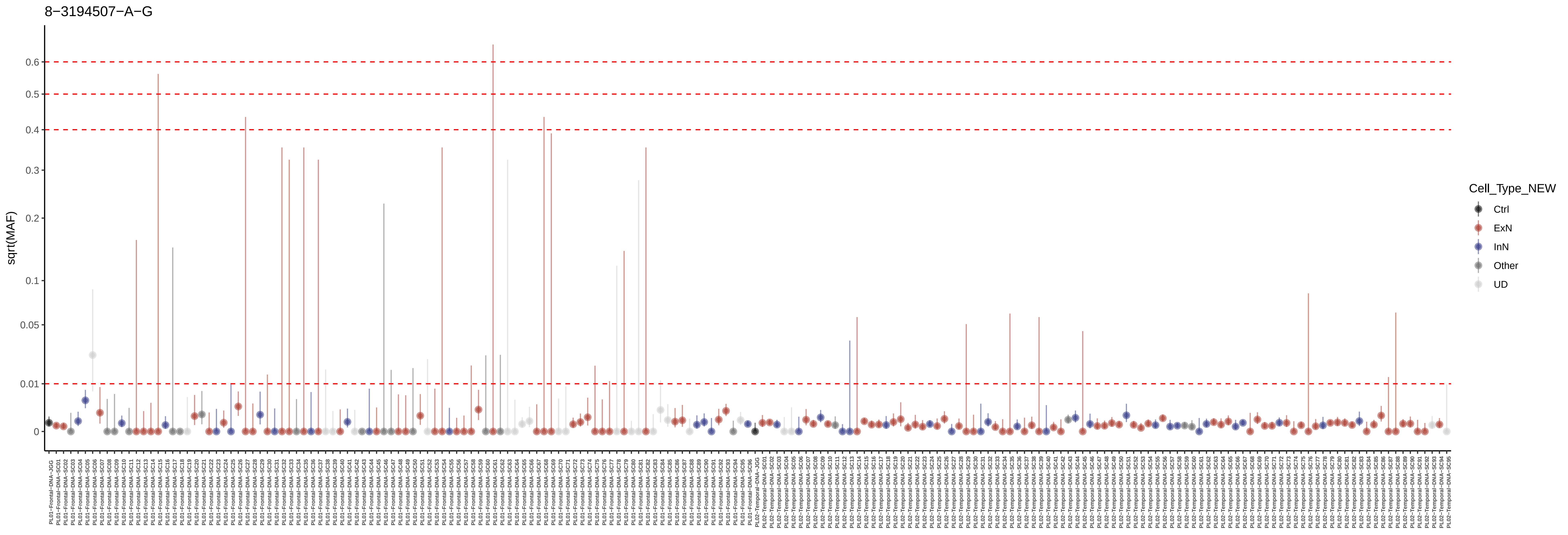
8−3194507−A−G
sqrt(MAF)
0.6
0.5
0.4
0.3
0.2
0.1
0.05
0.01
0
Cell_Type_NEW
Ctrl
ExN
InN
Other
UD

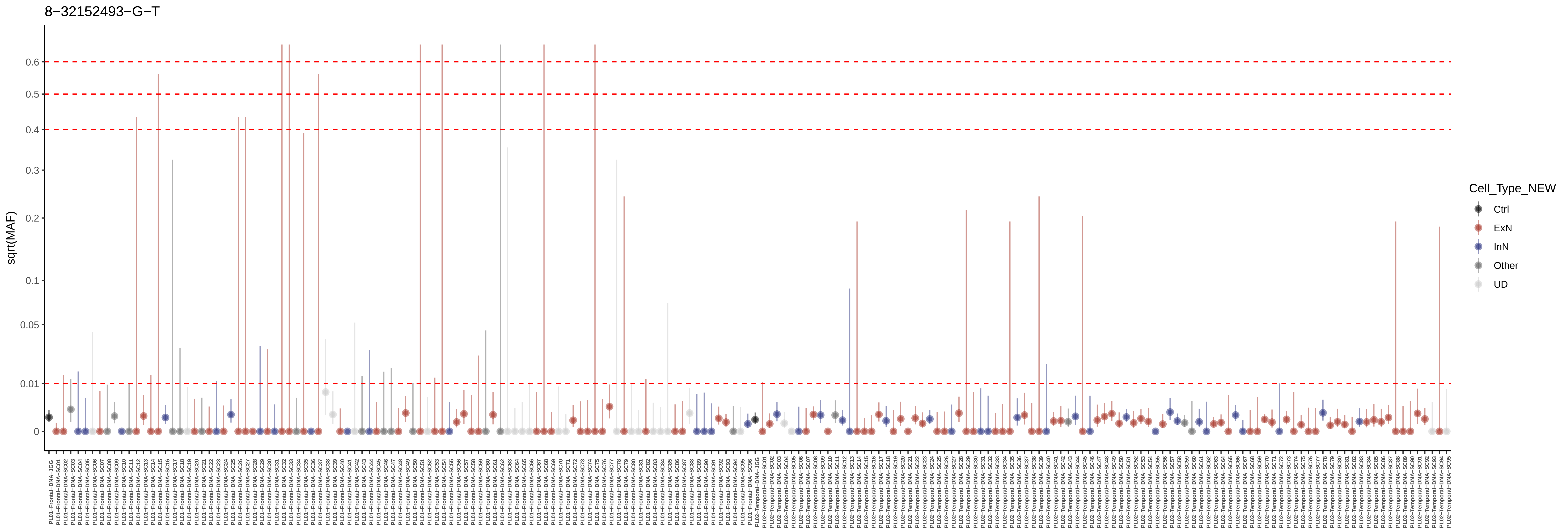
8−32152493−G−T
sqrt(MAF)
0.6
0.5
0.4
0.3
0.2
0.1
0.05
0.01
0
Cell_Type_NEW
Ctrl
ExN
InN
Other
UD

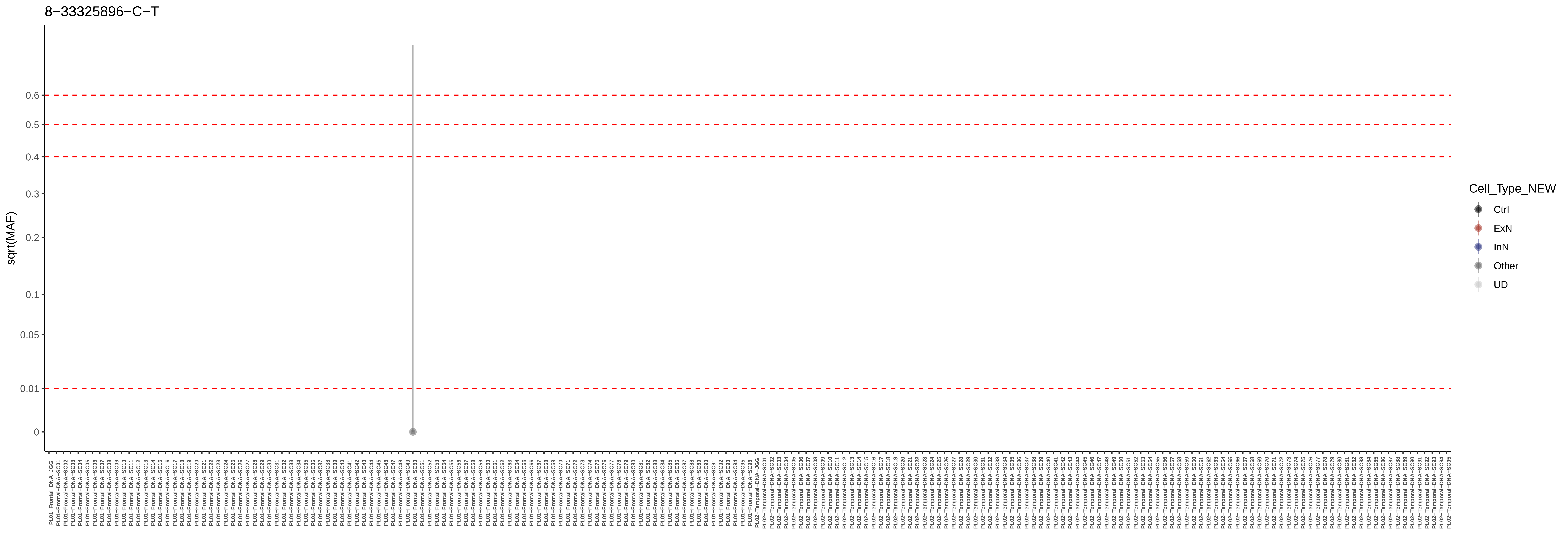
8−33325896−C−T
sqrt(MAF)
0.6
0.5
0.4
0.3
0.2
0.1
0.05
0.01
0
Cell_Type_NEW
Ctrl
ExN
InN
Other
UD

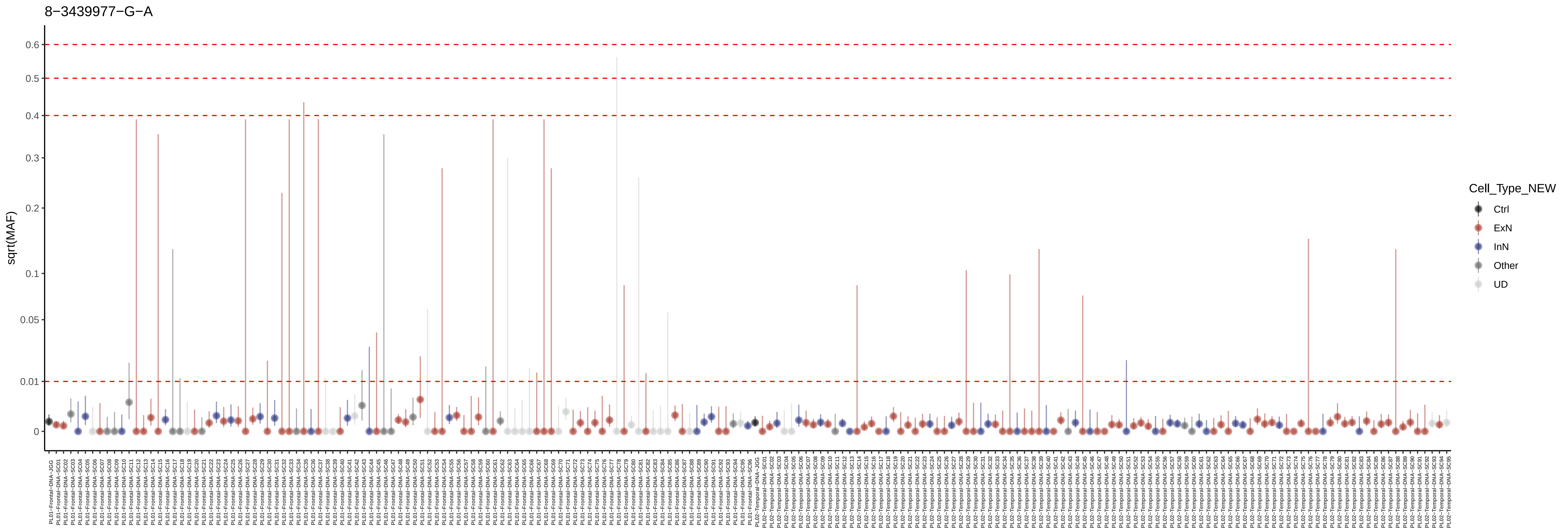
8−3439977−G−A
sqrt(MAF)
0.6
0.5
0.4
0.3
0.2
0.1
0.05
0.01
0
Cell_Type_NEW
Ctrl
ExN
InN
Other
UD

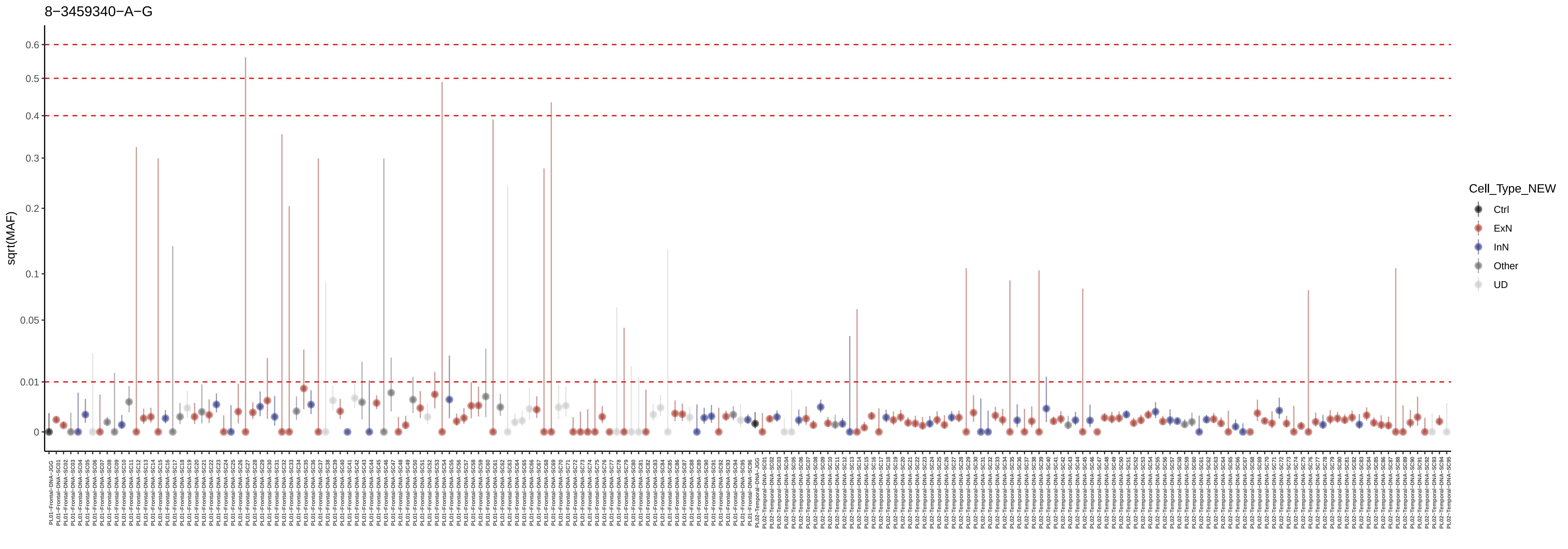

8−3459340−A−G
sqrt(MAF)
0.6
0.5
0.4
0.3
0.2
0.1
0.05
0.01
0
Cell_Type_NEW
Ctrl
ExN
InN
Other
UD

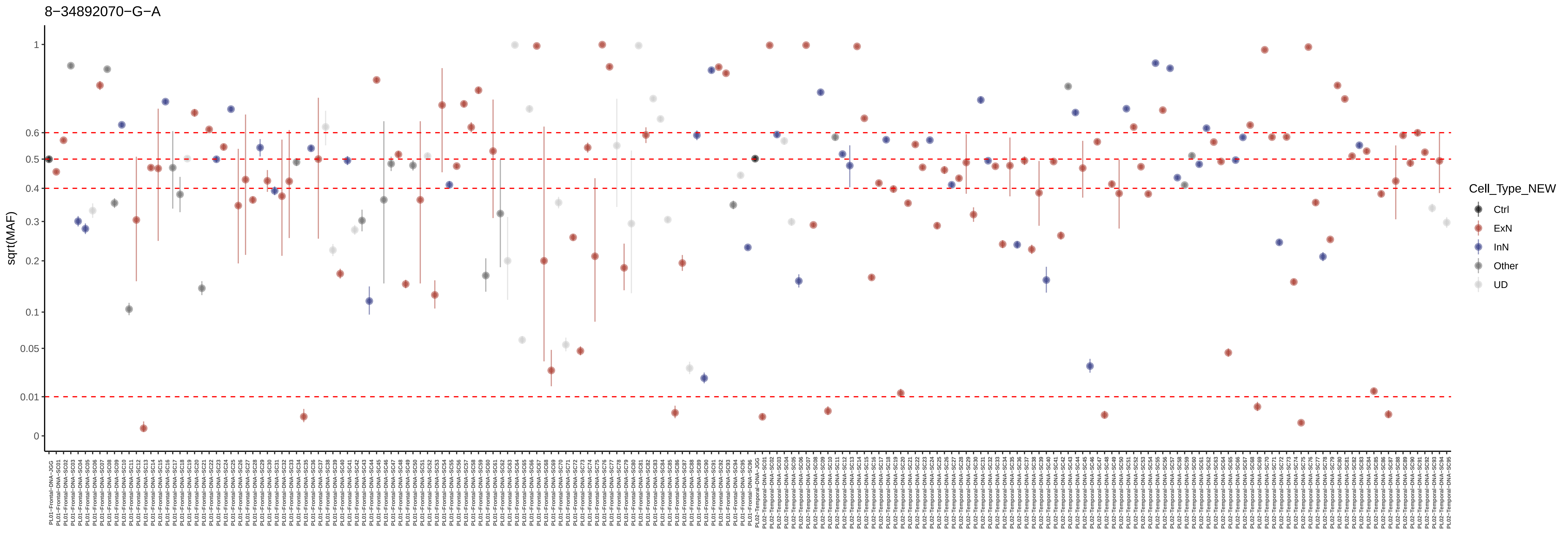
8−34892070−G−A
sqrt(MAF)
1
0.6
0.5
0.4
0.3
0.2
0.1
0.05
0.01
0
Cell_Type_NEW
Ctrl
ExN
InN
Other
UD

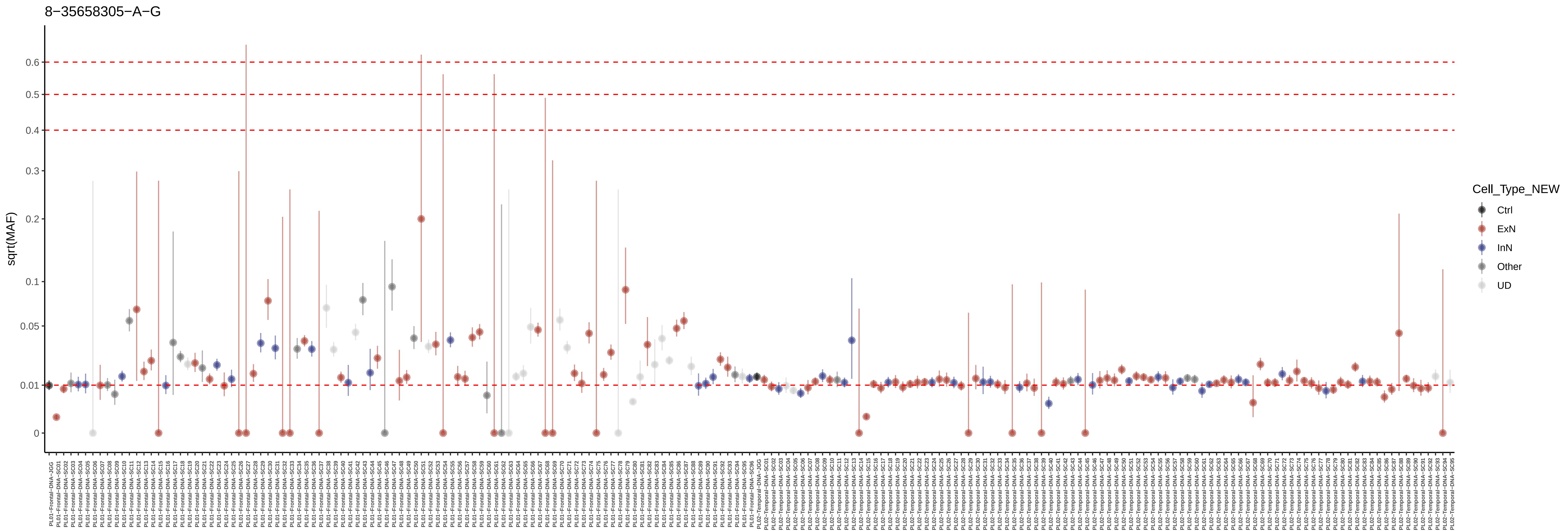
8−35658305−A−G
sqrt(MAF)
0.6
0.5
0.4
0.3
0.2
0.1
0.05
0.01
0
Cell_Type_NEW
Ctrl
ExN
InN
Other
UD

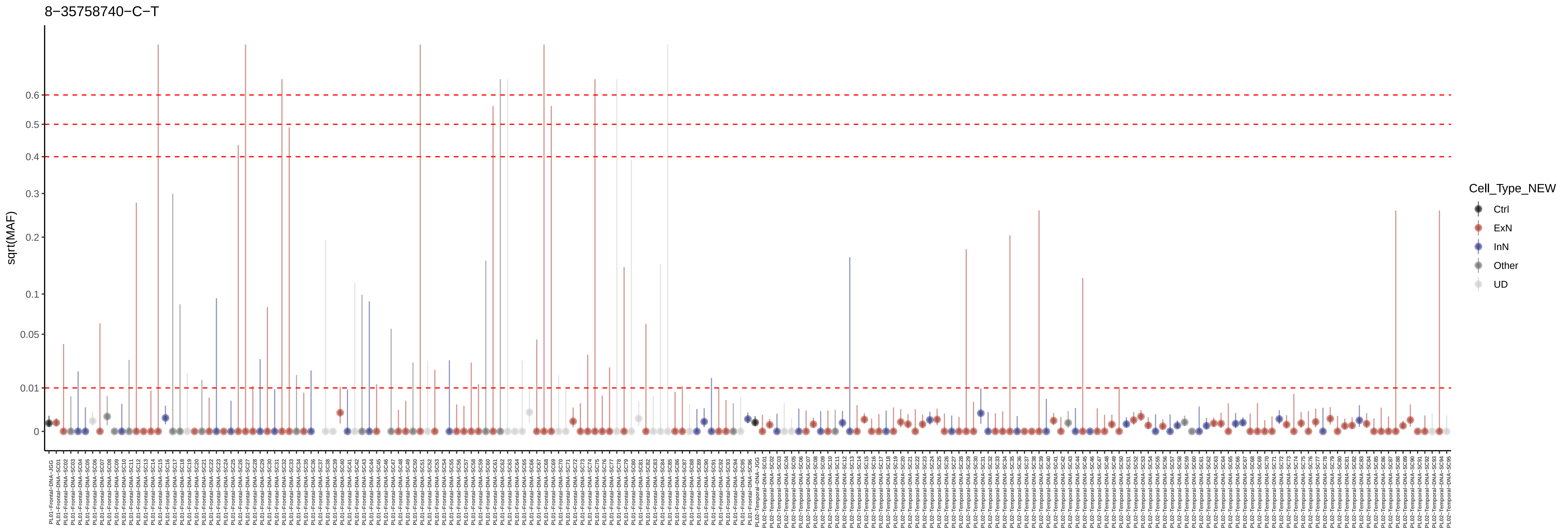
8−35758740−C−T
sqrt(MAF)
0.6
0.5
0.4
0.3
0.2
0.1
0.05
0.01
0
Cell_Type_NEW
Ctrl
ExN
InN
Other
UD

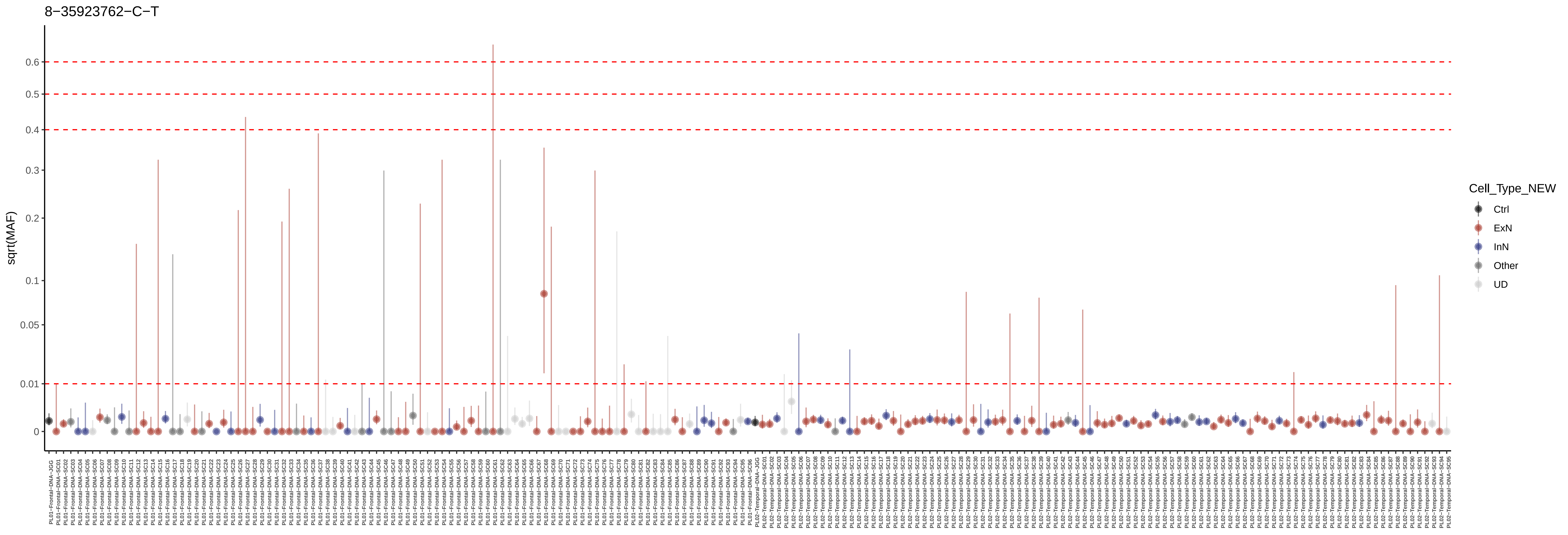
8−35923762−C−T
sqrt(MAF)
0.6
0.5
0.4
0.3
0.2
0.1
0.05
0.01
0
Cell_Type_NEW
Ctrl
ExN
InN
Other
UD

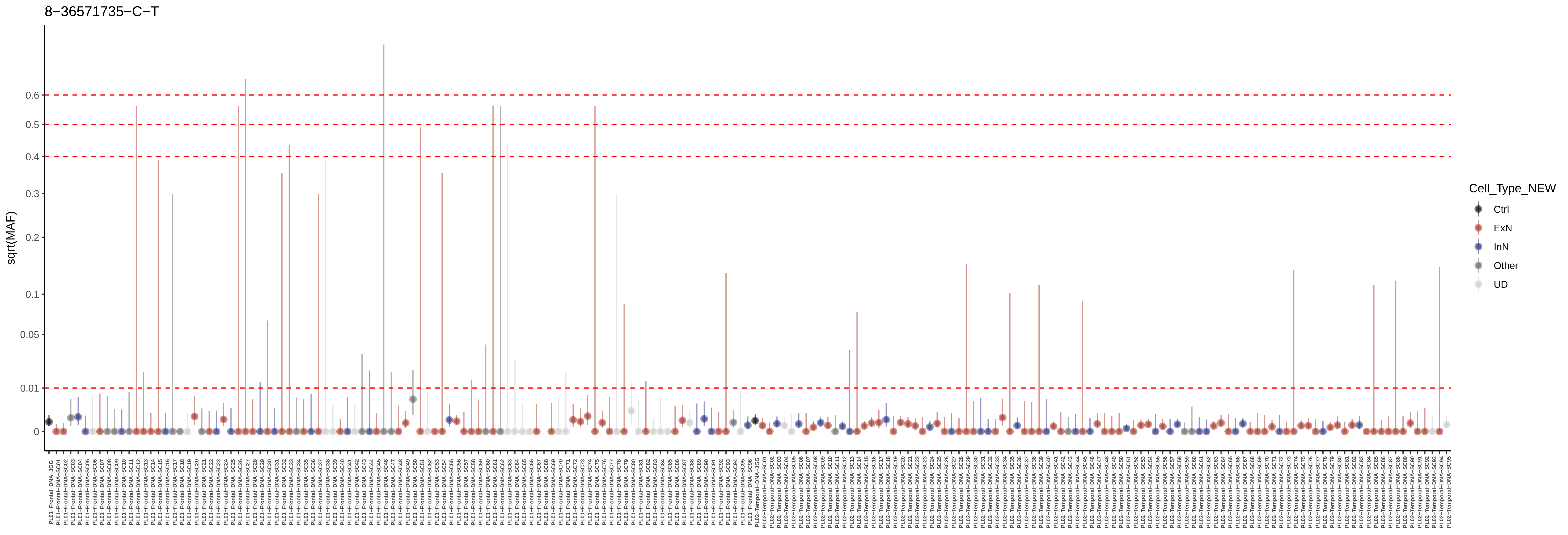
8−36571735−C−T
sqrt(MAF)
0.6
0.5
0.4
0.3
0.2
0.1
0.05
0.01
0
Cell_Type_NEW
Ctrl
ExN
InN
Other
UD

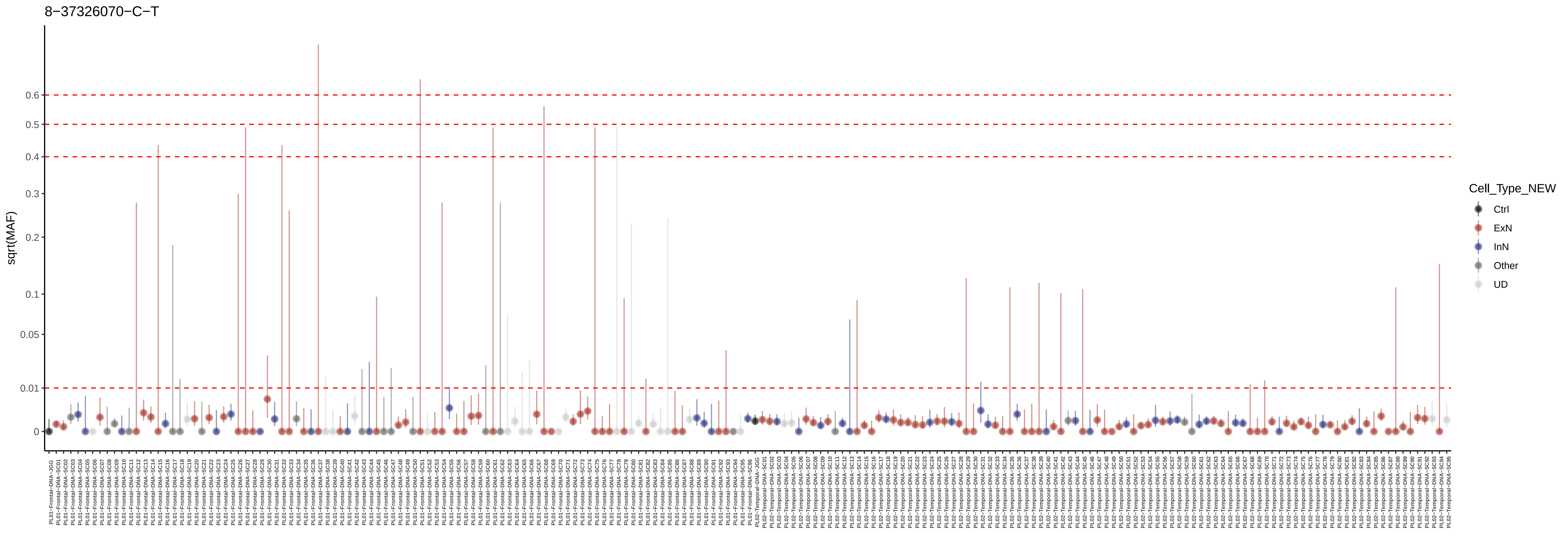
8−37326070−C−T
sqrt(MAF)
0.6
0.5
0.4
0.3
0.2
0.1
0.05
0.01
0
Cell_Type_NEW
Ctrl
ExN
InN
Other
UD

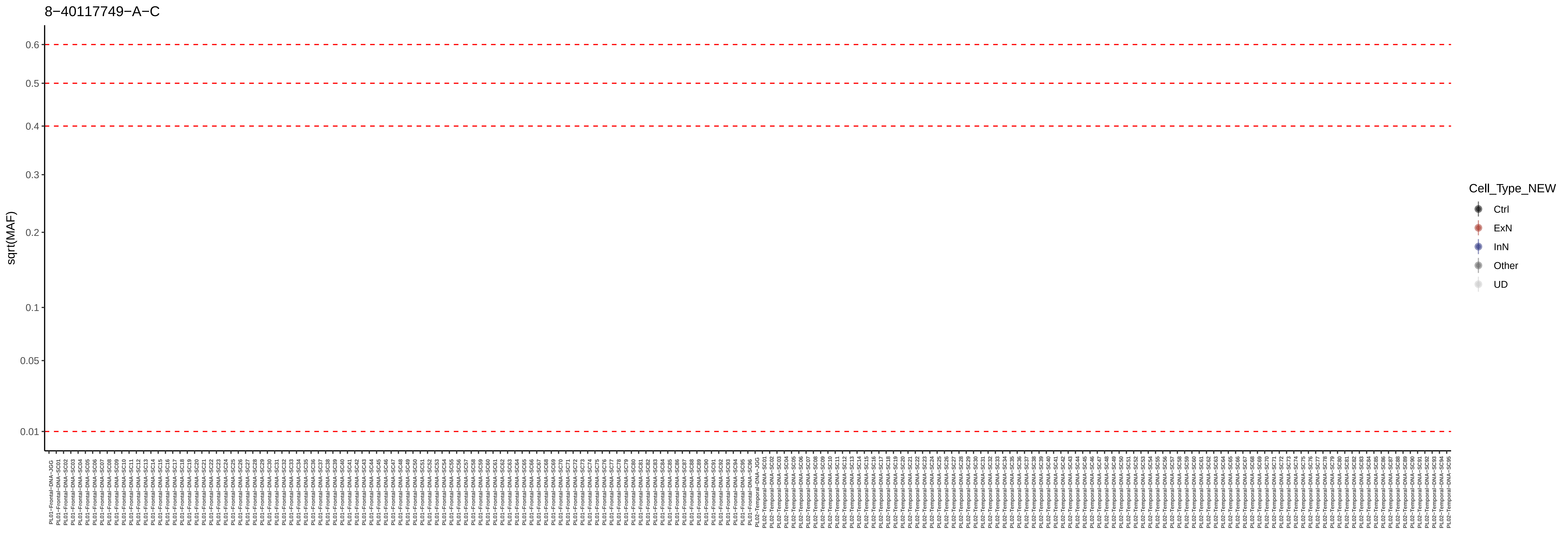
8−40117749−A−C
sqrt(MAF)
0.6
0.5
0.4
0.3
0.2
0.1
0.05
0.01
Cell_Type_NEW
Ctrl
ExN
InN
Other
UD

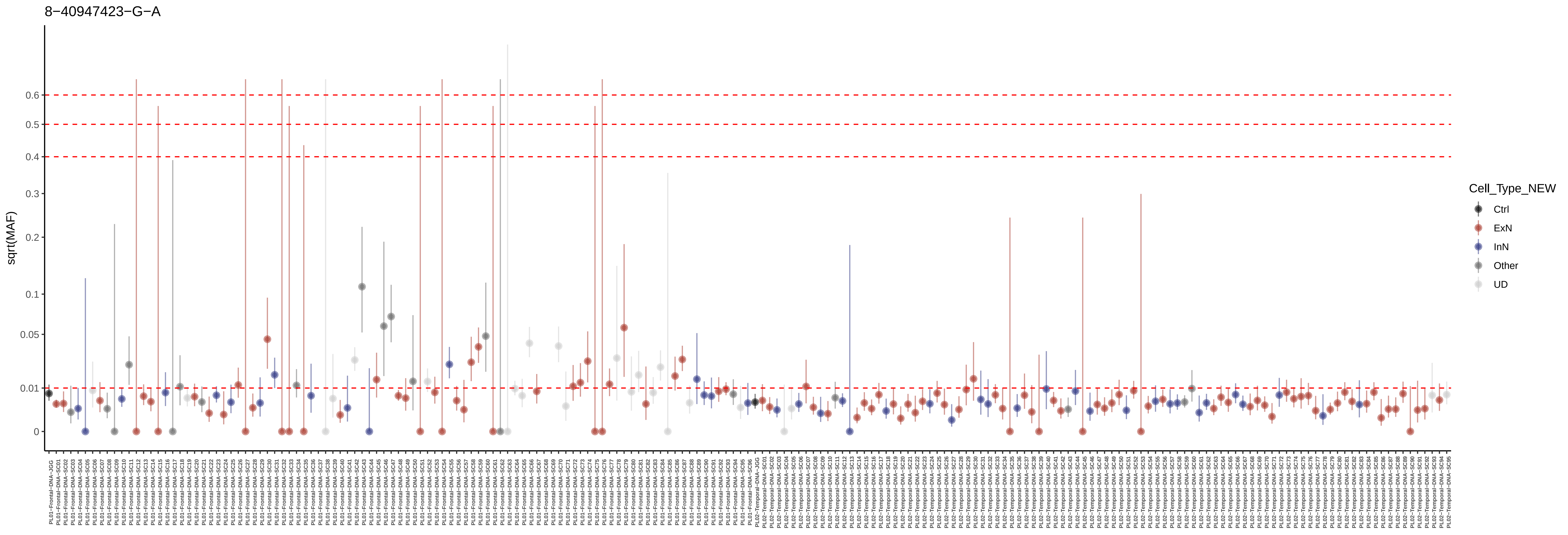

8−40947423−G−A
sqrt(MAF)
0.6
0.5
0.4
0.3
0.2
0.1
0.05
0.01
0
Cell_Type_NEW
Ctrl
ExN
InN
Other
UD

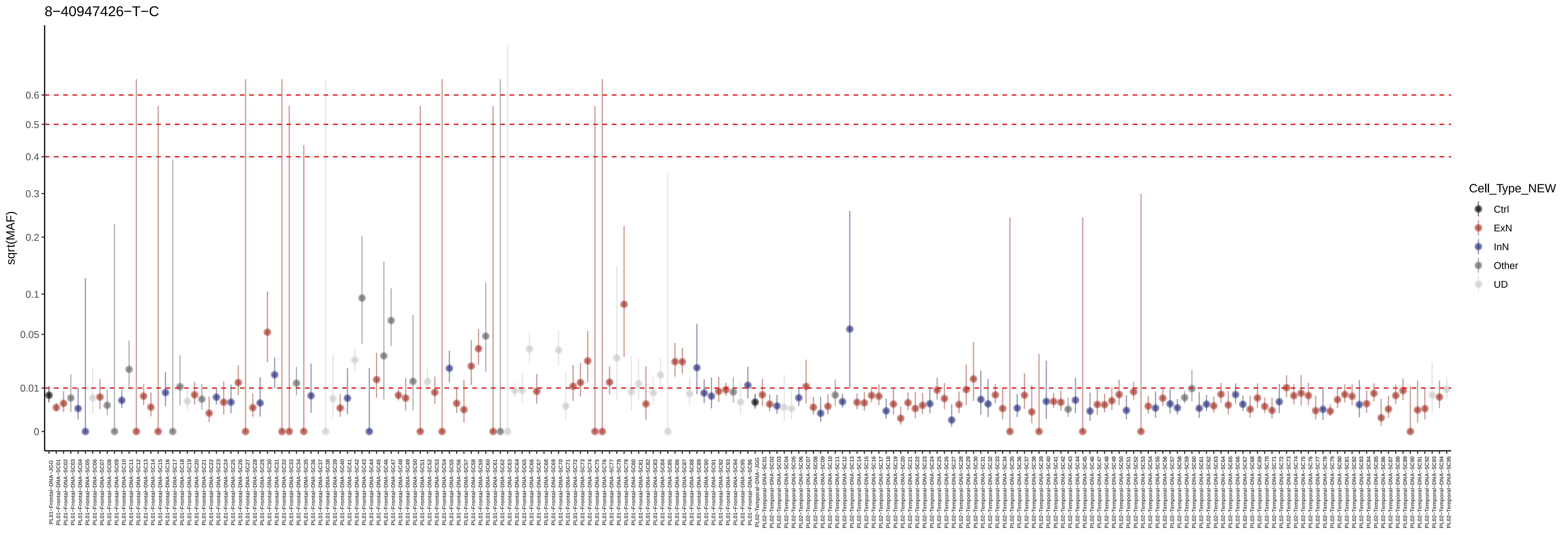
8−40947426−T−C
sqrt(MAF)
0.6
0.5
0.4
0.3
0.2
0.1
0.05
0.01
0
Cell_Type_NEW
Ctrl
ExN
InN
Other
UD

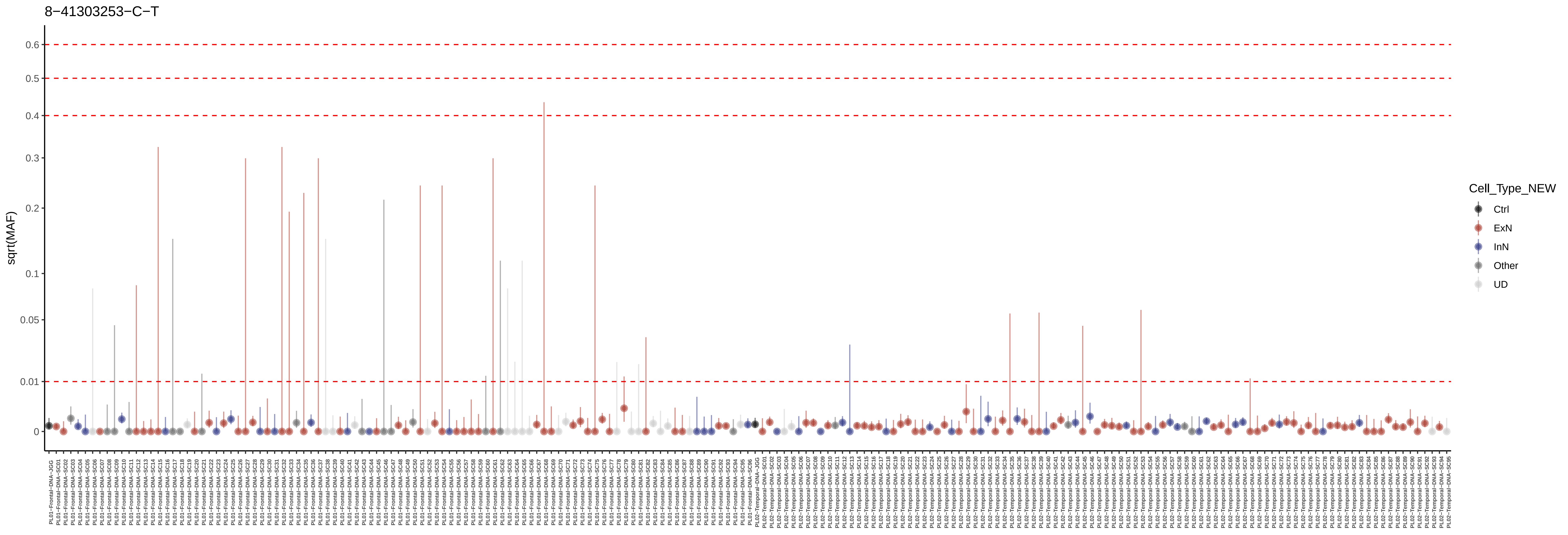
8−41303253−C−T
sqrt(MAF)
0.6
0.5
0.4
0.3
0.2
0.1
0.05
0.01
0
Cell_Type_NEW
Ctrl
ExN
InN
Other
UD

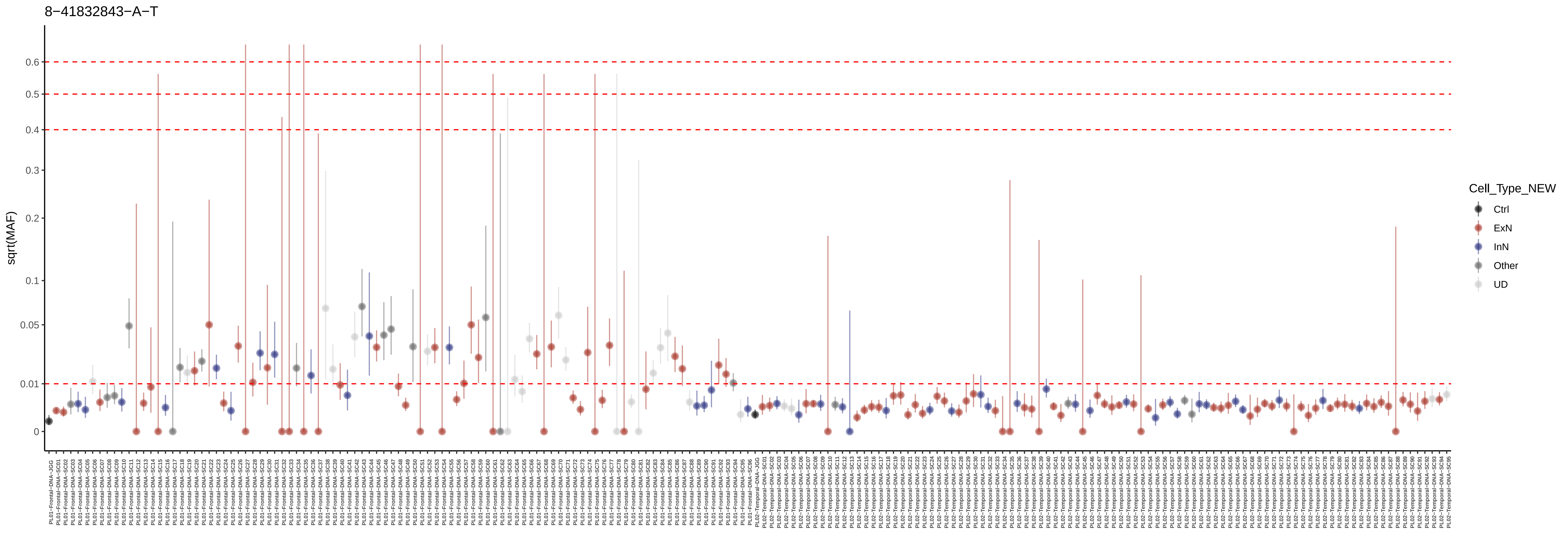
8−41832843−A−T
sqrt(MAF)
0.6
0.5
0.4
0.3
0.2
0.1
0.05
0.01
0
Cell_Type_NEW
Ctrl
ExN
InN
Other
UD

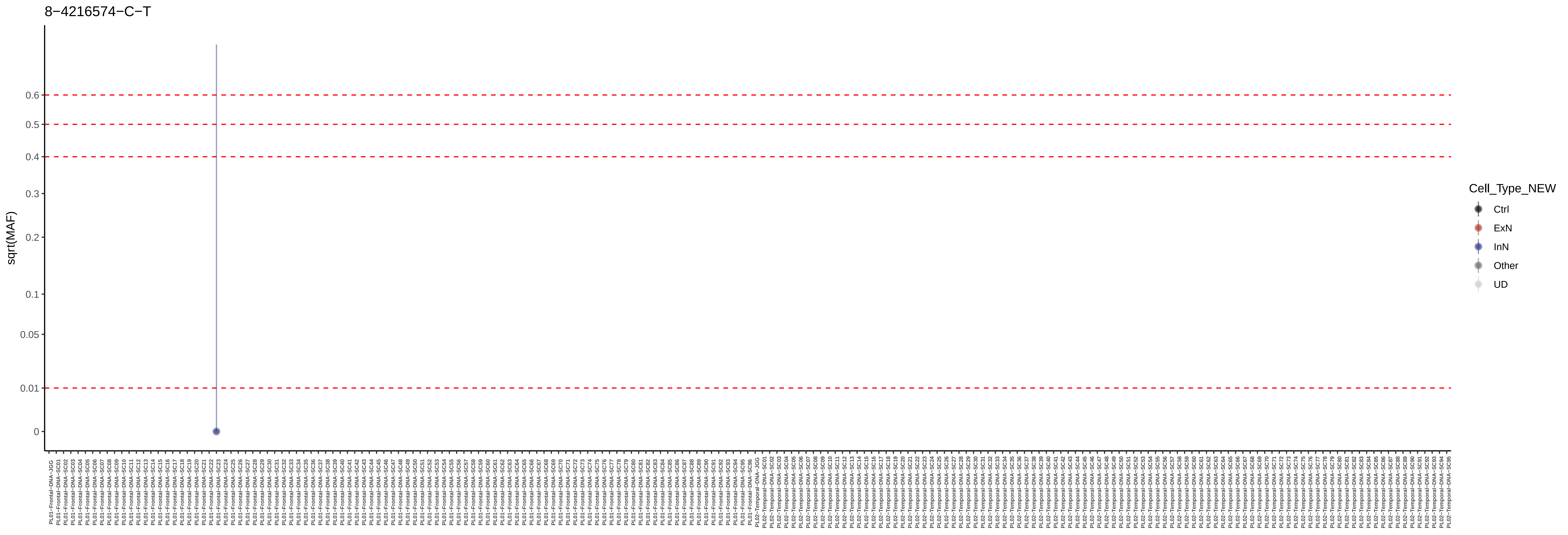

8−4216574−C−T
sqrt(MAF)
0.6
0.5
0.4
0.3
0.2
0.1
0.05
0.01
0
Cell_Type_NEW
Ctrl
ExN
InN
Other
UD

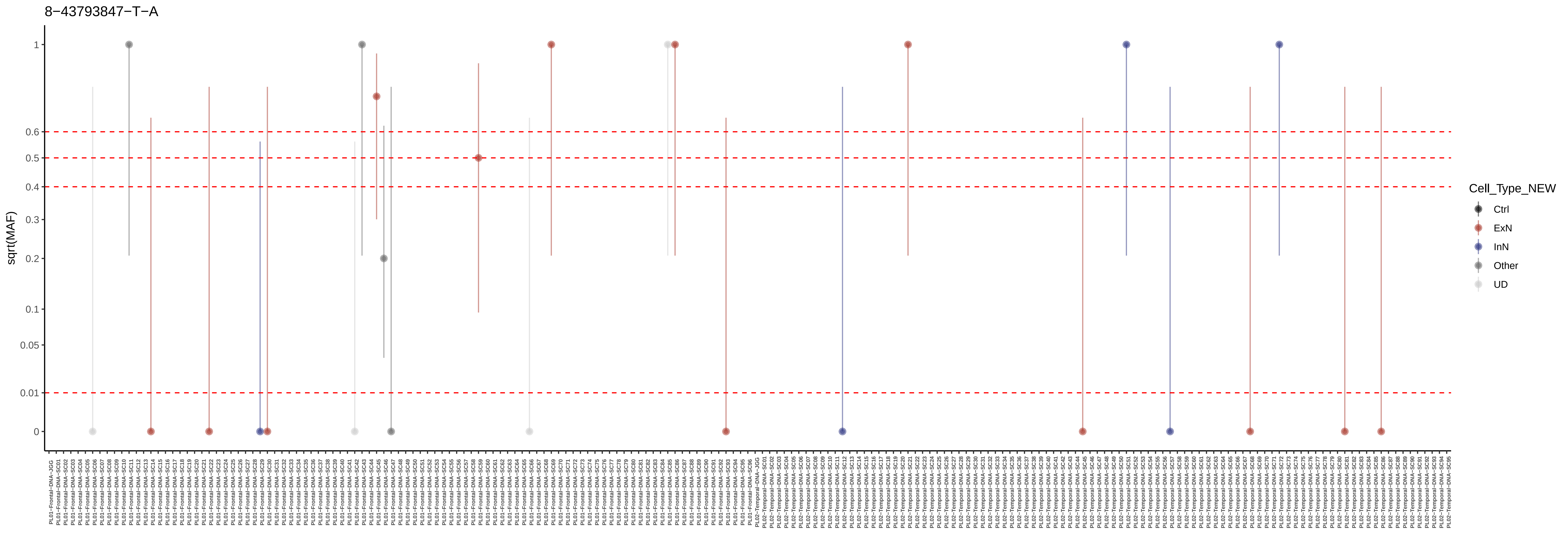
8−43793847−T−A
sqrt(MAF)
1
0.6
0.5
0.4
0.3
0.2
0.1
0.05
0.01
0
Cell_Type_NEW
Ctrl
ExN
InN
Other
UD

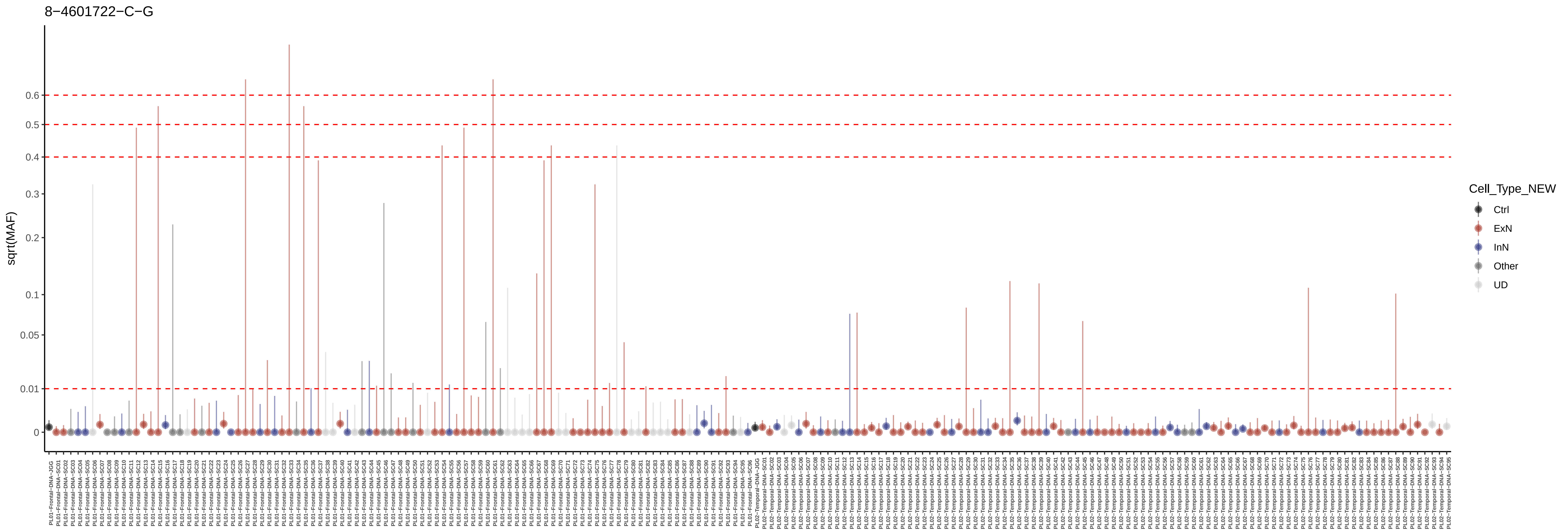

8−4601722−C−G
sqrt(MAF)
0.6
0.5
0.4
0.3
0.2
0.1
0.05
0.01
0
Cell_Type_NEW
Ctrl
ExN
InN
Other
UD

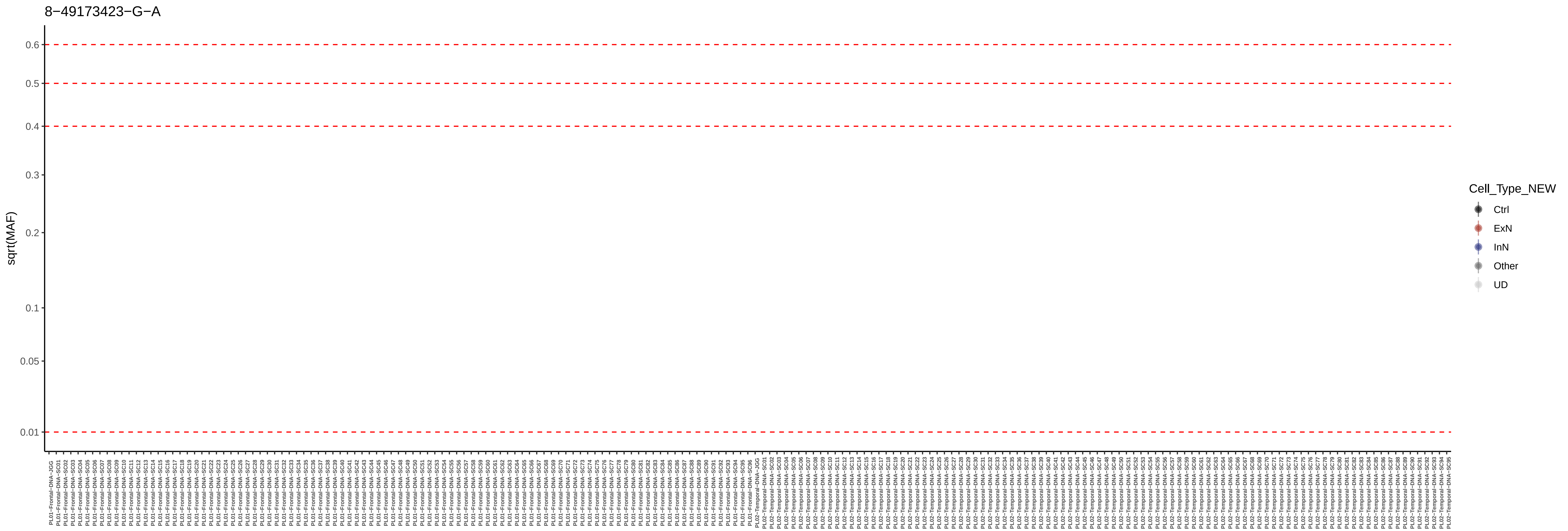

8−49173423−G−A
sqrt(MAF)
0.6
0.5
0.4
0.3
0.2
0.1
0.05
0.01
Cell_Type_NEW
Ctrl
ExN
InN
Other
UD

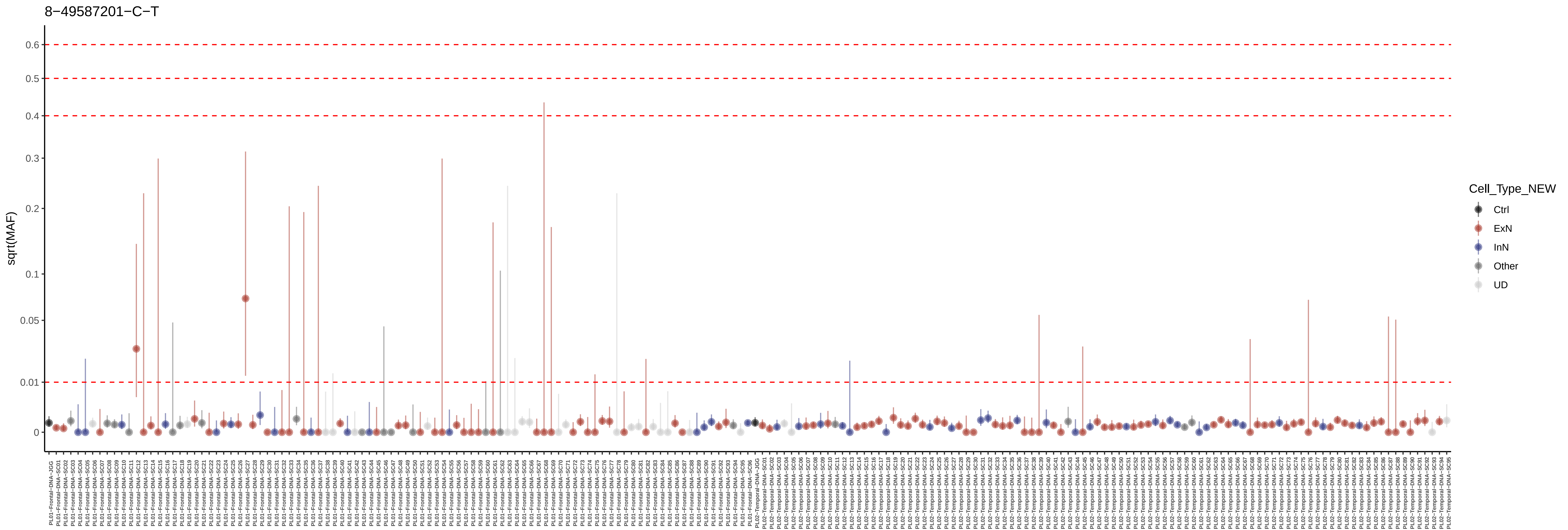

8−49587201−C−T
sqrt(MAF)
0.6
0.5
0.4
0.3
0.2
0.1
0.05
0.01
0
Cell_Type_NEW
Ctrl
ExN
InN
Other
UD

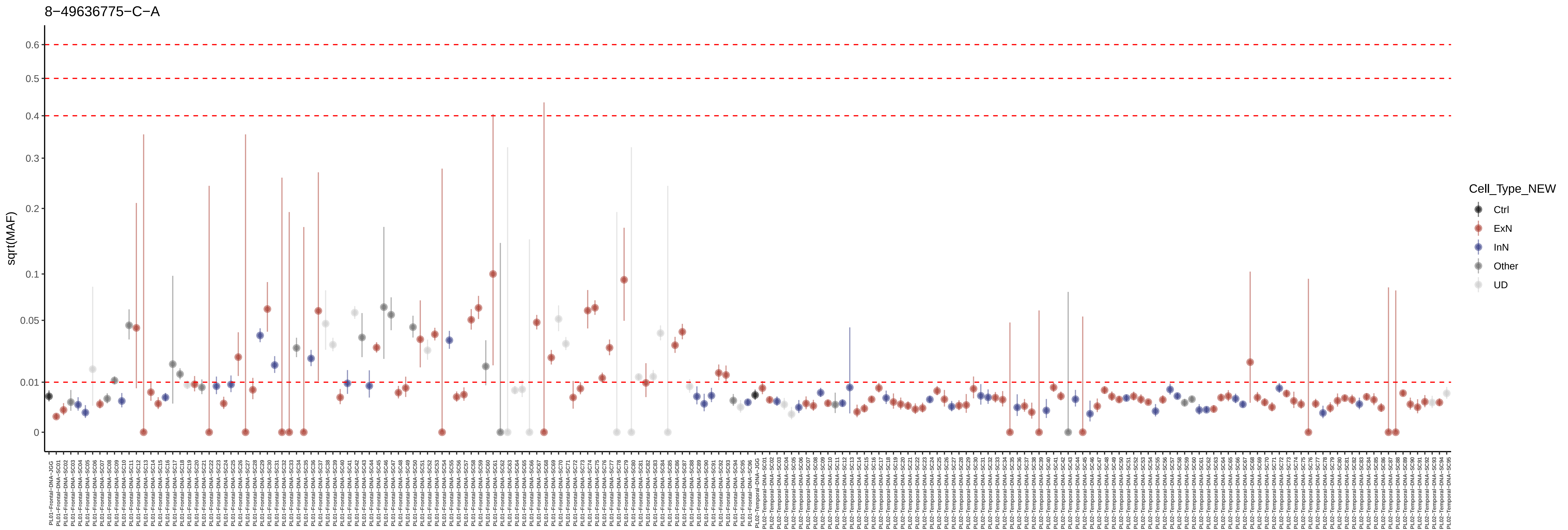
8−49636775−C−A
sqrt(MAF)
0.6
0.5
0.4
0.3
0.2
0.1
0.05
0.01
0
Cell_Type_NEW
Ctrl
ExN
InN
Other
UD

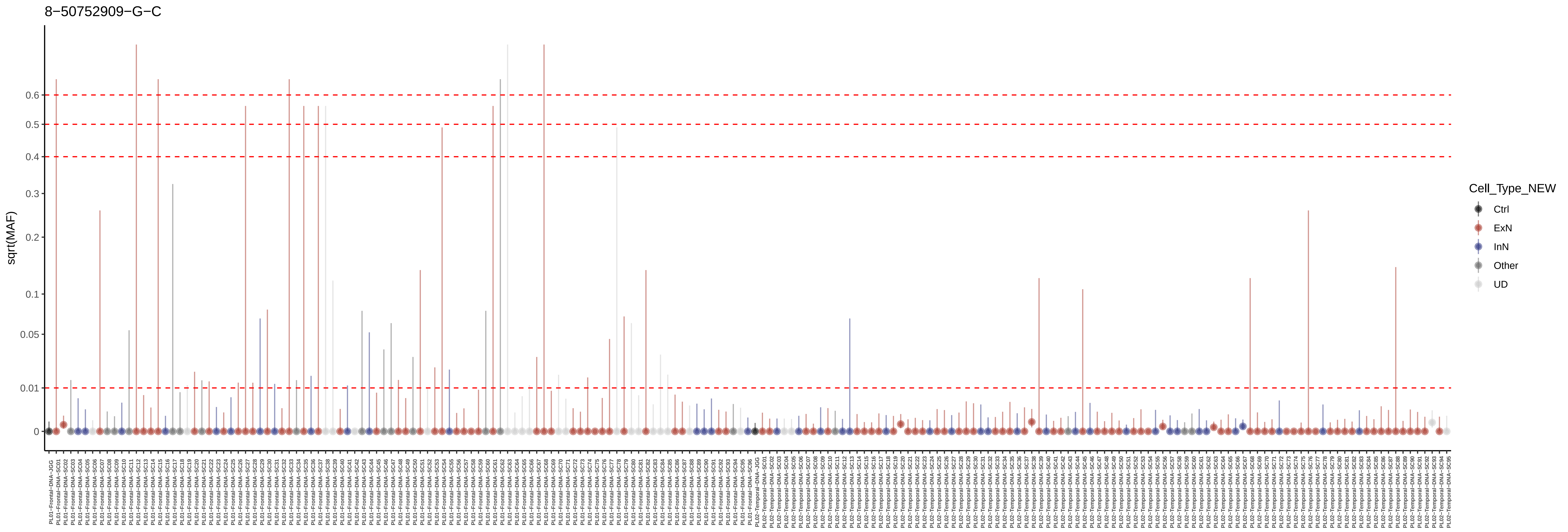
8−50752909−G−C
sqrt(MAF)
0.6
0.5
0.4
0.3
0.2
0.1
0.05
0.01
0
Cell_Type_NEW
Ctrl
ExN
InN
Other
UD

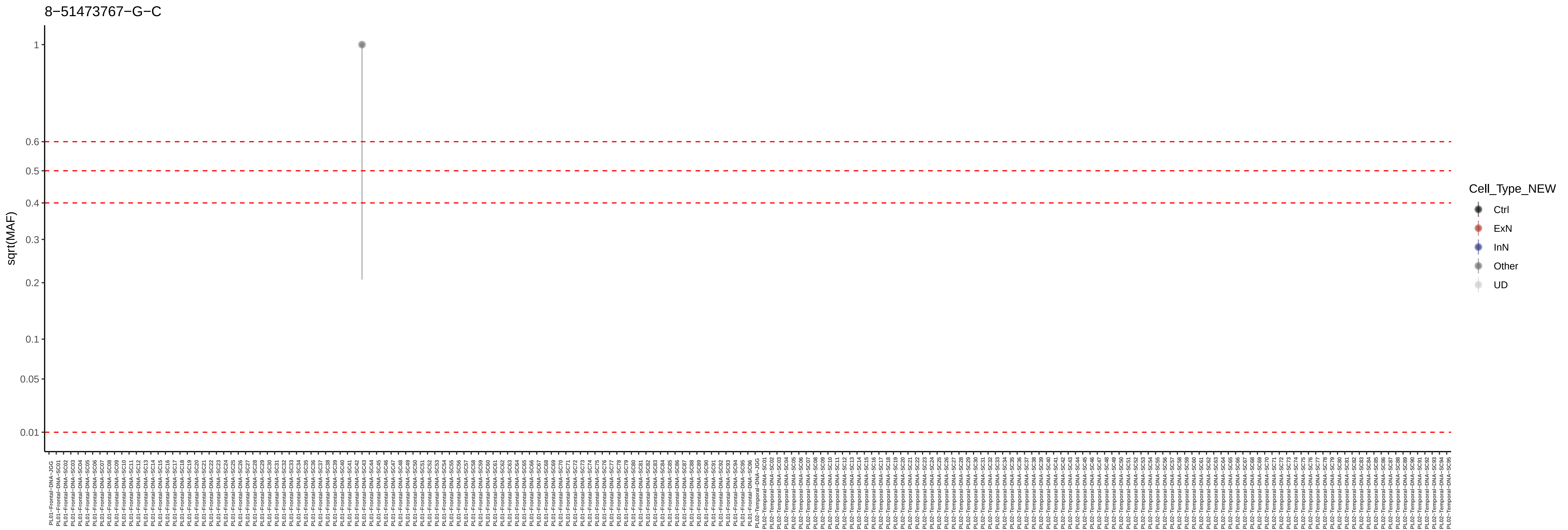
8−51473767−G−C
sqrt(MAF)
1
0.6
0.5
0.4
0.3
0.2
0.1
0.05
0.01
Cell_Type_NEW
Ctrl
ExN
InN
Other
UD

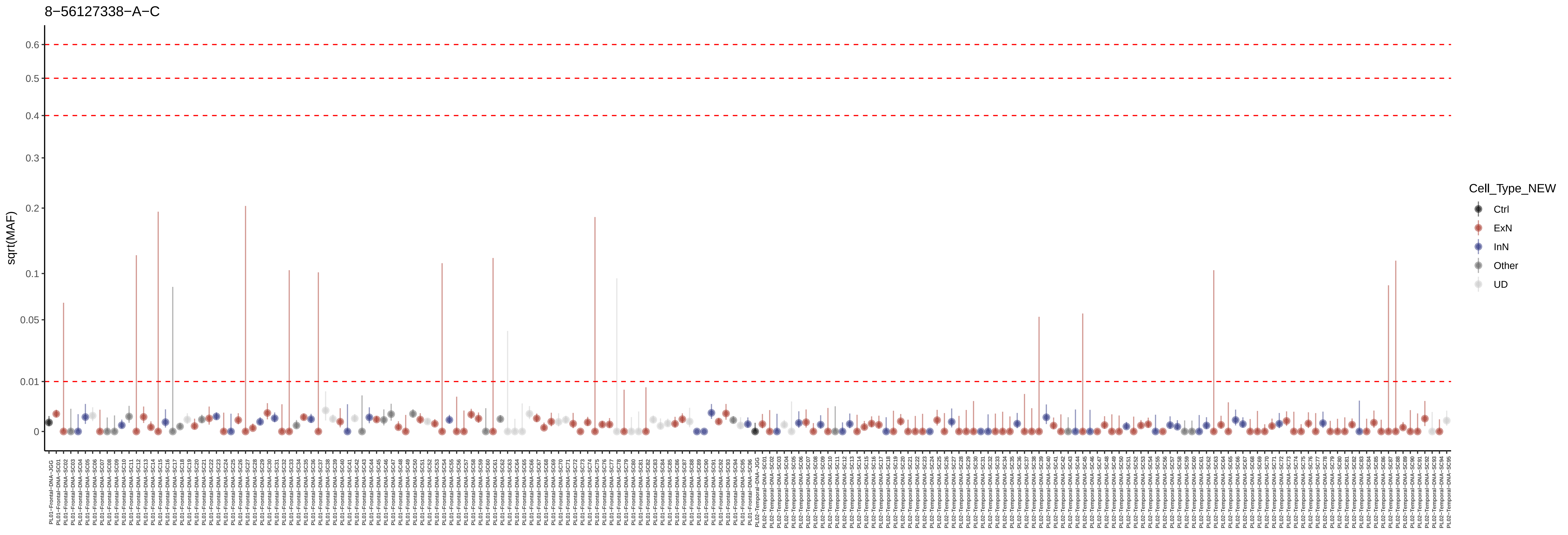
8−56127338−A−C
sqrt(MAF)
0.6
0.5
0.4
0.3
0.2
0.1
0.05
0.01
0
Cell_Type_NEW
Ctrl
ExN
InN
Other
UD

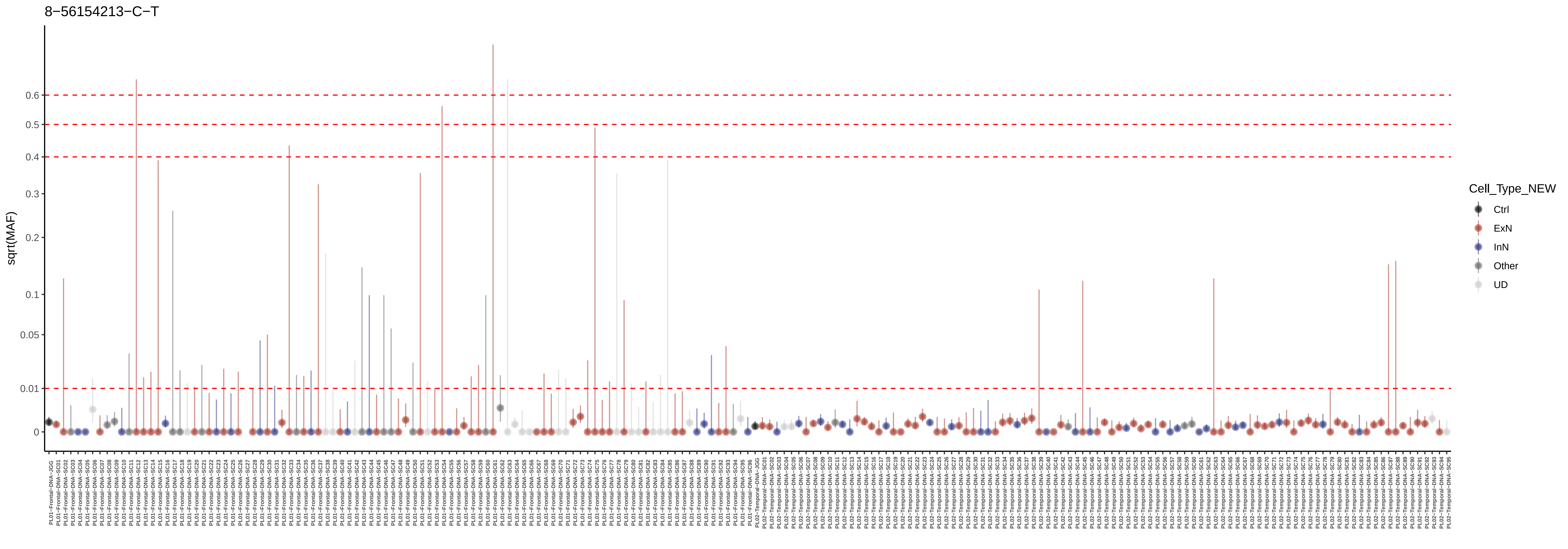
8−56154213−C−T
sqrt(MAF)
Cell_Type_NEW
Ctrl
ExN
InN
Other
UD

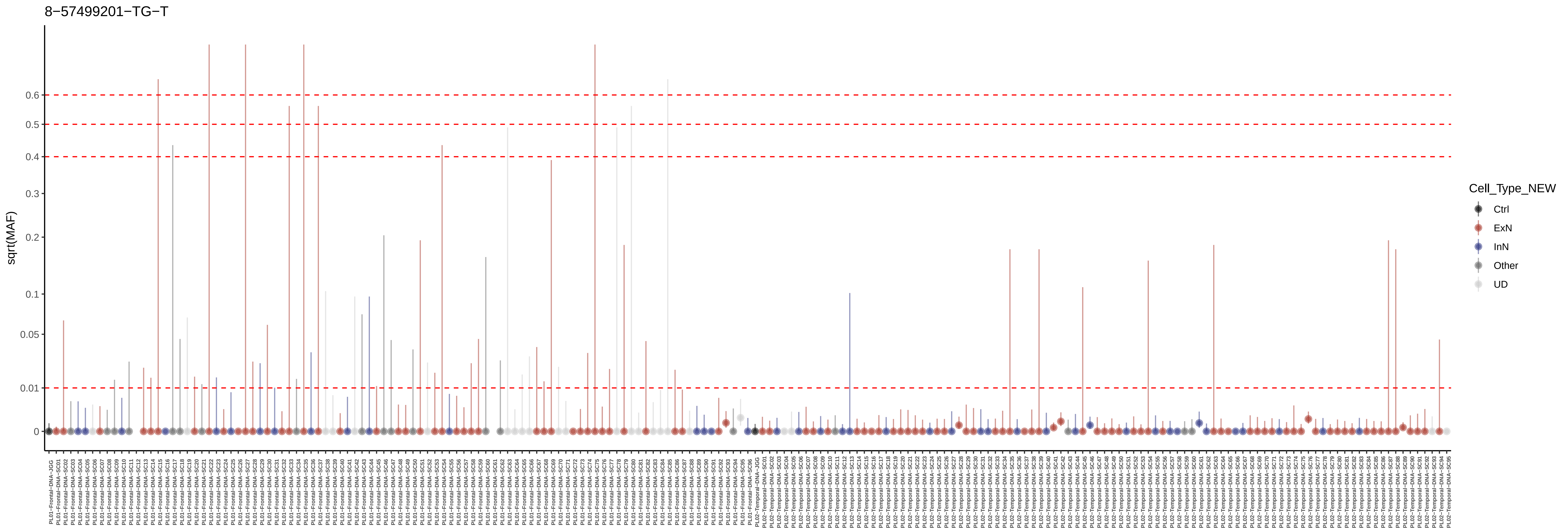
8−57499201−TG−T
sqrt(MAF)
0.6
0.5
0.4
0.3
0.2
0.1
0.05
0.01
0
Cell_Type_NEW
Ctrl
ExN
InN
Other
UD

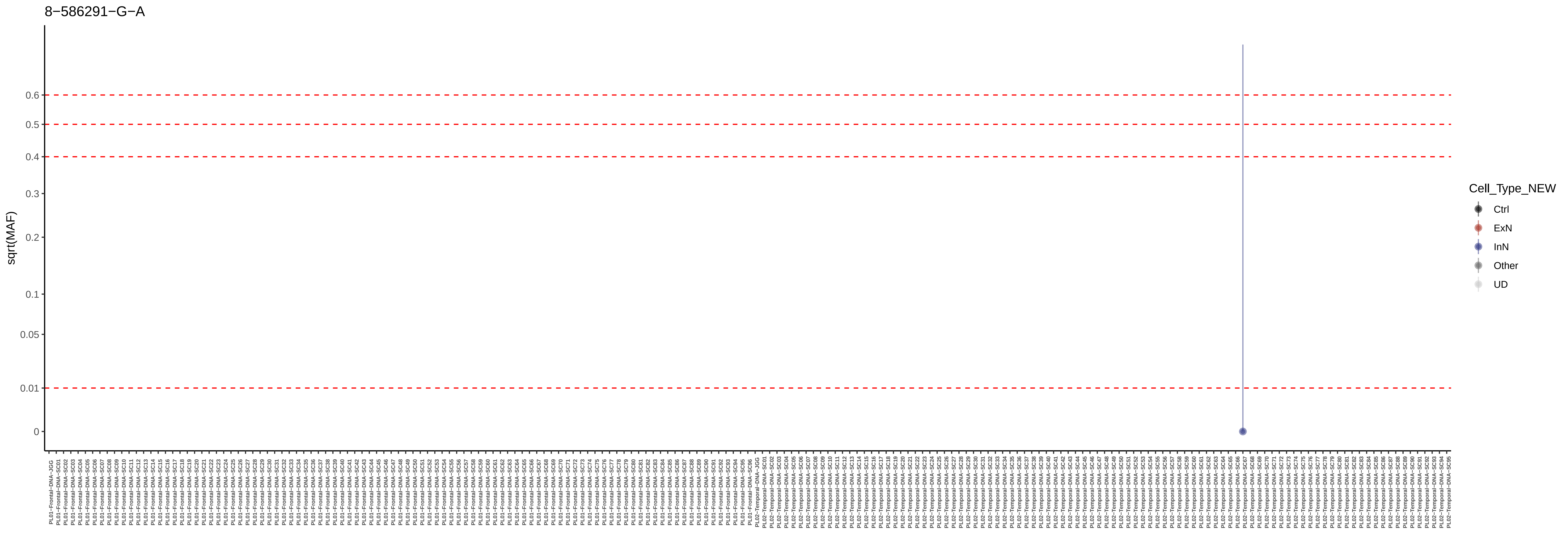
8−586291−G−A
sqrt(MAF)
0.6
0.5
0.4
0.3
0.2
0.1
0.05
0.01
0
Cell_Type_NEW
Ctrl
ExN
InN
Other
UD

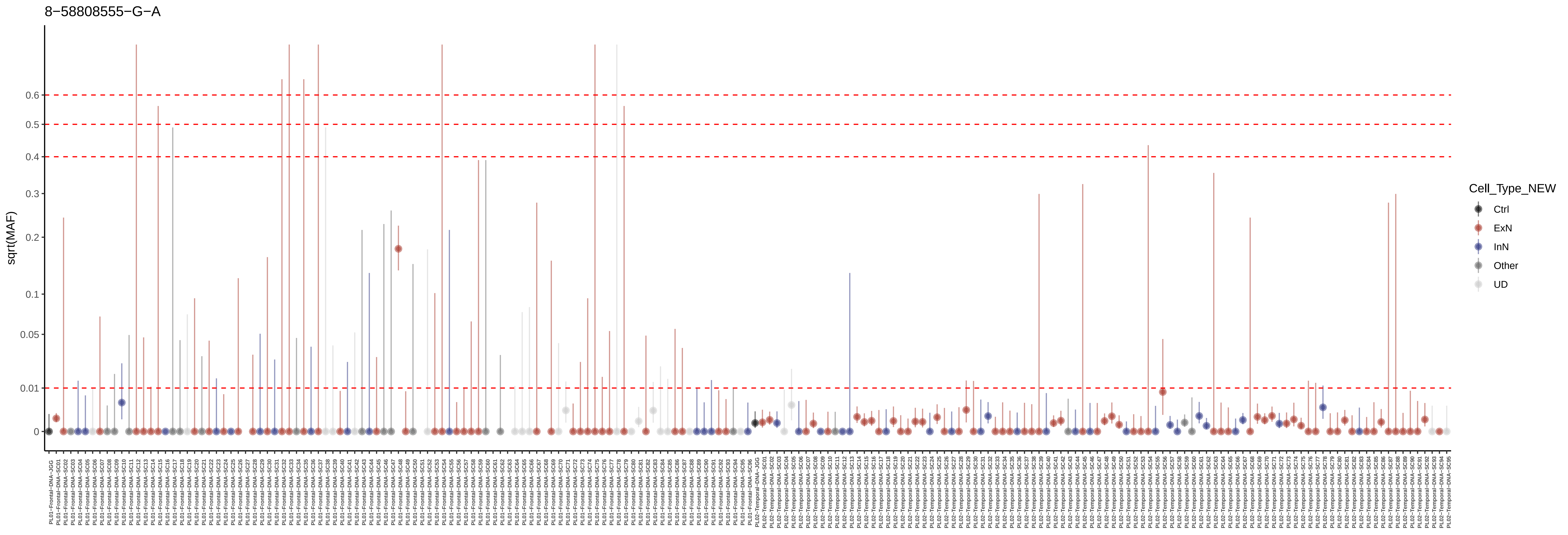
8−58808555−G−A
sqrt(MAF)
0.6
0.5
0.4
0.3
0.2
0.1
0.05
0.01
0
Cell_Type_NEW
Ctrl
ExN
InN
Other
UD

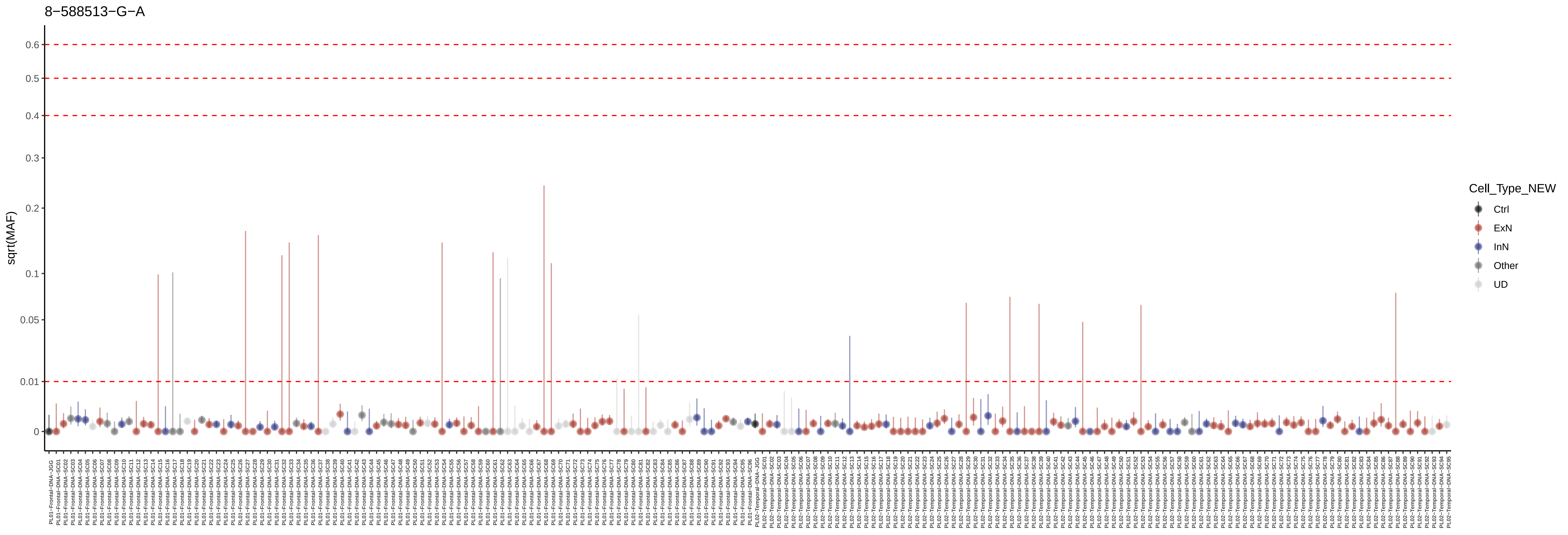
8−588513−G−A
sqrt(MAF)
0.6
0.5
0.4
0.3
0.2
0.1
0.05
0.01
0
Cell_Type_NEW
Ctrl
ExN
InN
Other
UD

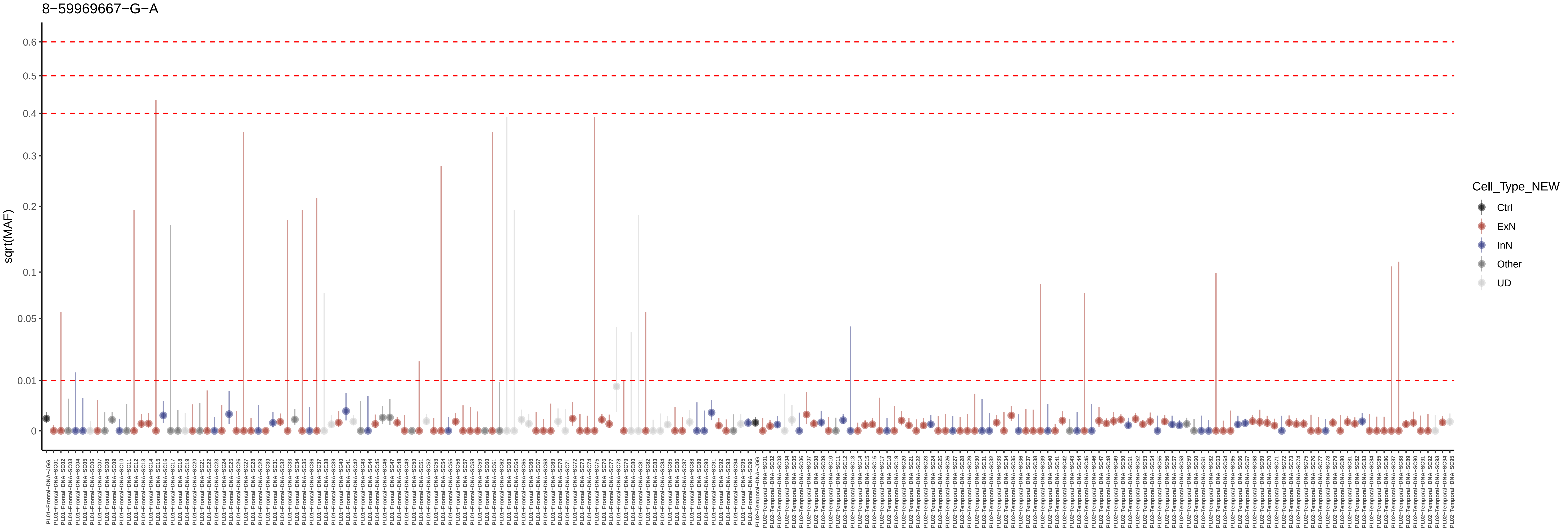

8−59969667−G−A
sqrt(MAF)
0.6
0.5
0.4
0.3
0.2
0.1
0.05
0.01
0
Cell_Type_NEW
Ctrl
ExN
InN
Other
UD

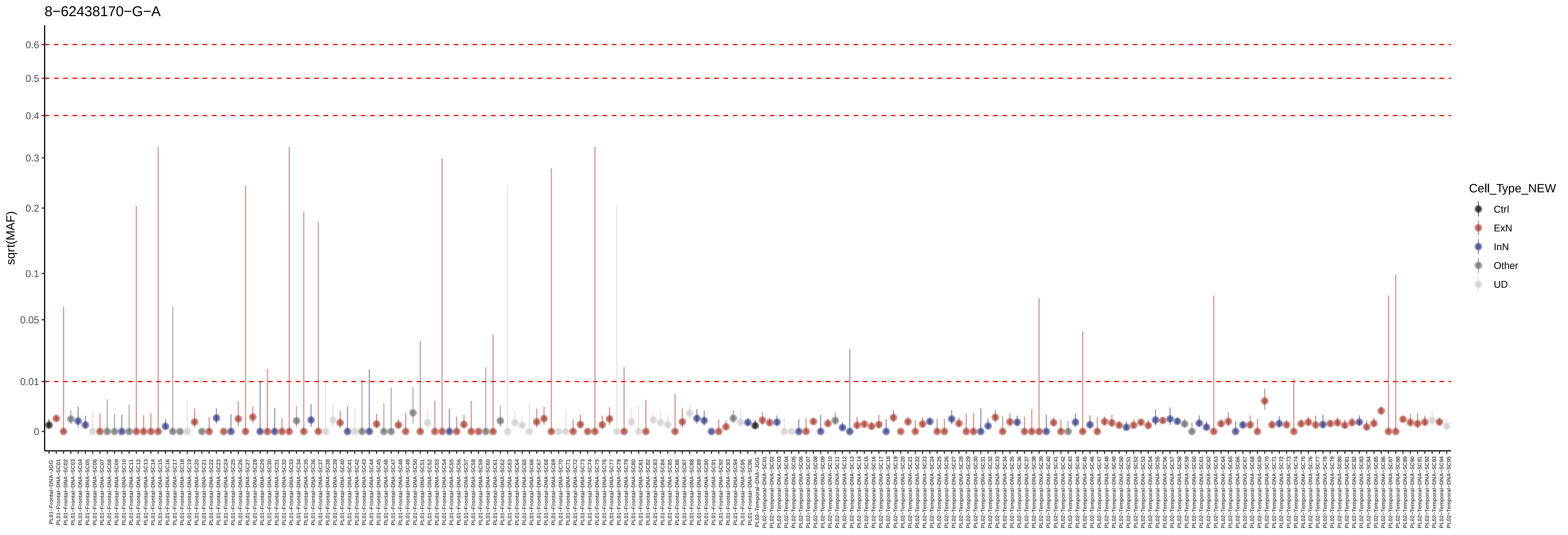
8−62438170−G−A
sqrt(MAF)
0.6
0.5
0.4
0.3
0.2
0.1
0.05
0.01
0
Cell_Type_NEW
Ctrl
ExN
InN
Other
UD

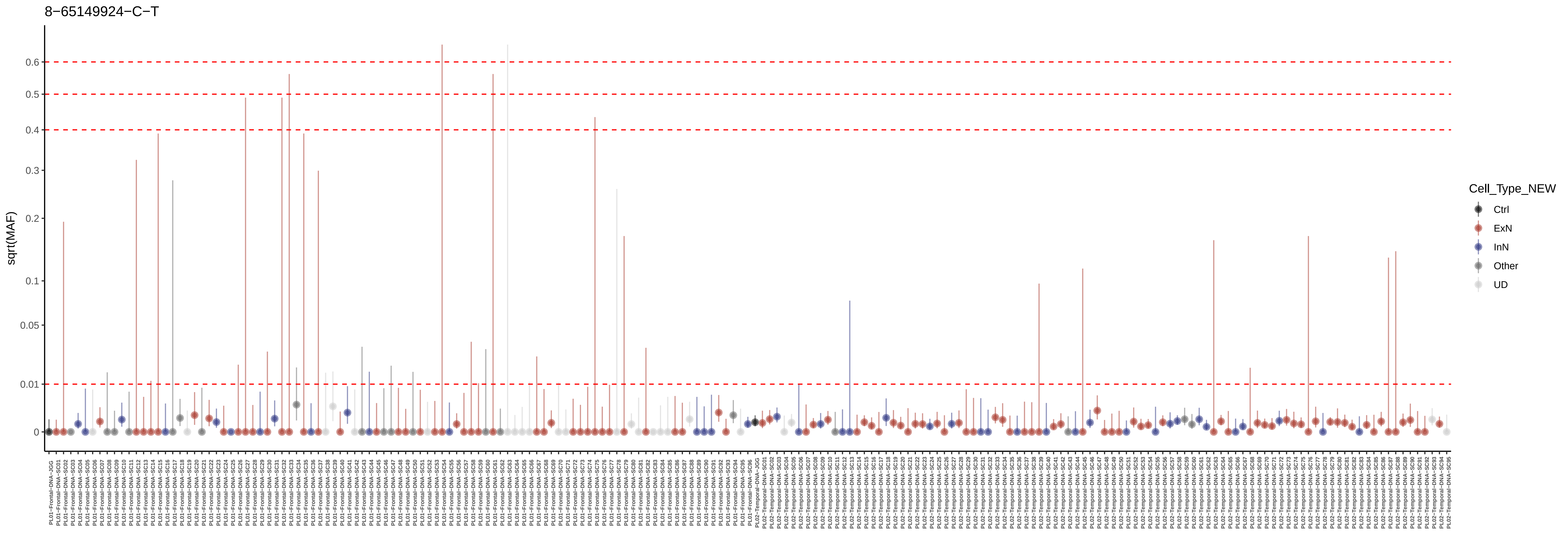
8−65149924−C−T
sqrt(MAF)
0.6
0.5
0.4
0.3
0.2
0.1
0.05
0.01
0
Cell_Type_NEW
Ctrl
ExN
InN
Other
UD

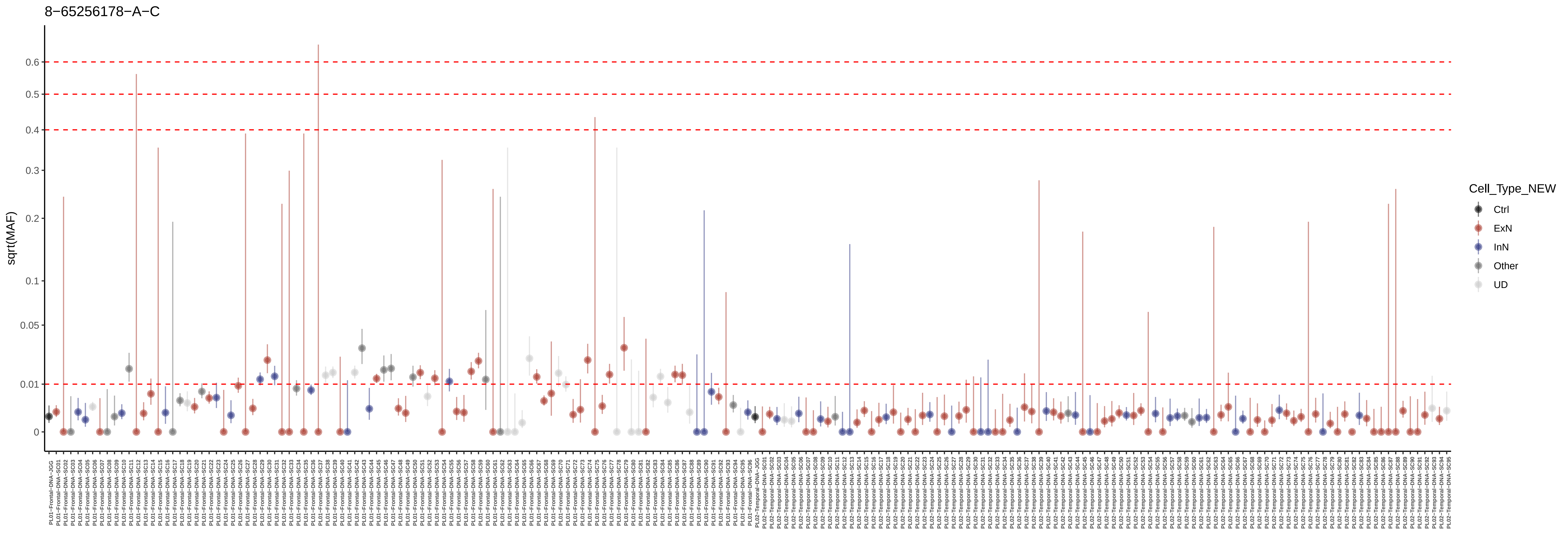

8−65256178−A−C
sqrt(MAF)
0.6
0.5
0.4
0.3
0.2
0.1
0.05
0.01
0
Cell_Type_NEW
Ctrl
ExN
InN
Other
UD

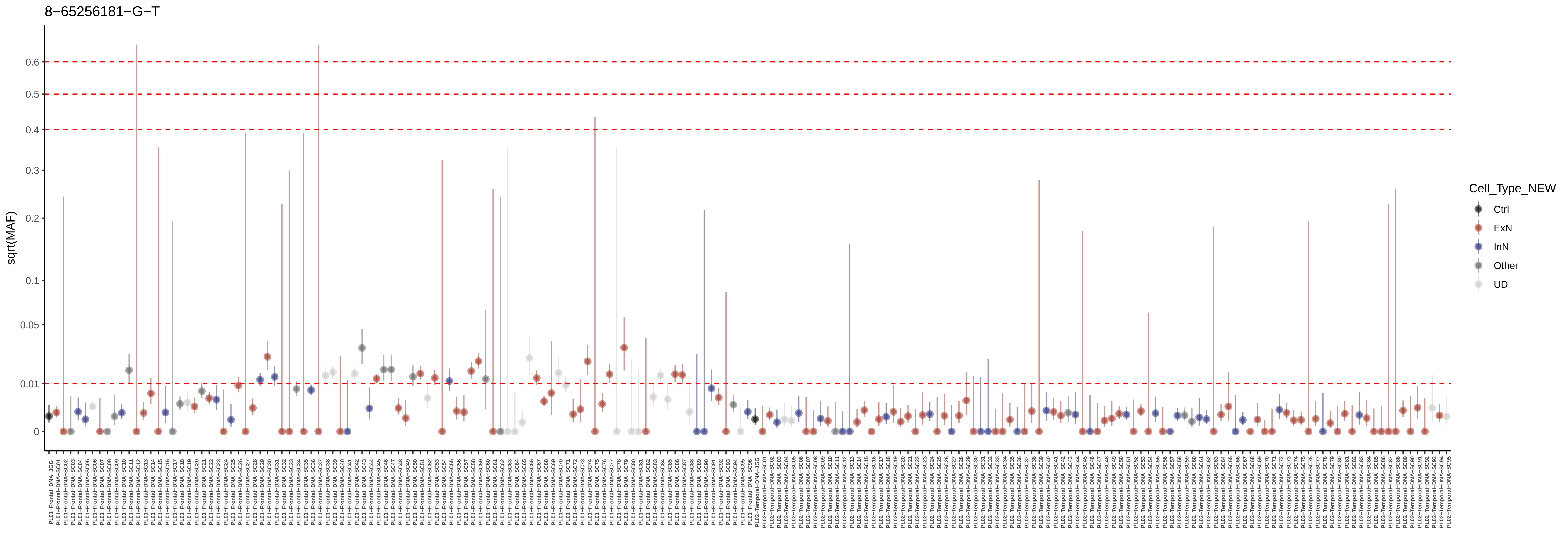
8−65256181−G−T
sqrt(MAF)
0.6
0.5
0.4
0.3
0.2
0.1
0.05
0.01
0
Cell_Type_NEW
Ctrl
ExN
InN
Other
UD

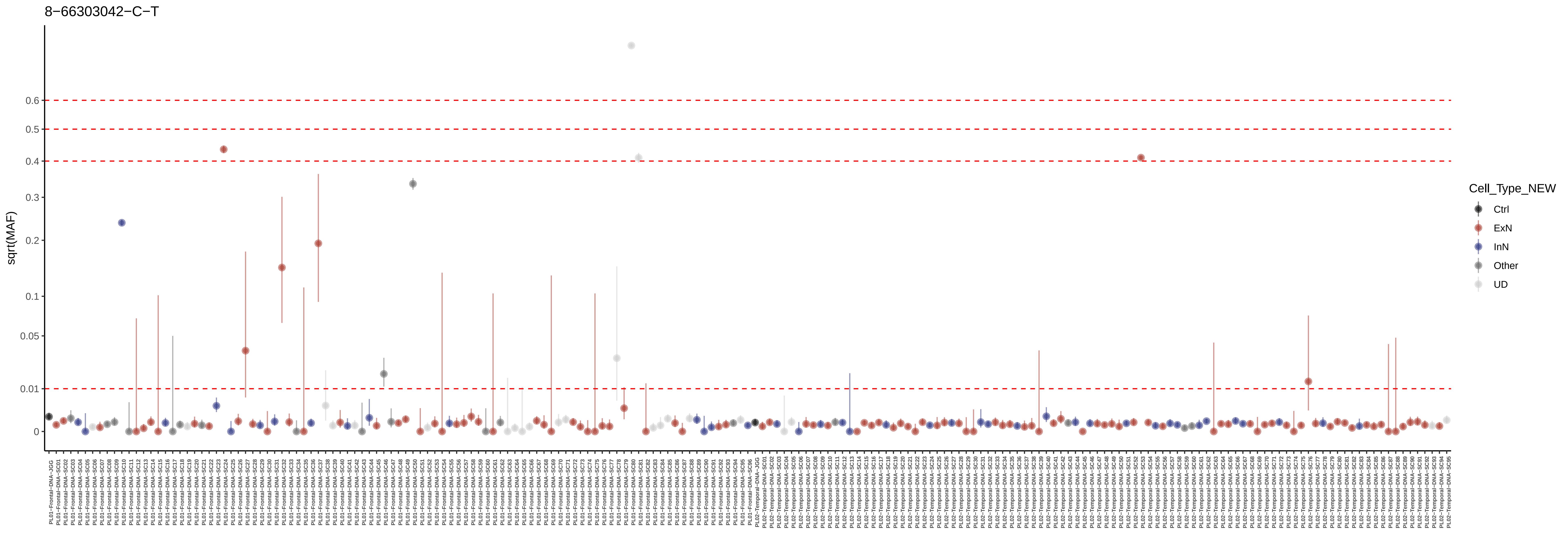
8−66303042−C−T
sqrt(MAF)
0.6
0.5
0.4
0.3
0.2
0.1
0.05
0.01
0
Cell_Type_NEW
Ctrl
ExN
InN
Other
UD

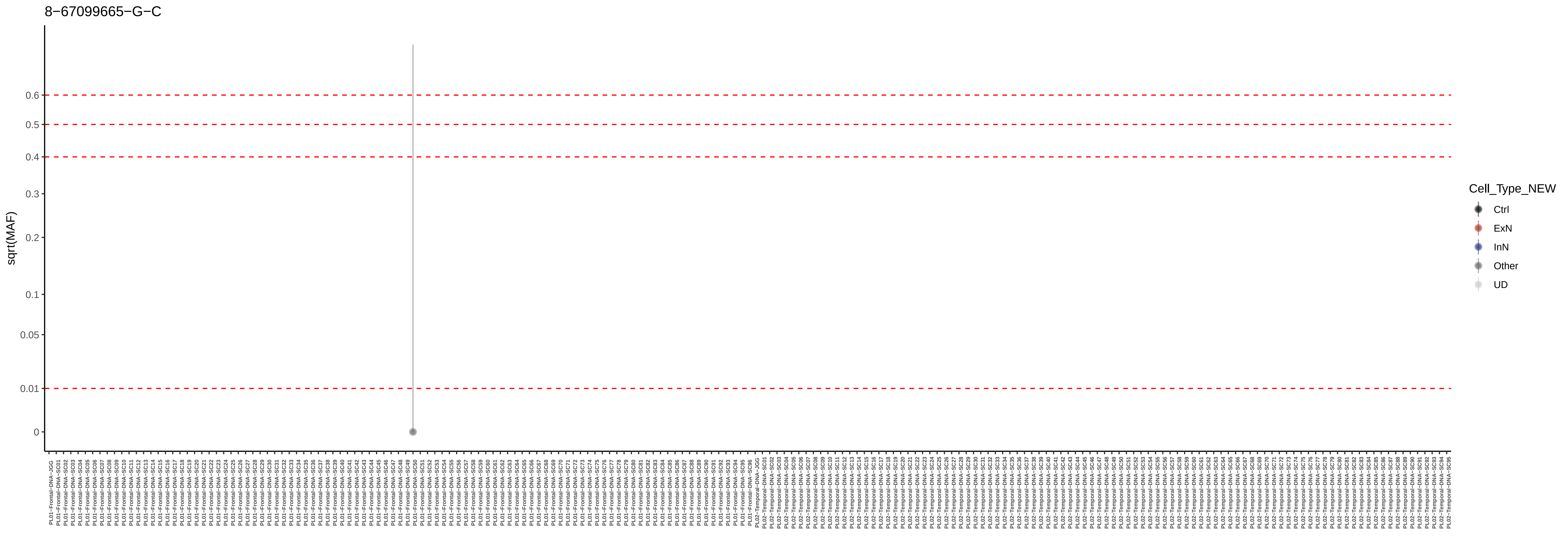
8−67099665−G−C
sqrt(MAF)
0.6
0.5
0.4
0.3
0.2
0.1
0.05
0.01
0
Cell_Type_NEW
Ctrl
ExN
InN
Other
UD

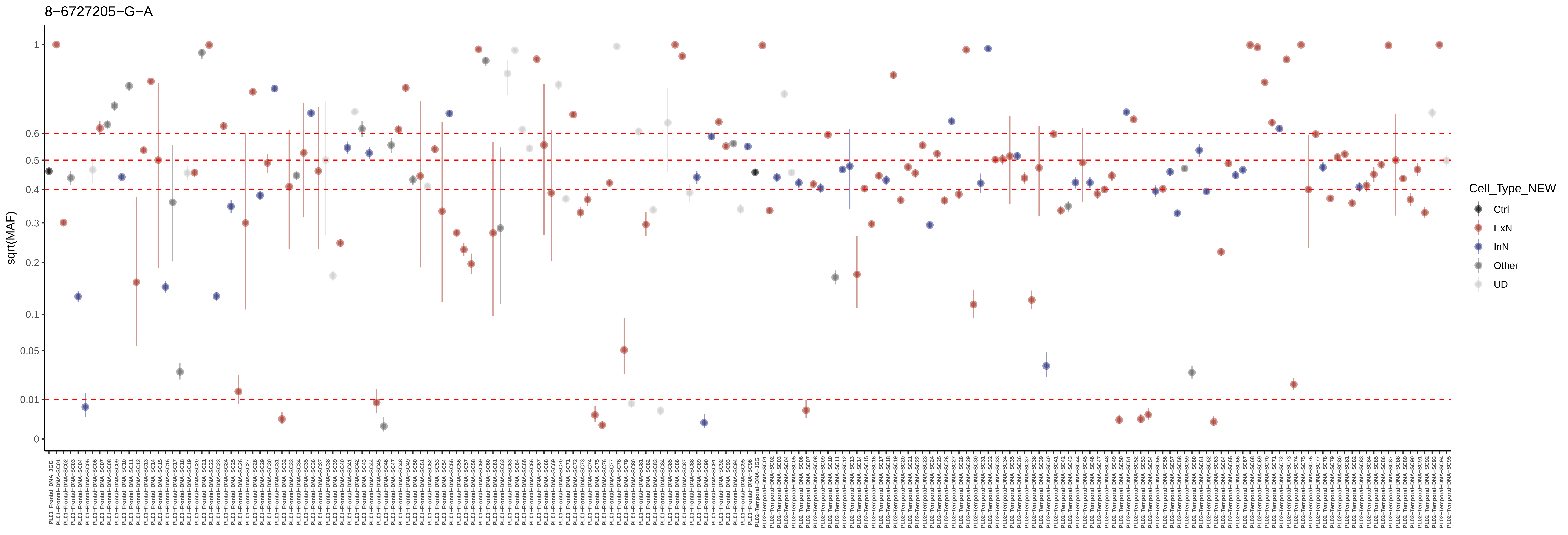

8−6727205−G−A
sqrt(MAF)
1
0.6
0.5
0.4
0.3
0.2
0.1
0.05
0.01
0
Cell_Type_NEW
Ctrl
ExN
InN
Other
UD

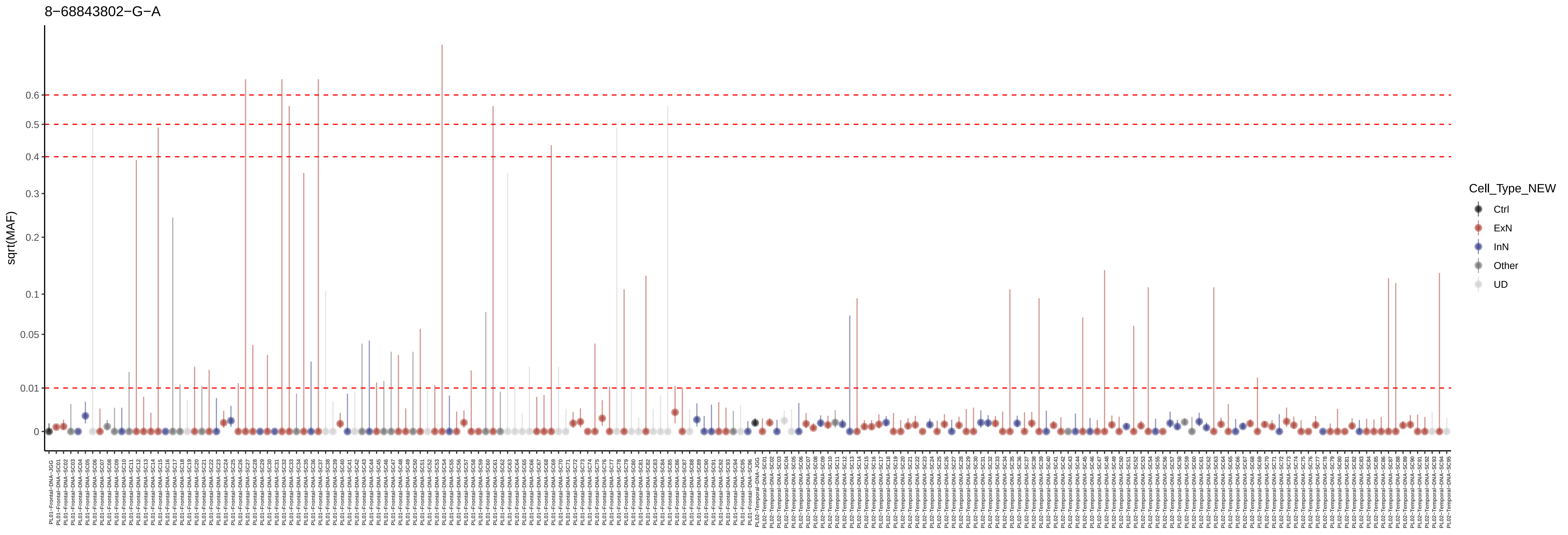

8-68843802-G-A
sqrt(MAF)
0.6
0.5
0.4
0.3
0.2
0.1
0.05
0.01
0
Cell_Type_NEW
Ctrl
ExN
InN
Other
UD

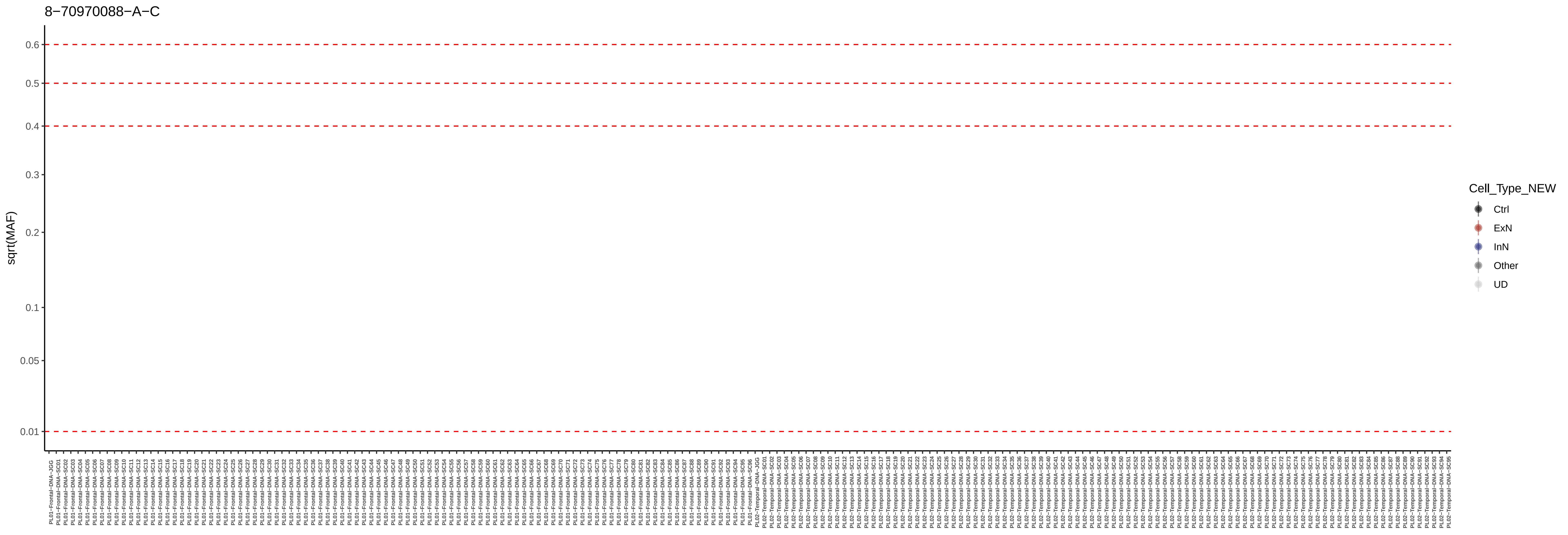
8−70970088−A−C
sqrt(MAF)
0.6
0.5
0.4
0.3
0.2
0.1
0.05
0.01
Cell_Type_NEW
Ctrl
ExN
InN
Other
UD

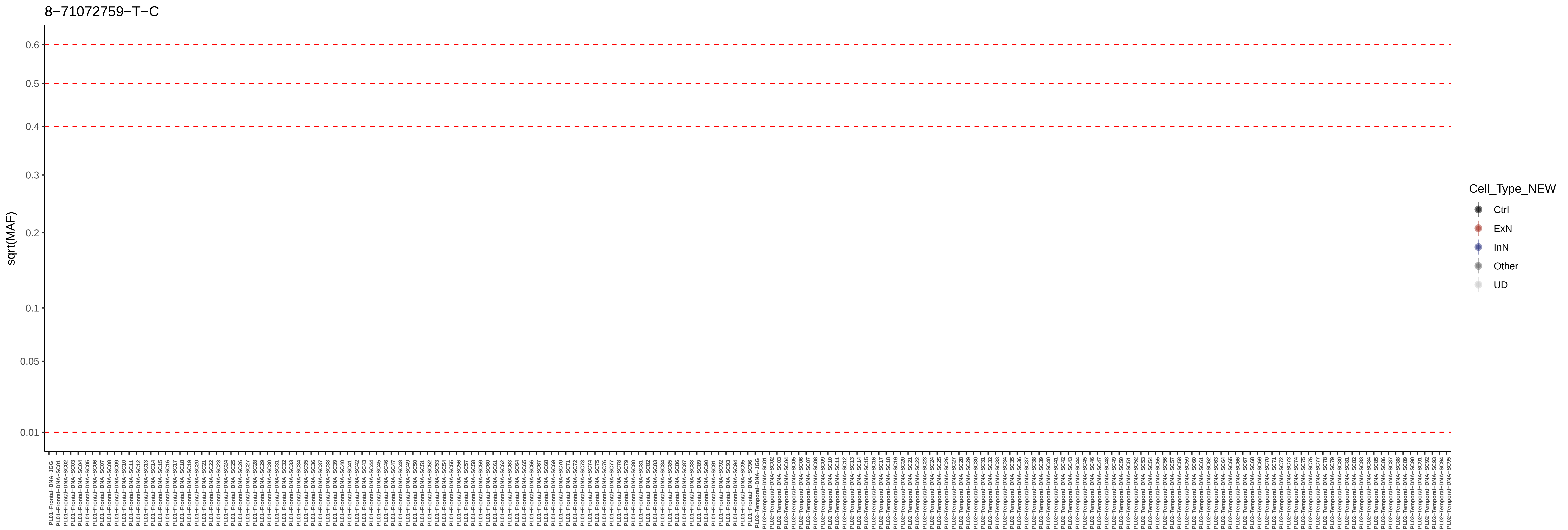

8−71072759−T−C
sqrt(MAF)
0.6
0.5
0.4
0.3
0.2
0.1
0.05
0.01
Cell_Type_NEW
Ctrl
ExN
InN
Other
UD

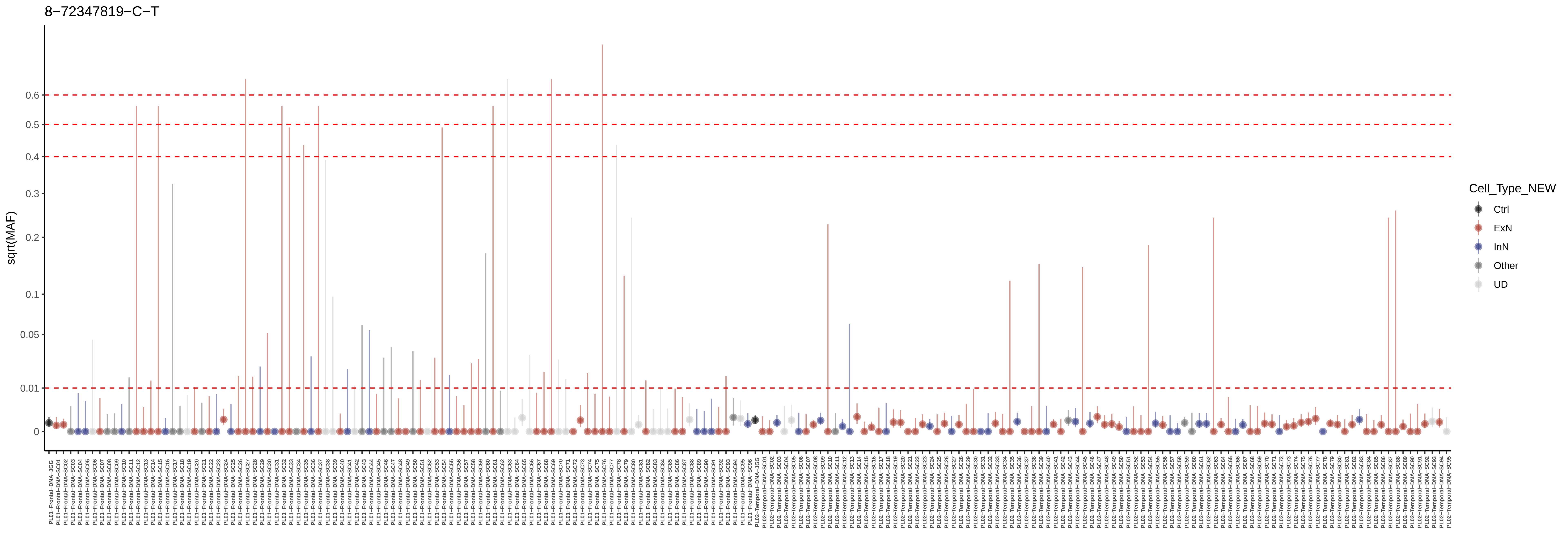
8−72347819−C−T
sqrt(MAF)
0.6
0.5
0.4
0.3
0.2
0.1
0.05
0.01
0
Cell_Type_NEW
Ctrl
ExN
InN
Other
UD

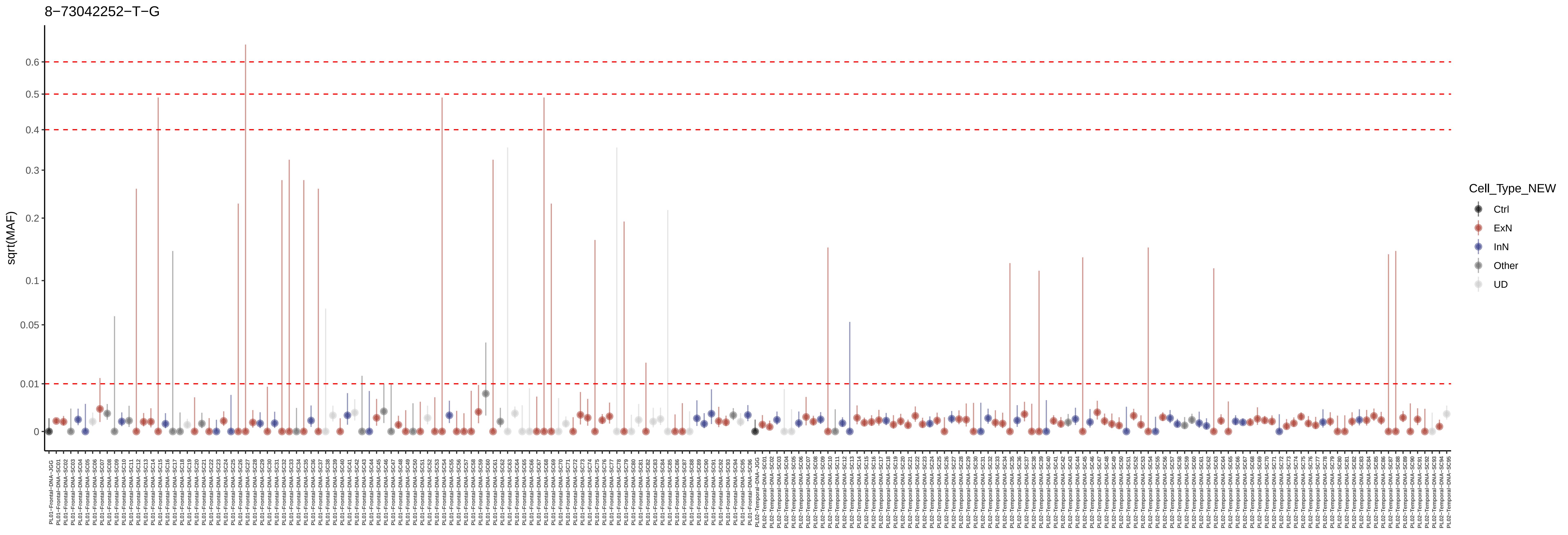
8−73042252−T−G
sqrt(MAF)
0.6
0.5
0.4
0.3
0.2
0.1
0.05
0.01
0
Cell_Type_NEW
Ctrl
ExN
InN
Other
UD

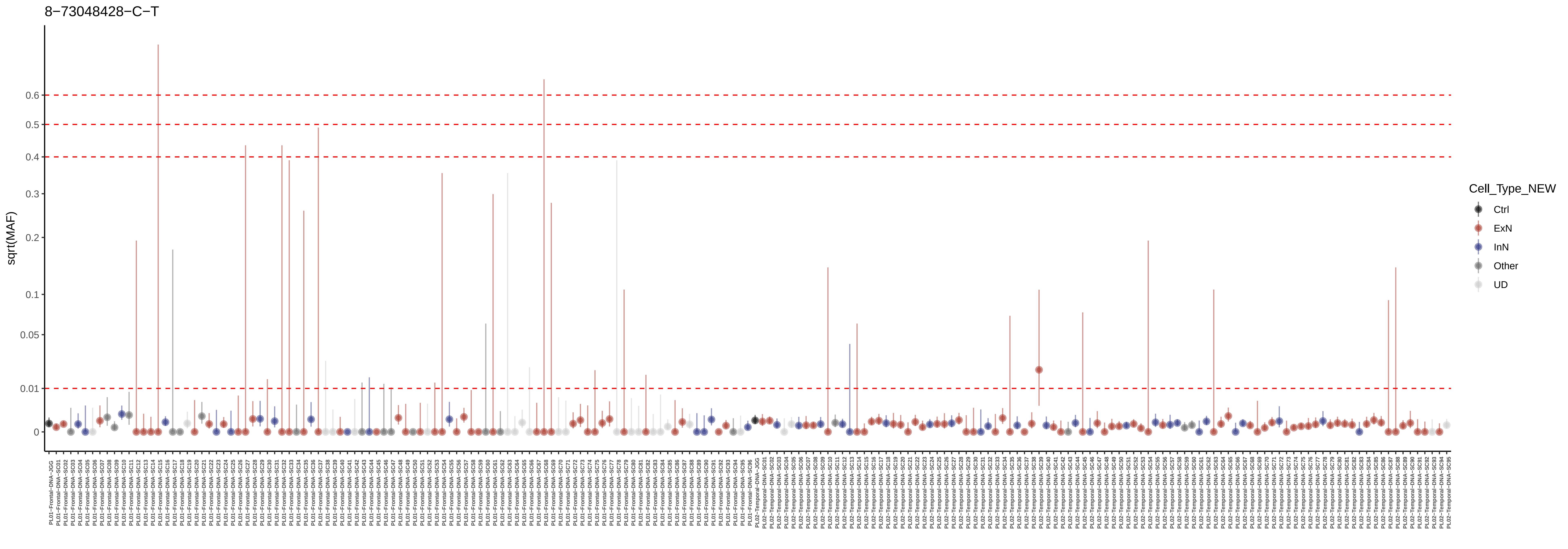
8−73048428−C−T
sqrt(MAF)
0.6
0.5
0.4
0.3
0.2
0.1
0.05
0.01
0
Cell_Type_NEW
Ctrl
ExN
InN
Other
UD

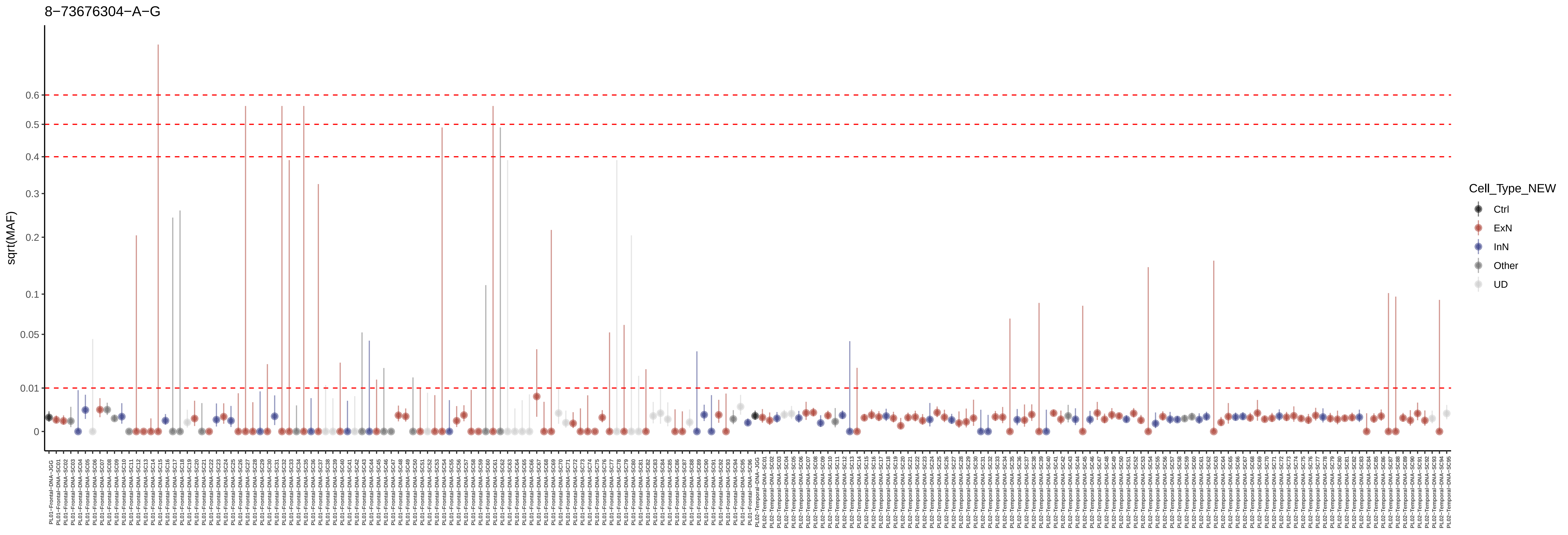

8−73676304−A−G
sqrt(MAF)
0.6
0.5
0.4
0.3
0.2
0.1
0.05
0.01
0
Cell_Type_NEW
Ctrl
ExN
InN
Other
UD

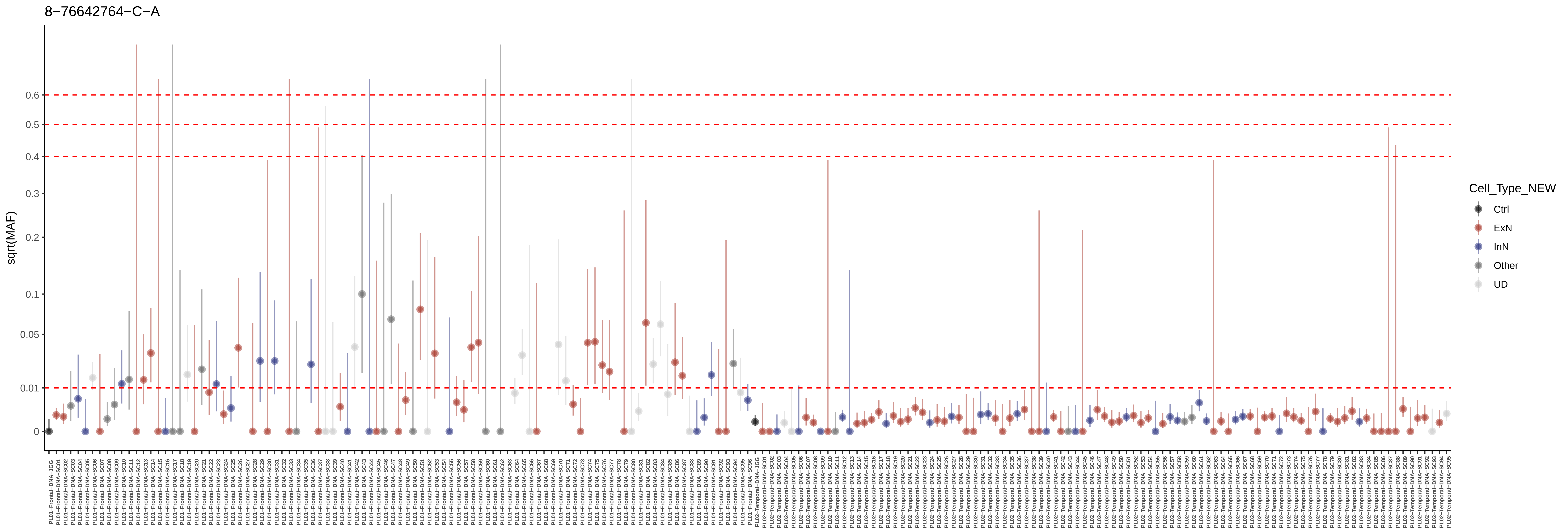
8−76642764−C−A
sqrt(MAF)
0.6
0.5
0.4
0.3
0.2
0.1
0.05
0.01
0
Cell_Type_NEW
Ctrl
ExN
InN
Other
UD

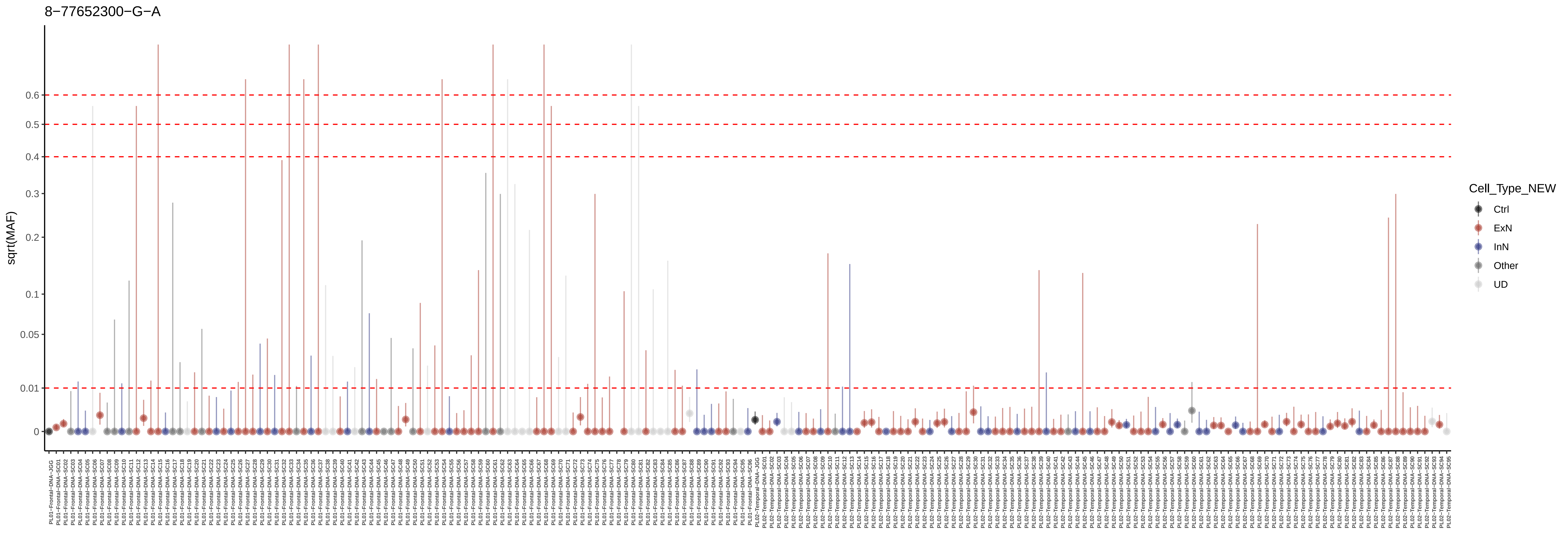

8−77652300−G−A
sqrt(MAF)
0.6
0.5
0.4
0.3
0.2
0.1
0.05
0.01
0
Cell_Type_NEW
Ctrl
ExN
InN
Other
UD

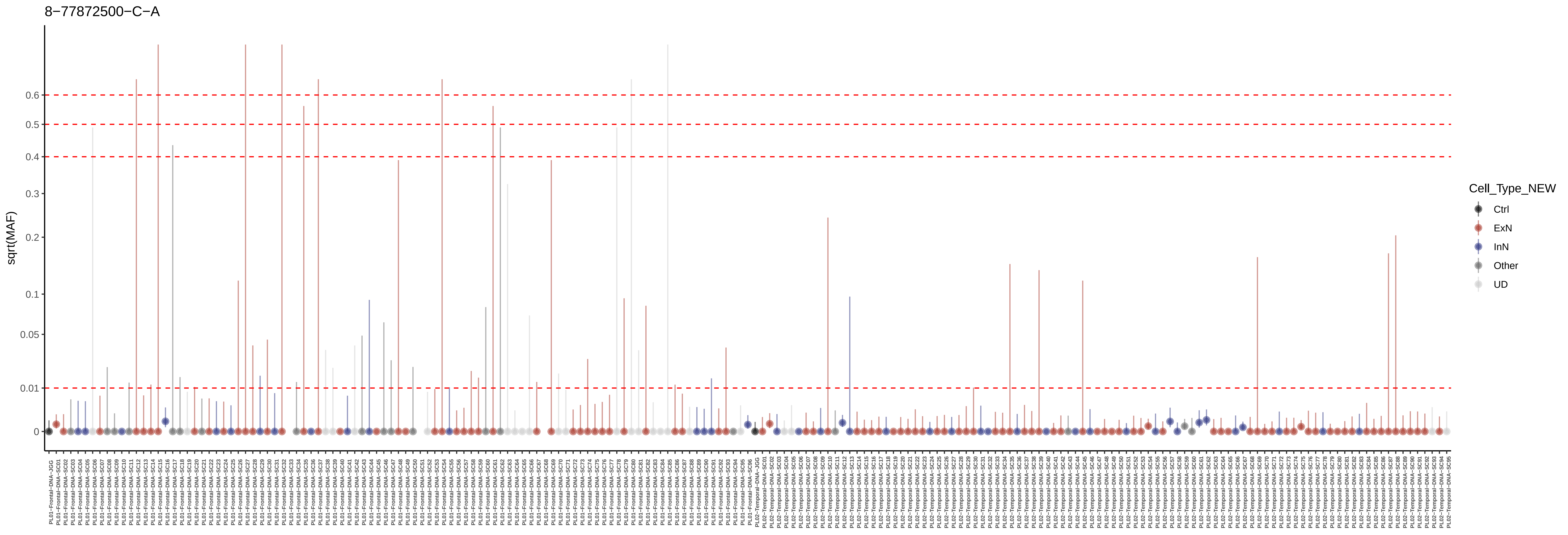

8−77872500−C−A
sqrt(MAF)
0.6
0.5
0.4
0.3
0.2
0.1
0.05
0.01
0
Cell_Type_NEW
Ctrl
ExN
InN
Other
UD

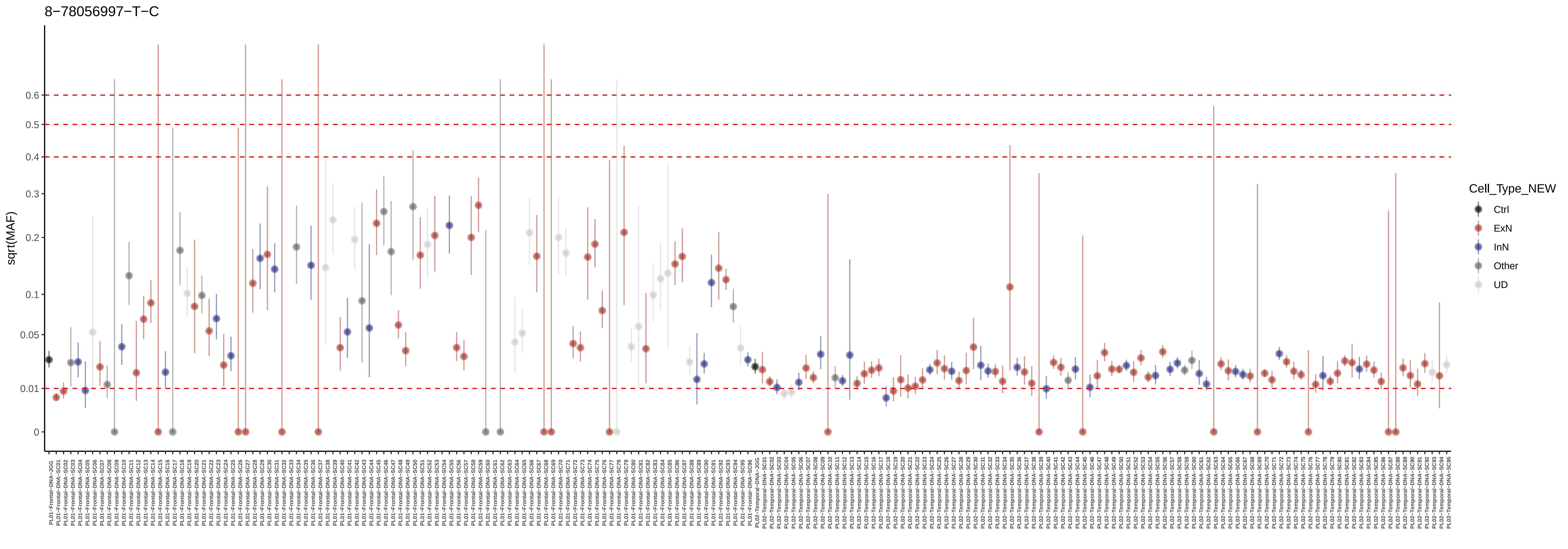

8−78056997−T−C
sqrt(MAF)
Cell_Type_NEW
Ctrl
ExN
InN
Other
UD

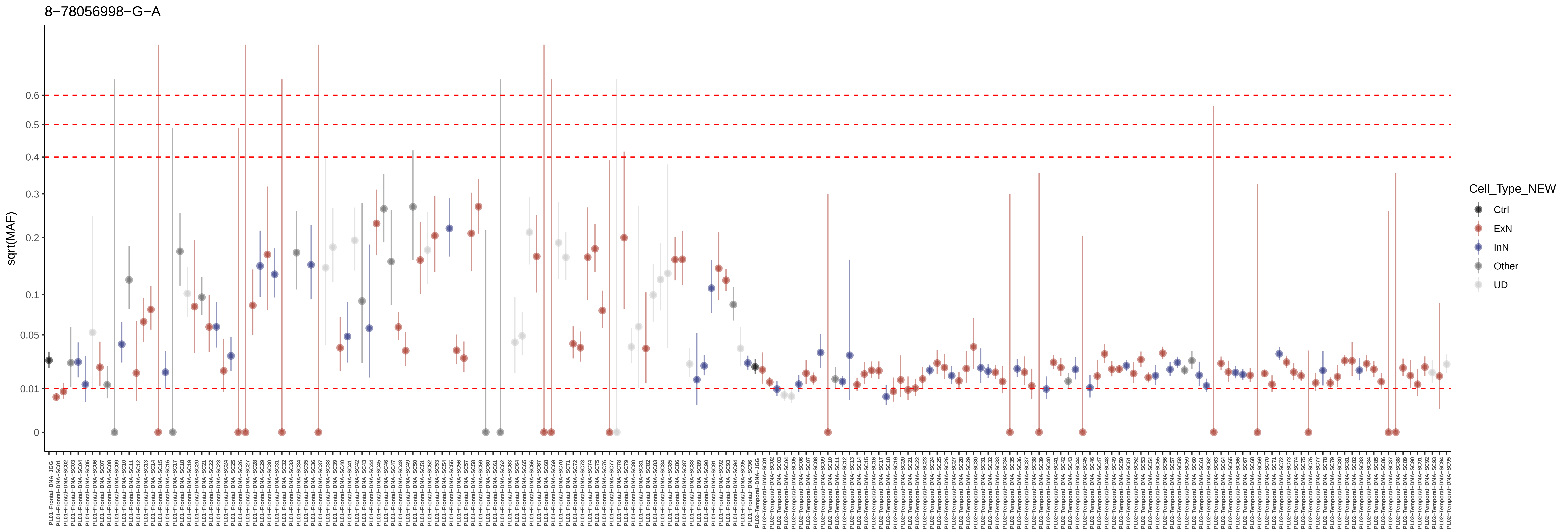

8−78056998−G−A
sqrt(MAF)
0
0.01
0.05
0.1
0.2
0.3
0.4
0.5
0.6
Cell_Type_NEW
Ctrl
ExN
InN
Other
UD

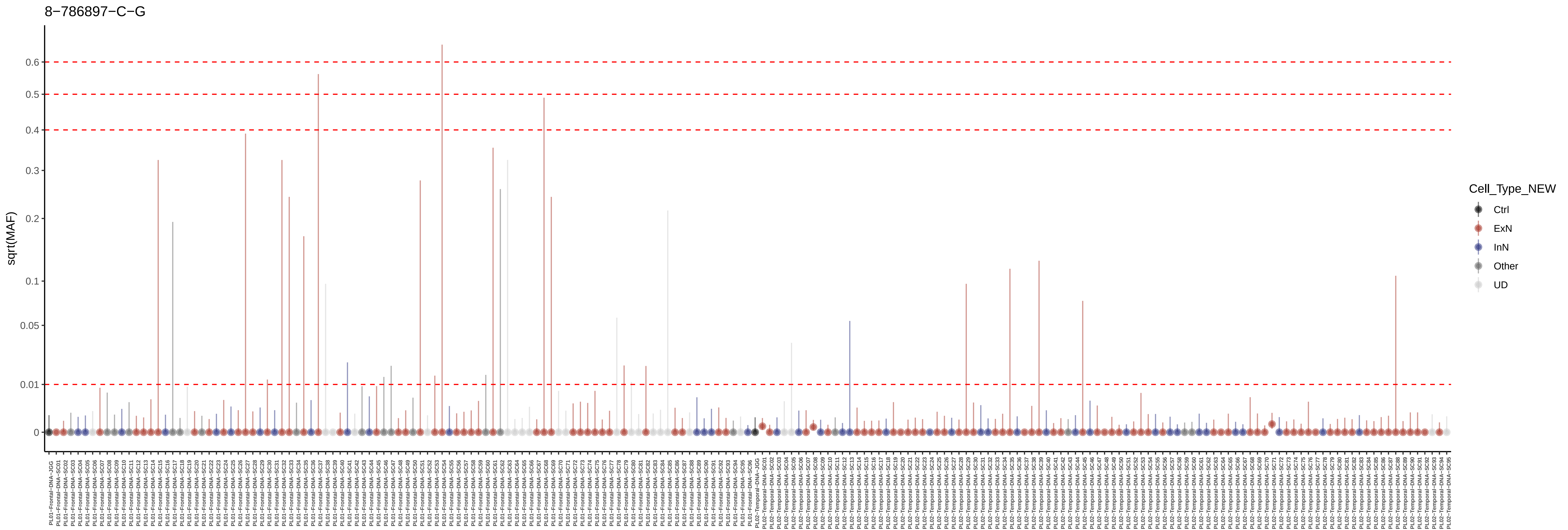
8−786897−C−G
sqrt(MAF)
0.6
0.5
0.4
0.3
0.2
0.1
0.05
0.01
0
Cell_Type_NEW
Ctrl
ExN
InN
Other
UD

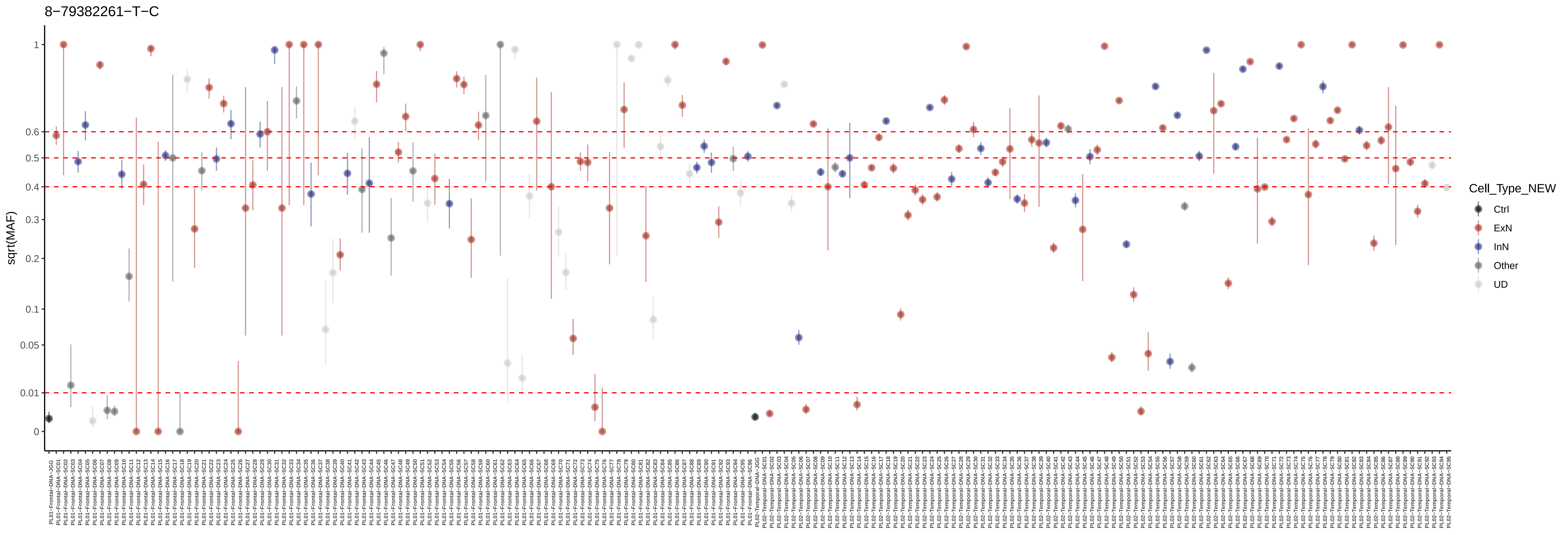
8−79382261−T−C
sqrt(MAF)
1
0.6
0.5
0.4
0.3
0.2
0.1
0.05
0.01
0
Cell_Type_NEW
Ctrl
ExN
InN
Other
UD

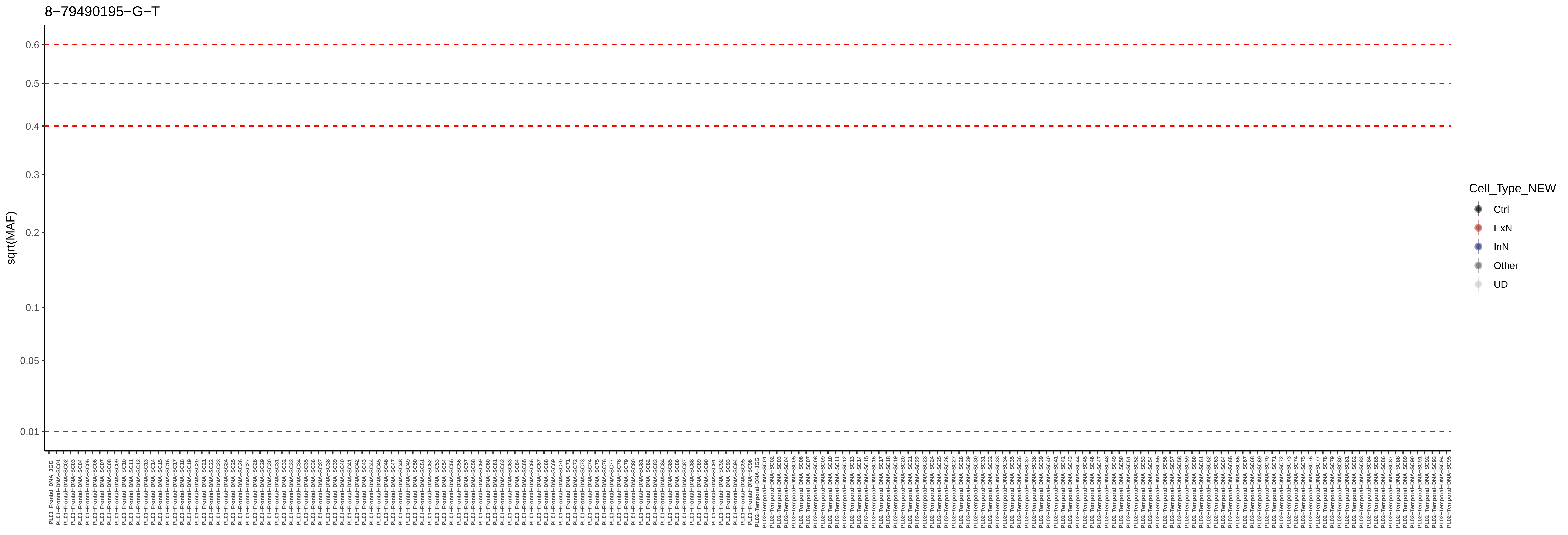
8−79490195−G−T
sqrt(MAF)
0.6
0.5
0.4
0.3
0.2
0.1
0.05
0.01
Cell_Type_NEW
Ctrl
ExN
InN
Other
UD

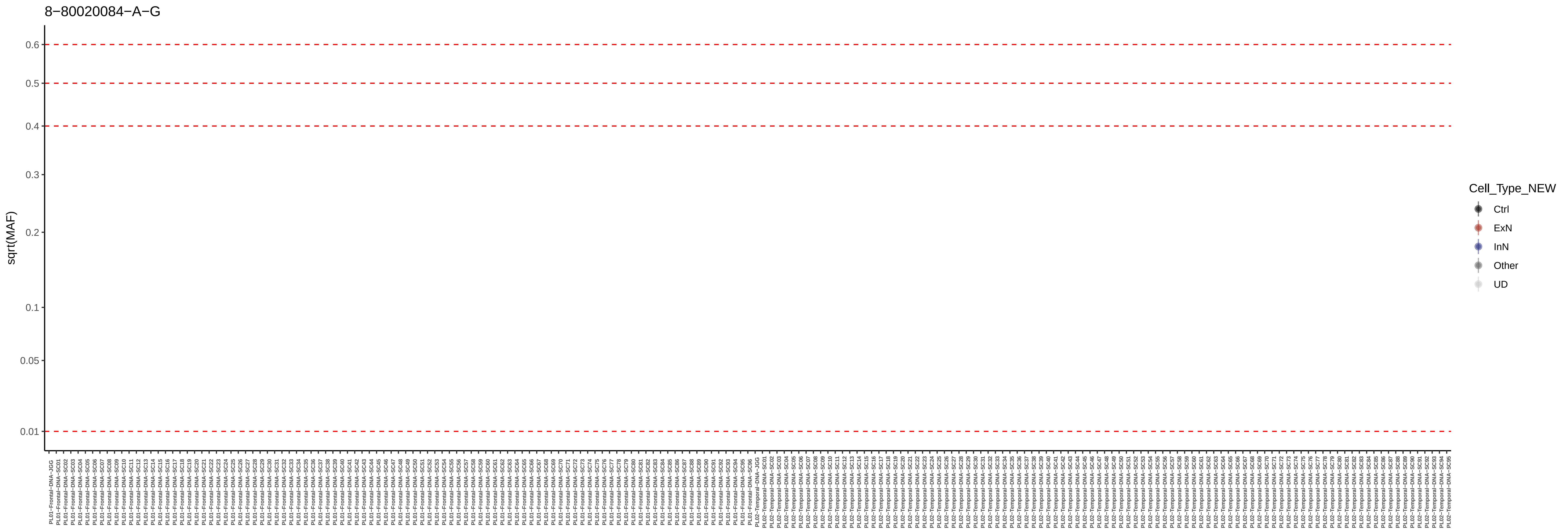
8−80020084−A−G
sqrt(MAF)
0.6
0.5
0.4
0.3
0.2
0.1
0.05
0.01
Cell_Type_NEW
Ctrl
ExN
InN
Other
UD

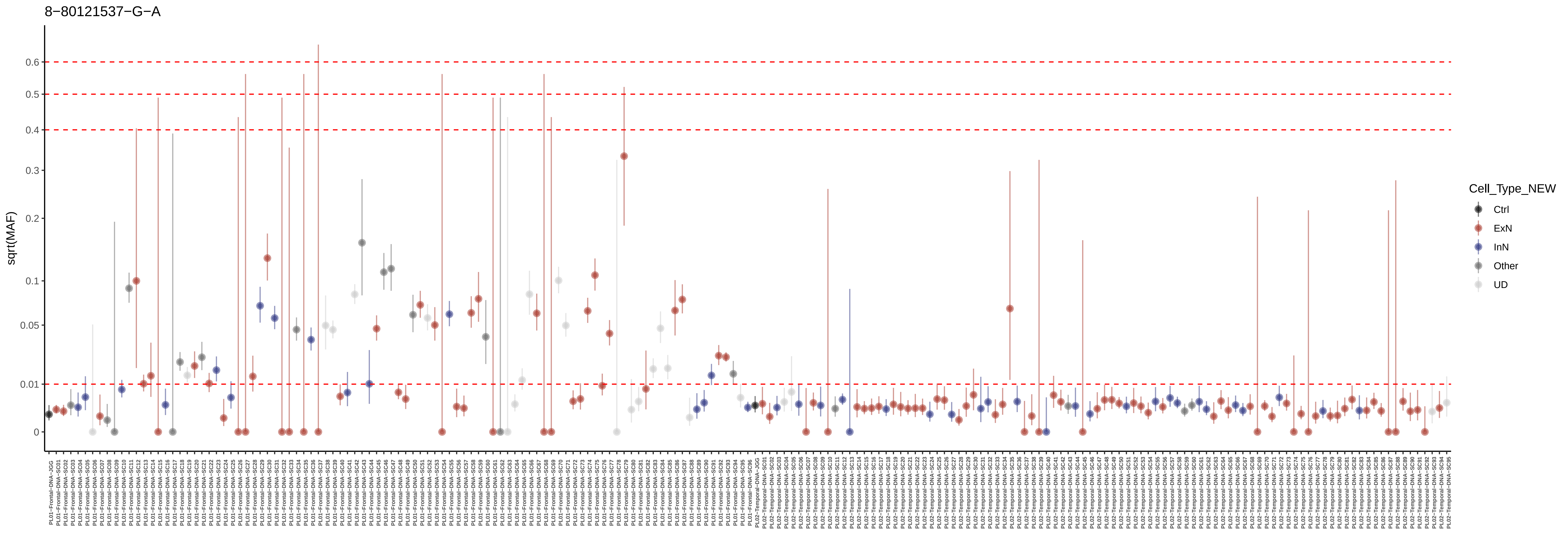

8−80121537−G−A
sqrt(MAF)
0
0.01
0.05
0.1
0.2
0.3
0.4
0.5
0.6
Cell_Type_NEW
Ctrl
ExN
InN
Other
UD

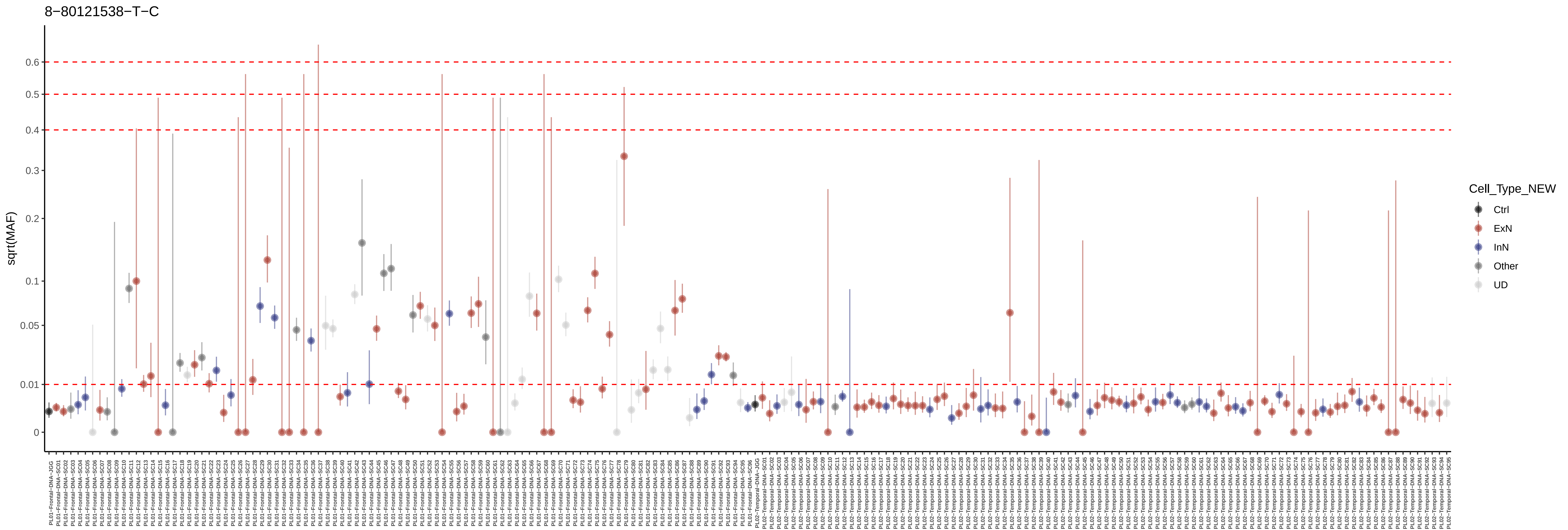
8−80121538−T−C
sqrt(MAF)
0.6
0.5
0.4
0.3
0.2
0.1
0.05
0.01
0
Cell_Type_NEW
Ctrl
ExN
InN
Other
UD

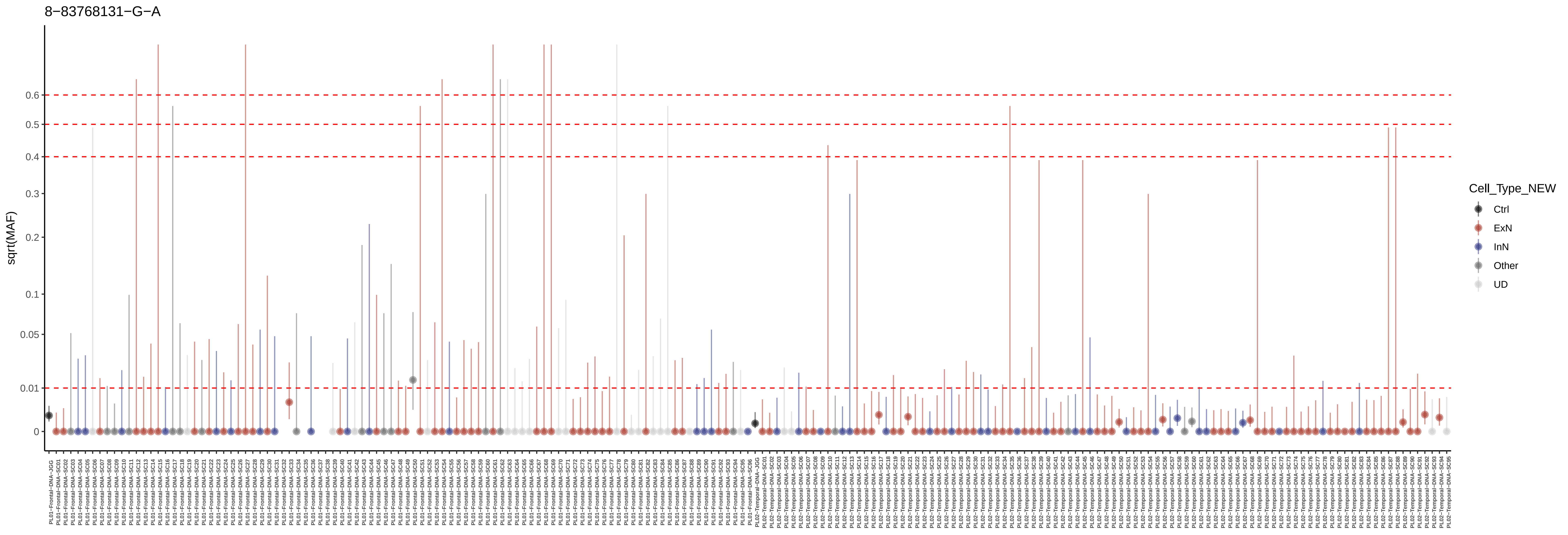
8−83768131−G−A
sqrt(MAF)
0.6
0.5
0.4
0.3
0.2
0.1
0.05
0.01
0
Cell_Type_NEW
Ctrl
ExN
InN
Other
UD

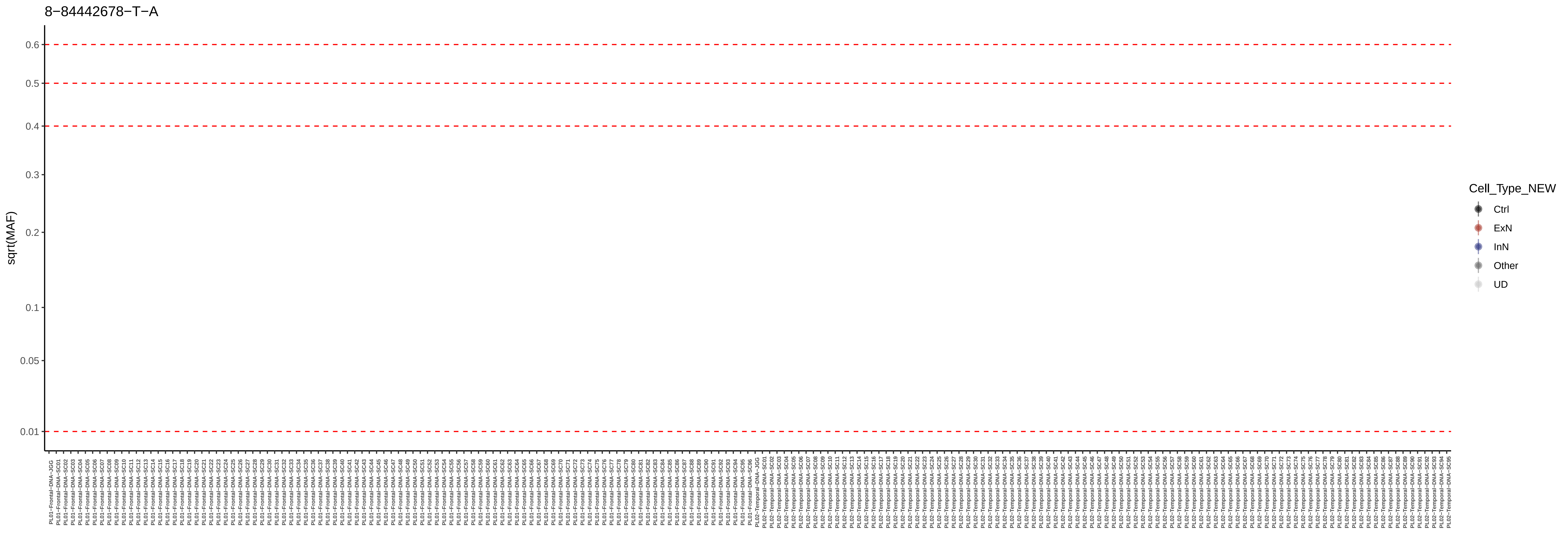

8−84442678−T−A
sqrt(MAF)
0.6
0.5
0.4
0.3
0.2
0.1
0.05
0.01
Cell_Type_NEW
Ctrl
ExN
InN
Other
UD

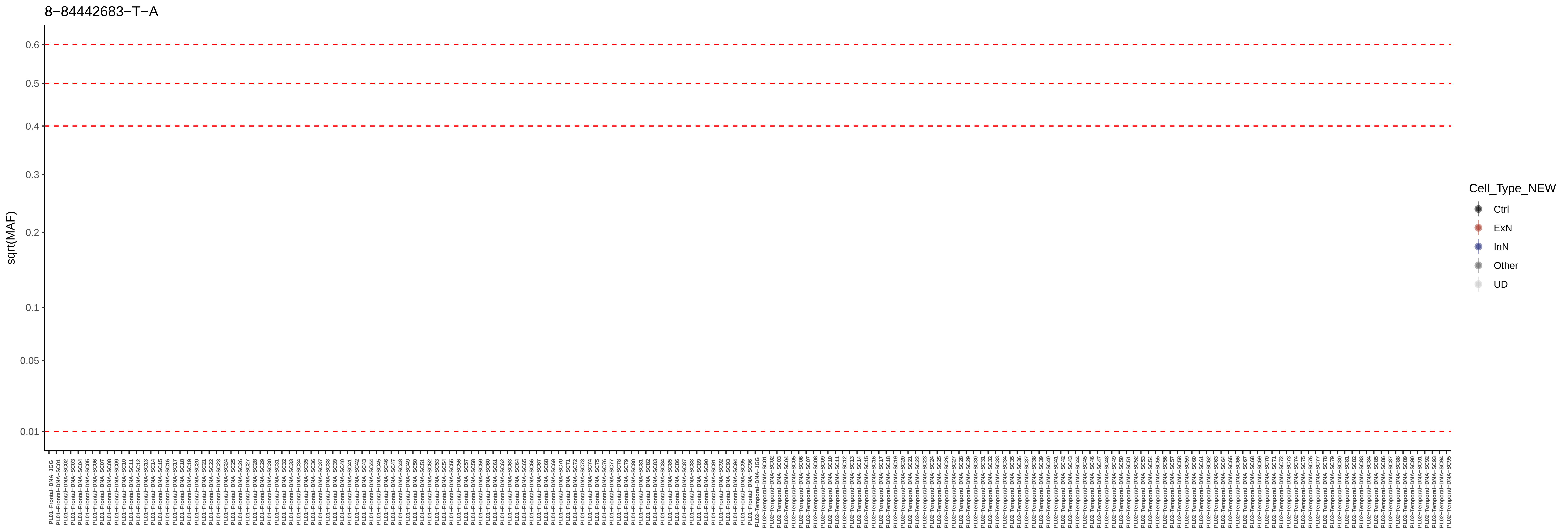
8−84442683−T−A
sqrt(MAF)
0.6
0.5
0.4
0.3
0.2
0.1
0.05
0.01
Cell_Type_NEW
Ctrl
ExN
InN
Other
UD

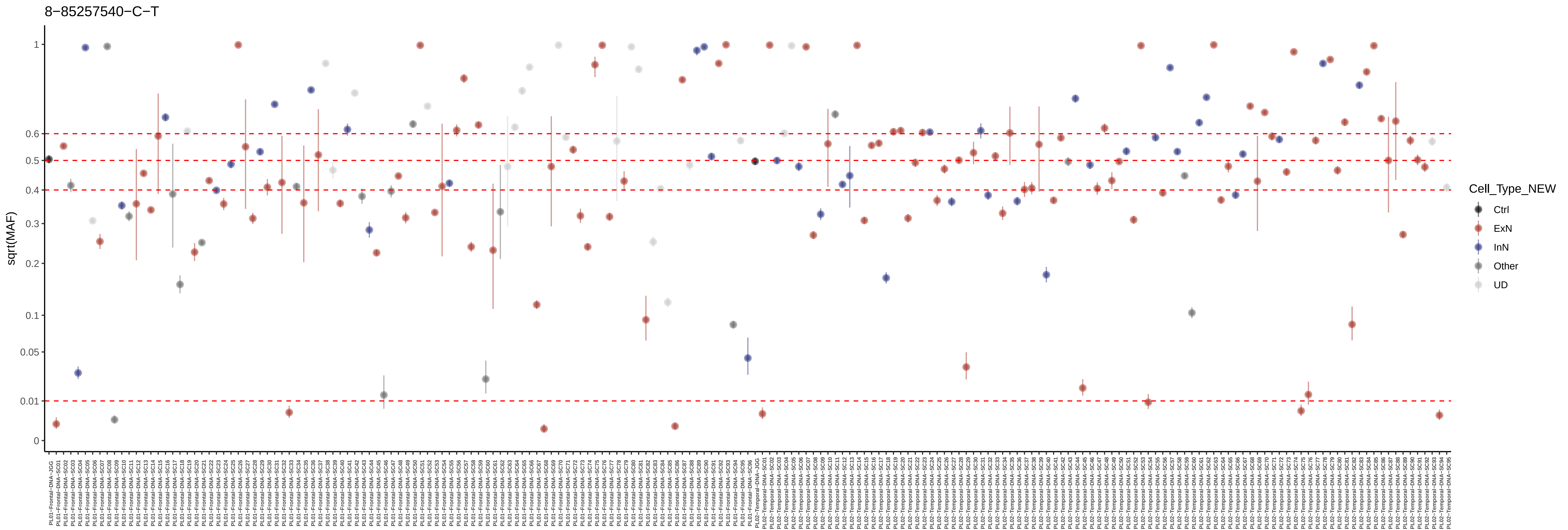

8−85257540−C−T
sqrt(MAF)
1
0.6
0.5
0.4
0.3
0.2
0.1
0.05
0.01
0
Cell_Type_NEW
Ctrl
ExN
InN
Other
UD

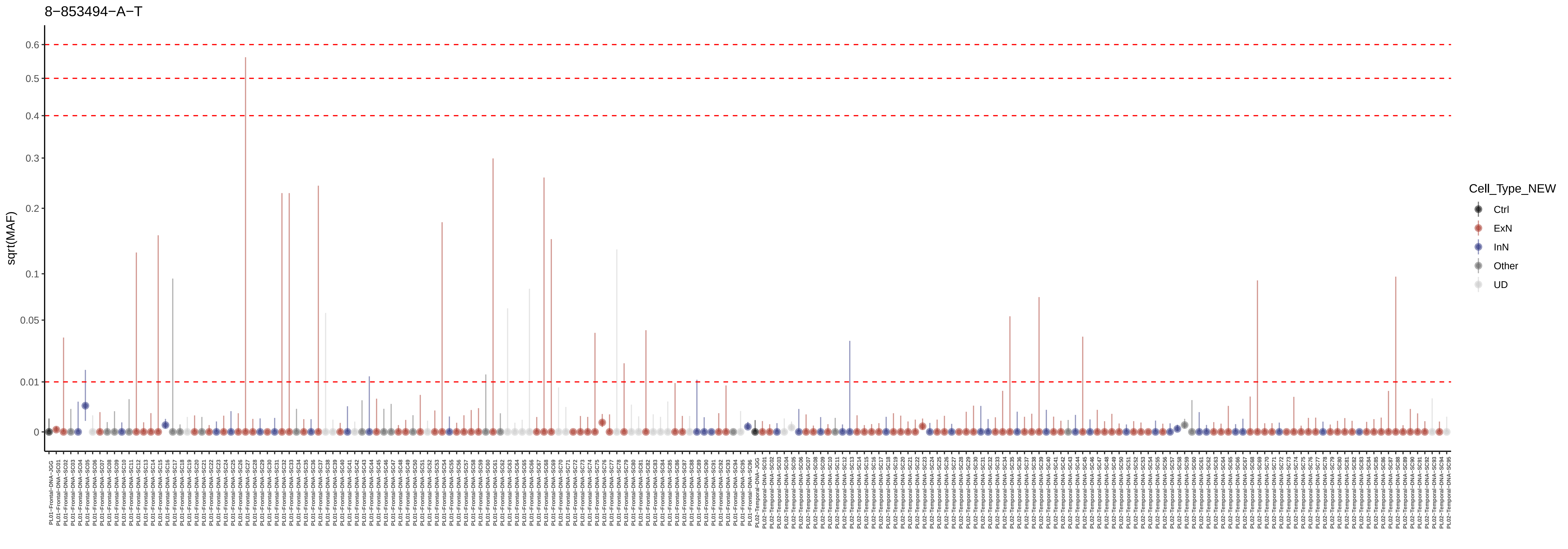

8−853494−A−T
sqrt(MAF)
Cell_Type_NEW
Ctrl
ExN
InN
Other
UD

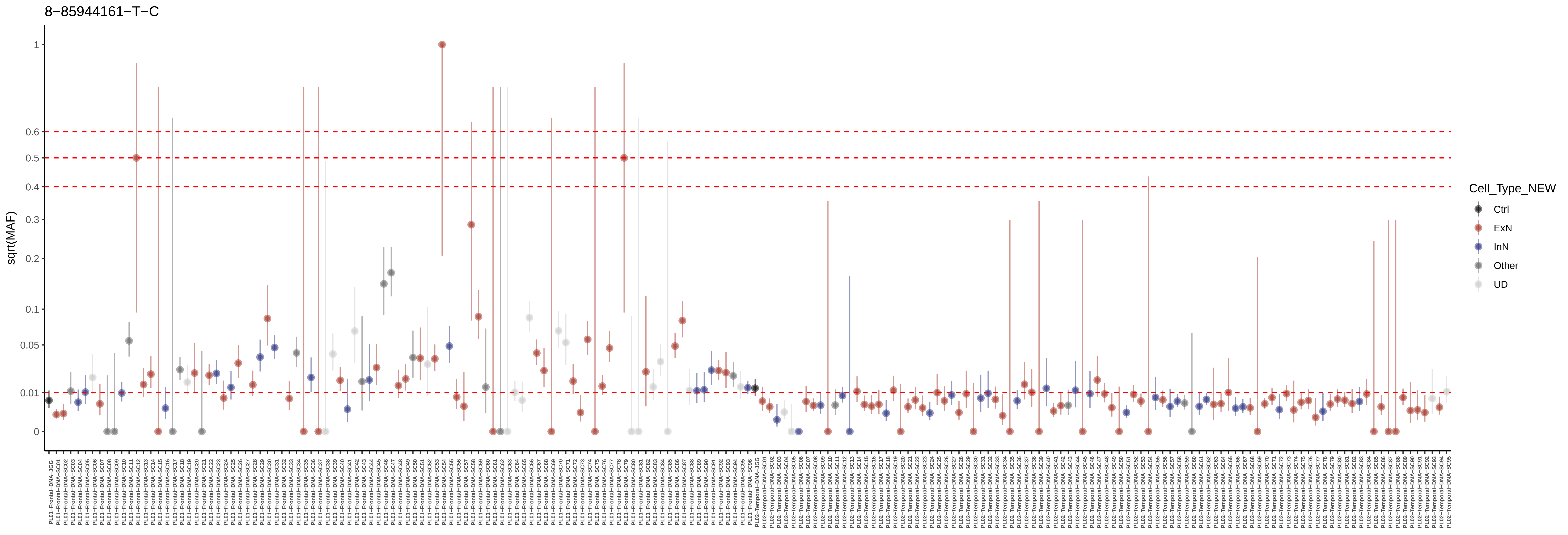

8−85944161−T−C
sqrt(MAF)
1
0.6
0.5
0.4
0.3
0.2
0.1
0.05
0.01
0
Cell_Type_NEW
Ctrl
ExN
InN
Other
UD

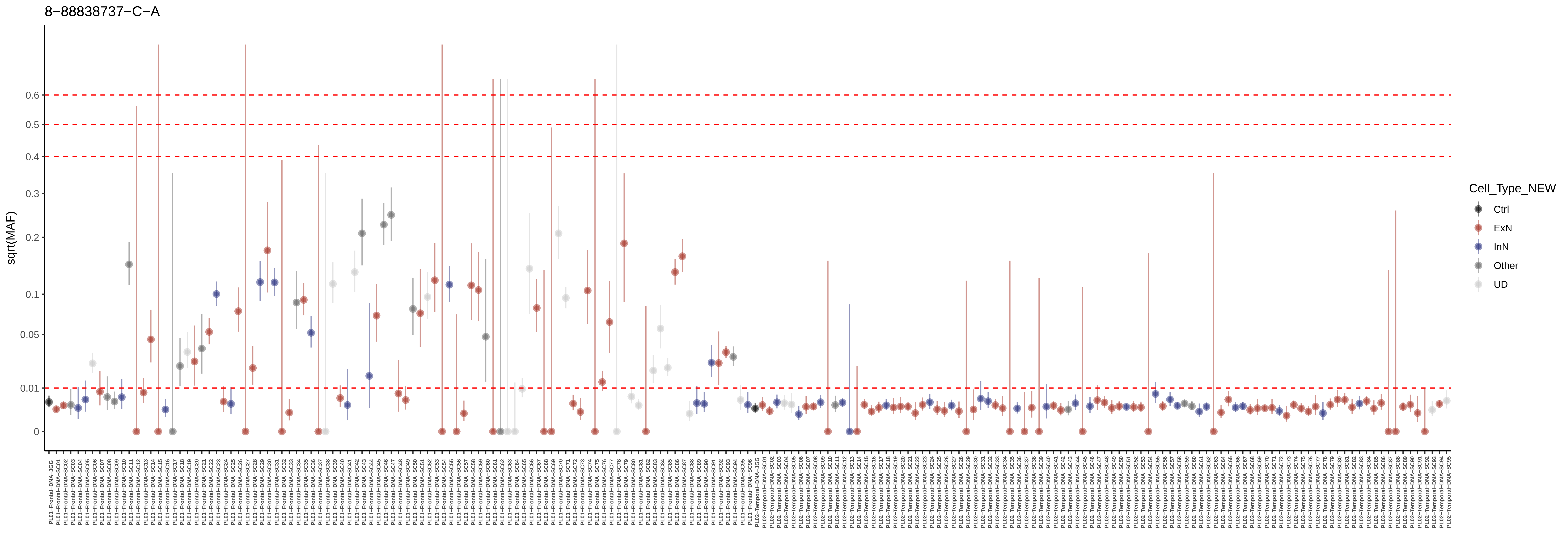
8−88838737−C−A
sqrt(MAF)
0.6
0.5
0.4
0.3
0.2
0.1
0.05
0.01
0
Cell_Type_NEW
Ctrl
ExN
InN
Other
UD

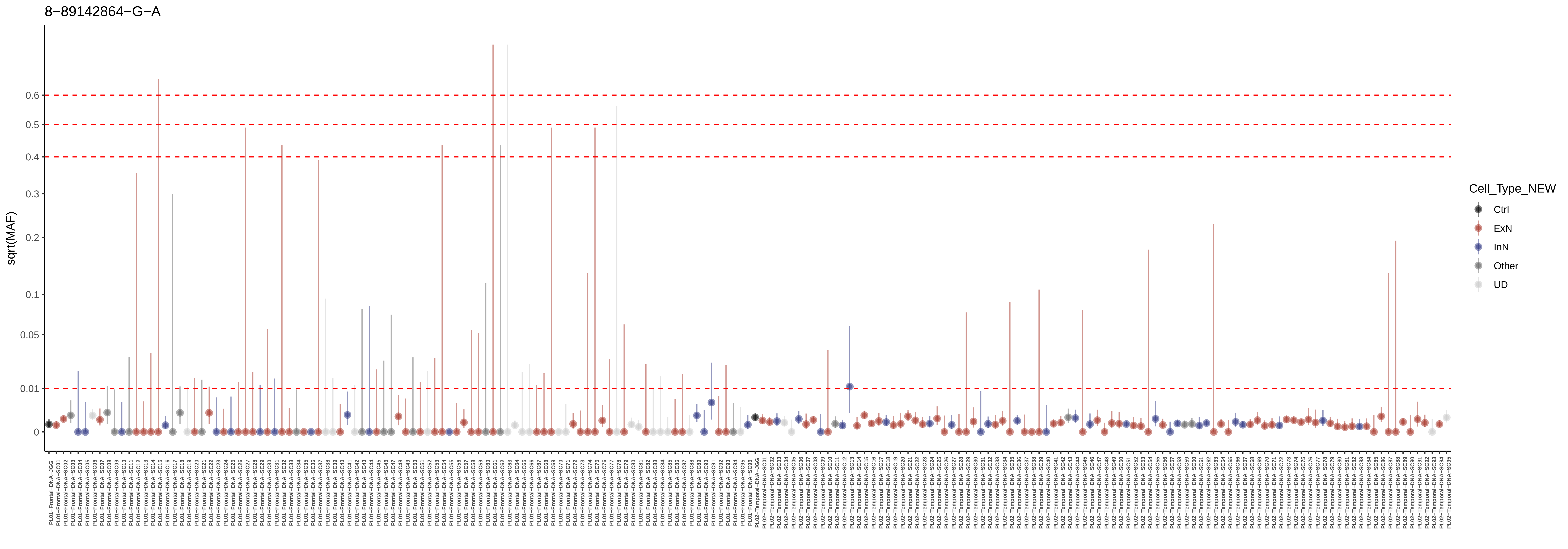
8−89142864−G−A
sqrt(MAF)
0.6
0.5
0.4
0.3
0.2
0.1
0.05
0.01
0
Cell_Type_NEW
Ctrl
ExN
InN
Other
UD

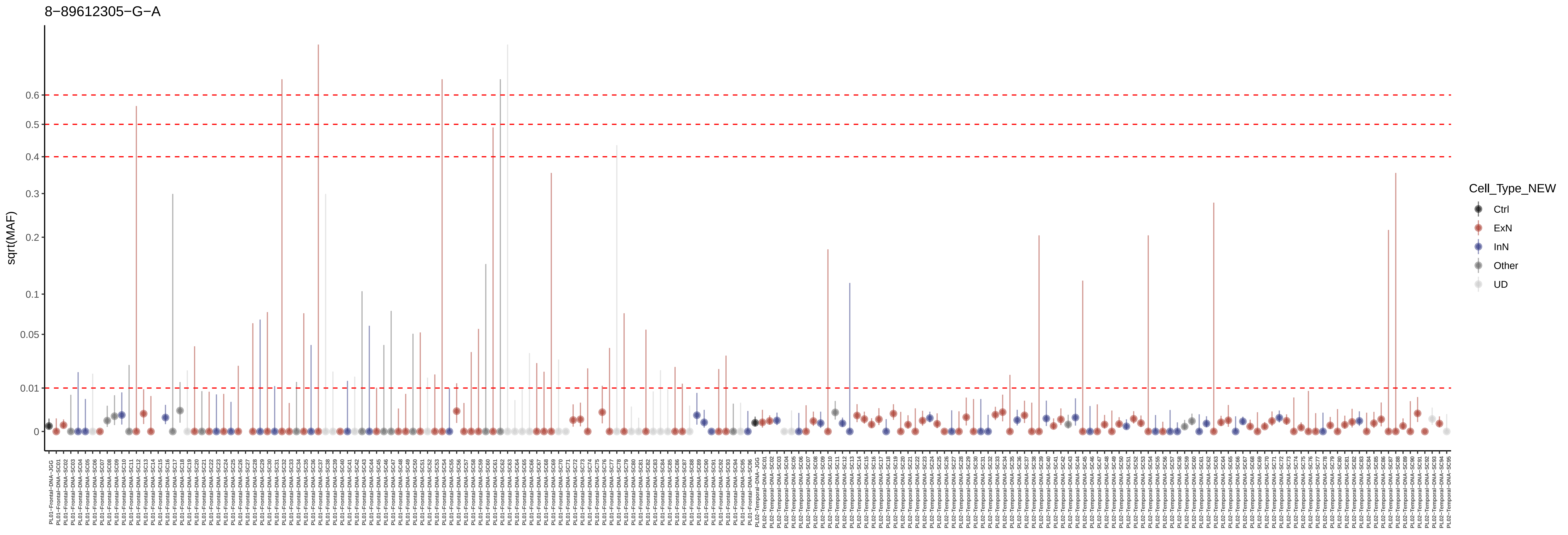
8−89612305−G−A
sqrt(MAF)
0.6
0.5
0.4
0.3
0.2
0.1
0.05
0.01
0
Cell_Type_NEW
Ctrl
ExN
InN
Other
UD

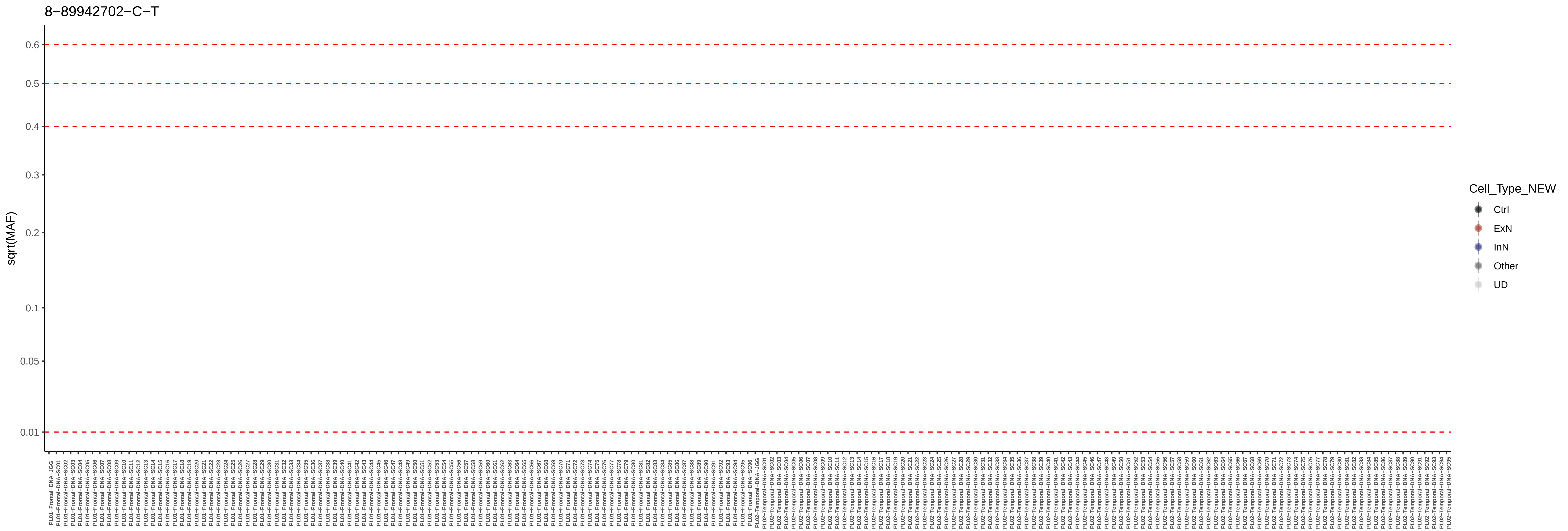
8−89942702−C−T
sqrt(MAF)
0.6
0.5
0.4
0.3
0.2
0.1
0.05
0.01
Cell_Type_NEW
Ctrl
ExN
InN
Other
UD

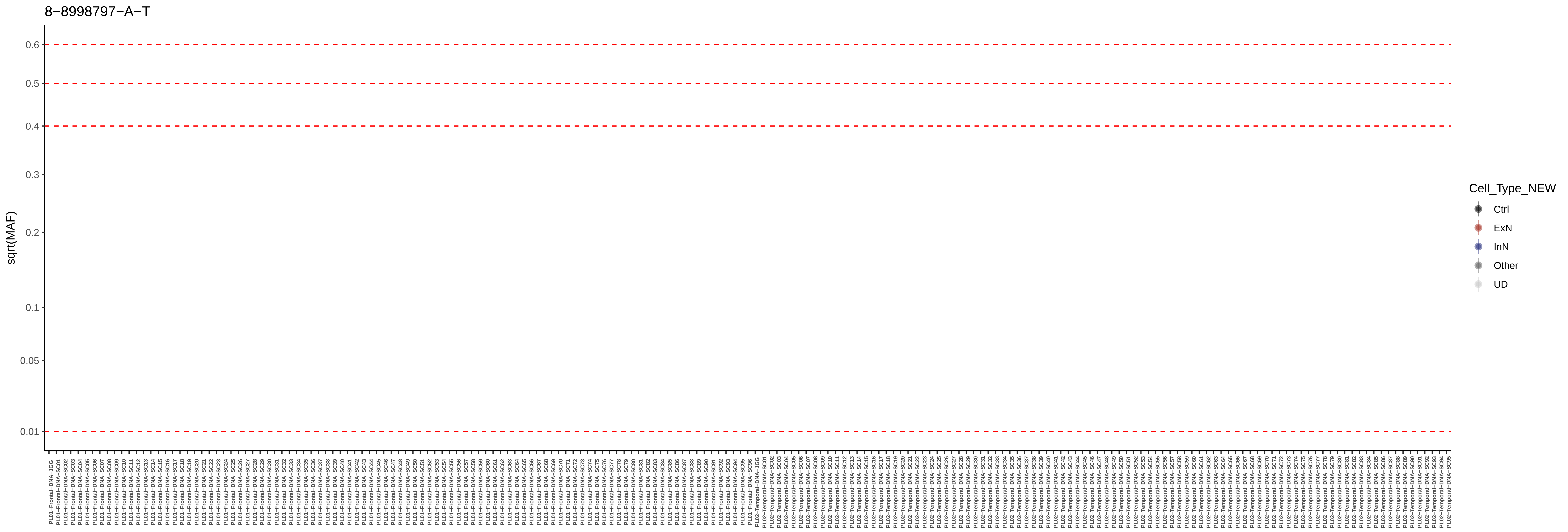
8−8998797−A−T
sqrt(MAF)
0.6
0.5
0.4
0.3
0.2
0.1
0.05
0.01
Cell_Type_NEW
Ctrl
ExN
InN
Other
UD

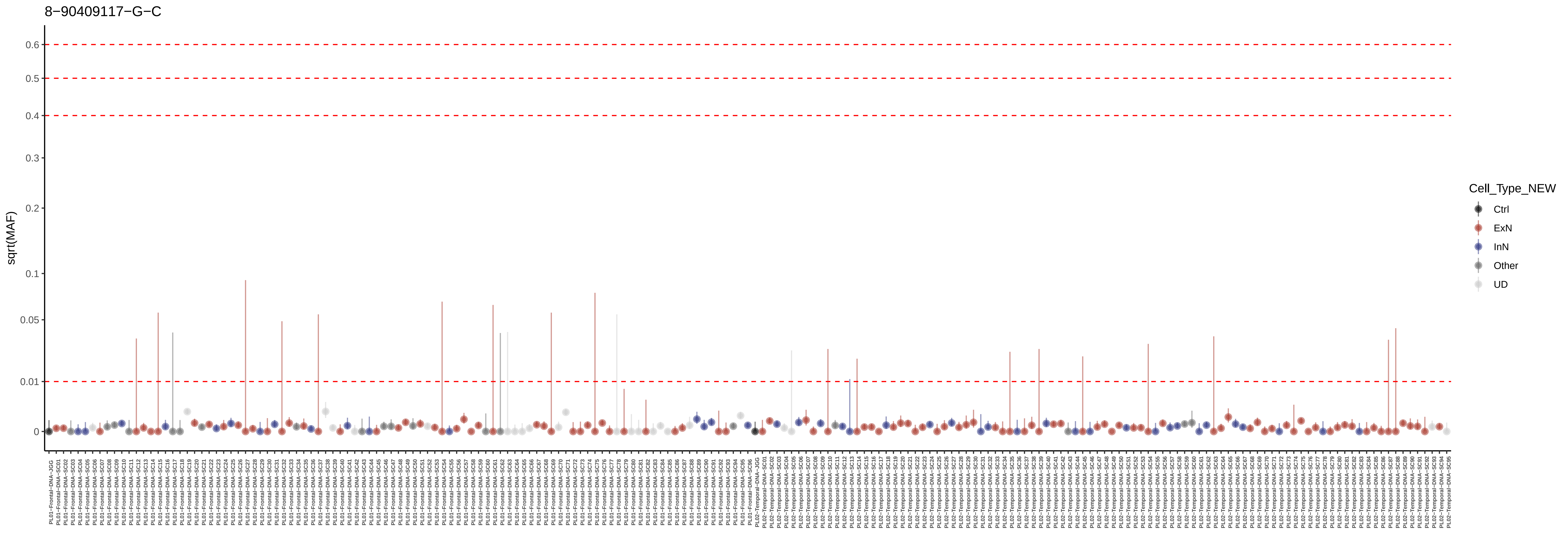

8−90409117−G−C
sqrt(MAF)
0.6
0.5
0.4
0.3
0.2
0.1
0.05
0.01
0
Cell_Type_NEW
Ctrl
ExN
InN
Other
UD

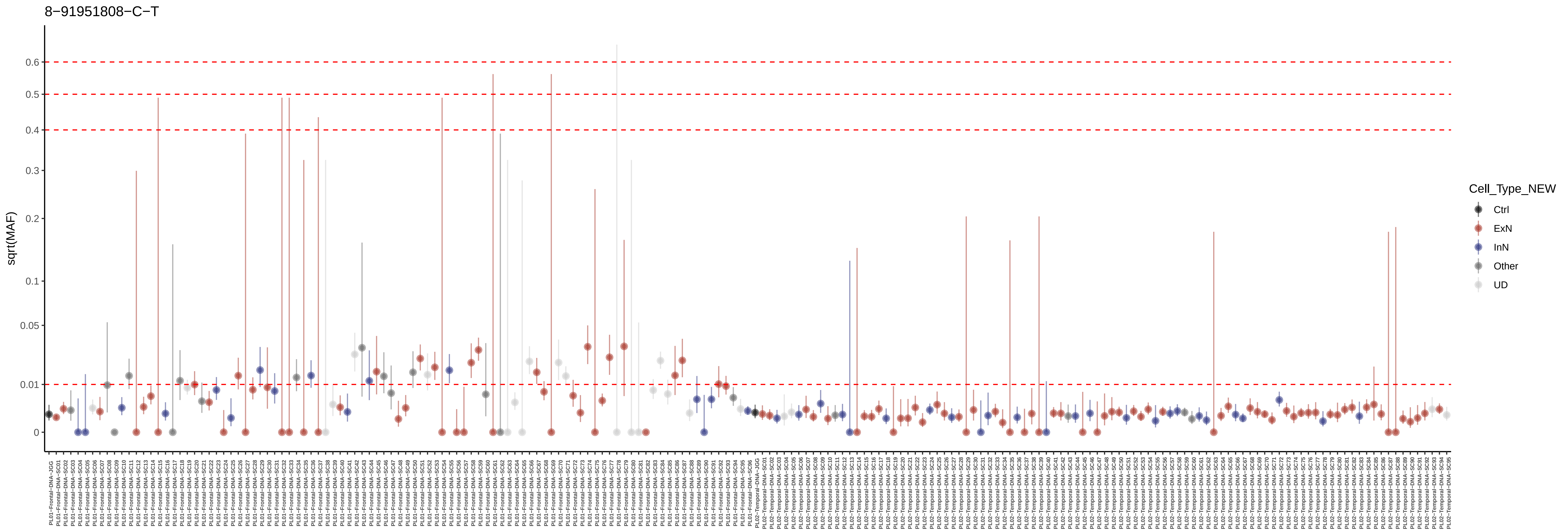
8−91951808−C−T
sqrt(MAF)
0.6
0.5
0.4
0.3
0.2
0.1
0.05
0.01
0
Cell_Type_NEW
Ctrl
ExN
InN
Other
UD

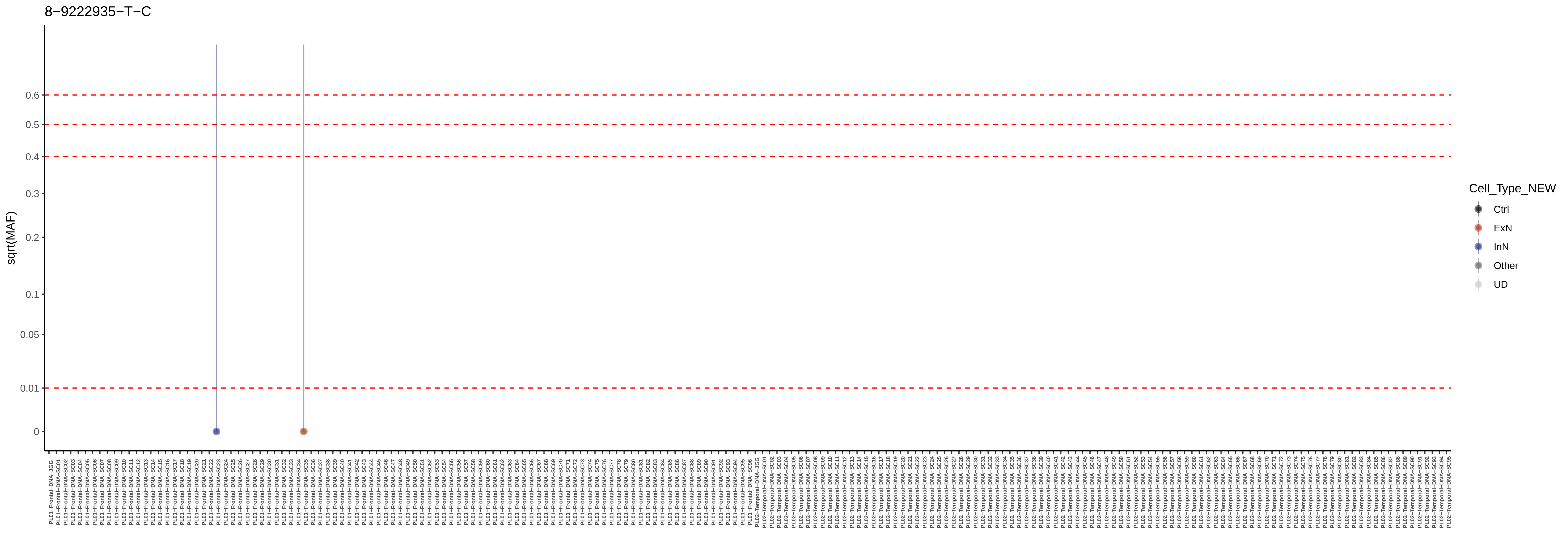
8−9222935−T−C
sqrt(MAF)
0.6
0.5
0.4
0.3
0.2
0.1
0.05
0.01
0
Cell_Type_NEW
Ctrl
ExN
InN
Other
UD

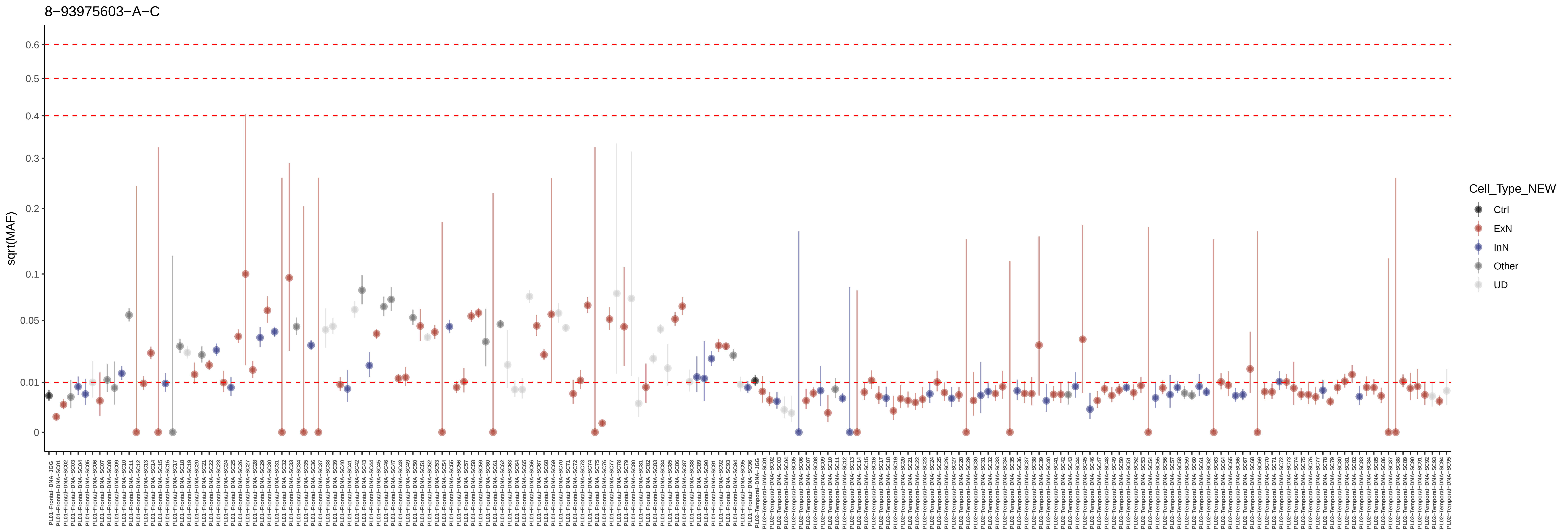

8−93975603−A−C
sqrt(MAF)
0.6
0.5
0.4
0.3
0.2
0.1
0.05
0.01
0
Cell_Type_NEW
Ctrl
ExN
InN
Other
UD

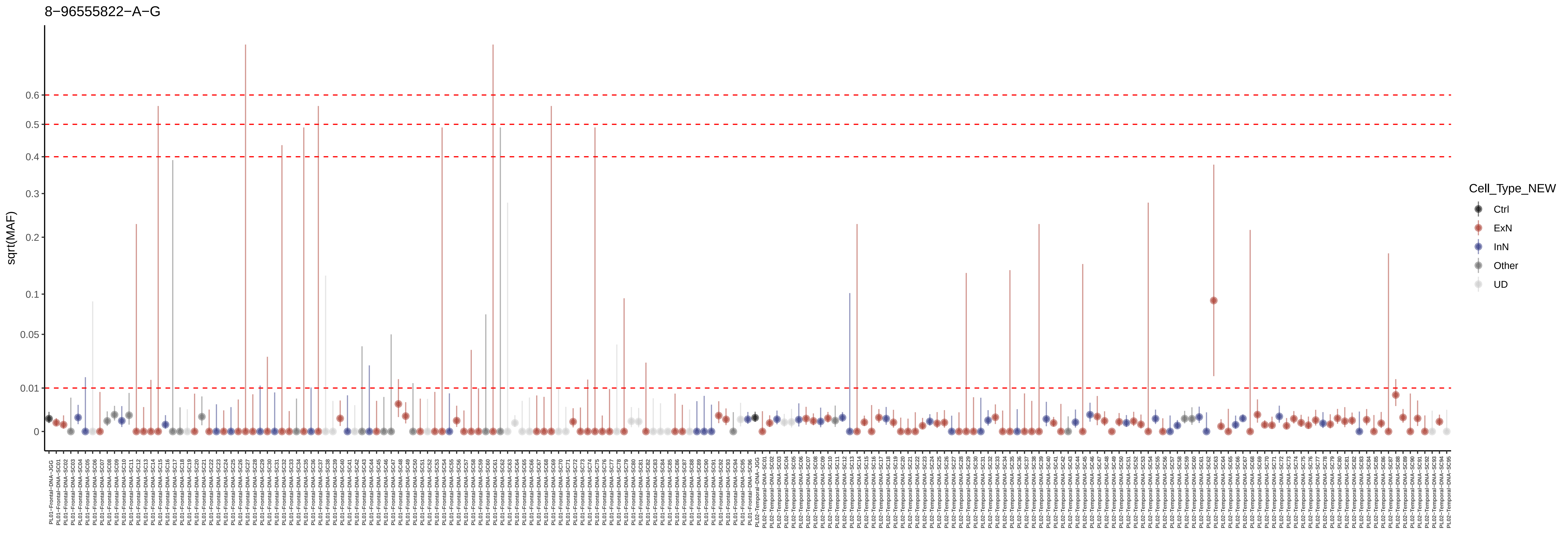
8−96555822−A−G
sqrt(MAF)
0.6
0.5
0.4
0.3
0.2
0.1
0.05
0.01
0
Cell_Type_NEW
Ctrl
ExN
InN
Other
UD

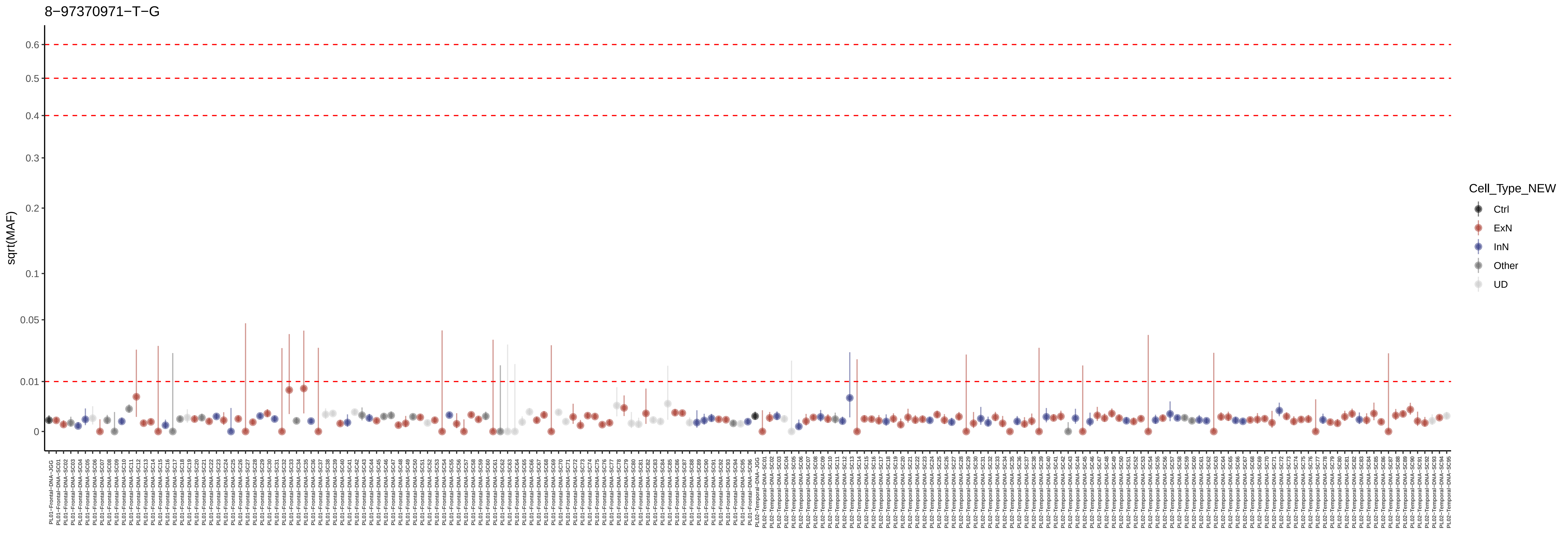

8−97370971−T−G
sqrt(MAF)
0.6
0.5
0.4
0.3
0.2
0.1
0.05
0.01
0
Cell_Type_NEW
Ctrl
ExN
InN
Other
UD

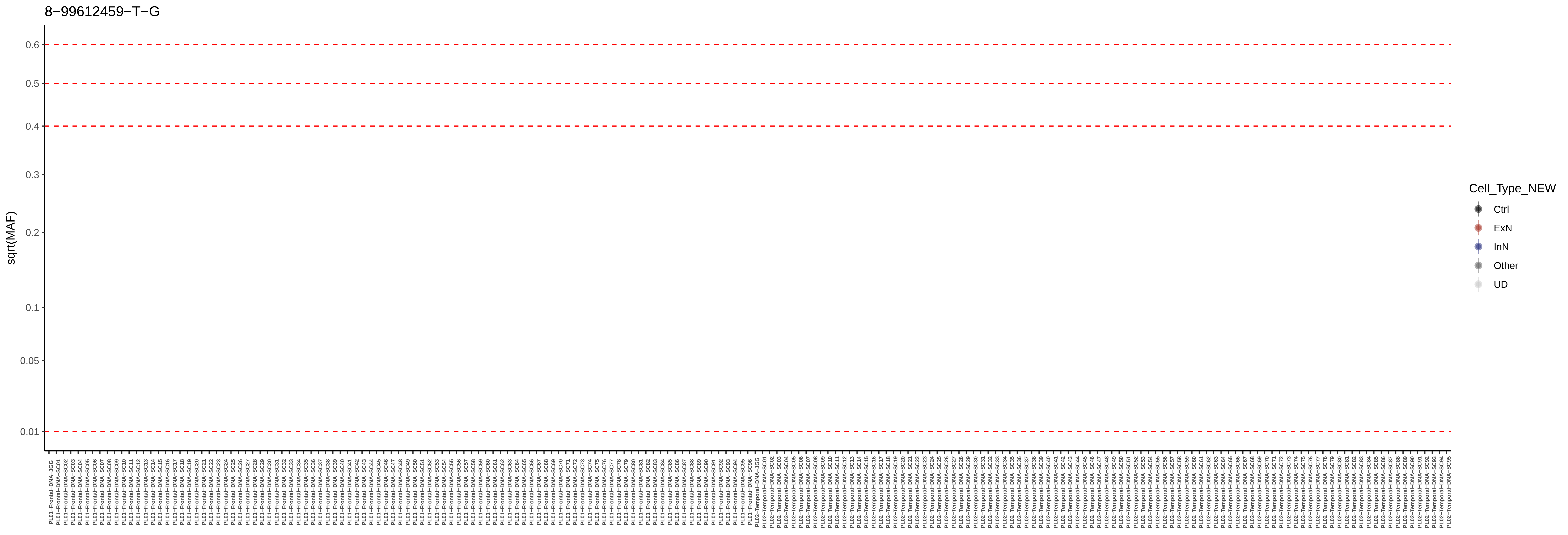
8−99612459−T−G
sqrt(MAF)
0.6
0.5
0.4
0.3
0.2
0.1
0.05
0.01
Cell_Type_NEW
Ctrl
ExN
InN
Other
UD

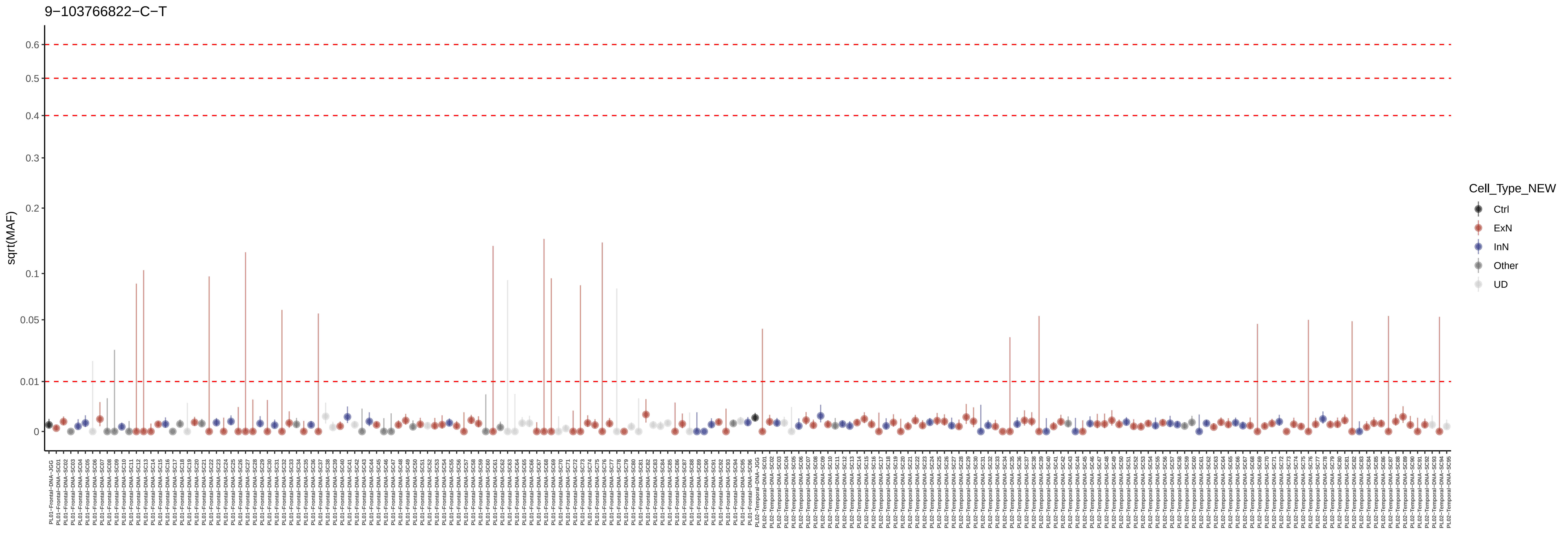

9−103766822−C−T
sqrt(MAF)
0.6
0.5
0.4
0.3
0.2
0.1
0.05
0.01
0
Cell_Type_NEW
Ctrl
ExN
InN
Other
UD

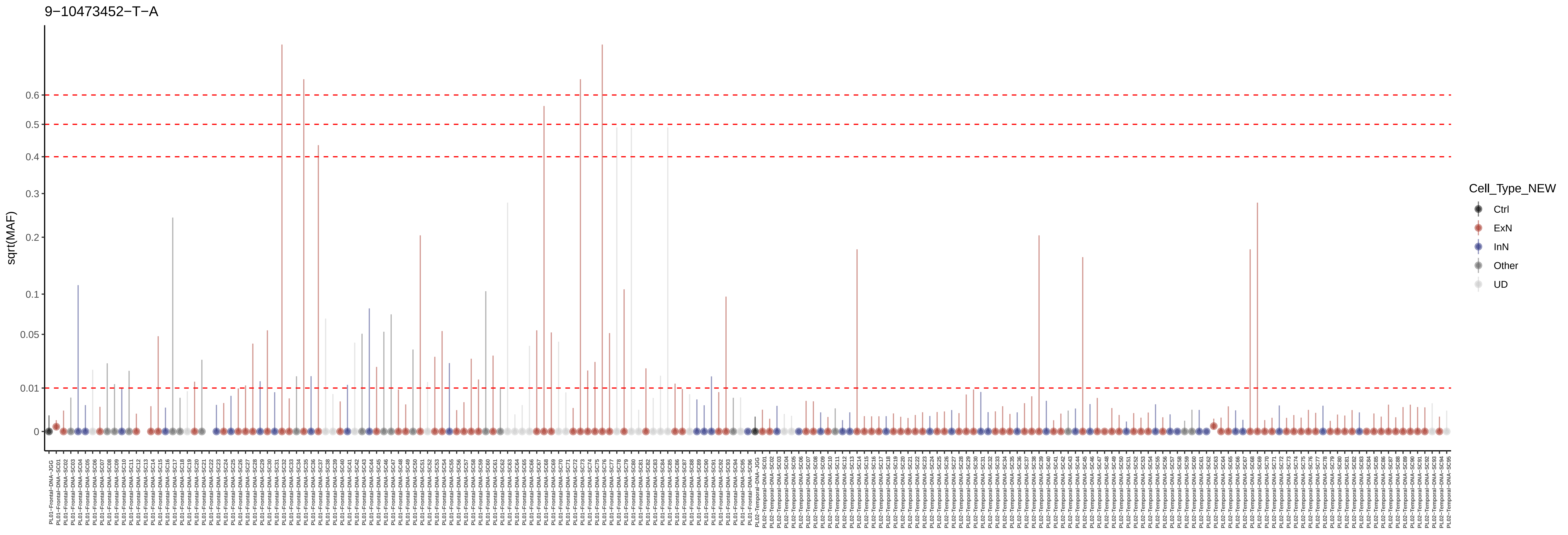
9−10473452−T−A
sqrt(MAF)
0.6
0.5
0.4
0.3
0.2
0.1
0.05
0.01
0
Cell_Type_NEW
Ctrl
ExN
InN
Other
UD

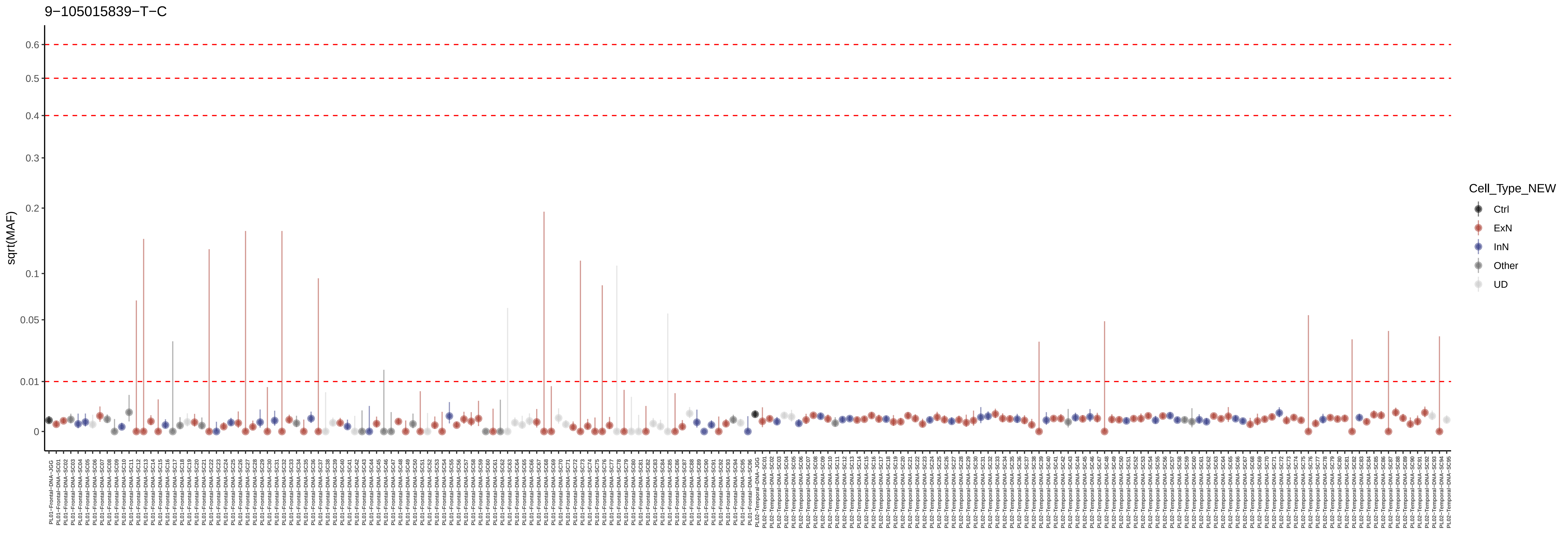
9−105015839−T−C
sqrt(MAF)
0.6
0.5
0.4
0.3
0.2
0.1
0.05
0.01
0
Cell_Type_NEW
Ctrl
ExN
InN
Other
UD

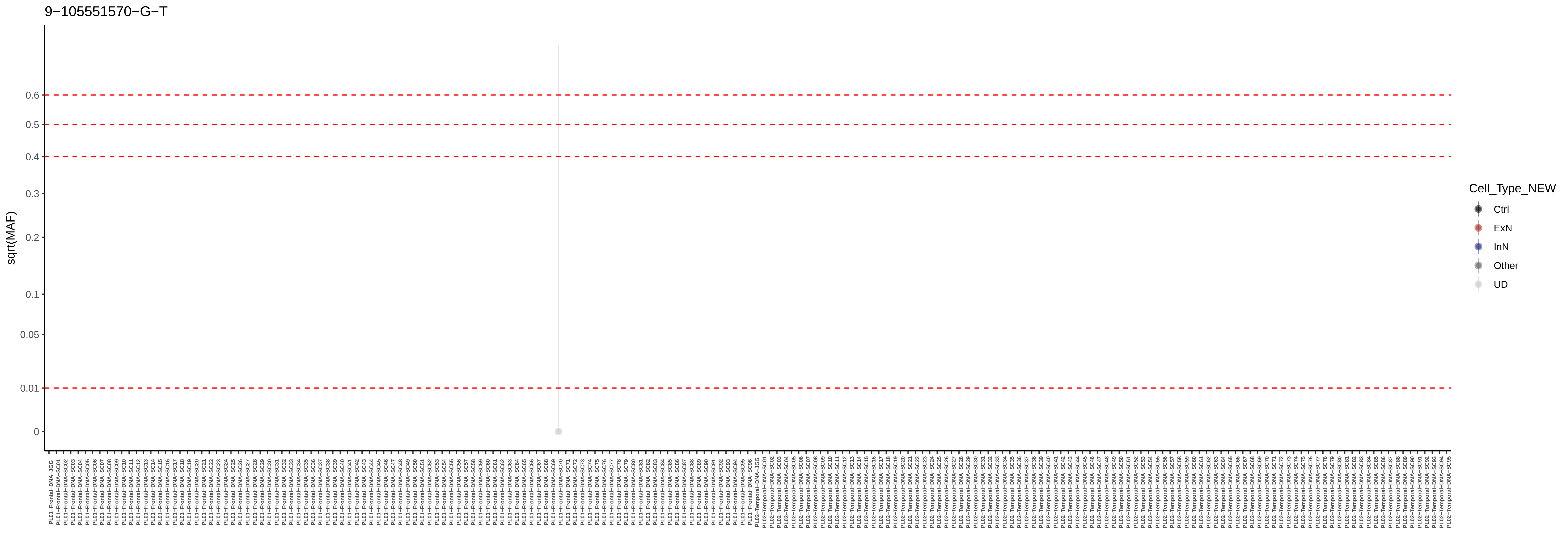
9−105551570−G−T
sqrt(MAF)
0.6
0.5
0.4
0.3
0.2
0.1
0.05
0.01
0
Cell_Type_NEW
Ctrl
ExN
InN
Other
UD

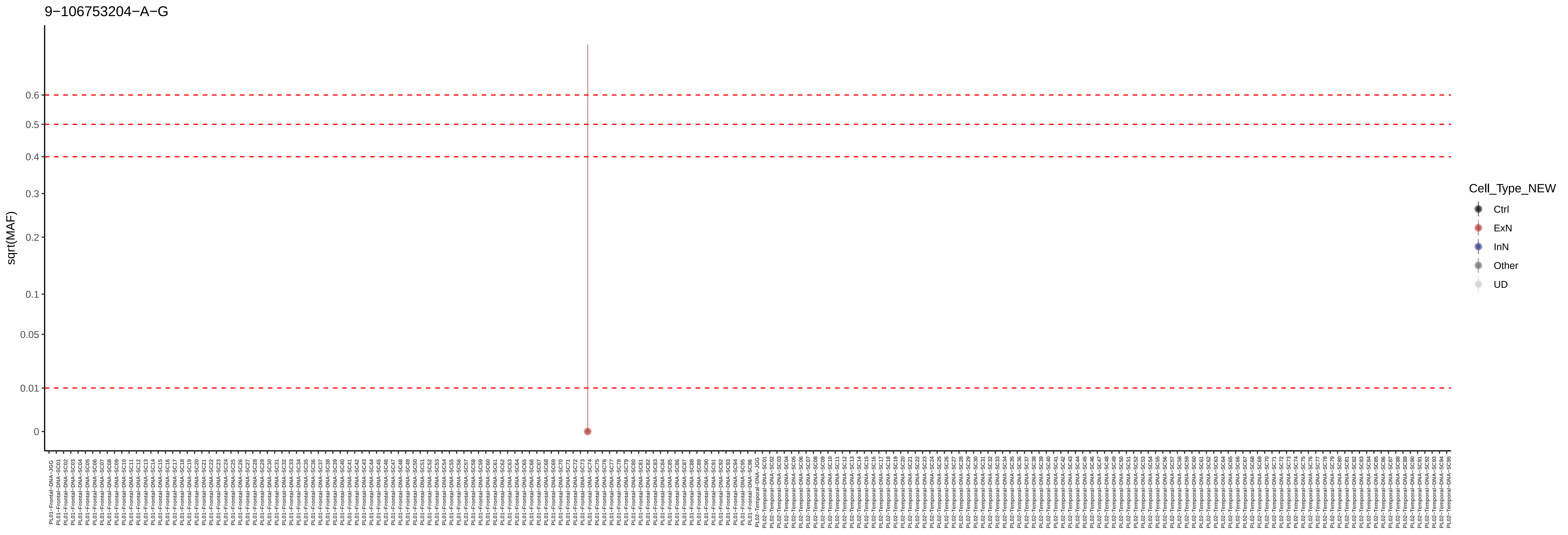

9−106753204−A−G
sqrt(MAF)
0.6
0.5
0.4
0.3
0.2
0.1
0.05
0.01
0
Cell_Type_NEW
Ctrl
ExN
InN
Other
UD

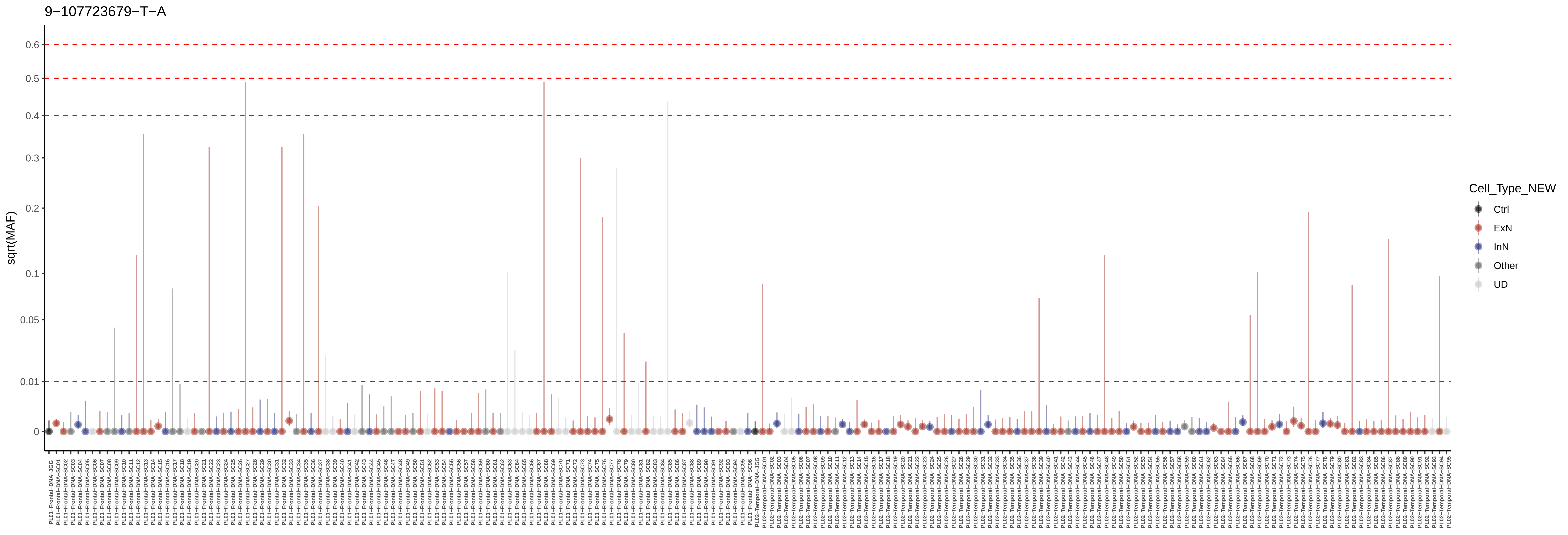

9−107723679−T−A
sqrt(MAF)
0.6
0.5
0.4
0.3
0.2
0.1
0.05
0.01
0
Cell_Type_NEW
Ctrl
ExN
InN
Other
UD

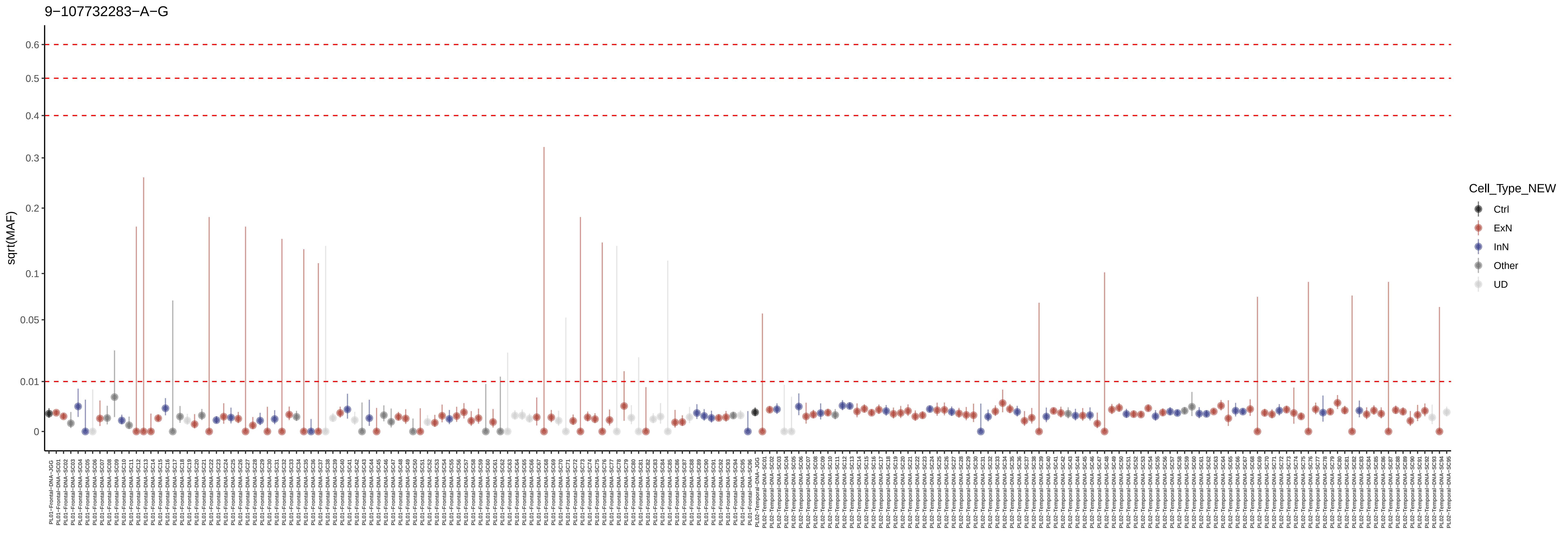
9−107732283−A−G
sqrt(MAF)
0.6
0.5
0.4
0.3
0.2
0.1
0.05
0.01
0
Cell_Type_NEW
Ctrl
ExN
InN
Other
UD

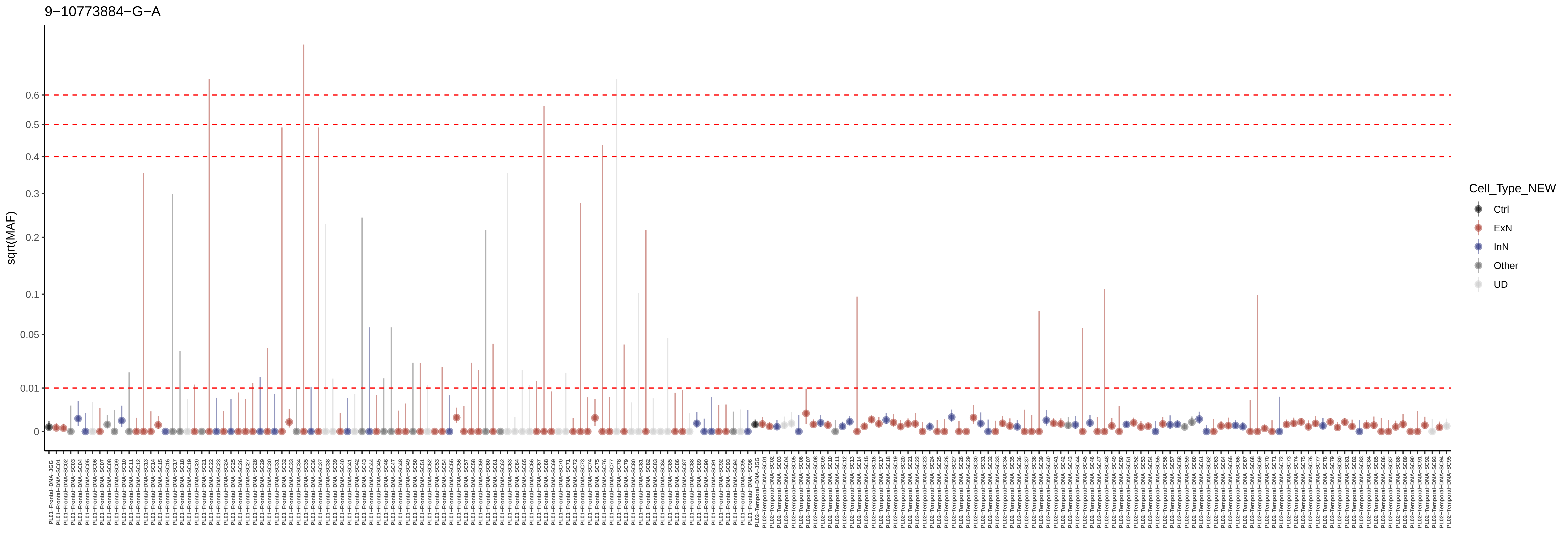

9−10773884−G−A
sqrt(MAF)
0.6
0.5
0.4
0.3
0.2
0.1
0.05
0.01
0
Cell_Type_NEW
Ctrl
ExN
InN
Other
UD

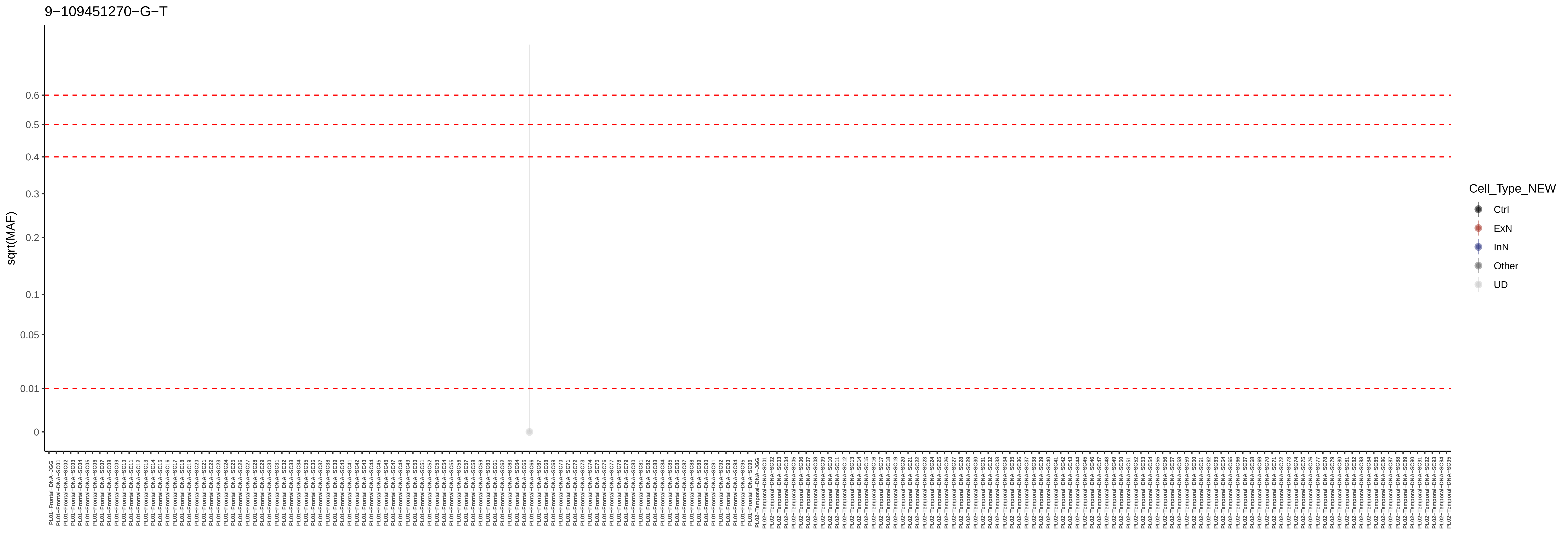
9−109451270−G−T
sqrt(MAF)
0.6
0.5
0.4
0.3
0.2
0.1
0.05
0.01
0
Cell_Type_NEW
Ctrl
ExN
InN
Other
UD

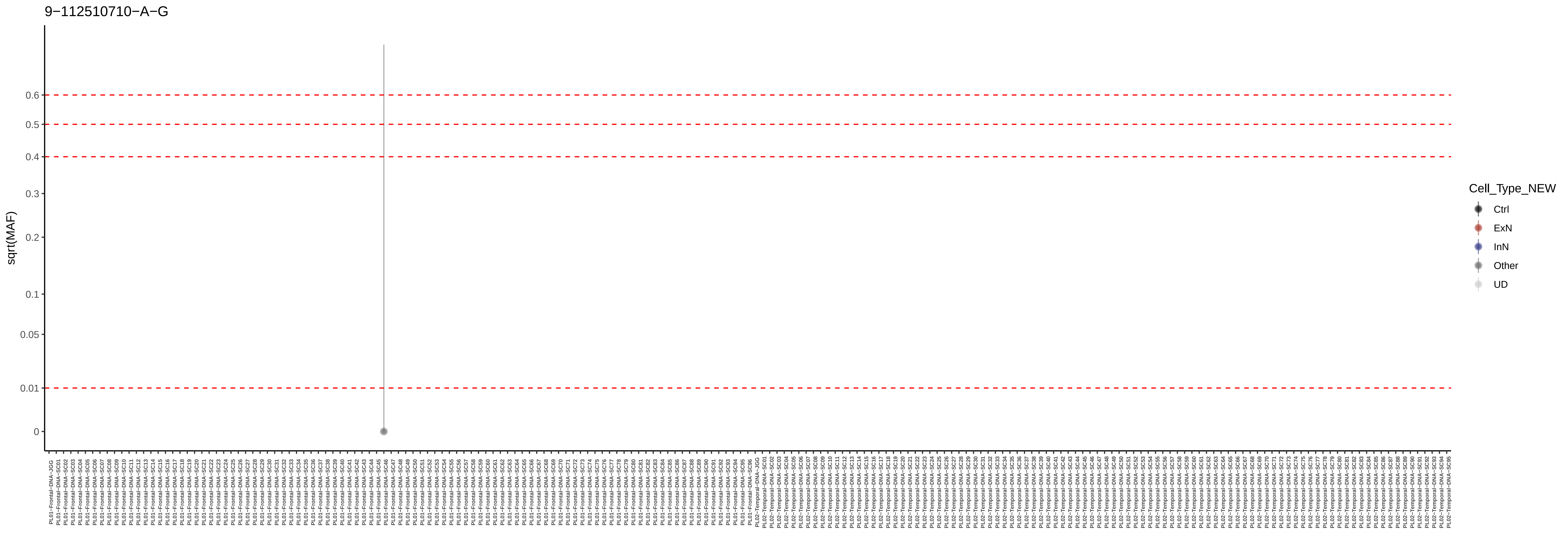
9−112510710−A−G
sqrt(MAF)
0.6
0.5
0.4
0.3
0.2
0.1
0.05
0.01
0
Cell_Type_NEW
Ctrl
ExN
InN
Other
UD

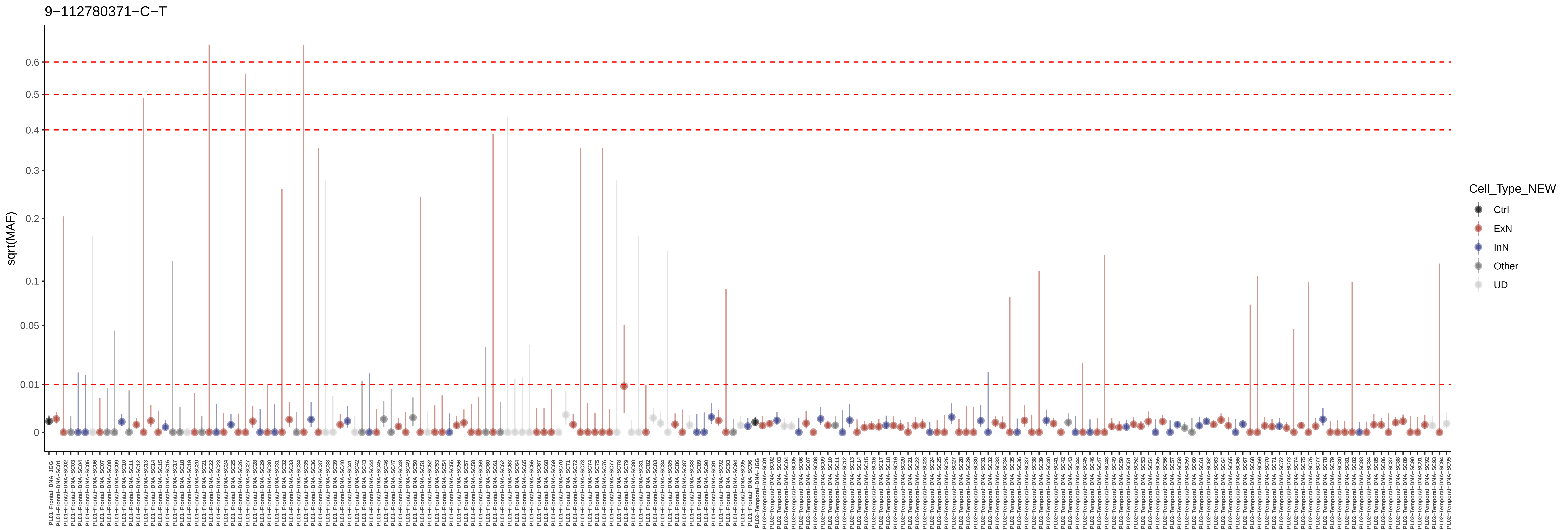
9−112780371−C−T
sqrt(MAF)
0.6
0.5
0.4
0.3
0.2
0.1
0.05
0.01
0
Cell_Type_NEW
Ctrl
ExN
InN
Other
UD

9−114921672−C−T

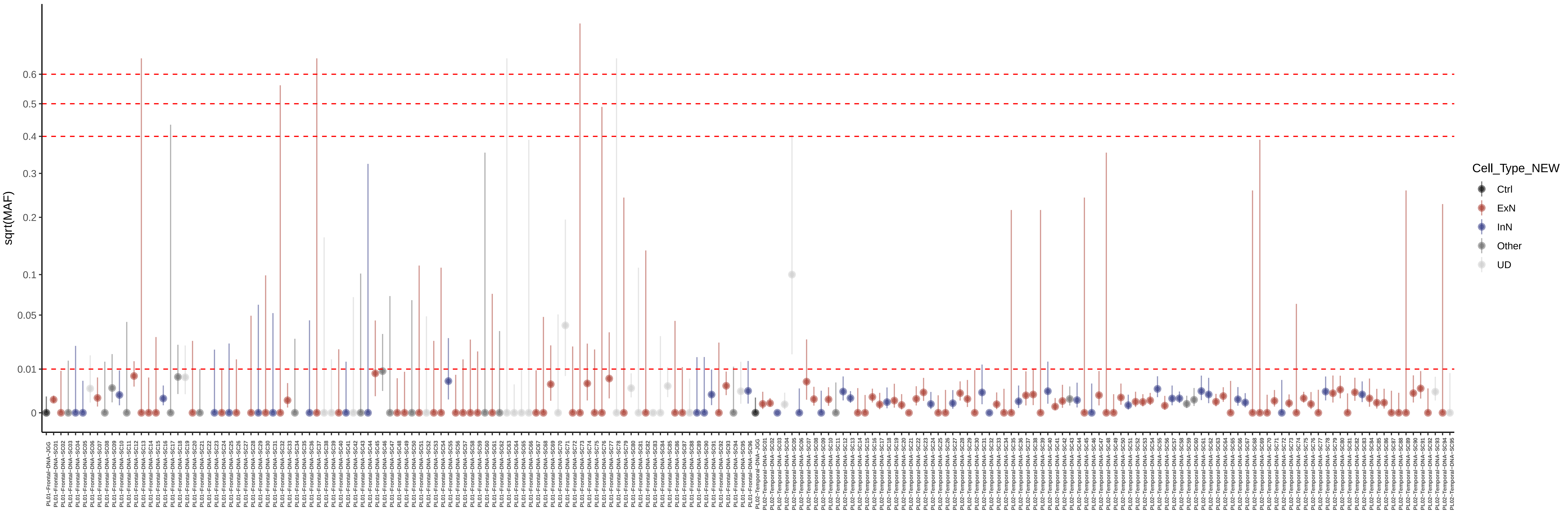

9−116754877−A−C

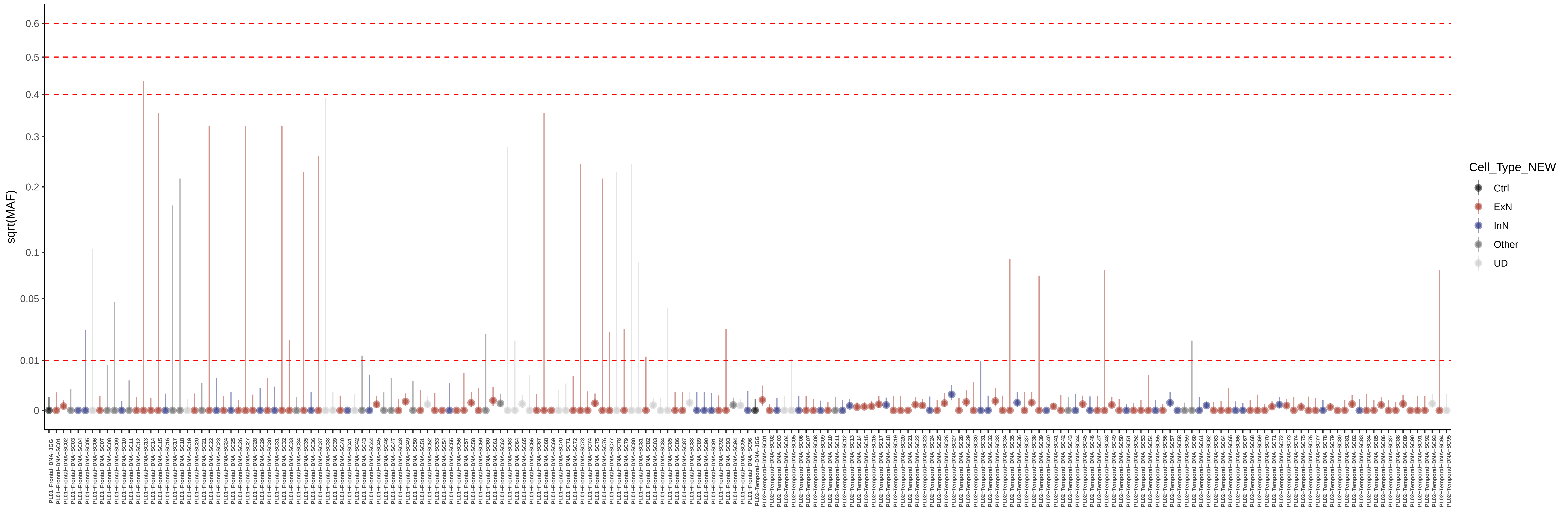

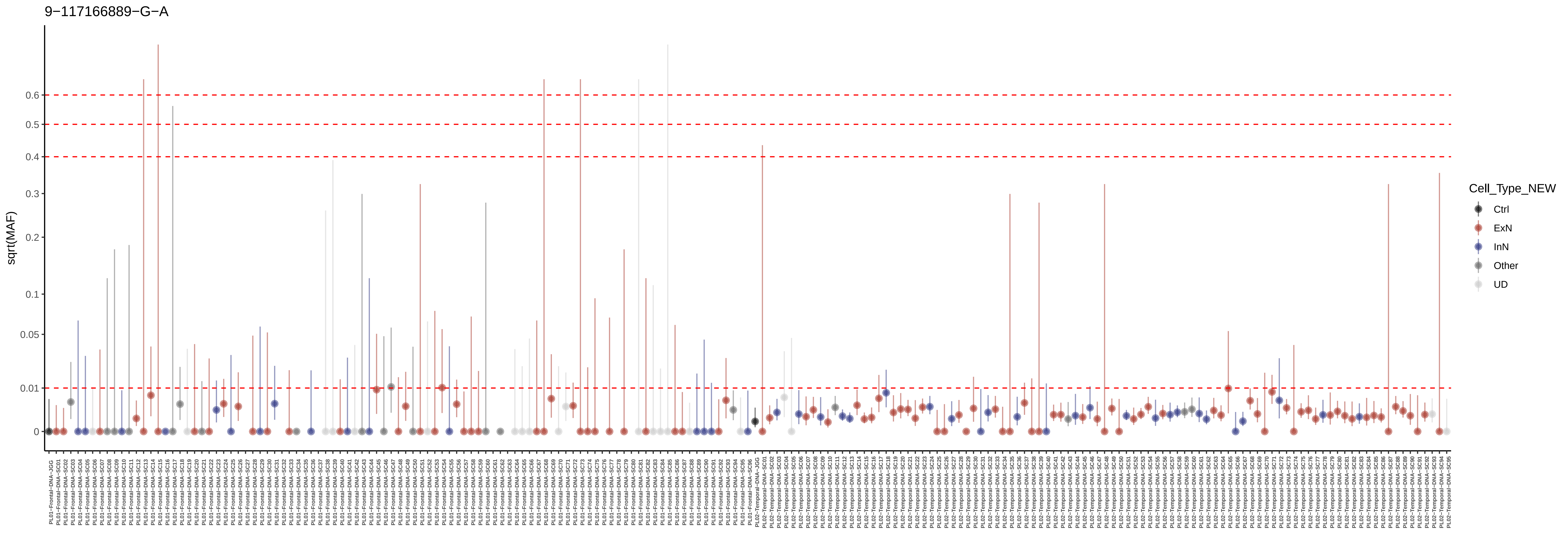
9−117166889−G−A
sqrt(MAF)
0.6
0.5
0.4
0.3
0.2
0.1
0.05
0.01
0
Cell_Type_NEW
Ctrl
ExN
InN
Other
UD

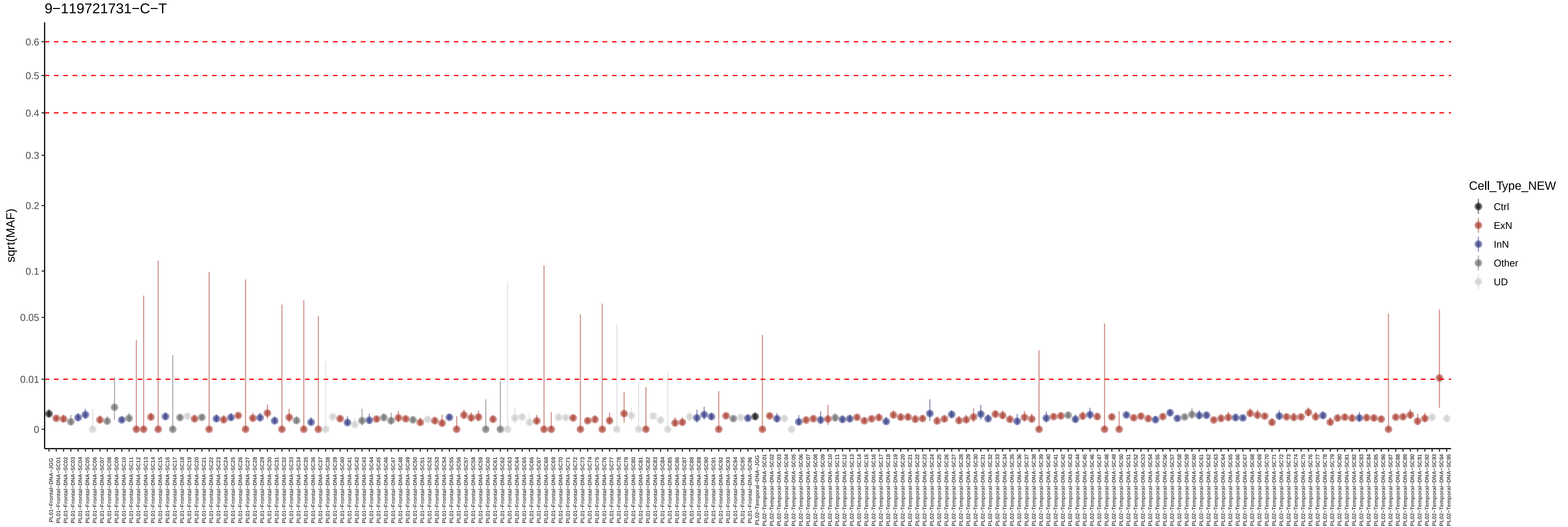
9−119721731−C−T
sqrt(MAF)
0.6
0.5
0.4
0.3
0.2
0.1
0.05
0.01
0
Cell_Type_NEW
Ctrl
ExN
InN
Other
UD

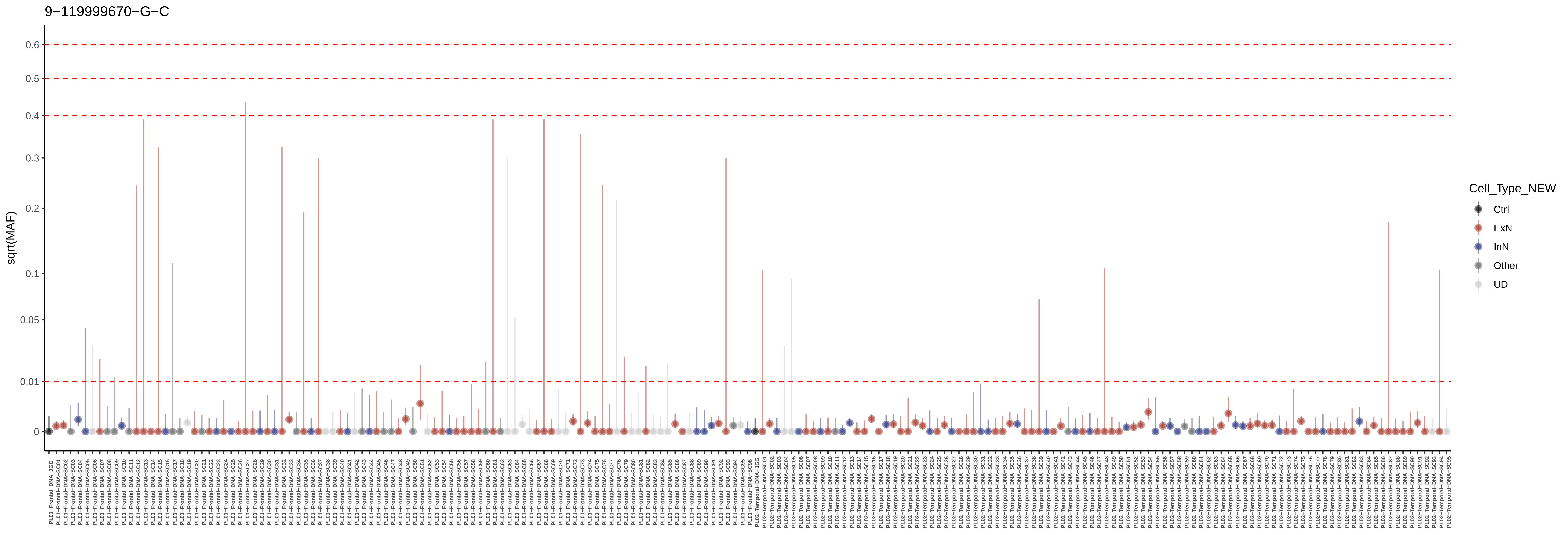

9−119999670−G−C
sqrt(MAF)
Cell_Type_NEW
Ctrl
ExN
InN
Other
UD

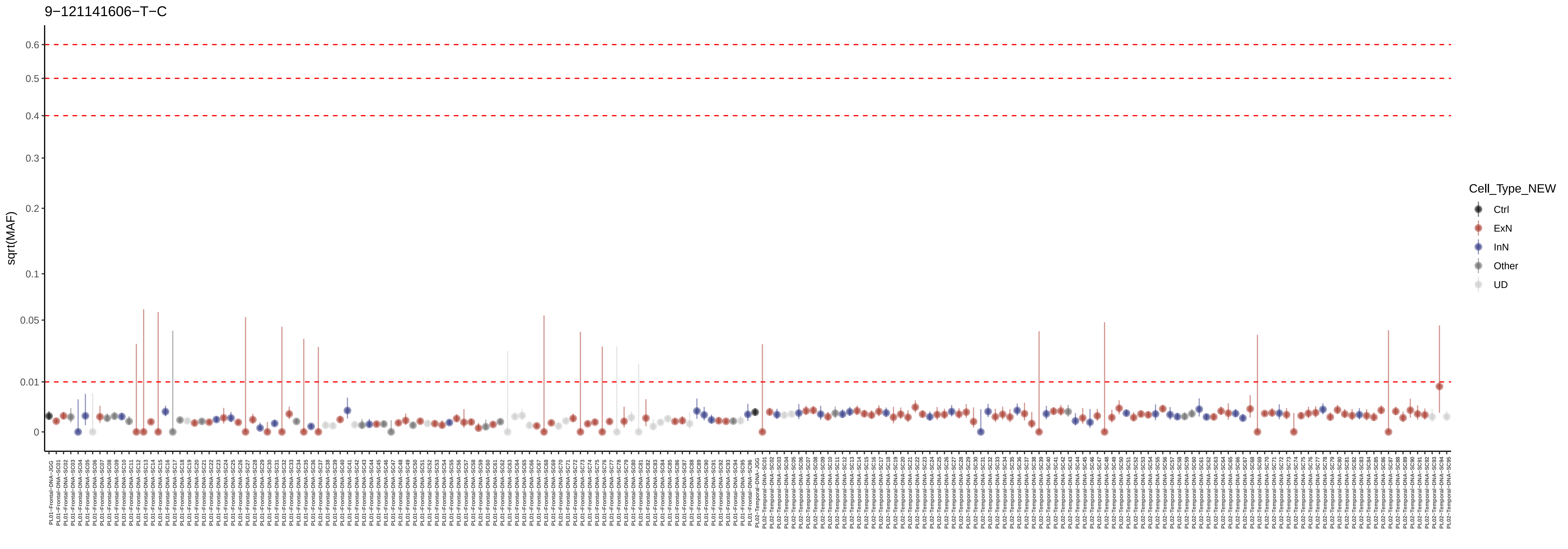
9−121141606−T−C
sqrt(MAF)
0.6
0.5
0.4
0.3
0.2
0.1
0.05
0.01
0
Cell_Type_NEW
Ctrl
ExN
InN
Other
UD

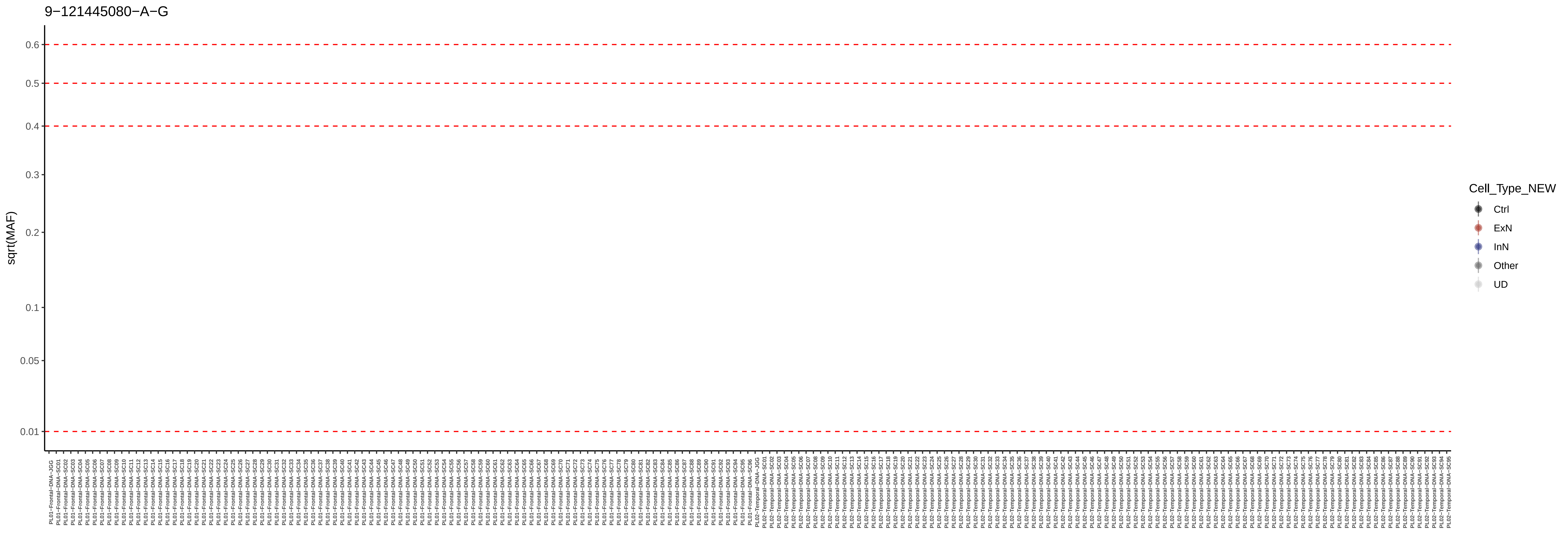
9−121445080−A−G
sqrt(MAF)
0.6
0.5
0.4
0.3
0.2
0.1
0.05
0.01
Cell_Type_NEW
Ctrl
ExN
InN
Other
UD

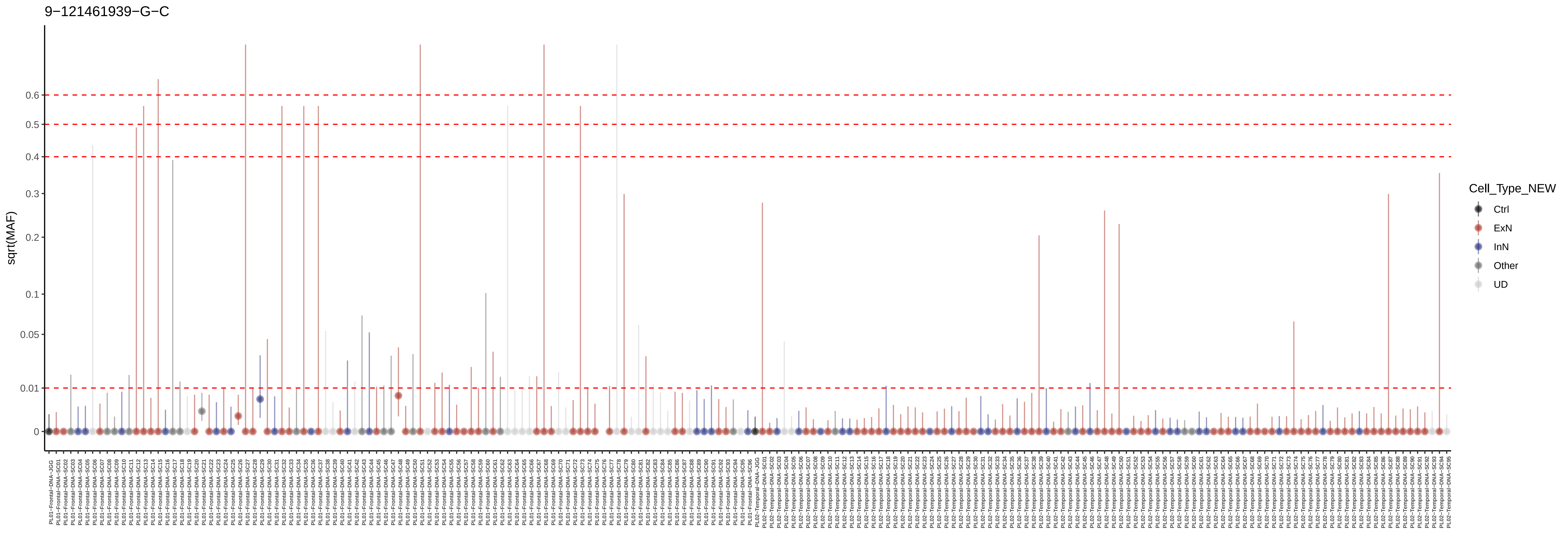

9−121461939−G−C
sqrt(MAF)
0.6
0.5
0.4
0.3
0.2
0.1
0.05
0.01
0
Cell_Type_NEW
Ctrl
ExN
InN
Other
UD

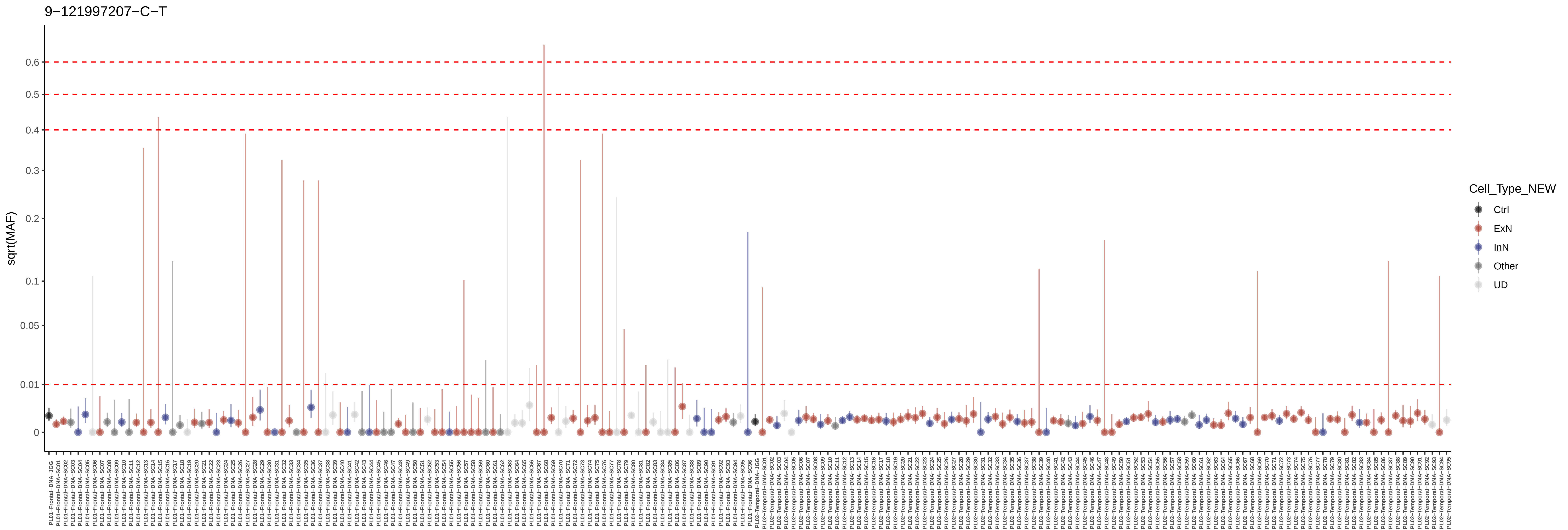

9−121997207−C−T
sqrt(MAF)
0.6
0.5
0.4
0.3
0.2
0.1
0.05
0.01
0
Cell_Type_NEW
Ctrl
ExN
InN
Other
UD

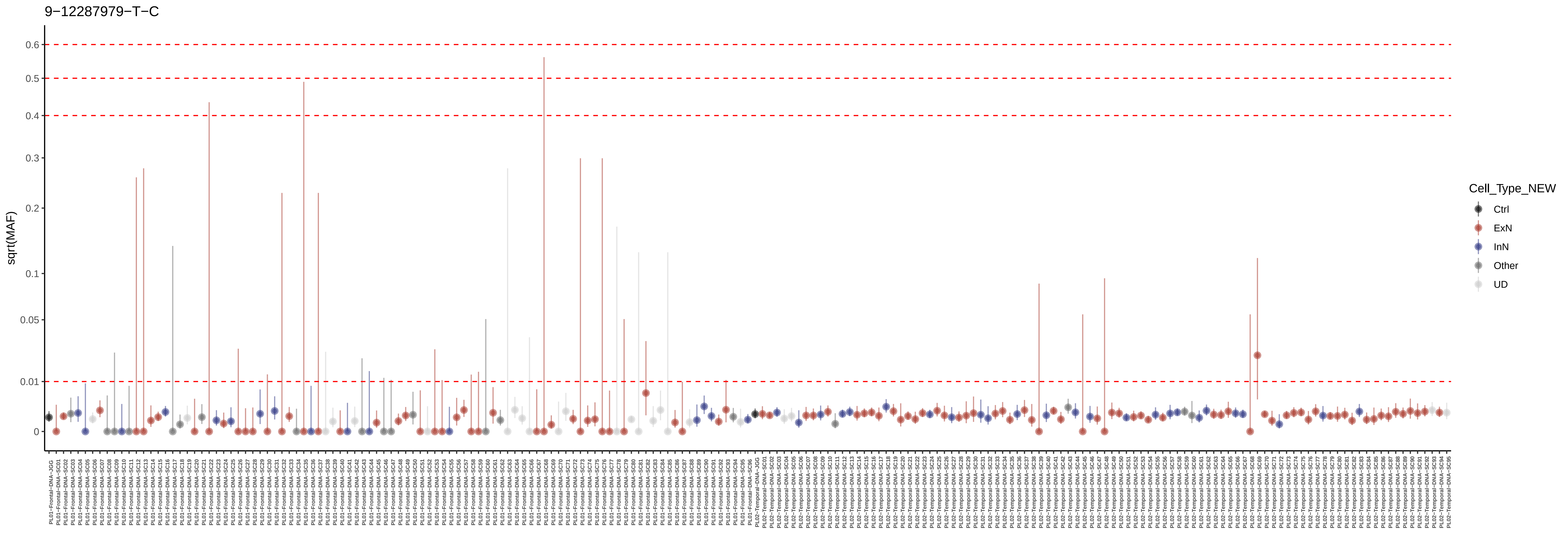

9−12287979−T−C
sqrt(MAF)
0.6
0.5
0.4
0.3
0.2
0.1
0.05
0.01
0
Cell_Type_NEW
Ctrl
ExN
InN
Other
UD

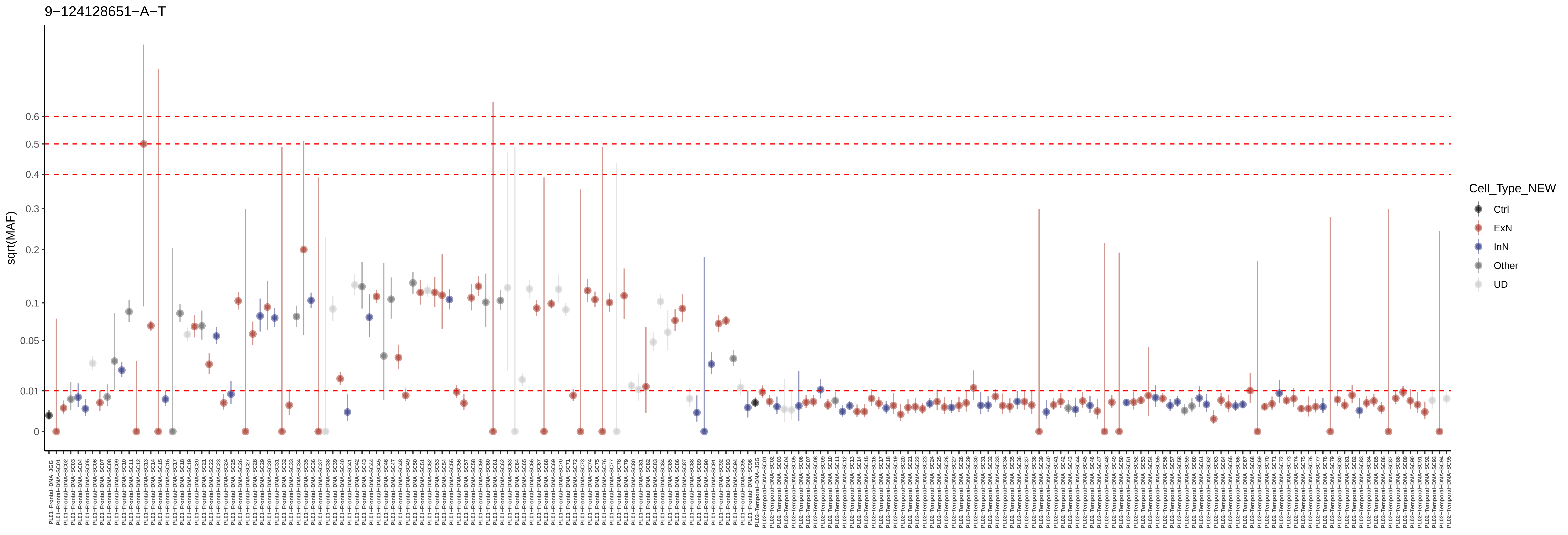
9−124128651−A−T
sqrt(MAF)
0.6
0.5
0.4
0.3
0.2
0.1
0.05
0.01
0
Cell_Type_NEW
Ctrl
ExN
InN
Other
UD

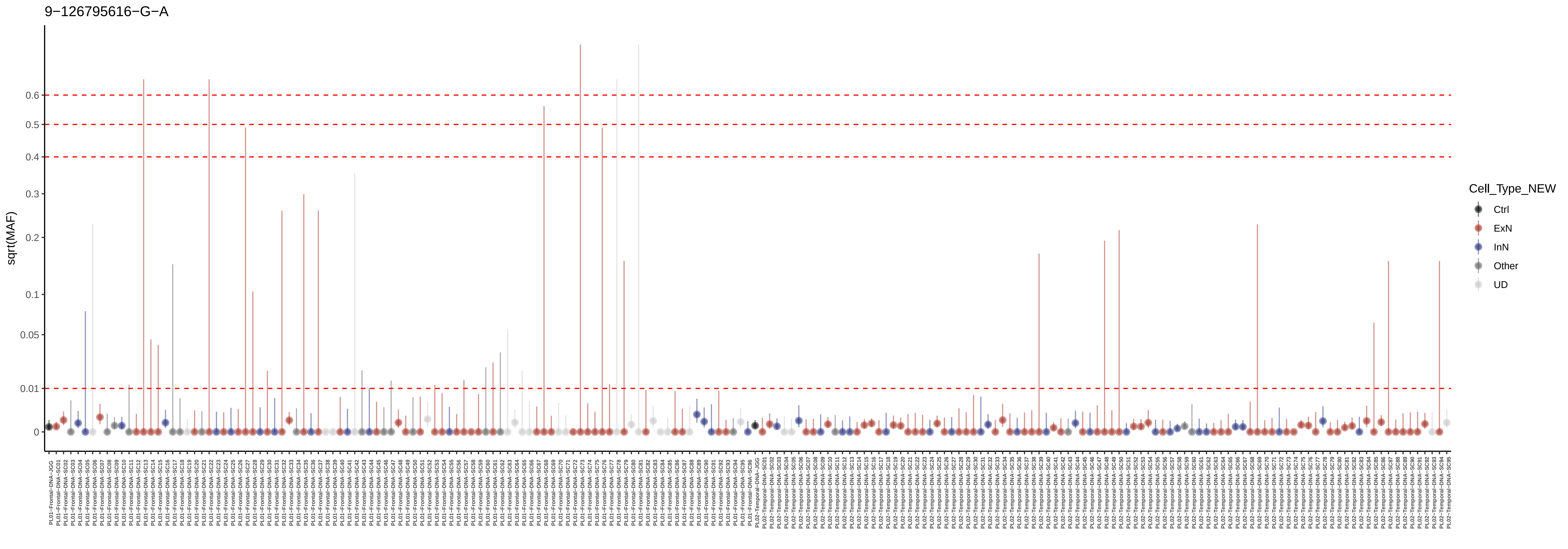
9−126795616−G−A
sqrt(MAF)
0.6
0.5
0.4
0.3
0.2
0.1
0.05
0.01
0
Cell_Type_NEW
Ctrl
ExN
InN
Other
UD

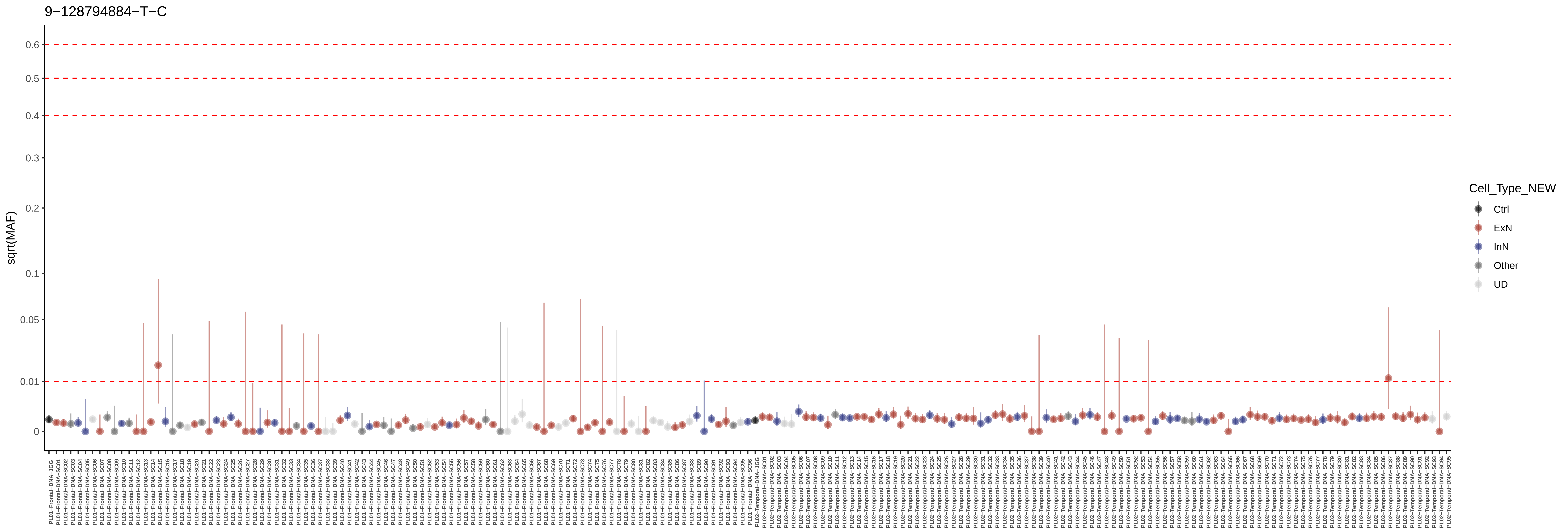
9−128794884−T−C
sqrt(MAF)
0.6
0.5
0.4
0.3
0.2
0.1
0.05
0.01
0
Cell_Type_NEW
Ctrl
ExN
InN
Other
UD

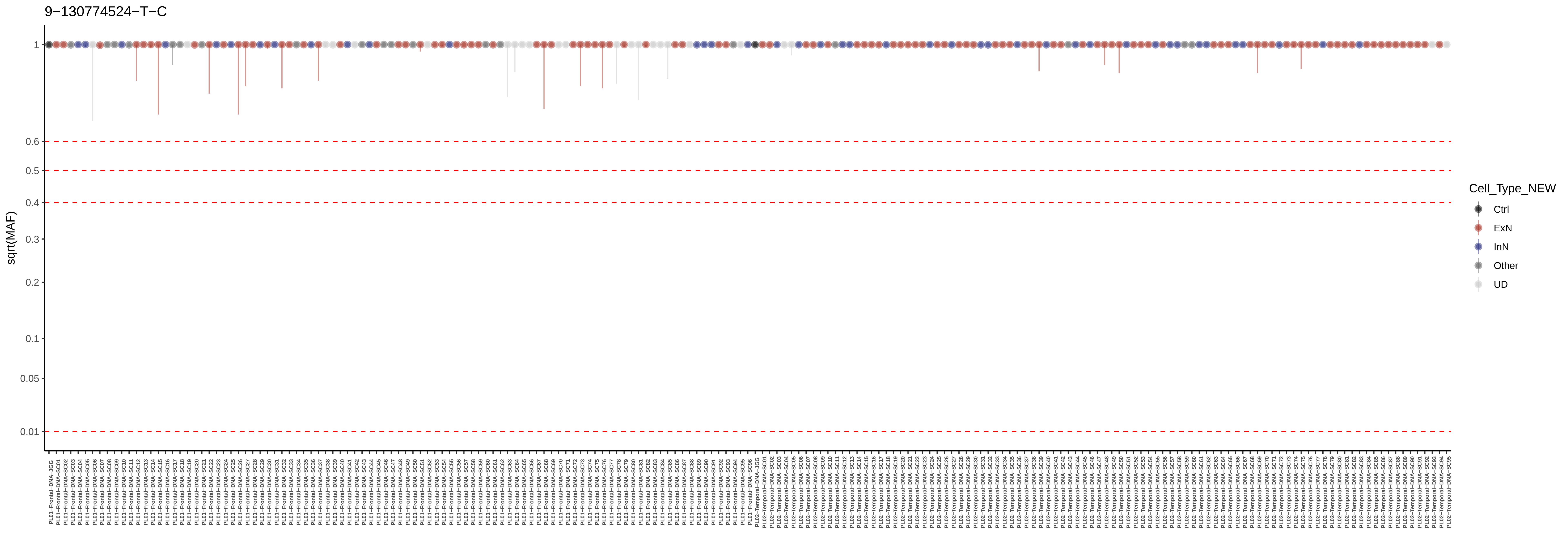
9−130774524−T−C
sqrt(MAF)
1
0.6
0.5
0.4
0.3
0.2
0.1
0.05
0.01
Cell_Type_NEW
Ctrl
ExN
InN
Other
UD

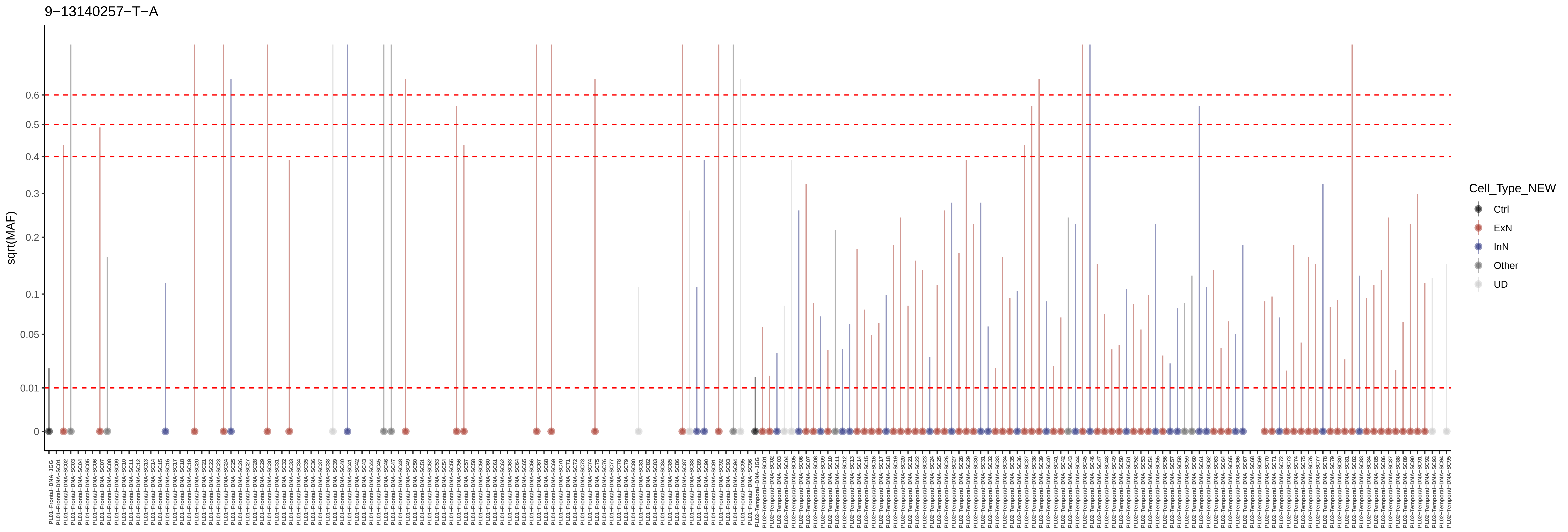
9−13140257−T−A
sqrt(MAF)
0.6
0.5
0.4
0.3
0.2
0.1
0.05
0.01
0
Cell_Type_NEW
Ctrl
ExN
InN
Other
UD

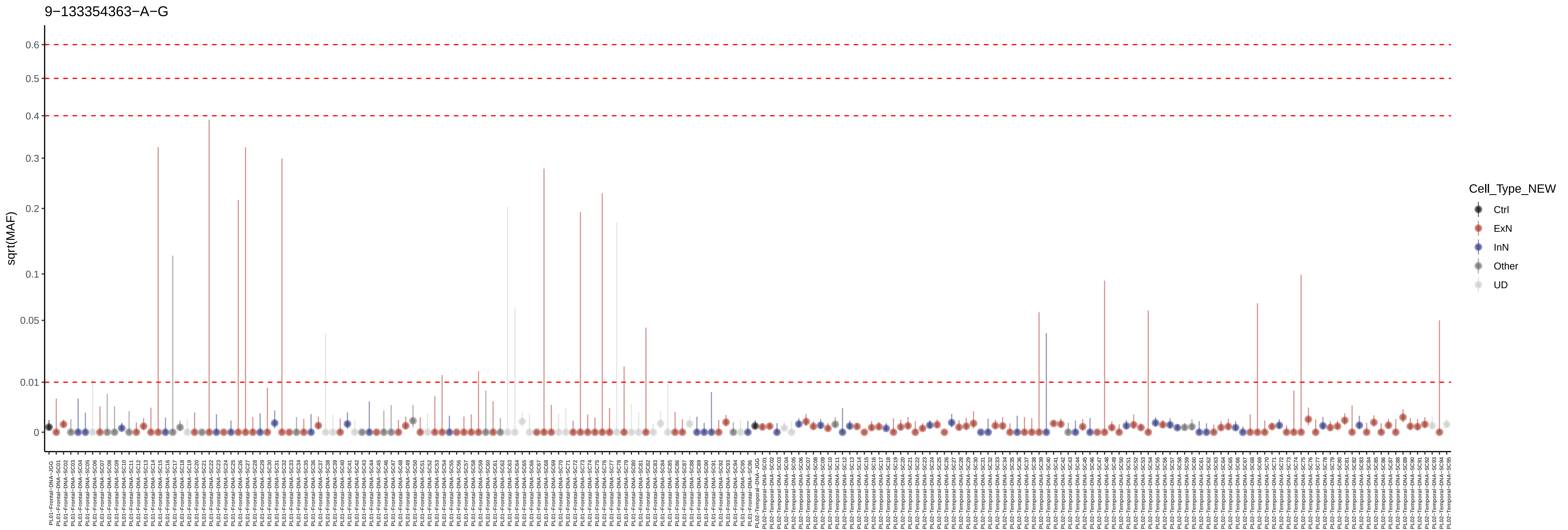
9−133354363−A−G
sqrt(MAF)
0.6
0.5
0.4
0.3
0.2
0.1
0.05
0.01
0
Cell_Type_NEW
Ctrl
ExN
InN
Other
UD

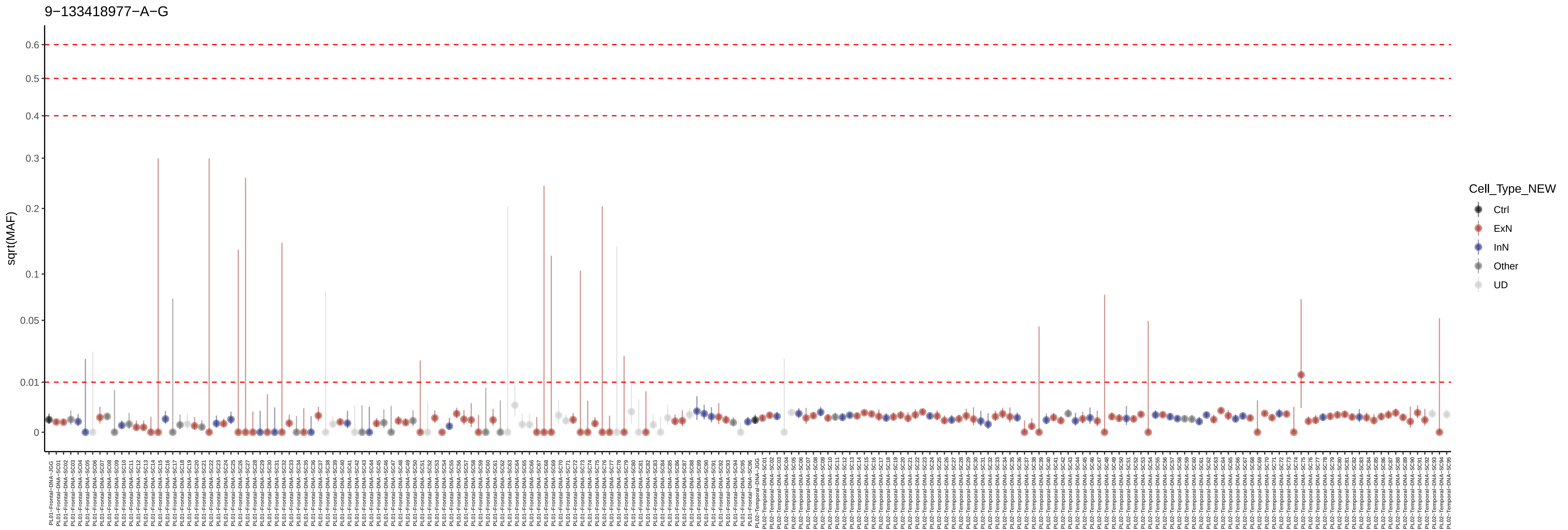
9−133418977−A−G
sqrt(MAF)
0.6
0.5
0.4
0.3
0.2
0.1
0.05
0.01
0
Cell_Type_NEW
Ctrl
ExN
InN
Other
UD

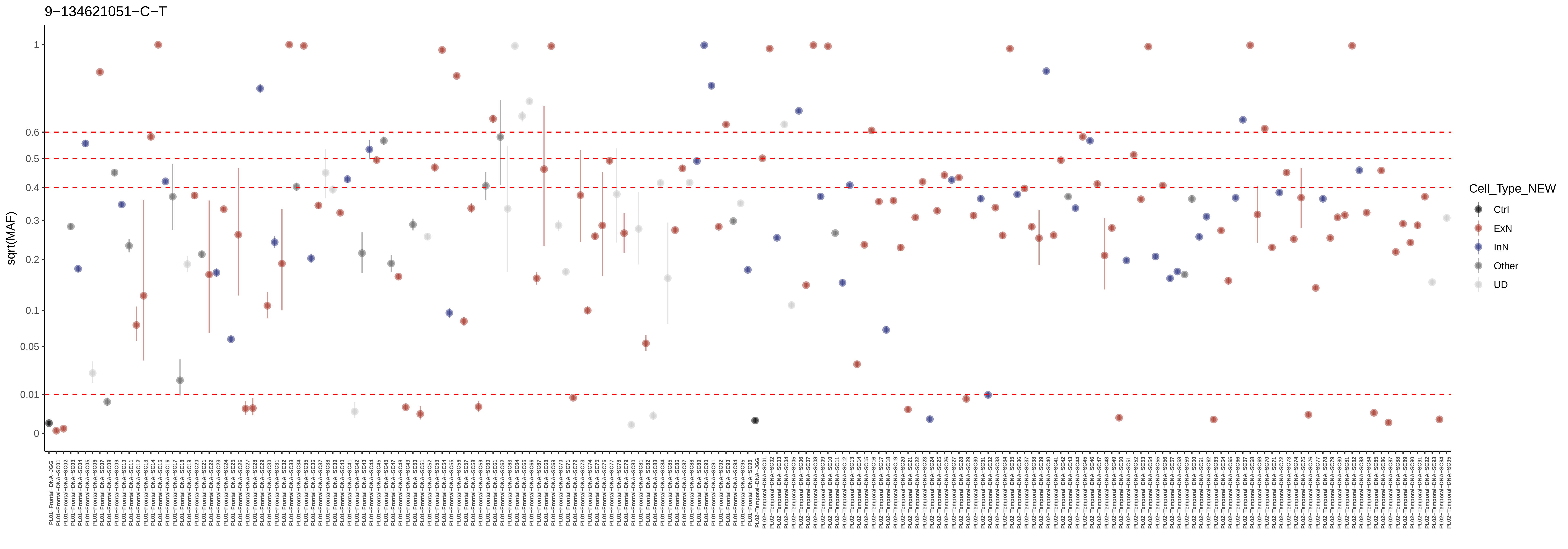
9−134621051−C−T
sqrt(MAF)
1
0.6
0.5
0.4
0.3
0.2
0.1
0.05
0.01
0
Cell_Type_NEW
Ctrl
ExN
InN
Other
UD

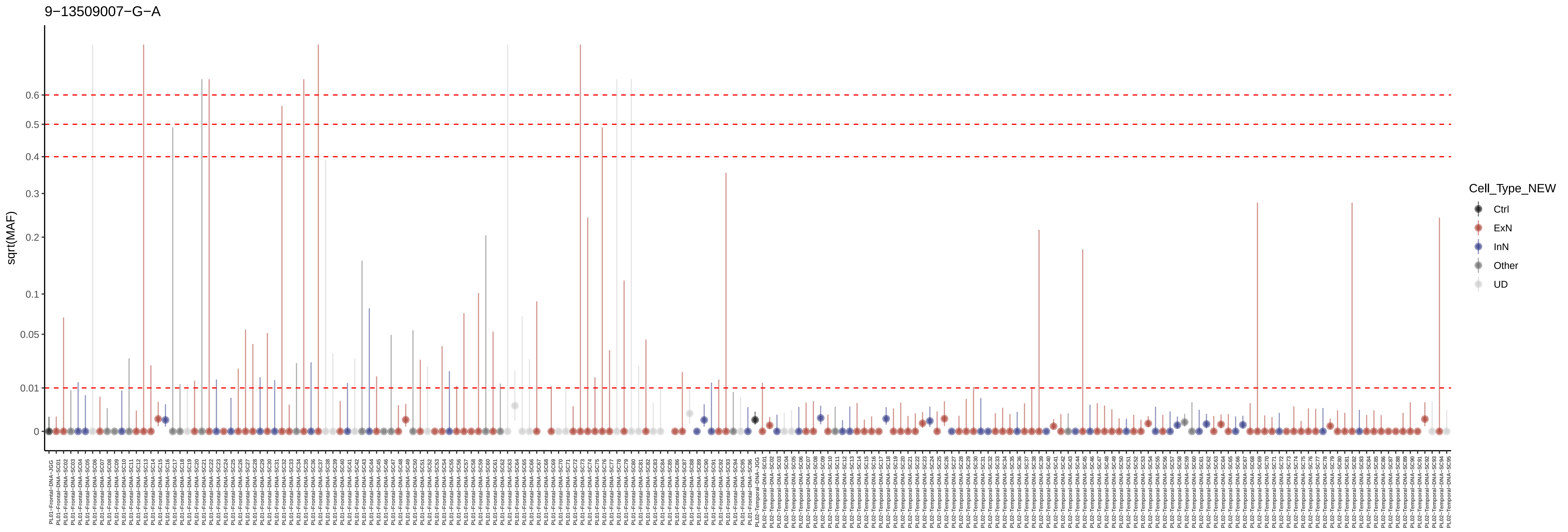
9−13509007−G−A
sqrt(MAF)
0.6
0.5
0.4
0.3
0.2
0.1
0.05
0.01
0
Cell_Type_NEW
Ctrl
ExN
InN
Other
UD

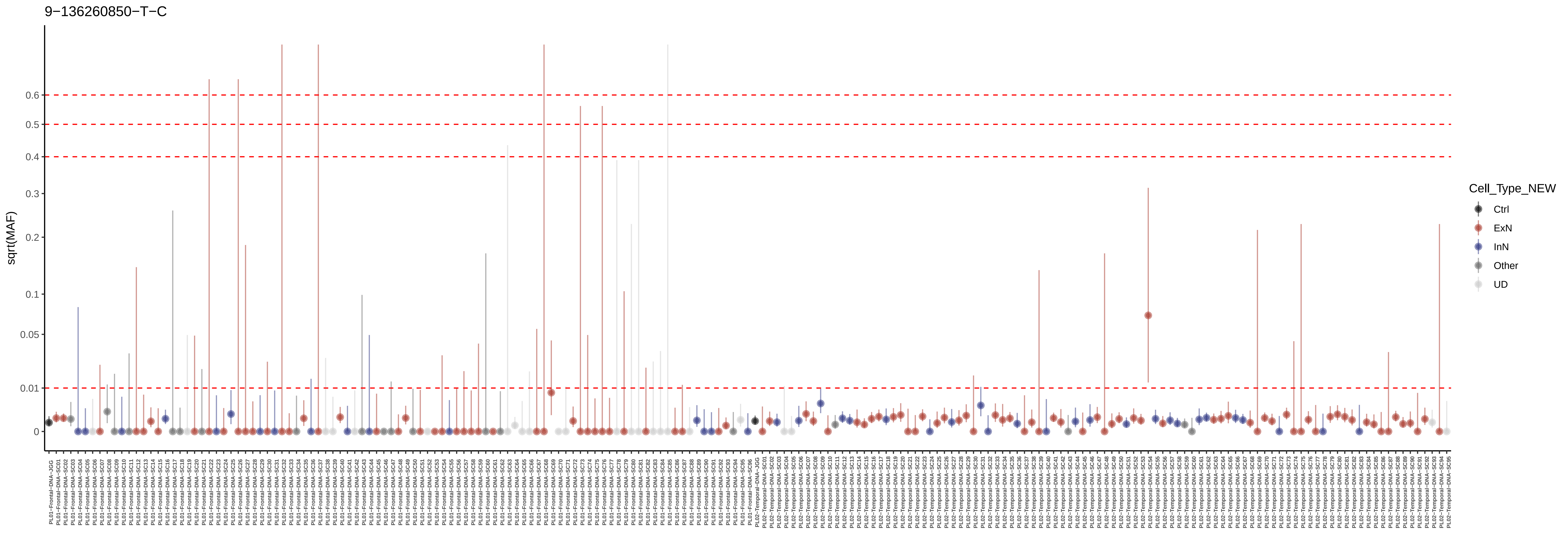
9−136260850−T−C
sqrt(MAF)
0.6
0.5
0.4
0.3
0.2
0.1
0.05
0.01
0
Cell_Type_NEW
Ctrl
ExN
InN
Other
UD

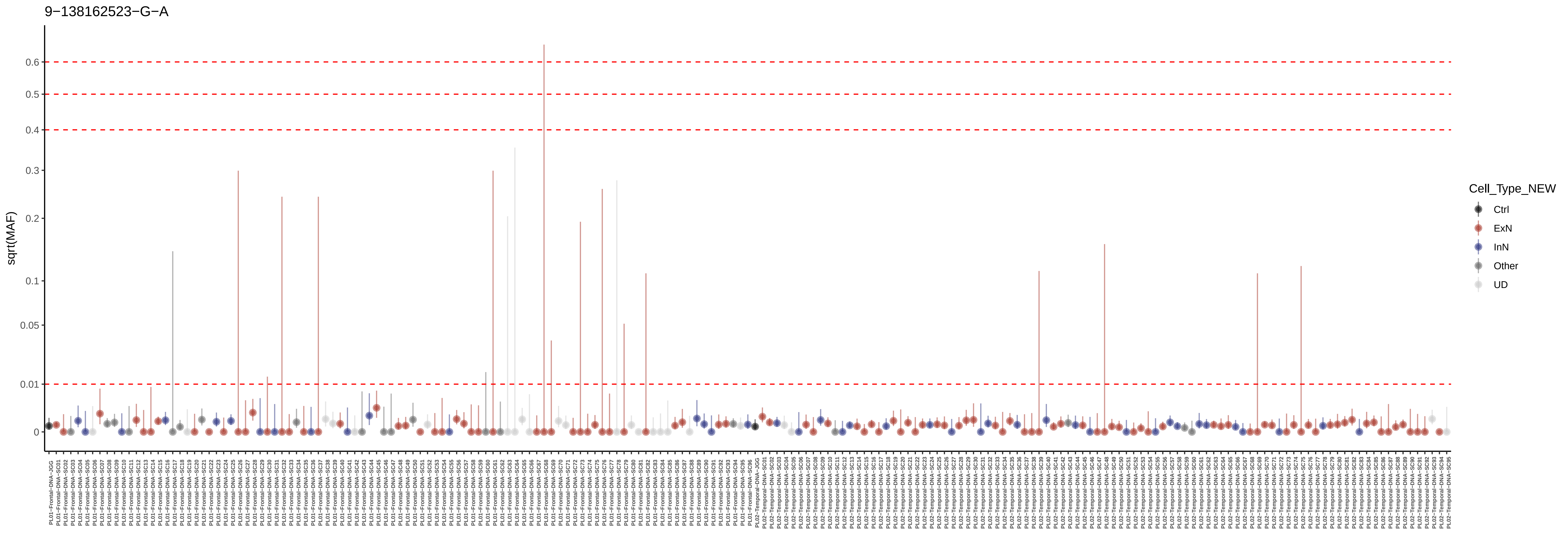

9−138162523−G−A
sqrt(MAF)
0.6
0.5
0.4
0.3
0.2
0.1
0.05
0.01
0
Cell_Type_NEW
Ctrl
ExN
InN
Other
UD

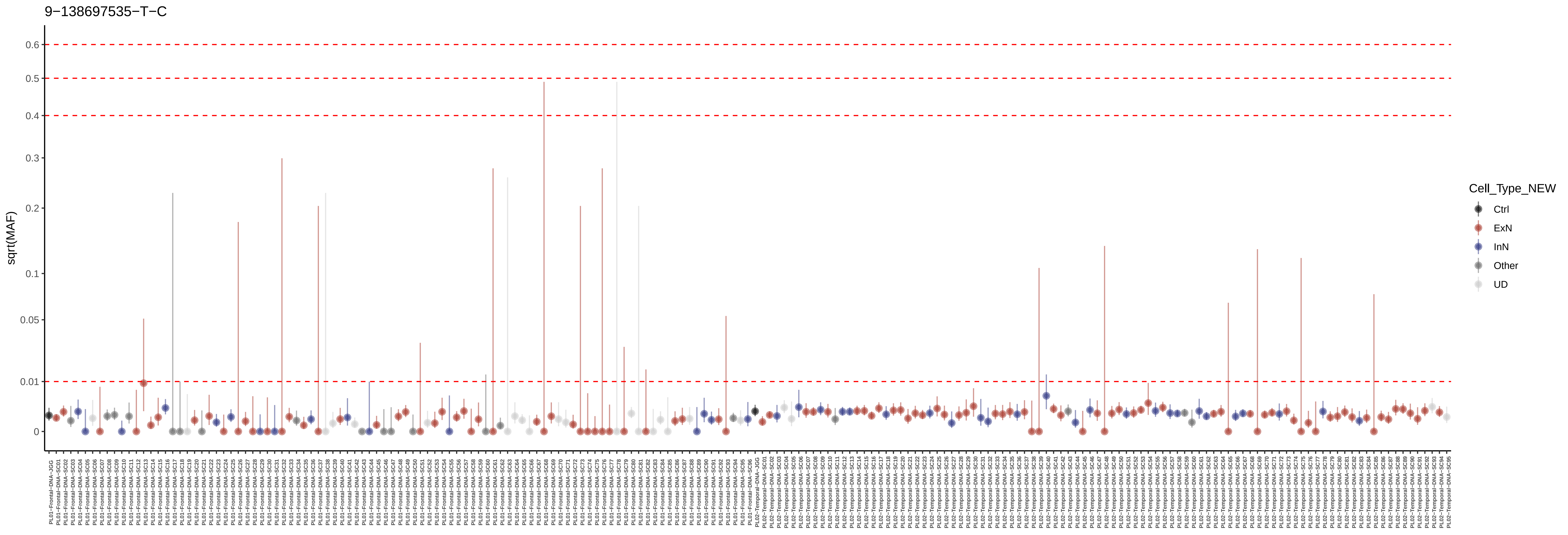

9−138697535−T−C
sqrt(MAF)
0.6
0.5
0.4
0.3
0.2
0.1
0.05
0.01
0
Cell_Type_NEW
Ctrl
ExN
InN
Other
UD

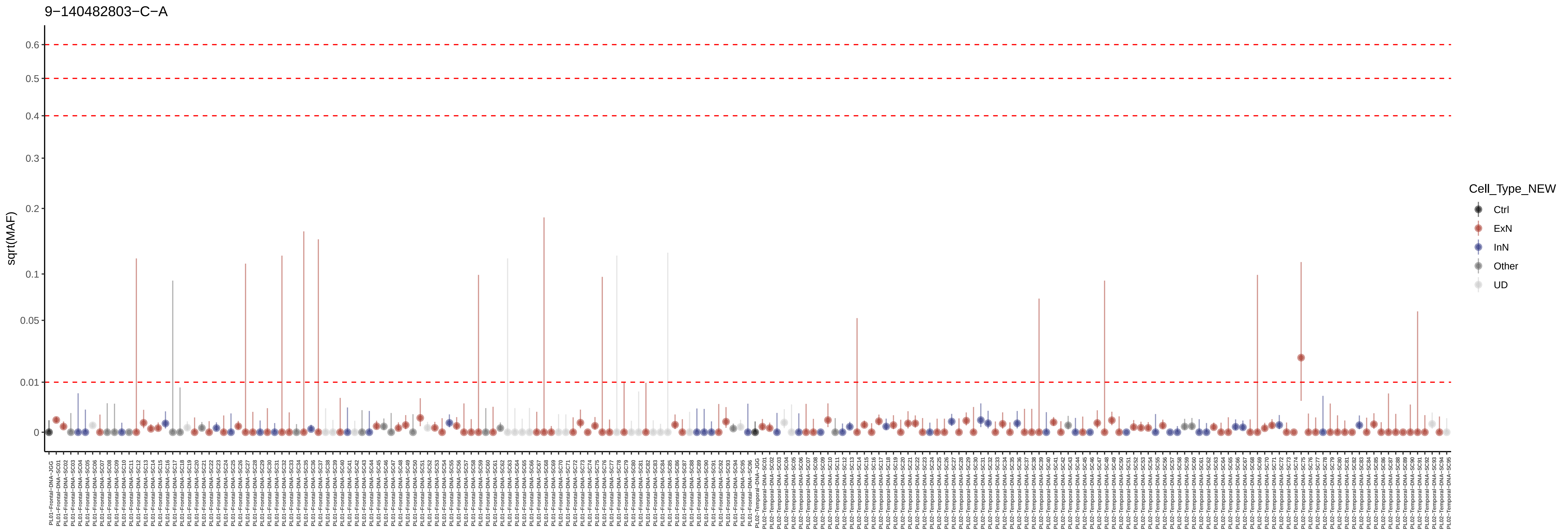
9−140482803−C−A
sqrt(MAF)
0.6
0.5
0.4
0.3
0.2
0.1
0.05
0.01
0
Cell_Type_NEW
Ctrl
ExN
InN
Other
UD

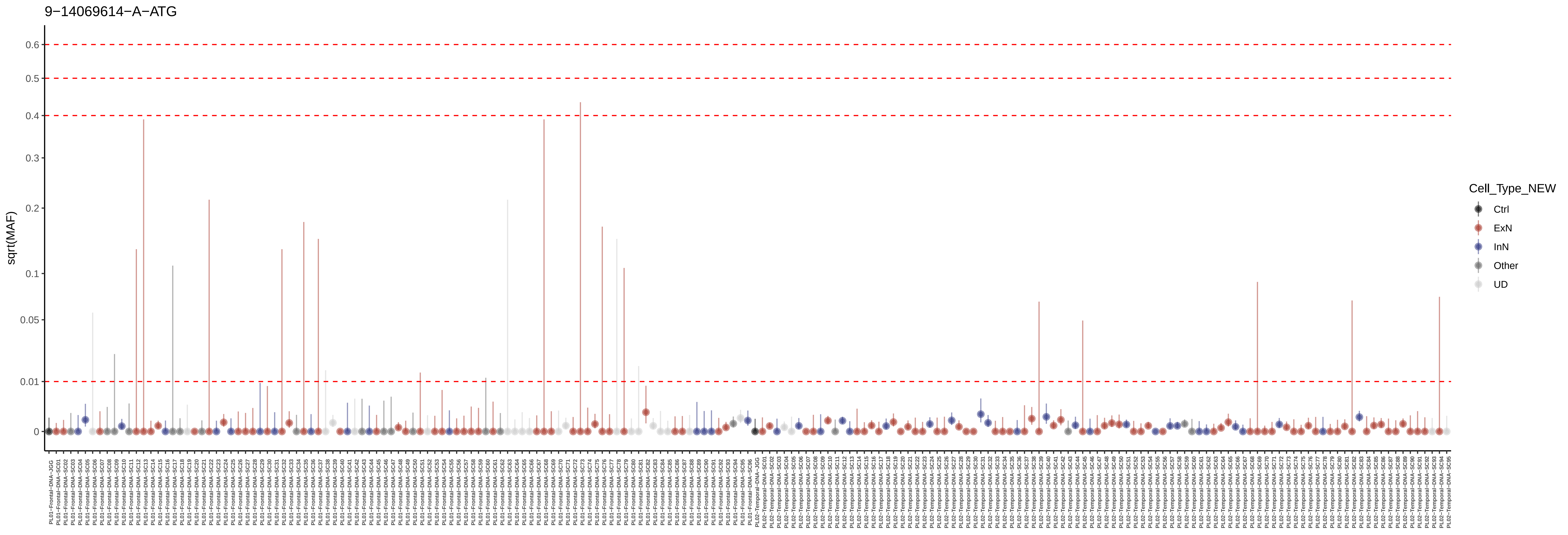
9−14069614−A−ATG
sqrt(MAF)
0.6
0.5
0.4
0.3
0.2
0.1
0.05
0.01
0
Cell_Type_NEW
Ctrl
ExN
InN
Other
UD

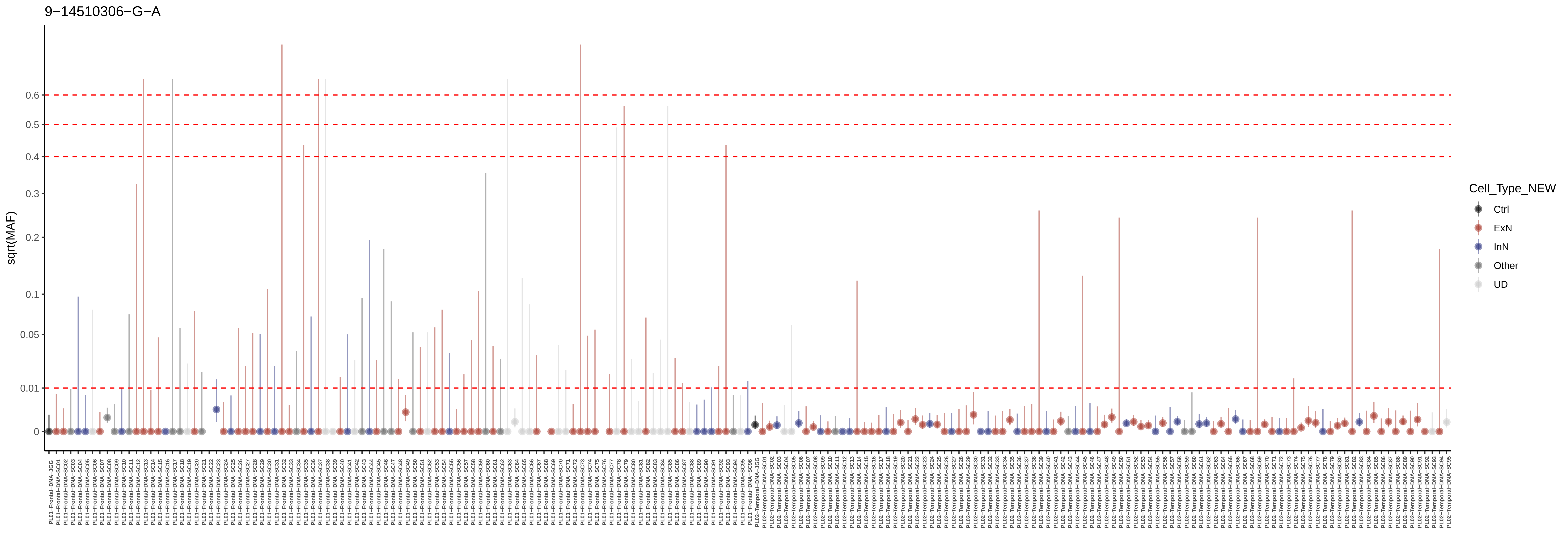
9−14510306−G−A
sqrt(MAF)
0.6
0.5
0.4
0.3
0.2
0.1
0.05
0.01
0
Cell_Type_NEW
Ctrl
ExN
InN
Other
UD

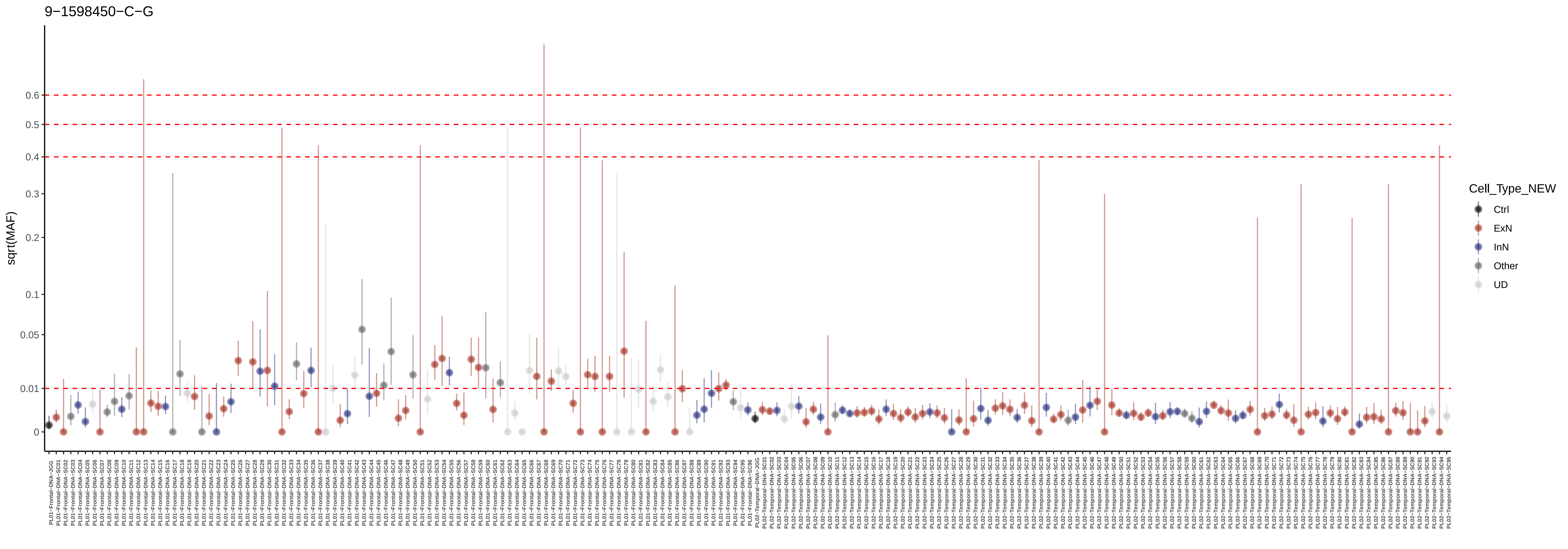

9−1598450−C−G
sqrt(MAF)
0.6
0.5
0.4
0.3
0.2
0.1
0.05
0.01
0
Cell_Type_NEW
Ctrl
ExN
InN
Other
UD

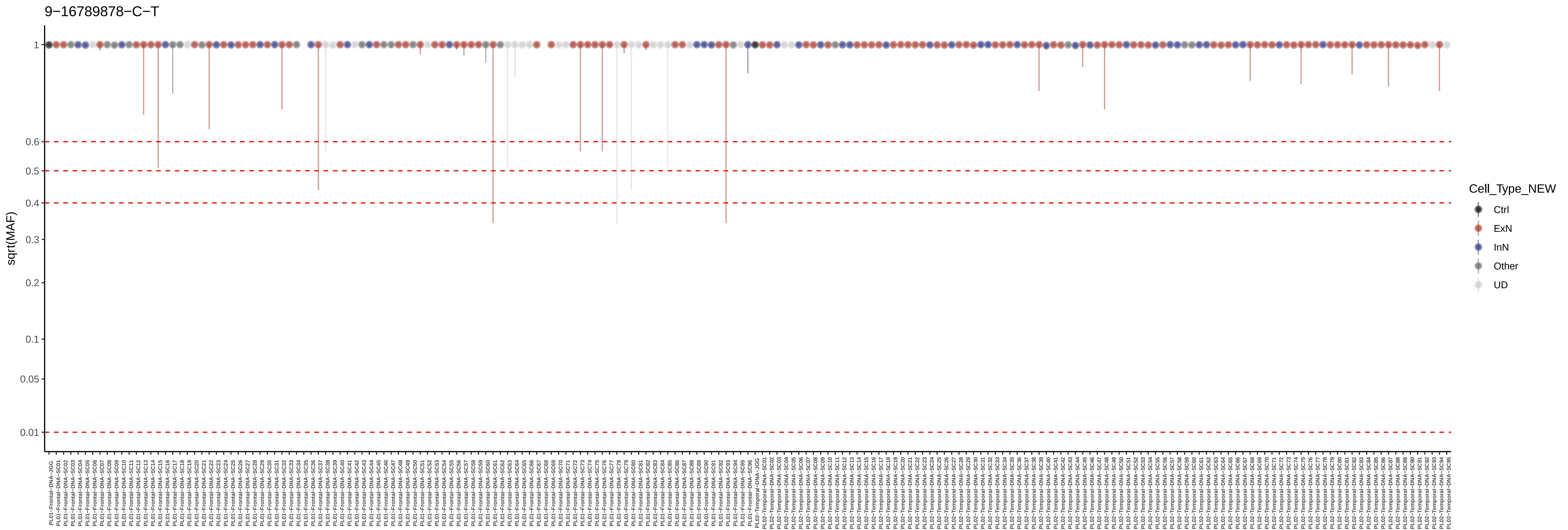

9−16789878−C−T
sqrt(MAF)
1
0.6
0.5
0.4
0.3
0.2
0.1
0.05
0.01
Cell_Type_NEW
Ctrl
ExN
InN
Other
UD

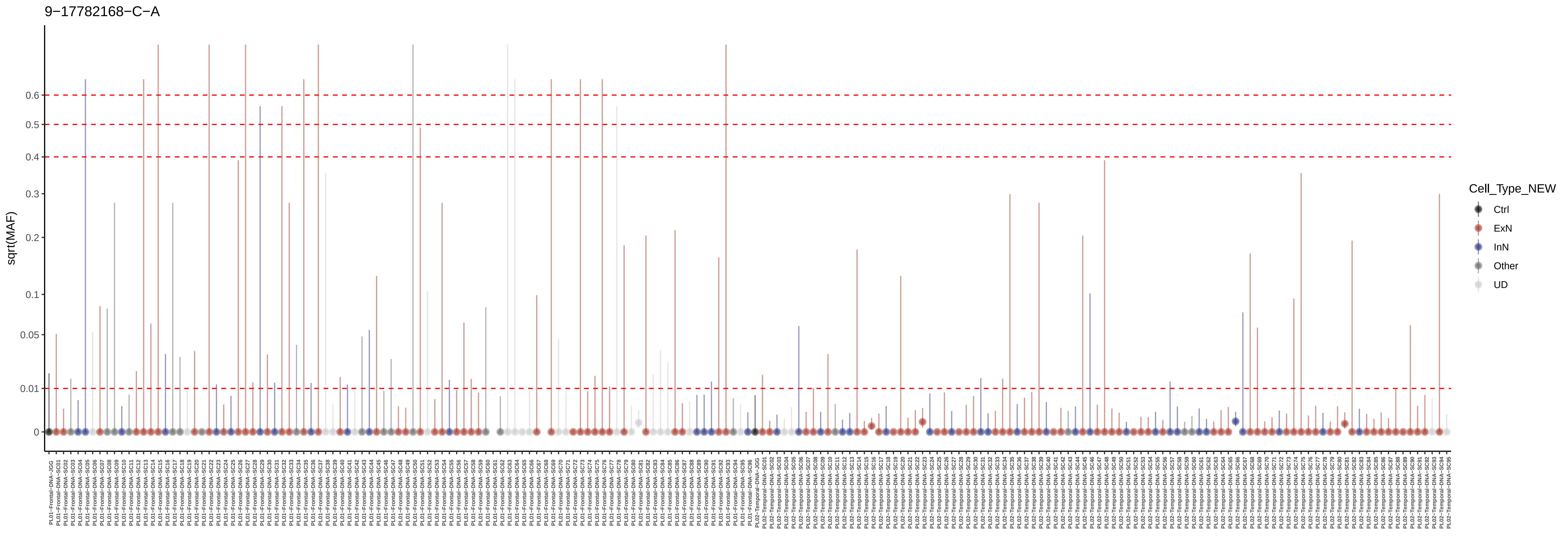
9−17782168−C−A
sqrt(MAF)
0.6
0.5
0.4
0.3
0.2
0.1
0.05
0.01
0
Cell_Type_NEW
Ctrl
ExN
InN
Other
UD

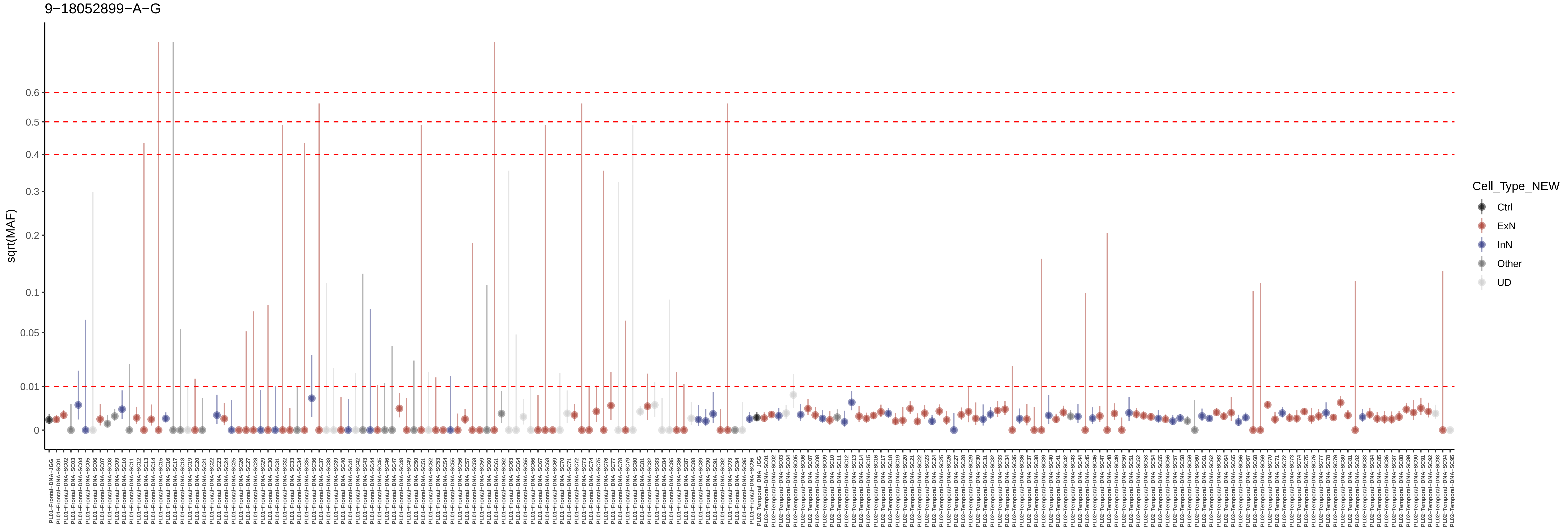
9−18052899−A−G
sqrt(MAF)
0.6
0.5
0.4
0.3
0.2
0.1
0.05
0.01
0
Cell_Type_NEW
Ctrl
ExN
InN
Other
UD

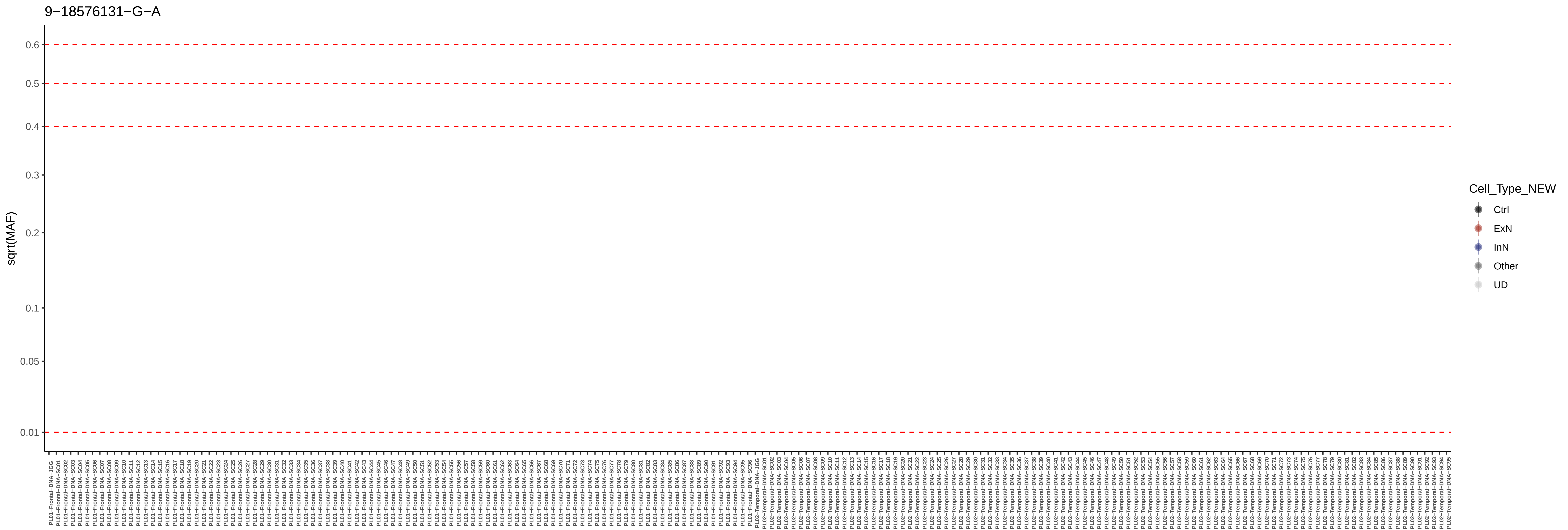
9−18576131−G−A
sqrt(MAF)
0.6
0.5
0.4
0.3
0.2
0.1
0.05
0.01
Cell_Type_NEW
Ctrl
ExN
InN
Other
UD

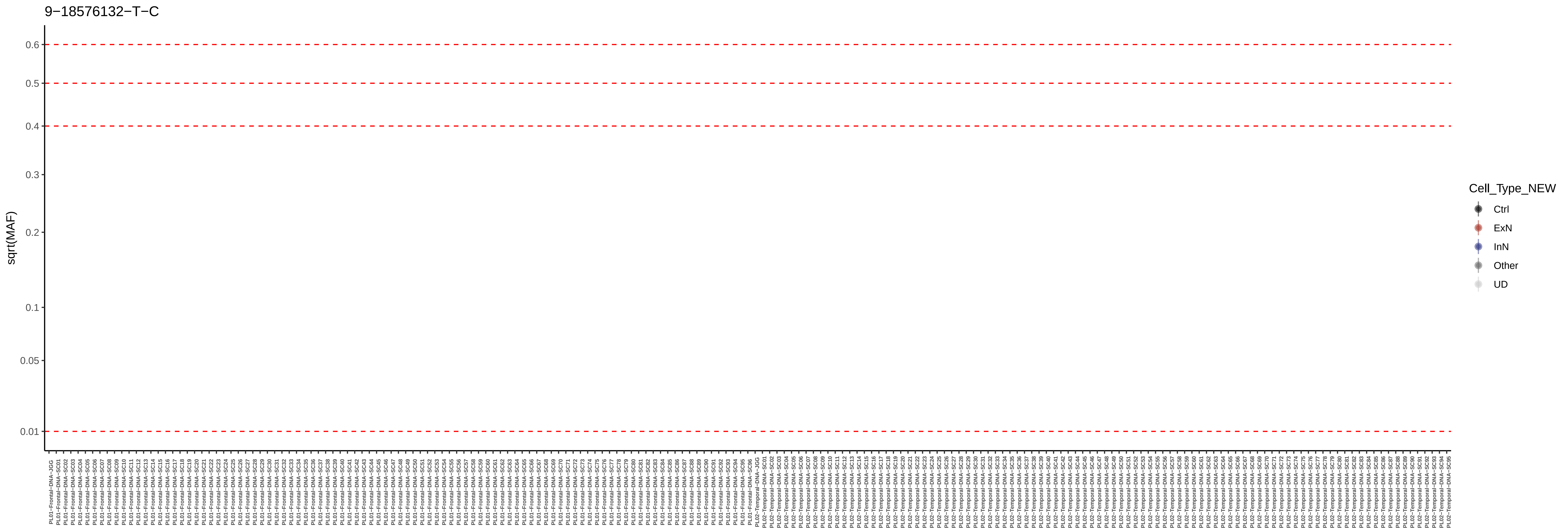

9−18576132−T−C
sqrt(MAF)
0.6
0.5
0.4
0.3
0.2
0.1
0.05
0.01
Cell_Type_NEW
Ctrl
ExN
InN
Other
UD

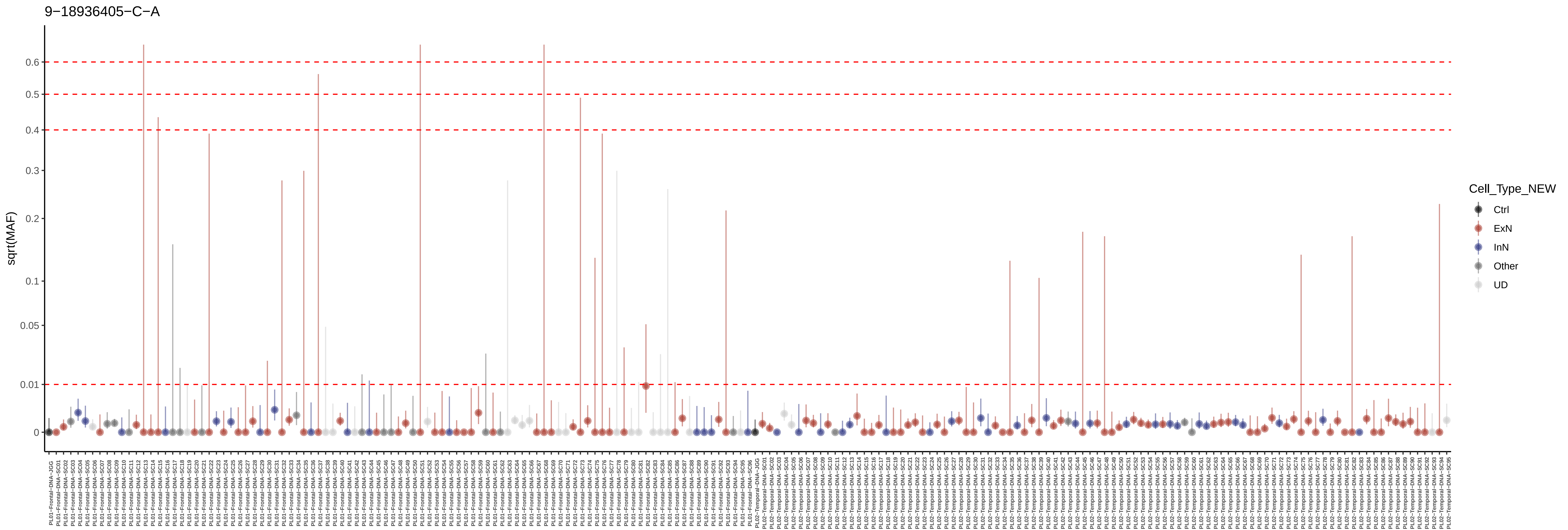
9−18936405−C−A
sqrt(MAF)
0.6
0.5
0.4
0.3
0.2
0.1
0.05
0.01
0
Cell_Type_NEW
Ctrl
ExN
InN
Other
UD

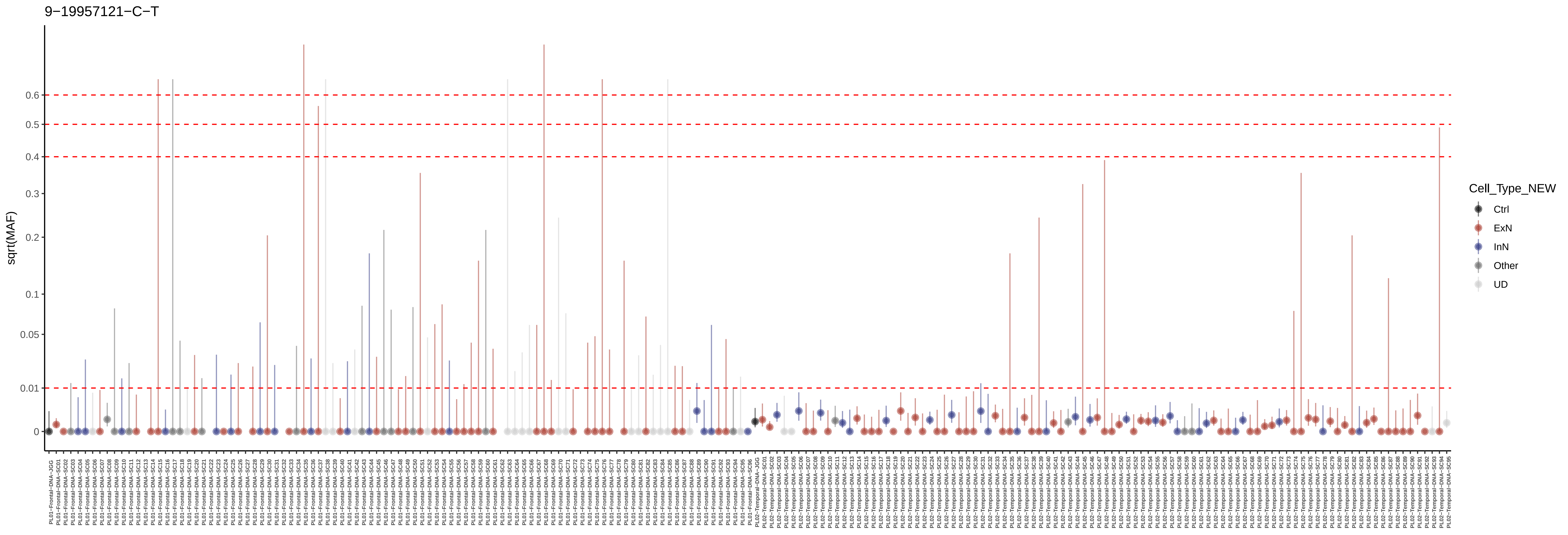
9−19957121−C−T
sqrt(MAF)
0.6
0.5
0.4
0.3
0.2
0.1
0.05
0.01
0
Cell_Type_NEW
Ctrl
ExN
InN
Other
UD

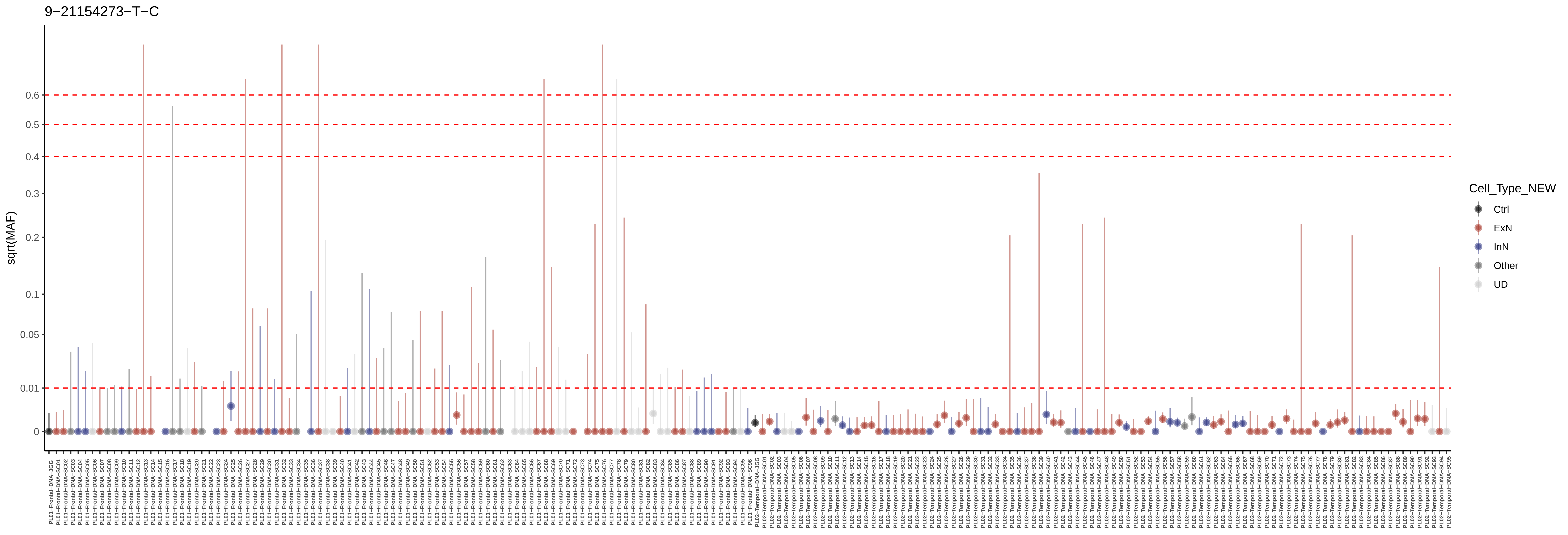
9−21154273−T−C
sqrt(MAF)
0.6
0.5
0.4
0.3
0.2
0.1
0.05
0.01
0
Cell_Type_NEW
Ctrl
ExN
InN
Other
UD

9−21506598−C−A

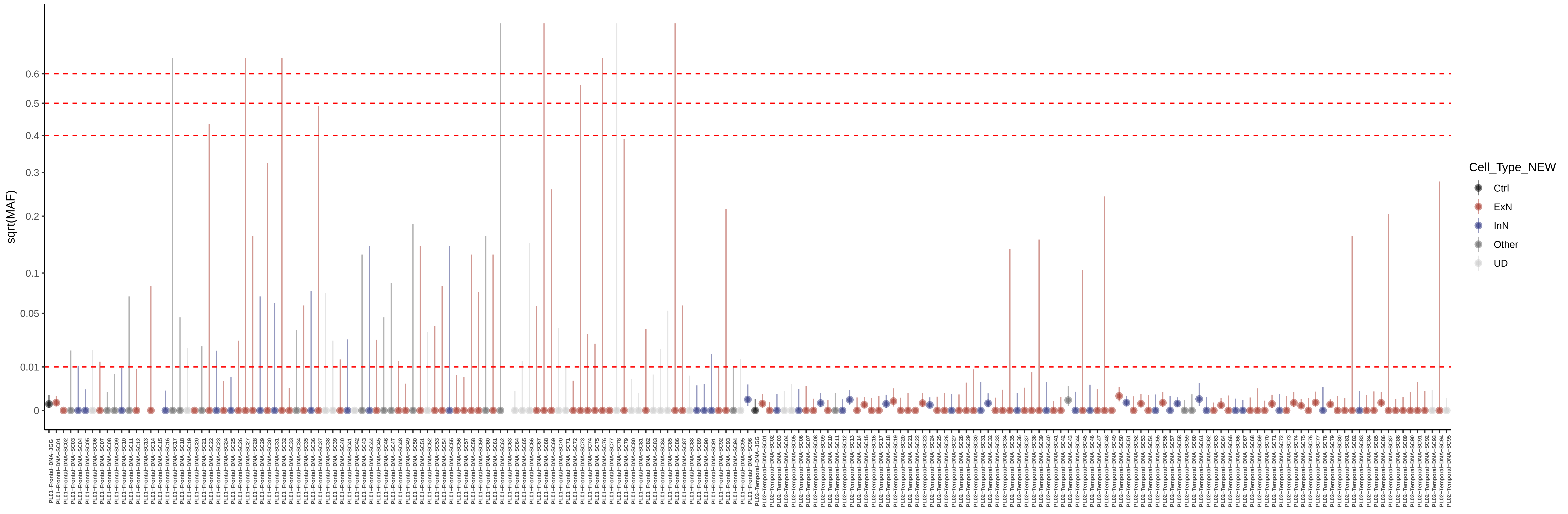

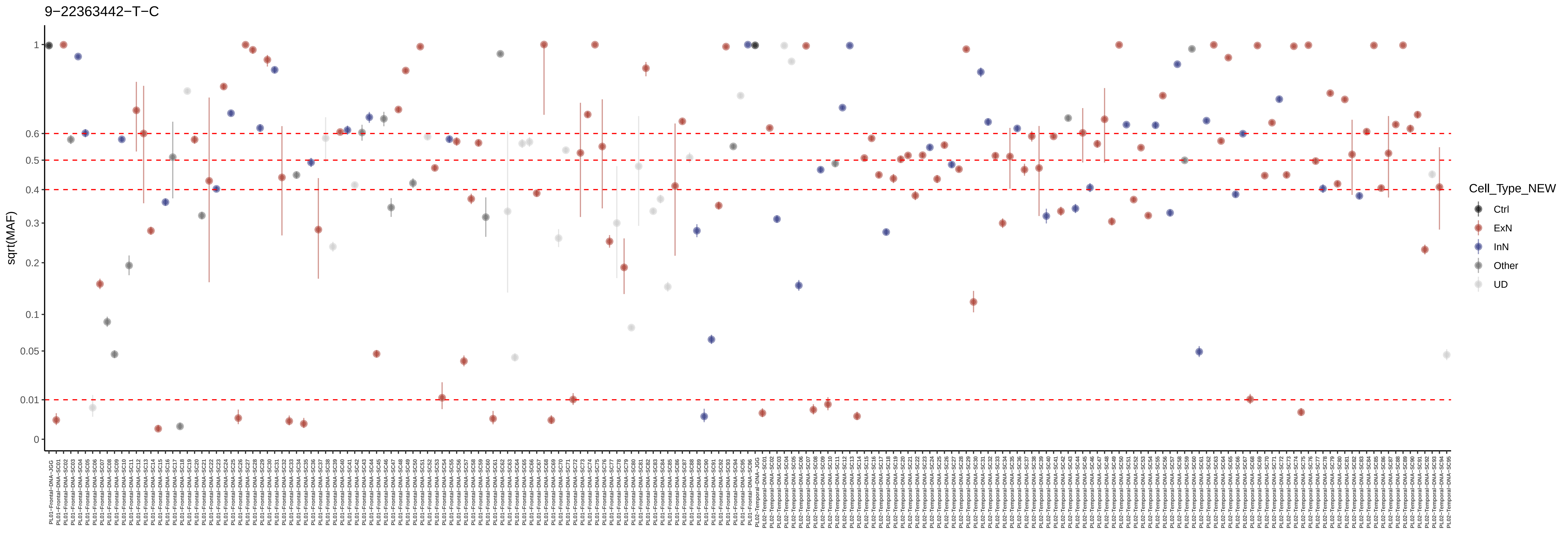
9−22363442−T−C
sqrt(MAF)
1
0.6
0.5
0.4
0.3
0.2
0.1
0.05
0.01
0
Cell_Type_NEW
Ctrl
ExN
InN
Other
UD

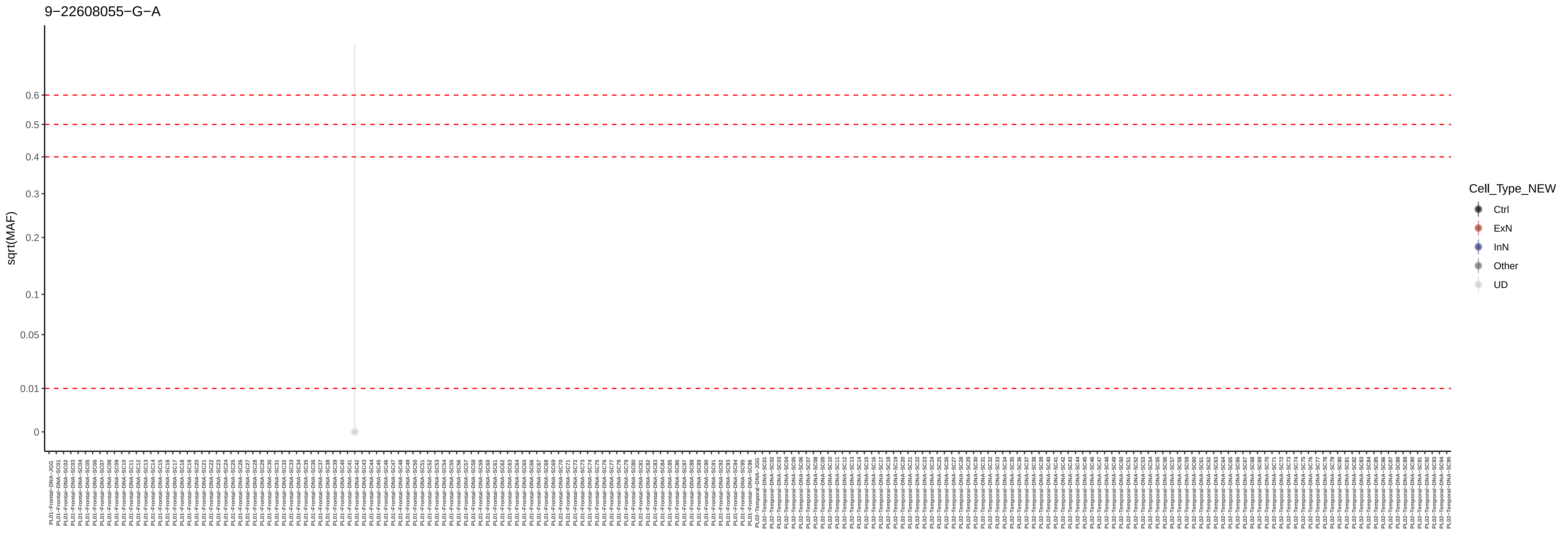

9−22608055−G−A
sqrt(MAF)
0.6
0.5
0.4
0.3
0.2
0.1
0.05
0.01
0
Cell_Type_NEW
Ctrl
ExN
InN
Other
UD

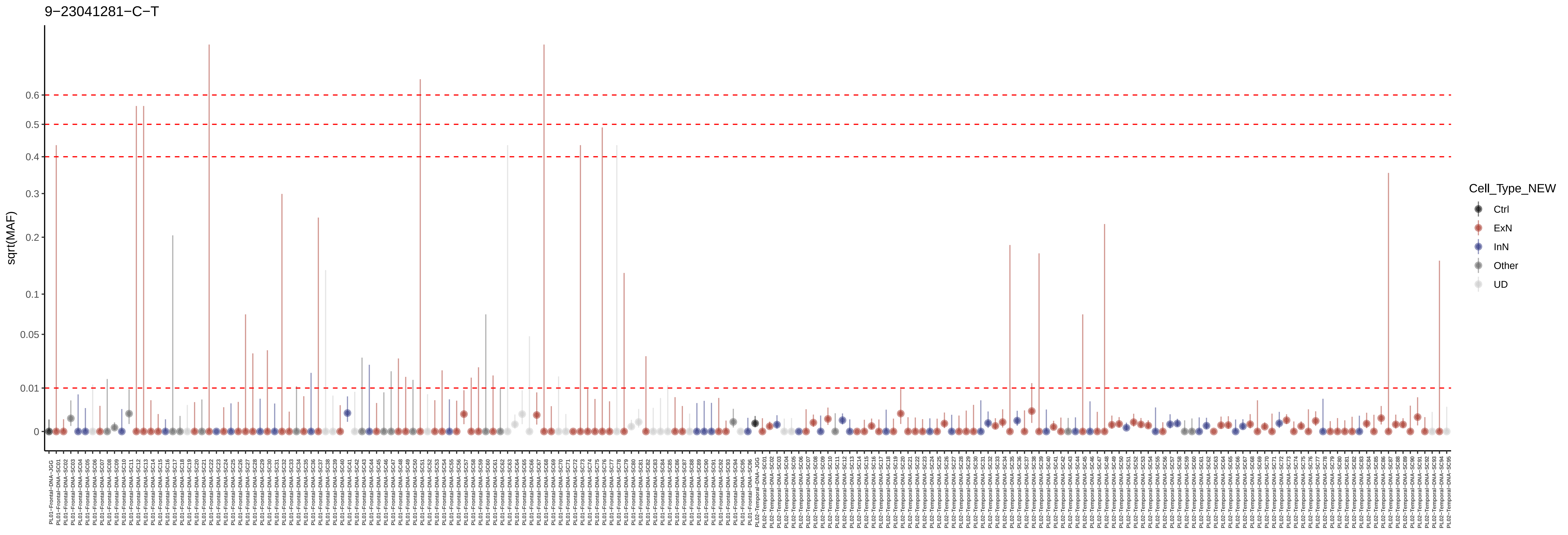
9−23041281−C−T
sqrt(MAF)
0.6
0.5
0.4
0.3
0.2
0.1
0.05
0.01
0
Cell_Type_NEW
Ctrl
ExN
InN
Other
UD

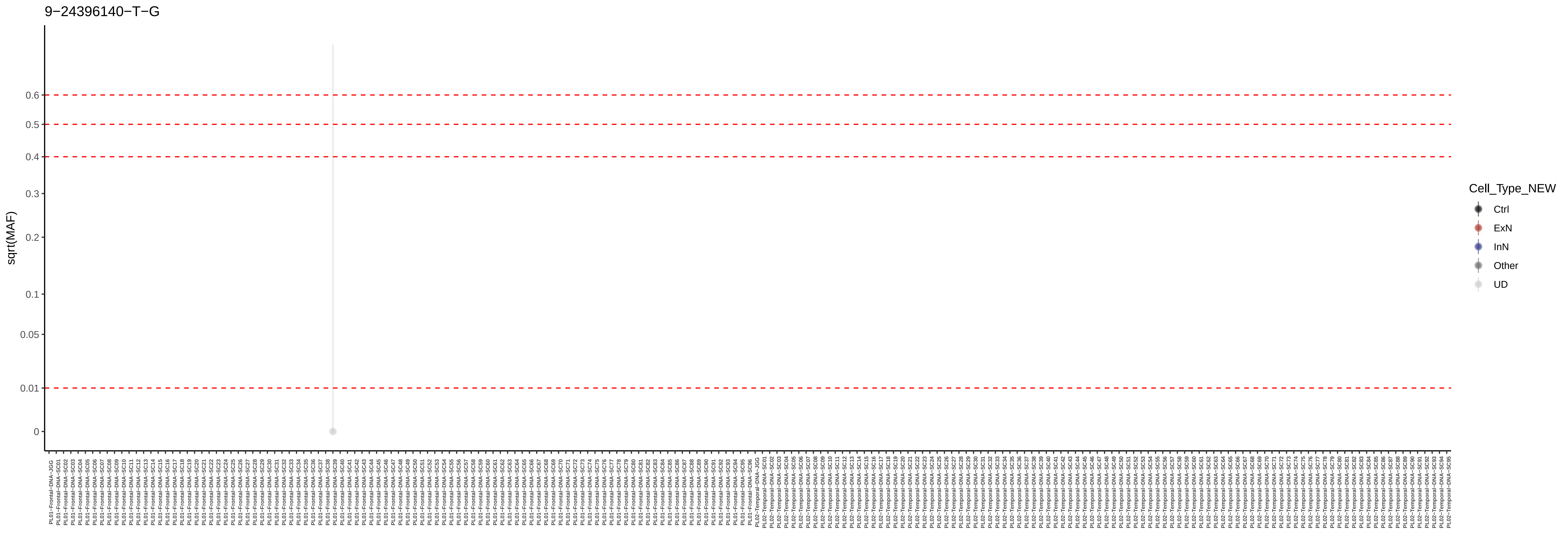
9−24396140−T−G
sqrt(MAF)
0.6
0.5
0.4
0.3
0.2
0.1
0.05
0.01
0
Cell_Type_NEW
Ctrl
ExN
InN
Other
UD

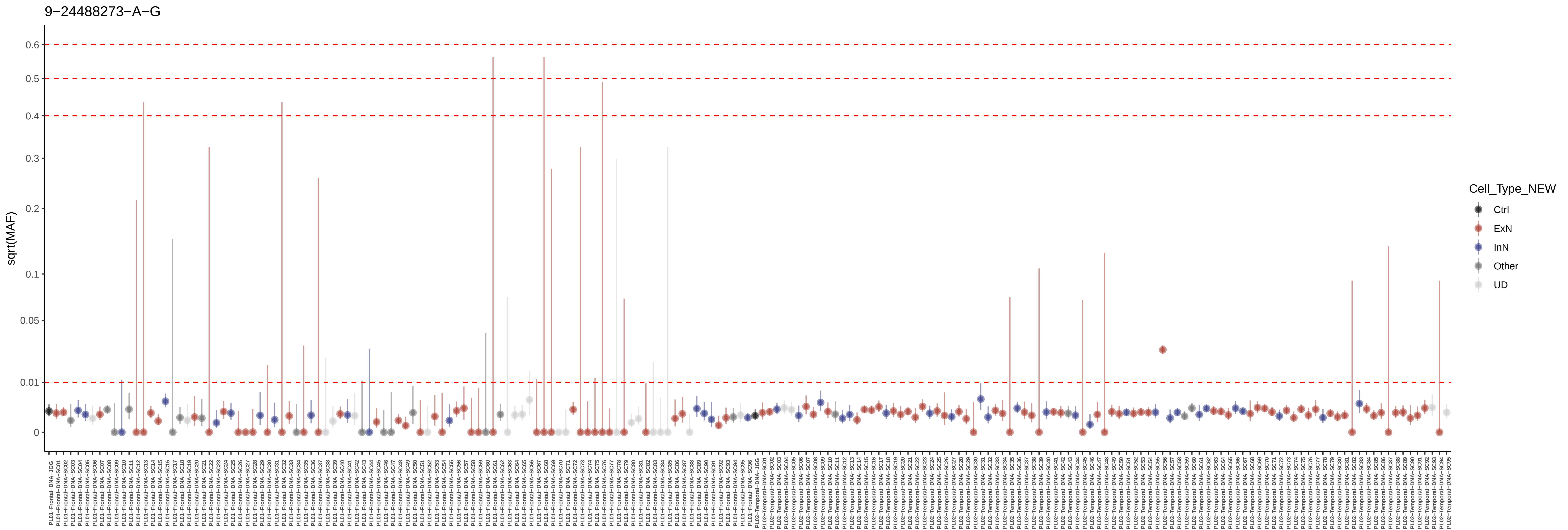
9−24488273−A−G
sqrt(MAF)
0.6
0.5
0.4
0.3
0.2
0.1
0.05
0.01
0
Cell_Type_NEW
Ctrl
ExN
InN
Other
UD

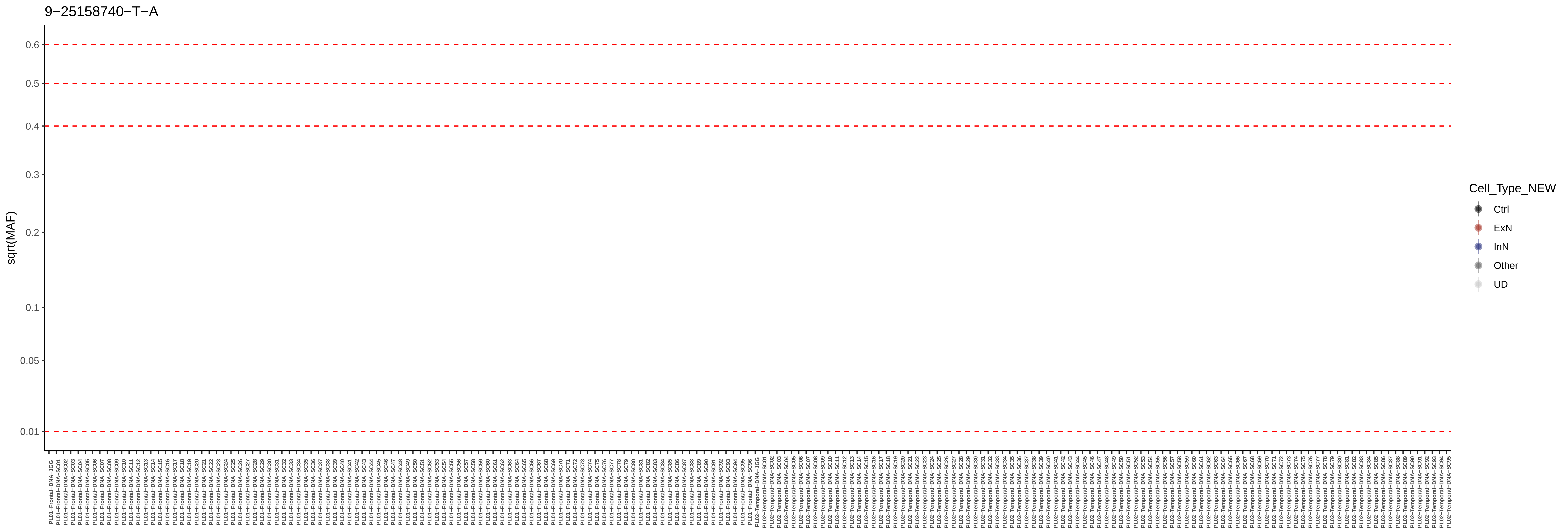
9−25158740−T−A
sqrt(MAF)
0.6
0.5
0.4
0.3
0.2
0.1
0.05
0.01
Cell_Type_NEW
Ctrl
ExN
InN
Other
UD

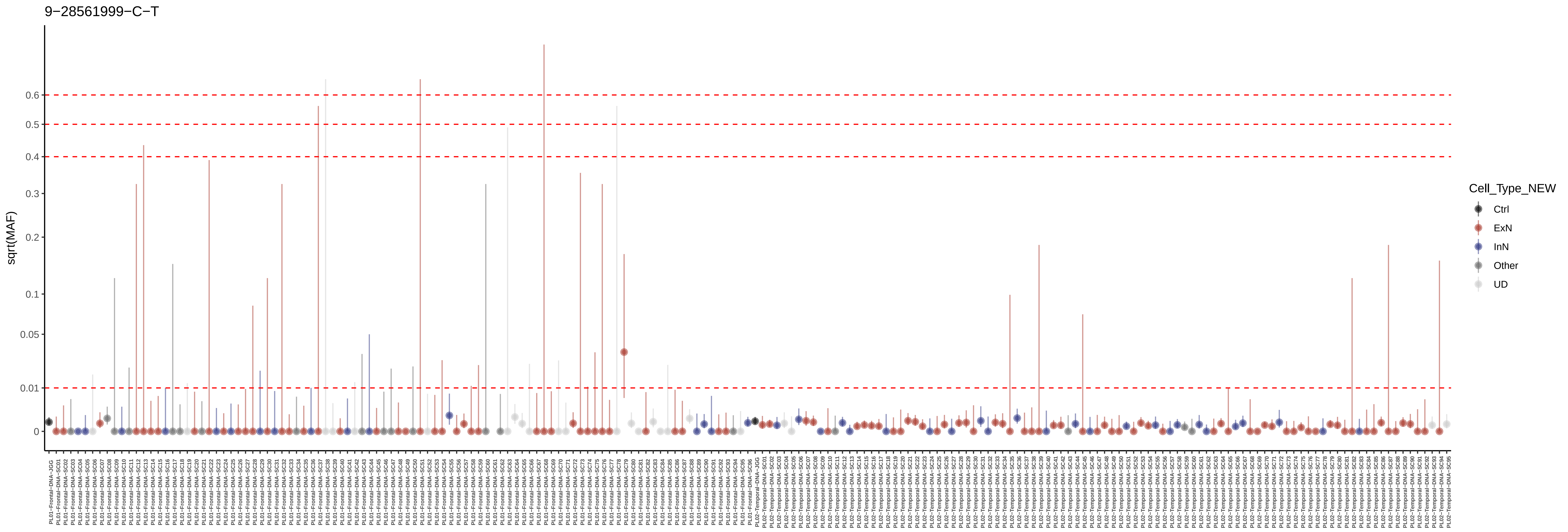
9−28561999−C−T
sqrt(MAF)
0.6
0.5
0.4
0.3
0.2
0.1
0.05
0.01
0
Cell_Type_NEW
Ctrl
ExN
InN
Other
UD

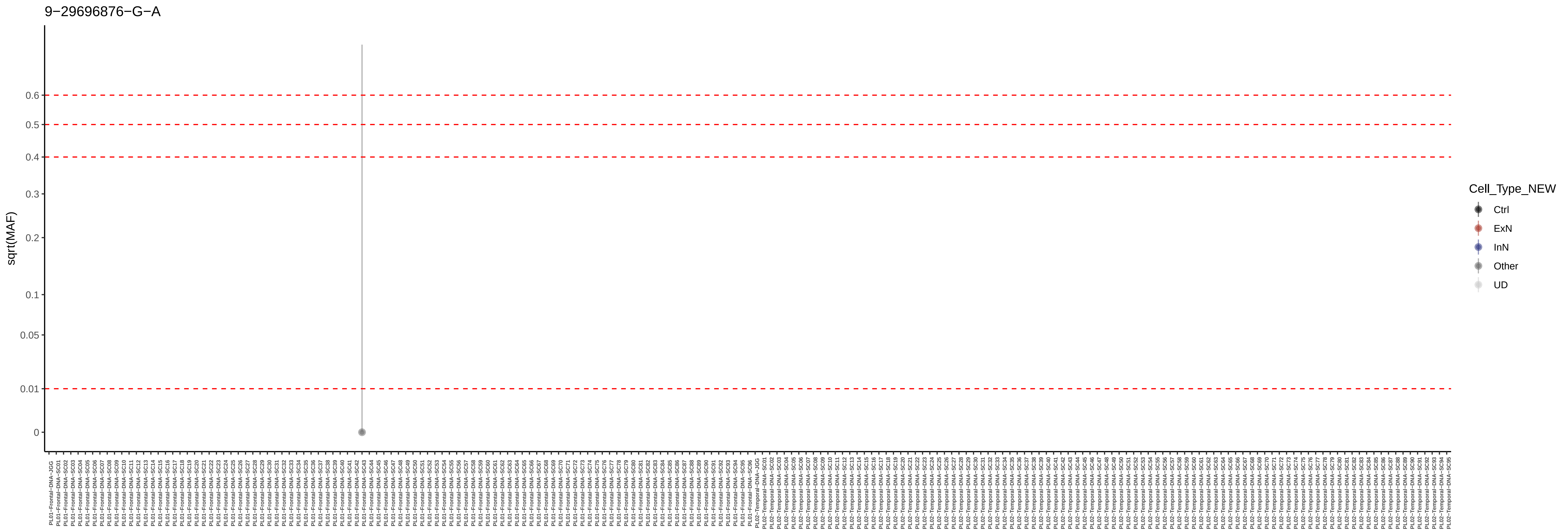
9−29696876−G−A
sqrt(MAF)
0.6
0.5
0.4
0.3
0.2
0.1
0.05
0.01
0
Cell_Type_NEW
Ctrl
ExN
InN
Other
UD

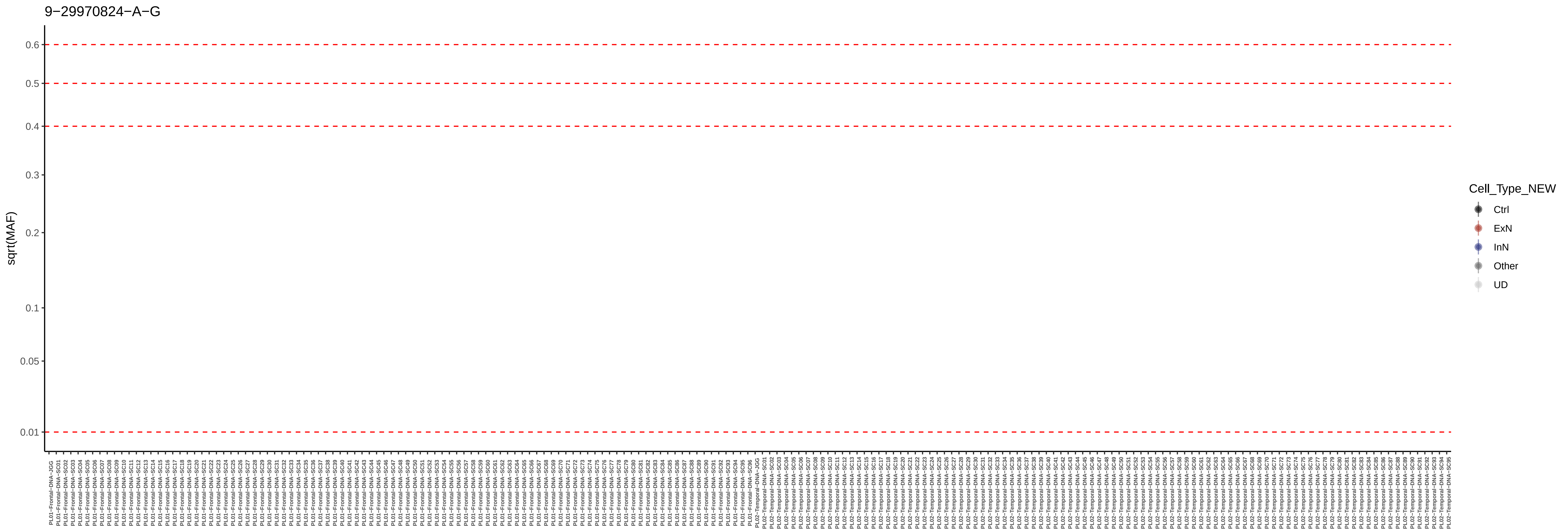
9−29970824−A−G
sqrt(MAF)
0.6
0.5
0.4
0.3
0.2
0.1
0.05
0.01
Cell_Type_NEW
Ctrl
ExN
InN
Other
UD

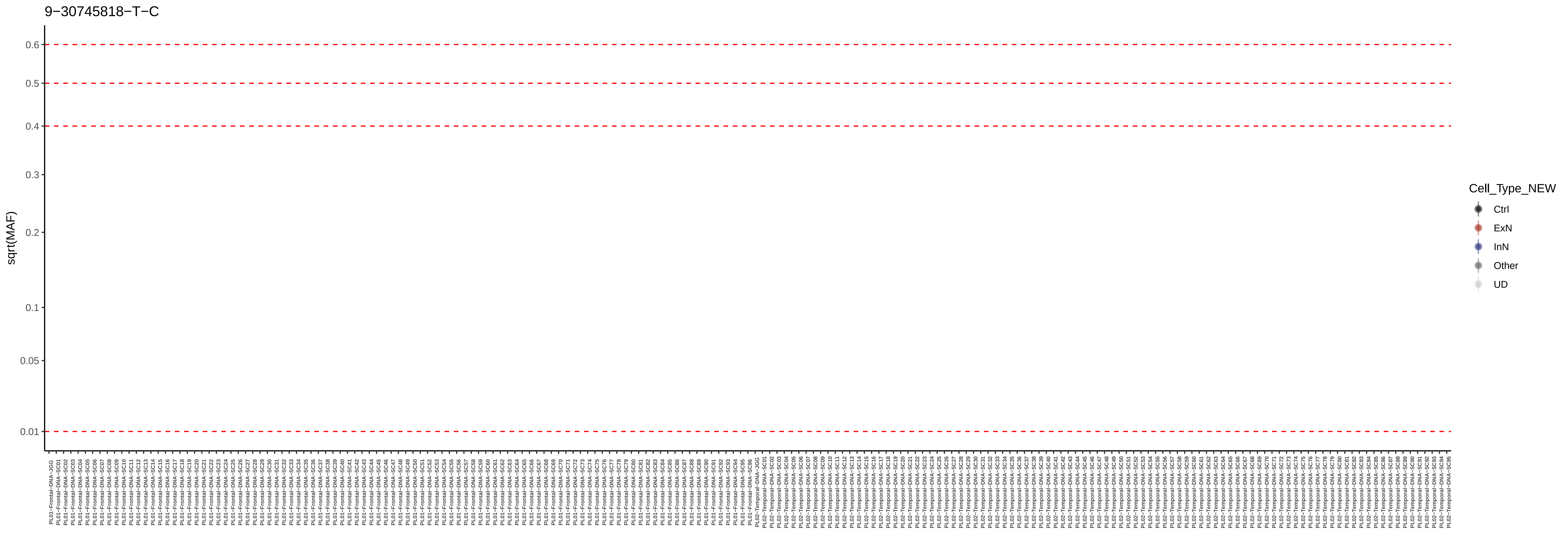
9−30745818−T−C
sqrt(MAF)
0.6
0.5
0.4
0.3
0.2
0.1
0.05
0.01
Cell_Type_NEW
Ctrl
ExN
InN
Other
UD

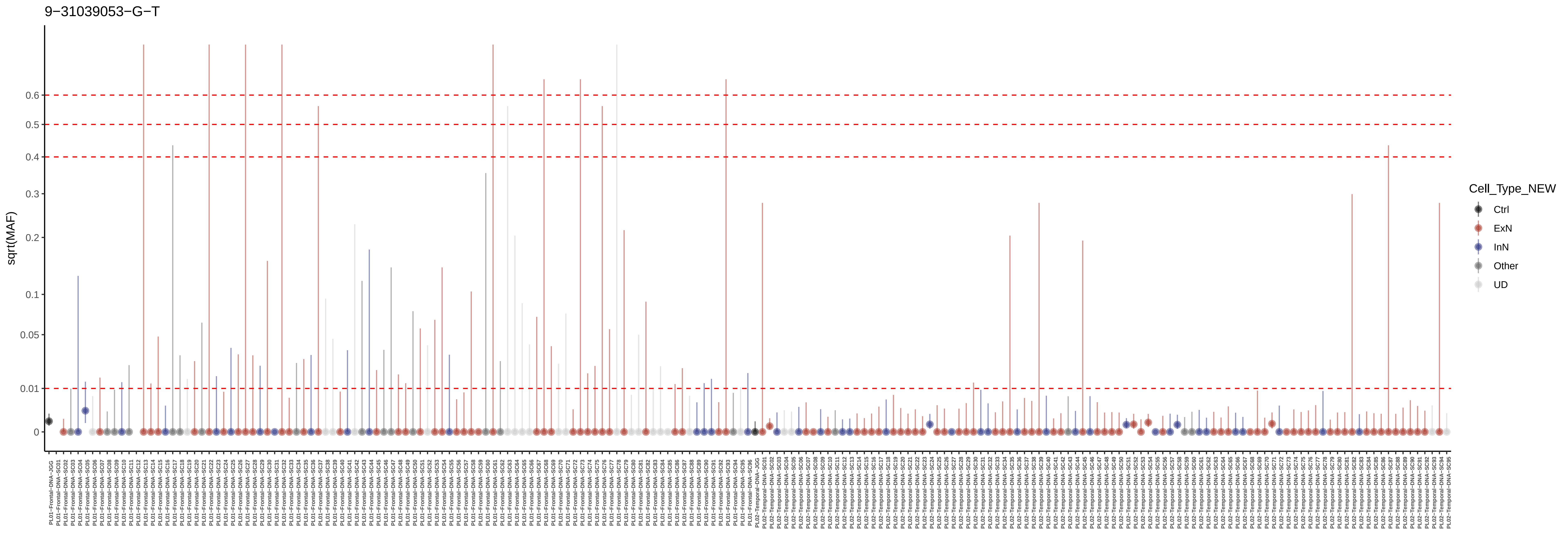
9−31039053−G−T
sqrt(MAF)
0.6
0.5
0.4
0.3
0.2
0.1
0.05
0.01
0
Cell_Type_NEW
Ctrl
ExN
InN
Other
UD

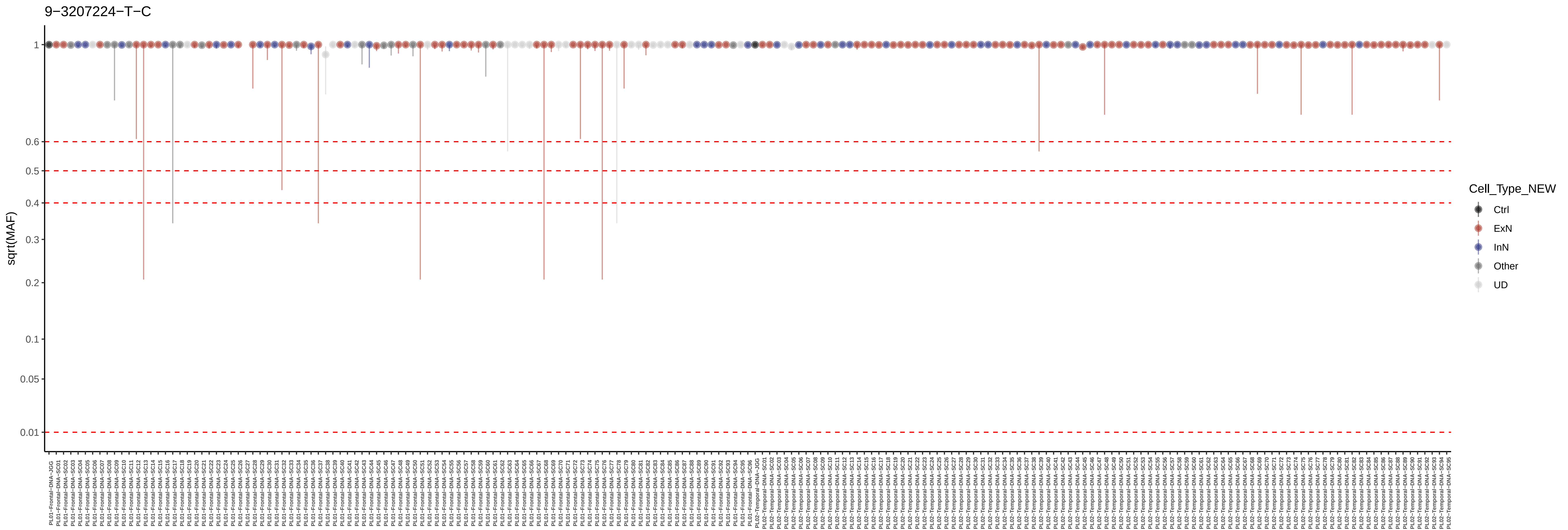
9−3207224−T−C
sqrt(MAF)
1
0.6
0.5
0.4
0.3
0.2
0.1
0.05
0.01
Cell_Type_NEW
Ctrl
ExN
InN
Other
UD

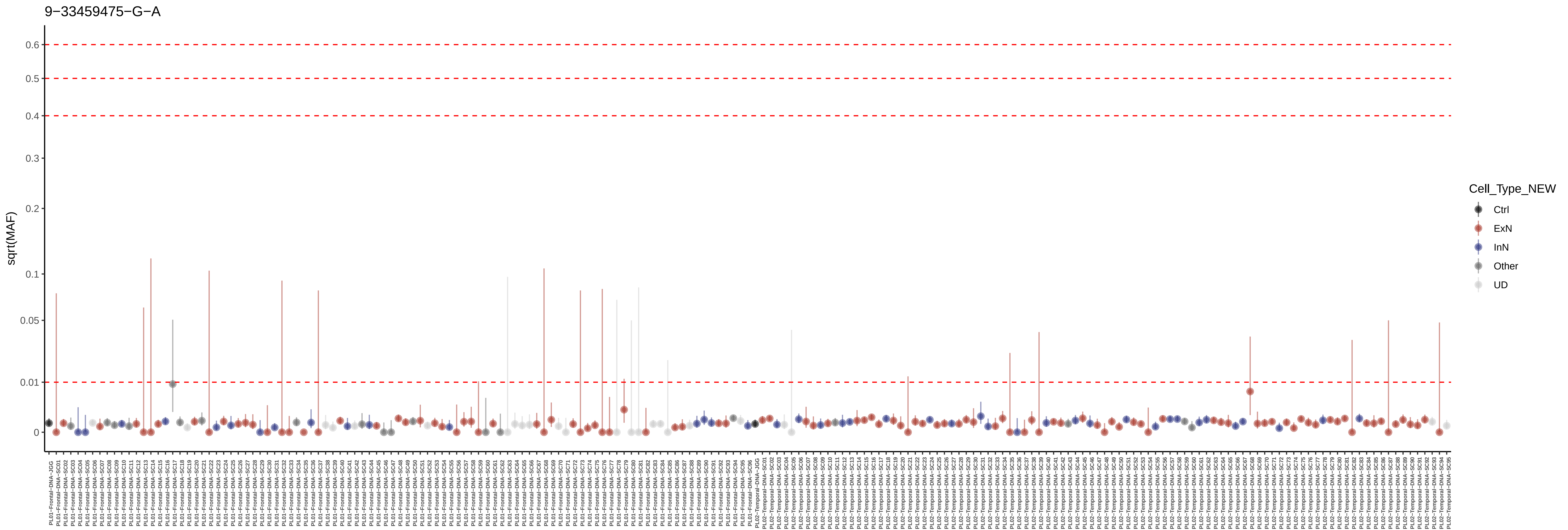
9−33459475−G−A
sqrt(MAF)
0.6
0.5
0.4
0.3
0.2
0.1
0.05
0.01
0
Cell_Type_NEW
Ctrl
ExN
InN
Other
UD

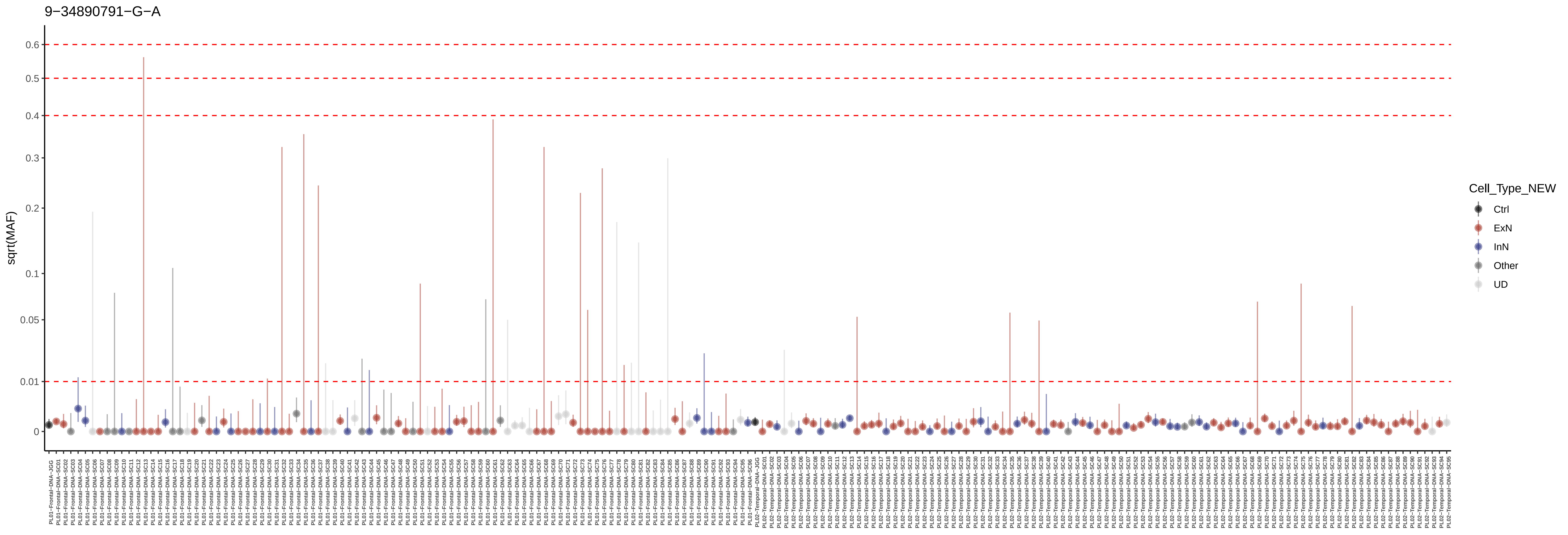

9−34890791−G−A
sqrt(MAF)
0.6
0.5
0.4
0.3
0.2
0.1
0.05
0.01
0
Cell_Type_NEW
Ctrl
ExN
InN
Other
UD

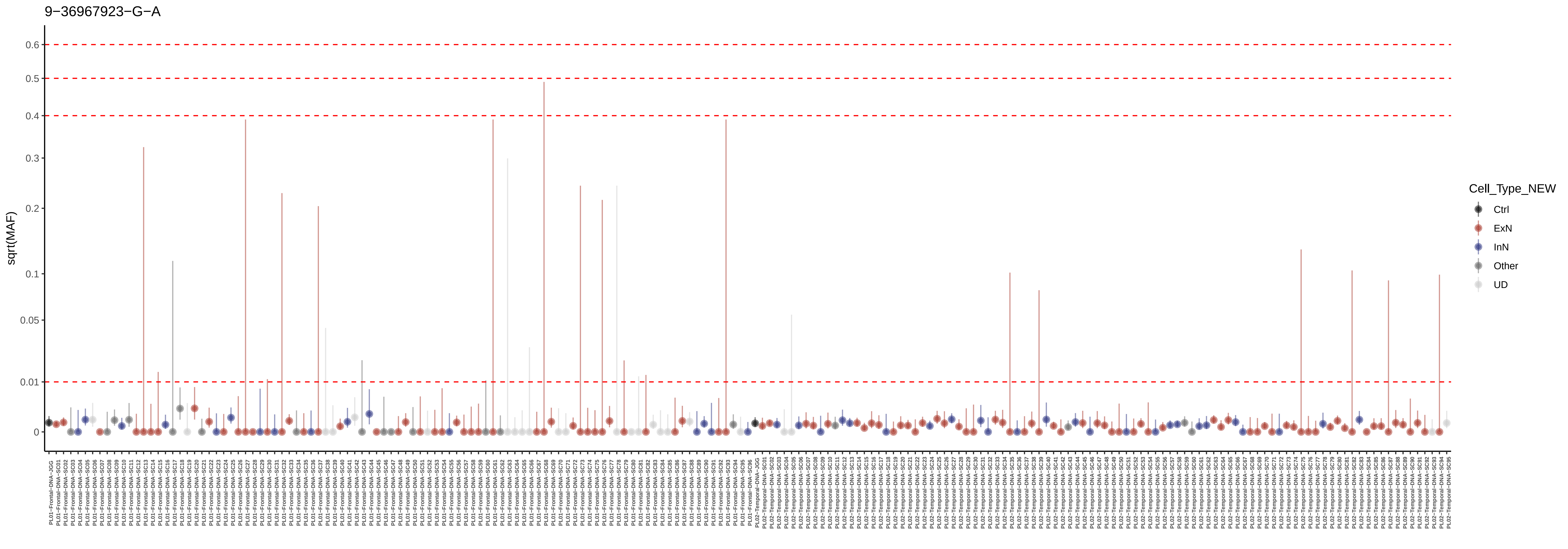

9−36967923−G−A
sqrt(MAF)
0.6
0.5
0.4
0.3
0.2
0.1
0.05
0.01
0
Cell_Type_NEW
Ctrl
ExN
InN
Other
UD

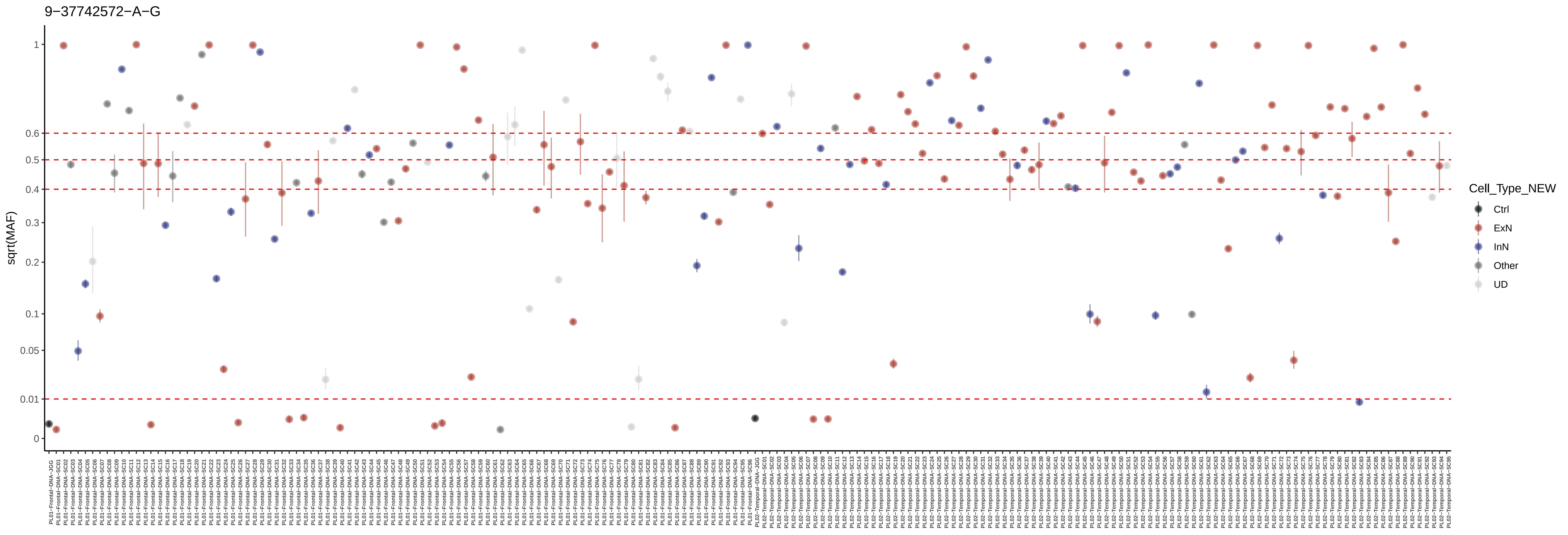

9−37742572−A−G
sqrt(MAF)
Cell_Type_NEW
Ctrl
ExN
InN
Other
UD

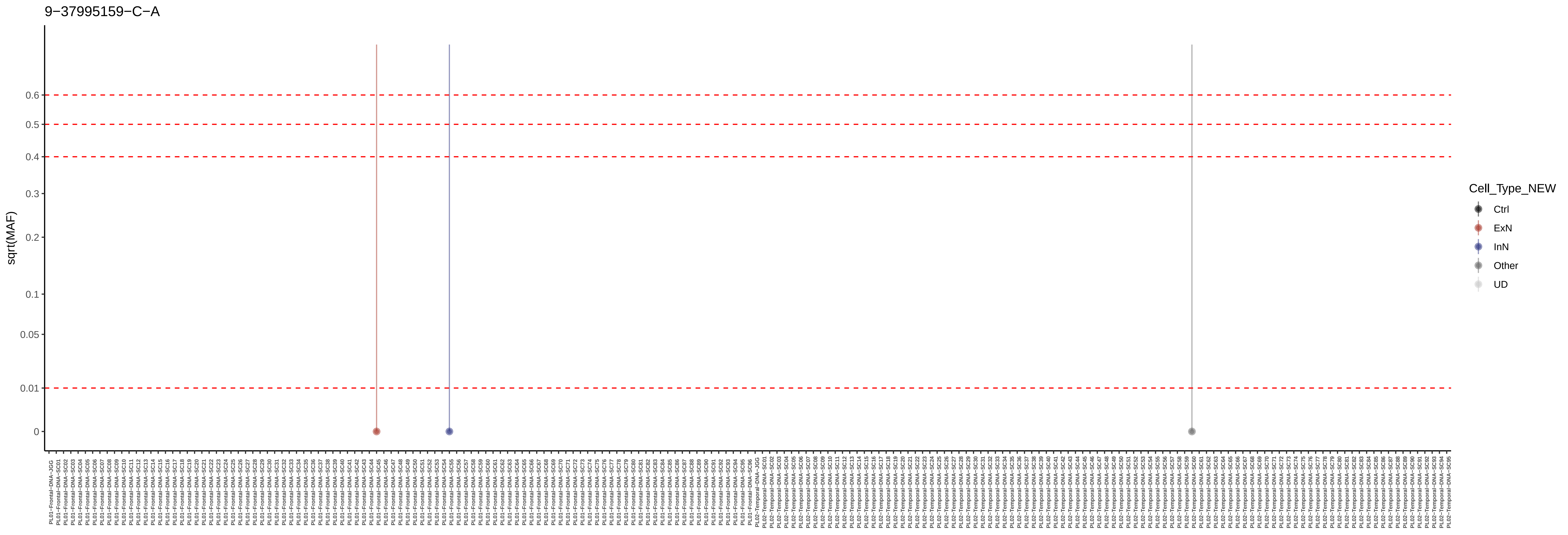

9−37995159−C−A
sqrt(MAF)
0.6
0.5
0.4
0.3
0.2
0.1
0.05
0.01
0
Cell_Type_NEW
Ctrl
ExN
InN
Other
UD

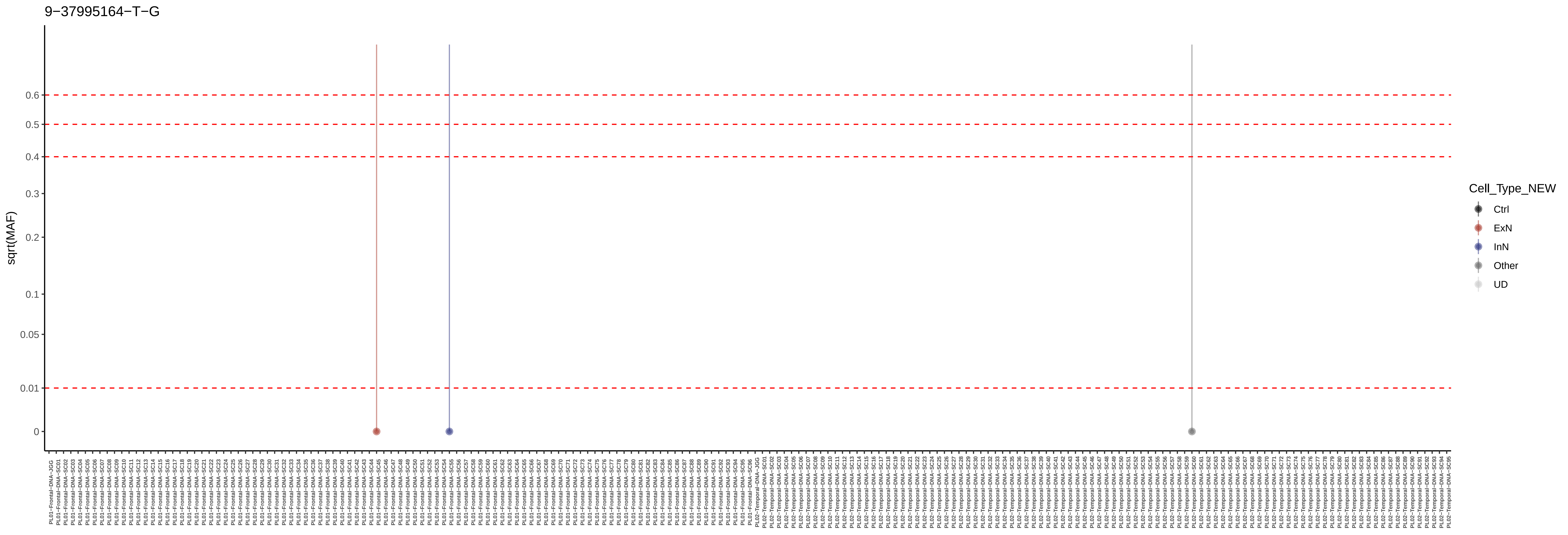

9−37995164−T−G
sqrt(MAF)
0.6
0.5
0.4
0.3
0.2
0.1
0.05
0.01
0
Cell_Type_NEW
Ctrl
ExN
InN
Other
UD

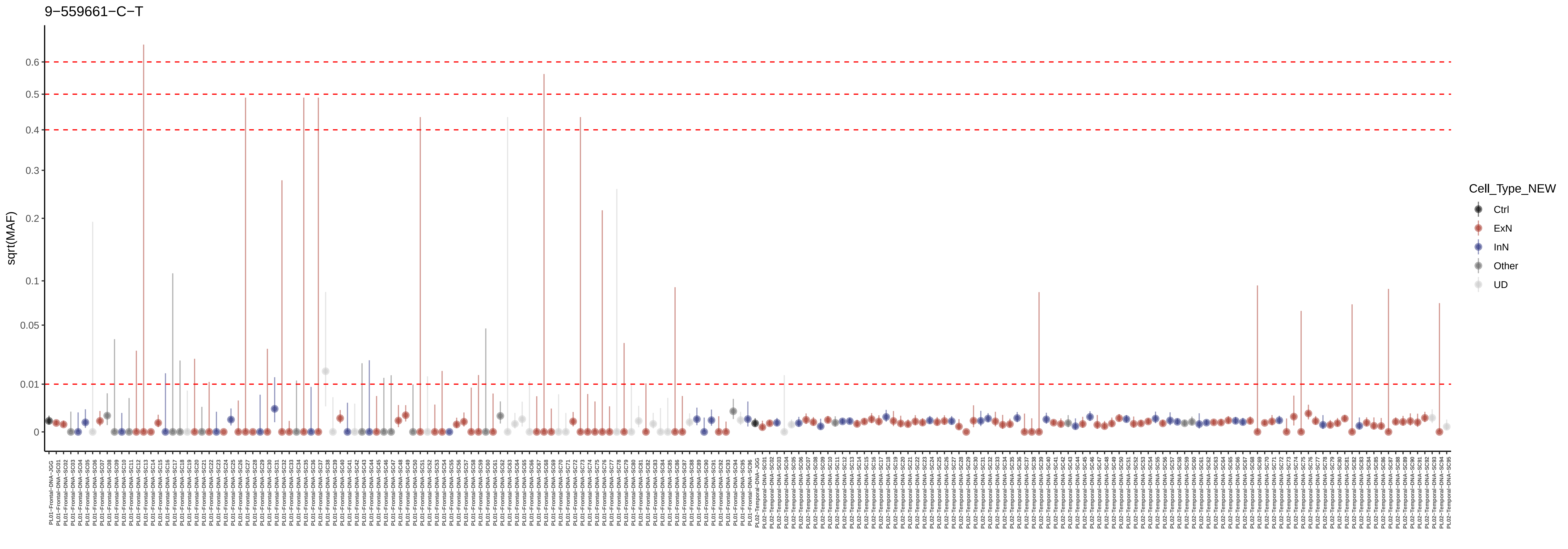
9−559661−C−T
sqrt(MAF)
0.6
0.5
0.4
0.3
0.2
0.1
0.05
0.01
0
Cell_Type_NEW
Ctrl
ExN
InN
Other
UD

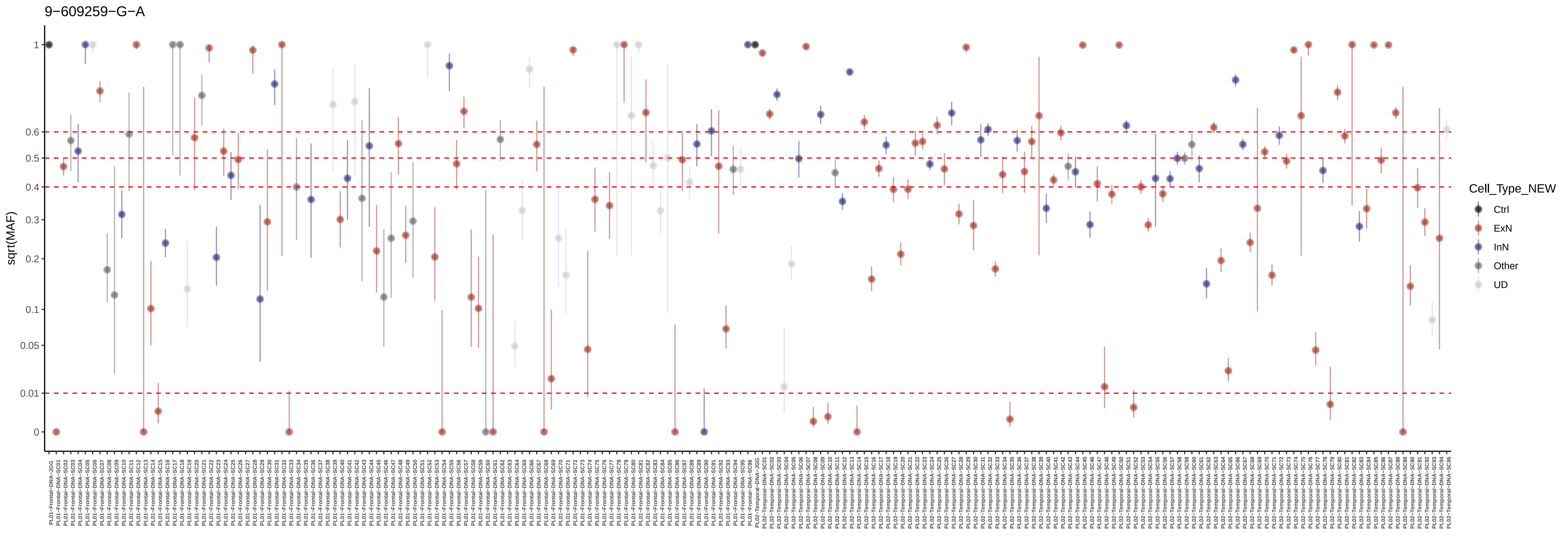

9−609259−G−A
sqrt(MAF)
1
0.6
0.5
0.4
0.3
0.2
0.1
0.05
0.01
0
Cell_Type_NEW
Ctrl
ExN
InN
Other
UD

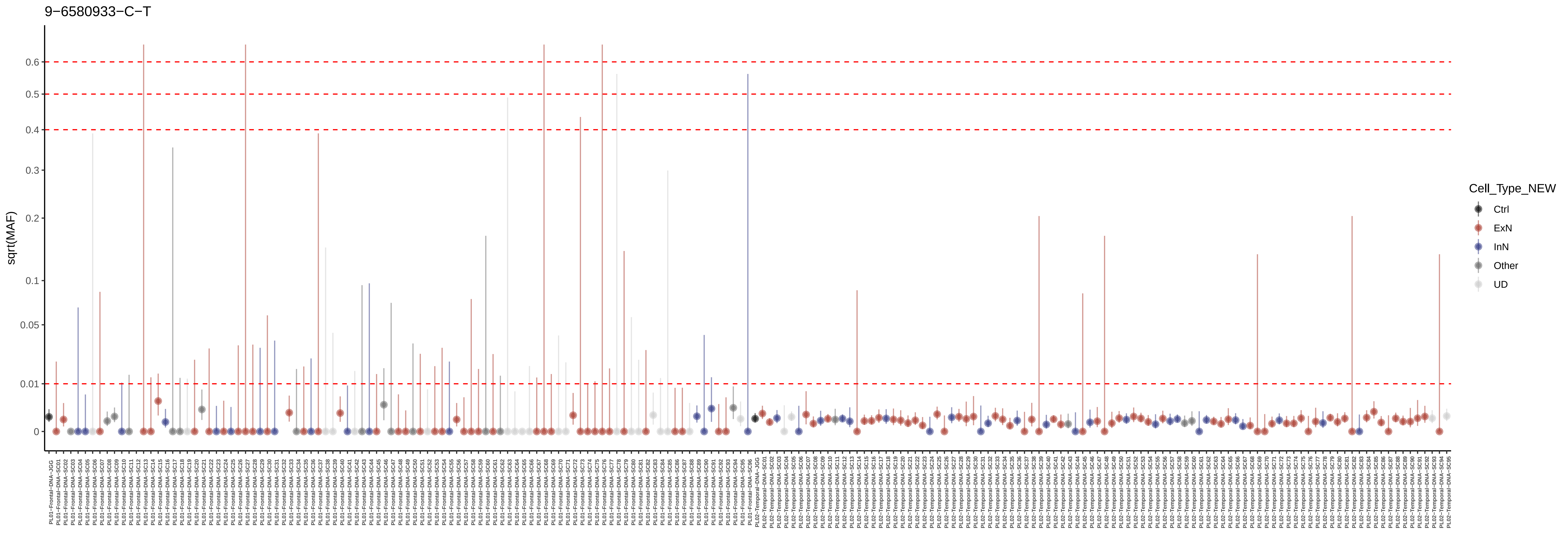
9−6580933−C−T
sqrt(MAF)
0
0.01
0.05
0.1
0.2
0.3
0.4
0.5
0.6
Cell_Type_NEW
Ctrl
ExN
InN
Other
UD

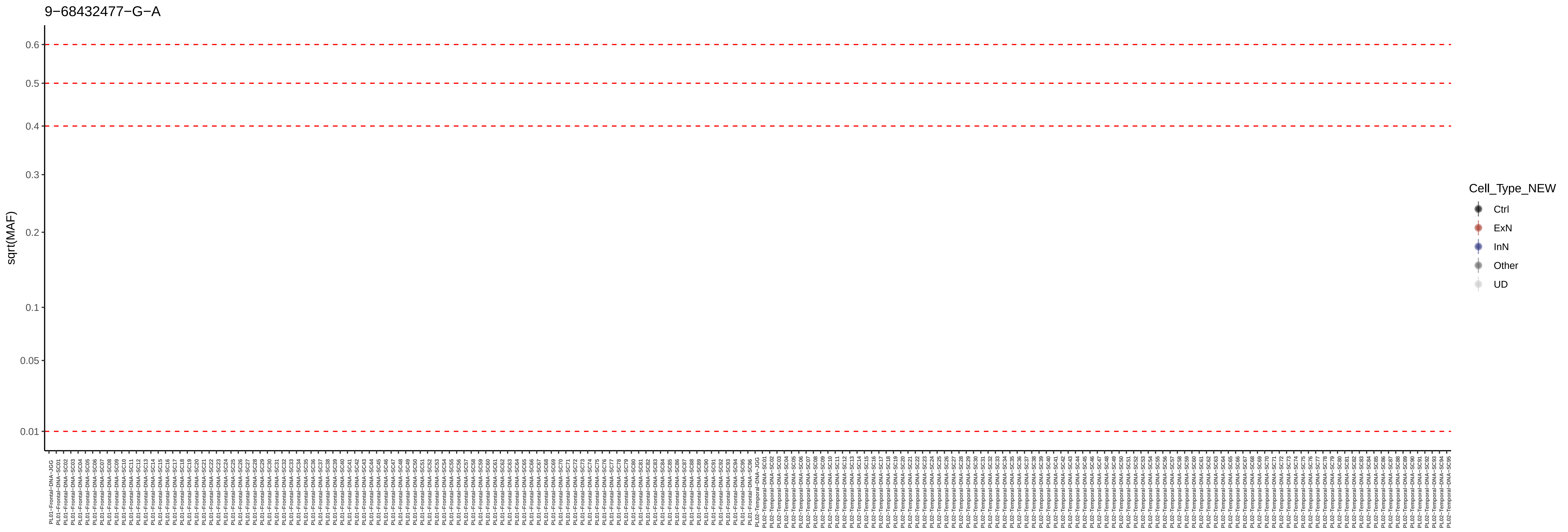
9−68432477−G−A
sqrt(MAF)
0.6
0.5
0.4
0.3
0.2
0.1
0.05
0.01
Cell_Type_NEW
Ctrl
ExN
InN
Other
UD

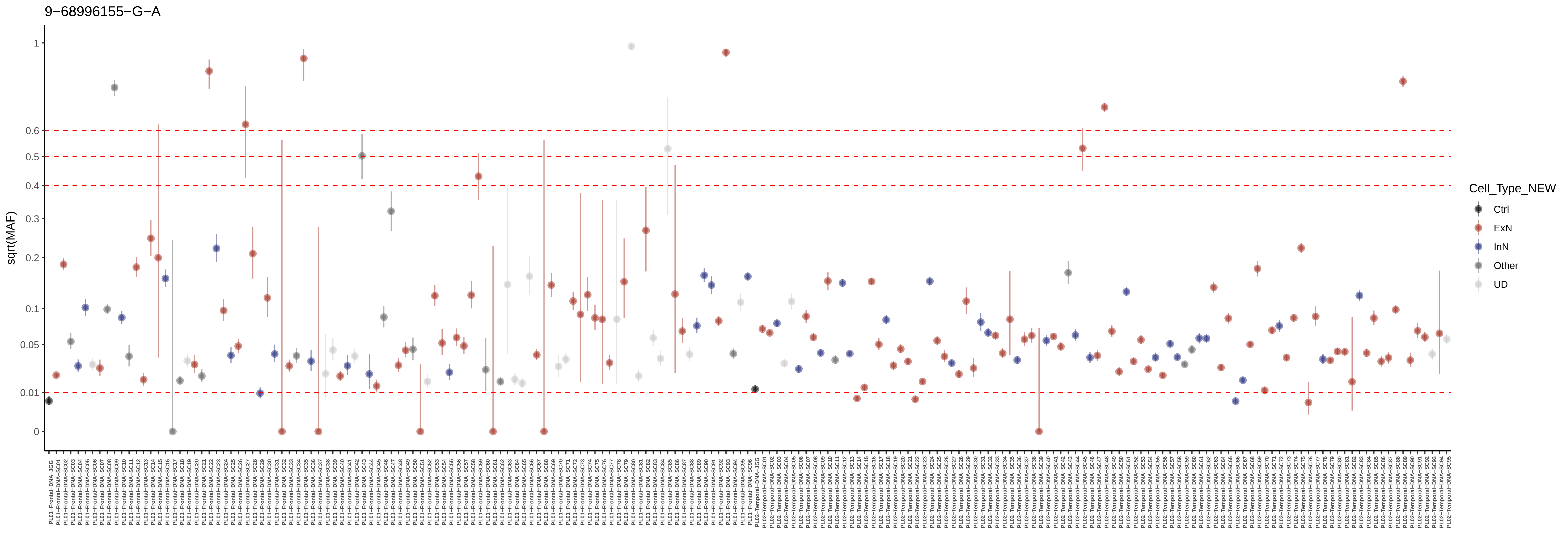
9−68996155−G−A
sqrt(MAF)
1
0.6
0.5
0.4
0.3
0.2
0.1
0.05
0.01
0
Cell_Type_NEW
Ctrl
ExN
InN
Other
UD

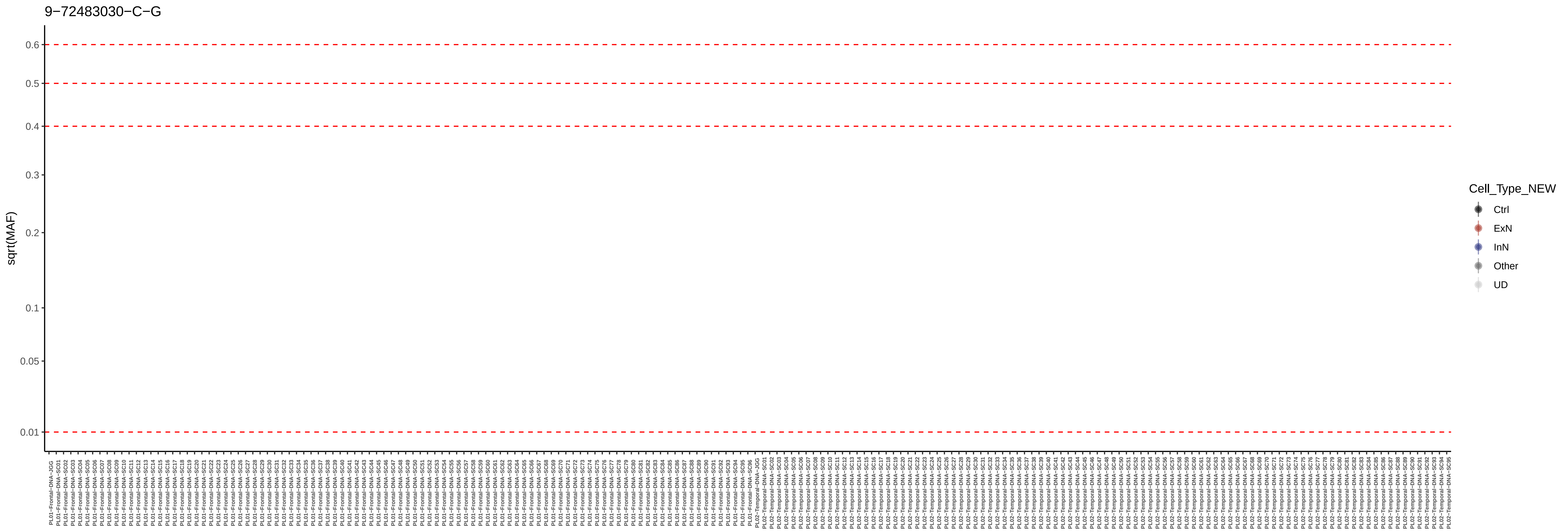

9−72483030−C−G
sqrt(MAF)
0.6
0.5
0.4
0.3
0.2
0.1
0.05
0.01
Cell_Type_NEW
Ctrl
ExN
InN
Other
UD

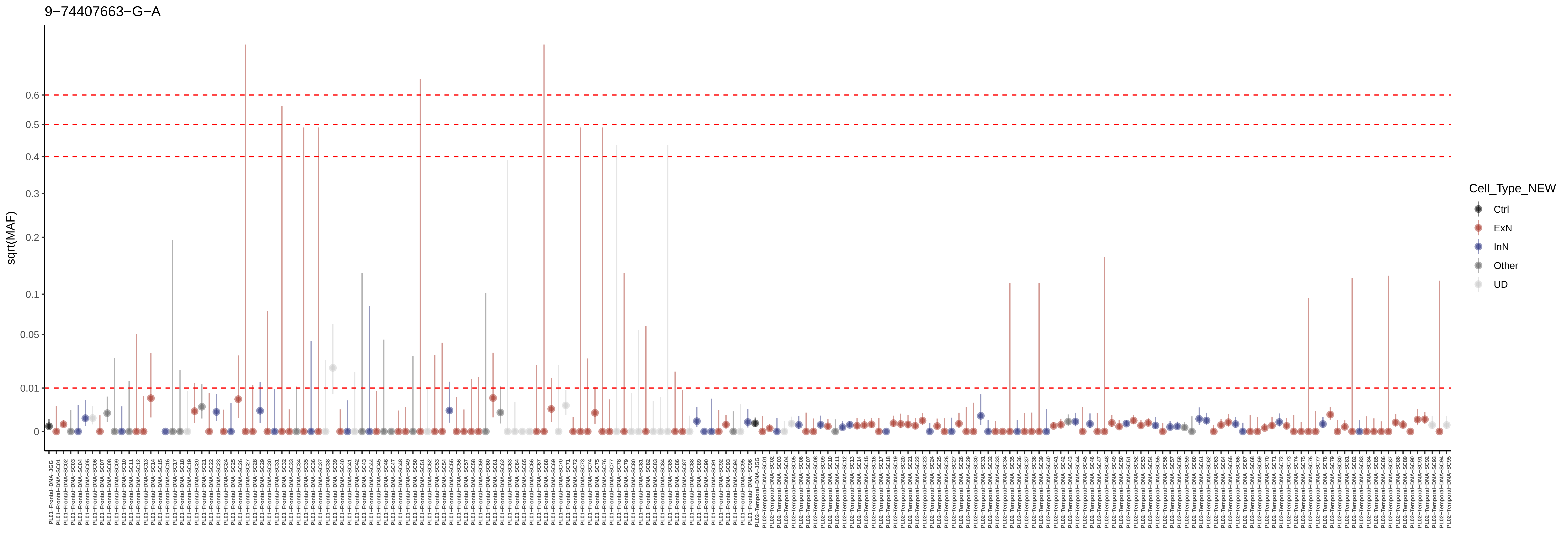
9−74407663−G−A
sqrt(MAF)
0.6
0.5
0.4
0.3
0.2
0.1
0.05
0.01
0
Cell_Type_NEW
Ctrl
ExN
InN
Other
UD

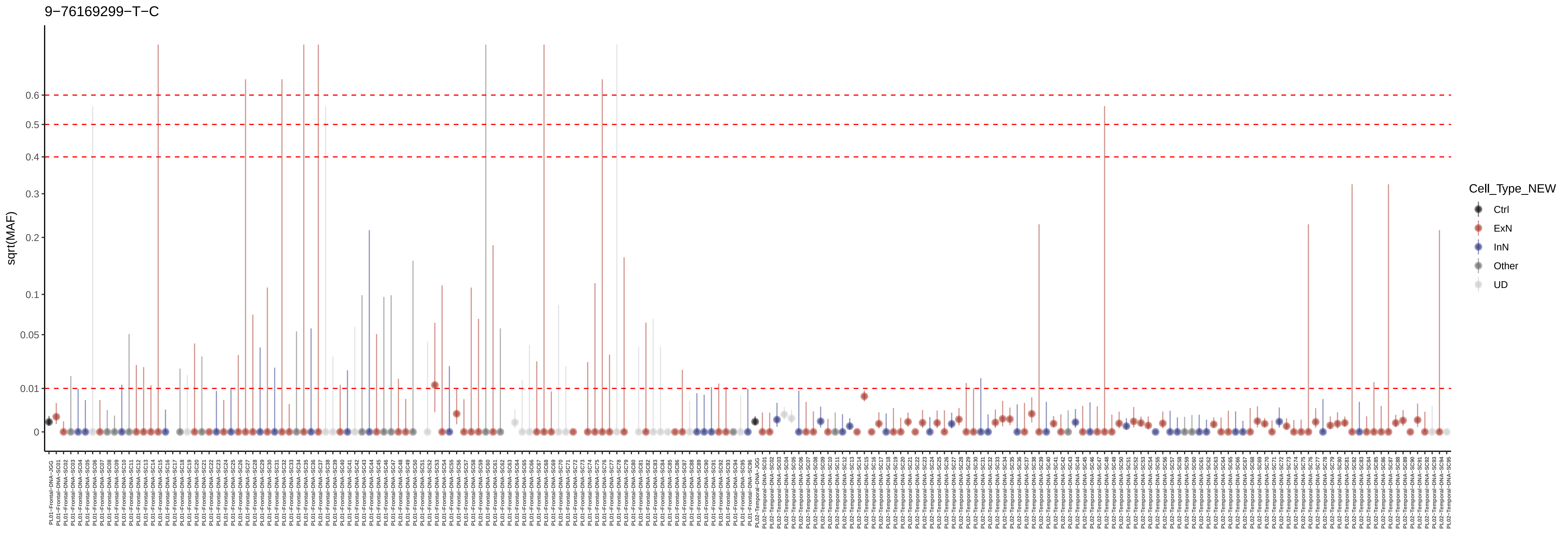
9−76169299−T−C
sqrt(MAF)
0.6
0.5
0.4
0.3
0.2
0.1
0.05
0.01
0
Cell_Type_NEW
Ctrl
ExN
InN
Other
UD

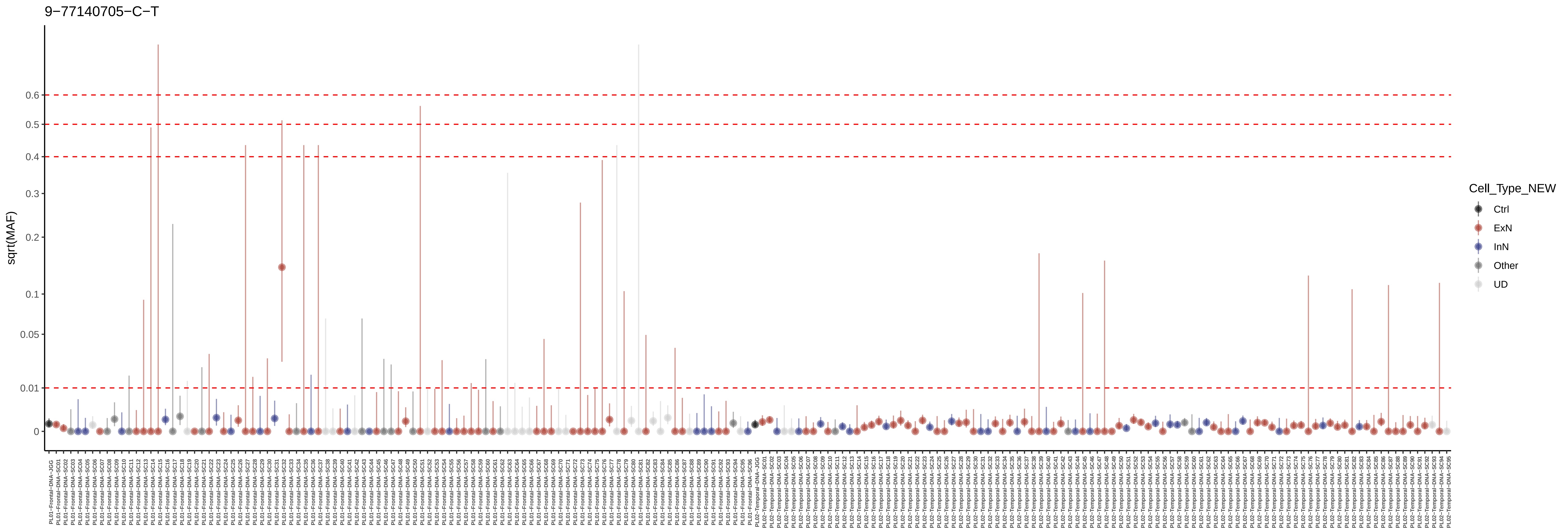
9−77140705−C−T
sqrt(MAF)
0.6
0.5
0.4
0.3
0.2
0.1
0.05
0.01
0
Cell_Type_NEW
Ctrl
ExN
InN
Other
UD

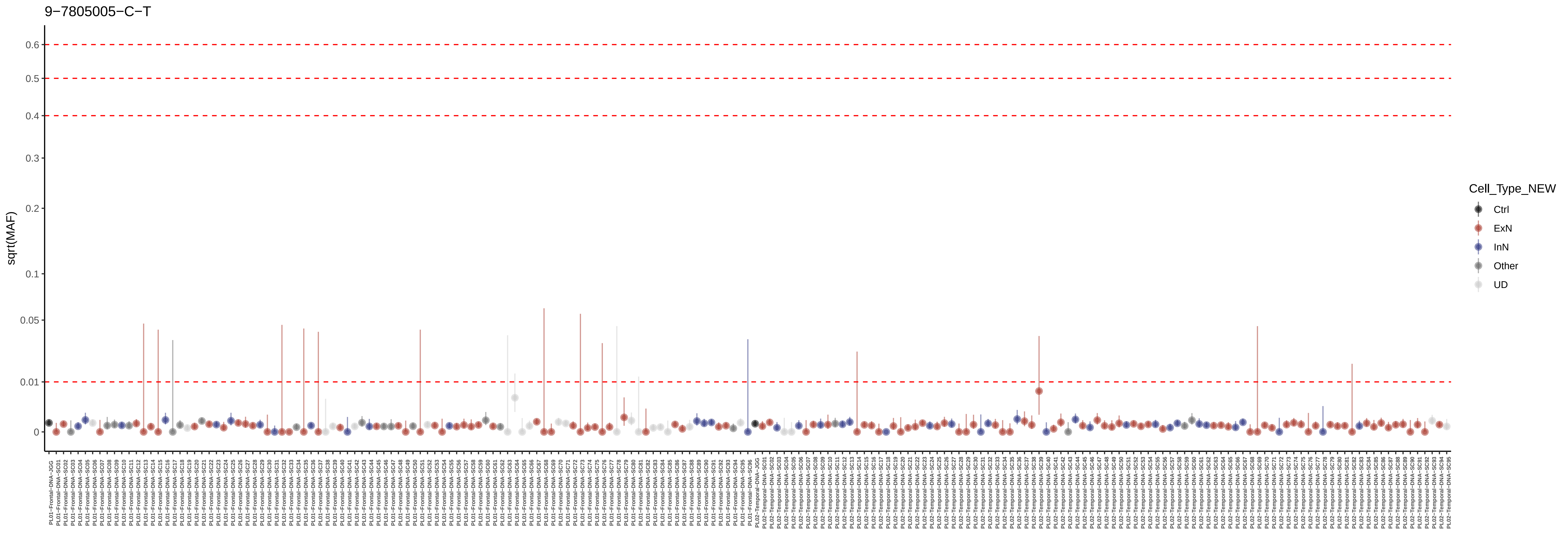

9−7805005−C−T
sqrt(MAF)
0.6
0.5
0.4
0.3
0.2
0.1
0.05
0.01
0
Cell_Type_NEW
Ctrl
ExN
InN
Other
UD

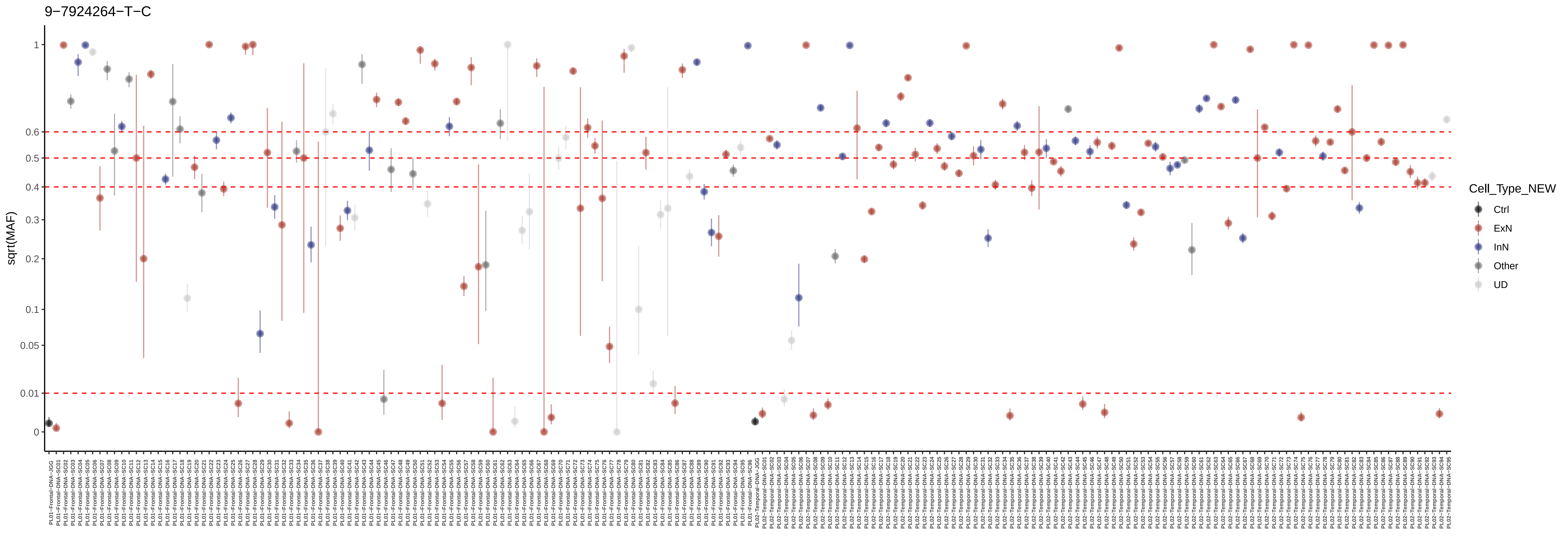
9−7924264−T−C
sqrt(MAF)
1
0.6
0.5
0.4
0.3
0.2
0.1
0.05
0.01
0
Cell_Type_NEW
Ctrl
ExN
InN
Other
UD

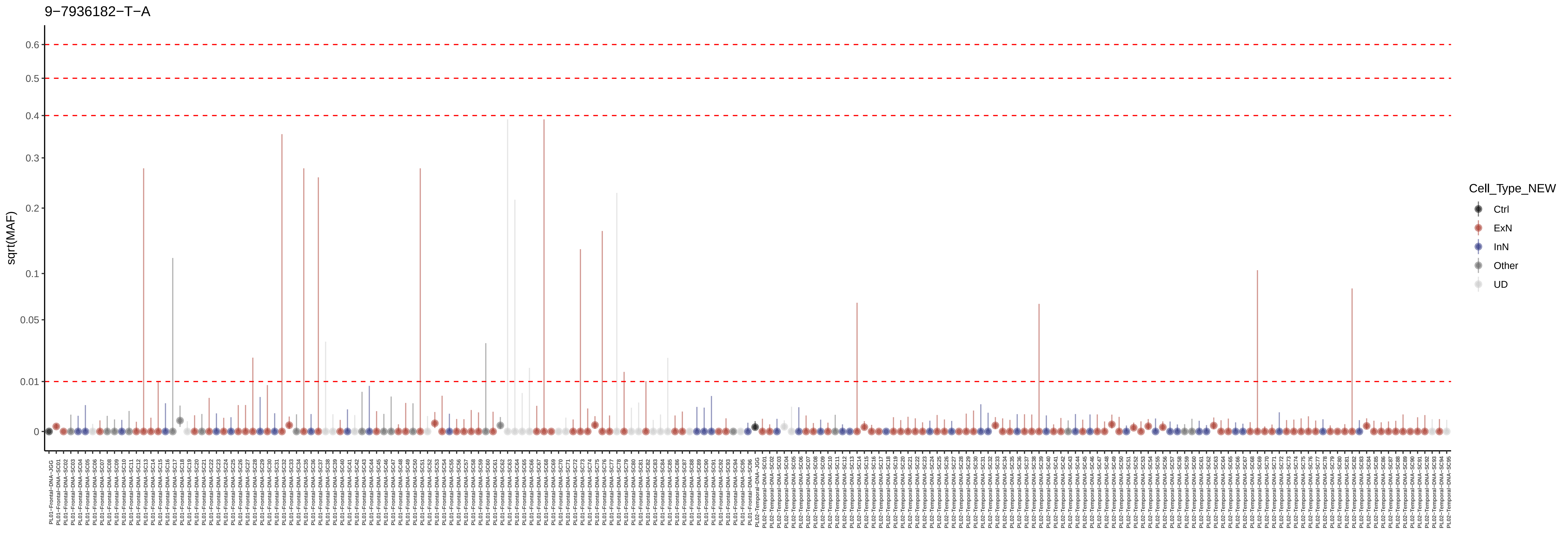

9−7936182−T−A
sqrt(MAF)
0.6
0.5
0.4
0.3
0.2
0.1
0.05
0.01
0
Cell_Type_NEW
Ctrl
ExN
InN
Other
UD

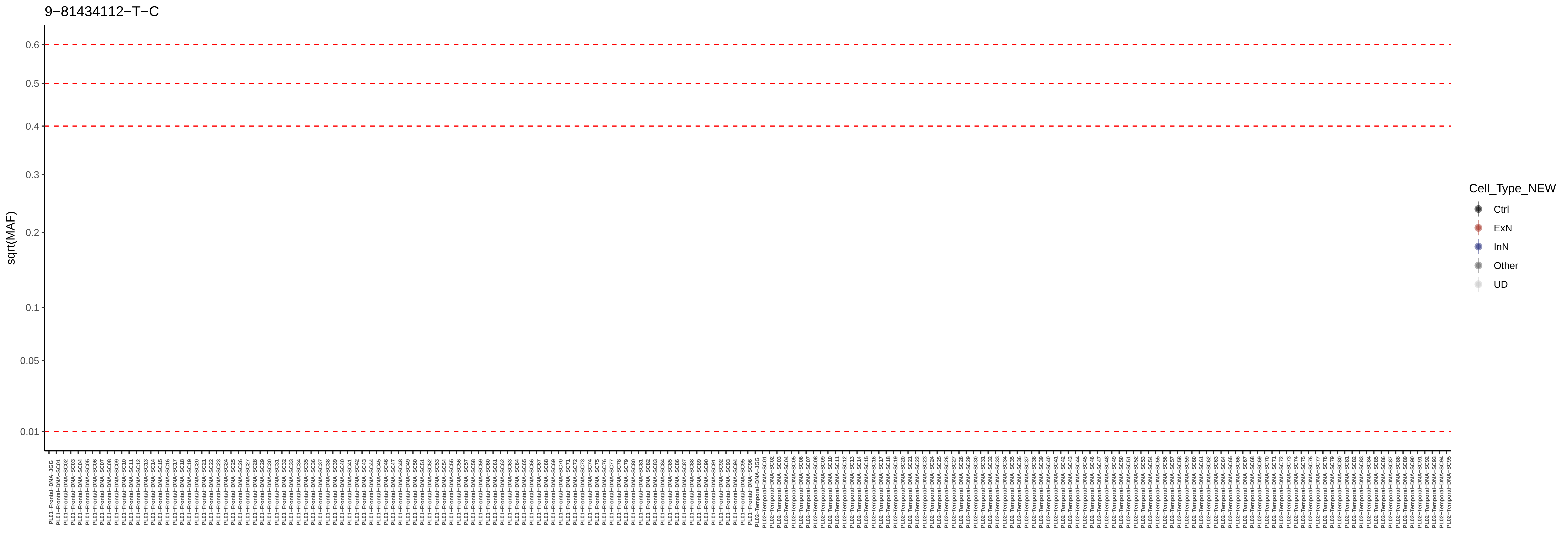

9−81434112−T−C
sqrt(MAF)
0.6
0.5
0.4
0.3
0.2
0.1
0.05
0.01
Cell_Type_NEW
Ctrl
ExN
InN
Other
UD

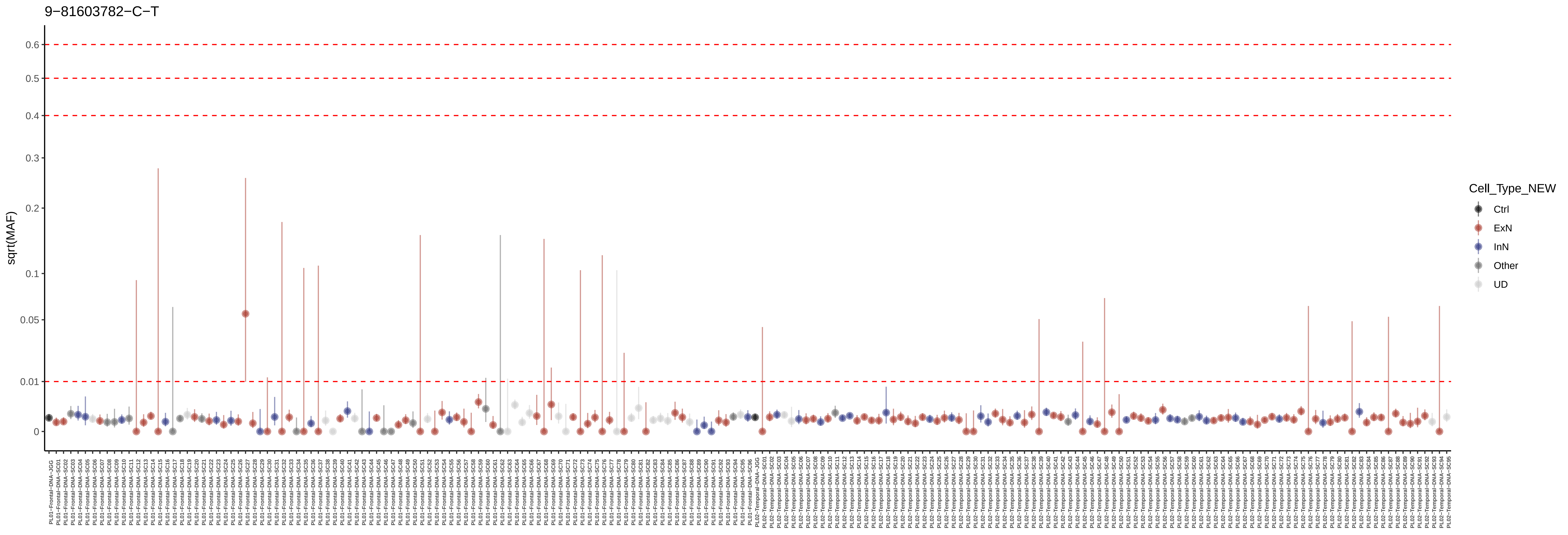
9−81603782−C−T
sqrt(MAF)
Cell_Type_NEW
Ctrl
ExN
InN
Other
UD

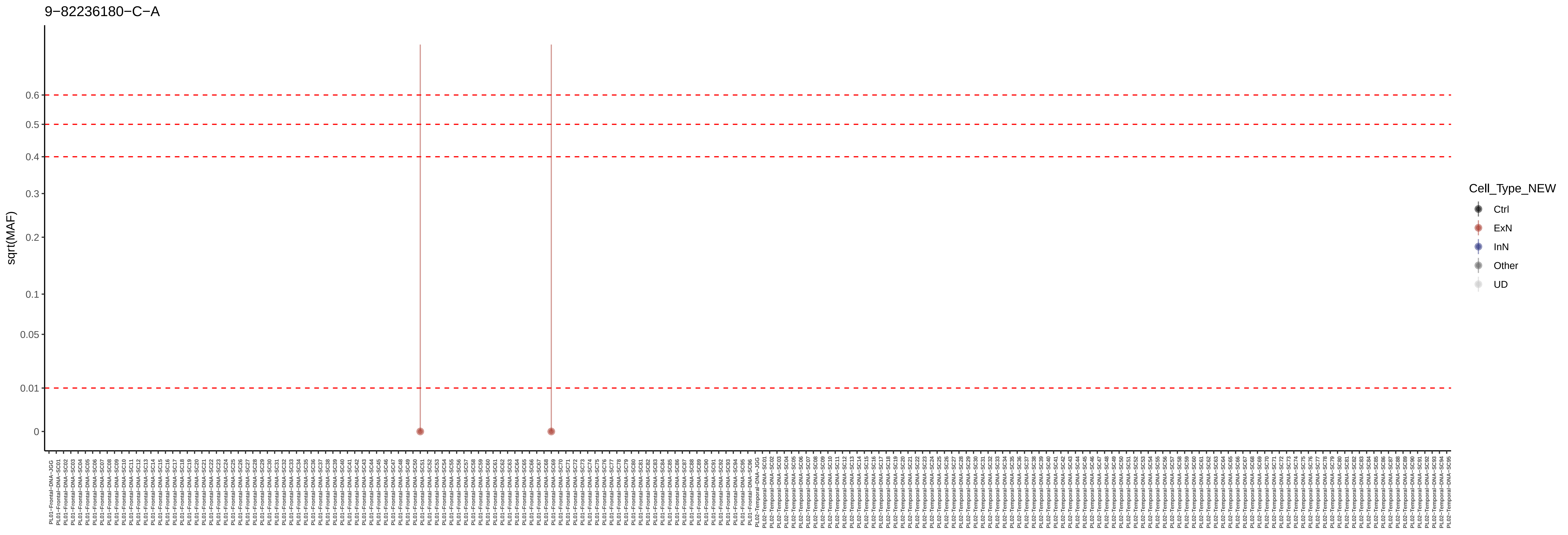

9−82236180−C−A
sqrt(MAF)
0.6
0.5
0.4
0.3
0.2
0.1
0.05
0.01
0
Cell_Type_NEW
Ctrl
ExN
InN
Other
UD

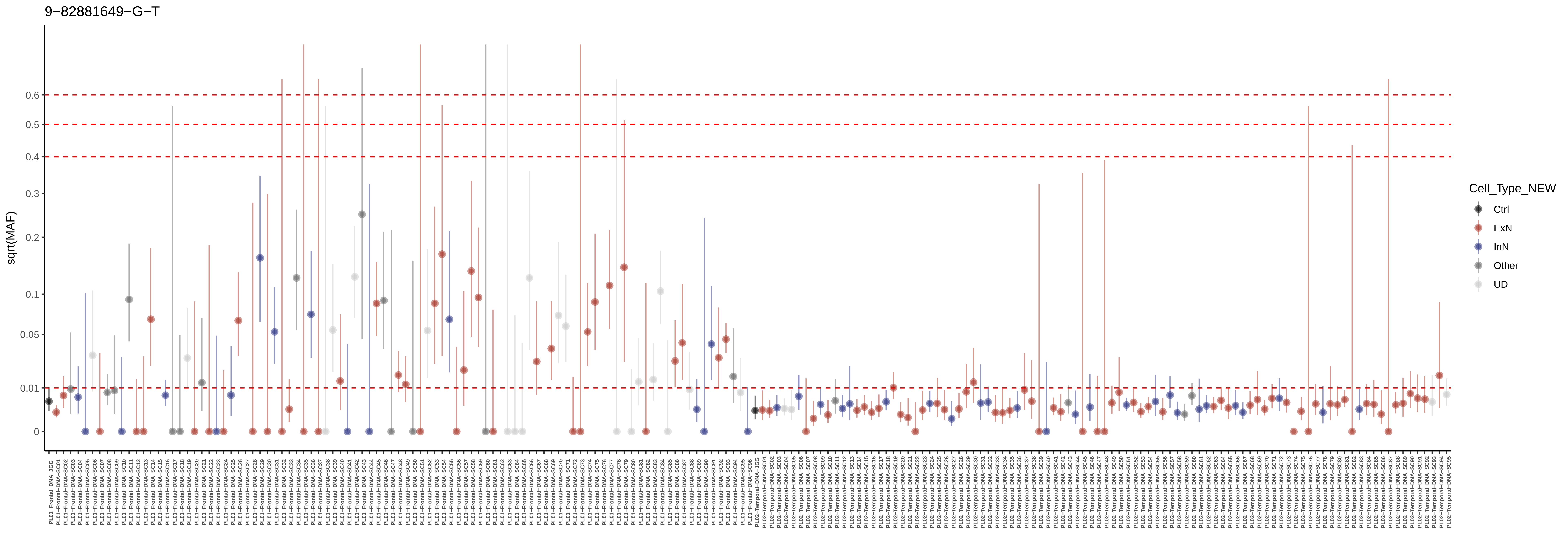
9−82881649−G−T
sqrt(MAF)
0.6
0.5
0.4
0.3
0.2
0.1
0.05
0.01
0
Cell_Type_NEW
Ctrl
ExN
InN
Other
UD

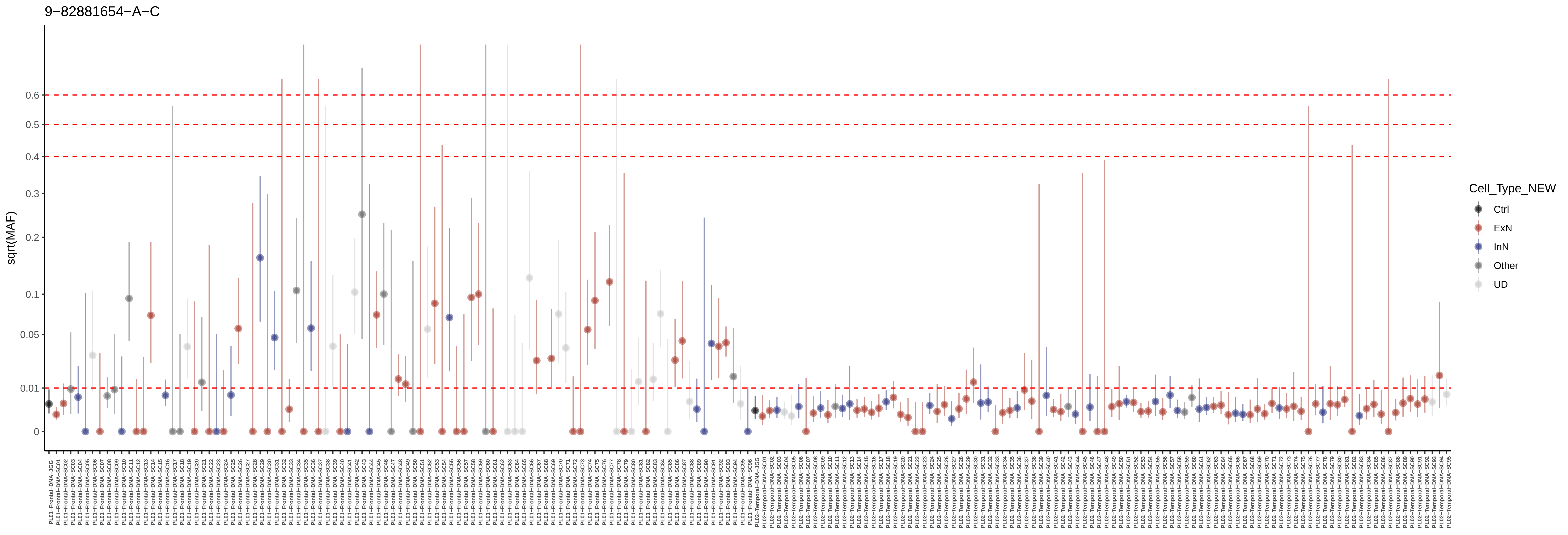

9−82881654−A−C
sqrt(MAF)
0.6
0.5
0.4
0.3
0.2
0.1
0.05
0.01
0
Cell_Type_NEW
Ctrl
ExN
InN
Other
UD

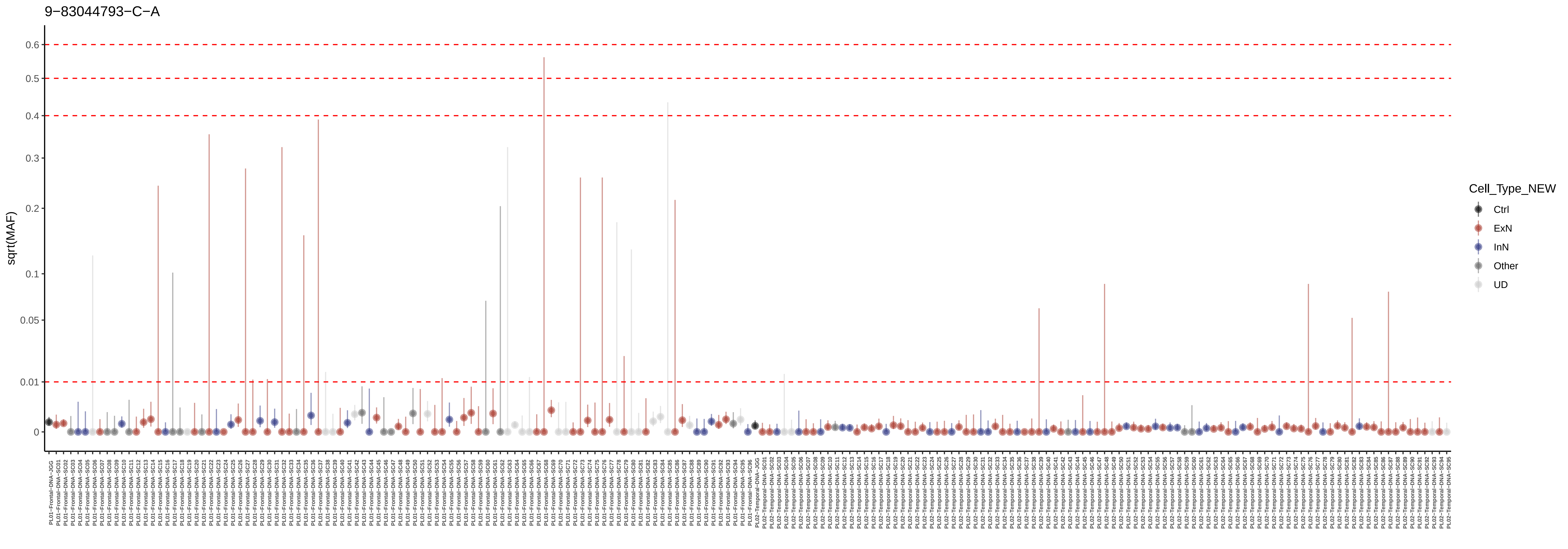
9−83044793−C−A
sqrt(MAF)
0.6
0.5
0.4
0.3
0.2
0.1
0.05
0.01
0
Cell_Type_NEW
Ctrl
ExN
InN
Other
UD

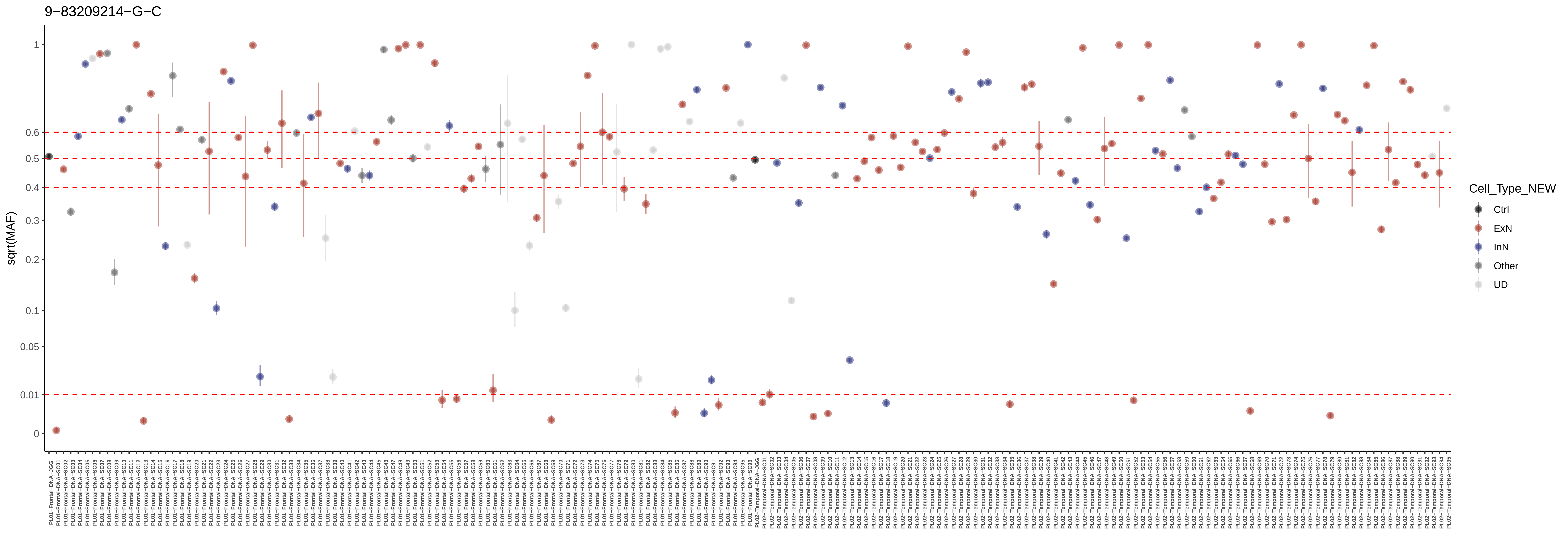
9−83209214−G−C
sqrt(MAF)
1
0.6
0.5
0.4
0.3
0.2
0.1
0.05
0.01
0
Cell_Type_NEW
Ctrl
ExN
InN
Other
UD

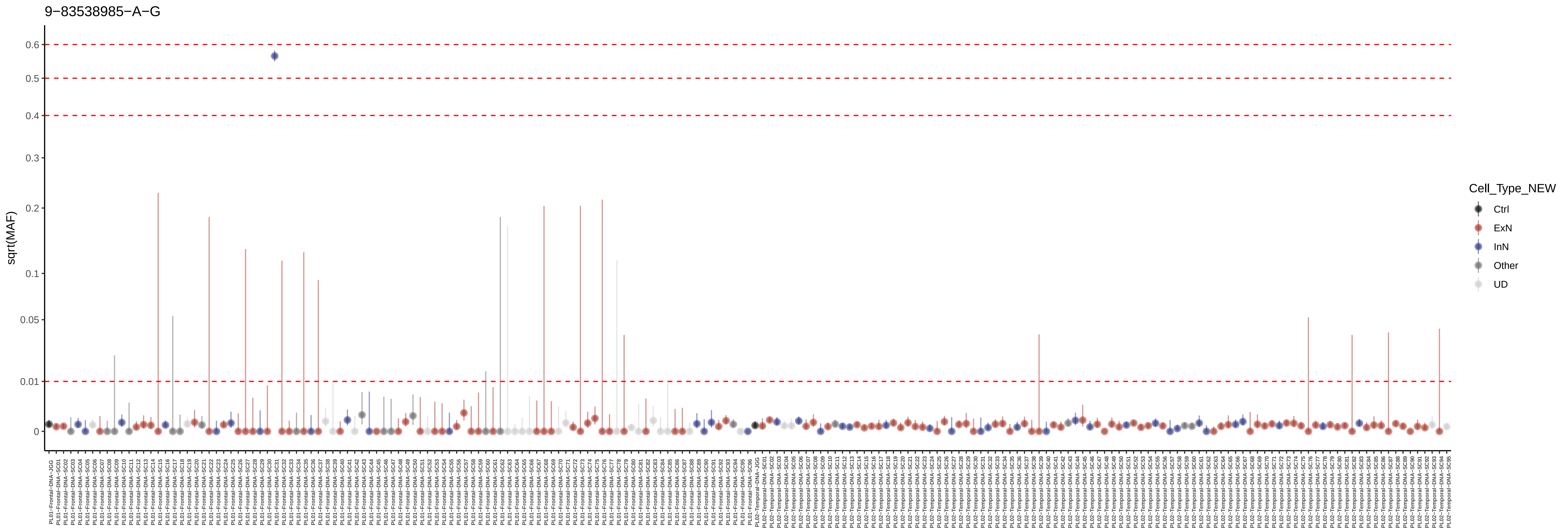

9−83538985−A−G
sqrt(MAF)
0.6
0.5
0.4
0.3
0.2
0.1
0.05
0.01
0
Cell_Type_NEW
Ctrl
ExN
InN
Other
UD

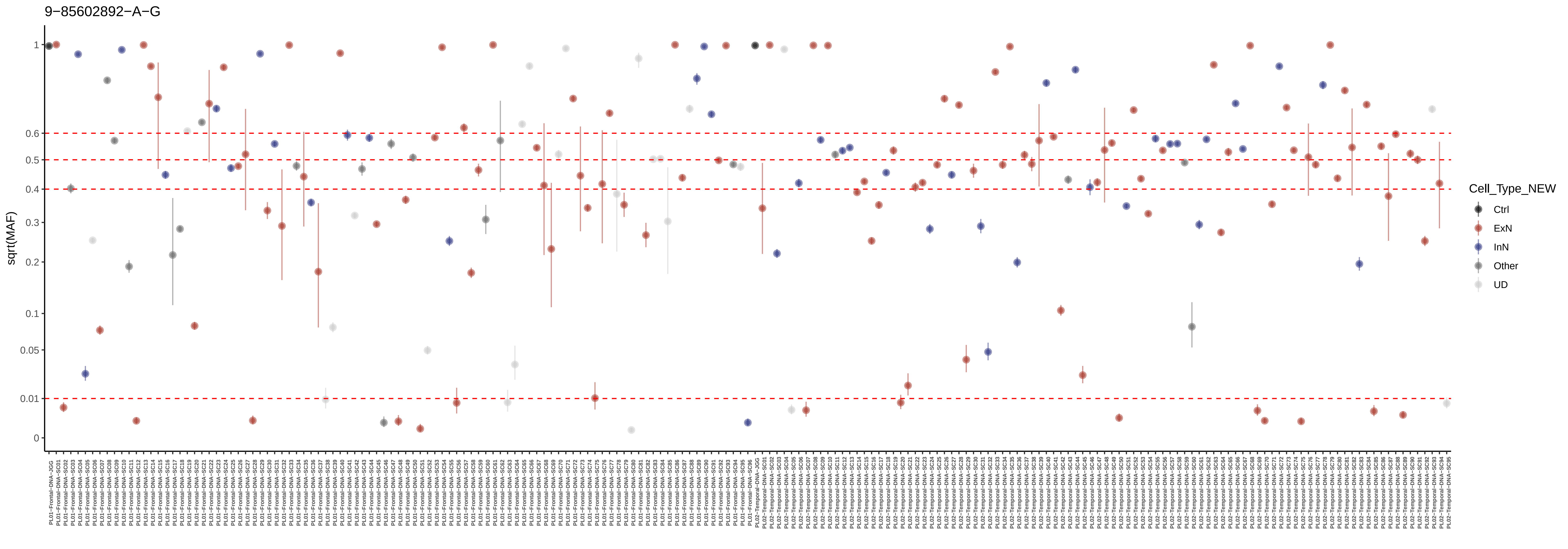
9−85602892−A−G
sqrt(MAF)
1
0.6
0.5
0.4
0.3
0.2
0.1
0.05
0.01
0
Cell_Type_NEW
Ctrl
ExN
InN
Other
UD

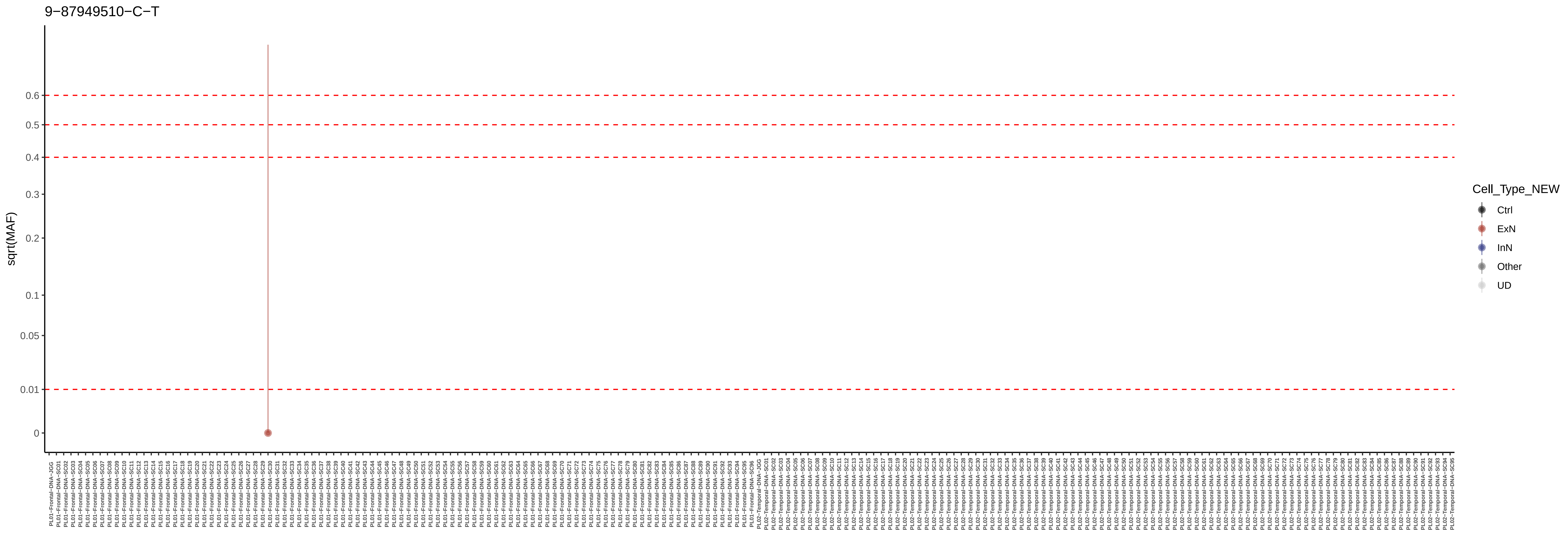

9−87949510−C−T
sqrt(MAF)
0.6
0.5
0.4
0.3
0.2
0.1
0.05
0.01
0
Cell_Type_NEW
Ctrl
ExN
InN
Other
UD

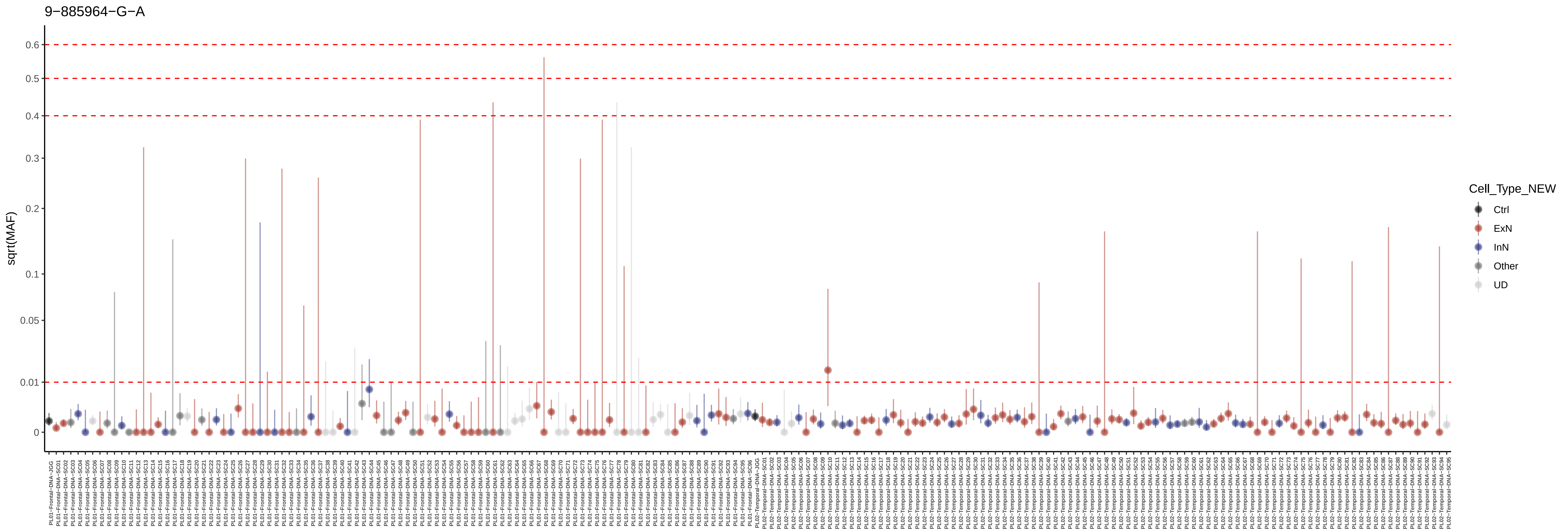

9−885964−G−A
sqrt(MAF)
0.6
0.5
0.4
0.3
0.2
0.1
0.05
0.01
0
Cell_Type_NEW
Ctrl
ExN
InN
Other
UD

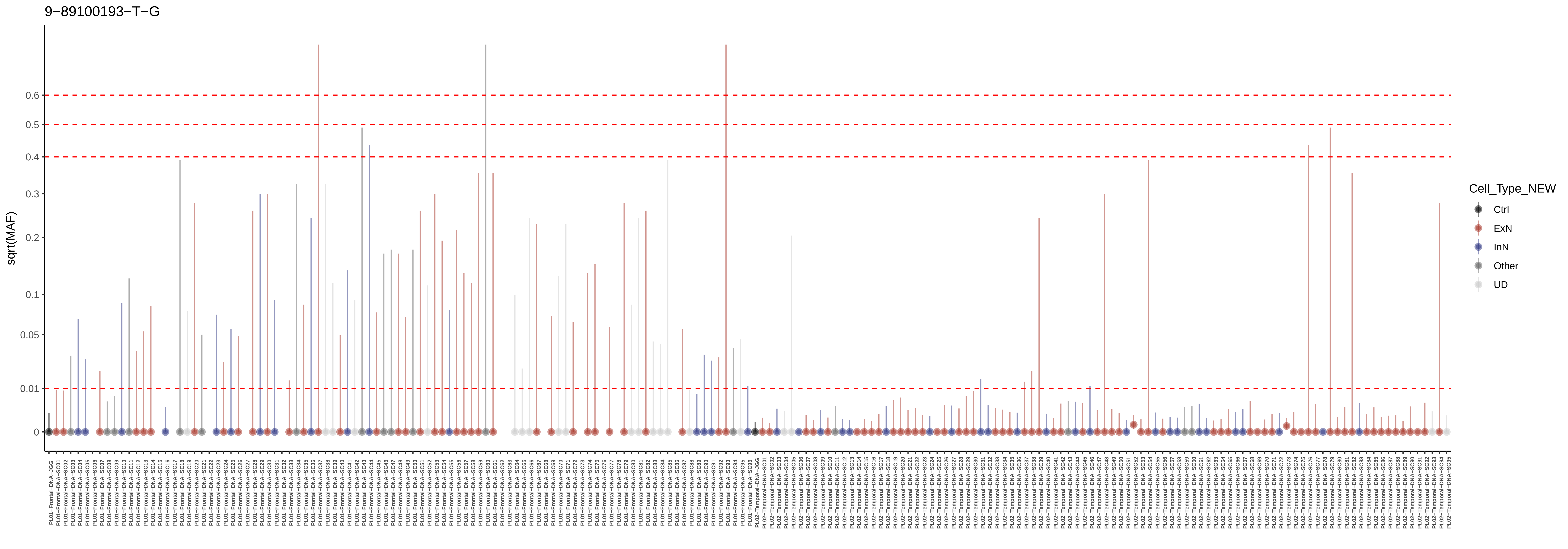
9−89100193−T−G
sqrt(MAF)
0.6
0.5
0.4
0.3
0.2
0.1
0.05
0.01
0
Cell_Type_NEW
Ctrl
ExN
InN
Other
UD

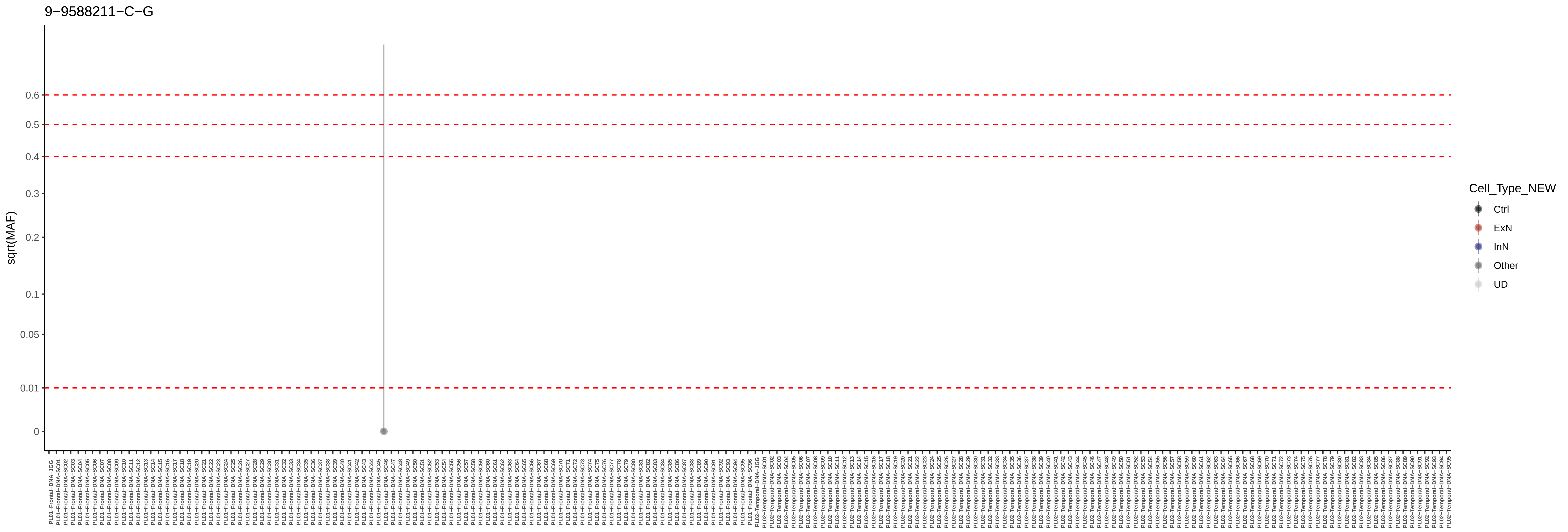
9−9588211−C−G
sqrt(MAF)
0.6
0.5
0.4
0.3
0.2
0.1
0.05
0.01
0
Cell_Type_NEW
Ctrl
ExN
InN
Other
UD

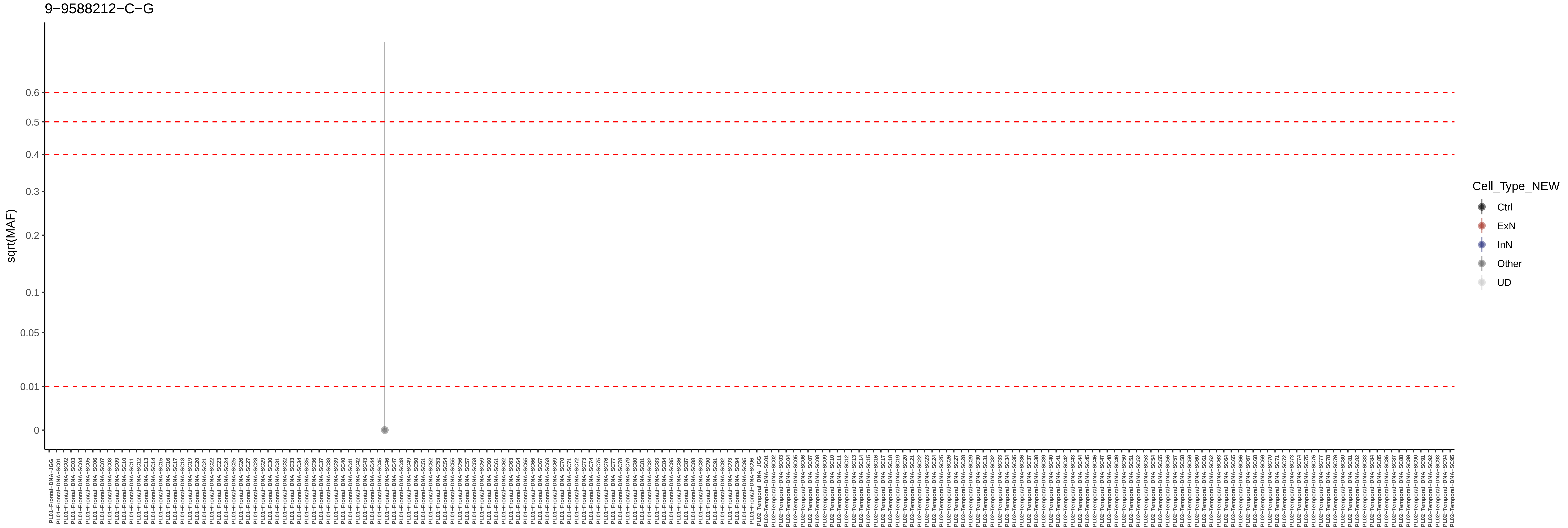

9−9588212−C−G
sqrt(MAF)
0.6
0.5
0.4
0.3
0.2
0.1
0.05
0.01
0
Cell_Type_NEW
Ctrl
ExN
InN
Other
UD

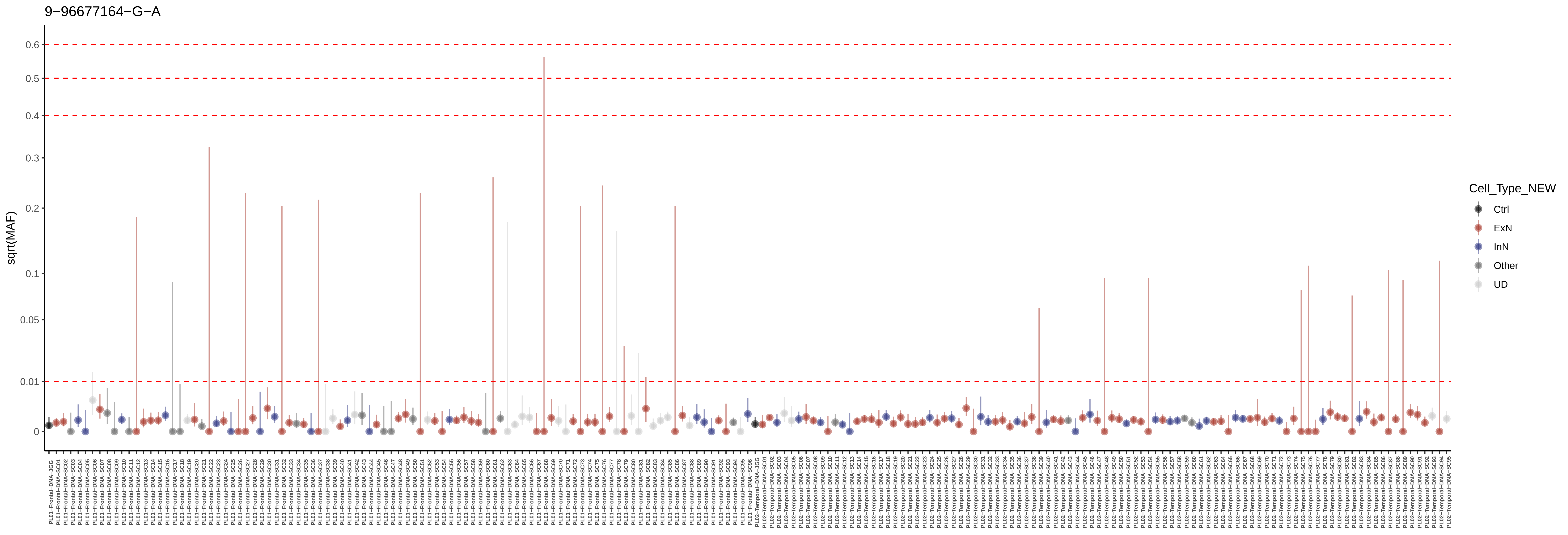

9−96677164−G−A
sqrt(MAF)
0.6
0.5
0.4
0.3
0.2
0.1
0.05
0.01
0
Cell_Type_NEW
Ctrl
ExN
InN
Other
UD

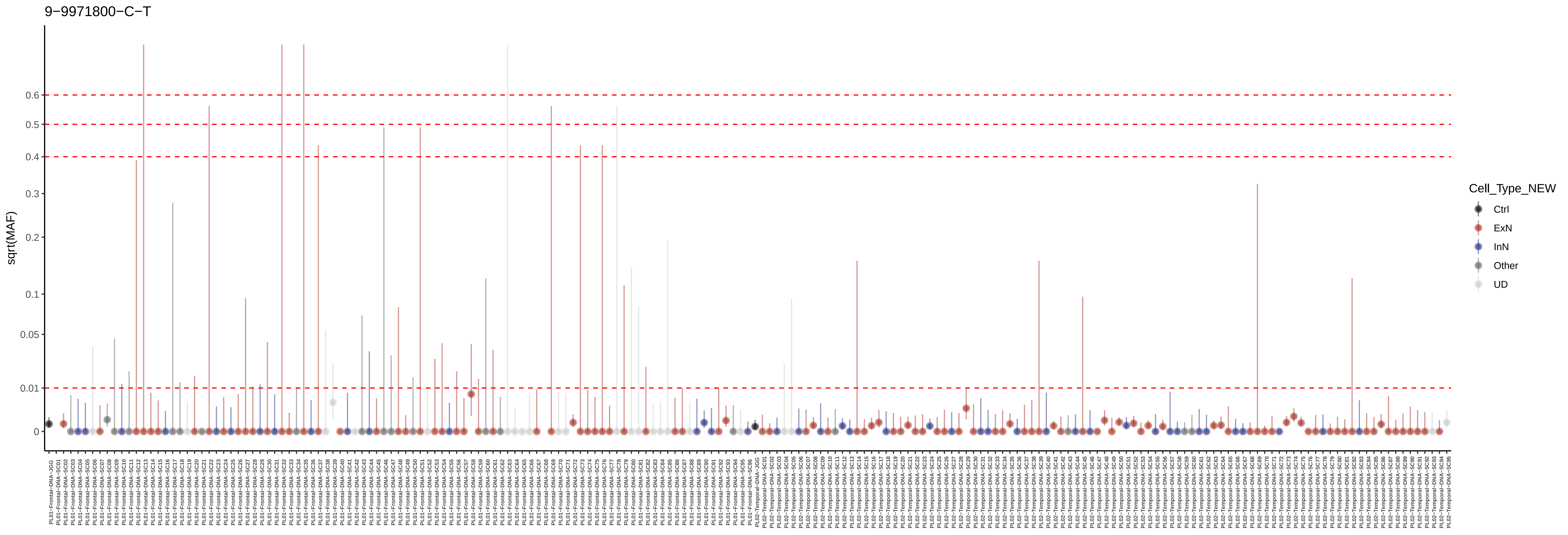
9−9971800−C−T
sqrt(MAF)
0.6
0.5
0.4
0.3
0.2
0.1
0.05
0.01
0
Cell_Type_NEW
Ctrl
ExN
InN
Other
UD

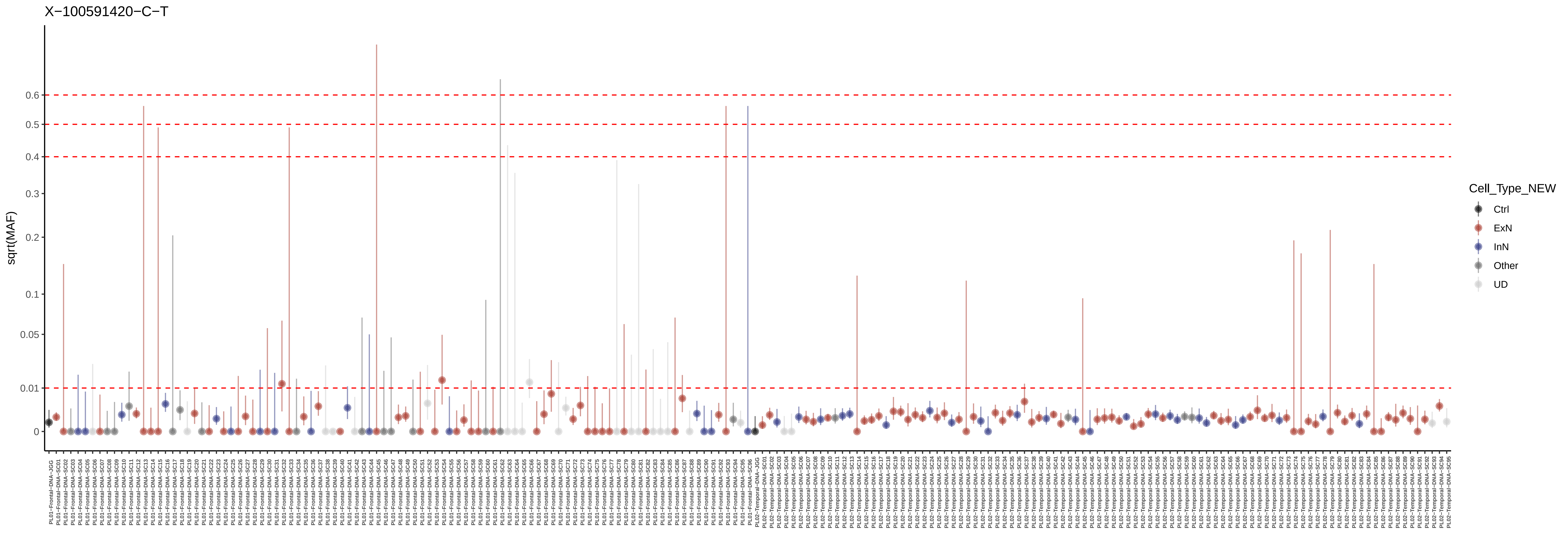
X−100591420−C−T
sqrt(MAF)
0.6
0.5
0.4
0.3
0.2
0.1
0.05
0.01
0
Cell_Type_NEW
Ctrl
ExN
InN
Other
UD

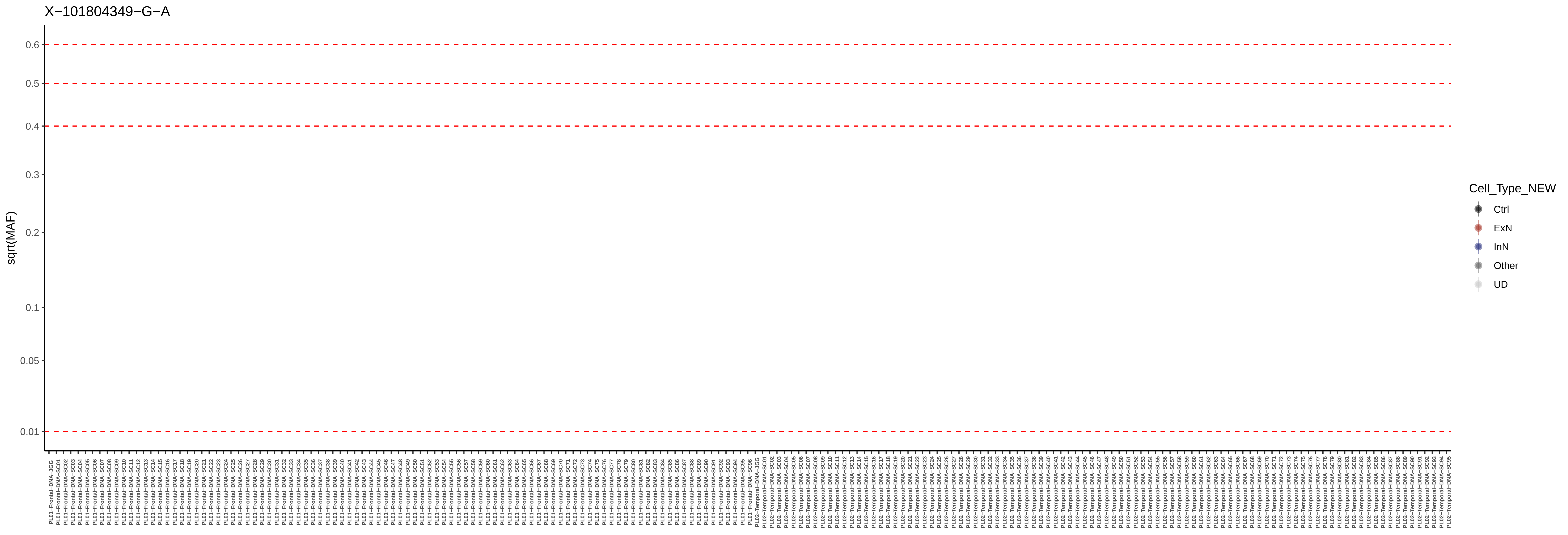

X−101804349−G−A
sqrt(MAF)
0.6
0.5
0.4
0.3
0.2
0.1
0.05
0.01
Cell_Type_NEW
Ctrl
ExN
InN
Other
UD

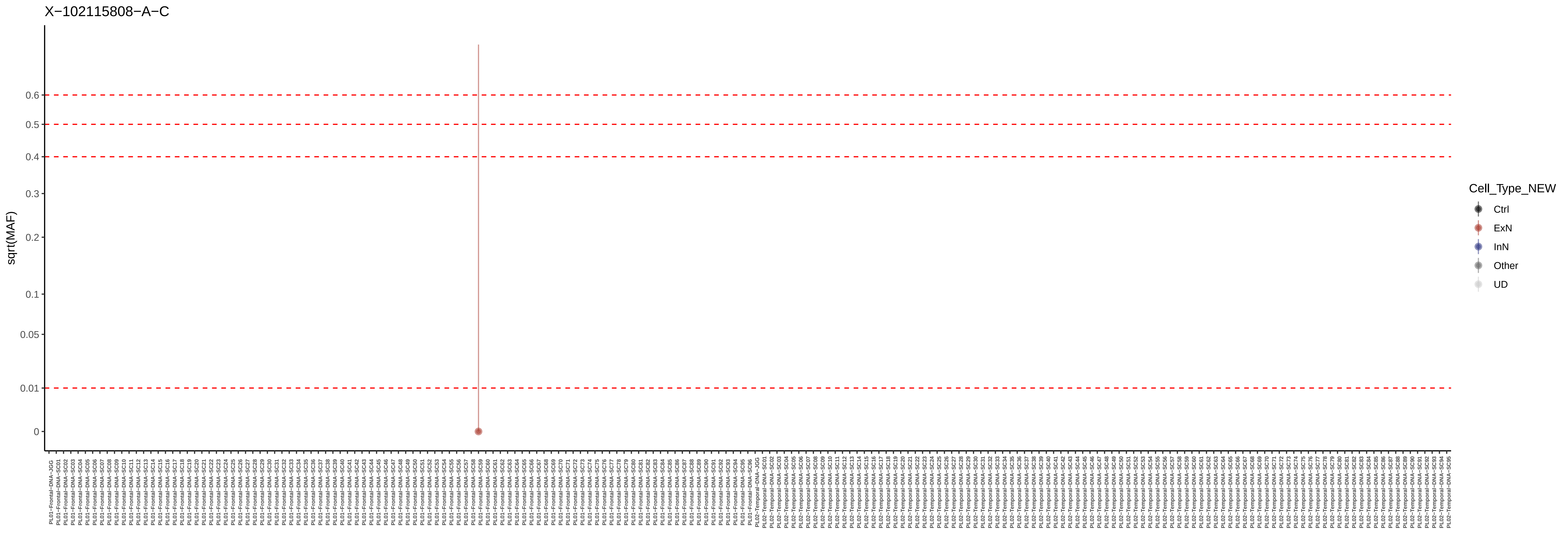
X−102115808−A−C
sqrt(MAF)
0.6
0.5
0.4
0.3
0.2
0.1
0.05
0.01
0
Cell_Type_NEW
Ctrl
ExN
InN
Other
UD

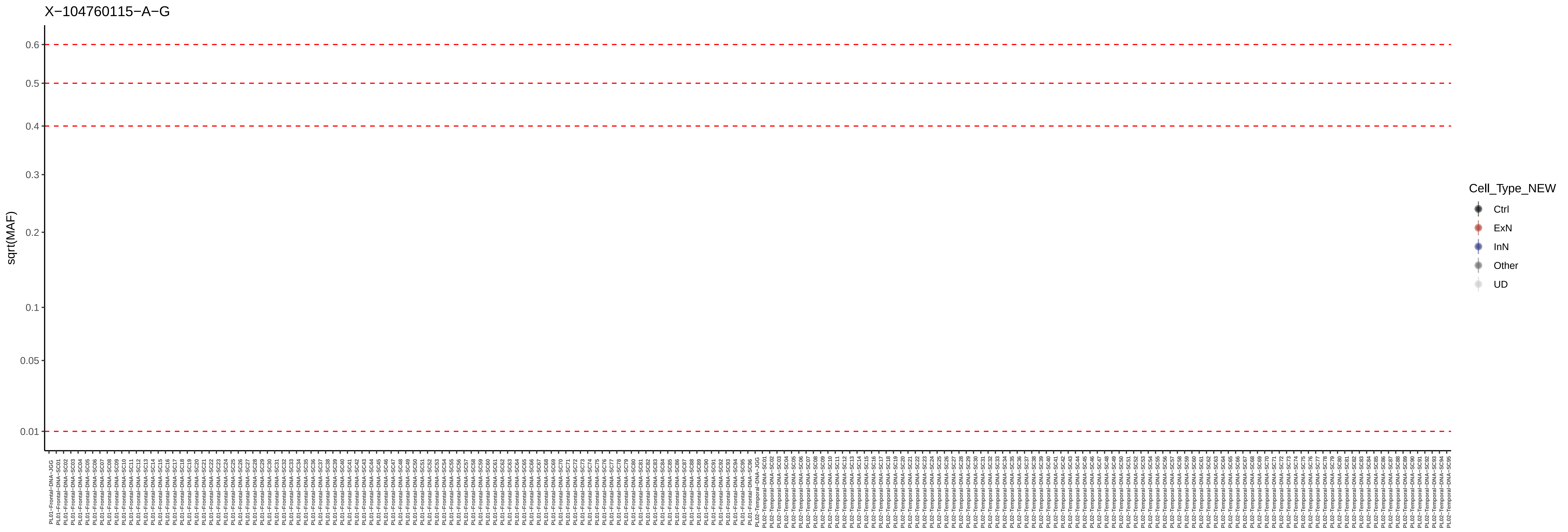
X−104760115−A−G
sqrt(MAF)
0.6
0.5
0.4
0.3
0.2
0.1
0.05
0.01
Cell_Type_NEW
Ctrl
ExN
InN
Other
UD

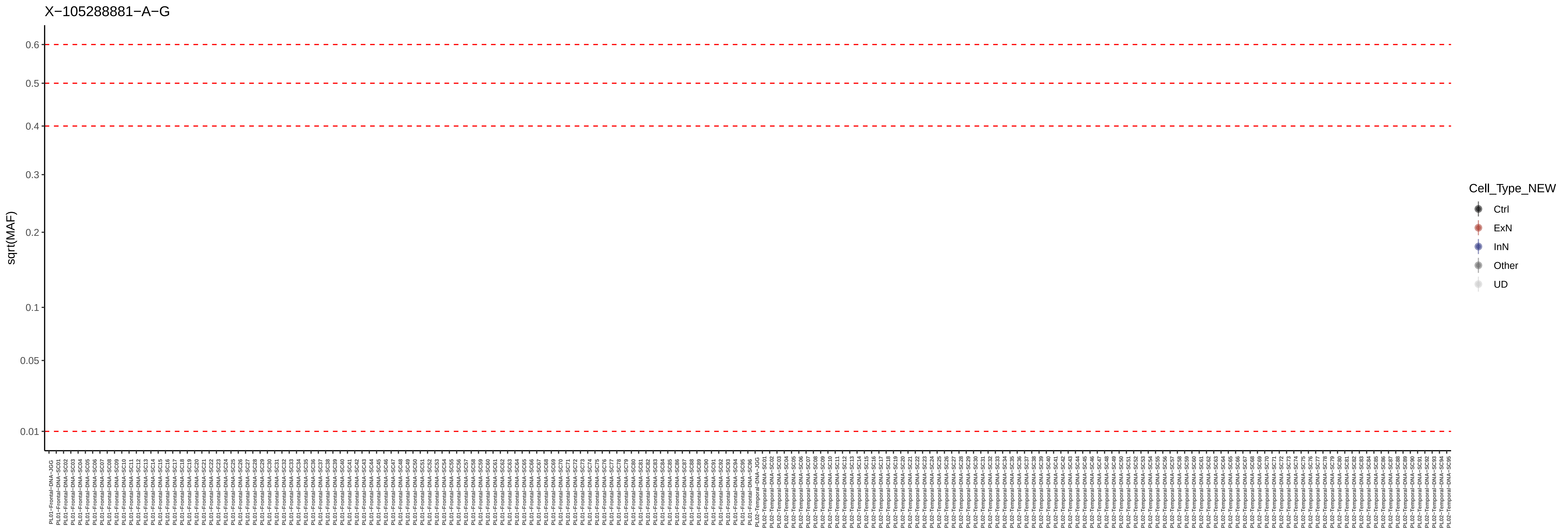
X−105288881−A−G
sqrt(MAF)
0.6
0.5
0.4
0.3
0.2
0.1
0.05
0.01
Cell_Type_NEW
Ctrl
ExN
InN
Other
UD

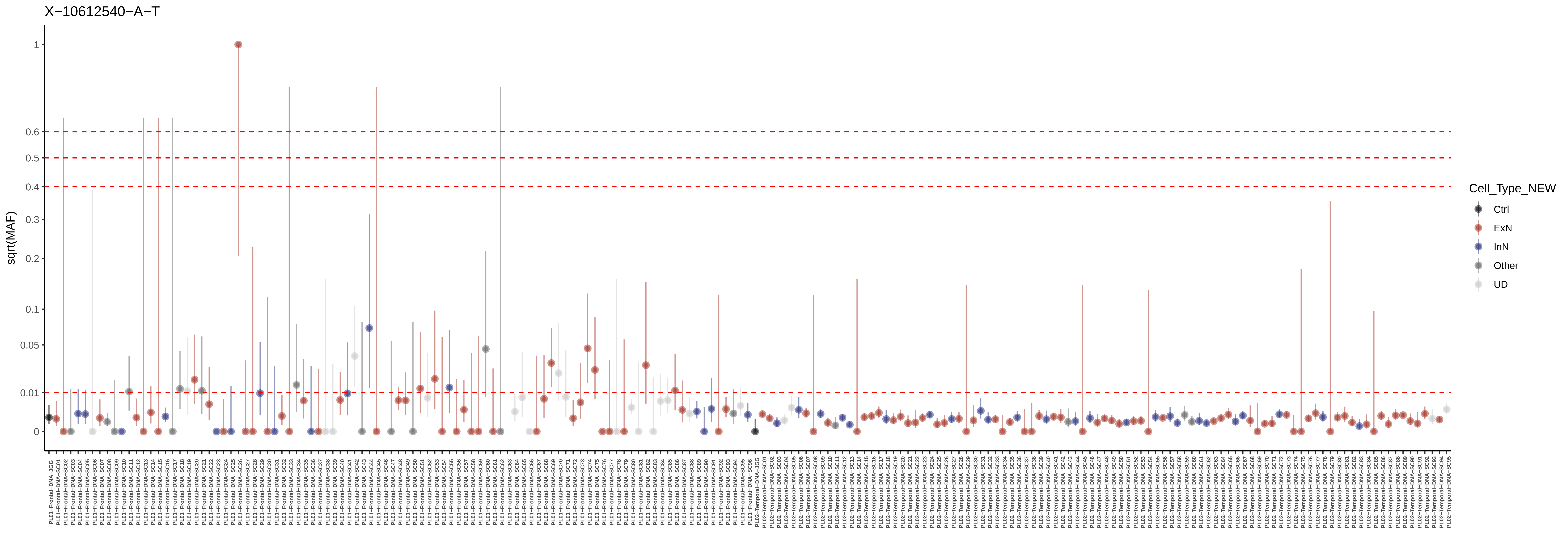
X−10612540−A−T
sqrt(MAF)
1
0.6
0.5
0.4
0.3
0.2
0.1
0.05
0.01
0
Cell_Type_NEW
Ctrl
ExN
InN
Other
UD

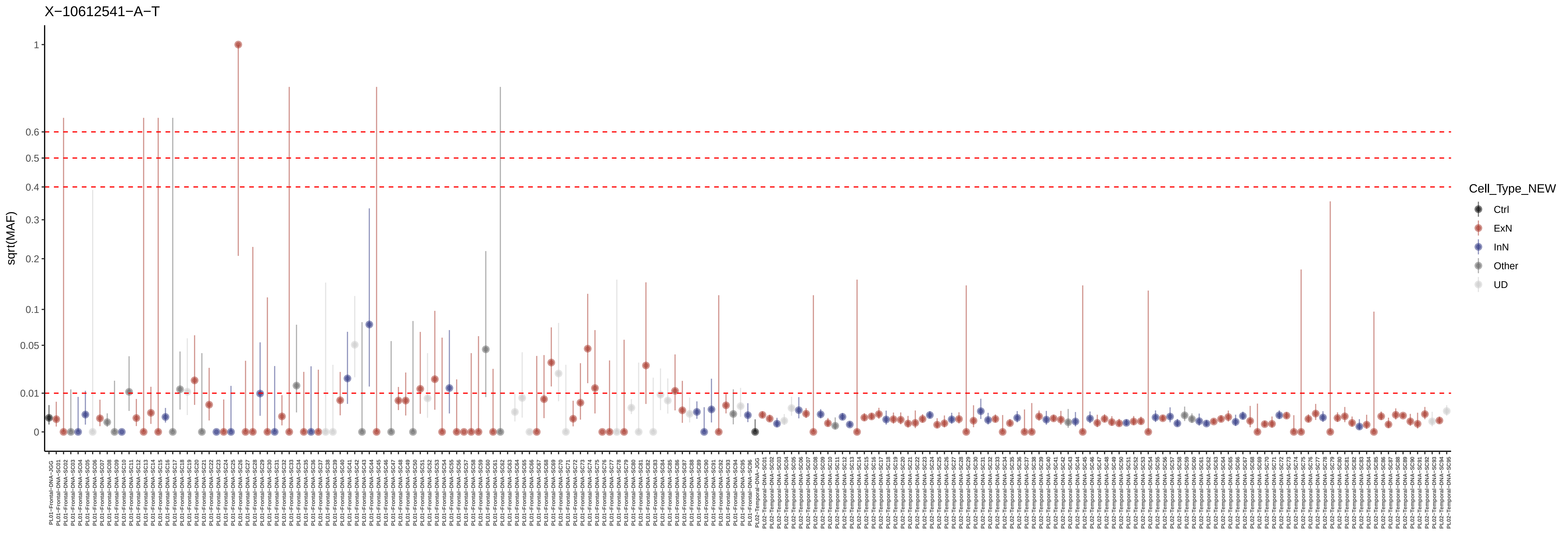
X−10612541−A−T
sqrt(MAF)
1
0.6
0.5
0.4
0.3
0.2
0.1
0.05
0.01
0
Cell_Type_NEW
Ctrl
ExN
InN
Other
UD

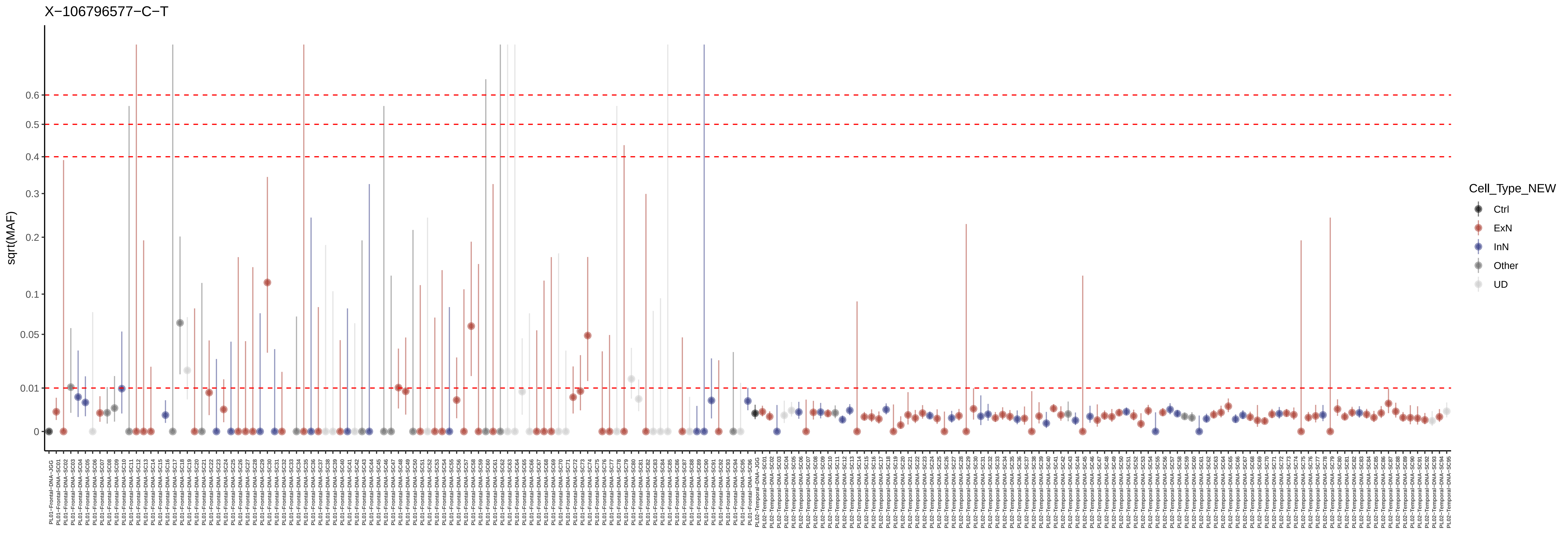
X−106796577−C−T
sqrt(MAF)
0.6
0.5
0.4
0.3
0.2
0.1
0.05
0.01
0
Cell_Type_NEW
Ctrl
ExN
InN
Other
UD

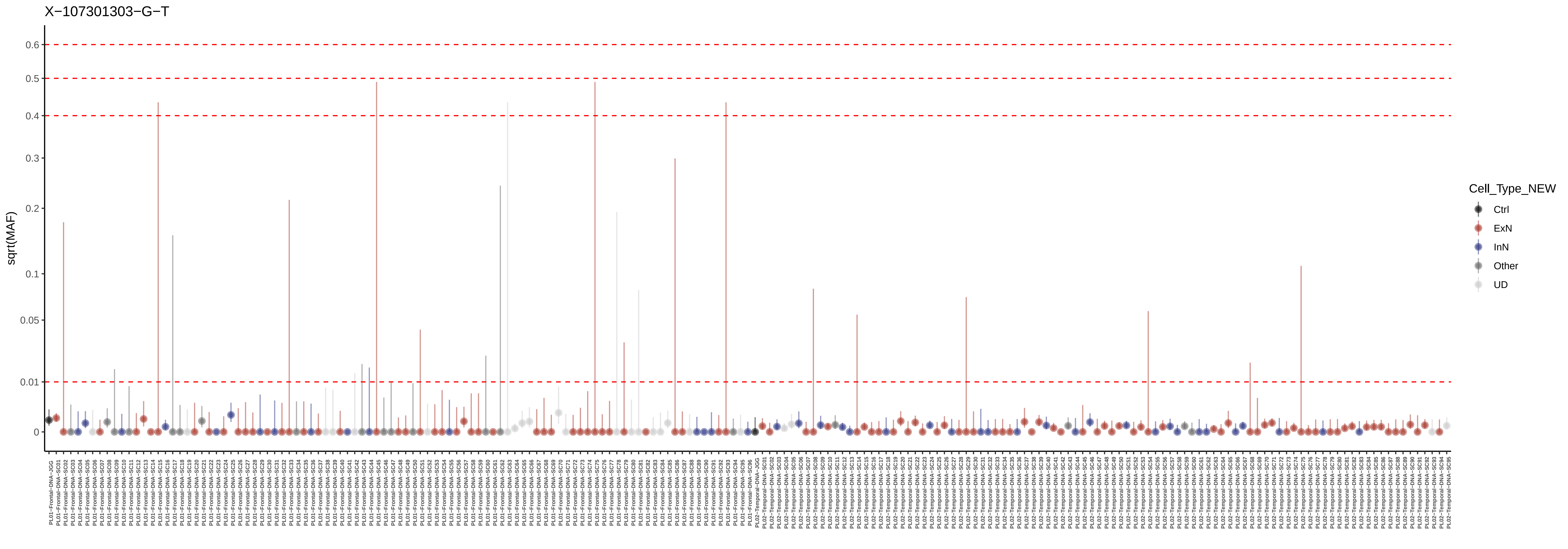

X−107301303−G−T
sqrt(MAF)
0.6
0.5
0.4
0.3
0.2
0.1
0.05
0.01
0
Cell_Type_NEW
Ctrl
ExN
InN
Other
UD

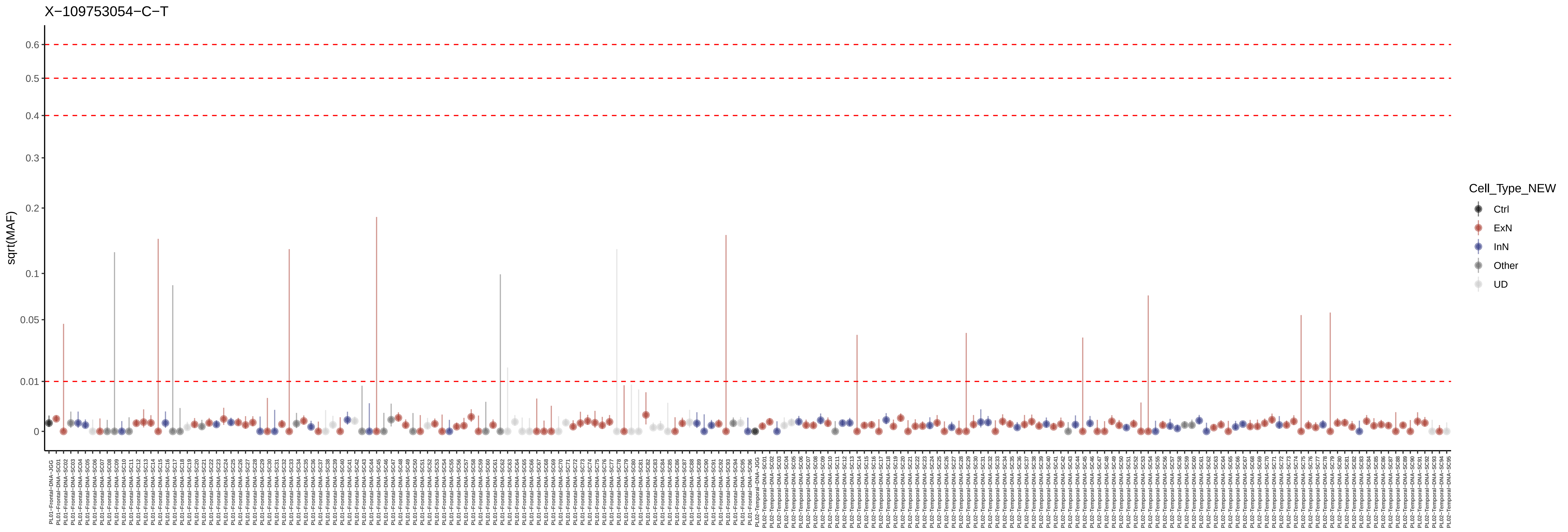
X−109753054−C−T
sqrt(MAF)
0.6
0.5
0.4
0.3
0.2
0.1
0.05
0.01
0
Cell_Type_NEW
Ctrl
ExN
InN
Other
UD

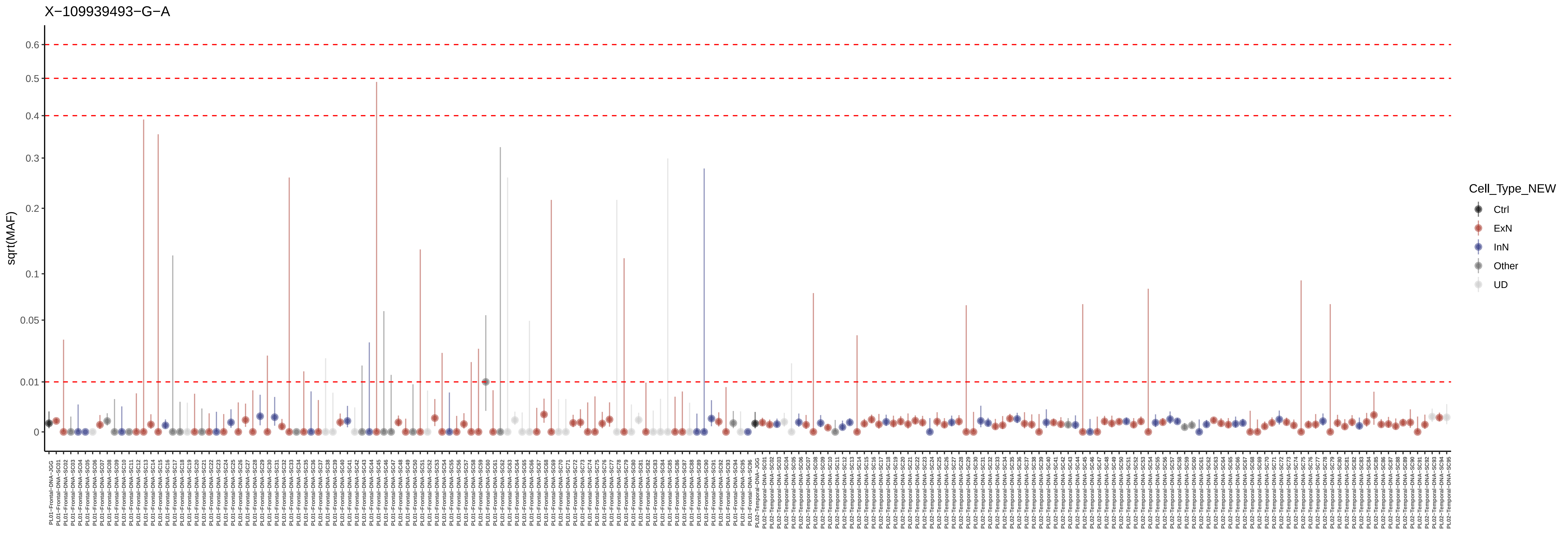
X−109939493−G−A
sqrt(MAF)
0.6
0.5
0.4
0.3
0.2
0.1
0.05
0.01
0
Cell_Type_NEW
Ctrl
ExN
InN
Other
UD

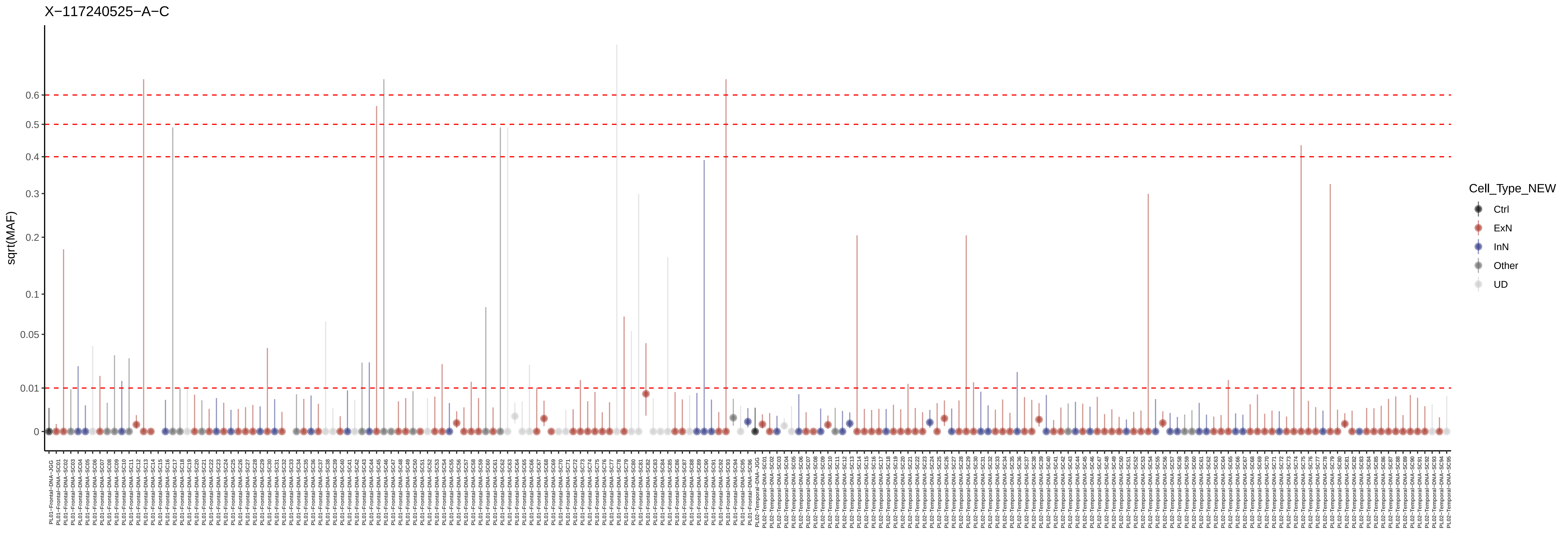
X−117240525−A−C
sqrt(MAF)
0.6
0.5
0.4
0.3
0.2
0.1
0.05
0.01
0
Cell_Type_NEW
Ctrl
ExN
InN
Other
UD

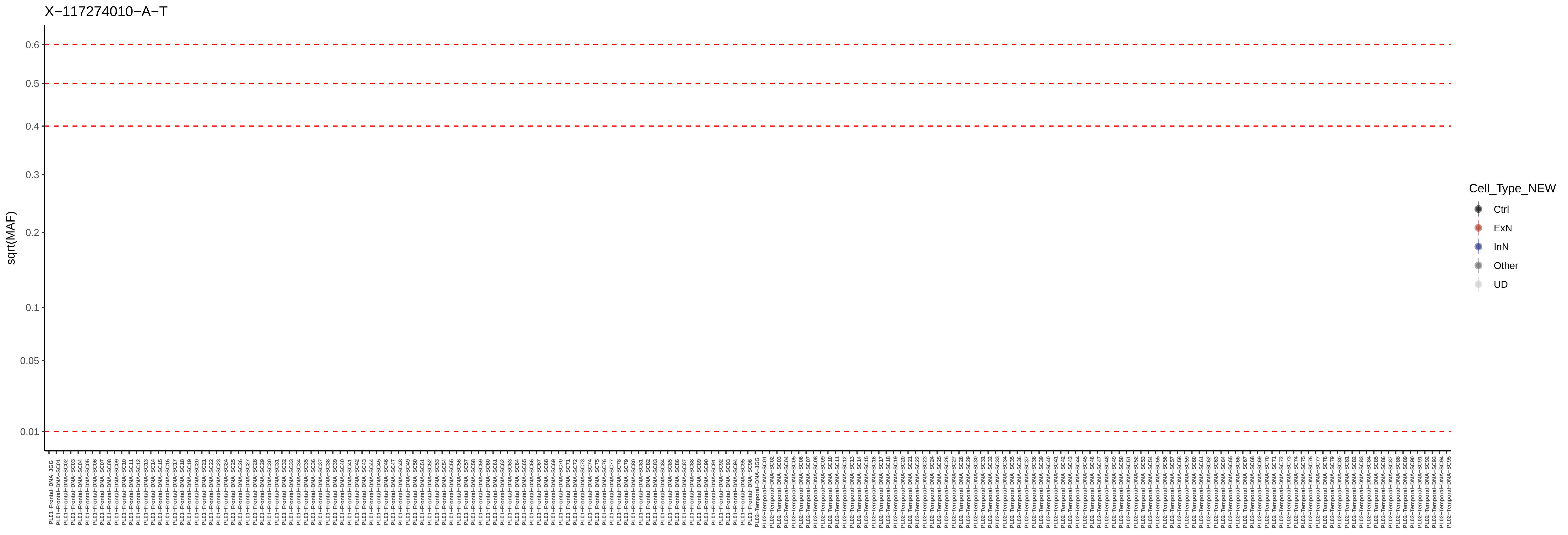

X−117274010−A−T
sqrt(MAF)
0.6
0.5
0.4
0.3
0.2
0.1
0.05
0.01
Cell_Type_NEW
Ctrl
ExN
InN
Other
UD

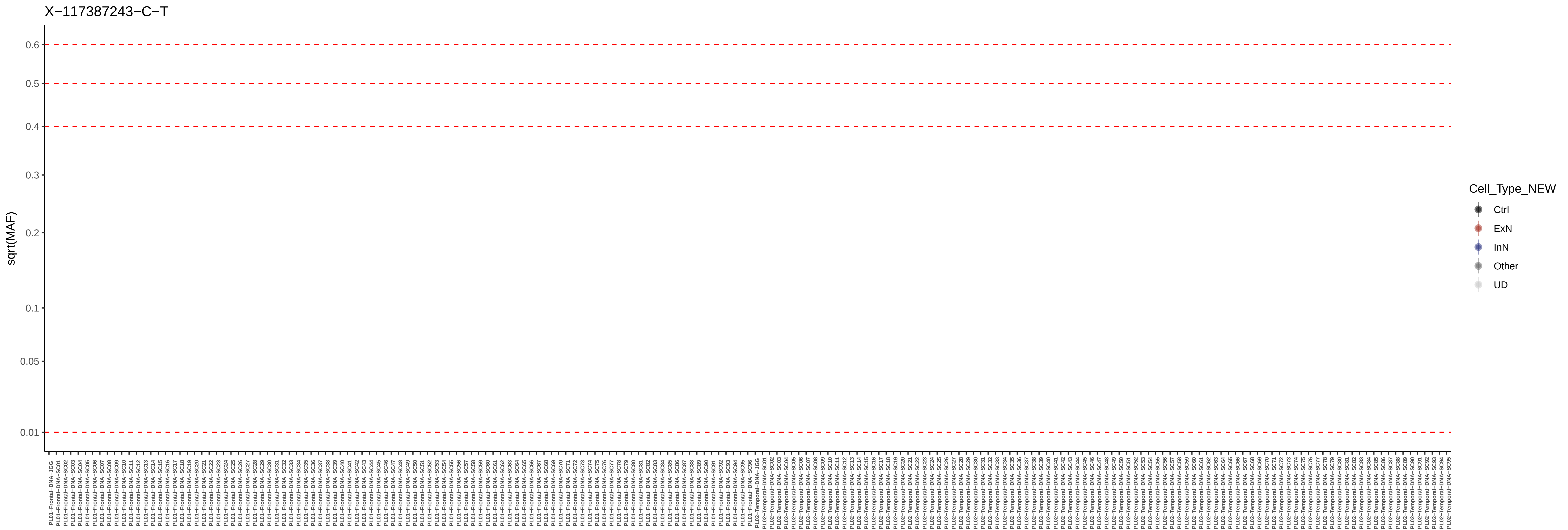
X−117387243−C−T
sqrt(MAF)
0.6
0.5
0.4
0.3
0.2
0.1
0.05
0.01
Cell_Type_NEW
Ctrl
ExN
InN
Other
UD

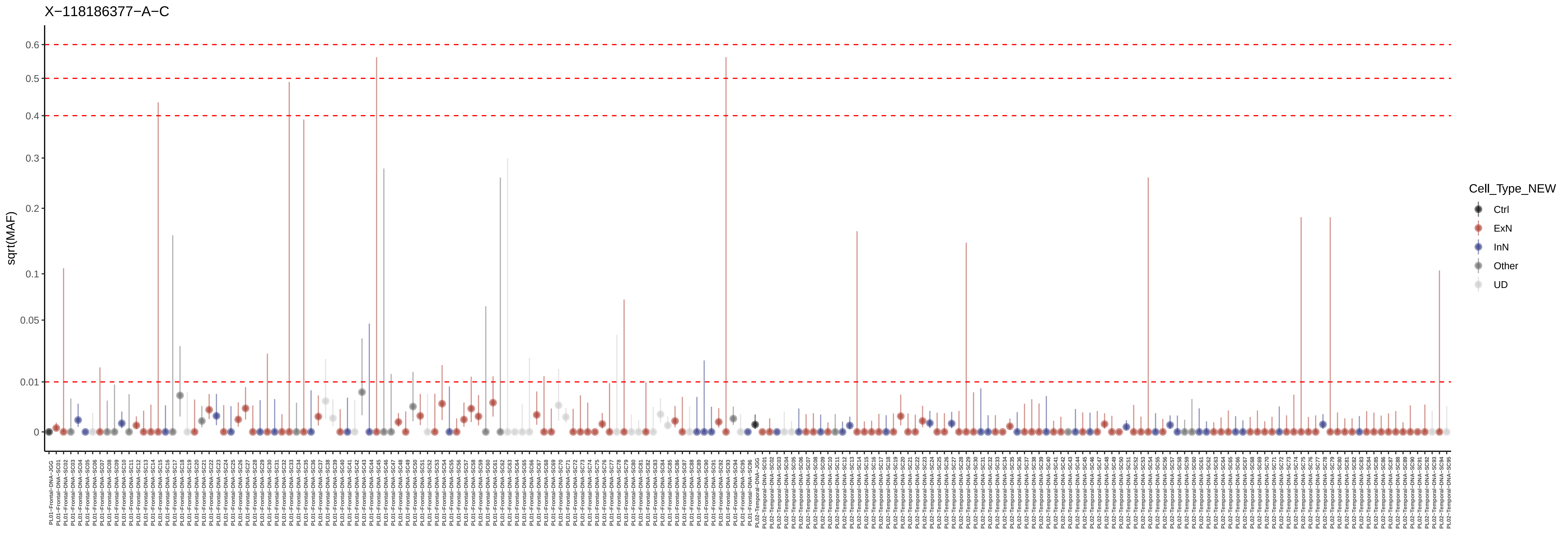
X−118186377−A−C
sqrt(MAF)
0.6
0.5
0.4
0.3
0.2
0.1
0.05
0.01
0
Cell_Type_NEW
Ctrl
ExN
InN
Other
UD

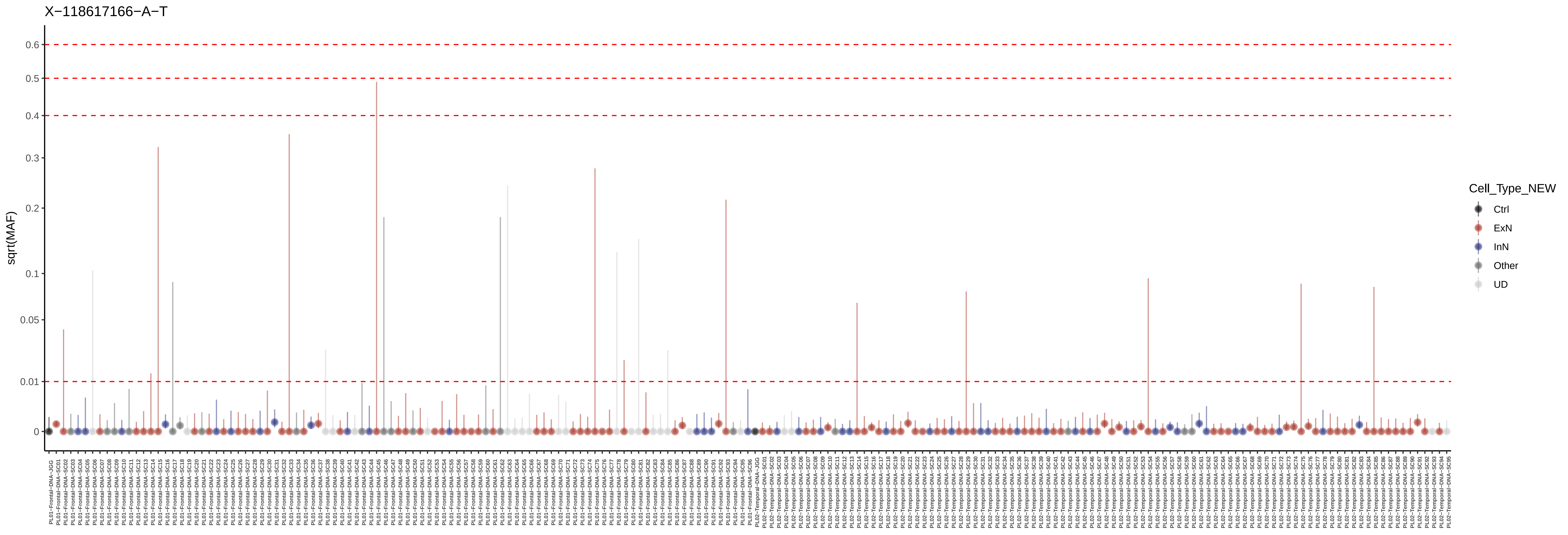
X−118617166−A−T
sqrt(MAF)
0.6
0.5
0.4
0.3
0.2
0.1
0.05
0.01
0
Cell_Type_NEW
Ctrl
ExN
InN
Other
UD

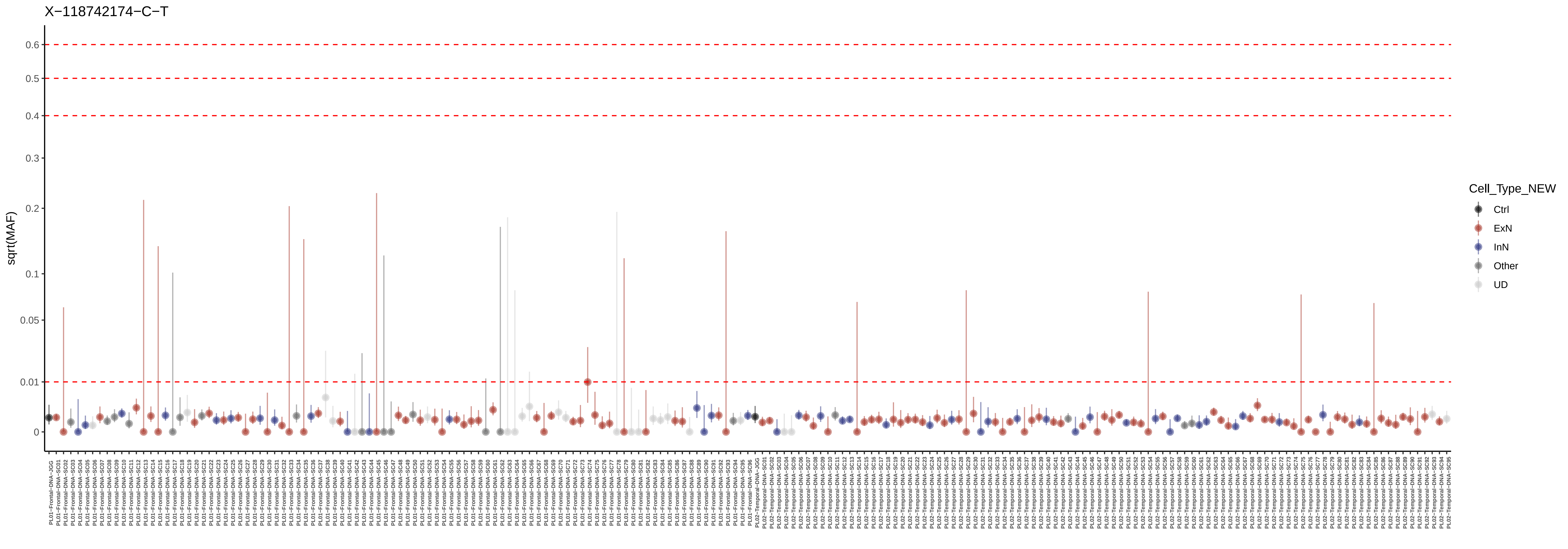
X−118742174−C−T
sqrt(MAF)
0.6
0.5
0.4
0.3
0.2
0.1
0.05
0.01
0
Cell_Type_NEW
Ctrl
ExN
InN
Other
UD

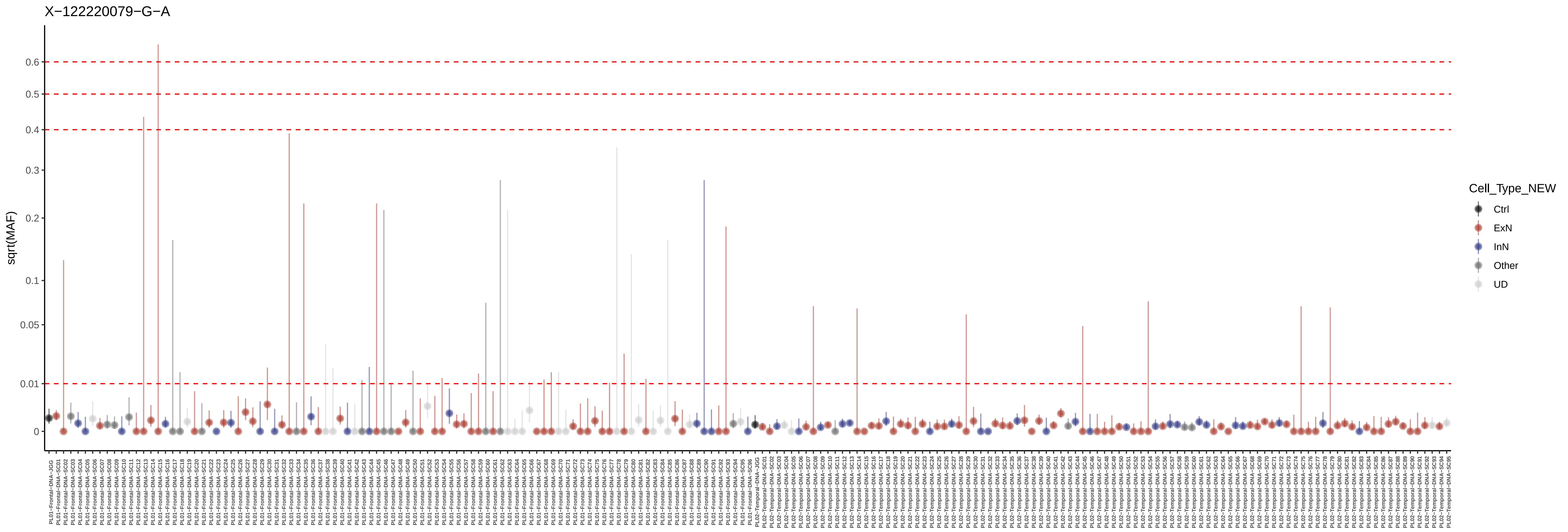
X−122220079−G−A
sqrt(MAF)
0.6
0.5
0.4
0.3
0.2
0.1
0.05
0.01
0
Cell_Type_NEW
Ctrl
ExN
InN
Other
UD

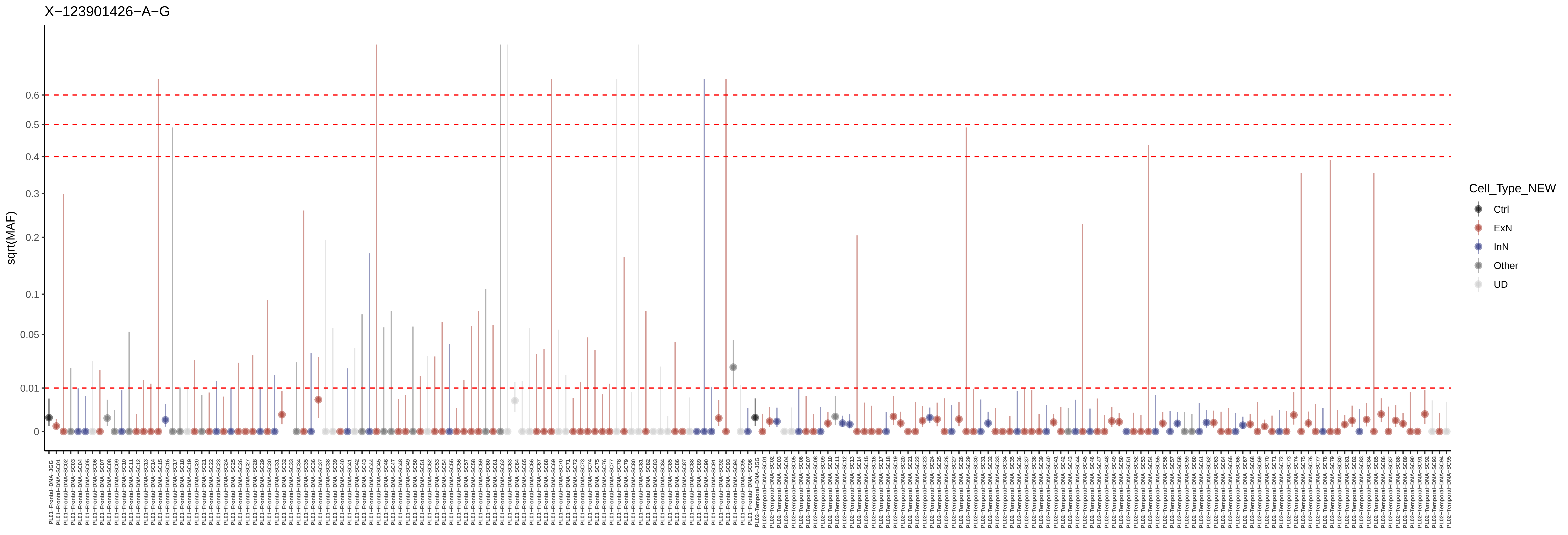
X−123901426−A−G
sqrt(MAF)
0.6
0.5
0.4
0.3
0.2
0.1
0.05
0.01
0
Cell_Type_NEW
Ctrl
ExN
InN
Other
UD

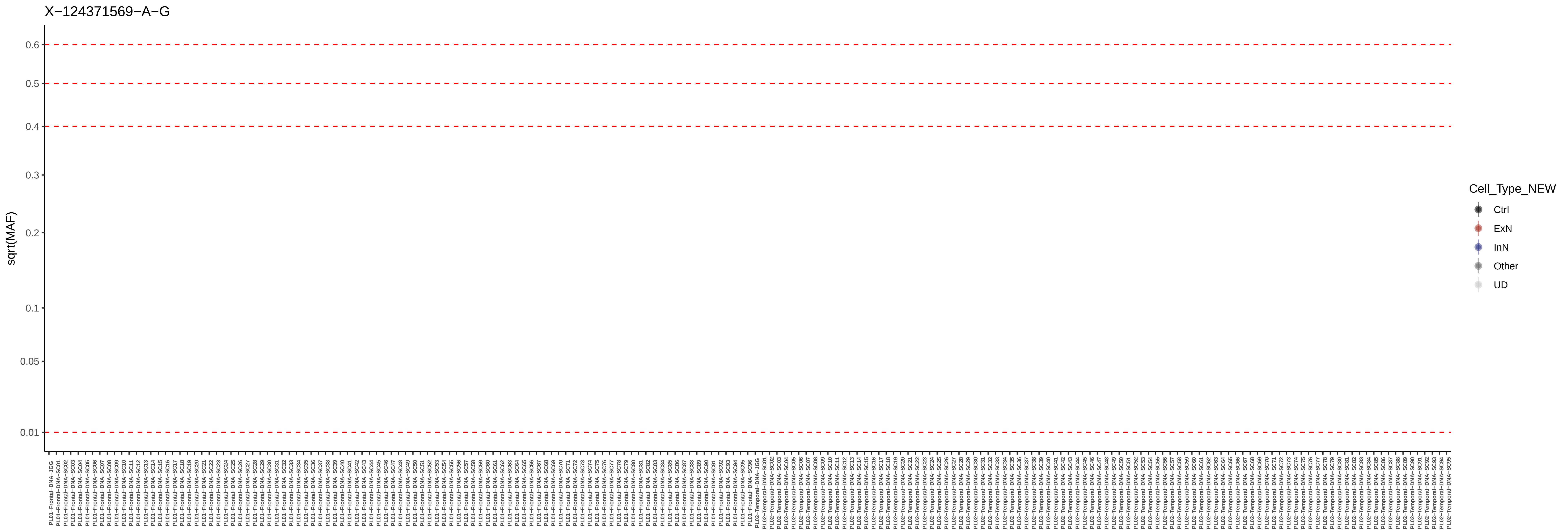
X−124371569−A−G
sqrt(MAF)
0.6
0.5
0.4
0.3
0.2
0.1
0.05
0.01
Cell_Type_NEW
Ctrl
ExN
InN
Other
UD

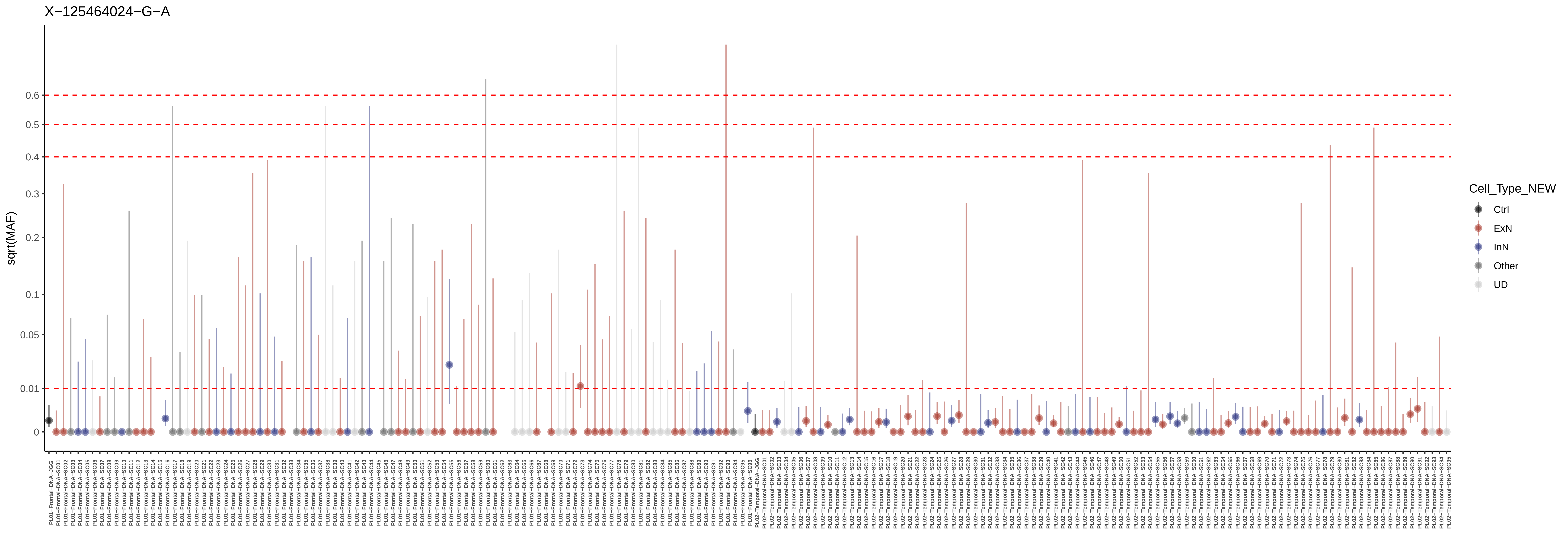

X−125464024−G−A
sqrt(MAF)
0.6
0.5
0.4
0.3
0.2
0.1
0.05
0.01
0
Cell_Type_NEW
Ctrl
ExN
InN
Other
UD

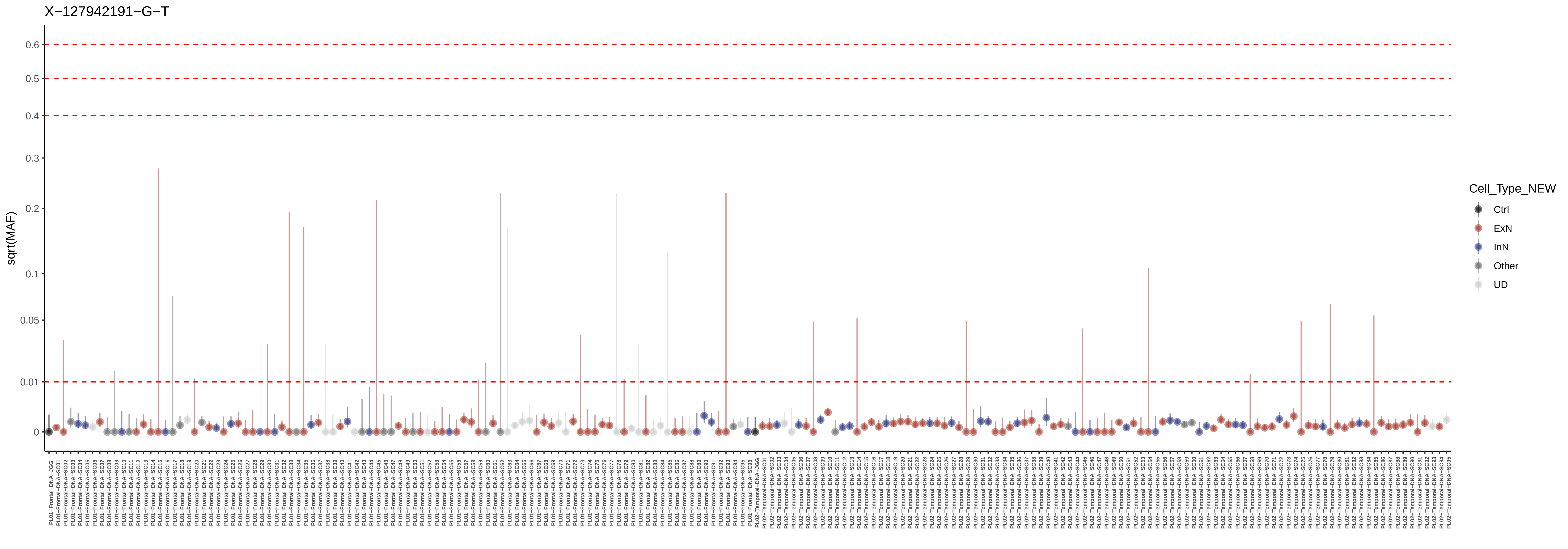
X−127942191−G−T
sqrt(MAF)
0.6
0.5
0.4
0.3
0.2
0.1
0.05
0.01
0
Cell_Type_NEW
Ctrl
ExN
InN
Other
UD

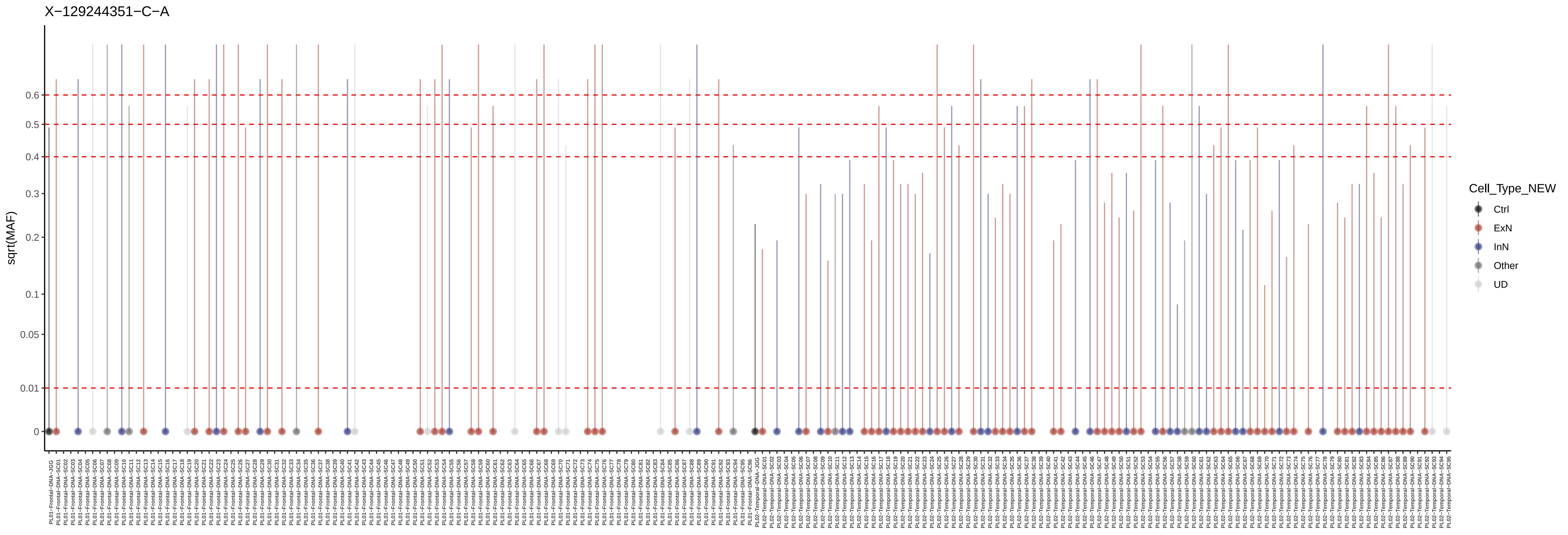

X−129244351−C−A
sqrt(MAF)
0.6
0.5
0.4
0.3
0.2
0.1
0.05
0.01
0
Cell_Type_NEW
Ctrl
ExN
InN
Other
UD

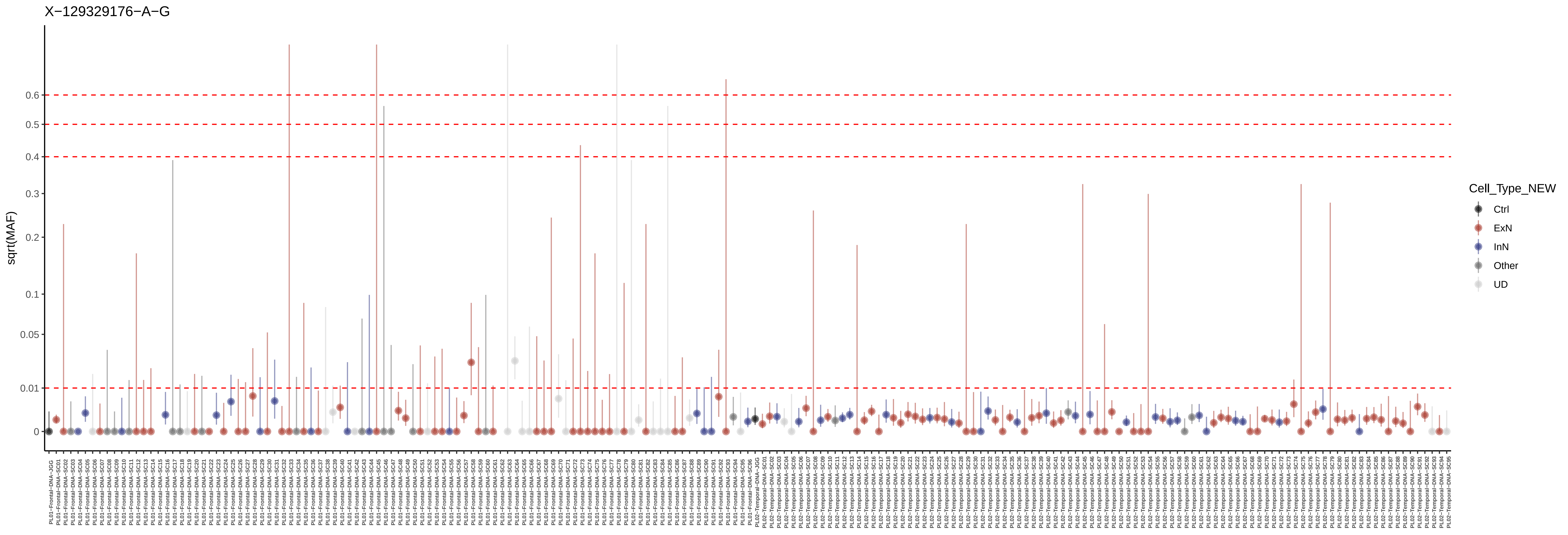
X−129329176−A−G
sqrt(MAF)
0.6
0.5
0.4
0.3
0.2
0.1
0.05
0.01
0
Cell_Type_NEW
Ctrl
ExN
InN
Other
UD

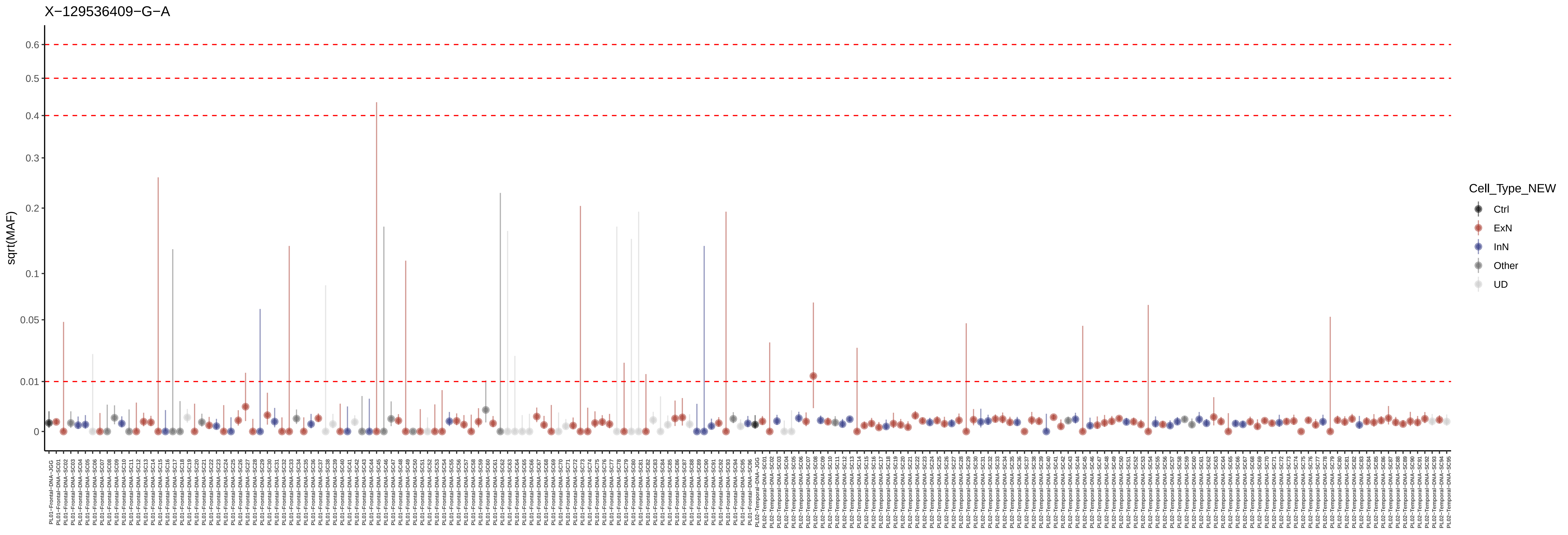

X−129536409−G−A
sqrt(MAF)
0.6
0.5
0.4
0.3
0.2
0.1
0.05
0.01
0
Cell_Type_NEW
Ctrl
ExN
InN
Other
UD

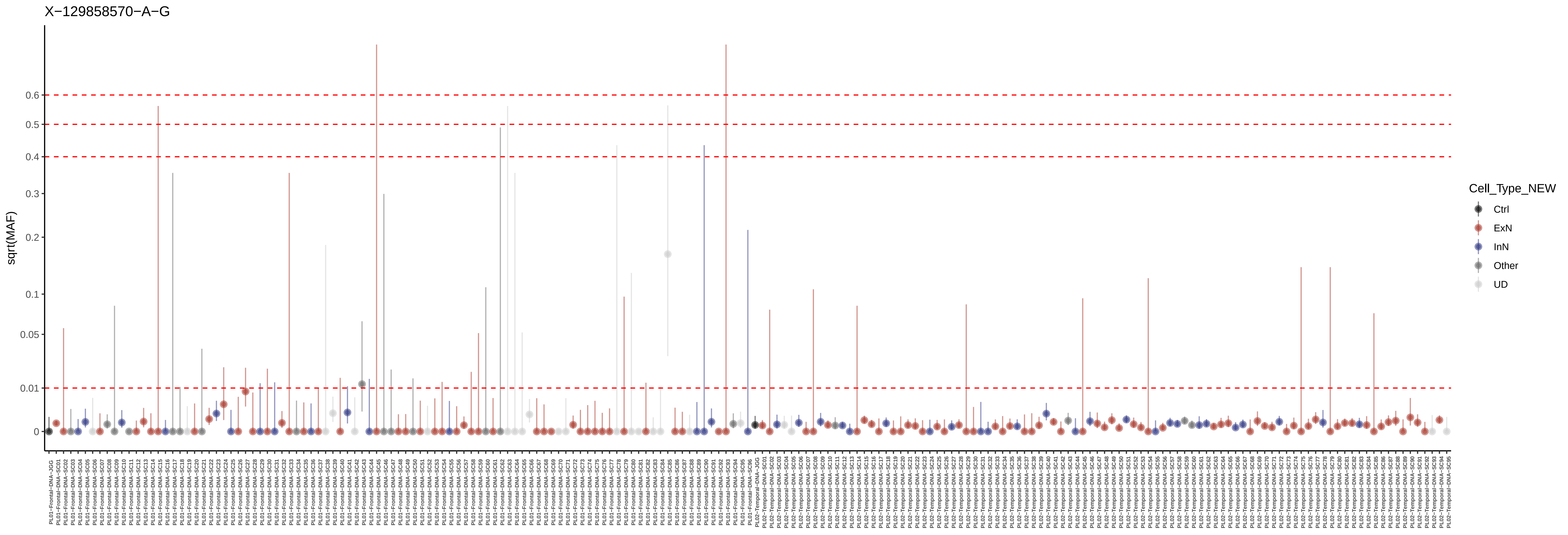

X−129858570−A−G
sqrt(MAF)
0.6
0.5
0.4
0.3
0.2
0.1
0.05
0.01
0
Cell_Type_NEW
Ctrl
ExN
InN
Other
UD

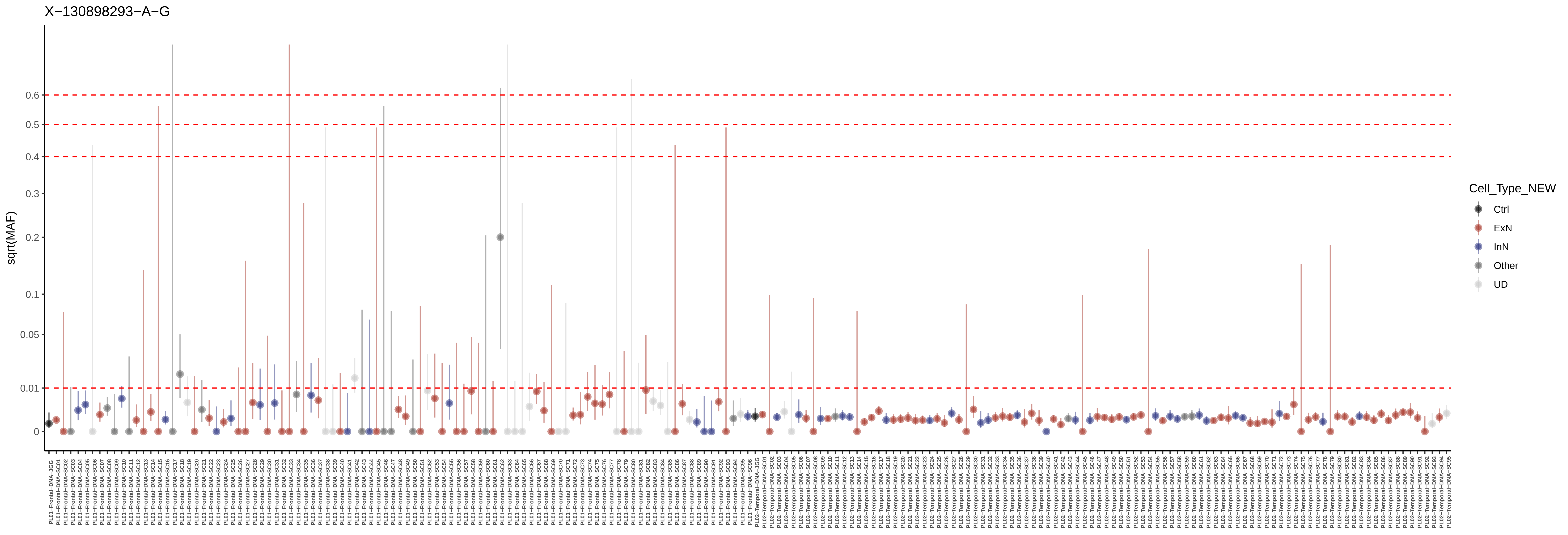
X−130898293−A−G
sqrt(MAF)
0.6
0.5
0.4
0.3
0.2
0.1
0.05
0.01
0
Cell_Type_NEW
Ctrl
ExN
InN
Other
UD

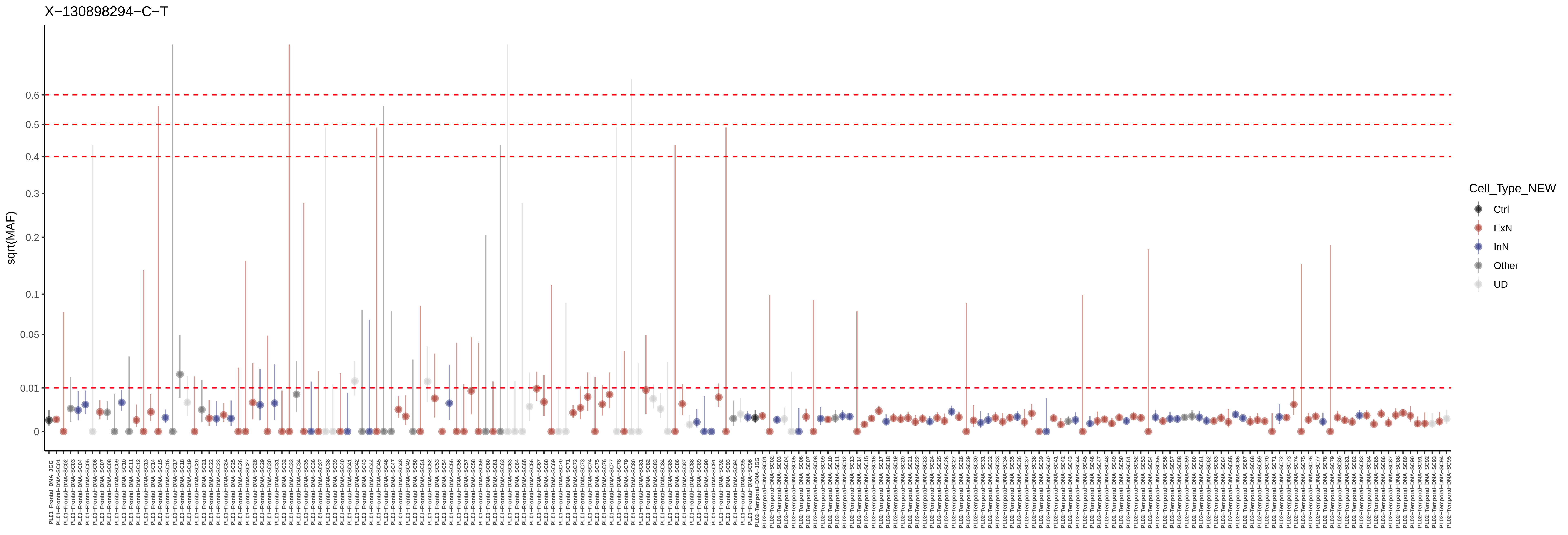

X−130898294−C−T
sqrt(MAF)
0.6
0.5
0.4
0.3
0.2
0.1
0.05
0.01
0
Cell_Type_NEW
Ctrl
ExN
InN
Other
UD

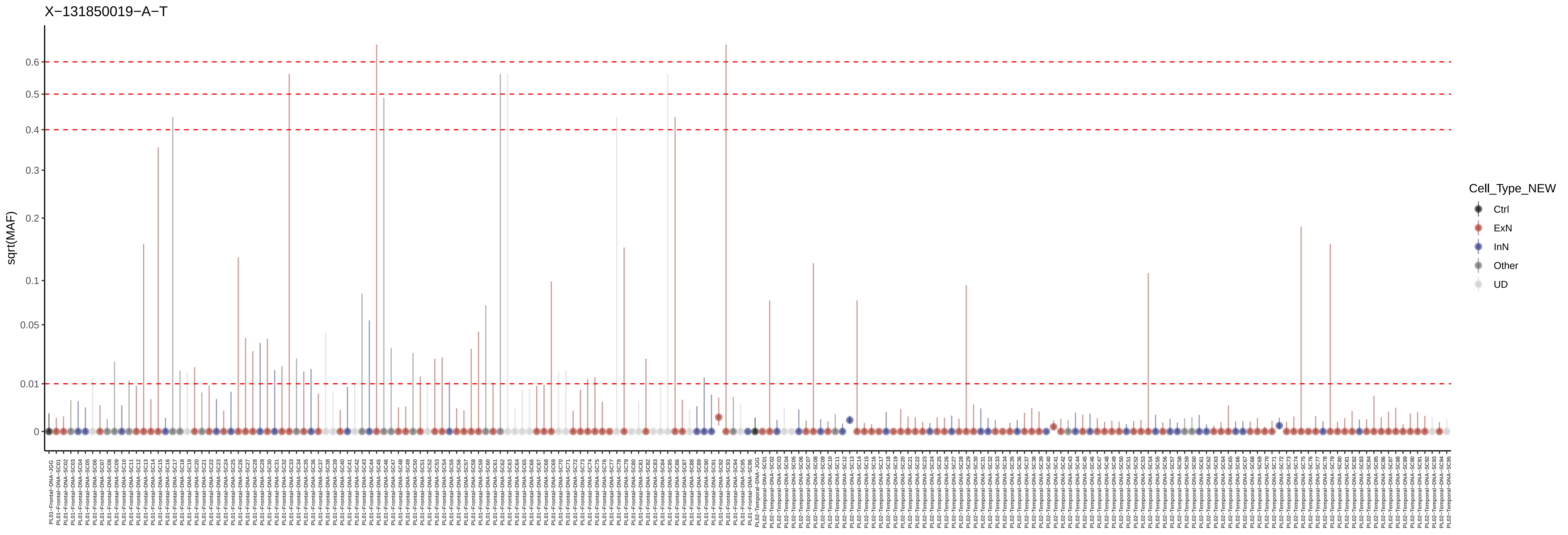
X−131850019−A−T
sqrt(MAF)
0.6
0.5
0.4
0.3
0.2
0.1
0.05
0.01
0
Cell_Type_NEW
Ctrl
ExN
InN
Other
UD

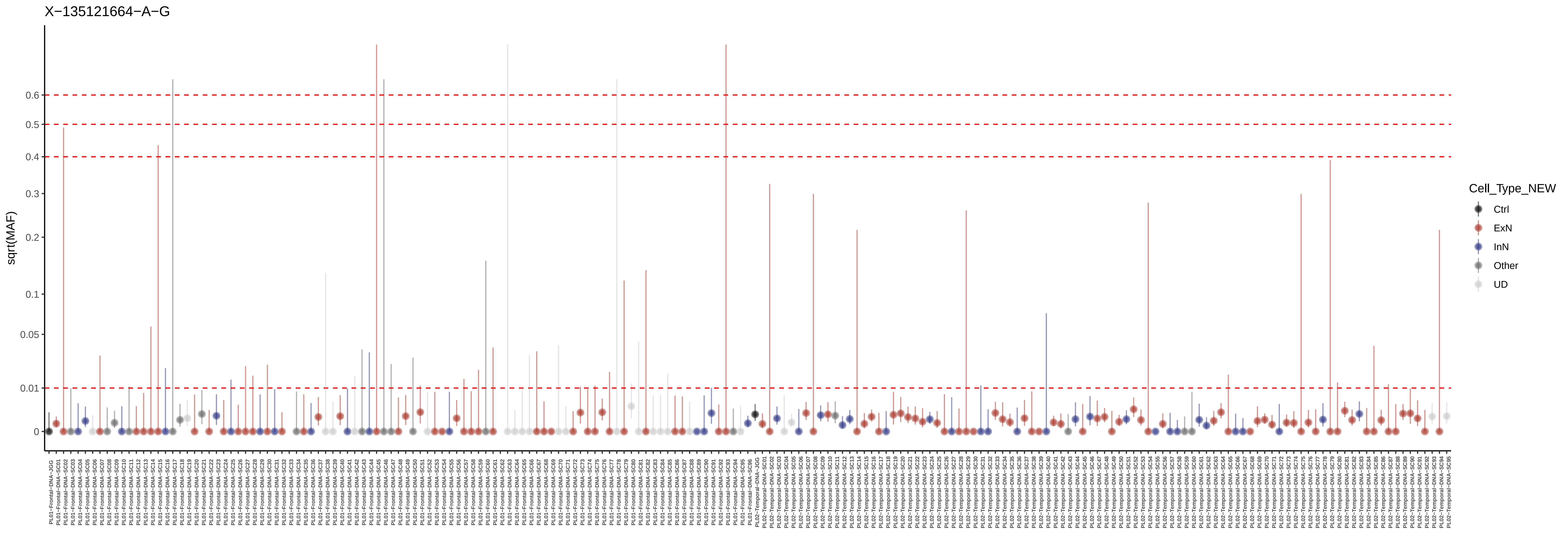
X−135121664−A−G
sqrt(MAF)
0.6
0.5
0.4
0.3
0.2
0.1
0.05
0.01
0
Cell_Type_NEW
Ctrl
ExN
InN
Other
UD

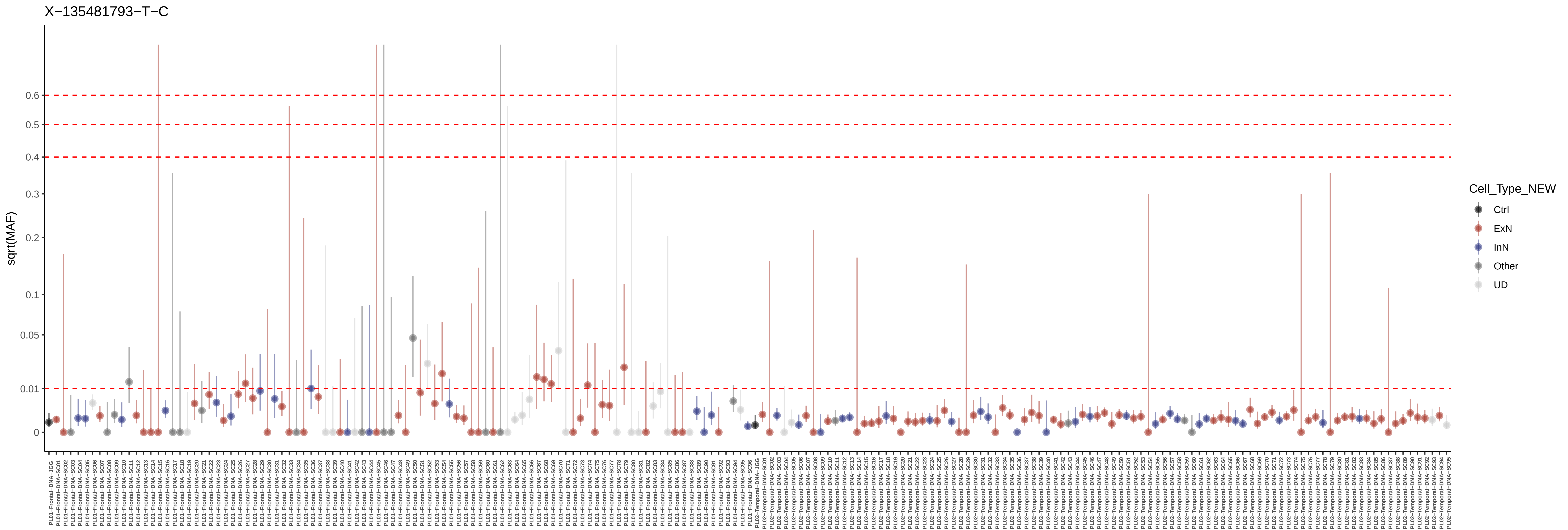
X−135481793−T−C
sqrt(MAF)
0.6
0.5
0.4
0.3
0.2
0.1
0.05
0.01
0
Cell_Type_NEW
Ctrl
ExN
InN
Other
UD

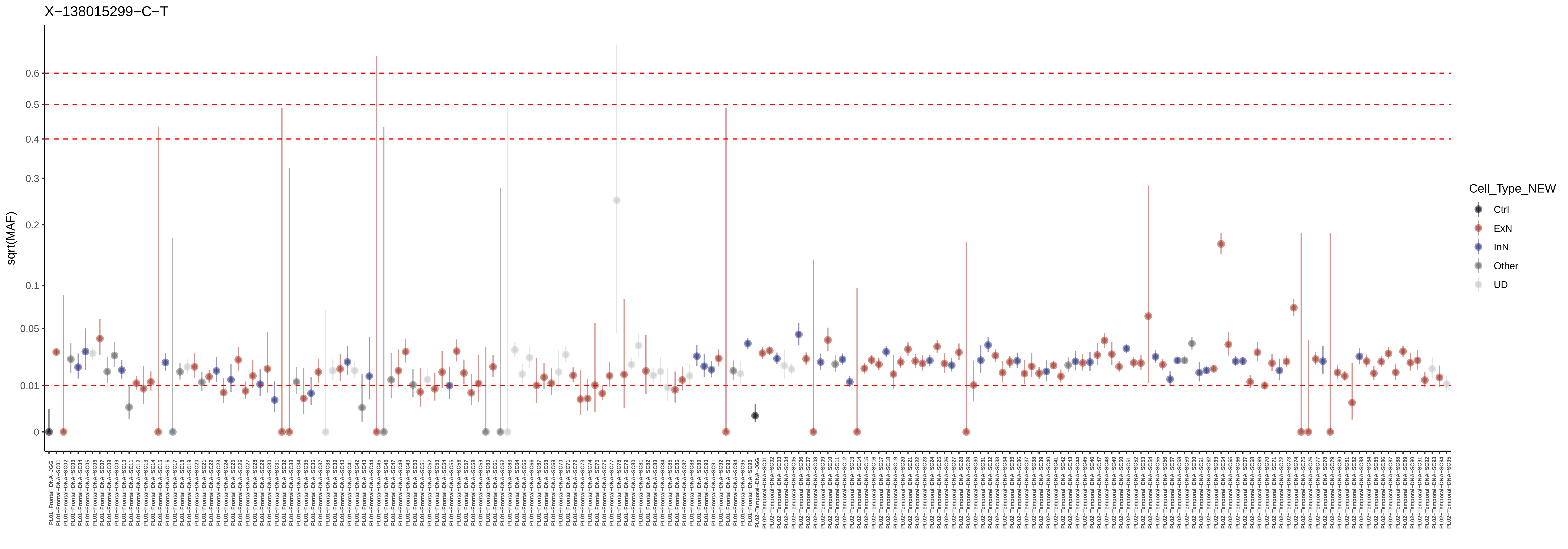

X−138015299−C−T
sqrt(MAF)
0.6
0.5
0.4
0.3
0.2
0.1
0.05
0.01
0
Cell_Type_NEW
Ctrl
ExN
InN
Other
UD

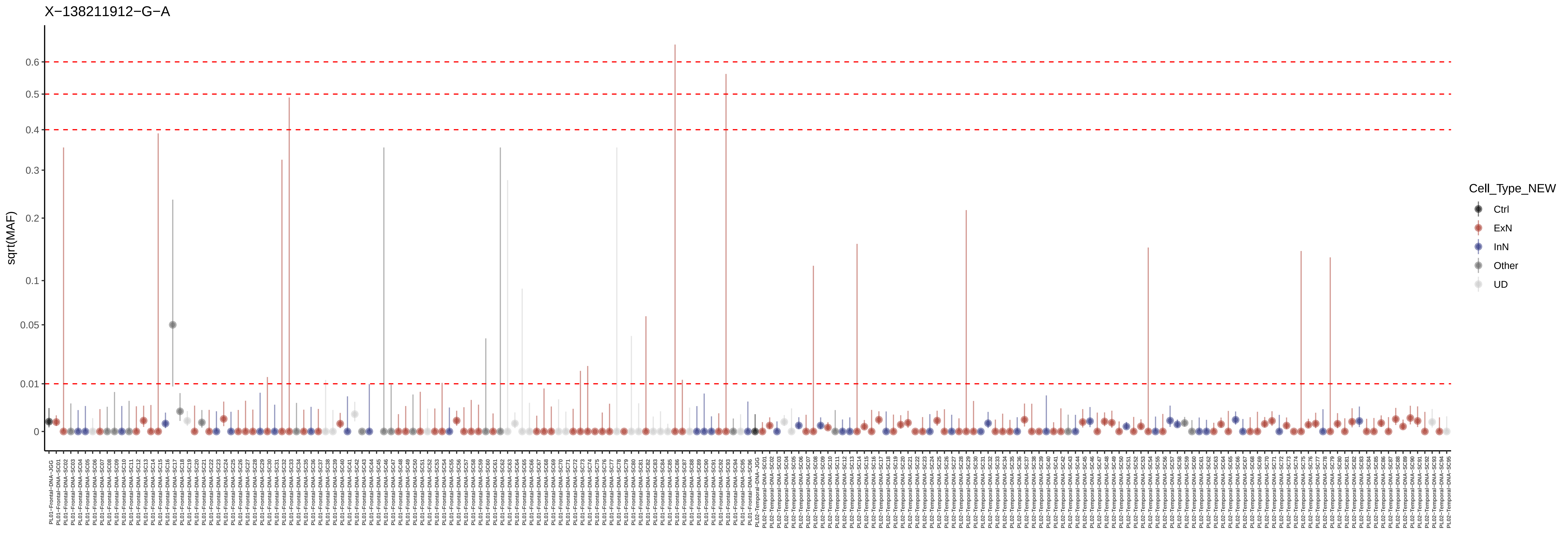
X−138211912−G−A
sqrt(MAF)
0.6
0.5
0.4
0.3
0.2
0.1
0.05
0.01
0
Cell_Type_NEW
Ctrl
ExN
InN
Other
UD

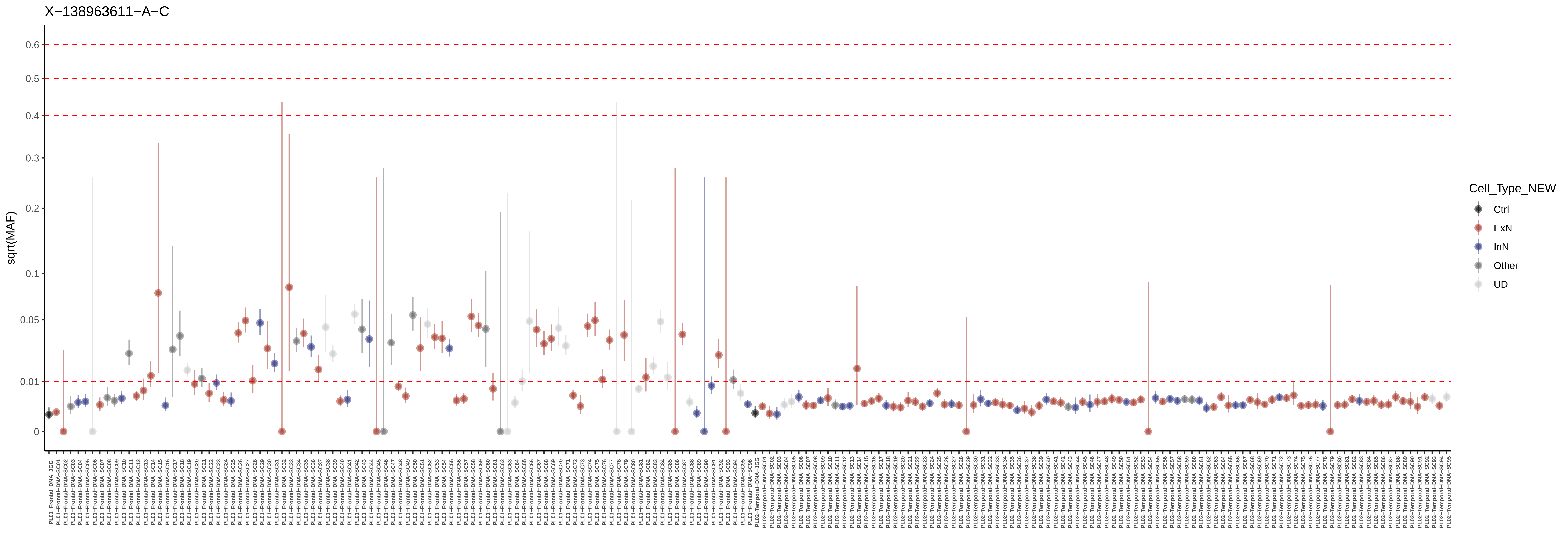

X−138963611−A−C
sqrt(MAF)
0.6
0.5
0.4
0.3
0.2
0.1
0.05
0.01
0
Cell_Type_NEW
Ctrl
ExN
InN
Other
UD

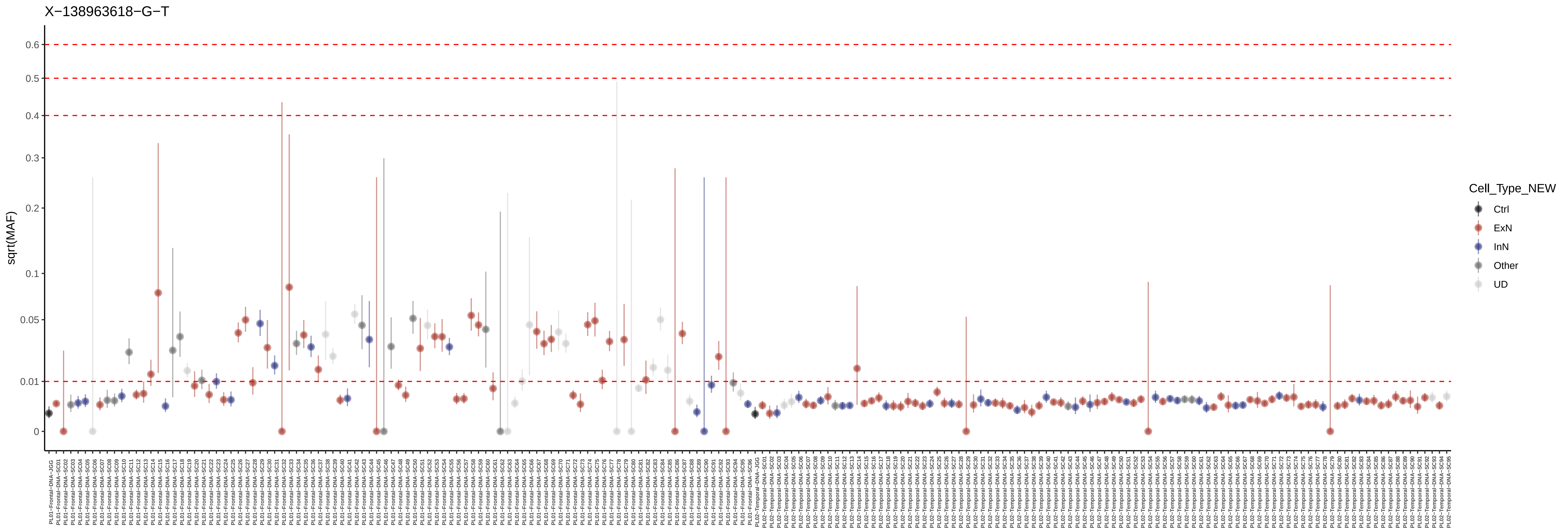
X−138963618−G−T
sqrt(MAF)
0.6
0.5
0.4
0.3
0.2
0.1
0.05
0.01
0
Cell_Type_NEW
Ctrl
ExN
InN
Other
UD

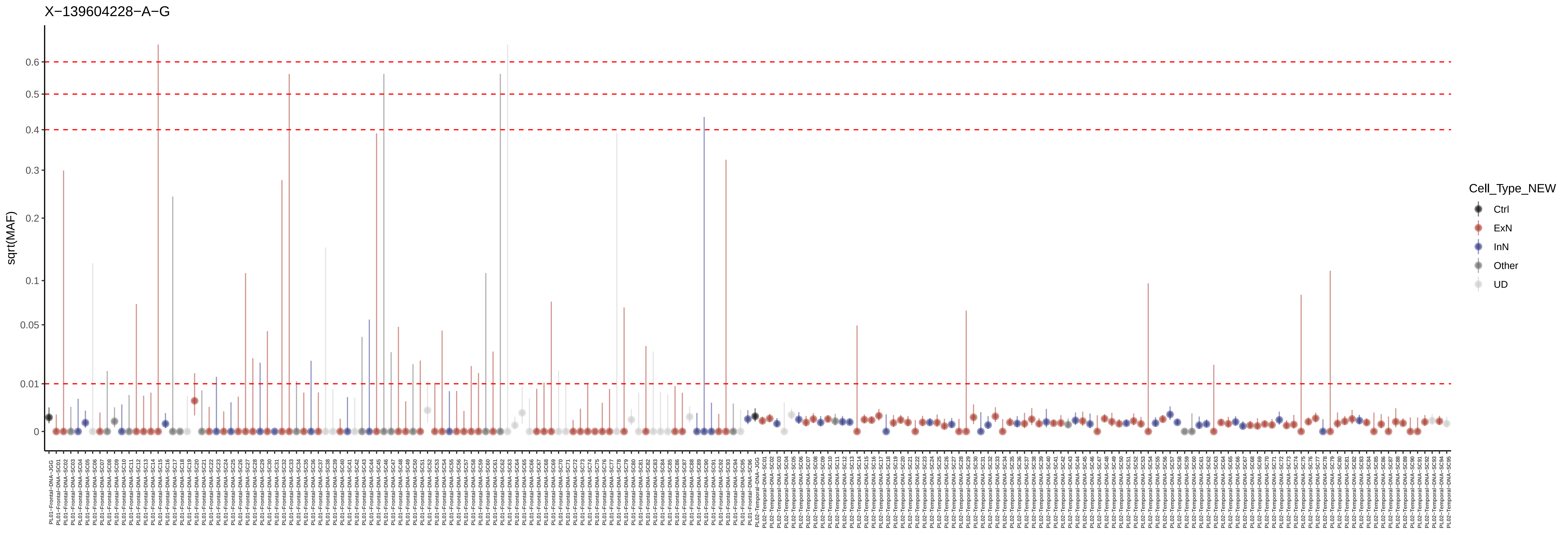
X−139604228−A−G
sqrt(MAF)
0.6
0.5
0.4
0.3
0.2
0.1
0.05
0.01
0
Cell_Type_NEW
Ctrl
ExN
InN
Other
UD

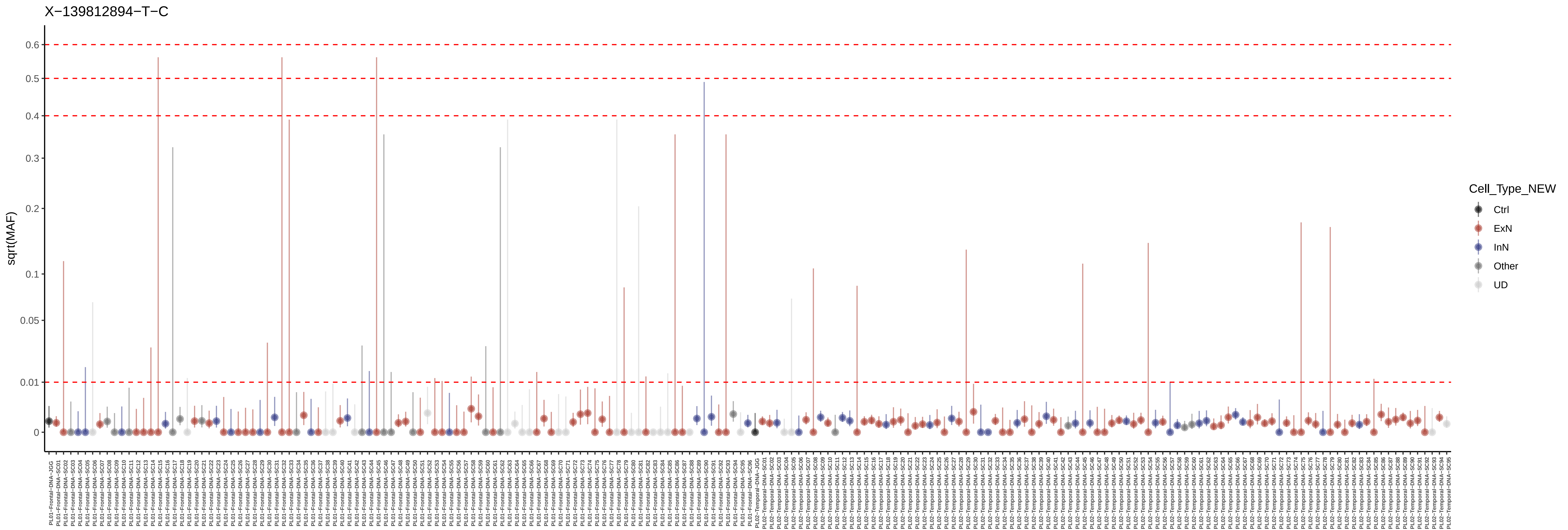

X−139812894−T−C
sqrt(MAF)
0.6
0.5
0.4
0.3
0.2
0.1
0.05
0.01
0
Cell_Type_NEW
Ctrl
ExN
InN
Other
UD

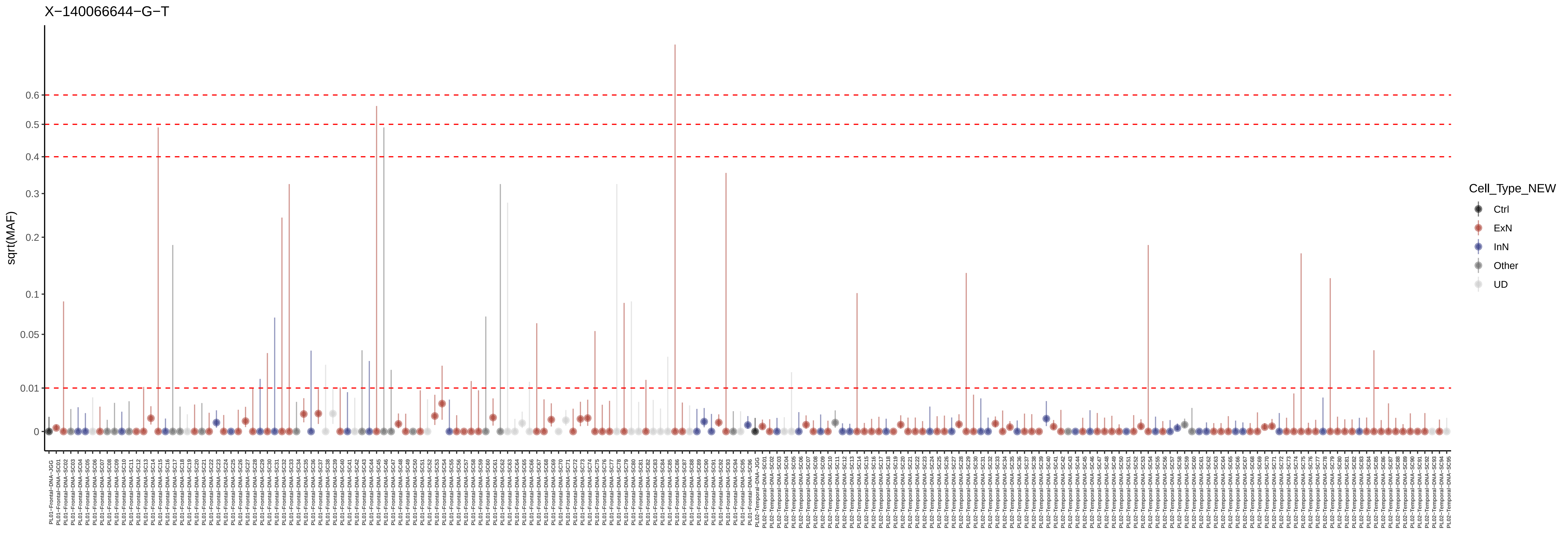

X−140066644−G−T
sqrt(MAF)
0.6
0.5
0.4
0.3
0.2
0.1
0.05
0.01
0
Cell_Type_NEW
Ctrl
ExN
InN
Other
UD

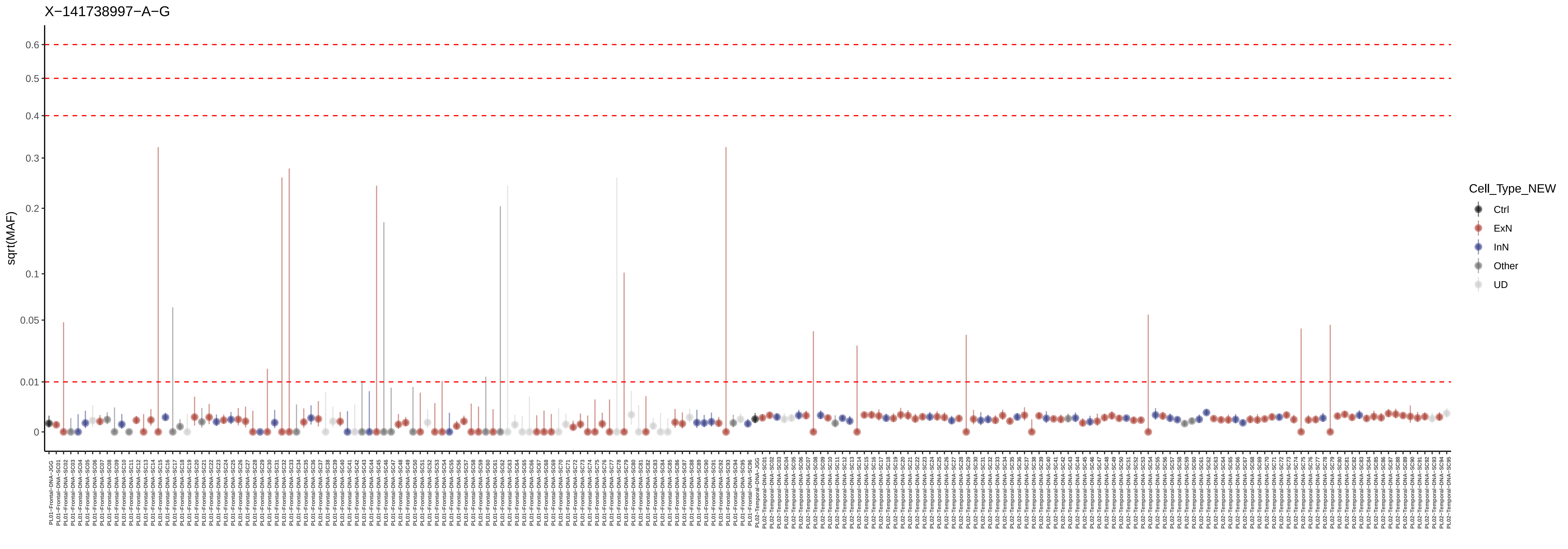
X−141738997−A−G
sqrt(MAF)
0.6
0.5
0.4
0.3
0.2
0.1
0.05
0.01
0
Cell_Type_NEW
Ctrl
ExN
InN
Other
UD

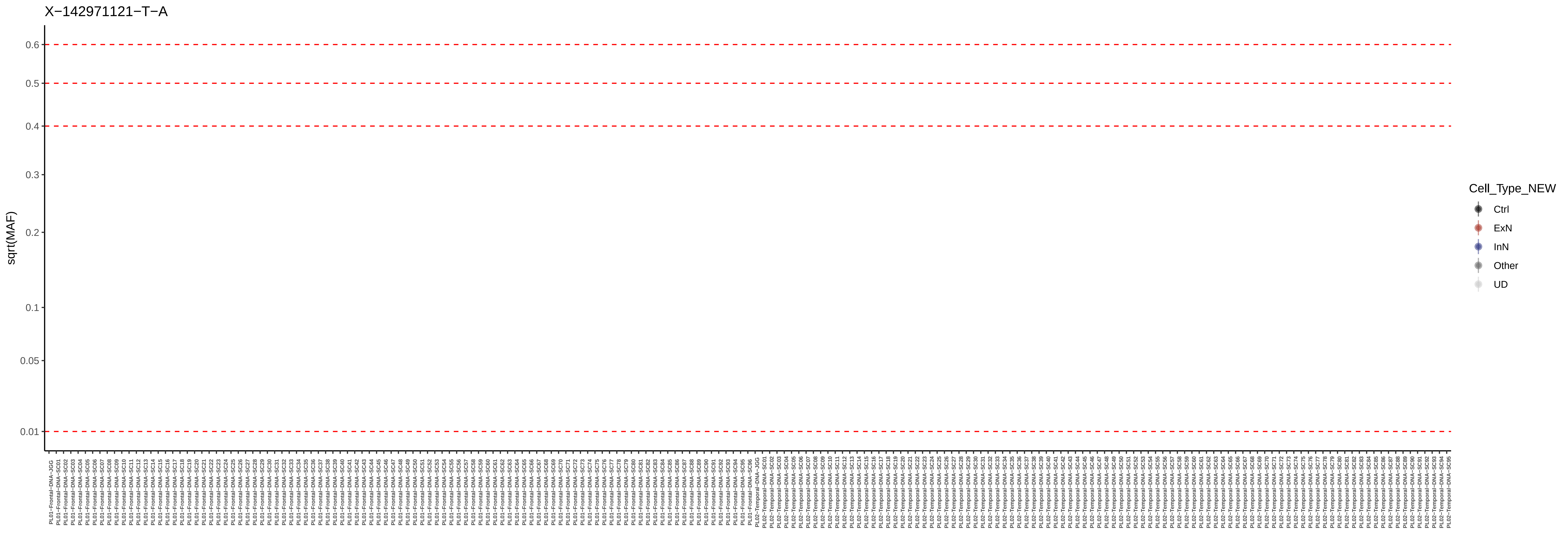
X−142971121−T−A
sqrt(MAF)
0.6
0.5
0.4
0.3
0.2
0.1
0.05
0.01
Cell_Type_NEW
Ctrl
ExN
InN
Other
UD

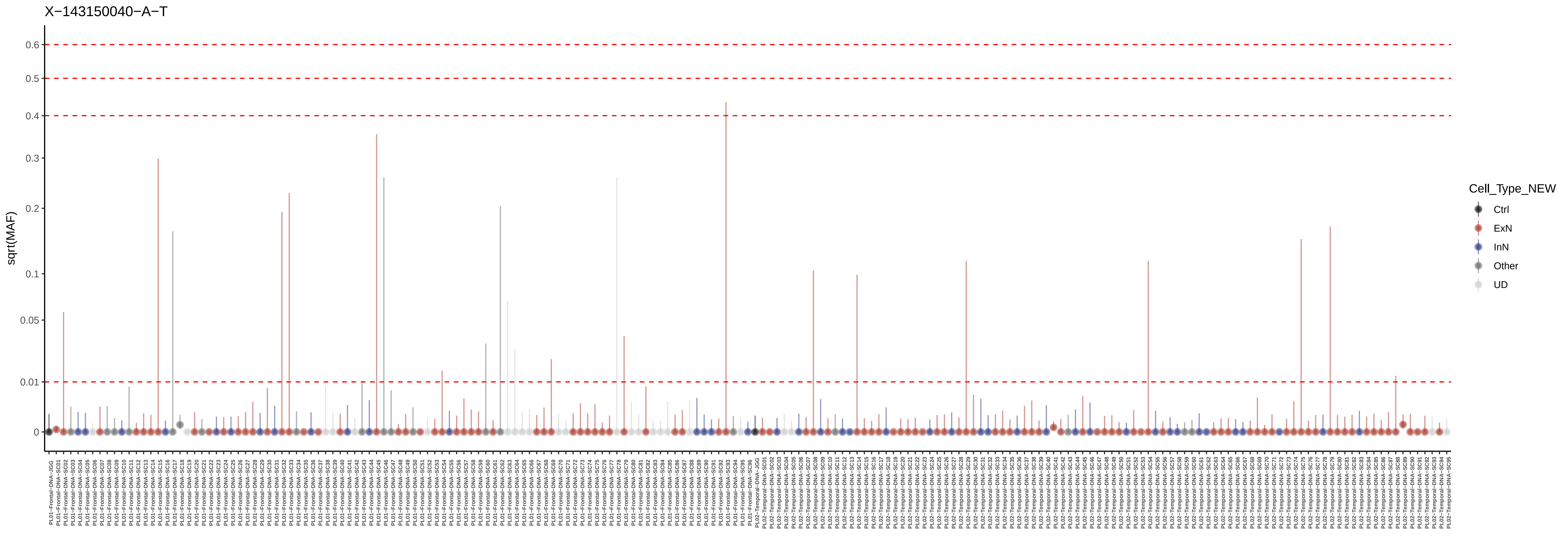
X−143150040−A−T
sqrt(MAF)
0.6
0.5
0.4
0.3
0.2
0.1
0.05
0.01
0
Cell_Type_NEW
Ctrl
ExN
InN
Other
UD

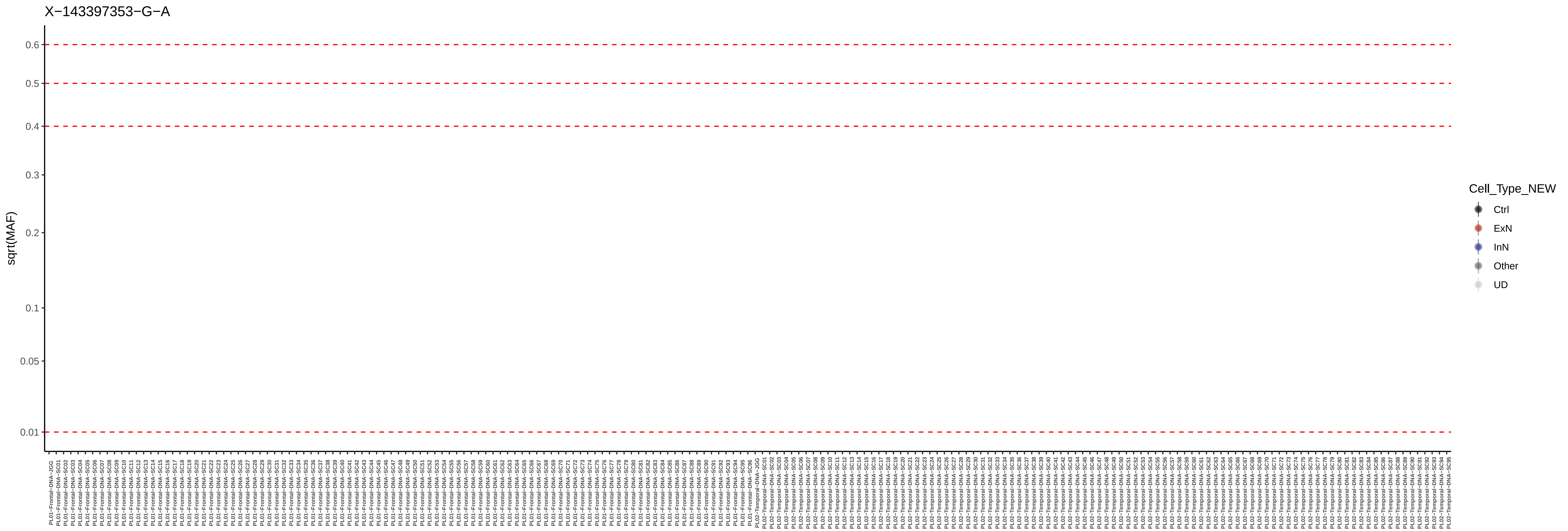

X−143397353−G−A
sqrt(MAF)
0.6
0.5
0.4
0.3
0.2
0.1
0.05
0.01
Cell_Type_NEW
Ctrl
ExN
InN
Other
UD

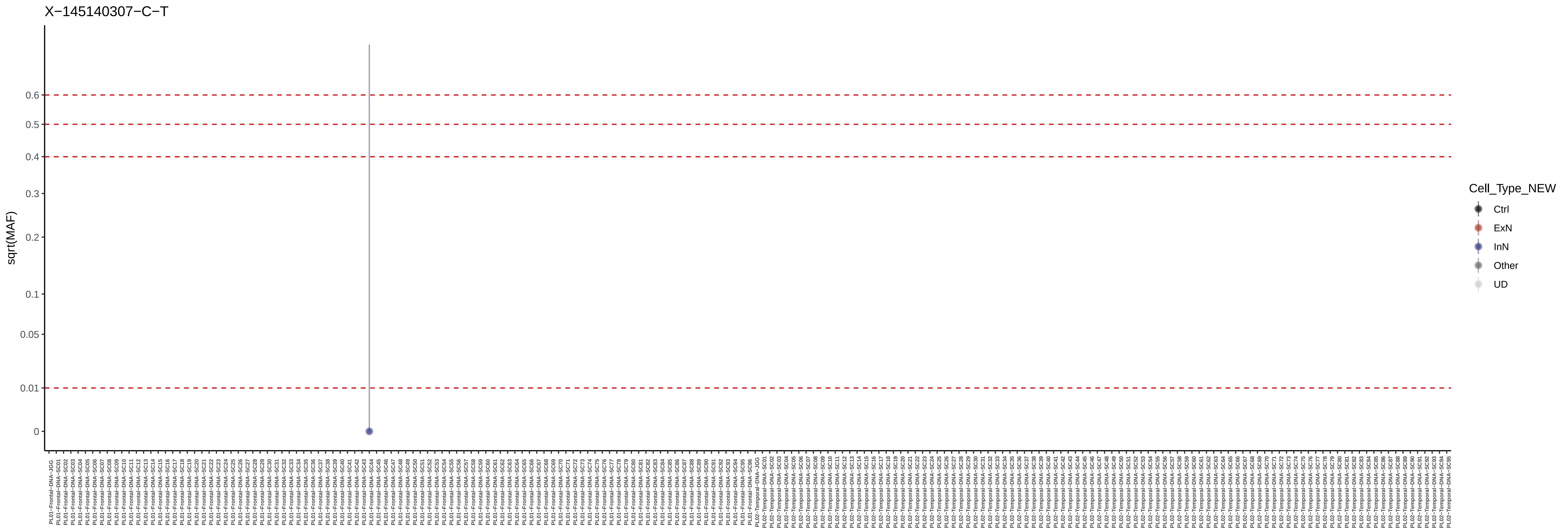
X−145140307−C−T
sqrt(MAF)
0.6
0.5
0.4
0.3
0.2
0.1
0.05
0.01
0
Cell_Type_NEW
Ctrl
ExN
InN
Other
UD

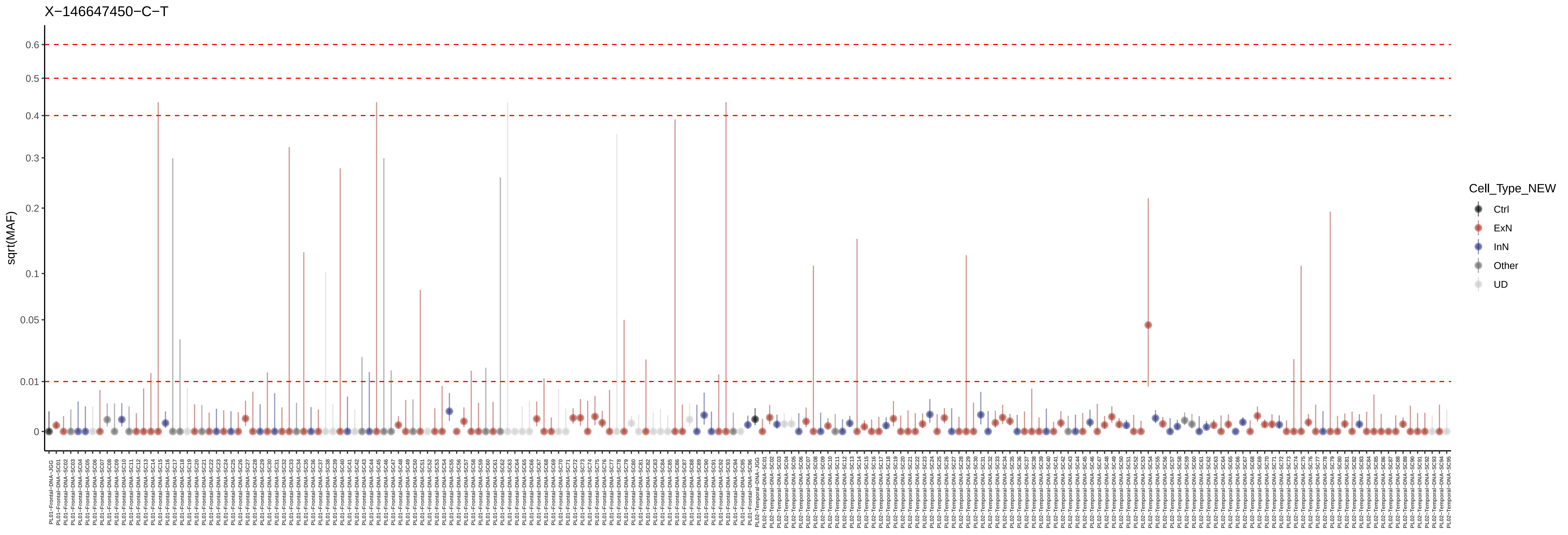
X−146647450−C−T
sqrt(MAF)
0.6
0.5
0.4
0.3
0.2
0.1
0.05
0.01
0
Cell_Type_NEW
Ctrl
ExN
InN
Other
UD

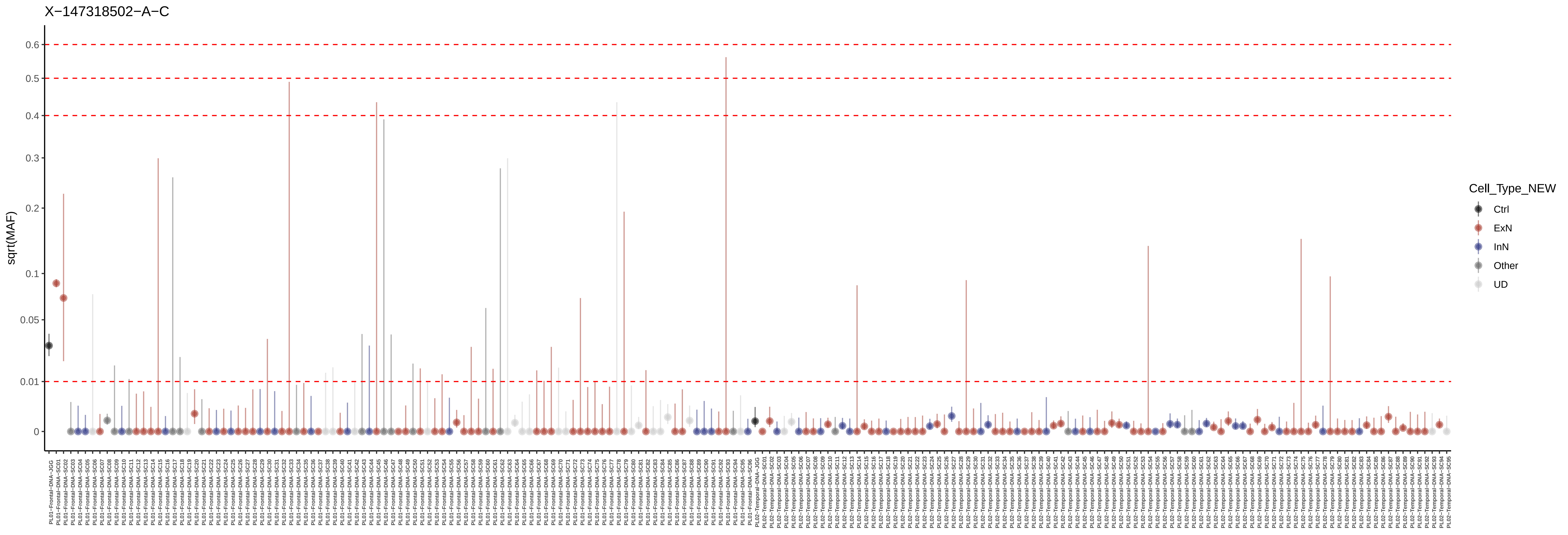

X−147318502−A−C
sqrt(MAF)
0.6
0.5
0.4
0.3
0.2
0.1
0.05
0.01
0
Cell_Type_NEW
Ctrl
ExN
InN
Other
UD

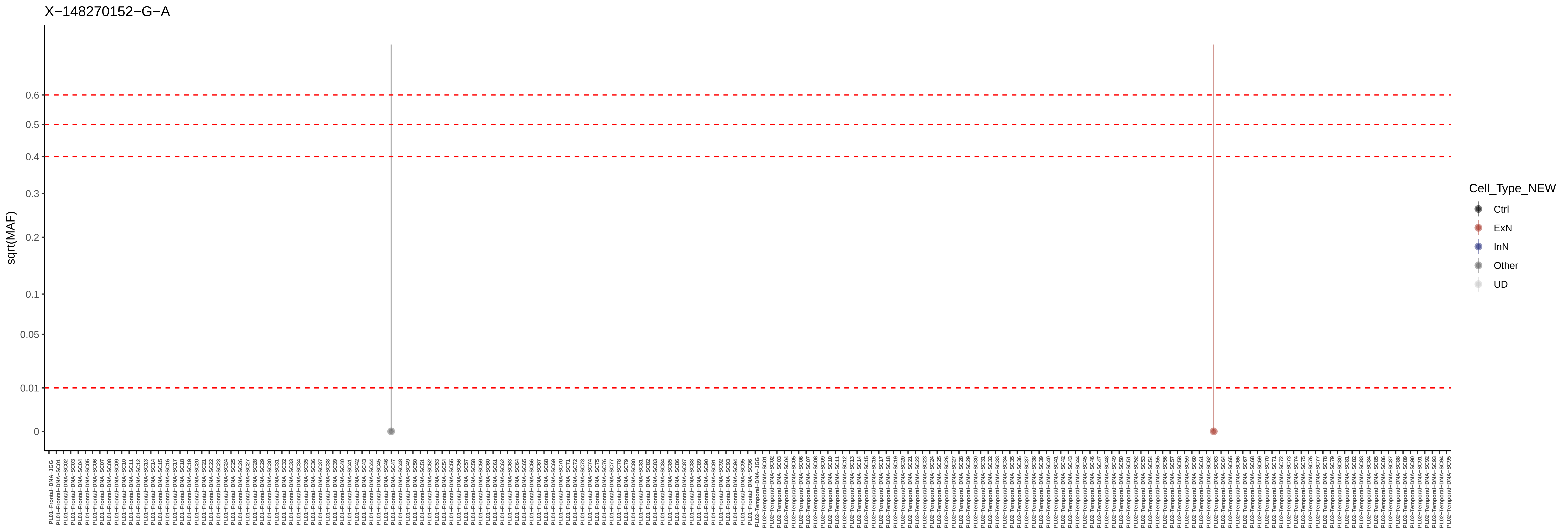
X−148270152−G−A
sqrt(MAF)
0.6
0.5
0.4
0.3
0.2
0.1
0.05
0.01
0
Cell_Type_NEW
Ctrl
ExN
InN
Other
UD

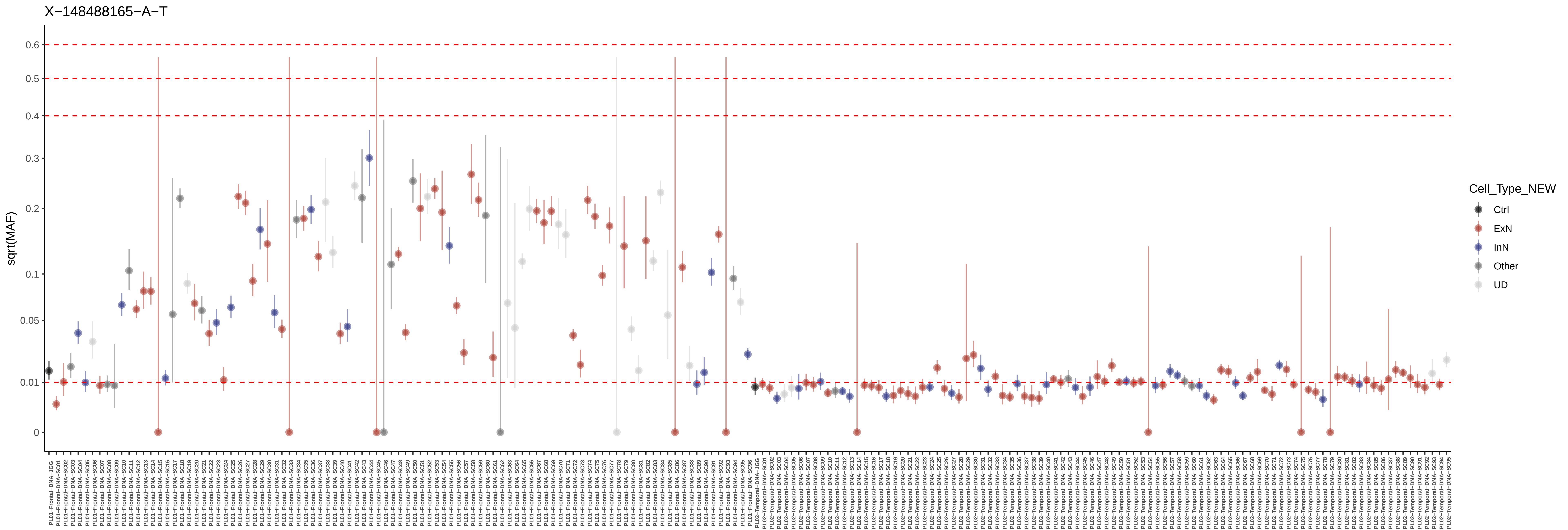

X−148488165−A−T
sqrt(MAF)
0.6
0.5
0.4
0.3
0.2
0.1
0.05
0.01
0
Cell_Type_NEW
Ctrl
ExN
InN
Other
UD

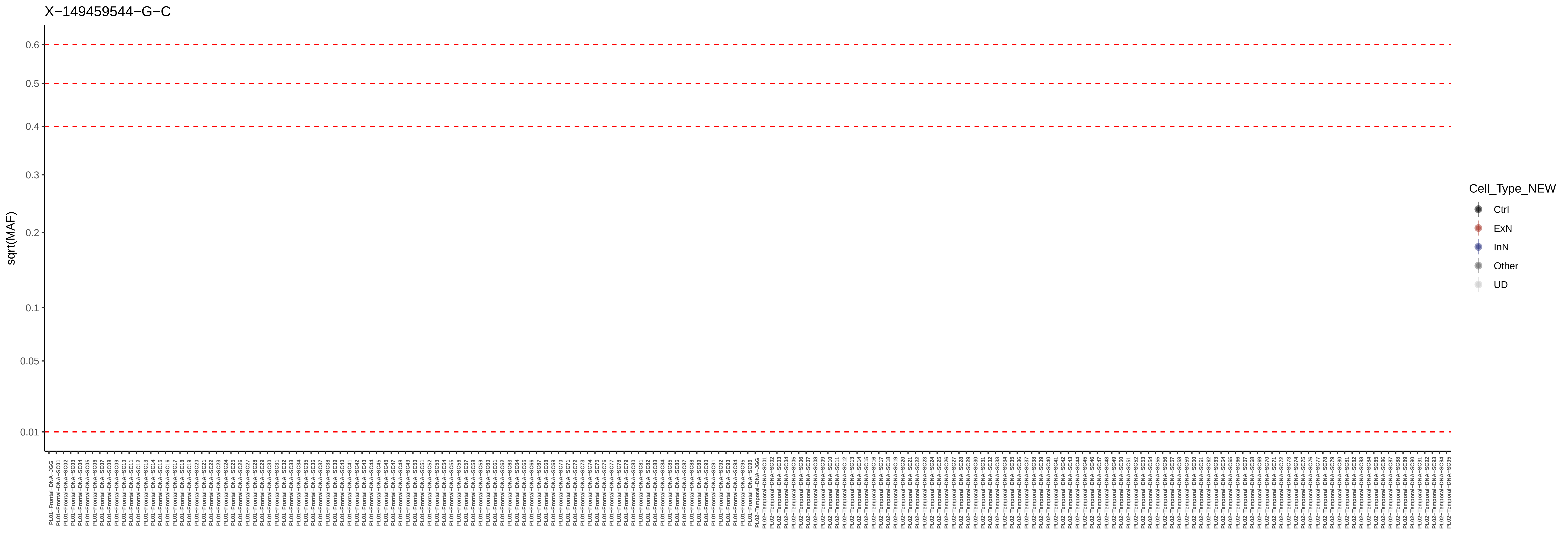
X−149459544−G−C
sqrt(MAF)
Cell_Type_NEW
Ctrl
ExN
InN
Other
UD

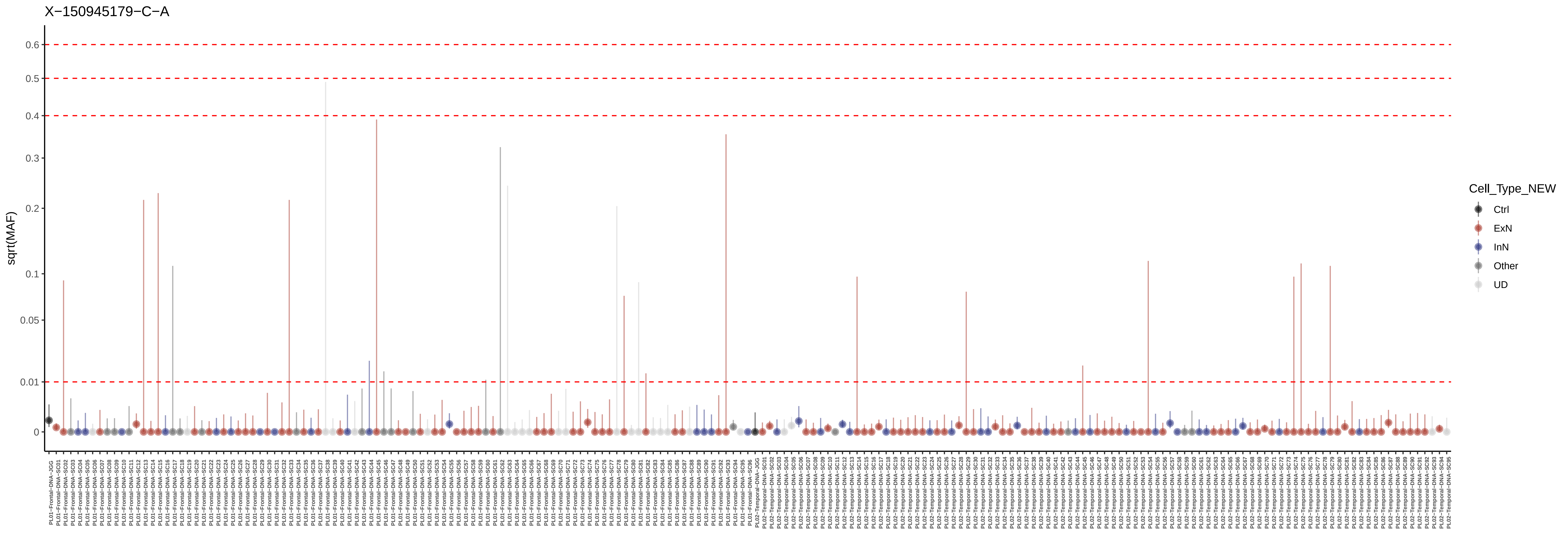
X−150945179−C−A
sqrt(MAF)
0.6
0.5
0.4
0.3
0.2
0.1
0.05
0.01
0
Cell_Type_NEW
Ctrl
ExN
InN
Other
UD

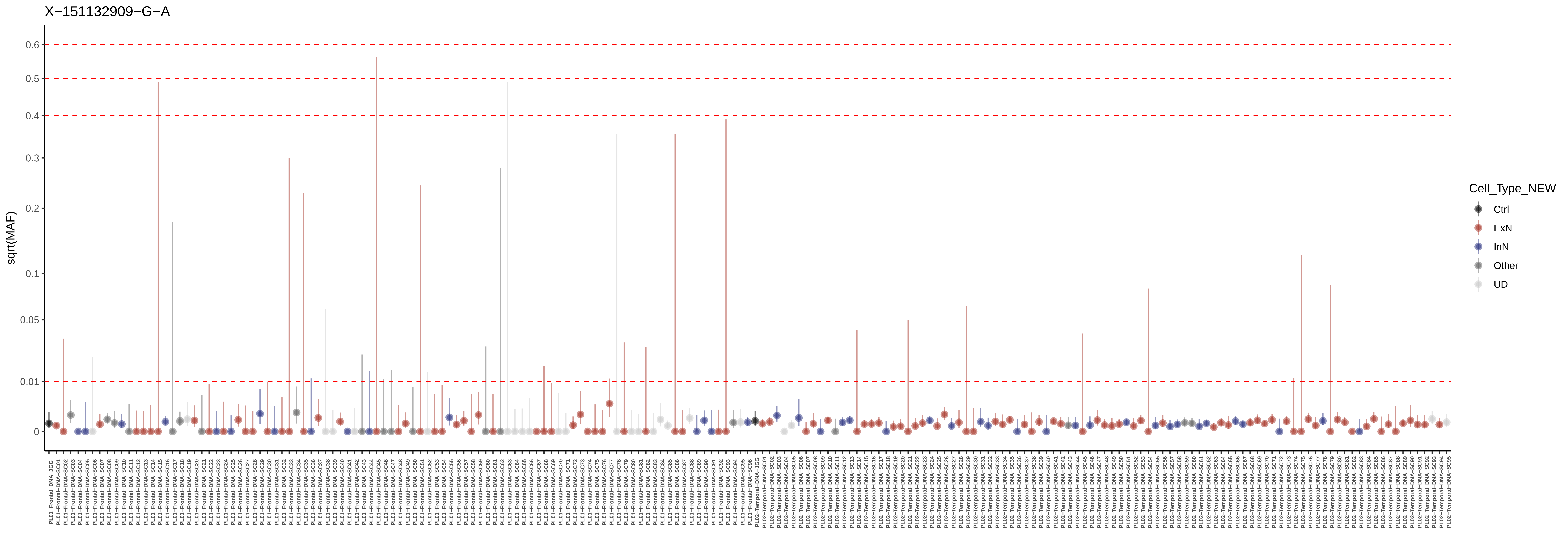
X−151132909−G−A
sqrt(MAF)
0.6
0.5
0.4
0.3
0.2
0.1
0.05
0.01
0
Cell_Type_NEW
Ctrl
ExN
InN
Other
UD

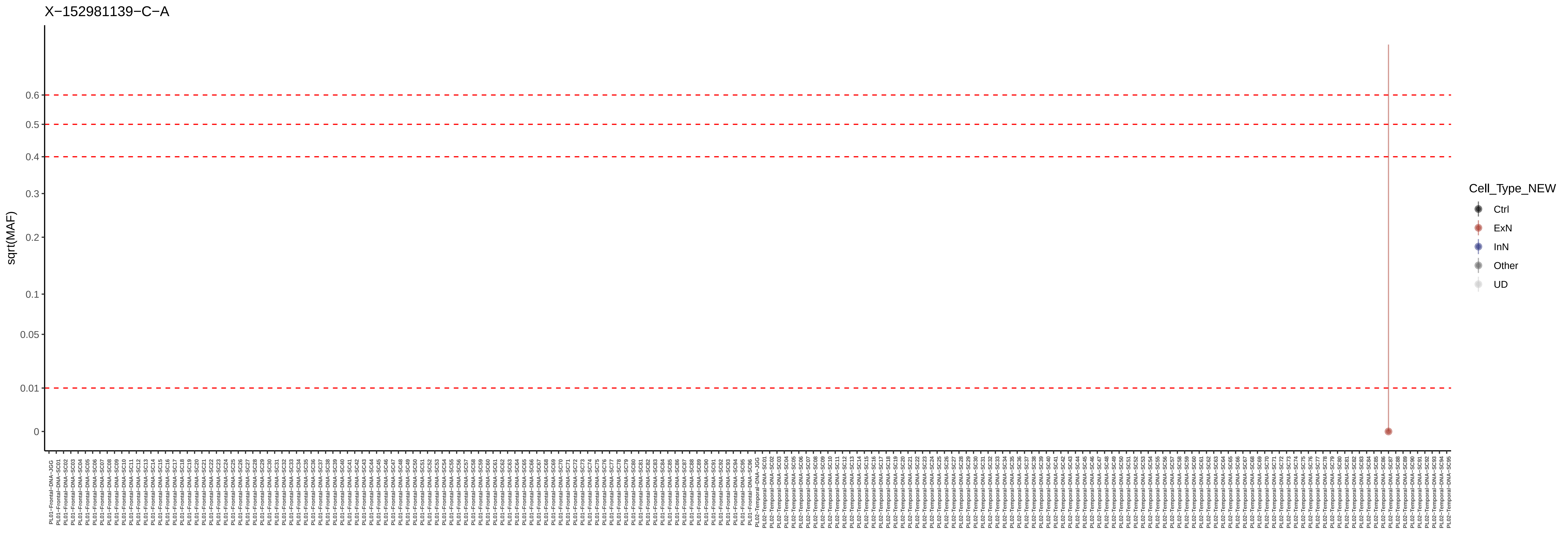
X−152981139−C−A
sqrt(MAF)
0.6
0.5
0.4
0.3
0.2
0.1
0.05
0.01
0
Cell_Type_NEW
Ctrl
ExN
InN
Other
UD

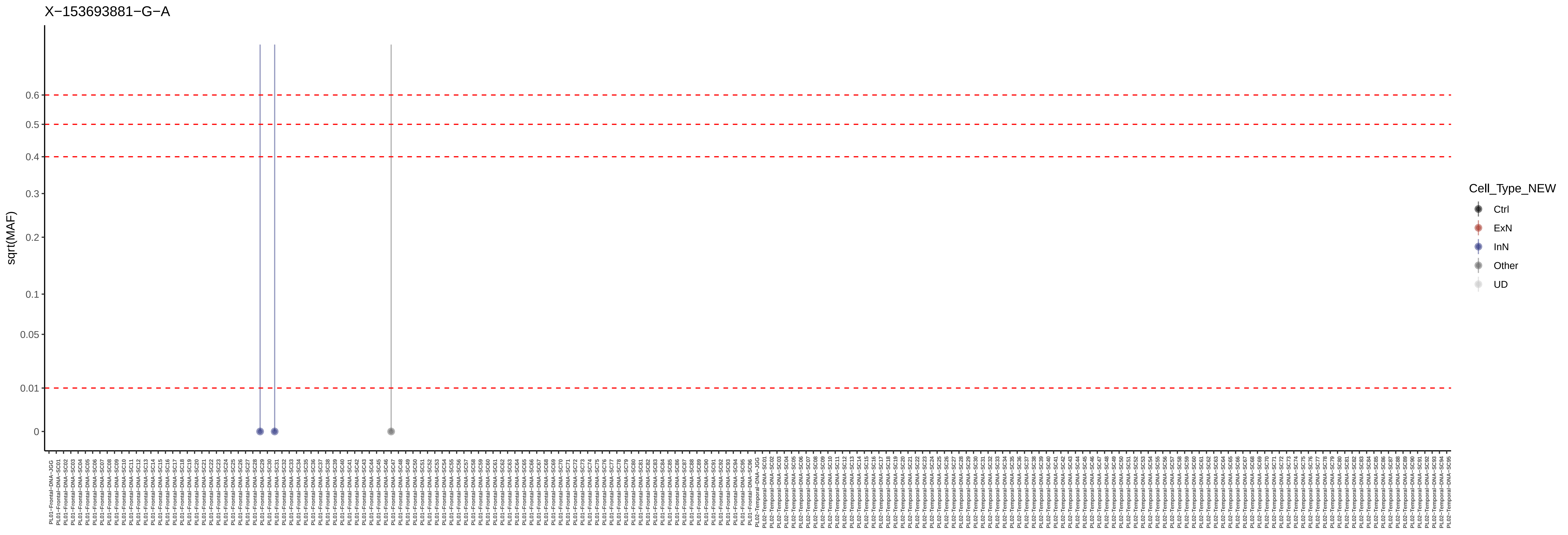

X−153693881−G−A
sqrt(MAF)
0.6
0.5
0.4
0.3
0.2
0.1
0.05
0.01
0
Cell_Type_NEW
Ctrl
ExN
InN
Other
UD

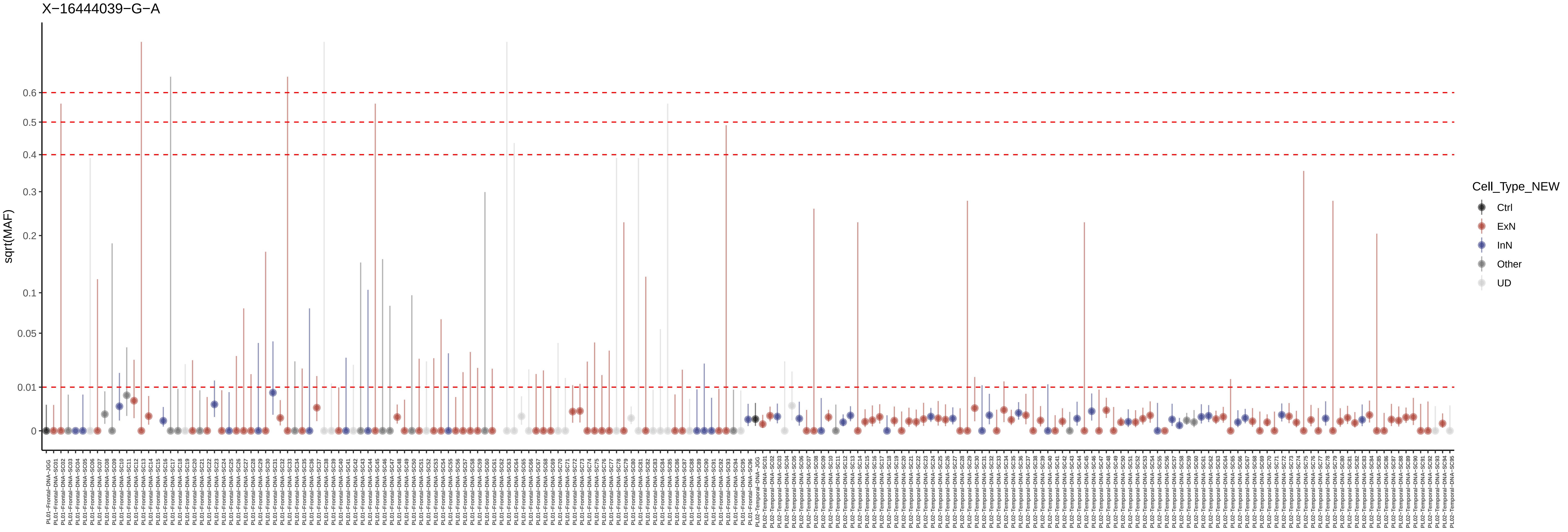
X−16444039−G−A
sqrt(MAF)
0.6
0.5
0.4
0.3
0.2
0.1
0.05
0.01
0
Cell_Type_NEW
Ctrl
ExN
InN
Other
UD

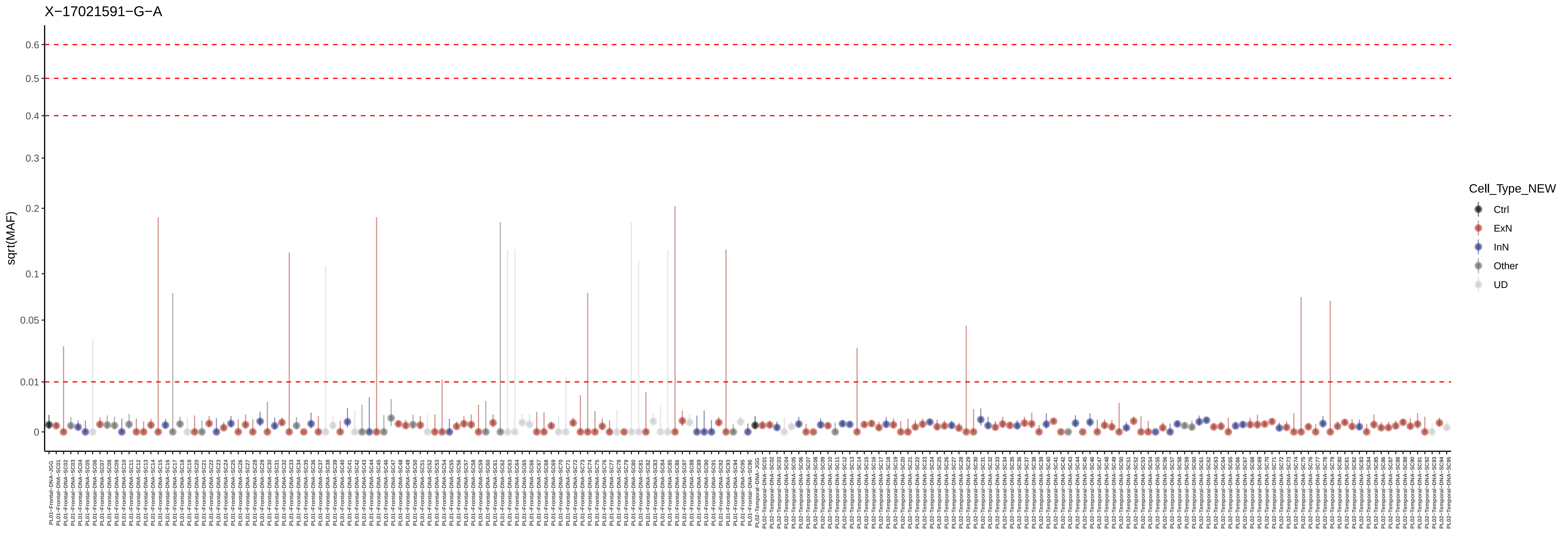
X−17021591−G−A
sqrt(MAF)
0.6
0.5
0.4
0.3
0.2
0.1
0.05
0.01
0
Cell_Type_NEW
Ctrl
ExN
InN
Other
UD
PL01−Frontal−DNA−JGG
PL02−Temporal−DNA−JGG

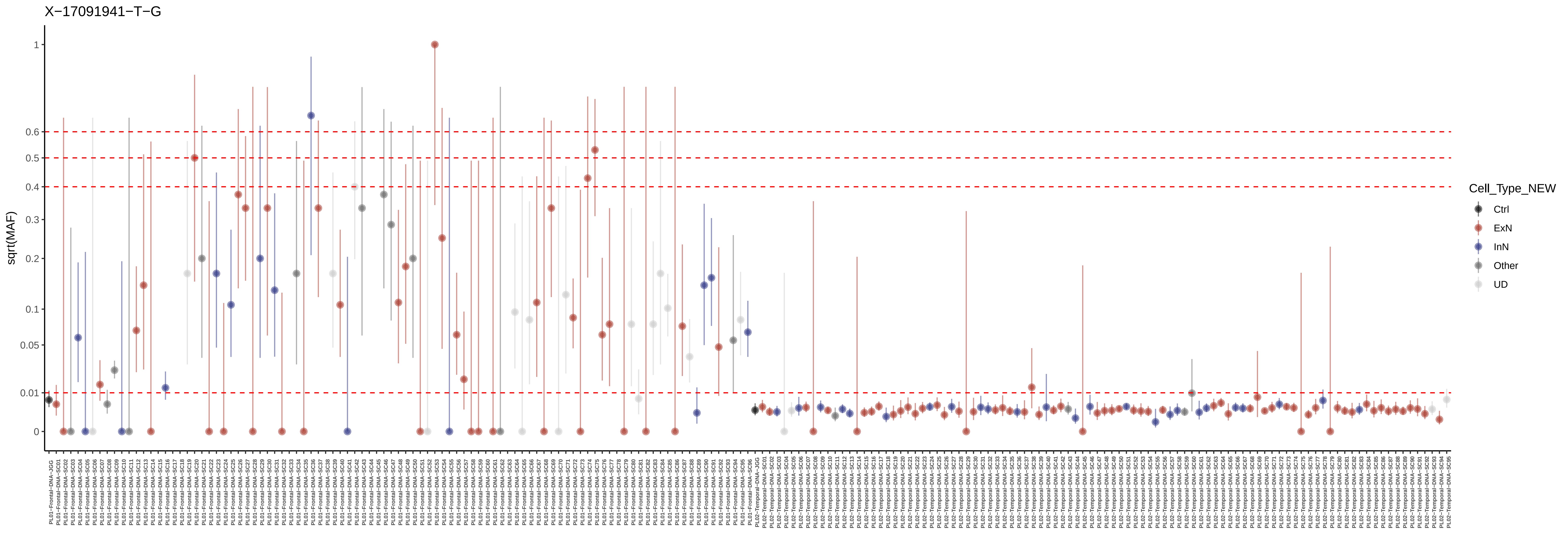
X−17091941−T−G
sqrt(MAF)
Cell_Type_NEW
Ctrl
ExN
InN
Other
UD

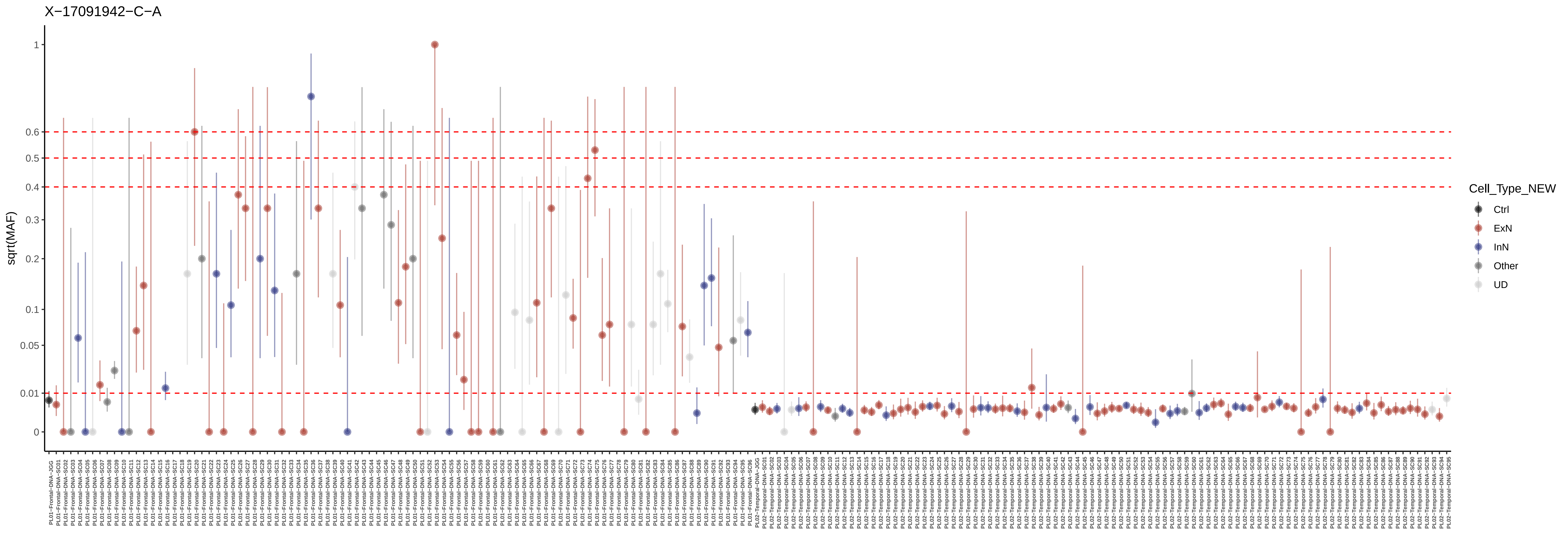
X−17091942−C−A
sqrt(MAF)
1
0.6
0.5
0.4
0.3
0.2
0.1
0.05
0.01
0
Cell_Type_NEW
Ctrl
ExN
InN
Other
UD

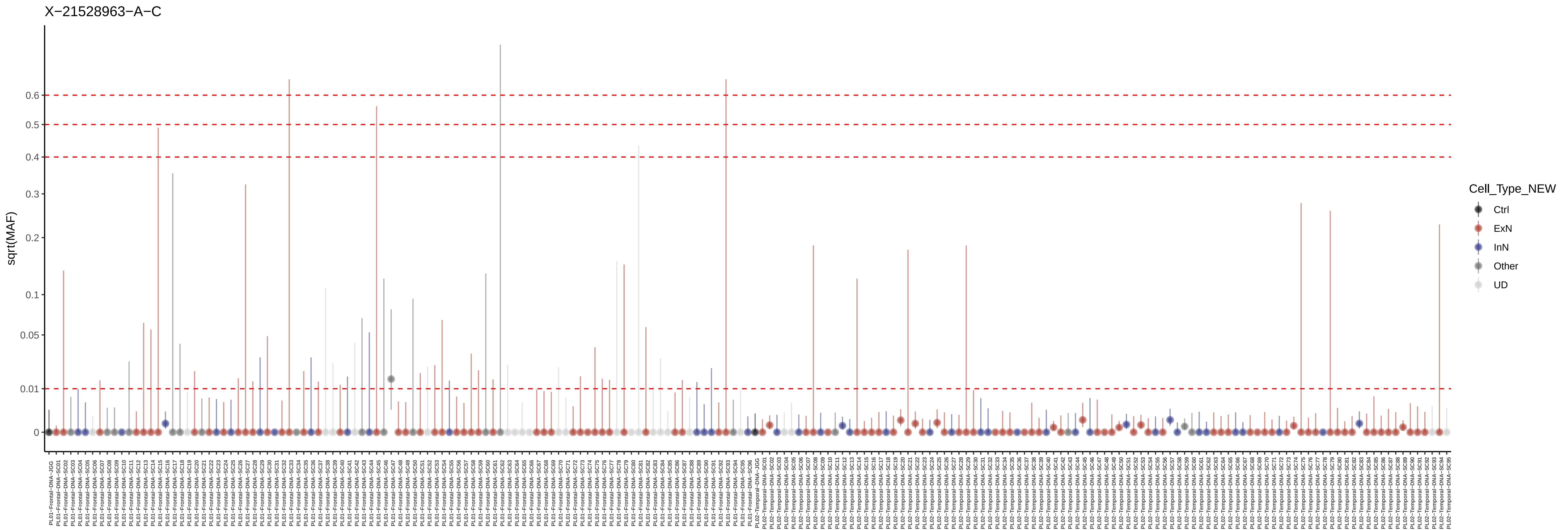
X−21528963−A−C
sqrt(MAF)
0.6
0.5
0.4
0.3
0.2
0.1
0.05
0.01
0
Cell_Type_NEW
Ctrl
ExN
InN
Other
UD

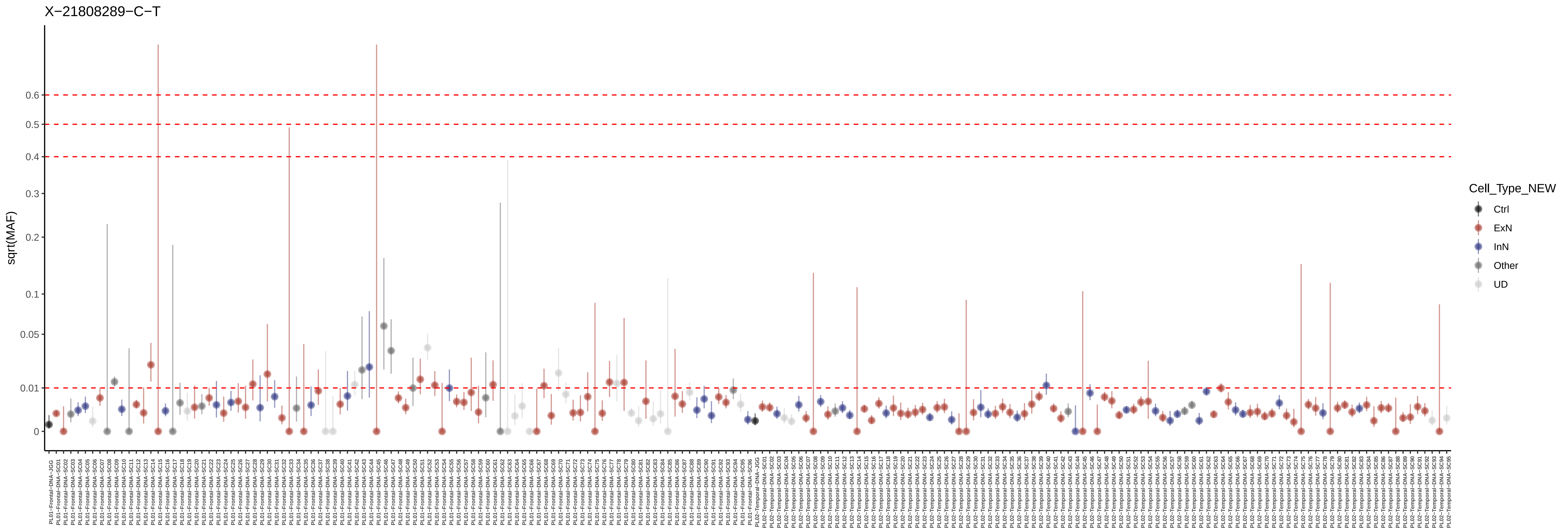
X−21808289−C−T
sqrt(MAF)
0.6
0.5
0.4
0.3
0.2
0.1
0.05
0.01
0
Cell_Type_NEW
Ctrl
ExN
InN
Other
UD

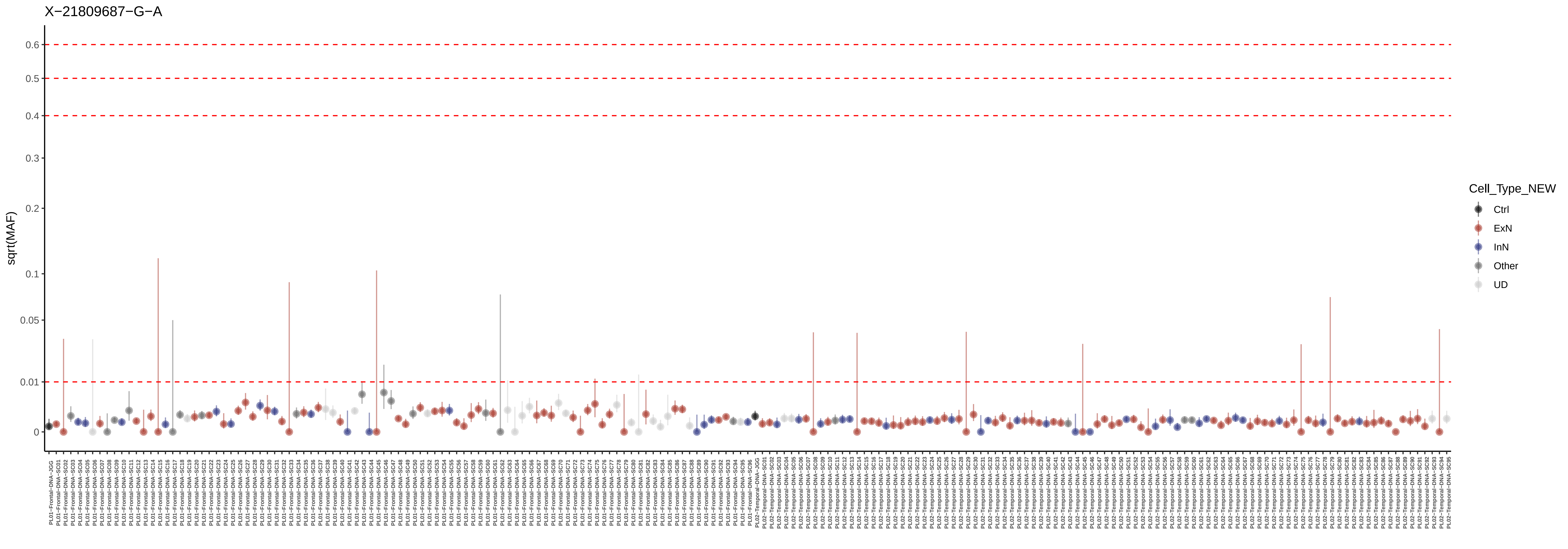

X−21809687−G−A
sqrt(MAF)
0.6
0.5
0.4
0.3
0.2
0.1
0.05
0.01
0
Cell_Type_NEW
Ctrl
ExN
InN
Other
UD

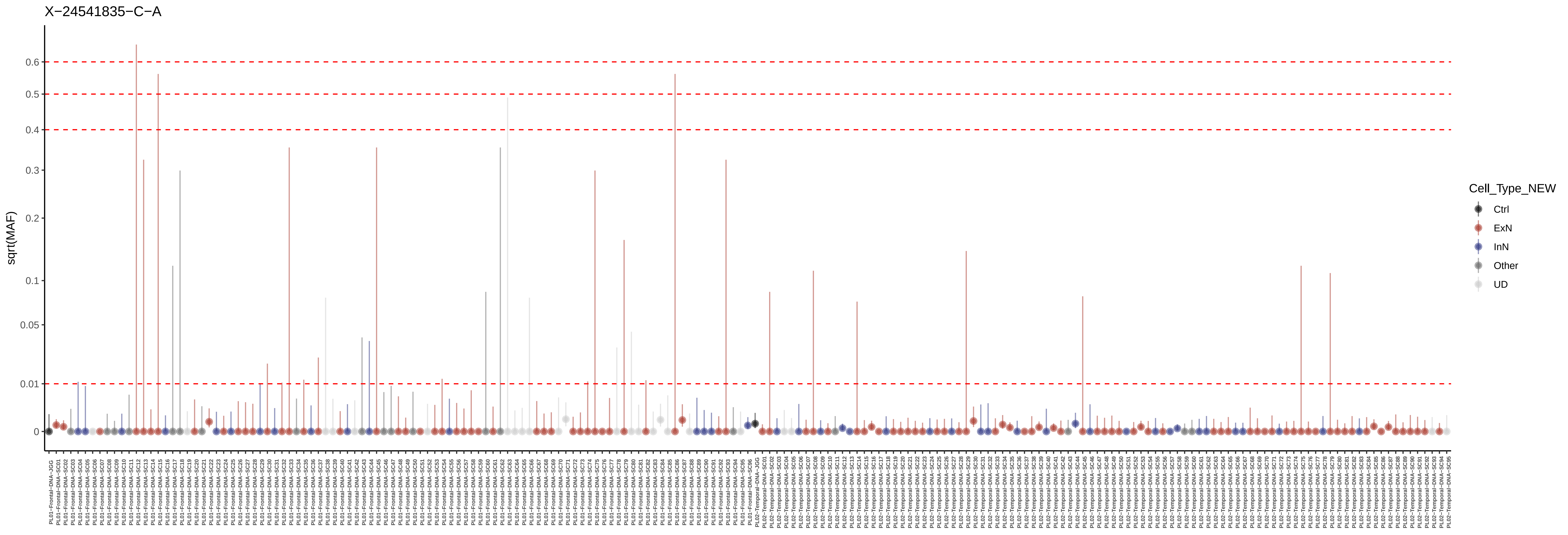
X−24541835−C−A
sqrt(MAF)
0.6
0.5
0.4
0.3
0.2
0.1
0.05
0.01
0
Cell_Type_NEW
Ctrl
ExN
InN
Other
UD

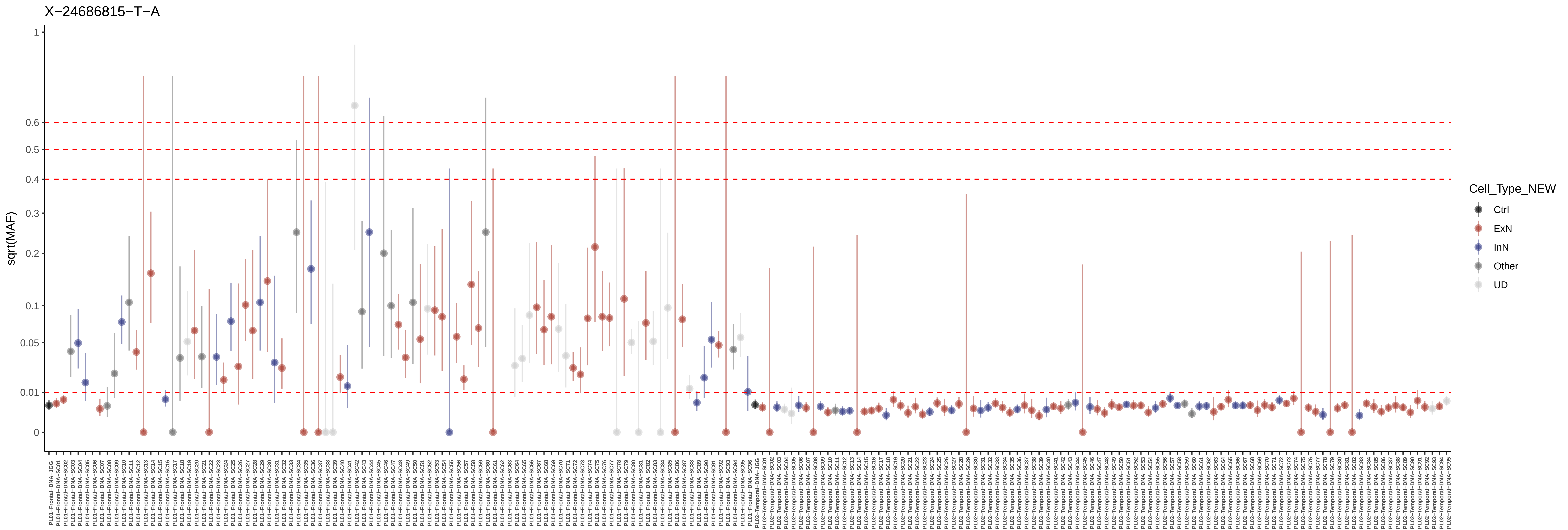
X−24686815−T−A
sqrt(MAF)
1
0.6
0.5
0.4
0.3
0.2
0.1
0.05
0.01
0
Cell_Type_NEW
Ctrl
ExN
InN
Other
UD

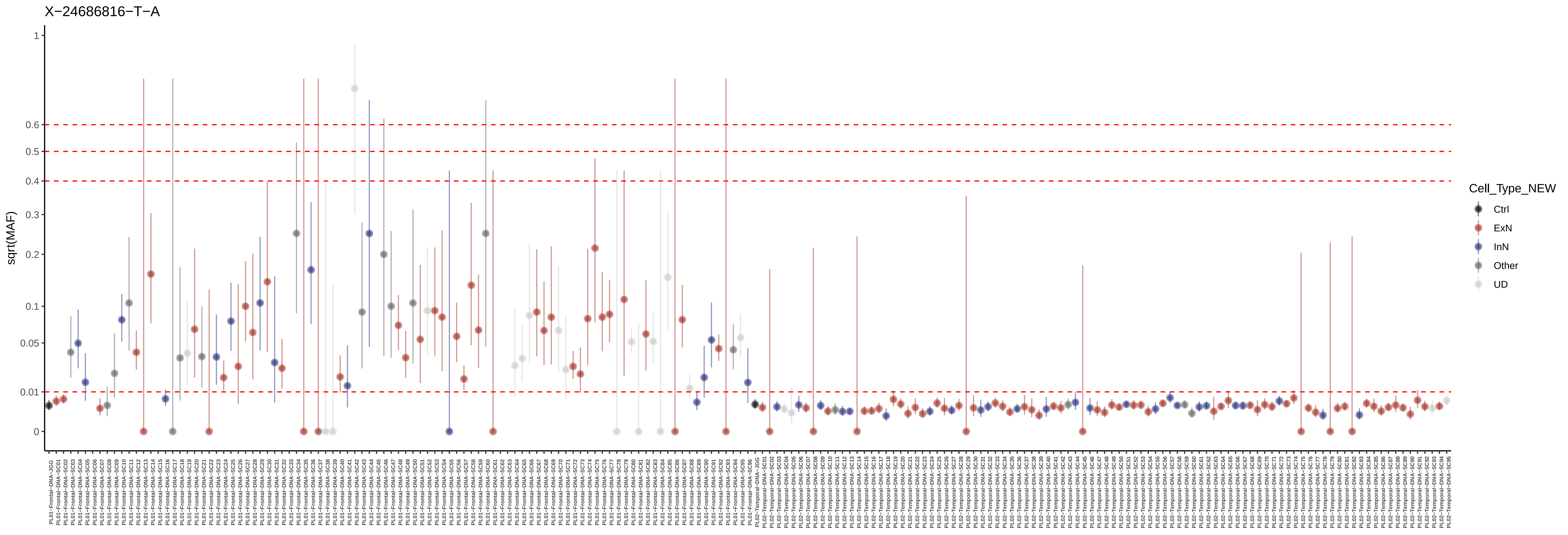
X−24686816−T−A
sqrt(MAF)
1
0.6
0.5
0.4
0.3
0.2
0.1
0.05
0.01
0
Cell_Type_NEW
Ctrl
ExN
InN
Other
UD

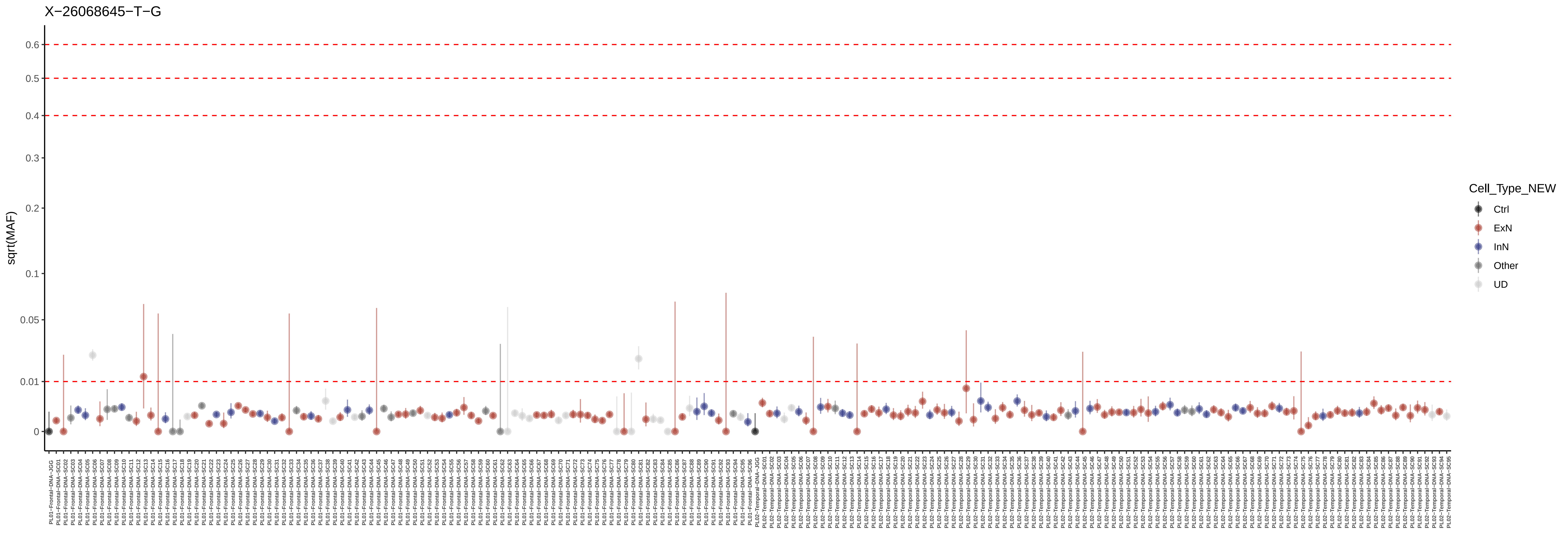
X−26068645−T−G
sqrt(MAF)
0.6
0.5
0.4
0.3
0.2
0.1
0.05
0.01
0
Cell_Type_NEW
Ctrl
ExN
InN
Other
UD

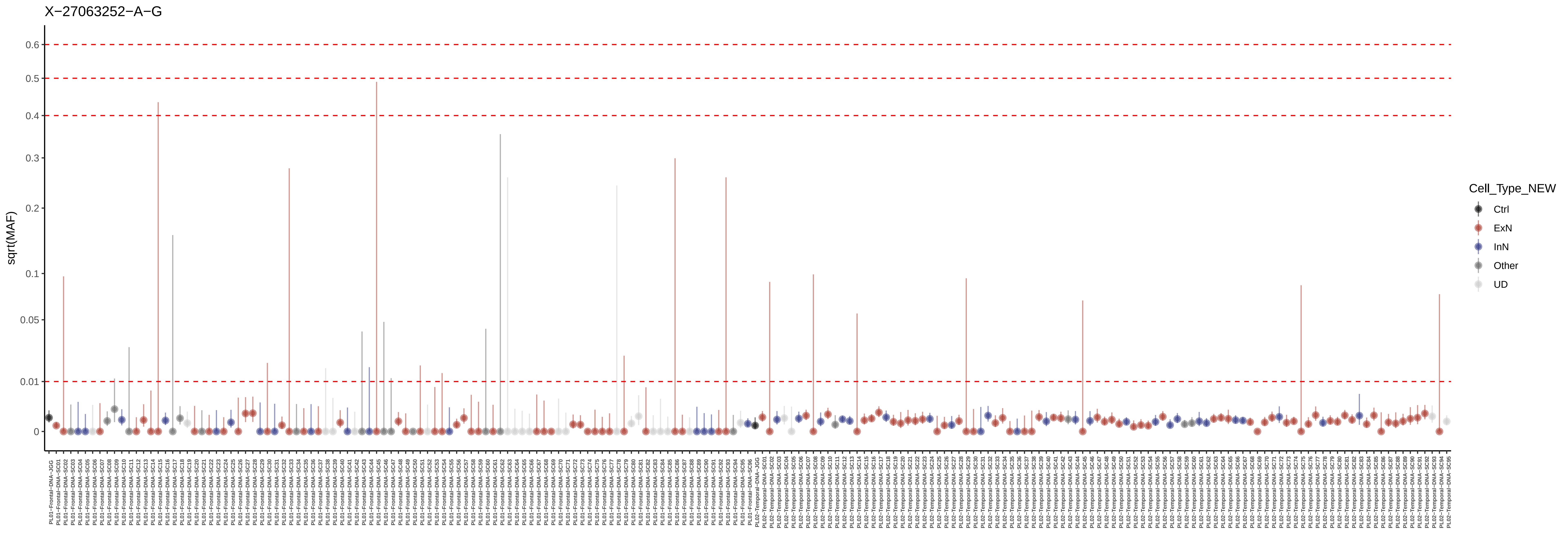
X−27063252−A−G
sqrt(MAF)
0.6
0.5
0.4
0.3
0.2
0.1
0.05
0.01
0
Cell_Type_NEW
Ctrl
ExN
InN
Other
UD

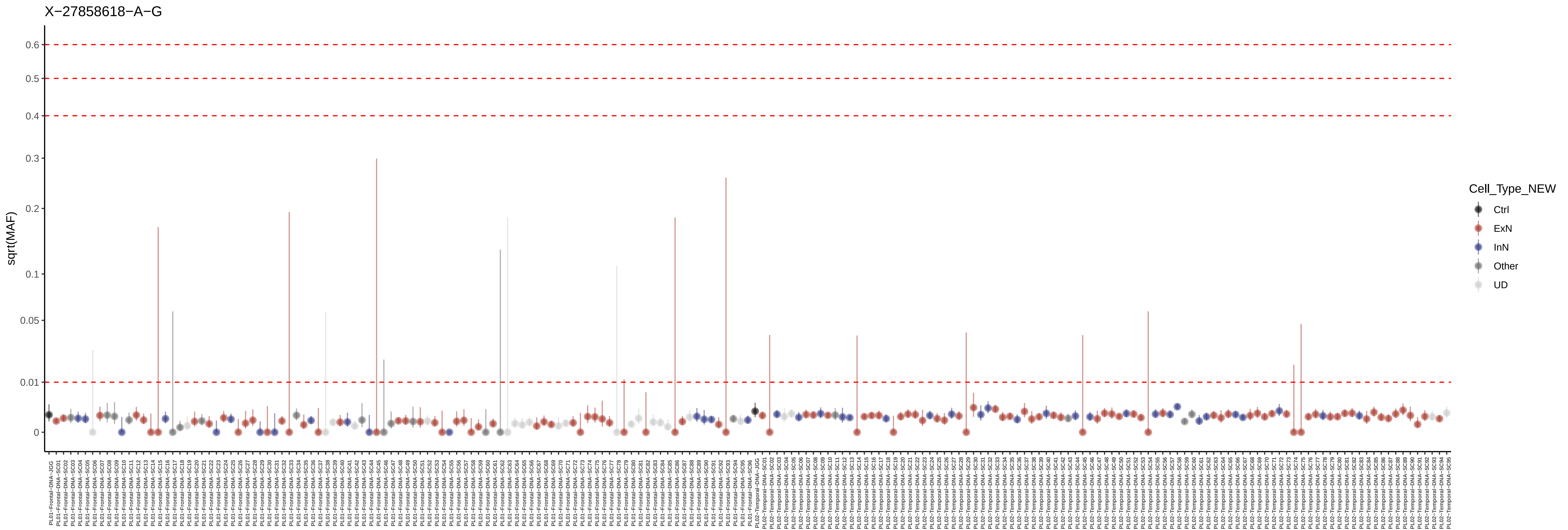
X−27858618−A−G
sqrt(MAF)
0.6
0.5
0.4
0.3
0.2
0.1
0.05
0.01
0
Cell_Type_NEW
Ctrl
ExN
InN
Other
UD

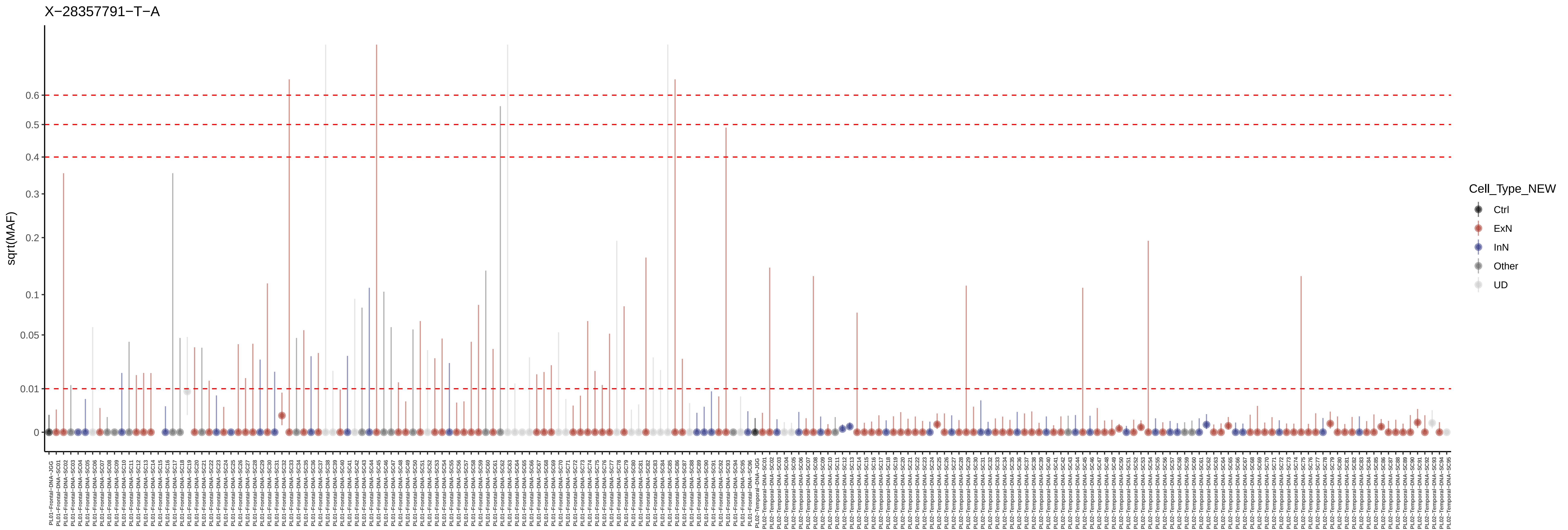

X−28357791−T−A
sqrt(MAF)
0.6
0.5
0.4
0.3
0.2
0.1
0.05
0.01
0
Cell_Type_NEW
Ctrl
ExN
InN
Other
UD

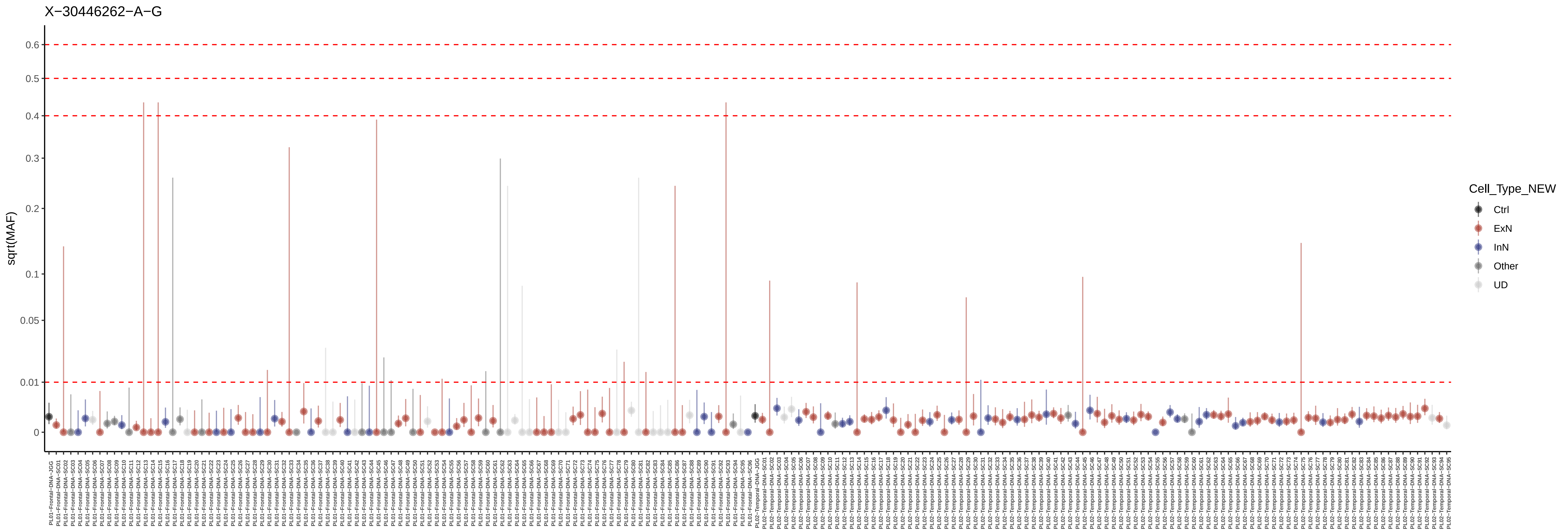
X−30446262−A−G
sqrt(MAF)
0.6
0.5
0.4
0.3
0.2
0.1
0.05
0.01
0
Cell_Type_NEW
Ctrl
ExN
InN
Other
UD

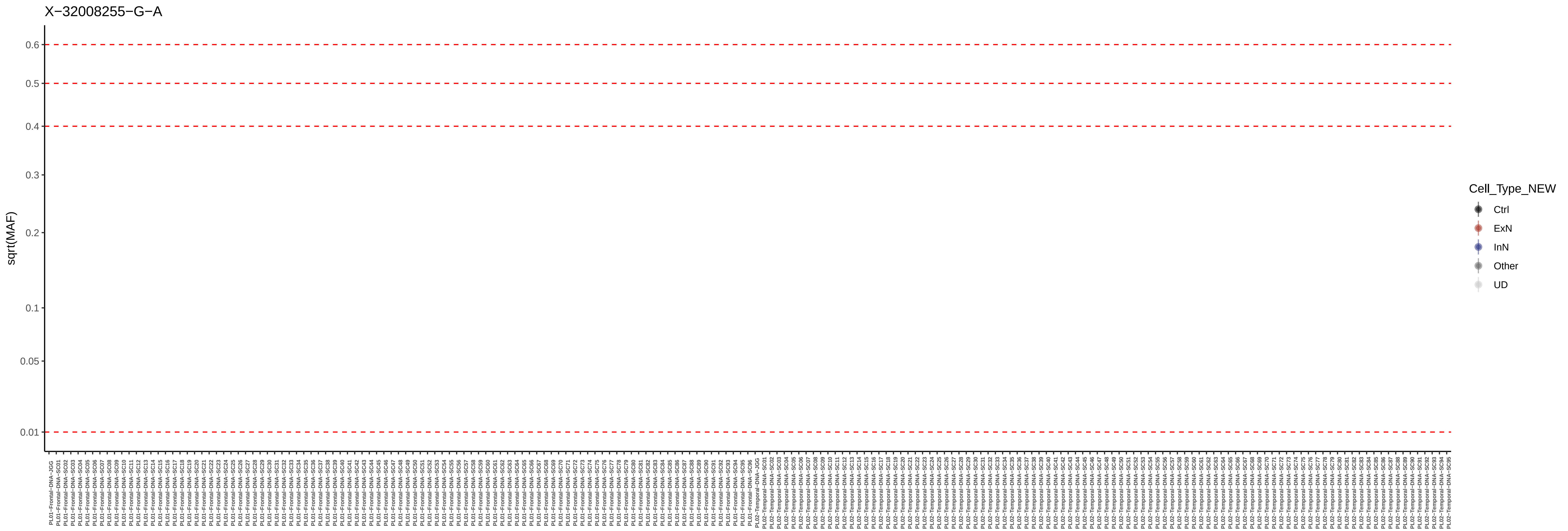

X−32008255−G−A
sqrt(MAF)
0.6
0.5
0.4
0.3
0.2
0.1
0.05
0.01
Cell_Type_NEW
Ctrl
ExN
InN
Other
UD

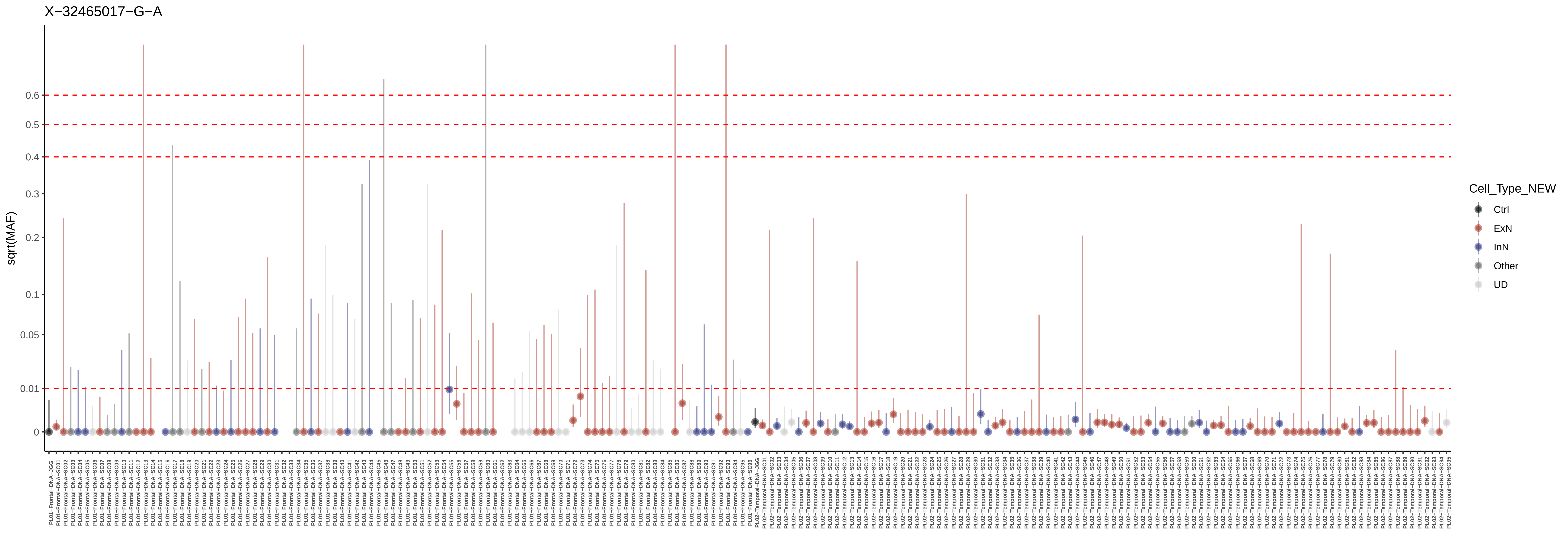

X−32465017−G−A
sqrt(MAF)
0
0.01
0.05
0.1
0.2
0.3
0.4
0.5
0.6
Cell_Type_NEW
Ctrl
ExN
InN
Other
UD

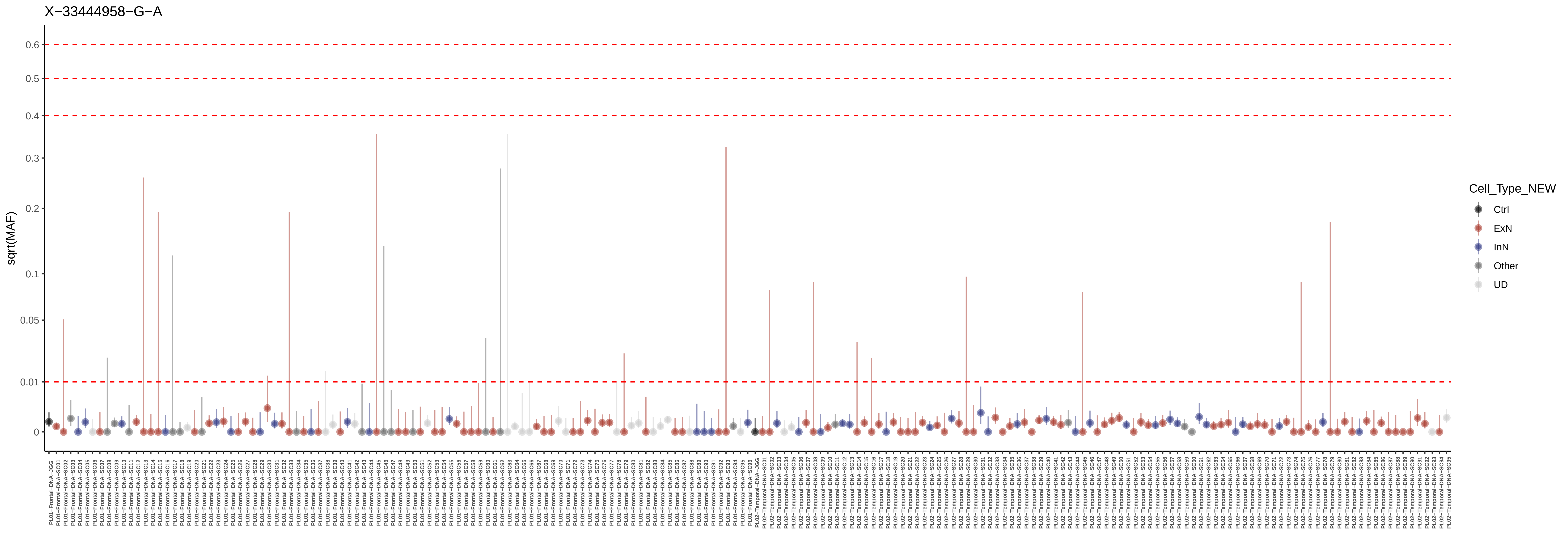
X−33444958−G−A
sqrt(MAF)
Cell_Type_NEW
Ctrl
ExN
InN
Other
UD

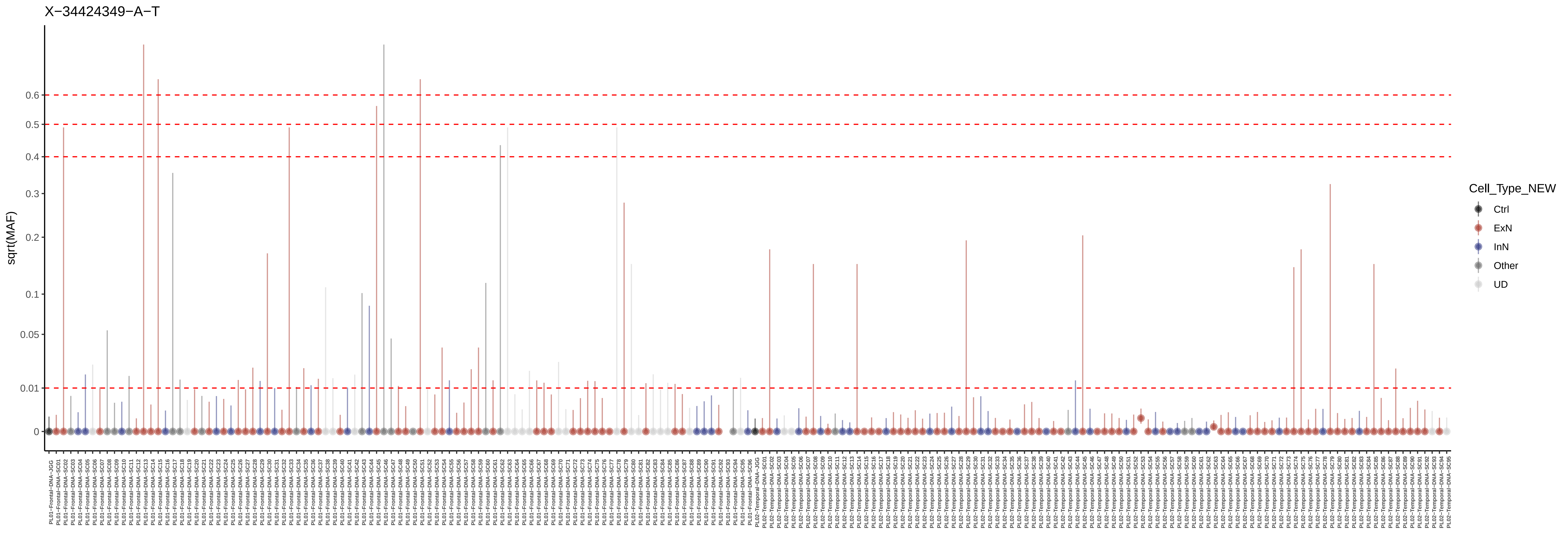
X−34424349−A−T
sqrt(MAF)
0.6
0.5
0.4
0.3
0.2
0.1
0.05
0.01
0
Cell_Type_NEW
Ctrl
ExN
InN
Other
UD

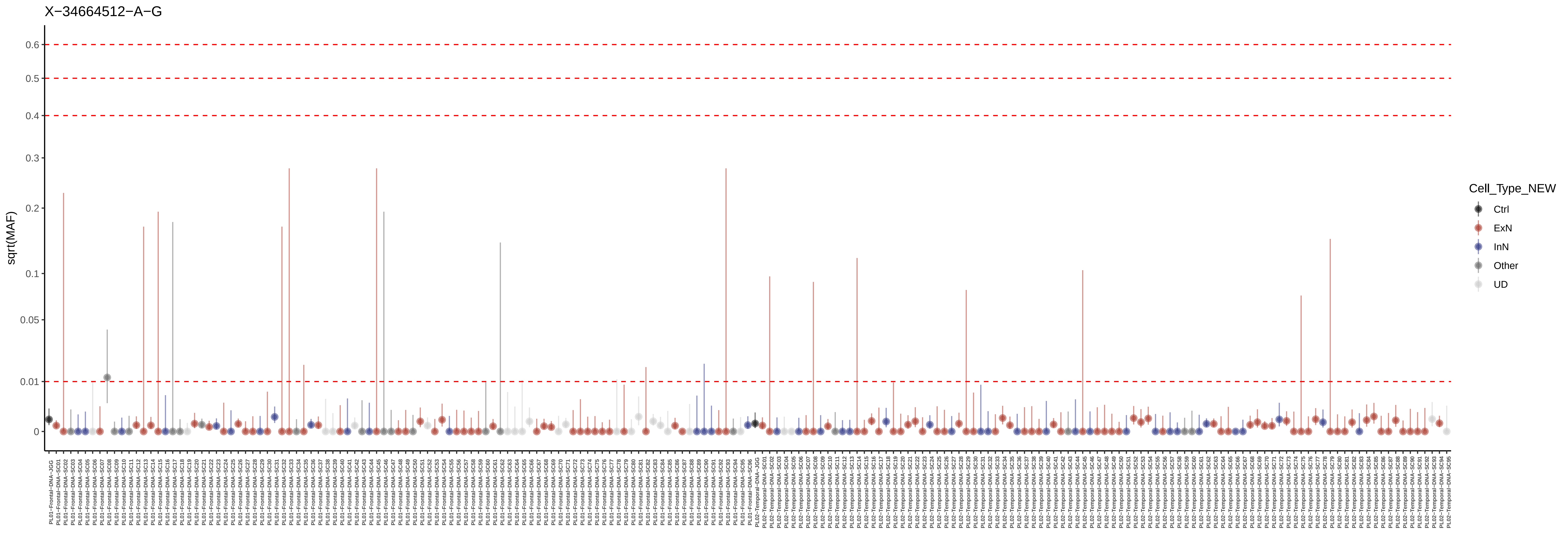
X−34664512−A−G
sqrt(MAF)
0.6
0.5
0.4
0.3
0.2
0.1
0.05
0.01
0
Cell_Type_NEW
Ctrl
ExN
InN
Other
UD

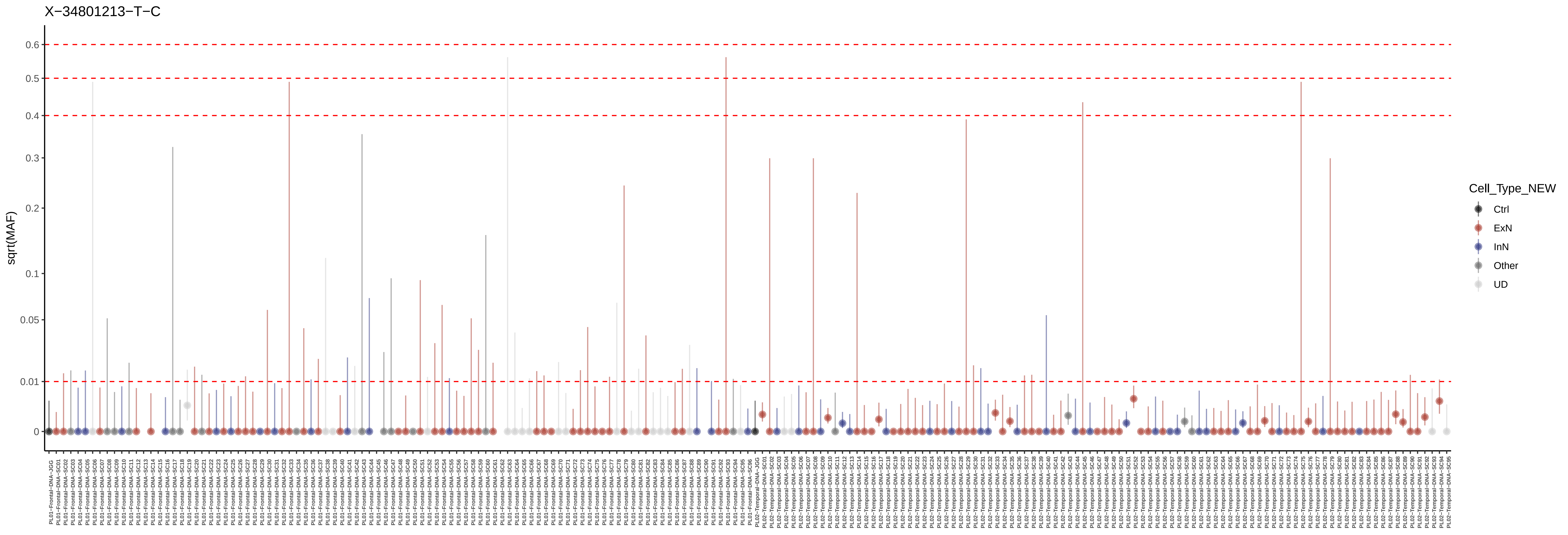

X−34801213−T−C
sqrt(MAF)
0.6
0.5
0.4
0.3
0.2
0.1
0.05
0.01
0
Cell_Type_NEW
Ctrl
ExN
InN
Other
UD

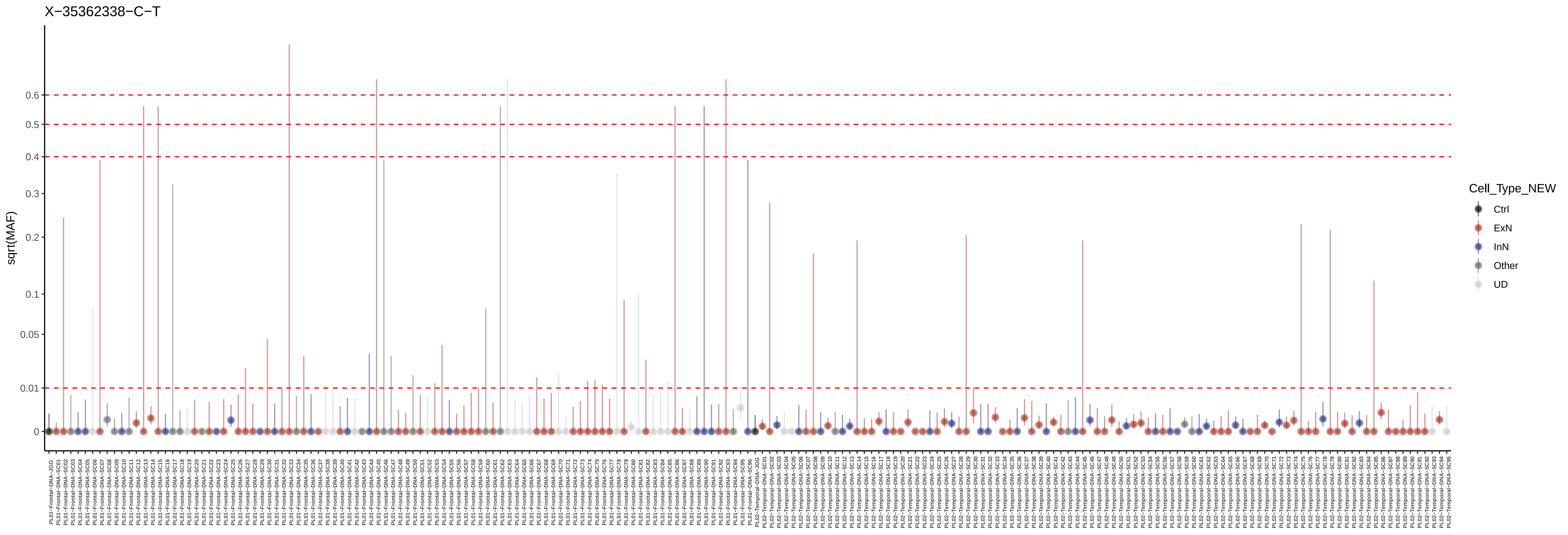

X−35362338−C−T
sqrt(MAF)
0.6
0.5
0.4
0.3
0.2
0.1
0.05
0.01
0
Cell_Type_NEW
Ctrl
ExN
InN
Other
UD

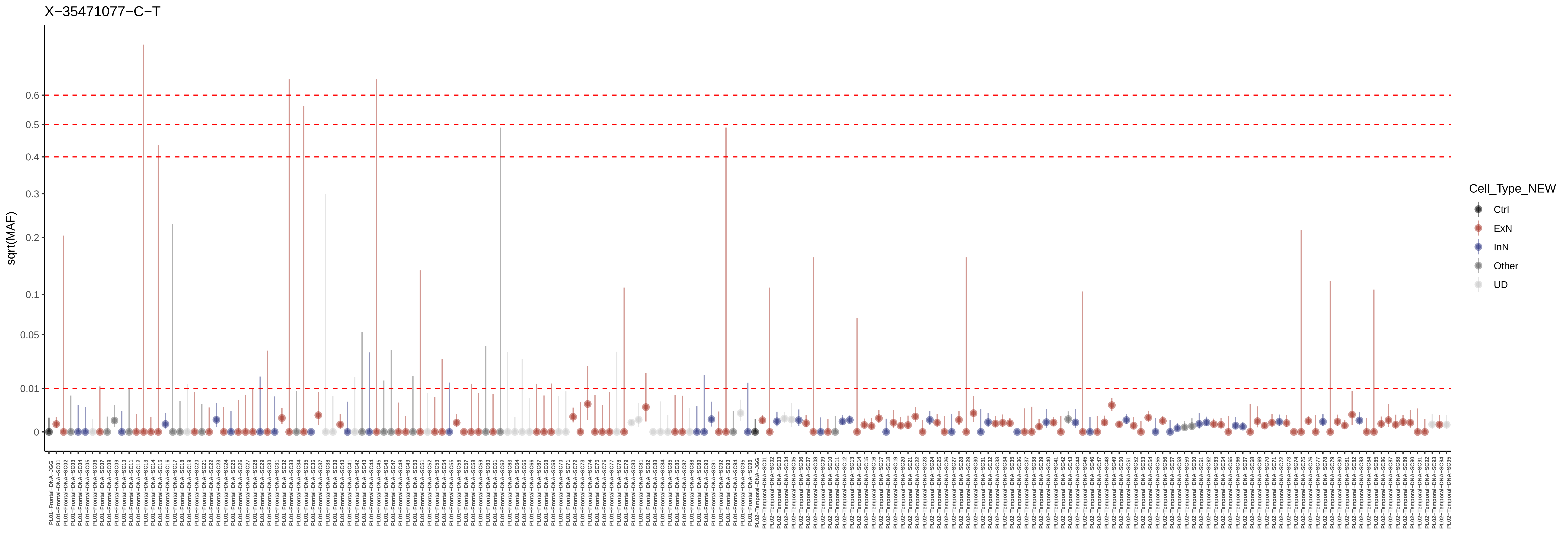

X−35471077−C−T
sqrt(MAF)
0.6
0.5
0.4
0.3
0.2
0.1
0.05
0.01
0
Cell_Type_NEW
Ctrl
ExN
InN
Other
UD

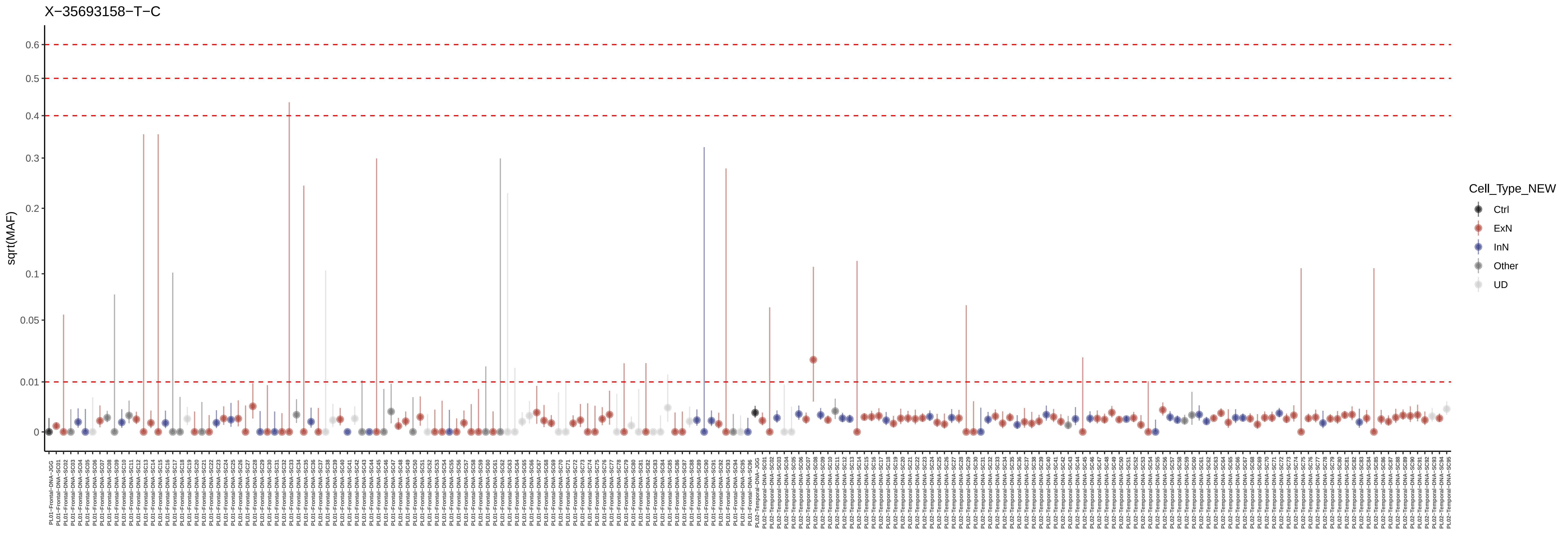
X−35693158−T−C
sqrt(MAF)
0.6
0.5
0.4
0.3
0.2
0.1
0.05
0.01
0
Cell_Type_NEW
Ctrl
ExN
InN
Other
UD

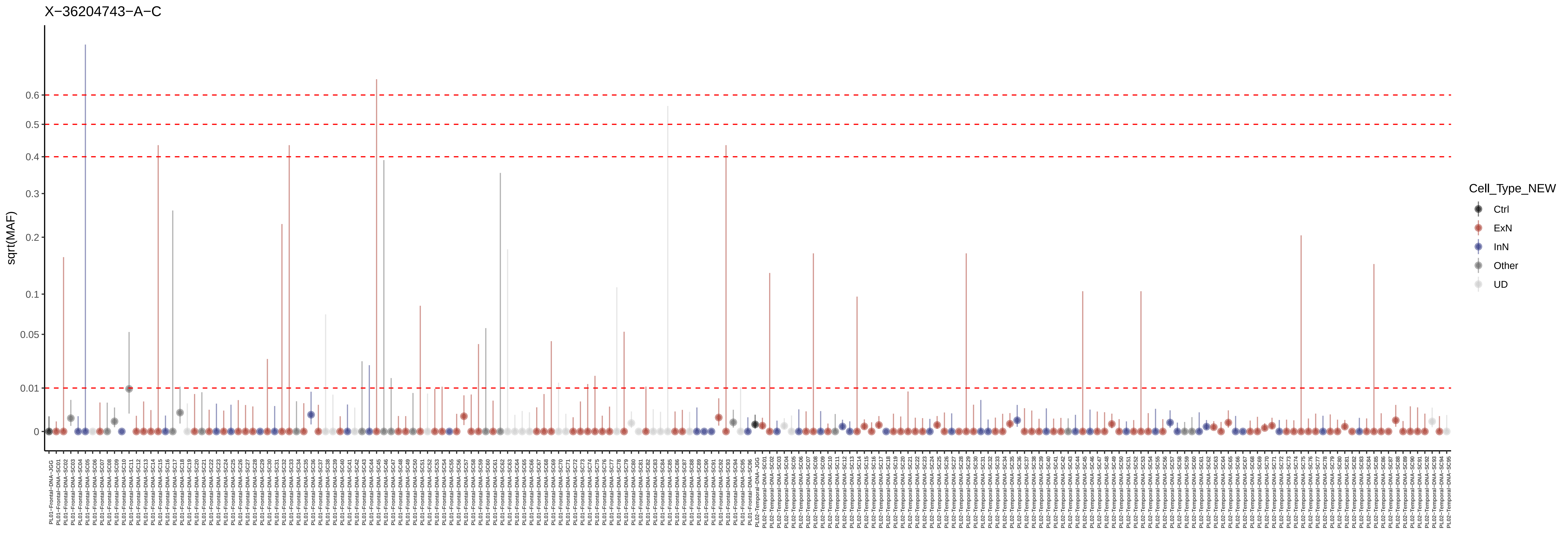
X−36204743−A−C
sqrt(MAF)
0.6
0.5
0.4
0.3
0.2
0.1
0.05
0.01
0
Cell_Type_NEW
Ctrl
ExN
InN
Other
UD

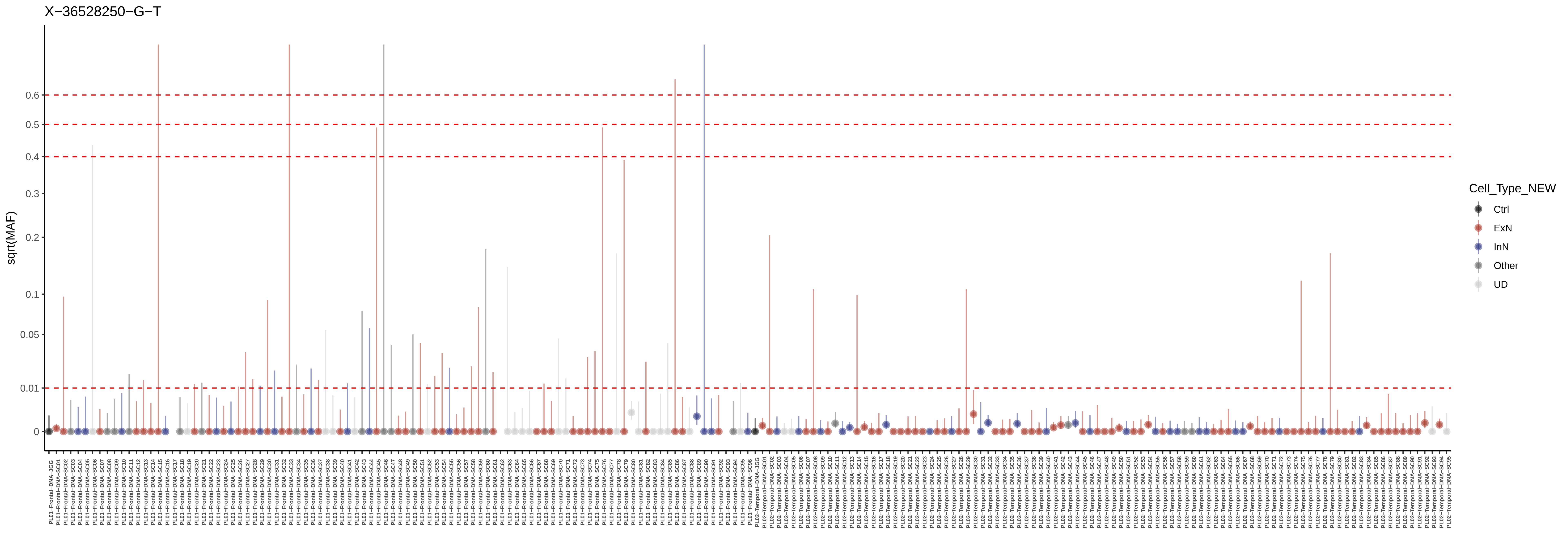

X−36528250−G−T
sqrt(MAF)
0.6
0.5
0.4
0.3
0.2
0.1
0.05
0.01
0
Cell_Type_NEW
Ctrl
ExN
InN
Other
UD

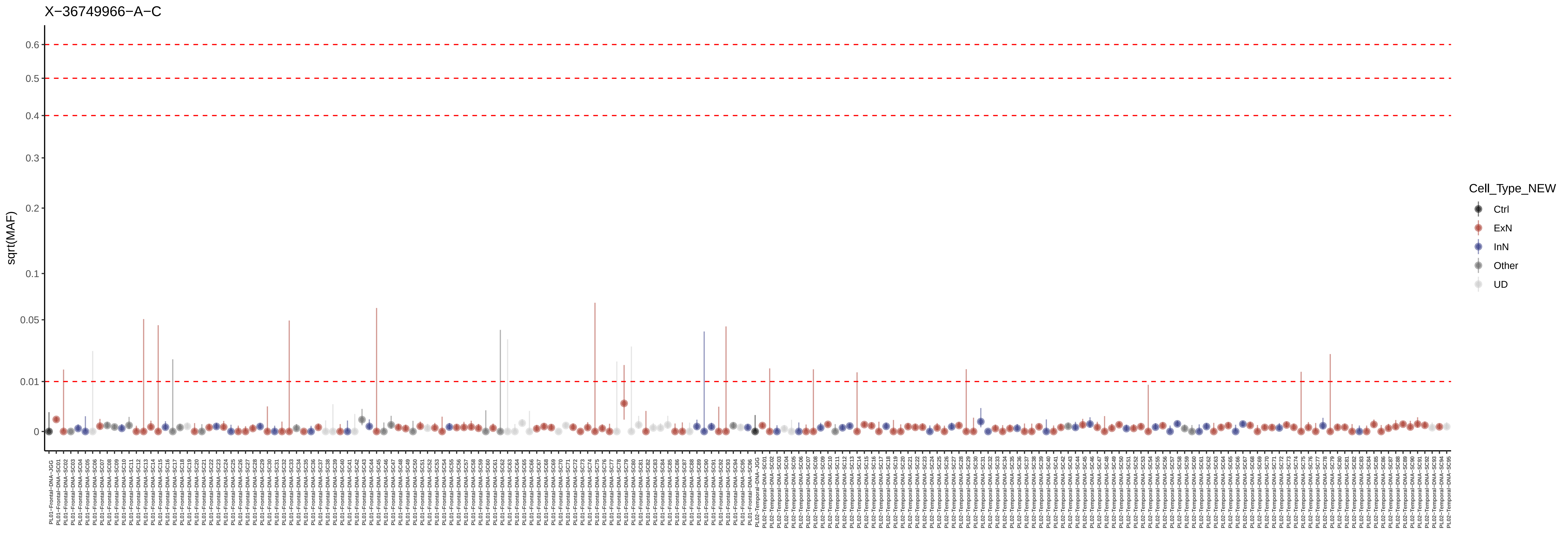
X−36749966−A−C
sqrt(MAF)
0.6
0.5
0.4
0.3
0.2
0.1
0.05
0.01
0
Cell_Type_NEW
Ctrl
ExN
InN
Other
UD

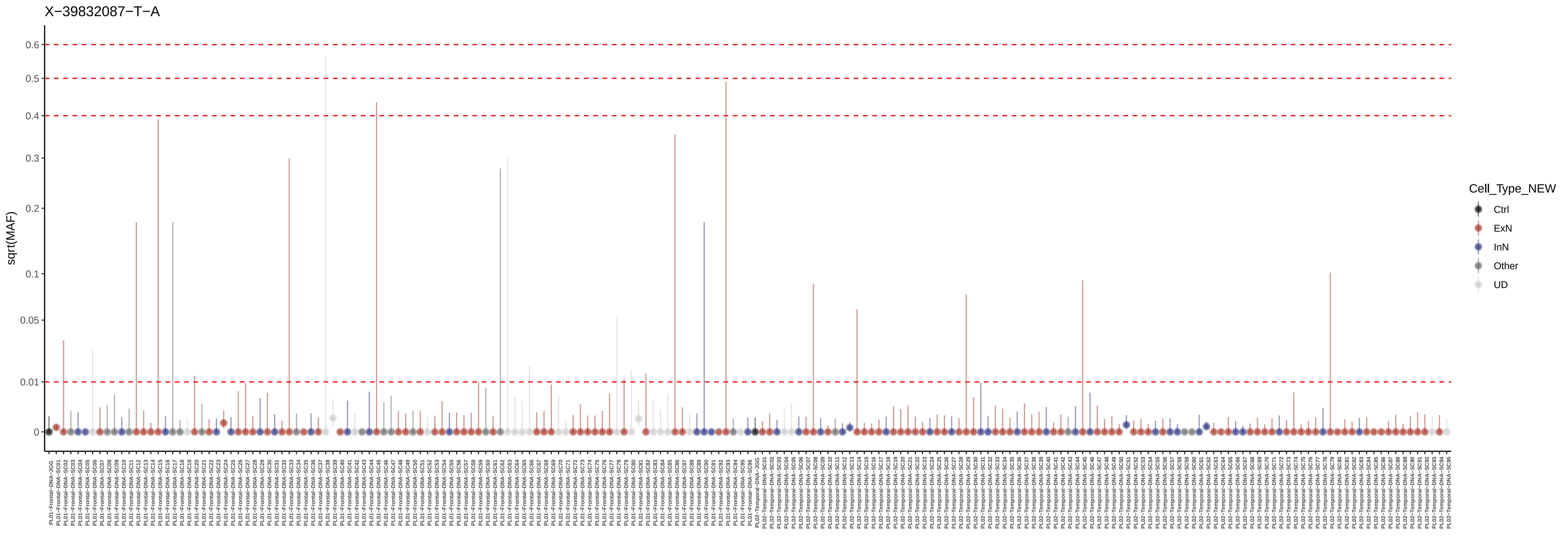
X−39832087−T−A
sqrt(MAF)
0.6
0.5
0.4
0.3
0.2
0.1
0.05
0.01
0
Cell_Type_NEW
Ctrl
ExN
InN
Other
UD

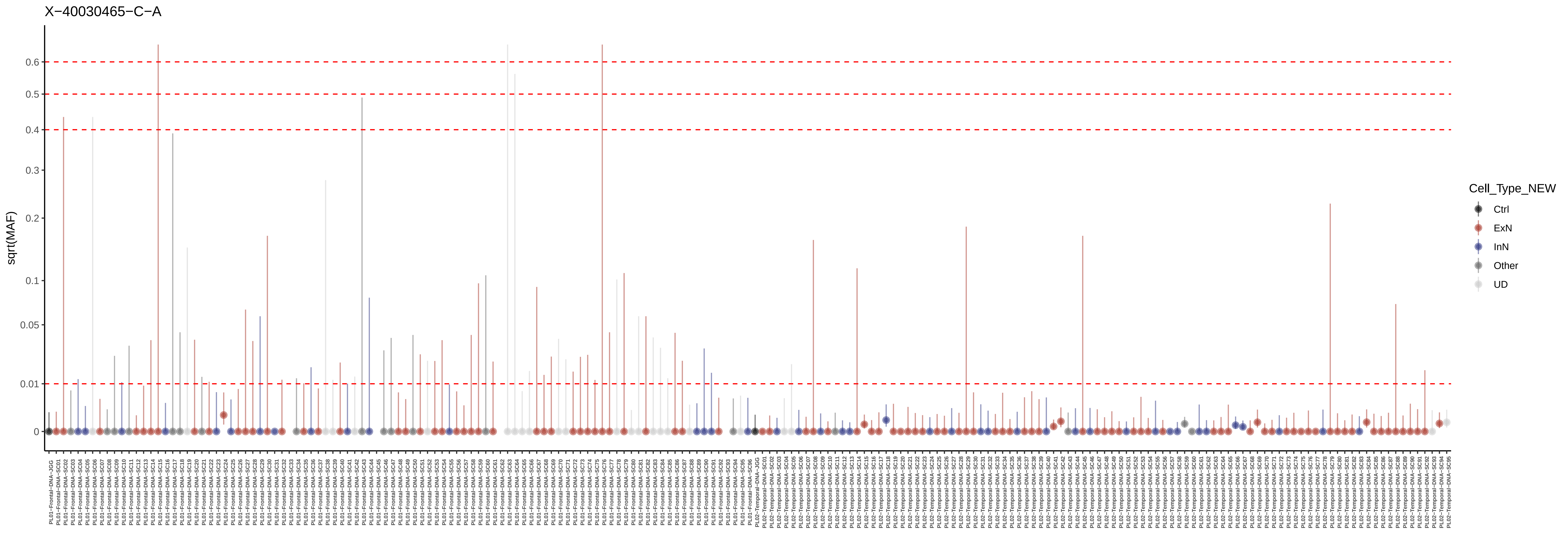

X−40030465−C−A
sqrt(MAF)
0.6
0.5
0.4
0.3
0.2
0.1
0.05
0.01
0
Cell_Type_NEW
Ctrl
ExN
InN
Other
UD

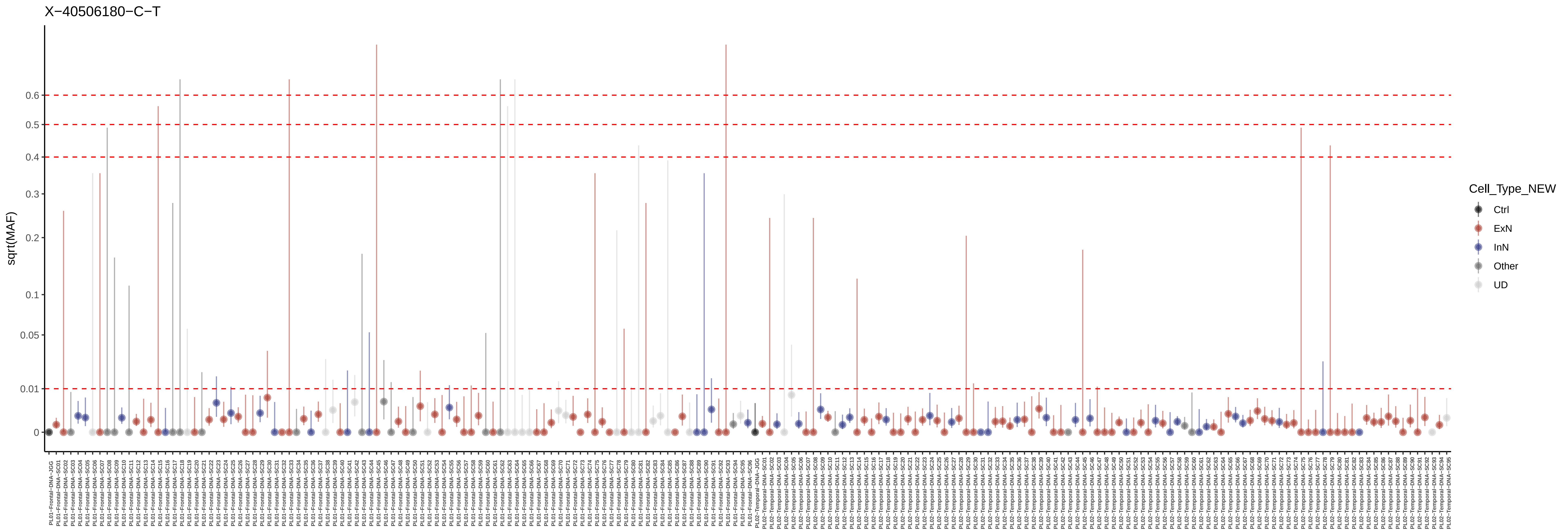
X−40506180−C−T
sqrt(MAF)
0.6
0.5
0.4
0.3
0.2
0.1
0.05
0.01
0
Cell_Type_NEW
Ctrl
ExN
InN
Other
UD

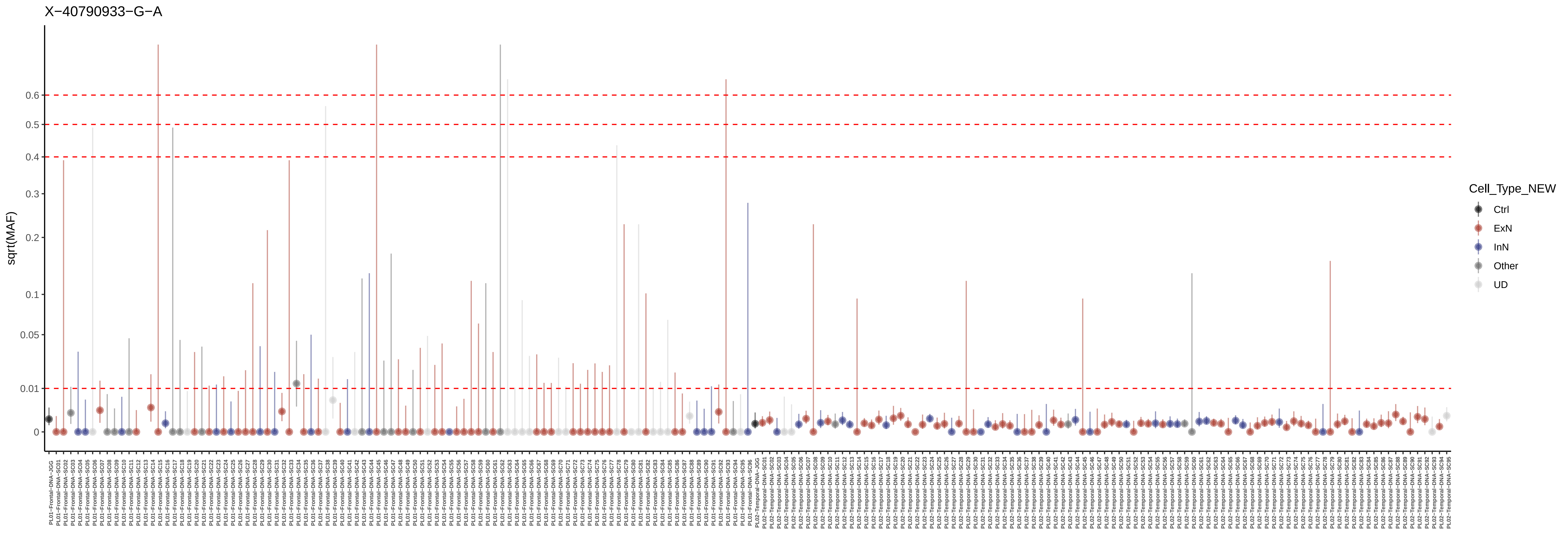
X−40790933−G−A
sqrt(MAF)
0.6
0.5
0.4
0.3
0.2
0.1
0.05
0.01
0
Cell_Type_NEW
Ctrl
ExN
InN
Other
UD

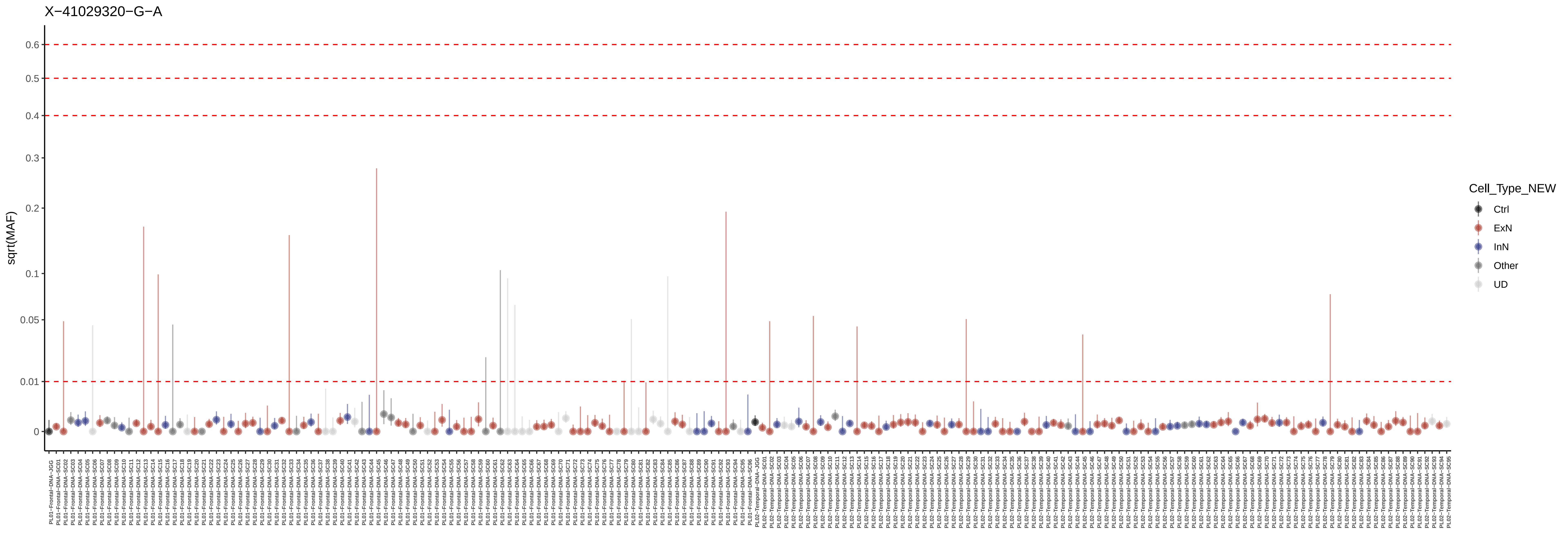
X−41029320−G−A
sqrt(MAF)
0.6
0.5
0.4
0.3
0.2
0.1
0.05
0.01
0
Cell_Type_NEW
Ctrl
ExN
InN
Other
UD

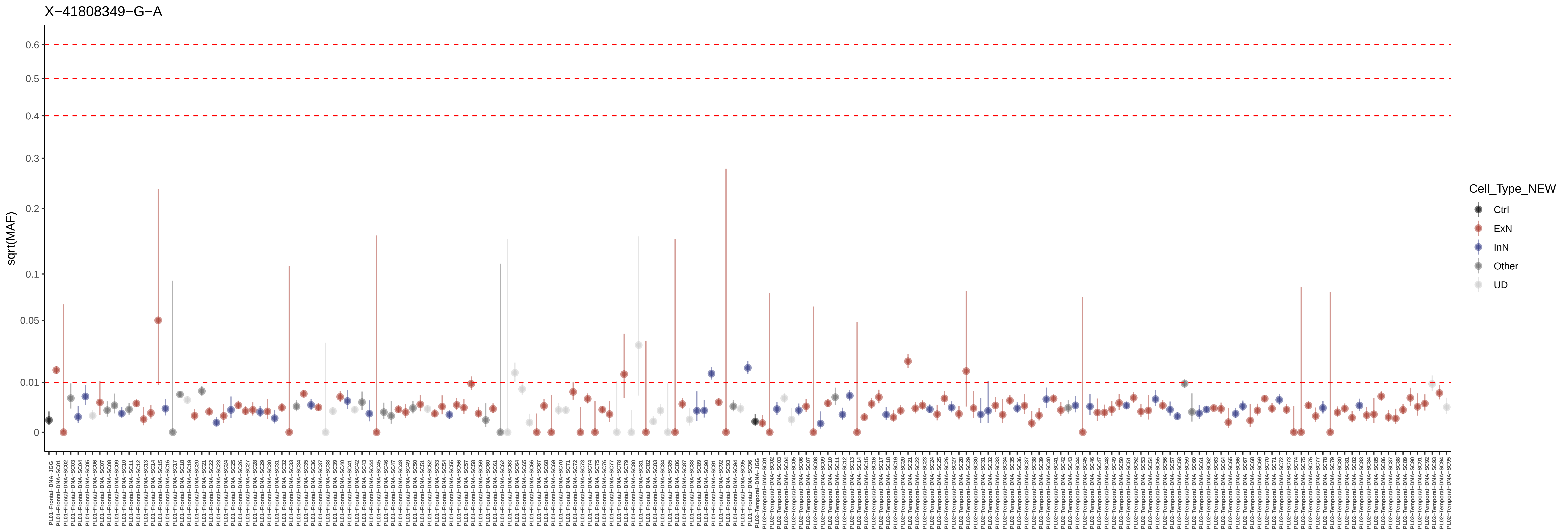
X−41808349−G−A
sqrt(MAF)
0.6
0.5
0.4
0.3
0.2
0.1
0.05
0.01
0
Cell_Type_NEW
Ctrl
ExN
InN
Other
UD

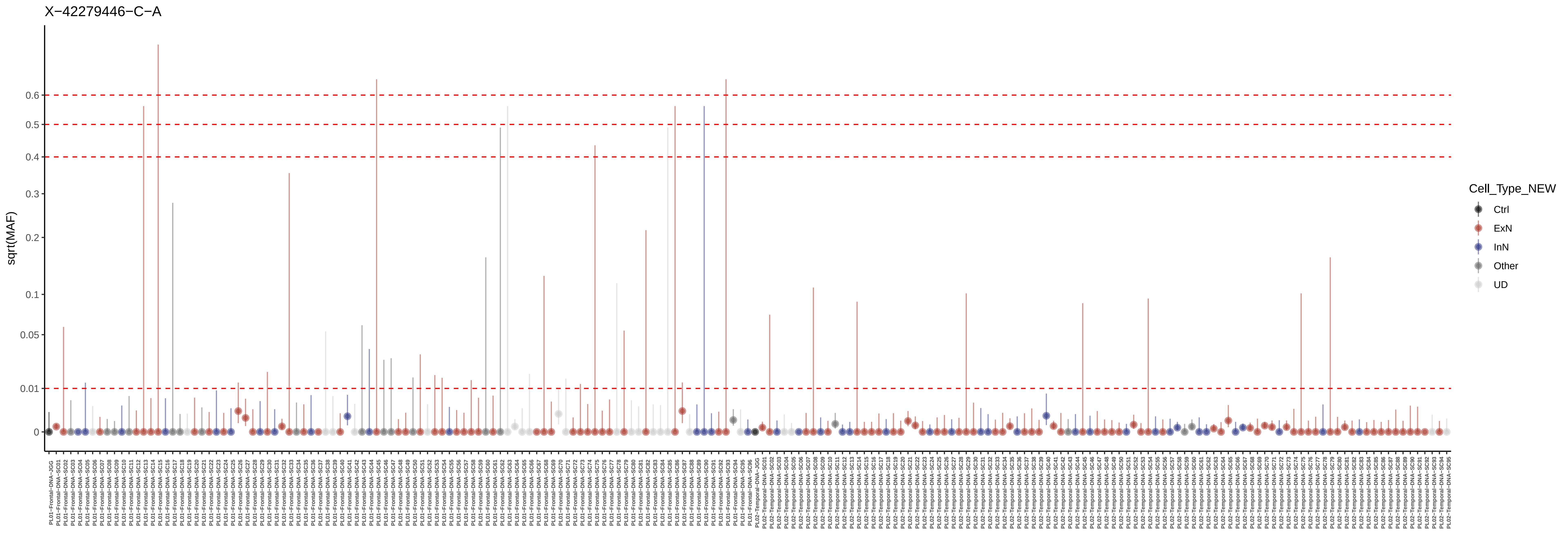

X−42279446−C−A
sqrt(MAF)
0.6
0.5
0.4
0.3
0.2
0.1
0.05
0.01
0
Cell_Type_NEW
Ctrl
ExN
InN
Other
UD

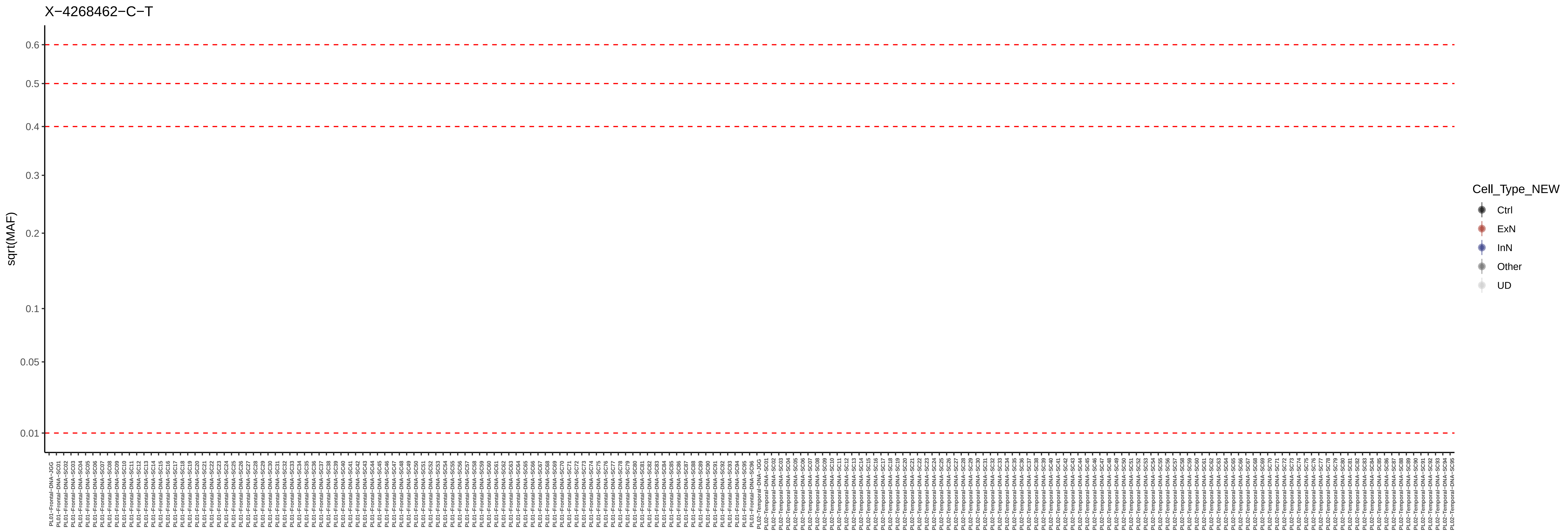
X−4268462−C−T
sqrt(MAF)
0.6
0.5
0.4
0.3
0.2
0.1
0.05
0.01
Cell_Type_NEW
Ctrl
ExN
InN
Other
UD

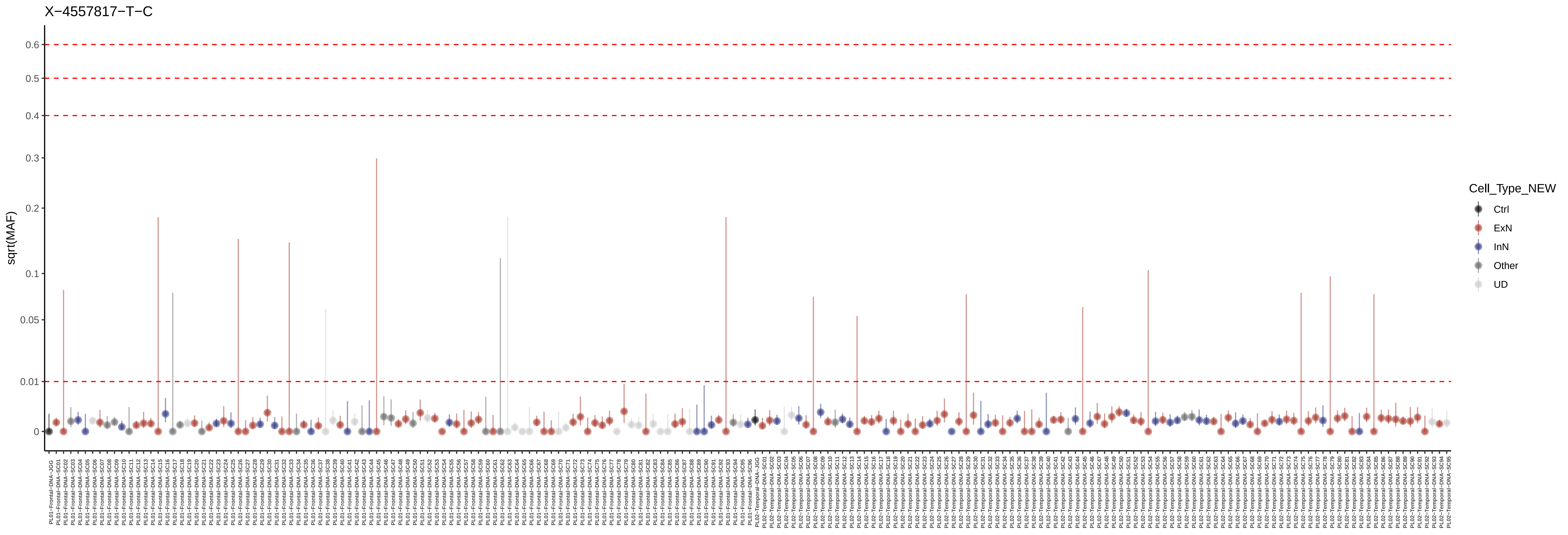
X−4557817−T−C
sqrt(MAF)
0.6
0.5
0.4
0.3
0.2
0.1
0.05
0.01
0
Cell_Type_NEW
Ctrl
ExN
InN
Other
UD

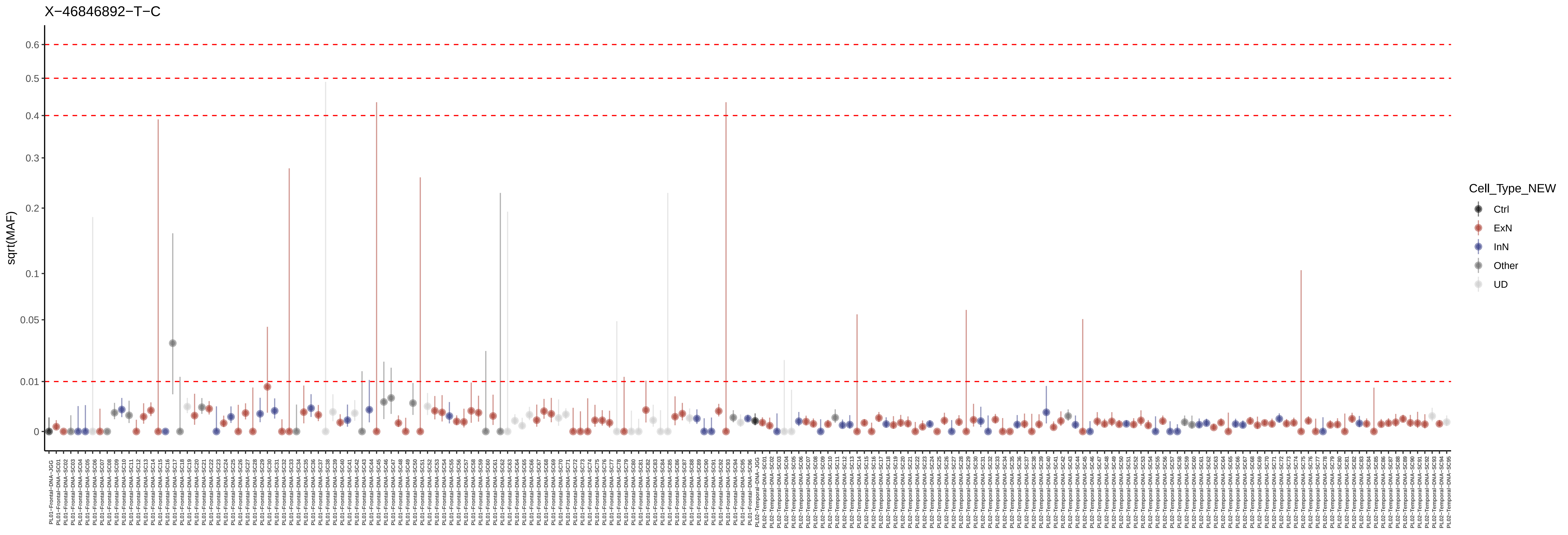
X−46846892−T−C
sqrt(MAF)
0.6
0.5
0.4
0.3
0.2
0.1
0.05
0.01
0
Cell_Type_NEW
Ctrl
ExN
InN
Other
UD

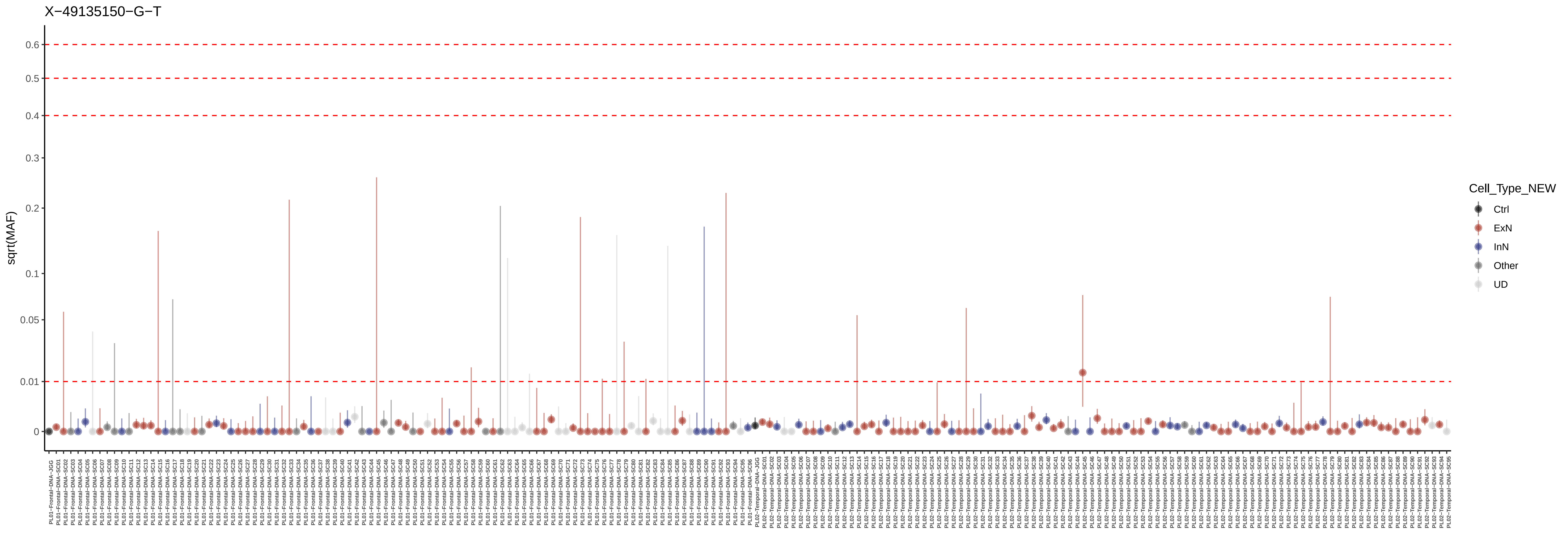

X−49135150−G−T
sqrt(MAF)
0.6
0.5
0.4
0.3
0.2
0.1
0.05
0.01
0
Cell_Type_NEW
Ctrl
ExN
InN
Other
UD

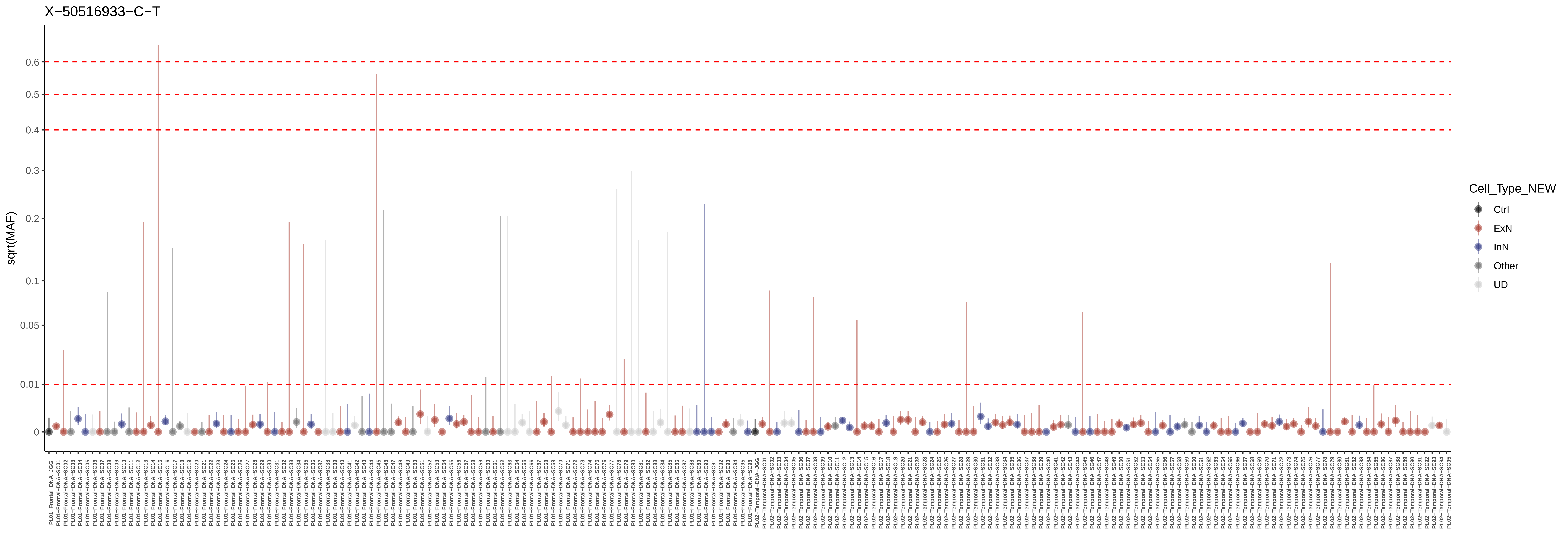
X−50516933−C−T
sqrt(MAF)
0.6
0.5
0.4
0.3
0.2
0.1
0.05
0.01
0
Cell_Type_NEW
Ctrl
ExN
InN
Other
UD

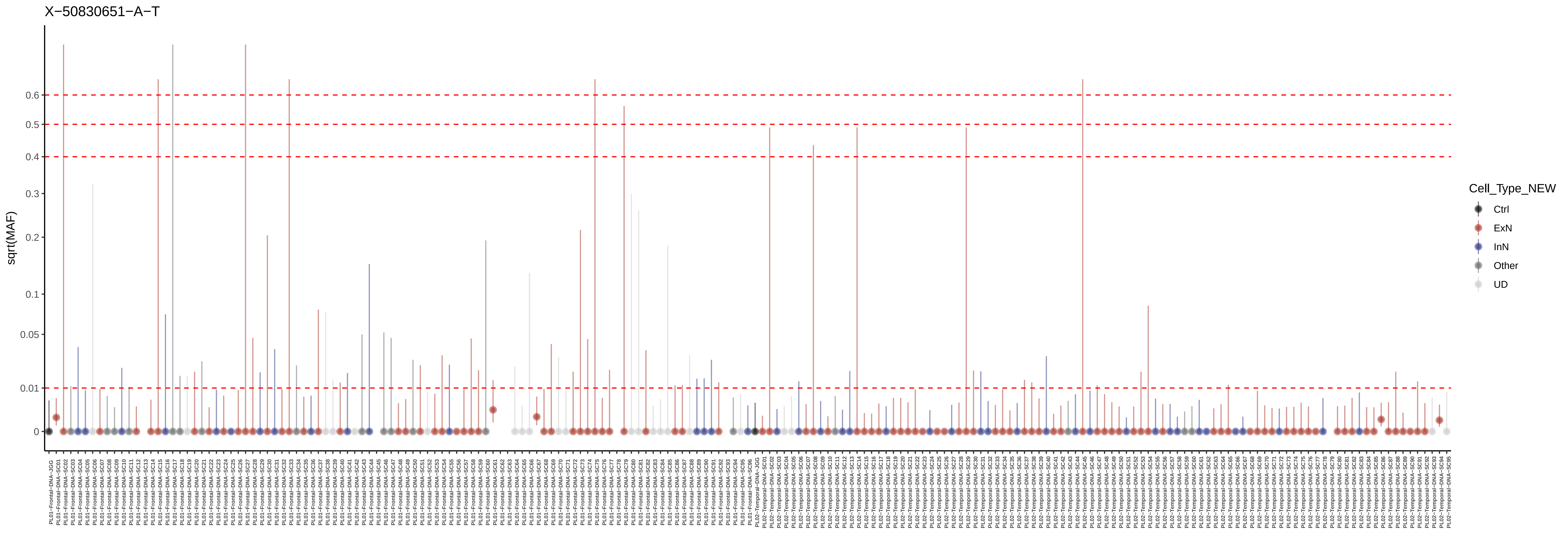

X−50830651−A−T
sqrt(MAF)
0.6
0.5
0.4
0.3
0.2
0.1
0.05
0.01
0
Cell_Type_NEW
Ctrl
ExN
InN
Other
UD

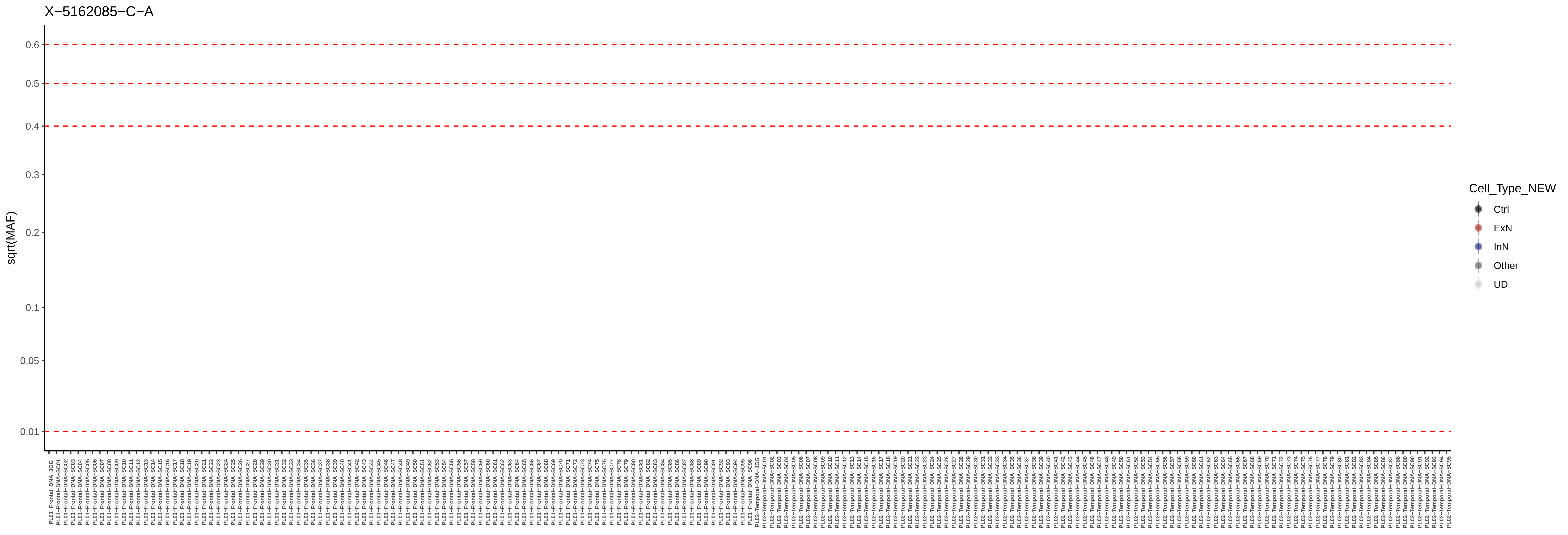

X−5162085−C−A
sqrt(MAF)
0.6
0.5
0.4
0.3
0.2
0.1
0.05
0.01
Cell_Type_NEW
Ctrl
ExN
InN
Other
UD

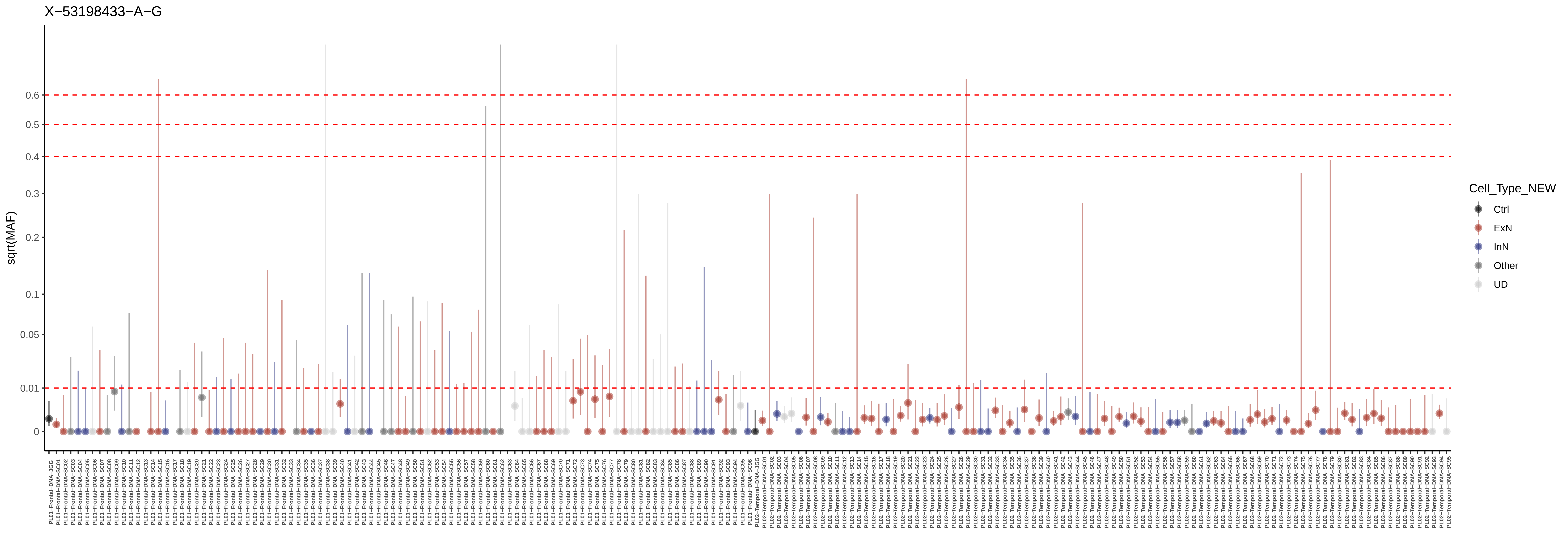

X−53198433−A−G
sqrt(MAF)
0.6
0.5
0.4
0.3
0.2
0.1
0.05
0.01
0
Cell_Type_NEW
Ctrl
ExN
InN
Other
UD

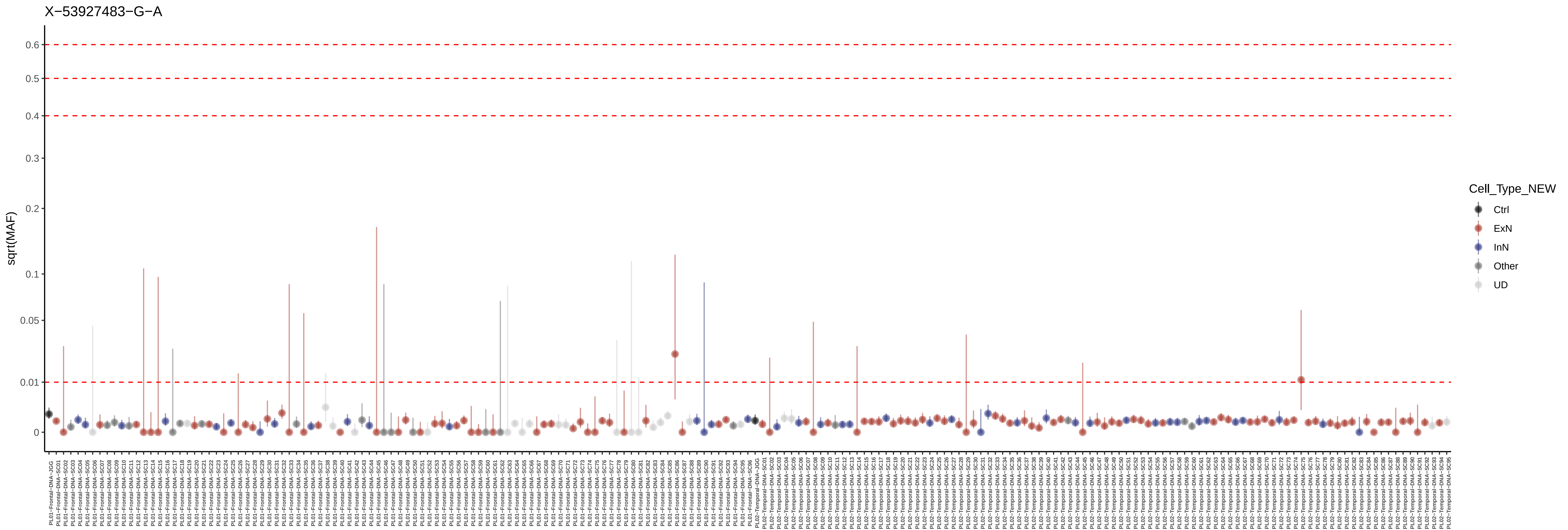
X−53927483−G−A
sqrt(MAF)
0.6
0.5
0.4
0.3
0.2
0.1
0.05
0.01
0
Cell_Type_NEW
Ctrl
ExN
InN
Other
UD

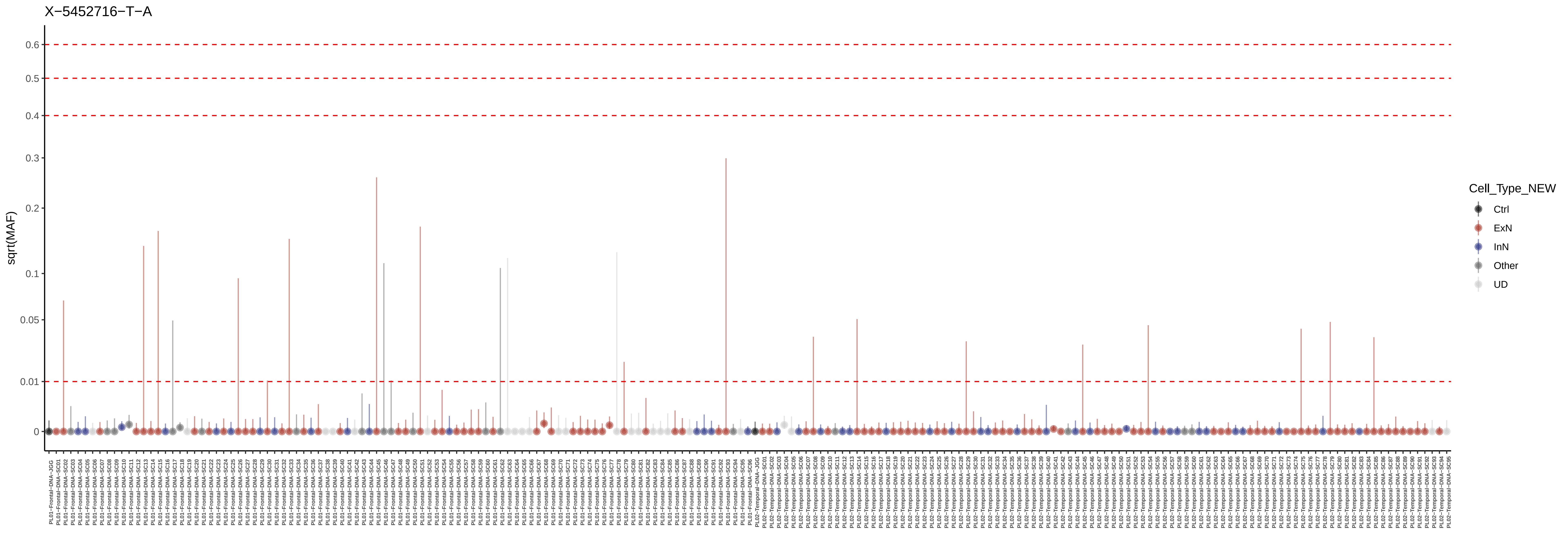
X−5452716−T−A
sqrt(MAF)
0.6
0.5
0.4
0.3
0.2
0.1
0.05
0.01
0
Cell_Type_NEW
Ctrl
ExN
InN
Other
UD

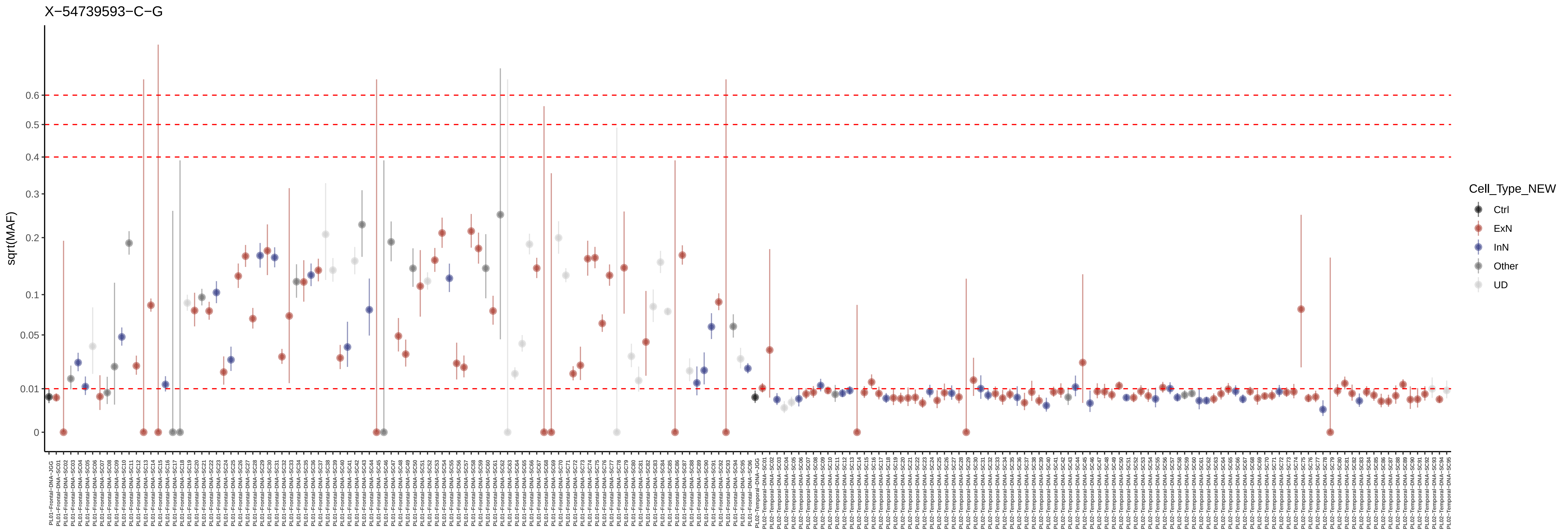
X−54739593−C−G
sqrt(MAF)
0.6
0.5
0.4
0.3
0.2
0.1
0.05
0.01
0
Cell_Type_NEW
Ctrl
ExN
InN
Other
UD

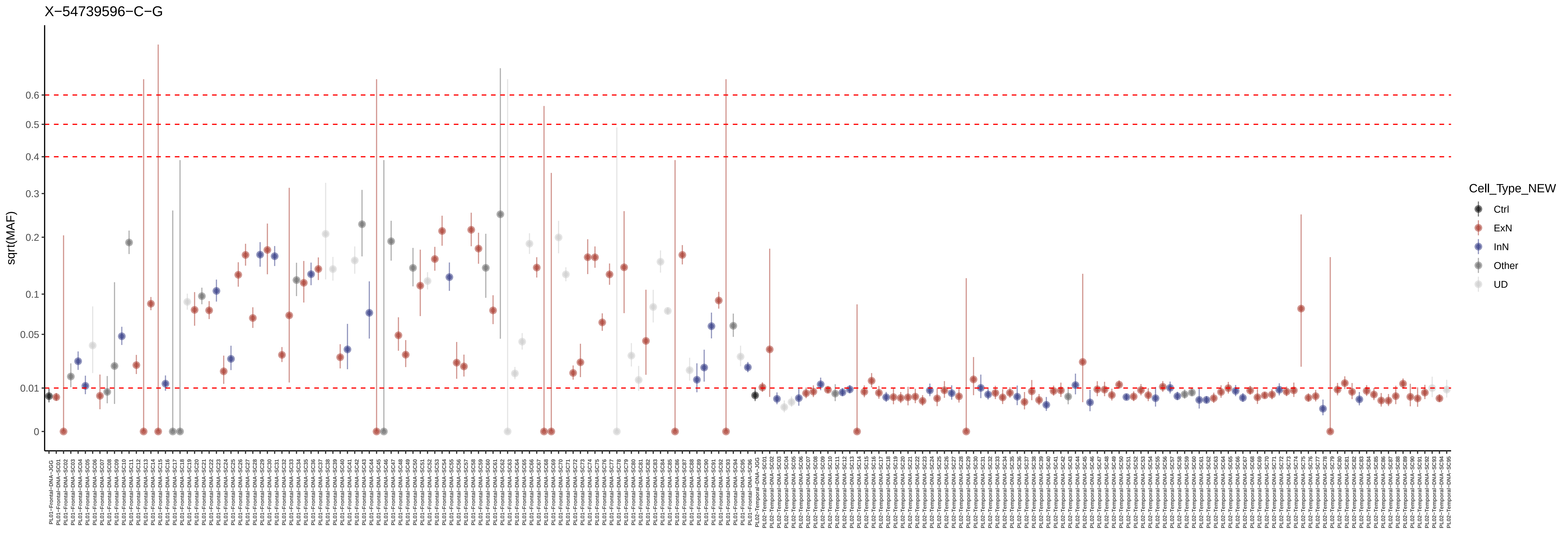
X−54739596−C−G
sqrt(MAF)
0.6
0.5
0.4
0.3
0.2
0.1
0.05
0.01
0
Cell_Type_NEW
Ctrl
ExN
InN
Other
UD

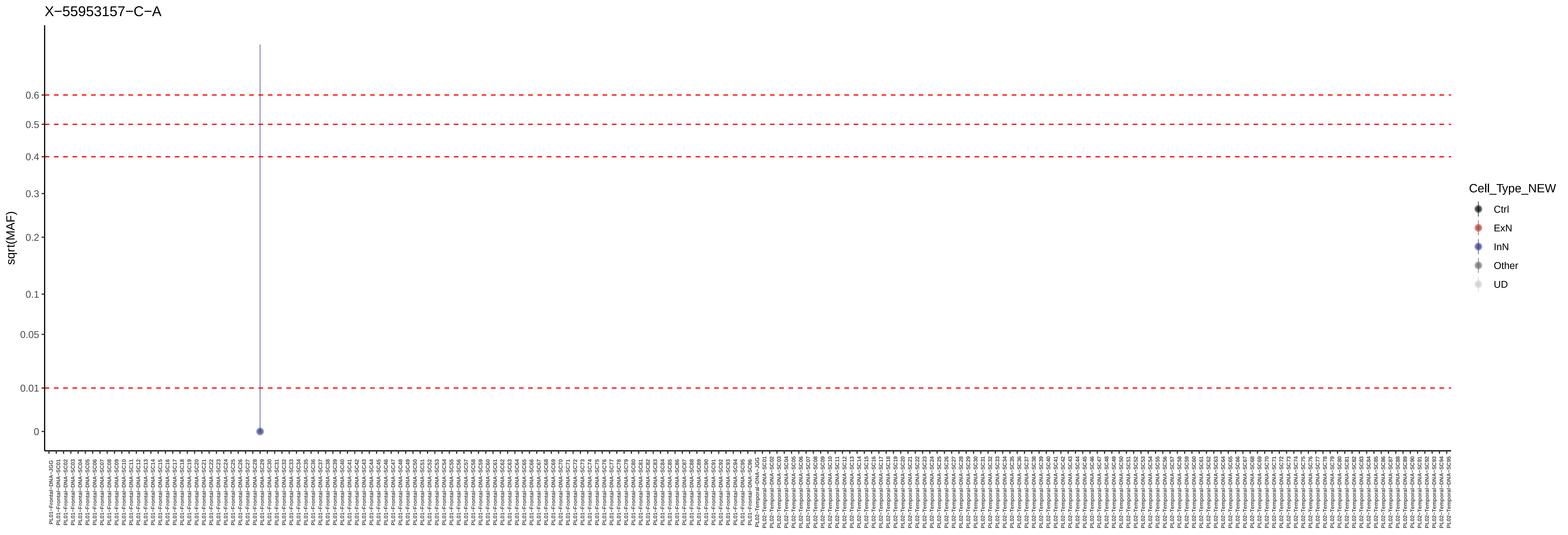

X−55953157−C−A
sqrt(MAF)
0.6
0.5
0.4
0.3
0.2
0.1
0.05
0.01
0
Cell_Type_NEW
Ctrl
ExN
InN
Other
UD

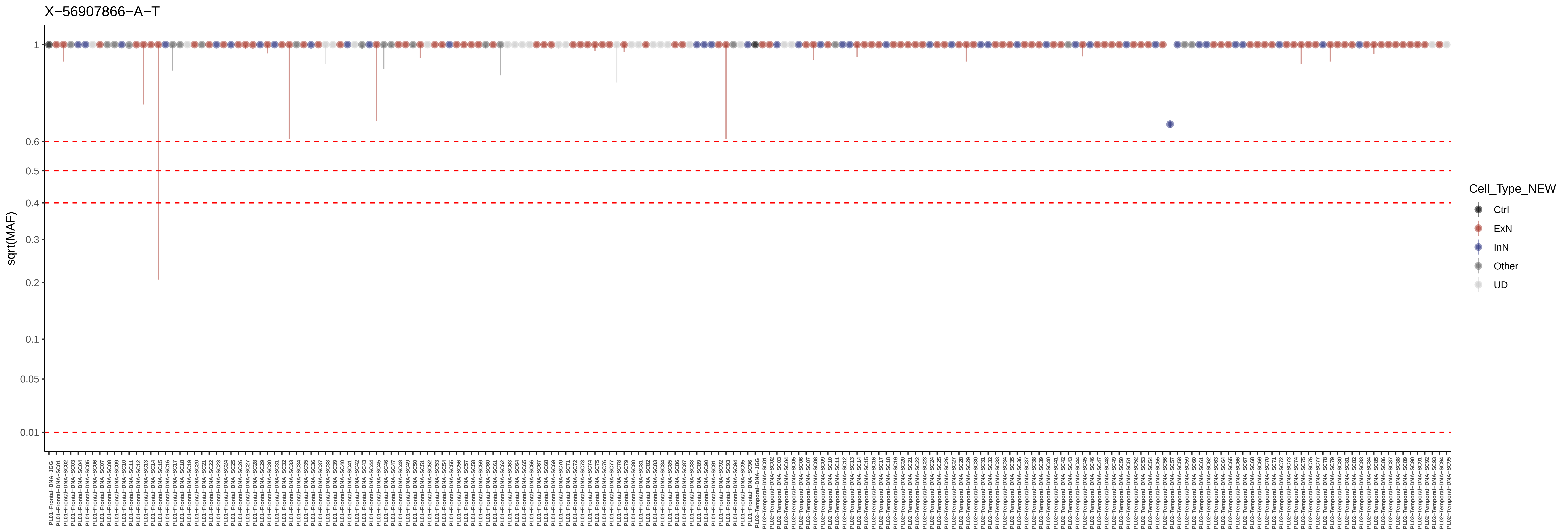

X−56907866−A−T
sqrt(MAF)
1
0.6
0.5
0.4
0.3
0.2
0.1
0.05
0.01
Cell_Type_NEW
Ctrl
ExN
InN
Other
UD

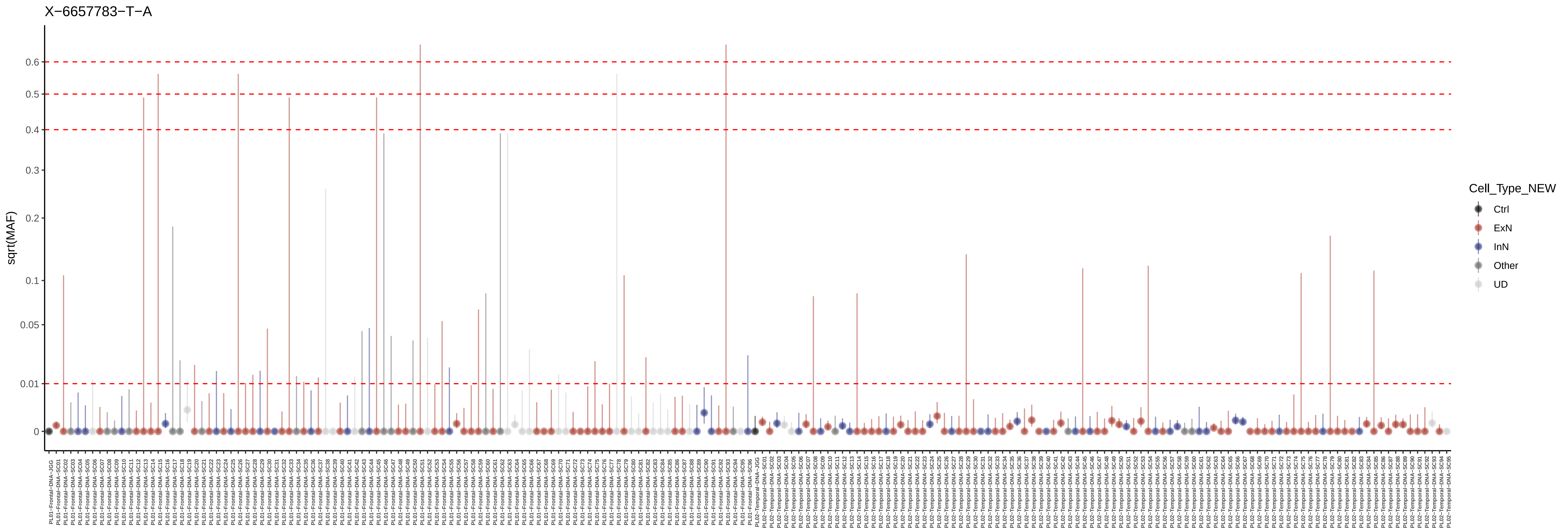
X−6657783−T−A
sqrt(MAF)
0.6
0.5
0.4
0.3
0.2
0.1
0.05
0.01
0
Cell_Type_NEW
Ctrl
ExN
InN
Other
UD

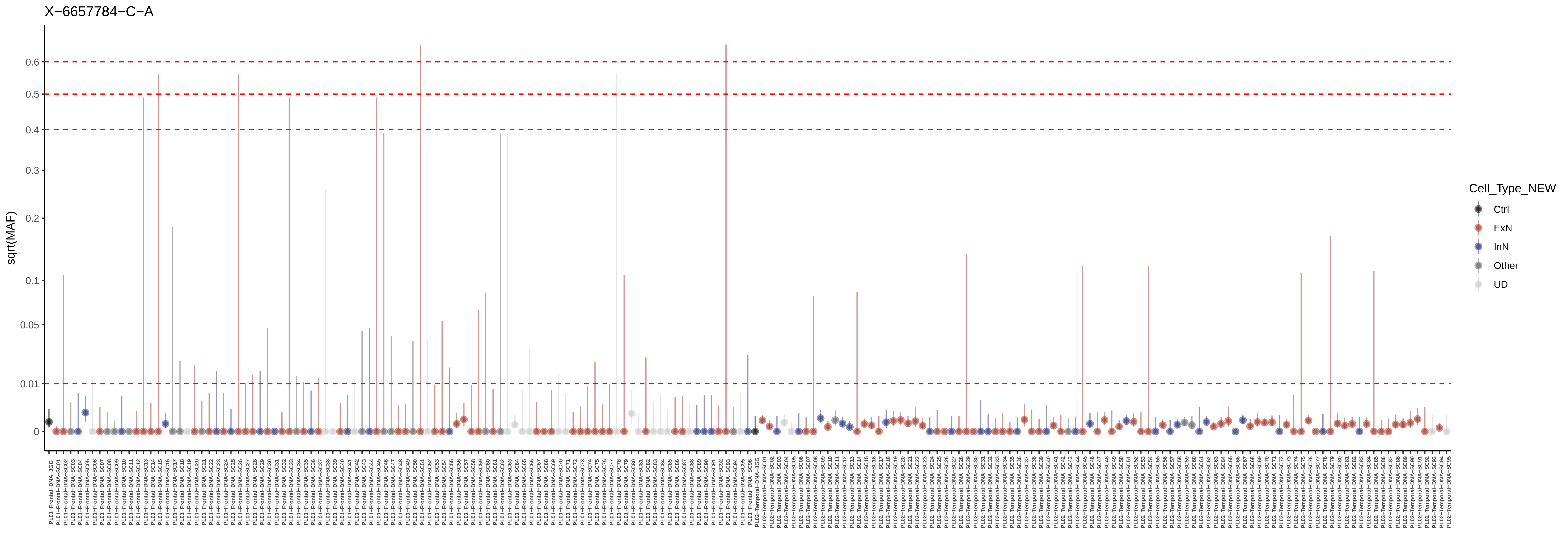

X−6657784−C−A
sqrt(MAF)
0.6
0.5
0.4
0.3
0.2
0.1
0.05
0.01
0
Cell_Type_NEW
Ctrl
ExN
InN
Other
UD

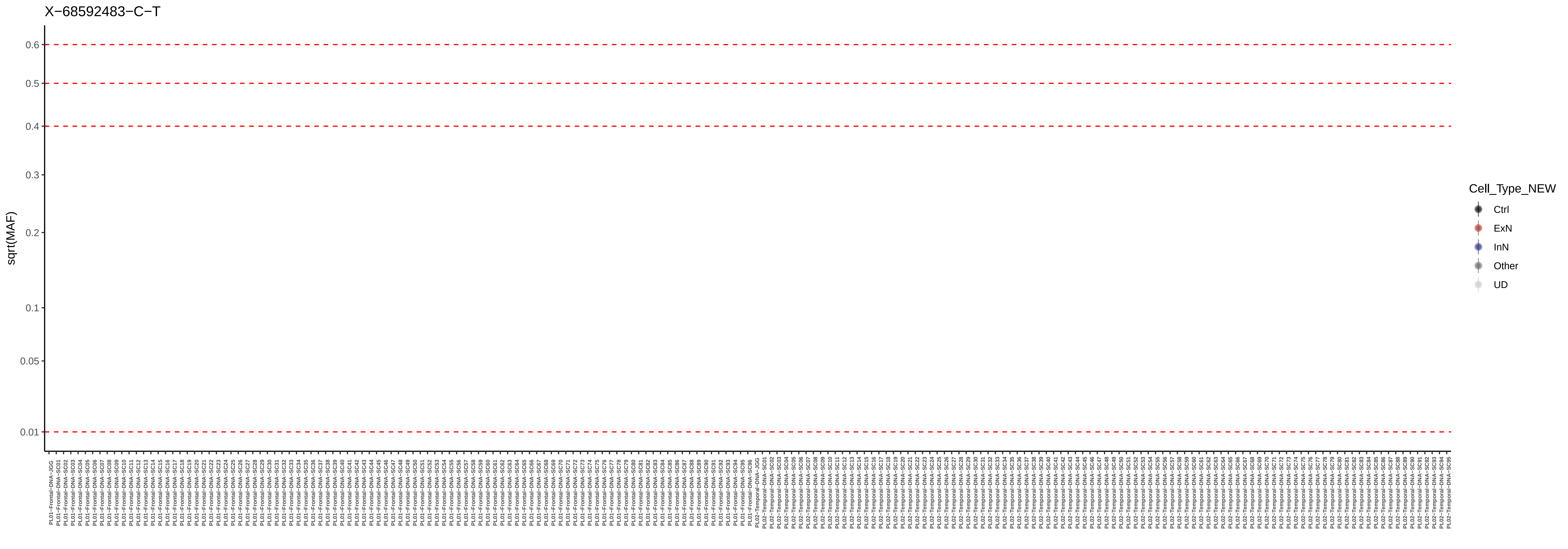
X−68592483−C−T
sqrt(MAF)
0.6
0.5
0.4
0.3
0.2
0.1
0.05
0.01
Cell_Type_NEW
Ctrl
ExN
InN
Other
UD

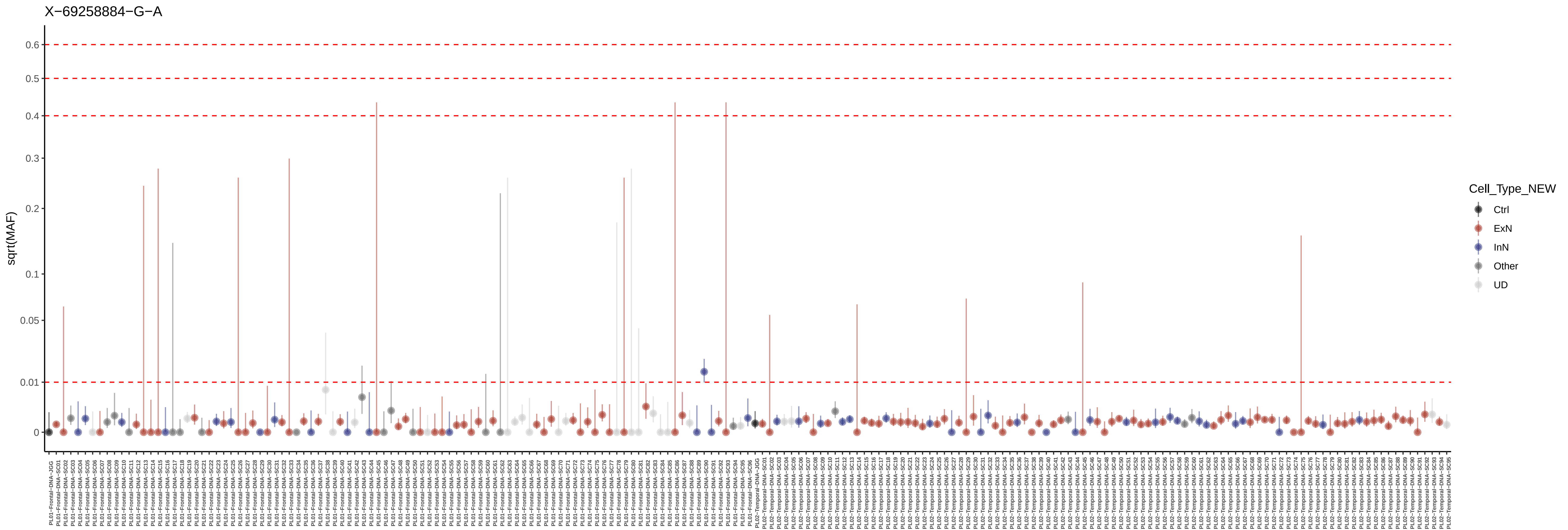
X−69258884−G−A
sqrt(MAF)
0.6
0.5
0.4
0.3
0.2
0.1
0.05
0.01
0
Cell_Type_NEW
Ctrl
ExN
InN
Other
UD

X−69794428−G−A

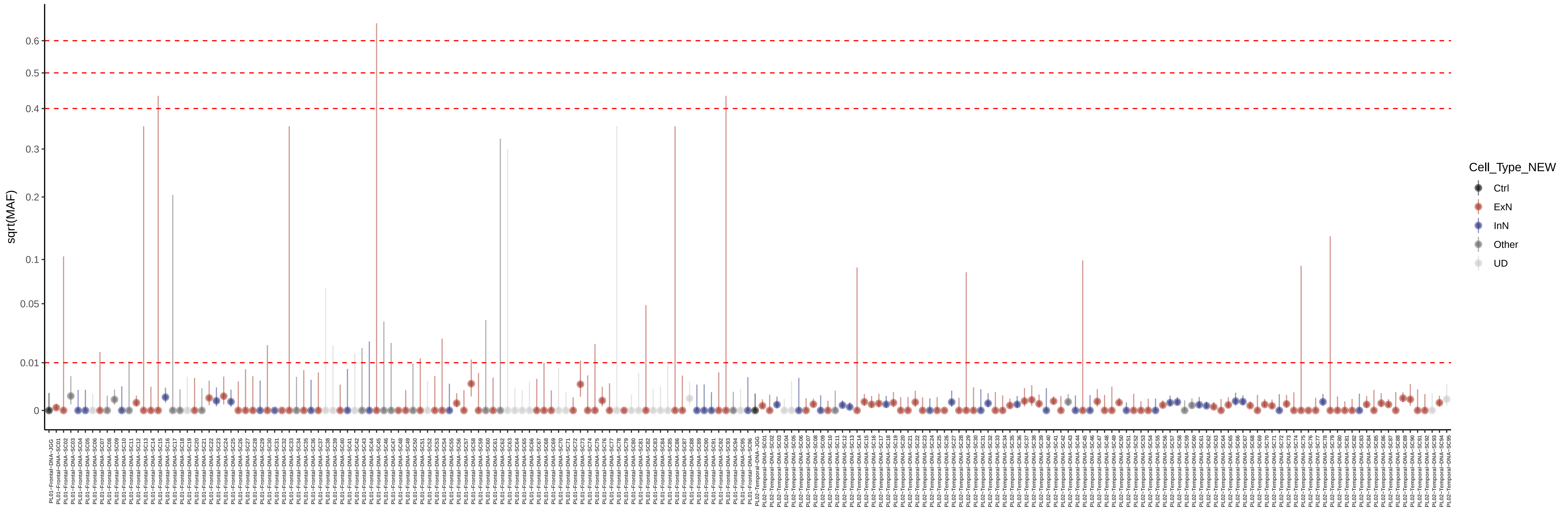

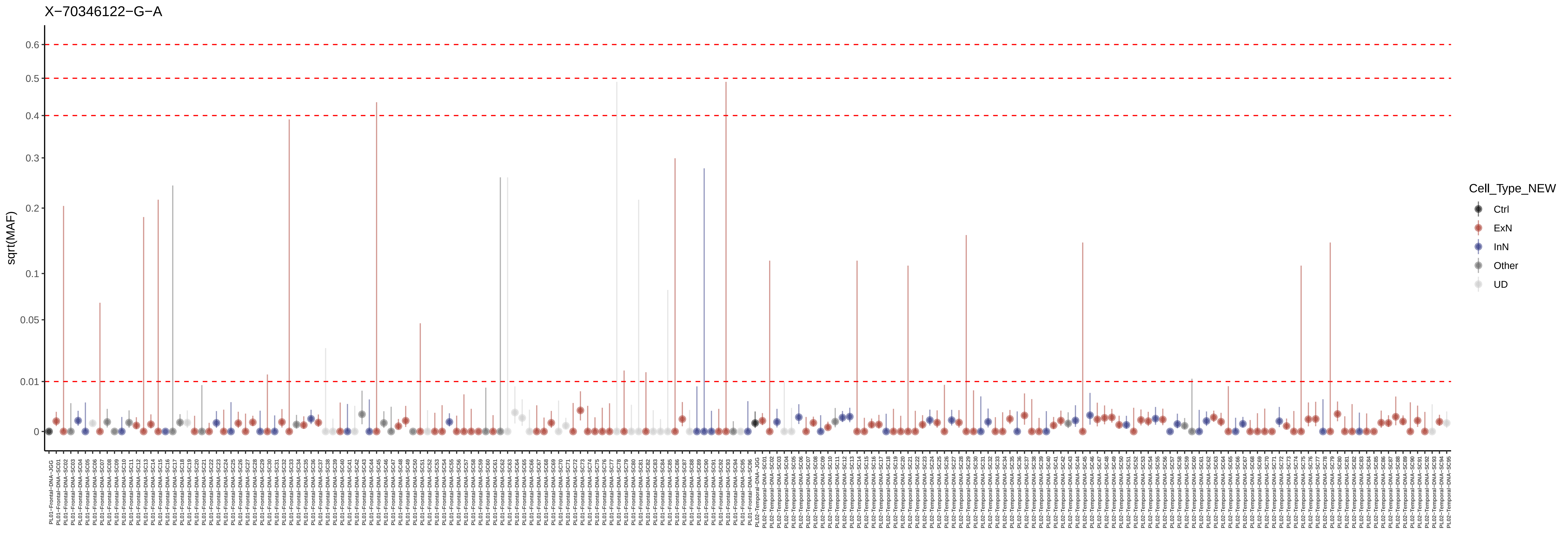
X−70346122−G−A
sqrt(MAF)
0.6
0.5
0.4
0.3
0.2
0.1
0.05
0.01
0
Cell_Type_NEW
Ctrl
ExN
InN
Other
UD

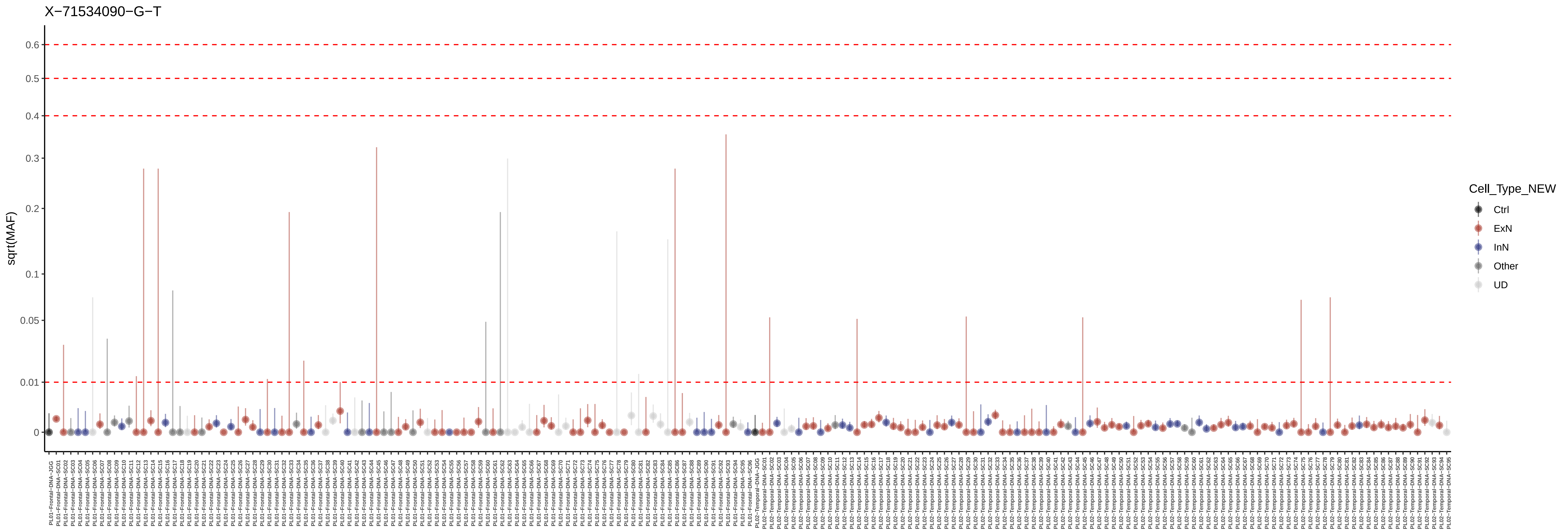
X−71534090−G−T
sqrt(MAF)
0.6
0.5
0.4
0.3
0.2
0.1
0.05
0.01
0
Cell_Type_NEW
Ctrl
ExN
InN
Other
UD

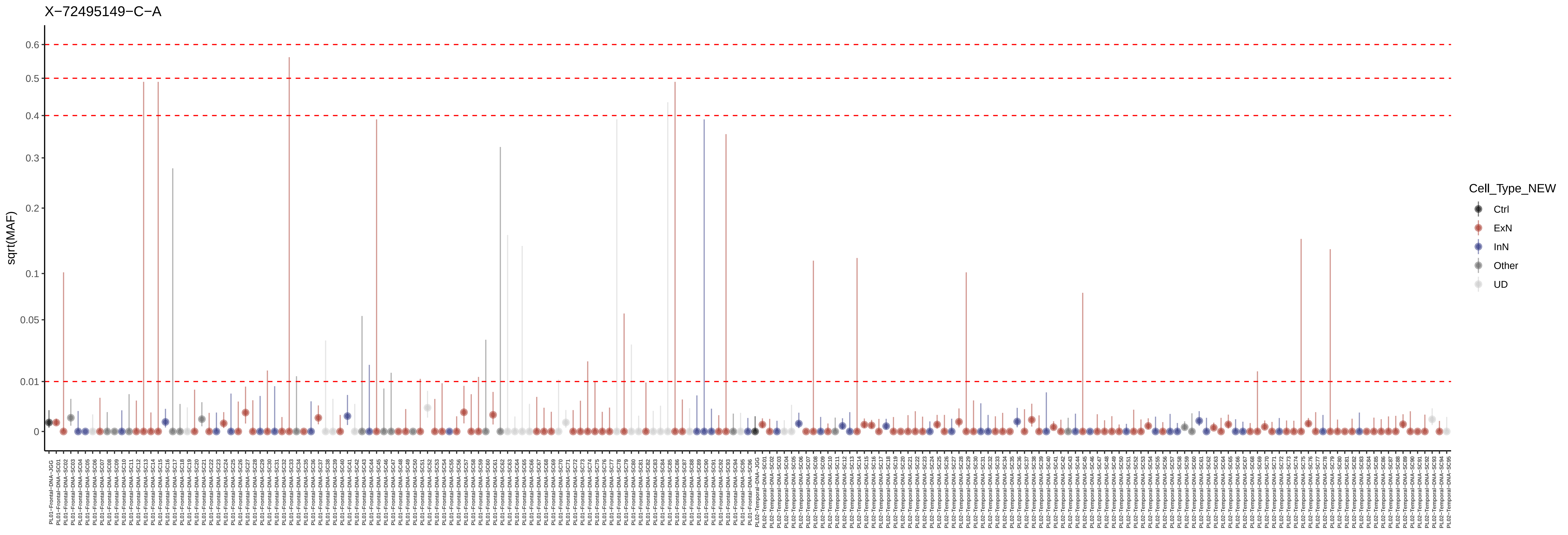
X−72495149−C−A
sqrt(MAF)
0.6
0.5
0.4
0.3
0.2
0.1
0.05
0.01
0
Cell_Type_NEW
Ctrl
ExN
InN
Other
UD

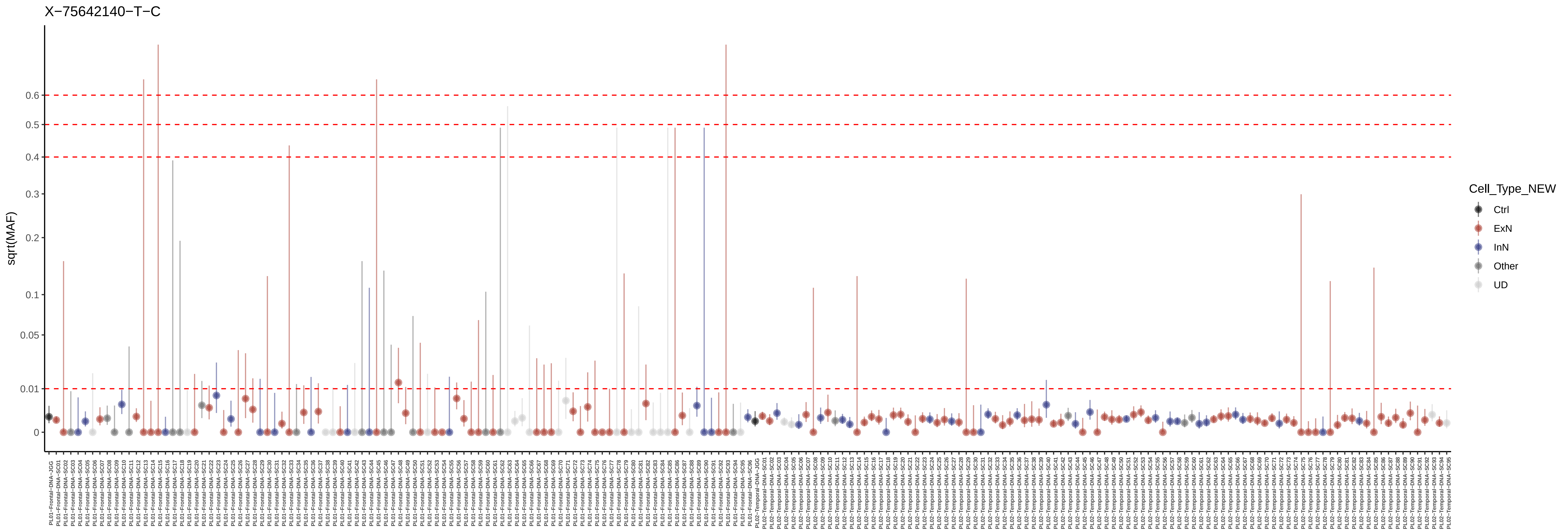

X−75642140−T−C
sqrt(MAF)
0.6
0.5
0.4
0.3
0.2
0.1
0.05
0.01
0
Cell_Type_NEW
Ctrl
ExN
InN
Other
UD

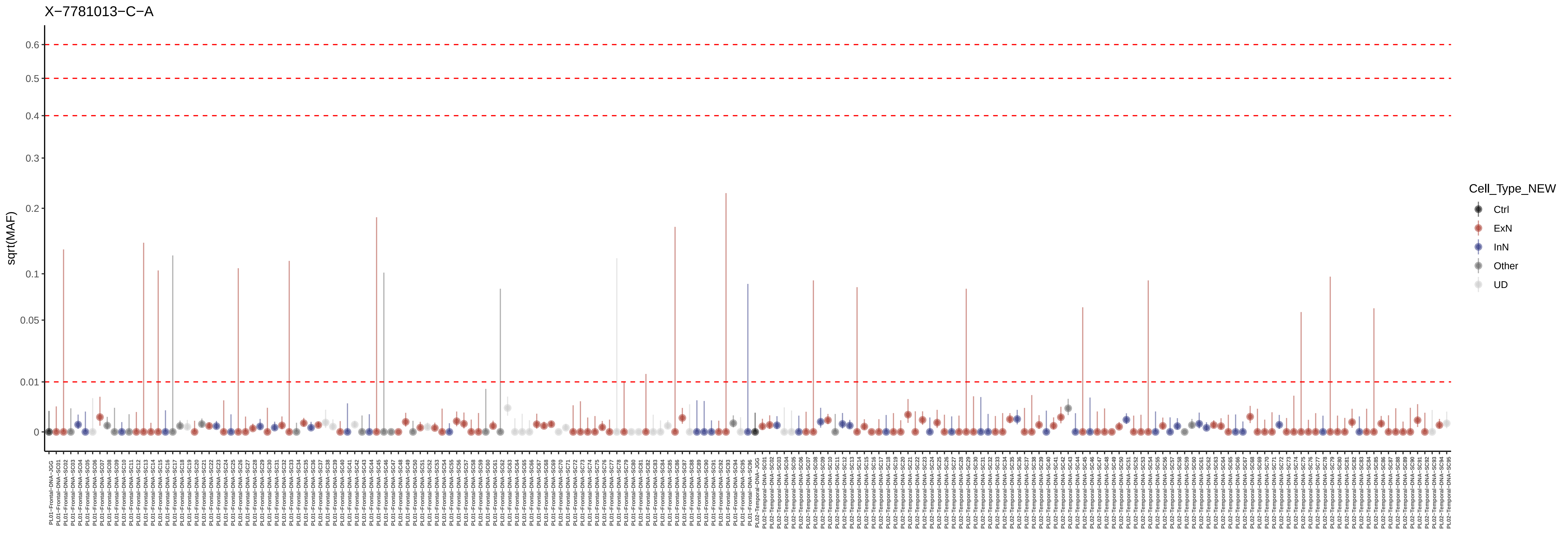
X−7781013−C−A
sqrt(MAF)
0.6
0.5
0.4
0.3
0.2
0.1
0.05
0.01
0
Cell_Type_NEW
Ctrl
ExN
InN
Other
UD

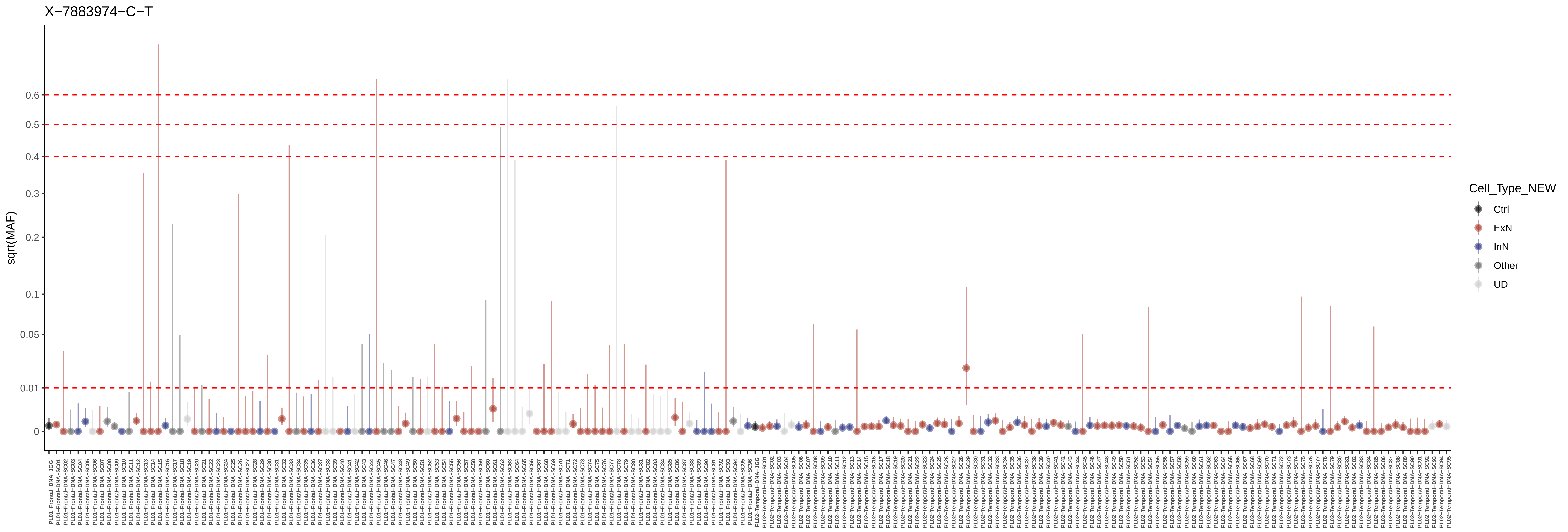
X−7883974−C−T
sqrt(MAF)
0.6
0.5
0.4
0.3
0.2
0.1
0.05
0.01
0
Cell_Type_NEW
Ctrl
ExN
InN
Other
UD

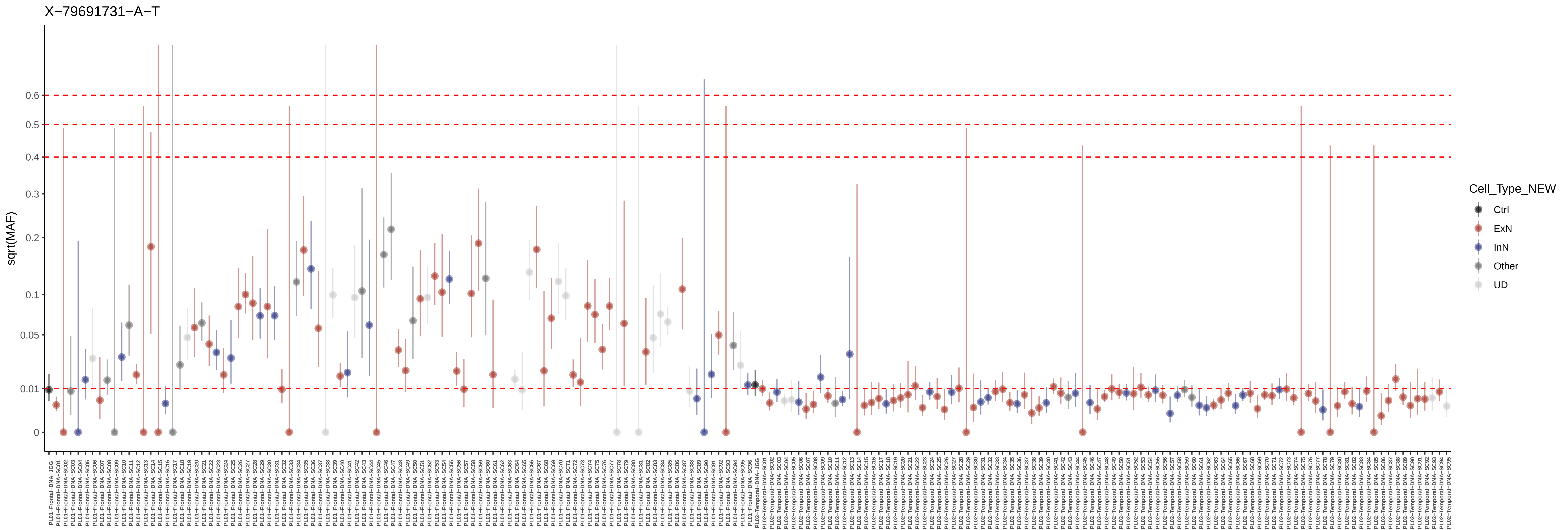

X−79691731−A−T
sqrt(MAF)
0.6
0.5
0.4
0.3
0.2
0.1
0.05
0.01
0
Cell_Type_NEW
Ctrl
ExN
InN
Other
UD

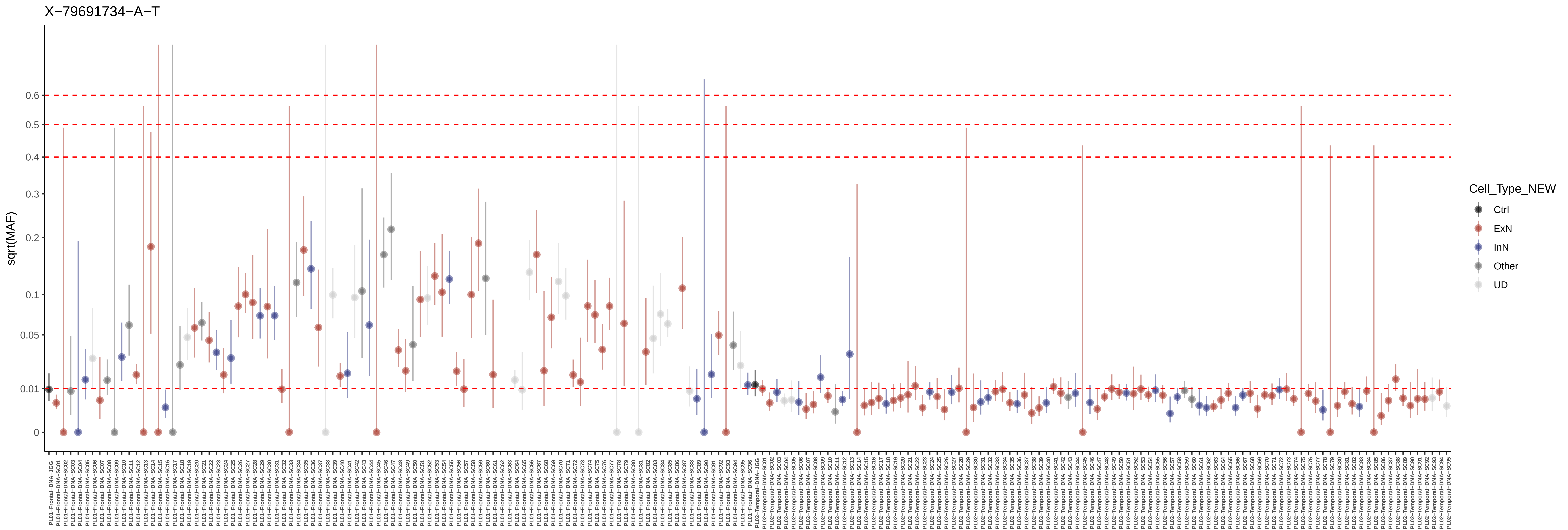
X−79691734−A−T
sqrt(MAF)
0.6
0.5
0.4
0.3
0.2
0.1
0.05
0.01
0
Cell_Type_NEW
Ctrl
ExN
InN
Other
UD

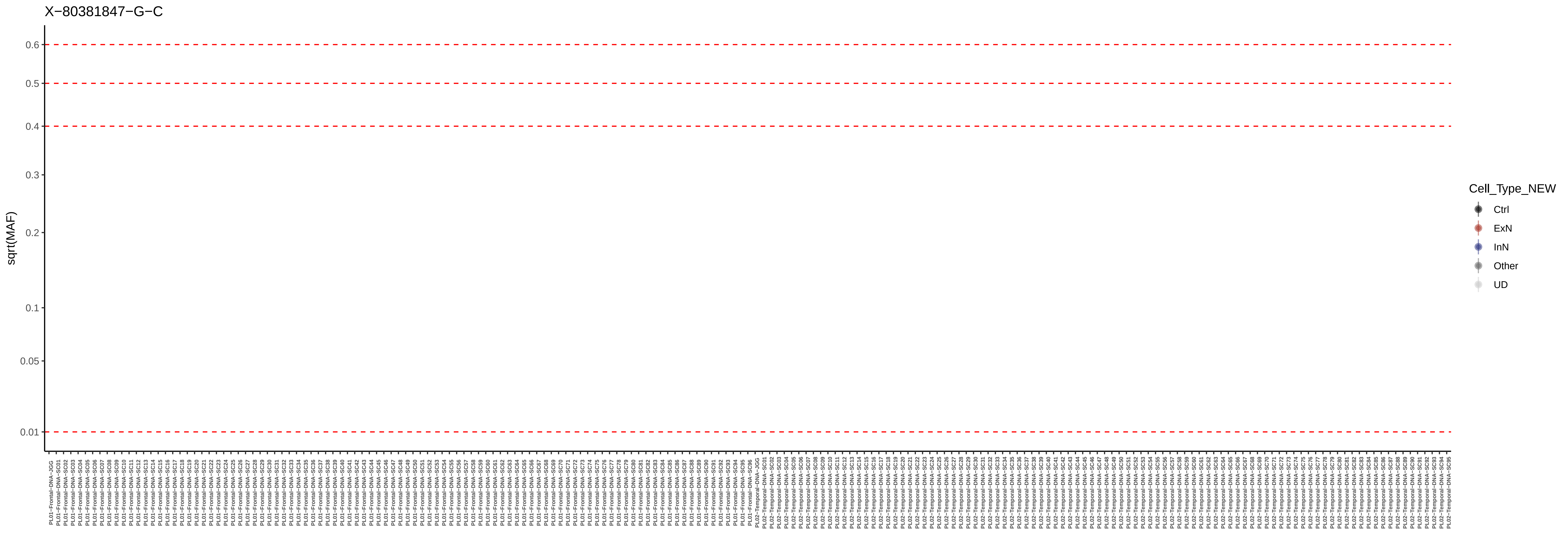
X−80381847−G−C
sqrt(MAF)
0.6
0.5
0.4
0.3
0.2
0.1
0.05
0.01
Cell_Type_NEW
Ctrl
ExN
InN
Other
UD

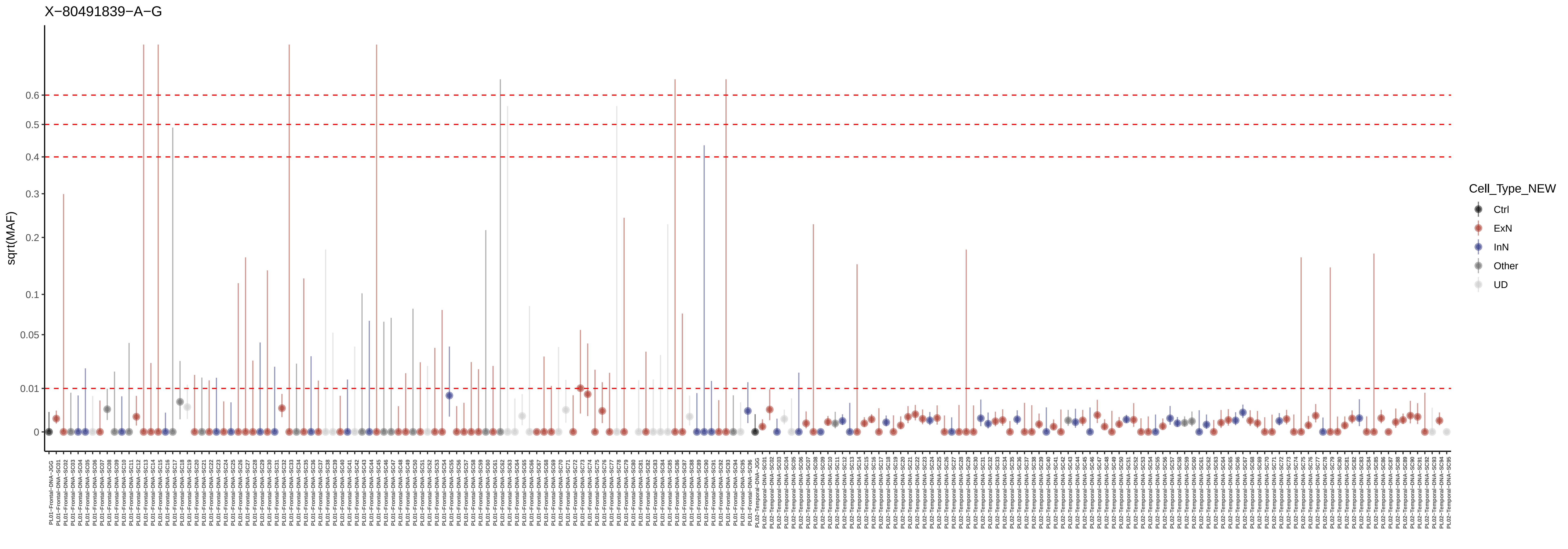

X−80491839−A−G
sqrt(MAF)
0.6
0.5
0.4
0.3
0.2
0.1
0.05
0.01
0
Cell_Type_NEW
Ctrl
ExN
InN
Other
UD

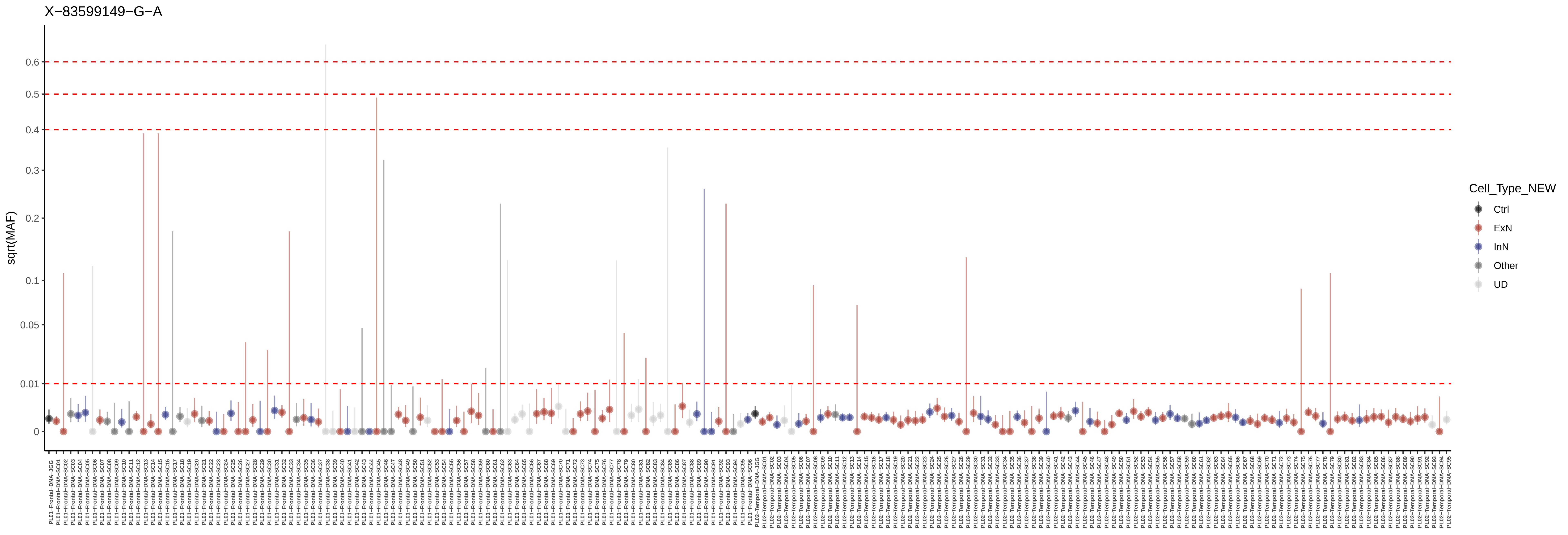

X−83599149−G−A
sqrt(MAF)
0.6
0.5
0.4
0.3
0.2
0.1
0.05
0.01
0
Cell_Type_NEW
Ctrl
ExN
InN
Other
UD

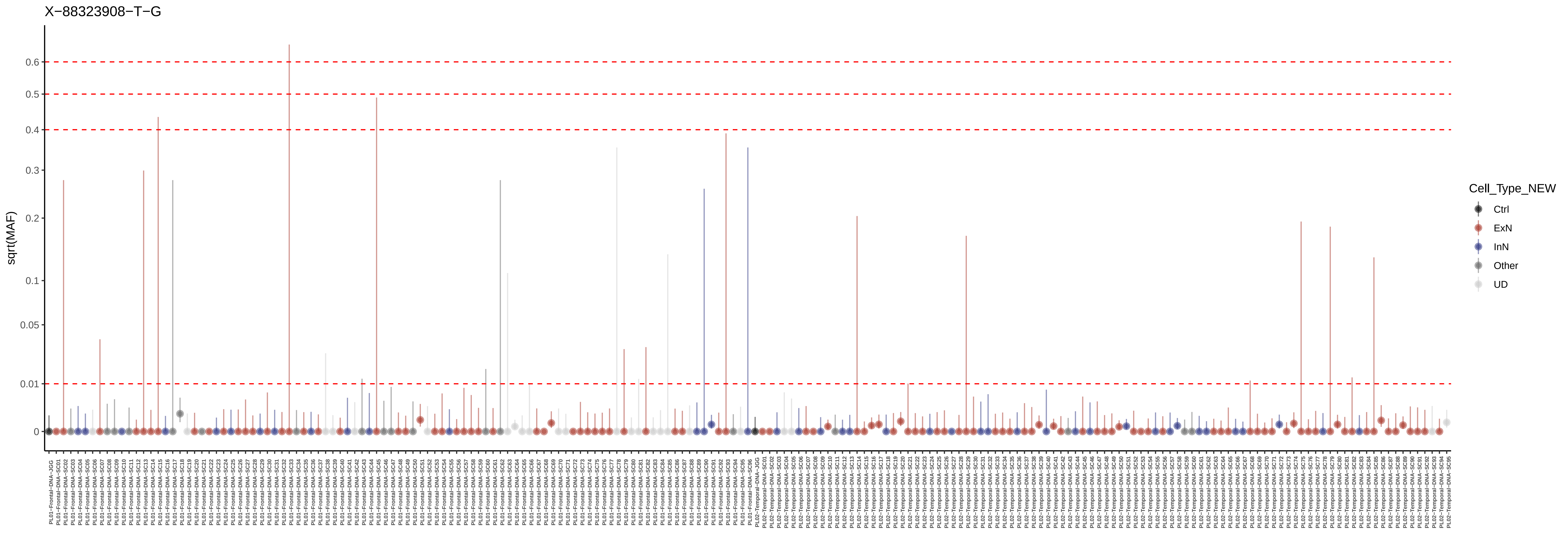
X−88323908−T−G
sqrt(MAF)
0.6
0.5
0.4
0.3
0.2
0.1
0.05
0.01
0
Cell_Type_NEW
Ctrl
ExN
InN
Other
UD

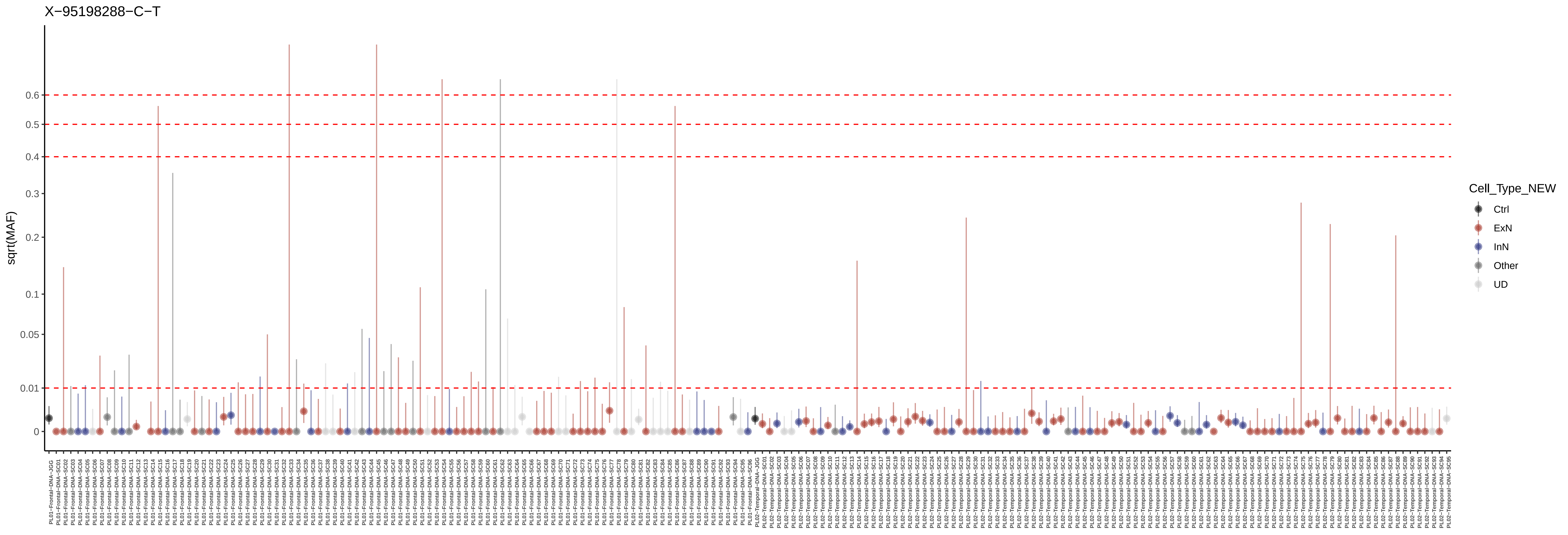
X−95198288−C−T
sqrt(MAF)
0.6
0.5
0.4
0.3
0.2
0.1
0.05
0.01
0
Cell_Type_NEW
Ctrl
ExN
InN
Other
UD

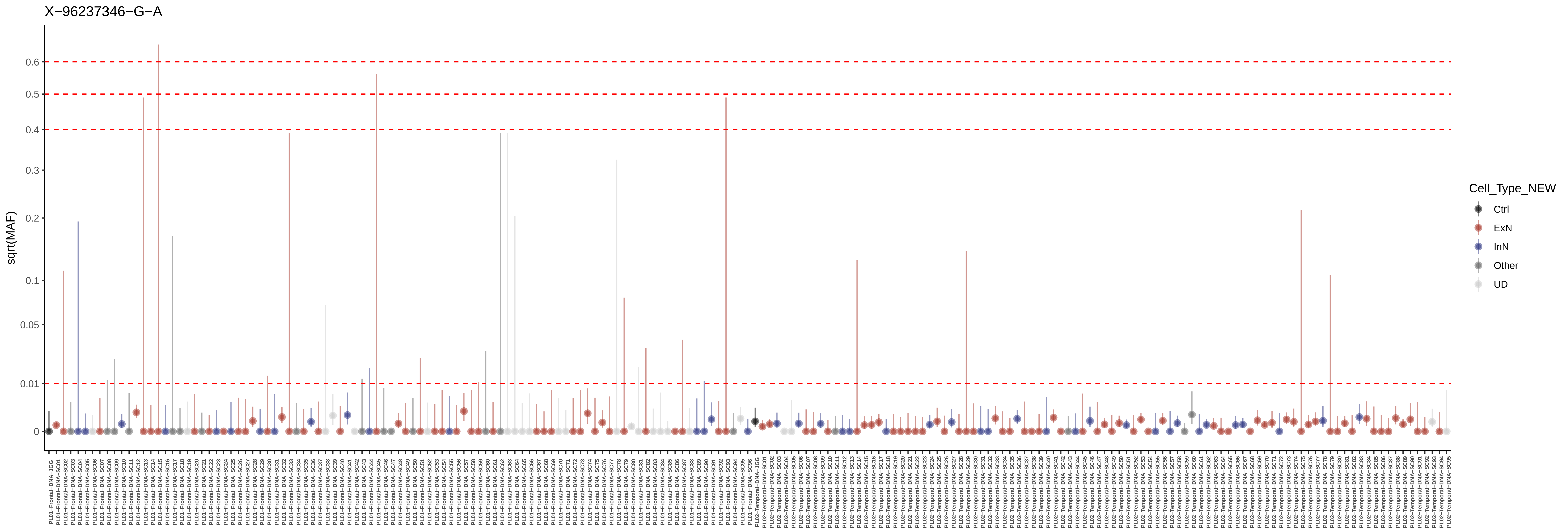
X−96237346−G−A
sqrt(MAF)
0.6
0.5
0.4
0.3
0.2
0.1
0.05
0.01
0
Cell_Type_NEW
Ctrl
ExN
InN
Other
UD

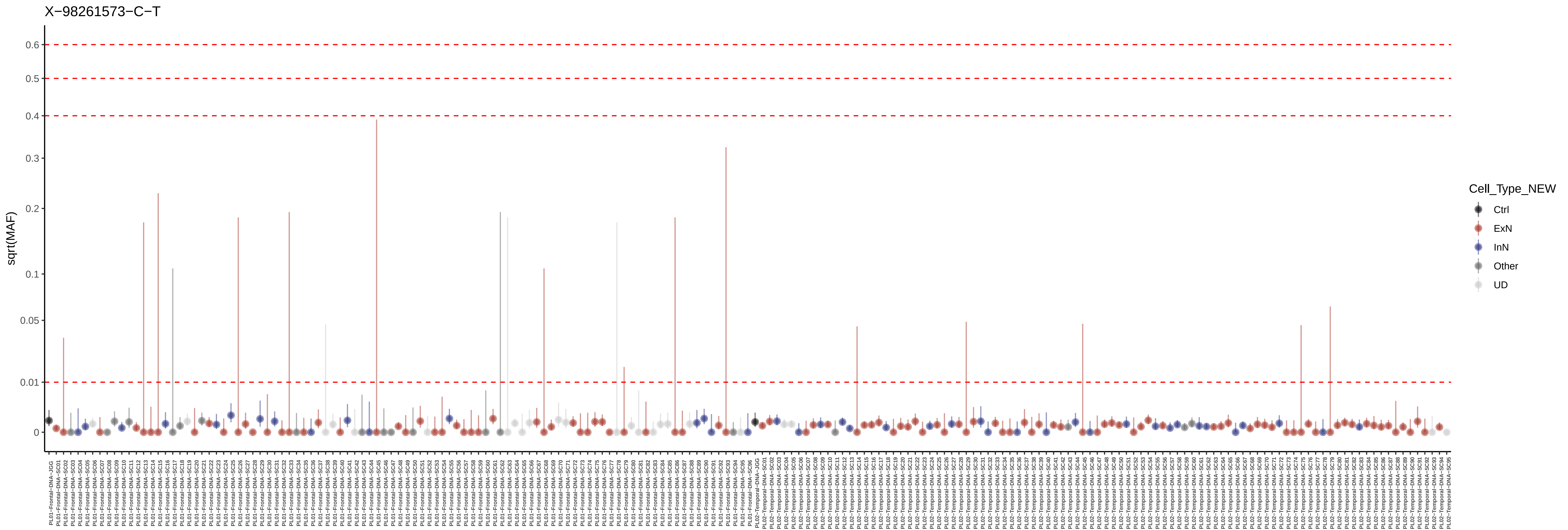

X−98261573−C−T
sqrt(MAF)
0.6
0.5
0.4
0.3
0.2
0.1
0.05
0.01
0
Cell_Type_NEW
Ctrl
ExN
InN
Other
UD

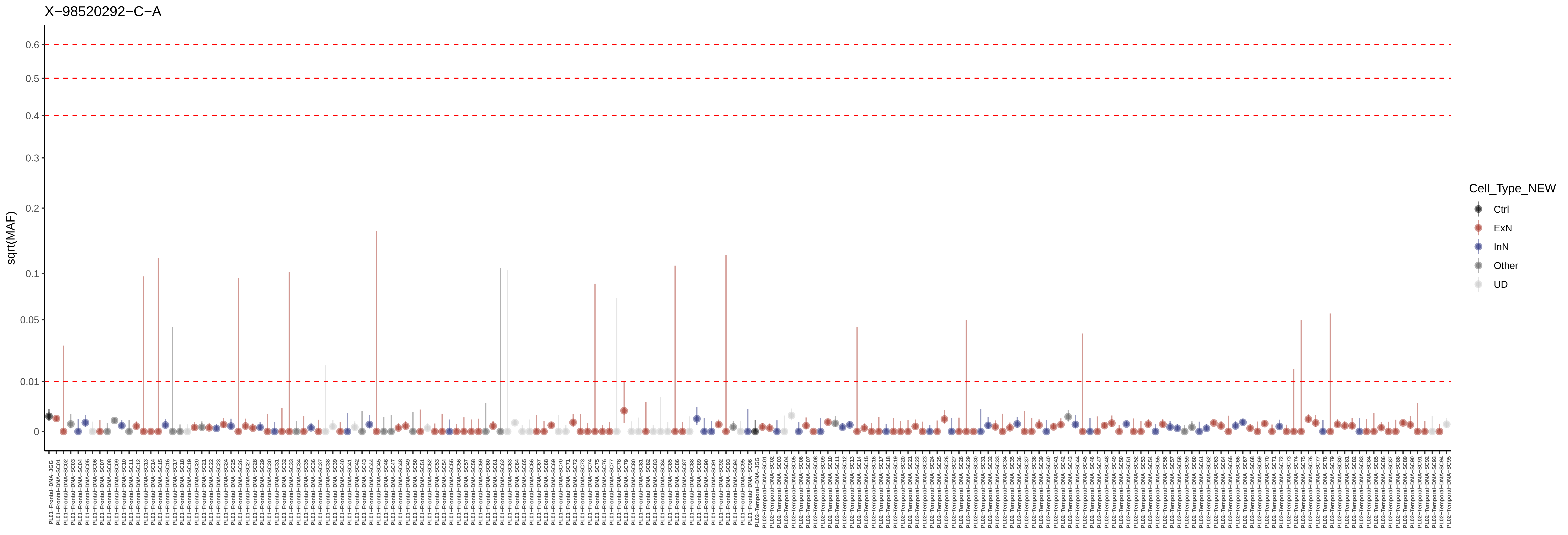
X−98520292−C−A
sqrt(MAF)
0.6
0.5
0.4
0.3
0.2
0.1
0.05
0.01
0
Cell_Type_NEW
Ctrl
ExN
InN
Other
UD

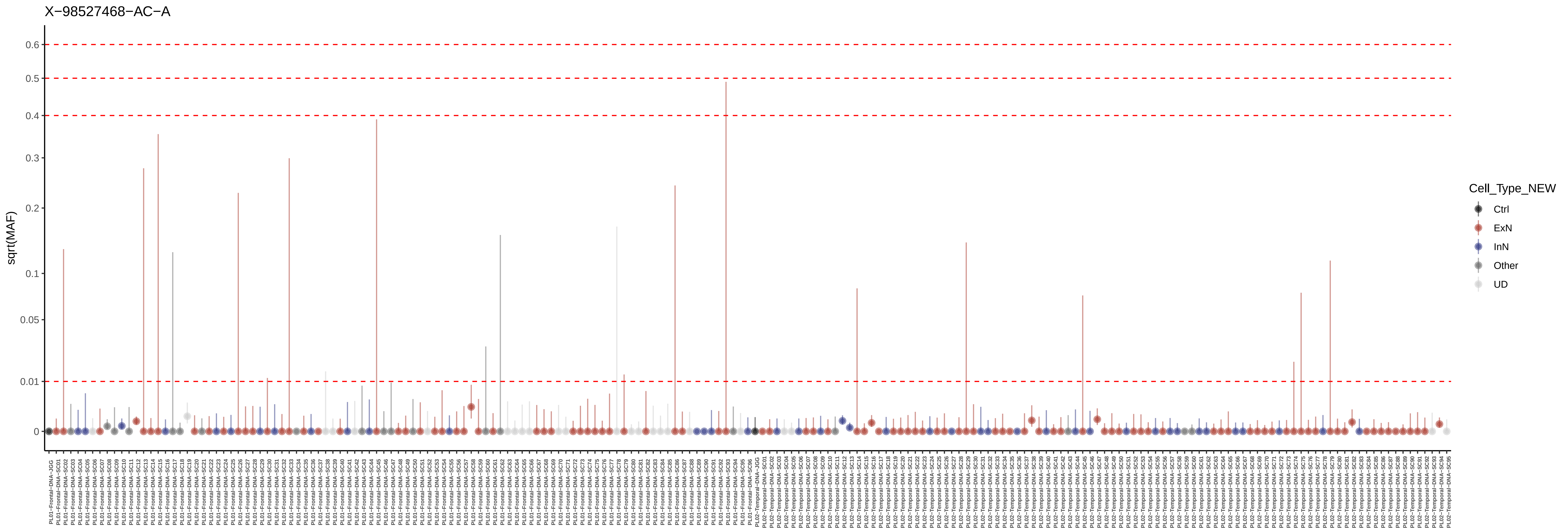
X−98527468−AC−A
sqrt(MAF)
0.6
0.5
0.4
0.3
0.2
0.1
0.05
0.01
0
Cell_Type_NEW
Ctrl
ExN
InN
Other
UD

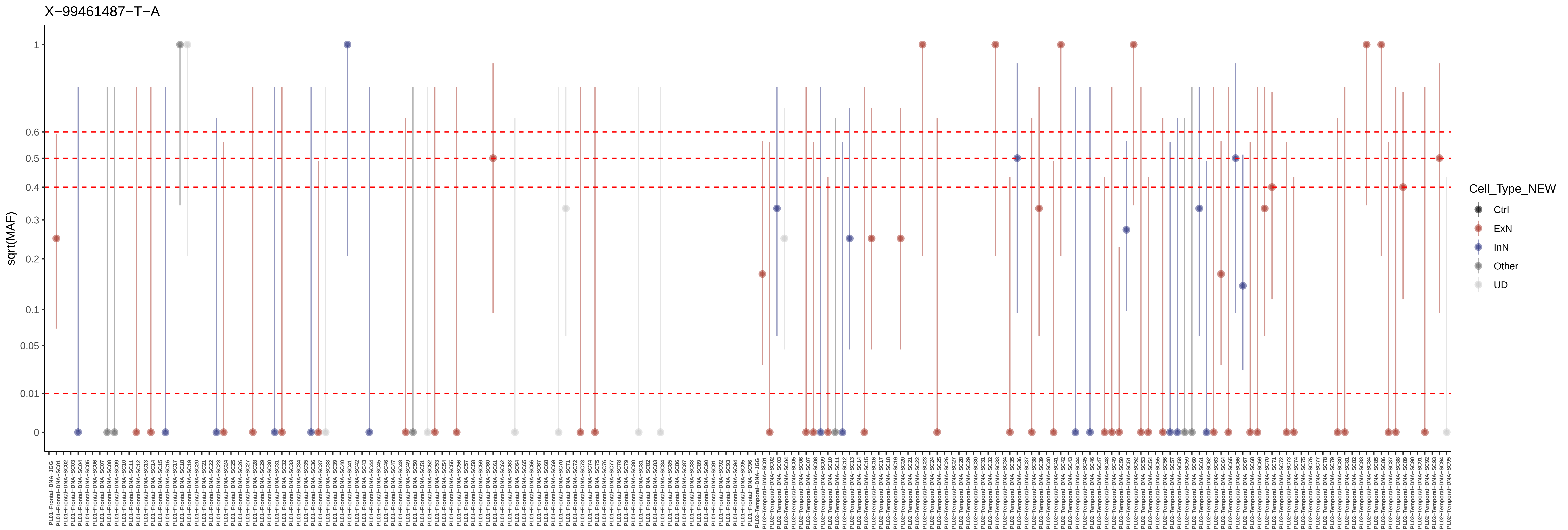

X−99461487−T−A
sqrt(MAF)
Cell_Type_NEW
Ctrl
ExN
InN
Other
UD

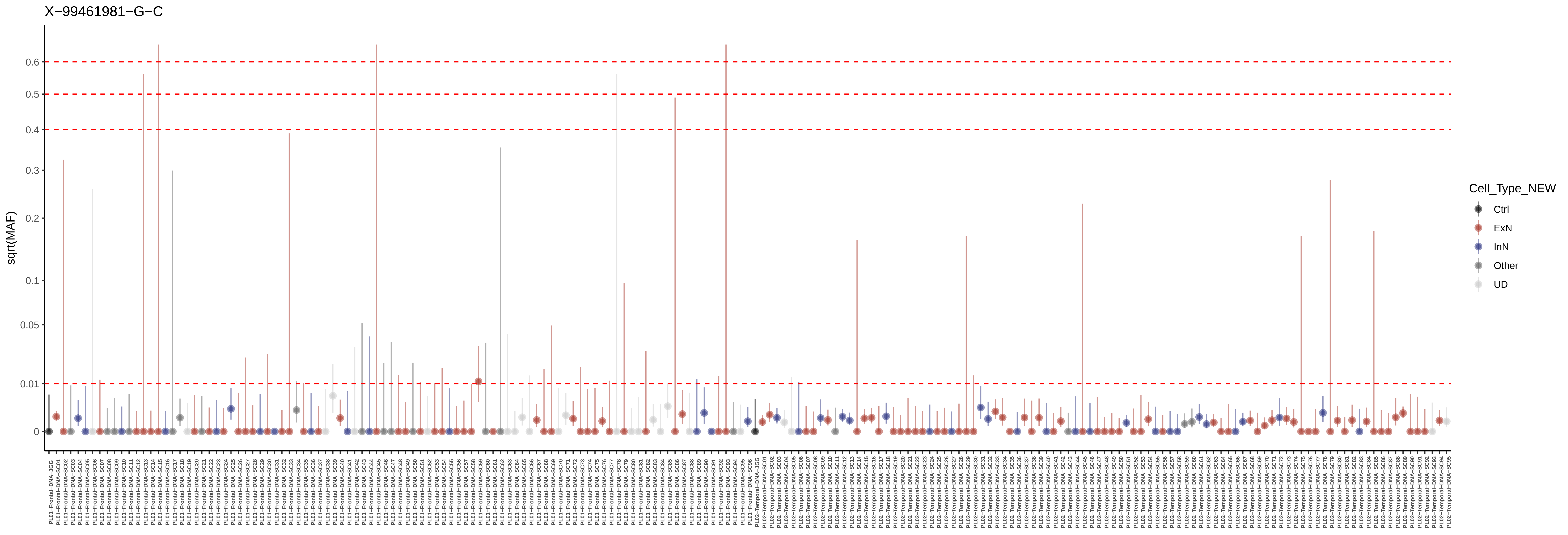

X−99461981−G−C
sqrt(MAF)
0.6
0.5
0.4
0.3
0.2
0.1
0.05
0.01
0
Cell_Type_NEW
Ctrl
ExN
InN
Other
UD

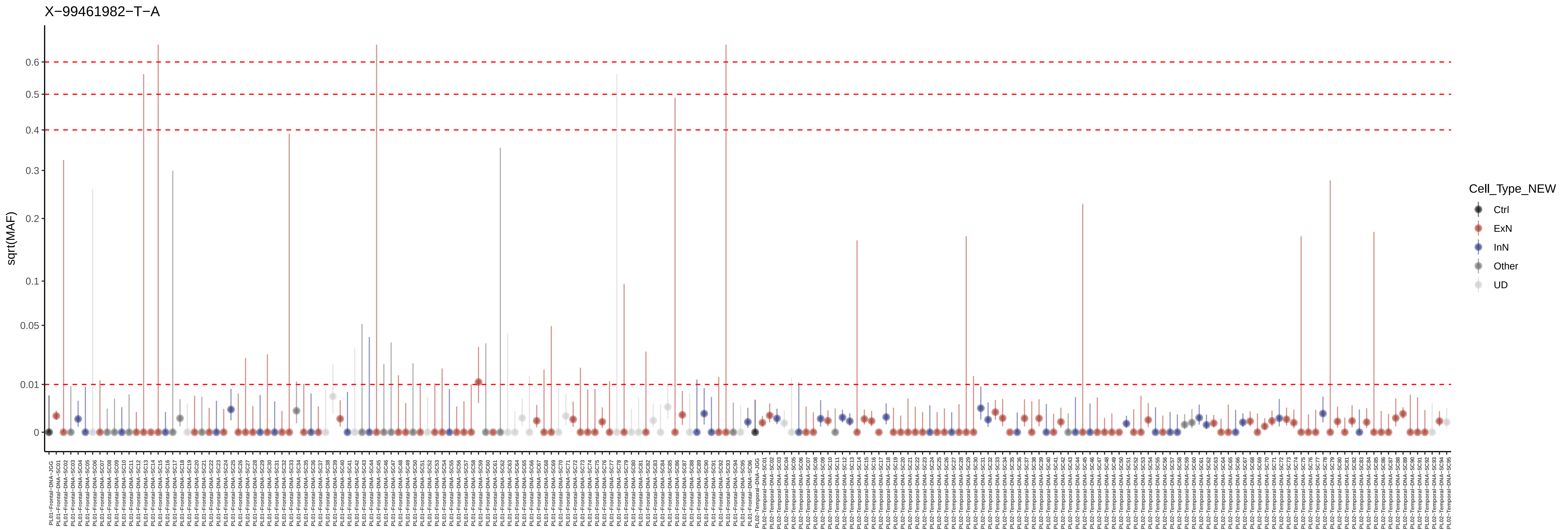
X−99461982−T−A
sqrt(MAF)
0.6
0.5
0.4
0.3
0.2
0.1
0.05
0.01
0
Cell_Type_NEW
Ctrl
ExN
InN
Other
UD

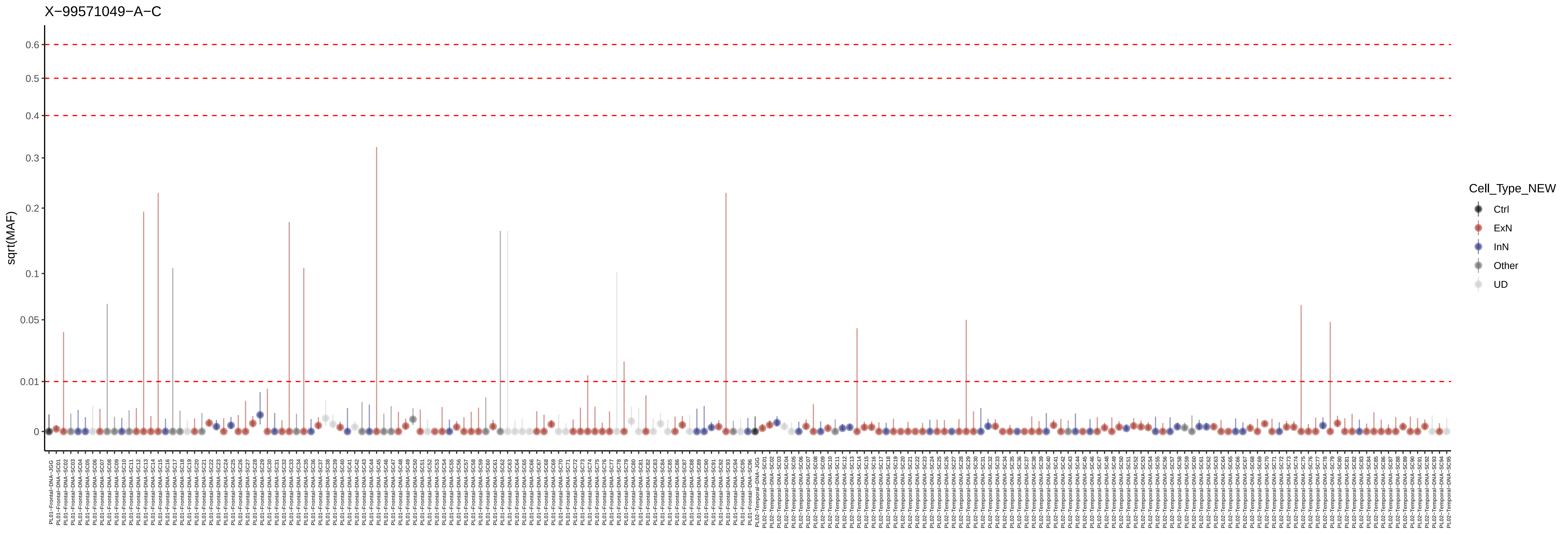
X−99571049−A−C
sqrt(MAF)
0.6
0.5
0.4
0.3
0.2
0.1
0.05
0.01
0
Cell_Type_NEW
Ctrl
ExN
InN
Other
UD

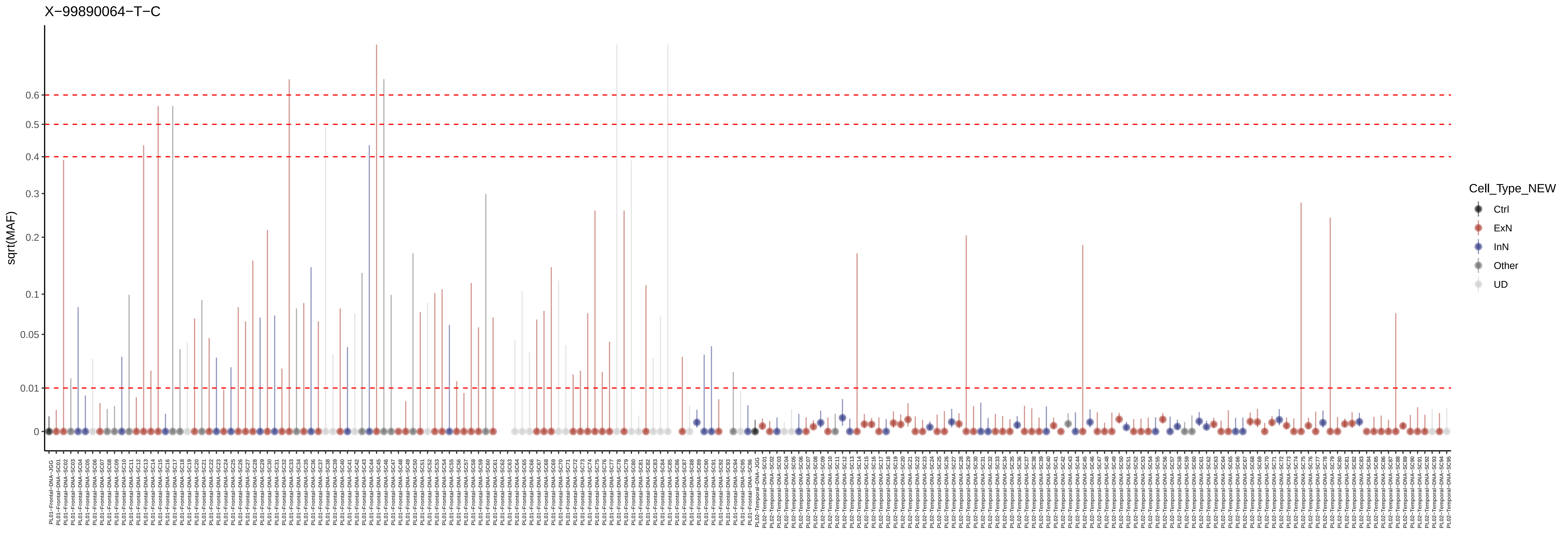
X−99890064−T−C
sqrt(MAF)
0.6
0.5
0.4
0.3
0.2
0.1
0.05
0.01
0
Cell_Type_NEW
Ctrl
ExN
InN
Other
UD
