## Supplementary Data 8 for "Cell-type-resolved somatic mosaicism reveals clonal dynamics of the human forebrain"

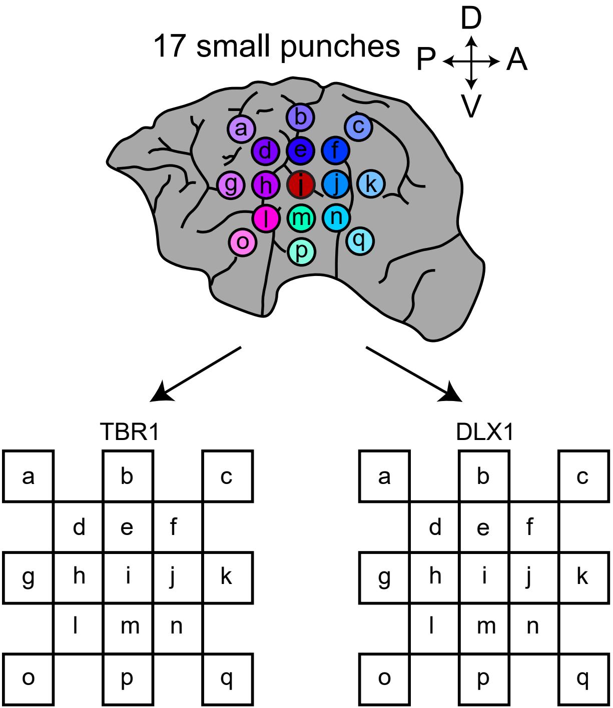
17 small punches
D
P
A
V
a
b
c
d
e
f
g
h
i
j
k
l
m
n
o
p
q
TBR1
DLX1

1−102152318−C−T

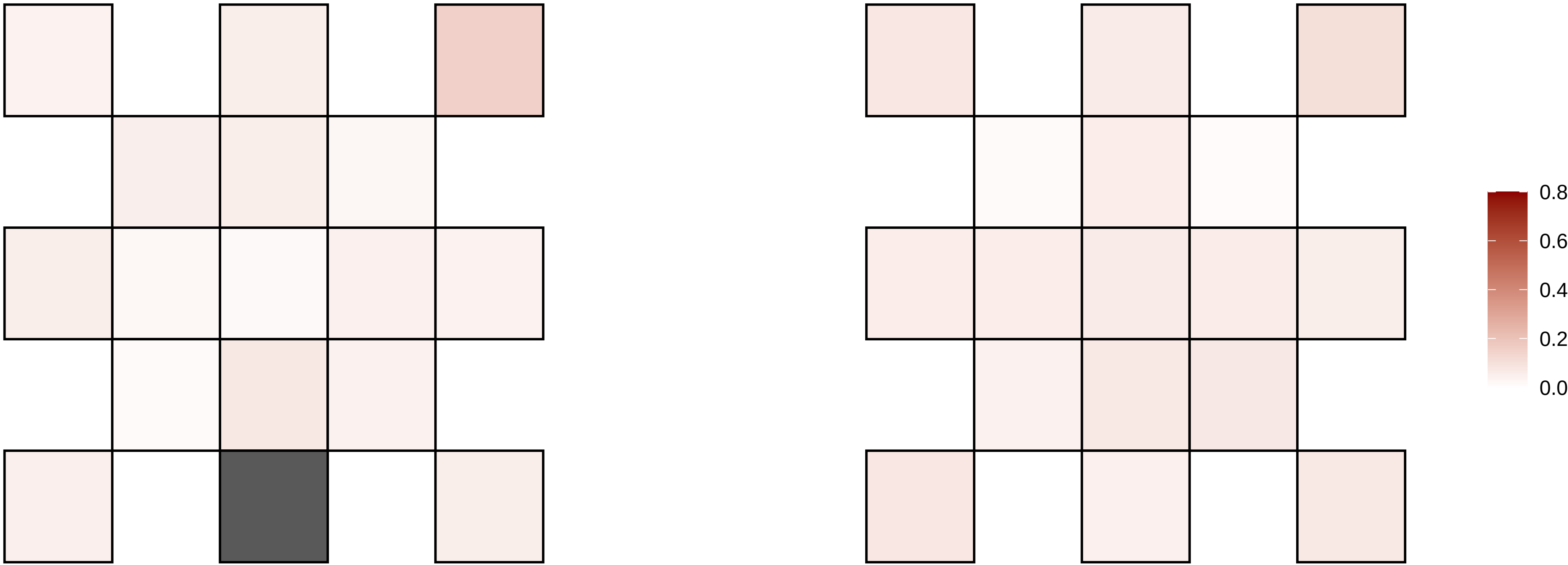

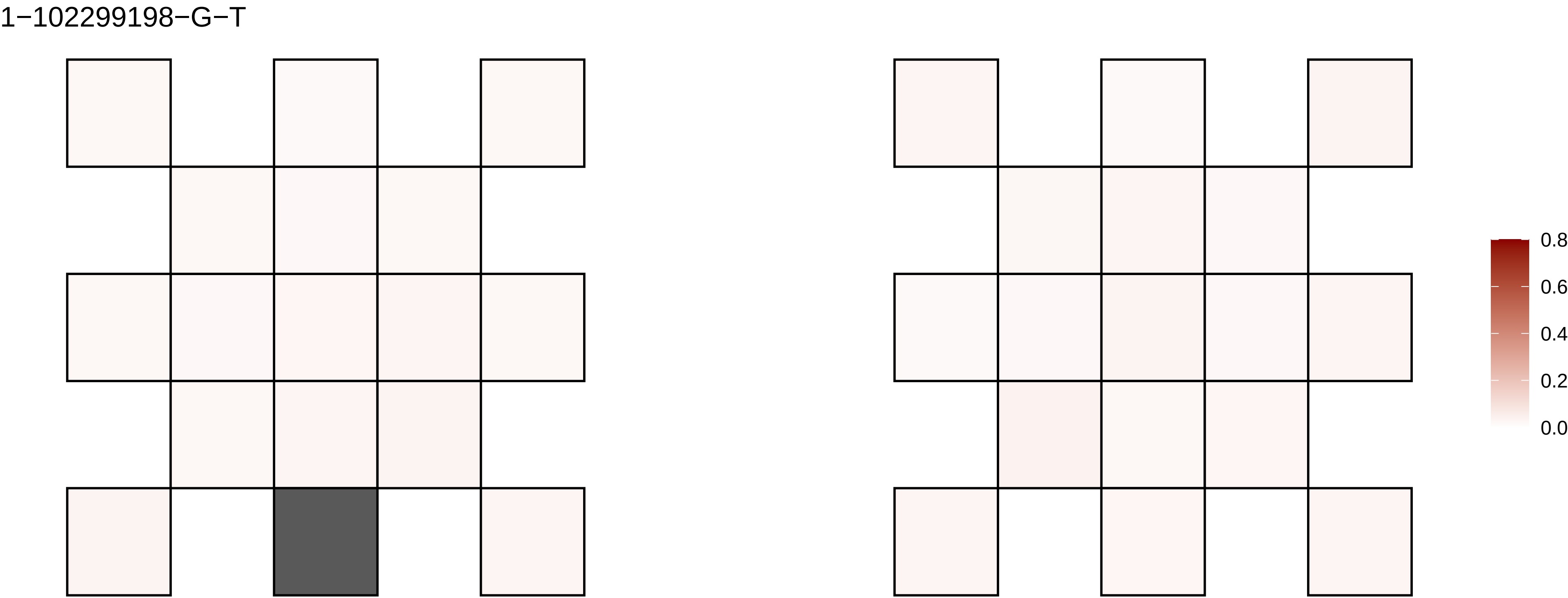
1−102299198−G−T
0.8
0.6
0.4
0.2
0.0

1−103746984−A−T

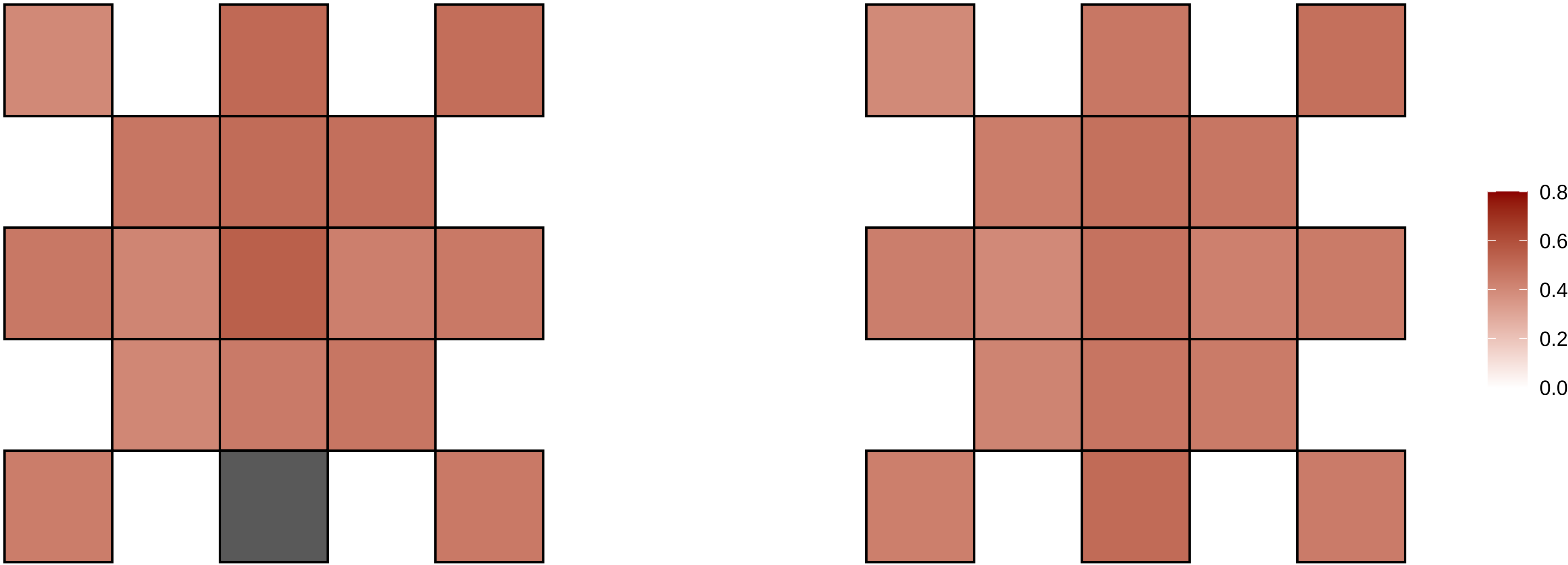

1−104816417−G−A

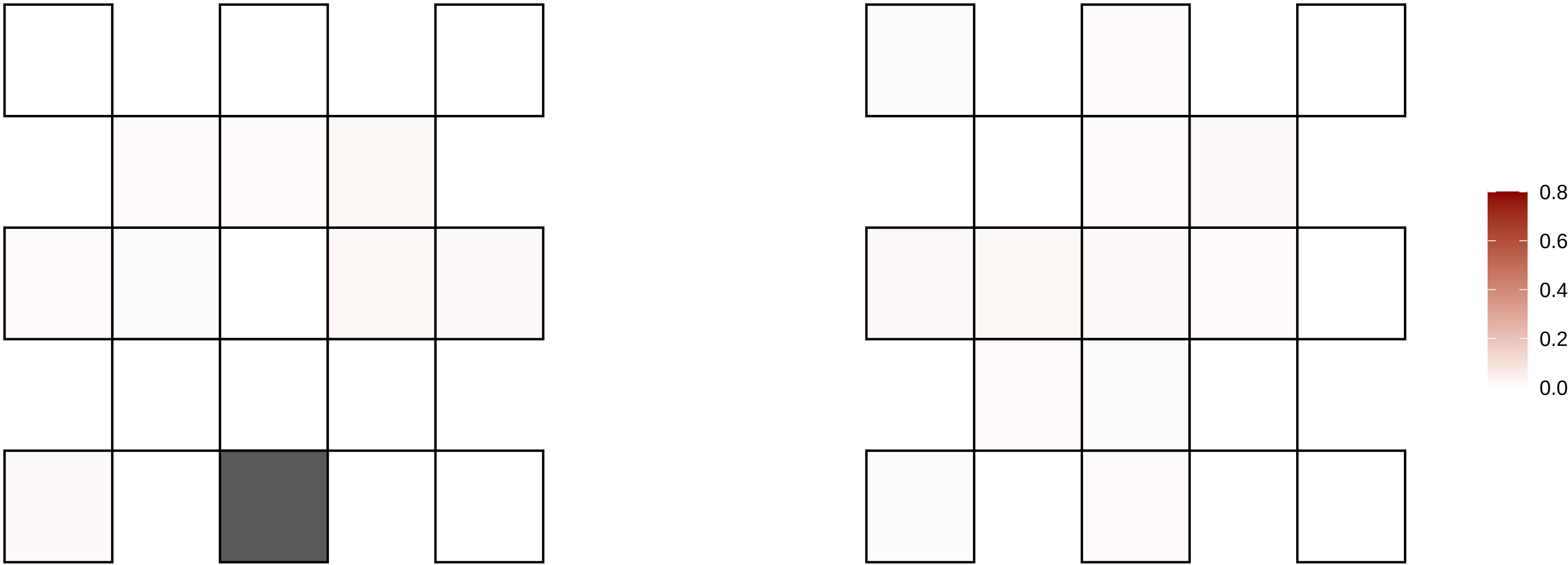

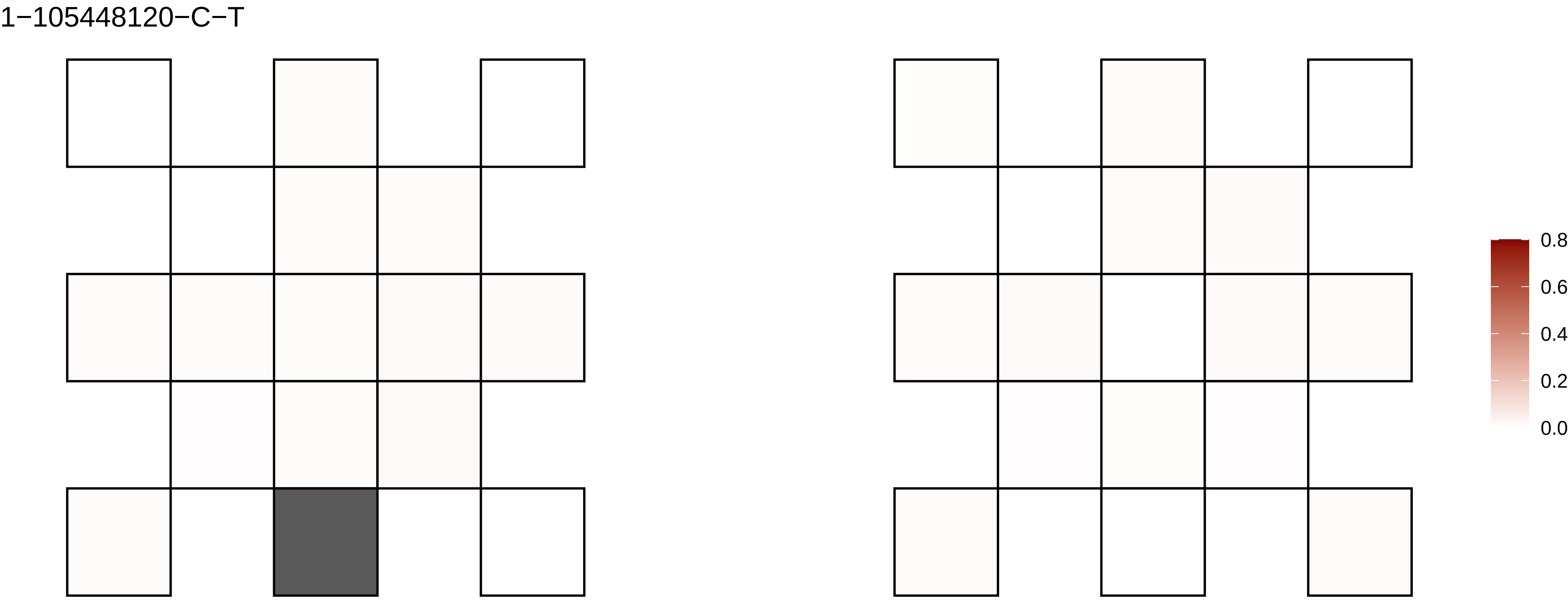
1−105448120−C−T
0.8
0.6
0.4
0.2
0.0

1−113168020−C−T

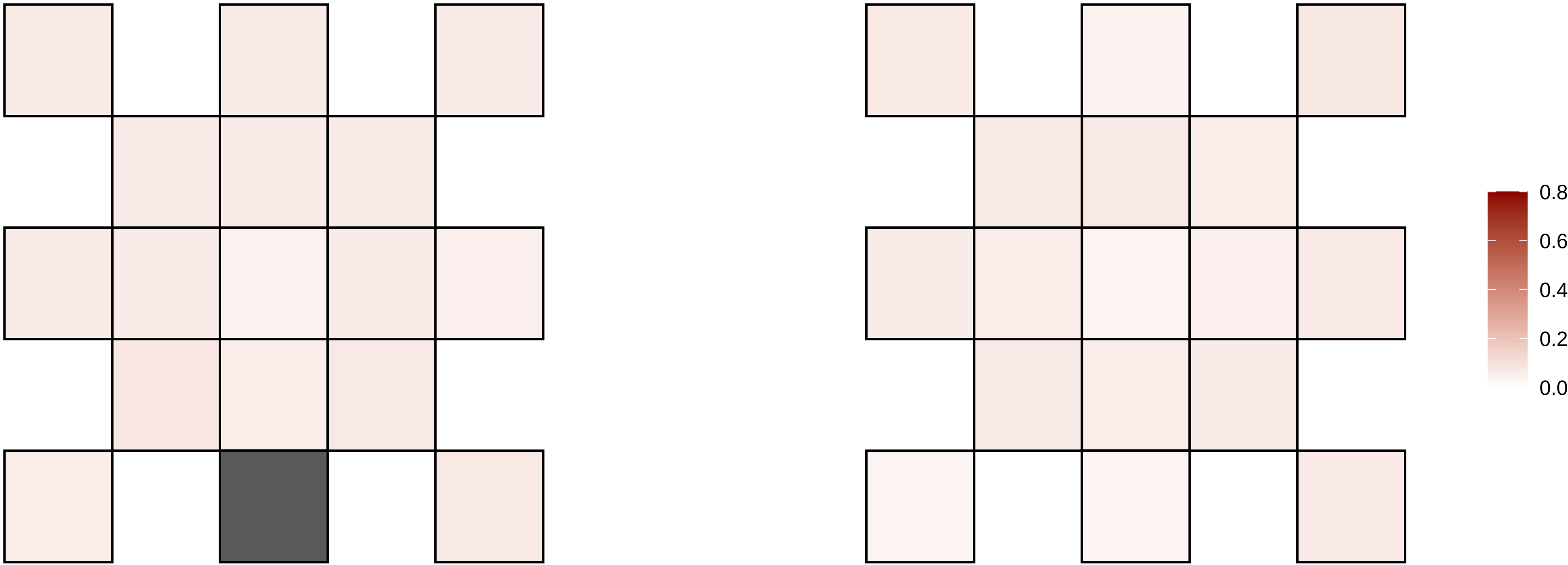

1−113168021−A−G

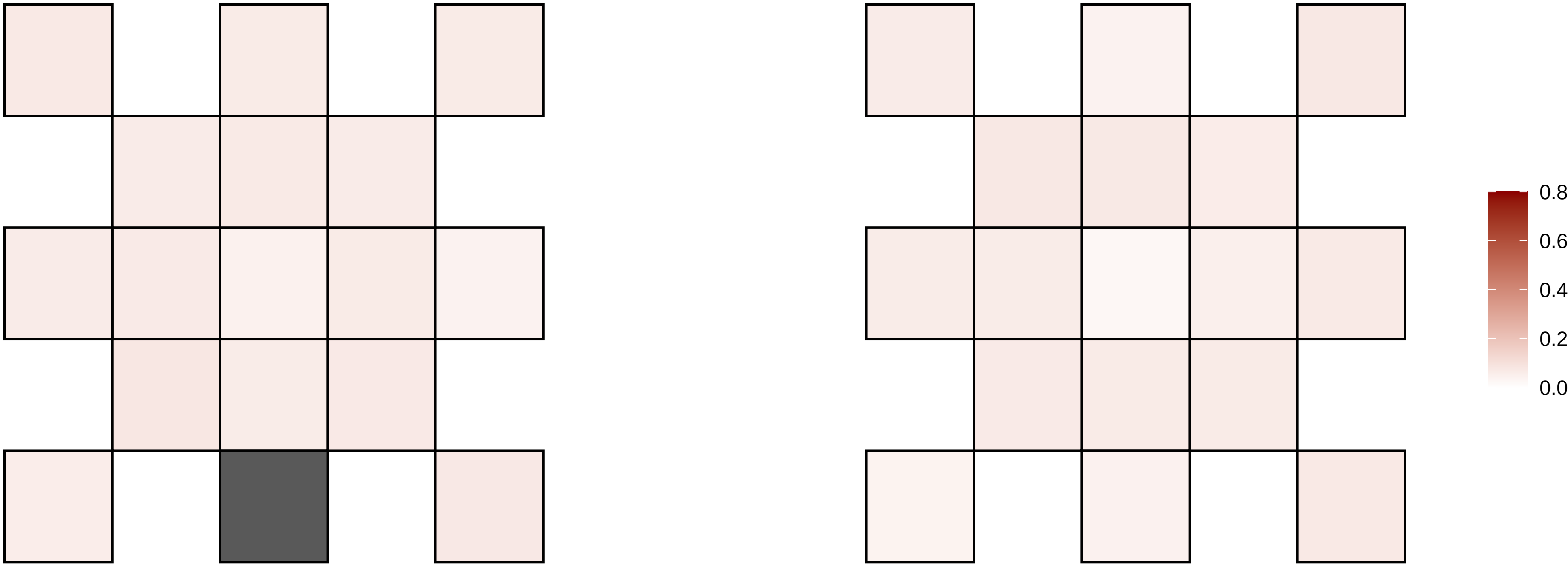

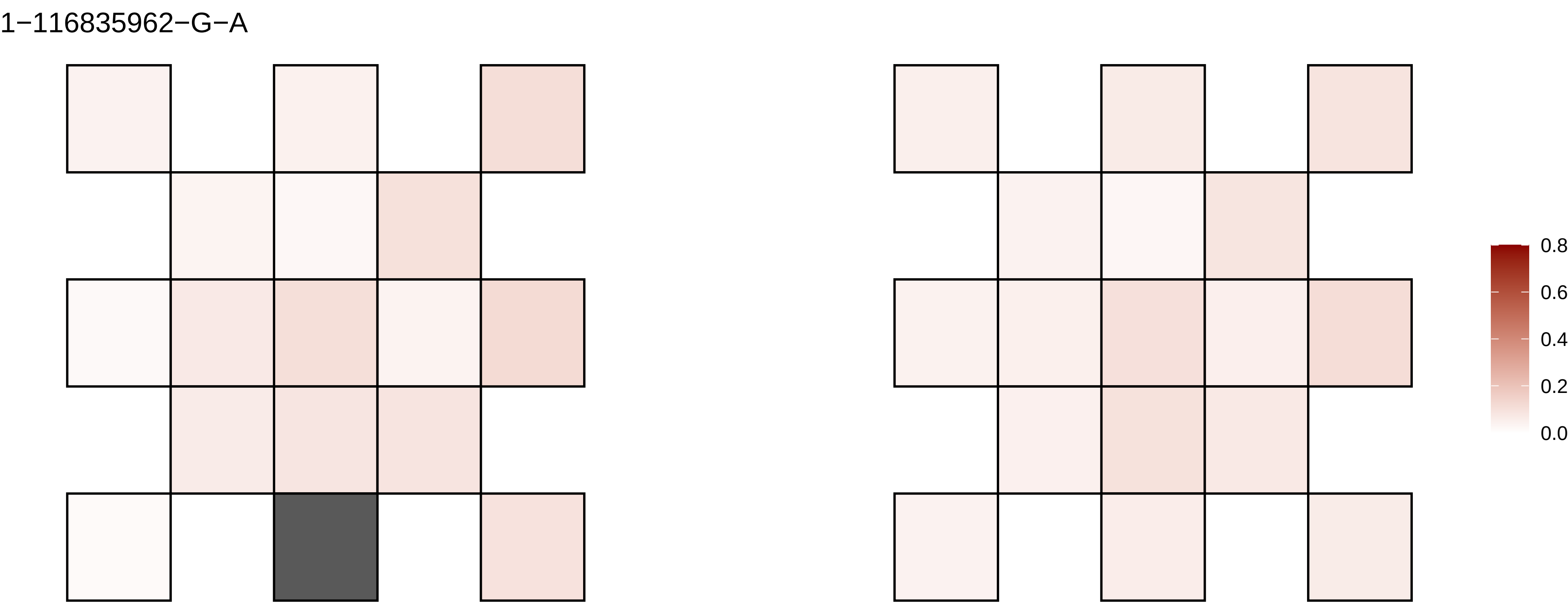

1−116835962−G−A
0.8
0.6
0.4
0.2
0.0

1−117129159−C−T

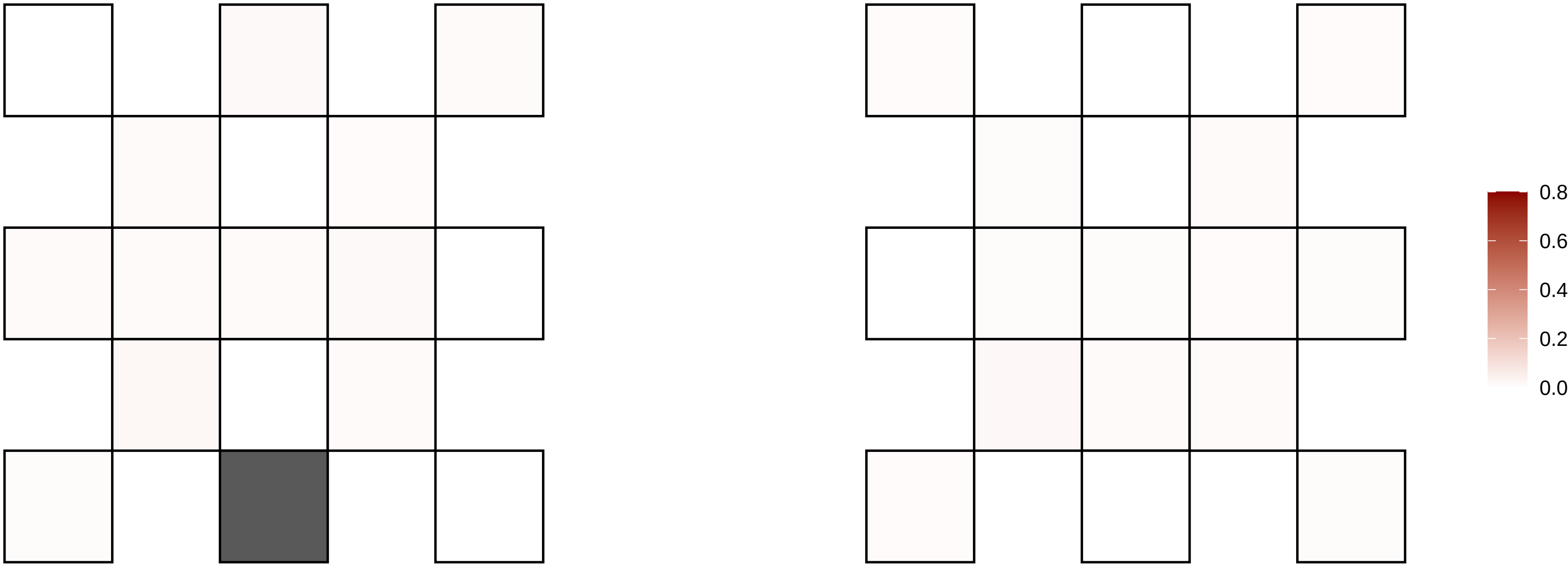

1−117326750−G−T

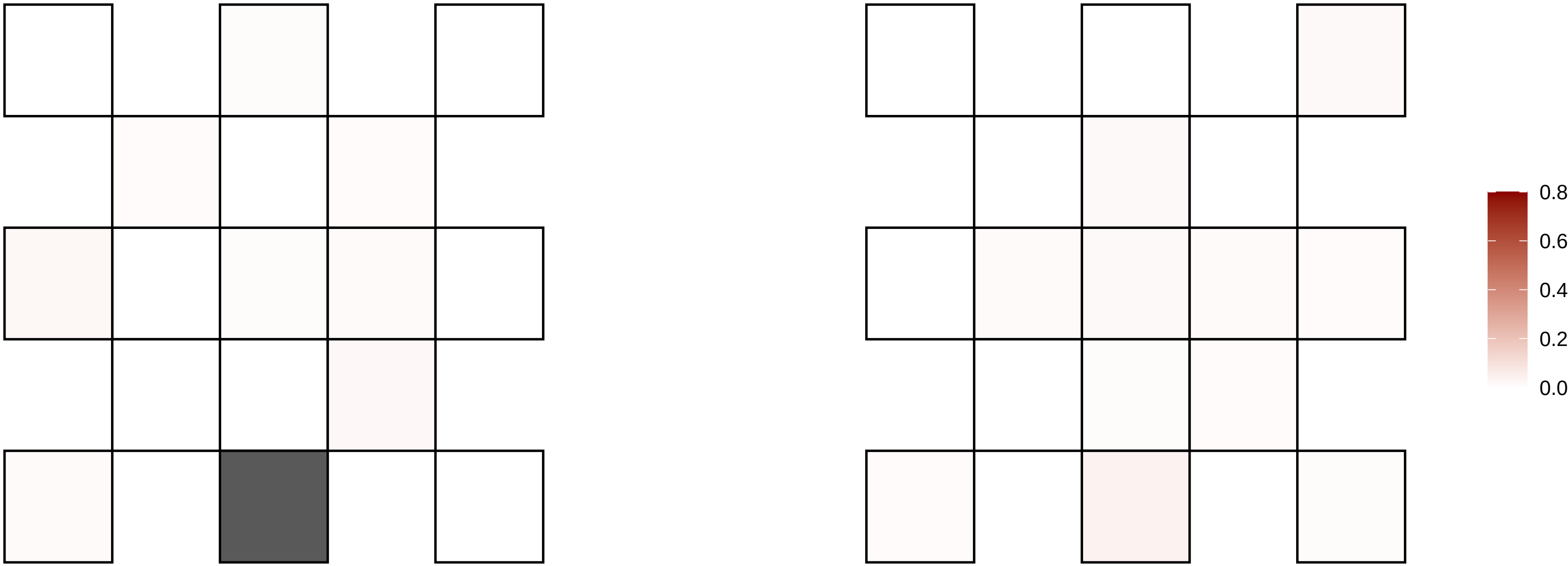

1−1196724−A−T

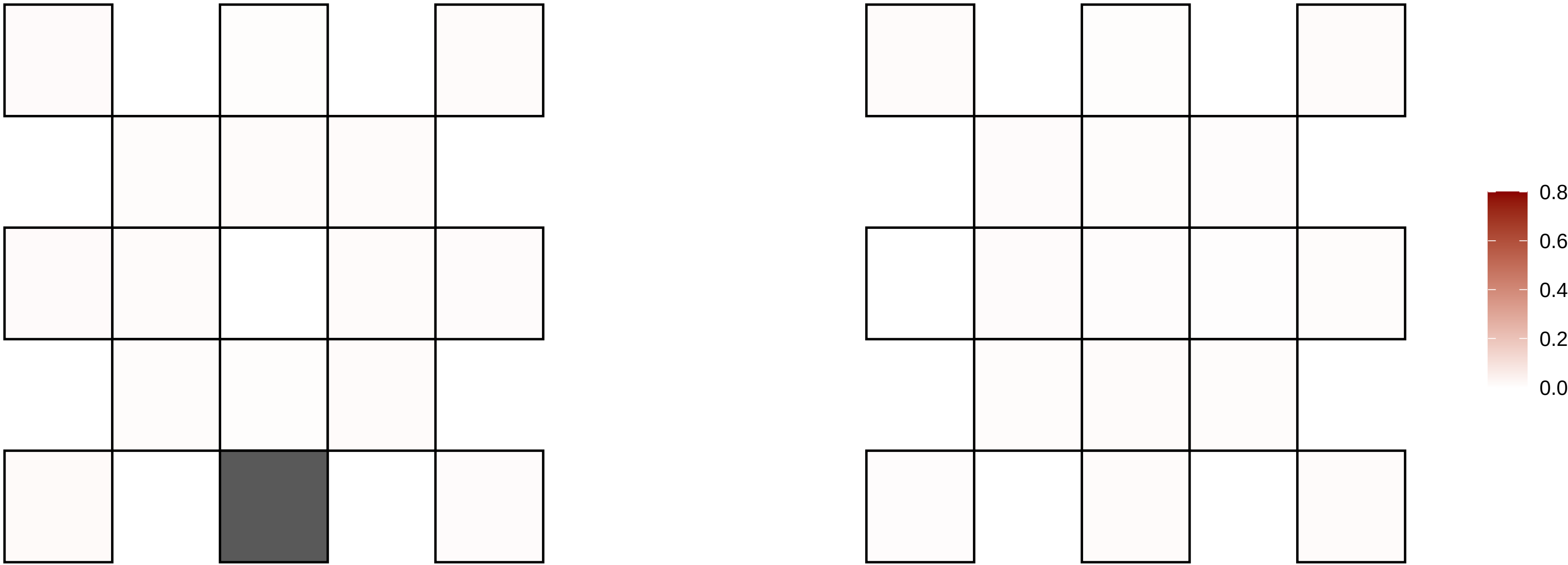

1−154786631−G−A

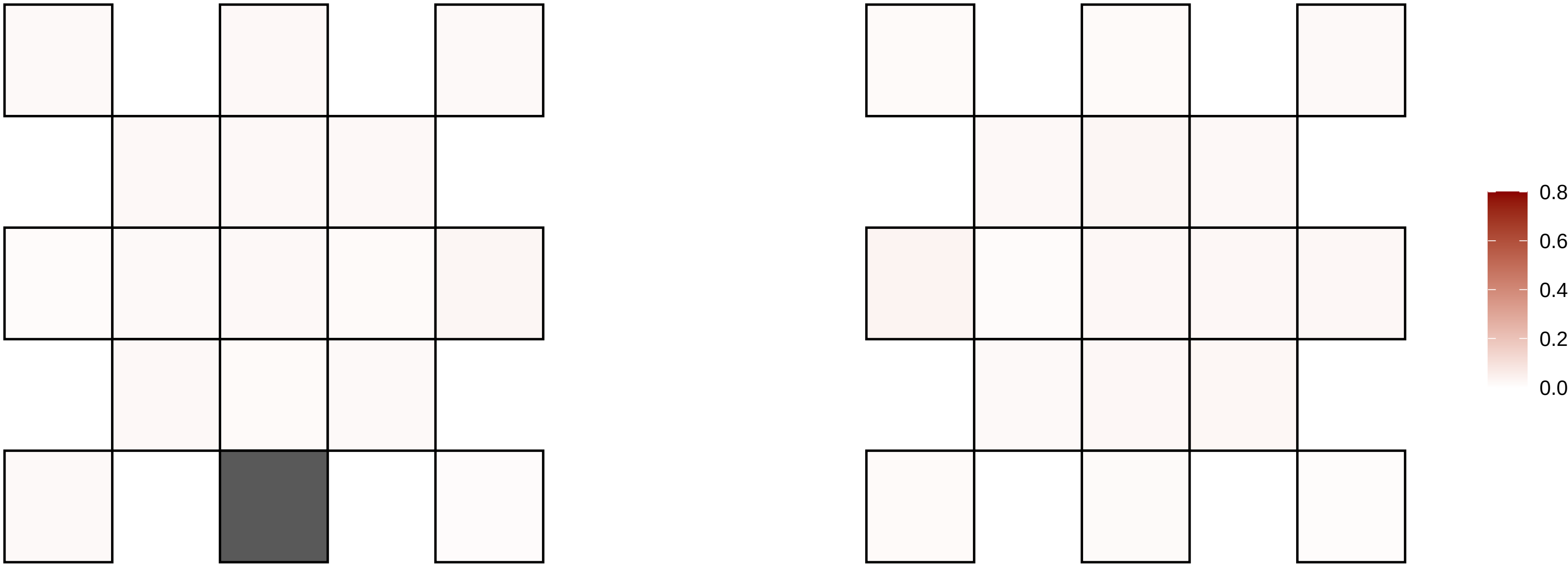

1−159646818−C−T

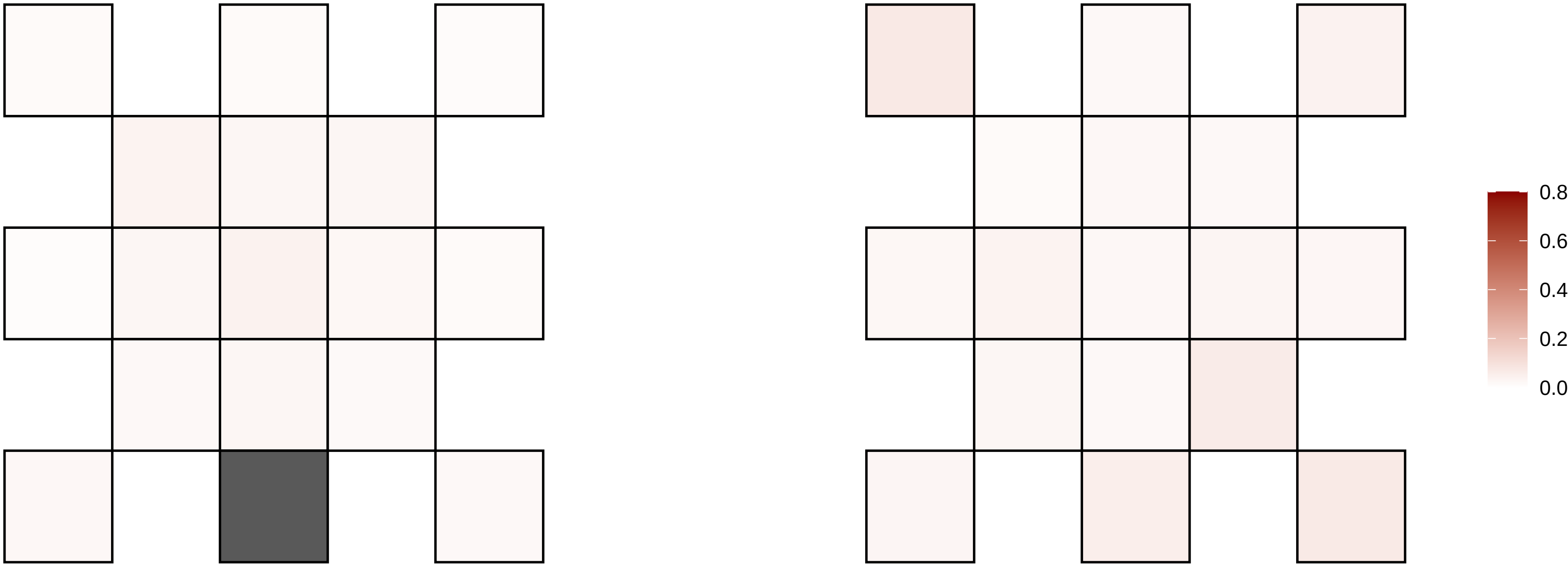

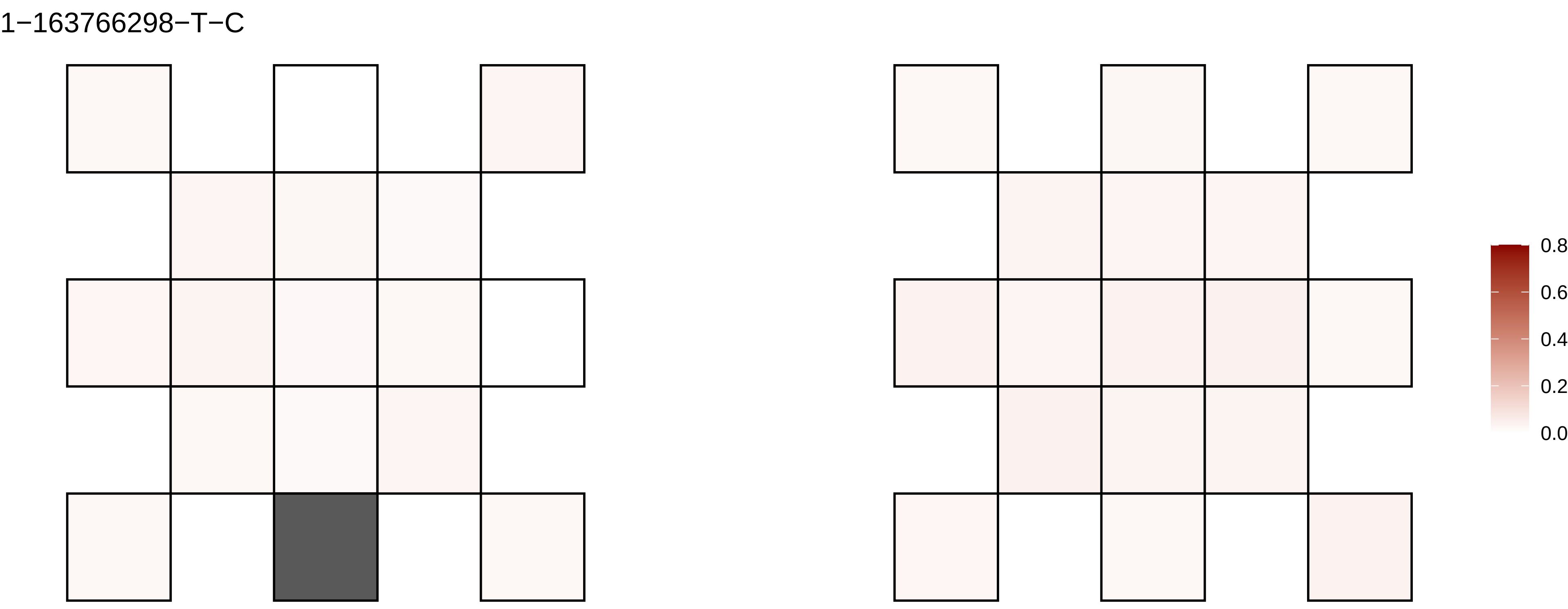
1−163766298−T−C
0.8
0.6
0.4
0.2
0.0

1−163770532−T−A

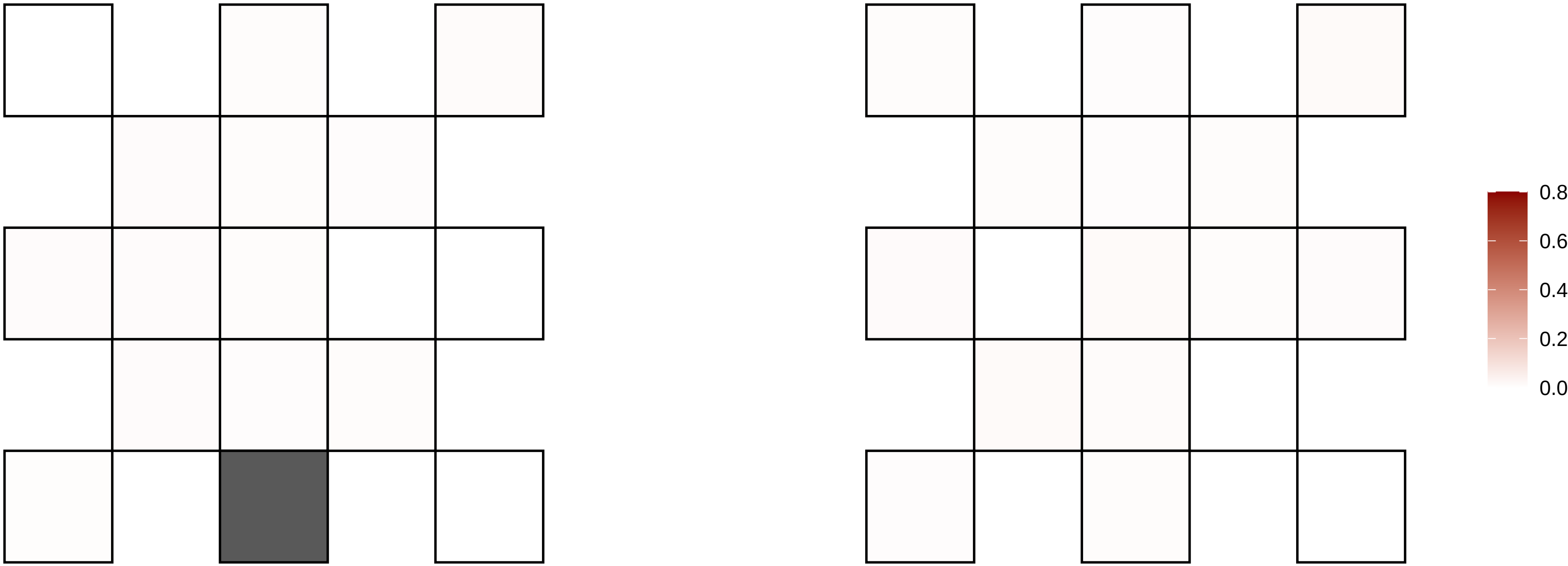

1−177430967−T−C

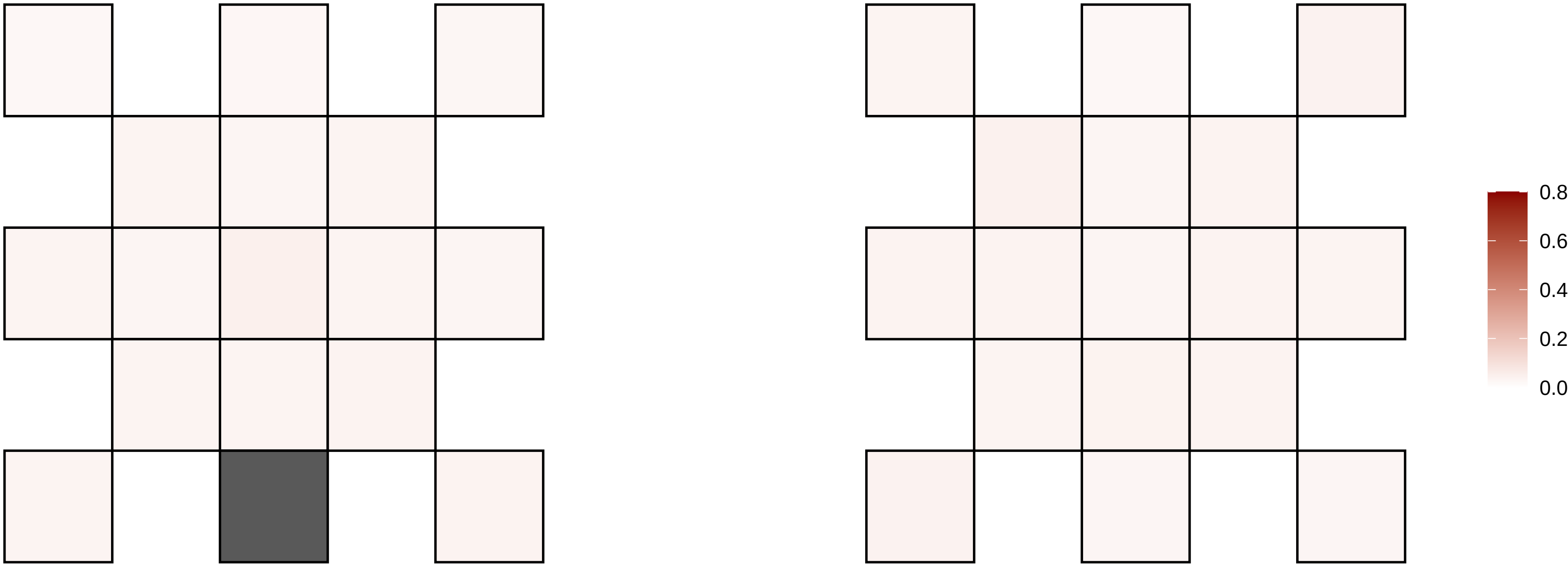

1−18387130−C−T

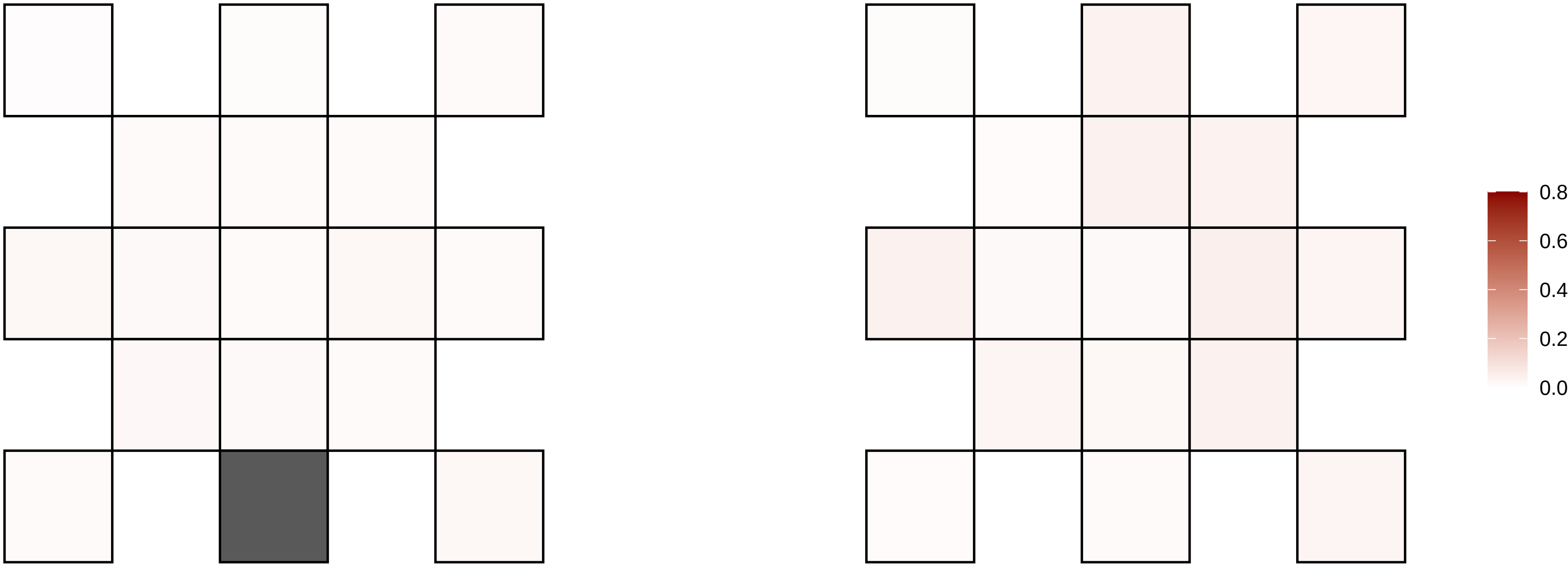

1−18400310−C−T

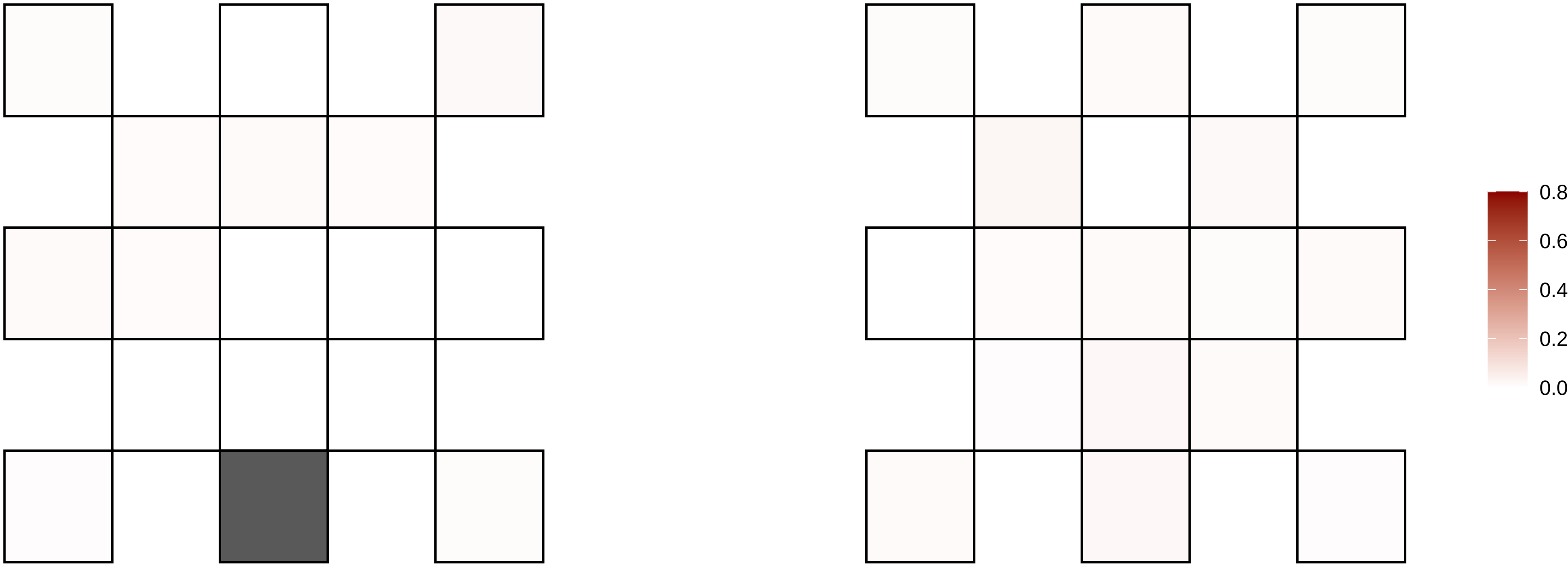

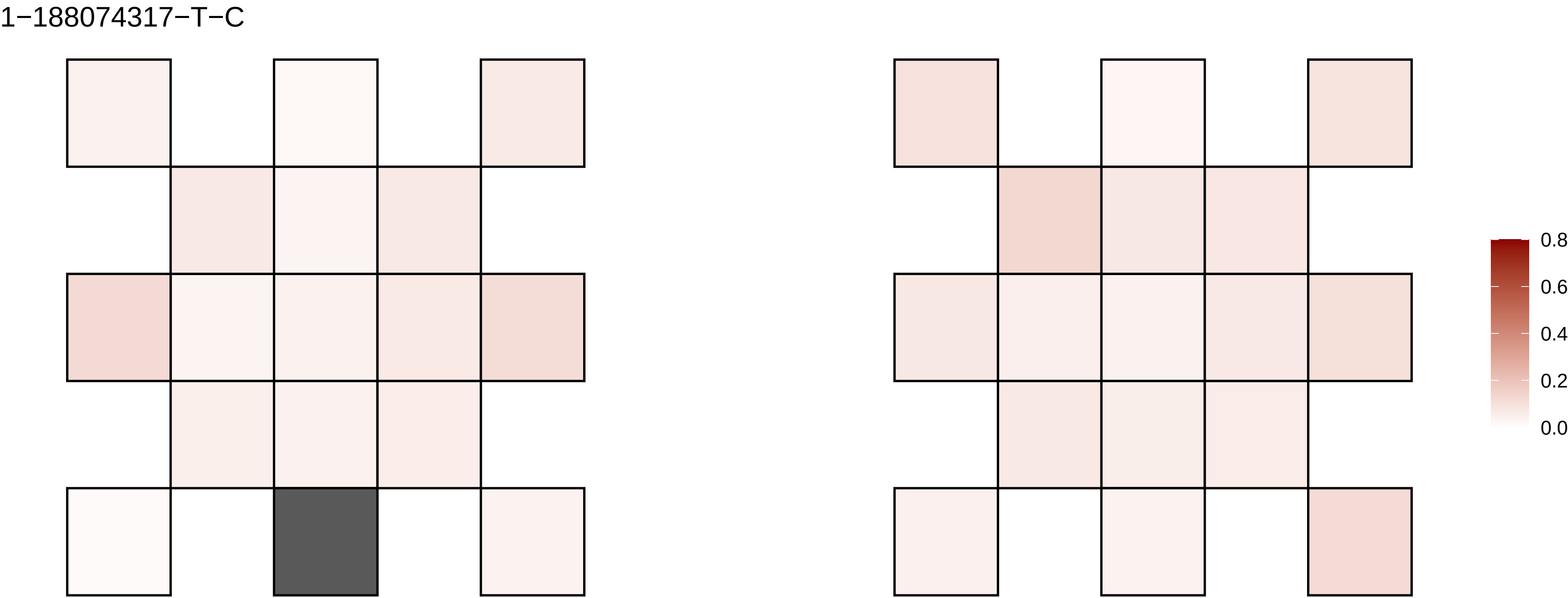

1−188074317−T−C
0.8
0.6
0.4
0.2
0.0

1−191248571−G−A

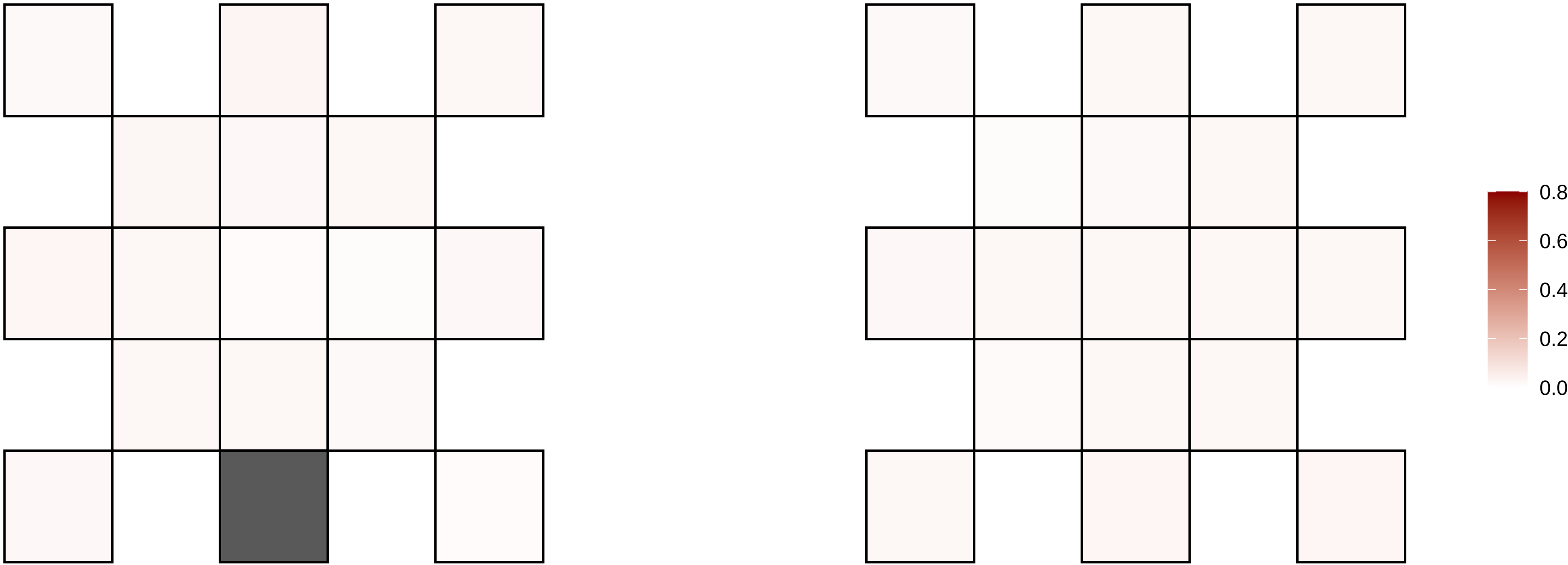

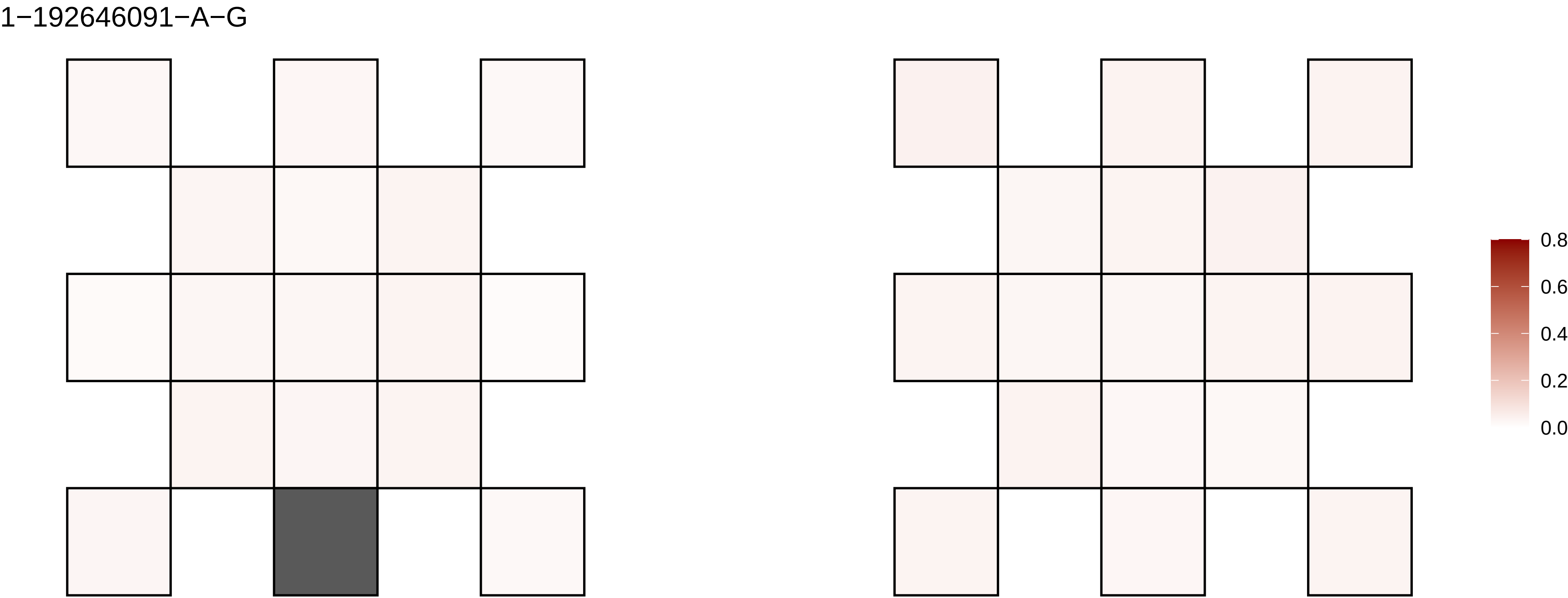
1−192646091−A−G
0.8
0.6
0.4
0.2
0.0

1−200010012−C−T

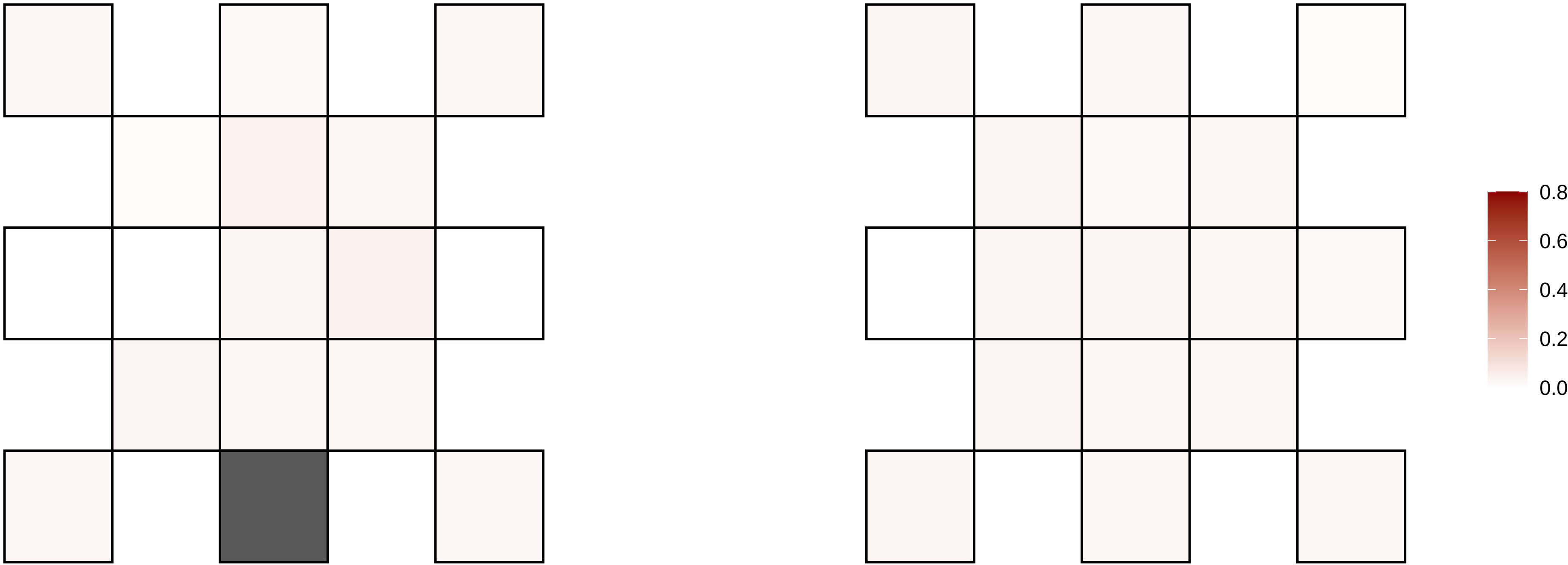

1−201266932−C−T

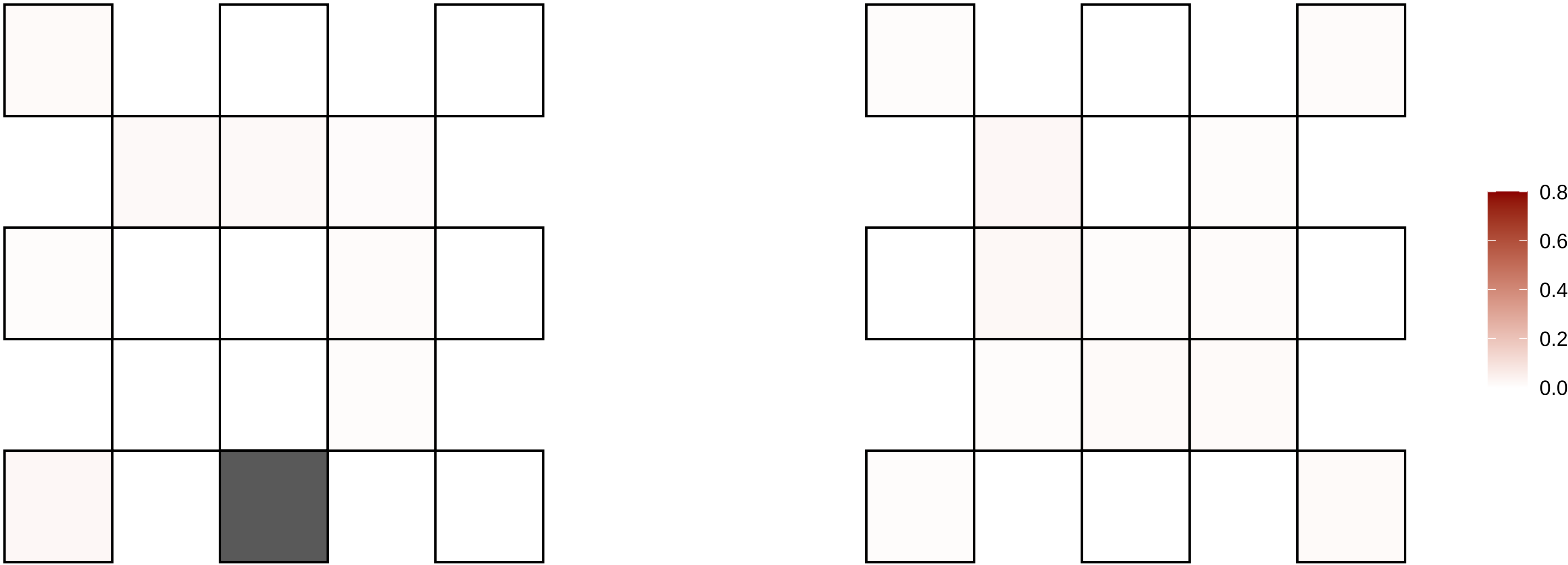

1−206627079−G−A

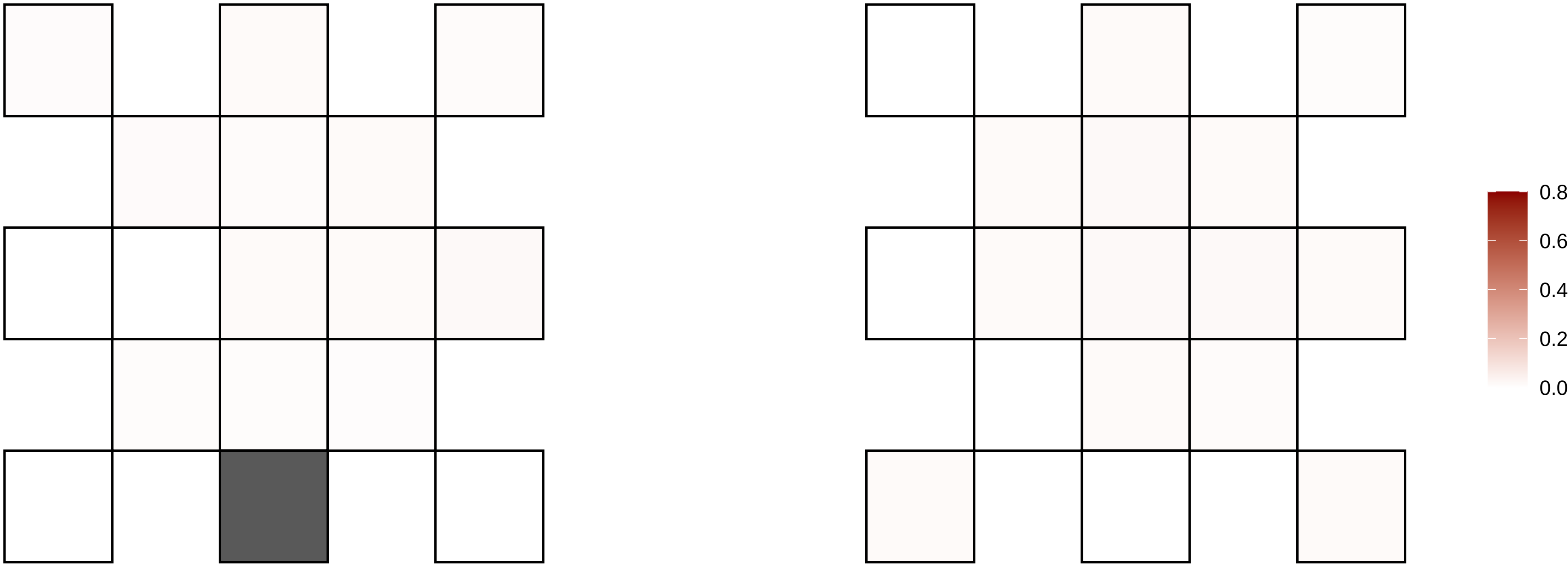

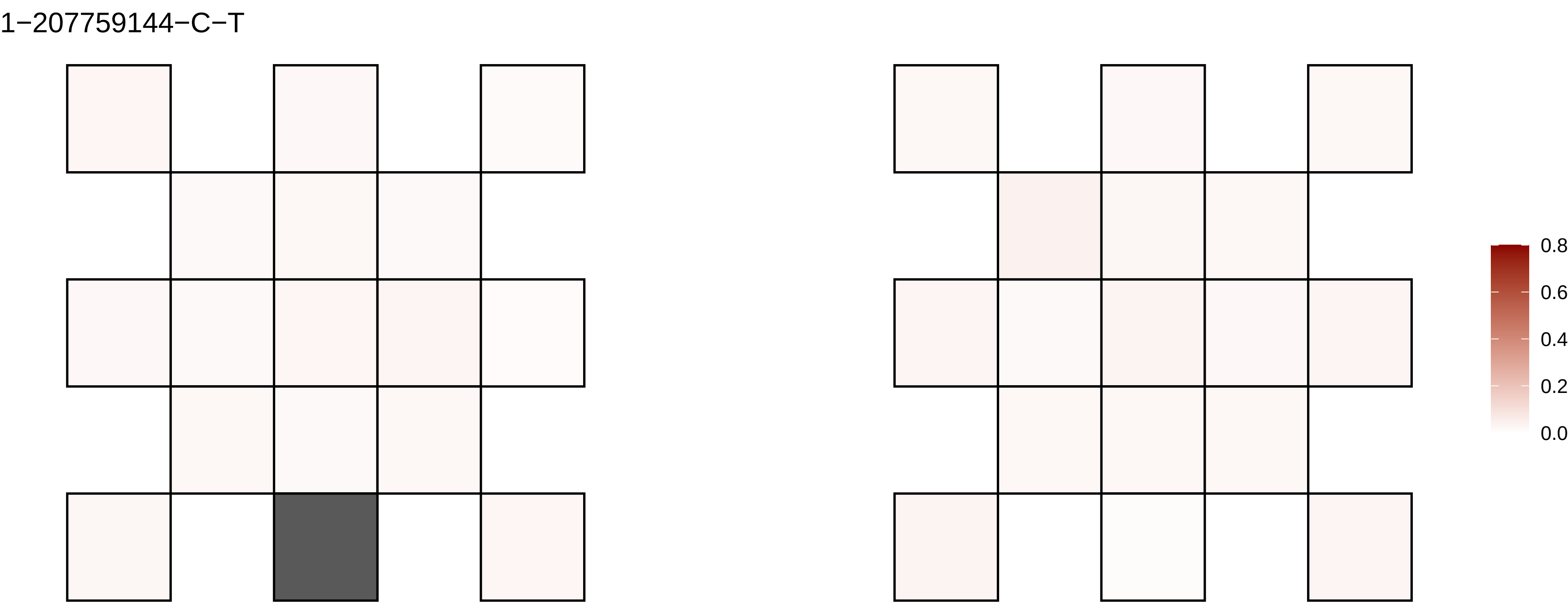

1−207759144−C−T
0.8
0.6
0.4
0.2
0.0

1−210808944−G−A

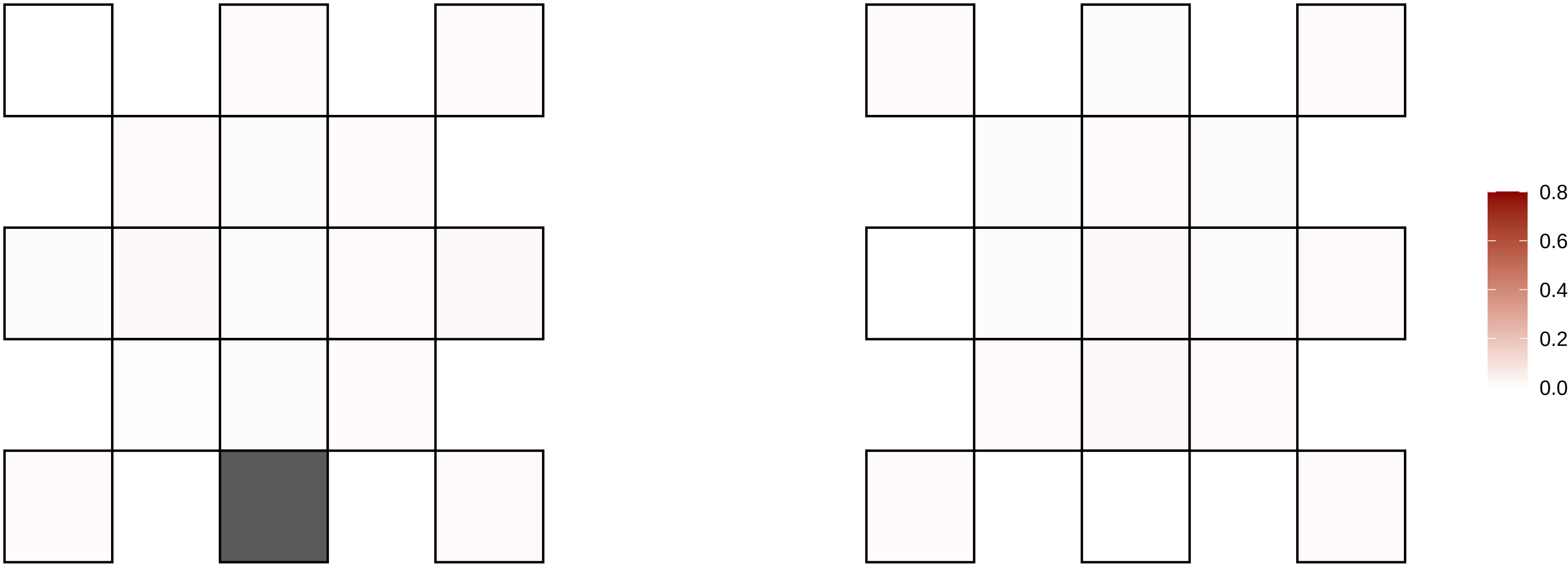

1−237812348−C−T

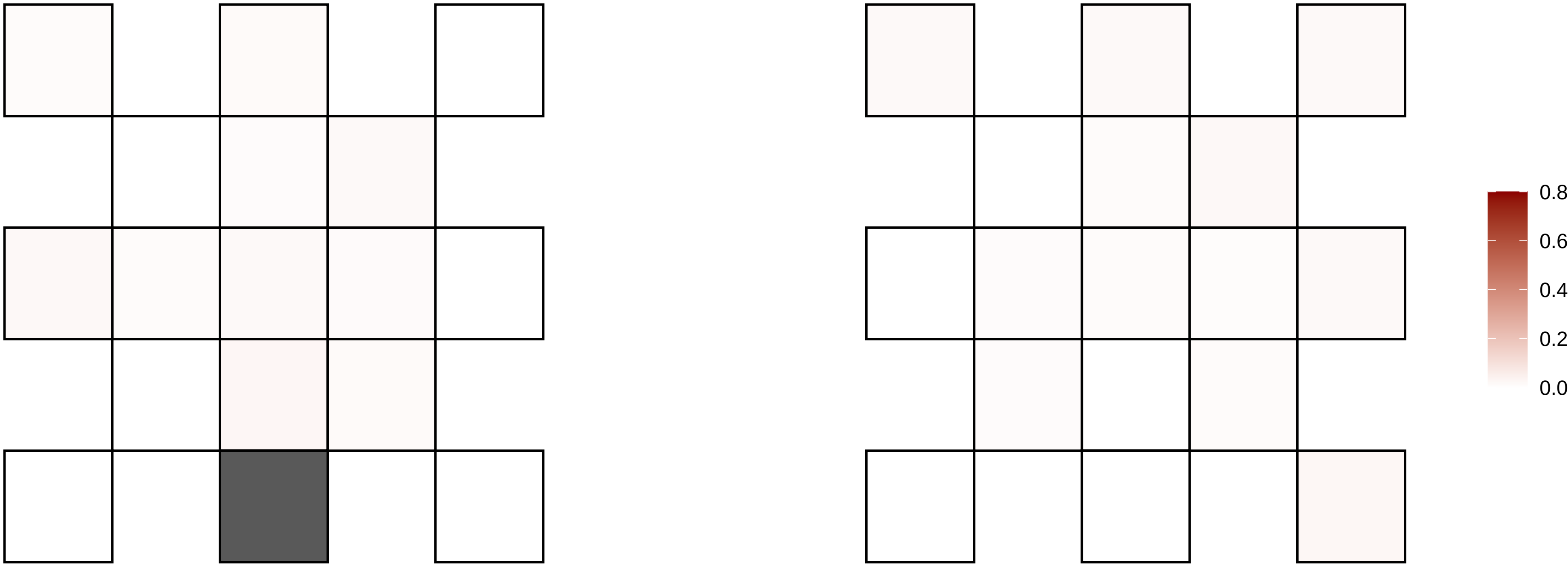

1−245457480−G−A

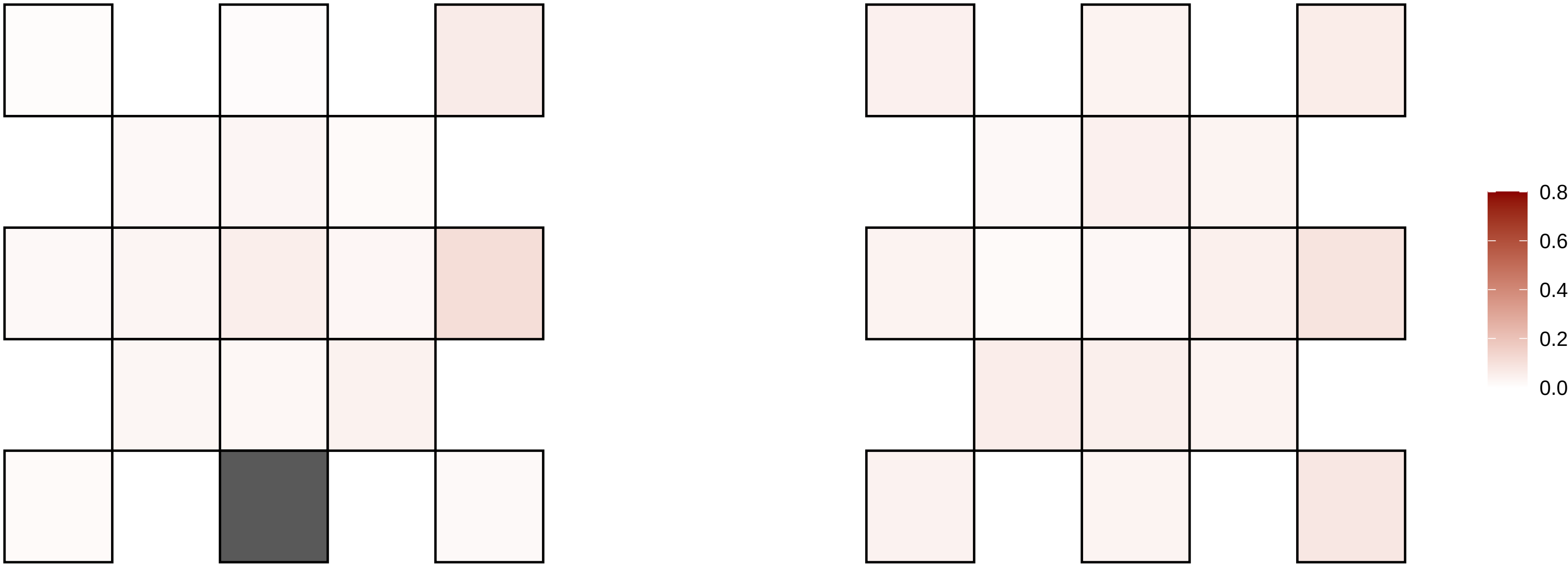

1−248261416−C−T

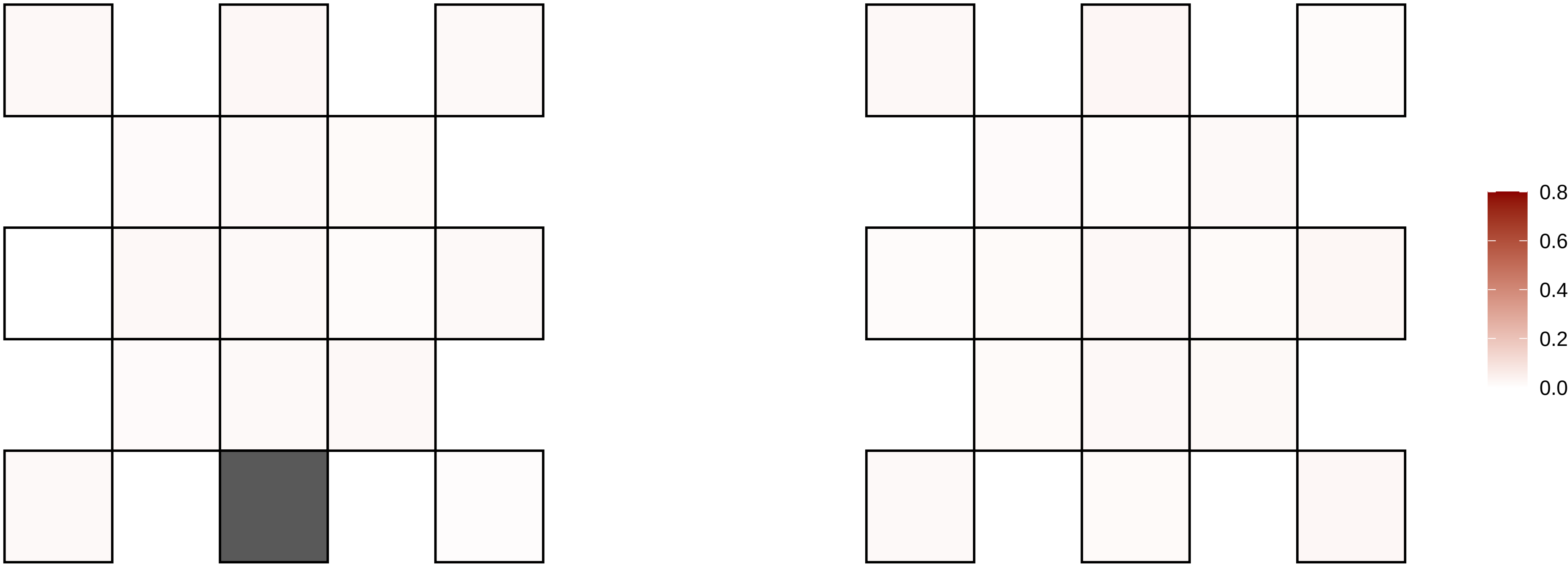

1−35110389−G−T

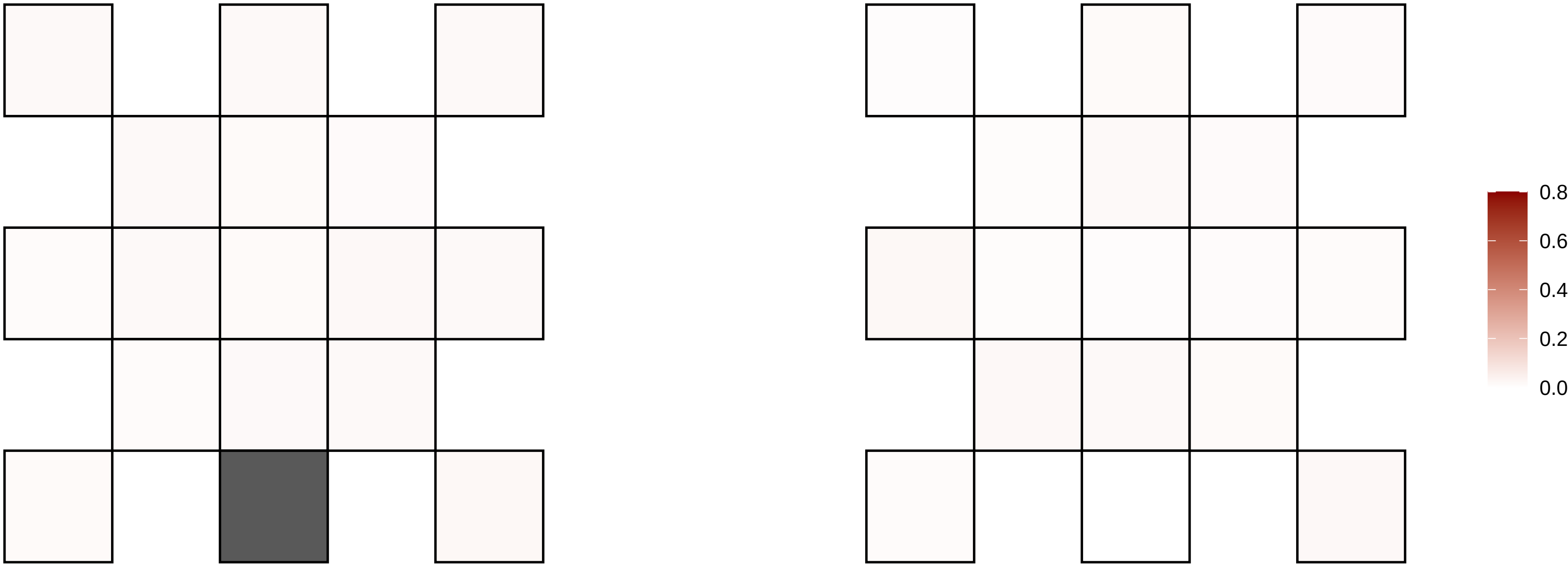

1−4450513−G−A

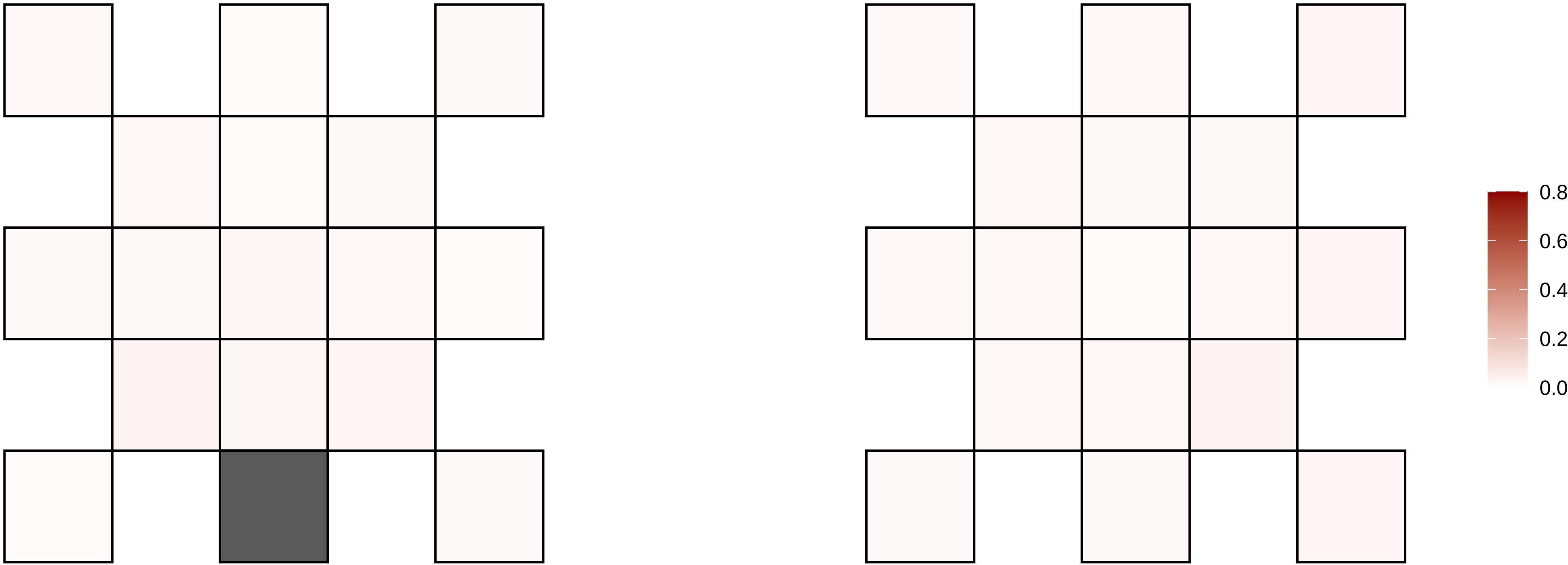

1−4465349−T−G

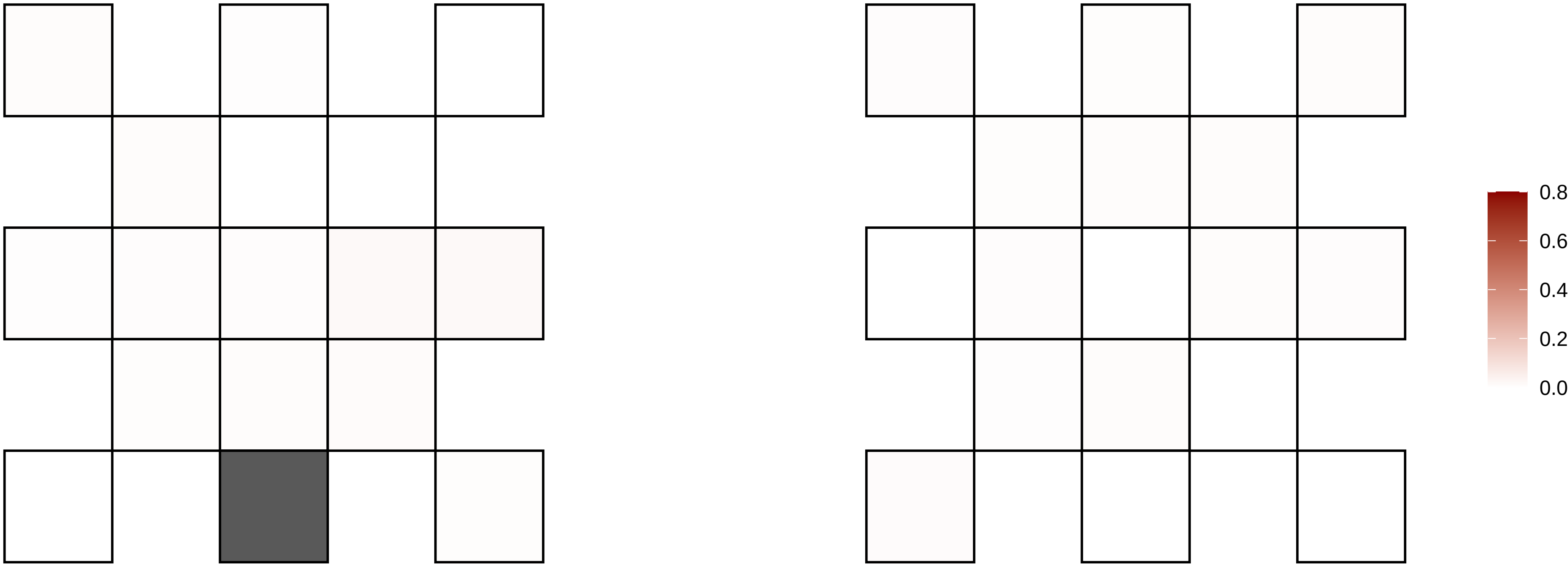

1−50289297−G−A

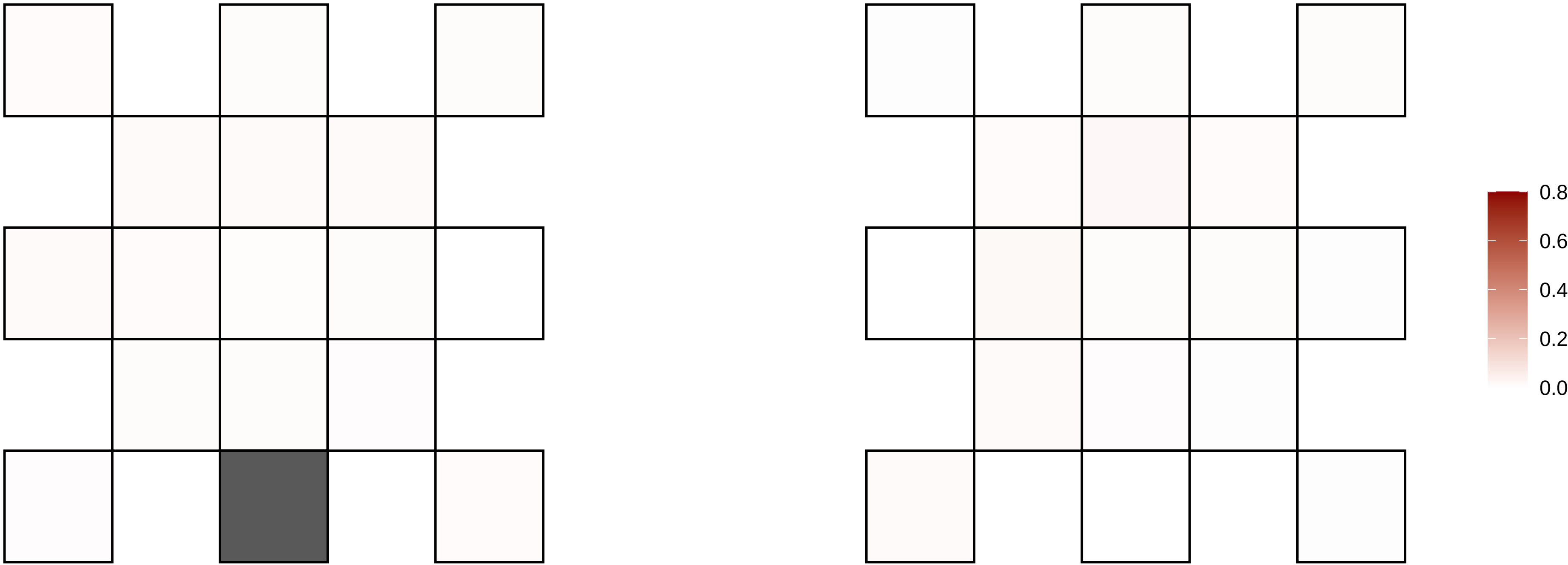

1−53912650−G−A

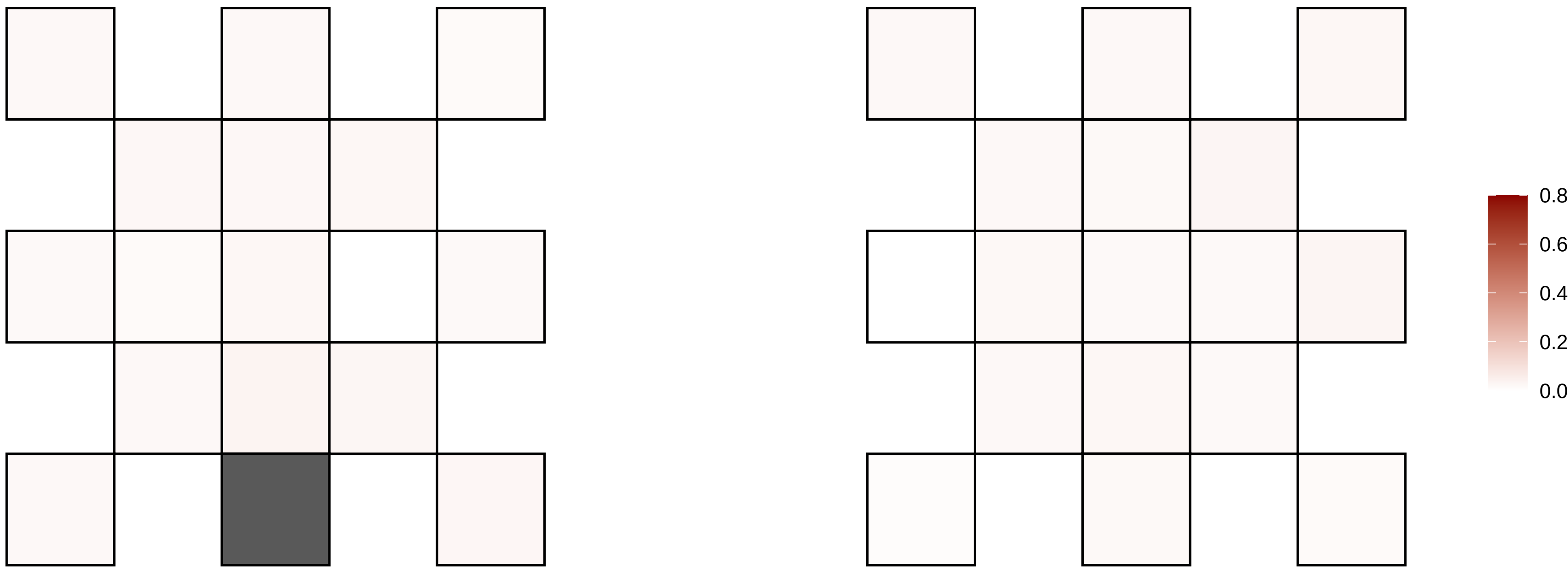

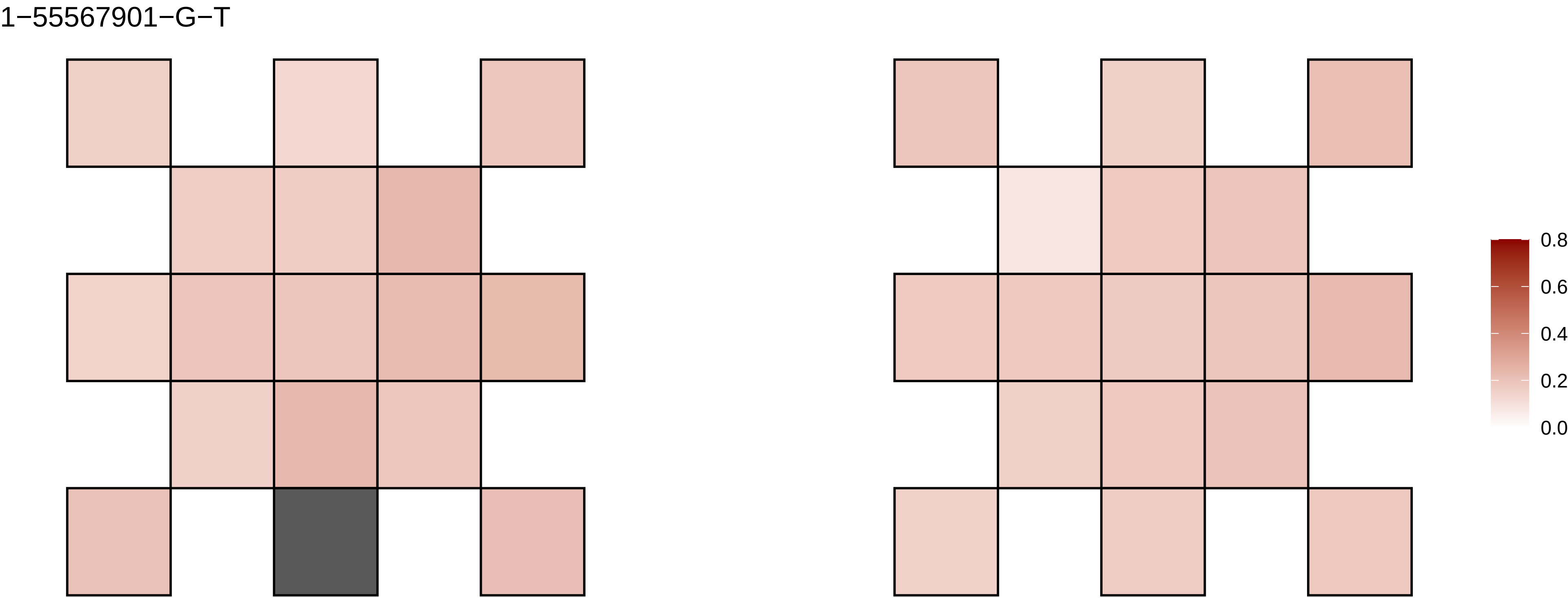

1−55567901−G−T
0.8
0.6
0.4
0.2
0.0

1−57886702−C−T

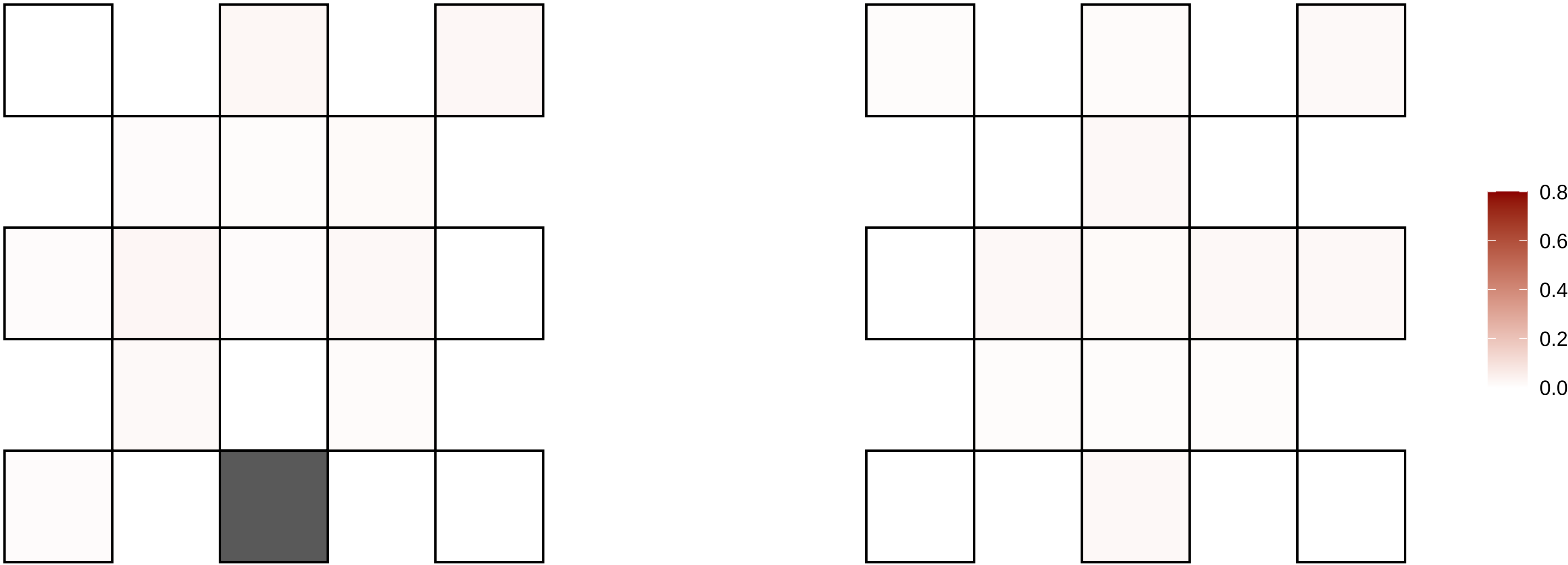

1−62675365−T−TTA

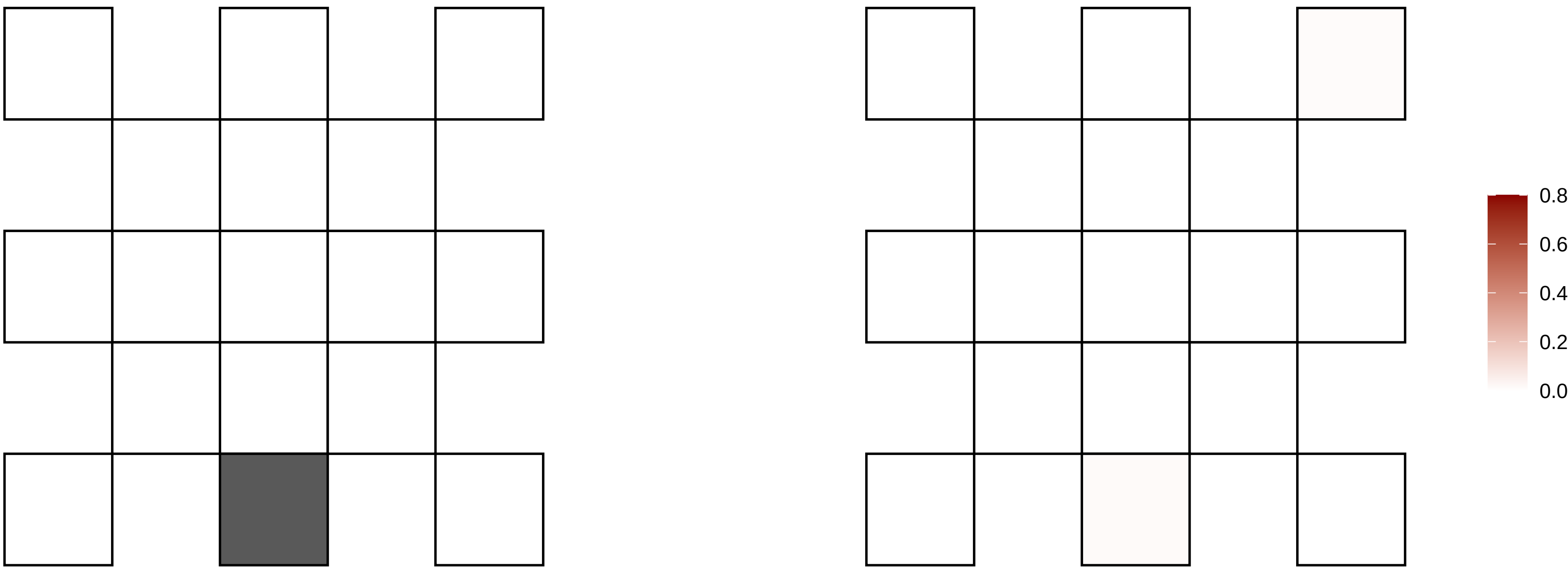

1−66719051−G−A

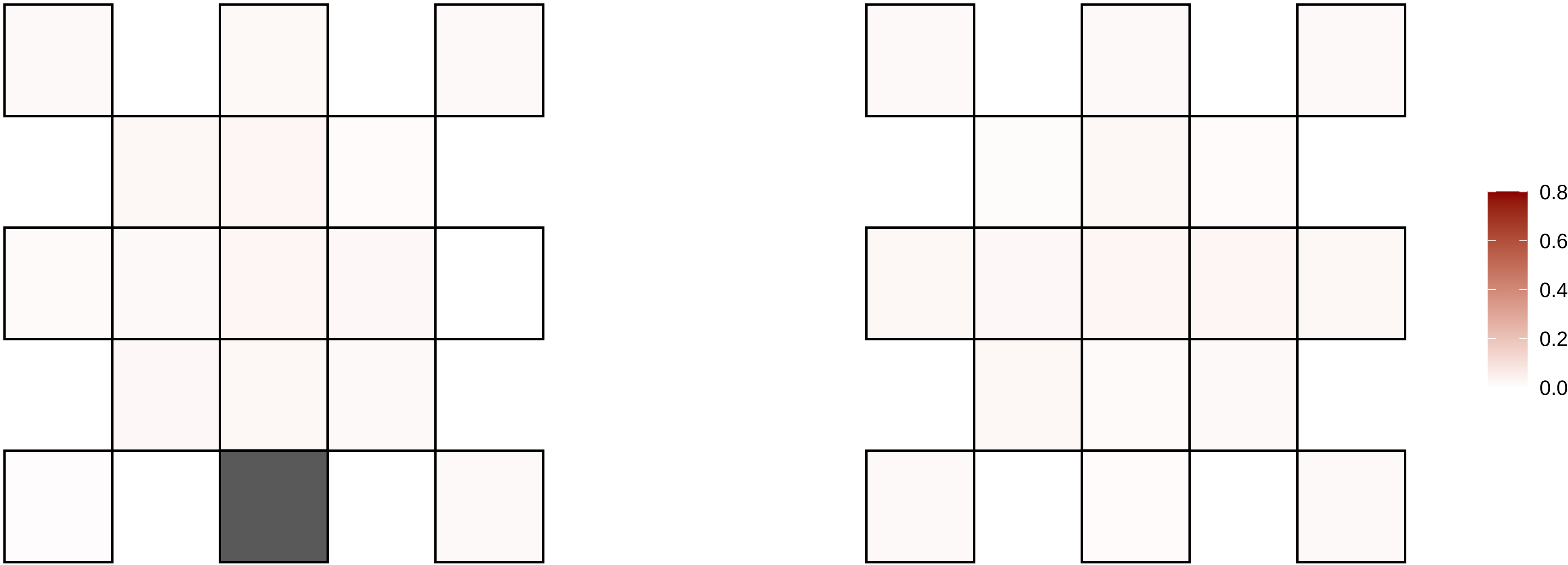

1−67722227−C−A

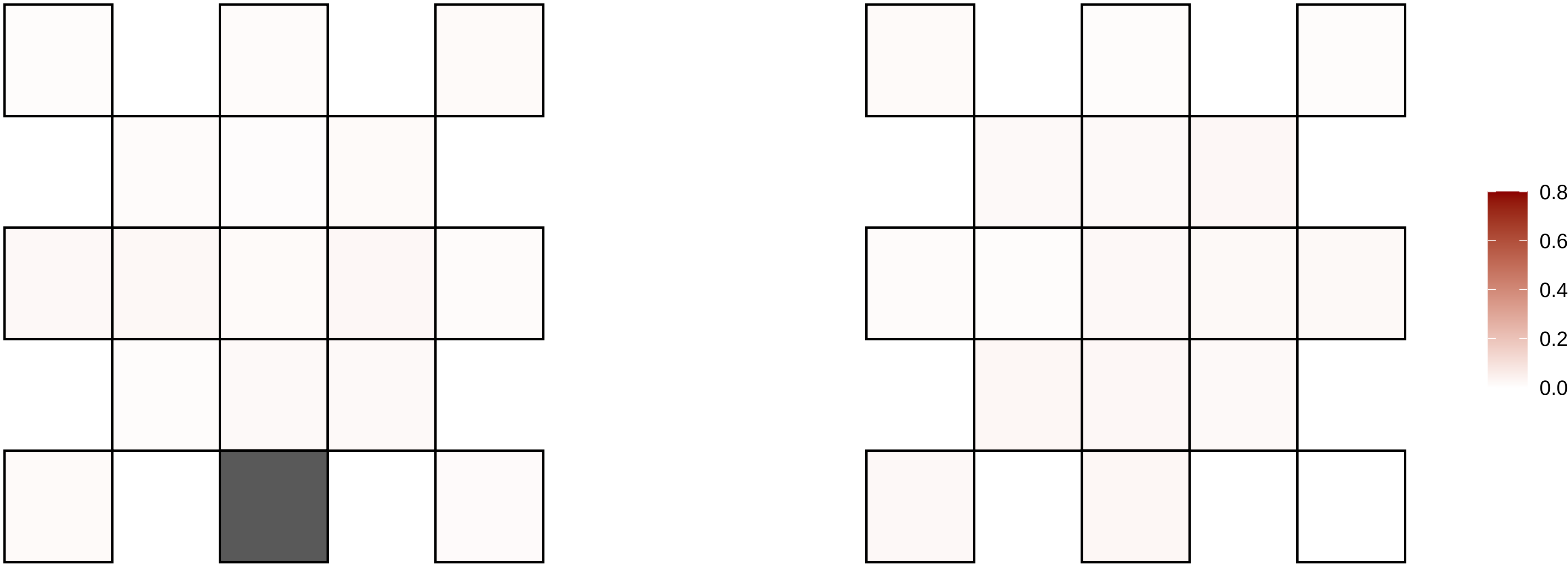

1−72003630−G−T

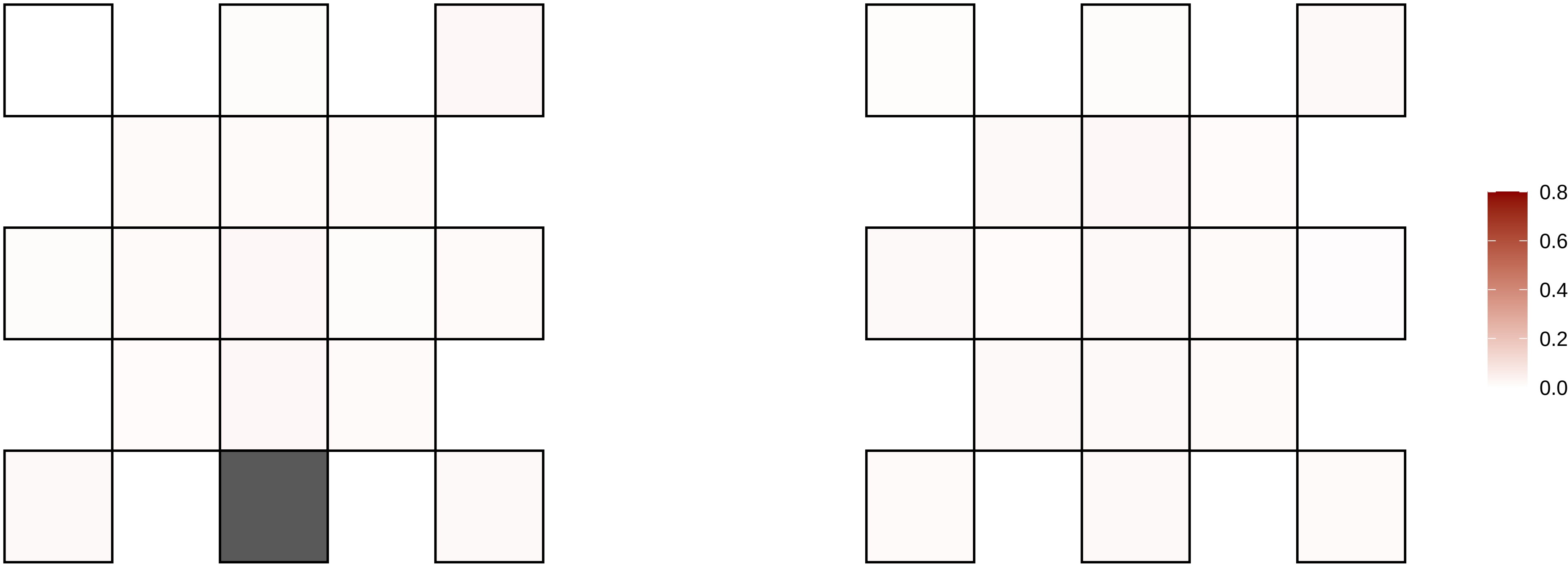

1−74501501−C−T

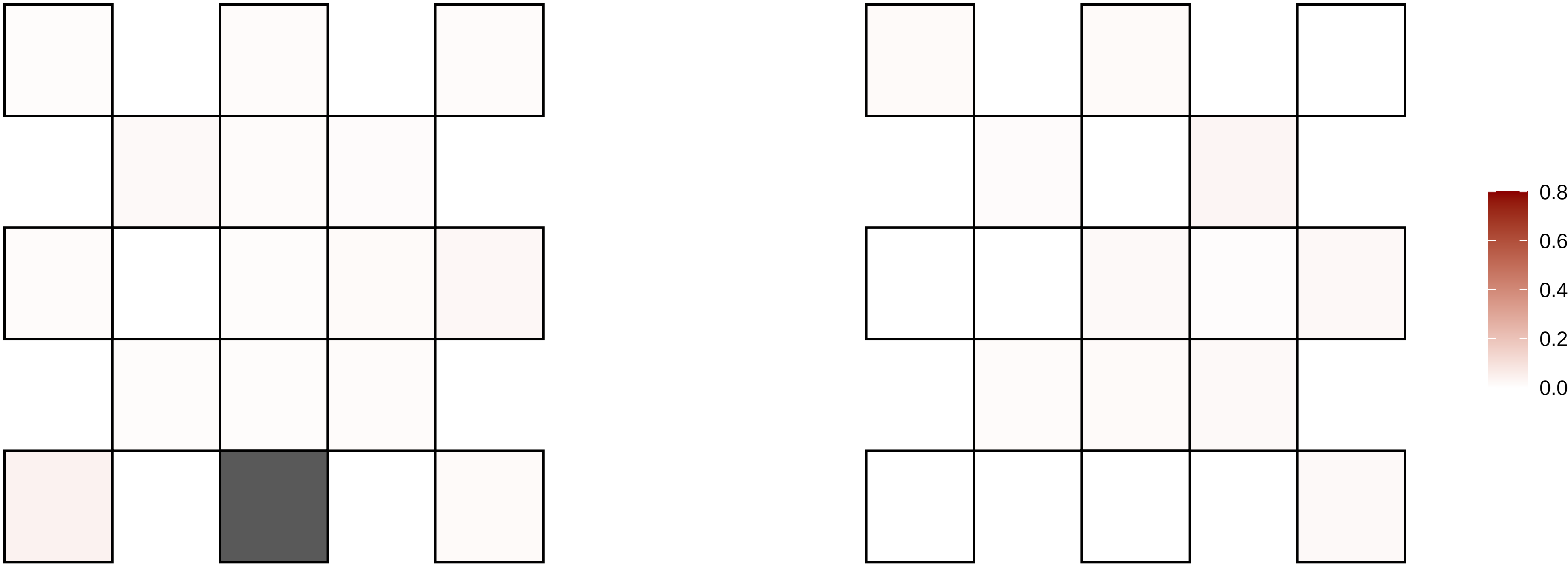

1−79102619−G−A

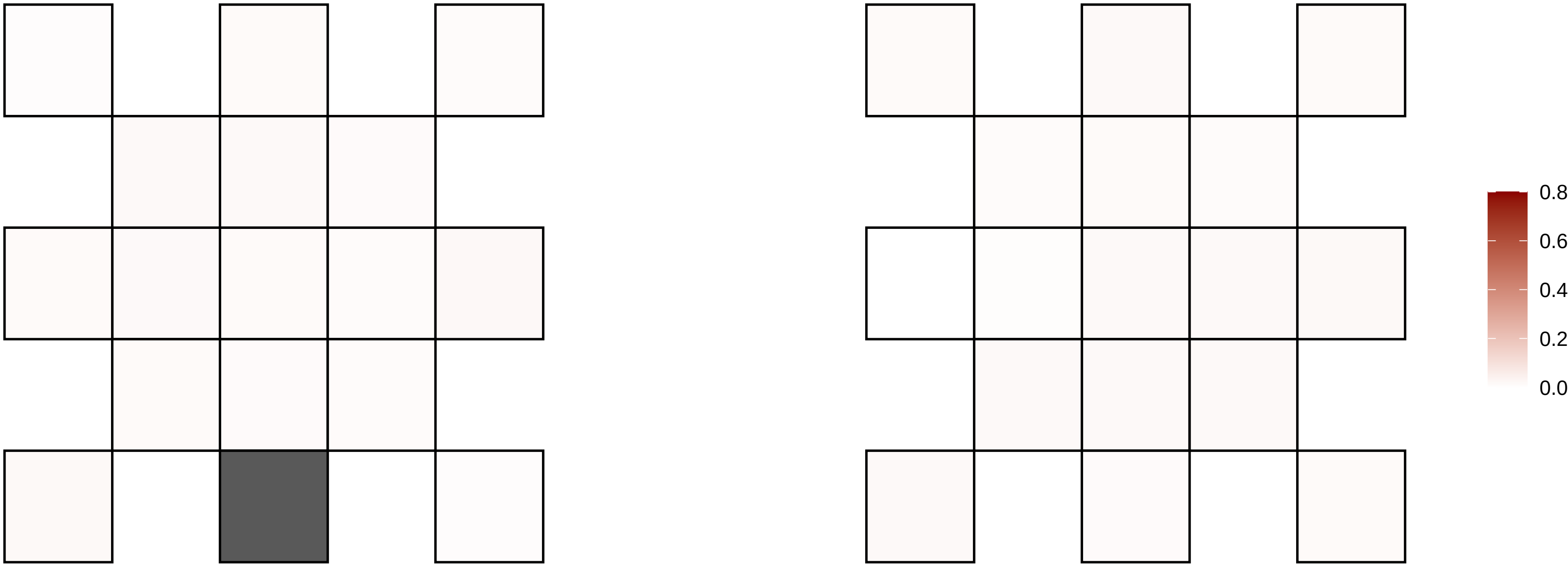

1−83361707−GC−G

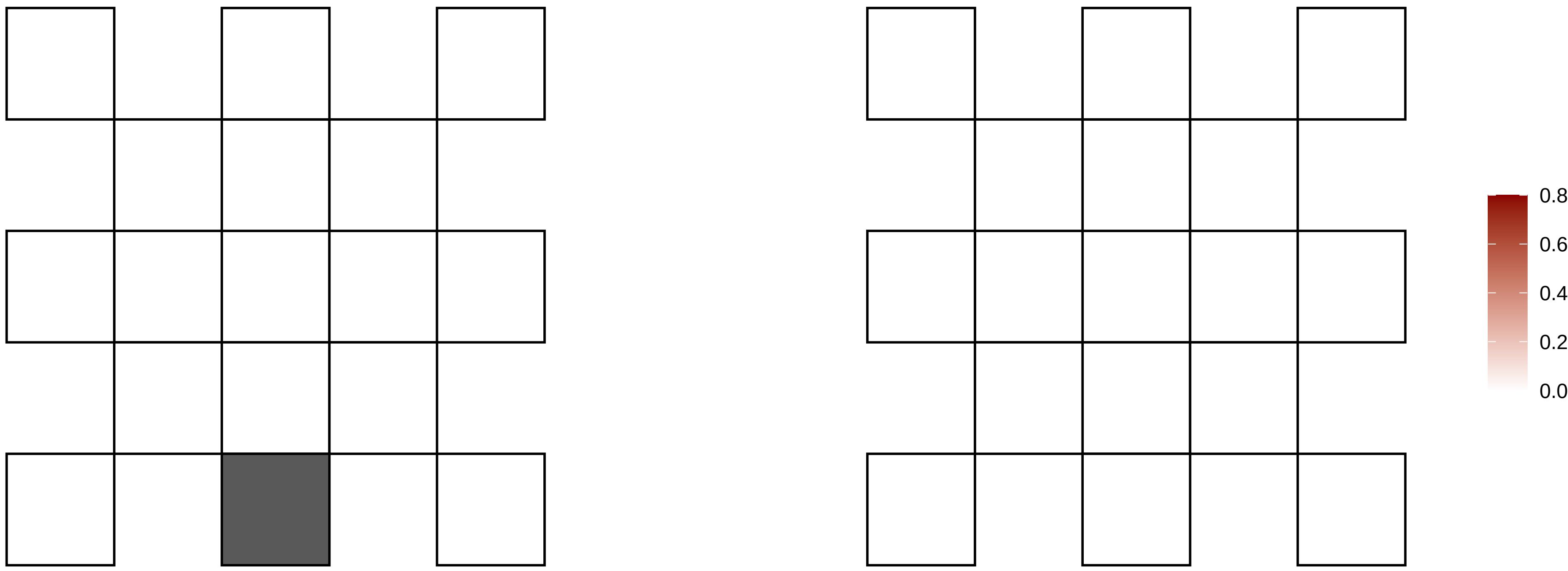

1−9049289−T−C

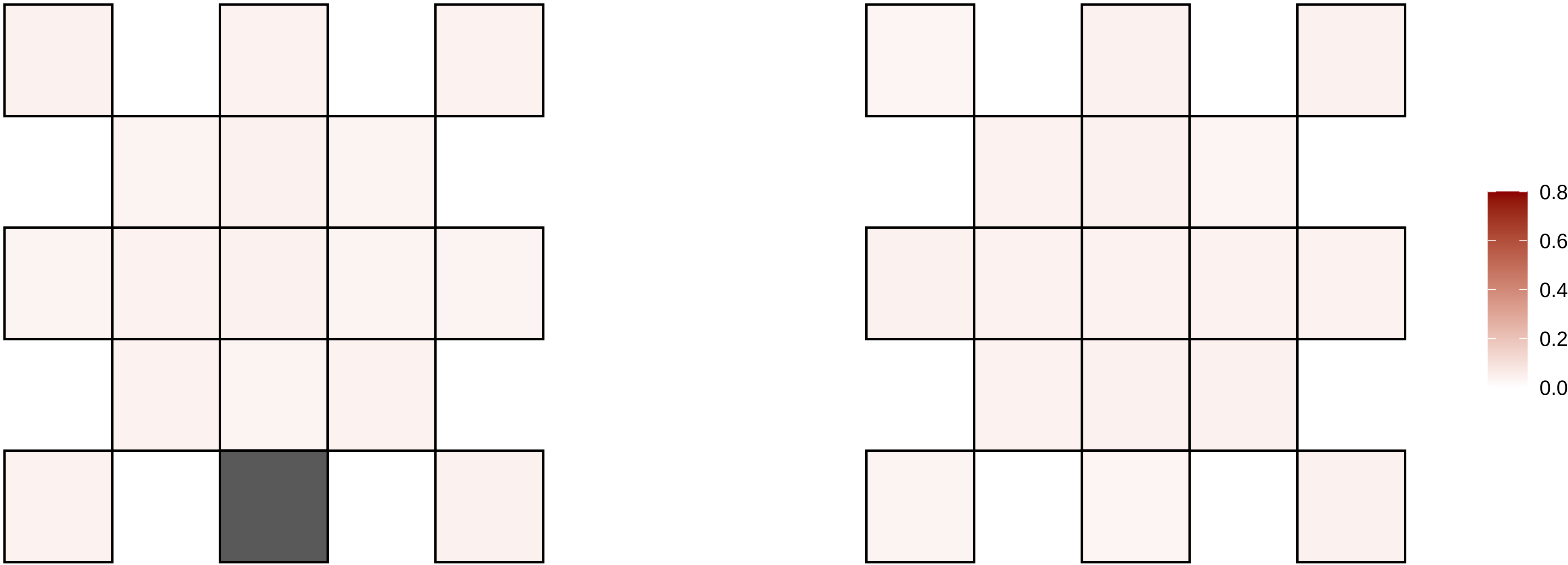

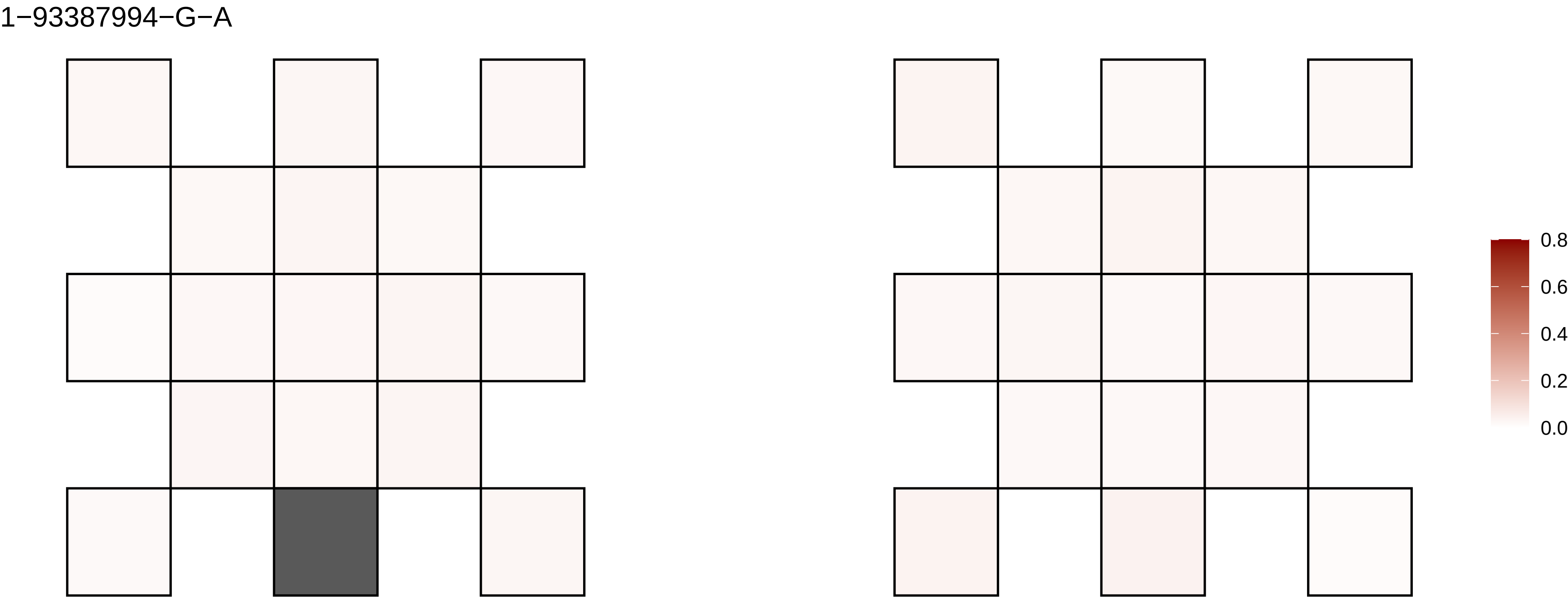
1−93387994−G−A
0.8
0.6
0.4
0.2
0.0

1−94219312−G−A

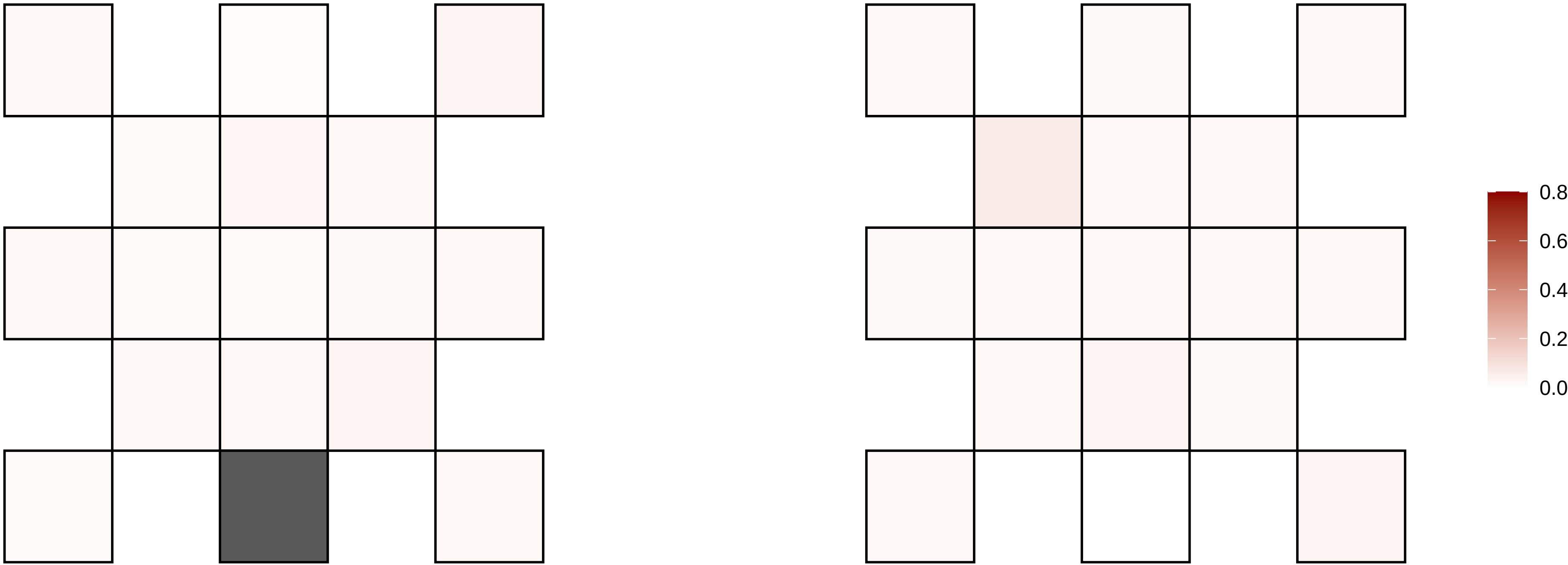

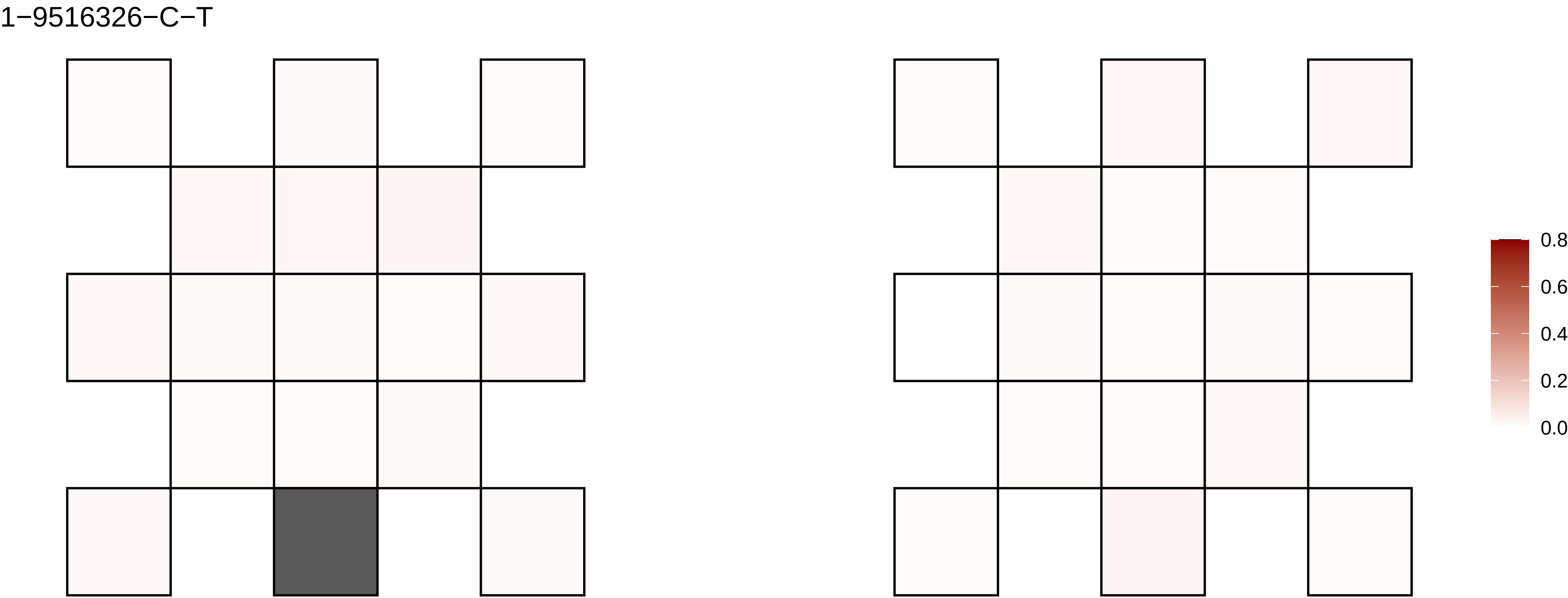
1−9516326−C−T
0.8
0.6
0.4
0.2
0.0

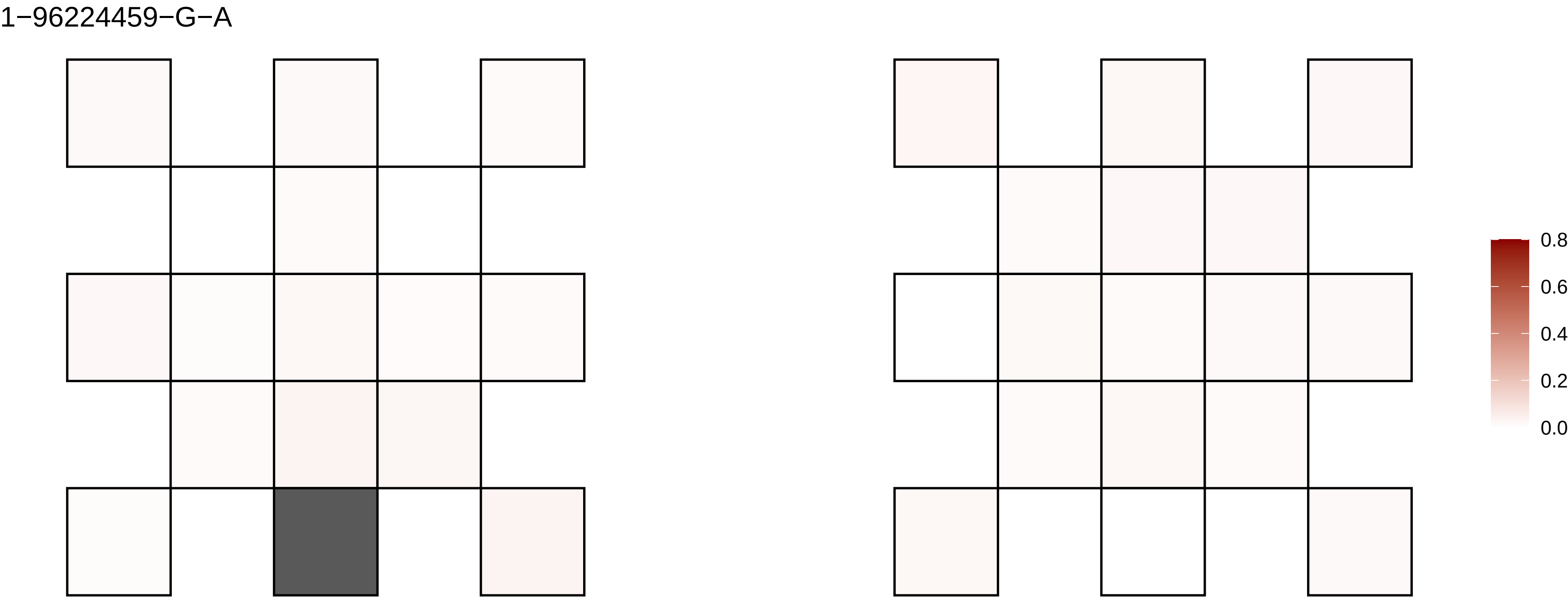
1−96224459−G−A
0.8
0.6
0.4
0.2
0.0

1−9672930−G−A

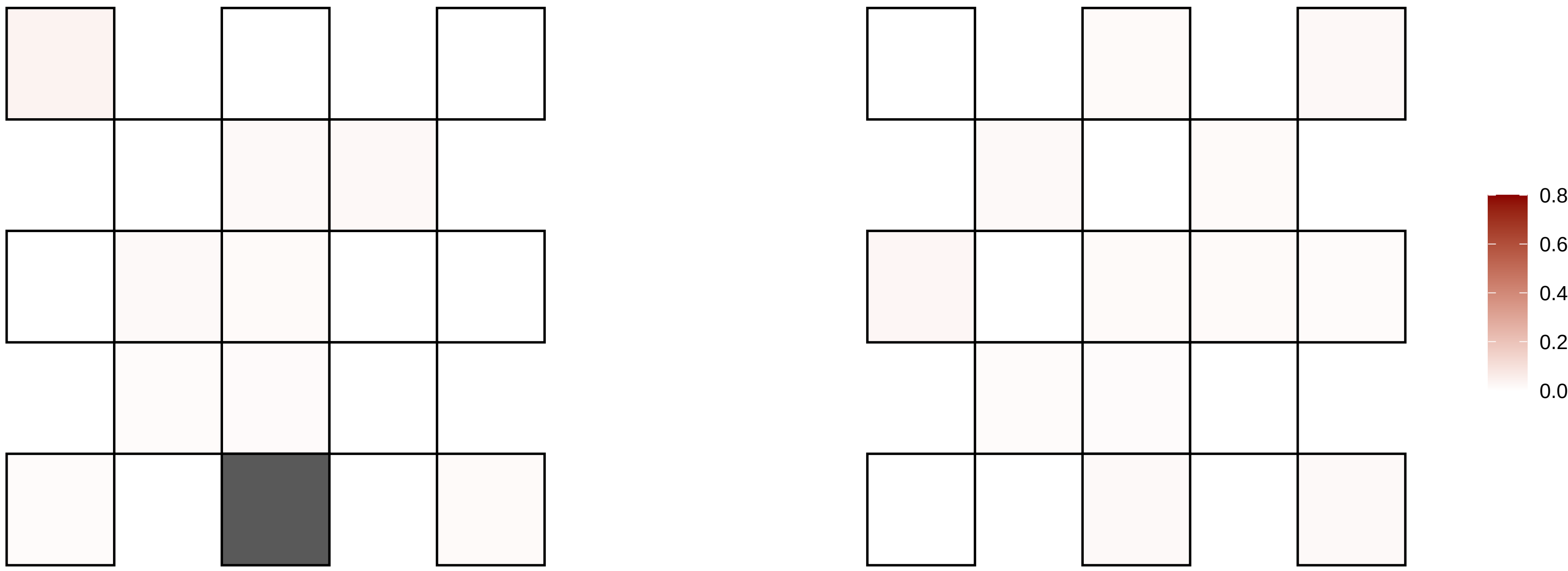

1−98301896−T−C

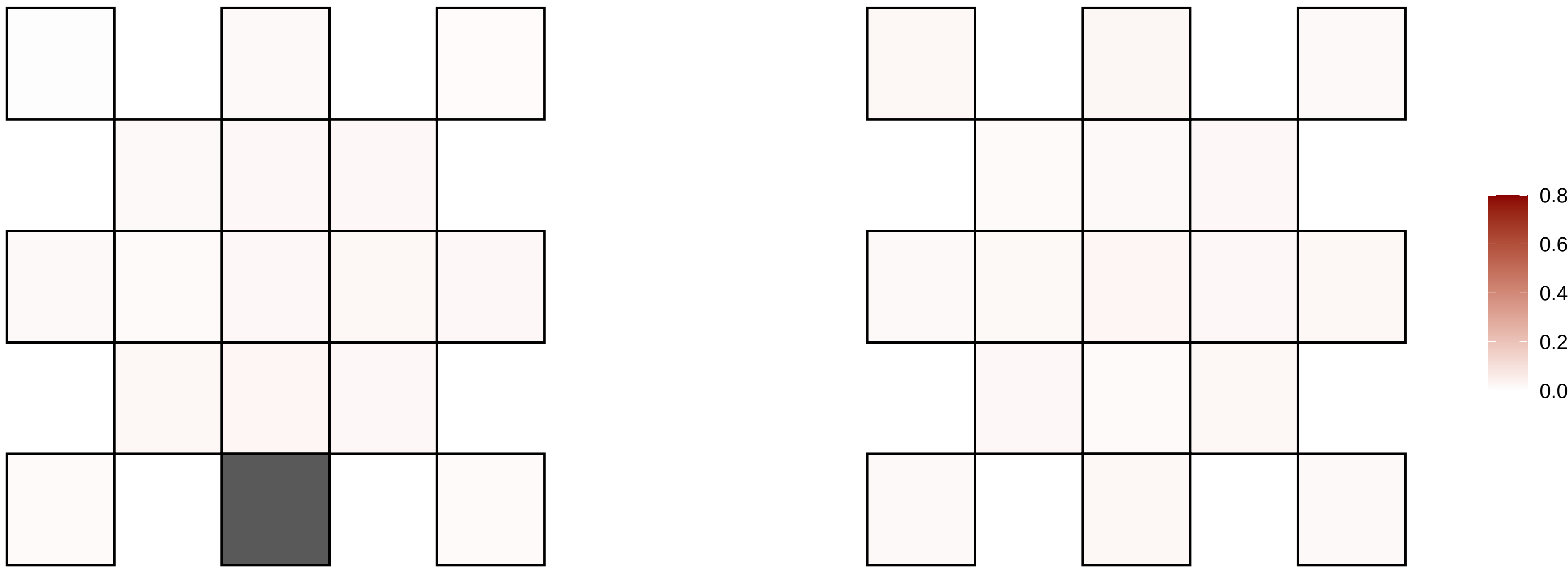

10−105764174−A−G

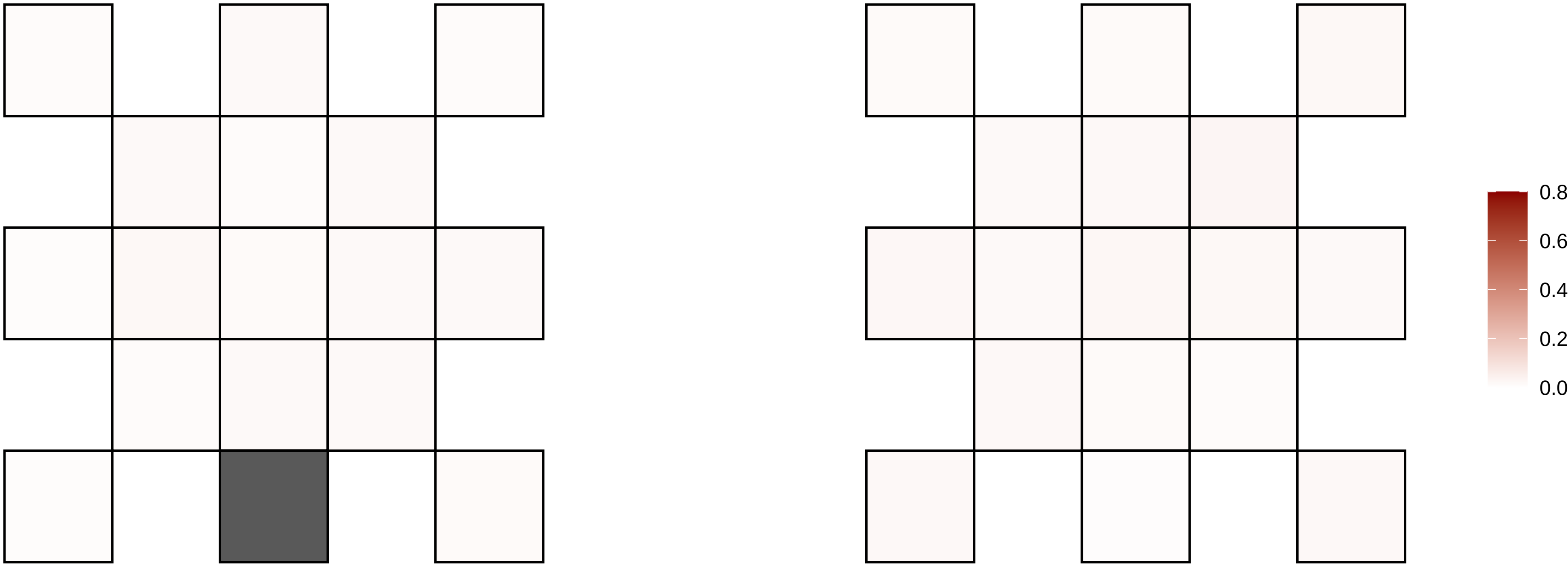

10−107355222−C−A

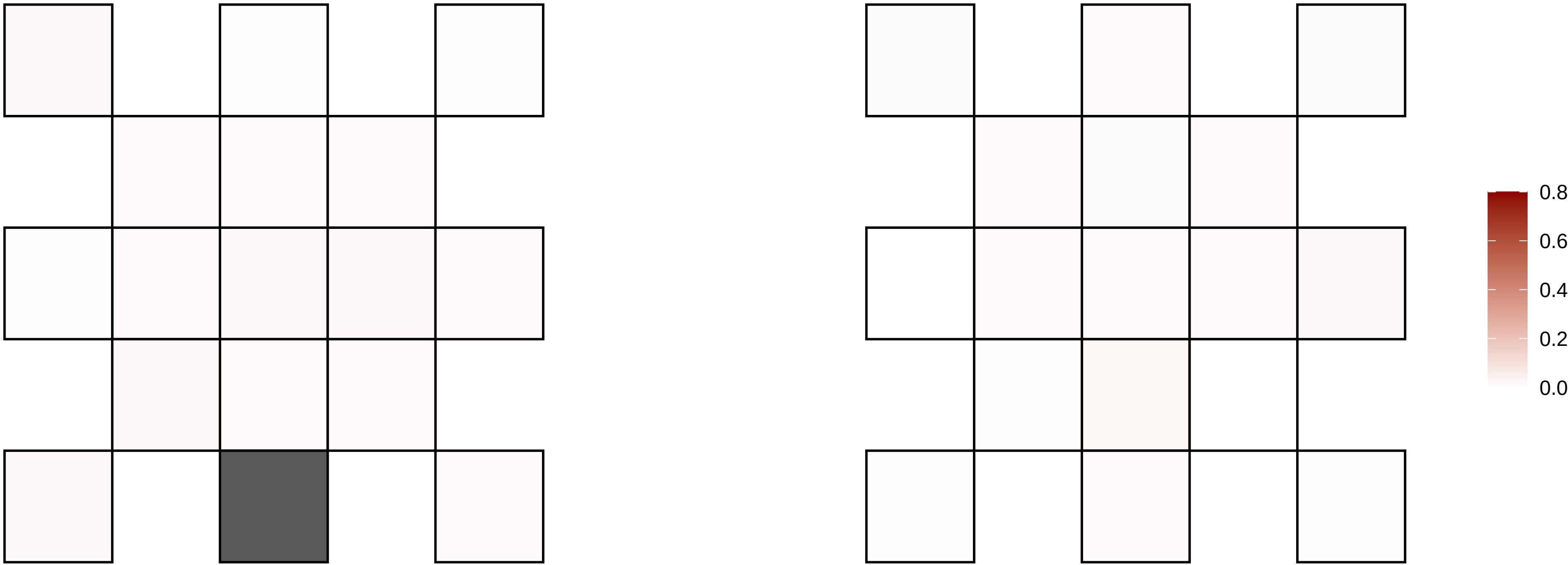

10−108671103−C−T

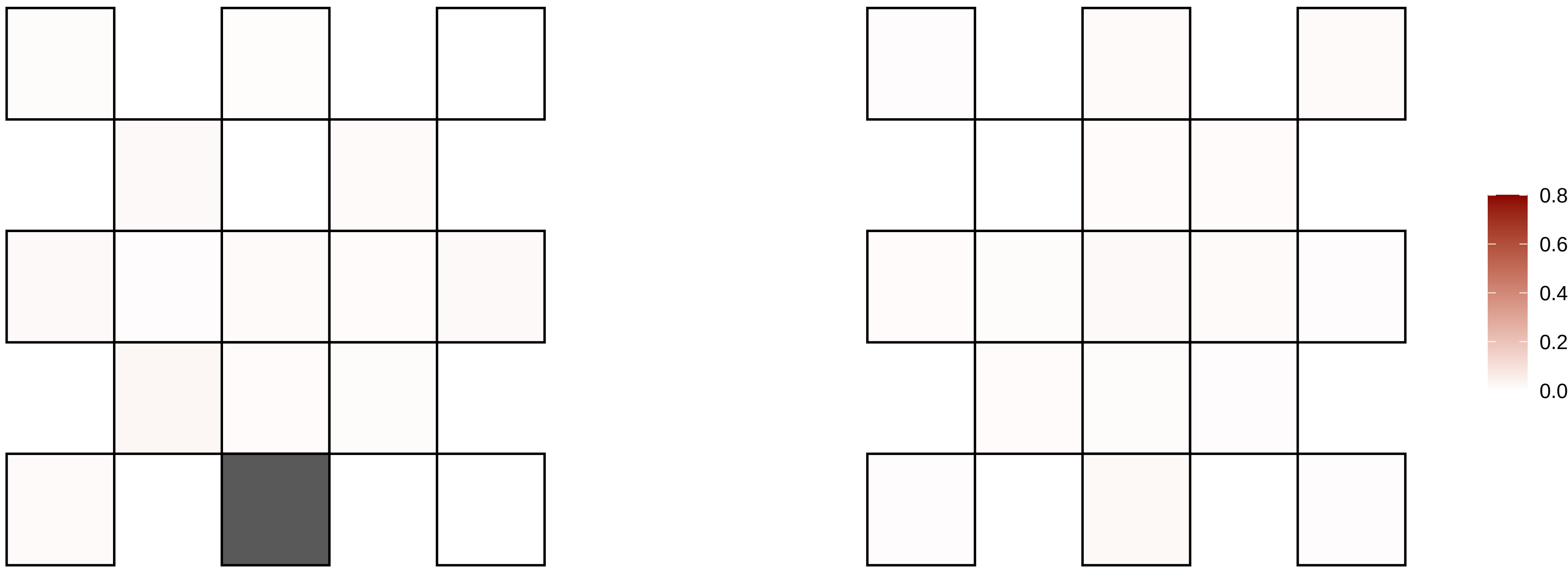

10−109795130−C−G

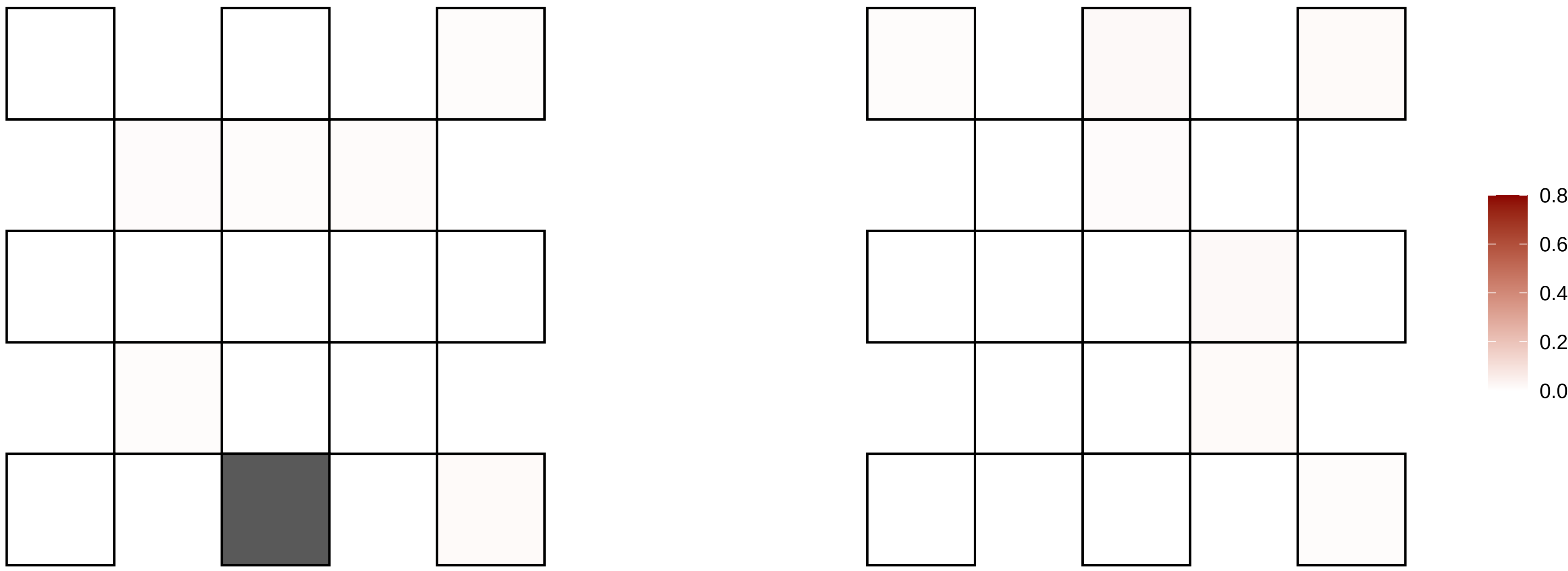

10−113404344−G−A

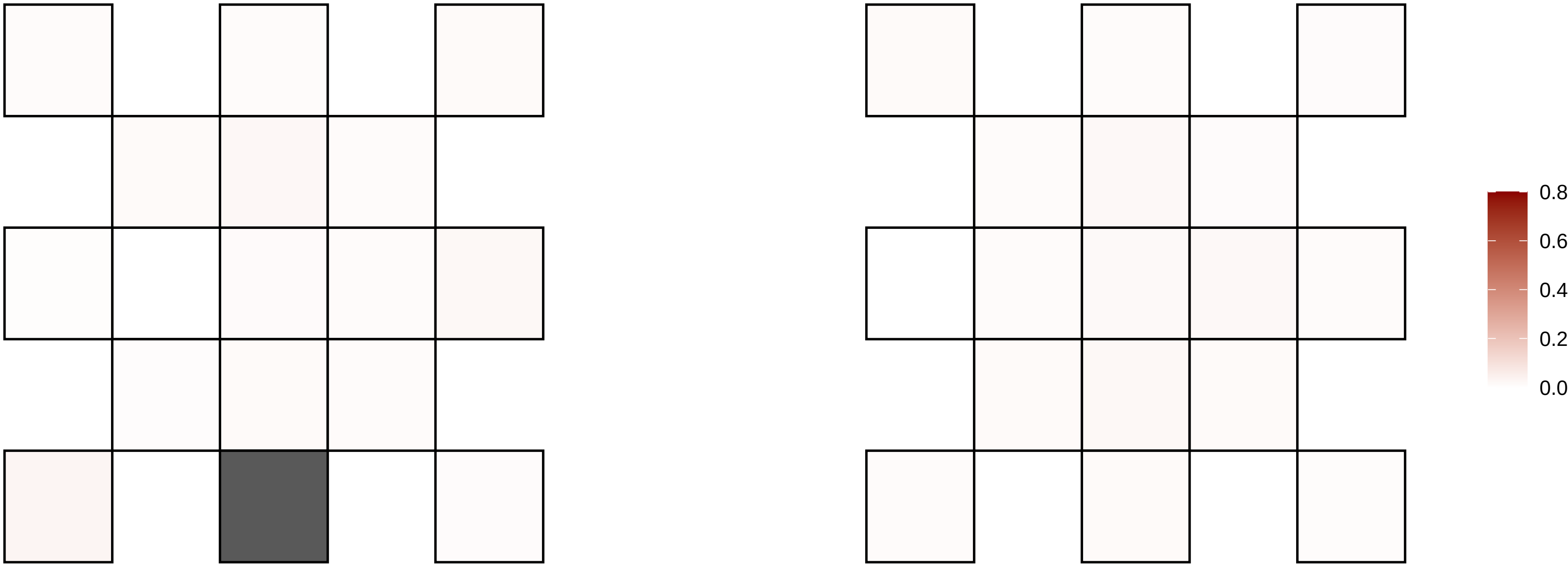

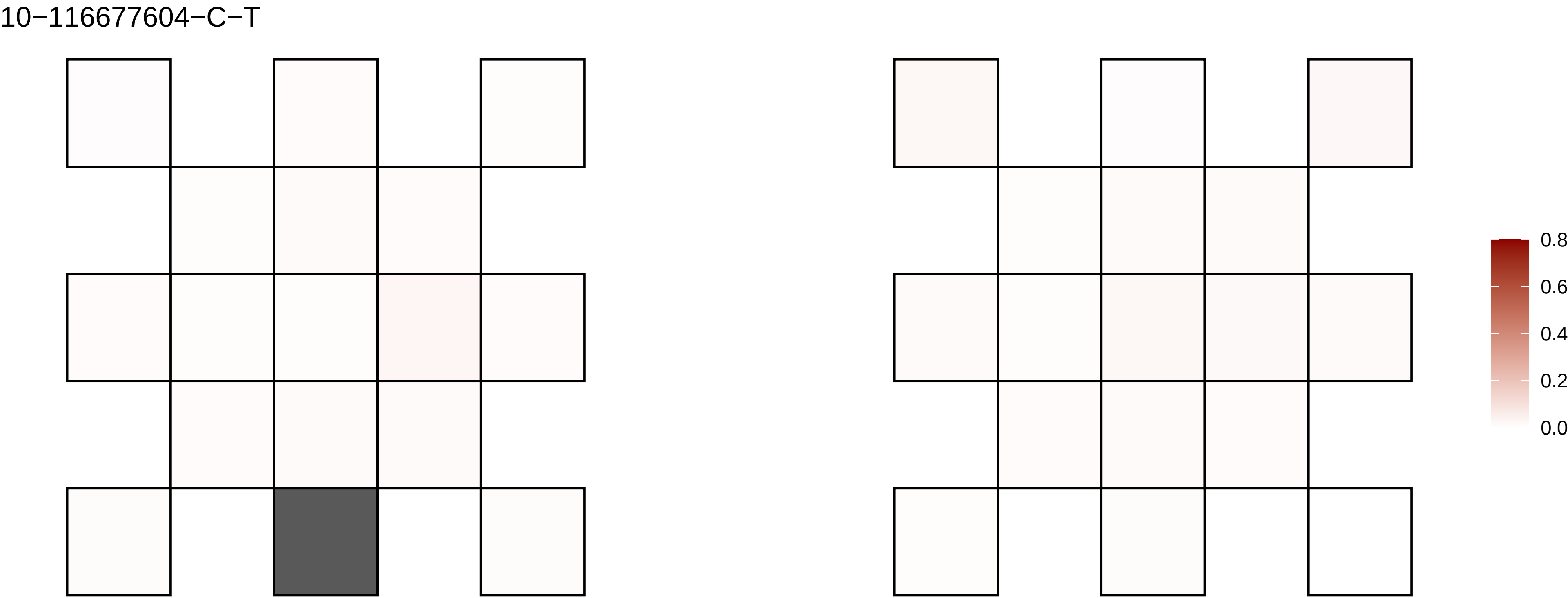
10−116677604−C−T
0.8
0.6
0.4
0.2
0.0

10−117540461−G−A

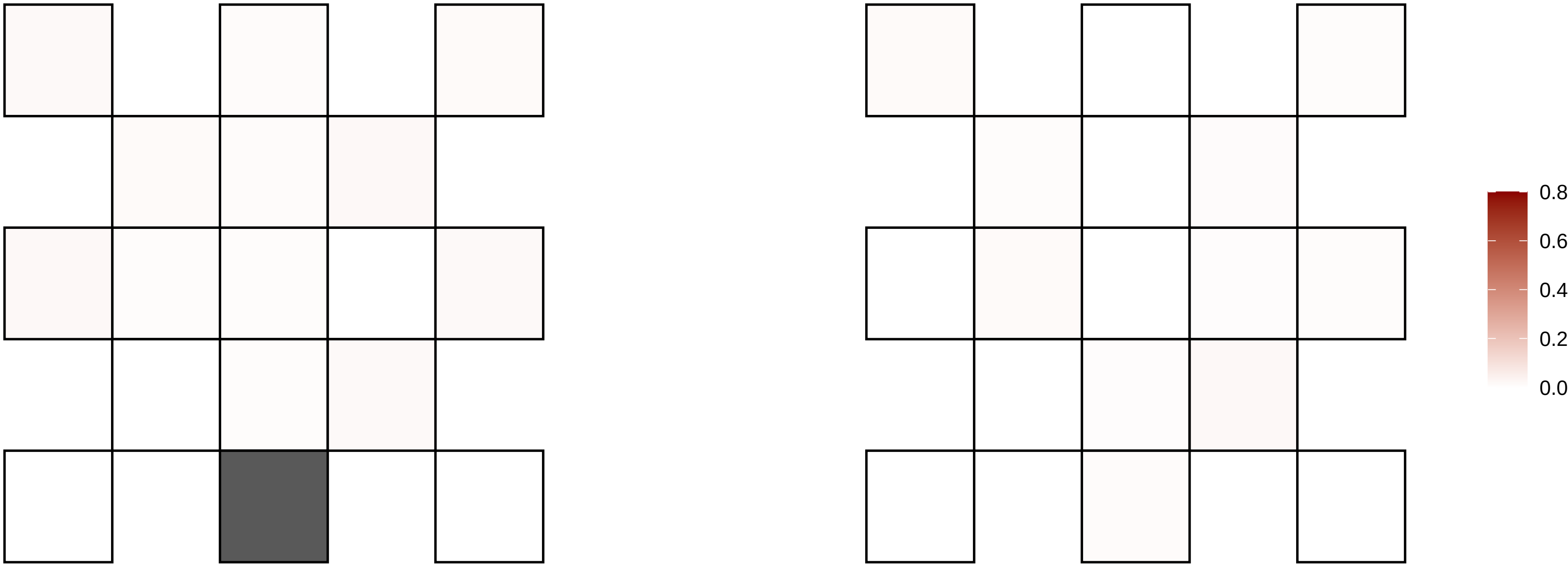

10−121073761−C−T

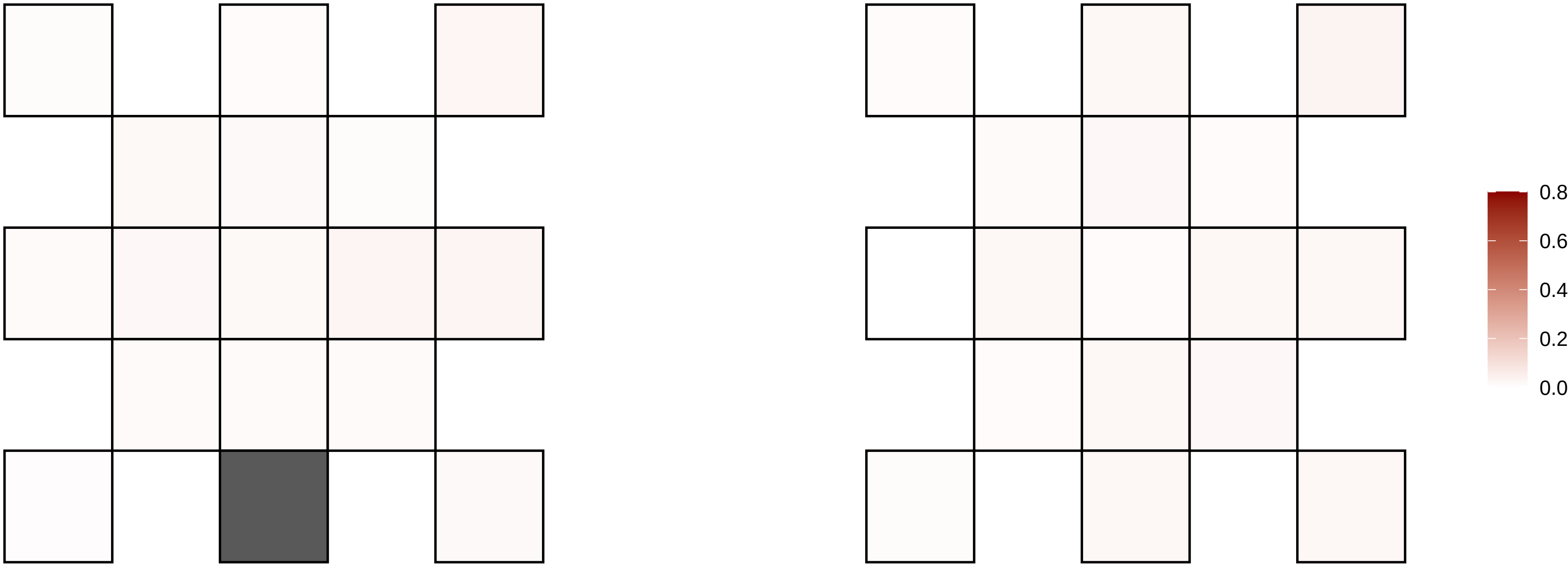

10−123873718−C−T

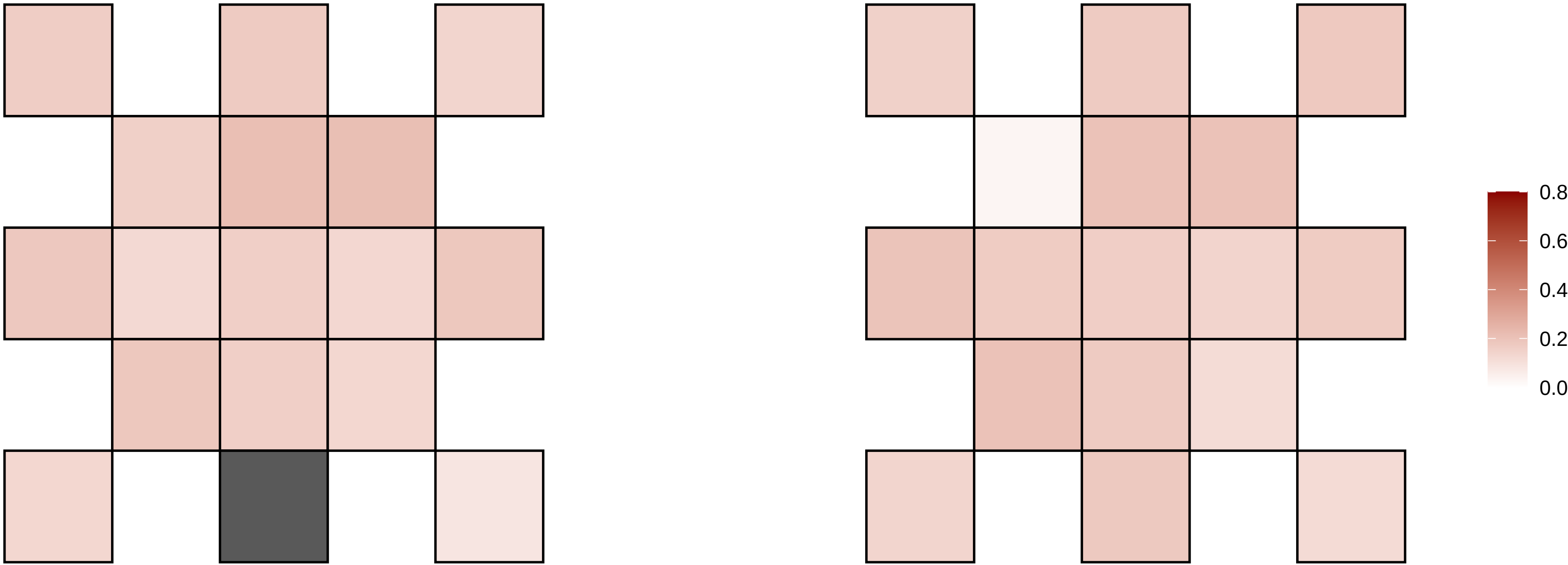

10−125075590−G−A

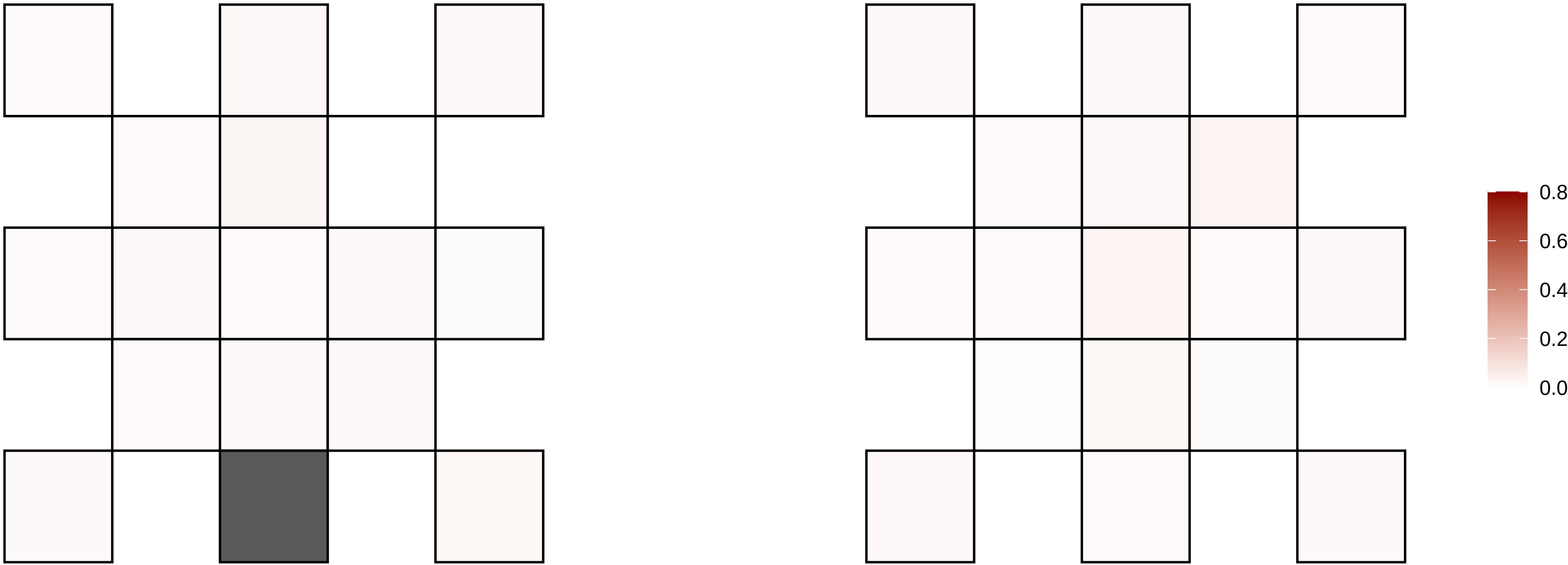

10−126824209−G−A

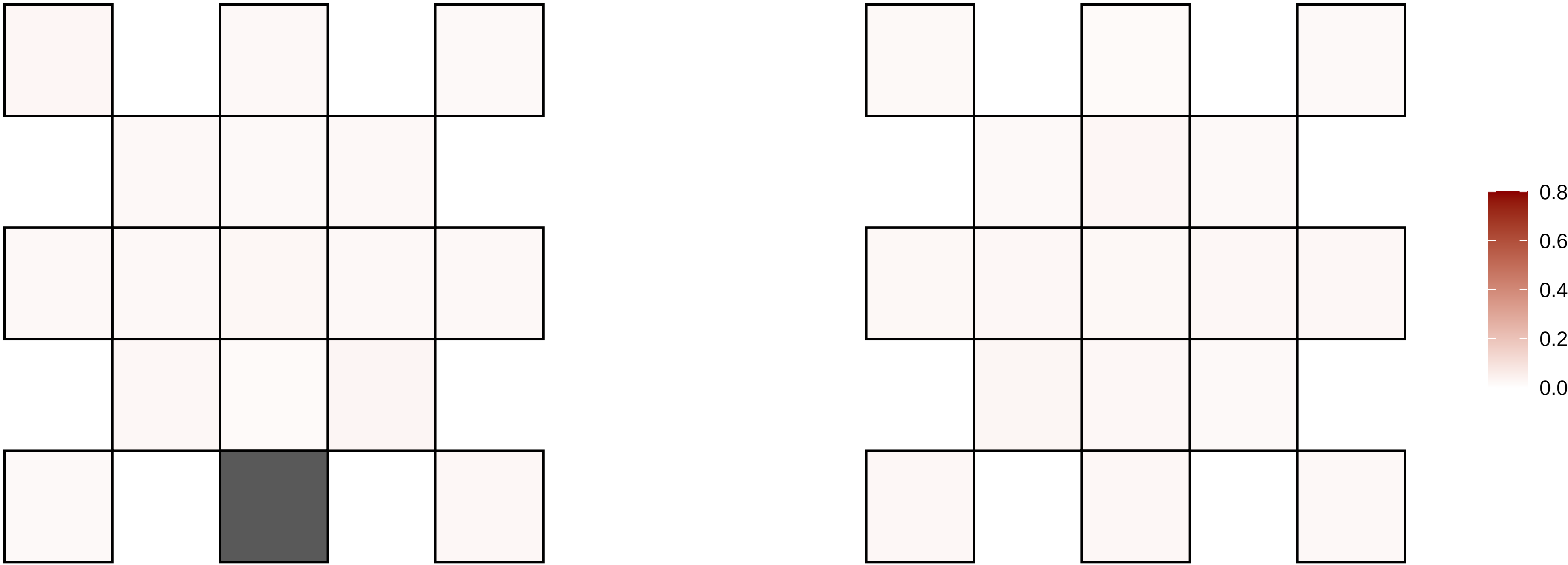

10−12696593−A−G

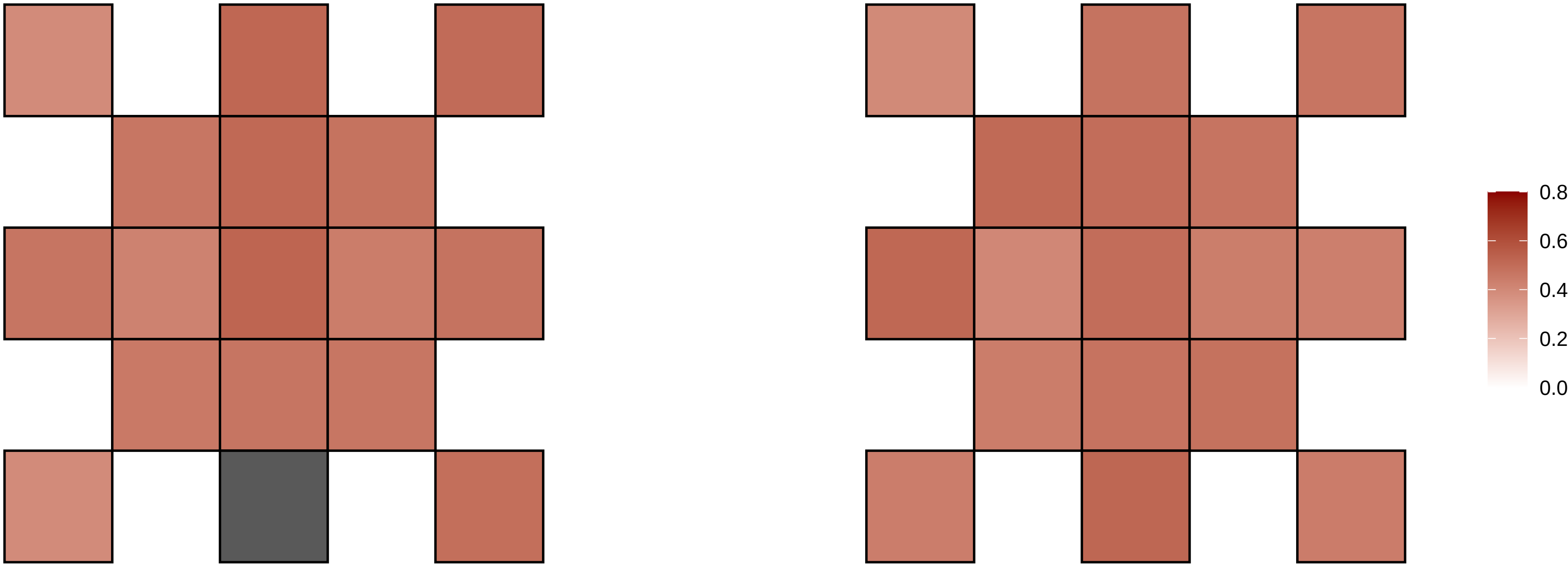

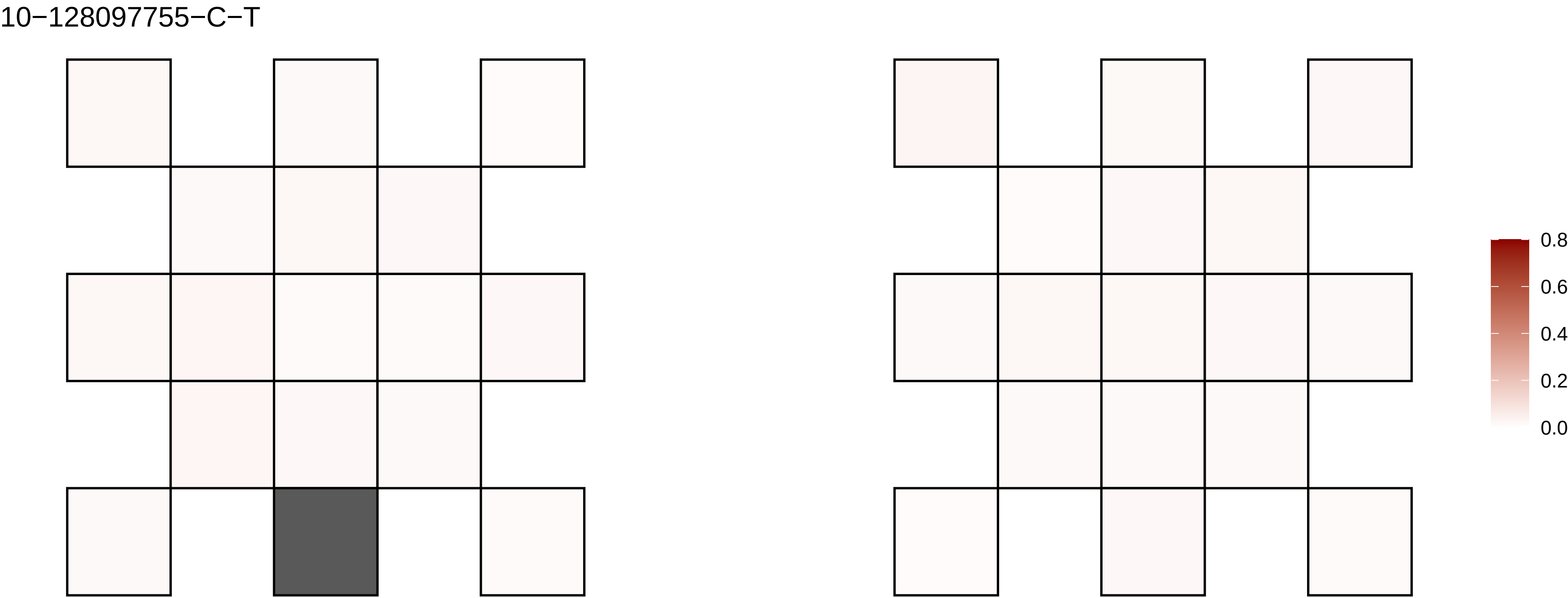
10−128097755−C−T
0.8
0.6
0.4
0.2
0.0

10−128228322−C−T

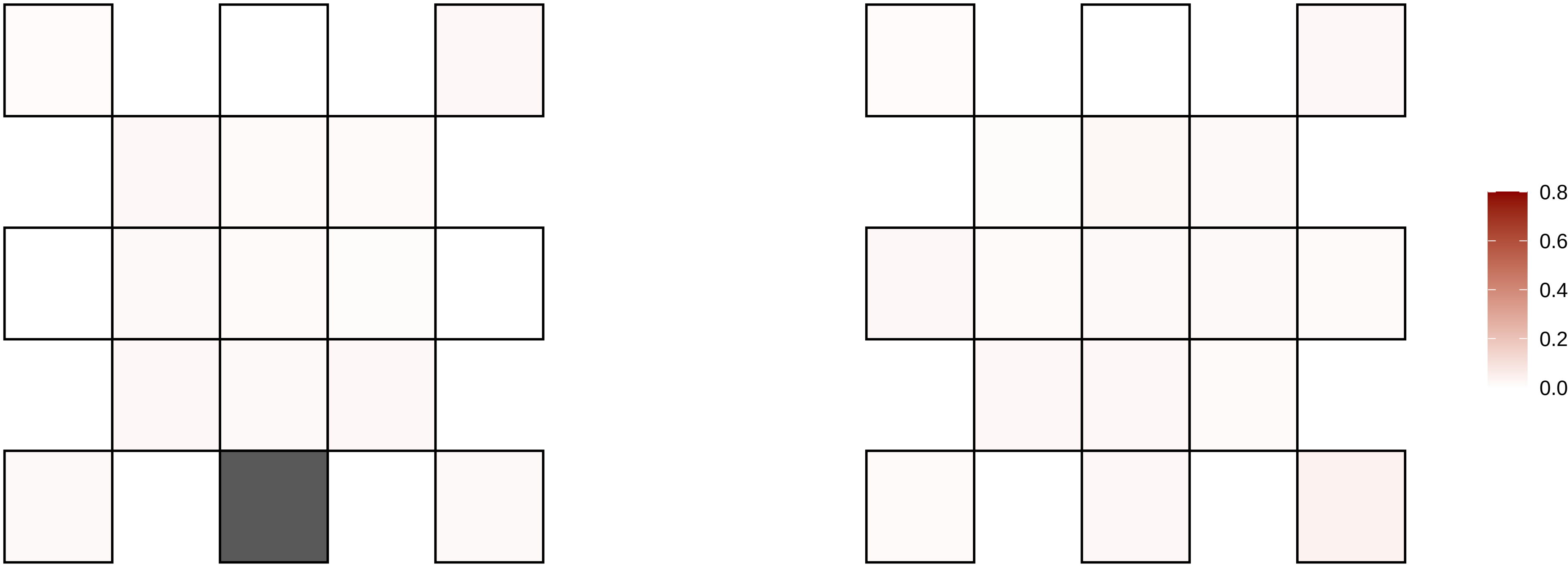

10−129376060−T−G

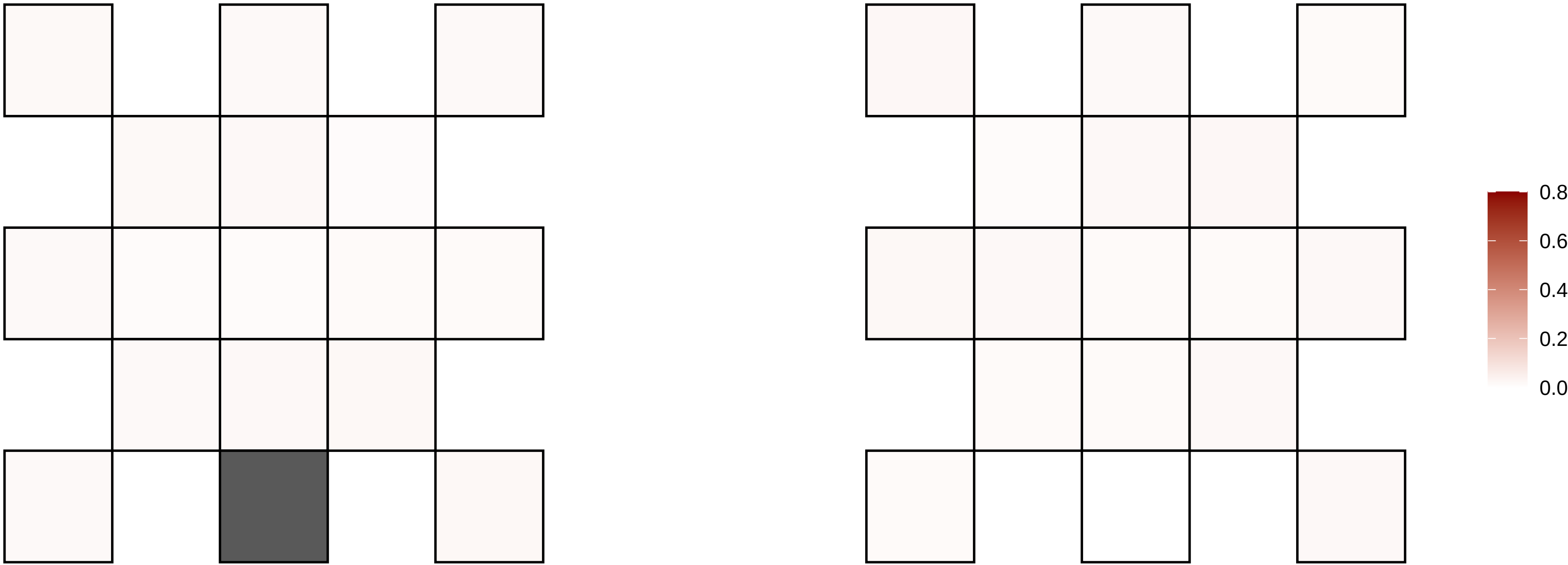

10−133221443−A−T

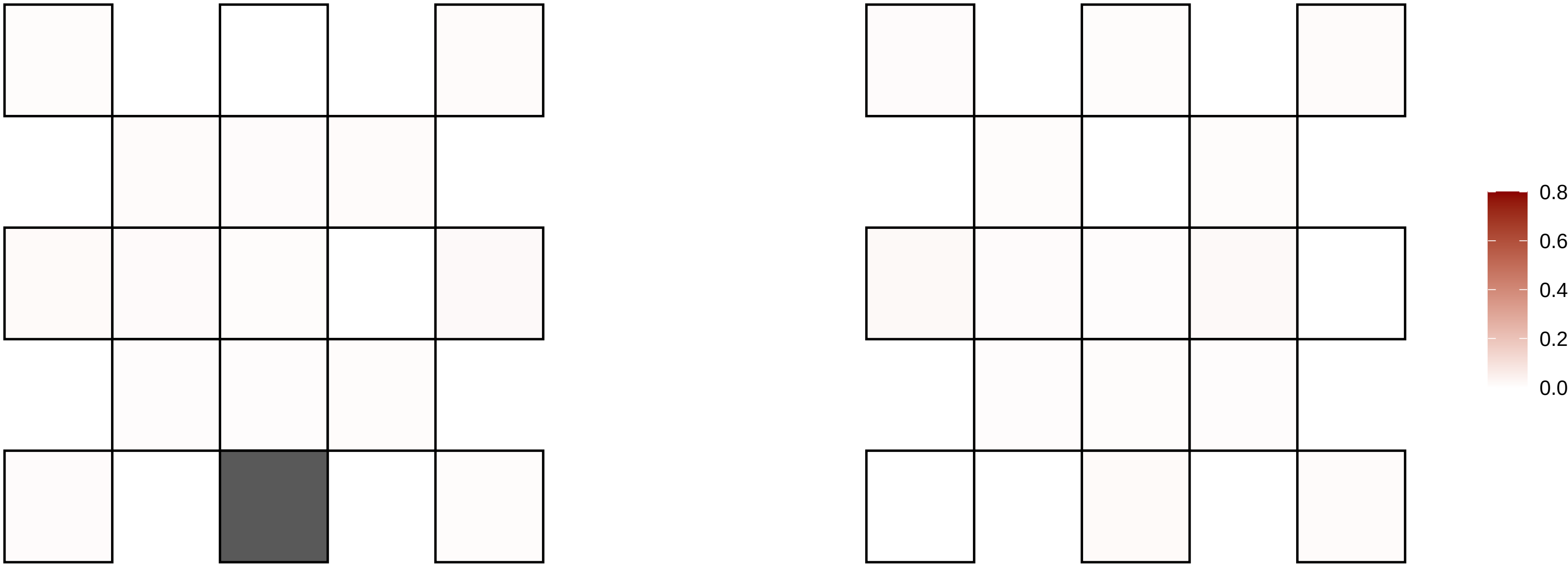

10−134211625−C−T

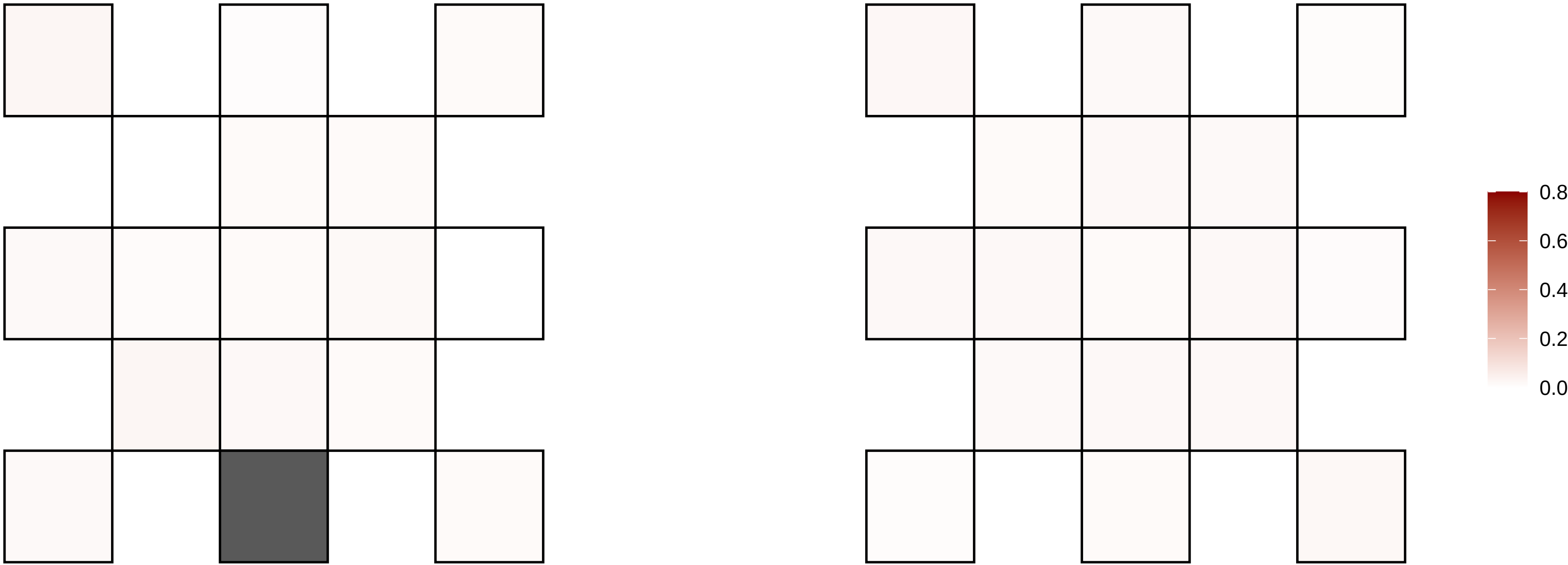

10−134280713−A−G

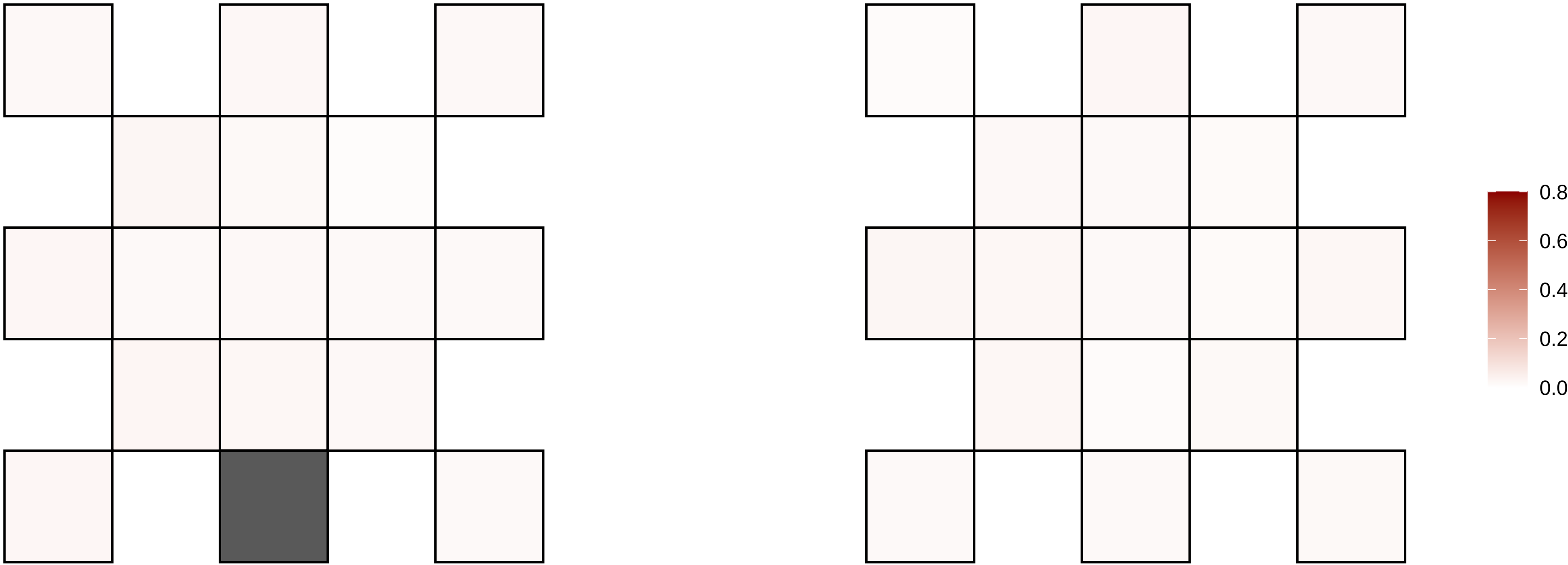

10−13796054−C−T

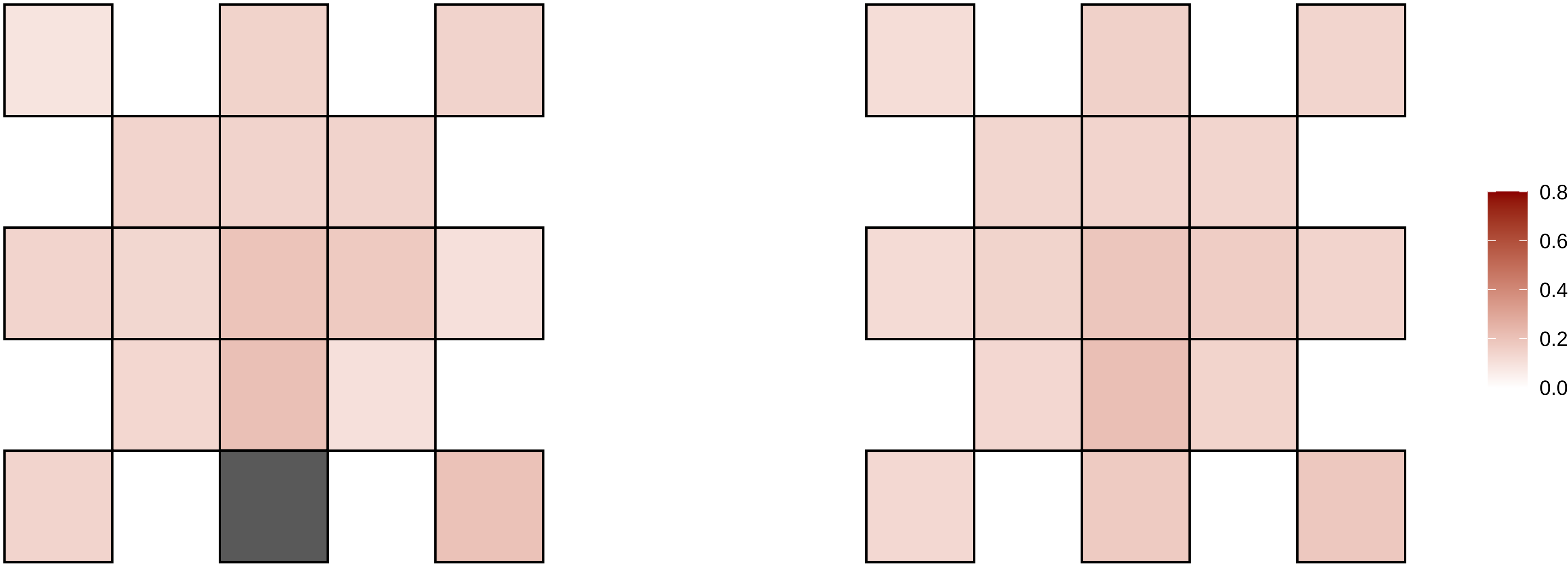

10−14490737−G−A

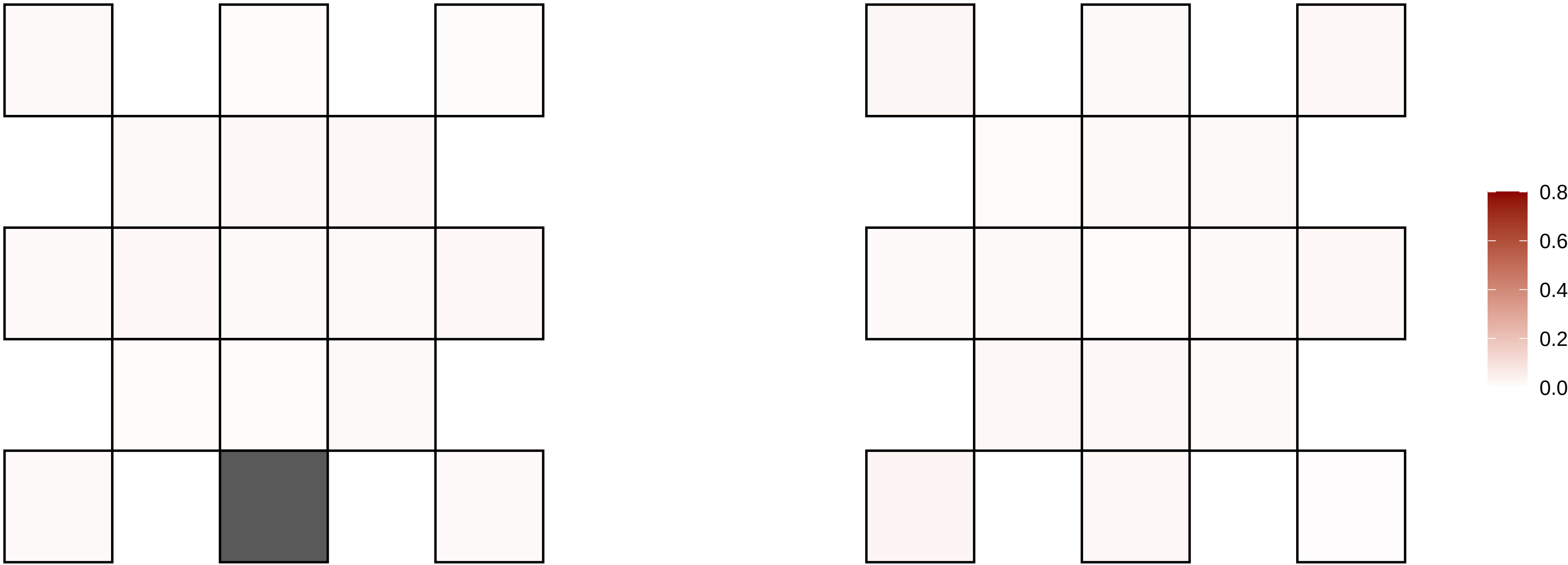

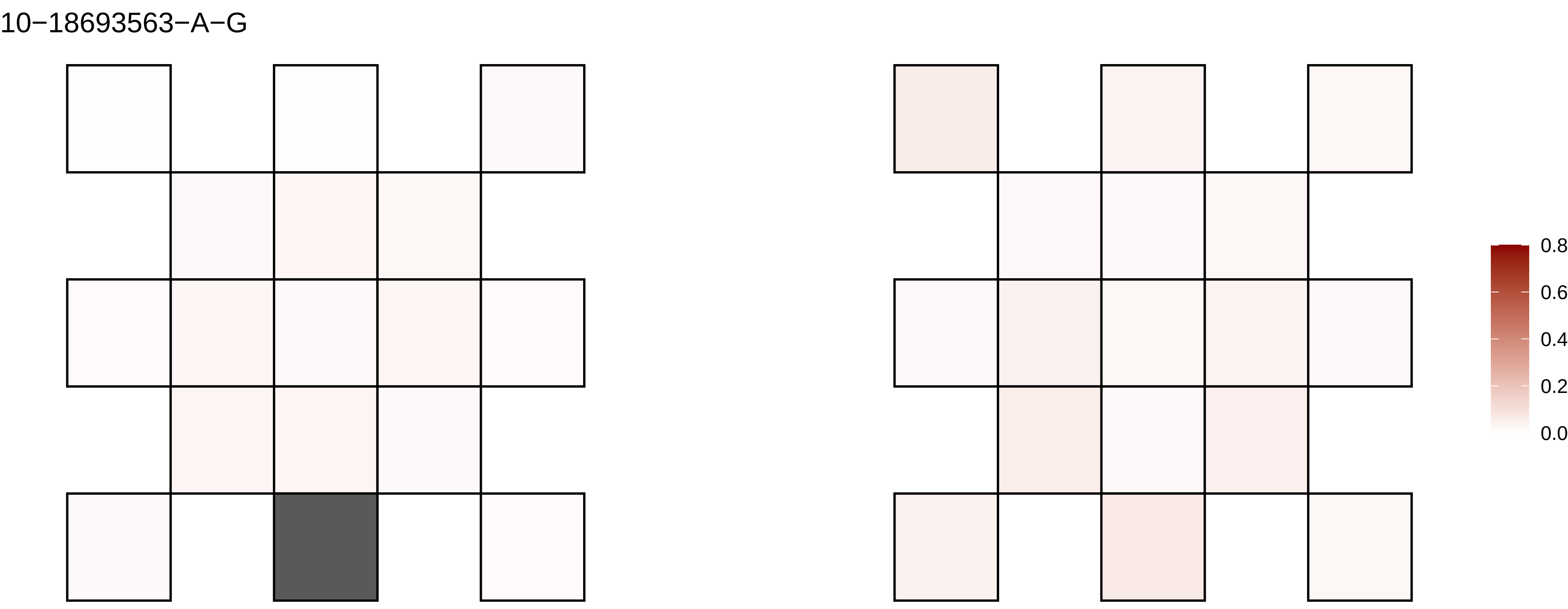
10−18693563−A−G
0.8
0.6
0.4
0.2
0.0

10−18760480−C−T

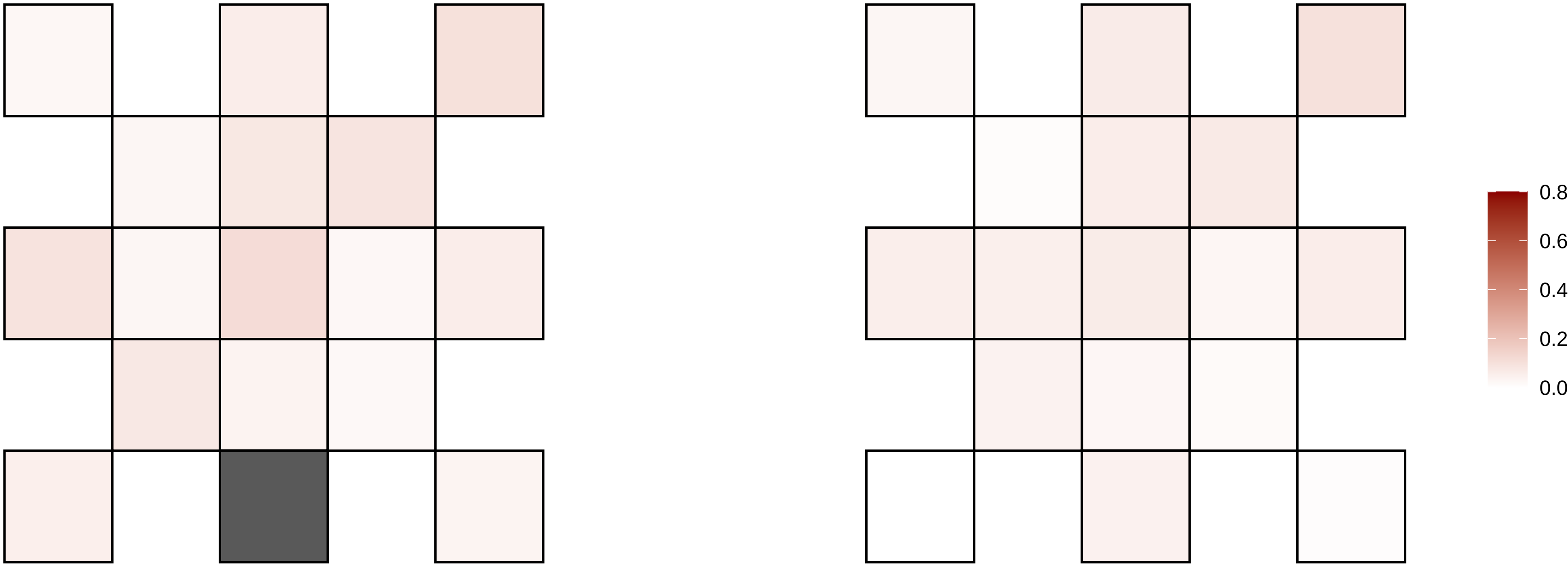

10−2261697−A−C

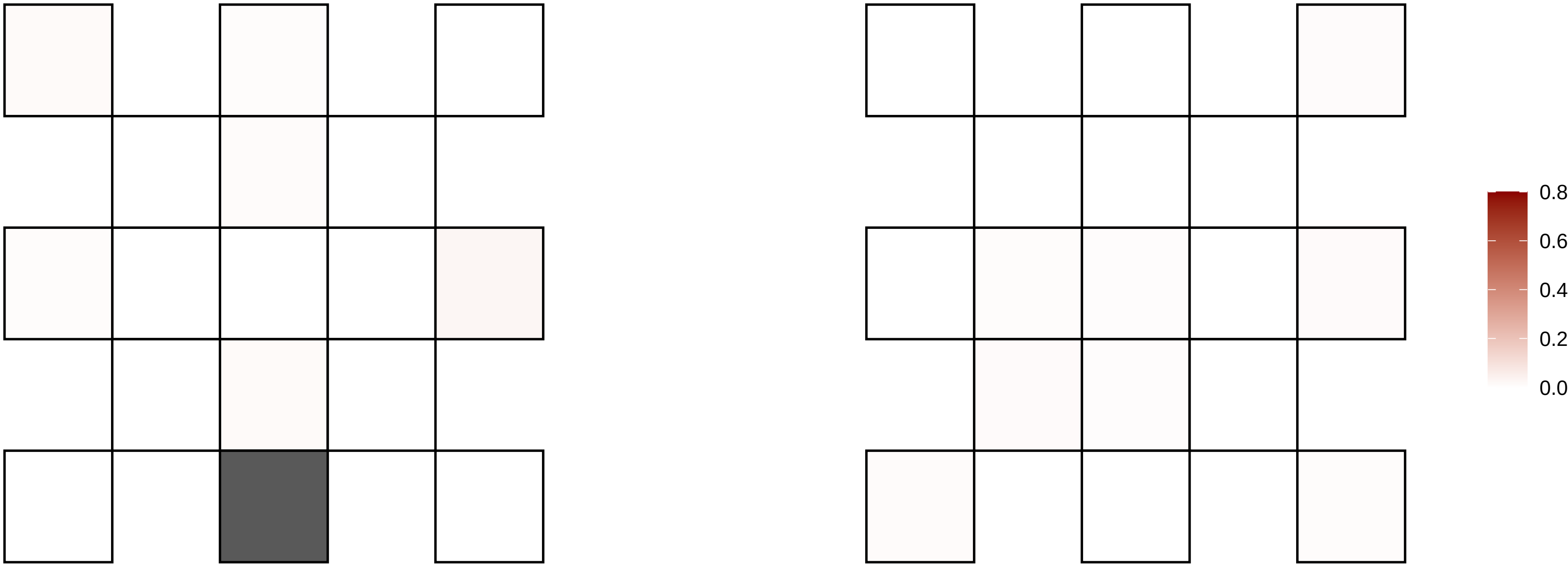

10−2261700−AT−A

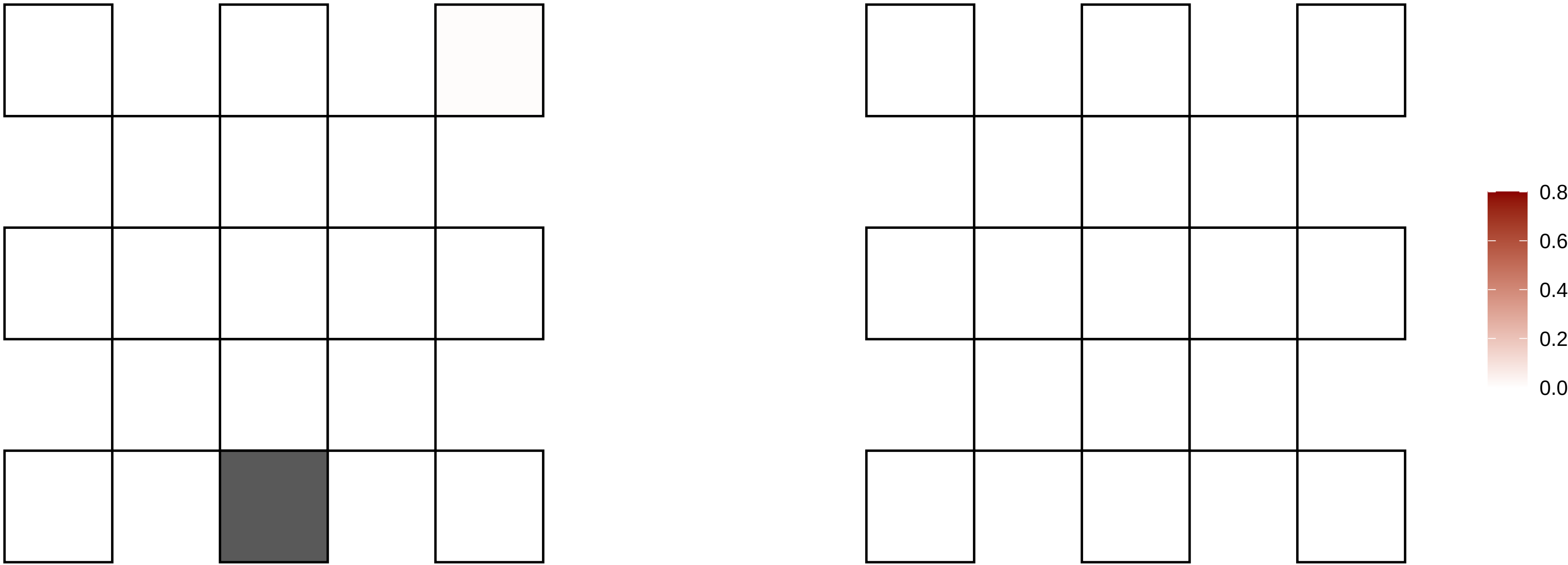

10−22765172−A−T

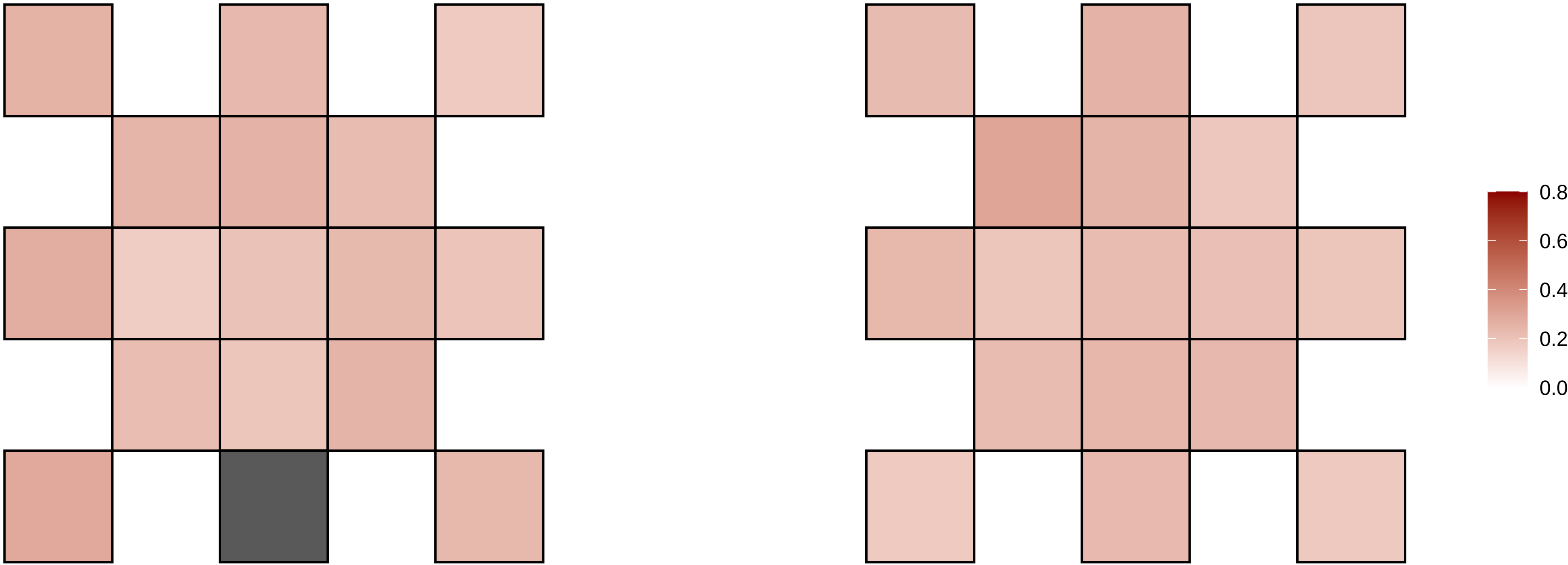

10−23976083−C−A

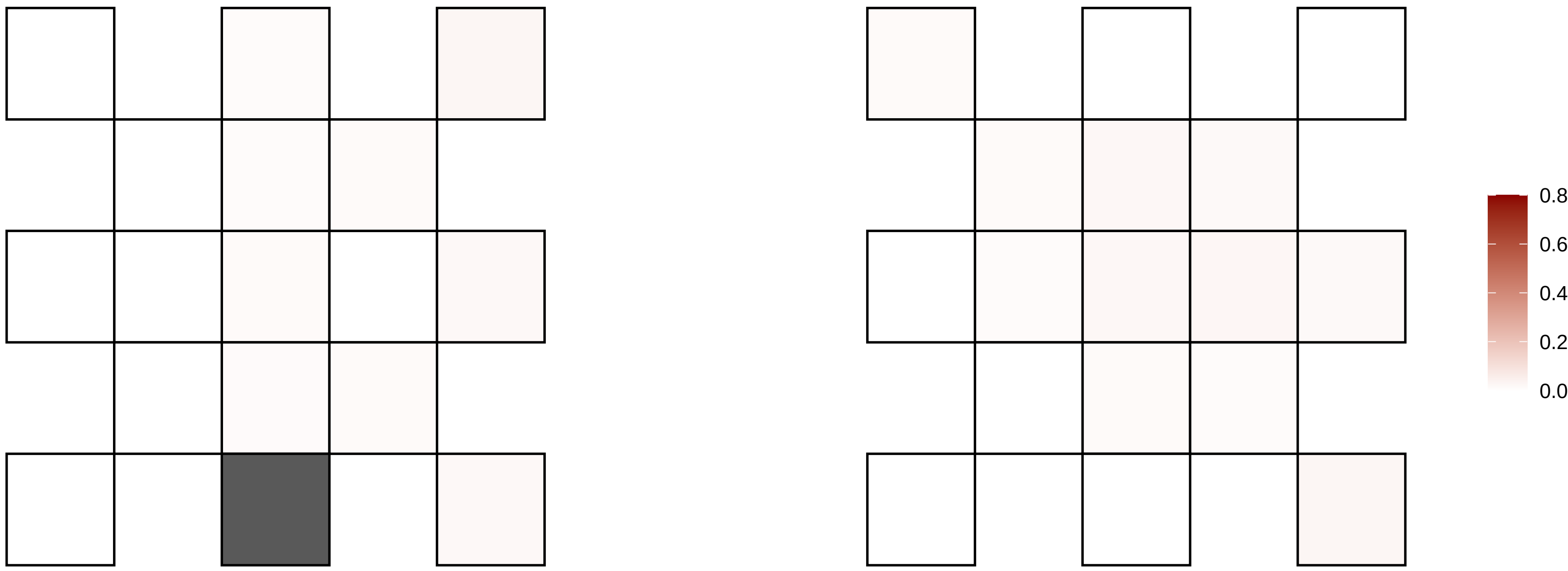

10−25457893−C−T

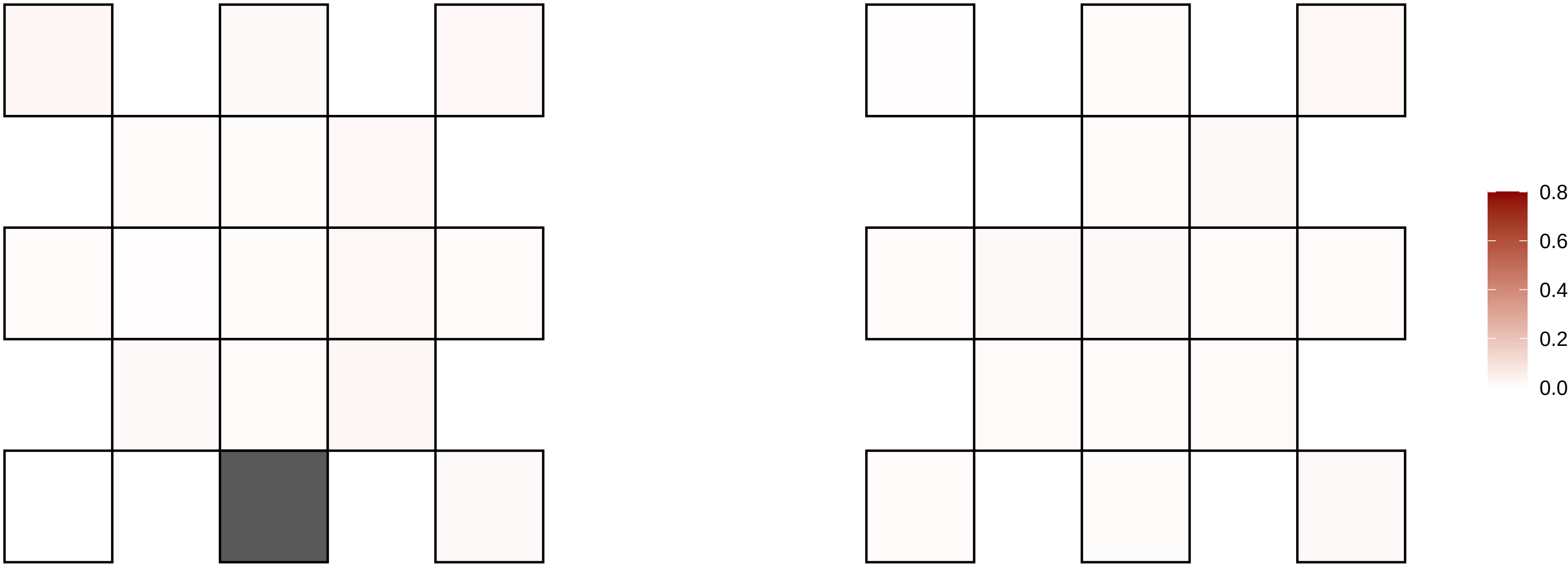

10−25610159−G−A

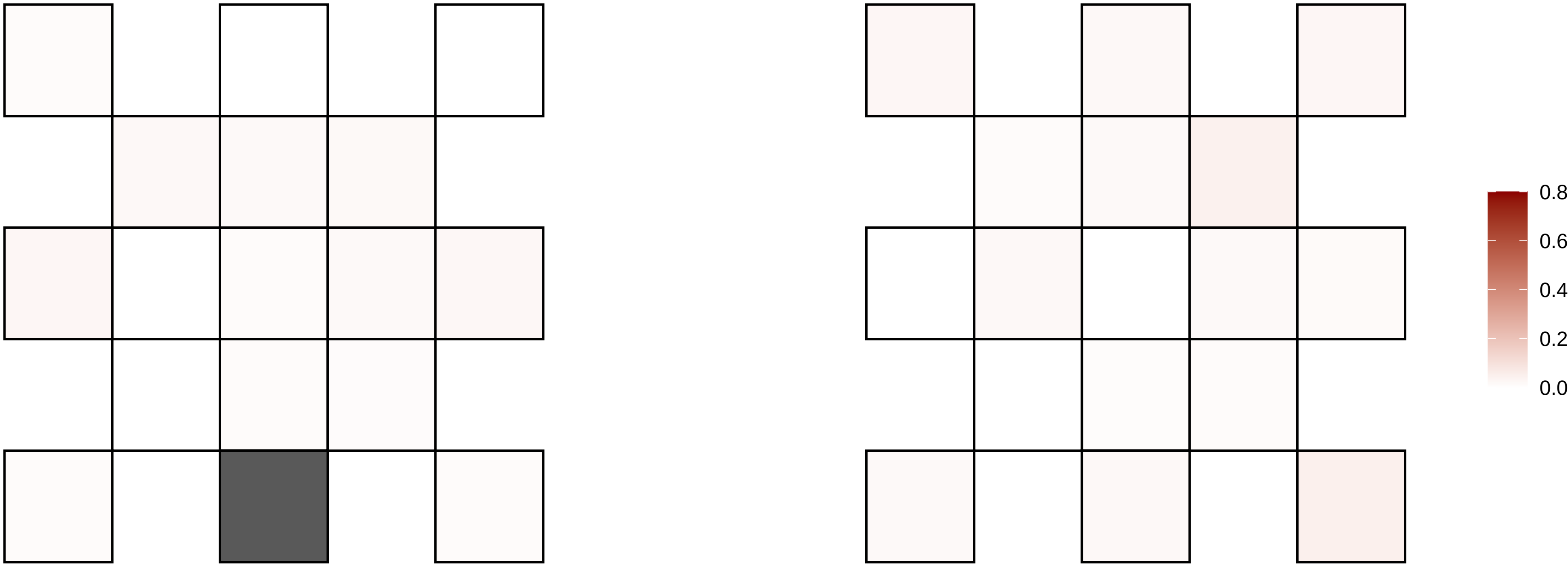

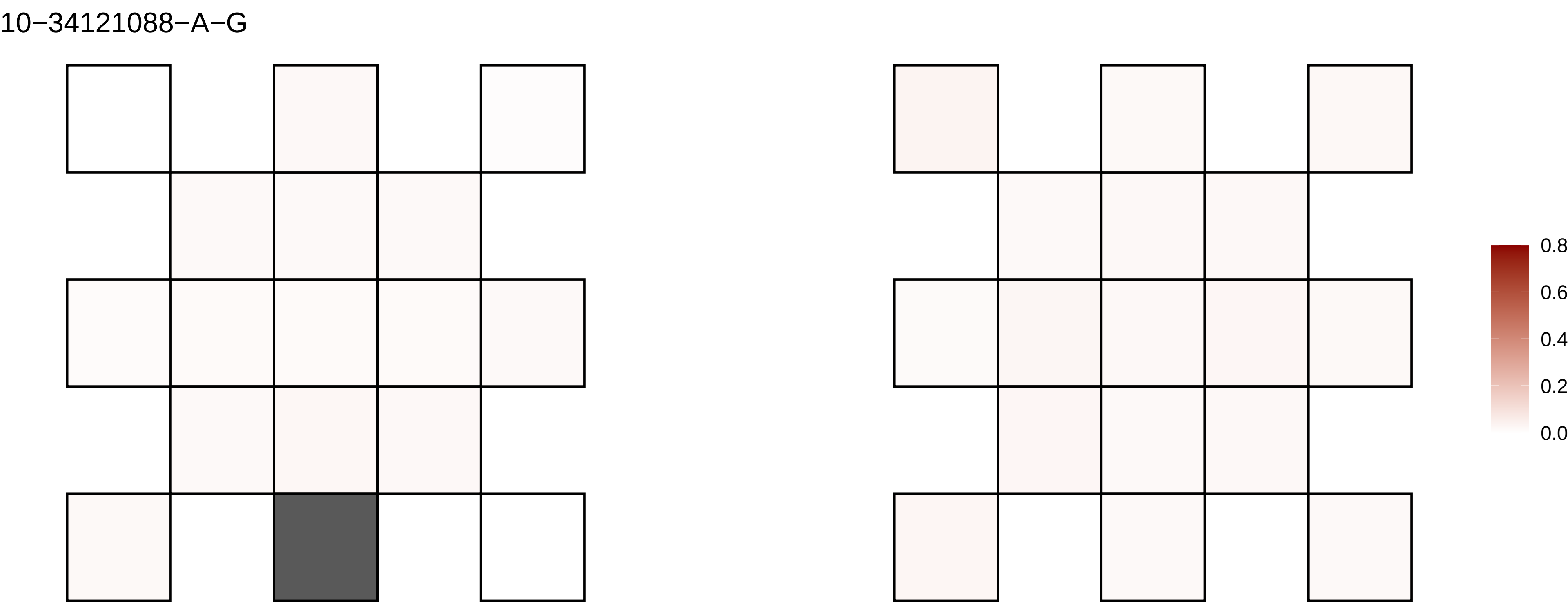

10−34121088−A−G
0.8
0.6
0.4
0.2
0.0

10−36039114−G−A

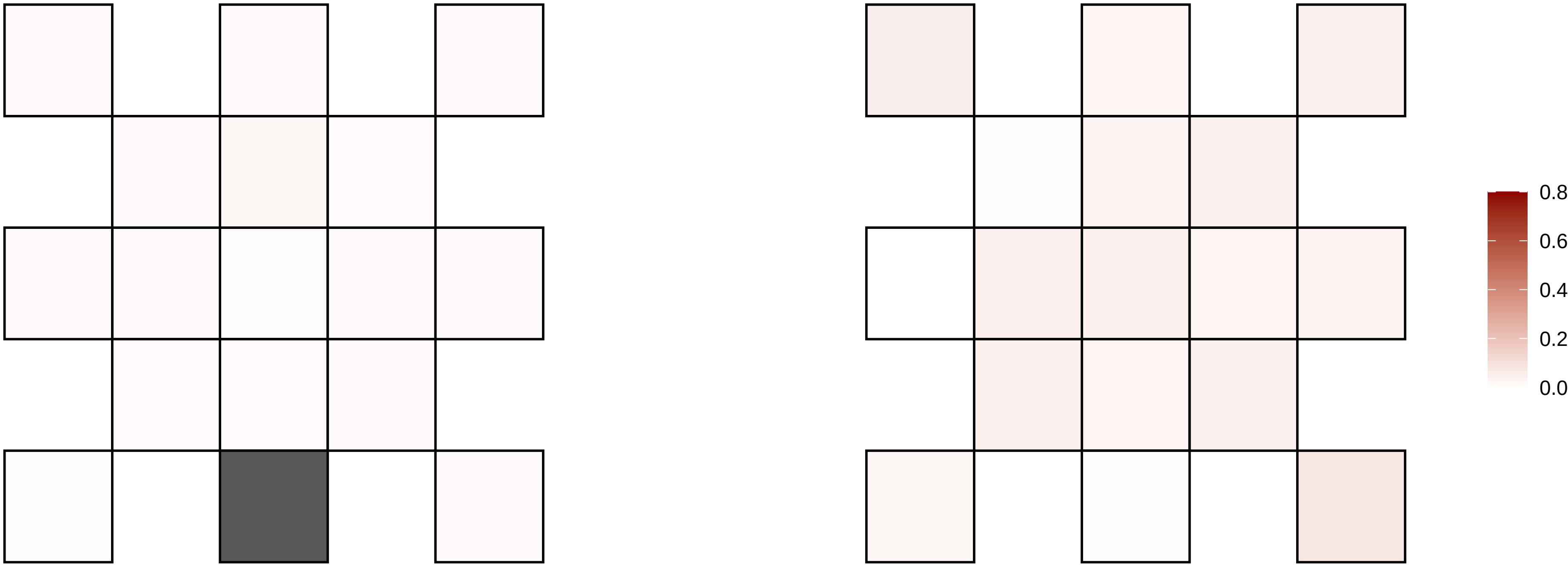

10−58152659−C−T

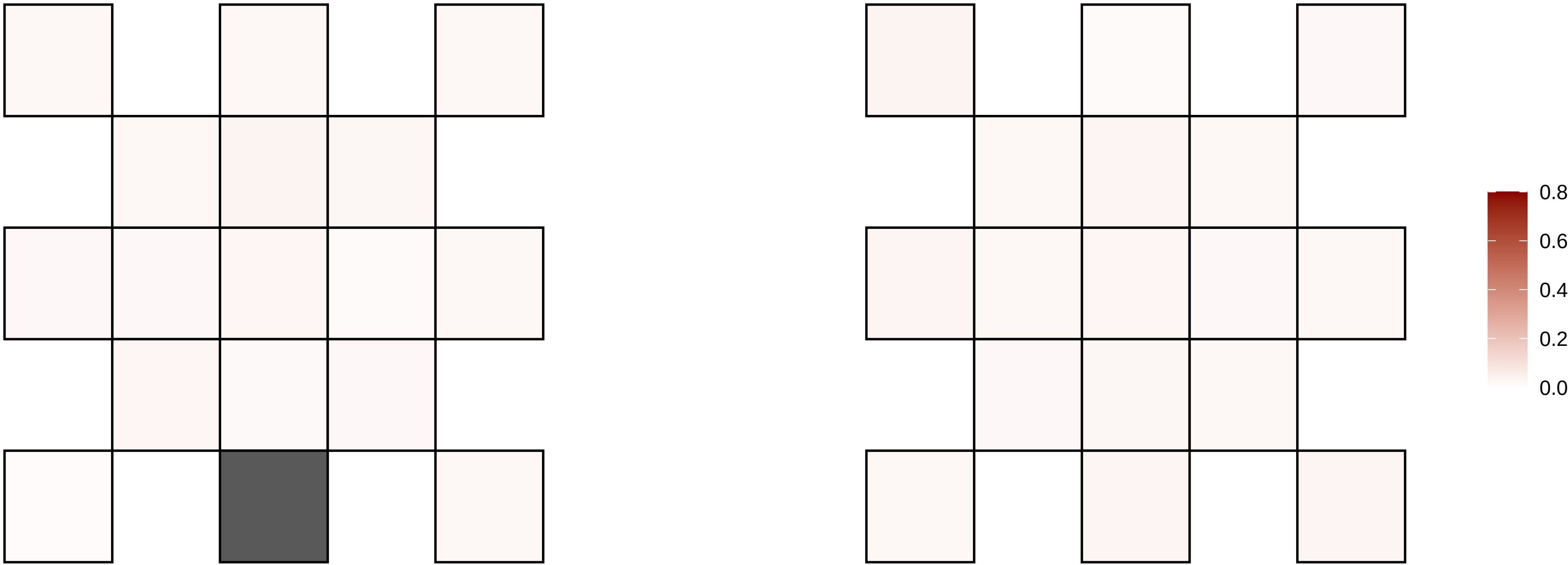

10−58457040−T−G

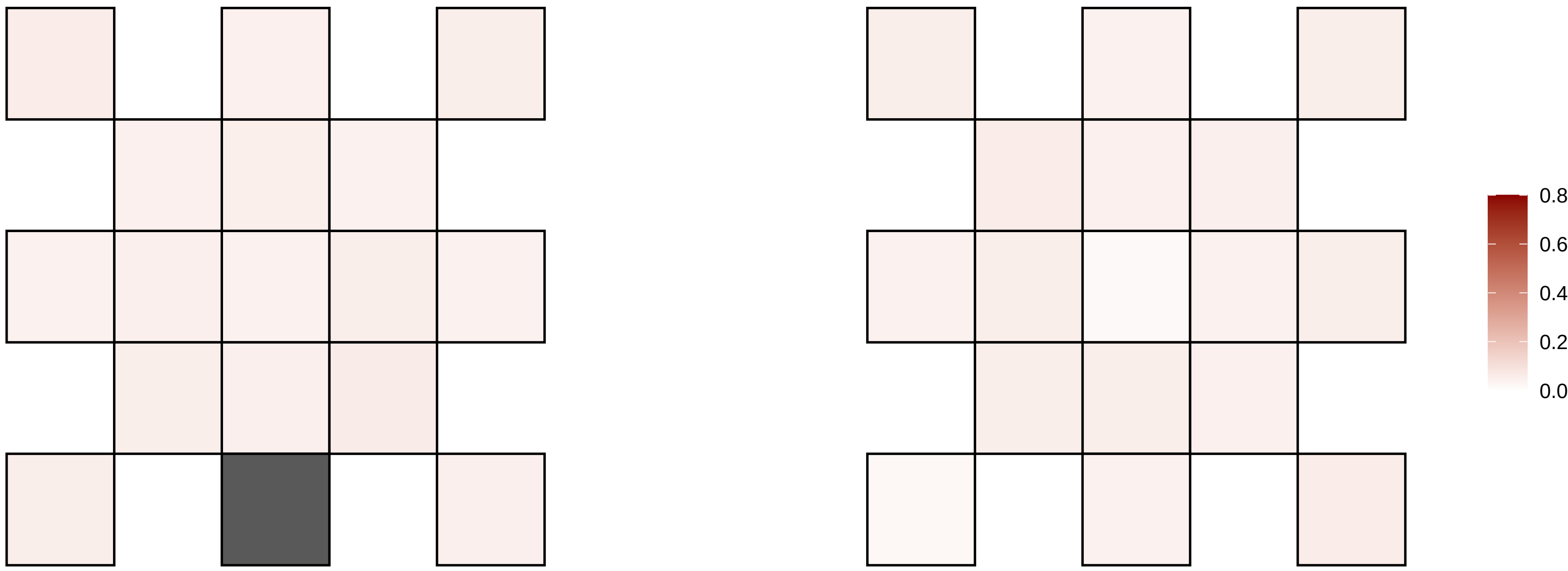

10−60559957−C−T

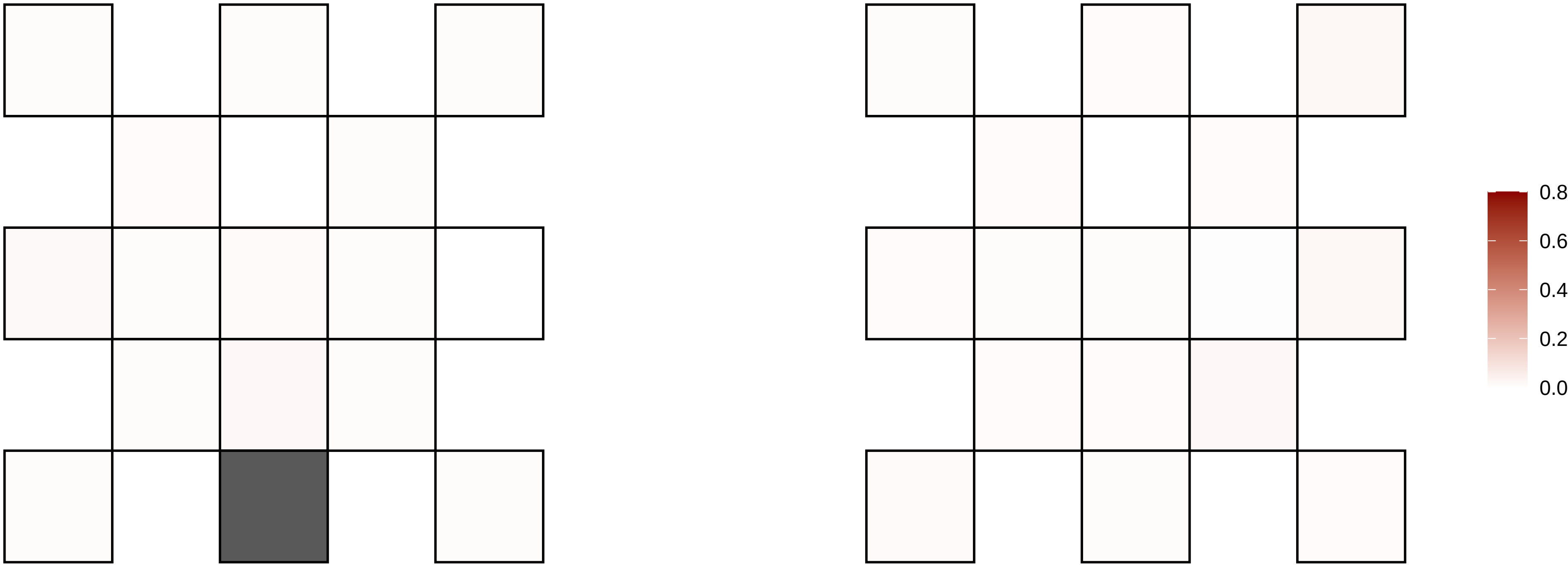

10−7459695−G−A

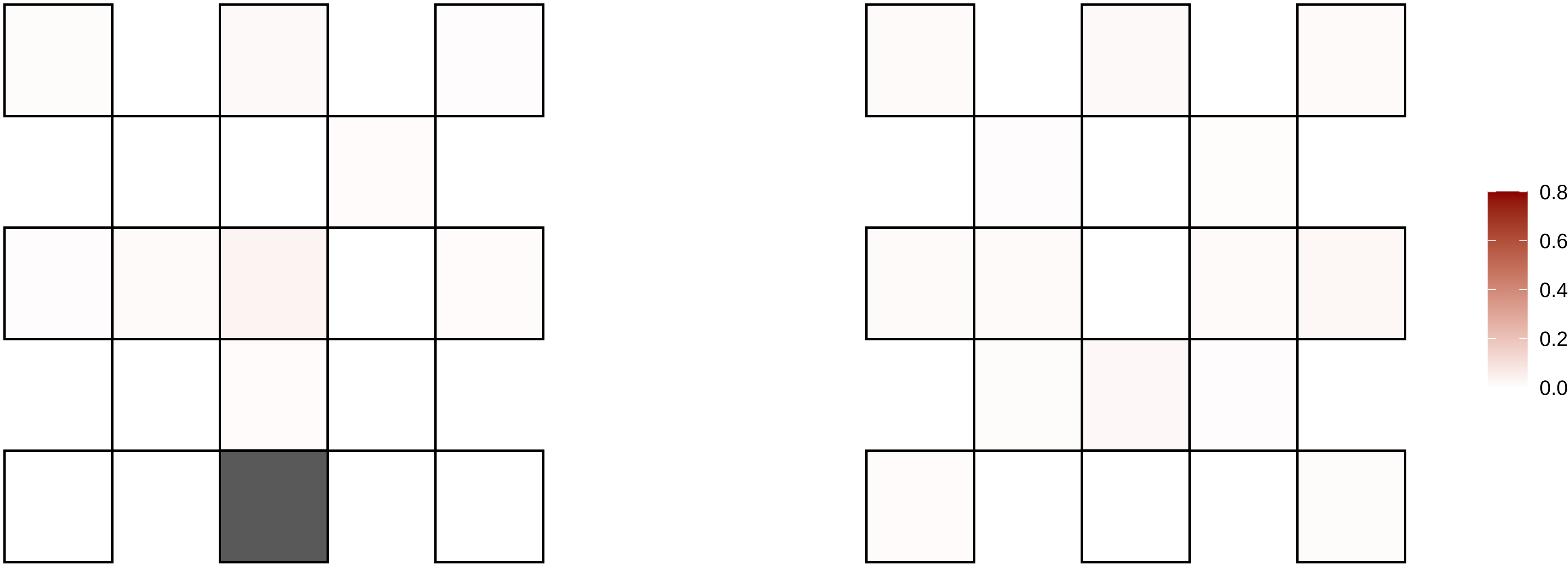

10−79843135−G−A

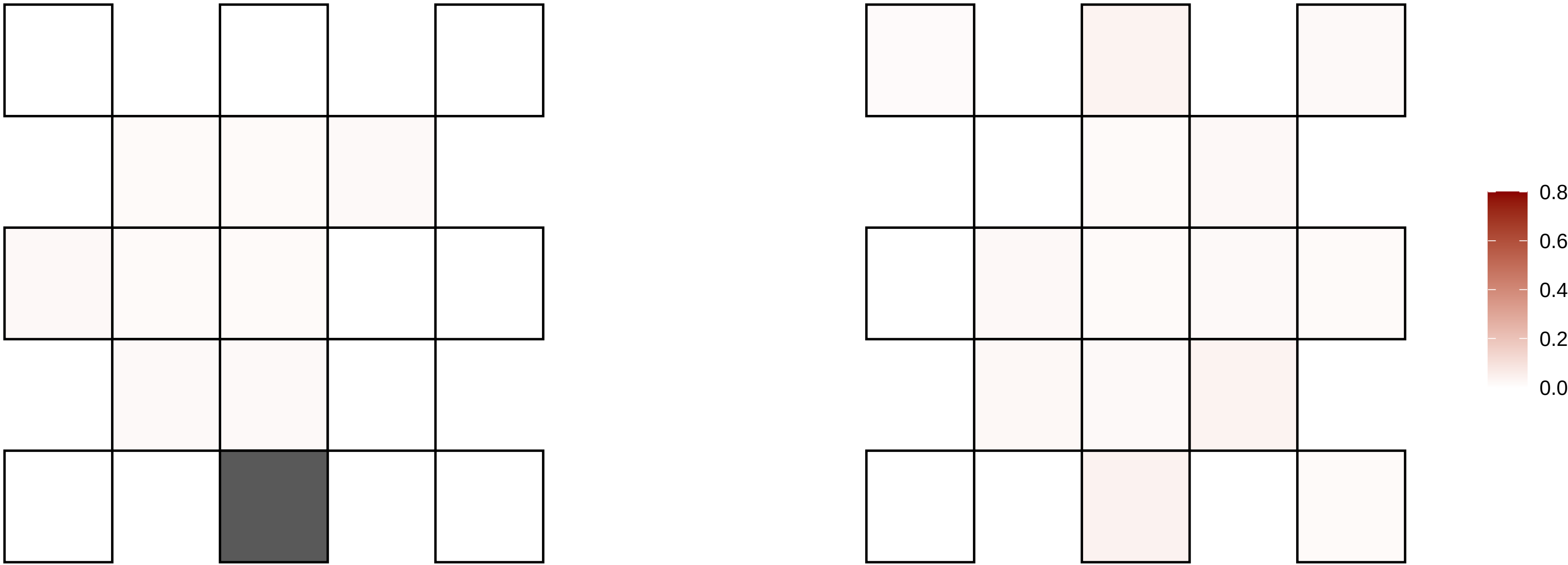

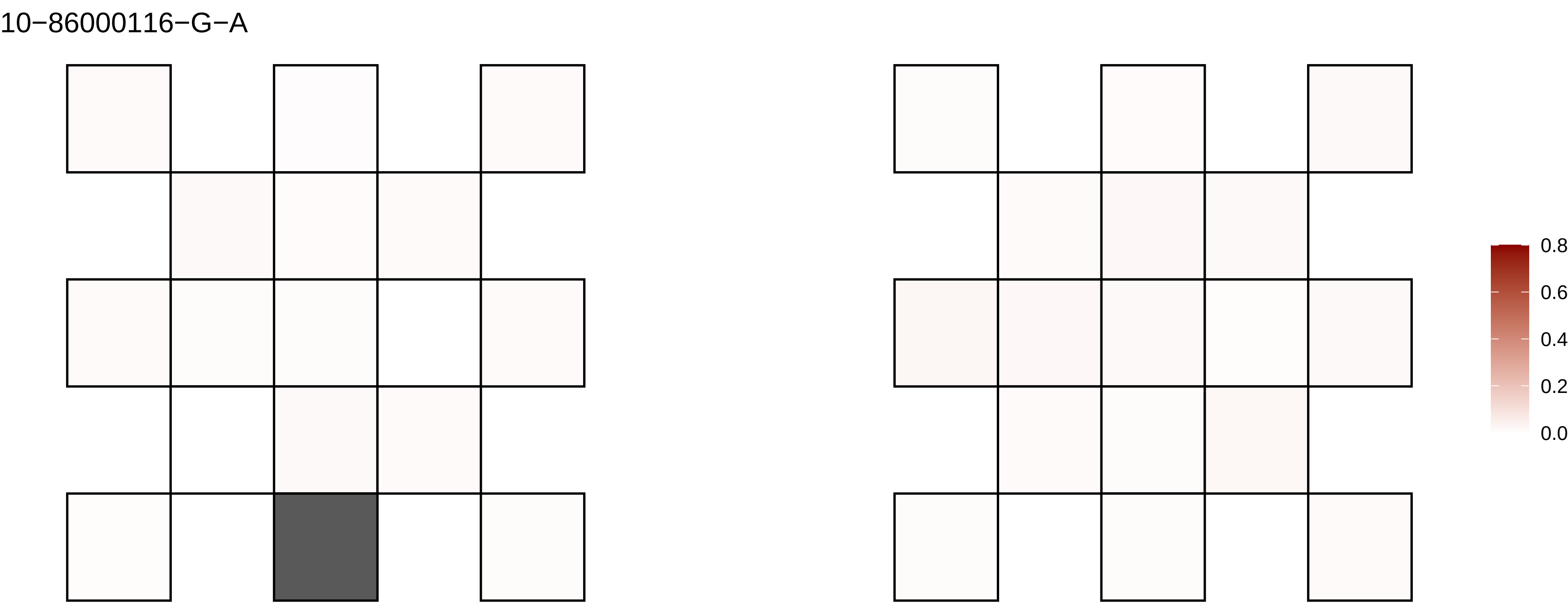
10−86000116−G−A
0.8
0.6
0.4
0.2
0.0

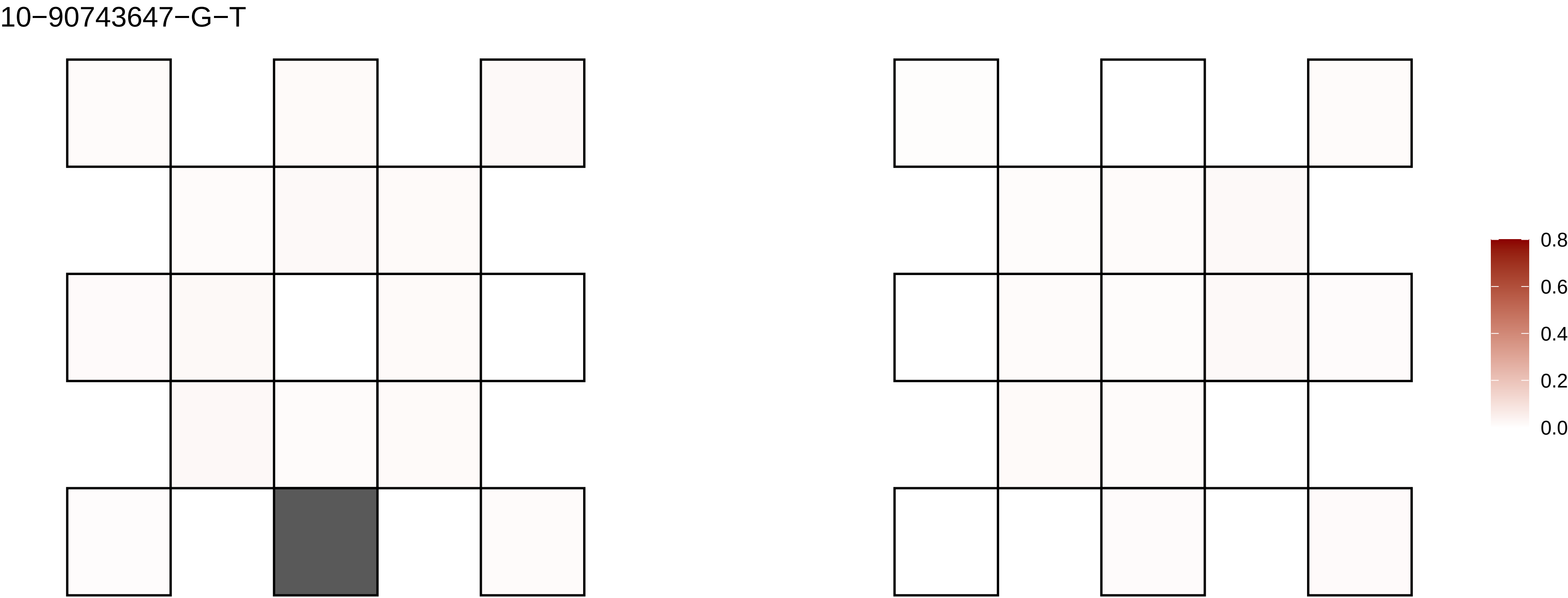
10−90743647−G−T
0.8
0.6
0.4
0.2
0.0

10−92052309−G−A

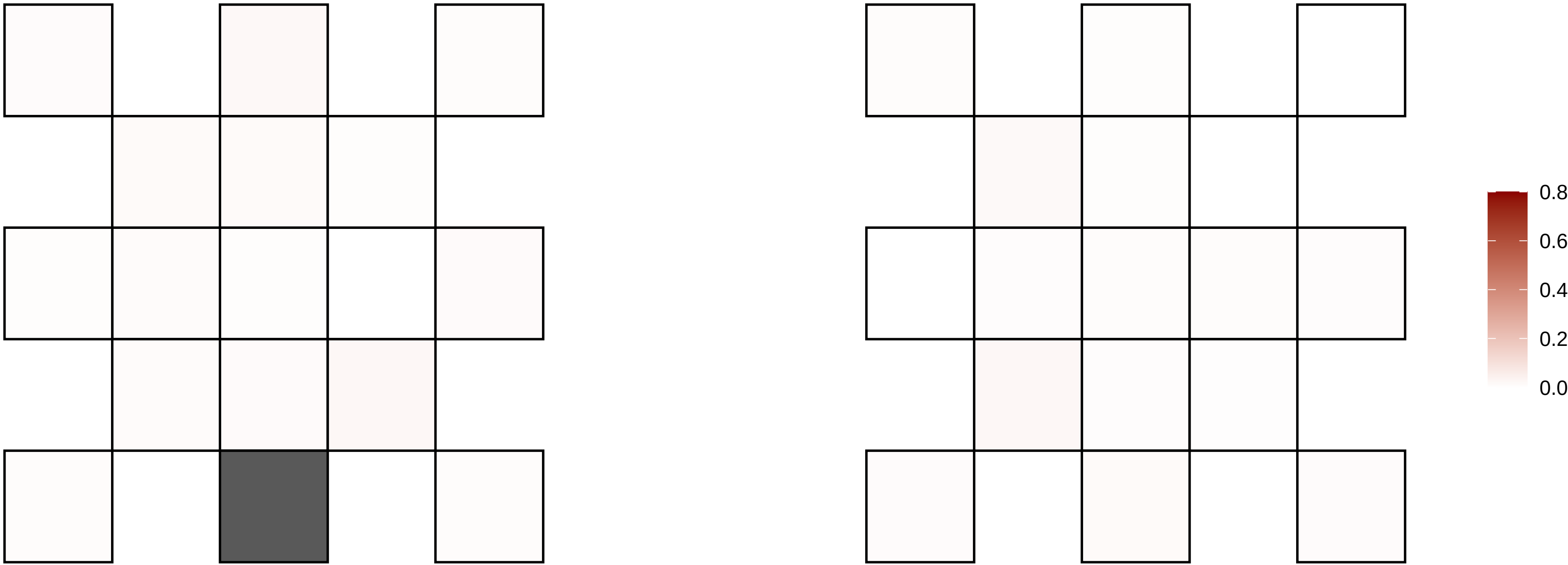

10−98832334−C−T

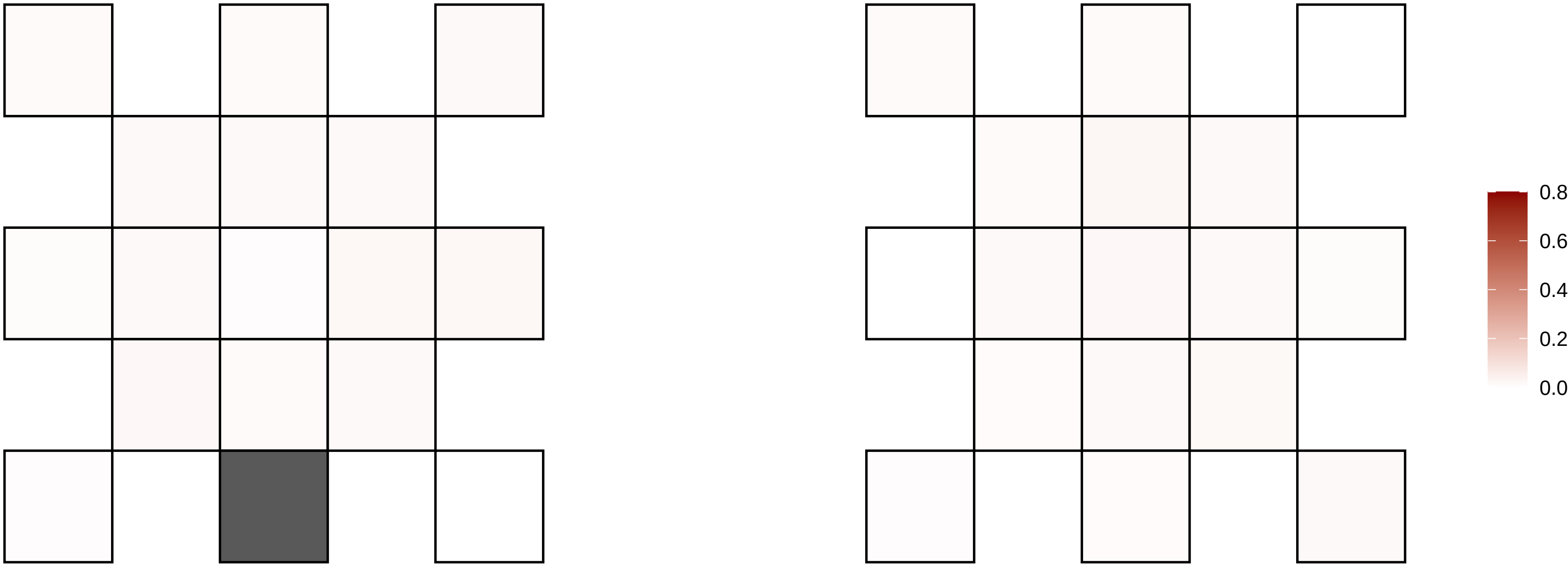

10−98832335−C−A

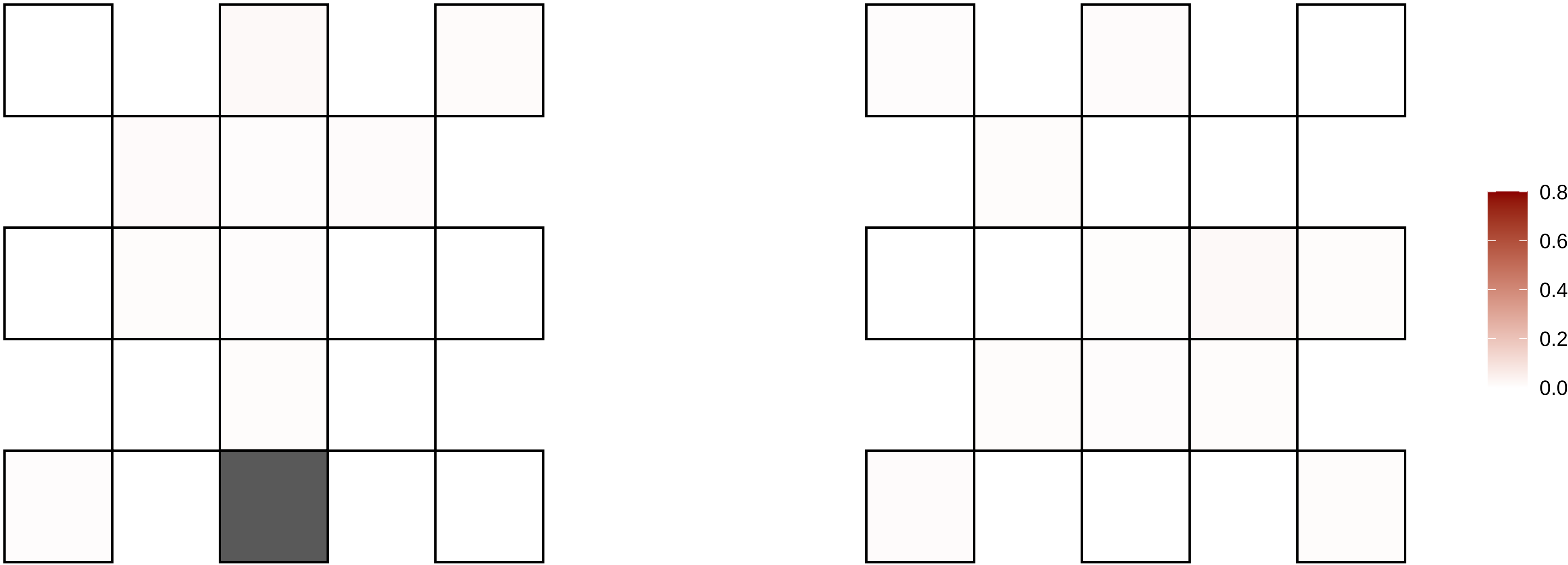

11−105769654−T−C

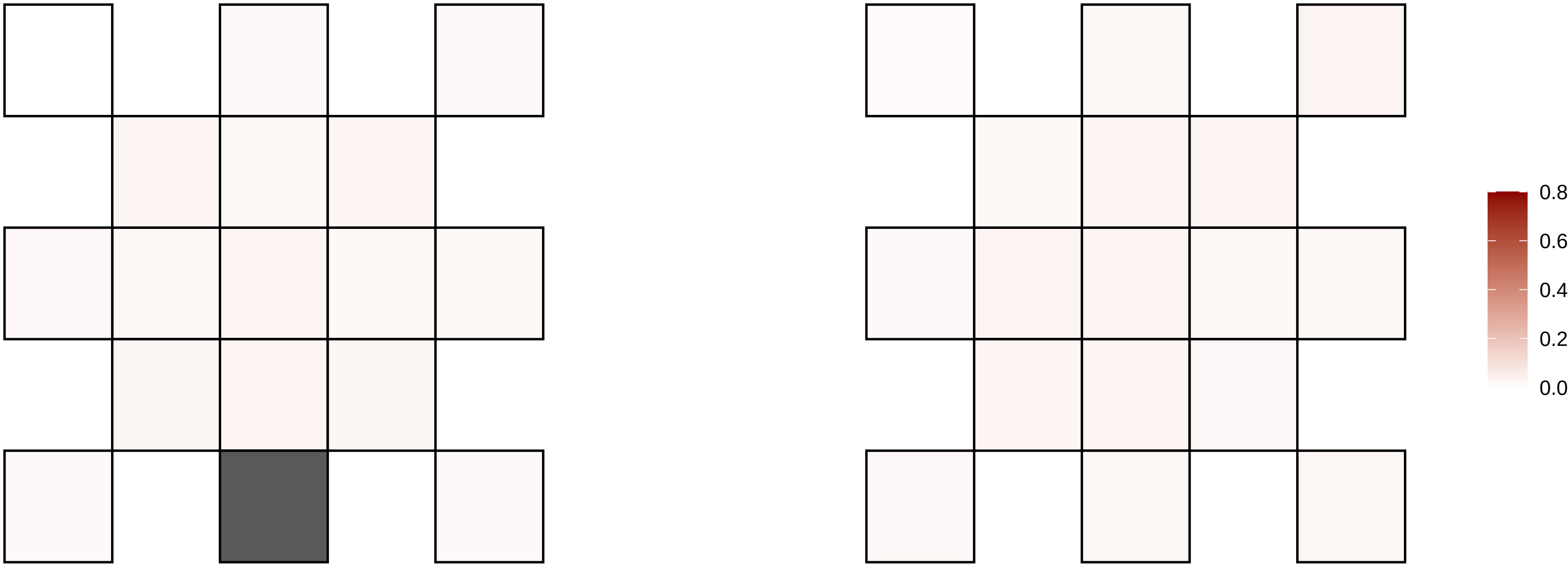

11−109546001−G−C

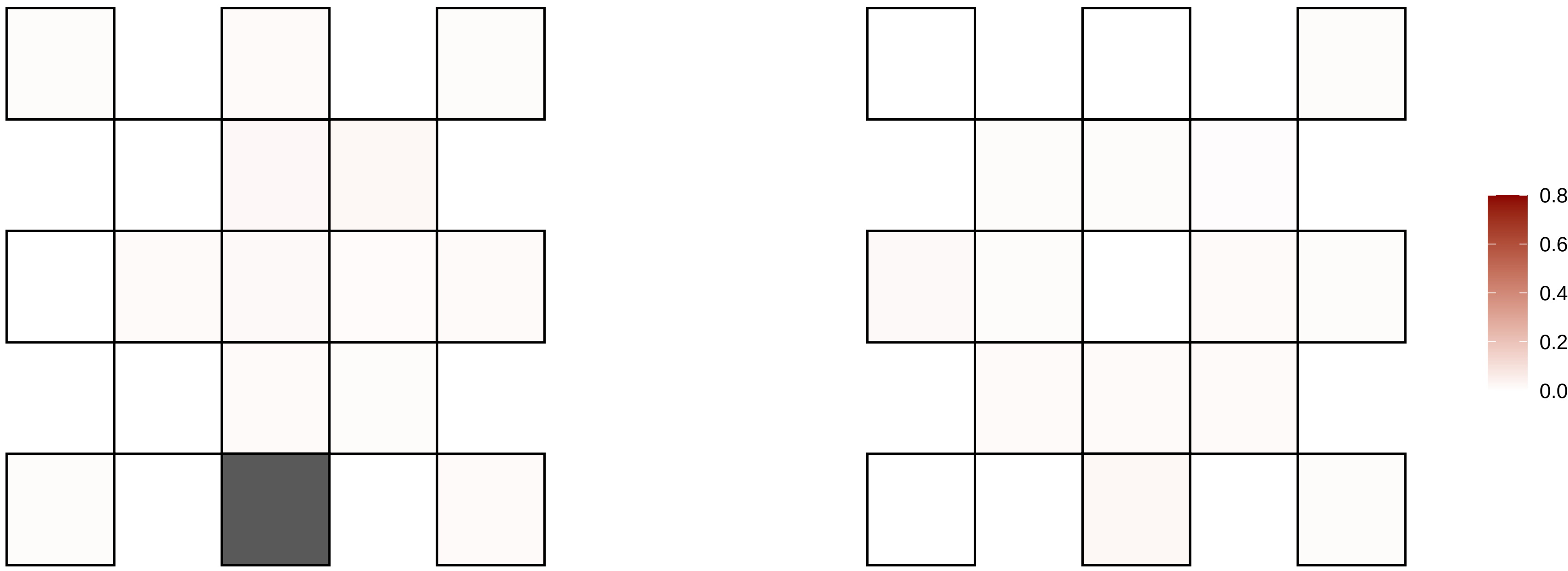

11−111580997−G−A

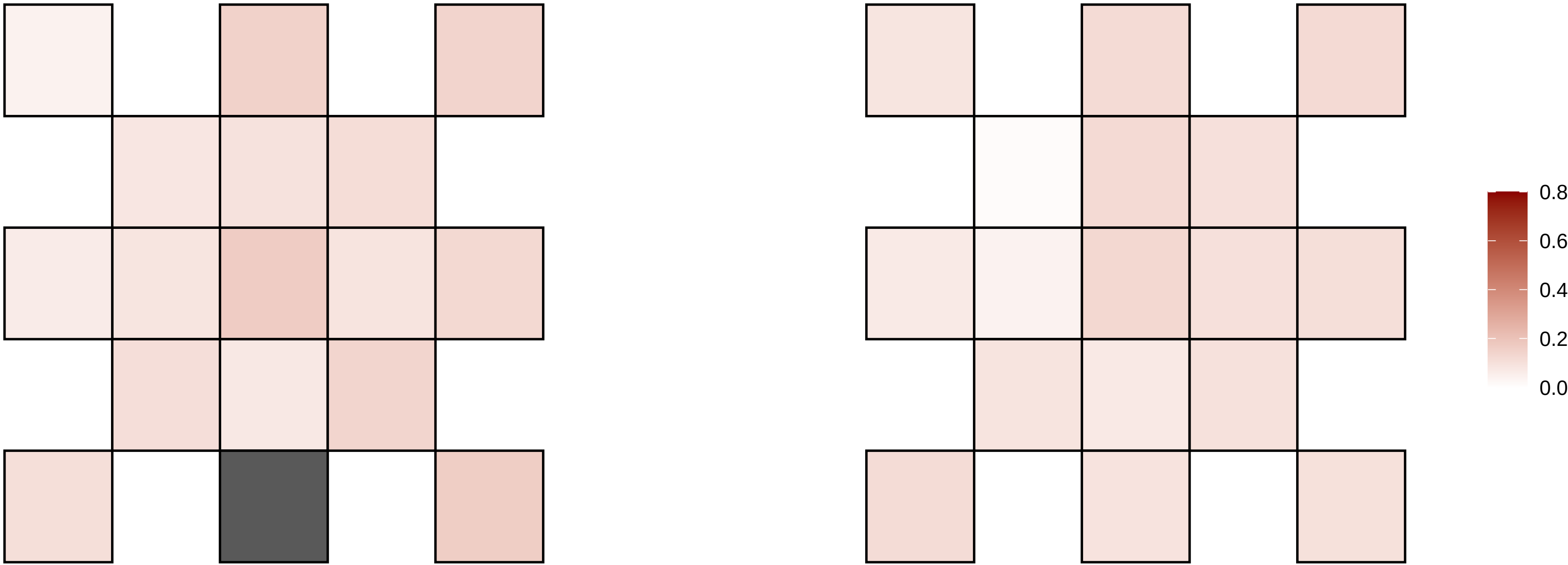

11−112581116−G−A

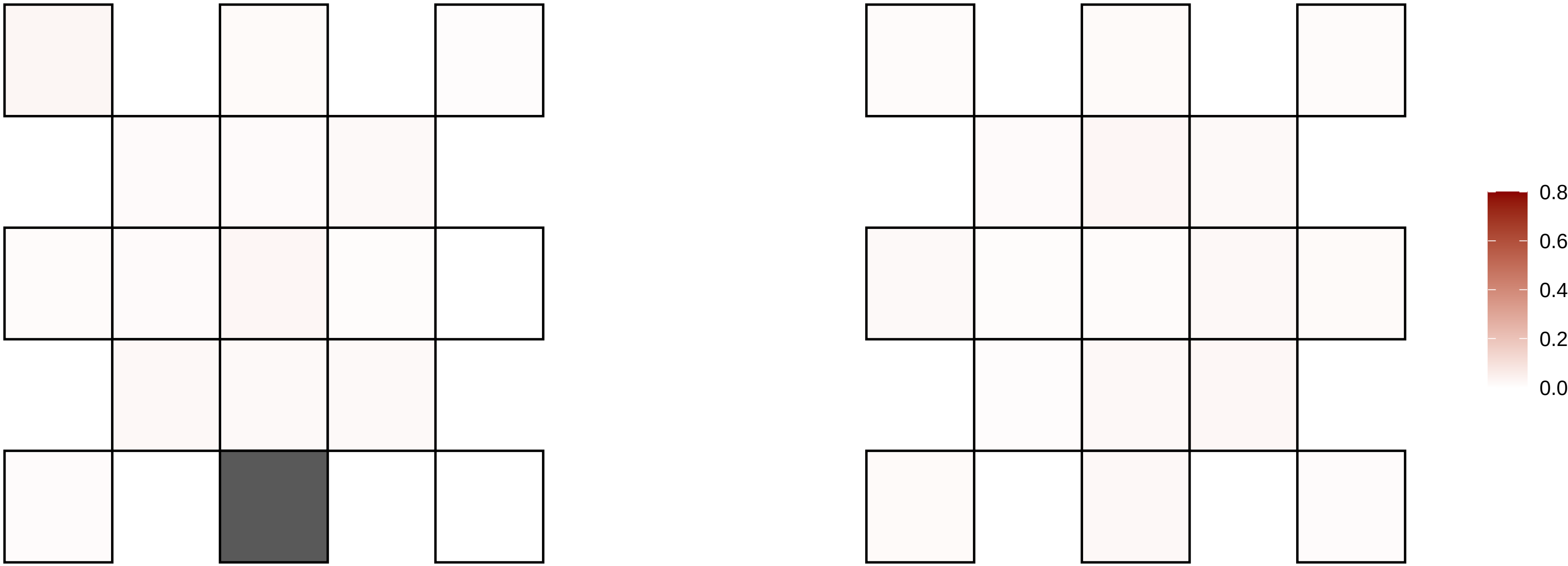

11−112824453−C−T

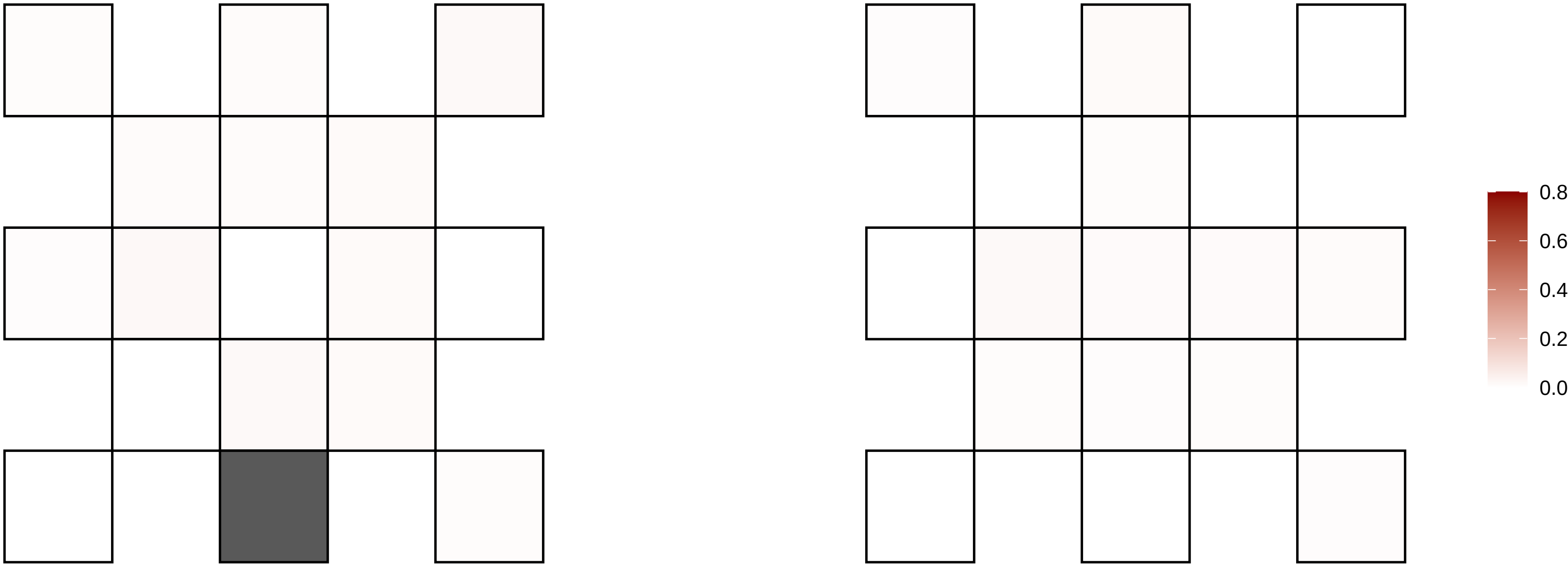

11−11681099−G−C

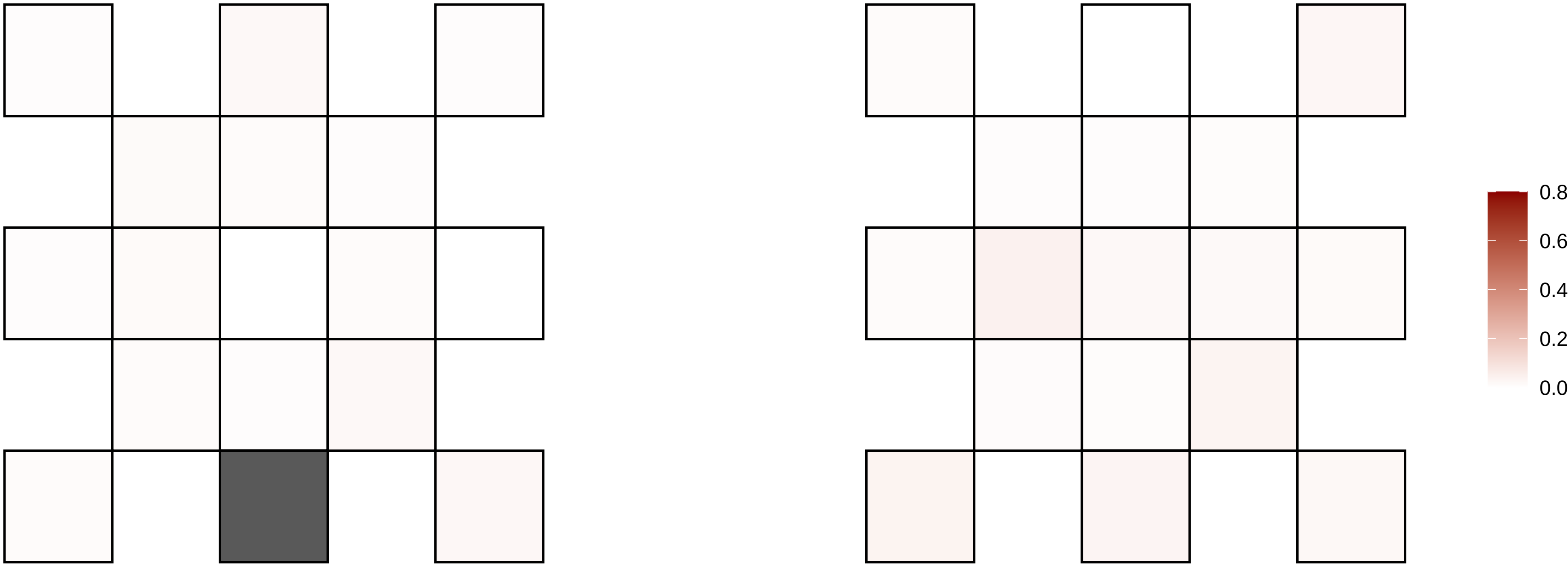

11−123817821−G−A

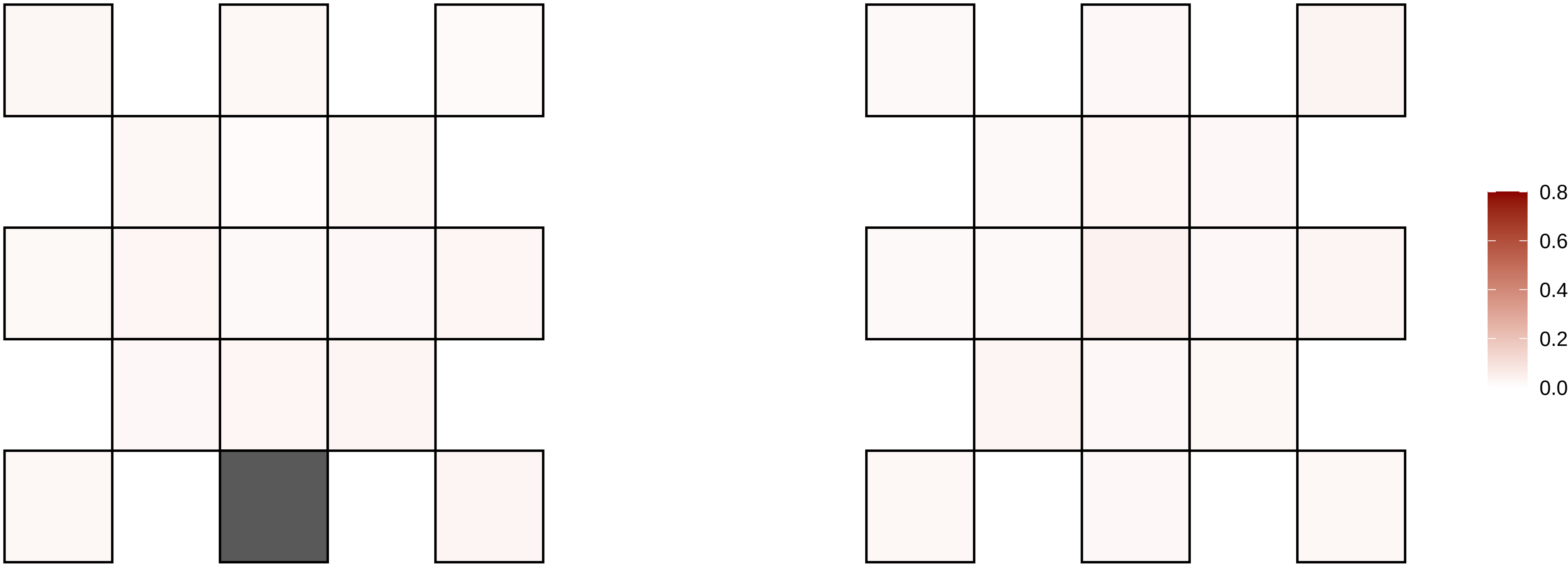

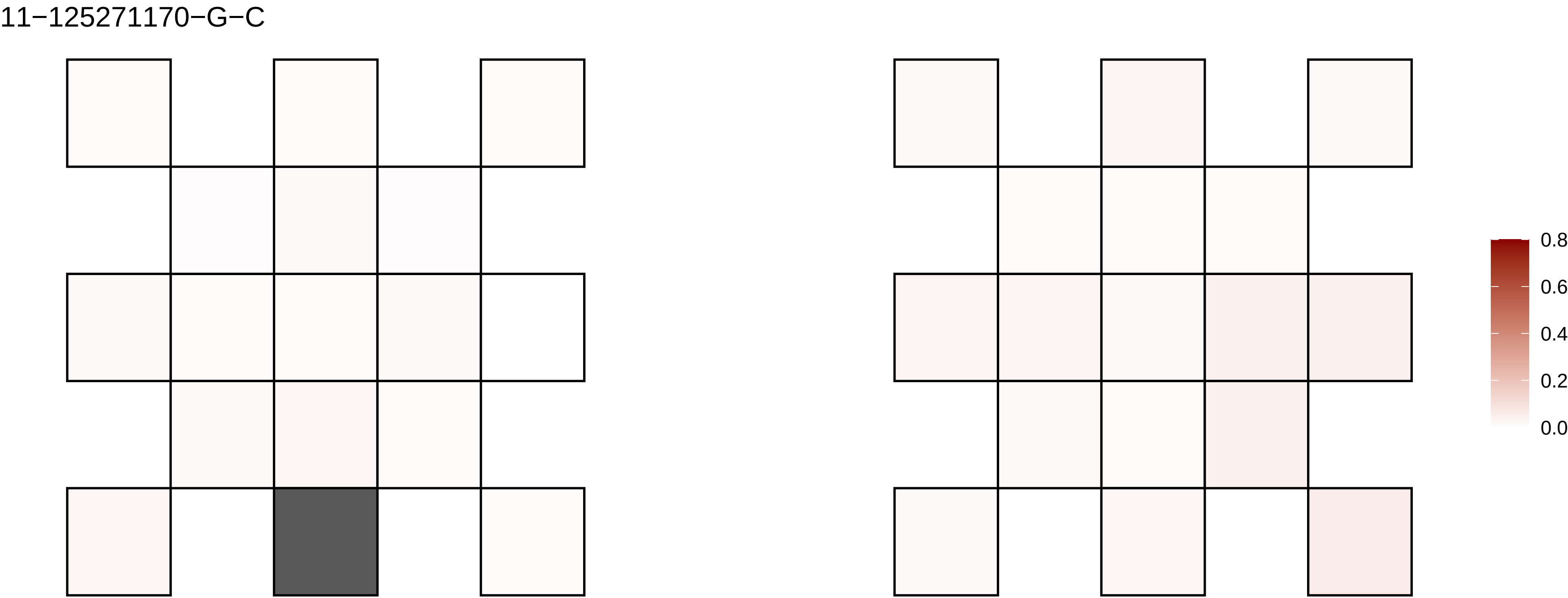
11−125271170−G−C
0.8
0.6
0.4
0.2
0.0

11−127350815−C−T

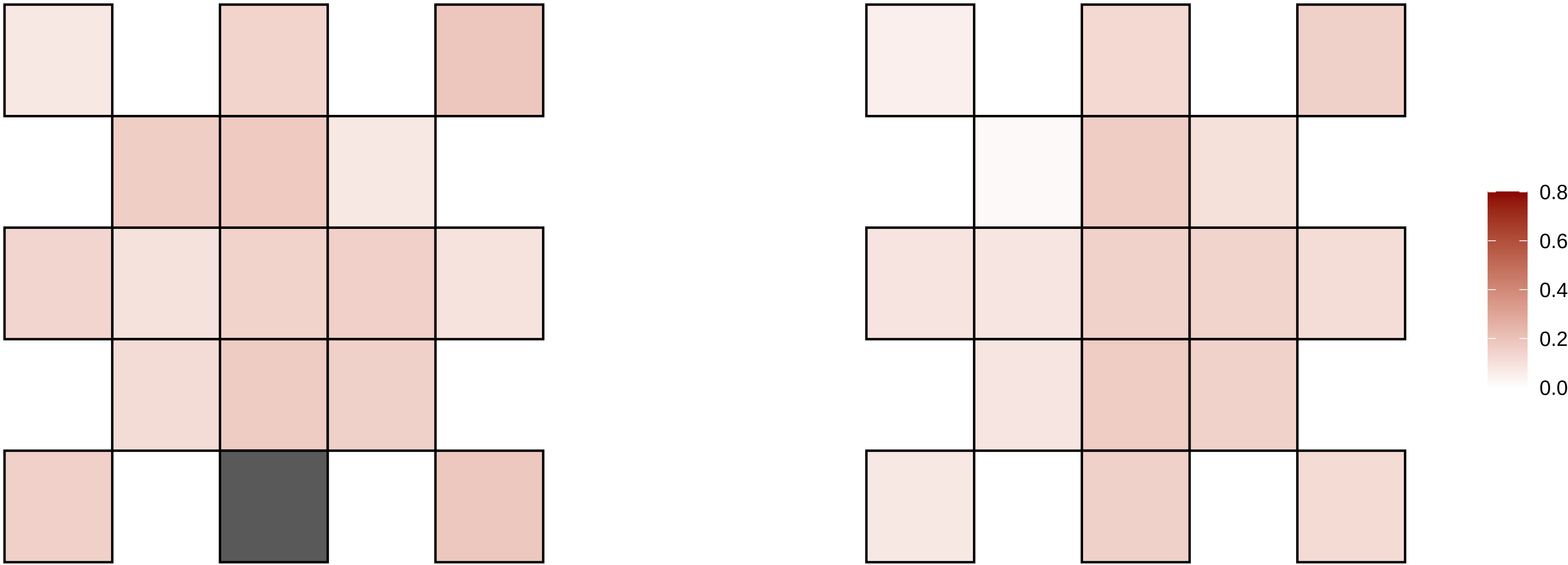

11−133014880−G−C

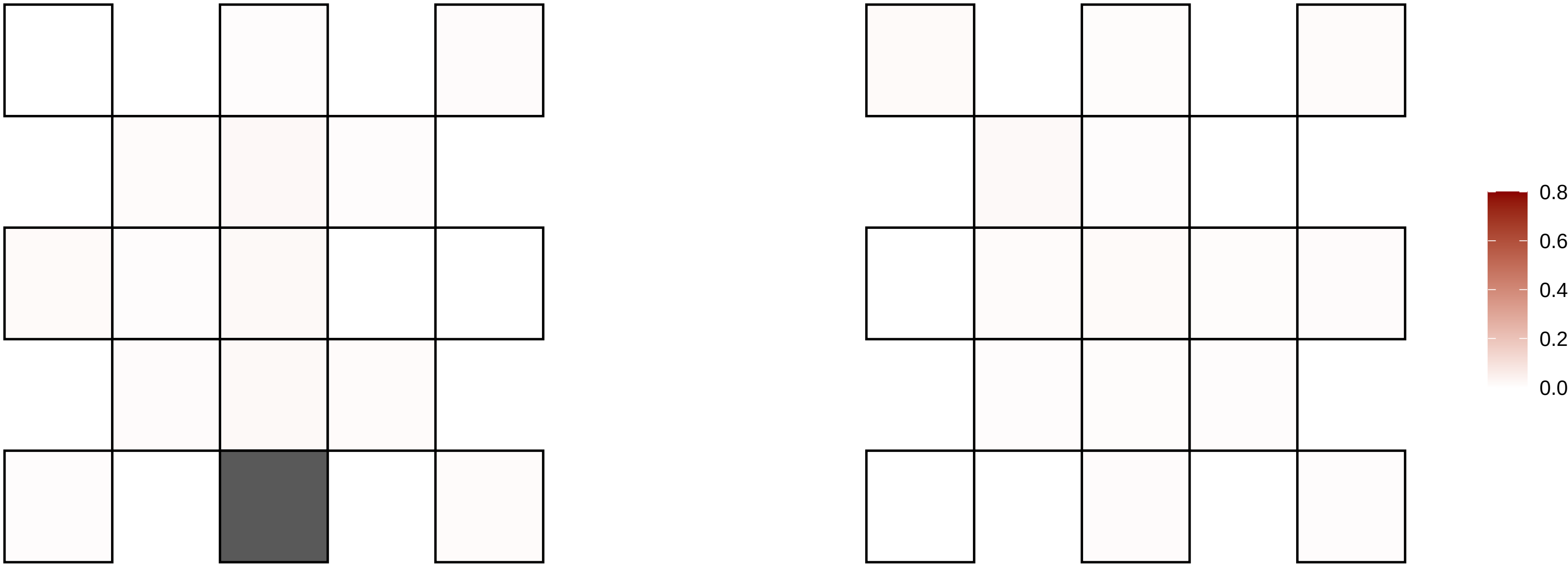

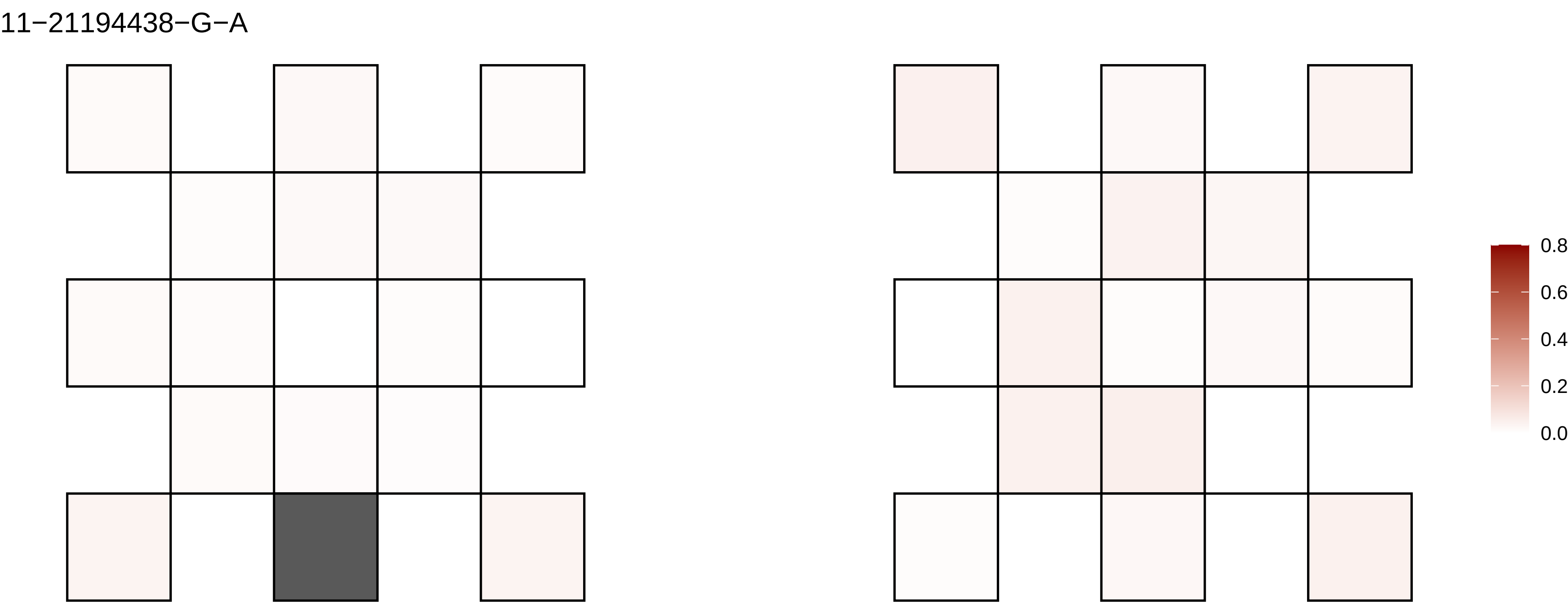
11−21194438−G−A
0.8
0.6
0.4
0.2
0.0

11−22211258−C−T

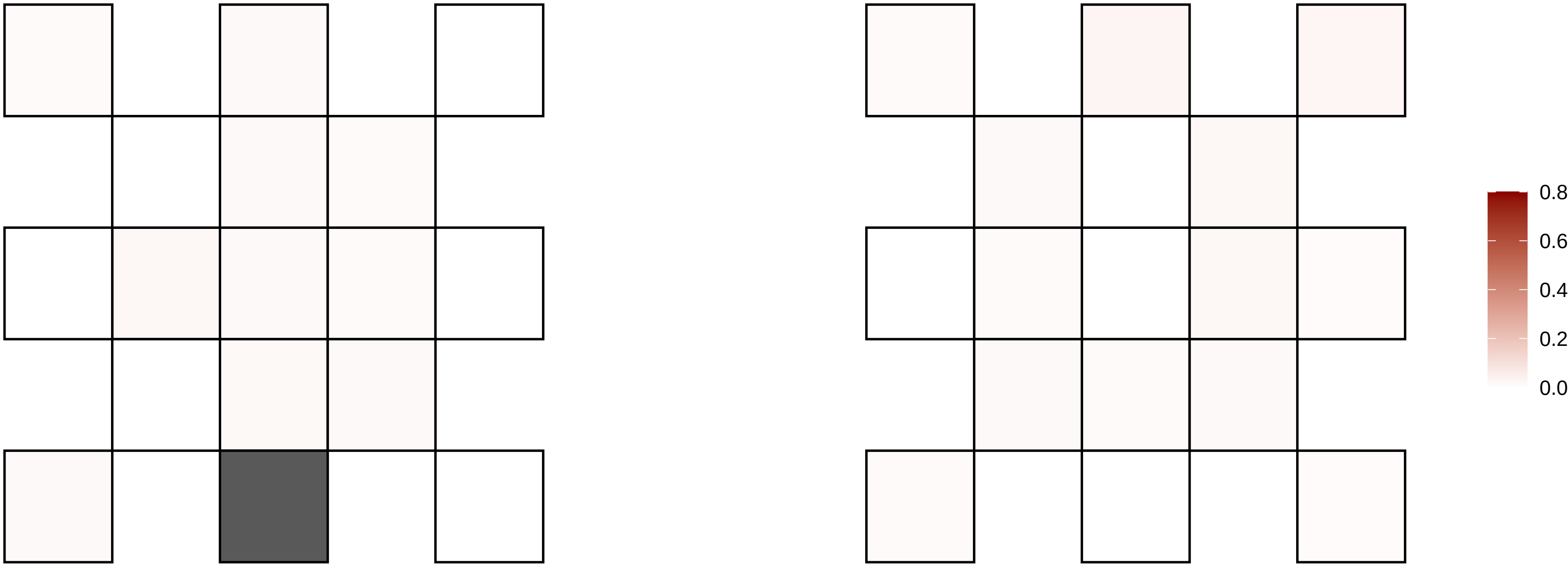

11−25958939−C−T

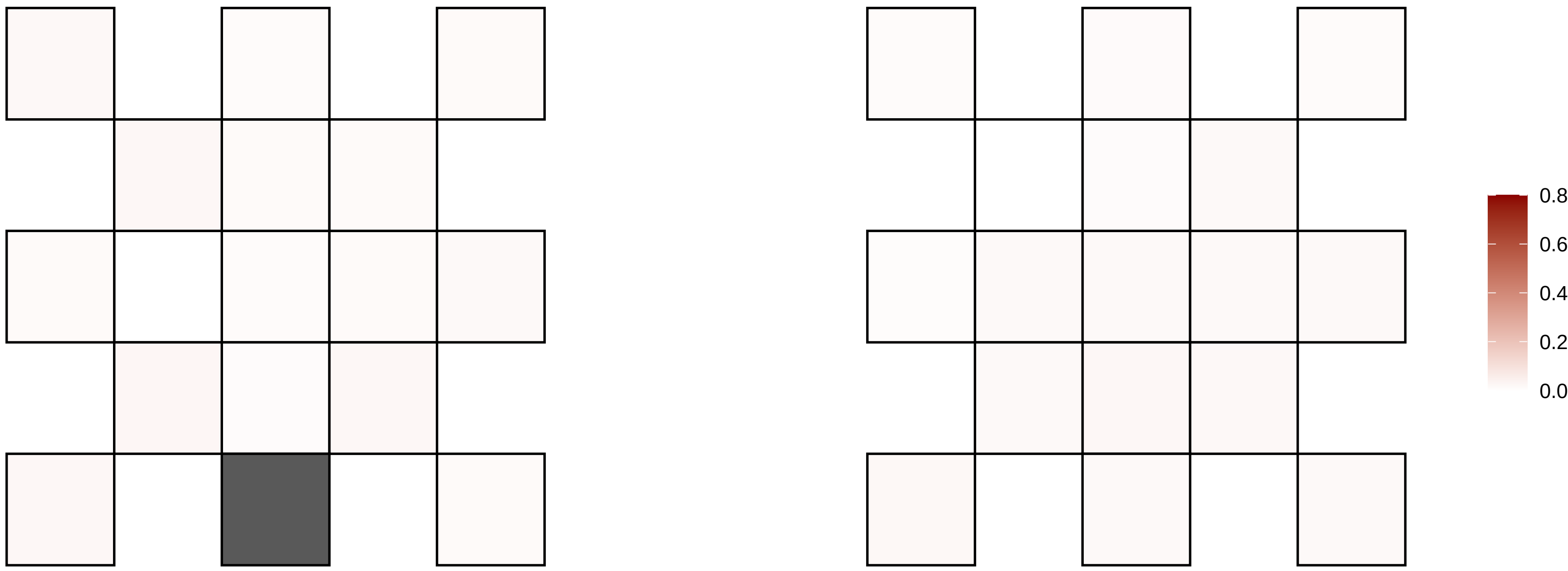

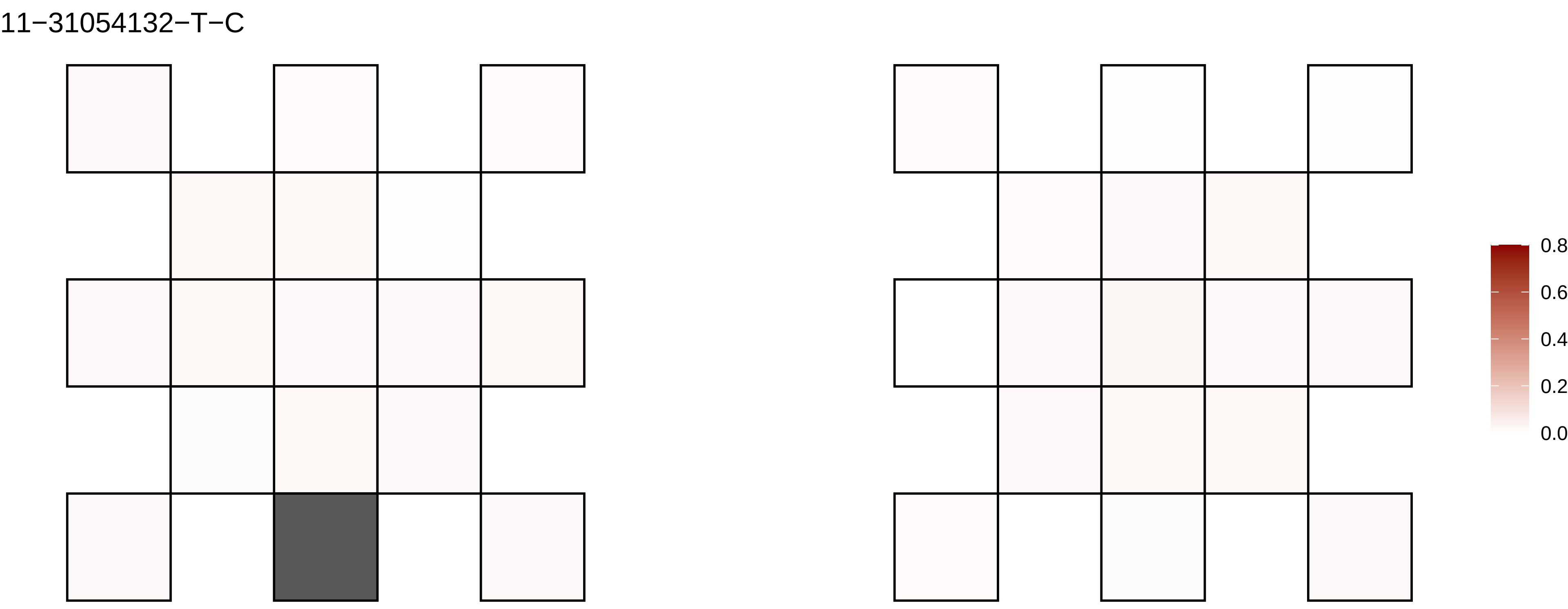
11−31054132−T−C
0.8
0.6
0.4
0.2
0.0

11−32349753−T−A

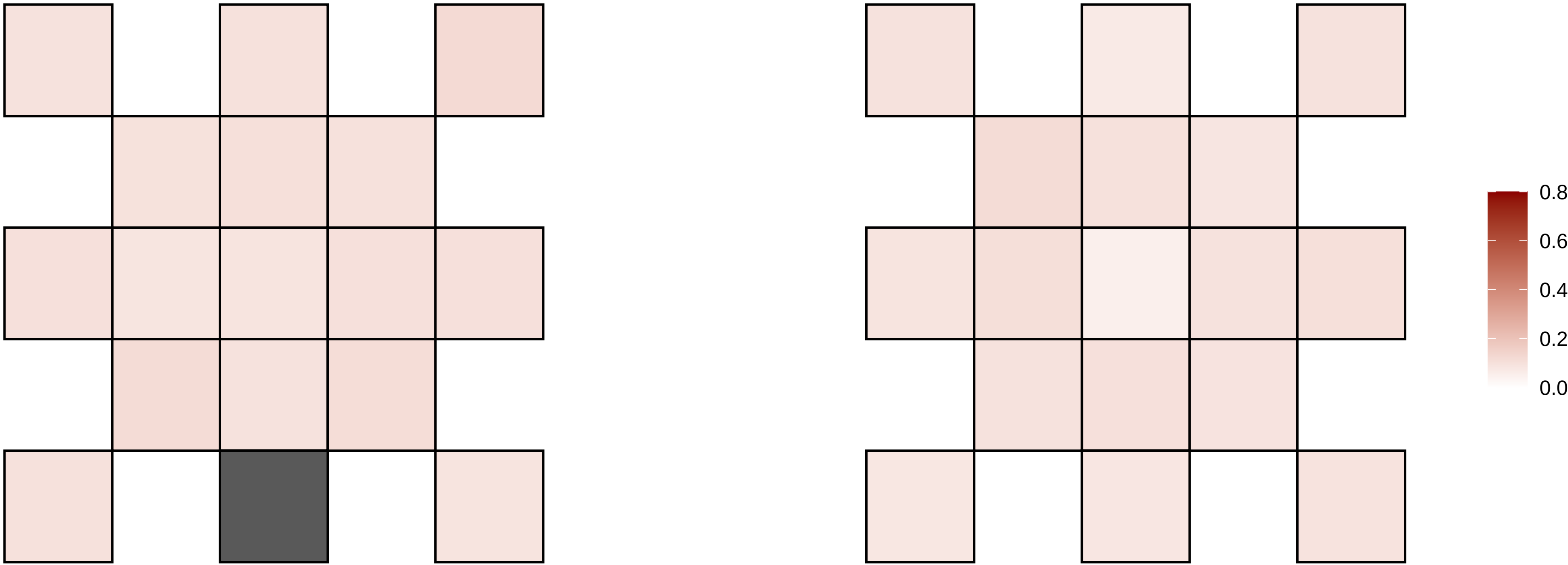

11−32349754−T−A

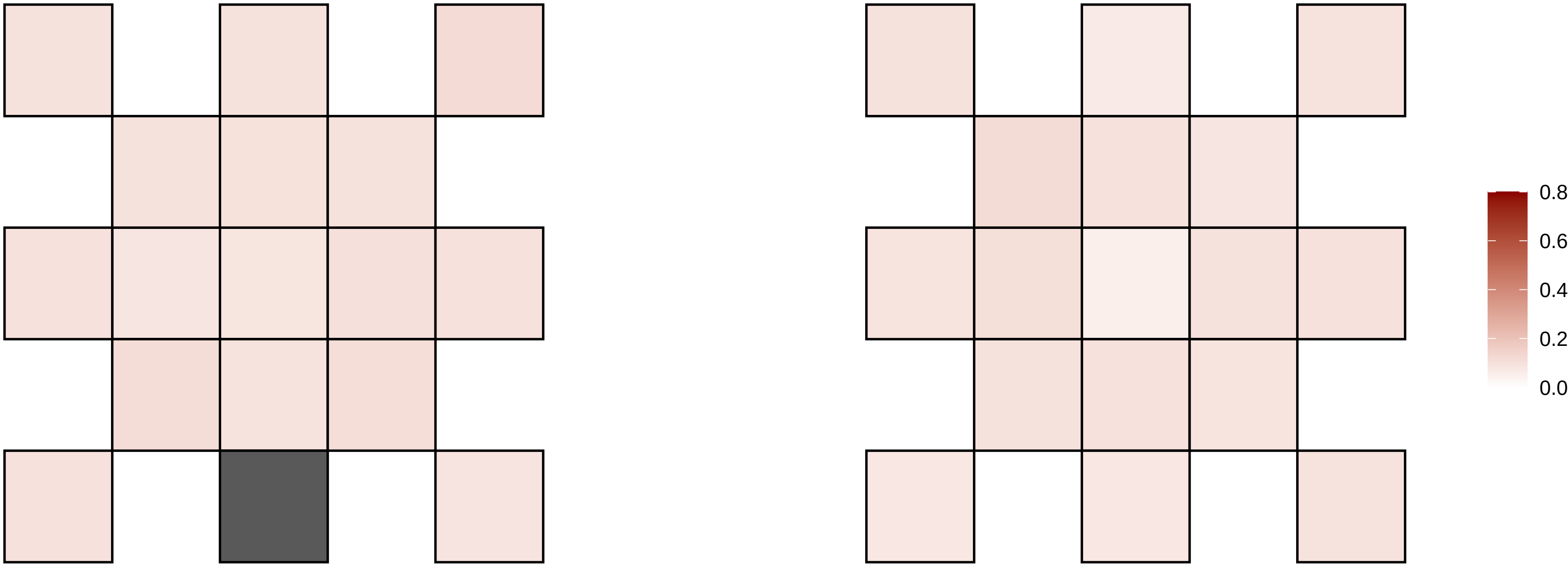

11−35512752−C−T

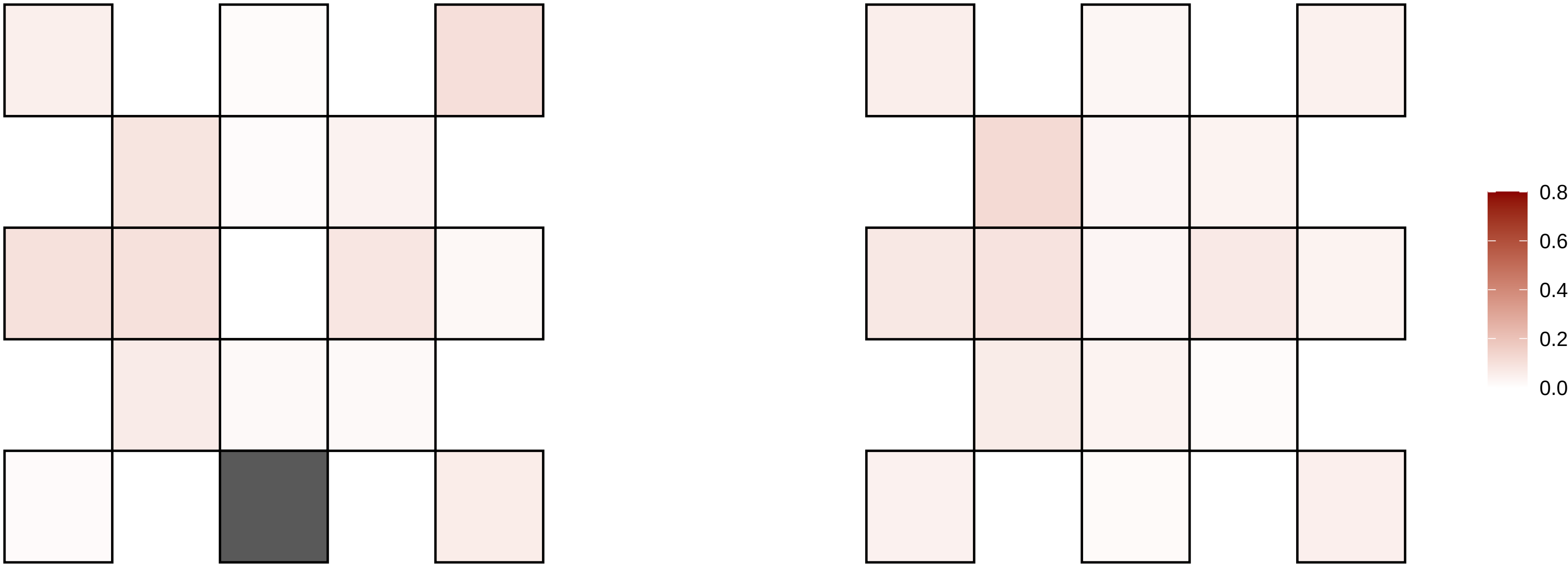

11−44221191−C−G

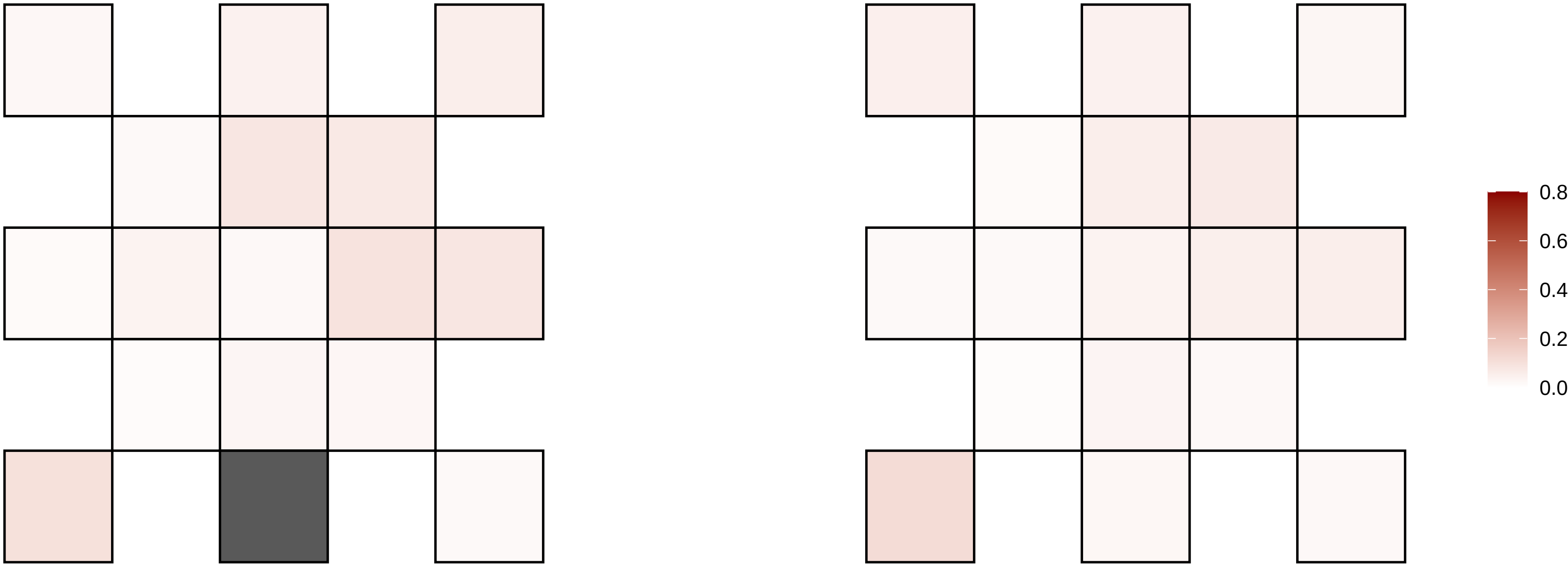

11−45873419−G−A

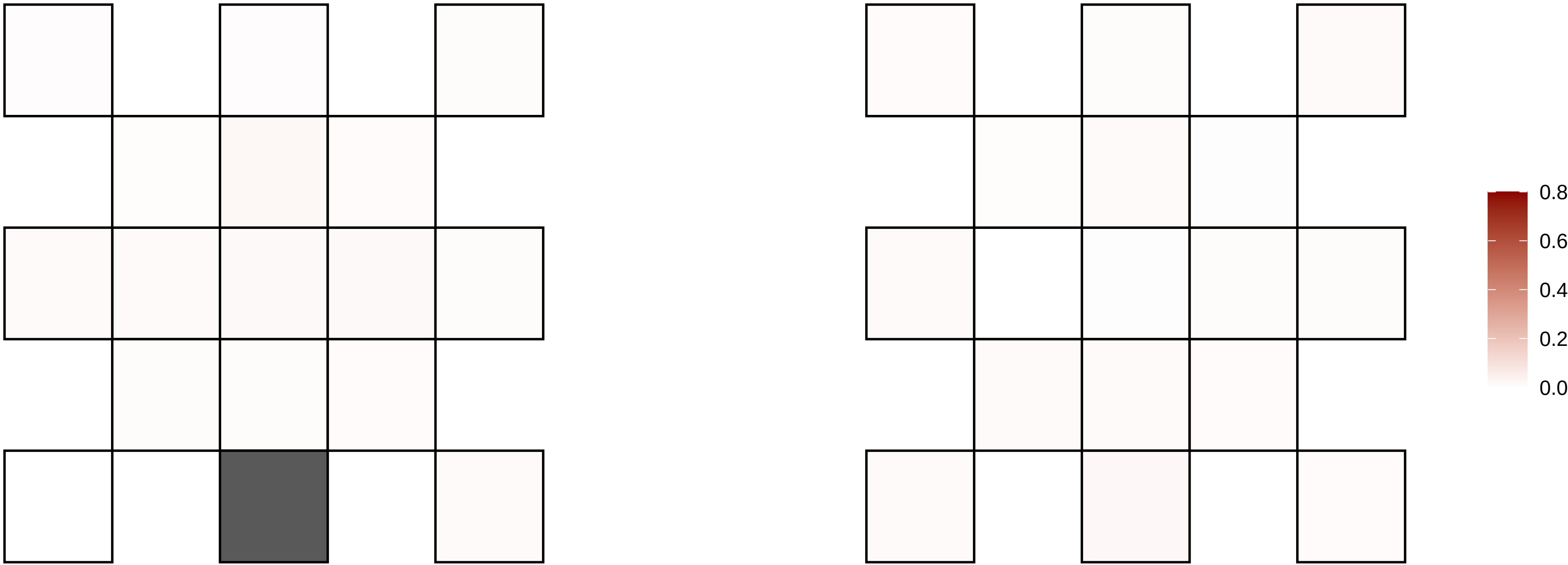

11−55838876−G−T

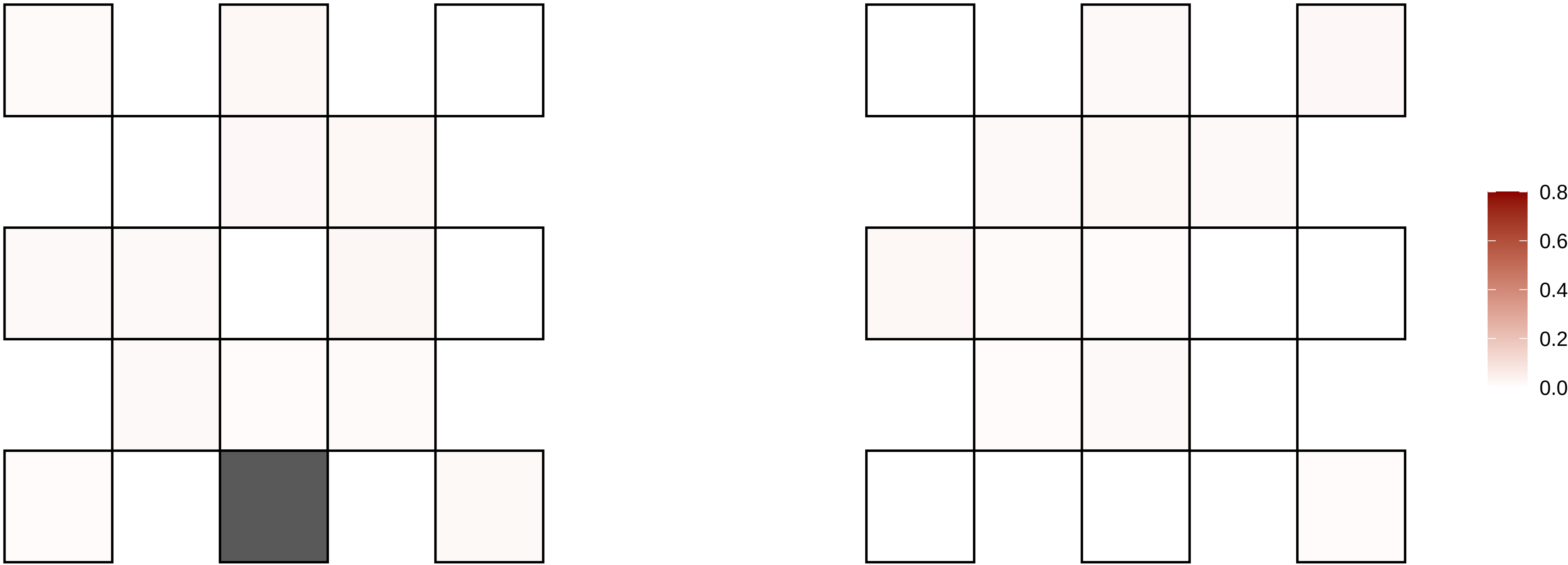

11−55997990−C−T

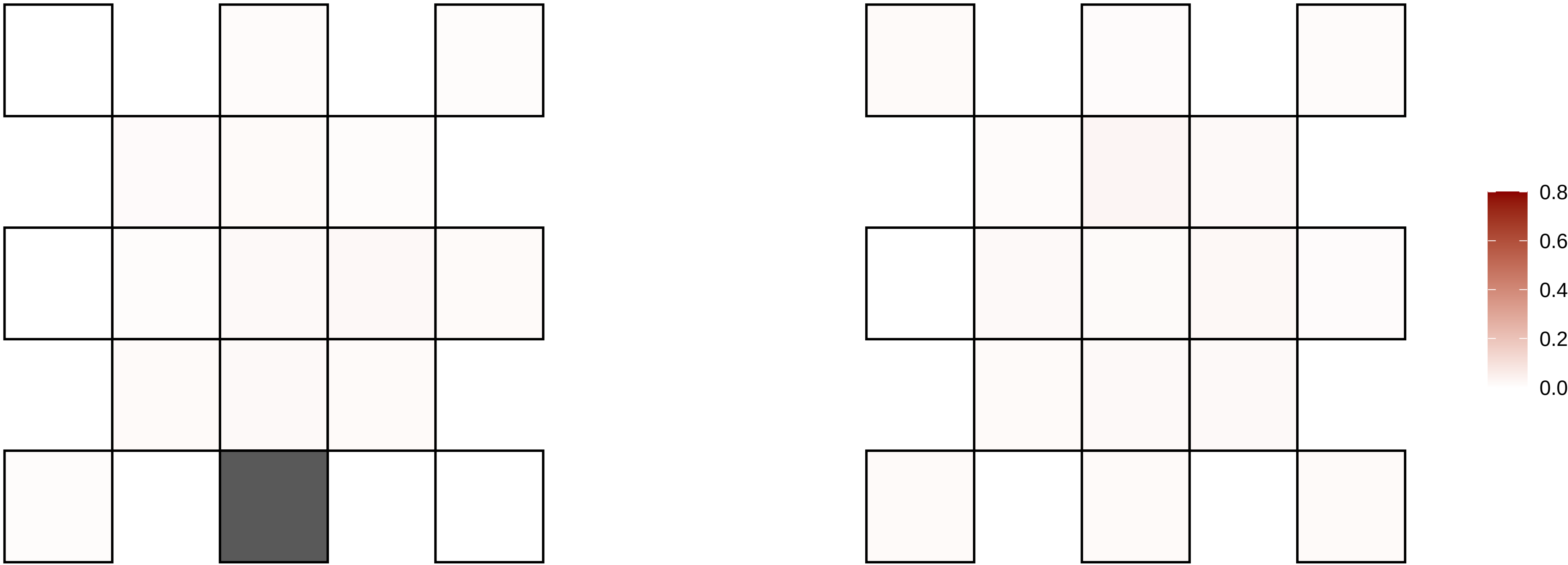

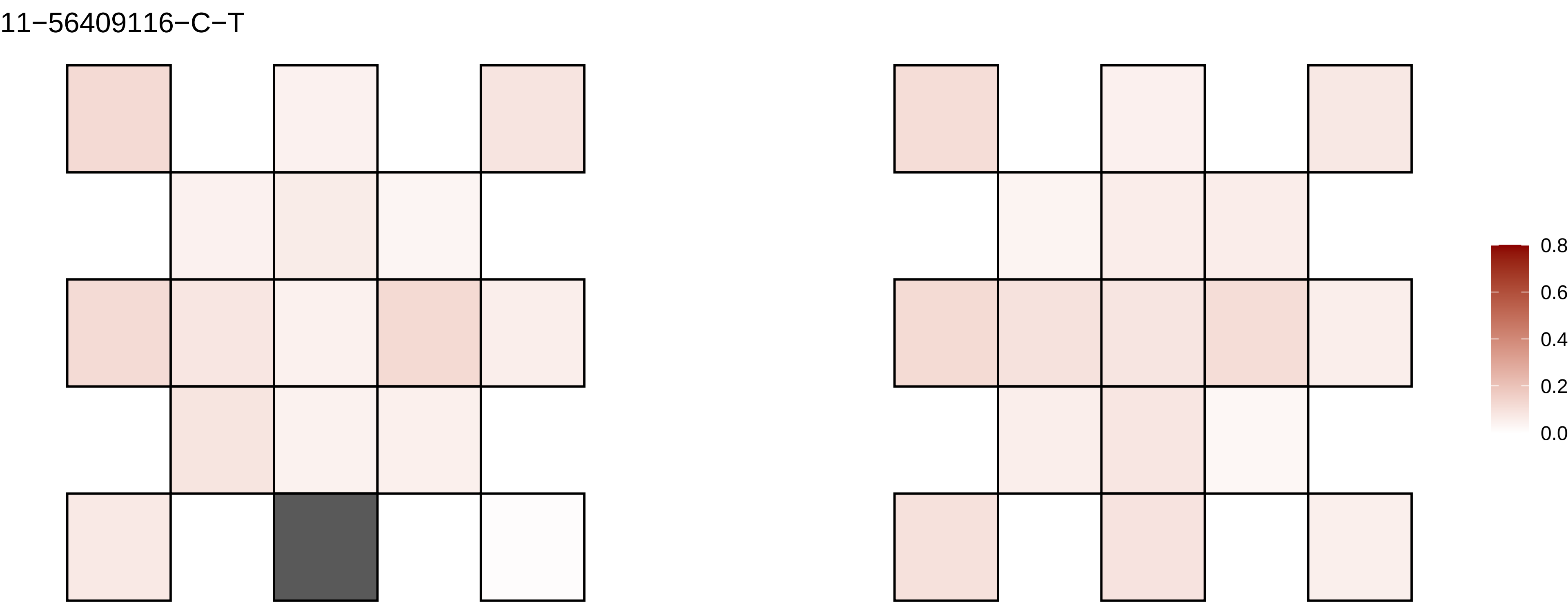
11−56409116−C−T
0.8
0.6
0.4
0.2
0.0

11−57985375−T−C

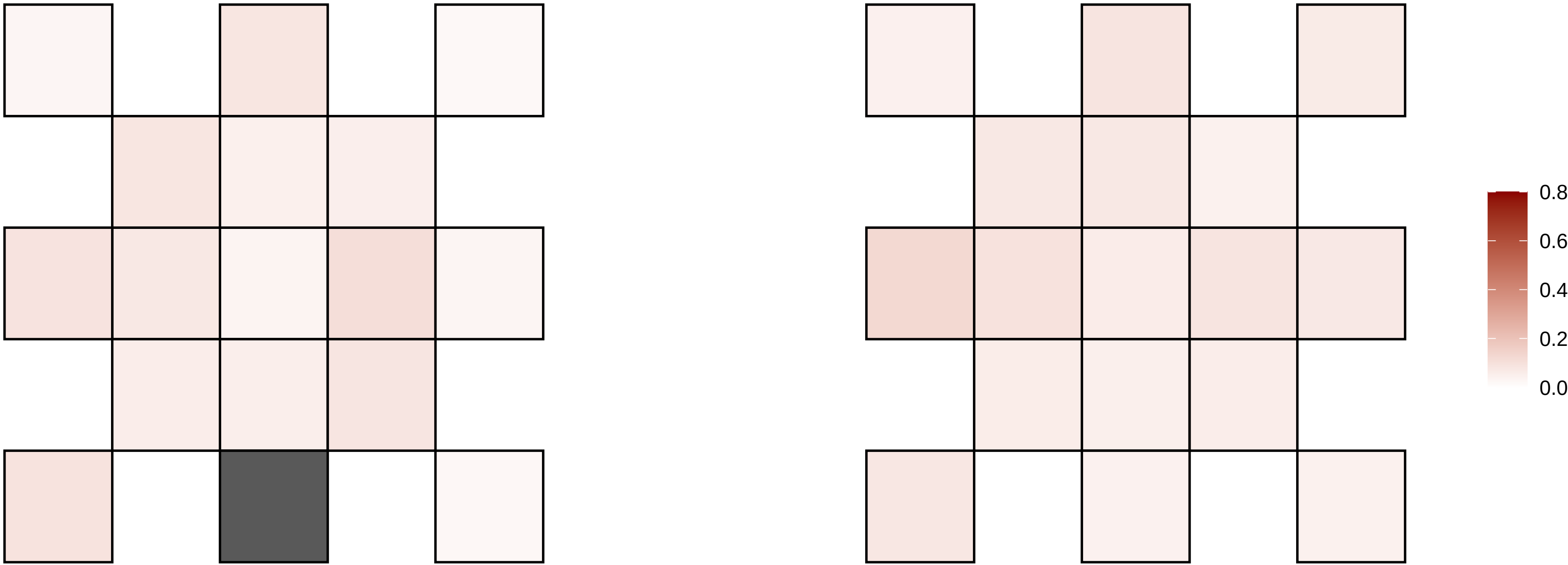

11−79785519−G−T

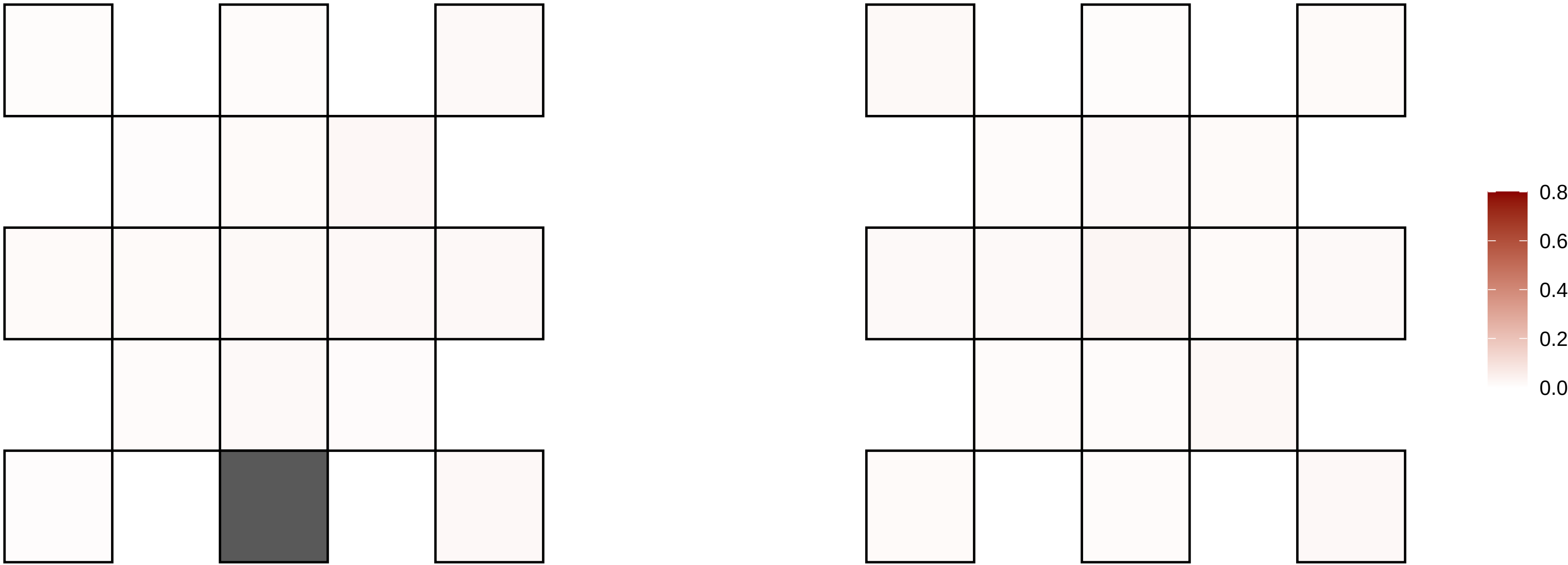

11−8331108−C−T

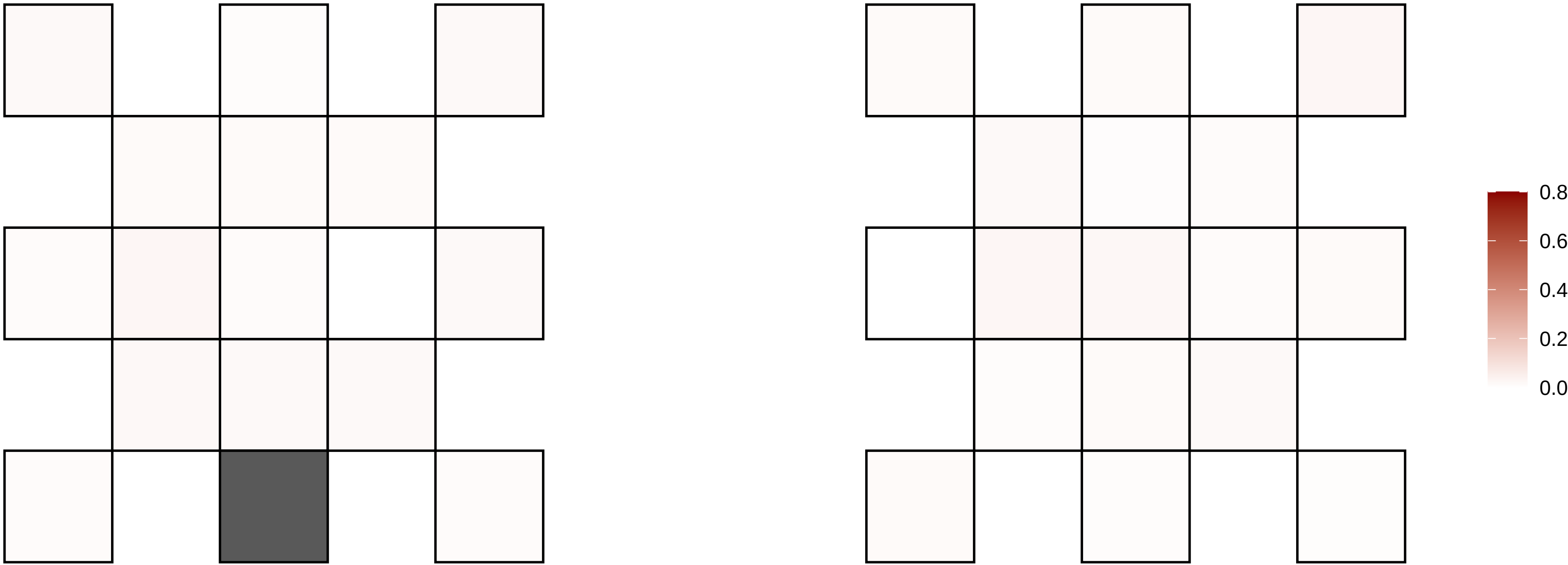

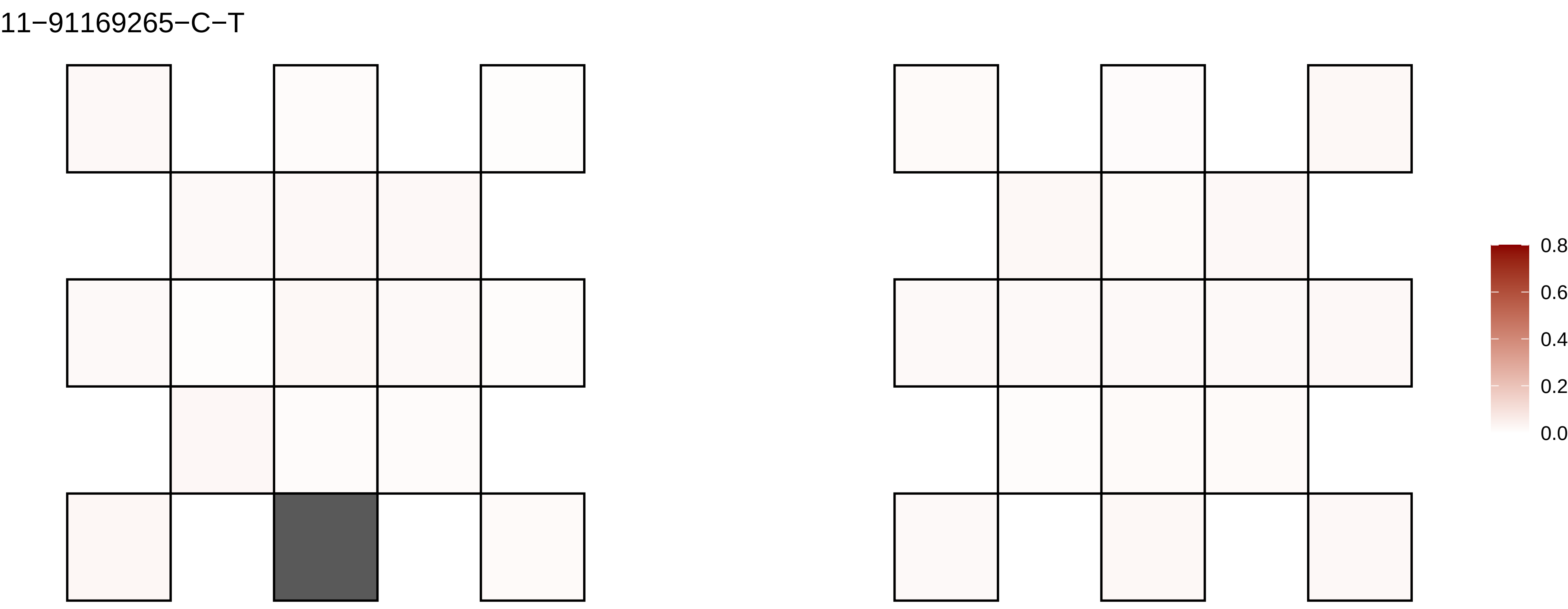
11−91169265−C−T
0.8
0.6
0.4
0.2
0.0

11−93547360−G−A

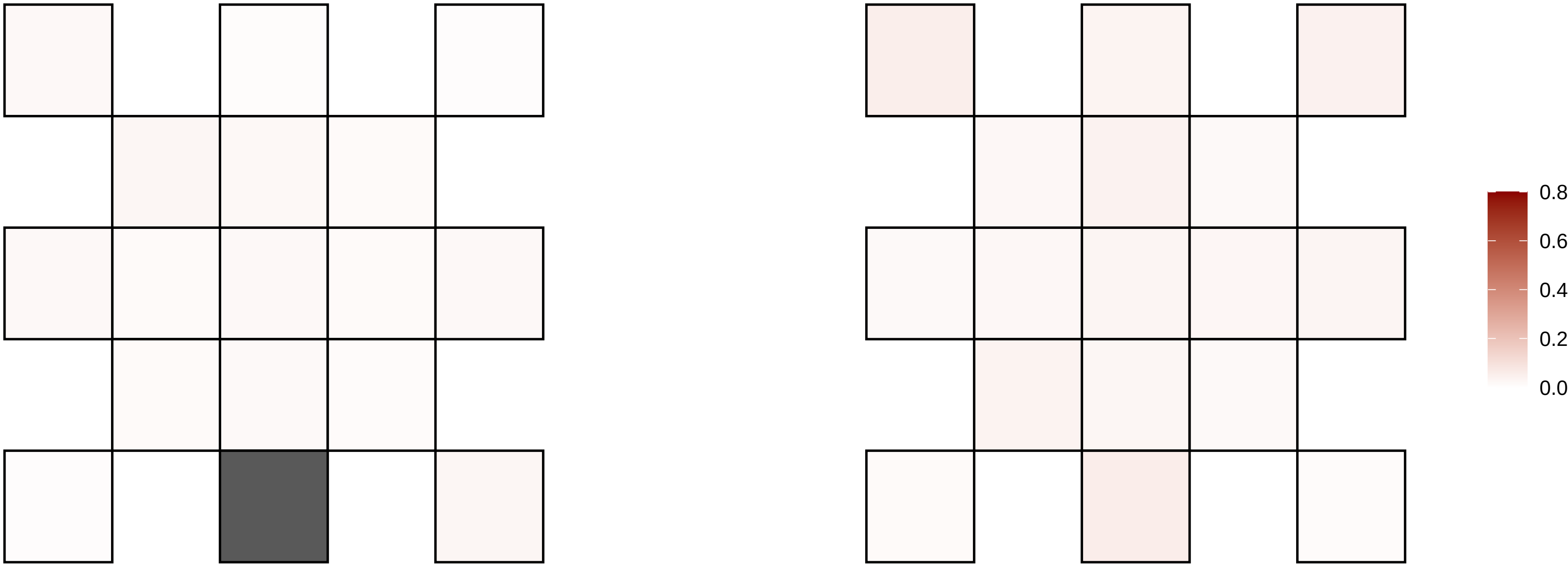

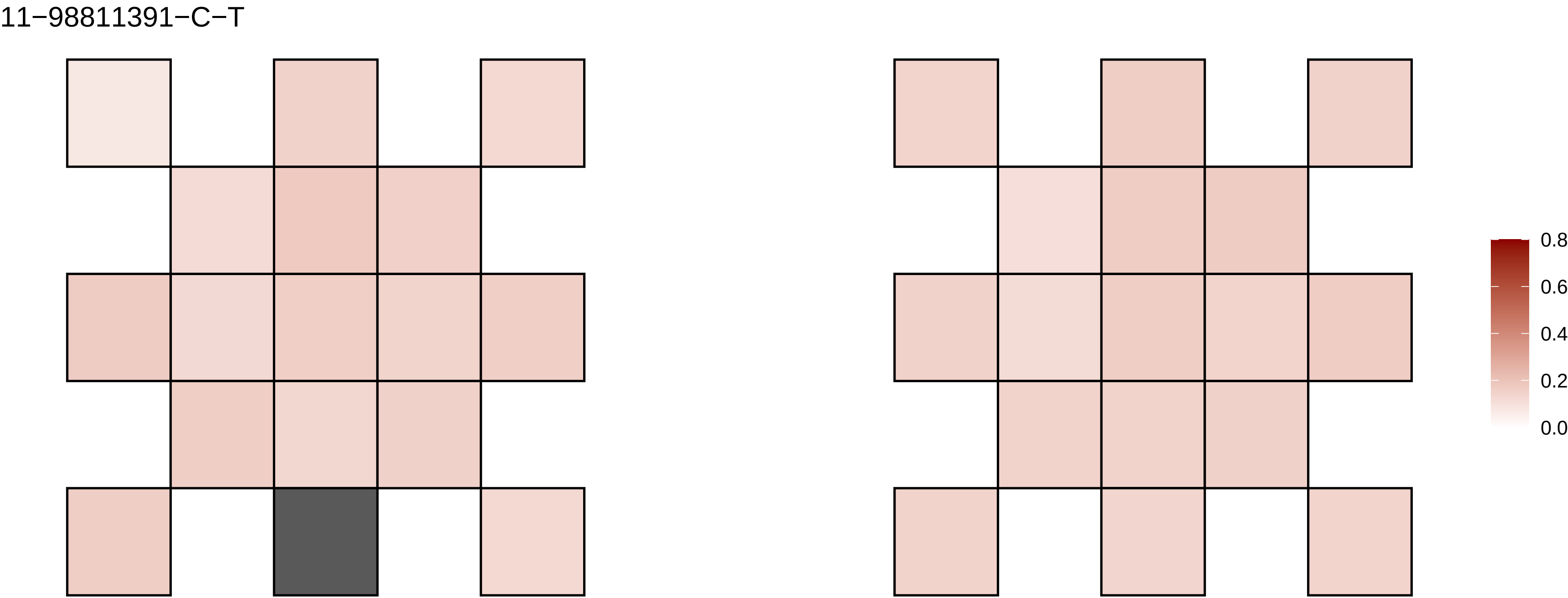
11−98811391−C−T
0.8
0.6
0.4
0.2
0.0

11−99288625−T−A

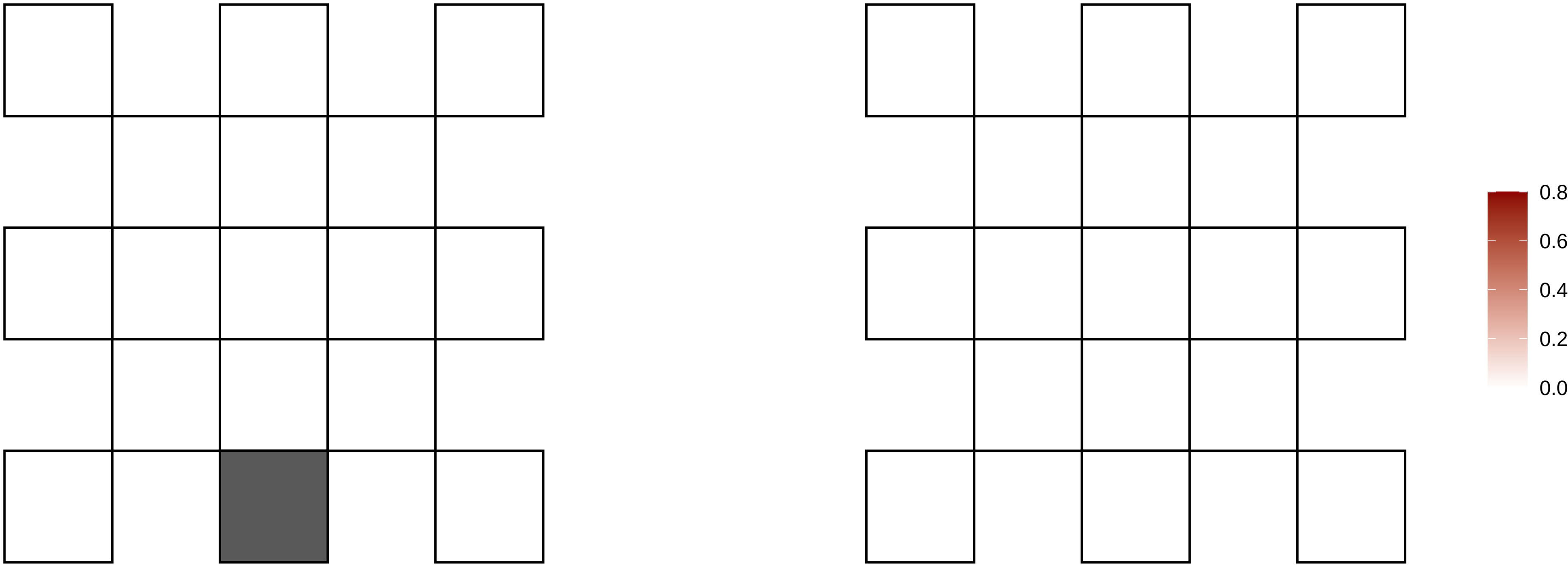

12−100202897−A−G

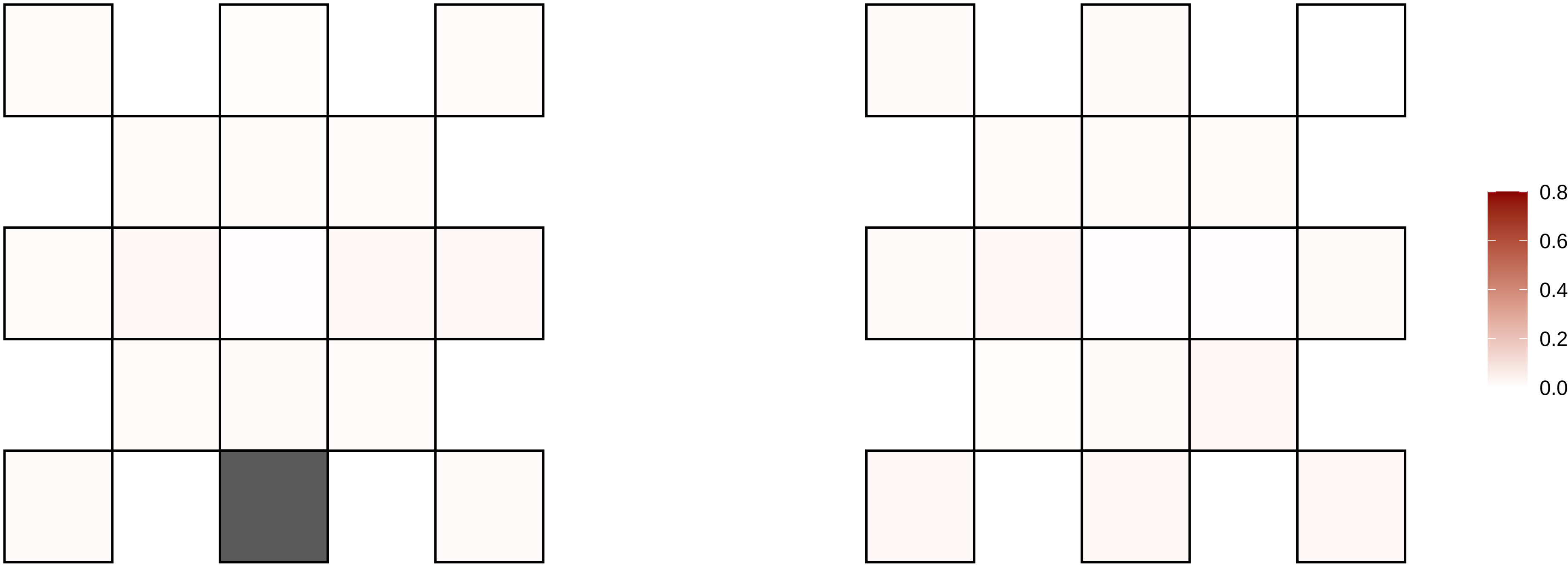

12−110209050−G−A

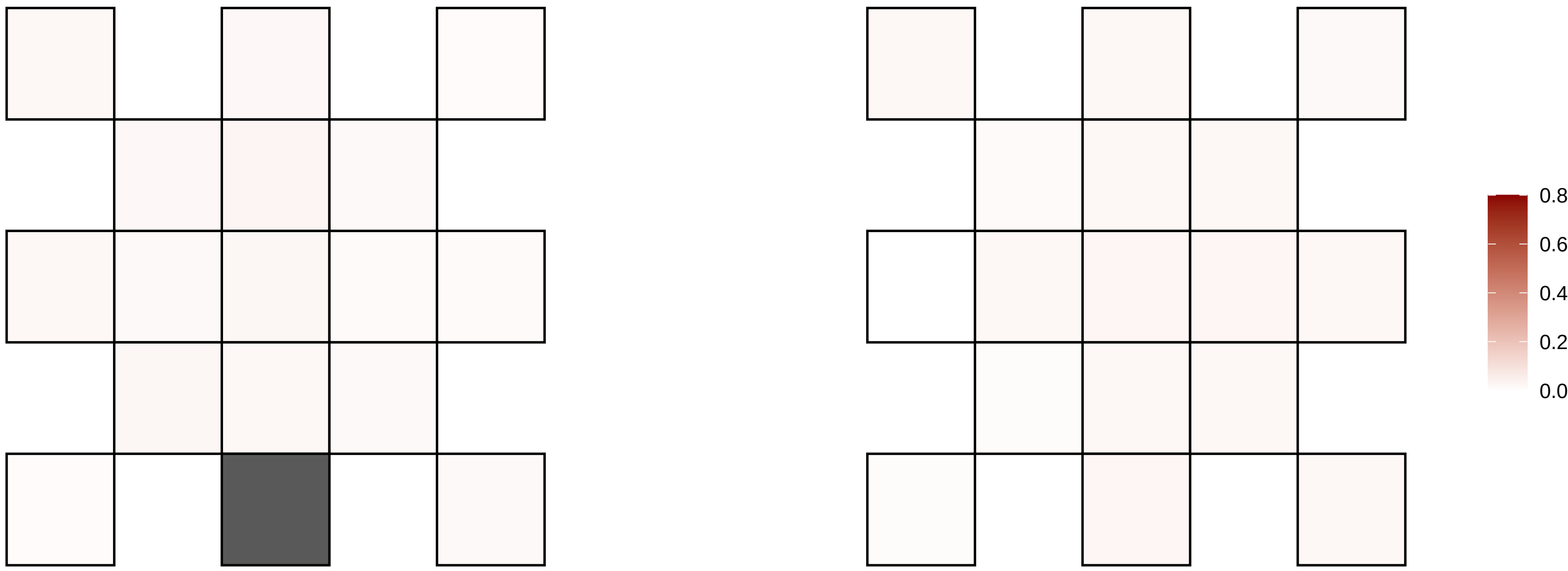

12−112926872−C−T

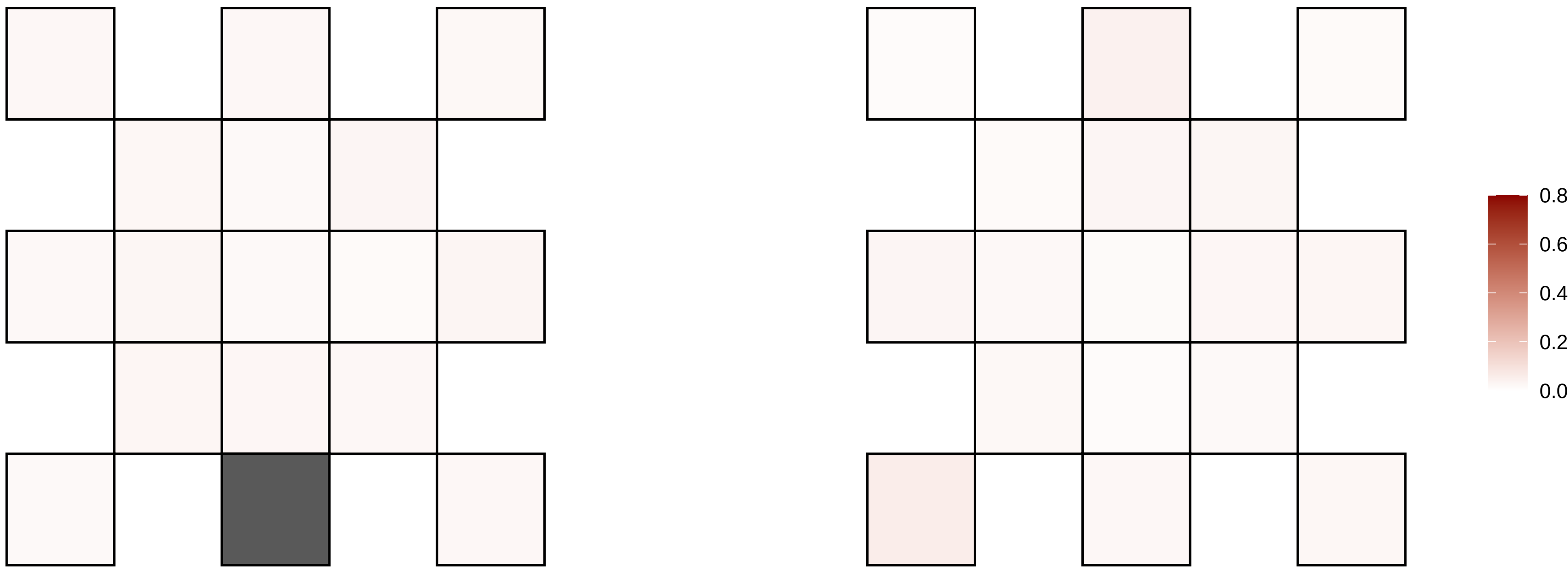

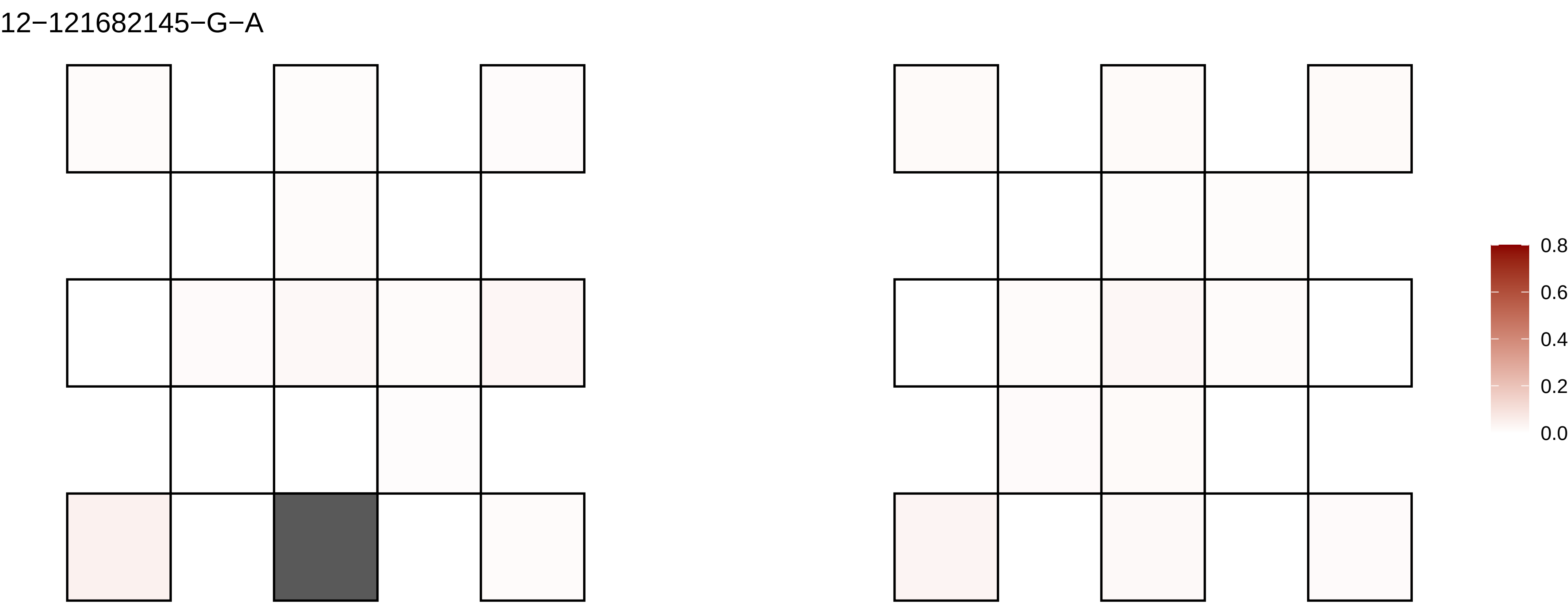
12−121682145−G−A
0.8
0.6
0.4
0.2
0.0

12−121707377−T−C

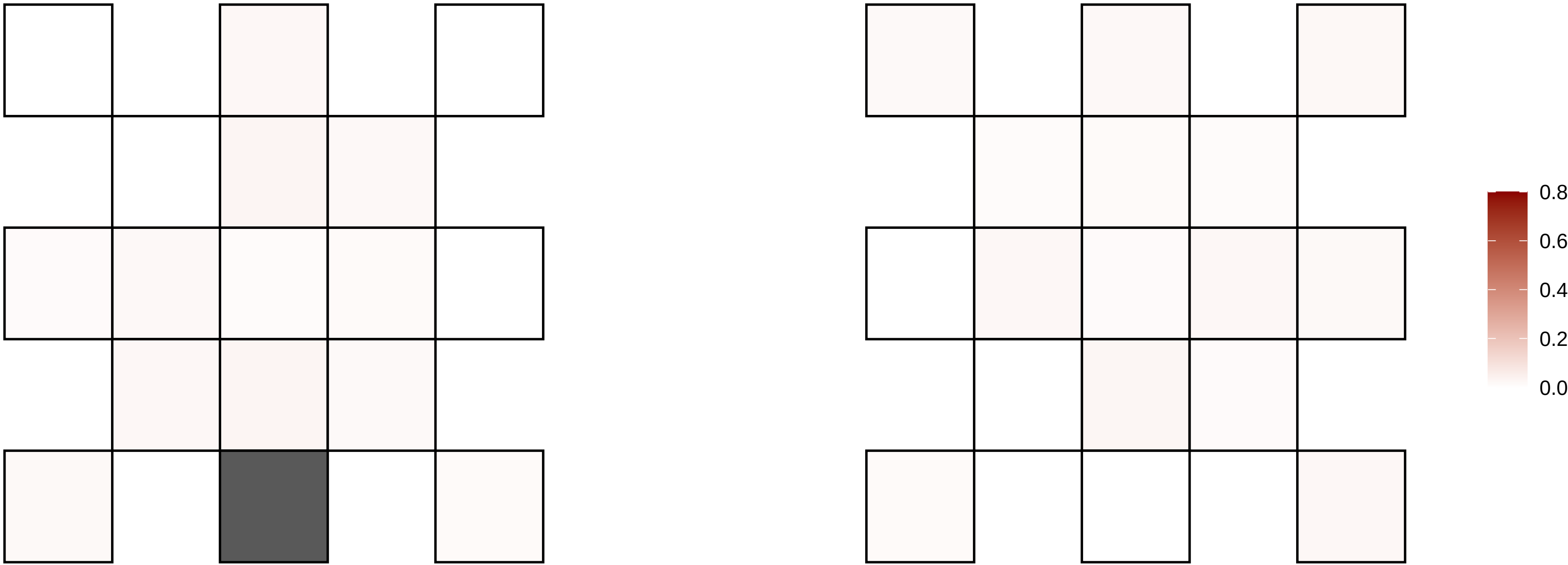

12−127258051−G−T

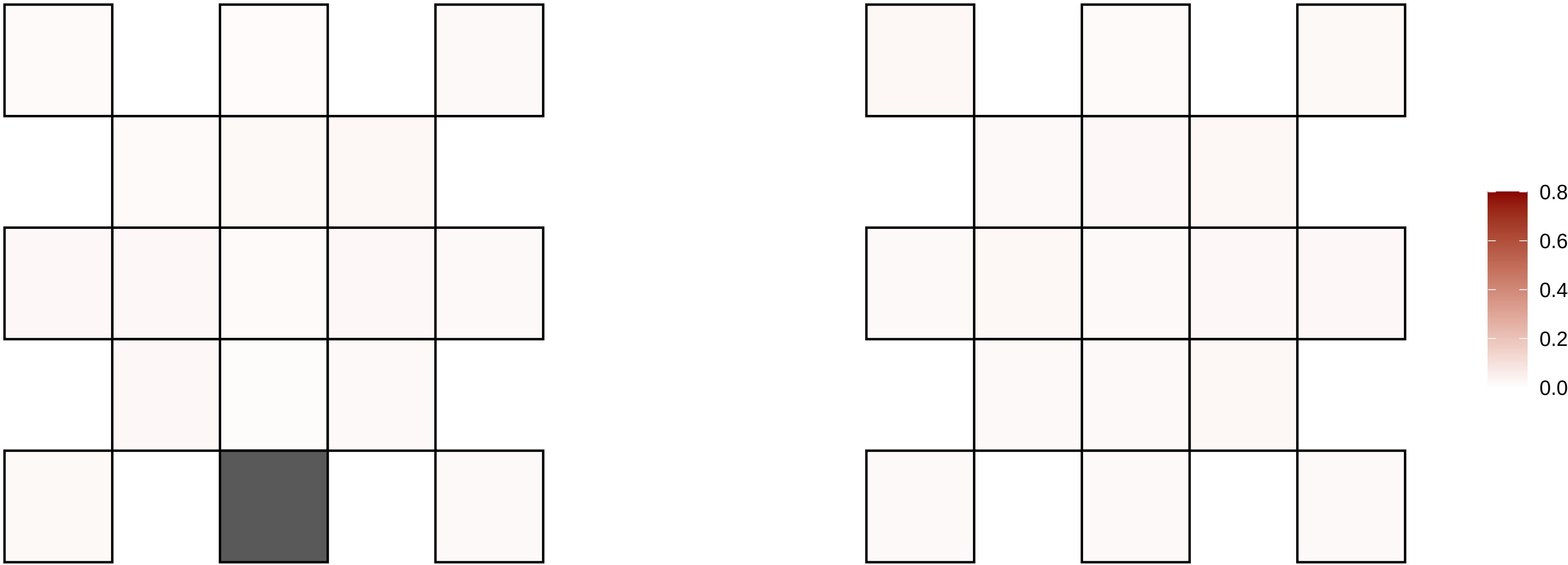

12−128703875−C−T

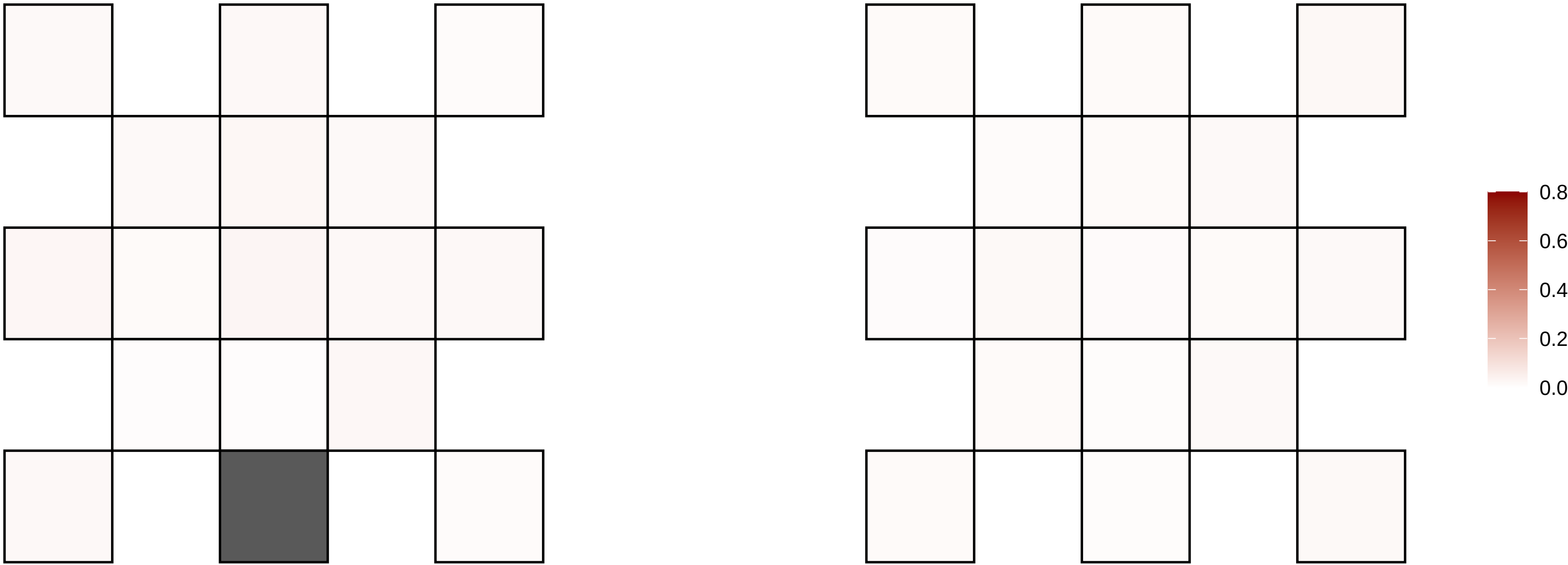

12−130892713−C−T

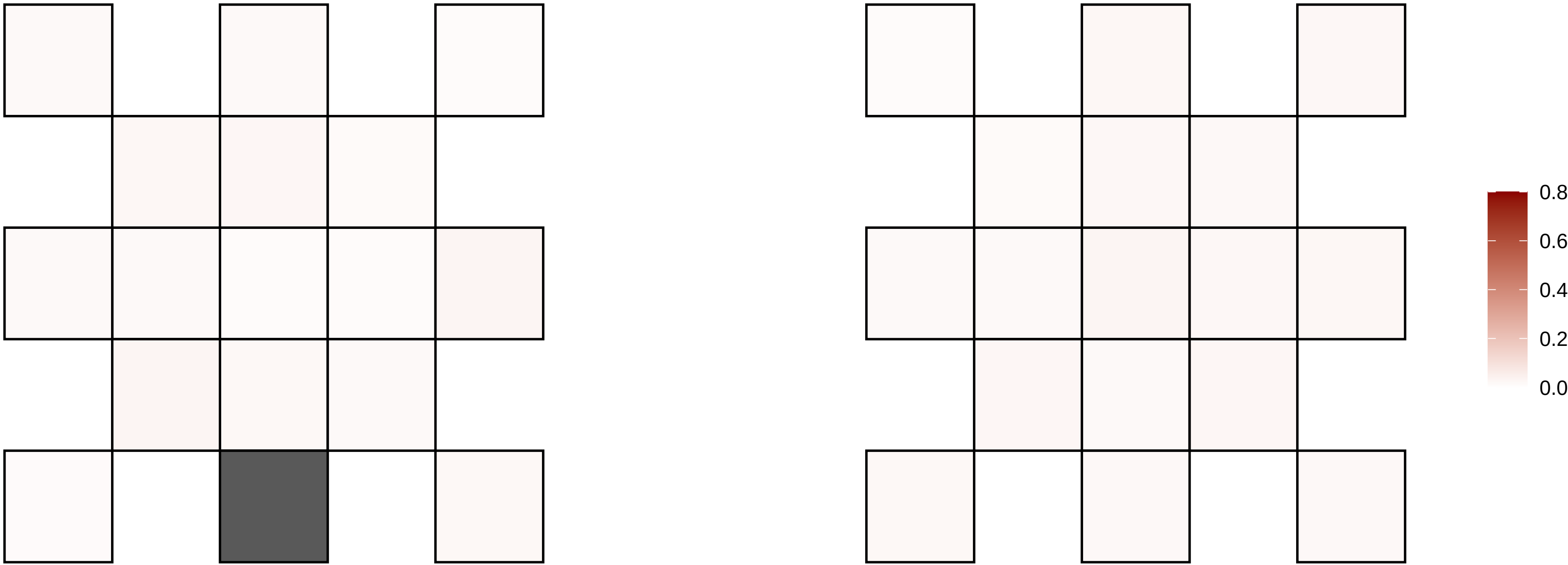

12−131906235−G−A

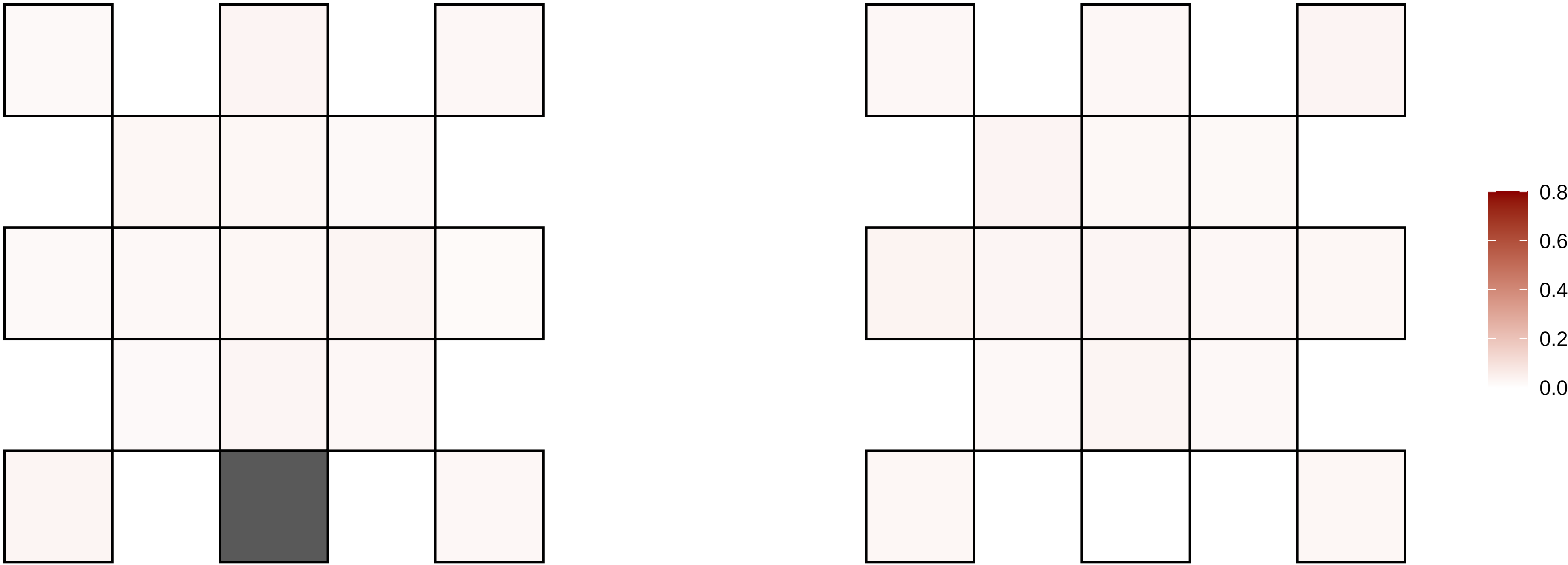

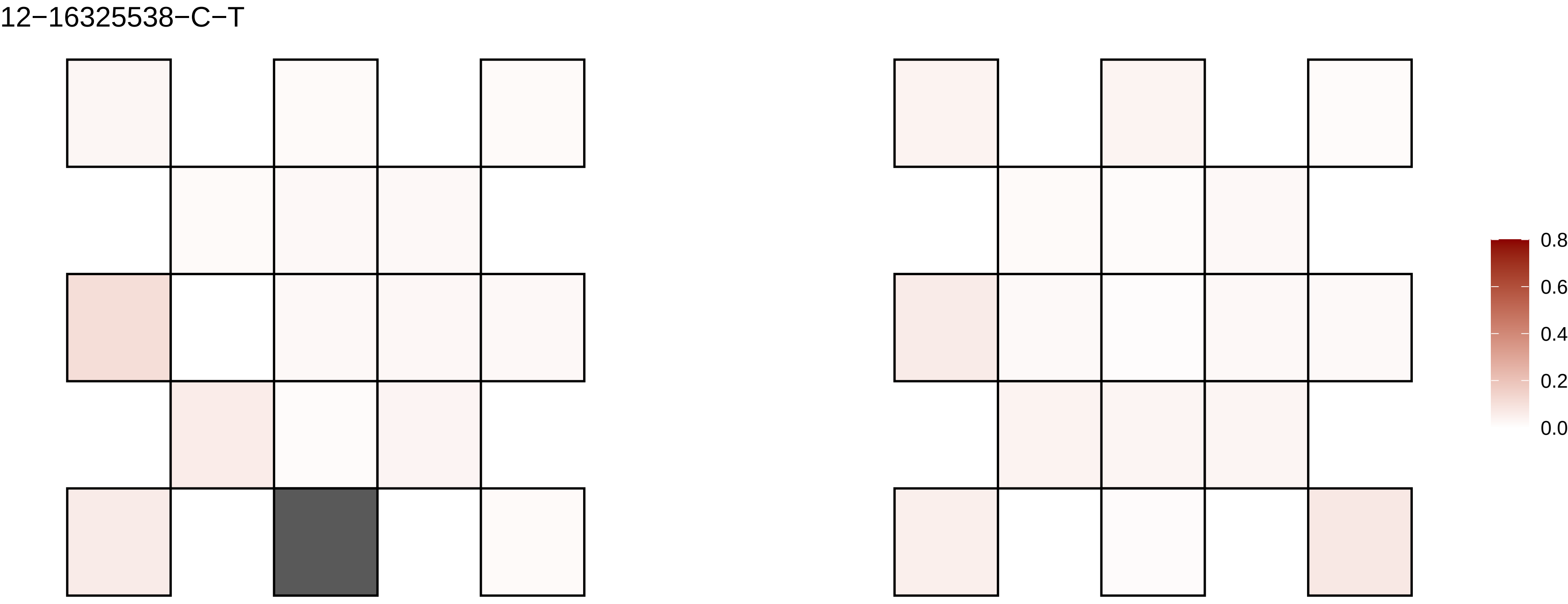
12−16325538−C−T
0.8
0.6
0.4
0.2
0.0

12−17467014−G−T

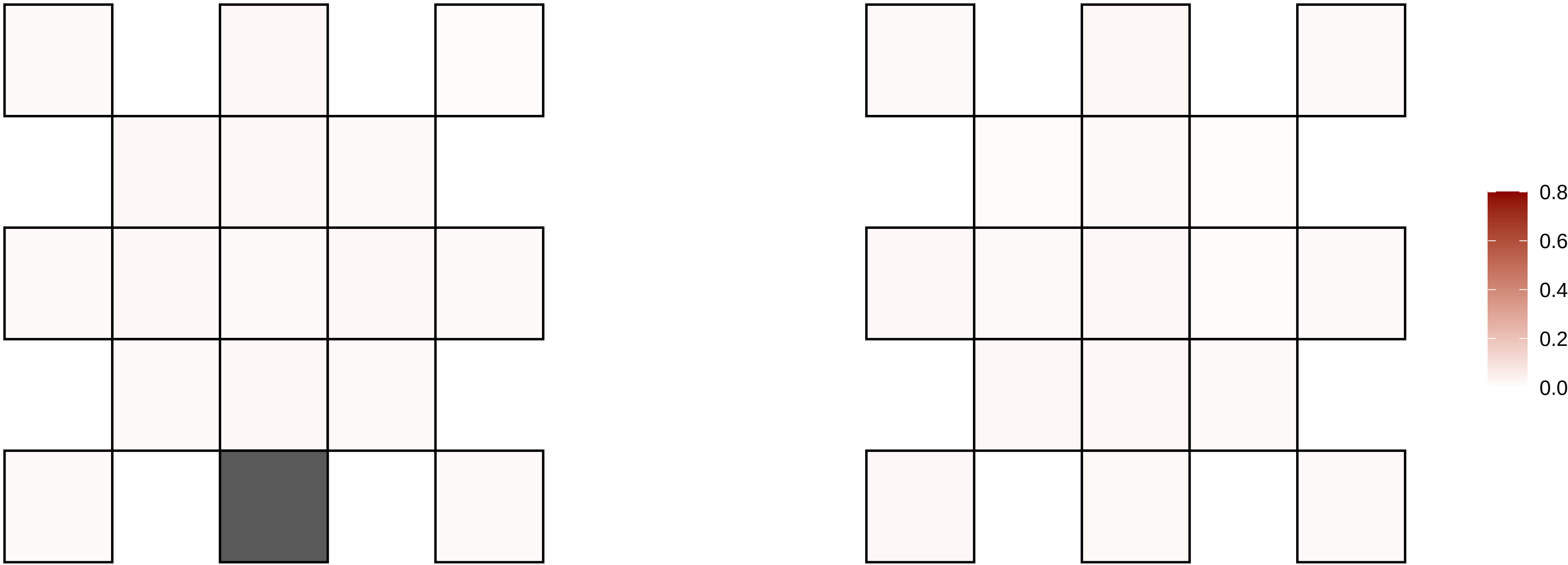

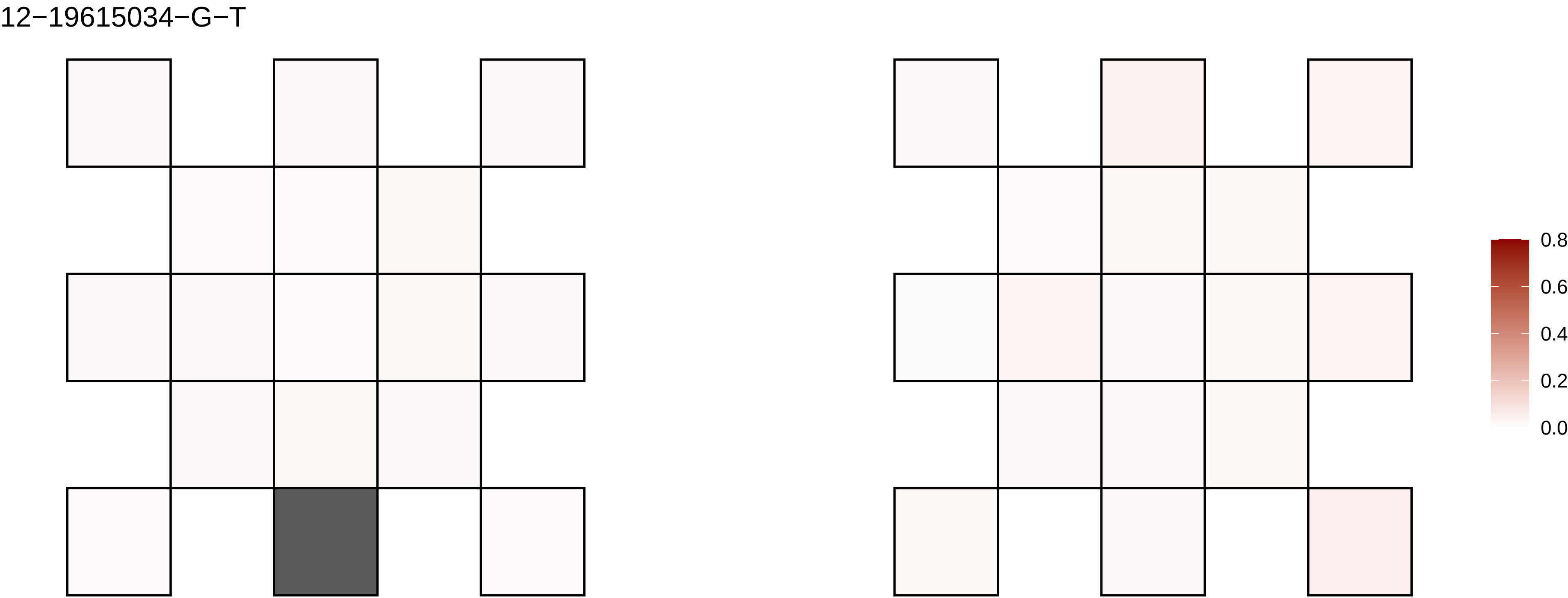

12−19615034−G−T
0.8
0.6
0.4
0.2
0.0

12−20421175−G−A

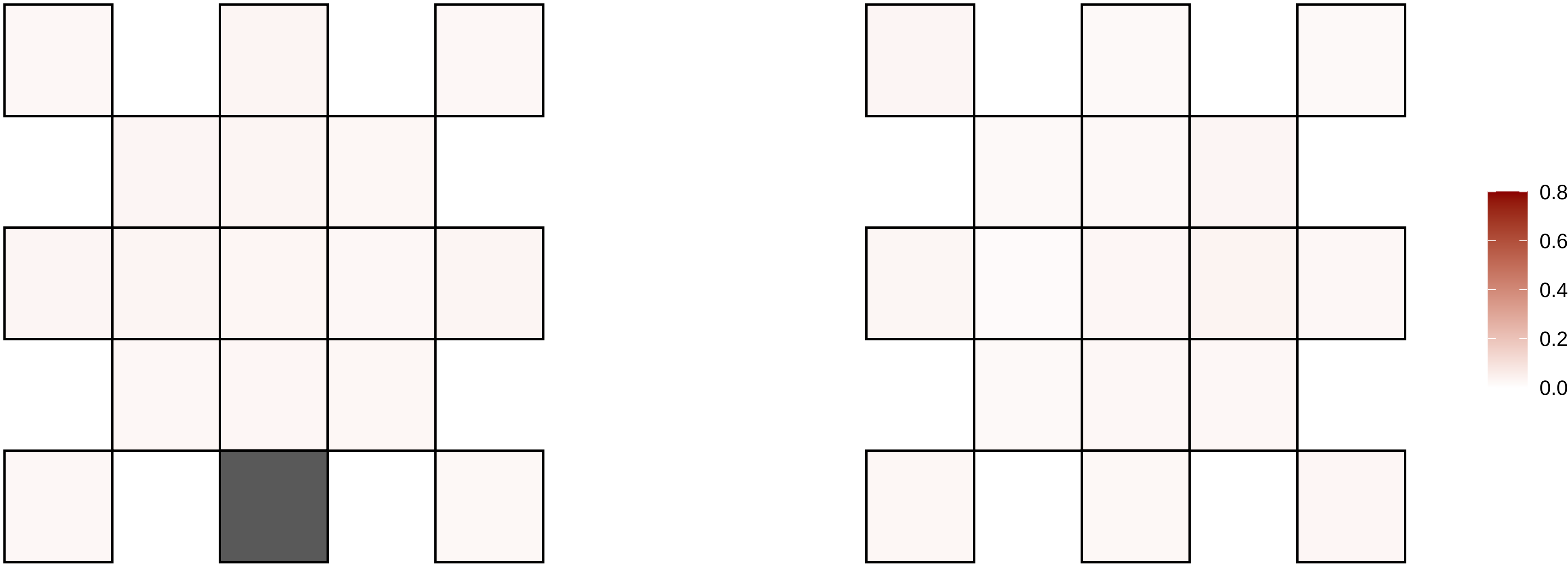

12−2204021−A−G

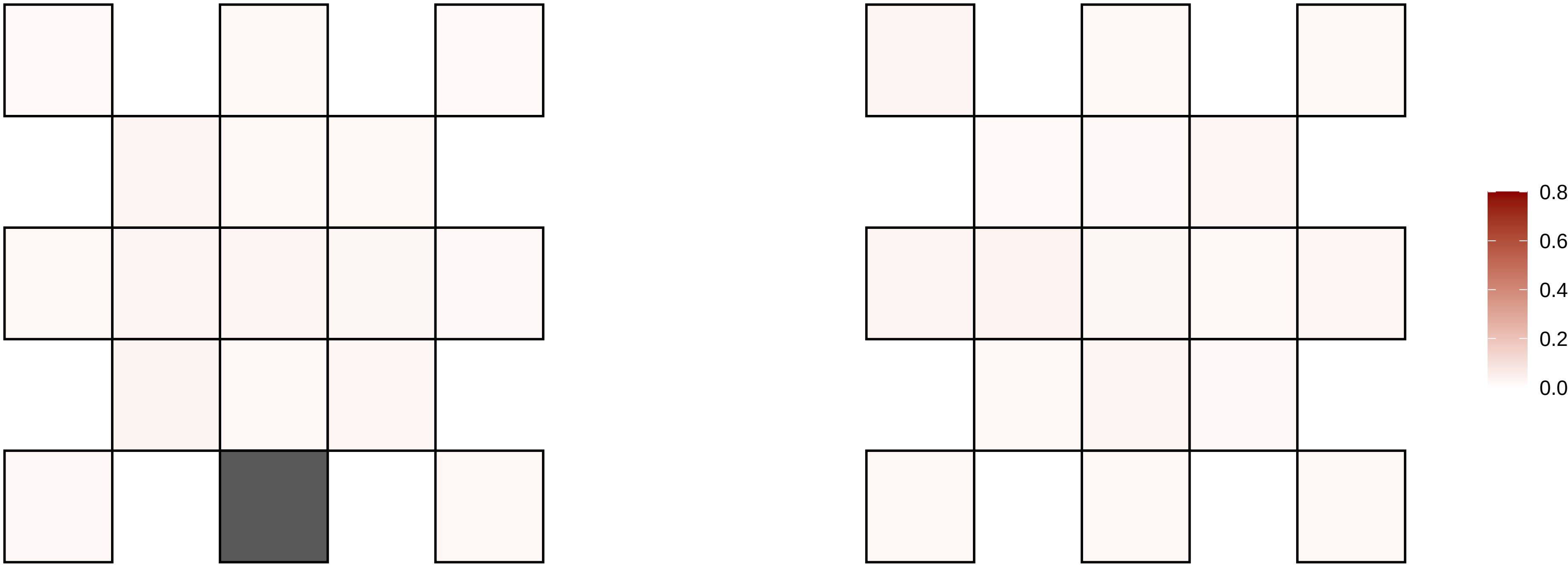

12−24436884−C−T

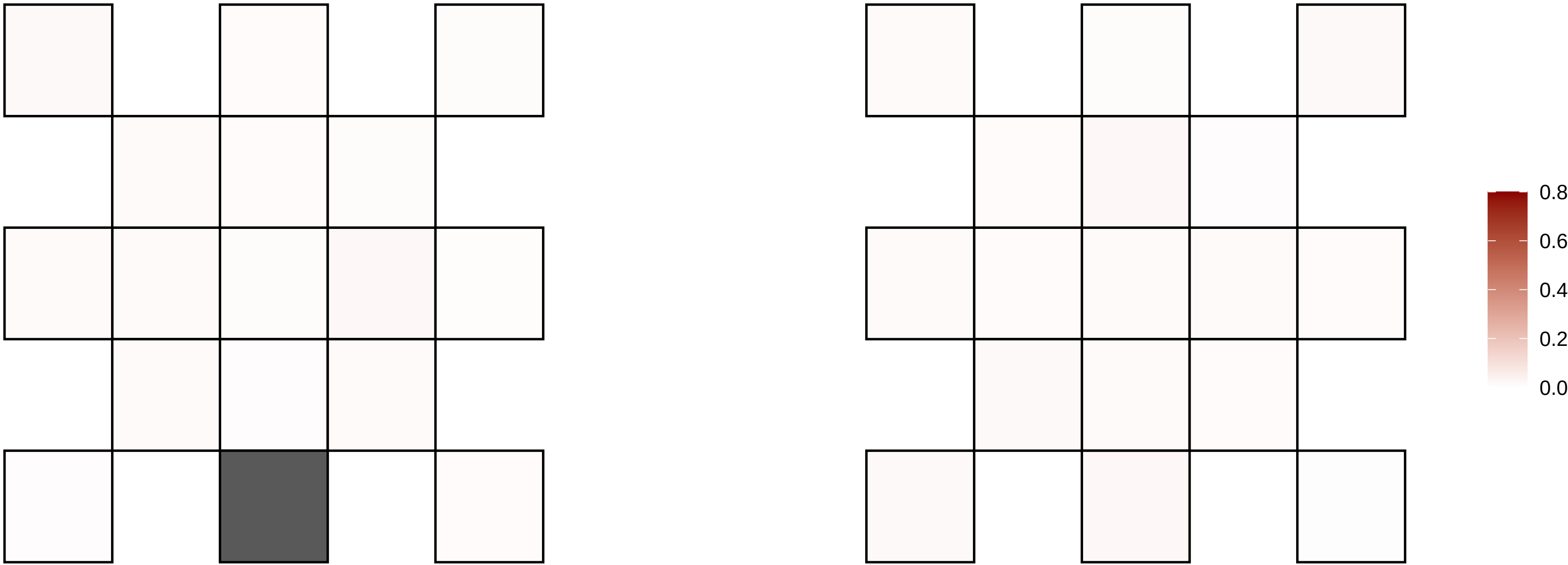

12−55677268−C−A

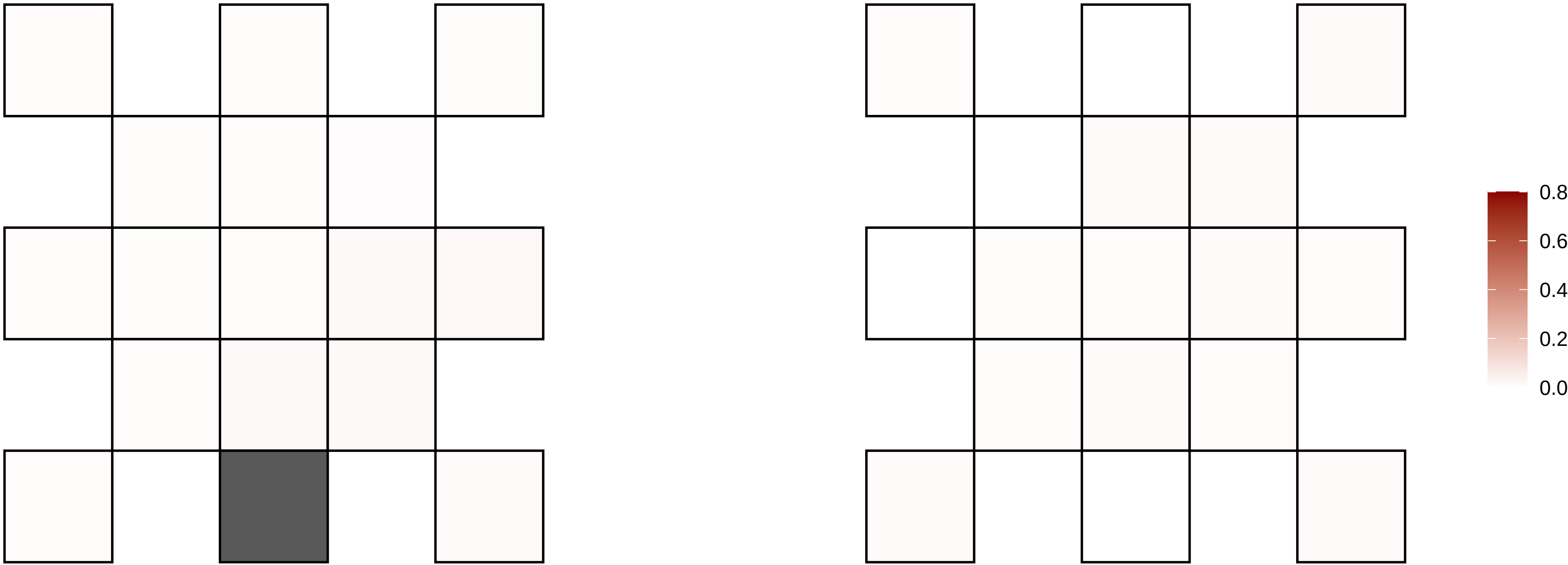

12−62486556−G−A

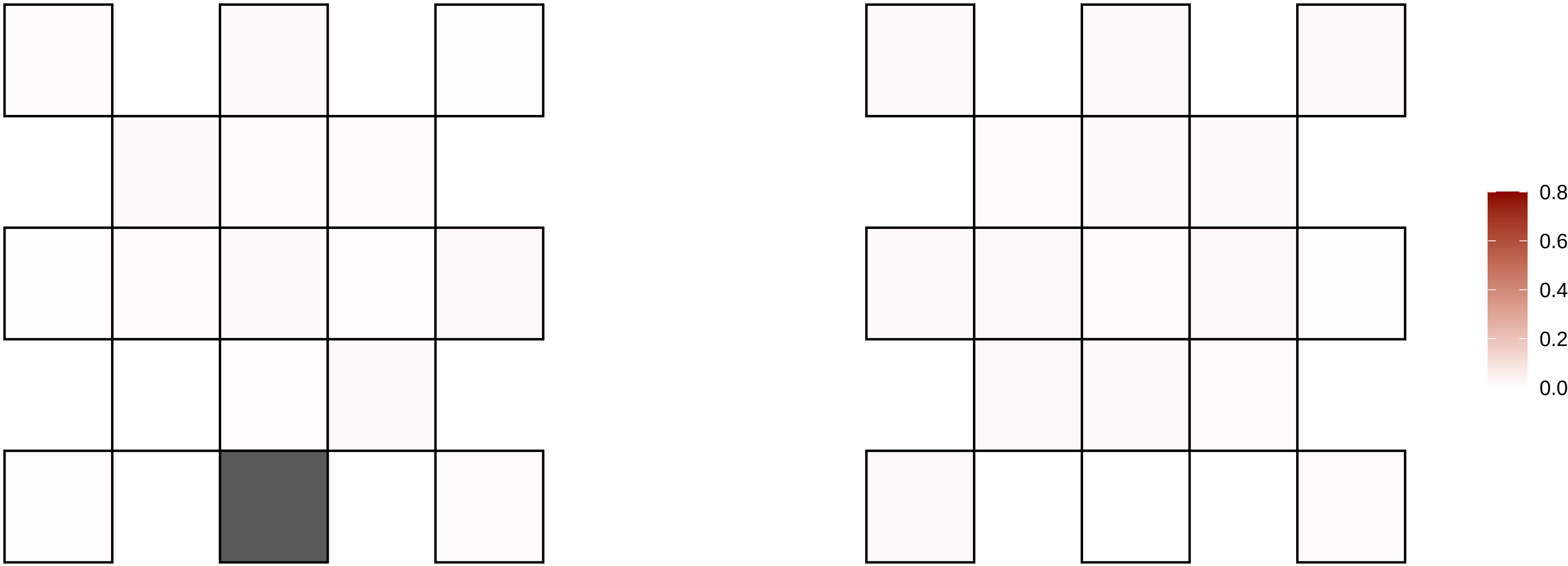

12−73783719−C−T

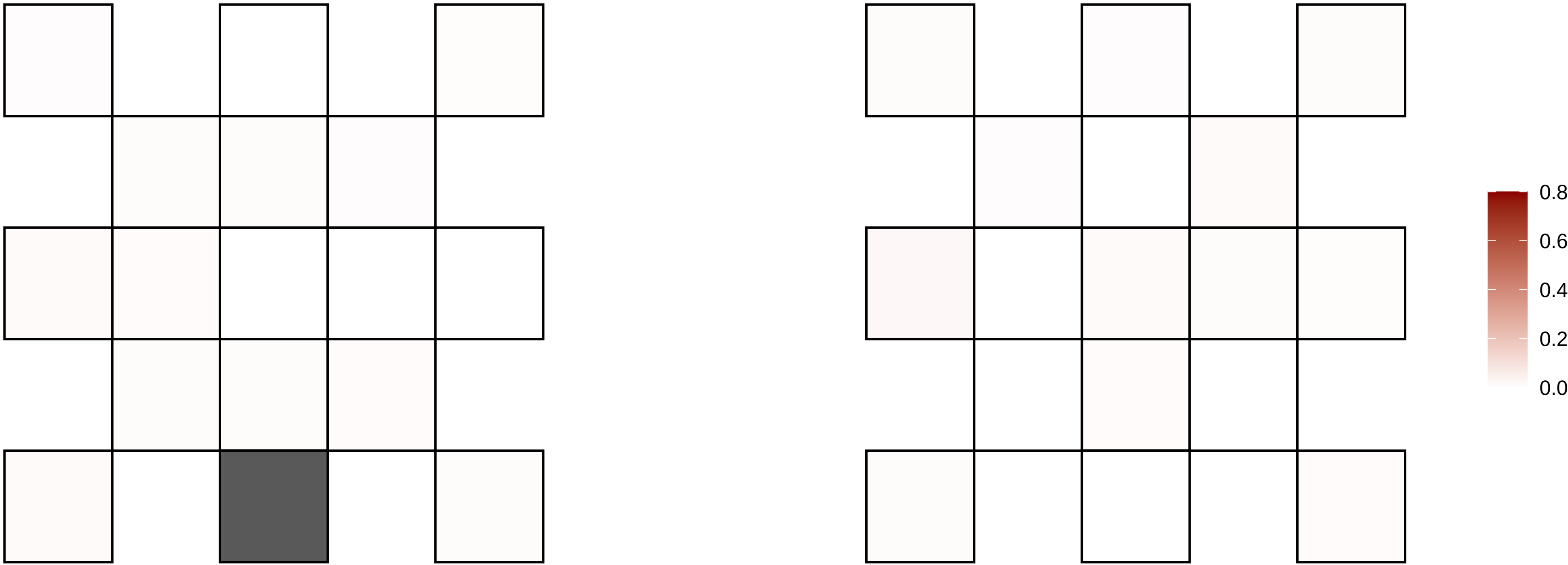

12−78761941−A−T

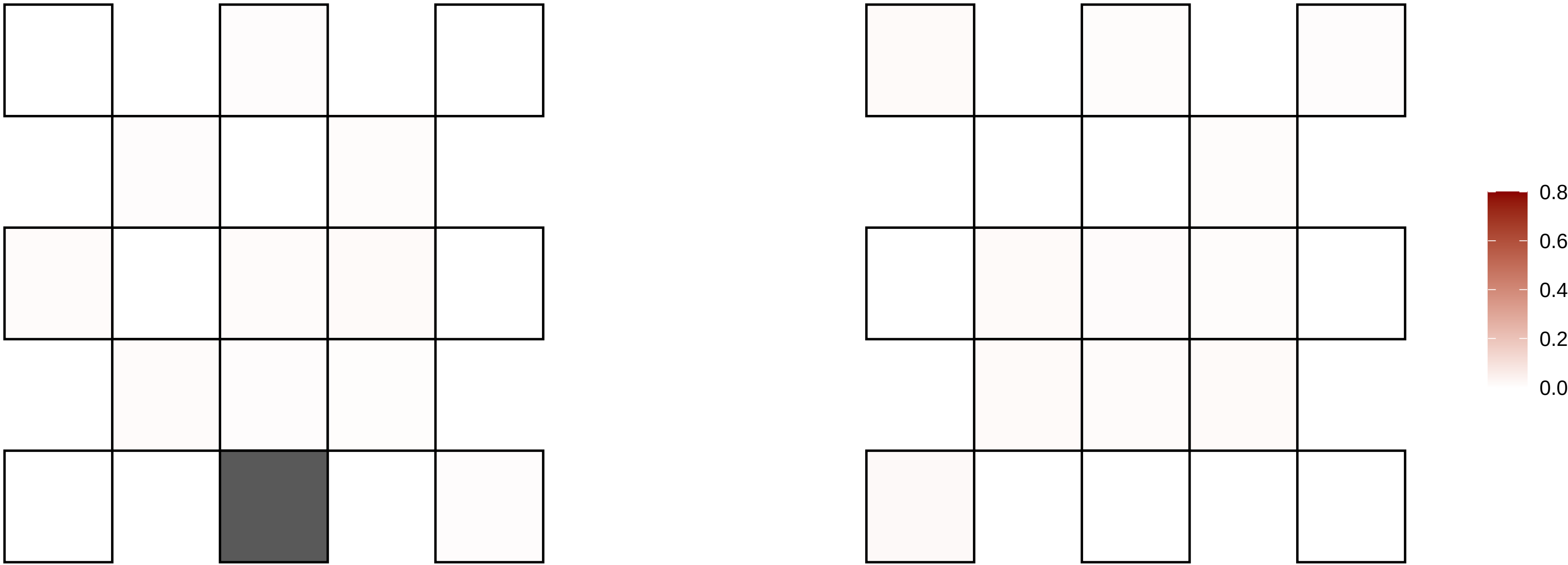

12−79536312−C−T

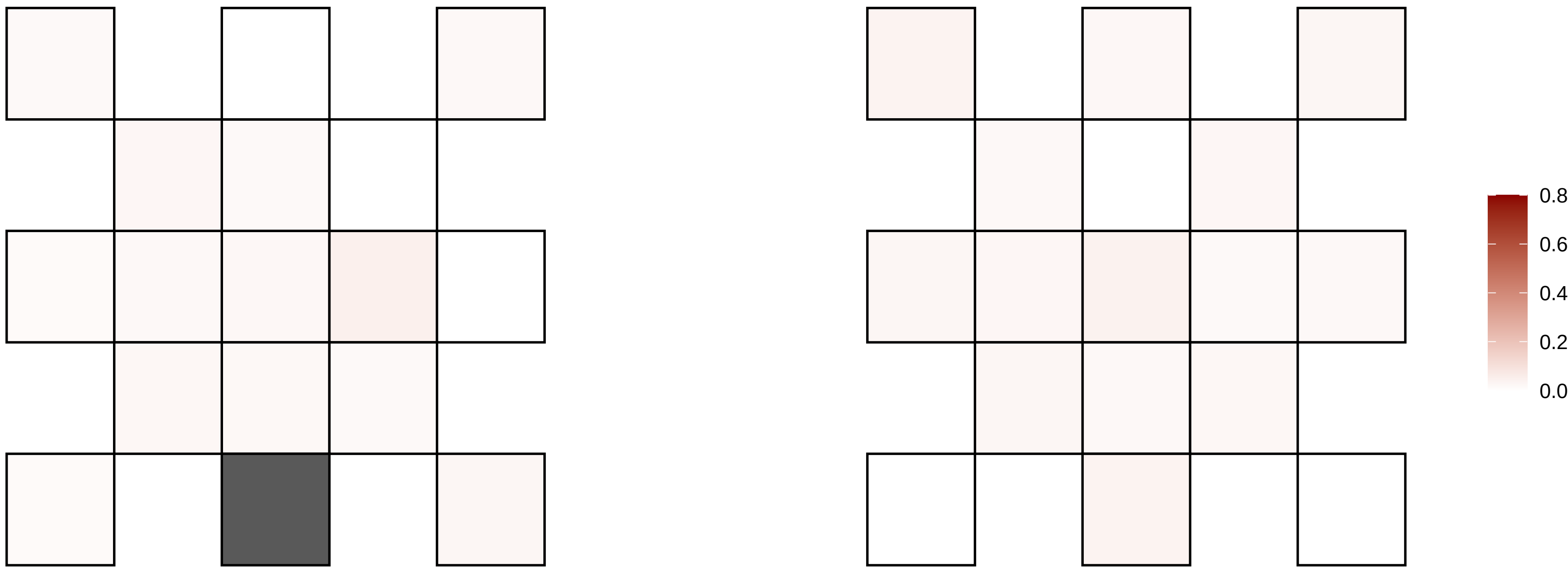

12−85014452−T−A

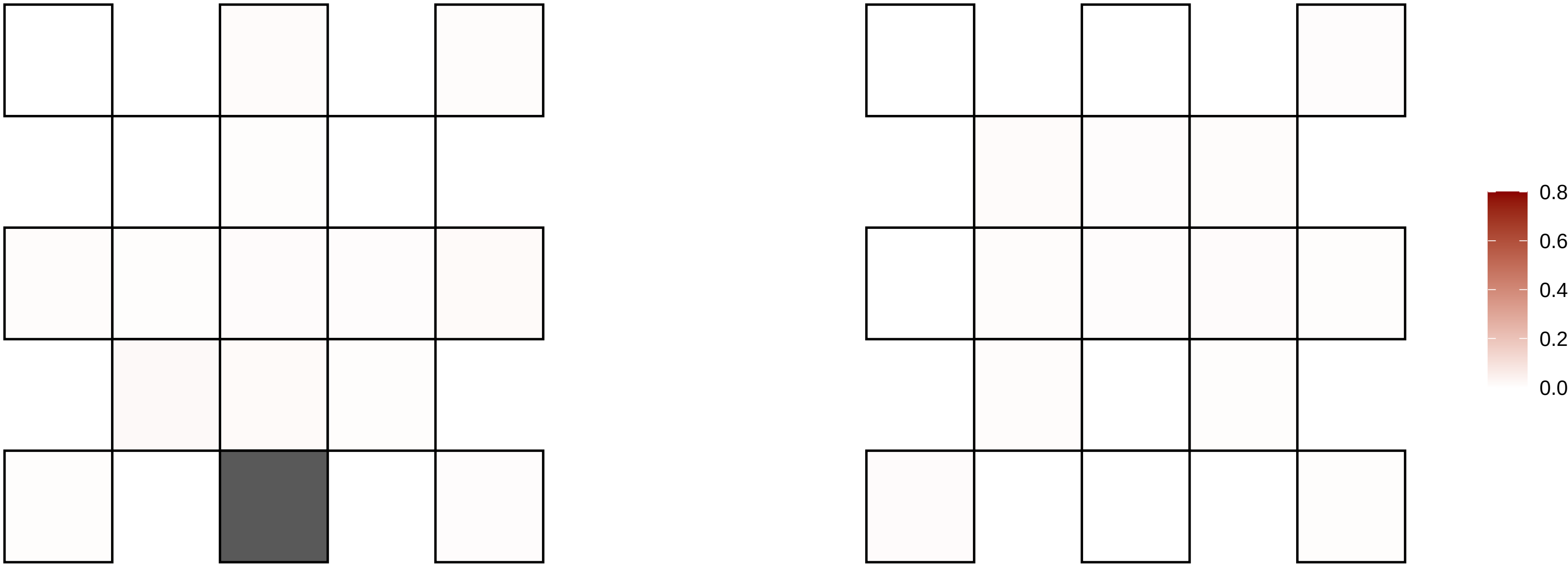

12−85787527−A−T

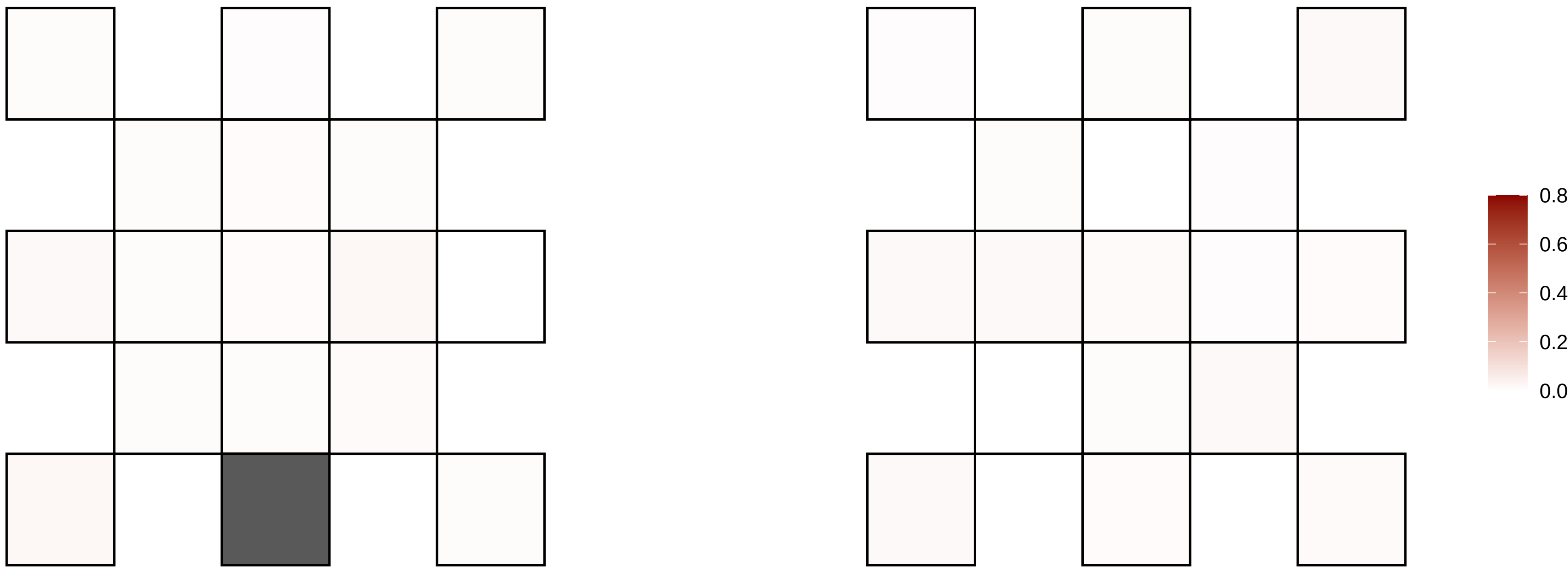

12−90774790−C−T

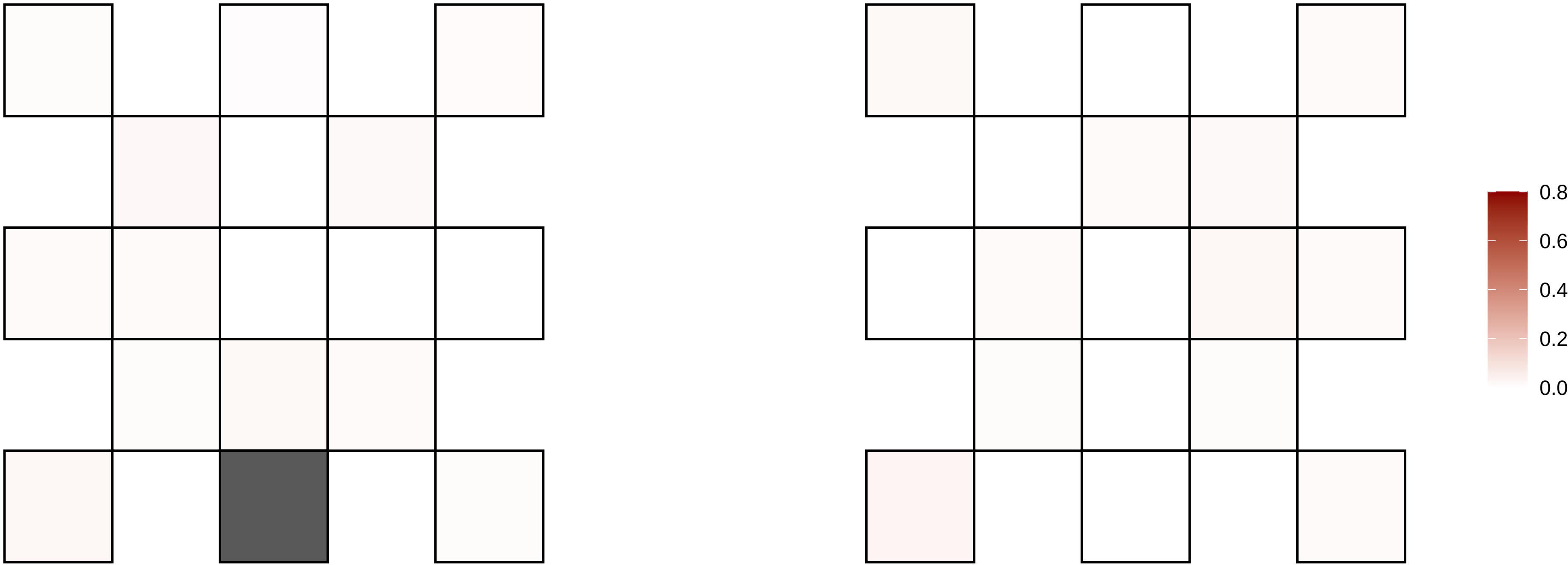

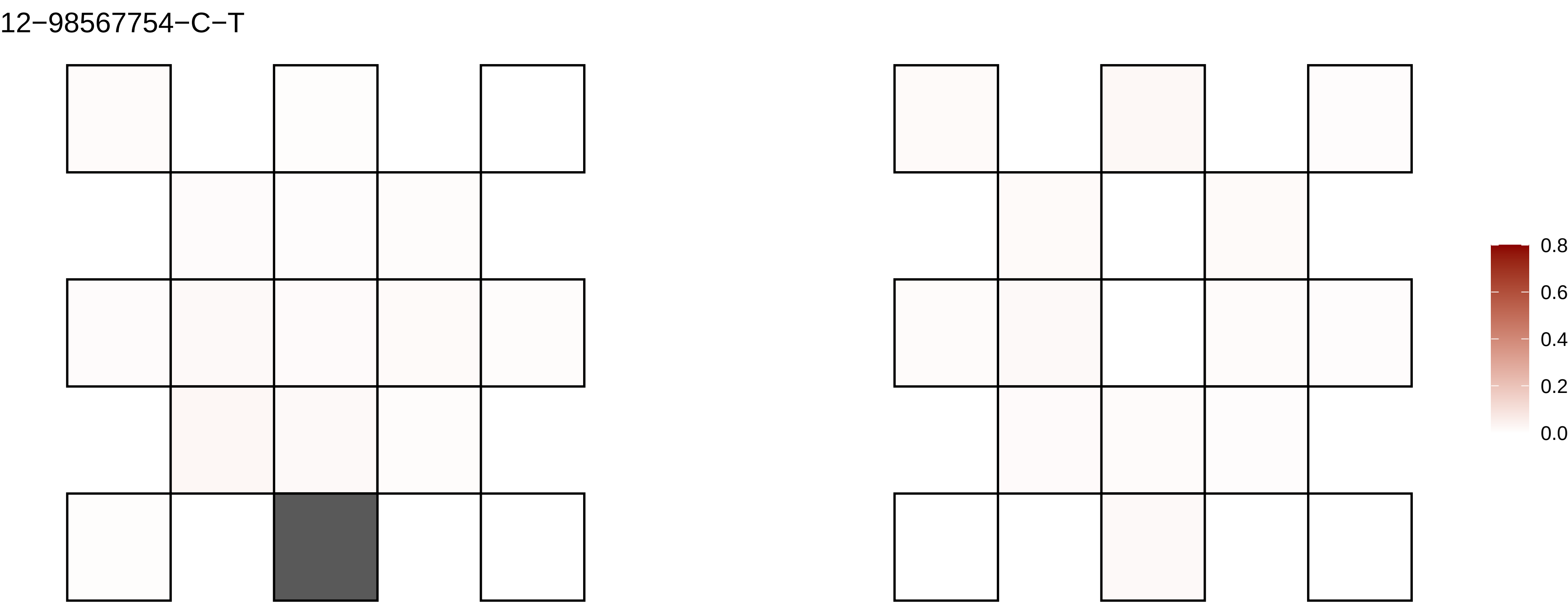
12−98567754−C−T
0.8
0.6
0.4
0.2
0.0

12−98674630−C−T

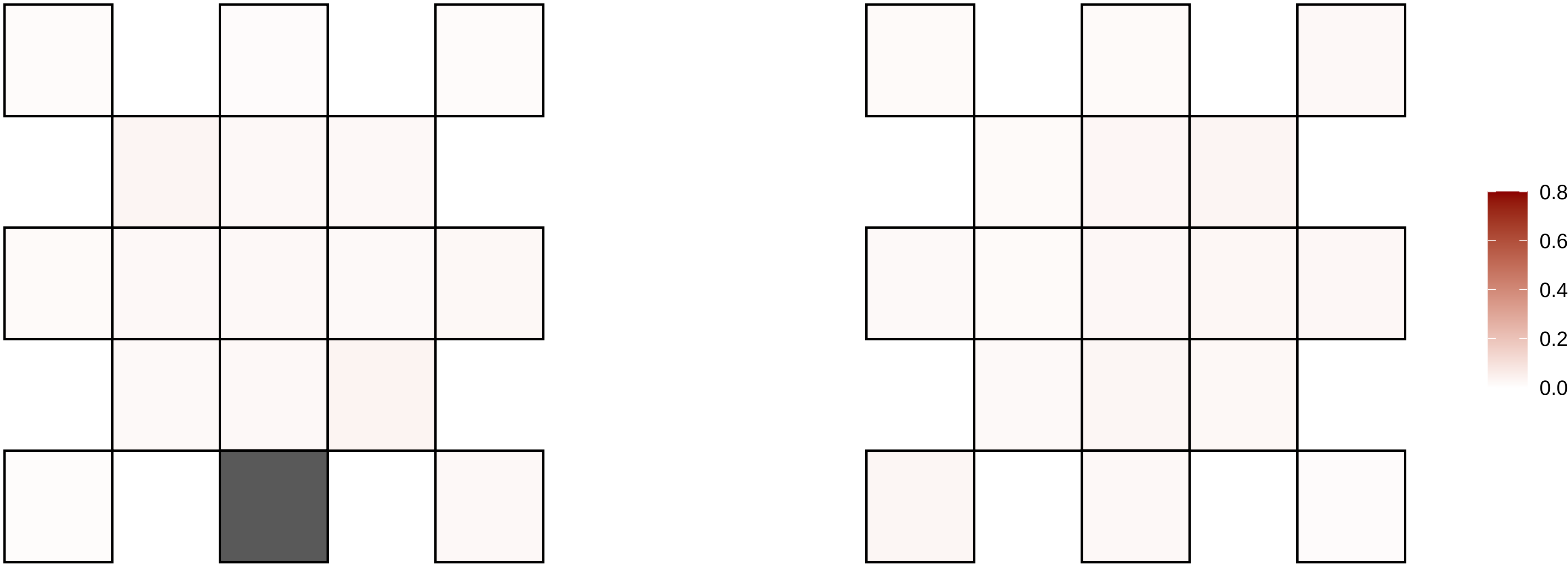

13−102240084−A−T

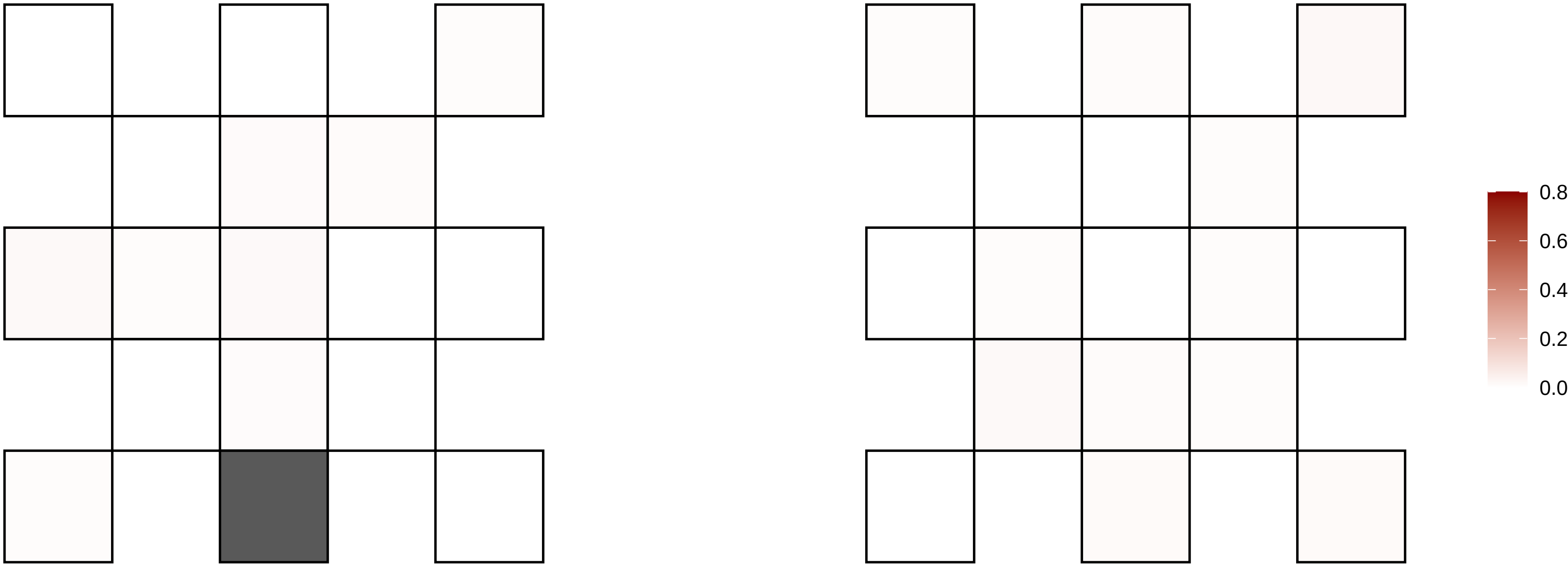

13−106141548−C−T

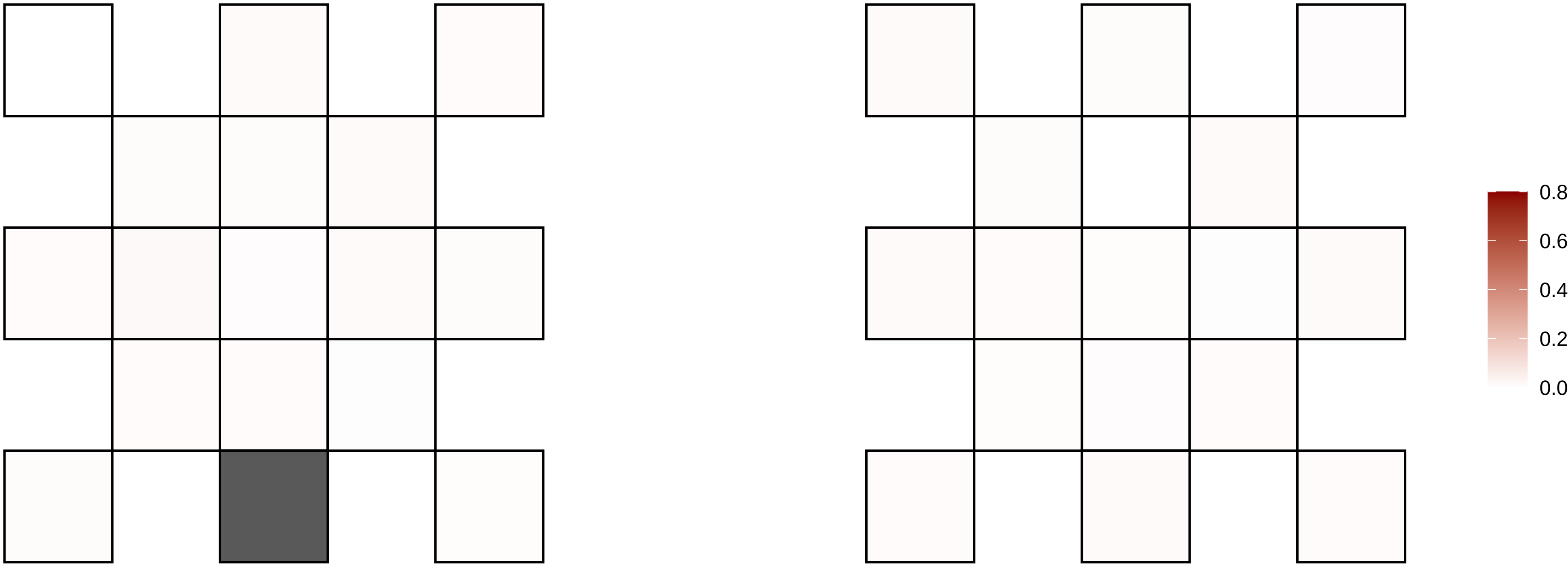

13−109148397−G−A

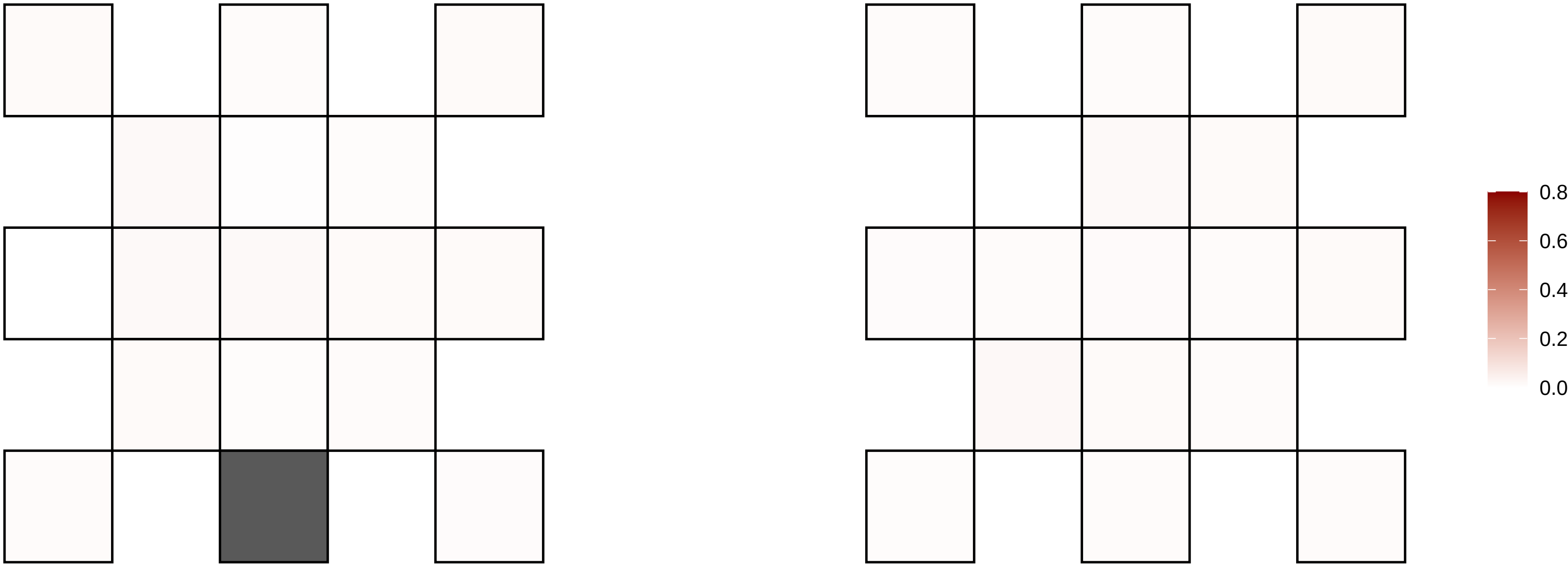

13−109292761−G−A

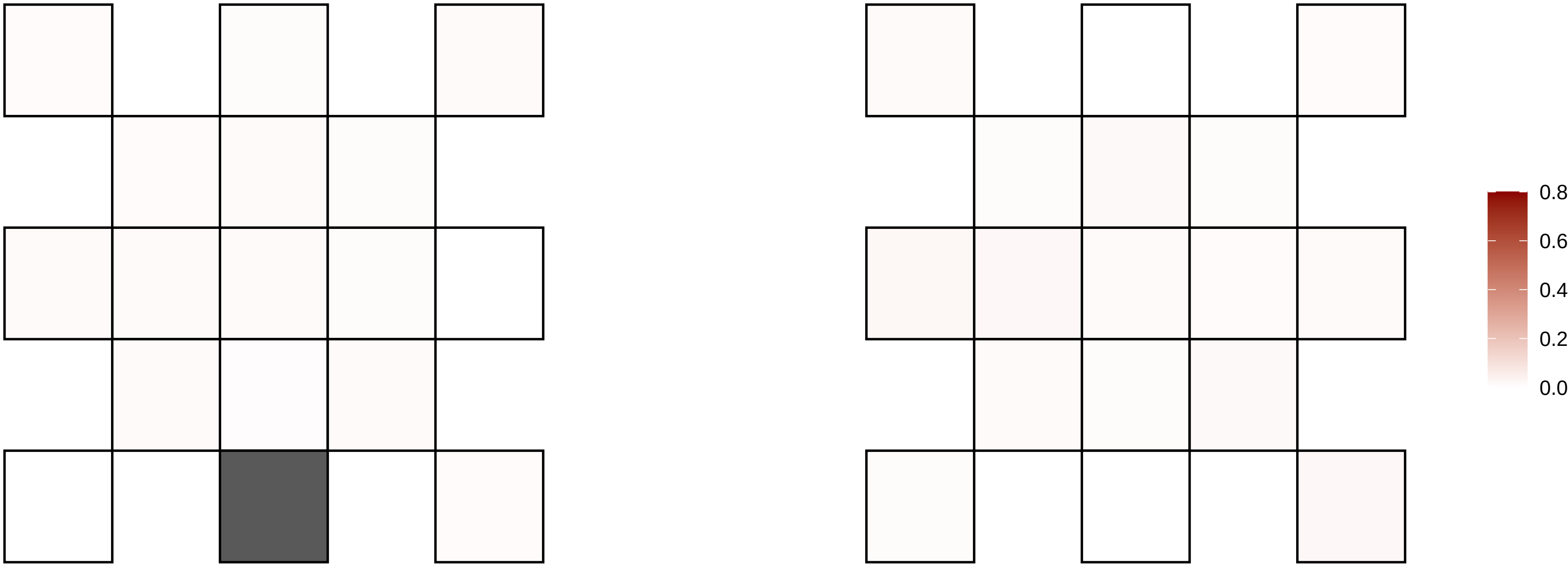

13−111450669−C−T

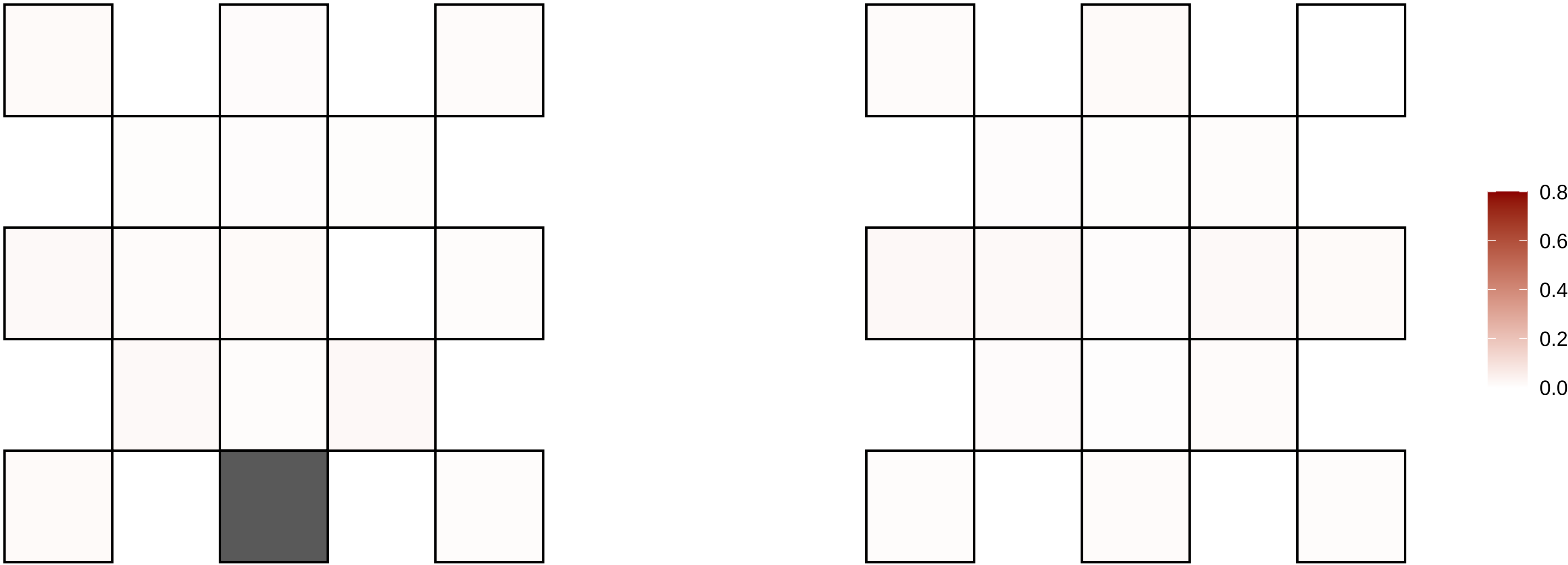

13−113879720−G−A

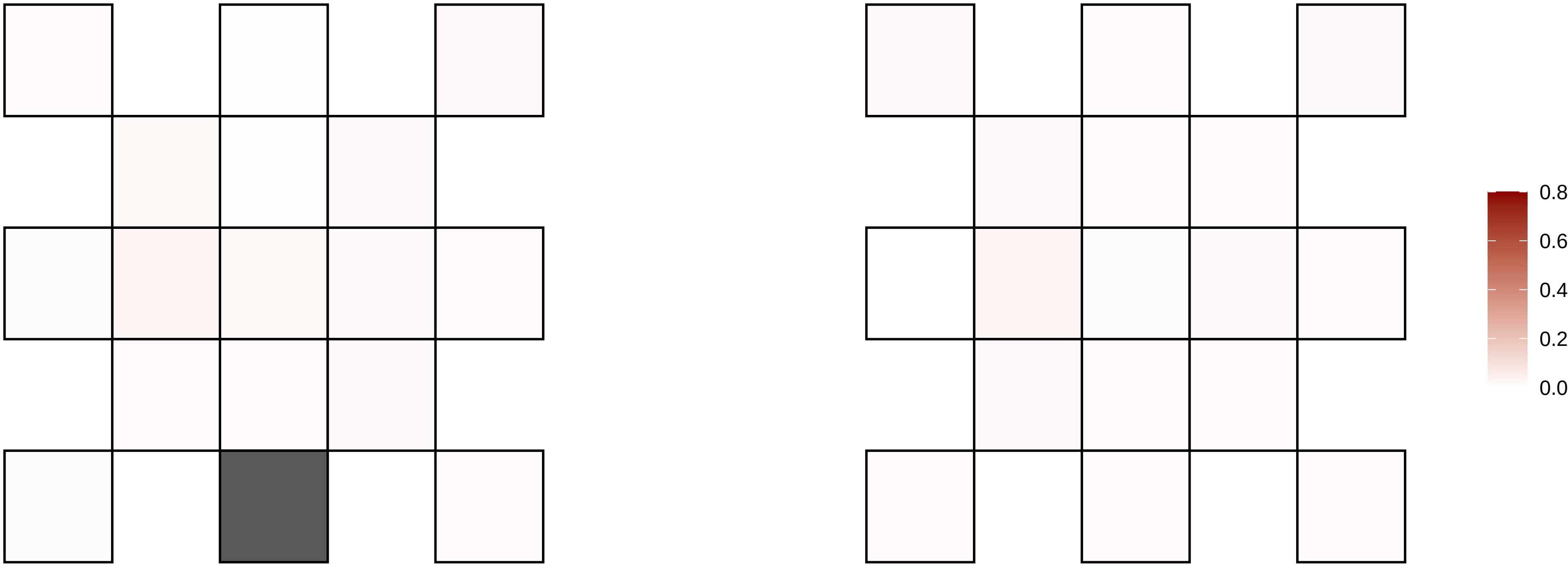

13−26972654−G−A

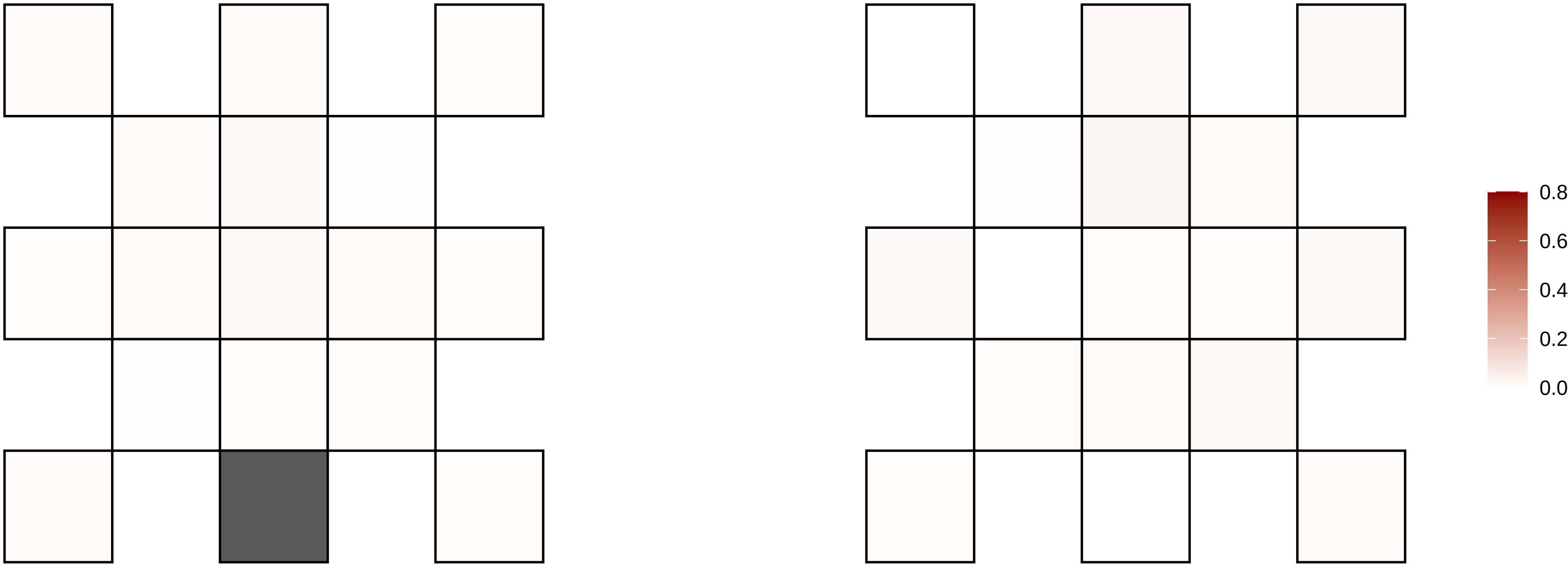

13−26972655−G−A

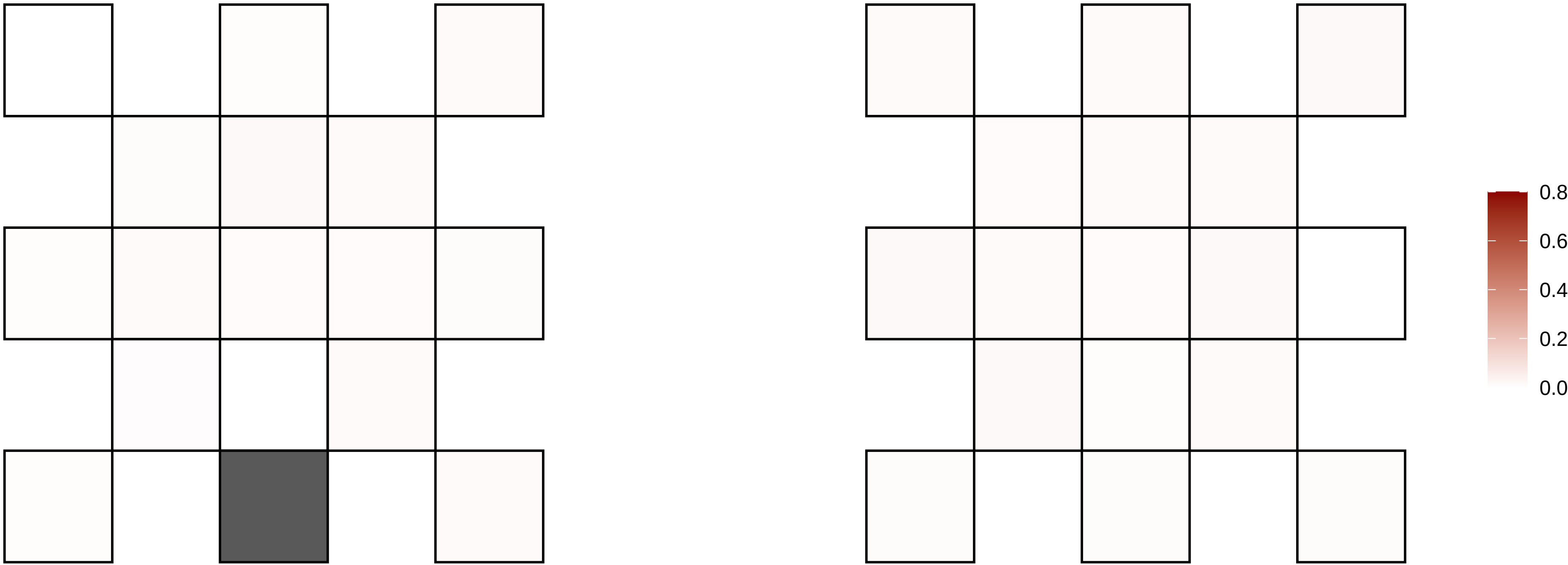

13−29040893−G−A

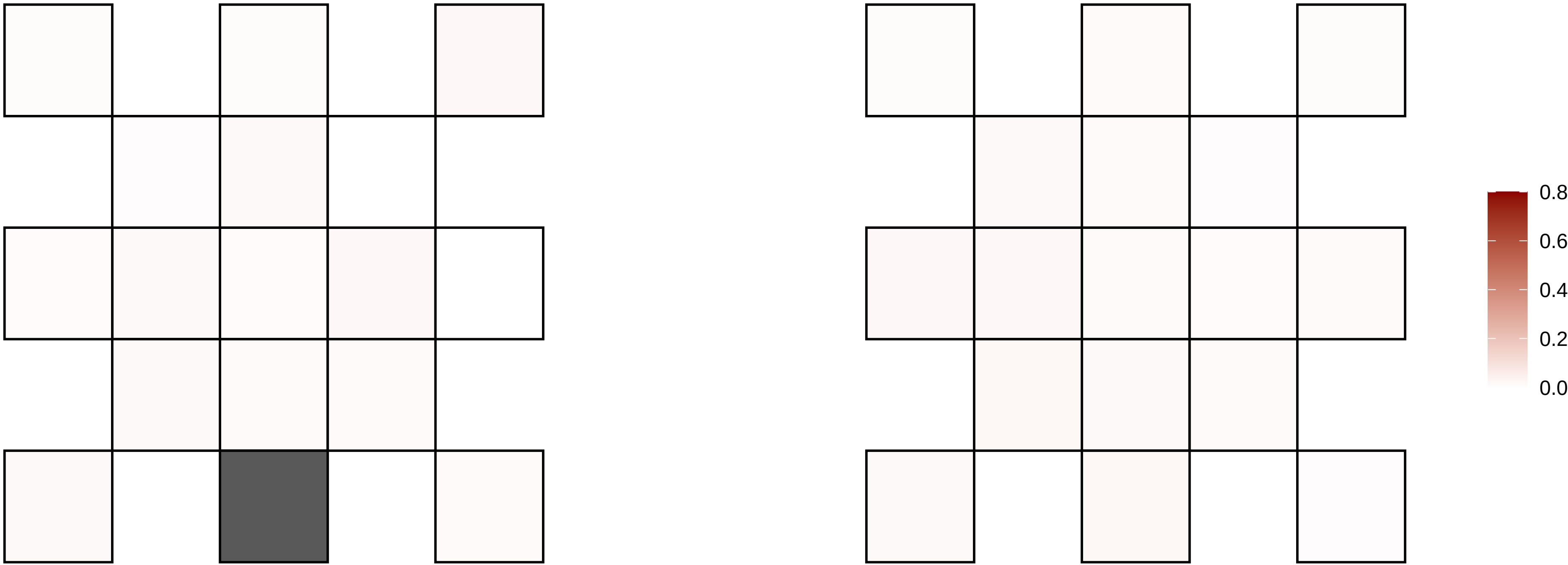

13−29577886−G−A

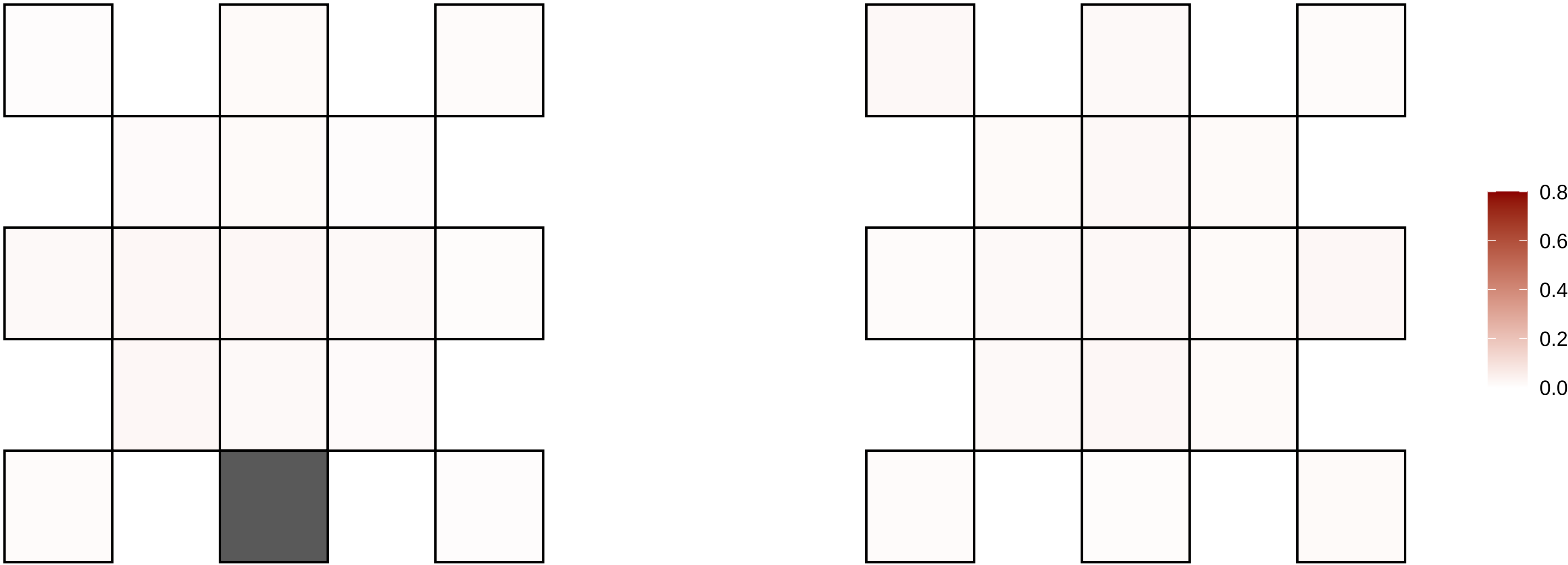

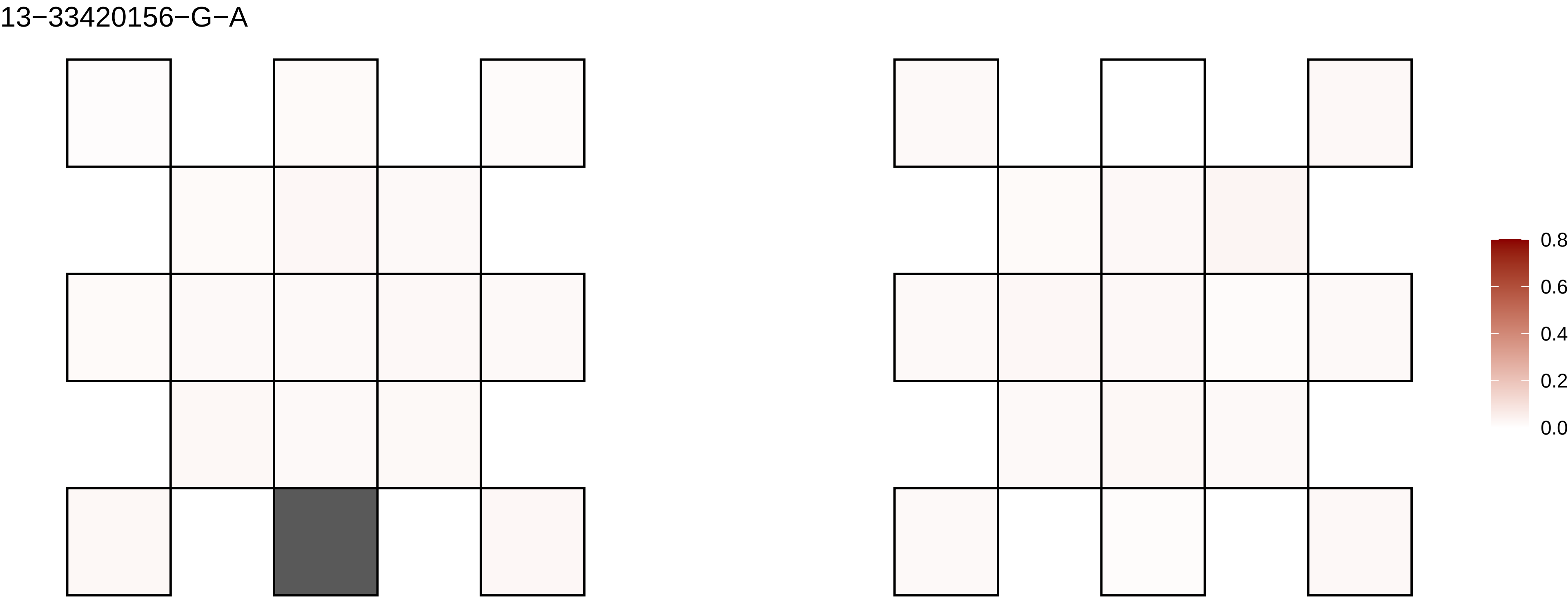
13−33420156−G−A
0.8
0.6
0.4
0.2
0.0

13−41556737−G−C

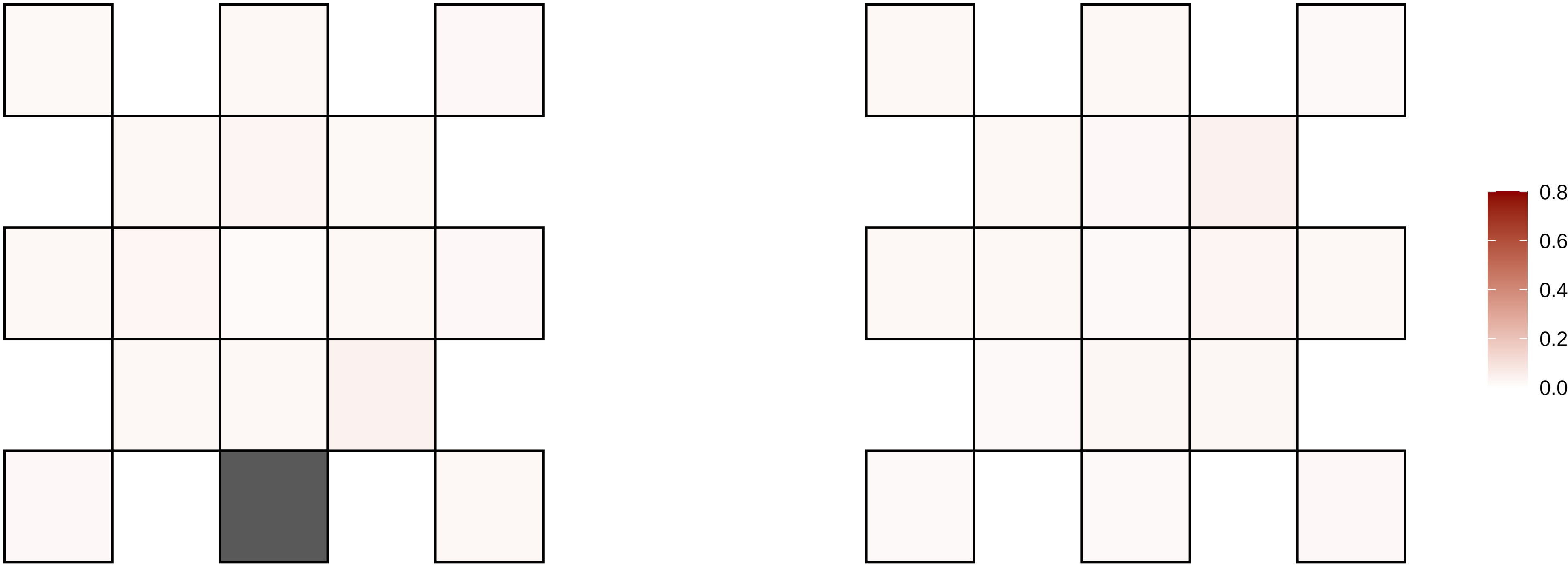

13−46288201−G−A

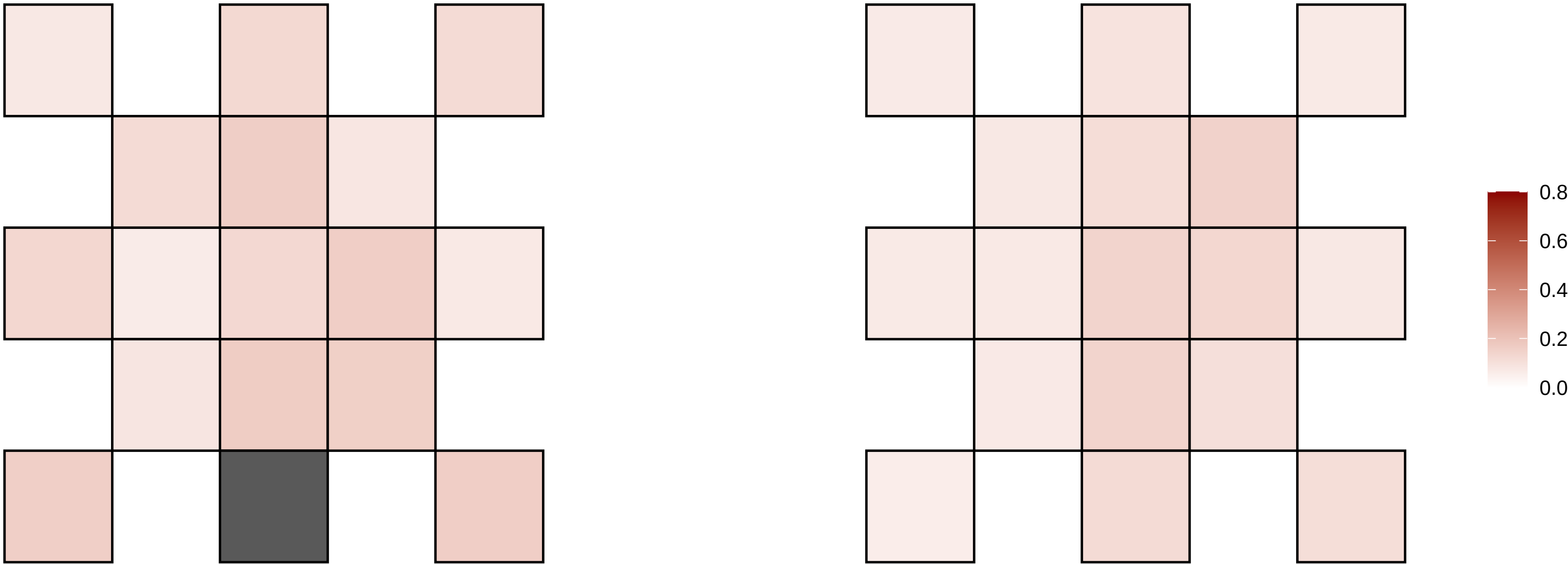

13−55140512−C−T

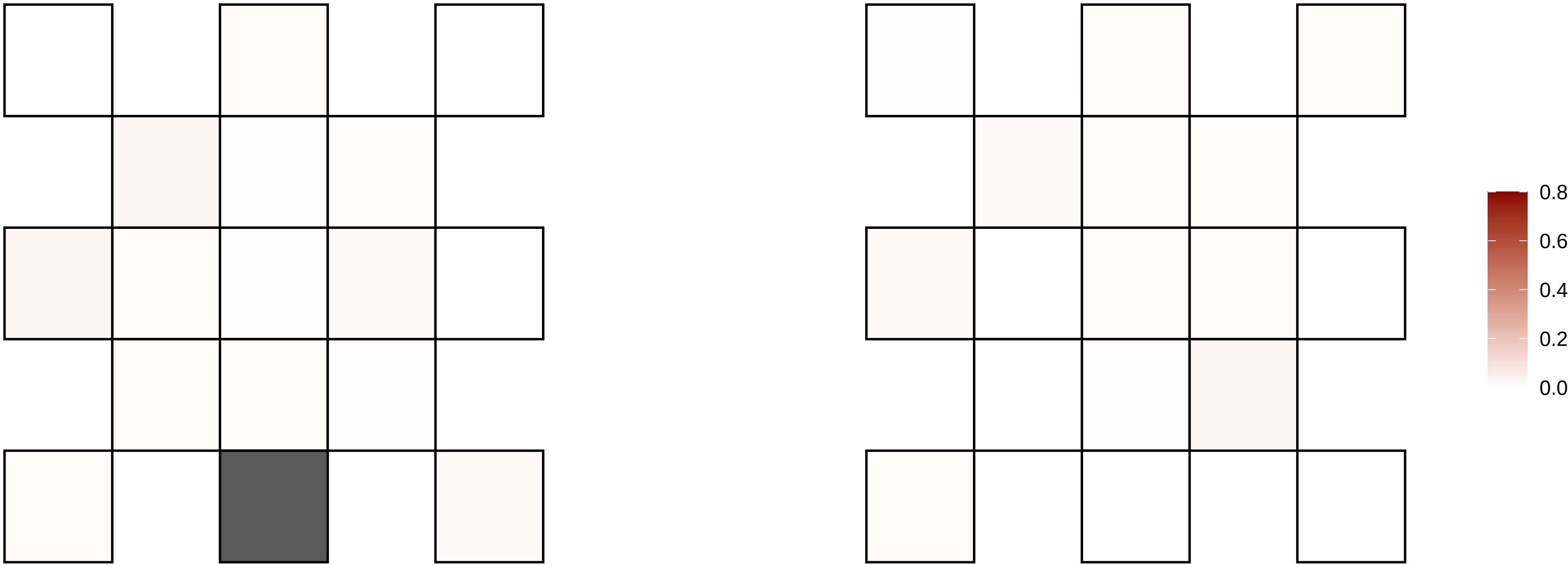

13−56431067−T−A

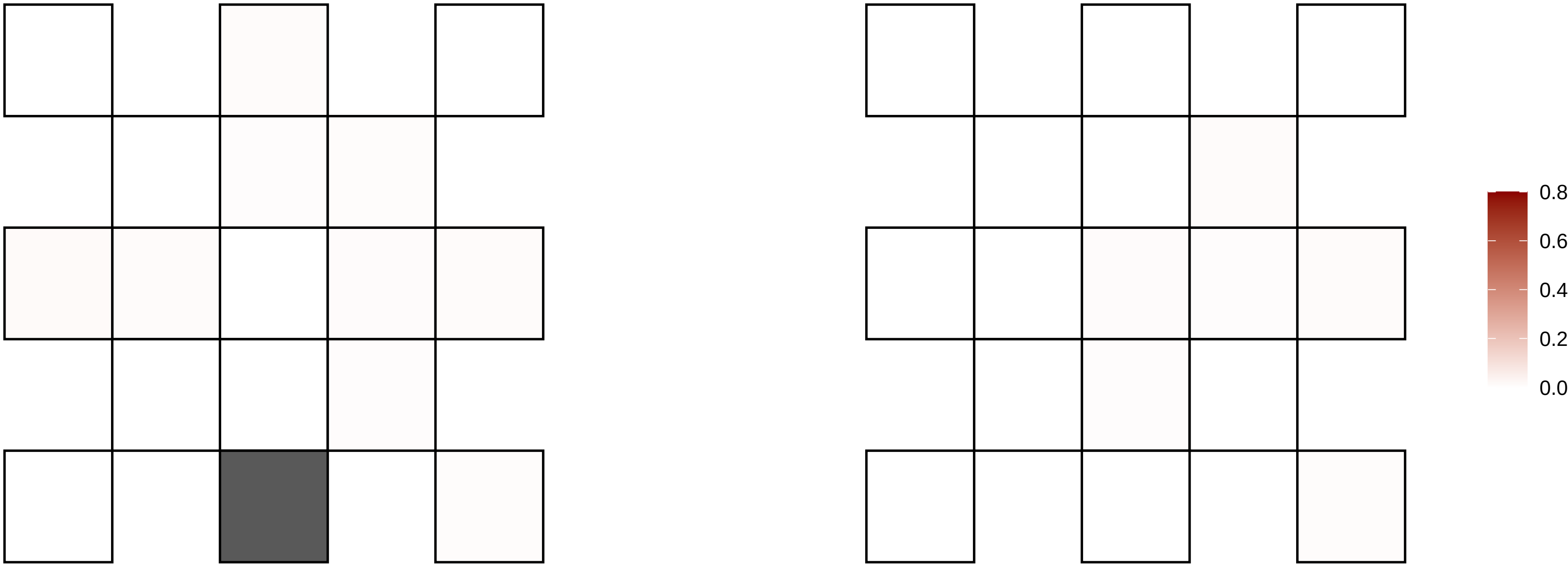

13−59159499−G−A

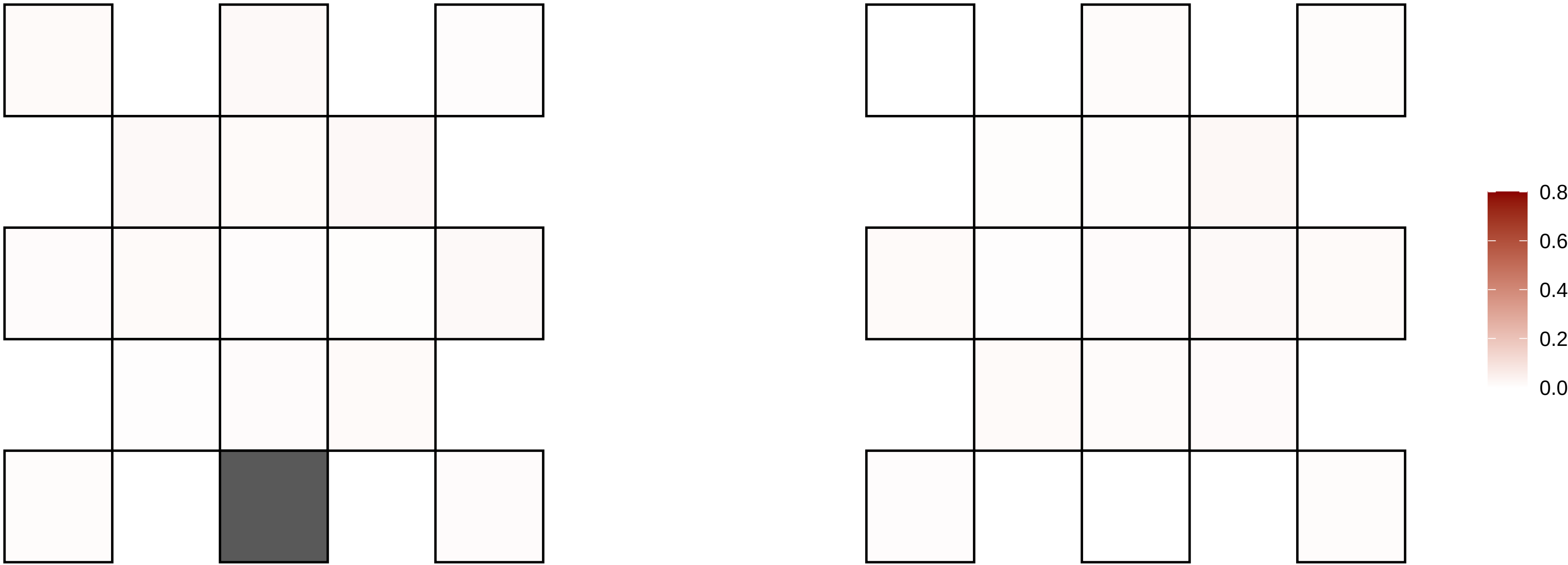

13−63432546−C−T

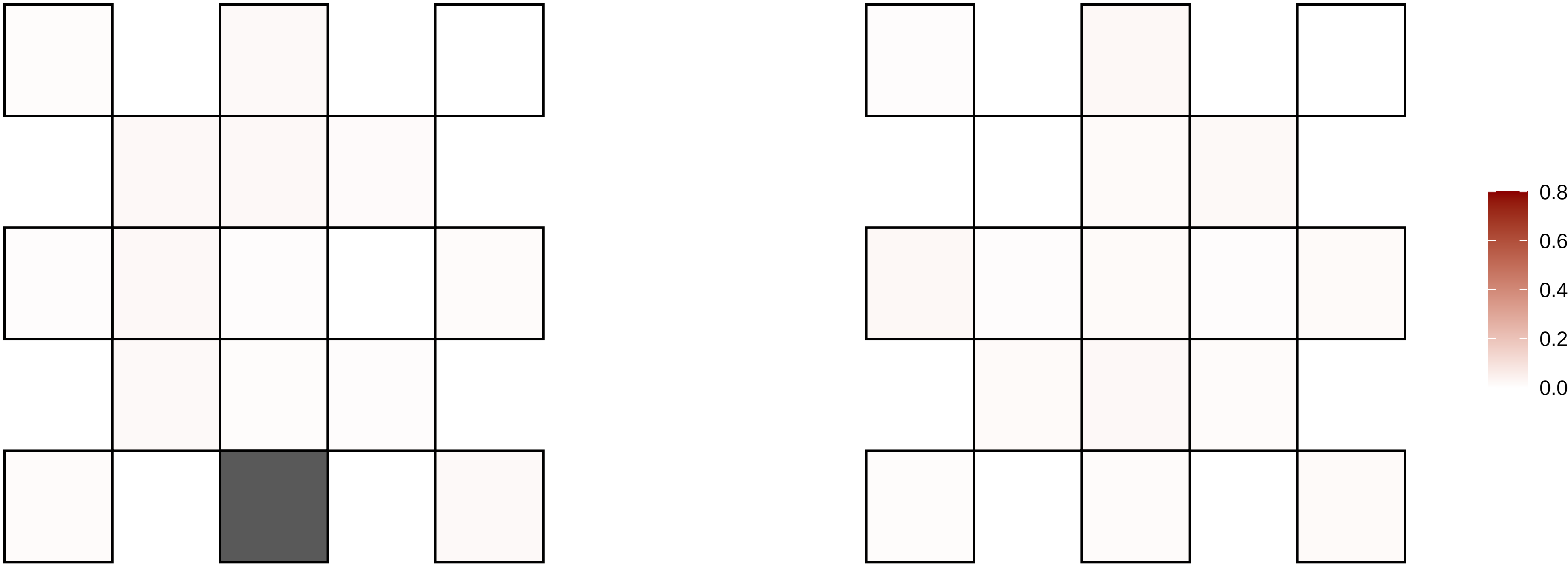

13−63856062−G−A

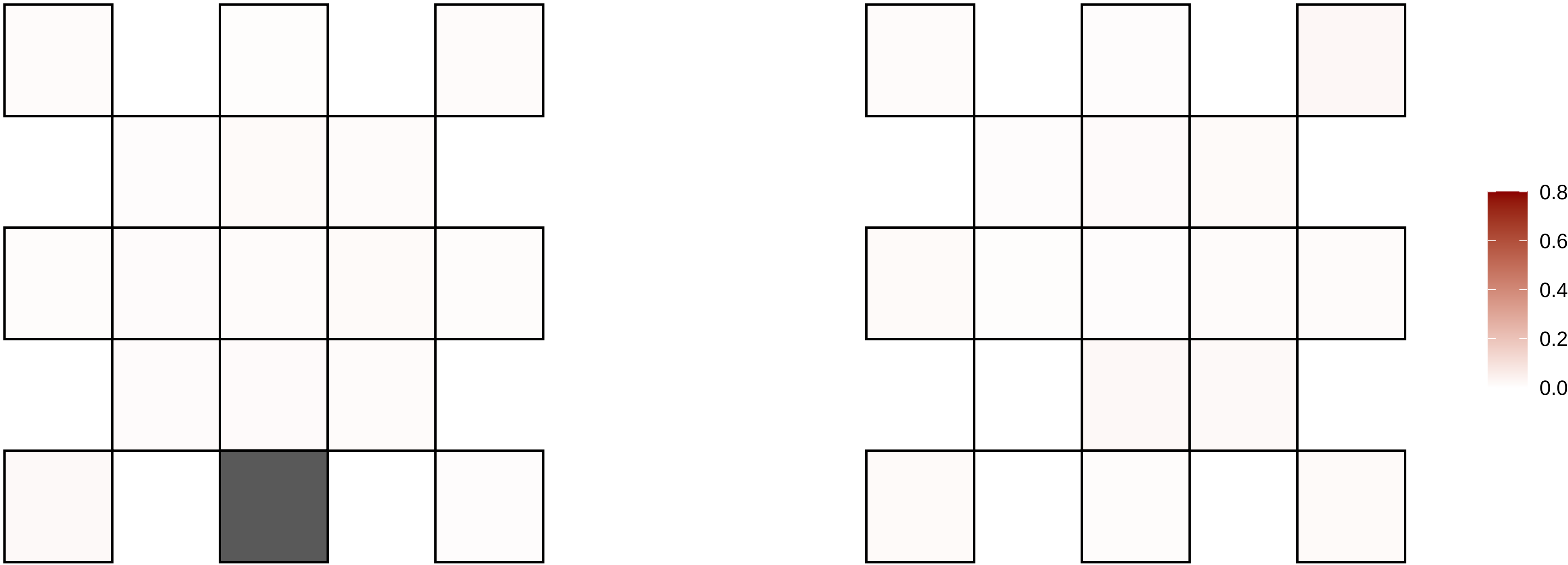

13−65586962−C−A

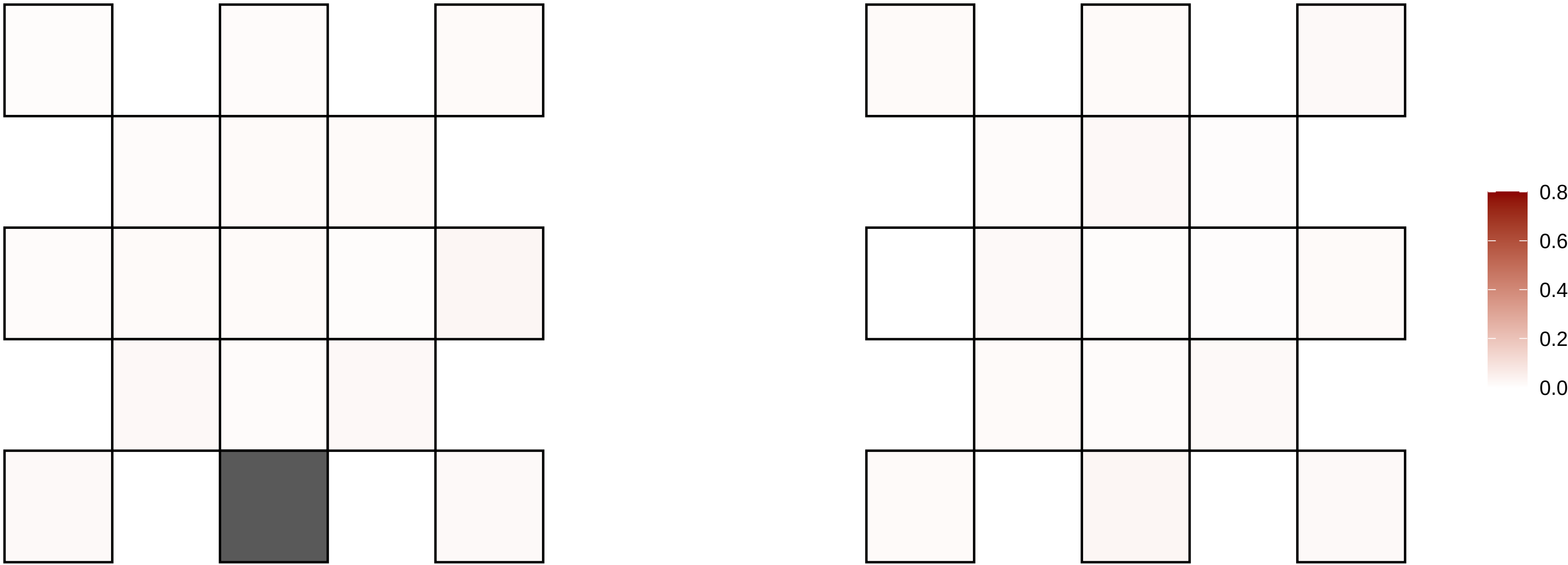

13−67322807−C−A

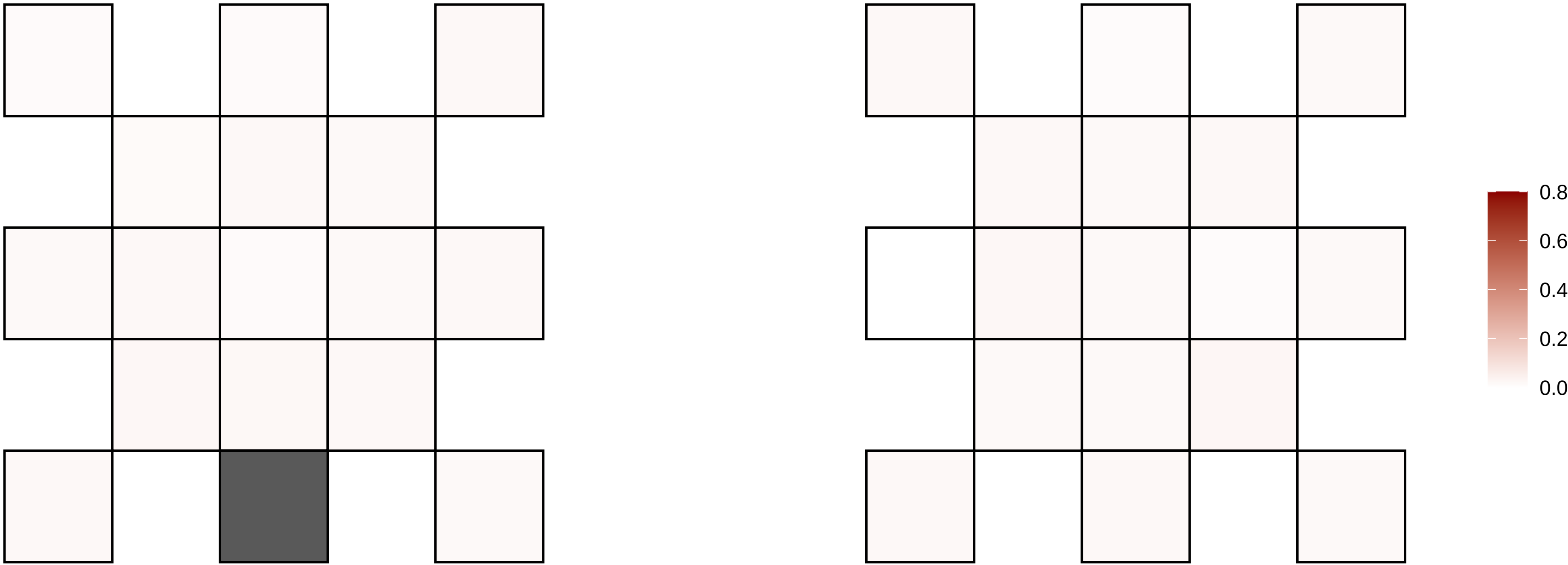

13−67852564−G−A

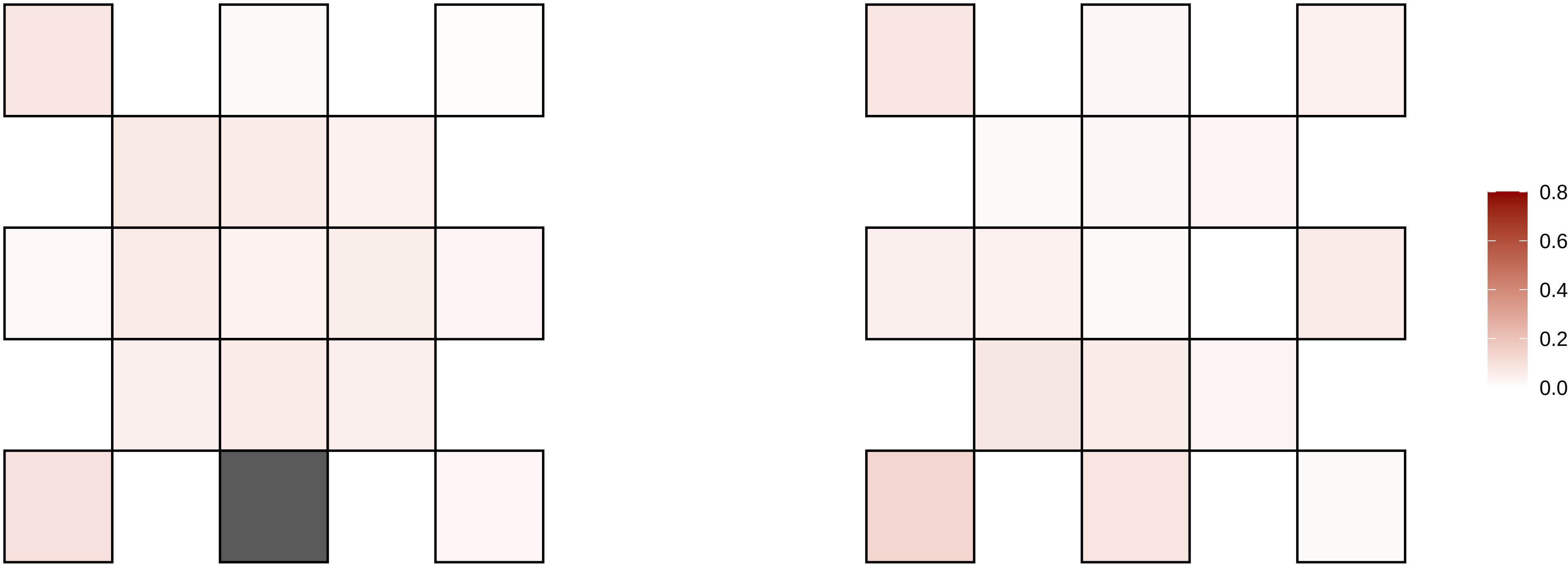

13−69093869−G−A

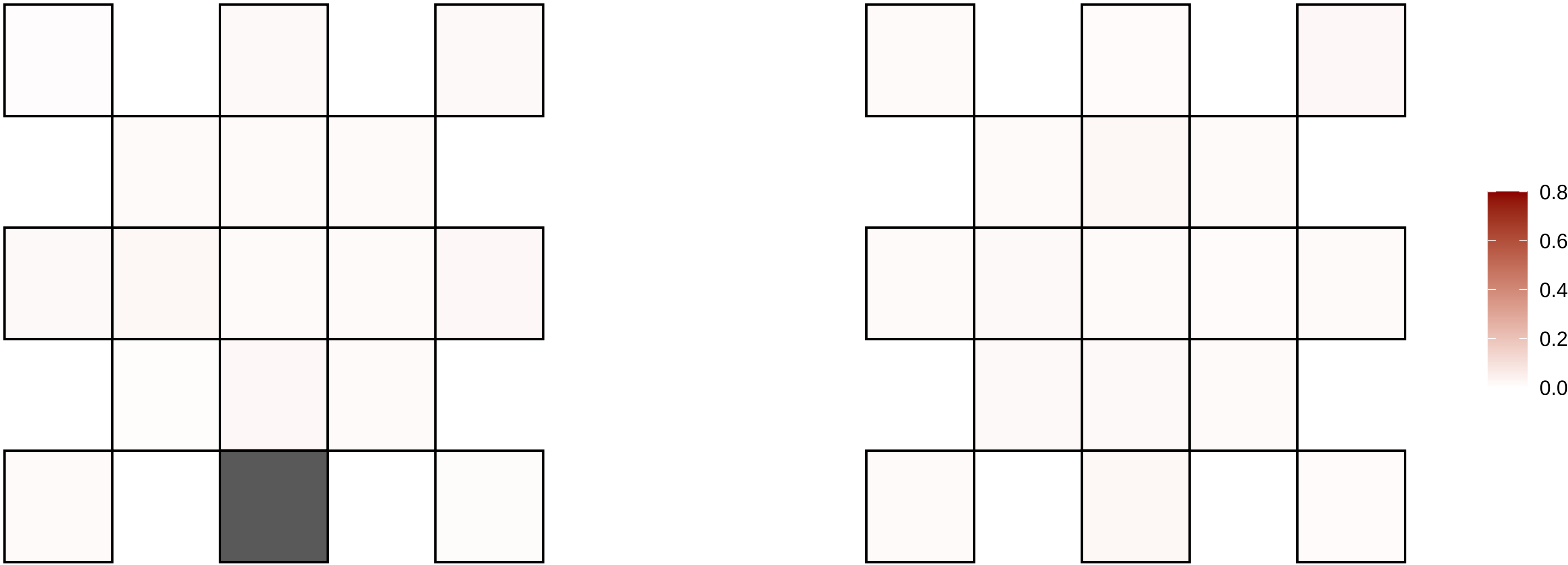

13−69308268−A−G

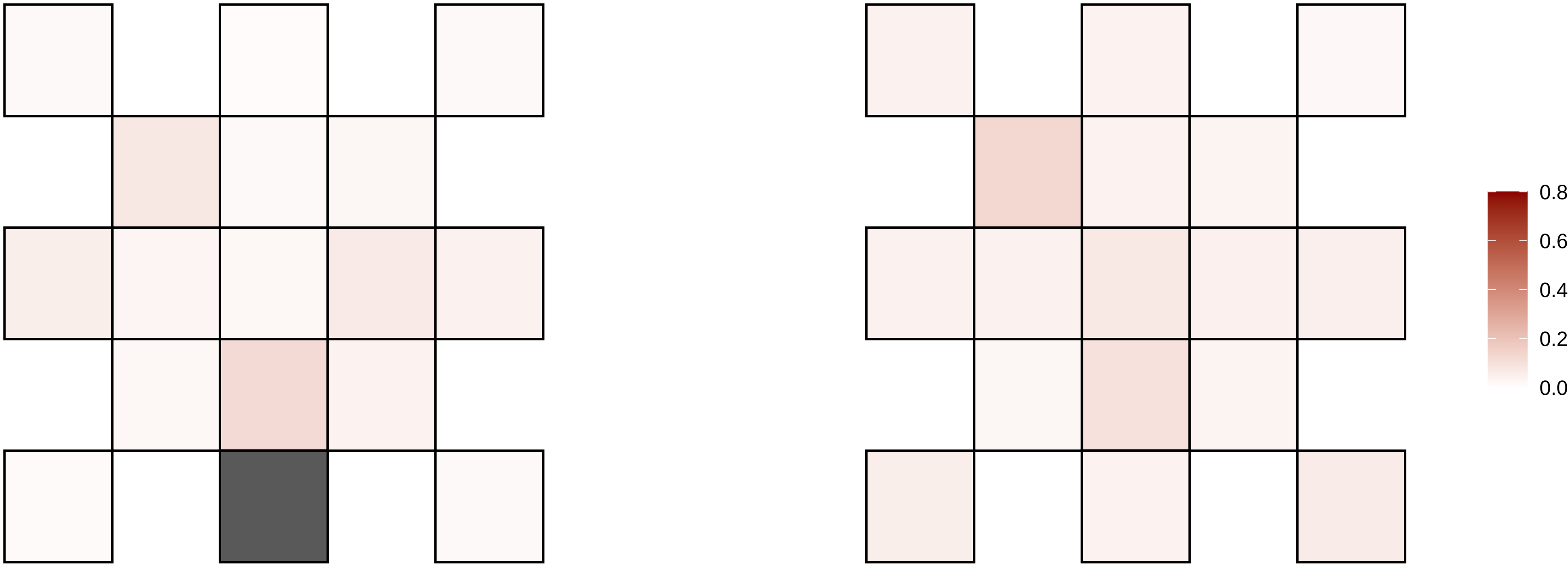

13−70279433−G−A

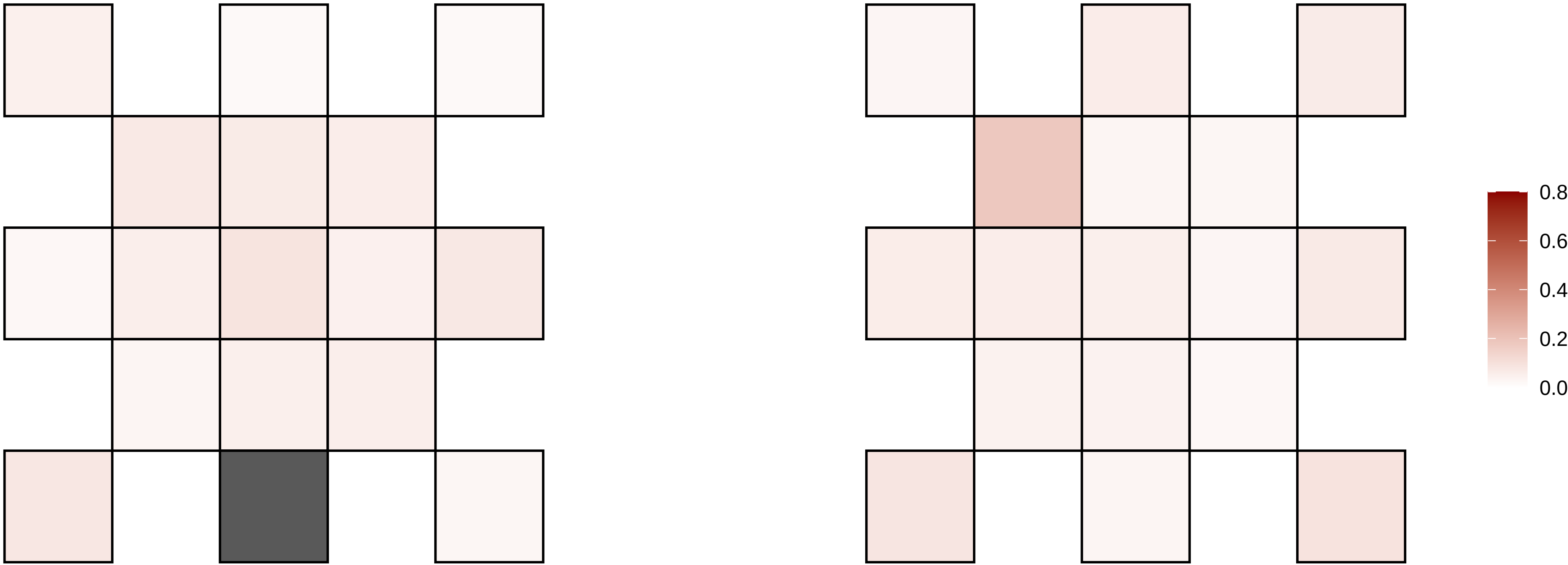

13−72218943−C−A

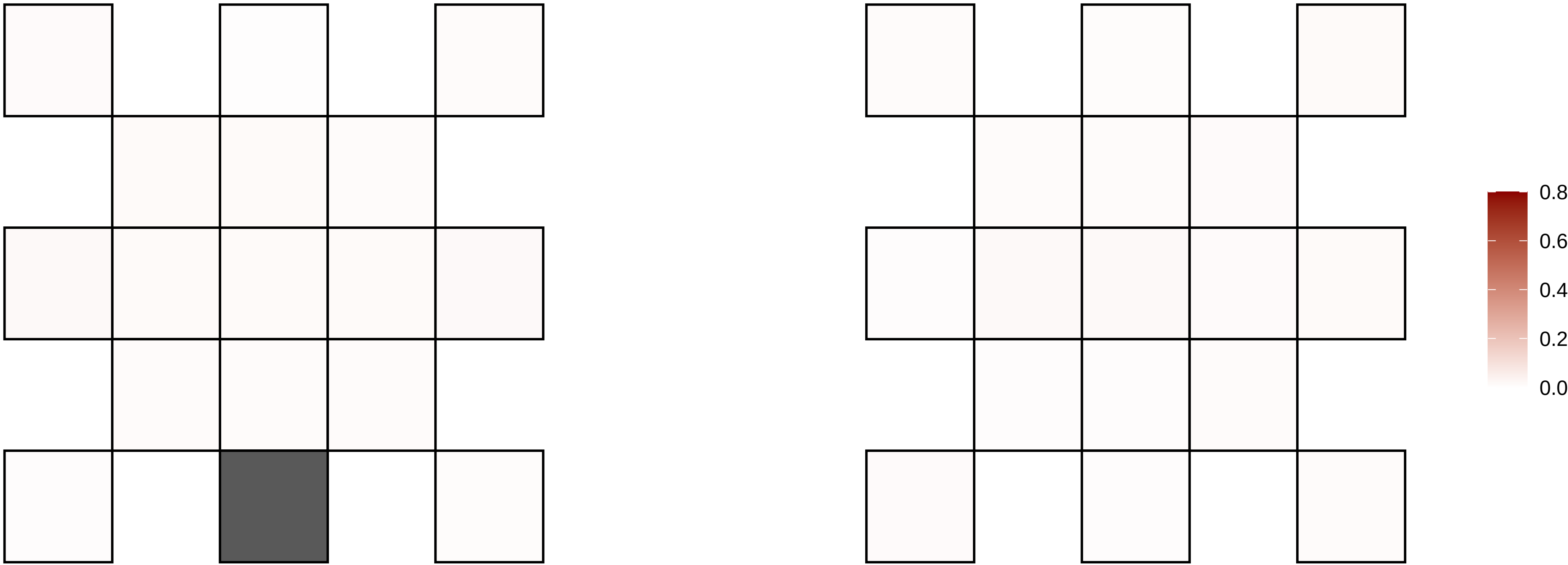

13−76279674−G−A

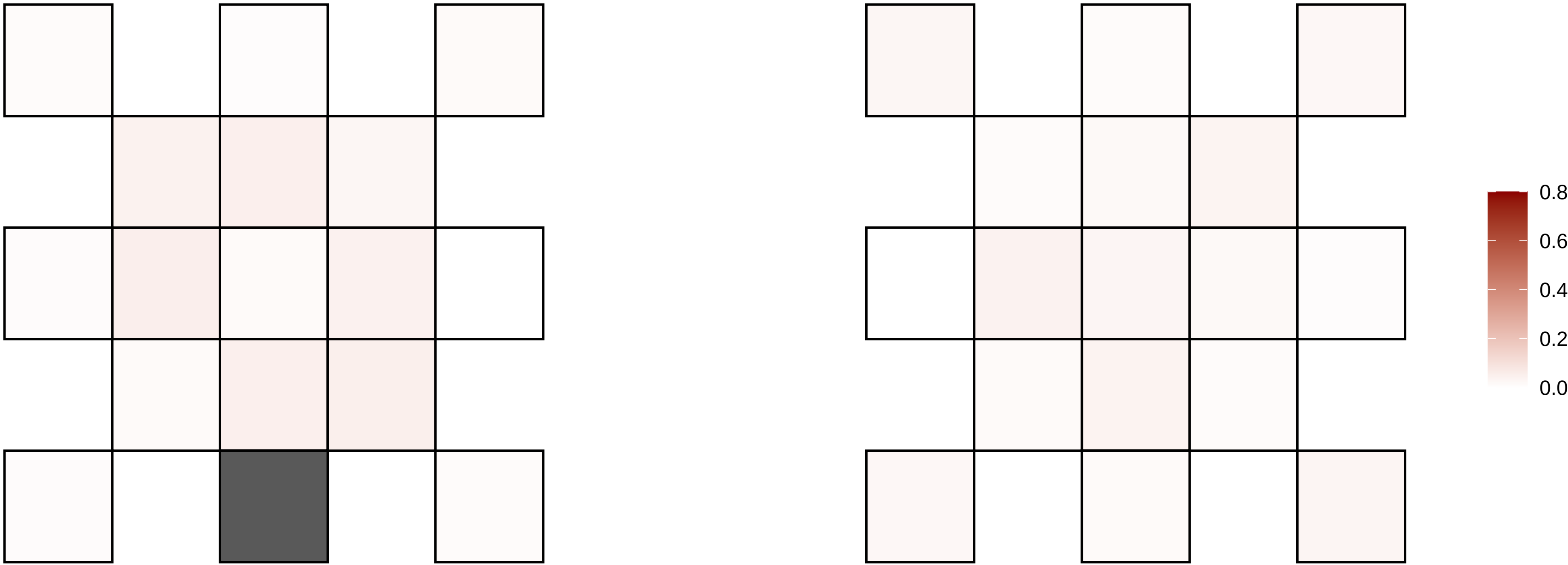

13−76874198−C−T

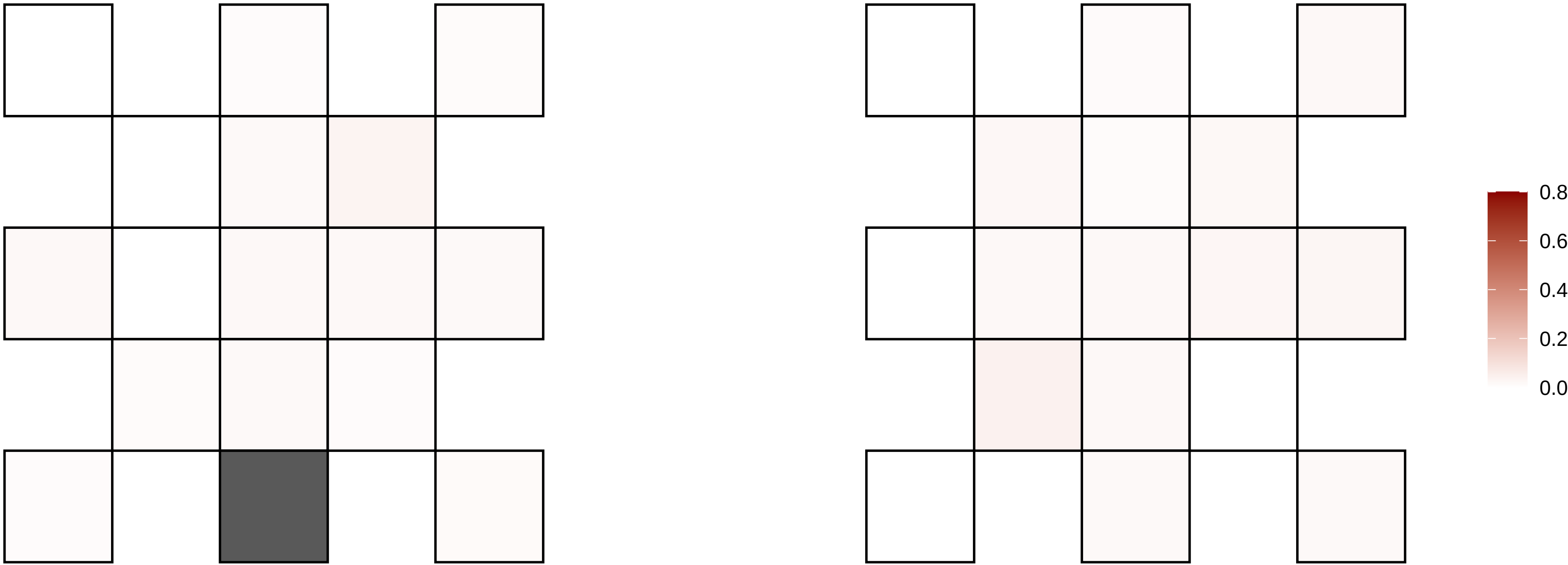

13−77509041−G−T

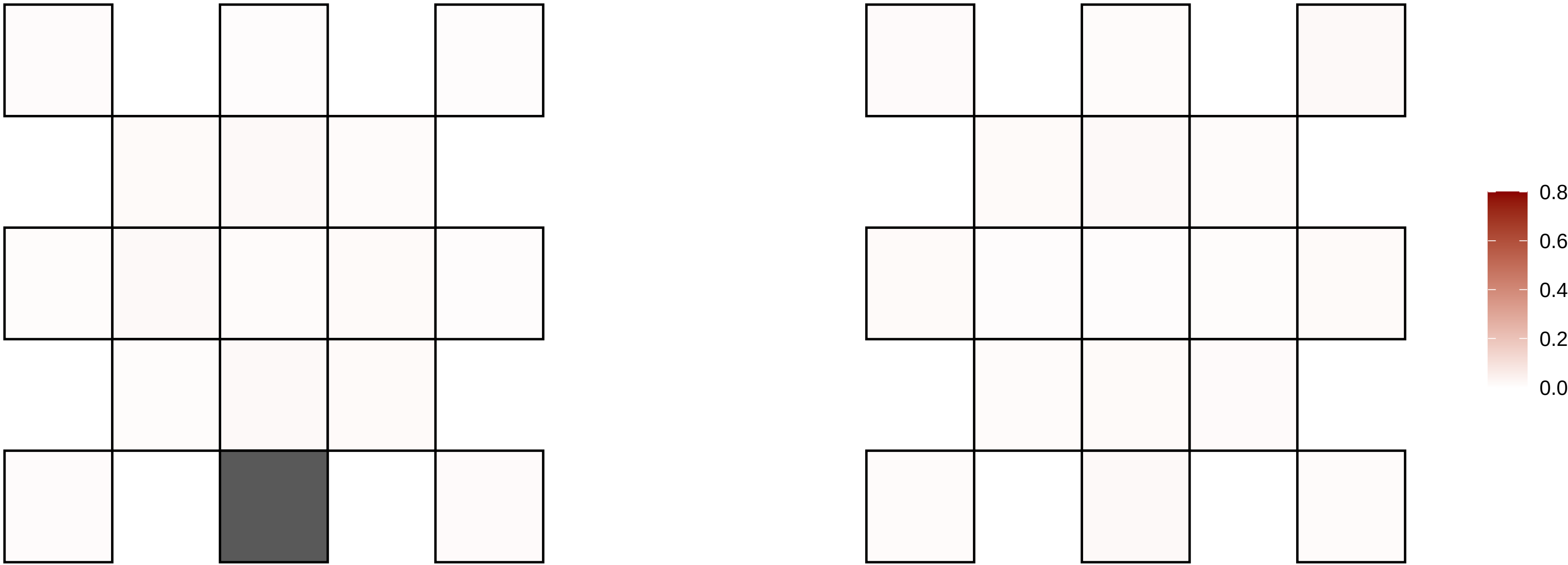

13−83875706−C−T

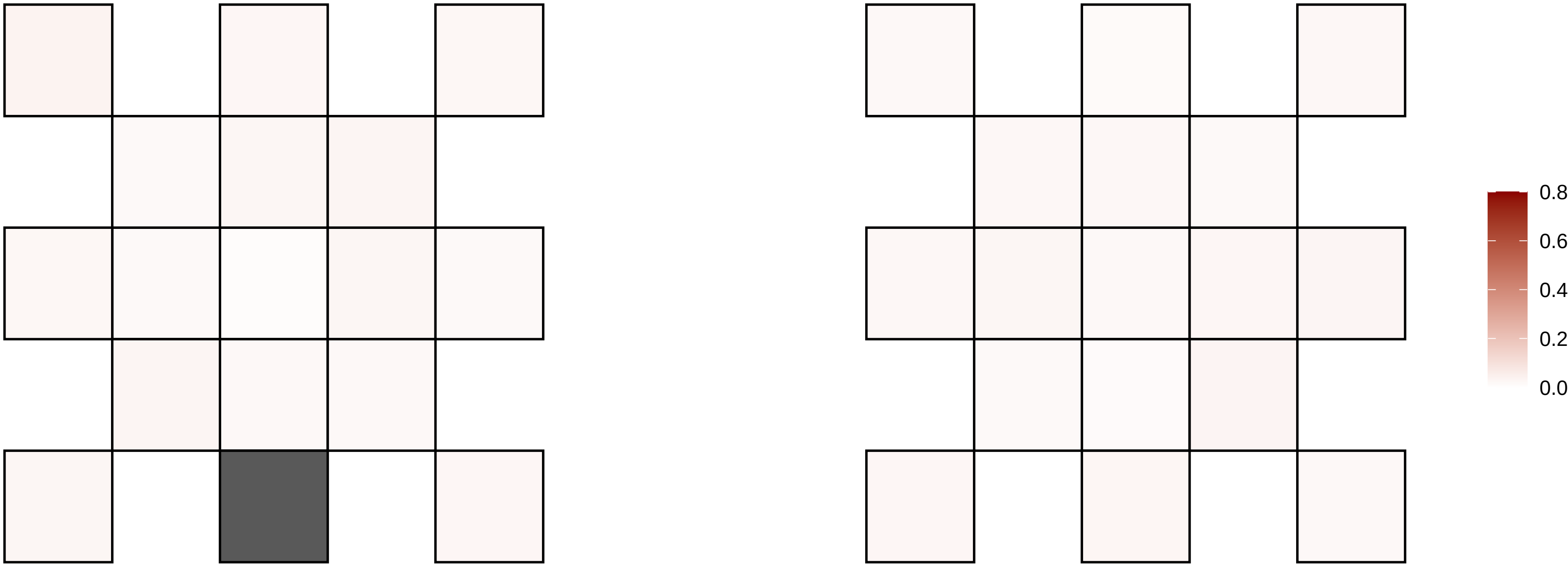

13−87055884−G−A

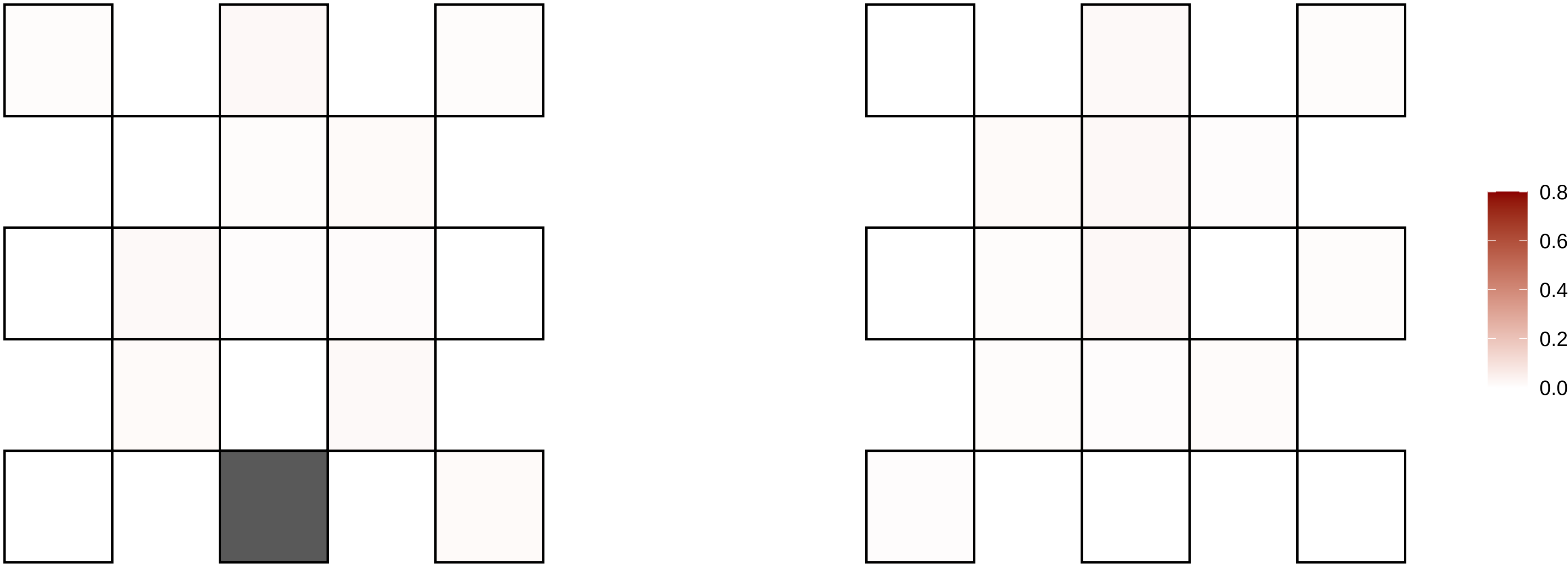

13−92867989−T−C

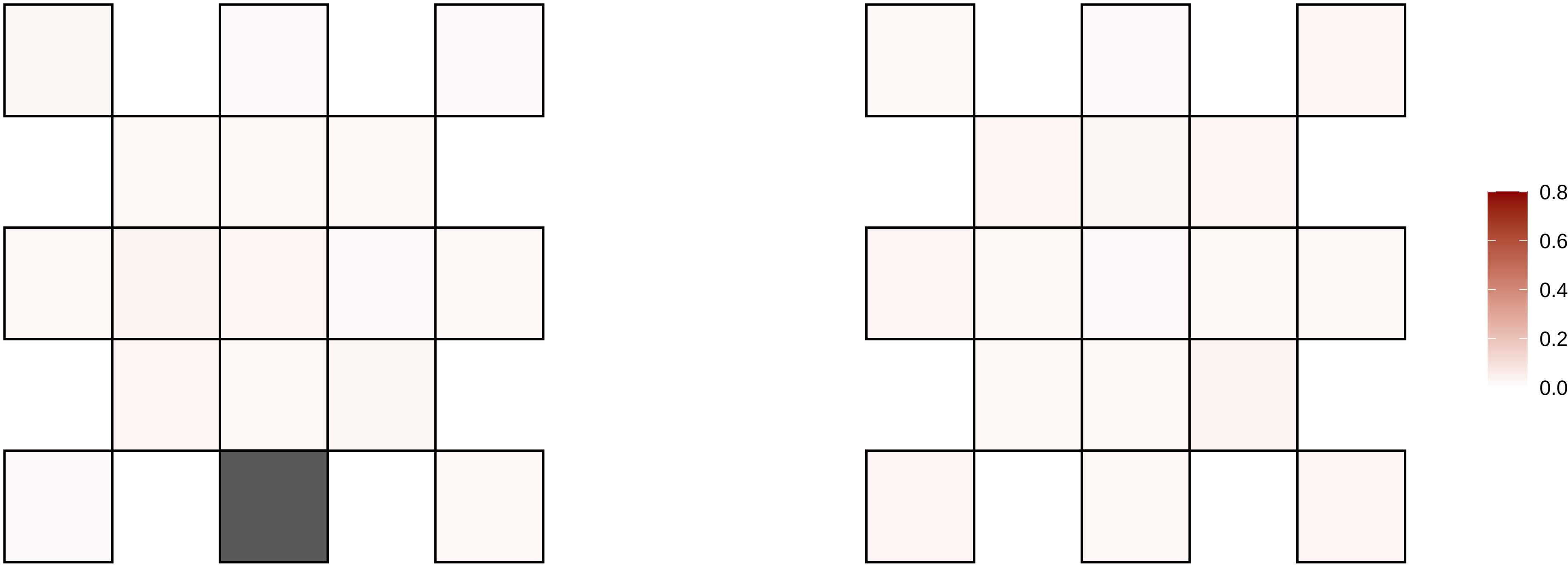

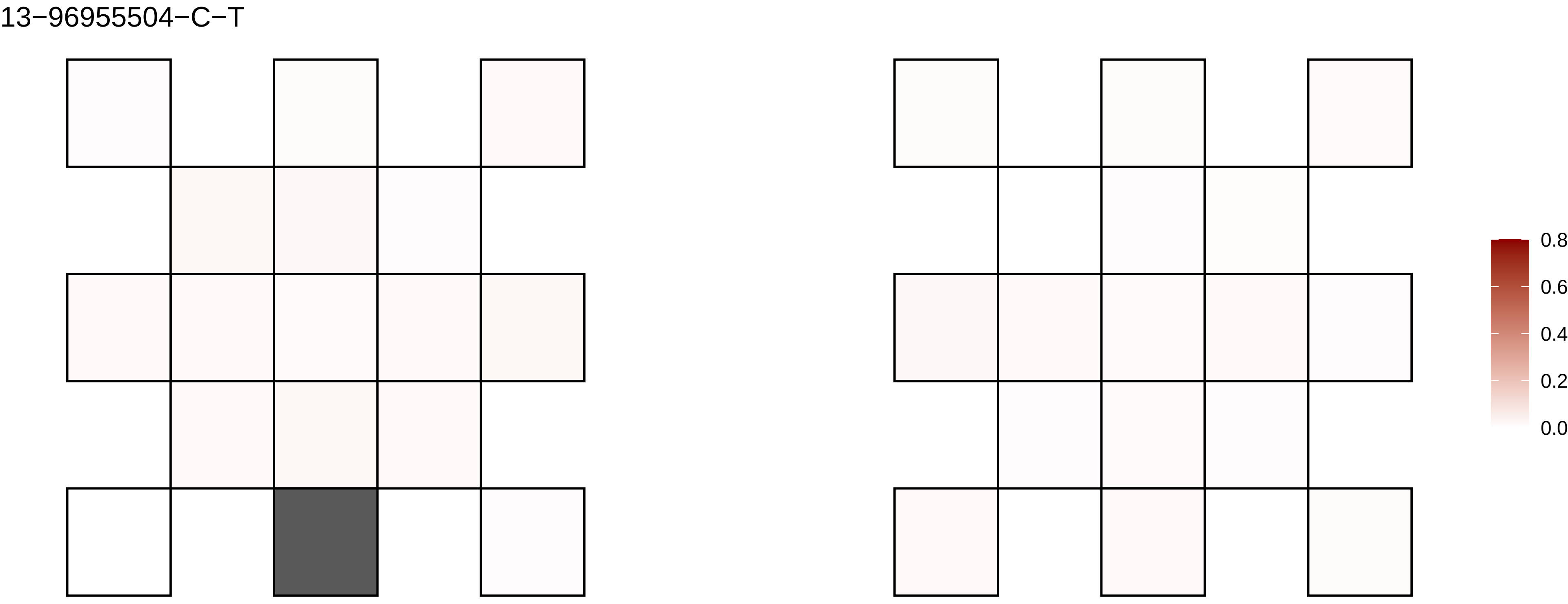
13−96955504−C−T
0.8
0.6
0.4
0.2
0.0

13−97072497−C−T

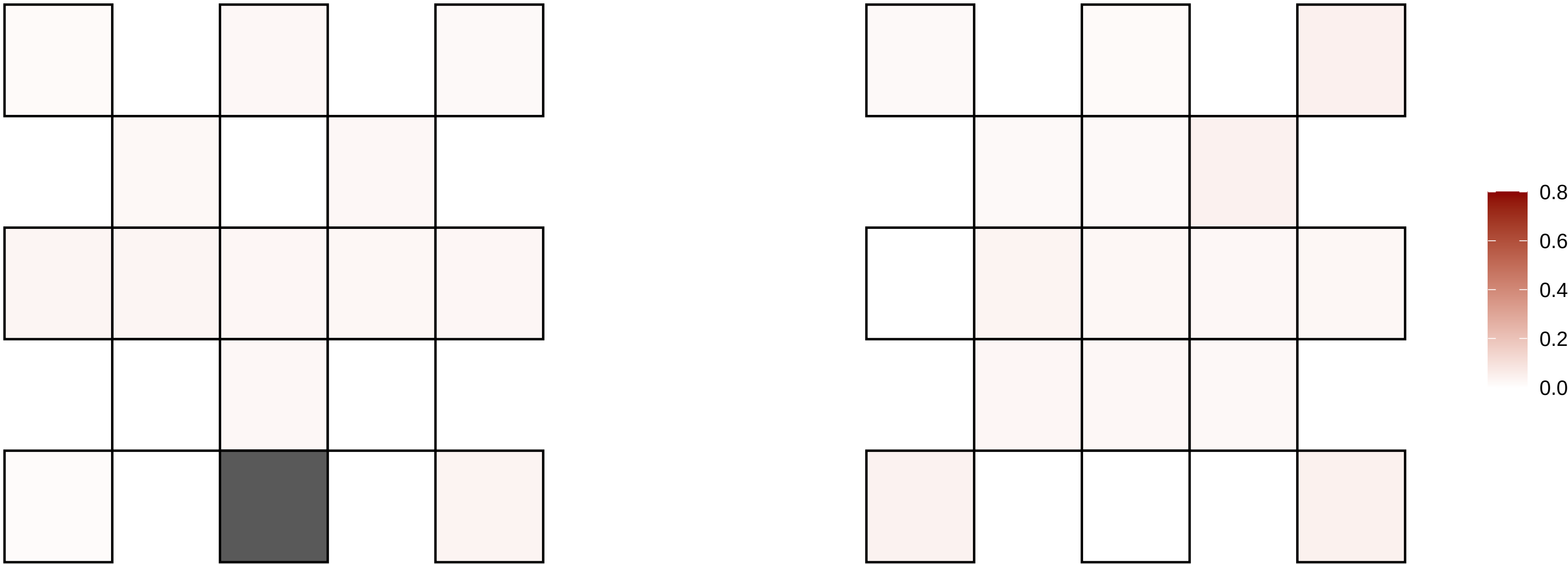

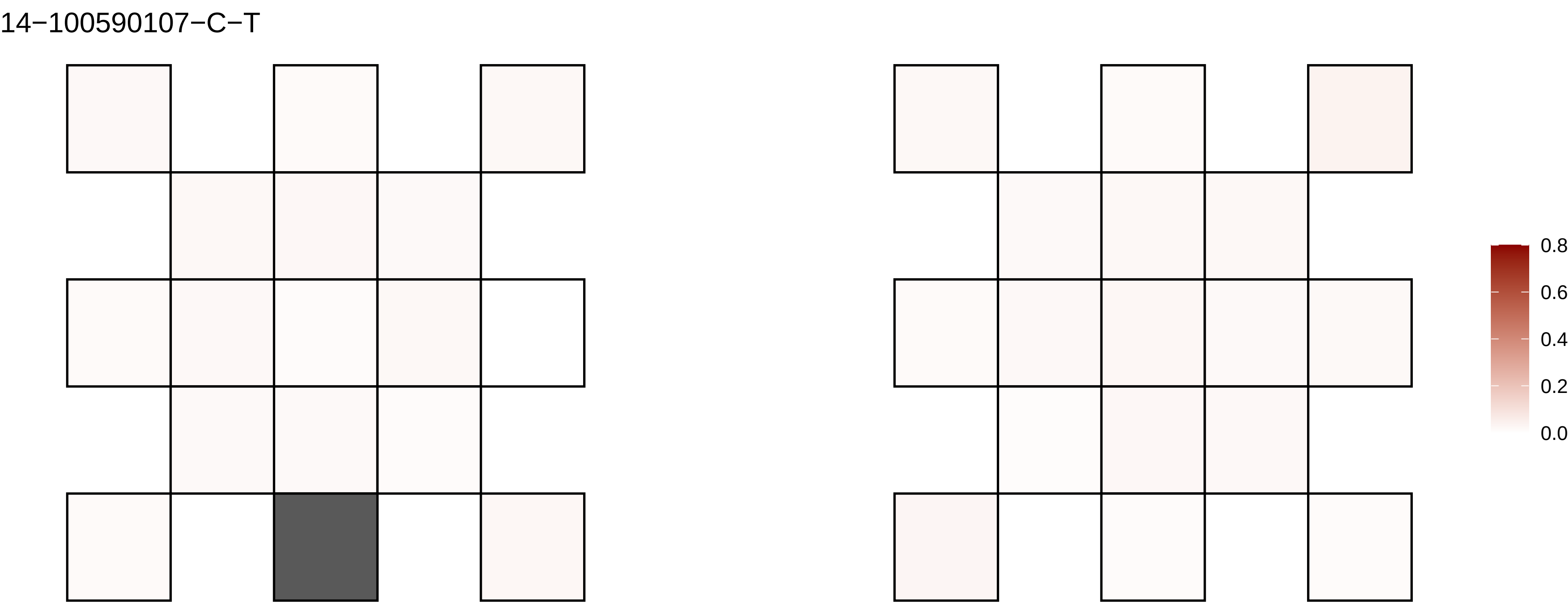
14−100590107−C−T
0.8
0.6
0.4
0.2
0.0

14−25316209−C−A

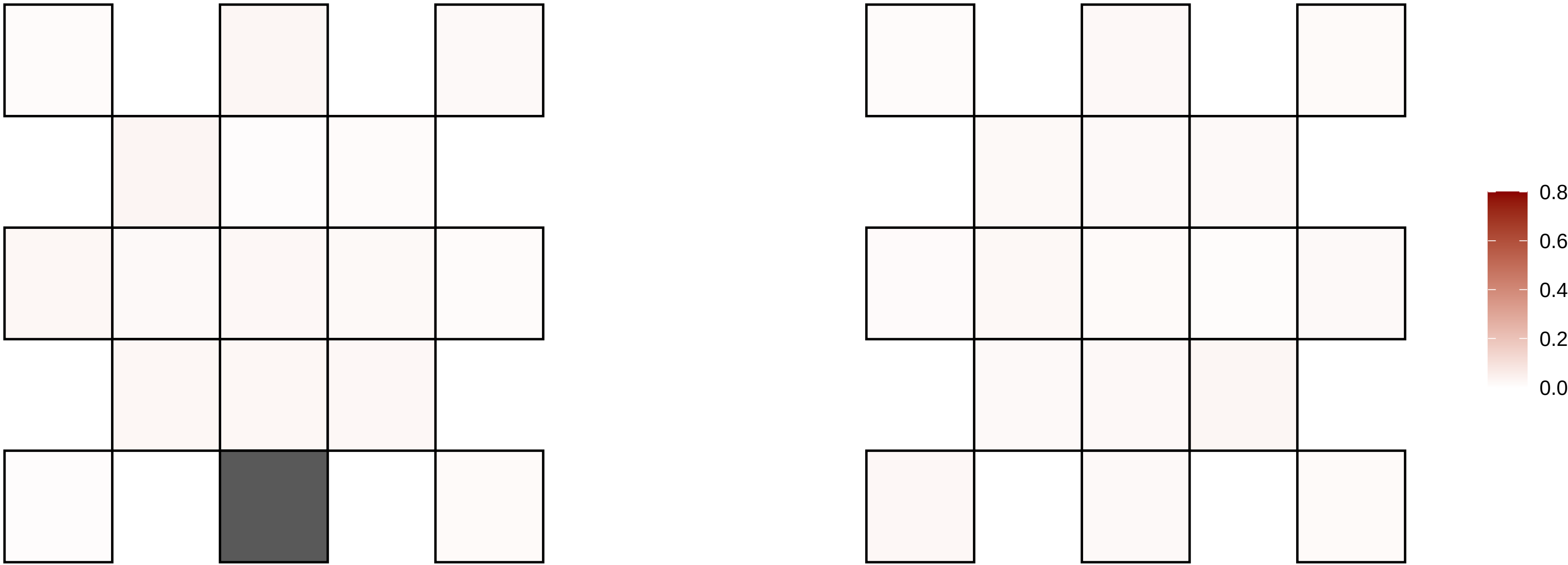

14−27412390−G−A

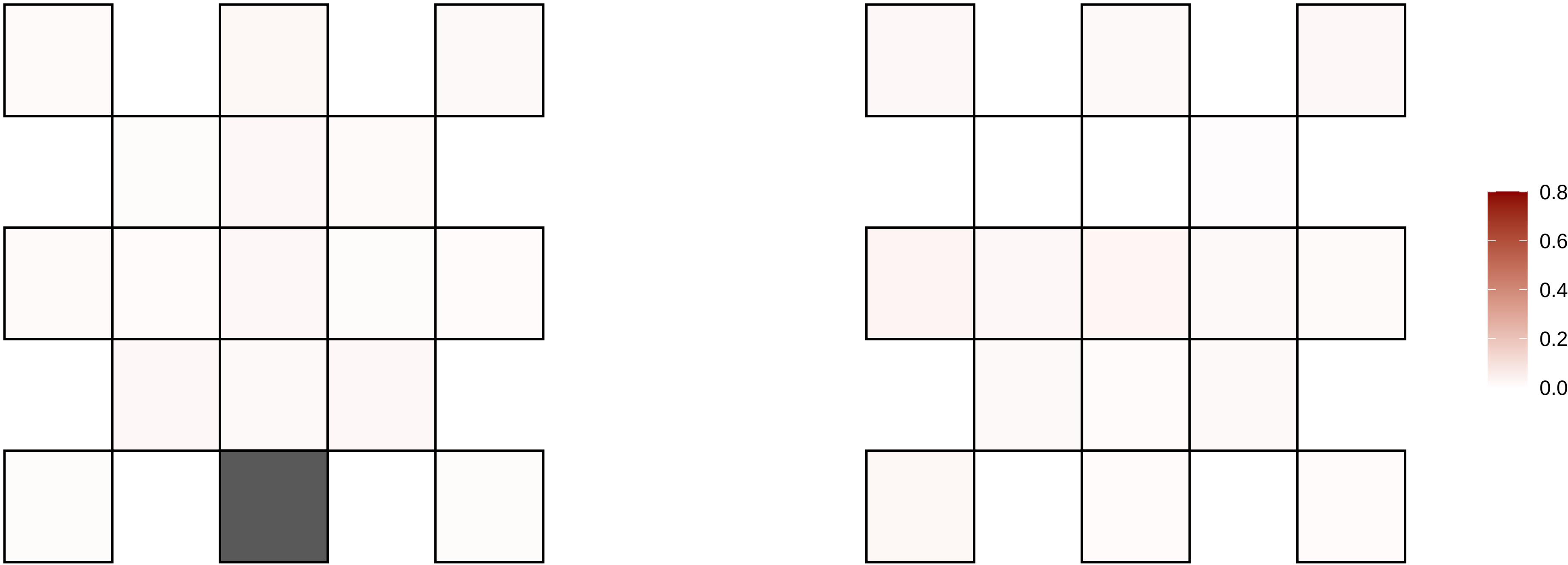

14−27649824−C−T

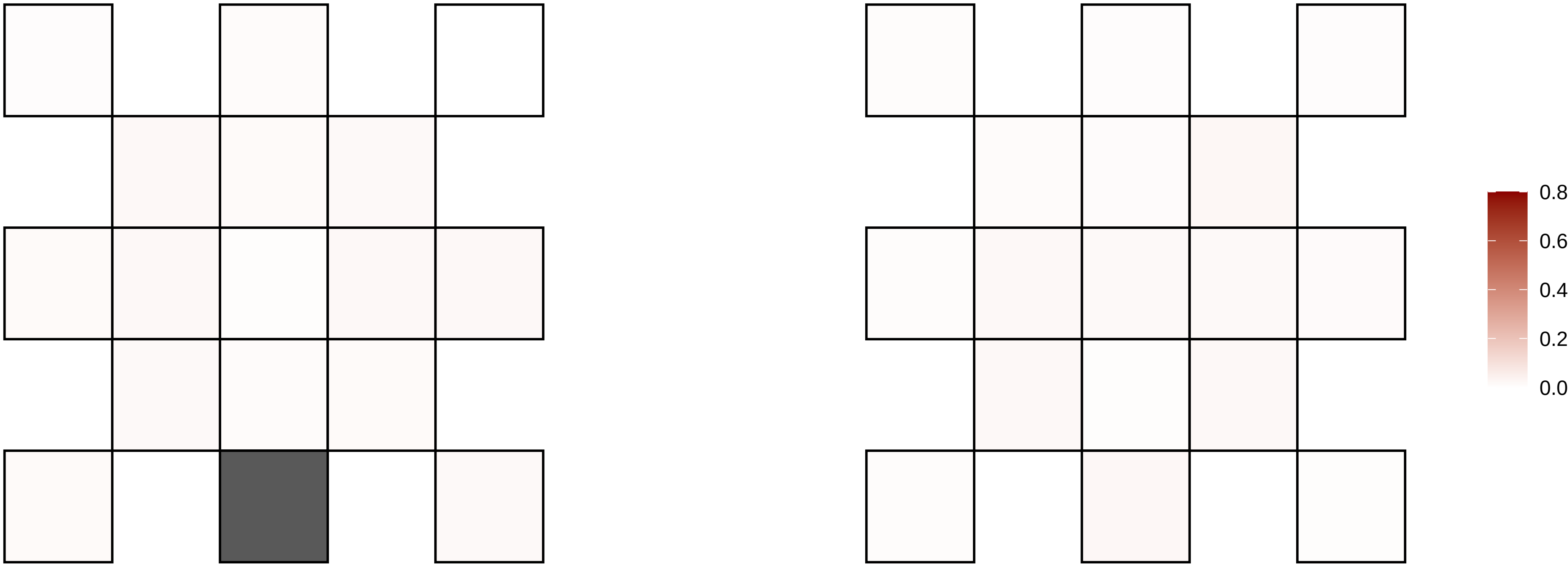

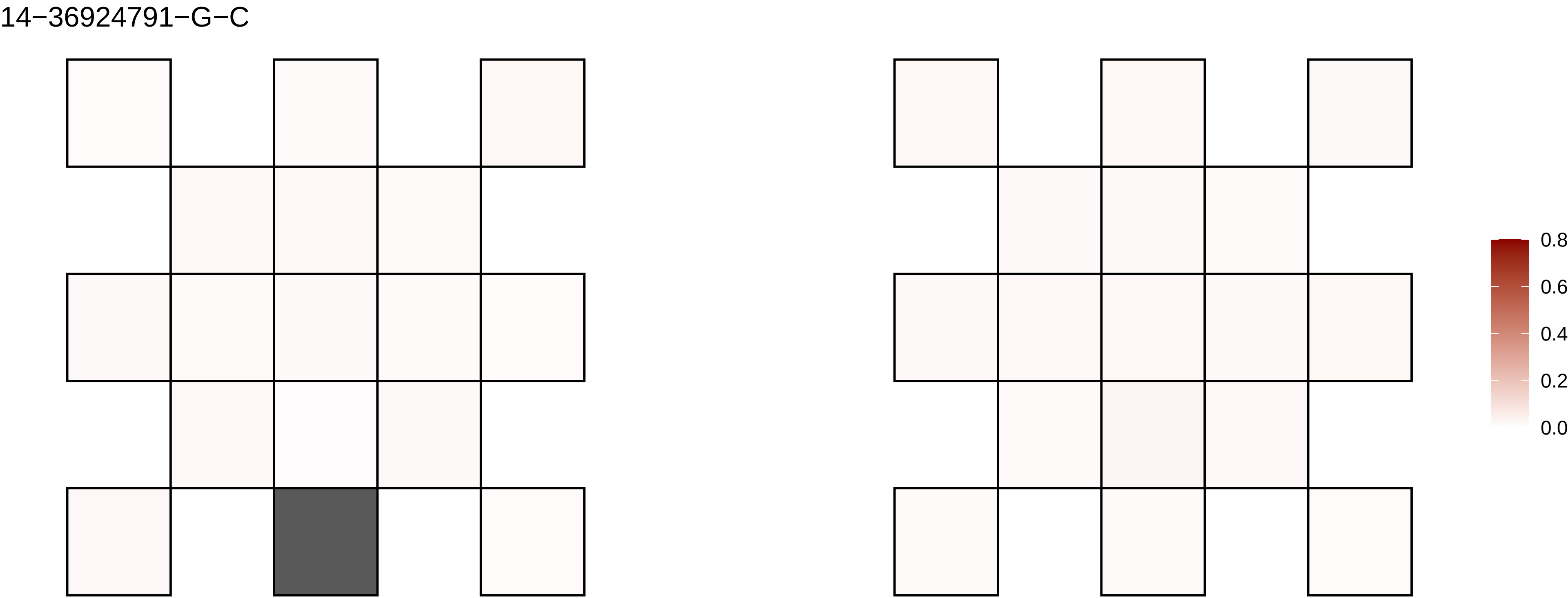
14−36924791−G−C
0.8
0.6
0.4
0.2
0.0

14−41027959−C−T

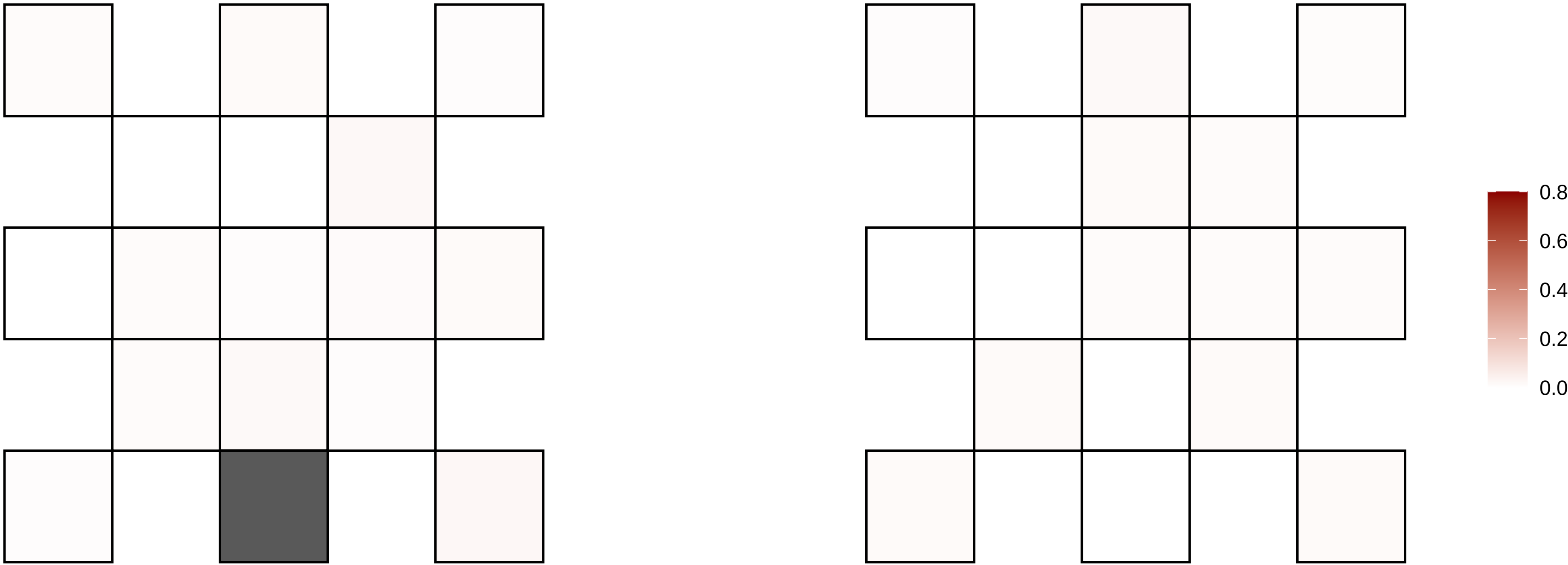

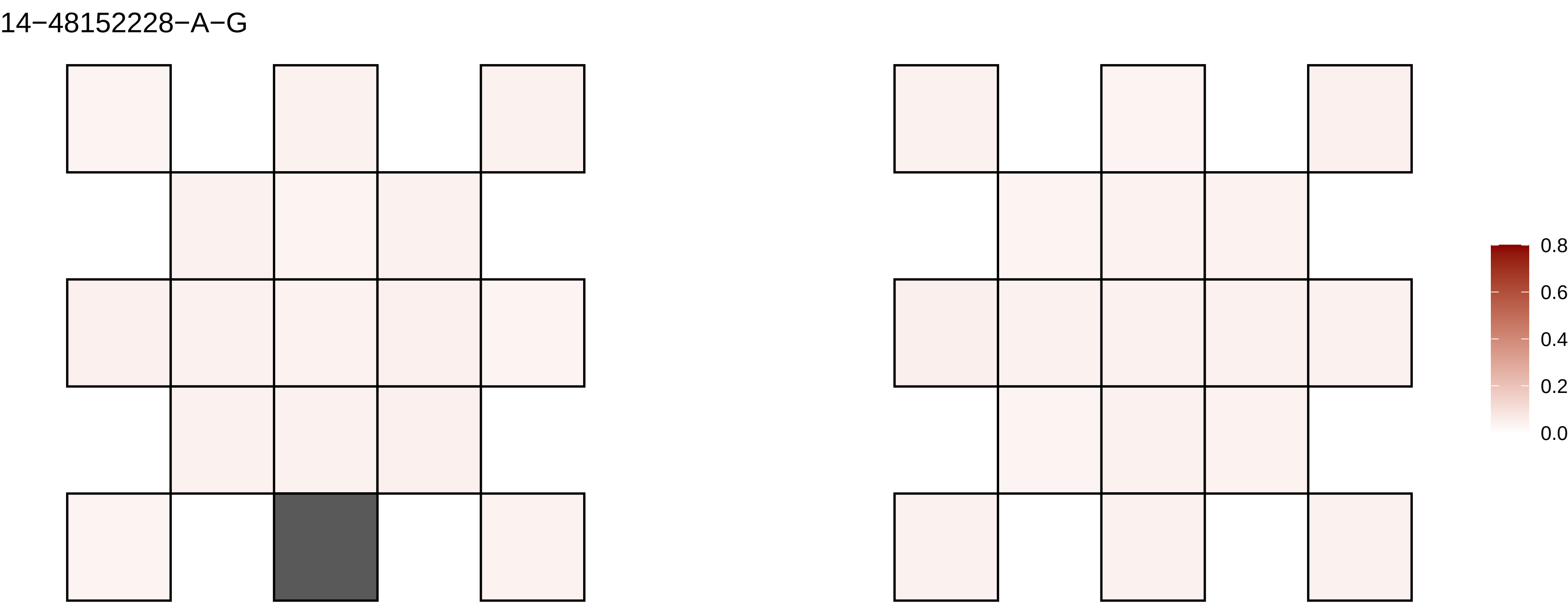
14−48152228−A−G
0.8
0.6
0.4
0.2
0.0

14−51478747−T−C

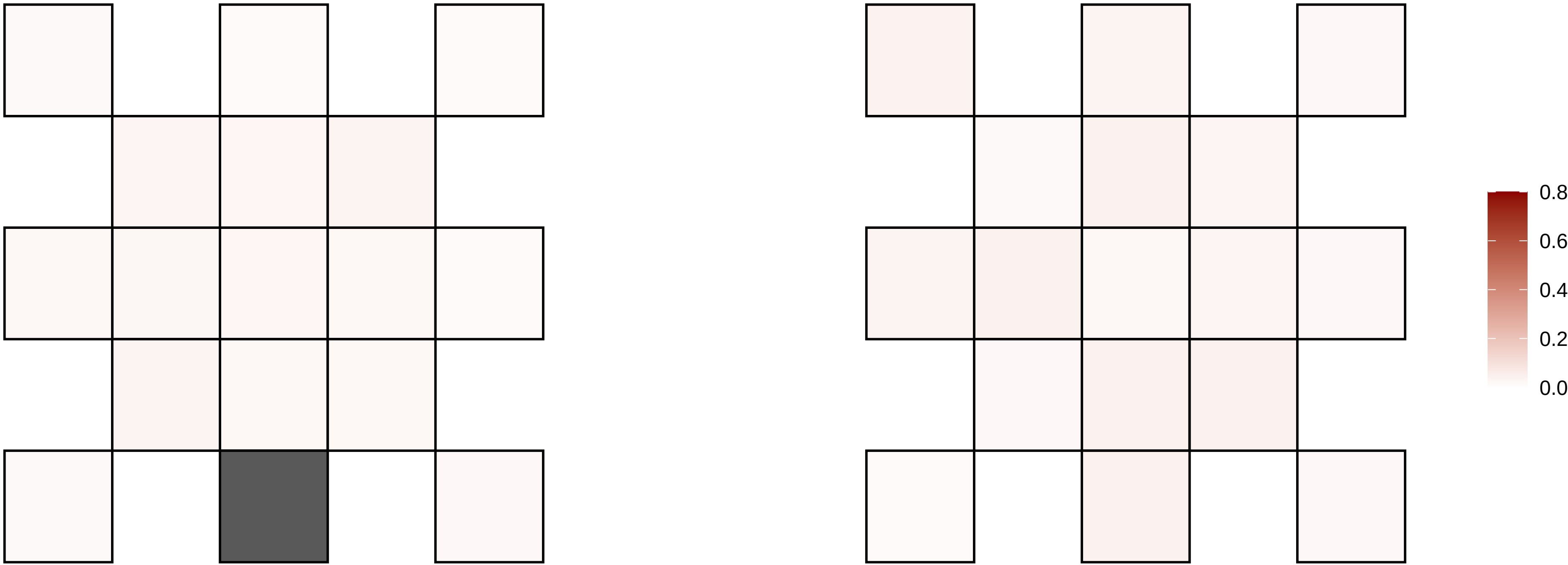

14−55887203−A−G

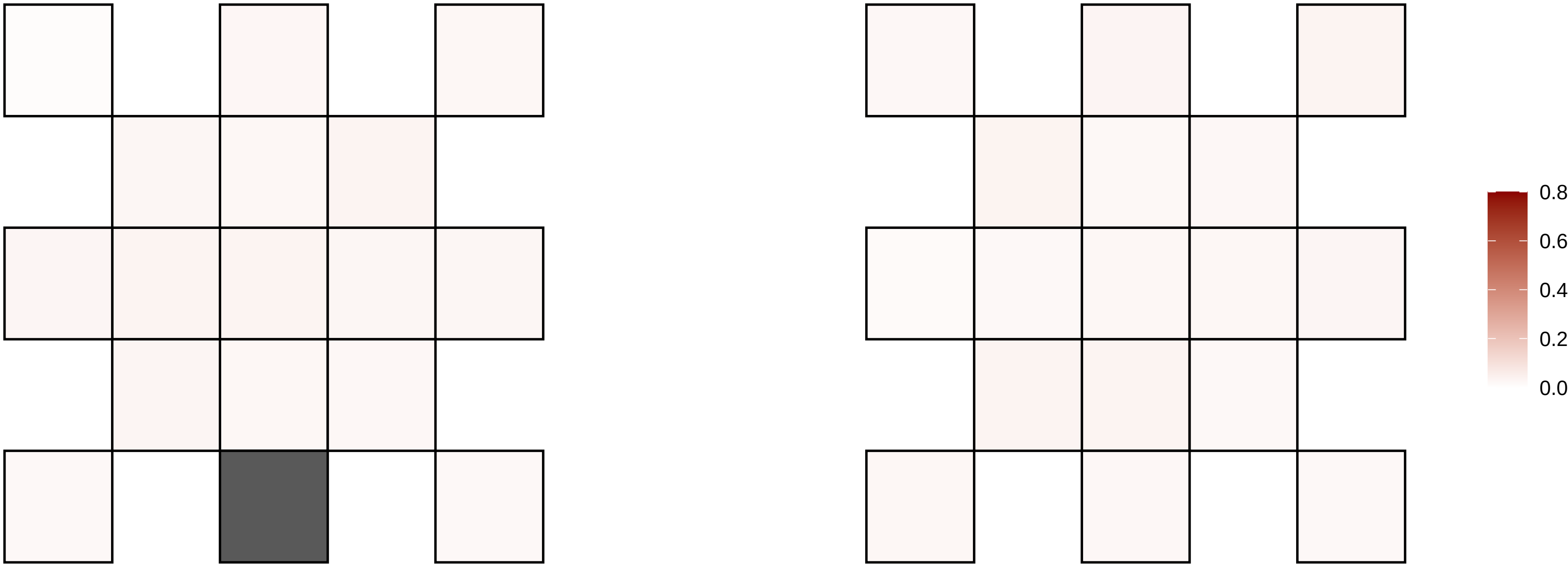

14−59827441−C−T

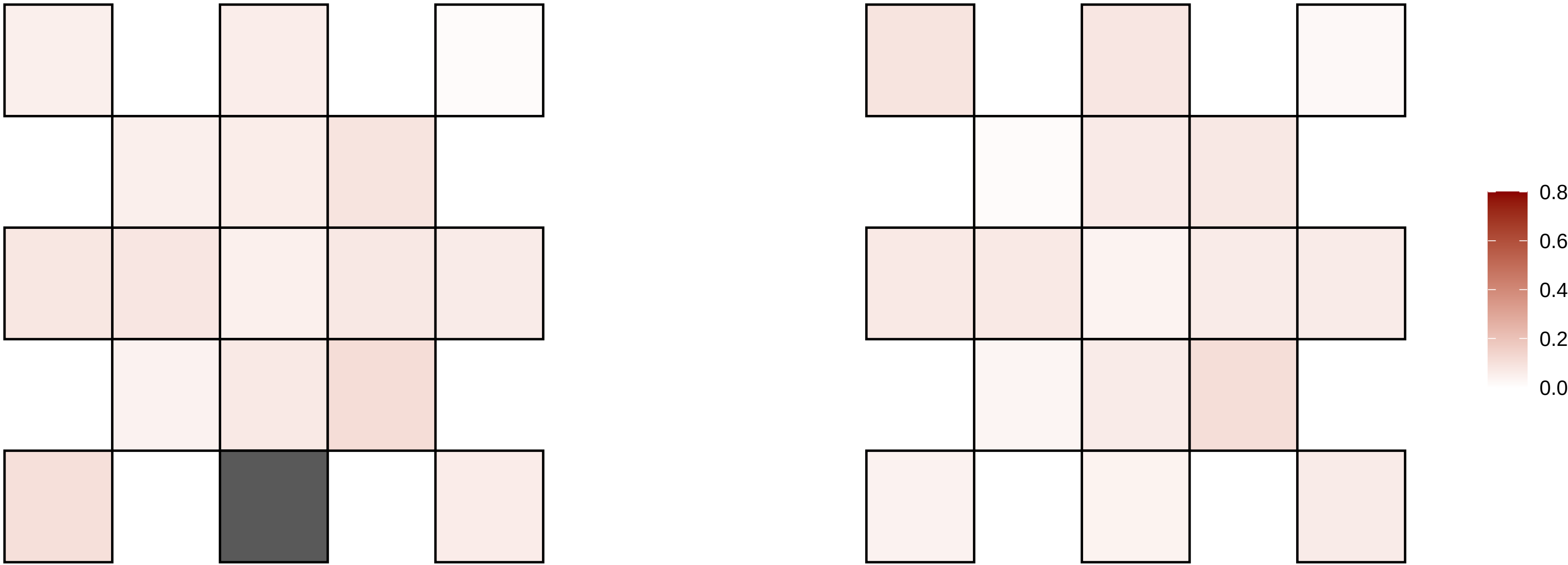

14−65264866−C−T

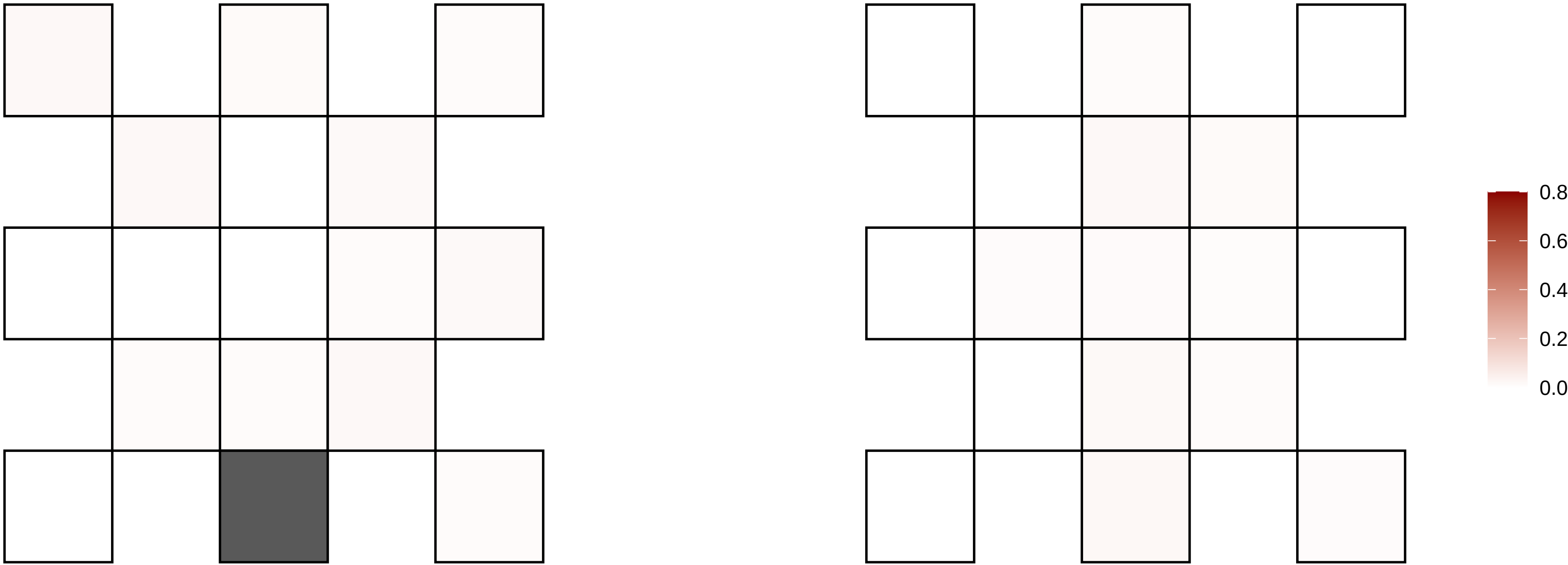

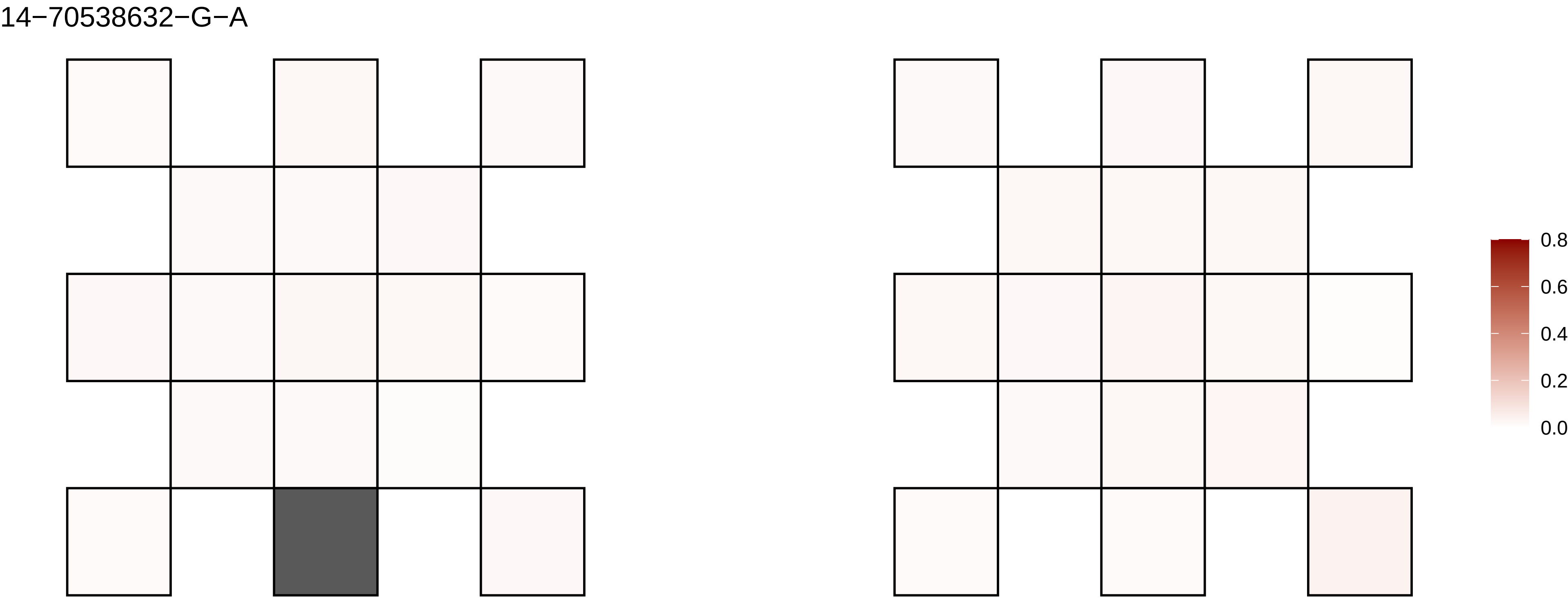
14−70538632−G−A
0.8
0.6
0.4
0.2
0.0

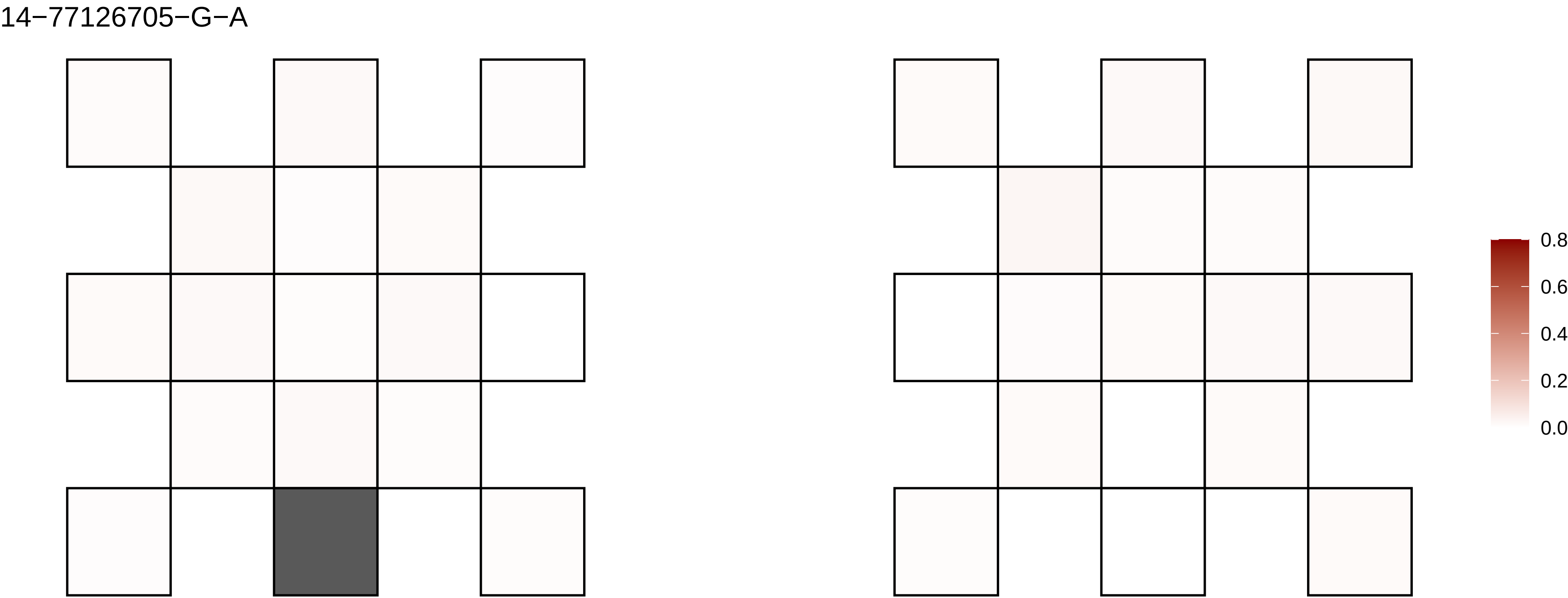
14−77126705−G−A
0.8
0.6
0.4
0.2
0.0

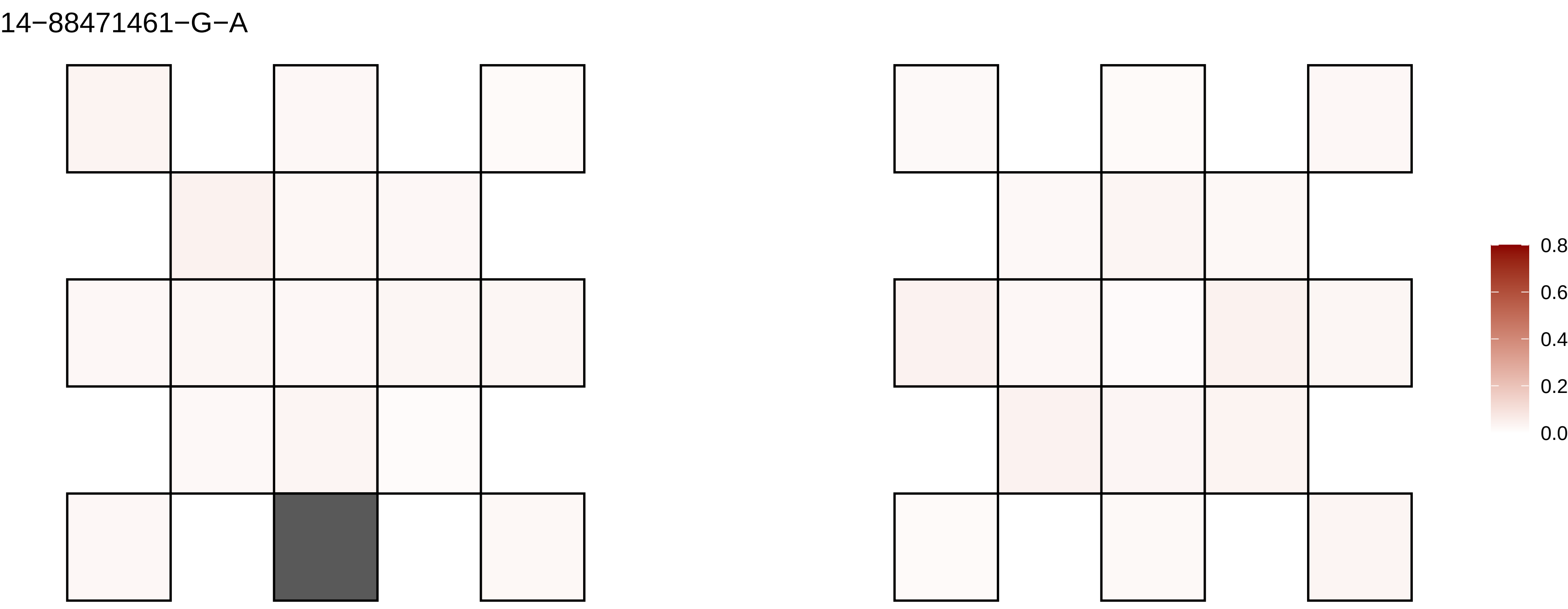

14−88471461−G−A
0.8
0.6
0.4
0.2
0.0

14−89375012−G−T

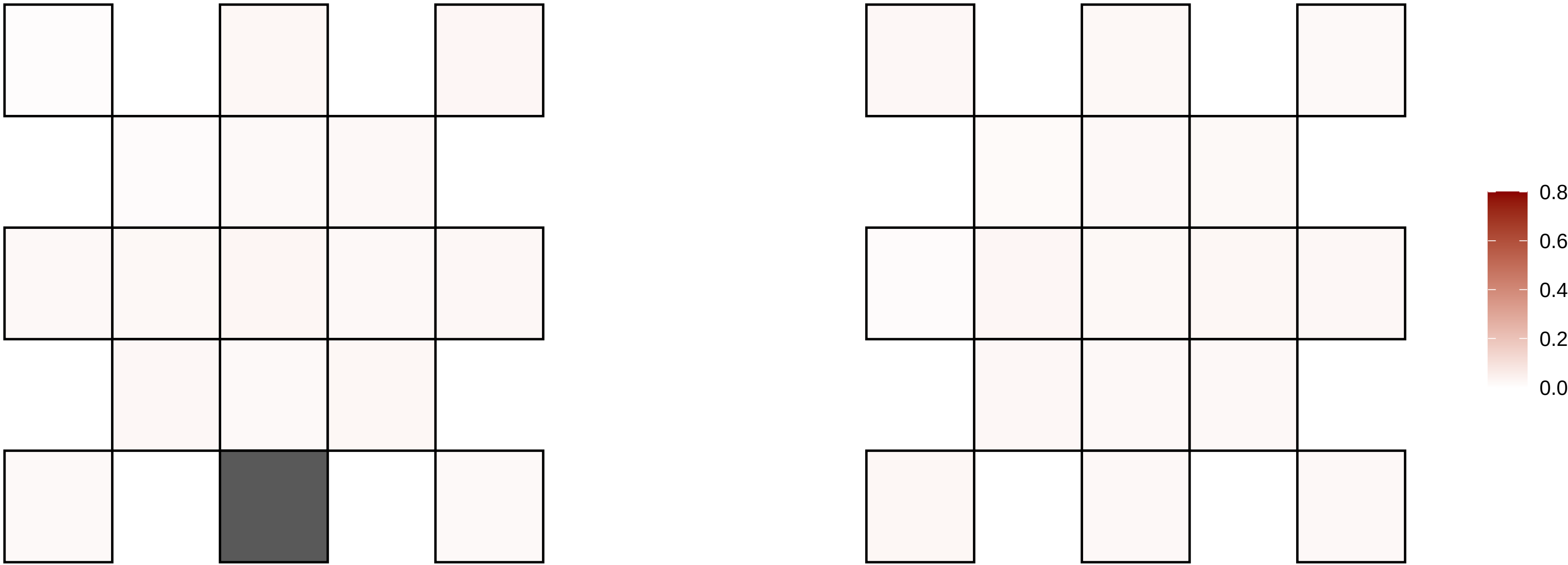

14−93813737−C−T

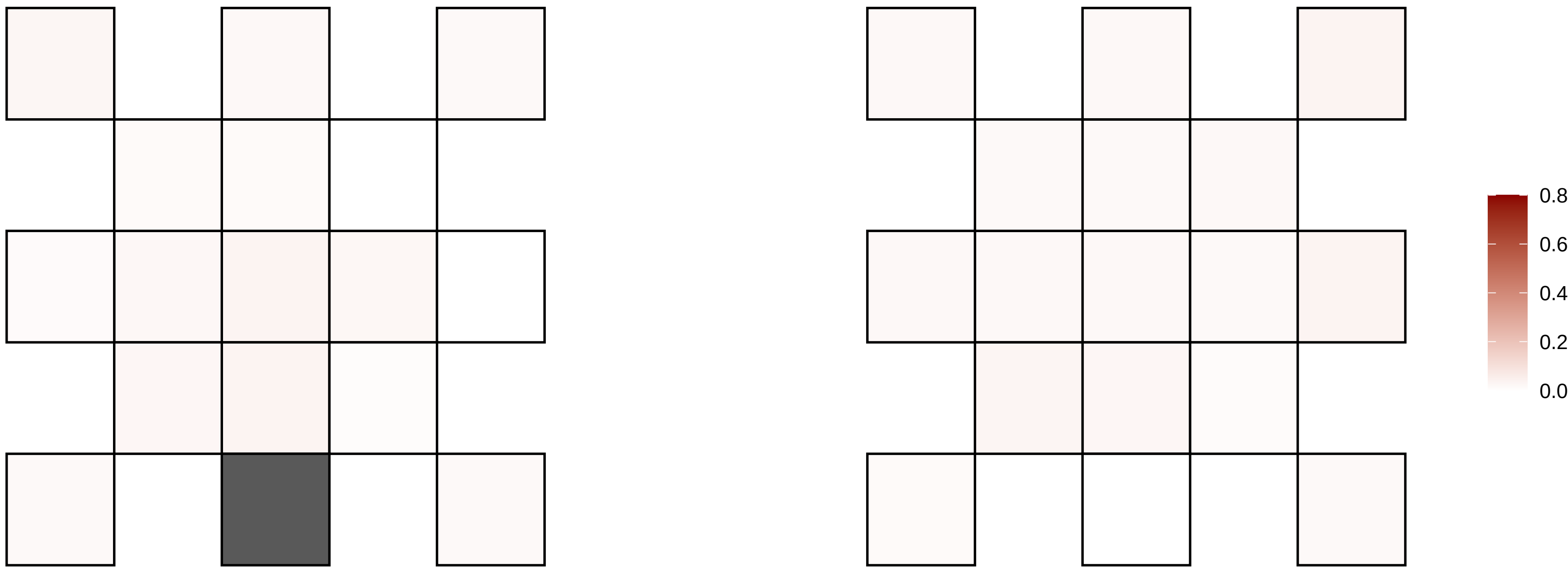

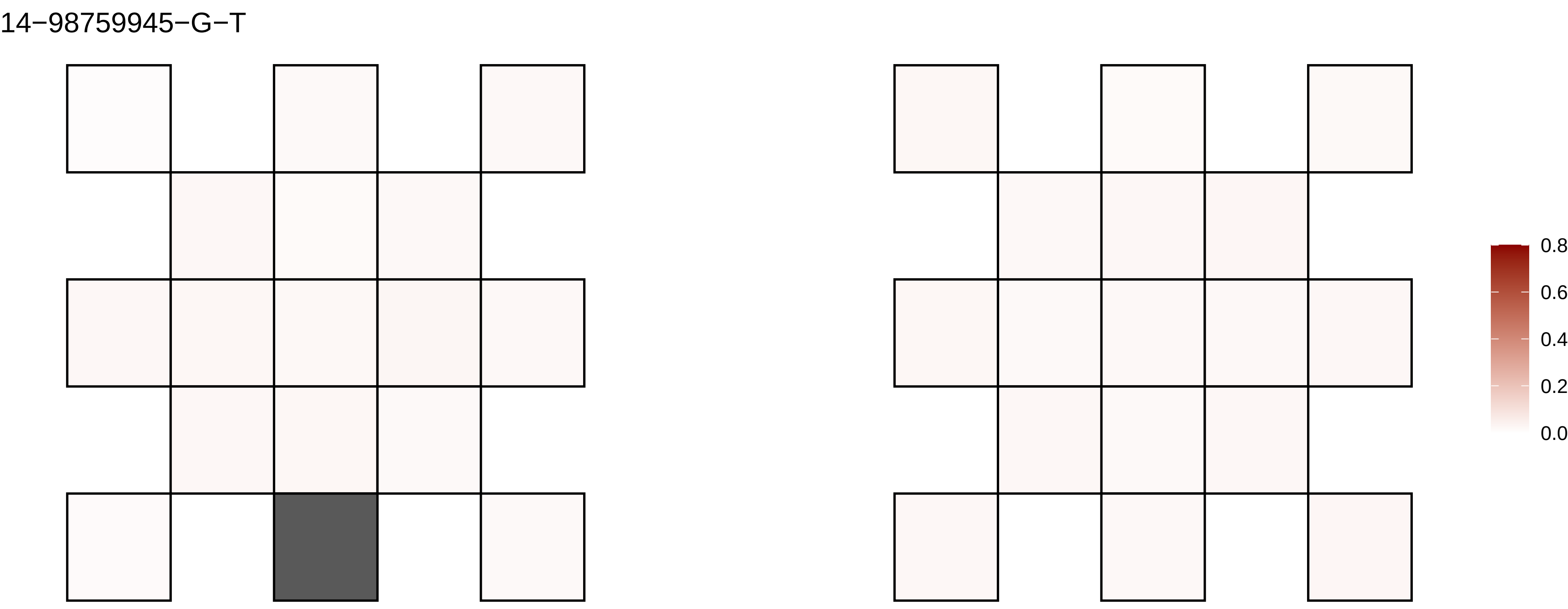
14−98759945−G−T
0.8
0.6
0.4
0.2
0.0

15−26356007−C−T

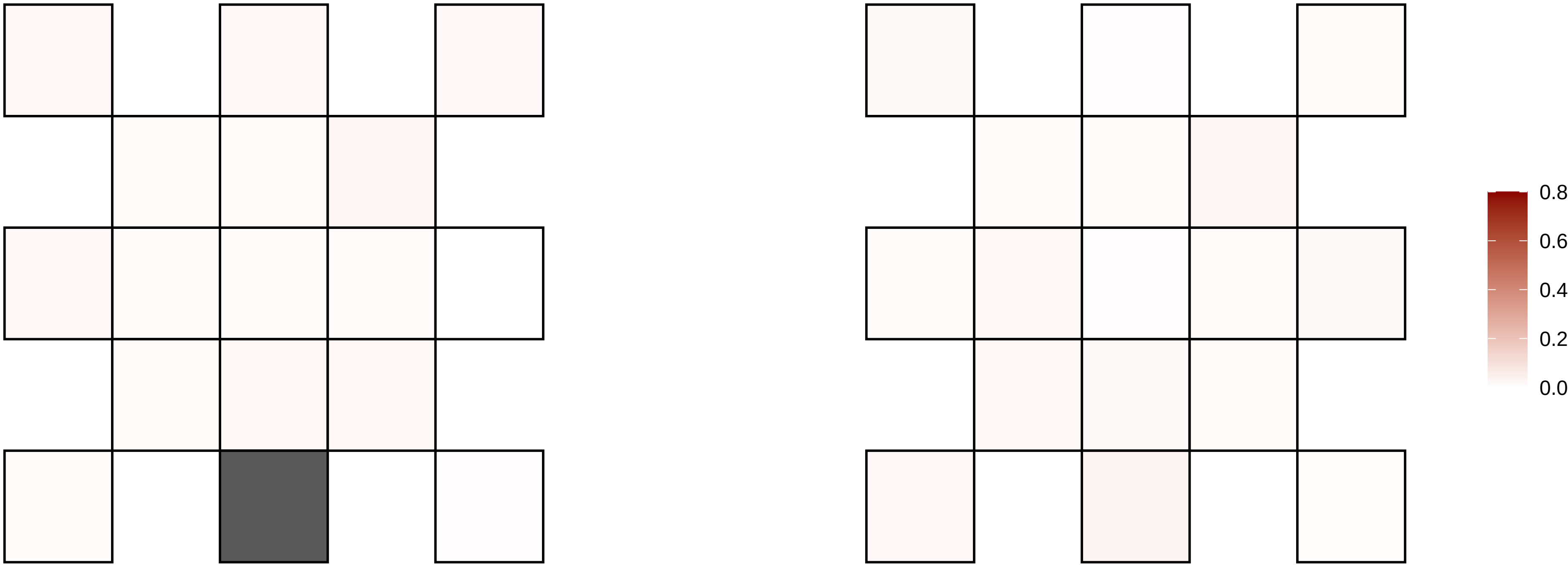

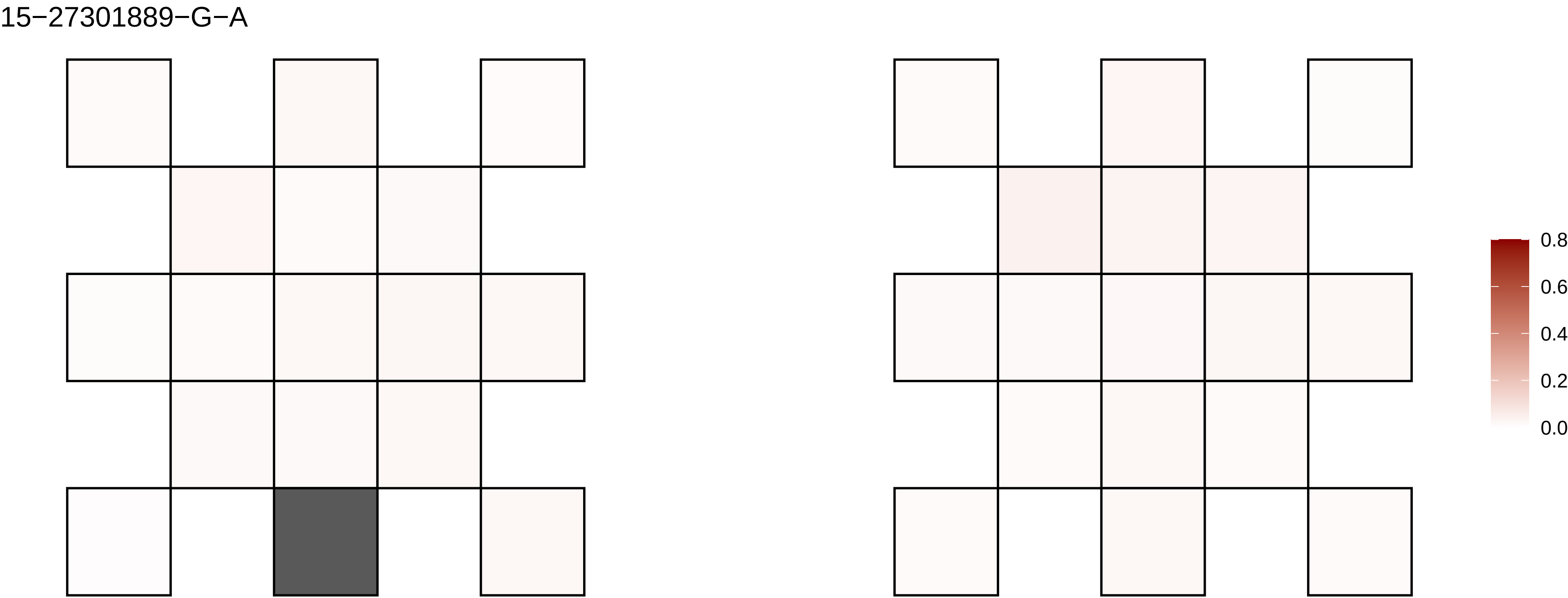
15−27301889−G−A
0.8
0.6
0.4
0.2
0.0

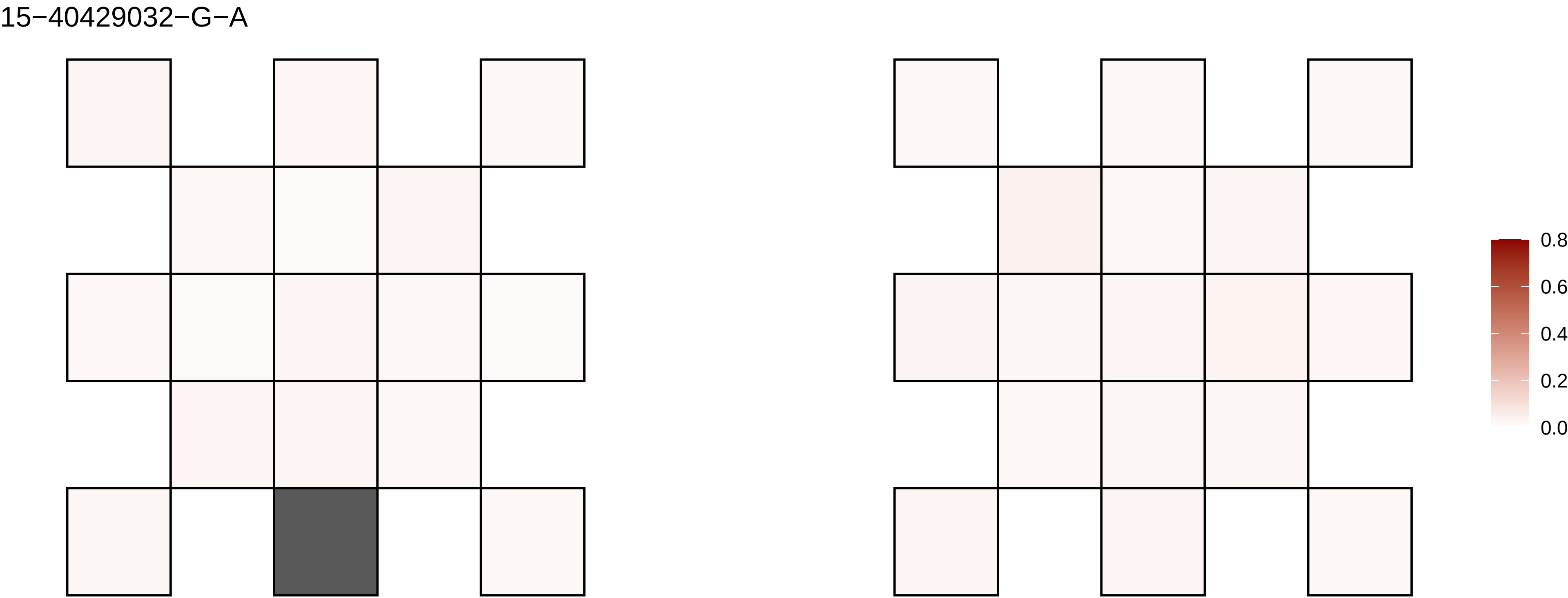
15−40429032−G−A
0.8
0.6
0.4
0.2
0.0

15−46932376−G−A

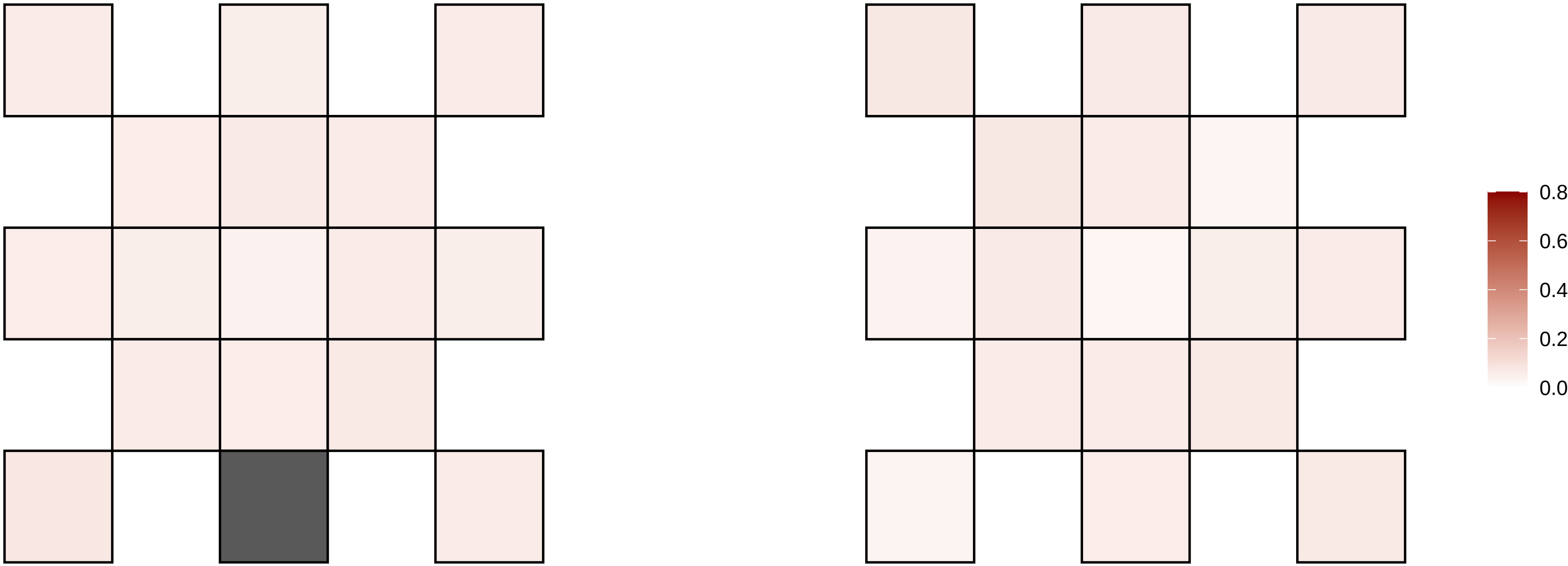

15−46962188−T−A

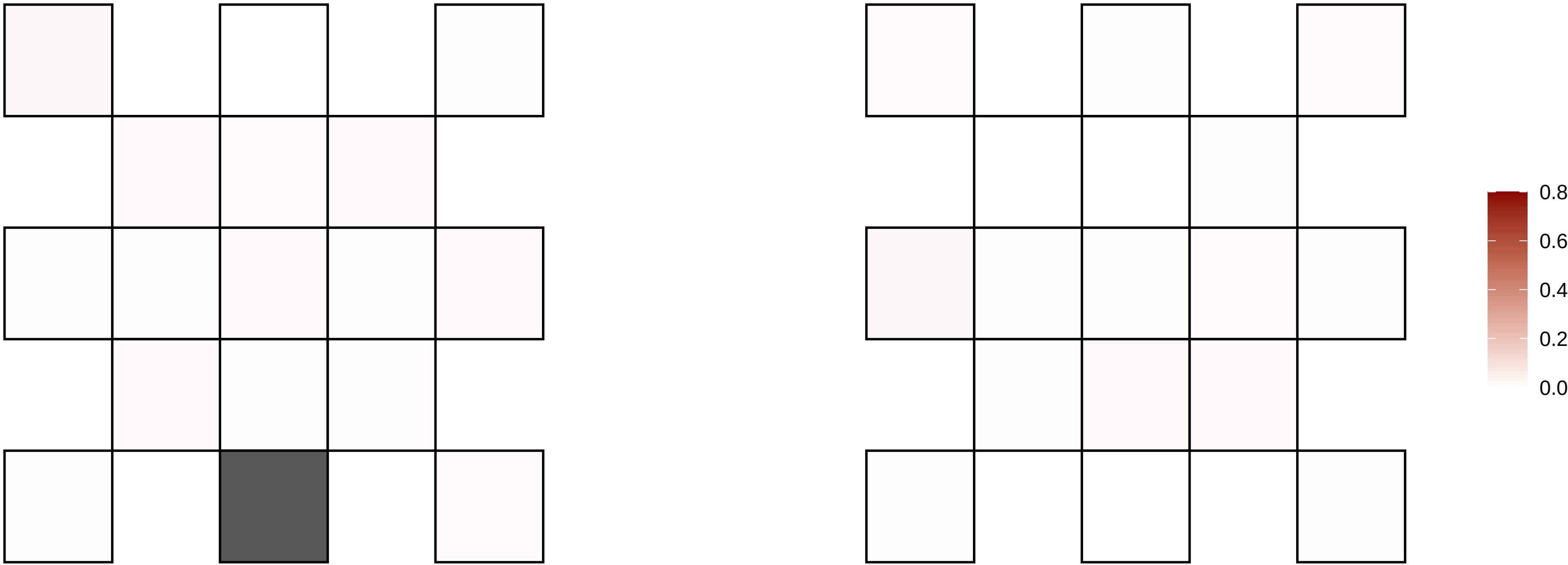

15−51489592−C−T

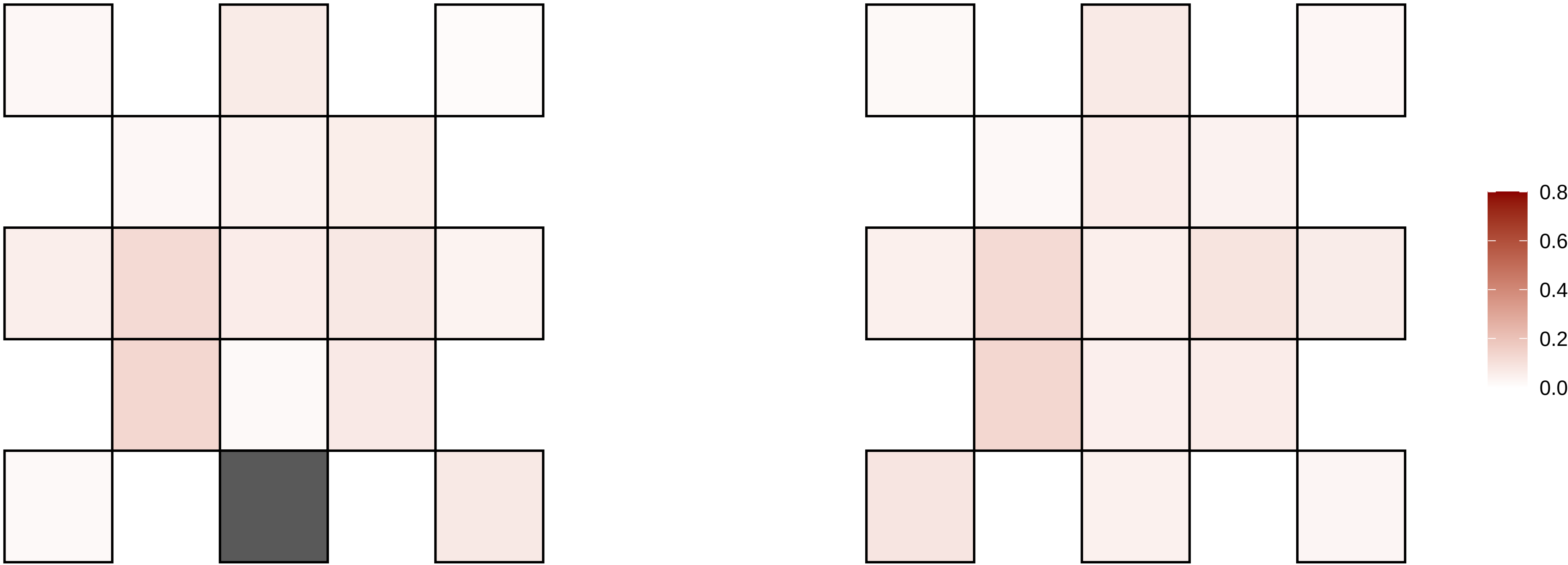

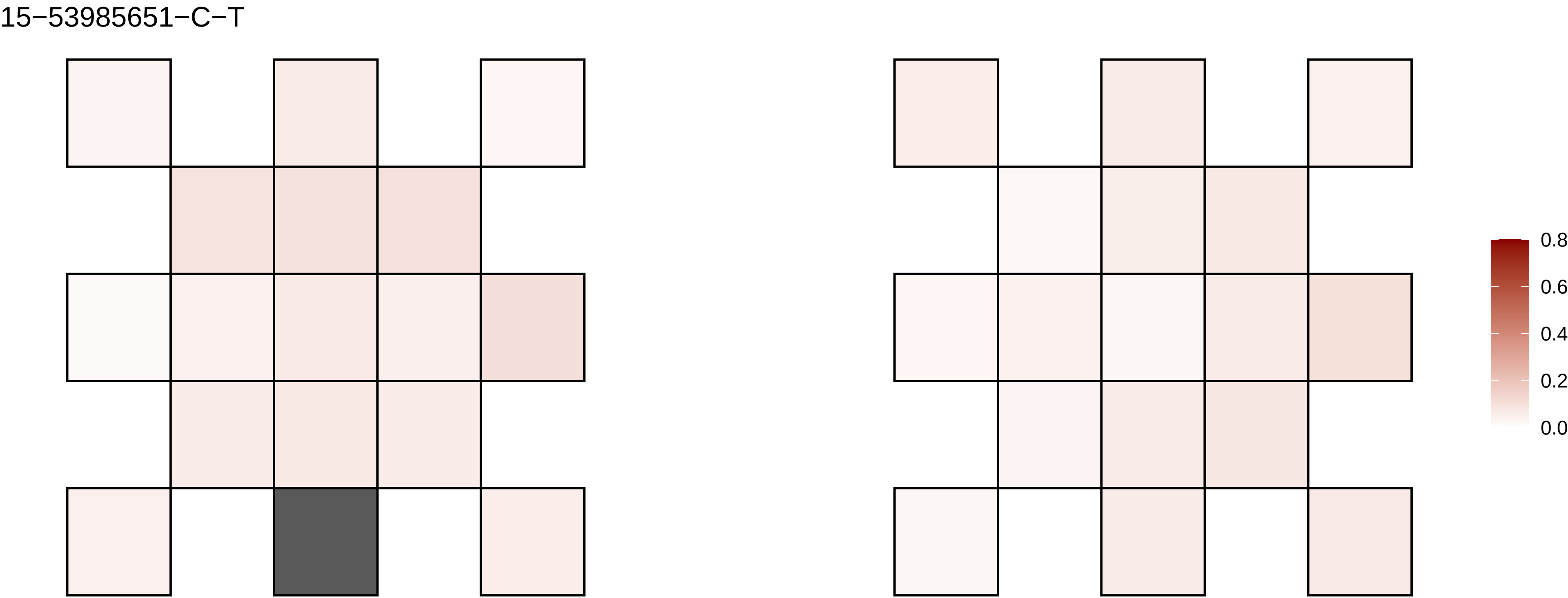

15−53985651−C−T
0.8
0.6
0.4
0.2
0.0

15−60164292−C−T

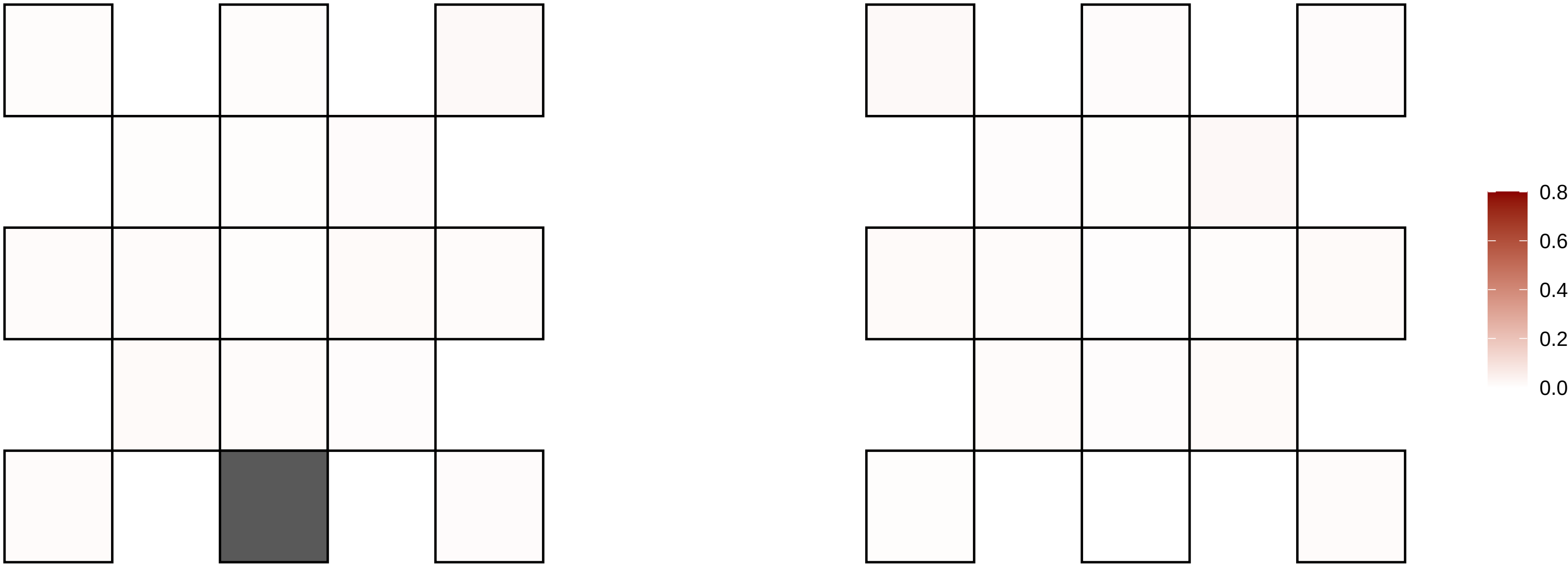

15−68132566−T−C

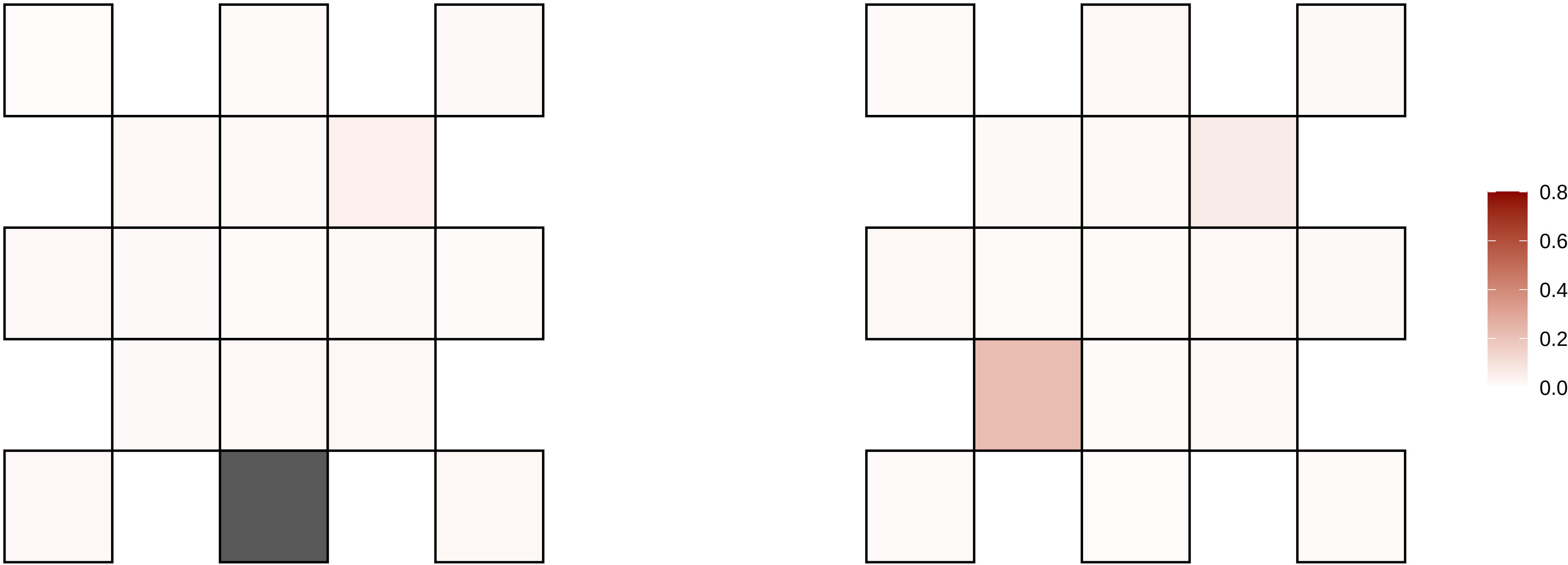

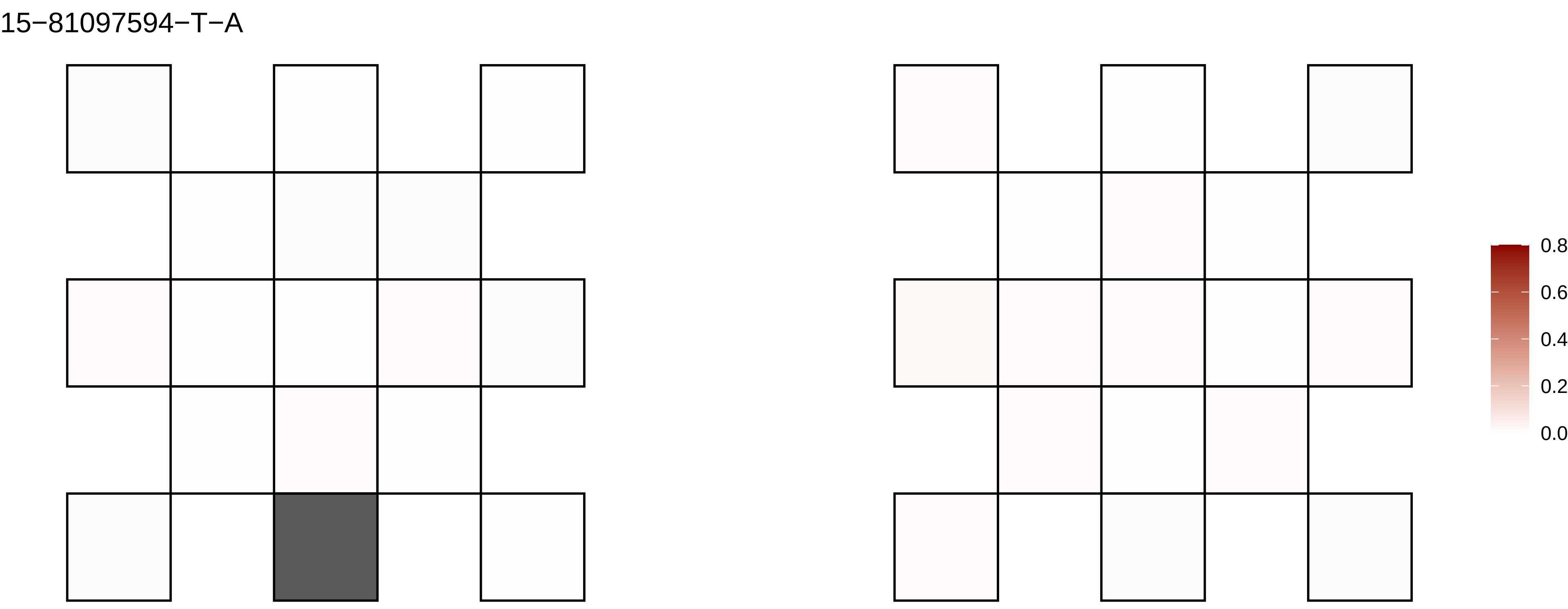
15−81097594−T−A
0.8
0.6
0.4
0.2
0.0

15−82103153−G−A

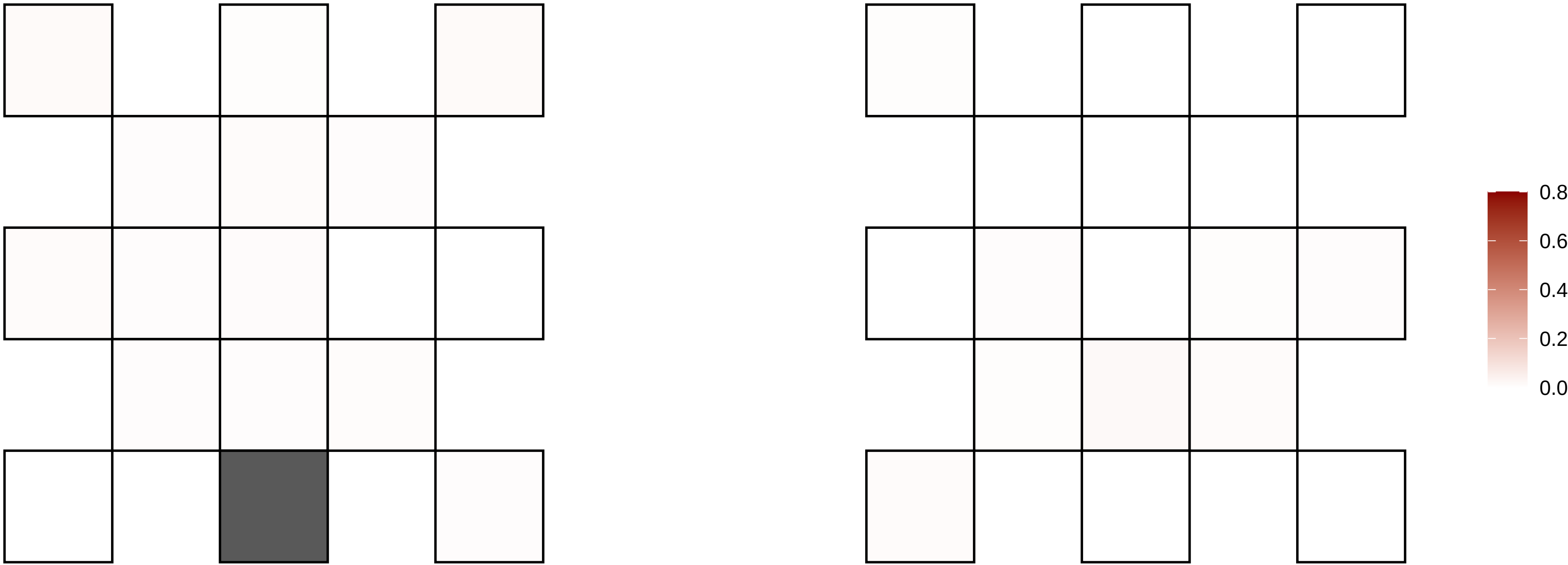

15−89449716−CT−C

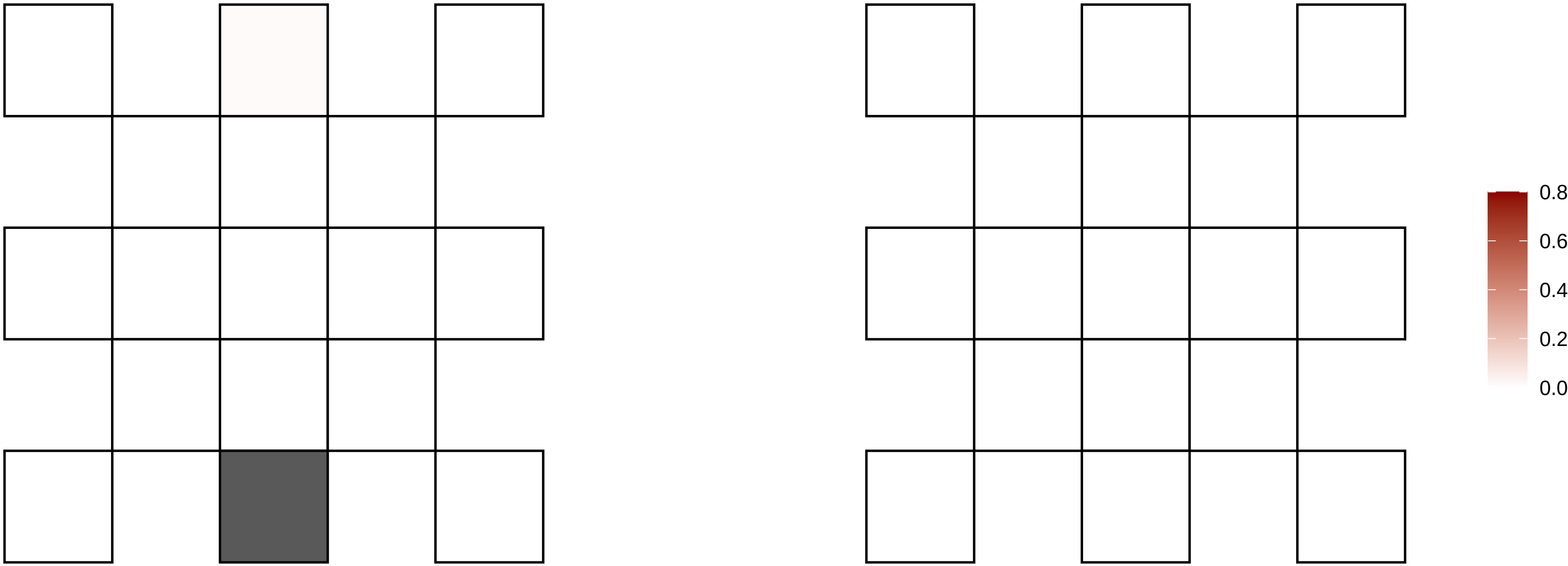

15−92865289−G−A

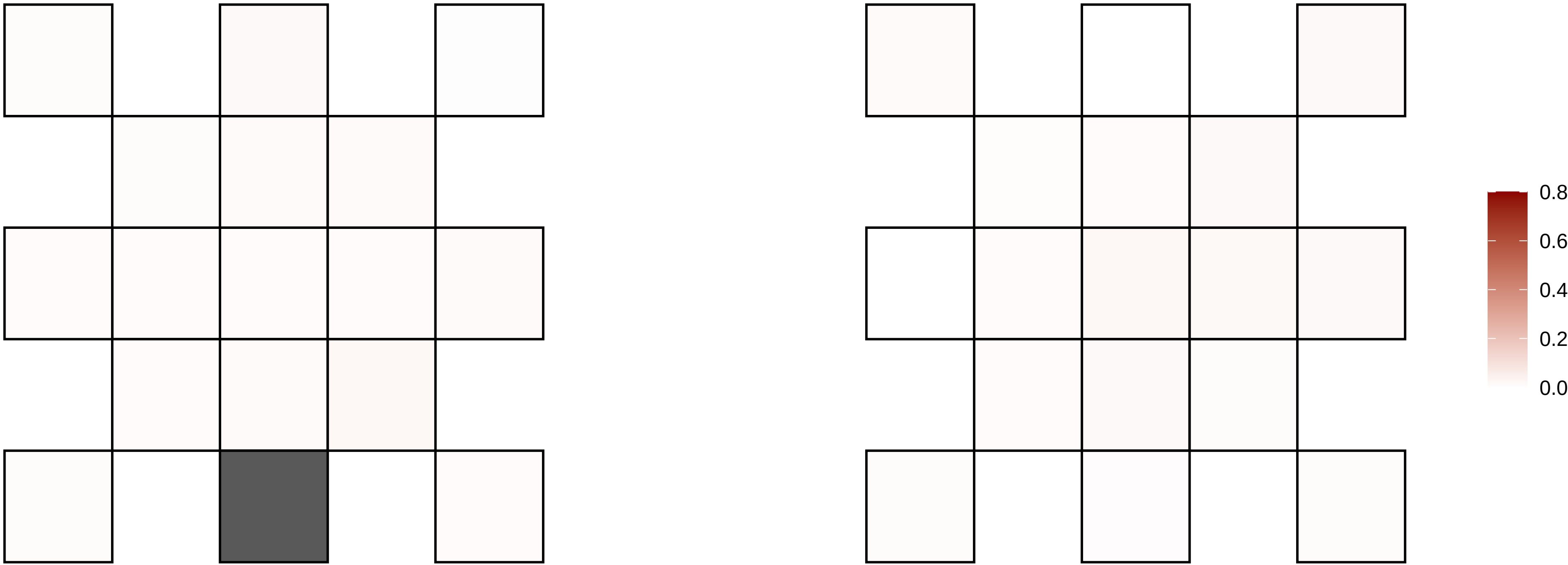

15−94903736−C−T

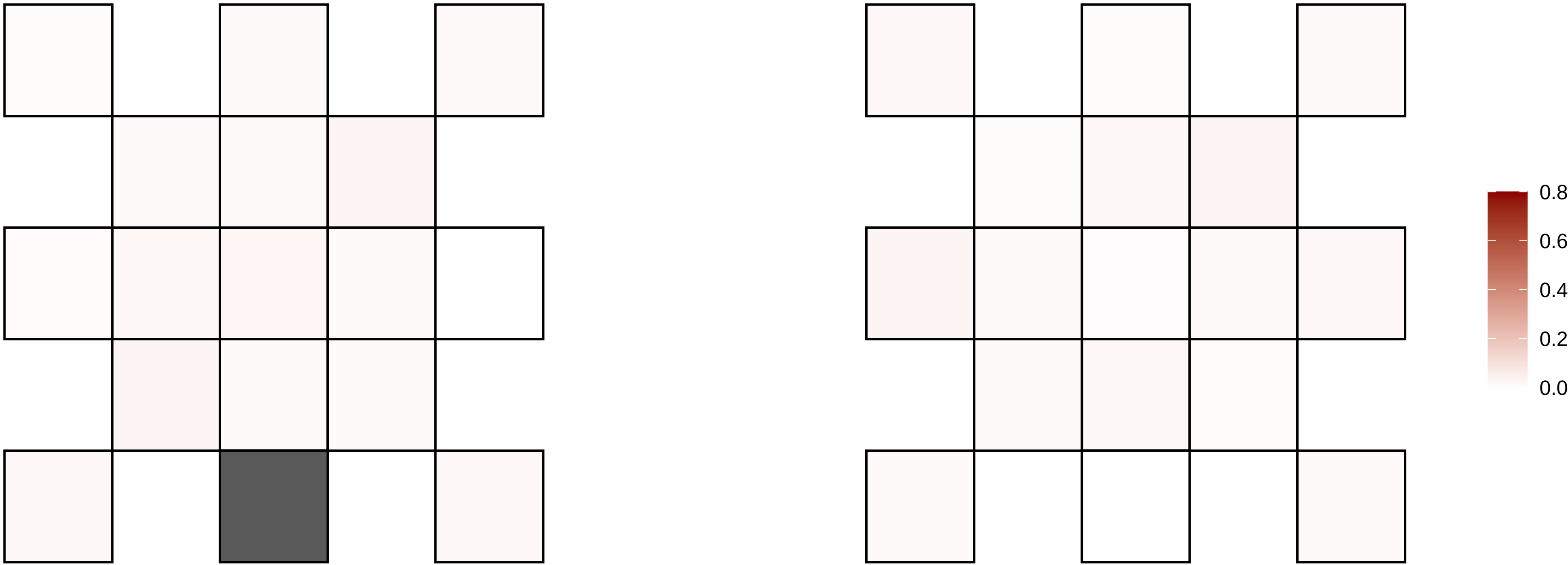

15−95374715−A−T

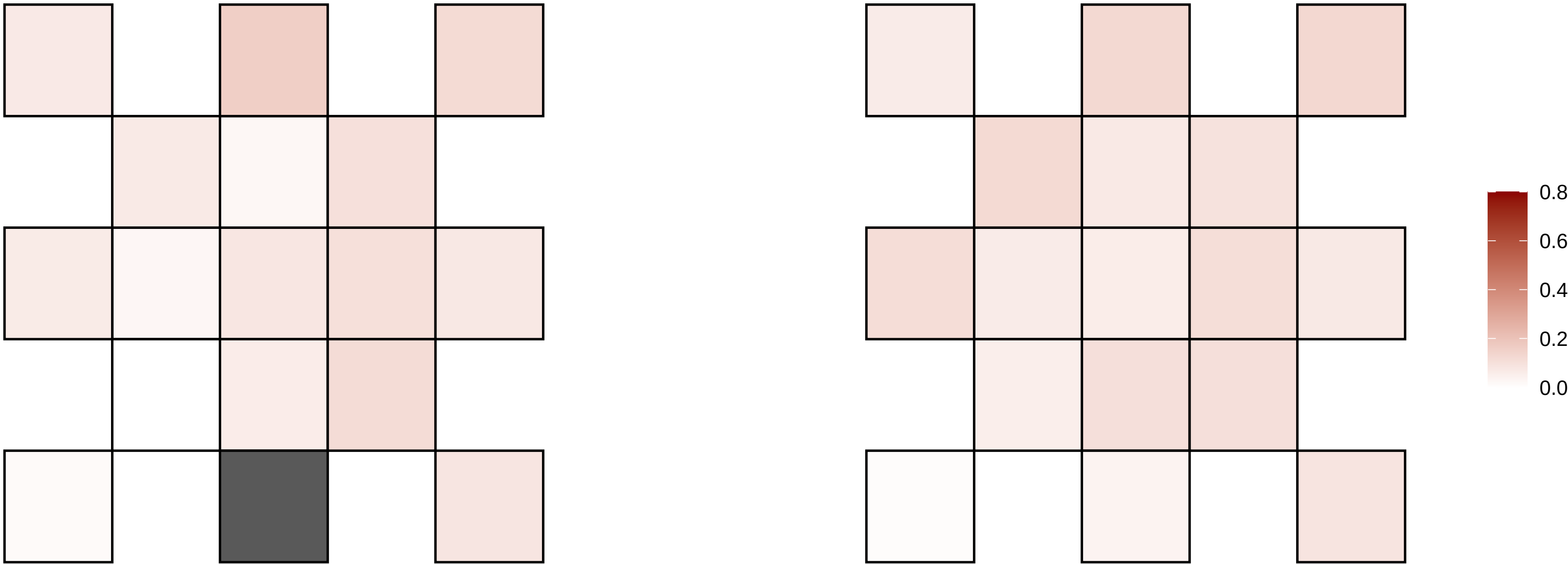

15−95789506−C−T

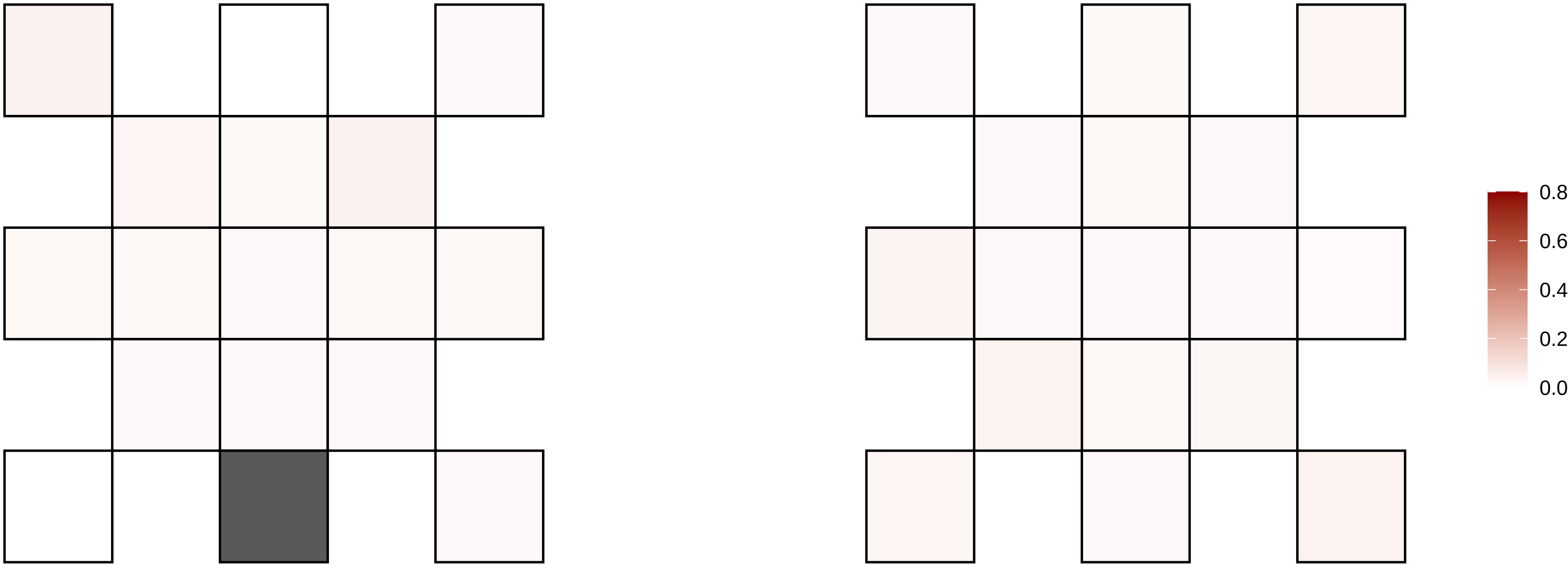

15−95947987−C−G

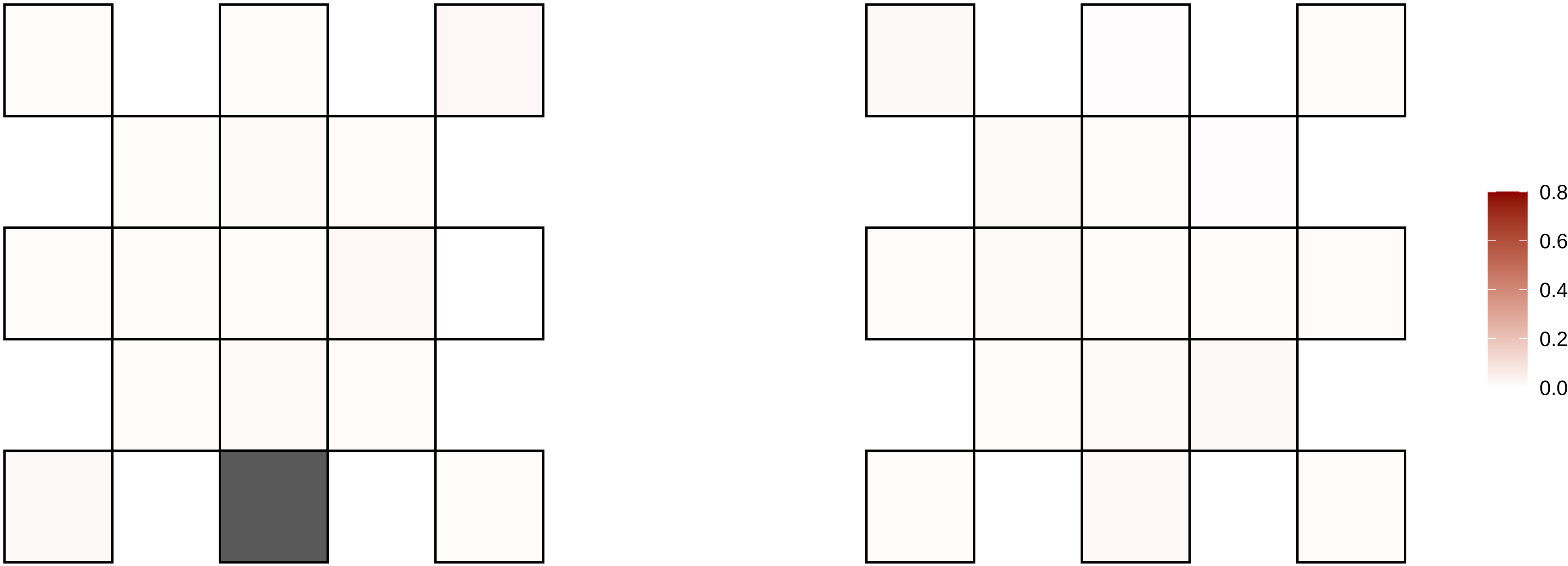

16−22921756−C−A

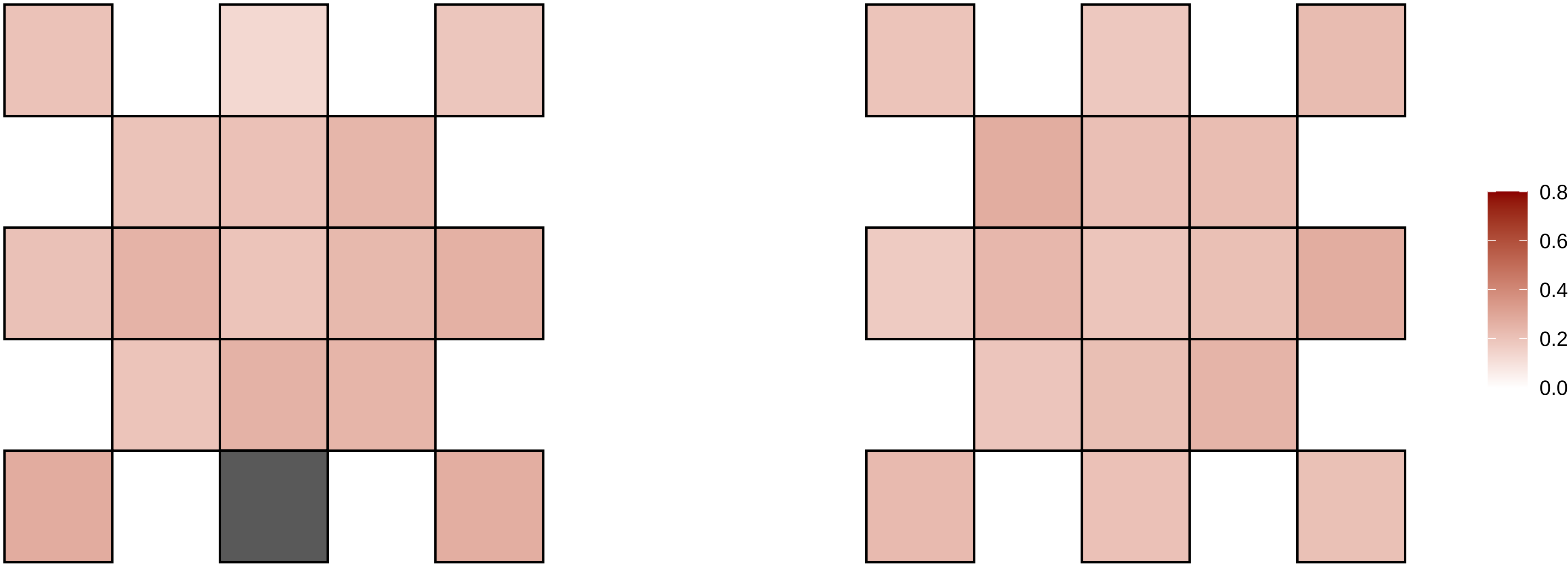

16−24164513−T−C

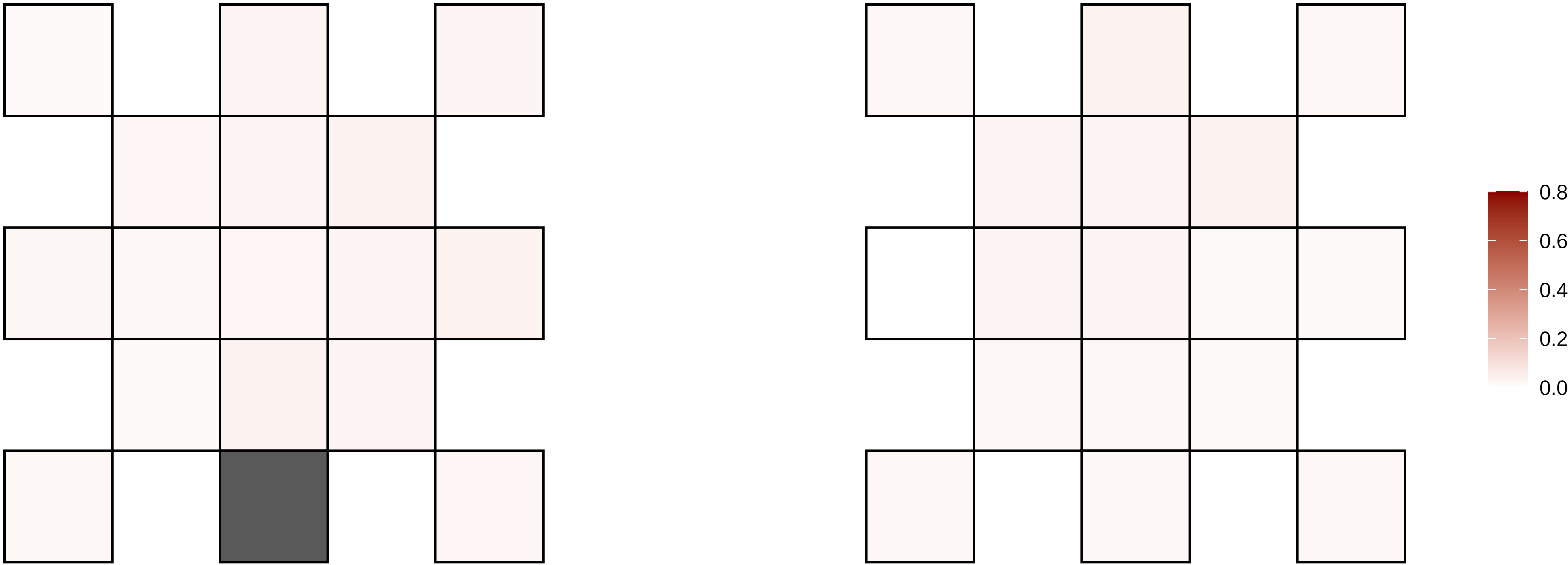

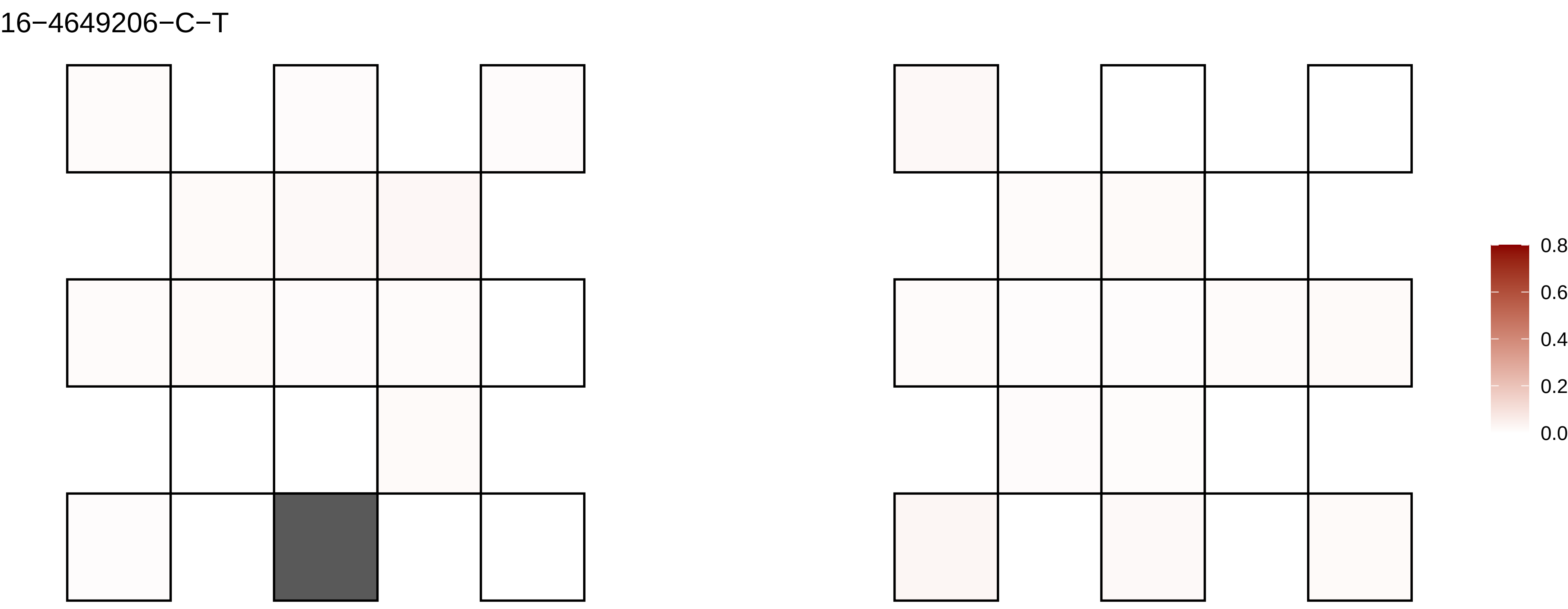
16−4649206−C−T
0.8
0.6
0.4
0.2
0.0

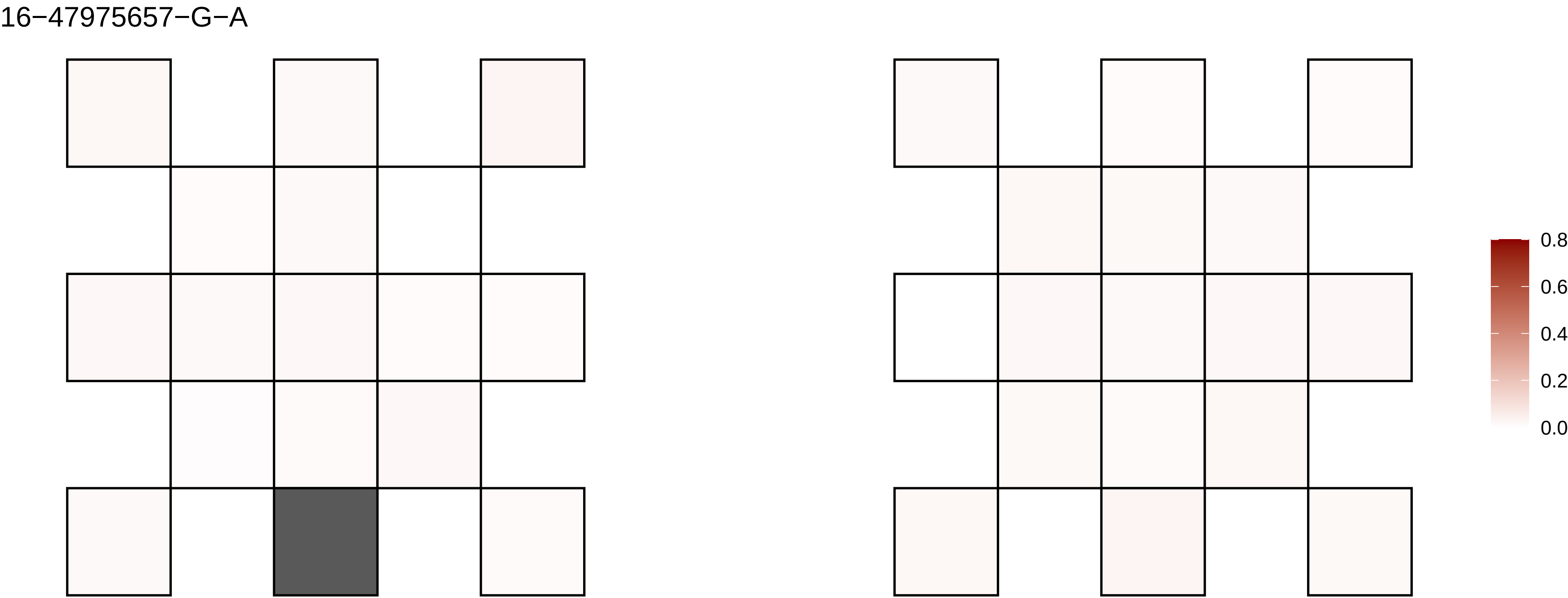
16−47975657−G−A
0.8
0.6
0.4
0.2
0.0

16−53451083−T−A

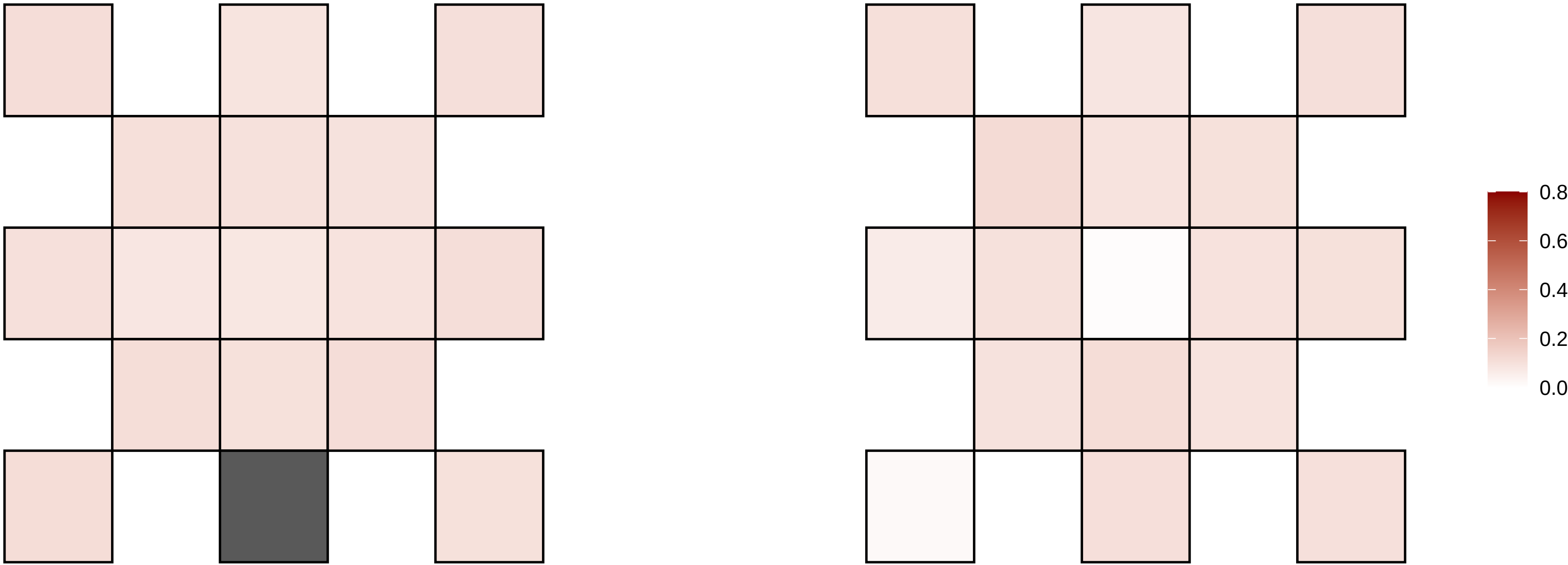

16−53451084−T−A

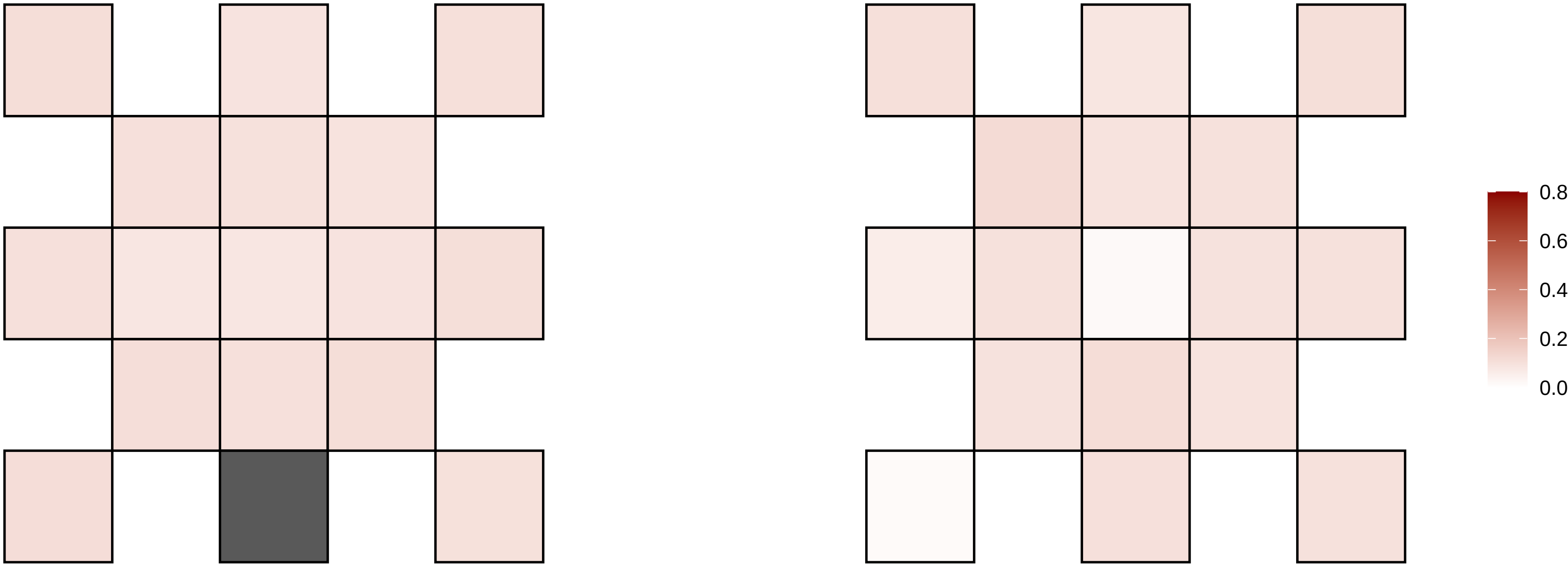

16−5864193−G−T

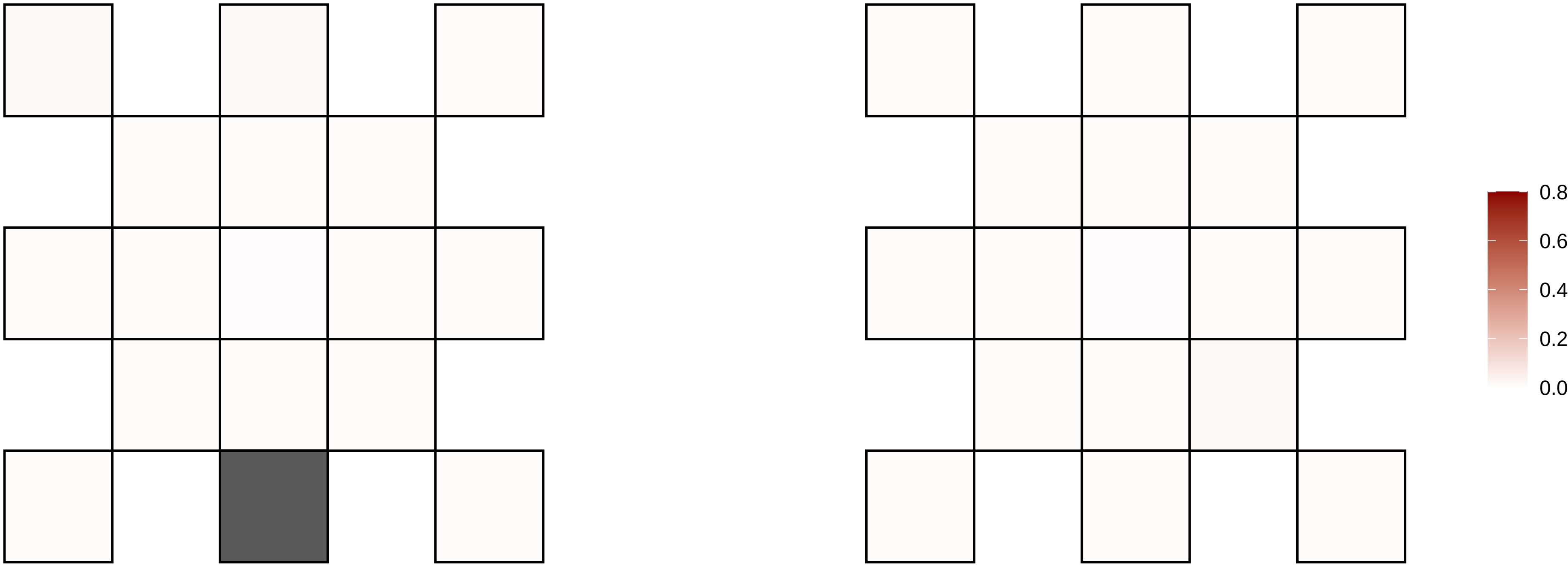

16−59987114−A−G

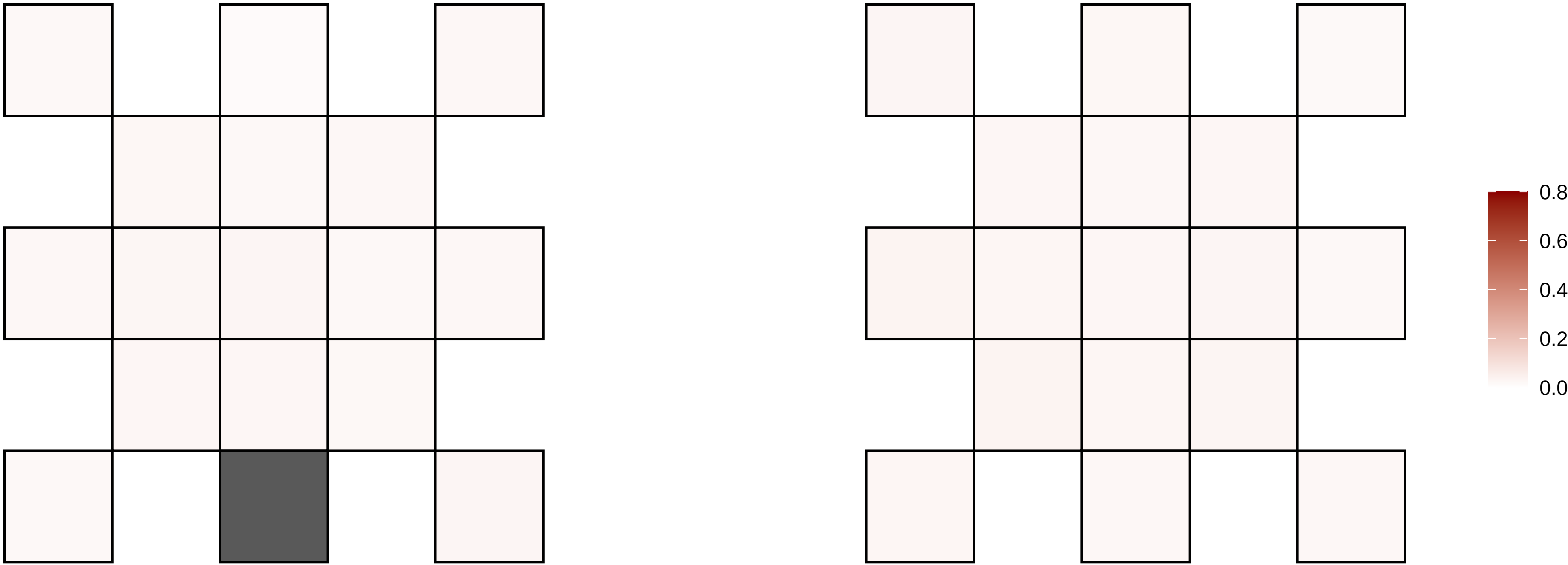

16−60721807−G−T

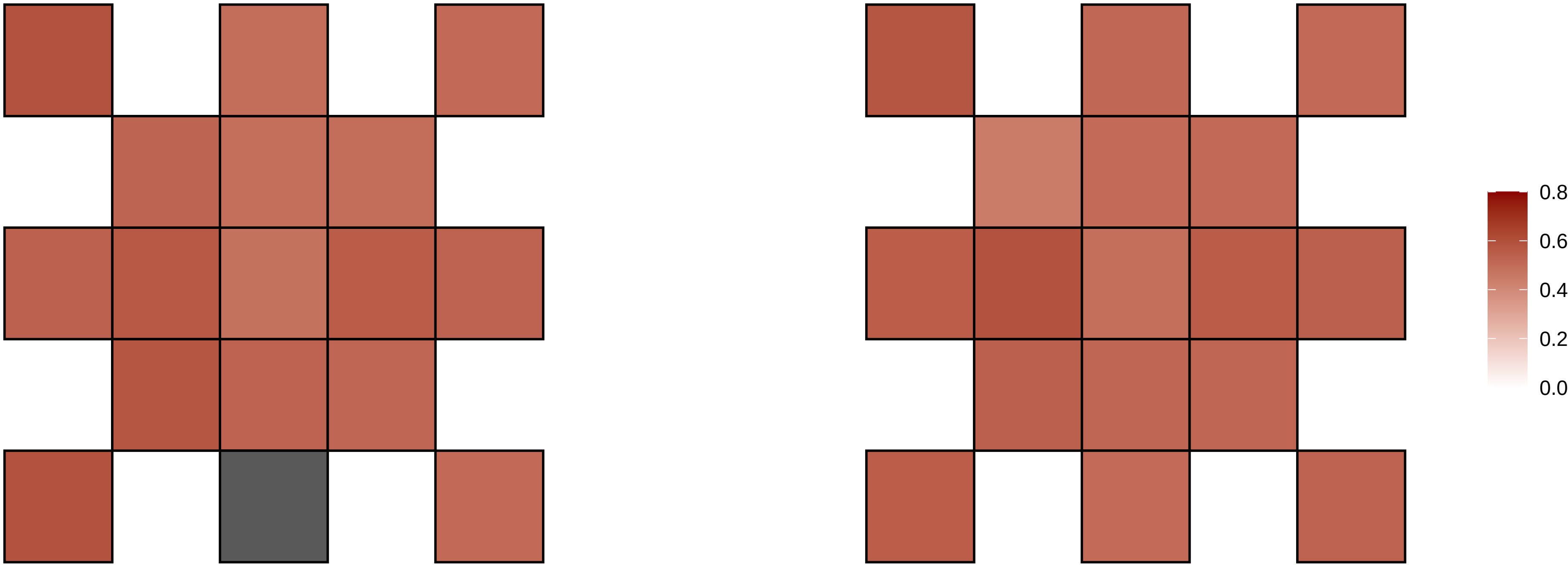

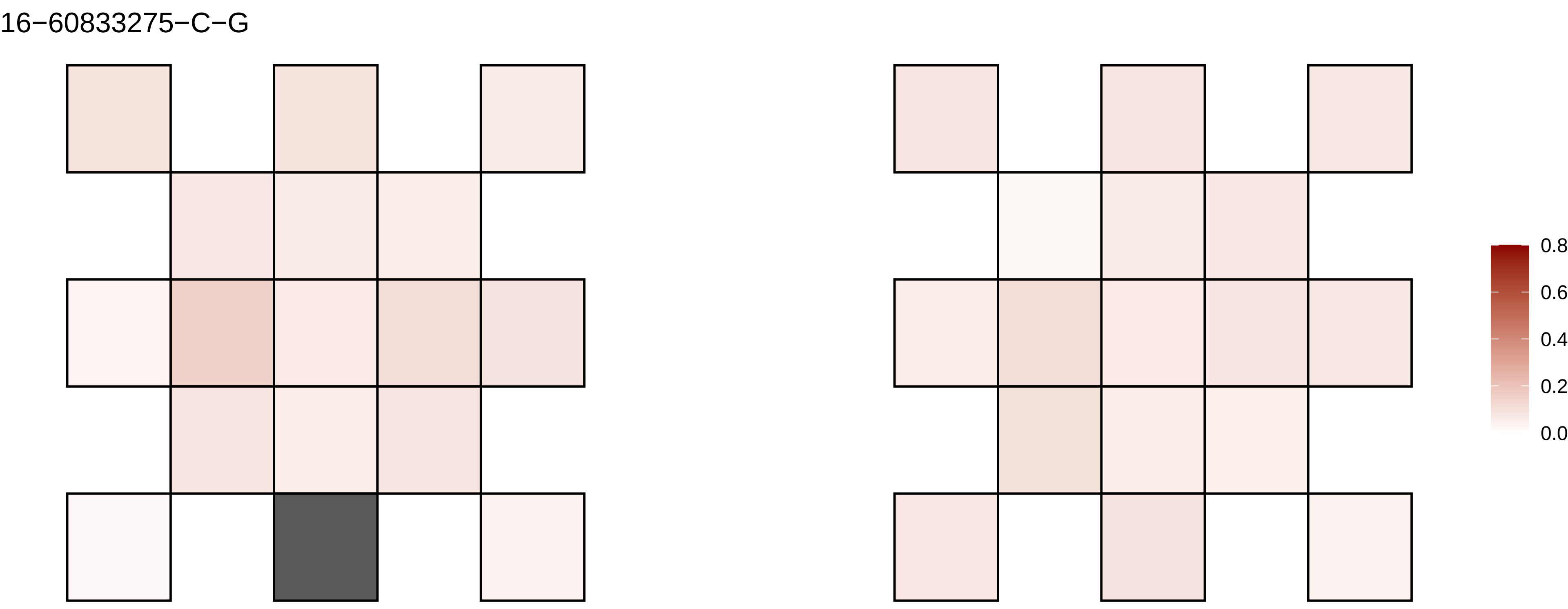
16−60833275−C−G
0.8
0.6
0.4
0.2
0.0

16−63481903−A−T

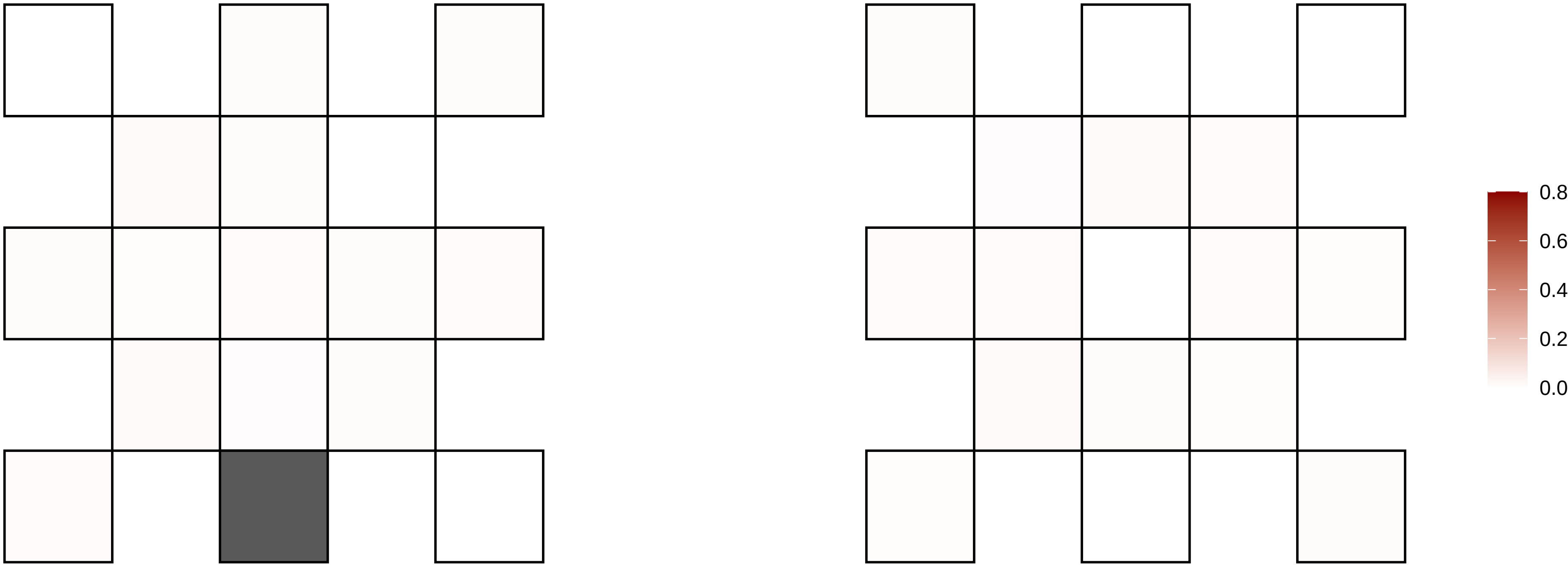

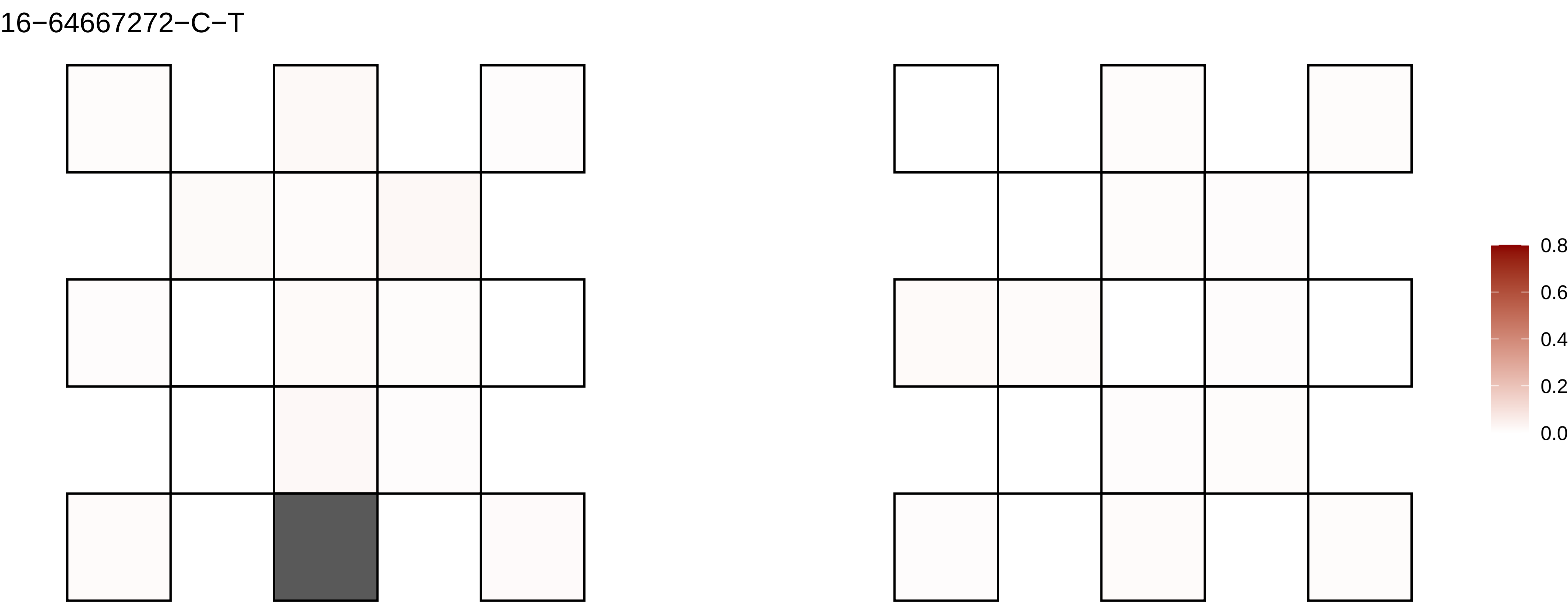
16−64667272−C−T
0.8
0.6
0.4
0.2
0.0

16−64780863−G−A

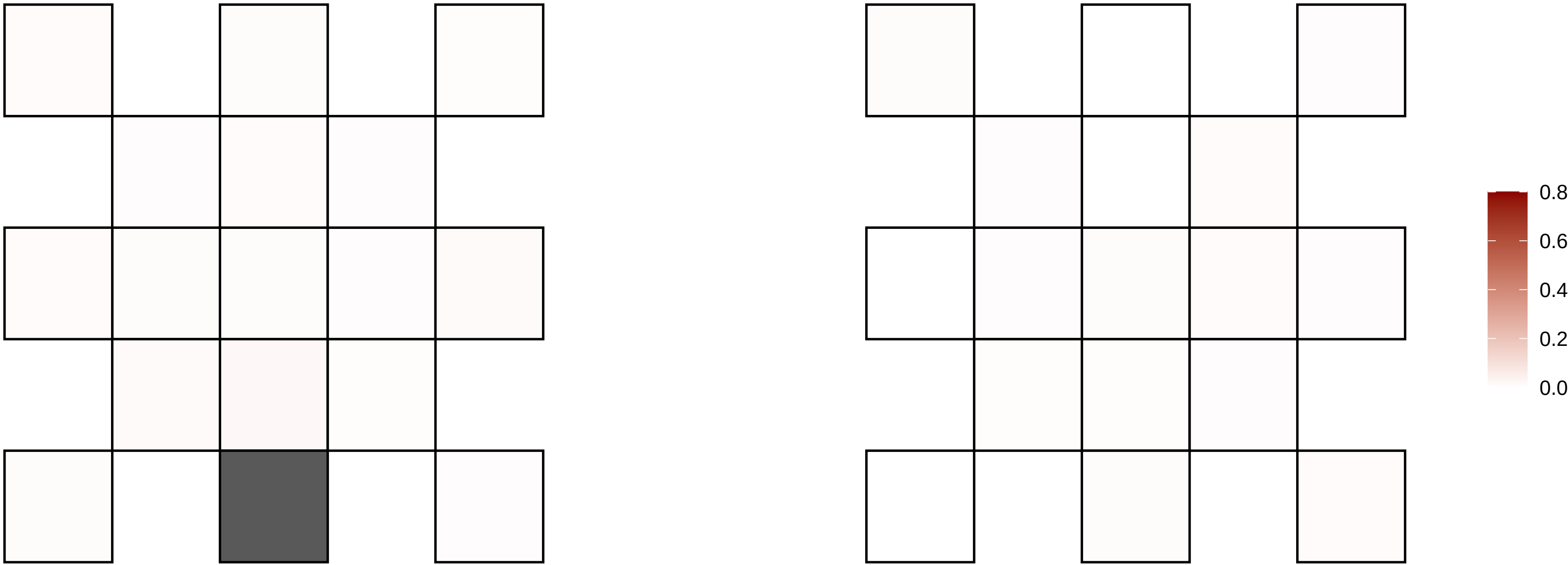

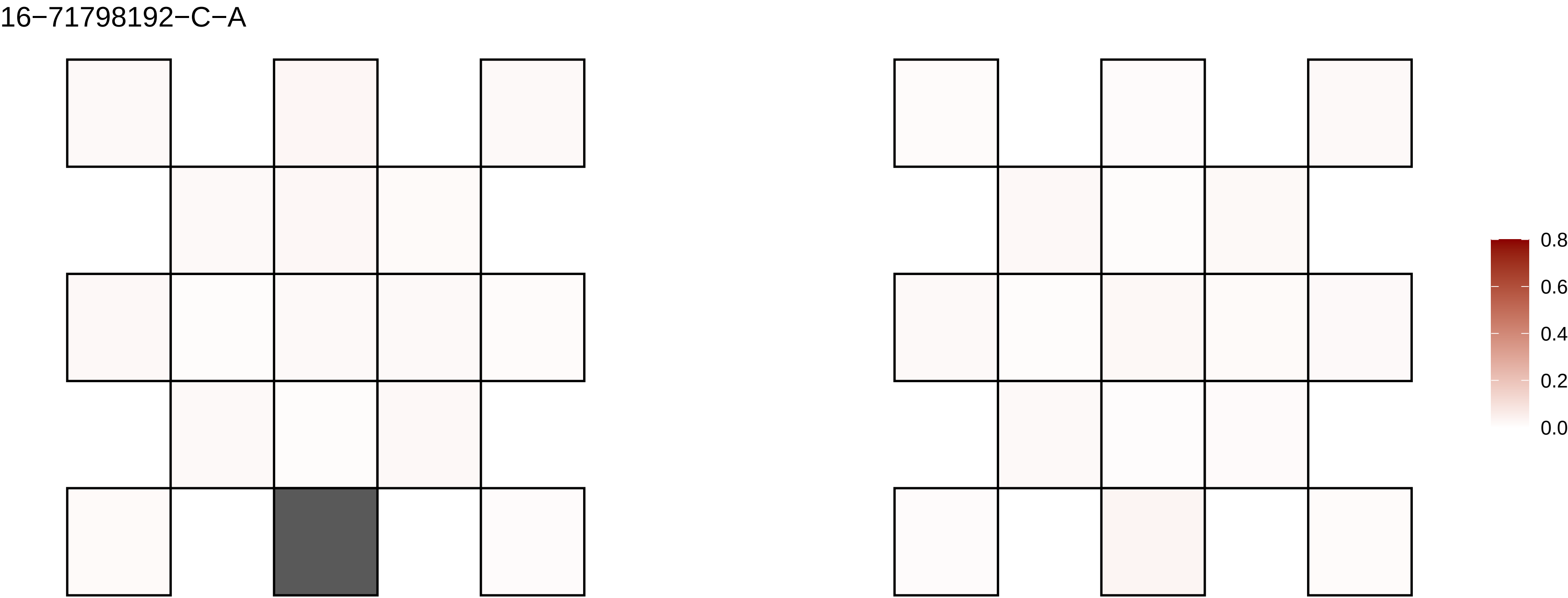
16−71798192−C−A
0.8
0.6
0.4
0.2
0.0

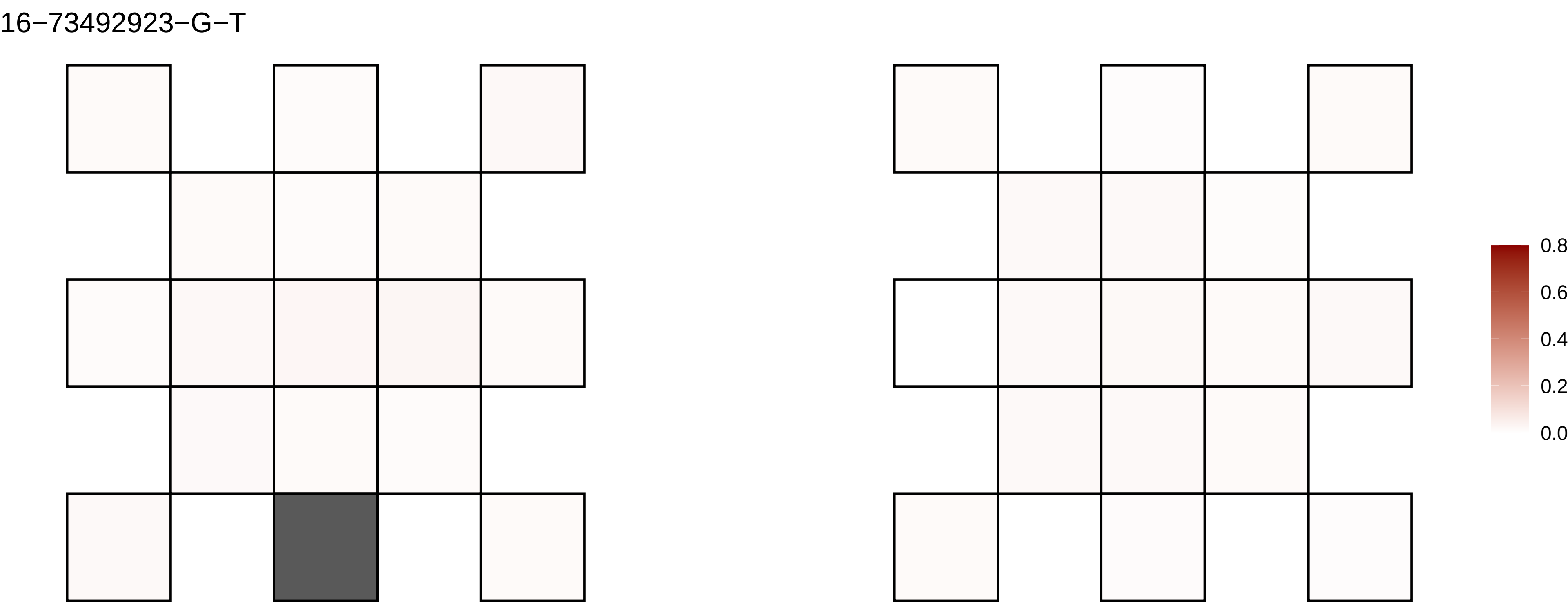
16−73492923−G−T
0.8
0.6
0.4
0.2
0.0

16−77437520−G−T

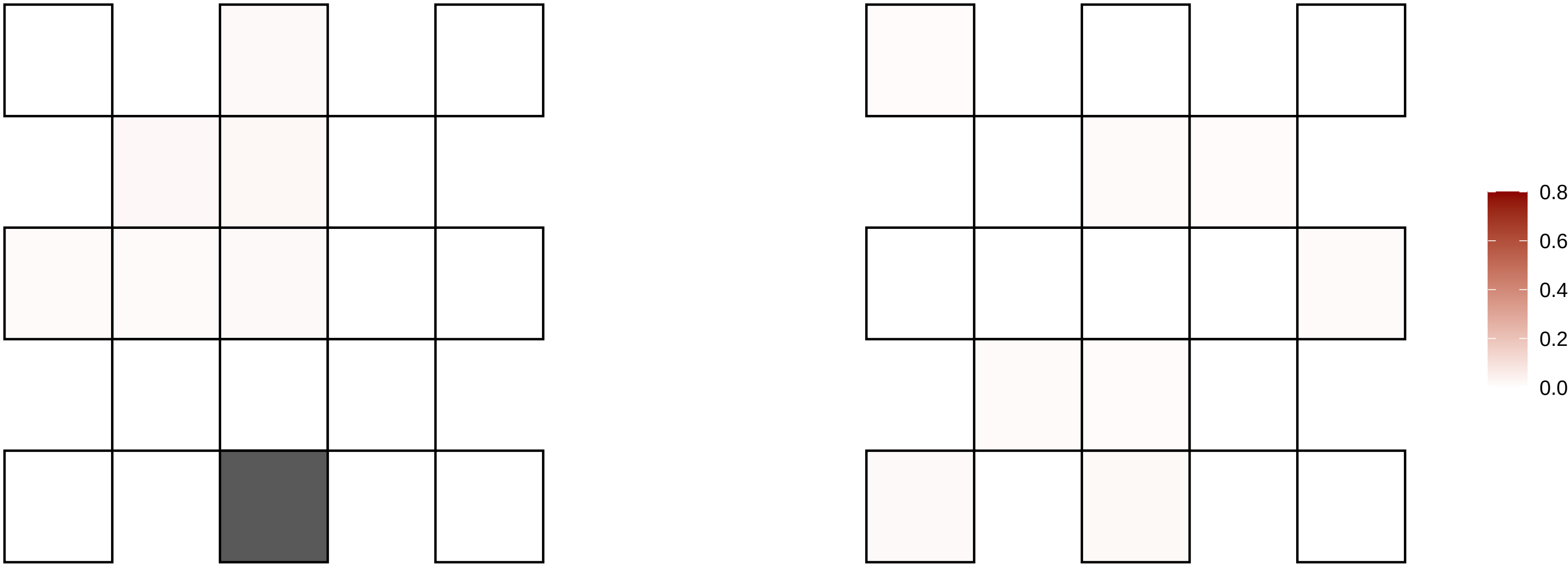

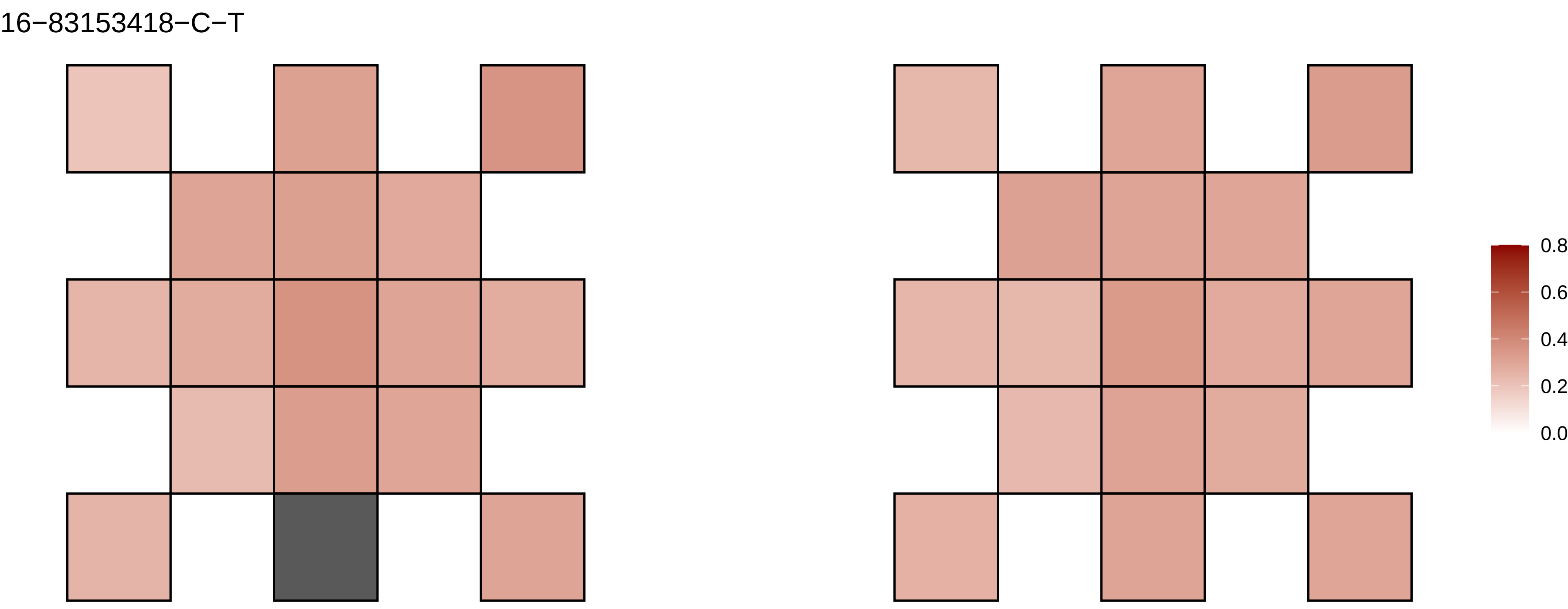
16−83153418−C−T
0.8
0.6
0.4
0.2
0.0

16−86500807−C−T

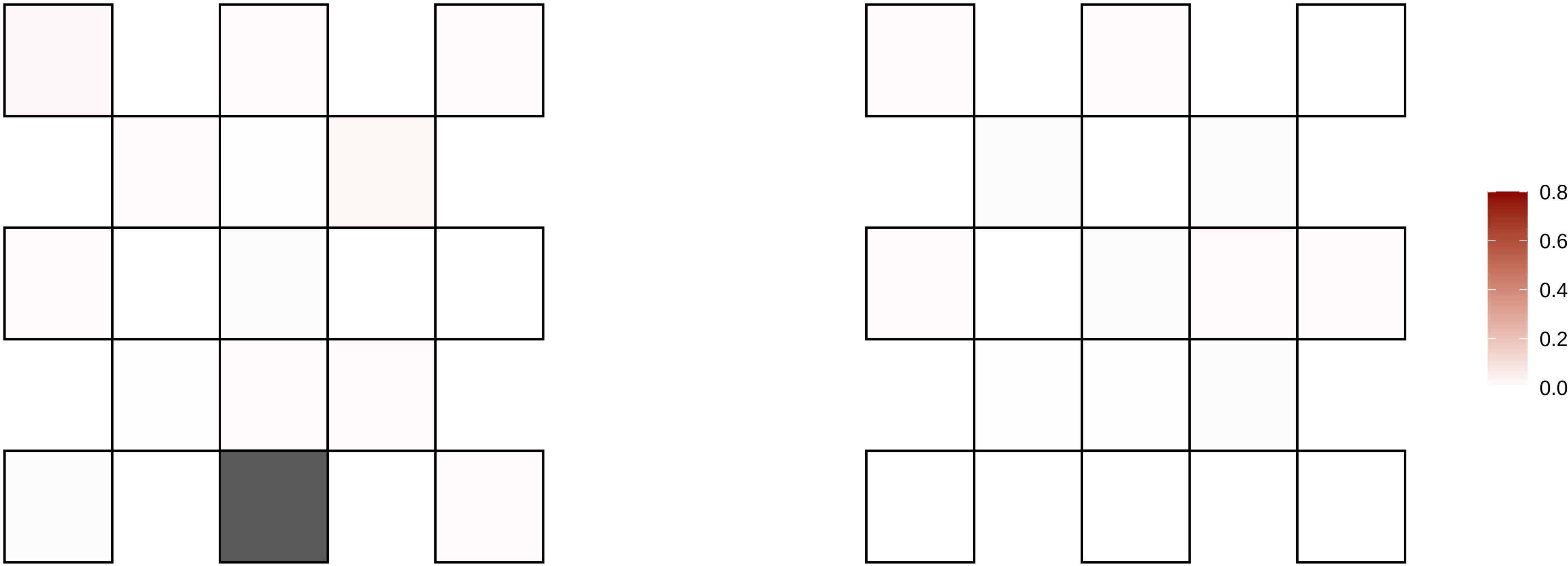

16−8802550−C−T

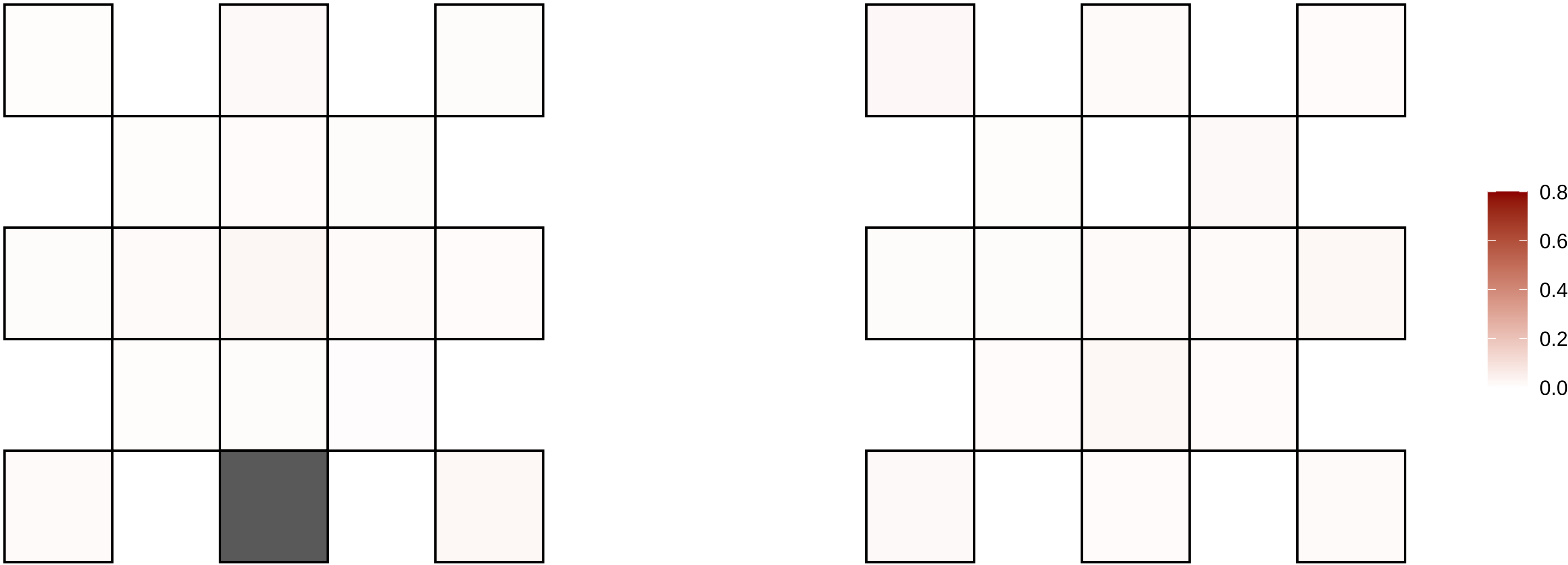

16−925921−C−T

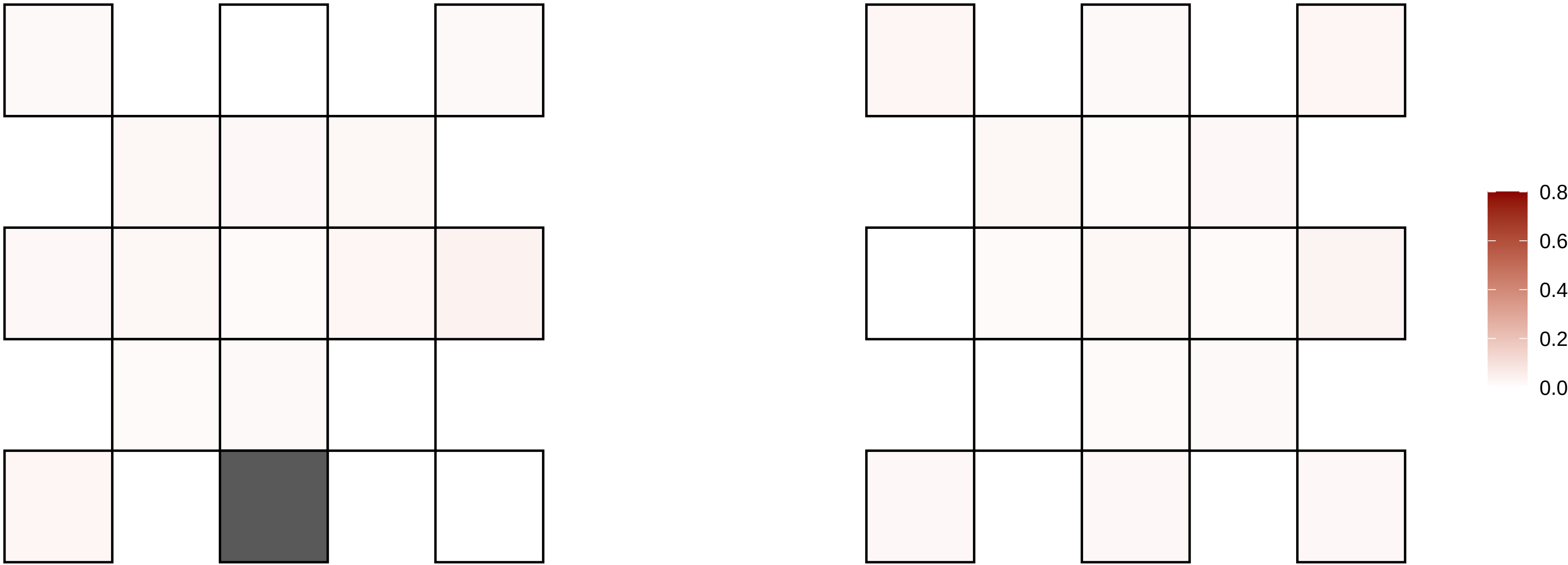

16−9382504−C−T

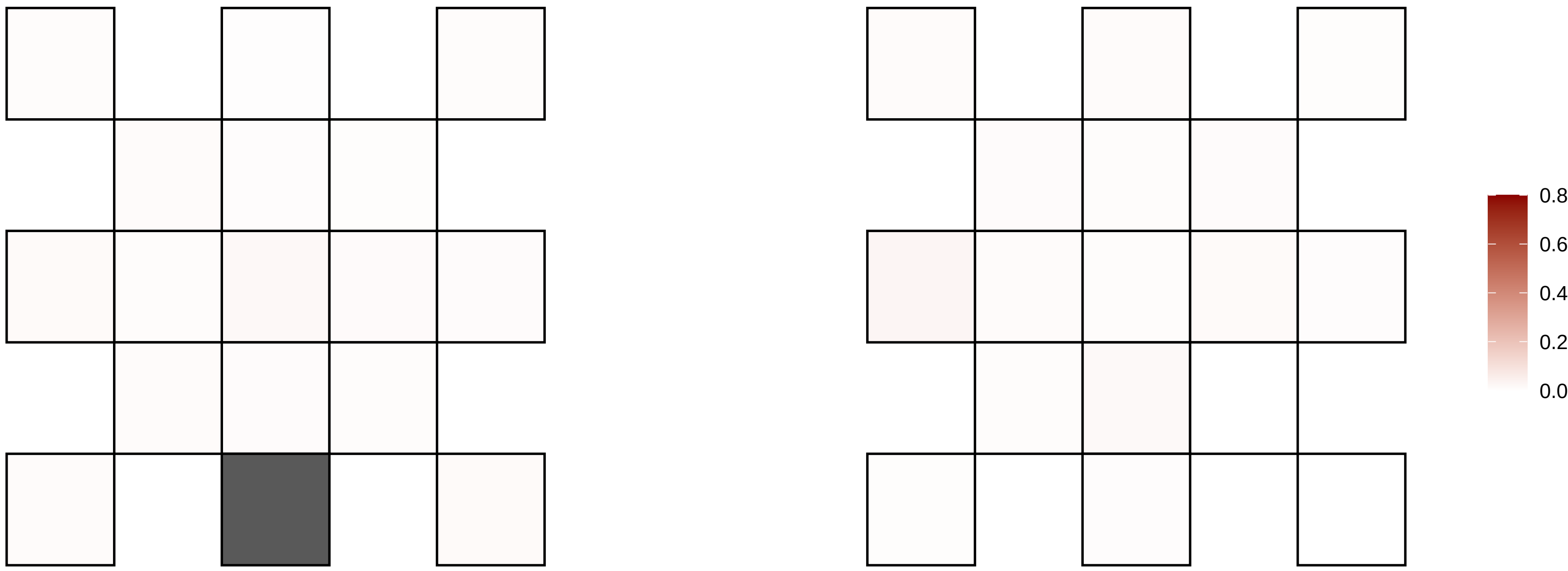

17−10283611−C−T

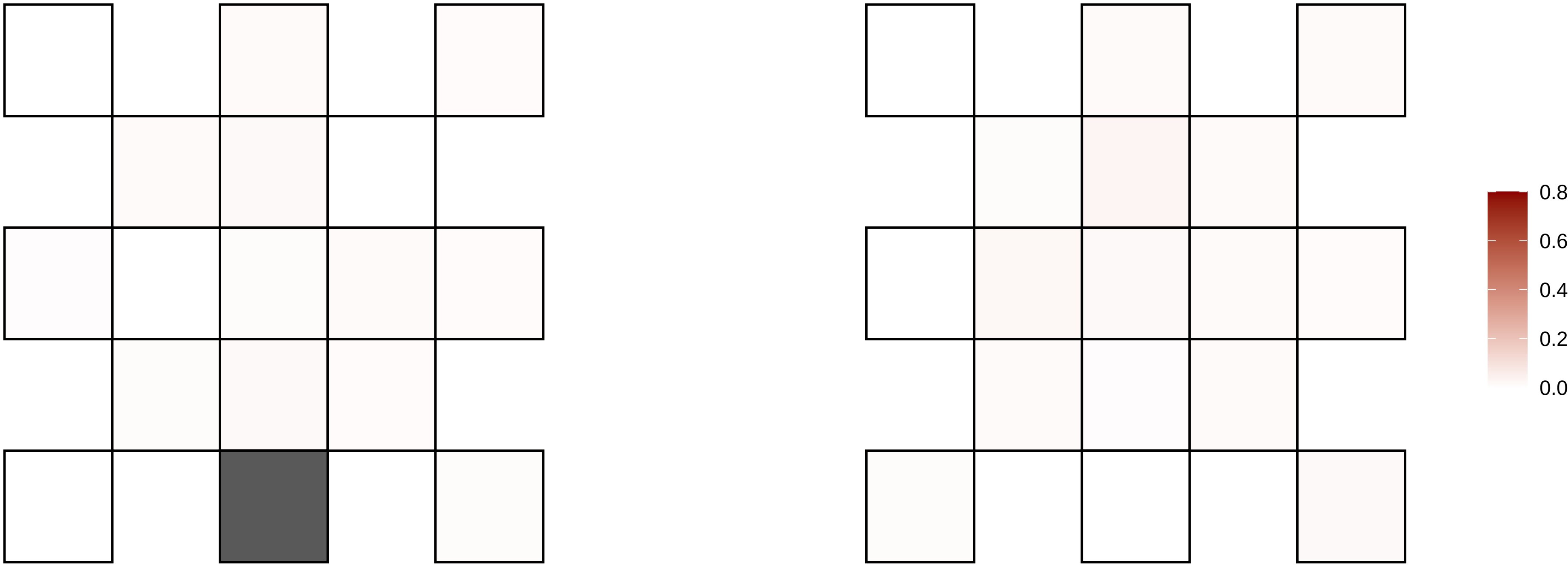

17−12534415−G−T

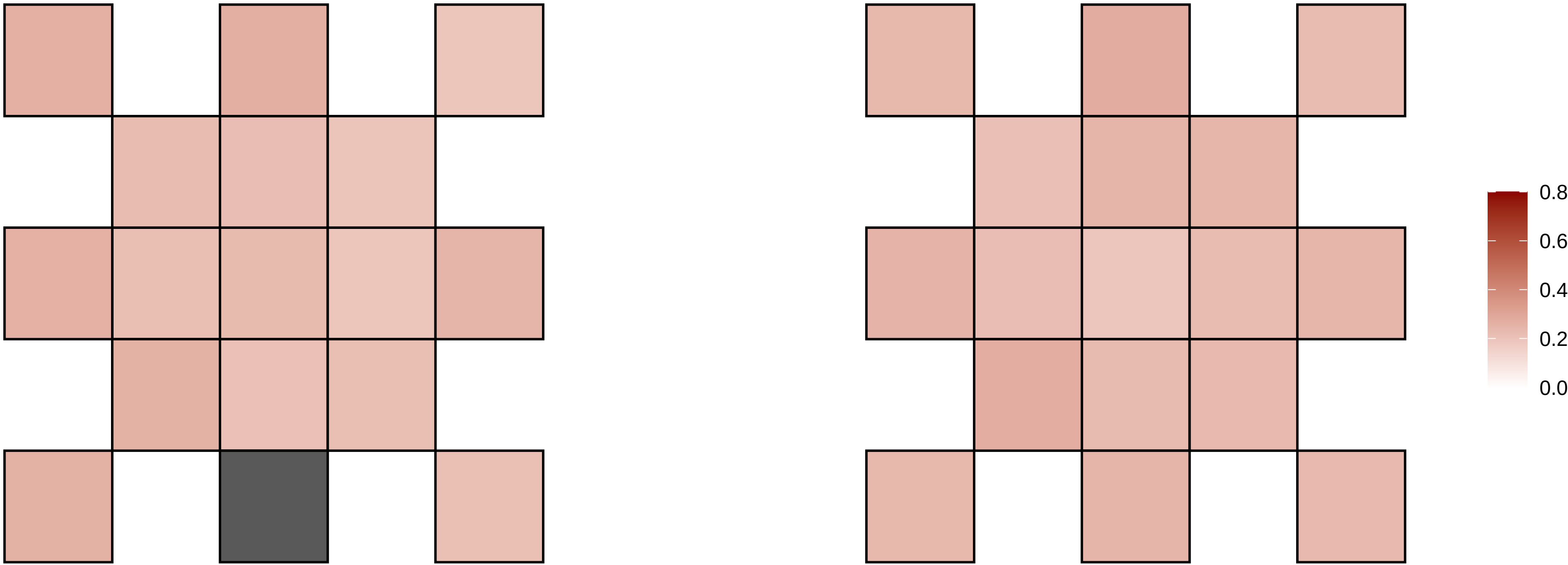

17−1478793−G−A

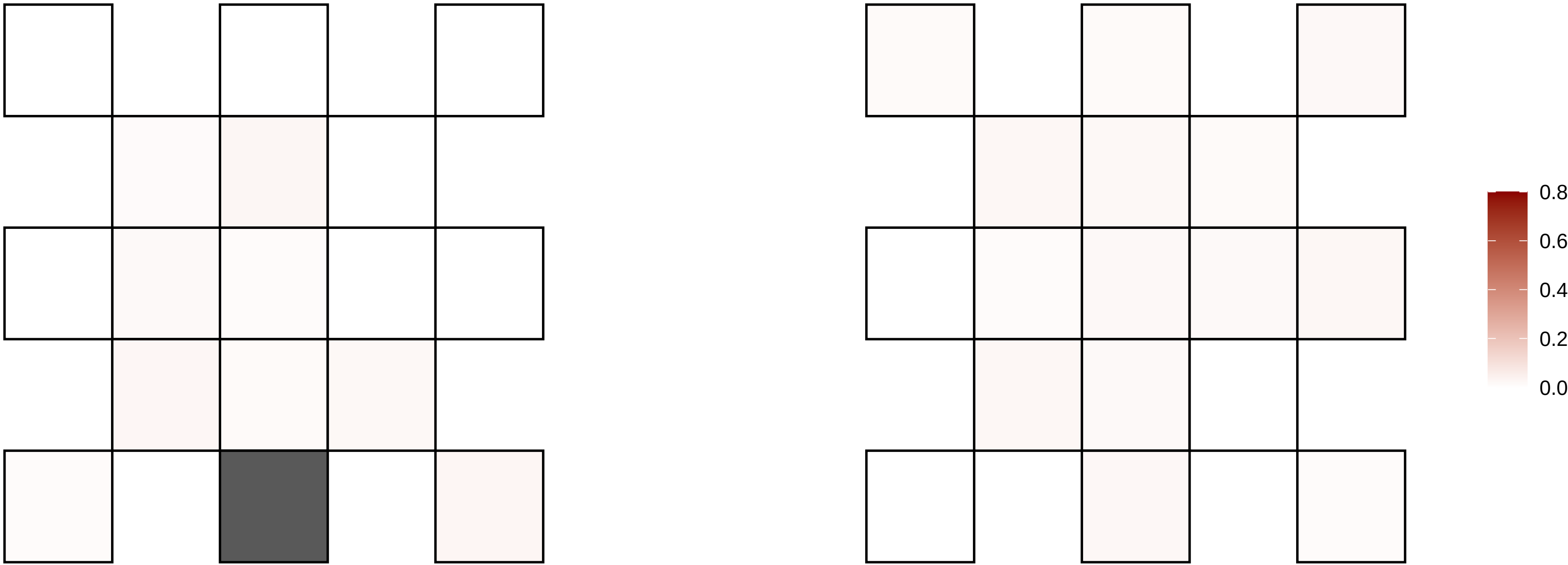

17−1620800−G−T

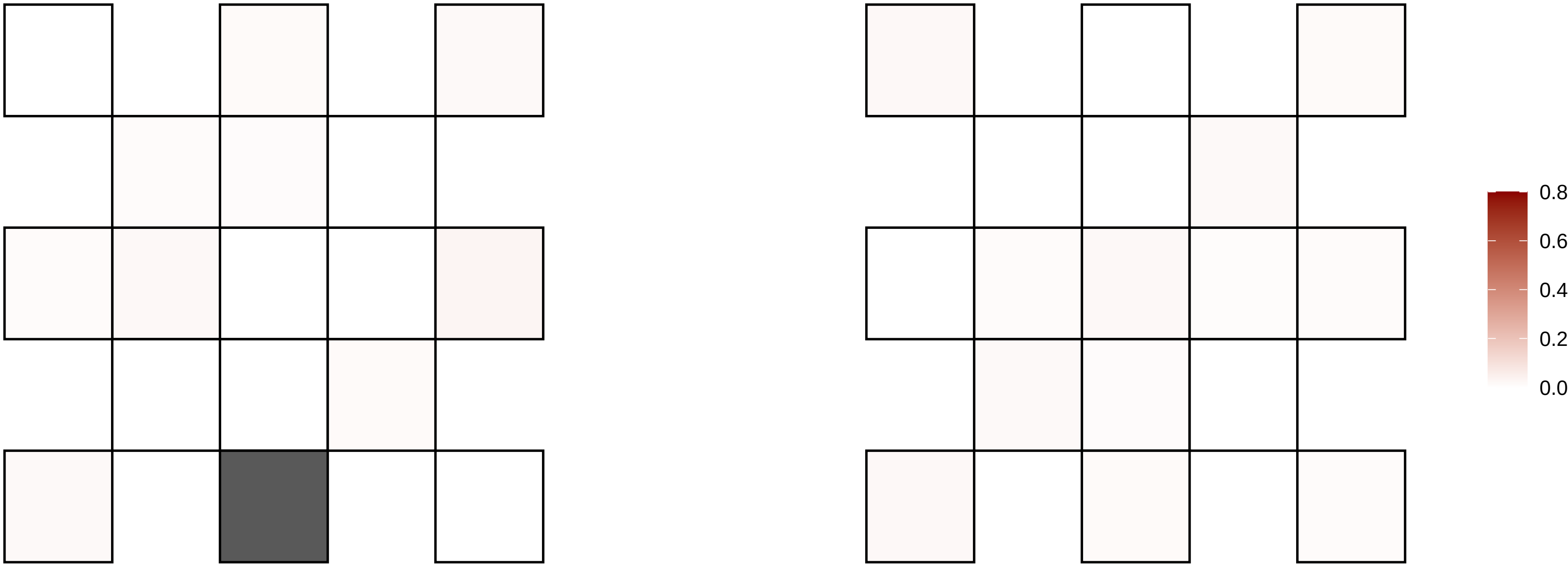

17−17411221−G−A

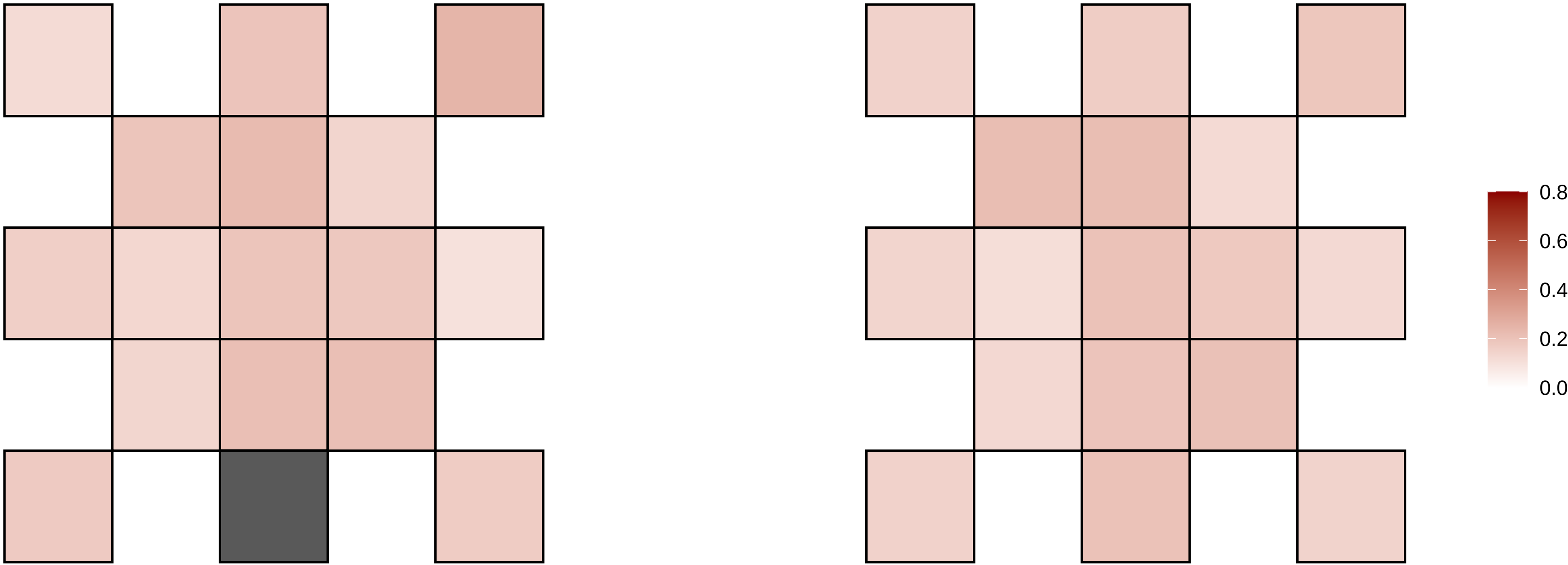

17−184881−G−A

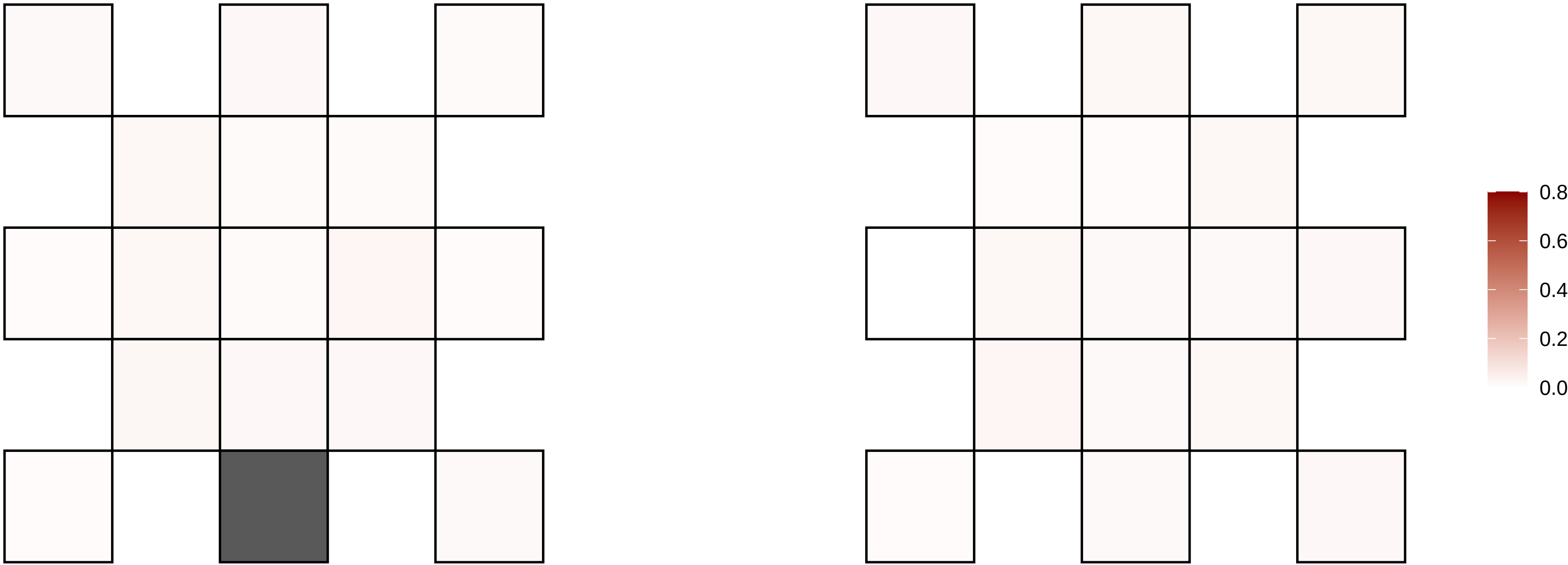

17−20005504−G−A

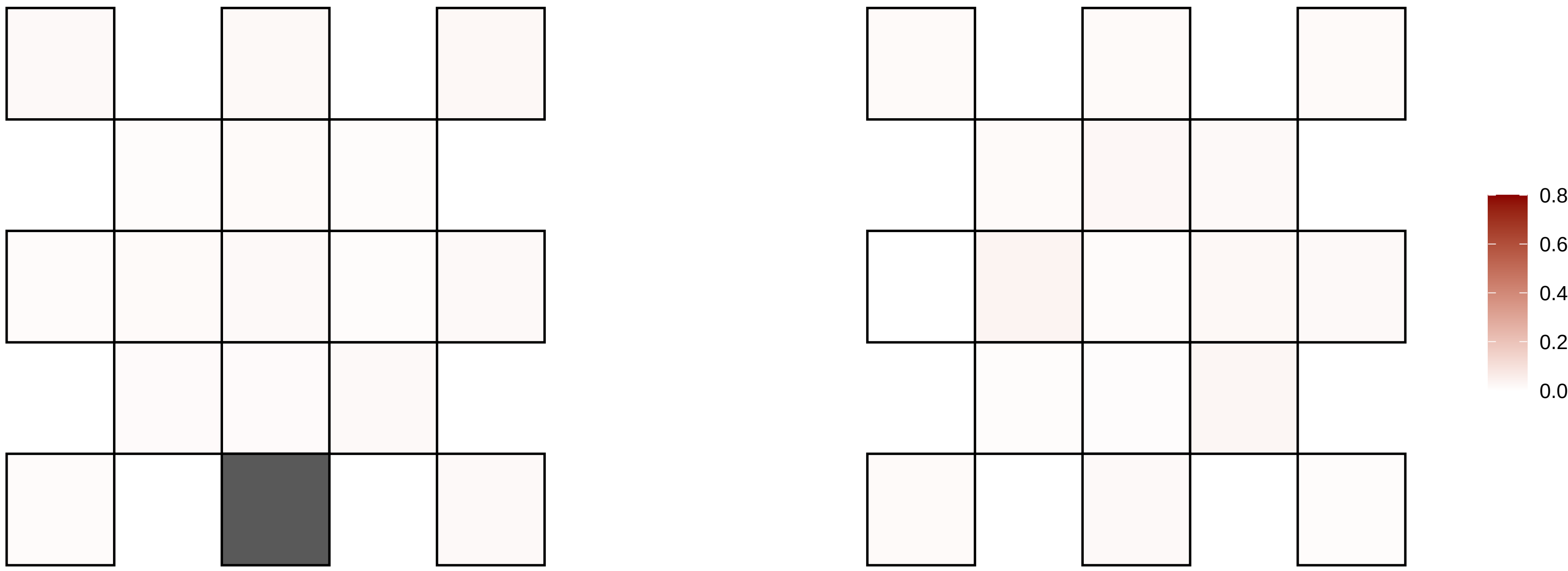

17−20005505−G−A

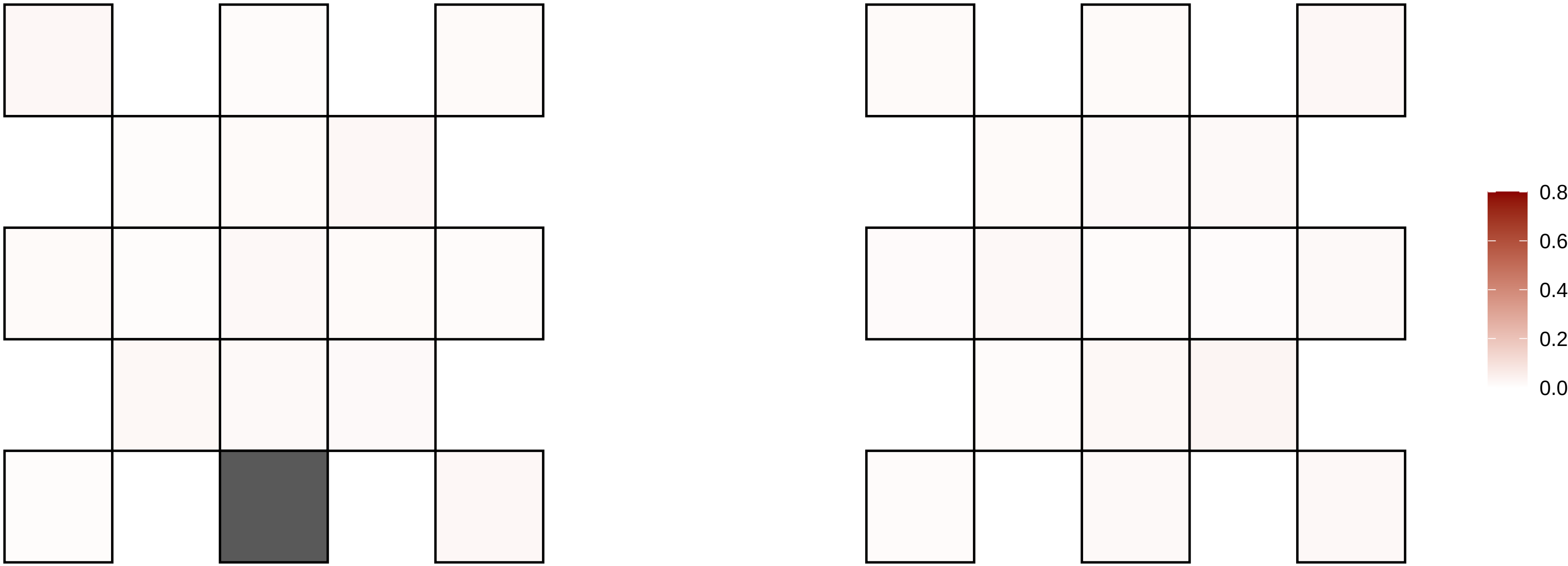

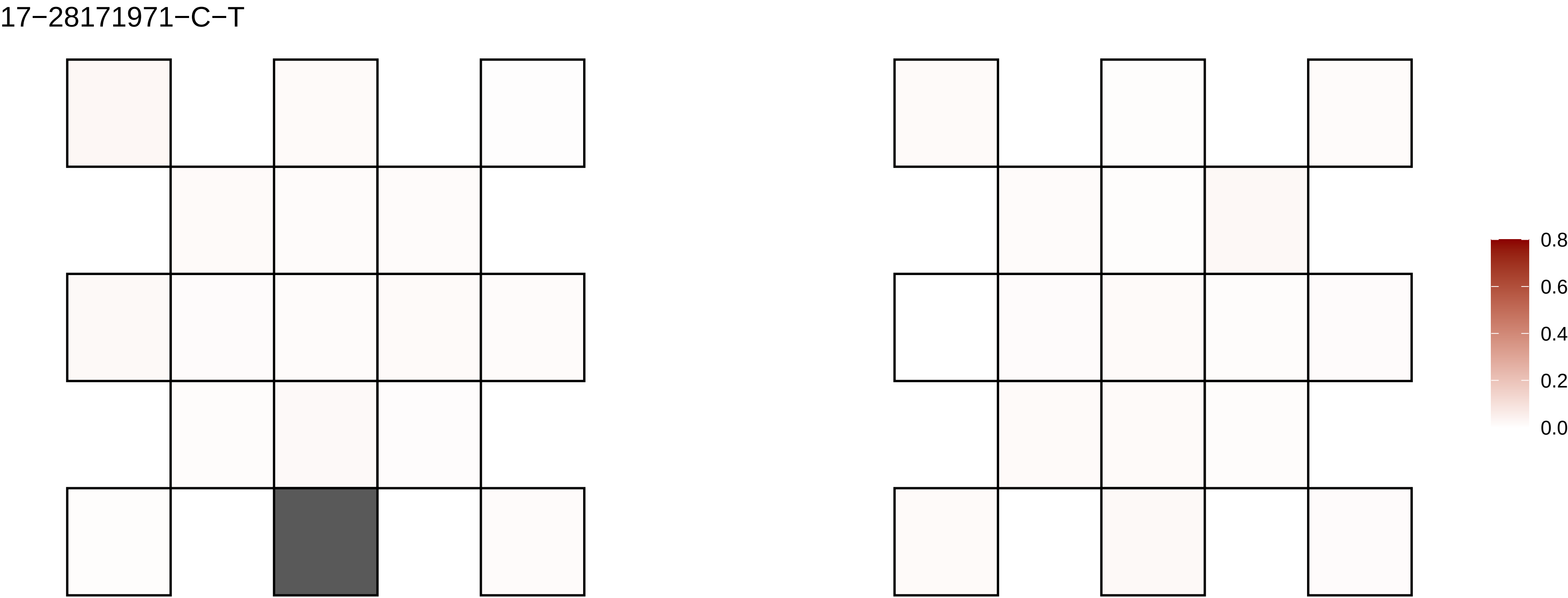

17−28171971−C−T
0.8
0.6
0.4
0.2
0.0

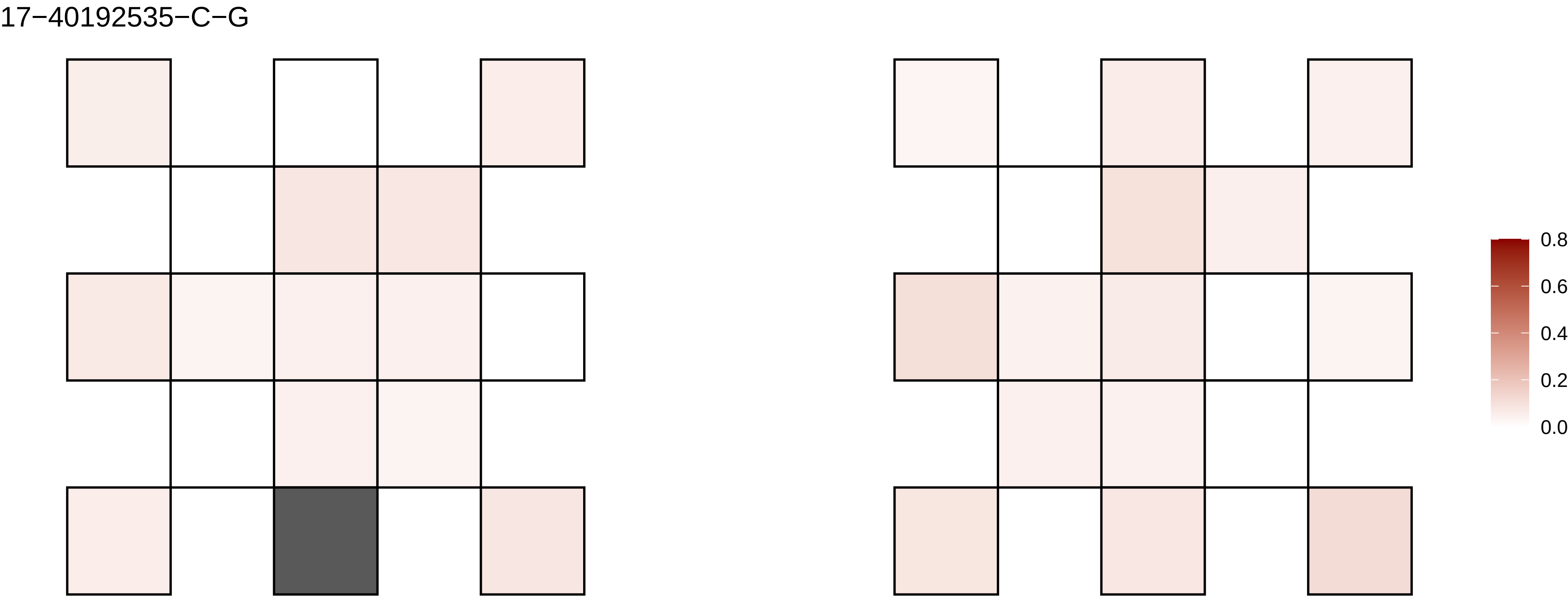

17−40192535−C−G
0.8
0.6
0.4
0.2
0.0

17−43076022−G−A

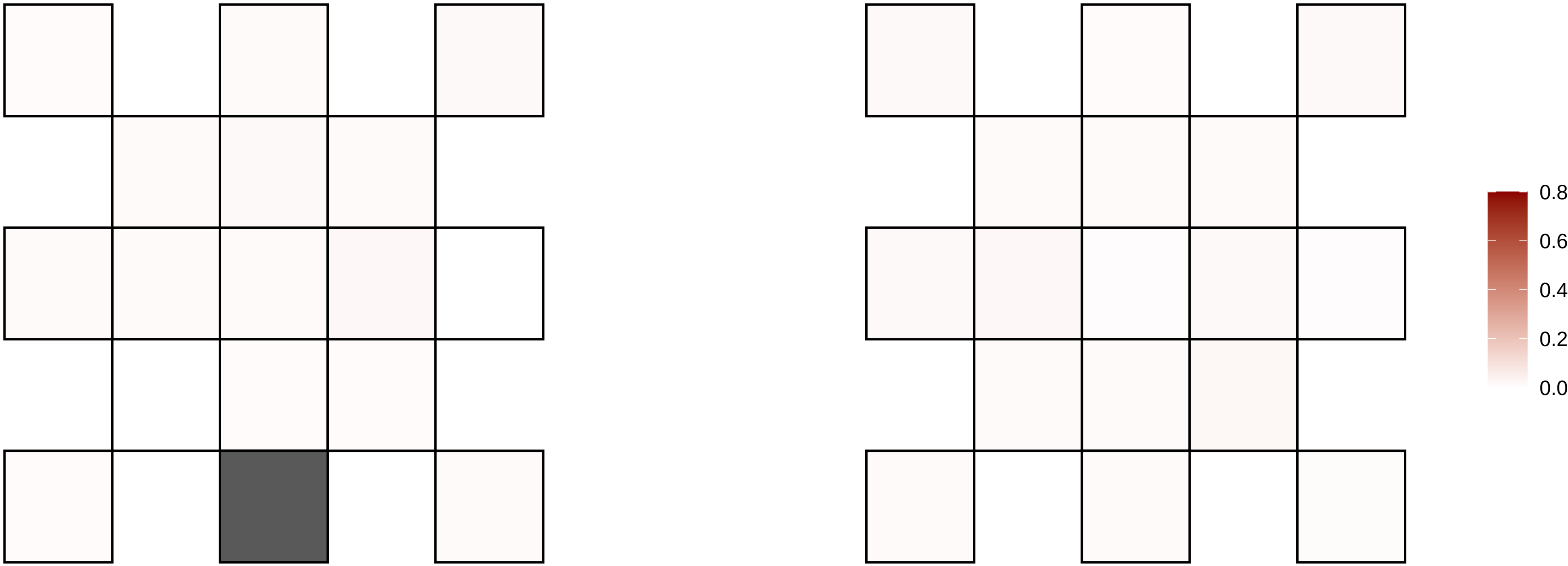

17−45401979−G−A

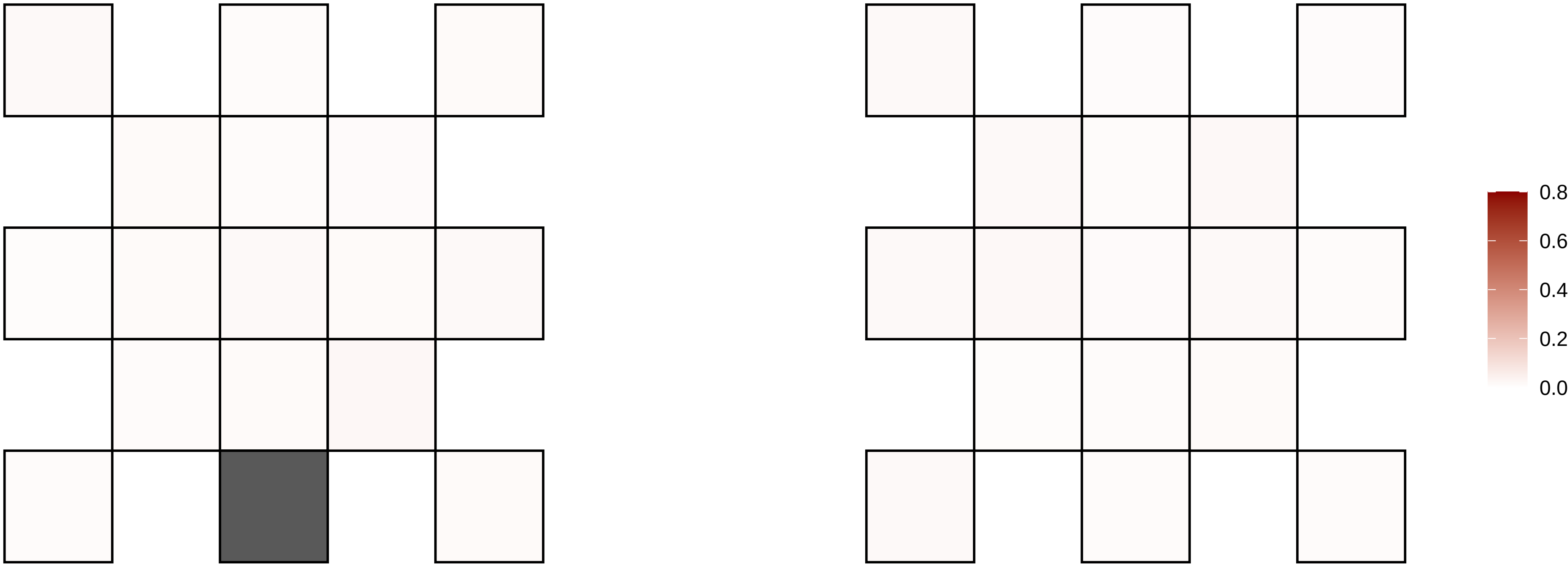

17−50011156−A−T

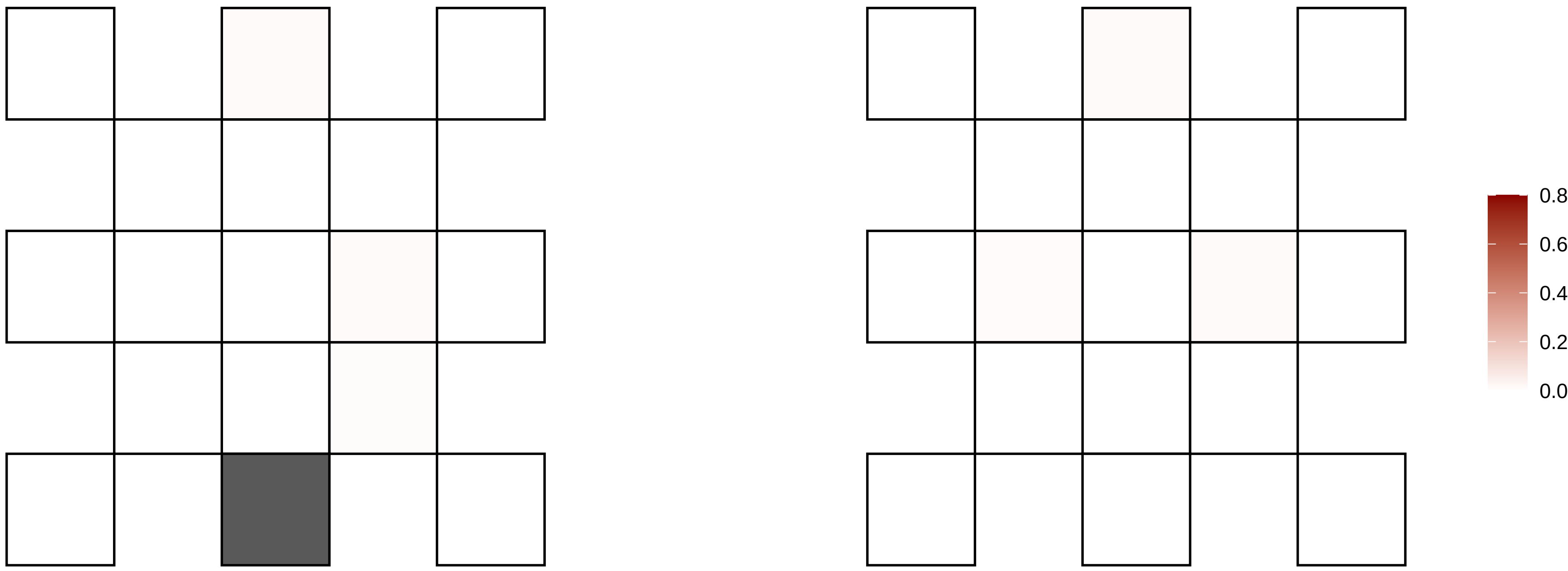

17−59031927−C−T

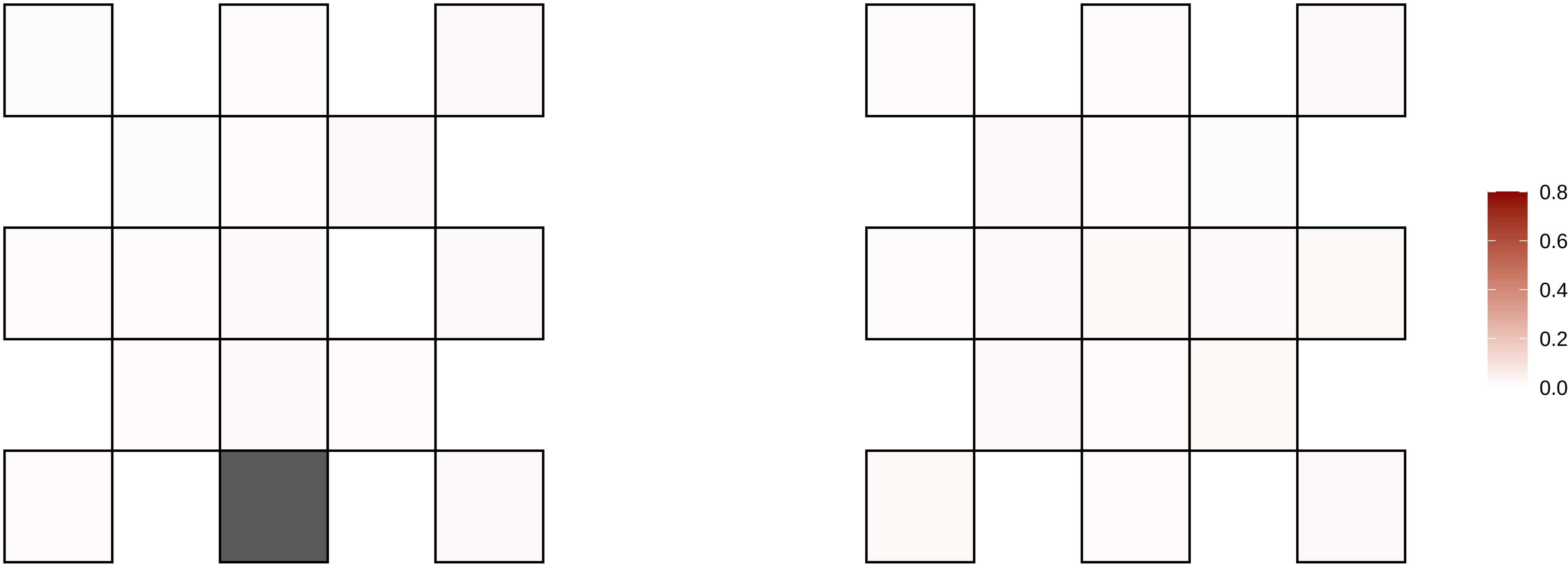

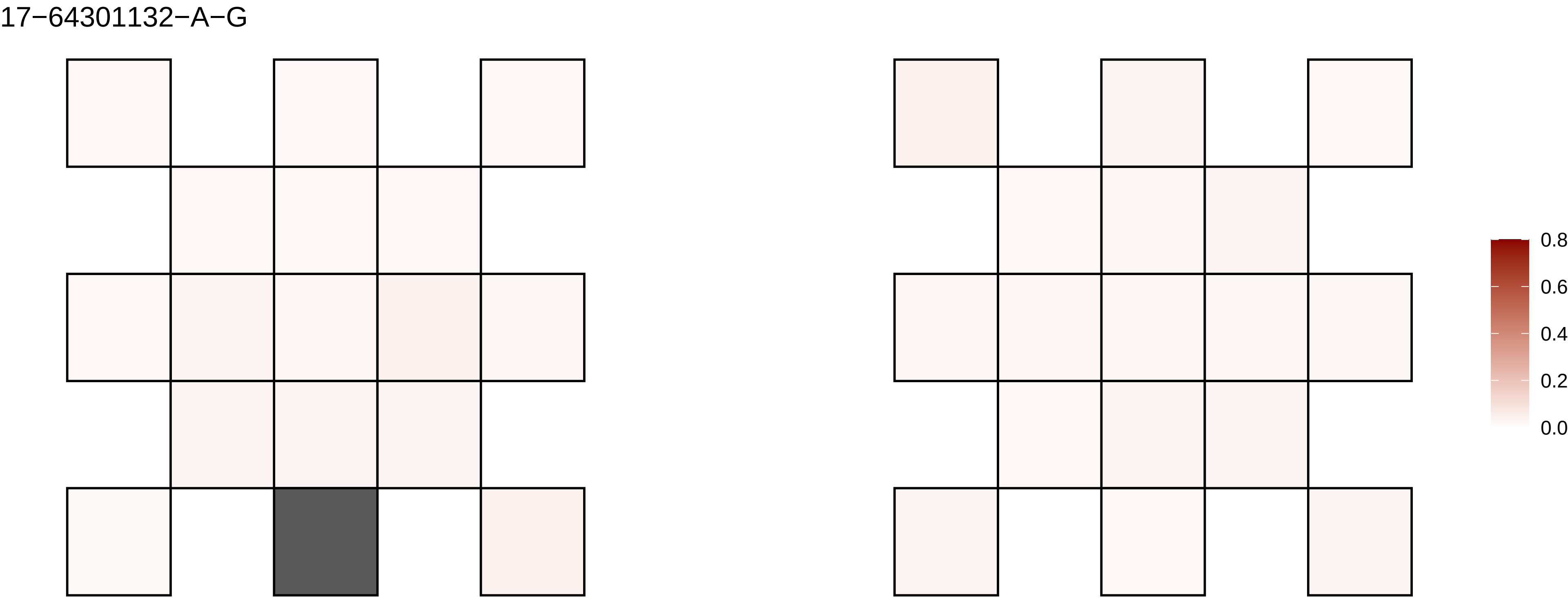
17−64301132−A−G
0.8
0.6
0.4
0.2
0.0

17−66742030−T−G

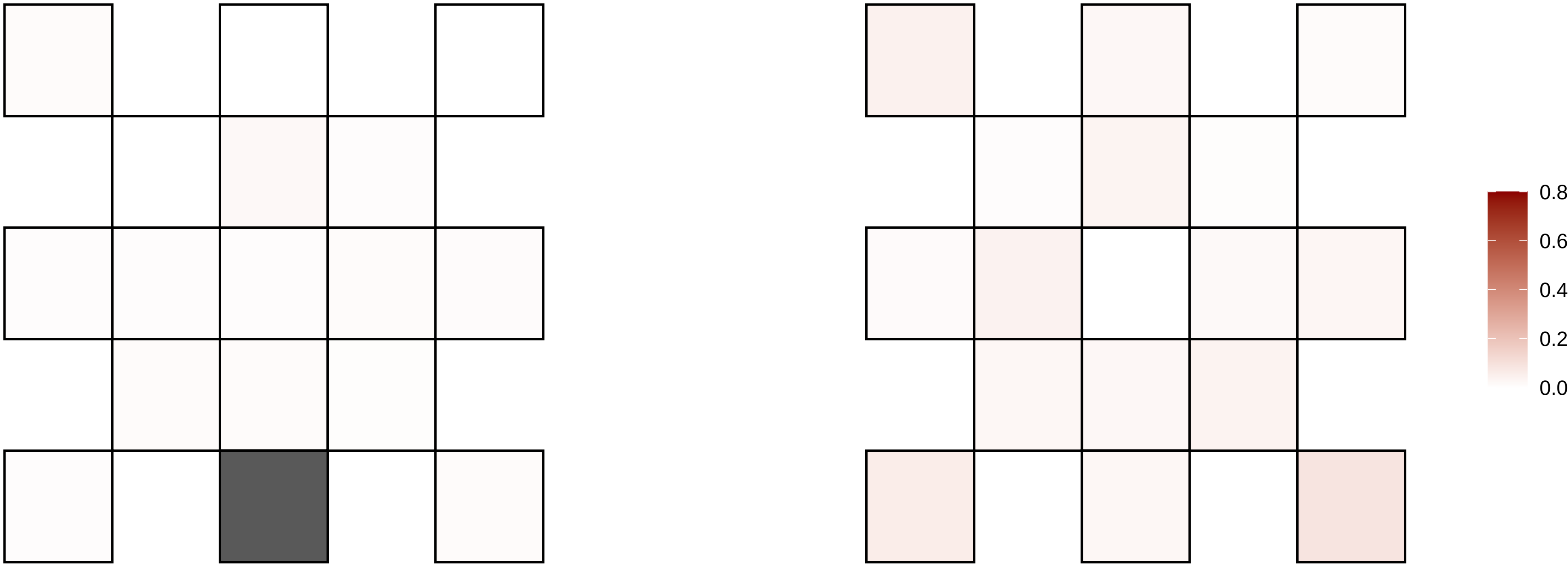

17−73327453−G−A

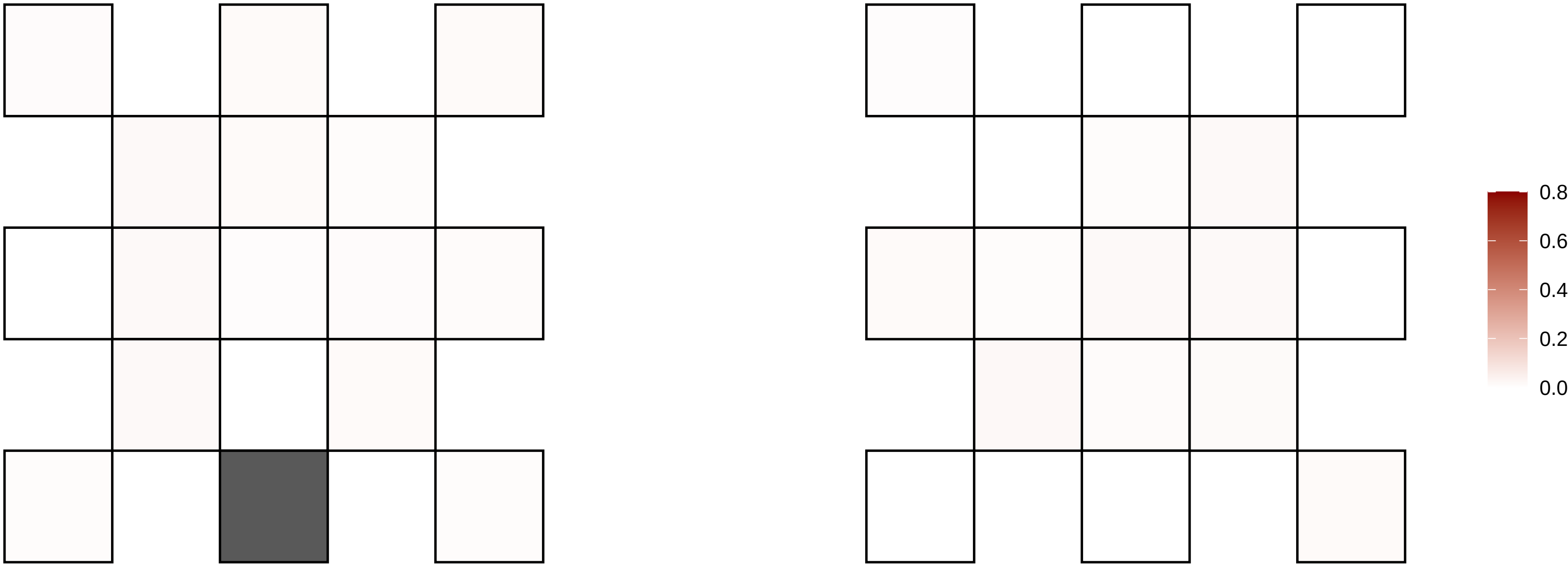

17−77323400−C−T

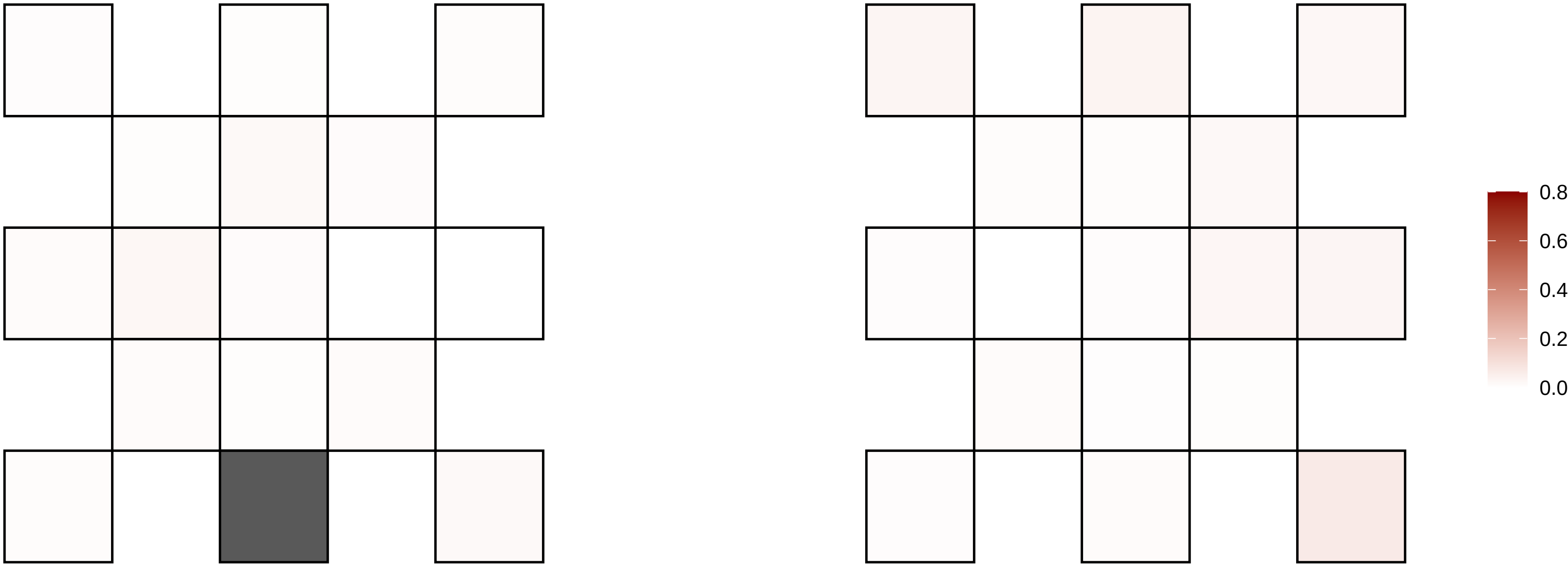

17−79305344−G−A

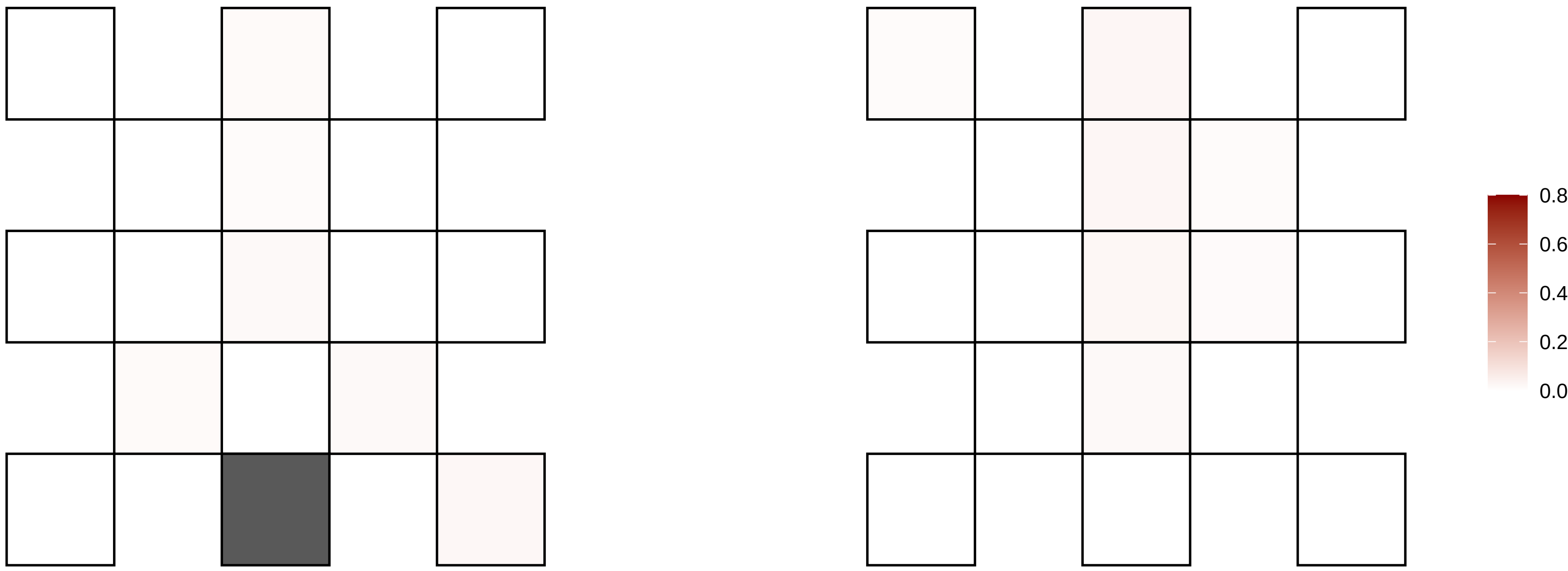

17−80207448−G−A

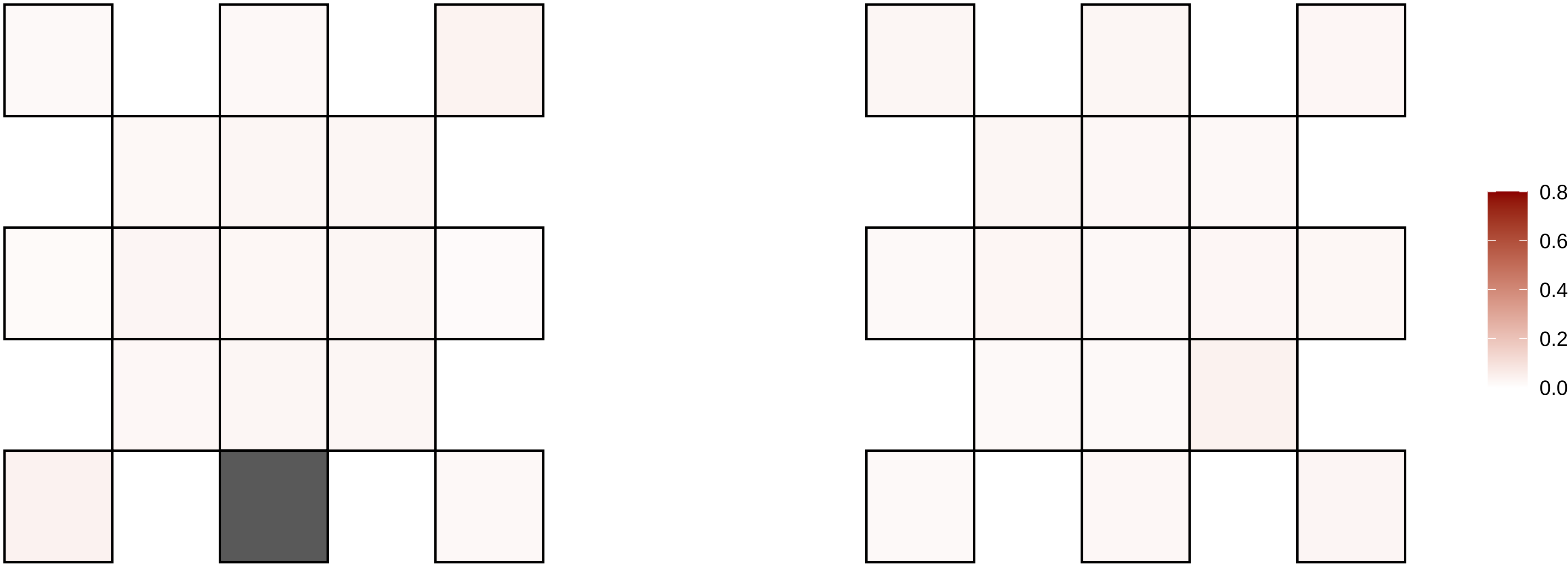

17−80643417−G−A

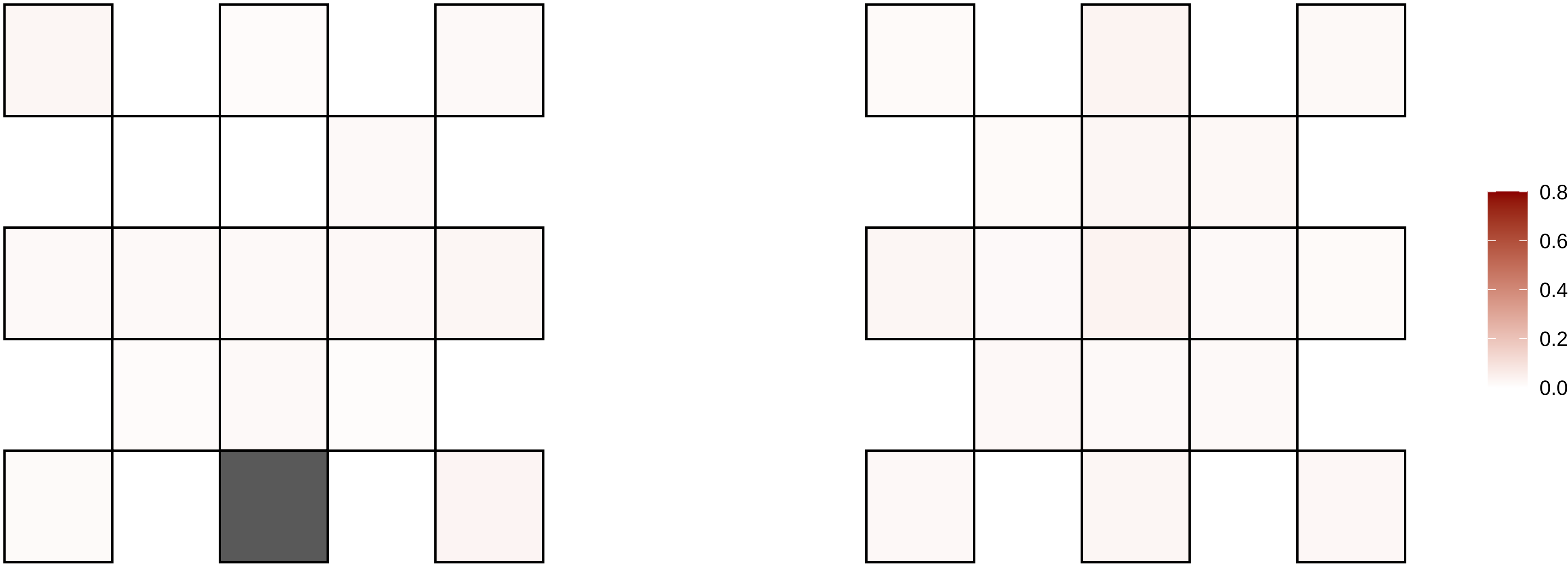

17−8815877−G−A

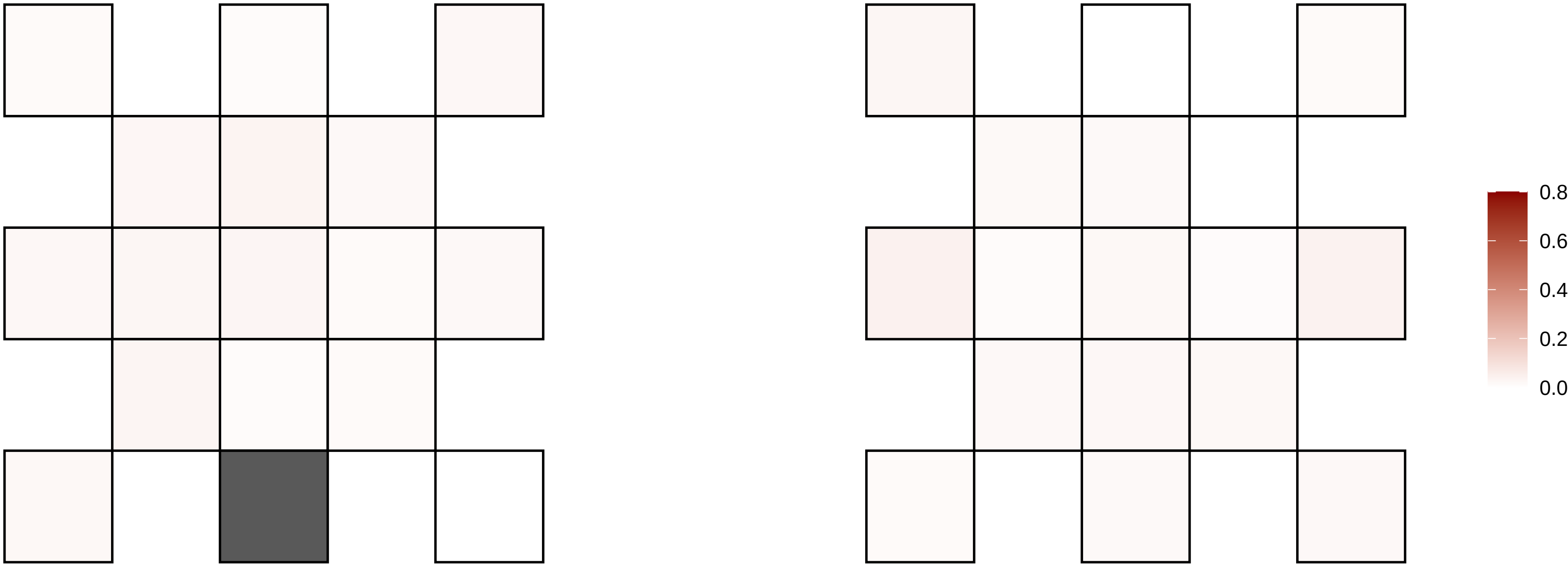

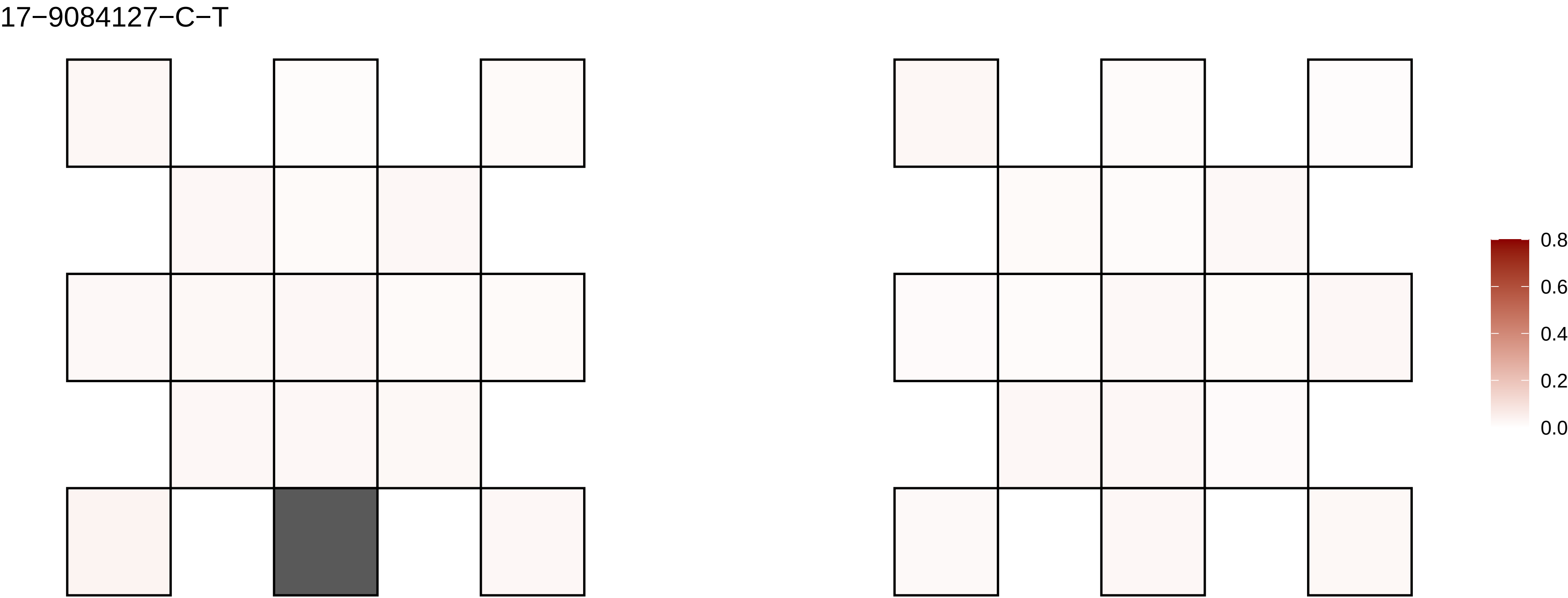
17−9084127−C−T
0.8
0.6
0.4
0.2
0.0

18−1735890−A−C

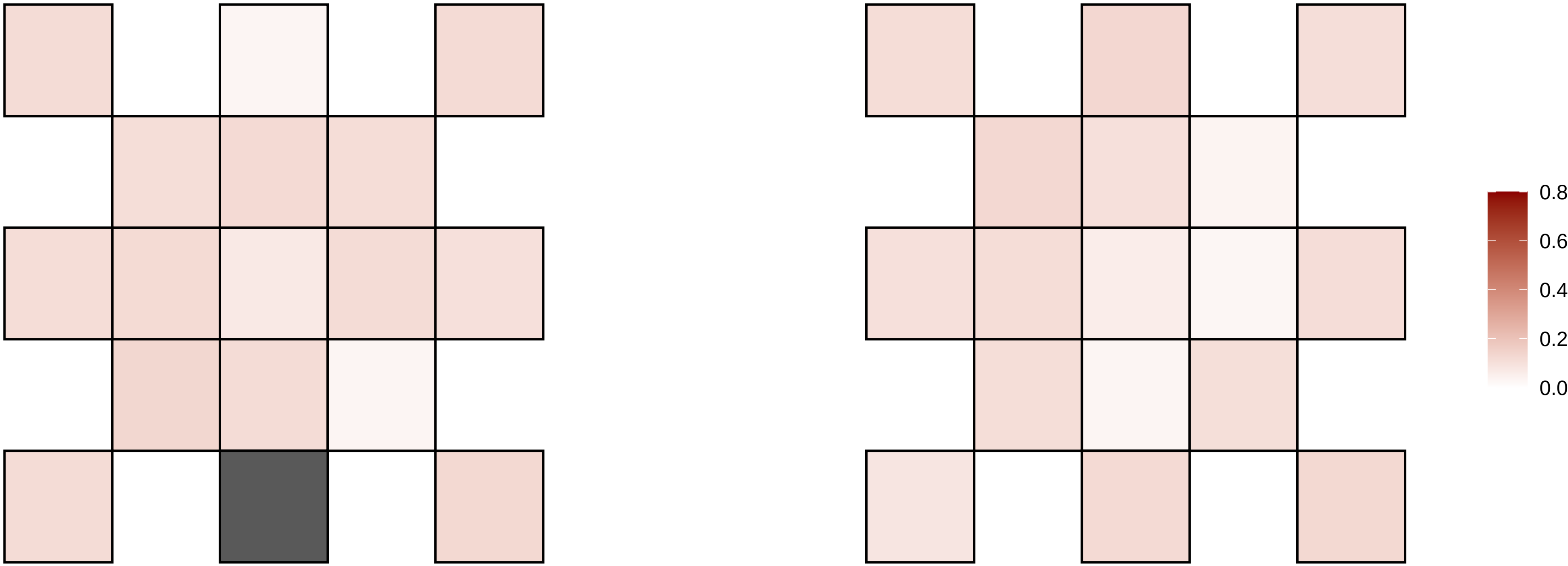

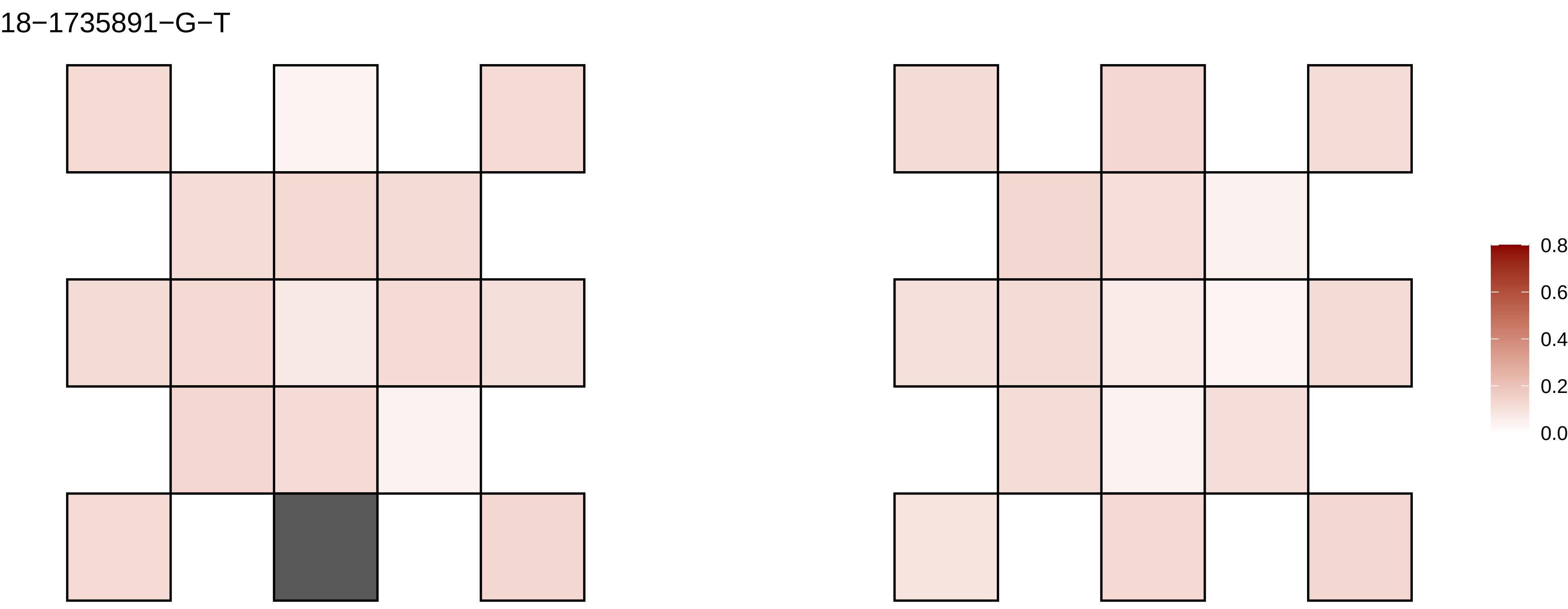
18−1735891−G−T
0.8
0.6
0.4
0.2
0.0

18−24782762−T−A

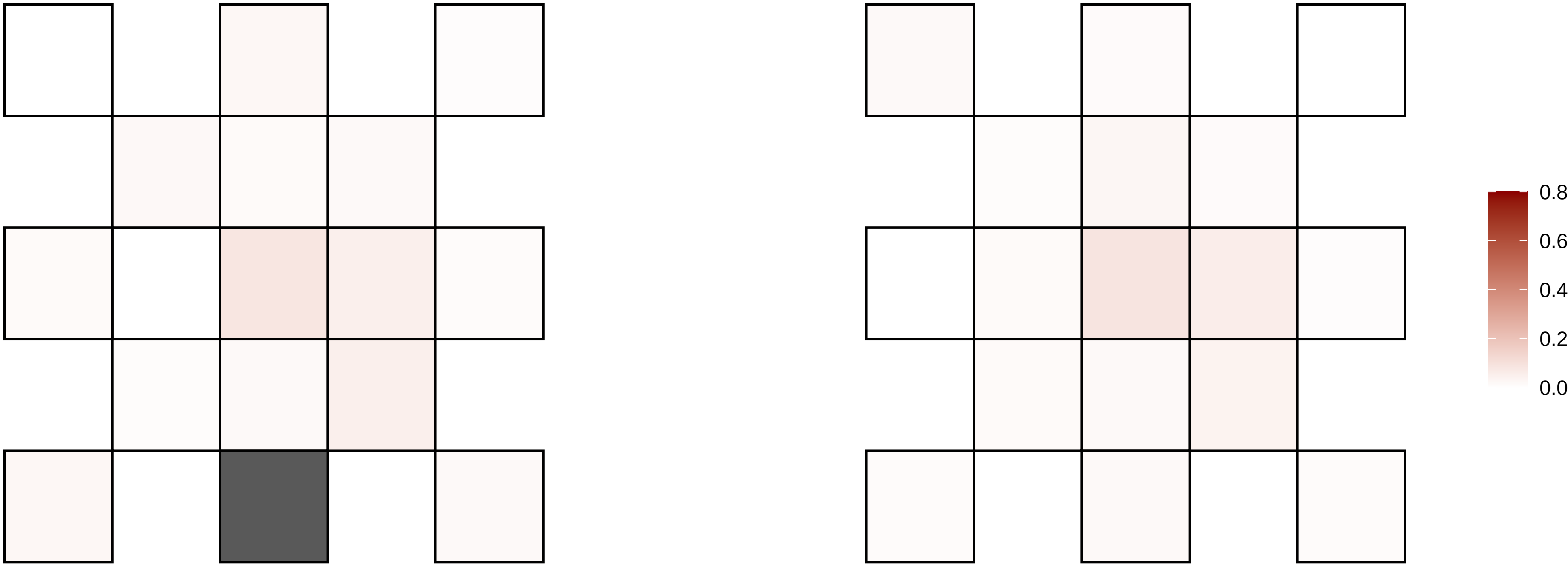

18−25171573−C−T

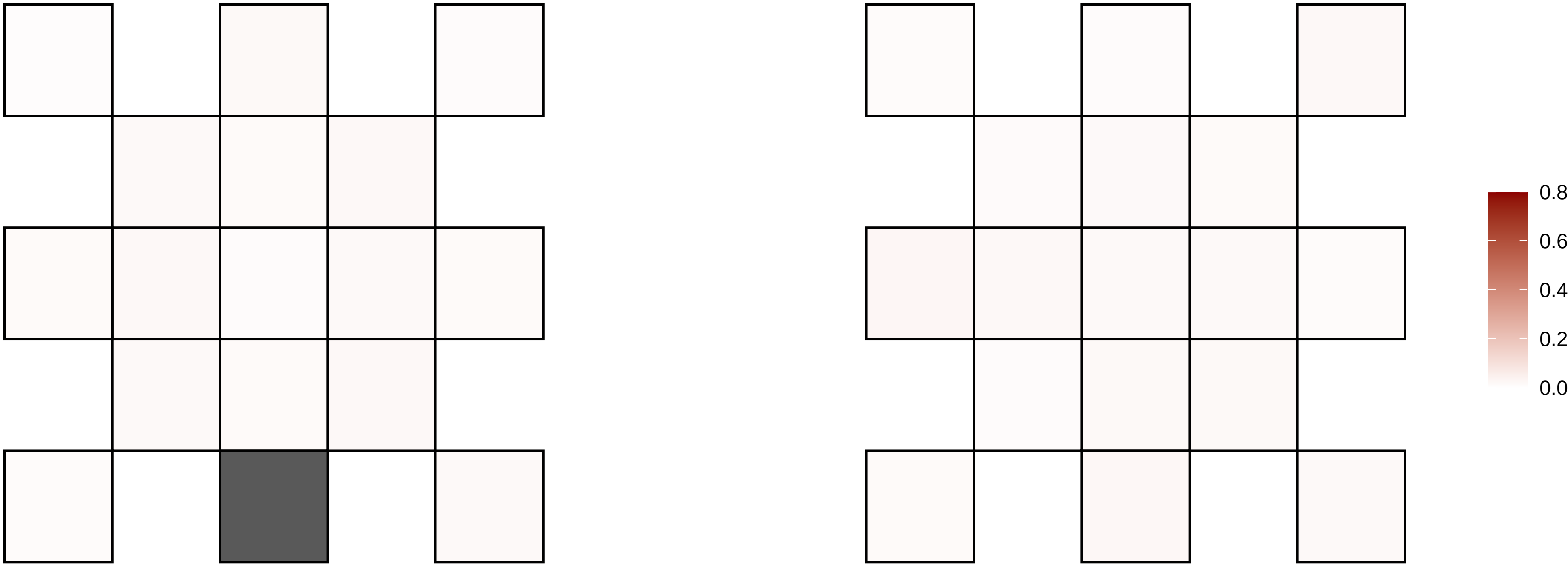

18−33999883−C−T

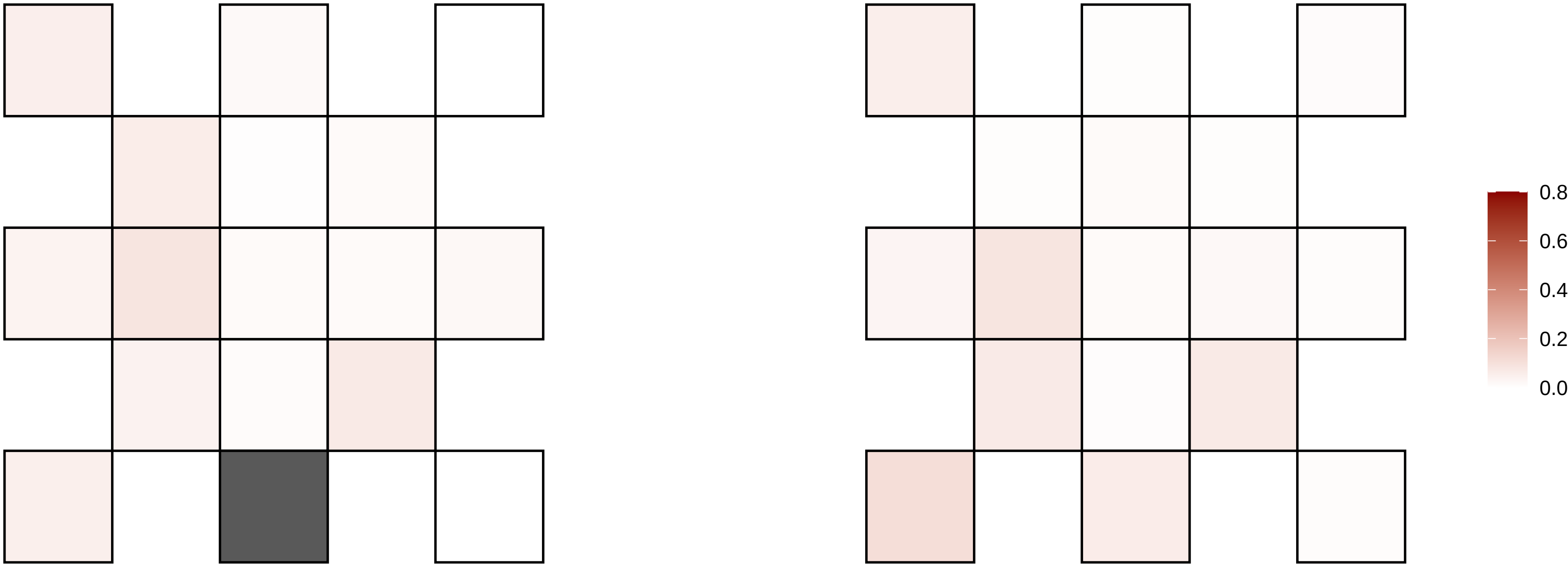

18−36090085−A−C

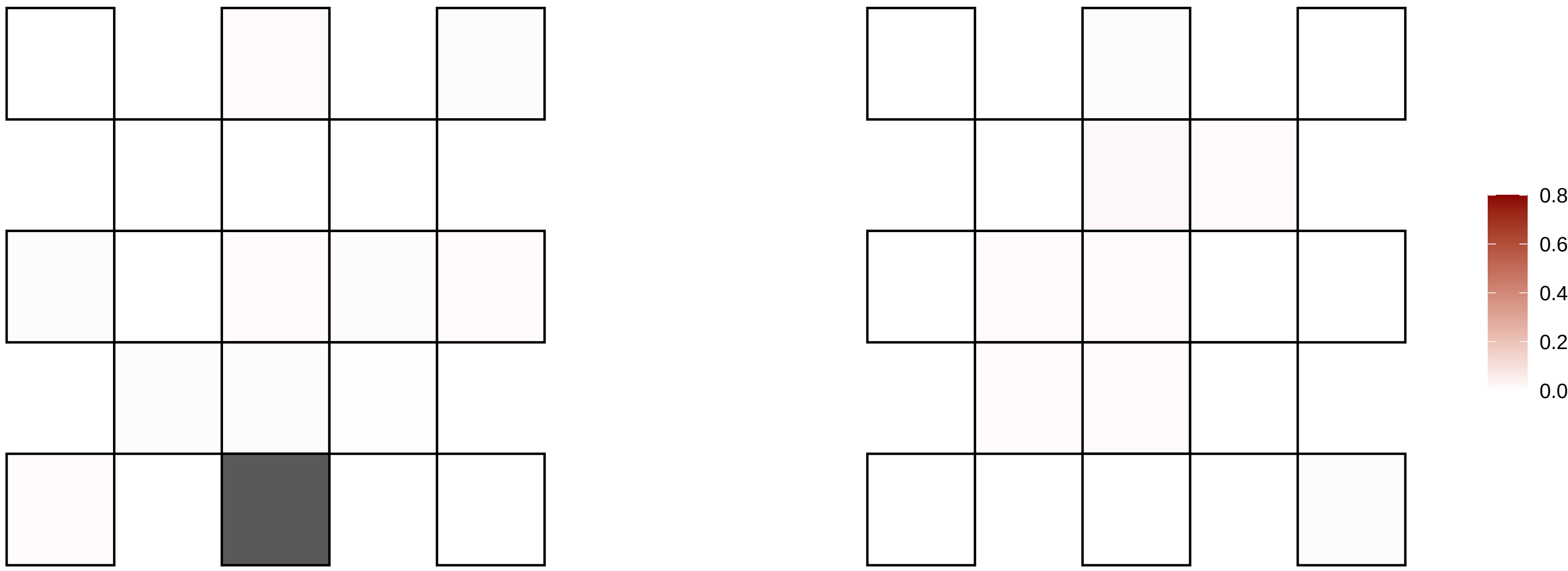

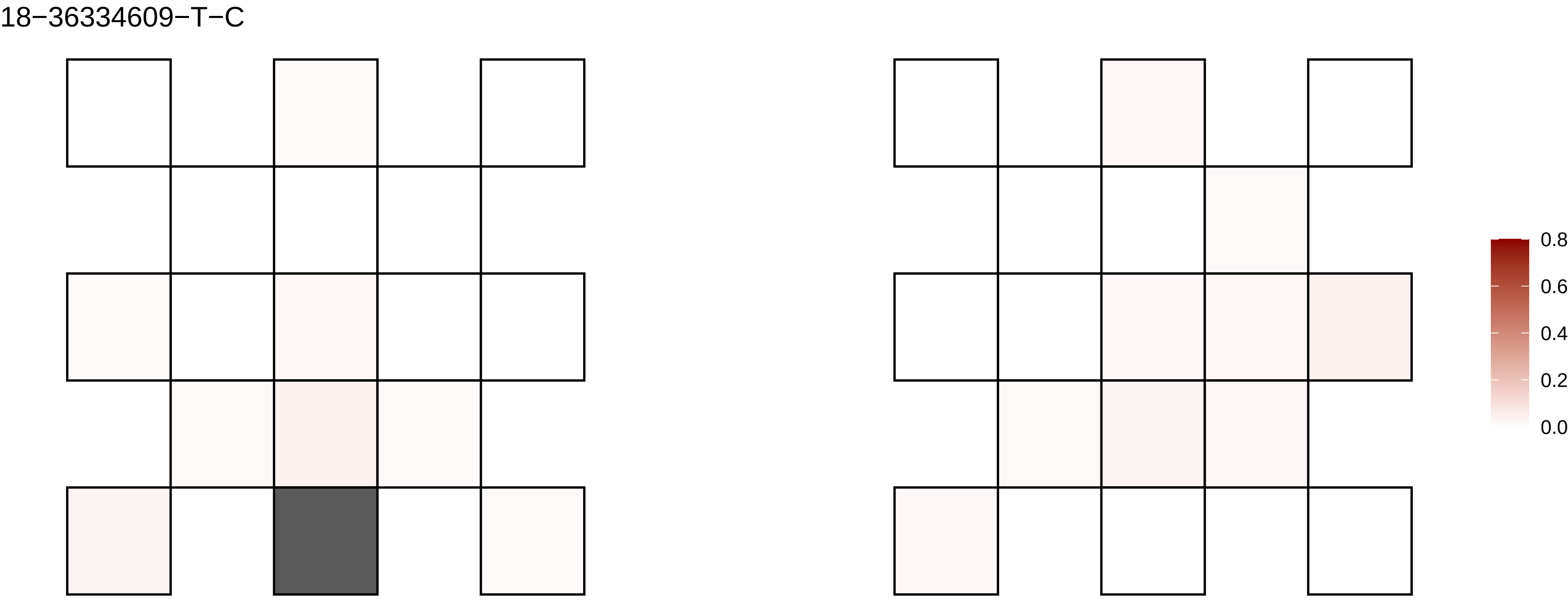

18−36334609−T−C
0.8
0.6
0.4
0.2
0.0

18−37021461−T−C

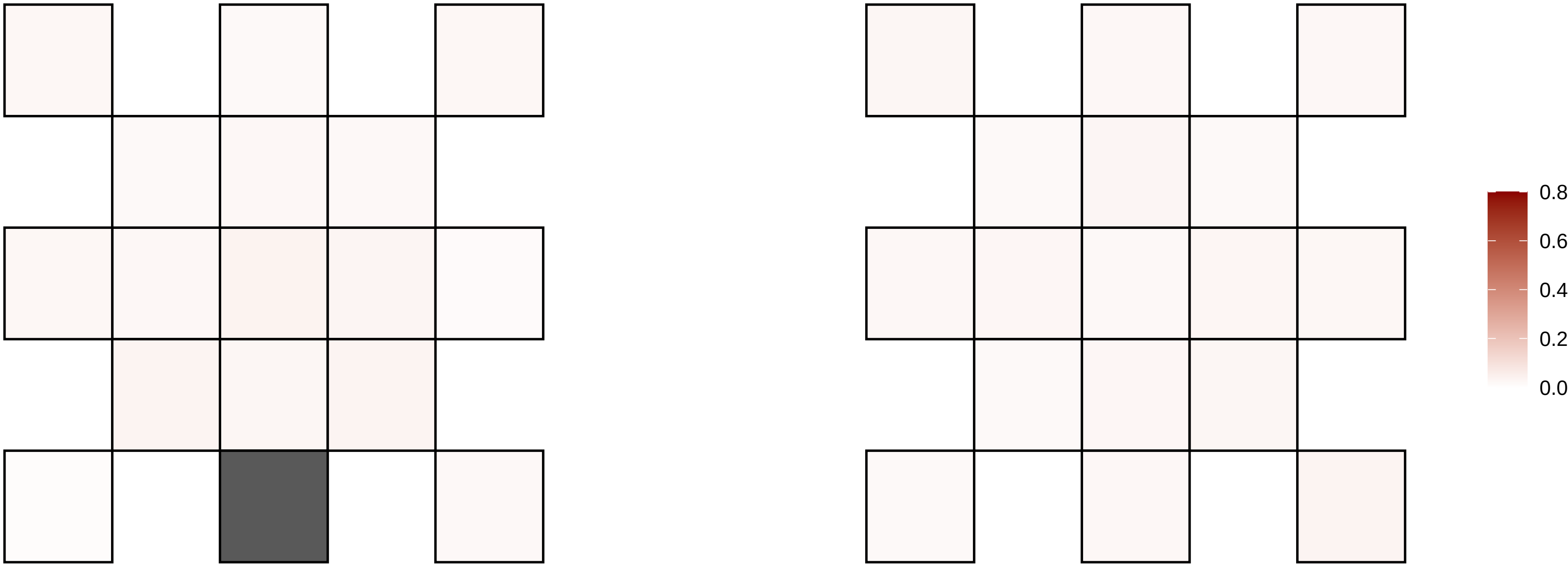

18−42932839−C−T

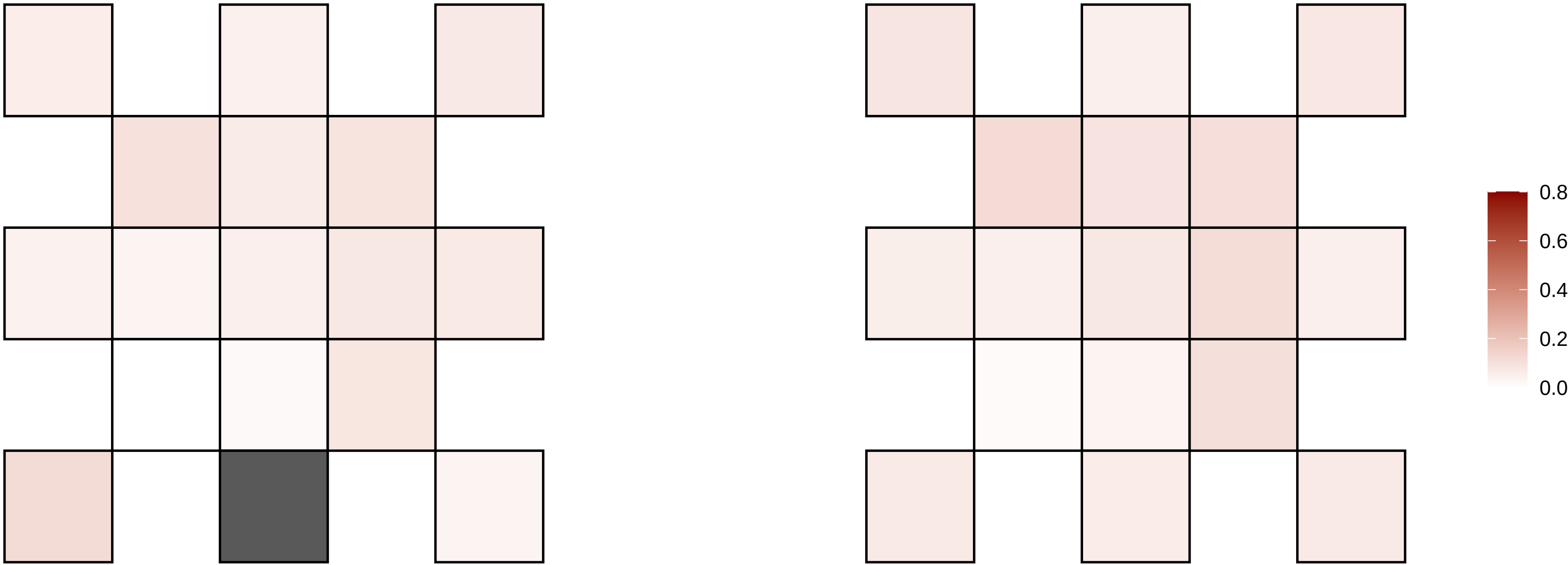

18−49194474−G−A

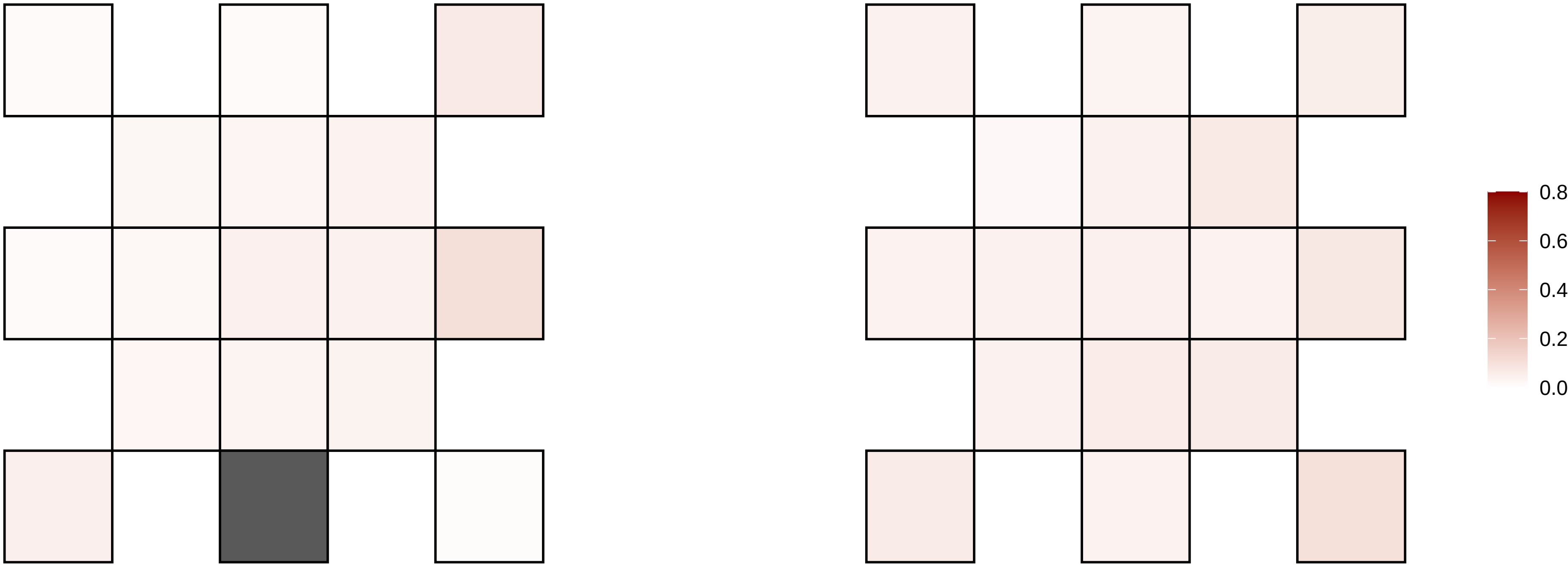

18−49557610−G−T

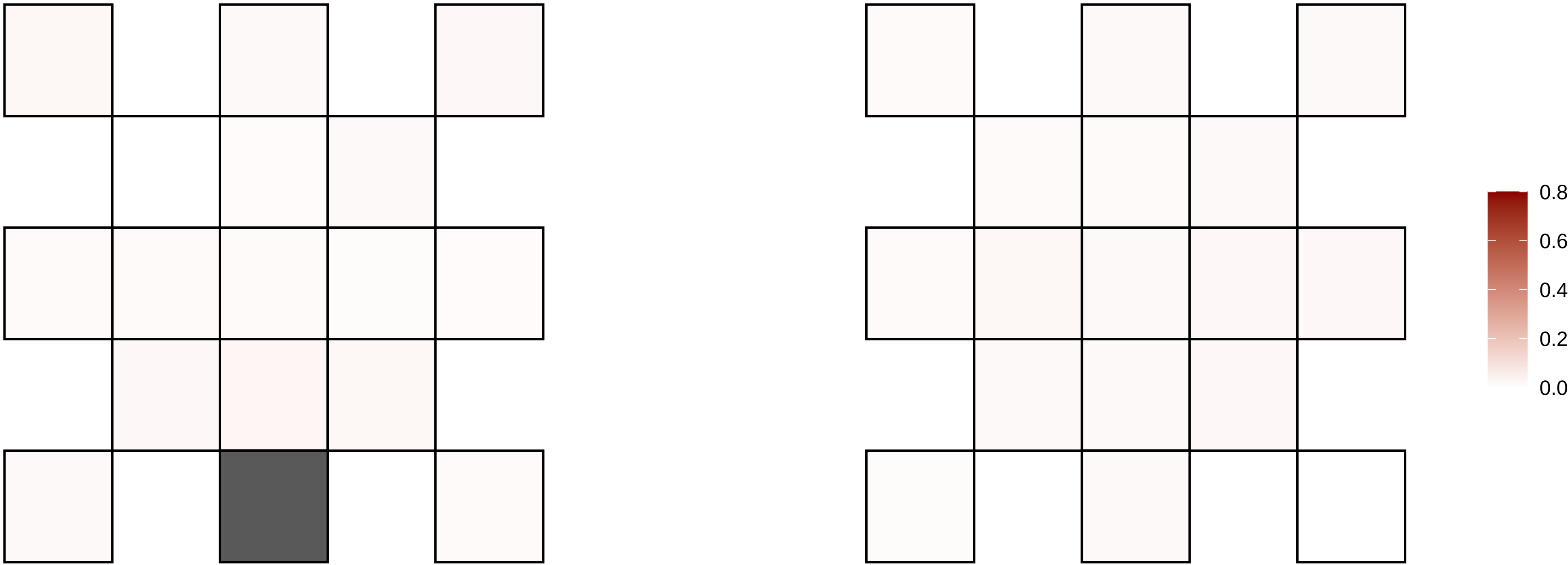

18−60849046−C−T

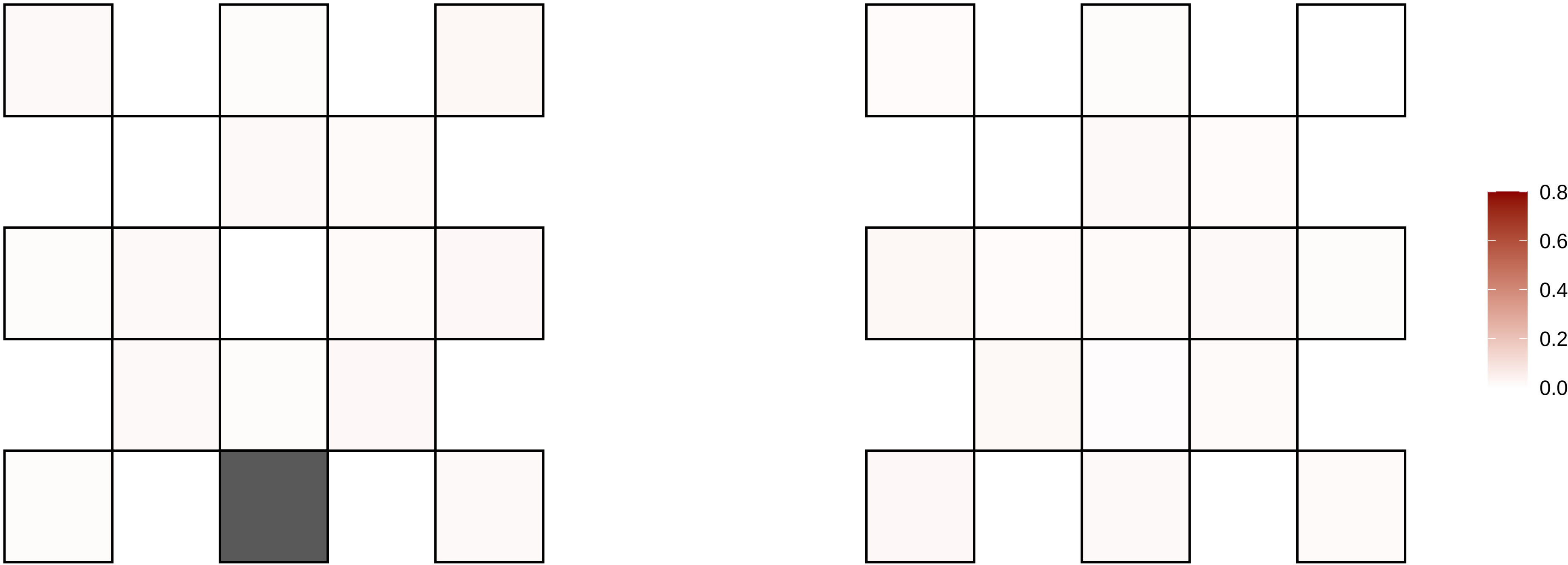

18−64016149−C−A

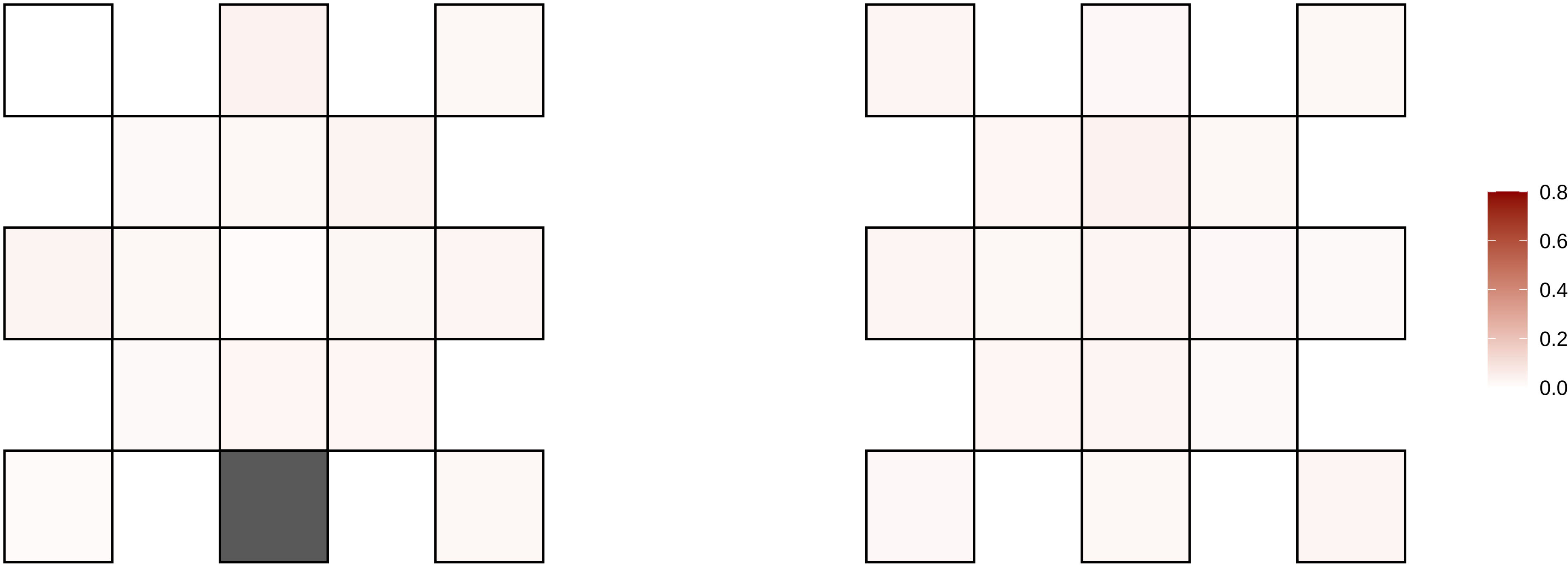

18−64125536−C−G

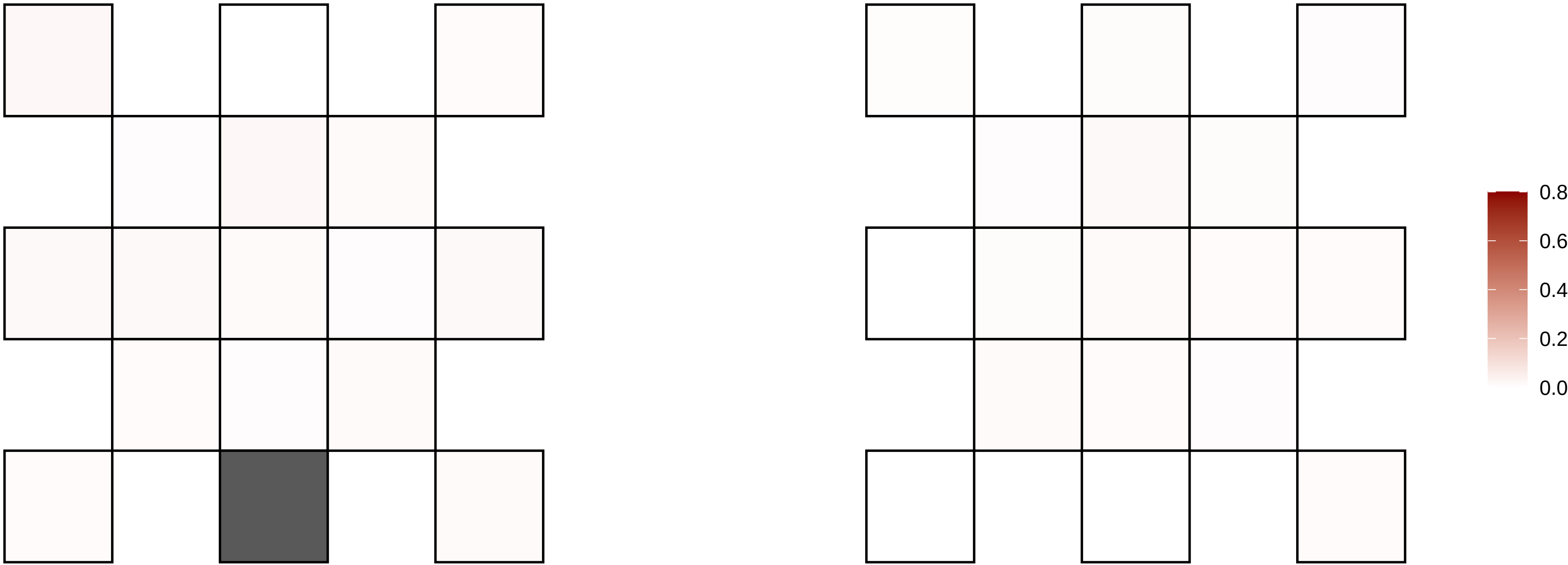

18−66103772−A−T

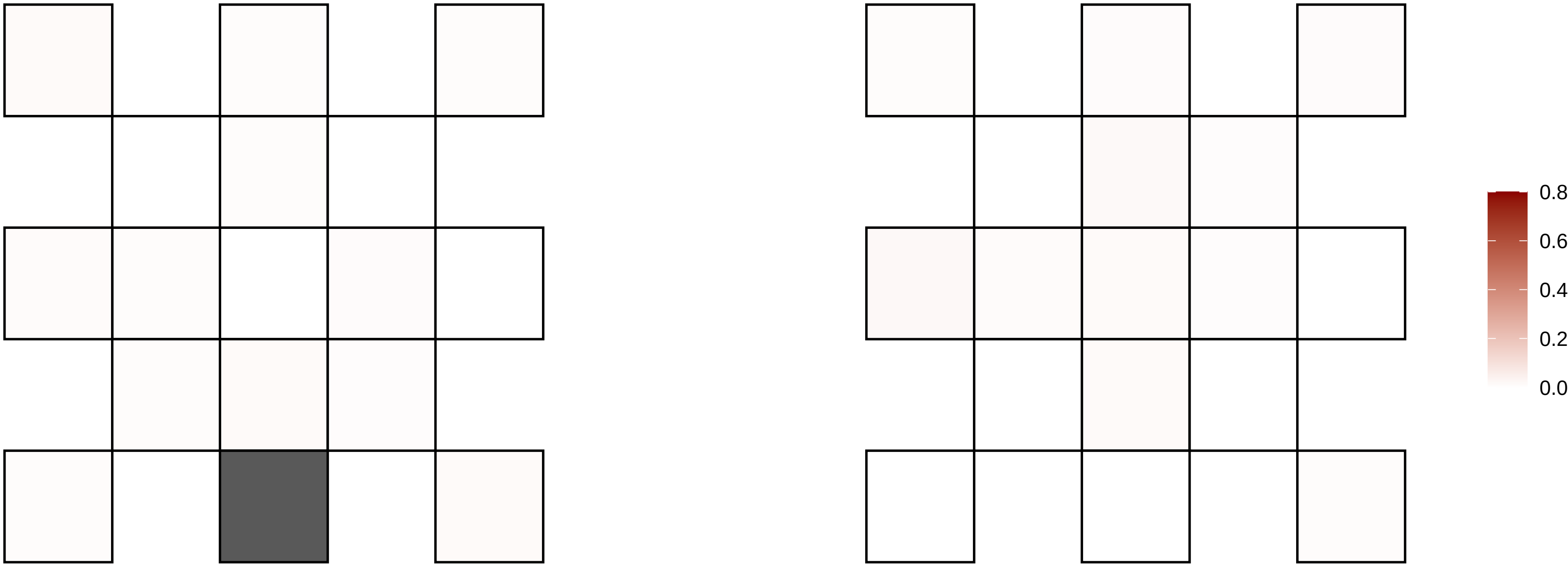

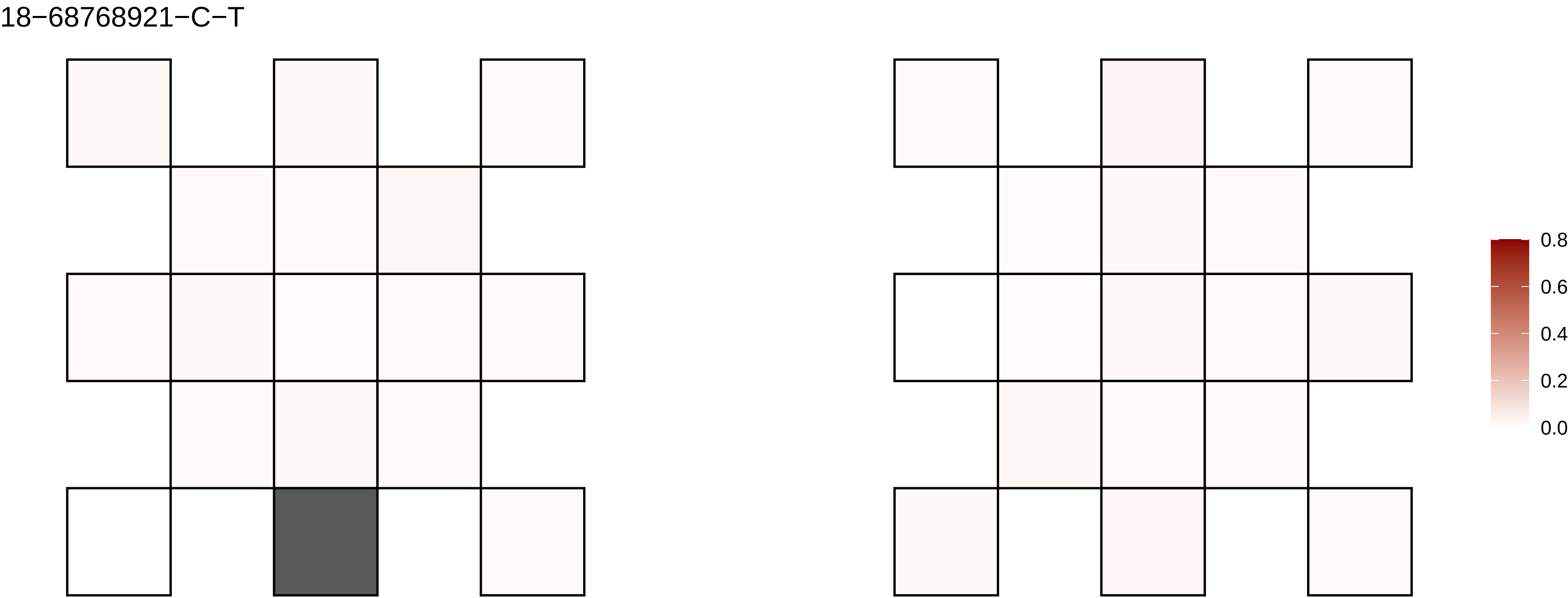
18−68768921−C−T
0.8
0.6
0.4
0.2
0.0

19−1124954−C−T

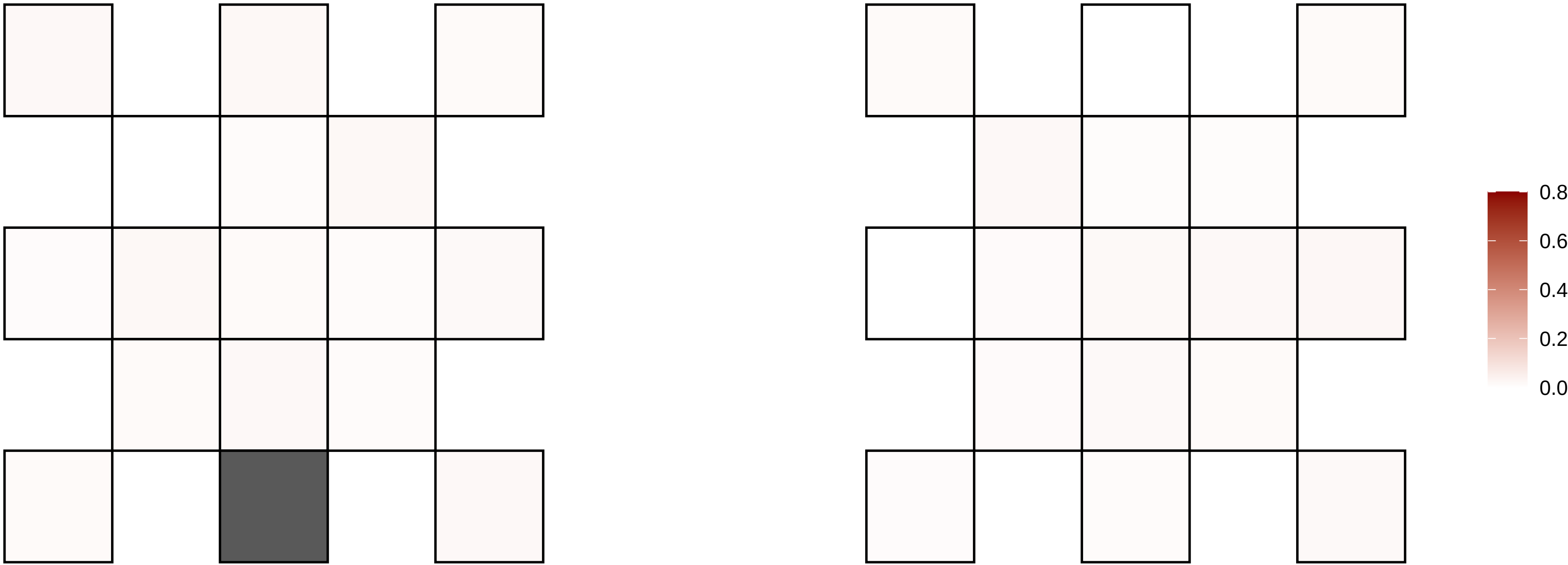

19−13941229−G−A

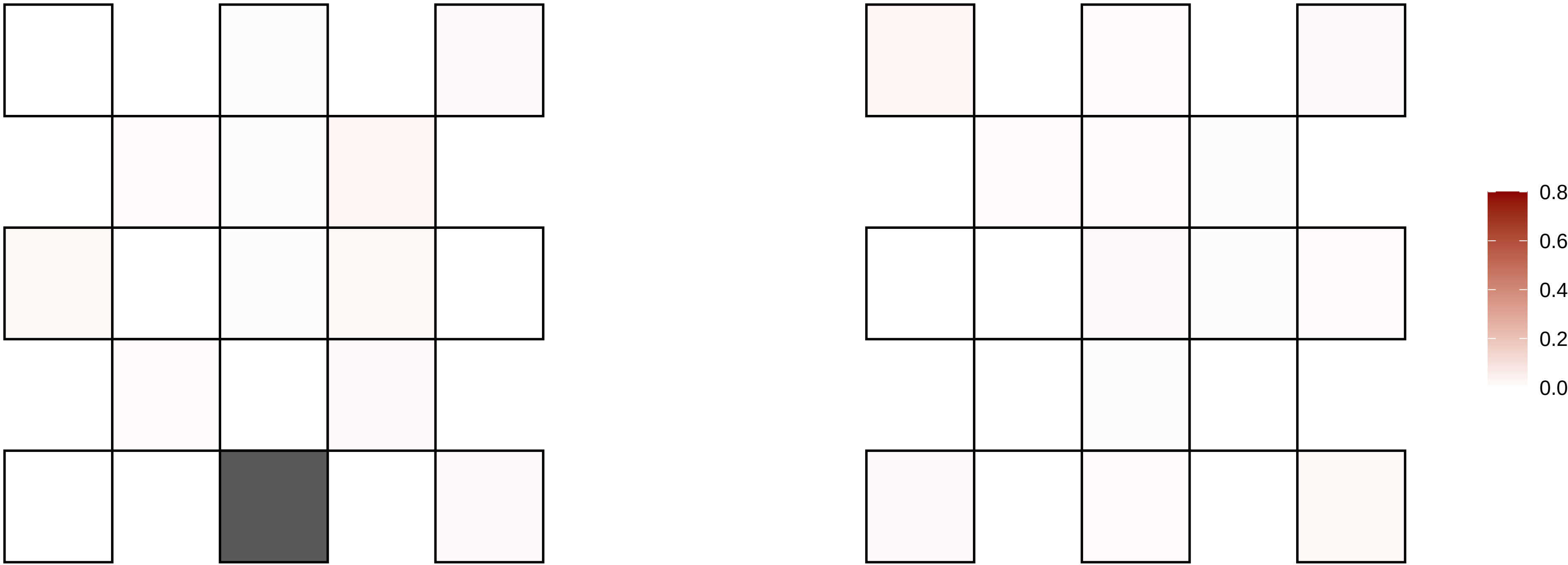

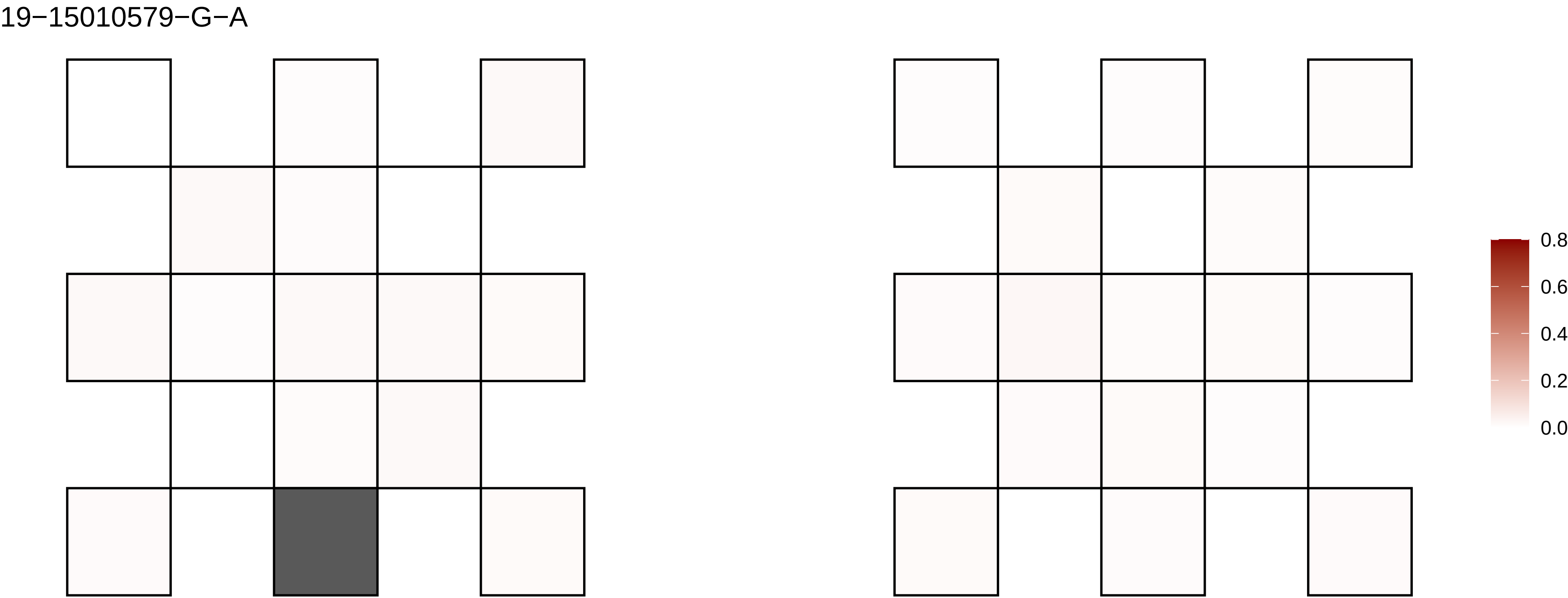
19−15010579−G−A
0.8
0.6
0.4
0.2
0.0

19−1621289−C−T

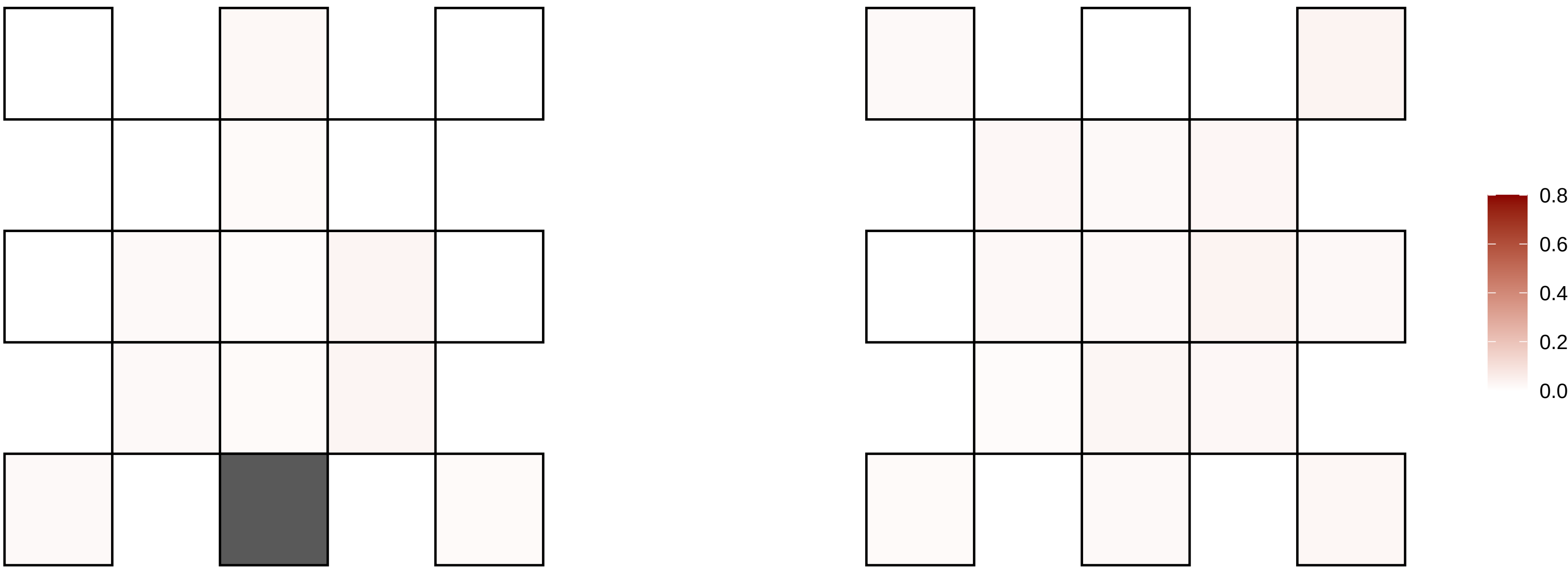

19−30808081−C−T

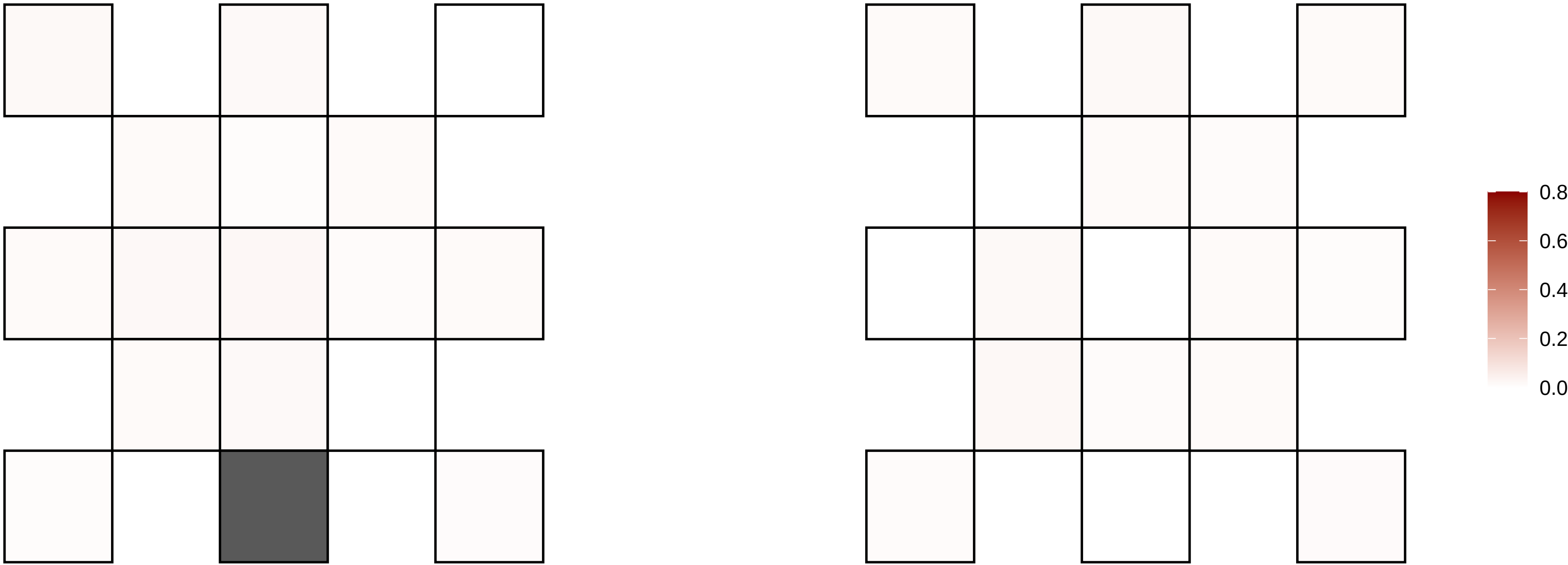

19−35551269−T−A

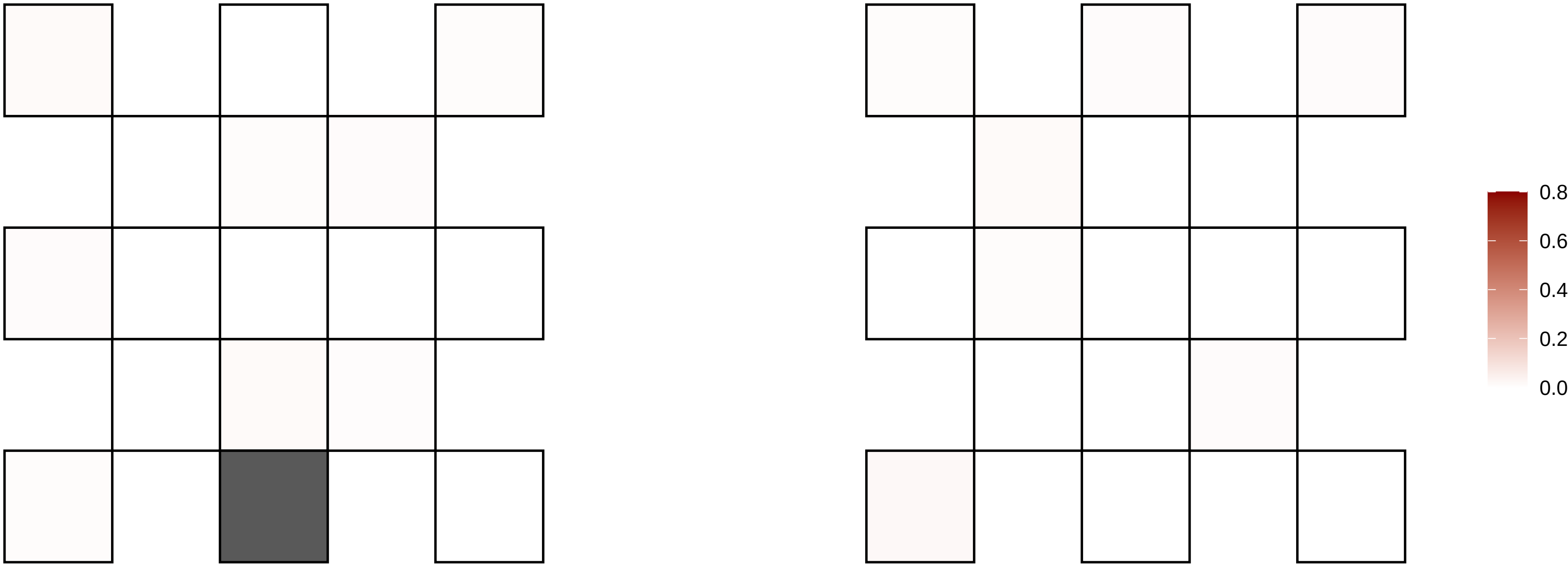

19−36516973−C−T

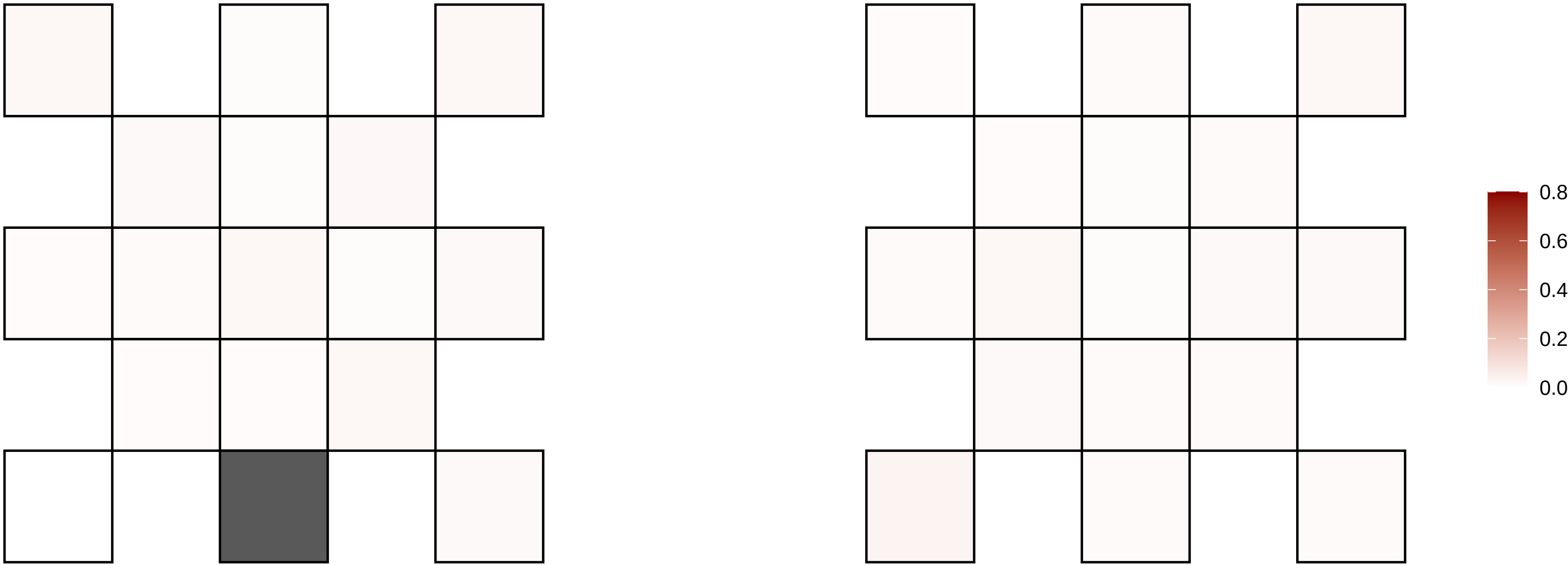

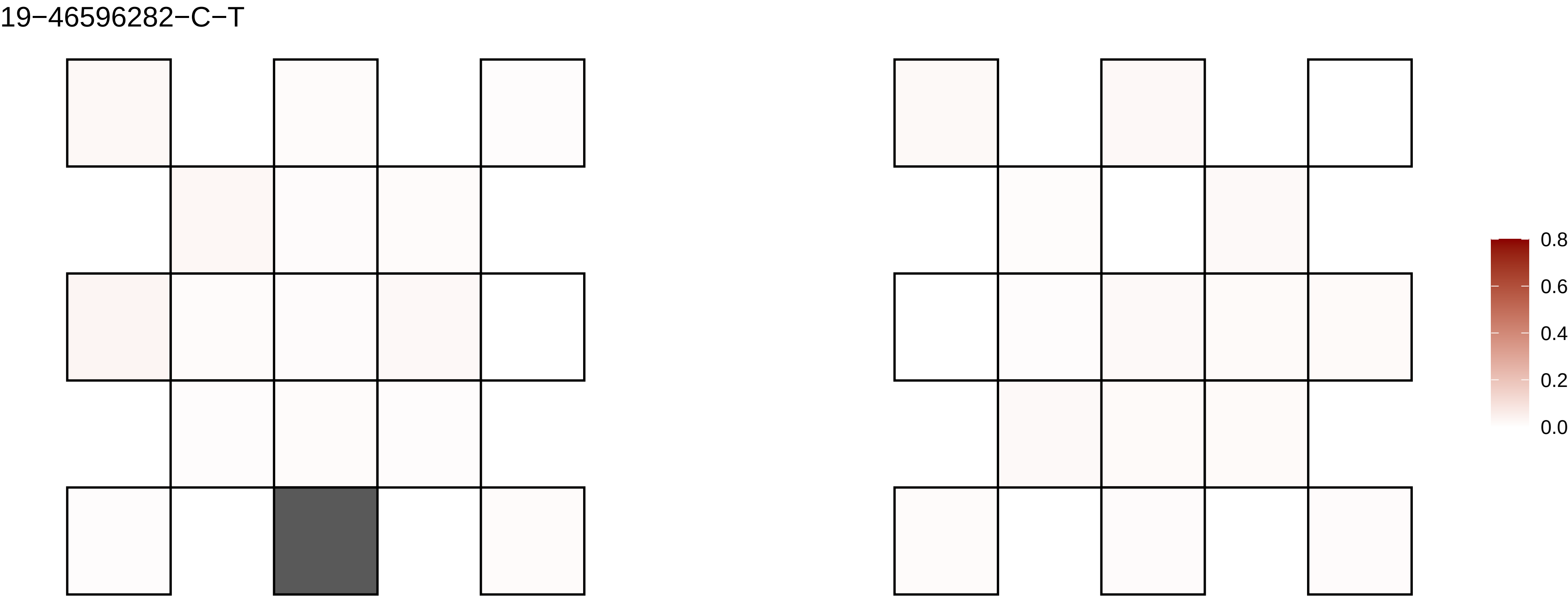
19−46596282−C−T
0.8
0.6
0.4
0.2
0.0

19−53880285−C−T

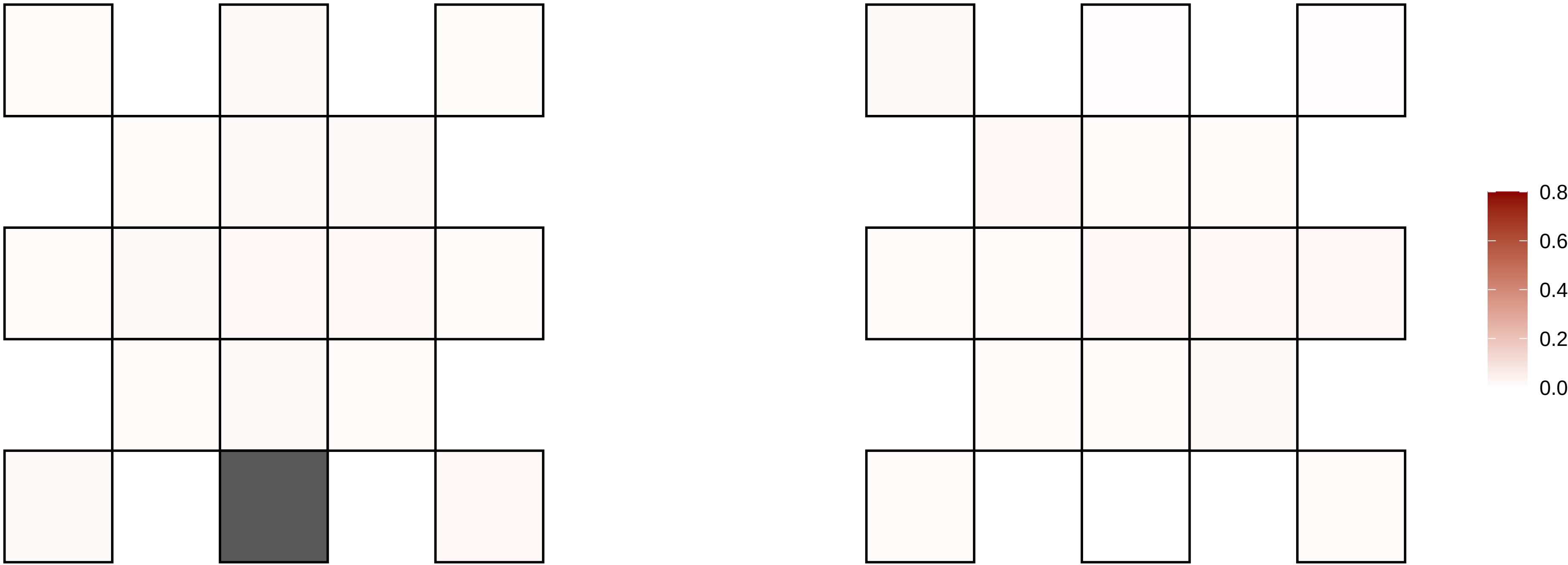

19−5499386−G−C

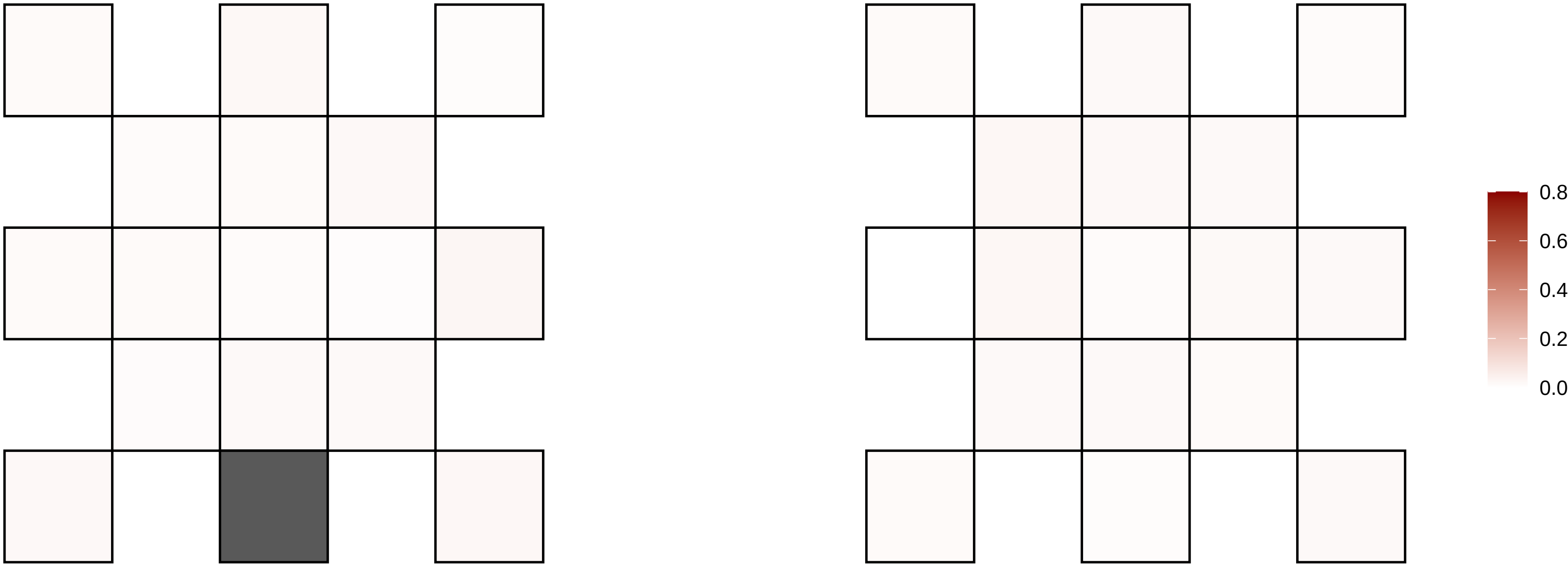

19−55665618−G−A

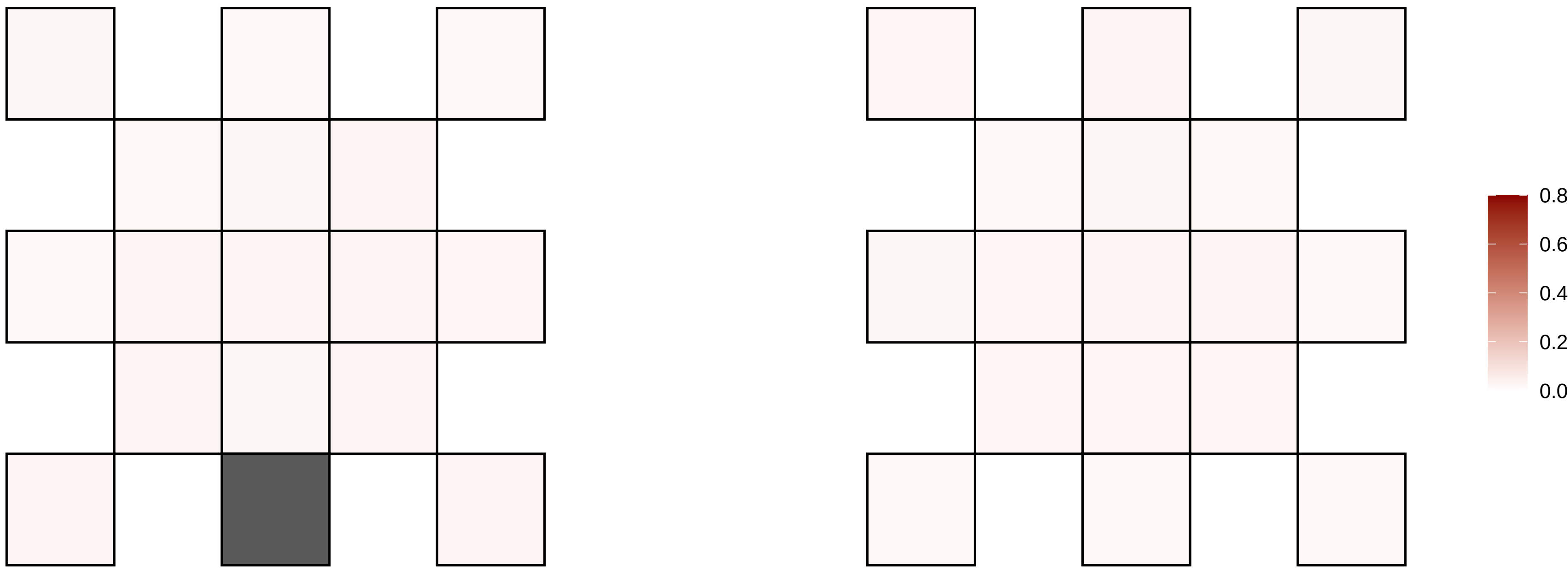

19−57342142−G−A

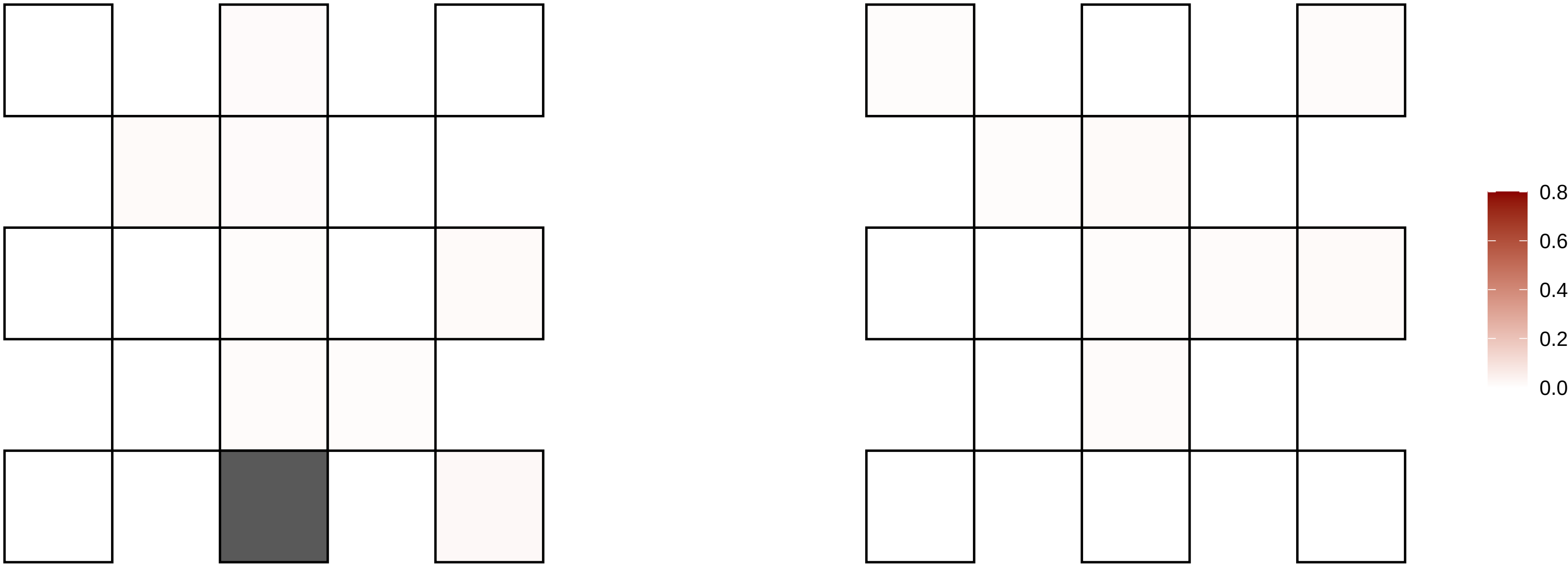

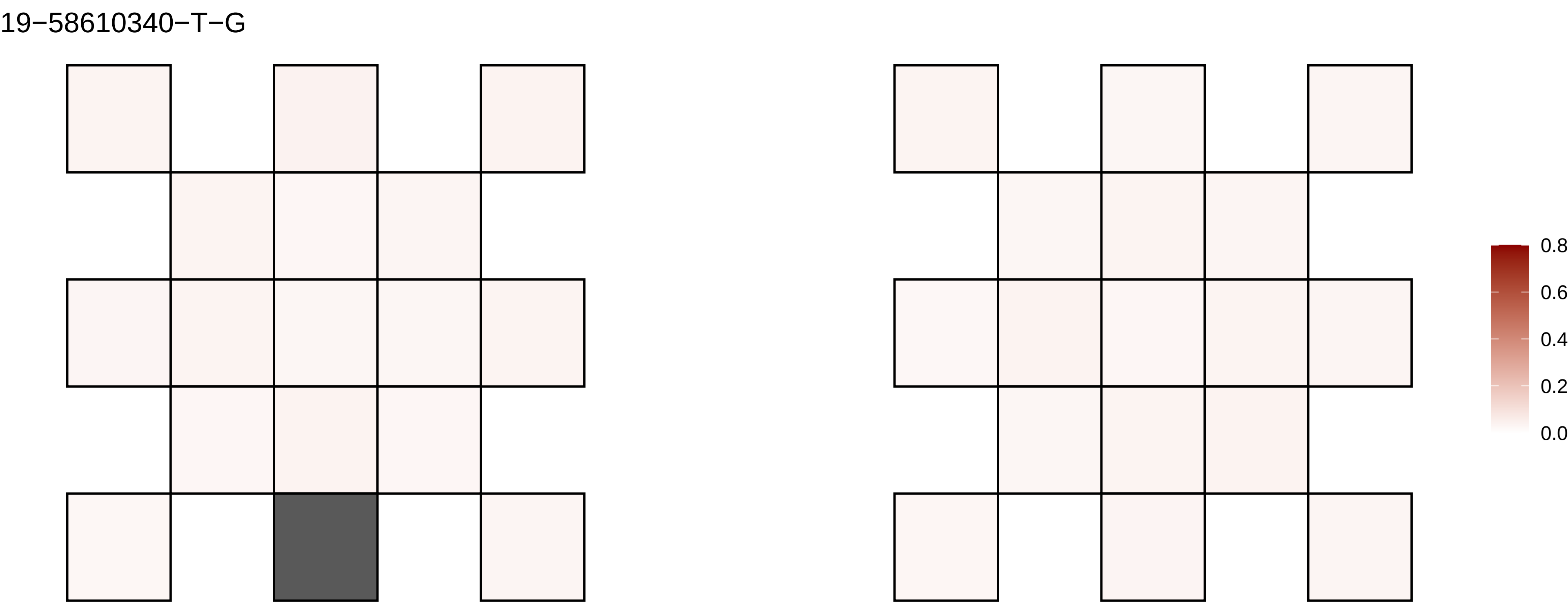
19−58610340−T−G
0.8
0.6
0.4
0.2
0.0

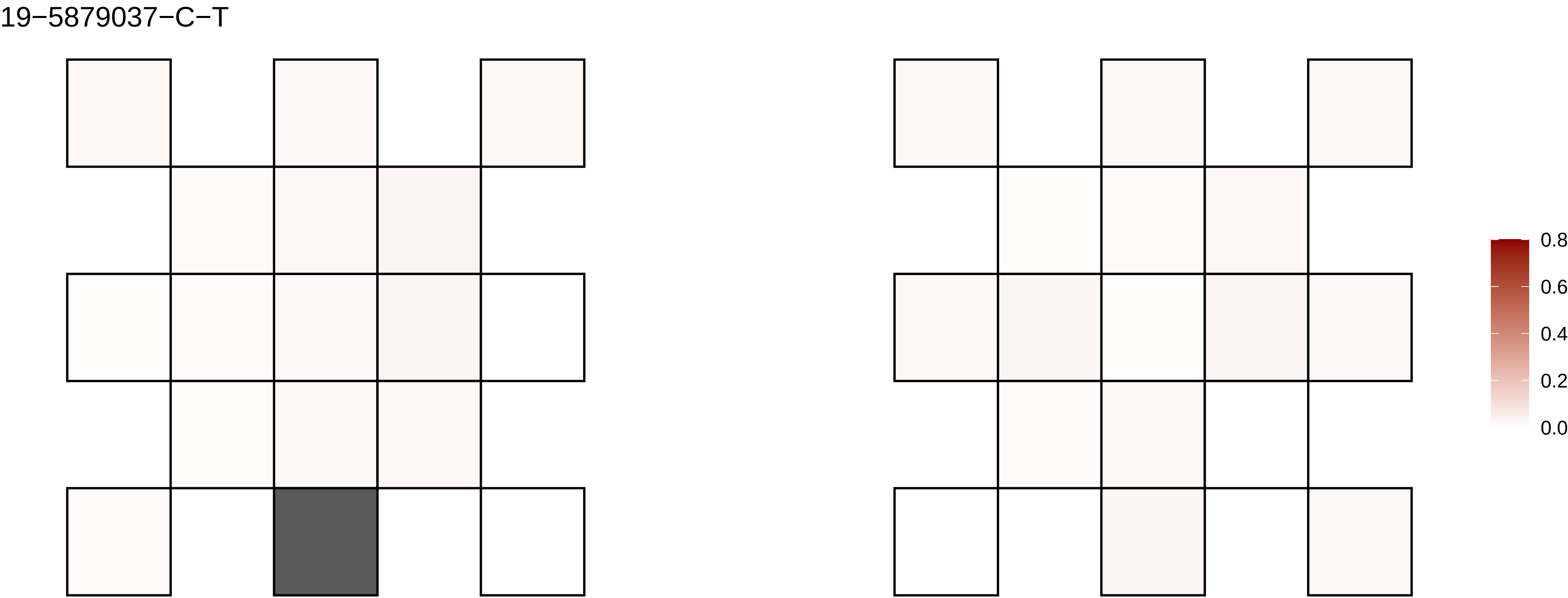

19−5879037−C−T
0.8
0.6
0.4
0.2
0.0

2−104851040−C−T

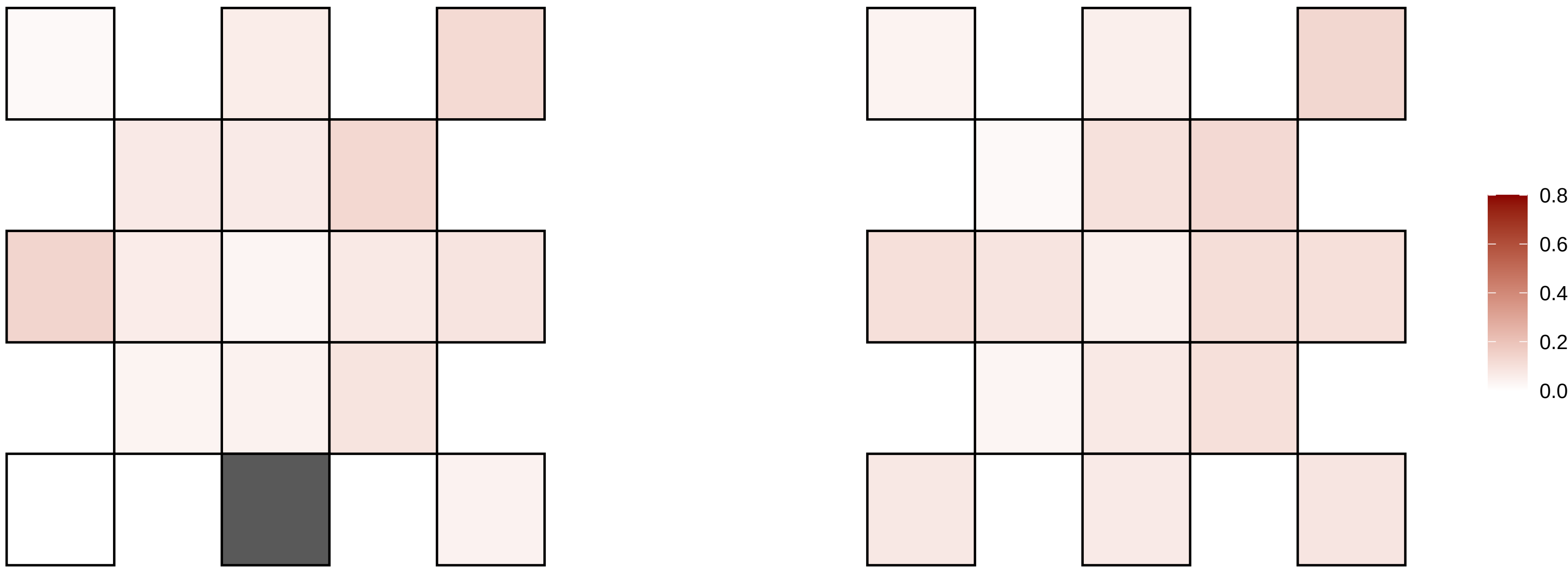

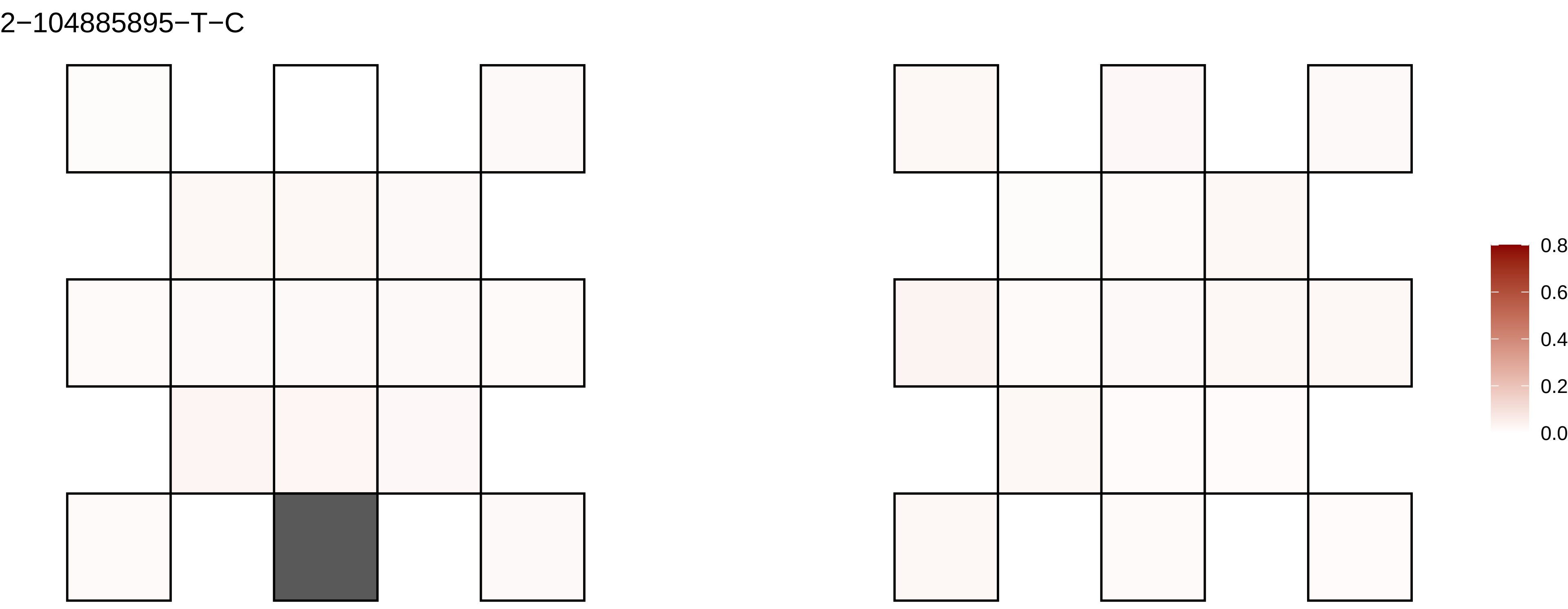
2−104885895−T−C
0.8
0.6
0.4
0.2
0.0

2−105083647−G−A

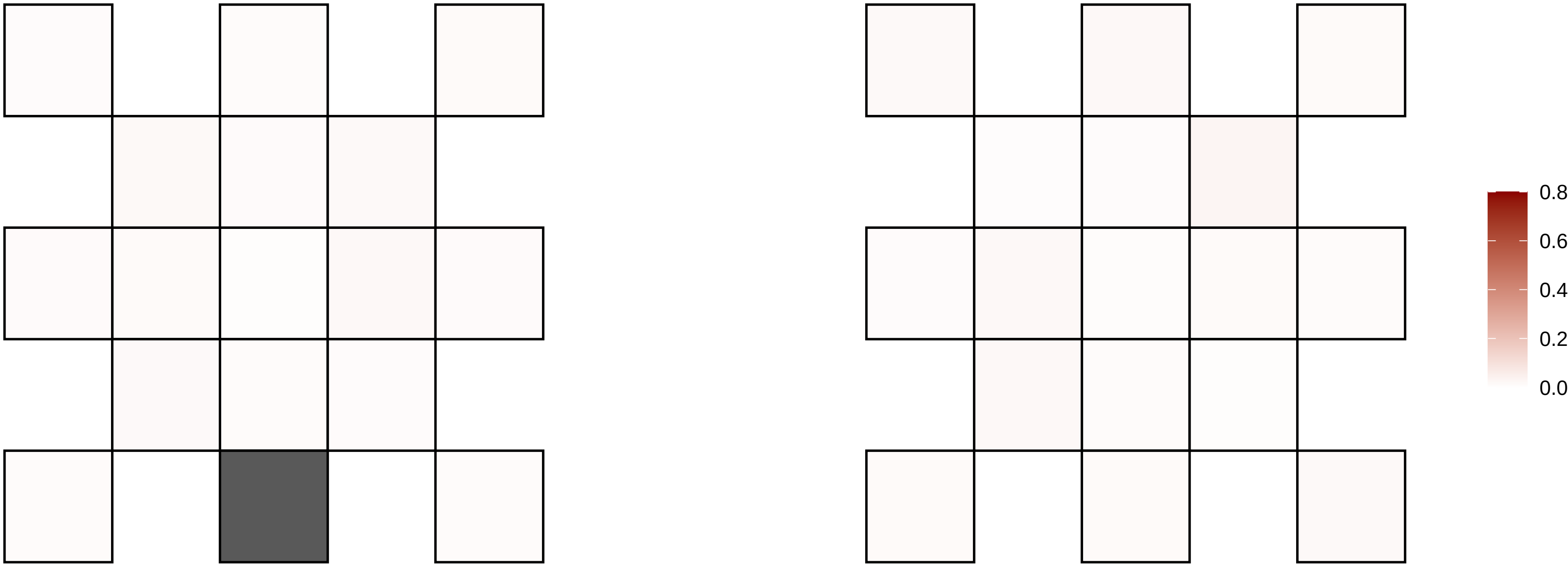

2−110272685−G−A

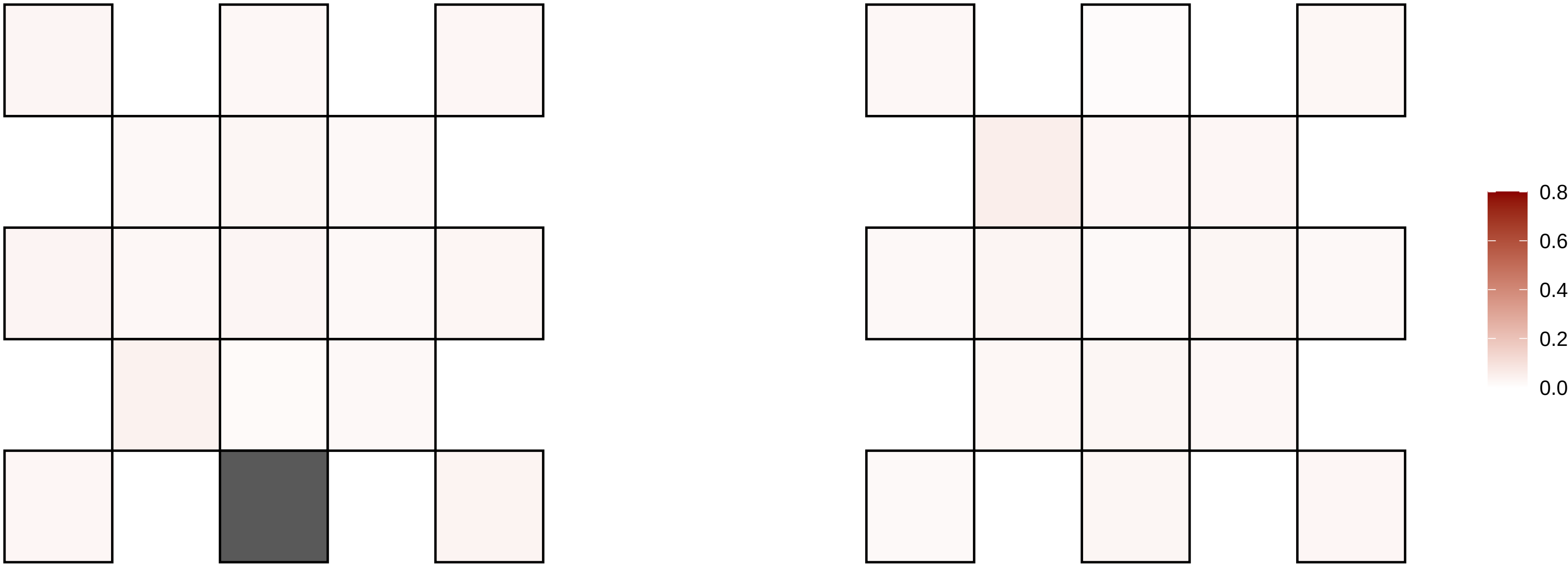

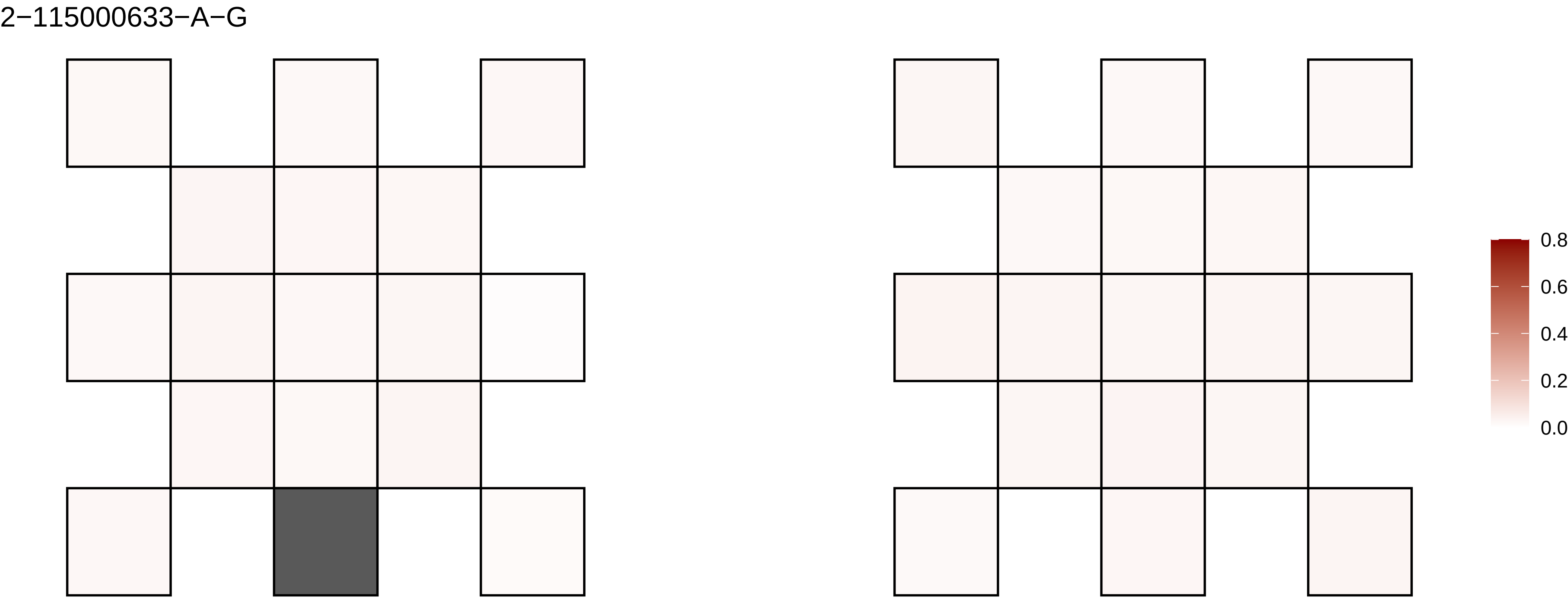
2−115000633−A−G
0.8
0.6
0.4
0.2
0.0

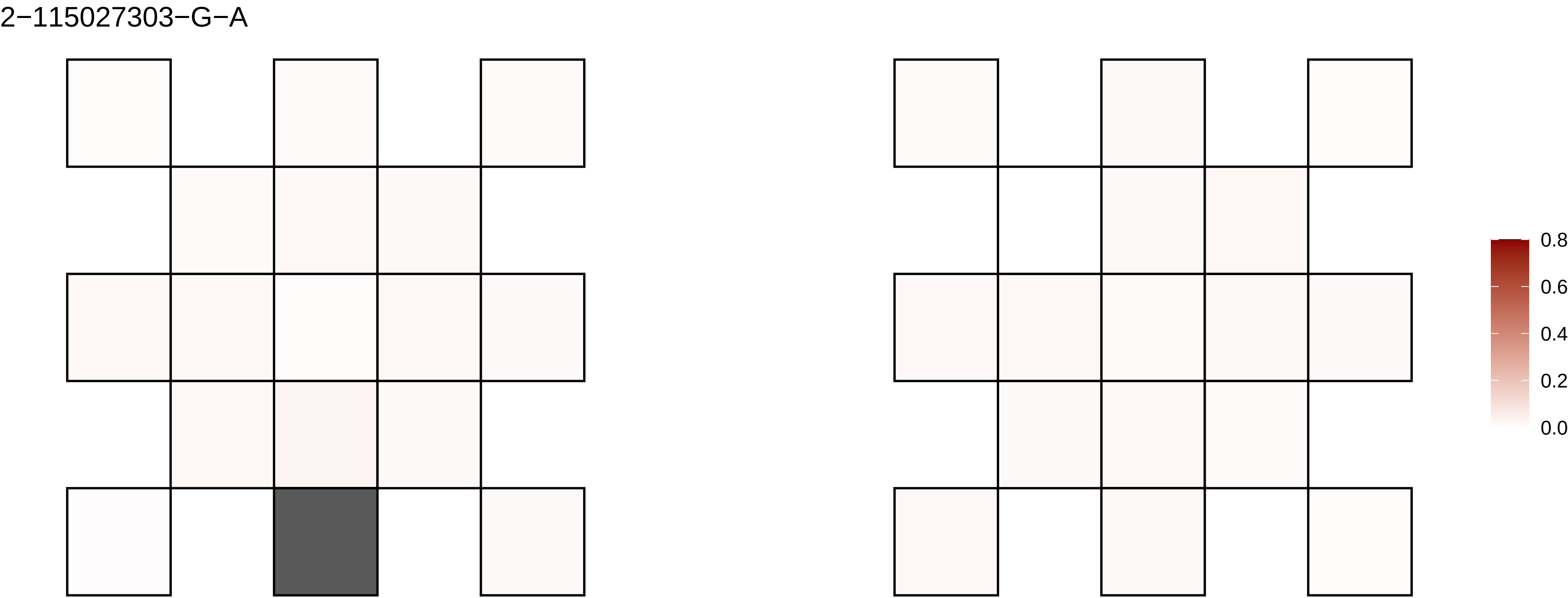

2−115027303−G−A
0.8
0.6
0.4
0.2
0.0

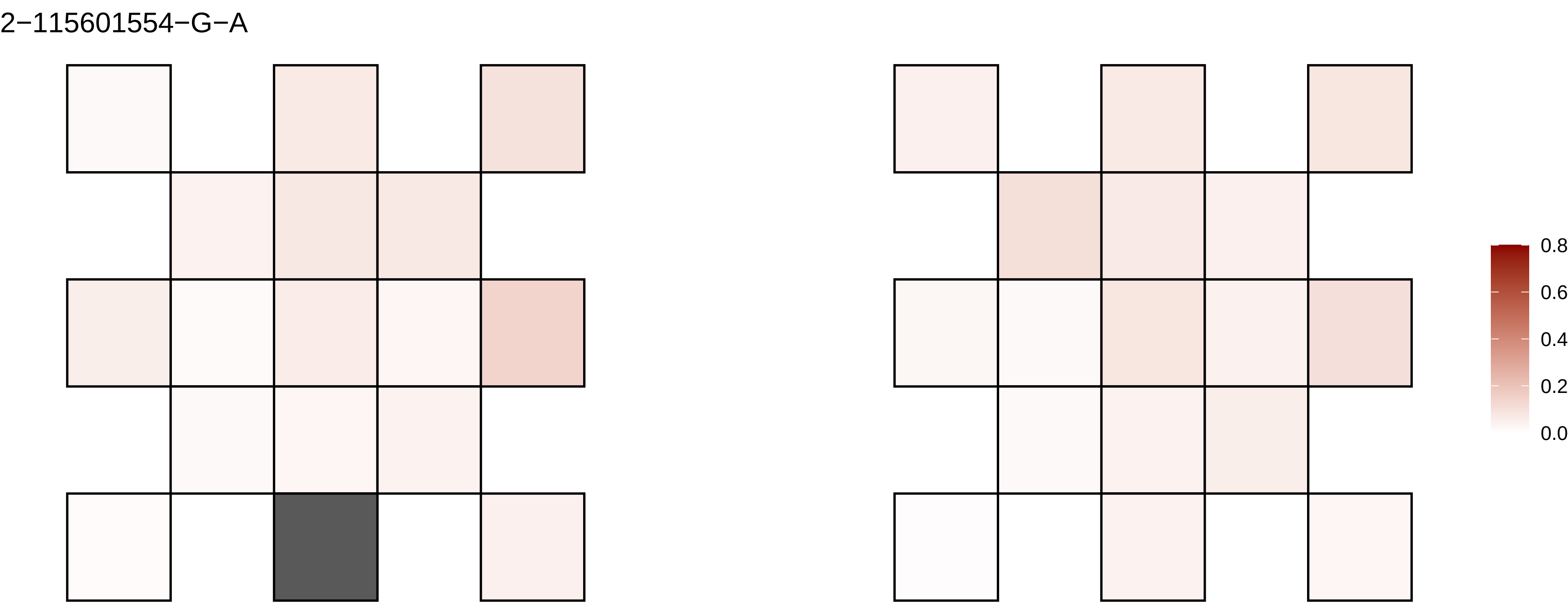
2−115601554−G−A
0.8
0.6
0.4
0.2
0.0

2−115708162−C−T

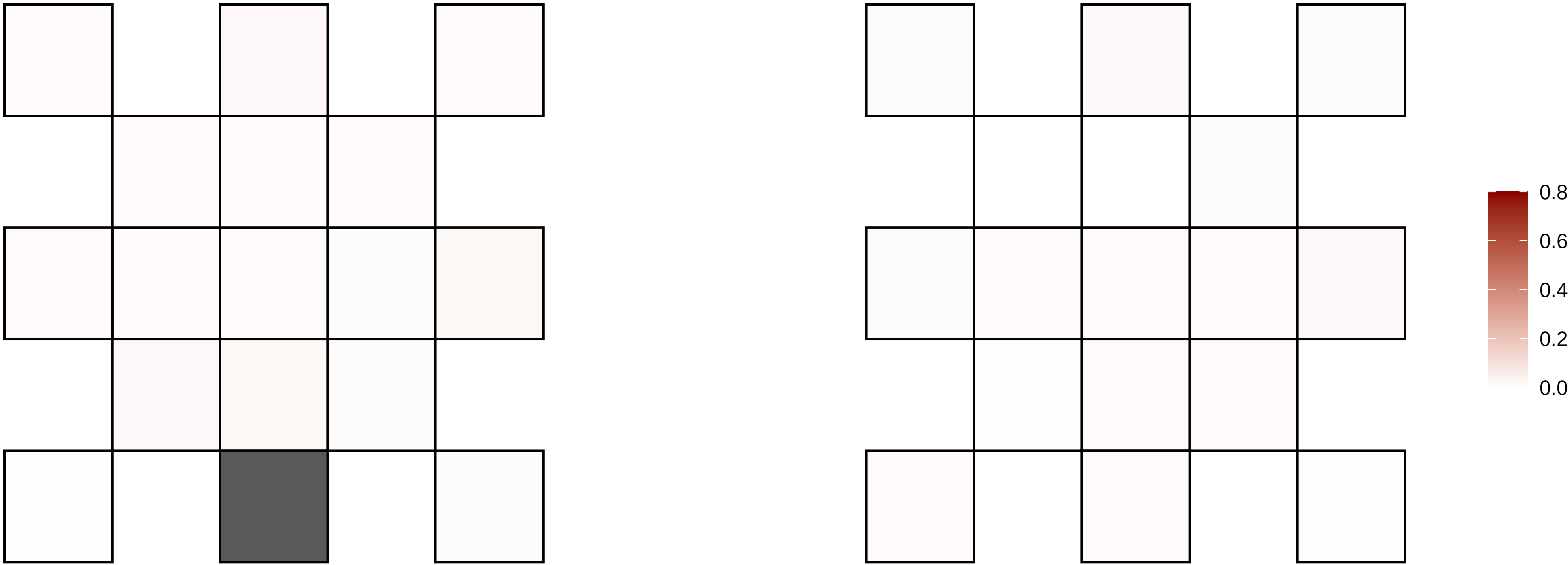

2−116395374−G−T

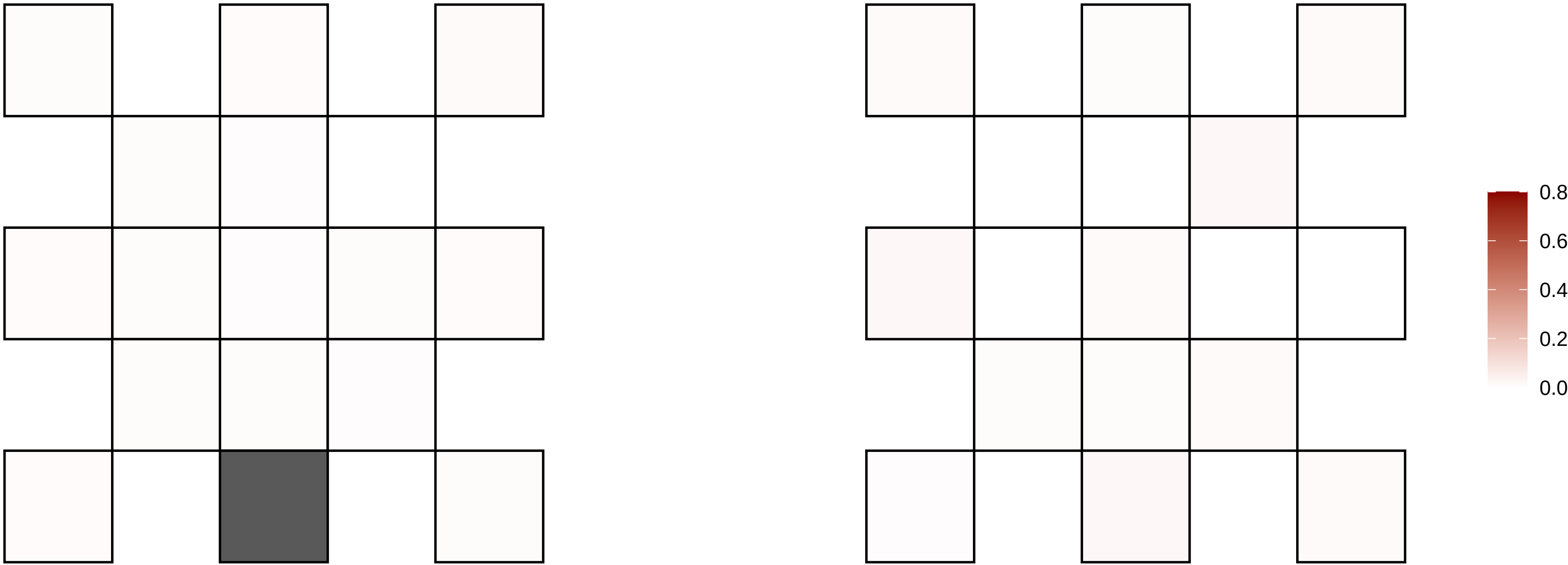

2−118425584−C−T

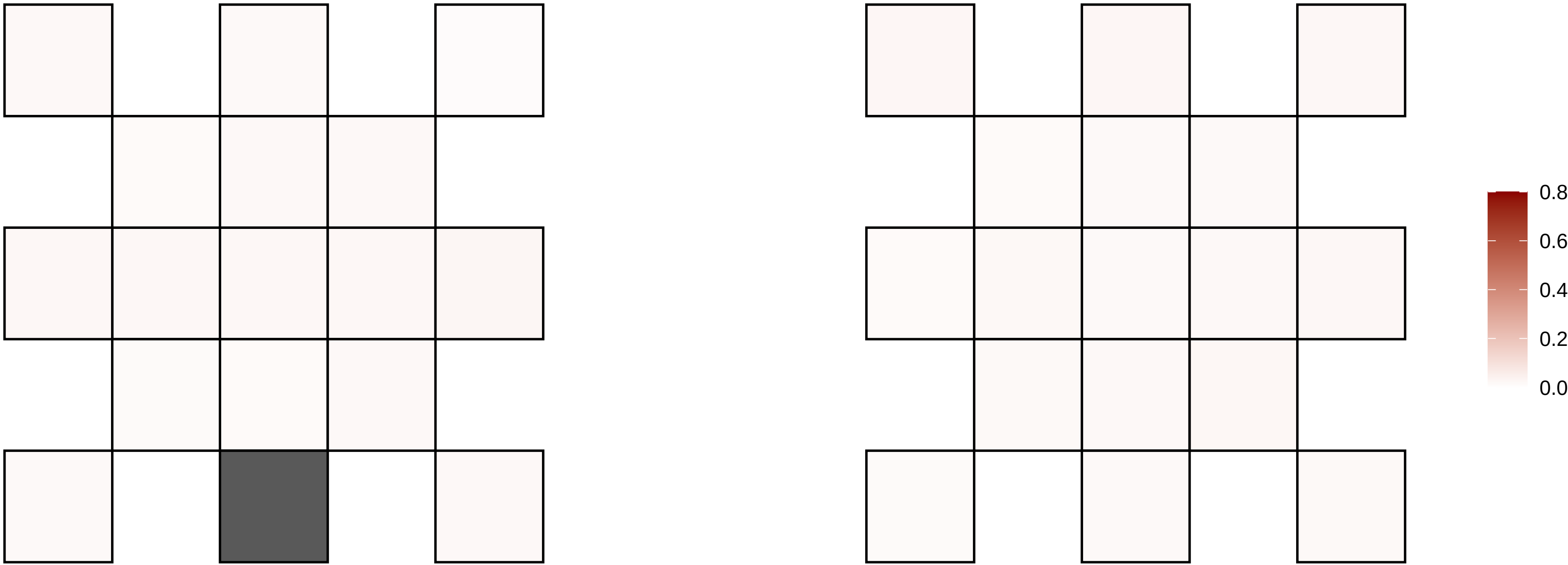

2−129713862−G−A

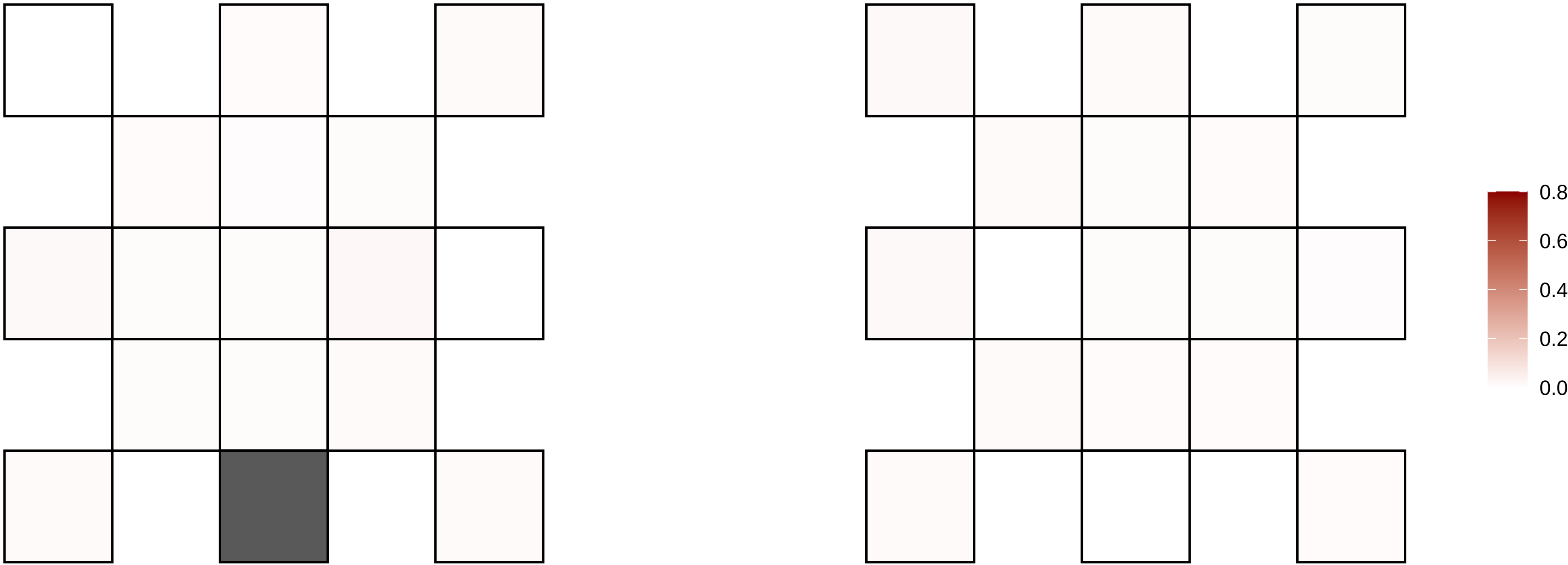

2−130483876−C−T

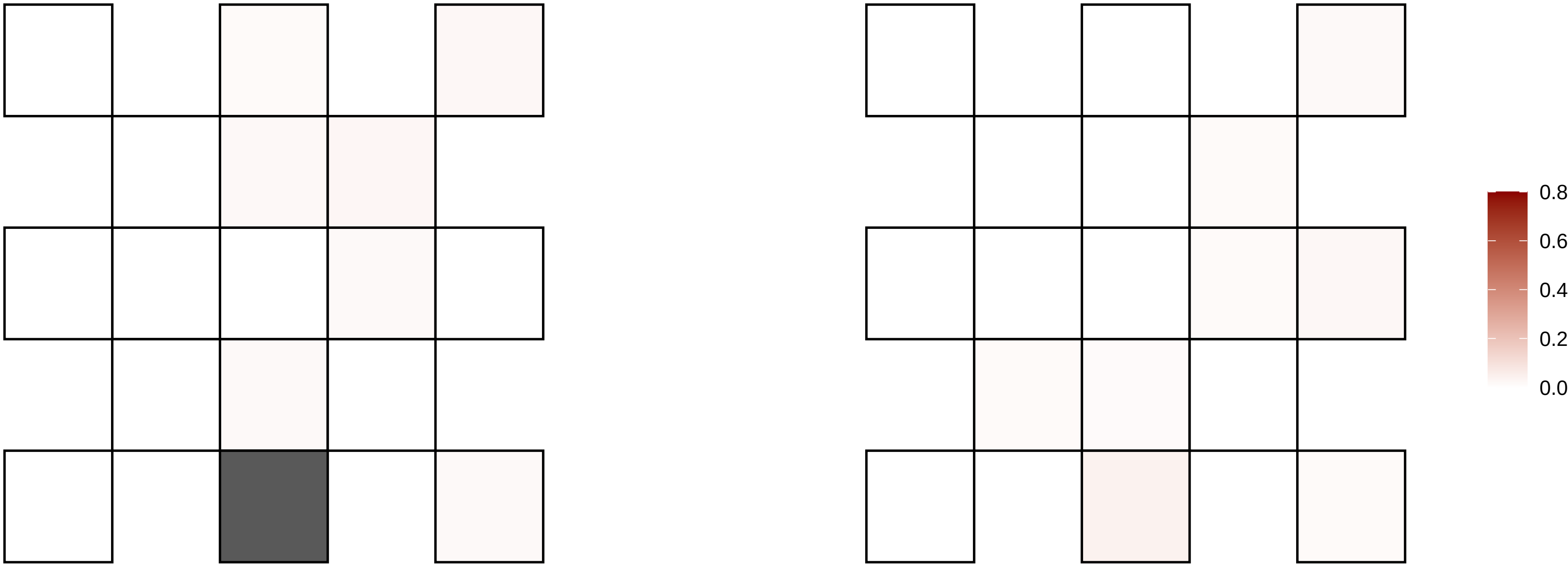

2−137813385−AT−A

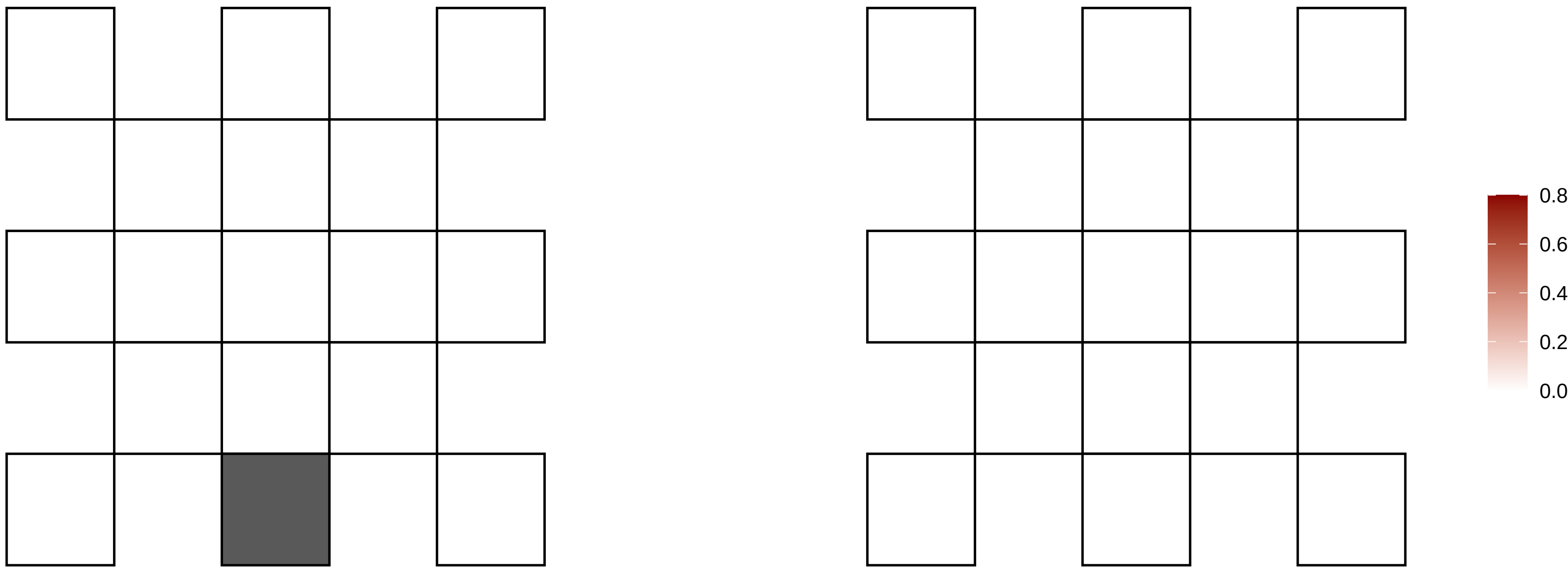

2−139934078−G−A

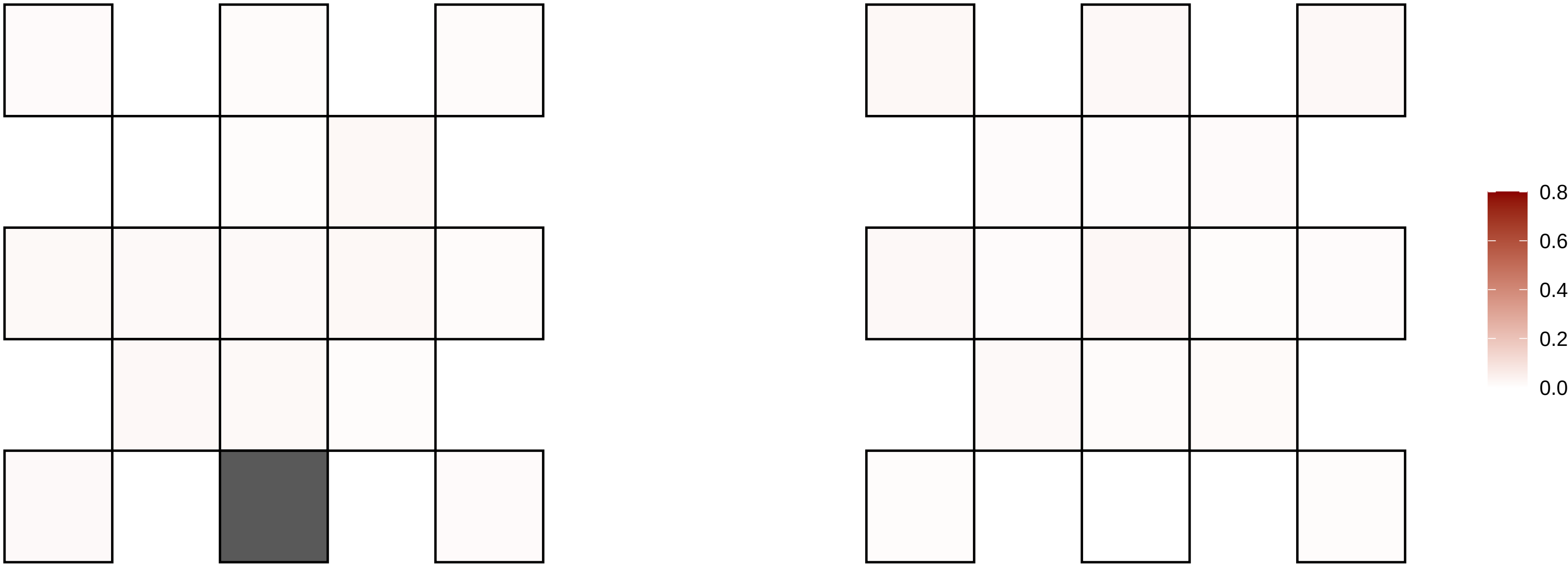

2−141914766−A−T

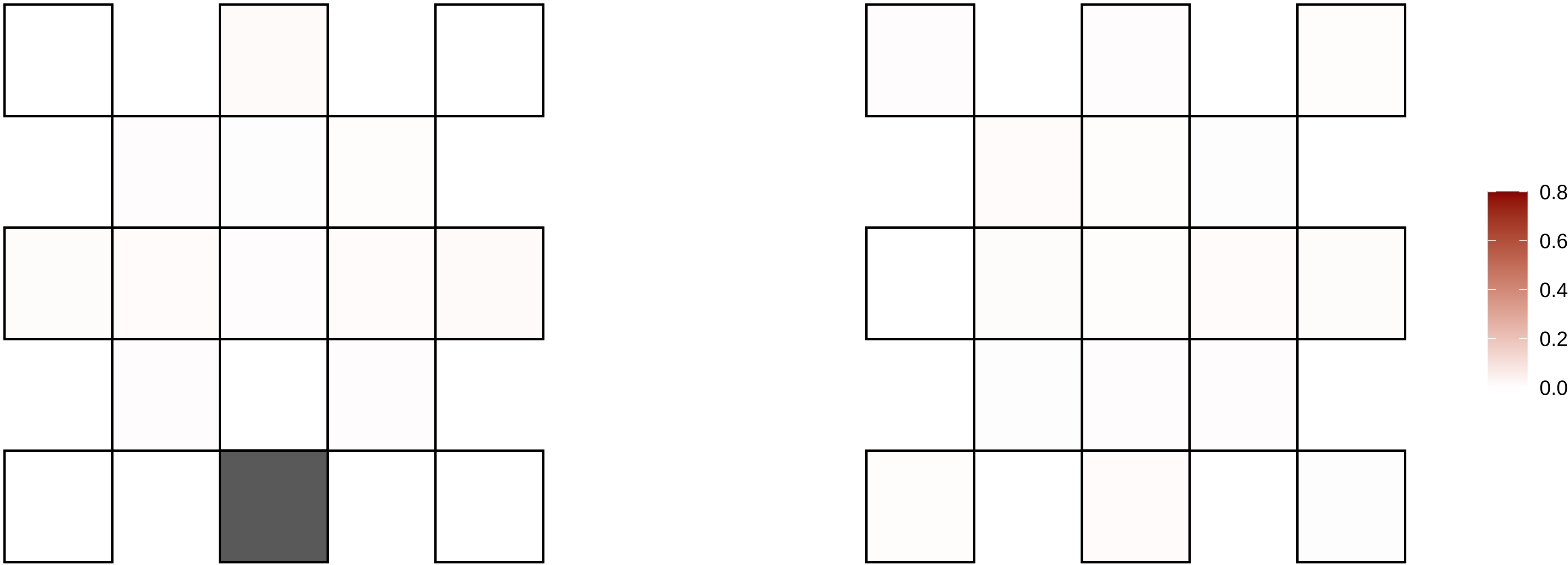

2−143167083−A−G

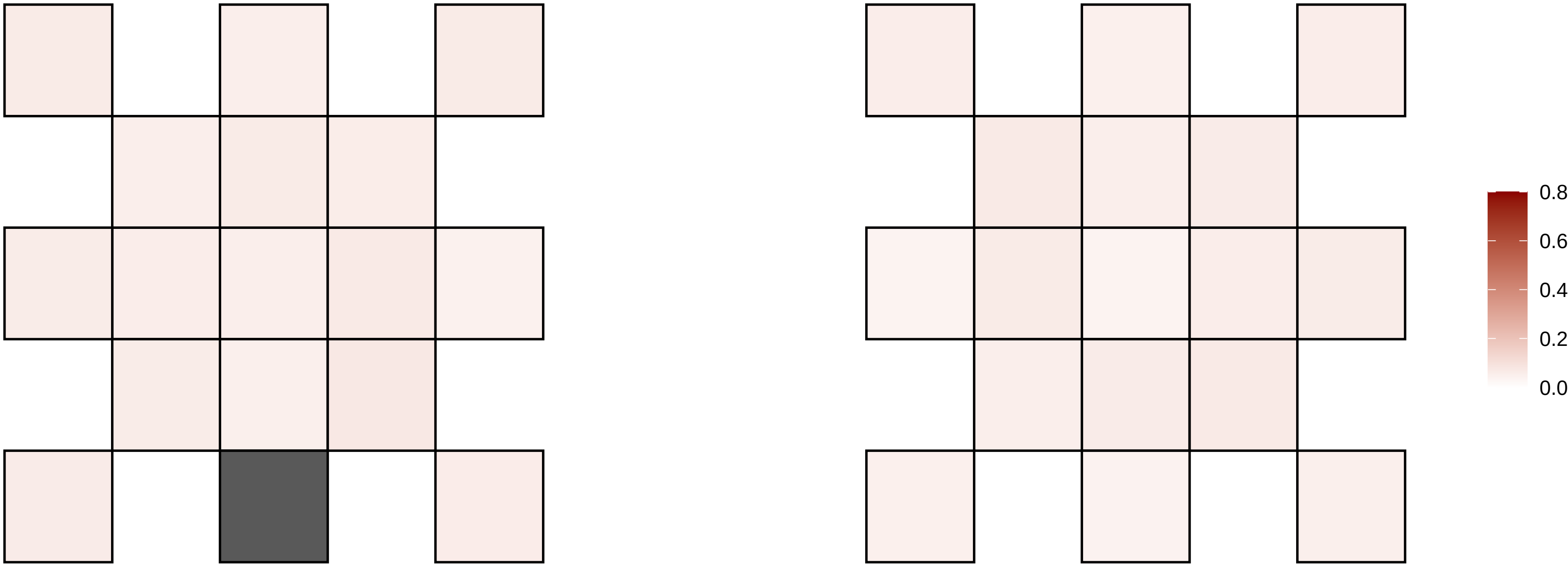

2−14466159−G−A

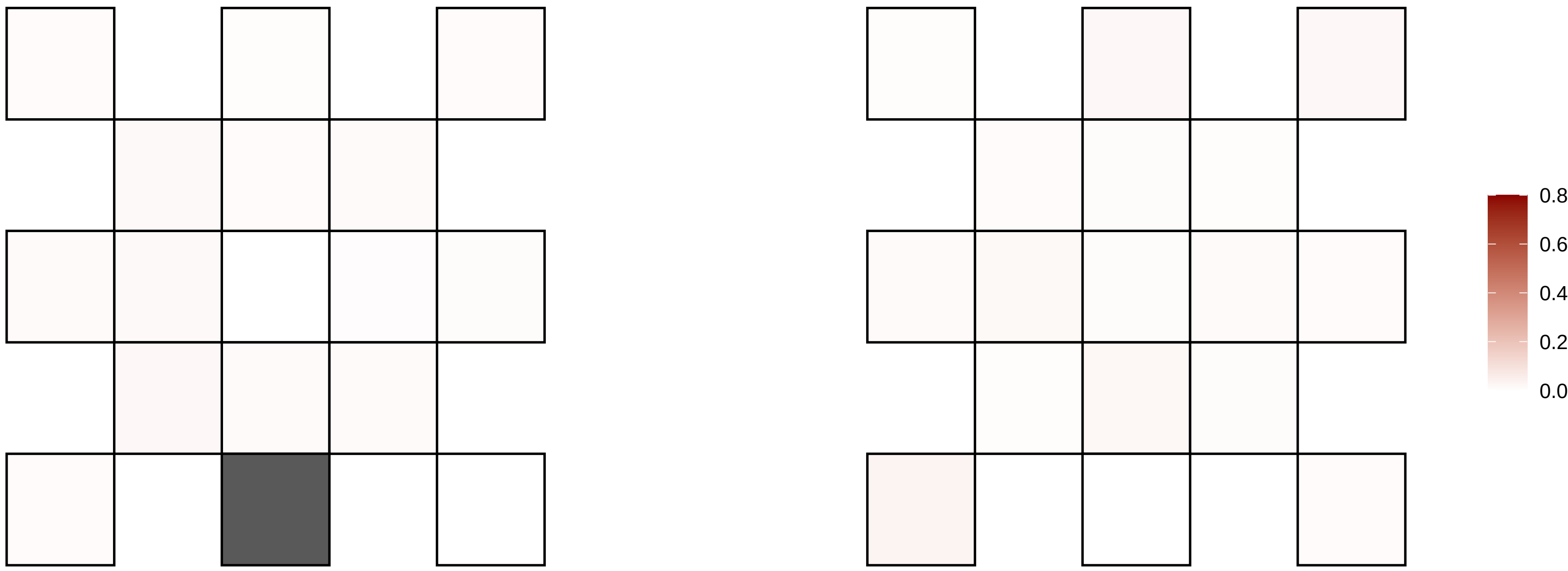

2−159760148−C−T

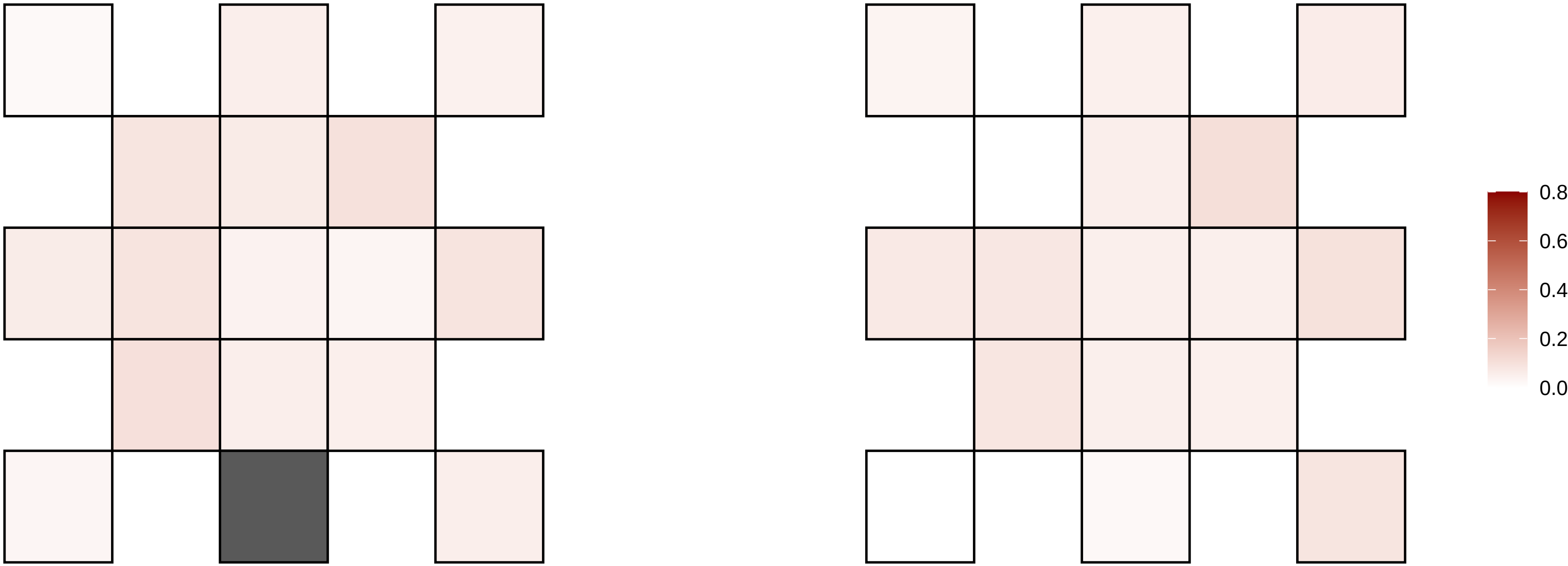

2−163073318−G−A

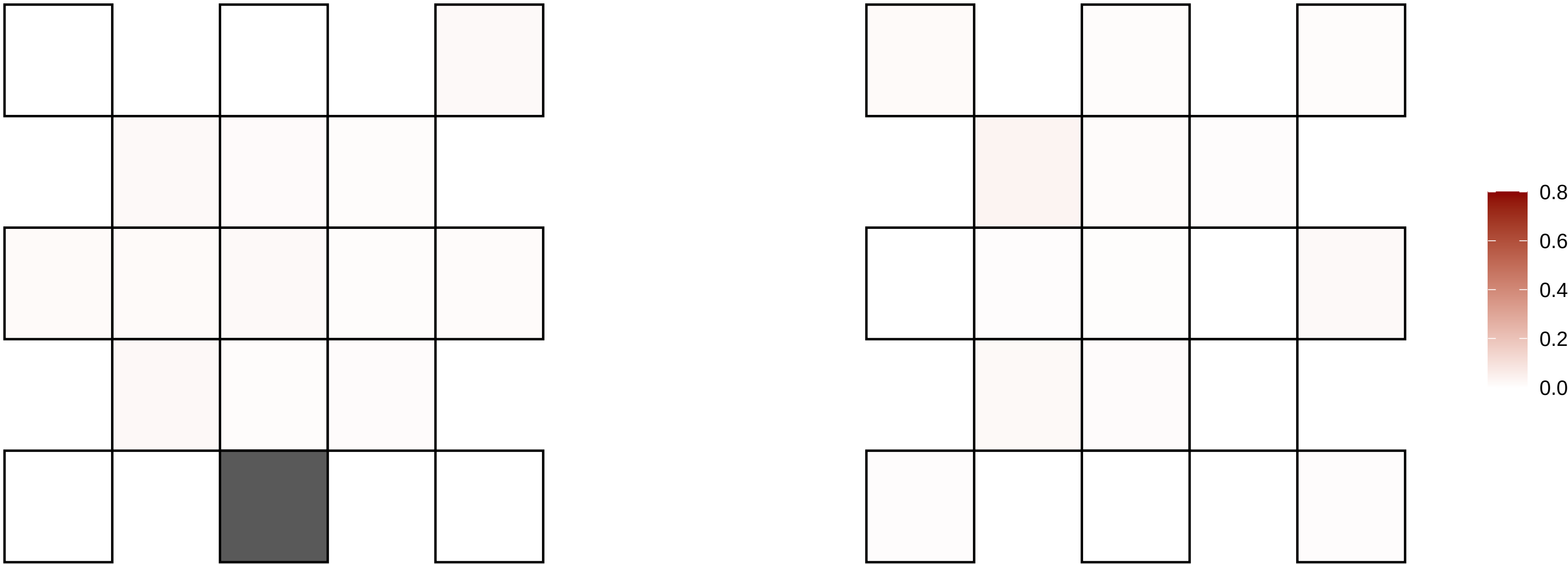

2−164210723−G−A

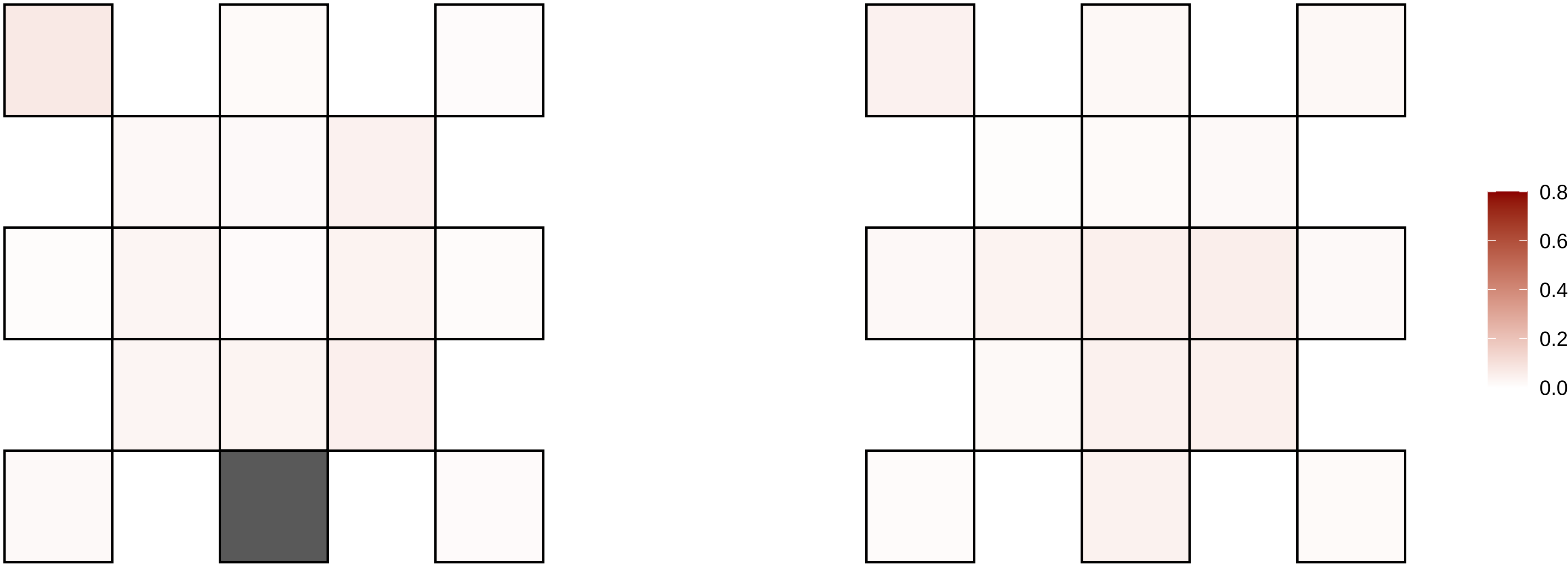

2−167087829−G−A

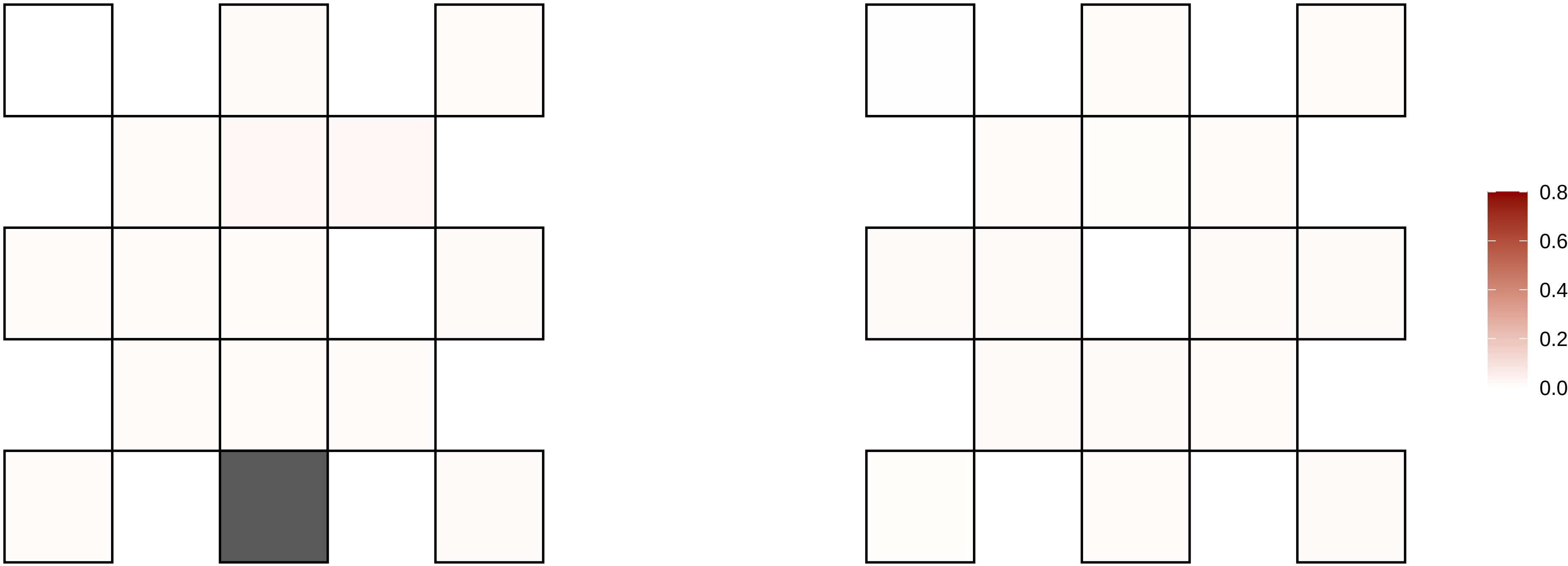

2−167342653−C−T

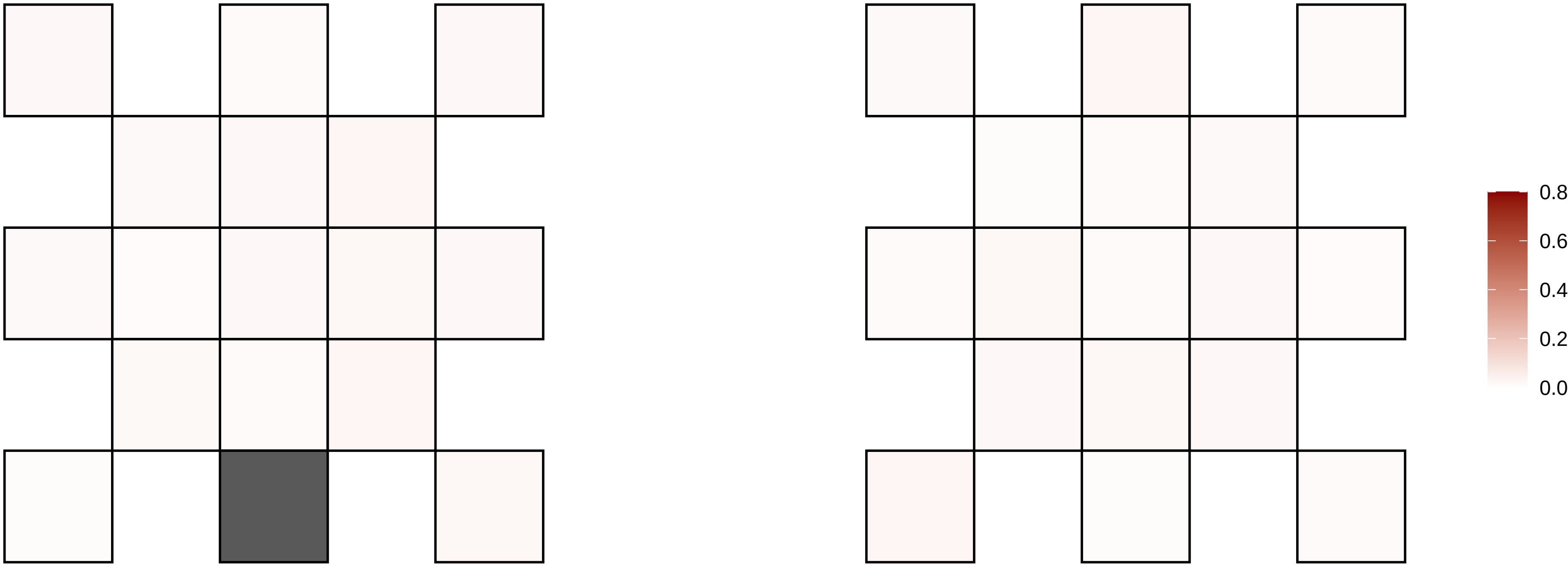

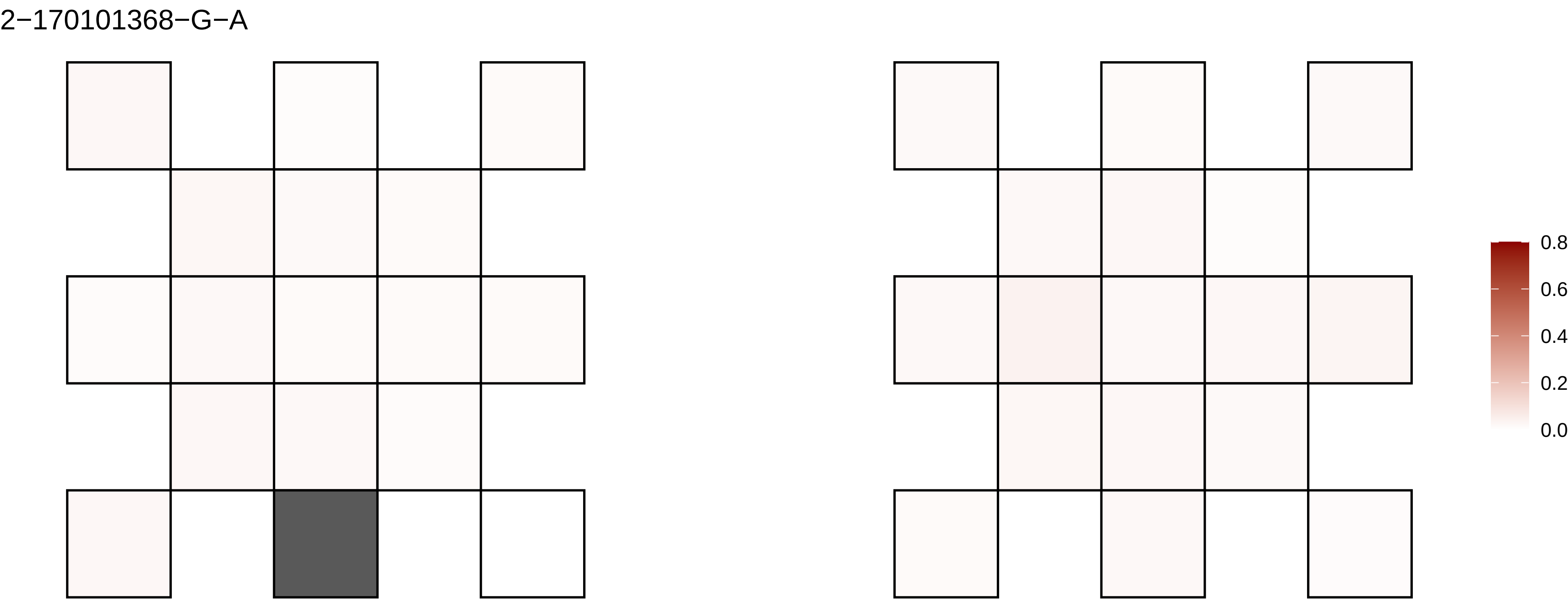

2−170101368−G−A
0.8
0.6
0.4
0.2
0.0

2−172334983−T−G

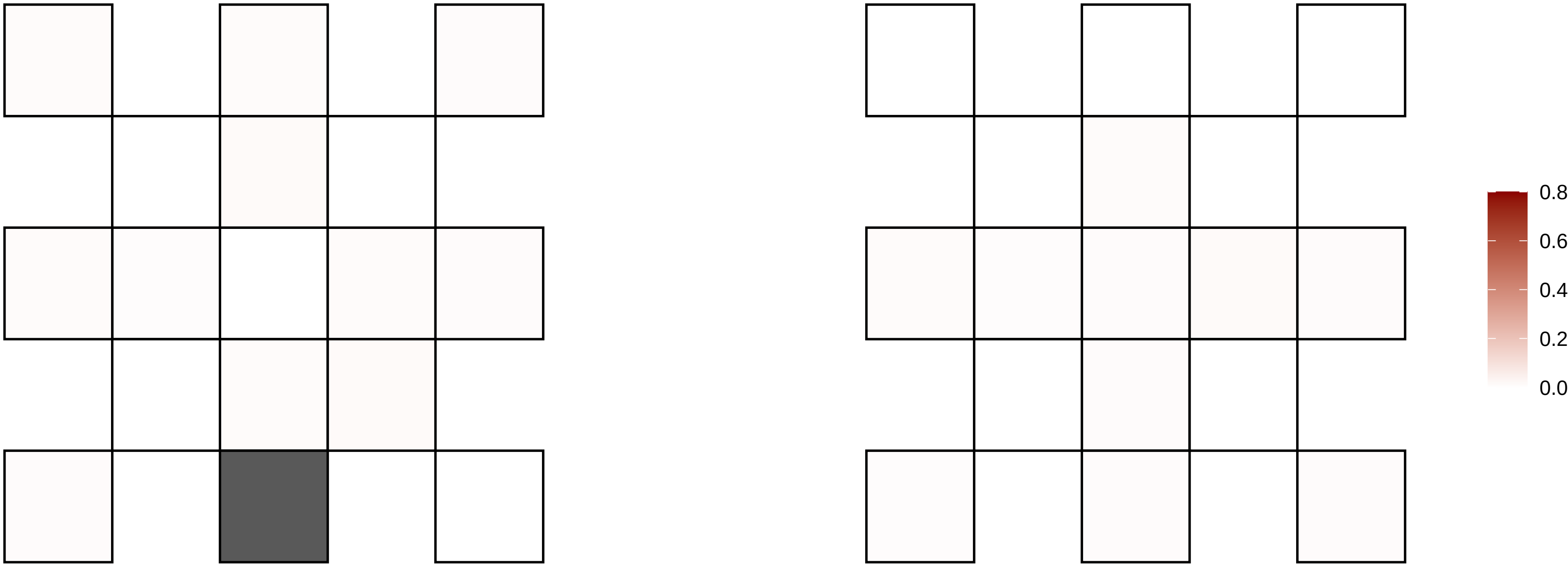

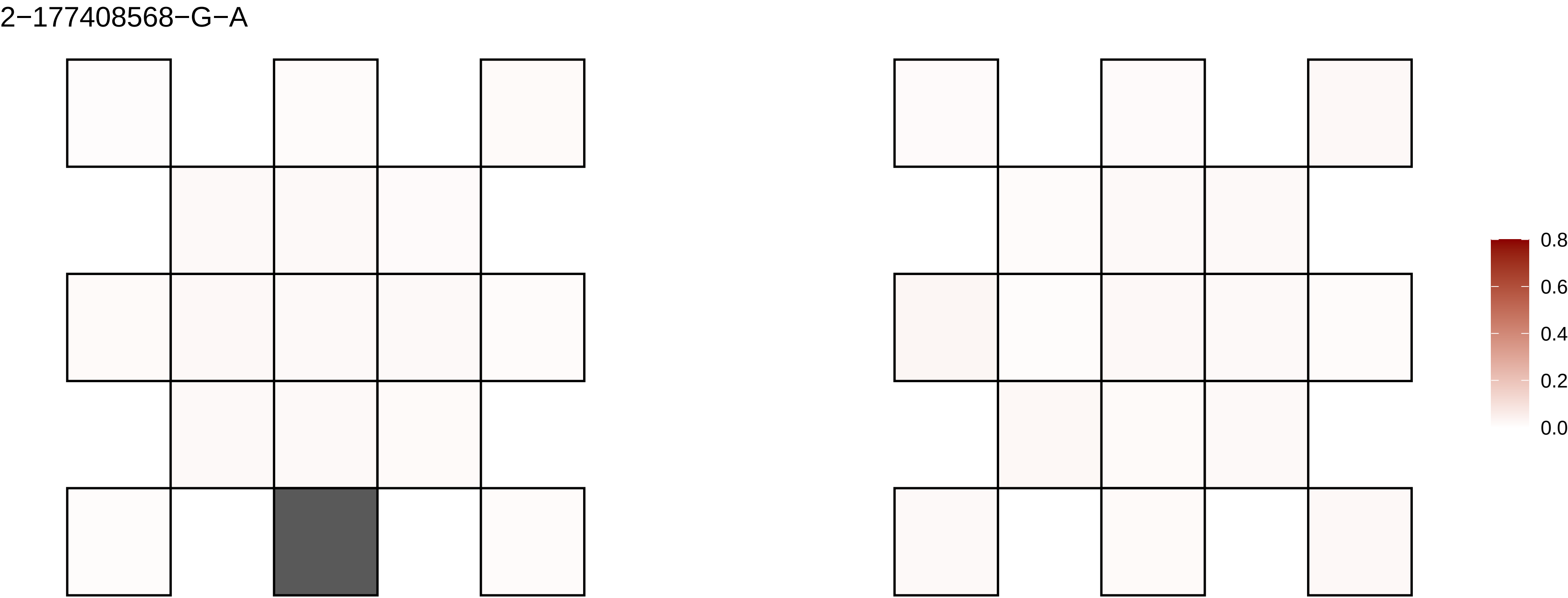
2−177408568−G−A
0.8
0.6
0.4
0.2
0.0

2−178026730−A−T

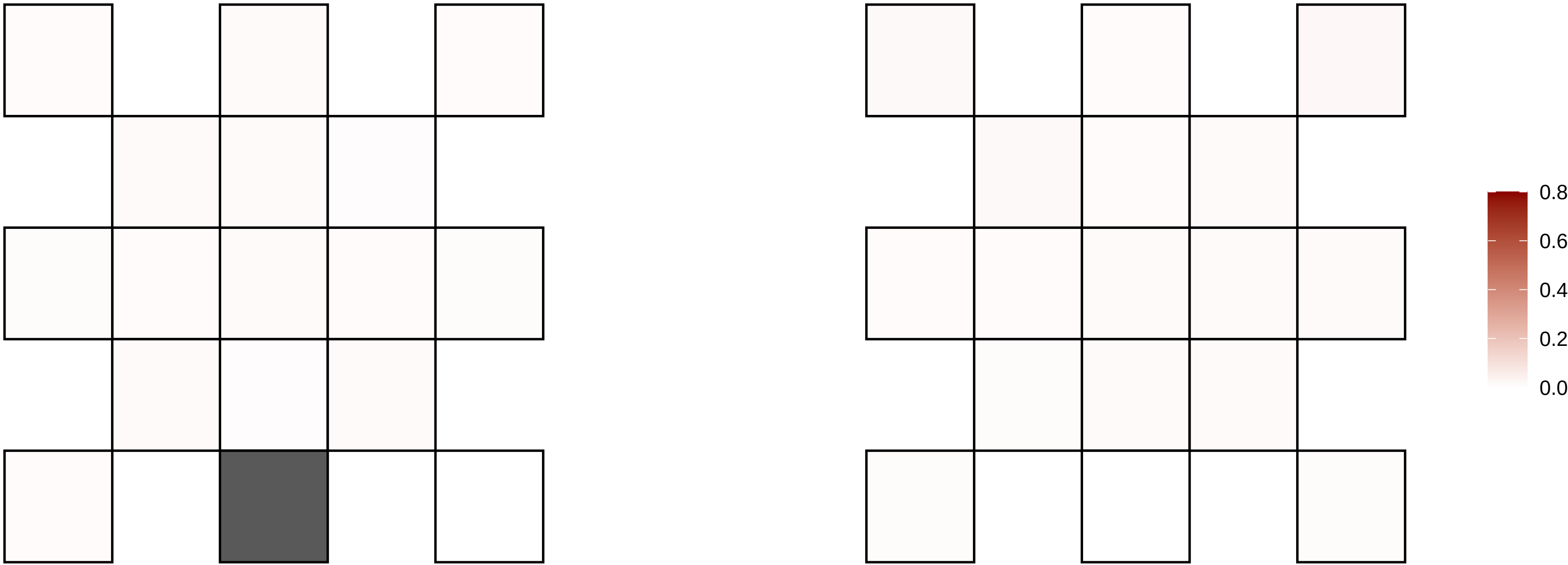

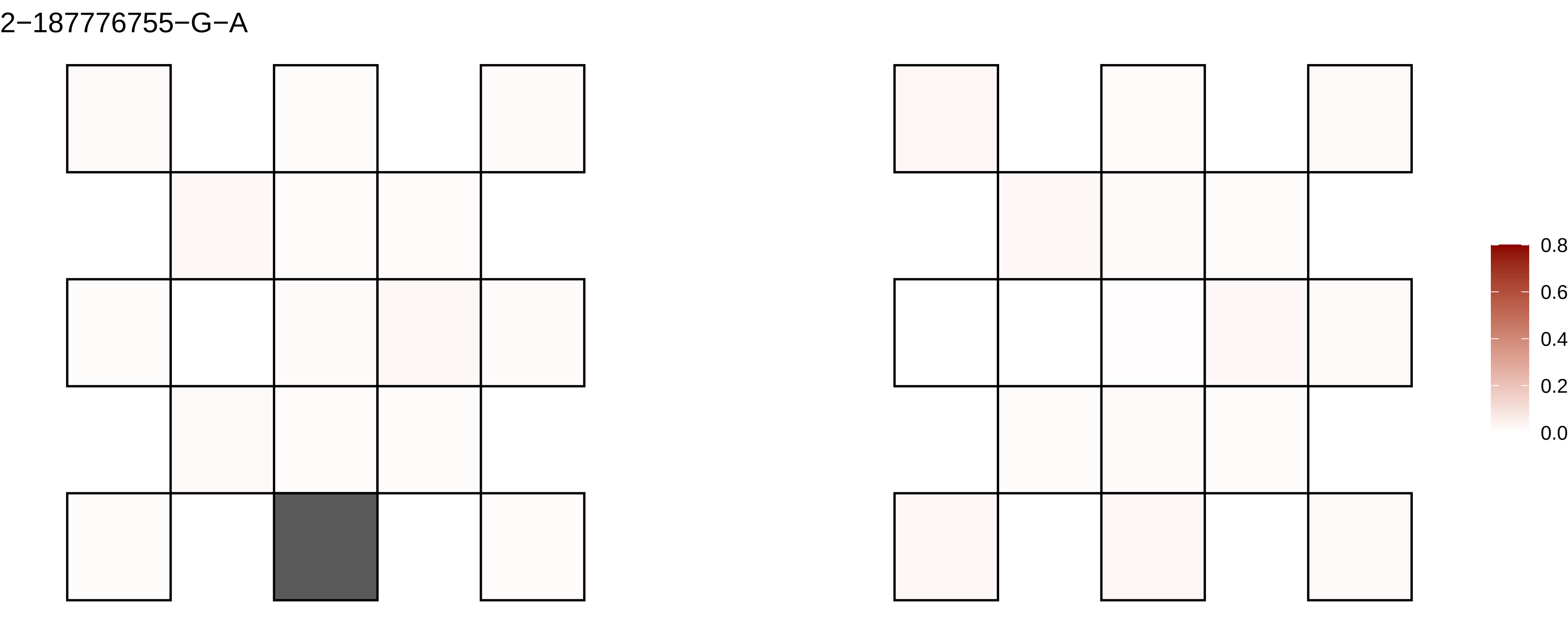
2−187776755−G−A
0.8
0.6
0.4
0.2
0.0

2−190501942−A−G

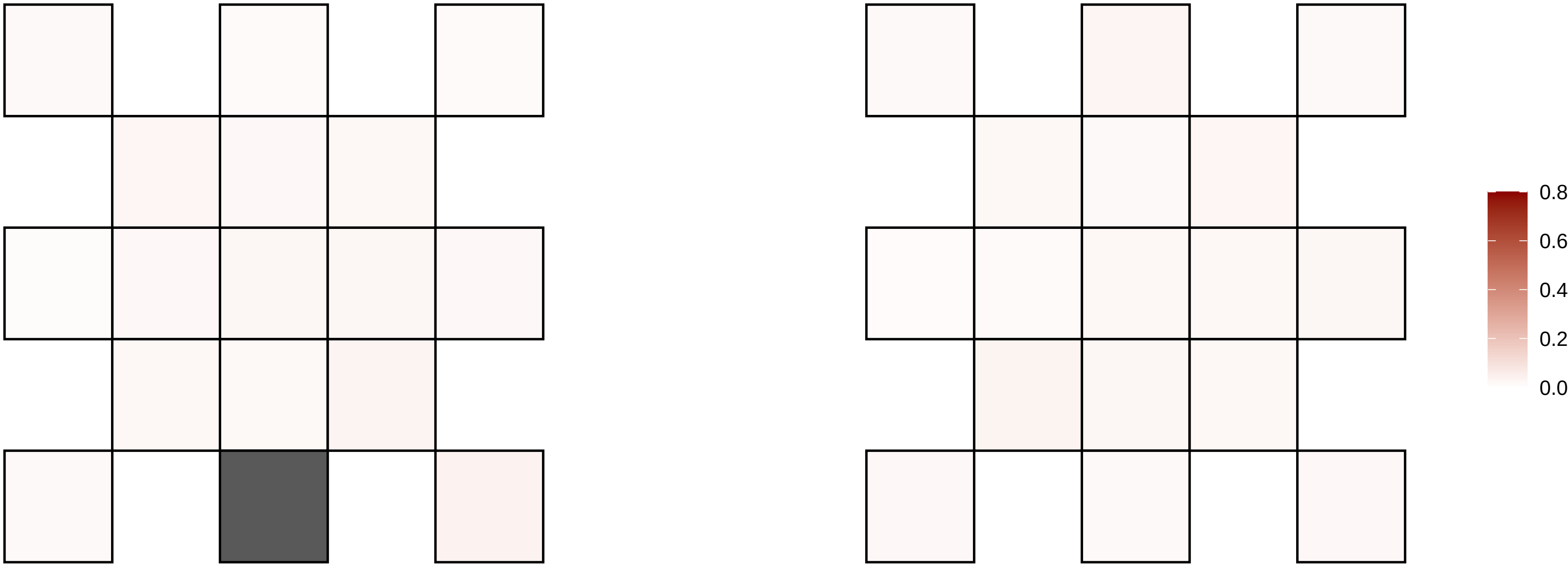

2−193626603−C−A

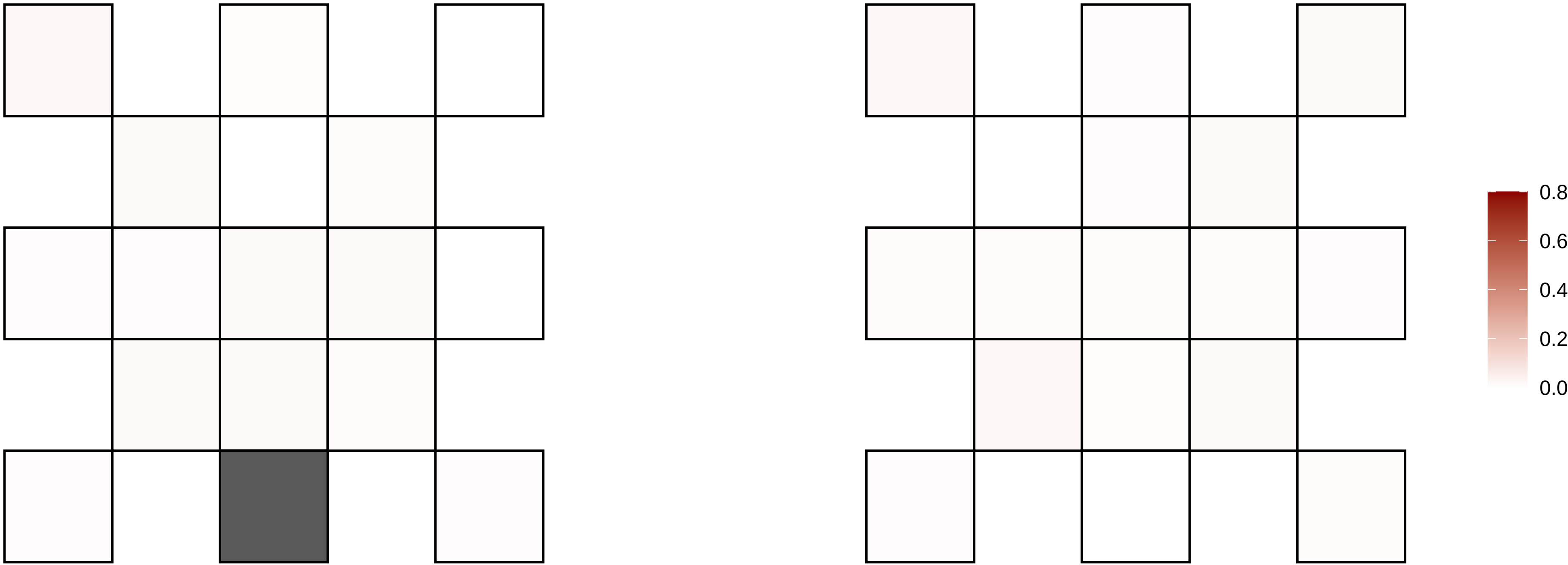

2−193831190−T−A

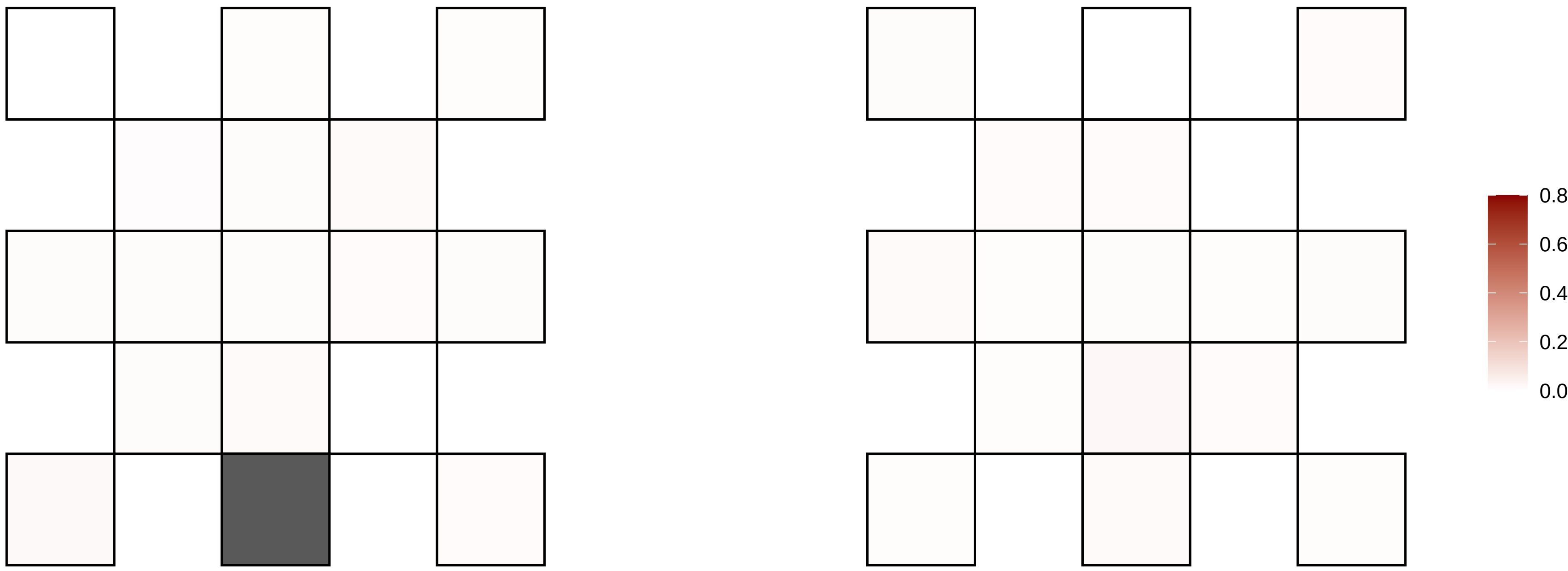

2−198773820−A−G

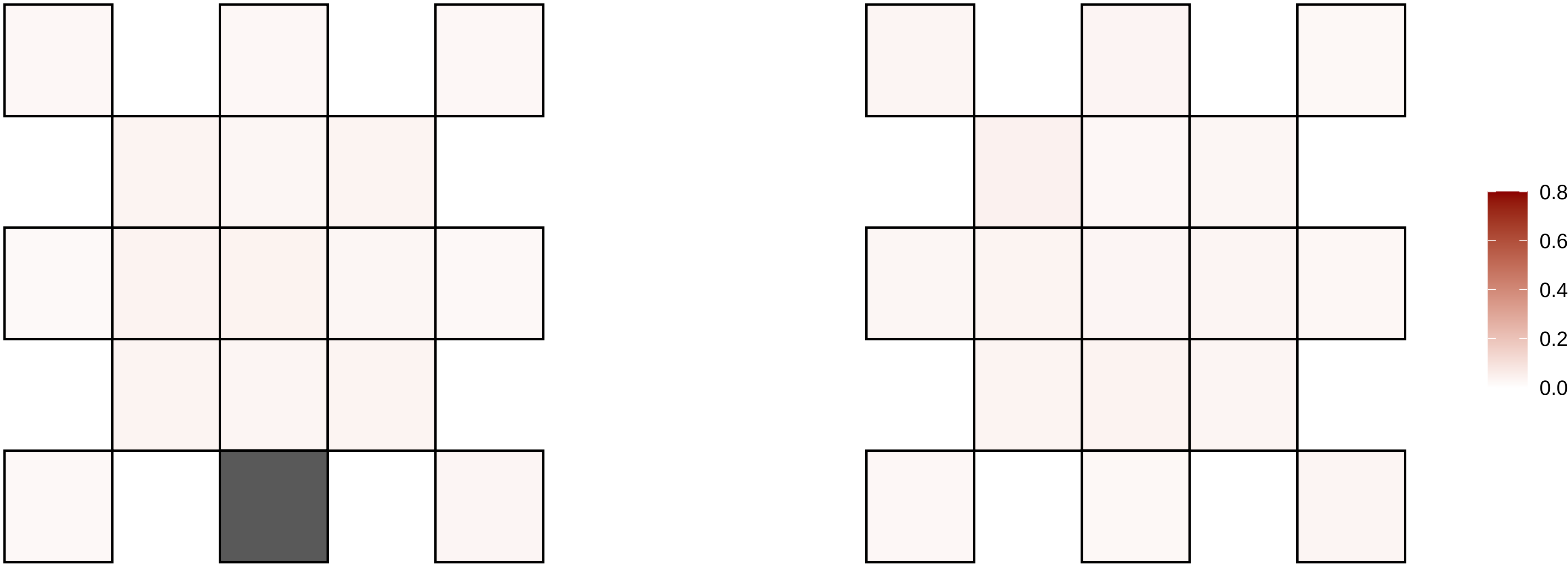

2−203467456−T−C

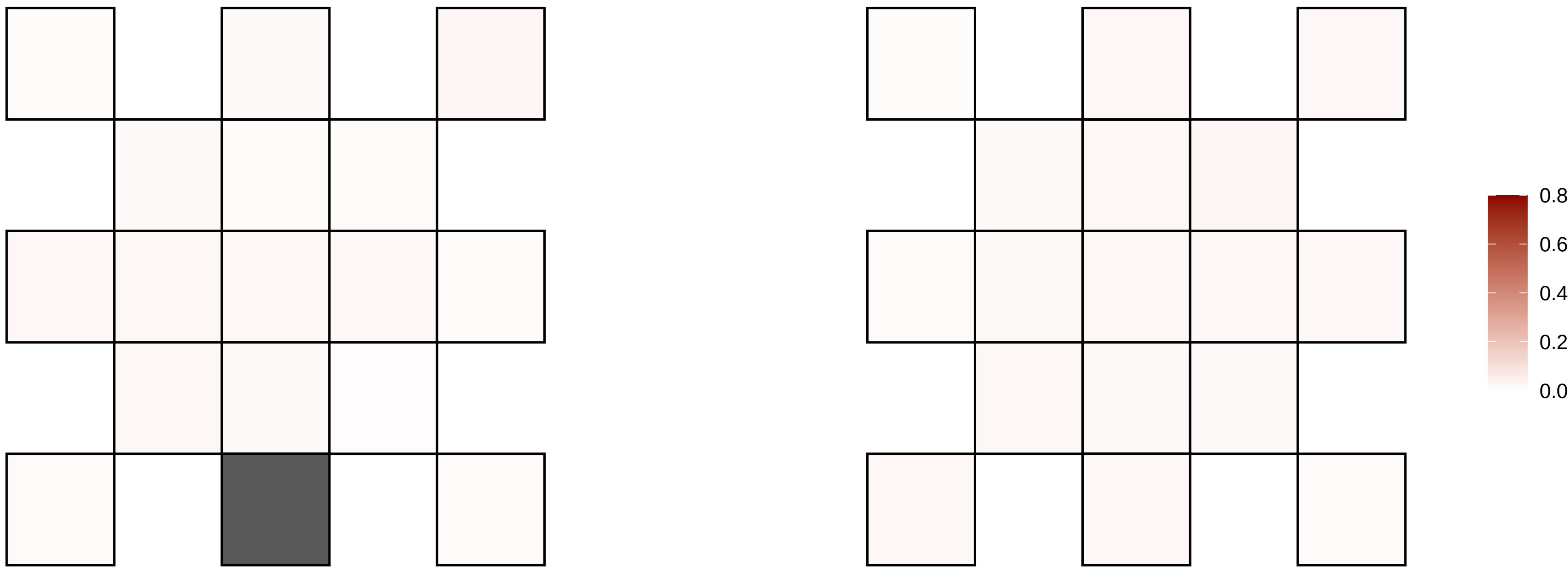

2−204722237−C−T

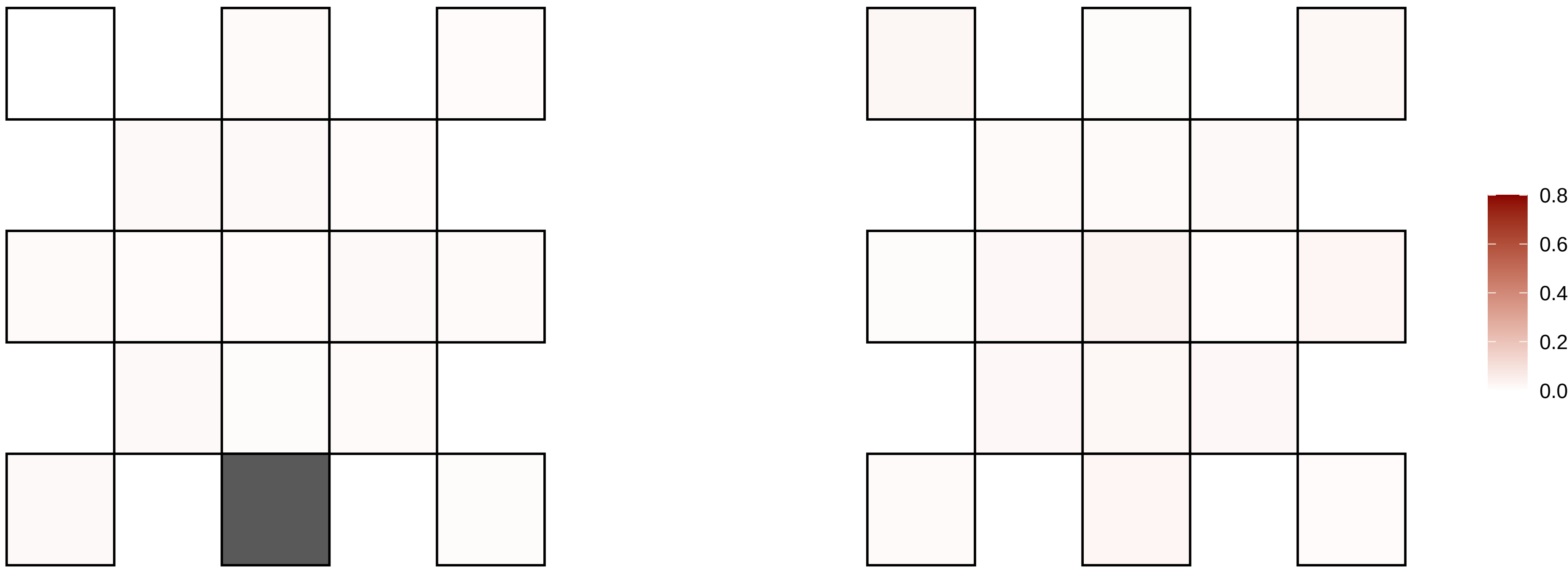

2−209569113−C−T

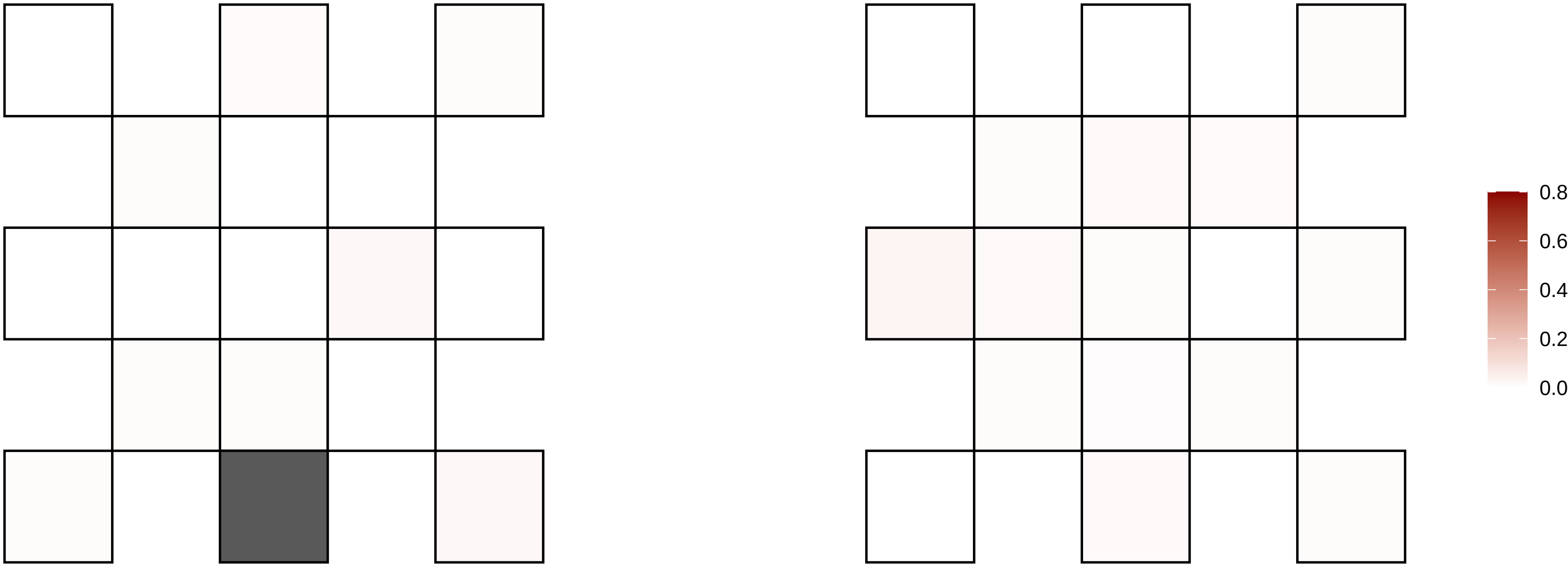

2−213063972−C−T

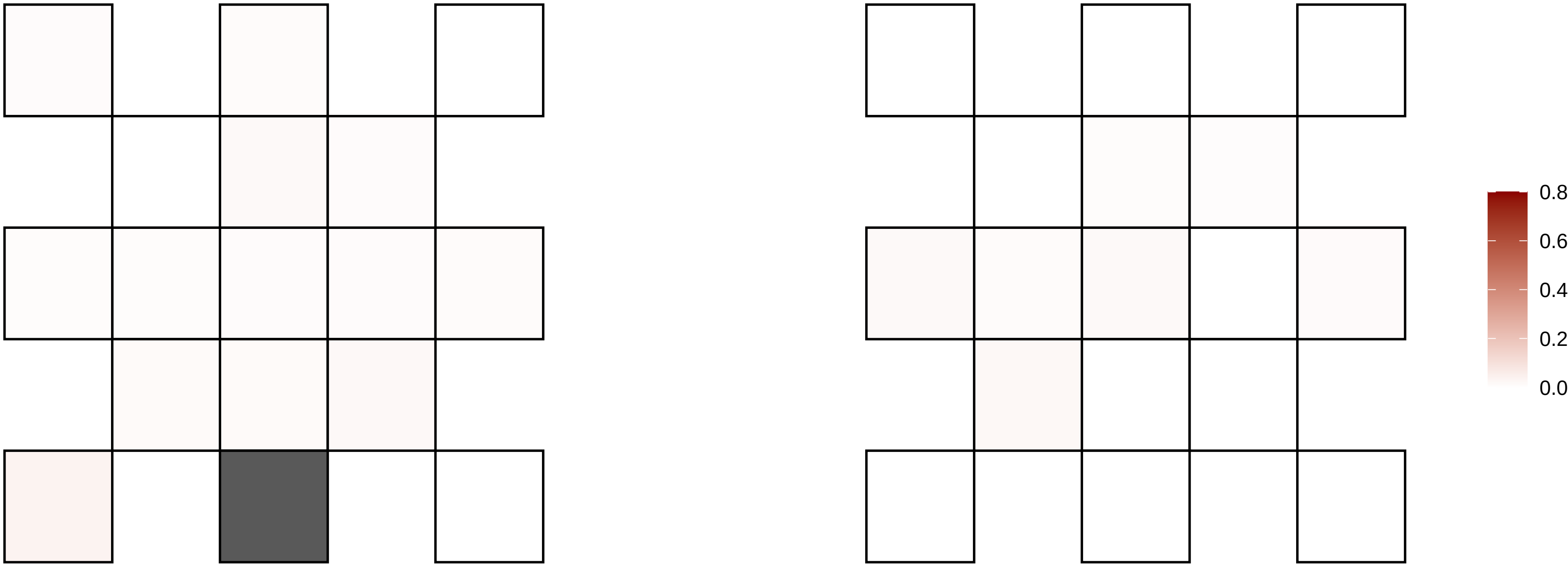

2−22209916−G−A

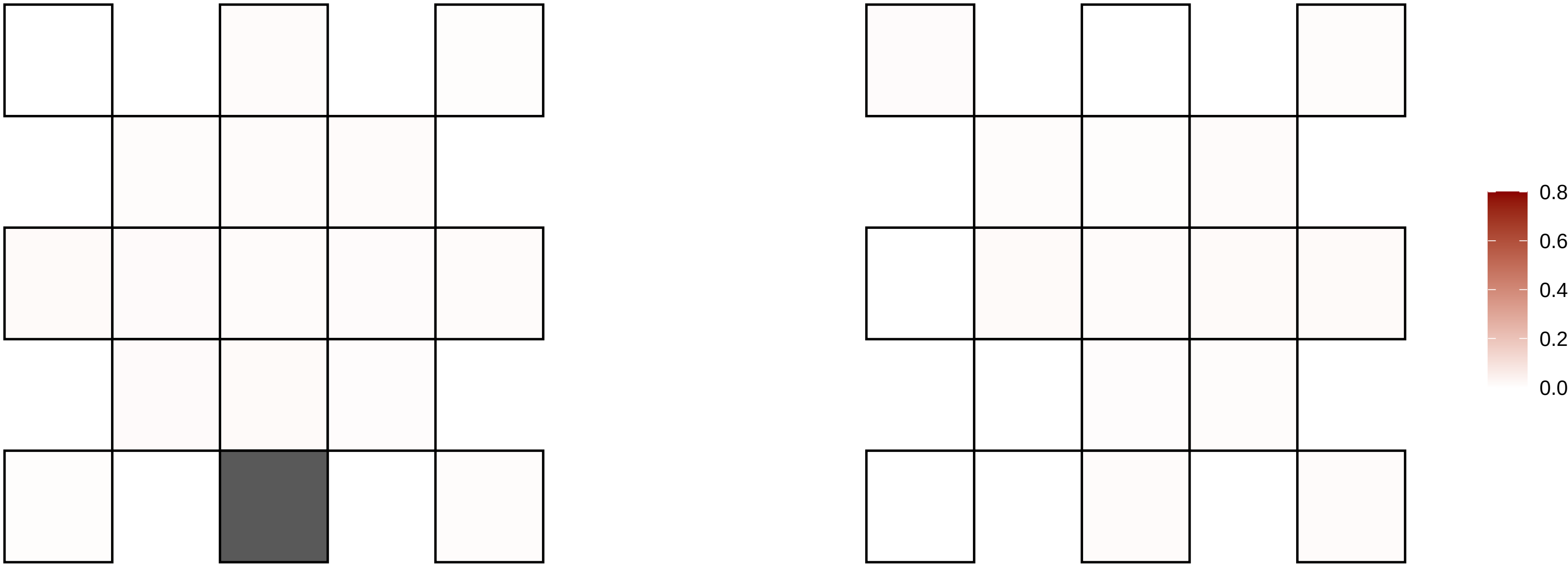

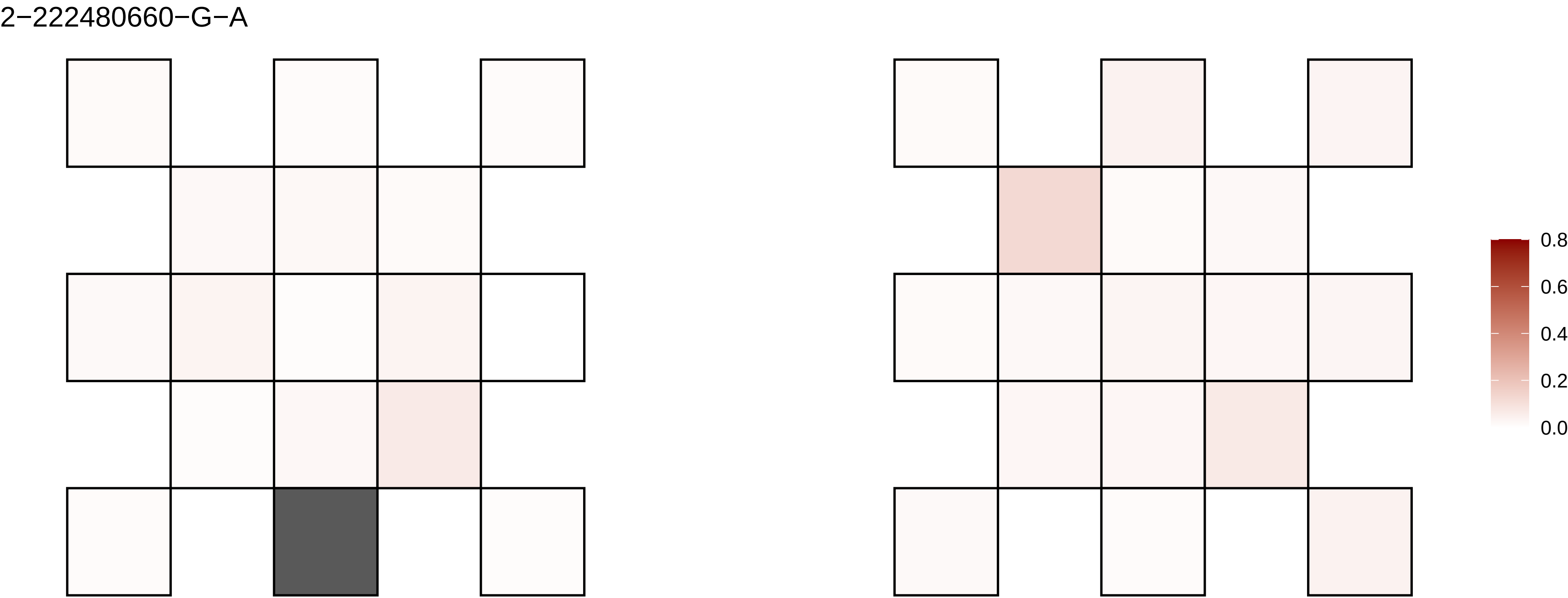
2−222480660−G−A
0.8
0.6
0.4
0.2
0.0

2−225875953−A−G

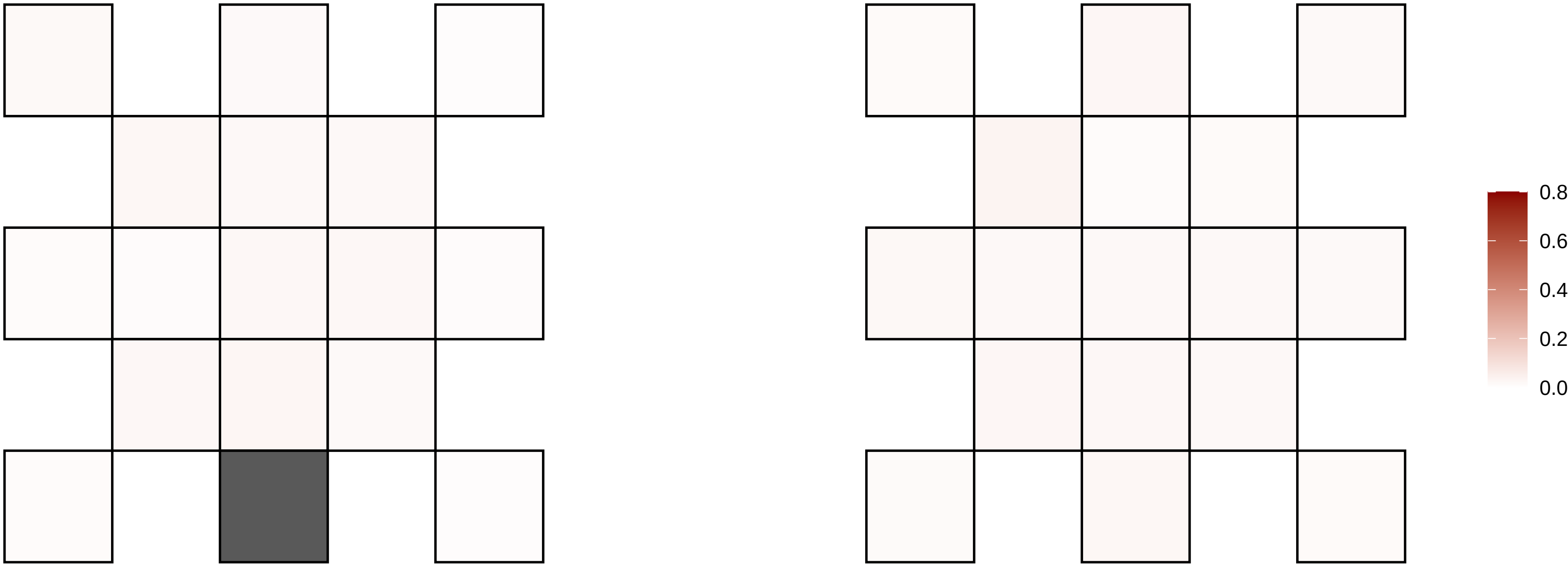

2−226074050−C−A

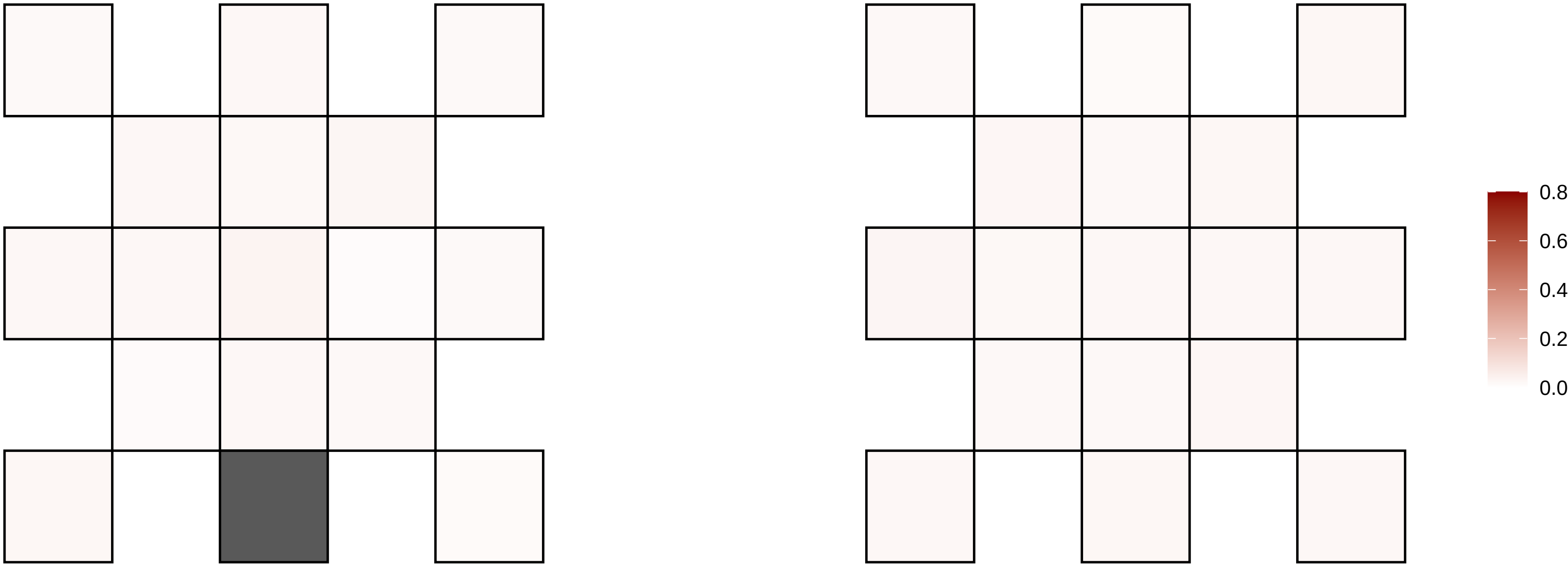

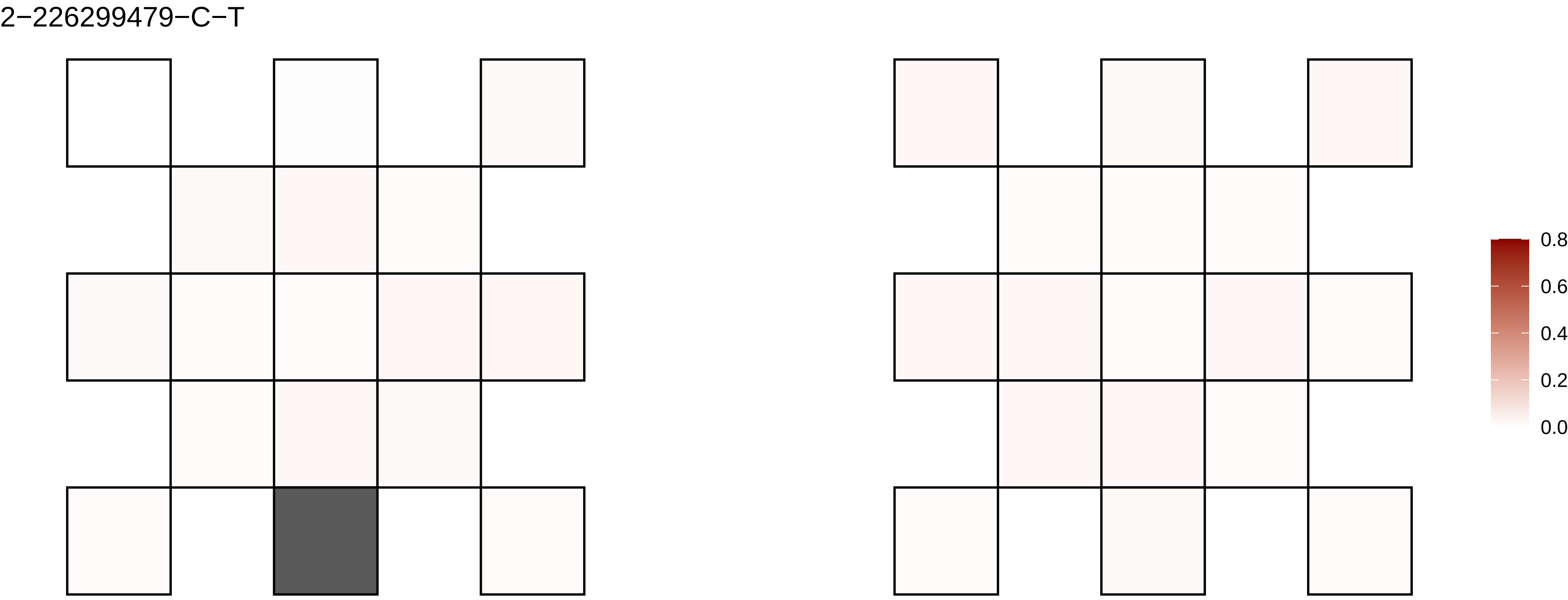
2−226299479−C−T
0.8
0.6
0.4
0.2
0.0

2−229992854−G−A

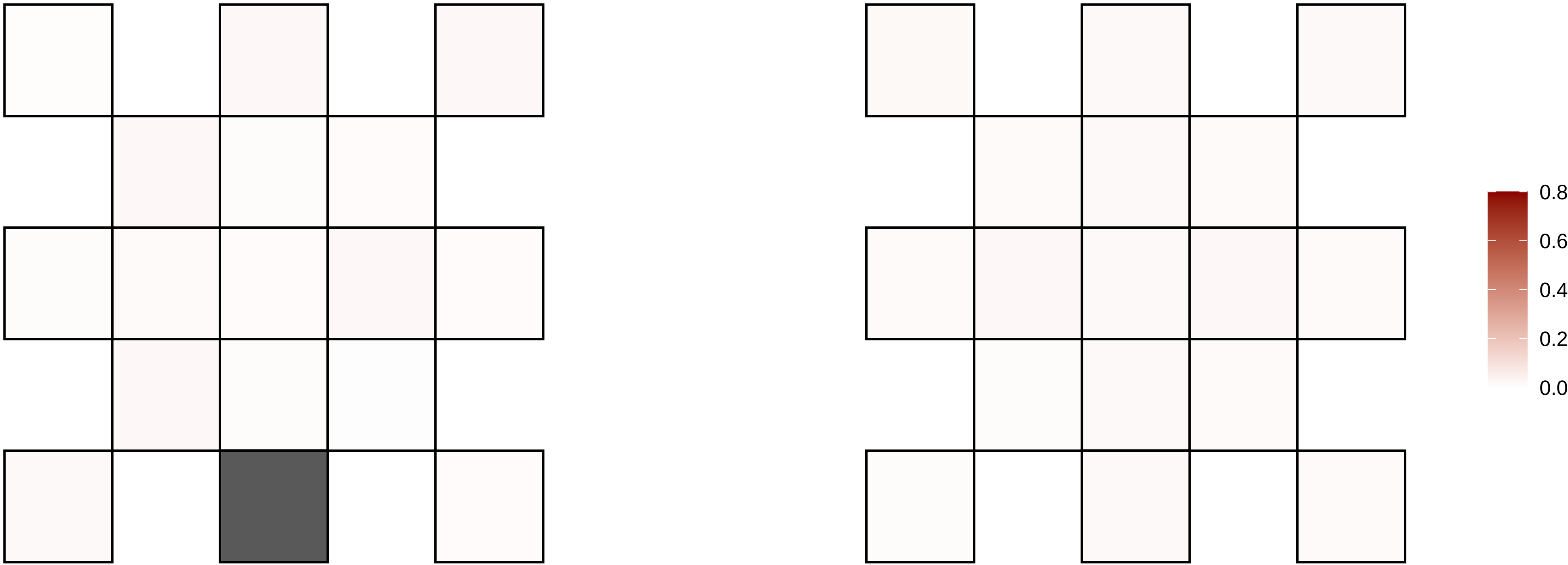

2−234349292−A−G

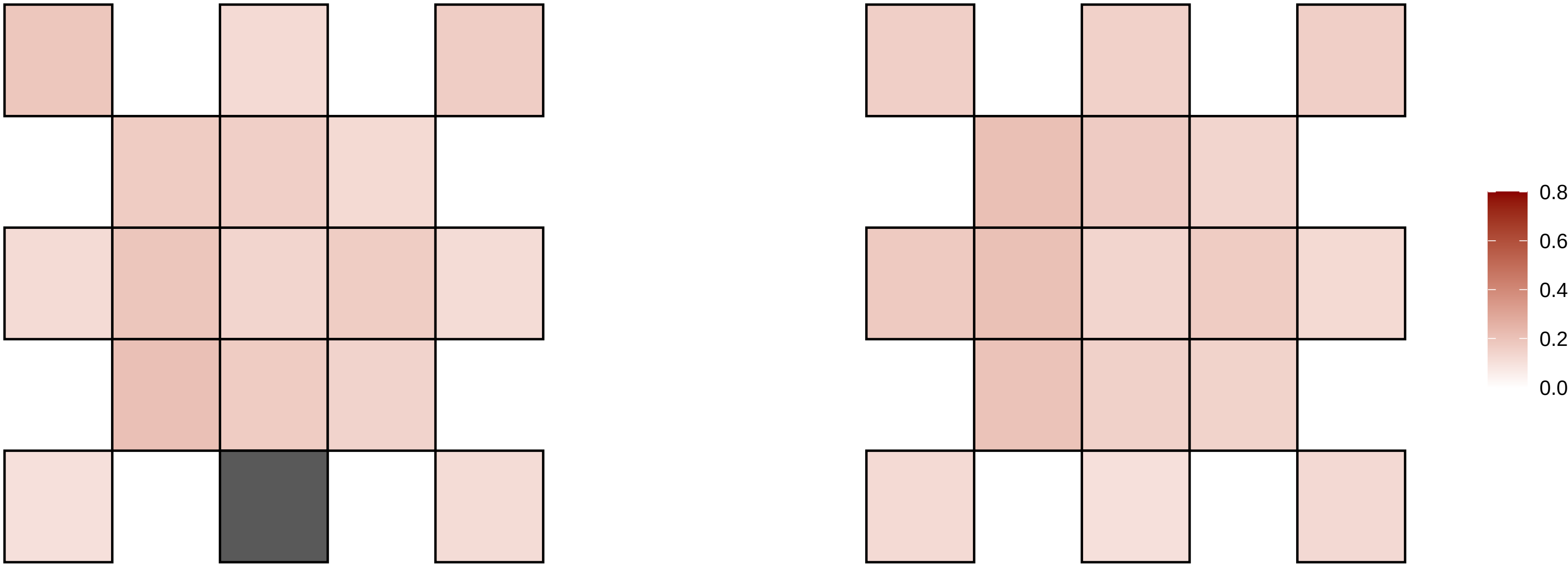

2−240582791−G−A

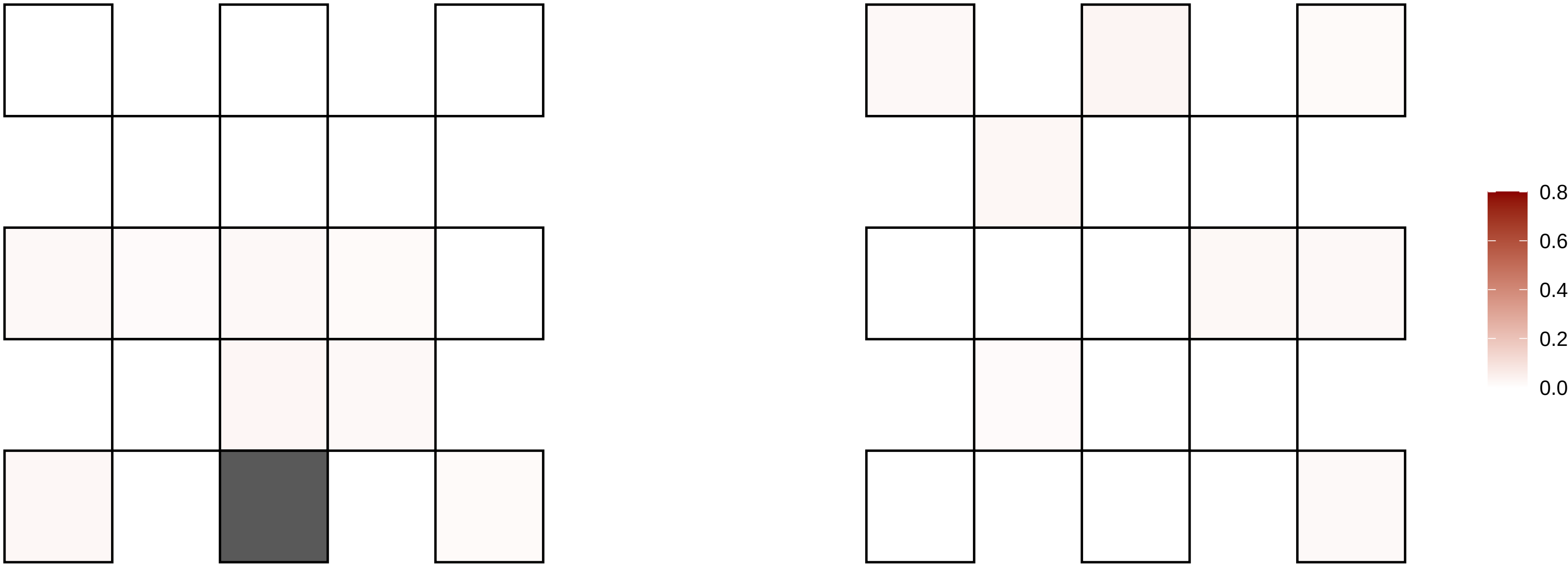

2−241015446−G−A

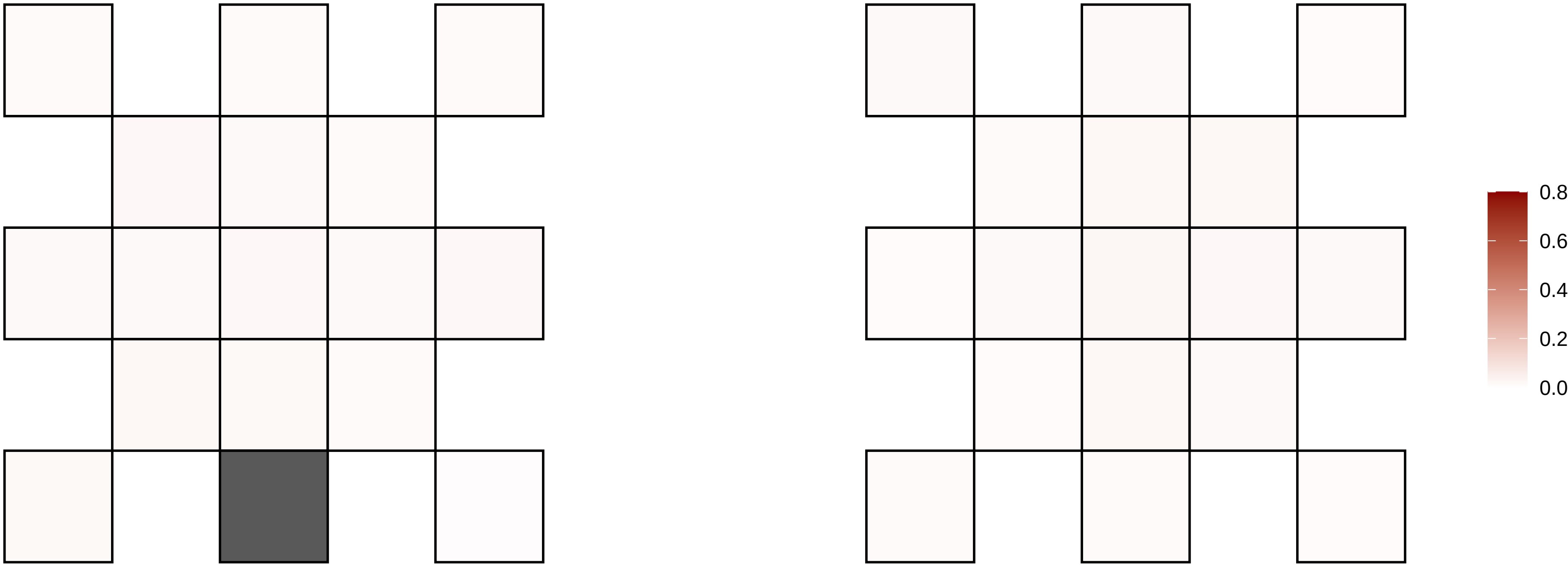

2−241258643−G−A

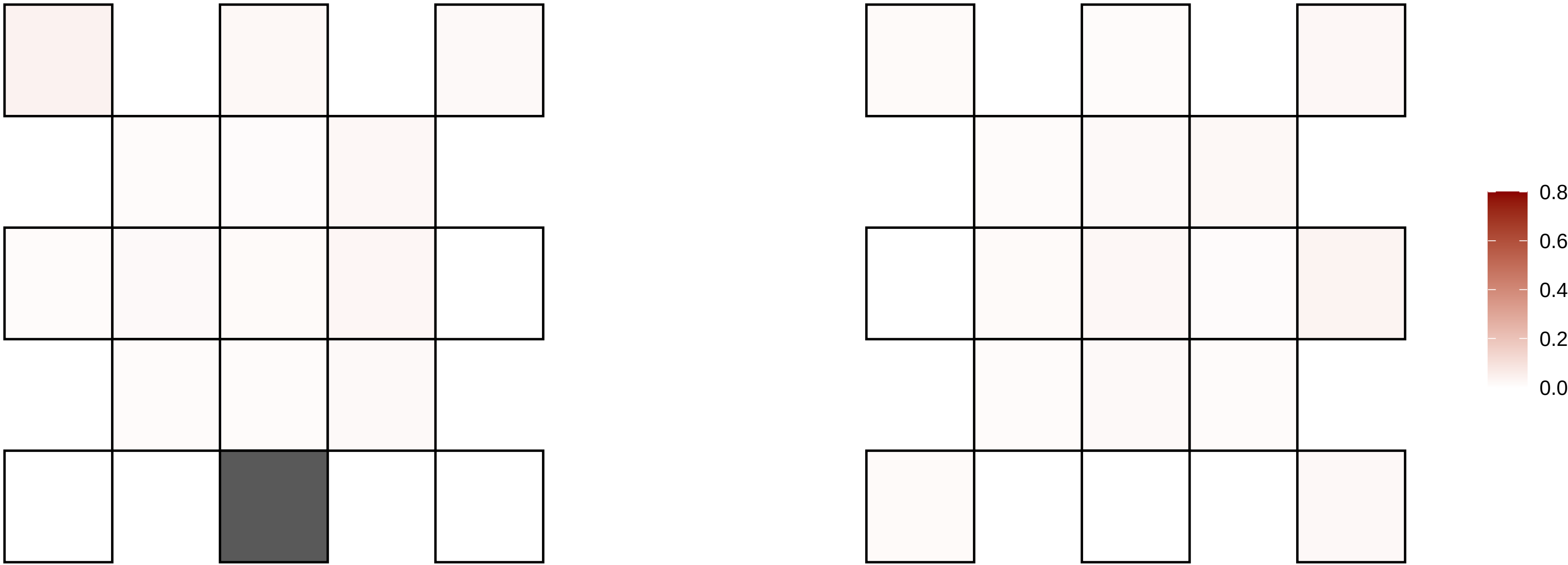

2−27102490−G−A

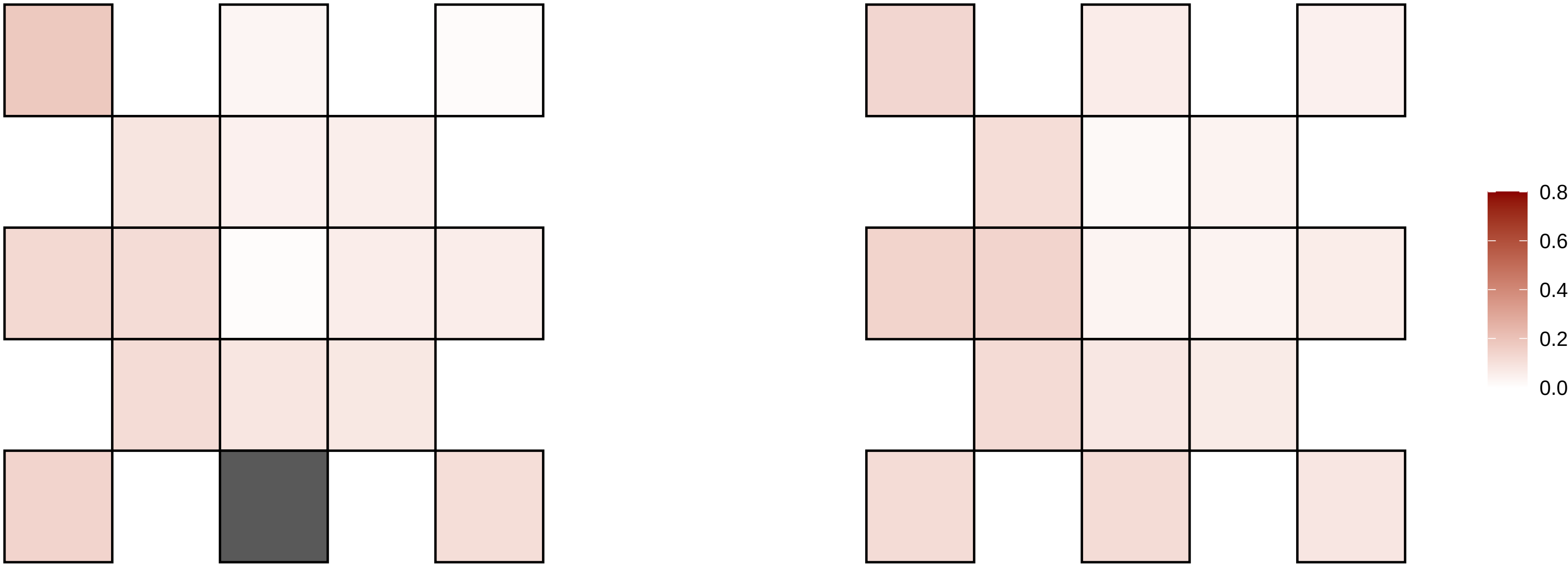

2−3282940−C−T

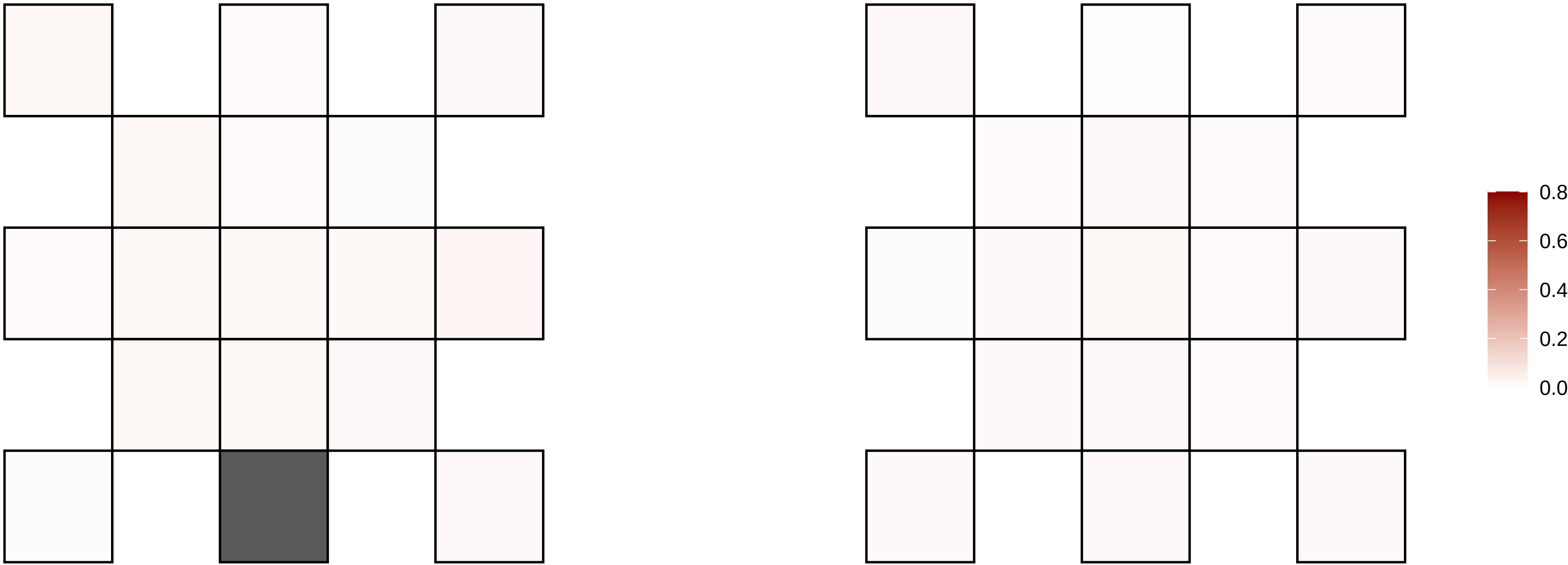

2−34945794−T−C

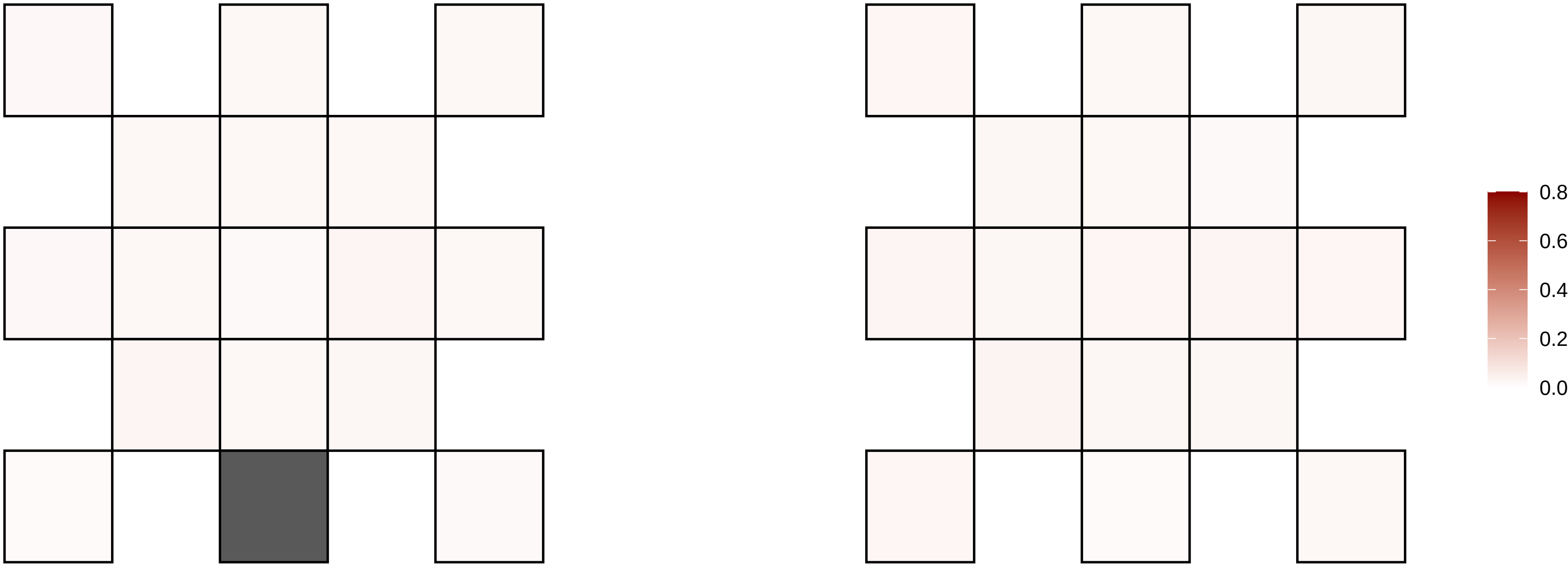

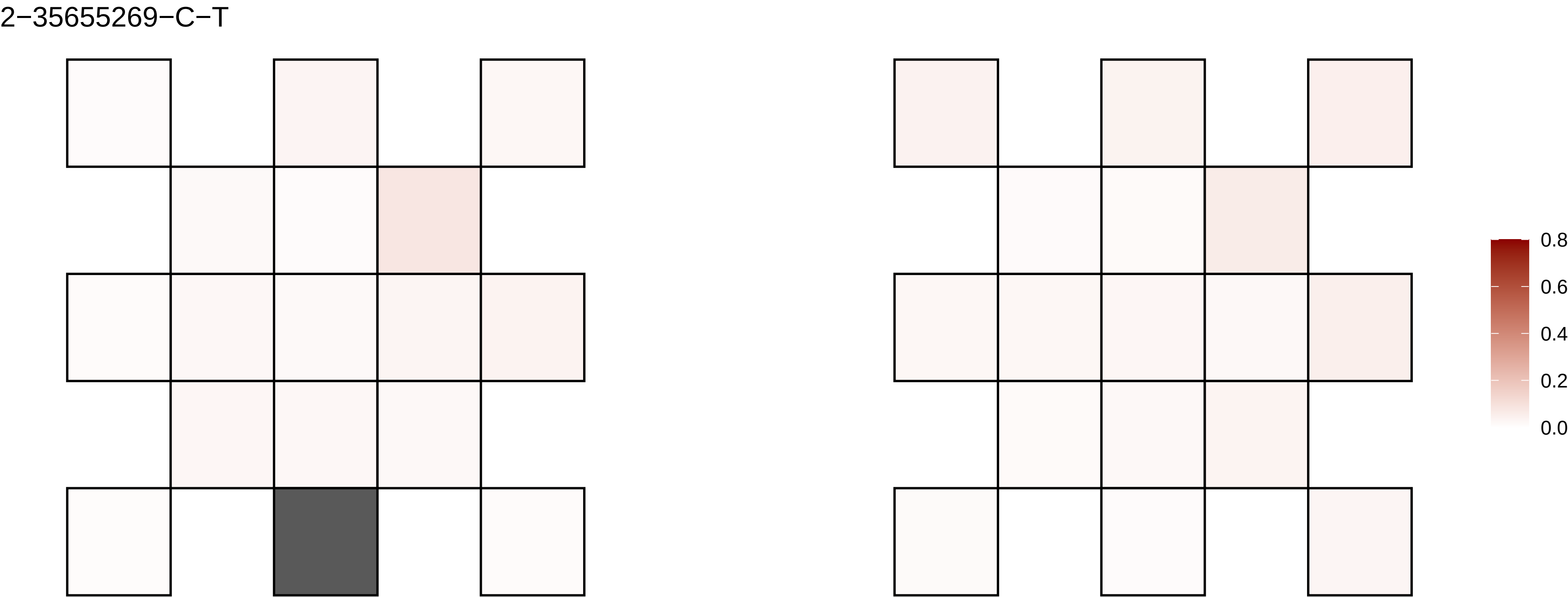
2−35655269−C−T
0.8
0.6
0.4
0.2
0.0

2−41714937−C−A

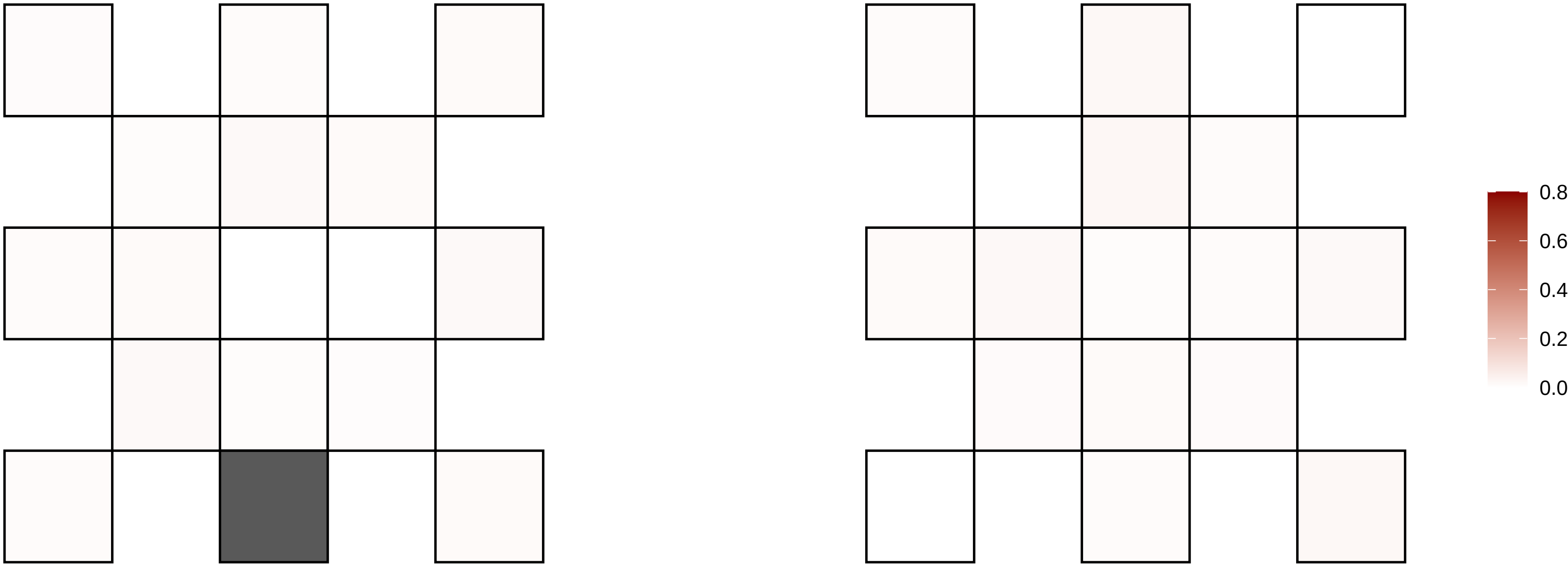

2−424135−C−T

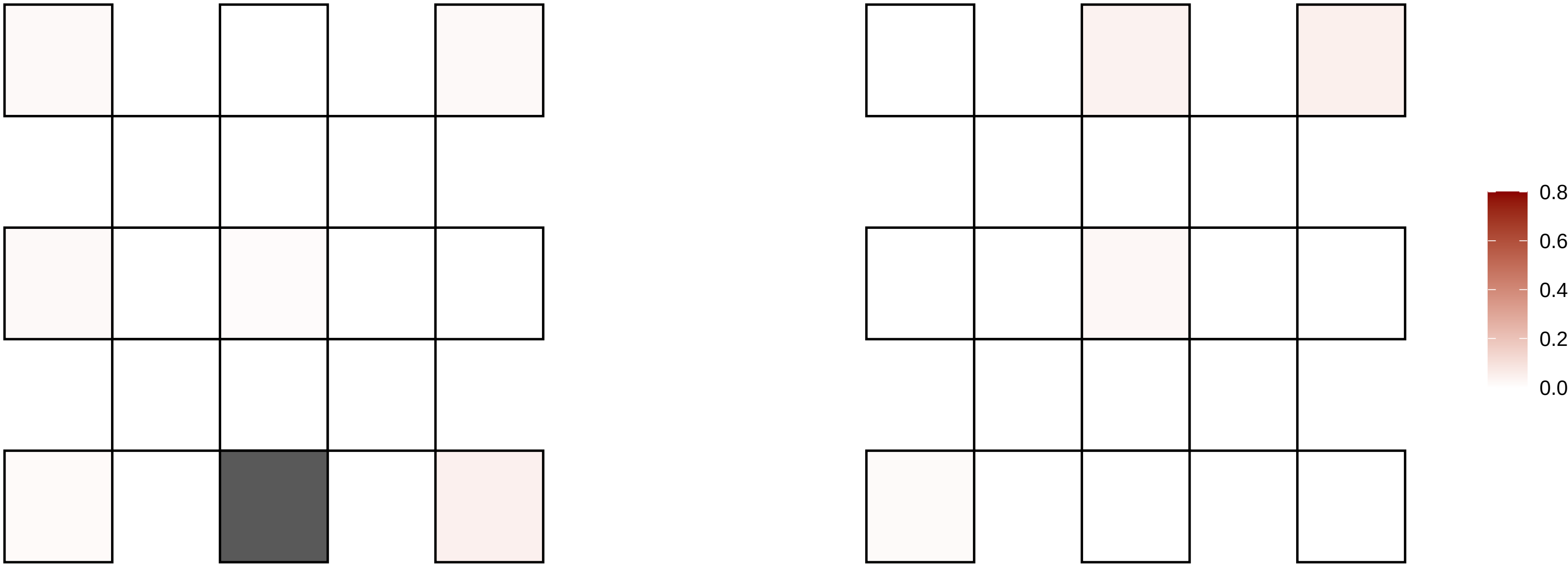

2−45142013−G−A

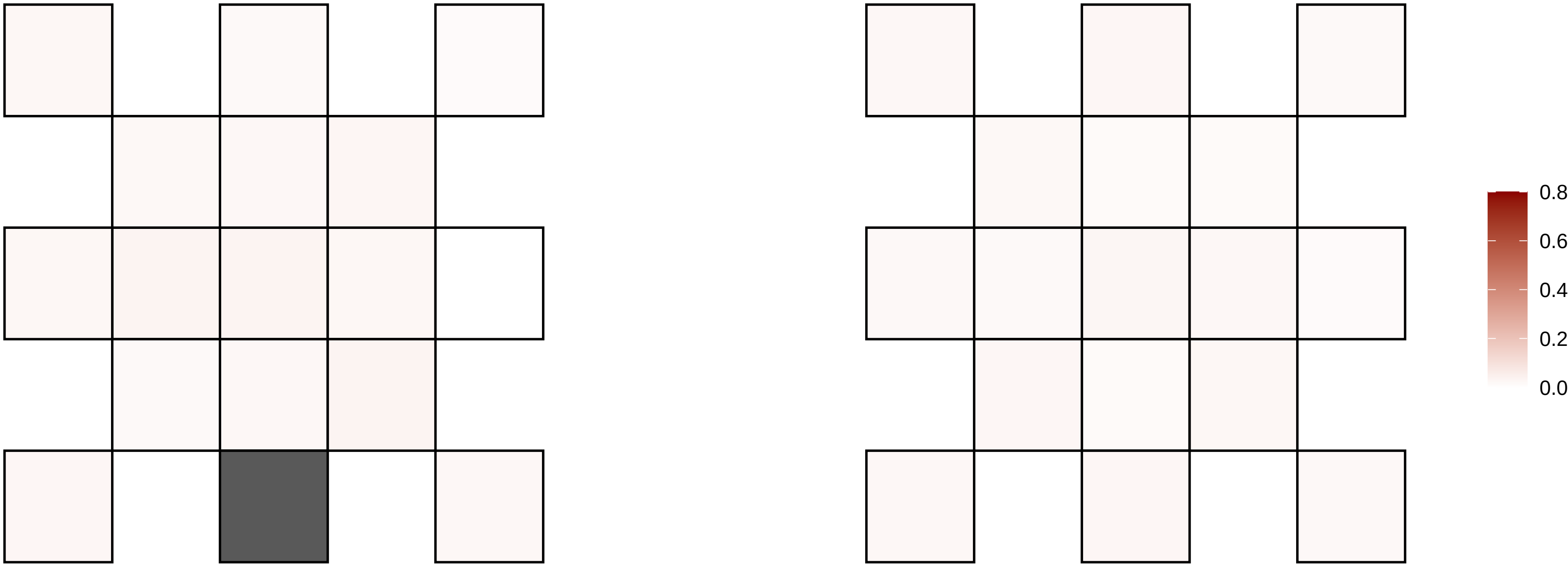

2−48643619−C−T

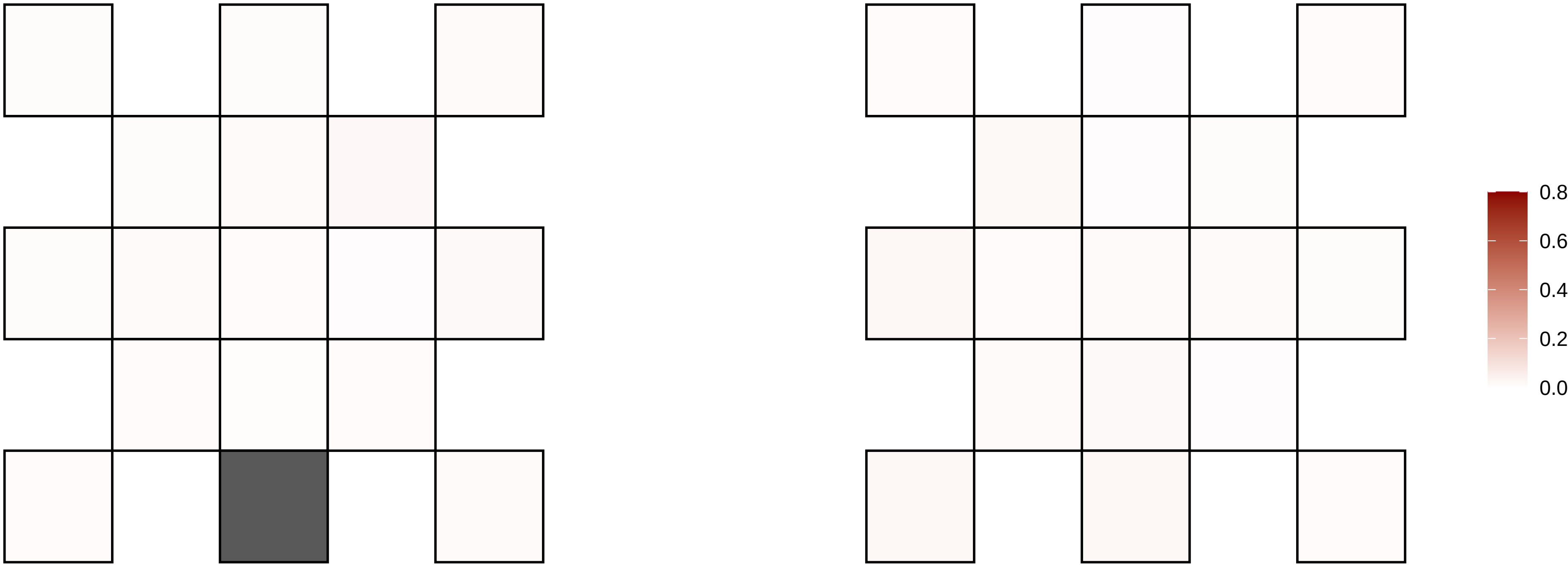

2−52452769−G−A

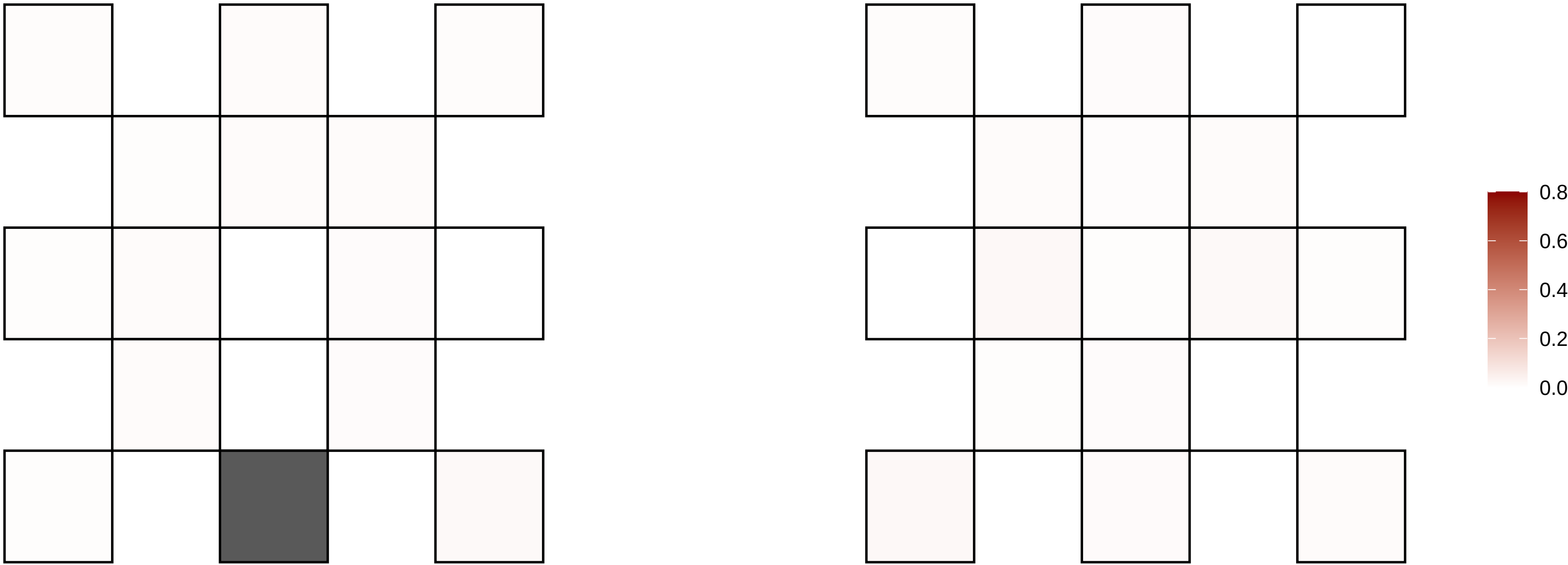

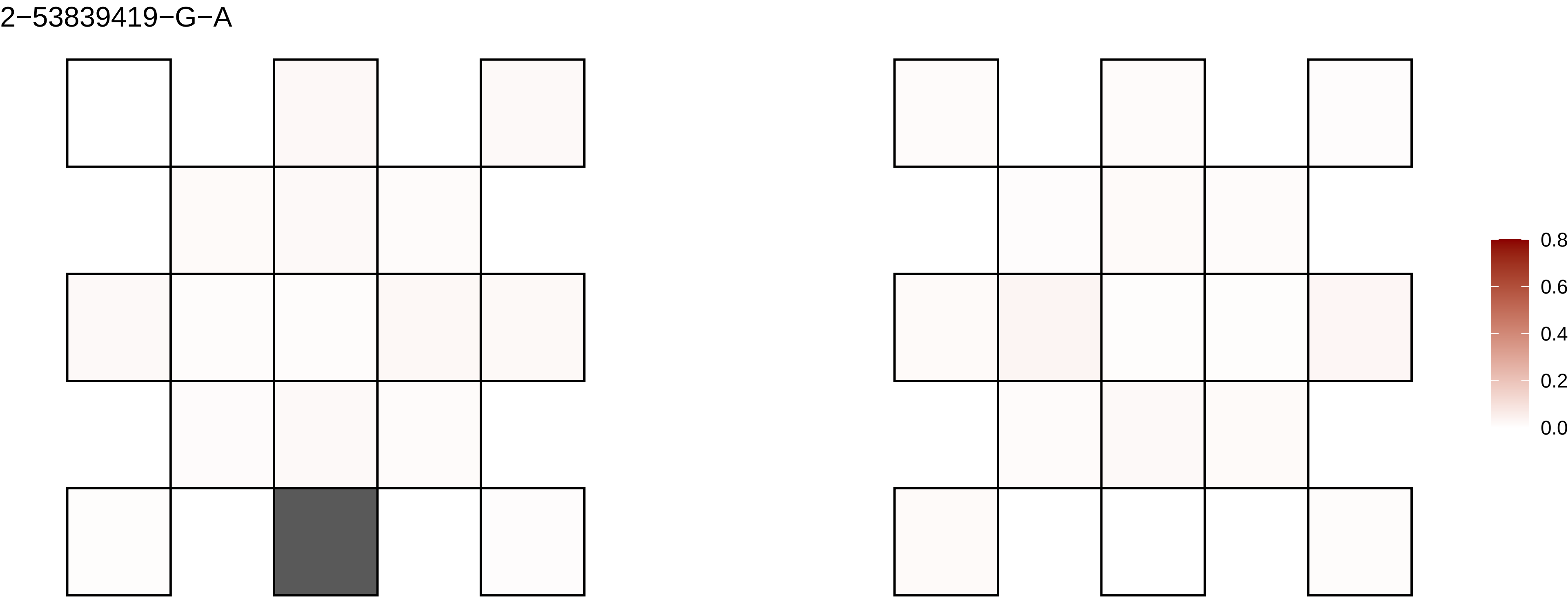
2−53839419−G−A
0.8
0.6
0.4
0.2
0.0

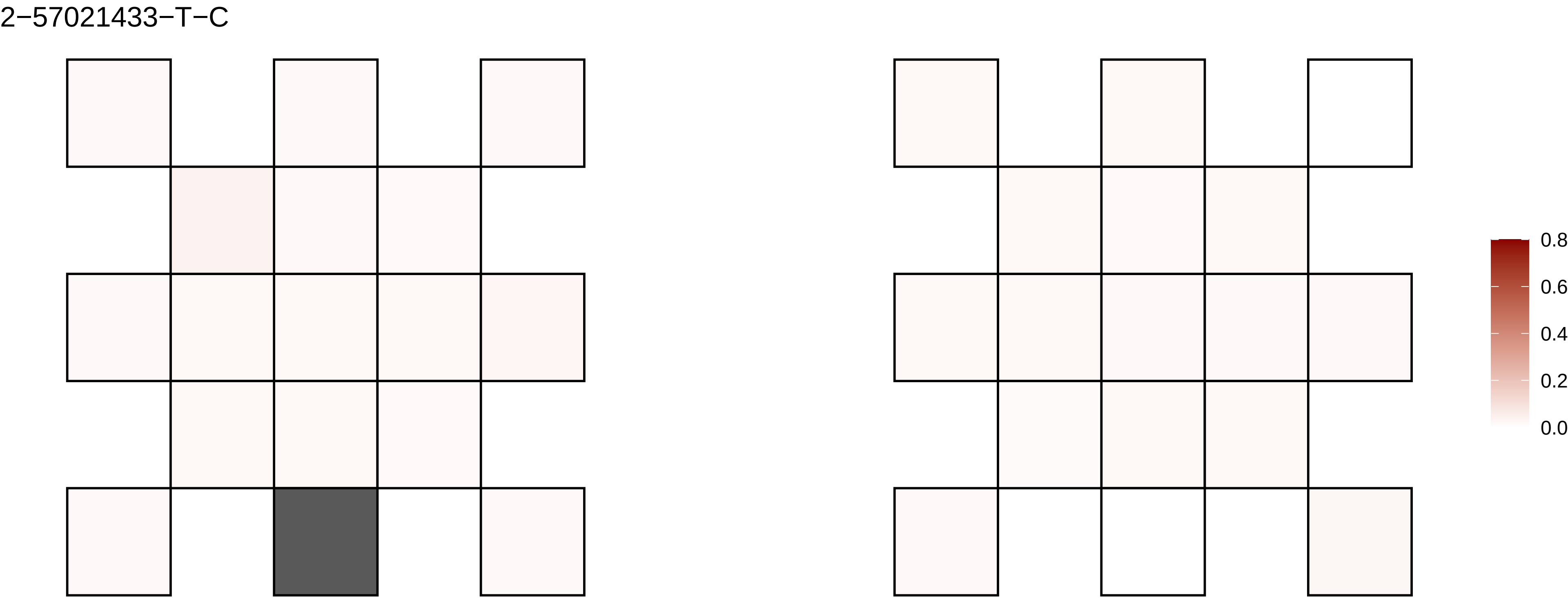

2−57021433−T−C
0.8
0.6
0.4
0.2
0.0

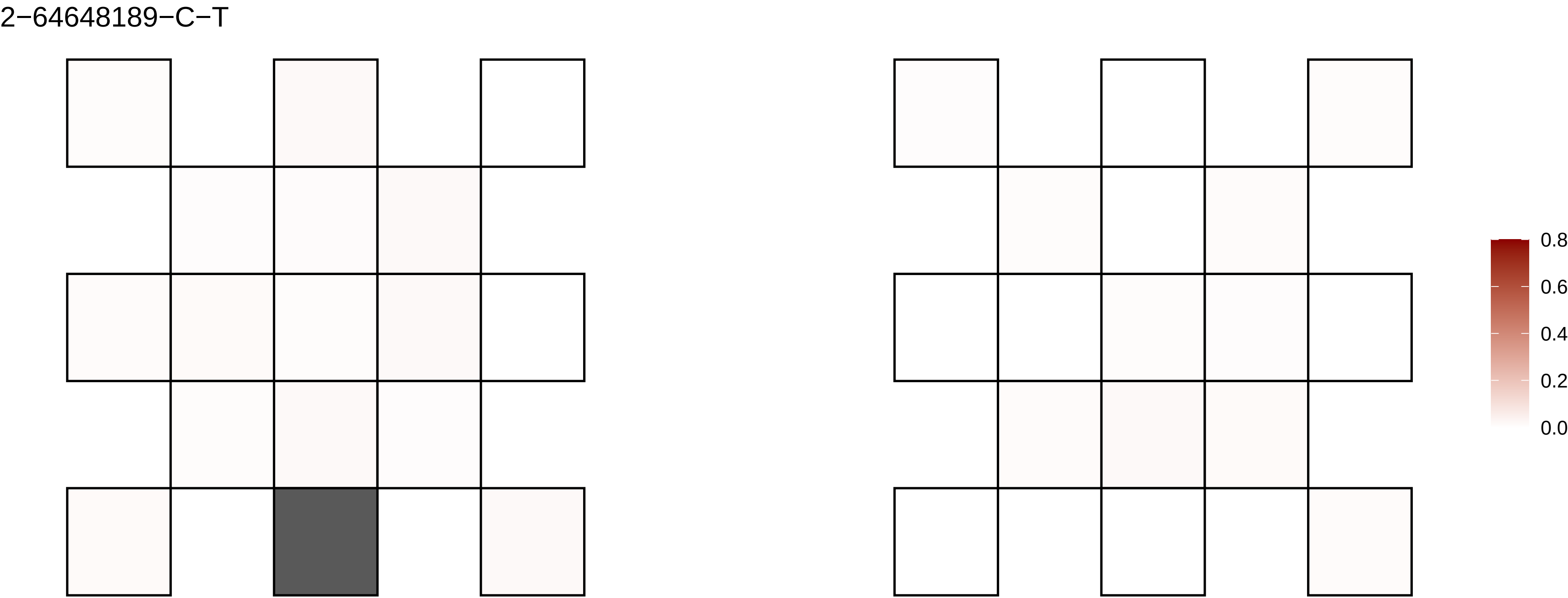

2−64648189−C−T
0.8
0.6
0.4
0.2
0.0

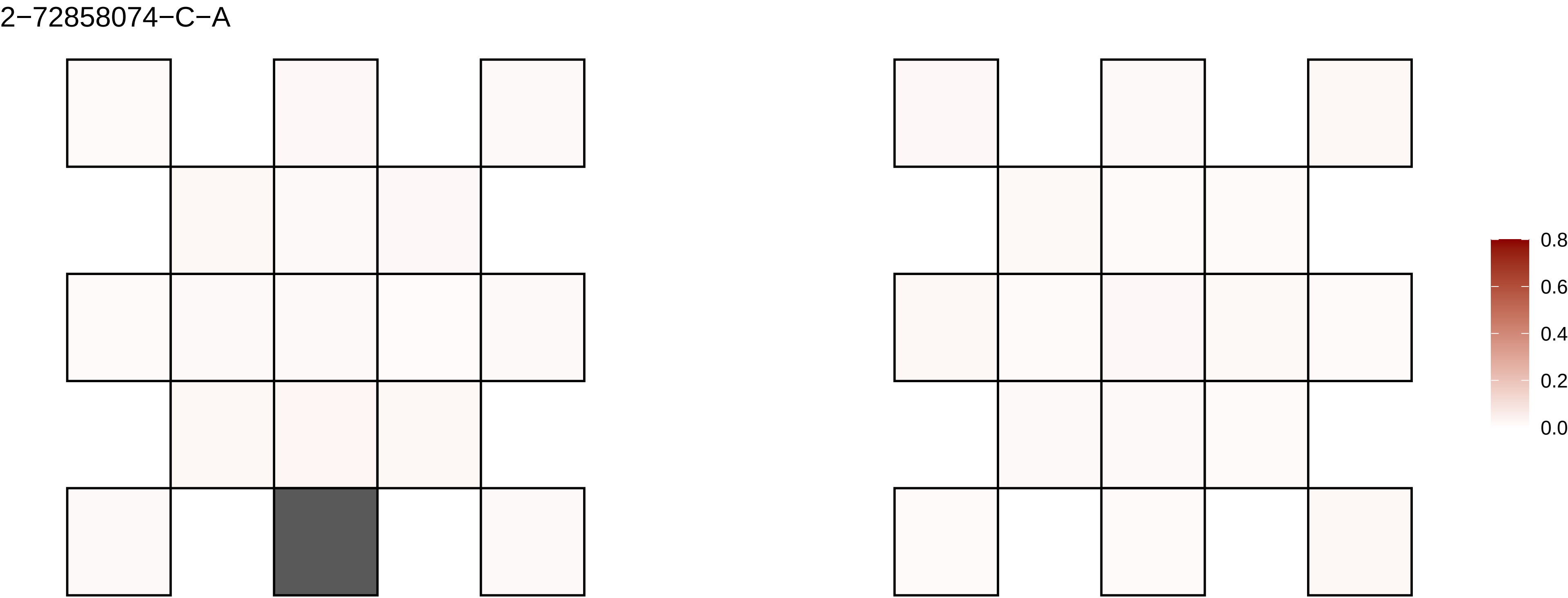
2−72858074−C−A
0.8
0.6
0.4
0.2
0.0

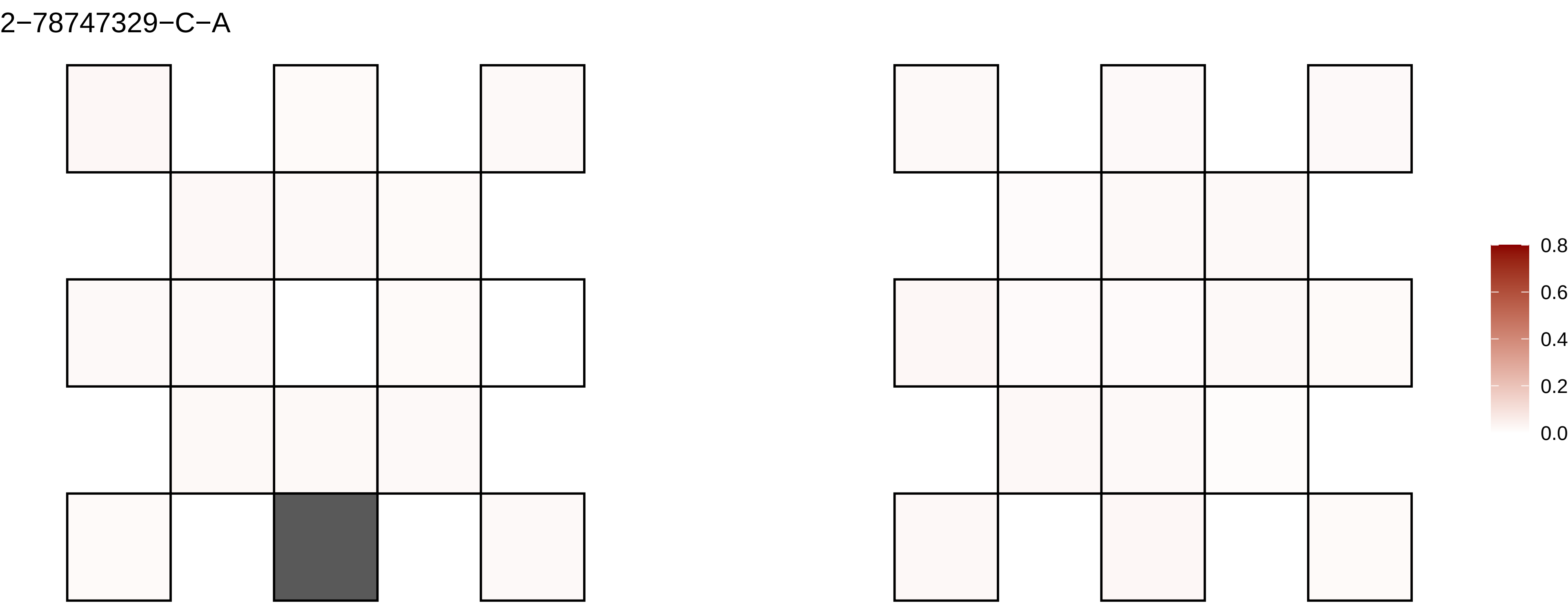
2−78747329−C−A
0.8
0.6
0.4
0.2
0.0

2−82189643−C−A

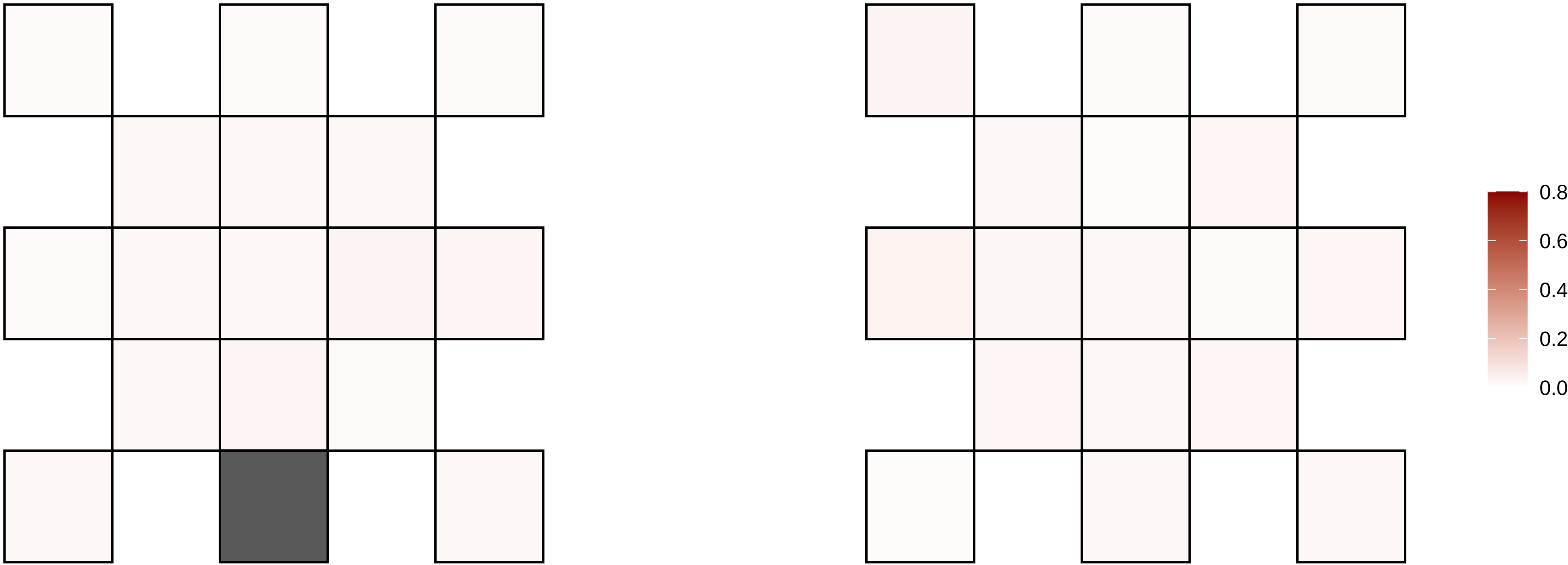

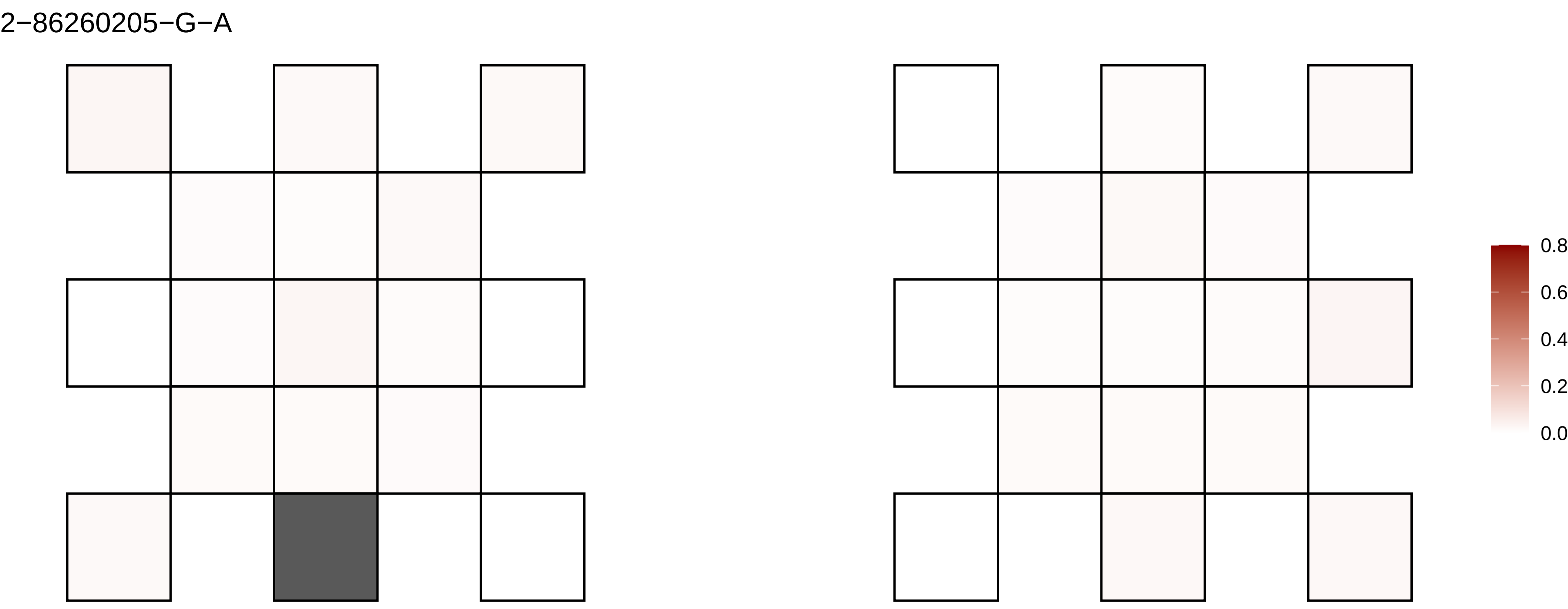

2−86260205−G−A
0.8
0.6
0.4
0.2
0.0

2−88993566−C−T

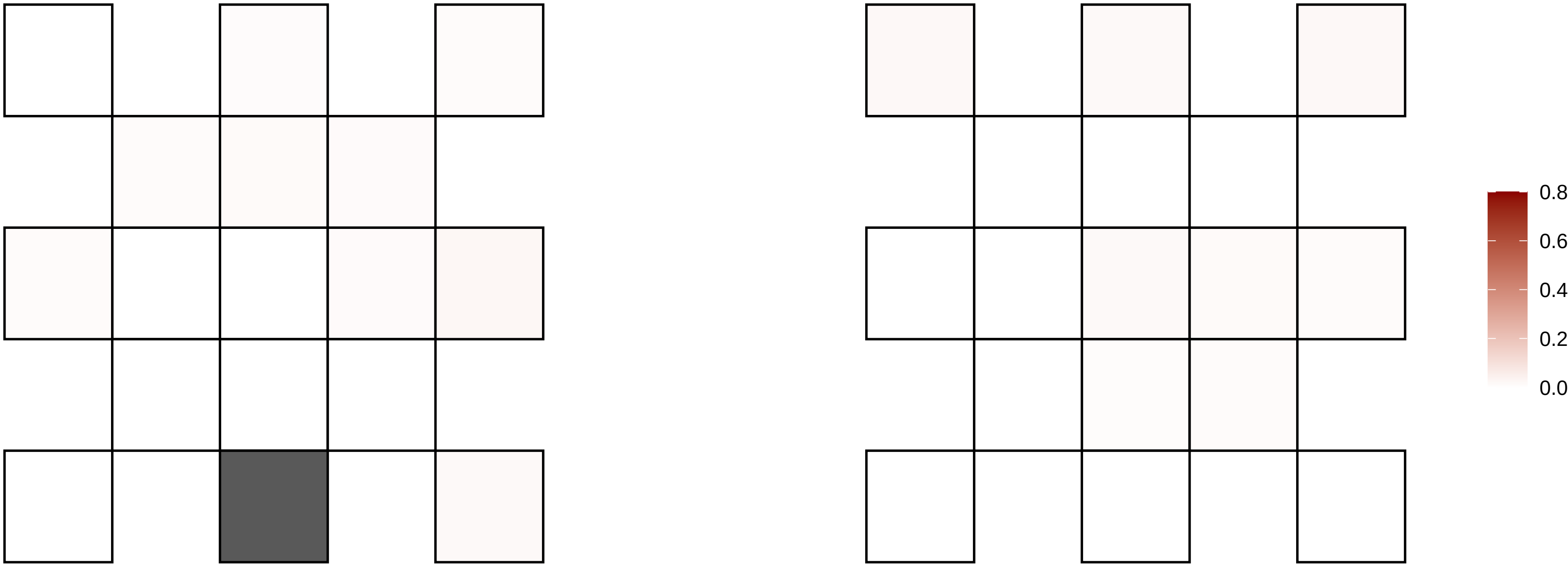

2−97528065−C−T

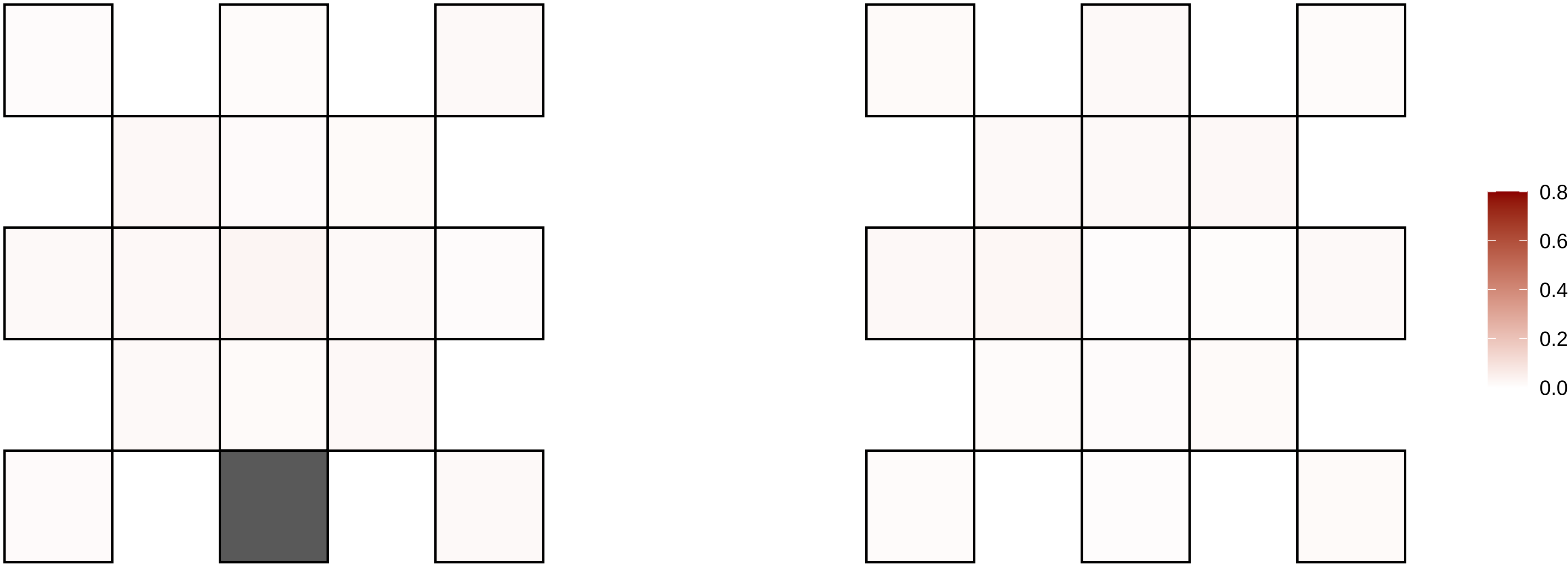

20−10701398−A−G

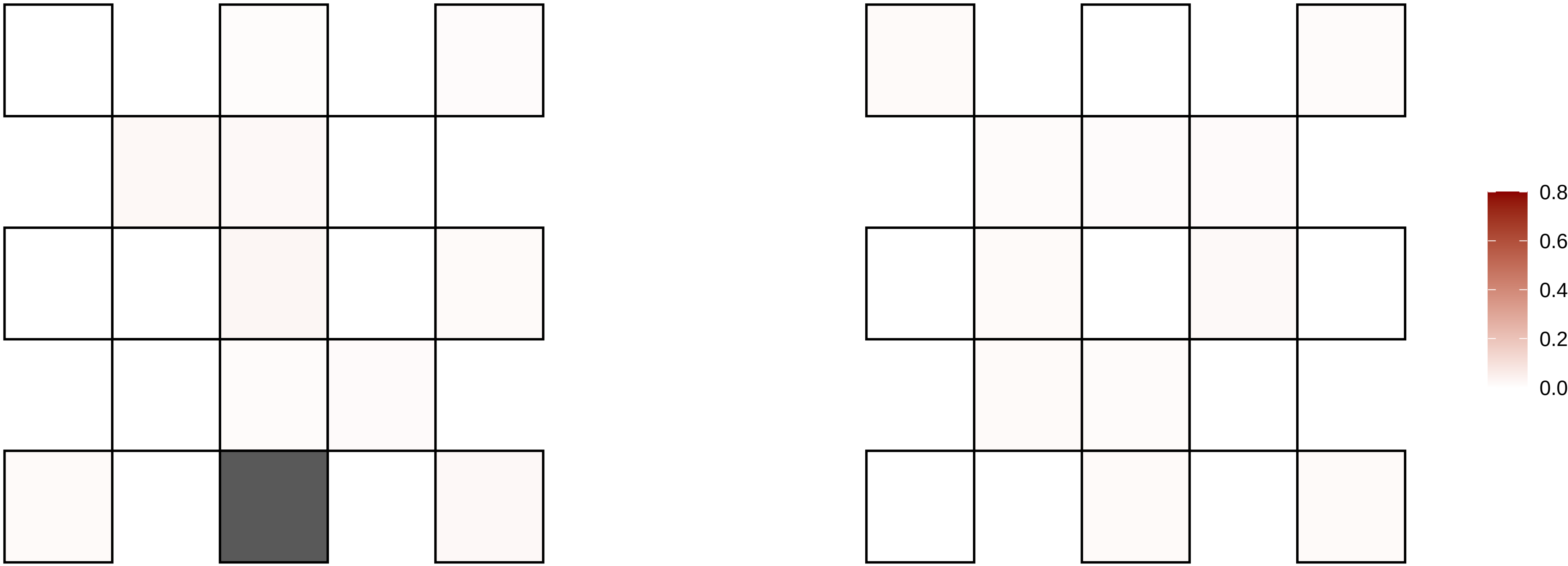

20−10742088−G−A

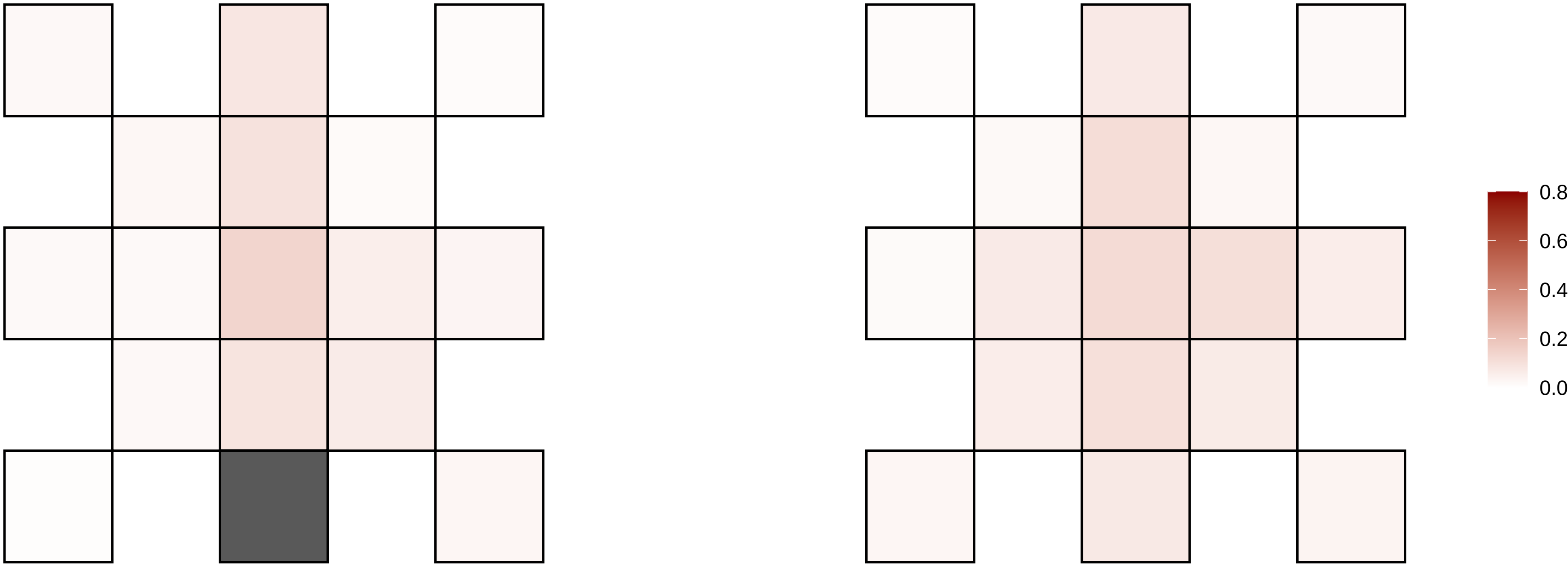

20−11798381−C−G

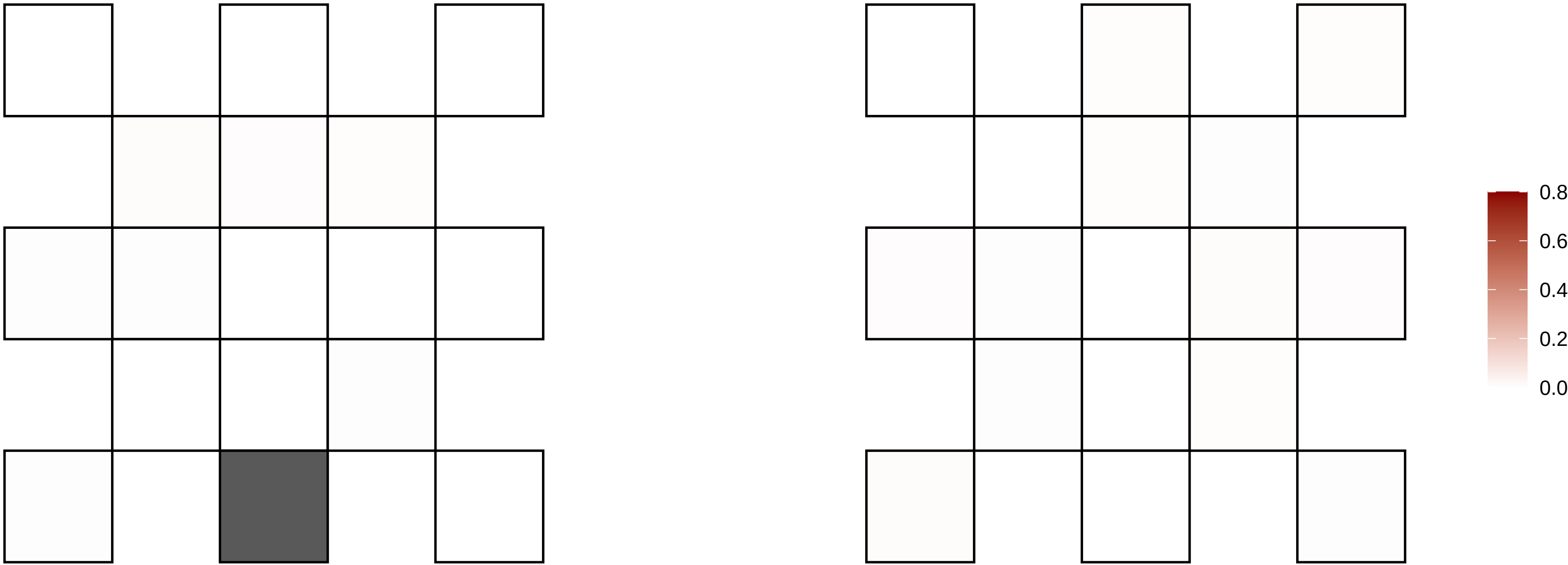

20−12514966−T−G

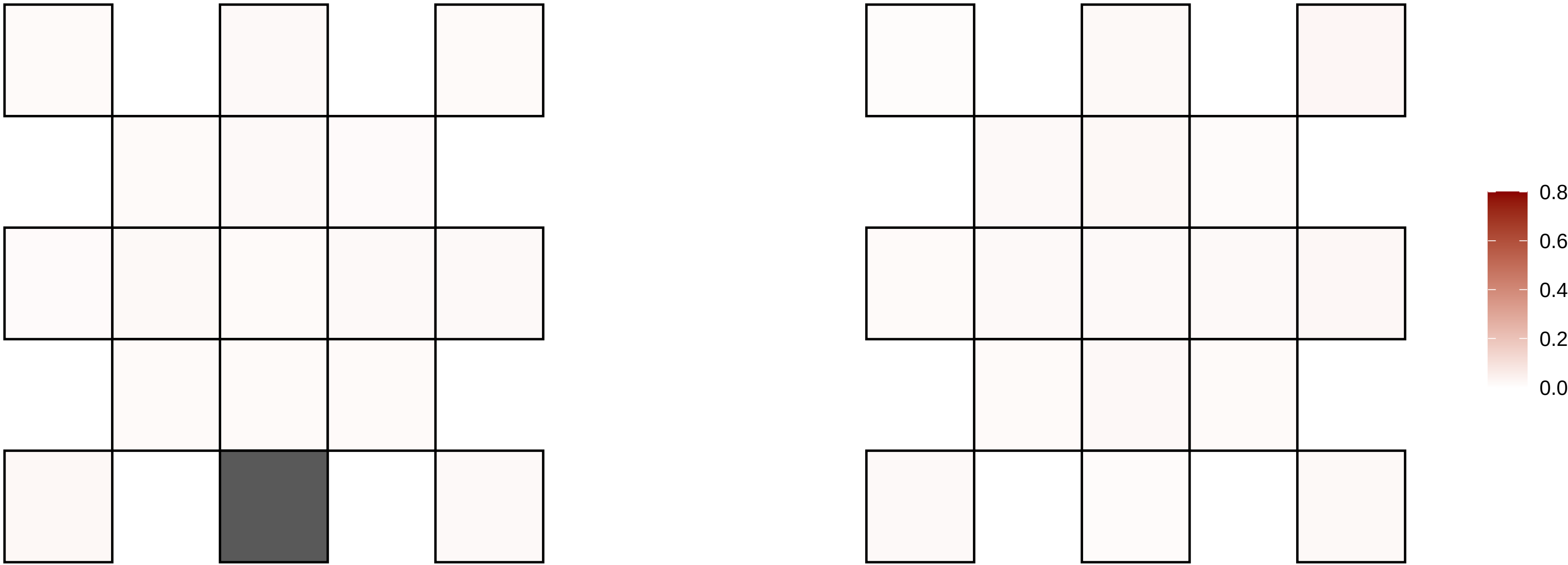

20−15344074−C−T

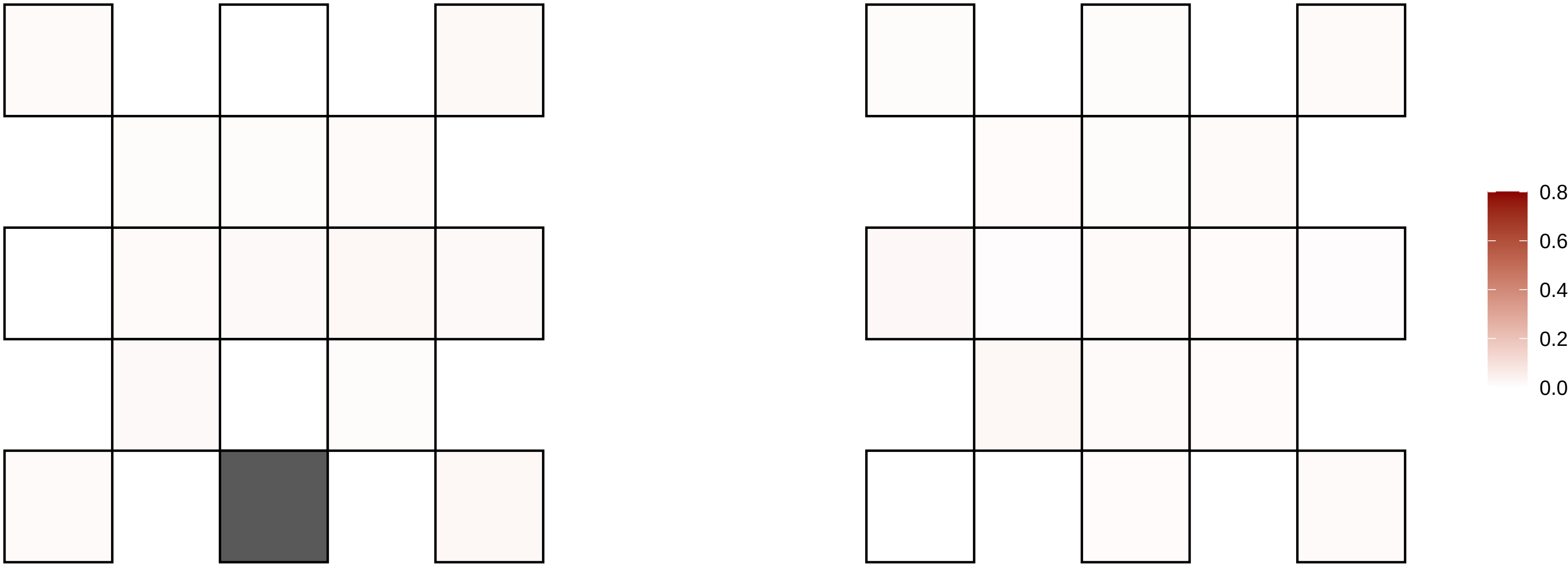

20−16783172−C−T

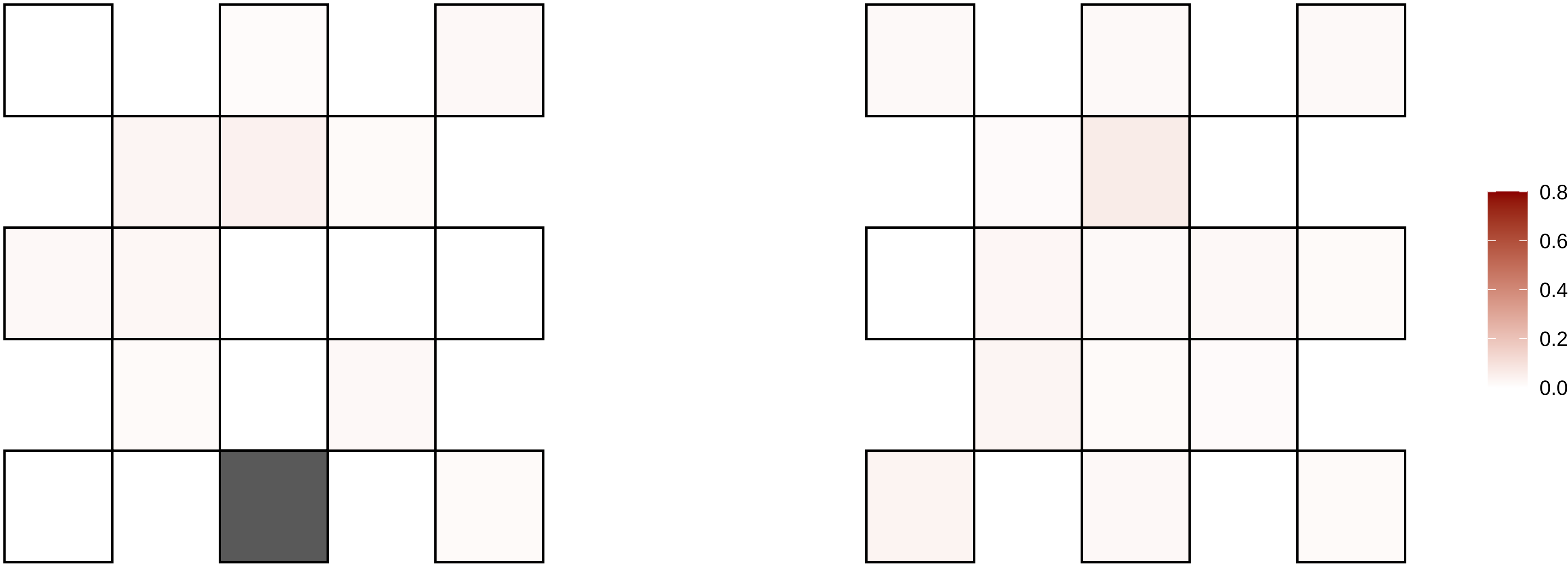

20−18482771−G−T

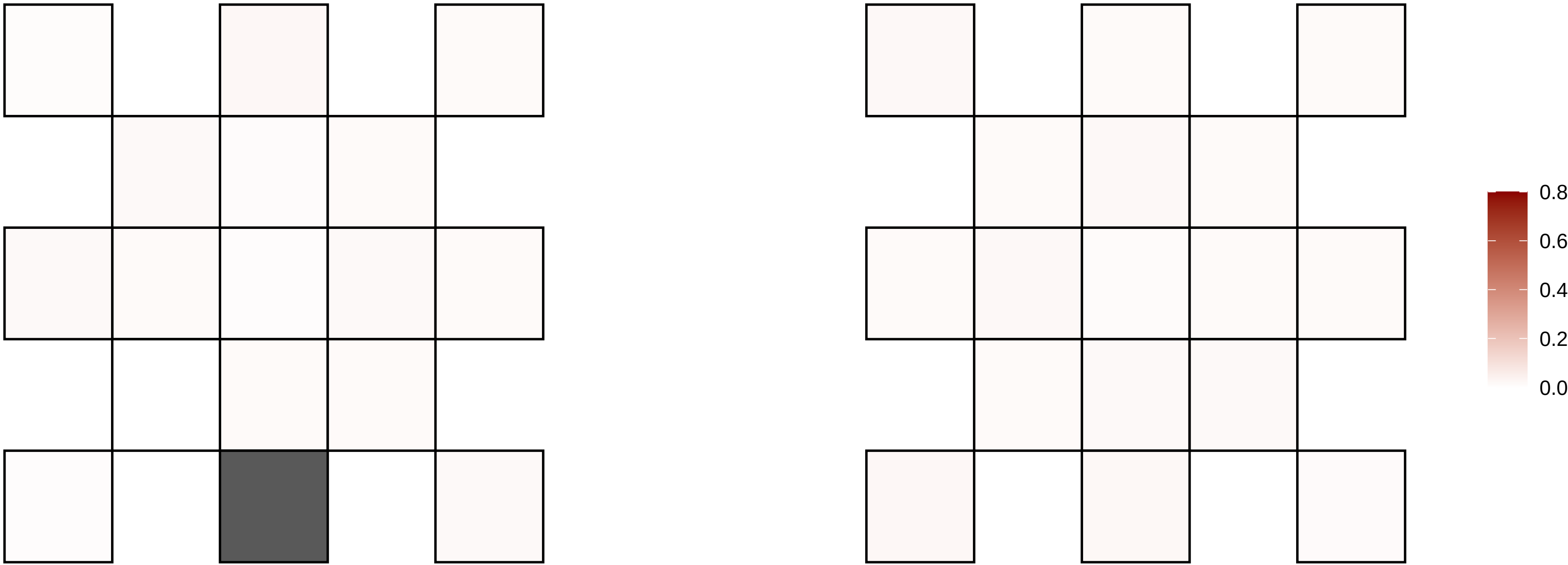

20−21698027−C−T

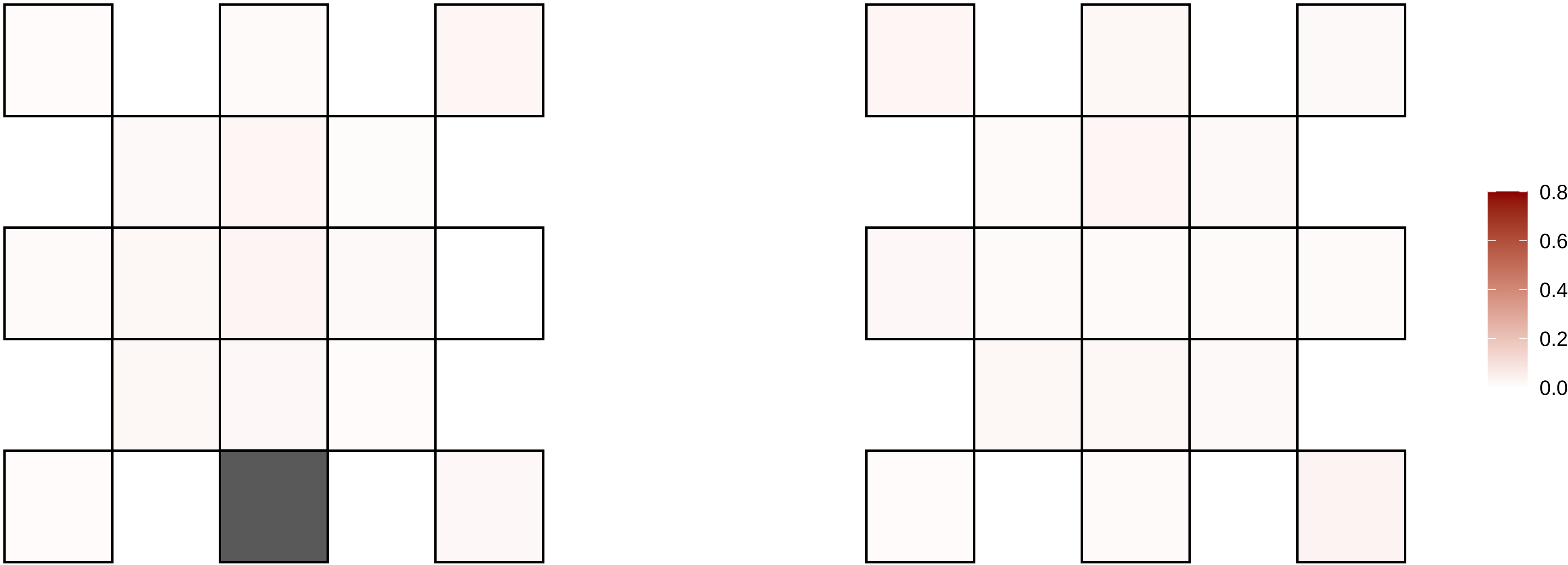

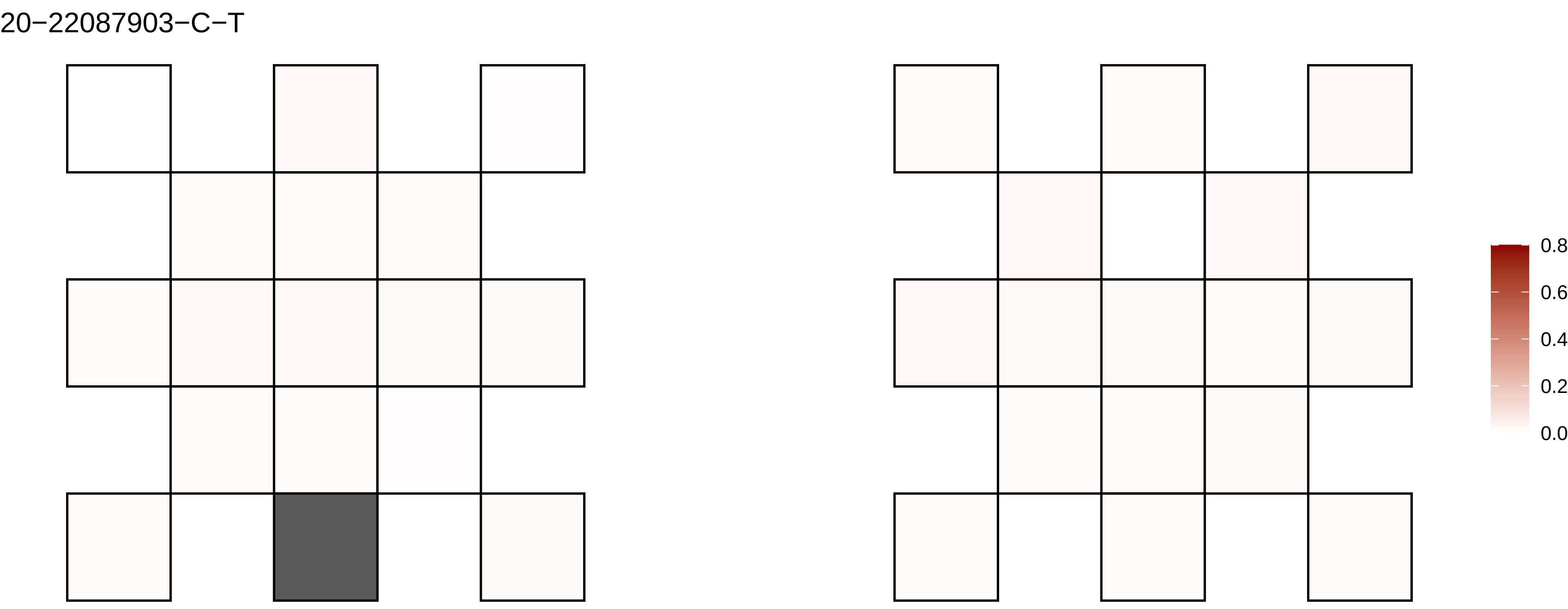
20−22087903−C−T
0.8
0.6
0.4
0.2
0.0

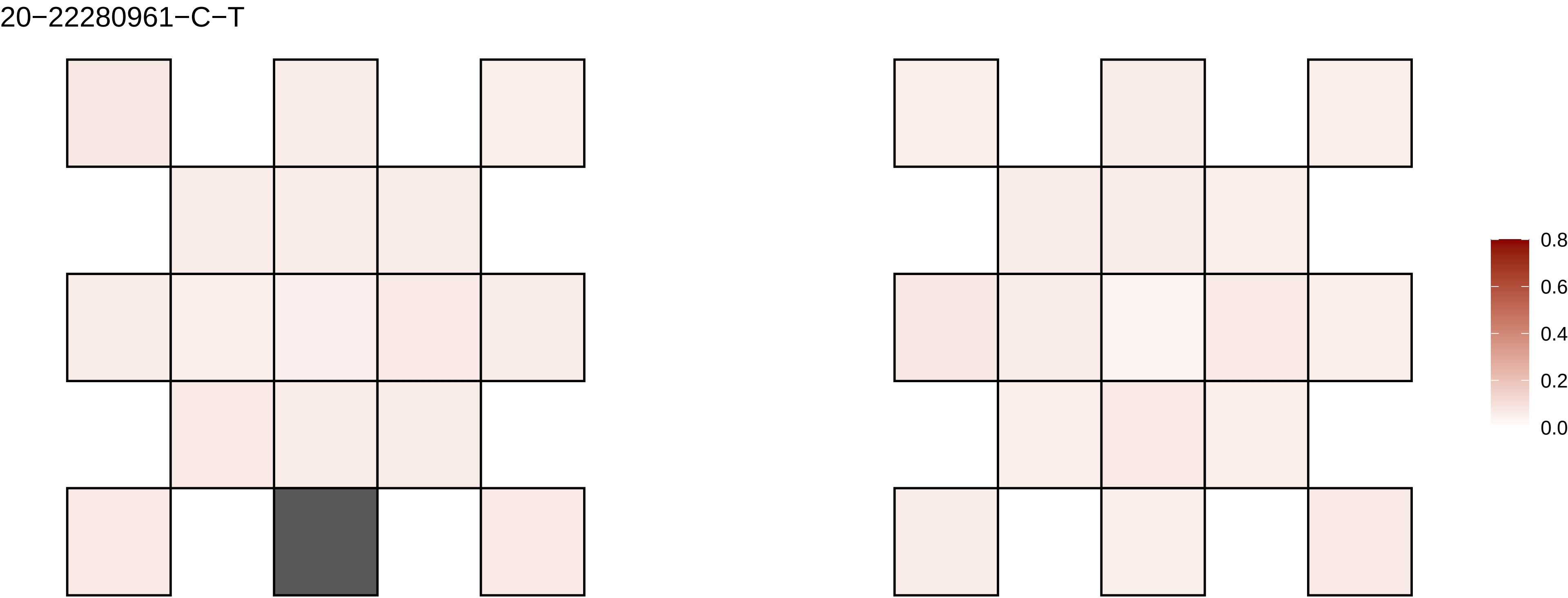
20−22280961−C−T
0.8
0.6
0.4
0.2
0.0

20−22383923−C−A

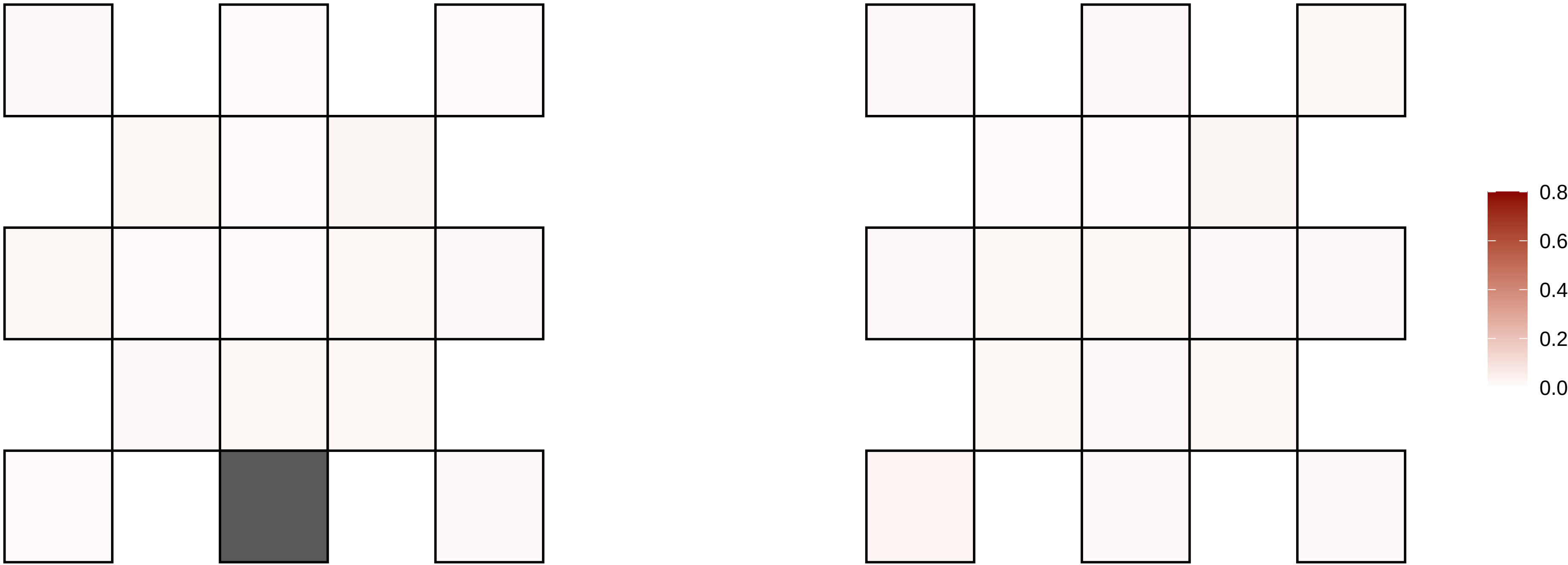

20−24978911−G−A

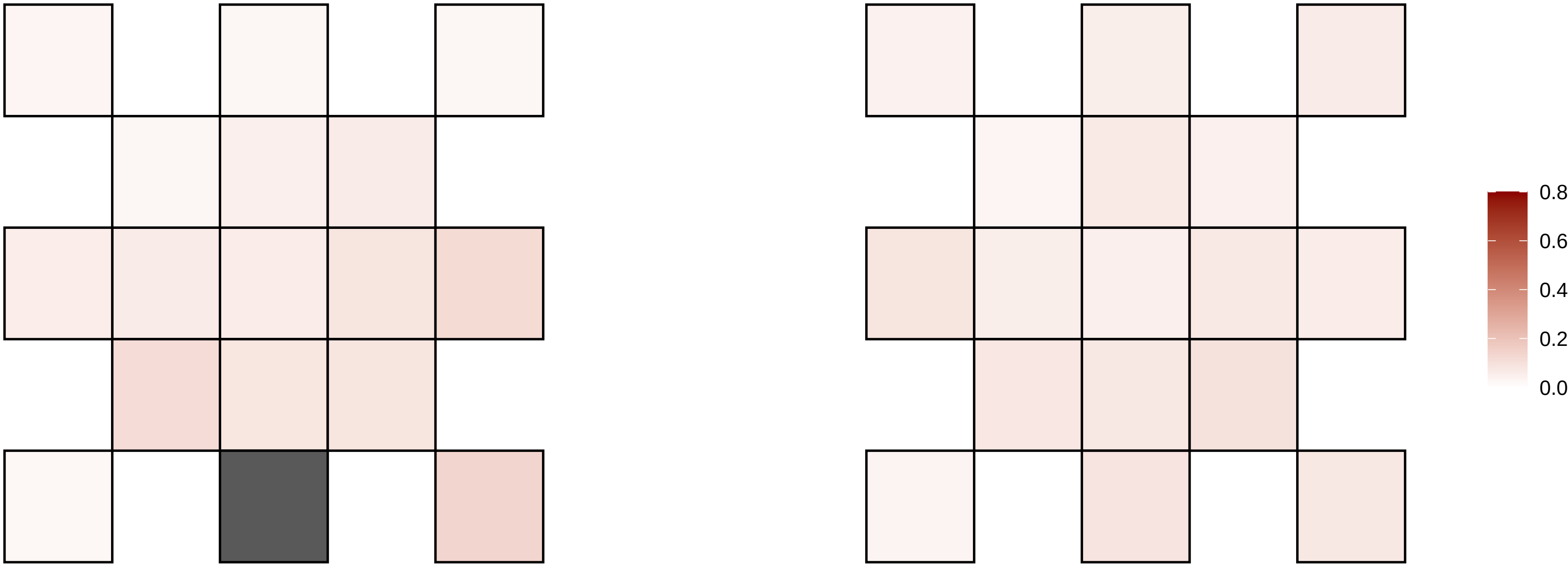

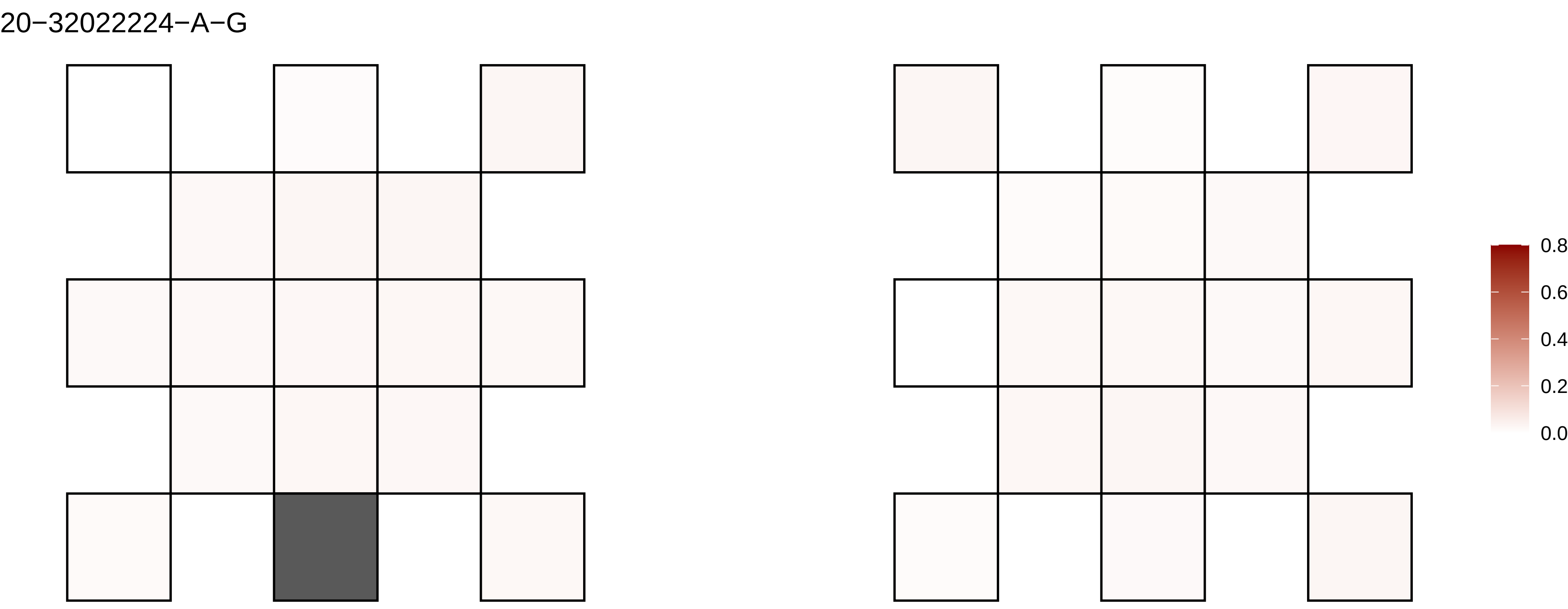
20−32022224−A−G
0.8
0.6
0.4
0.2
0.0

20−34064358−C−T

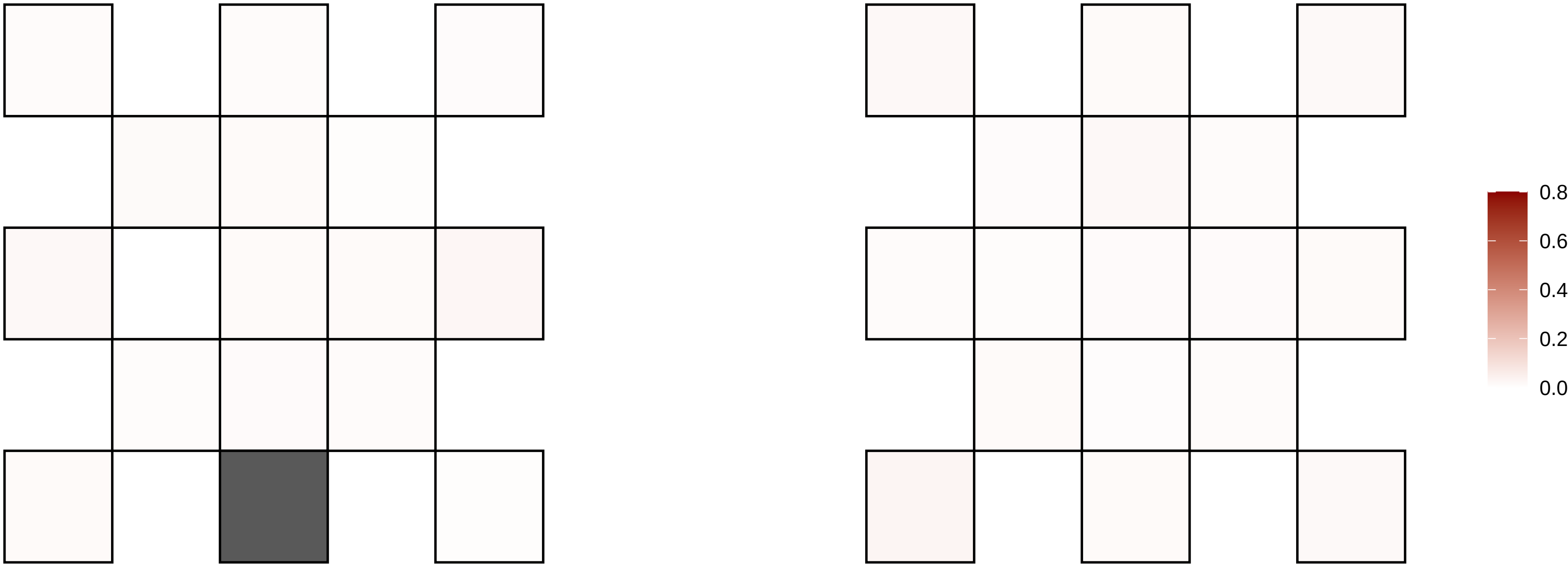

20−38271172−C−T

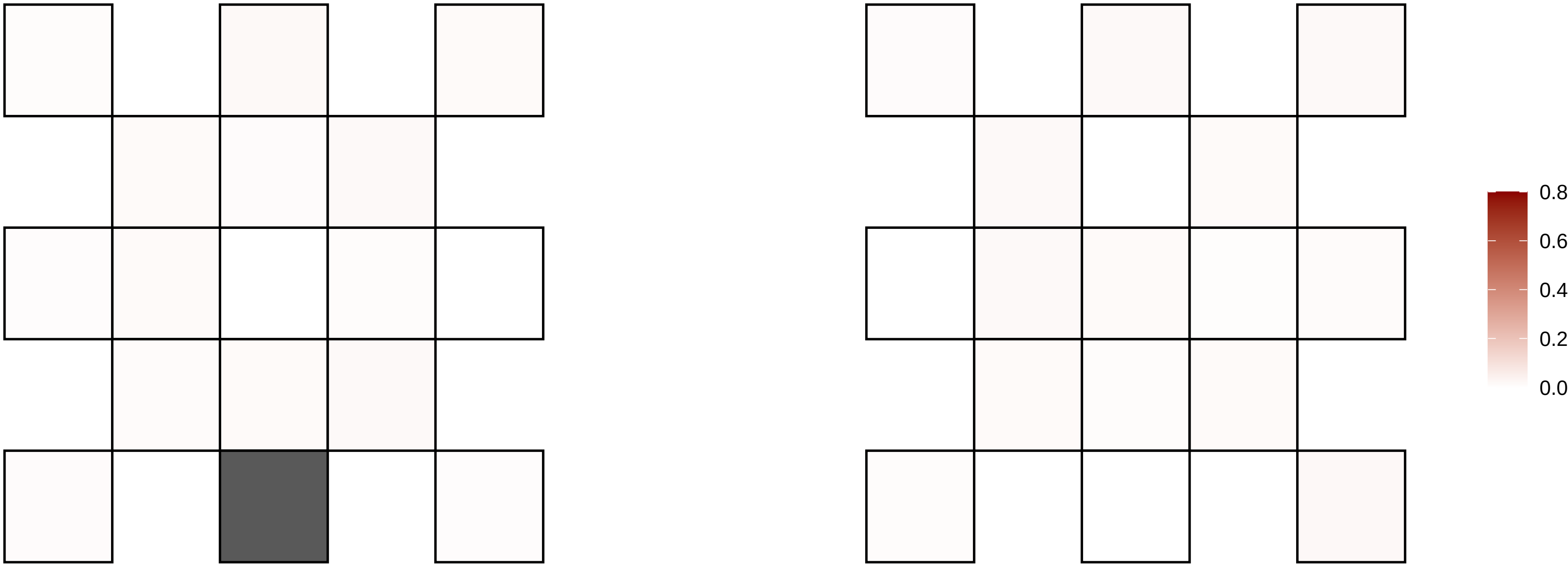

20−39222576−G−T

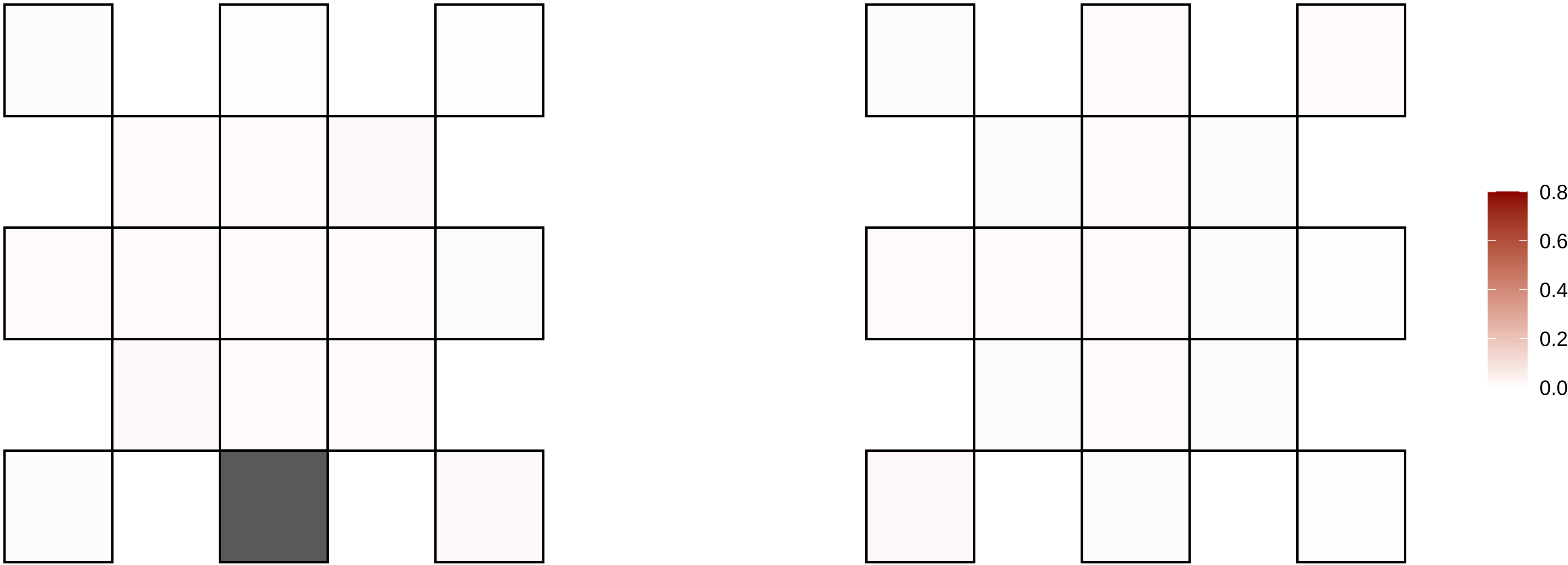

20−39714075−G−A

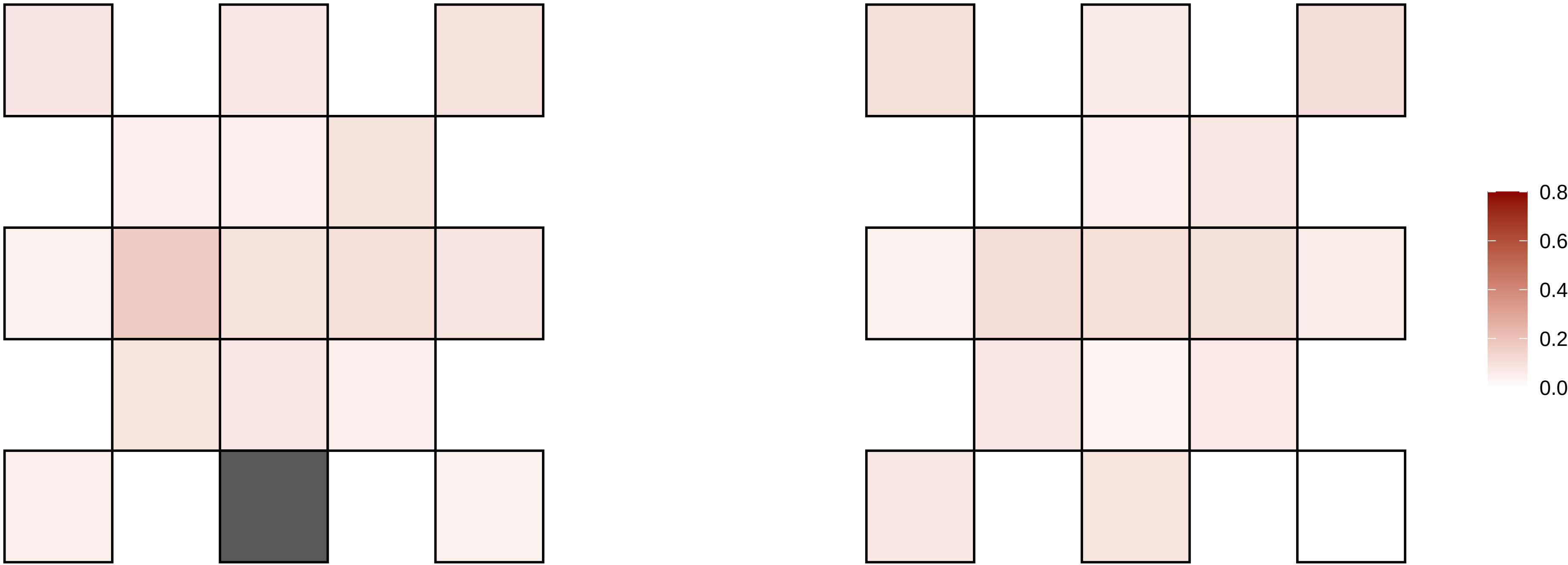

20−41427297−T−C

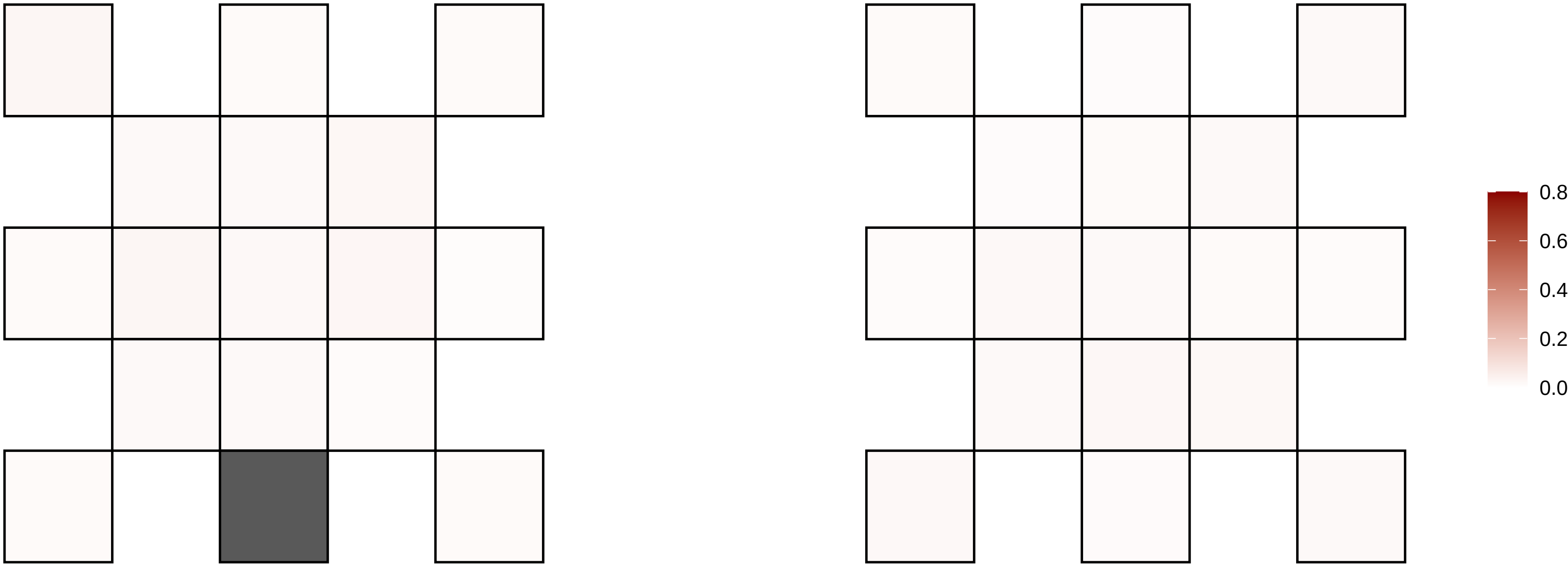

20−52196787−G−A

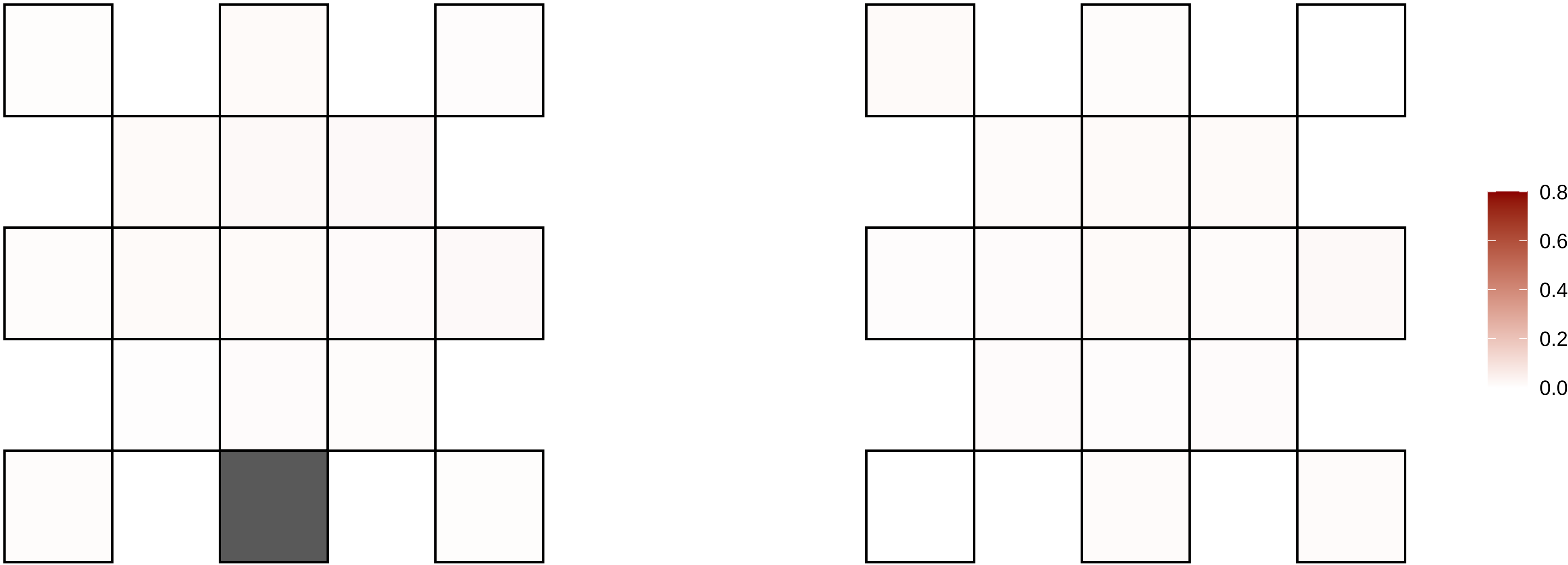

20−5456475−C−T

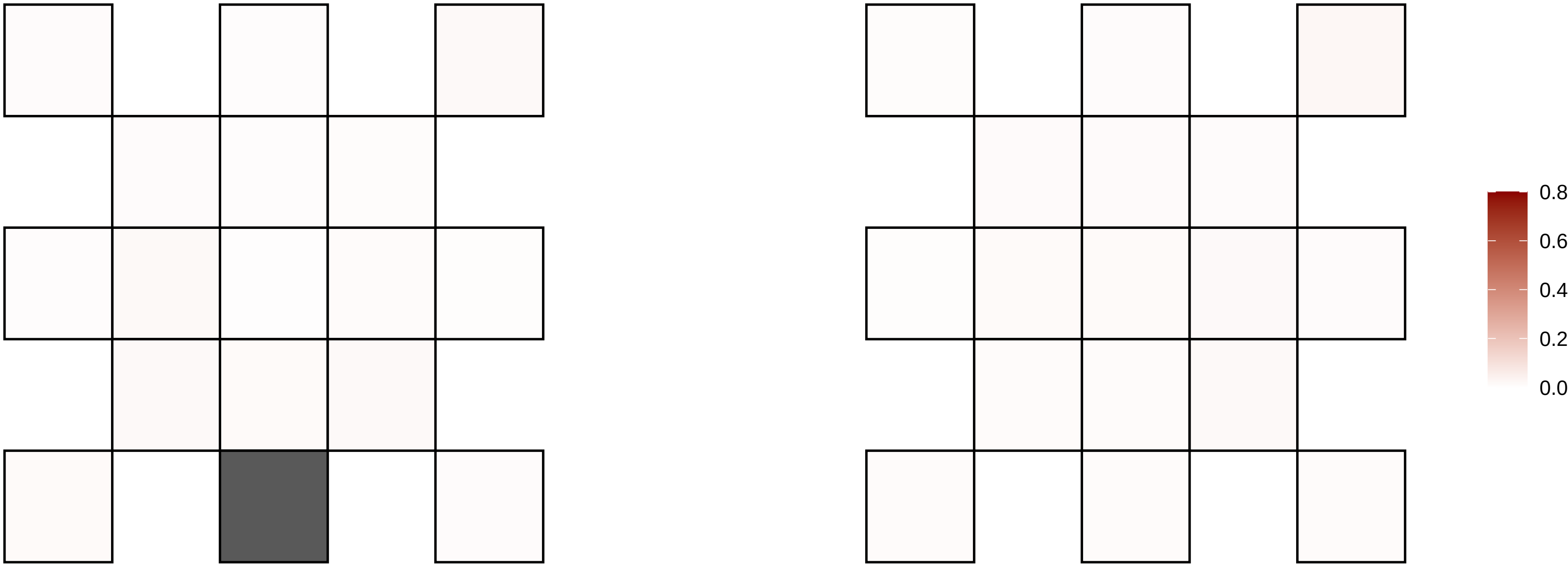

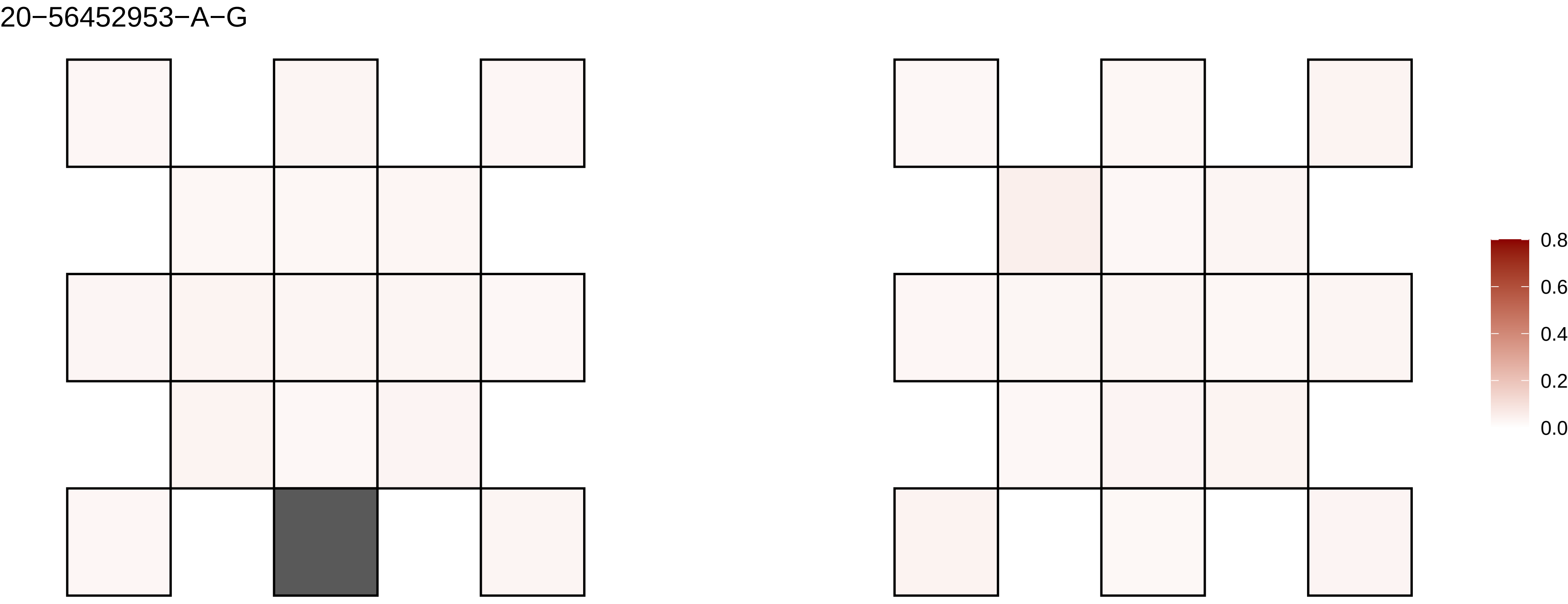
20−56452953−A−G
0.8
0.6
0.4
0.2
0.0

20−61014143−G−A

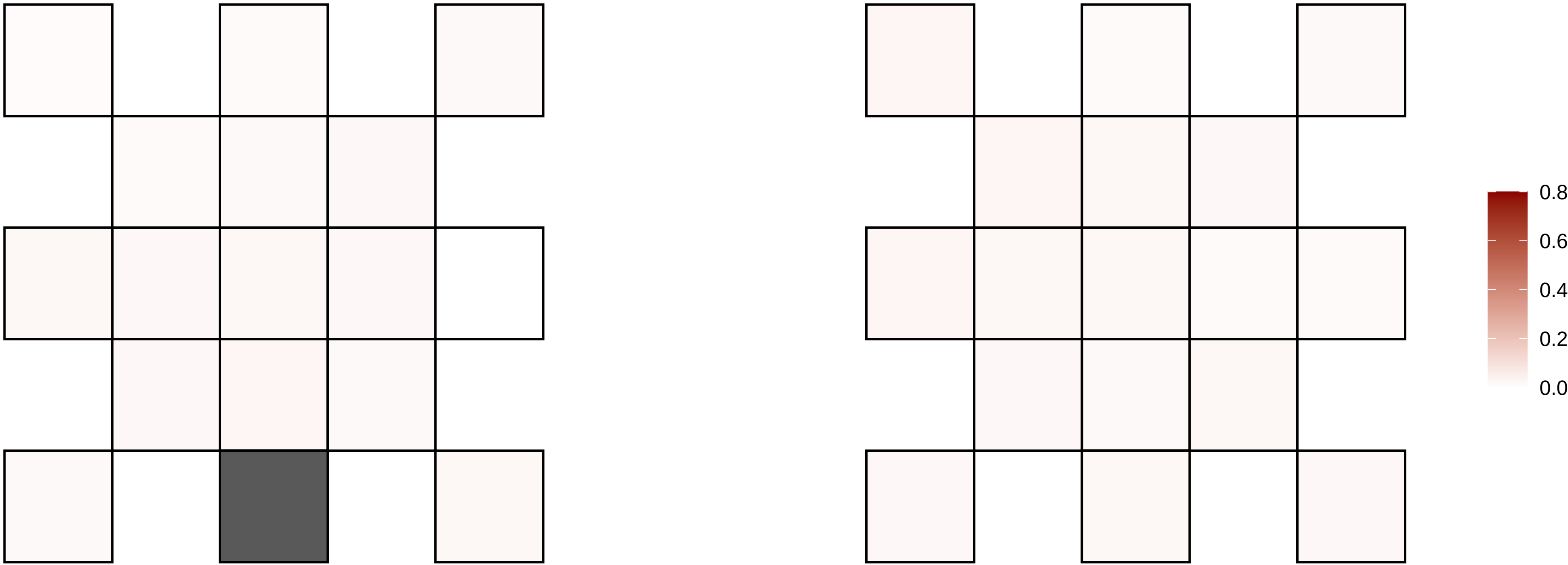

20−9698158−A−G

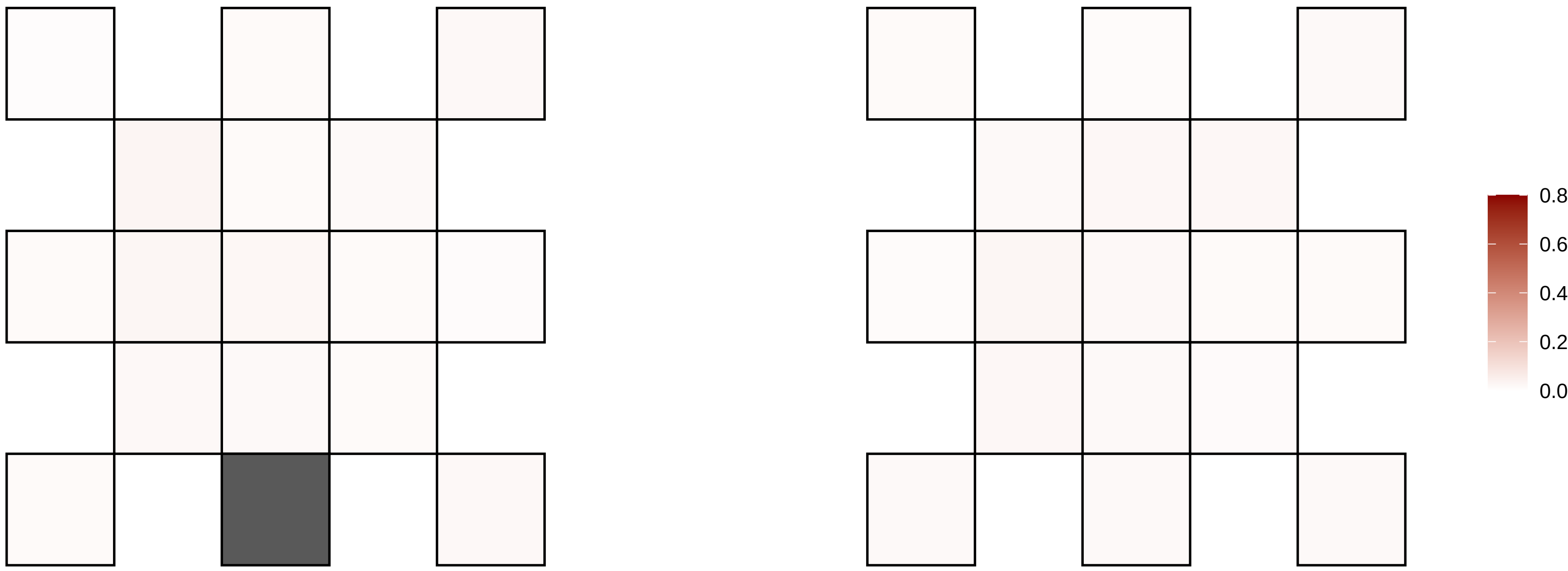

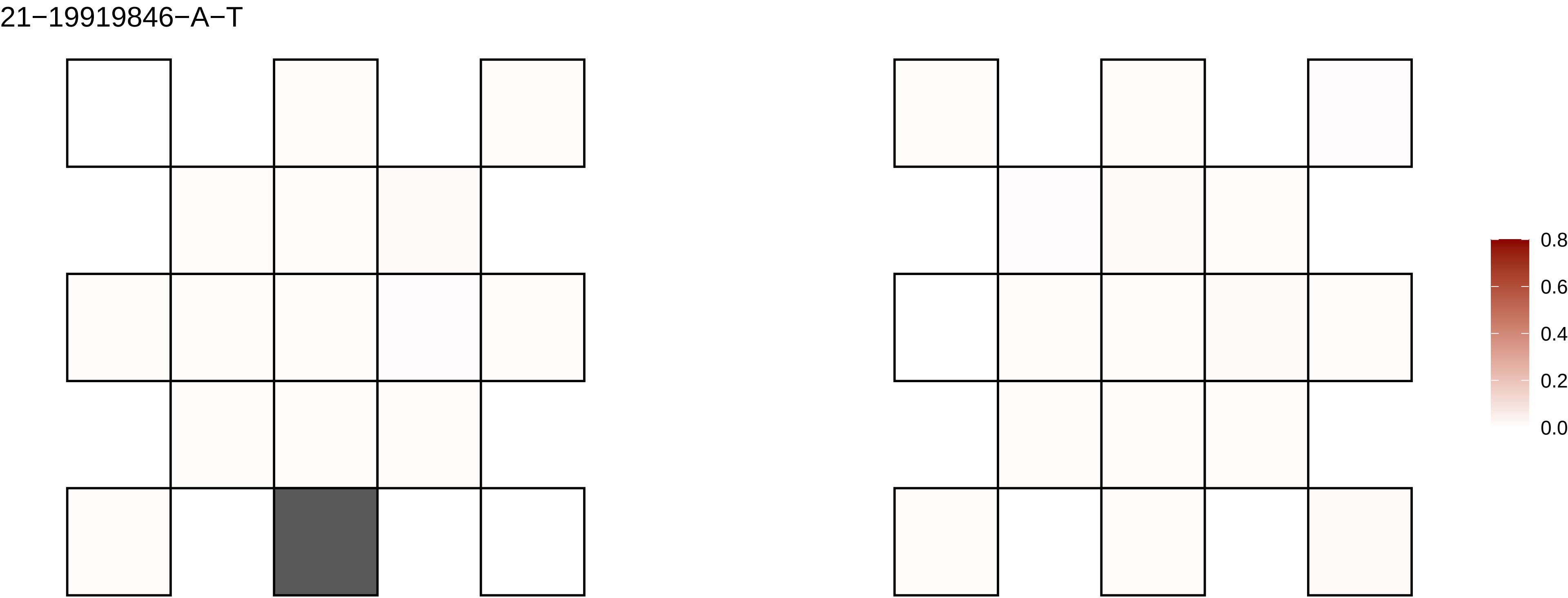

21−19919846−A−T
0.8
0.6
0.4
0.2
0.0

21−21779235−G−T

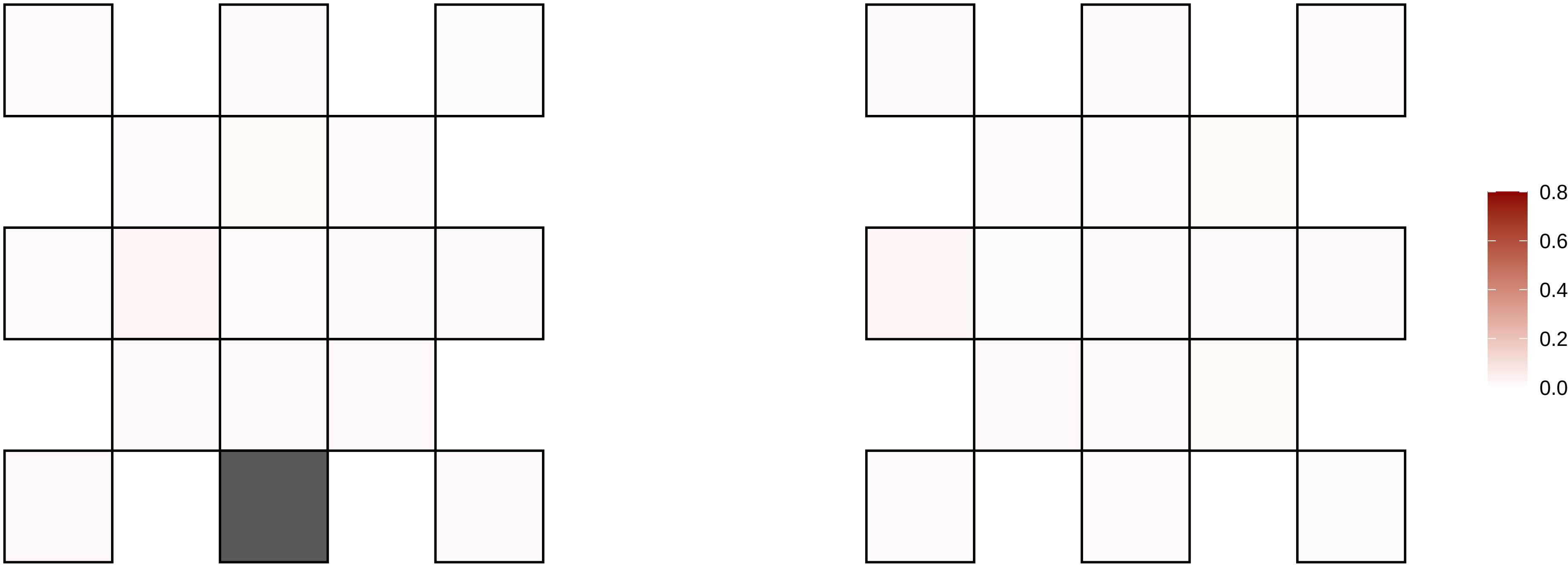

21−24508933−T−C

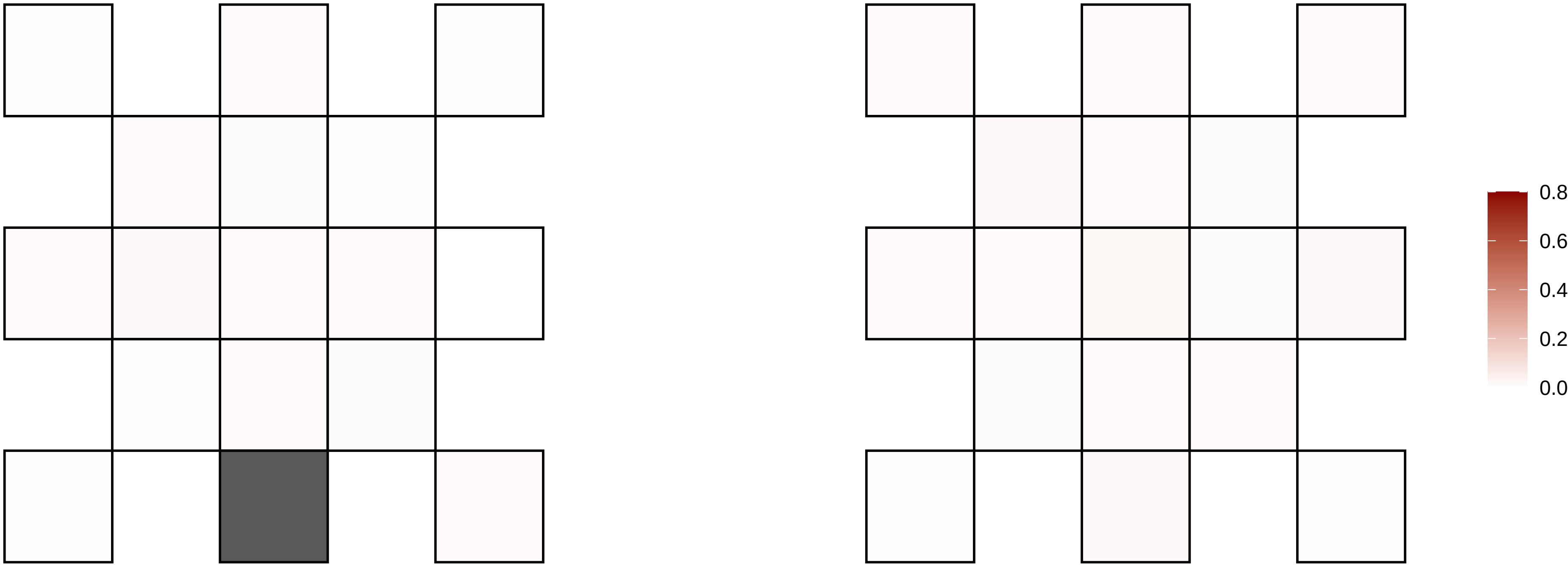

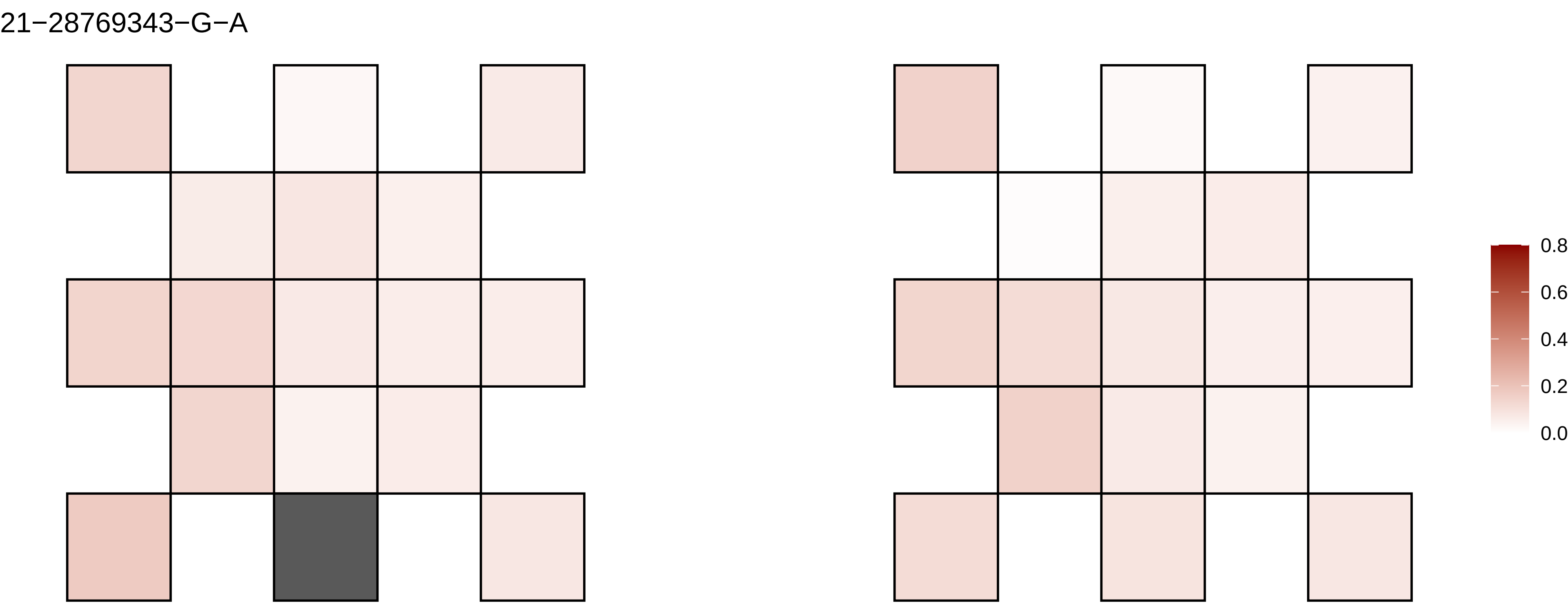

21−28769343−G−A
0.8
0.6
0.4
0.2
0.0

21−32316236−C−T

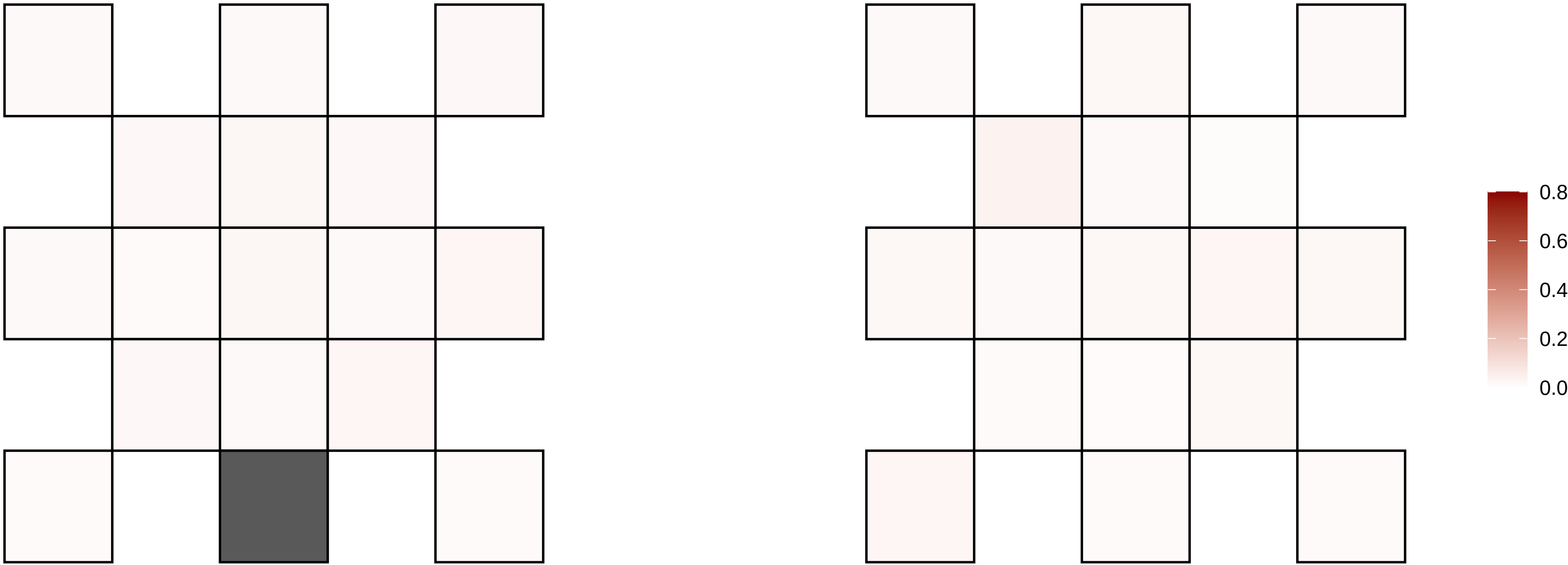

21−32682912−C−T

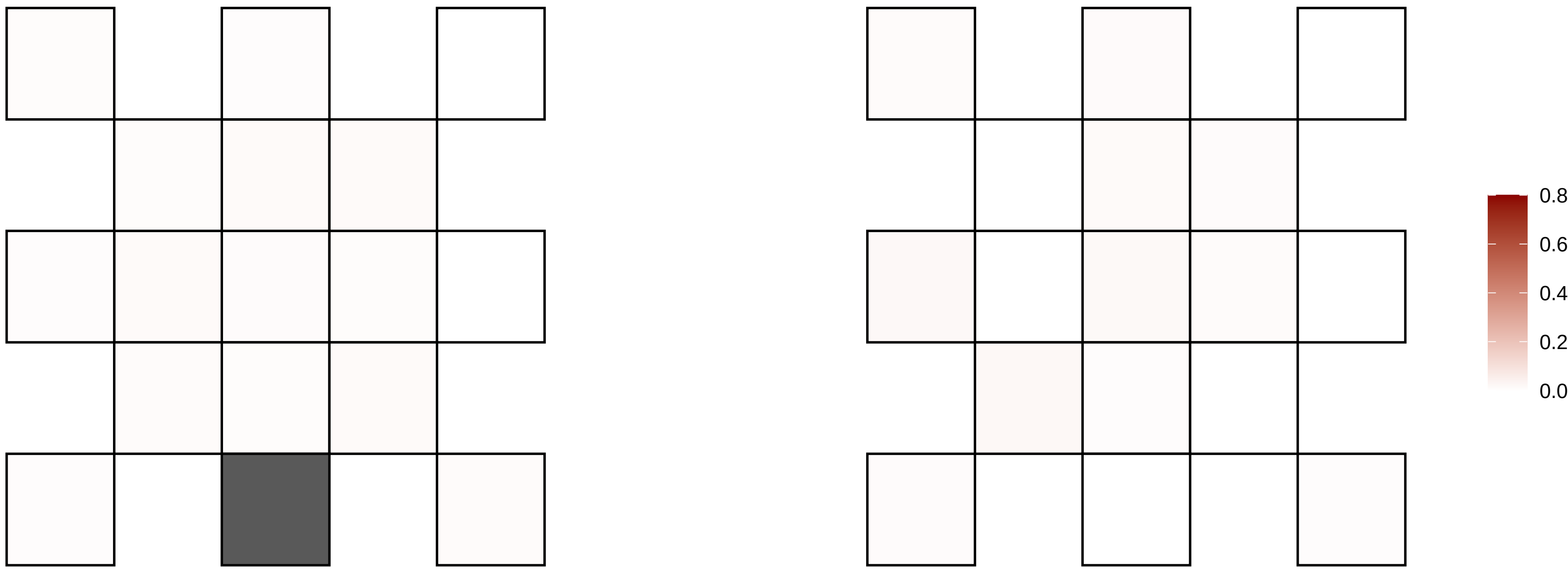

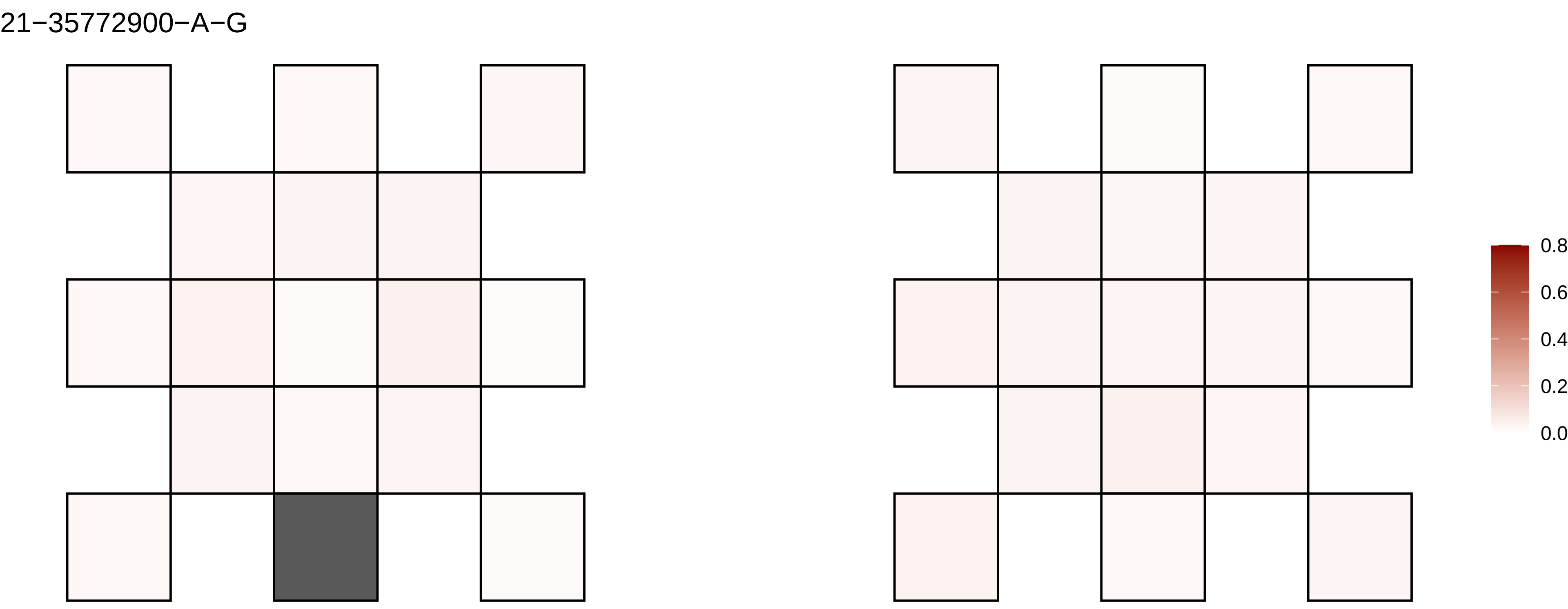
21−35772900−A−G
0.8
0.6
0.4
0.2
0.0

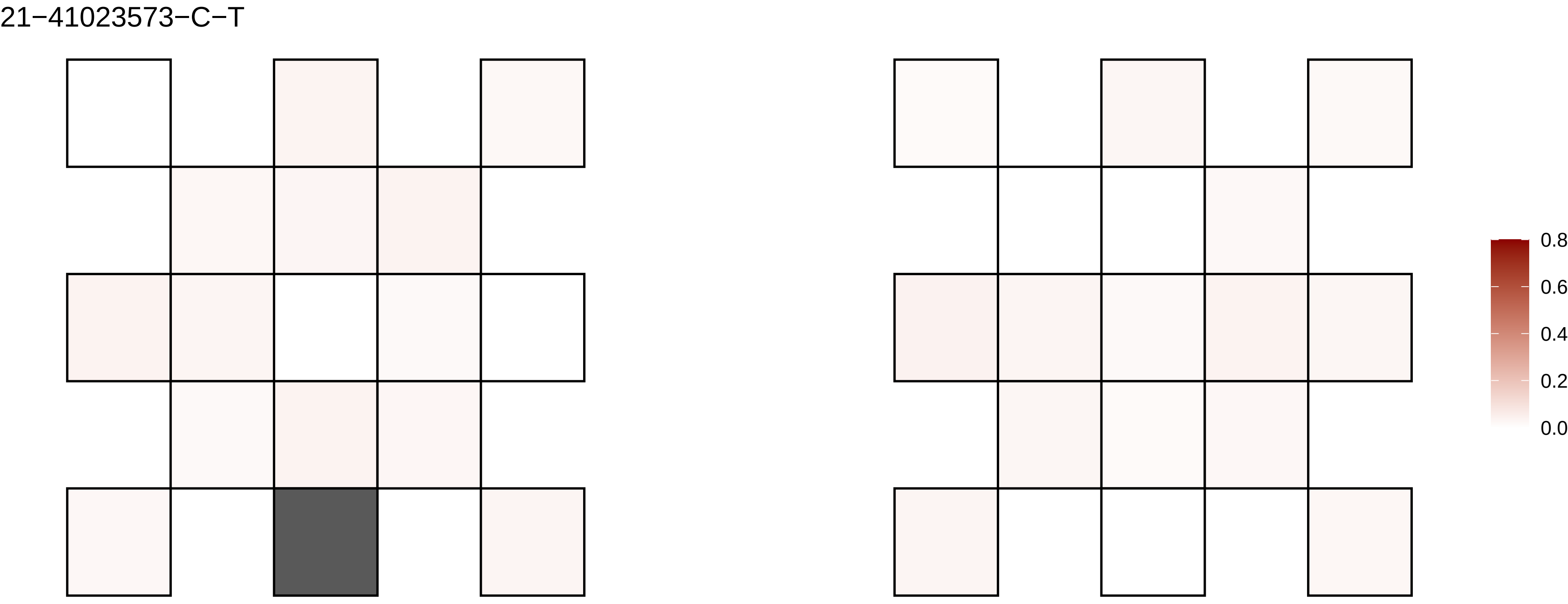
21−41023573−C−T
0.8
0.6
0.4
0.2
0.0

21−41236582−C−T

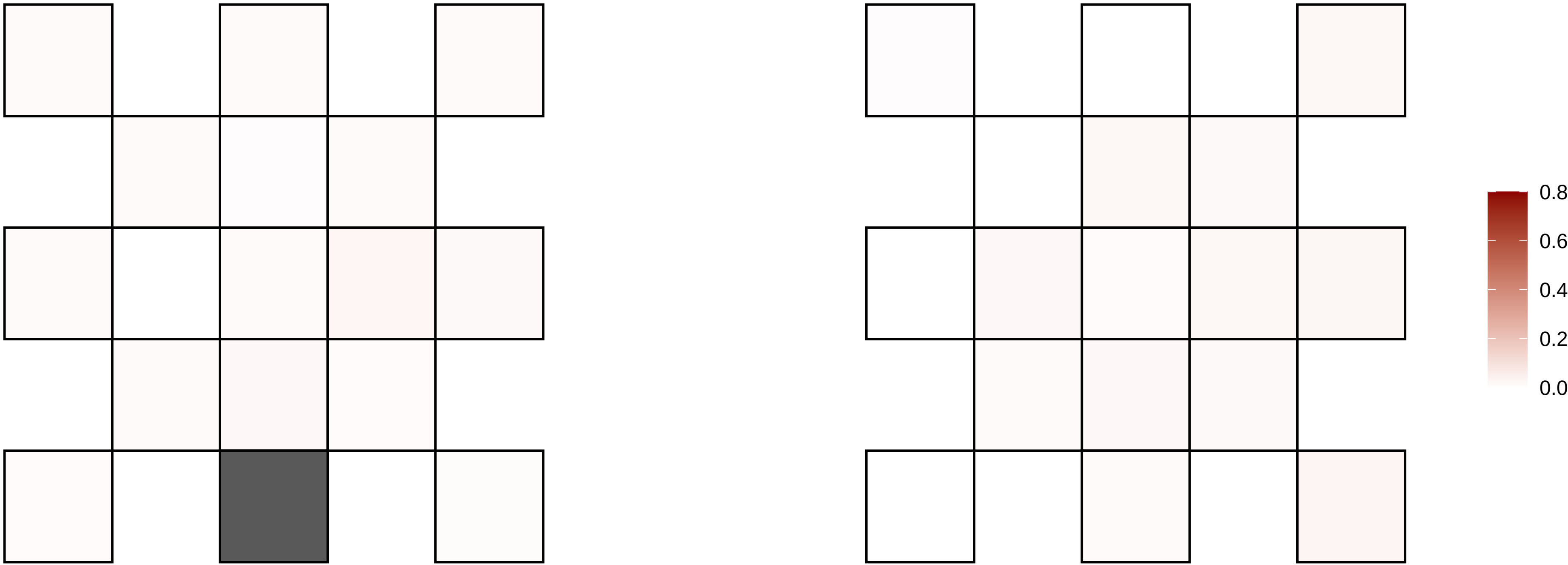

21−41370607−T−C

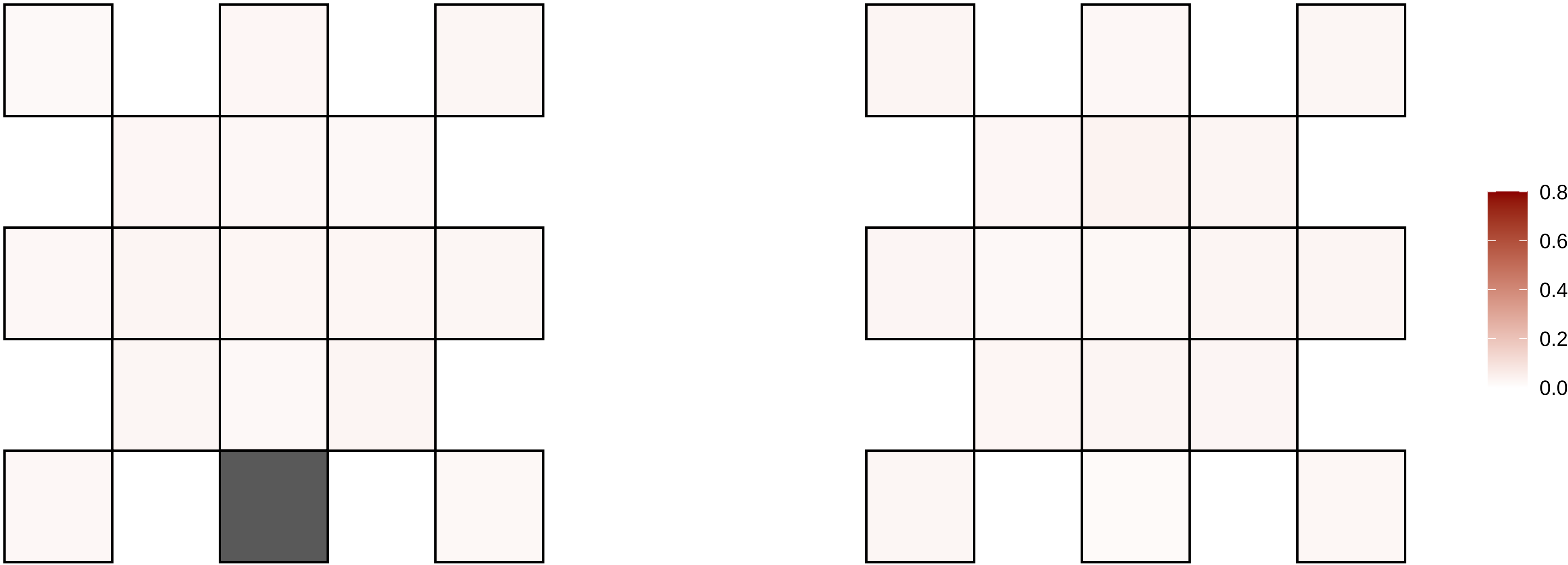

21−42343147−G−A

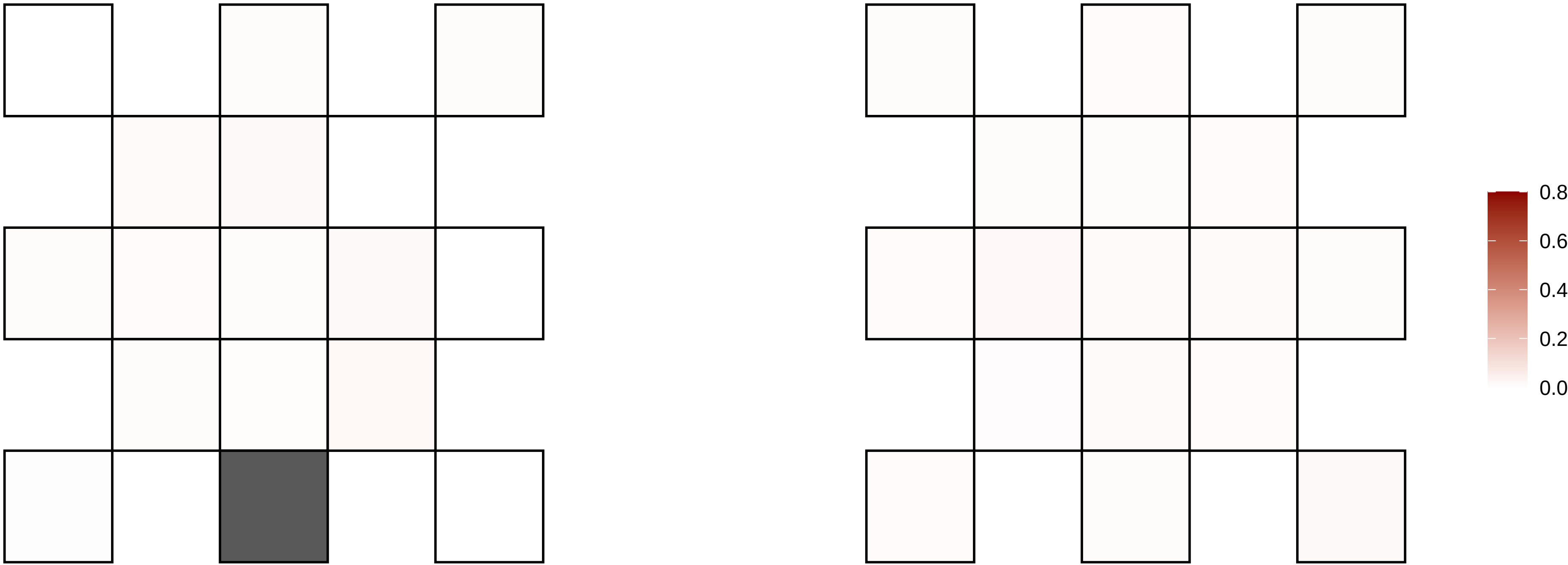

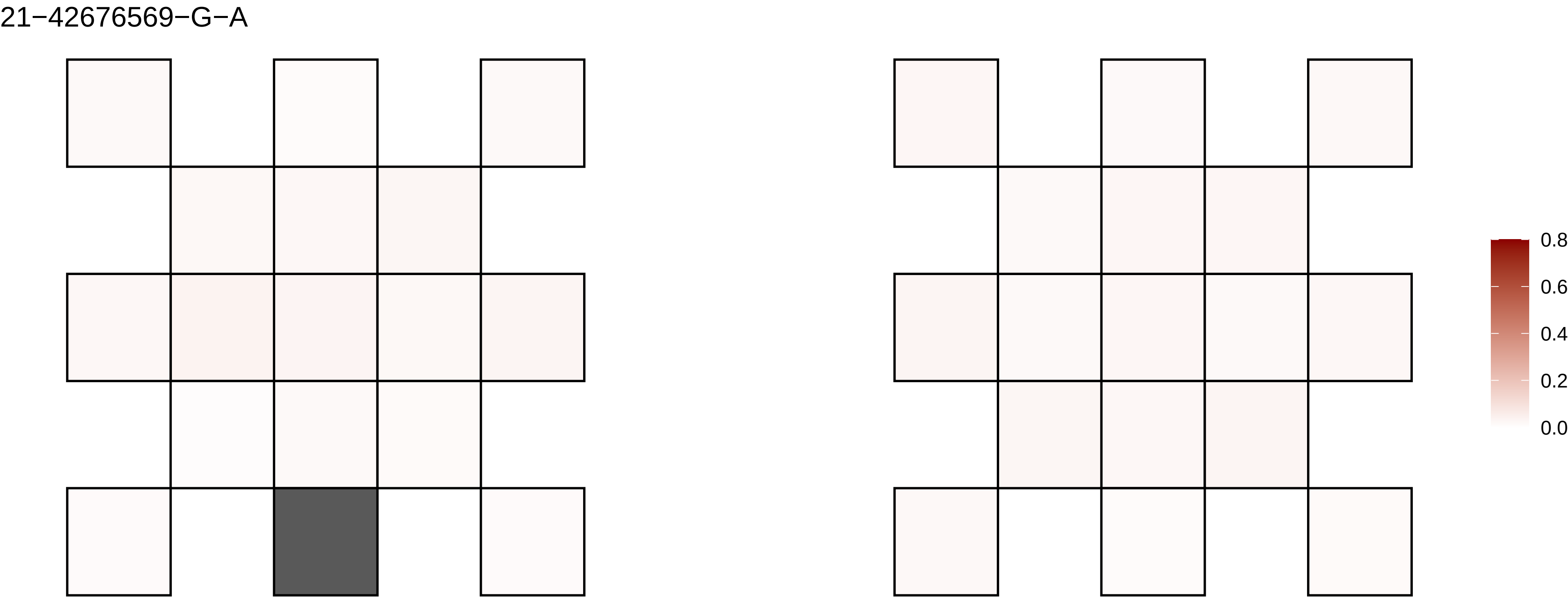
21−42676569−G−A
0.8
0.6
0.4
0.2
0.0

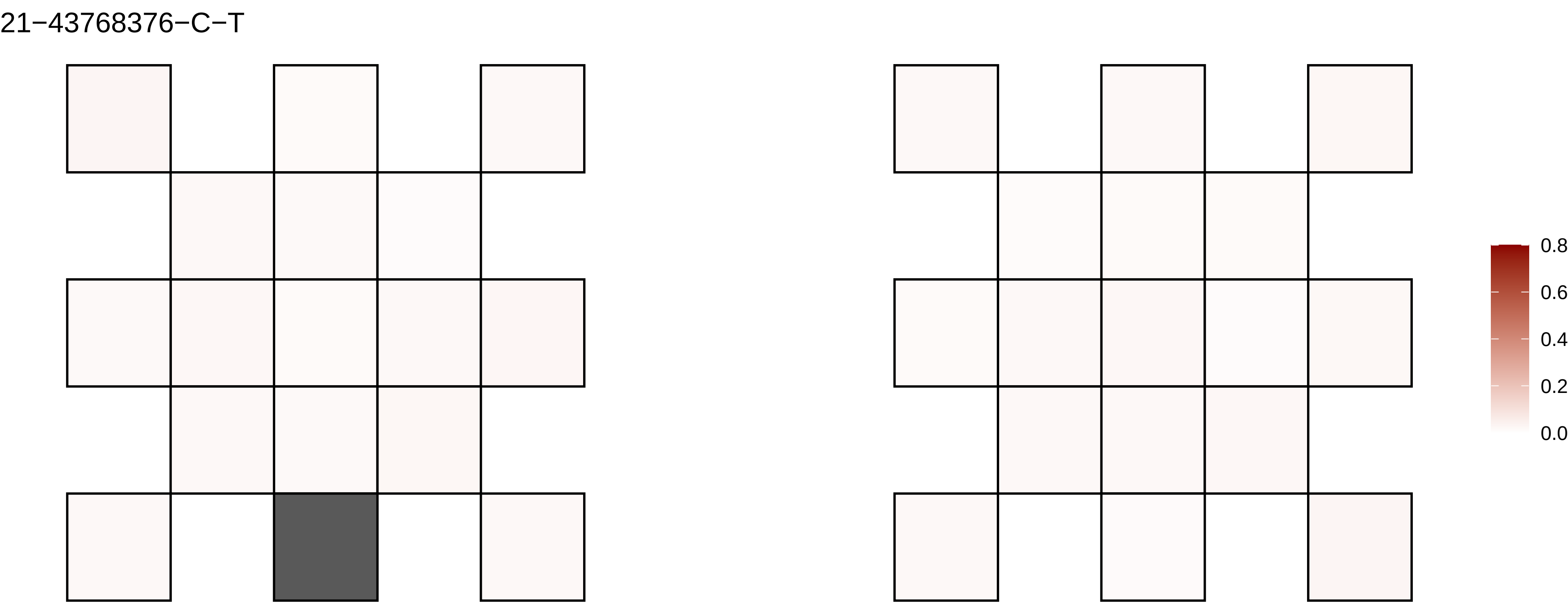
21−43768376−C−T
0.8
0.6
0.4
0.2
0.0

21−47553269−G−A

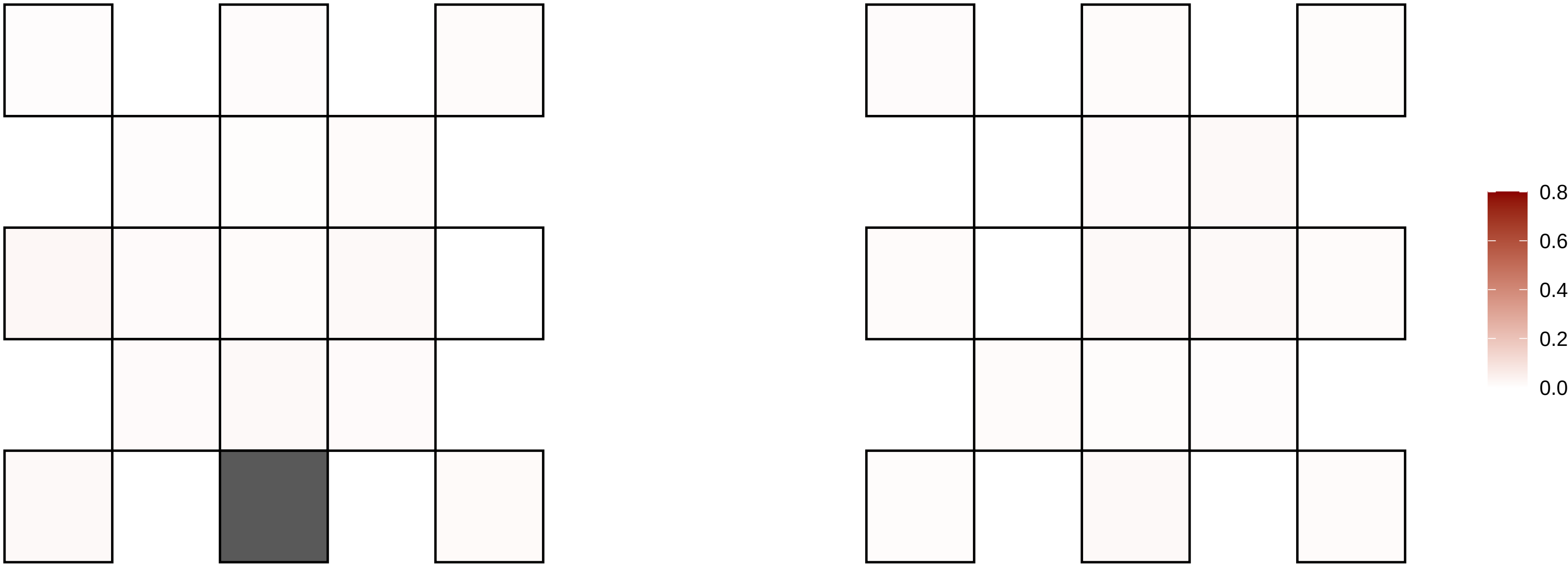

22−23315541−G−A

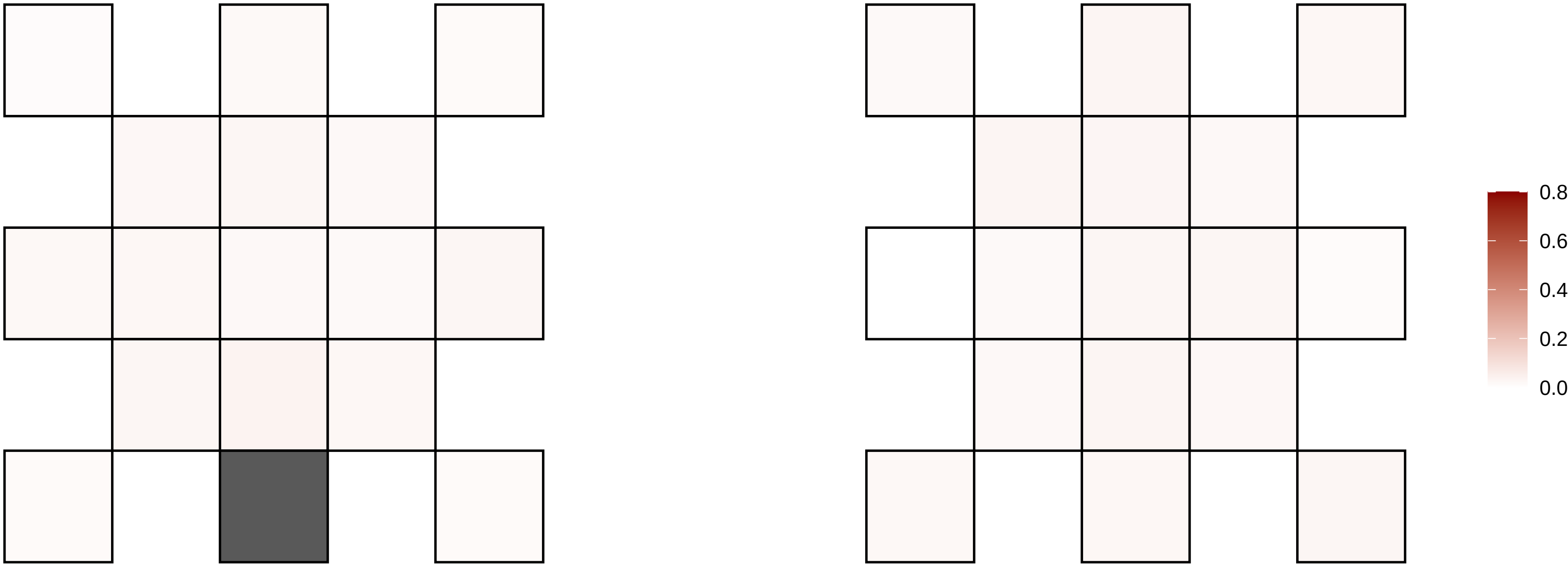

22−46301414−G−T

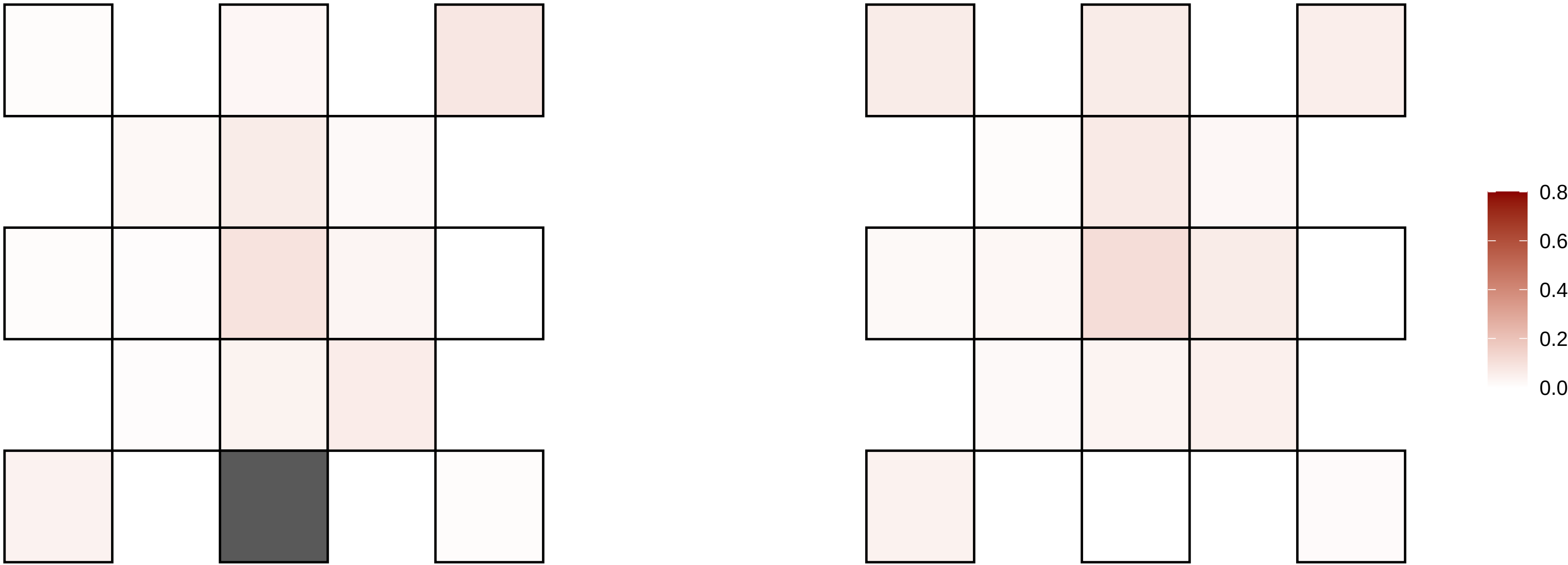

22−47149979−G−A

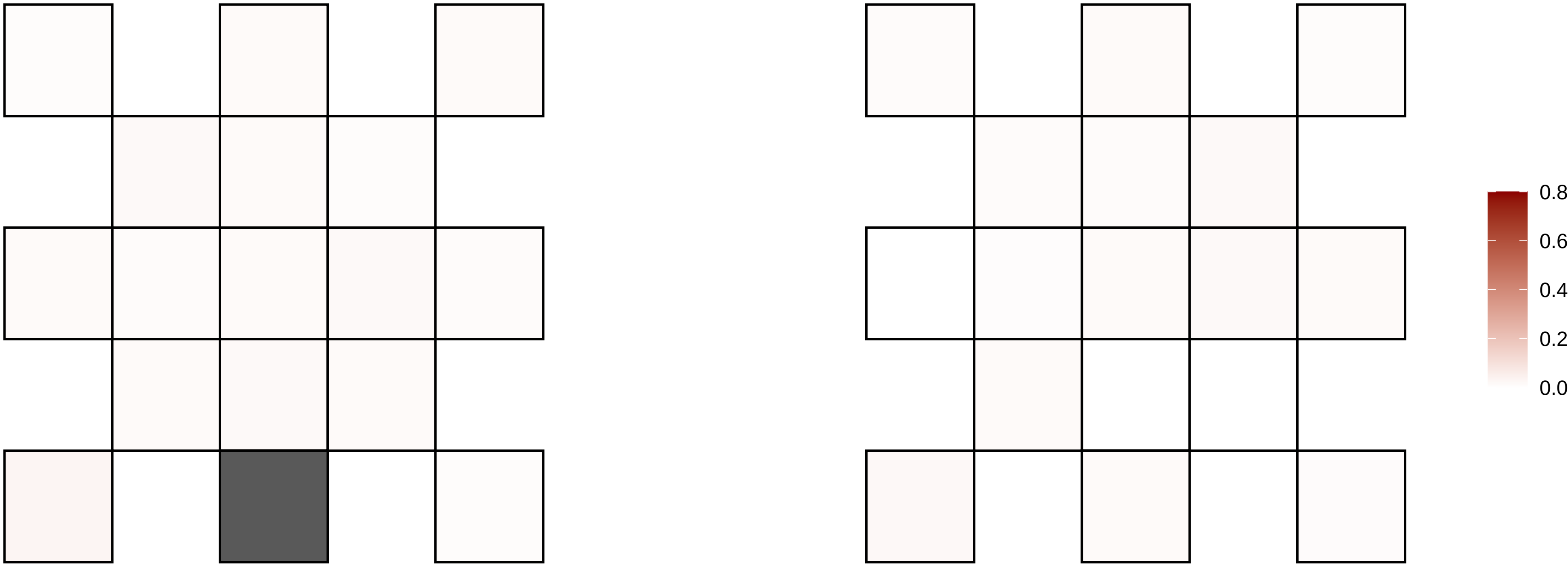

22−49202968−G−A

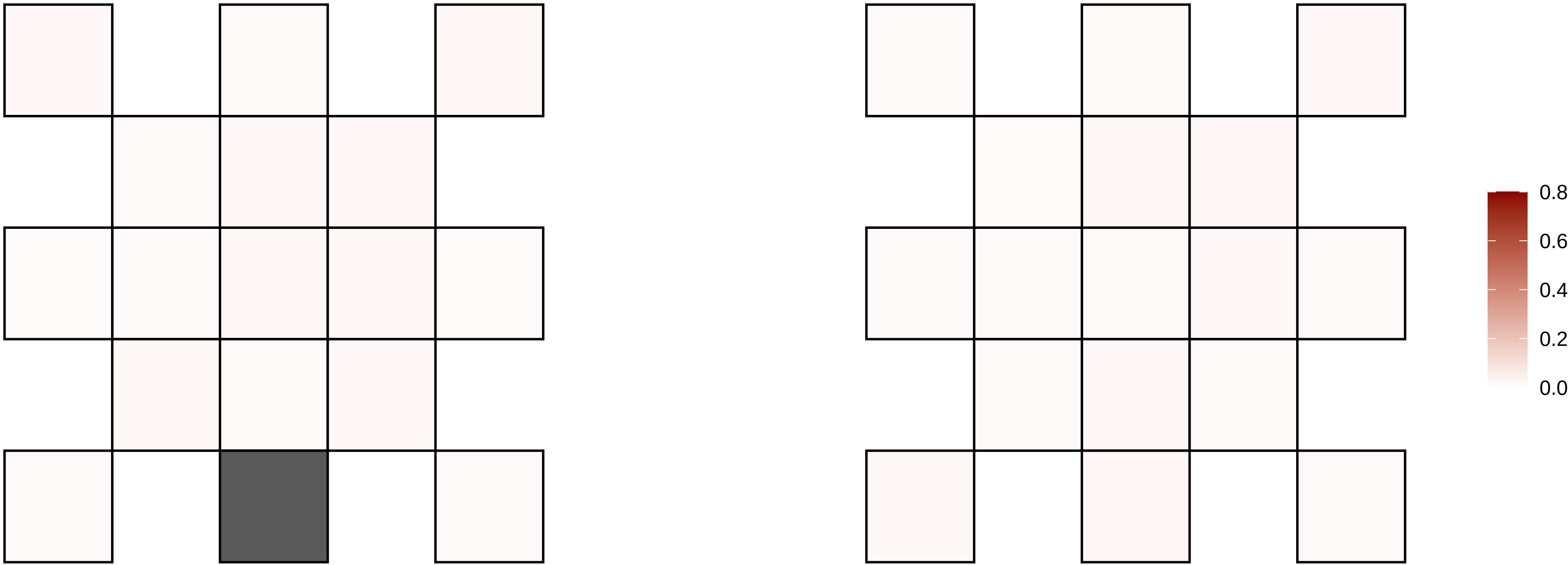

22−49376519−G−A

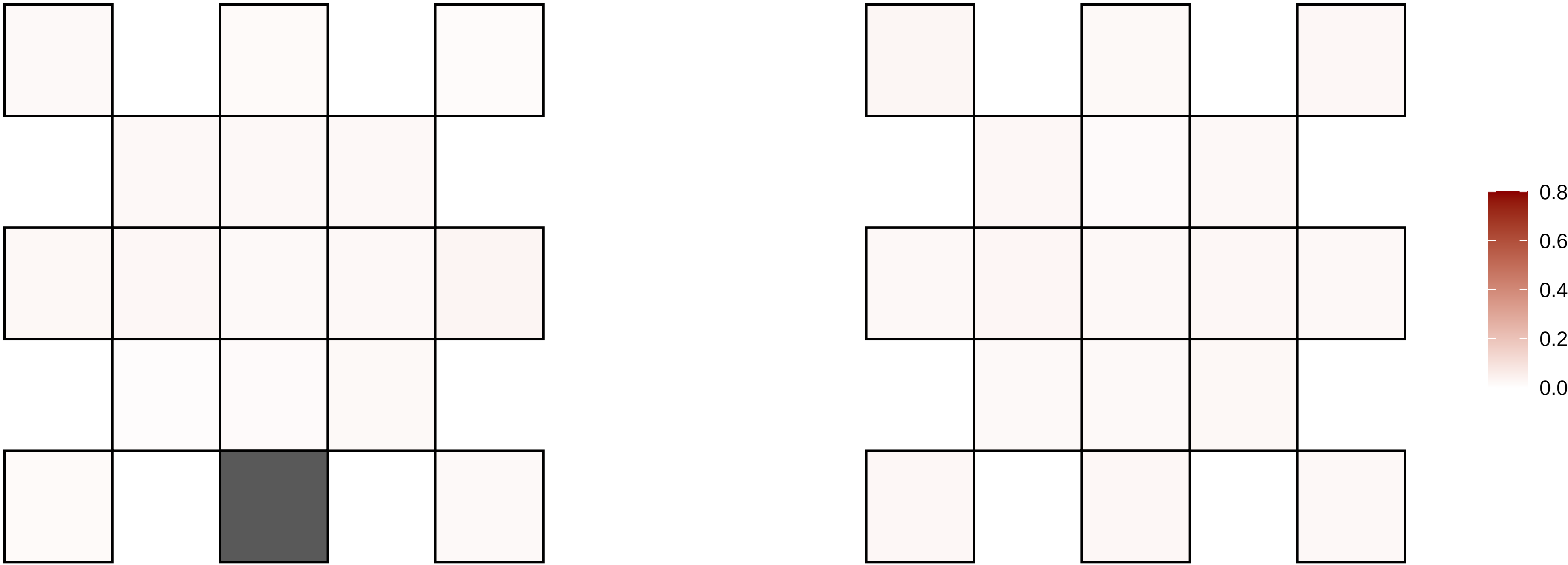

22−49717540−G−A

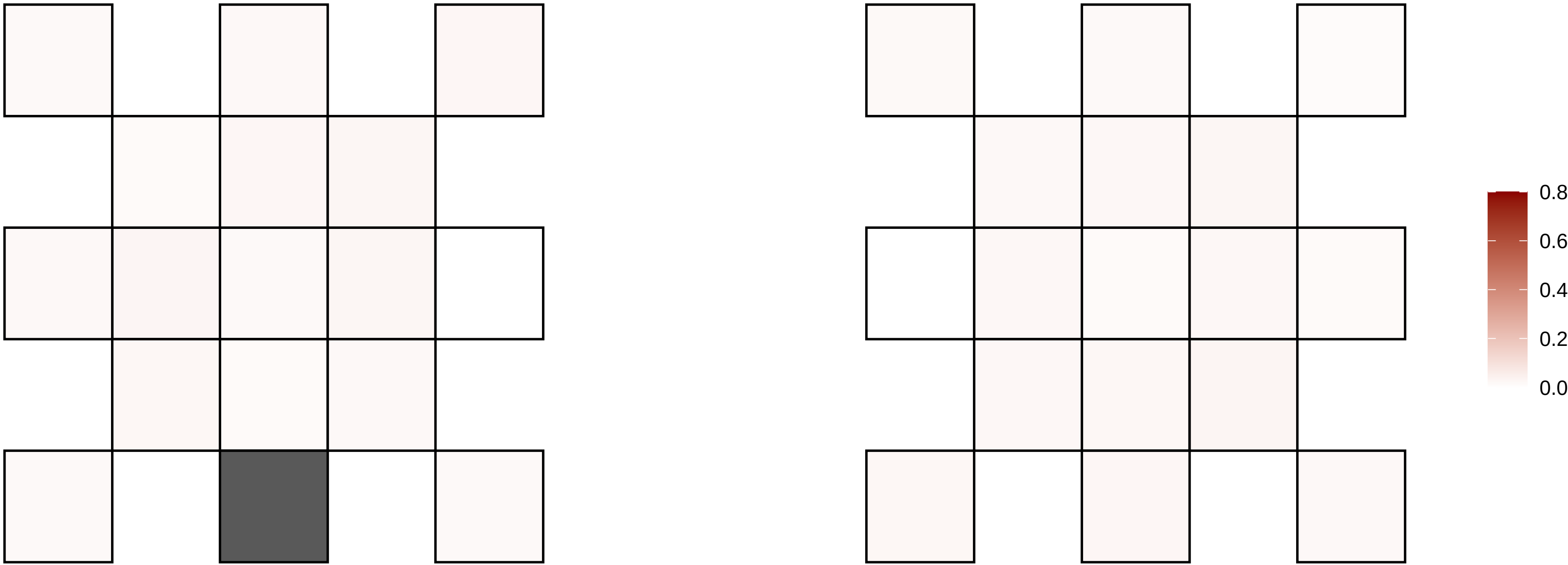

22−49847904−G−A

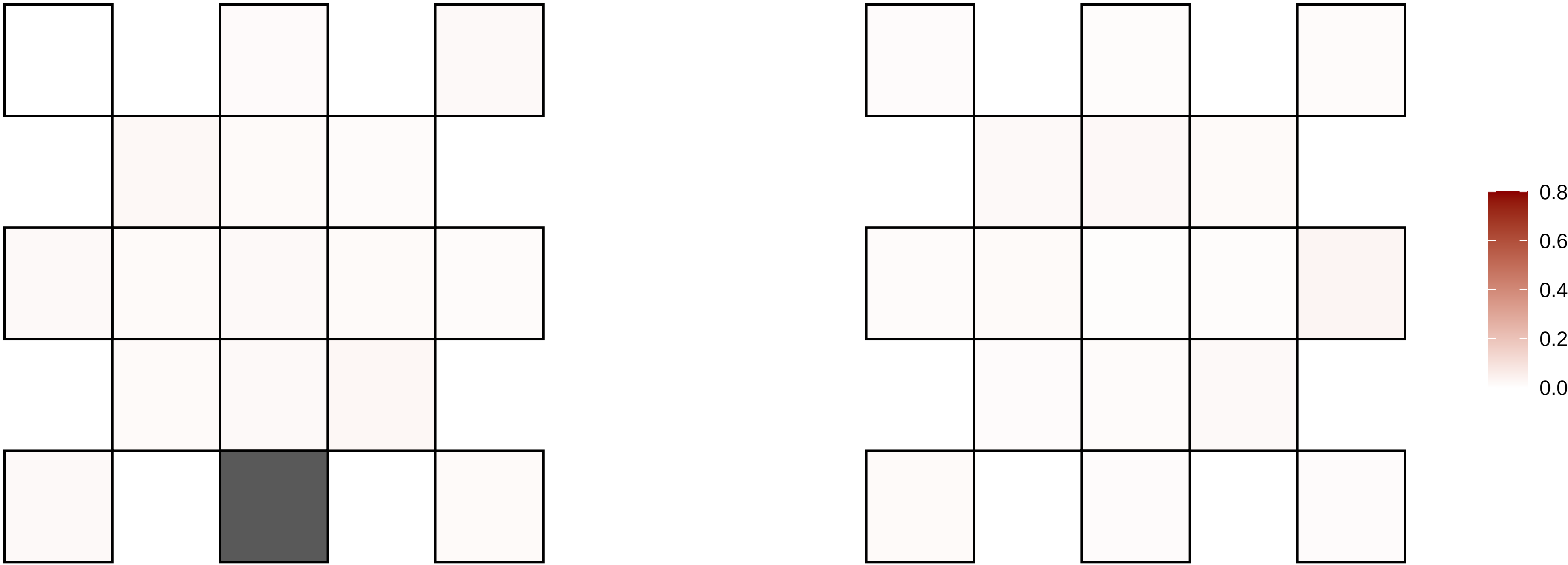

22−51170068−G−A

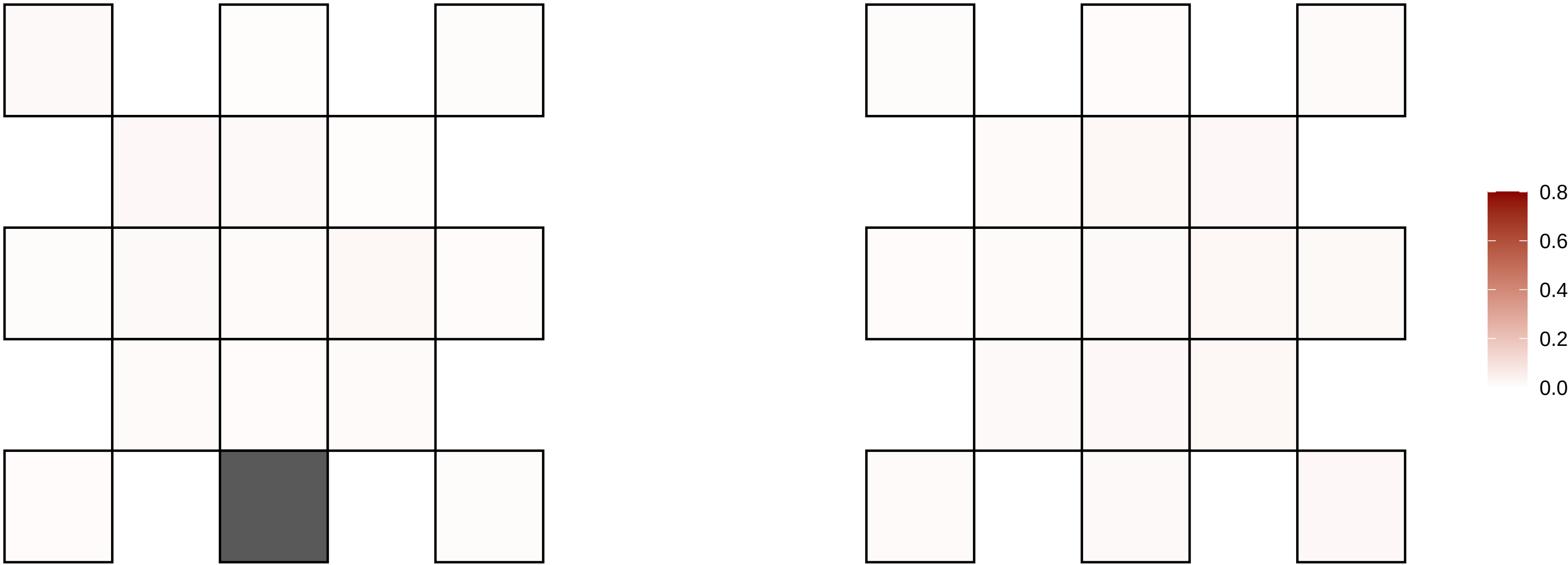

3−101156372−T−C

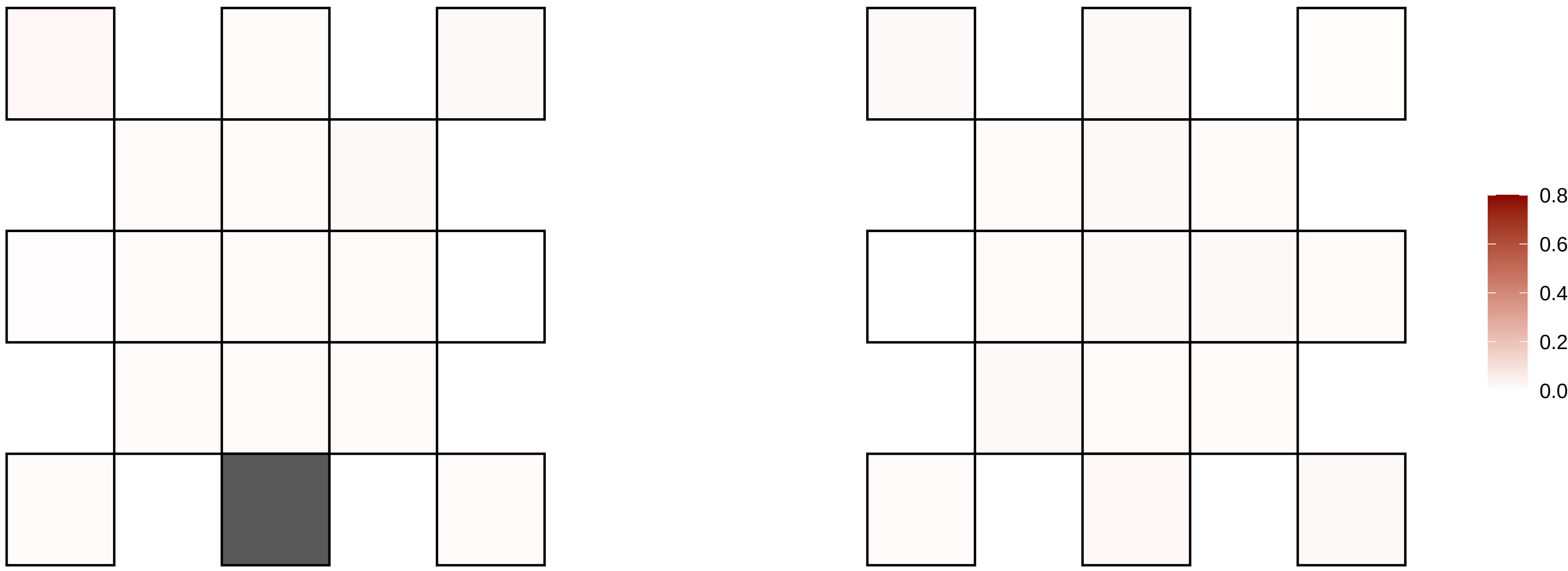

3−103294509−T−G

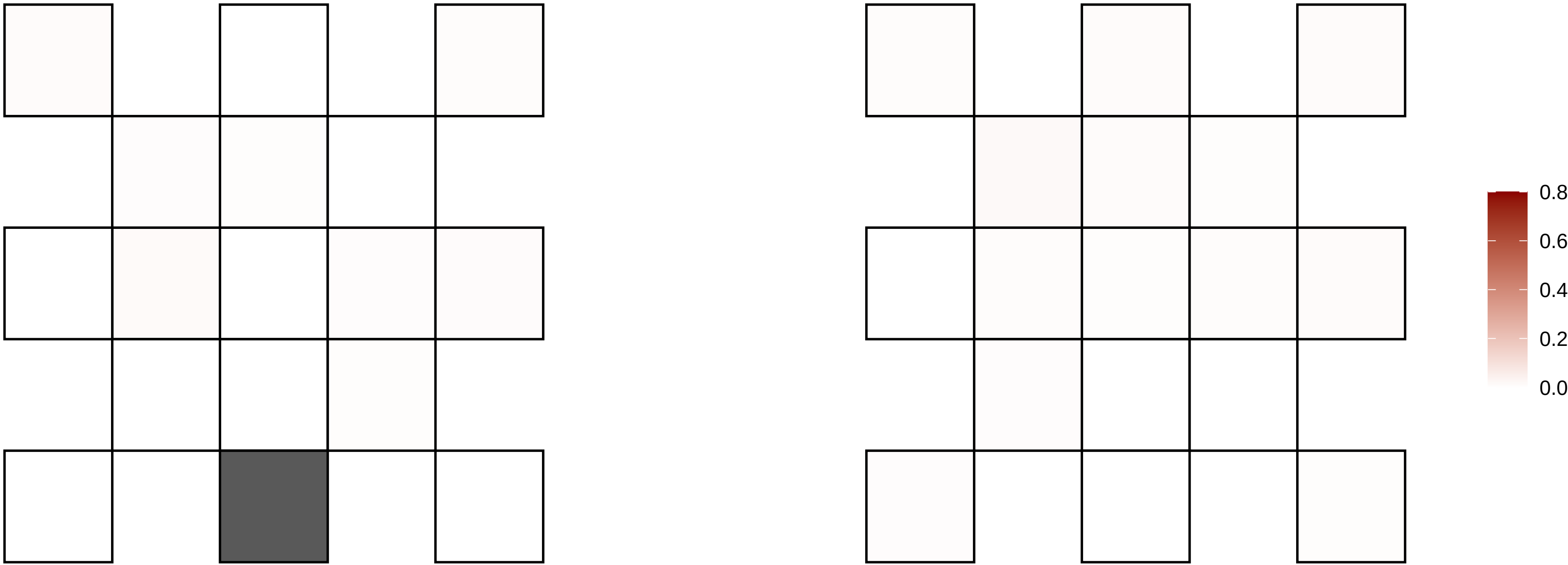

3−106102898−G−T

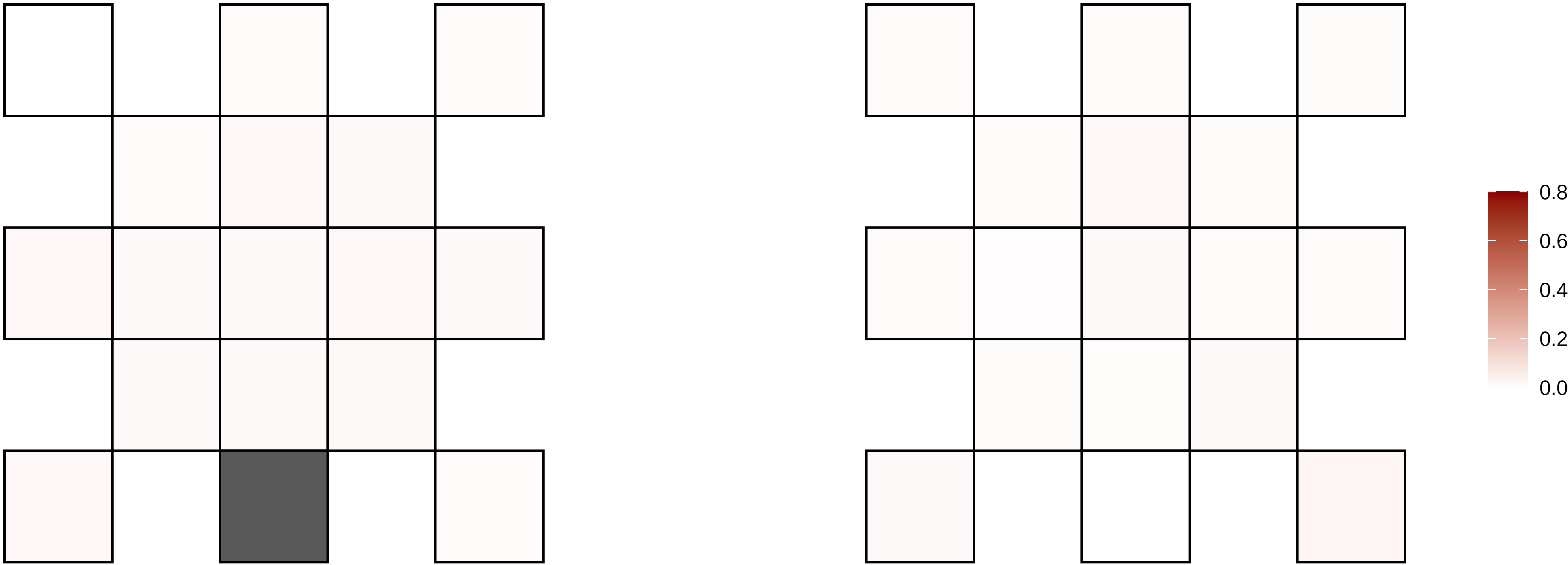

3−107821326−C−T

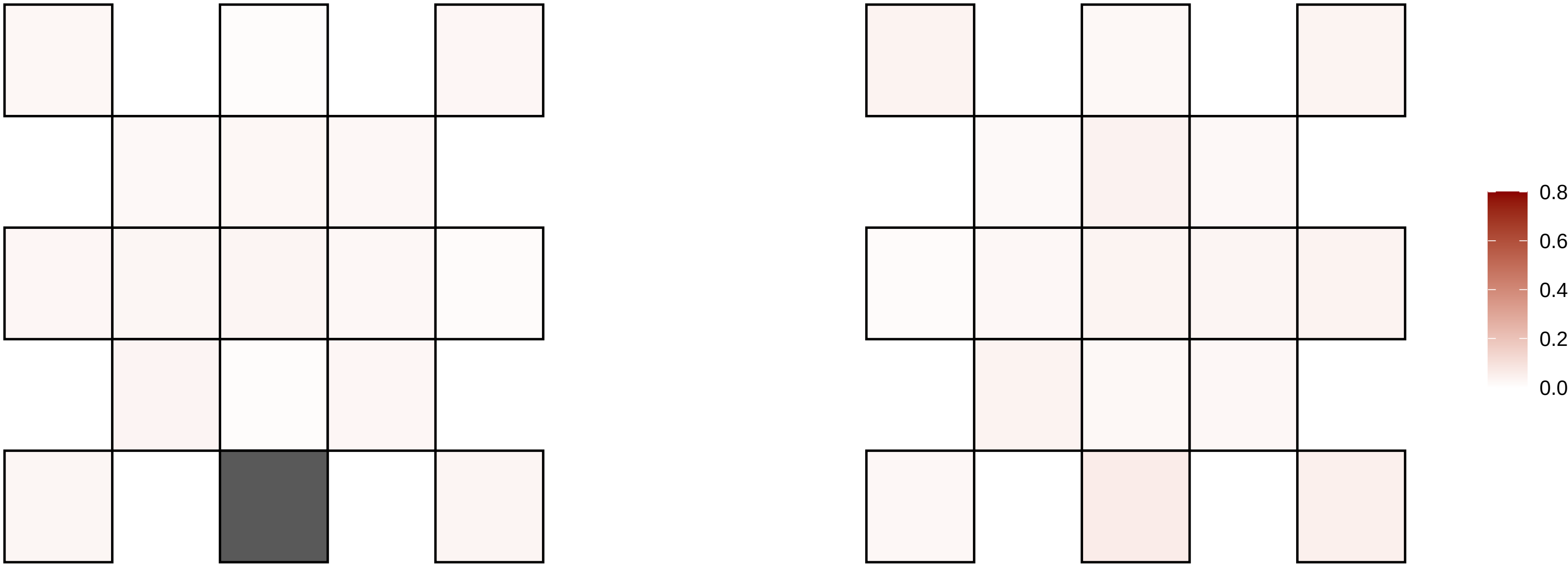

3−108008662−A−T

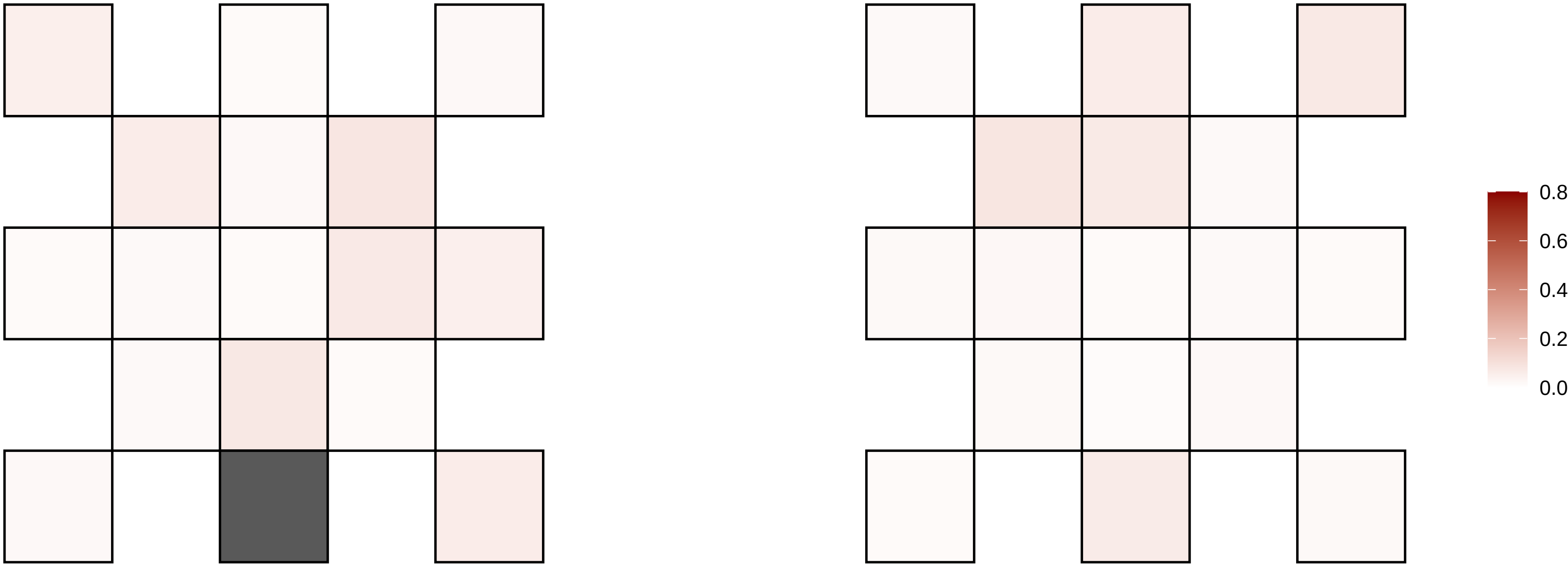

3−109900234−C−T

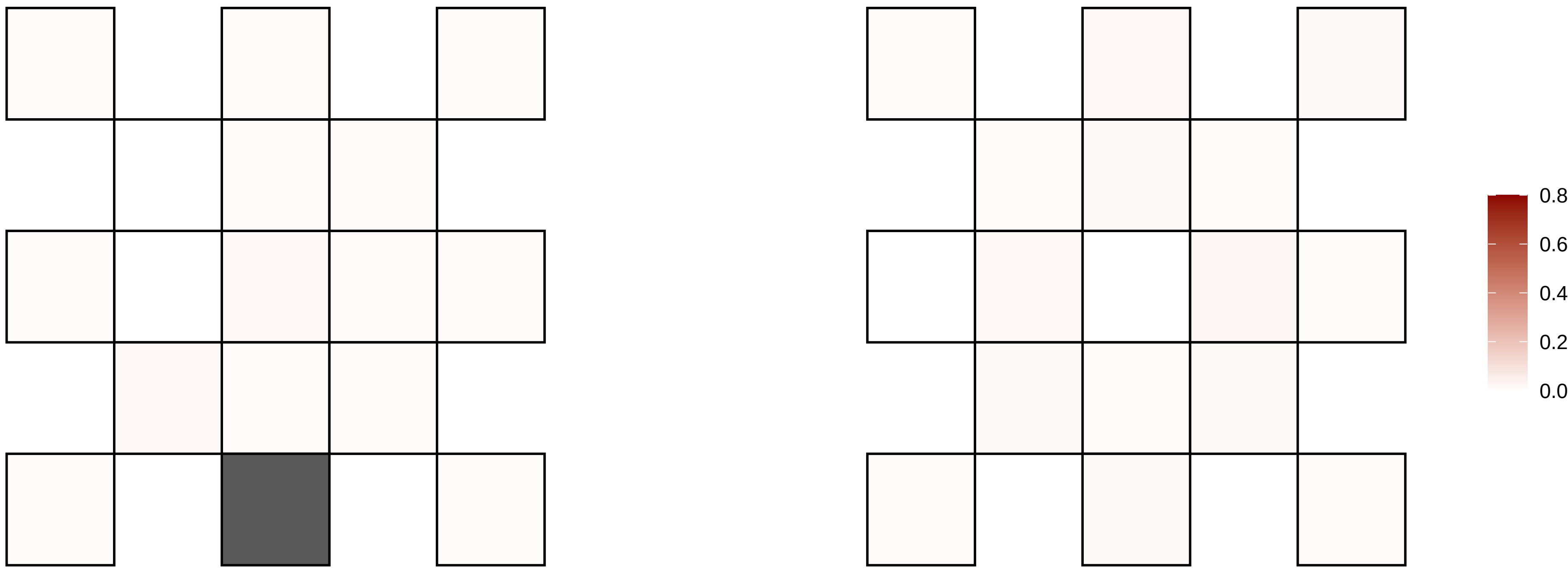

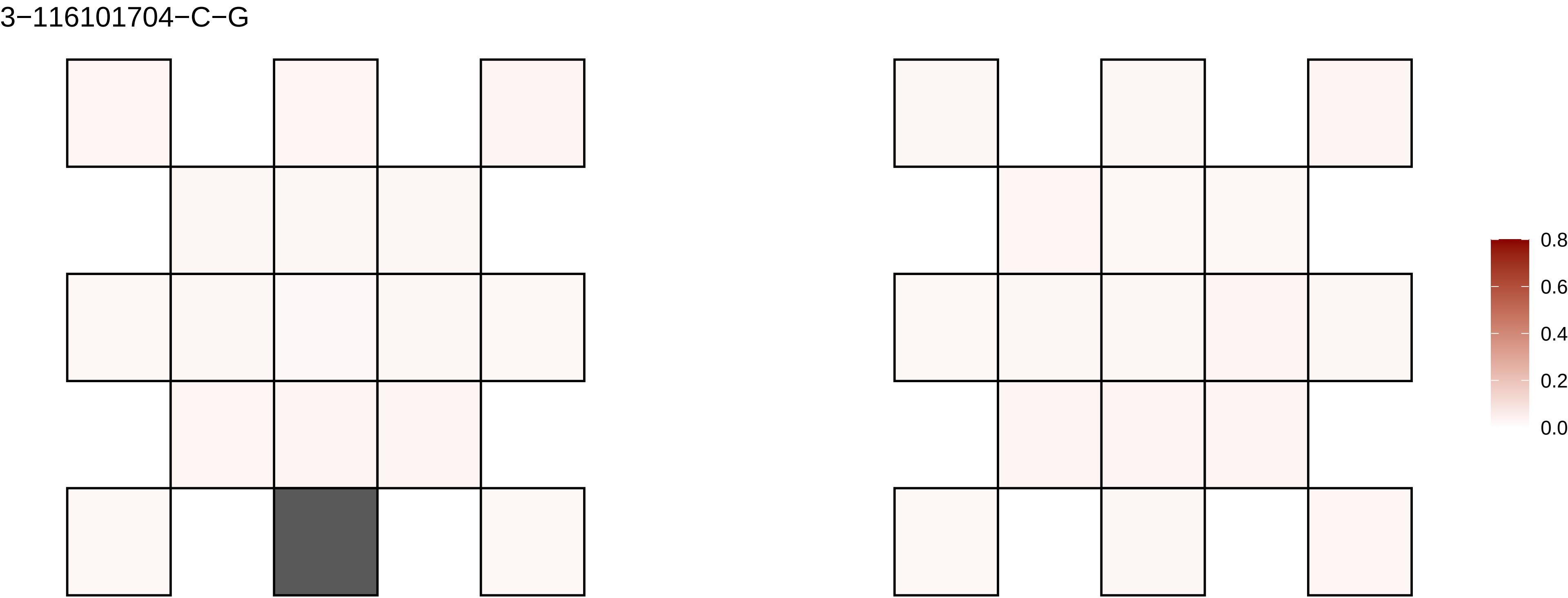
3−116101704−C−G
0.8
0.6
0.4
0.2
0.0

3−116224135−C−T

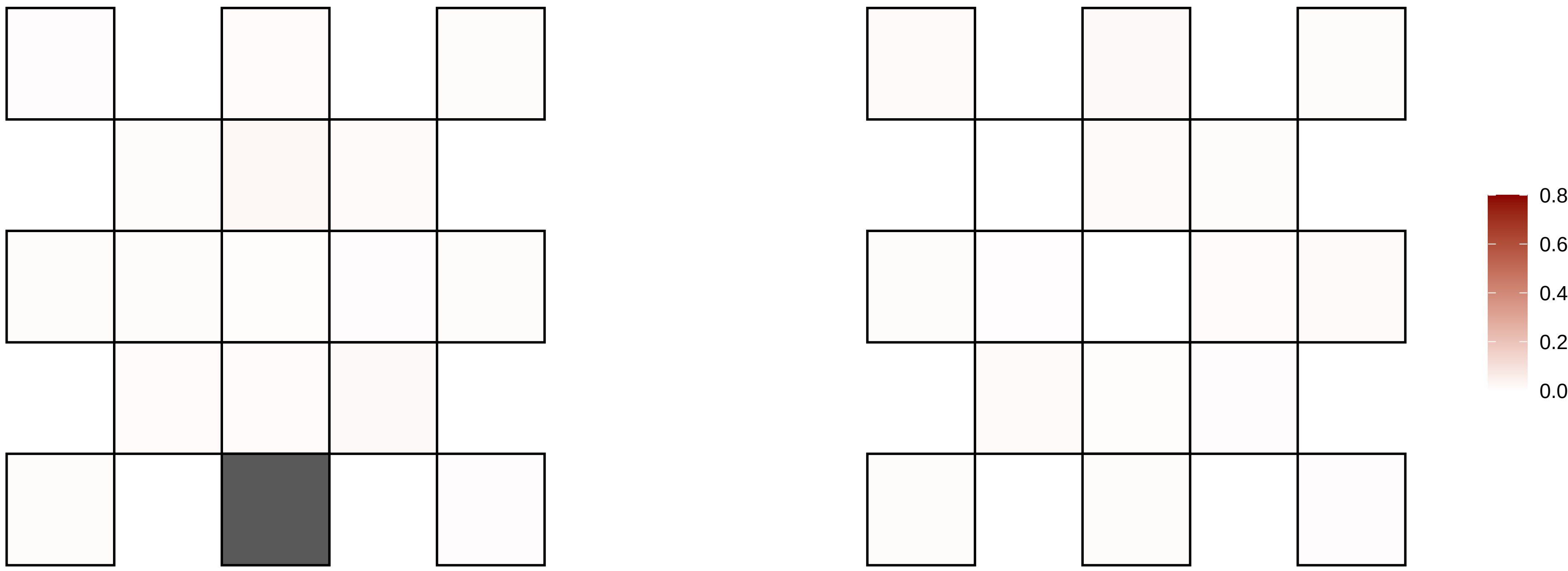

3−119866148−A−C

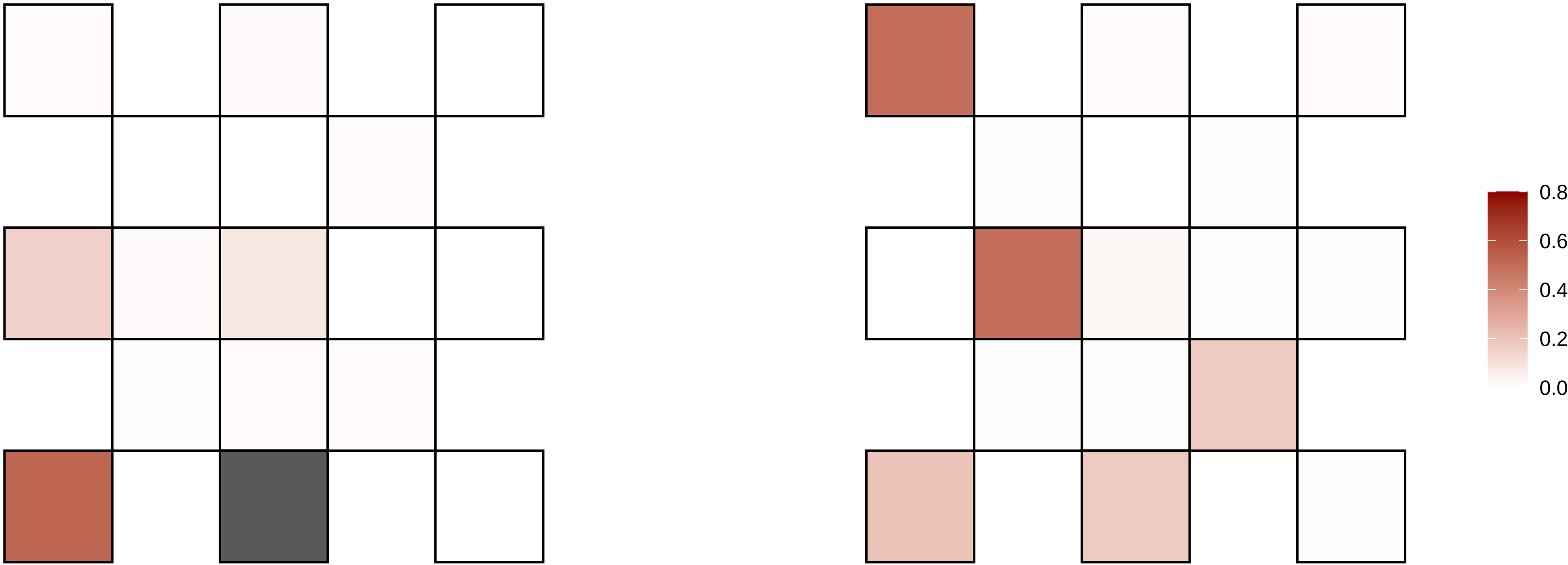

3−124402098−G−A

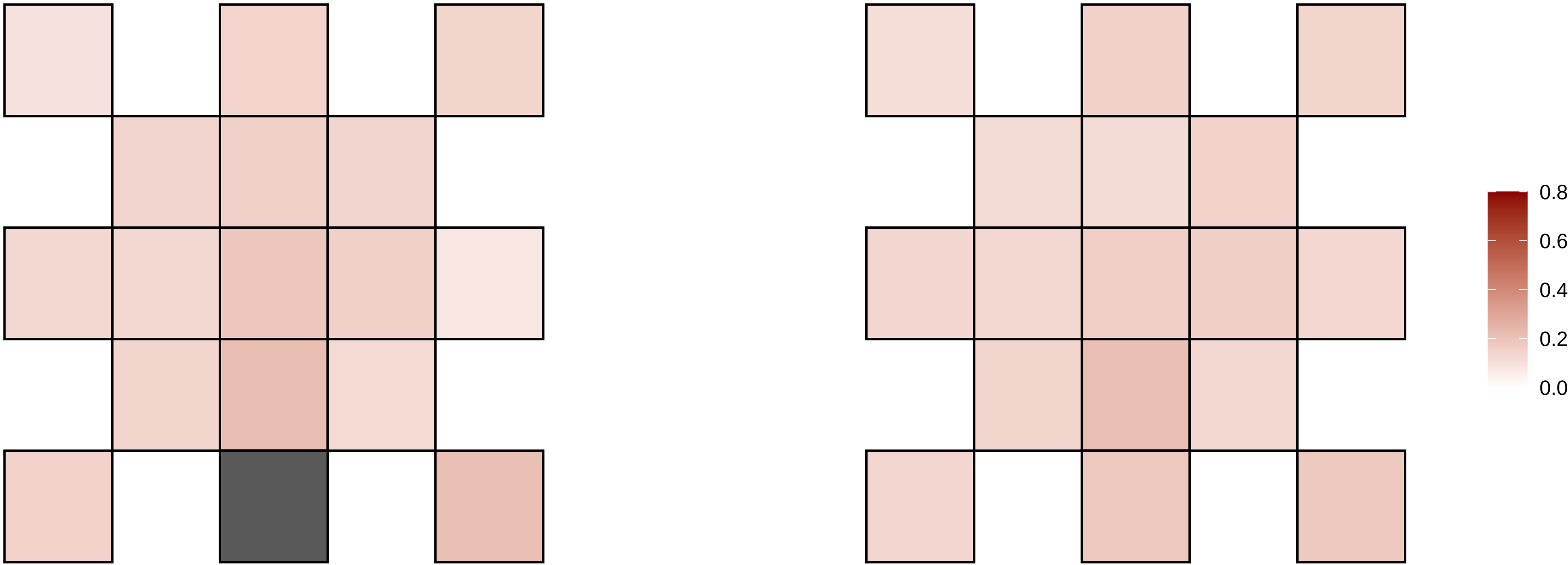

3−130175696−A−G

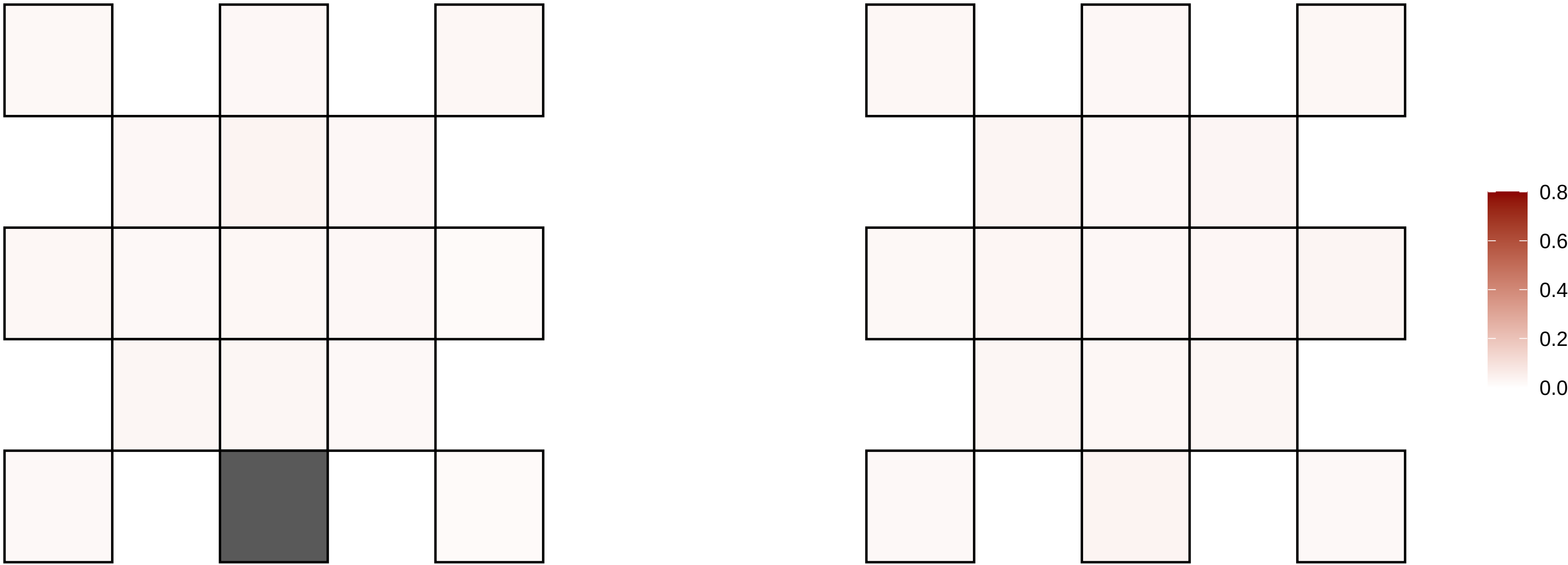

3−137569791−G−A

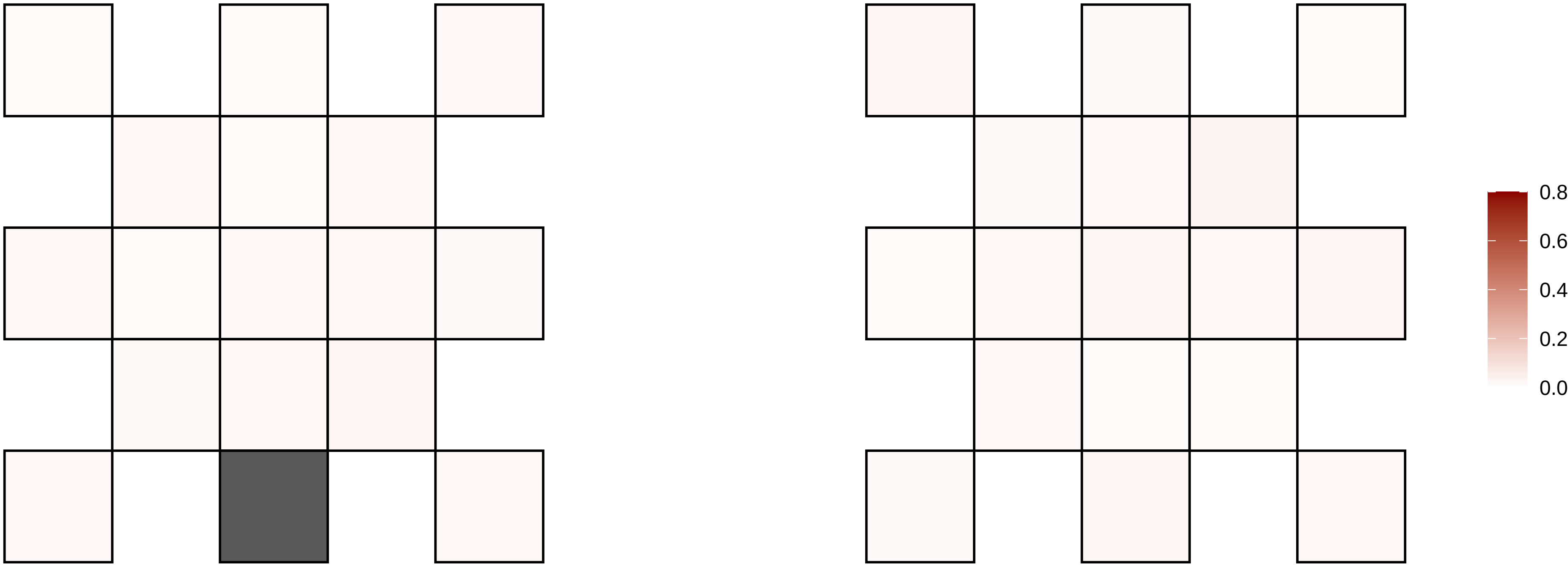

3−145391171−A−G

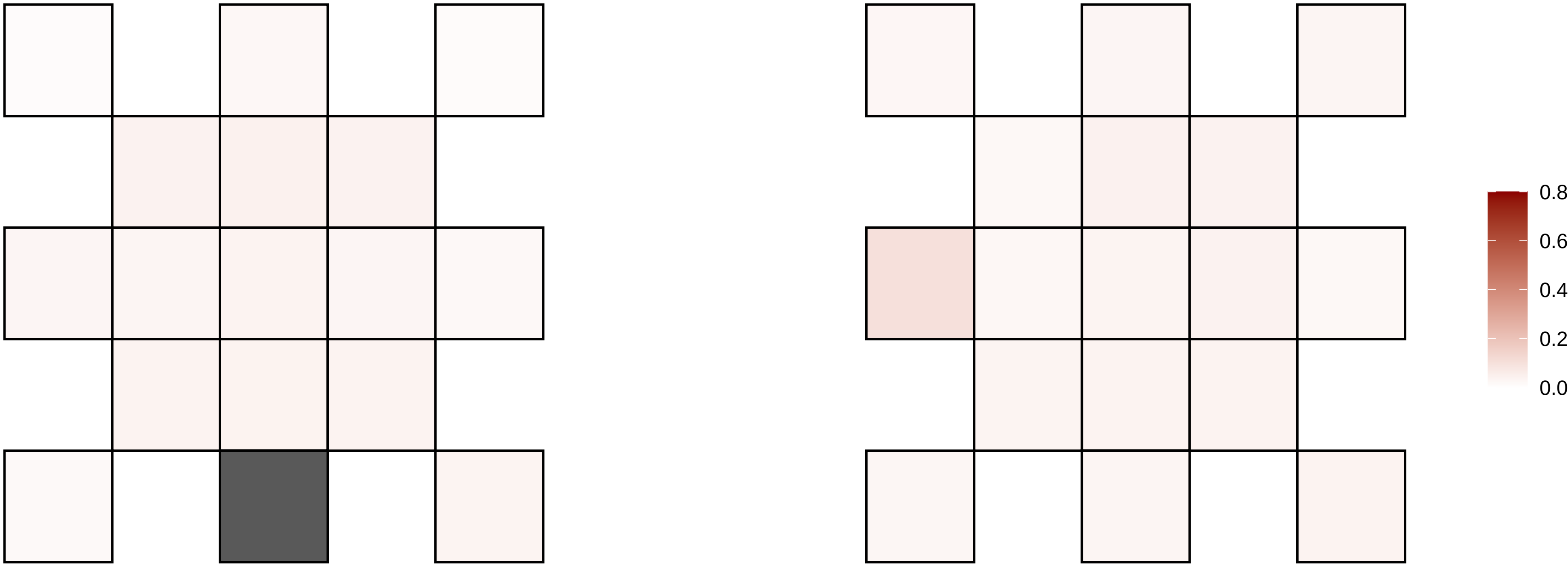

3−147144597−G−A

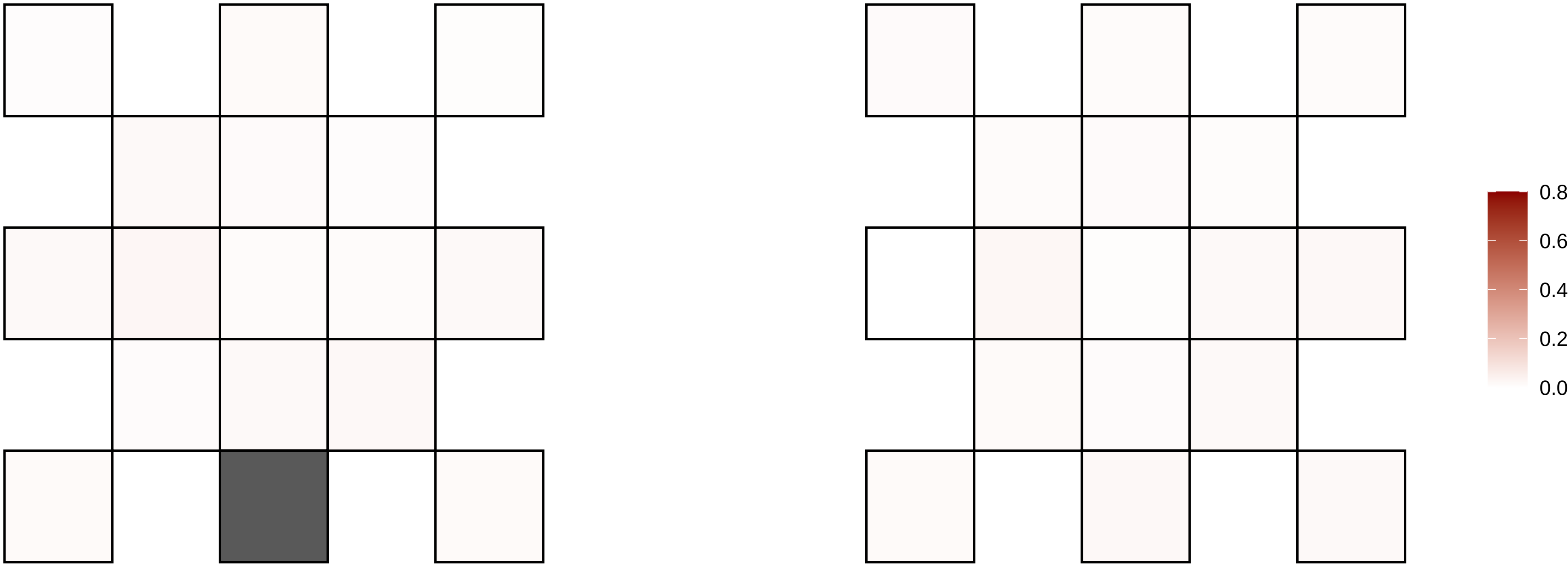

3−154786585−A−G

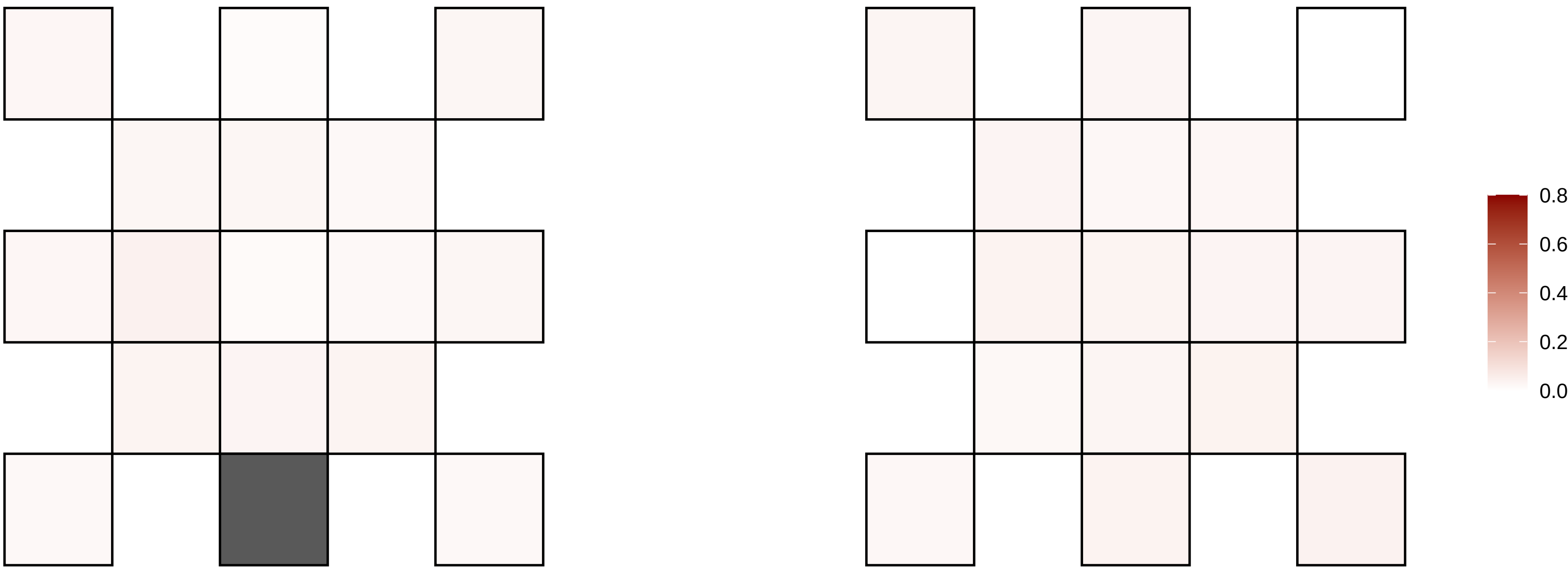

3−155866578−C−T

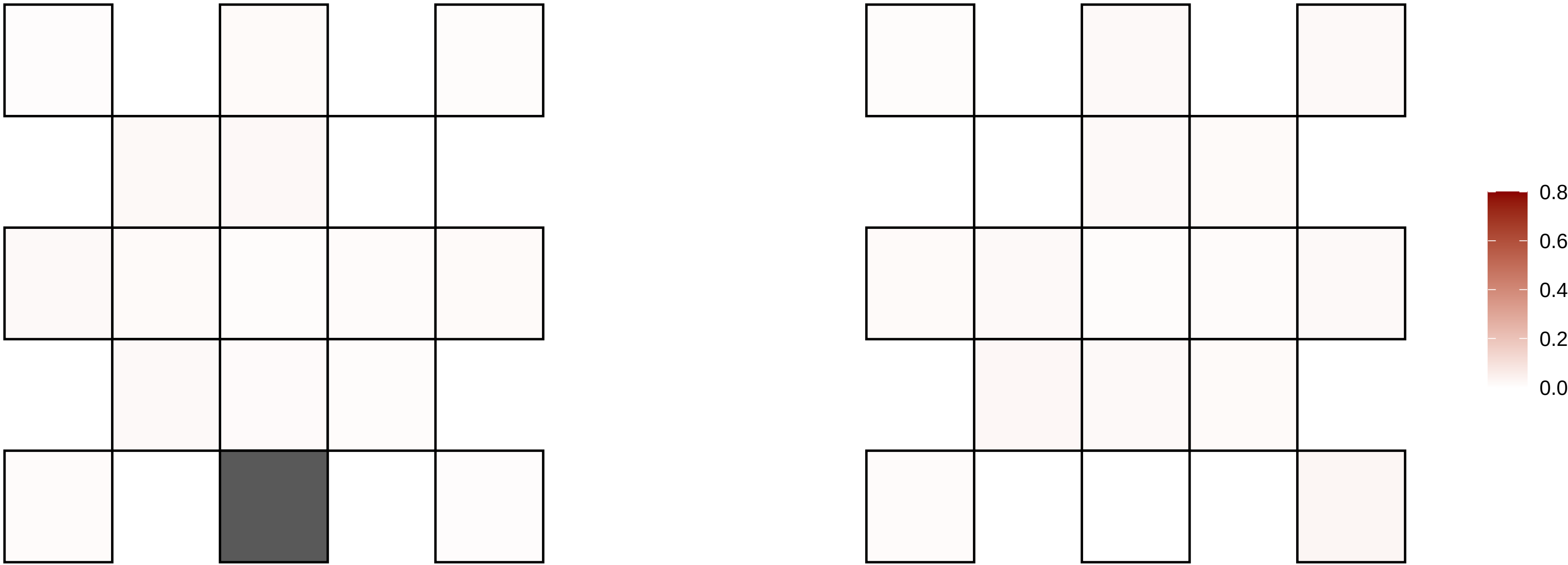

3−156768561−C−T

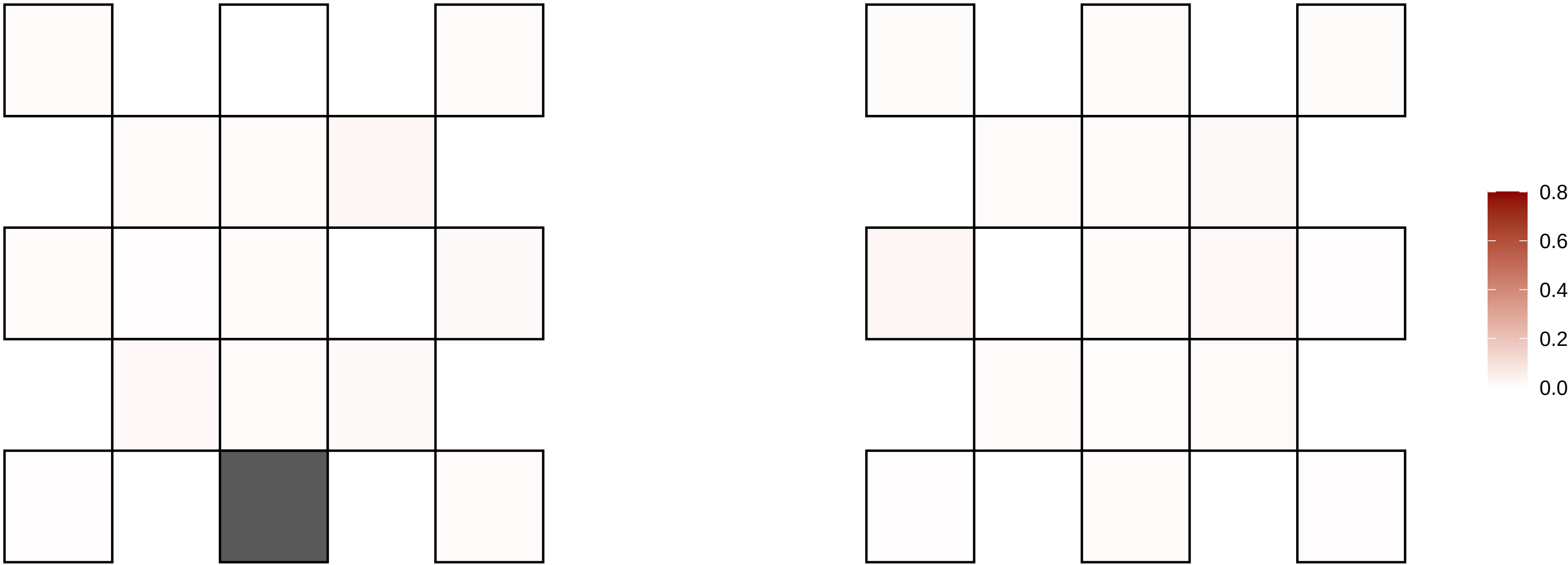

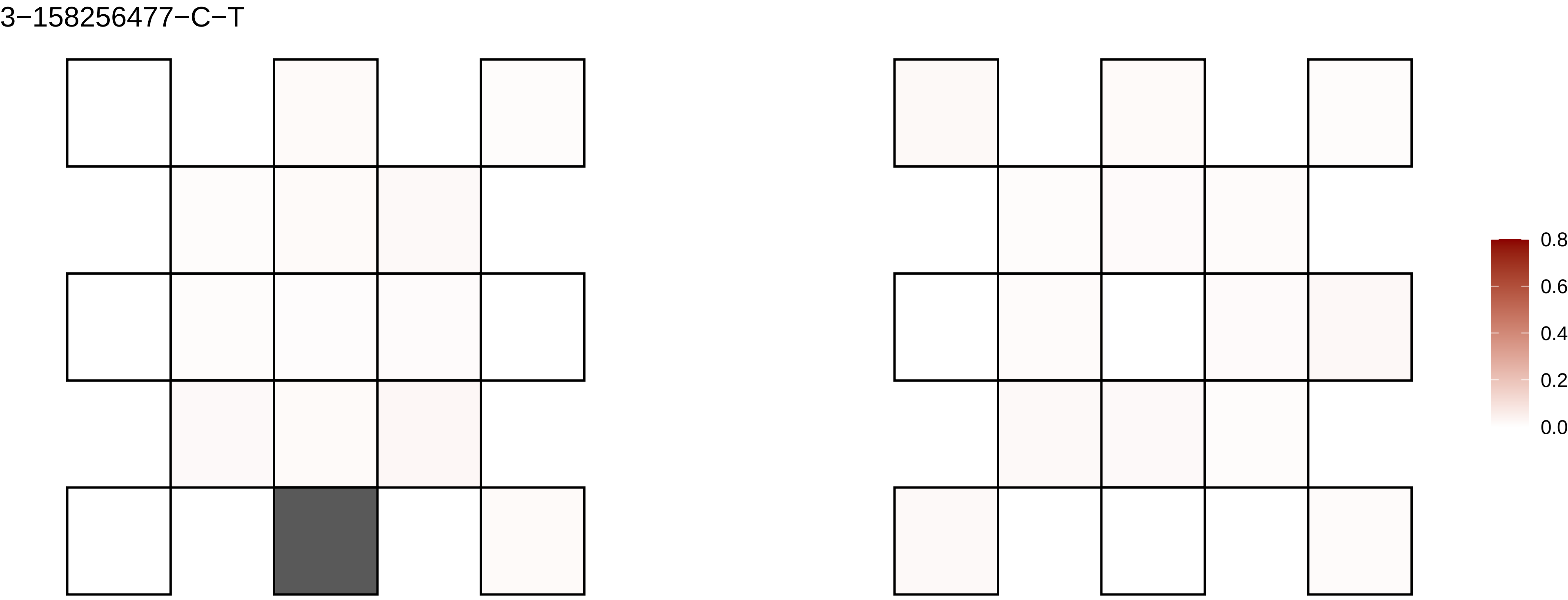

3−158256477−C−T
0.8
0.6
0.4
0.2
0.0

3−159483100−C−T

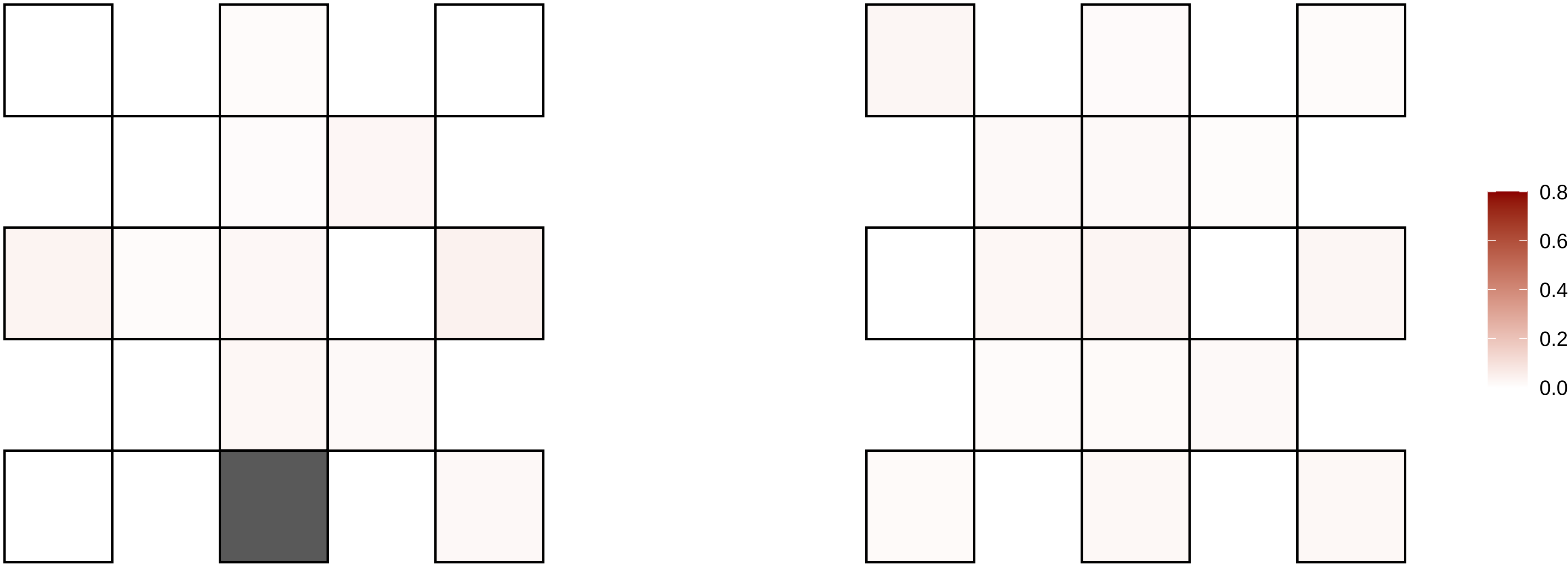

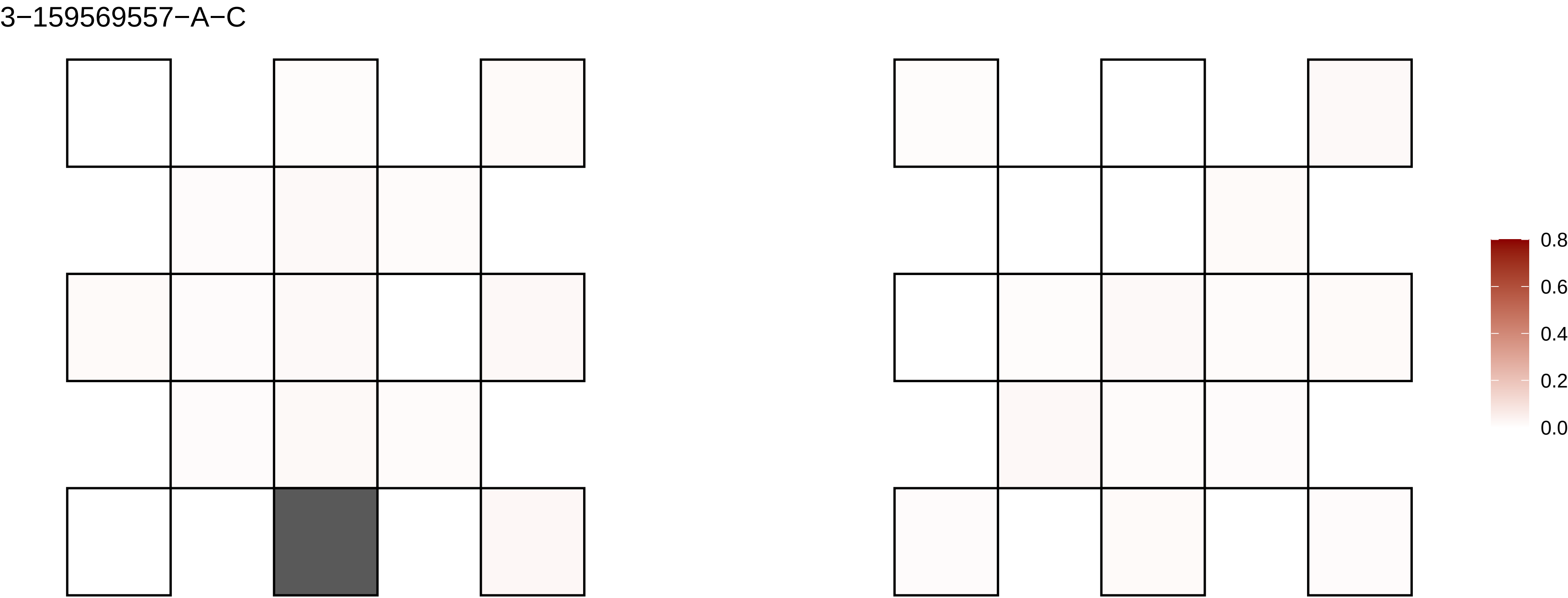

3−159569557−A−C
0.8
0.6
0.4
0.2
0.0

3−164879470−C−T

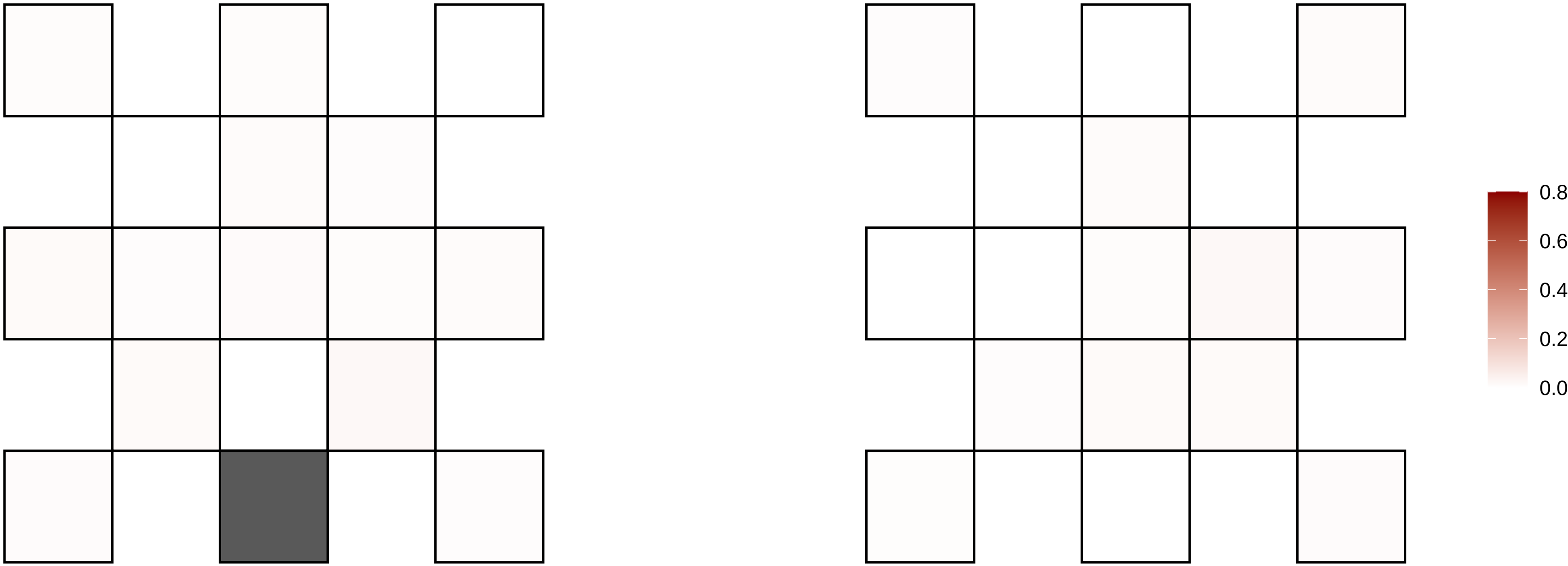

3−16511350−A−C

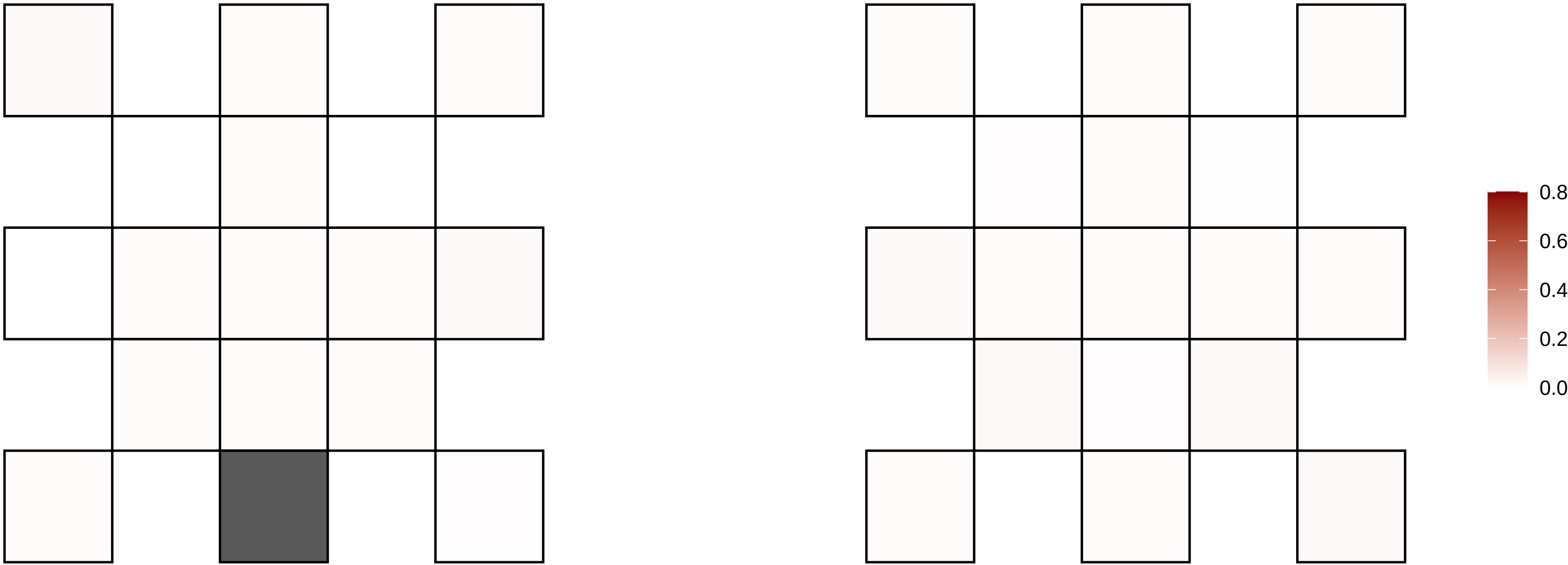

3−16568702−C−T

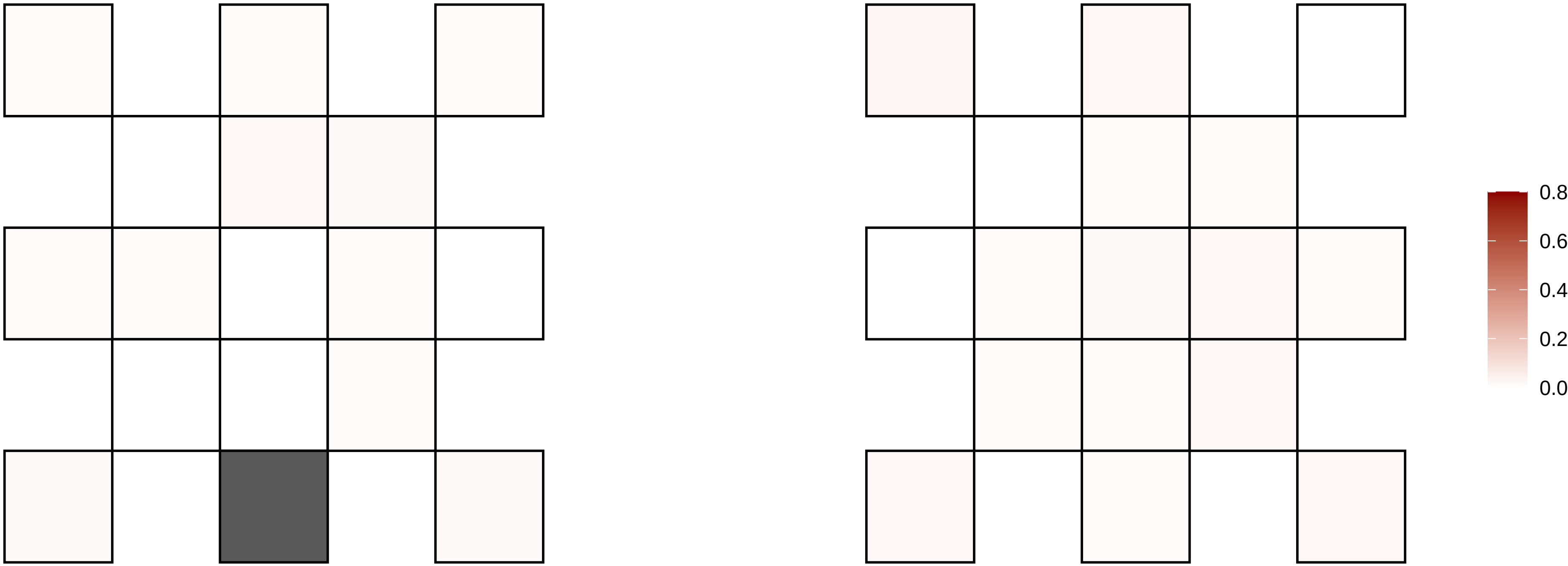

3−169833345−G−A

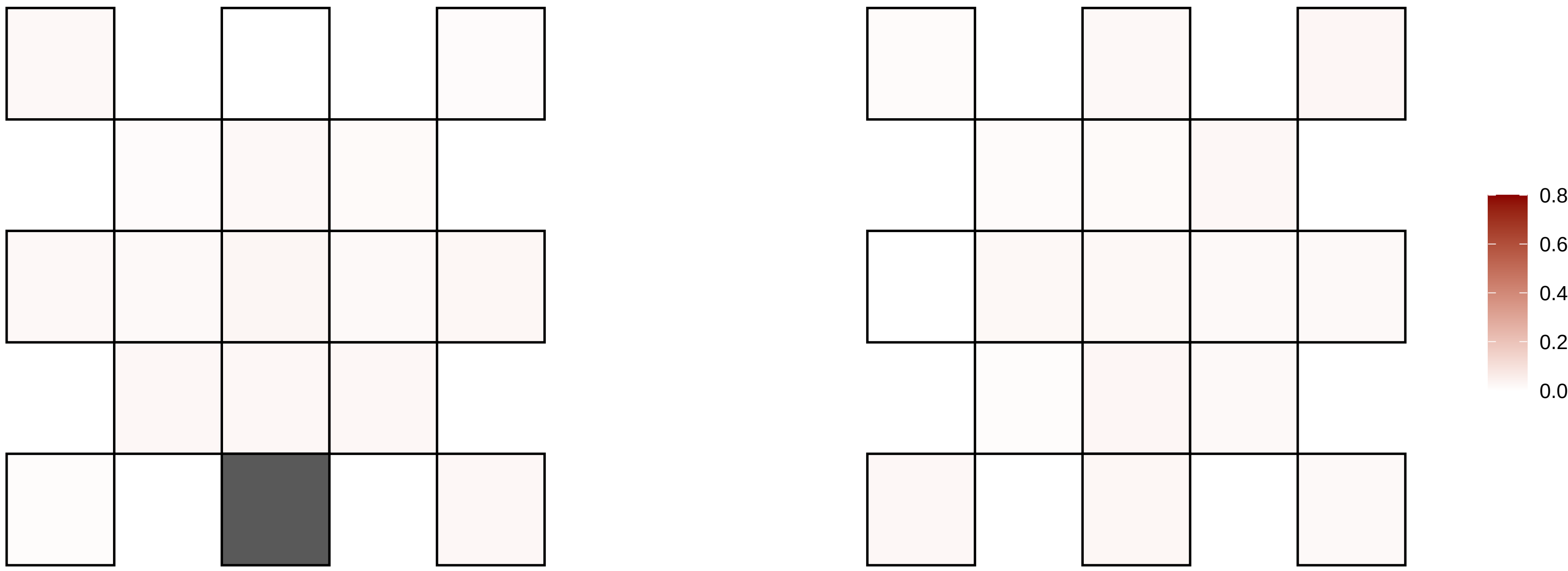

3−172635725−G−A

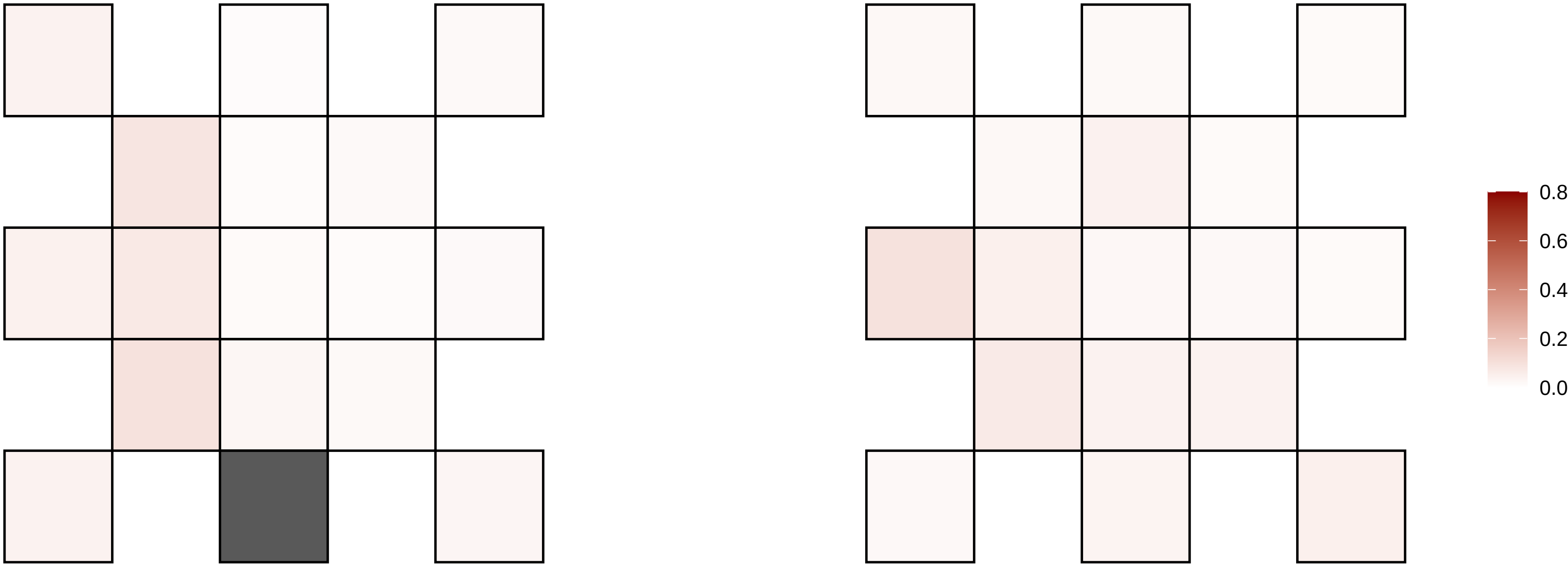

3−173423567−C−A

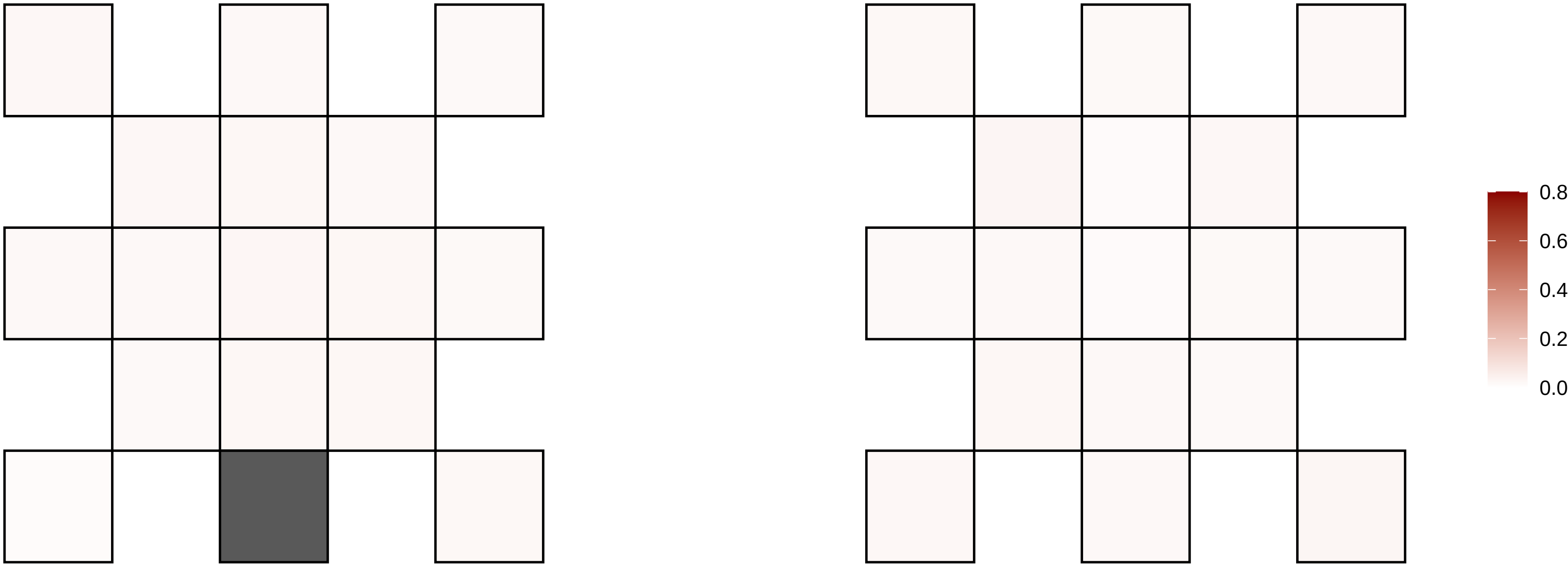

3−181259475−C−T

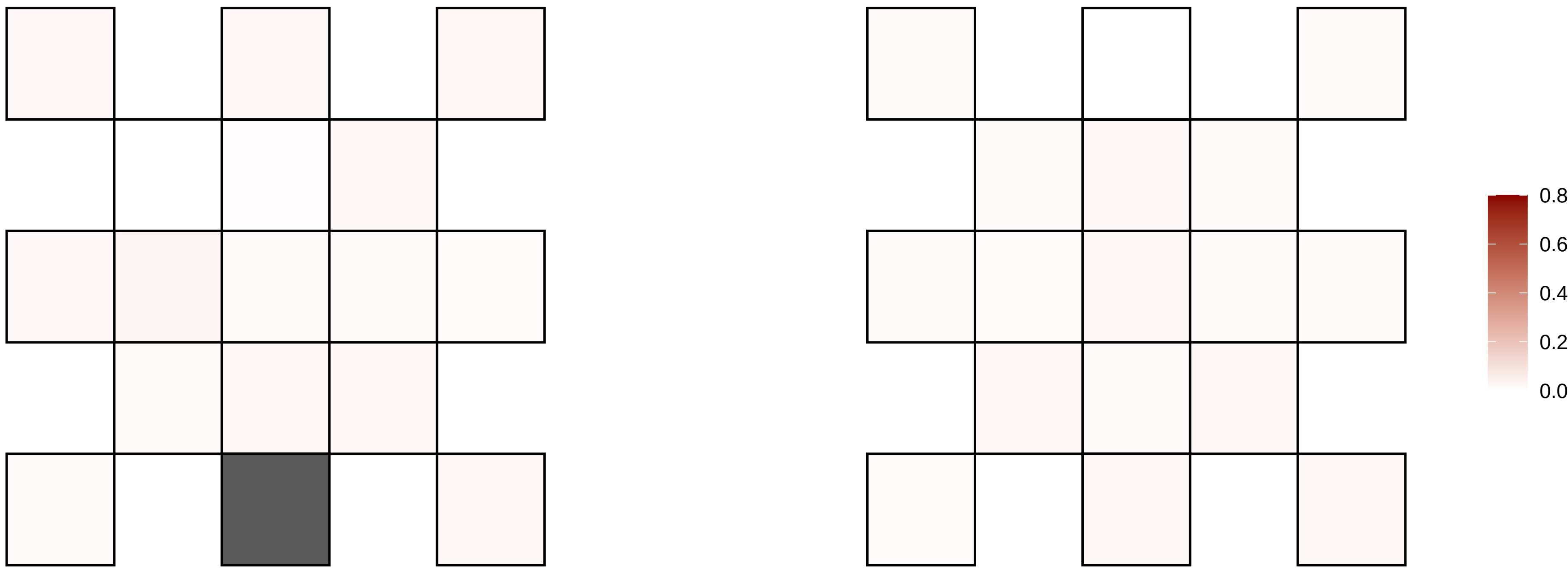

3−185077743−C−T

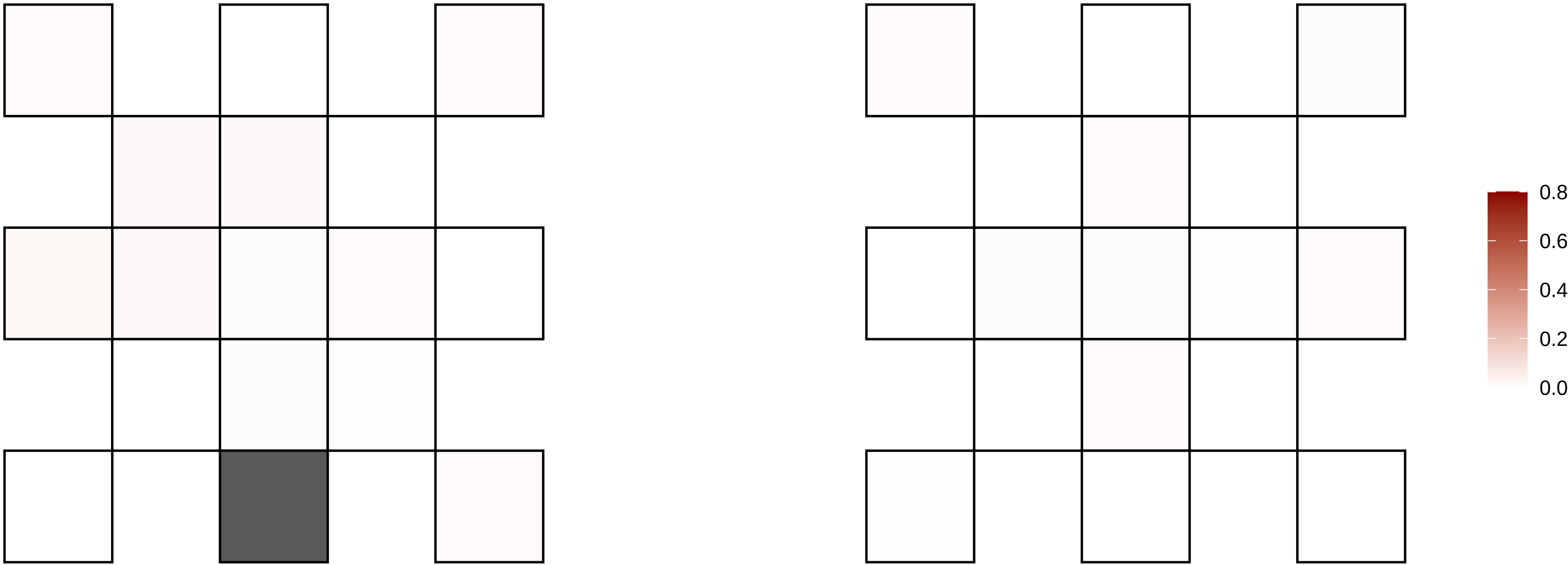

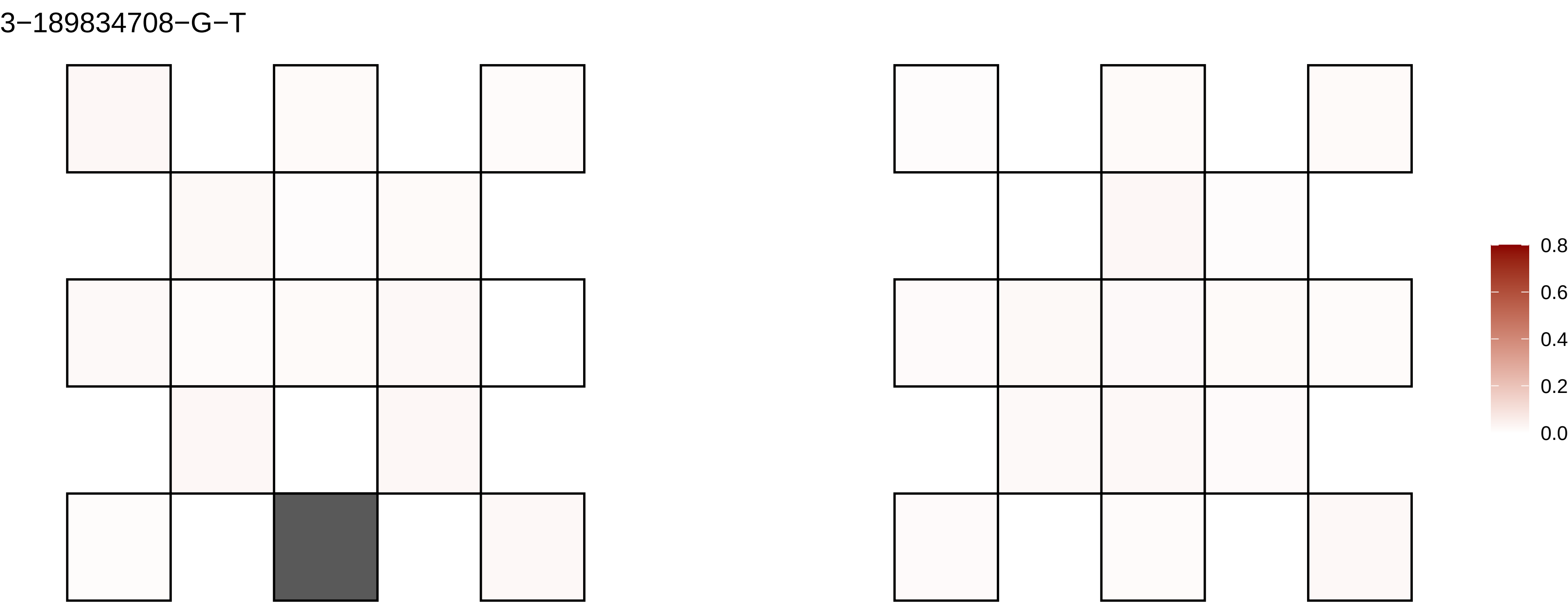
3−189834708−G−T
0.8
0.6
0.4
0.2
0.0

3−190125961−G−A

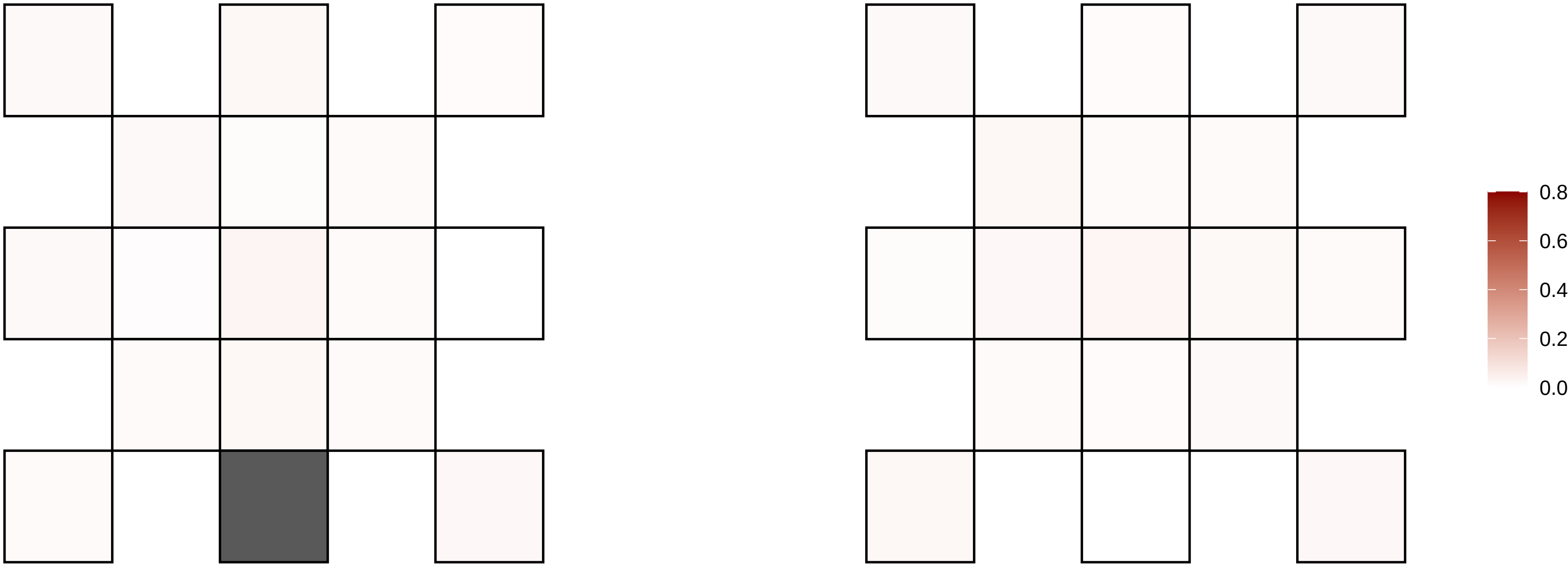

3−193720234−C−T

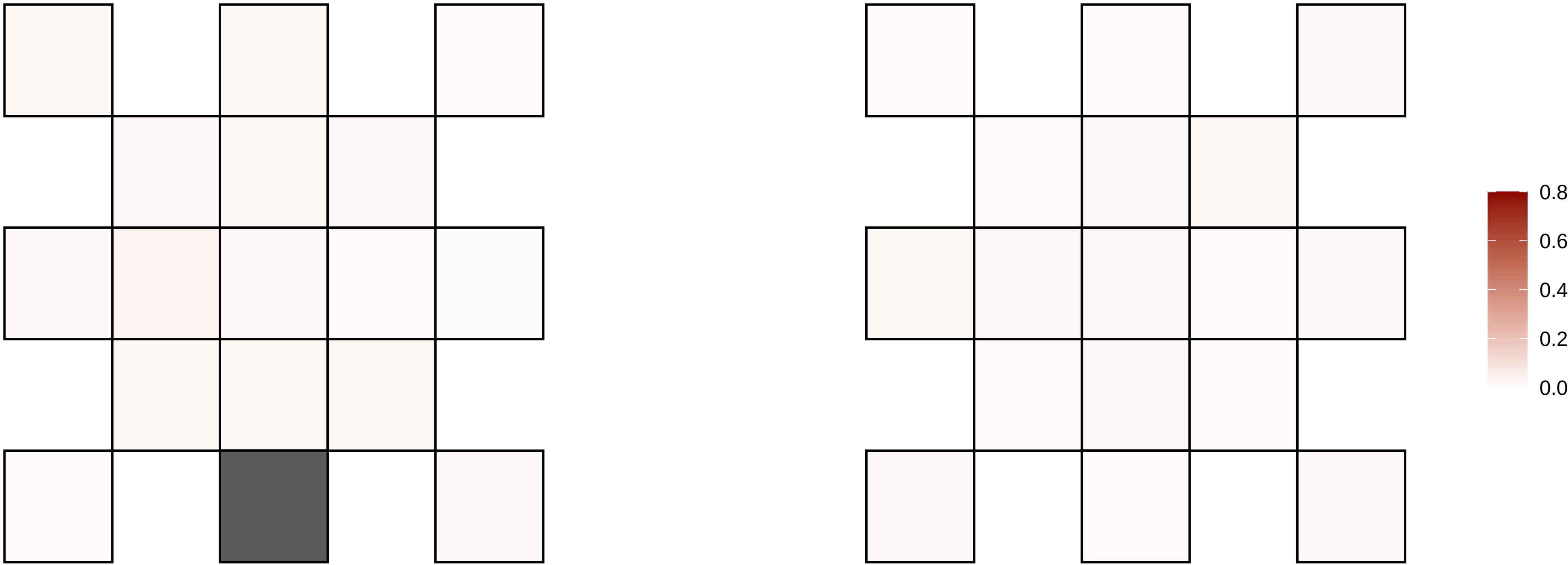

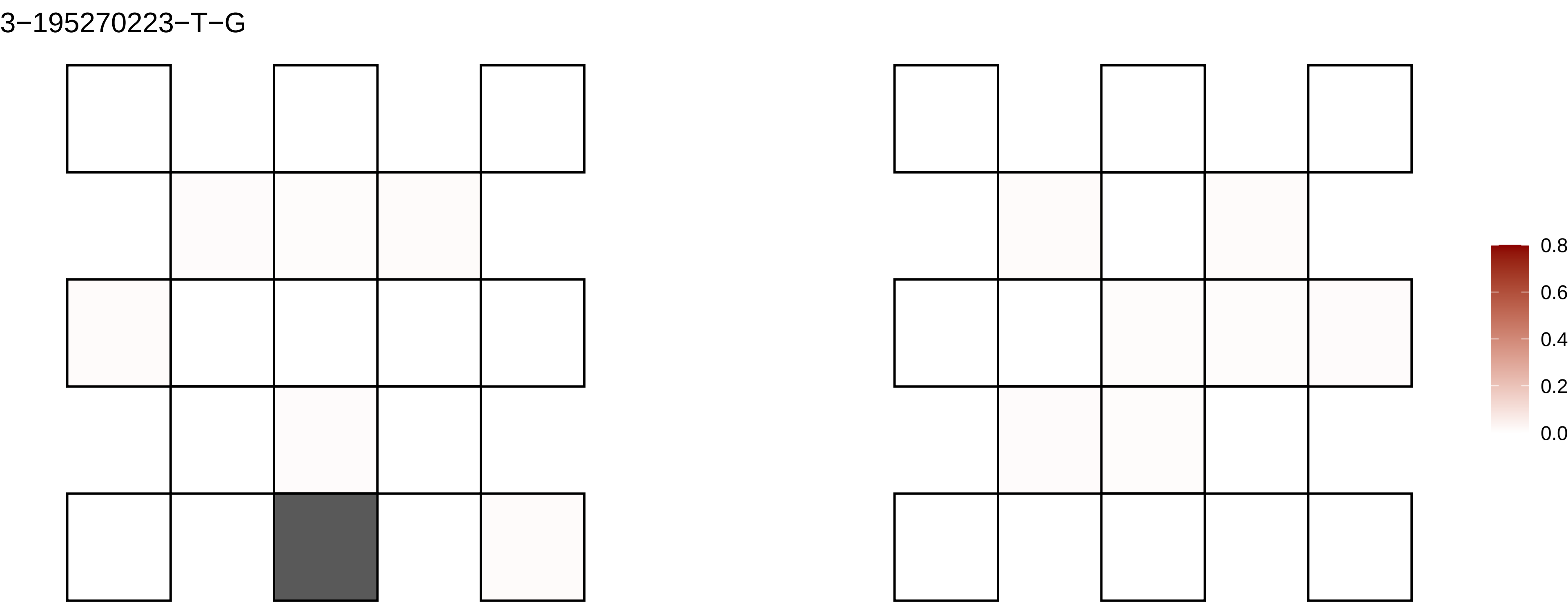
3−195270223−T−G
0.8
0.6
0.4
0.2
0.0

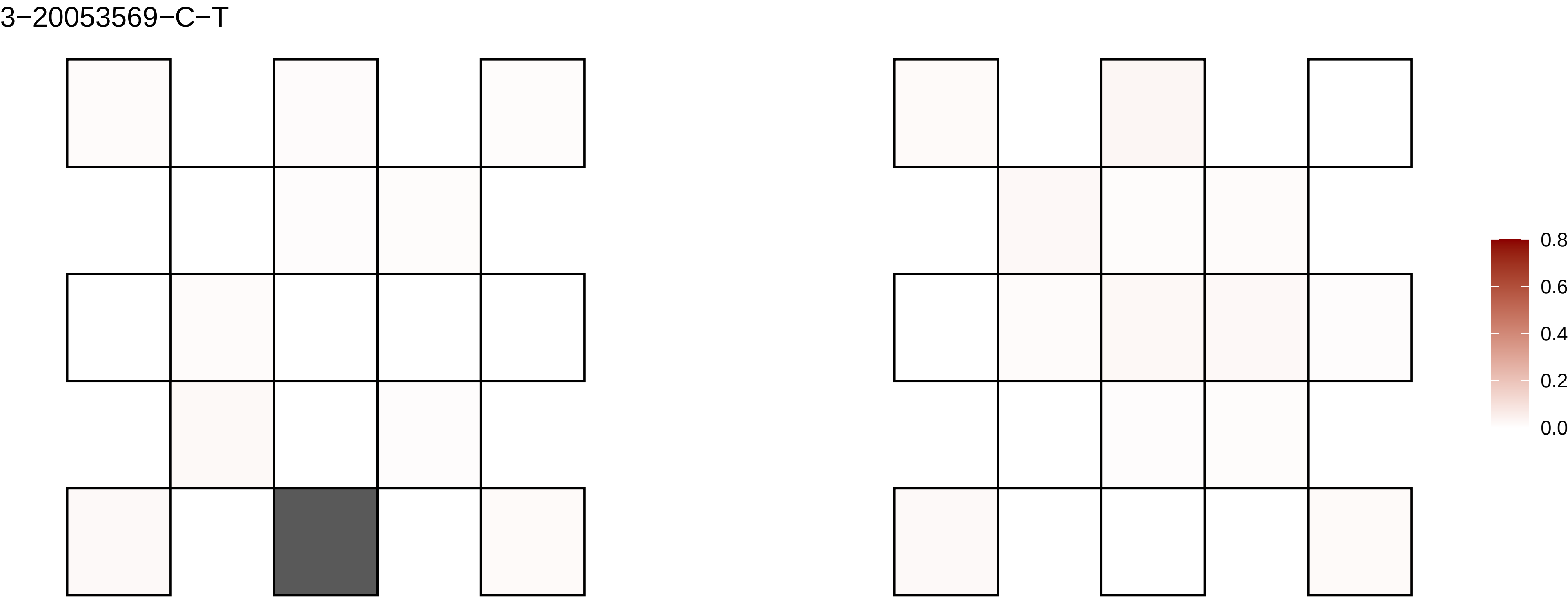
3−20053569−C−T
0.8
0.6
0.4
0.2
0.0

3−20145262−G−C

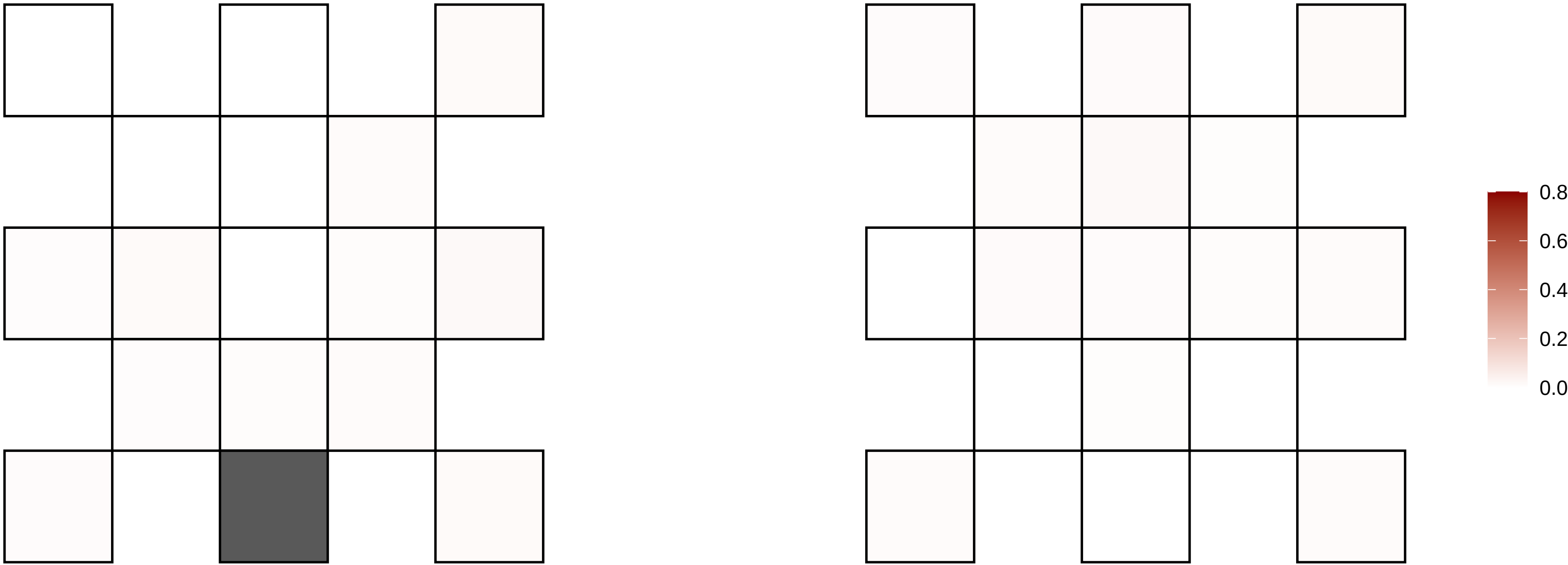

3−20222654−C−T

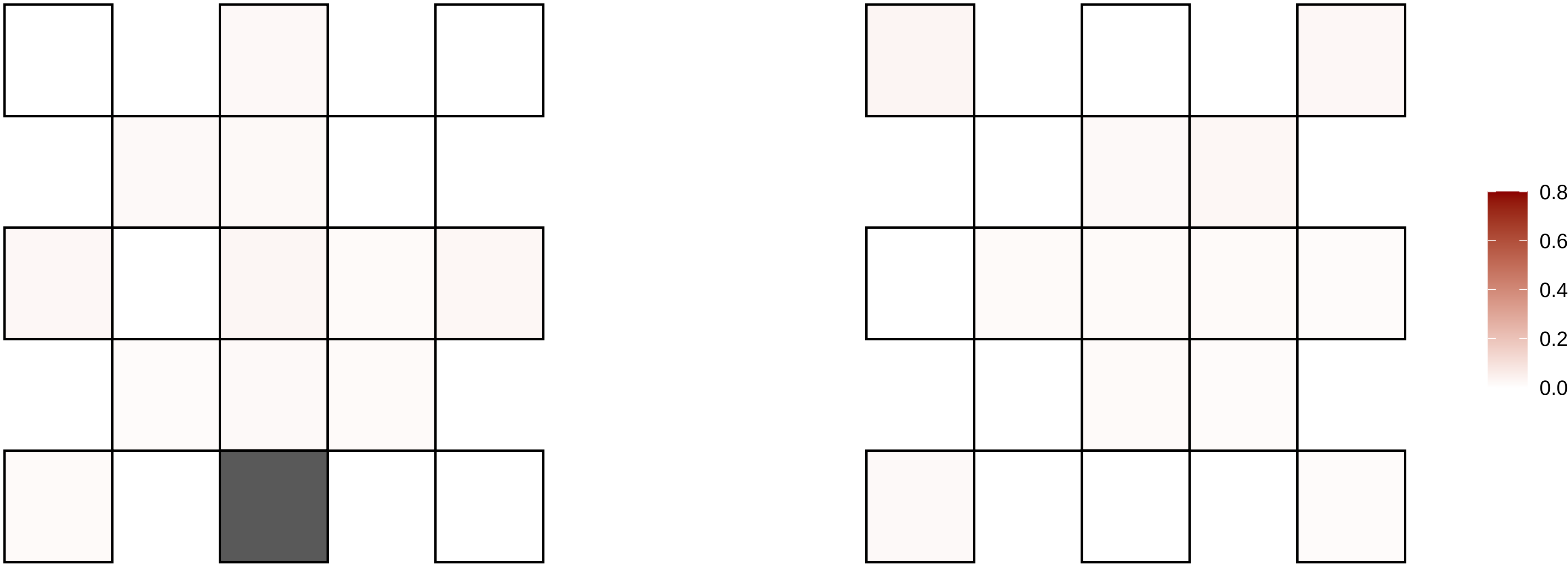

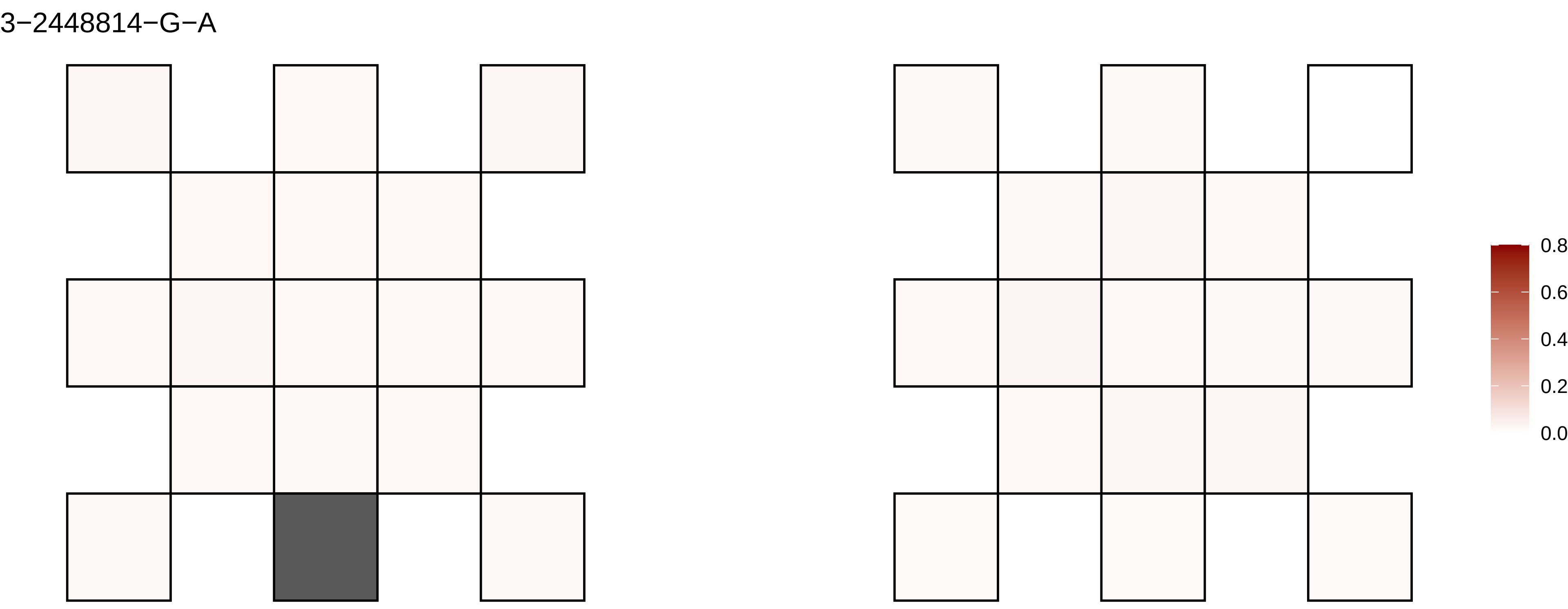

3−2448814−G−A
0.8
0.6
0.4
0.2
0.0

3−29362488−G−A

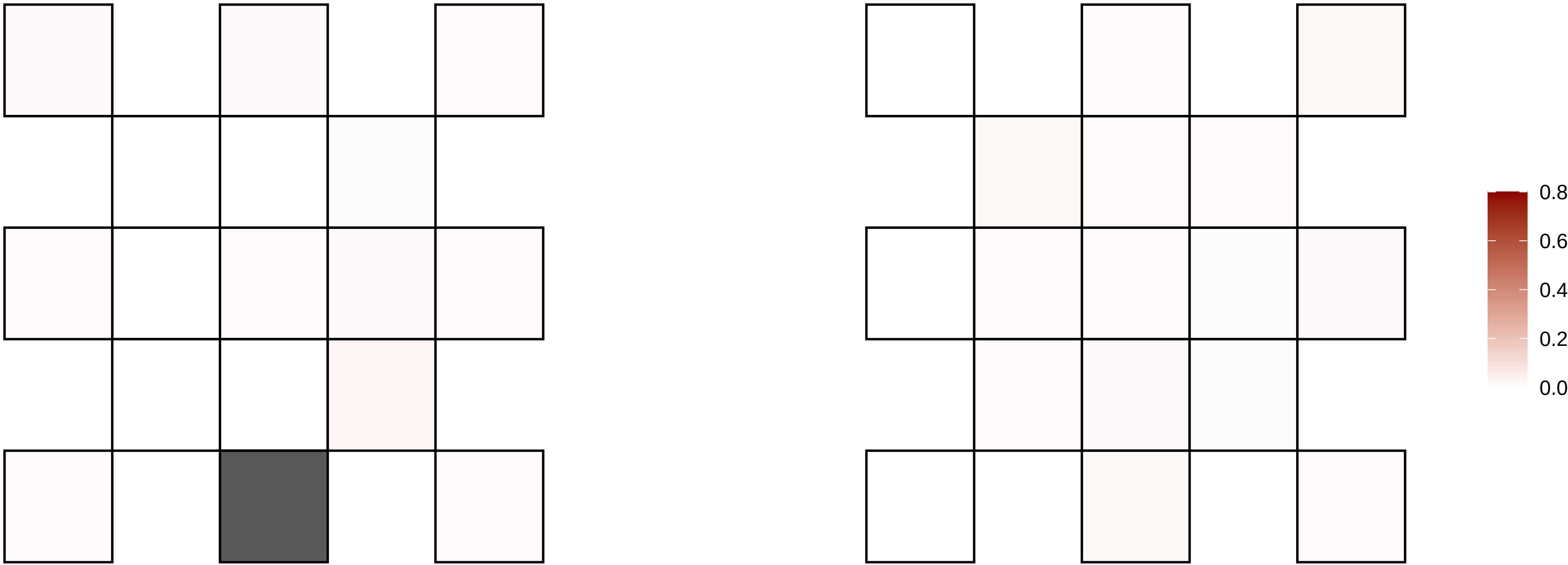

3−29763350−T−C

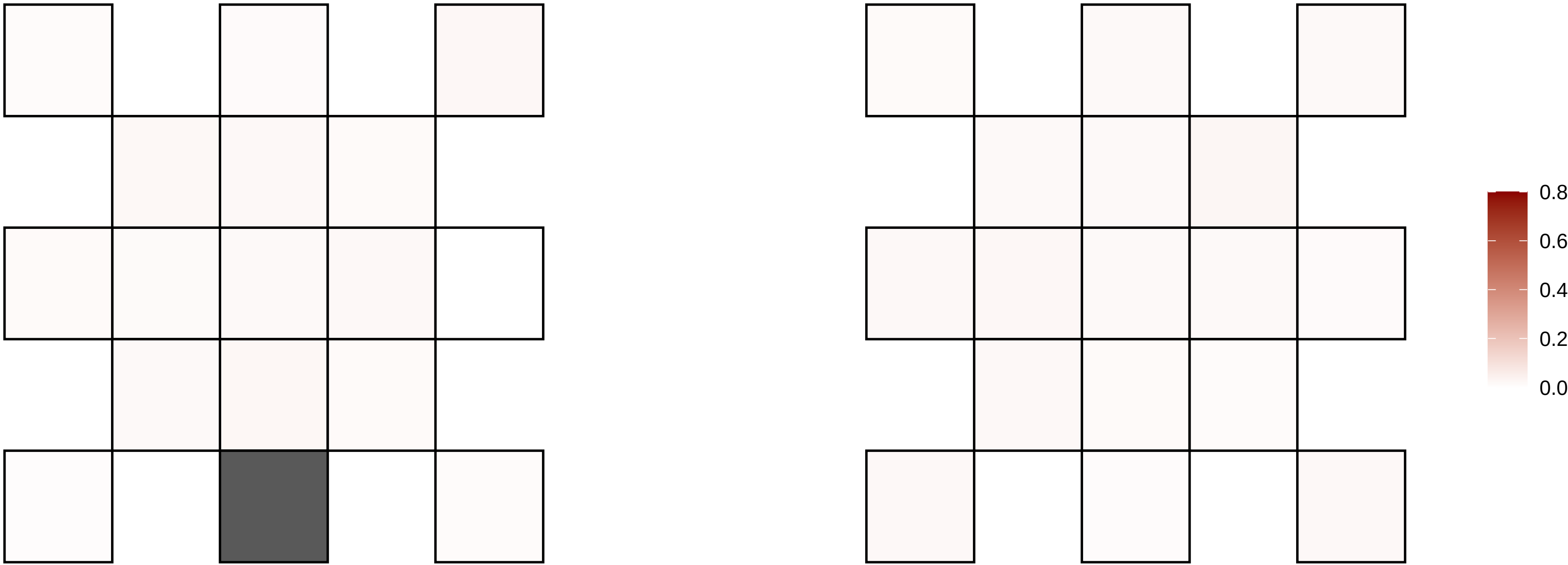

3−32863852−G−A

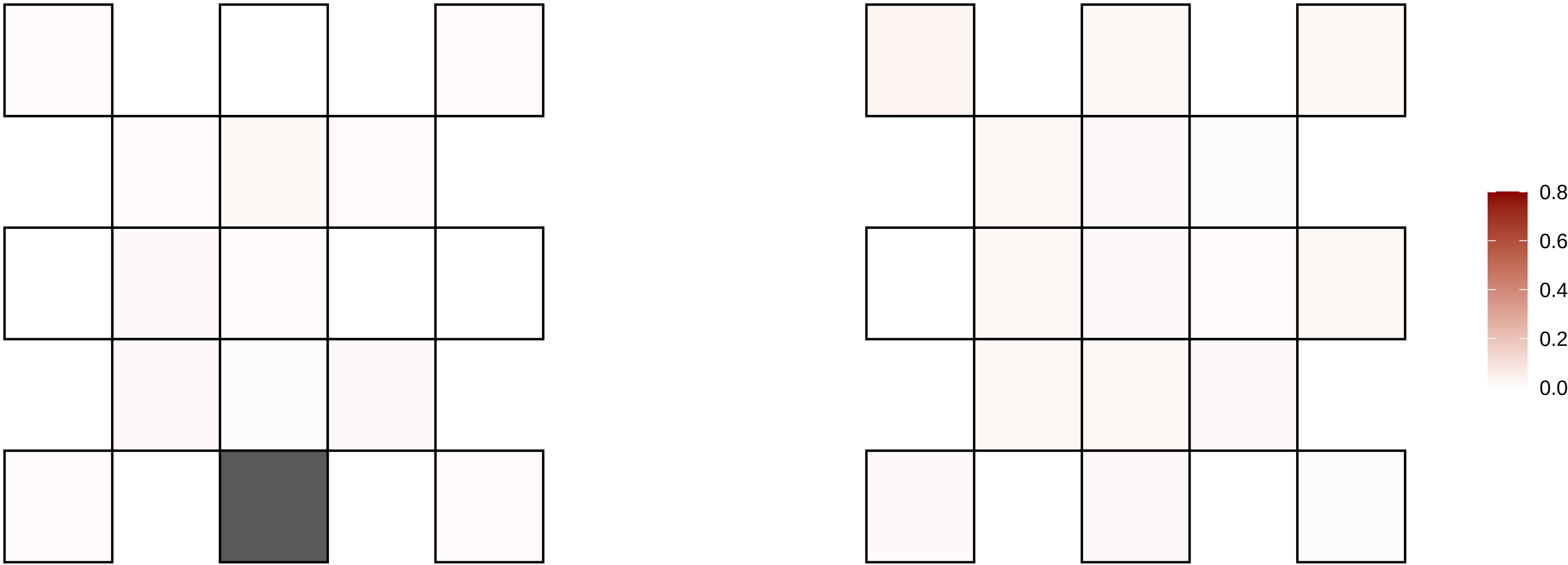

3−3458468−TG−T

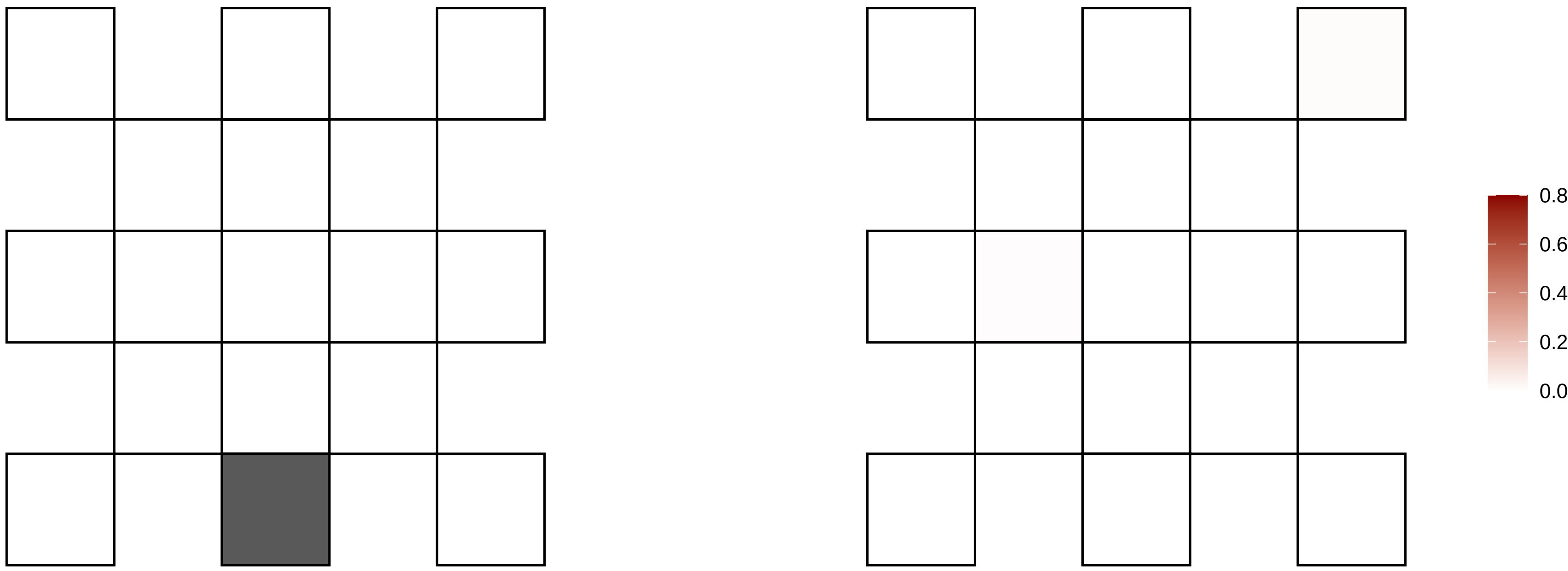

3−36389918−C−T

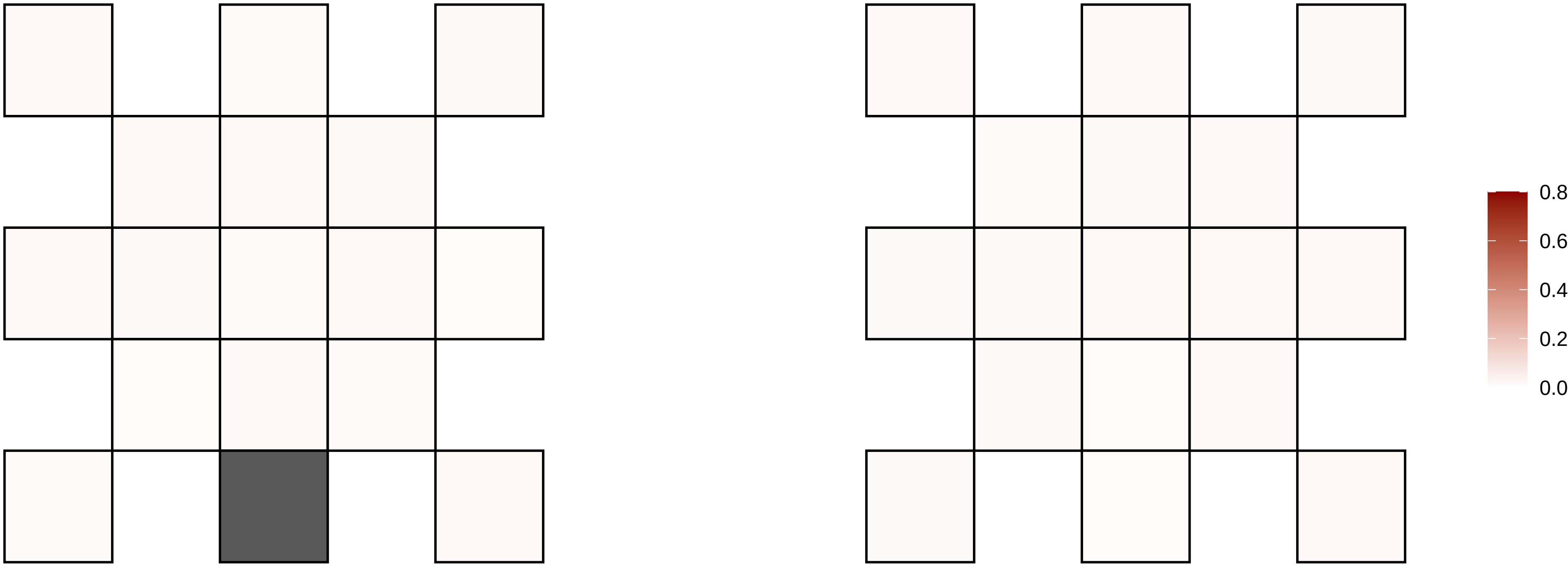

3−36414768−C−G

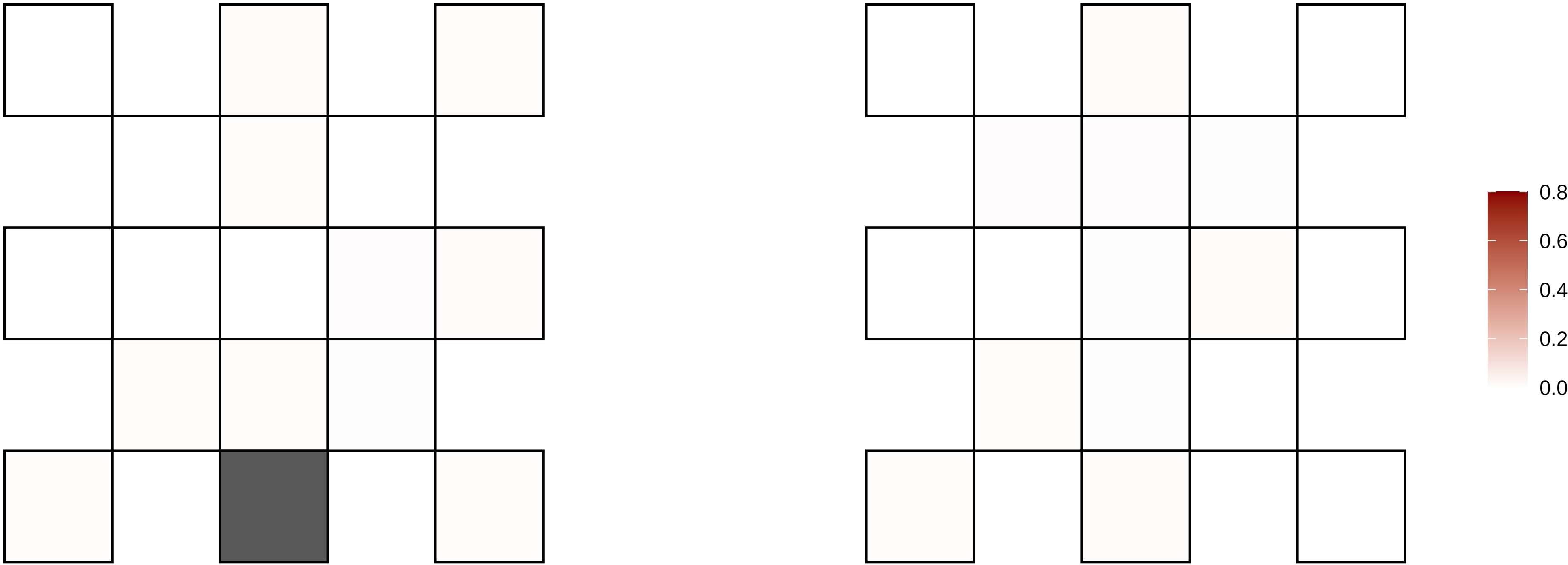

3−40086182−G−T

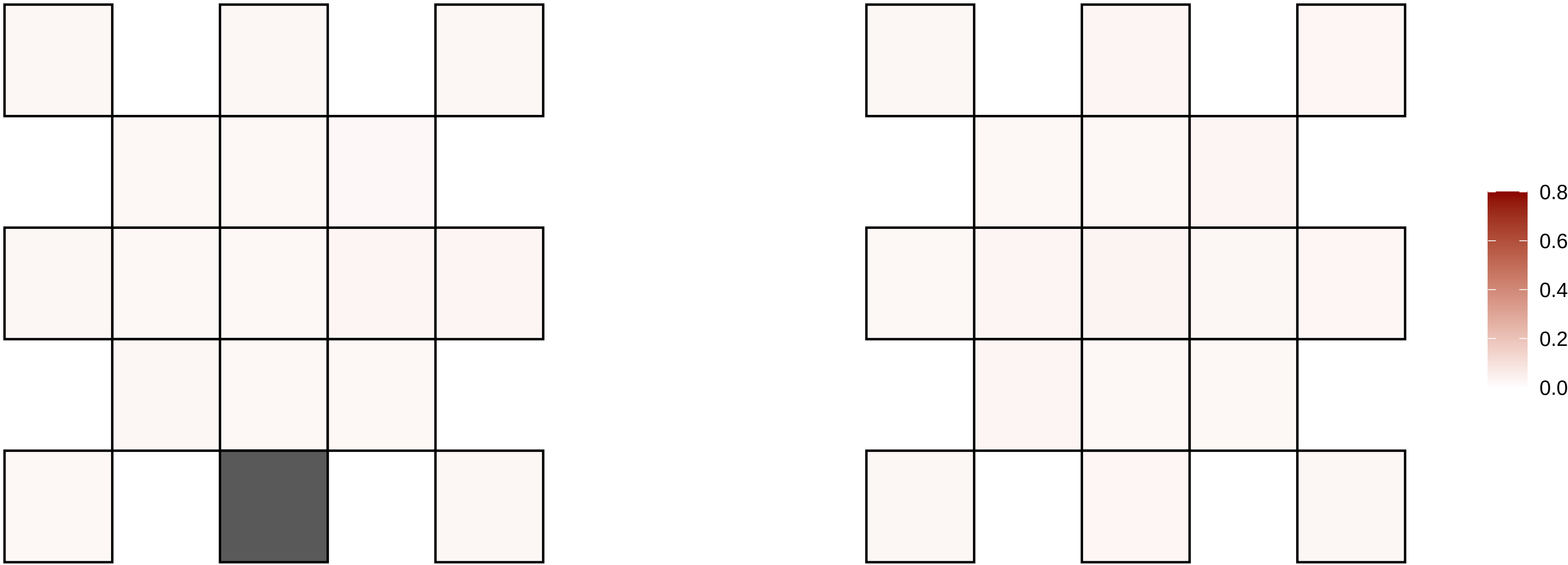

3−44078495−G−A

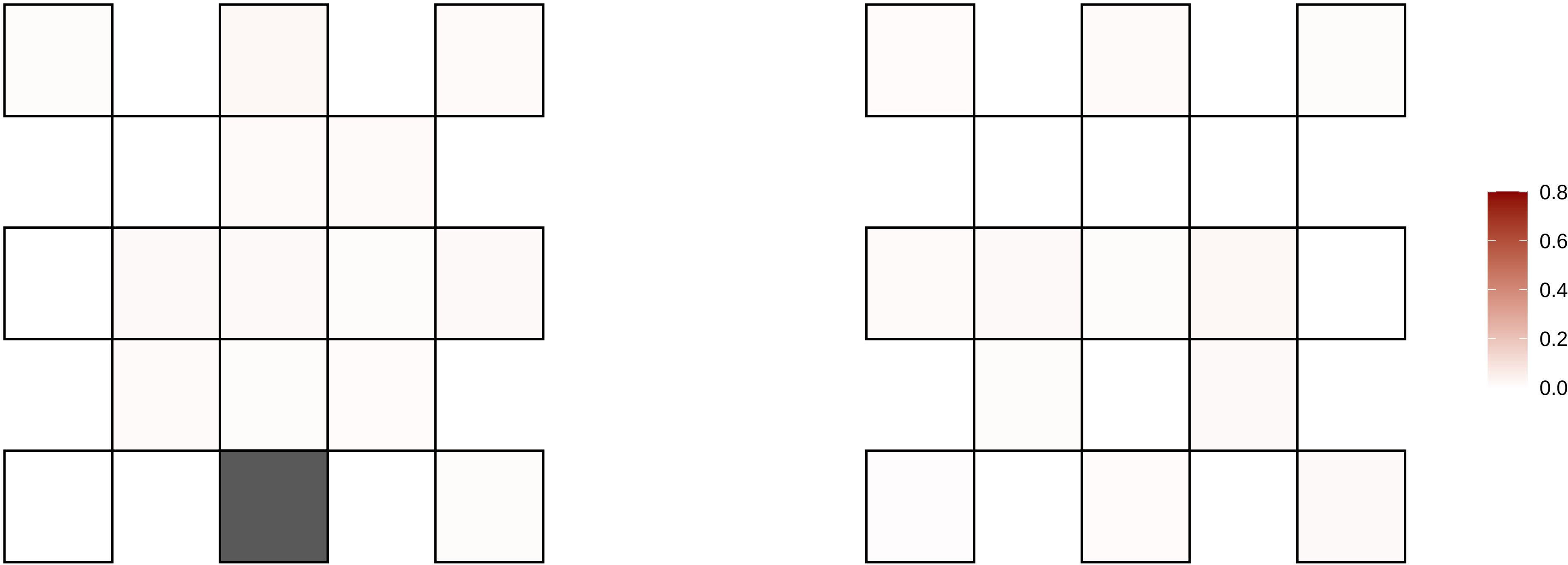

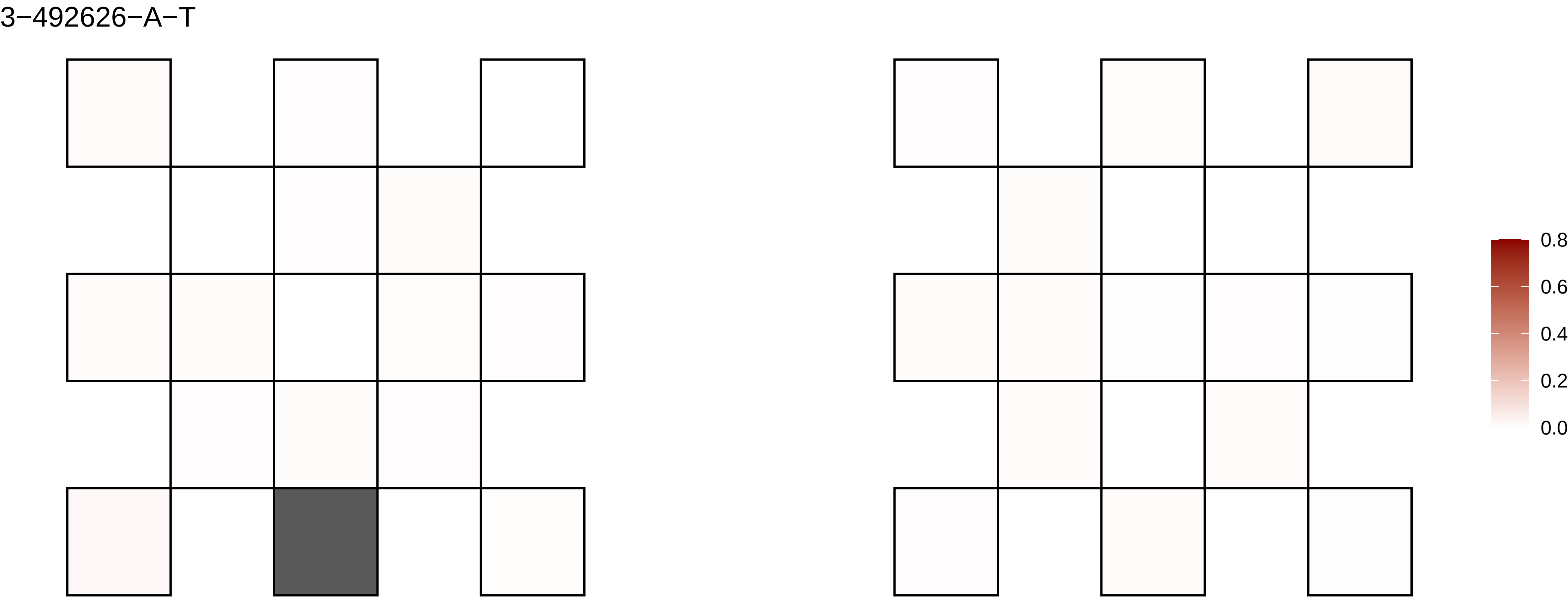
3−492626−A−T
0.8
0.6
0.4
0.2
0.0

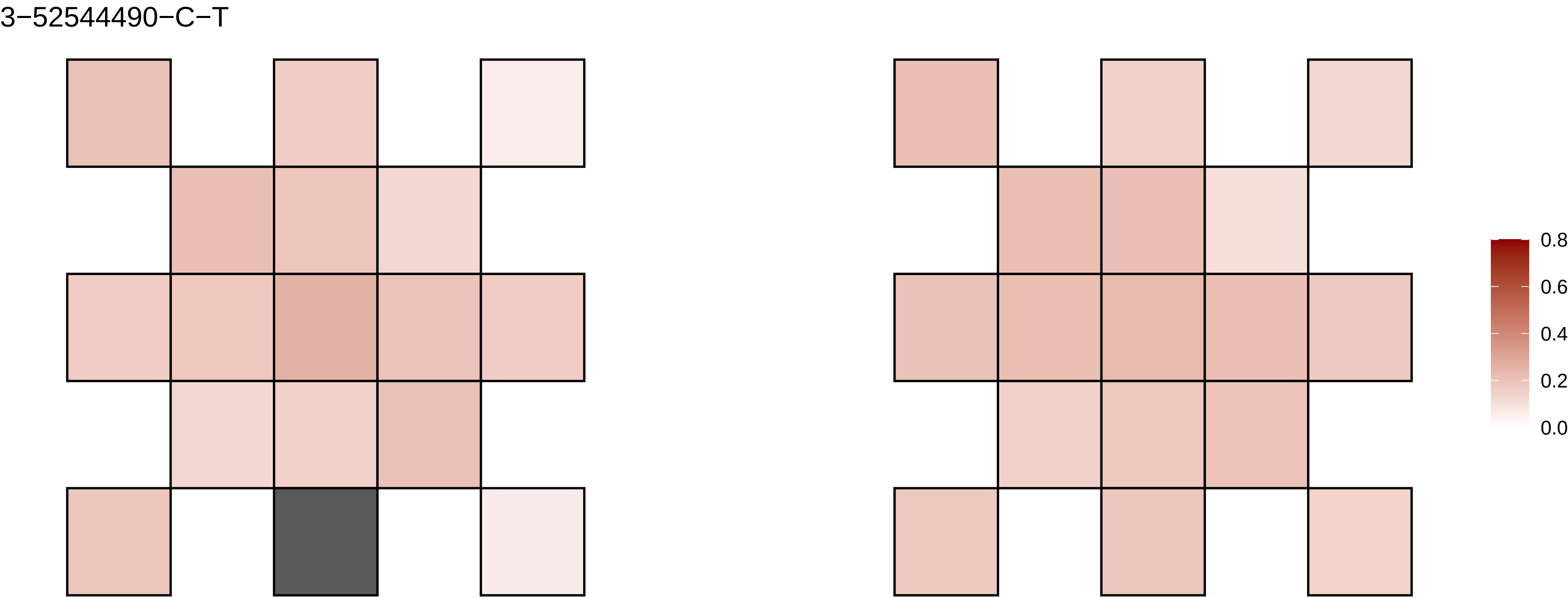
3−52544490−C−T
0.8
0.6
0.4
0.2
0.0

3−5352719−C−T

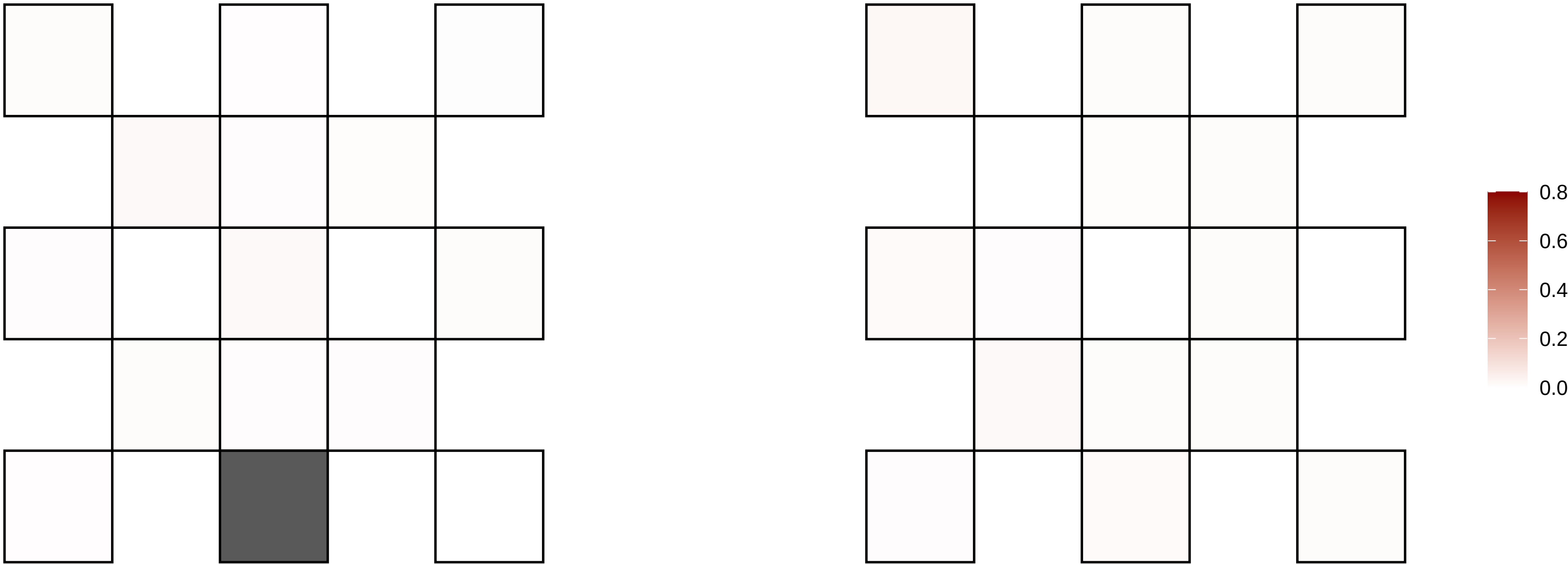

3−54233044−C−T

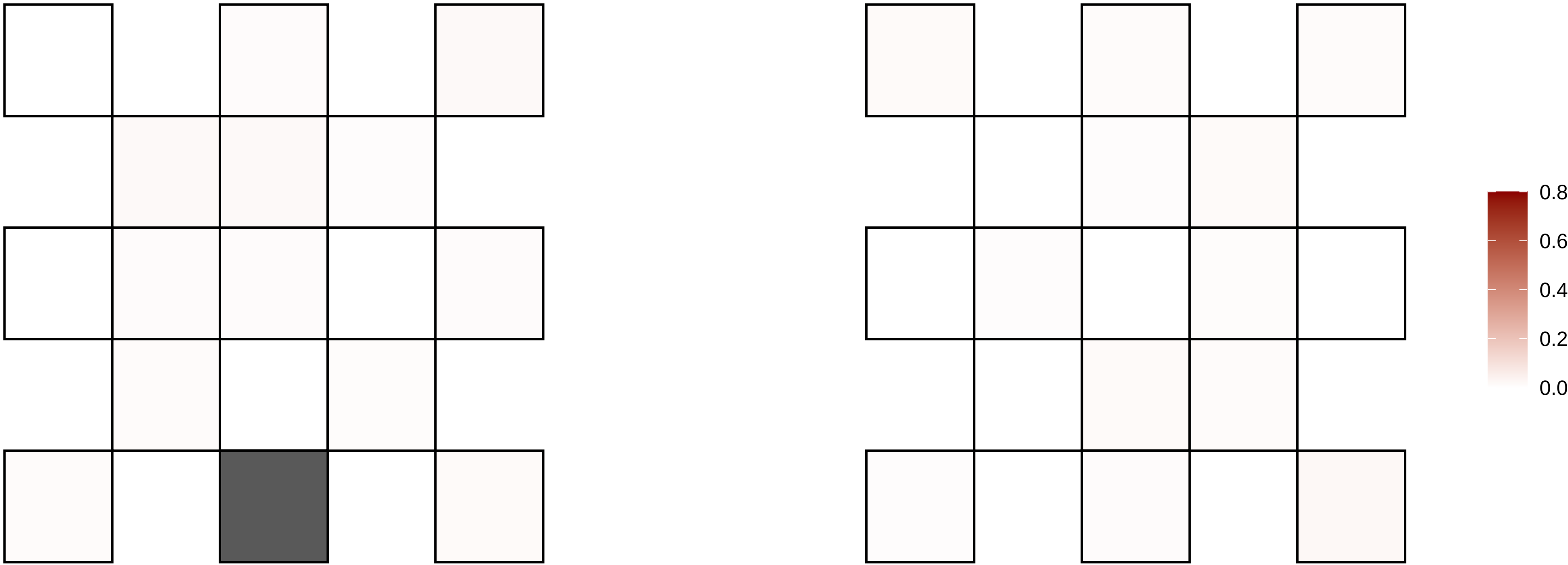

3−57851309−G−A

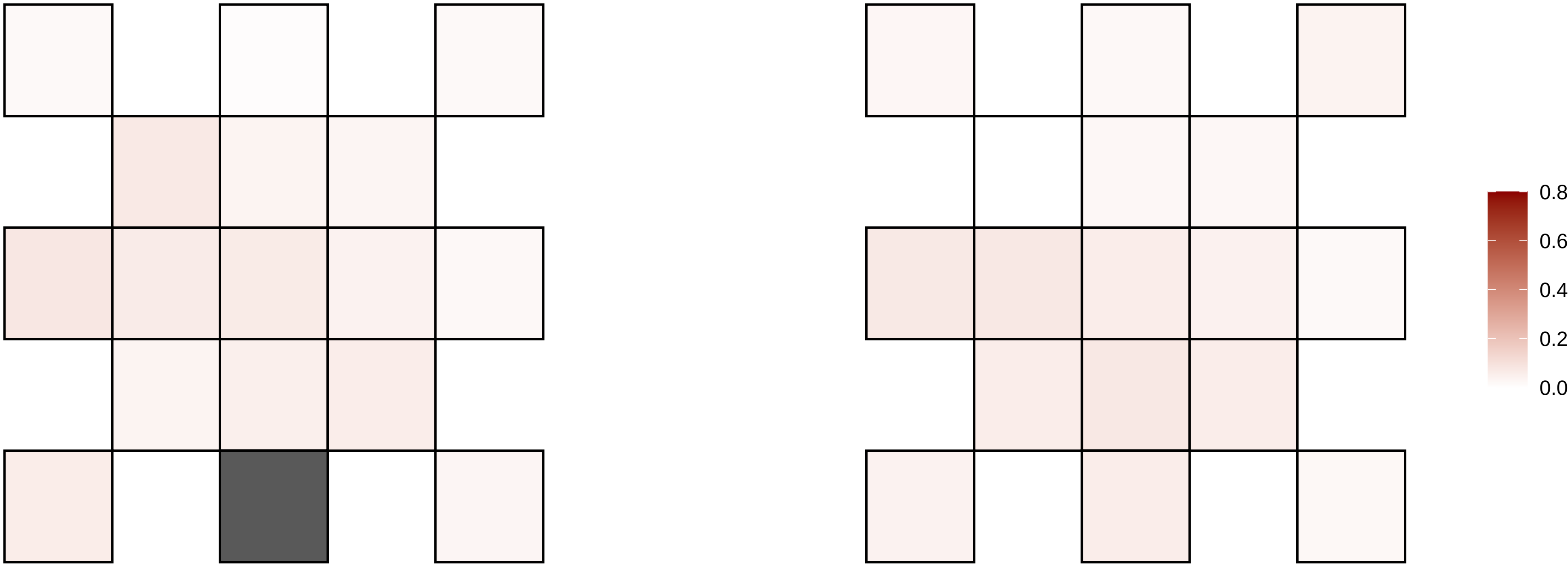

3−58915984−C−A

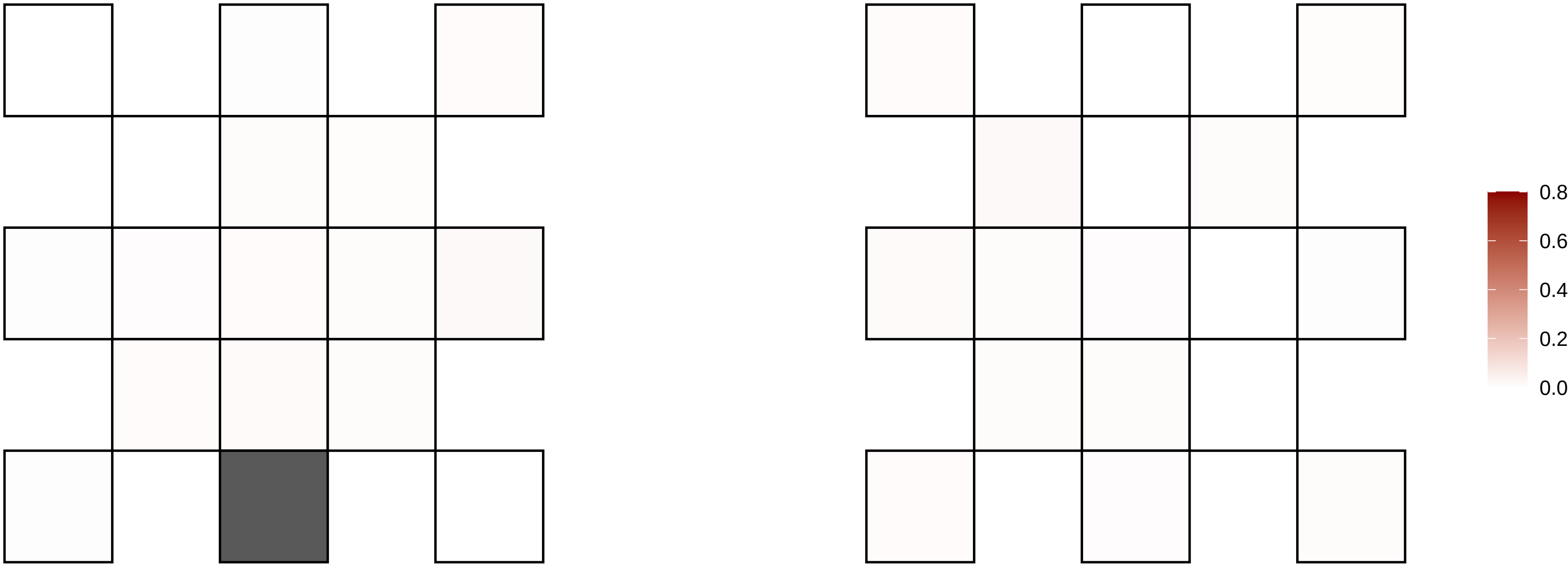

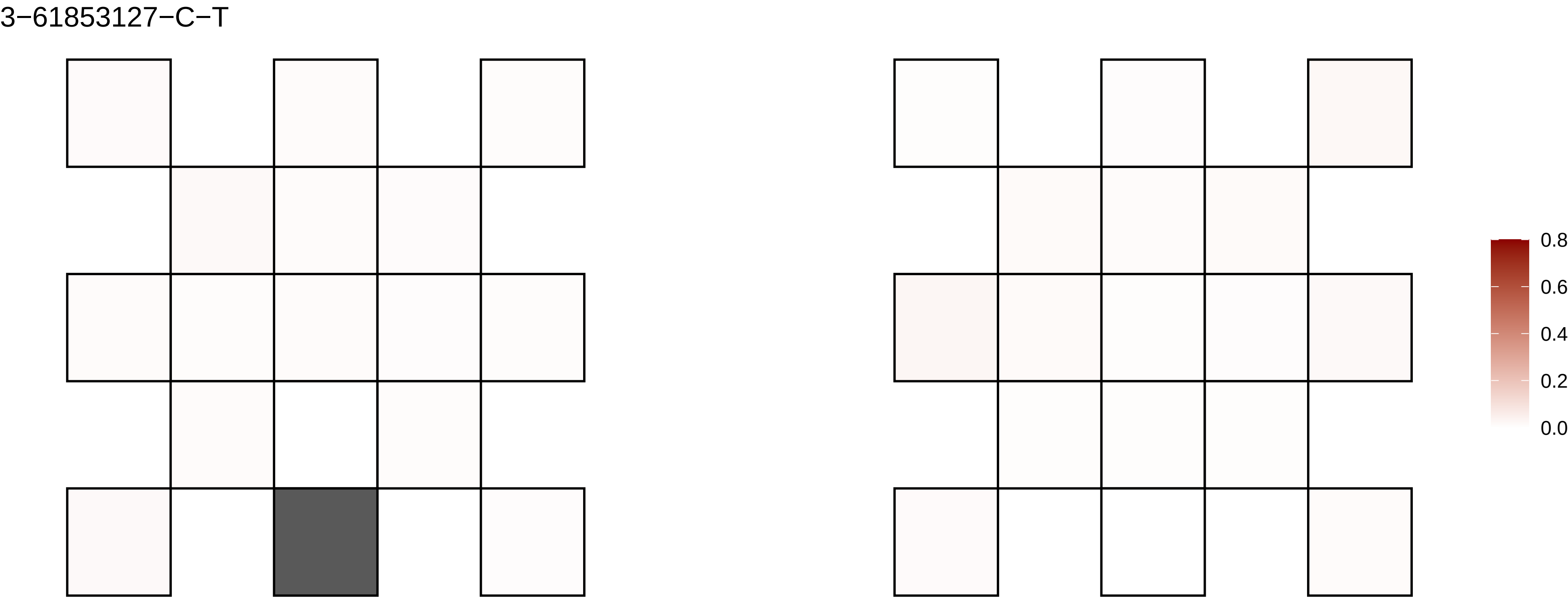
3−61853127−C−T
0.8
0.6
0.4
0.2
0.0

3−69001163−C−T

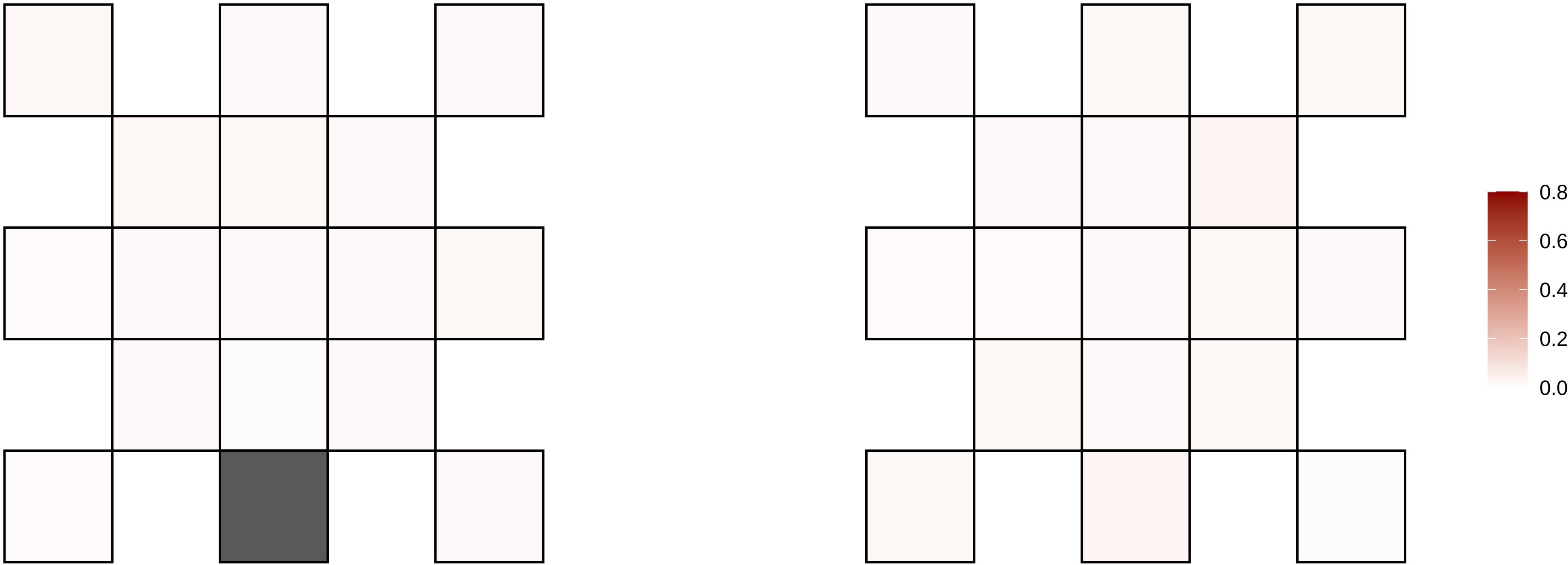

3−69456970−G−A

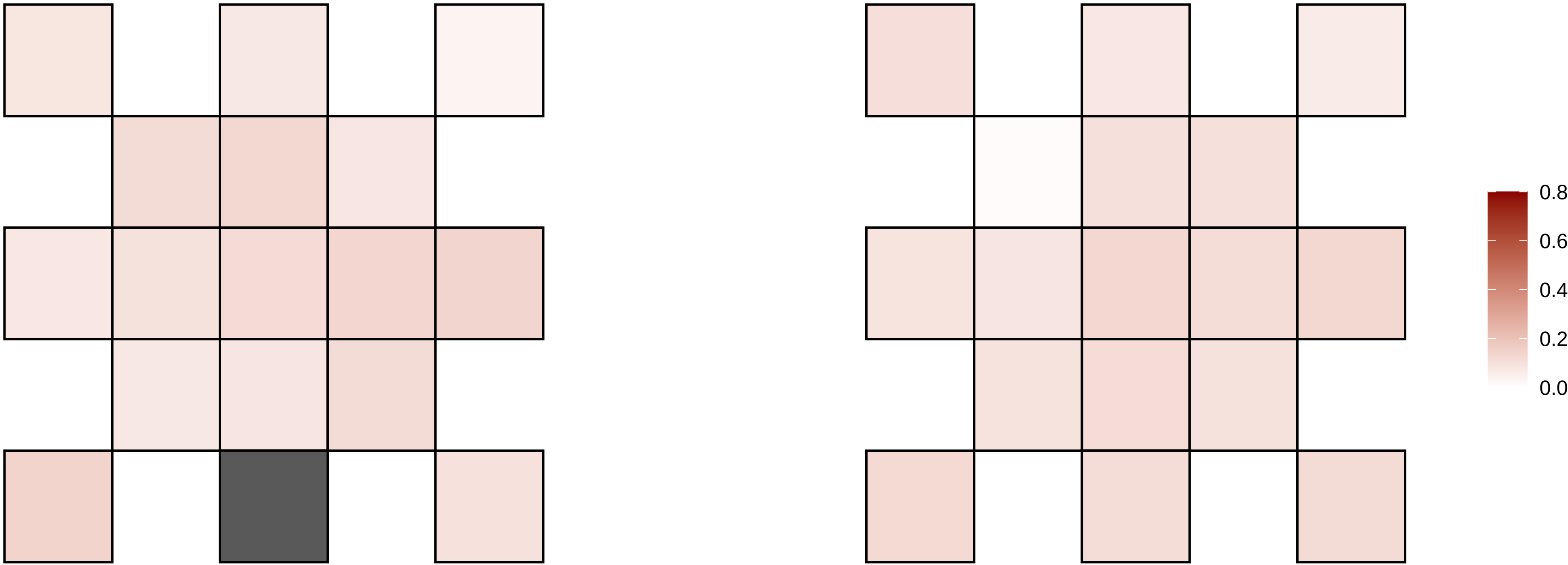

3−78738978−T−C

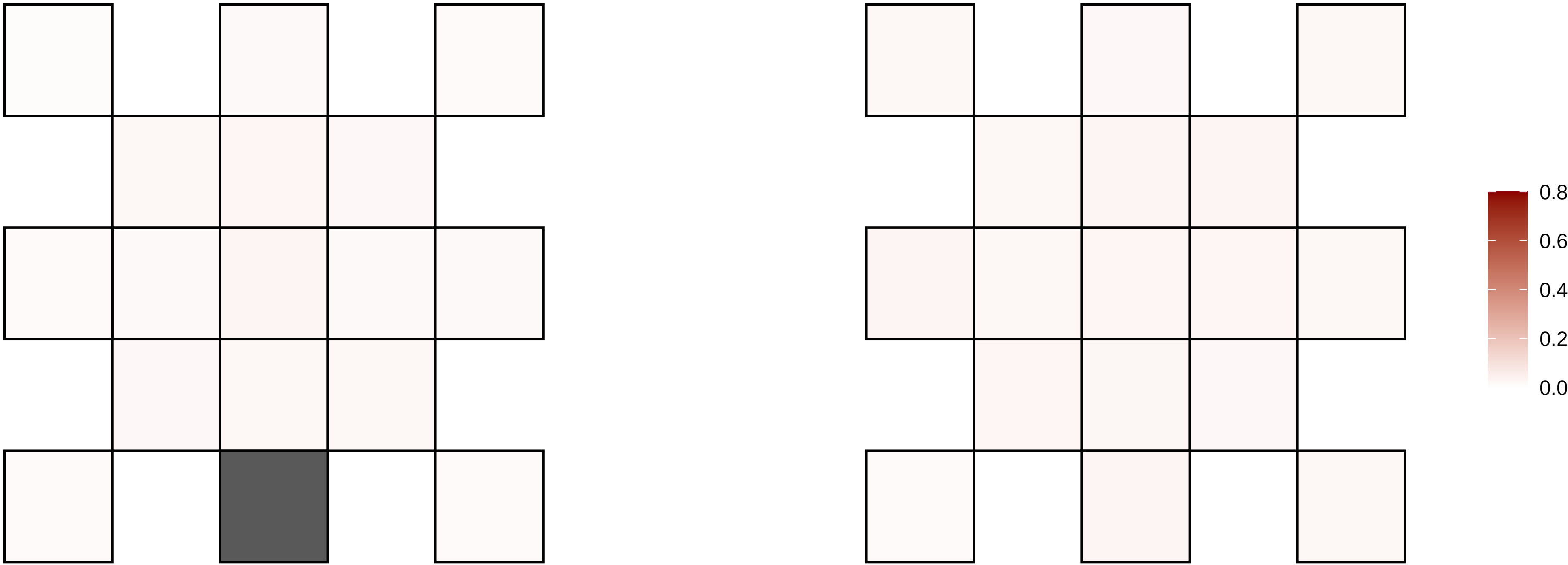

3−79357944−G−C

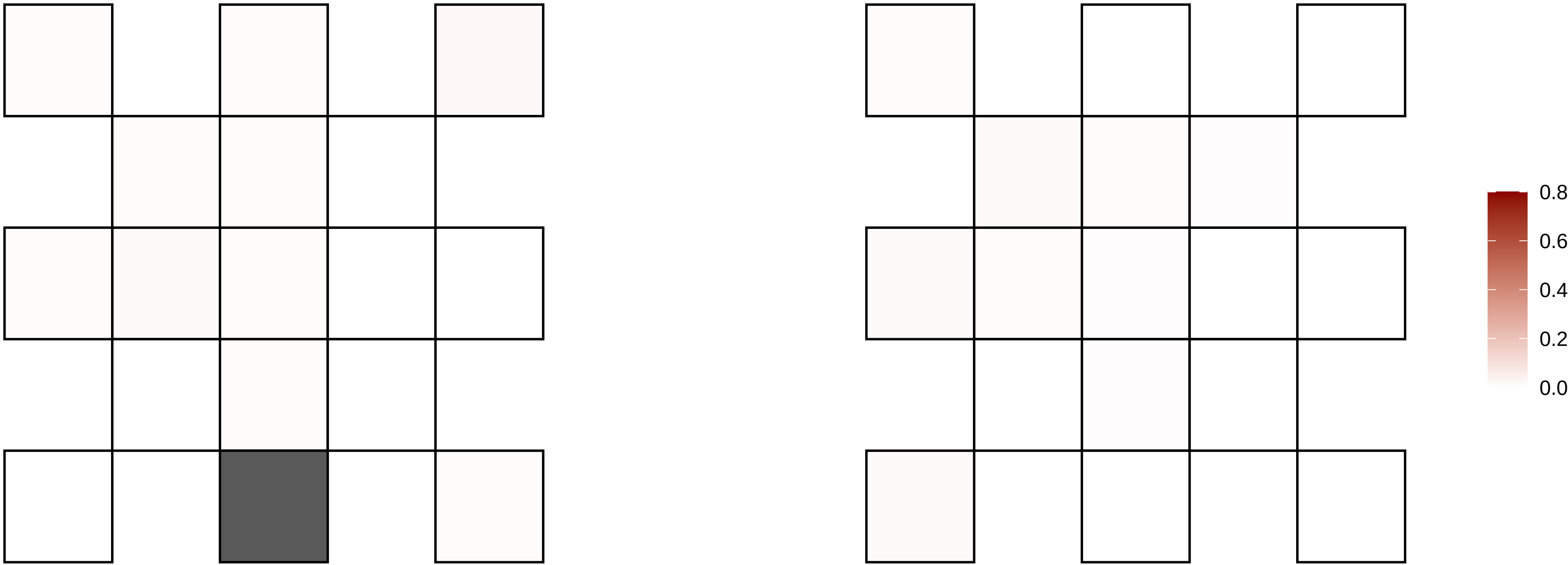

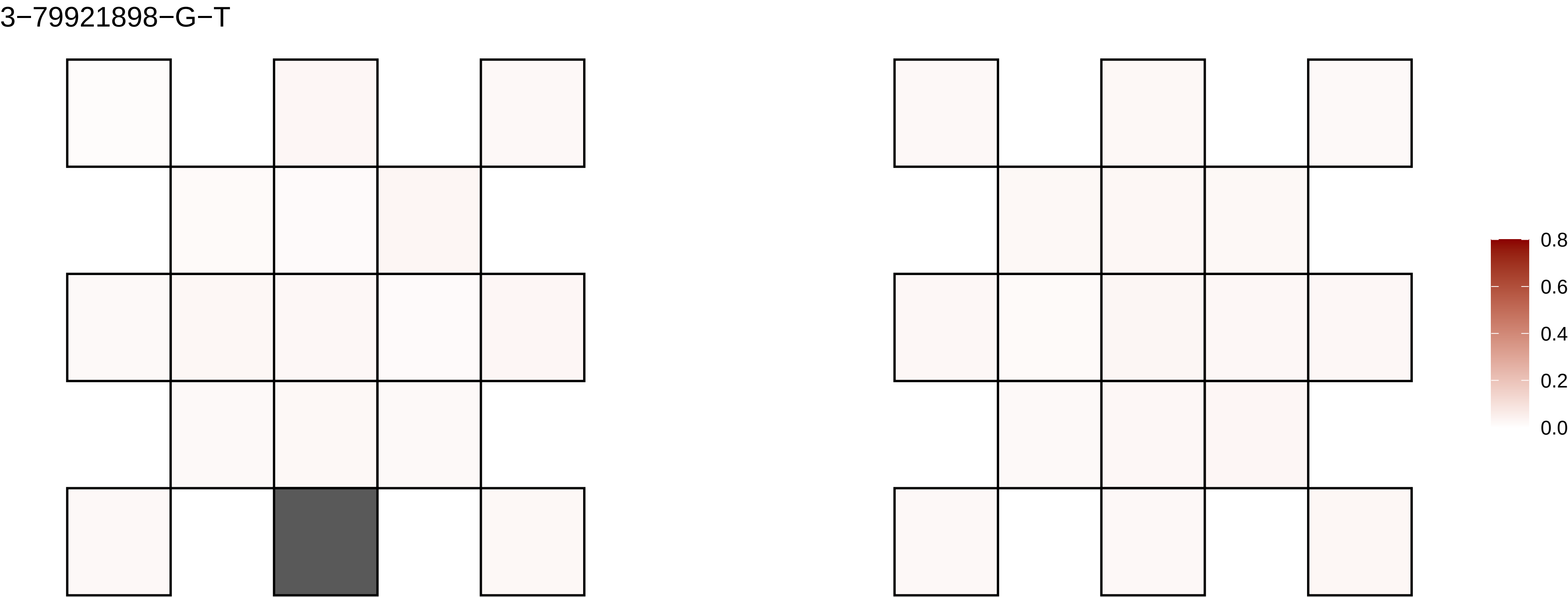

3−79921898−G−T
0.8
0.6
0.4
0.2
0.0

3−82072417−C−T

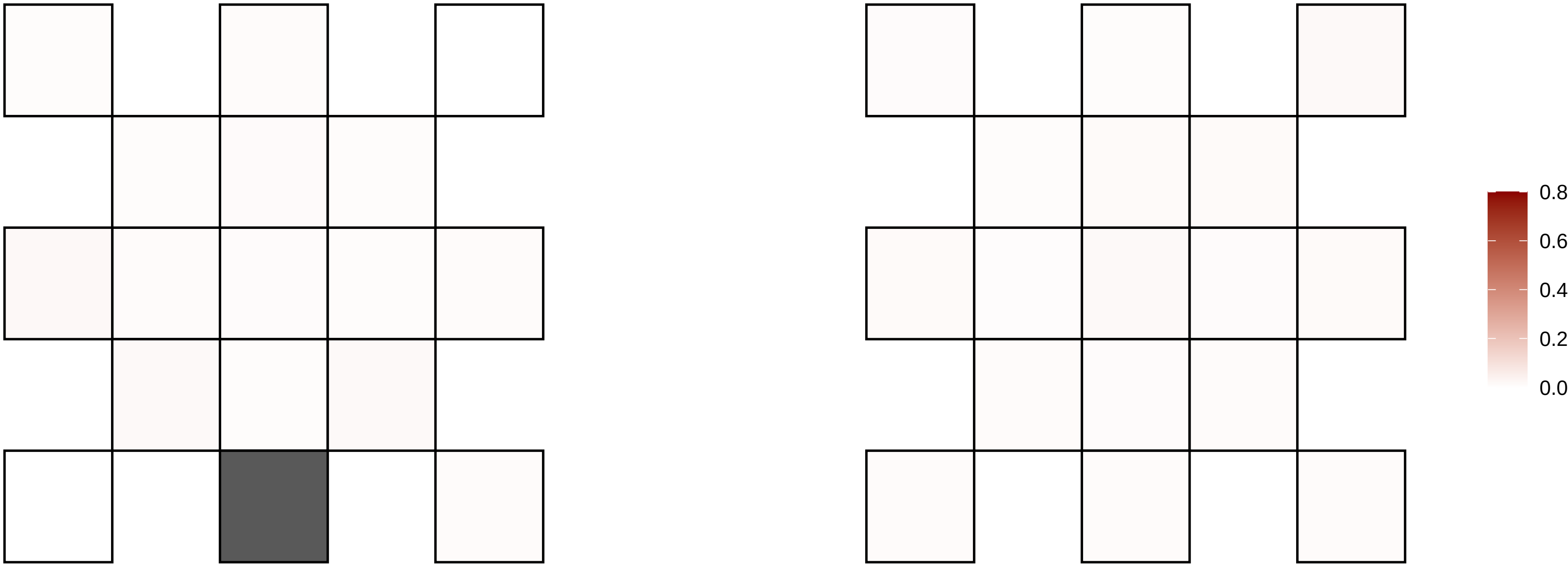

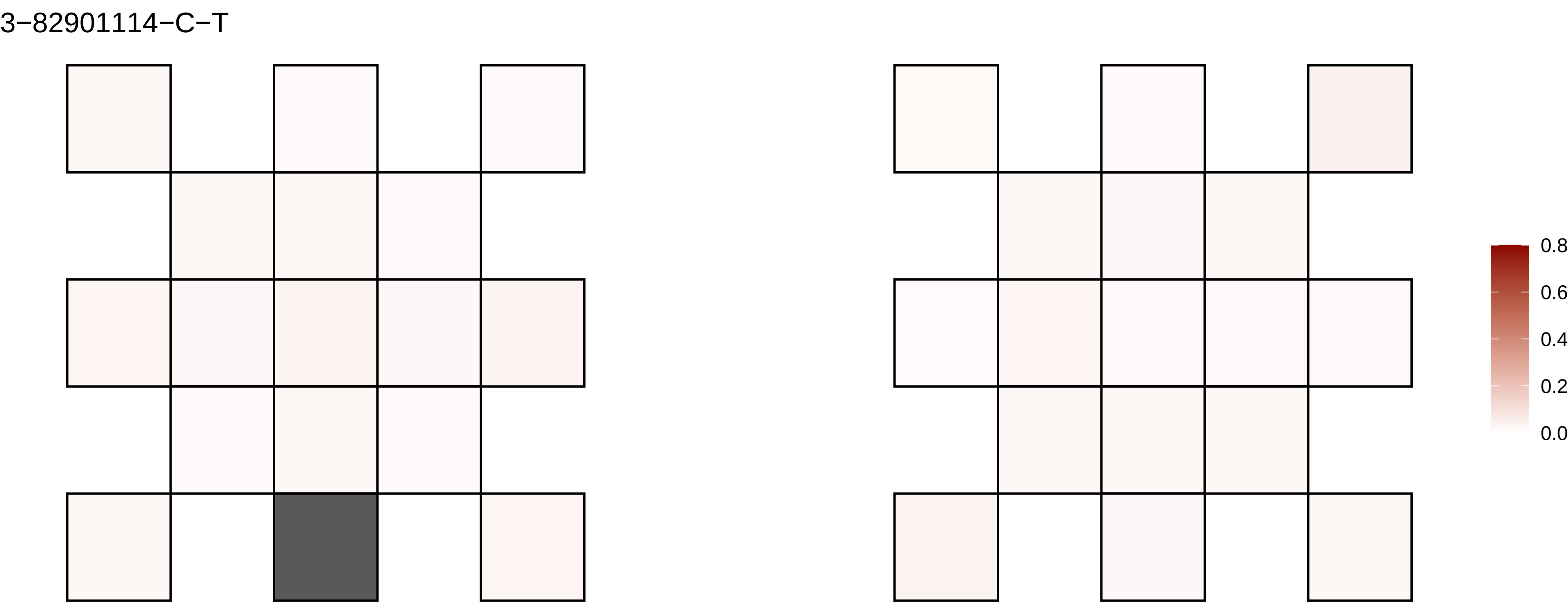
3−82901114−C−T
0.8
0.6
0.4
0.2
0.0

3−83516288−A−C

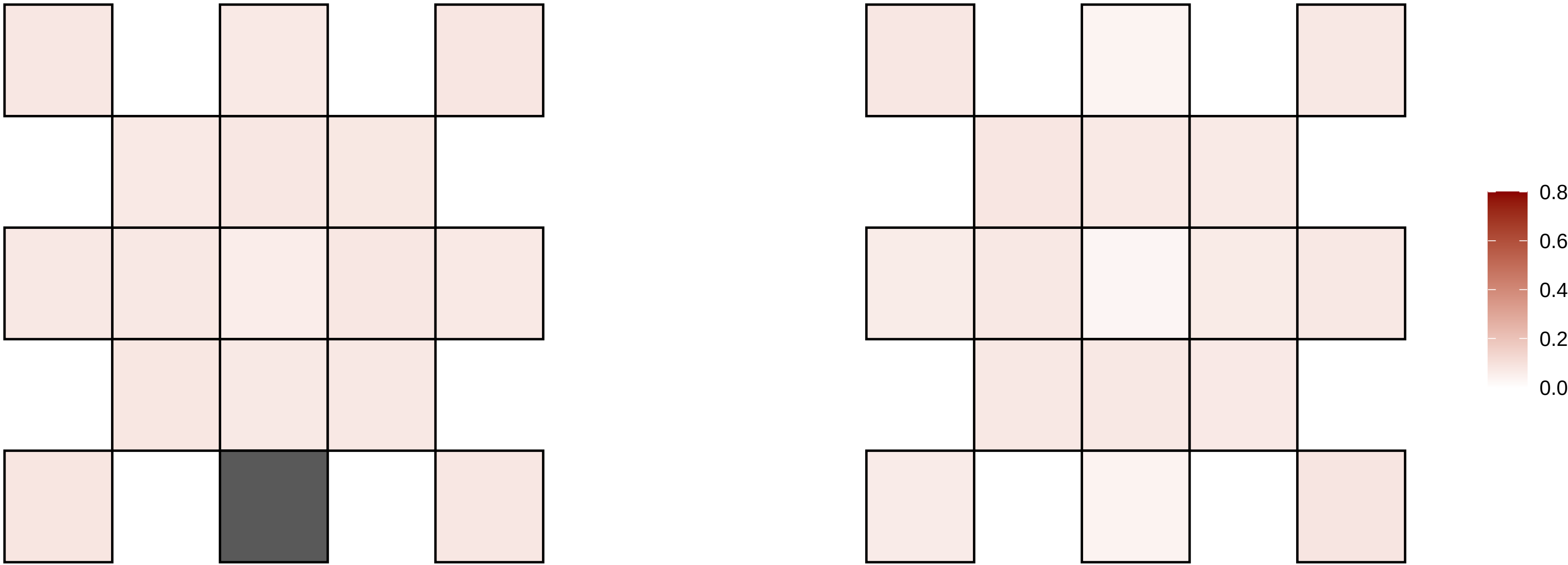

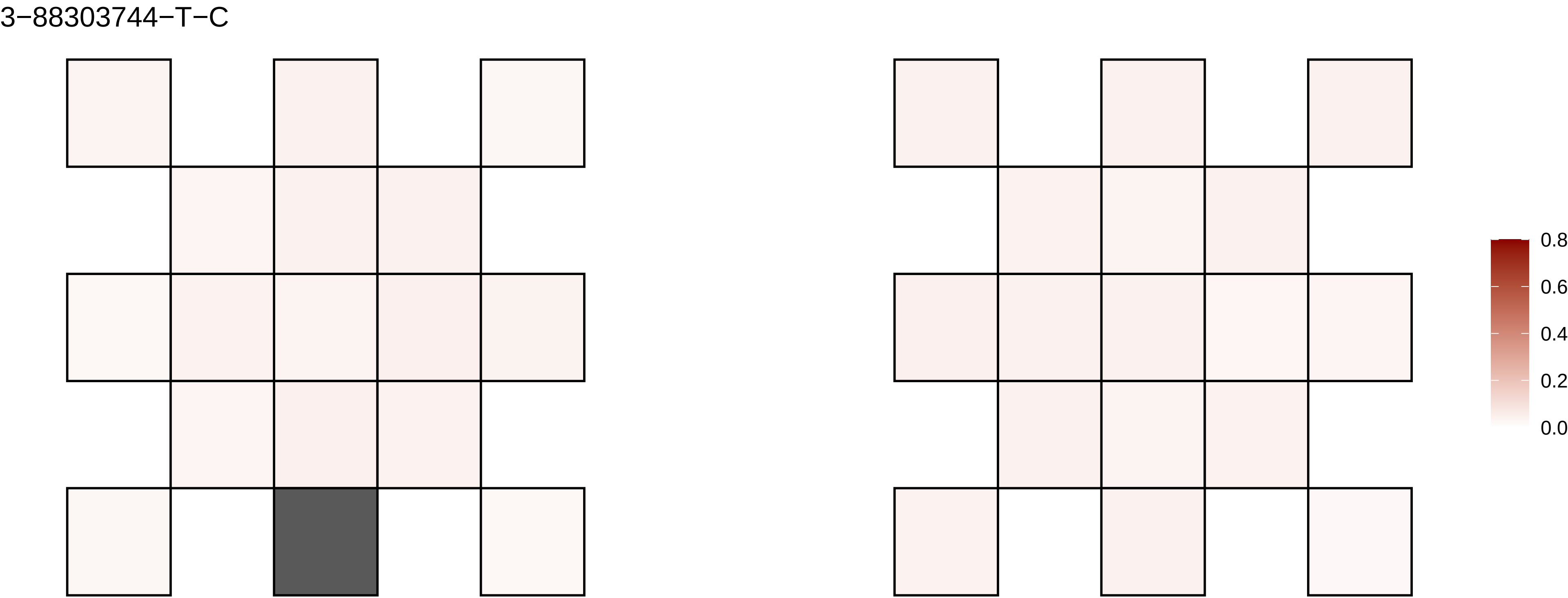
3−88303744−T−C
0.8
0.6
0.4
0.2
0.0

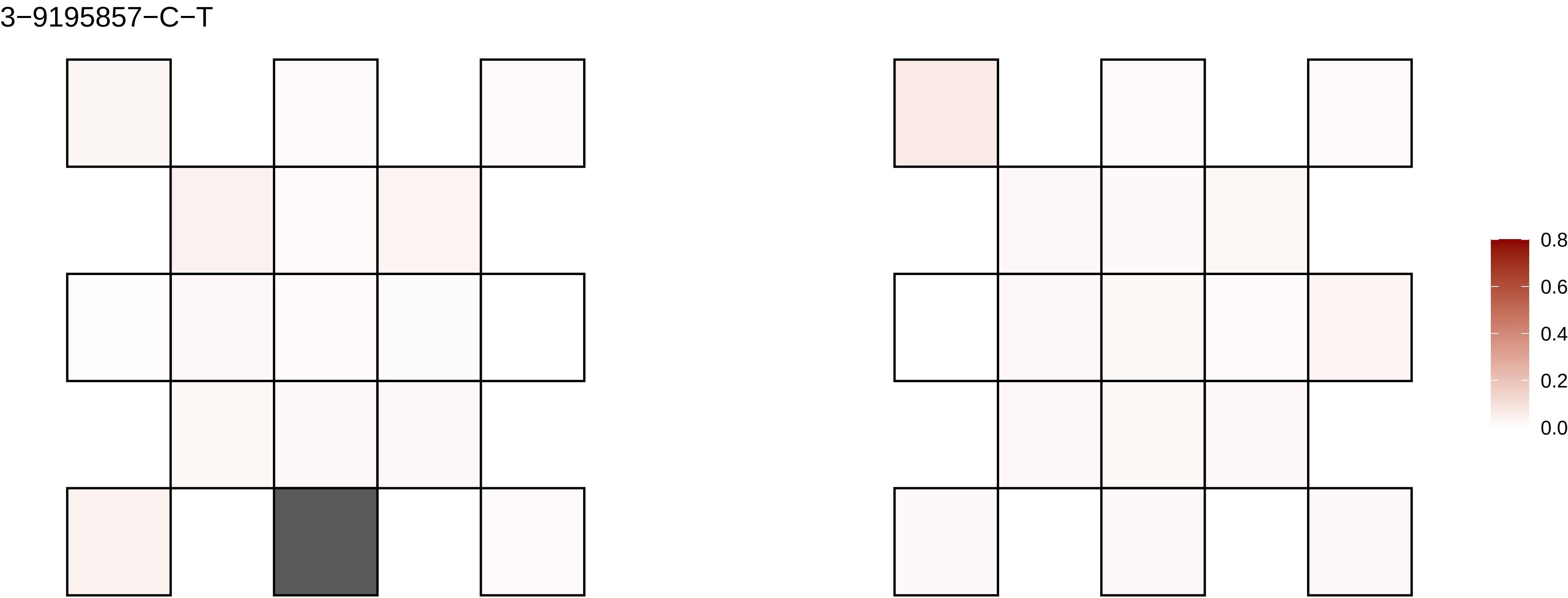
3−9195857−C−T
0.8
0.6
0.4
0.2
0.0

3−9291135−G−A

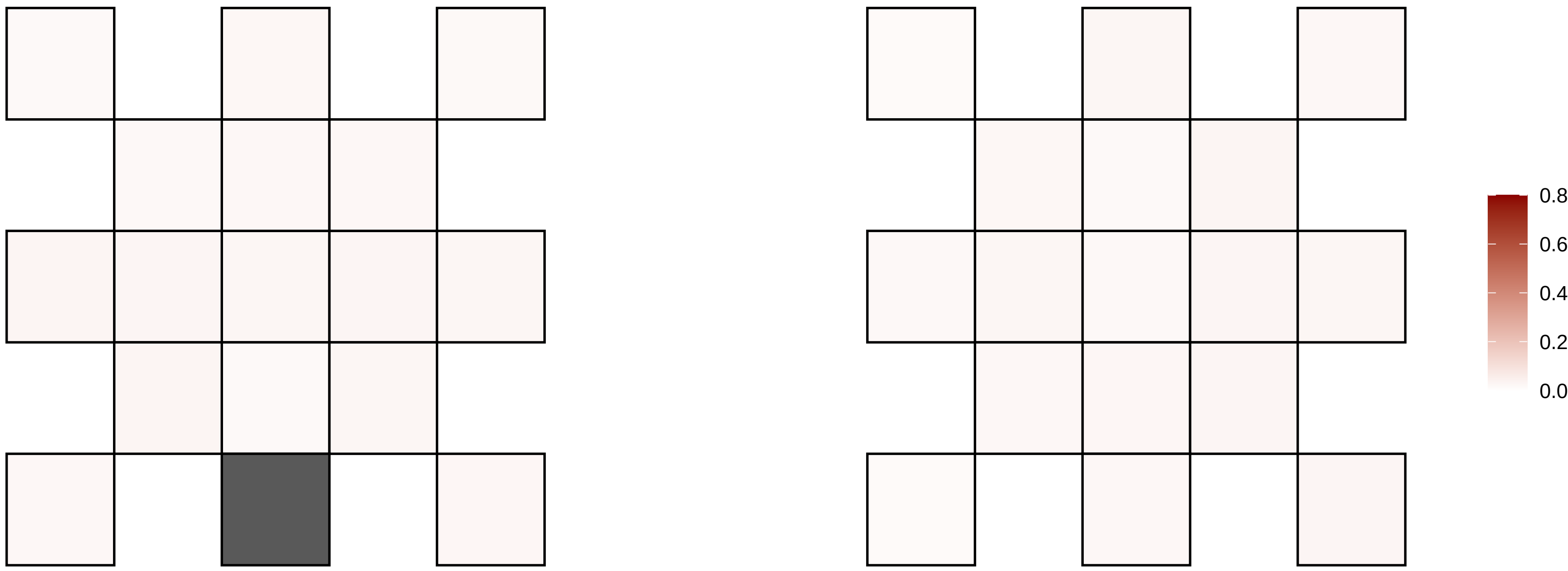

3−9859360−C−T

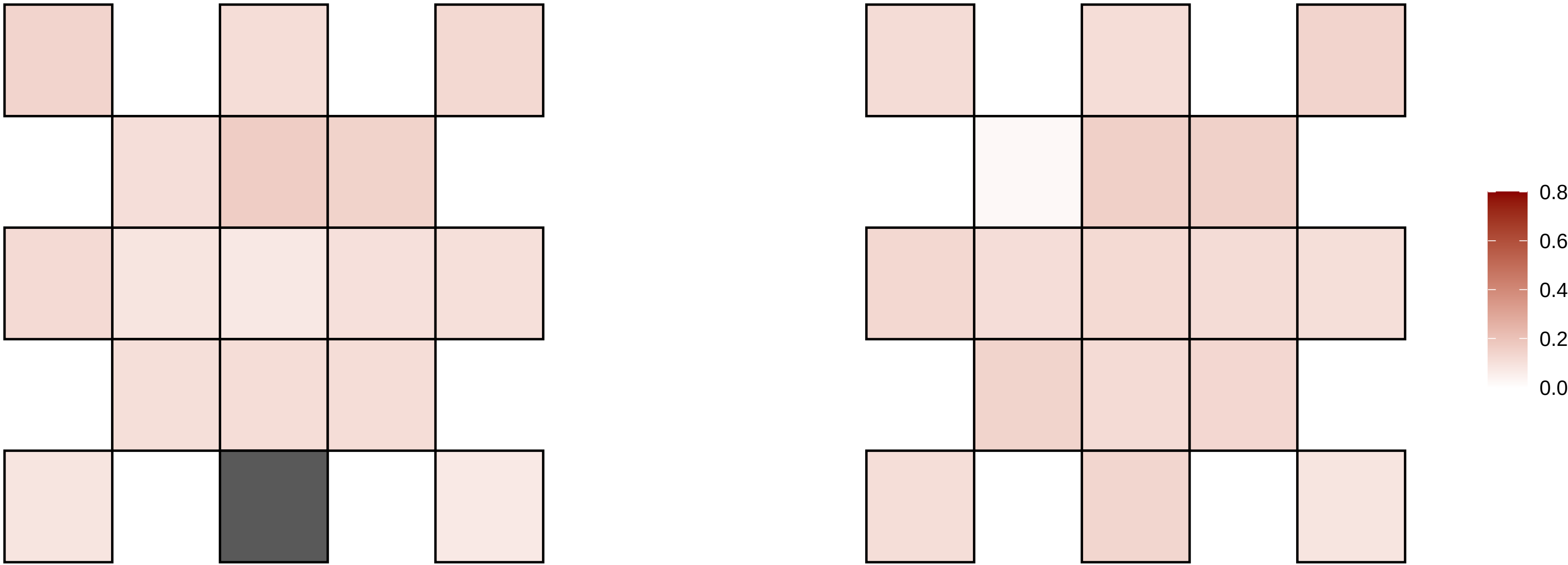

3−99869280−C−T

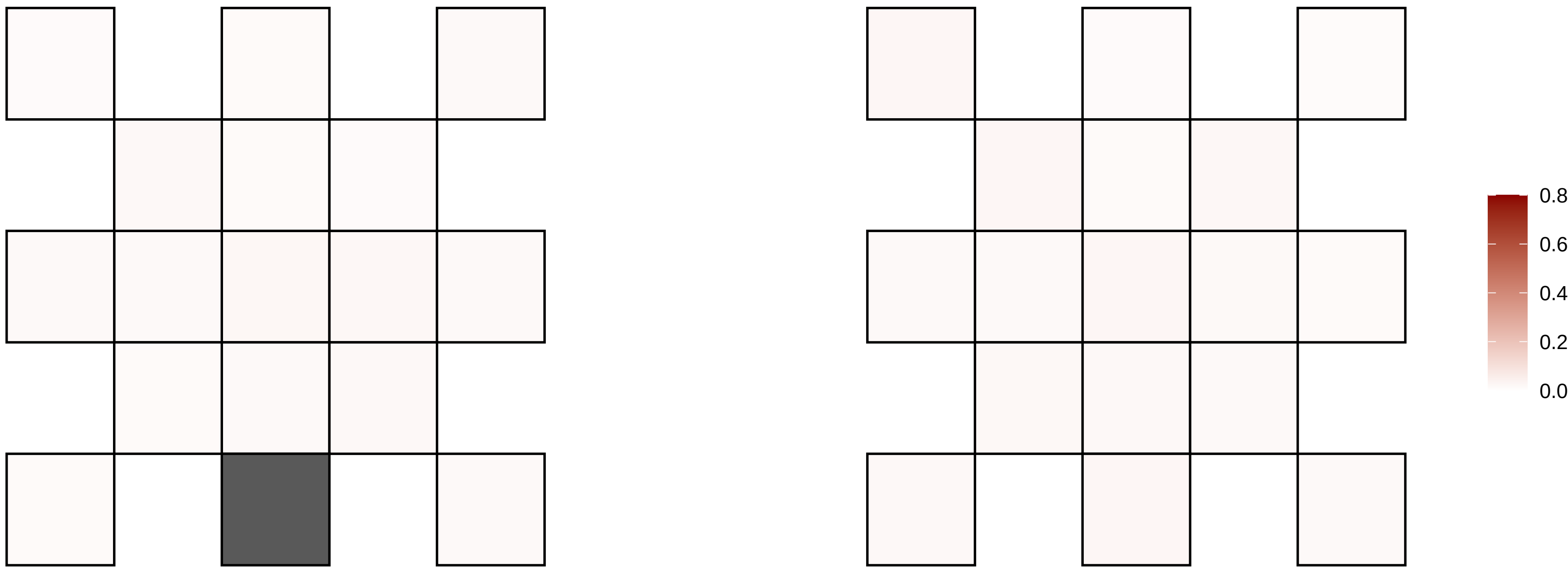

4−100236559−G−A

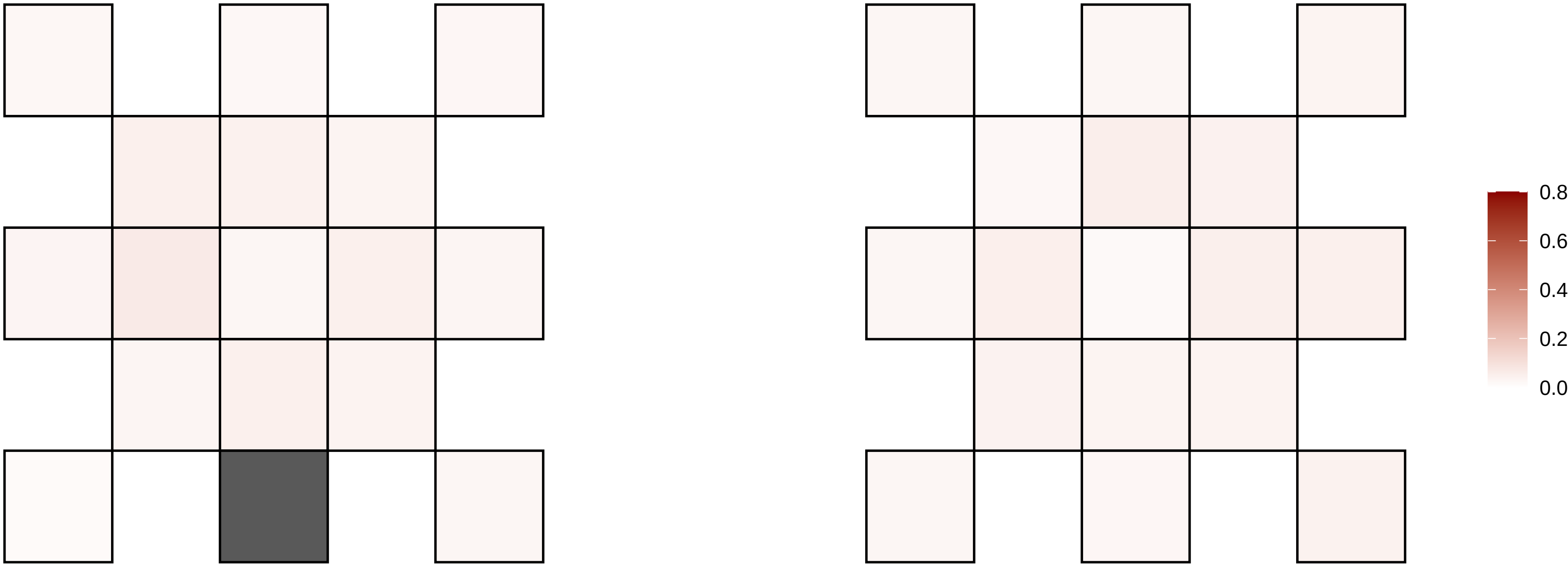

4−100484738−G−A

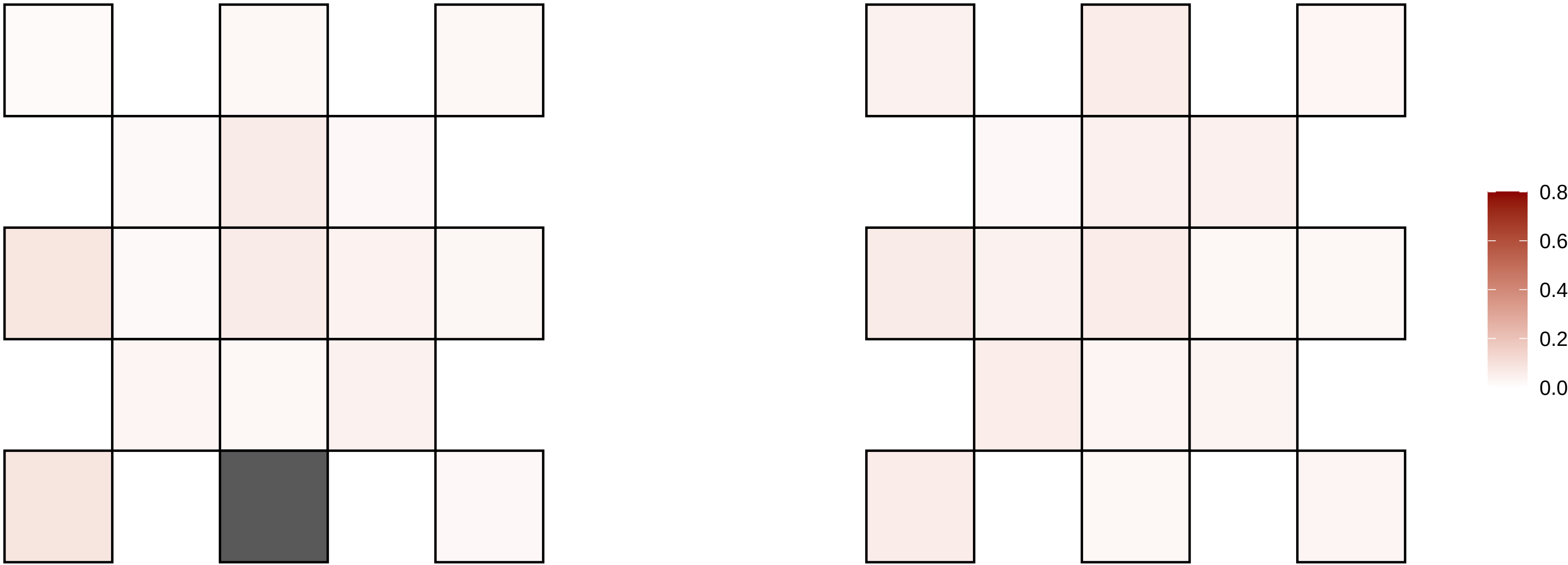

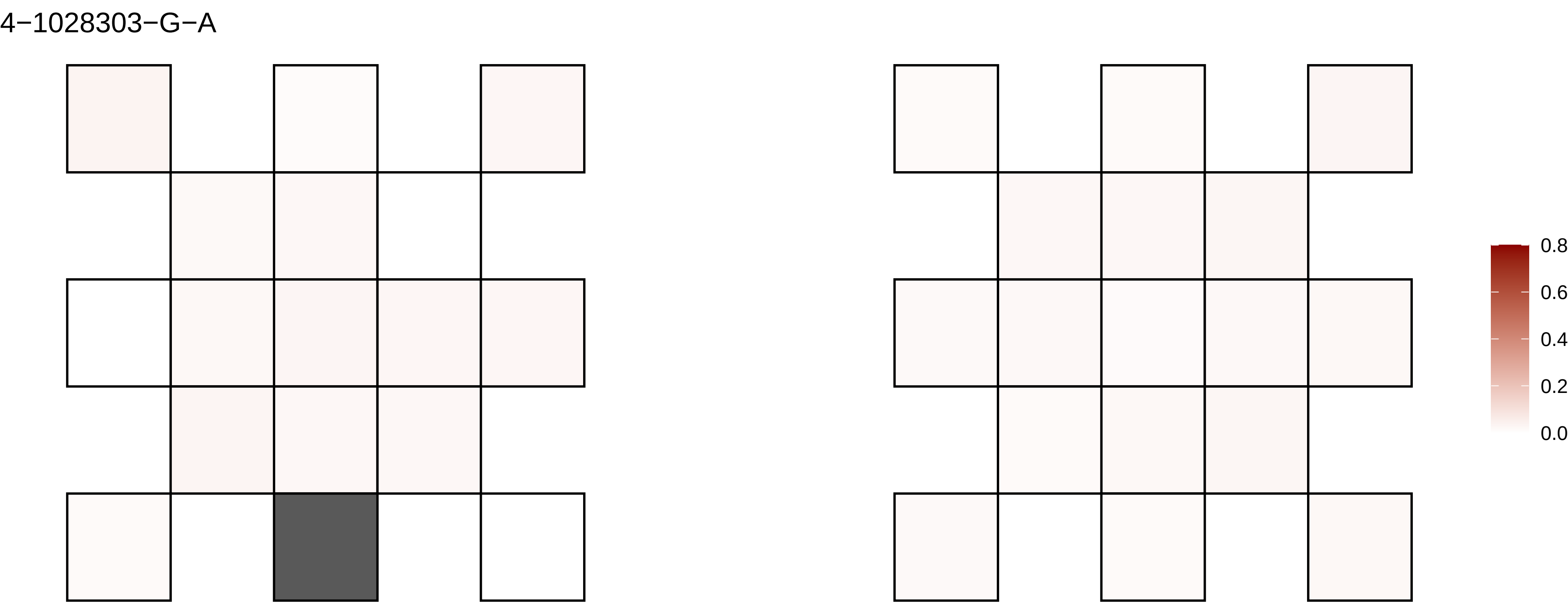
4−1028303−G−A
0.8
0.6
0.4
0.2
0.0

4−1028304−G−A

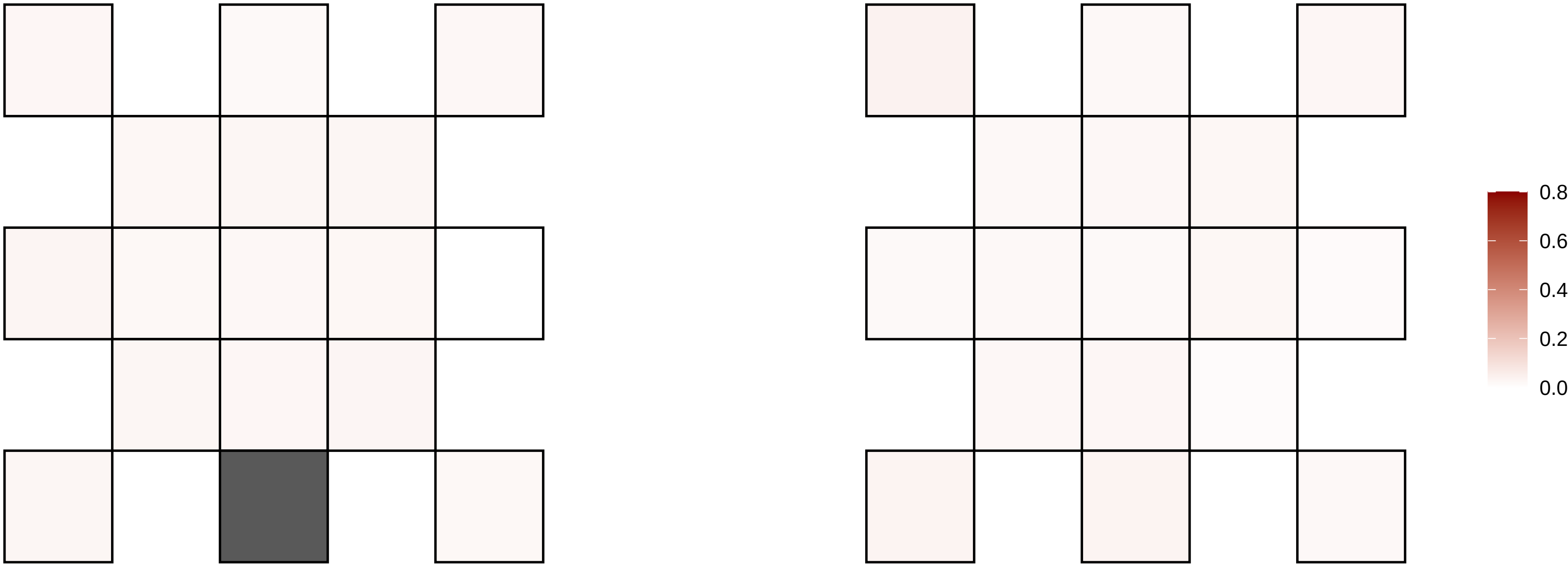

4−103751165−A−G

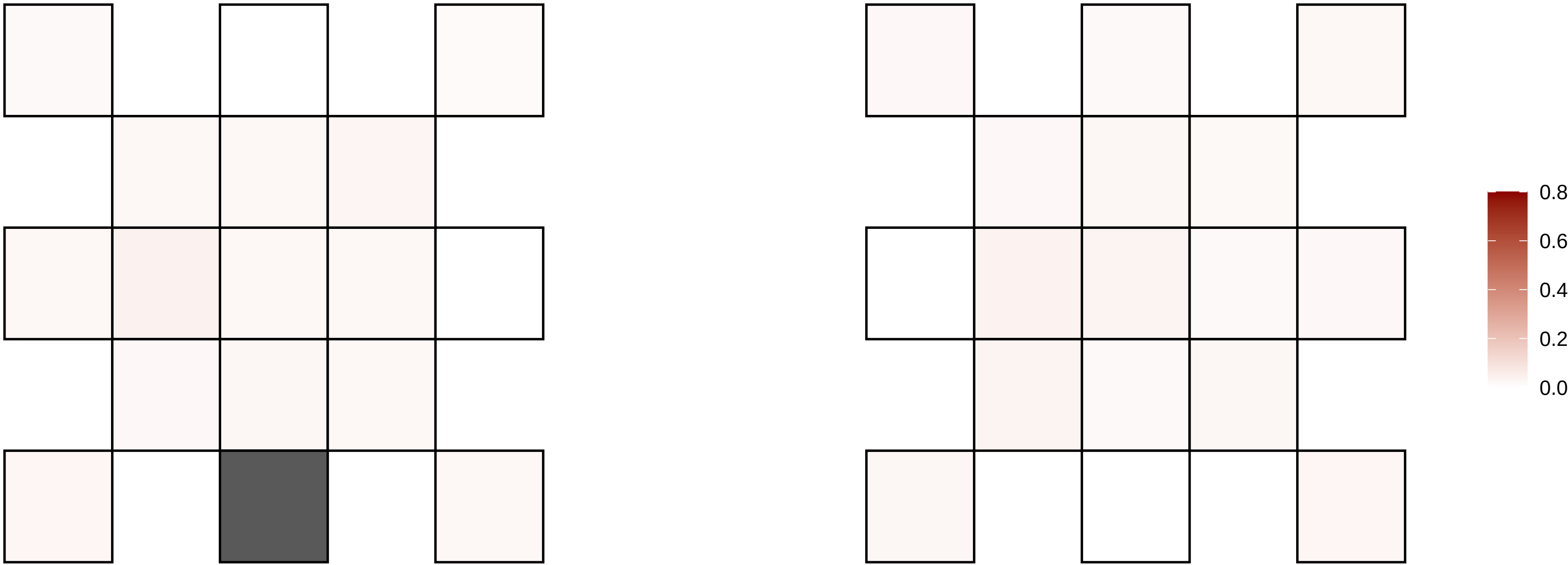

4−104214112−C−T

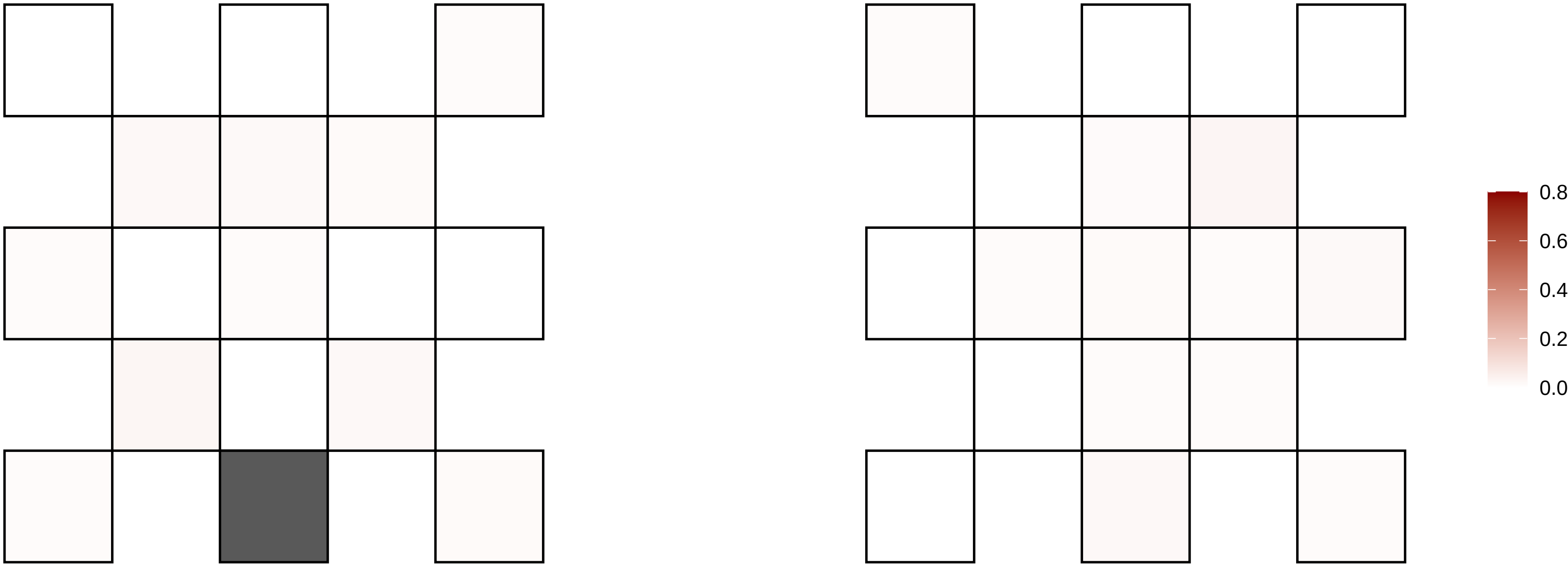

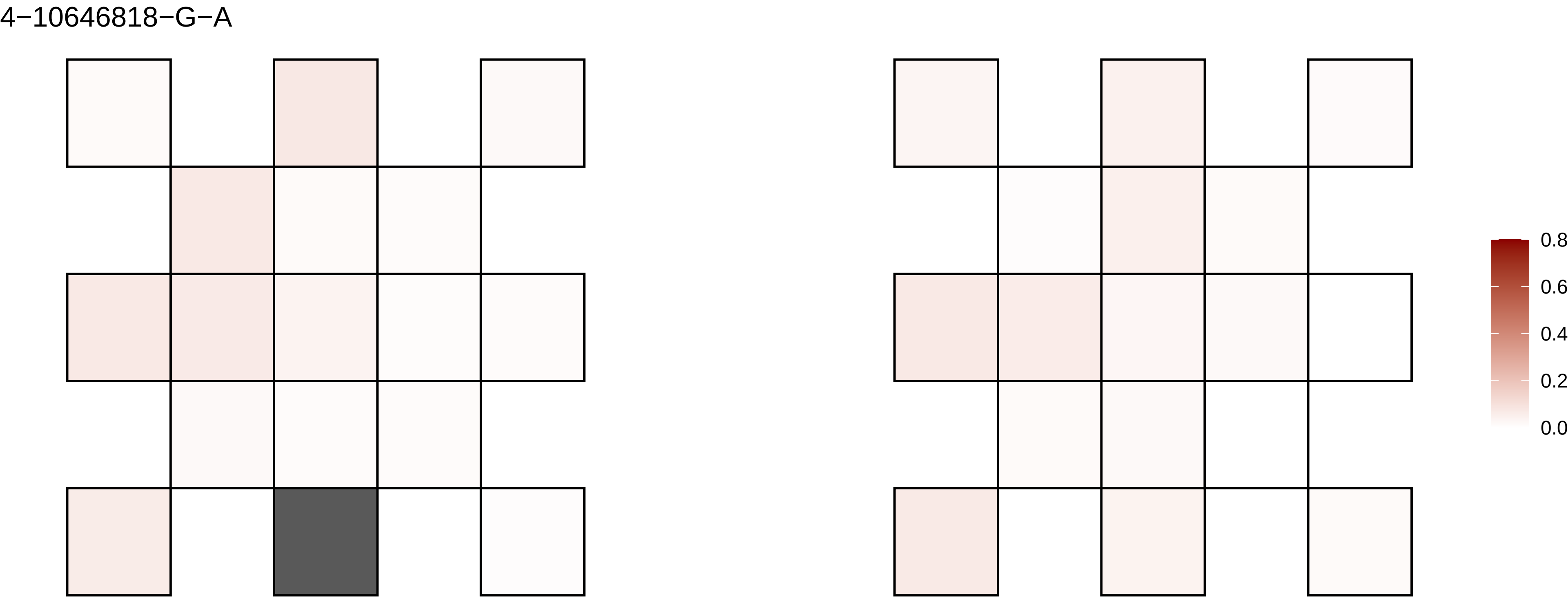
4−10646818−G−A
0.8
0.6
0.4
0.2
0.0

4−109178107−T−A

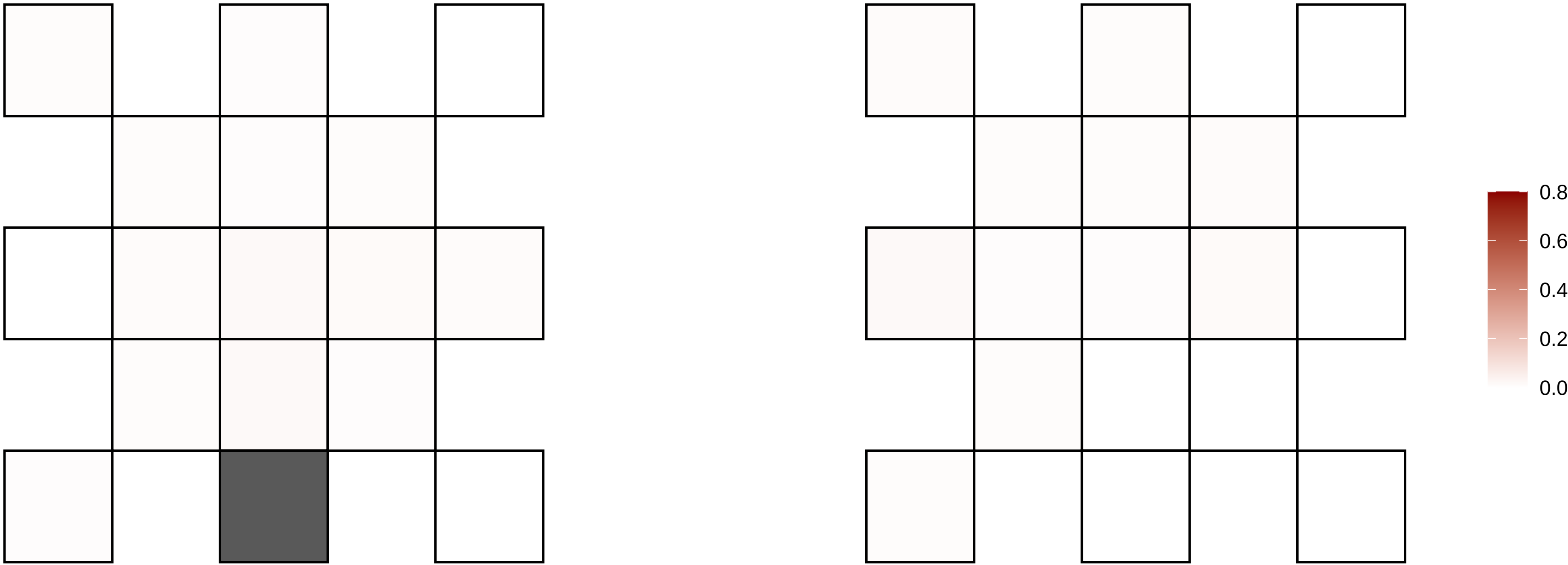

4−111676042−A−T

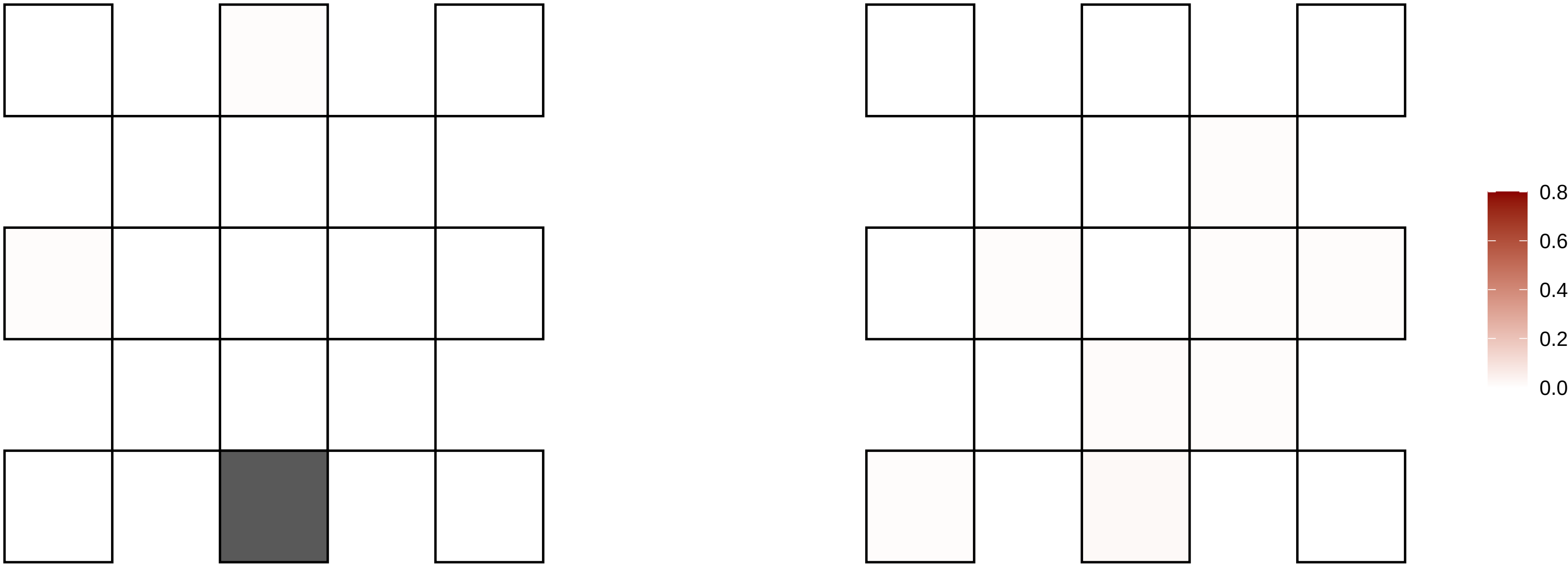

4−111915329−C−T

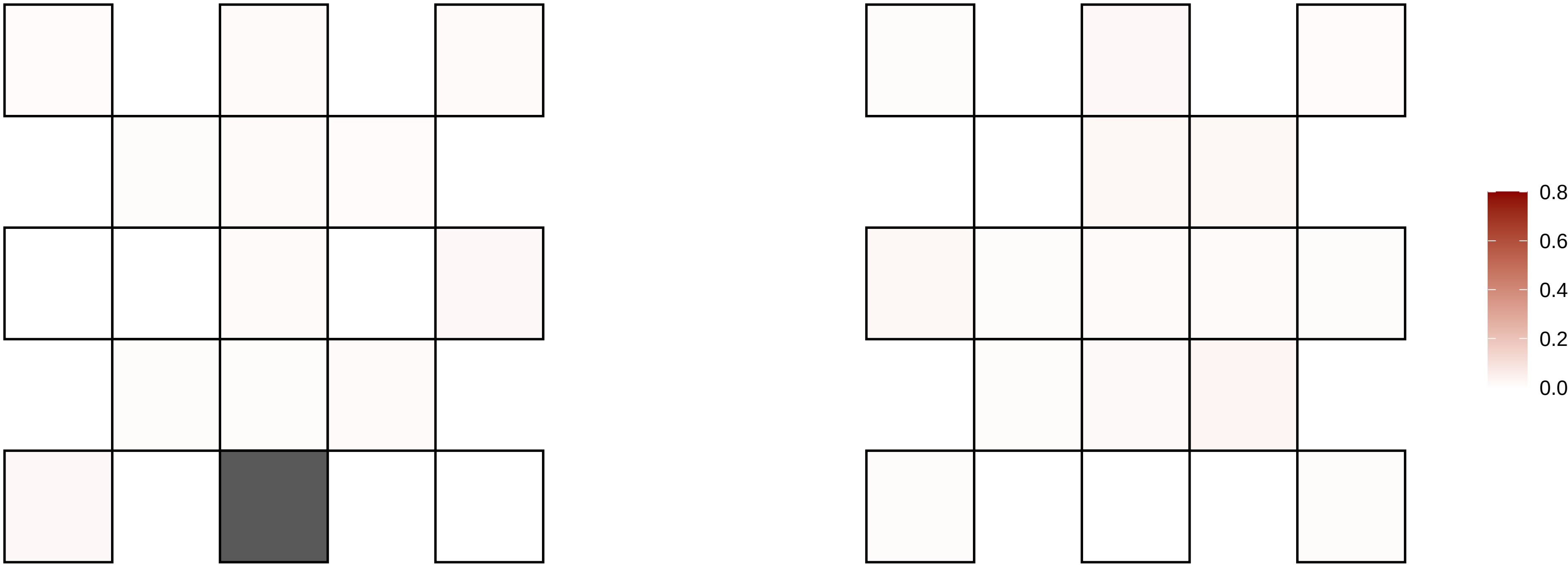

4−112309482−G−A

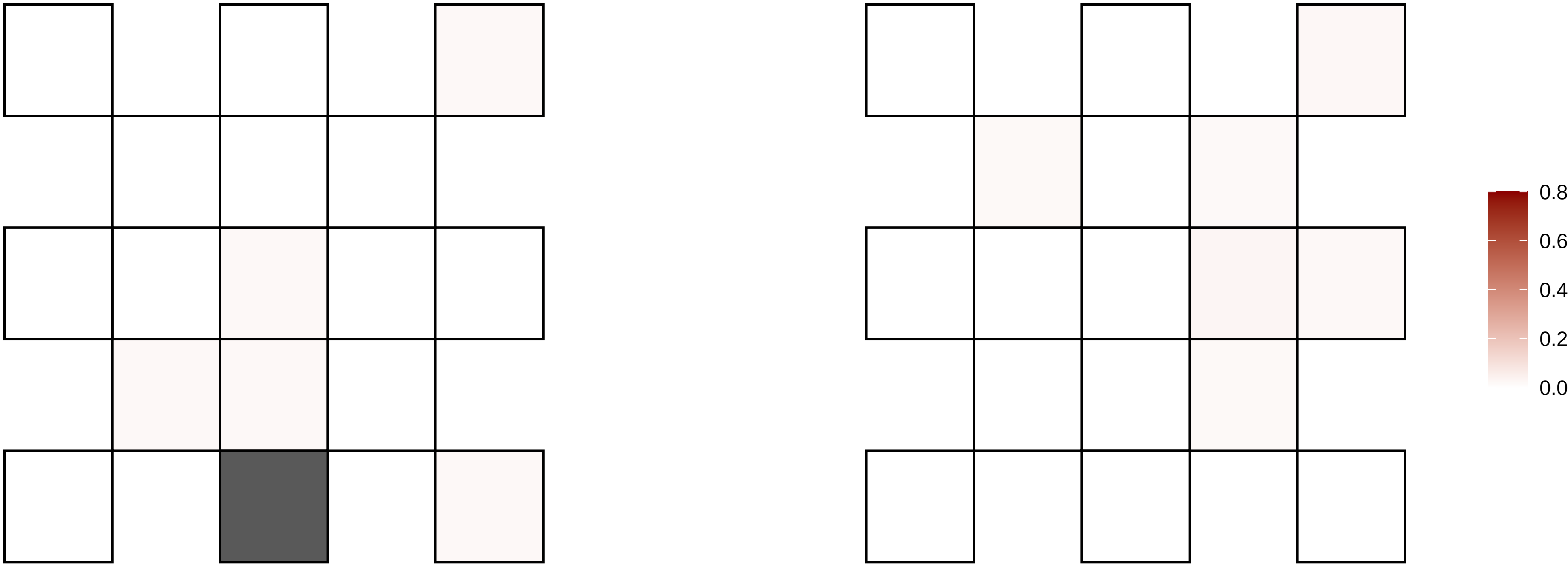

4−116228416−A−G

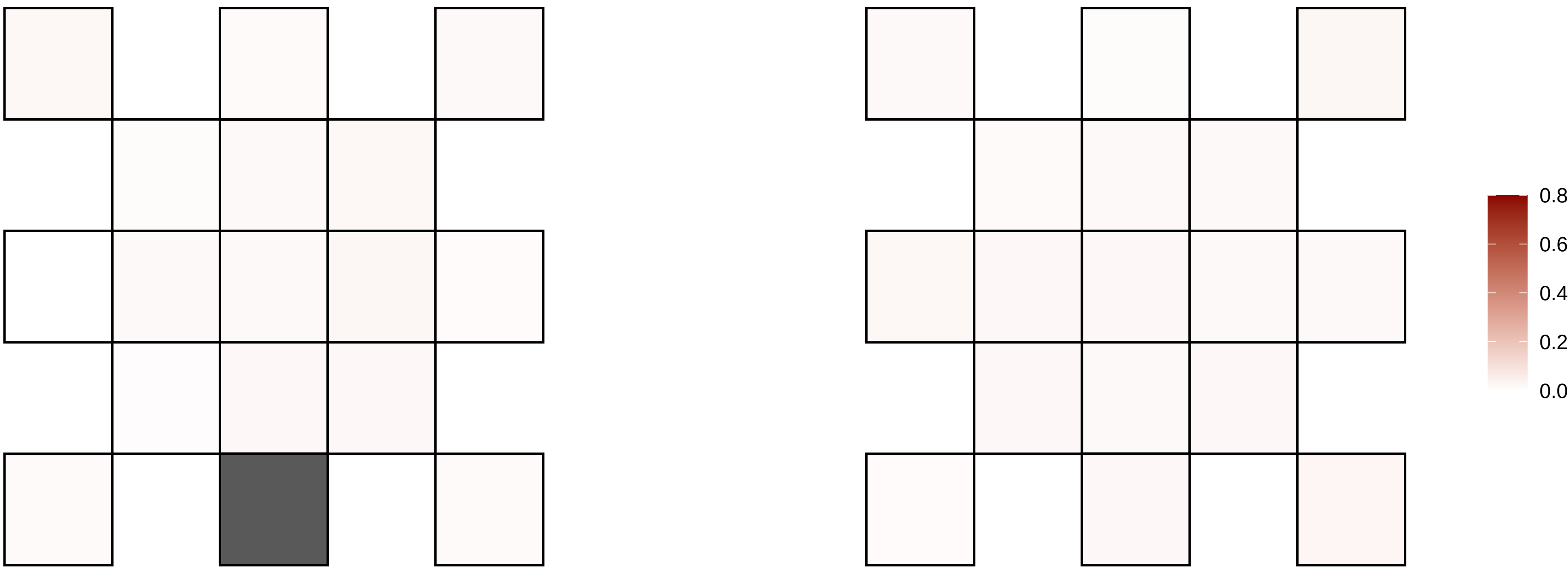

4−120426084−C−T

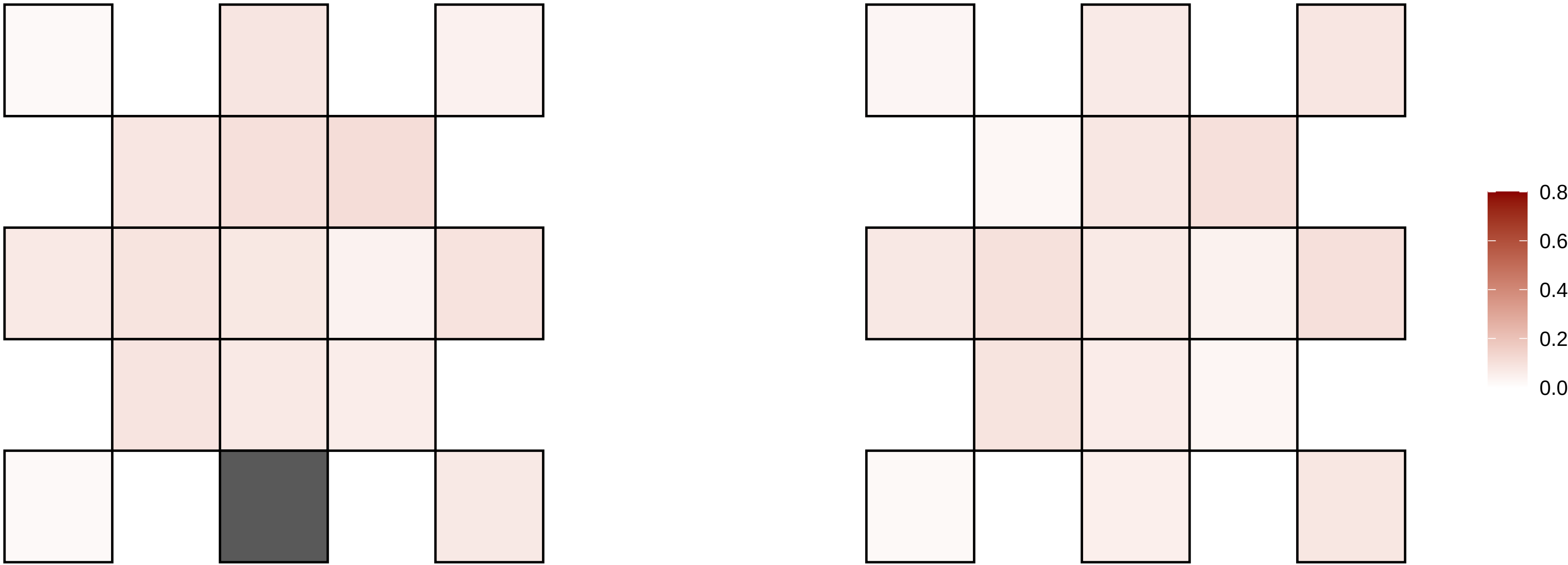

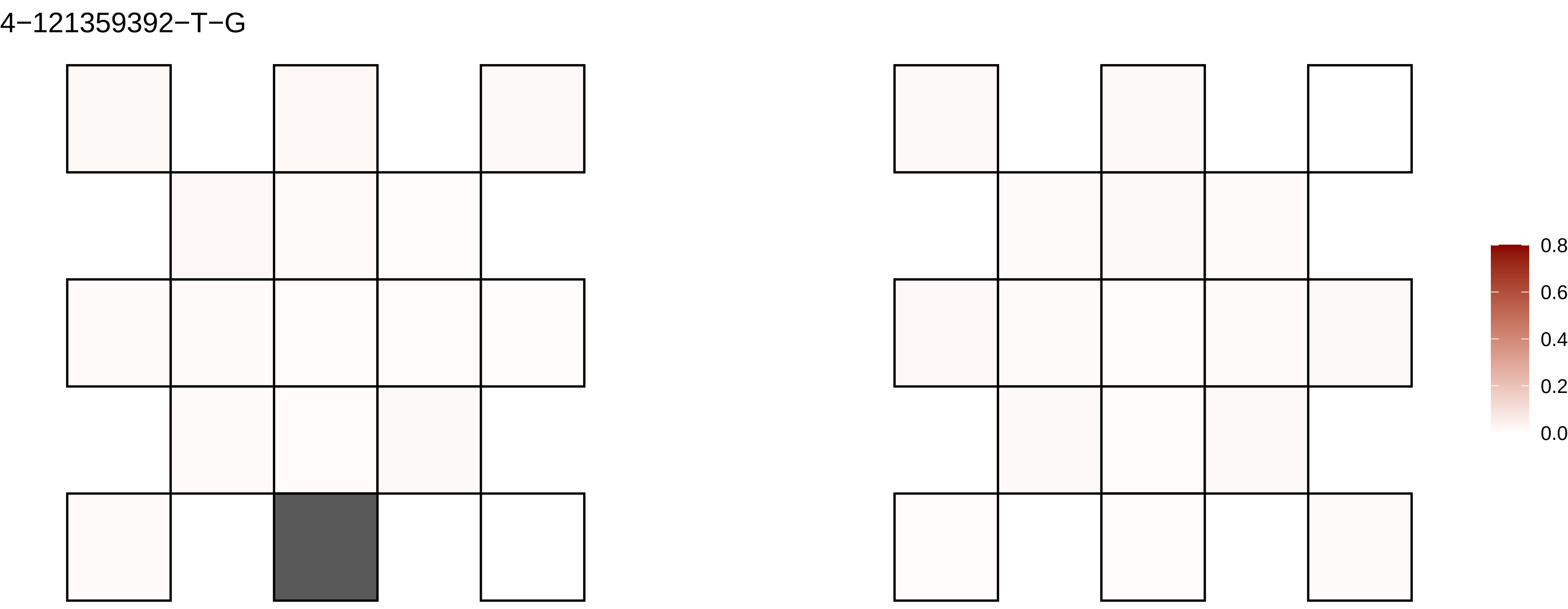
4−121359392−T−G
0.8
0.6
0.4
0.2
0.0

4−122936525−G−A

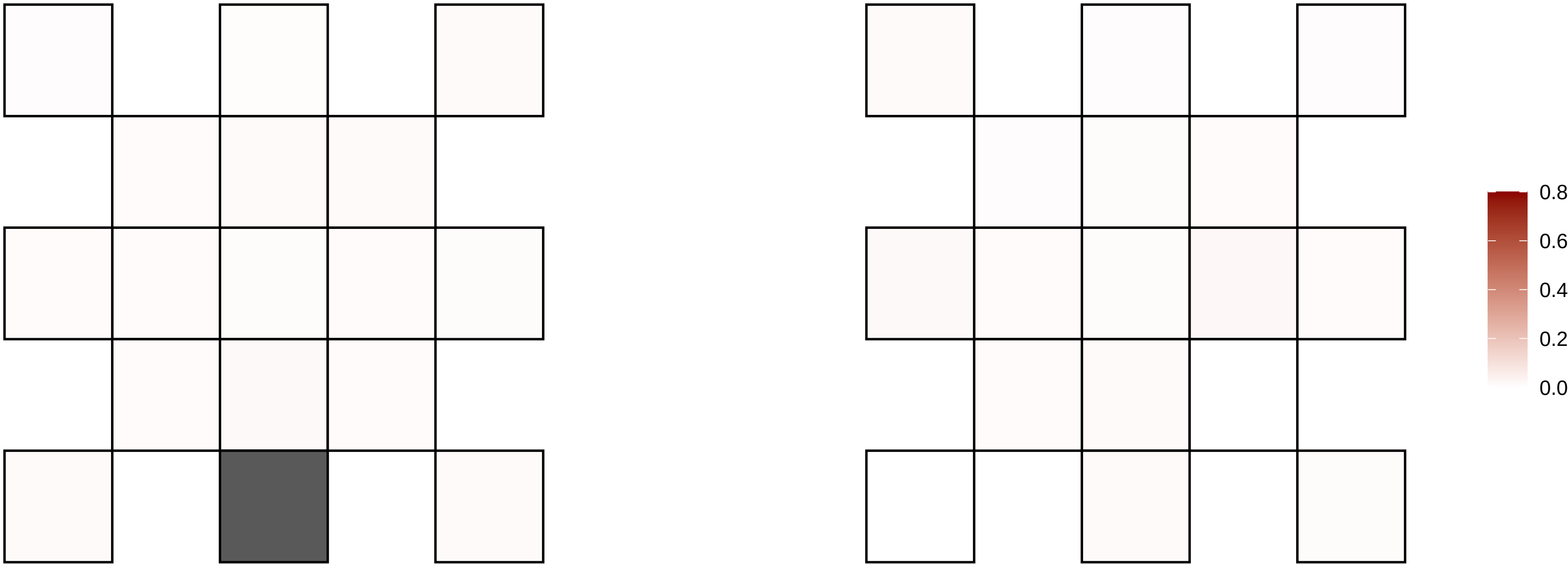

4−123000327−T−G

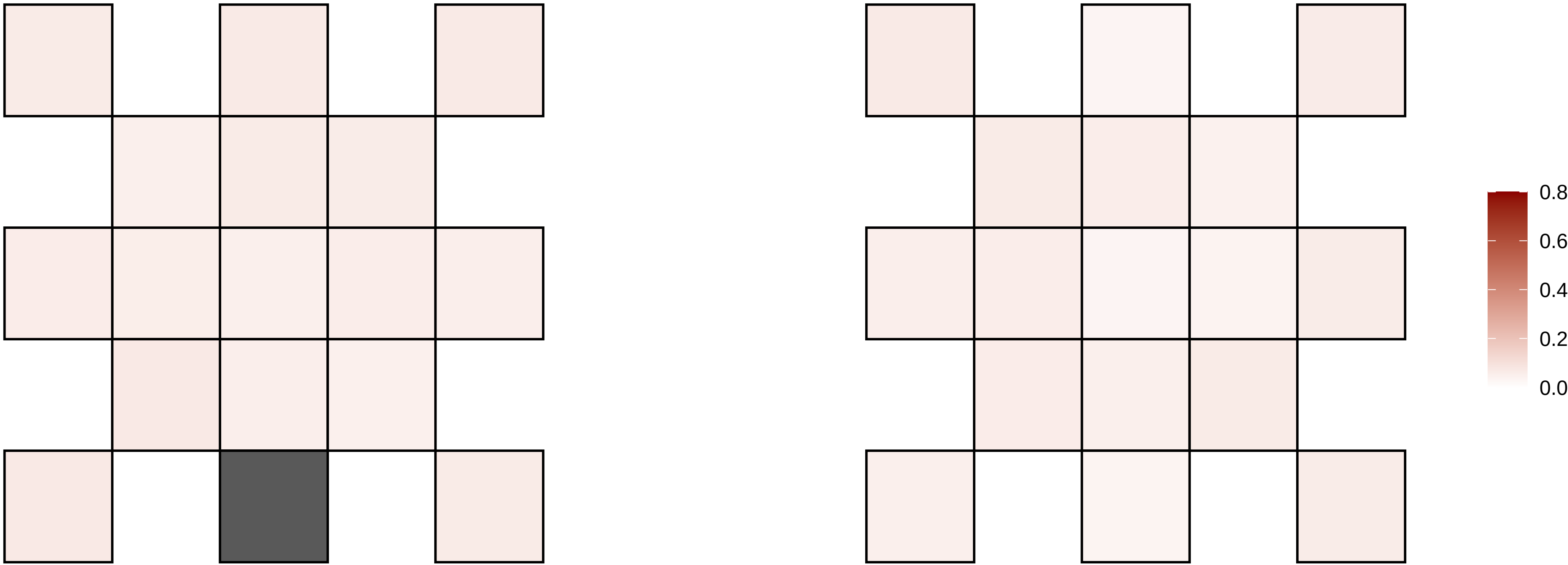

4−123000328−C−A

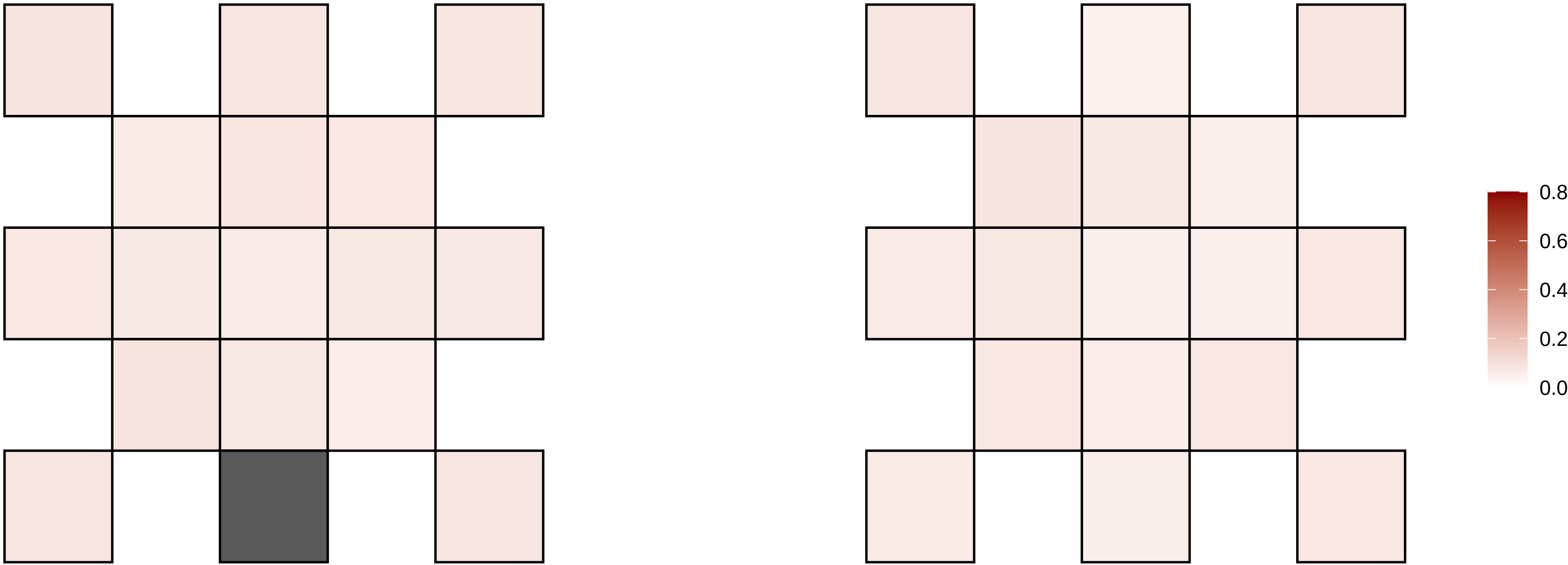

4−125806400−T−C

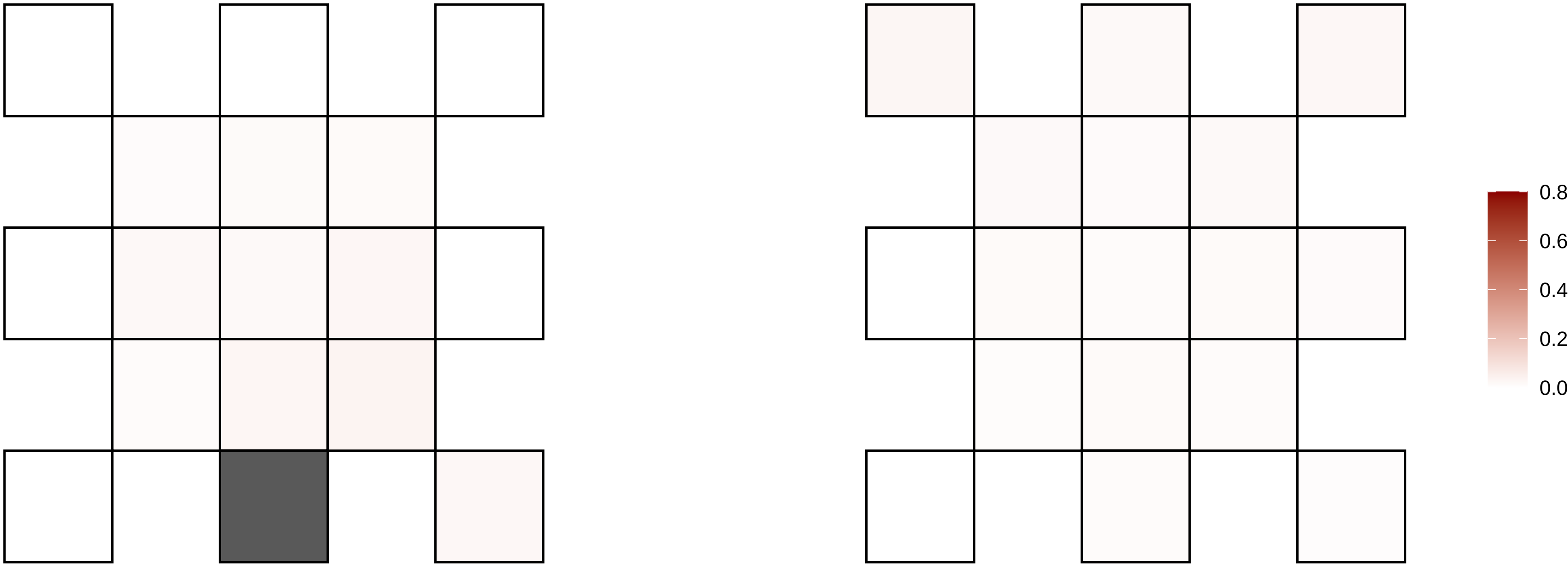

4−135038063−C−T

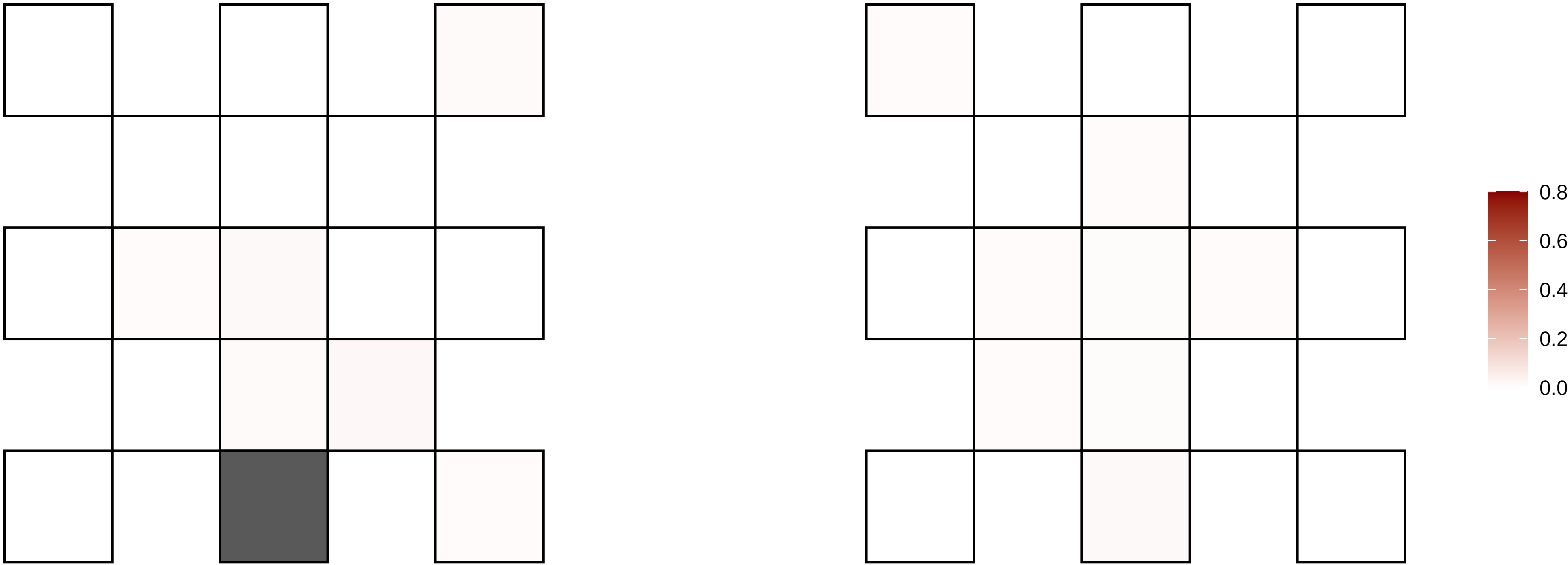

4−137937295−G−T

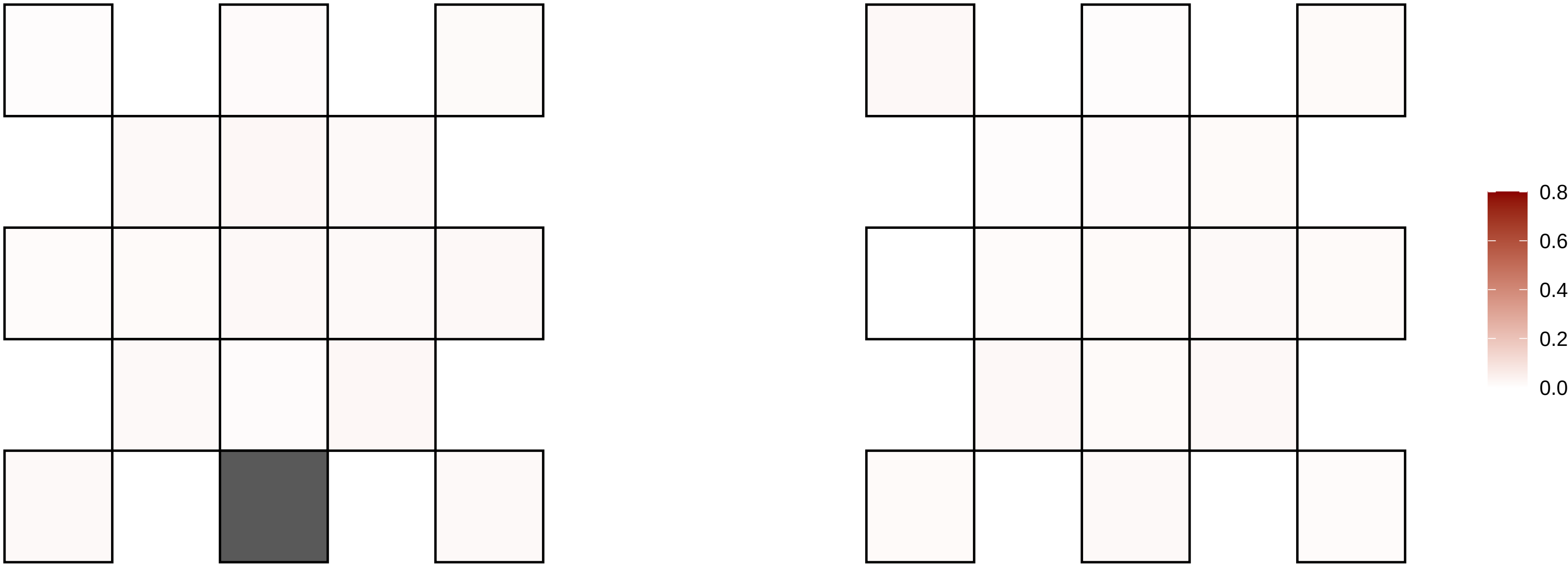

4−139482903−G−A

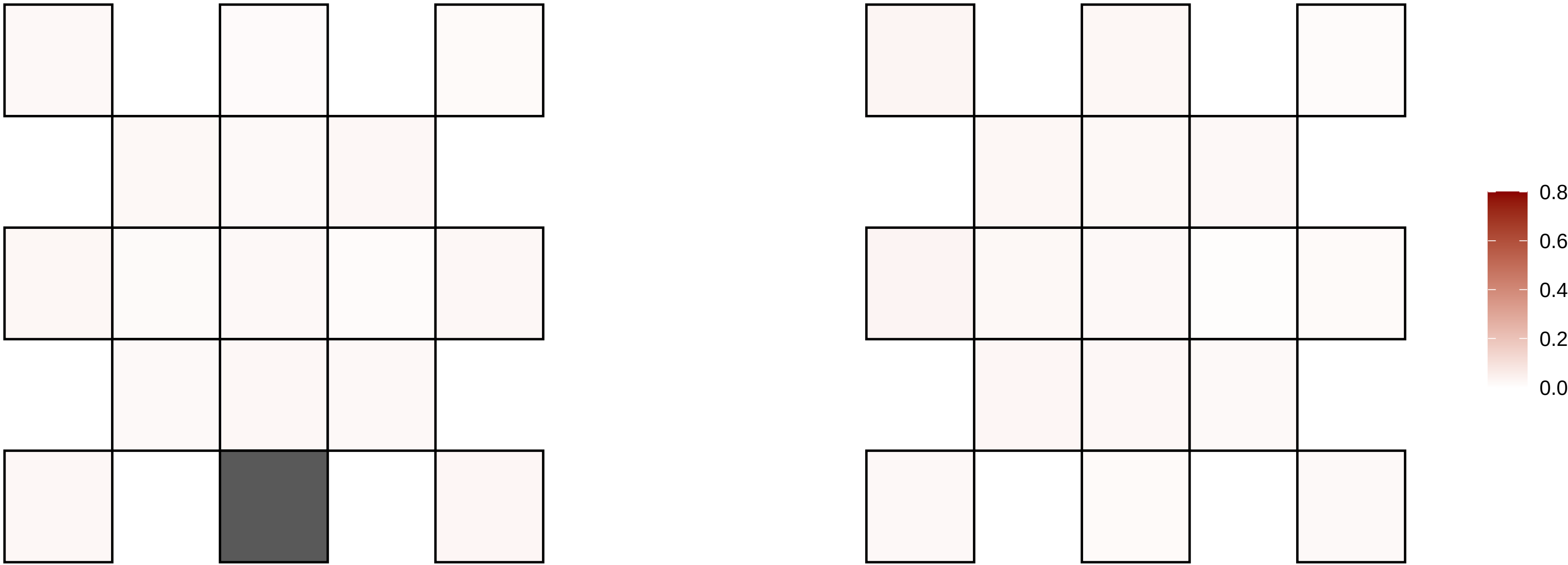

4−146067239−G−A

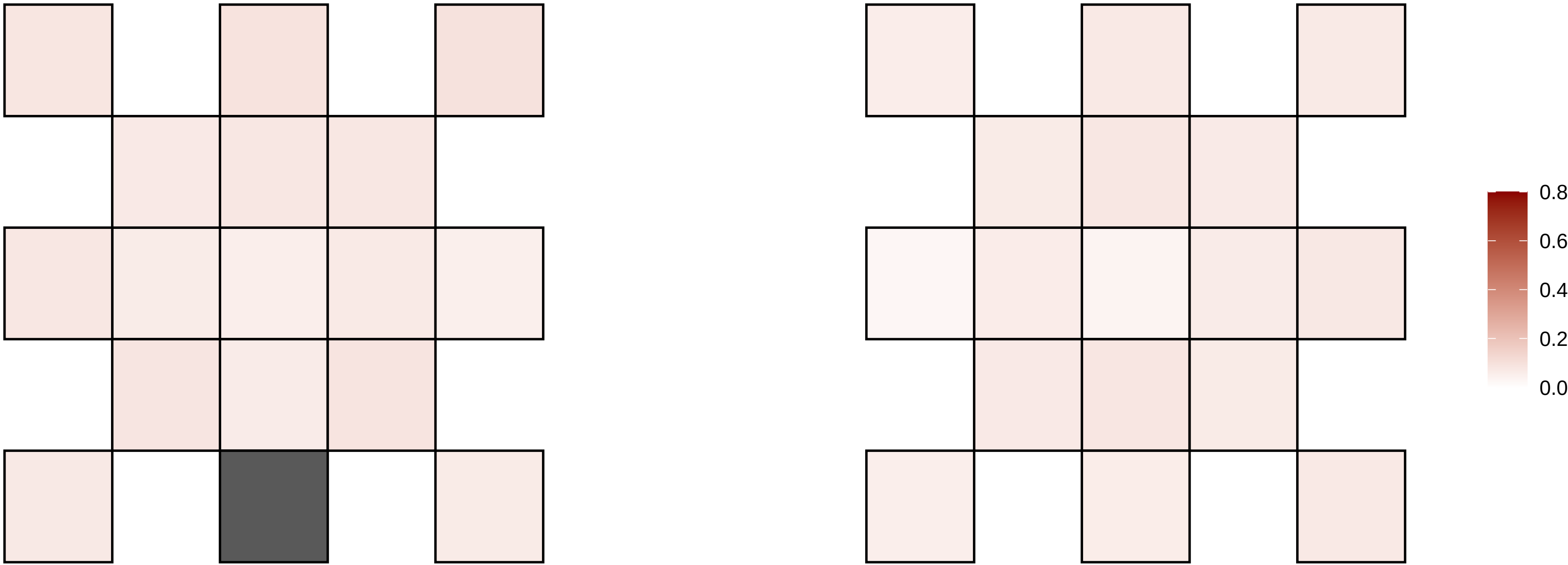

4−152509643−C−A

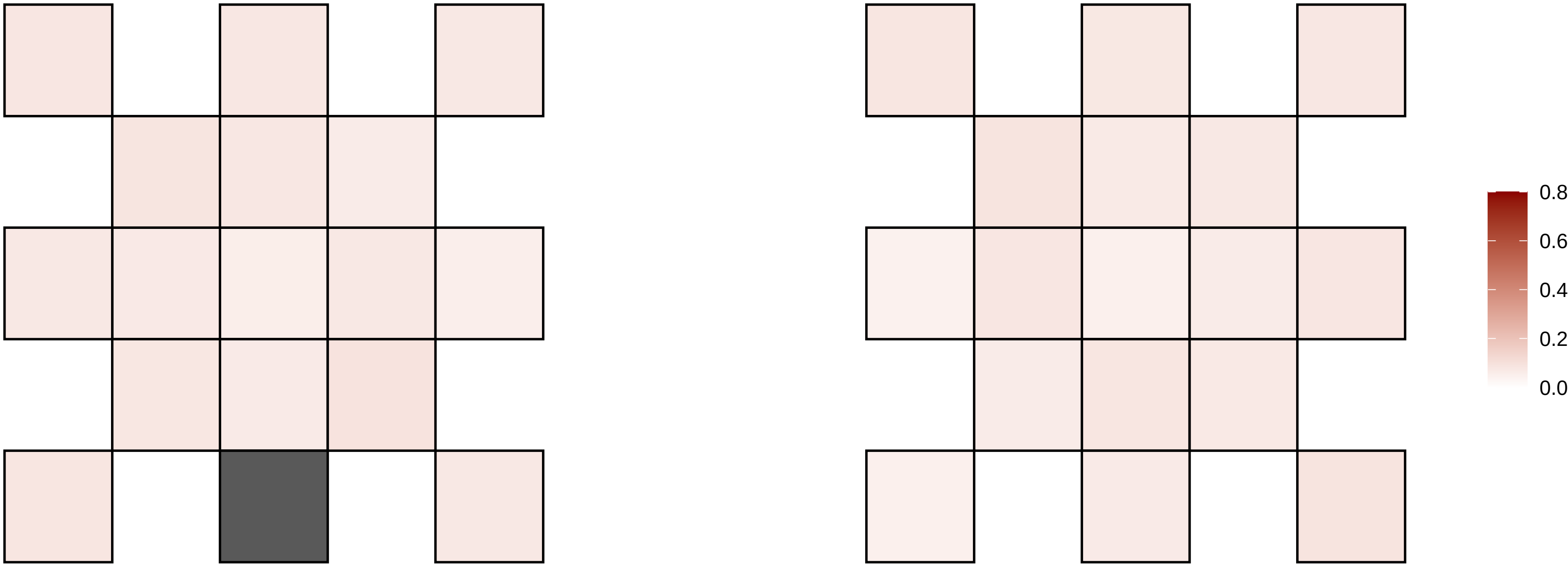

4−153029643−C−T

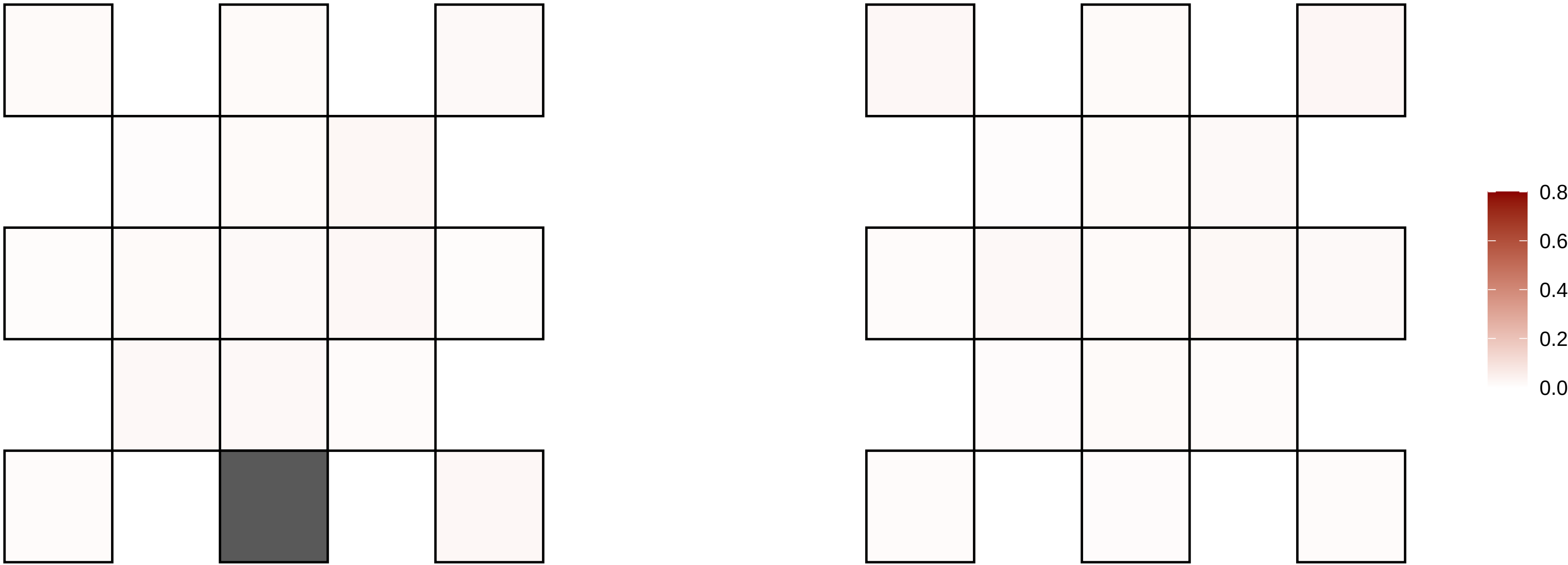

4−155190936−C−T

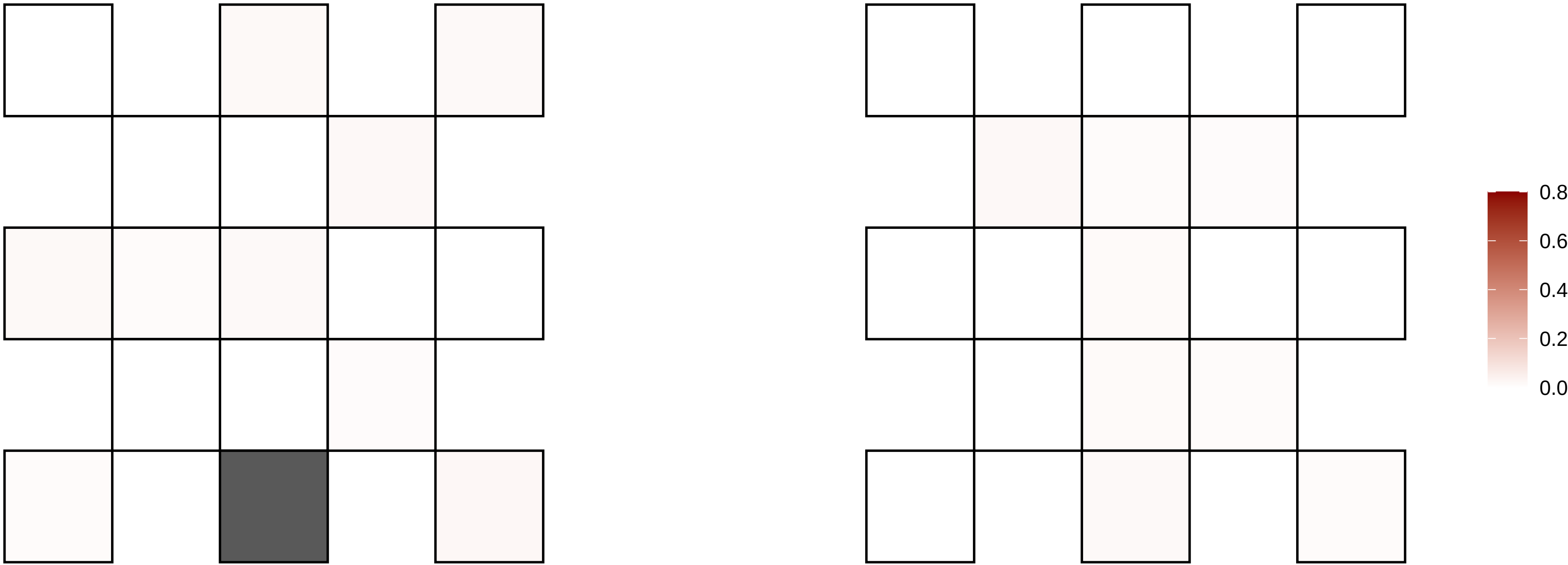

4−155889870−C−T

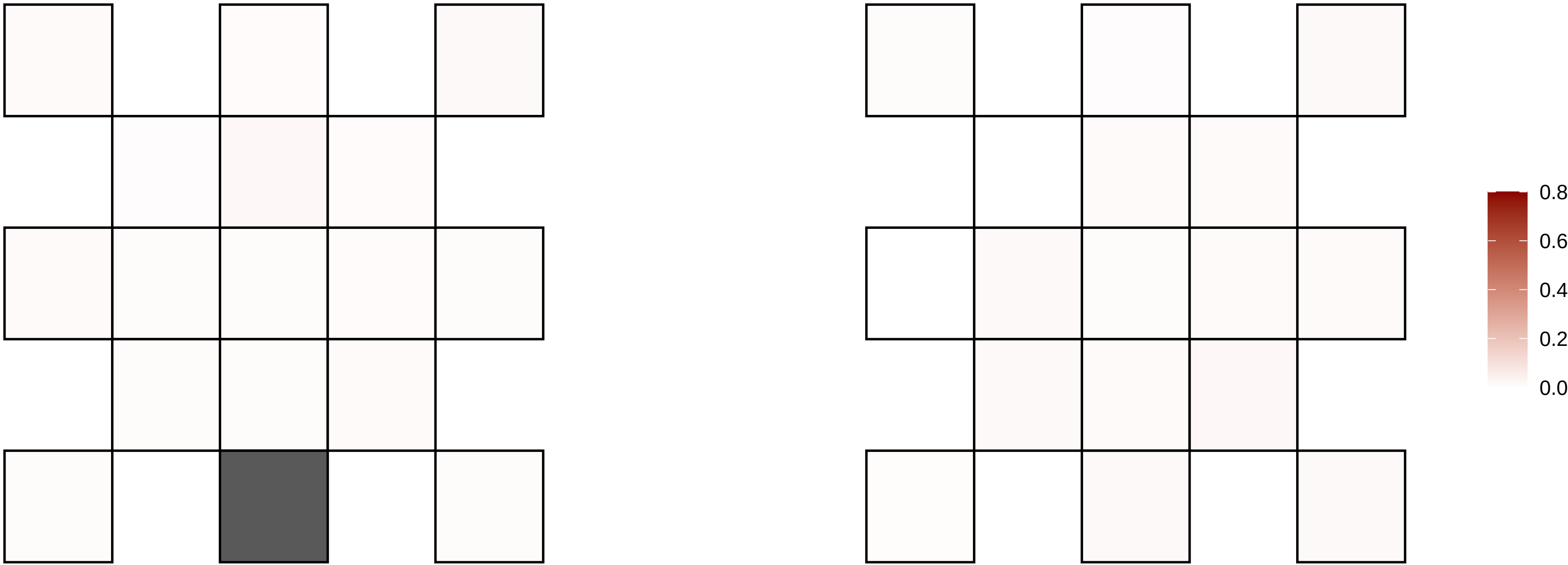

4−155904014−T−G

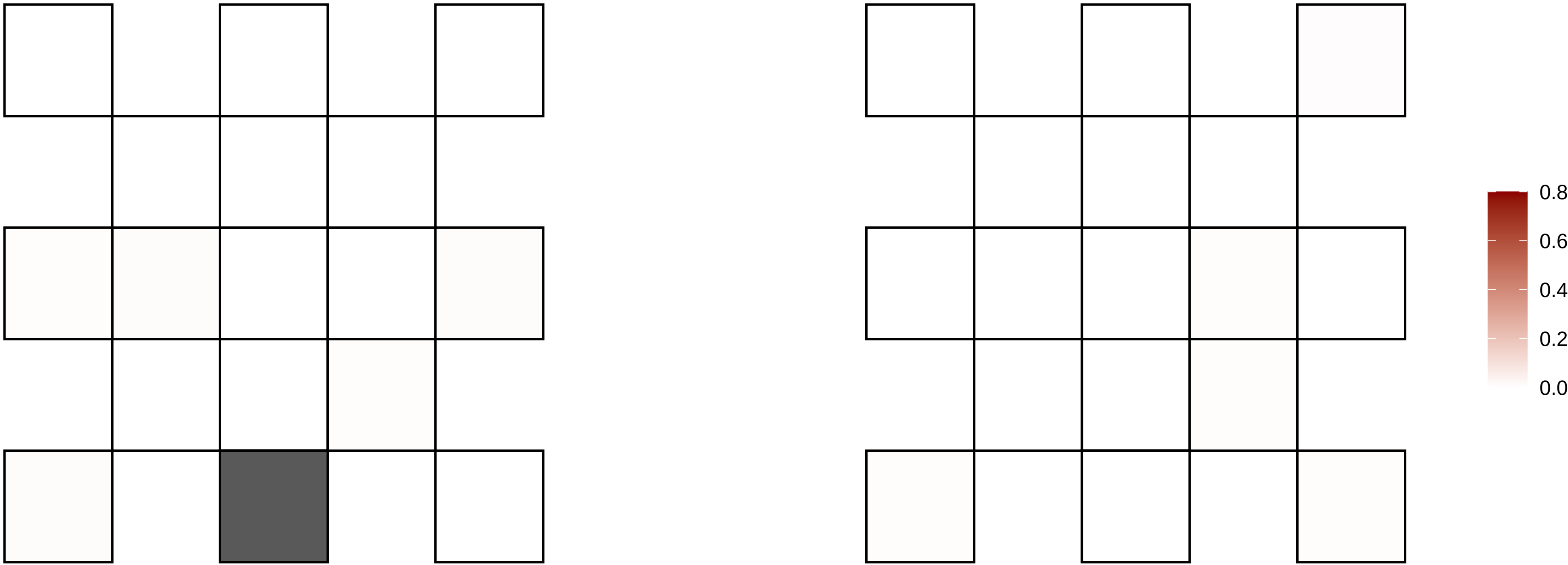

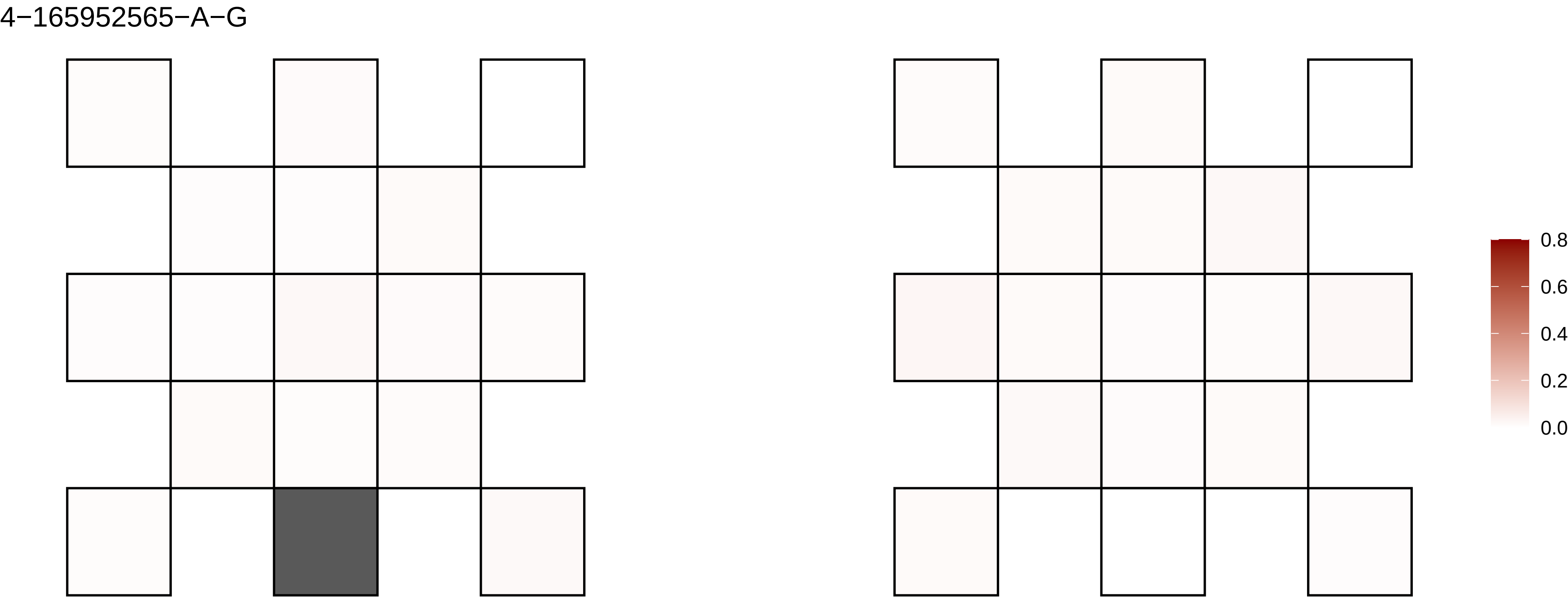

4−165952565−A−G
0.8
0.6
0.4
0.2
0.0

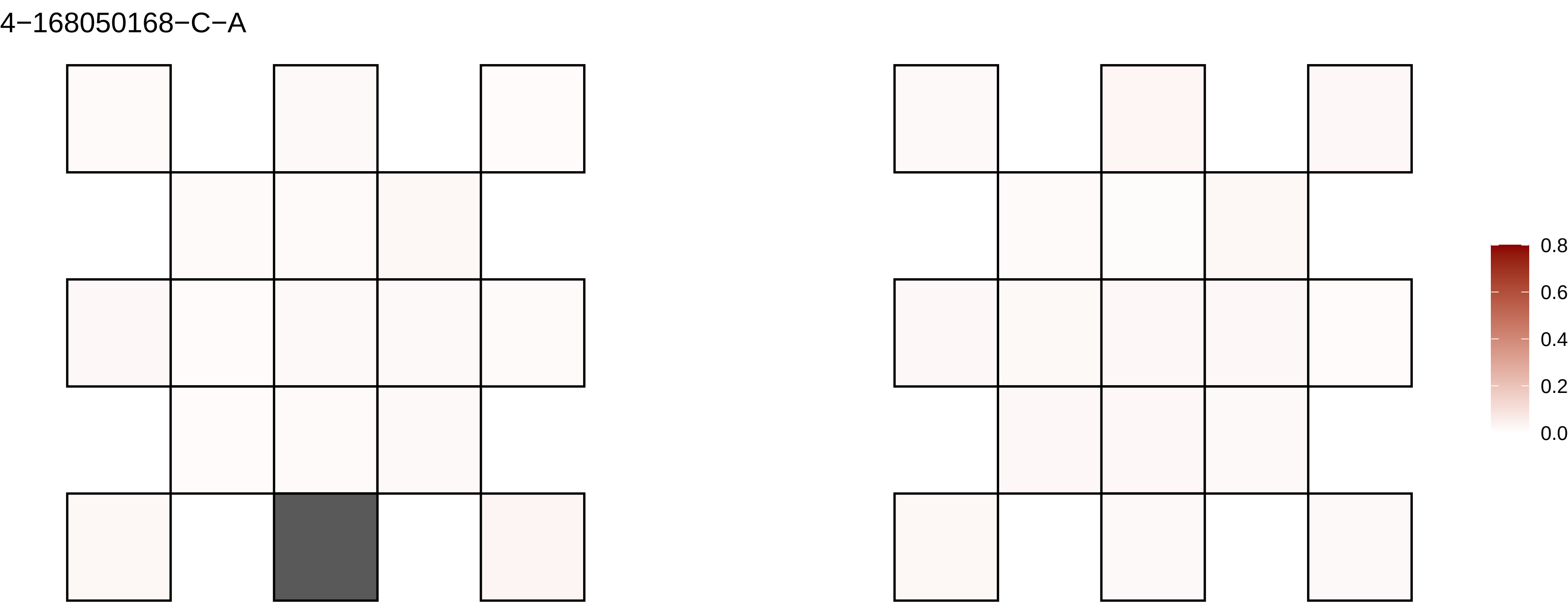
4−168050168−C−A
0.8
0.6
0.4
0.2
0.0

4−183244212−C−T

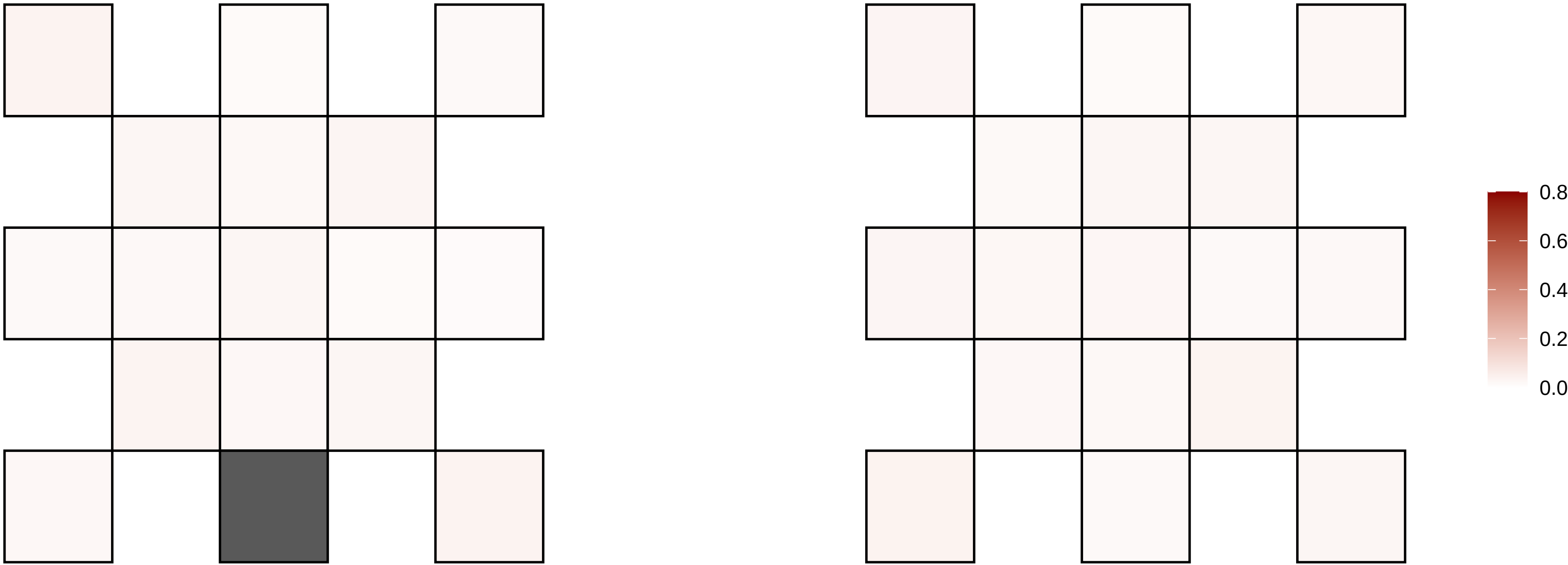

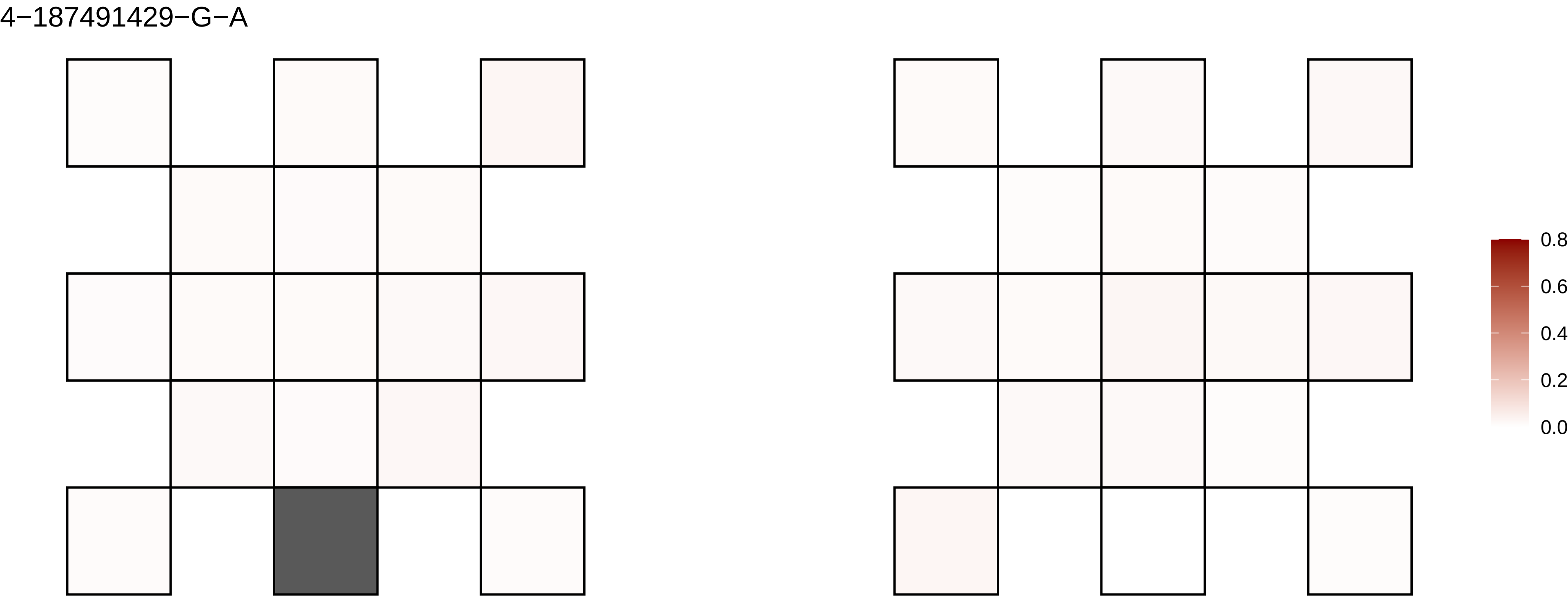

4−187491429−G−A
0.8
0.6
0.4
0.2
0.0

4−187539951−G−A

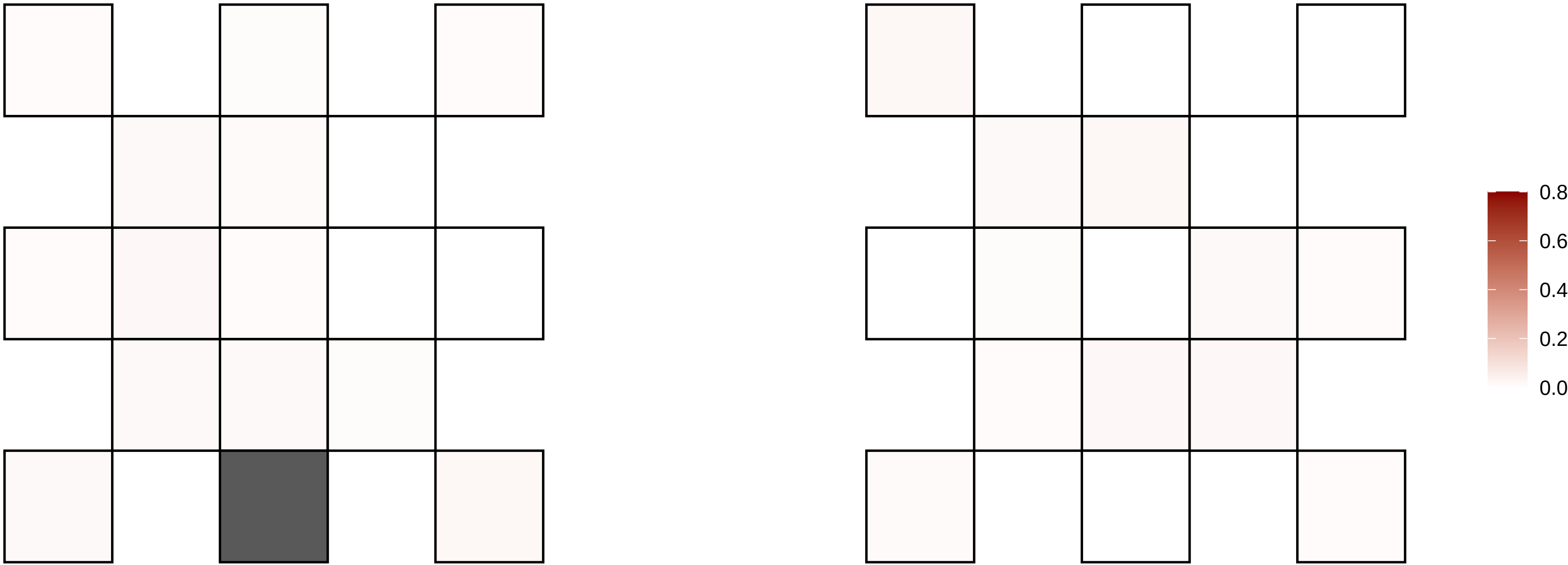

4−188115218−G−T

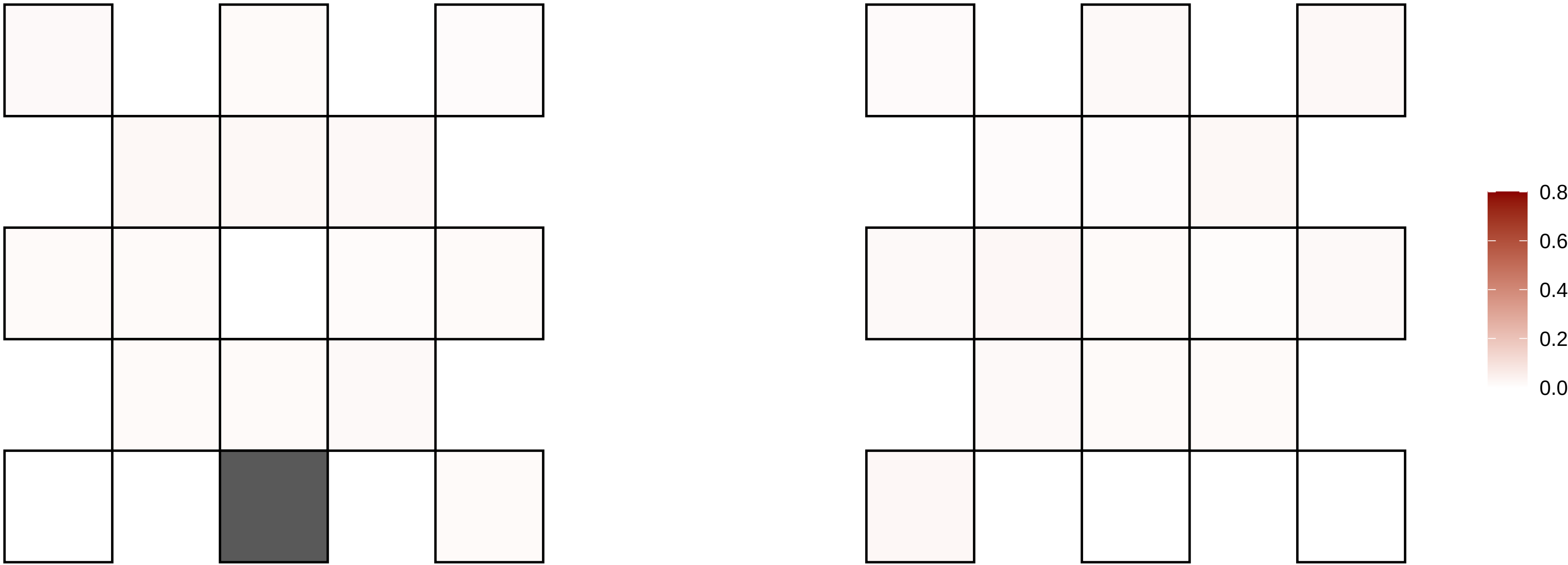

4−21301357−T−G

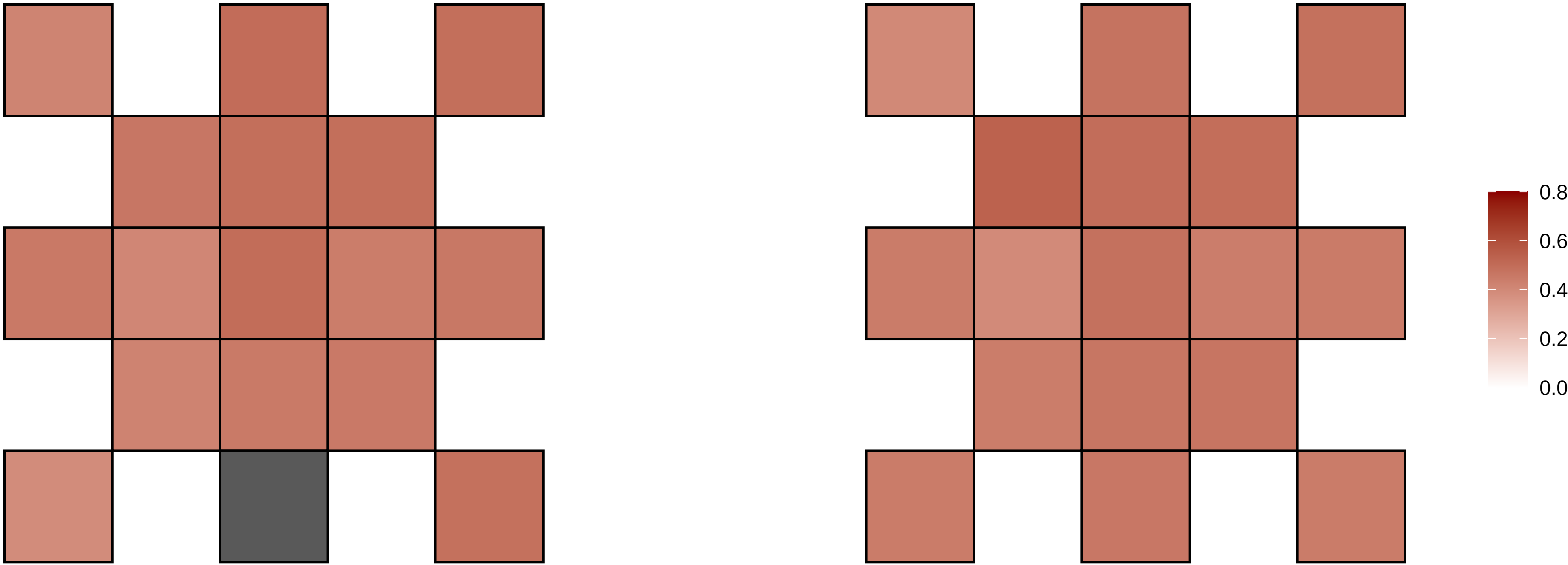

4−22428363−C−T

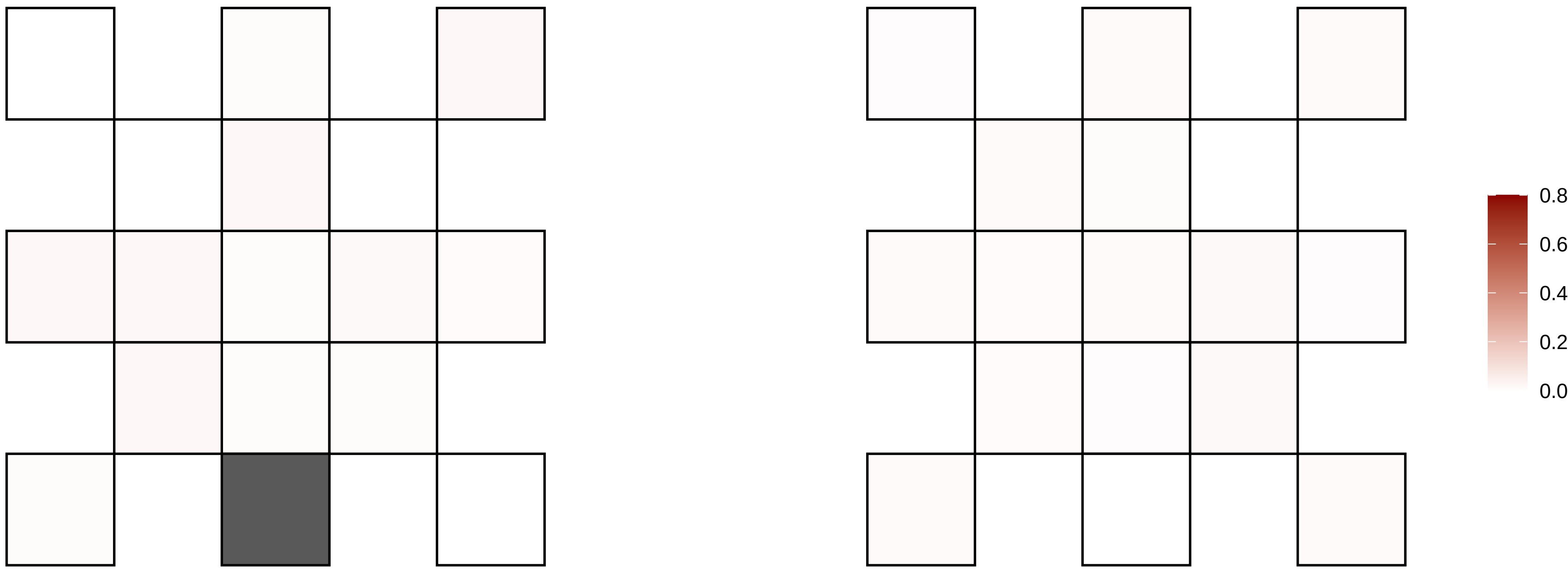

4−23552226−G−A

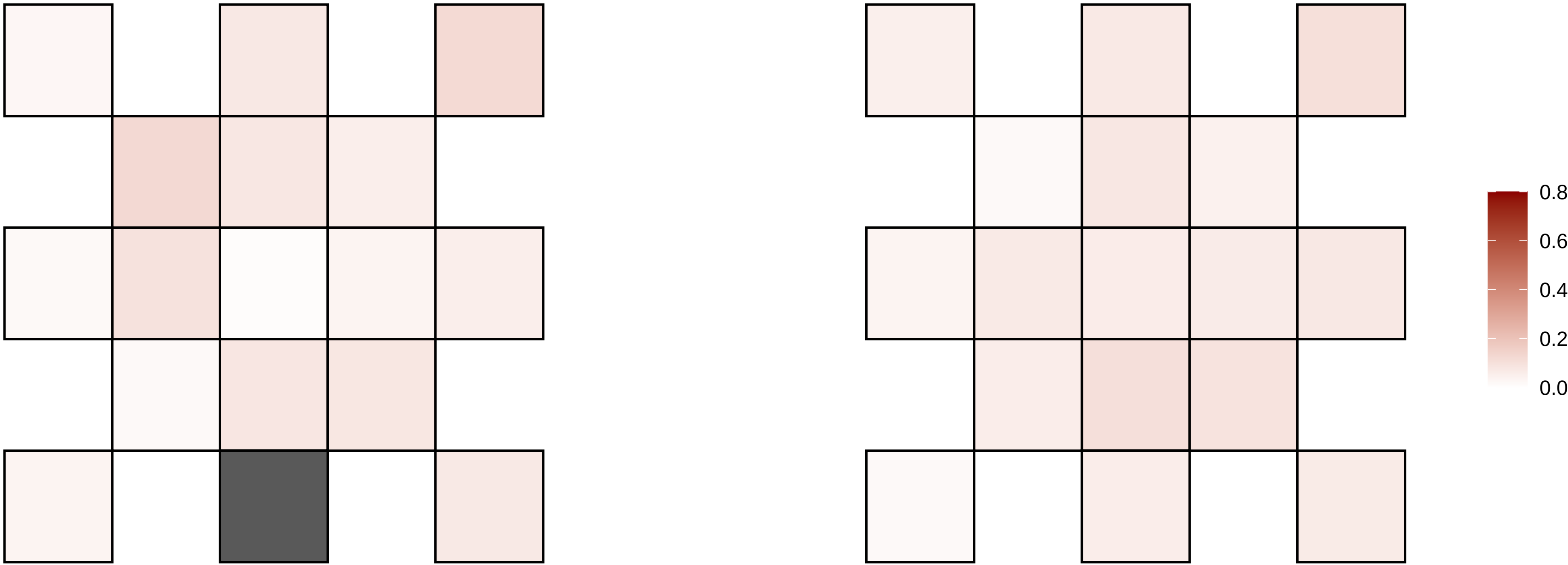

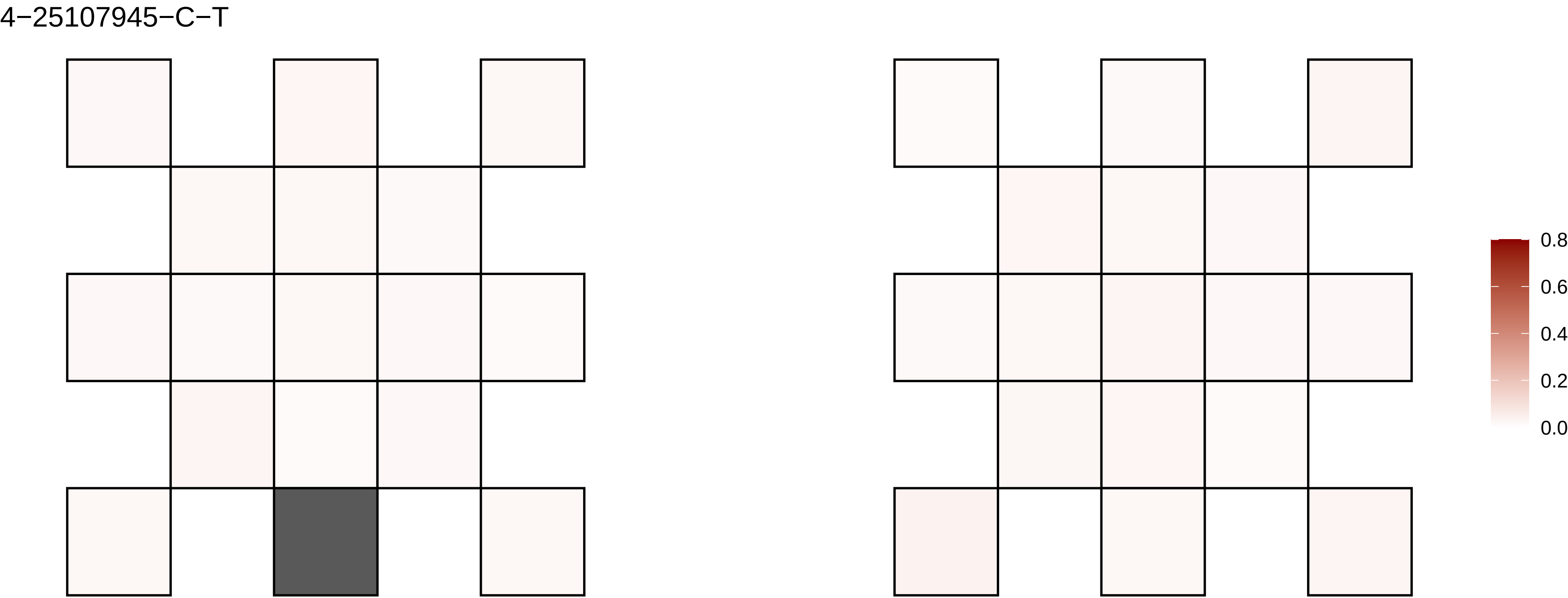
4−25107945−C−T
0.8
0.6
0.4
0.2
0.0

4−26808694−C−T

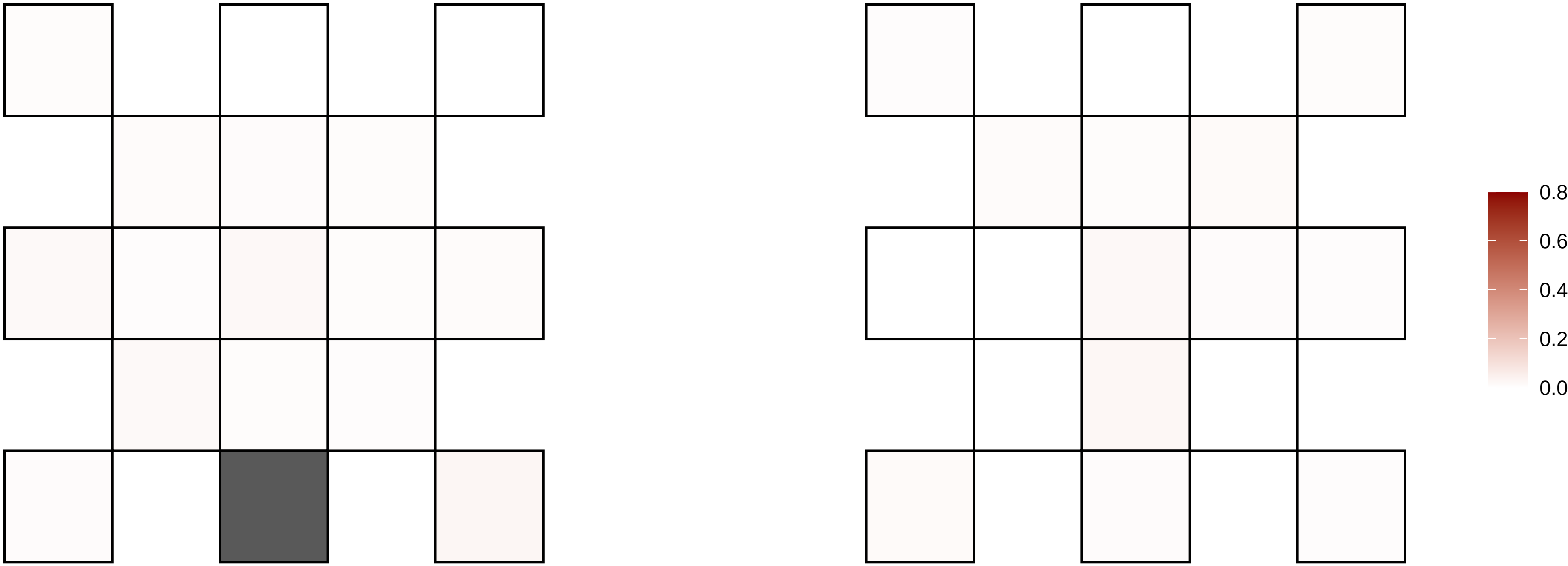

4−30123079−T−G

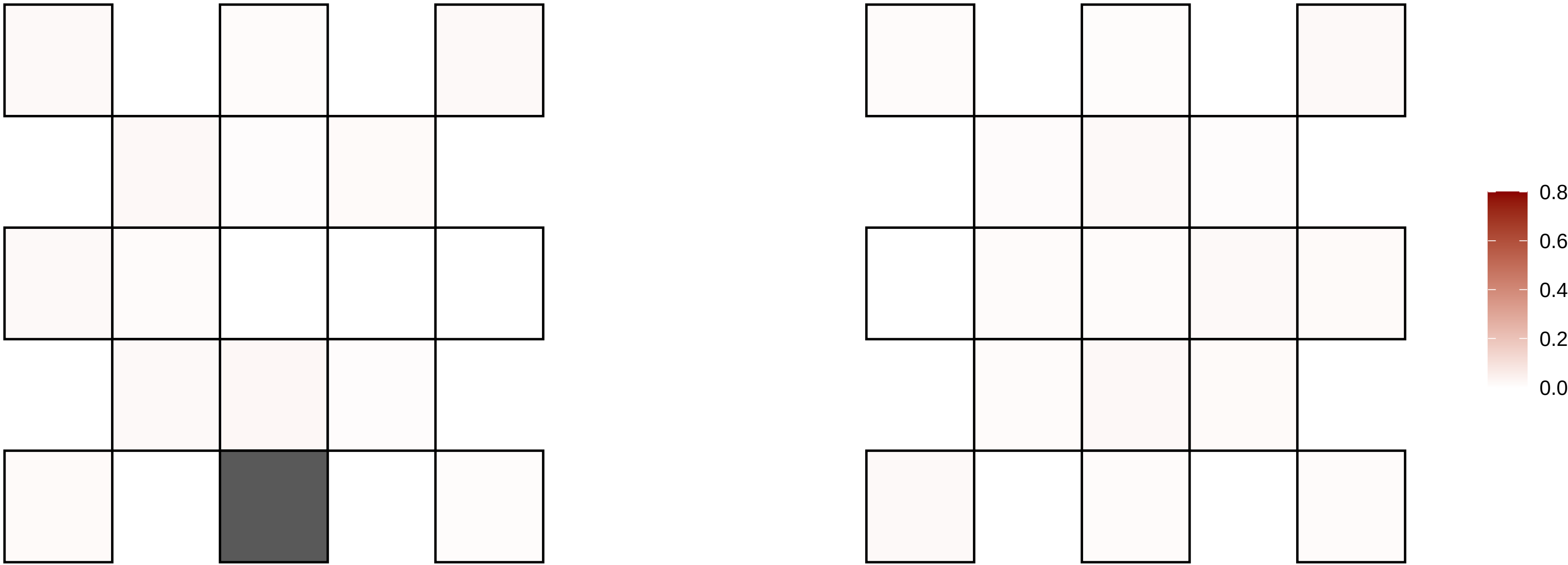

4−32745748−A−T

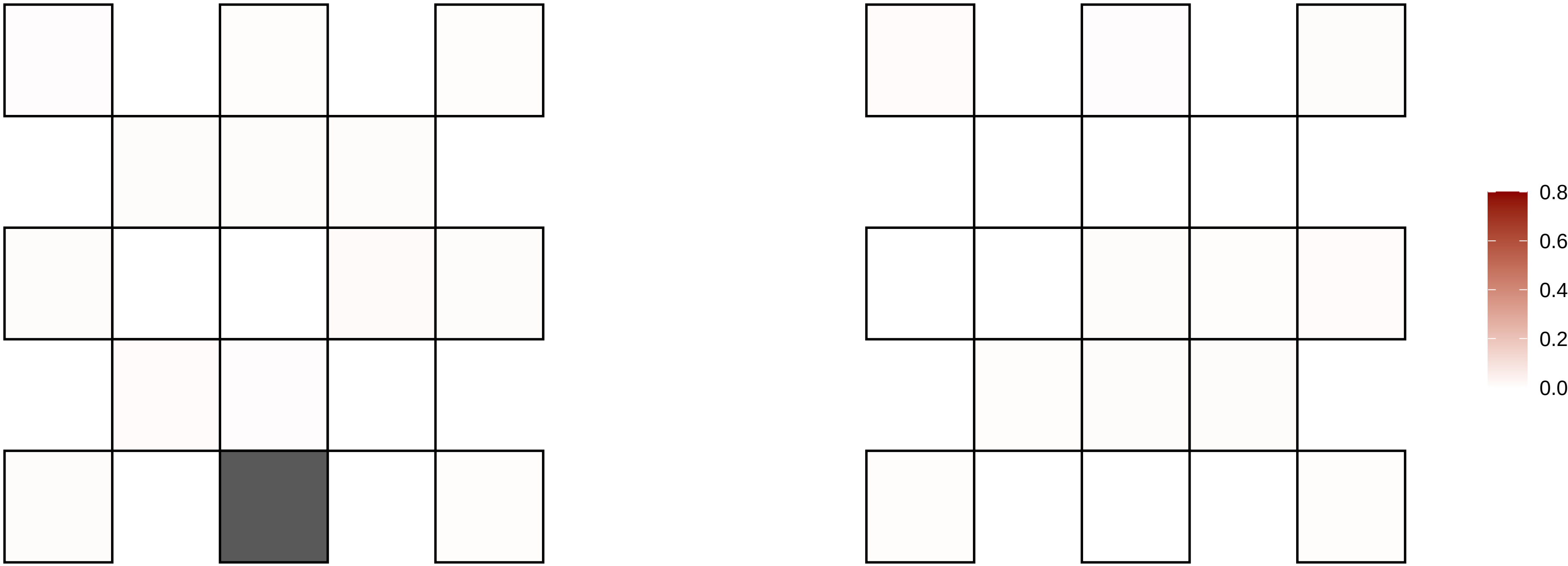

4−34386270−C−T

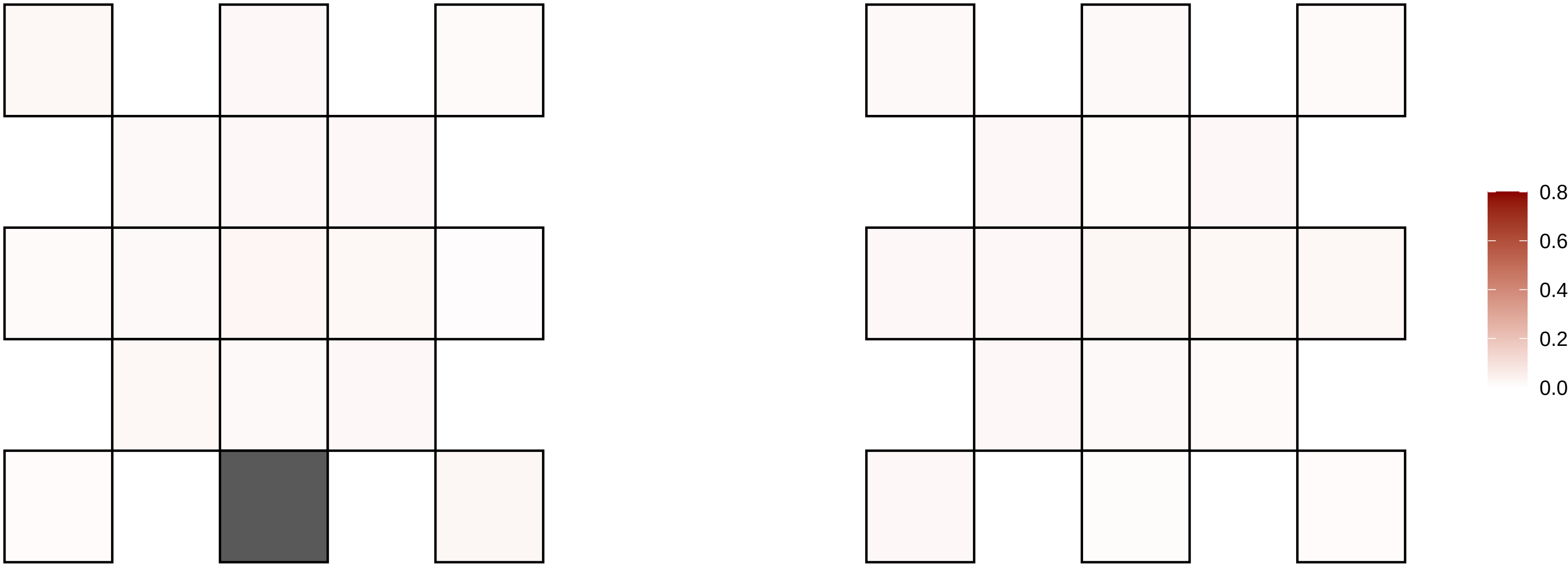

4−34586097−A−T

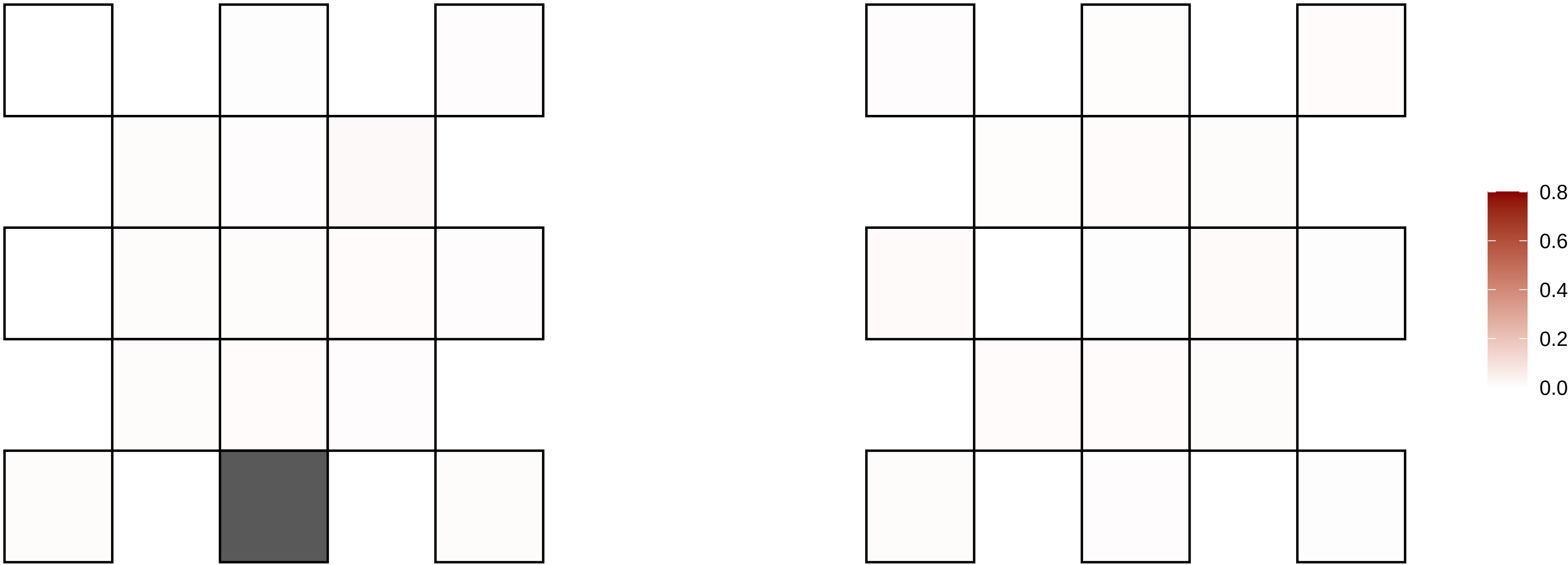

4−36163098−C−T

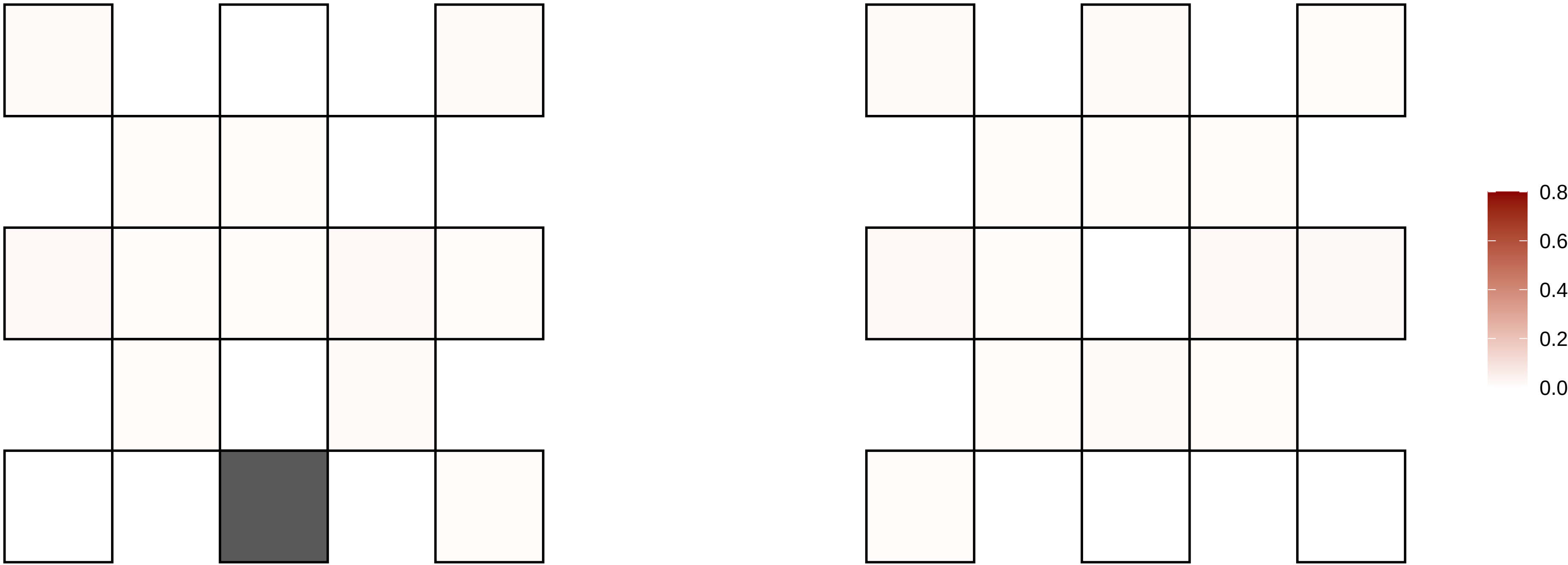

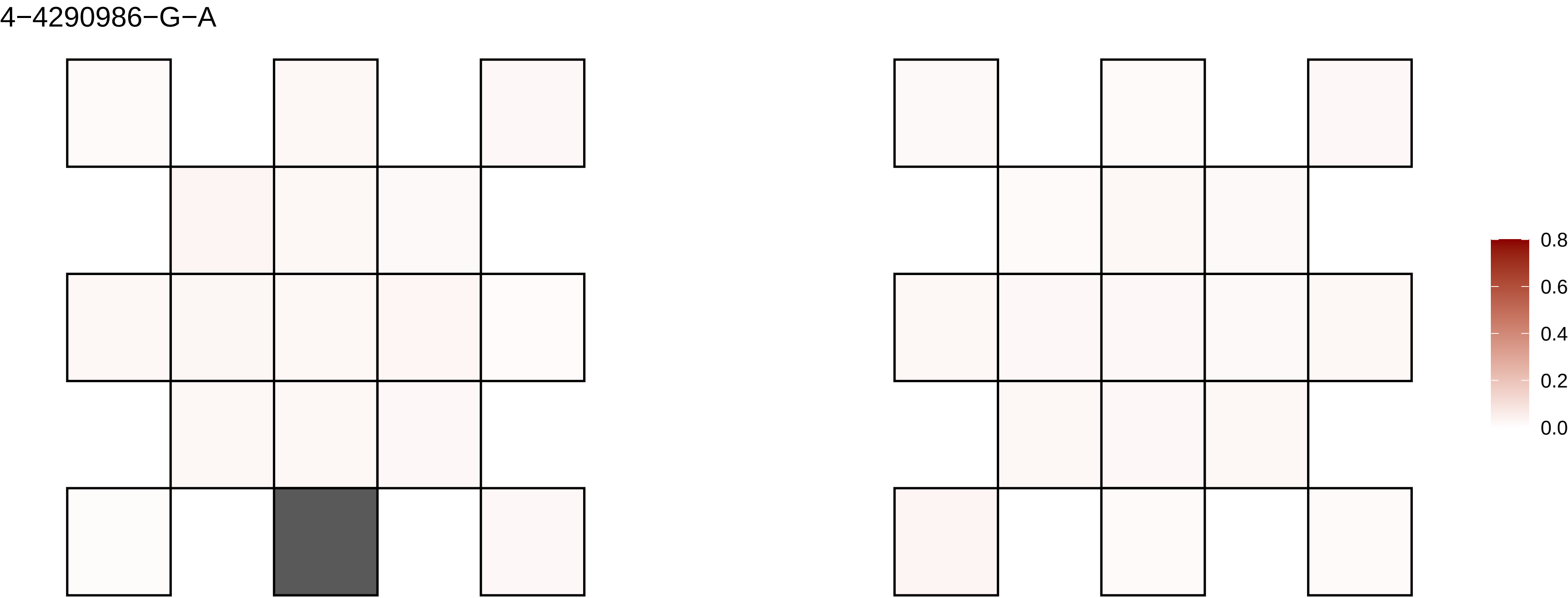
4−4290986−G−A
0.8
0.6
0.4
0.2
0.0

4−44006646−G−A

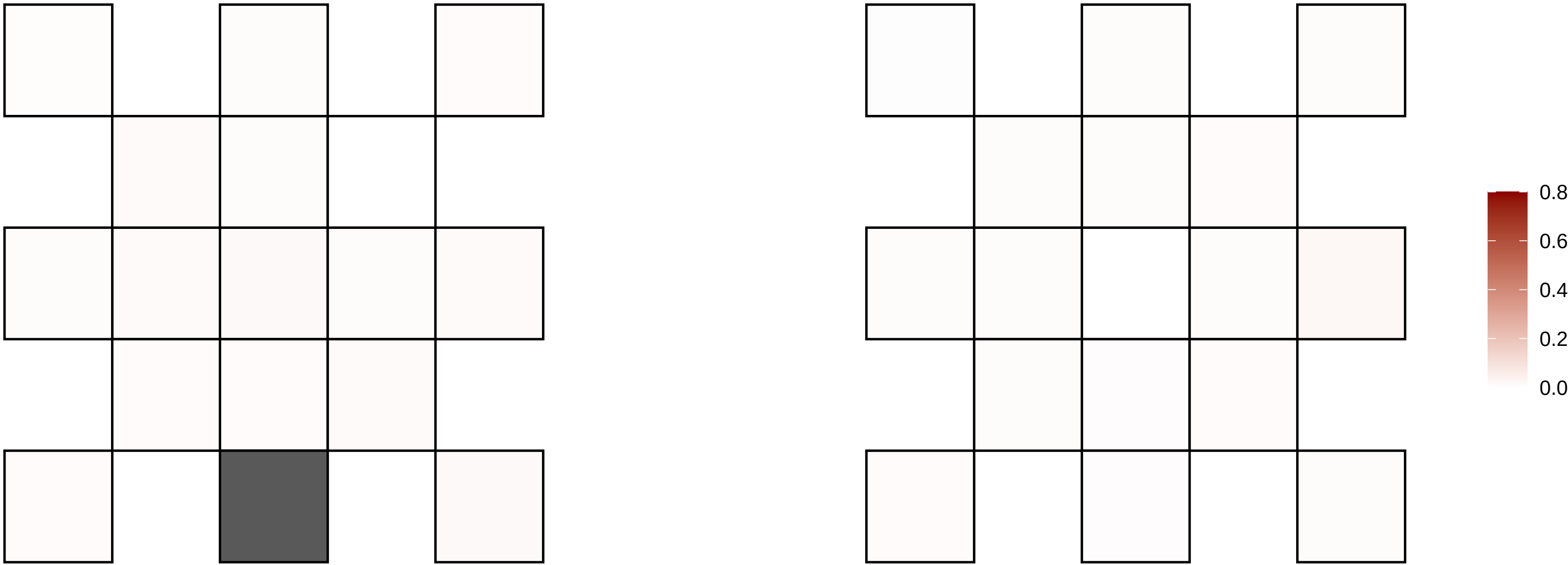

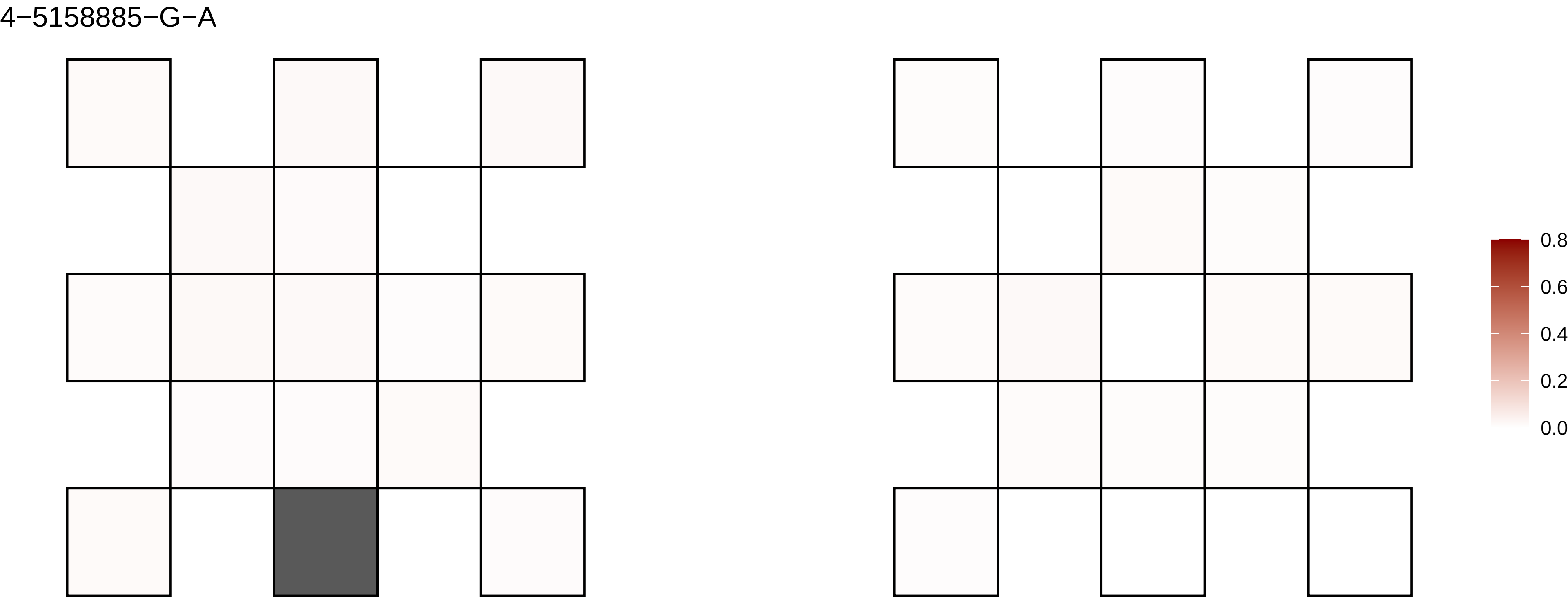
4−5158885−G−A
0.8
0.6
0.4
0.2
0.0

4−52773−C−T

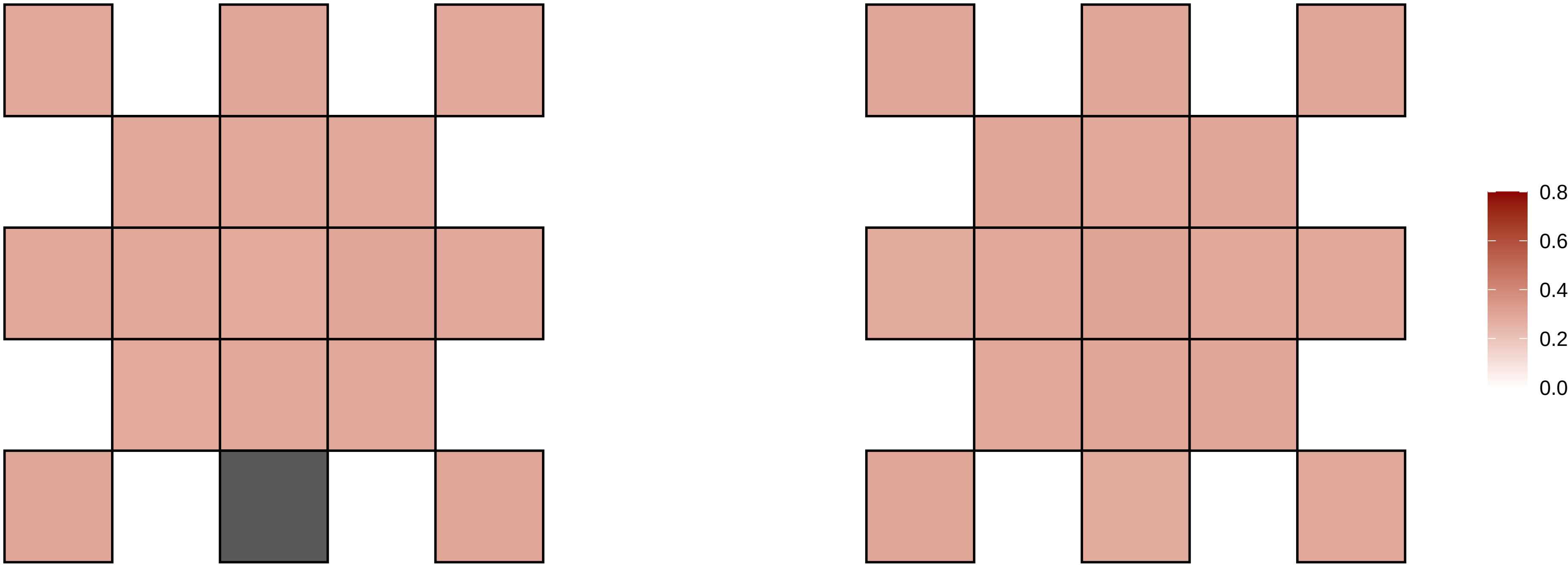

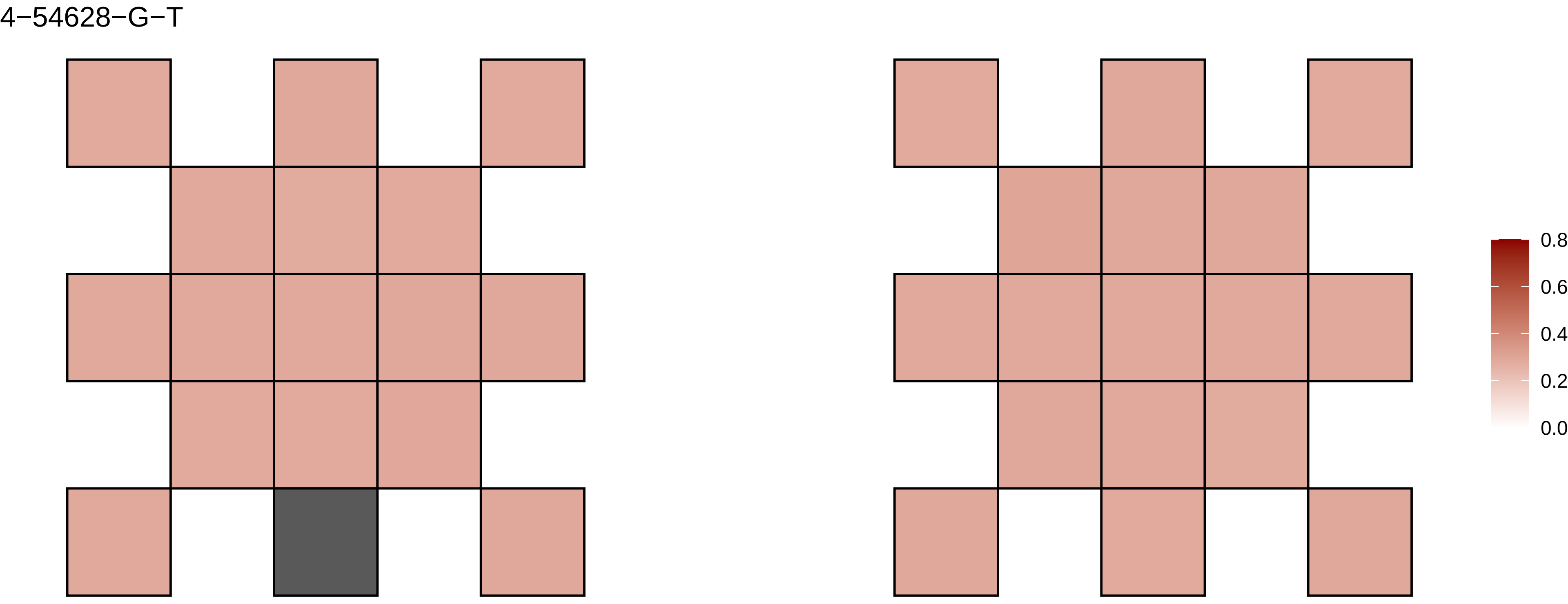
4−54628−G−T
0.8
0.6
0.4
0.2
0.0

4−57773855−A−T

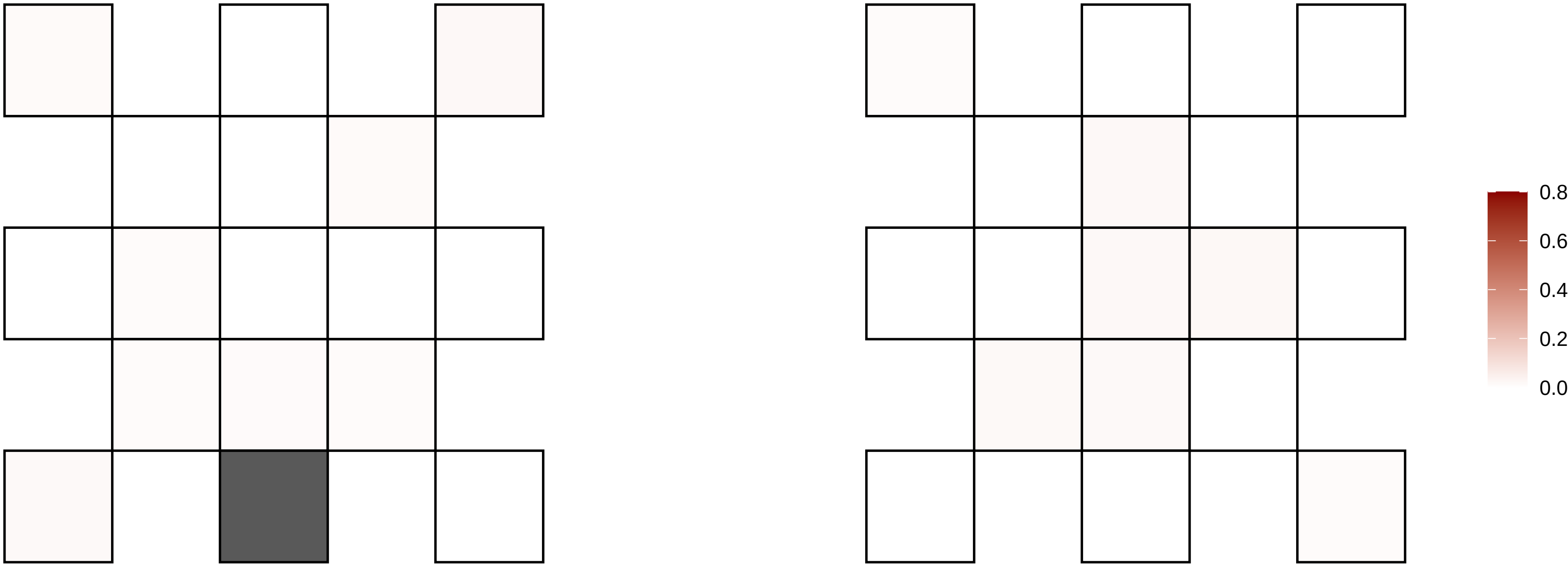

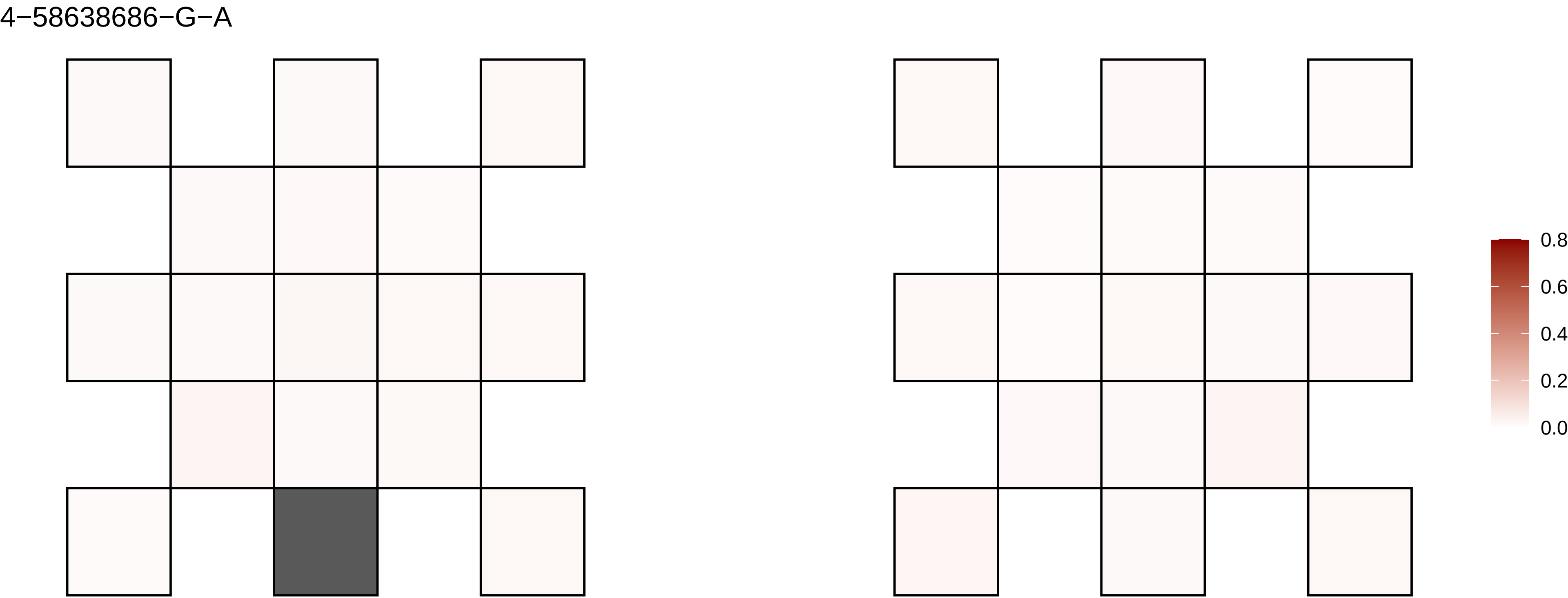

4−58638686−G−A
0.8
0.6
0.4
0.2
0.0

4−59641558−C−T

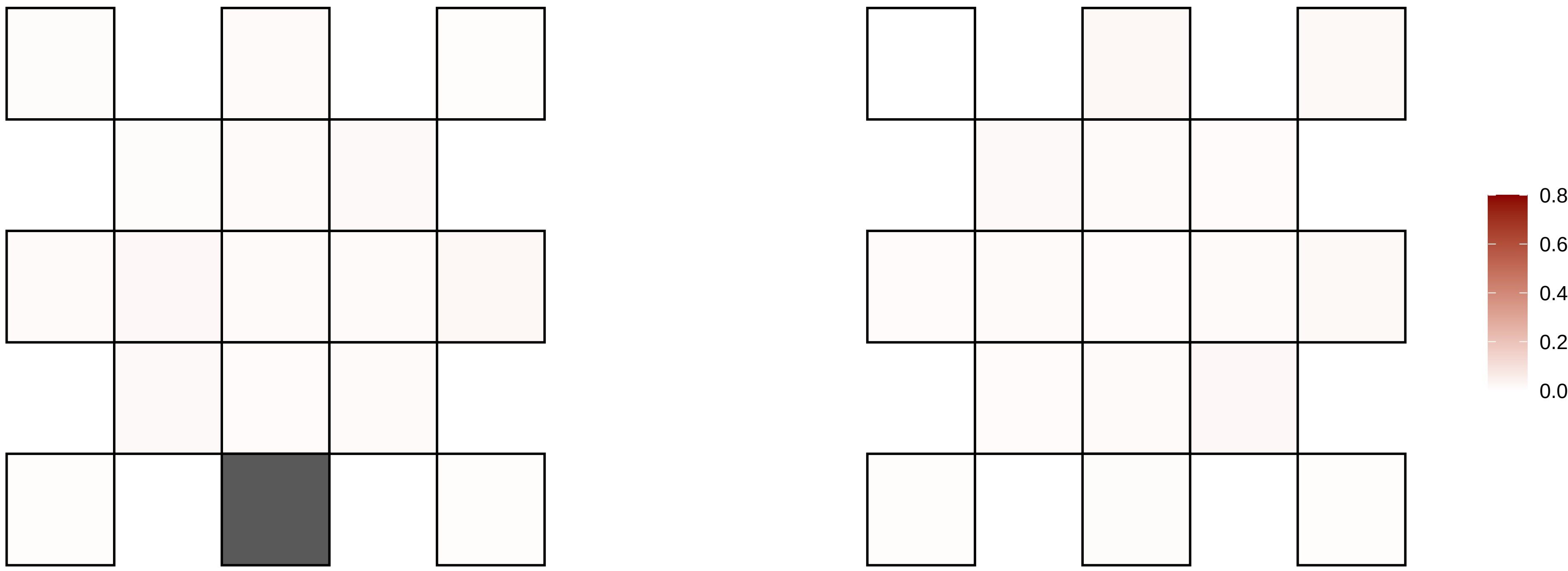

4−60968373−G−A

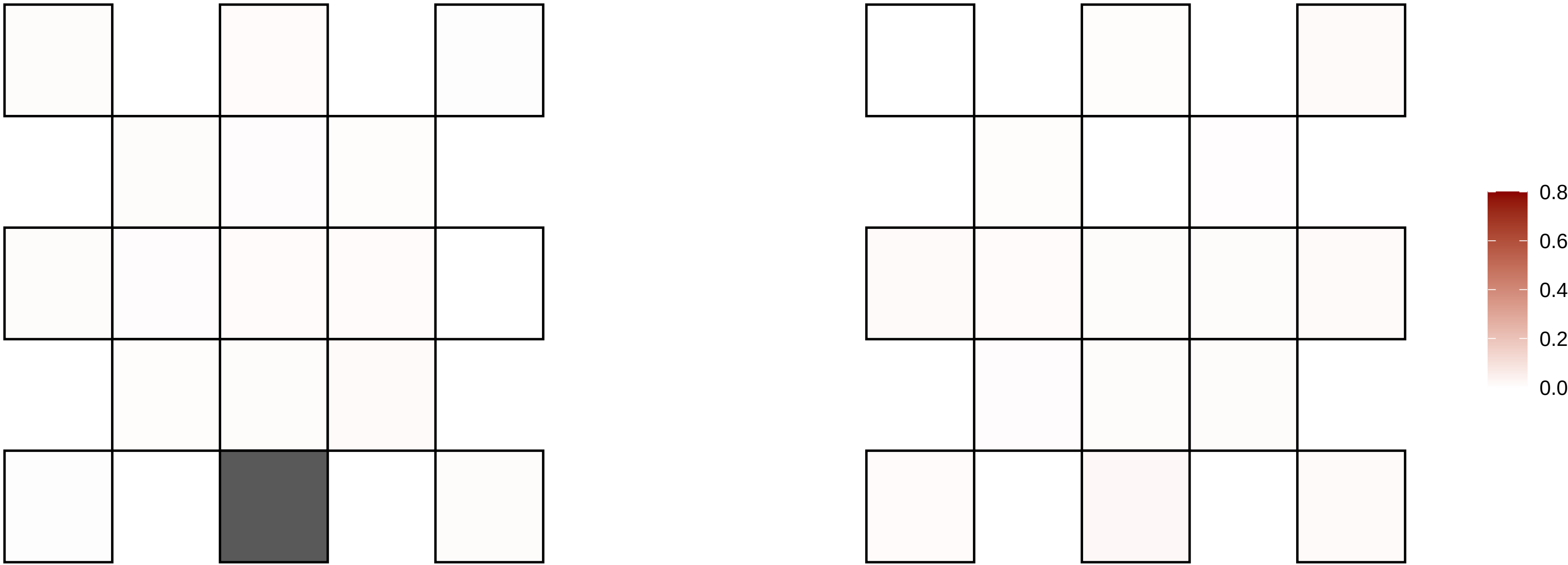

4−63640827−A−T

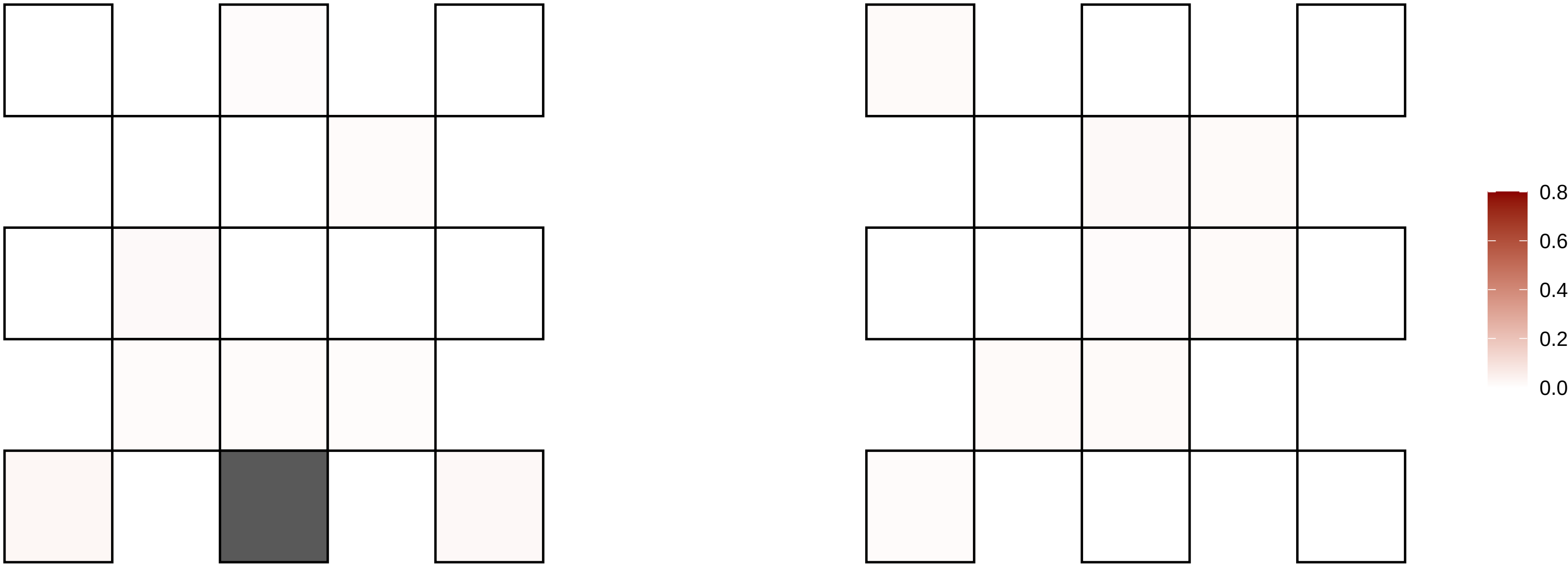

4−64742736−G−A

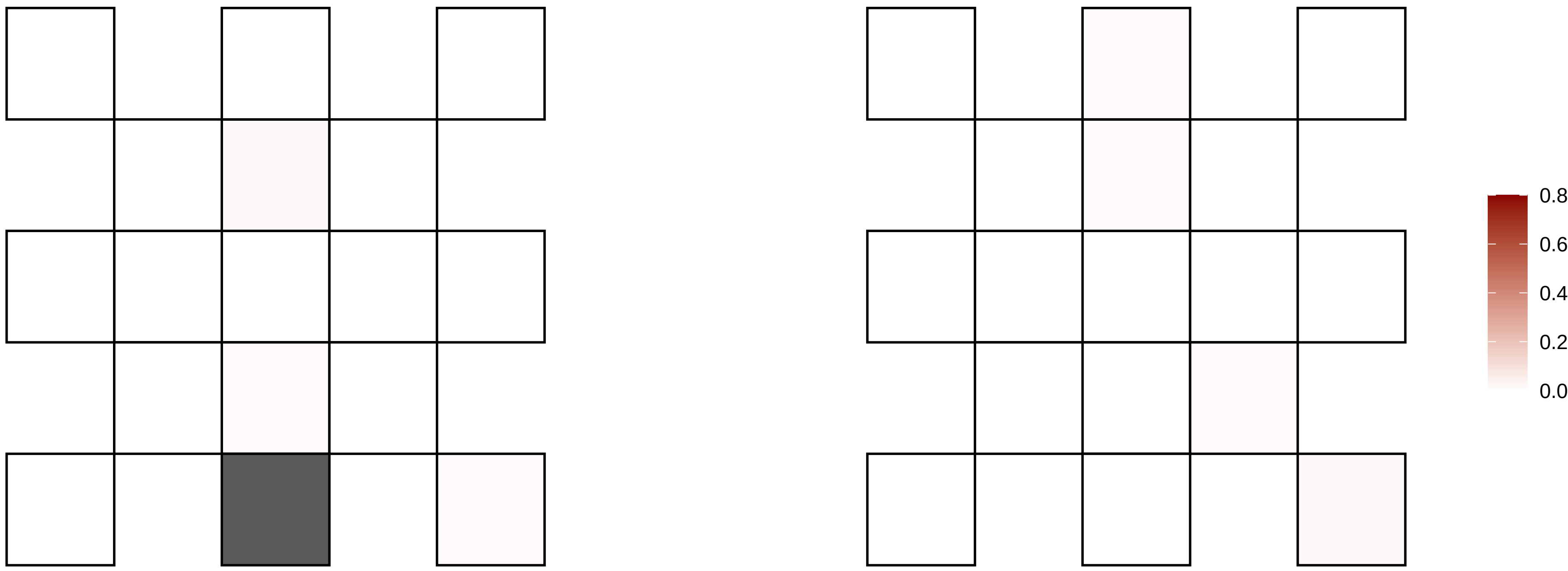

4−66798015−A−T

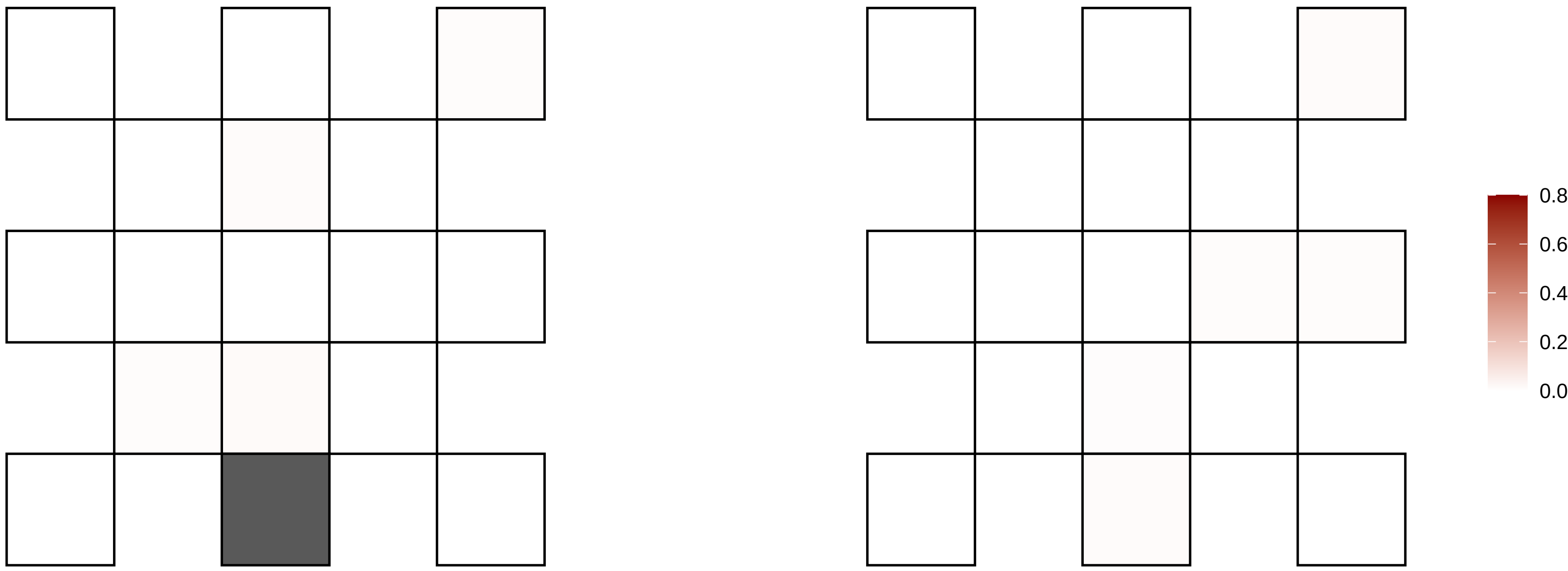

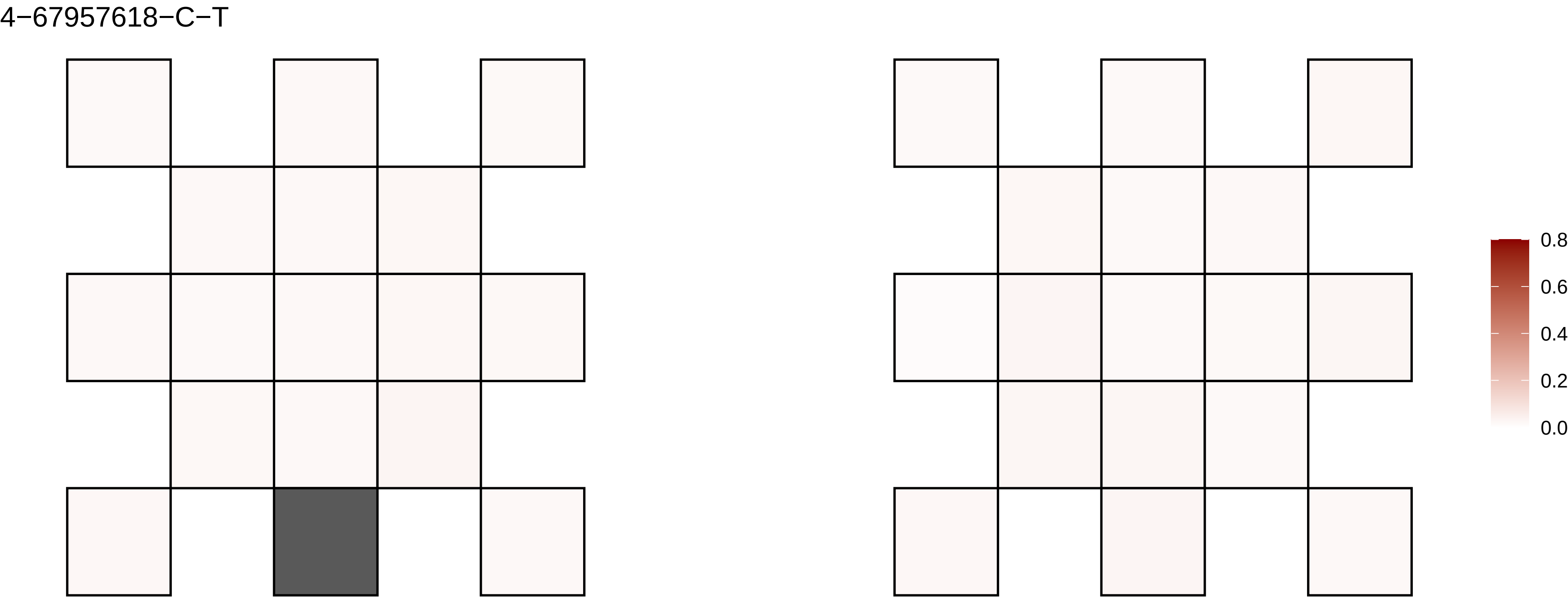
4−67957618−C−T
0.8
0.6
0.4
0.2
0.0

4−72321186−G−A

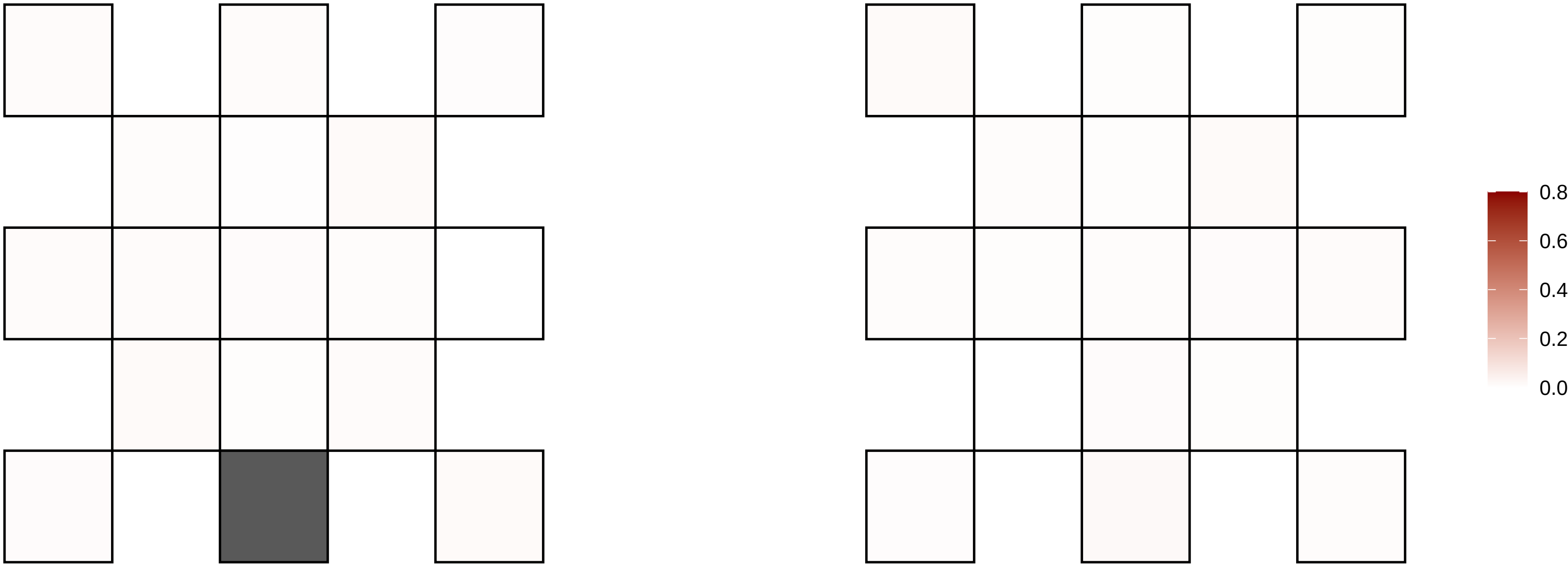

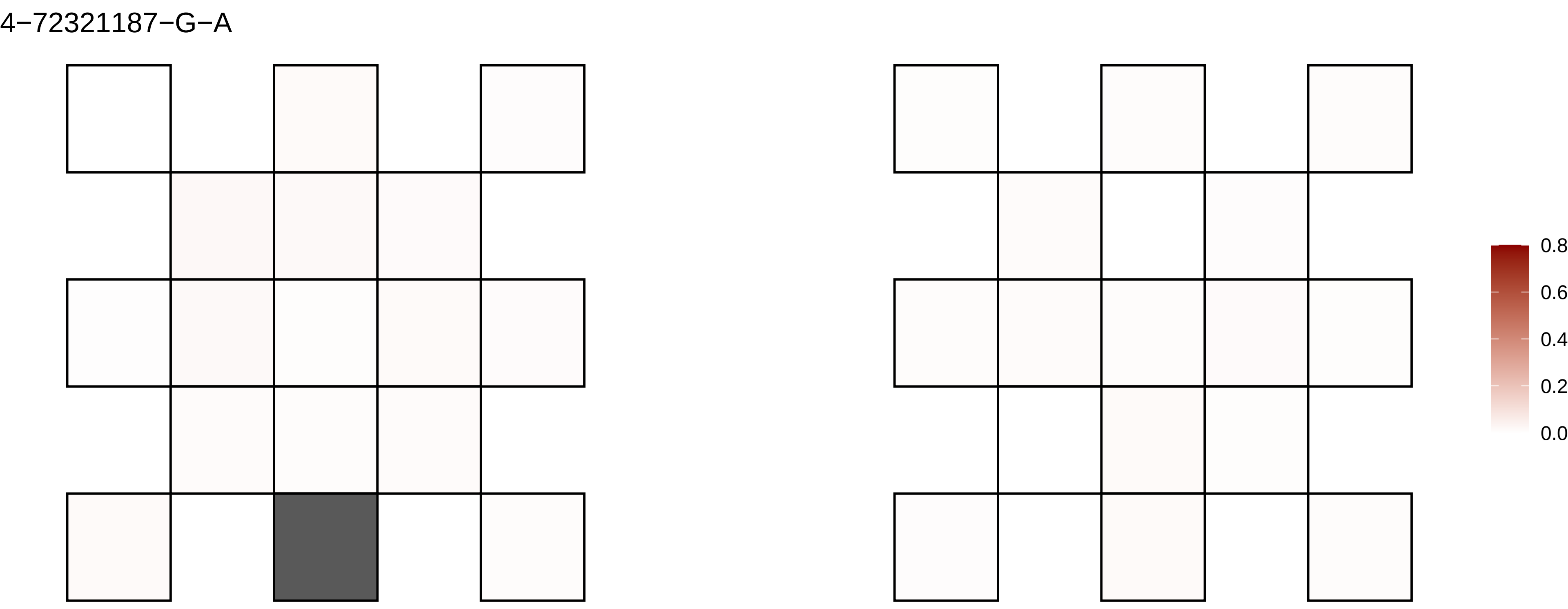
4−72321187−G−A
0.8
0.6
0.4
0.2
0.0

4−73709991−A−T

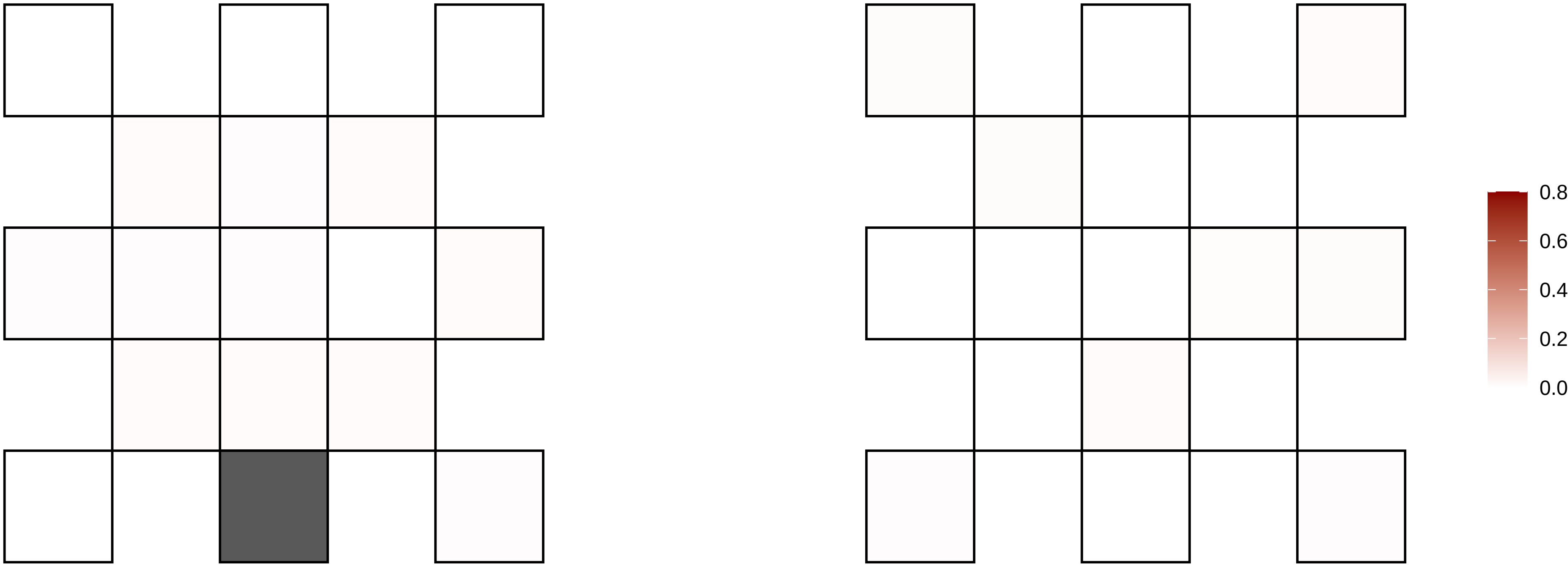

4−74784894−G−A

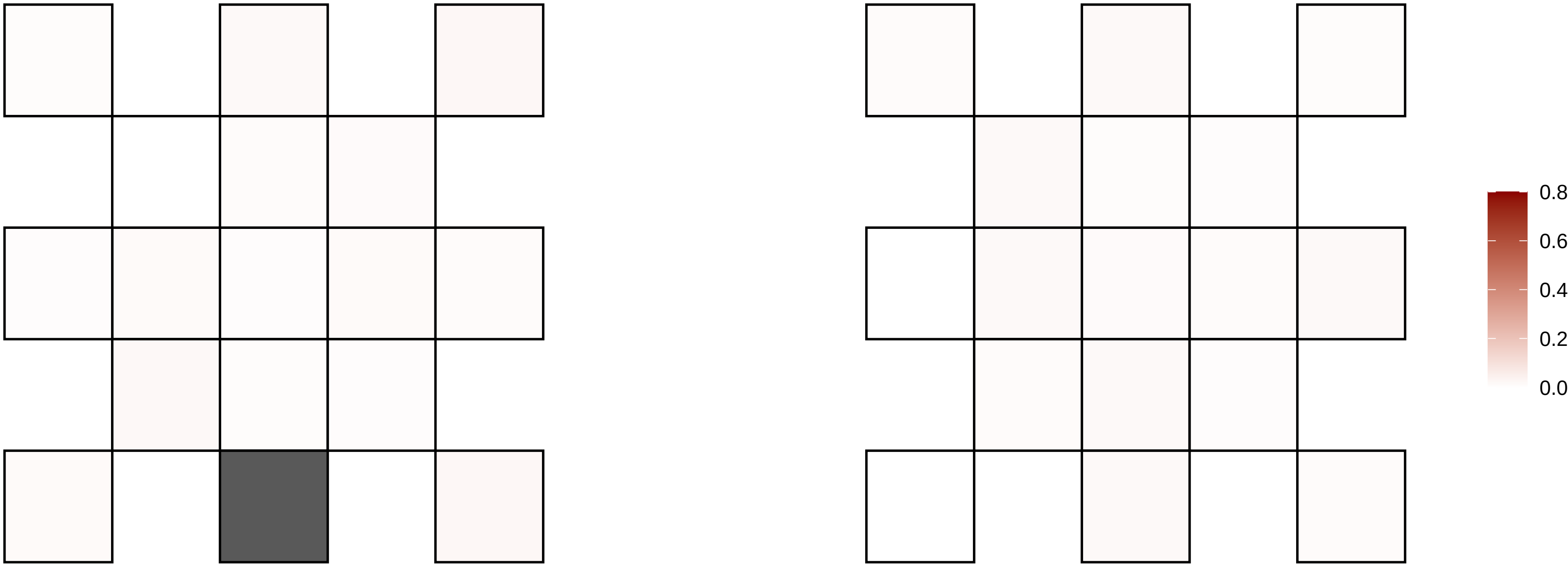

4−85164675−G−A

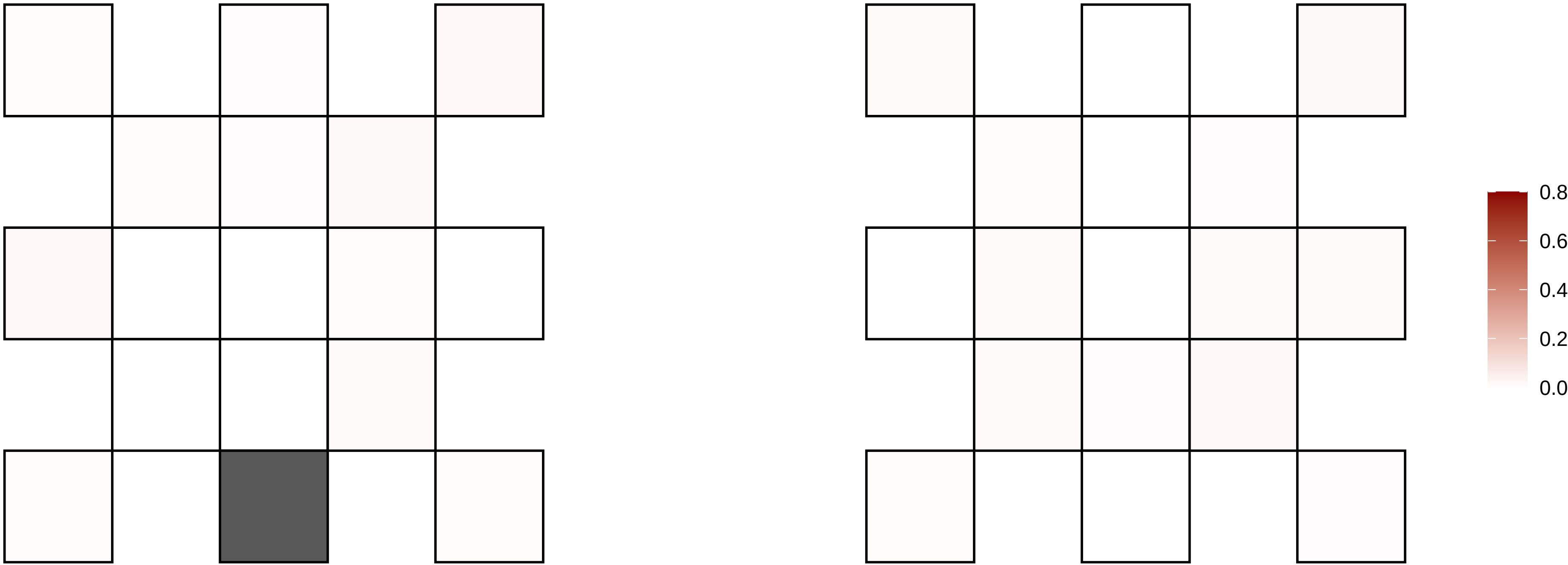

4−91993313−G−A

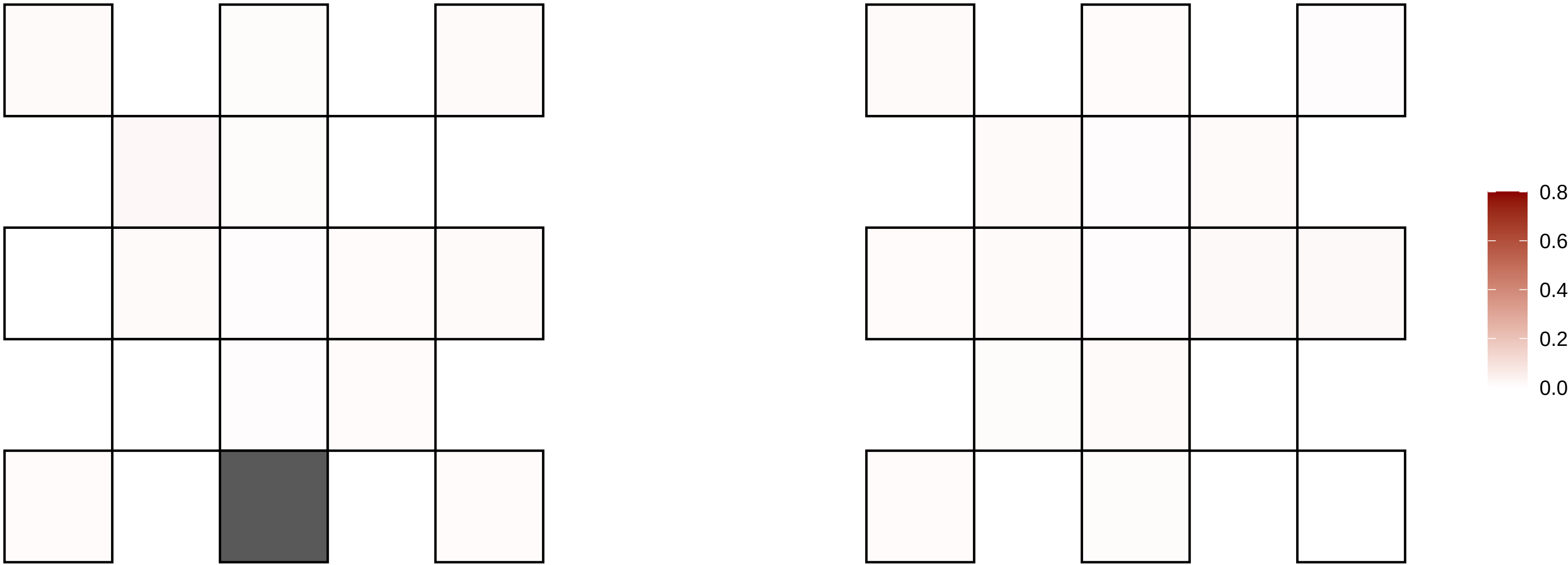

4−92330313−C−G

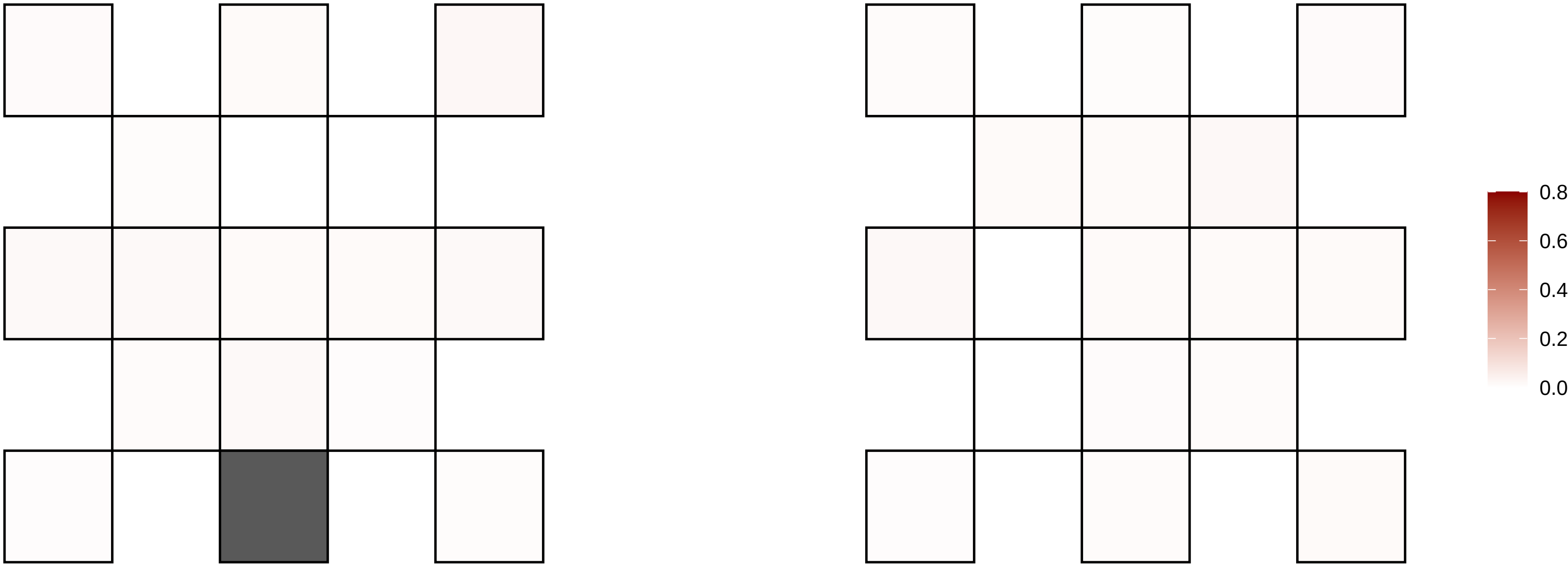

4−92660887−G−A

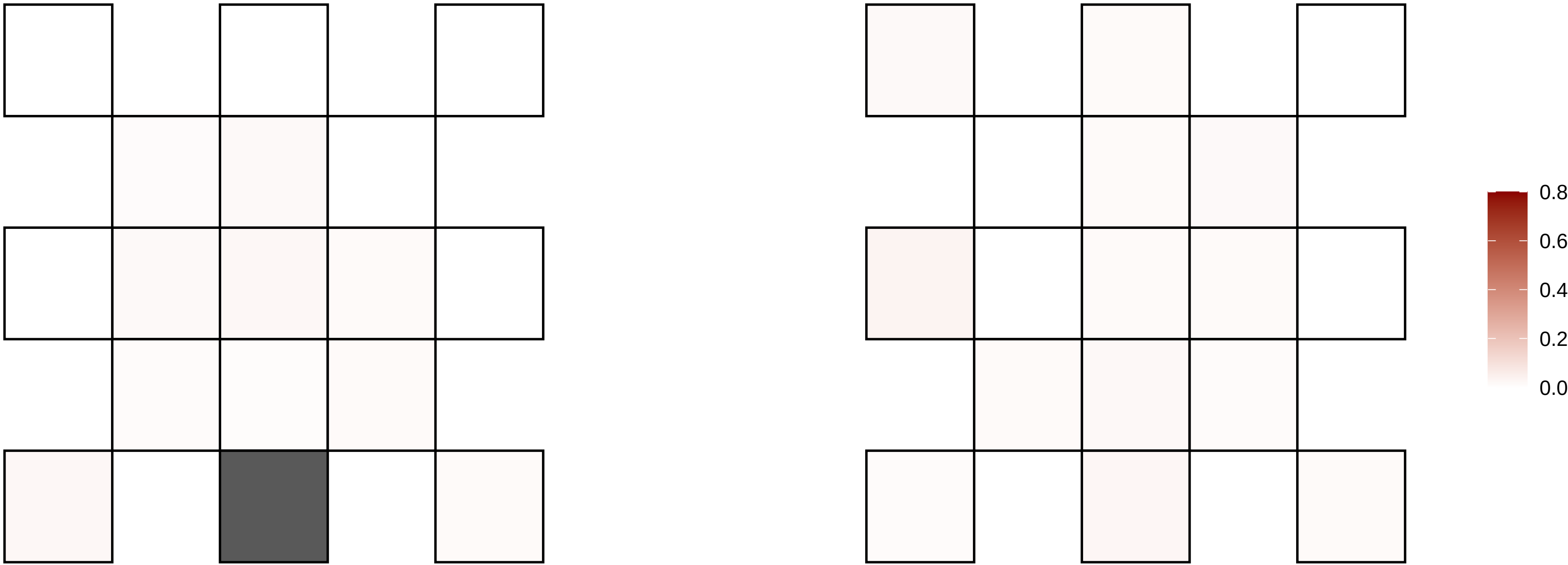

4−93137328−T−A

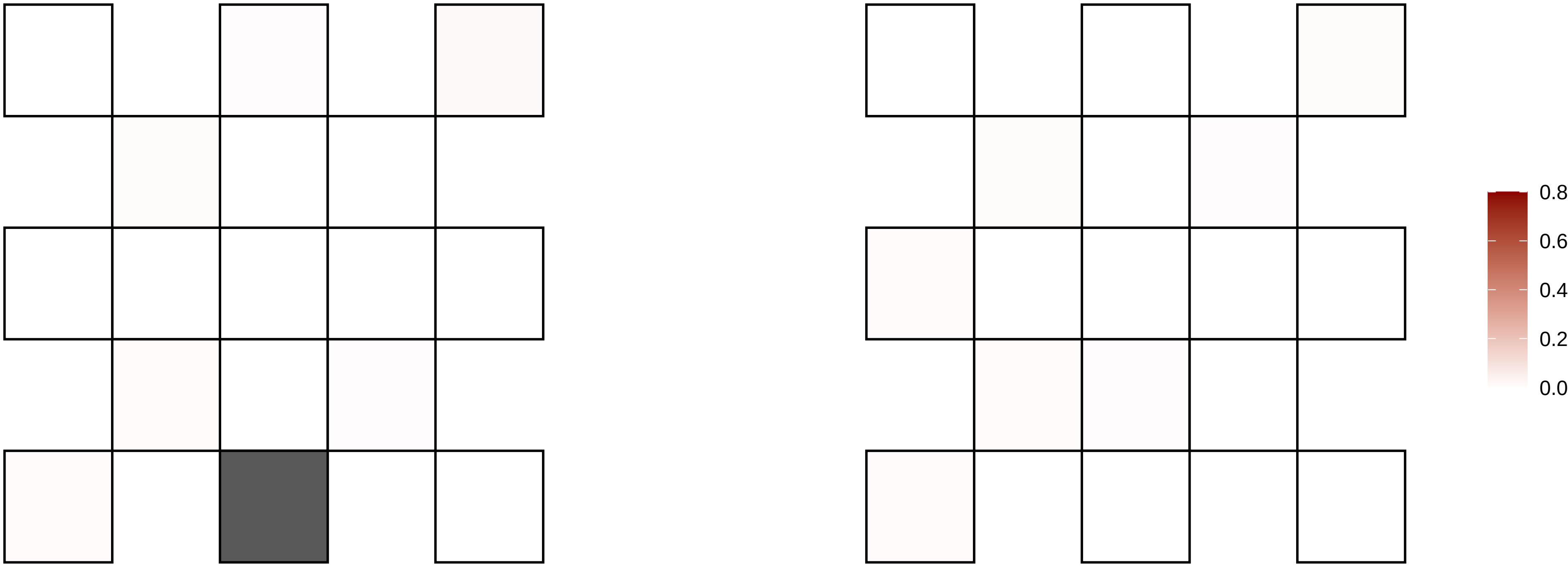

4−94277150−C−T

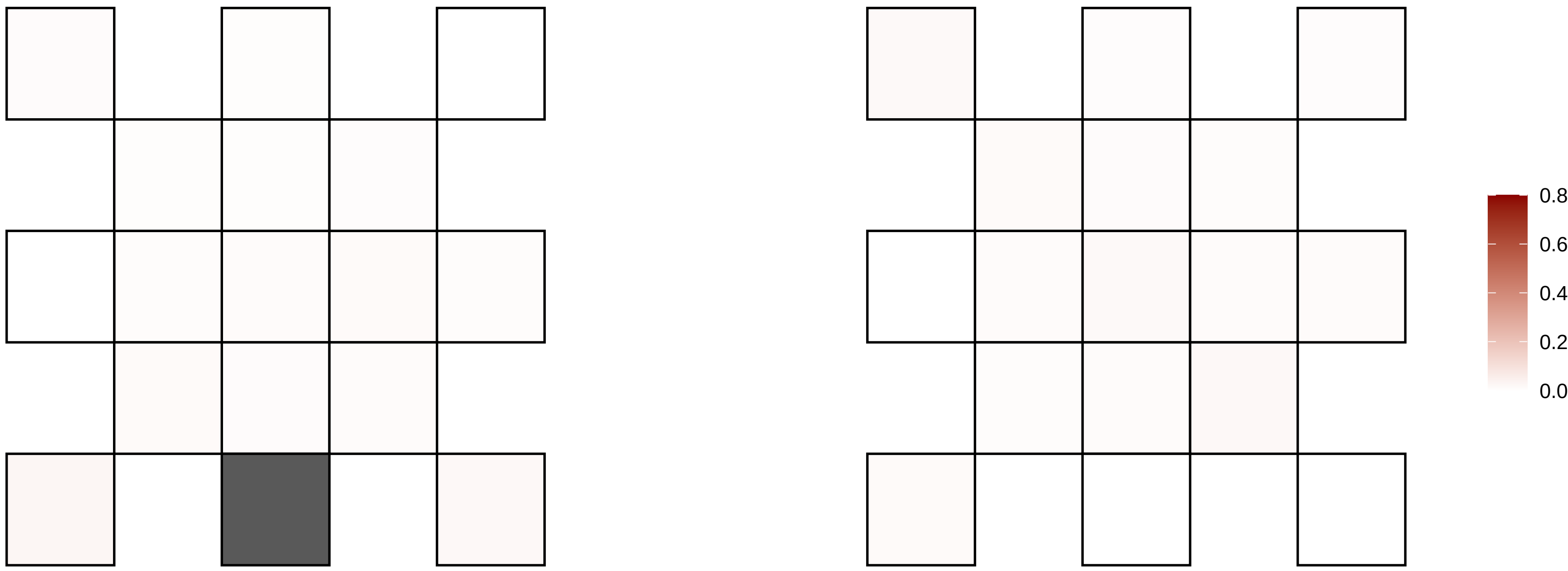

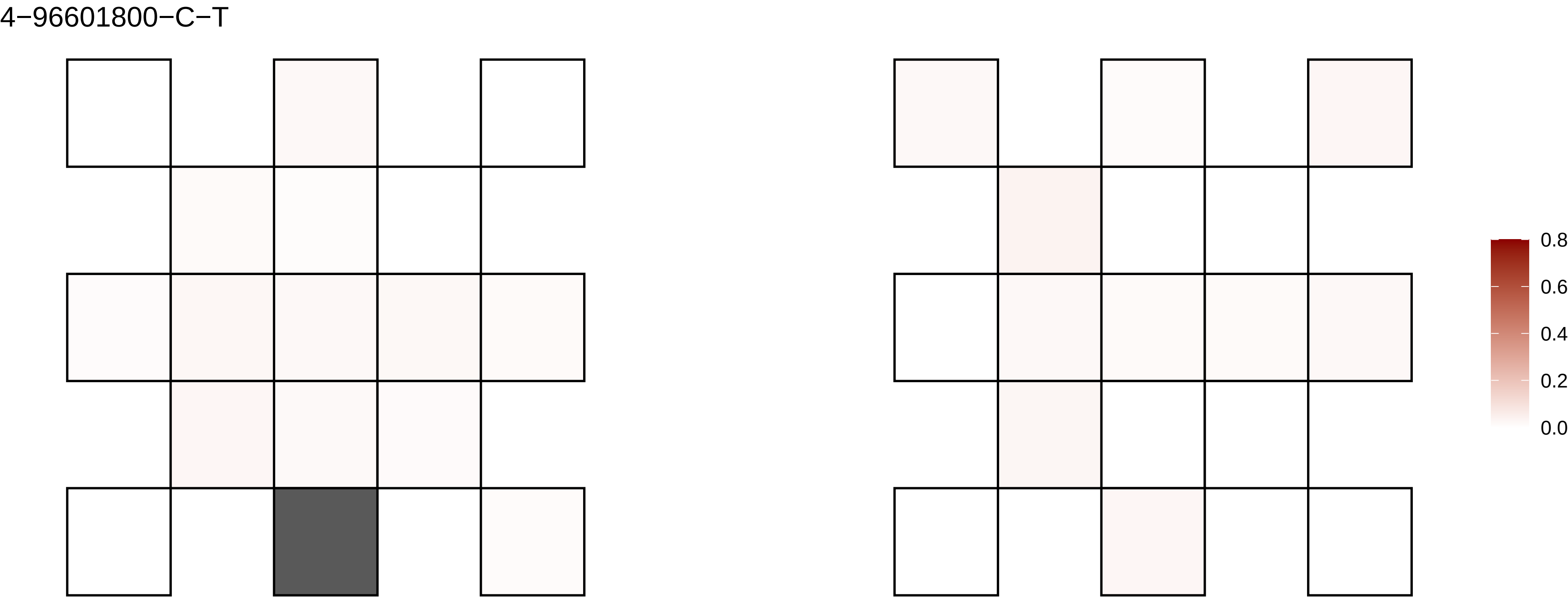
4−96601800−C−T
0.8
0.6
0.4
0.2
0.0

5−10808399−G−A

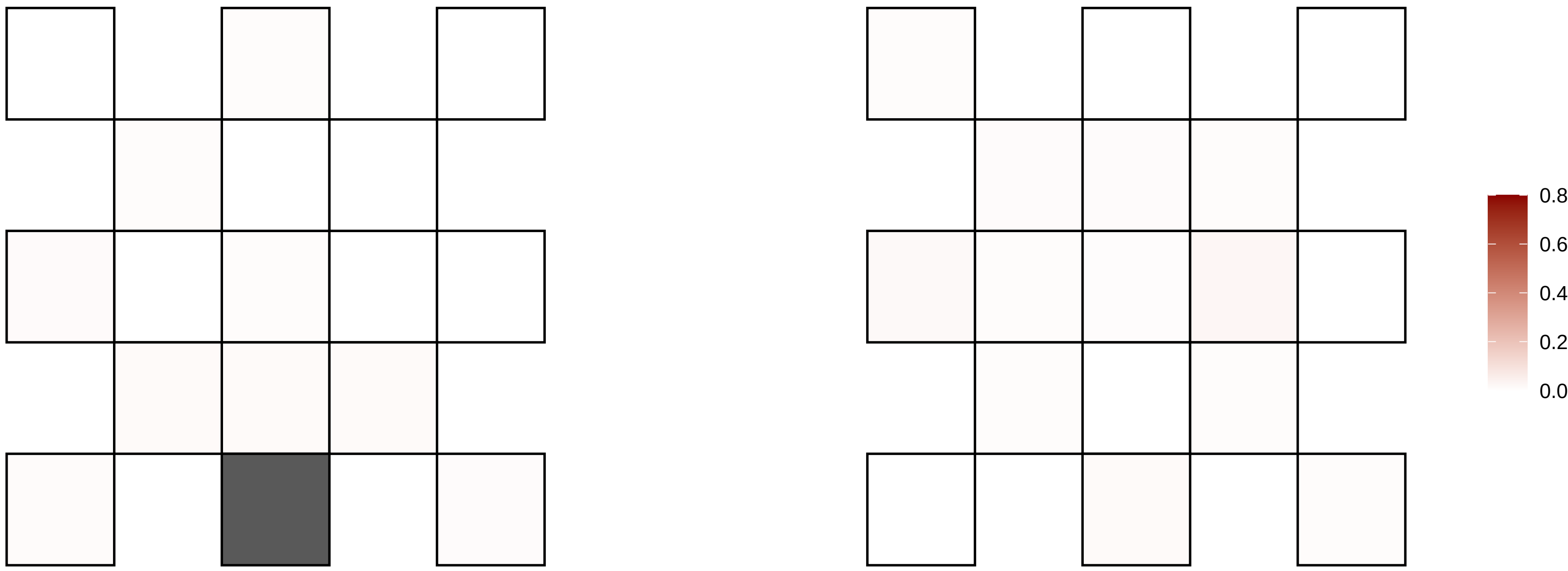

5−115045074−T−C

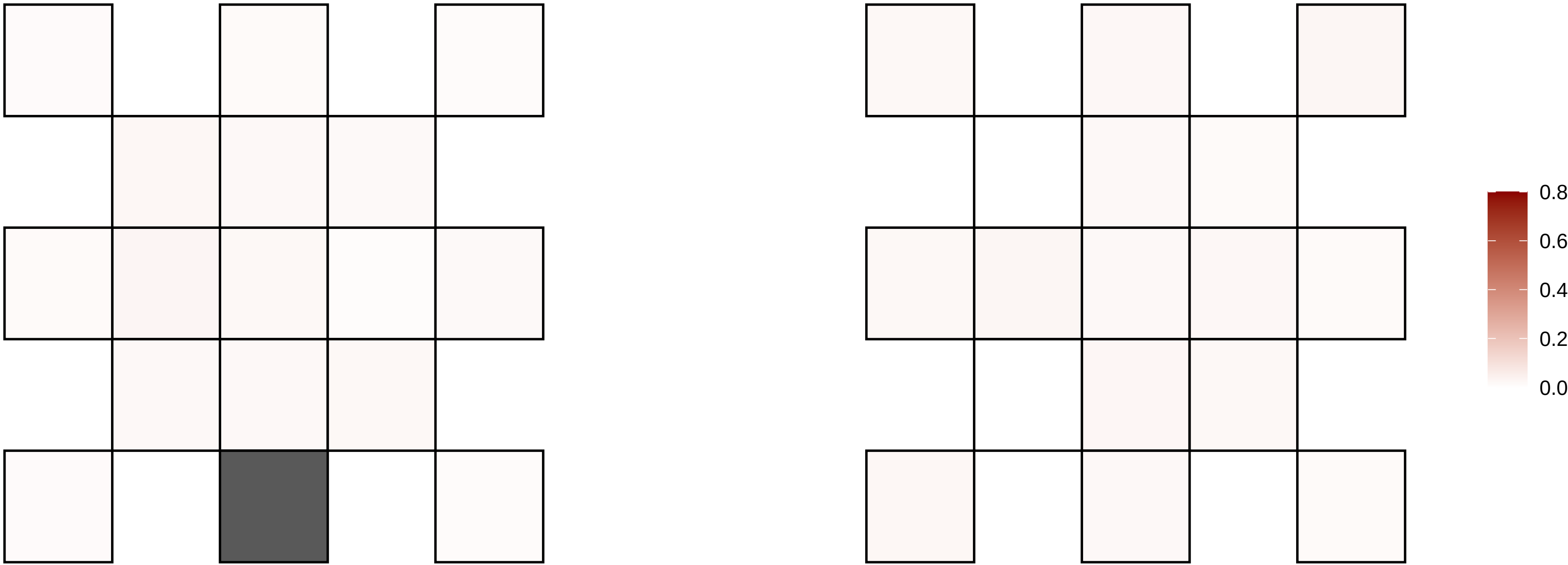

5−115316467−A−C

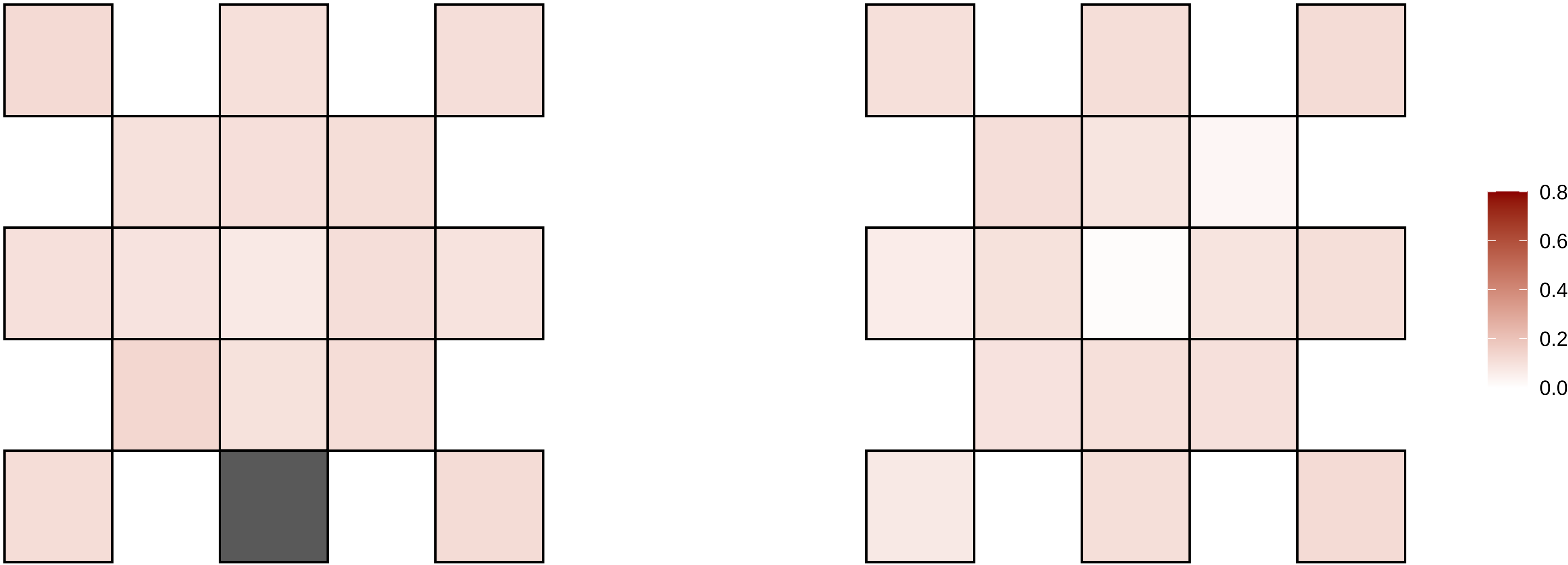

5−115316468−G−T

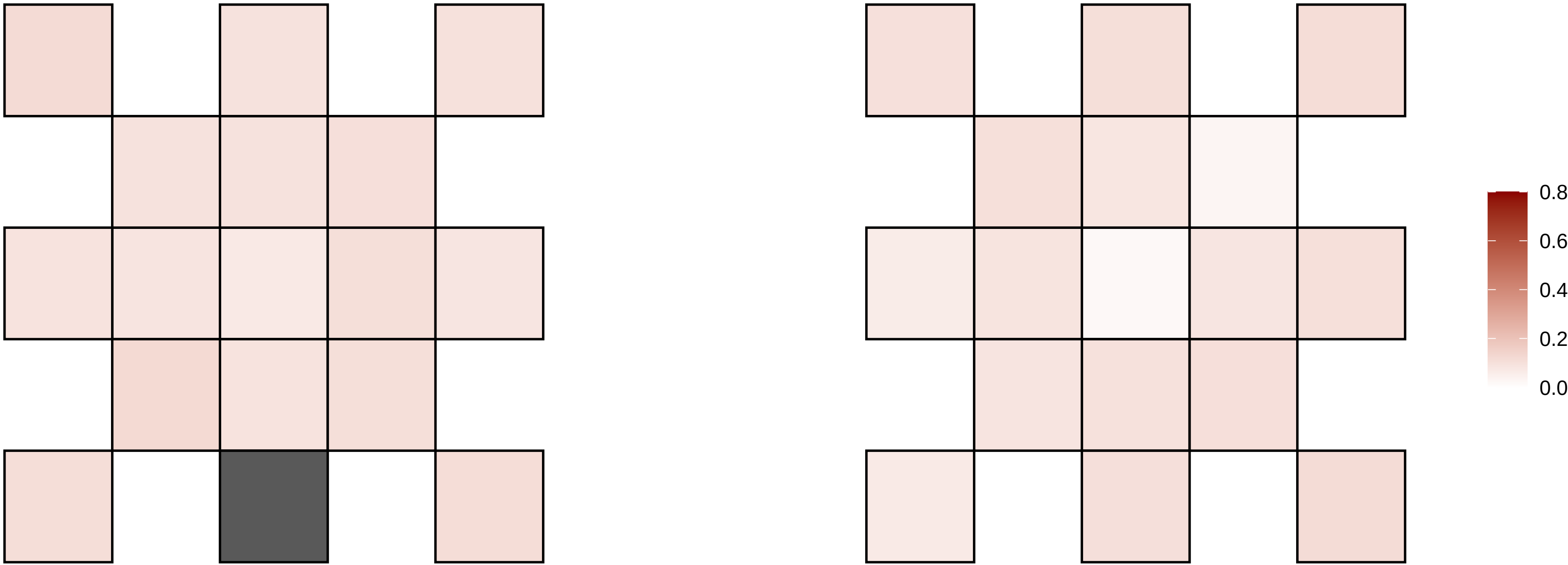

5−117219941−G−T

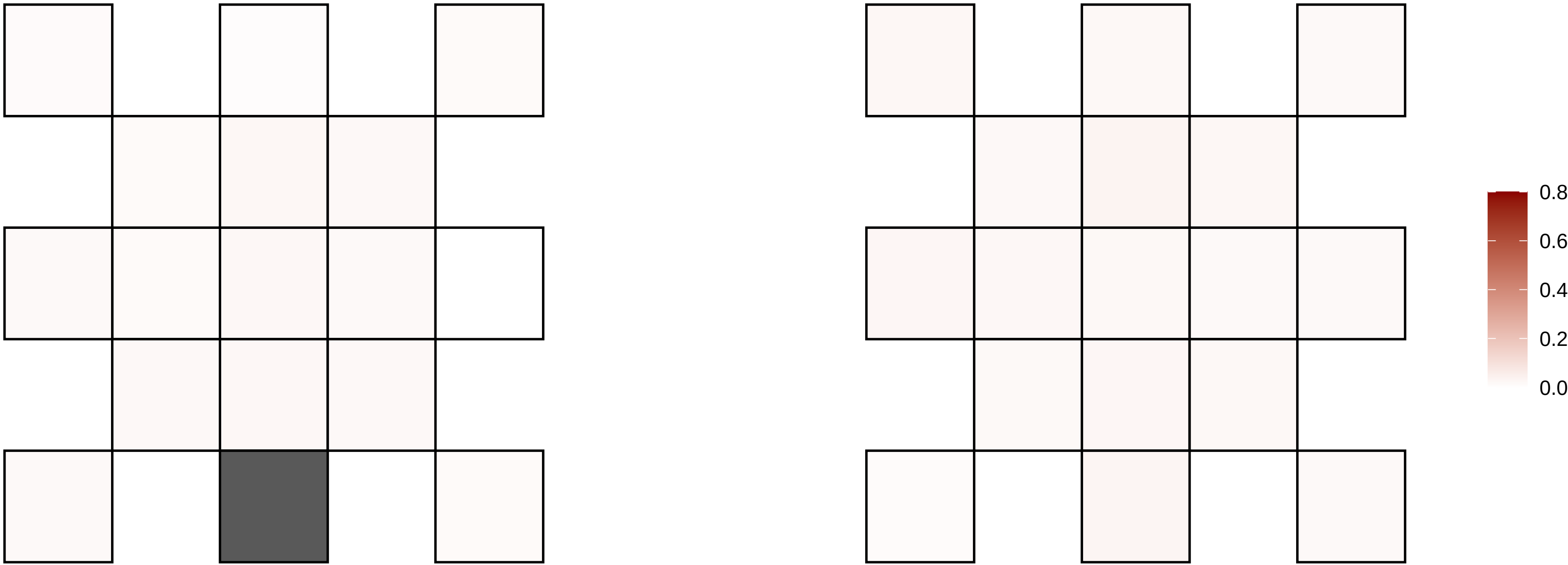

5−117725395−G−A

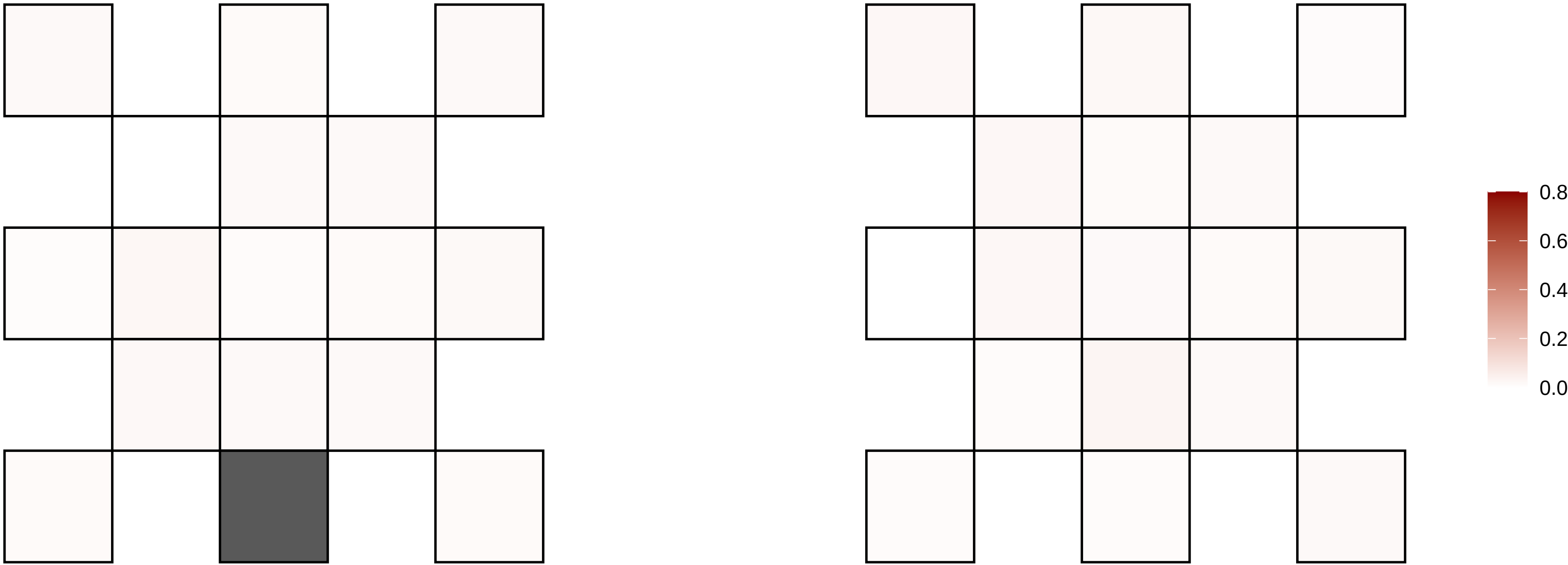

5−121446225−C−T

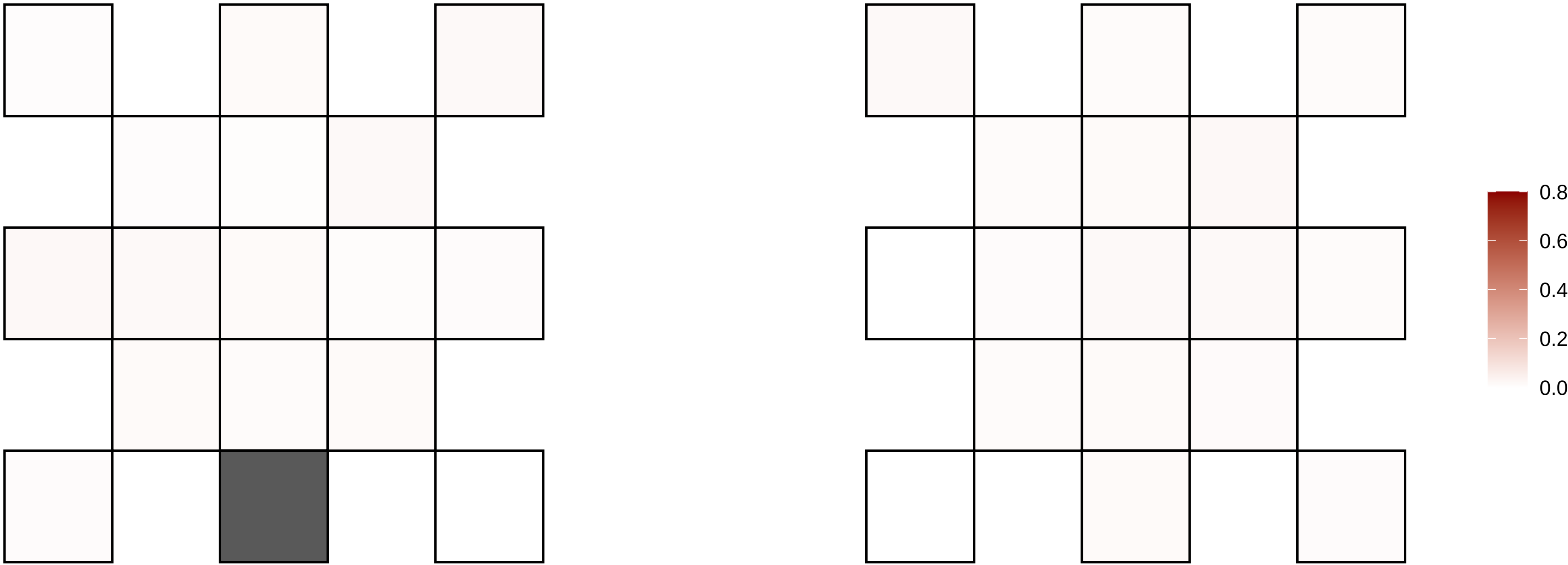

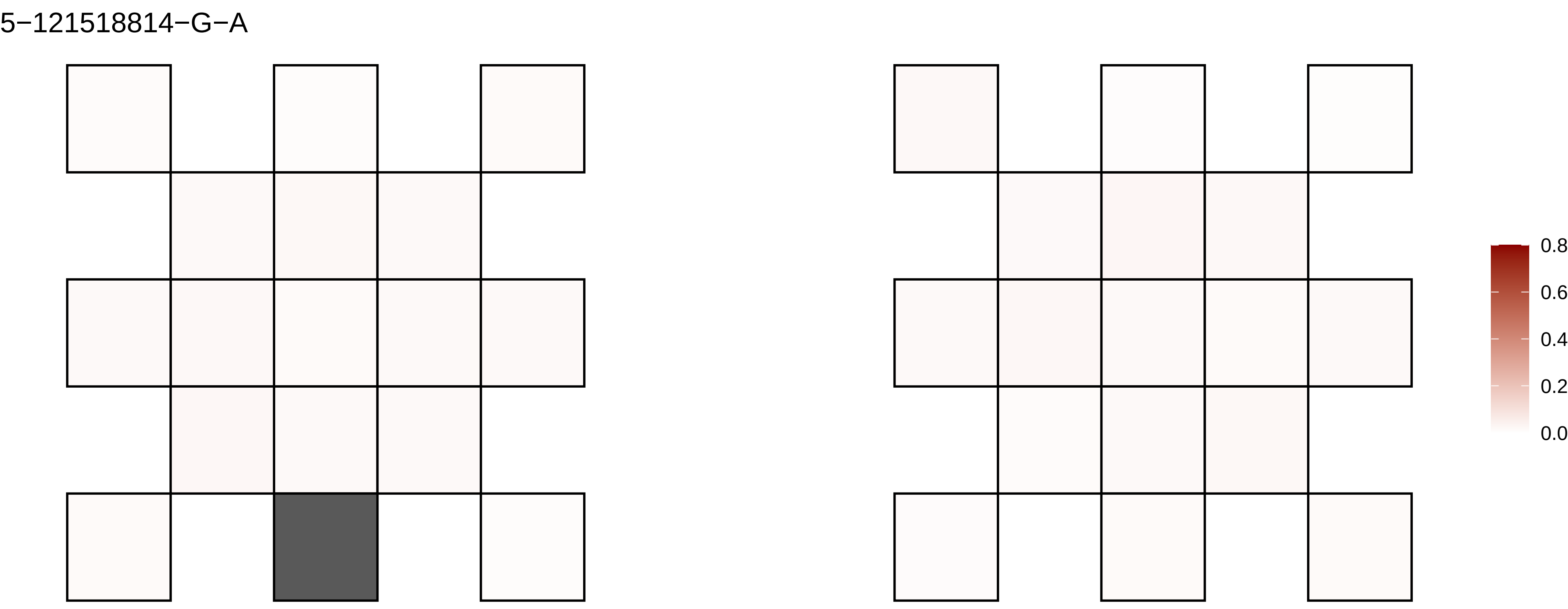
5−121518814−G−A
0.8
0.6
0.4
0.2
0.0

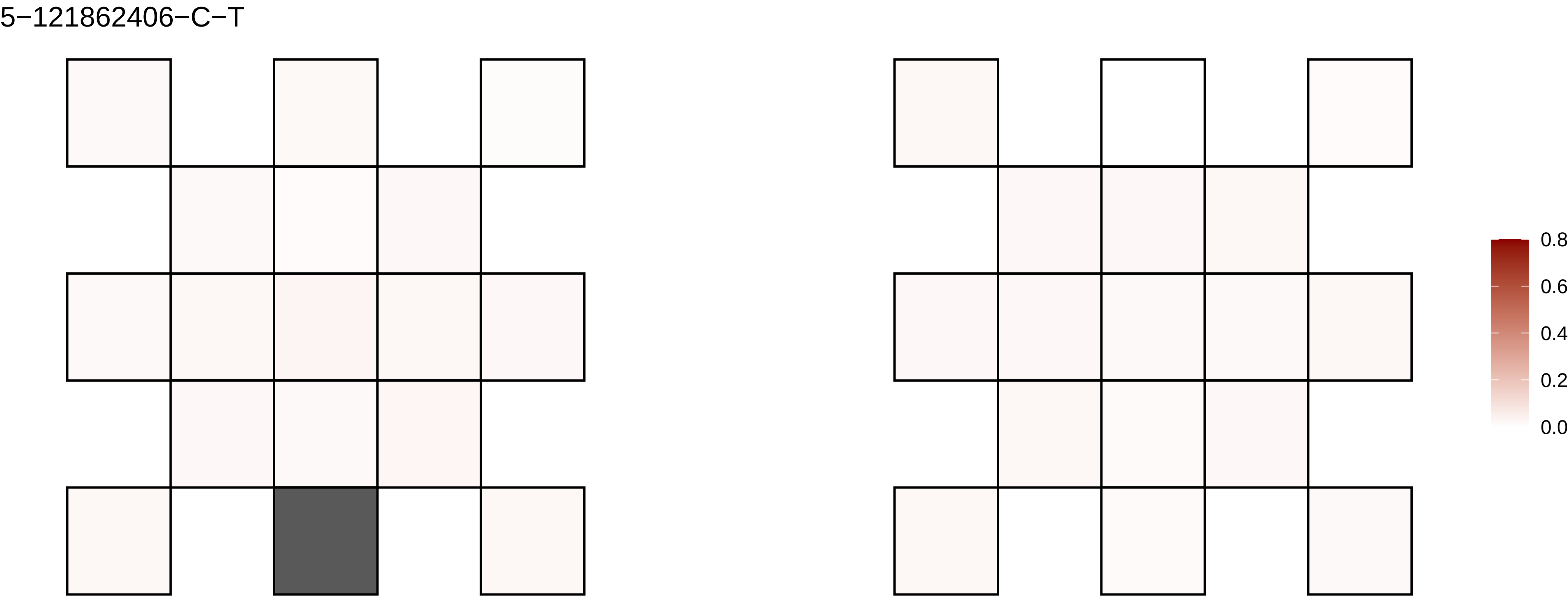
5−121862406−C−T
0.8
0.6
0.4
0.2
0.0

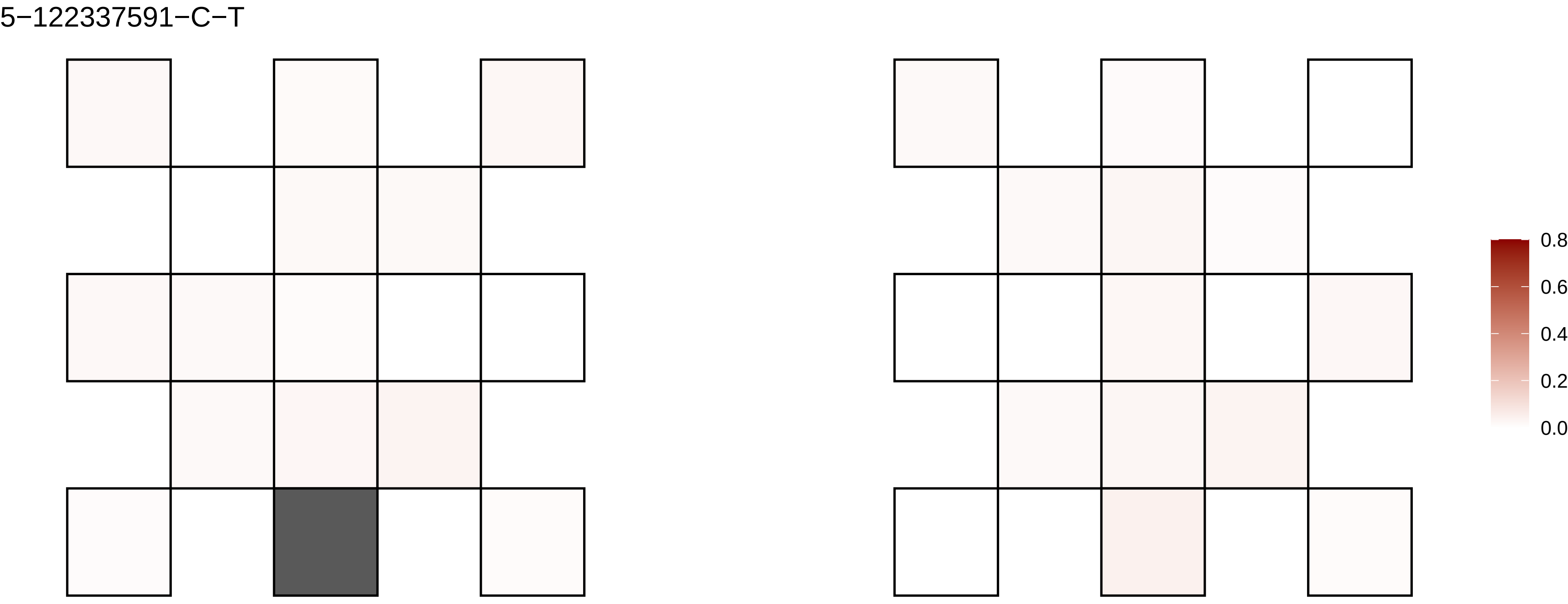
5−122337591−C−T
0.8
0.6
0.4
0.2
0.0

5−13021088−C−T

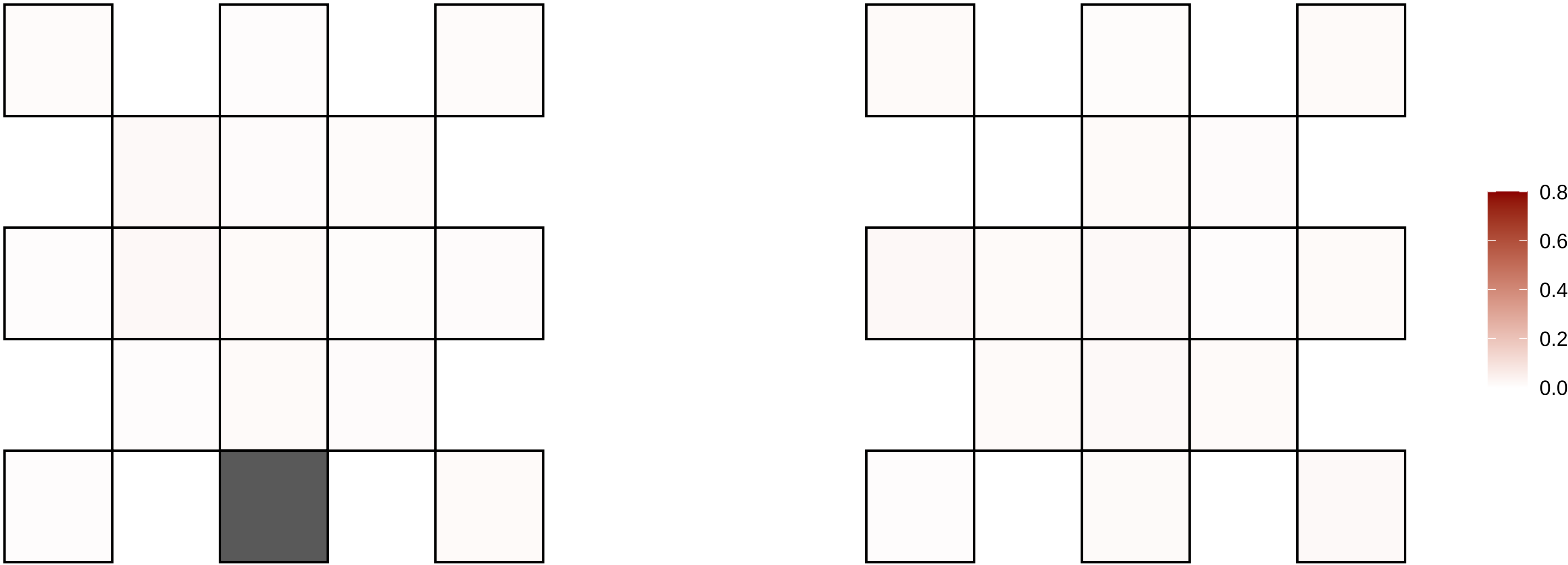

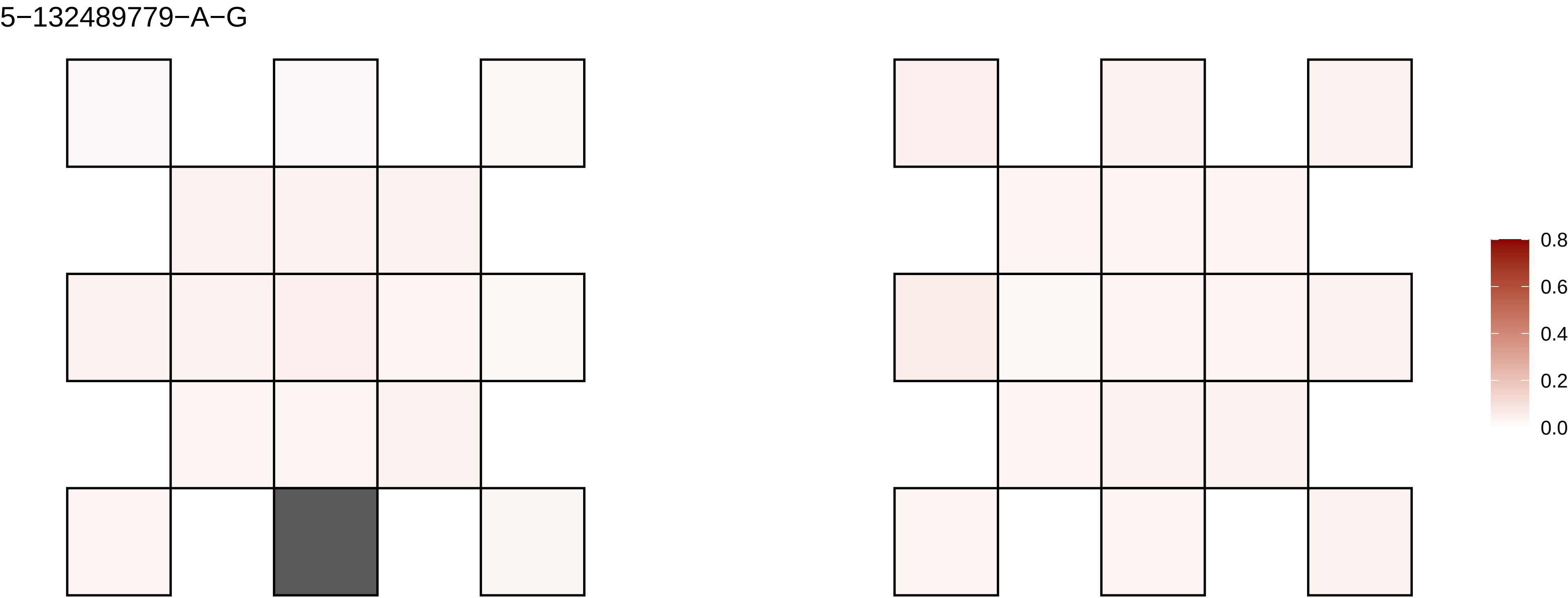
5−132489779−A−G
0.8
0.6
0.4
0.2
0.0

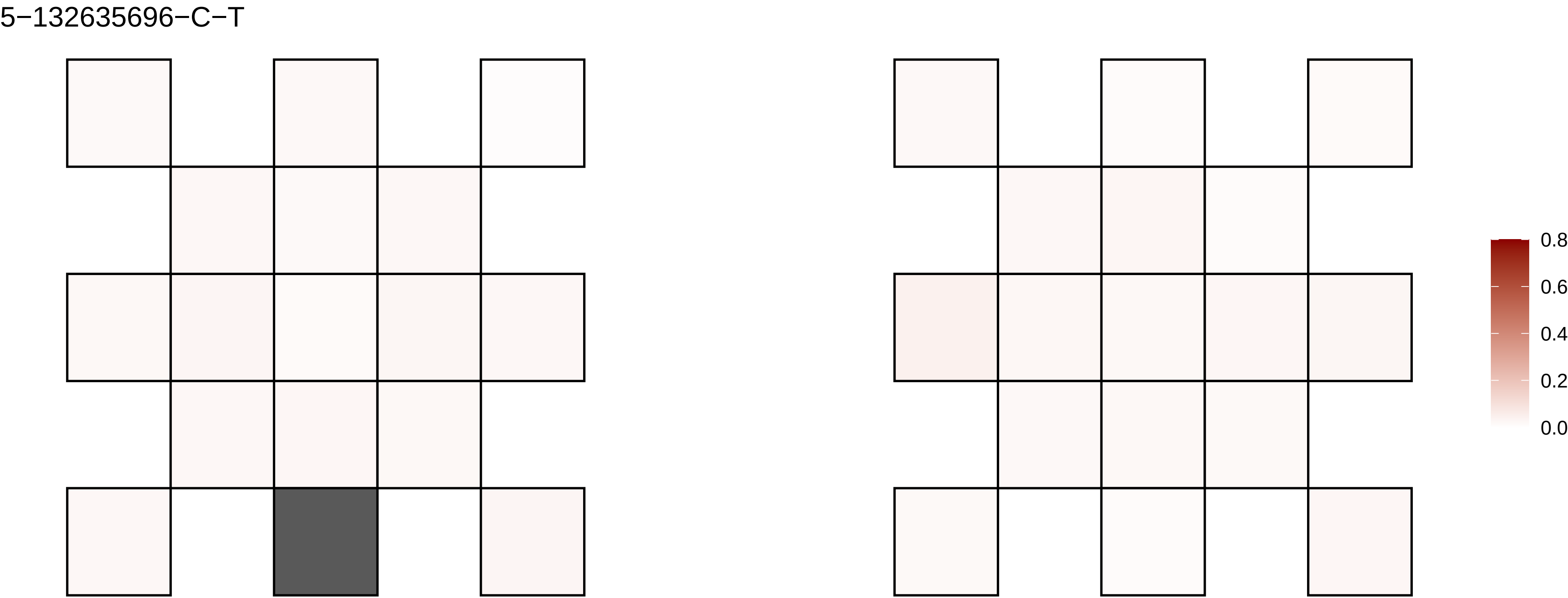
5−132635696−C−T
0.8
0.6
0.4
0.2
0.0

5−133636296−C−A

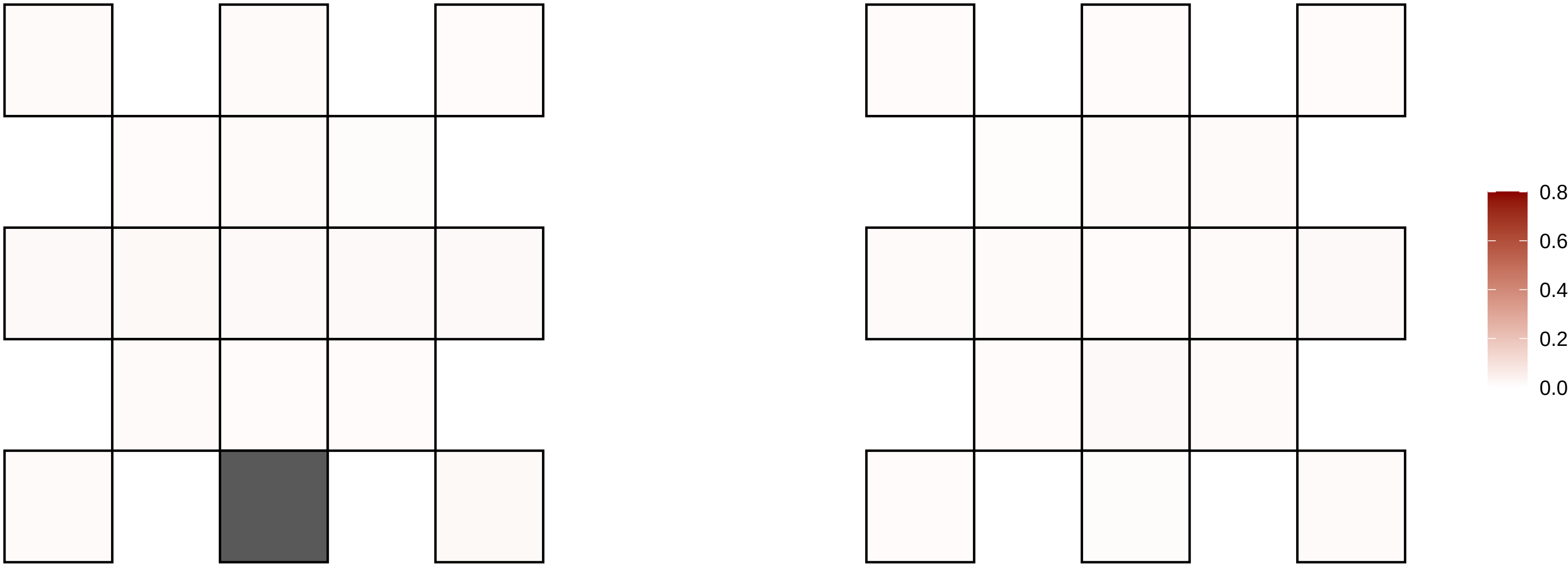

5−142247168−G−A

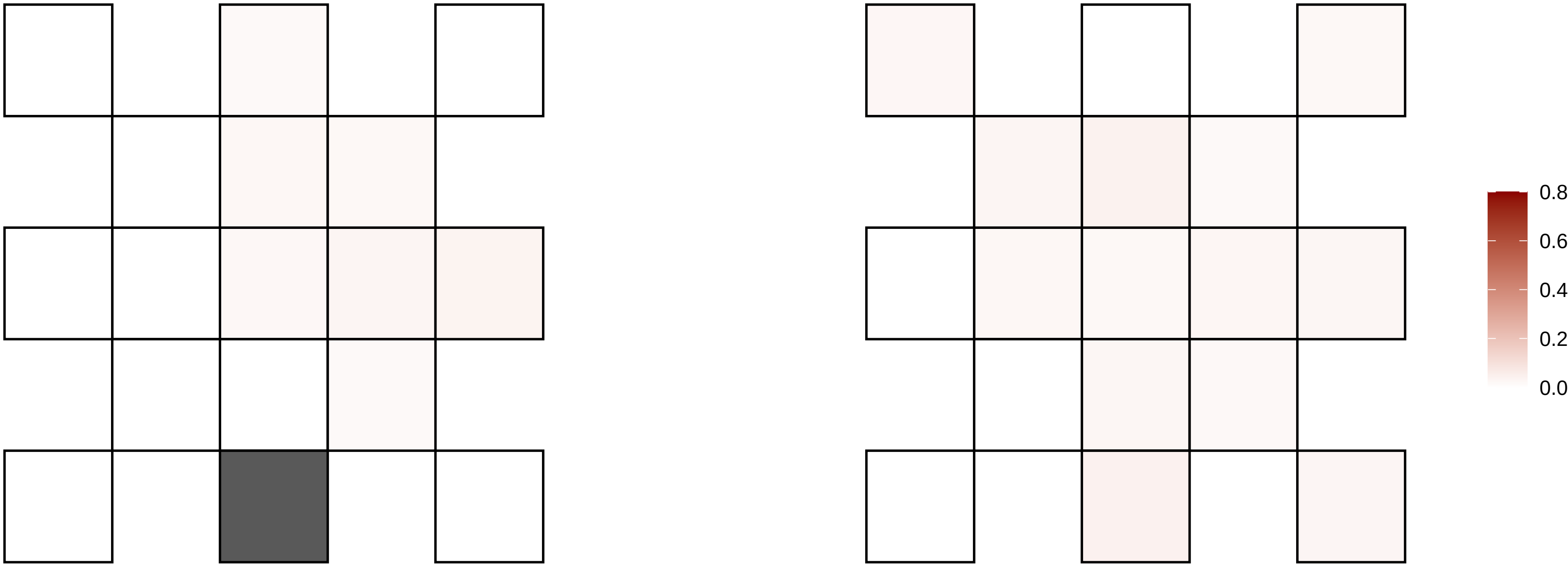

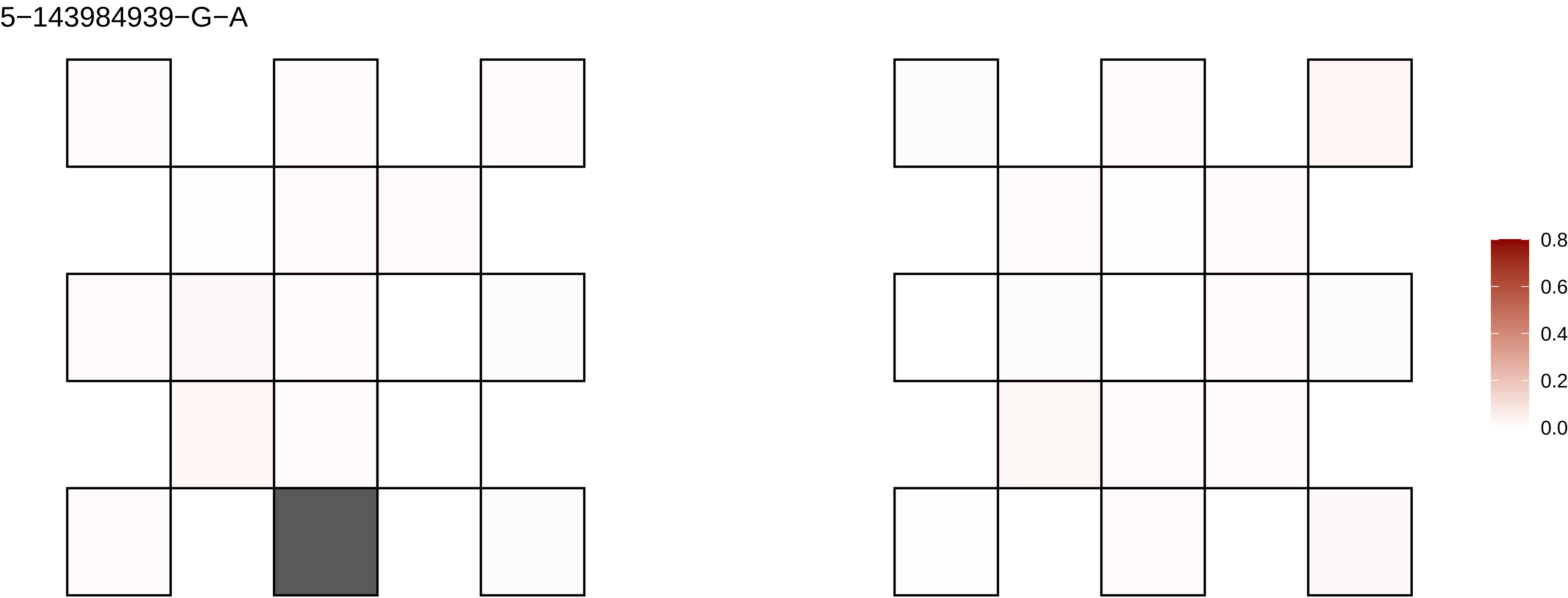

5−143984939−G−A
0.8
0.6
0.4
0.2
0.0

5−155350078−C−T

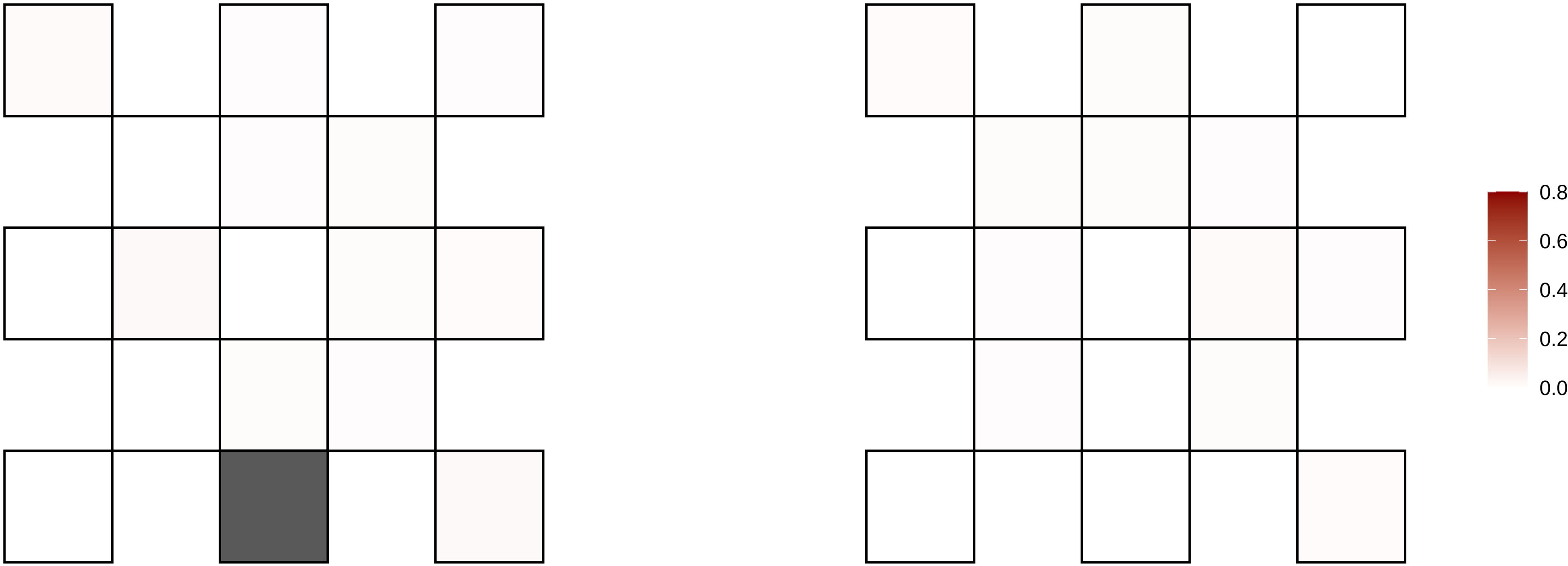

5−155351365−C−A

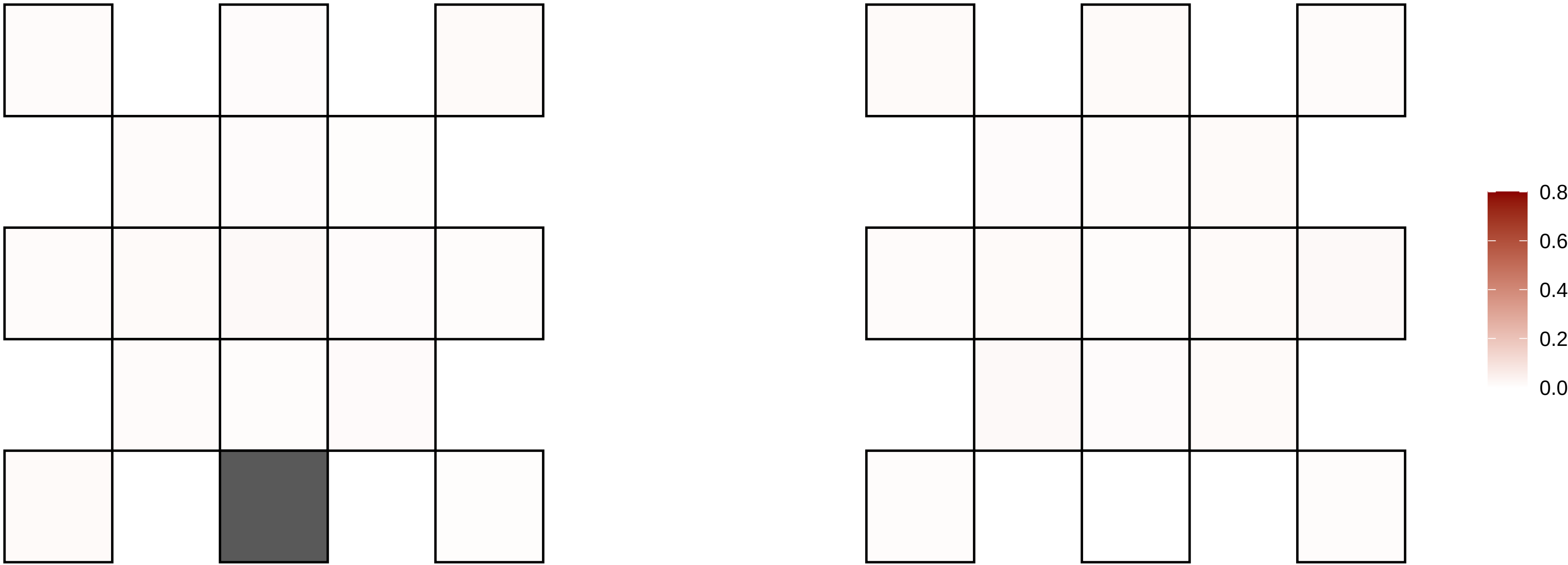

5−158180215−G−A

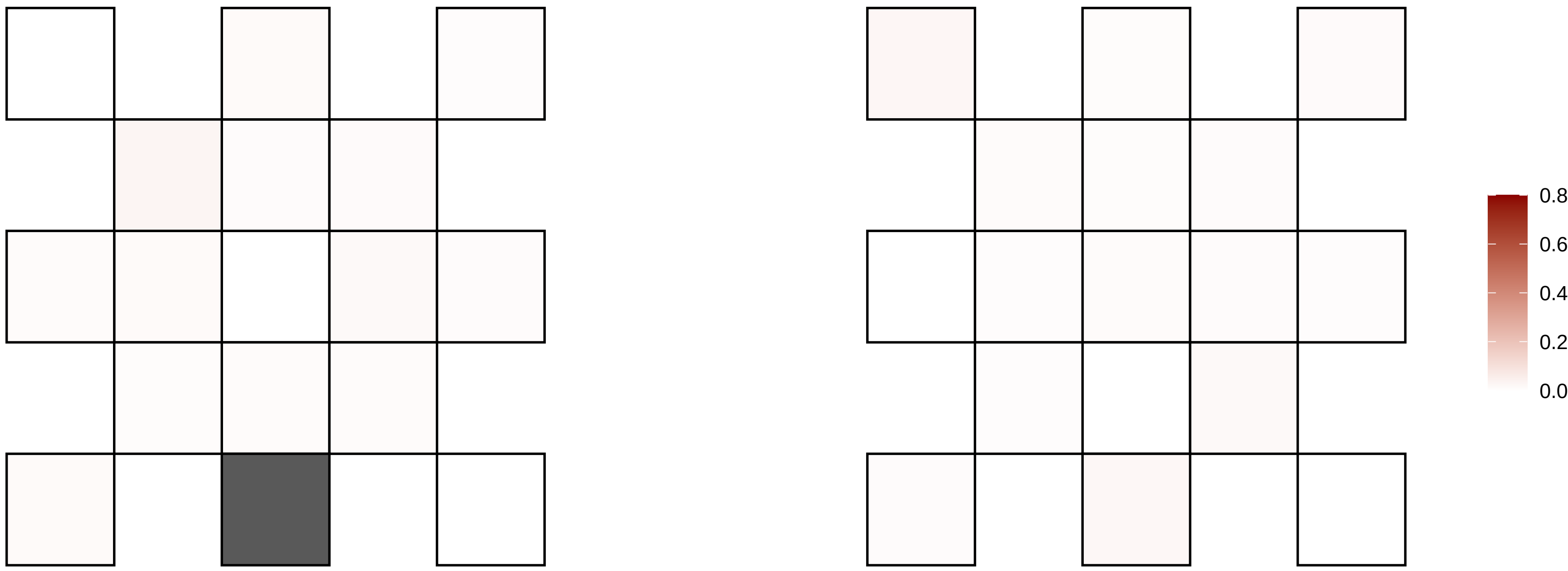

5−15936810−G−A

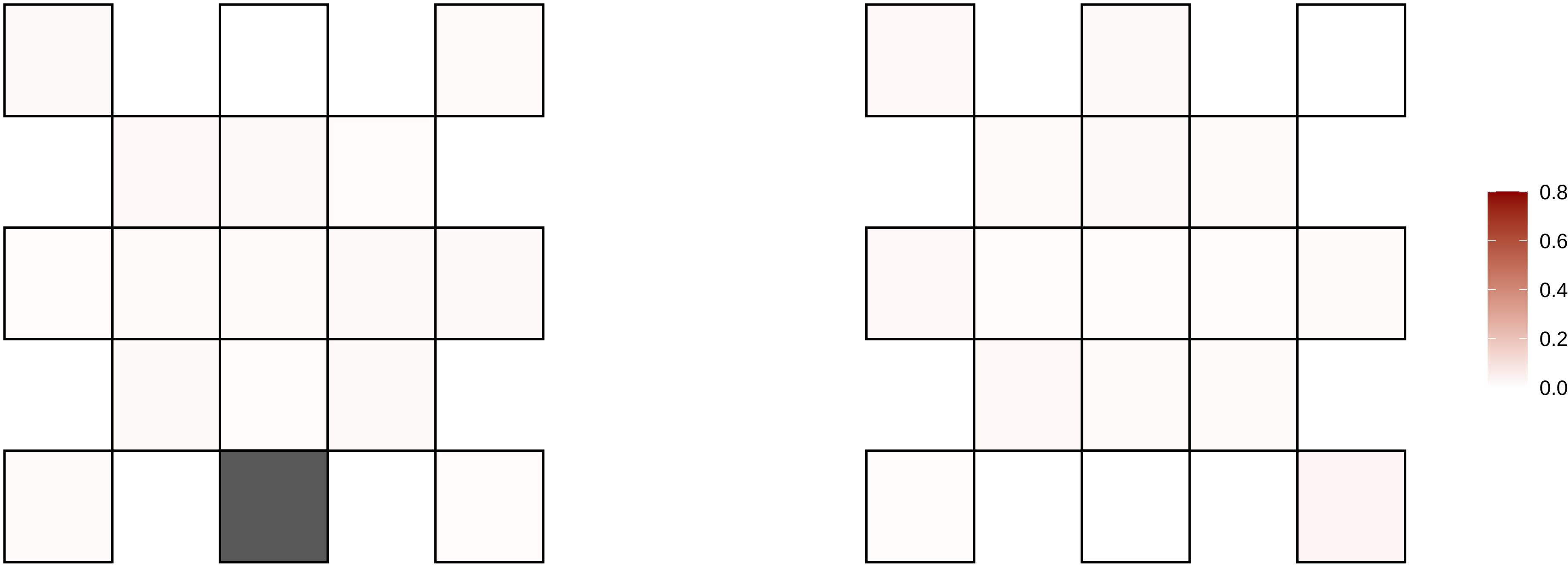

5−160583527−G−A

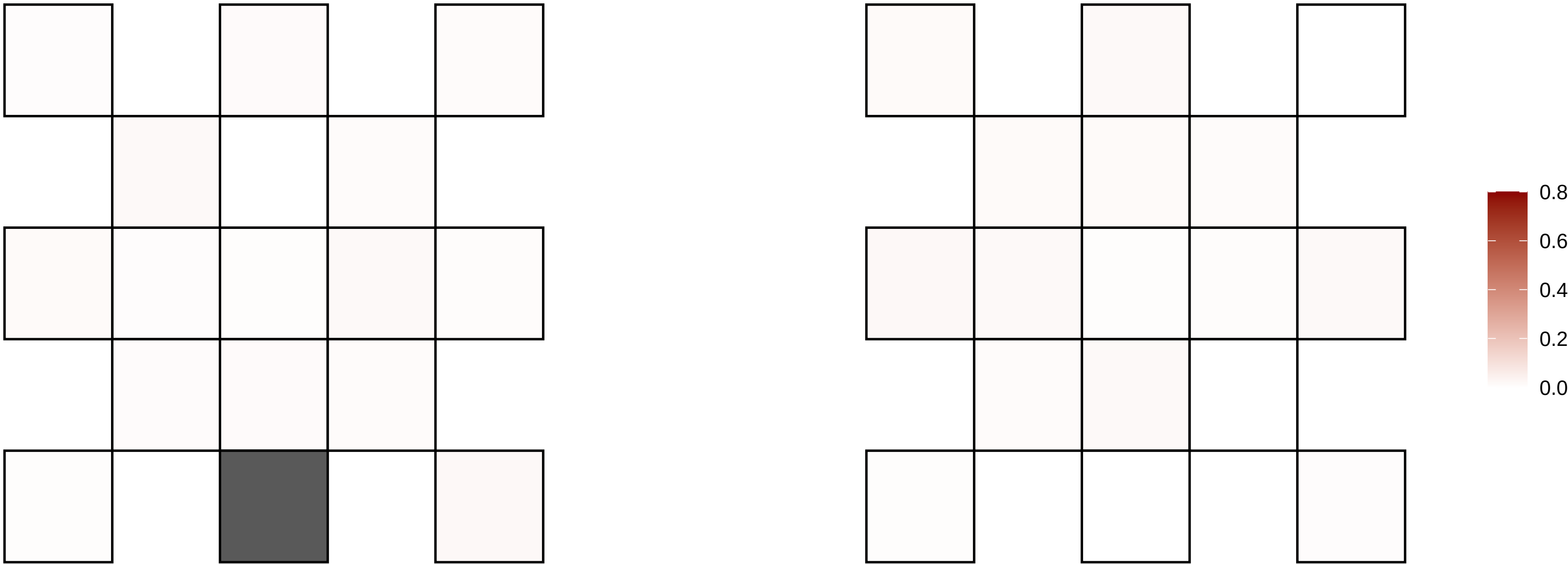

5−165691201−A−G

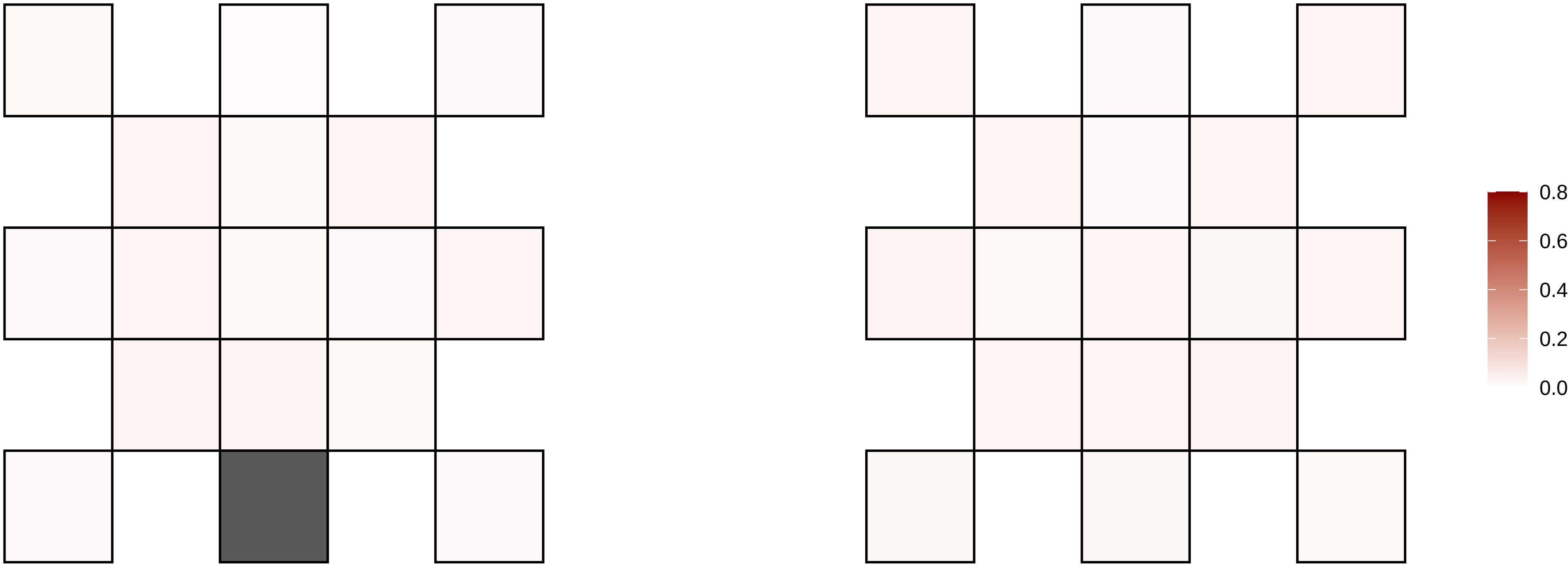

5−167949614−A−G

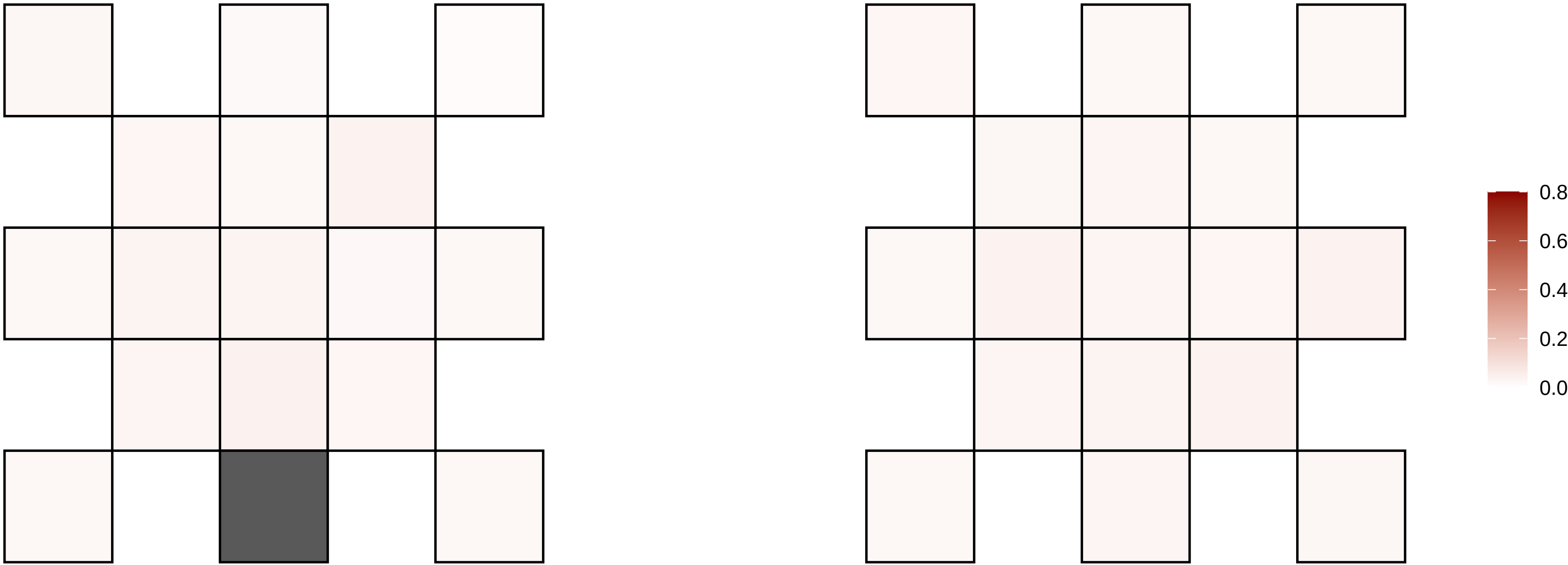

5−179554887−C−T

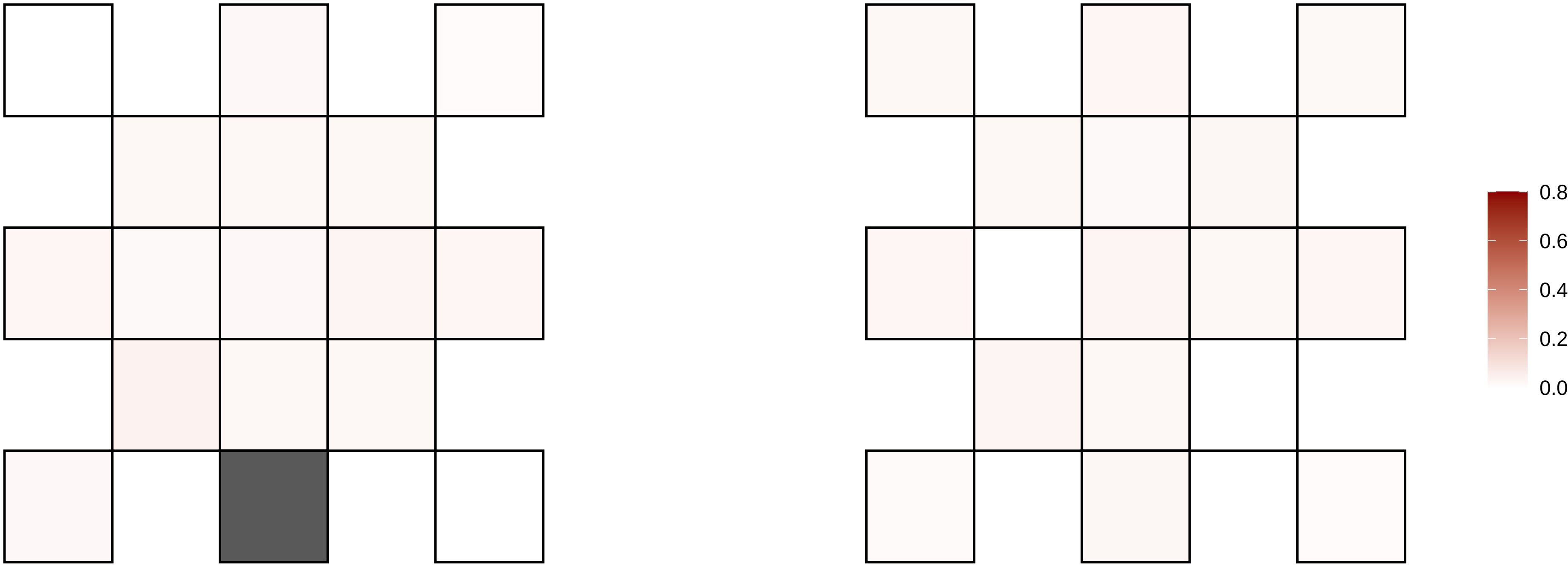

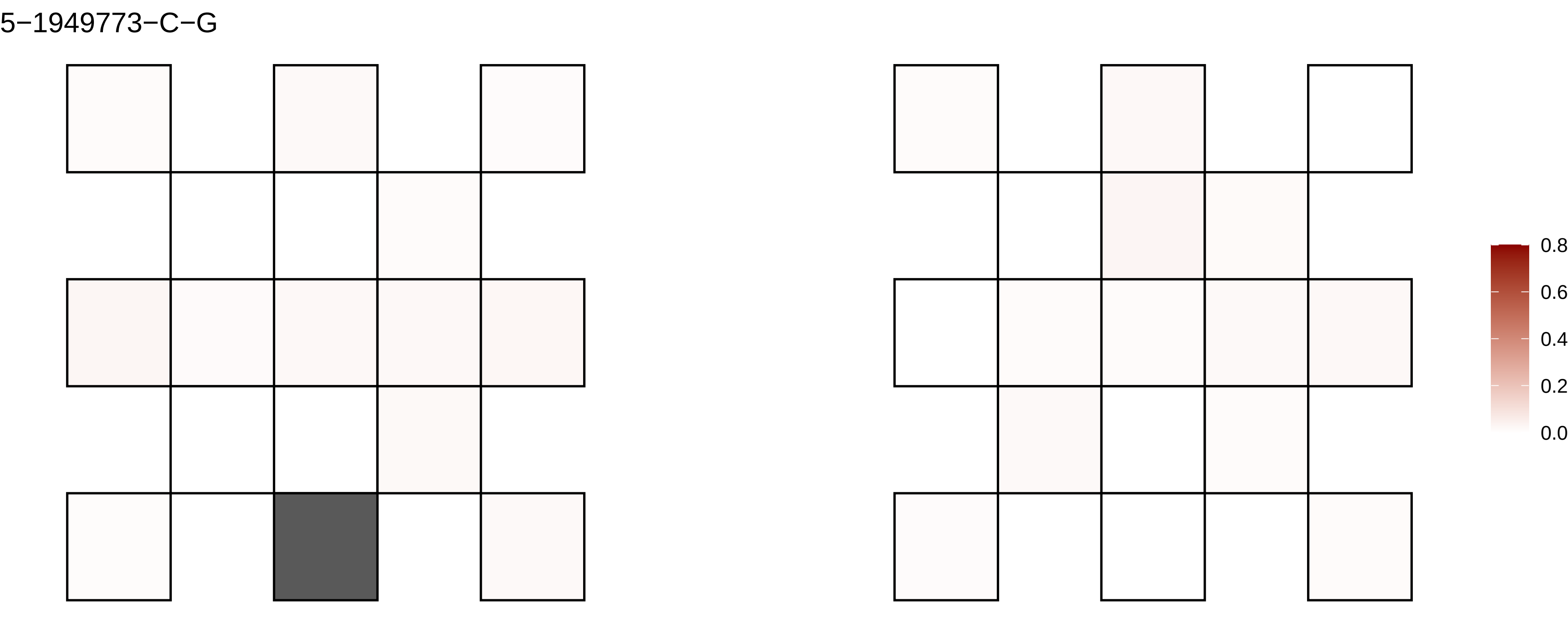
5−1949773−C−G
0.8
0.6
0.4
0.2
0.0

5−26502402−A−G

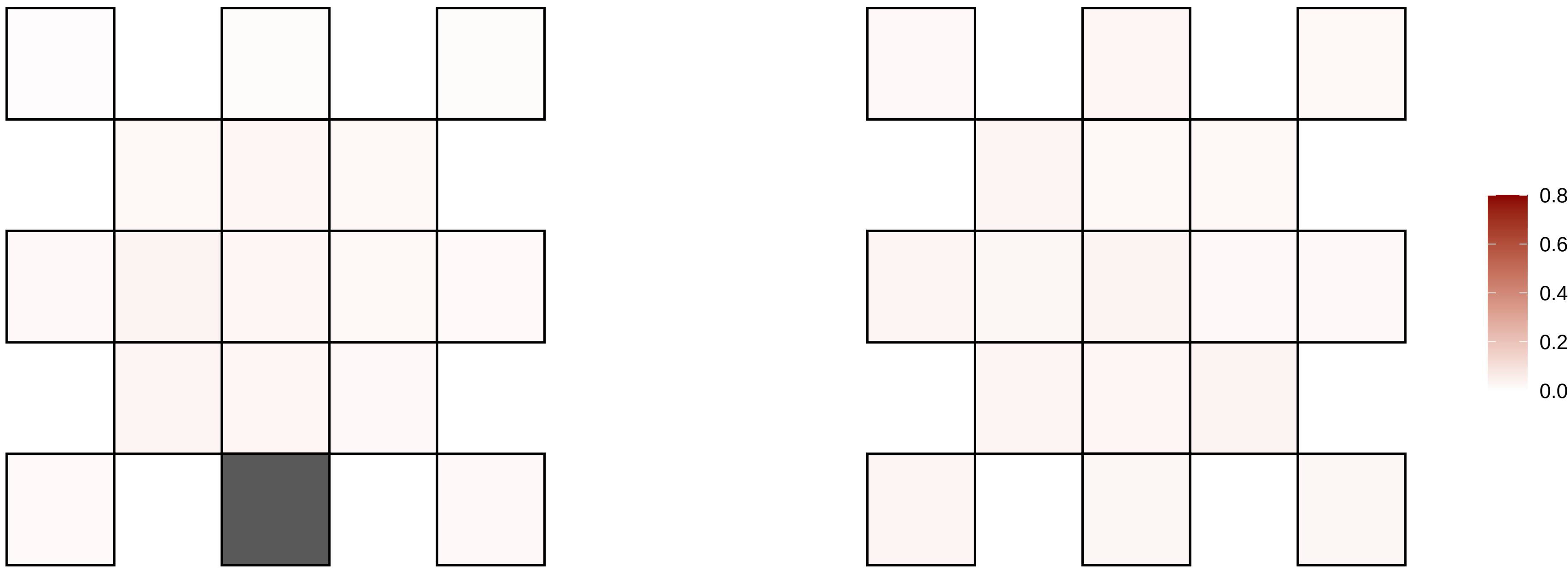

5−28689952−A−G

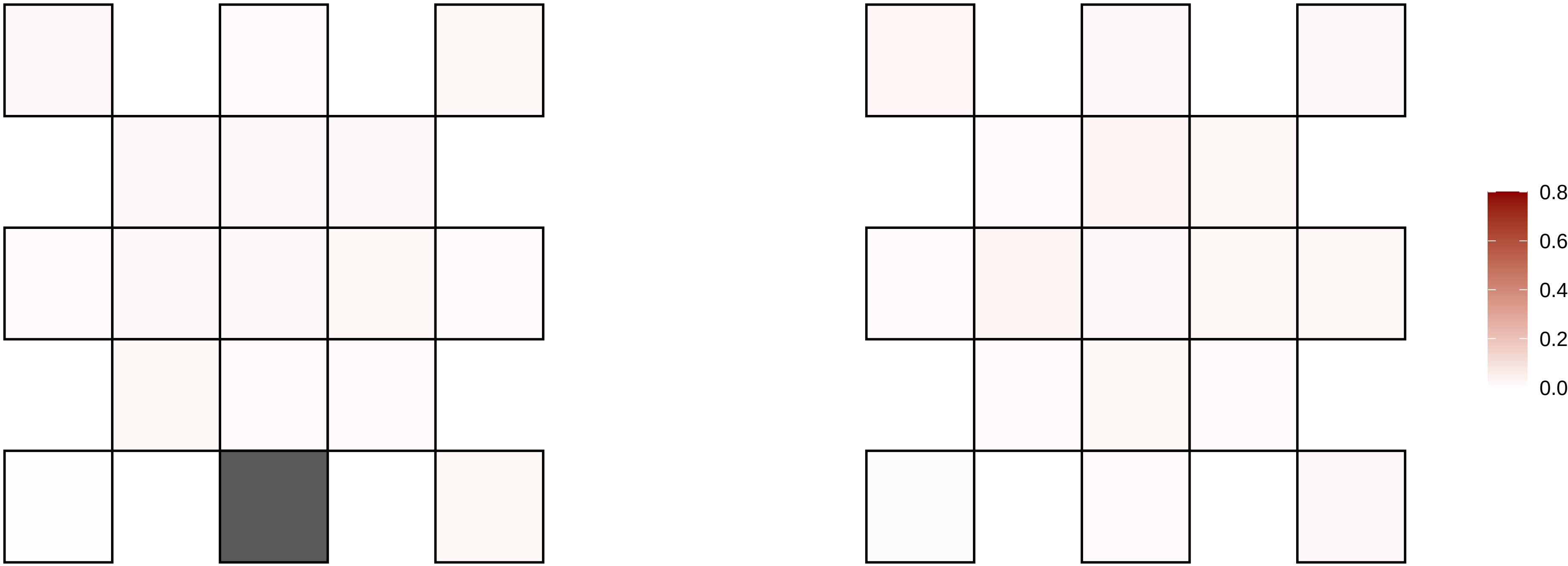

5−30650591−C−A

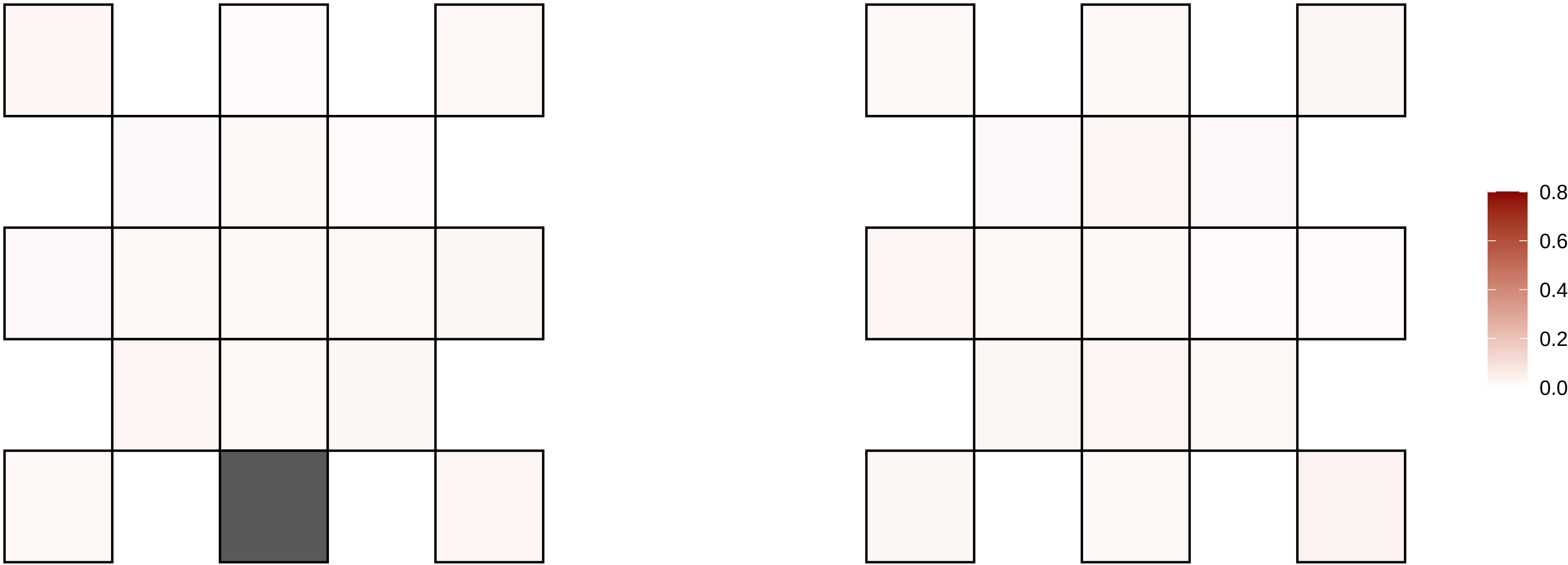

5−30923233−C−T

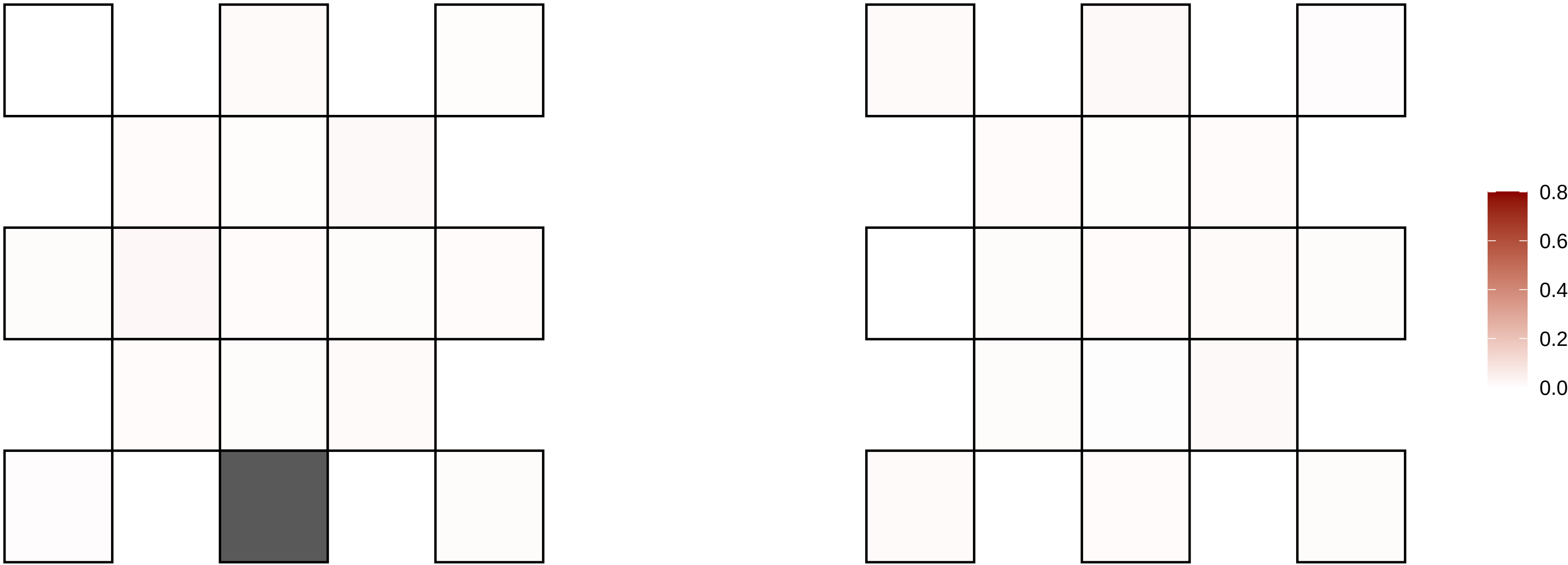

5−42394330−C−T

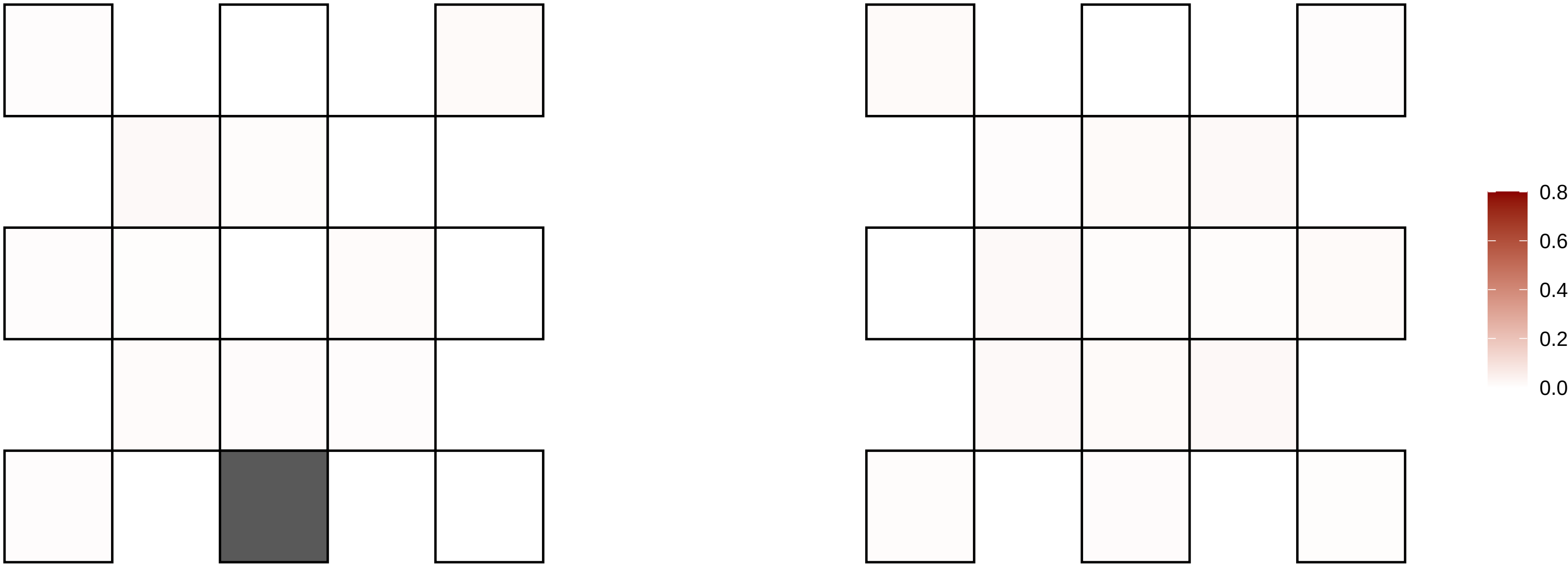

5−42474677−G−T

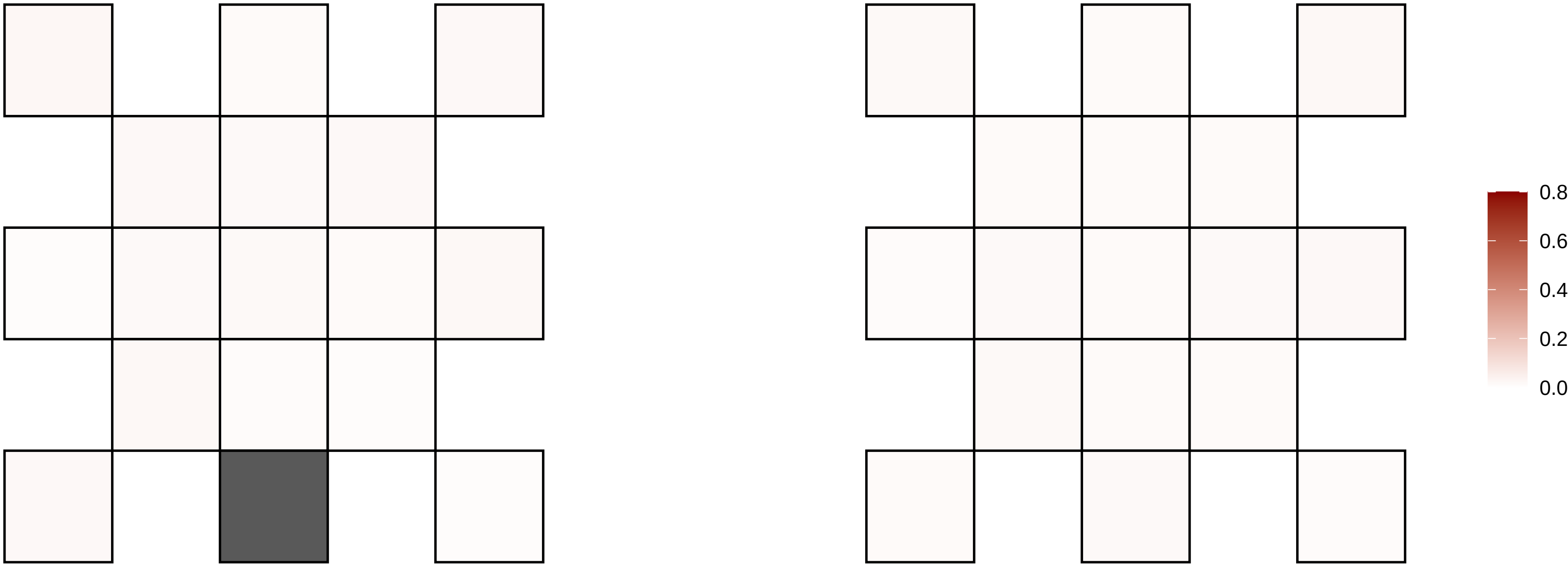

5−50102837−T−C

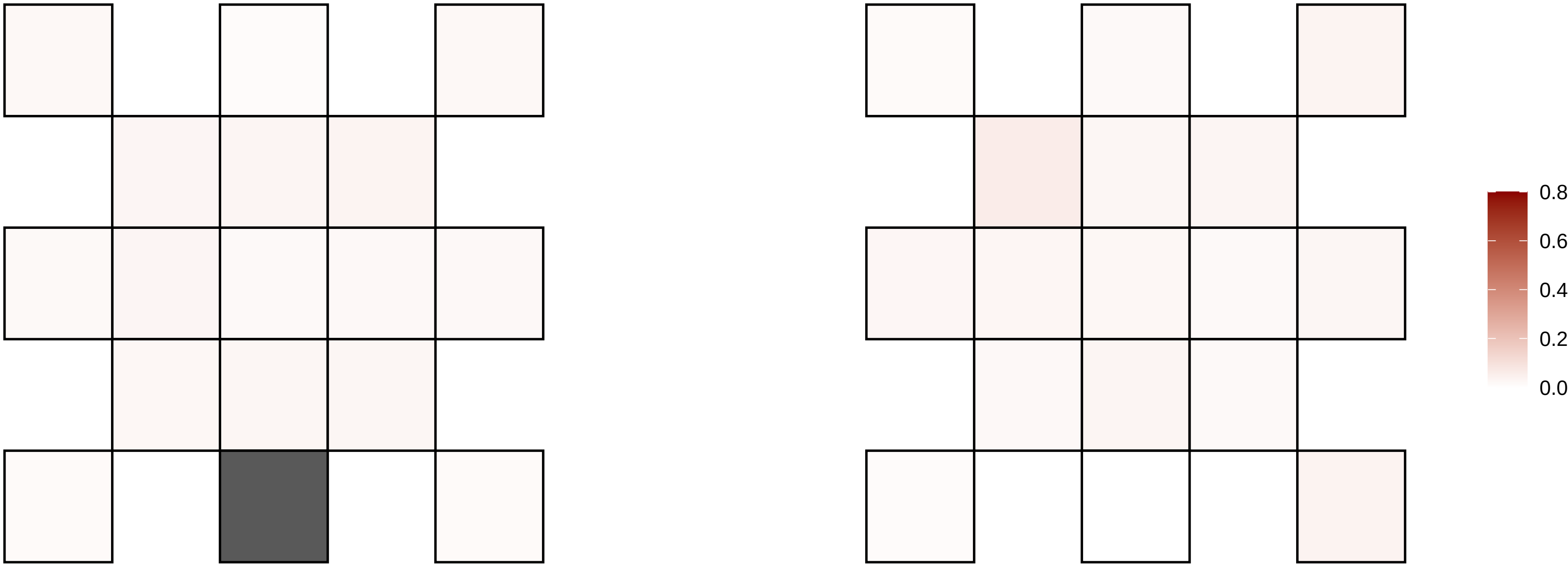

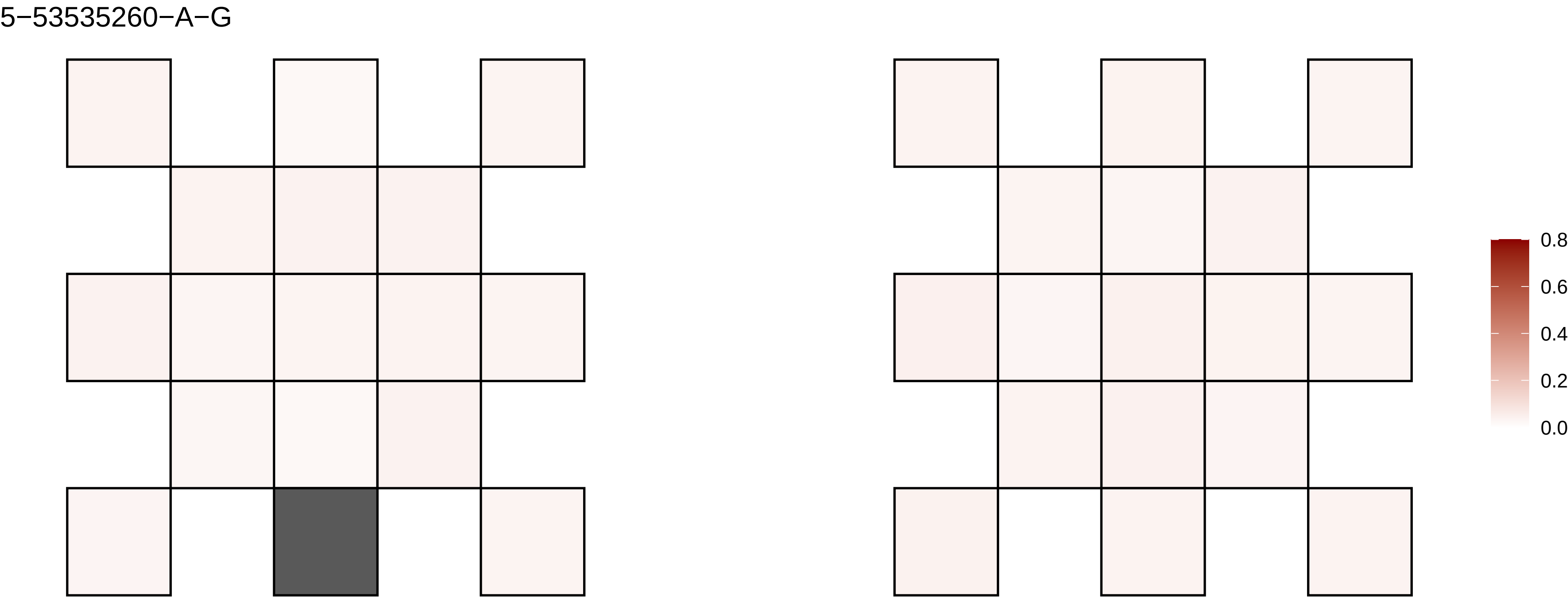

5−53535260−A−G
0.8
0.6
0.4
0.2
0.0

5−57965108−C−T

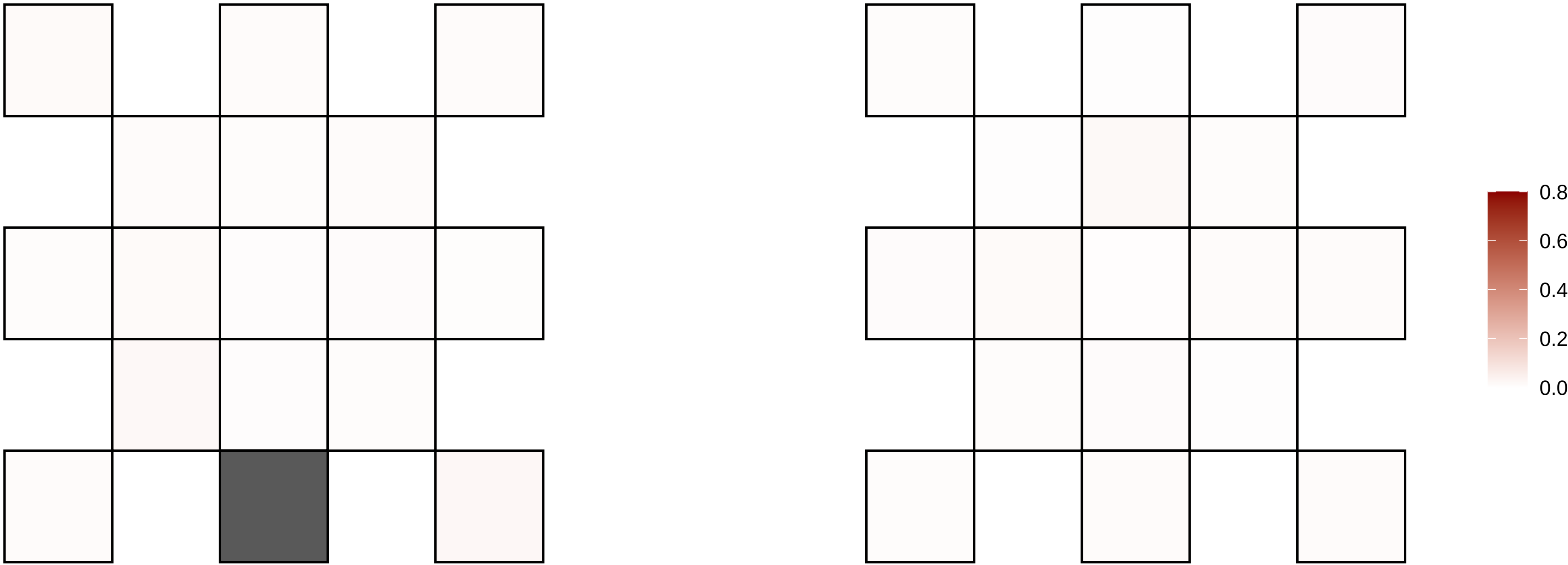

5−62110494−G−A

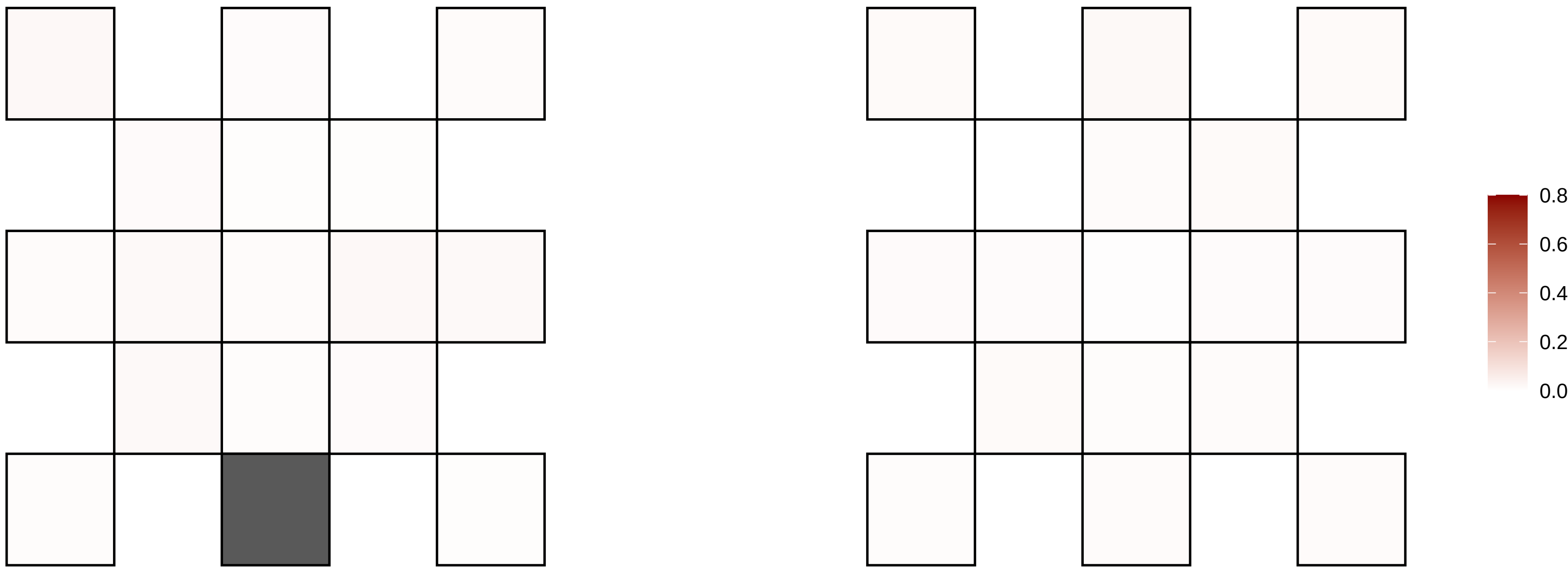

5−64328747−C−T

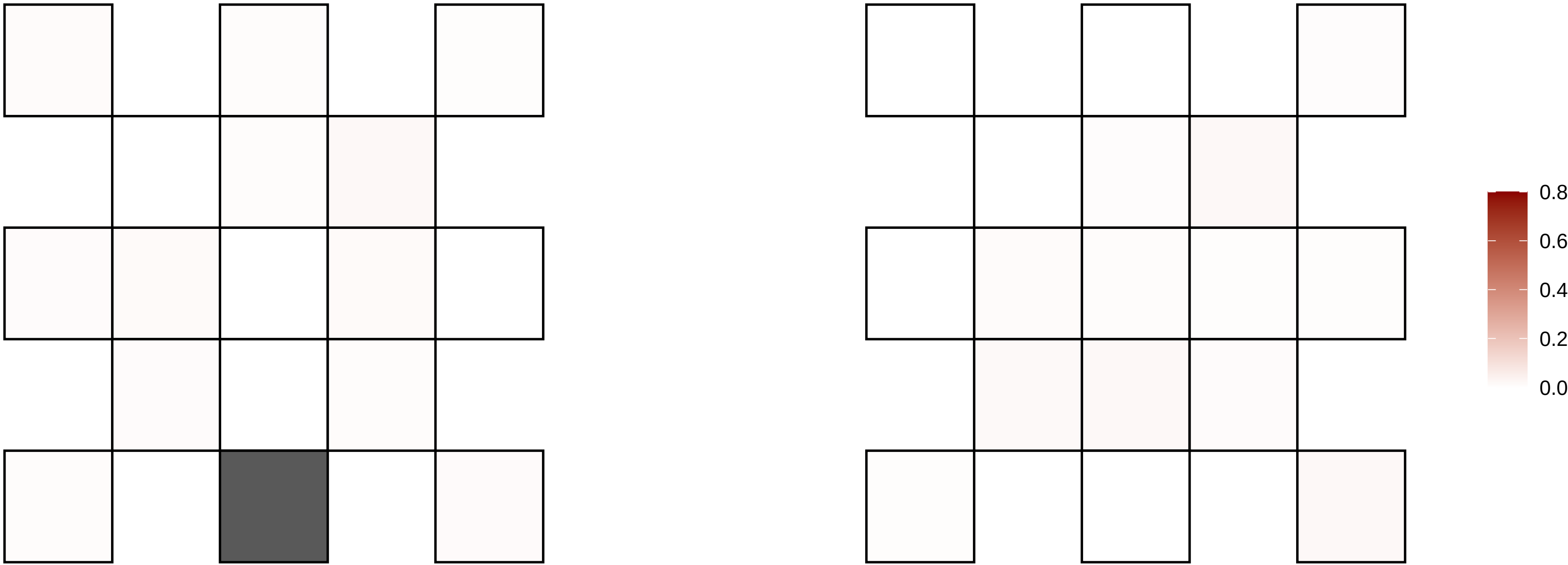

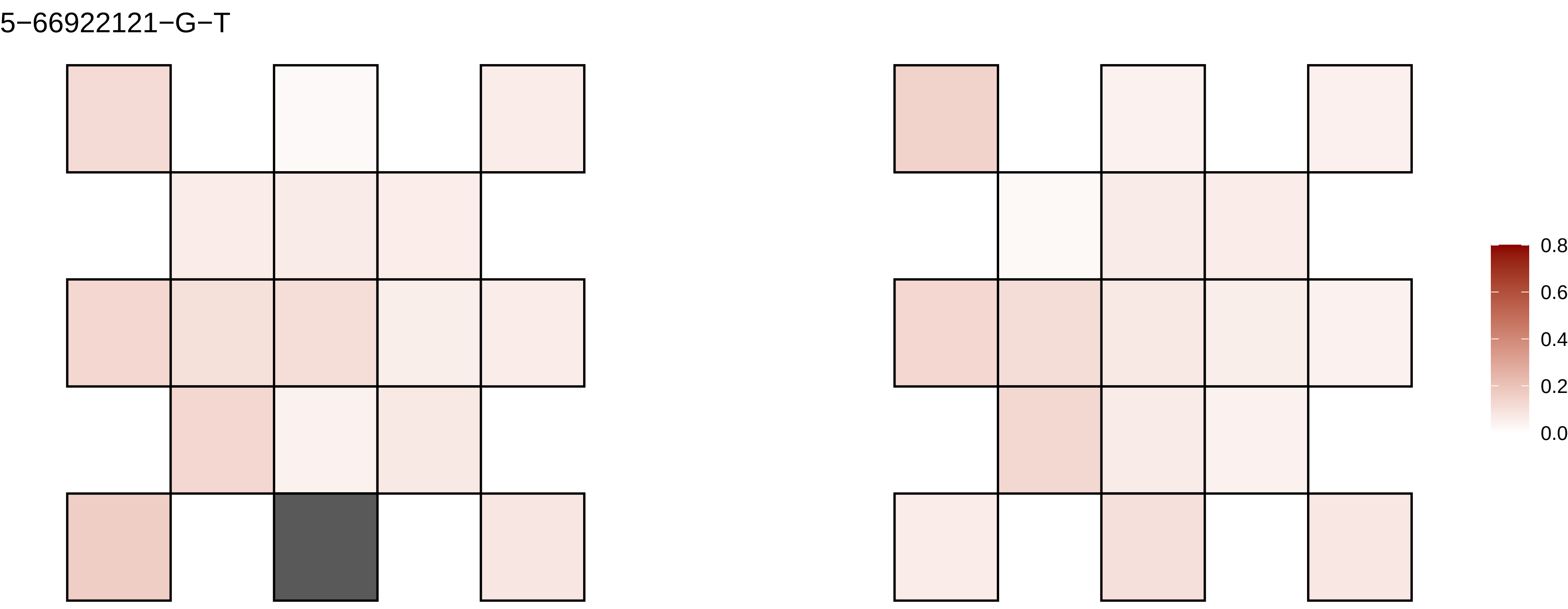
5−66922121−G−T
0.8
0.6
0.4
0.2
0.0

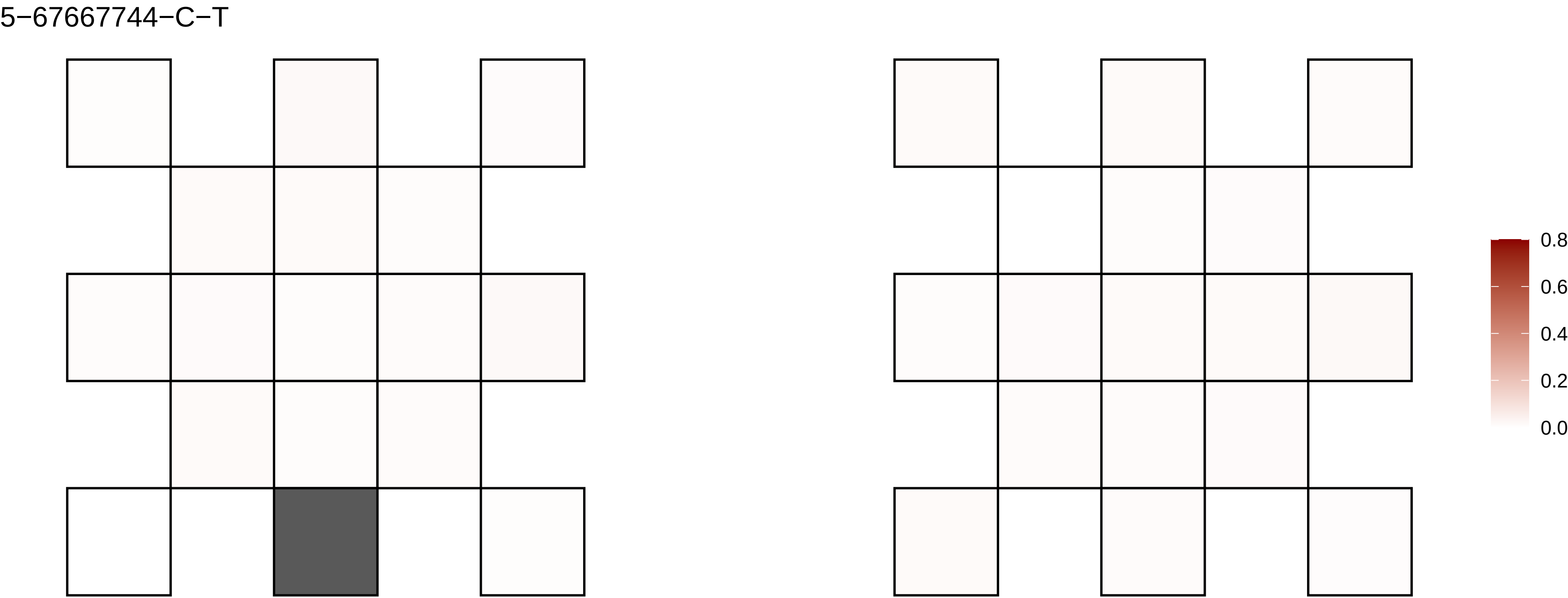
5−67667744−C−T
0.8
0.6
0.4
0.2
0.0

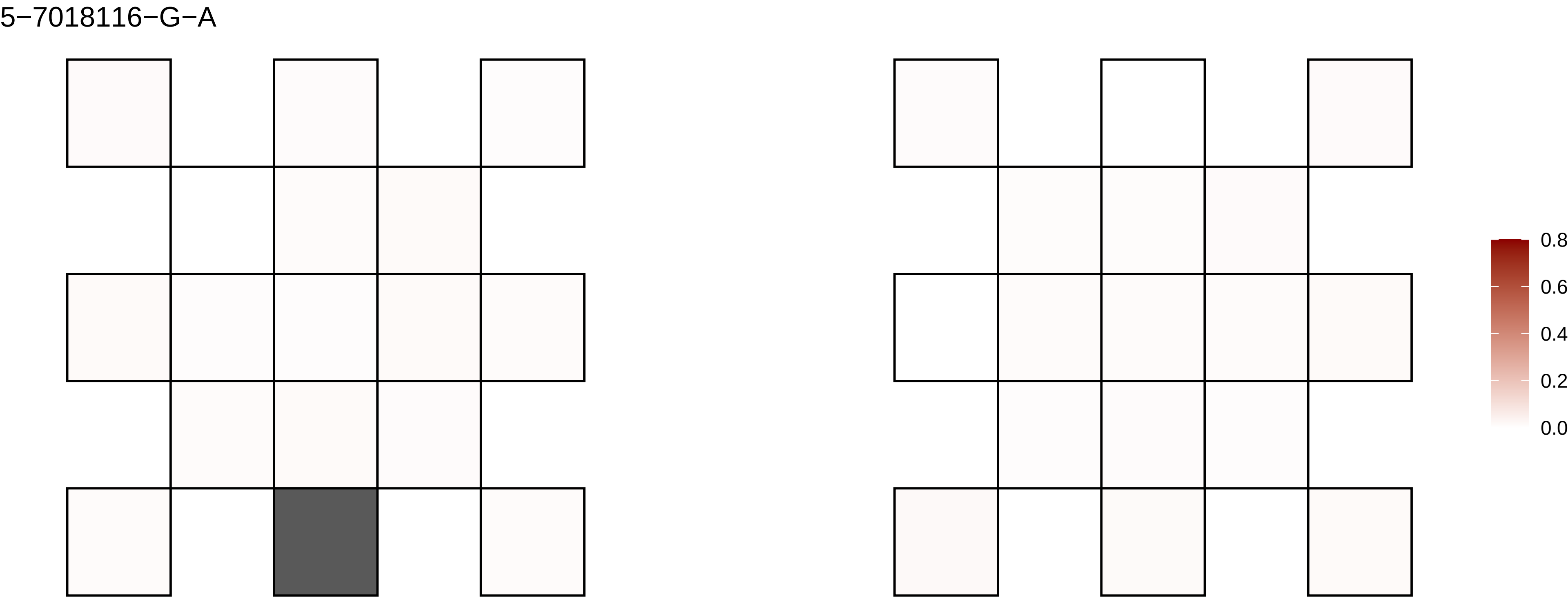
5−7018116−G−A
0.8
0.6
0.4
0.2
0.0

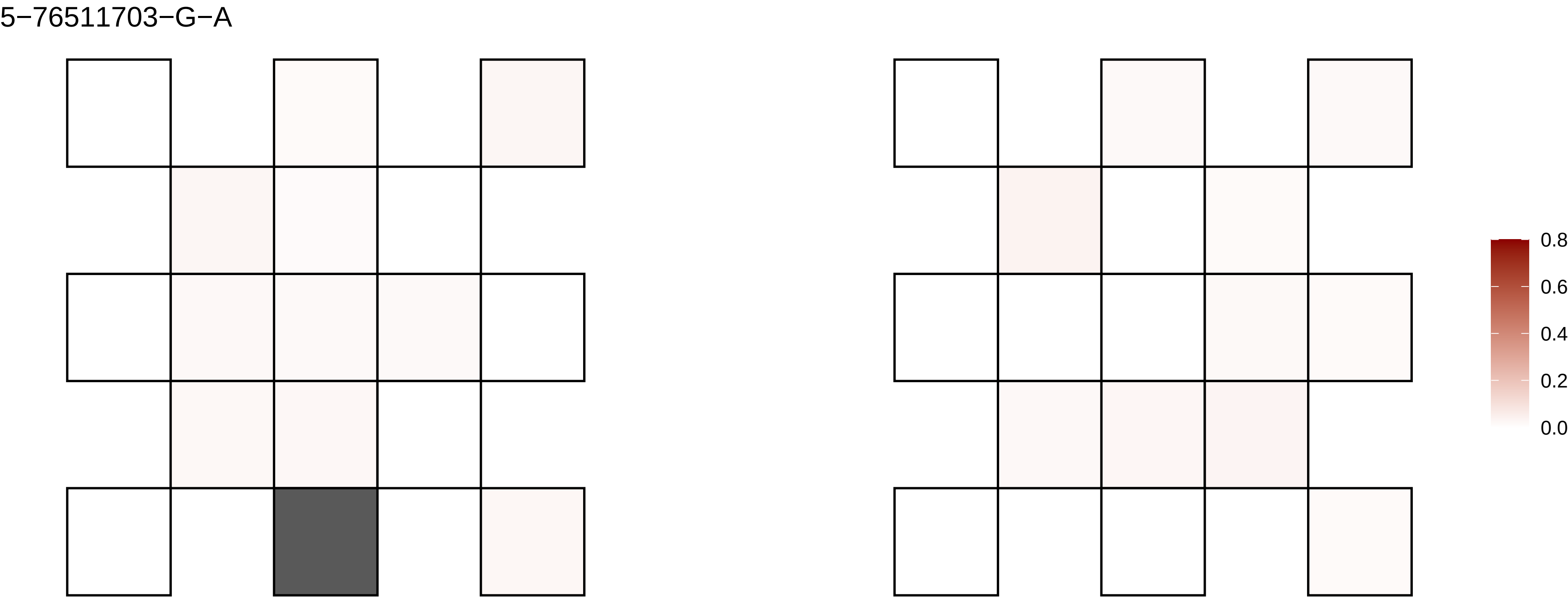
5−76511703−G−A
0.8
0.6
0.4
0.2
0.0

5−7667409−C−G

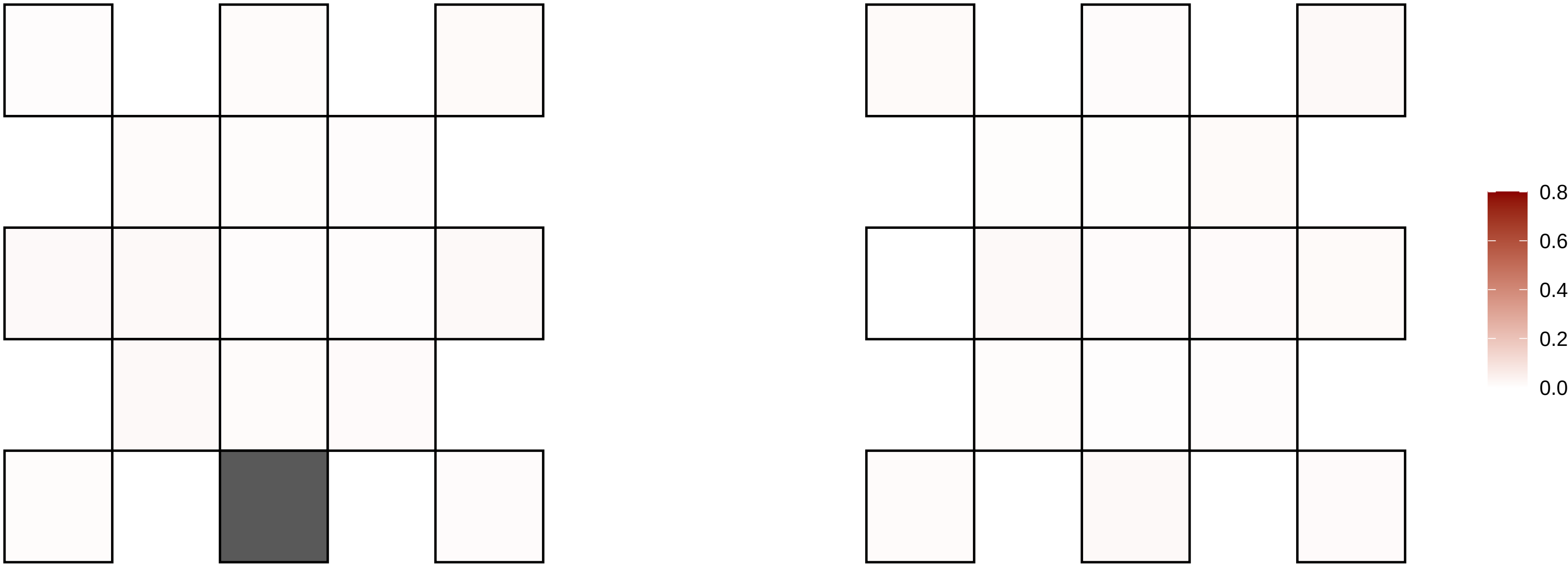

5−77299567−G−A

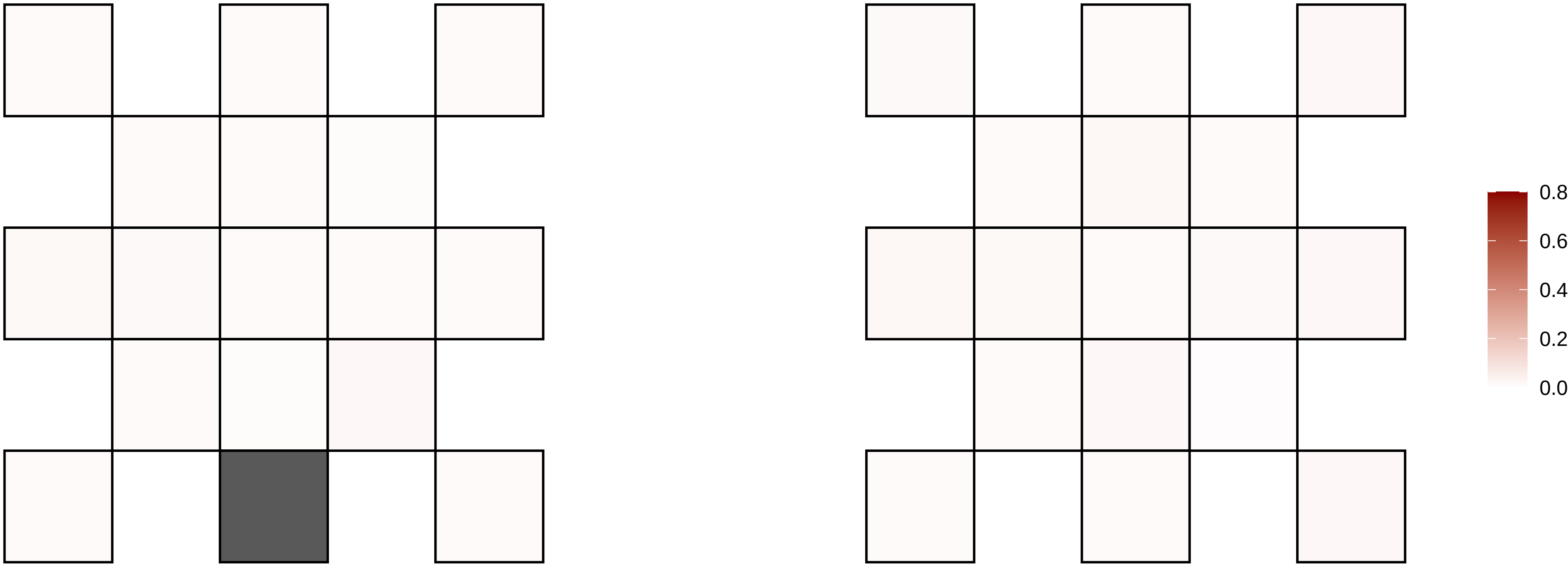

5−7771752−C−T

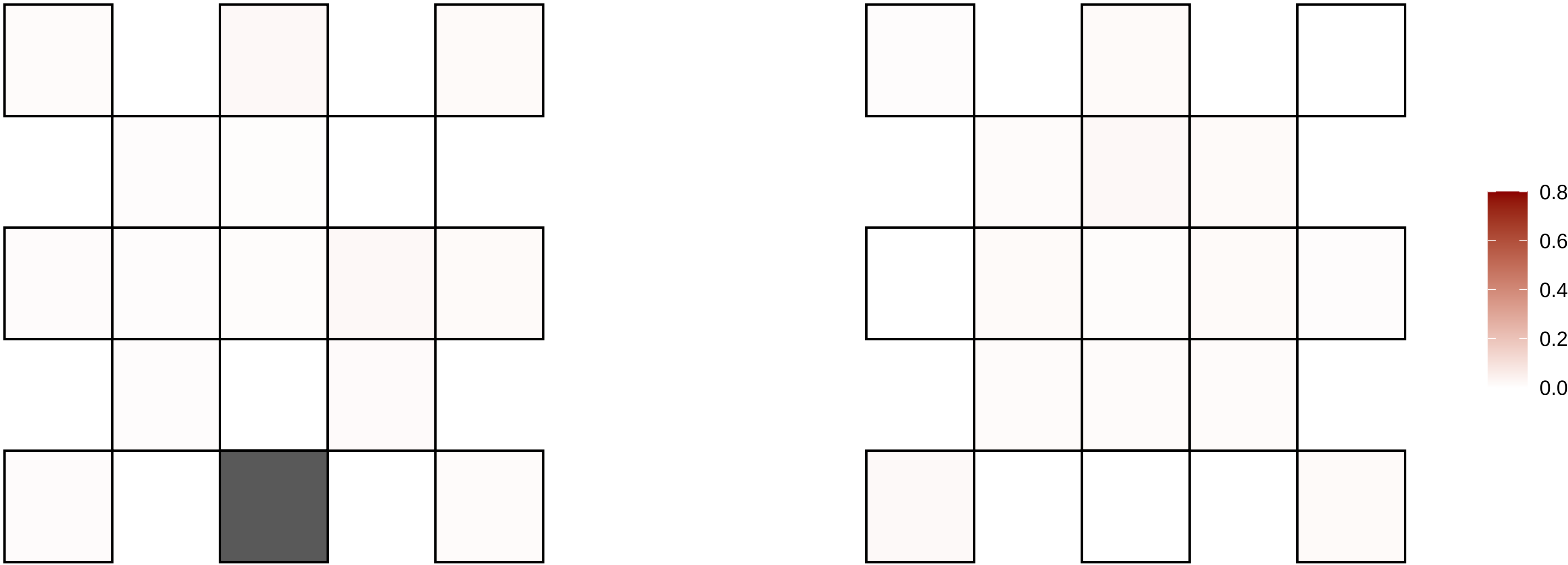

5−96456832−G−A

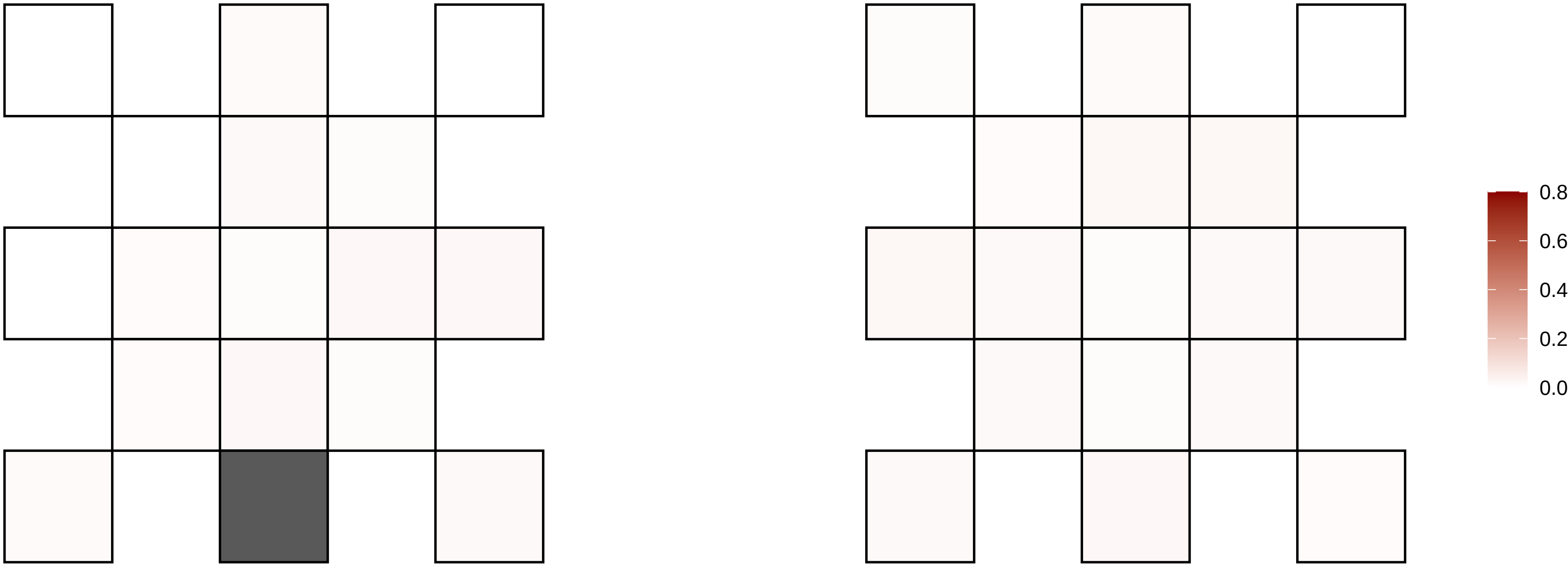

5−99372353−G−A

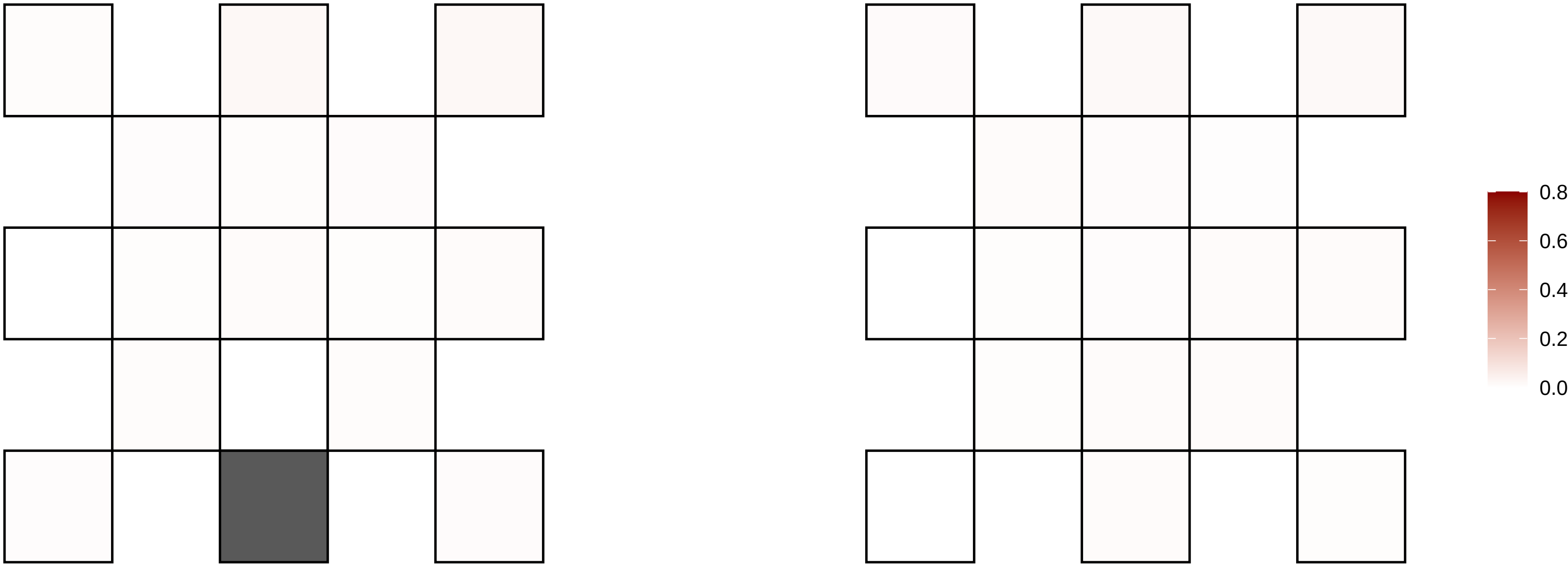

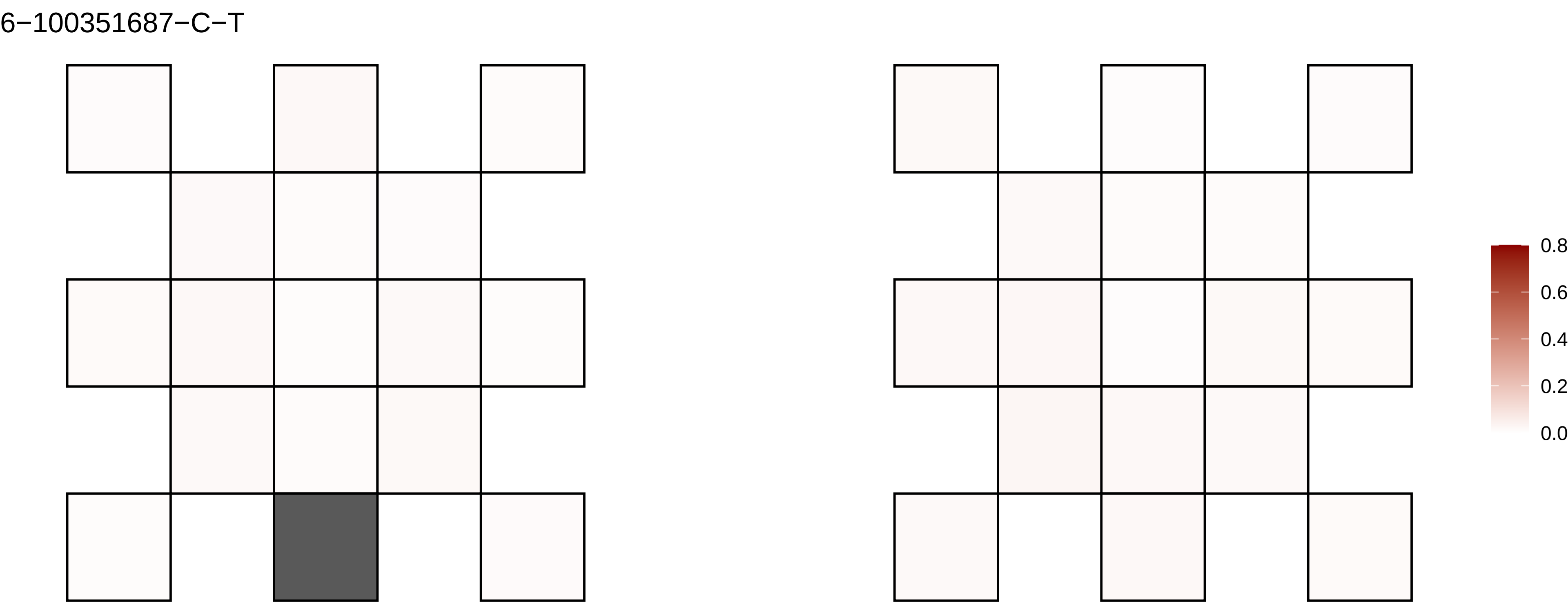
6−100351687−C−T
0.8
0.6
0.4
0.2
0.0

6−102878754−T−A

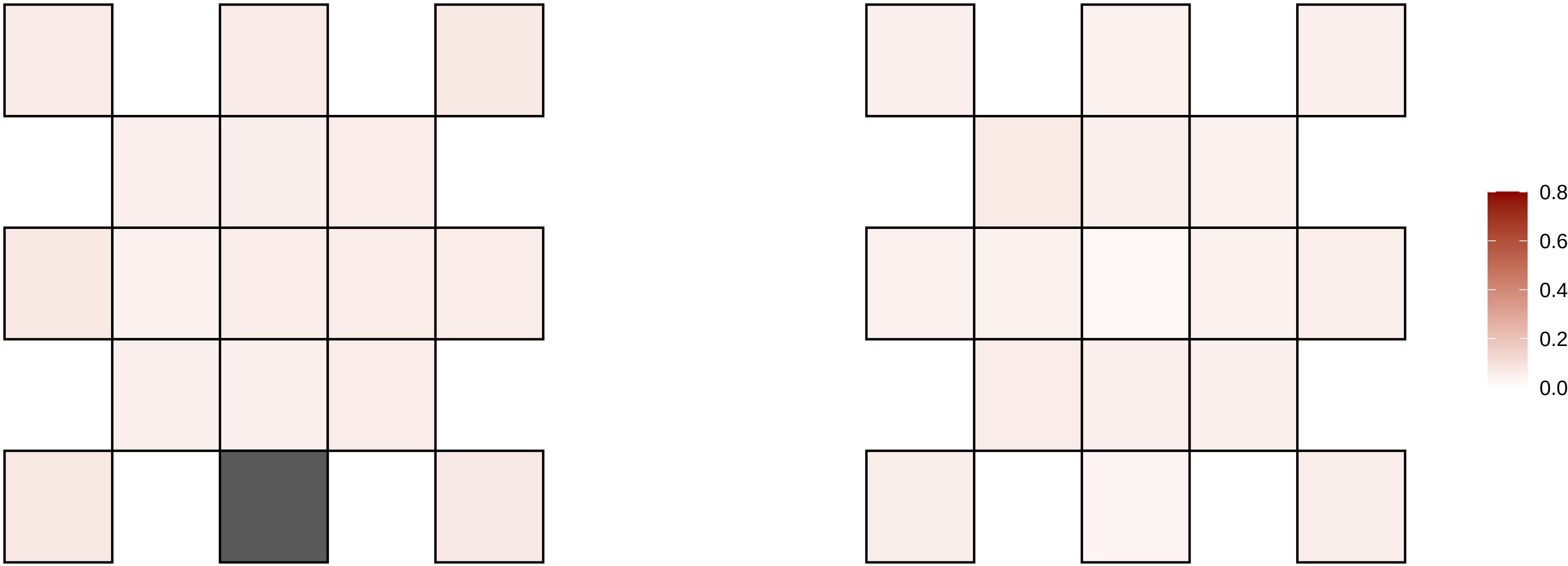

6−10399242−G−A

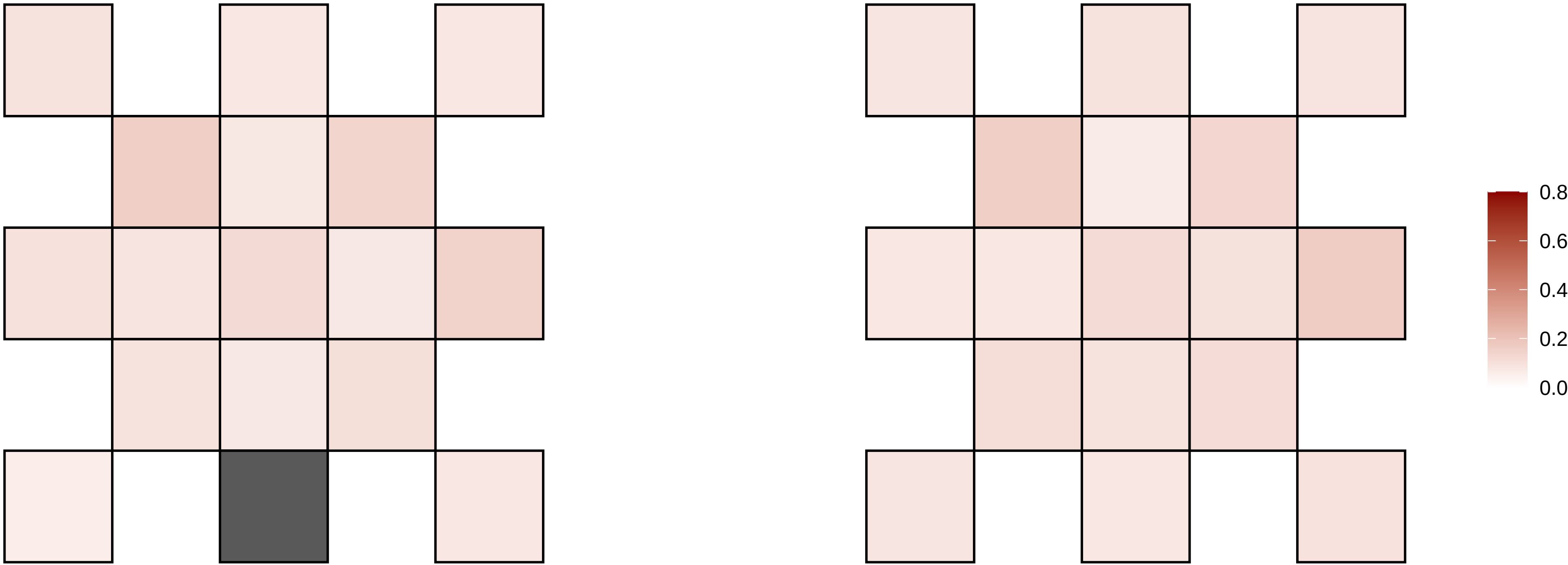

6−108325070−A−G

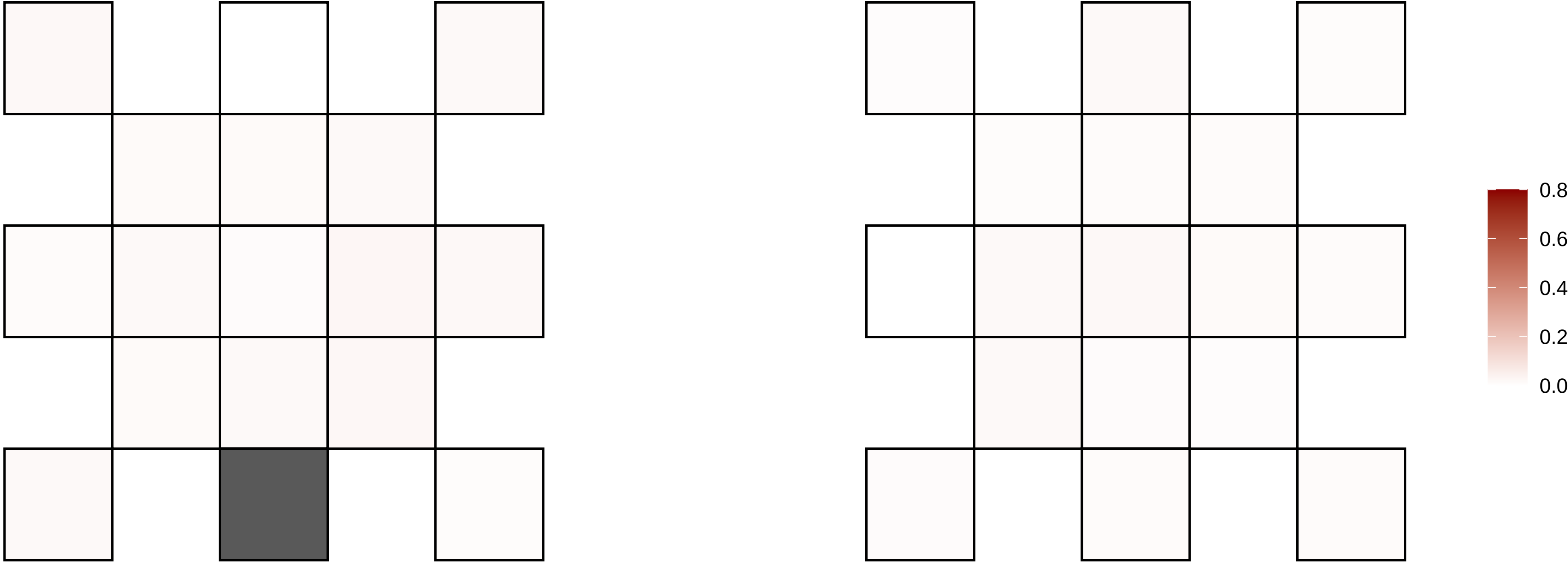

6−114240573−A−T

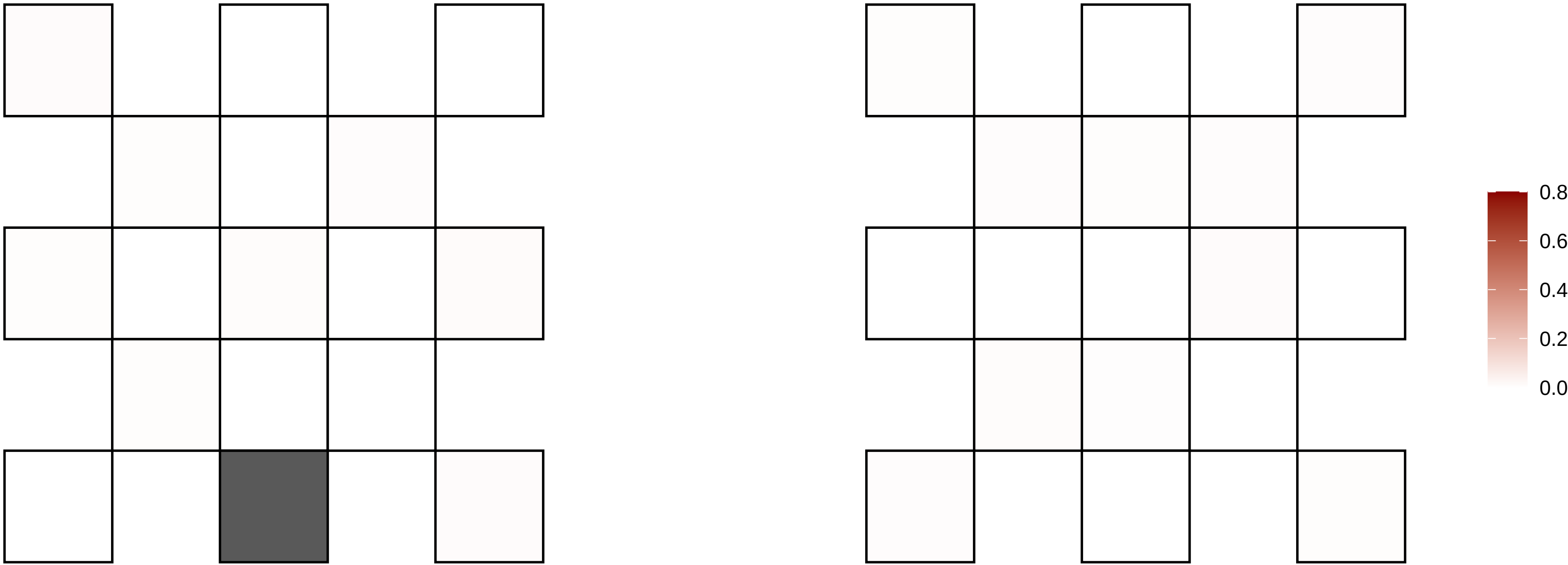

6−115212381−G−A

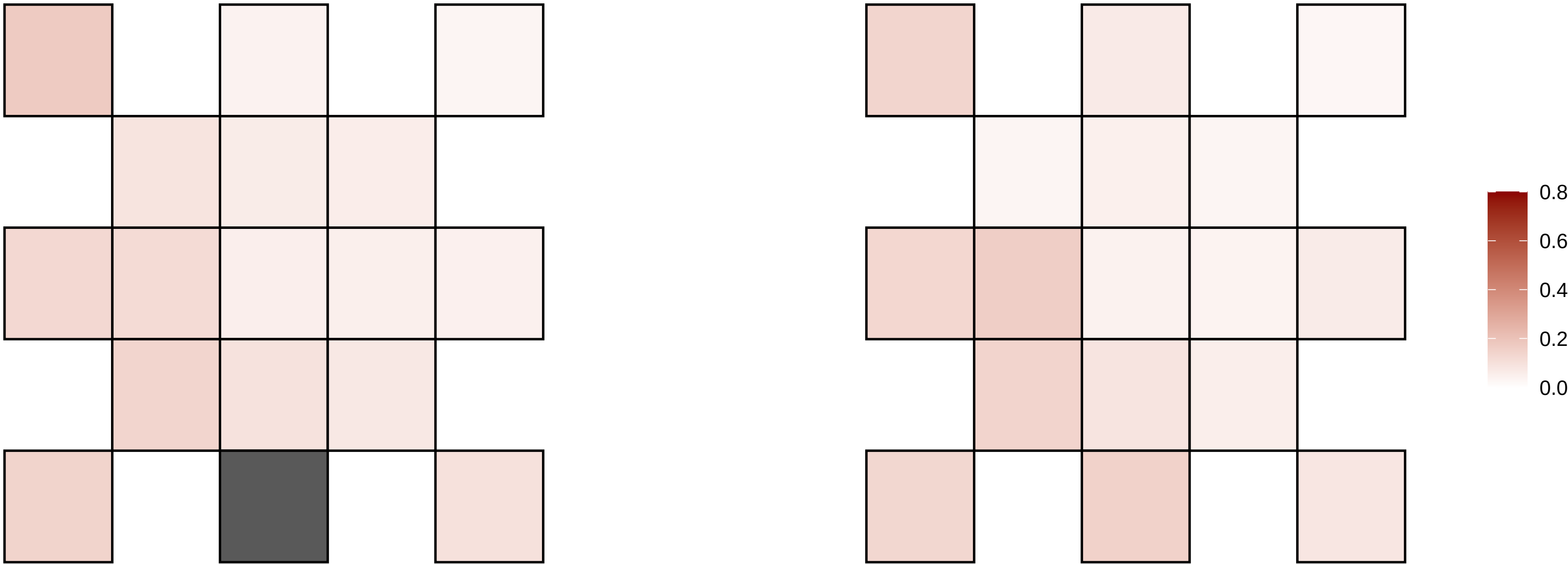

6−115259744−C−T

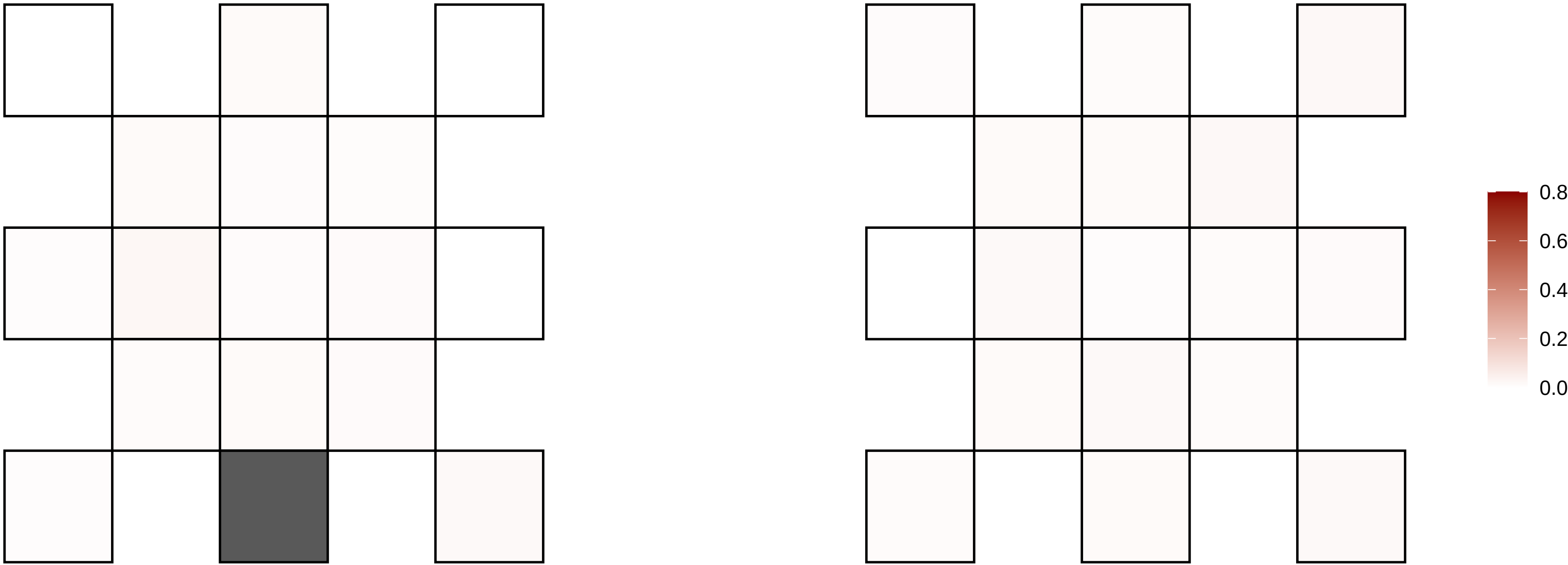

6−124787238−C−T

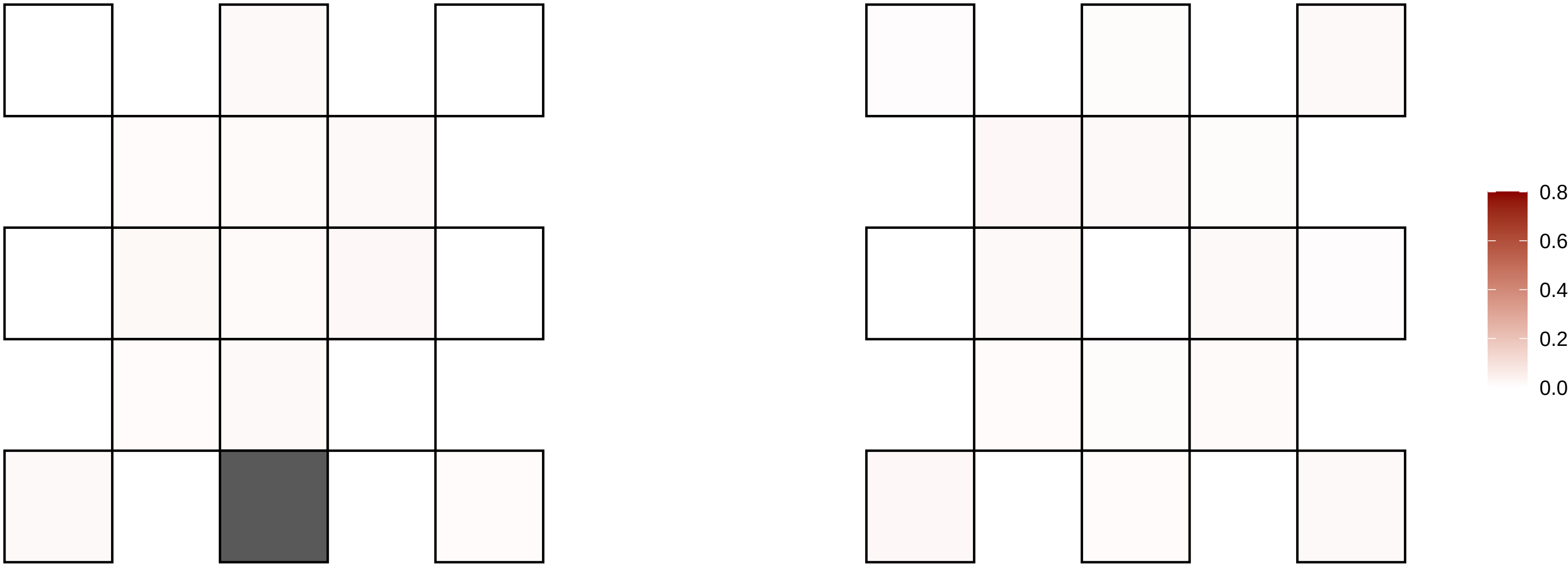

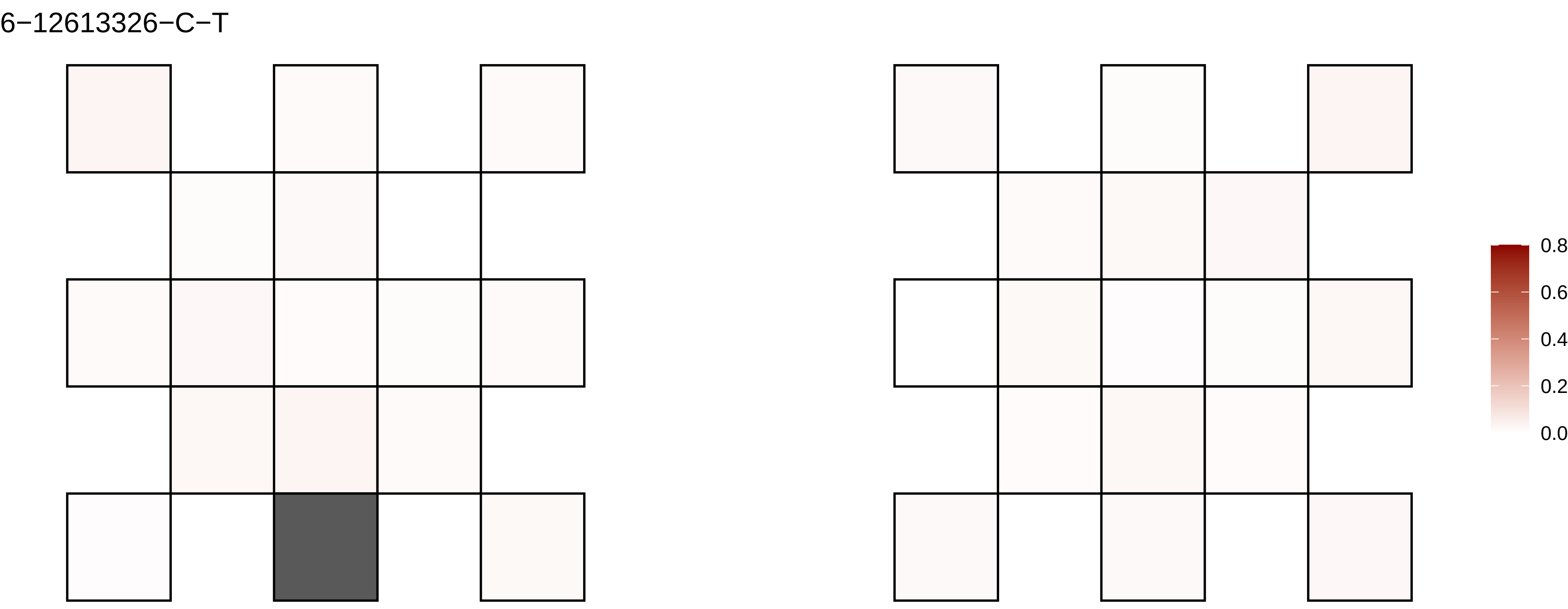
6−12613326−C−T
0.8
0.6
0.4
0.2
0.0

6−132170864−G−A

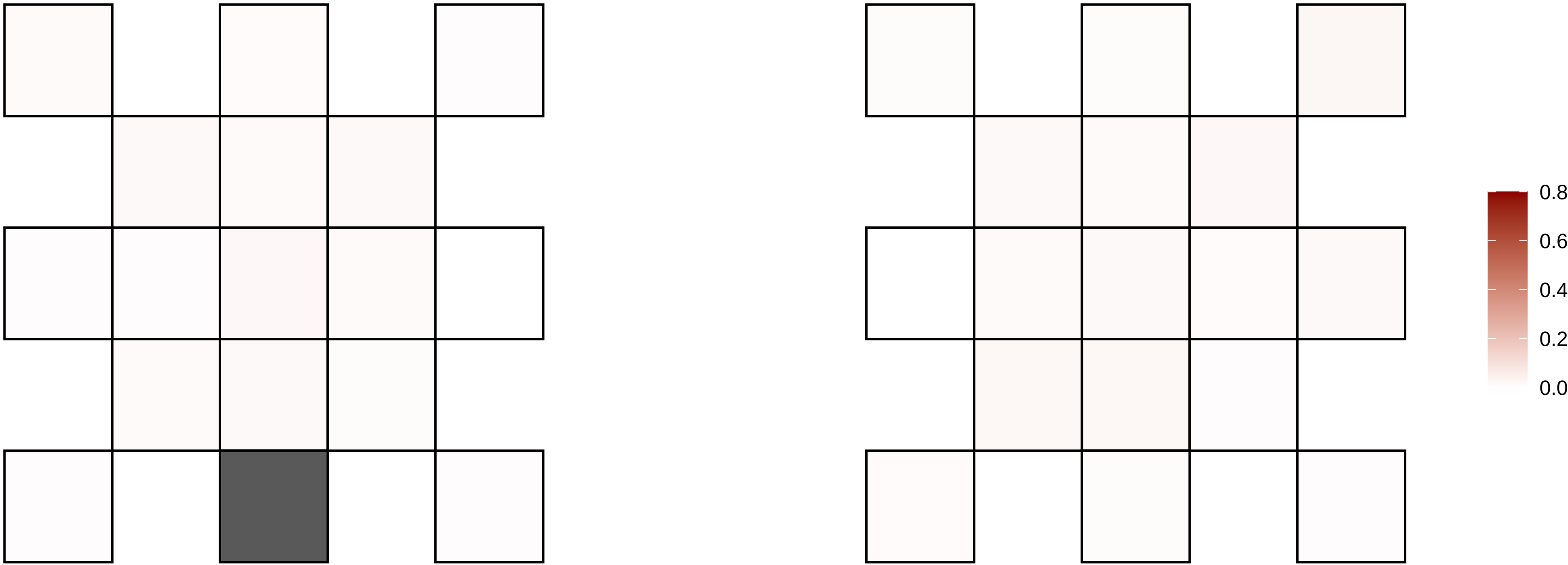

6−138352132−G−A

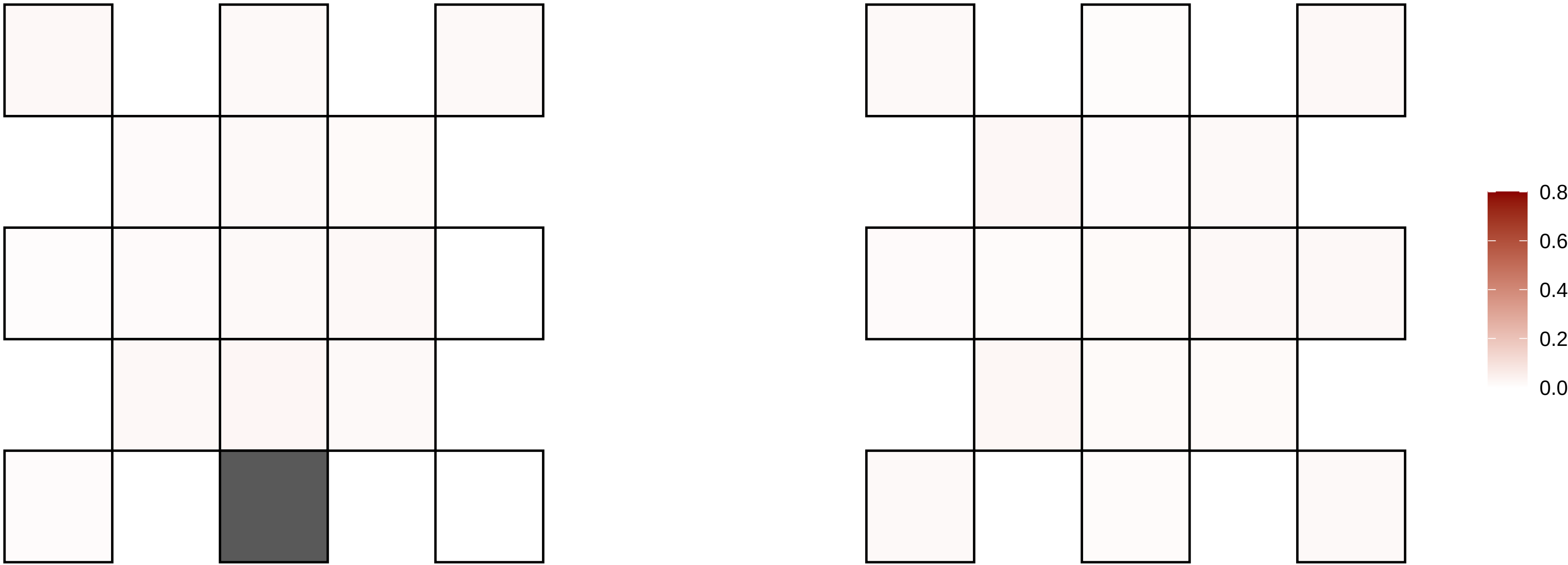

6−139194275−C−T

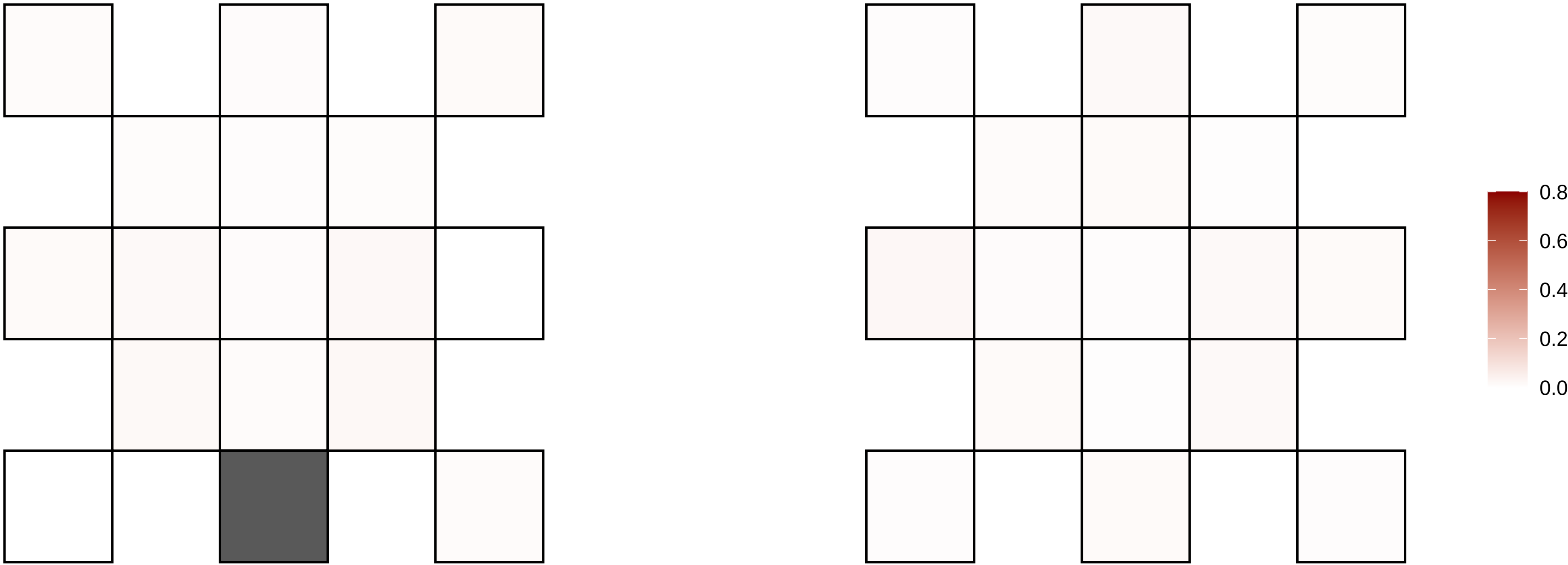

6−142200306−C−T

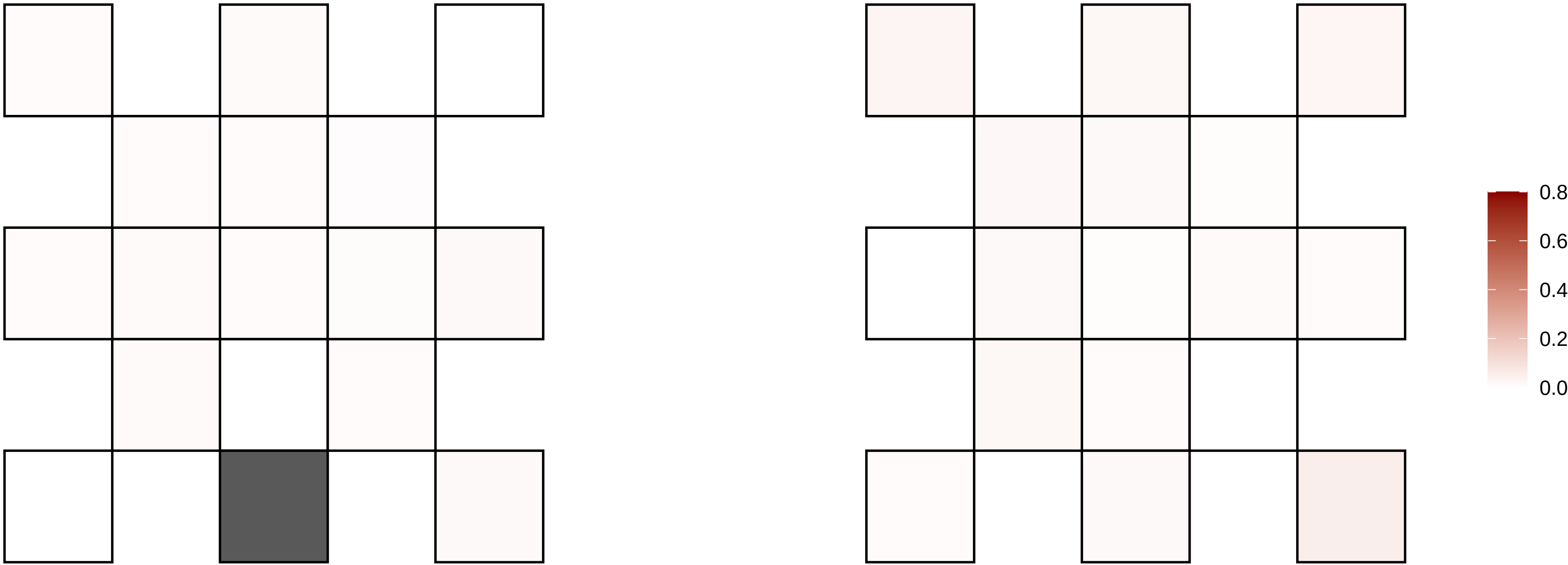

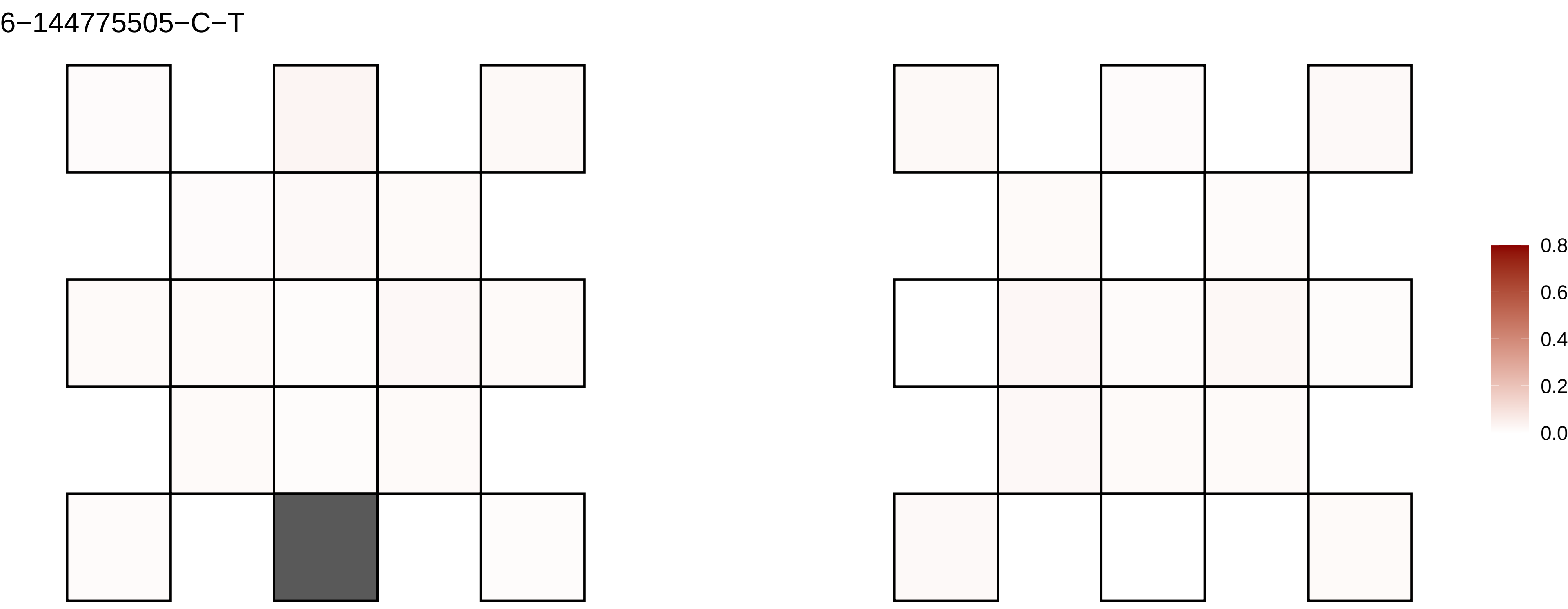

6−144775505−C−T
0.8
0.6
0.4
0.2
0.0

6−145560357−G−T

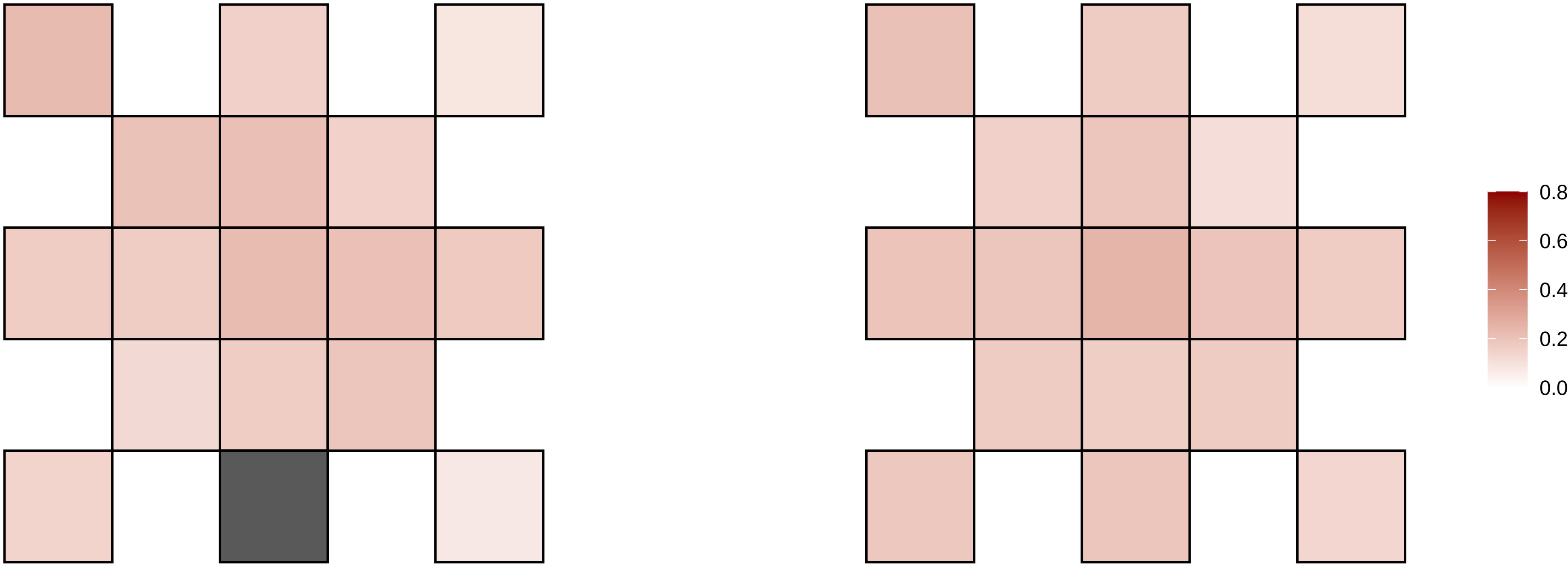

6−146019805−C−T

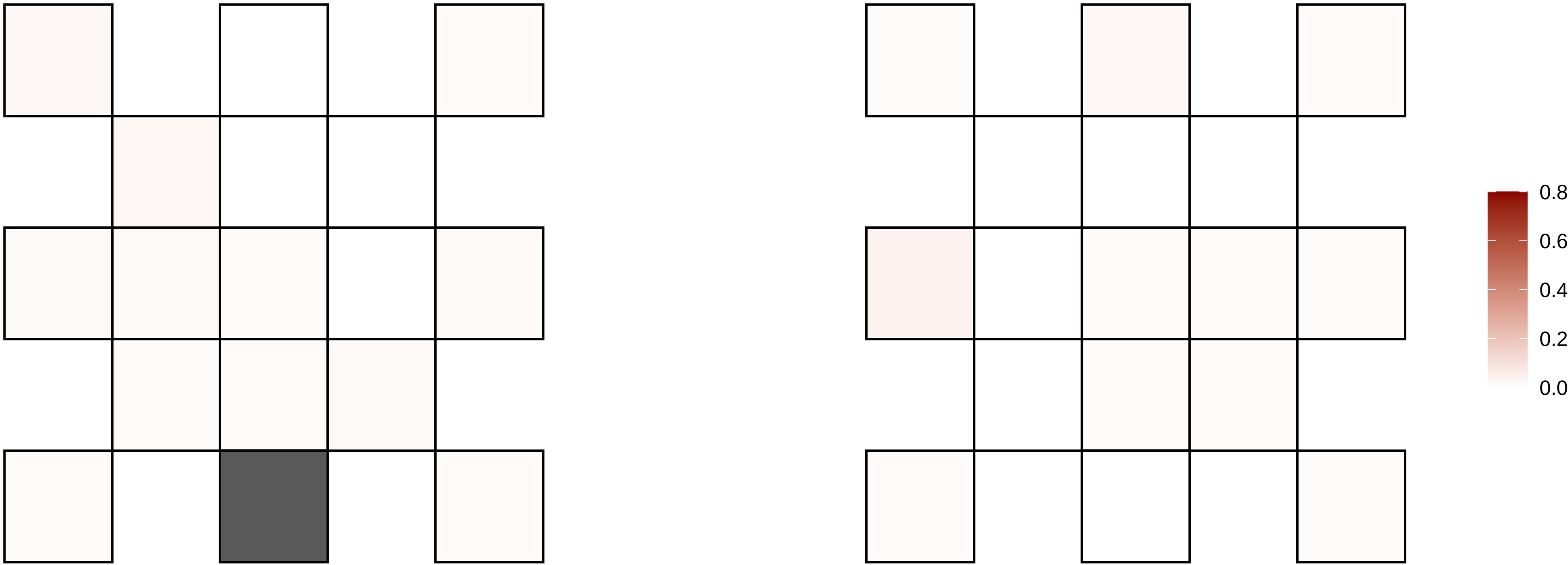

6−151519709−G−T

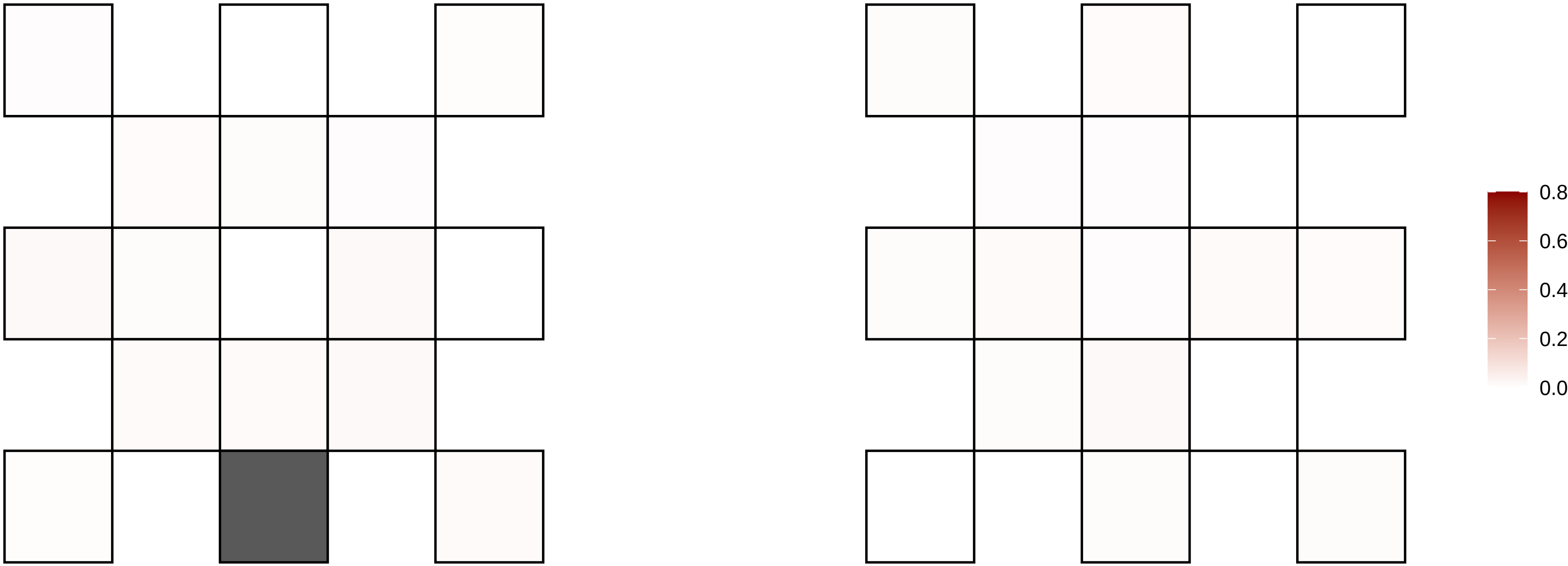

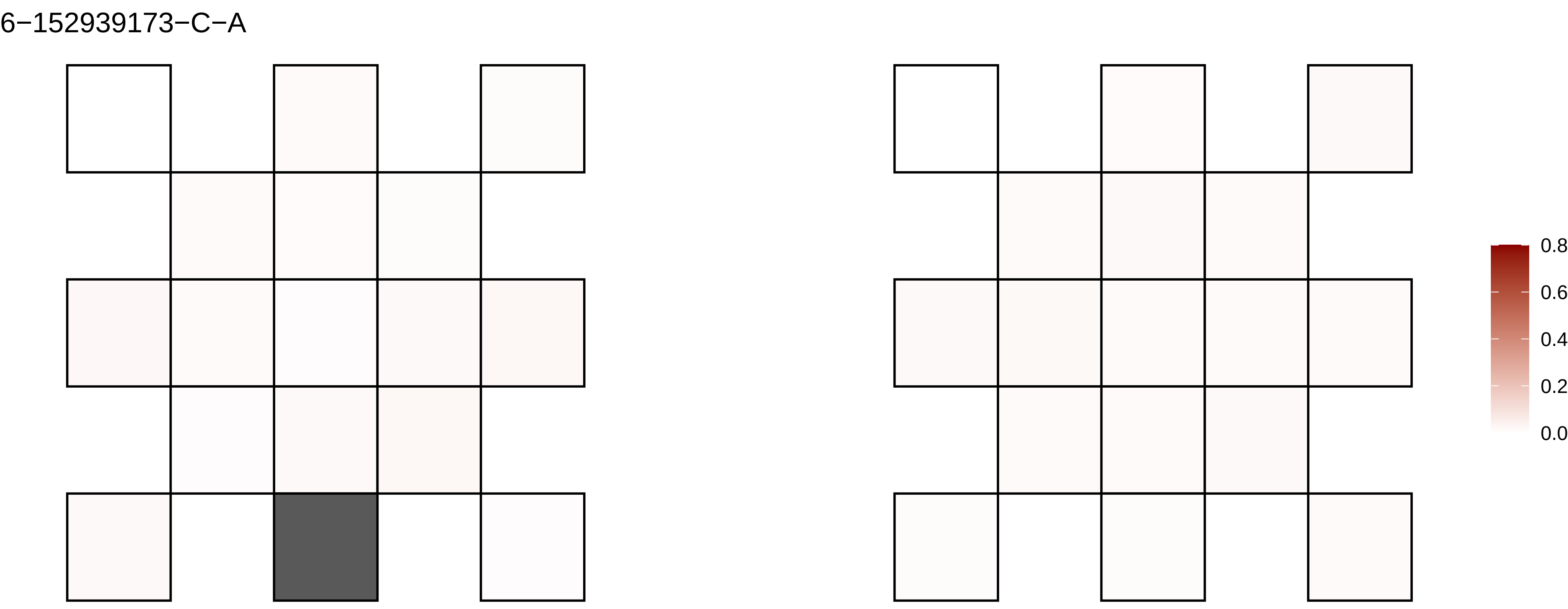
6−152939173−C−A
0.8
0.6
0.4
0.2
0.0

6−153112948−G−T

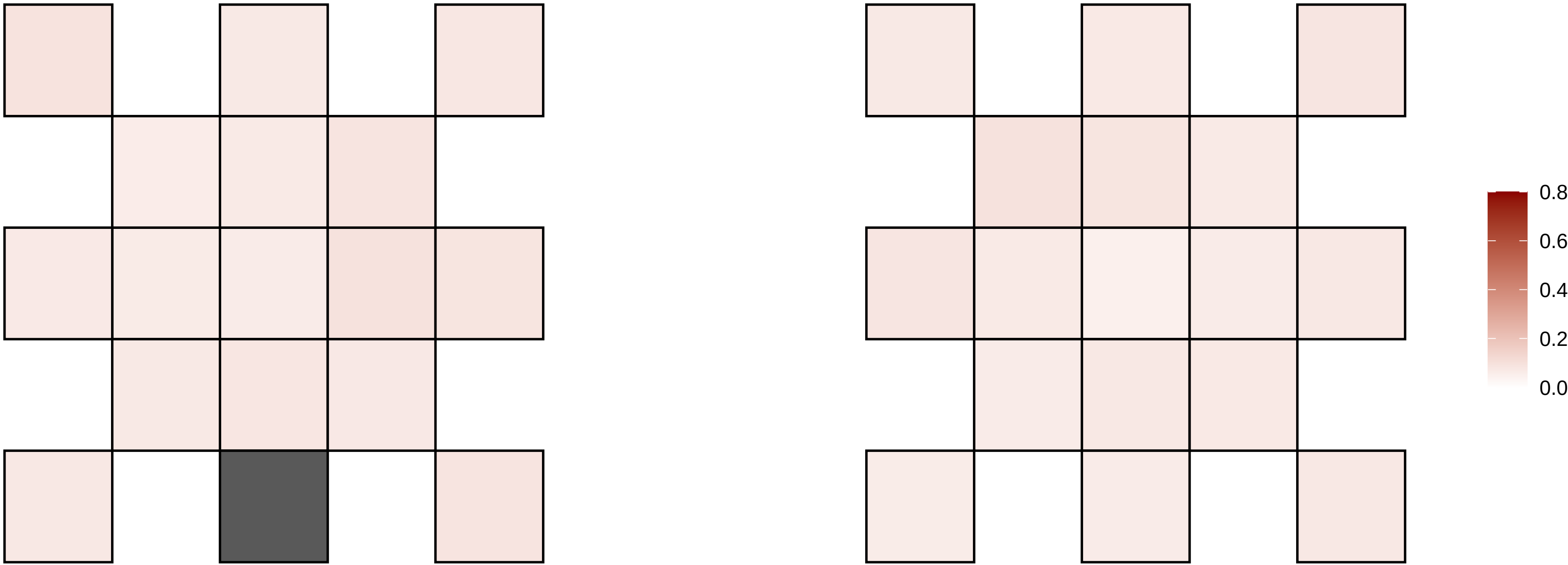

6−153564460−T−G

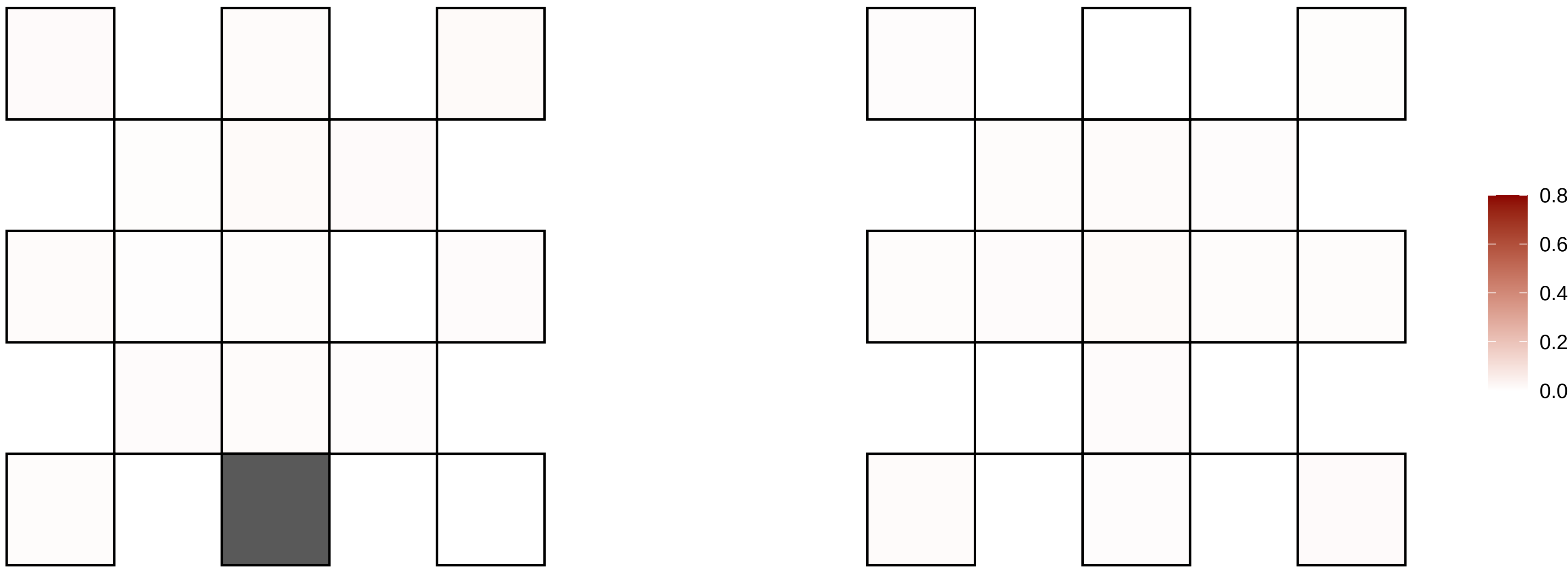

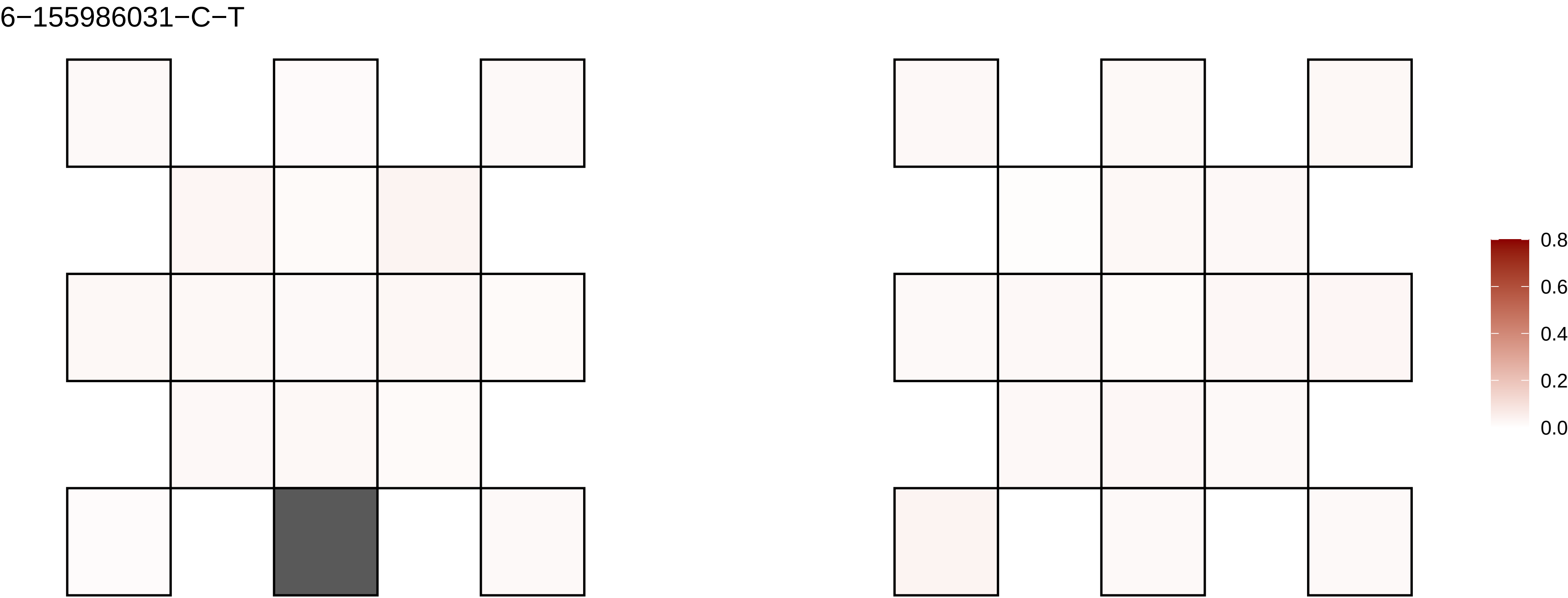
6−155986031−C−T
0.8
0.6
0.4
0.2
0.0

6−156635821−G−A

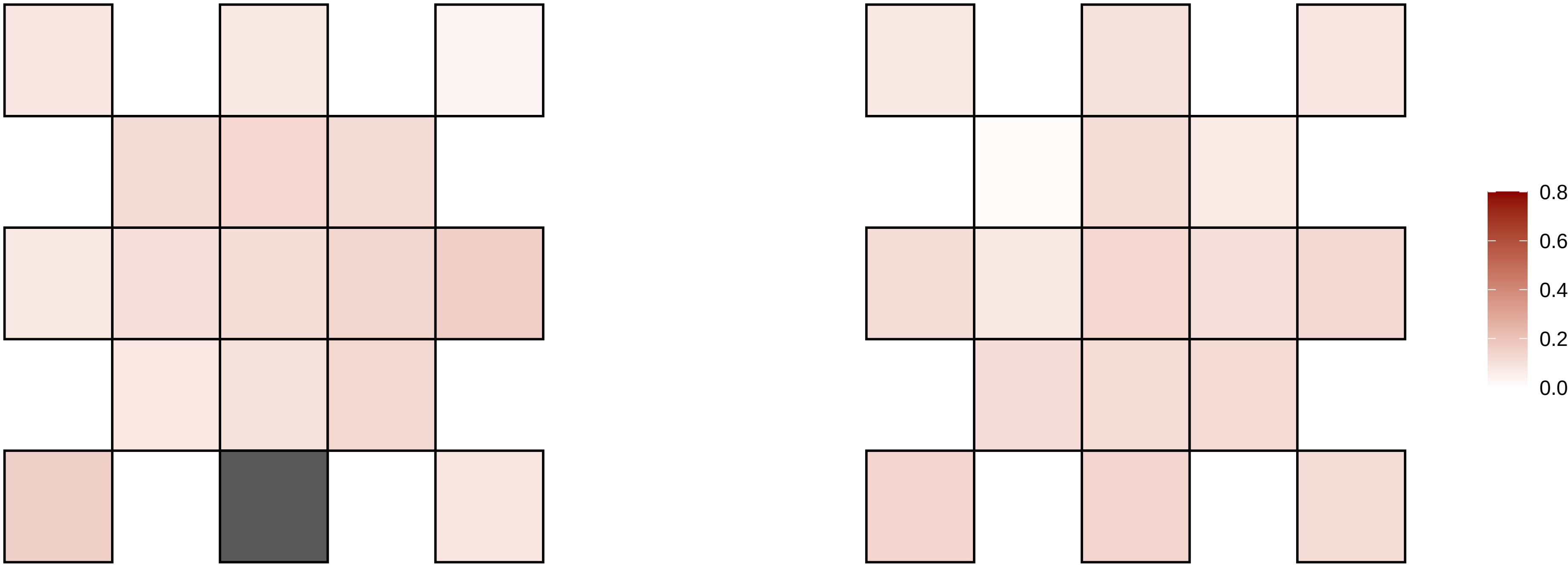

6−156792200−G−A

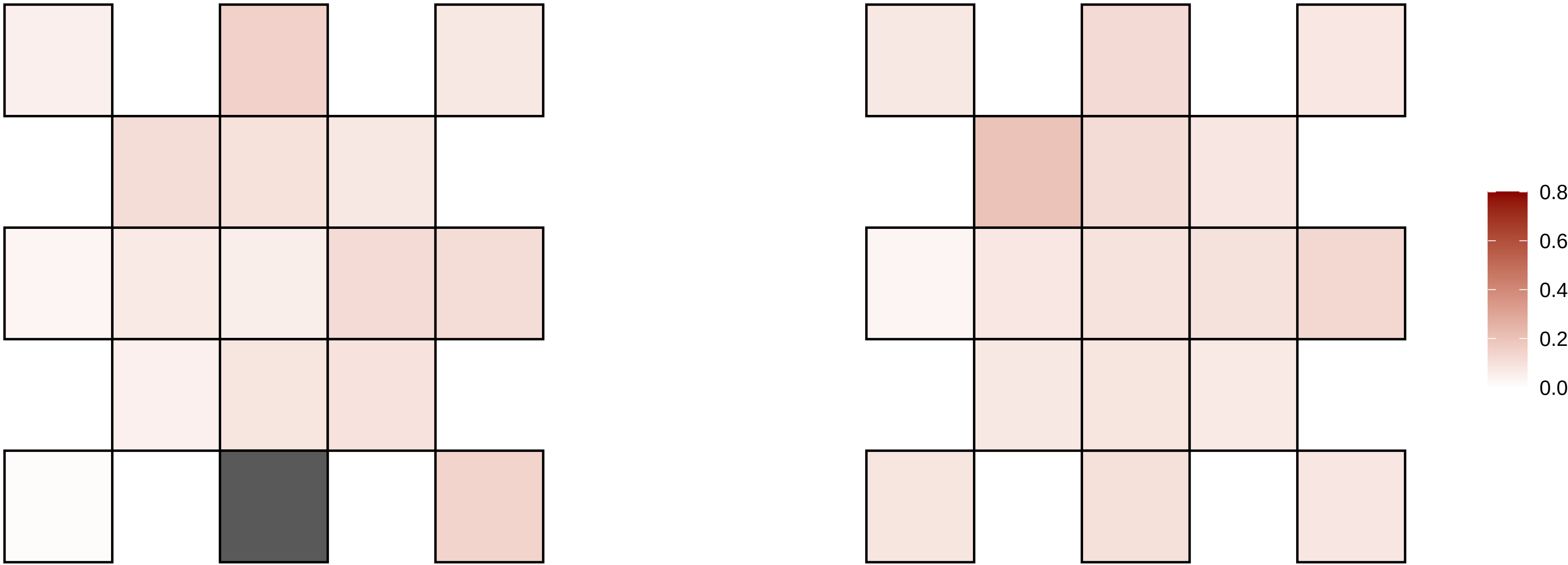

6−159935200−C−T

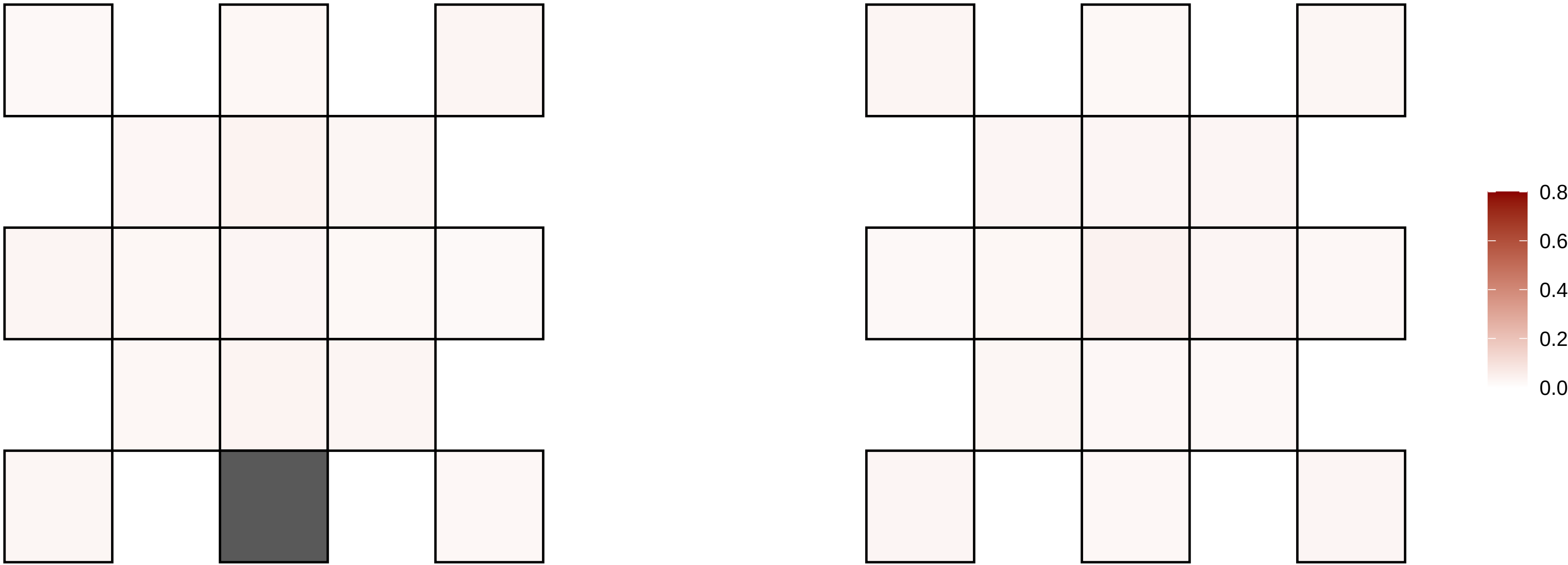

6−161158616−T−G

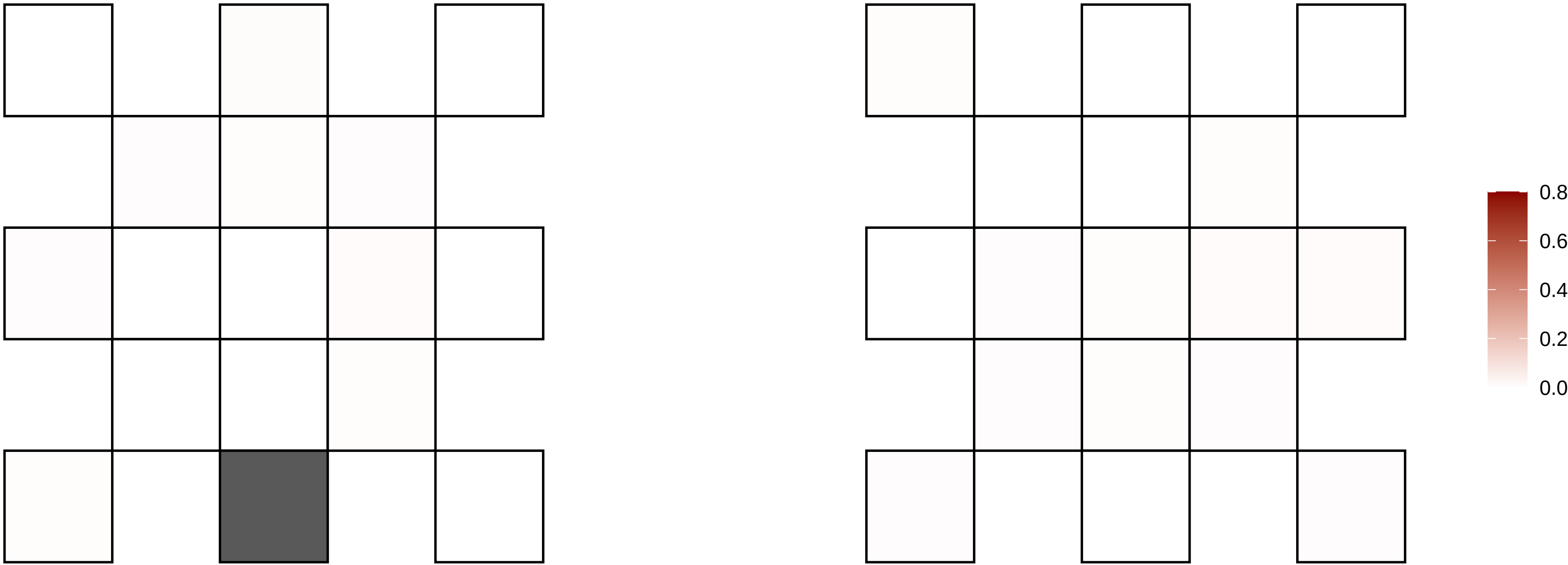

6−162806617−C−T

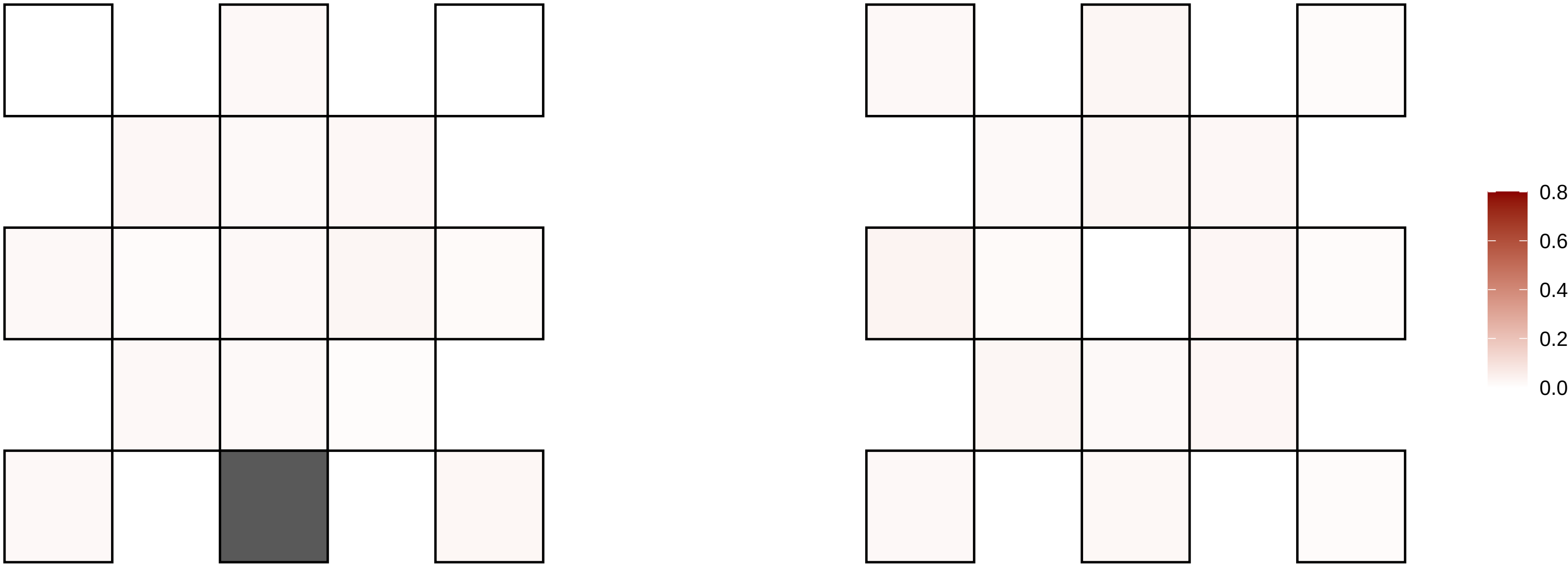

6−1661827−C−T

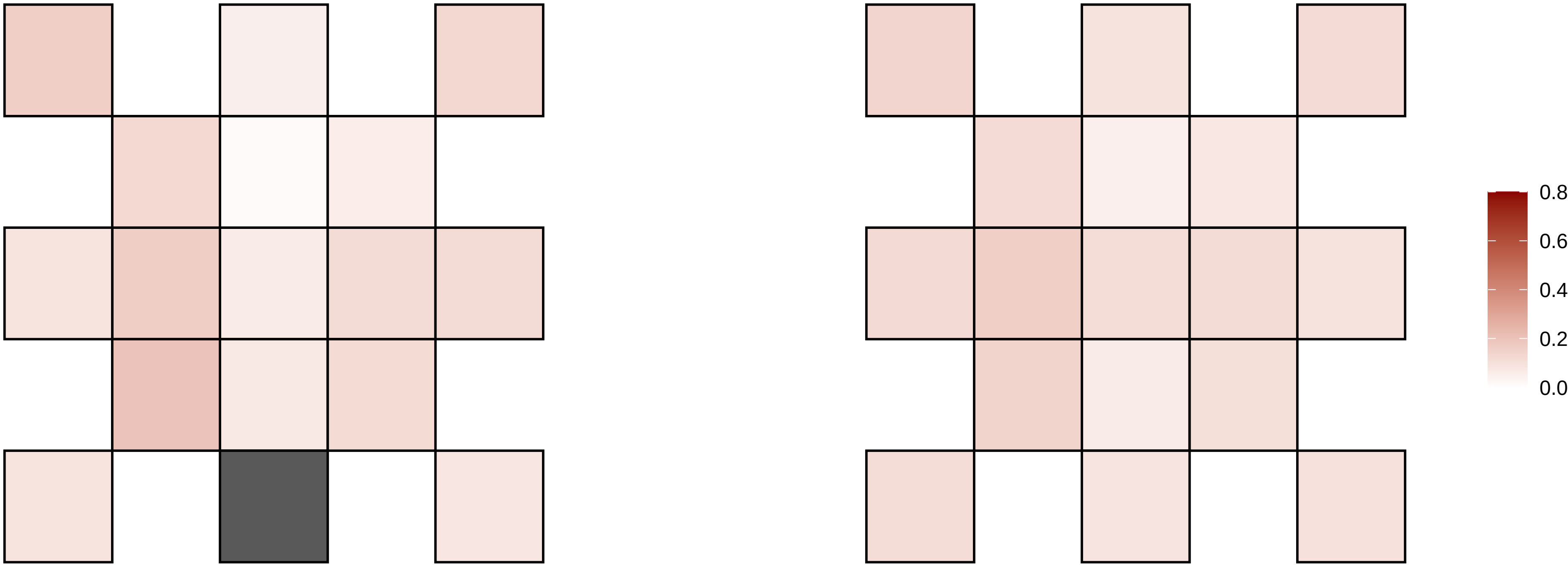

6−166482454−T−C

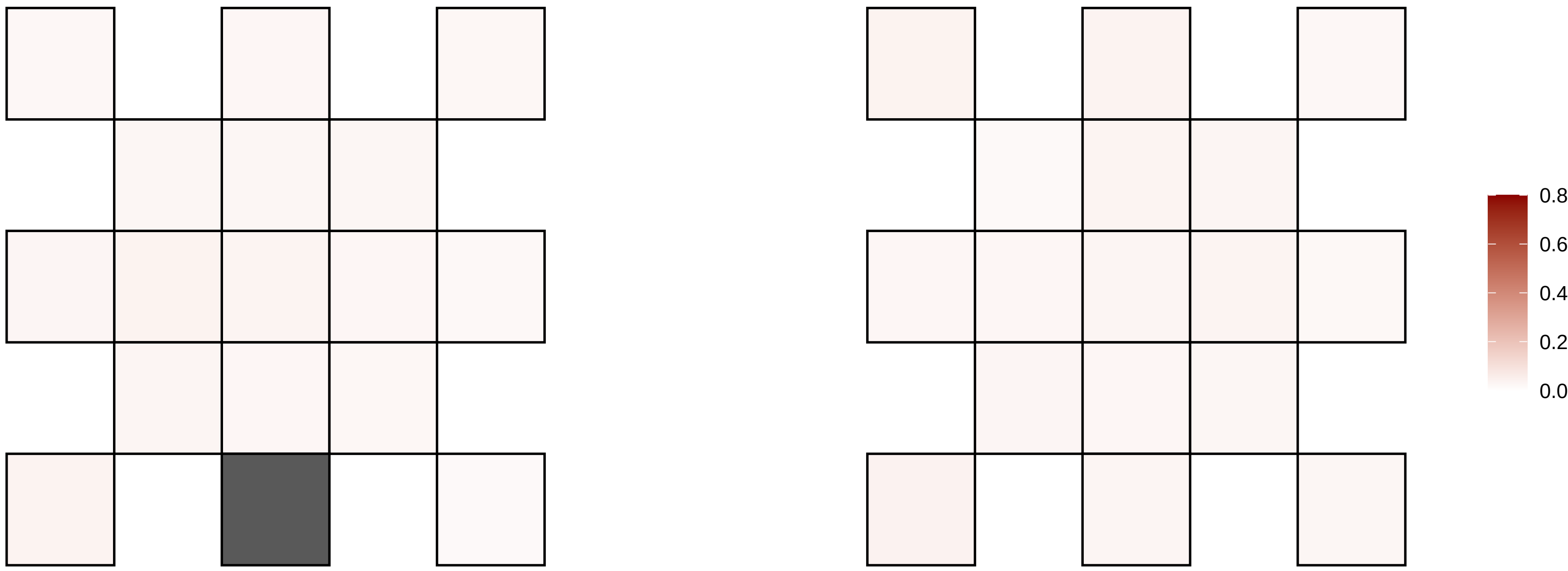

6−169577791−G−A

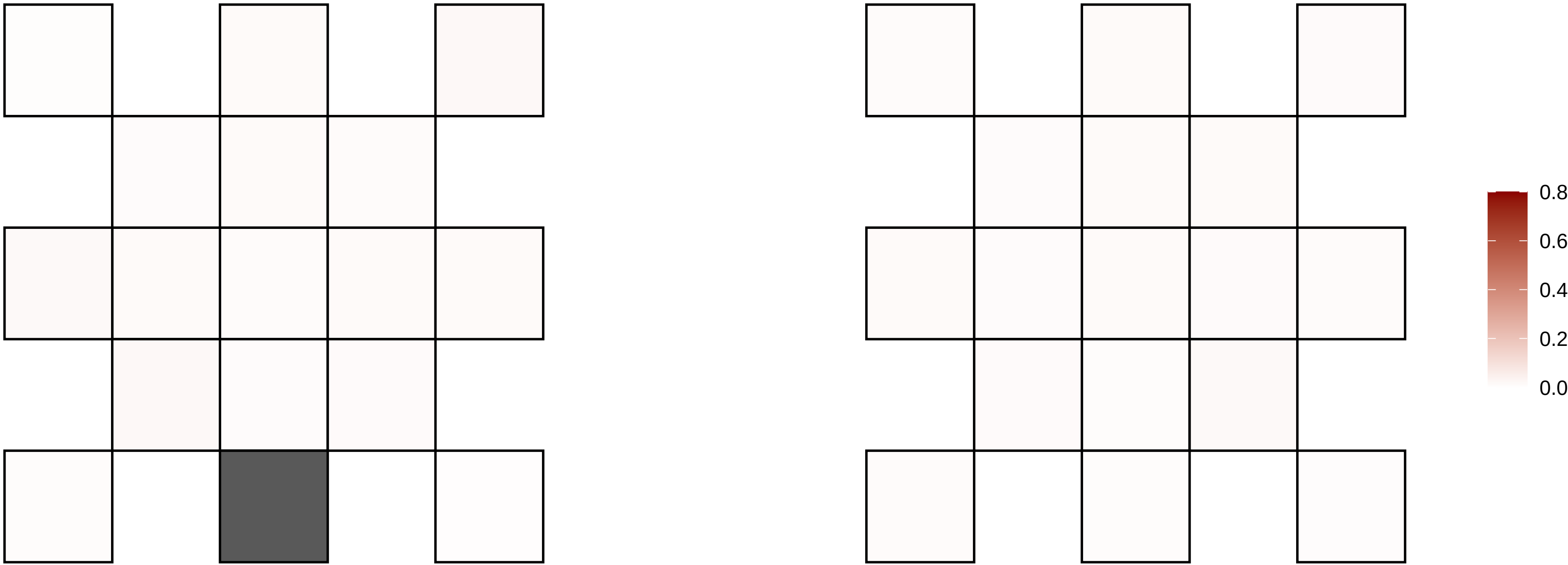

6−18819400−C−T

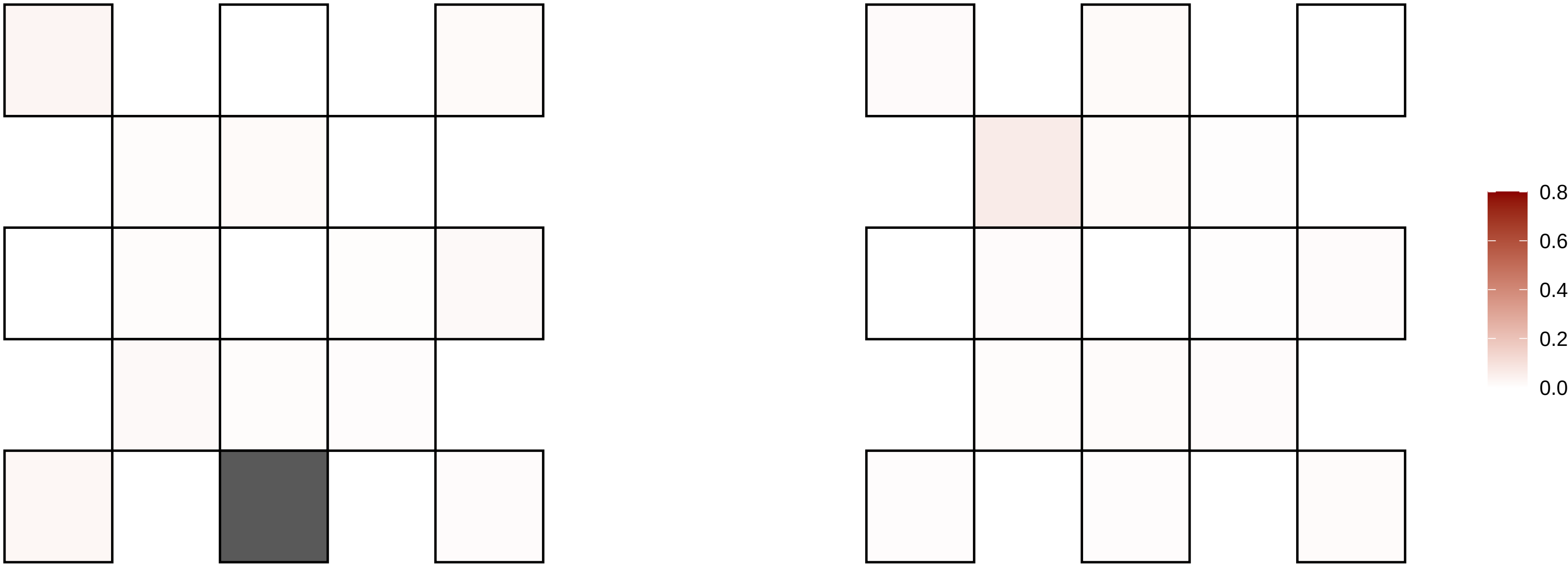

6−18903997−G−A

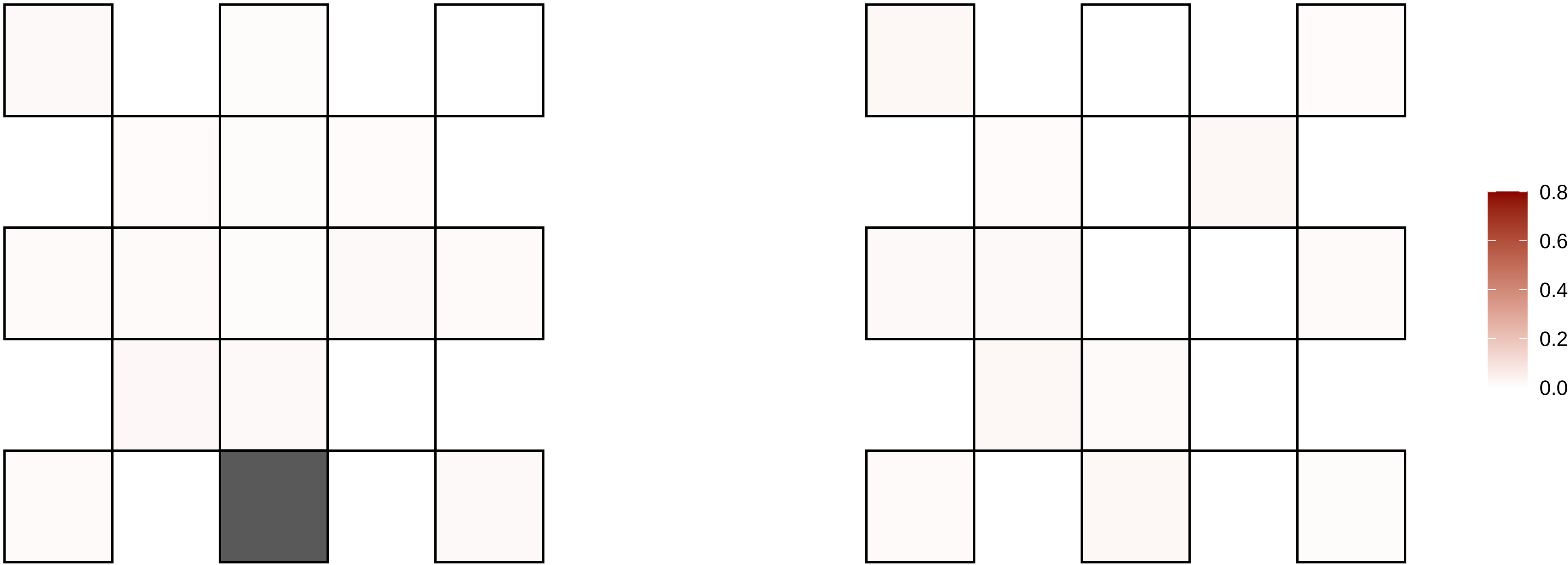

6−24537268−G−A

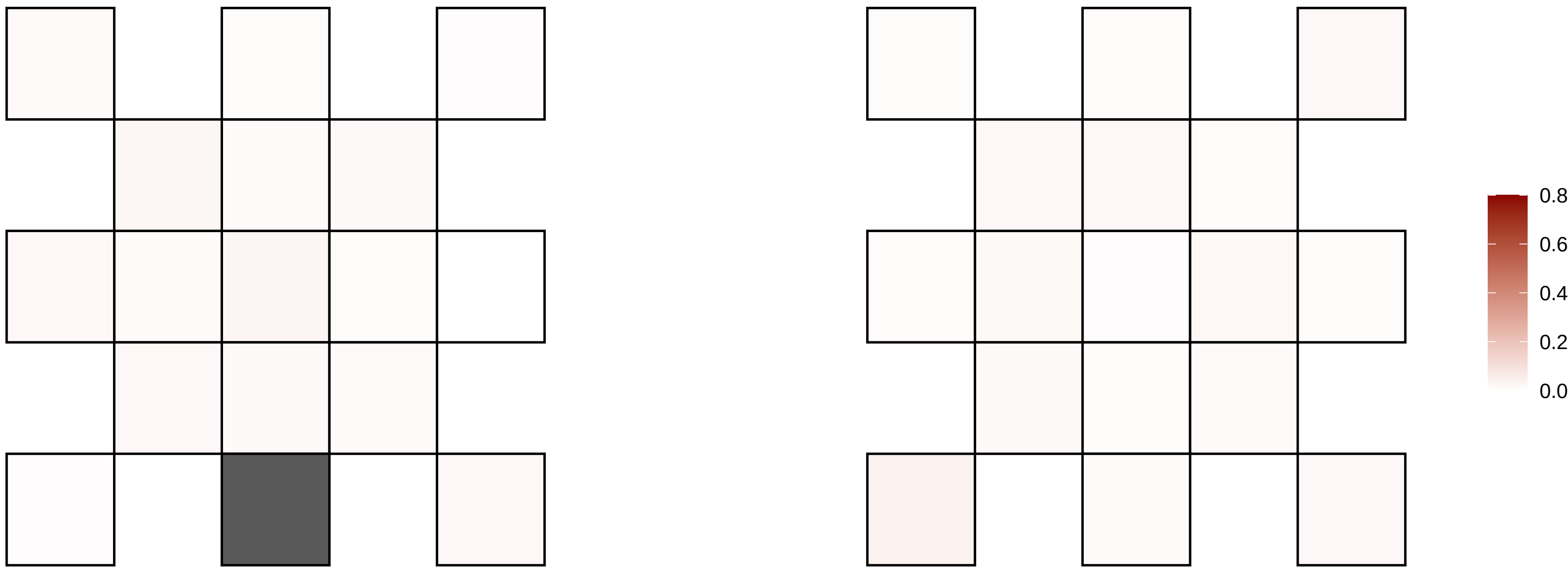

6−28244213−C−T

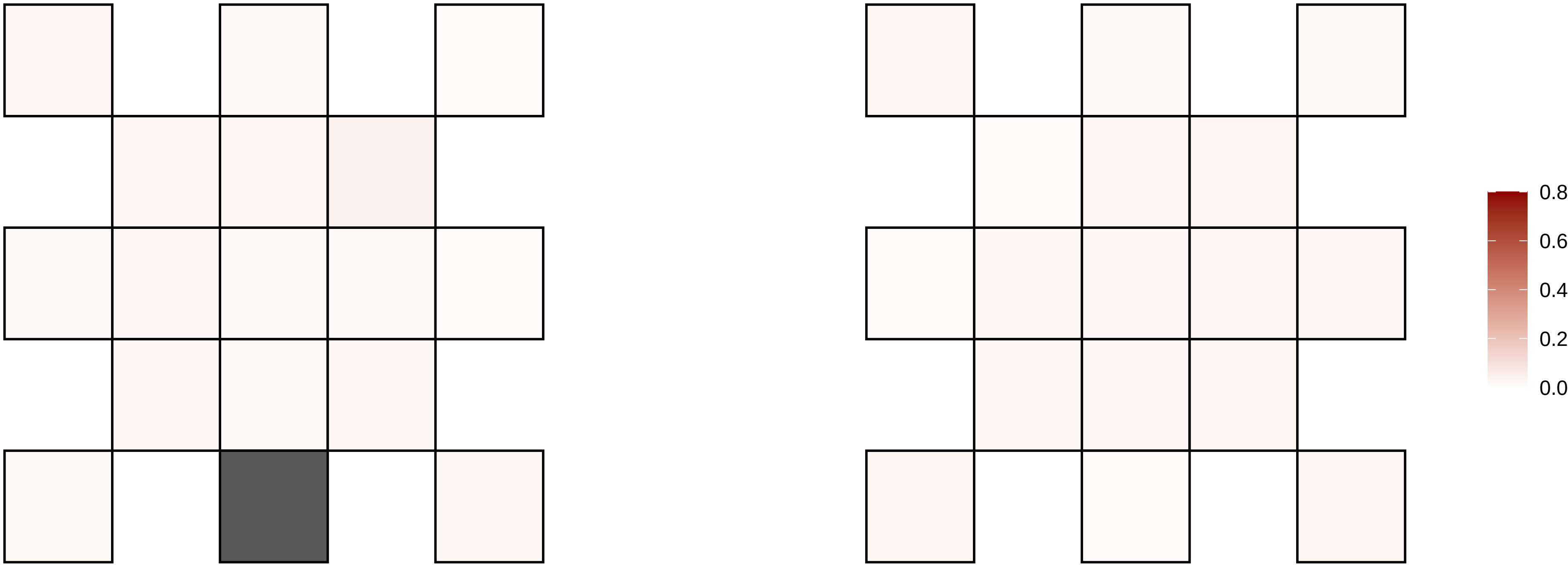

6−3287273−C−T

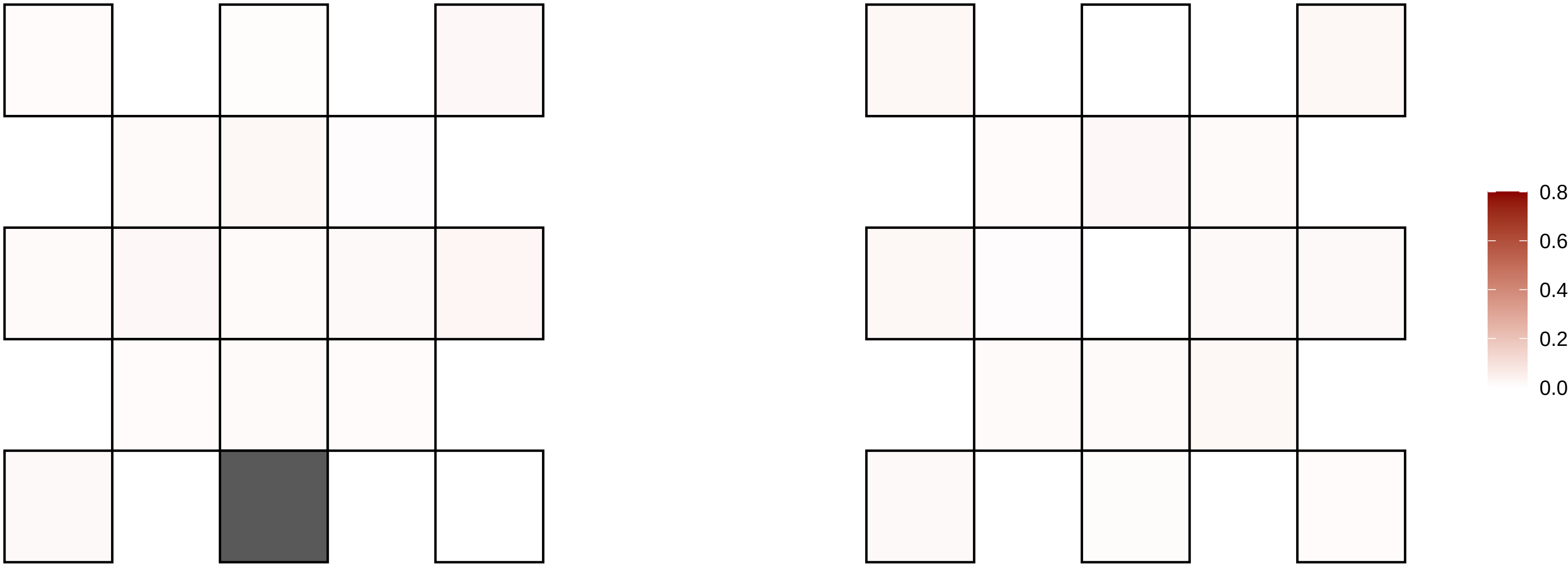

6−36931639−G−C

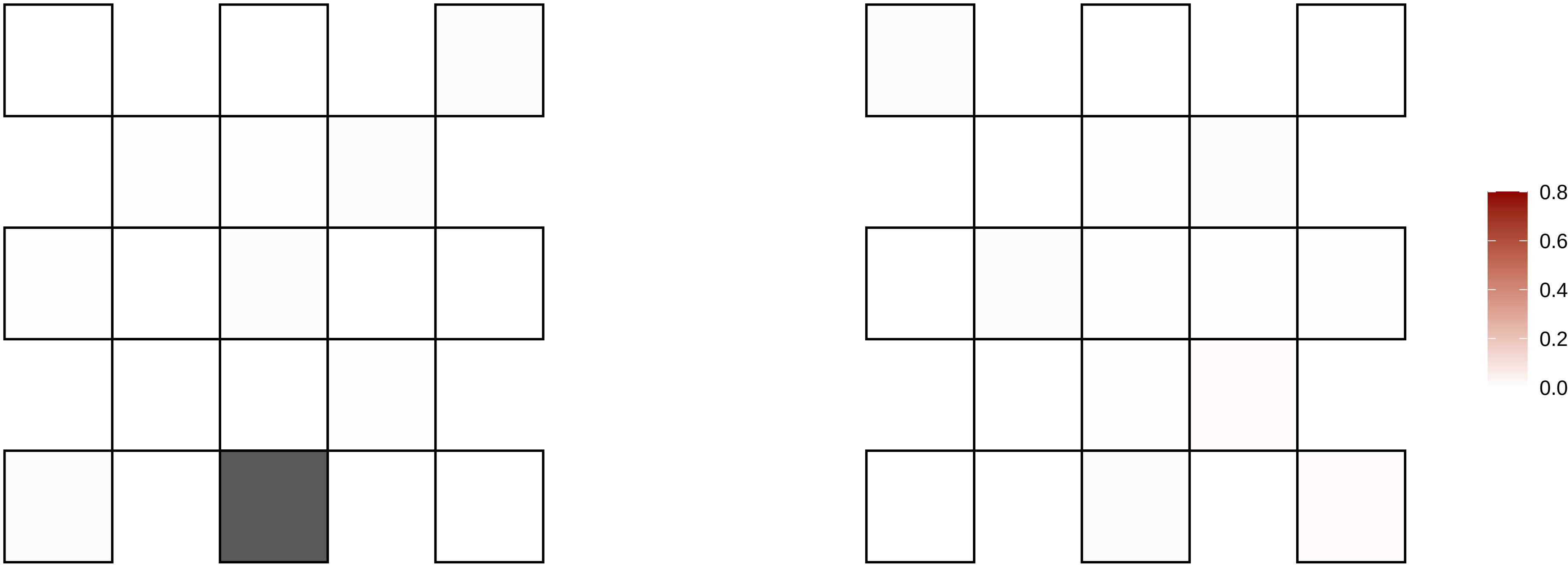

6−41864337−C−T

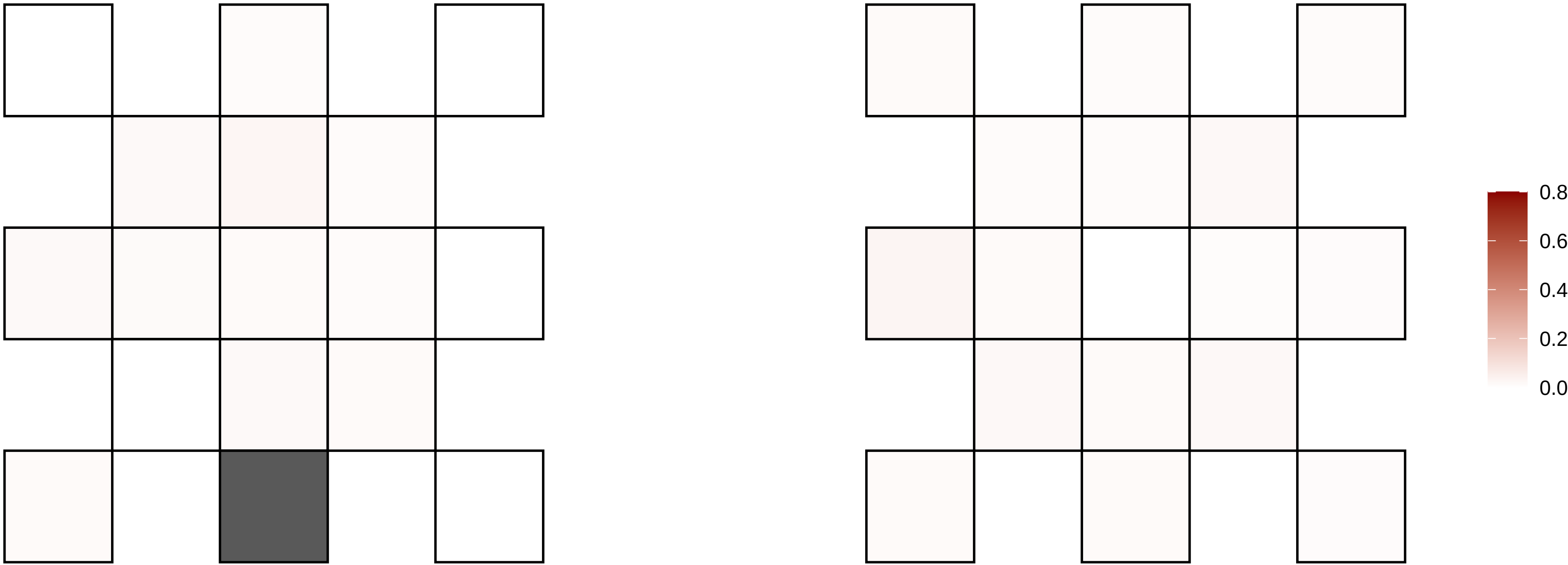

6−47367037−G−A

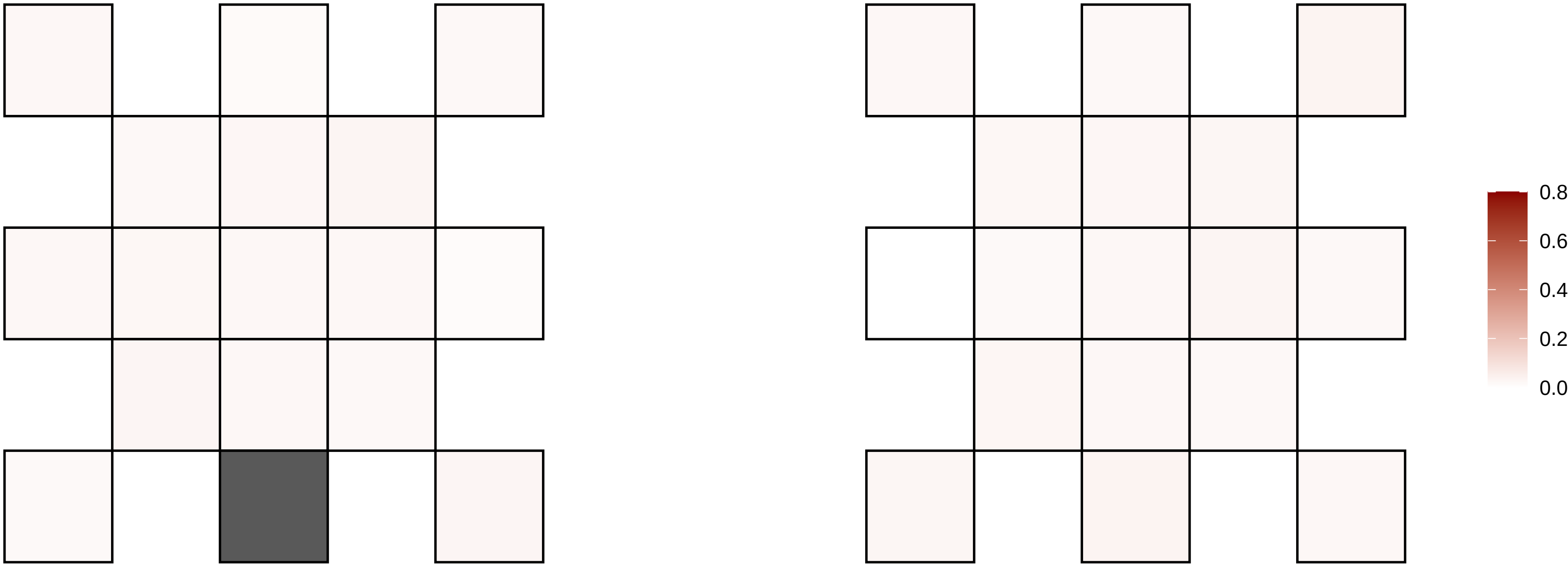

6−49937325−C−G

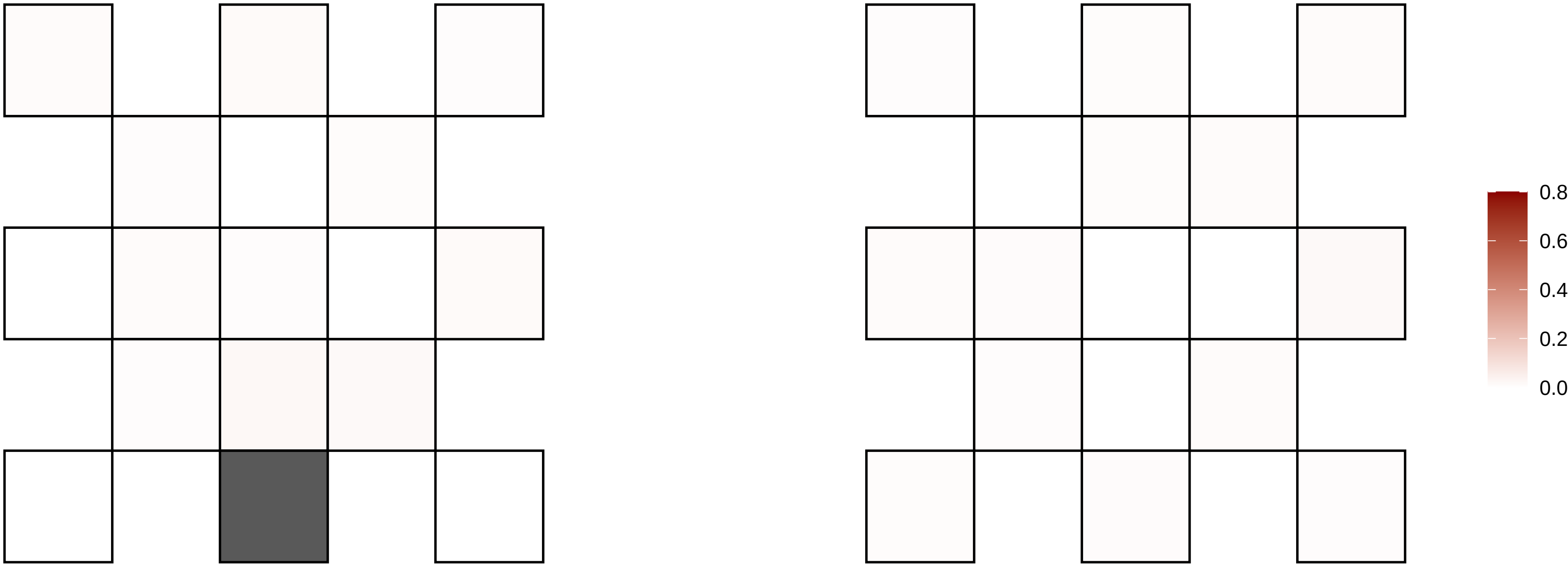

6−53413261−G−A

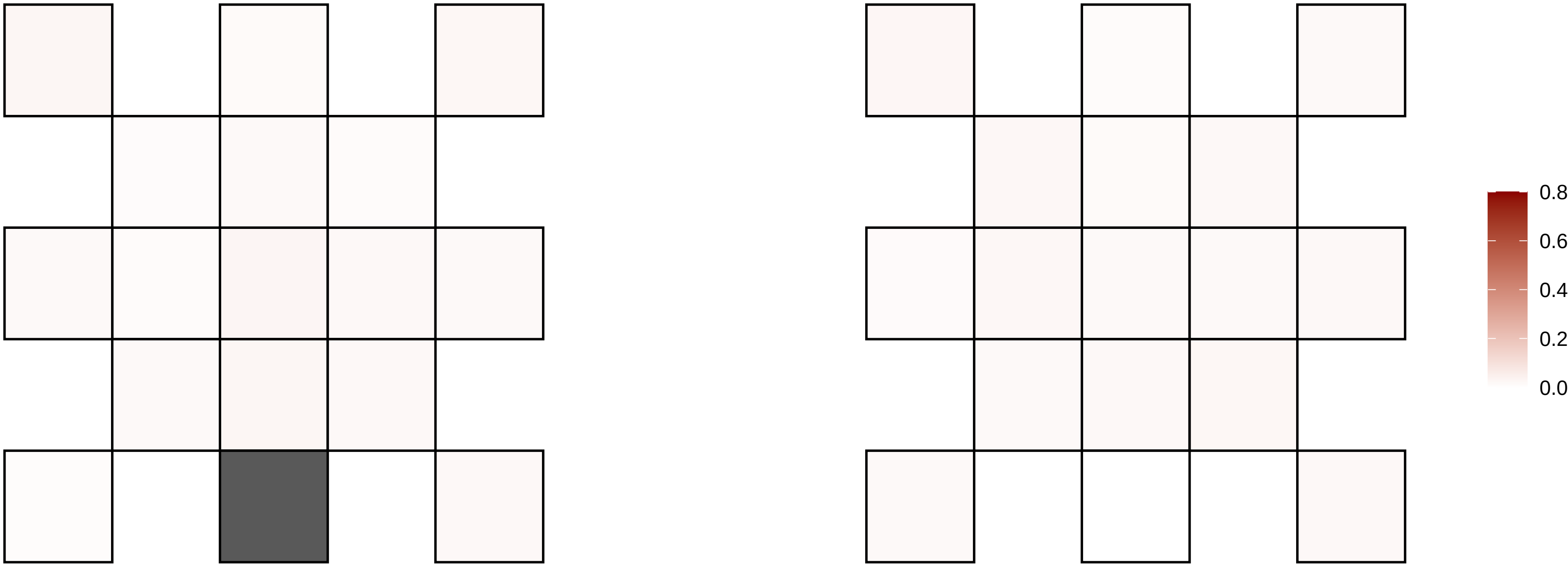

6−53918630−A−G

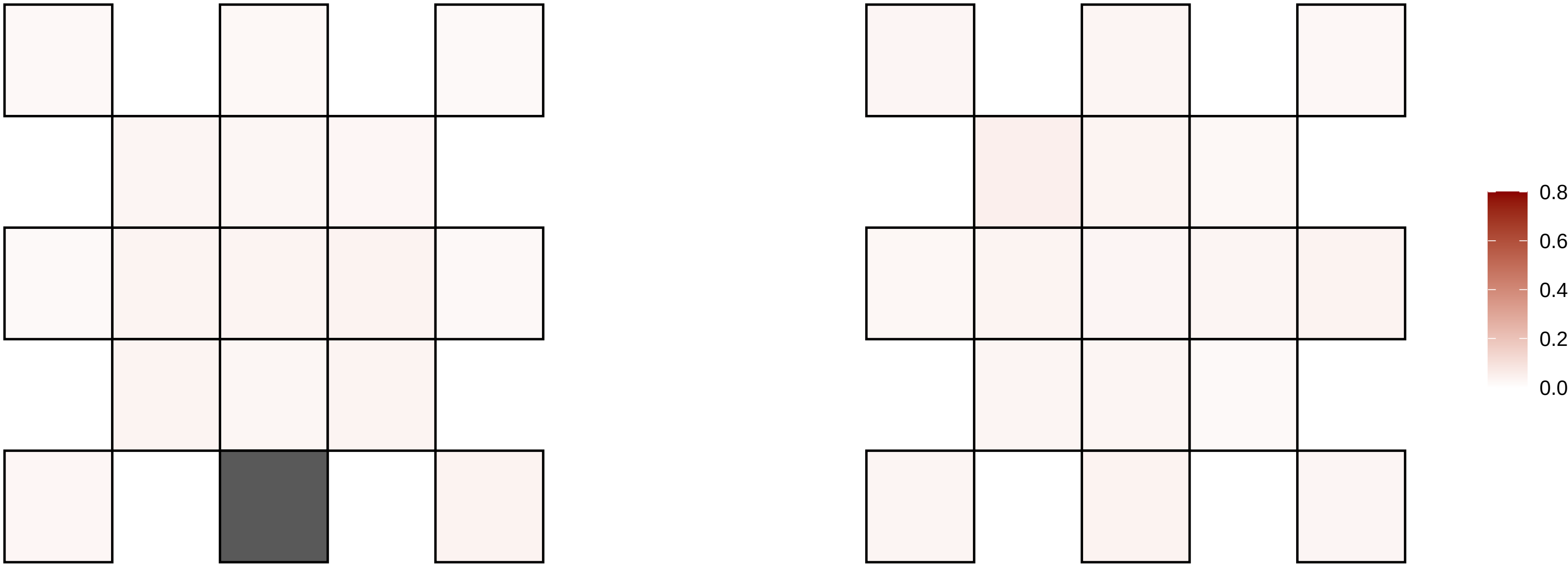

6−56684720−G−A

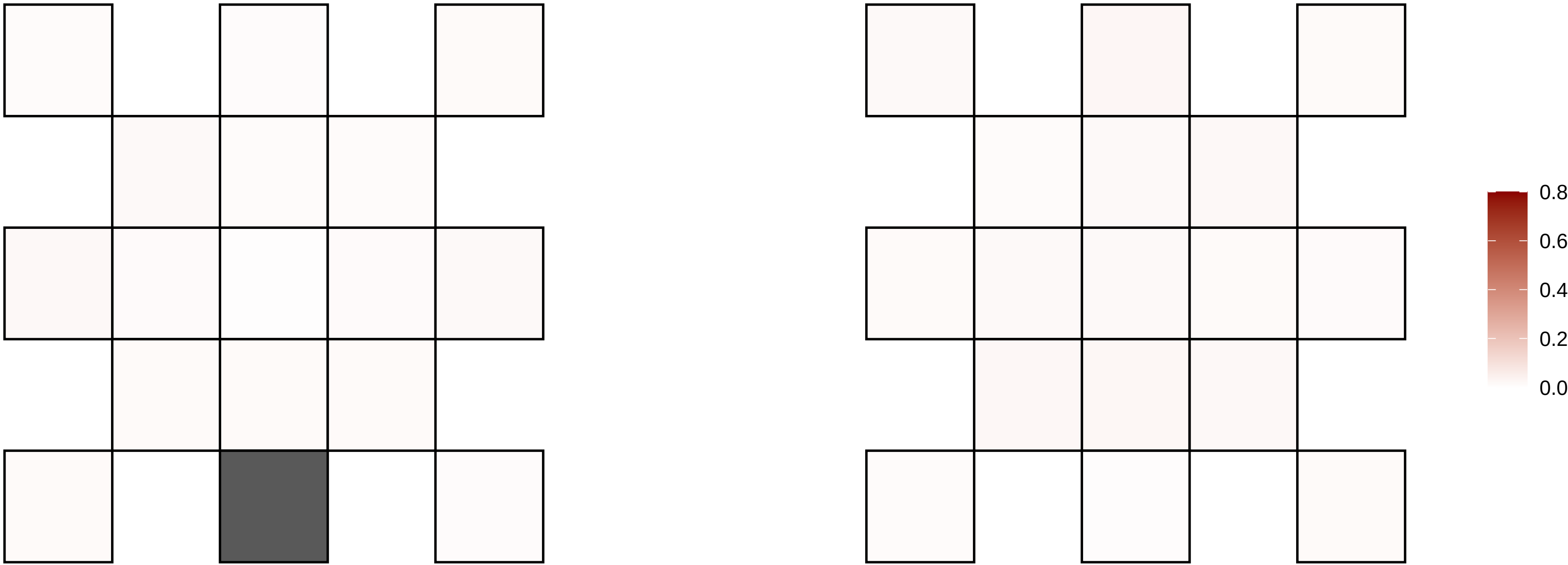

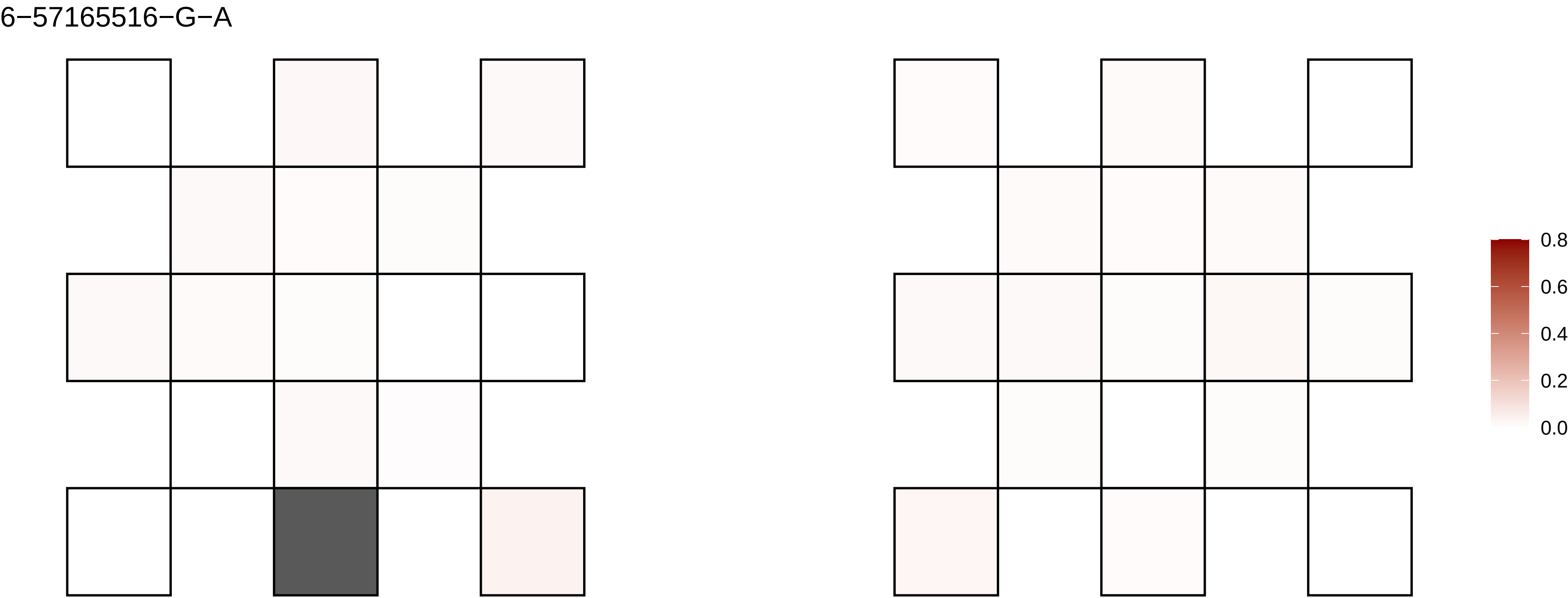
6−57165516−G−A
0.8
0.6
0.4
0.2
0.0

6−66934852−G−A

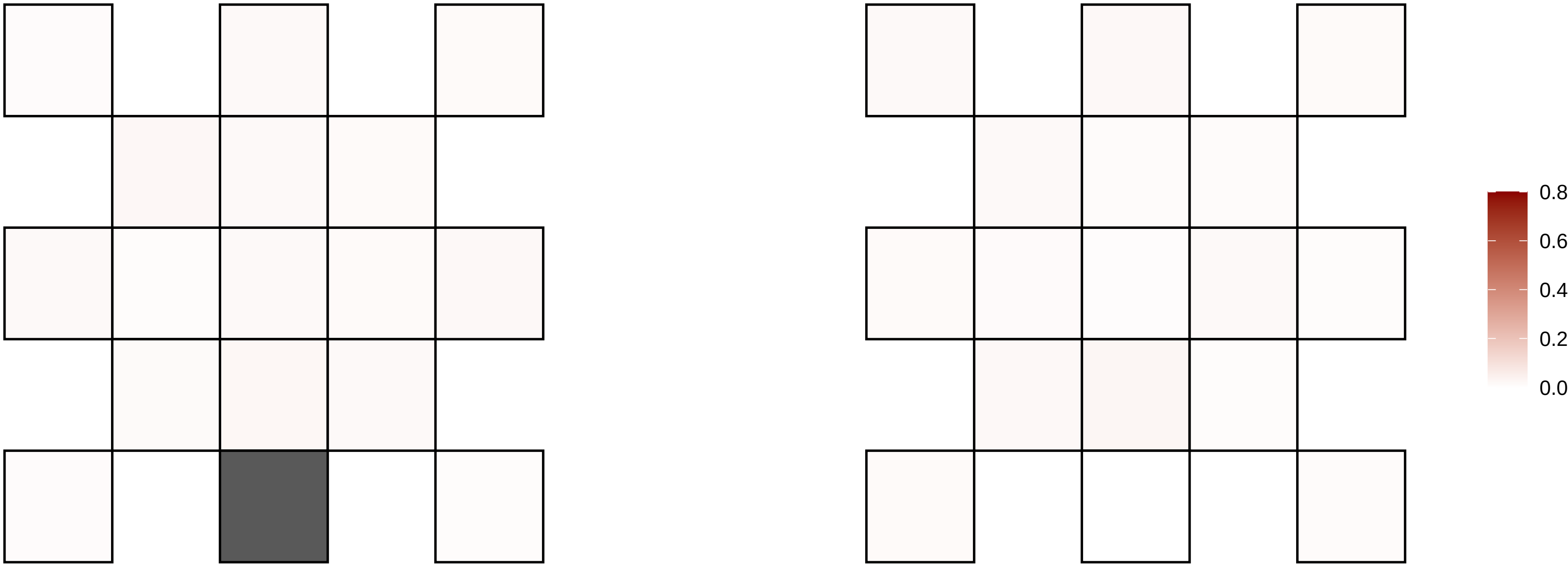

6−68312890−A−T

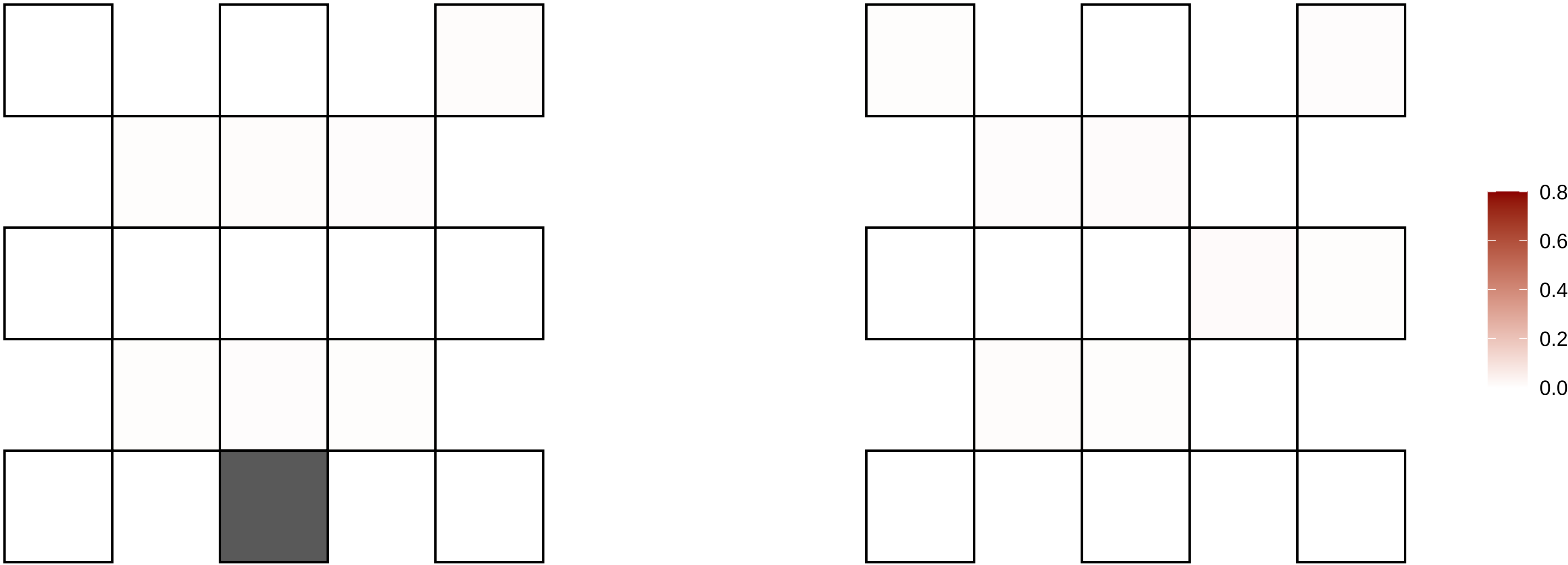

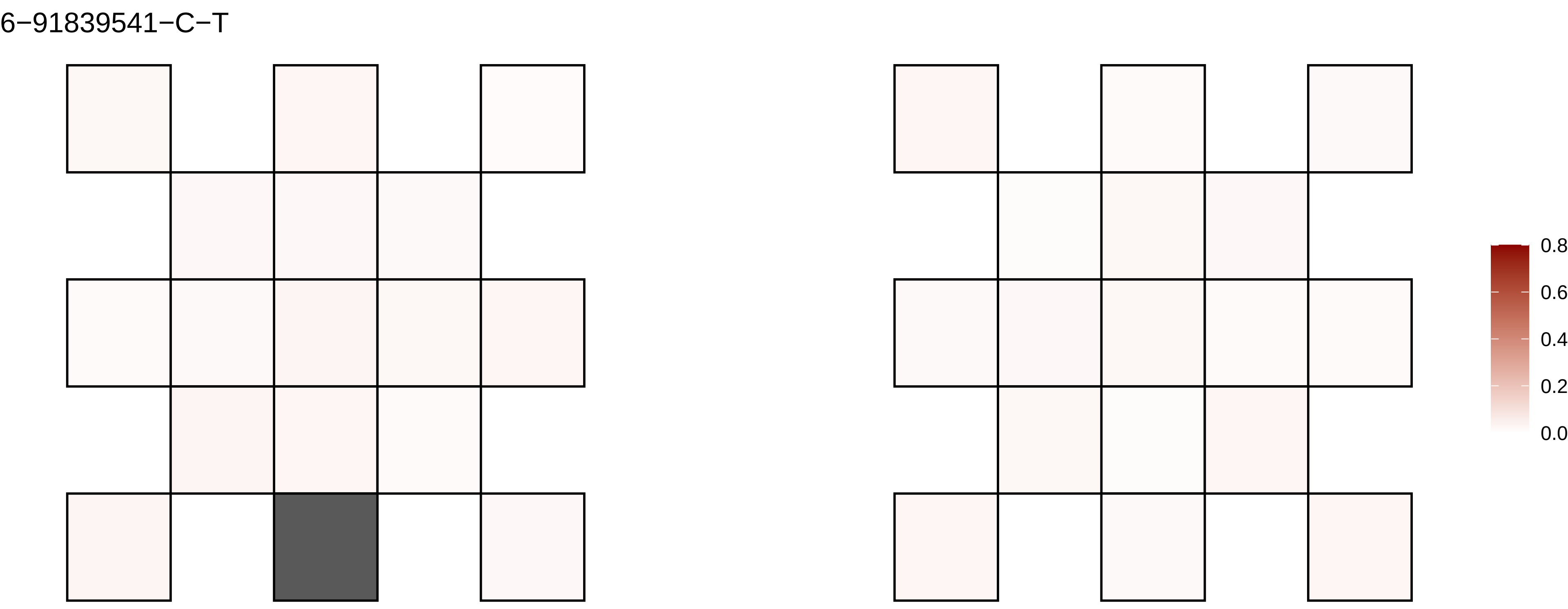

6−91839541−C−T
0.8
0.6
0.4
0.2
0.0

6−92211371−C−T

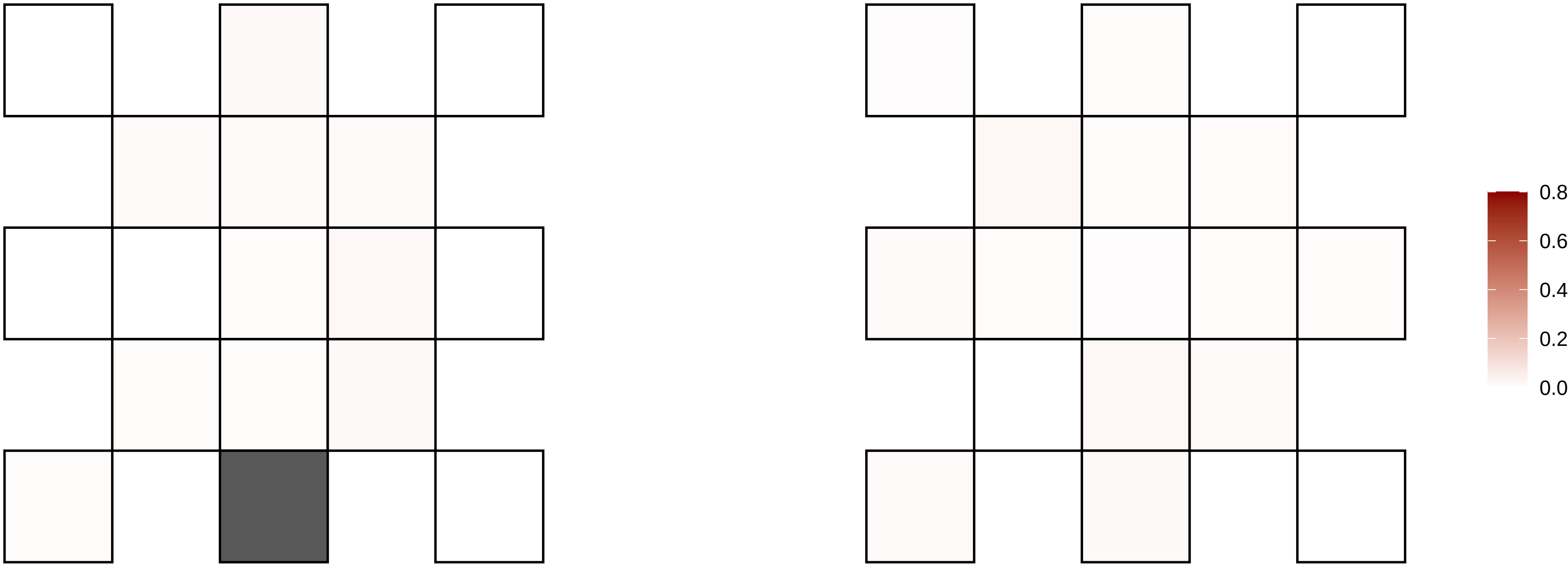

6−94855972−G−A

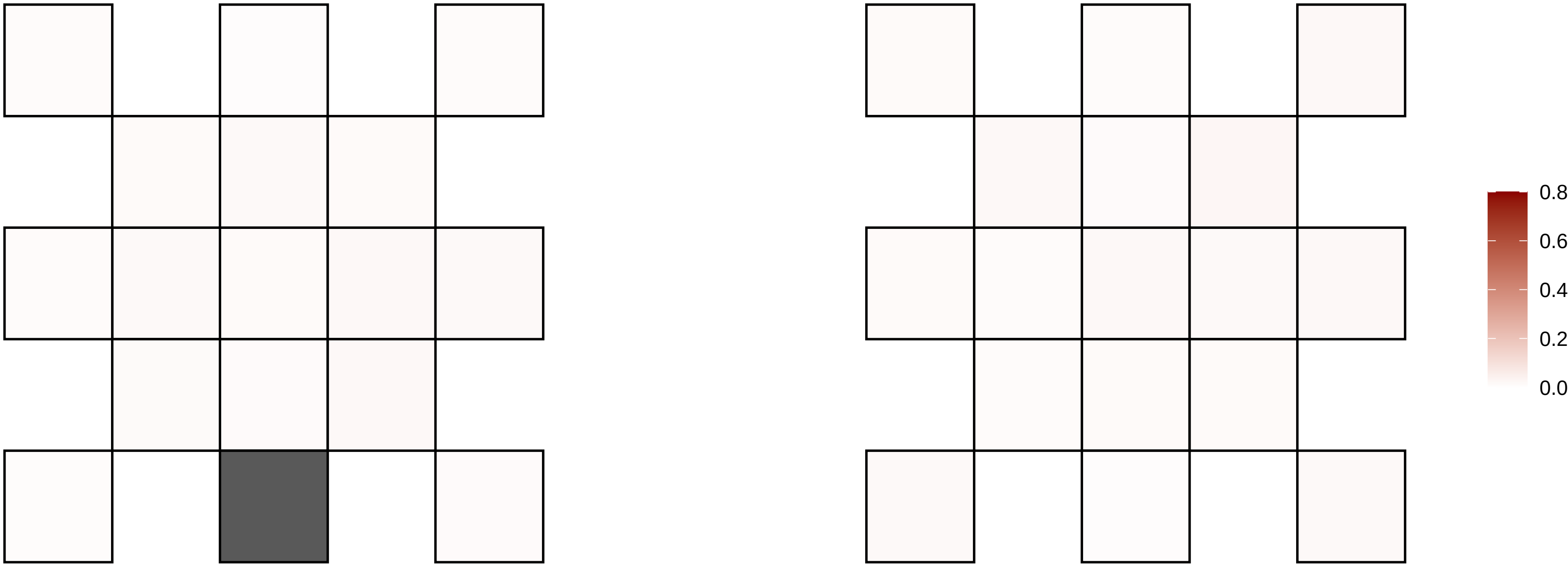

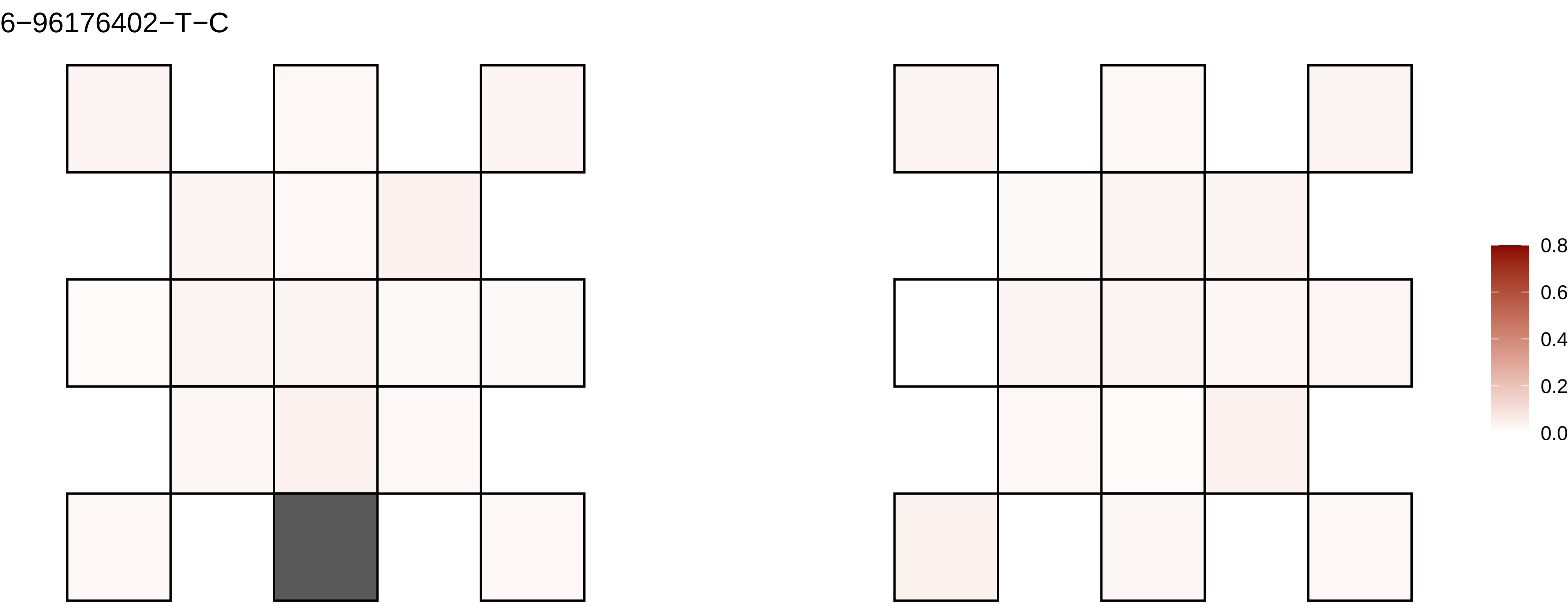

6−96176402−T−C
0.8
0.6
0.4
0.2
0.0

6−98372583−C−T

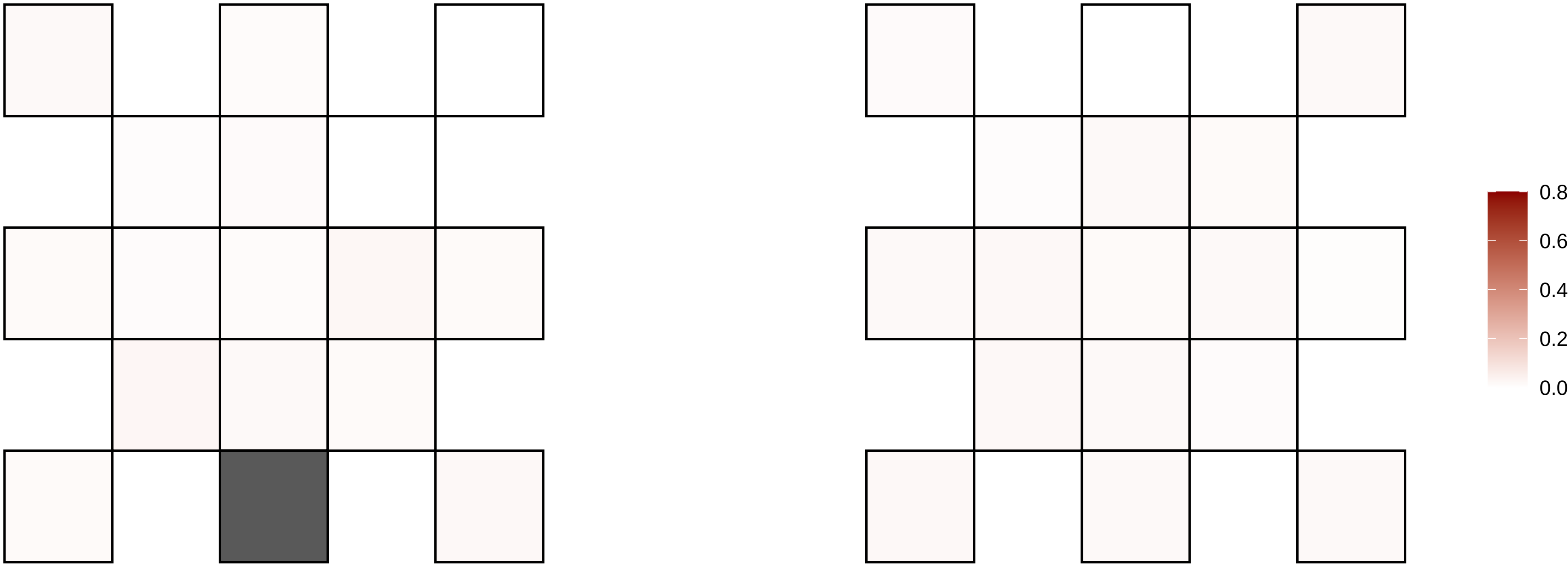

6−99090478−G−T

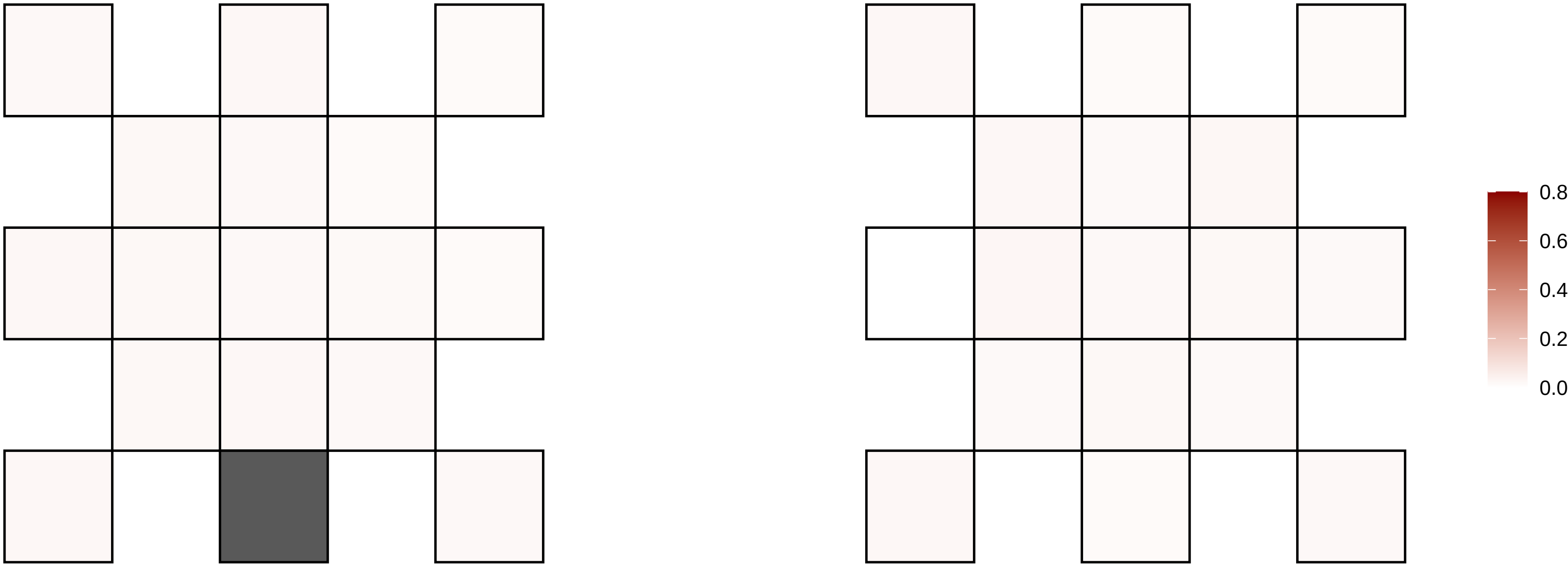

6−99303058−CCT−C

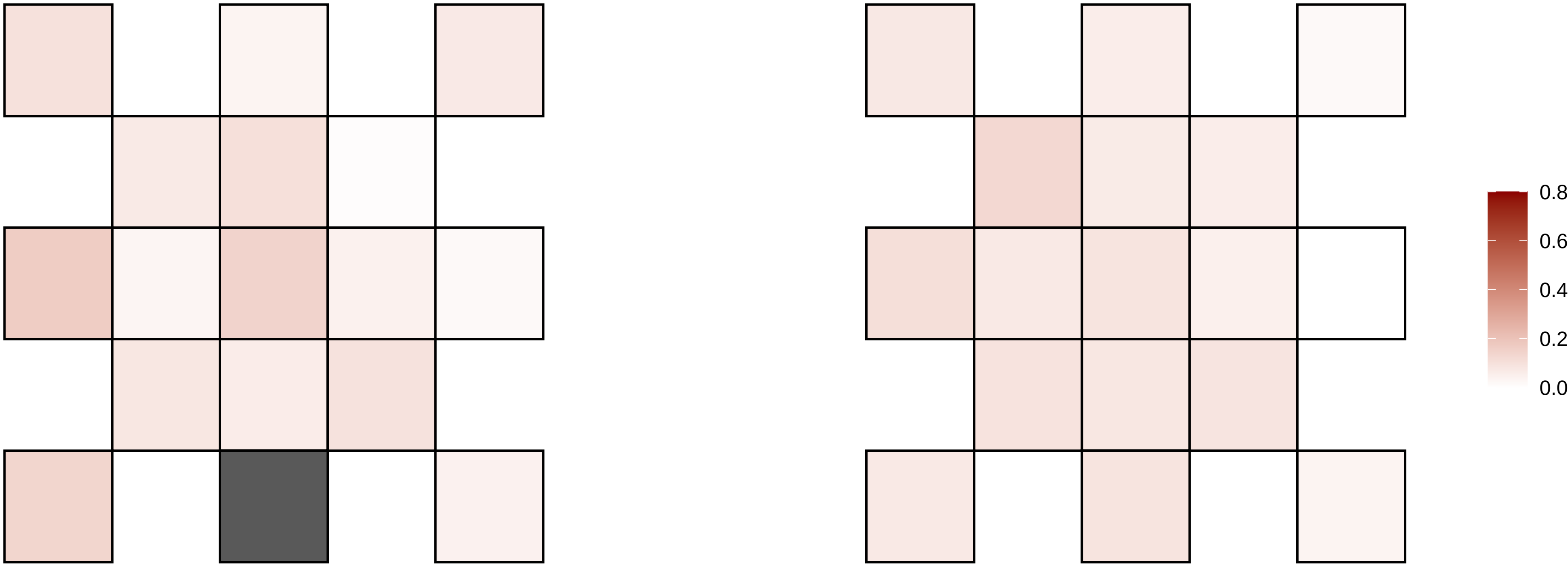

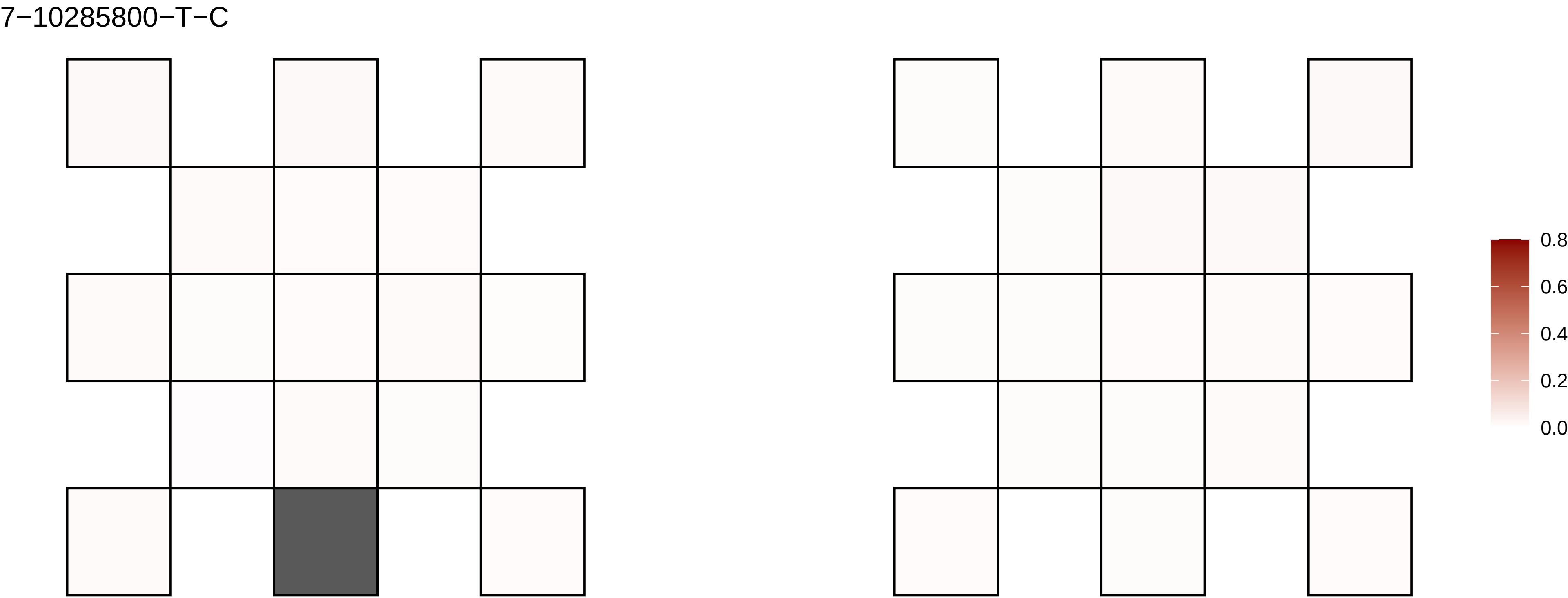

7−10285800−T−C
0.8
0.6
0.4
0.2
0.0

7−109736410−A−T

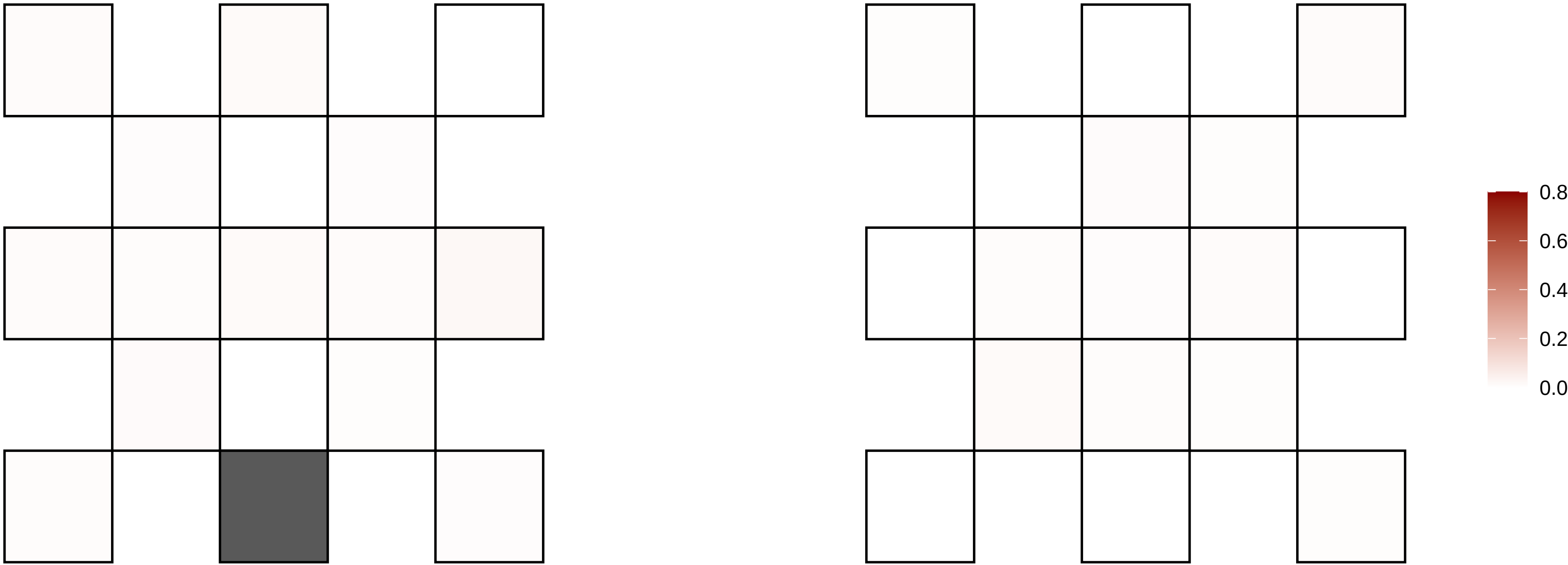

7−114017234−C−T

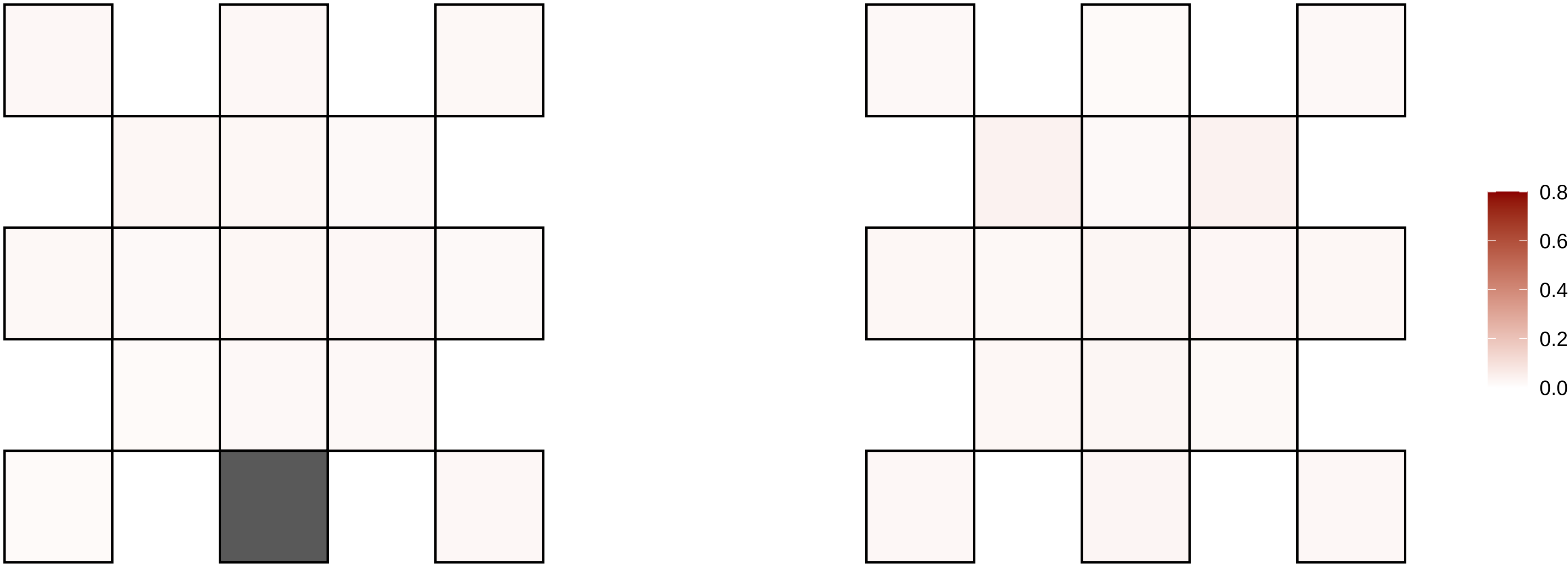

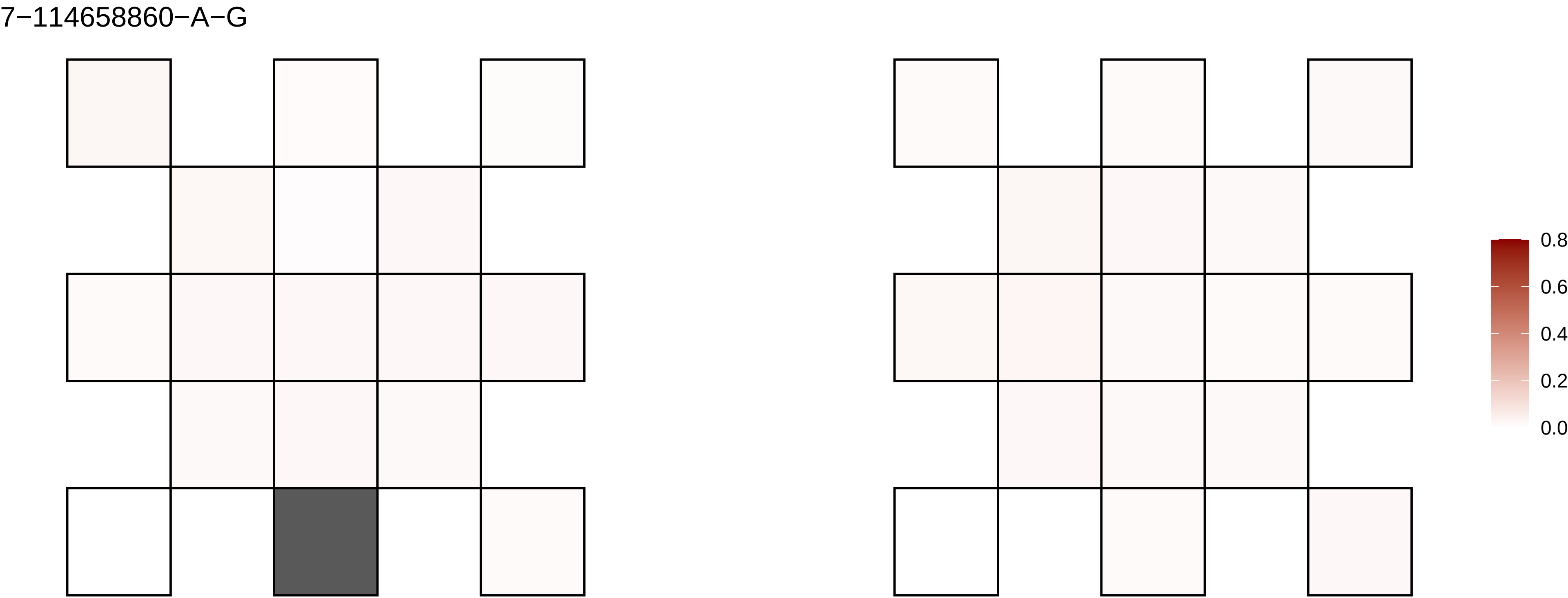
7−114658860−A−G
0.8
0.6
0.4
0.2
0.0

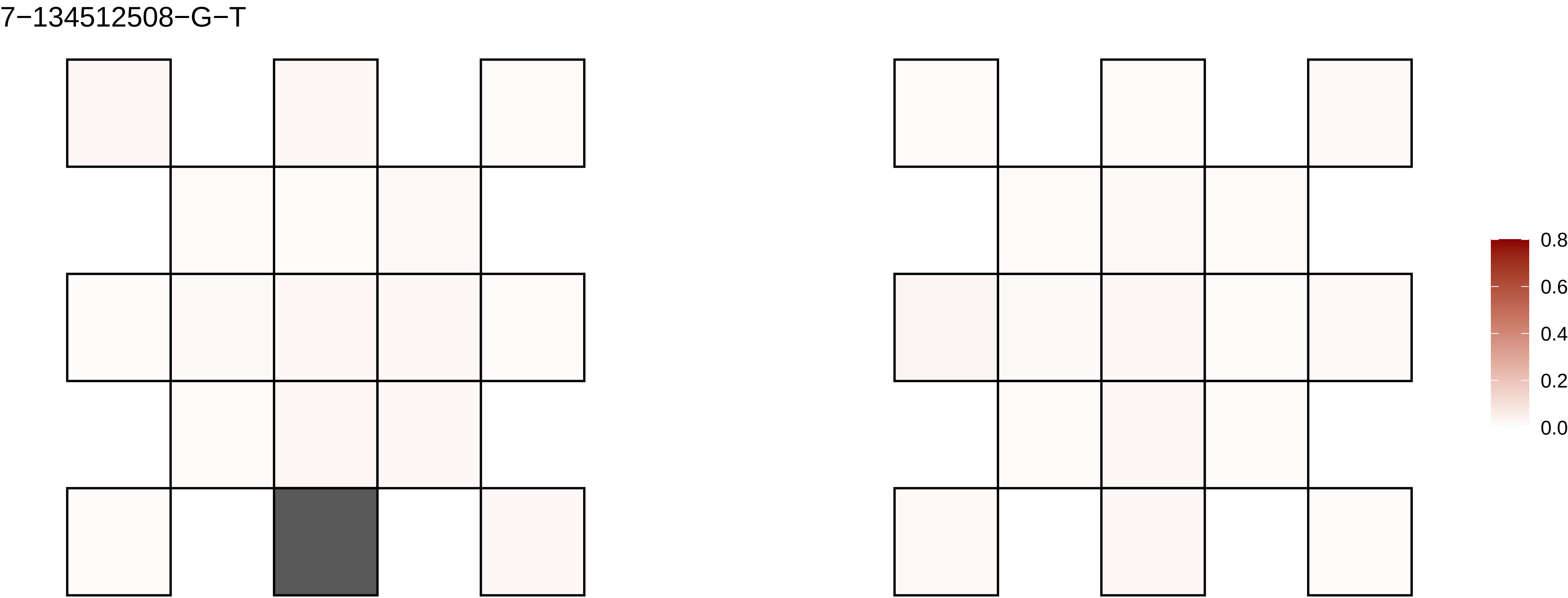
7−134512508−G−T
0.8
0.6
0.4
0.2
0.0

7−137220703−G−A

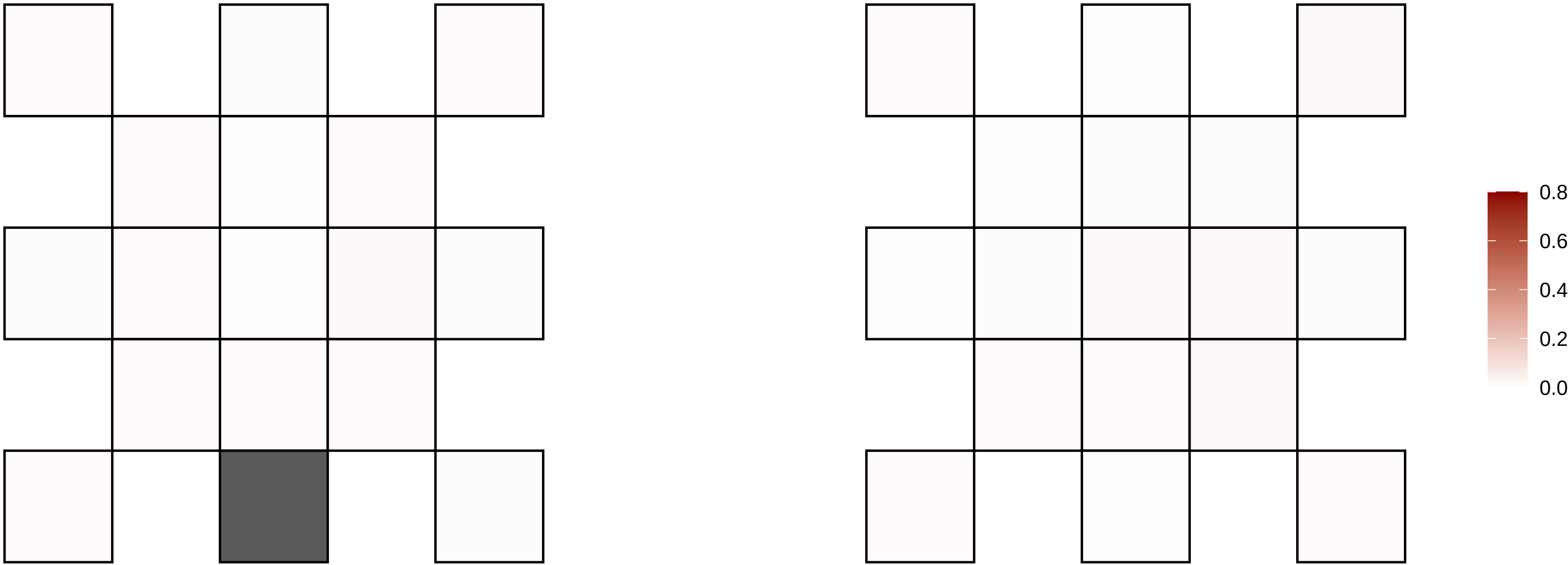

7−139192862−C−T

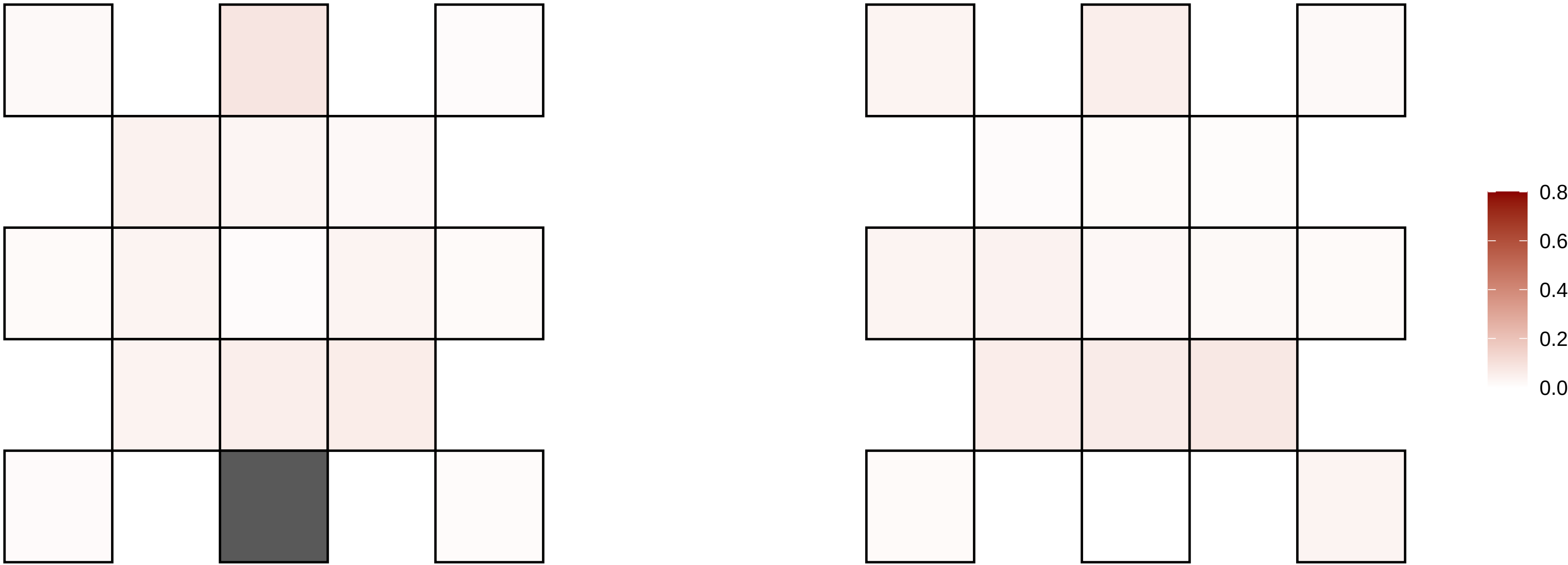

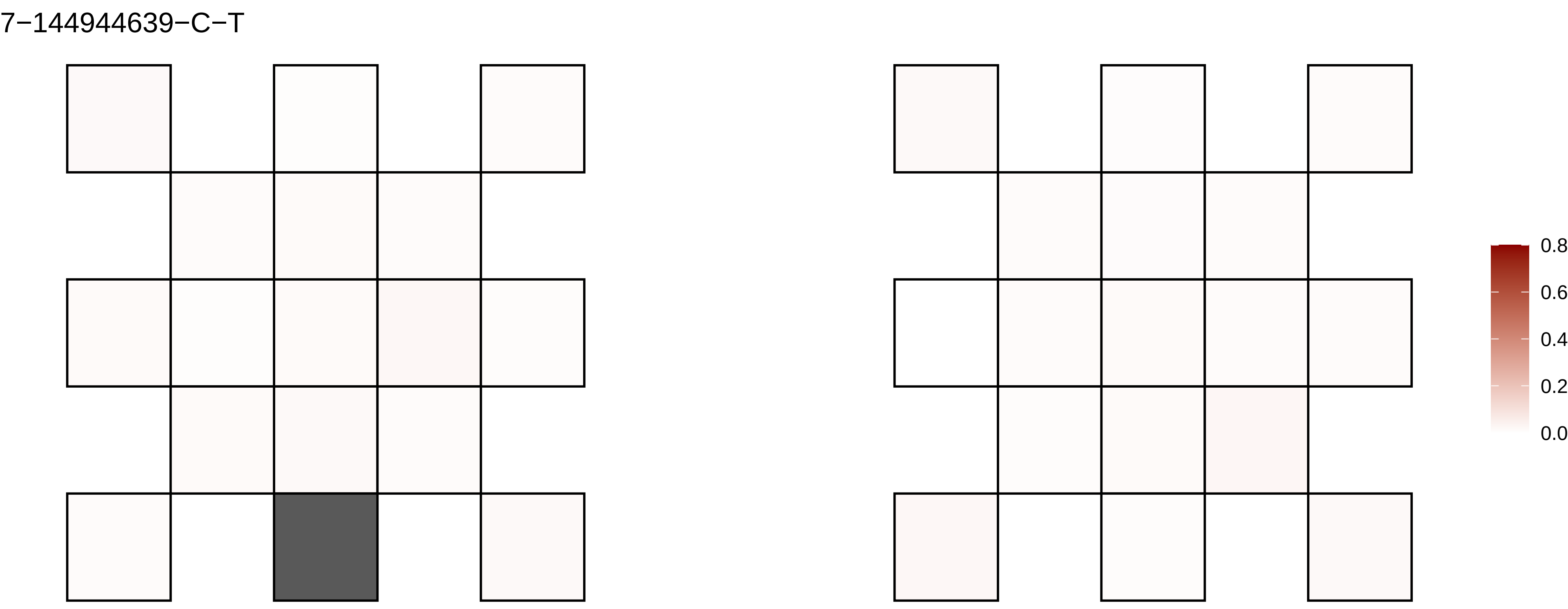
7−144944639−C−T
0.8
0.6
0.4
0.2
0.0

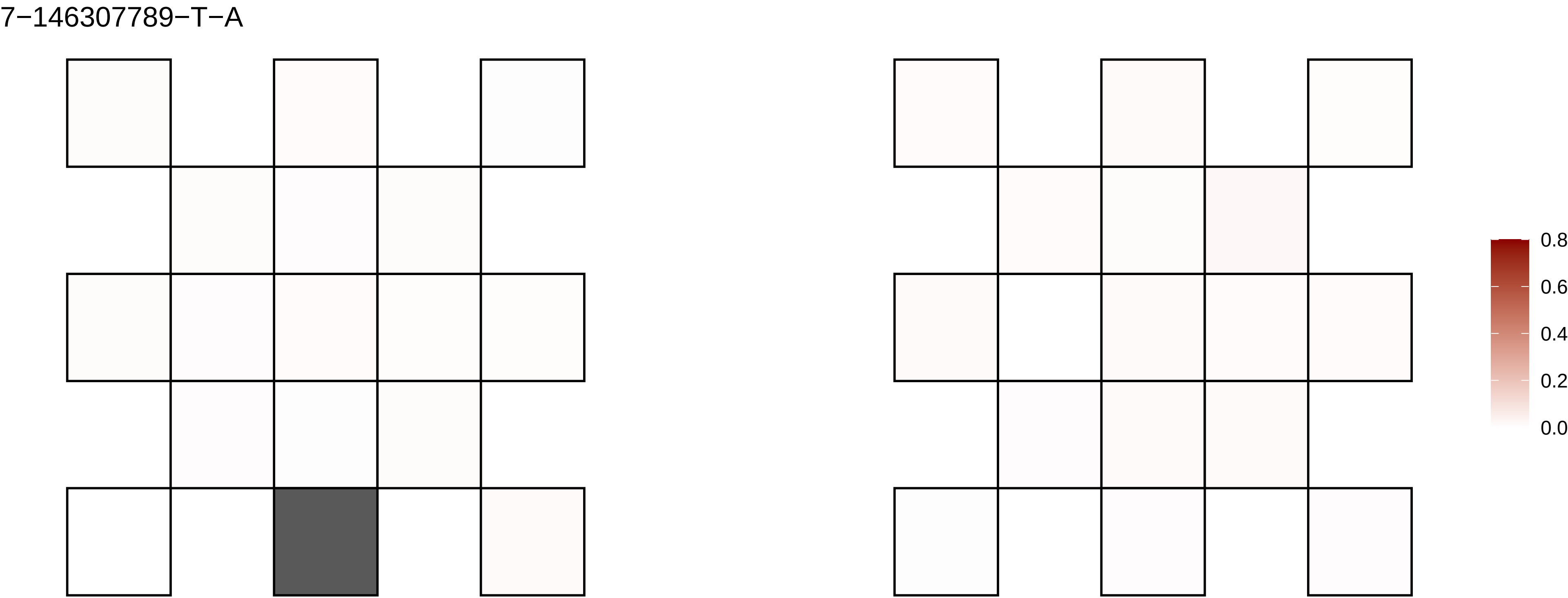
7−146307789−T−A
0.8
0.6
0.4
0.2
0.0

7−147380993−G−A

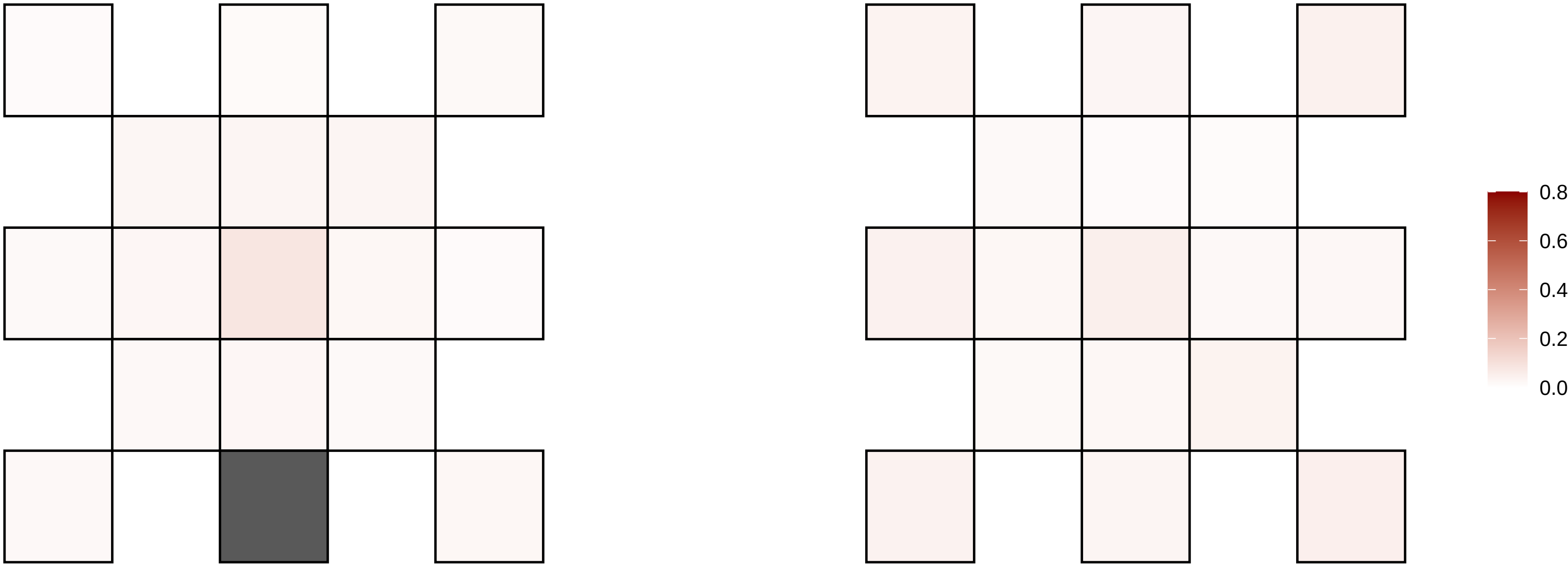

7−151491278−C−T

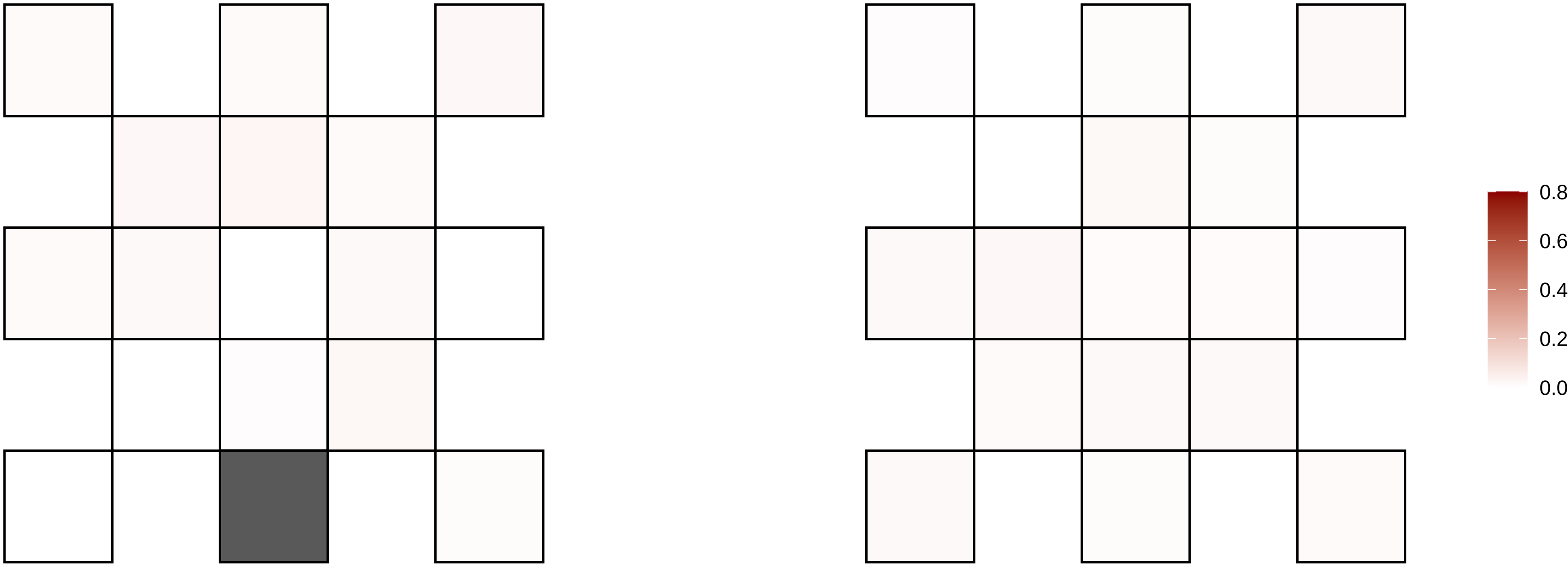

7−154551172−G−A

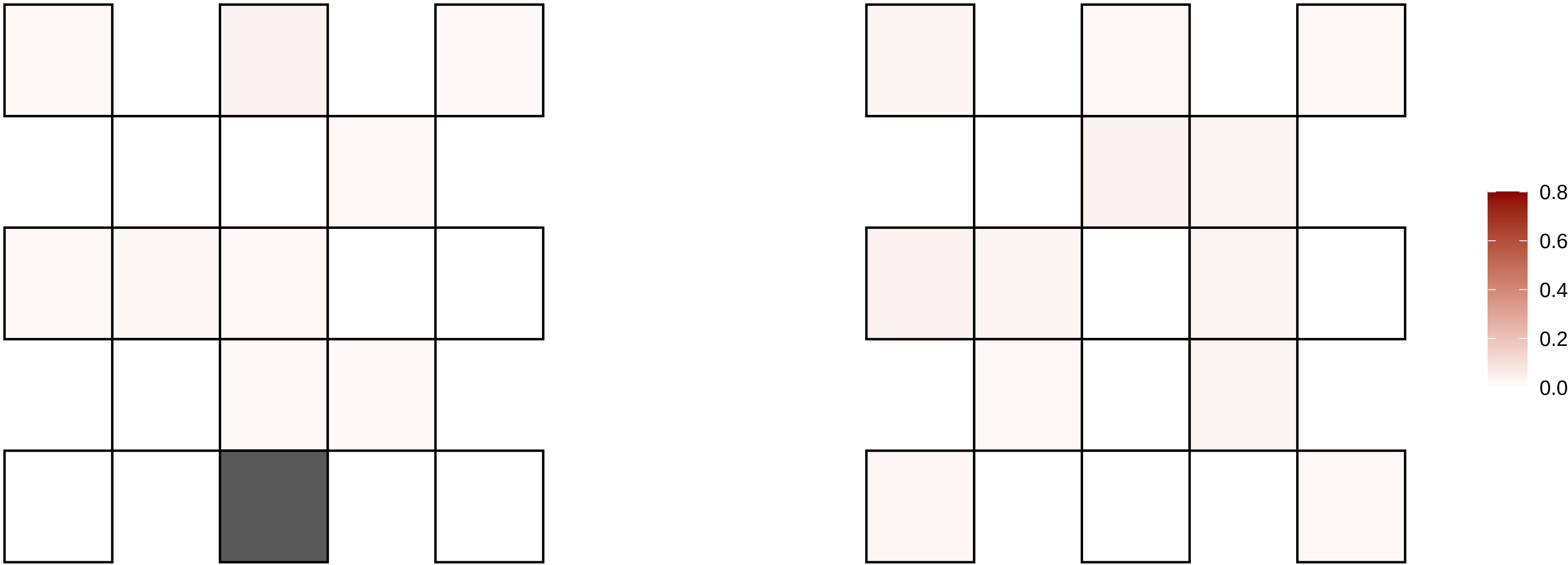

7−155855403−T−C

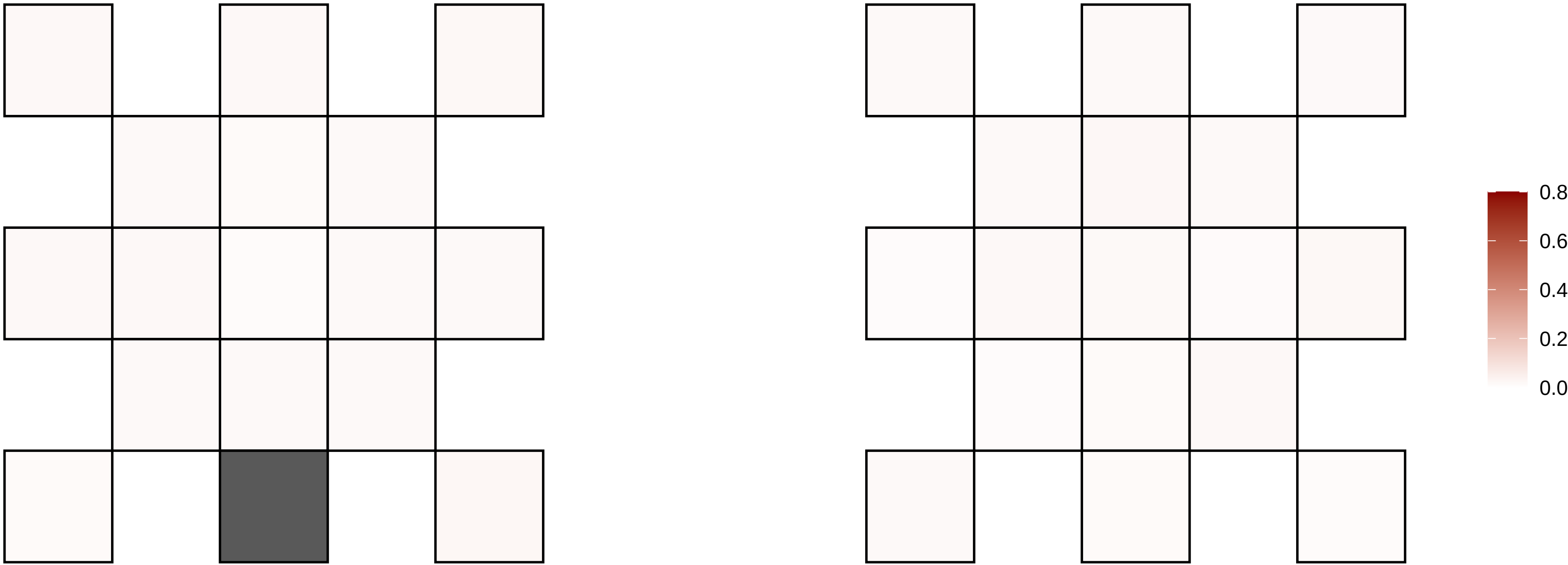

7−16742061−C−T

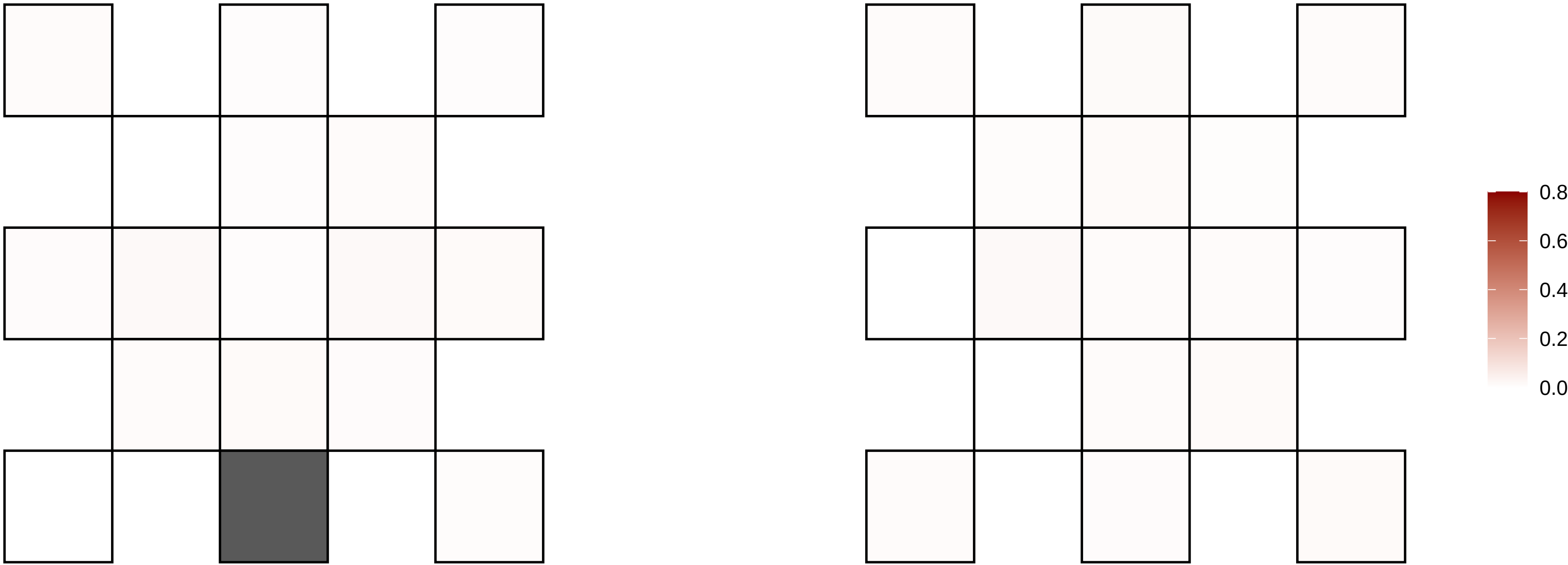

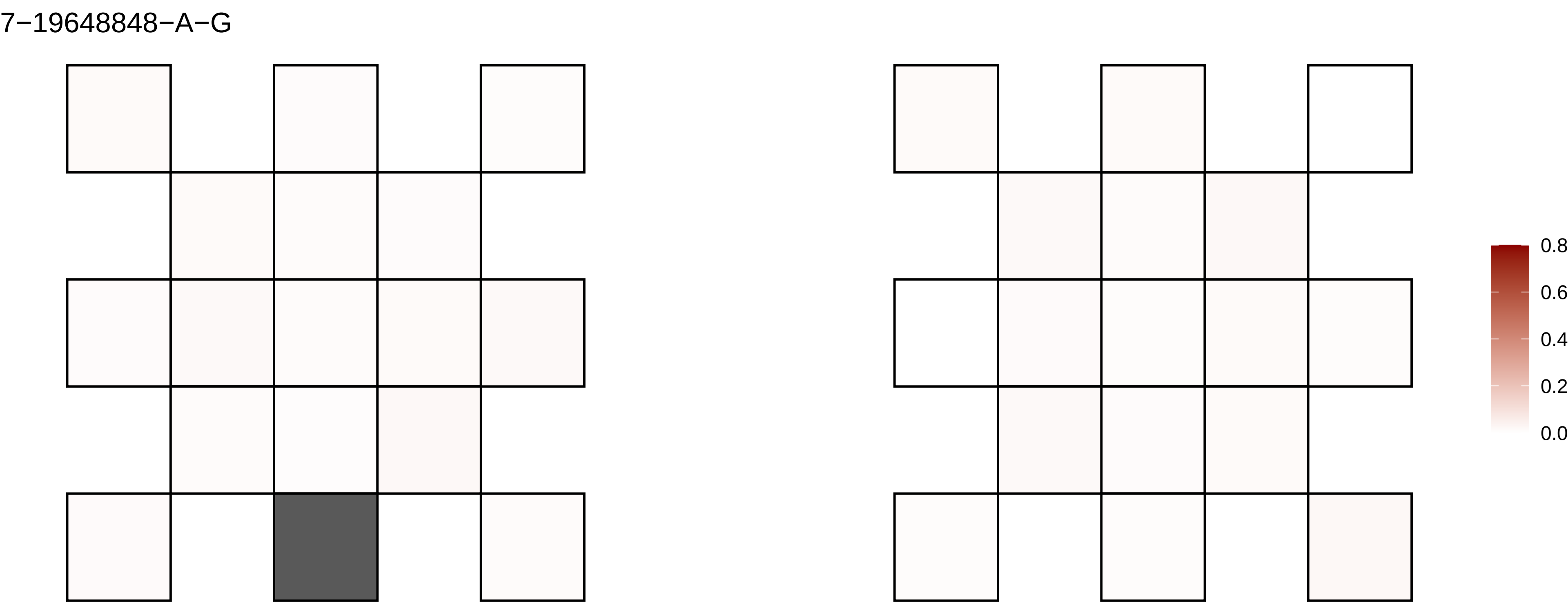
7−19648848−A−G
0.8
0.6
0.4
0.2
0.0

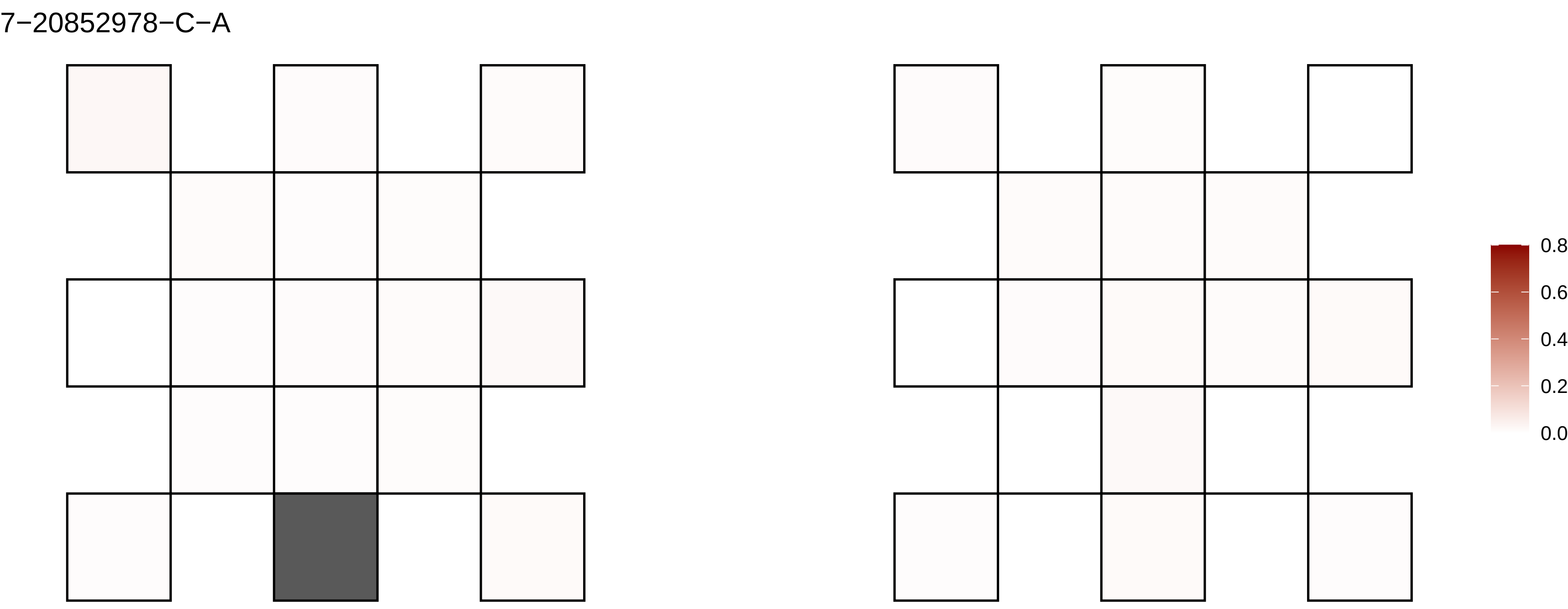
7−20852978−C−A
0.8
0.6
0.4
0.2
0.0

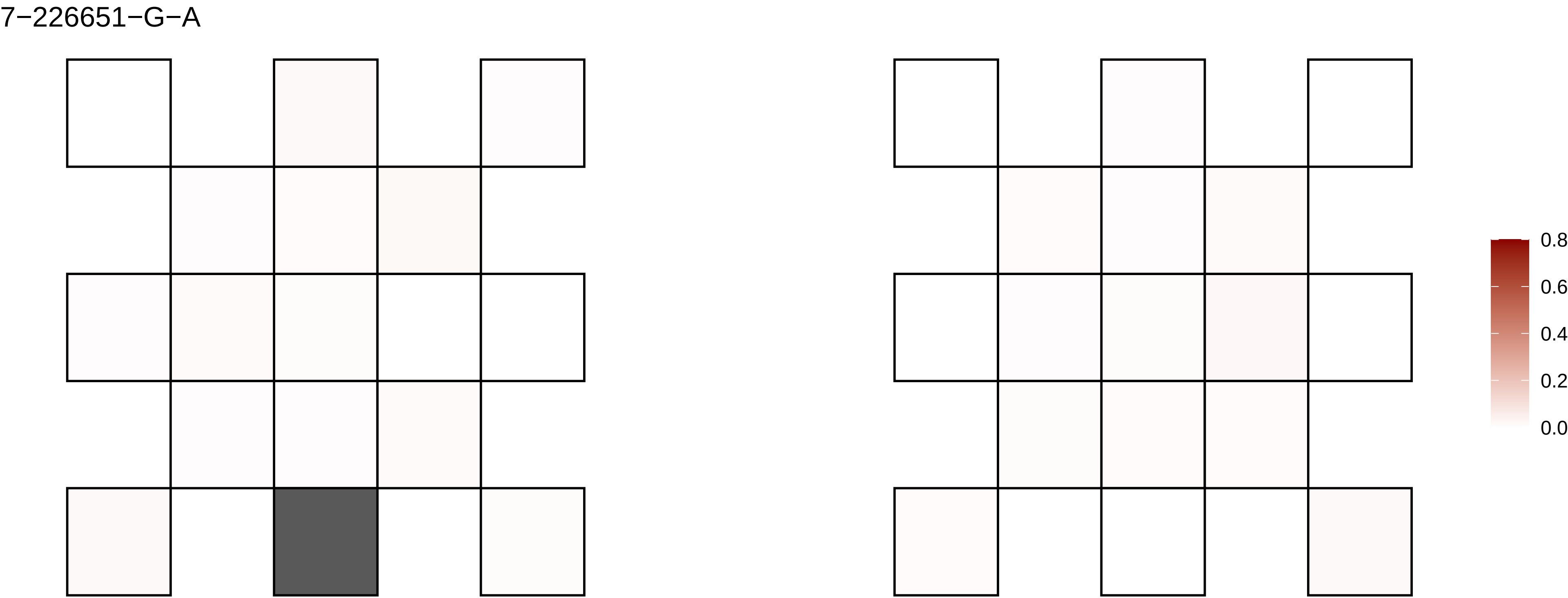

7−226651−G−A
0.8
0.6
0.4
0.2
0.0

7−24504428−A−G

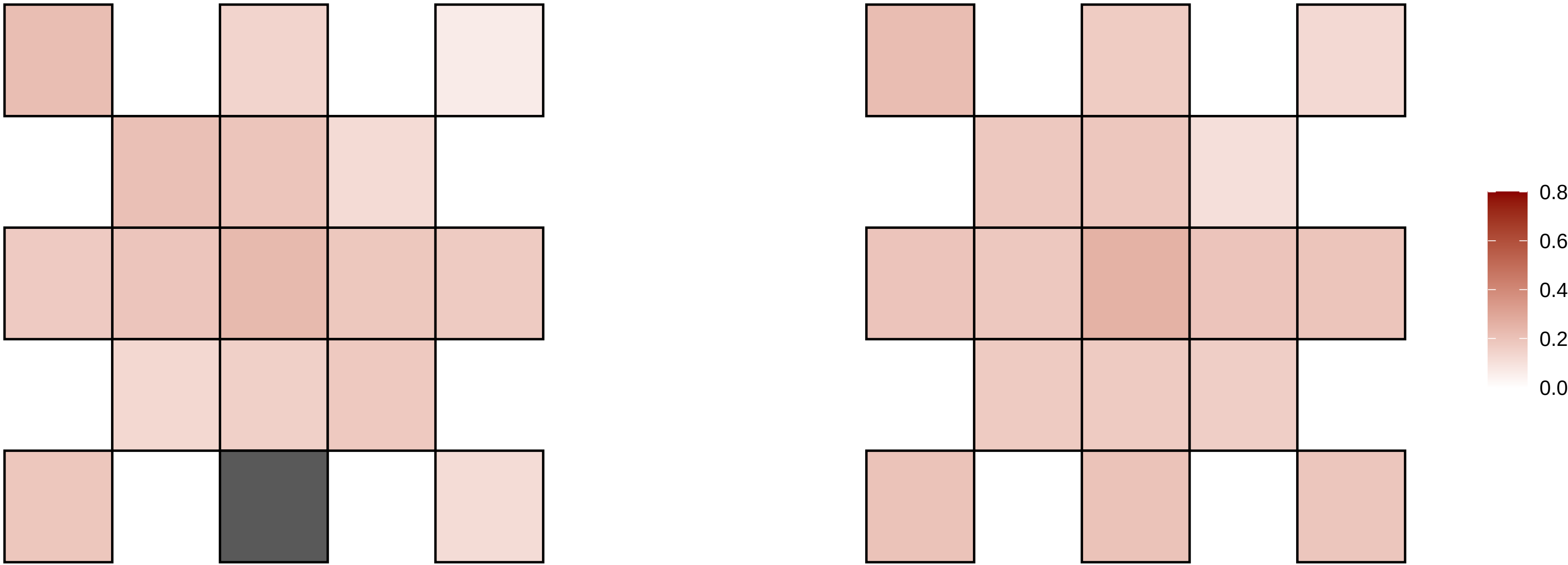

7−26490020−G−A

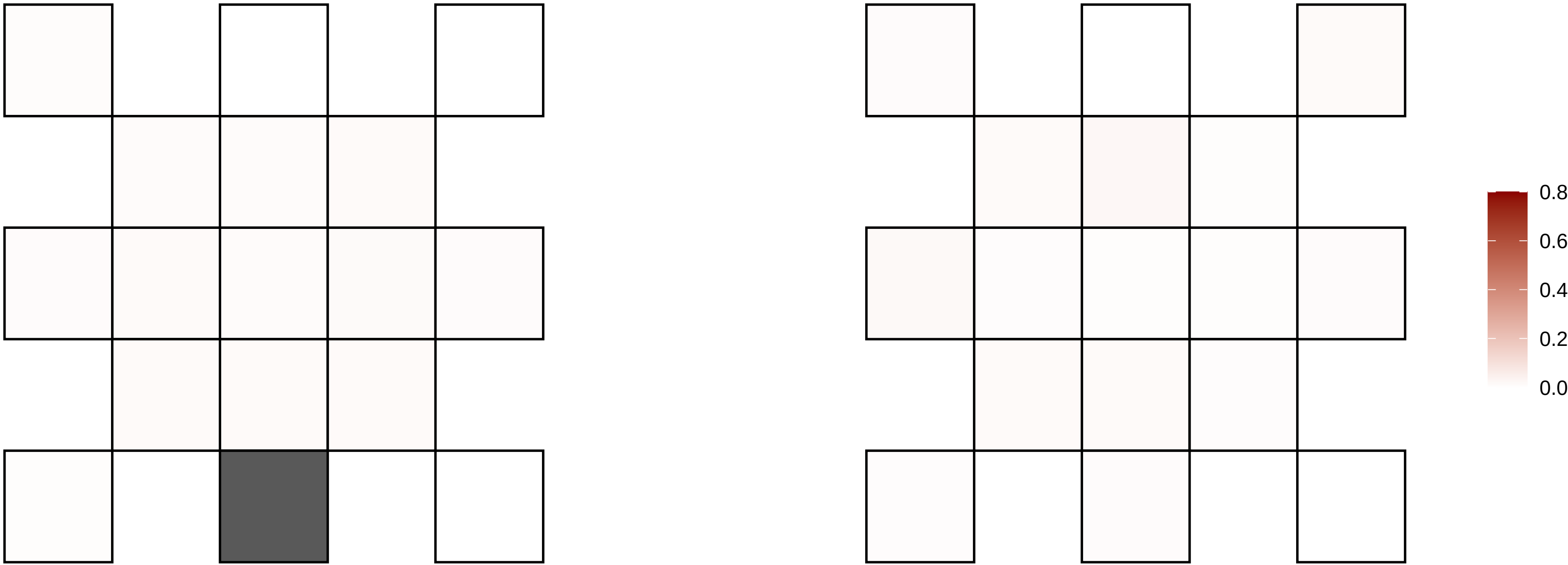

7−30962886−G−A

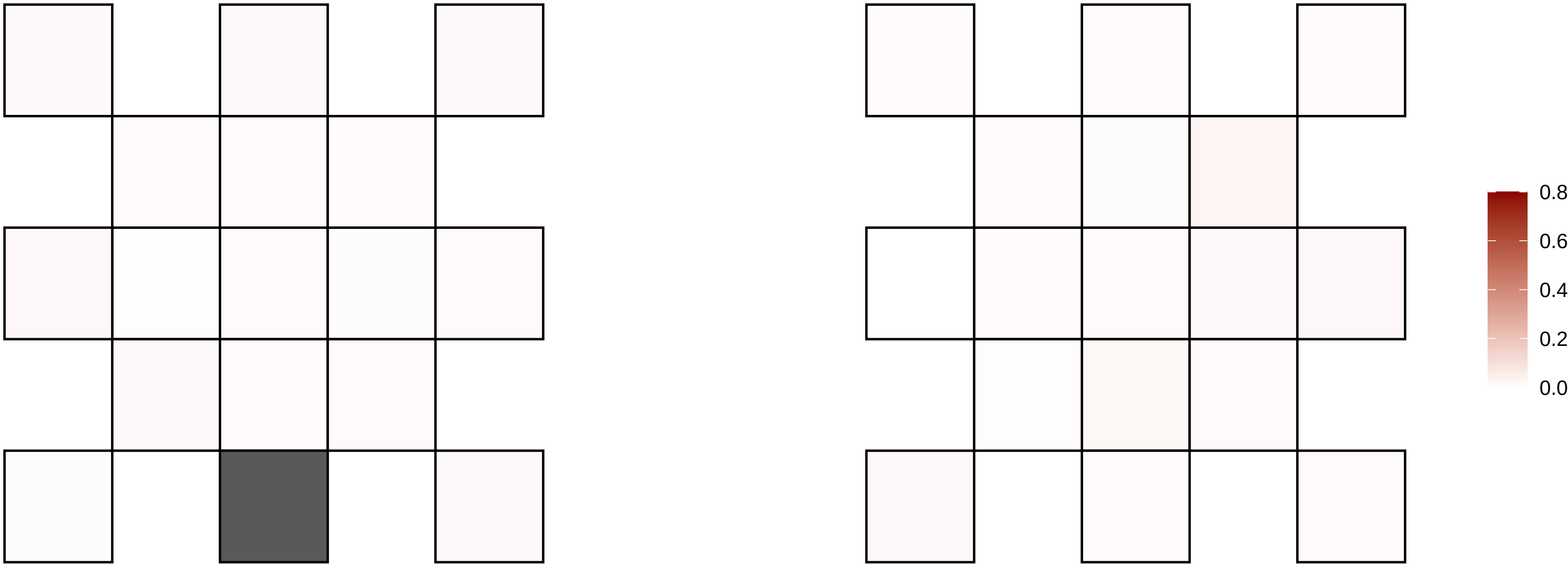

7−31462228−G−A

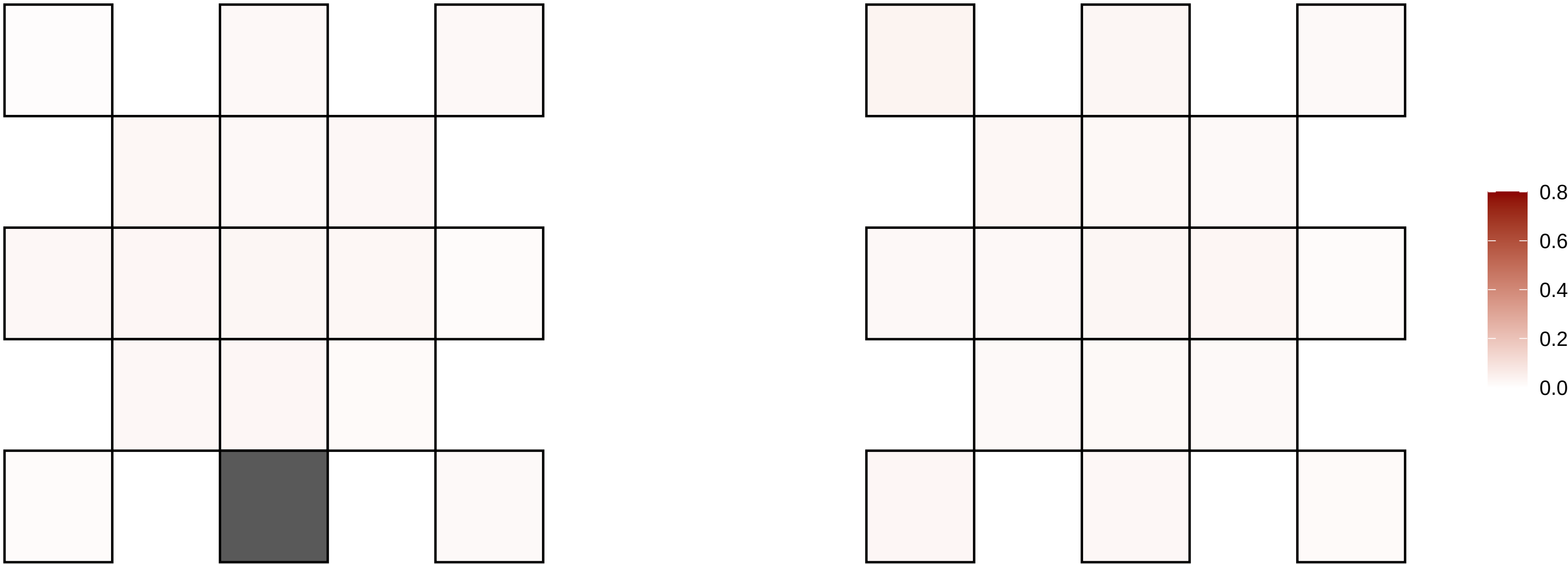

7−35583026−G−C

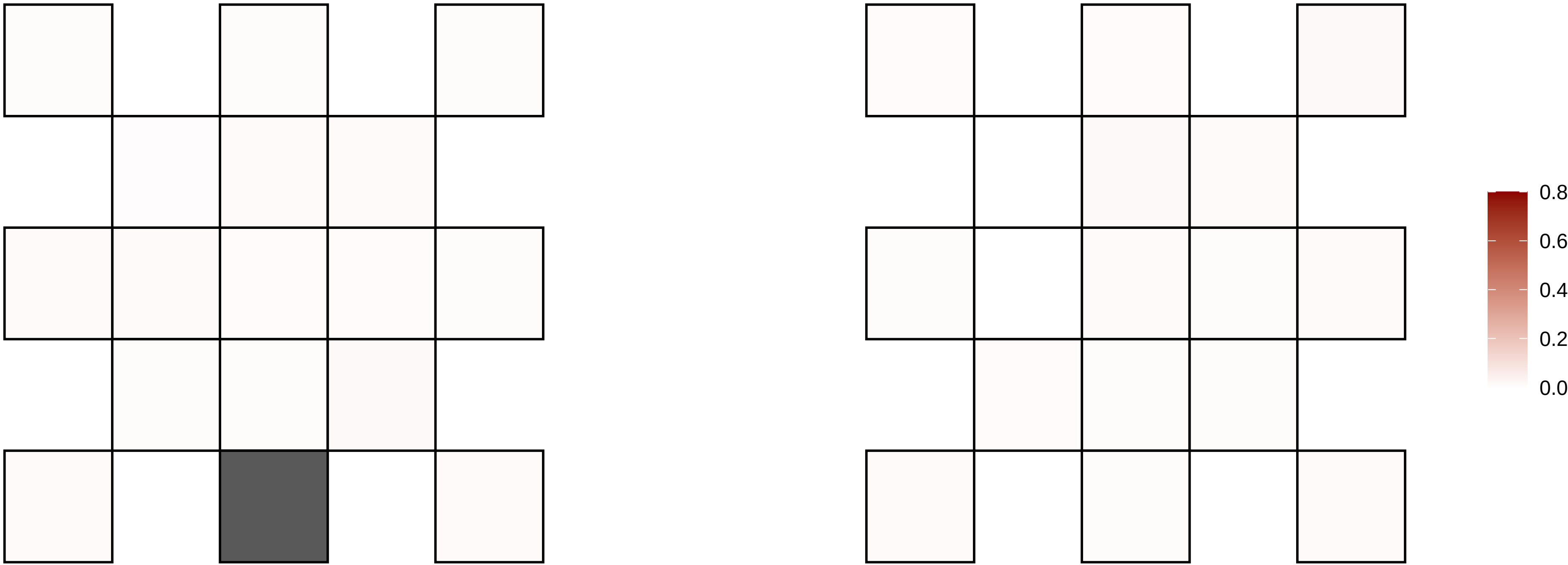

7−41735414−A−G

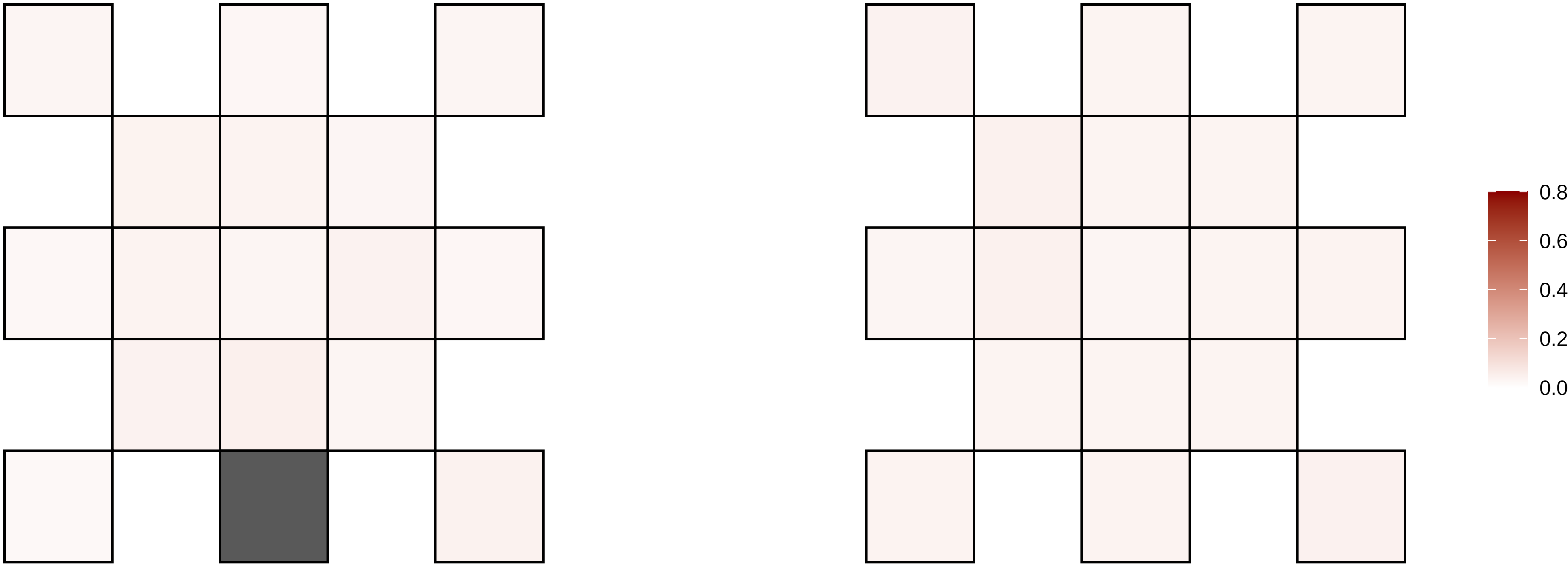

7−46623362−G−C

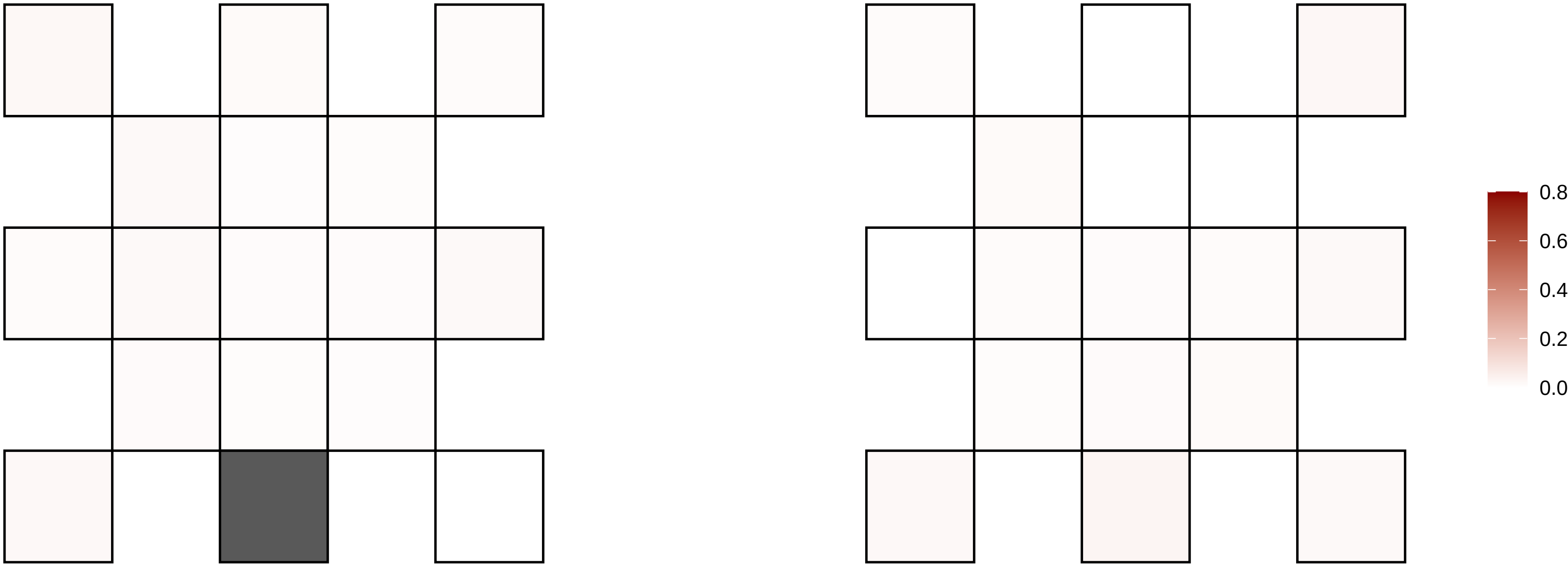

7−54130903−T−G

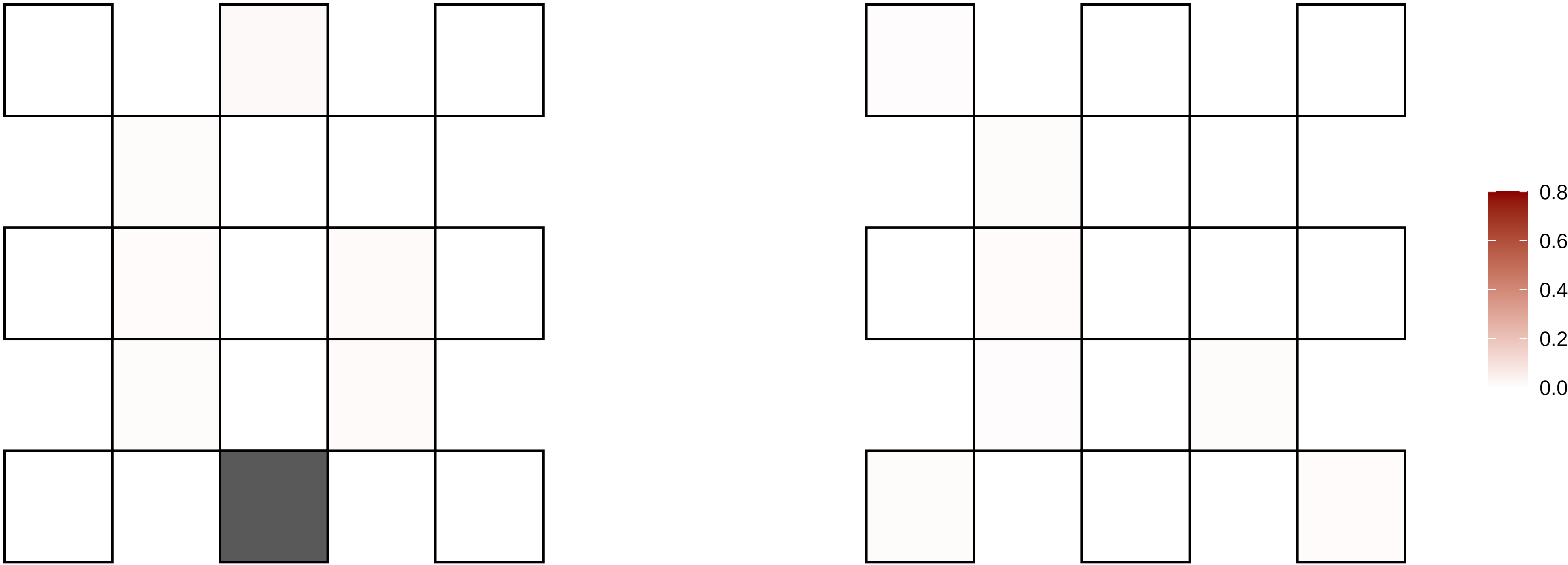

7−75695715−A−G

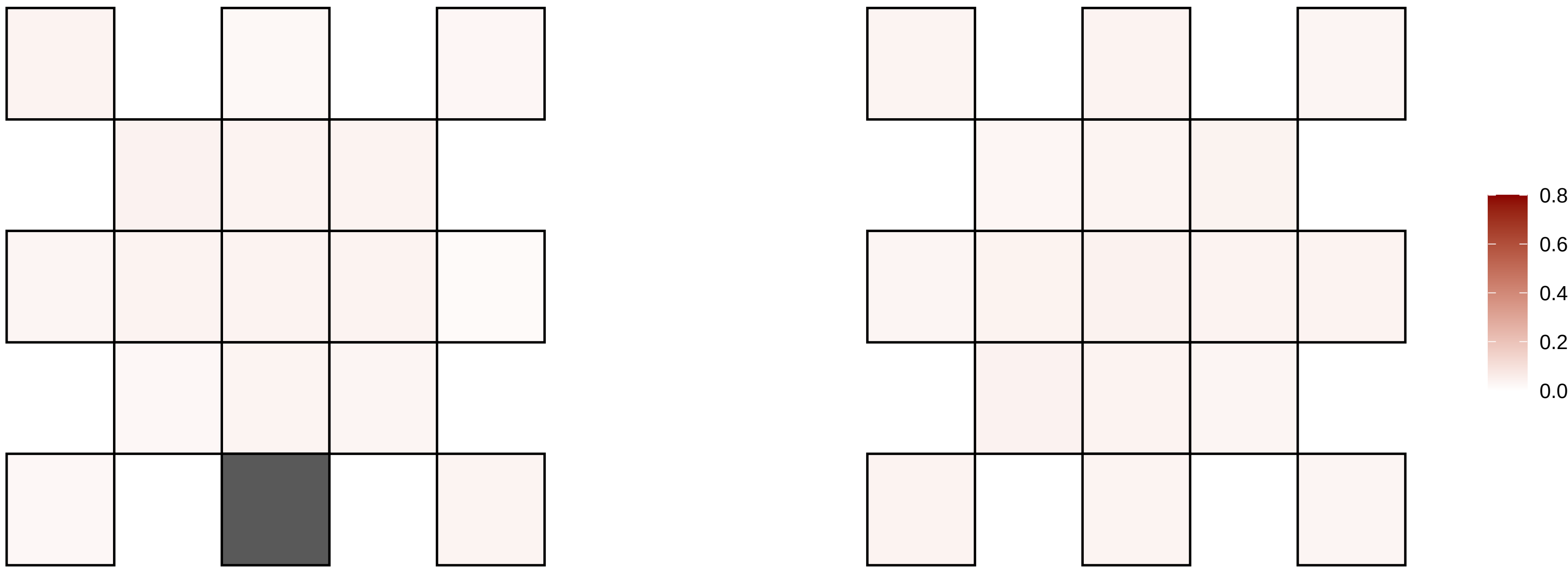

7−78830807−C−T

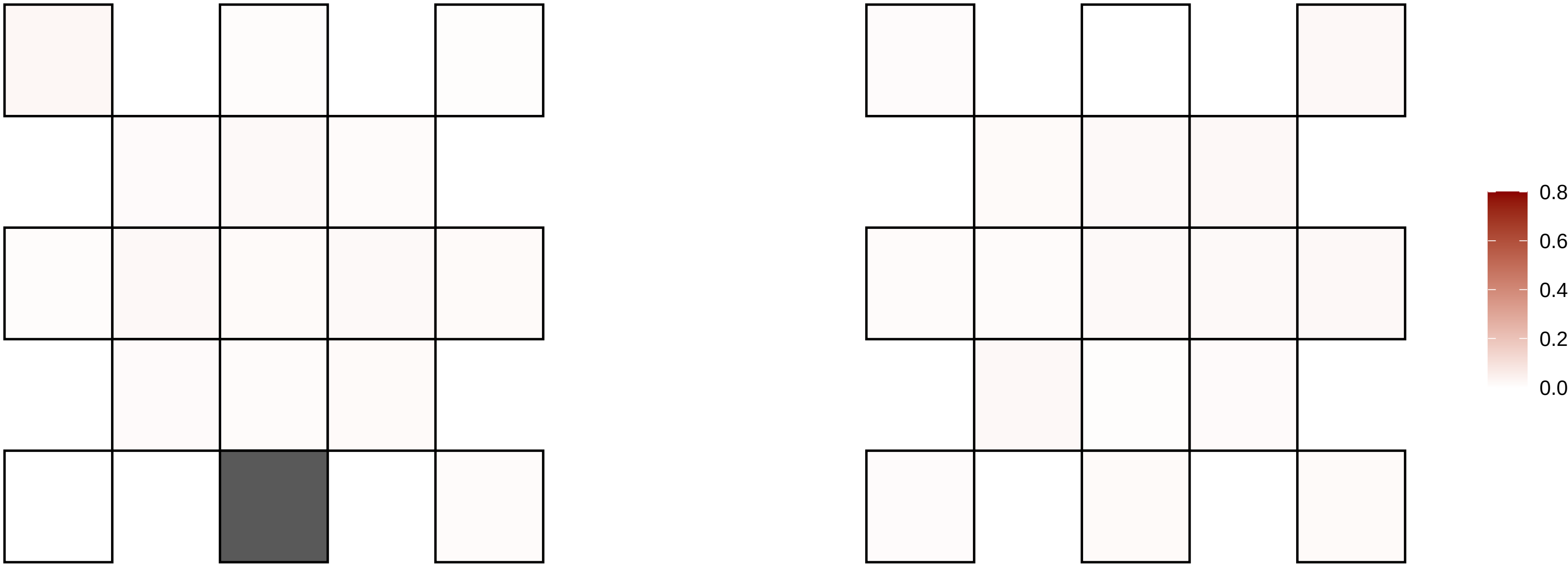

7−87643499−T−A

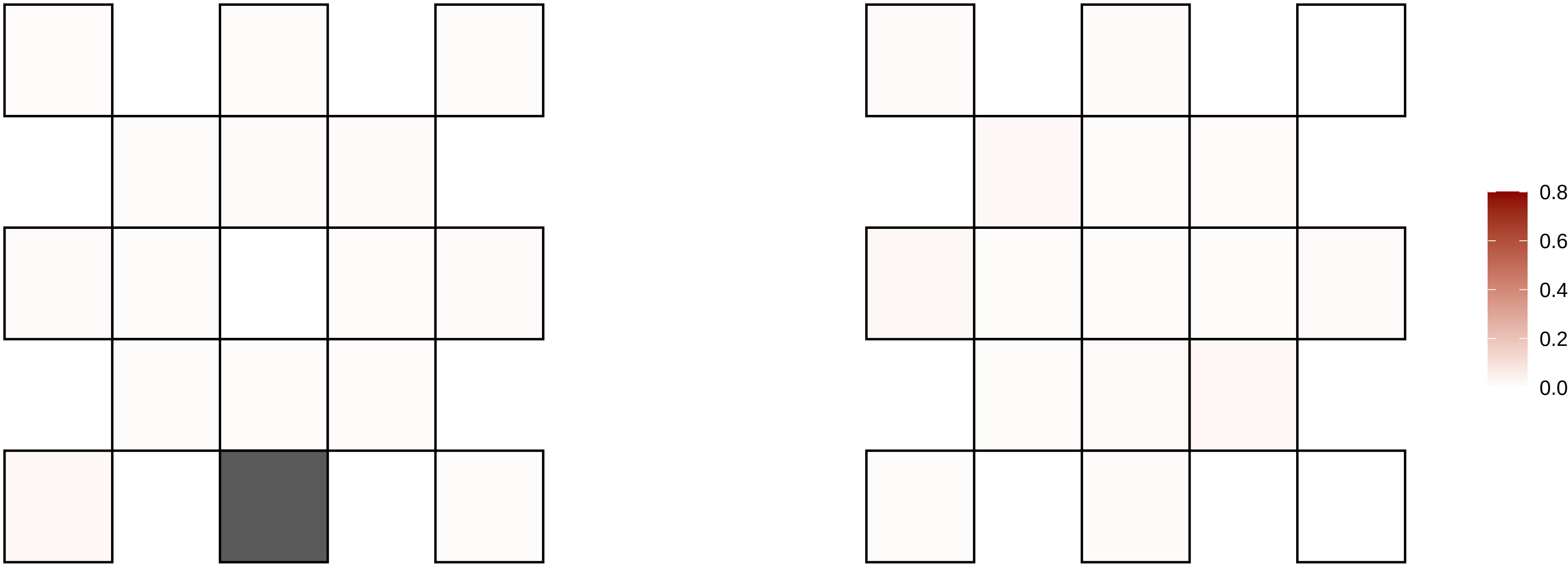

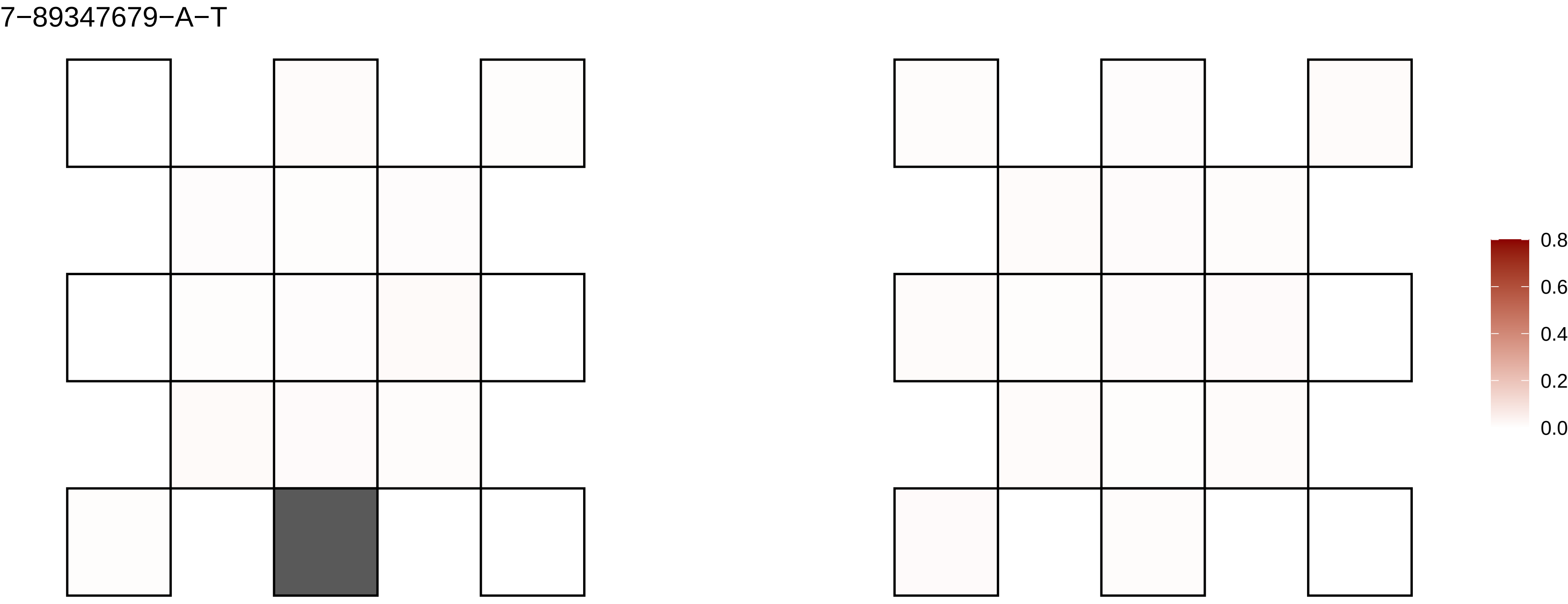
7−89347679−A−T
0.8
0.6
0.4
0.2
0.0

7−9192631−C−T

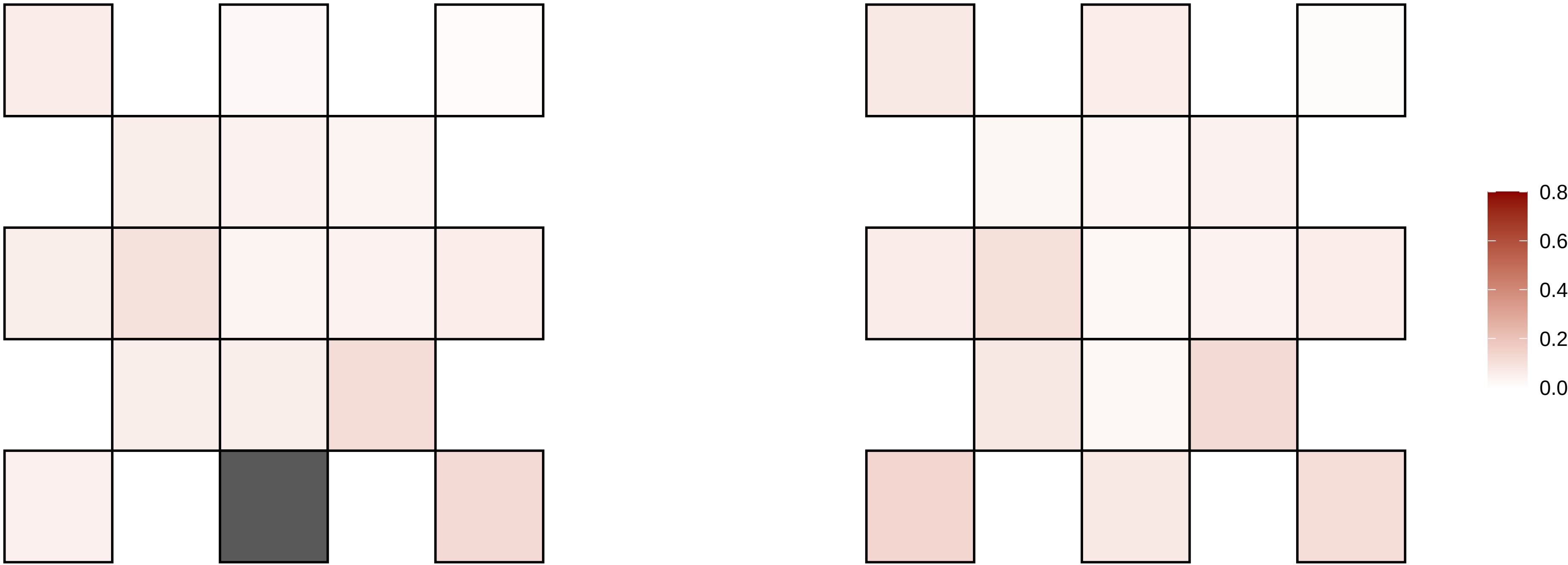

7−94919994−G−C

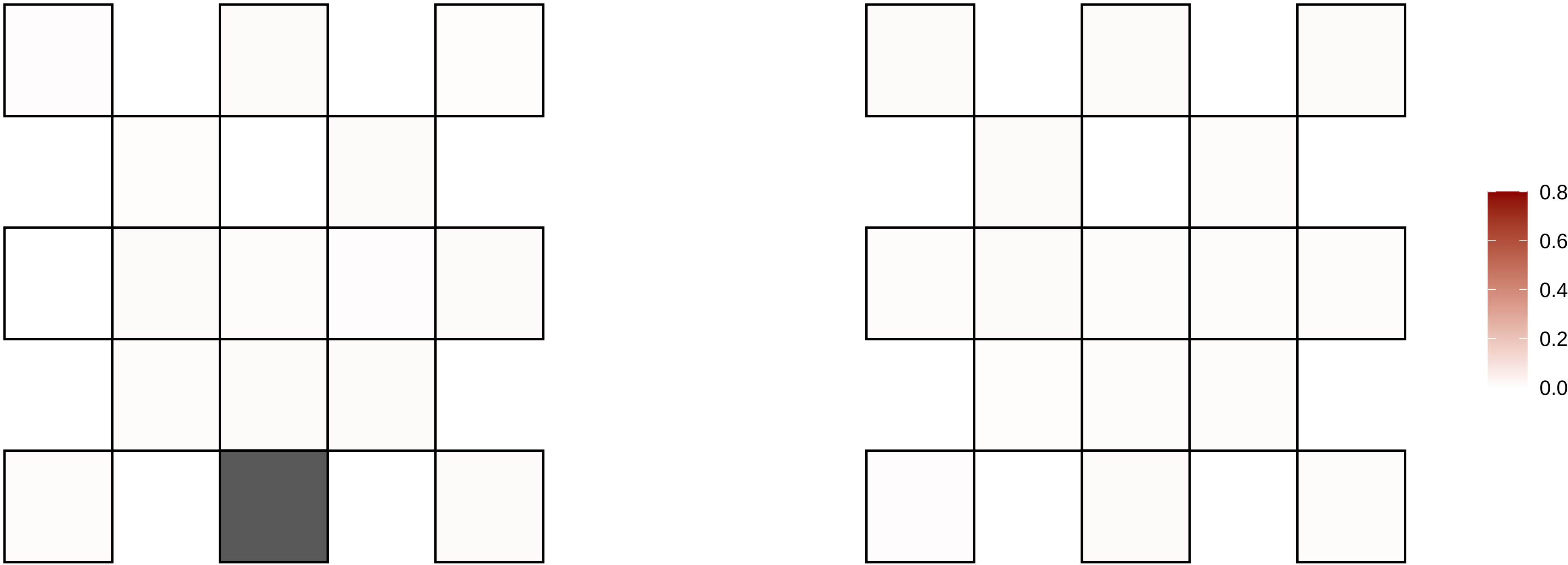

7−98311420−C−T

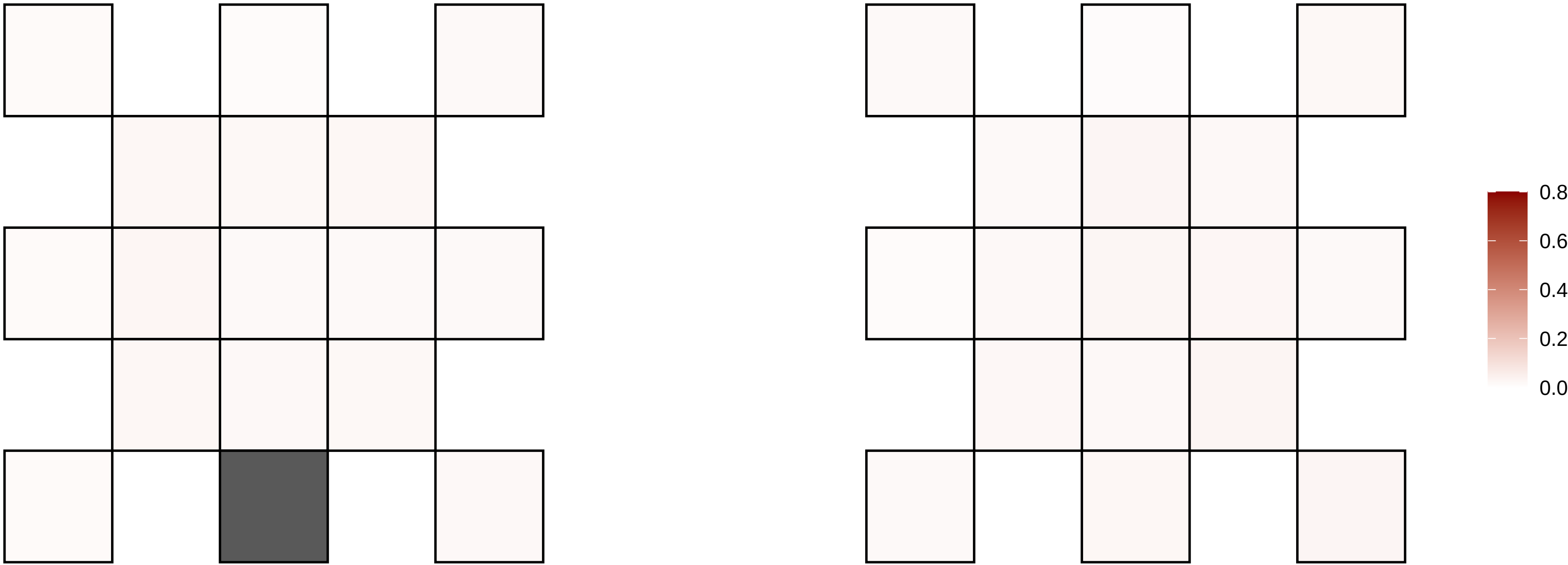

8−101369438−G−A

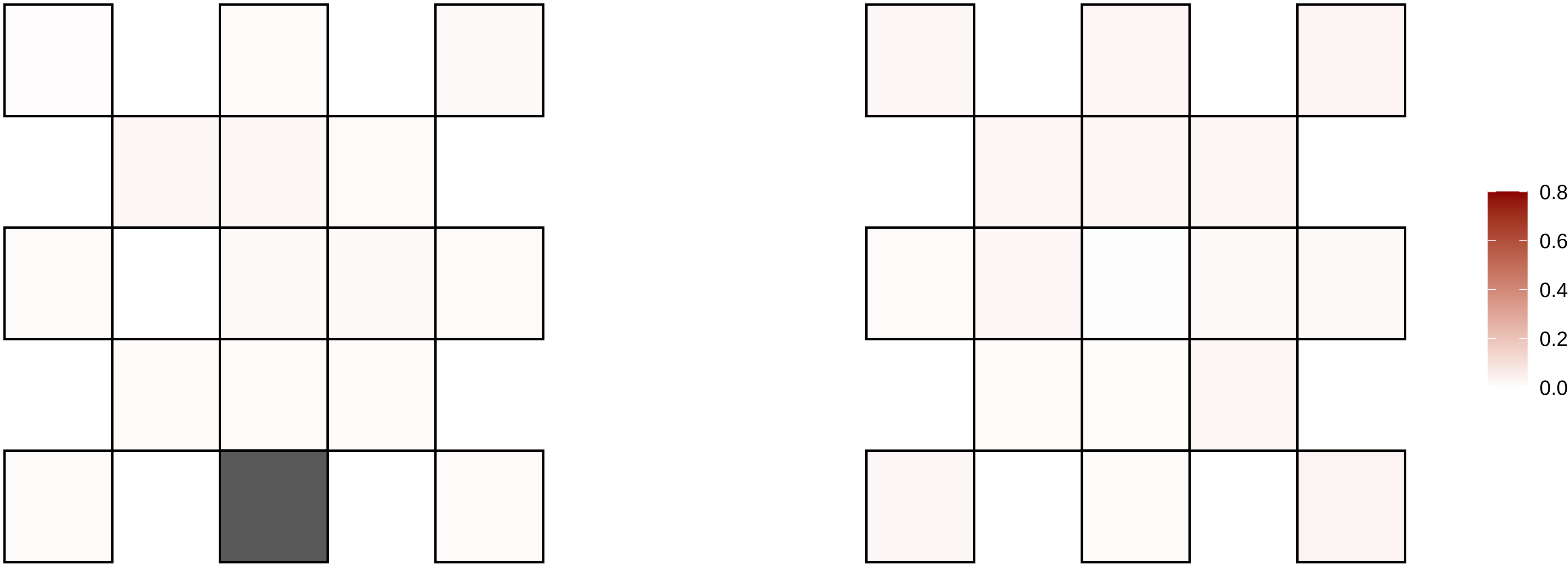

8−101774660−C−A

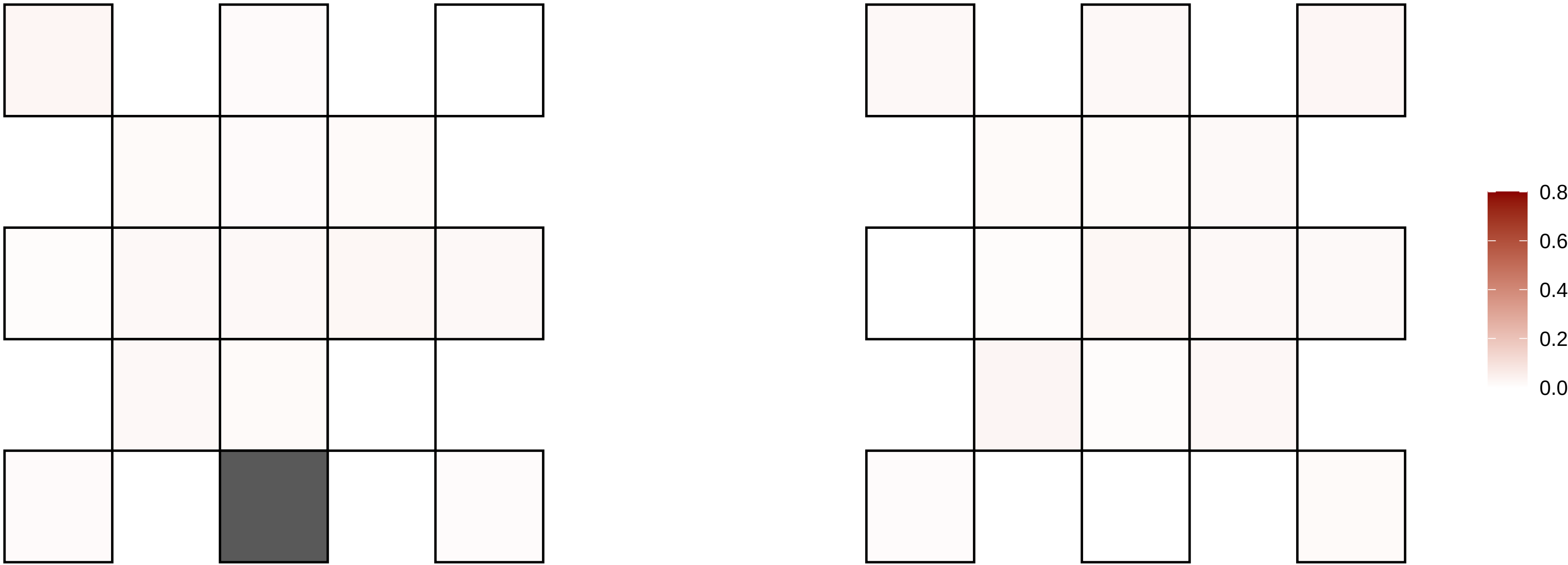

8−110907861−T−A

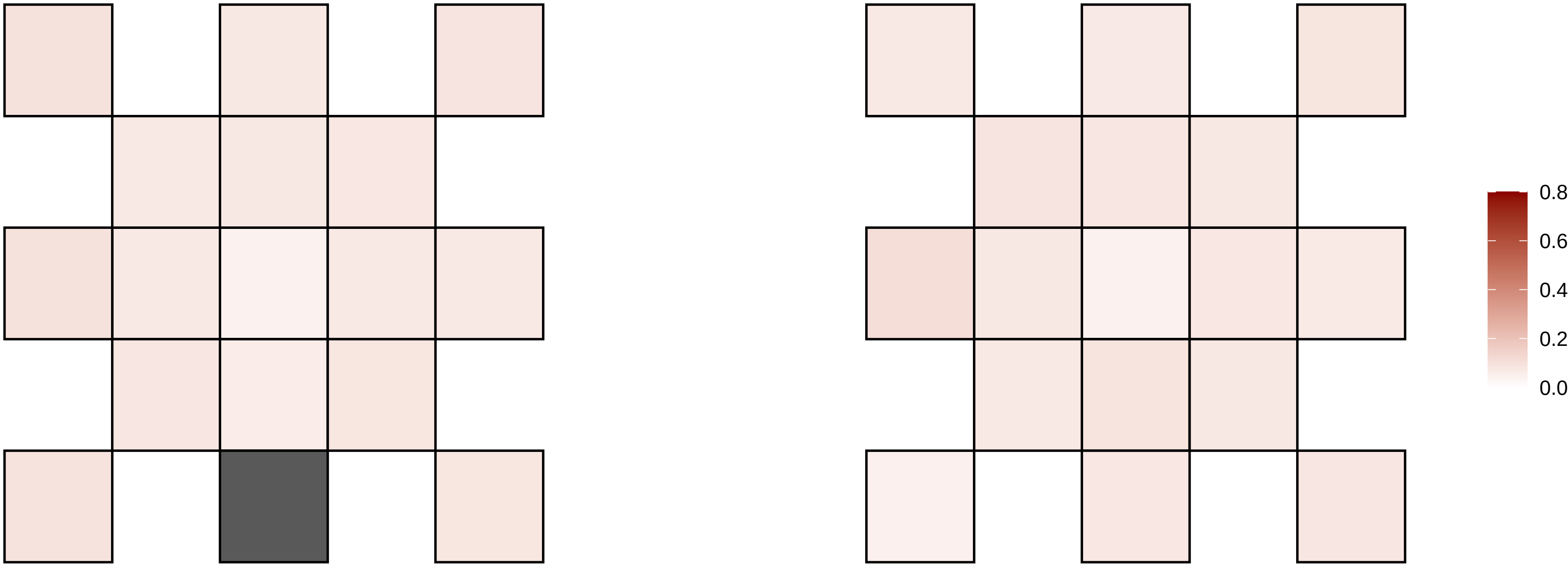

8−117647581−C−T

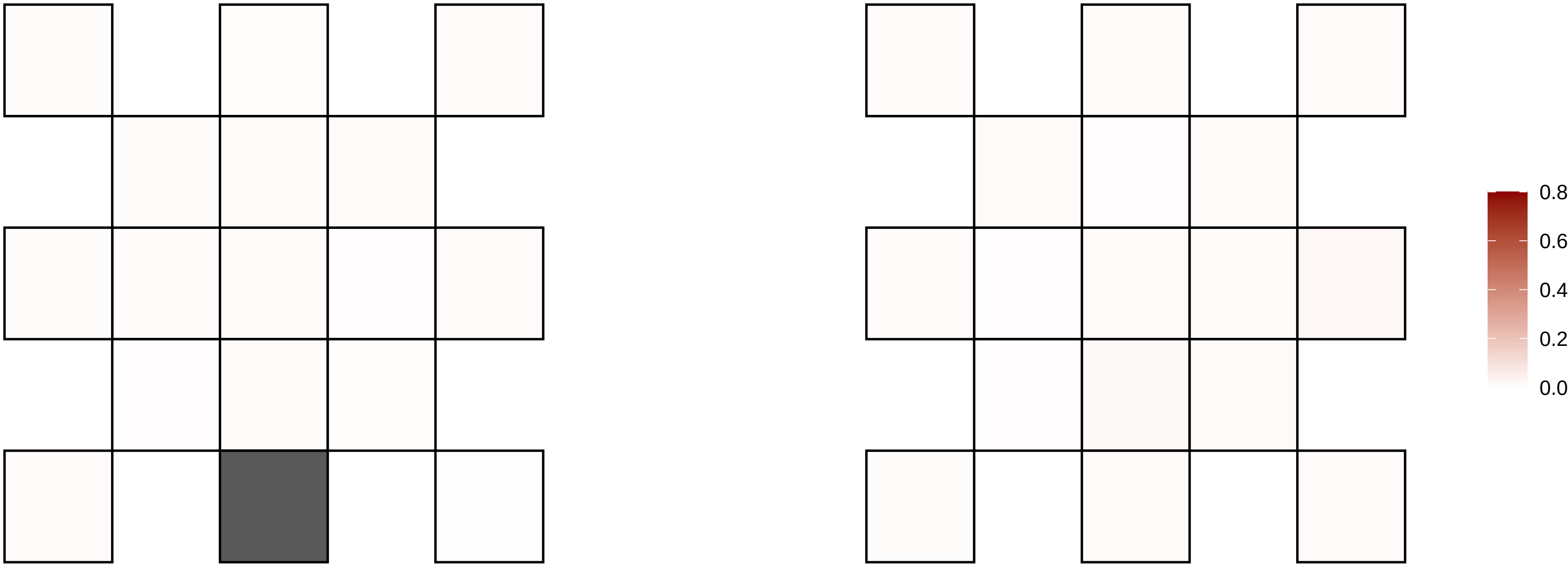

8−122970954−A−T

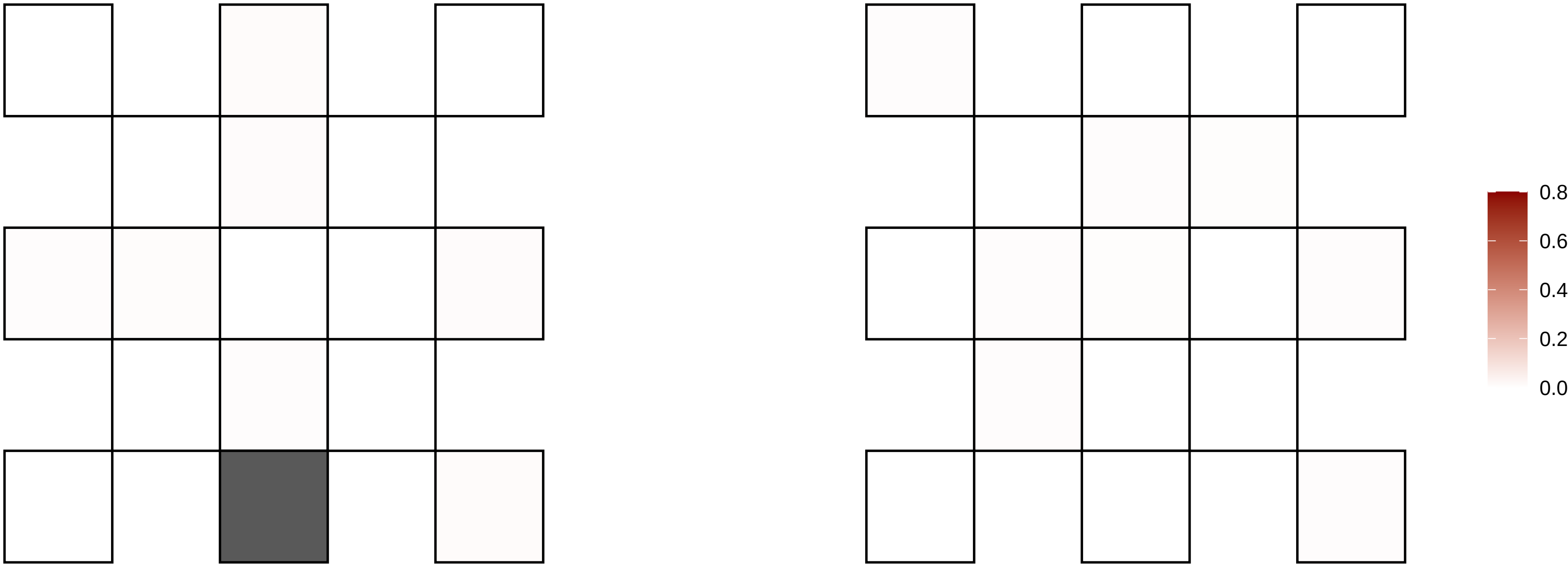

8−125891540−G−A

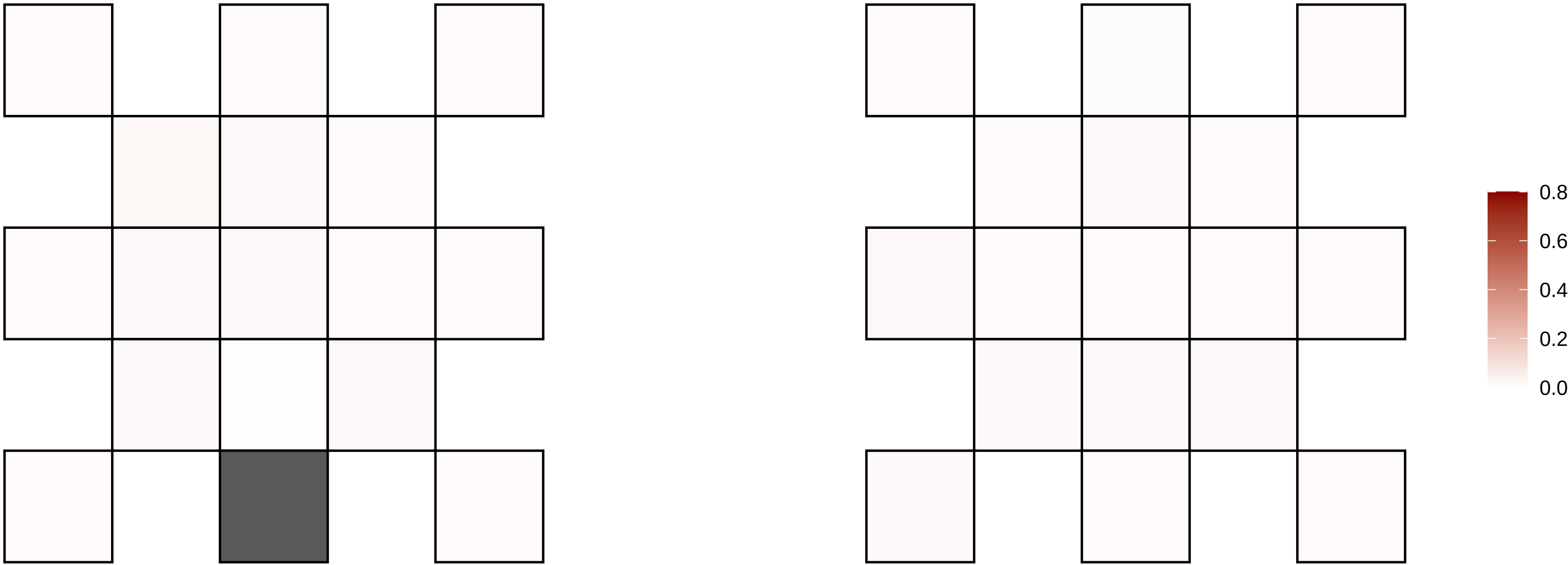

8−135596207−G−A

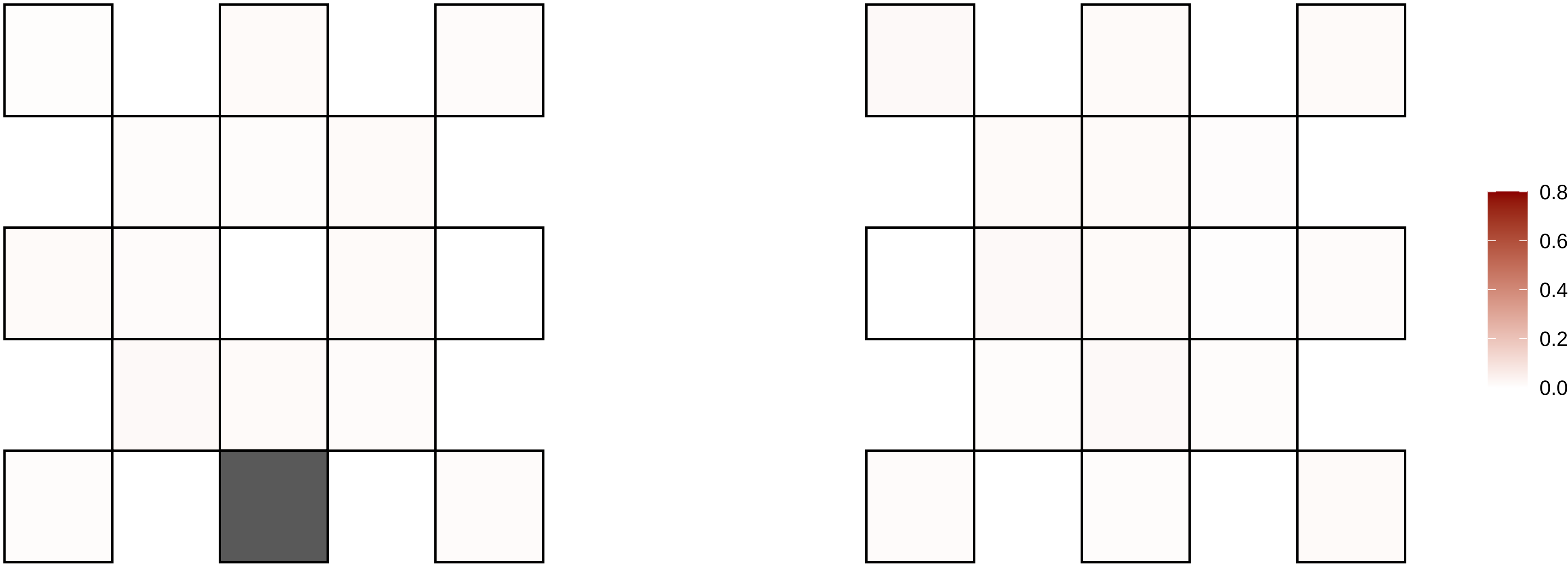

8−140637848−C−A

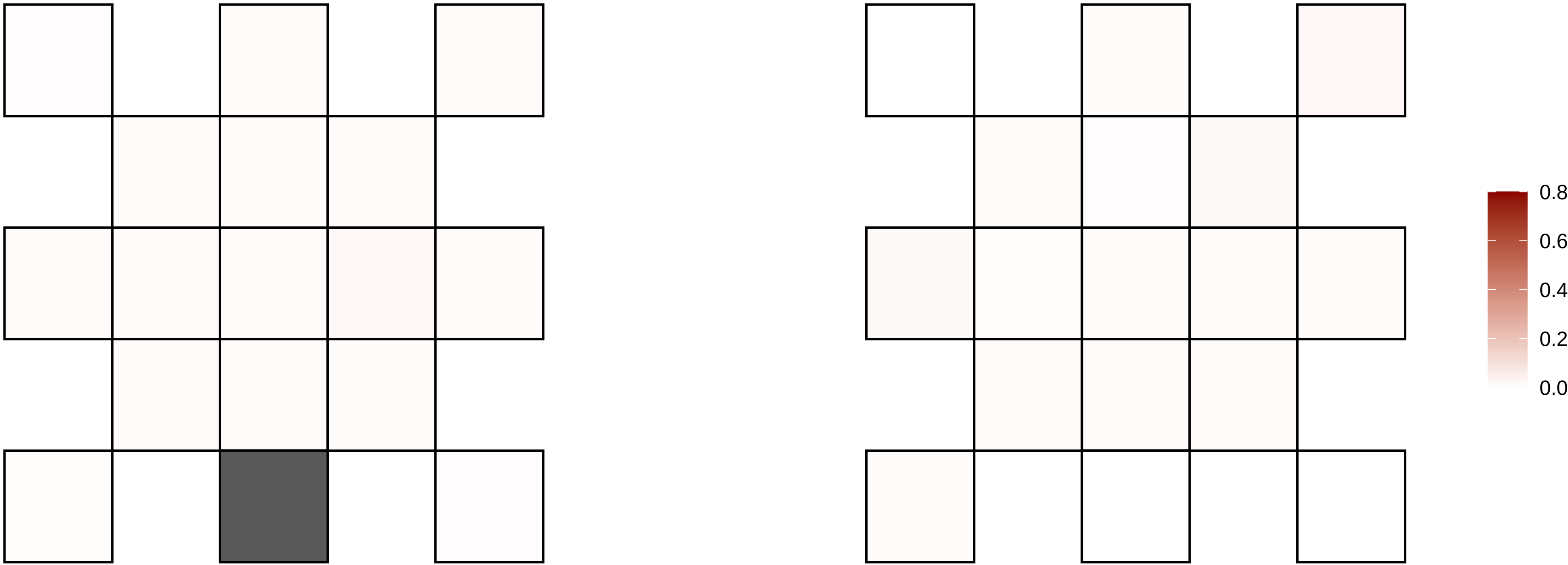

8−20681102−G−A

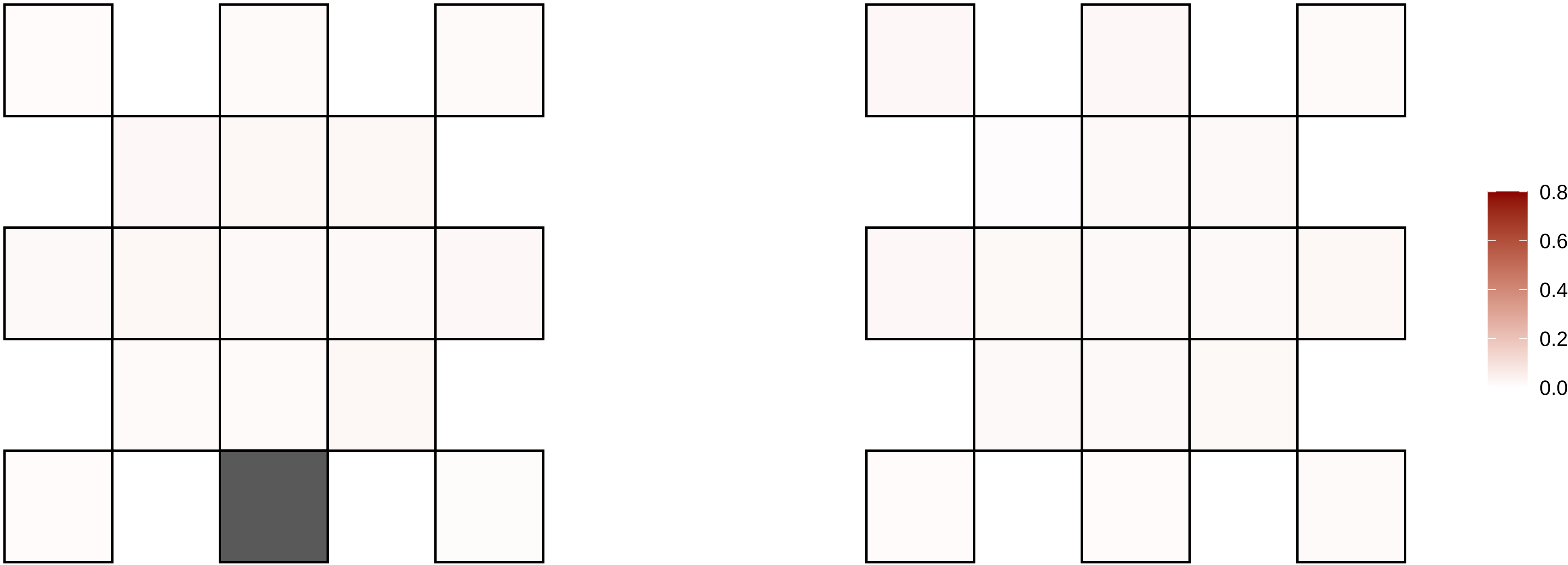

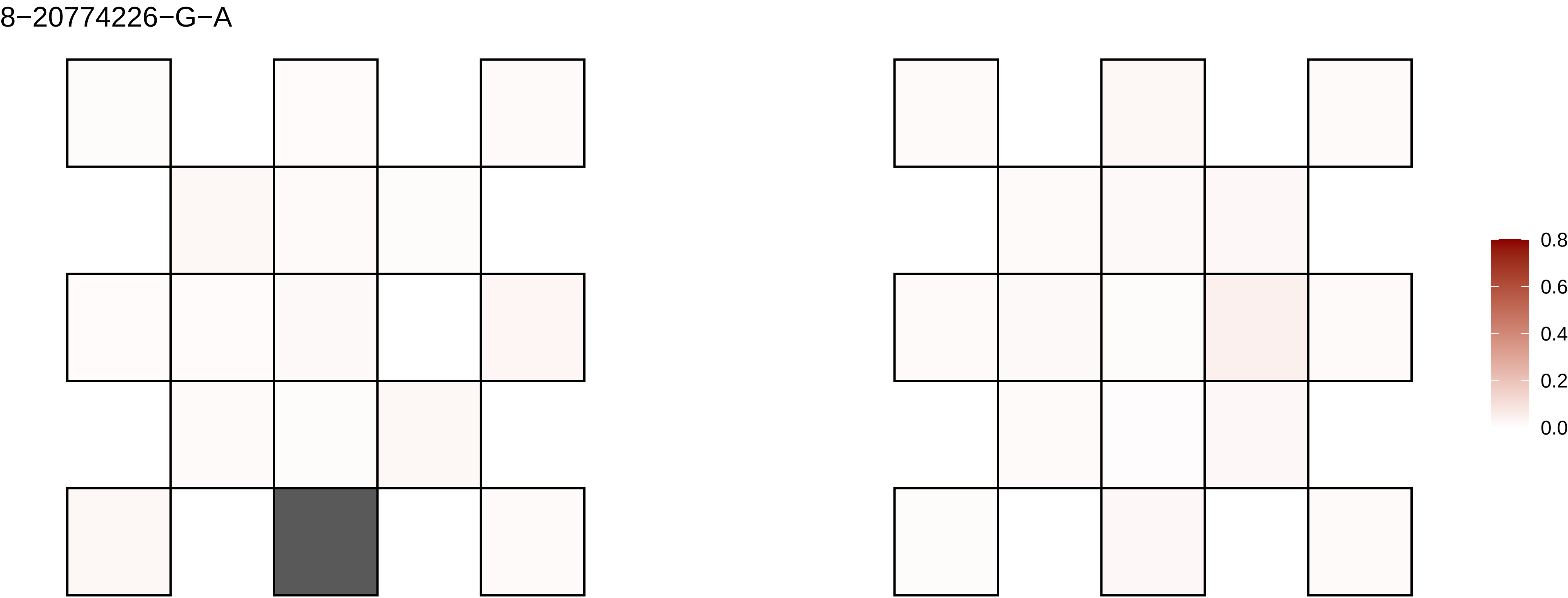
8−20774226−G−A
0.8
0.6
0.4
0.2
0.0

8−24188792−G−A

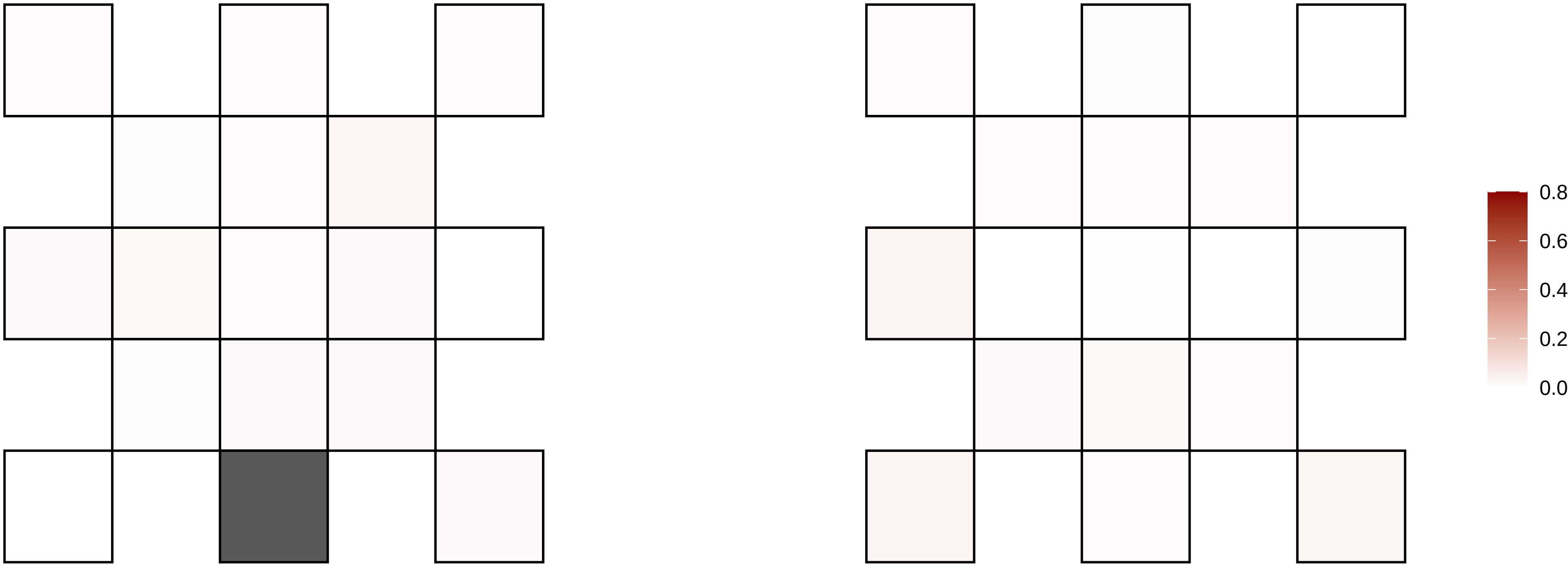

8−29866666−C−T

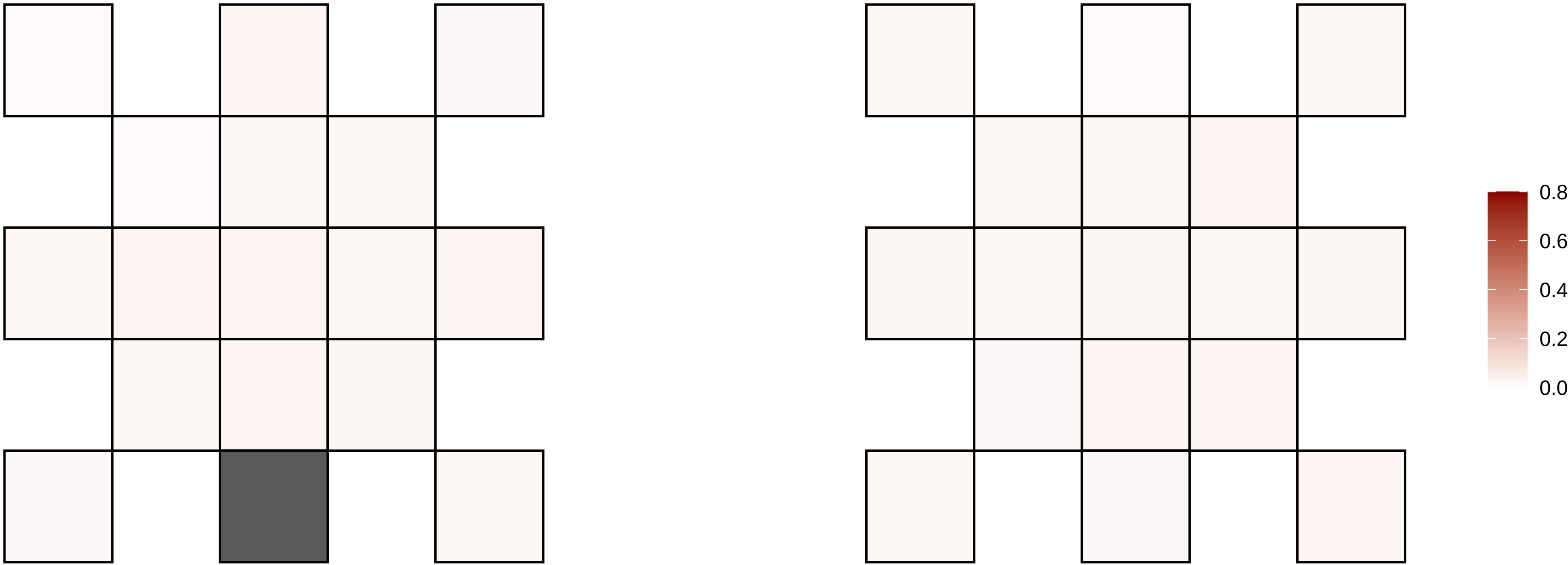

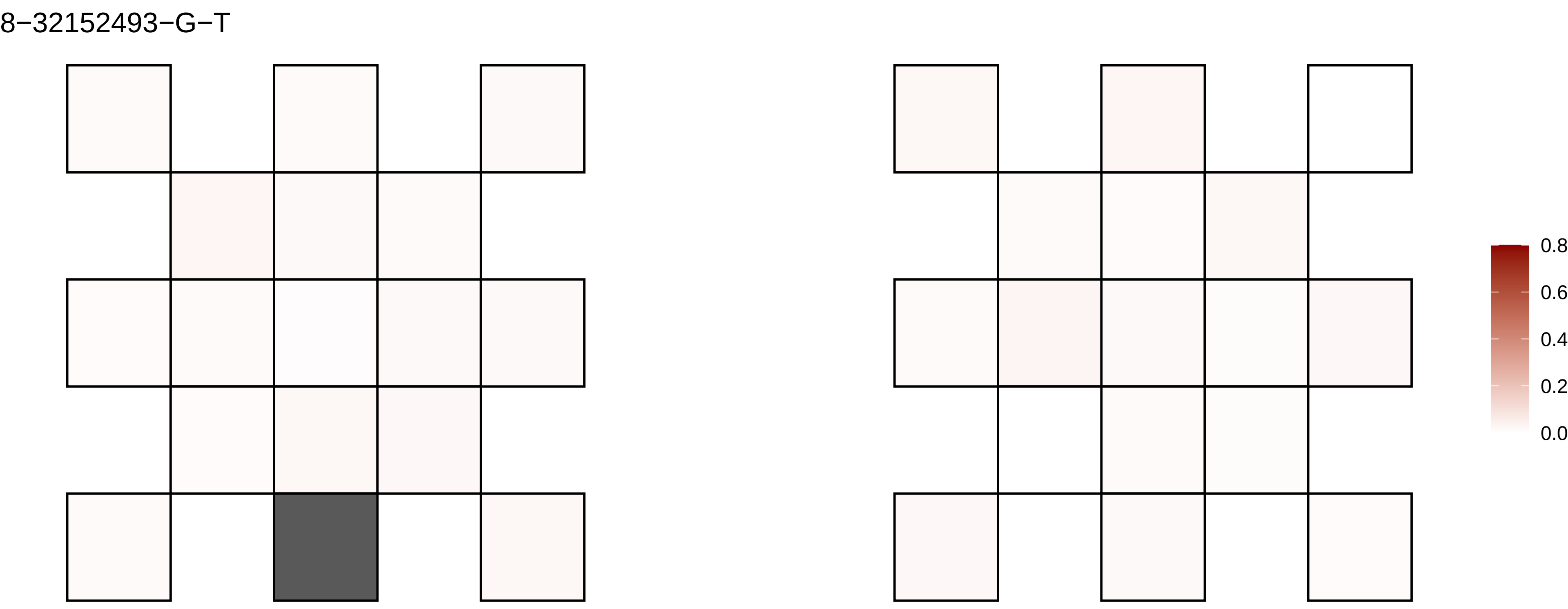
8−32152493−G−T
0.8
0.6
0.4
0.2
0.0

8−3439977−G−A

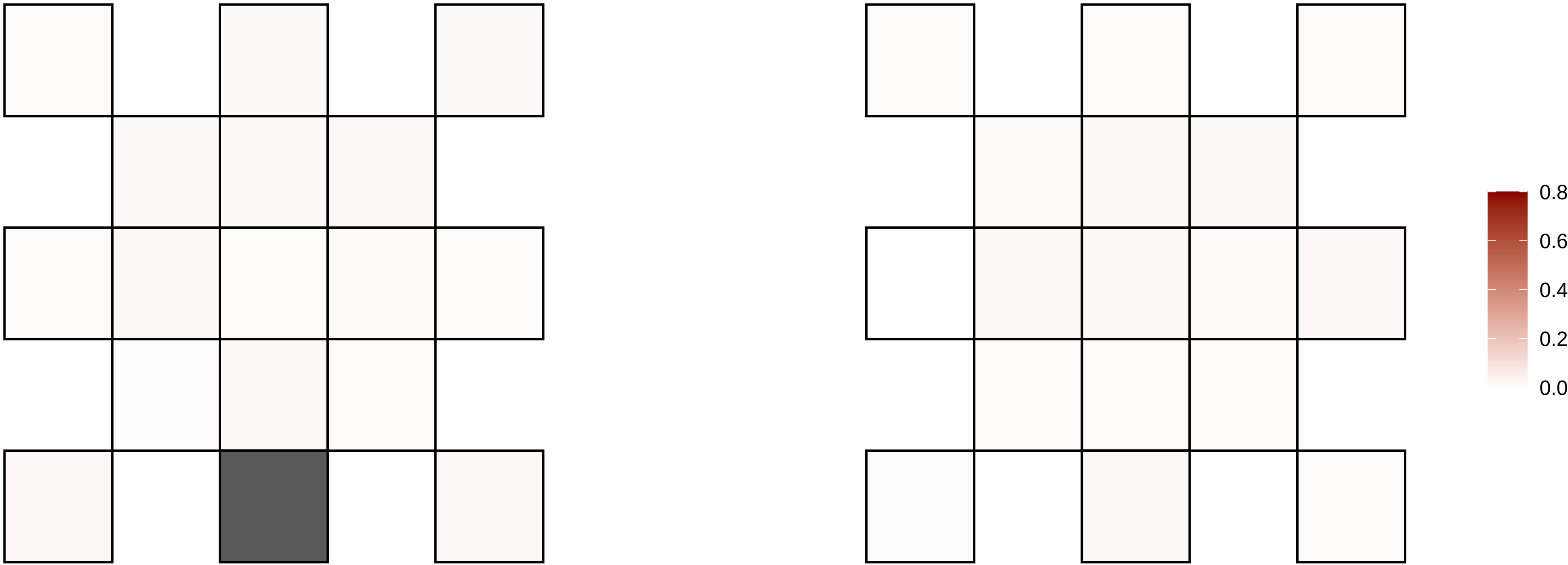

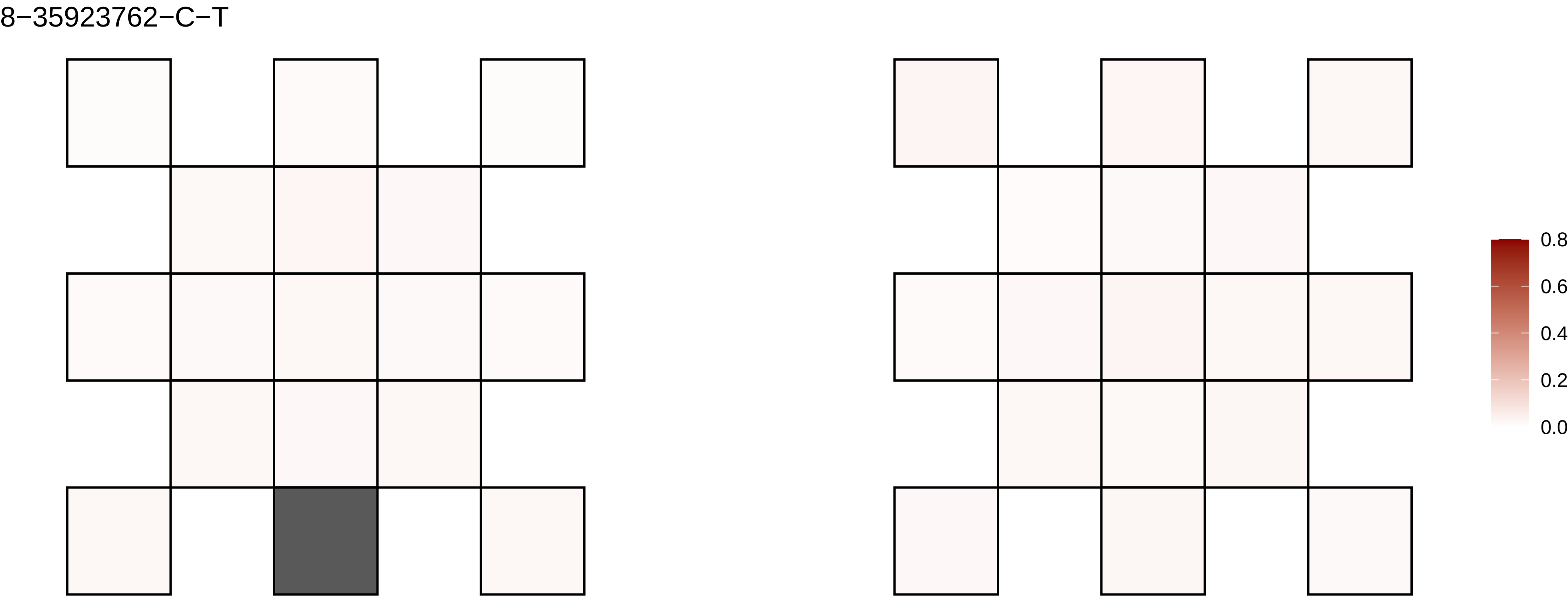
8−35923762−C−T
0.8
0.6
0.4
0.2
0.0

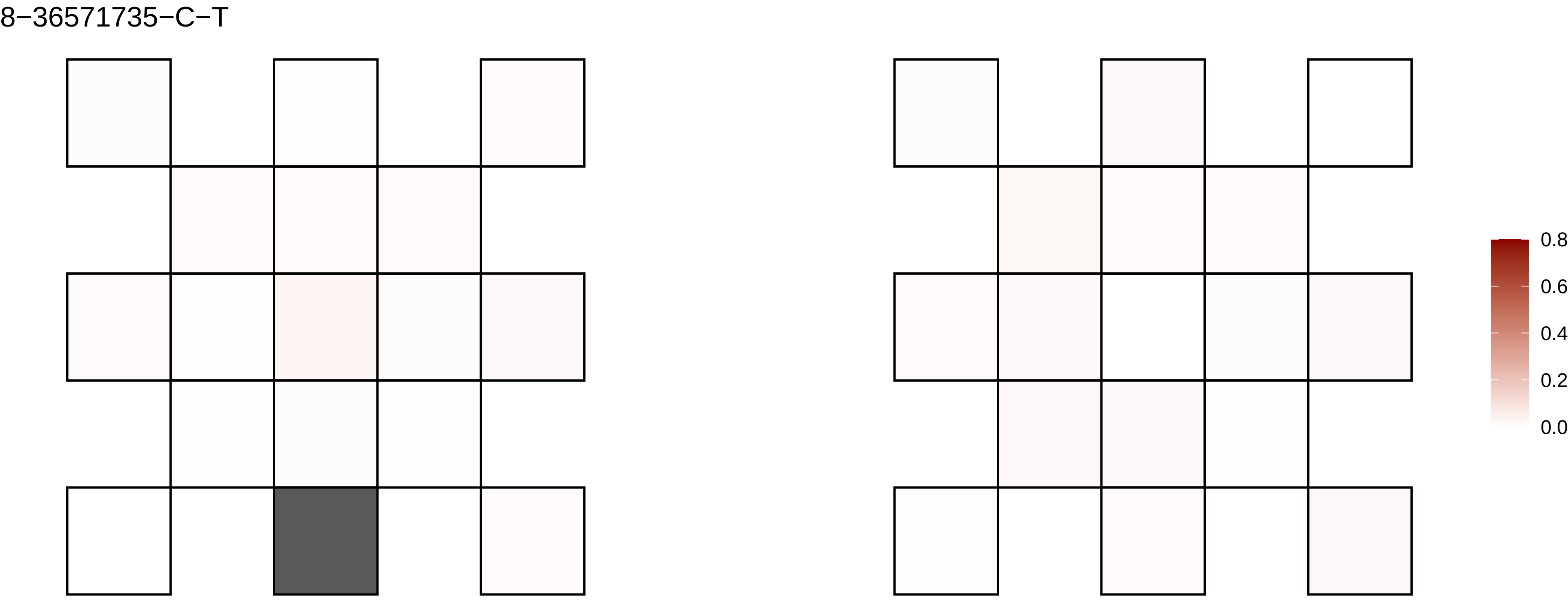

8−36571735−C−T
0.8
0.6
0.4
0.2
0.0

8−41303253−C−T

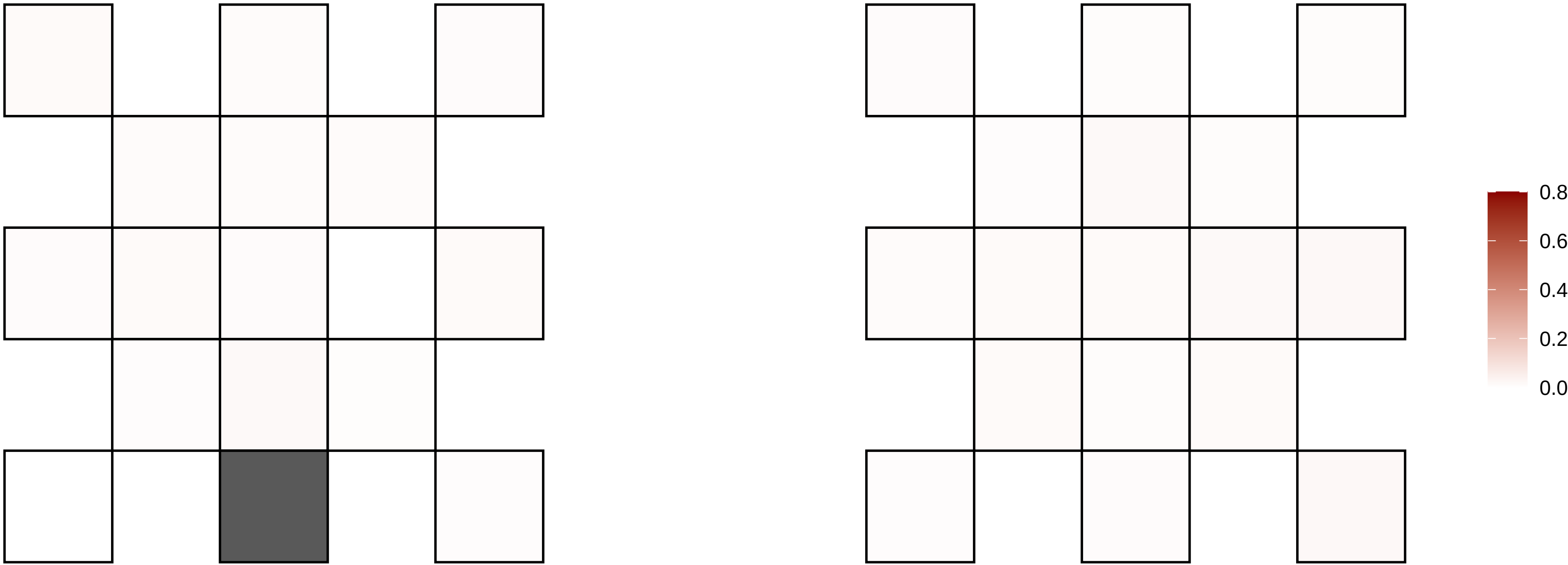

8−4601722−C−G

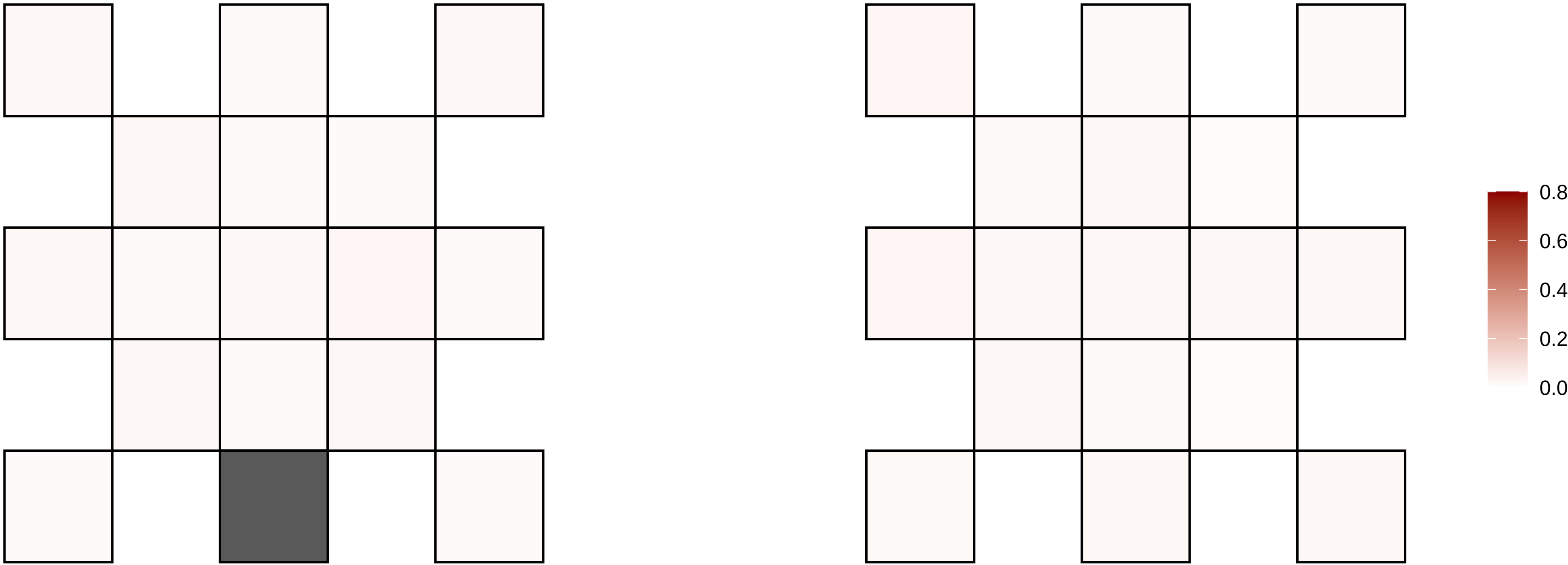

8−49587201−C−T

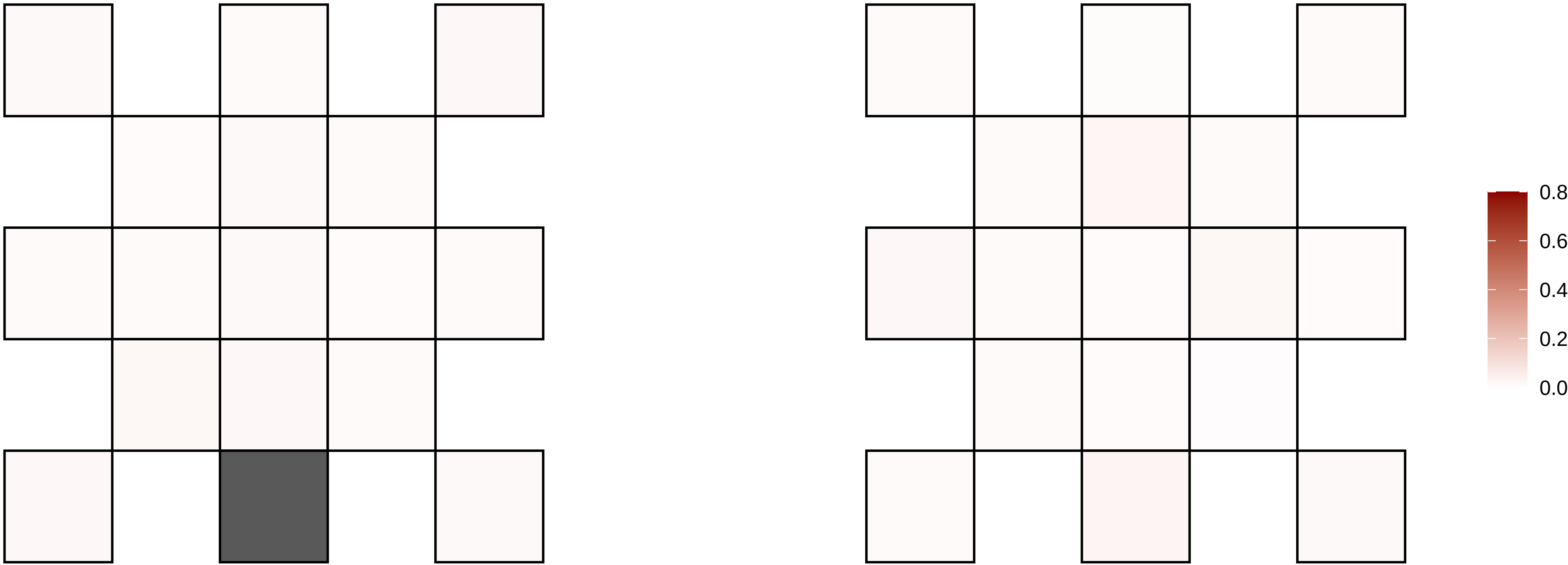

8−49636775−C−A

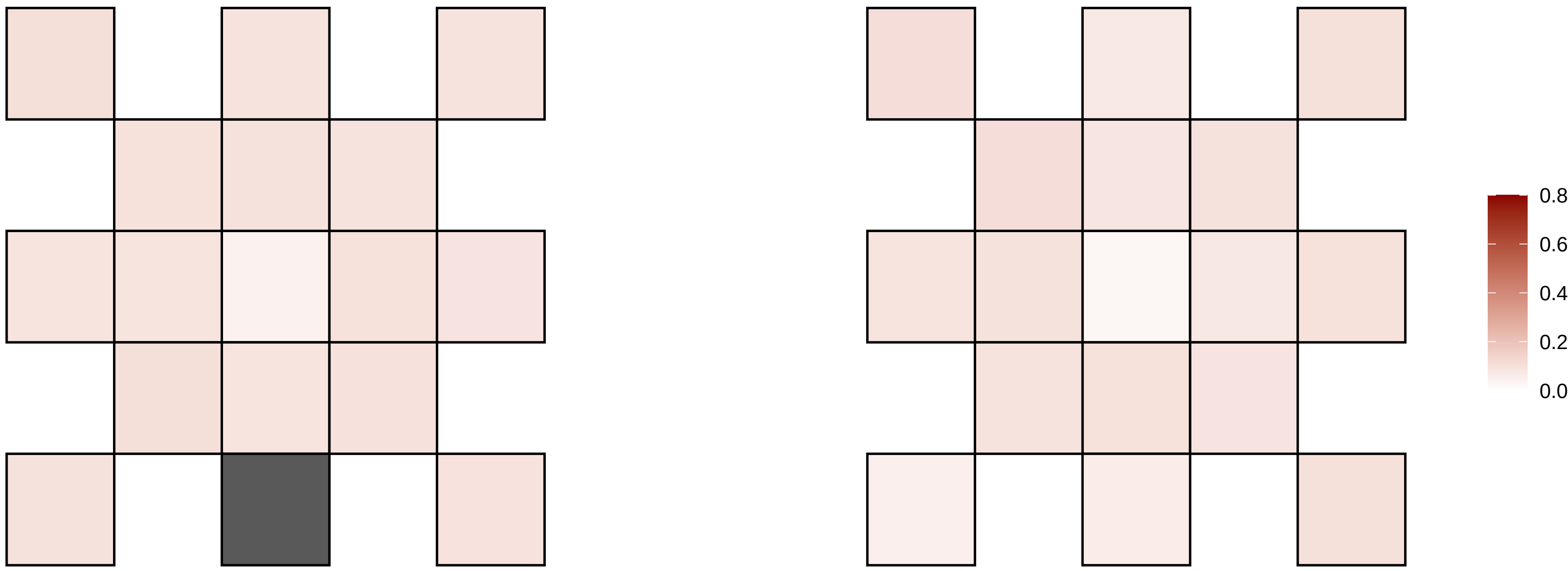

8−50752909−G−C

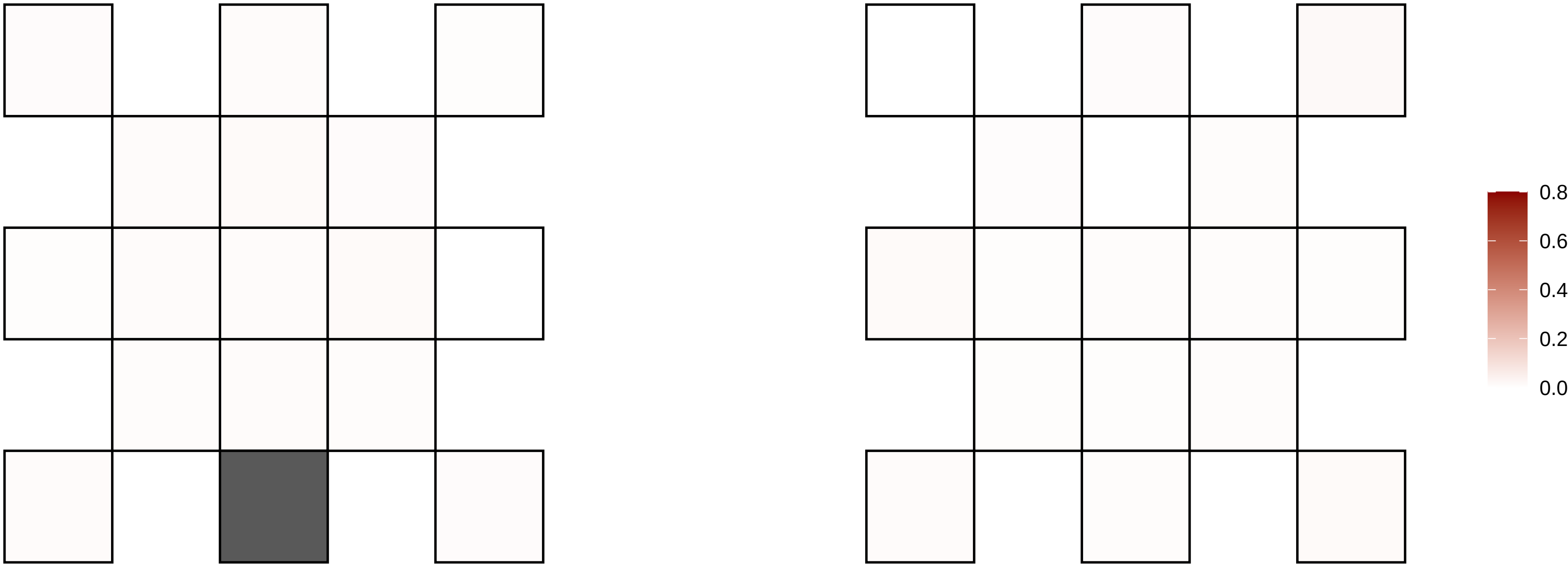

8−56154213−C−T

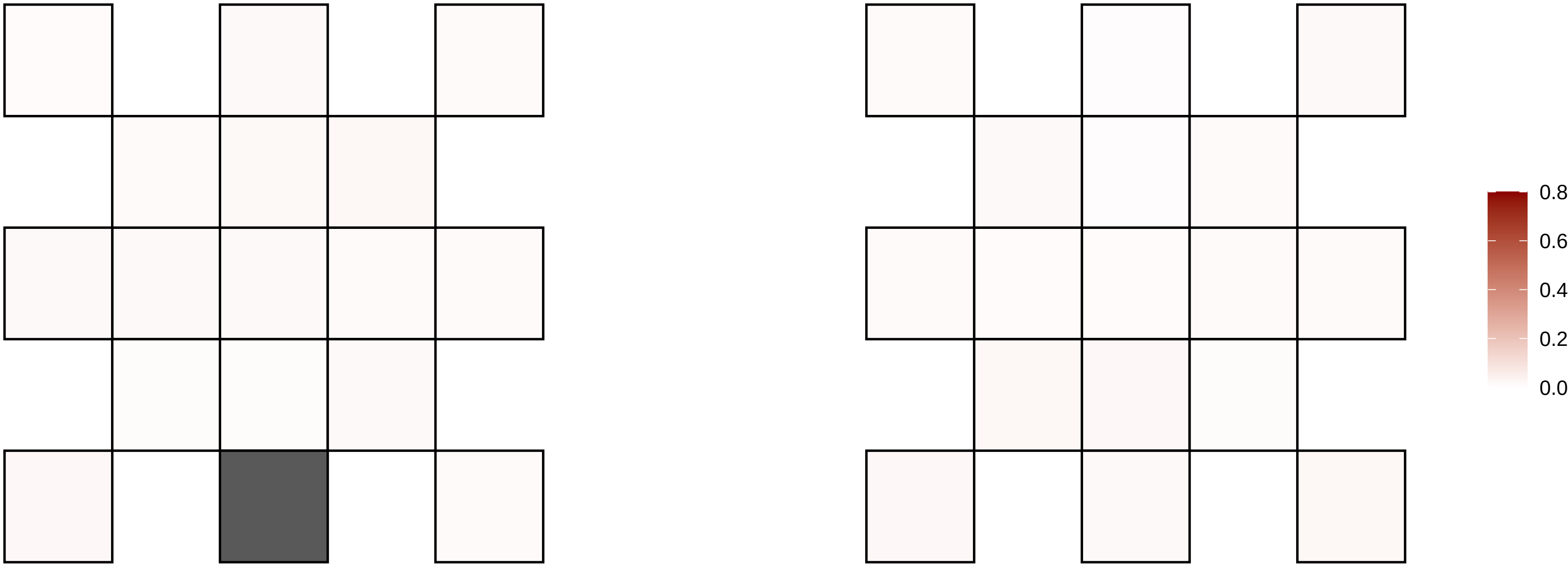

8−57499201−TG−T

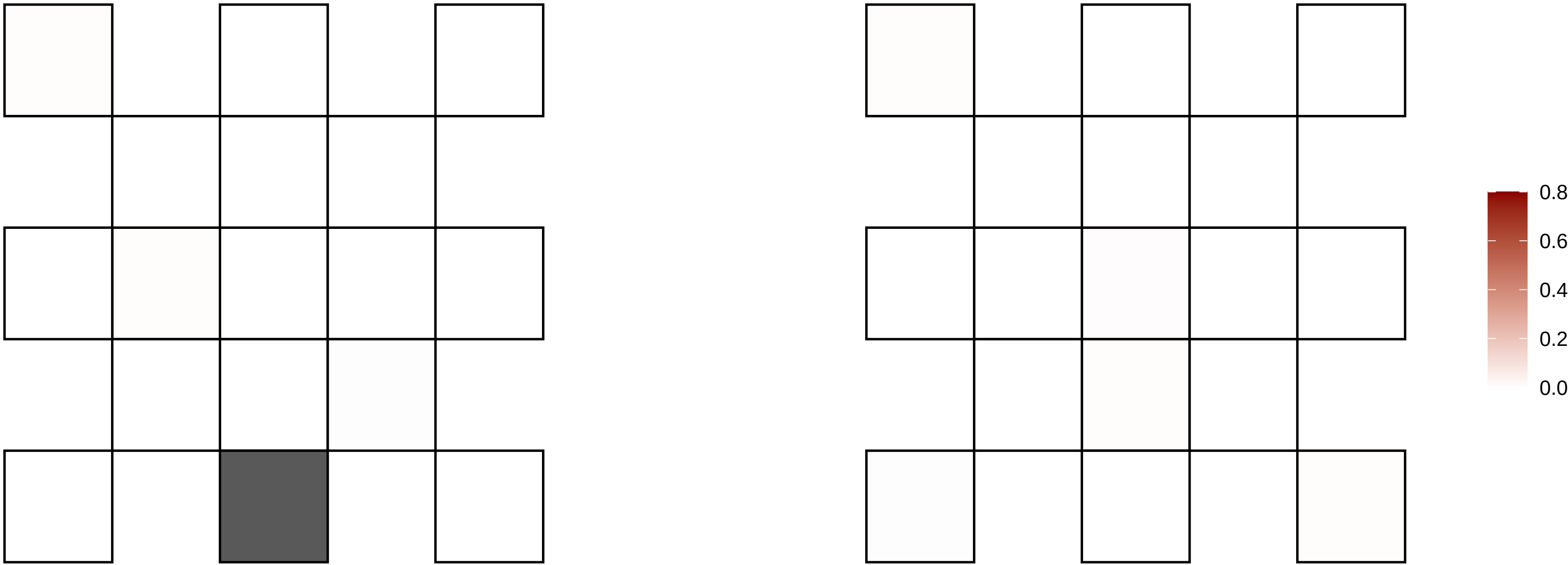

8−58808555−G−A

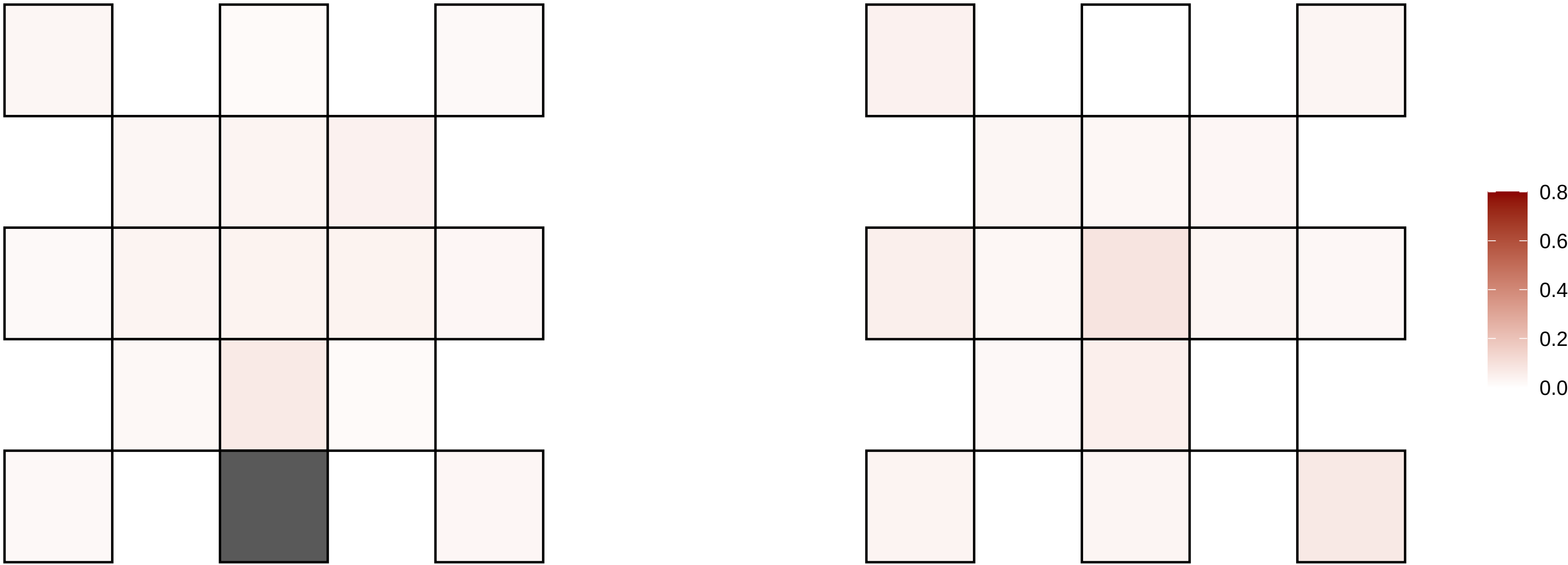

8−62438170−G−A

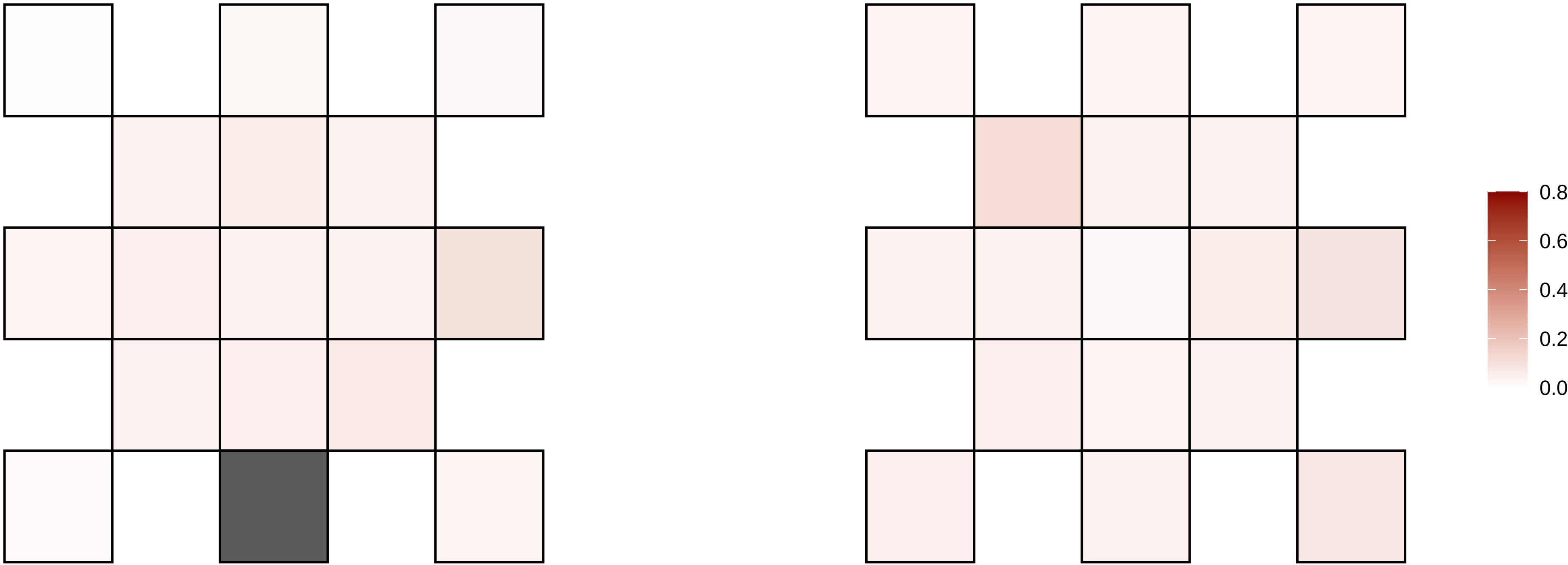

8−65149924−C−T

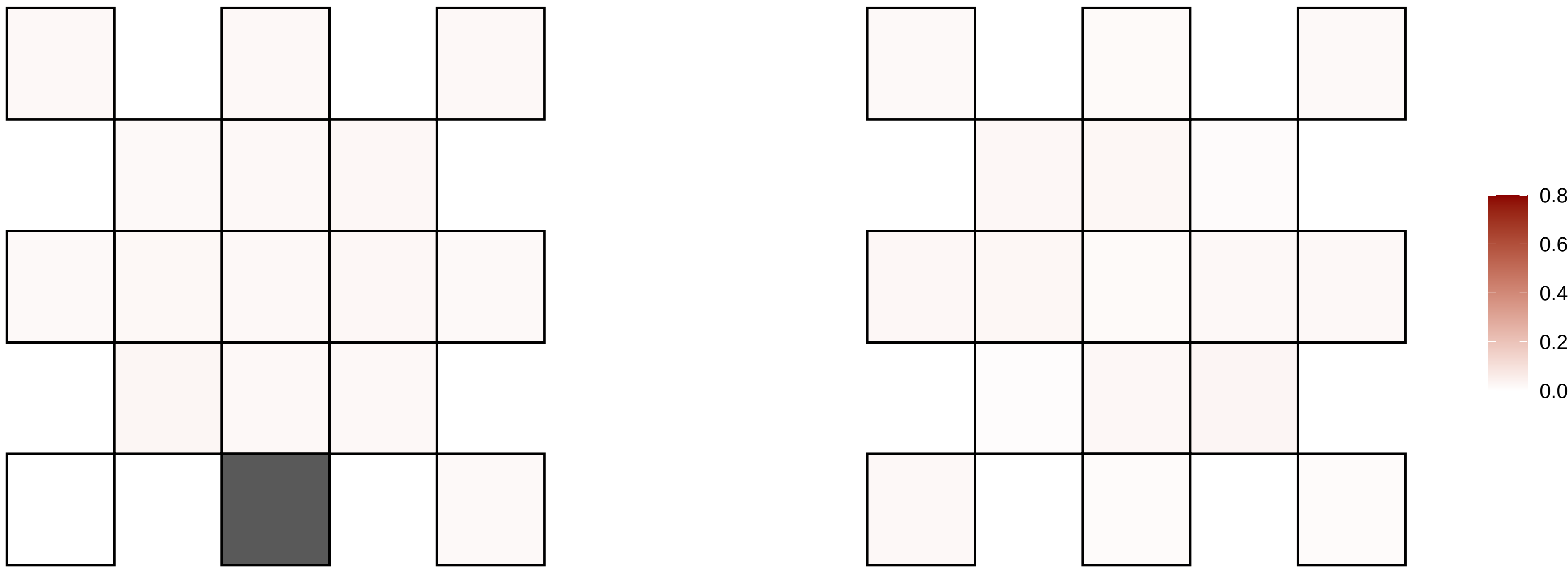

8−66303042−C−T

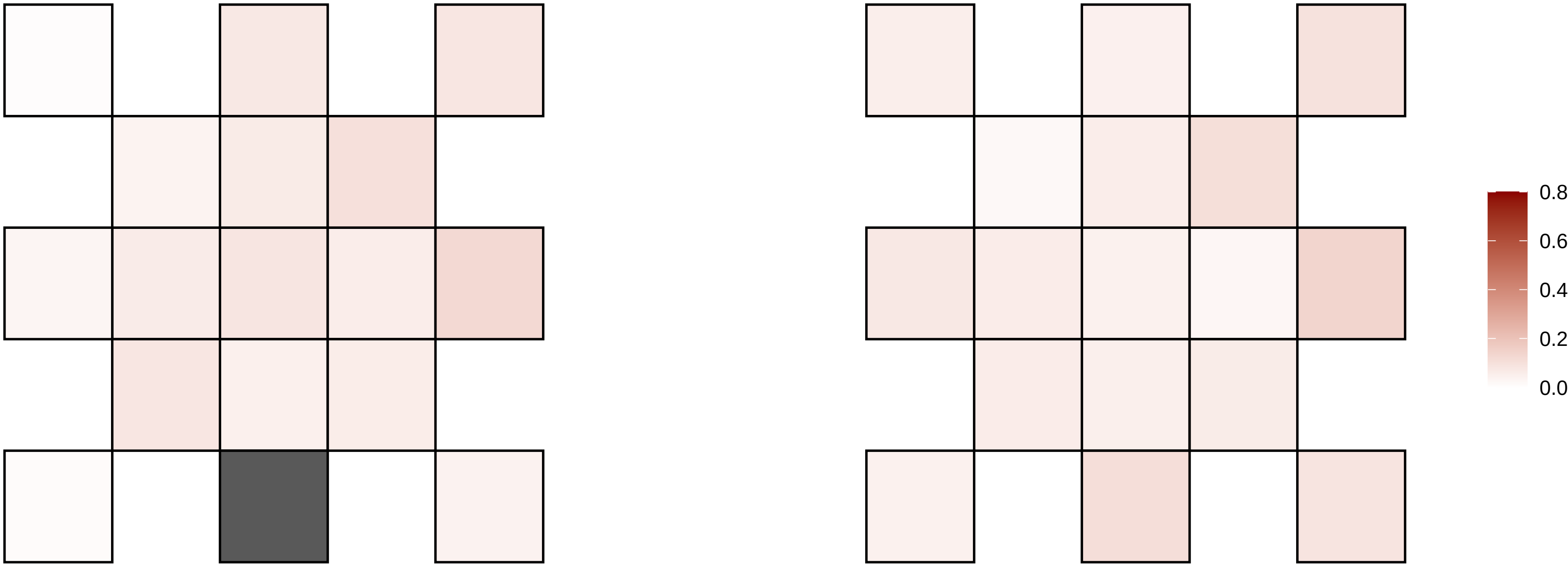

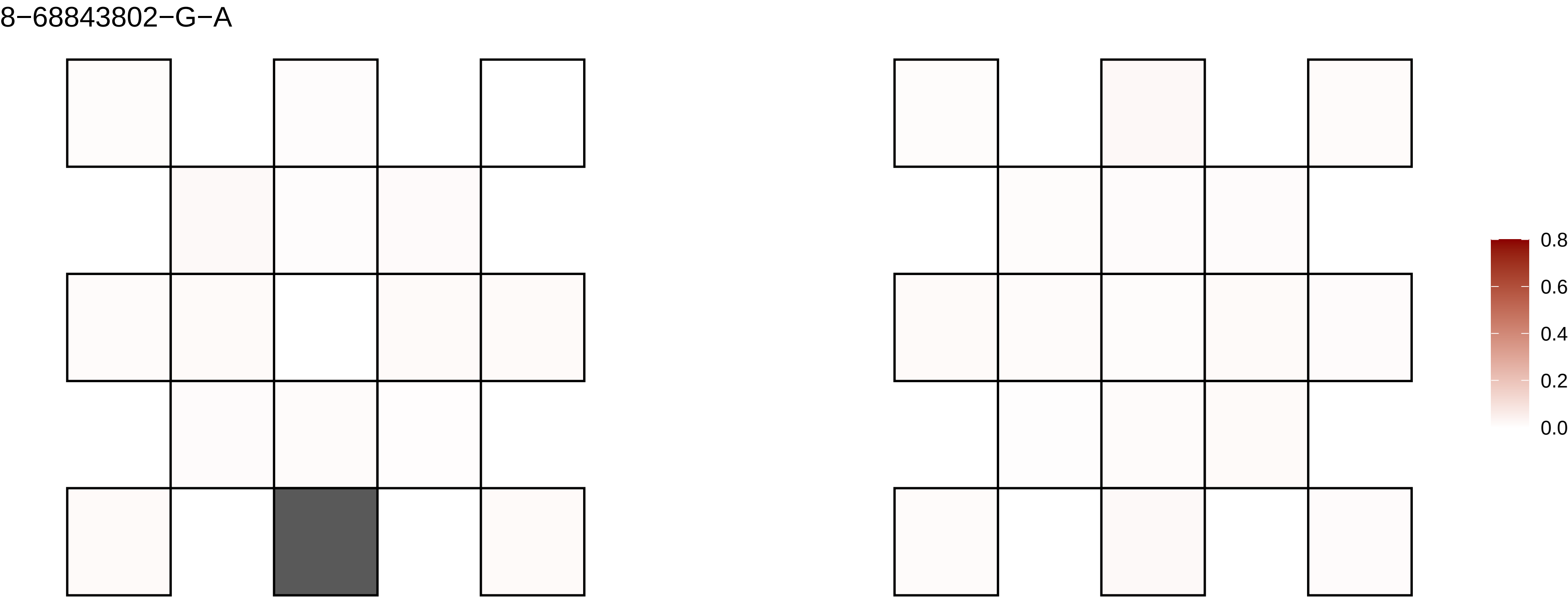
8−68843802−G−A
0.8
0.6
0.4
0.2
0.0

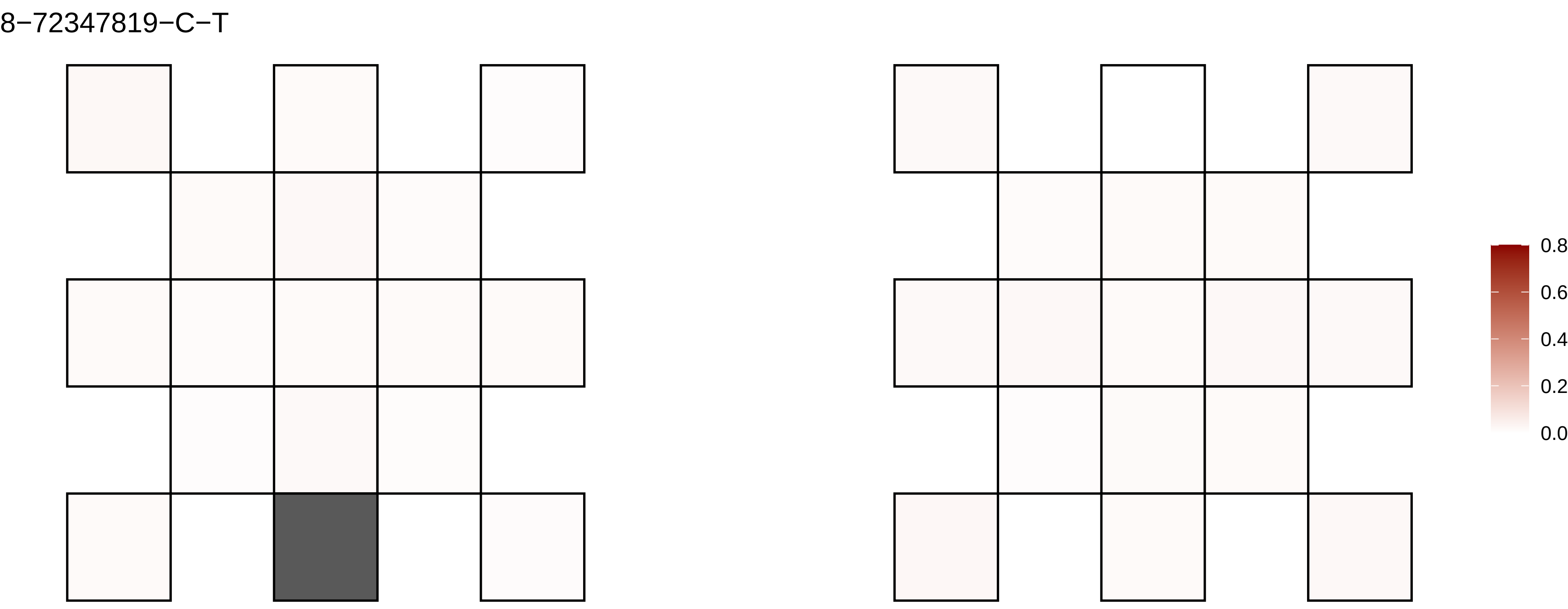

8−72347819−C−T
0.8
0.6
0.4
0.2
0.0

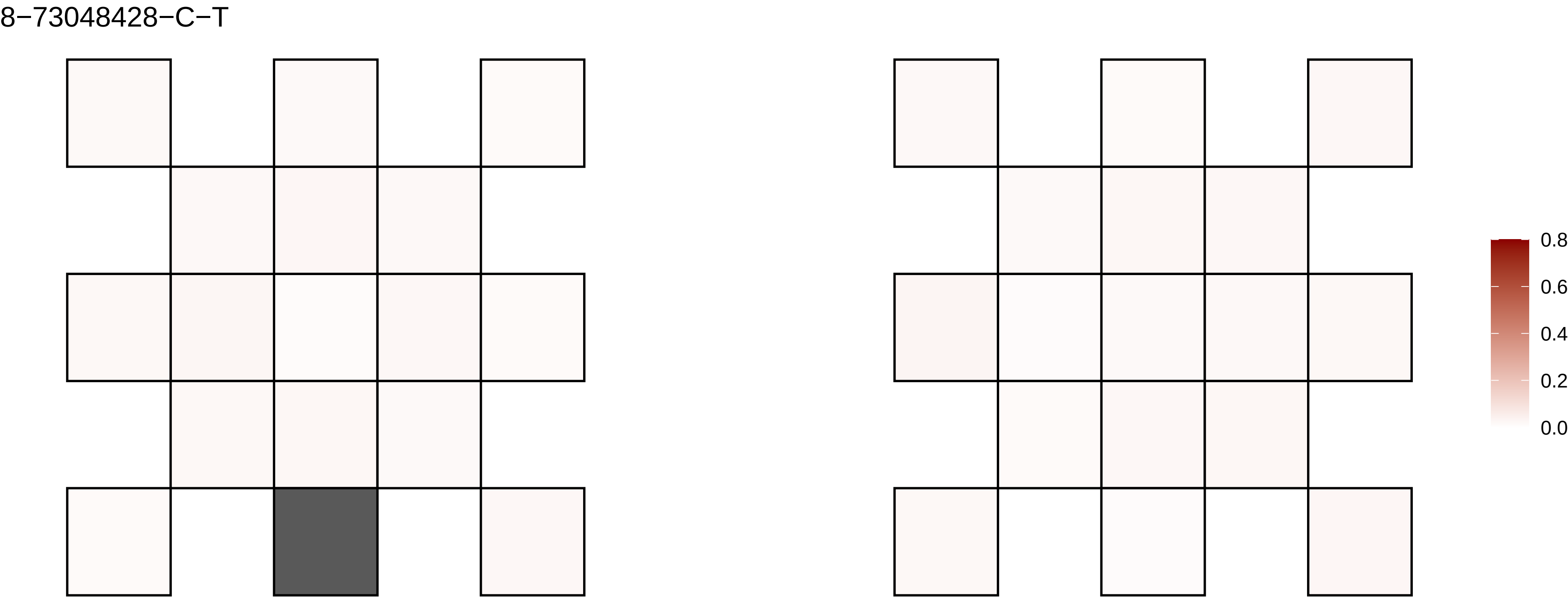

8−73048428−C−T
0.8
0.6
0.4
0.2
0.0

8−77652300−G−A

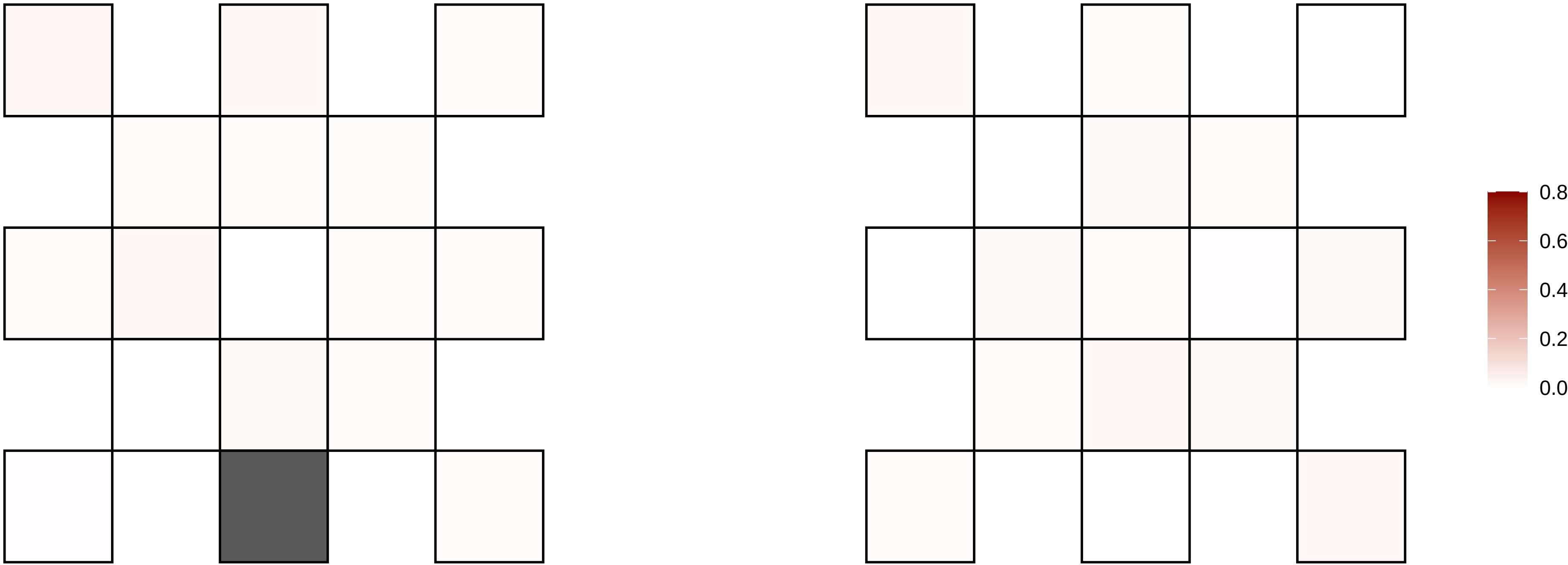

8−786897−C−G

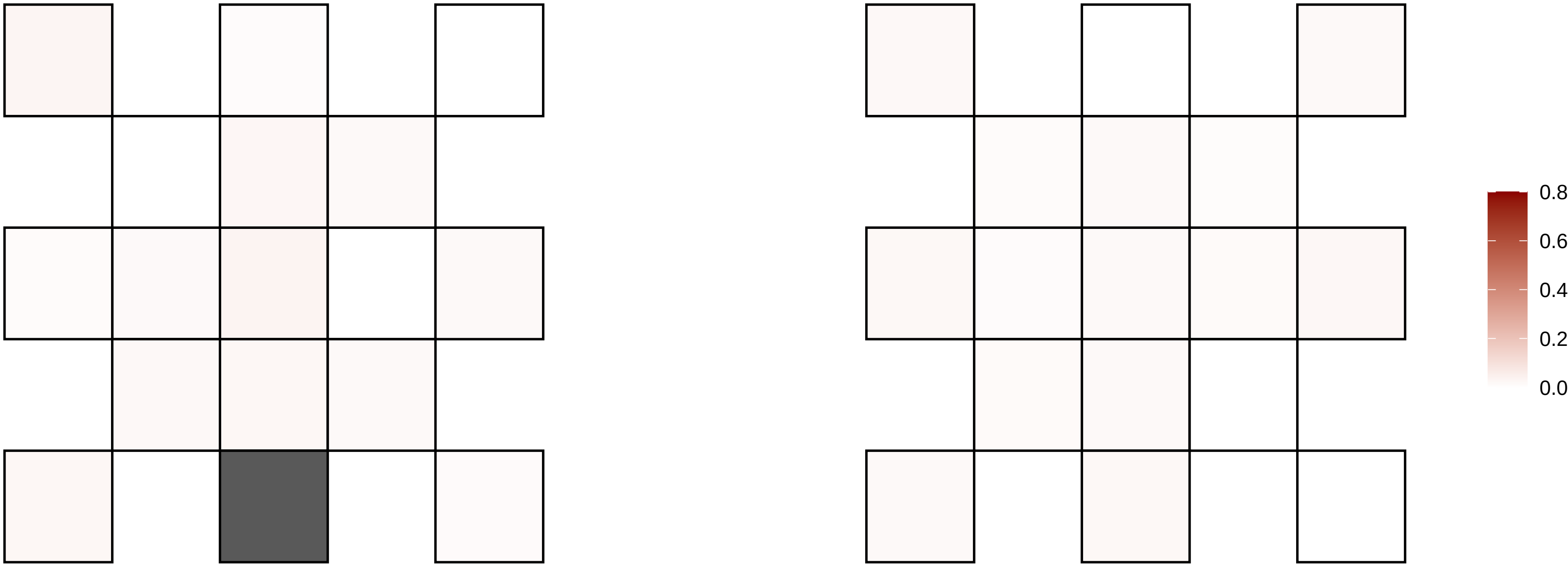

8−80121537−G−A

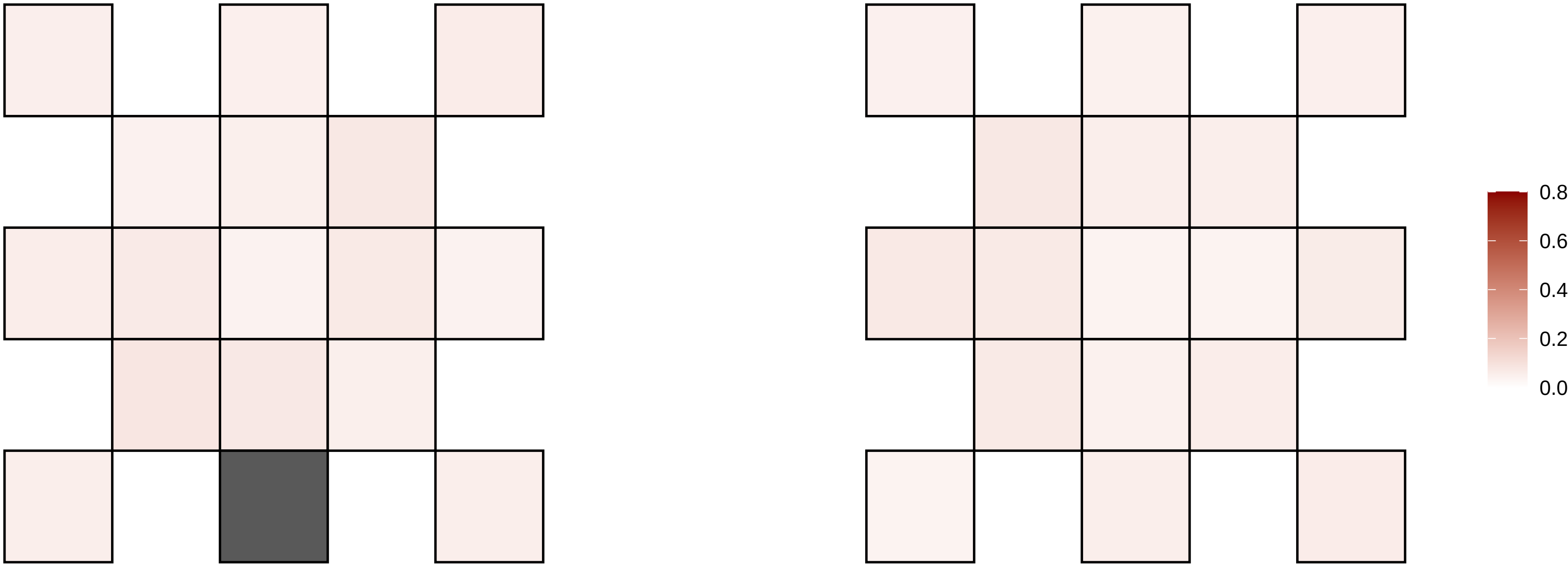

8−80121538−T−C

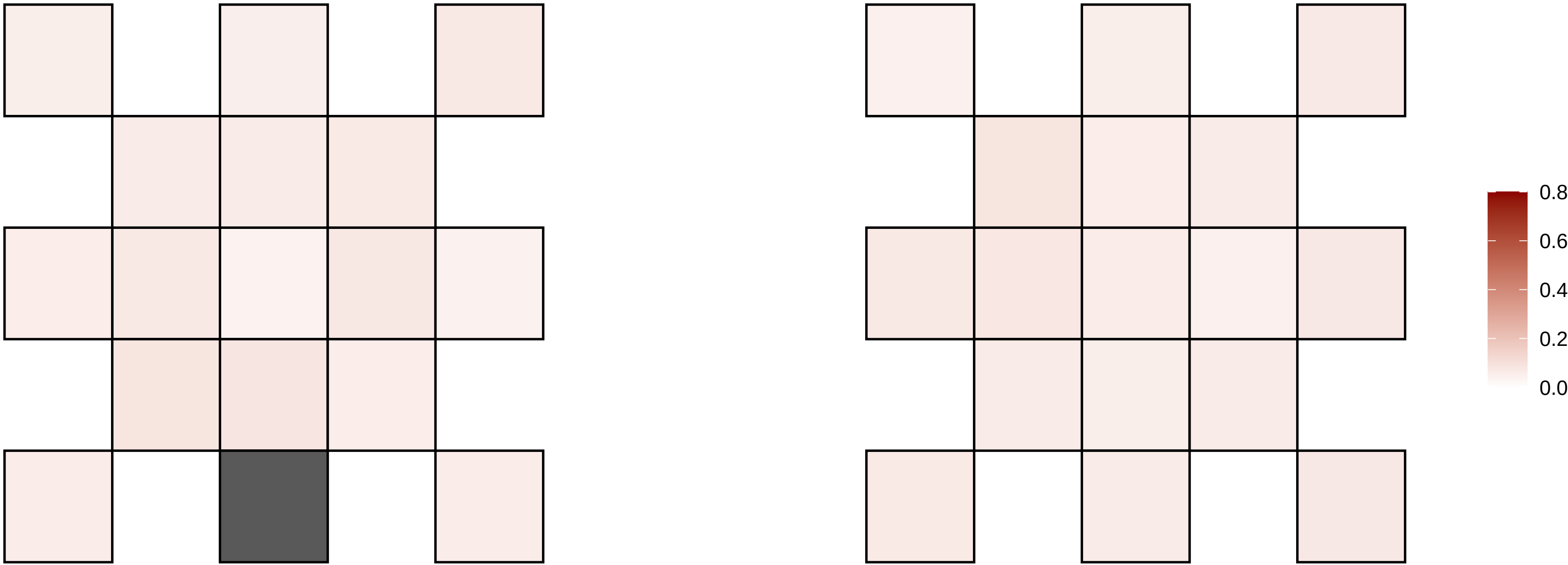

8−83768131−G−A

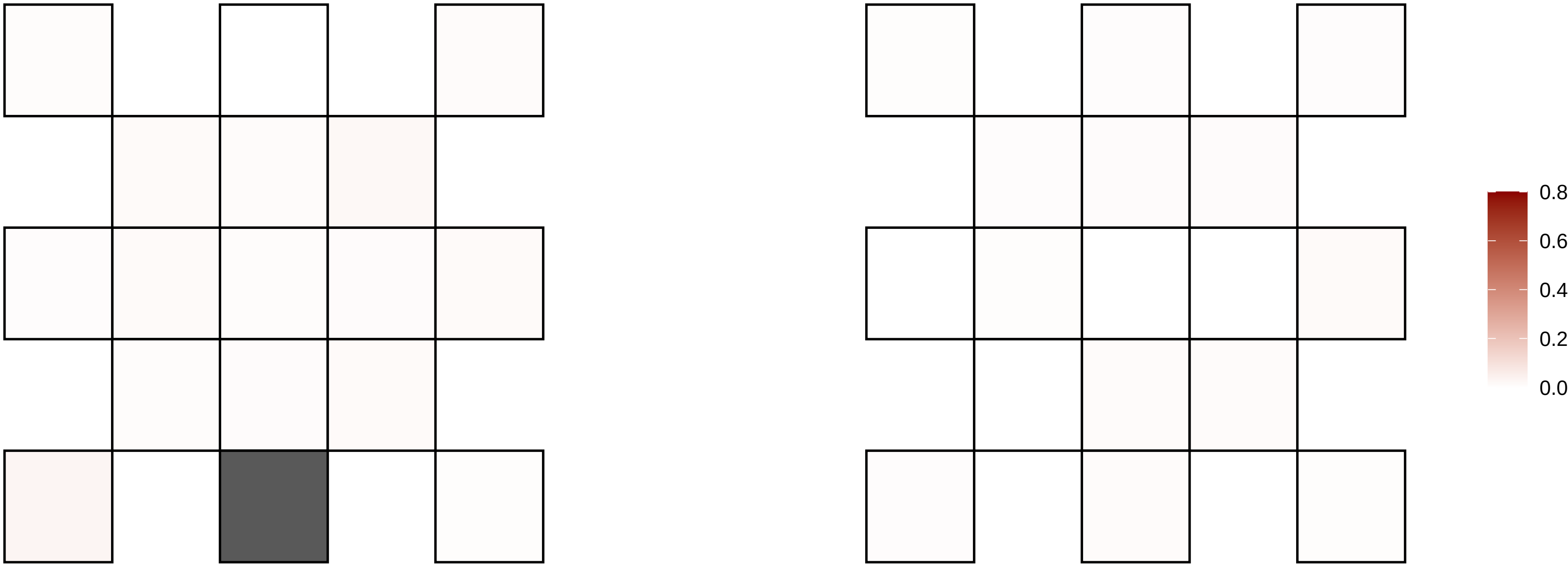

8−89142864−G−A

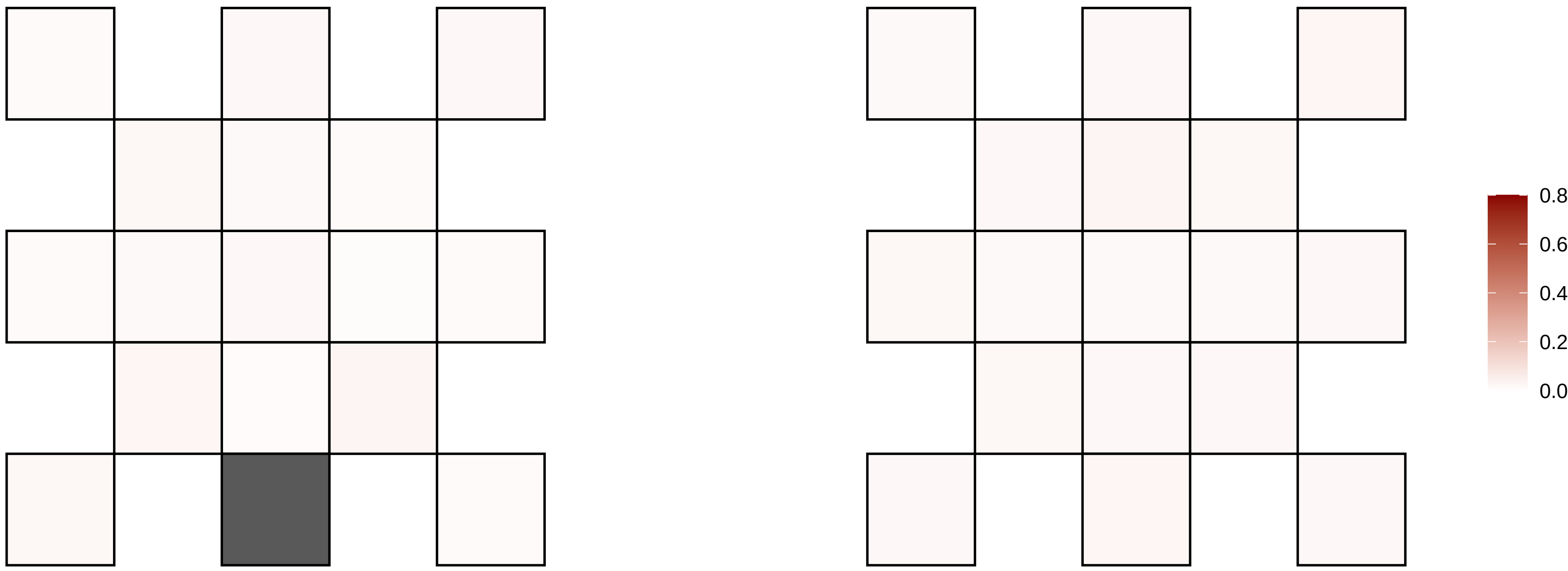

8−89612305−G−A

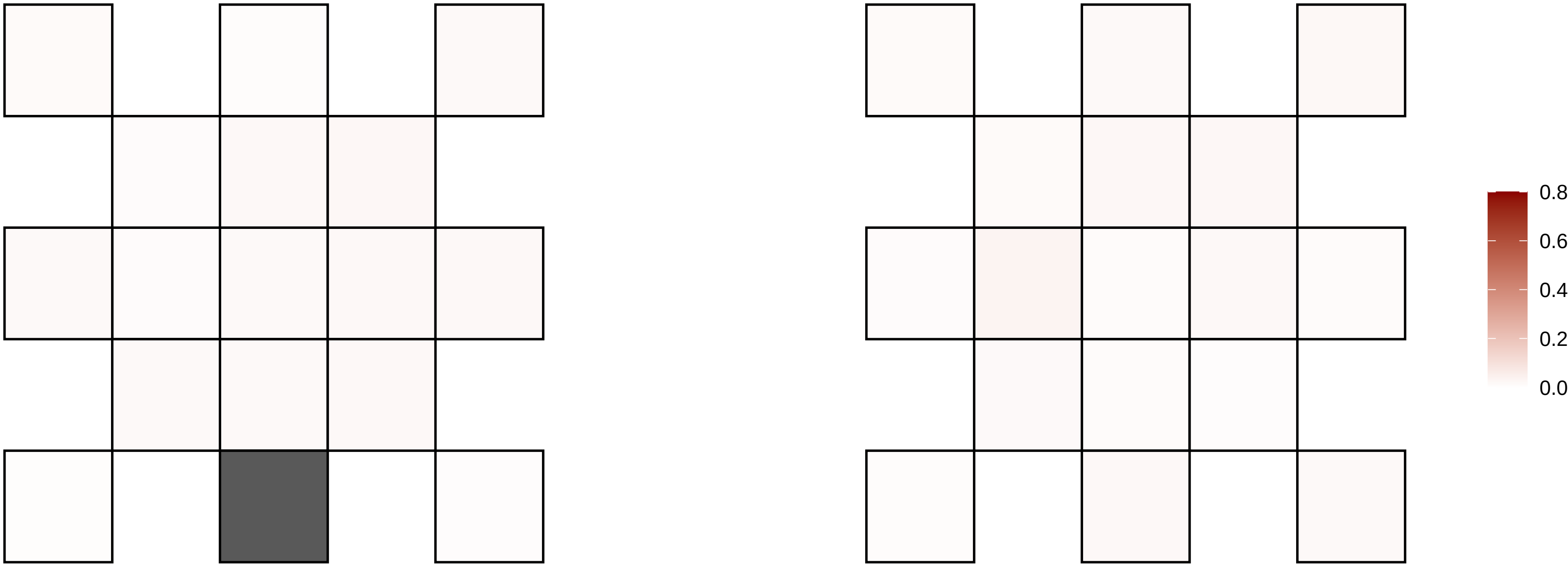

8−91951808−C−T

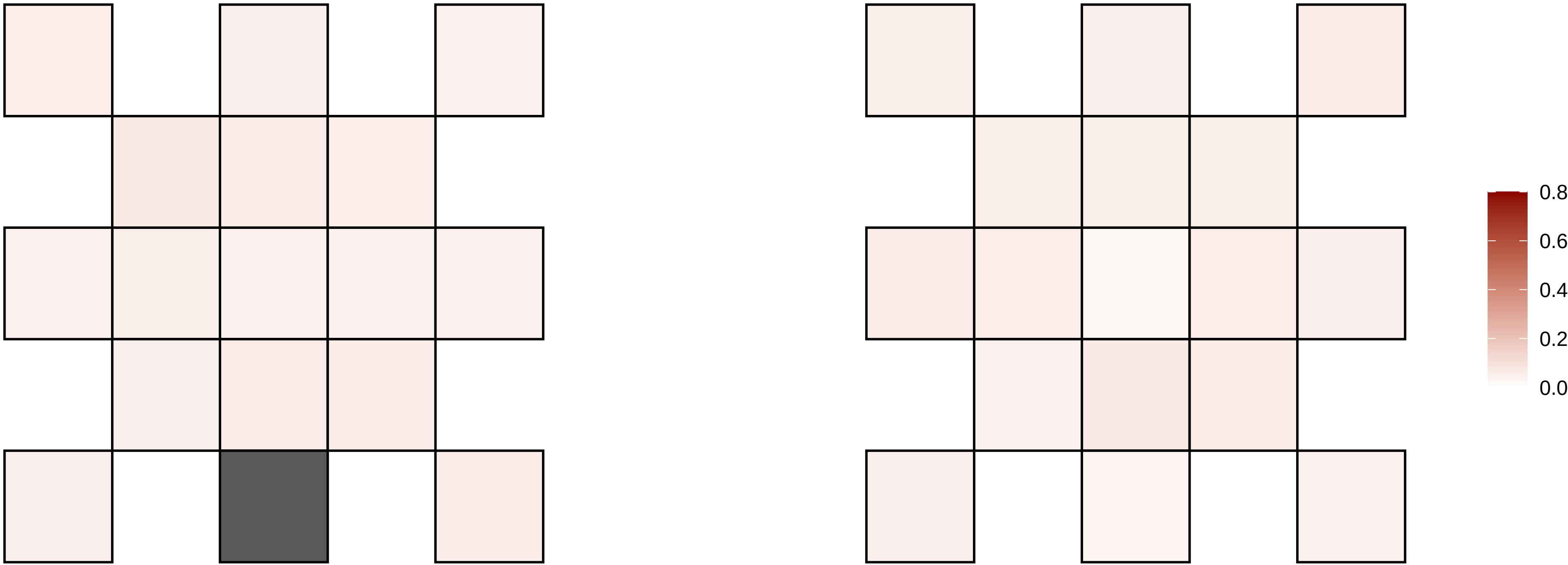

9−103766822−C−T

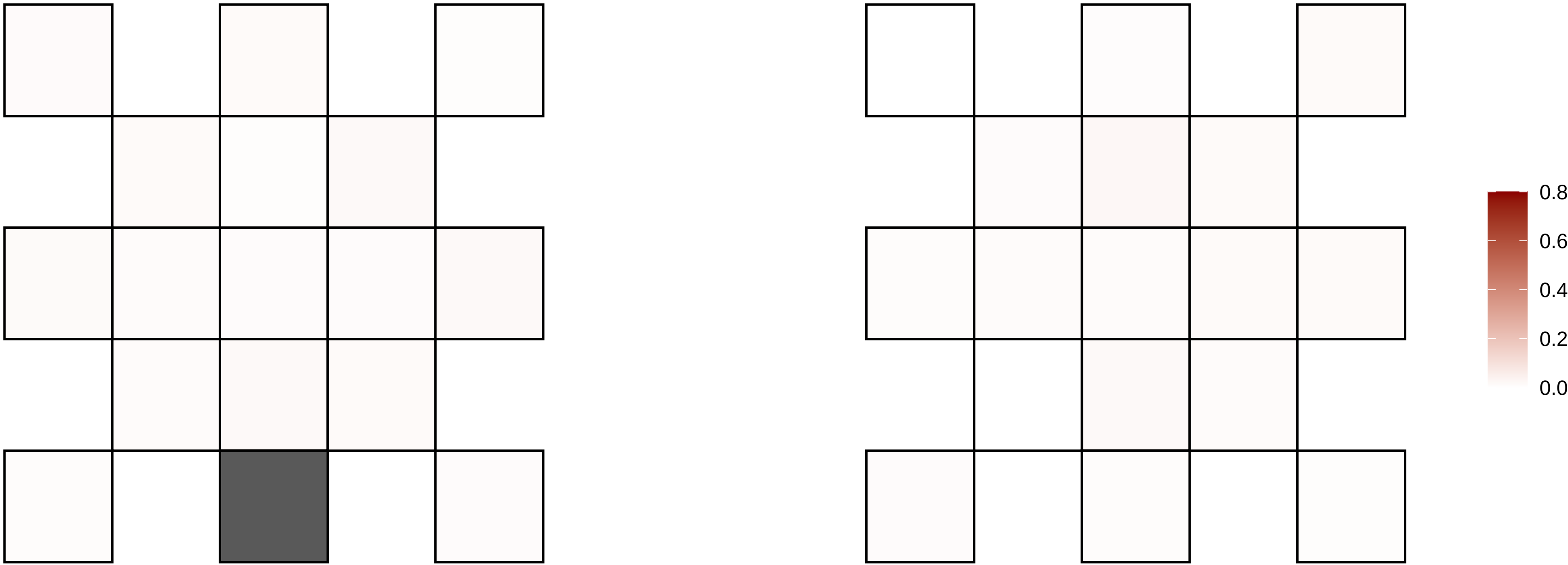

9−10473452−T−A

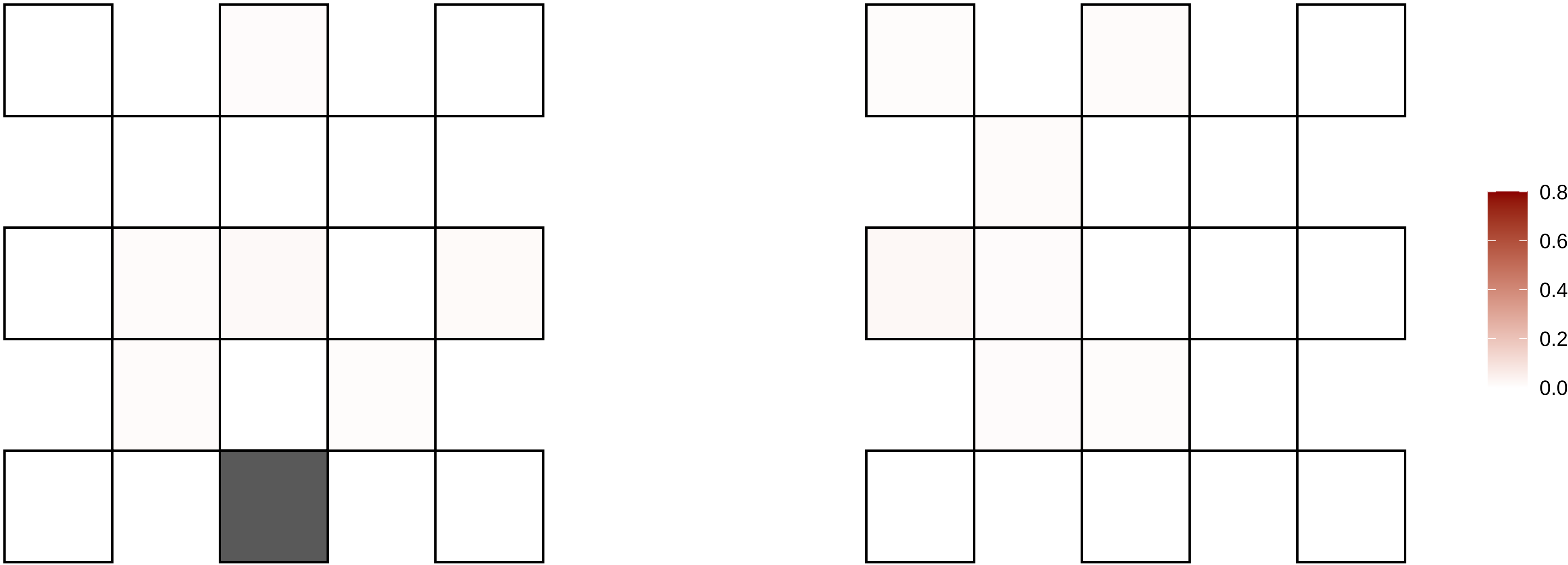

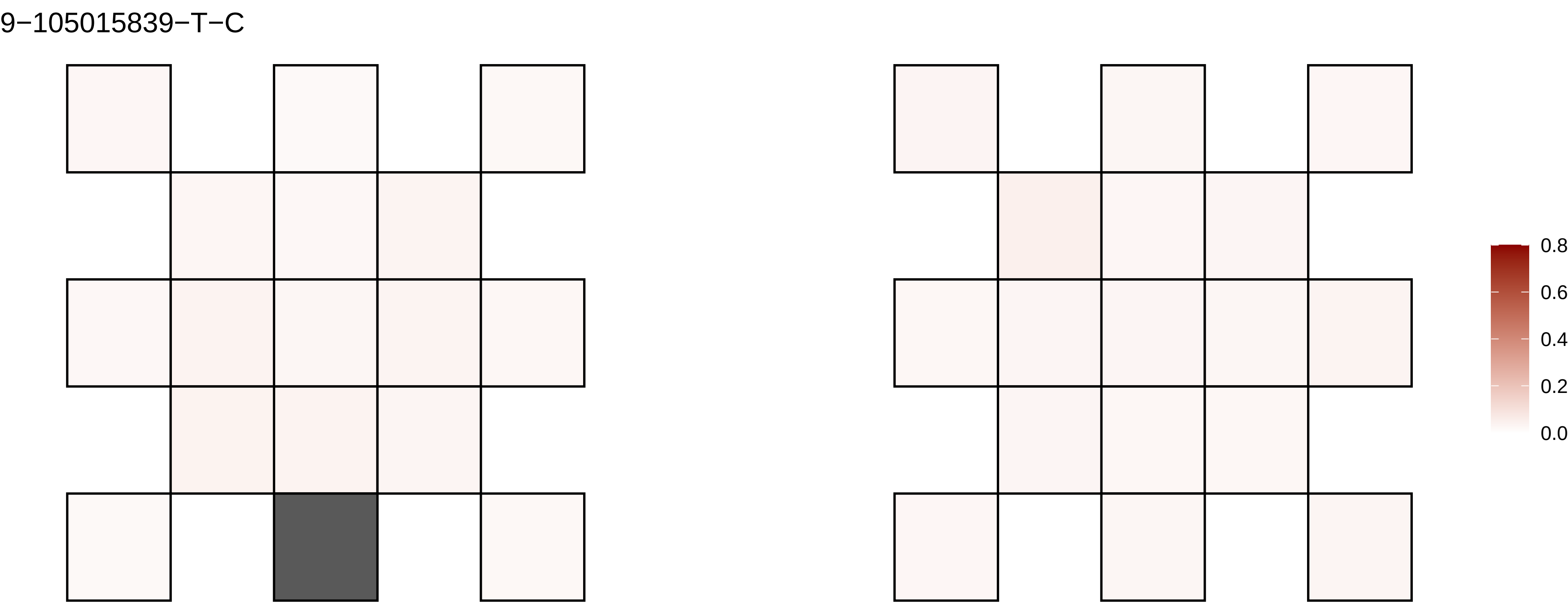

9−105015839−T−C
0.8
0.6
0.4
0.2
0.0

9−107723679−T−A

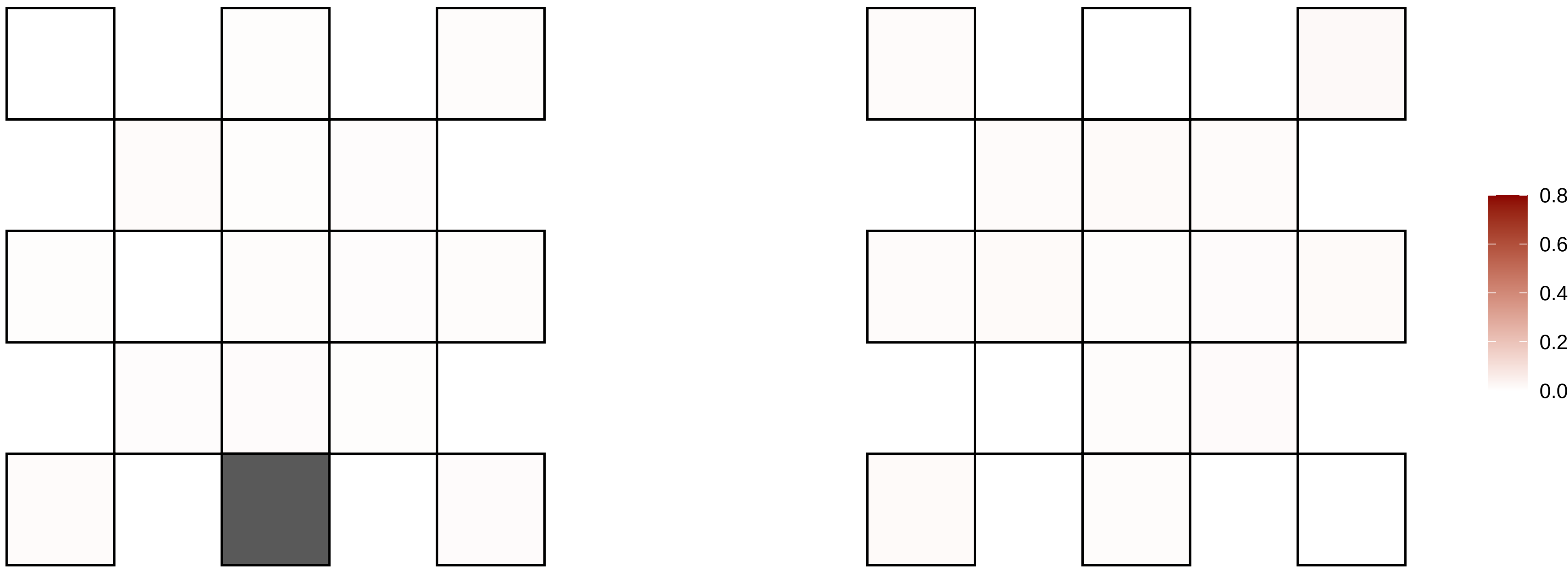

9−10773884−G−A

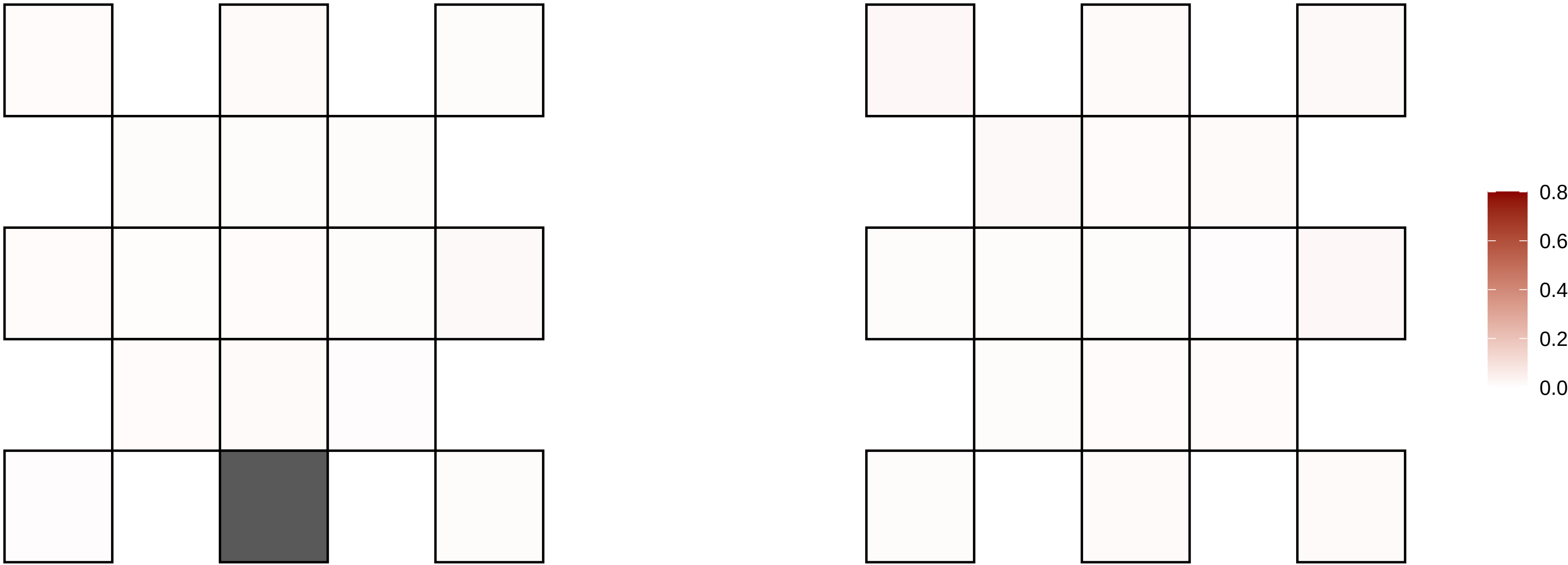

9−119721731−C−T

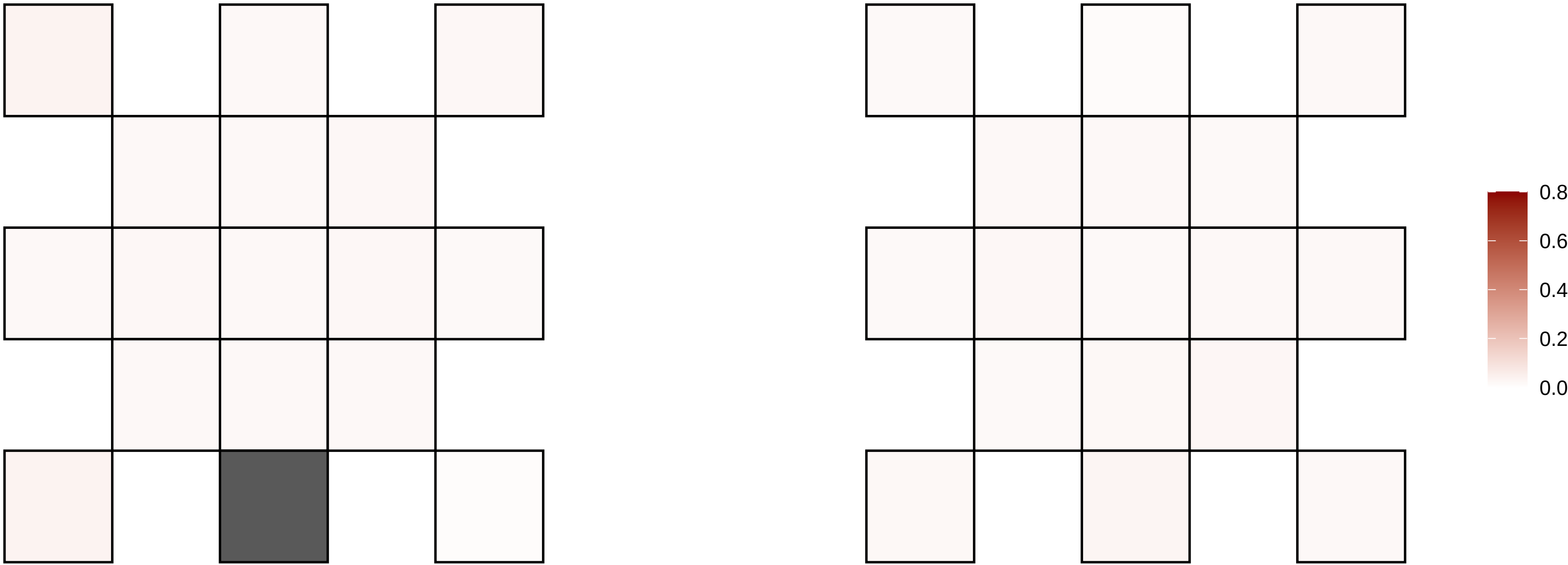

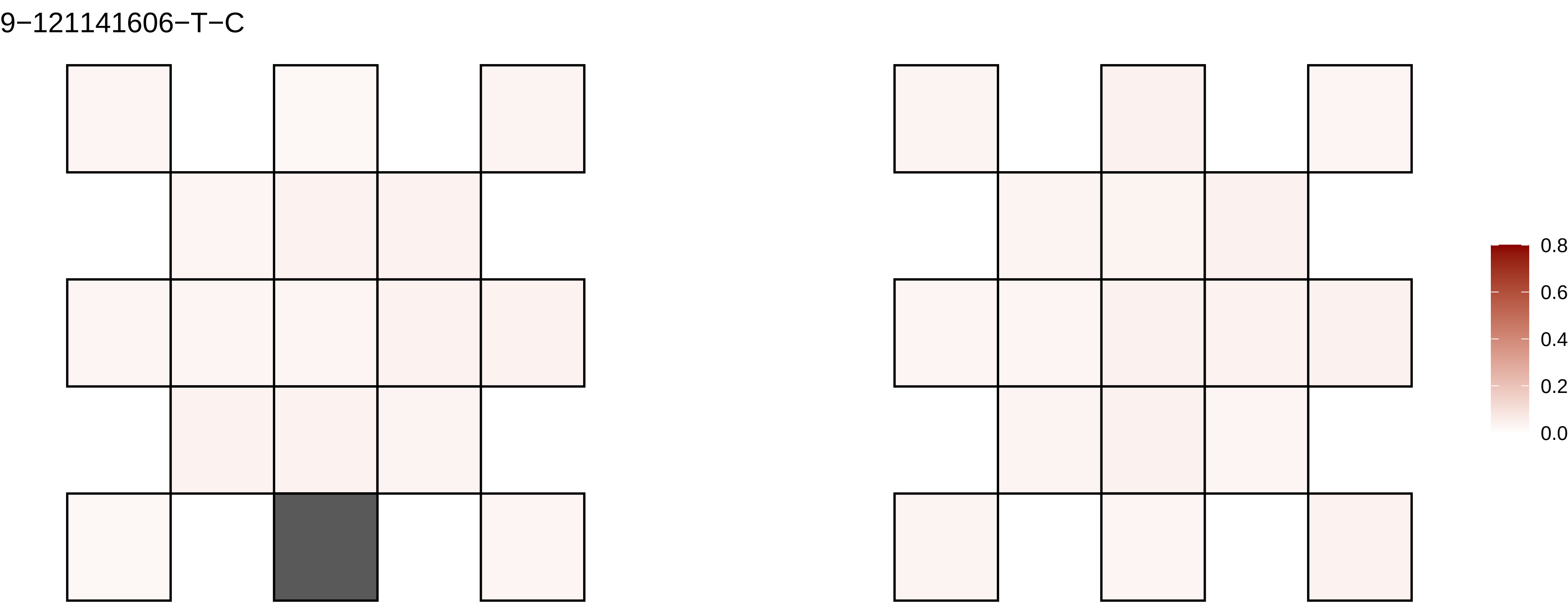
9−121141606−T−C
0.8
0.6
0.4
0.2
0.0

9−121997207−C−T

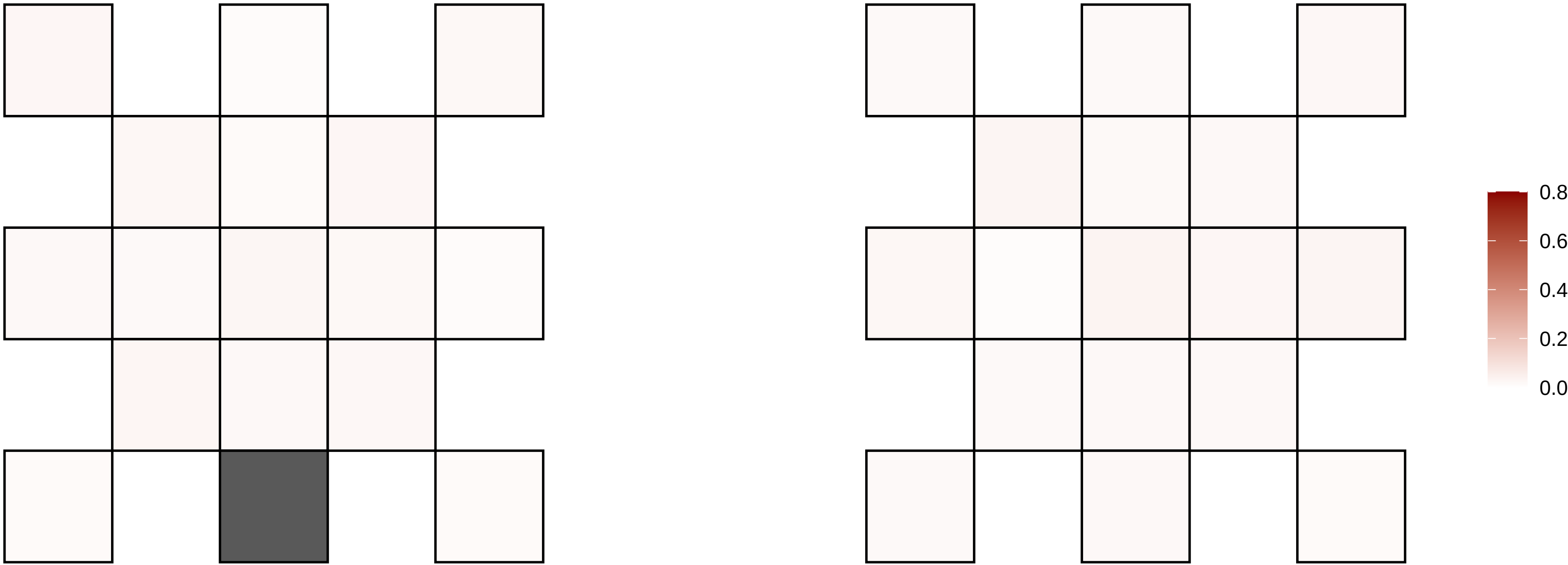

9−12287979−T−C

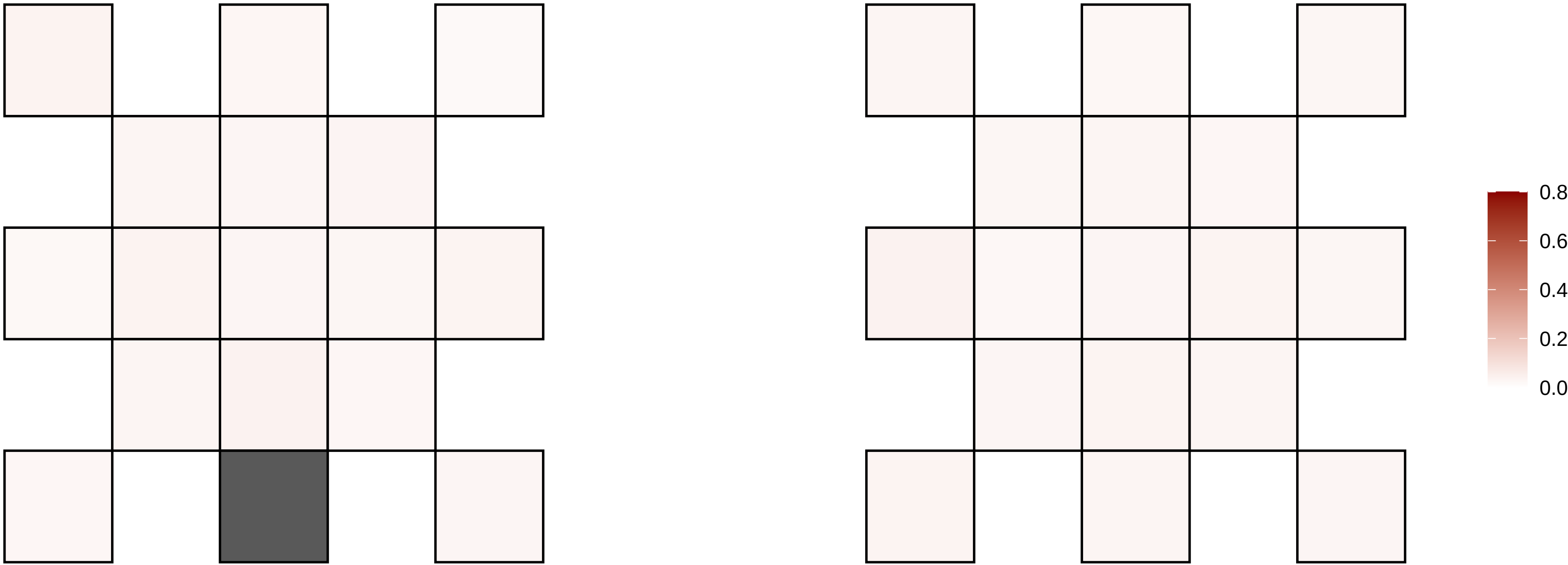

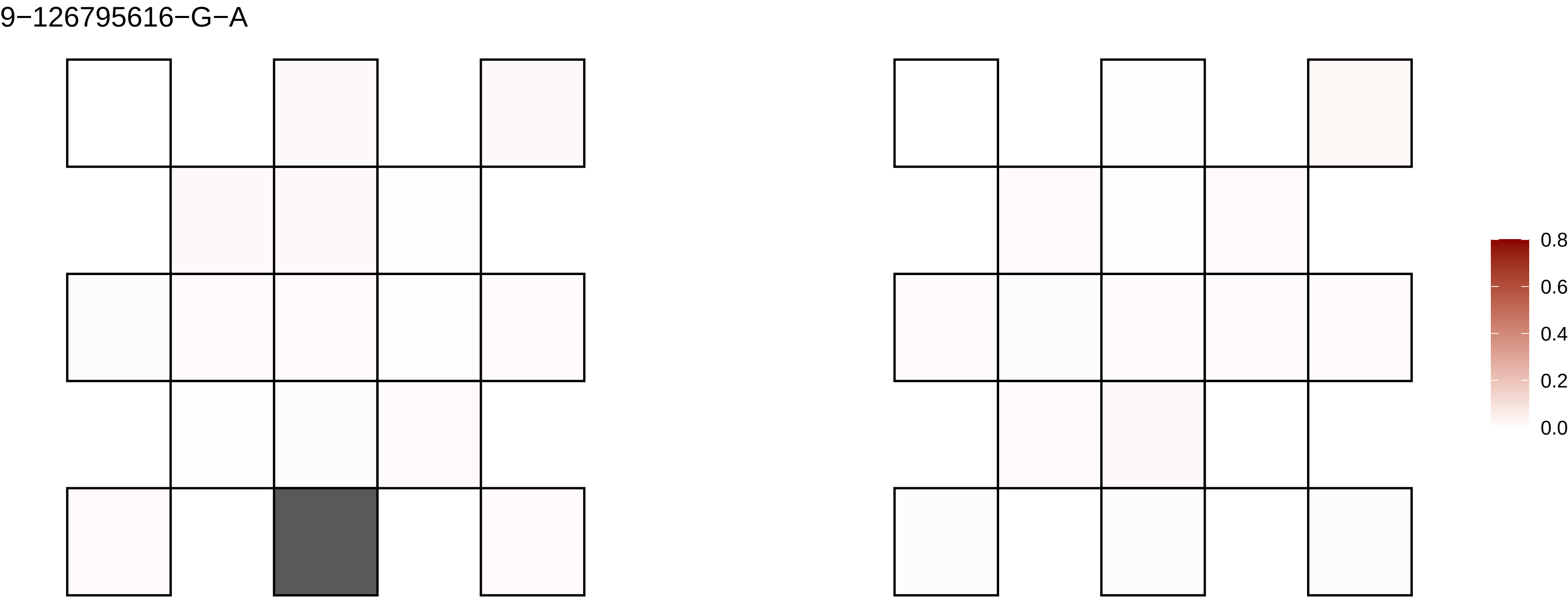
9−126795616−G−A
0.8
0.6
0.4
0.2
0.0

9−134621051−C−T

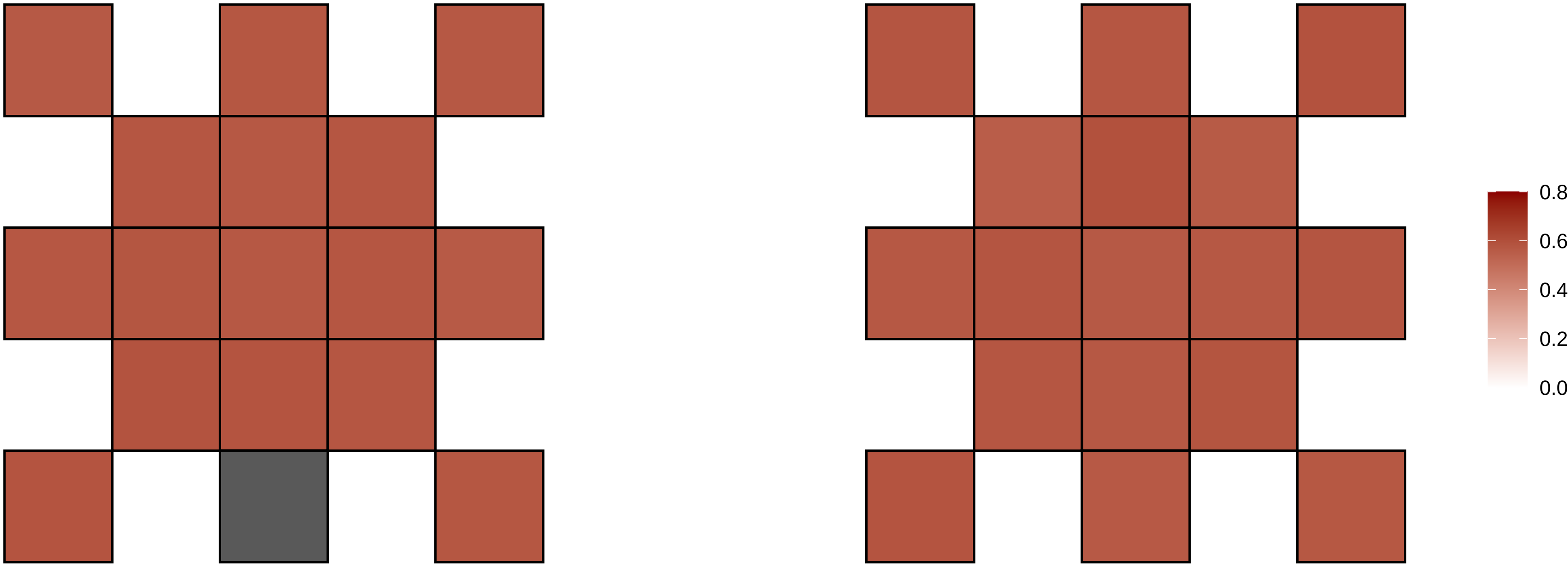

9−13509007−G−A

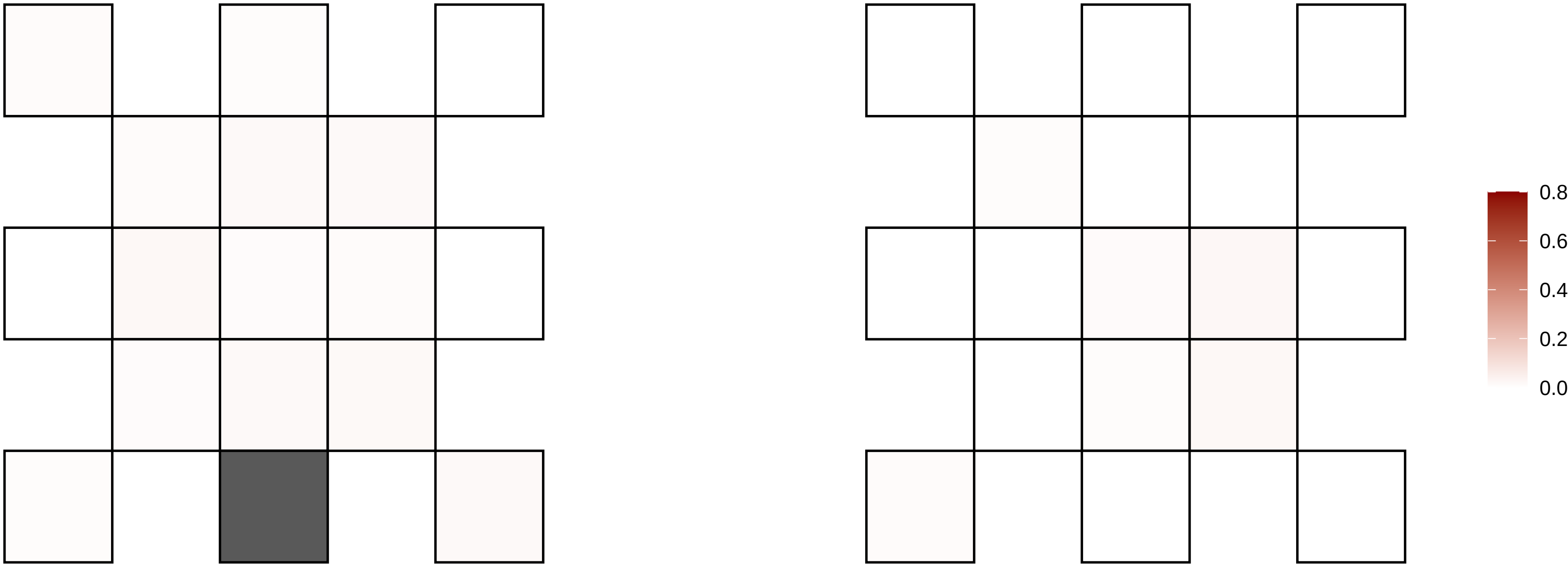

9−138162523−G−A

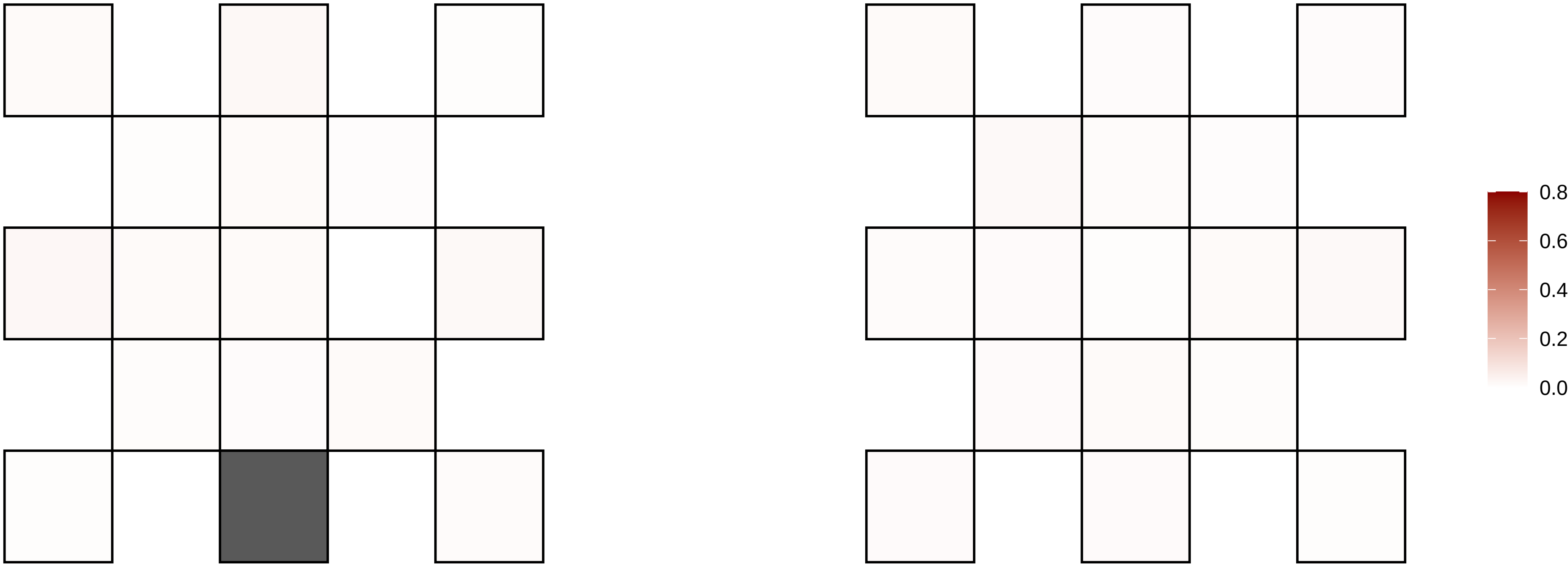

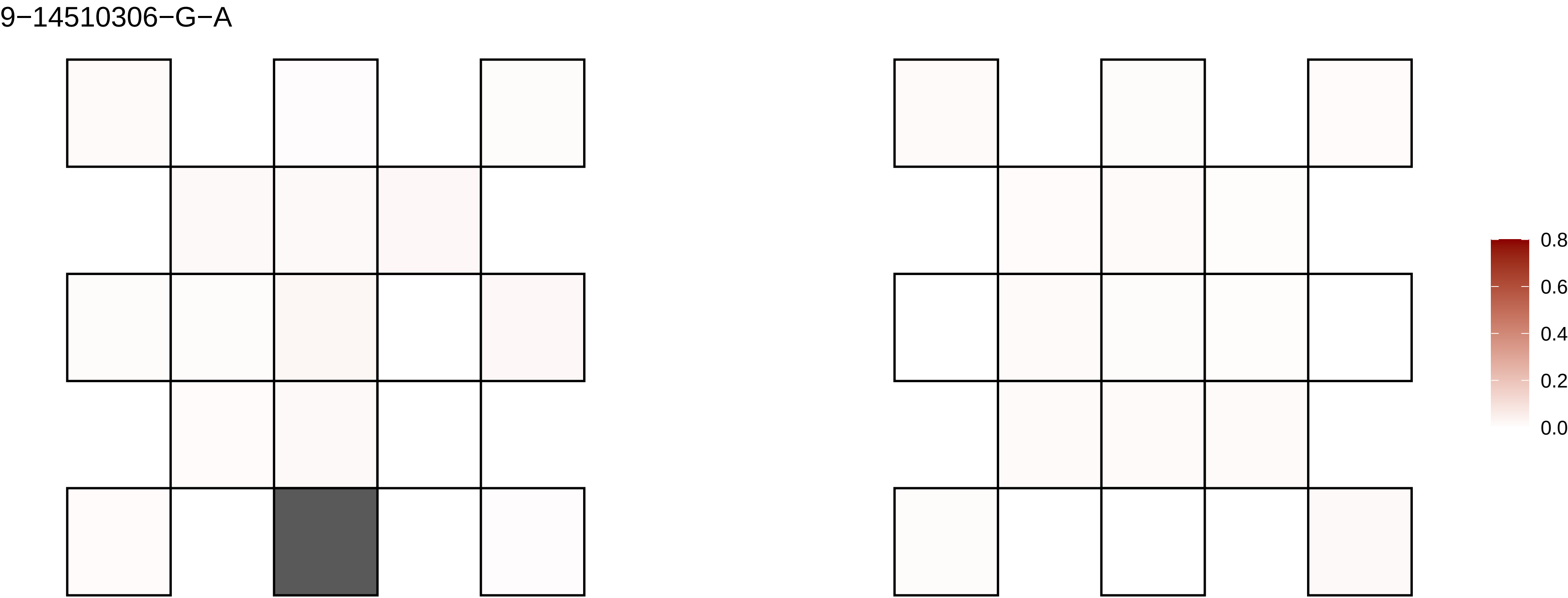
9−14510306−G−A
0.8
0.6
0.4
0.2
0.0

9−19957121−C−T

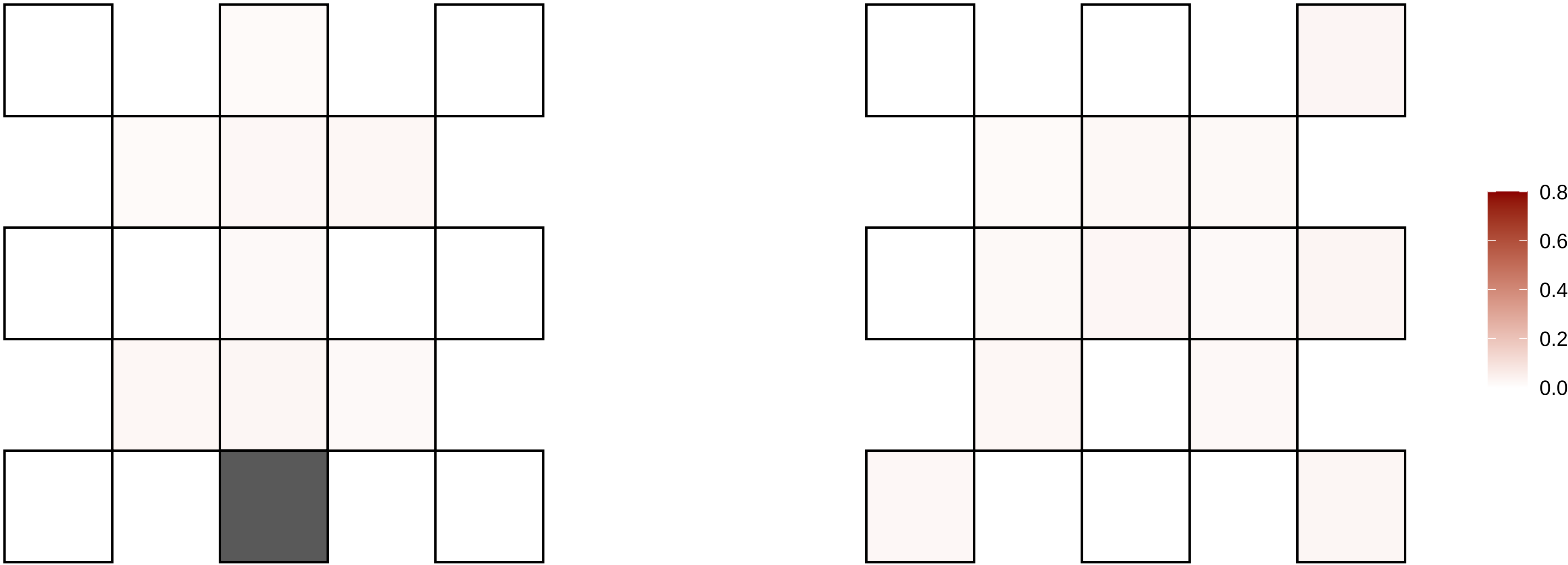

9−21154273−T−C

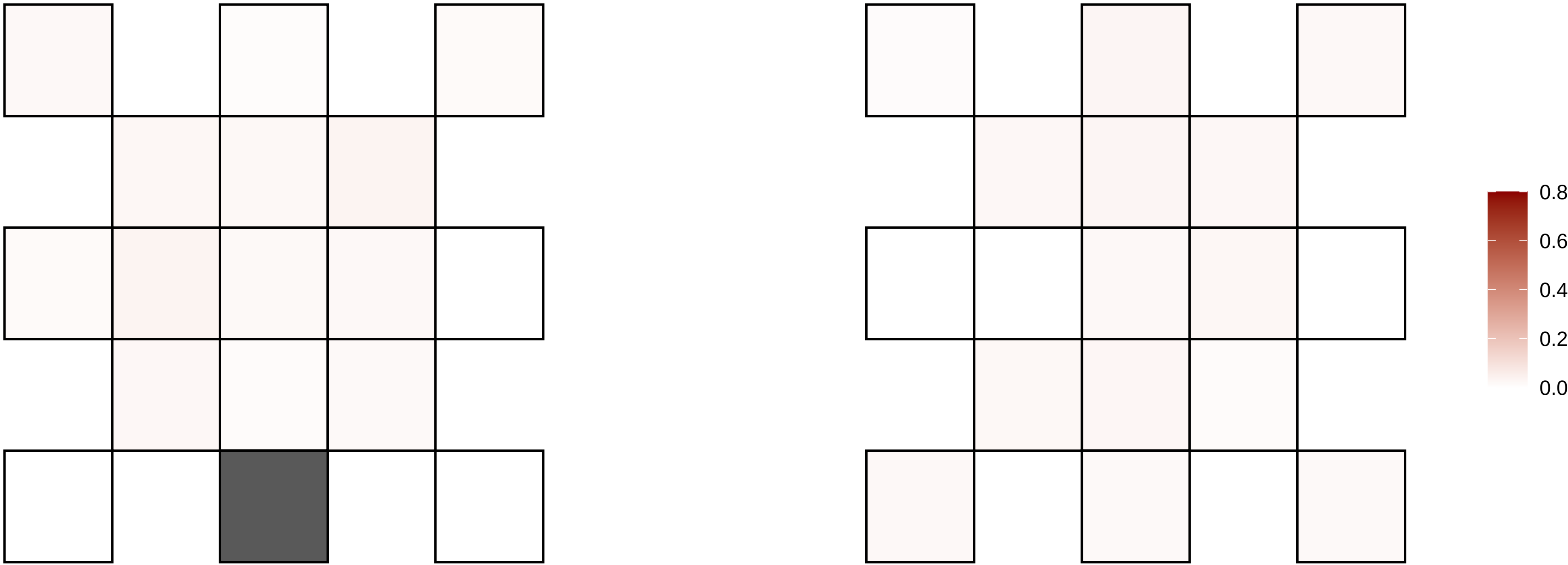

9−21506598−C−A

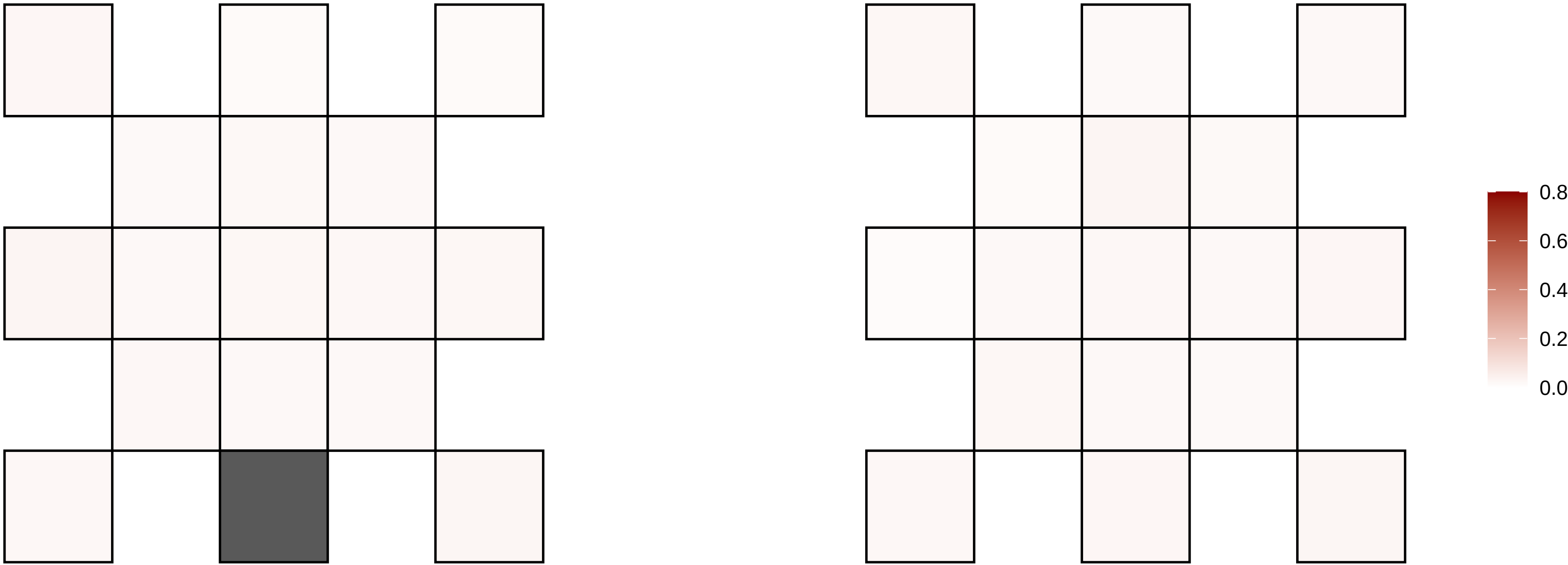

9−23041281−C−T

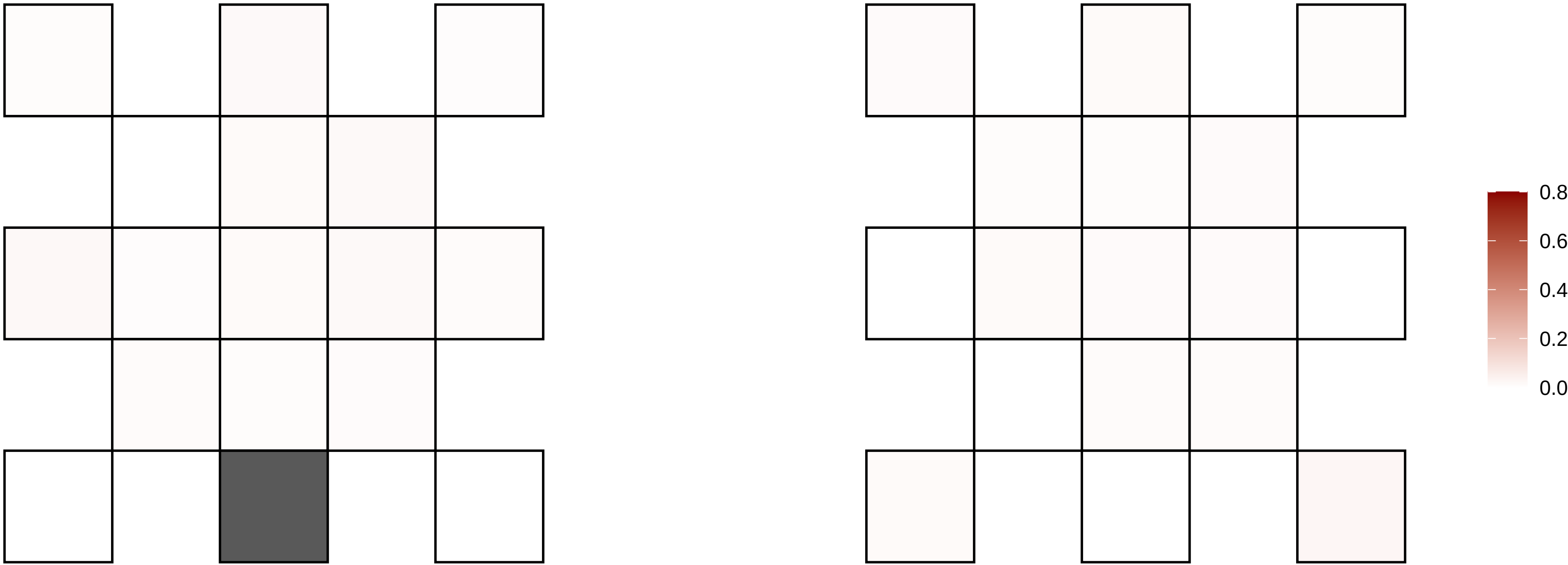

9−28561999−C−T

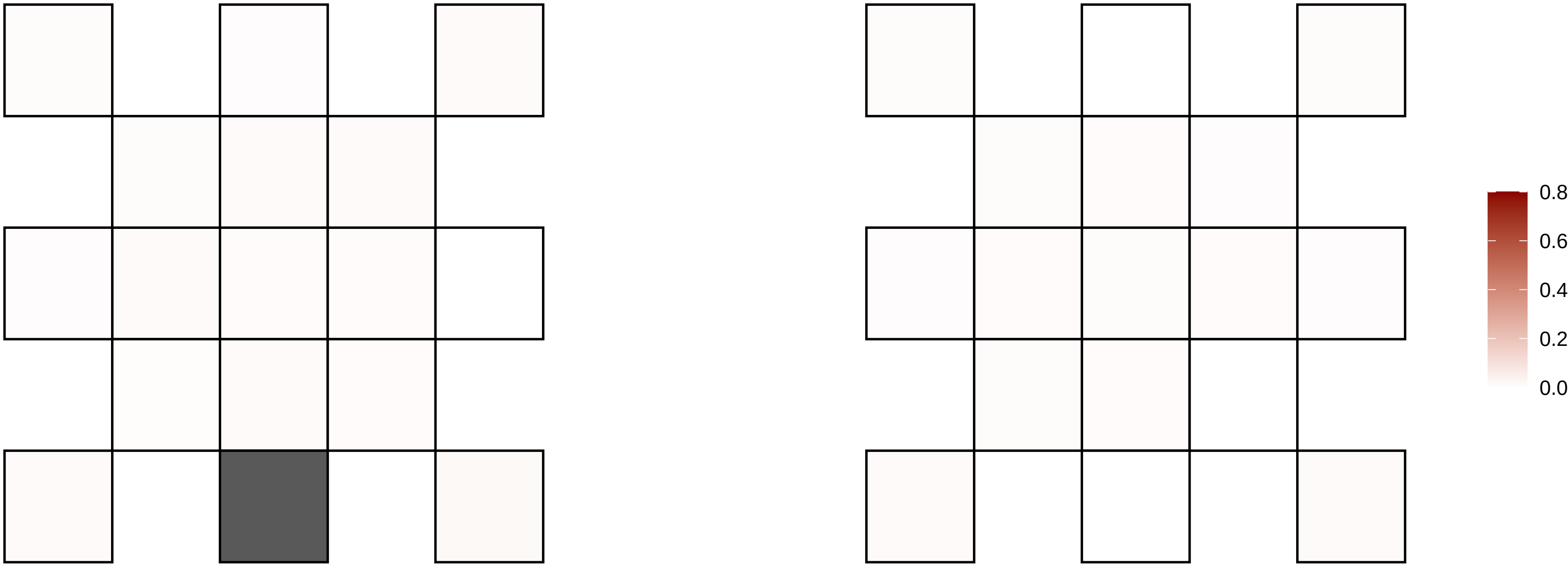

9−31039053−G−T

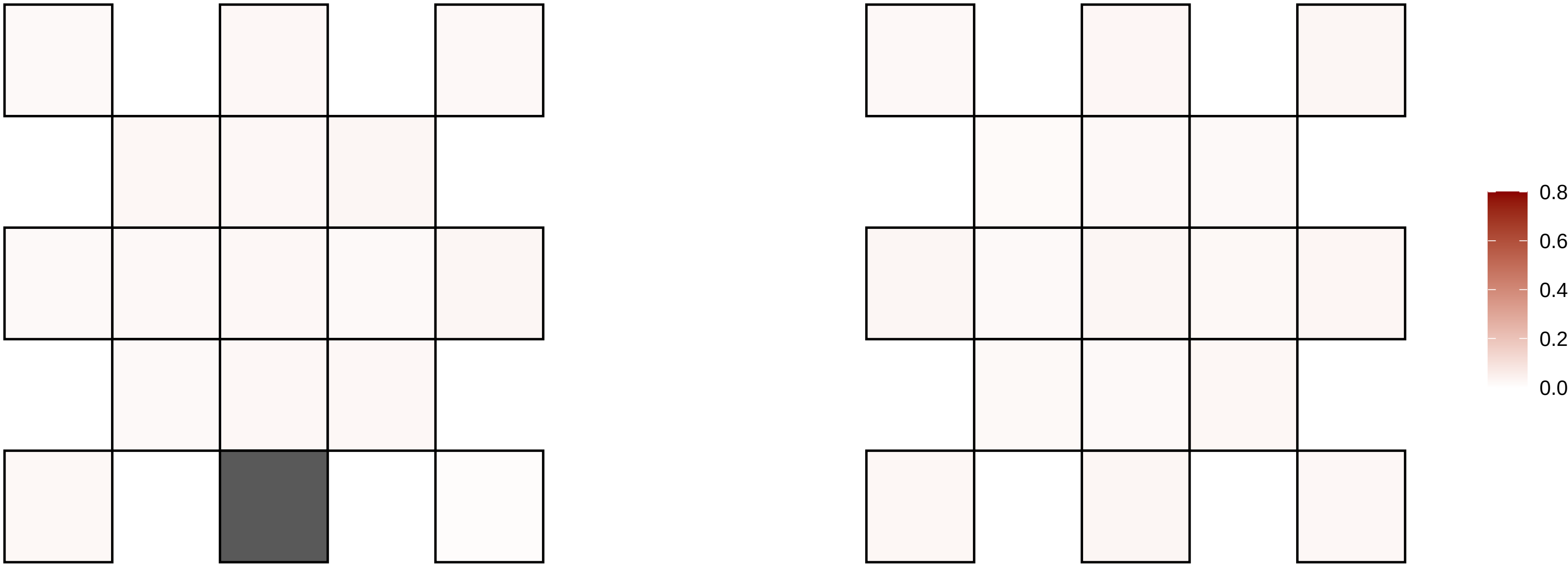

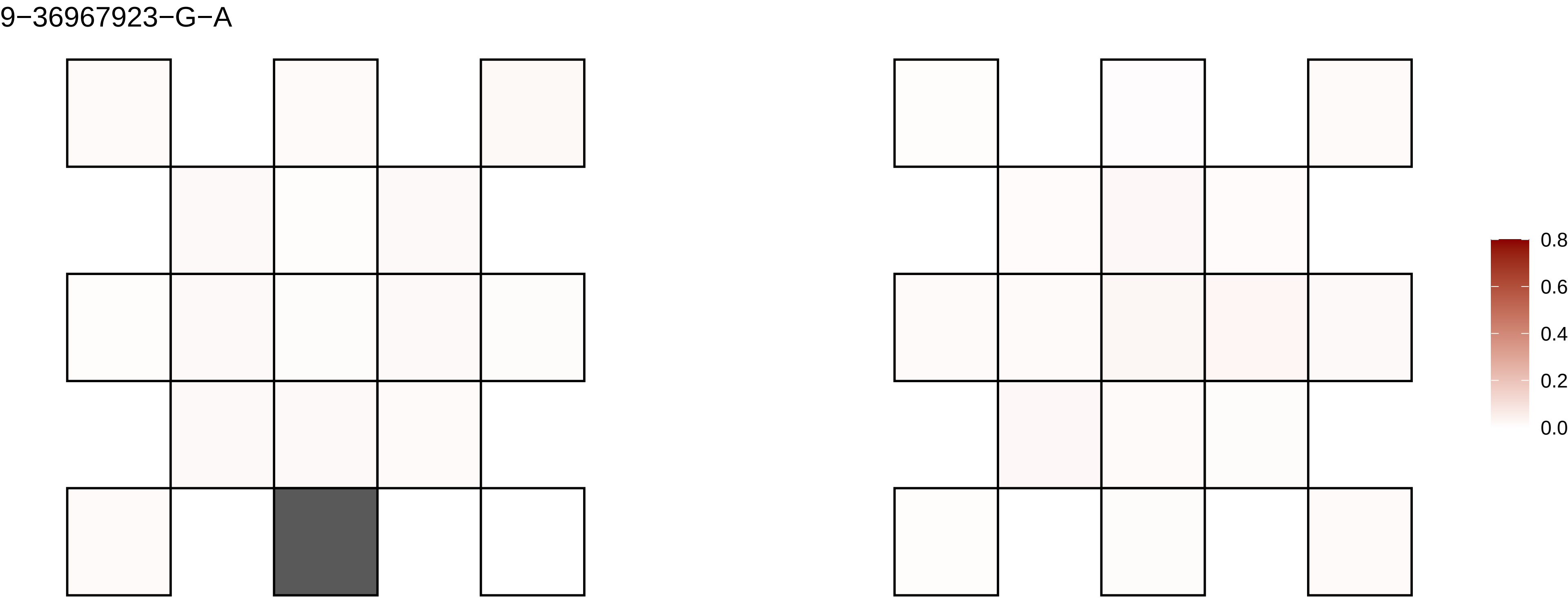
9−36967923−G−A
0.8
0.6
0.4
0.2
0.0

9−559661−C−T

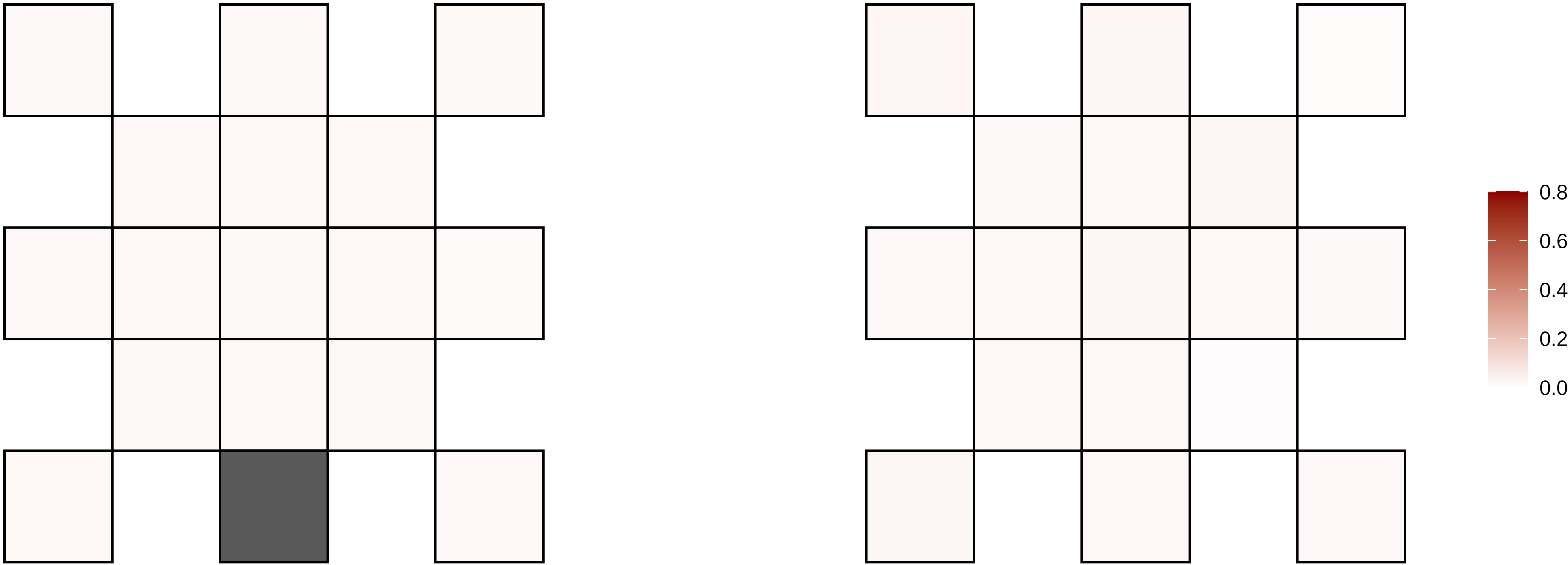

9−74407663−G−A

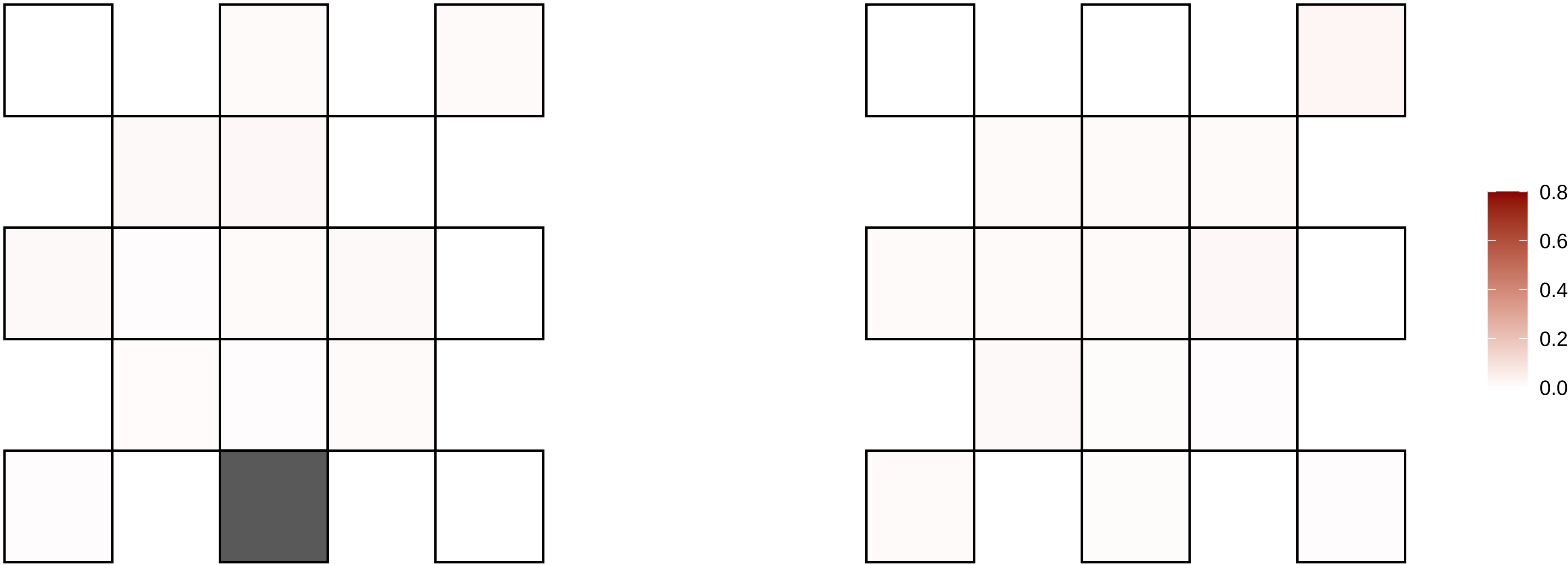

9−76169299−T−C

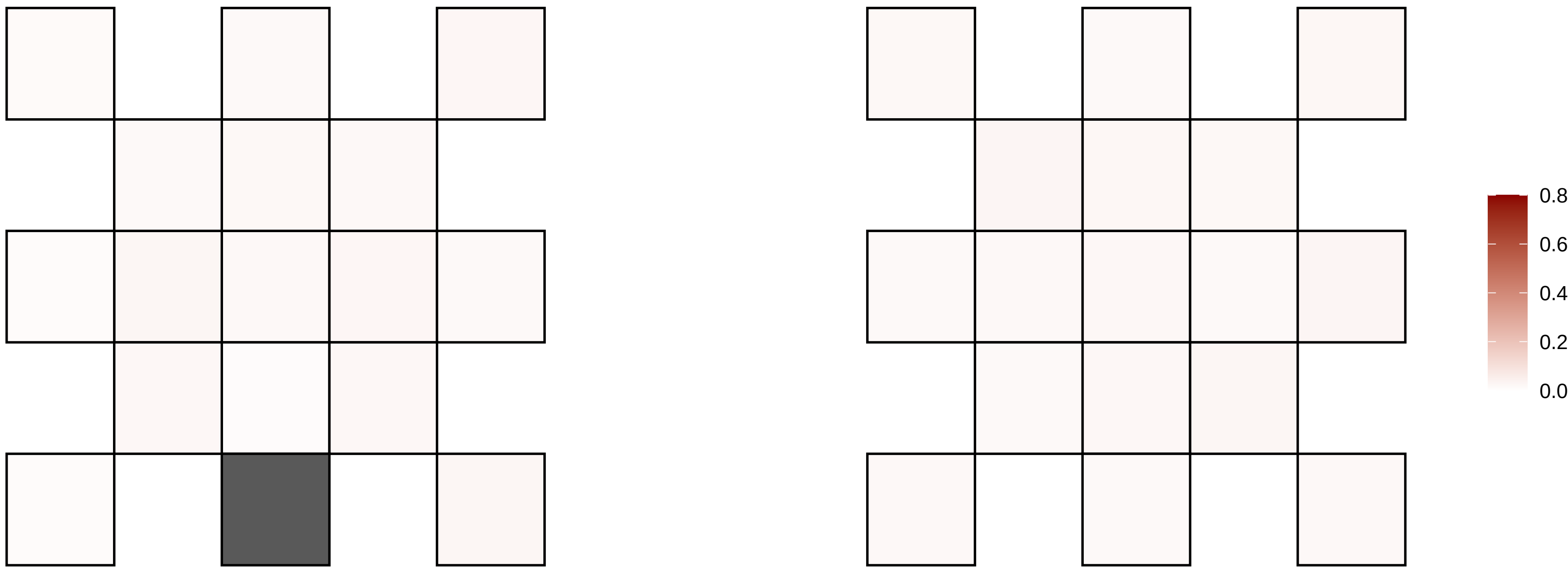

9−77140705−C−T

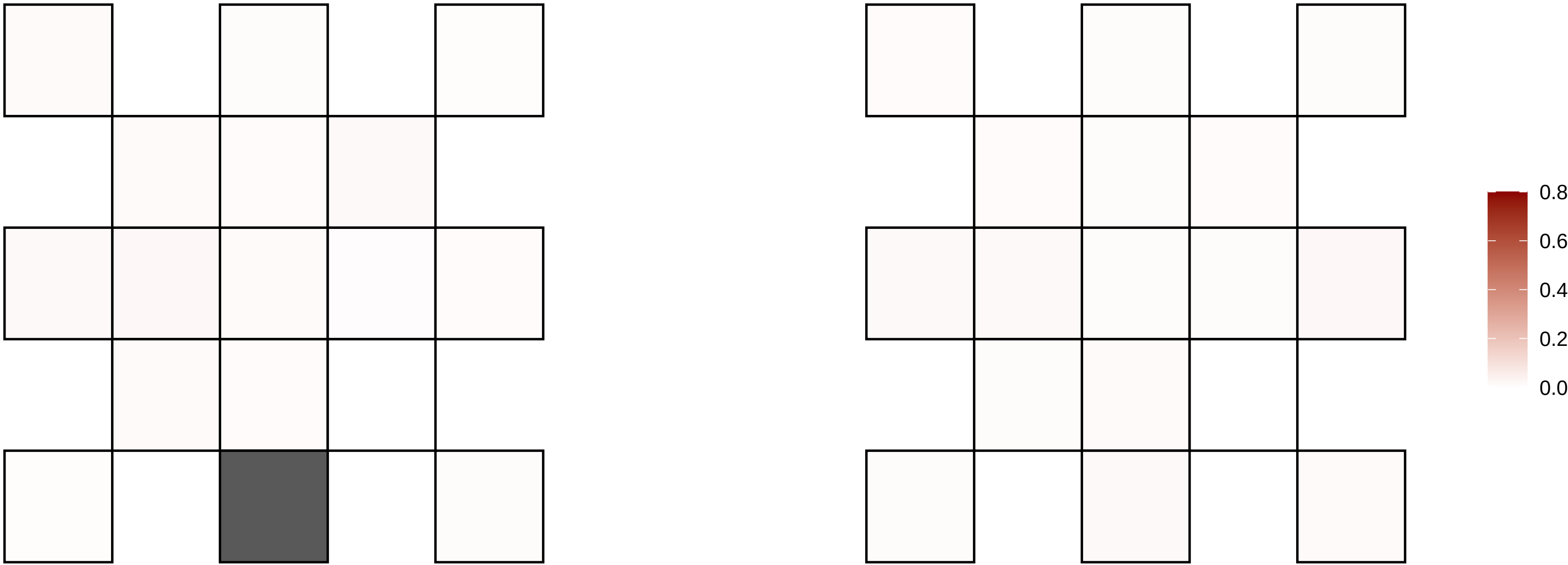

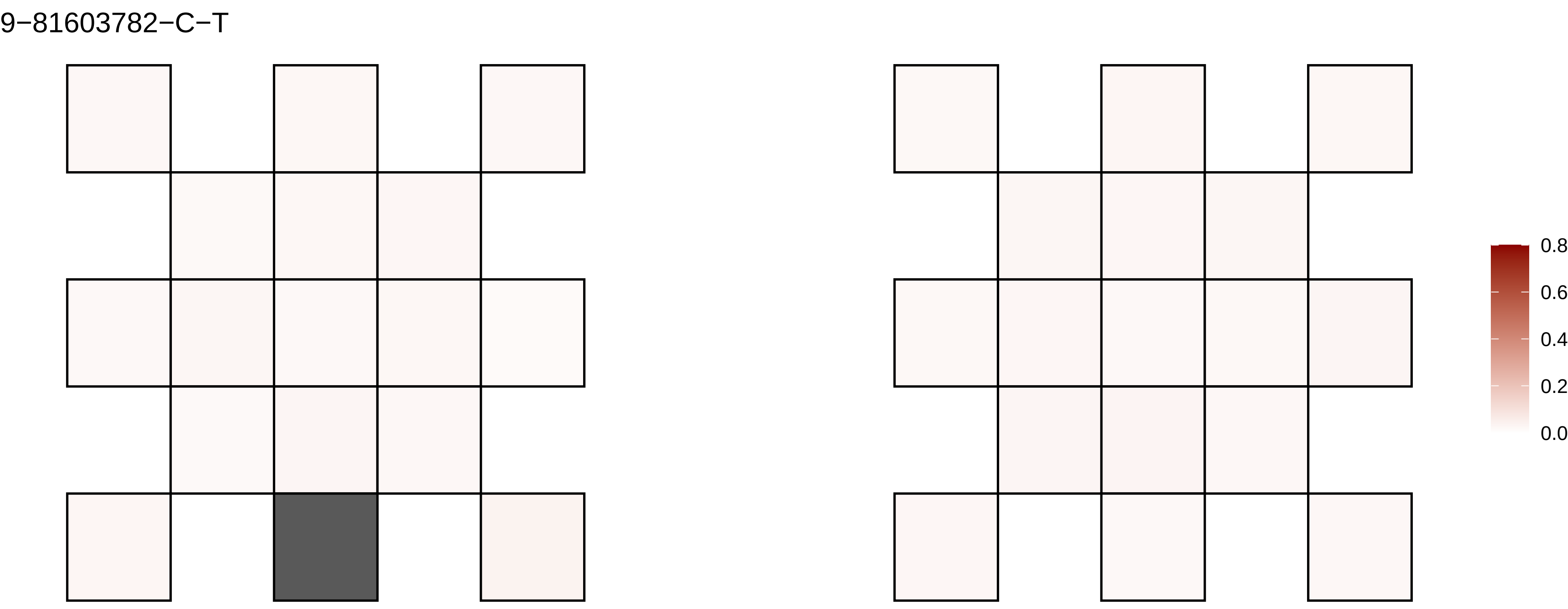
9−81603782−C−T
0.8
0.6
0.4
0.2
0.0

9−82881649−G−T

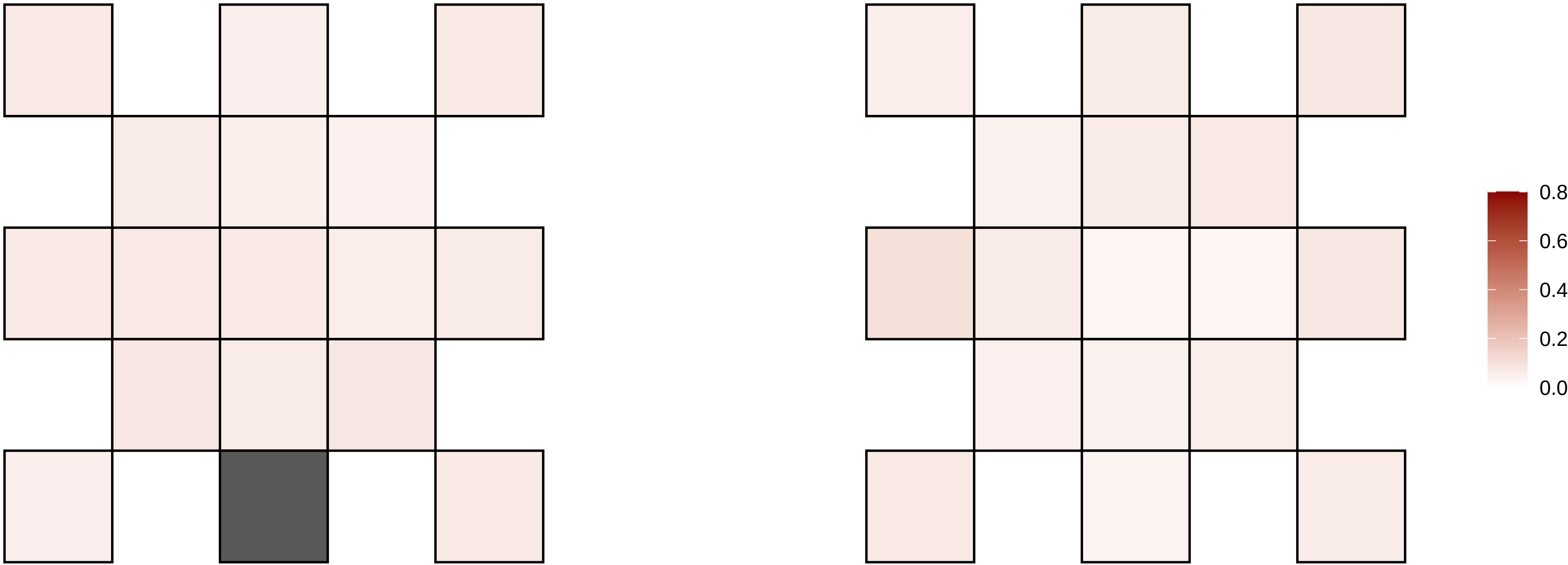

9−82881654−A−C

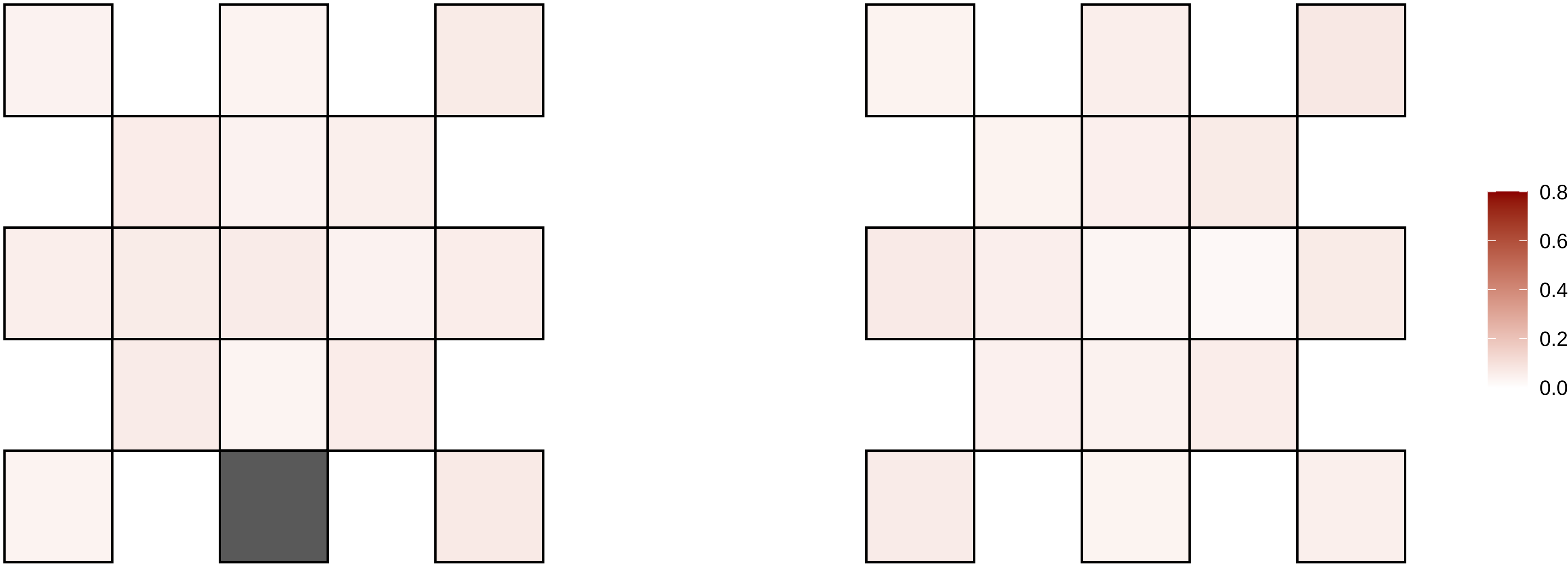

9−83044793−C−A

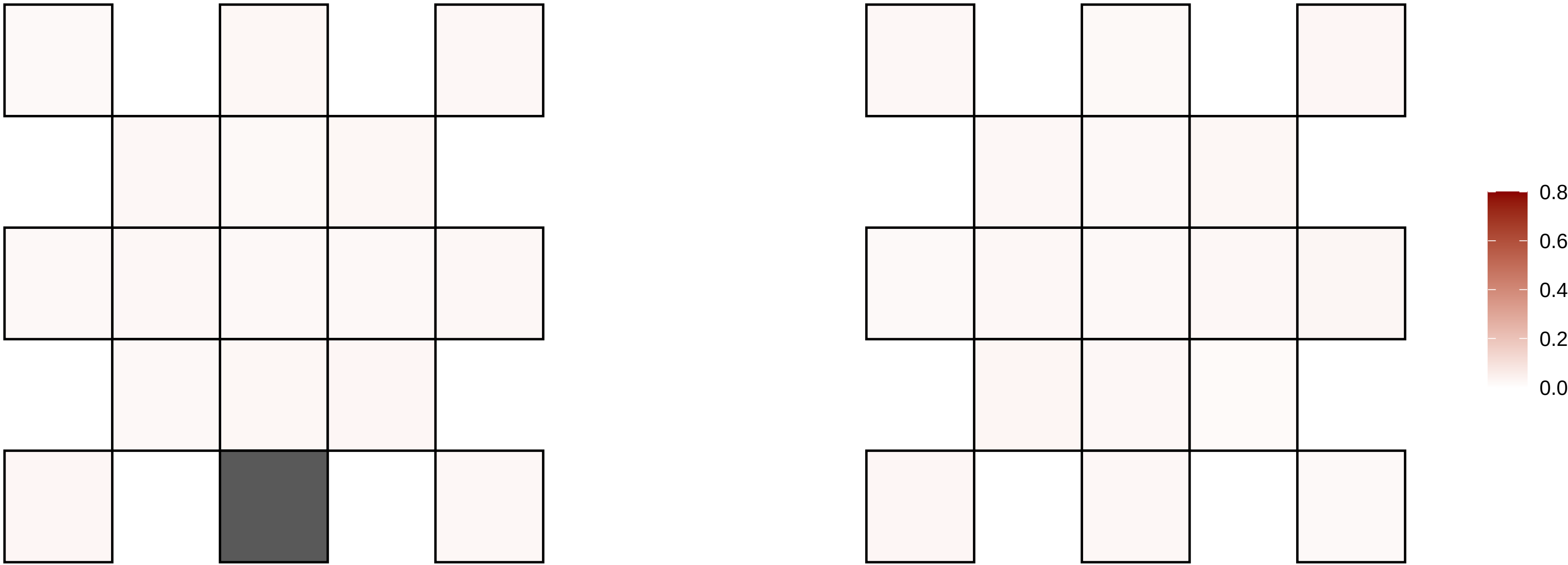

9−83538985−A−G

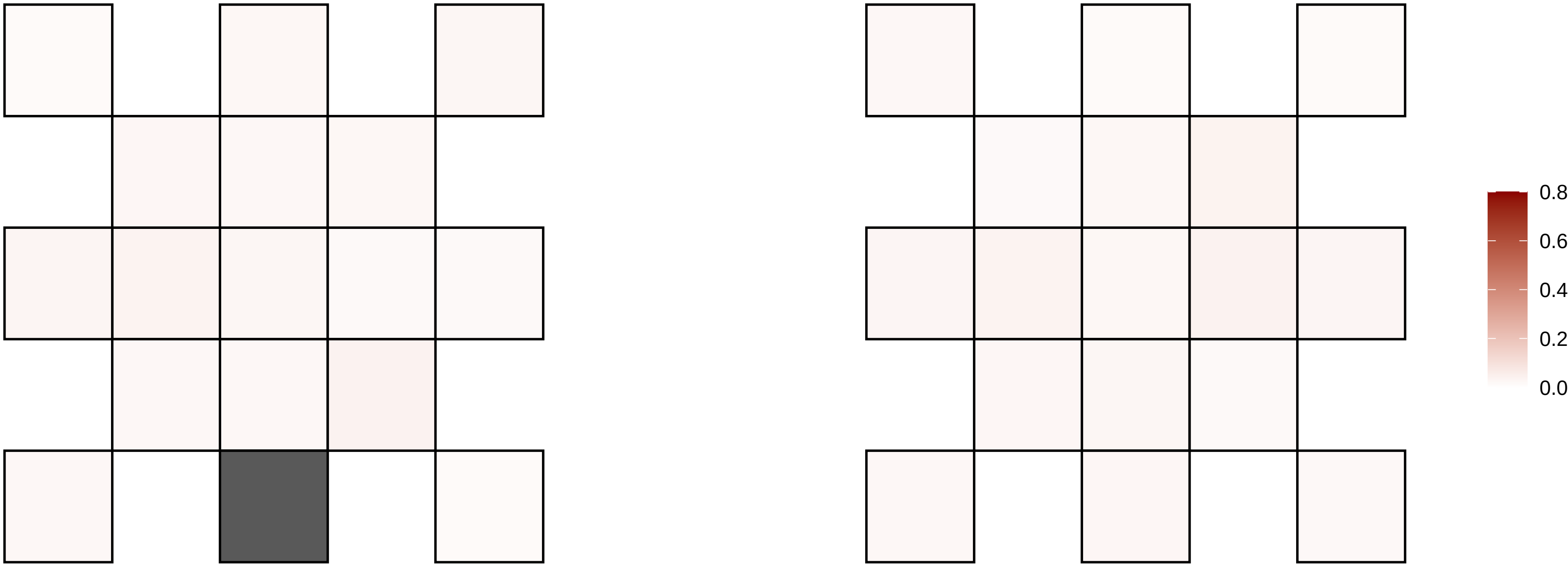

9−96677164−G−A

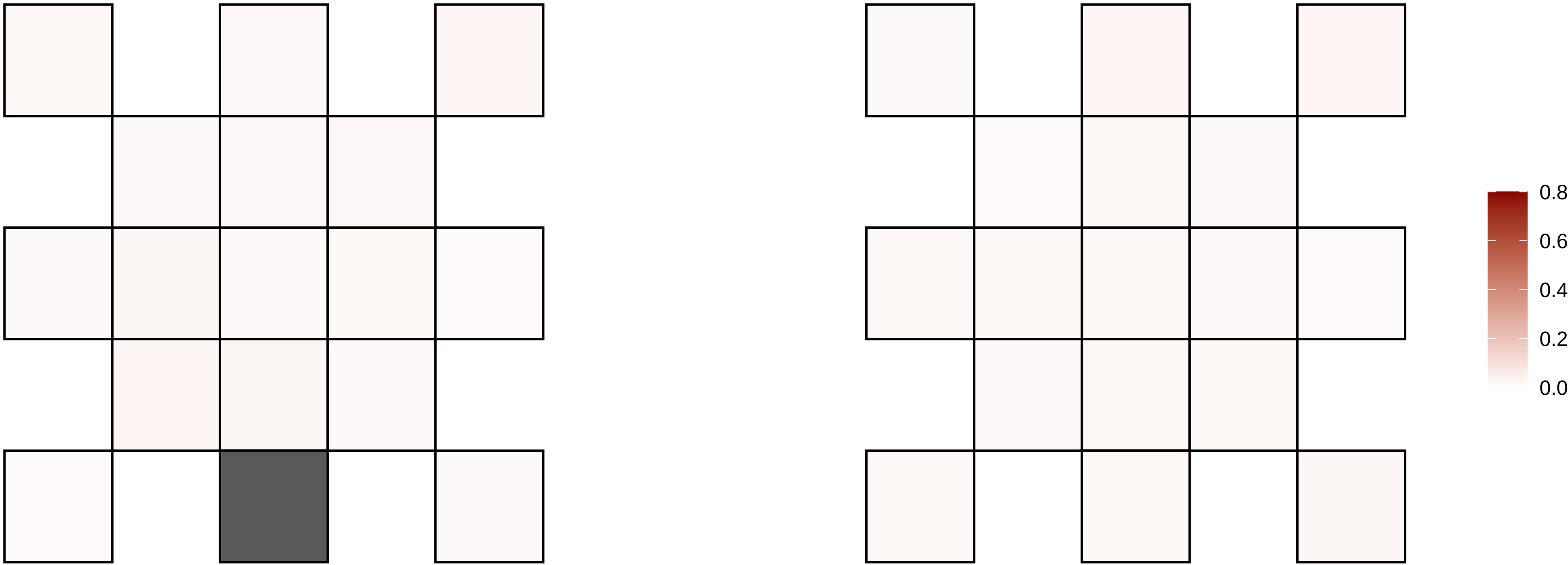

9−9971800−C−T

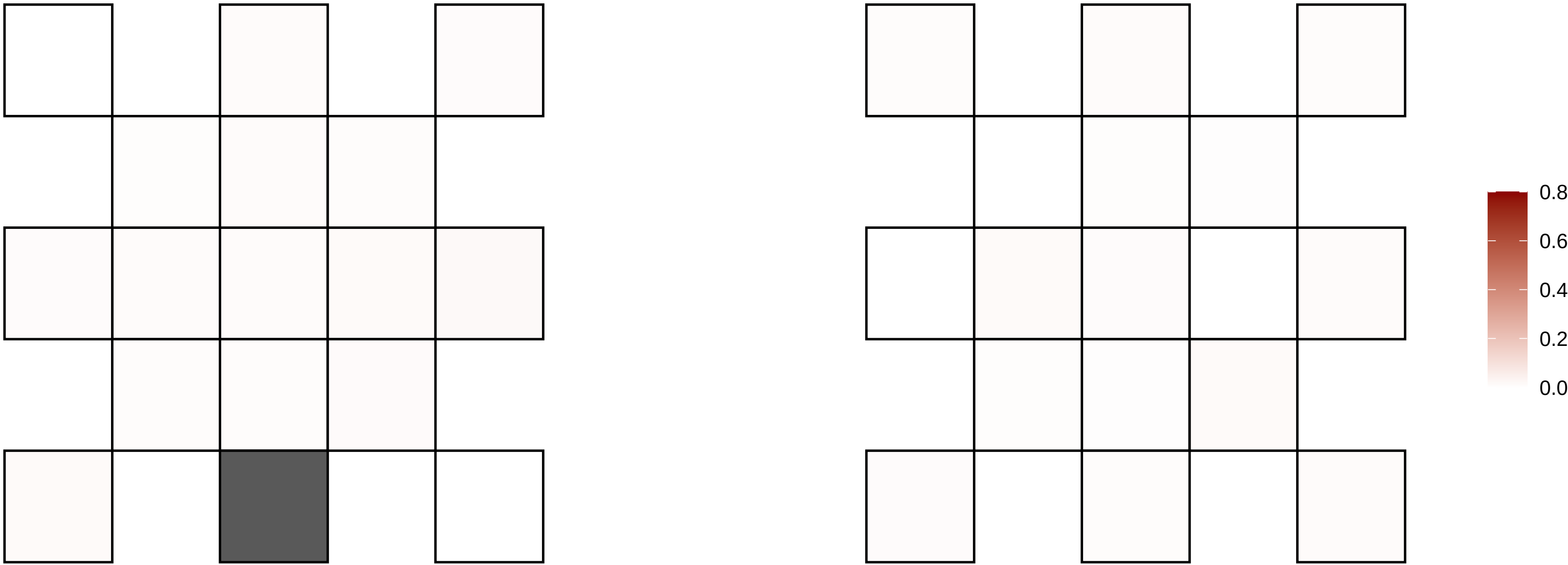

X−107301303−G−T

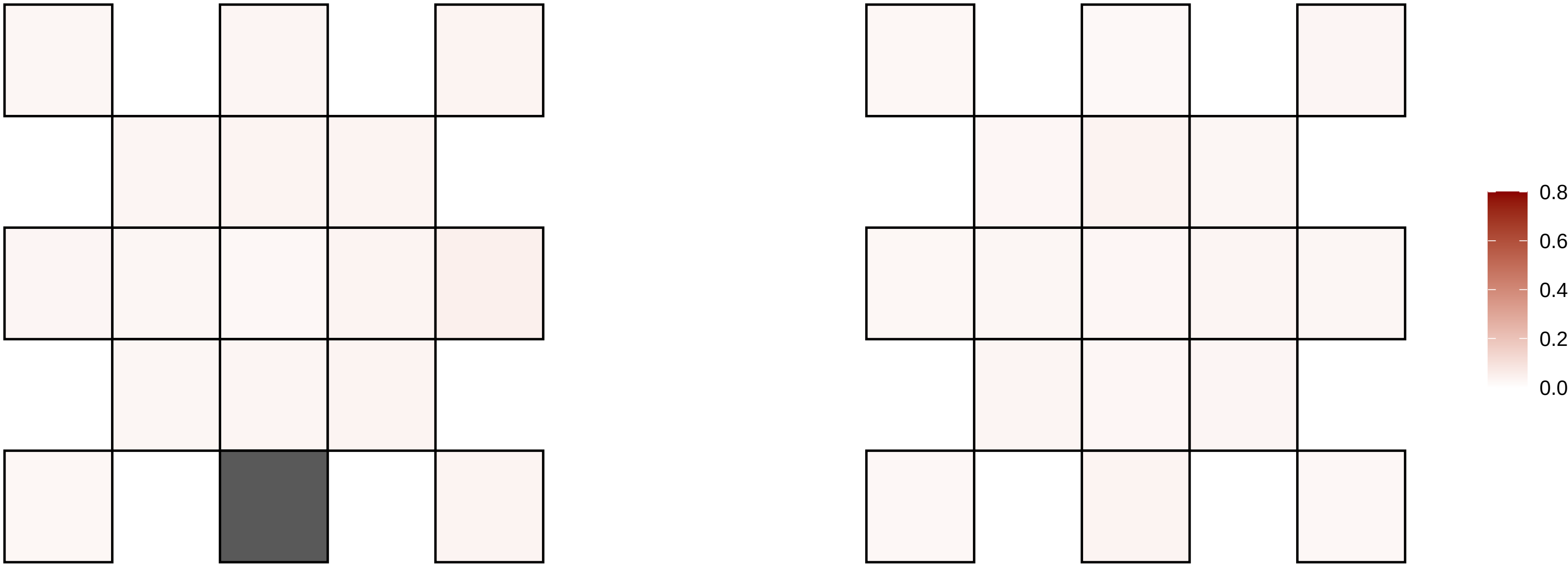

X−109753054−C−T

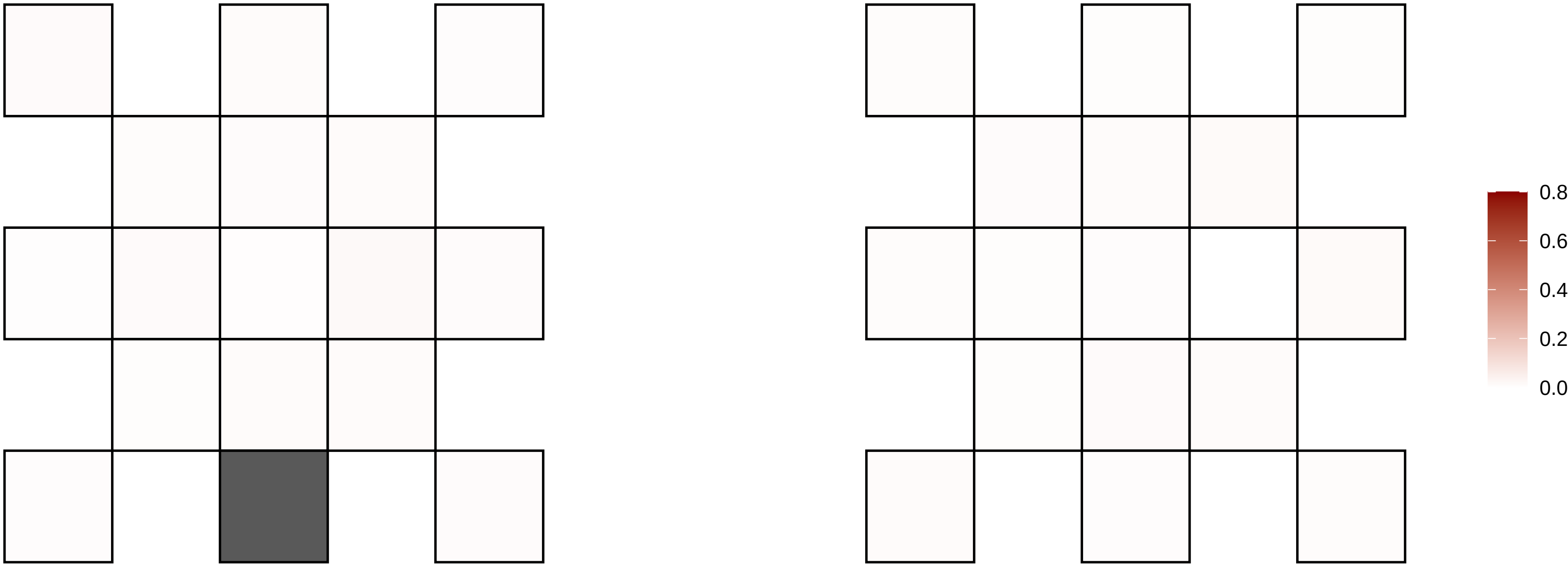

X−109939493−G−A

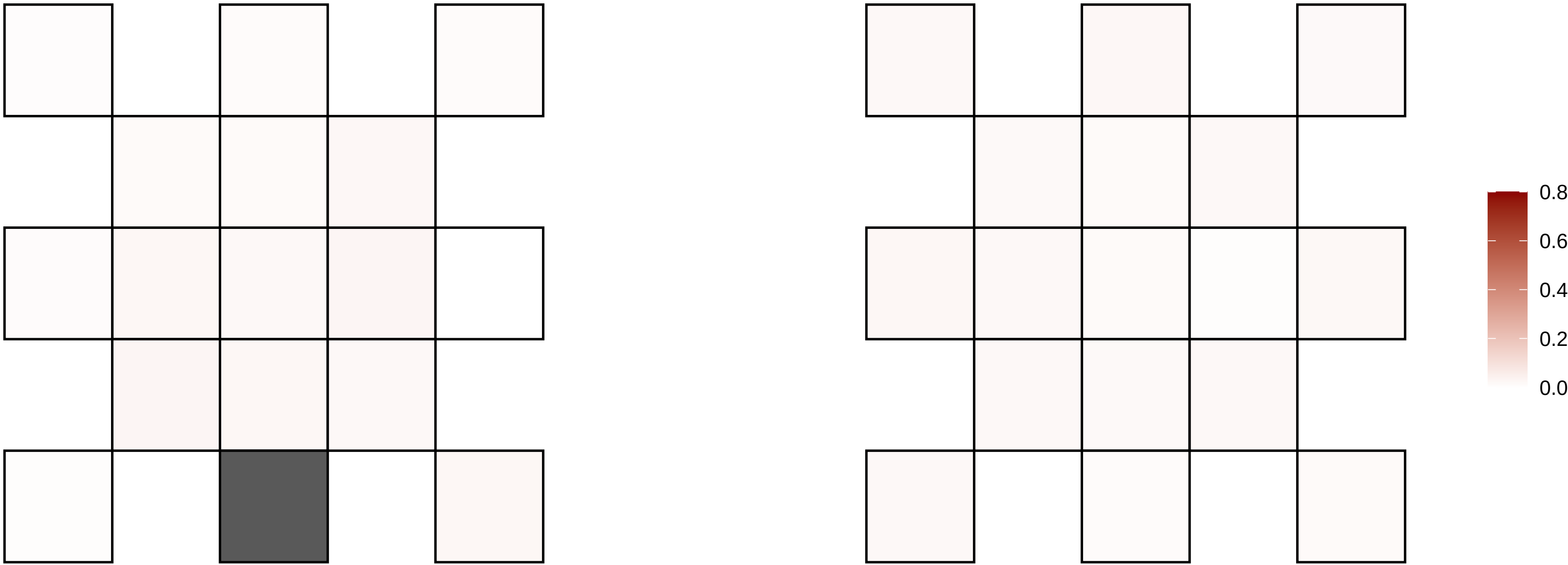

X−117240525−A−C

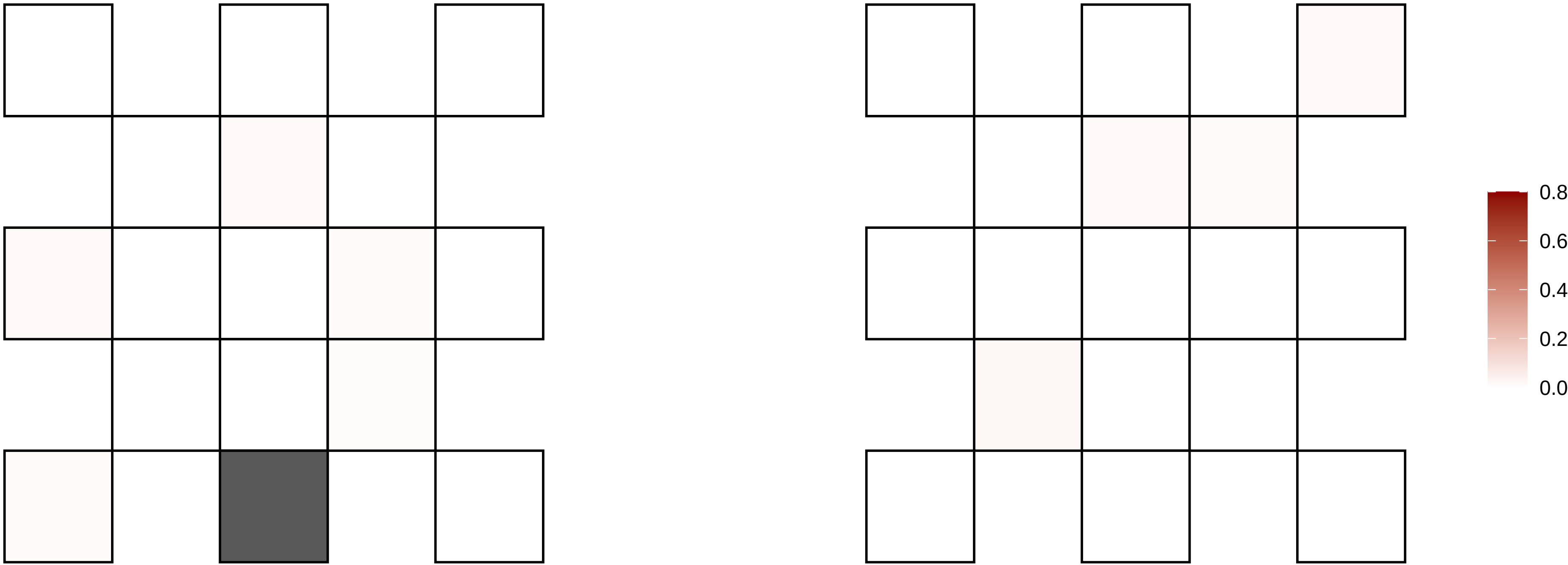

X−122220079−G−A

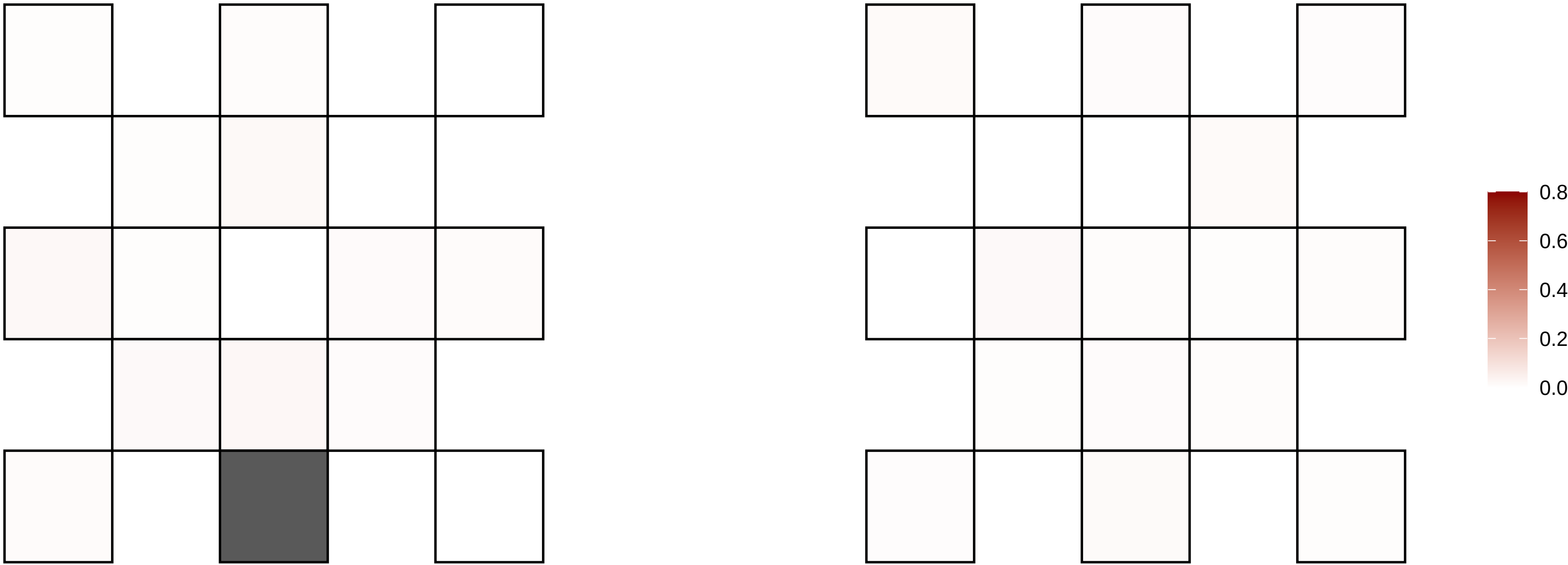

X−125464024−G−A

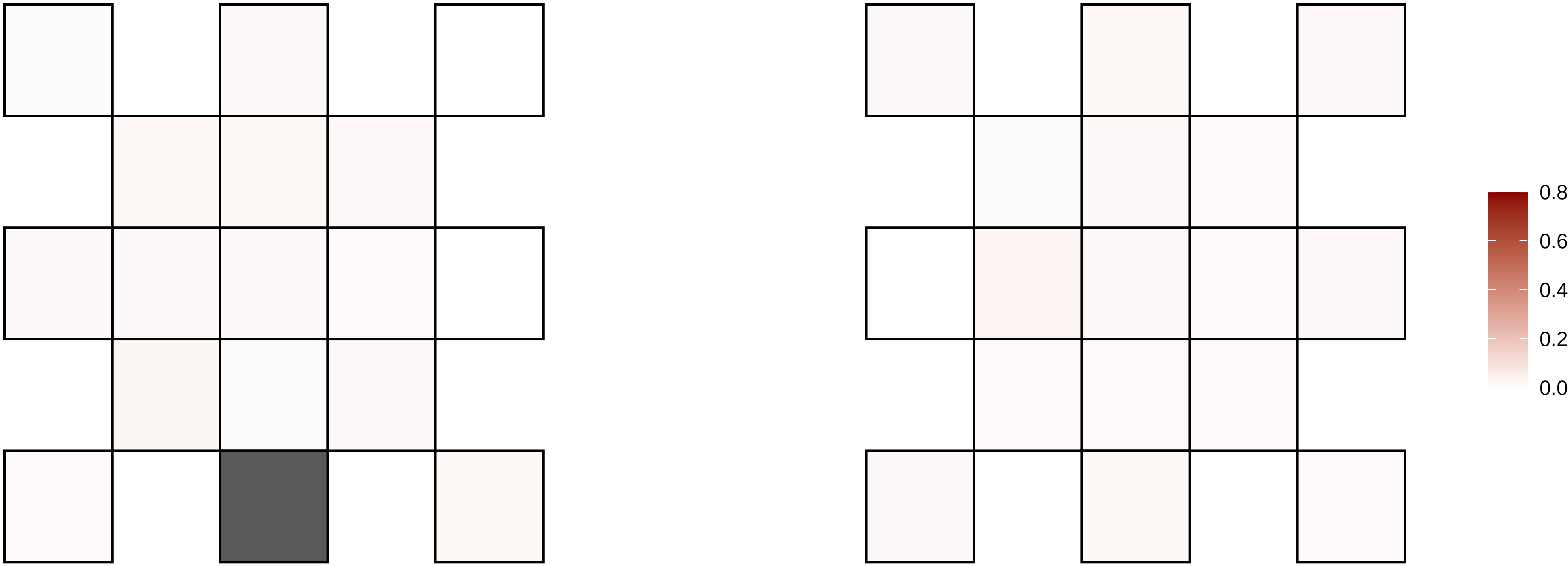

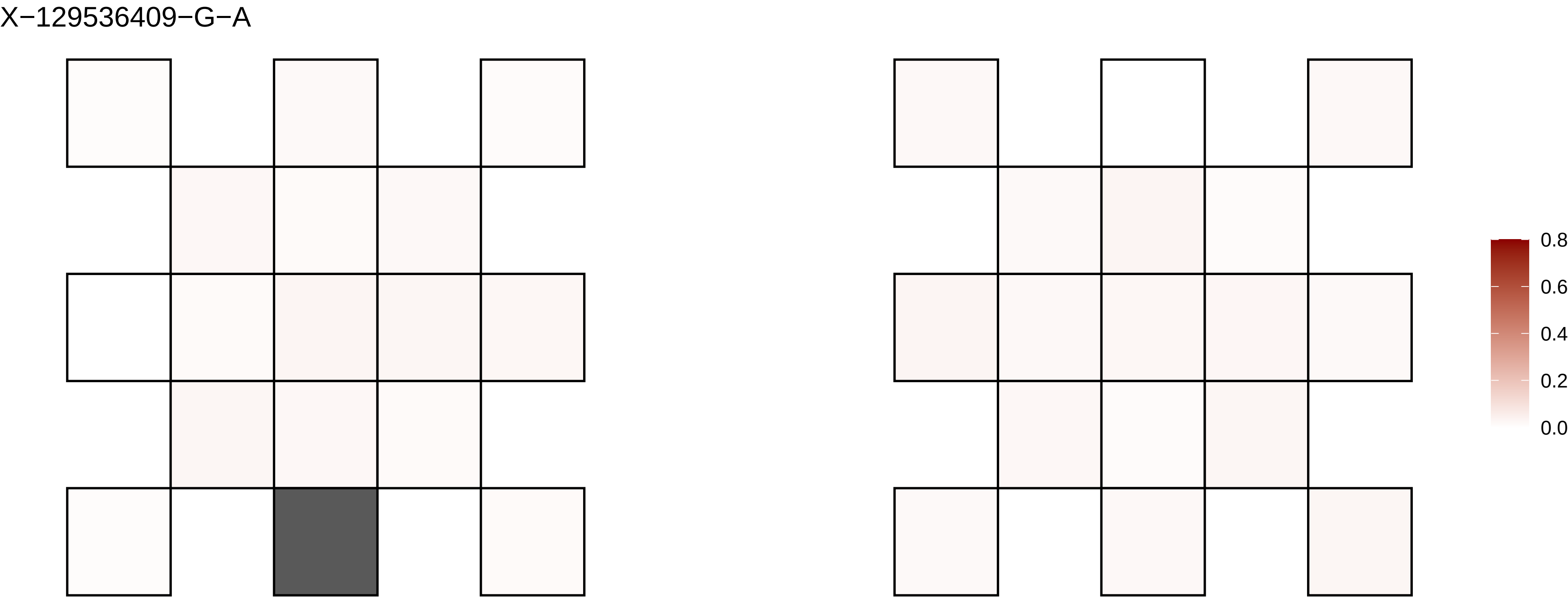
X−129536409−G−A
0.8
0.6
0.4
0.2
0.0

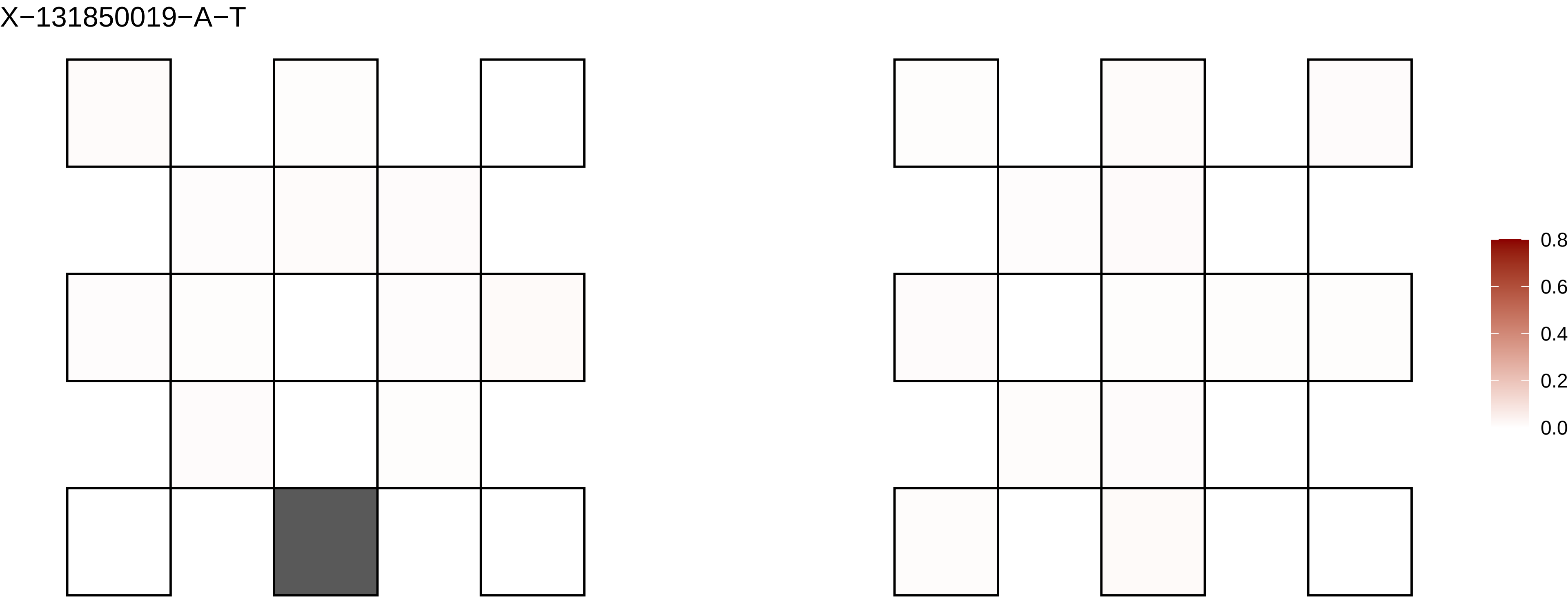
X−131850019−A−T
0.8
0.6
0.4
0.2
0.0

X−138211912−G−A

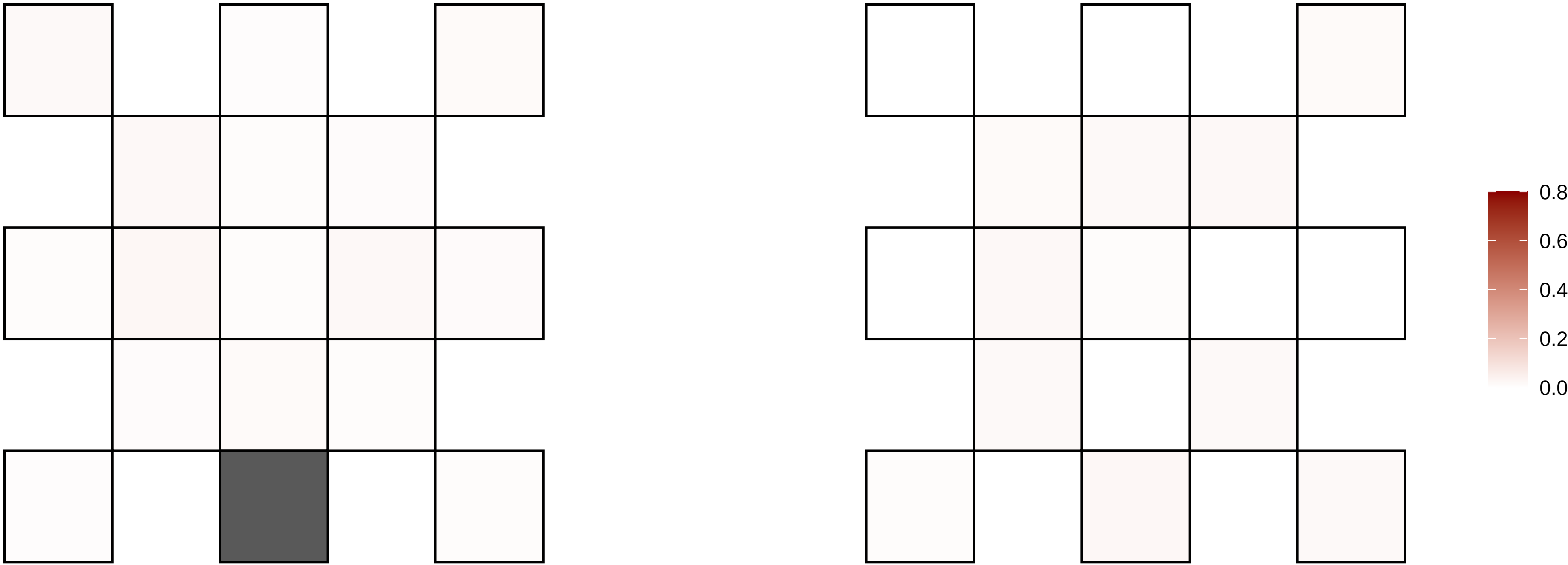

X−139604228−A−G

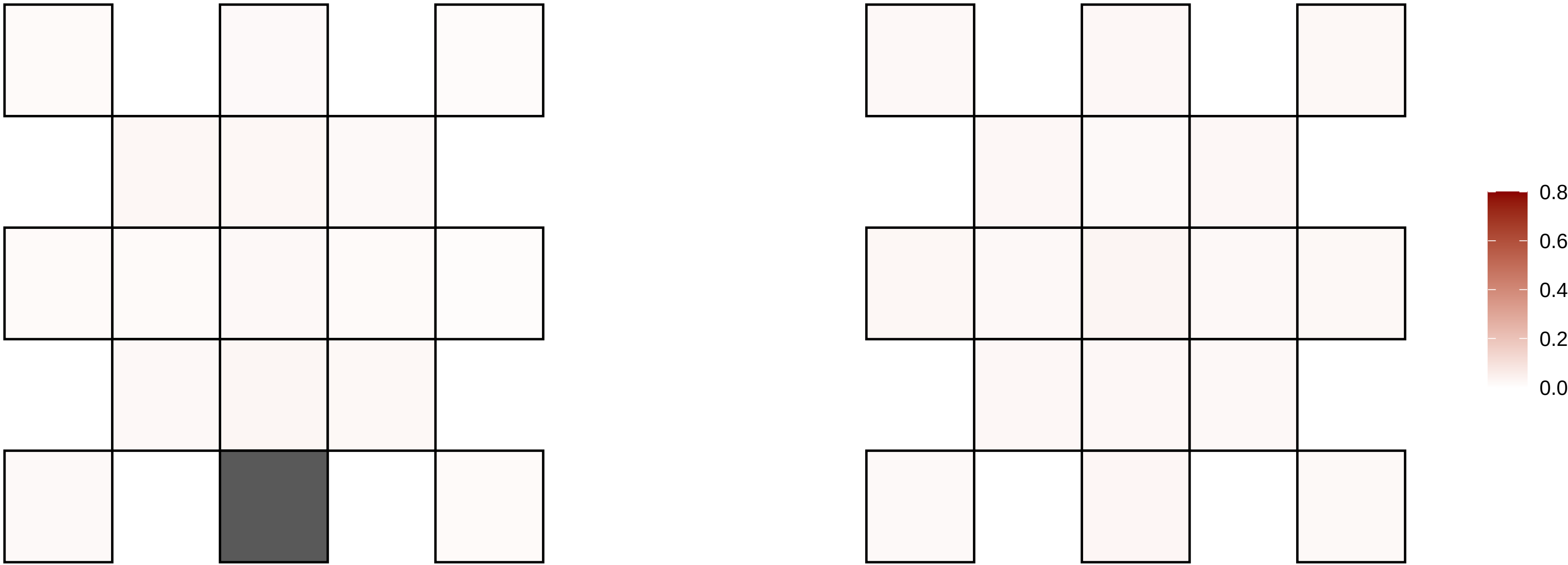

X−140066644−G−T

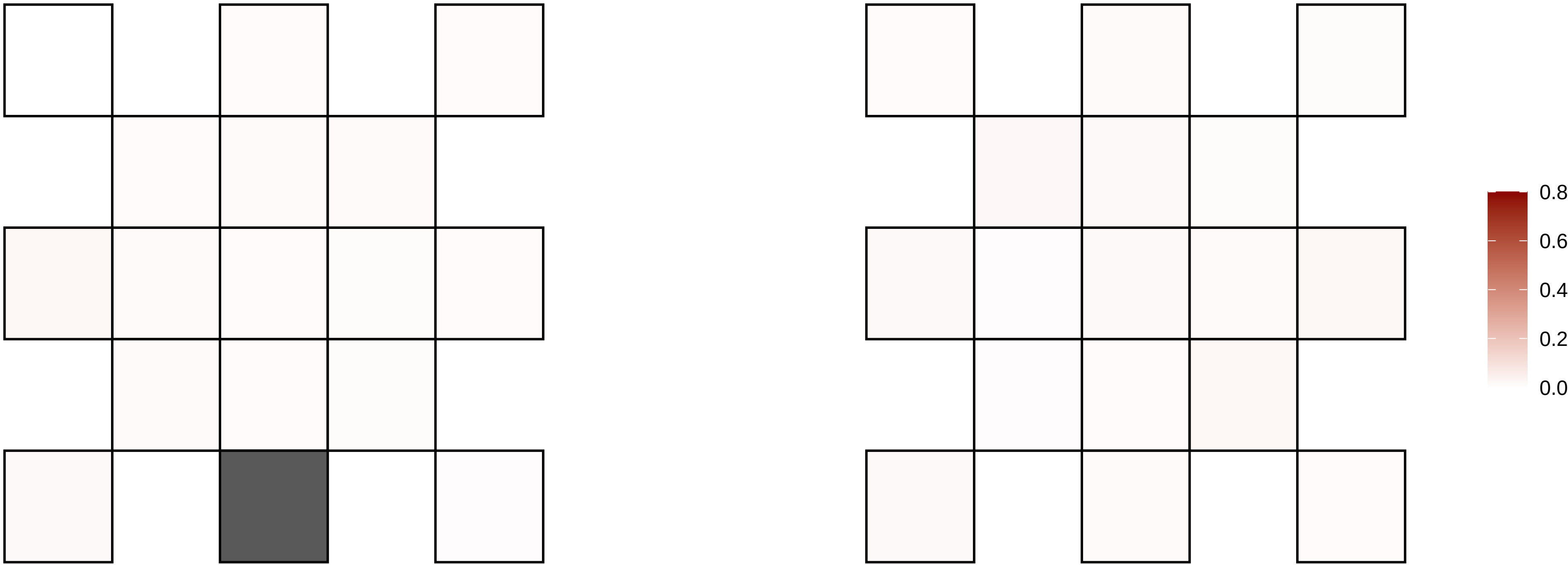

X−141738997−A−G

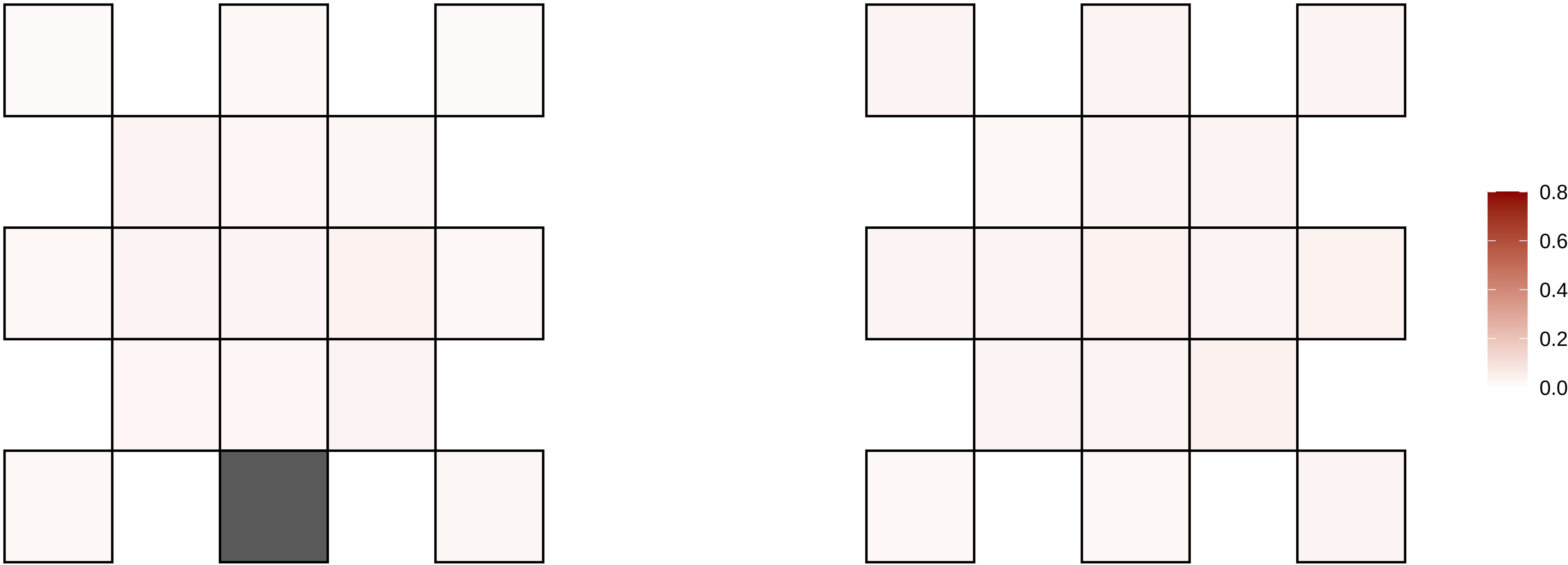

X−143150040−A−T

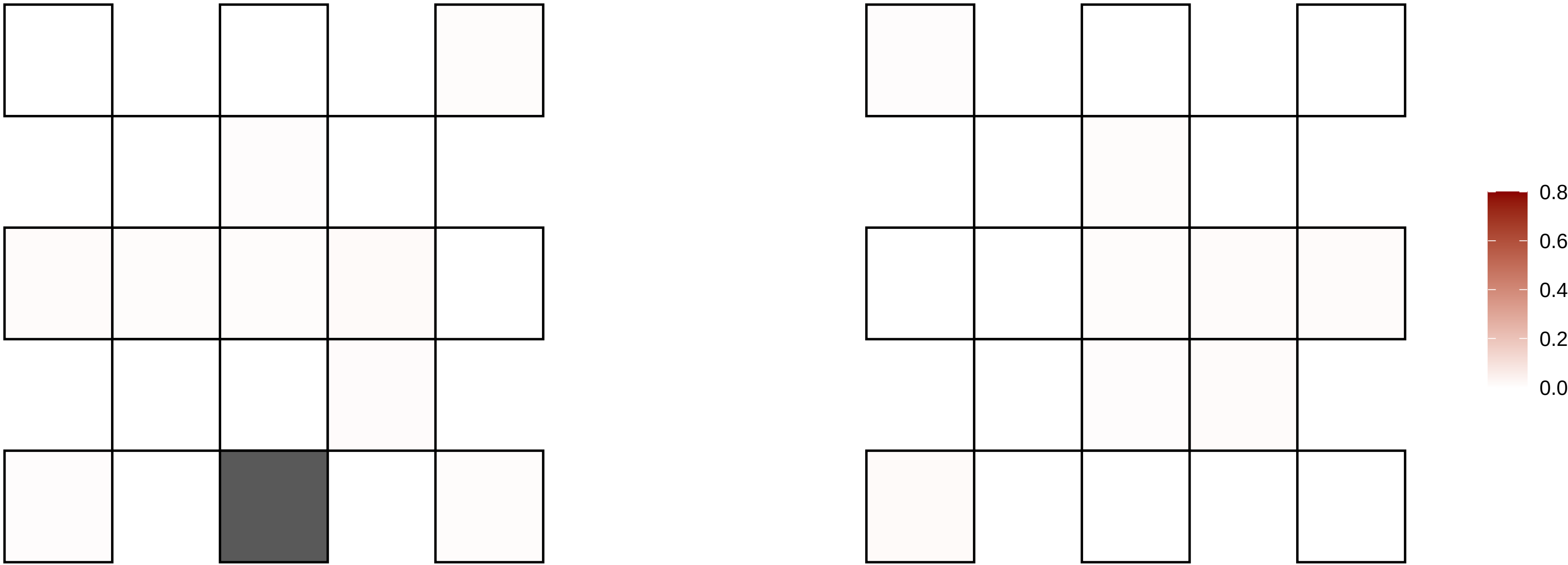

X−146647450−C−T

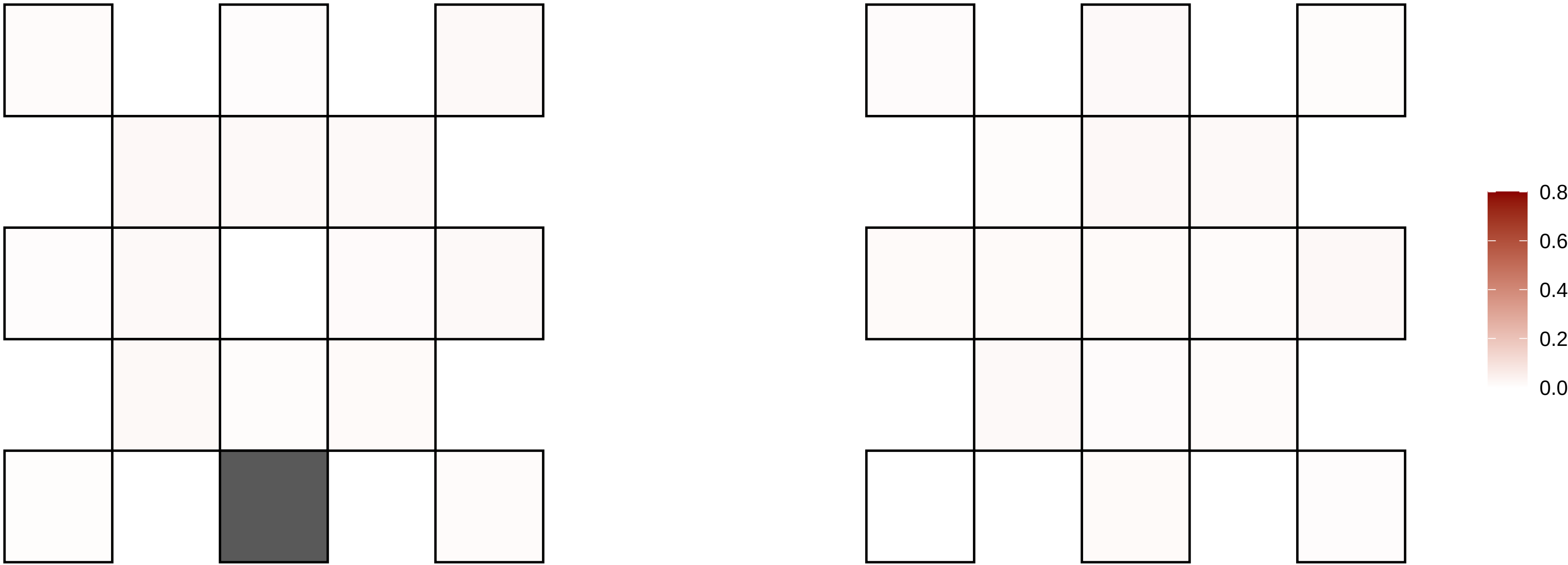

X−150945179−C−A

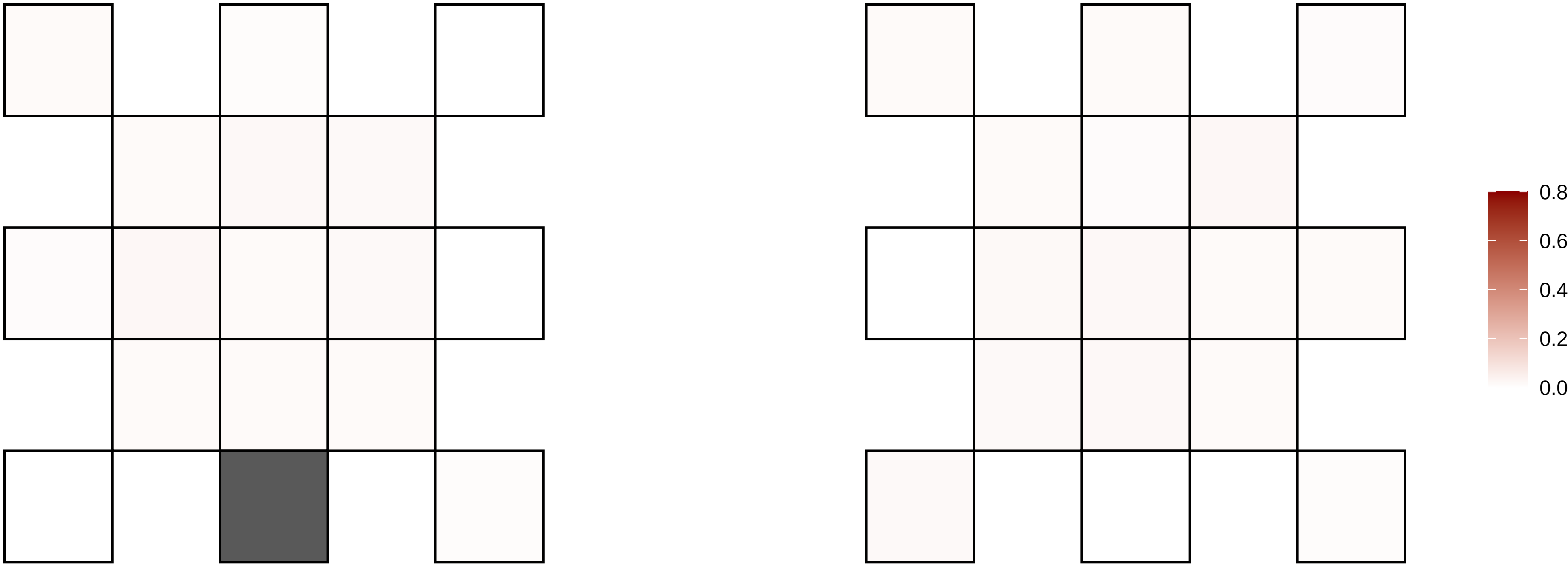

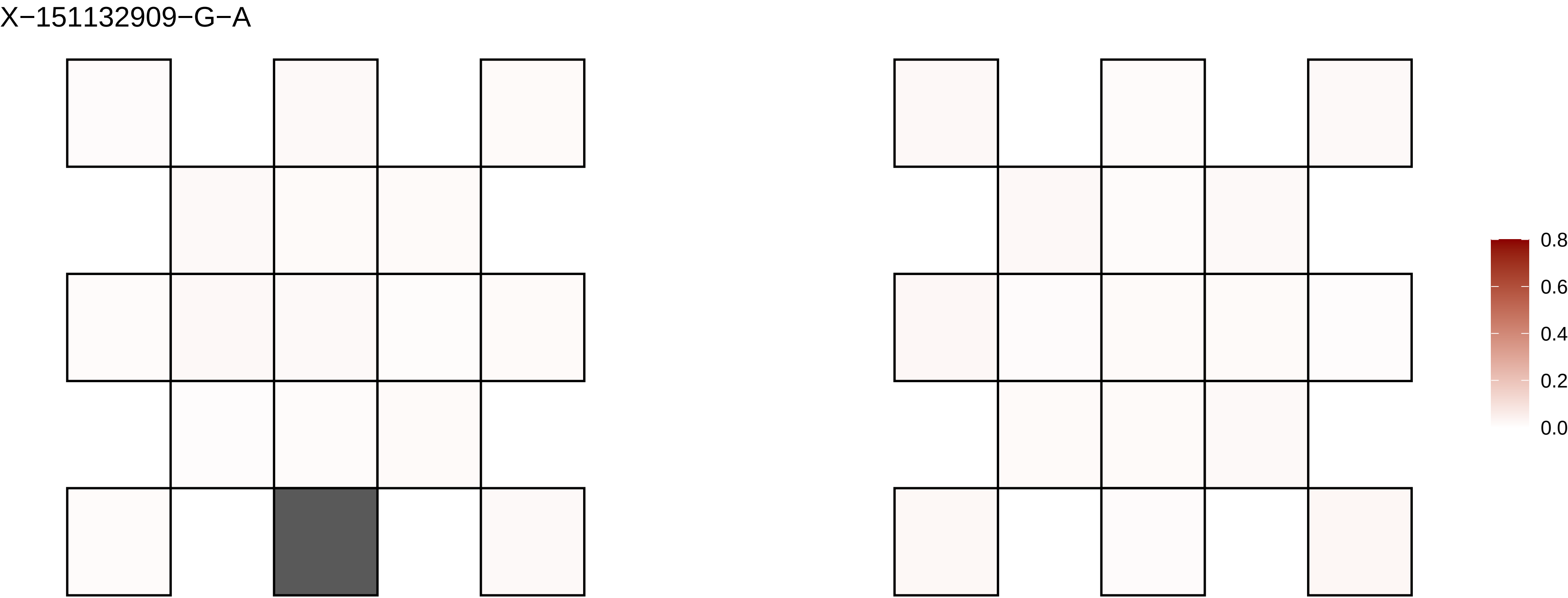
X−151132909−G−A
0.8
0.6
0.4
0.2
0.0

X−16444039−G−A

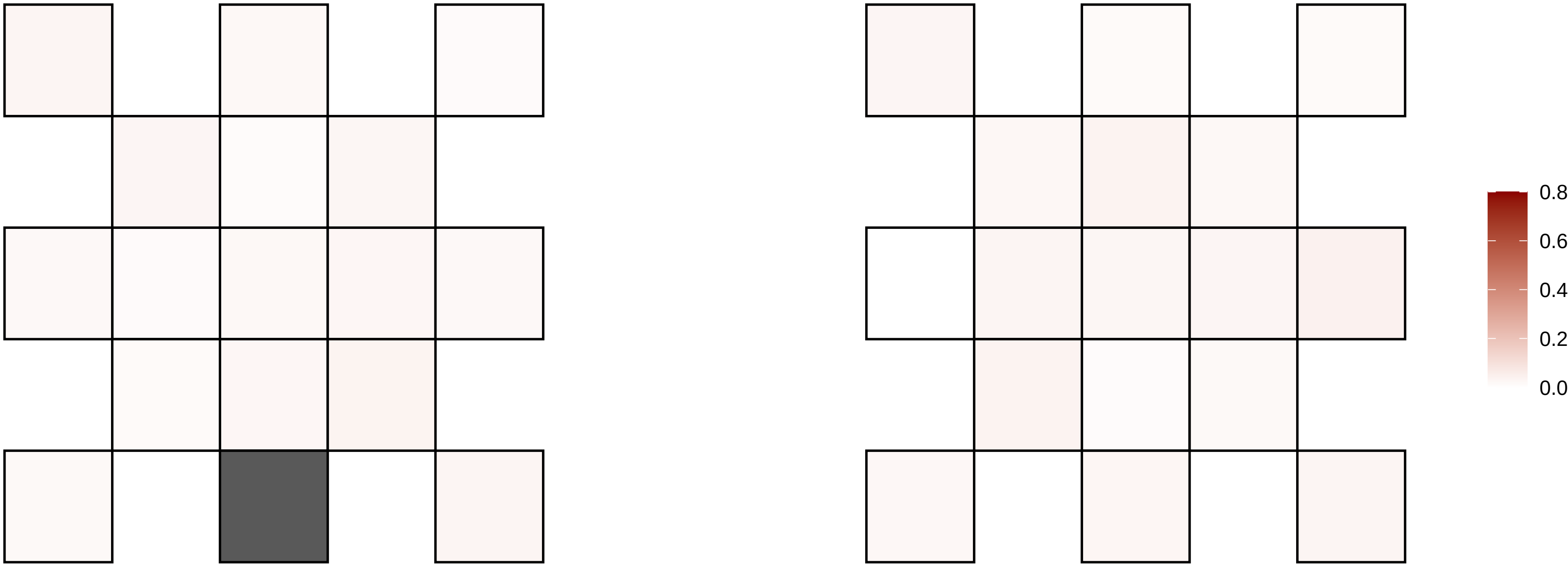

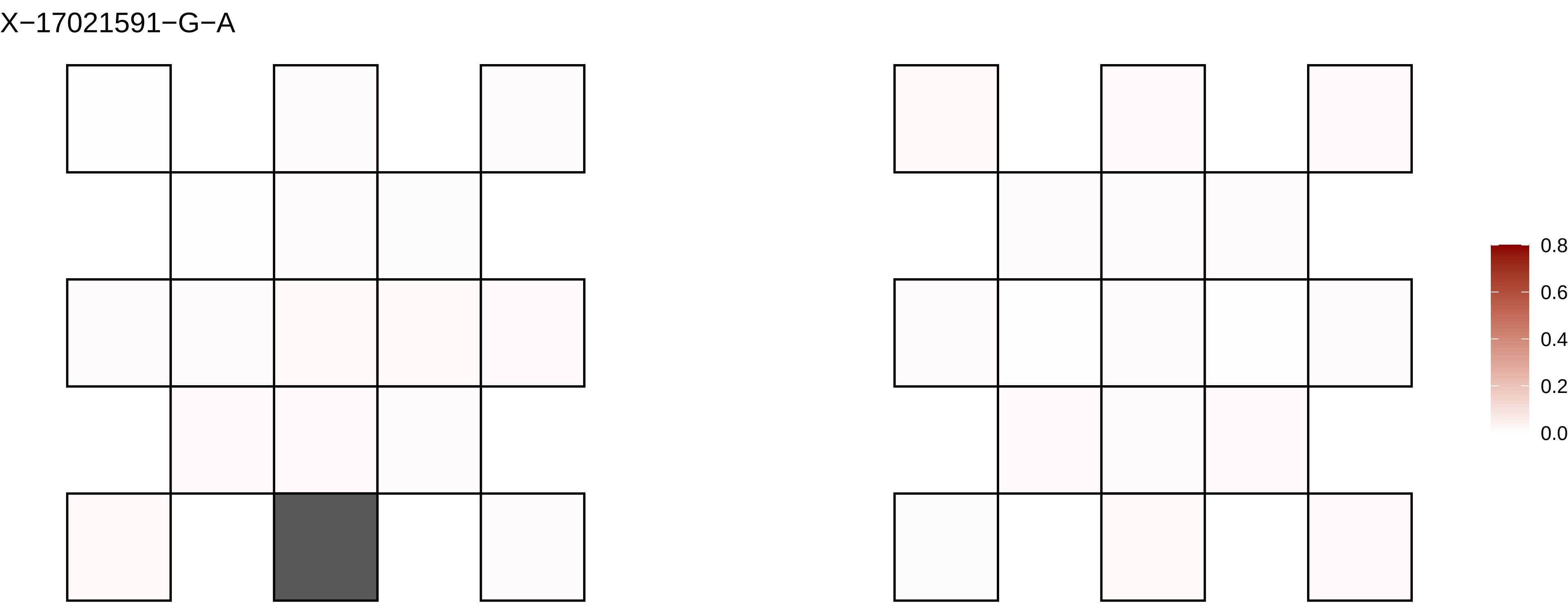
X−17021591−G−A
0.8
0.6
0.4
0.2
0.0

X−17091941−T−G

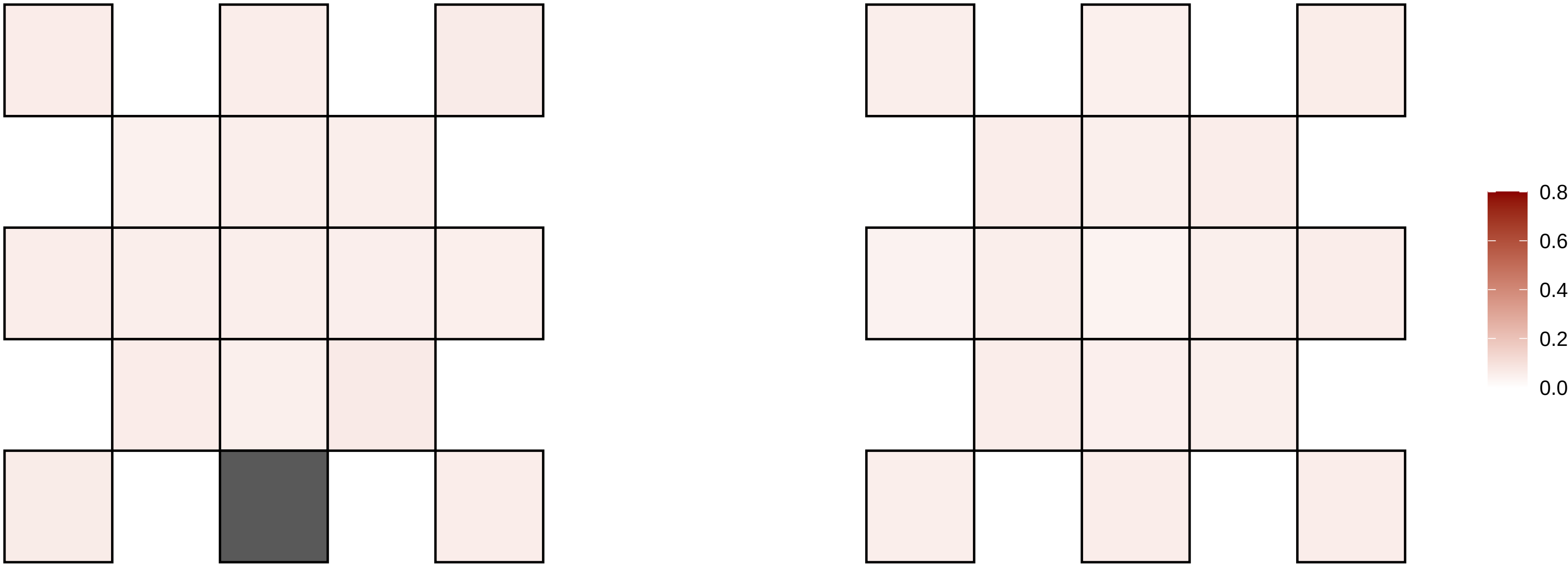

X−17091942−C−A

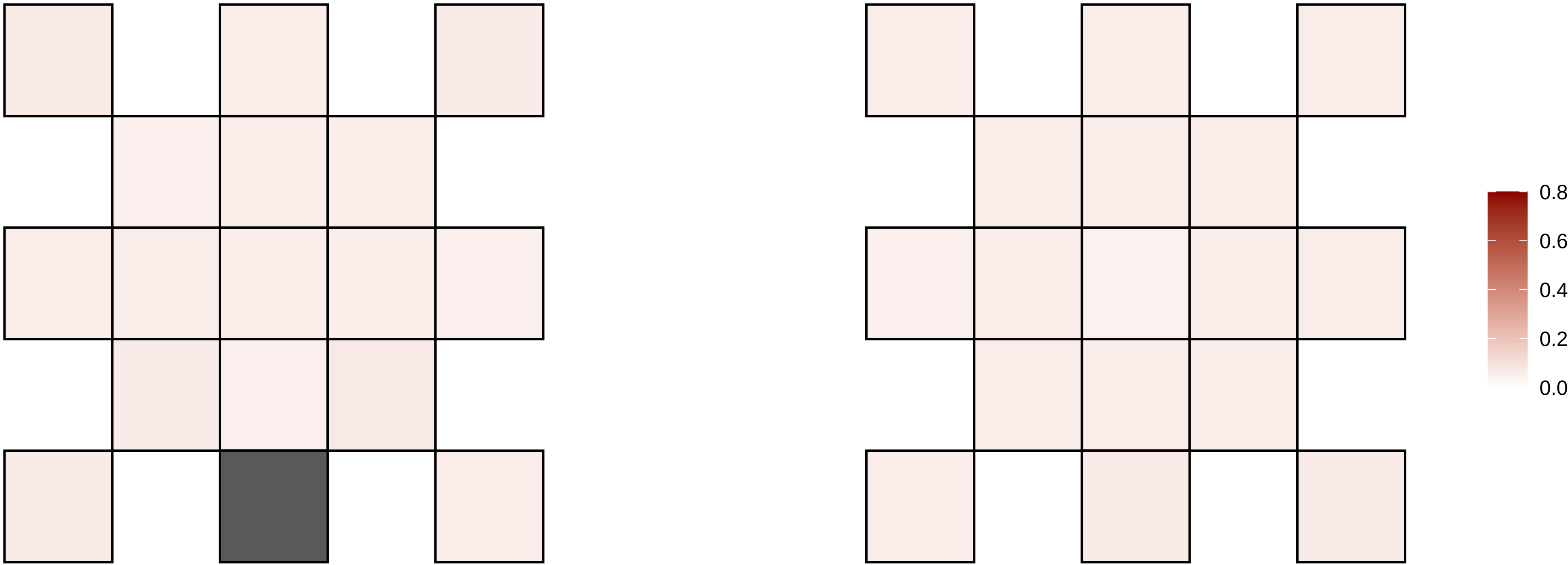

X−21528963−A−C

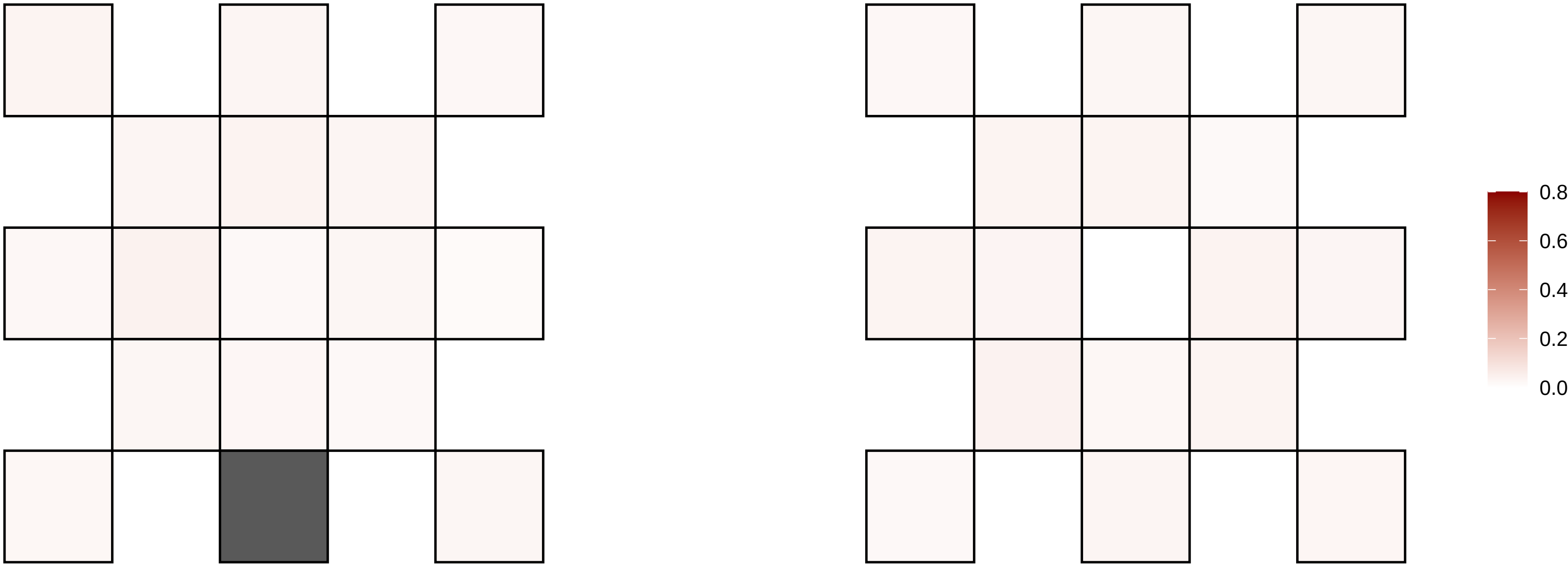

X−24541835−C−A

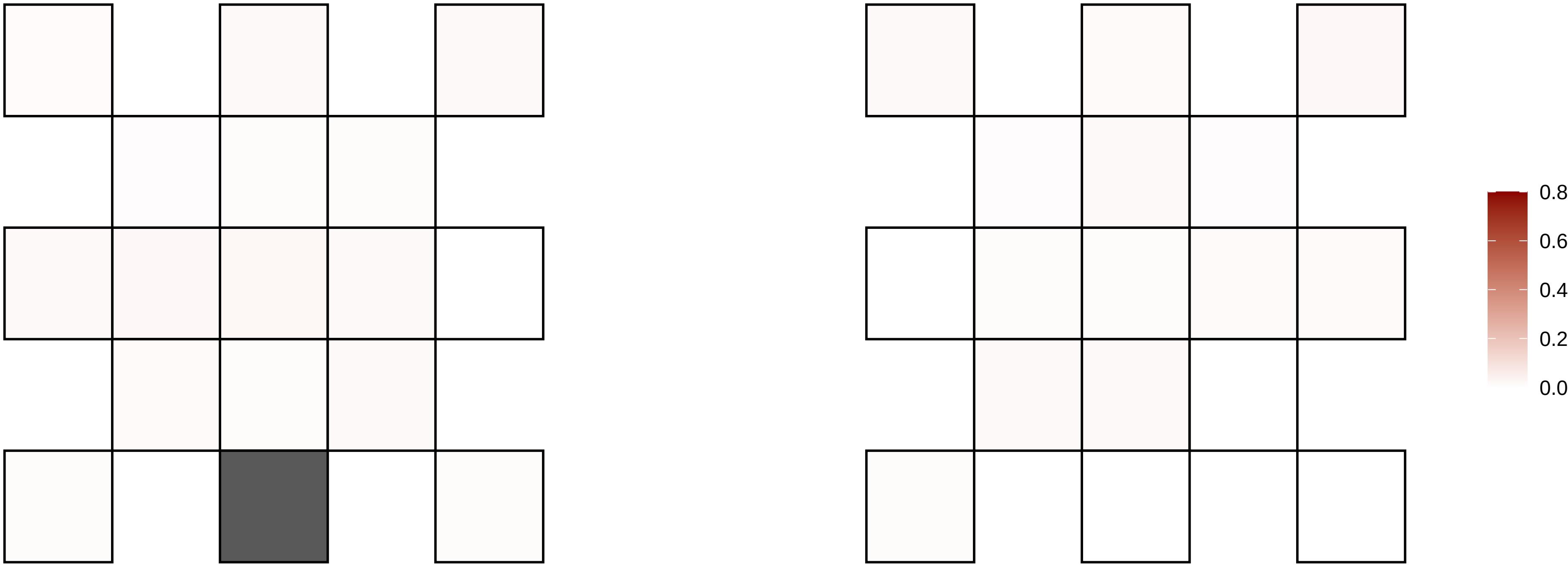

X−27063252−A−G

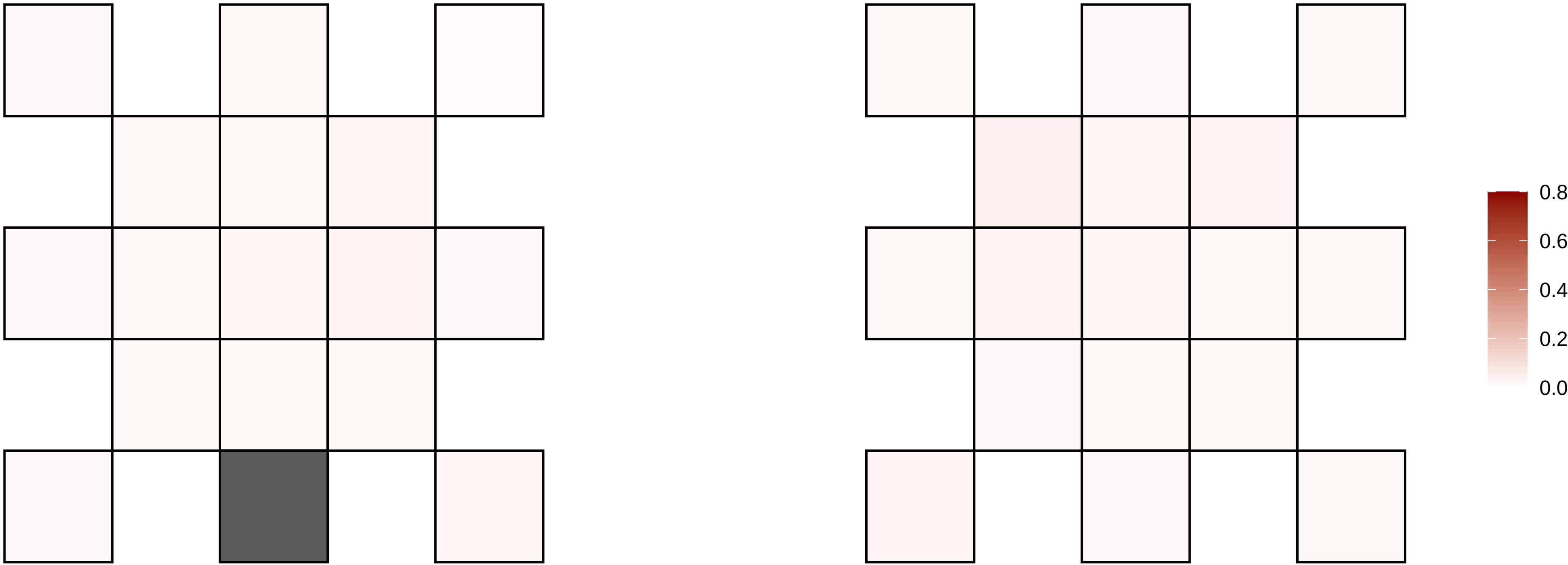

X−28357791−T−A

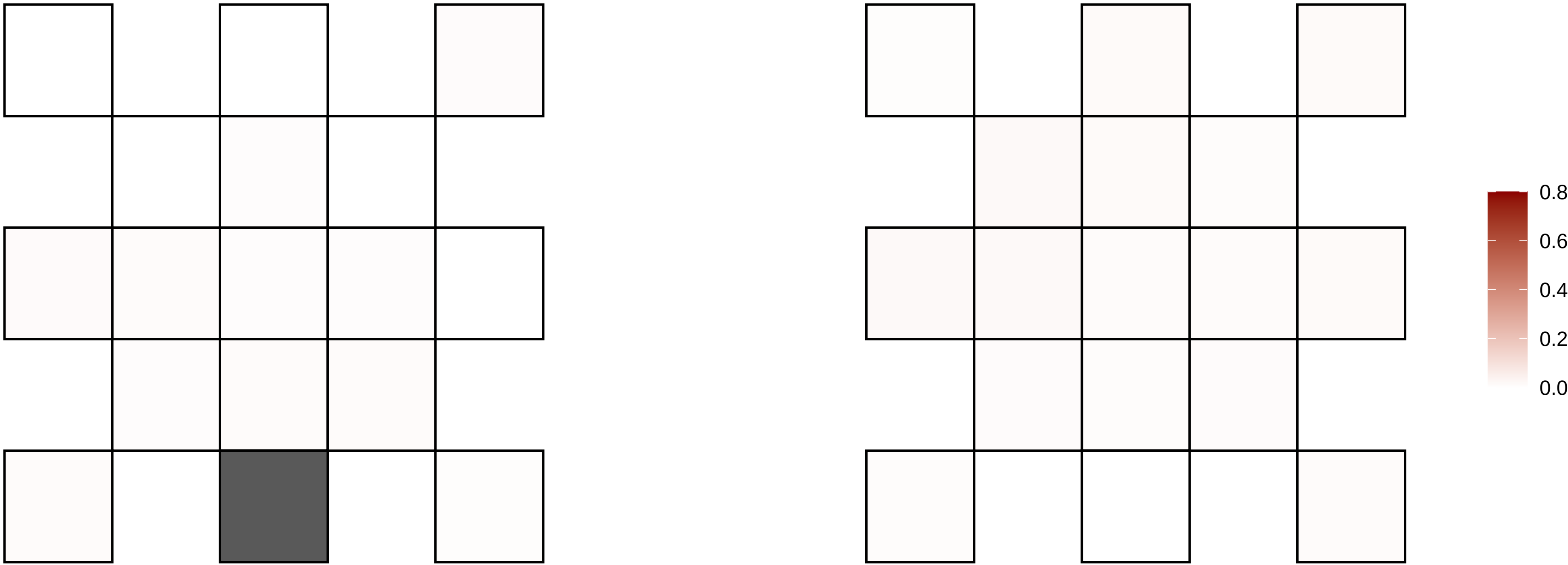

X−32465017−G−A

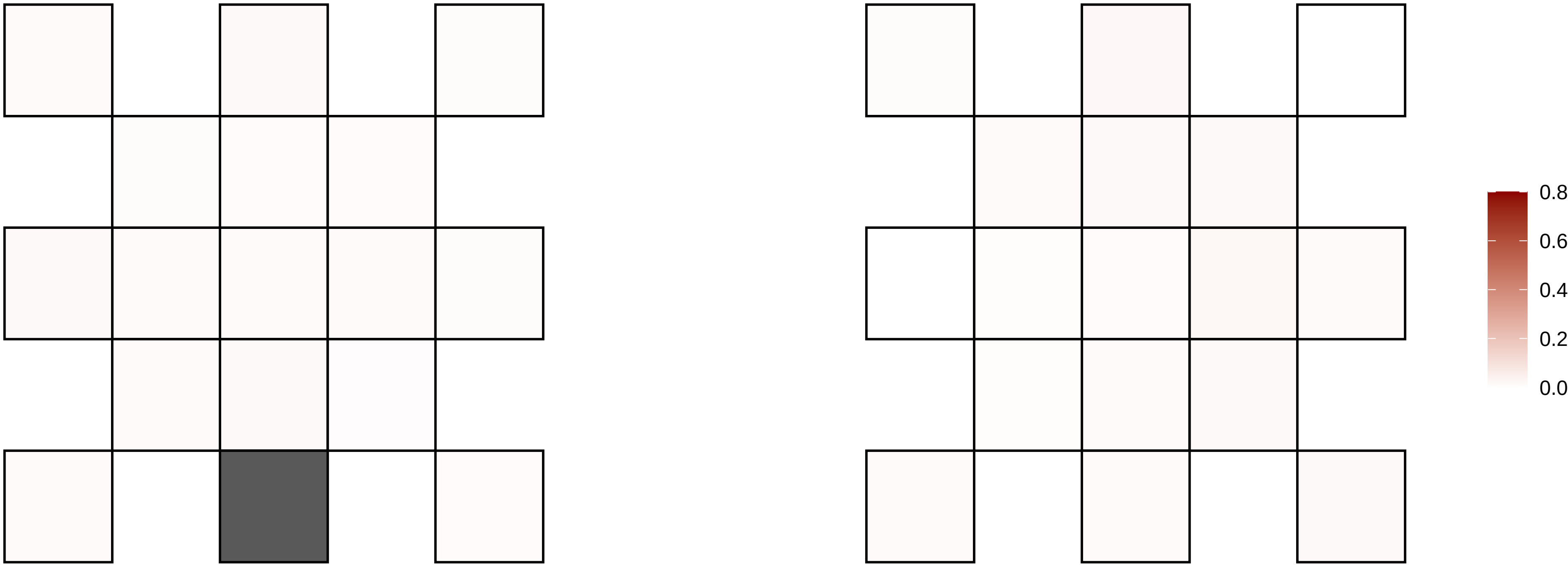

X−33444958−G−A

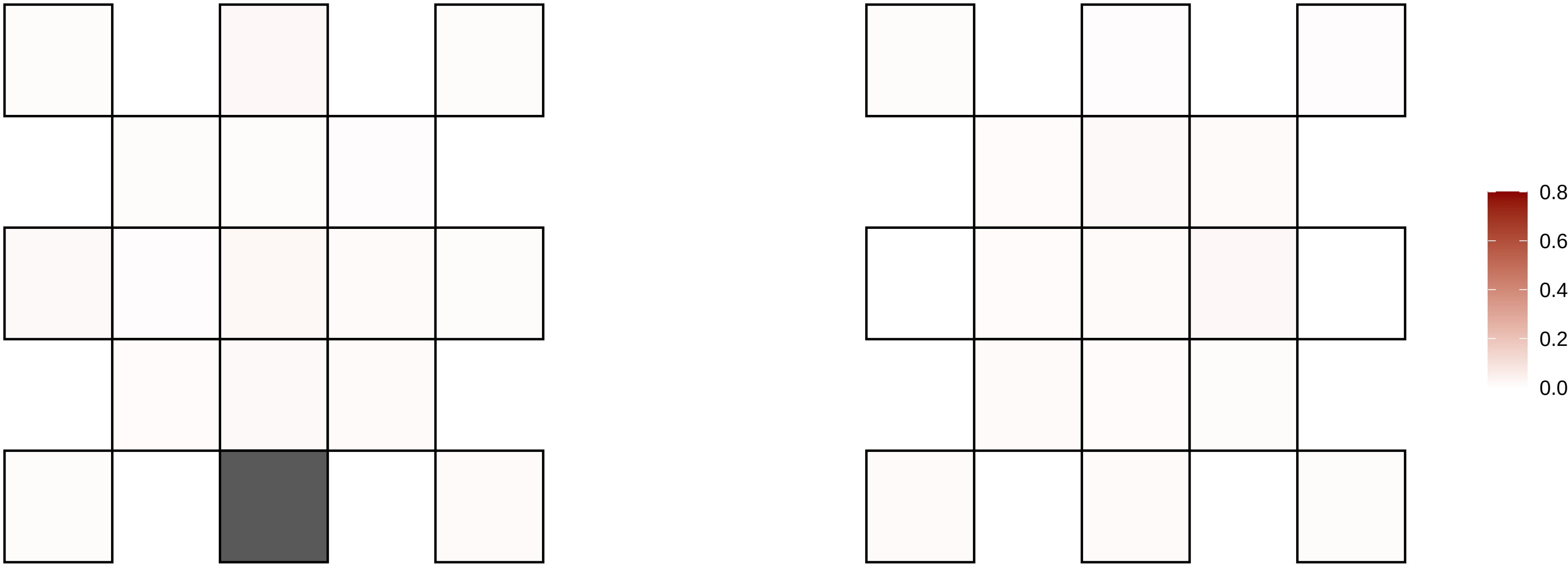

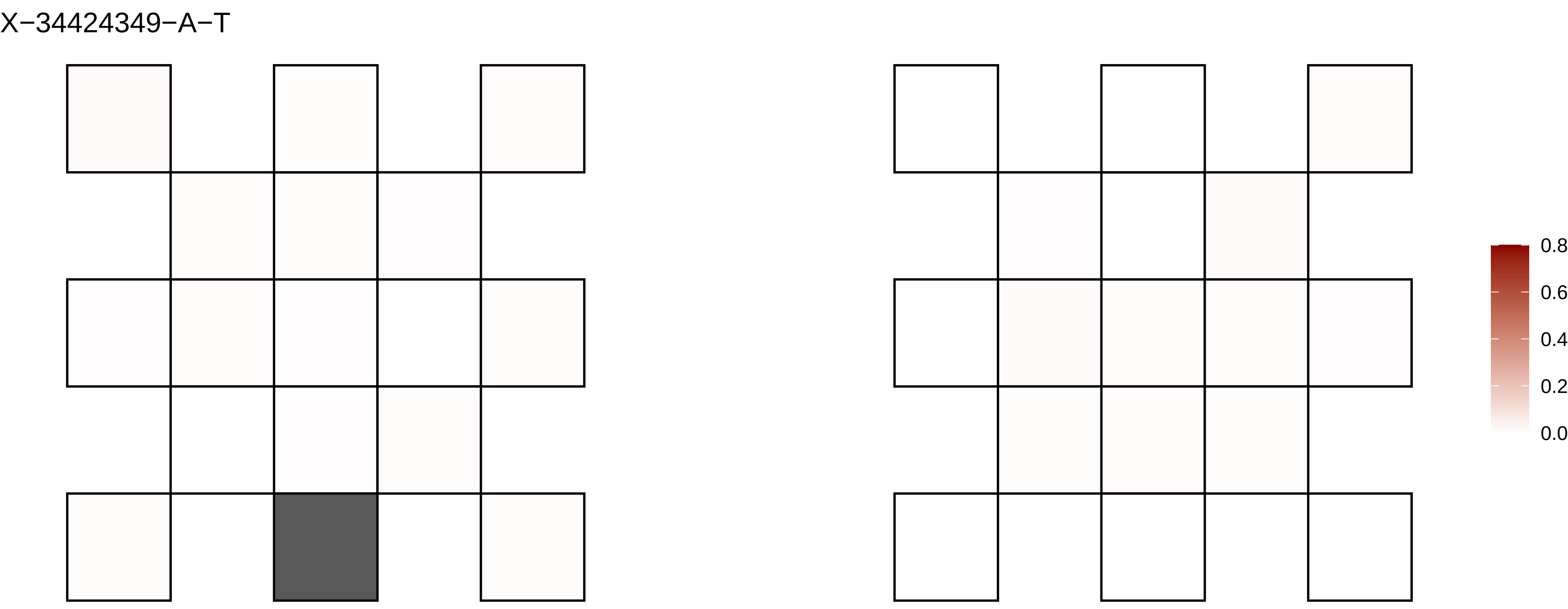

X−34424349−A−T
0.8
0.6
0.4
0.2
0.0

X−34664512−A−G

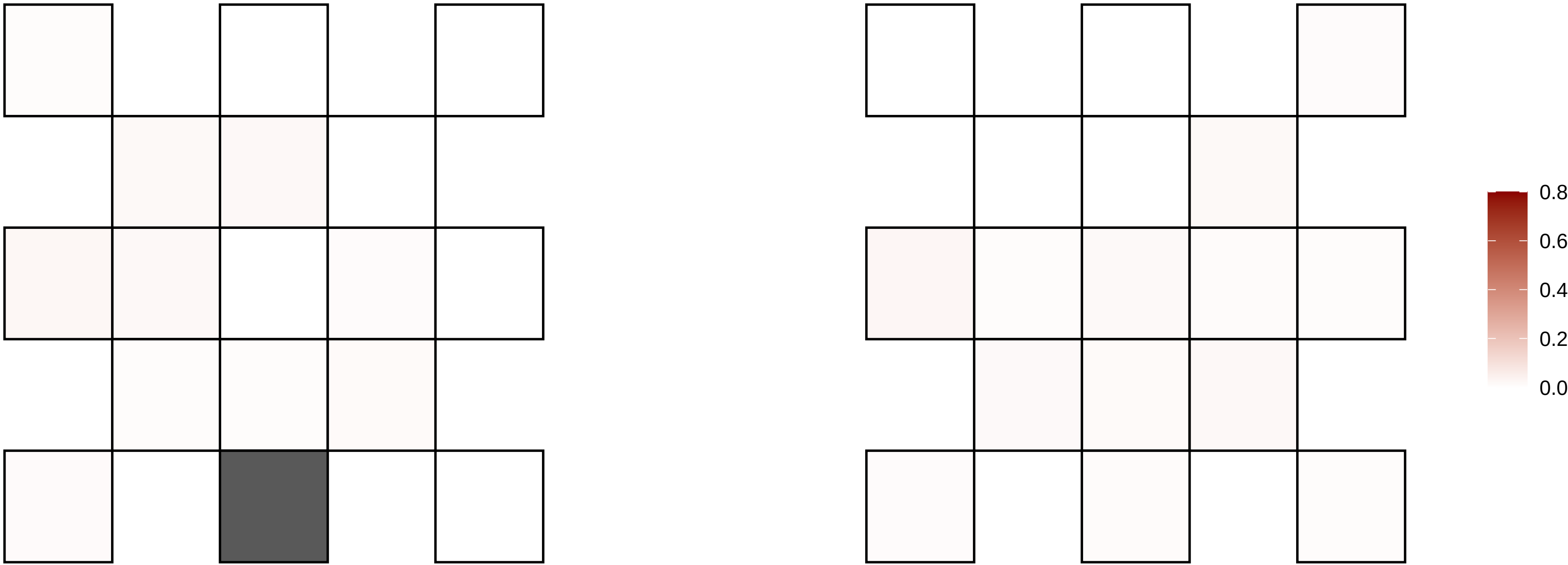

X−34801213−T−C

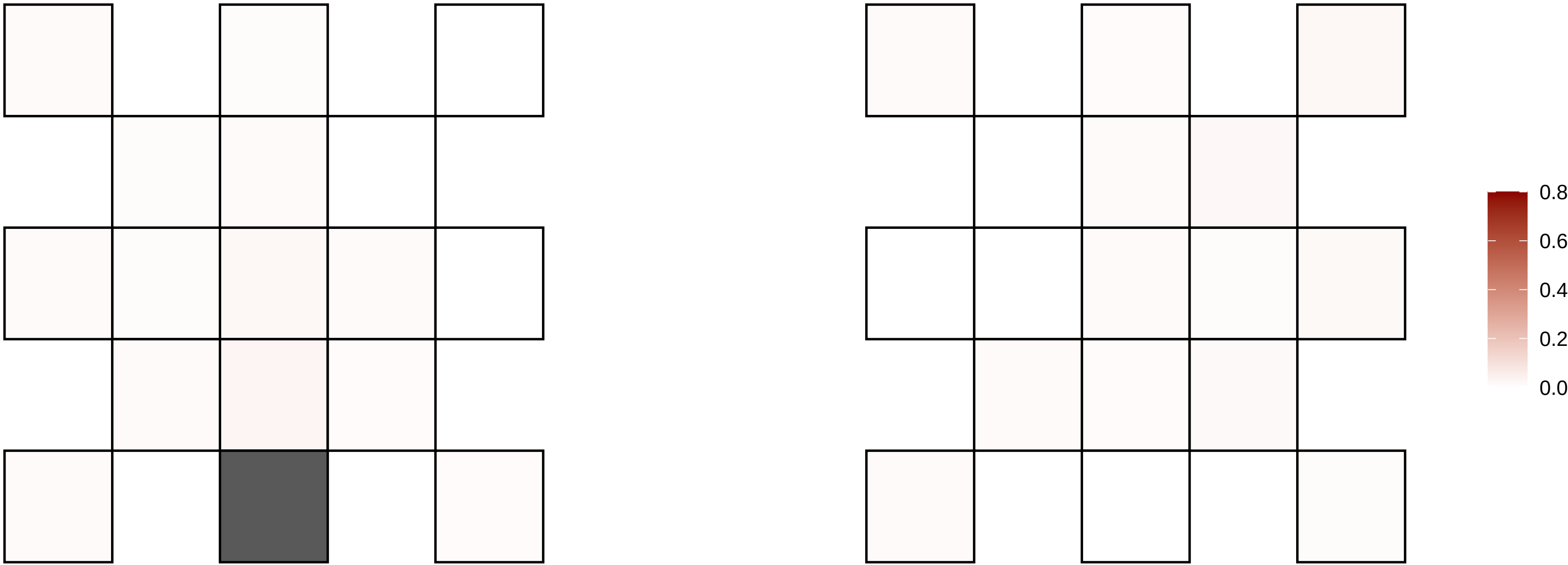

X−35362338−C−T

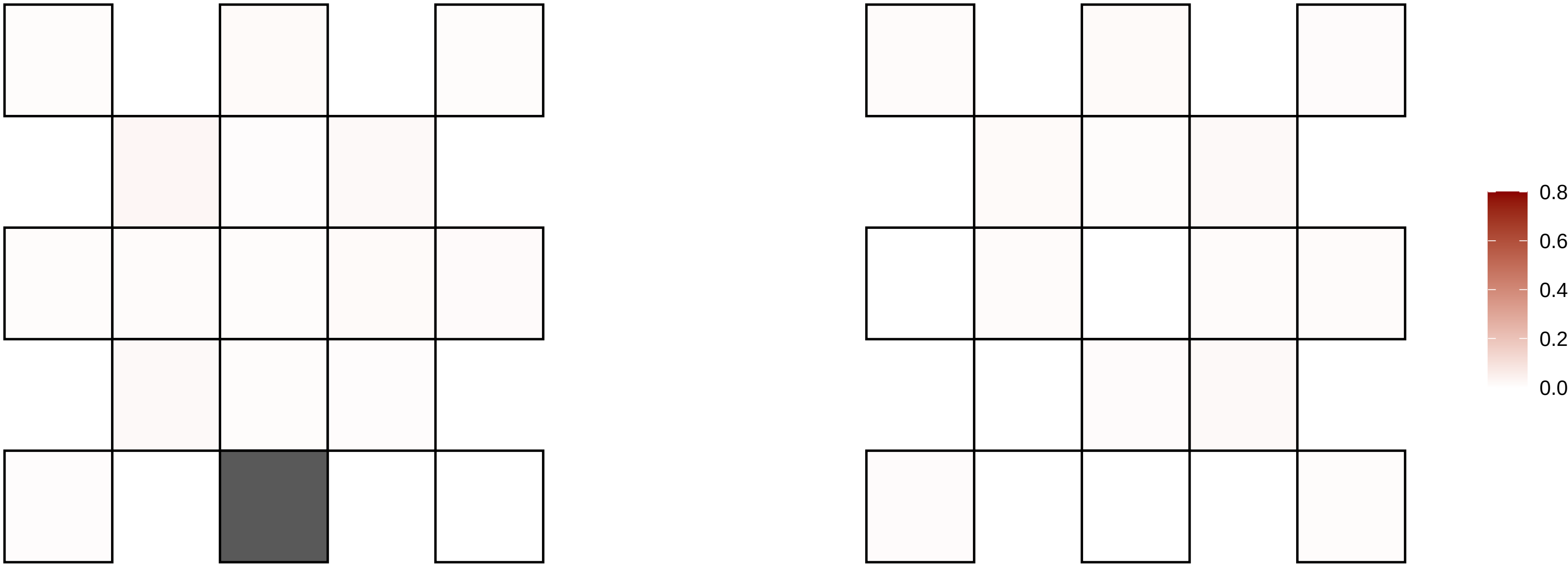

X−35471077−C−T

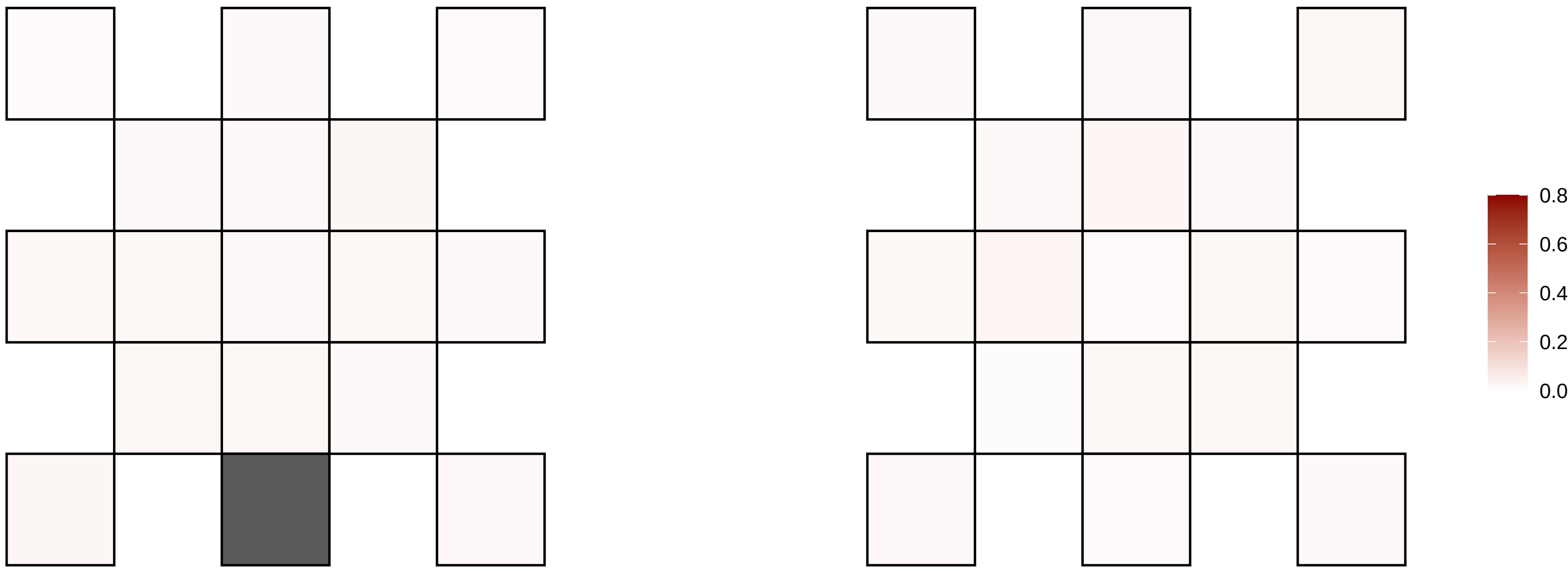

X−36528250−G−T

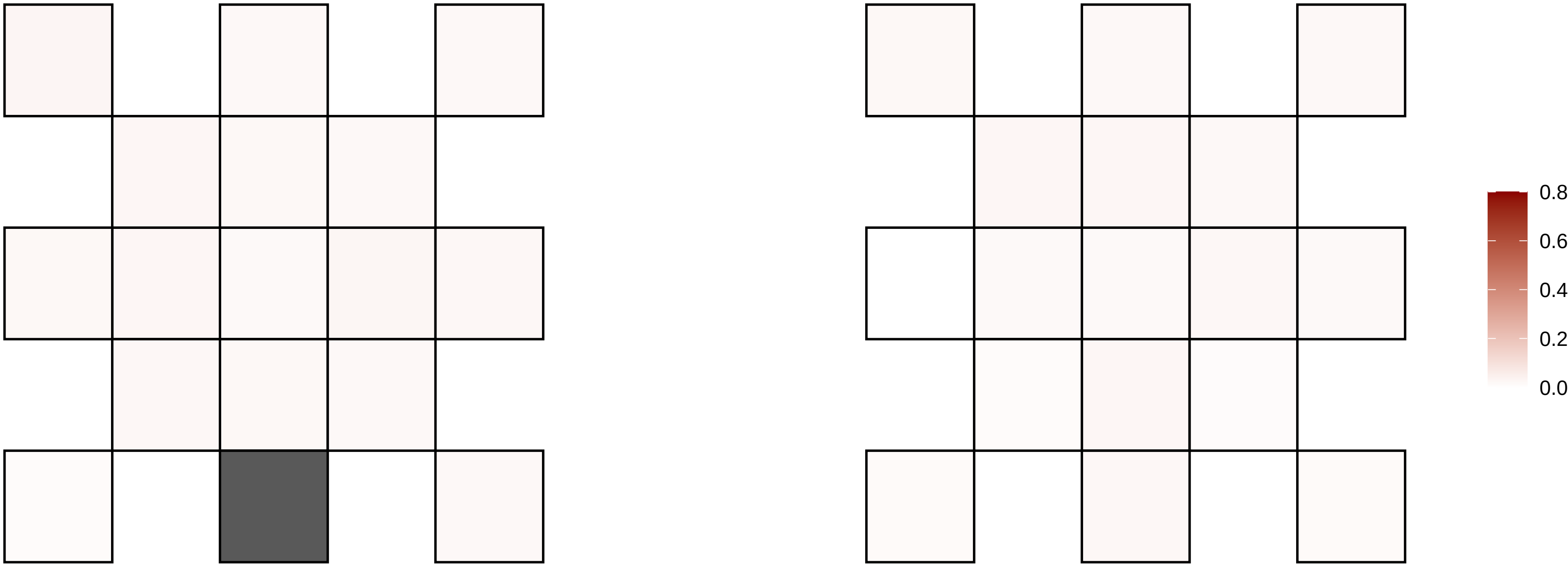

X−39832087−T−A

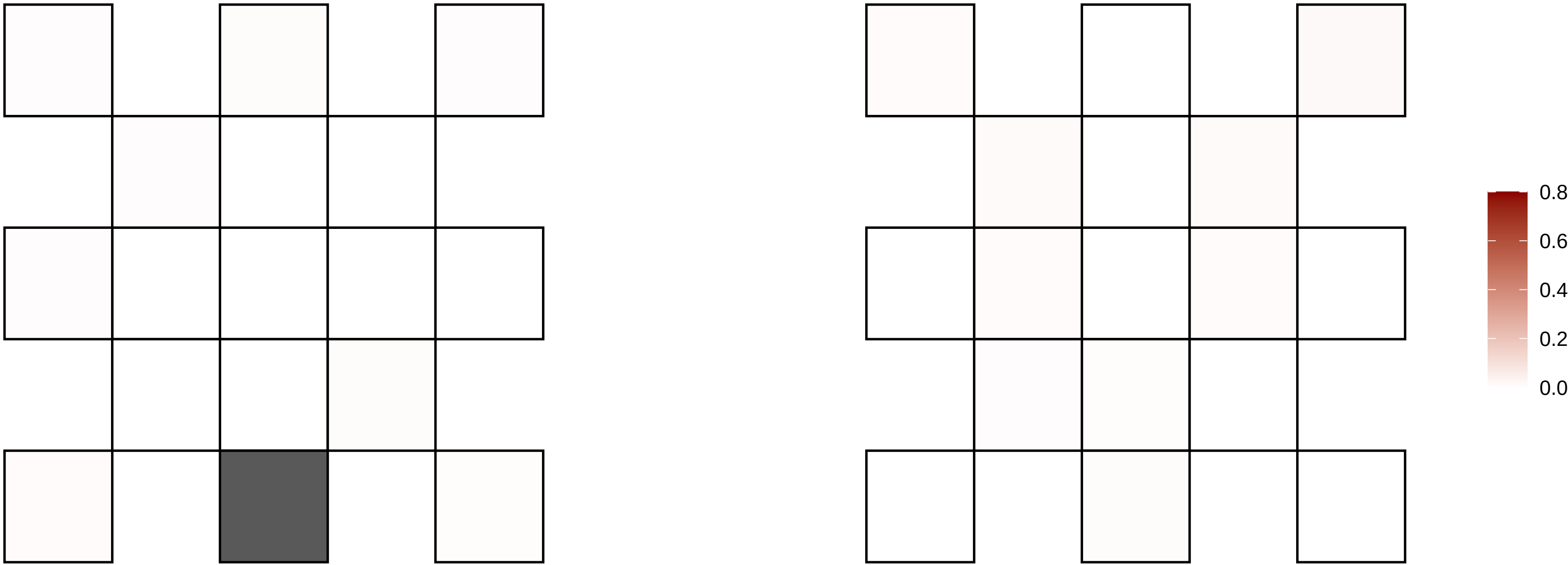

X−40030465−C−A

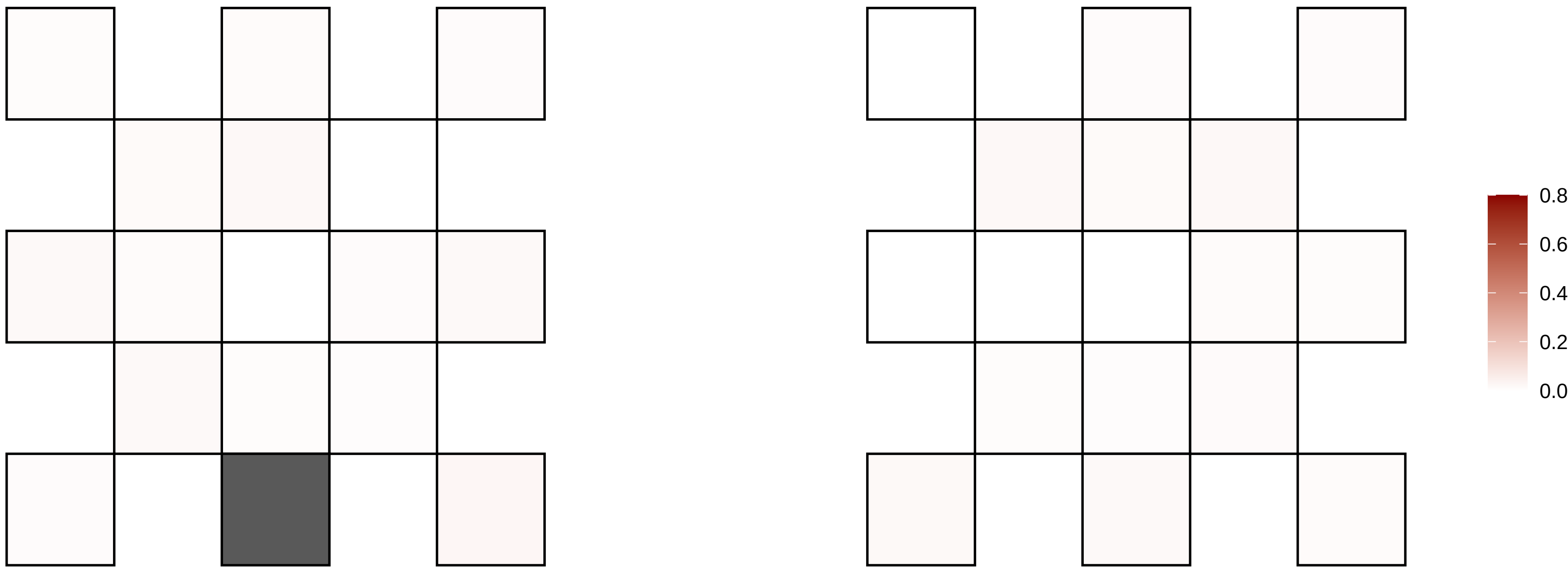

X−40790933−G−A

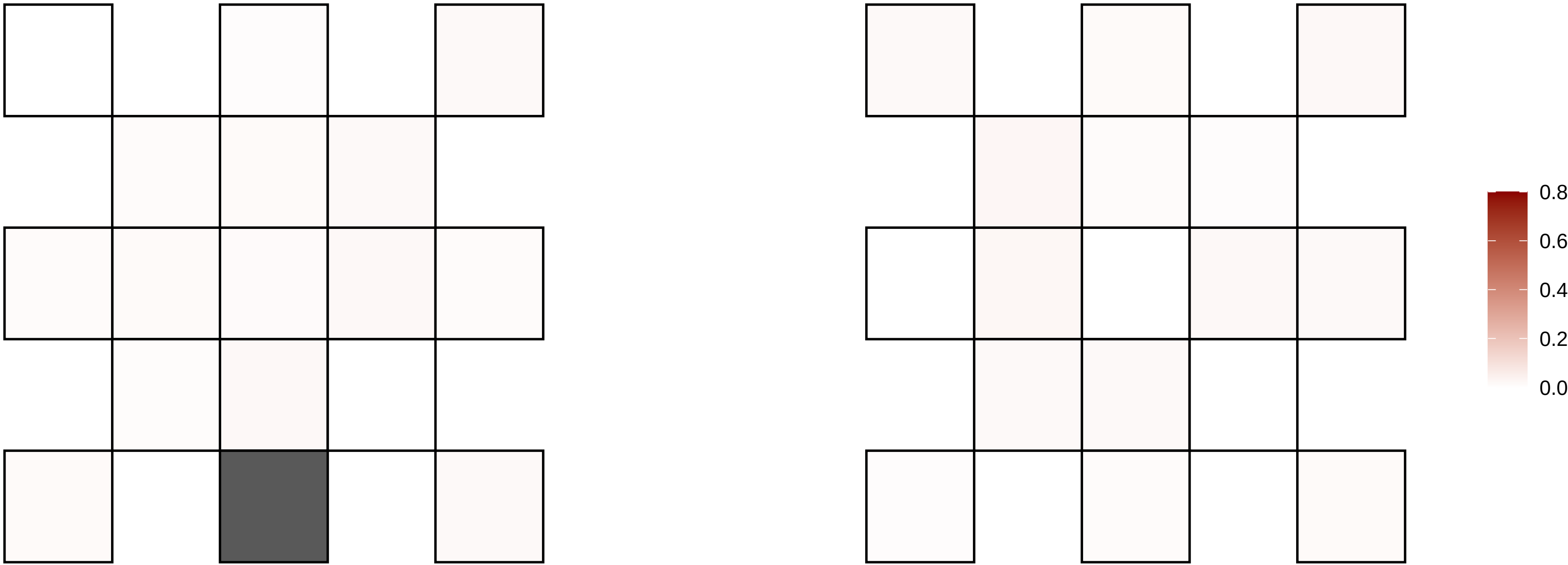

X−41029320−G−A

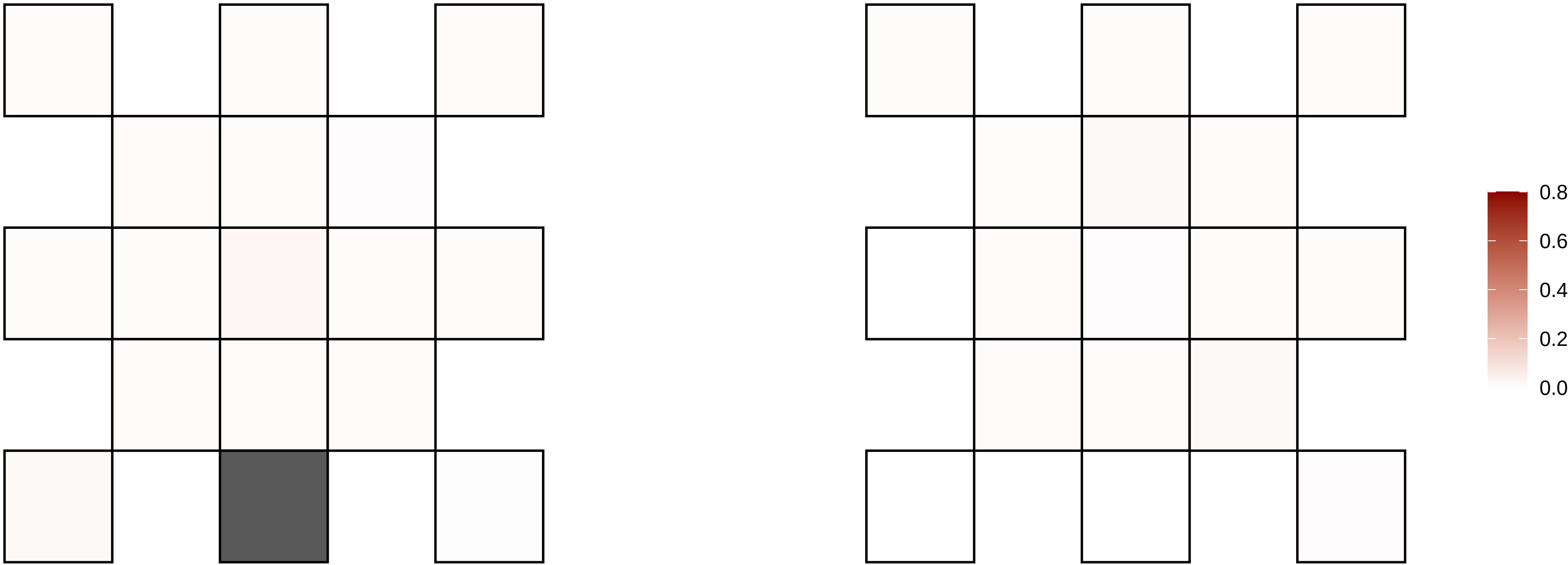

X−4557817−T−C

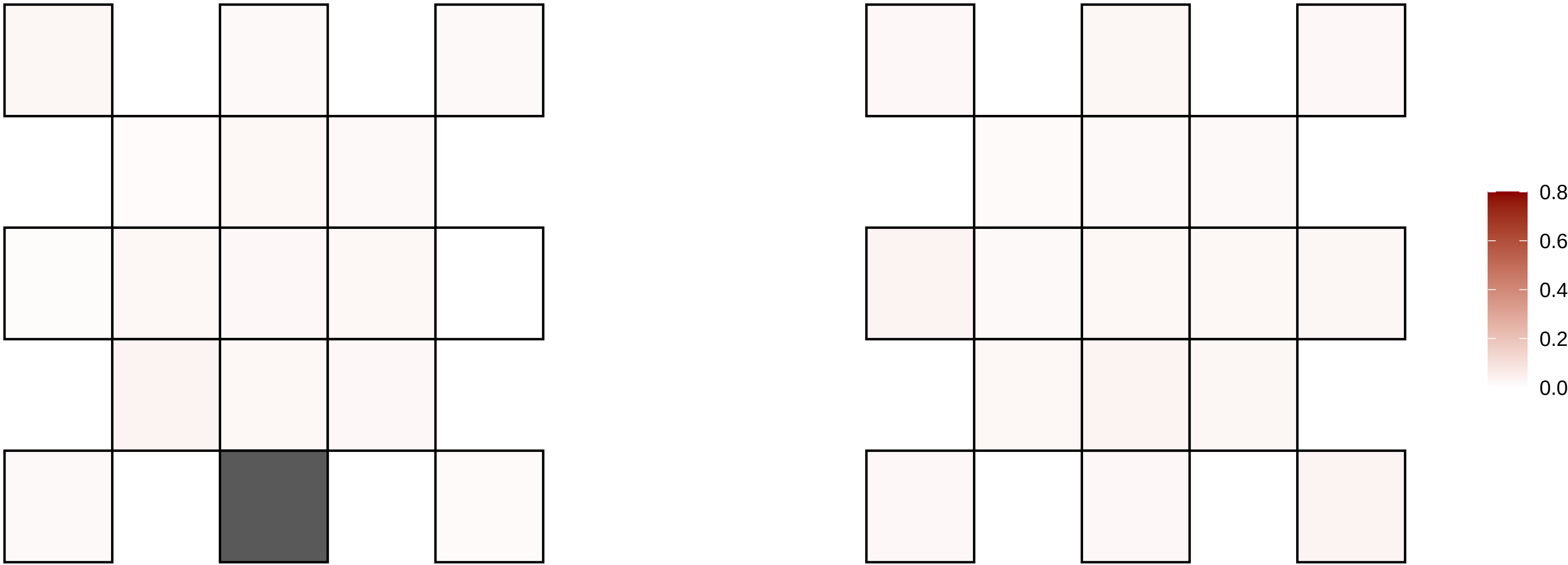

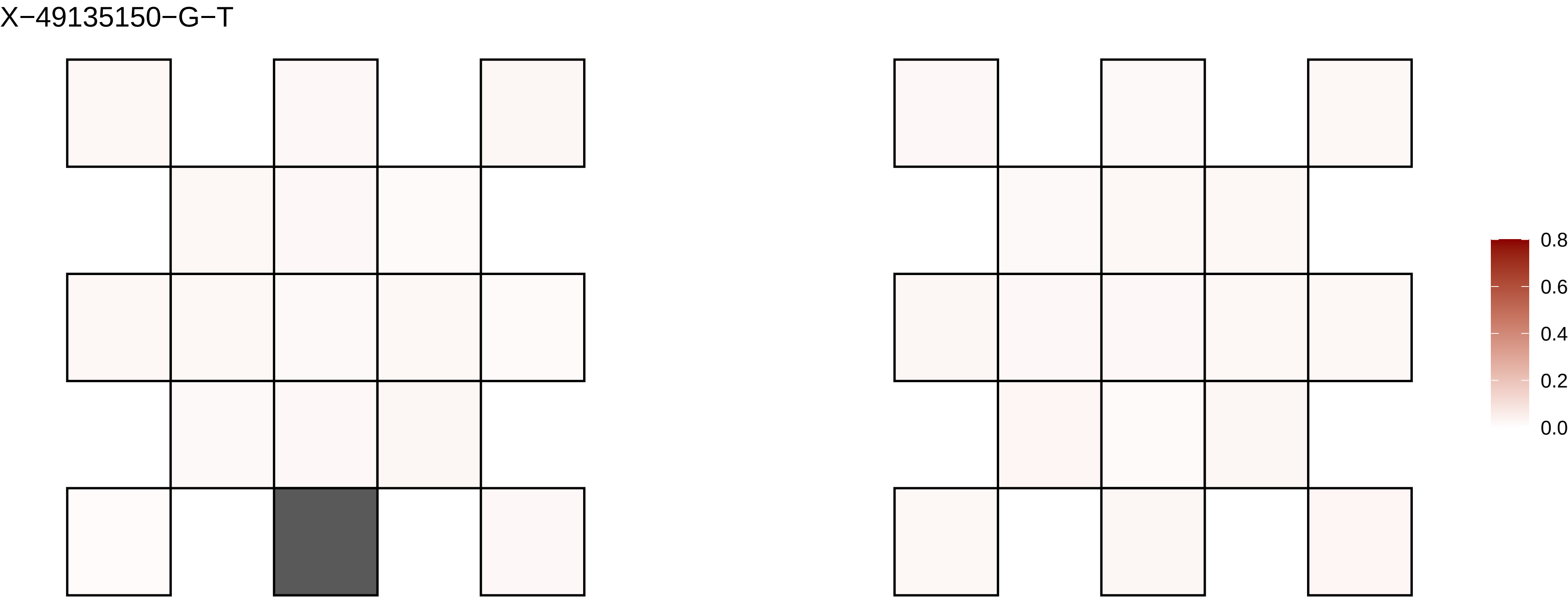
X−49135150−G−T
0.8
0.6
0.4
0.2
0.0

X−53927483−G−A

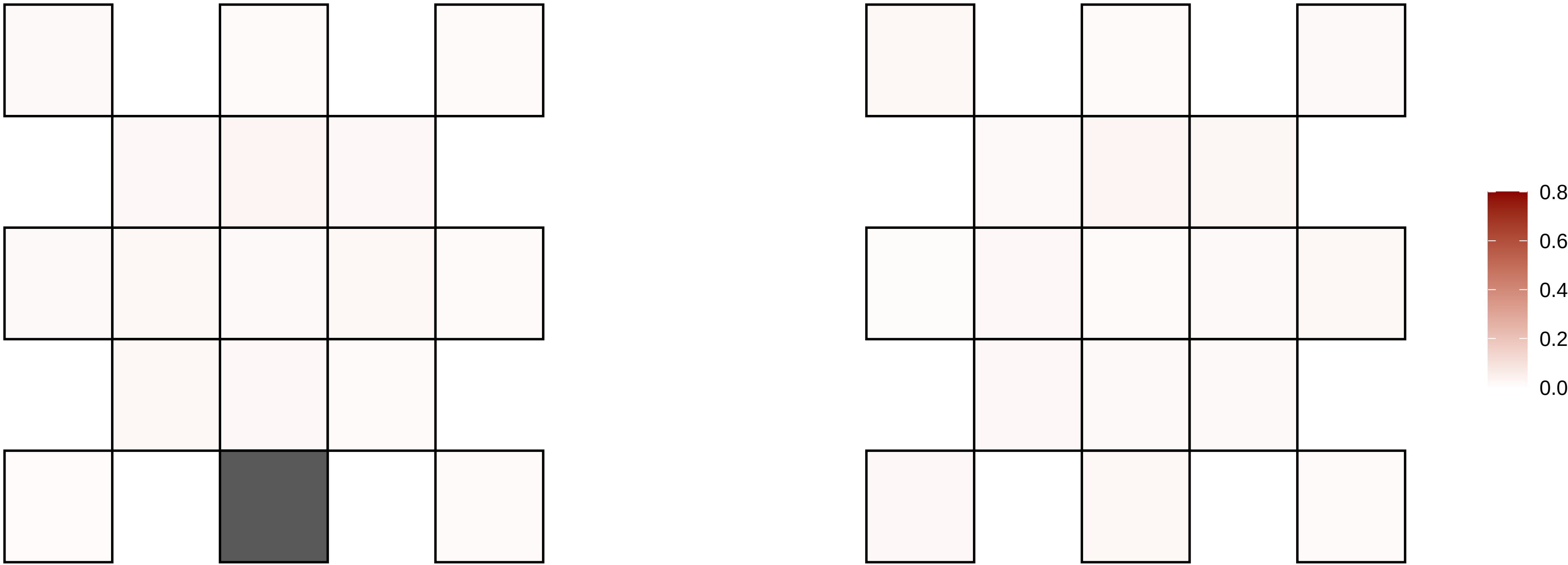

X−5452716−T−A

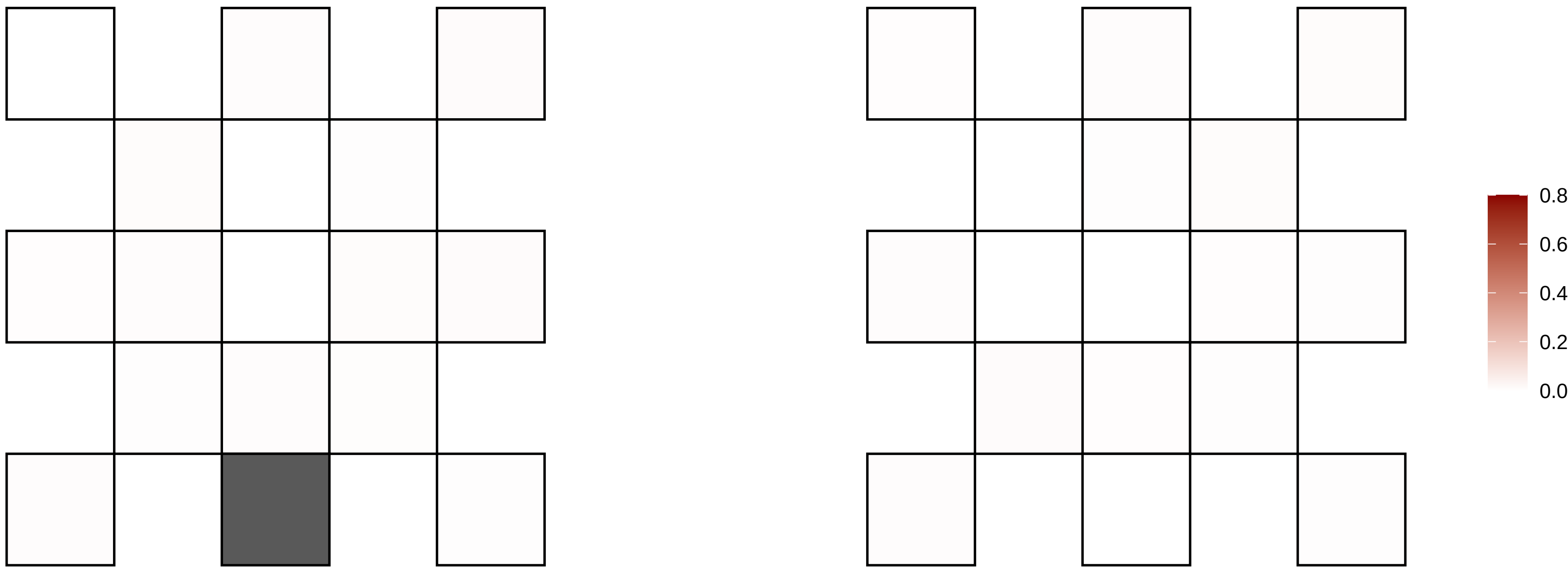

X−6657783−T−A

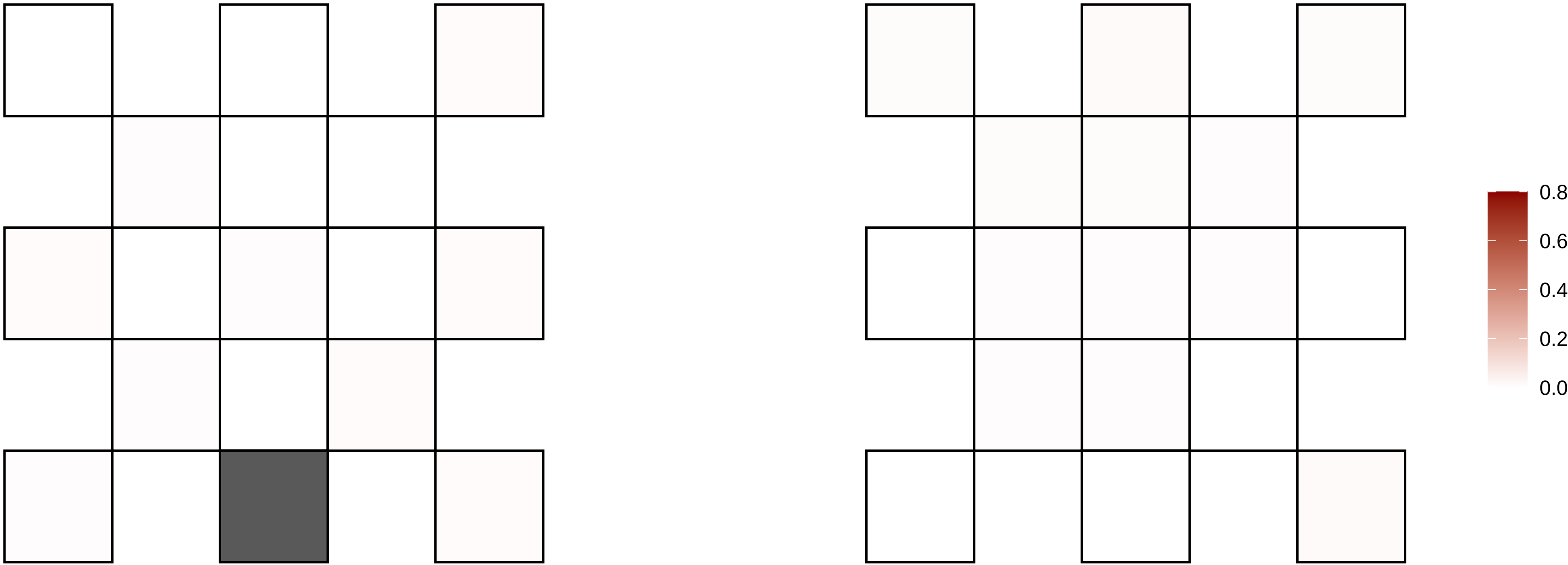

X−6657784−C−A

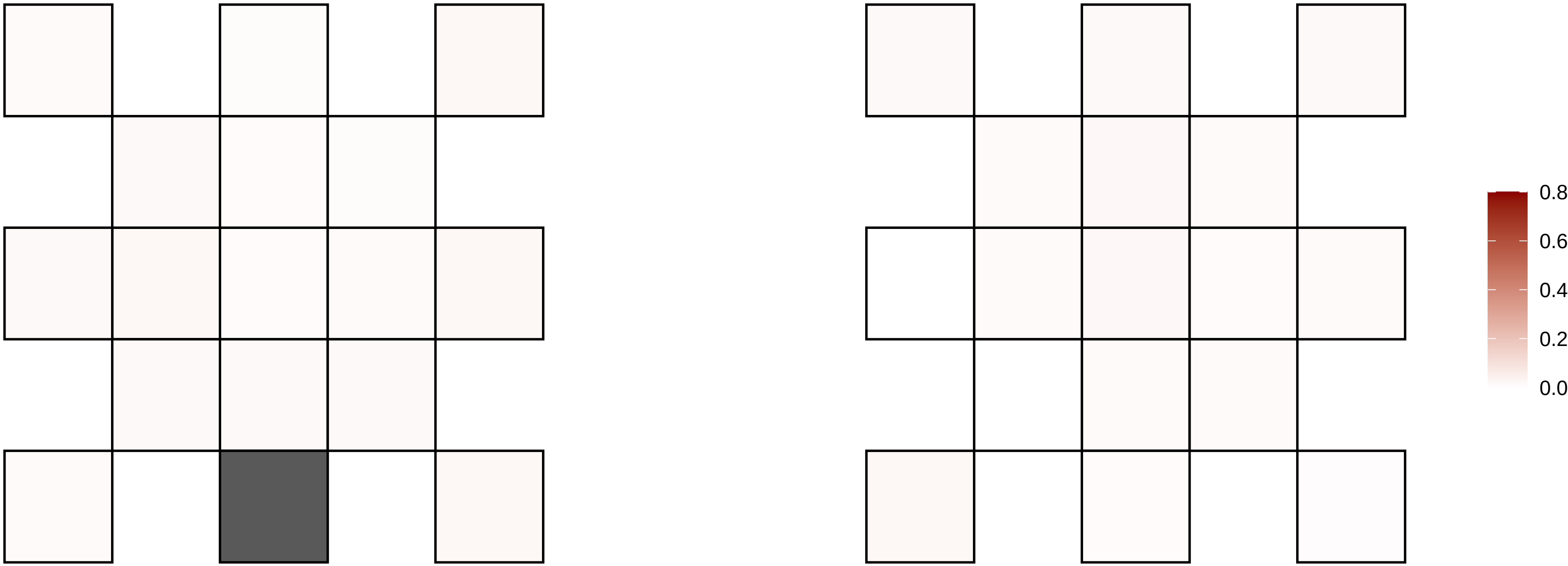

X−69258884−G−A

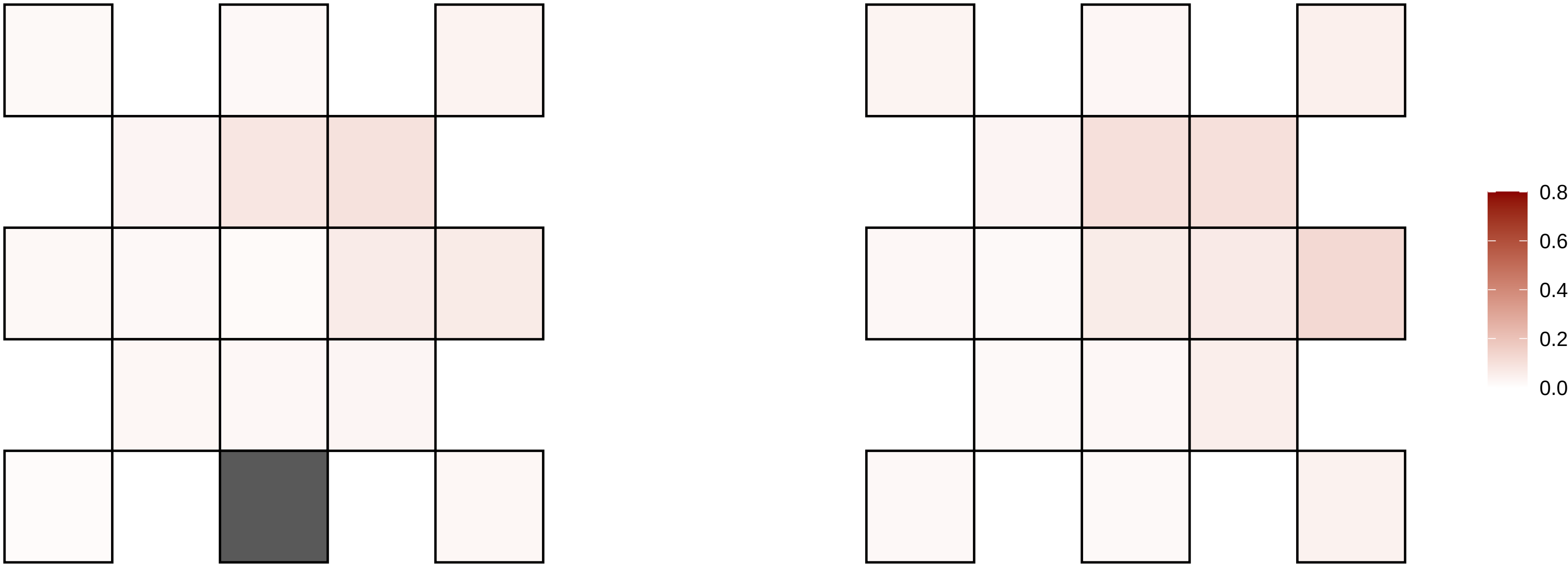

X−69794428−G−A

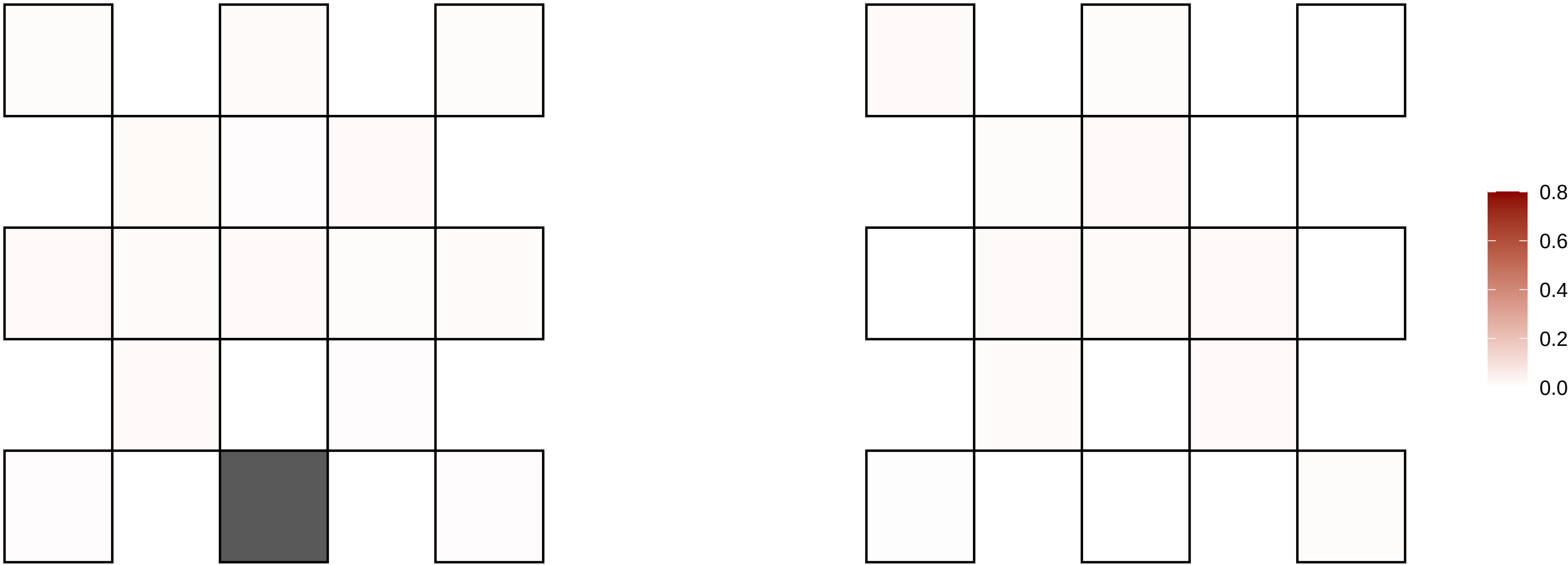

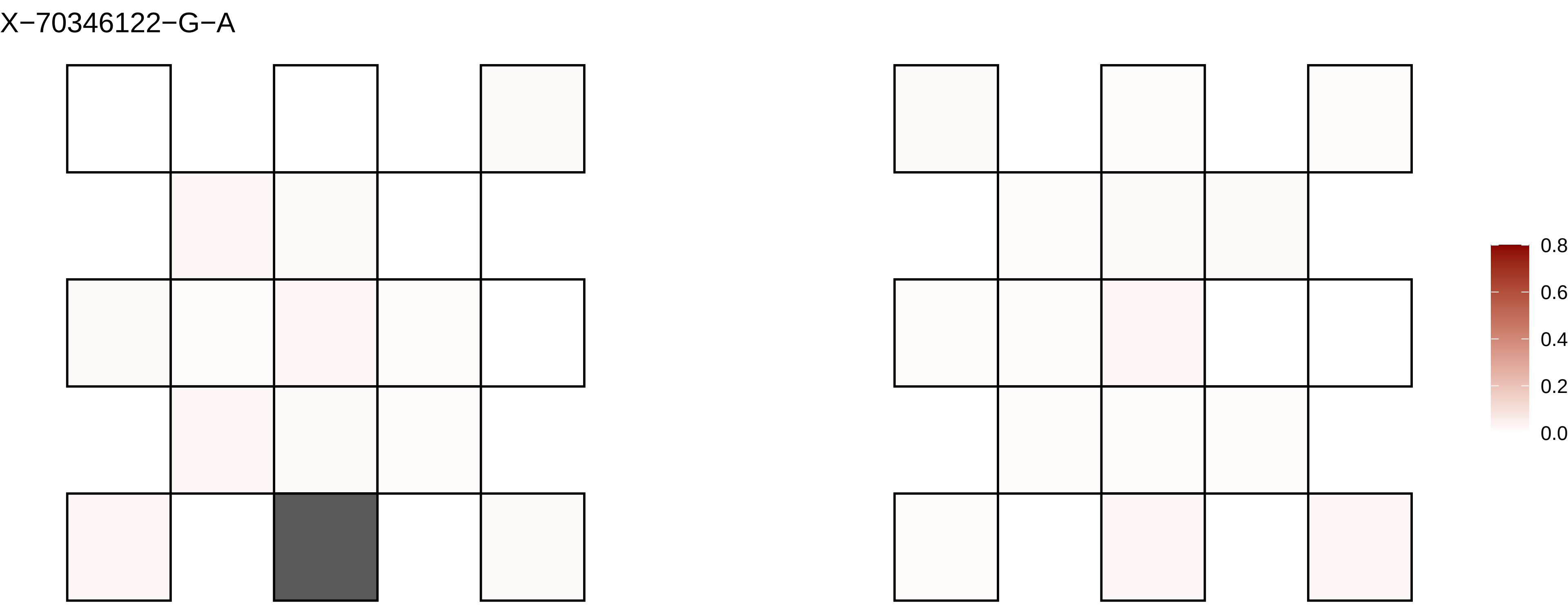
X−70346122−G−A
0.8
0.6
0.4
0.2
0.0

X−72495149−C−A

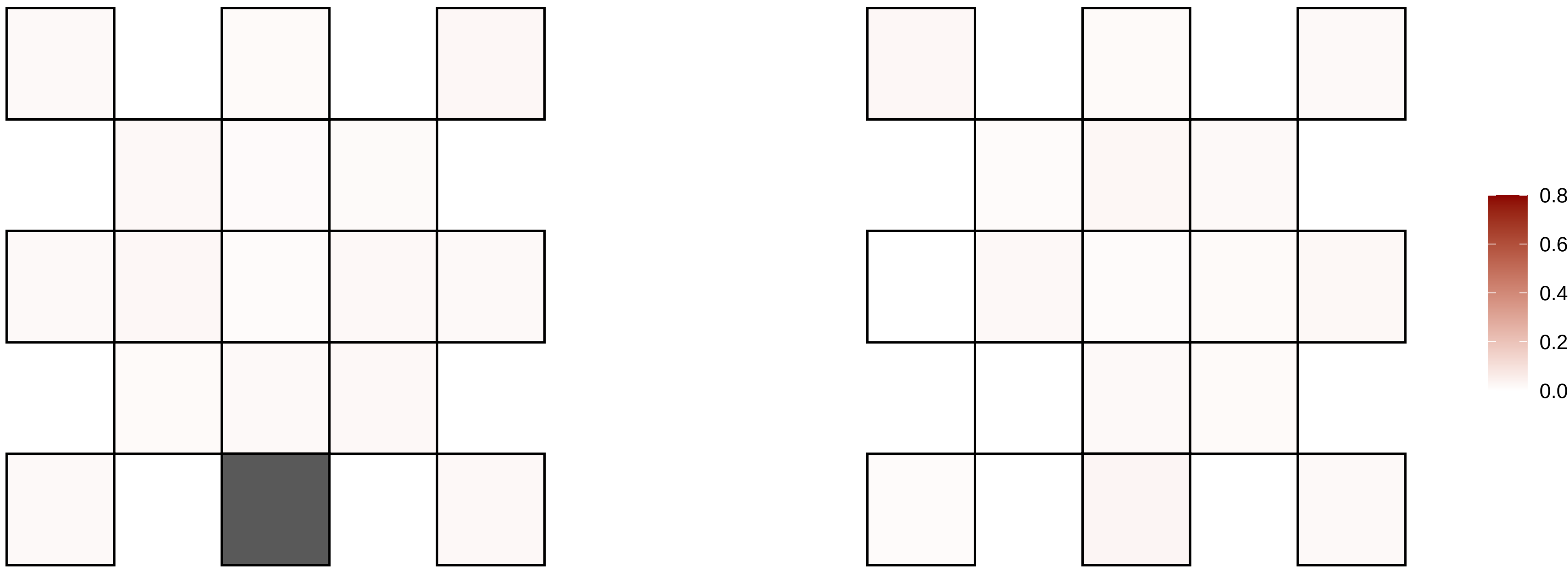

X−7883974−C−T

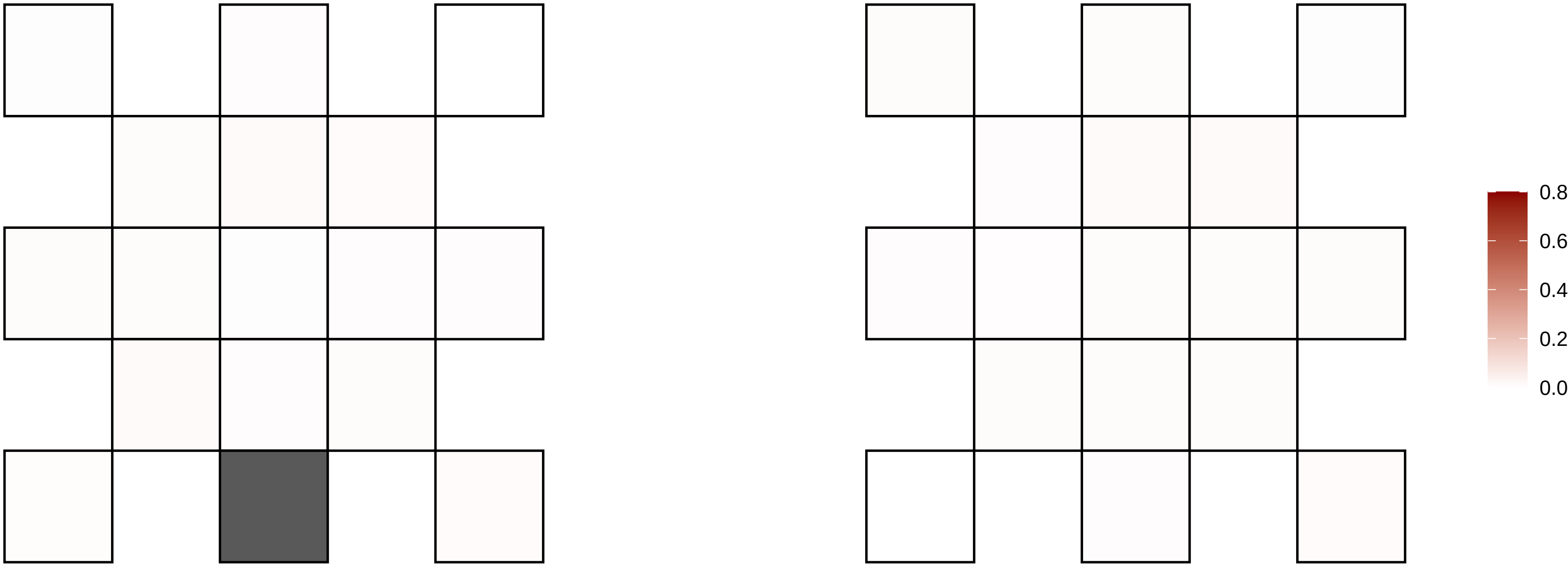

X−79691731−A−T

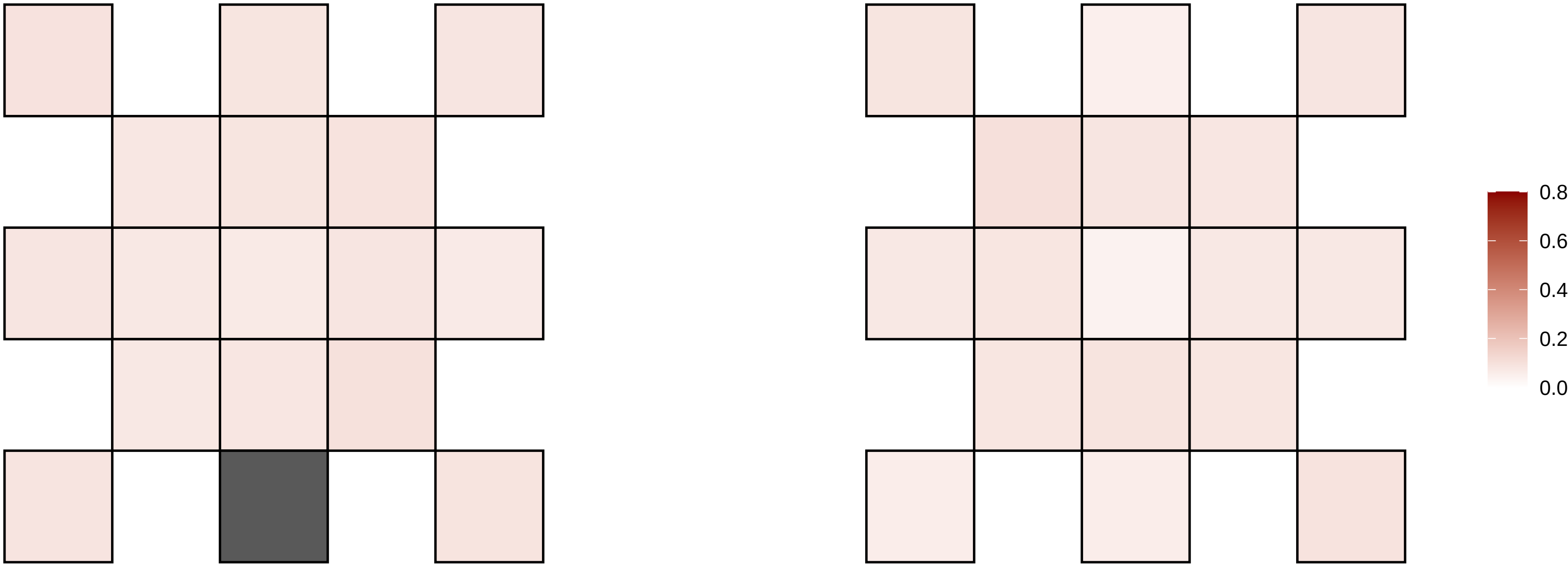

X−79691734−A−T

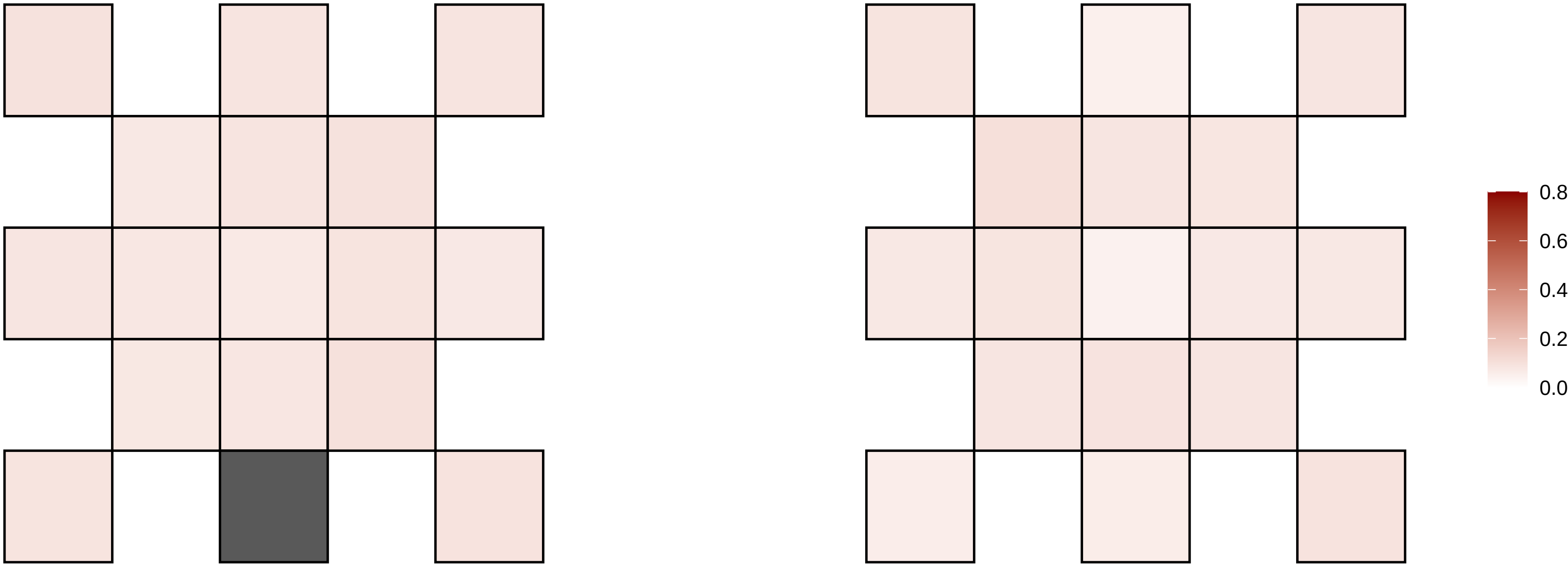

X−83599149−G−A

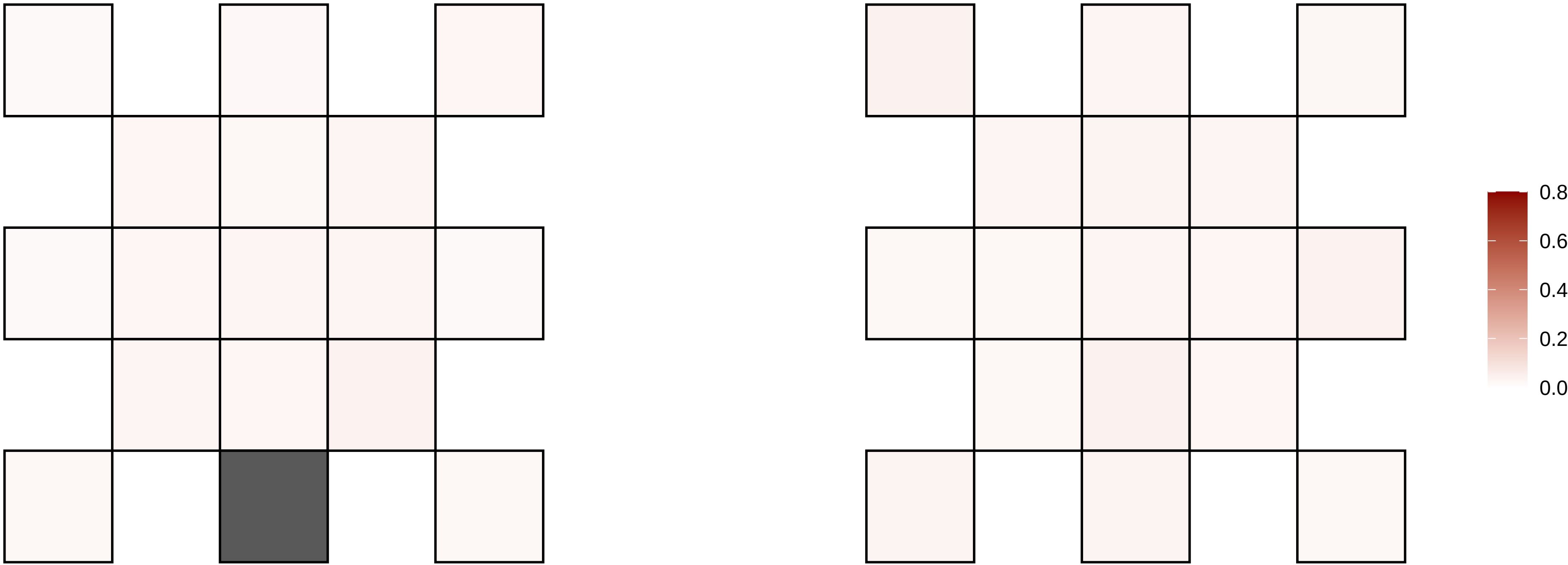

X−88323908−T−G

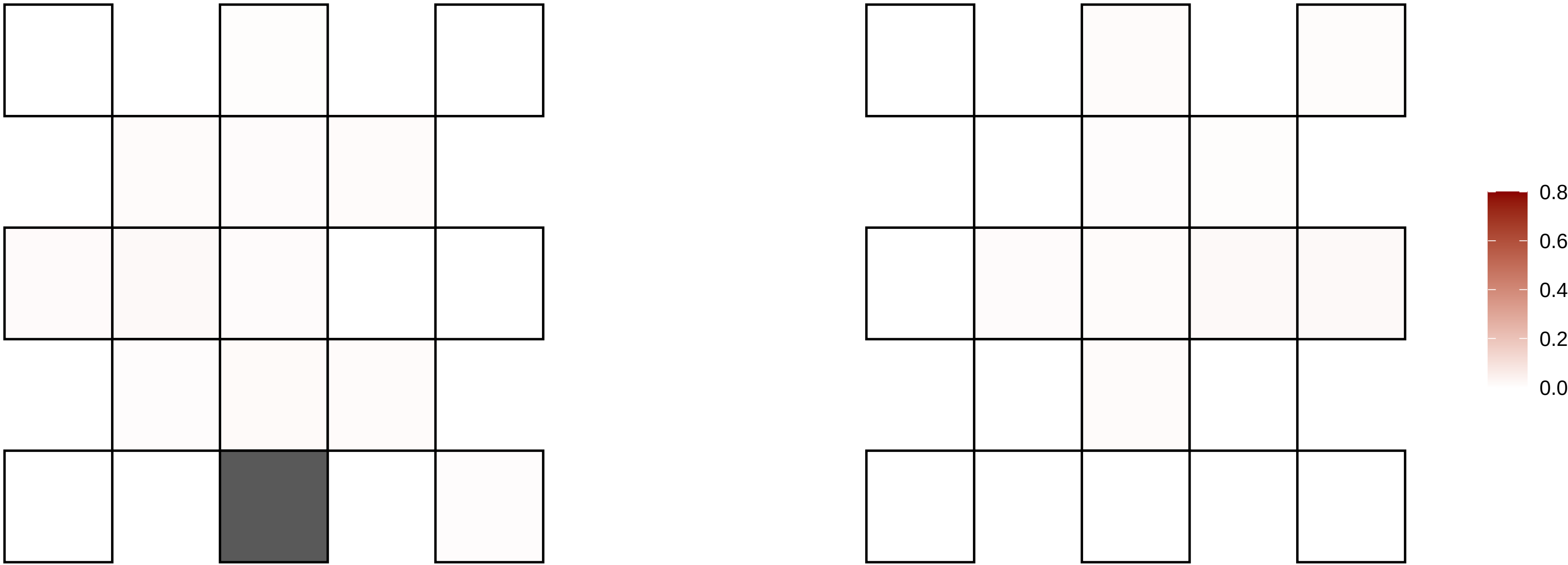

X−95198288−C−T

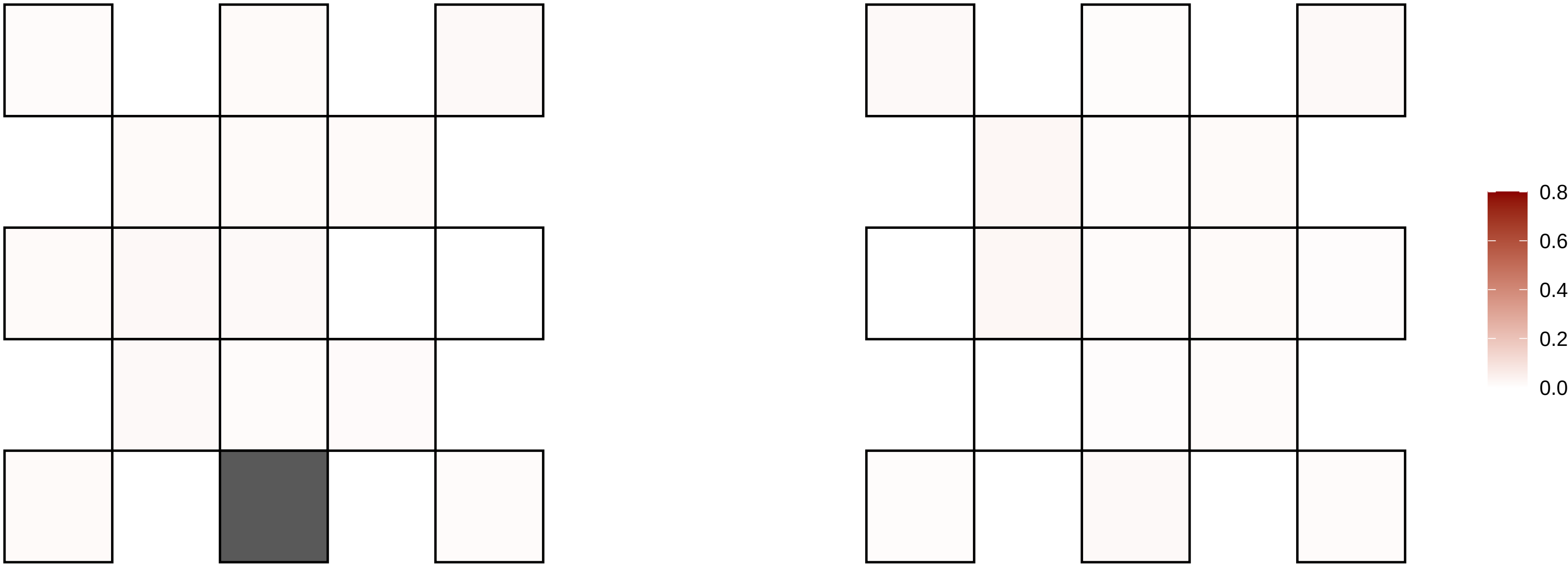

X−96237346−G−A

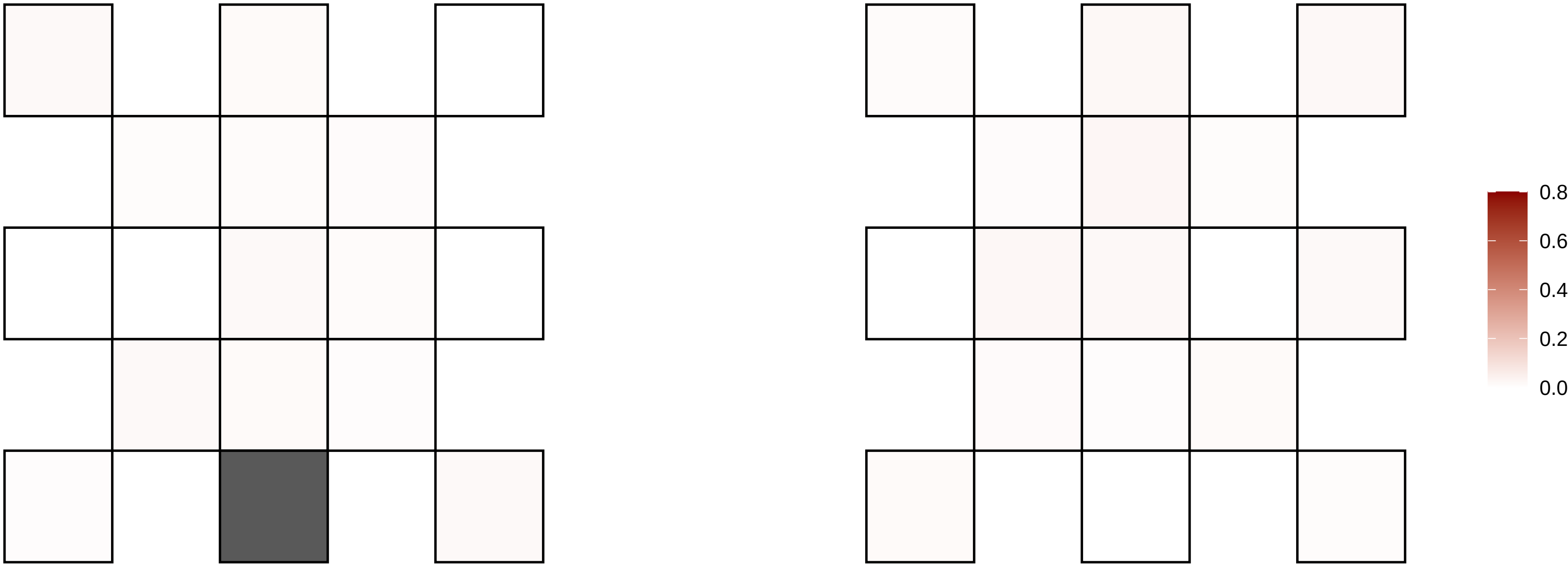

X−98261573−C−T

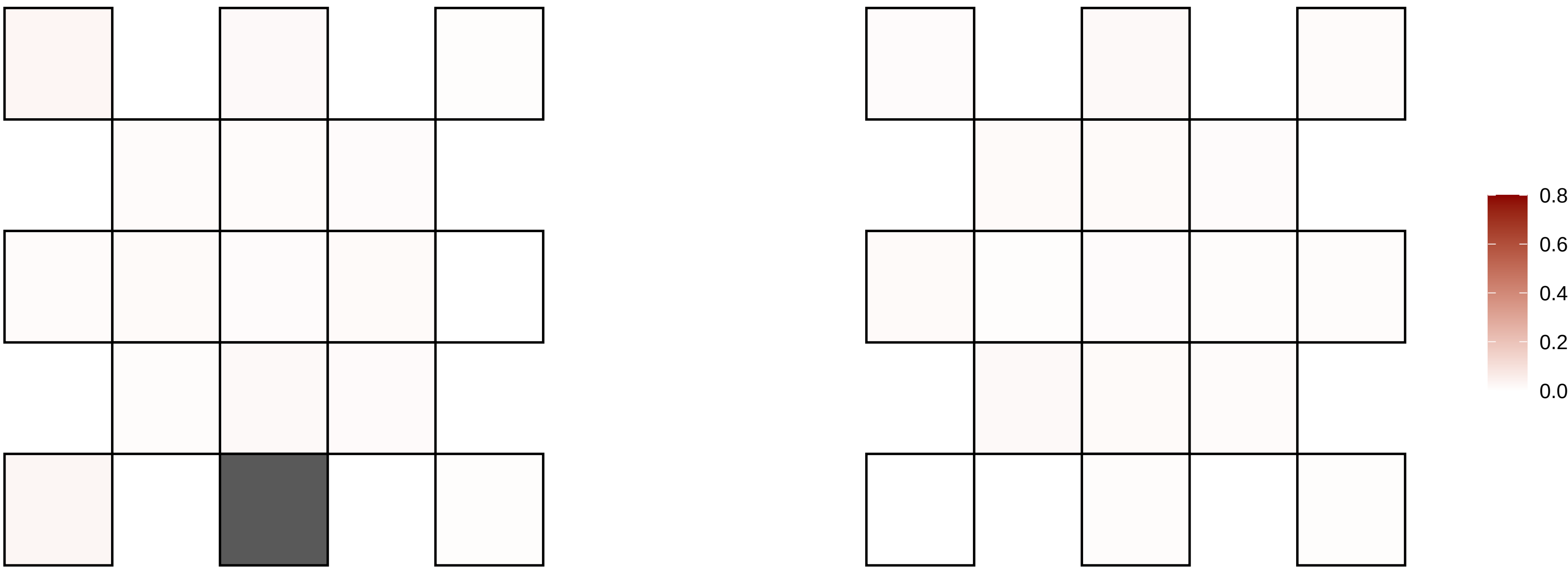

X−98520292−C−A

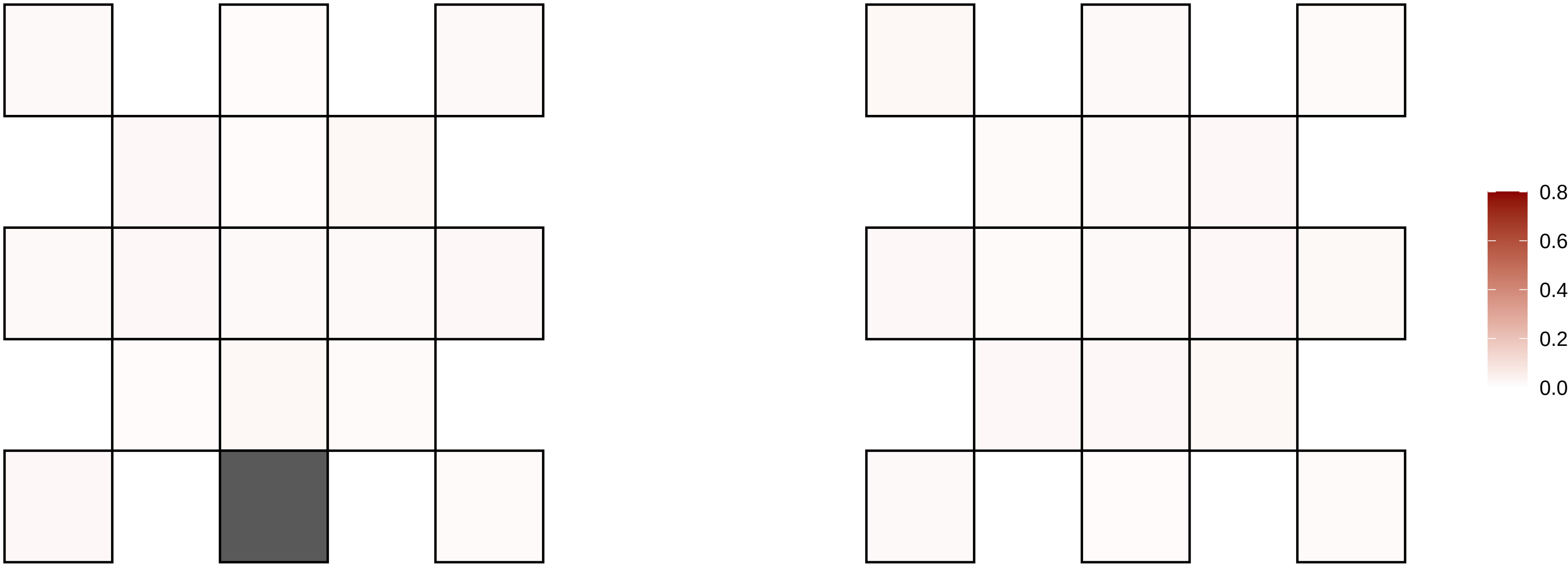

X−98527468−AC−A

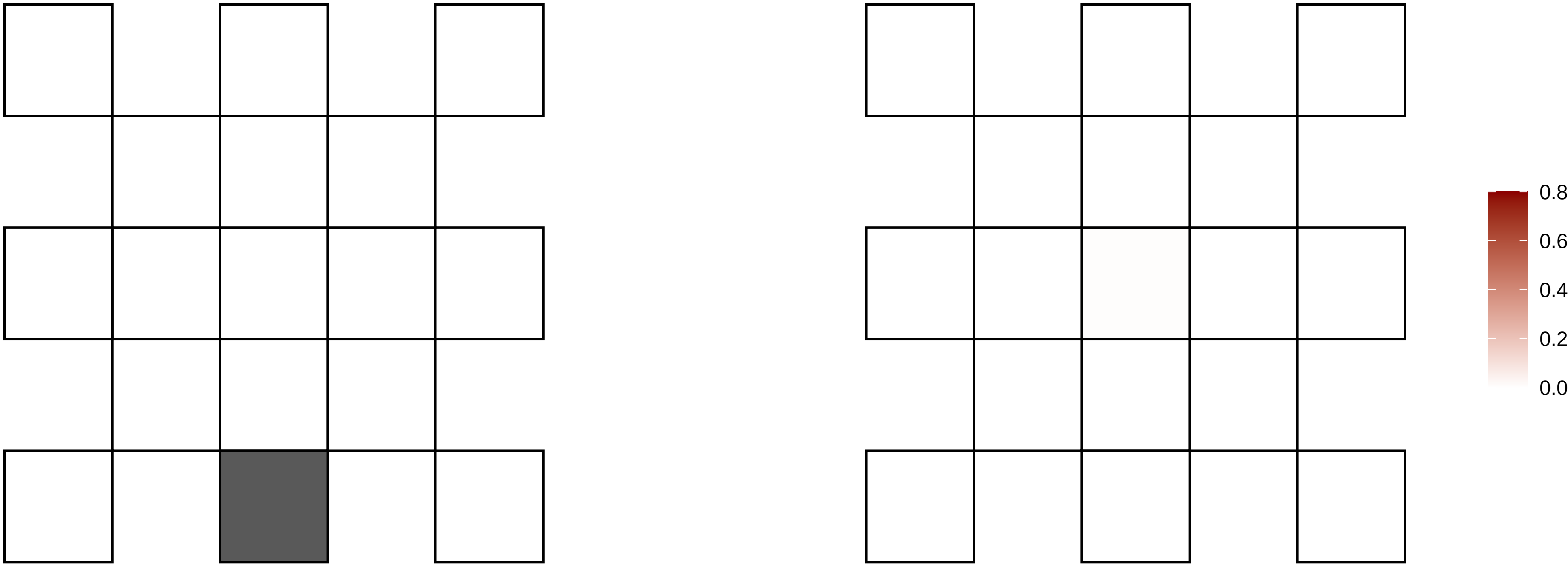

X−99571049−A−C

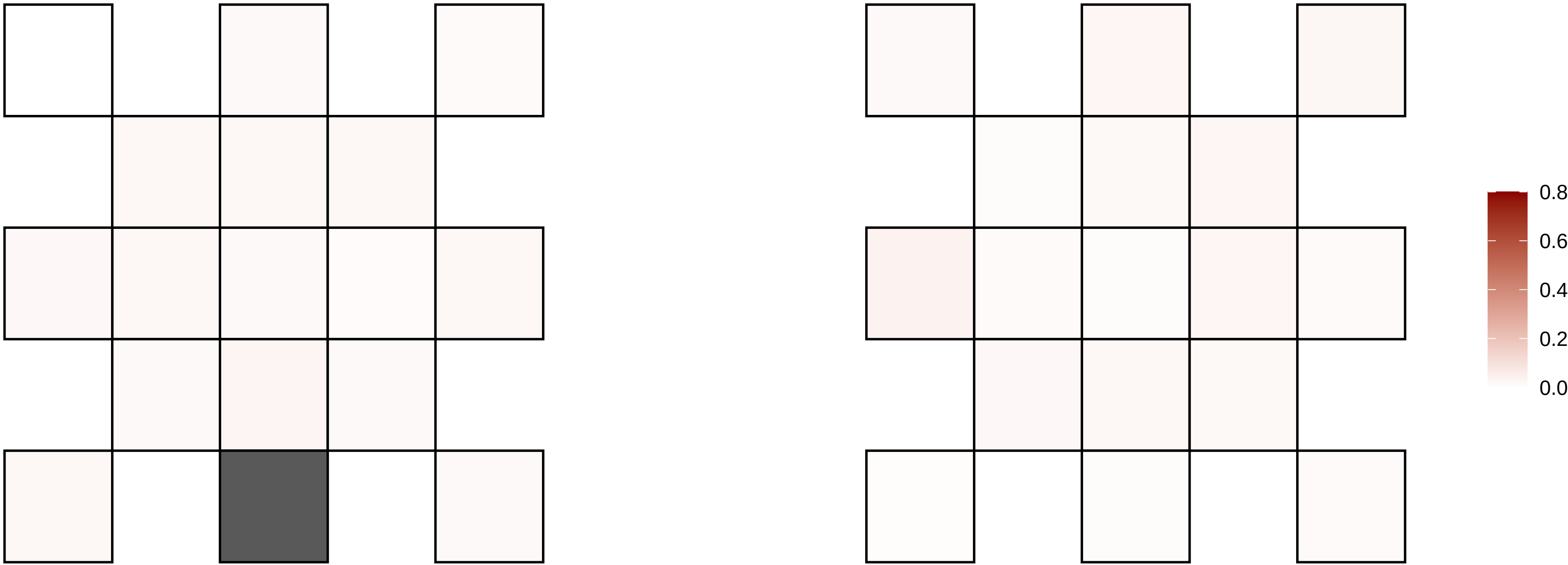

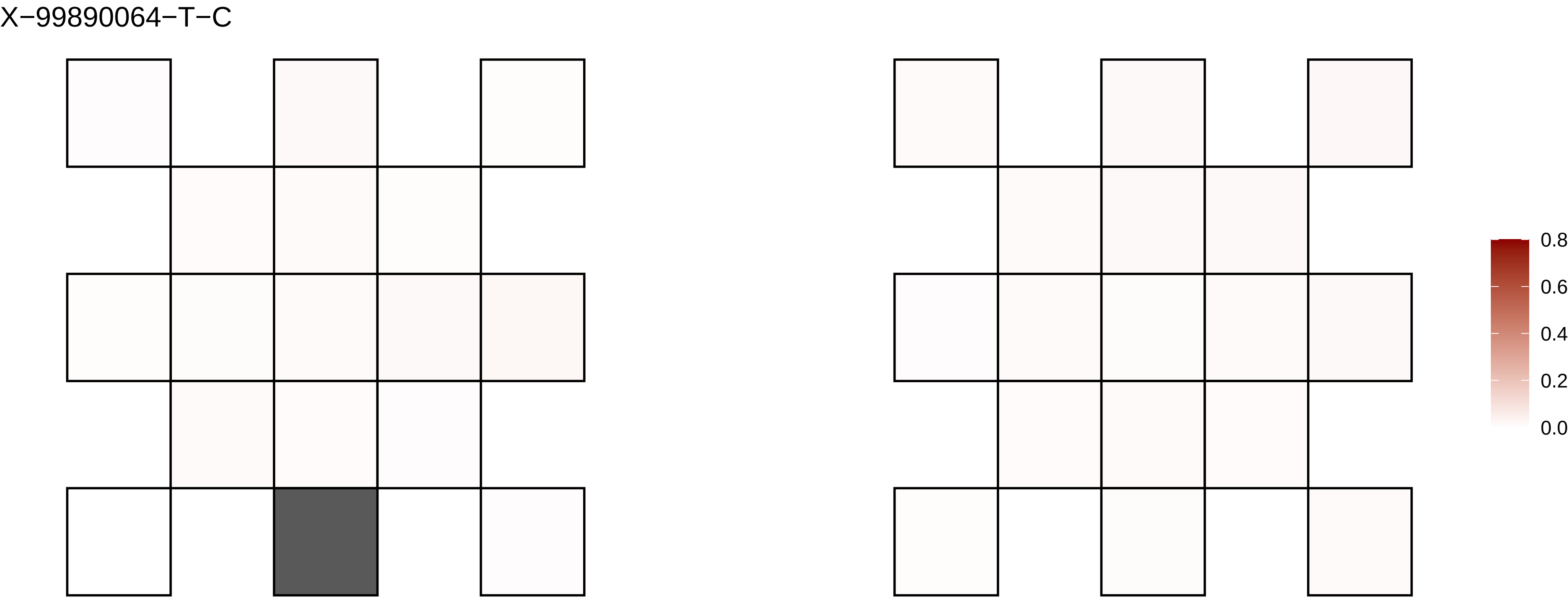
X−99890064−T−C
0.8
0.6
0.4
0.2
0.0
