## Supplementary Data 3 for "Cell-type-resolved somatic mosaicism reveals clonal dynamics of the human forebrain"

ID01 1−106934533−C−T

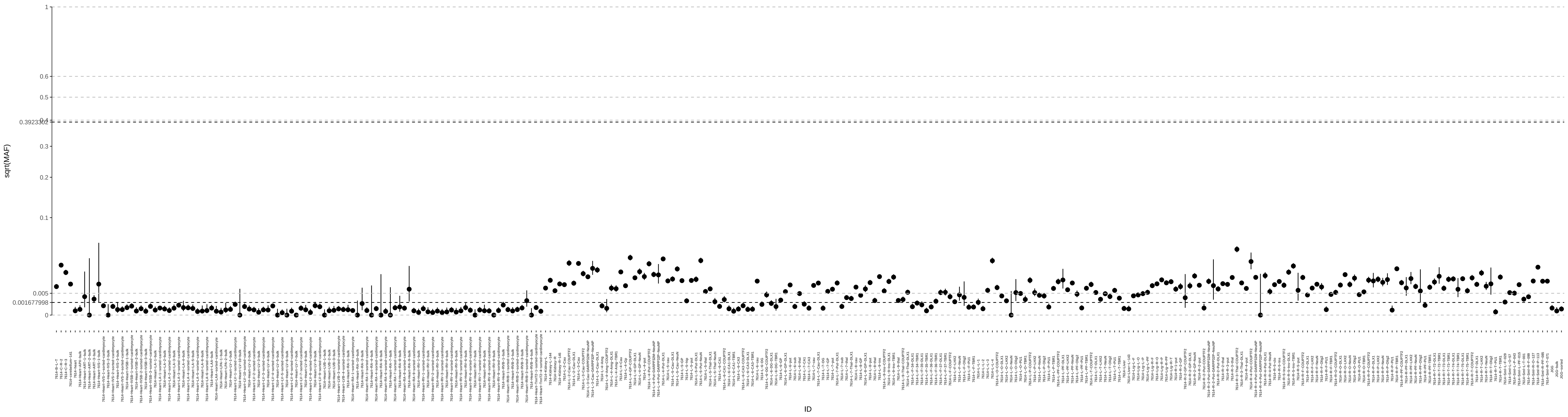

ID01 1−168467950−G−A

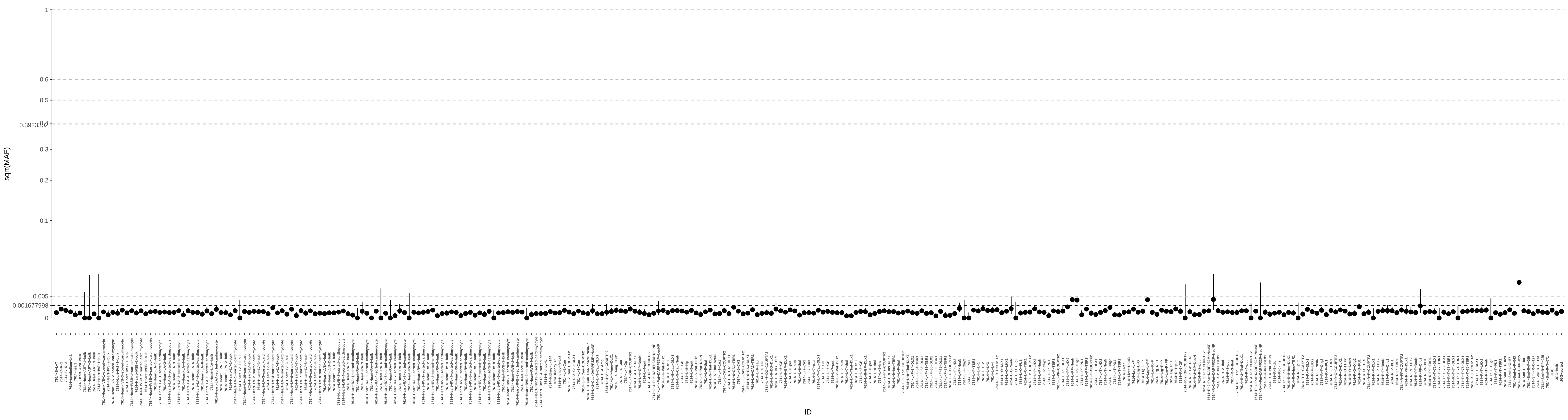

ID01 1−169329191−G−A

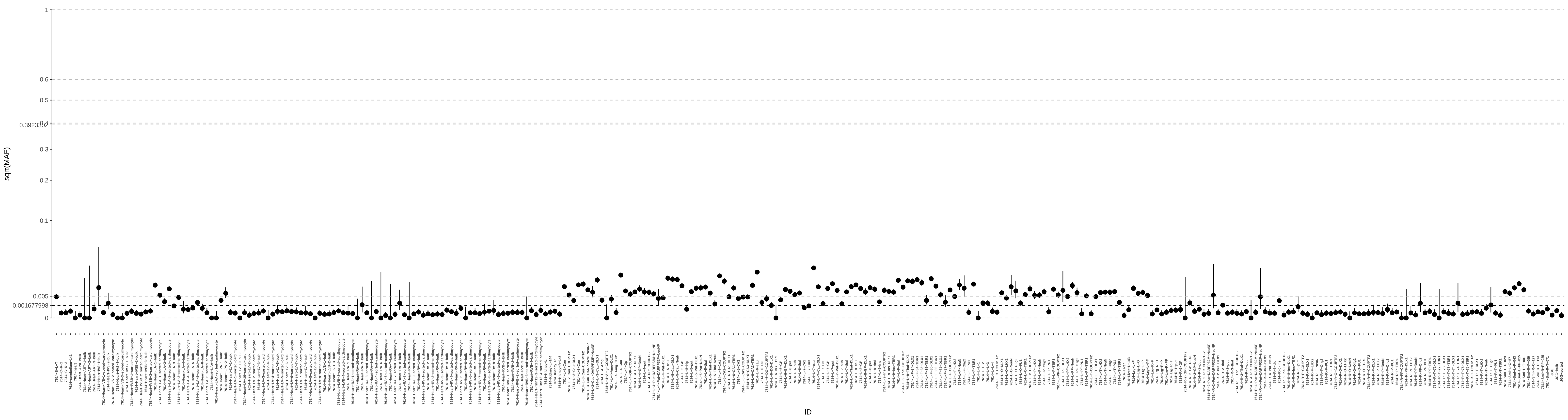

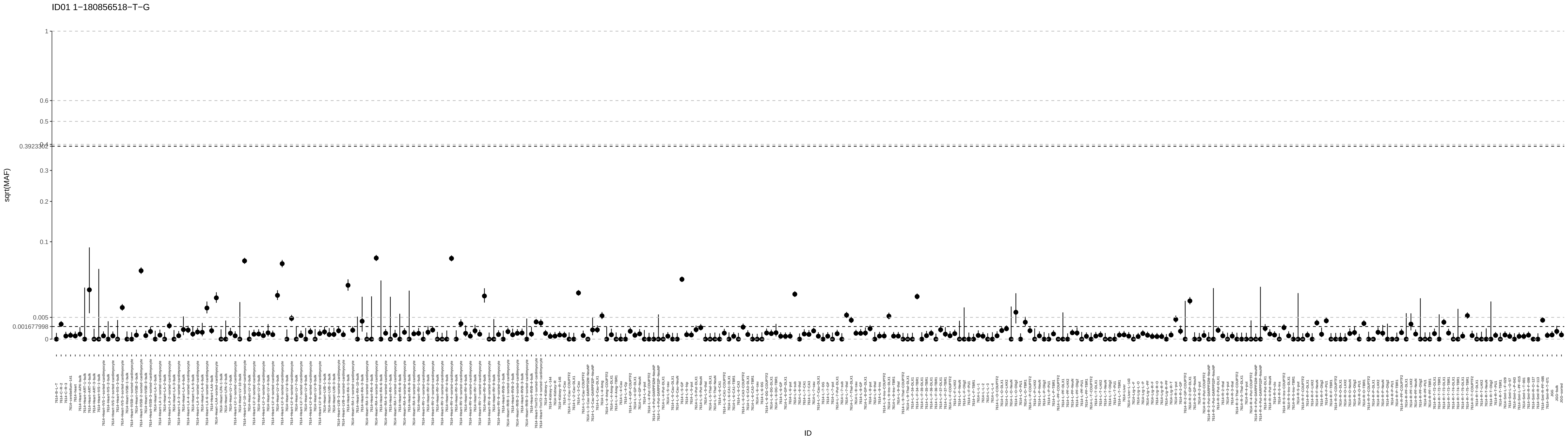
ID01 1−180856518−T−G
sqrt(MAF)
1
0.6
0.5
0.3923302
0.3
0.2
0.1
0.005
0.001677998
0
ID

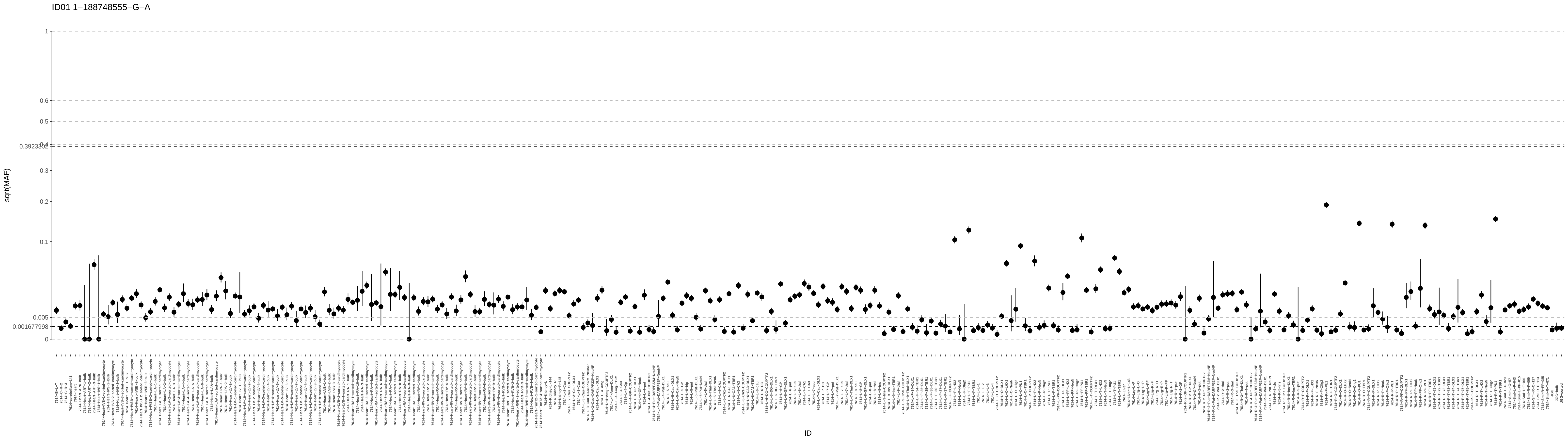
ID01 1−188748555−G−A
sqrt(MAF)
1
0.6
0.5
0.4
0.3923302
0.3
0.2
0.1
0.005
0.001677998
0
ID

ID01 1−212352377−G−T

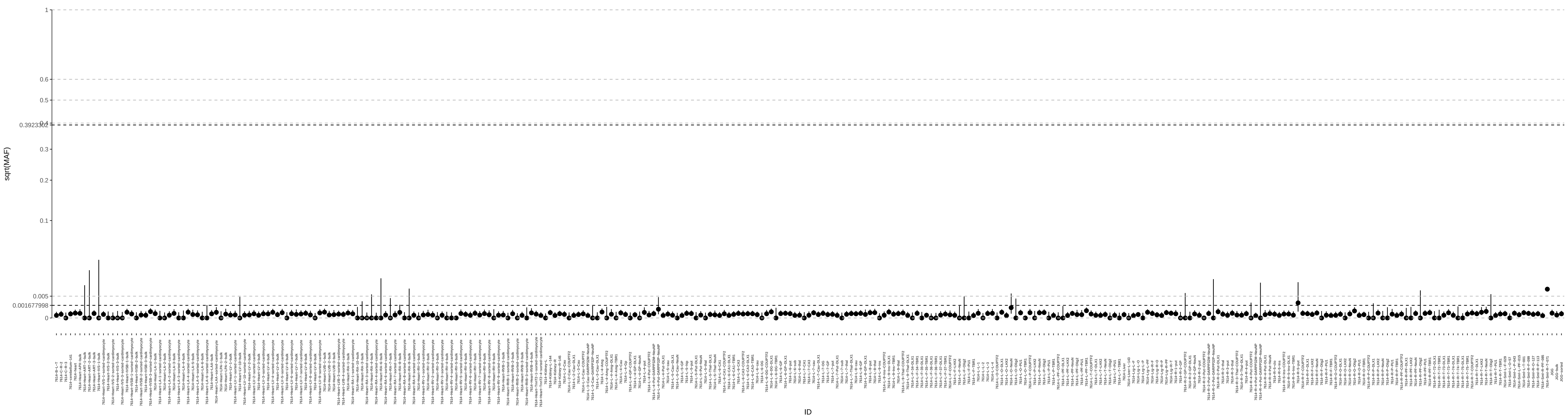

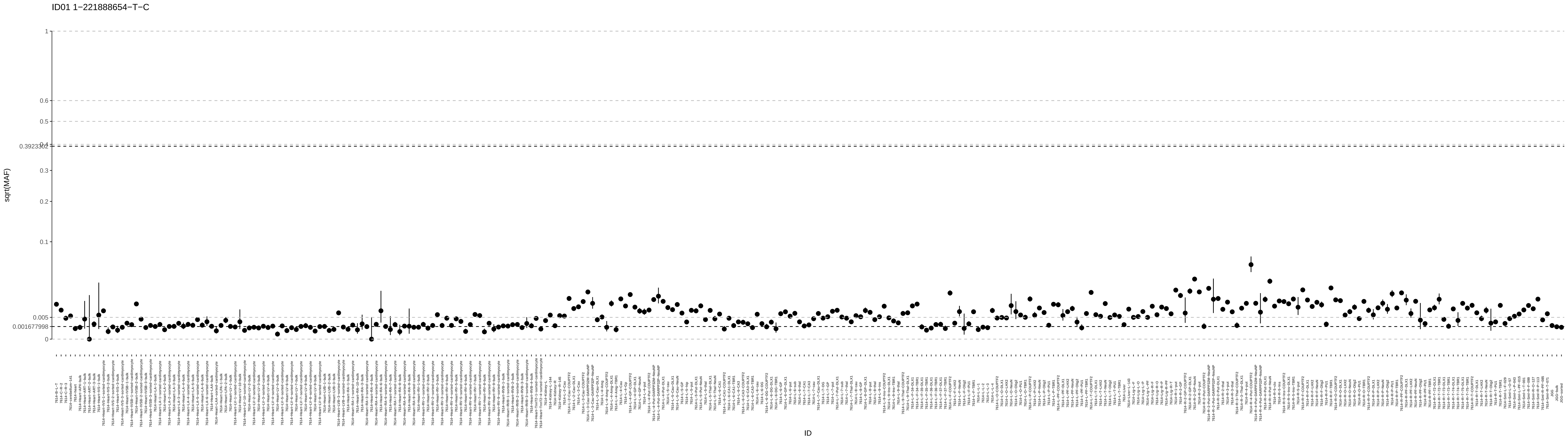
ID01 1−221888654−T−C
sqrt(MAF)
1
0.6
0.5
0.3923302
0.3
0.2
0.1
0.005
0.001677998
0
ID

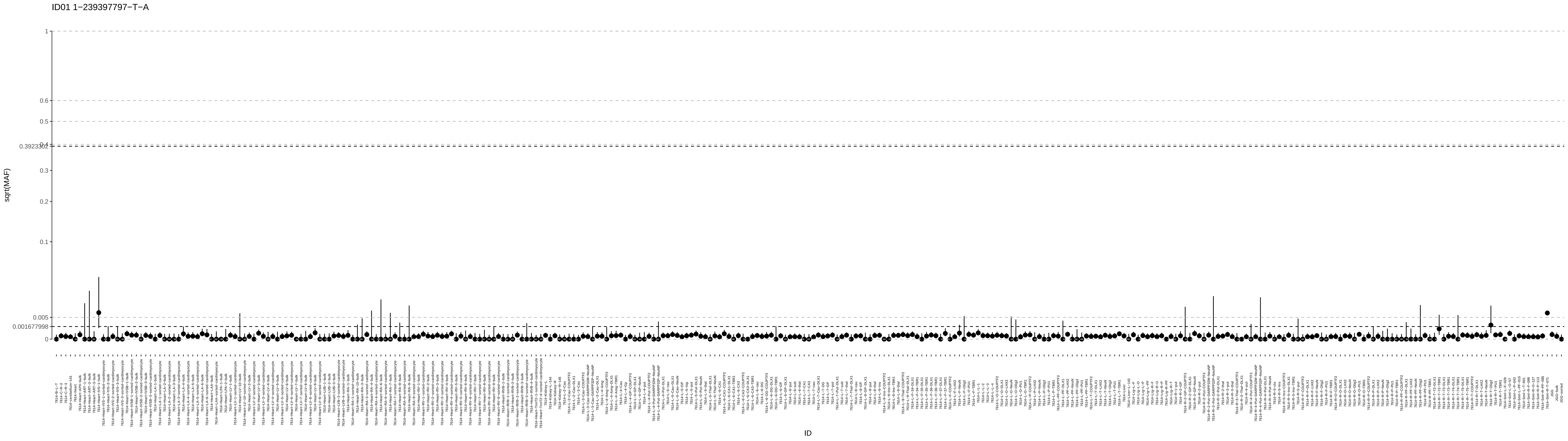
ID01 1−239397797−T−A
sqrt(MAF)
1
0.6
0.5
0.4
0.3923302
0.3
0.2
0.1
0.005
0.001677998
0
ID

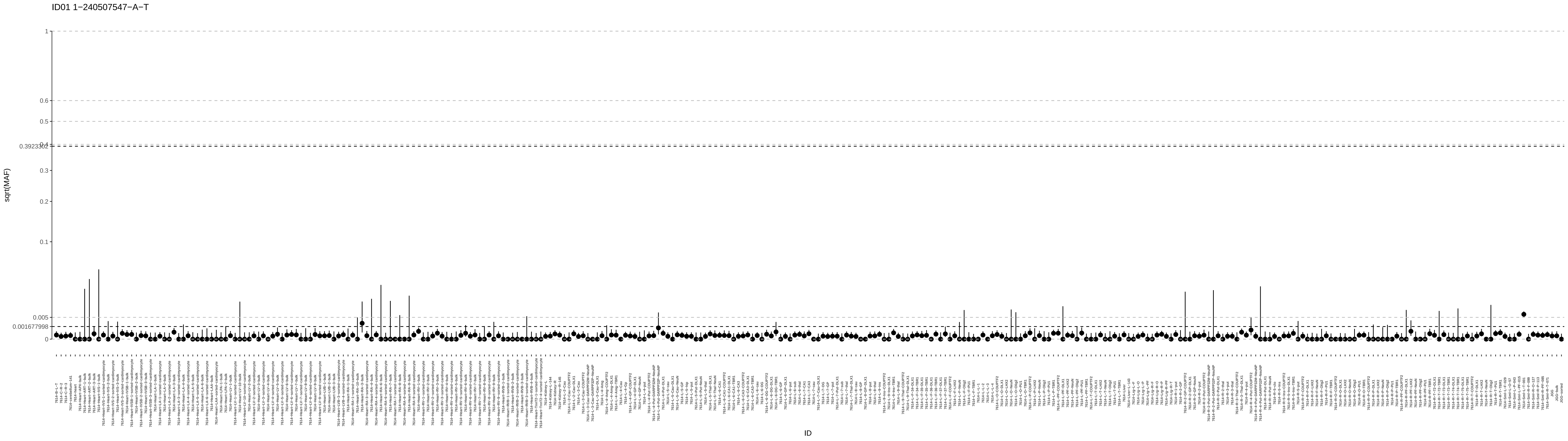
ID01 1−240507547−A−T
sqrt(MAF)
ID

ID01 1−241434408−C−T

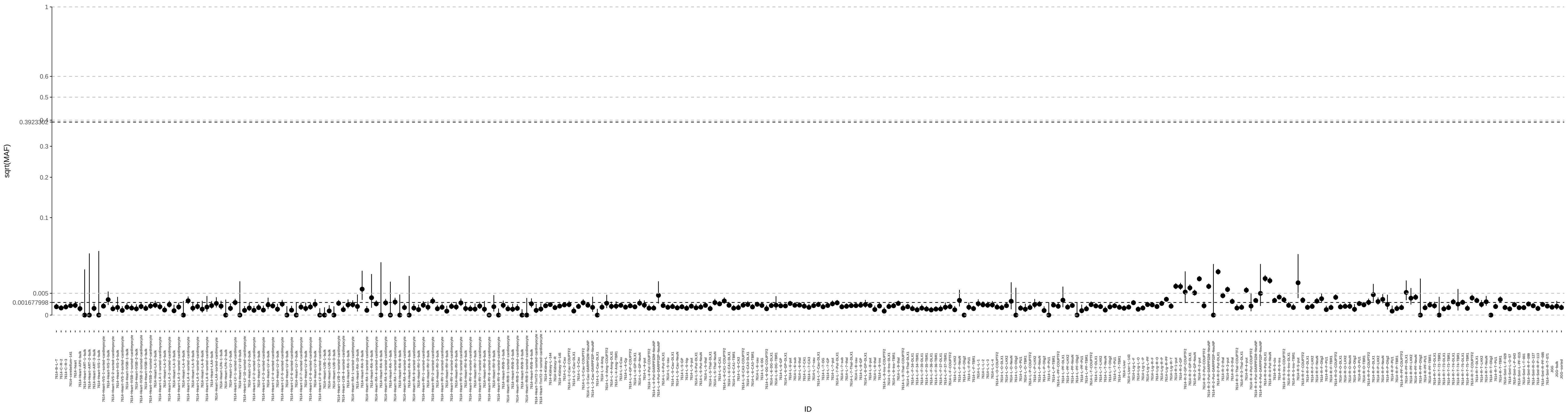

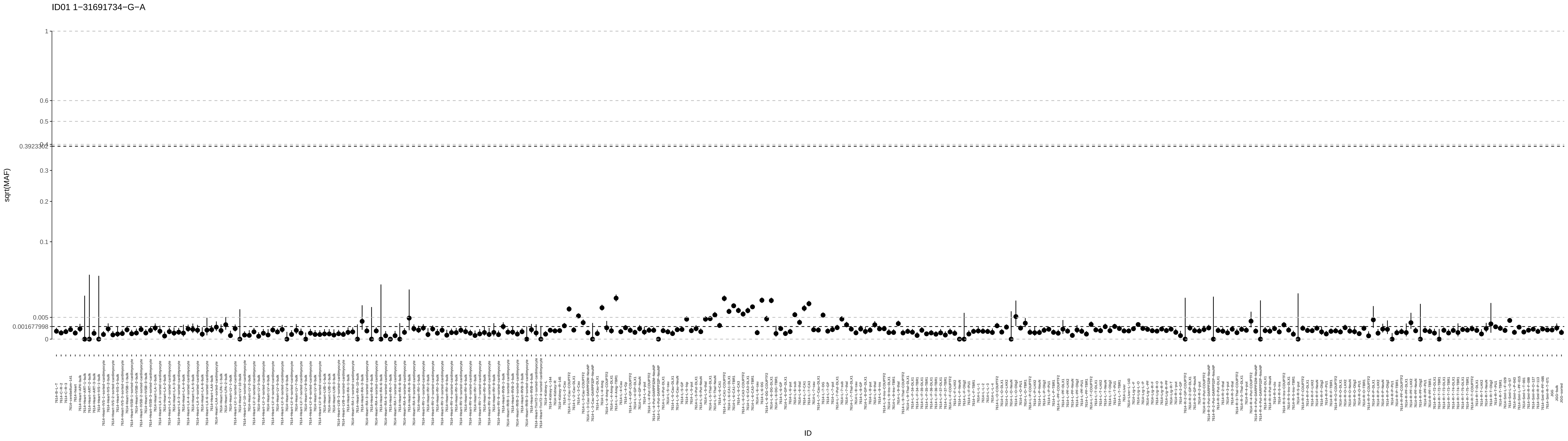
ID01 1−31691734−G−A
sqrt(MAF)
1
0.6
0.5
0.4
0.3923302
0.3
0.2
0.1
0.005
0.001677998
0
ID

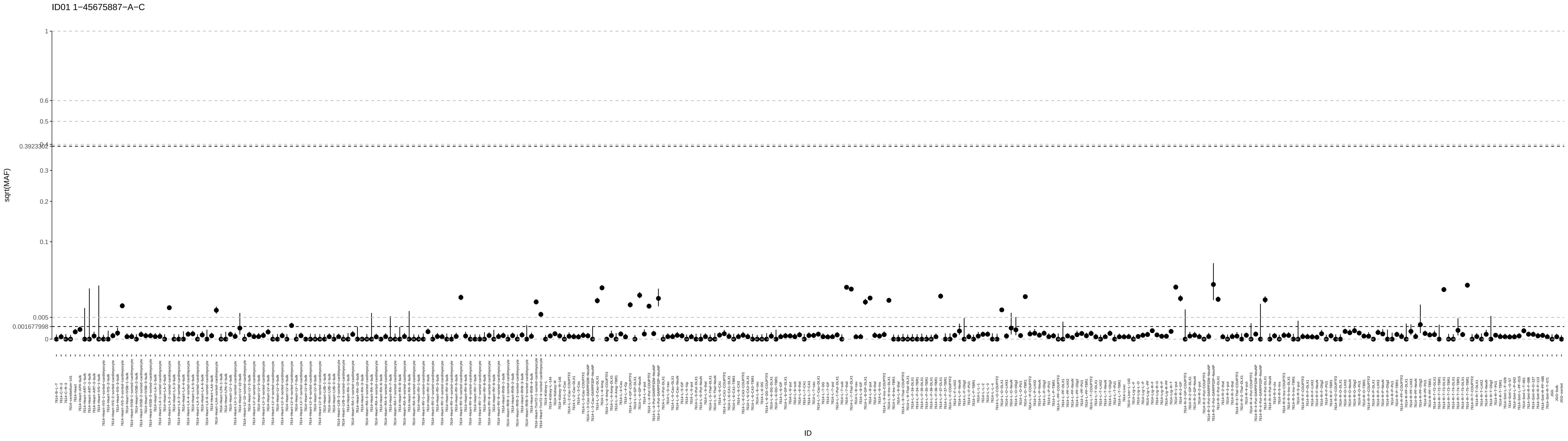
ID01 1−45675887−A−C
sqrt(MAF)
1
0.6
0.5
0.3923302
0.3
0.2
0.1
0.005
0.001677998
0
ID

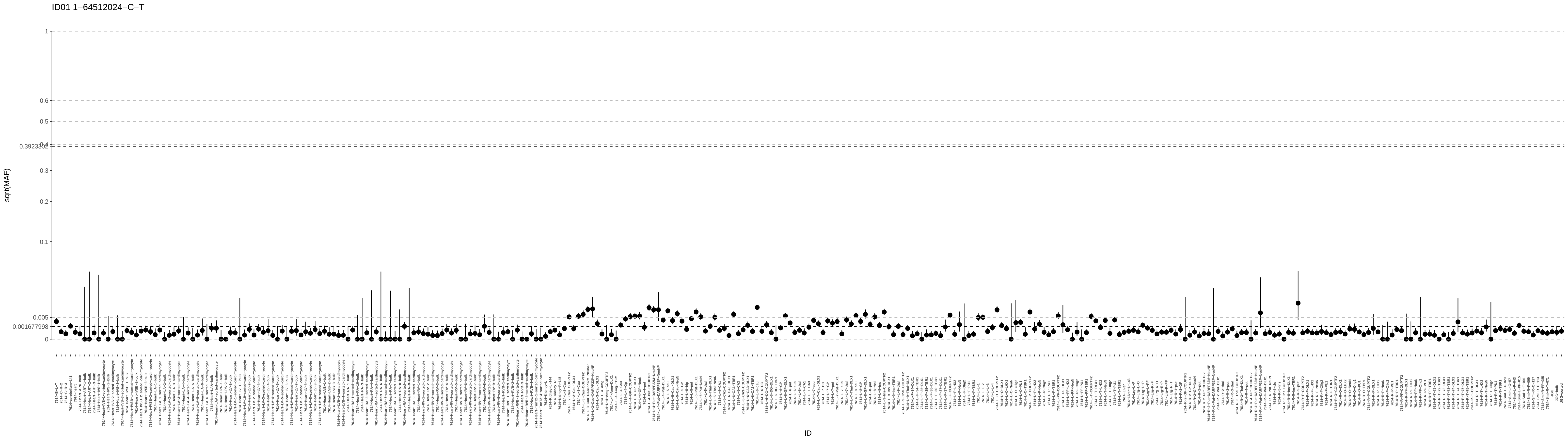
ID01 1−64512024−C−T
sqrt(MAF)
1
0.6
0.5
0.3923302
0.3
0.2
0.1
0.005
0.001677998
0
ID

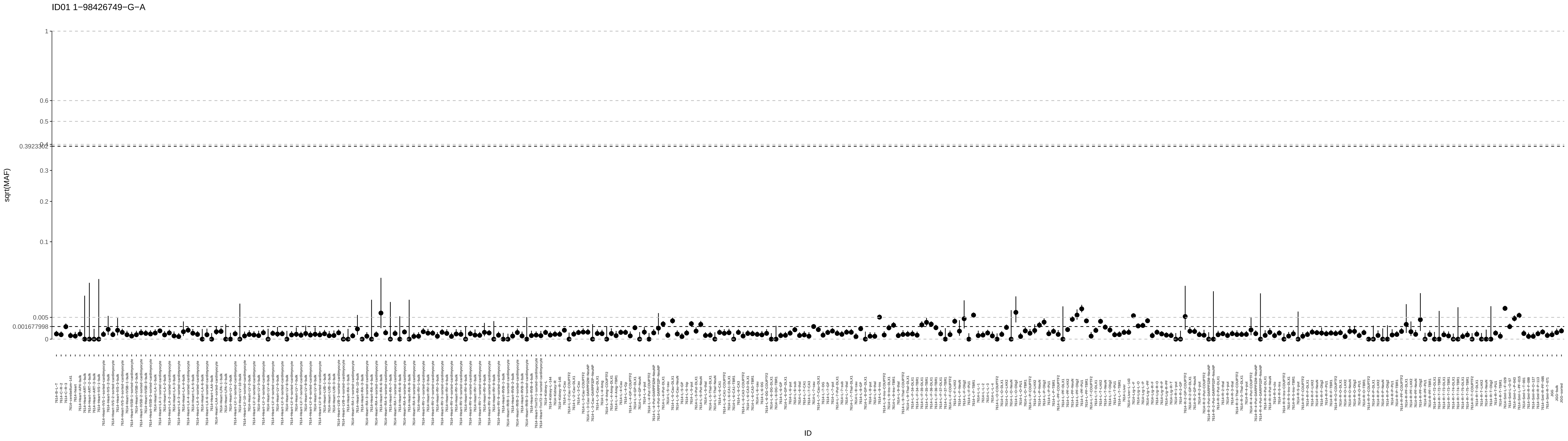
ID01 1−98426749−G−A
sqrt(MAF)
1
0.6
0.5
0.4
0.3923302
0.3
0.2
0.1
0.005
0.001677998
0
ID

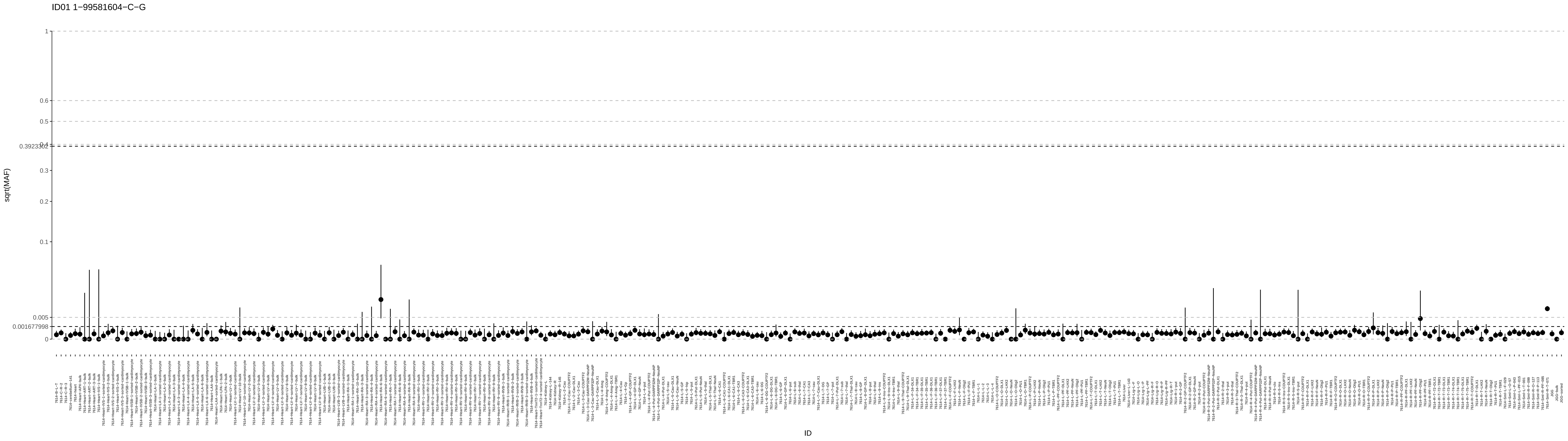
ID01 1−99581604−C−G
sqrt(MAF)
ID
1
0.6
0.5
0.3923302
0.3
0.2
0.1
0.005
0.001677998
0

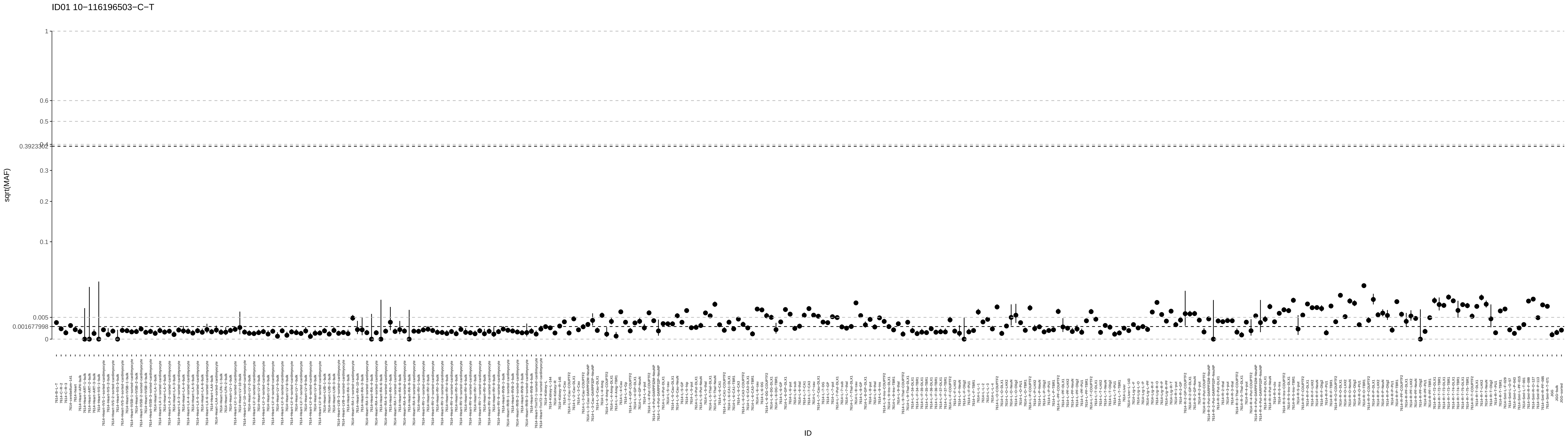

ID01 10−116196503−C−T
sqrt(MAF)
1
0.6
0.5
0.3923302
0.3
0.2
0.1
0.005
0.001677998
0
ID

ID01 10−127139902−G−A

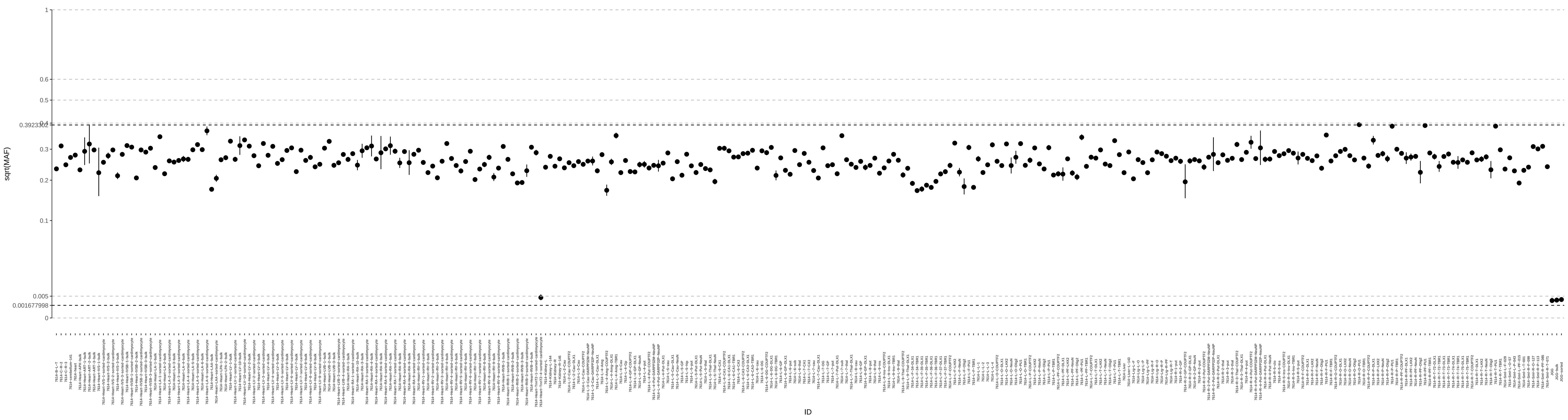

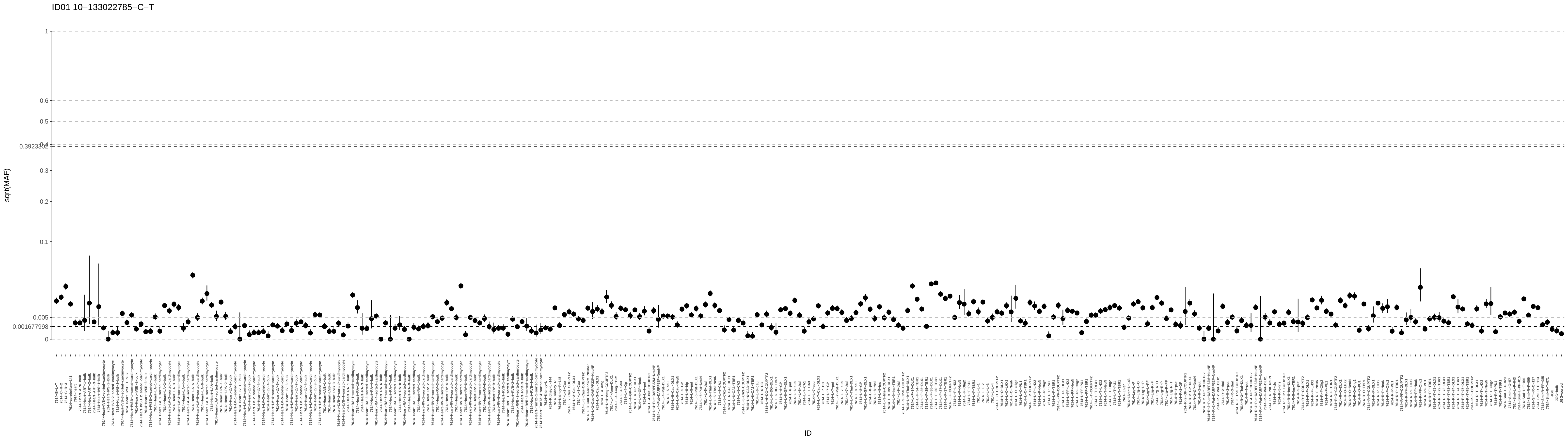
ID01 10−133022785−C−T
1
0.6
0.5
0.4
0.3923302
0.3
0.2
0.1
0.005
0.001677998
0
sqrt(MAF)
ID

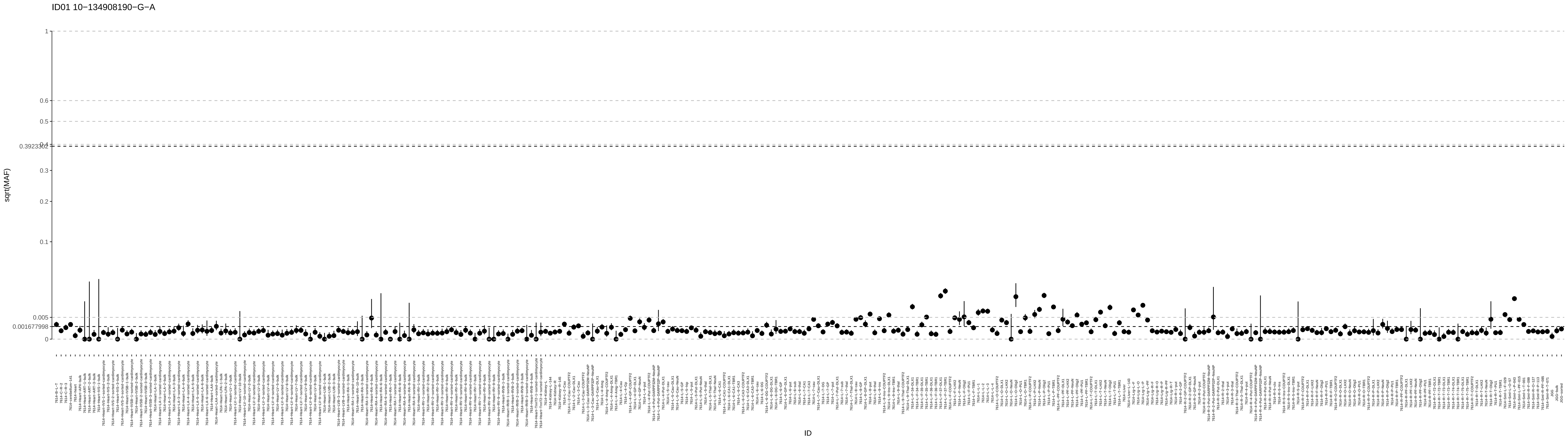
ID01 10−134908190−G−A
1
0.6
0.5
0.3923302
0.3
0.2
0.1
0.005
0.001677998
0
sqrt(MAF)
ID

ID01 10−23042694−GAGTT−G

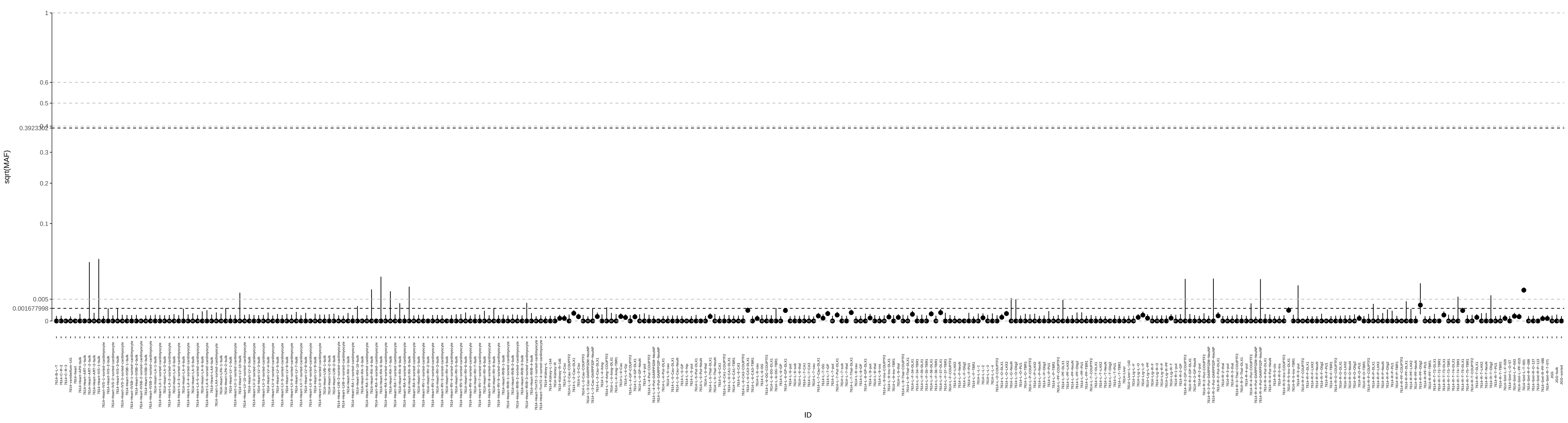

ID01 10−71631216−G−A

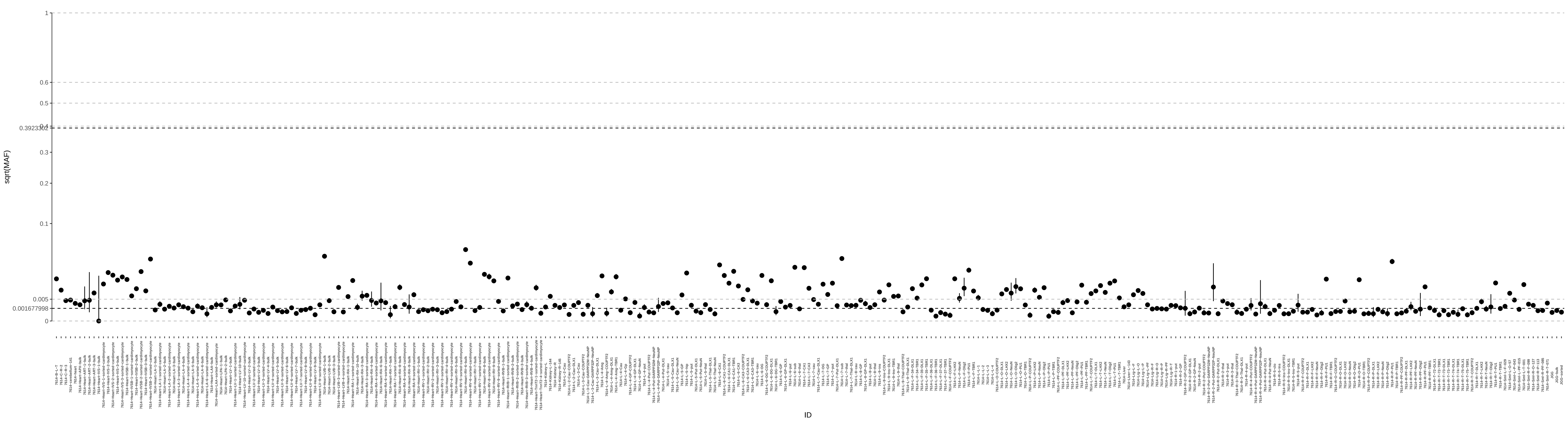

ID01 10−84641815−C−T

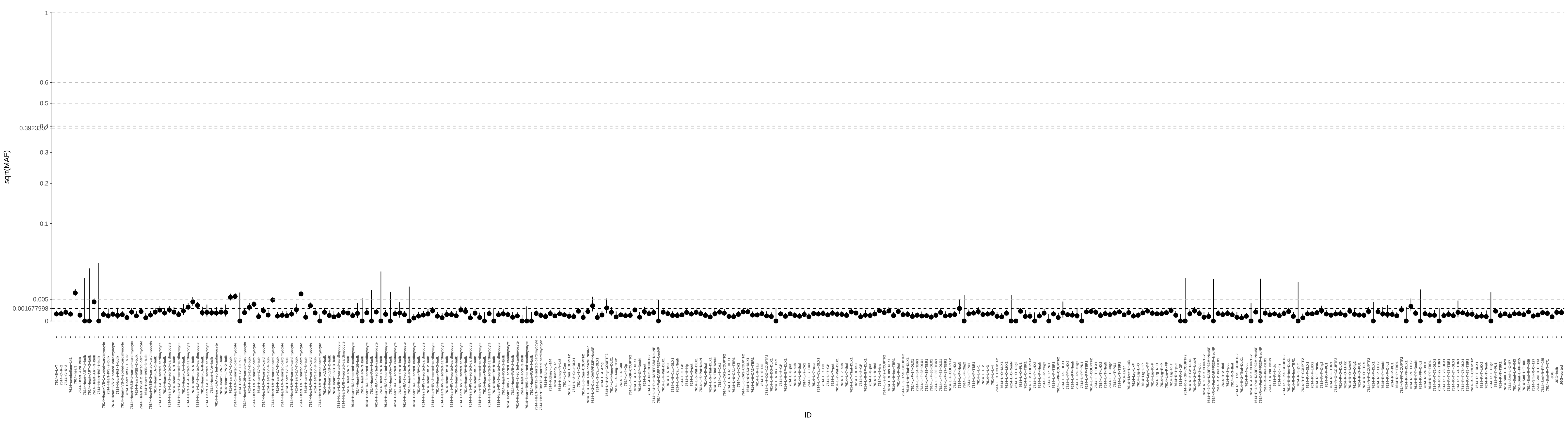

ID01 10−97951160−C−T

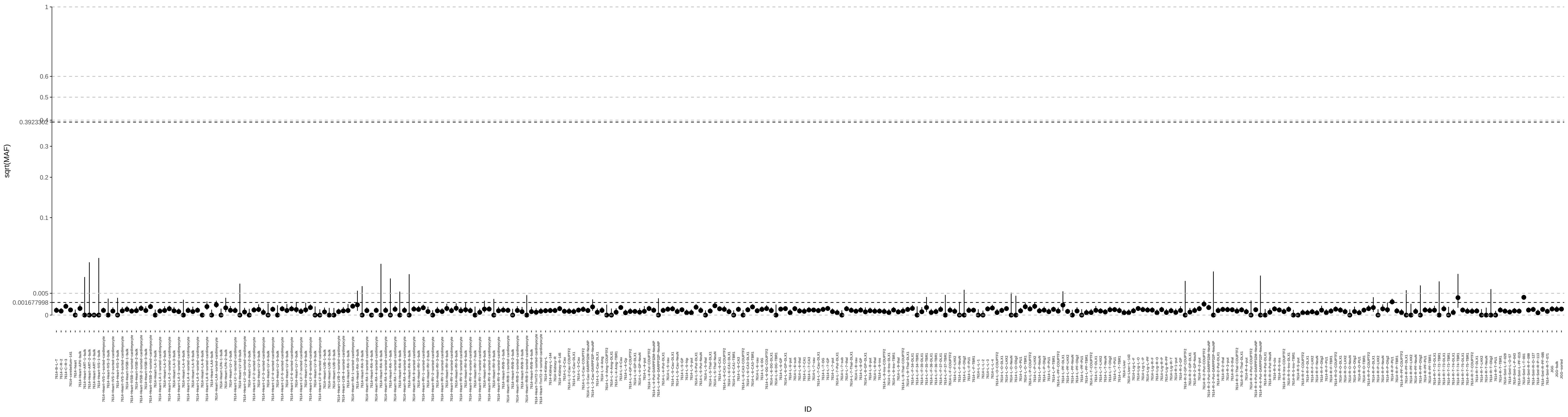

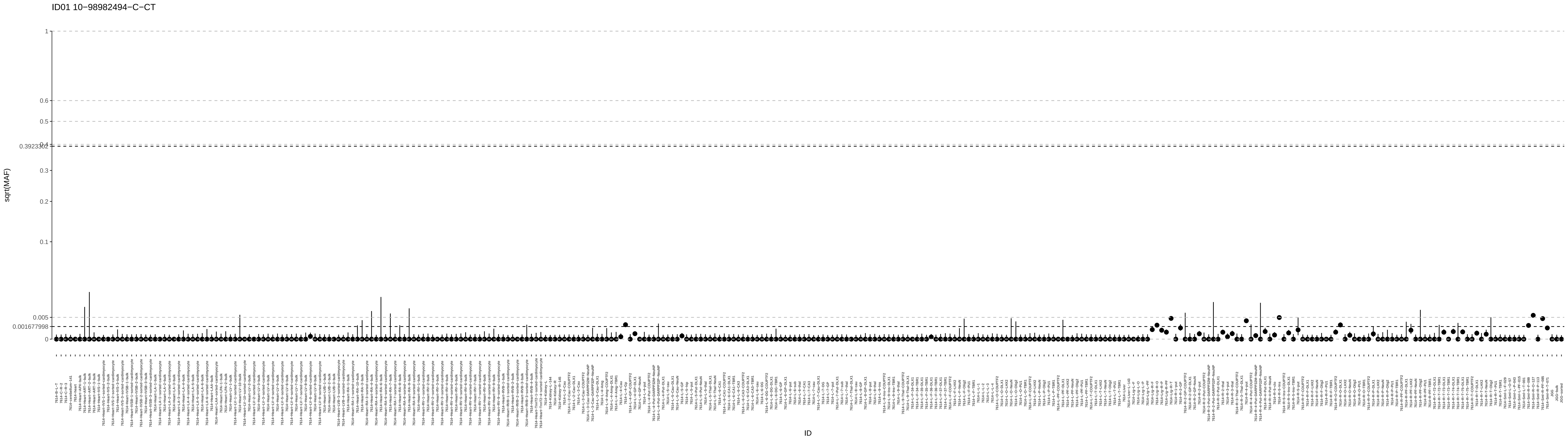

ID01 10−98982494−C−CT
sqrt(MAF)
1
0.6
0.5
0.3923302
0.3
0.2
0.1
0.005
0.001677998
0
ID

ID01 11−110591774−A−C

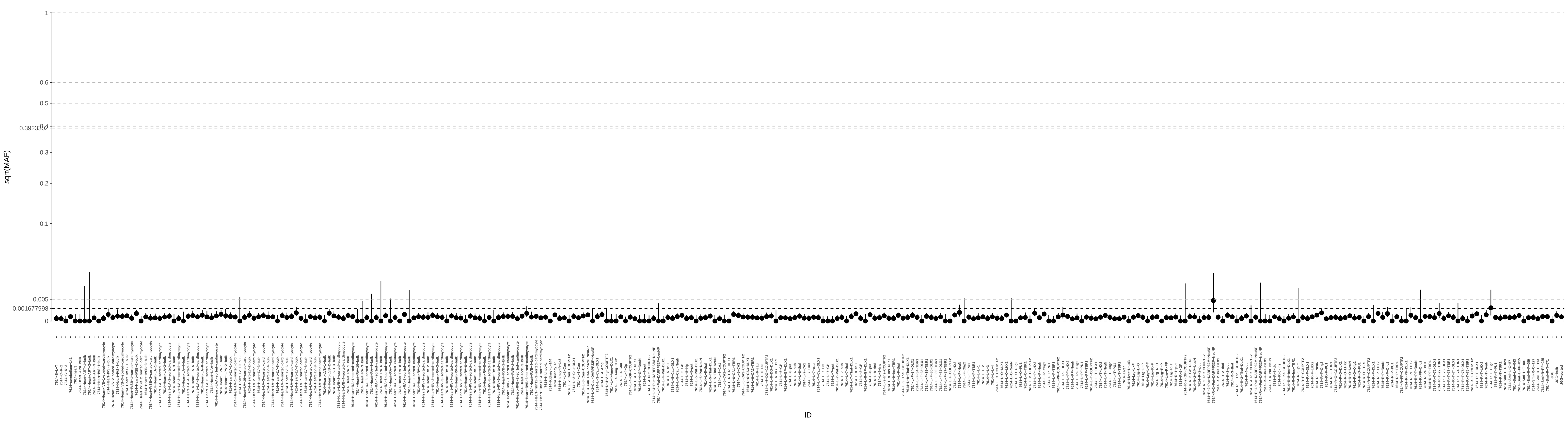

ID01 11−114745430−C−A

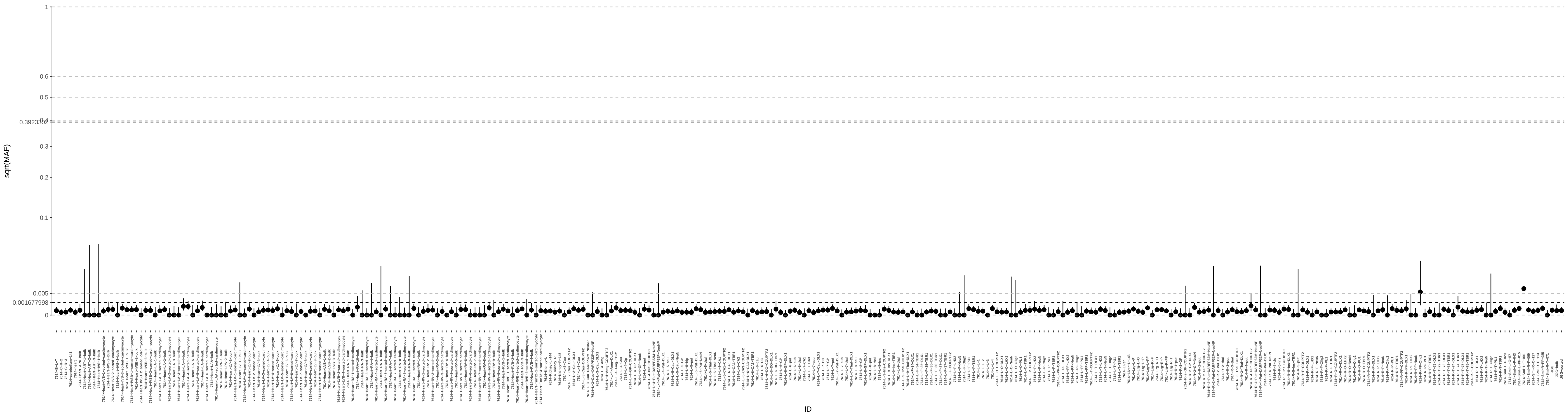

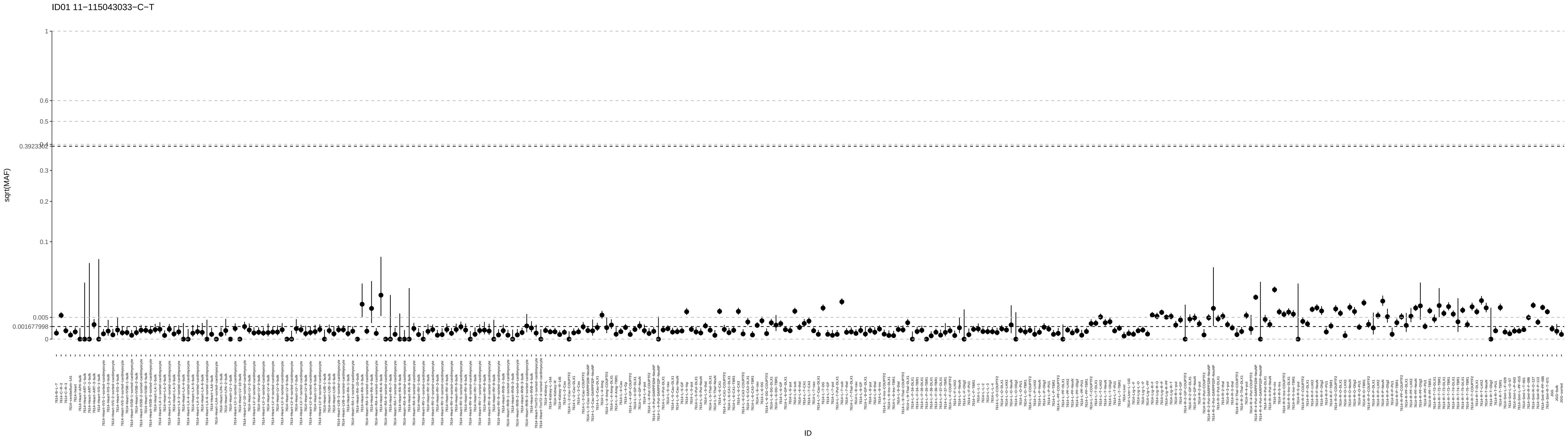

ID01 11−115043033−C−T
sqrt(MAF)
1
0.6
0.5
0.3923302
0.3
0.2
0.1
0.005
0.001677998
0
ID

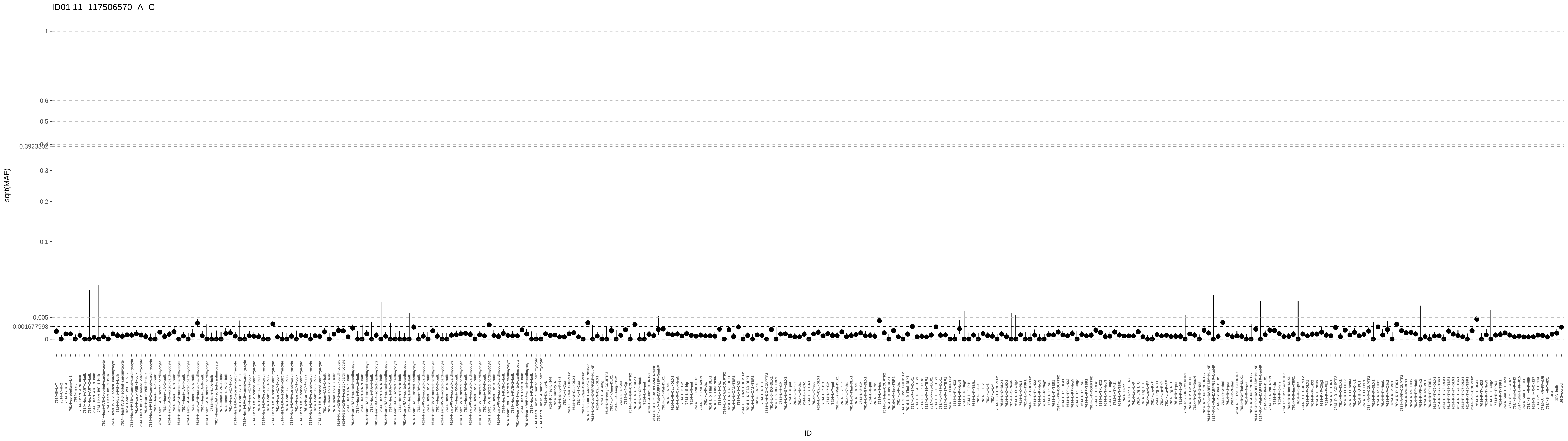
ID01 11−117506570−A−C
sqrt(MAF)
1
0.6
0.5
0.3923302
0.3
0.2
0.1
0.005
0.001677998
0
ID

ID01 11−123516528−G−A

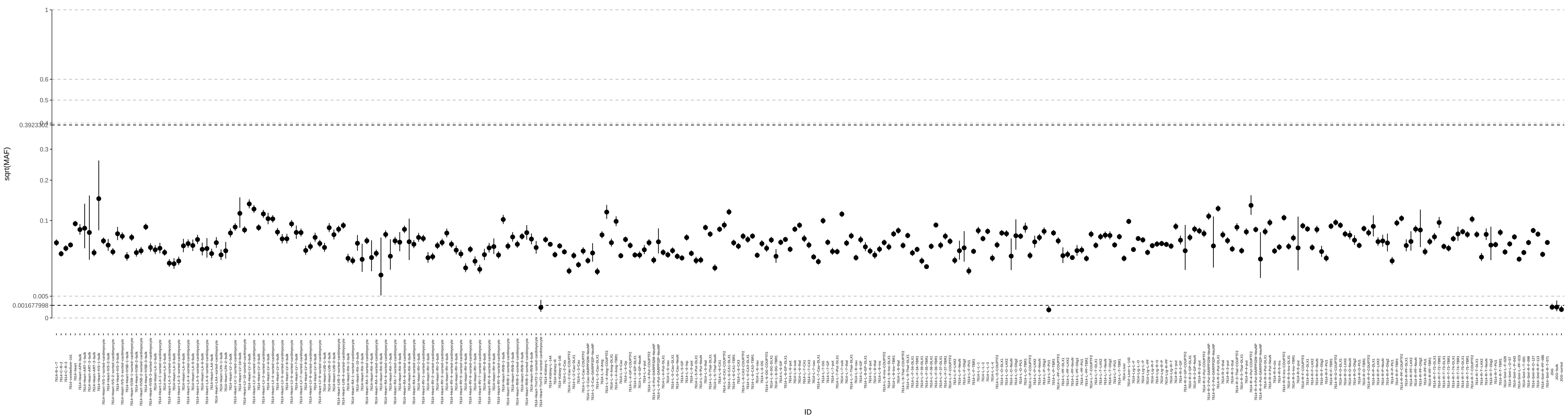

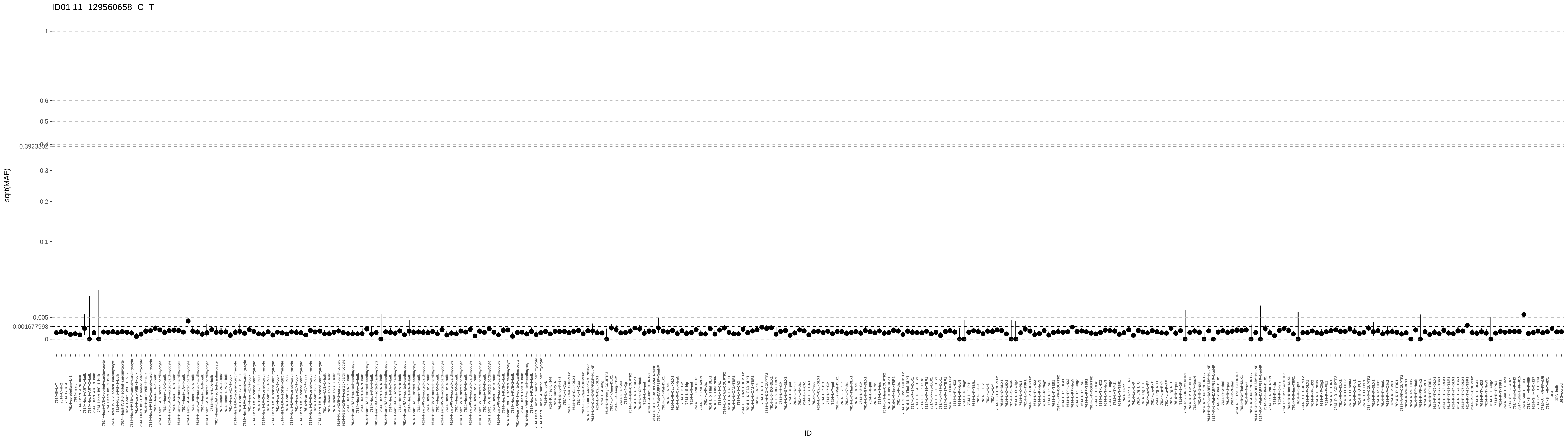
ID01 11−129560658−C−T
sqrt(MAF)
1
0.6
0.5
0.3923302
0.3
0.2
0.1
0.005
0.001677998
0
ID

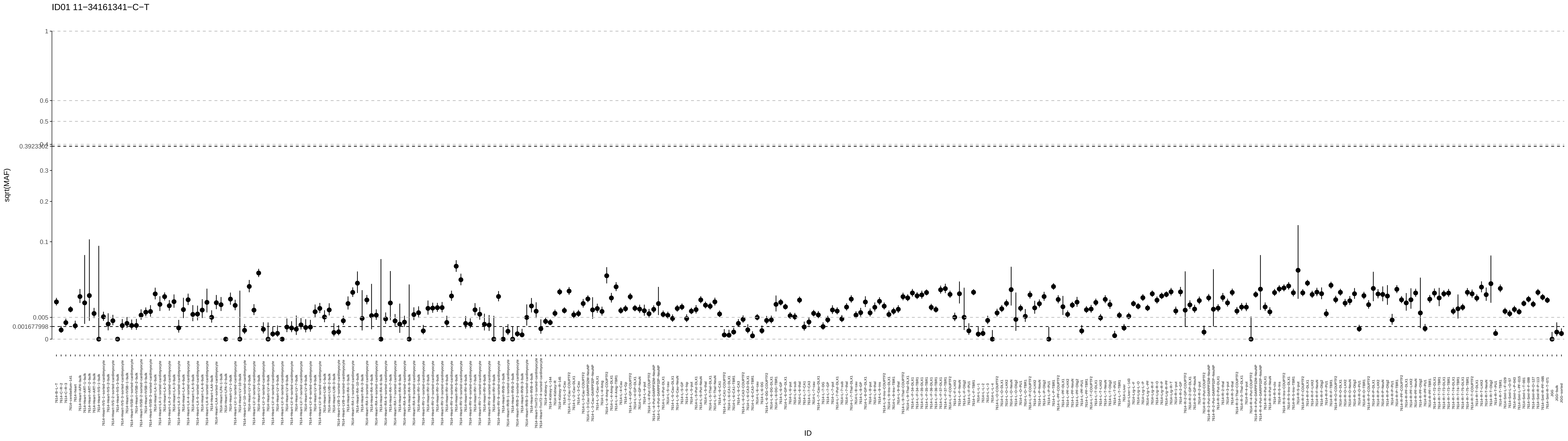

ID01 11−34161341−C−T
sqrt(MAF)
1
0.6
0.5
0.3923302
0.3
0.2
0.1
0.005
0.001677998
0
ID

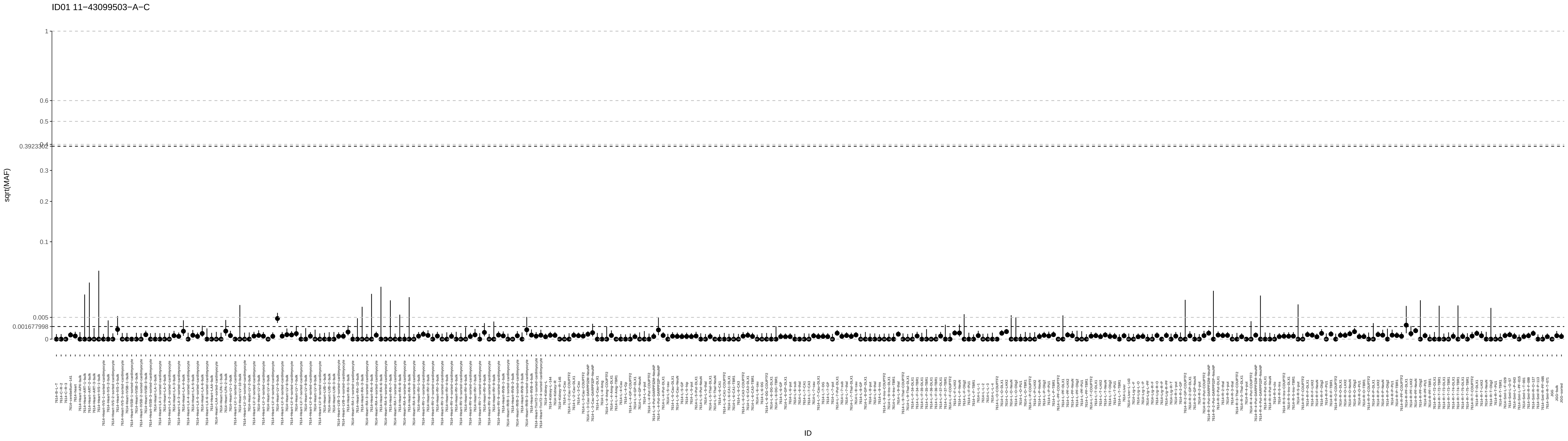
ID01 11−43099503−A−C
sqrt(MAF)
1
0.6
0.5
0.3923302
0.3
0.2
0.1
0.005
0.001677998
0
ID

ID01 11−5470408−T−C

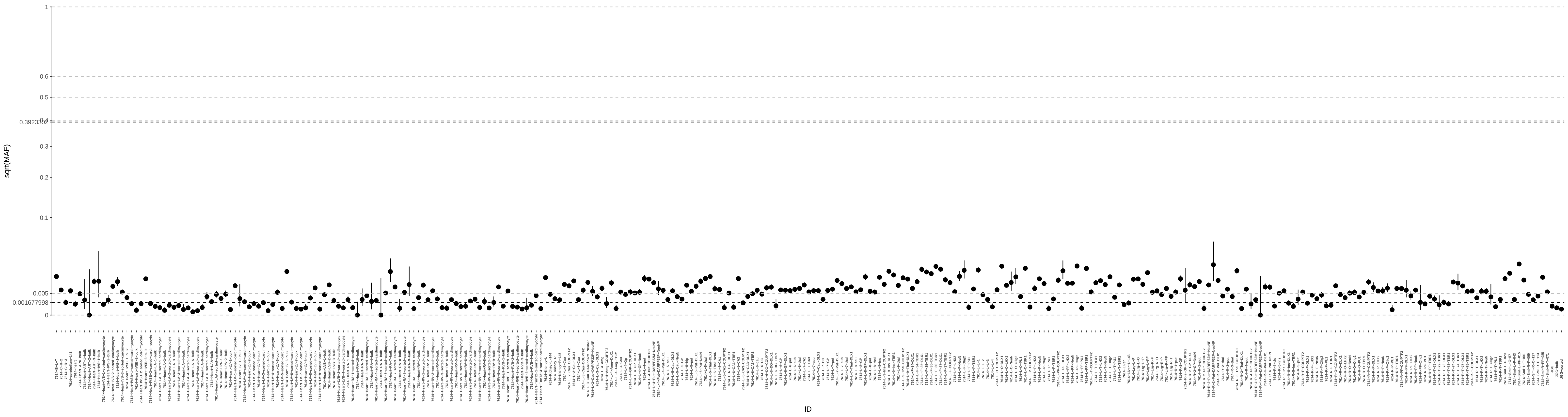

ID01 11−8350555−G−A

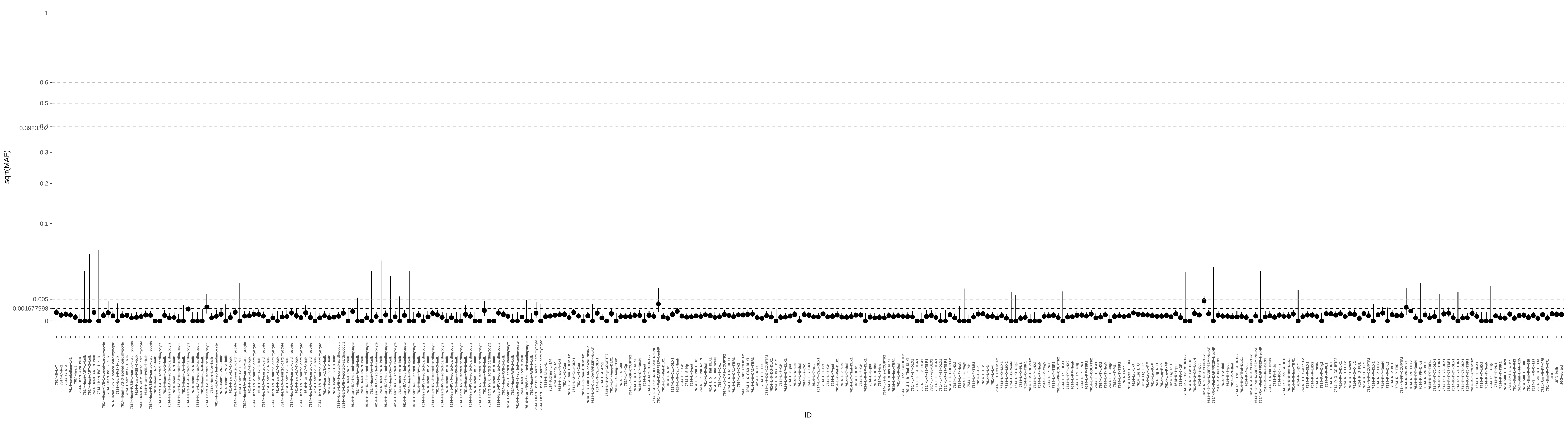

ID01 11−84980734−G−A

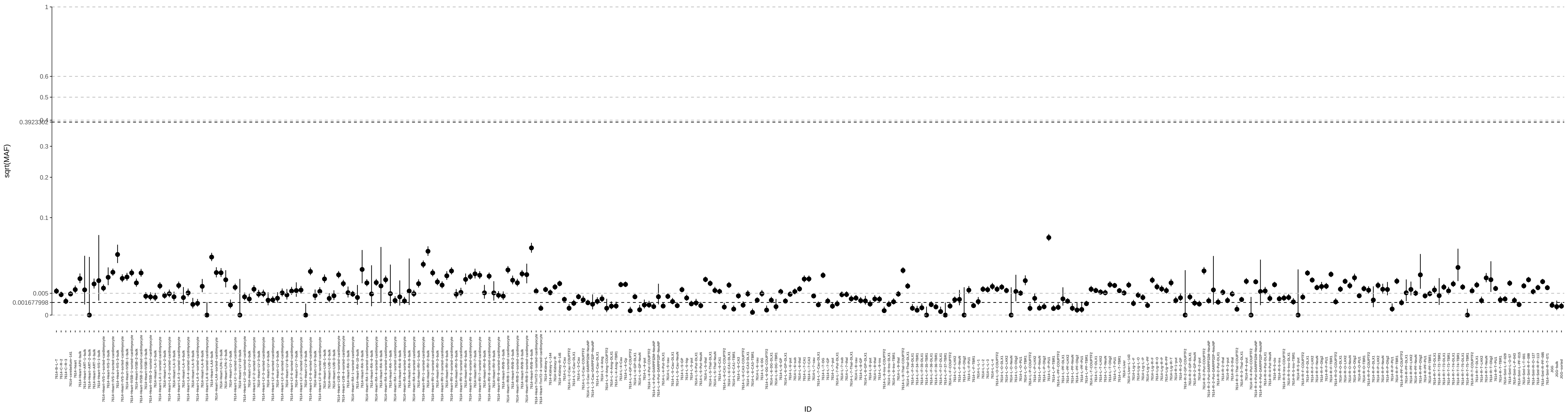

ID01 11−88087255−A−C

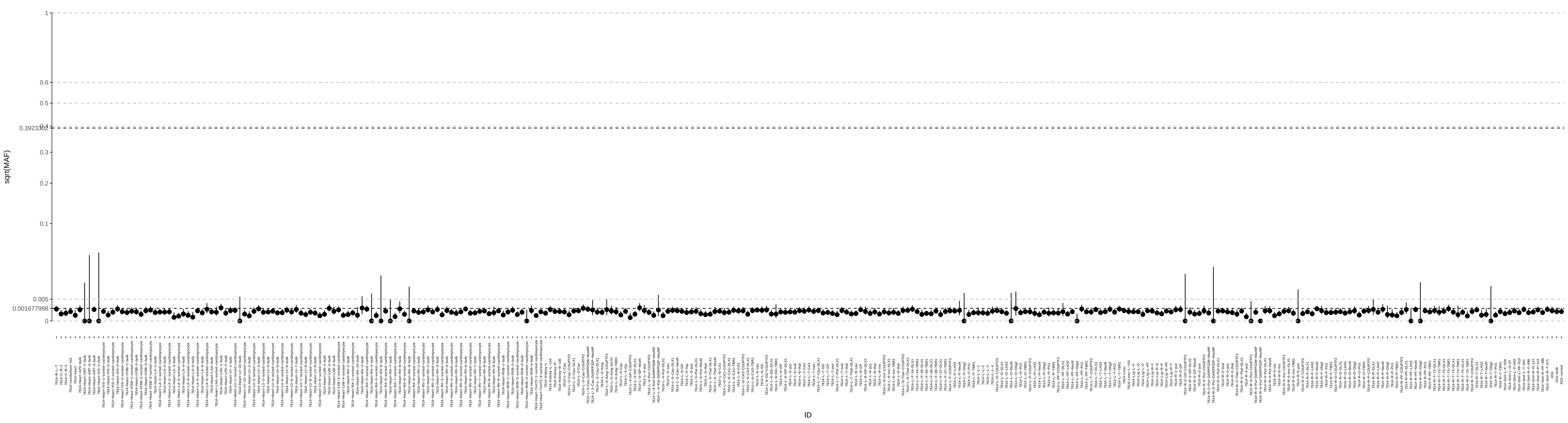

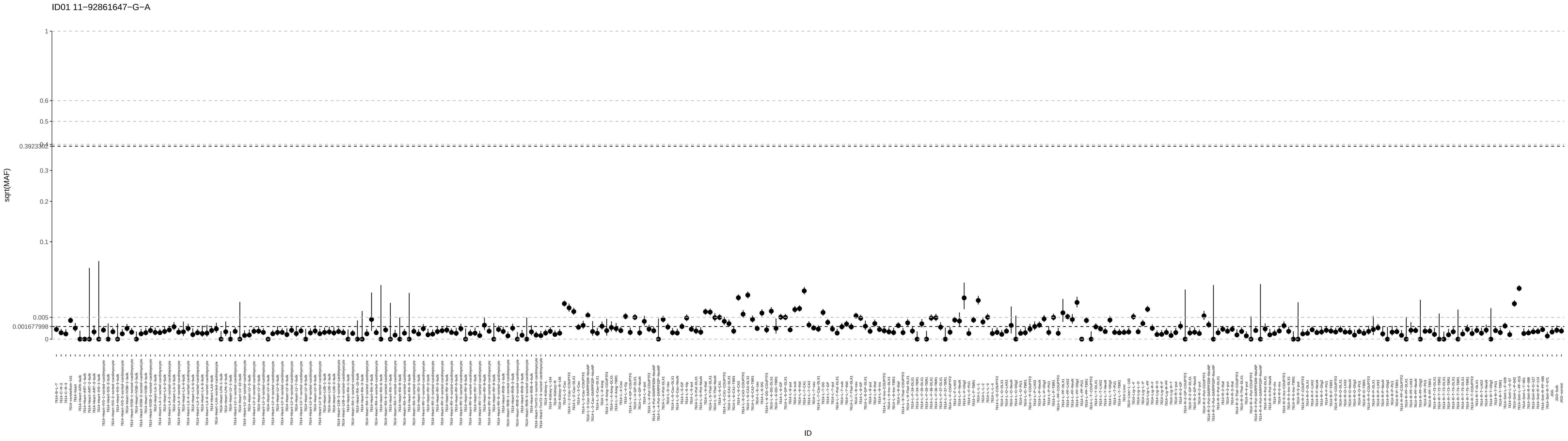

ID01 11−92861647−G−A
sqrt(MAF)
1
0.6
0.5
0.3923302
0.3
0.2
0.1
0.005
0.001677998
0
ID

ID01 11−94045731−T−C

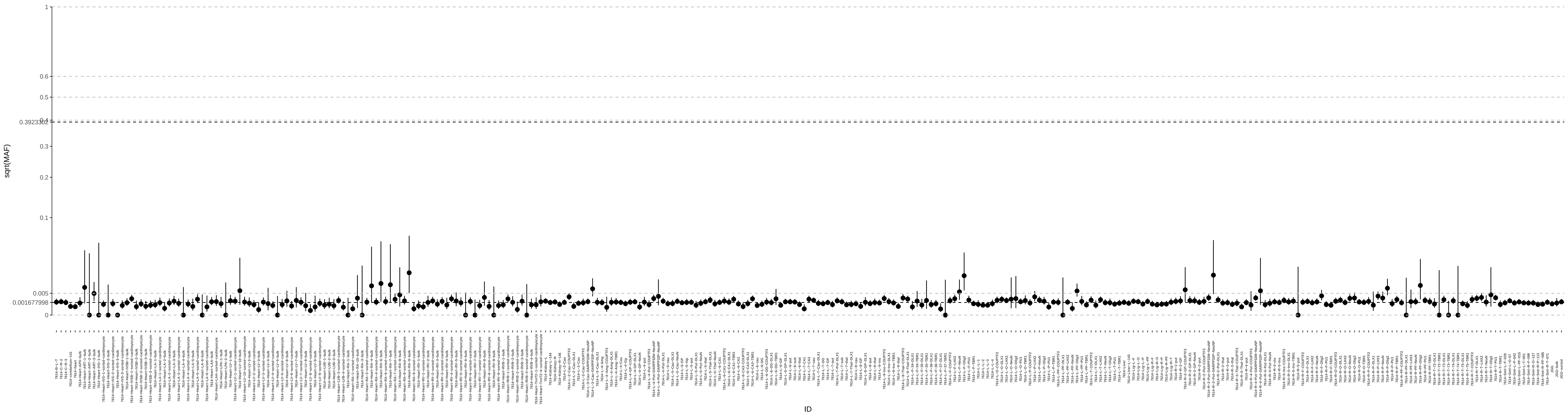

ID01 11−94048896−G−A

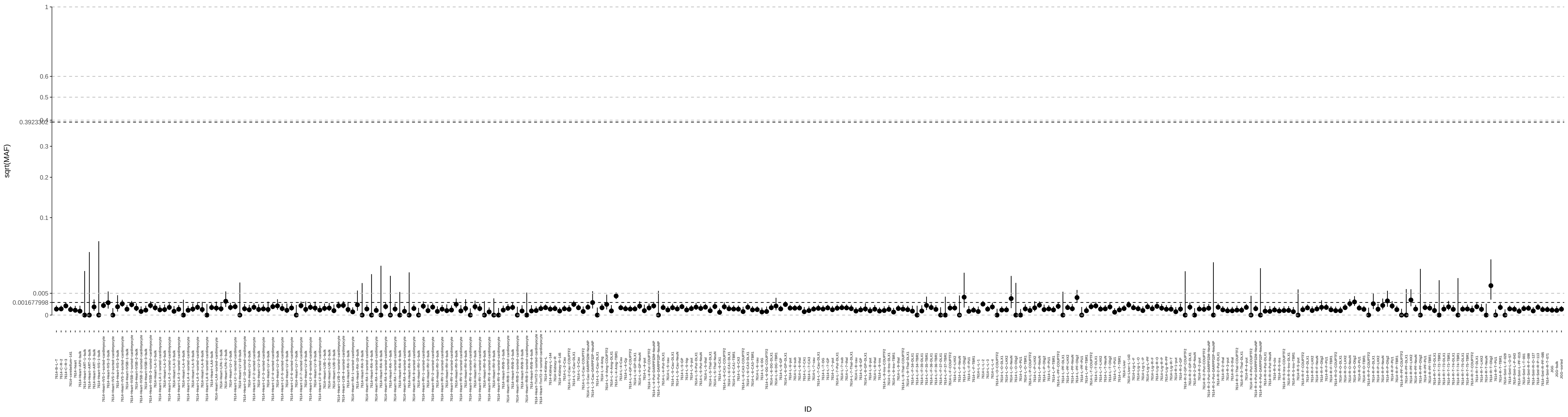

ID01 11−97326785−G−A

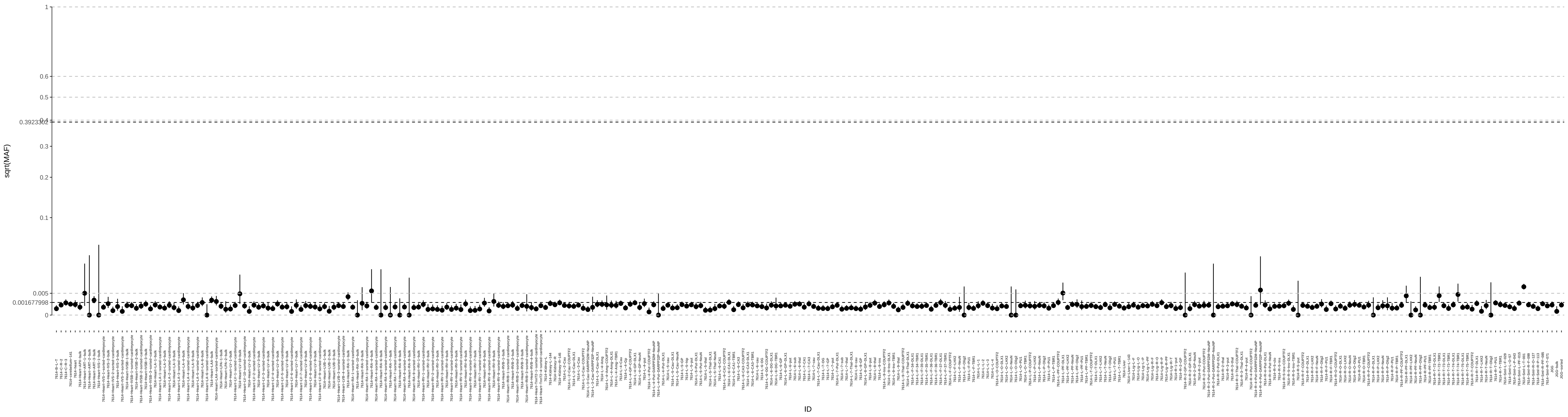

ID01 11−99258149−G−A

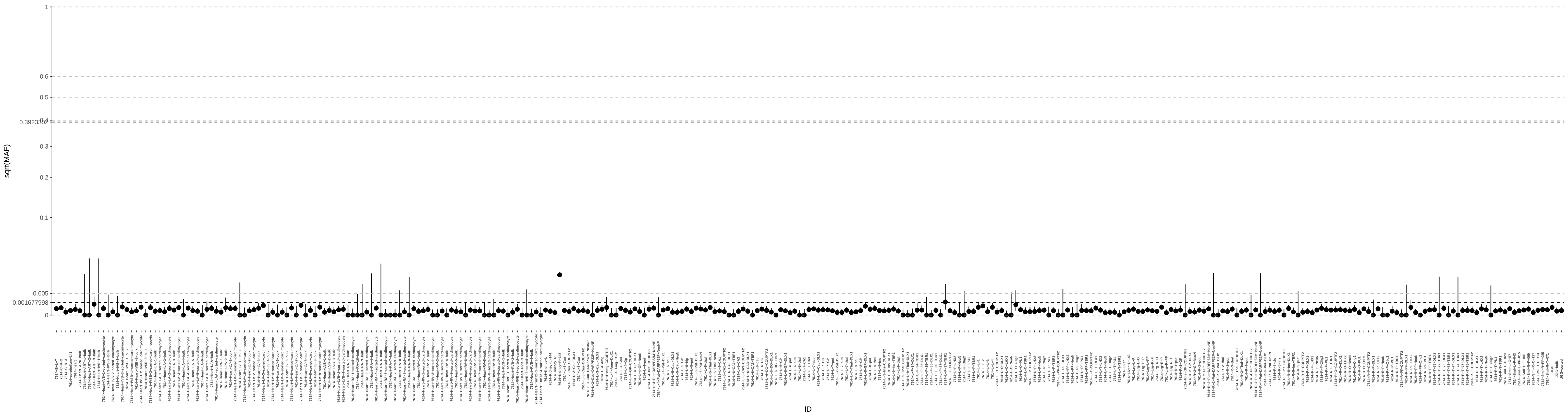

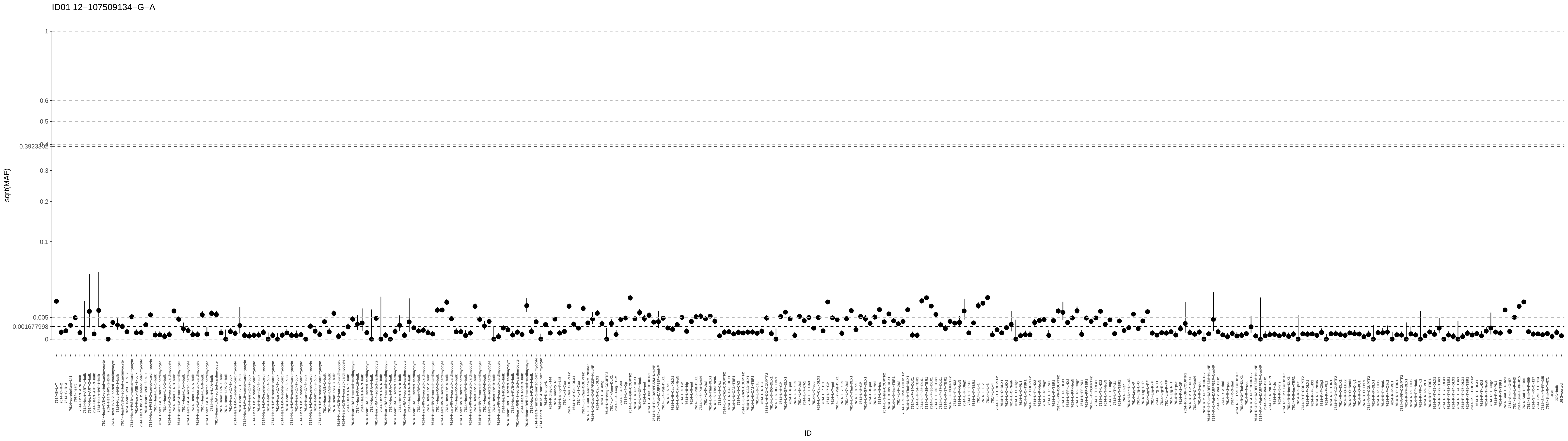
ID01 12−107509134−G−A
sqrt(MAF)
1
0.6
0.5
0.3923302
0.3
0.2
0.1
0.005
0.001677998
0
ID

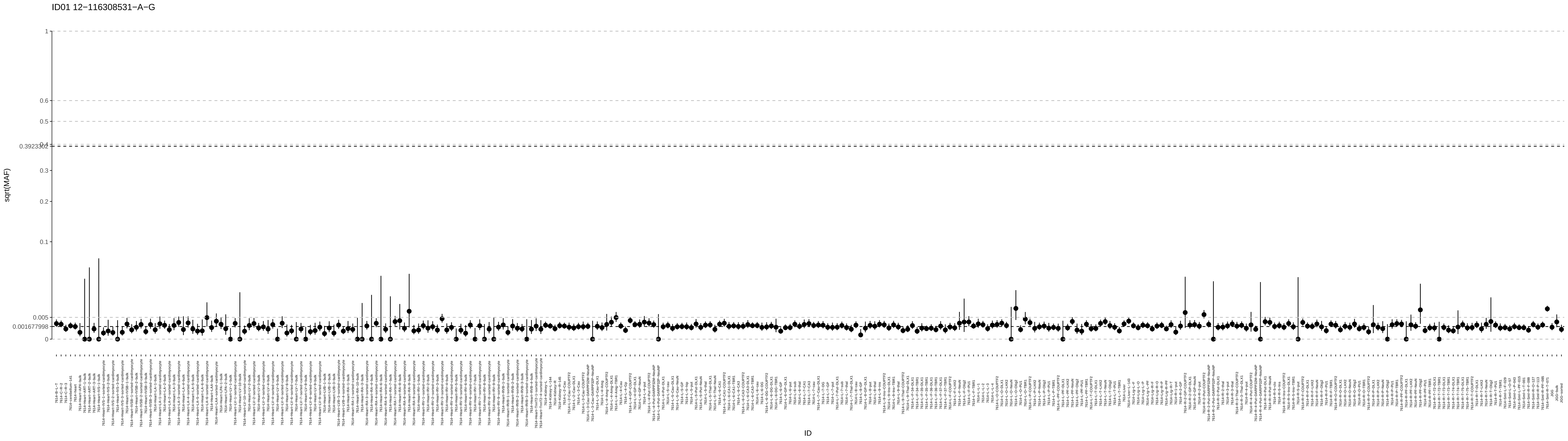
ID01 12−116308531−A−G
sqrt(MAF)
1
0.6
0.5
0.3923302
0.3
0.2
0.1
0.005
0.001677998
0
ID

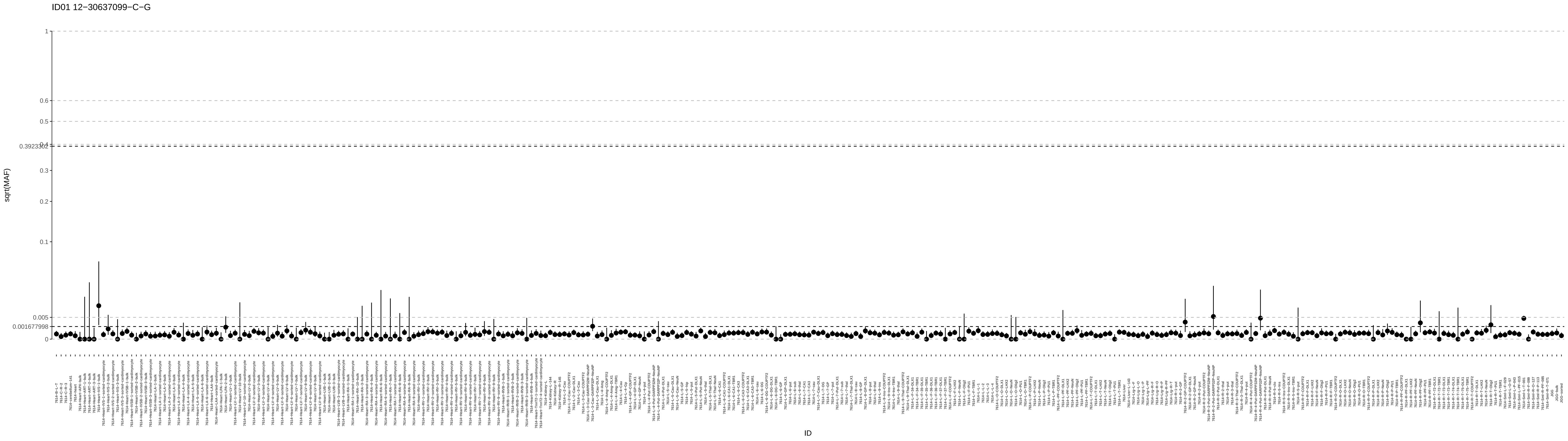
ID01 12−30637099−C−G
1
0.6
0.5
0.3923302
0.3
0.2
0.1
0.005
0.001677998
0
sqrt(MAF)
ID

ID01 12−30749782−G−T

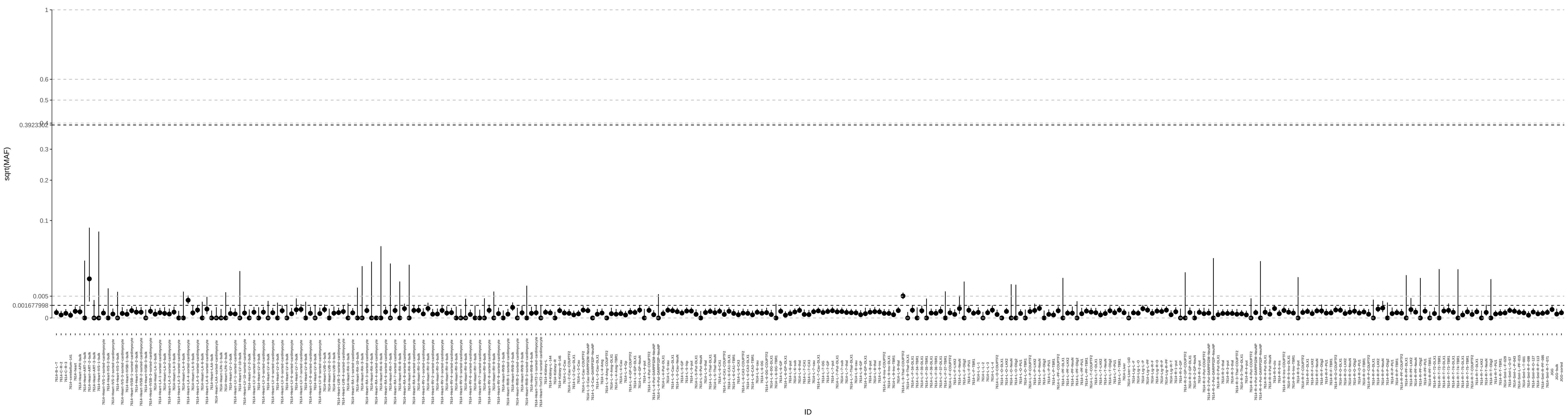

ID01 12−40717911−G−A

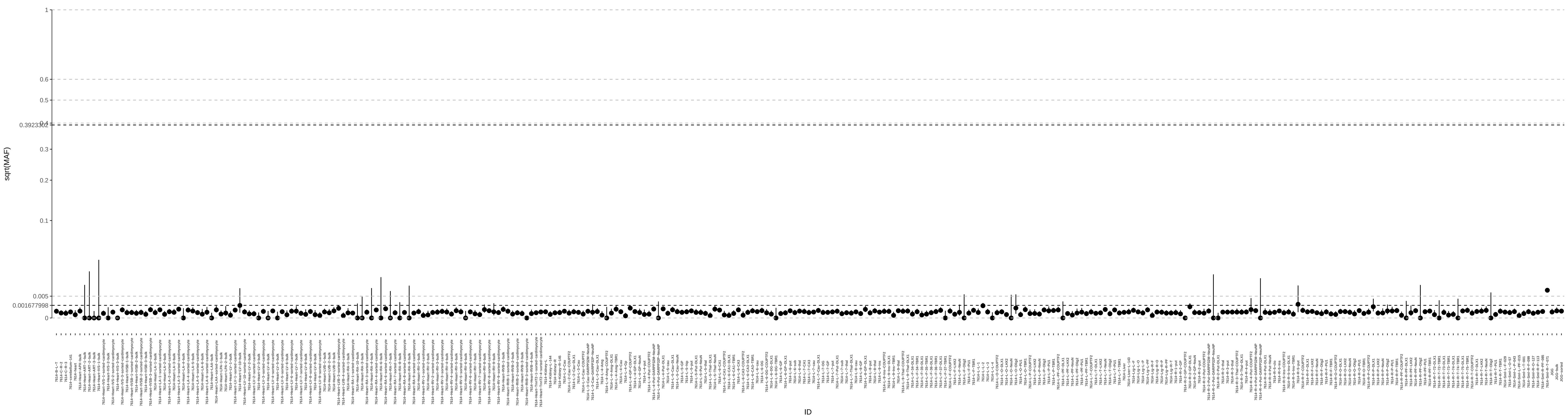

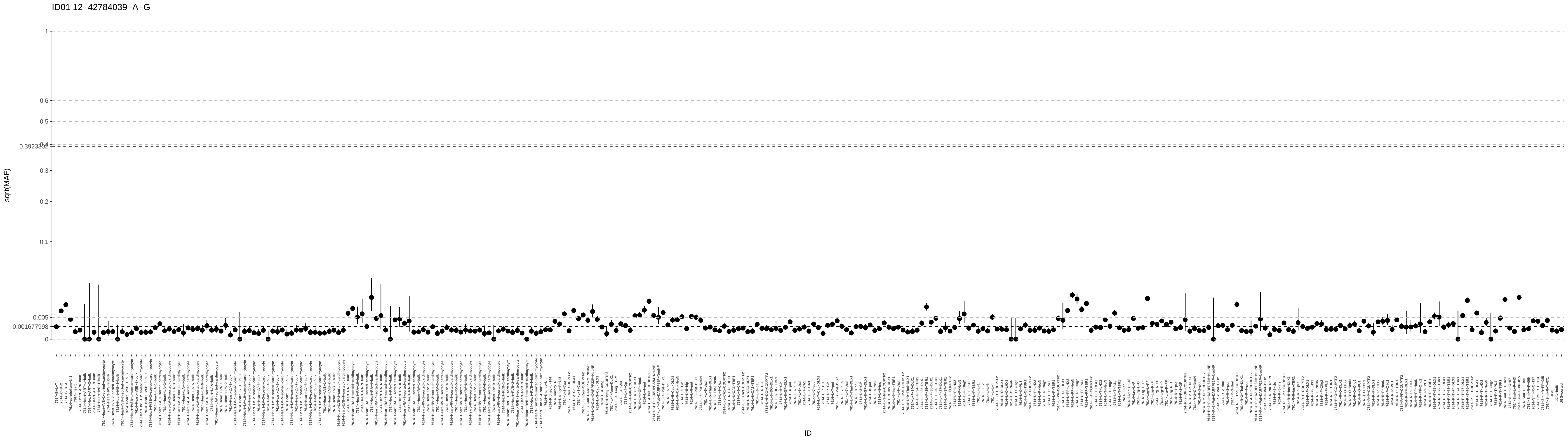
ID01 12−42784039−A−G
sqrt(MAF)
1
0.6
0.5
0.4
0.3923302
0.3
0.2
0.1
0.005
0.001677998
0
ID

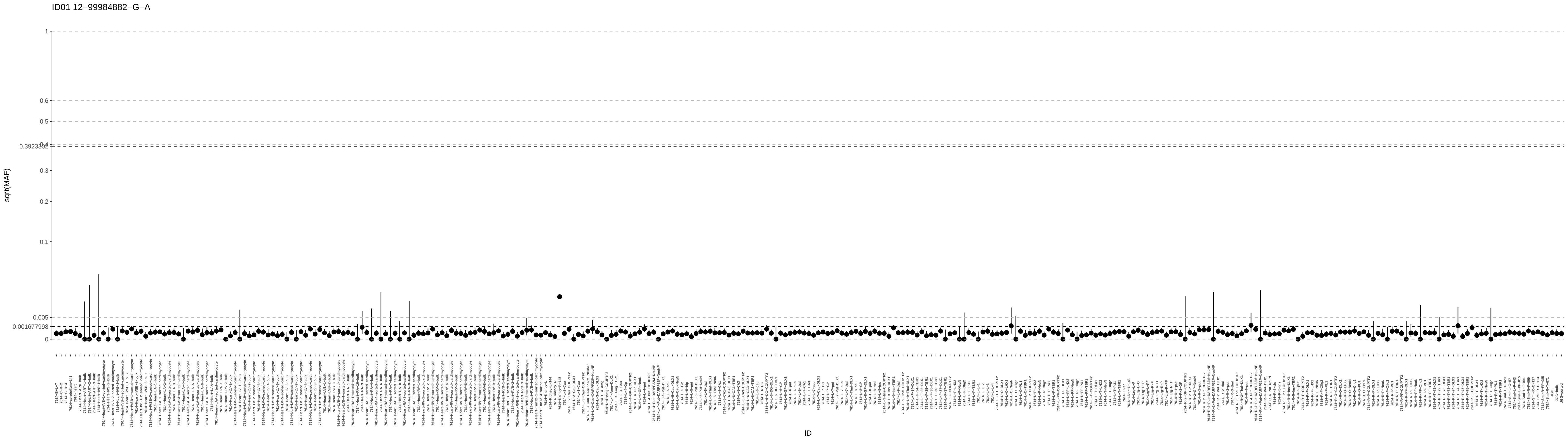
ID01 12−99984882−G−A
sqrt(MAF)
1
0.6
0.5
0.3923302
0.3
0.2
0.1
0.005
0.001677998
0
ID

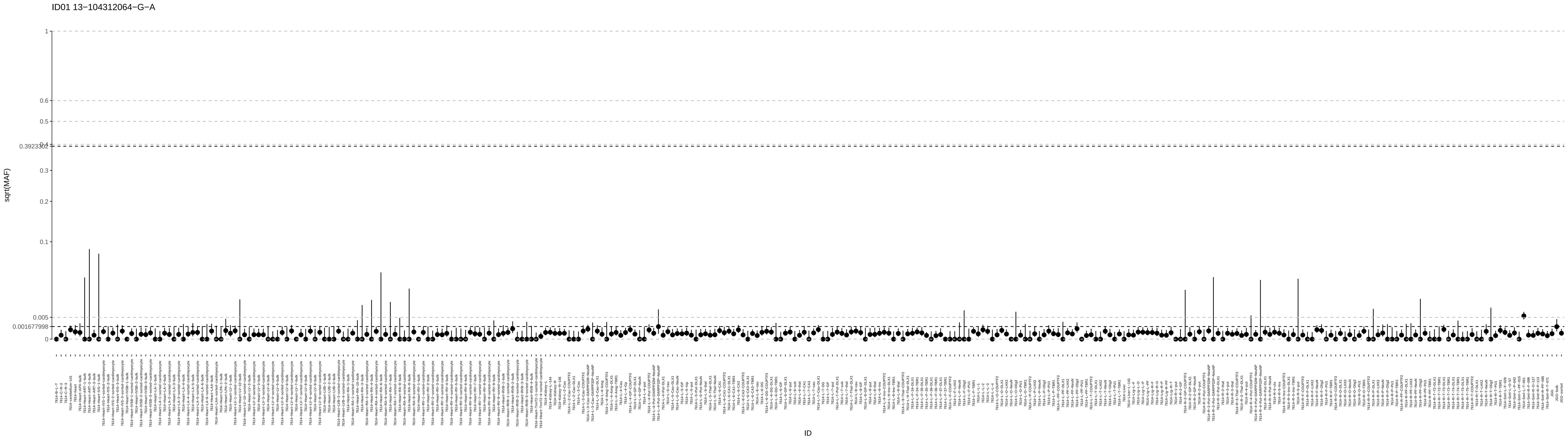
ID01 13−104312064−G−A
sqrt(MAF)
1
0.6
0.5
0.3923302
0.3
0.2
0.1
0.005
0.001677998
0
ID

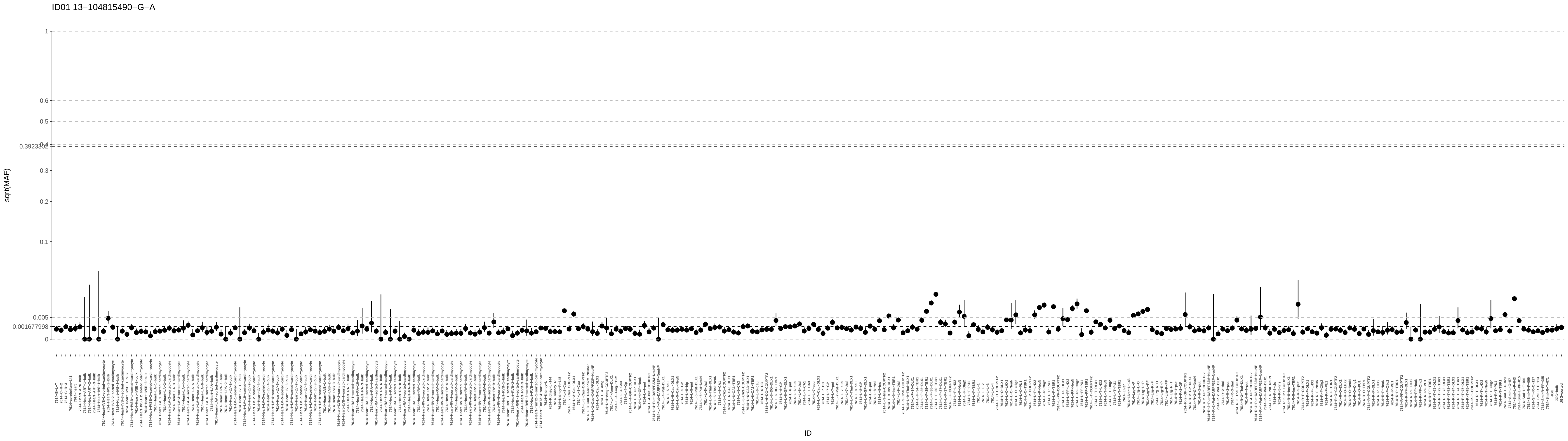
ID01 13−104815490−G−A
sqrt(MAF)
1
0.6
0.5
0.3923302
0.3
0.2
0.1
0.005
0.001677998
0
ID

ID01 13-113887705-CCAG-C

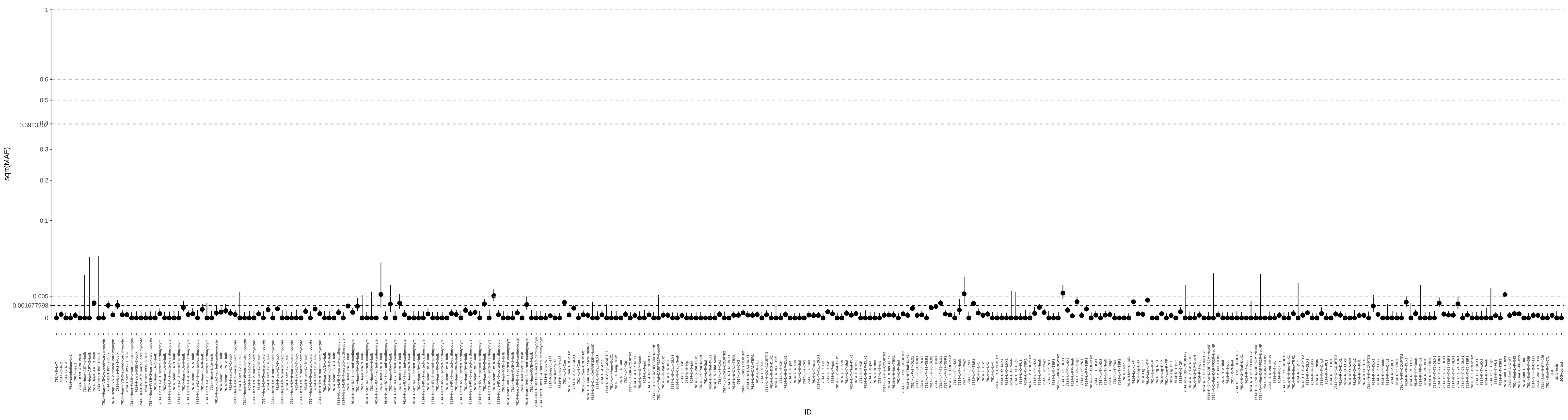

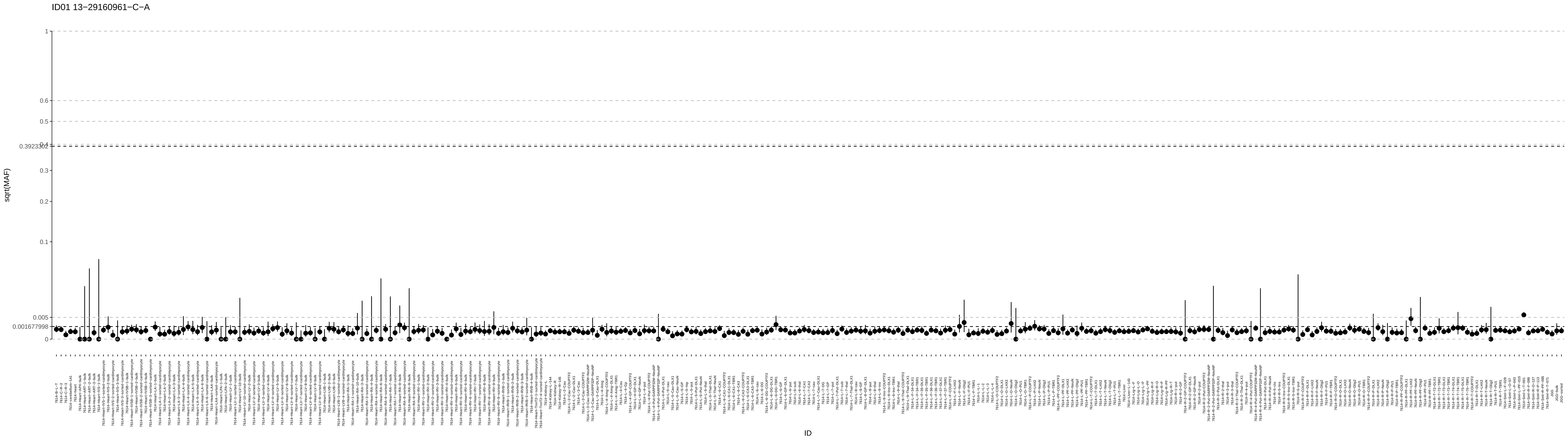
ID01 13−29160961−C−A
sqrt(MAF)
1
0.6
0.5
0.3923302
0.3
0.2
0.1
0.005
0.001677998
0
ID

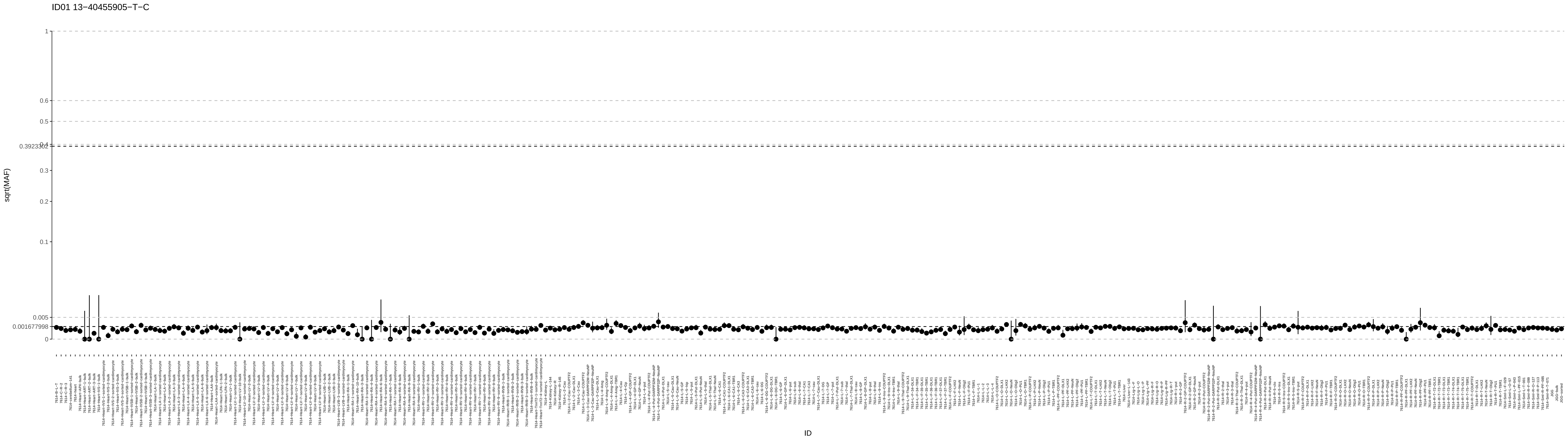
ID01 13−40455905−T−C
sqrt(MAF)
1
0.6
0.5
0.4
0.3923302
0.3
0.2
0.1
0.005
0.001677998
0
ID

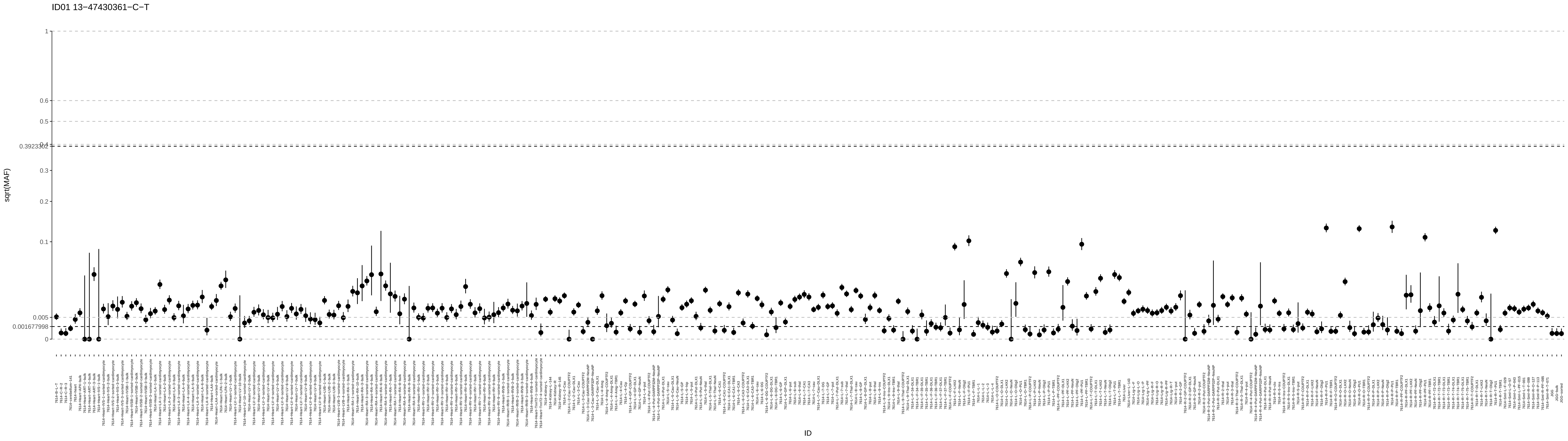
ID01 13−47430361−C−T
sqrt(MAF)
1
0.6
0.5
0.3923302
0.3
0.2
0.1
0.005
0.001677998
0
ID

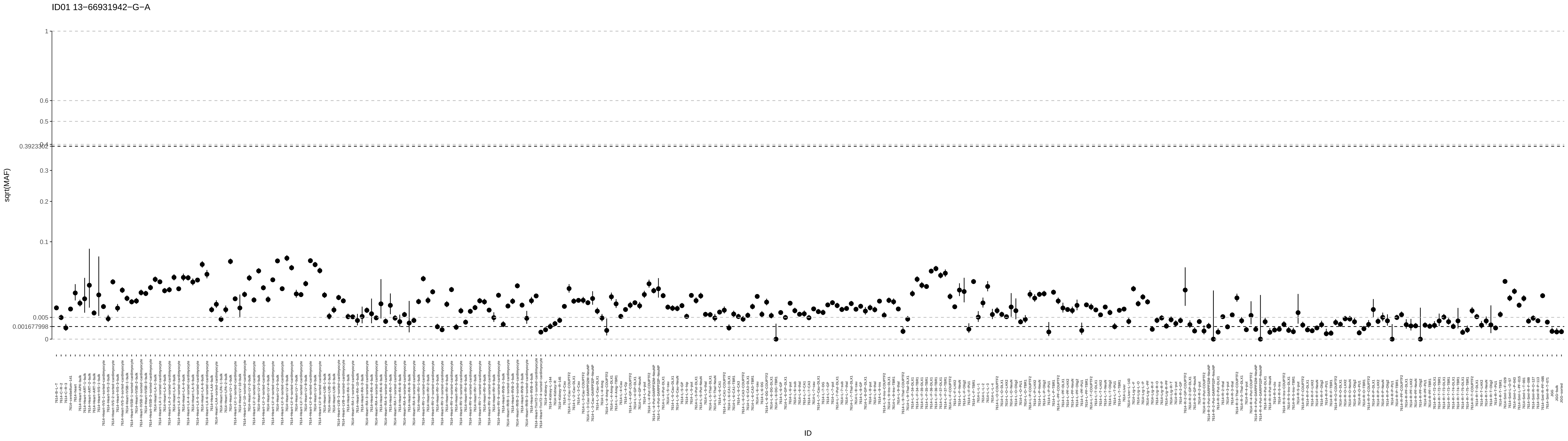
ID01 13−66931942−G−A
sqrt(MAF)
ID
1
0.6
0.5
0.3923302
0.3
0.2
0.1
0.005
0.001677998
0

ID01 13−79159985−T−C

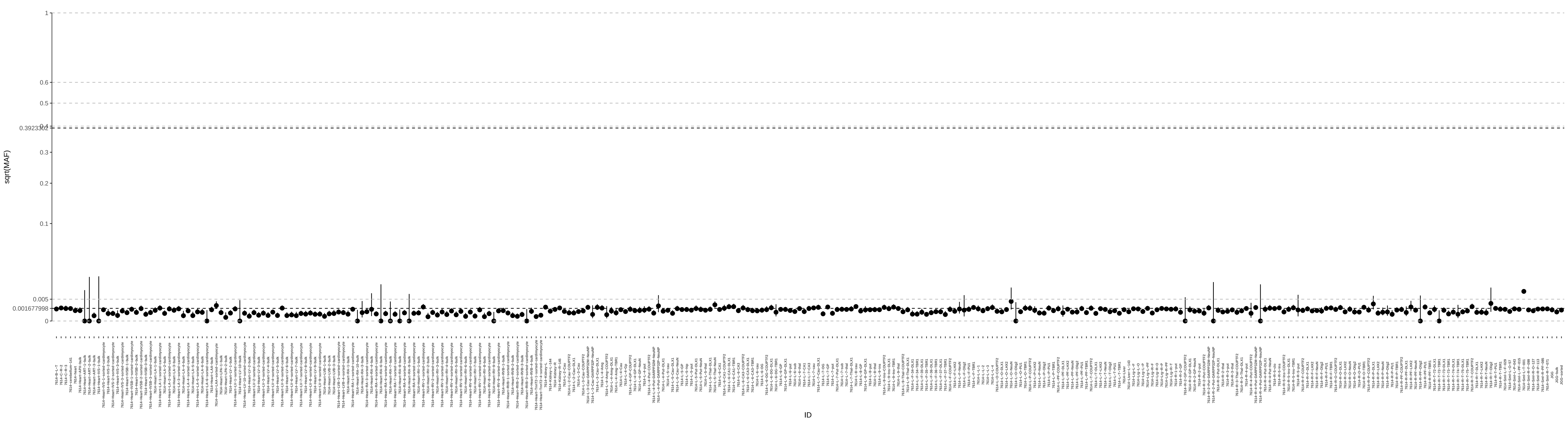

ID01 13−84595018−A−G

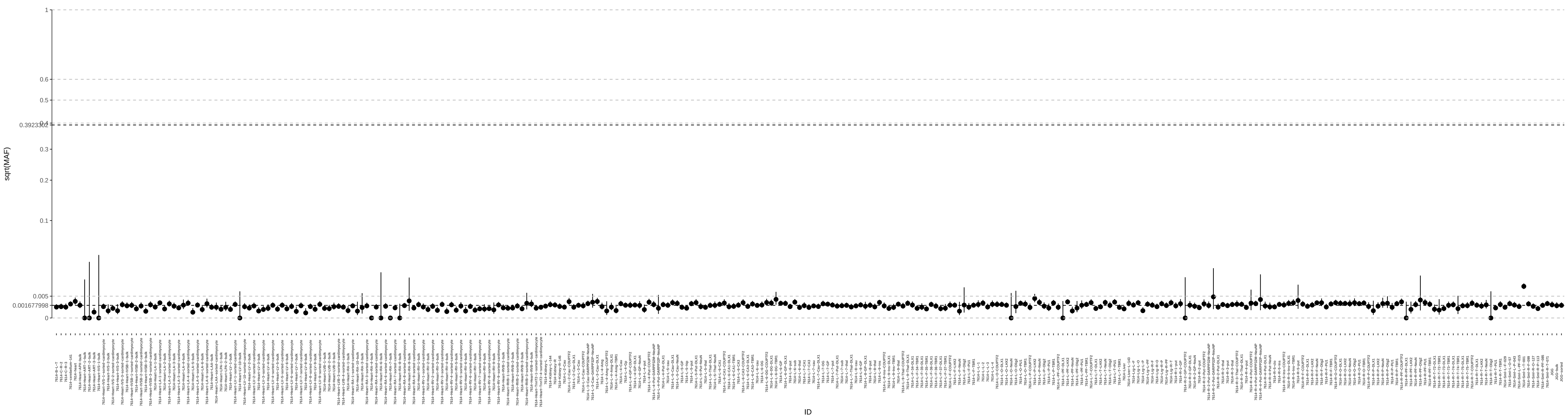

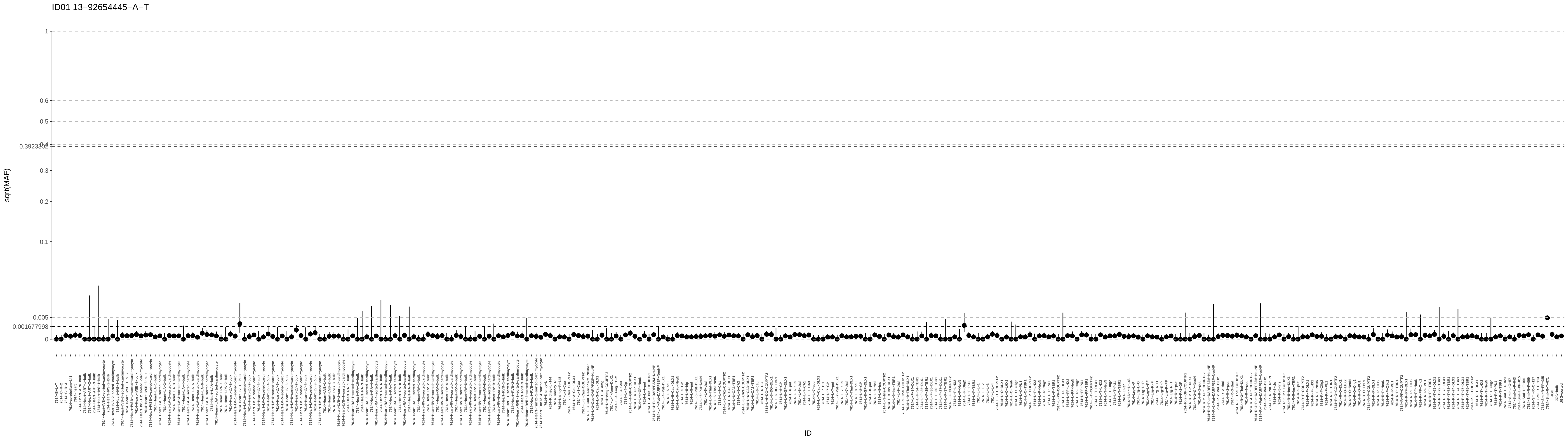
ID01 13−92654445−A−T
sqrt(MAF)
1
0.6
0.5
0.3923302
0.3
0.2
0.1
0.005
0.001677998
0
ID

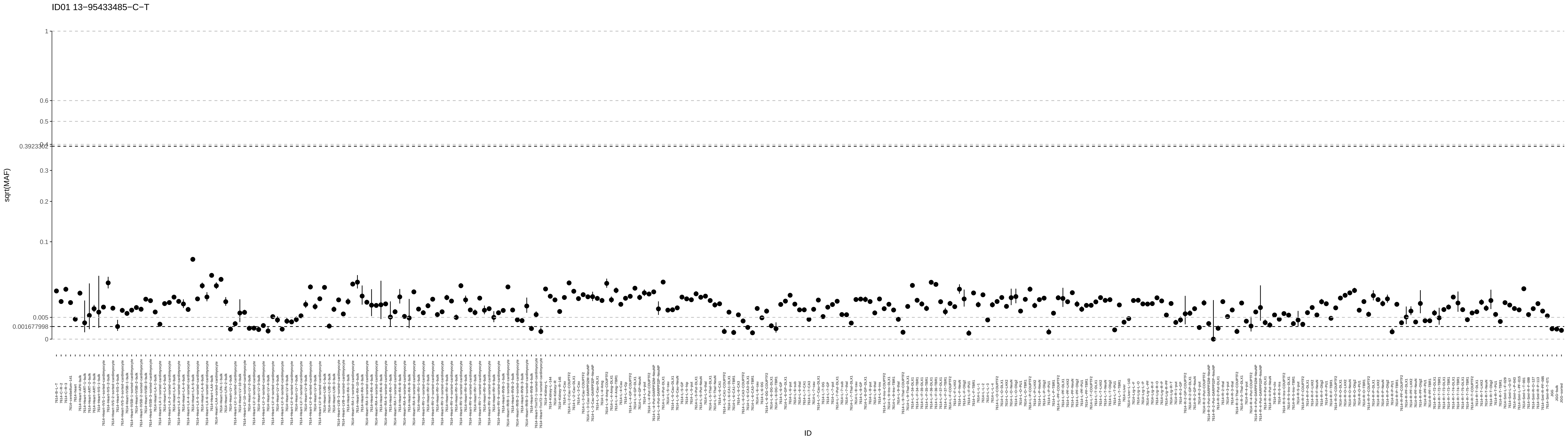
ID01 13−95433485−C−T
sqrt(MAF)
1
0.6
0.5
0.3923302
0.3
0.2
0.1
0.005
0.001677998
0
ID

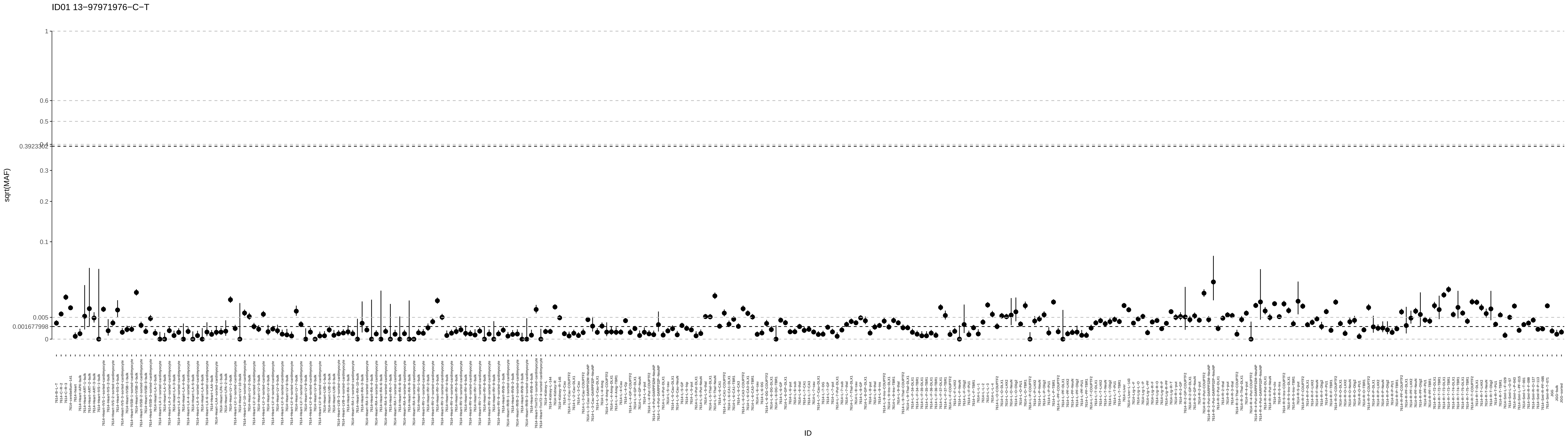
ID01 13−97971976−C−T
sqrt(MAF)
1
0.6
0.5
0.3923302
0.3
0.2
0.1
0.005
0.001677998
0
ID

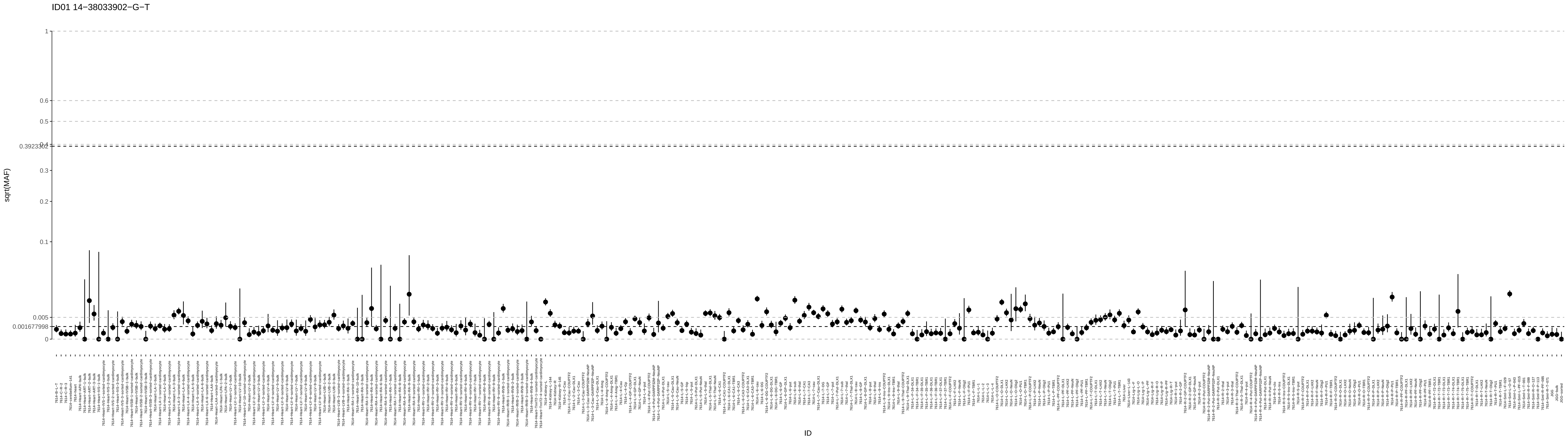
ID01 14−38033902−G−T
sqrt(MAF)
1
0.6
0.5
0.3923302
0.3
0.2
0.1
0.005
0.001677998
0
ID

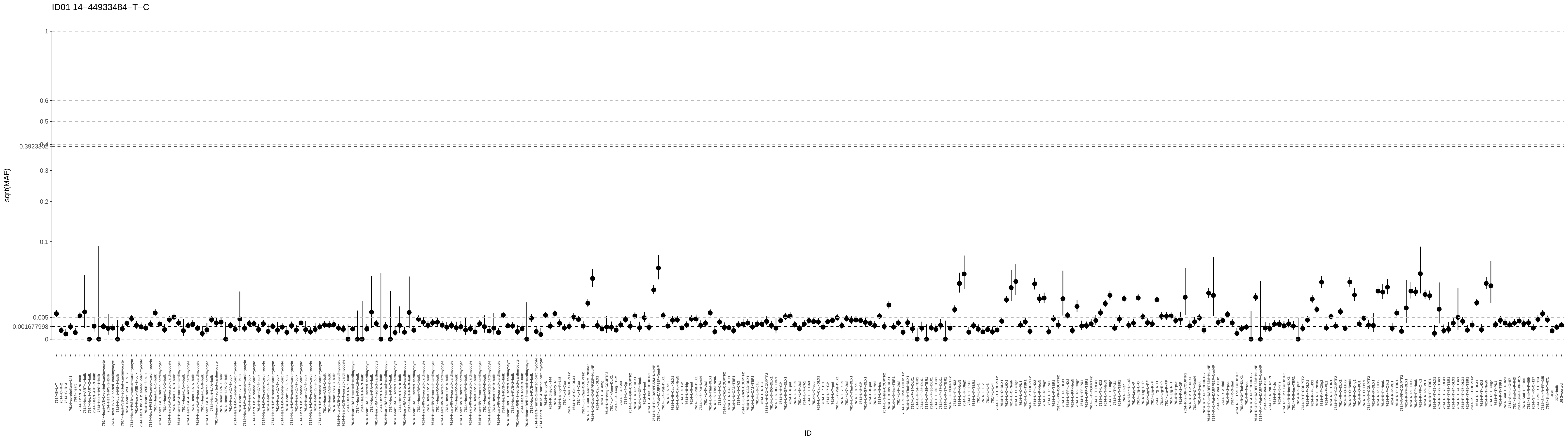
ID01 14−44933484−T−C
sqrt(MAF)
1
0.6
0.5
0.4
0.3923302
0.3
0.2
0.1
0.005
0.001677998
0
ID

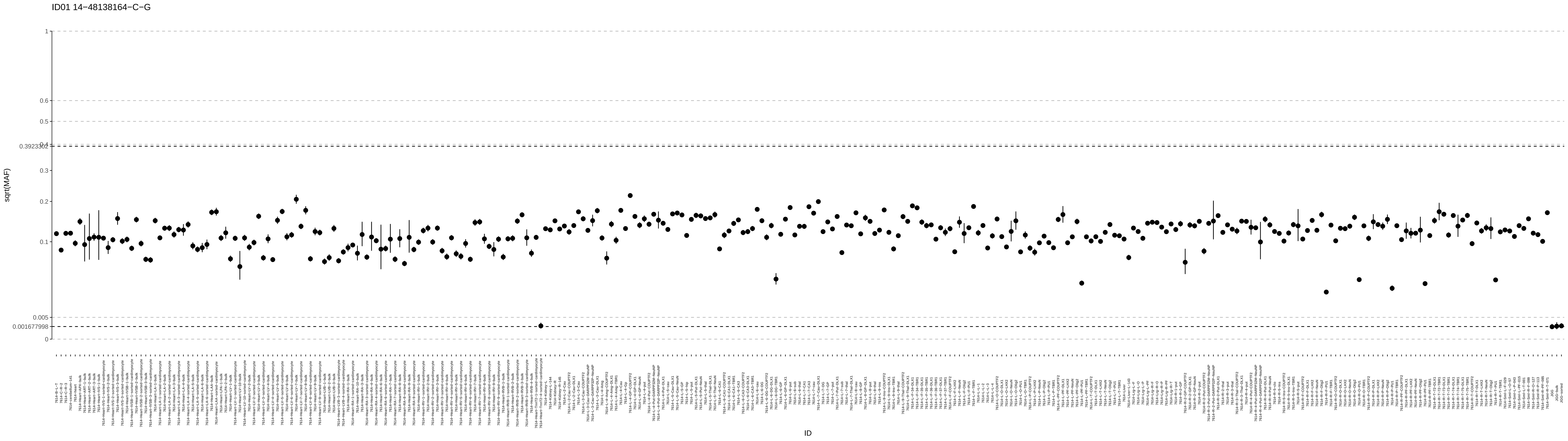
ID01 14−48138164−C−G
sqrt(MAF)
1
0.6
0.5
0.4
0.3923302
0.3
0.2
0.1
0.005
0.001677998
0
ID

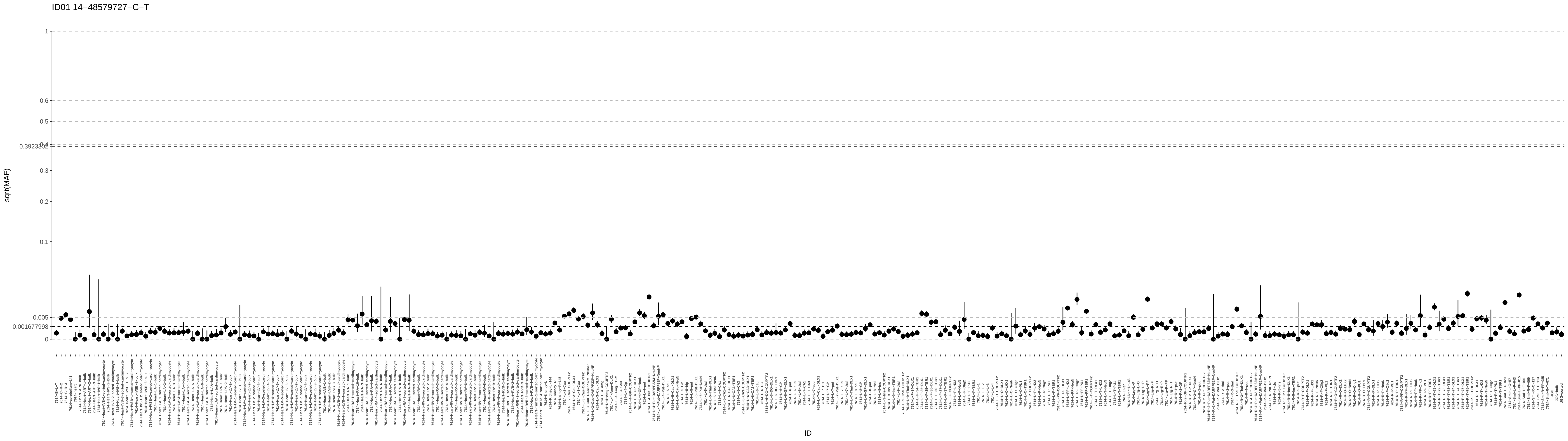
ID01 14−48579727−C−T
sqrt(MAF)
1
0.6
0.5
0.3923302
0.3
0.2
0.1
0.005
0.001677998
0
ID

ID01 14−54432218−C−T

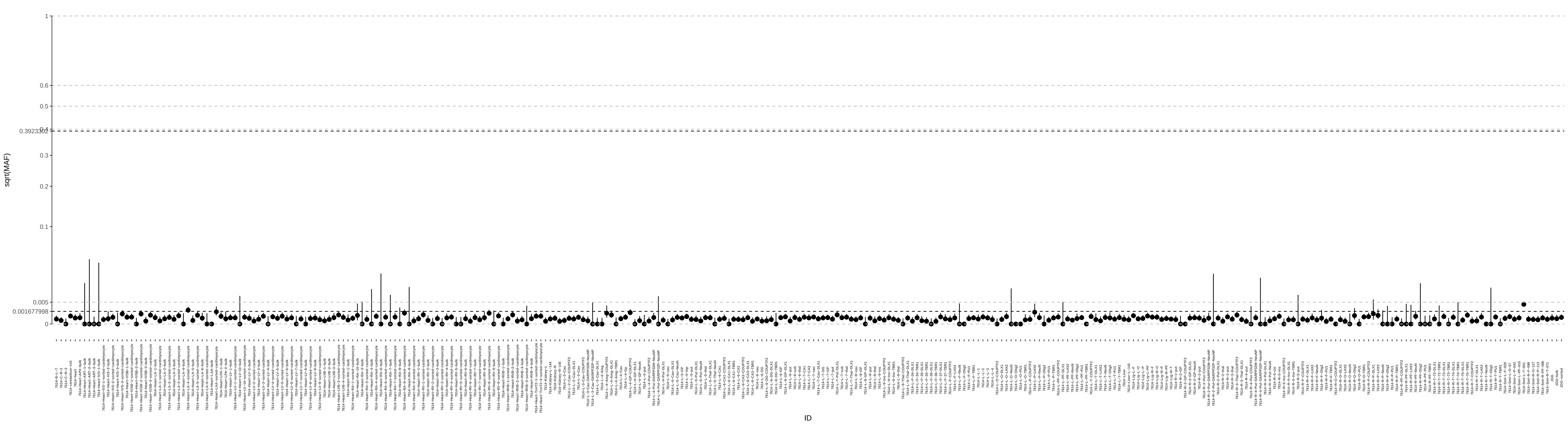

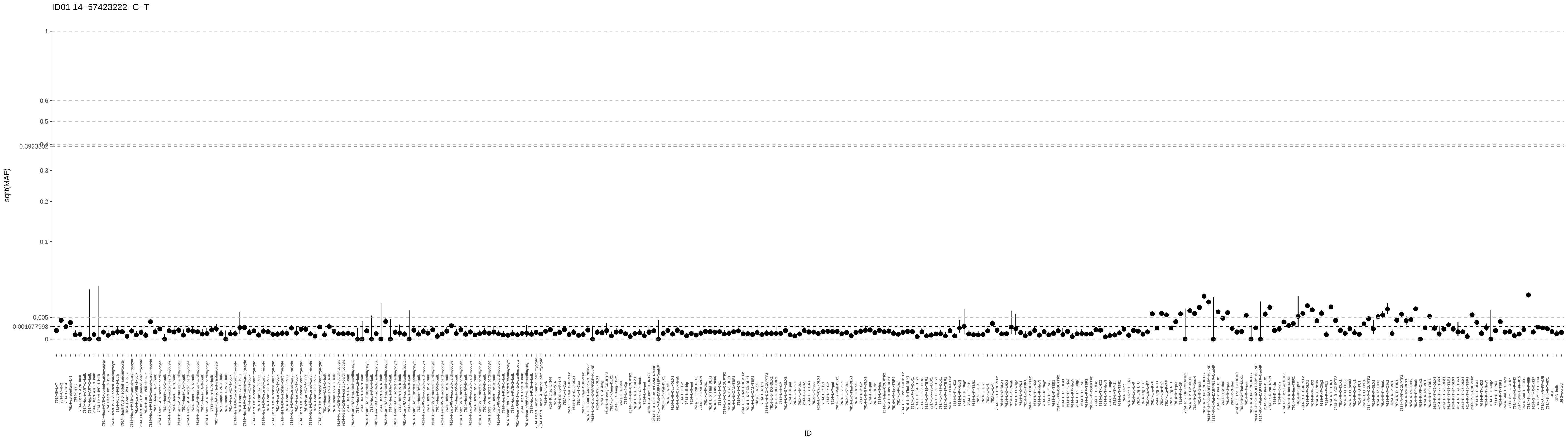

ID01 14−57423222−C−T
sqrt(MAF)
1
0.6
0.5
0.3923302
0.3
0.2
0.1
0.005
0.001677998
0
ID

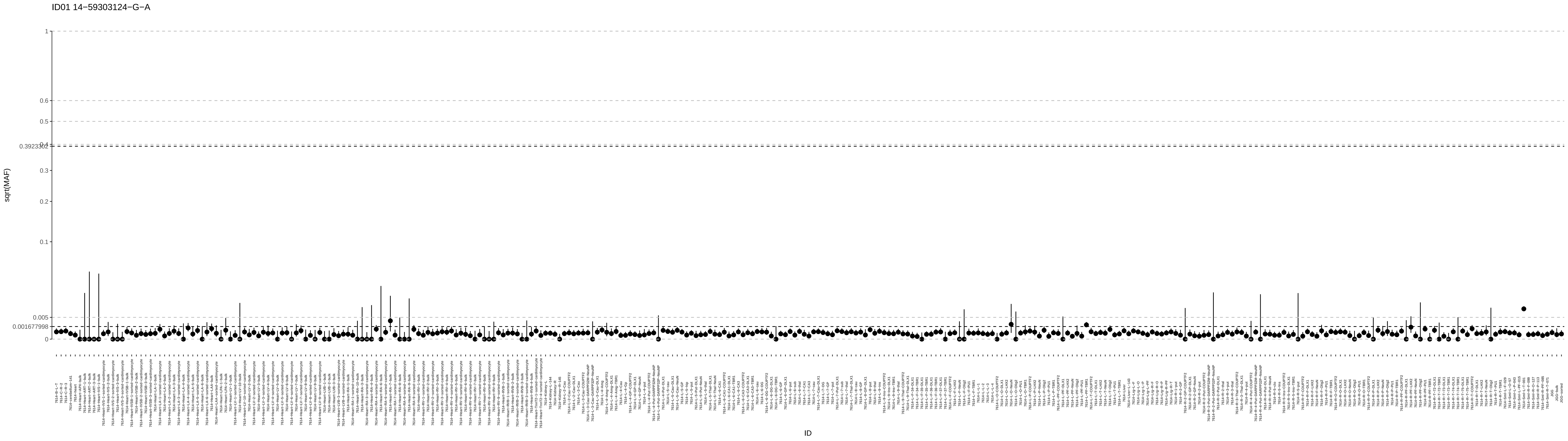
ID01 14−59303124−G−A
sqrt(MAF)
1
0.6
0.5
0.3923302
0.3
0.2
0.1
0.005
0.001677998
0
ID

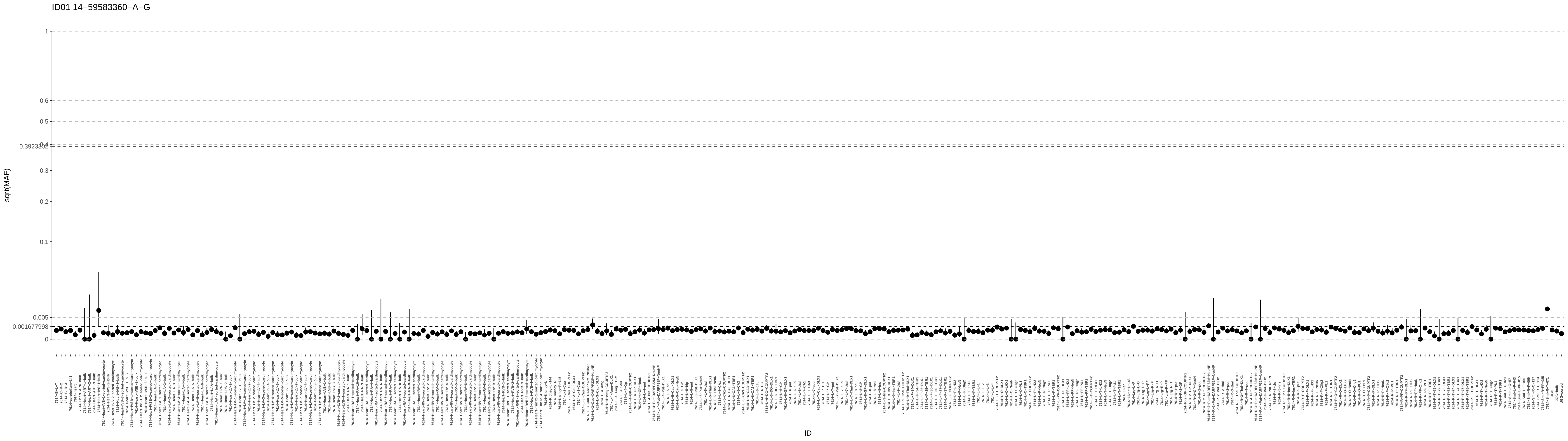

ID01 14−59583360−A−G
sqrt(MAF)
1
0.6
0.5
0.3923302
0.3
0.2
0.1
0.005
0.001677998
0
ID

ID01 14−65962309−A−C

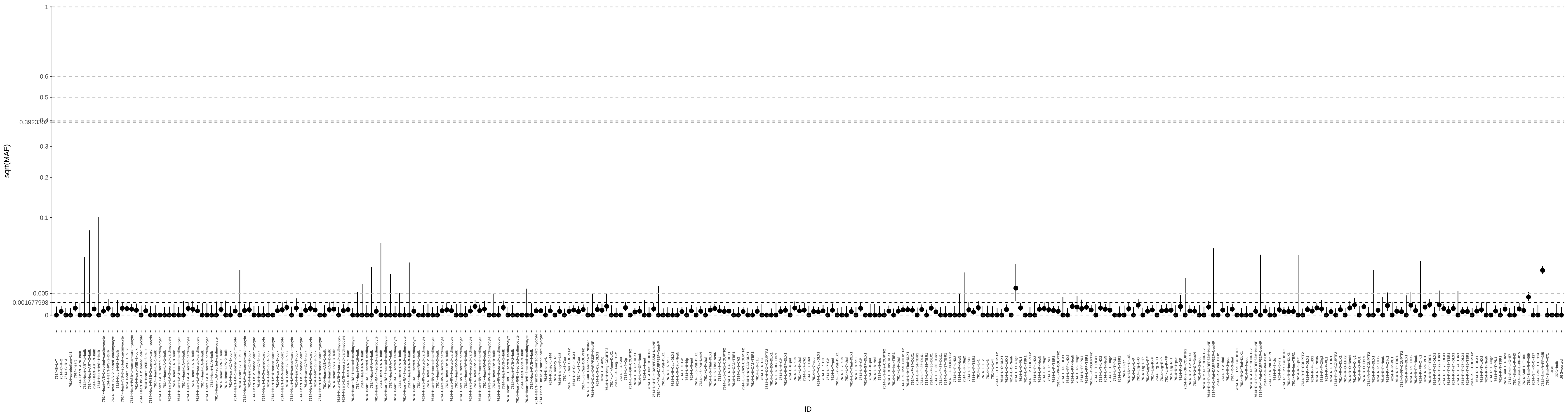

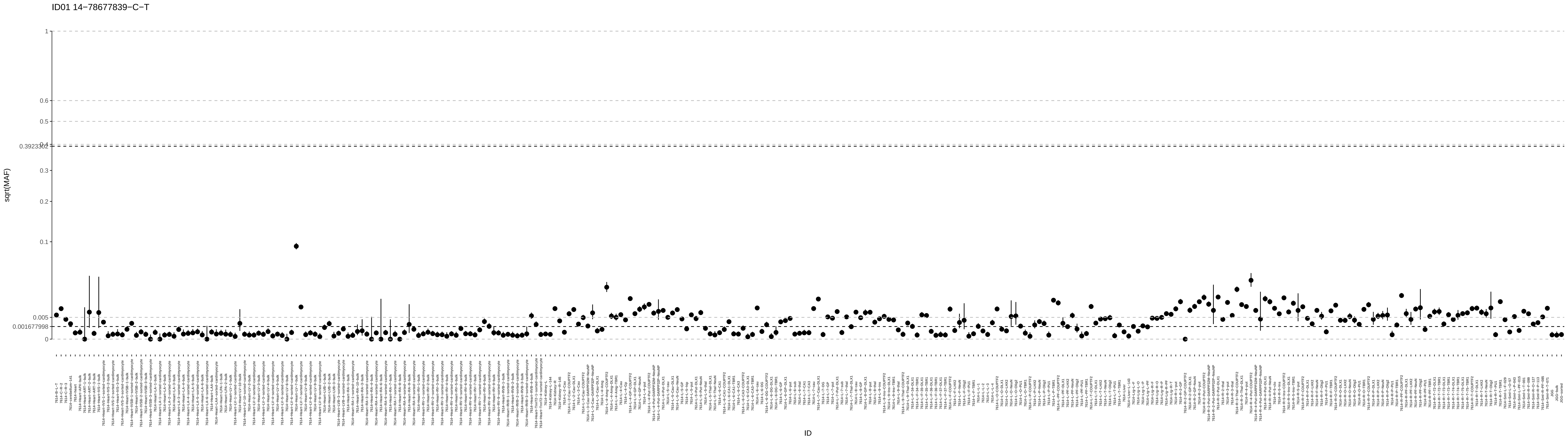
ID01 14−78677839−C−T
sqrt(MAF)
1
0.6
0.5
0.4
0.3923302
0.3
0.2
0.1
0.005
0.001677998
0
ID

ID01 14−82666170−G−A

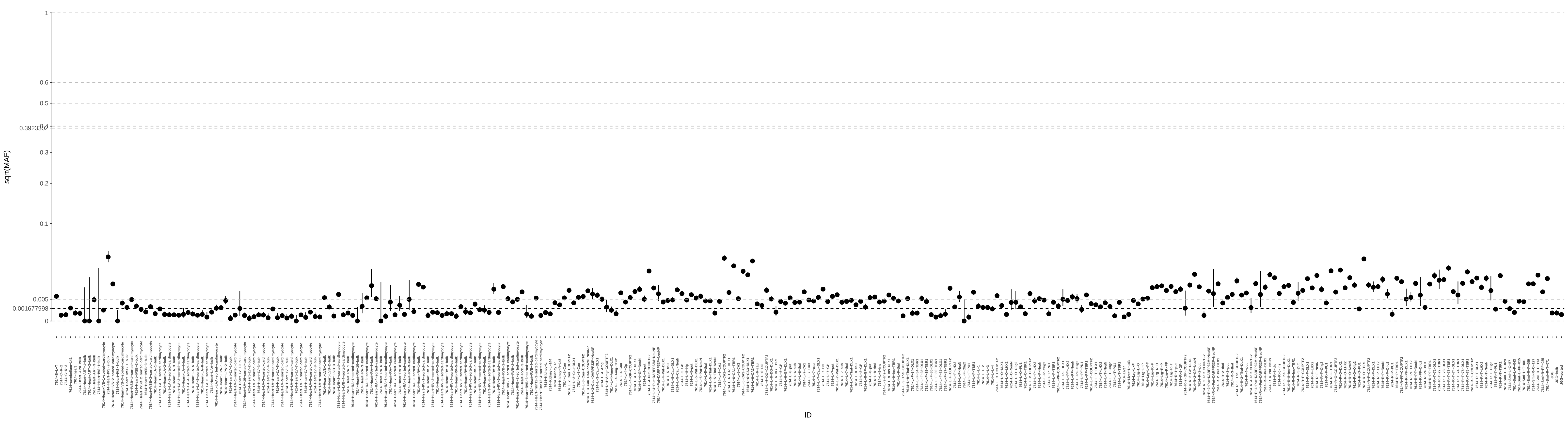

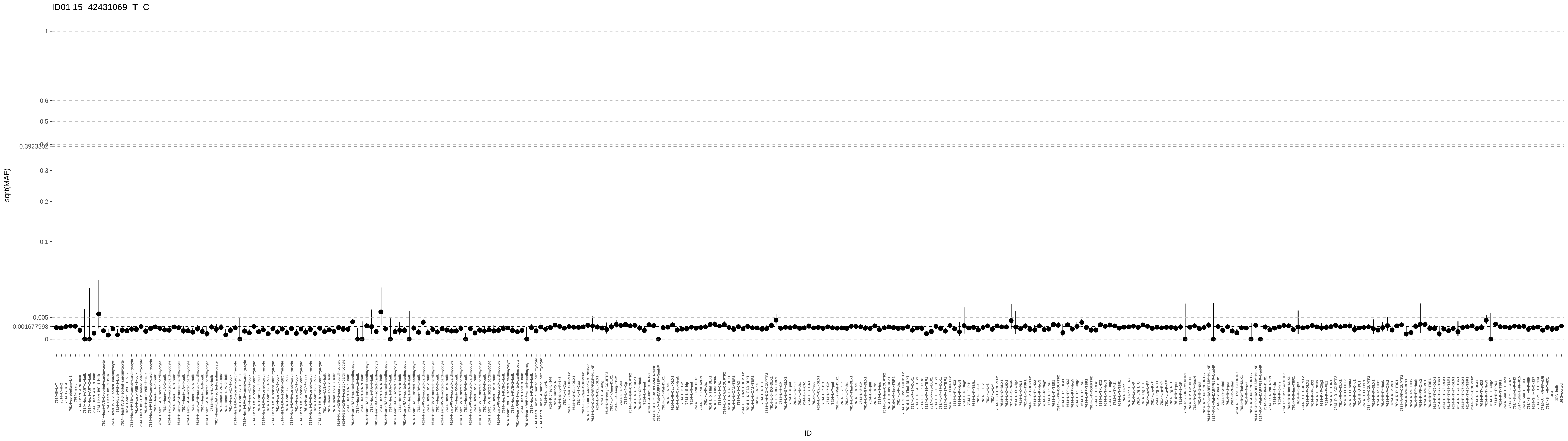
ID01 15−42431069−T−C
sqrt(MAF)
0.3923302
0.001677998
ID

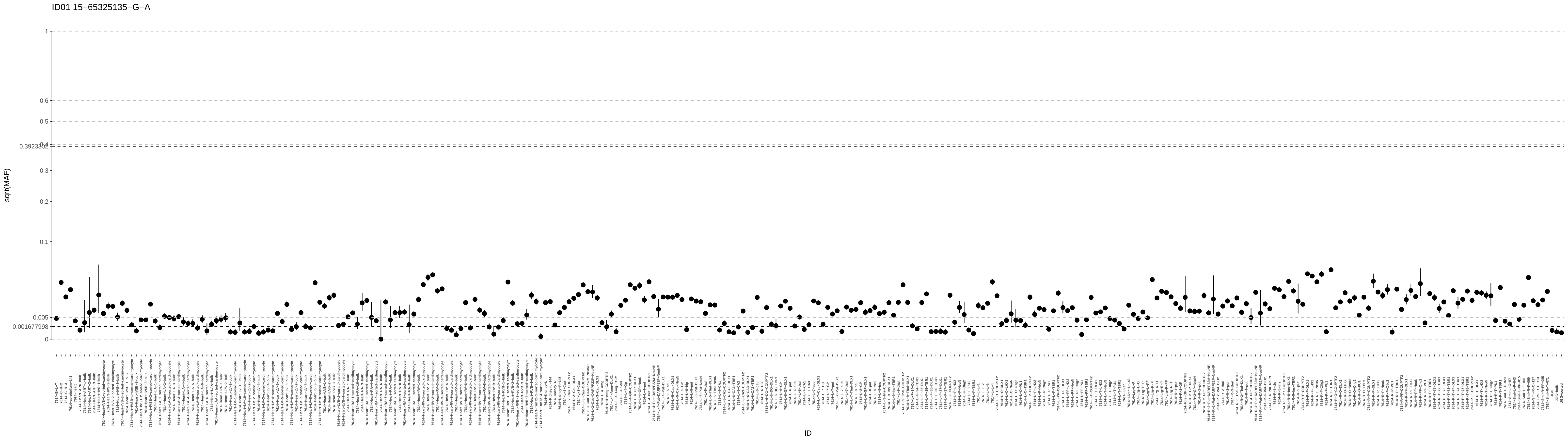
ID01 15−65325135−G−A
sqrt(MAF)
1
0.6
0.5
0.3923302
0.3
0.2
0.1
0.005
0.001677998
0
ID

ID01 15−89933068−T−G

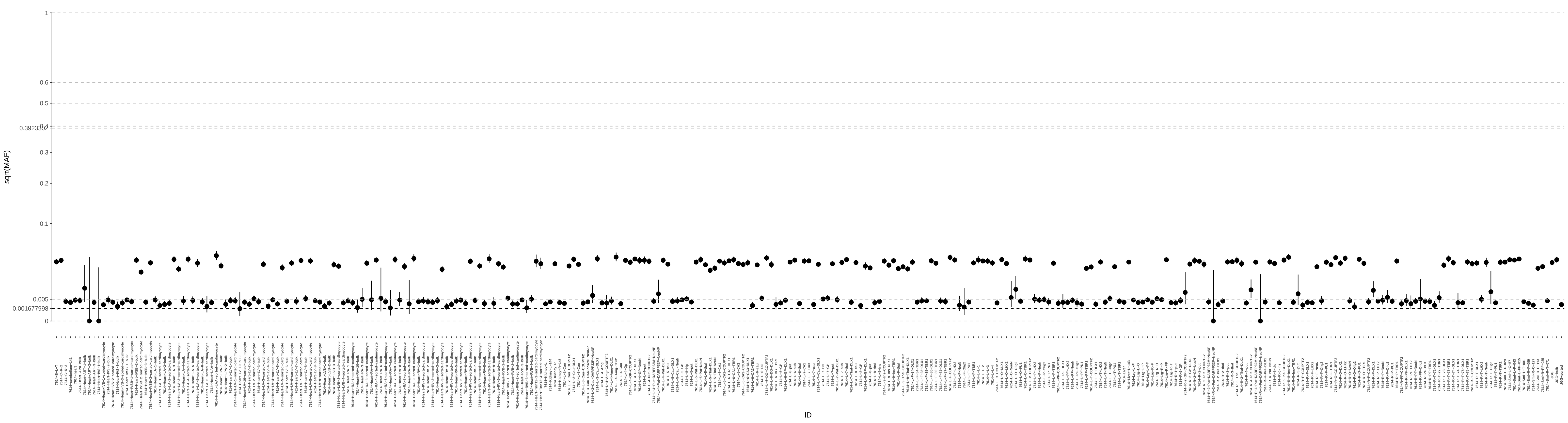

ID01 15−93474236−T−G

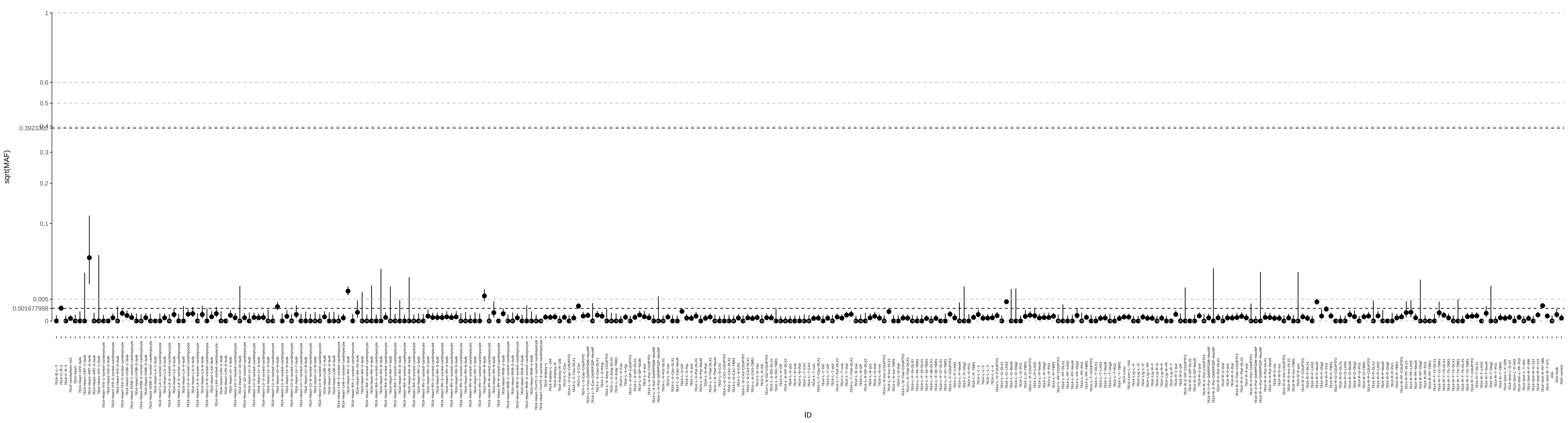

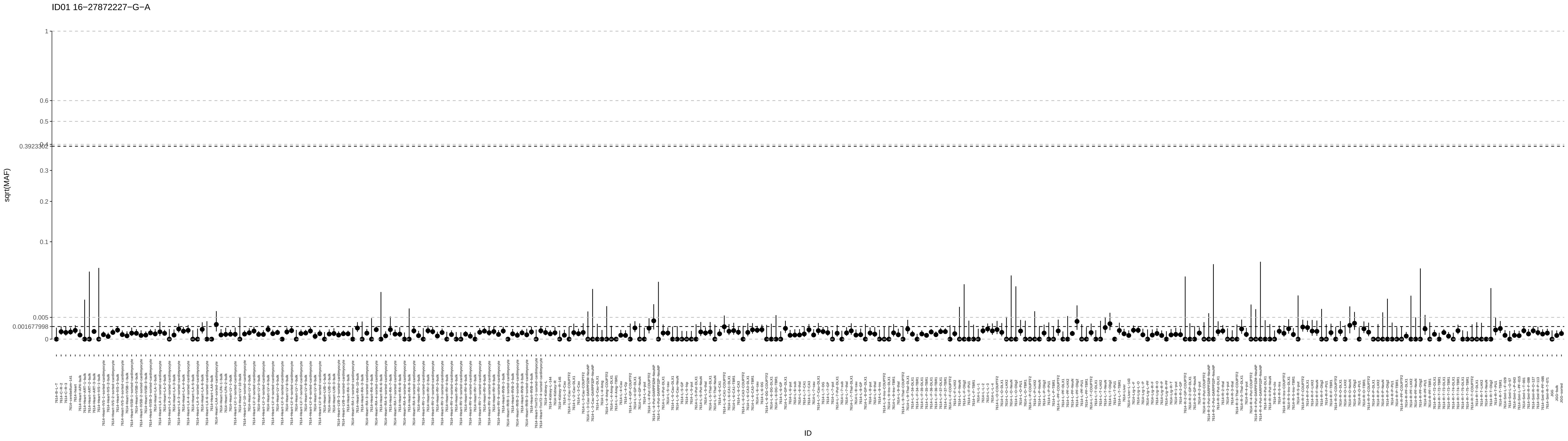
ID01 16−27872227−G−A
sqrt(MAF)
1
0.6
0.5
0.3923302
0.3
0.2
0.1
0.005
0.001677998
0
ID

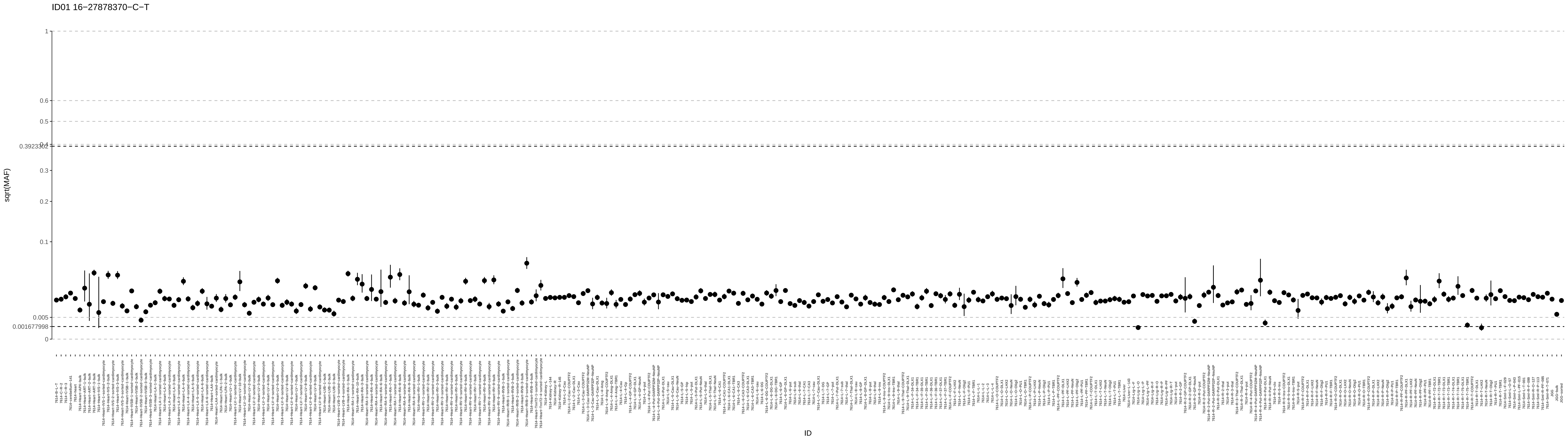
ID01 16−27878370−C−T
sqrt(MAF)
1
0.6
0.5
0.3923302
0.3
0.2
0.1
0.005
0.001677998
0
ID

ID01 16−27878380−T−C

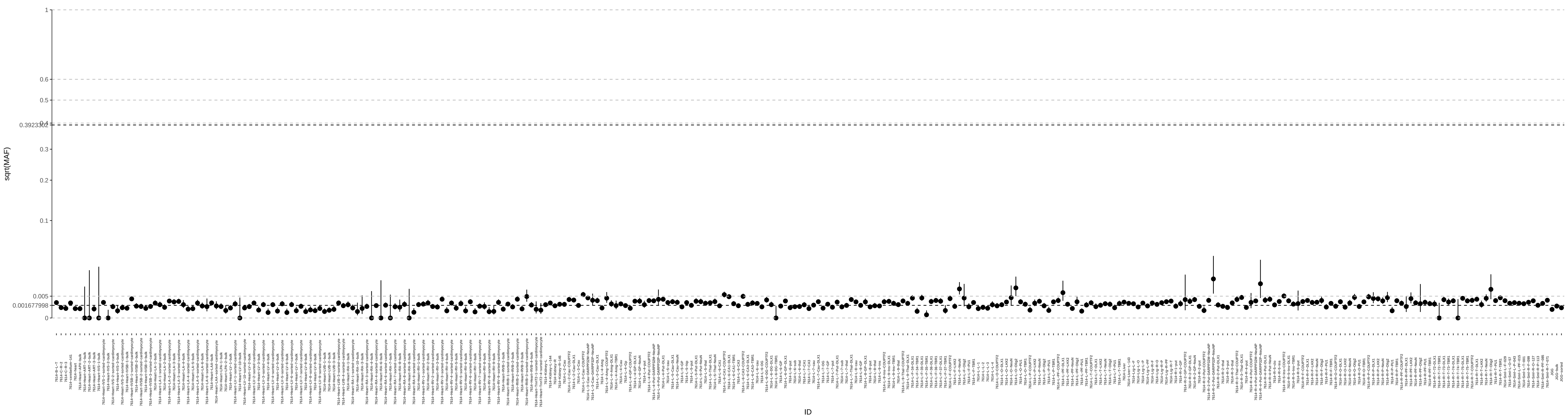

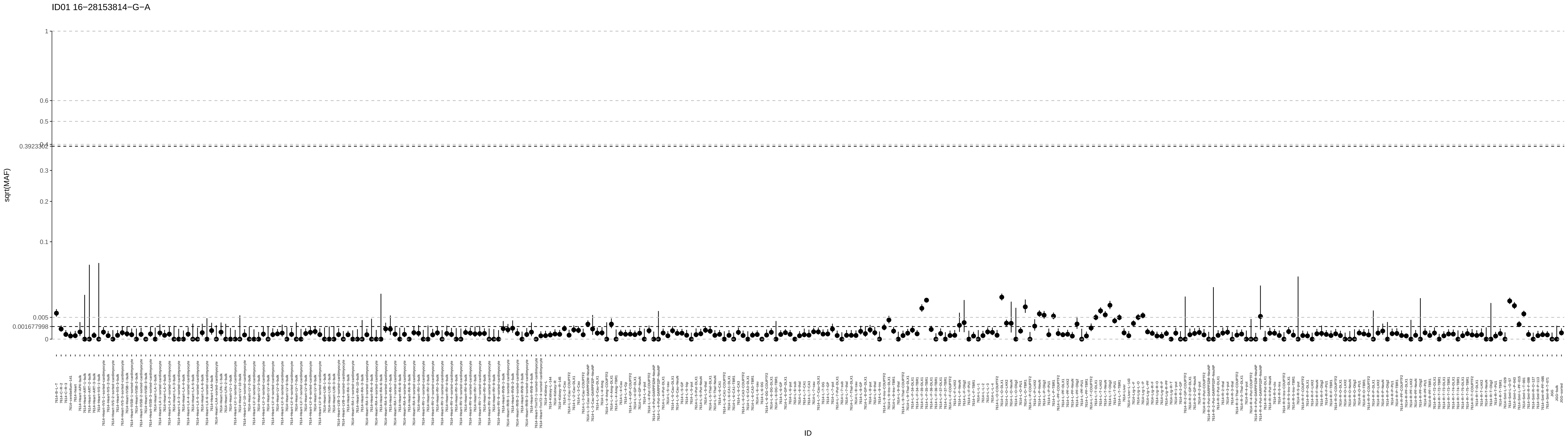

ID01 16−28153814−G−A
1
0.6
0.5
0.3923302
0.3
0.2
0.1
0.005
0.001677998
0
sqrt(MAF)
ID

ID01 16−58426889−C−A

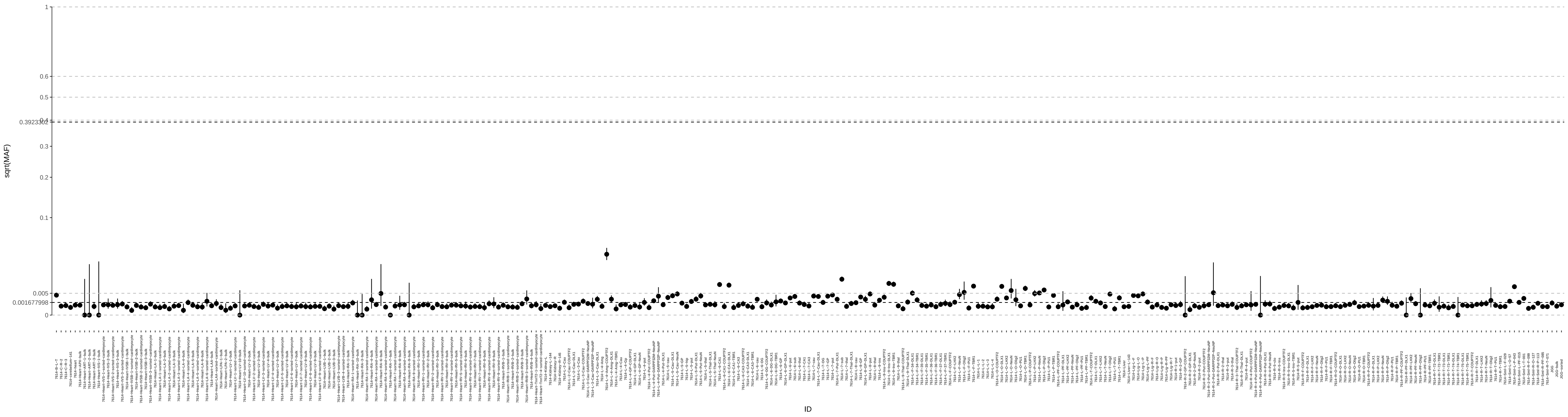

ID01 16−69679204−G−C

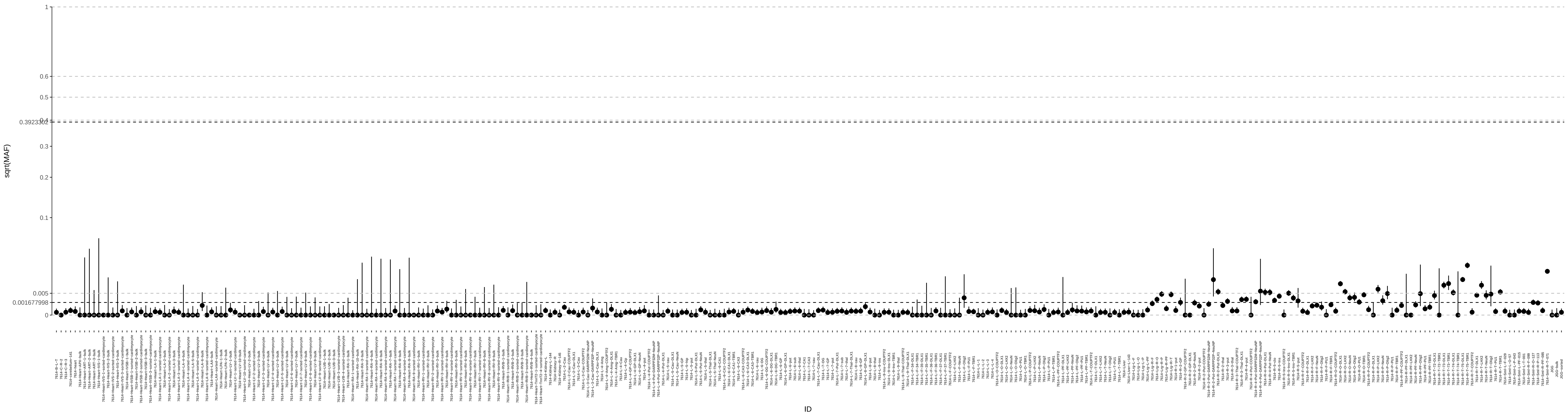

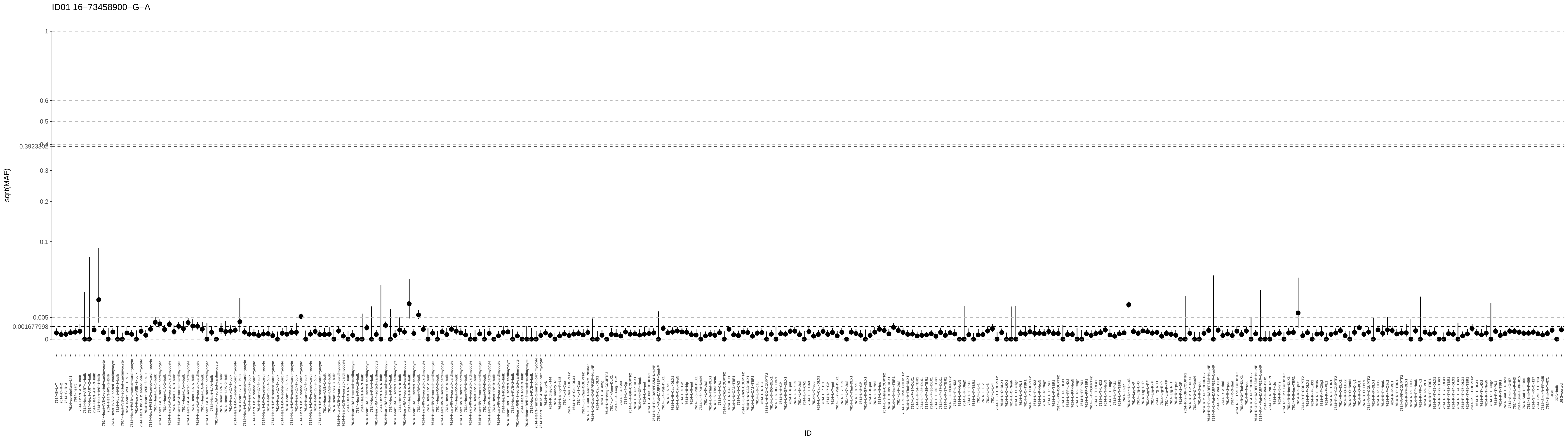
ID01 16−73458900−G−A
sqrt(MAF)
1
0.6
0.5
0.3923302
0.3
0.2
0.1
0.005
0.001677998
0
ID

ID01 16−73751747−A−G

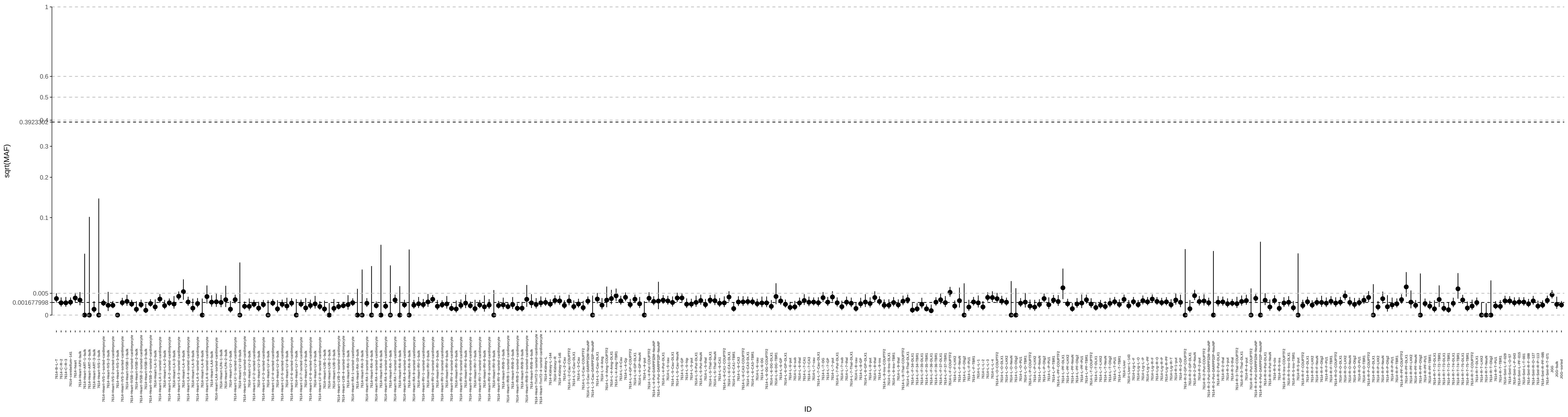

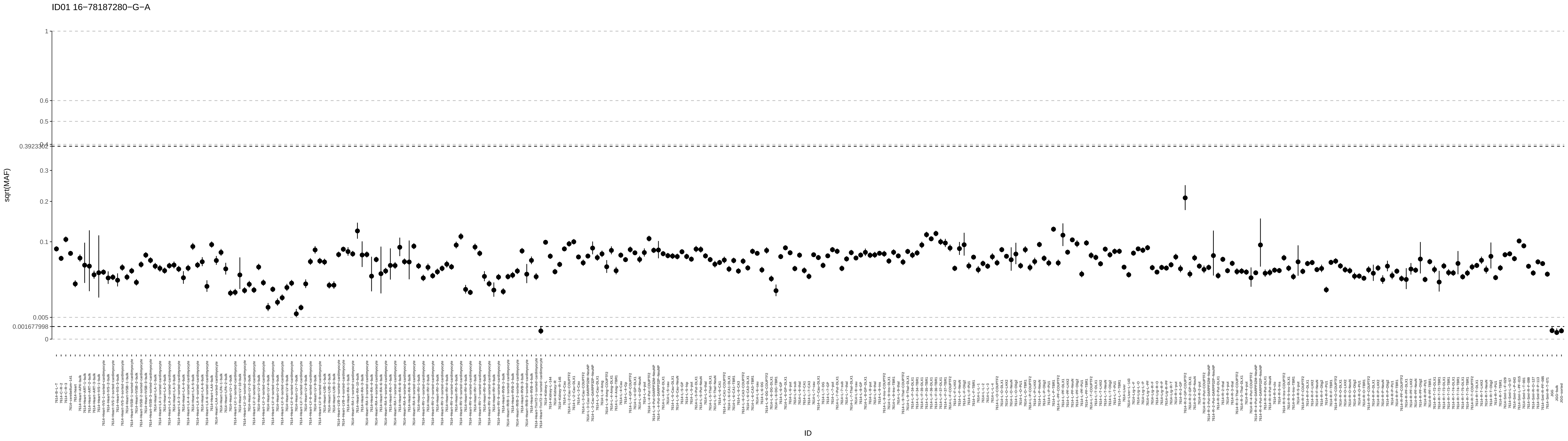
ID01 16−78187280−G−A
sqrt(MAF)
1
0.6
0.5
0.4
0.3923302
0.3
0.2
0.1
0.005
0.001677998
0
ID

ID01 16−78774167−C−T

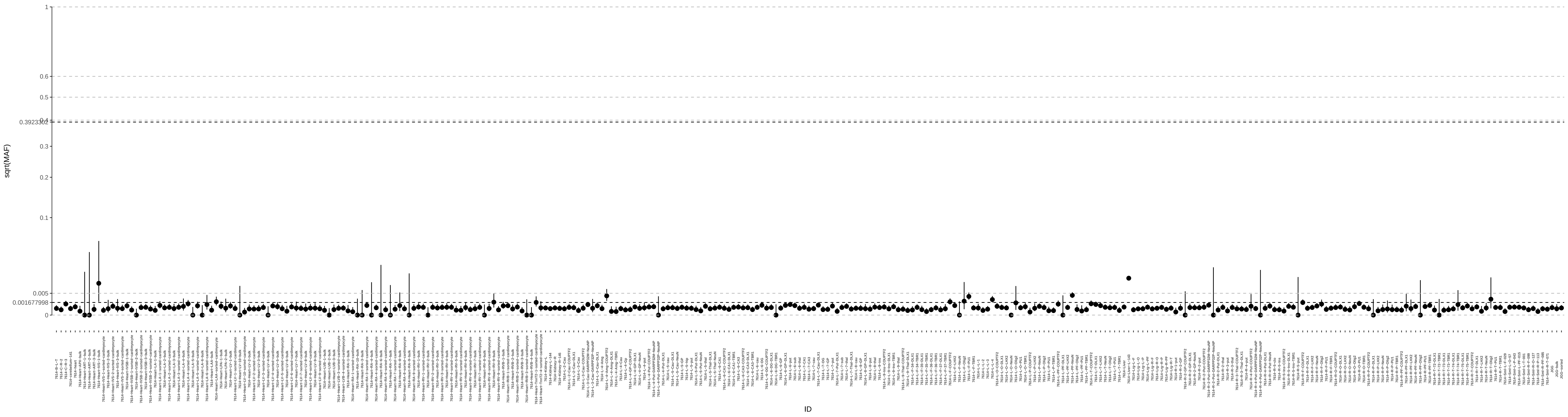

ID01 16−80604095−C−T

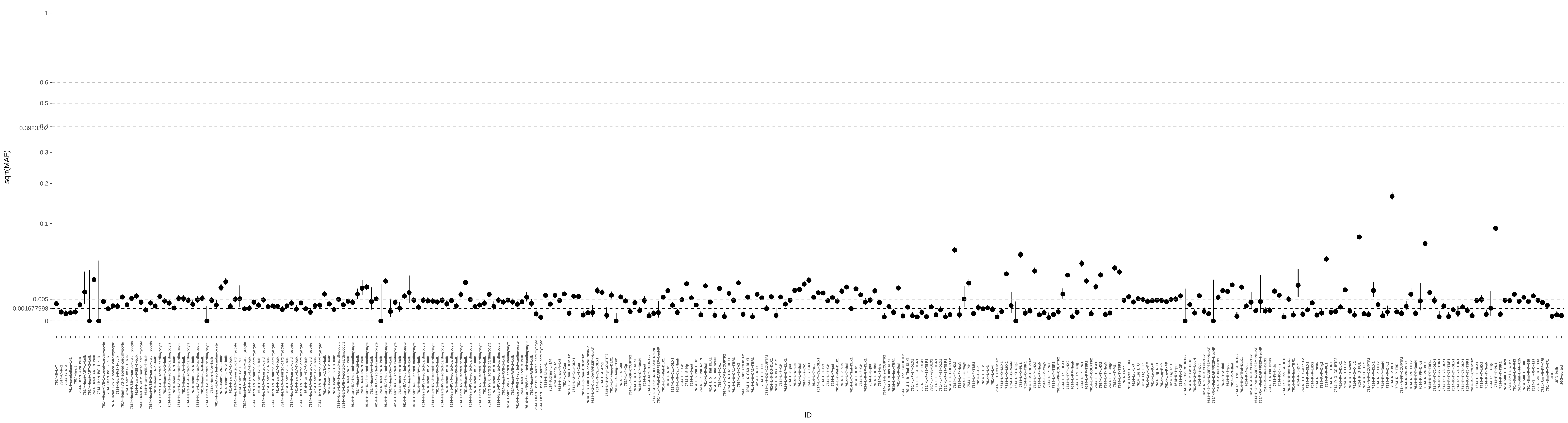

ID01 16−83174915−T−C

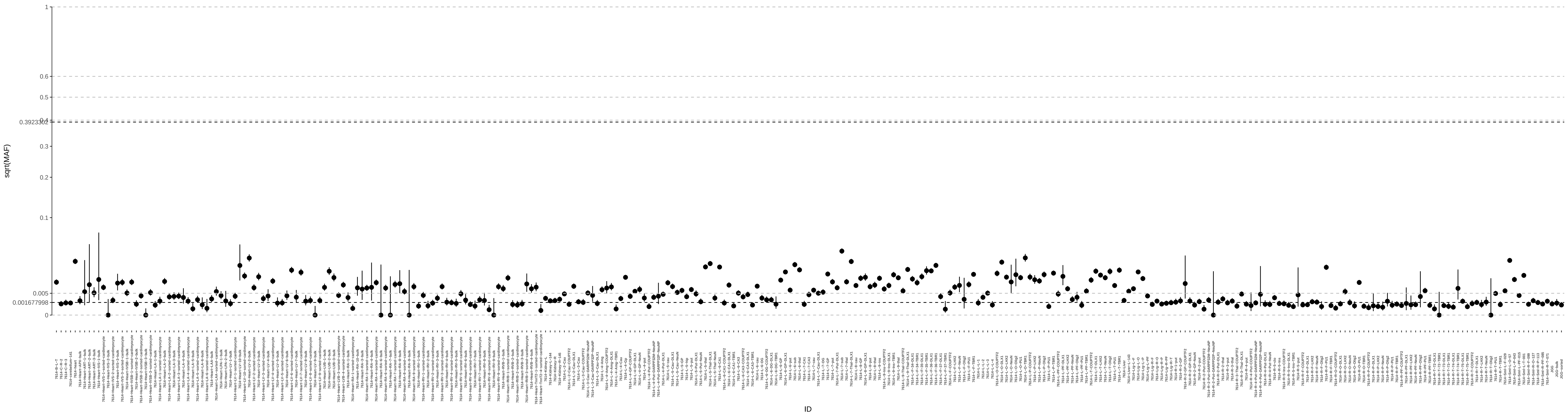

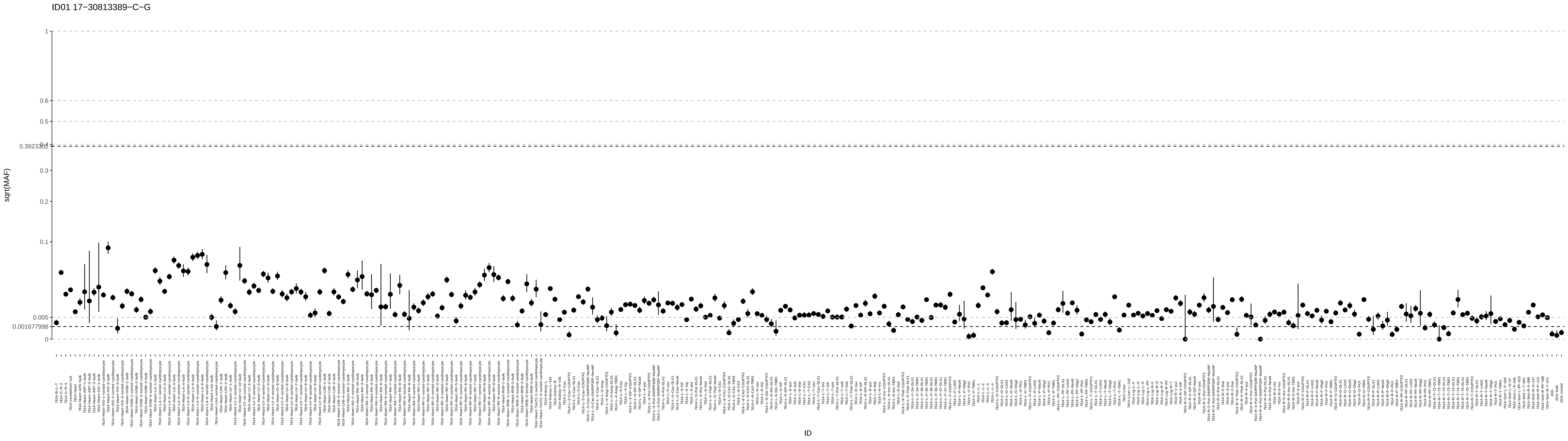
ID01 17−30813389−C−G
sqrt(MAF)
1
0.6
0.5
0.3923302
0.3
0.2
0.1
0.005
0.001677998
0
ID

ID01 17−31854088−G−A

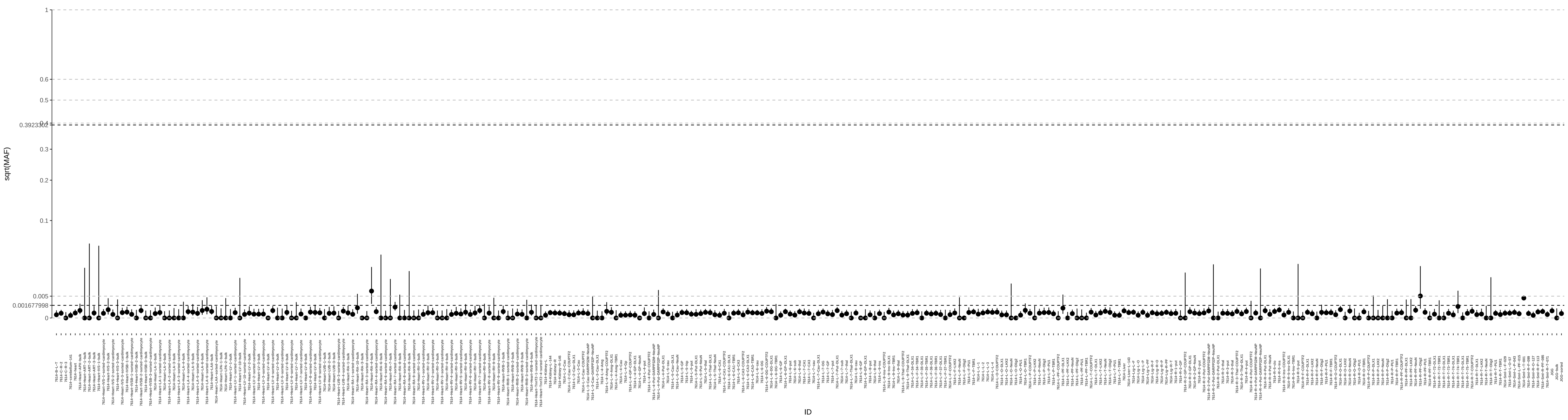

ID01 17−32373696−G−A

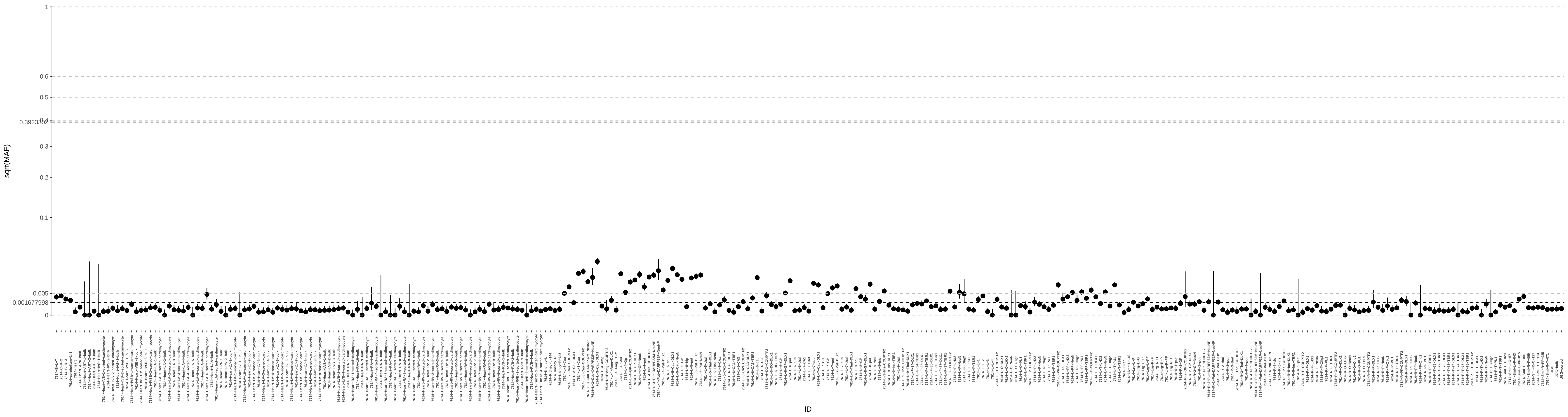

ID01 17−5657235−A−C

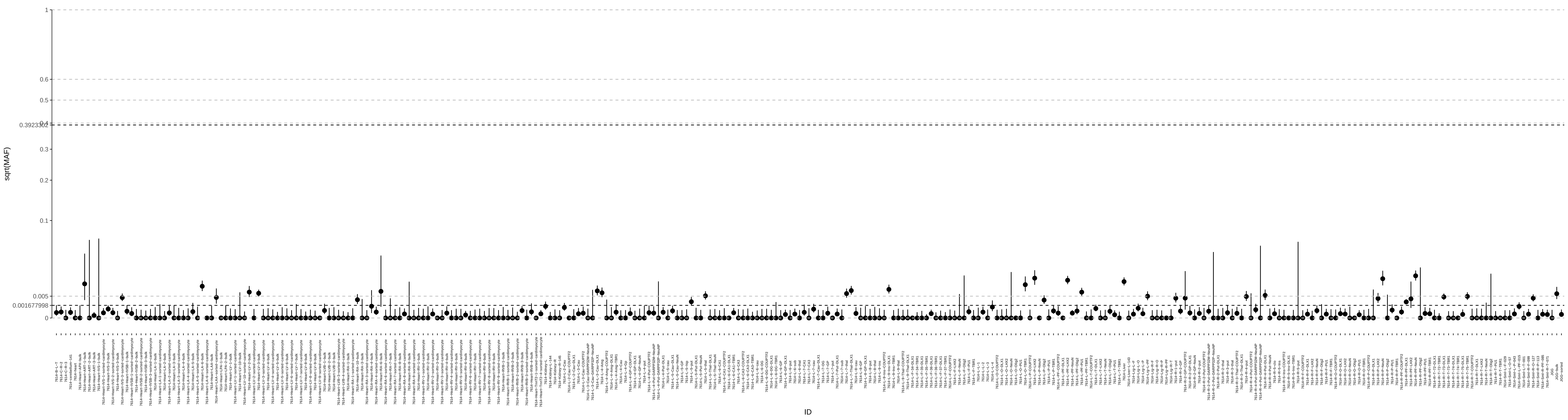

ID01 17−56786985−T−C

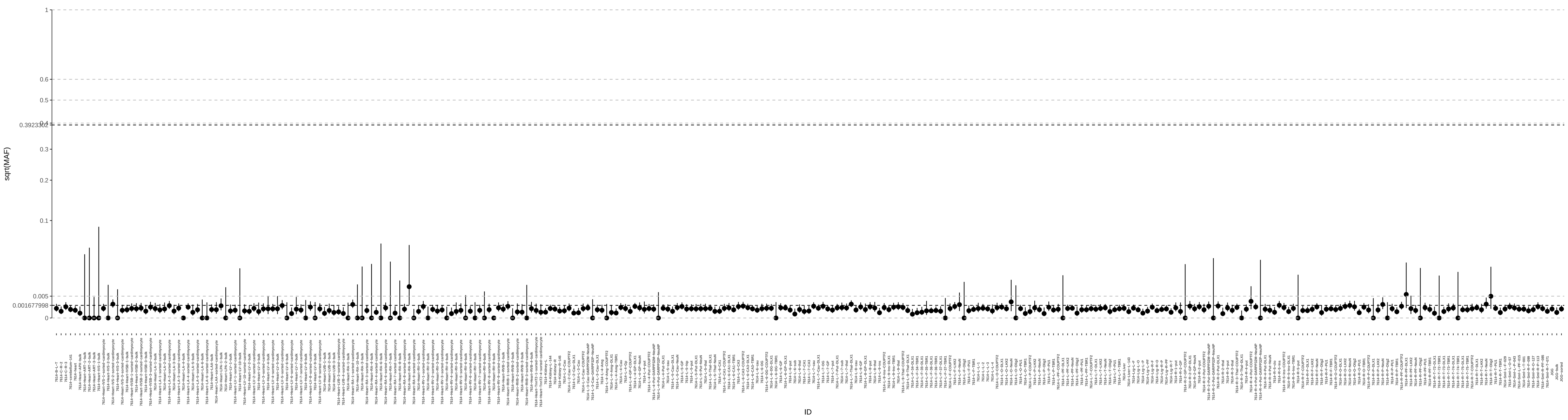

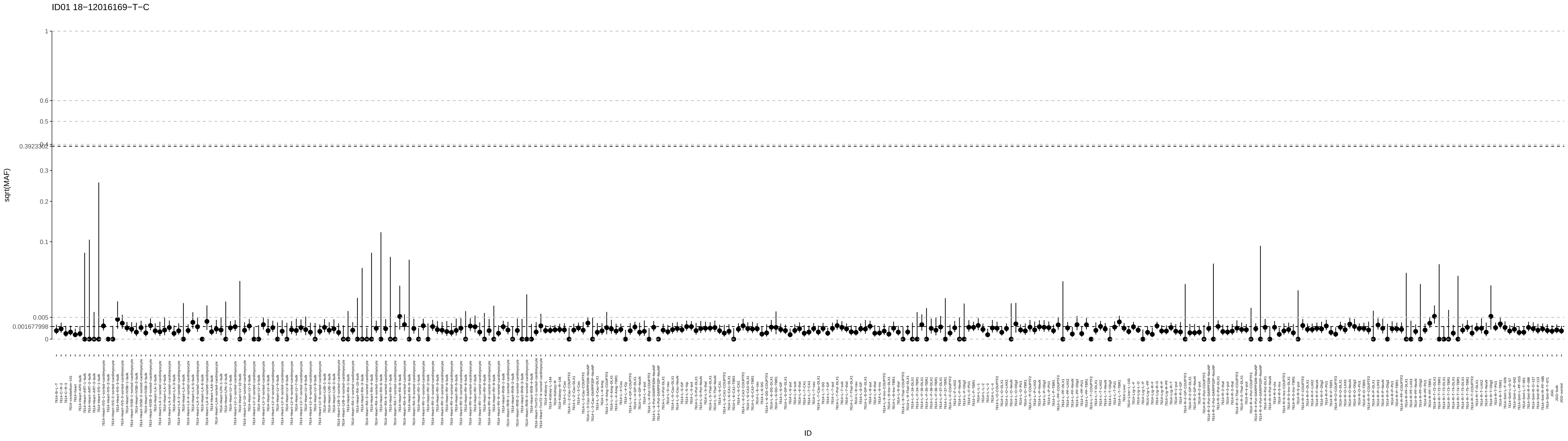
ID01 18-12016169-T-C
sqrt(MAF)
1
0.6
0.5
0.3923302
0.3
0.2
0.1
0.005
0.001677998
0
ID

ID01 18−27189441−TA−T

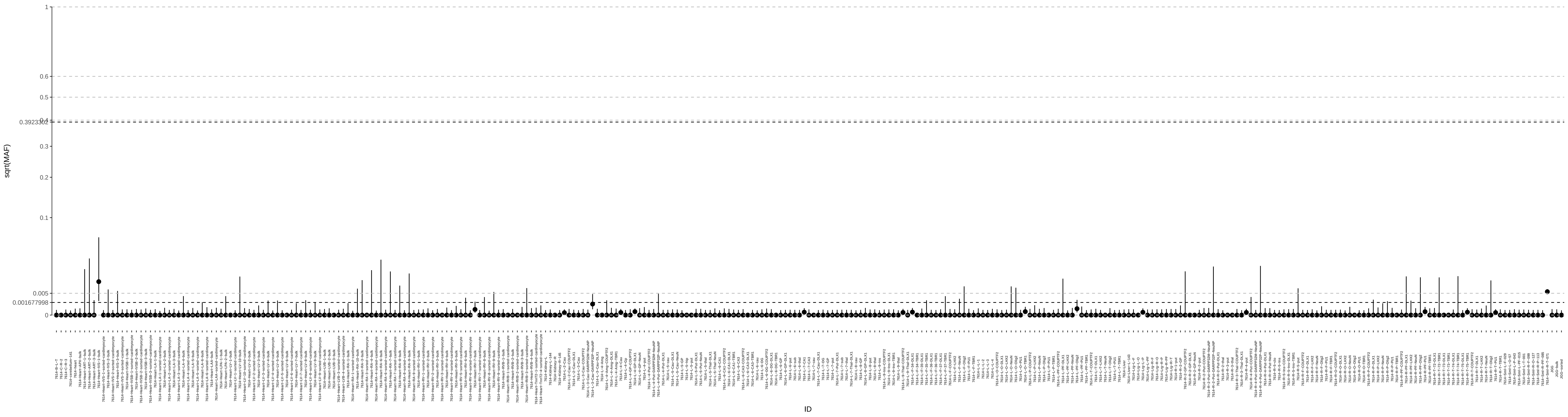

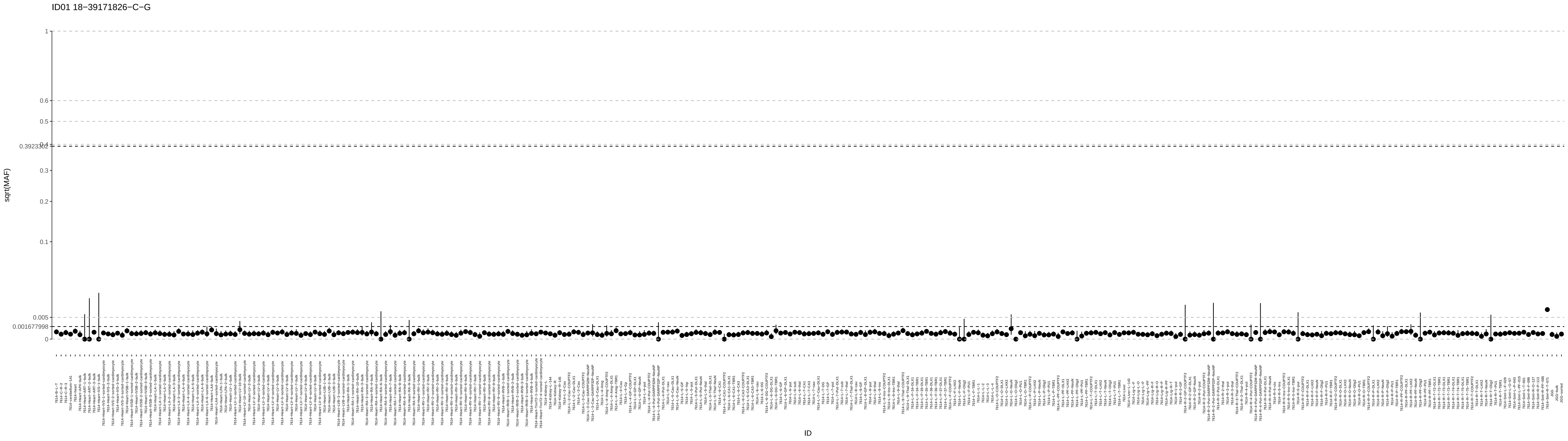
ID01 18−39171826−C−G
sqrt(MAF)
1
0.6
0.5
0.3923302
0.3
0.2
0.1
0.005
0.001677998
0
ID

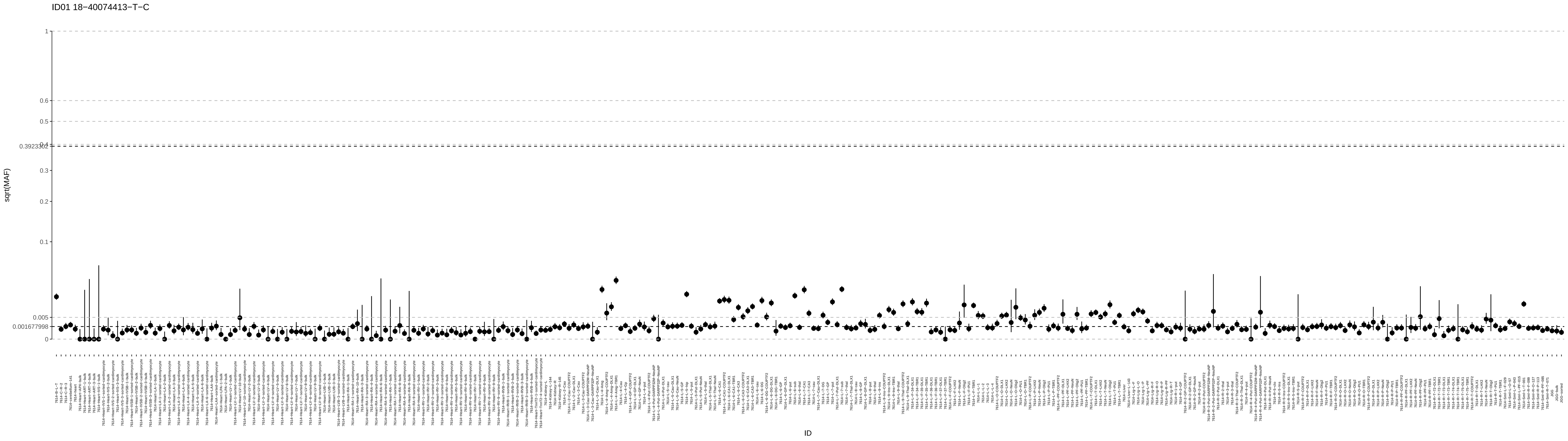
ID01 18−40074413−T−C
sqrt(MAF)
1
0.6
0.5
0.3923302
0.3
0.2
0.1
0.005
0.001677998
0
ID

ID01 18−56480516−G−A

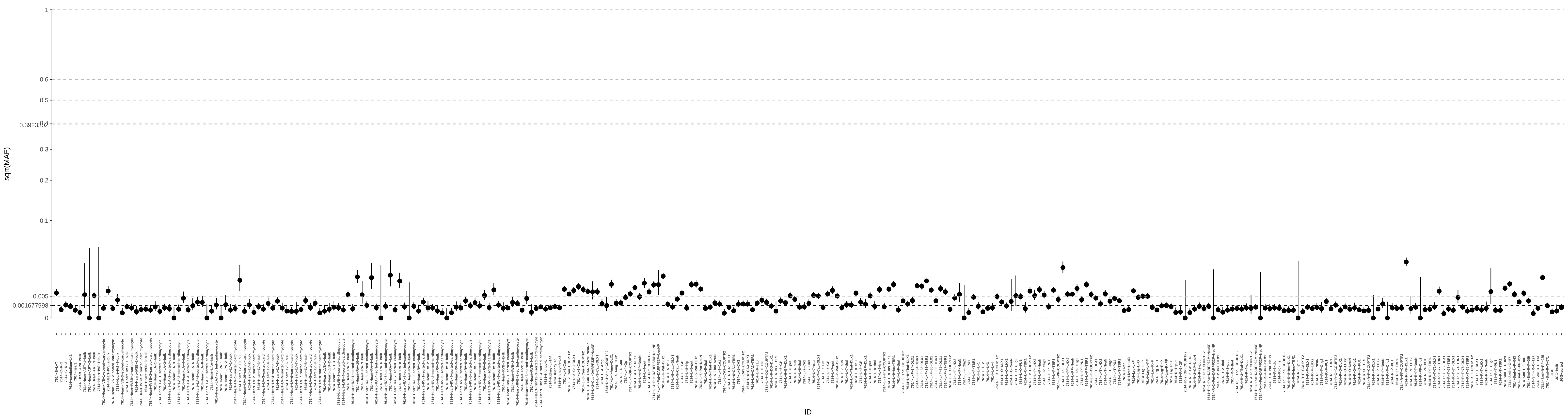

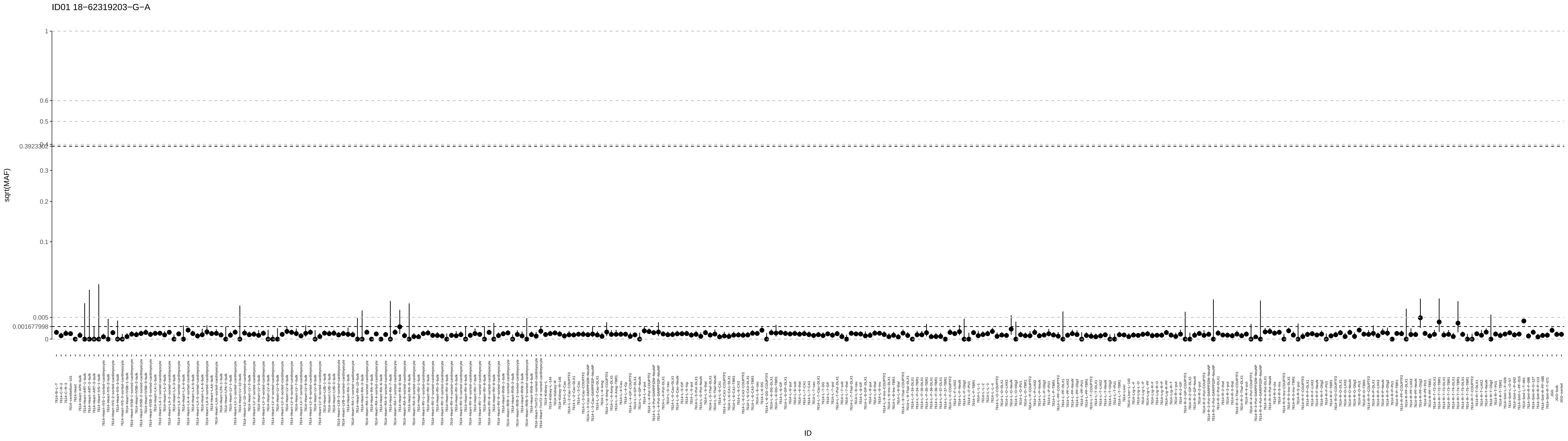
ID01 18−62319203−G−A
1
0.6
0.5
0.4
0.3923302
0.3
0.2
0.1
0.005
0.001677998
0
sqrt(MAF)
ID

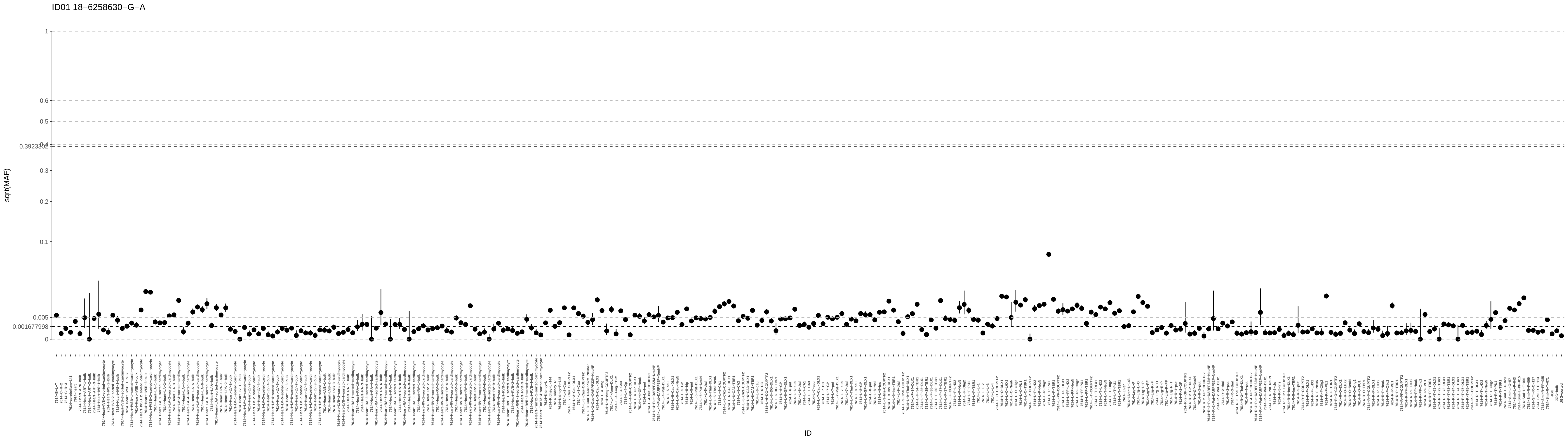
ID01 18−6258630−G−A
sqrt(MAF)
1
0.6
0.5
0.3923302
0.3
0.2
0.1
0.005
0.001677998
0
ID

ID01 18−65335071−G−A

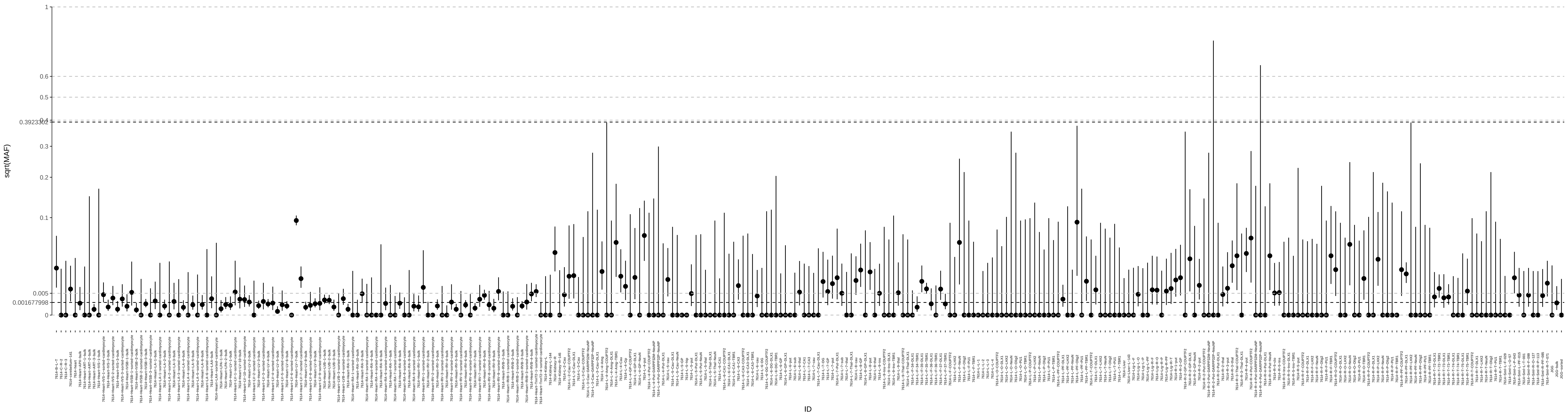

ID01 18−72031228−C−T

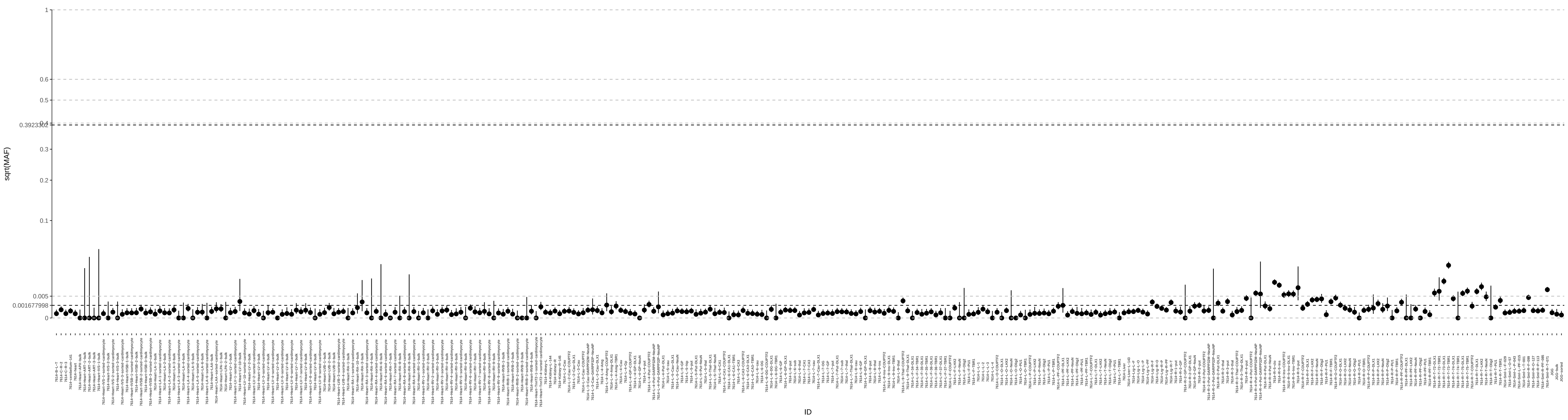

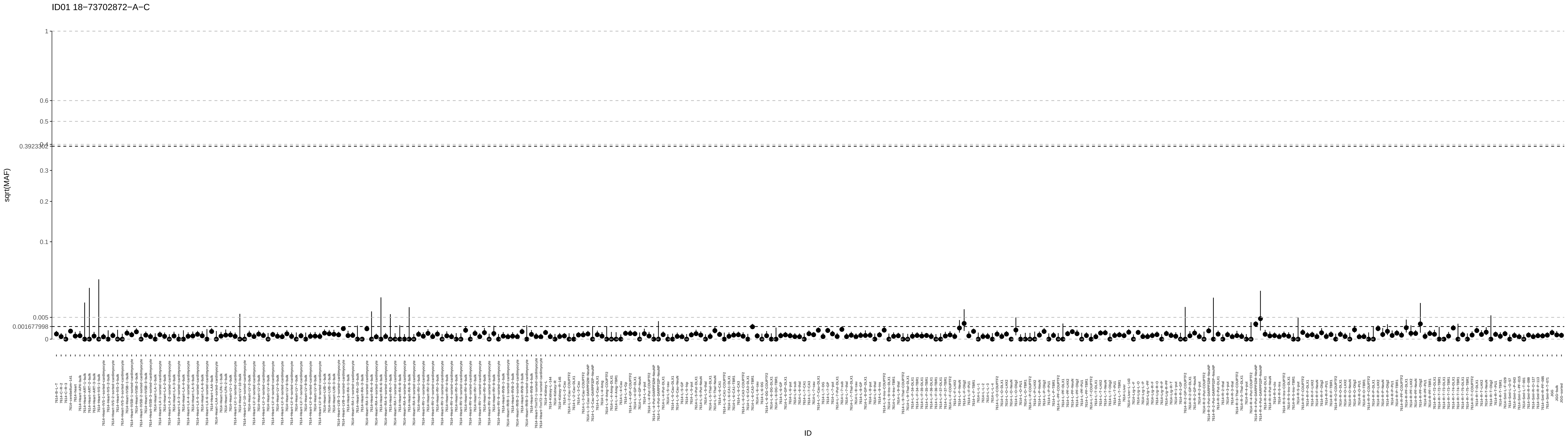
ID01 18−73702872−A−C
sqrt(MAF)
1
0.6
0.5
0.3923302
0.3
0.2
0.1
0.005
0.001677998
0
ID

ID01 19−23463606−T−C

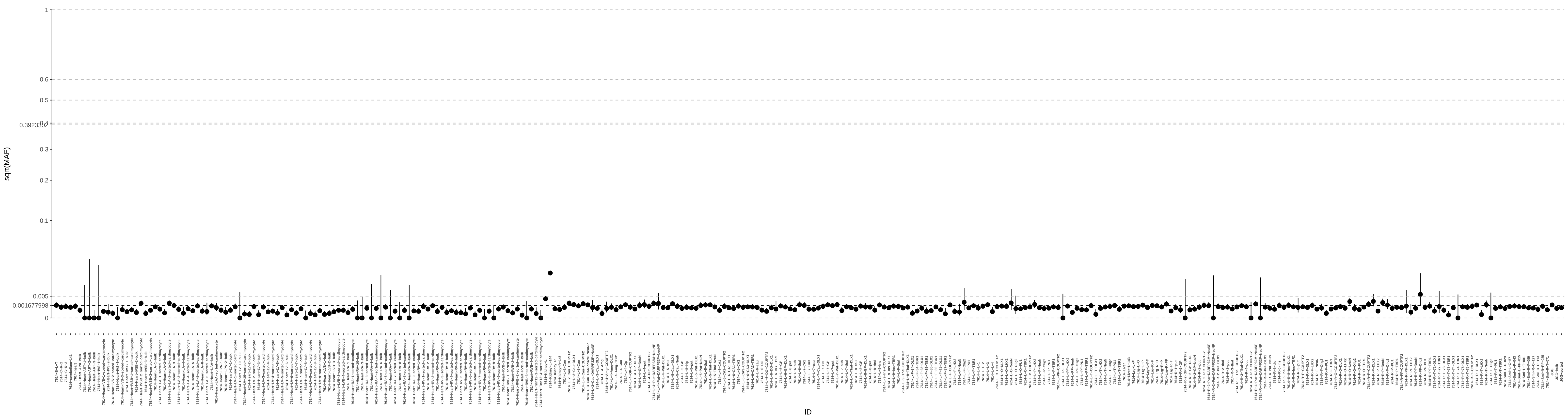

ID01 19−29424630−G−T

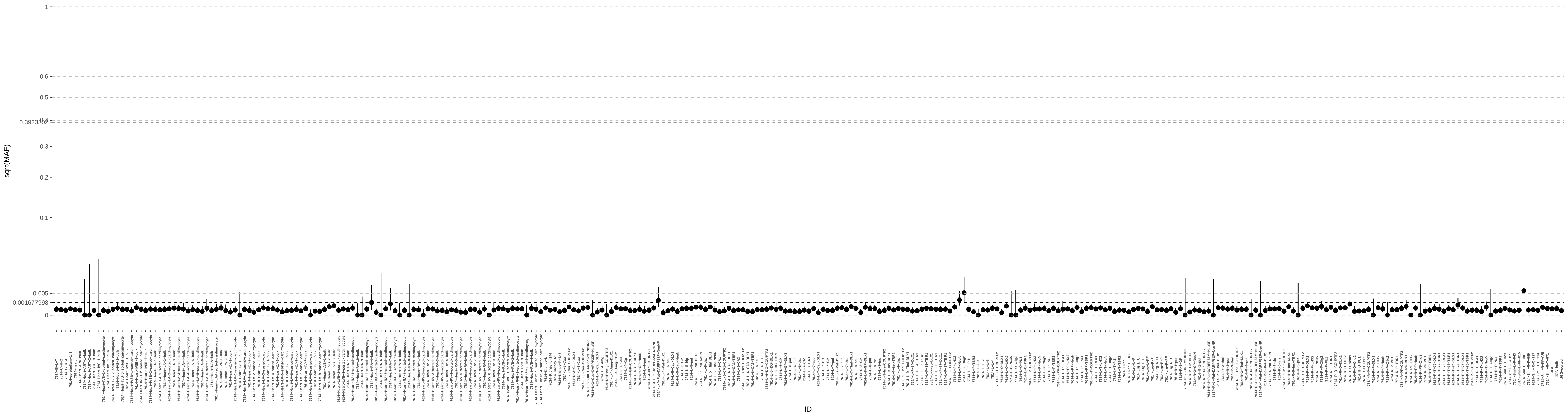

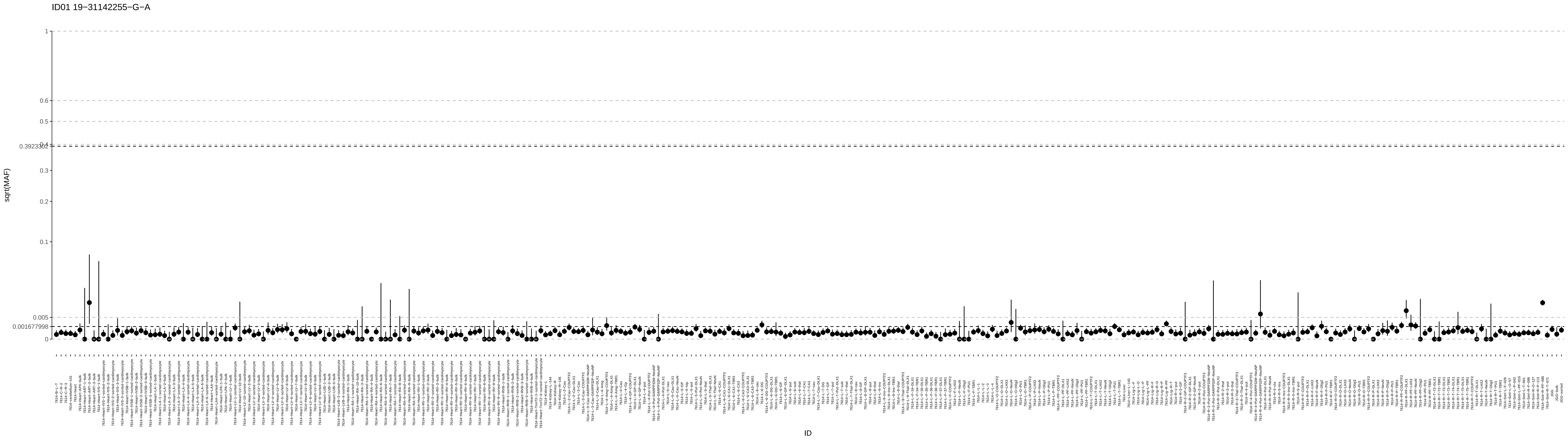
ID01 19−31142255−G−A
sqrt(MAF)
1
0.6
0.5
0.4
0.3923302
0.3
0.2
0.1
0.005
0.001677998
0
ID

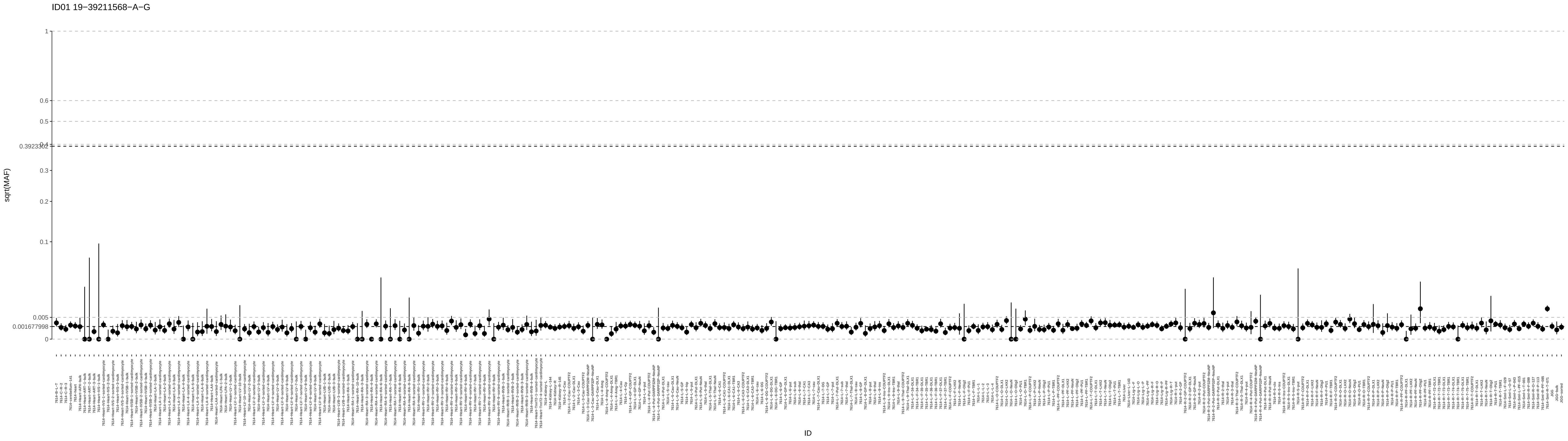
ID01 19−39211568−A−G
sqrt(MAF)
1
0.6
0.5
0.3923302
0.3
0.2
0.1
0.005
0.001677998
0
ID

ID01 19−47833717−G−A

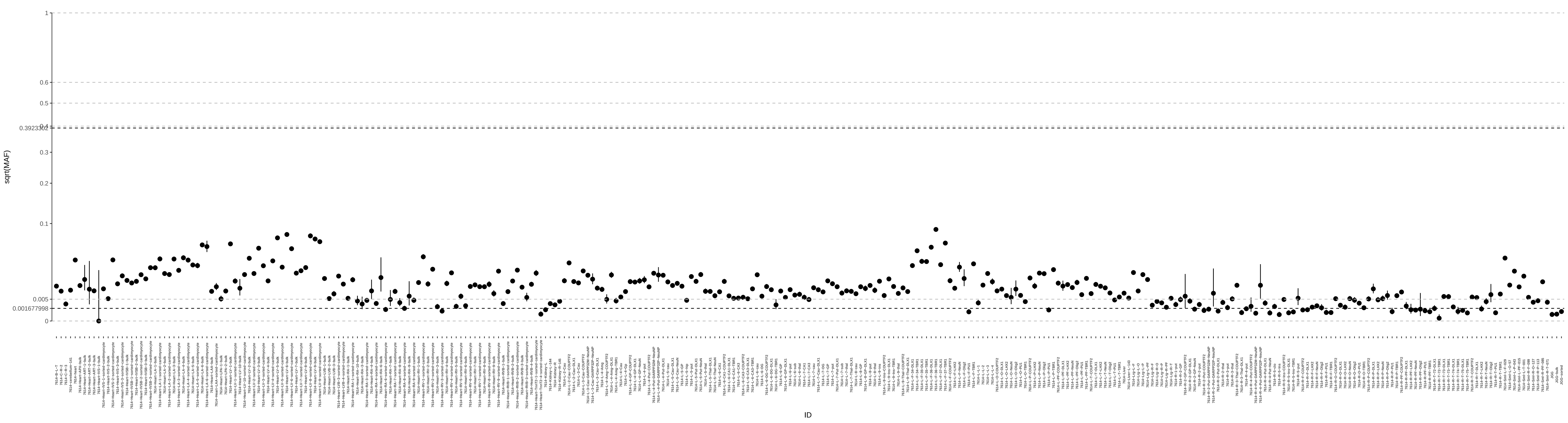

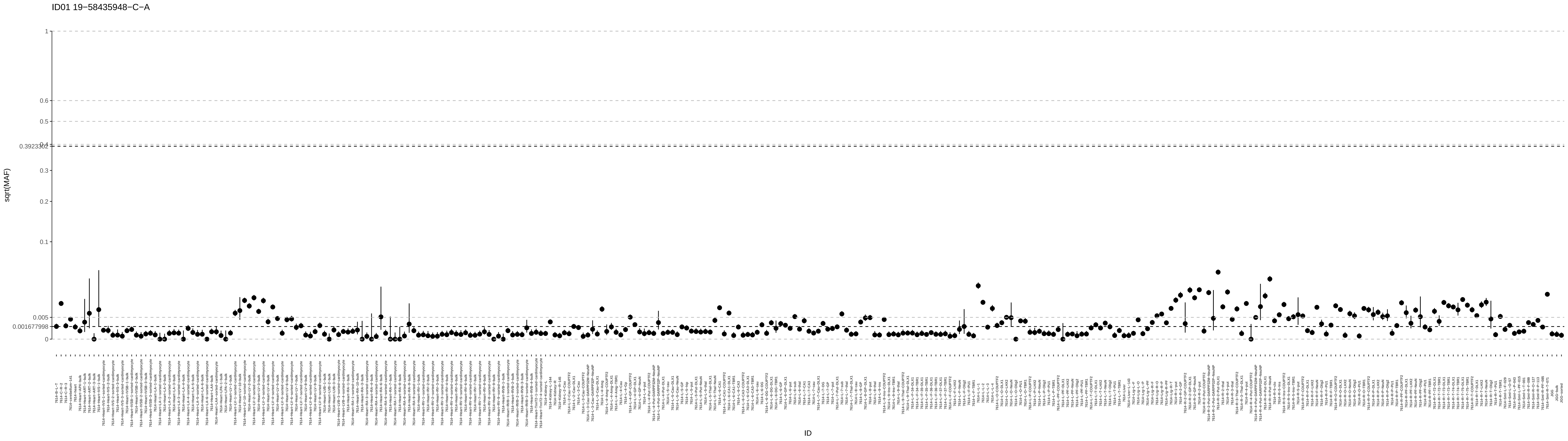
ID01 19−58435948−C−A
sqrt(MAF)
1
0.6
0.5
0.3923302
0.3
0.2
0.1
0.005
0.001677998
0
ID

ID01 2−101097017−C−T

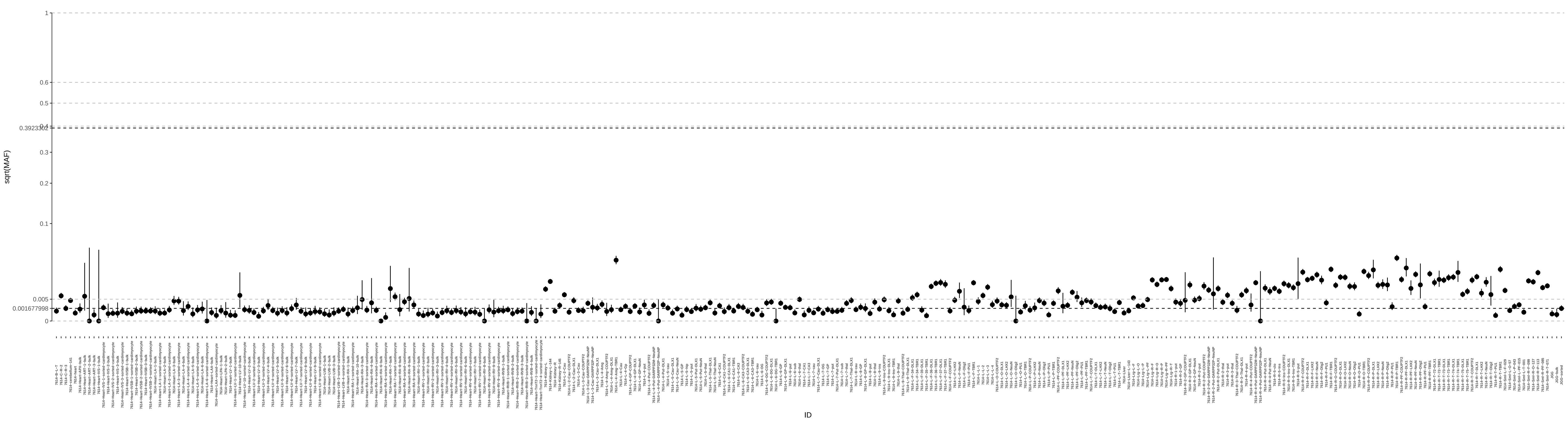

ID01 2−119890951−G−A

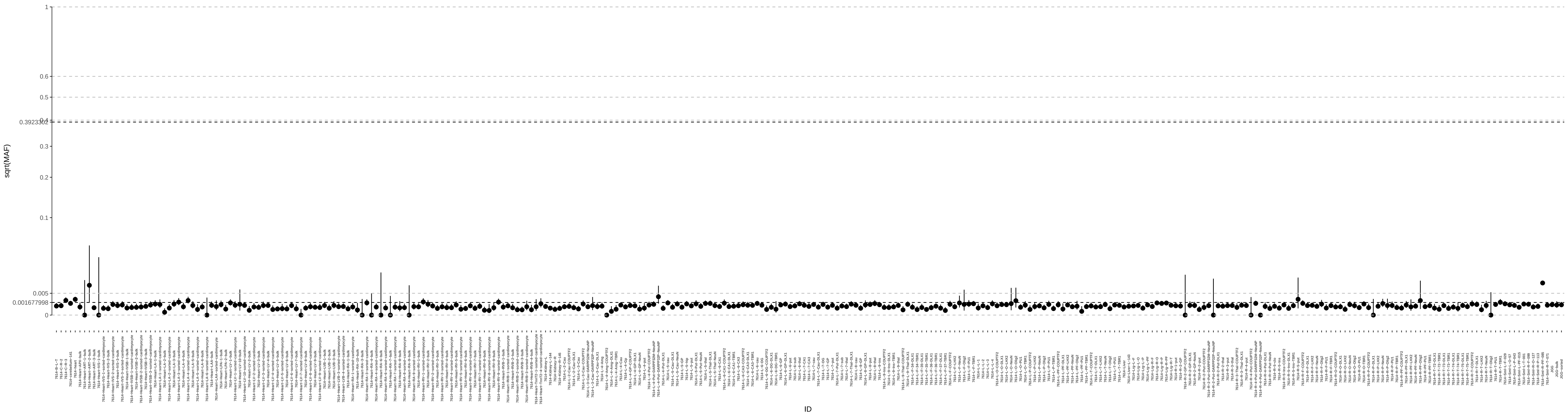

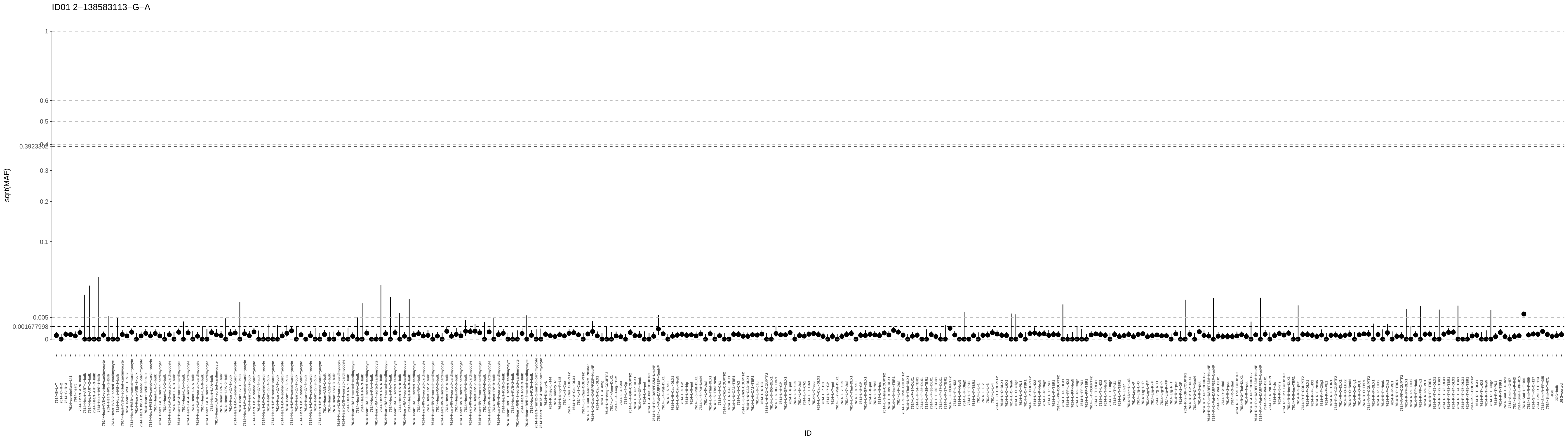
ID01 2−138583113−G−A
sqrt(MAF)
1
0.6
0.5
0.3923302
0.3
0.2
0.1
0.005
0.001677998
0
ID

ID01 2-139753954-C-T

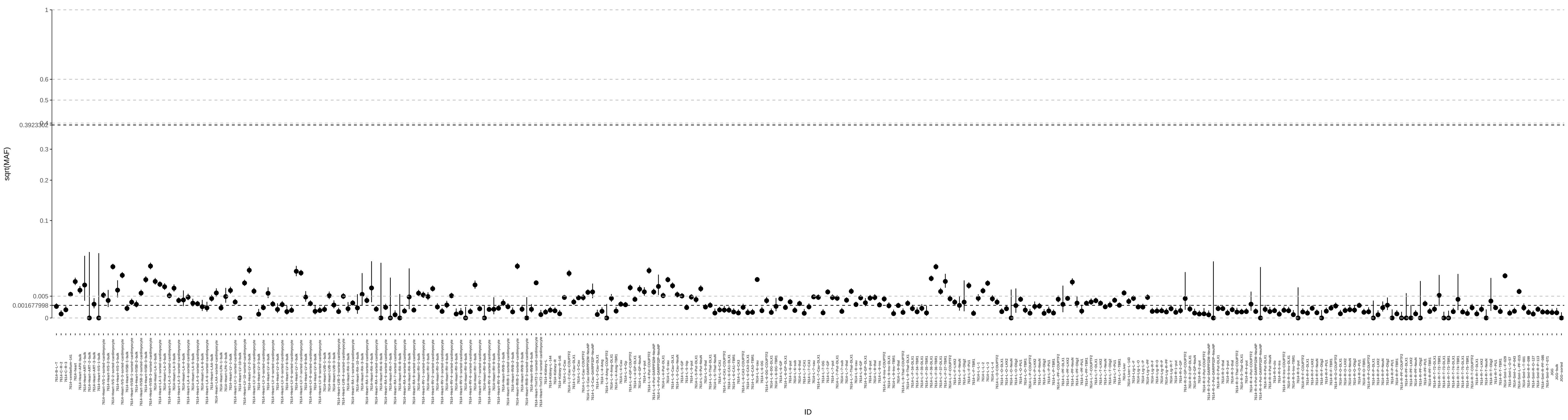

ID01 2−142968696−G−T

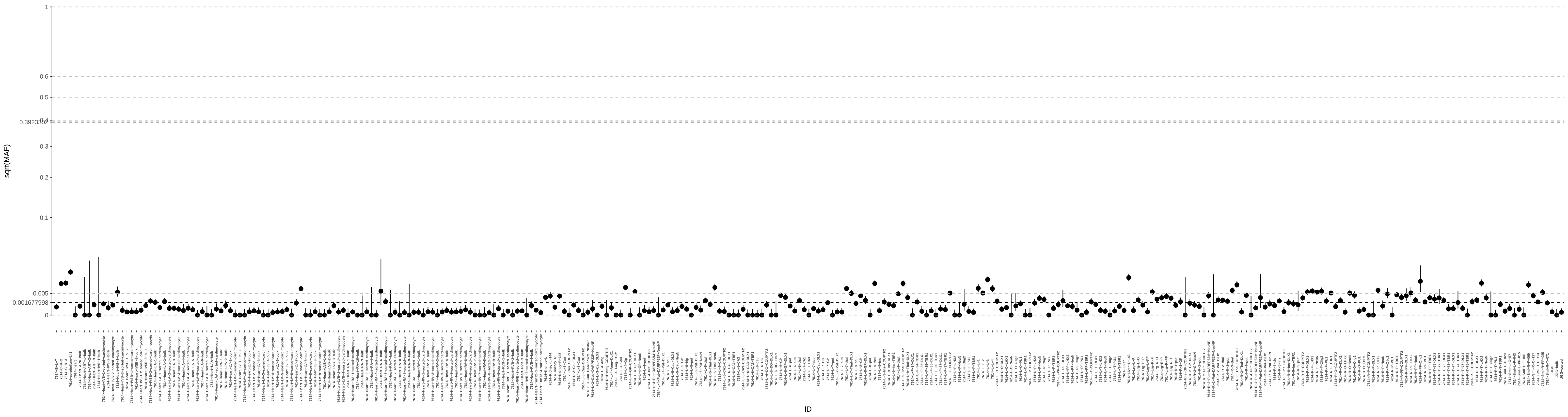

ID01 2−143727379−C−T

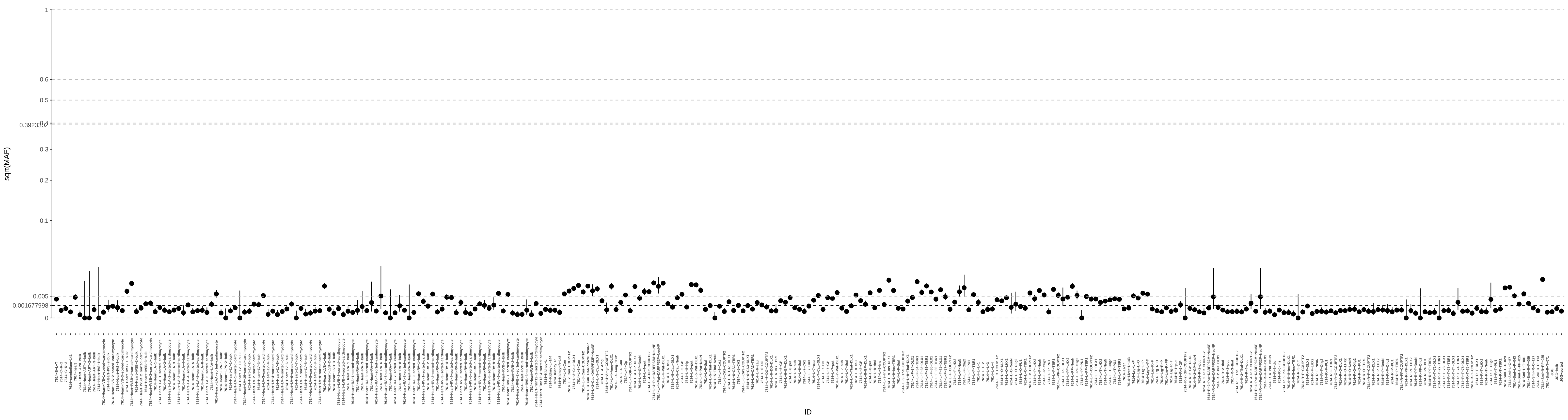

ID01 2−1519264−AAC−A

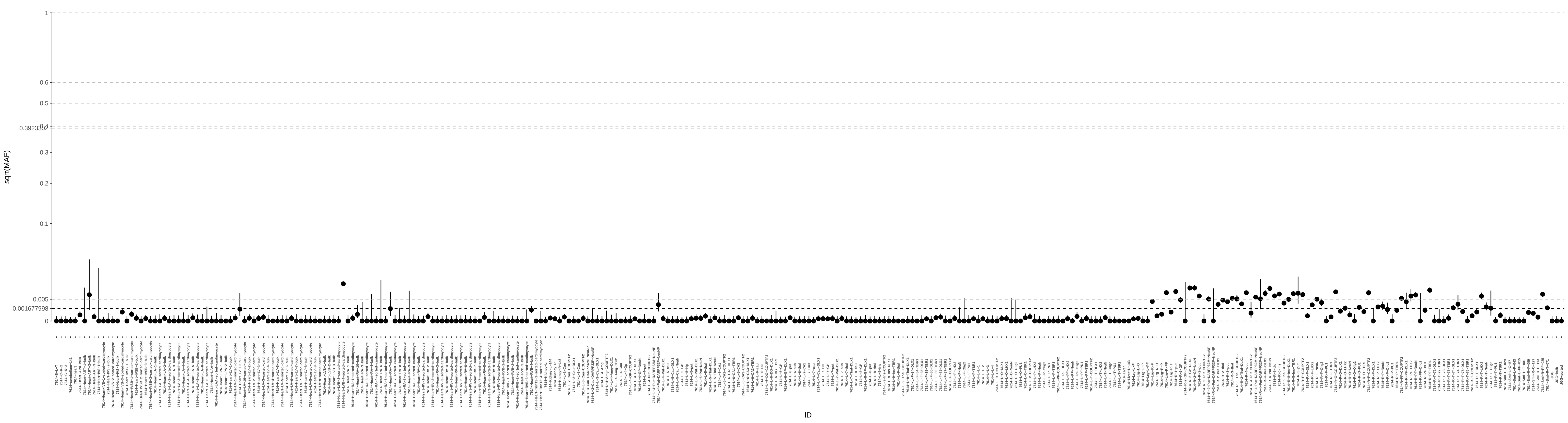

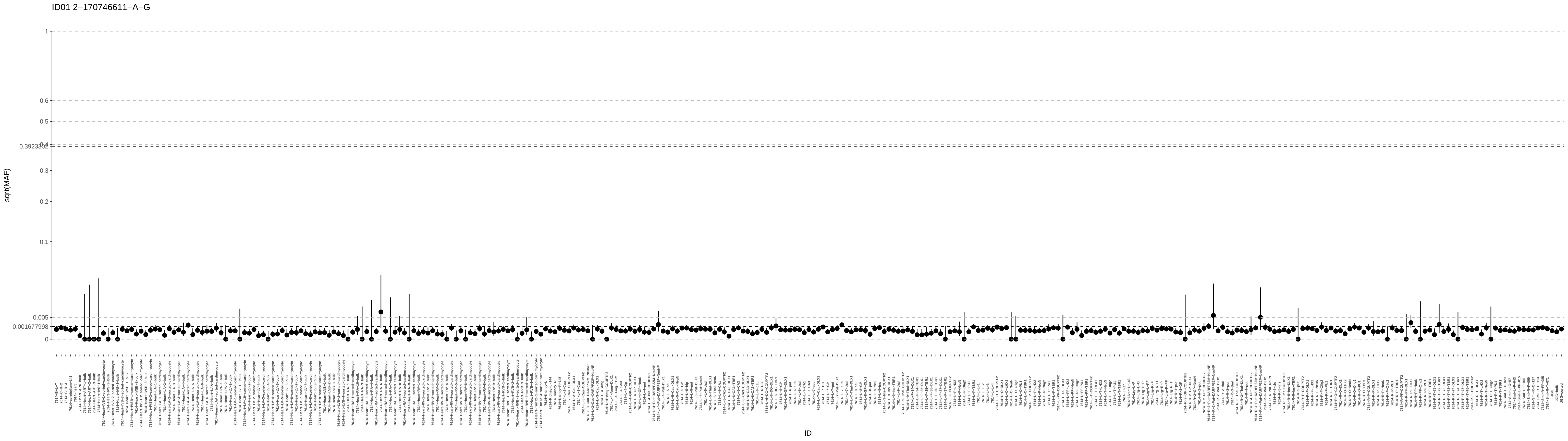
ID01 2−170746611−A−G
sqrt(MAF)
1
0.6
0.5
0.3923302
0.3
0.2
0.1
0.005
0.001677998
0
ID

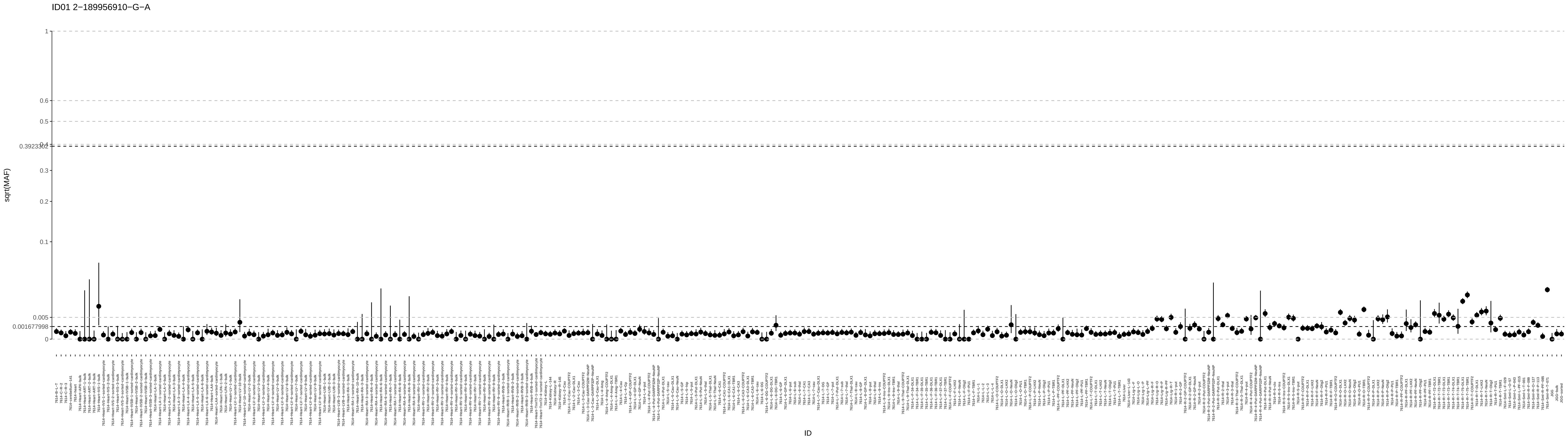
ID01 2−189956910−G−A
sqrt(MAF)
1
0.6
0.5
0.3923302
0.3
0.2
0.1
0.005
0.001677998
0
ID

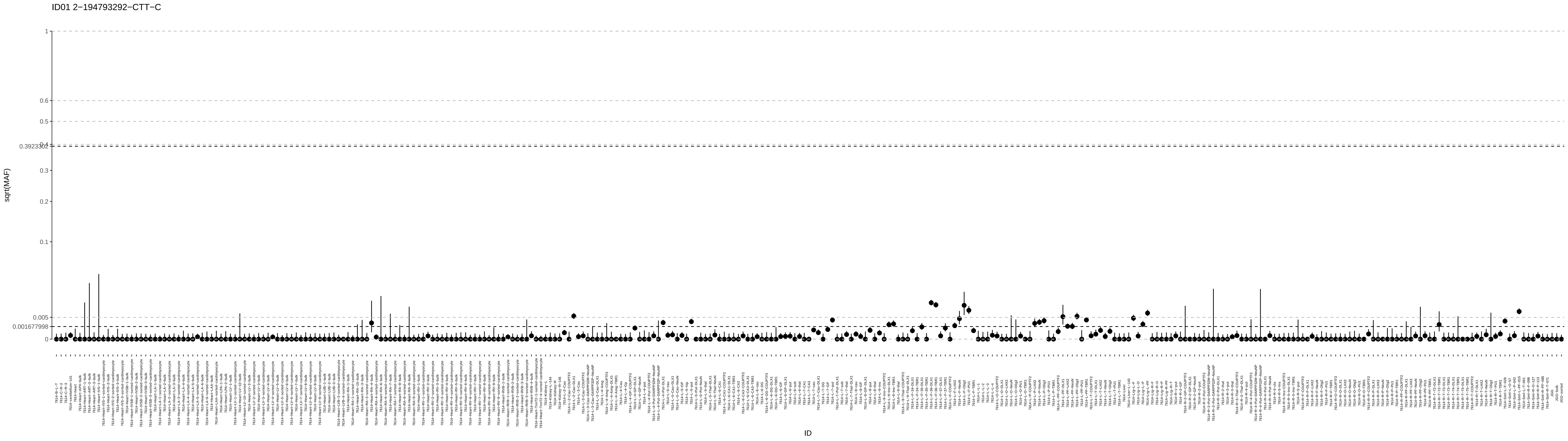
ID01 2−194793292−CTT−C
sqrt(MAF)
1
0.6
0.5
0.3923302
0.3
0.2
0.1
0.005
0.001677998
0
ID

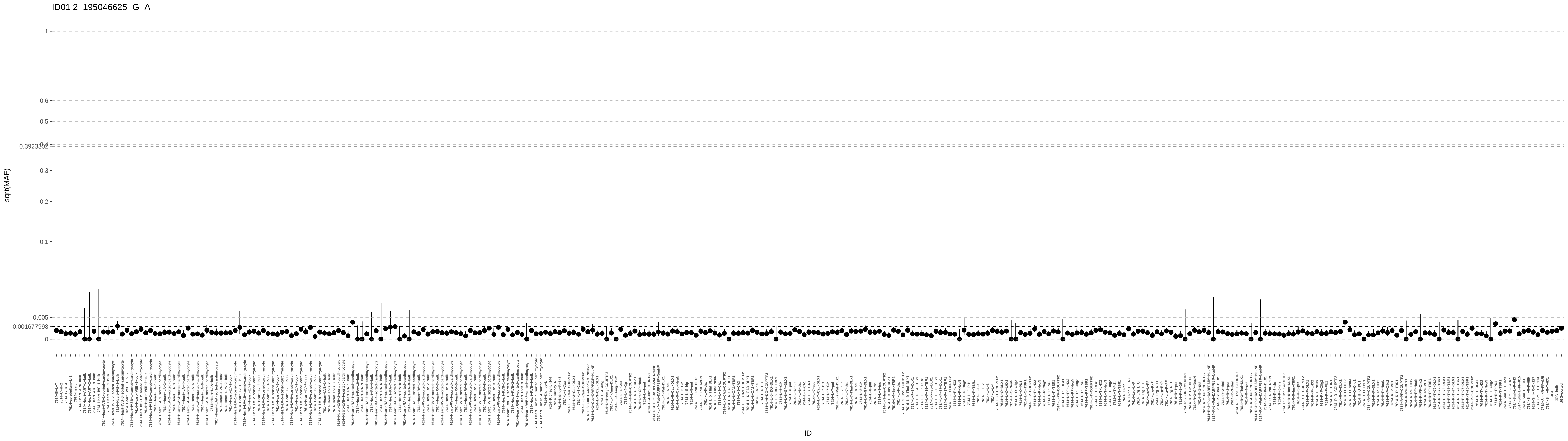

ID01 2−195046625−G−A
sqrt(MAF)
1
0.6
0.5
0.3923302
0.3
0.2
0.1
0.005
0.001677998
0
ID

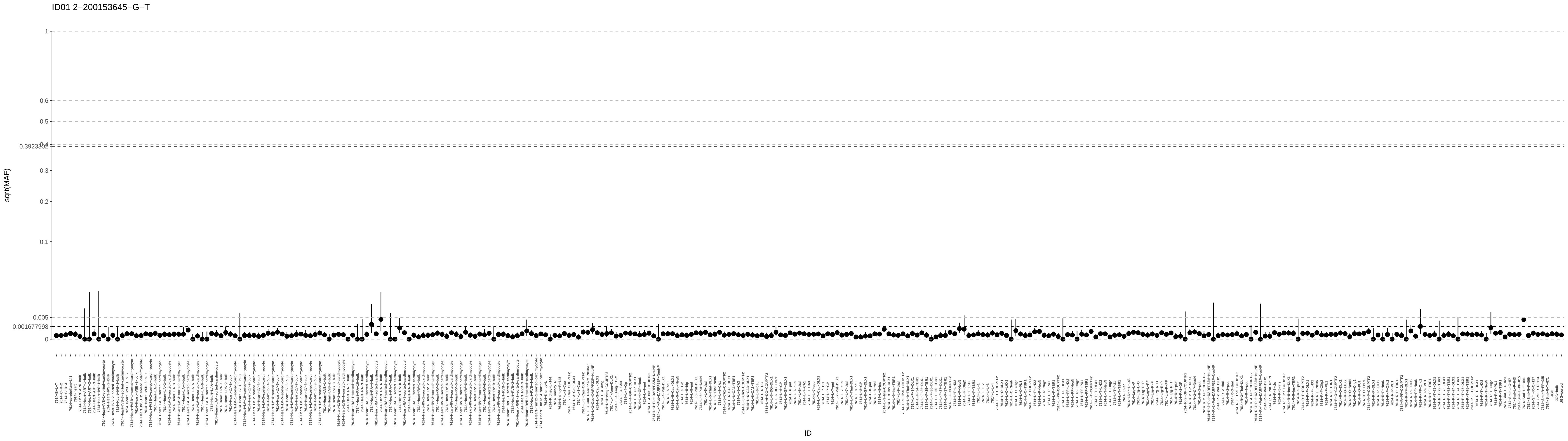
ID01 2−200153645−G−T
sqrt(MAF)
1
0.6
0.5
0.4
0.3923302
0.3
0.2
0.1
0.005
0.001677998
0
ID

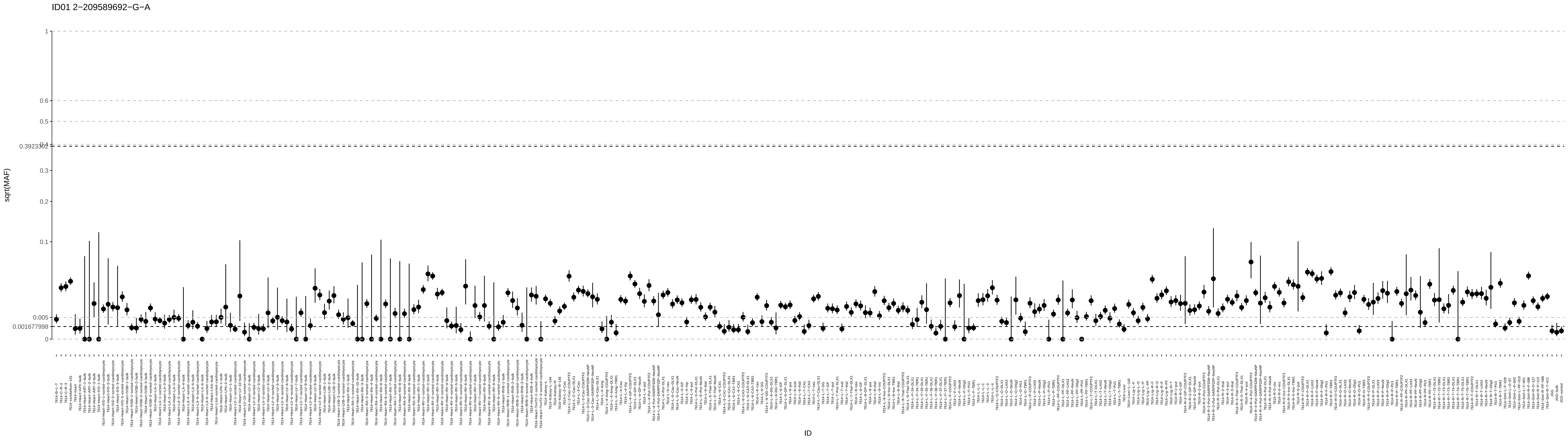
ID01 2−209589692−G−A
sqrt(MAF)
1
0.6
0.5
0.3923302
0.3
0.2
0.1
0.005
0.001677998
0
ID

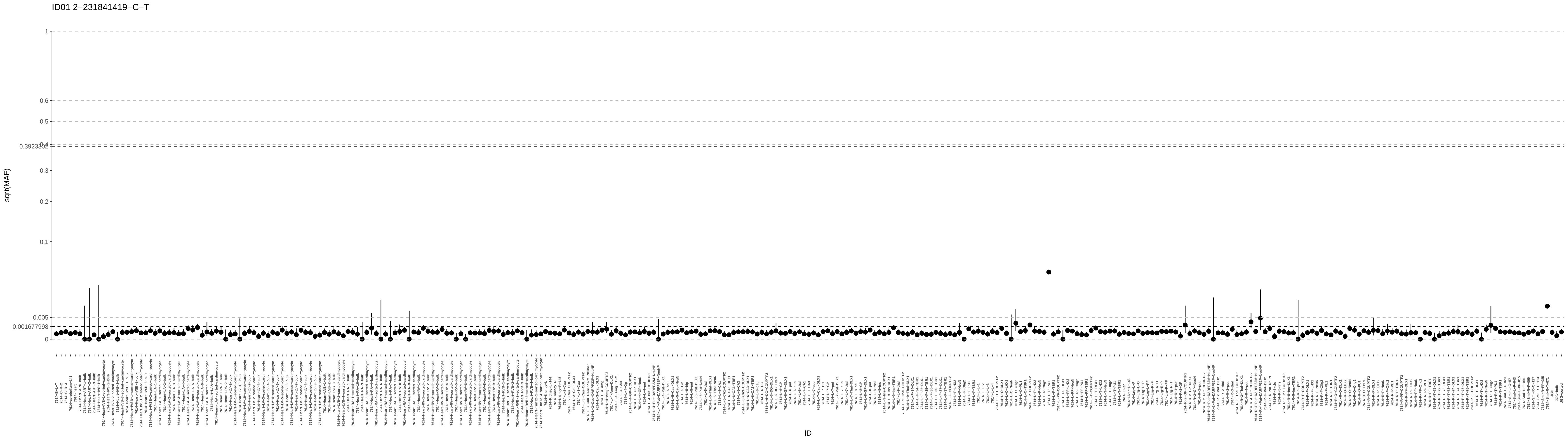
ID01 2−231841419−C−T
sqrt(MAF)
1
0.6
0.5
0.4
0.3923302
0.3
0.2
0.1
0.005
0.001677998
0
ID

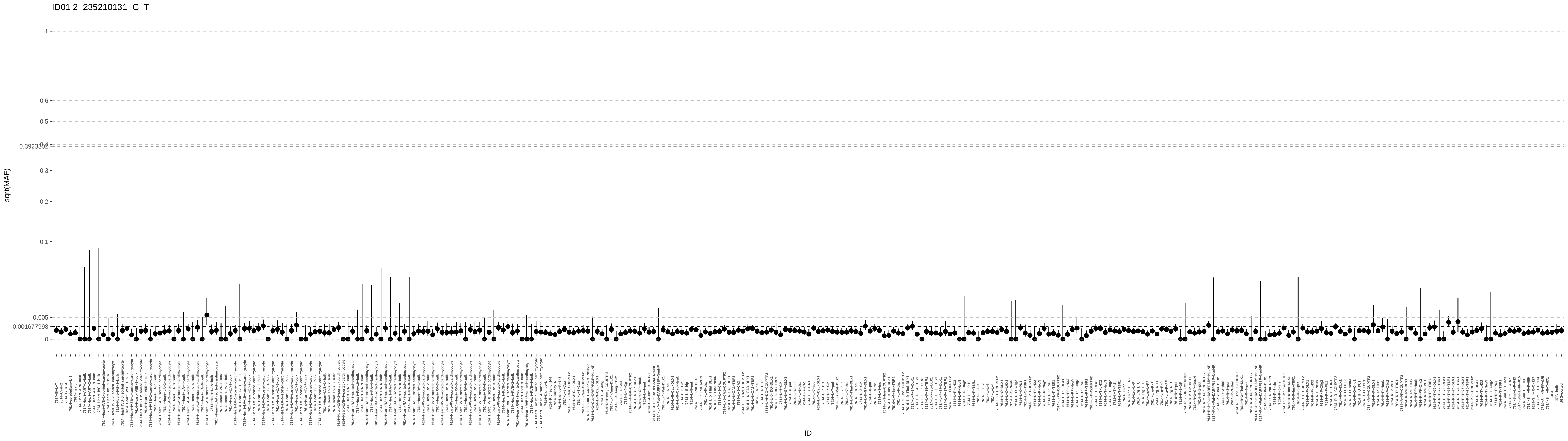

ID01 2−235210131−C−T
sqrt(MAF)
1
0.6
0.5
0.3923302
0.3
0.2
0.1
0.005
0.001677998
0
ID

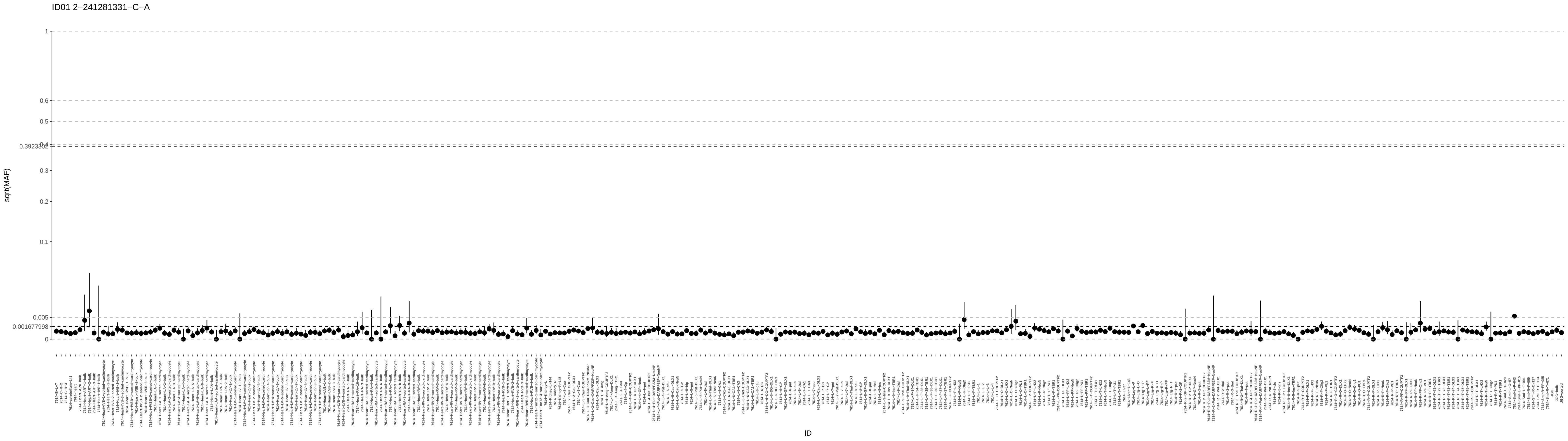
ID01 2−241281331−C−A
sqrt(MAF)
1
0.6
0.5
0.3923302
0.3
0.2
0.1
0.005
0.001677998
0
ID

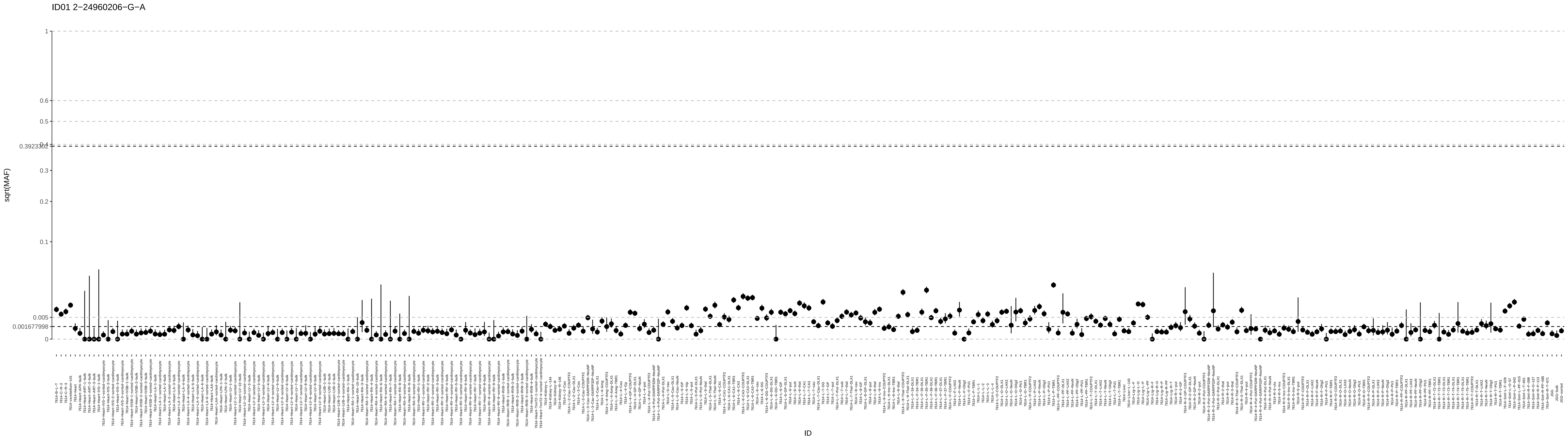

ID01 2−24960206−G−A
sqrt(MAF)
1
0.6
0.5
0.3923302
0.3
0.2
0.1
0.005
0.001677998
0
ID

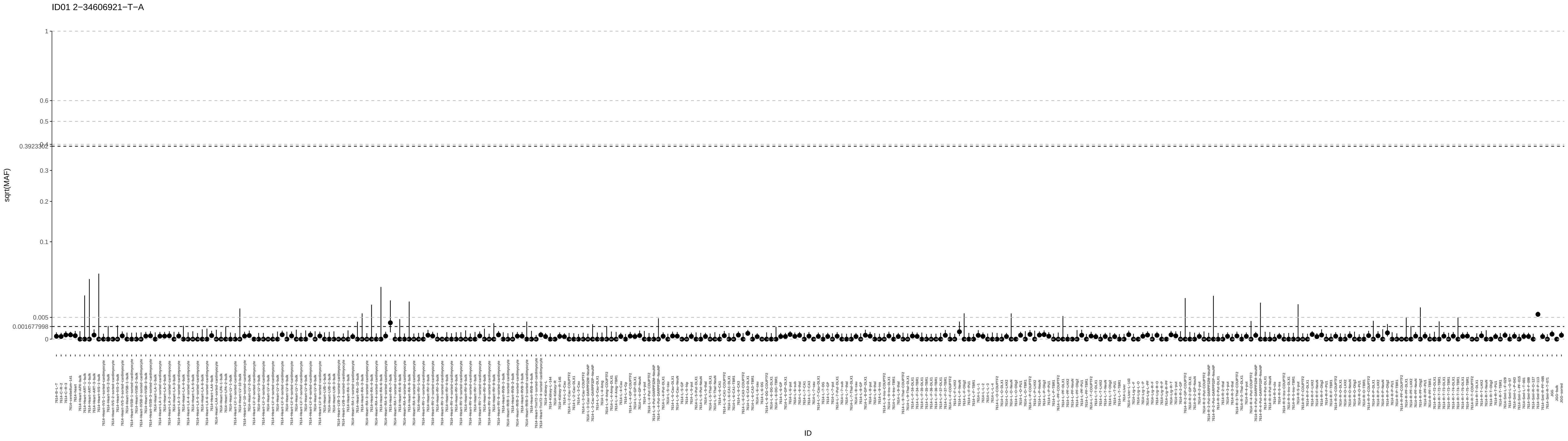
ID01 2−34606921−T−A
sqrt(MAF)
1
0.6
0.5
0.3923302
0.3
0.2
0.1
0.005
0.001677998
0
ID

ID01 2−49268814−A−C

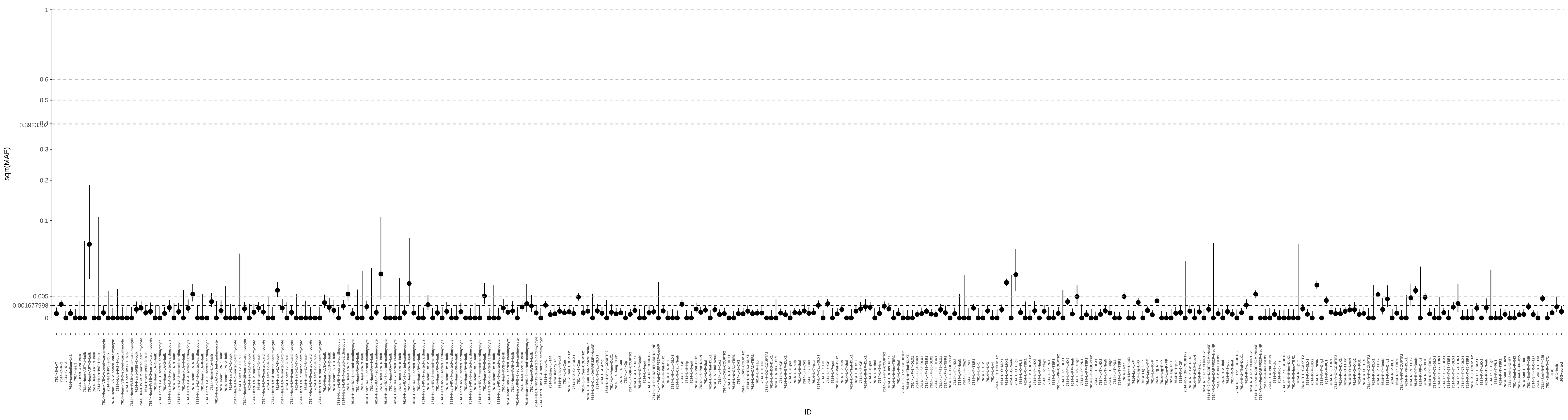

ID01 2−53799113−A−T

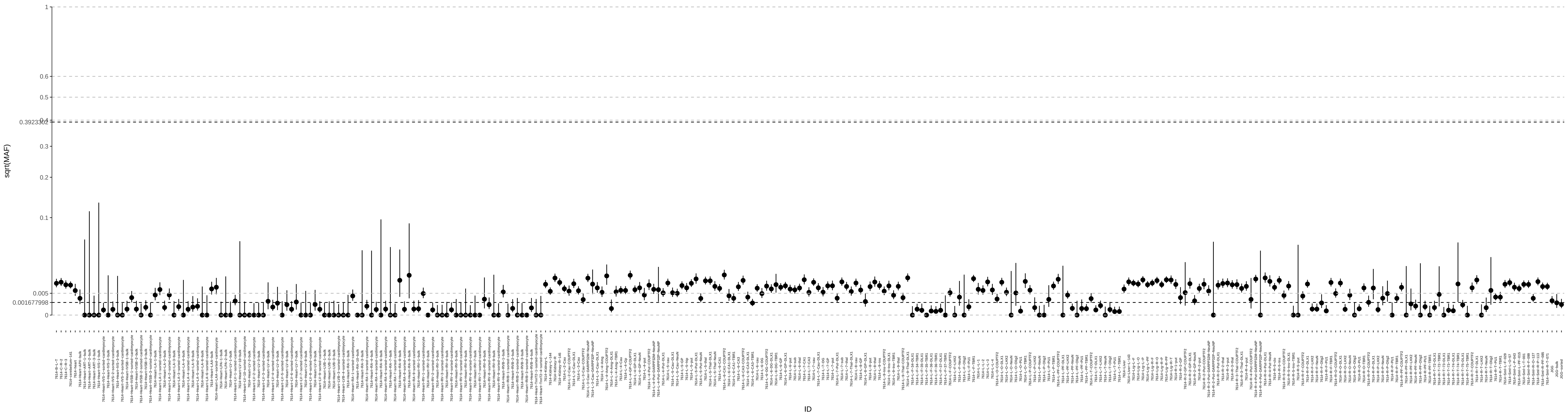

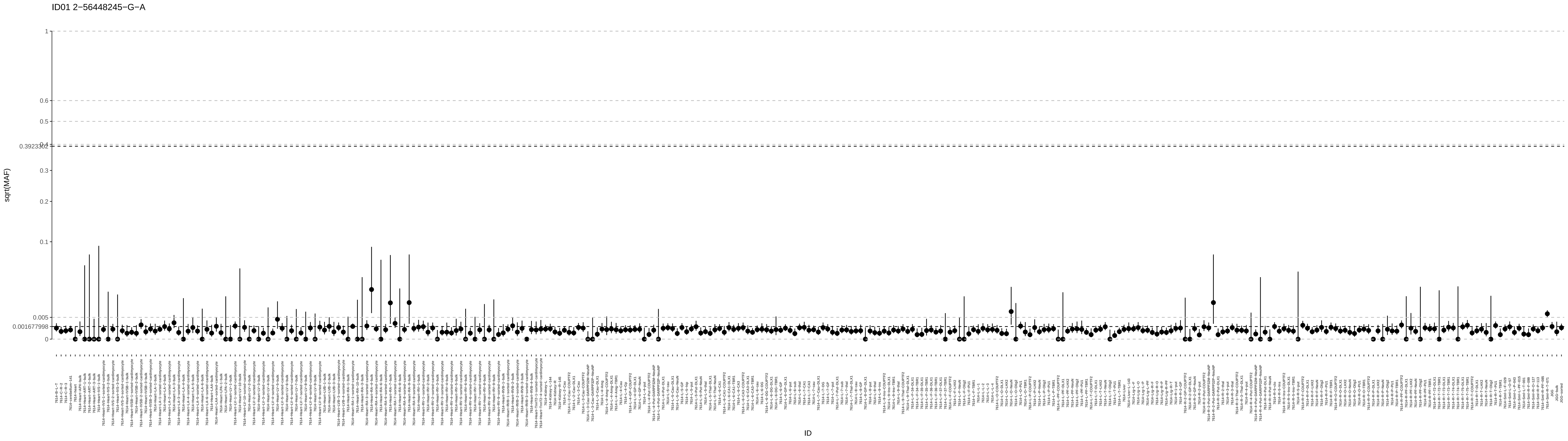

ID01 2−56448245−G−A
sqrt(MAF)
1
0.6
0.5
0.3923302
0.3
0.2
0.1
0.005
0.001677998
0
ID

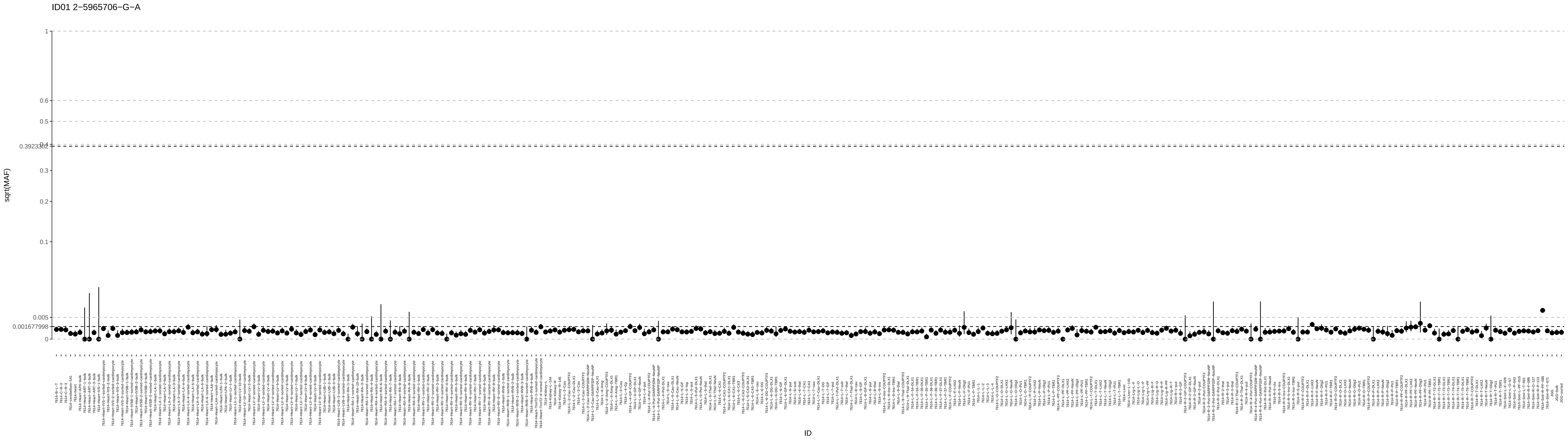
ID01 2−5965706−G−A
sqrt(MAF)
1
0.6
0.5
0.3923302
0.3
0.2
0.1
0.005
0.001677998
0
ID

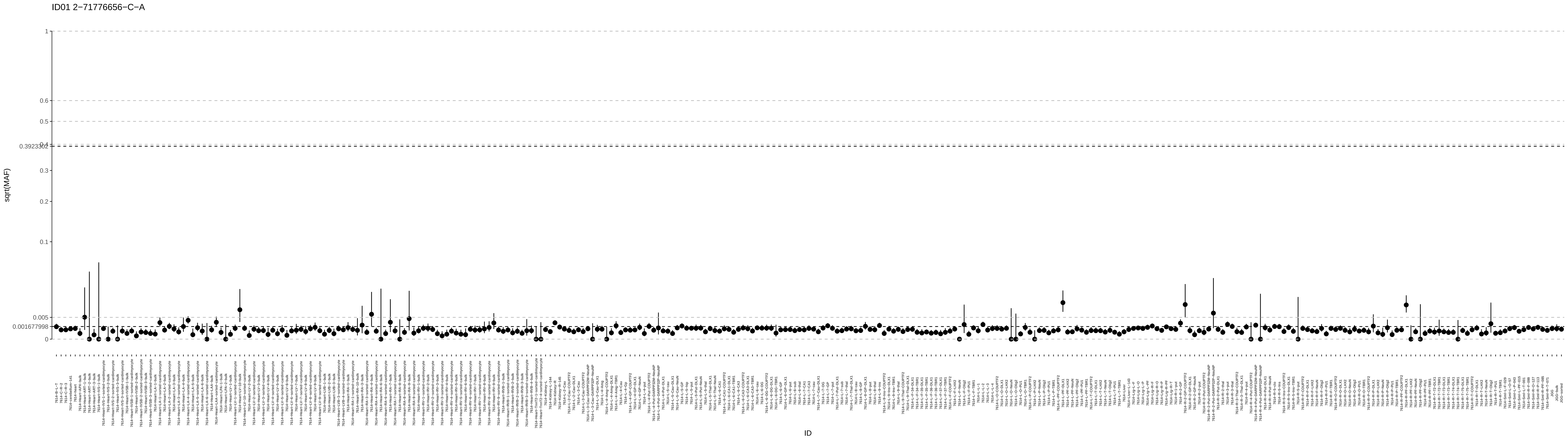
ID01 2-71776656-C-A
sqrt(MAF)
1
0.6
0.5
0.3923302
0.3
0.2
0.1
0.005
0.001677998
0
ID

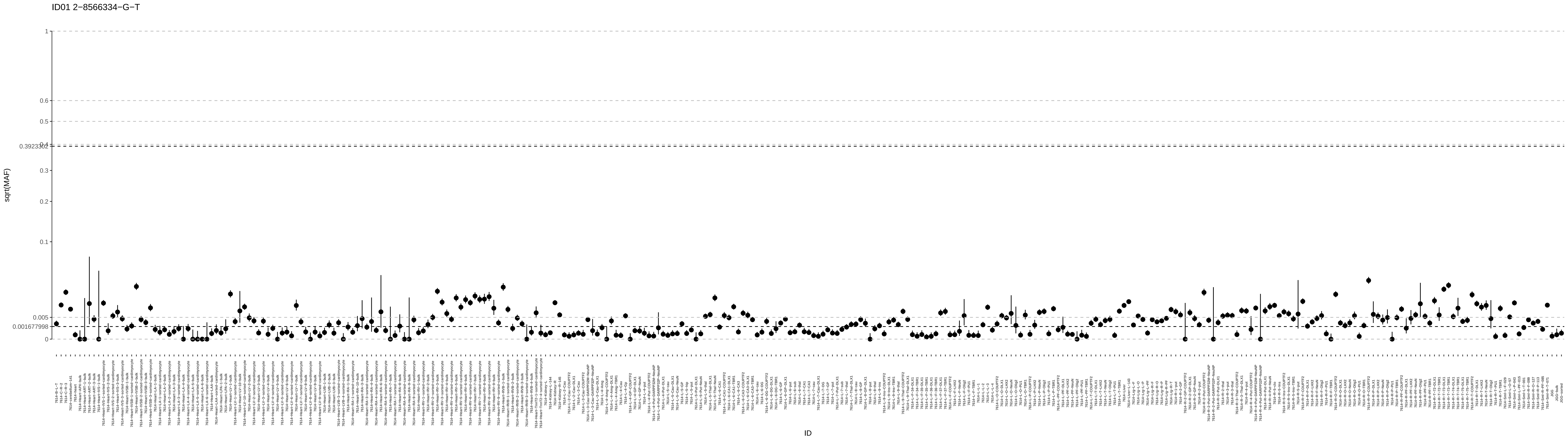
ID01 2-8566334-G-T
sqrt(MAF)
1
0.6
0.5
0.3923302
0.3
0.2
0.1
0.005
0.001677998
0
ID

ID01 2−90231584−G−T

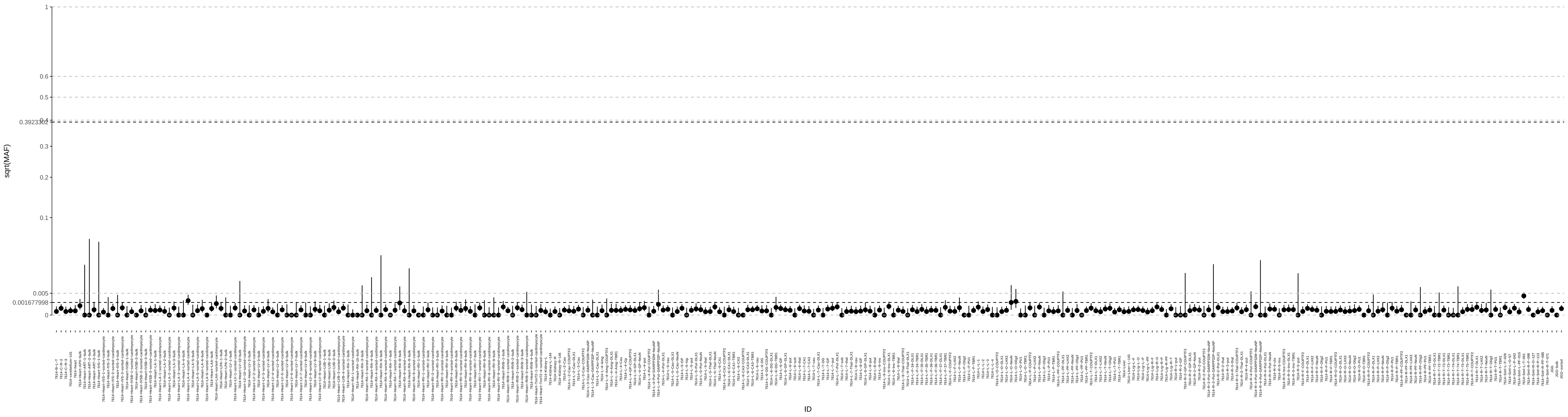

ID01 2−97358394−C−T

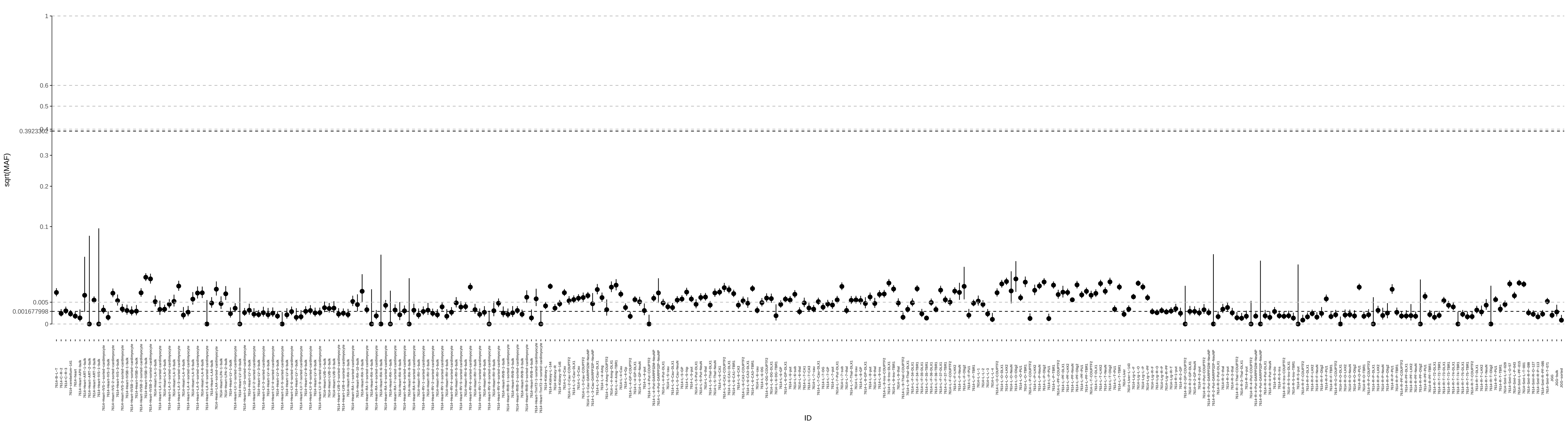

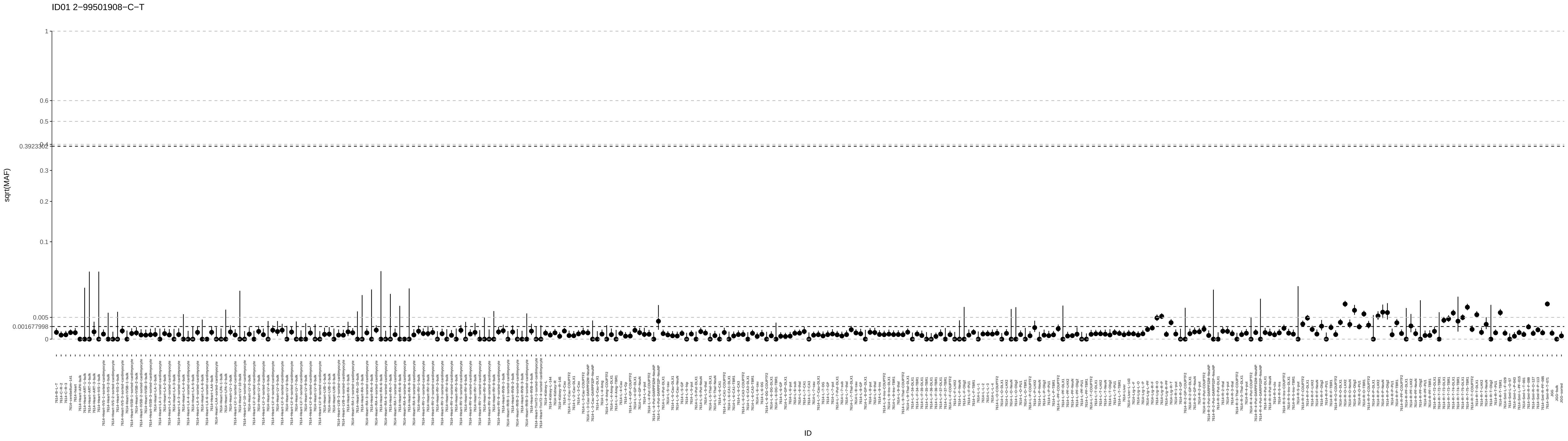
ID01 2−99501908−C−T
sqrt(MAF)
1
0.6
0.5
0.3923302
0.3
0.2
0.1
0.005
0.001677998
0
ID

ID01 20−13255185−C−A

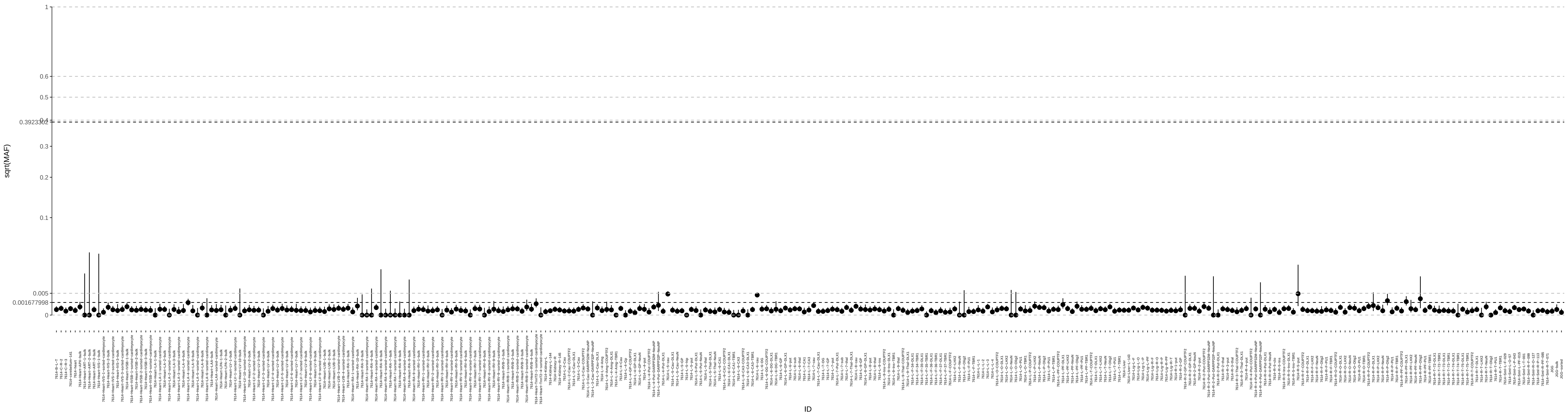

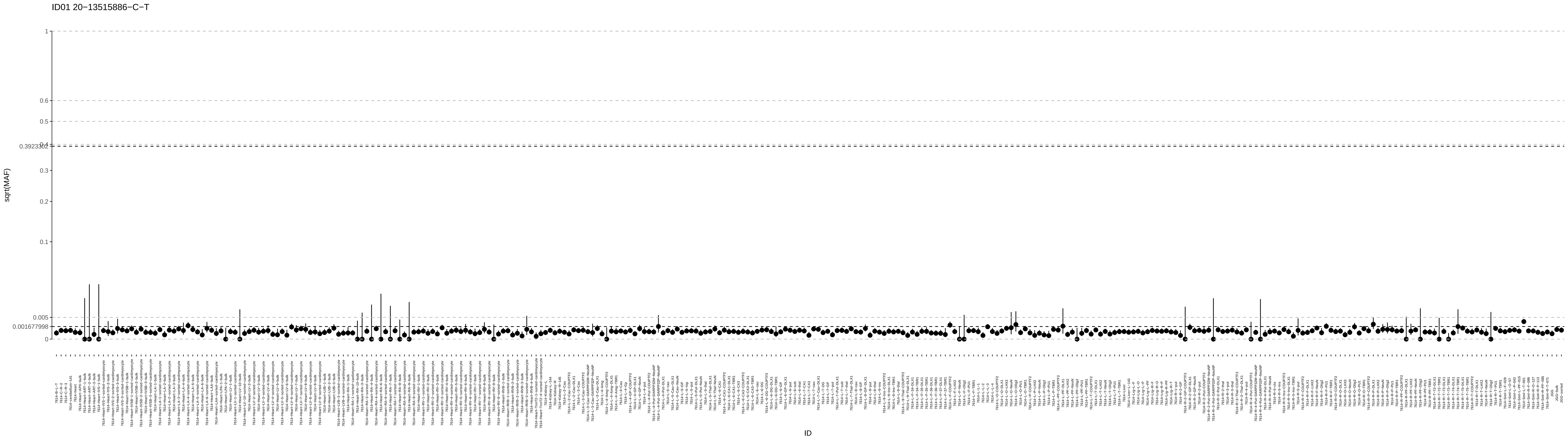
ID01 20−13515886−C−T
sqrt(MAF)
1
0.6
0.5
0.4
0.3923302
0.3
0.2
0.1
0.005
0.001677998
0
ID

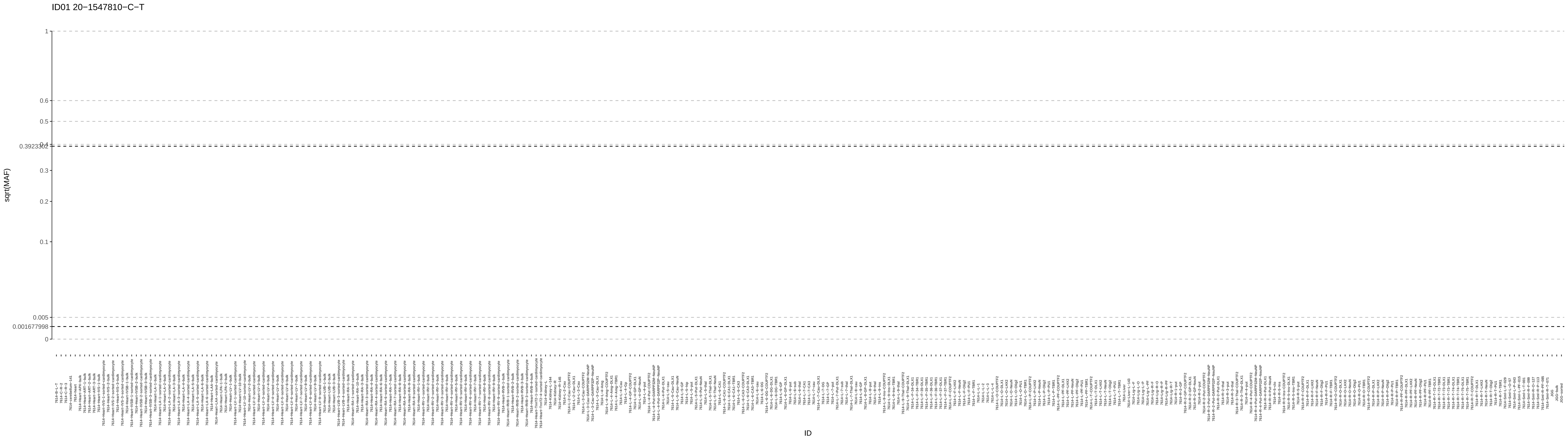

ID01 20−1547810−C−T
sqrt(MAF)
ID

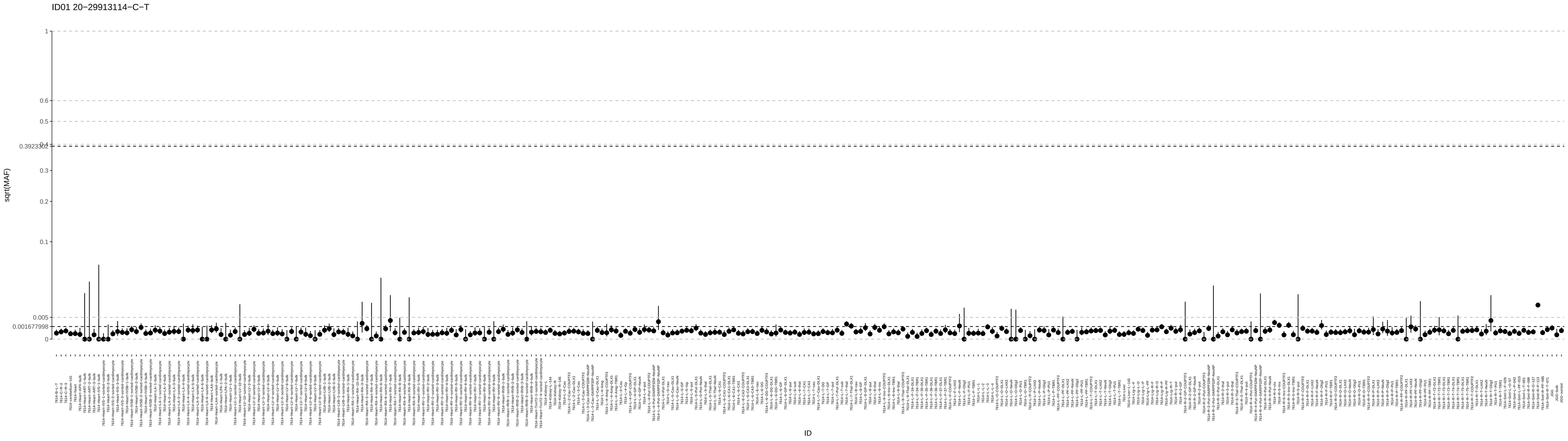
ID01 20−29913114−C−T
sqrt(MAF)
1
0.6
0.5
0.3923302
0.3
0.2
0.1
0.005
0.001677998
0
ID

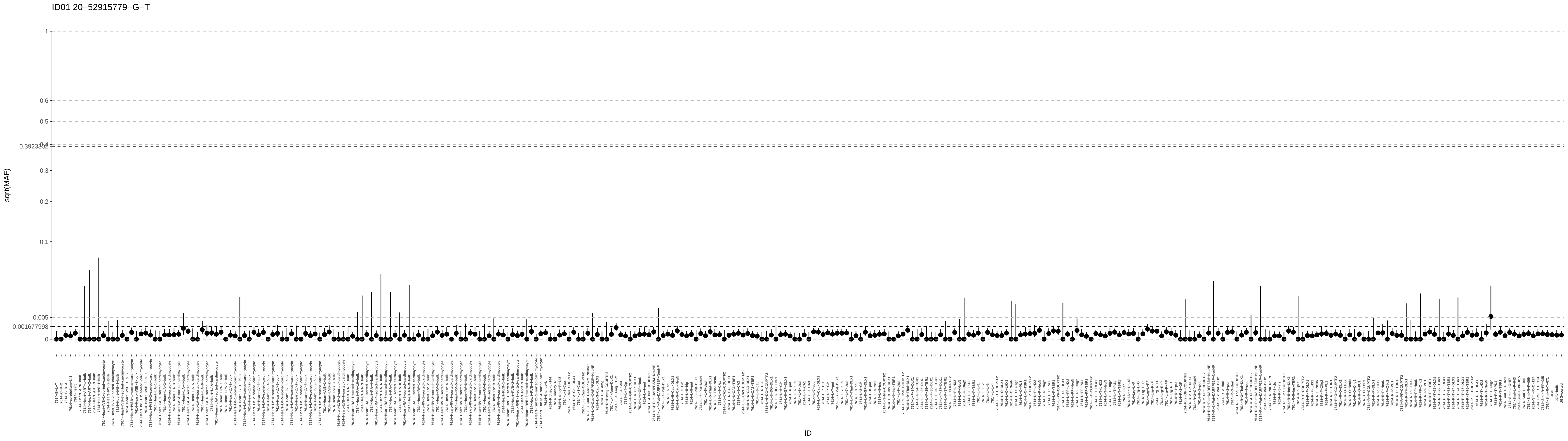
ID01 20−52915779−G−T
sqrt(MAF)
1
0.6
0.5
0.3923302
0.3
0.2
0.1
0.005
0.001677998
0
ID

ID01 20−52916352−A−C

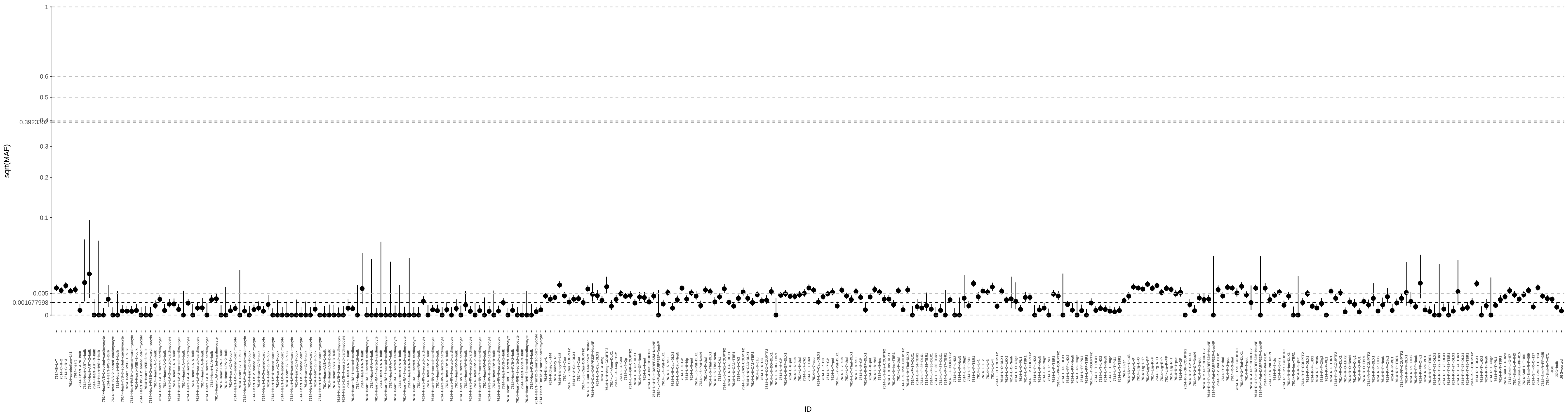

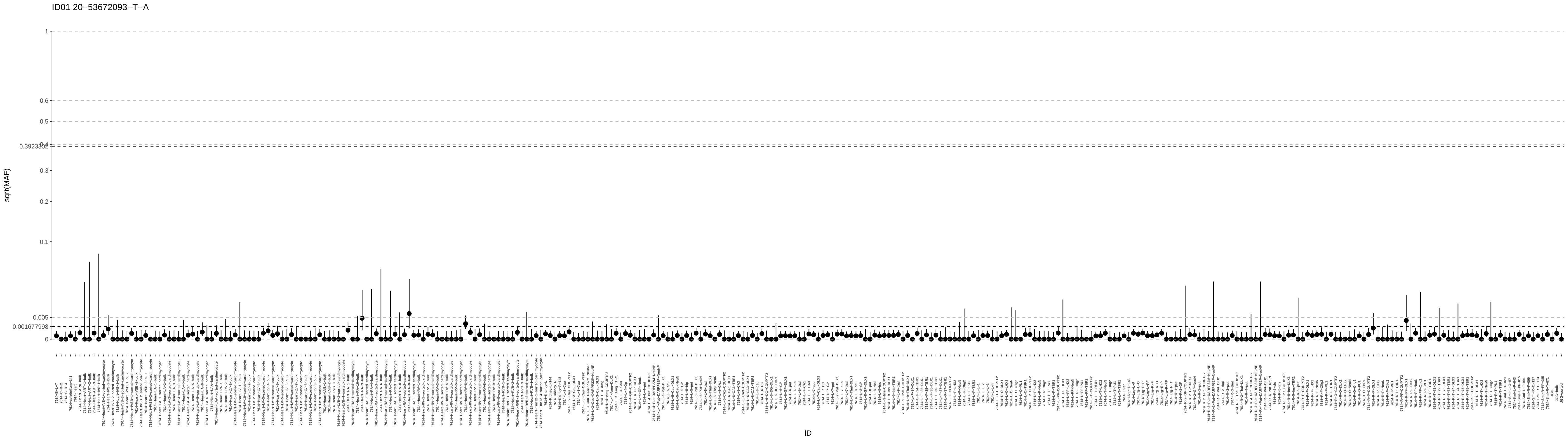
ID01 20−53672093−T−A
sqrt(MAF)
1
0.6
0.5
0.3923302
0.4
0.3
0.2
0.1
0.005
0.001677998
0
ID

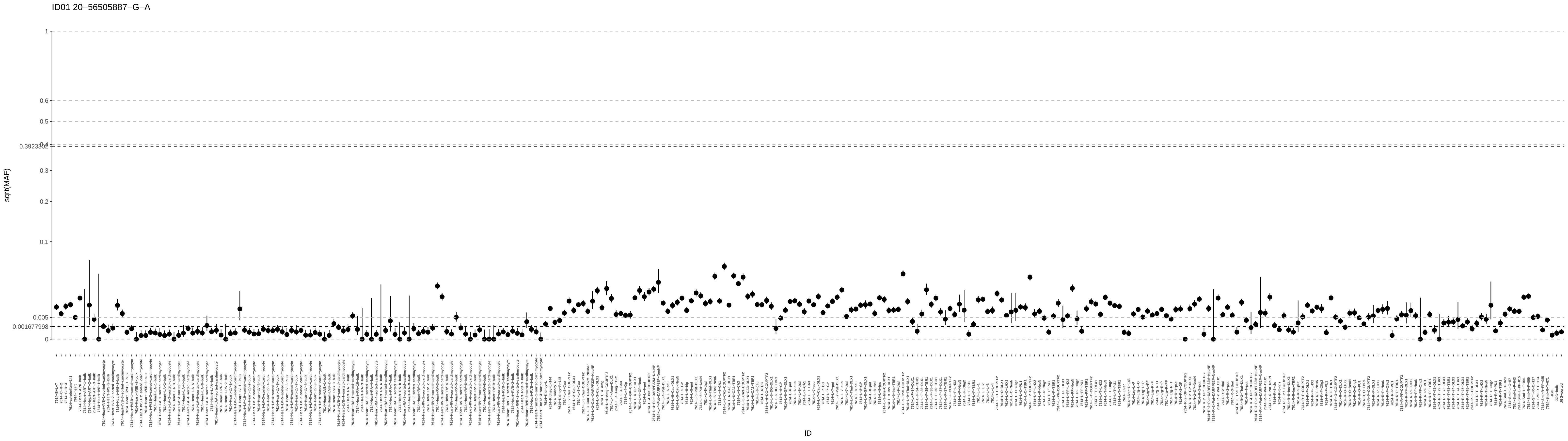
ID01 20−56505887−G−A
1
0.6
0.5
0.4
0.3923302
0.3
0.2
0.1
0.005
0.001677998
0
sqrt(MAF)
ID

ID01 20−61748407−G−A

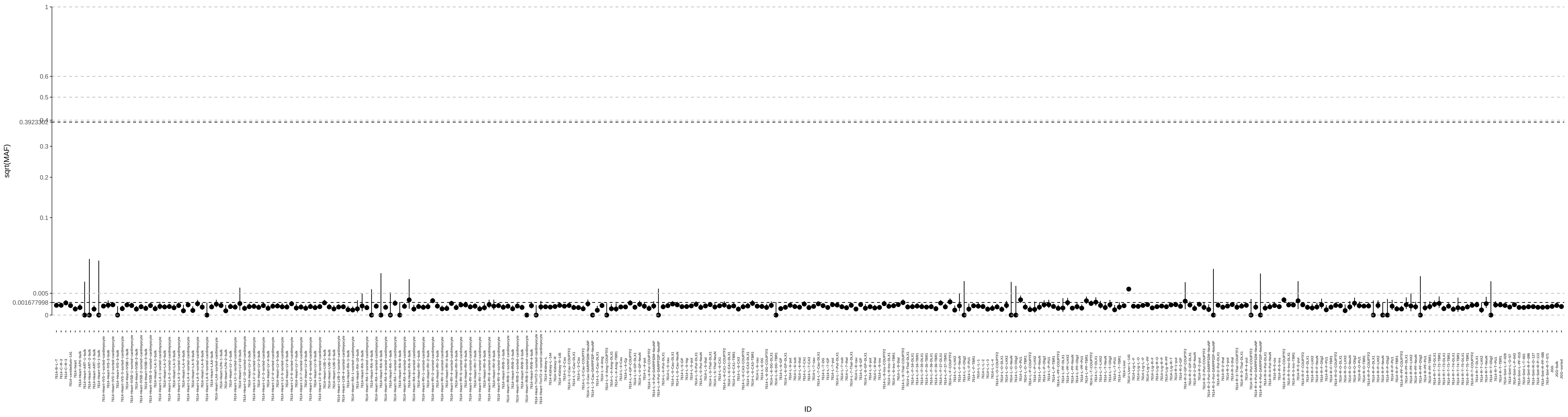

ID01 21−17683648−TTA−T

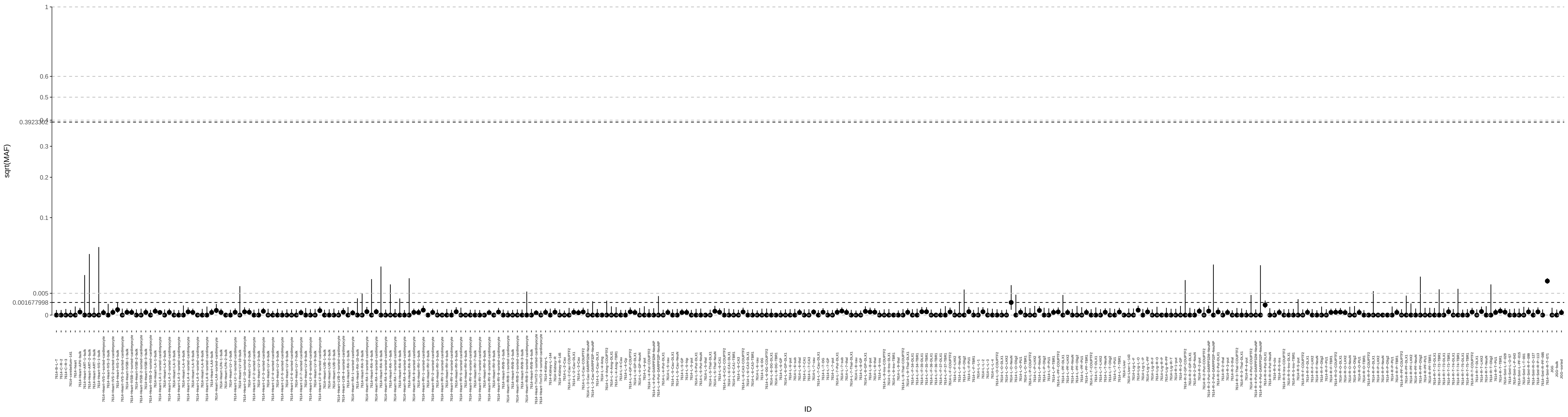

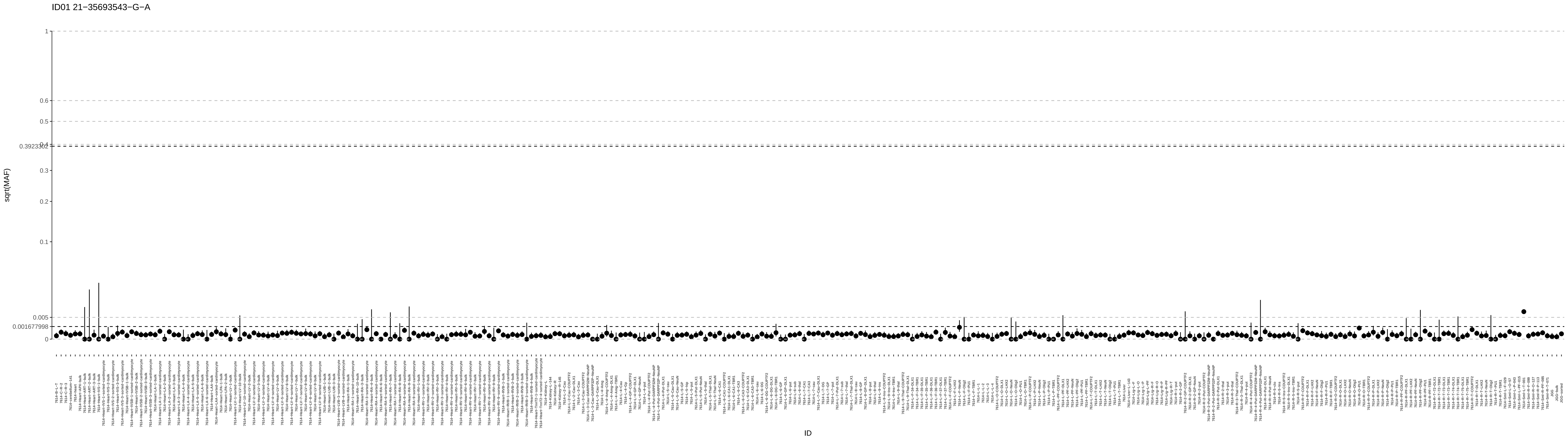
ID01 21−35693543−G−A
sqrt(MAF)
1
0.6
0.5
0.4
0.3923302
0.3
0.2
0.1
0.005
0.001677998
0
ID

ID01 21−42801460−G−A

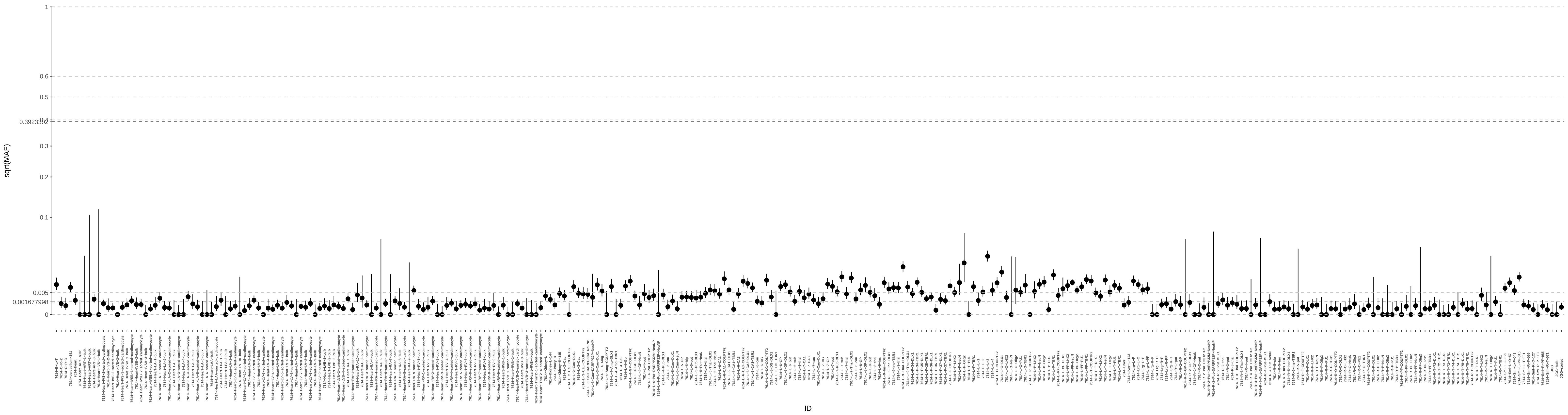

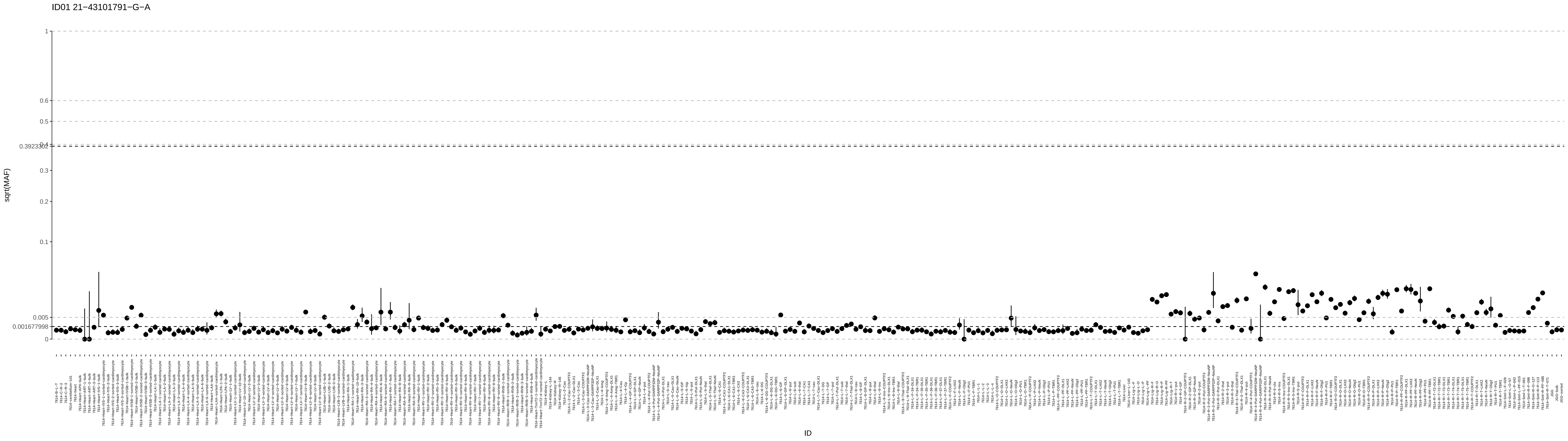
ID01 21−43101791−G−A
sqrt(MAF)
1
0.6
0.5
0.4
0.3923302
0.3
0.2
0.1
0.005
0.001677998
0
ID

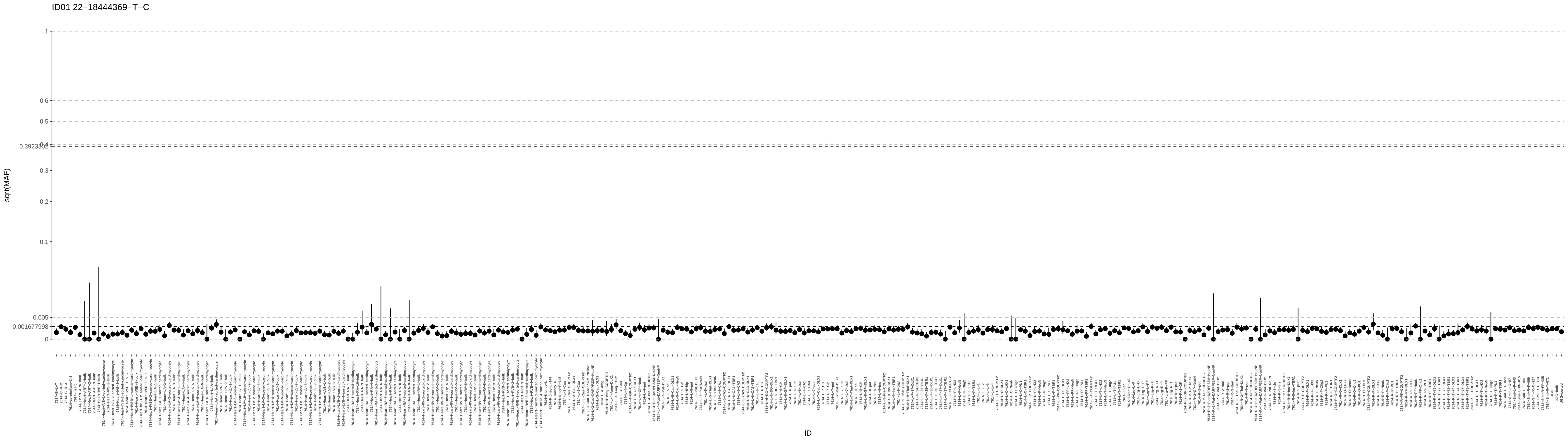
ID01 22-18444369-T-C
sqrt(MAF)
ID

ID01 22−19049947−T−TCCTC

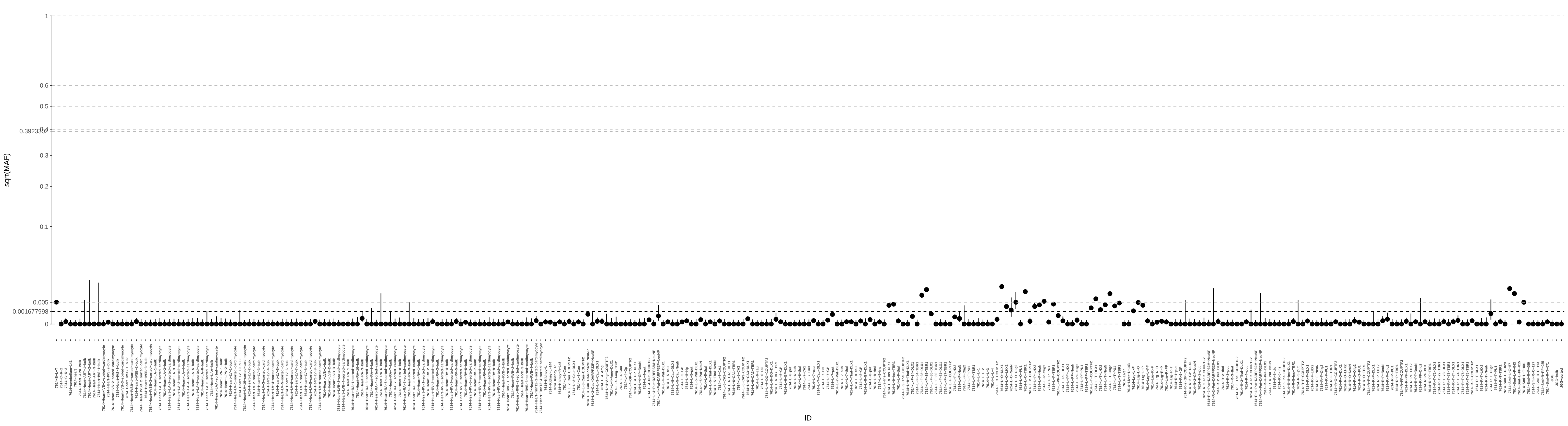

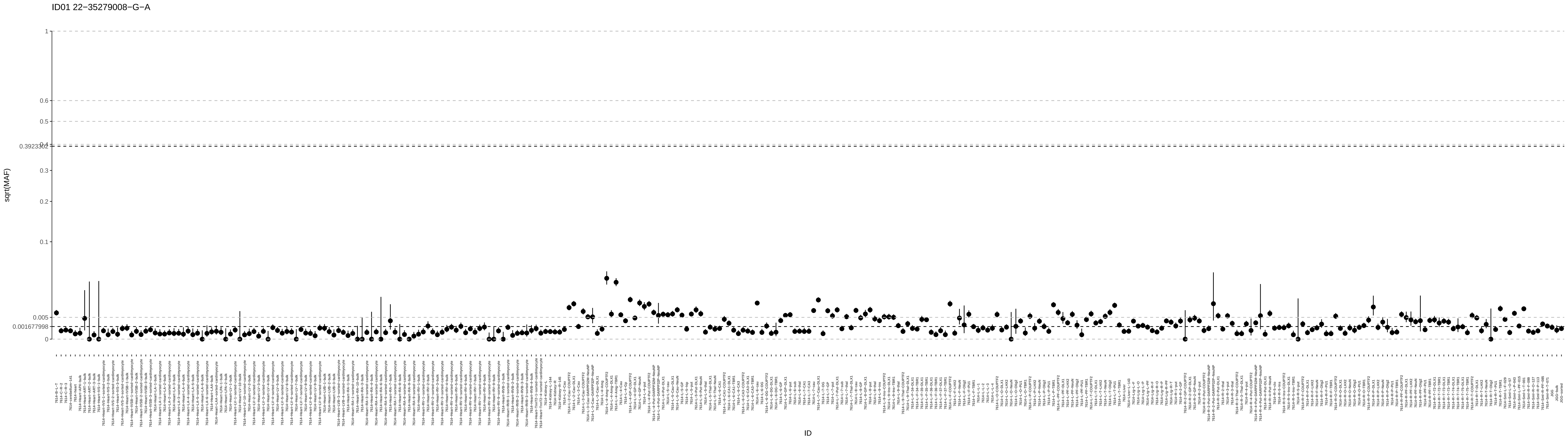
ID01 22−35279008−G−A
sqrt(MAF)
1
0.6
0.5
0.3923302
0.3
0.2
0.1
0.005
0.001677998
0
ID

ID01 22−46947808−G−A

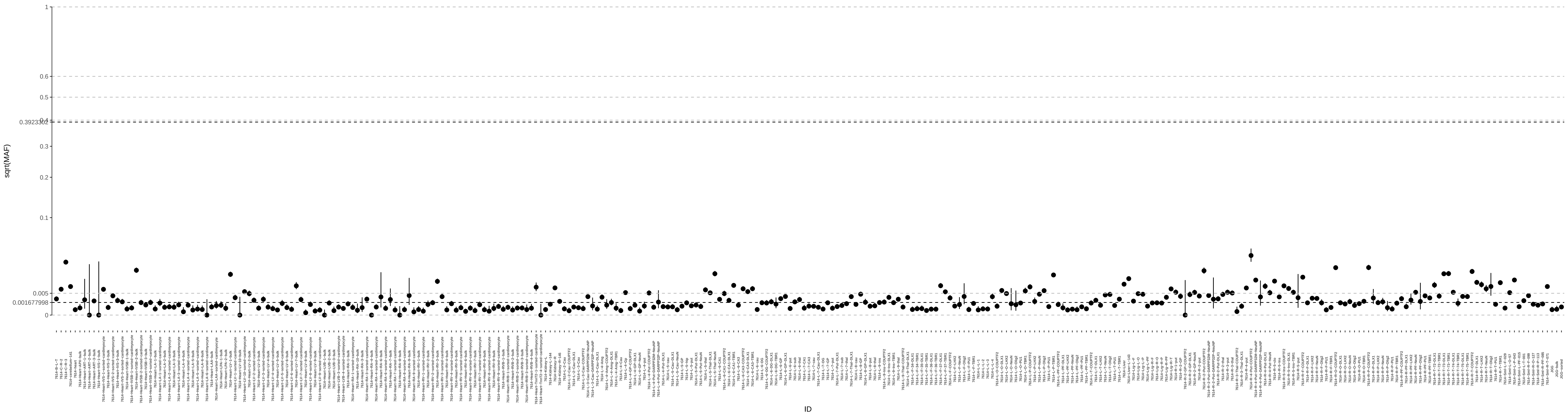

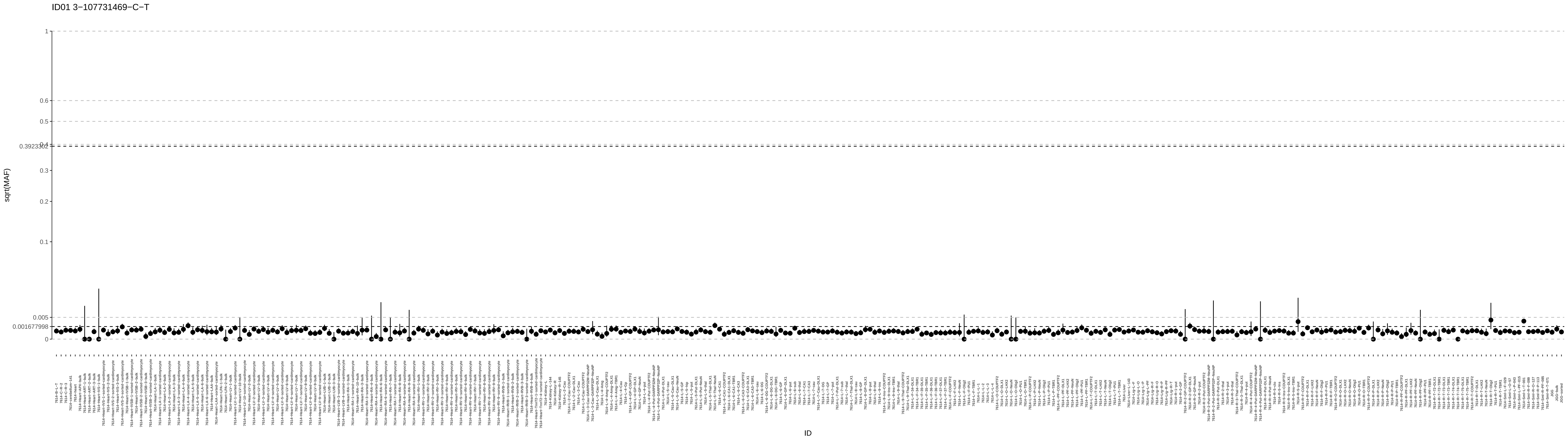
ID01 3−107731469−C−T
sqrt(MAF)
1
0.6
0.5
0.3923302
0.3
0.2
0.1
0.005
0.001677998
0
ID

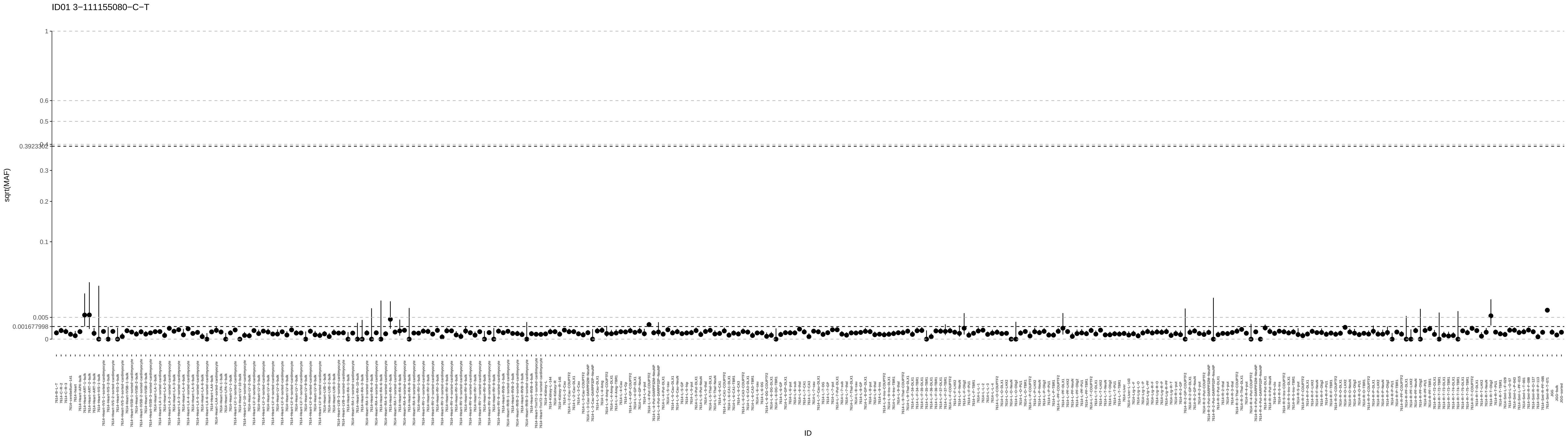
ID01 3−111155080−C−T
sqrt(MAF)
1
0.6
0.5
0.3923302
0.3
0.2
0.1
0.005
0.001677998
0
ID

ID01 3−112163628−C−T

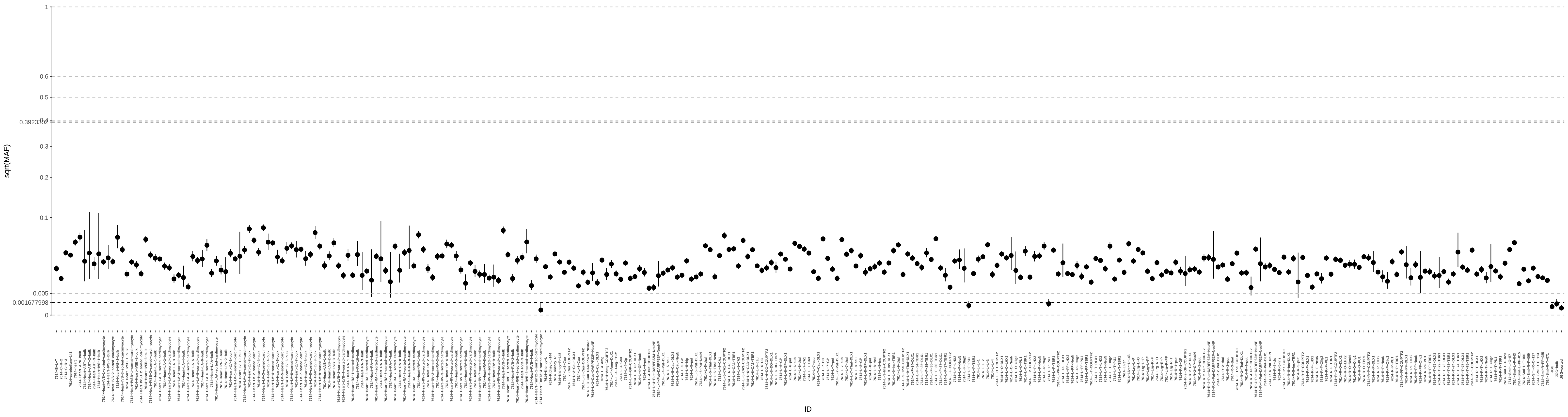

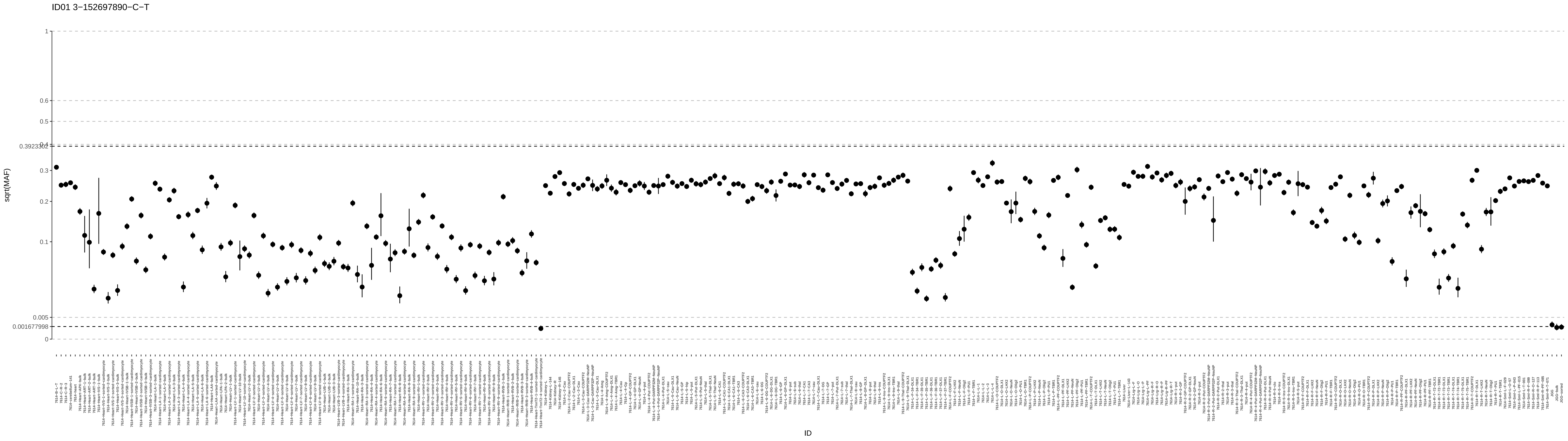
ID01 3−152697890−C−T
1
0.6
0.5
0.3923302
0.3
0.2
0.1
0.005
0.001677998
0
sqrt(MAF)
ID

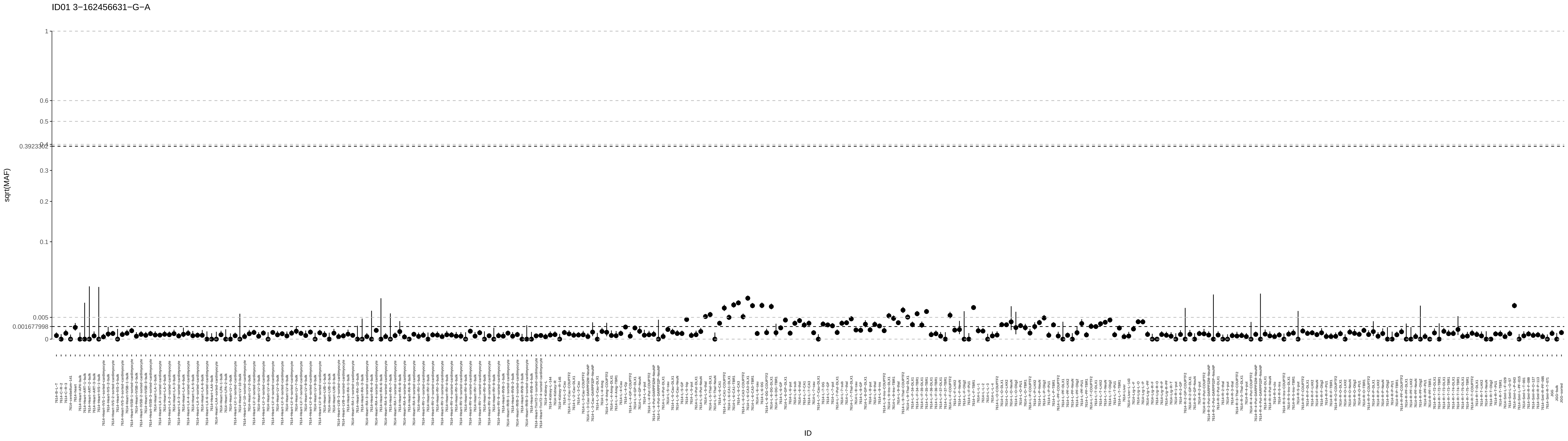

ID01 3−162456631−G−A
sqrt(MAF)
1
0.6
0.5
0.4
0.3923302
0.3
0.2
0.1
0.005
0.001677998
0
ID

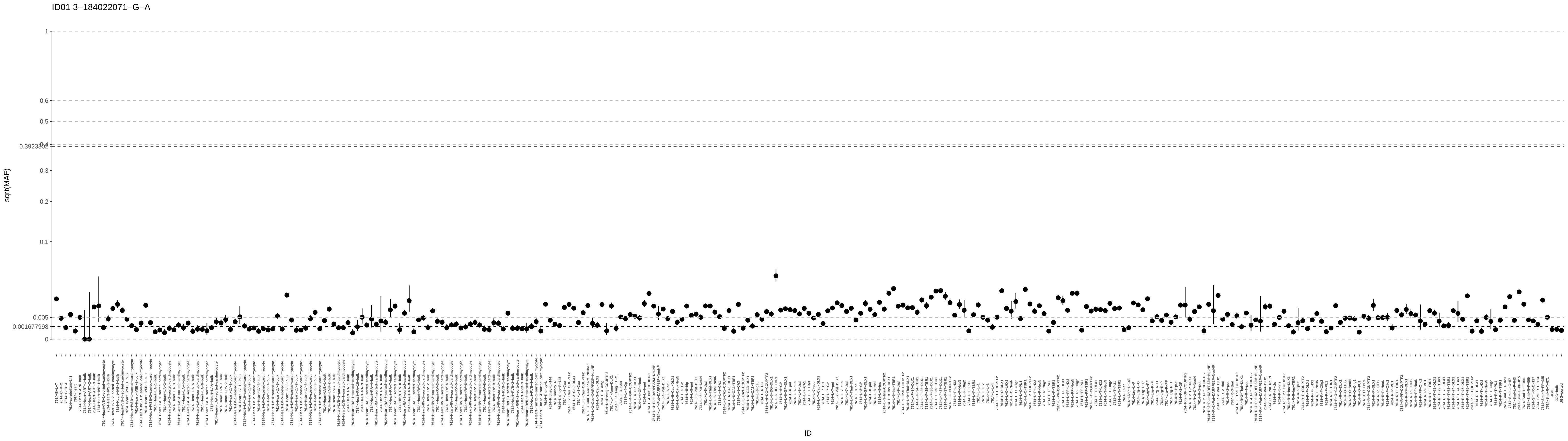
ID01 3−184022071−G−A
sqrt(MAF)
1
0.6
0.5
0.3923302
0.3
0.2
0.1
0.005
0.001677998
0
ID

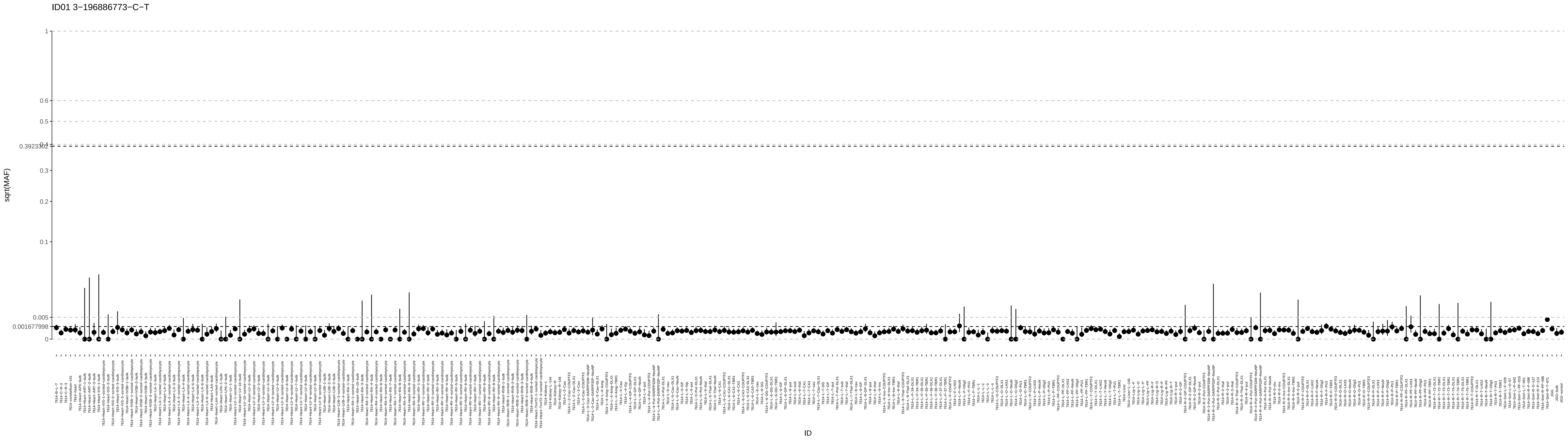

ID01 3−196886773−C−T
sqrt(MAF)
1
0.6
0.5
0.4
0.3923302
0.3
0.2
0.1
0.005
0.001677998
0
ID

ID01 3−23502134−AGAAAG−A

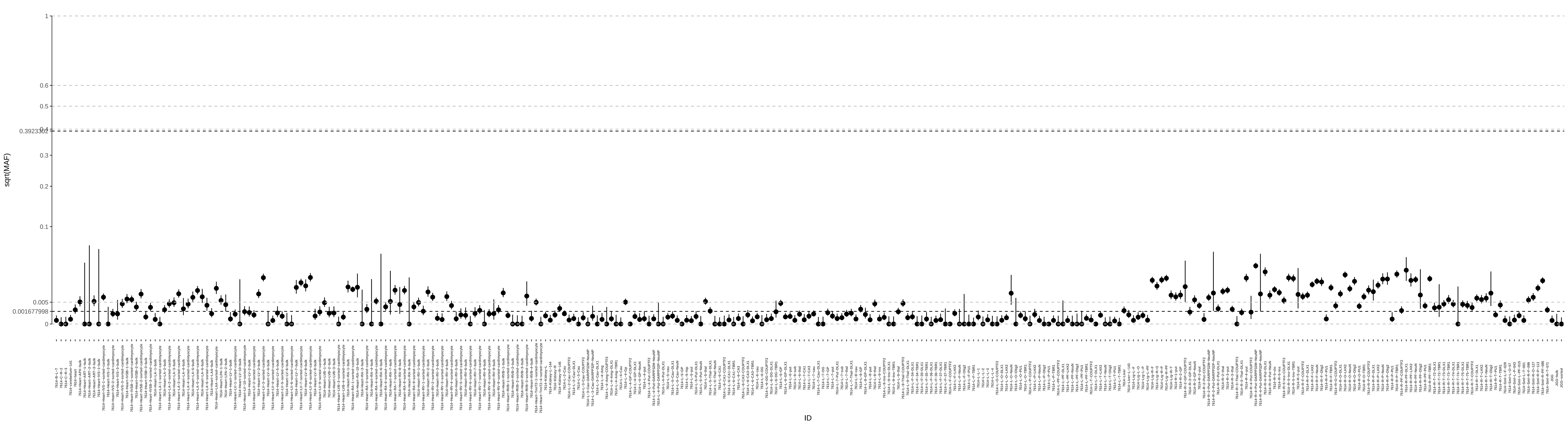

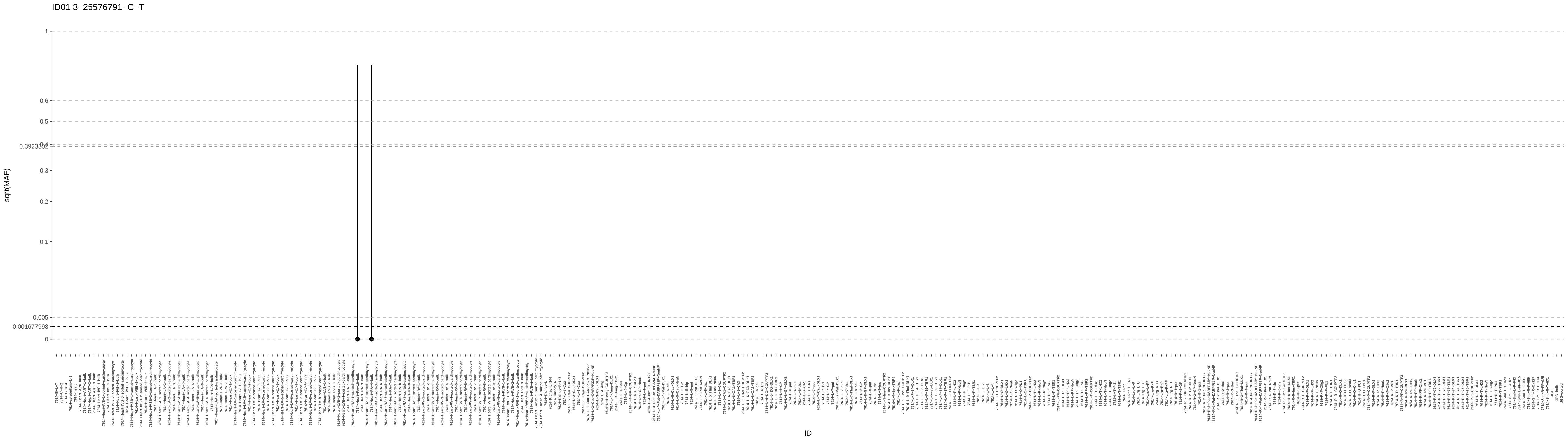
ID01 3−25576791−C−T
sqrt(MAF)
1
0.6
0.5
0.3923302
0.3
0.2
0.1
0.005
0.001677998
0
ID

ID01 3−31854915−A−C

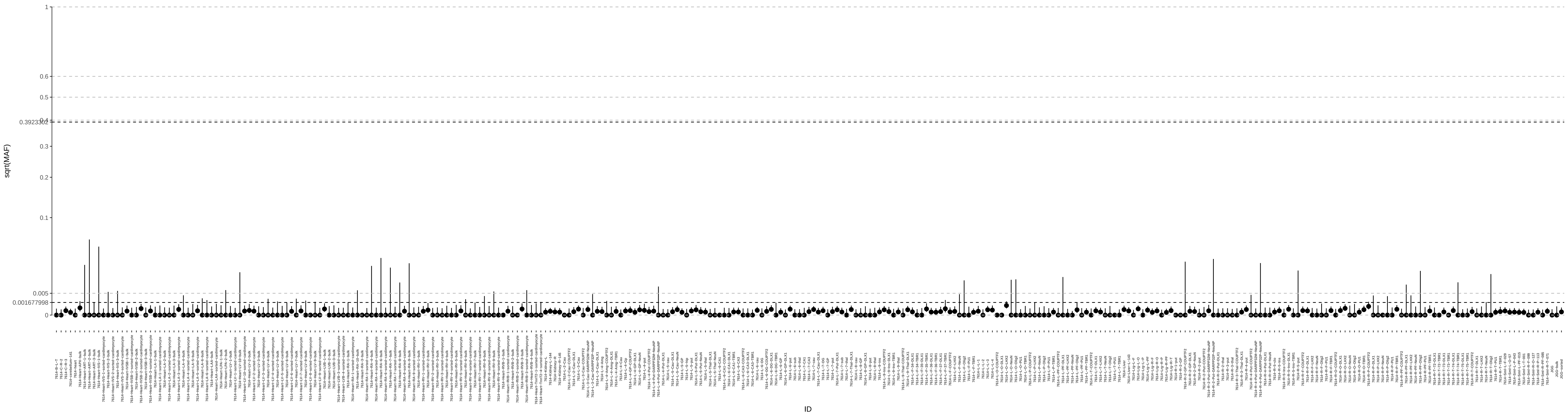

ID01 3-34583283-C-T

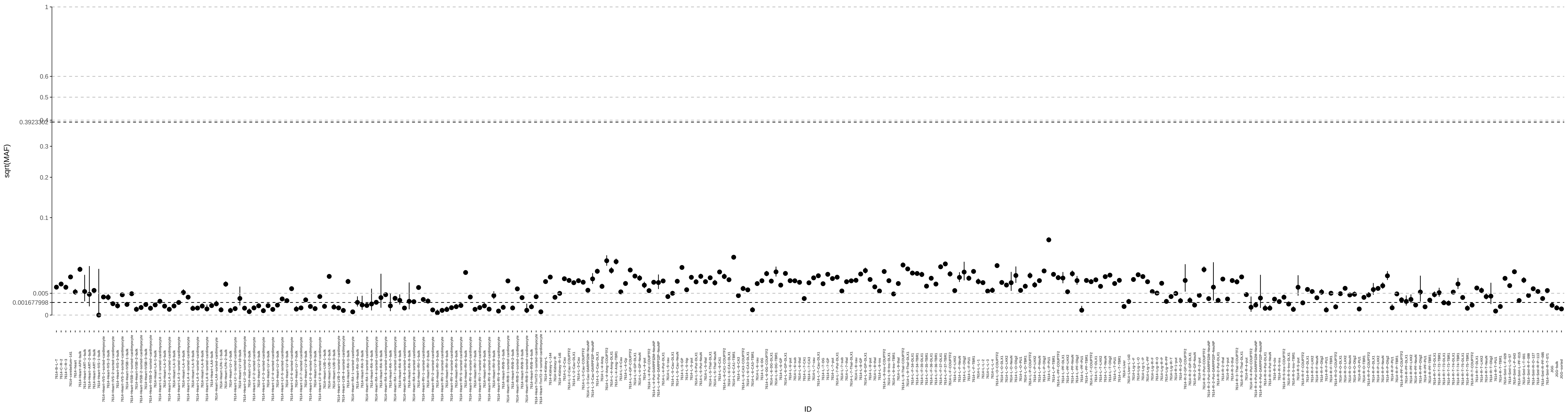

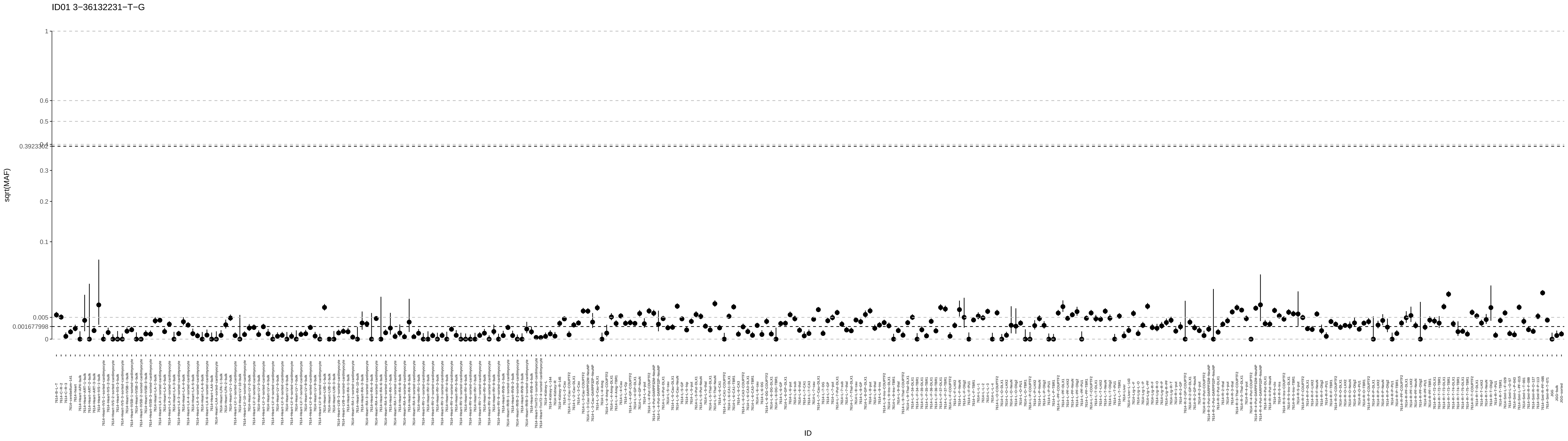

ID01 3−36132231−T−G
sqrt(MAF)
1
0.6
0.5
0.3923302
0.3
0.2
0.1
0.005
0.001677998
0
ID

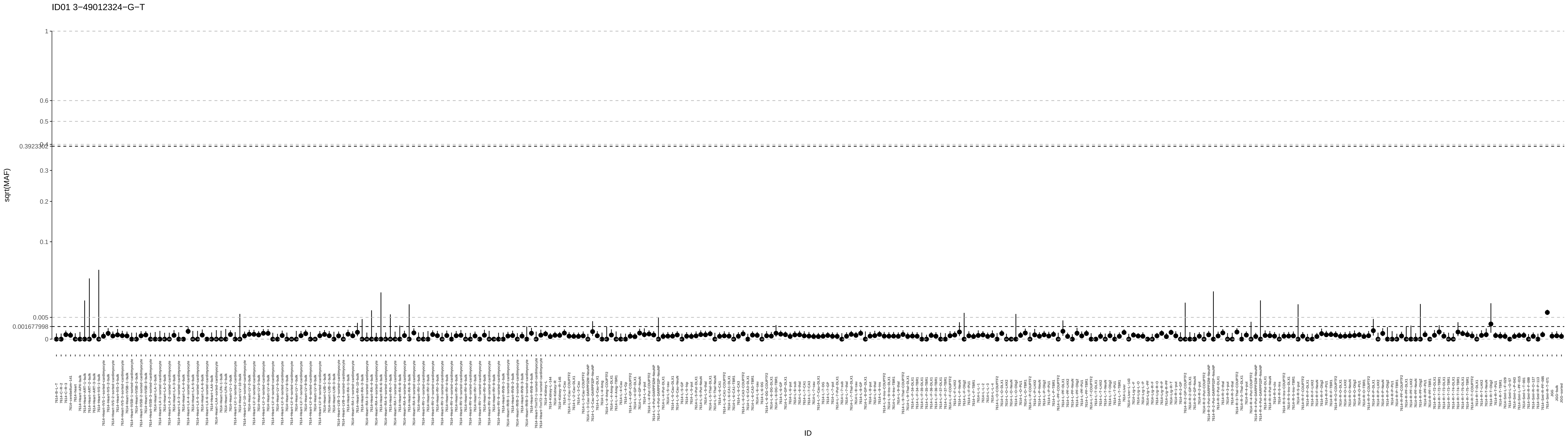
ID01 3−49012324−G−T
sqrt(MAF)
ID

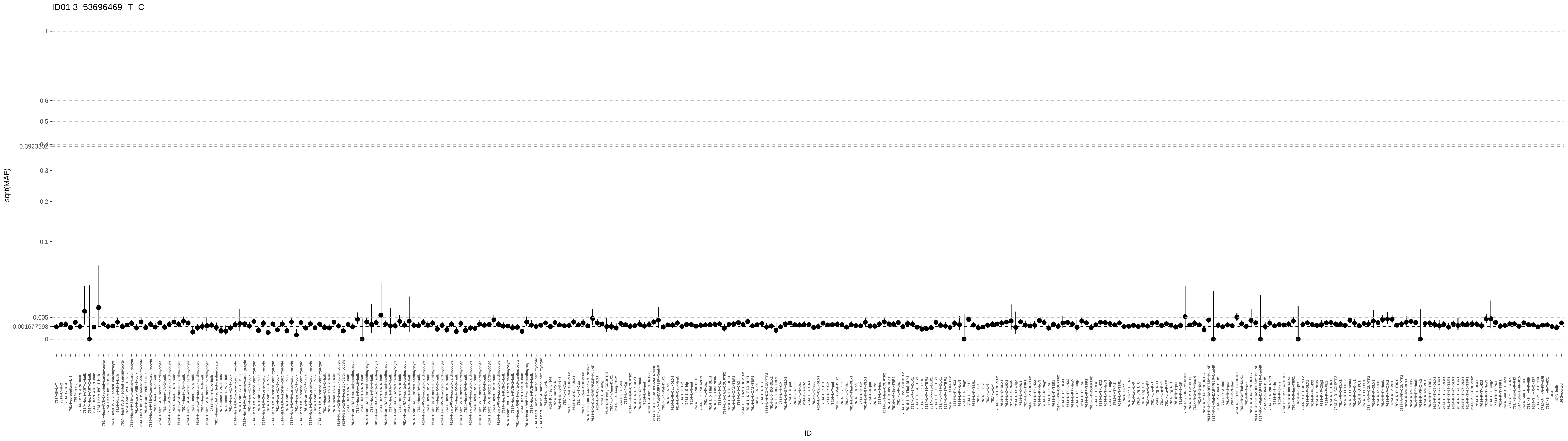
ID01 3−53696469−T−C
1
0.6
0.5
0.3923302
0.3
0.2
0.1
0.005
0.001677998
0
sqrt(MAF)
ID

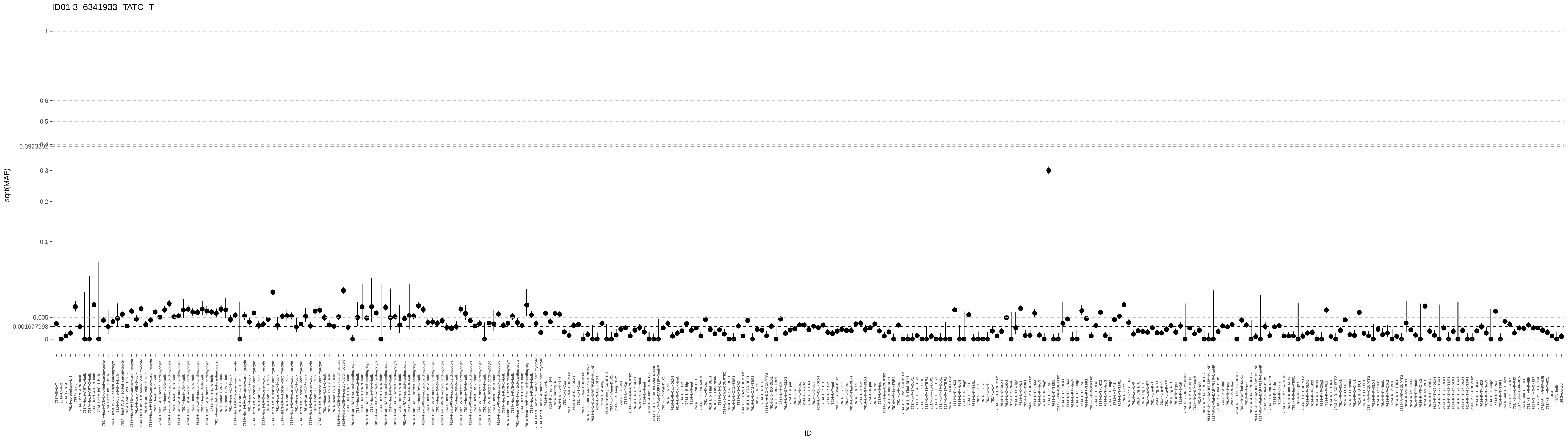

ID01 3−6341933−TATC−T
sqrt(MAF)
ID

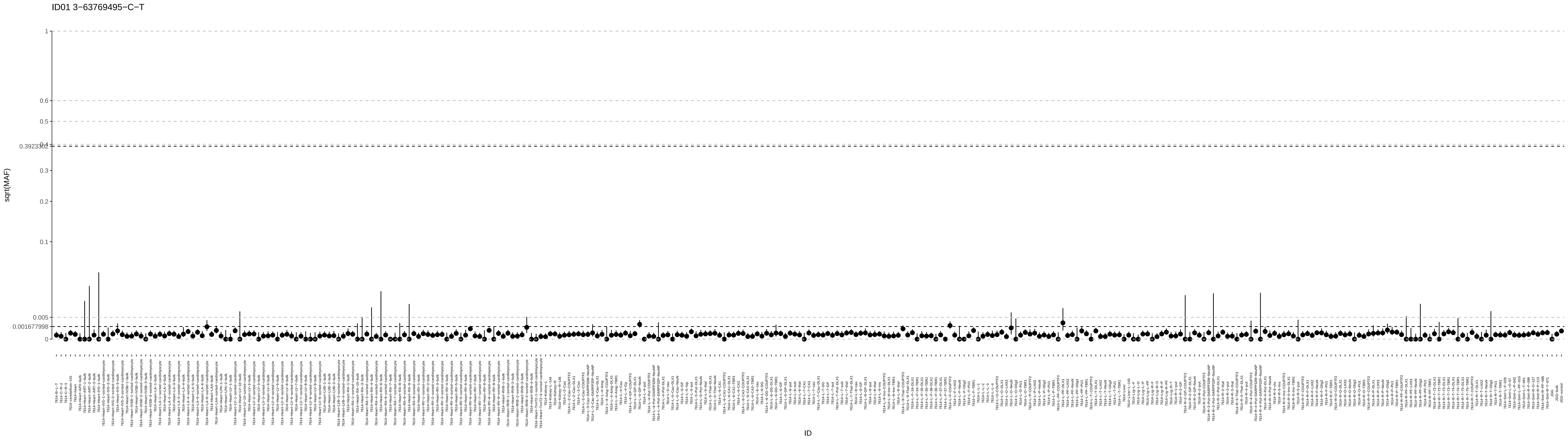

ID01 3-63769495-C-T
sqrt(MAF)
1
0.6
0.5
0.3923302
0.3
0.2
0.1
0.005
0.001677998
0
ID

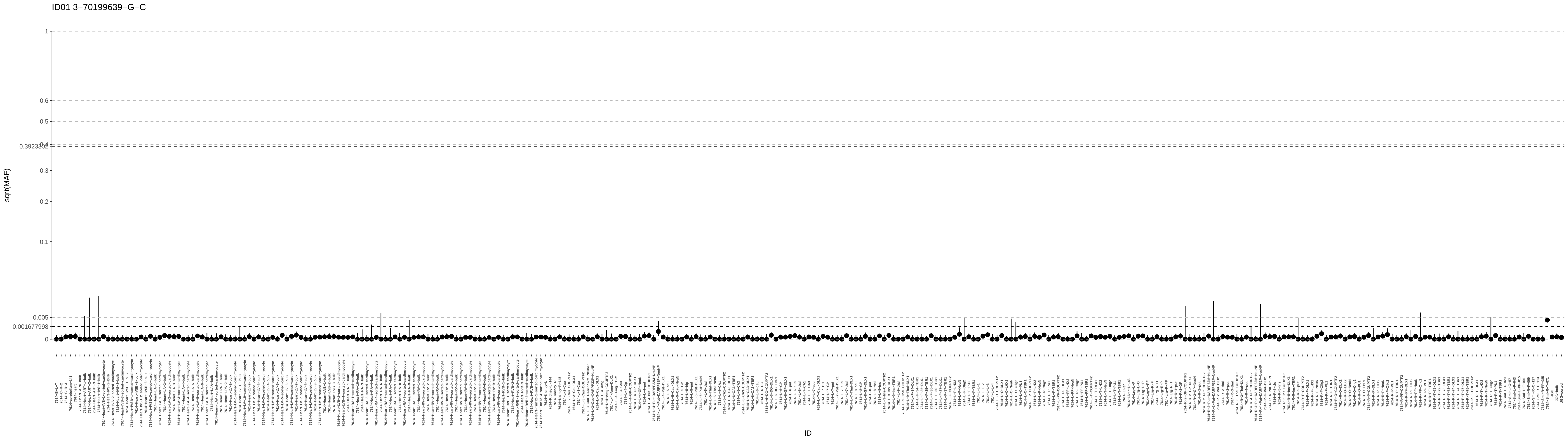
ID01 3−70199639−G−C
sqrt(MAF)
1
0.6
0.5
0.3923302
0.3
0.2
0.1
0.005
0.001677998
0
ID

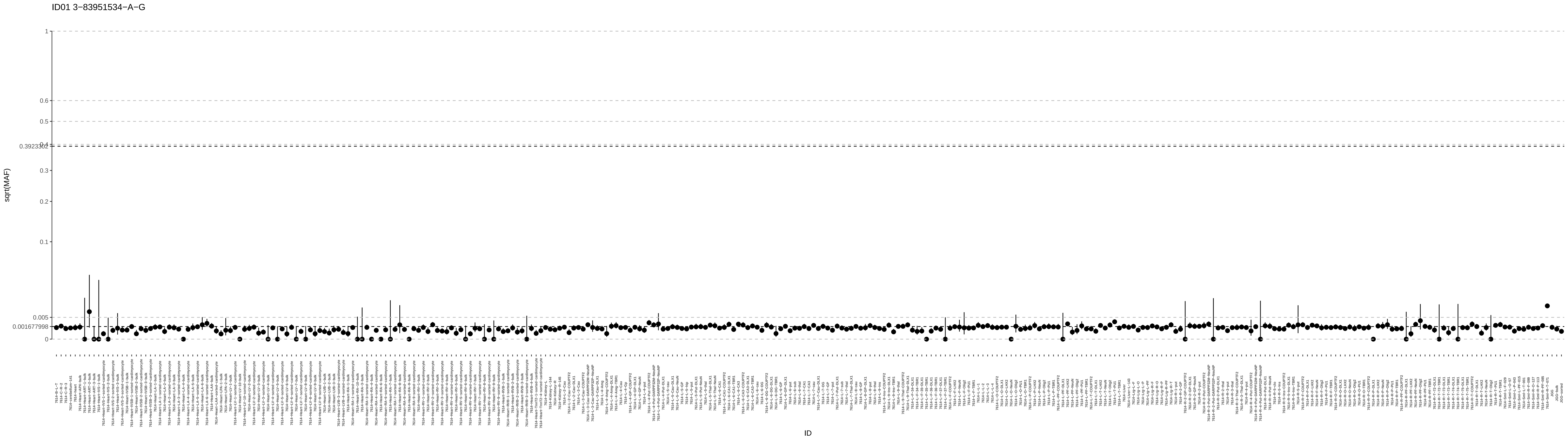
ID01 3−83951534−A−G
sqrt(MAF)
1
0.6
0.5
0.3923302
0.3
0.2
0.1
0.005
0.001677998
0
ID

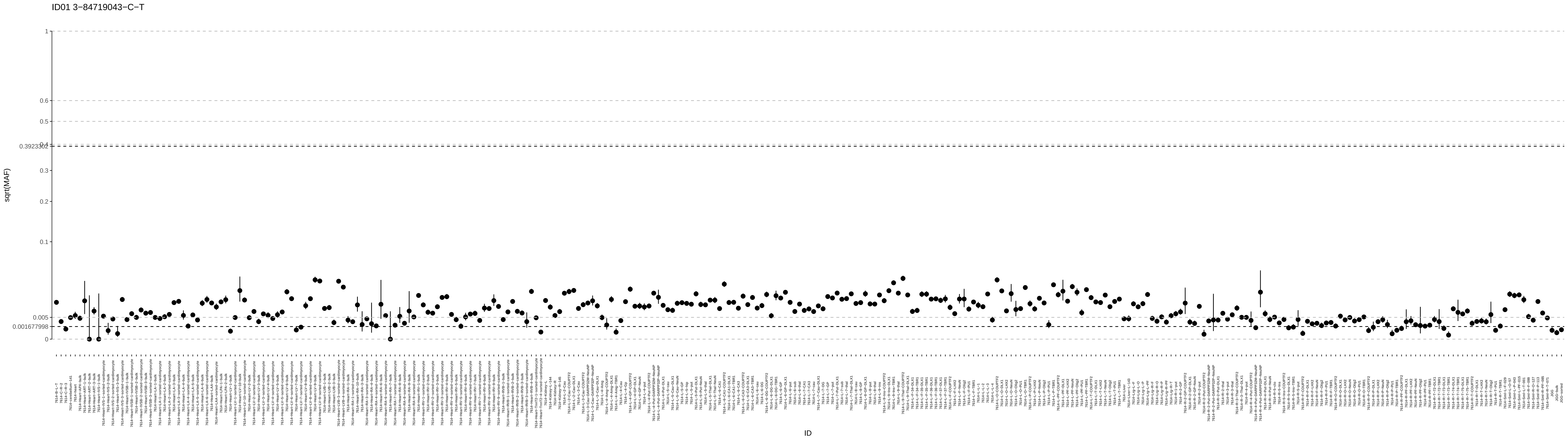
ID01 3−84719043−C−T
sqrt(MAF)
1
0.6
0.5
0.4
0.3923302
0.3
0.2
0.1
0.005
0.001677998
0
ID

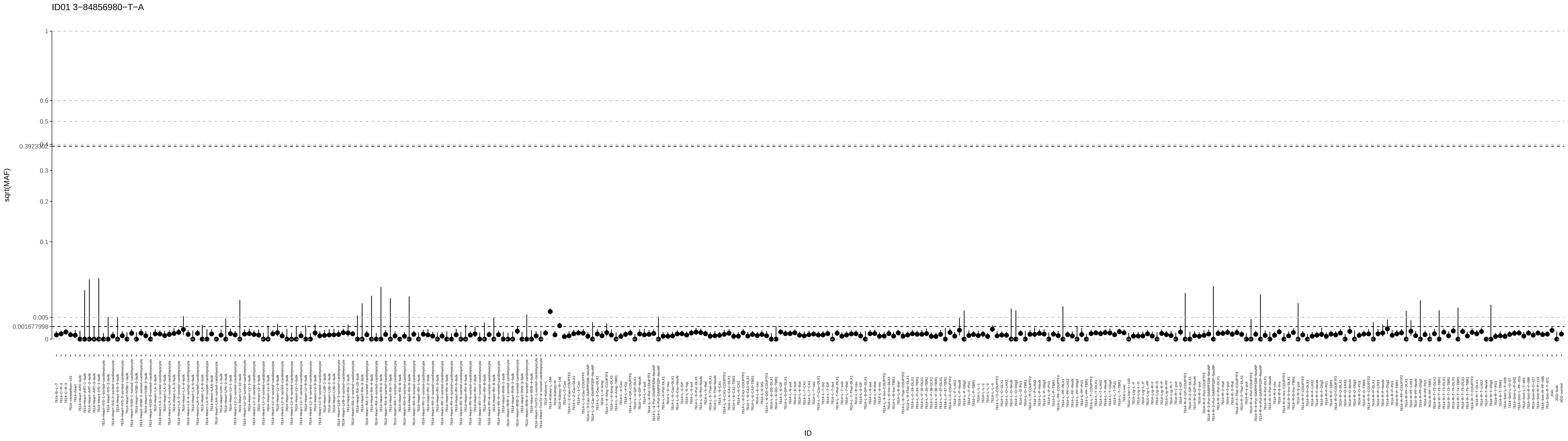
ID01 3−84856980−T−A
sqrt(MAF)
1
0.6
0.5
0.4
0.3923302
0.3
0.2
0.1
0.005
0.001677998
0
ID

ID01 3−85794290−A−C

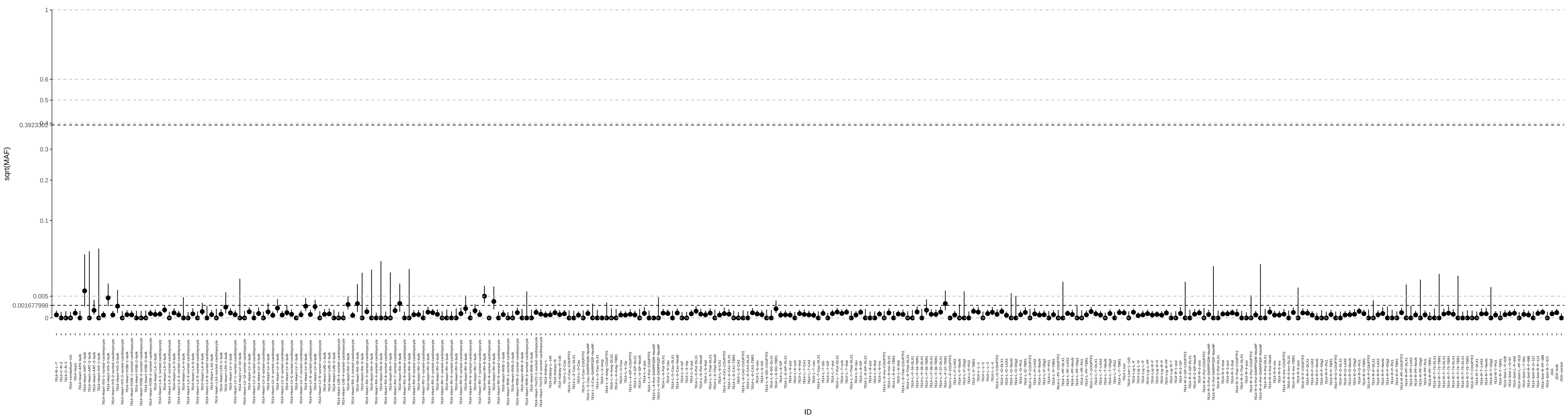

ID01 3-94668508-A-C

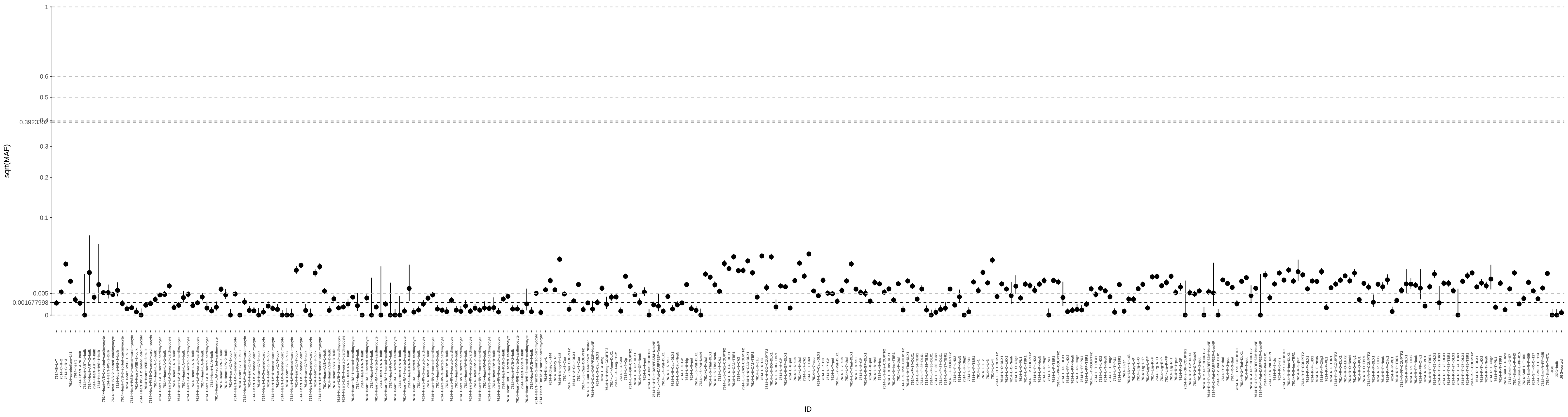

ID01 3−96090255−C−T

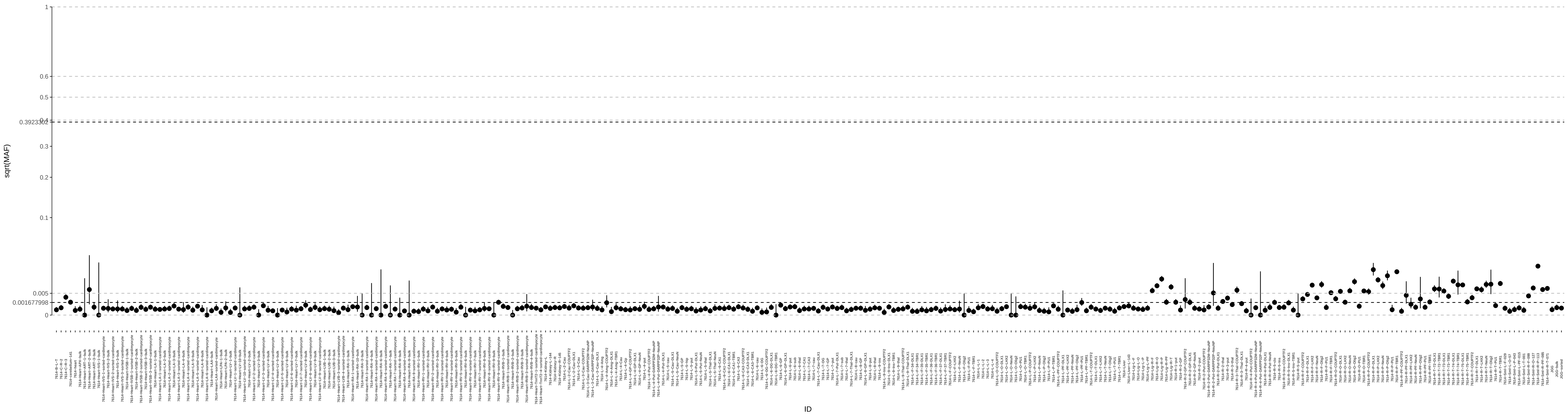

ID01 4−130387276−A−G

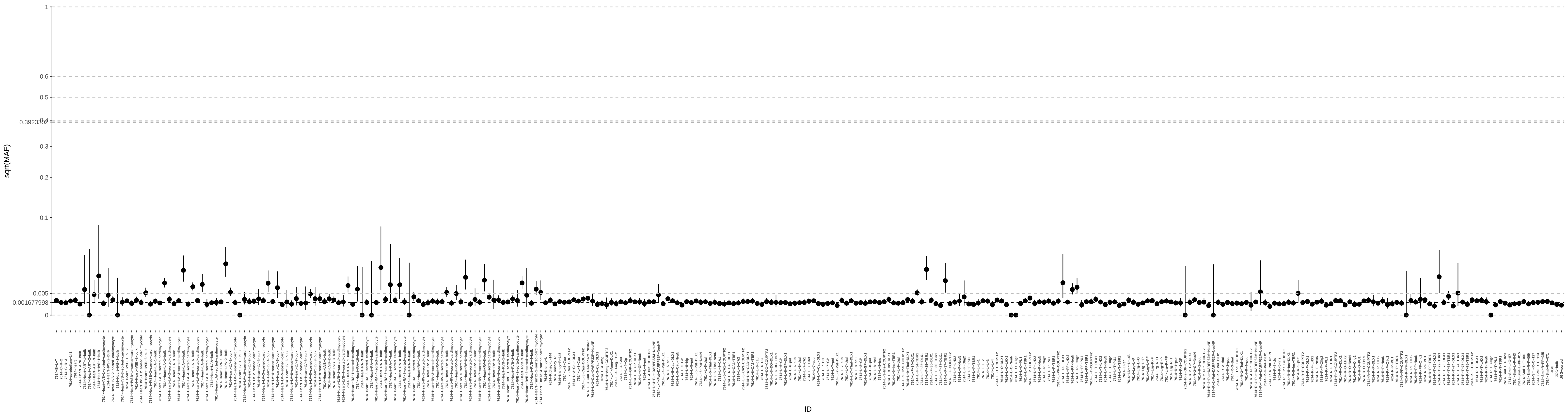

ID01 4−149096018−G−A

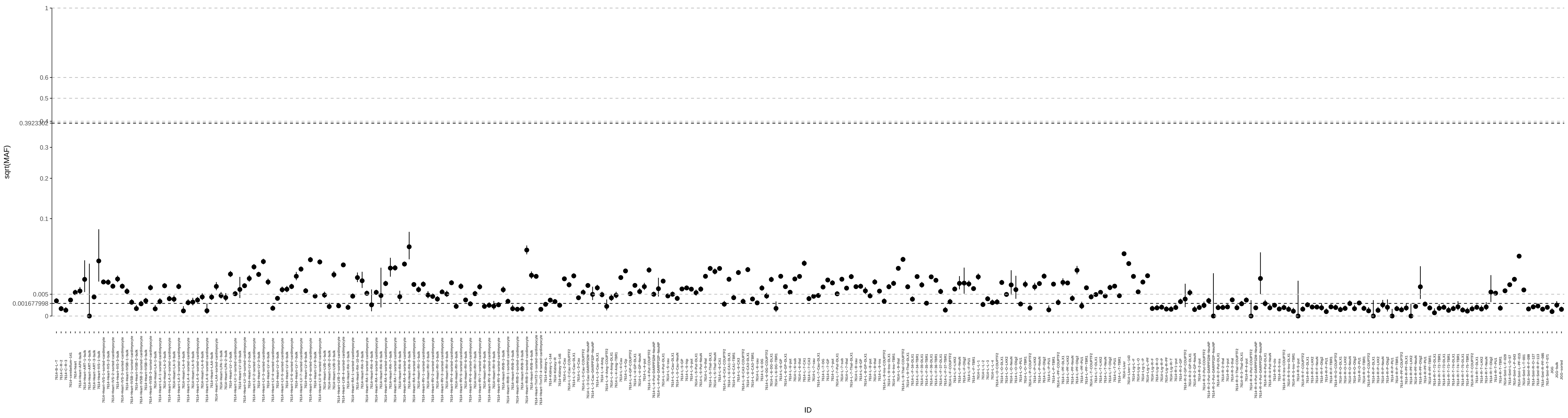

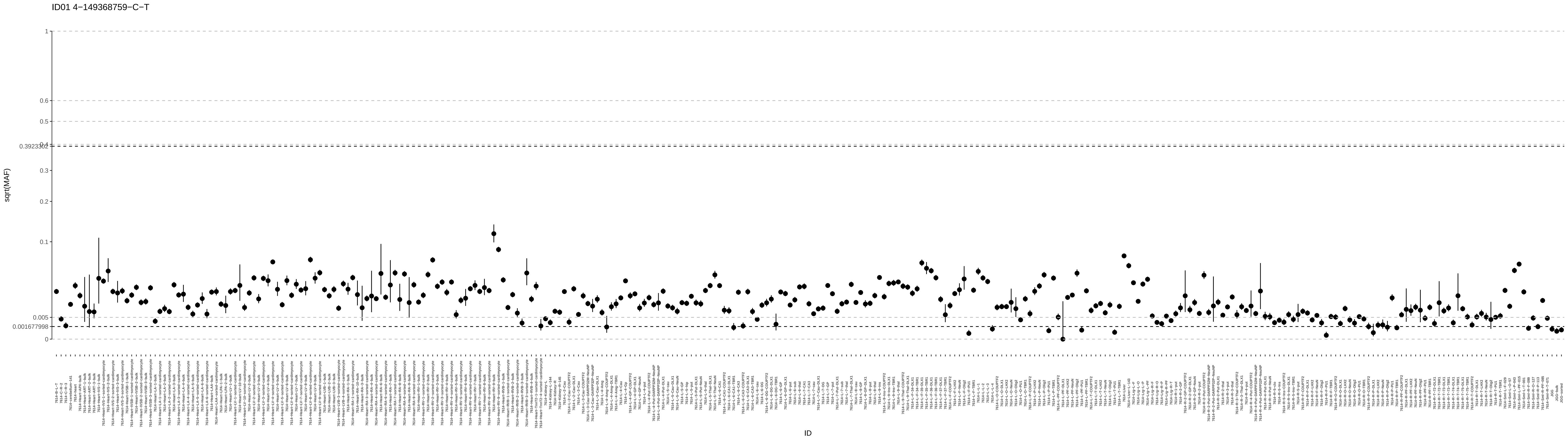

ID01 4−149368759−C−T
sqrt(MAF)
1
0.6
0.5
0.3923302
0.3
0.2
0.1
0.005
0.001677998
0
ID

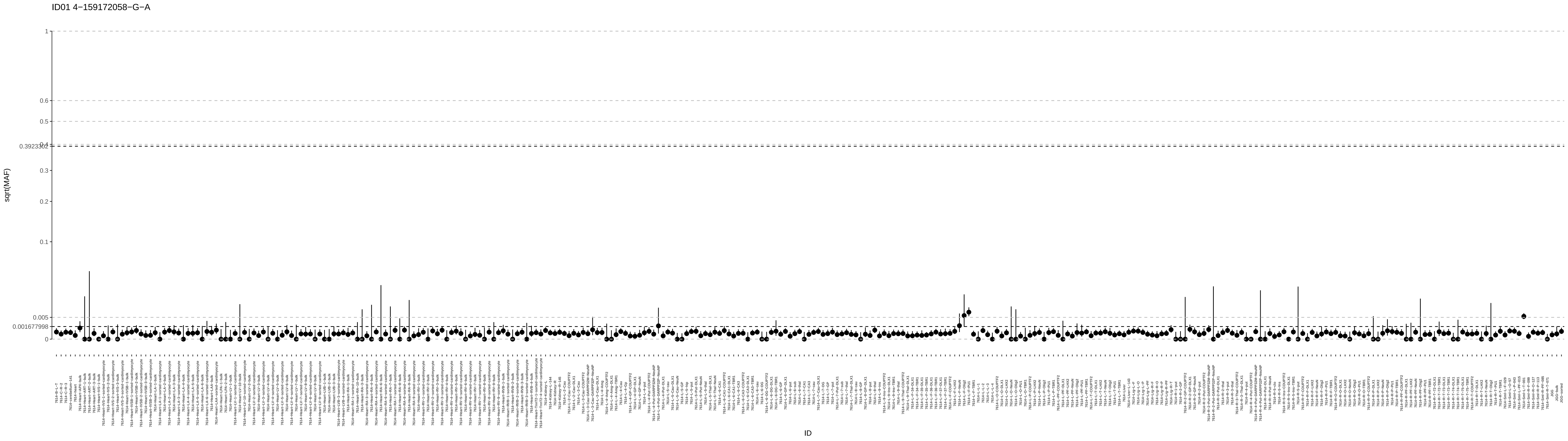

ID01 4−159172058−G−A
sqrt(MAF)
1
0.6
0.5
0.4
0.3923302
0.3
0.2
0.1
0.005
0.001677998
0
ID

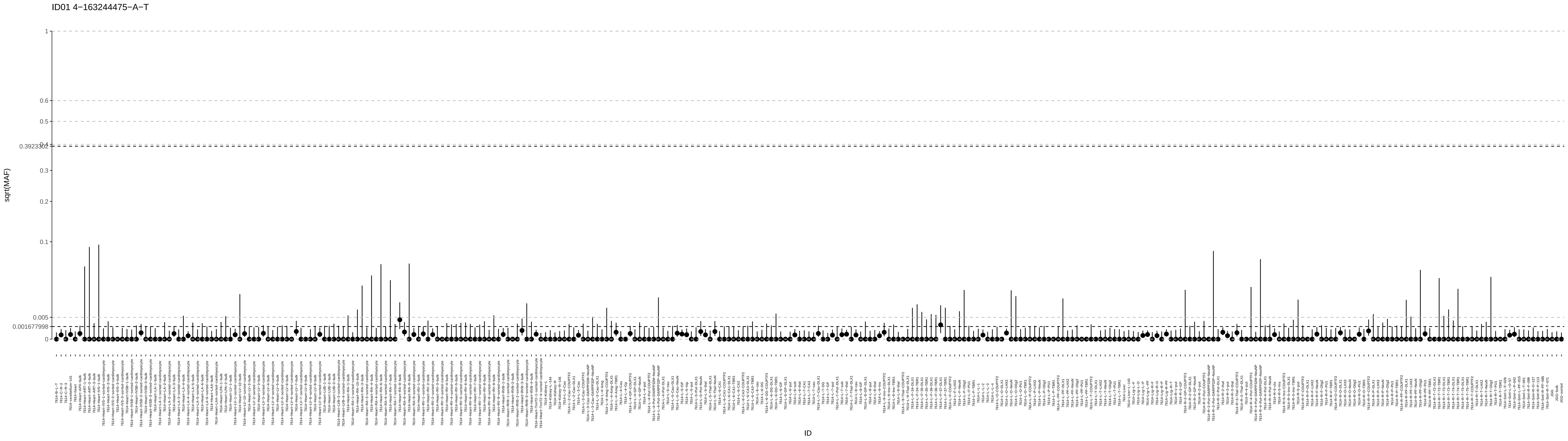

ID01 4−163244475−A−T
sqrt(MAF)
1
0.6
0.5
0.3923302
0.3
0.2
0.1
0.005
0.001677998
0
ID

ID01 4−16800988−T−A

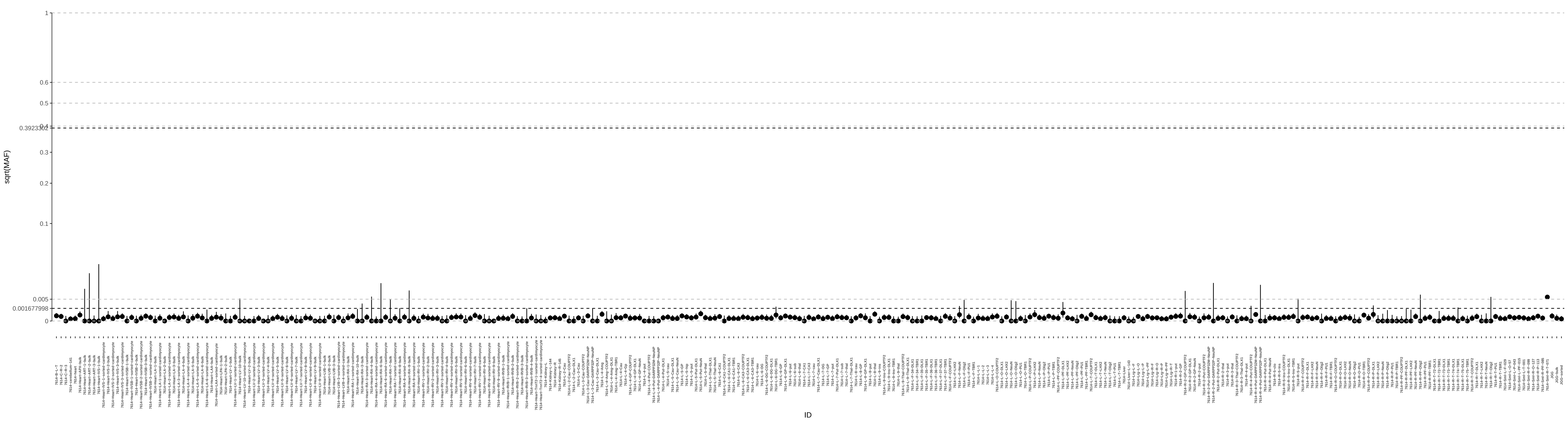

ID01 4−176659497−G−A

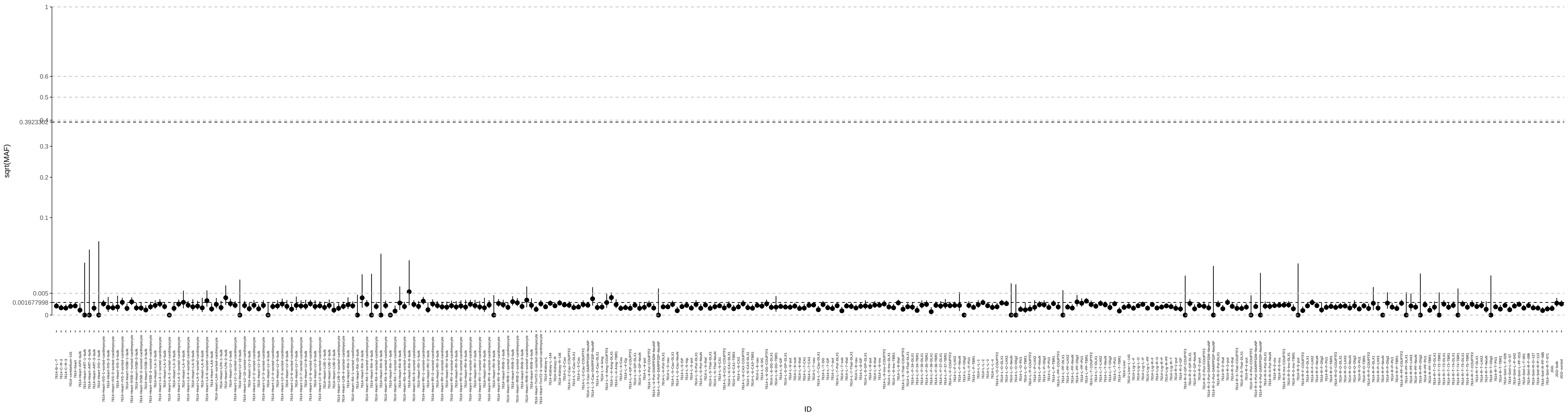

ID01 4−181698552−T−C

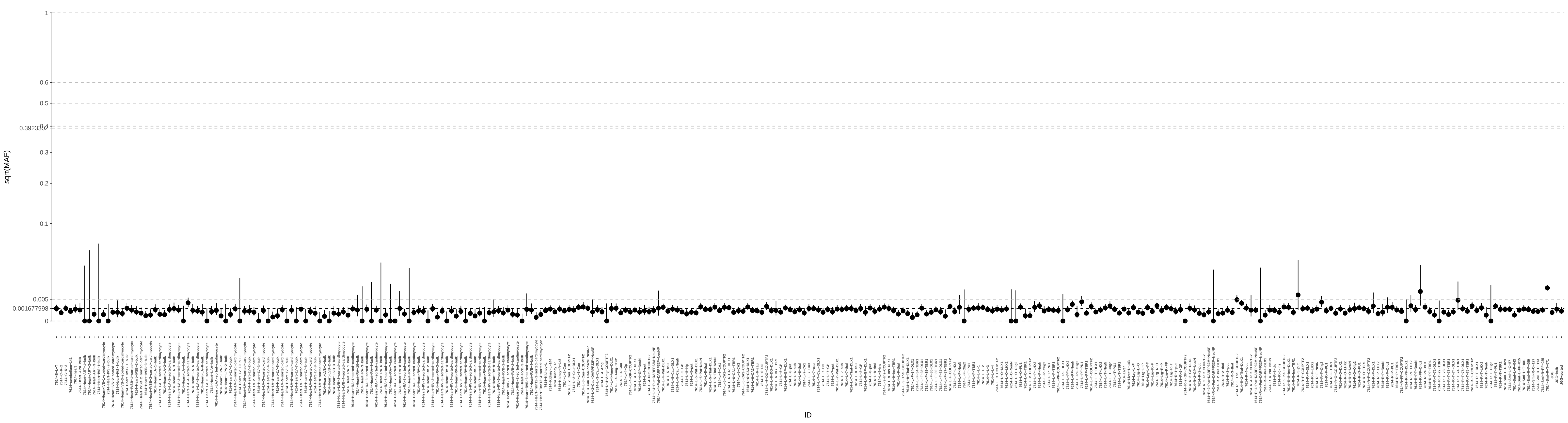

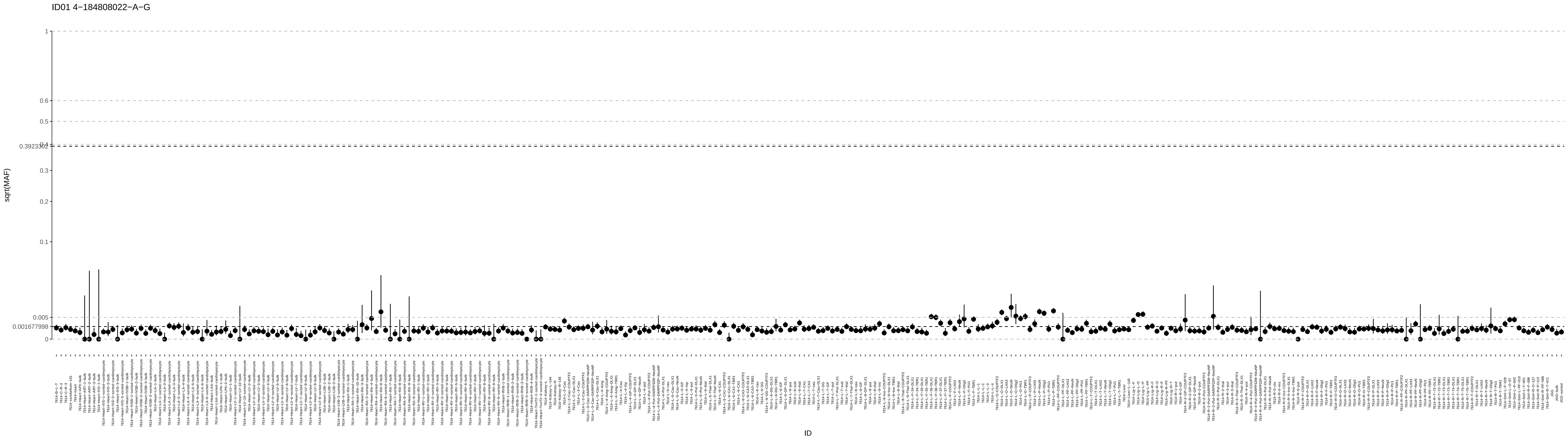
ID01 4−184808022−A−G
sqrt(MAF)
1
0.6
0.5
0.3923302
0.3
0.2
0.1
0.005
0.001677998
0
ID

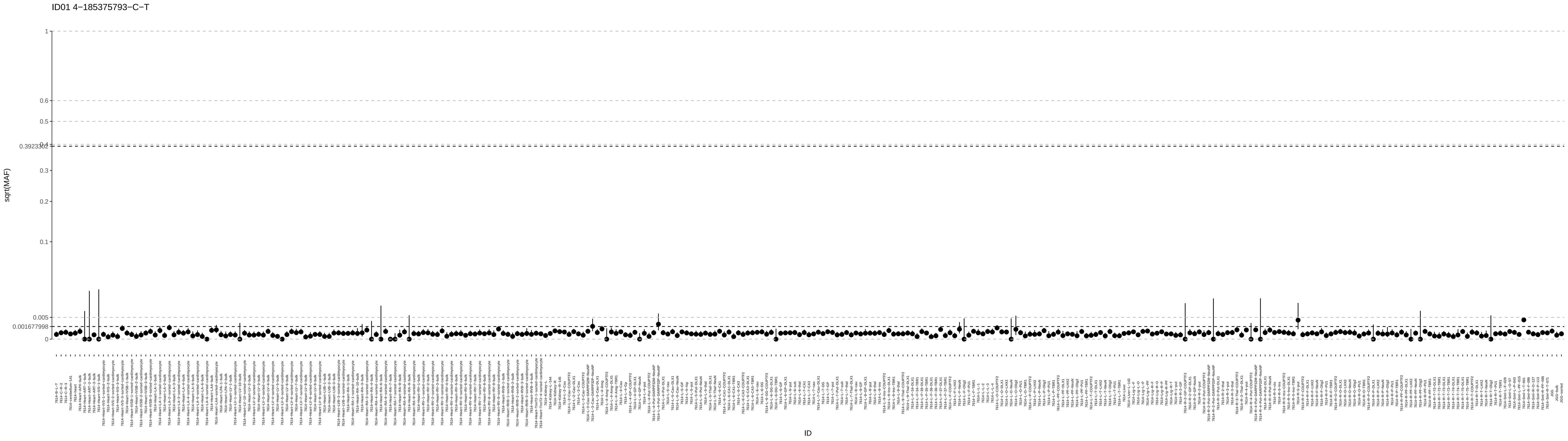
ID01 4−185375793−C−T
sqrt(MAF)
1
0.6
0.5
0.3923302
0.3
0.2
0.1
0.005
0.001677998
0
ID

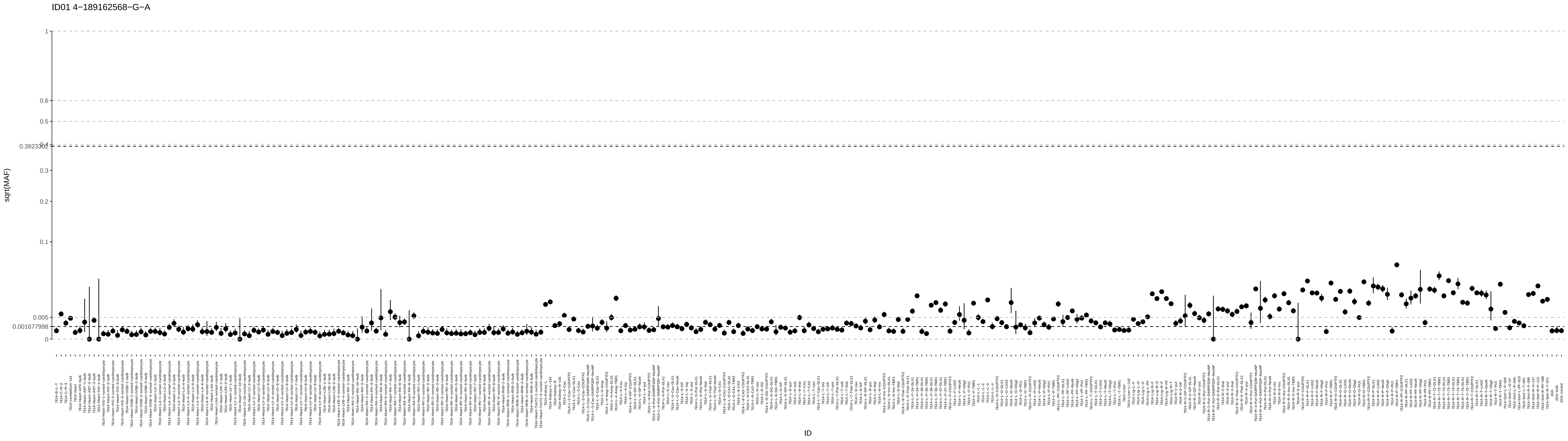

ID01 4−189162568−G−A
sqrt(MAF)
1
0.6
0.5
0.3923302
0.3
0.2
0.1
0.005
0.001677998
0
ID

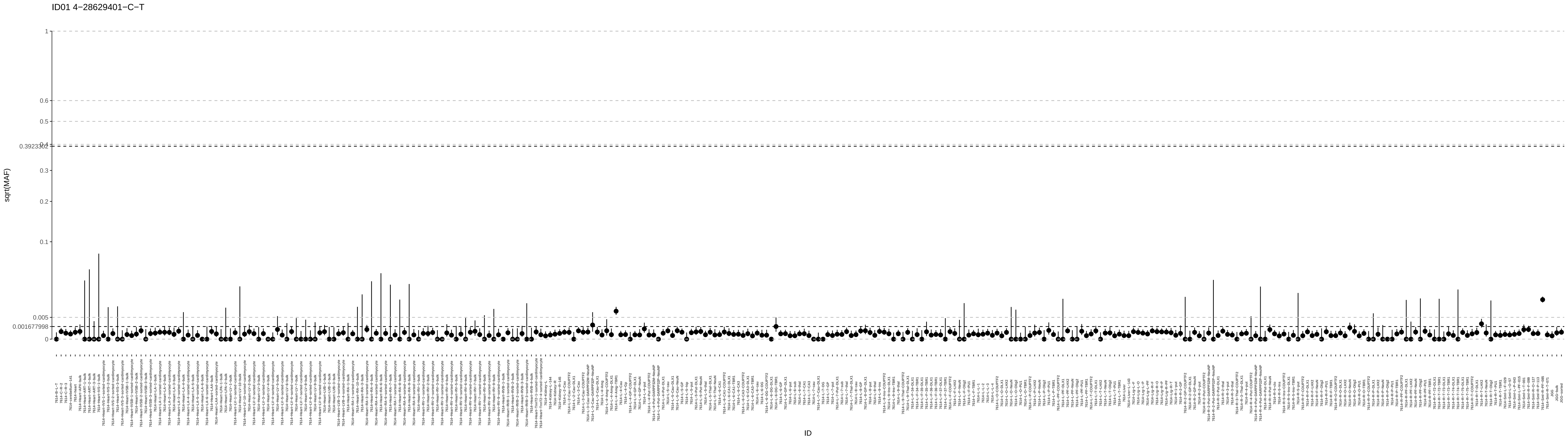
ID01 4−28629401−C−T
sqrt(MAF)
1
0.6
0.5
0.3923302
0.3
0.2
0.1
0.005
0.001677998
0
ID

ID01 4-33192479-C-A

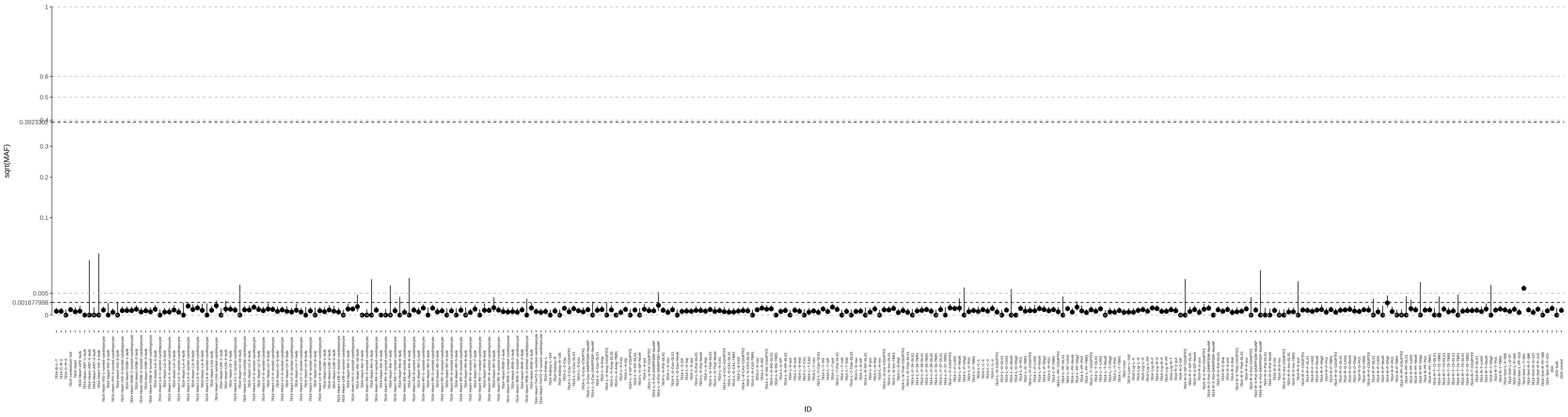

ID01 4−55310609−G−A

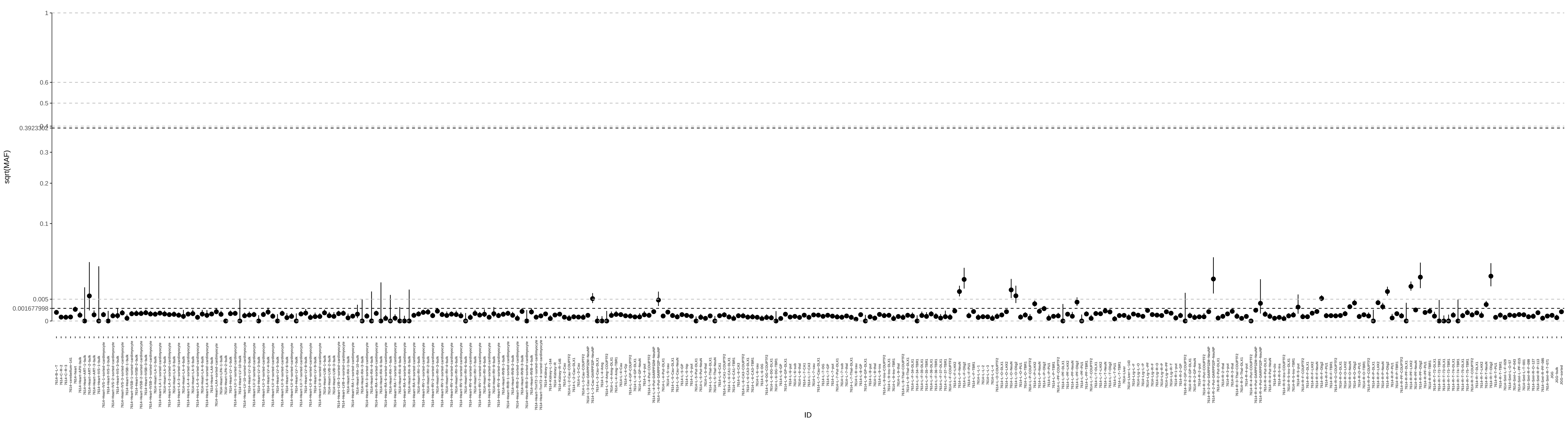

ID01 4−55312148−T−C

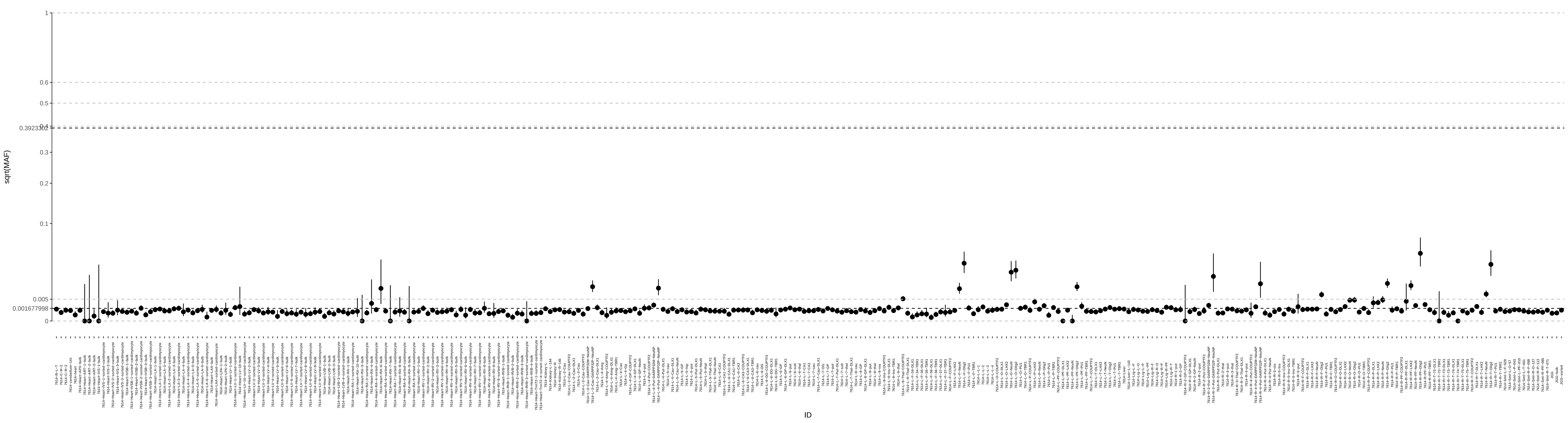

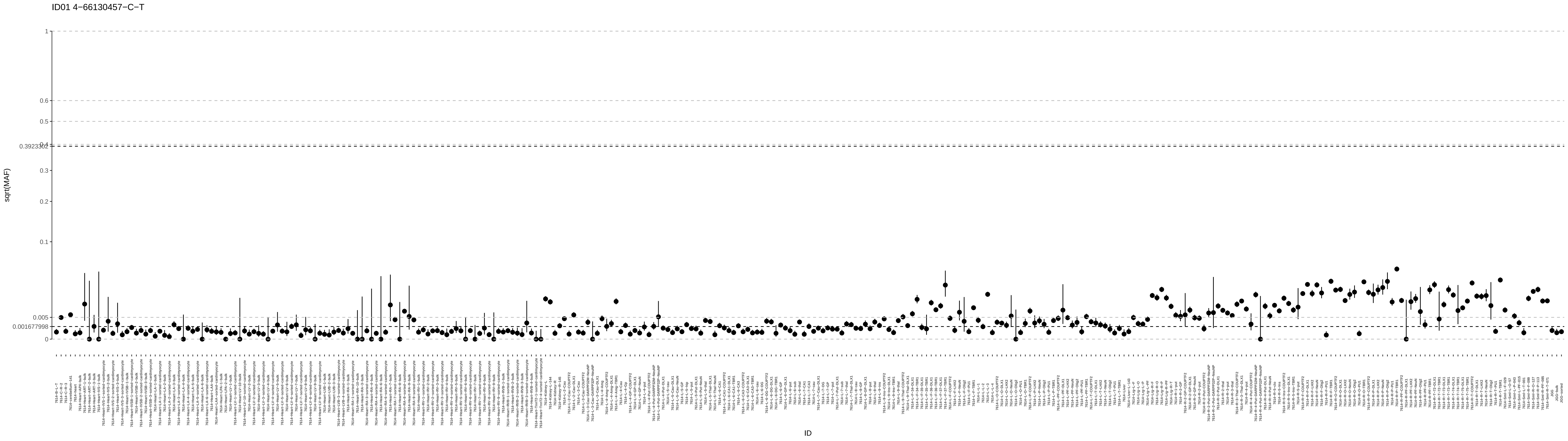
ID01 4−66130457−C−T
sqrt(MAF)
1
0.6
0.5
0.3923302
0.3
0.2
0.1
0.005
0.001677998
0
ID

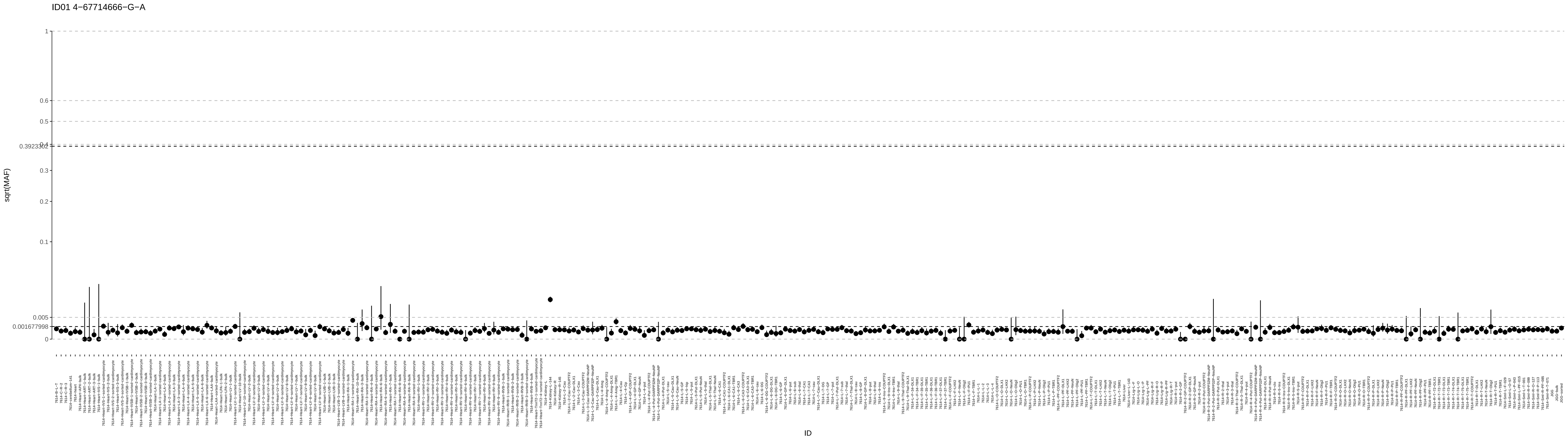
ID01 4−67714666−G−A
sqrt(MAF)
ID
1
0.6
0.5
0.3923302
0.3
0.2
0.1
0.005
0.001677998
0

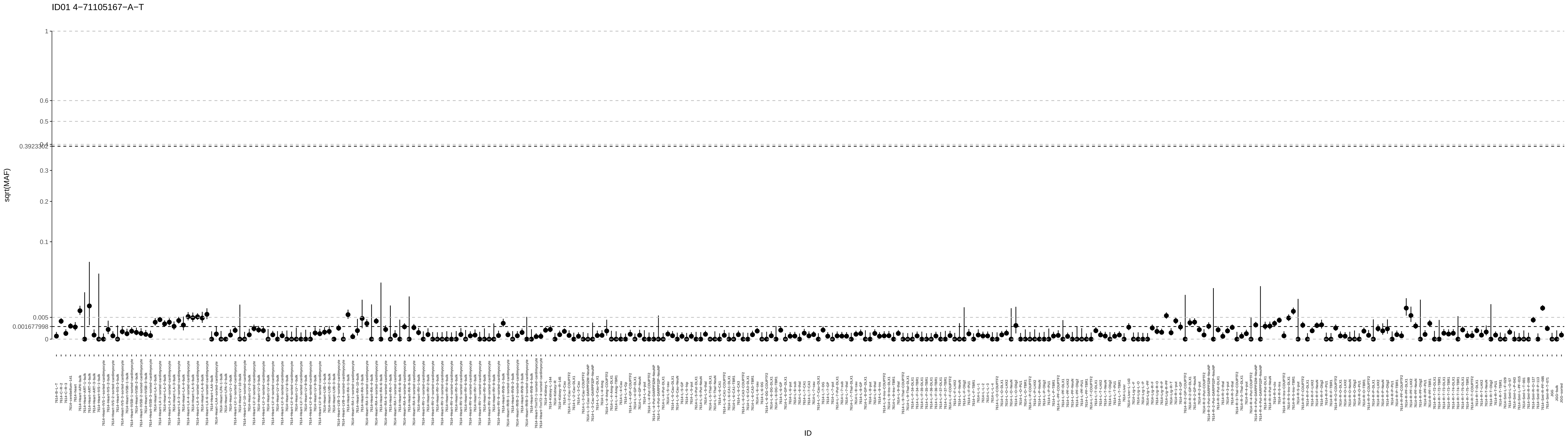
ID01 4−71105167−A−T
sqrt(MAF)
1
0.6
0.5
0.3923302
0.3
0.2
0.1
0.005
0.001677998
0
ID

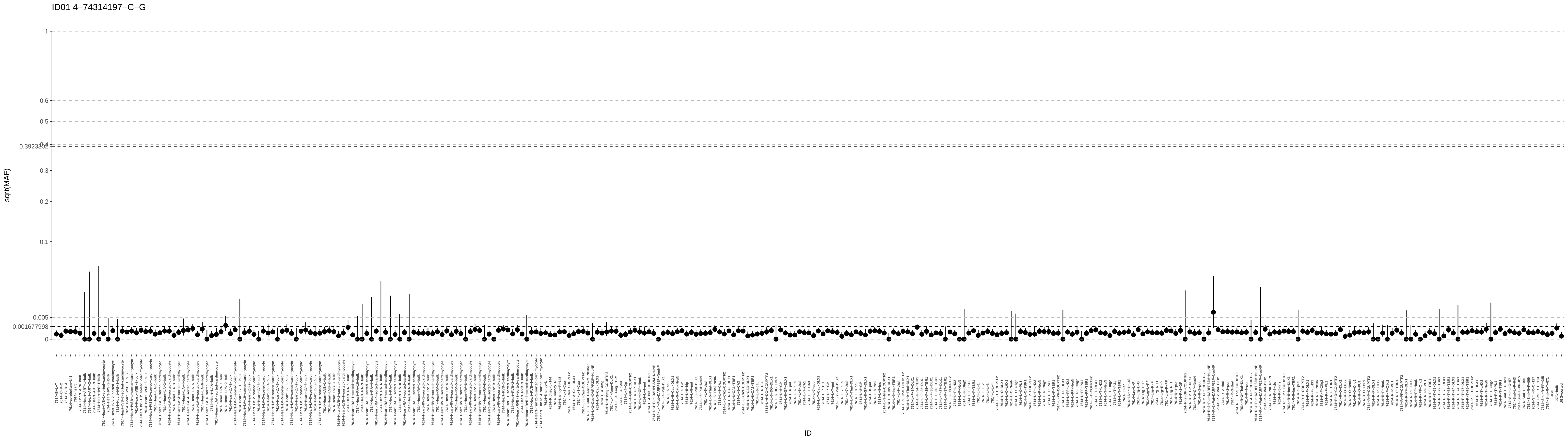
ID01 4−74314197−C−G
sqrt(MAF)
1
0.6
0.5
0.3923302
0.3
0.2
0.1
0.005
0.001677998
0
ID

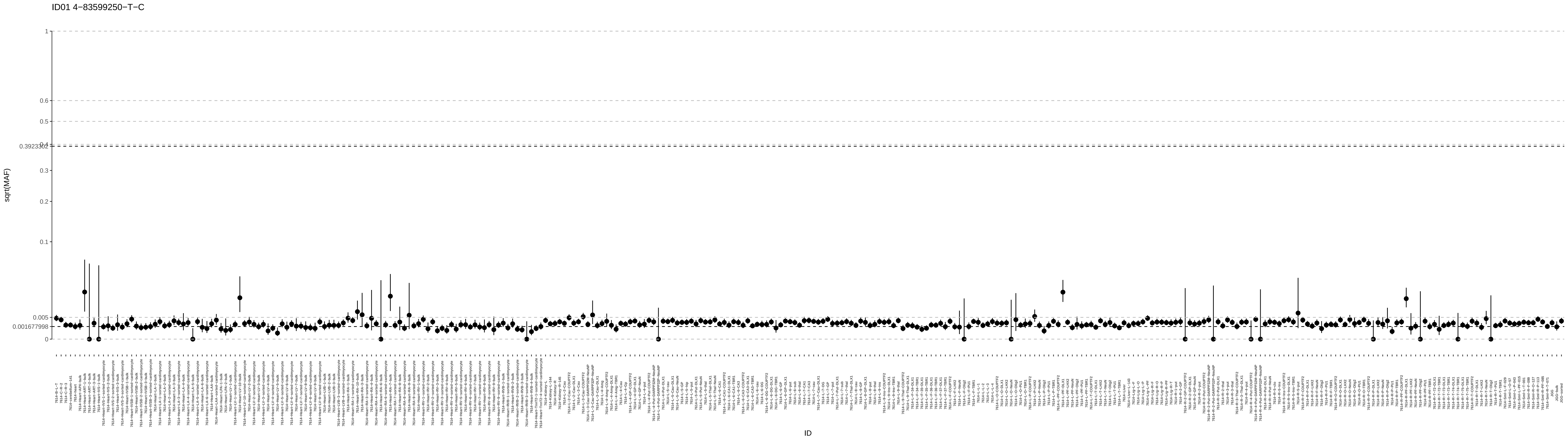
ID01 4−83599250−T−C
sqrt(MAF)
1
0.6
0.5
0.3923302
0.3
0.2
0.1
0.005
0.001677998
0
ID

ID01 4−93868224−A−C

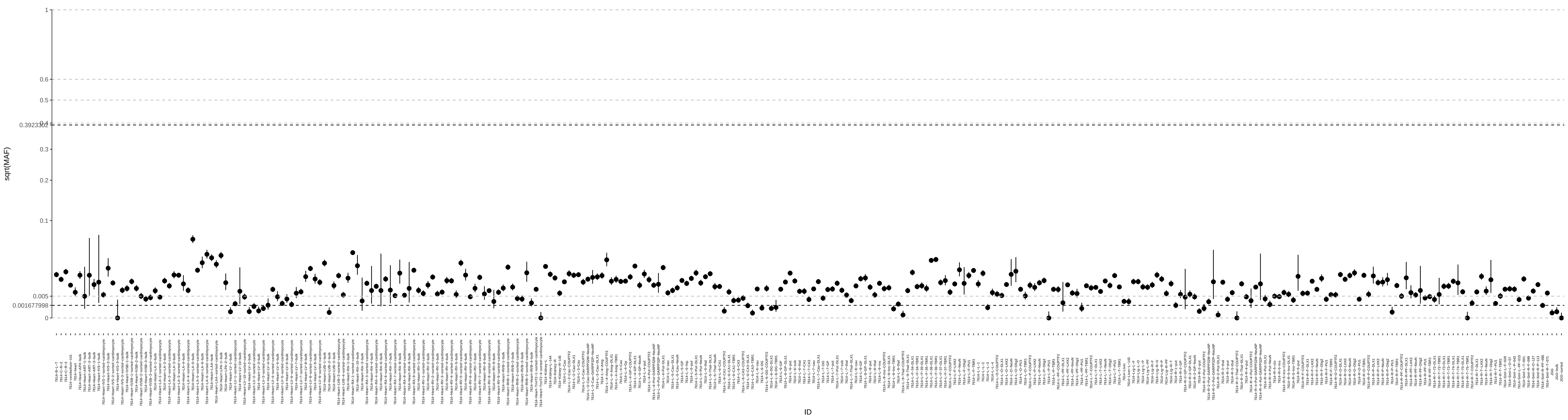

ID01 5−109537368−T−C

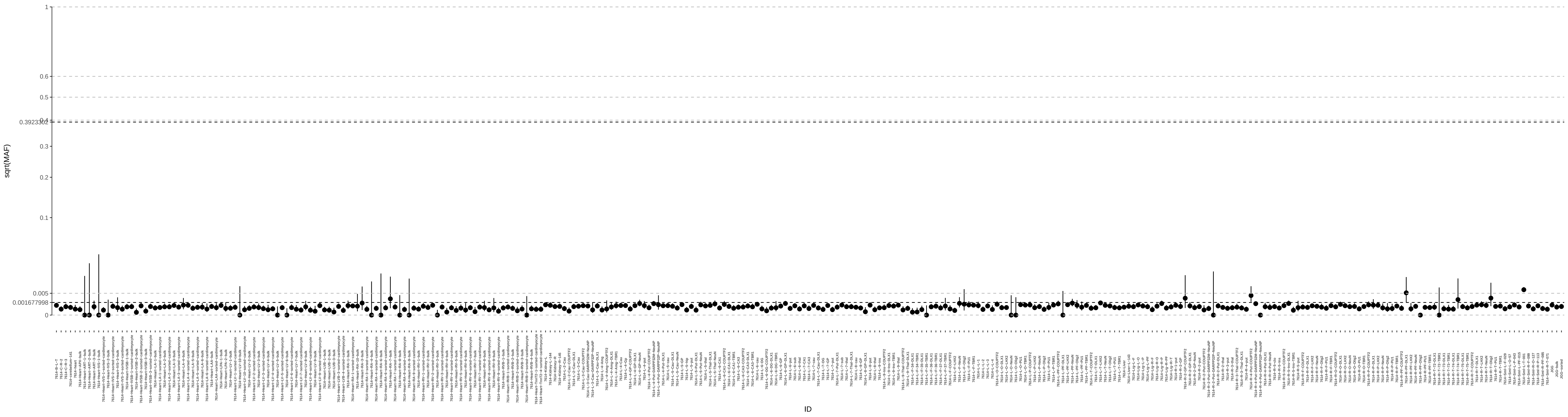

ID01 5−132659937−G−A

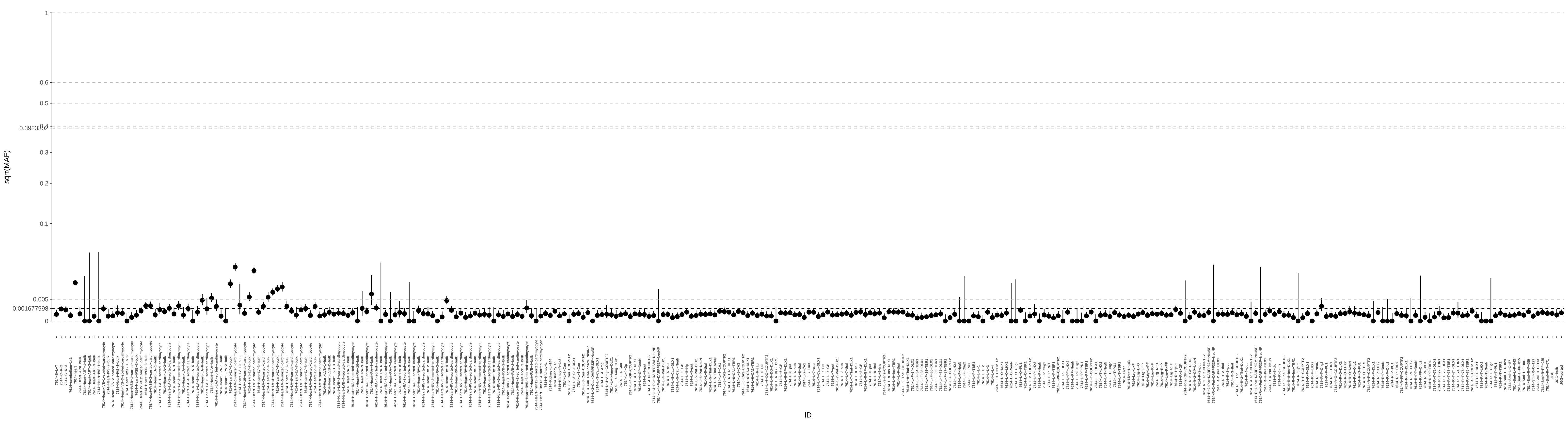

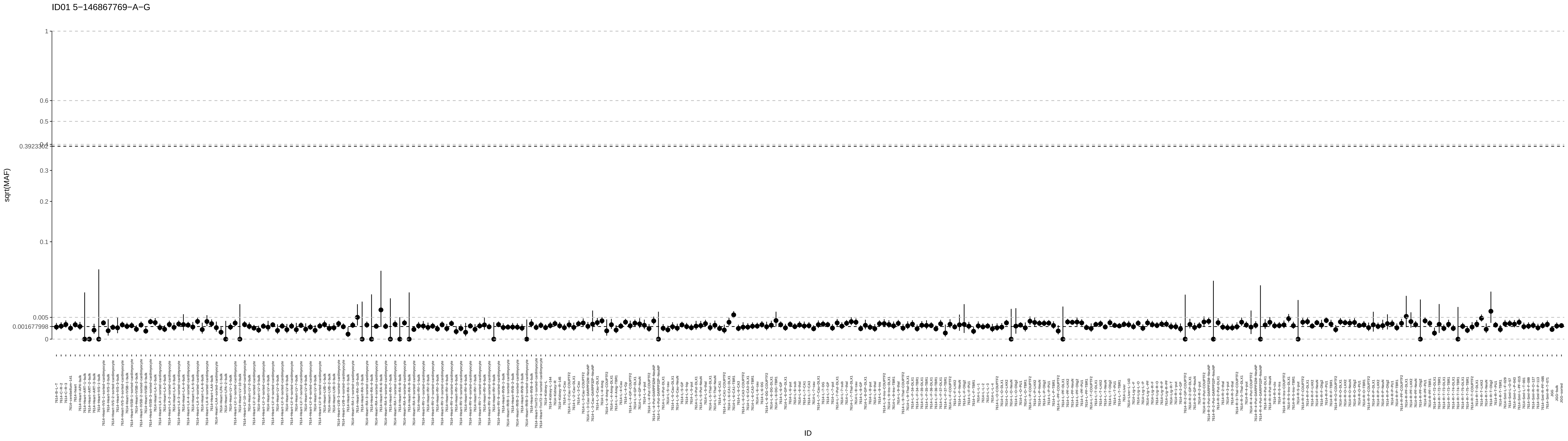
ID01 5−146867769−A−G
sqrt(MAF)
ID
1
0.6
0.5
0.4
0.3923302
0.3
0.2
0.1
0.005
0.001677998
0

ID01 5−160411647−T−G

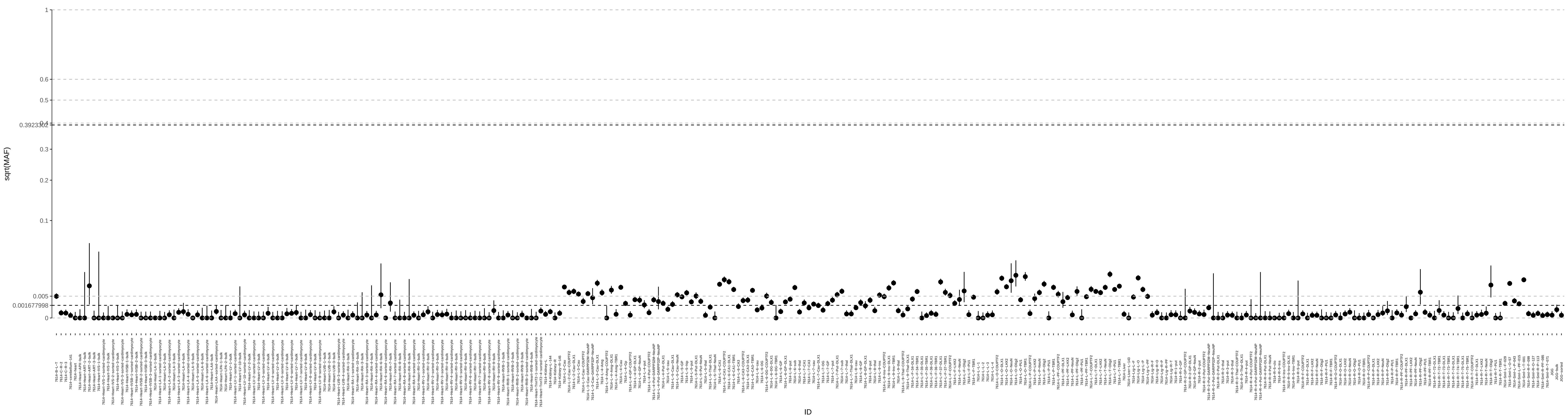

ID01 5−17226551−A−G

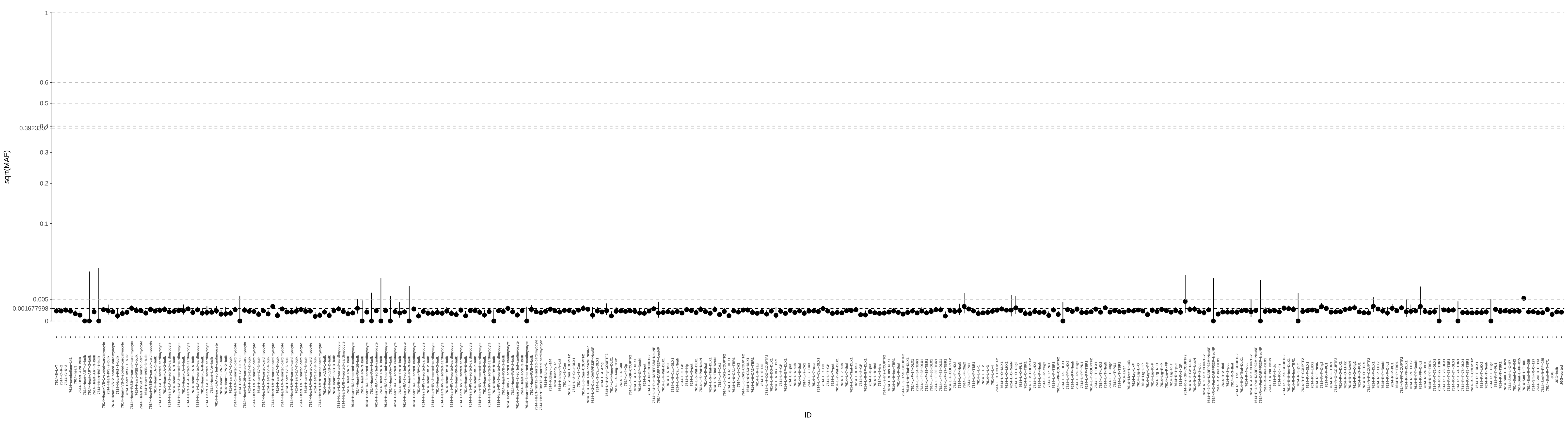

ID01 5−172837236−C−T

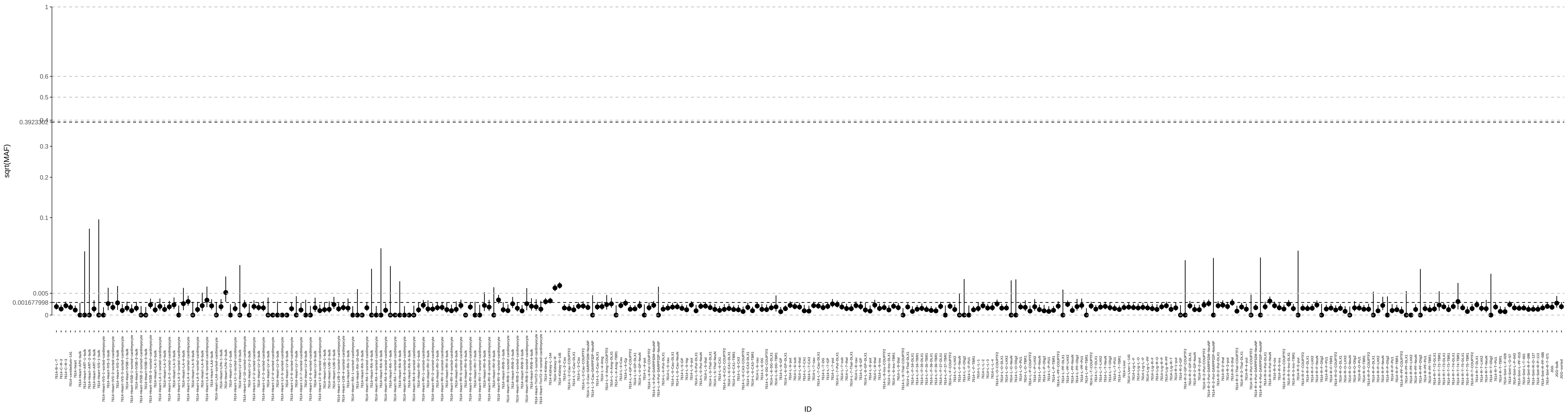

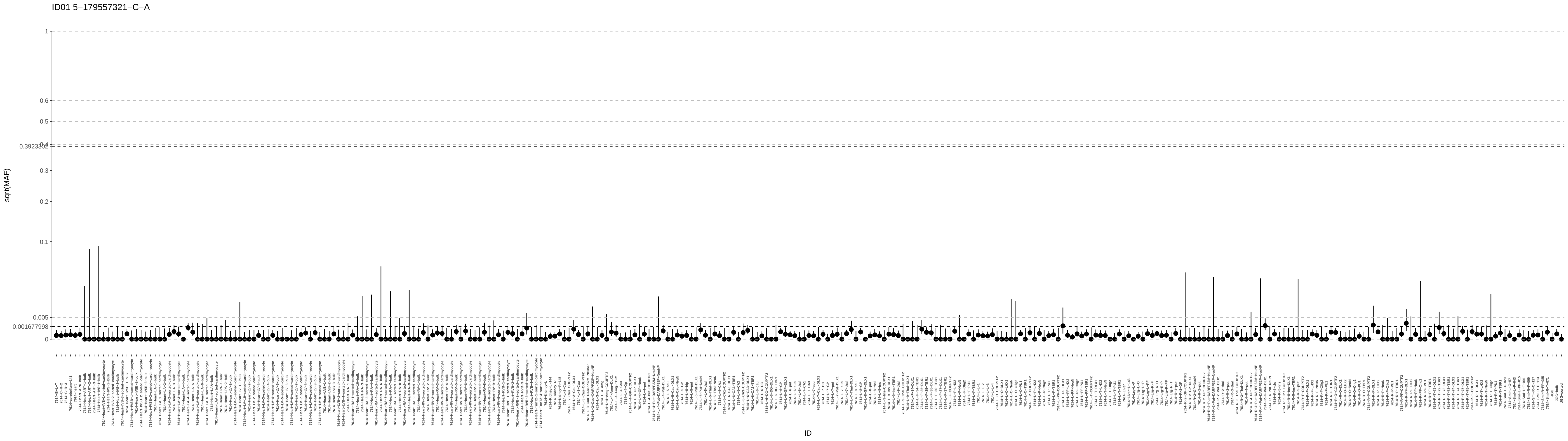
ID01 5−179557321−C−A
sqrt(MAF)
1
0.6
0.5
0.3923302
0.3
0.2
0.1
0.005
0.001677998
0
ID

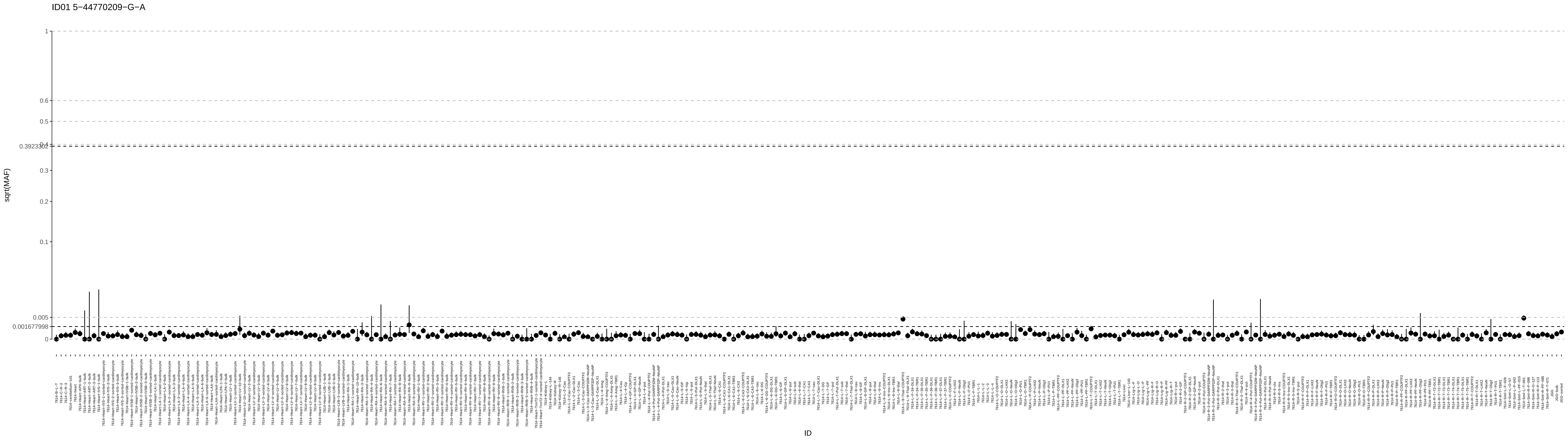
ID01 5−44770209−G−A
sqrt(MAF)
1
0.6
0.5
0.3923302
0.3
0.2
0.1
0.005
0.001677998
0
ID

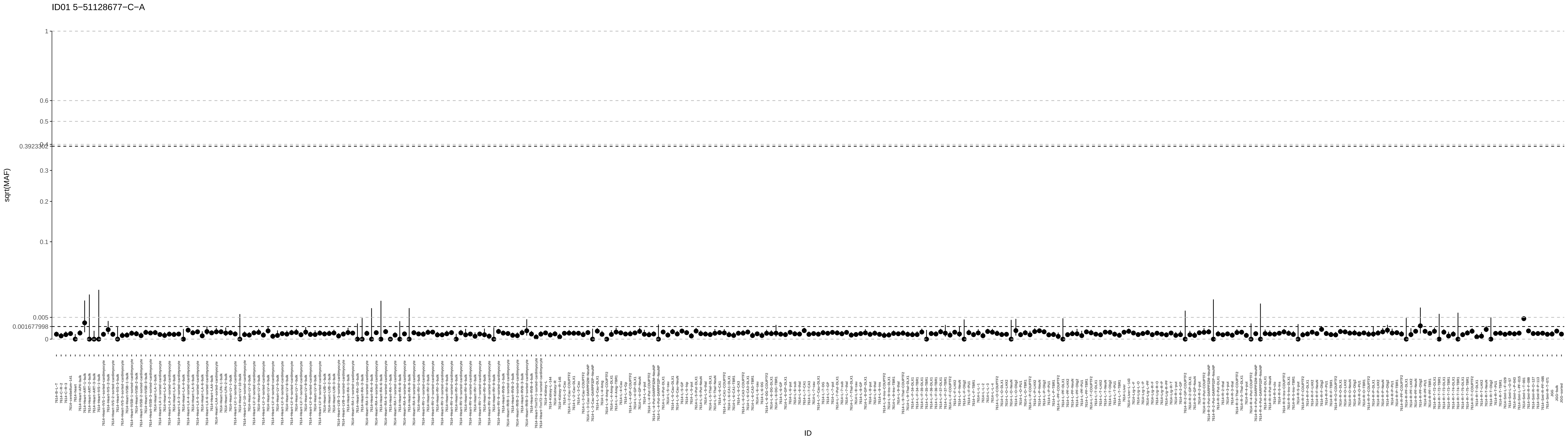
ID01 5−51128677−C−A
1
0.6
0.5
0.3923302
0.3
0.2
0.1
0.005
0.001677998
0
sqrt(MAF)
ID

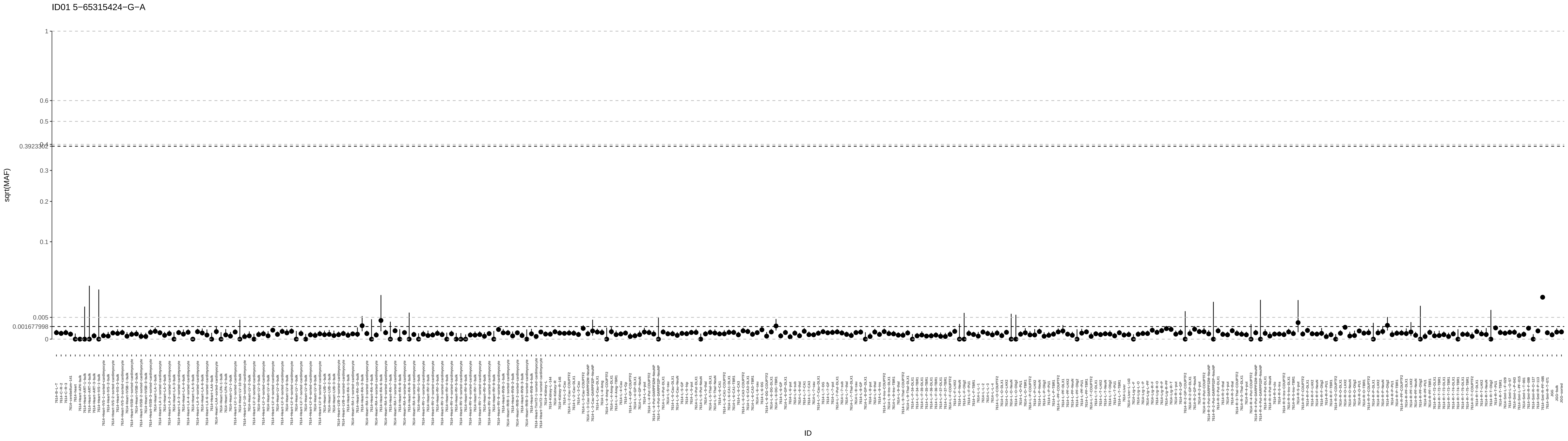
ID01 5−65315424−G−A
sqrt(MAF)
ID
1
0.6
0.5
0.3923302
0.3
0.2
0.1
0.005
0.001677998
0

ID01 5−72178944−ACG−A

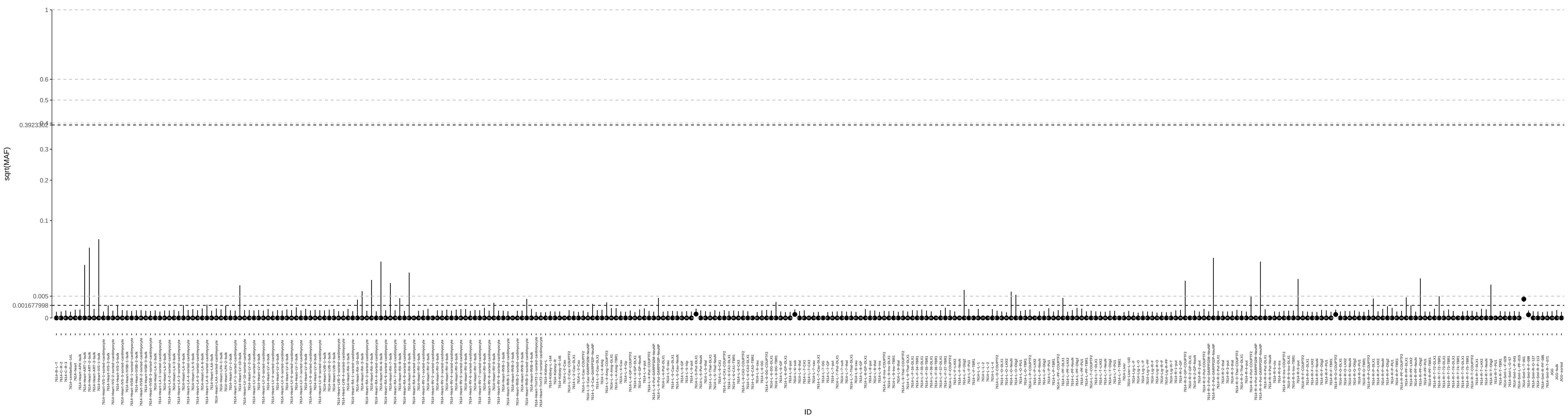

ID01 5−72392587−A−G

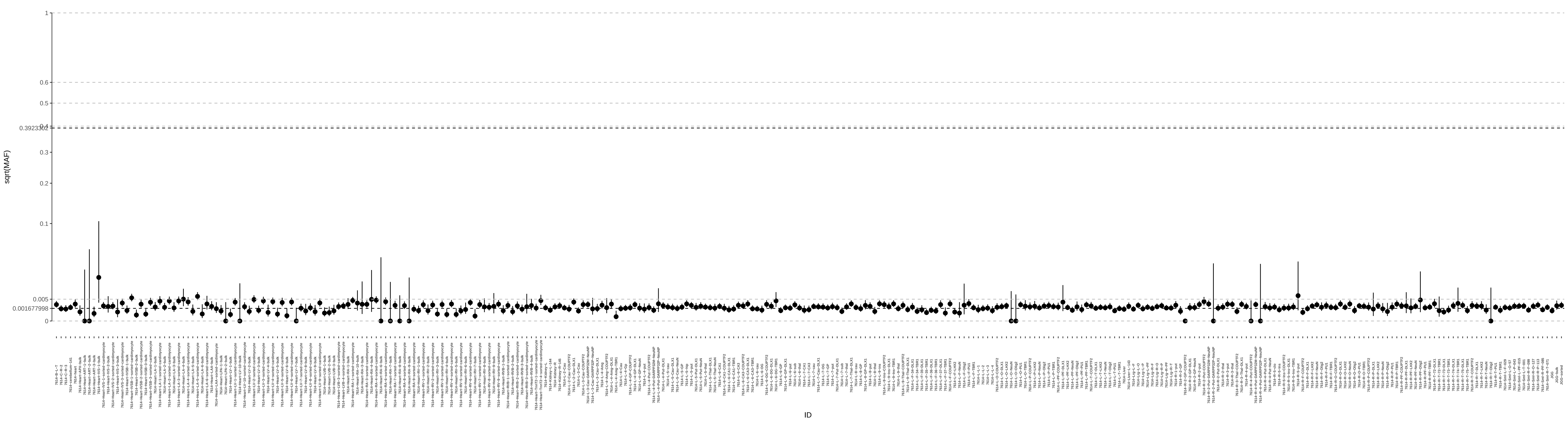

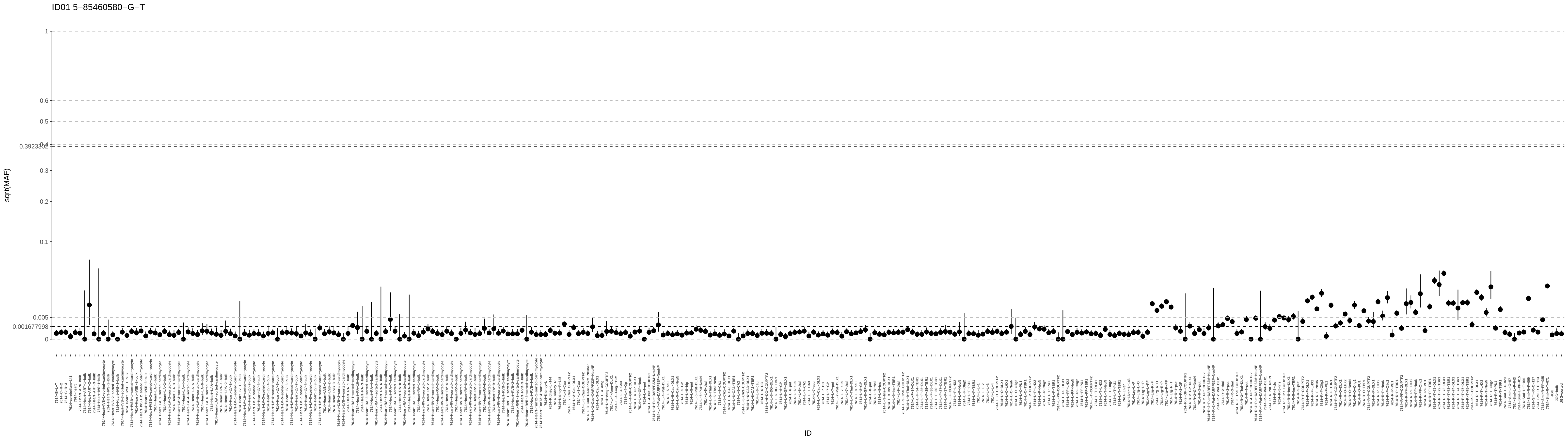
ID01 5−85460580−G−T
sqrt(MAF)
ID
1
0.6
0.5
0.3923302
0.3
0.2
0.1
0.005
0.001677998
0

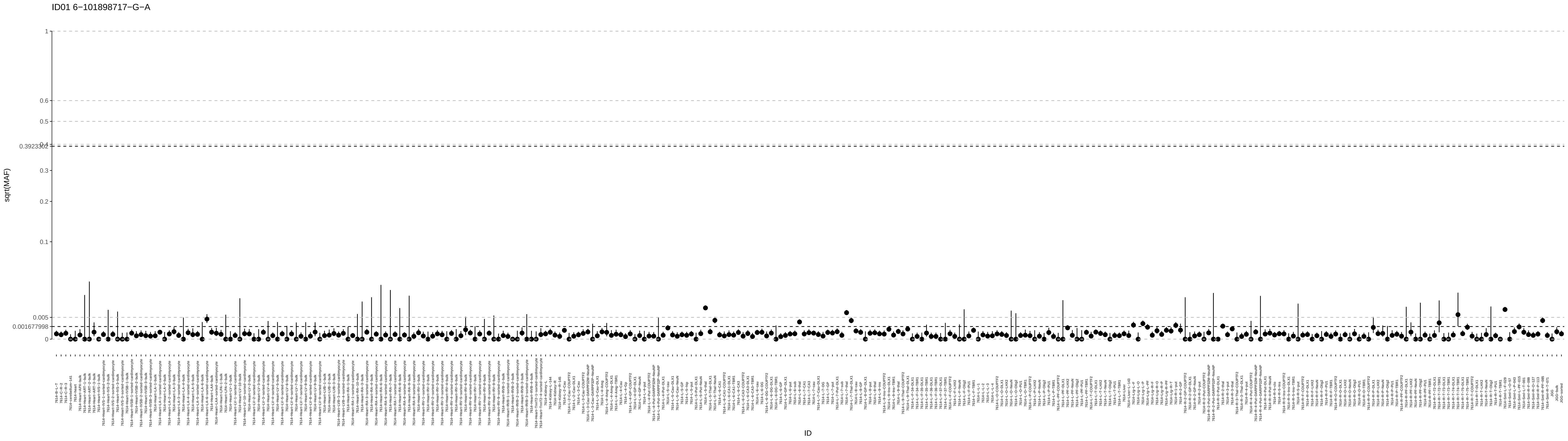
ID01 6−101898717−G−A
sqrt(MAF)
1
0.6
0.5
0.3923302
0.3
0.2
0.1
0.005
0.001677998
0
ID

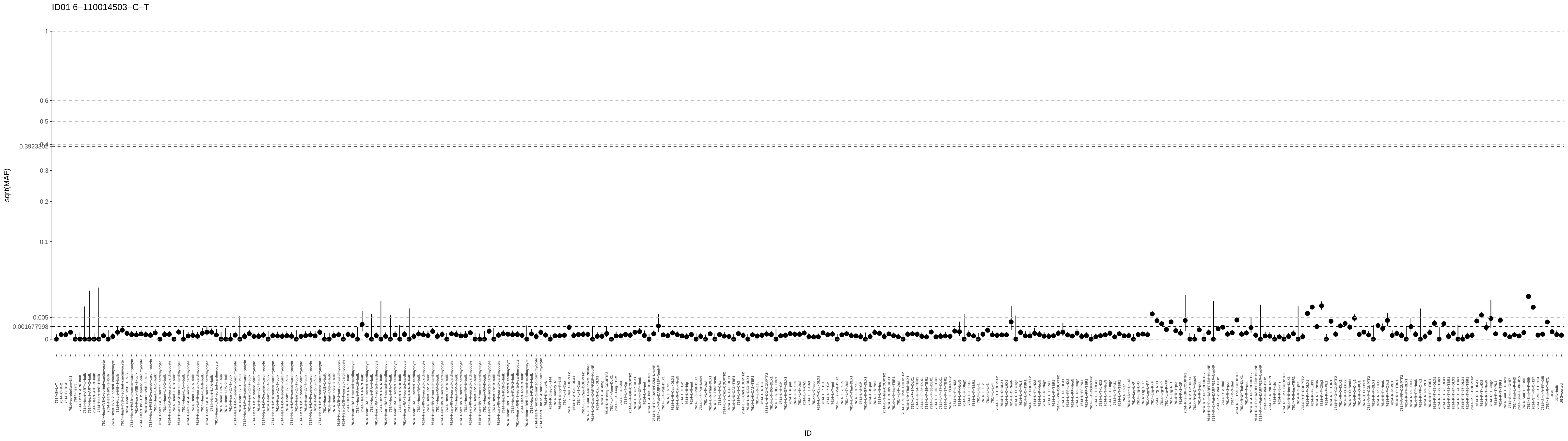

ID01 6−110014503−C−T
sqrt(MAF)
1
0.6
0.5
0.3923302
0.3
0.2
0.1
0.005
0.001677998
0
ID

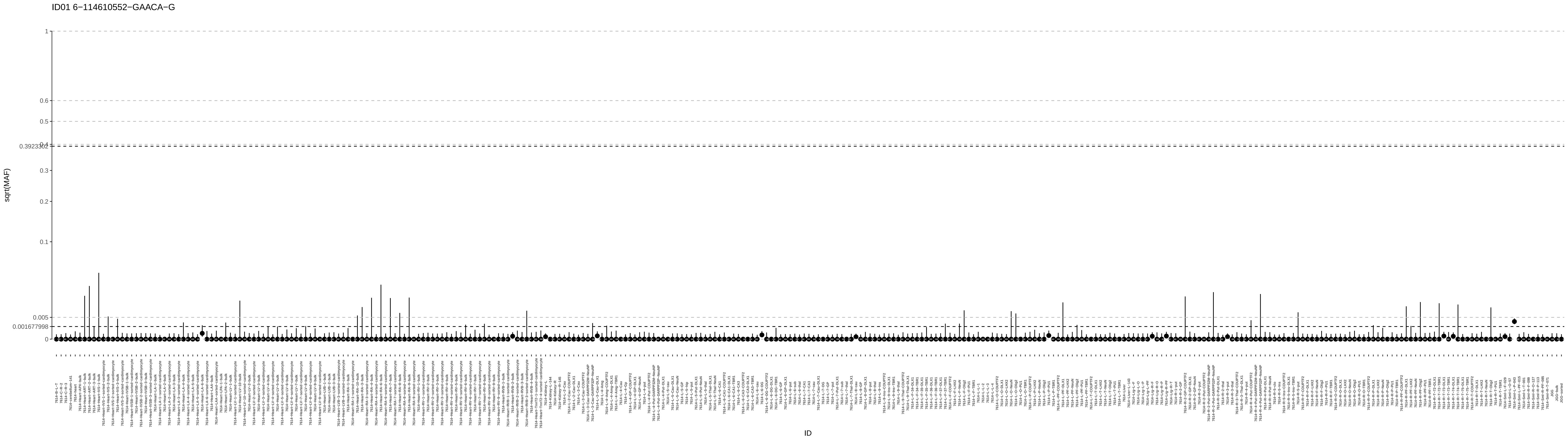
ID01 6−114610552−GAACA−G
sqrt(MAF)
1
0.6
0.5
0.4
0.3923302
0.3
0.2
0.1
0.005
0.001677998
0
ID

ID01 6−121420086−C−T

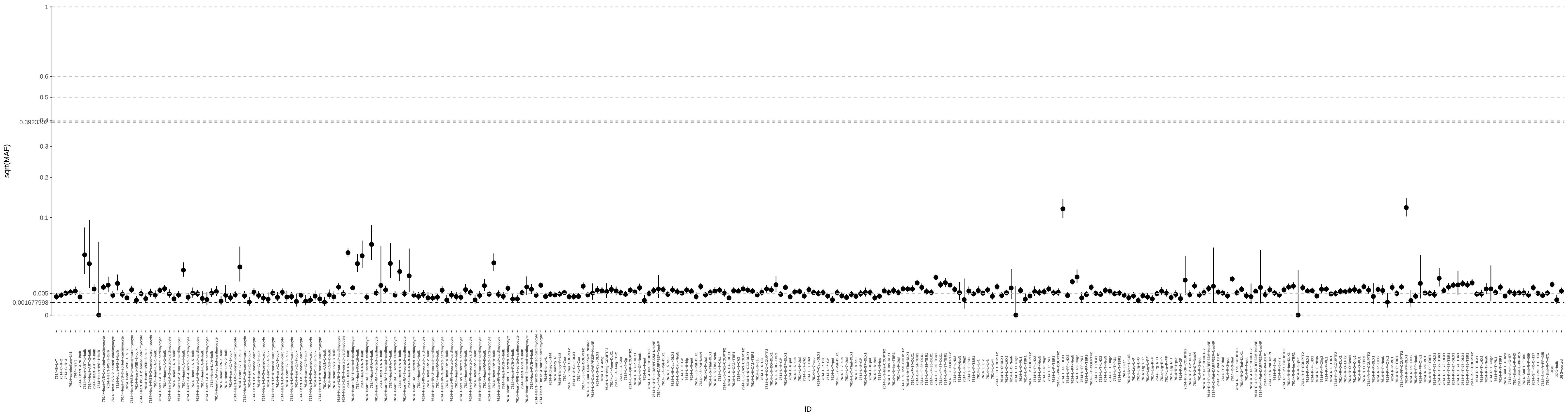

ID01 6−122610884−C−T

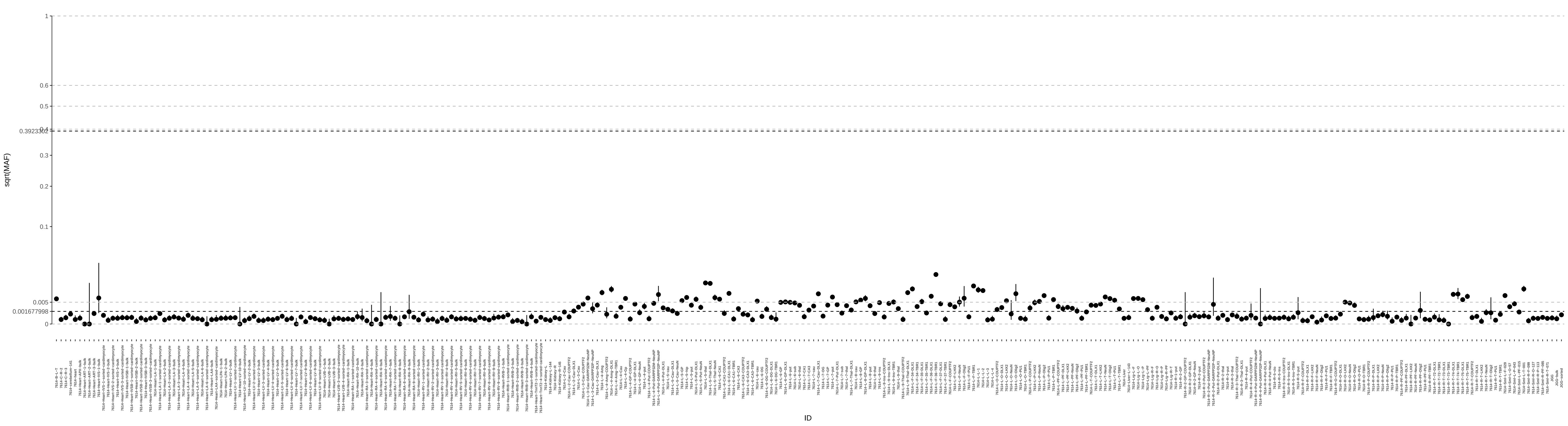

ID01 6−125153224−A−C

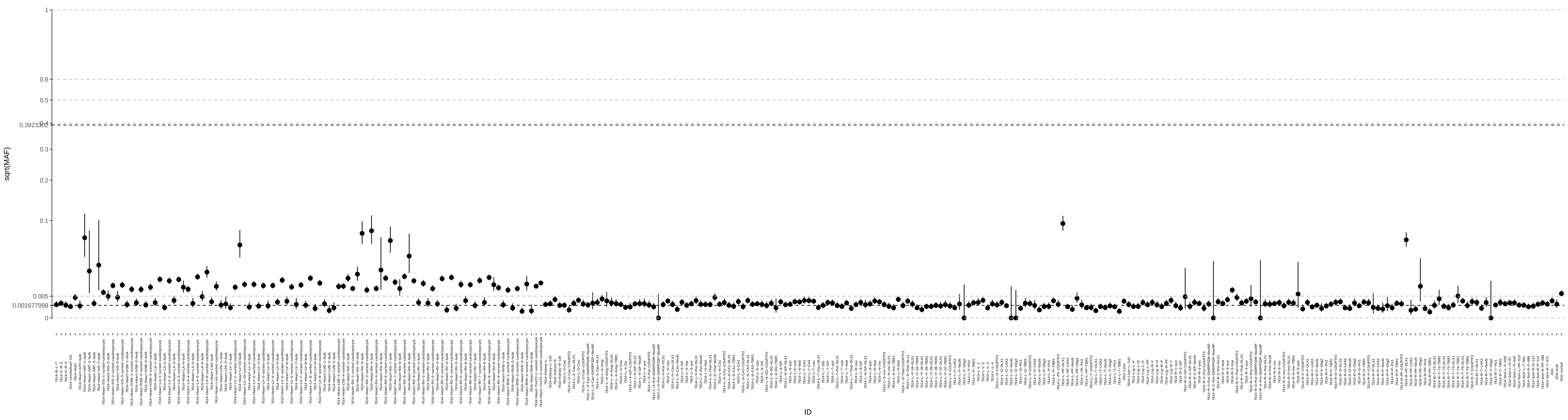

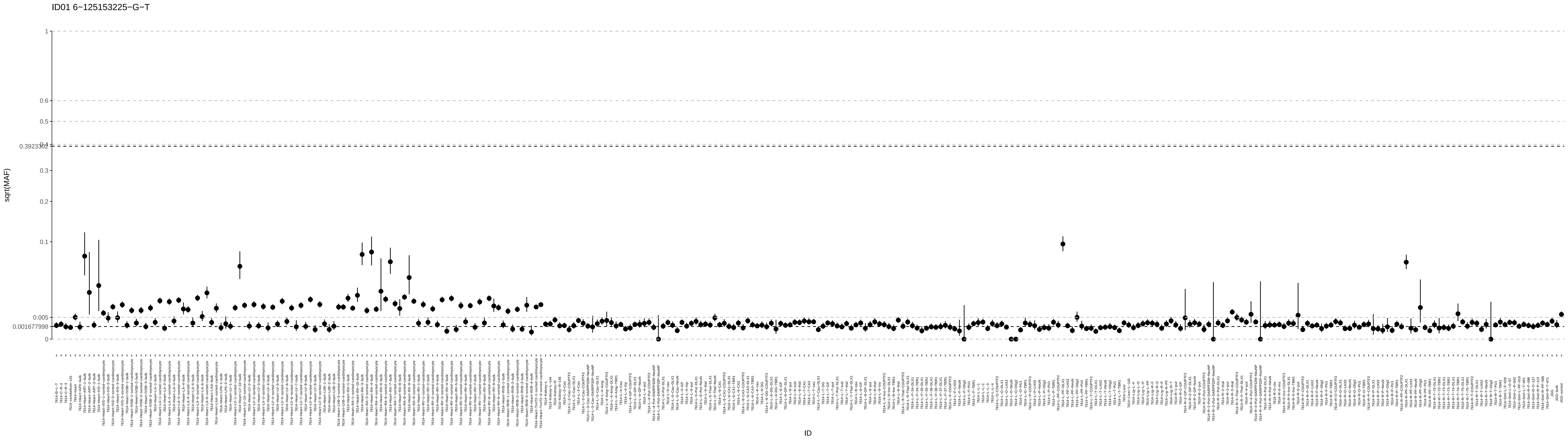
ID01 6−125153225−G−T
sqrt(MAF)
1
0.6
0.5
0.4
0.3923302
0.3
0.2
0.1
0.005
0.001677998
0
ID

ID01 6−131367555−G−A

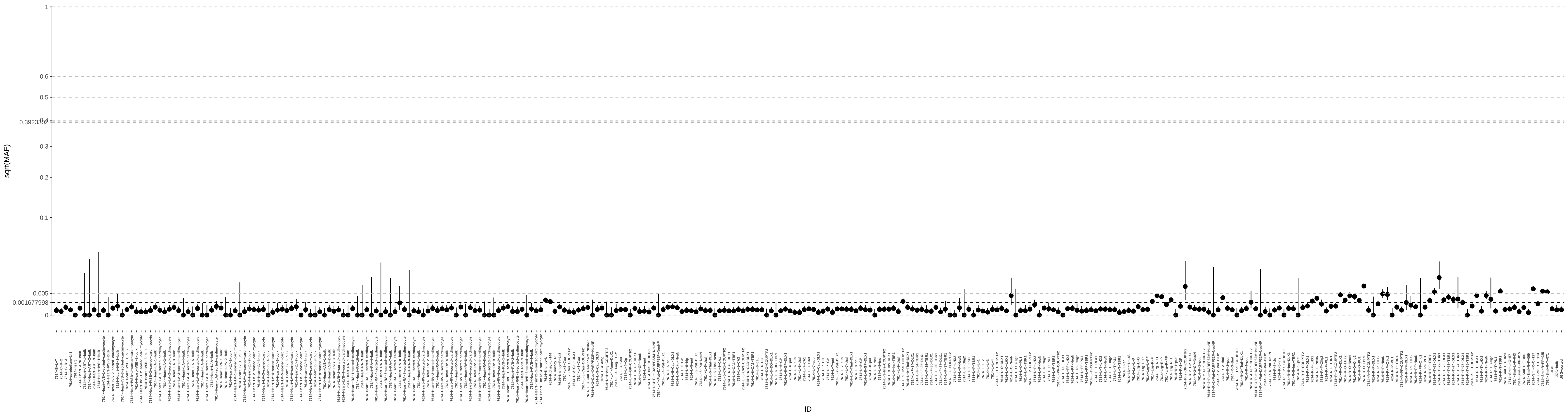

ID01 6−154659392−C−T

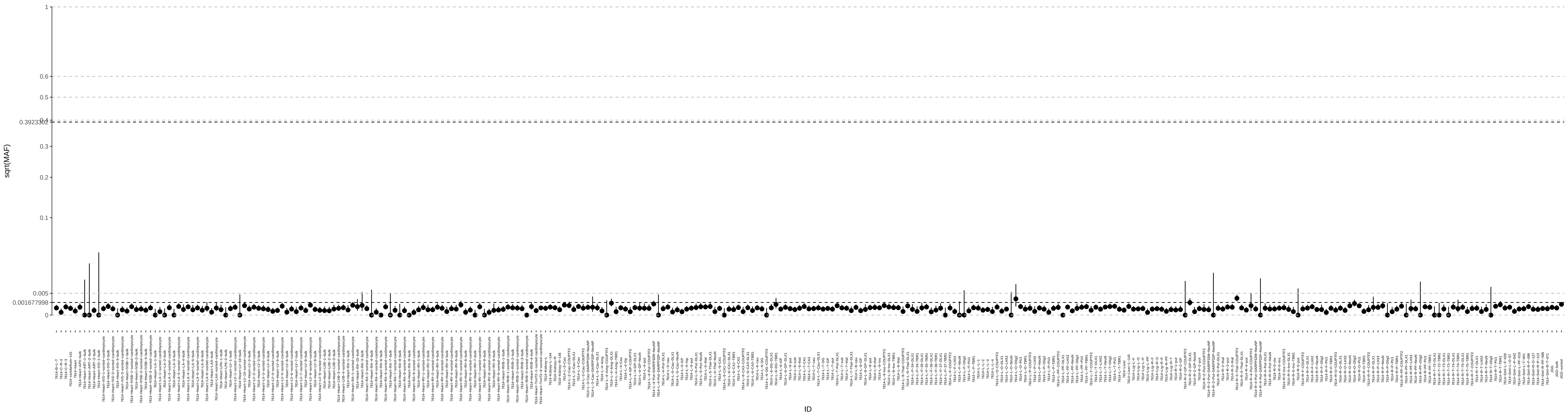

ID01 6−156279892−A−G

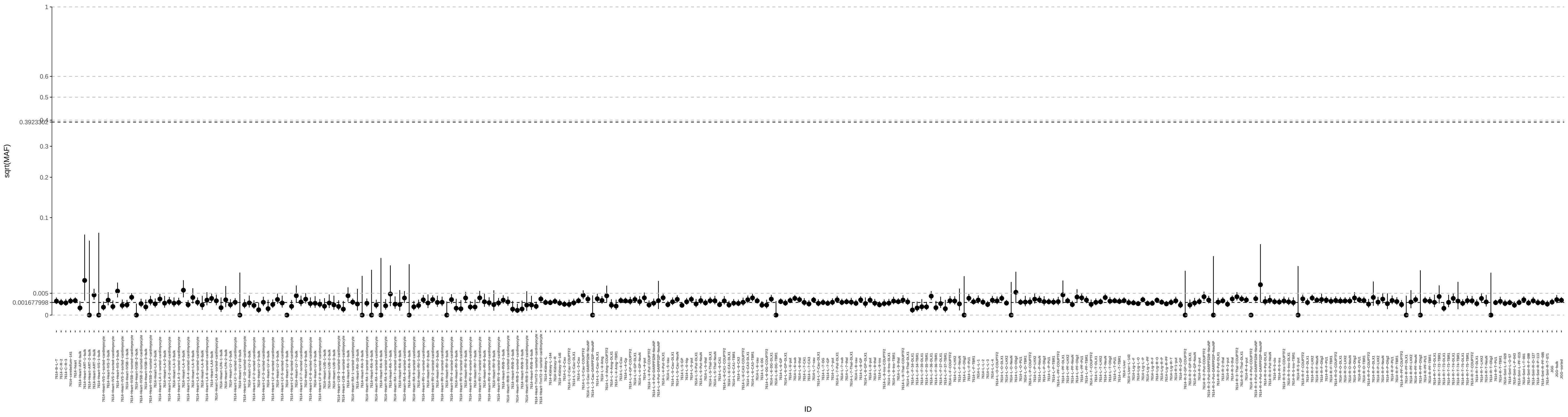

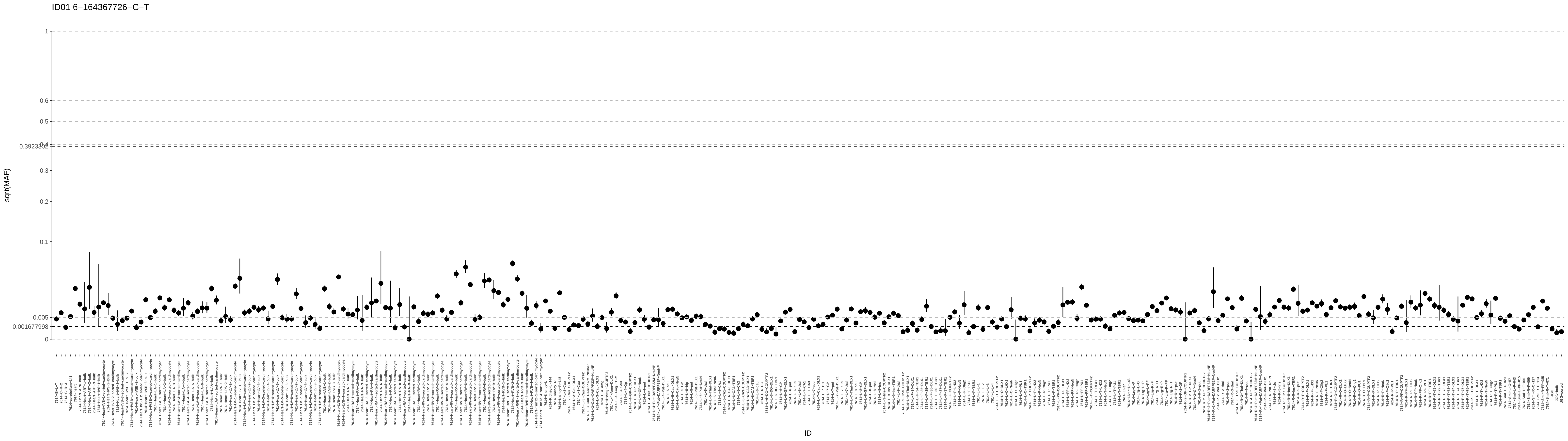
ID01 6−164367726−C−T
sqrt(MAF)
1
0.6
0.5
0.3923302
0.3
0.2
0.1
0.005
0.001677998
0
ID

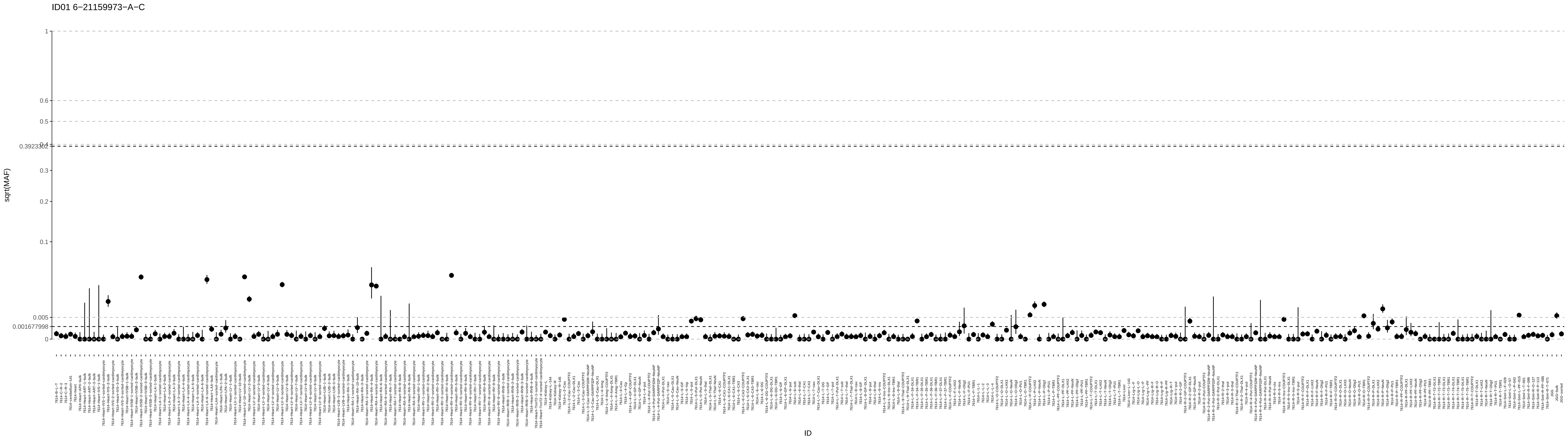

ID01 6−21159973−A−C
sqrt(MAF)
1
0.6
0.5
0.4
0.3923302
0.3
0.2
0.1
0.005
0.001677998
0
ID

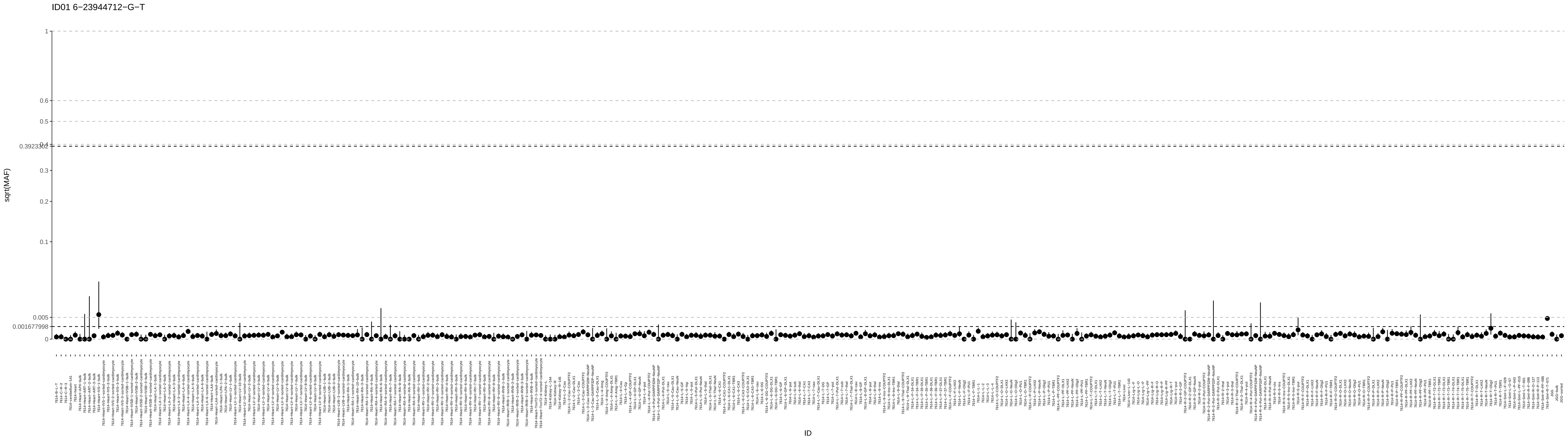
ID01 6−23944712−G−T
1
0.6
0.5
0.3923302
0.3
0.2
0.1
0.005
0.001677998
0
sqrt(MAF)
ID

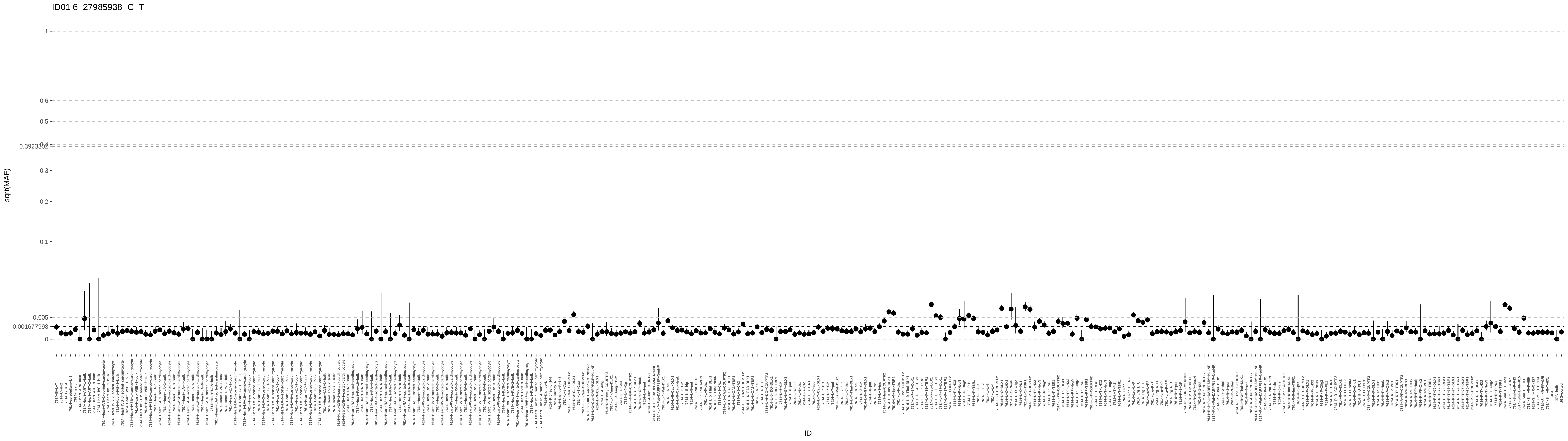
ID01 6−27985938−C−T
sqrt(MAF)
1
0.6
0.5
0.4
0.3923302
0.3
0.2
0.1
0.005
0.001677998
0
ID

ID01 6-40371400-C-T

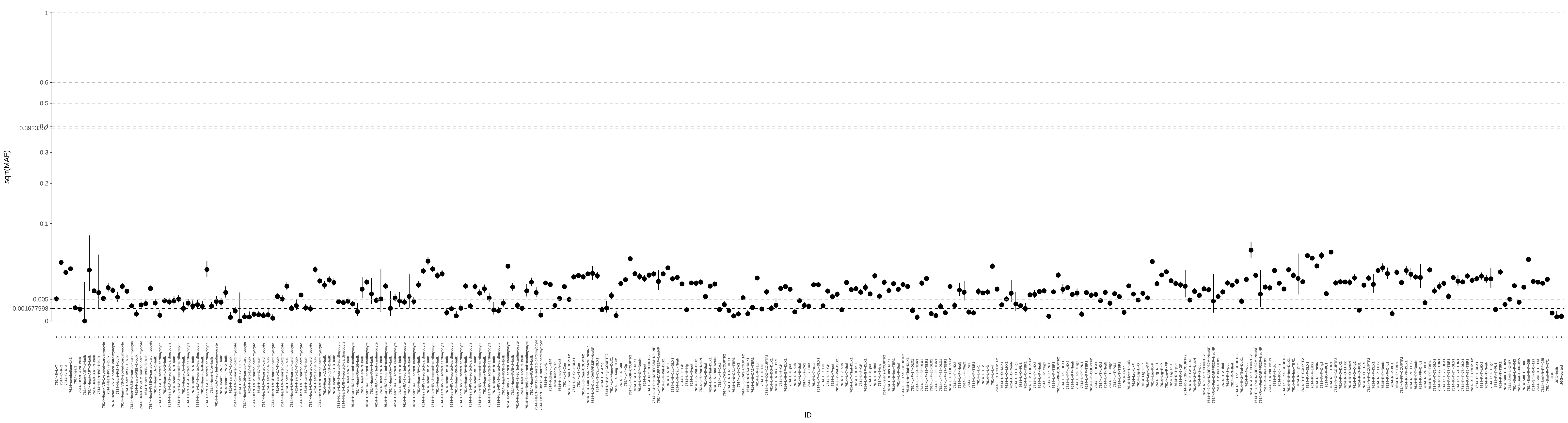

ID01 6−45630553−G−A

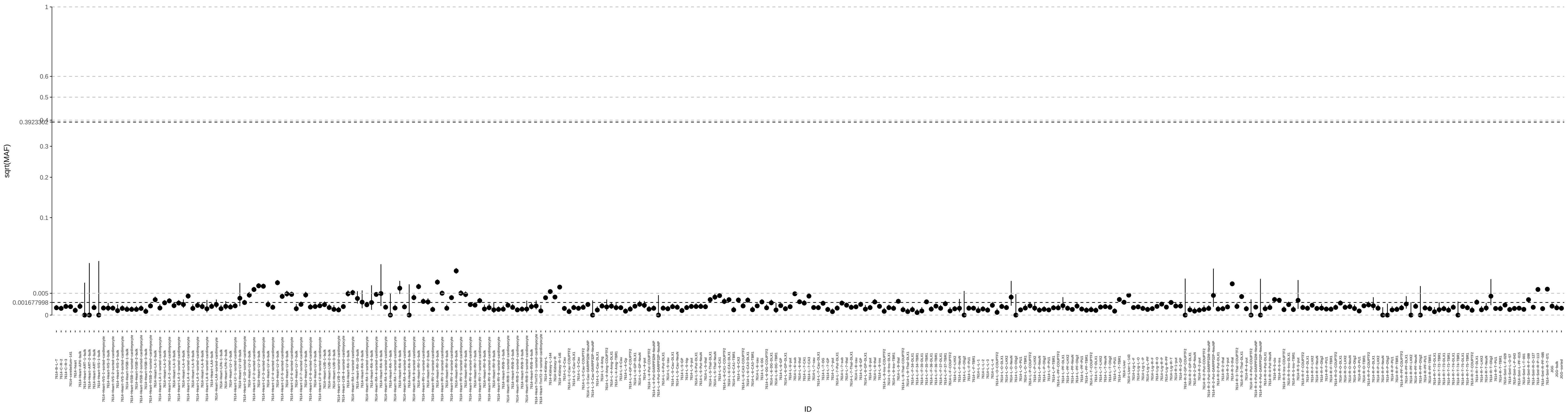

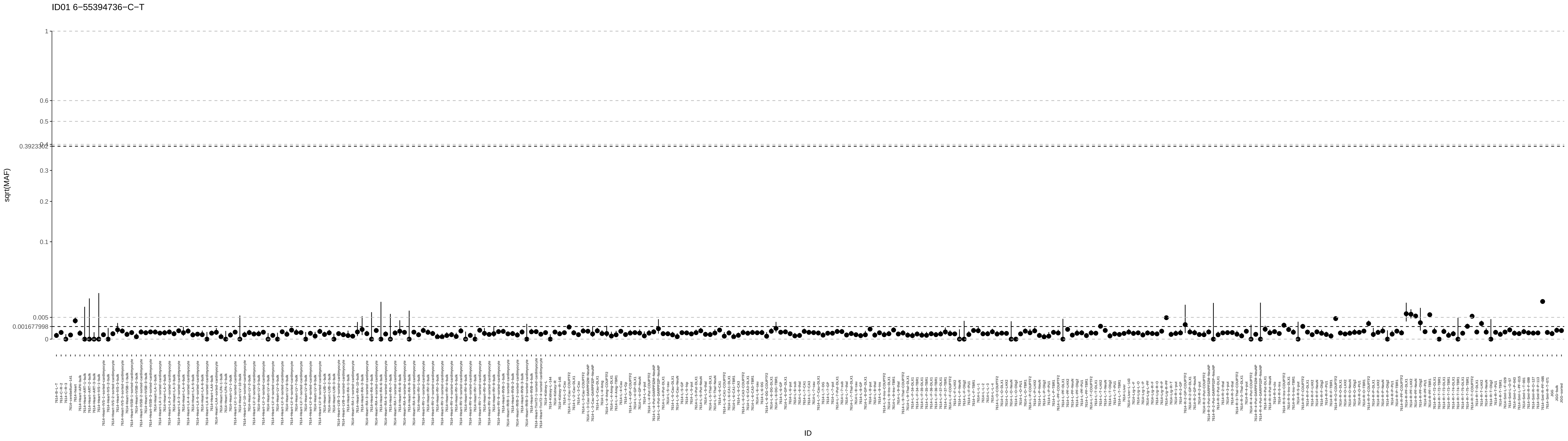
ID01 6−55394736−C−T
sqrt(MAF)
1
0.6
0.5
0.4
0.3923302
0.3
0.2
0.1
0.005
0.001677998
0
ID

ID01 6−57218601−A−G

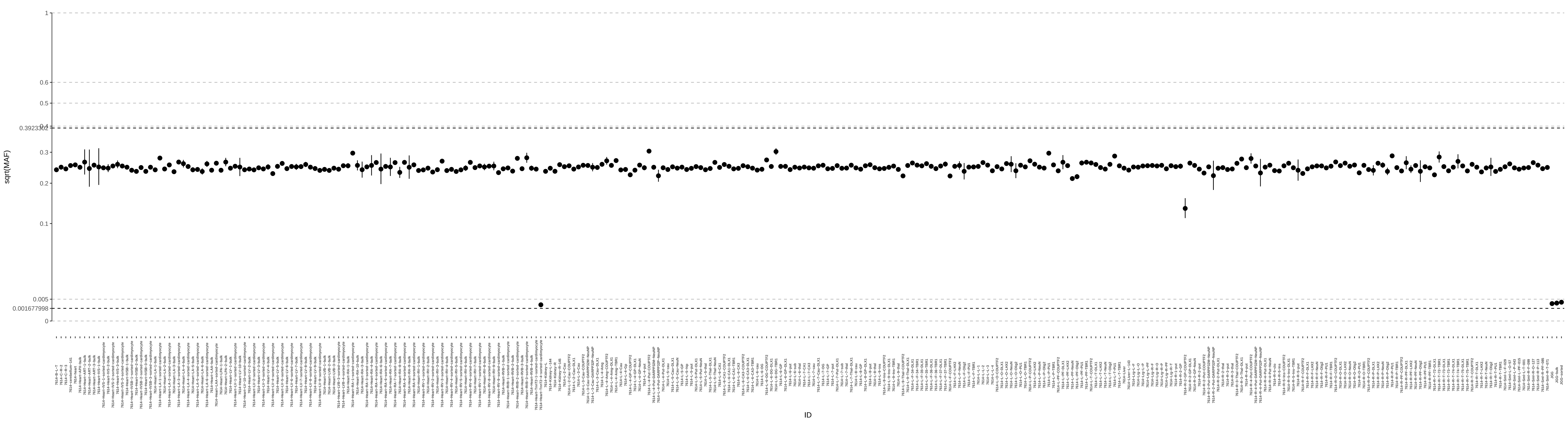

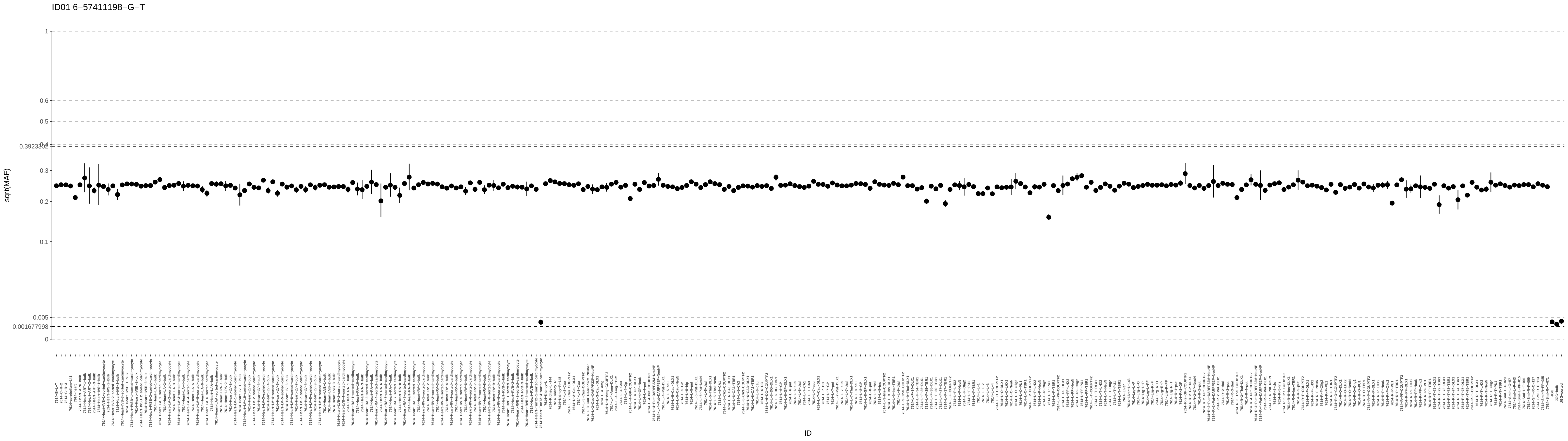
ID01 6−57411198−G−T
sqrt(MAF)
1
0.6
0.5
0.3923302
0.3
0.2
0.1
0.005
0.001677998
0
ID

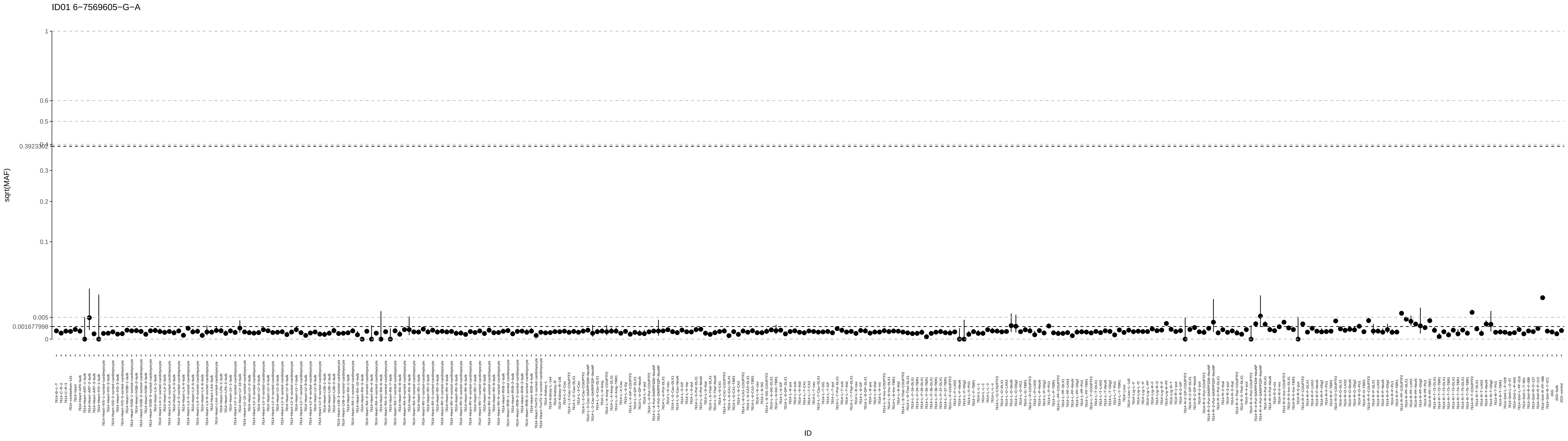

ID01 6−7569605−G−A
sqrt(MAF)
1
0.6
0.5
0.3923302
0.3
0.2
0.1
0.005
0.001677998
0
ID

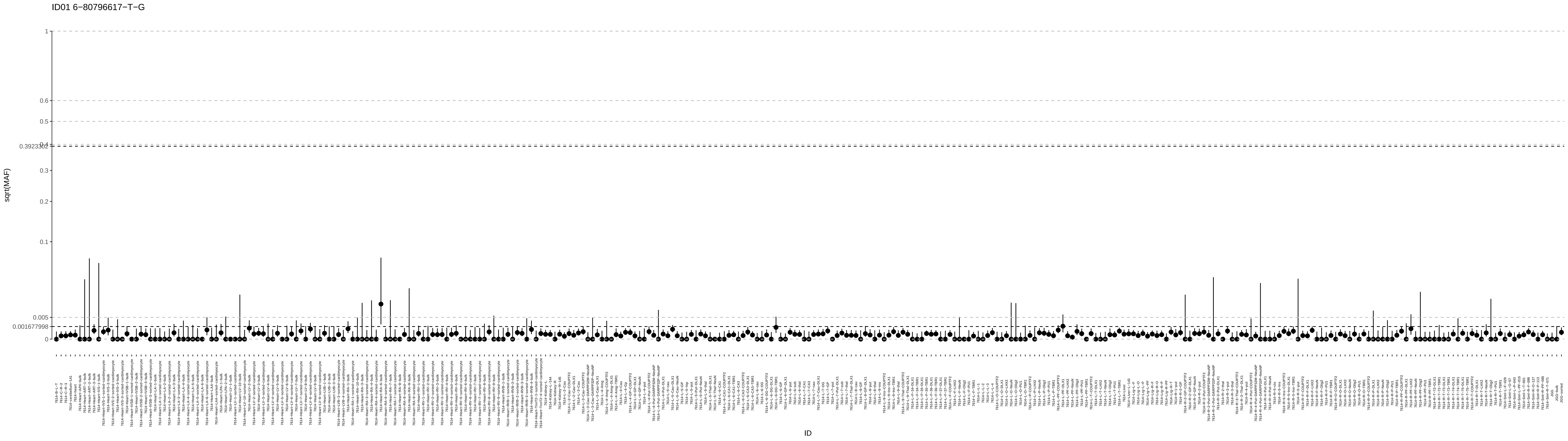

ID01 6−80796617−T−G
sqrt(MAF)
1
0.6
0.5
0.3923302
0.3
0.2
0.1
0.005
0.001677998
0
ID

ID01 7-121347585-T-G

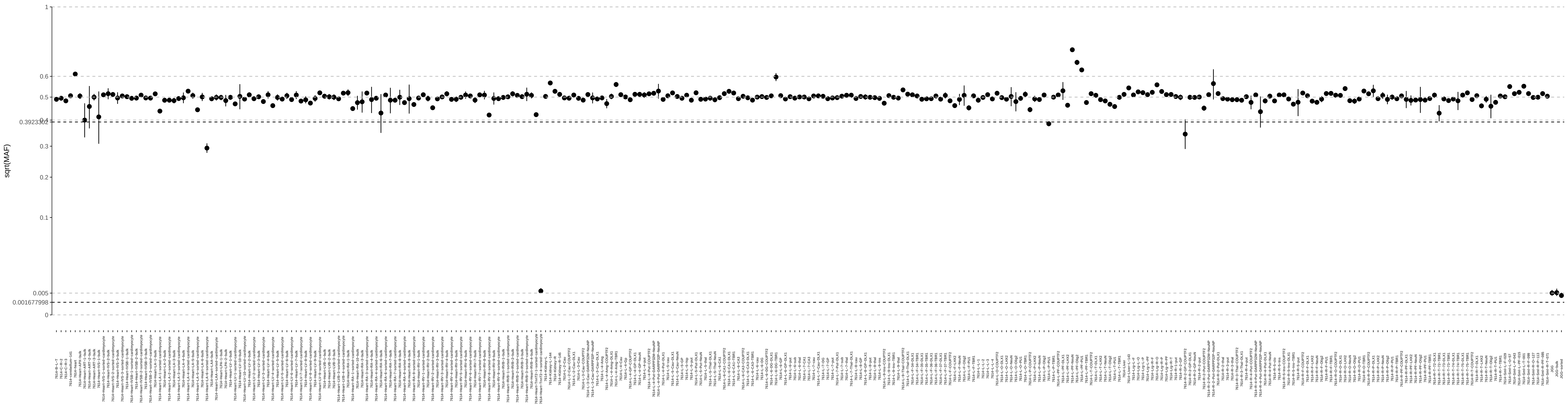

ID01 7−121766164−G−A

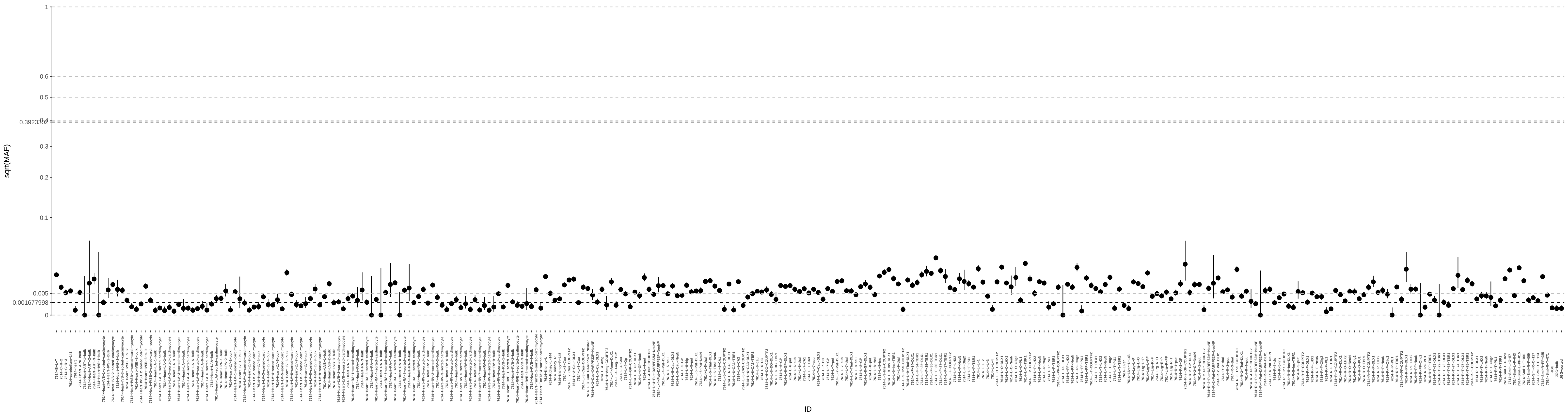

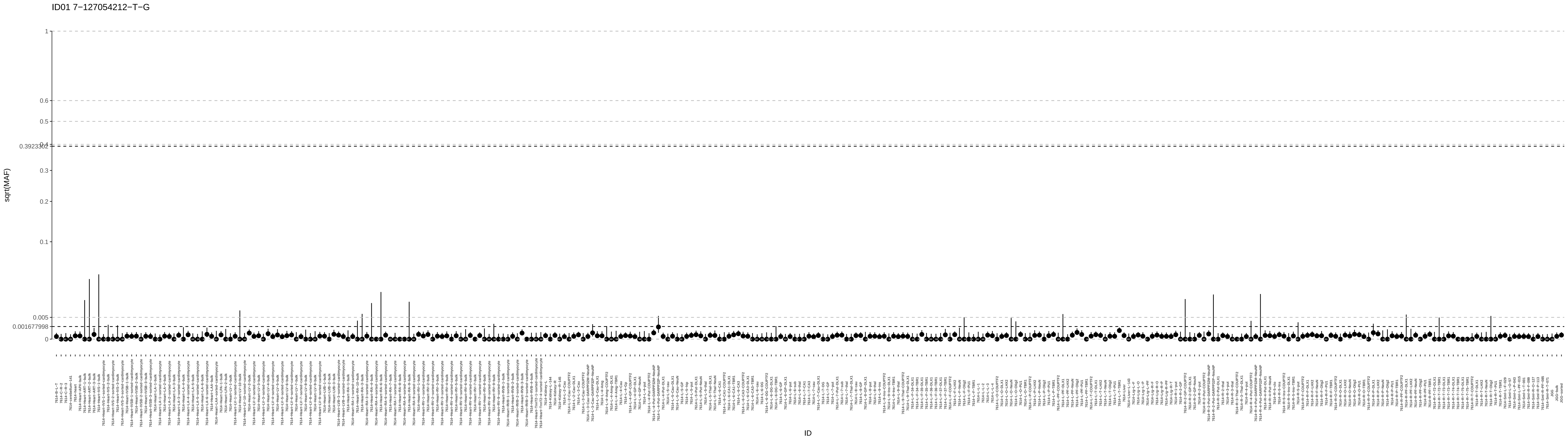

ID01 7−127054212−T−G
sqrt(MAF)
1
0.6
0.5
0.3923302
0.3
0.2
0.1
0.005
0.001677998
0
ID

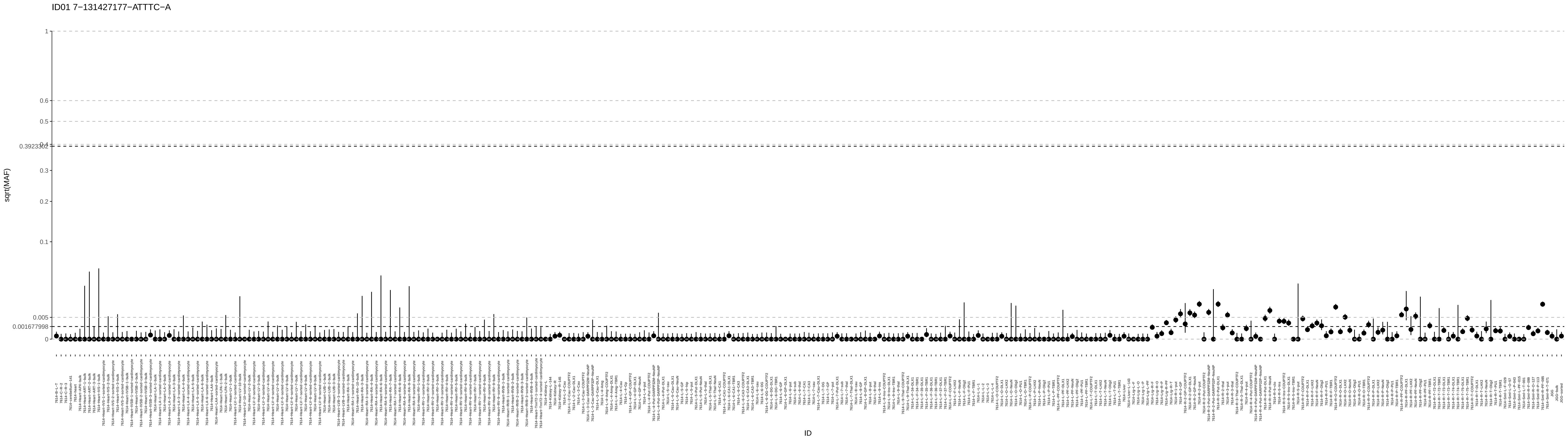
ID01 7−131427177−ATTTC−A
sqrt(MAF)
1
0.6
0.5
0.3923302
0.3
0.2
0.1
0.005
0.001677998
0
ID

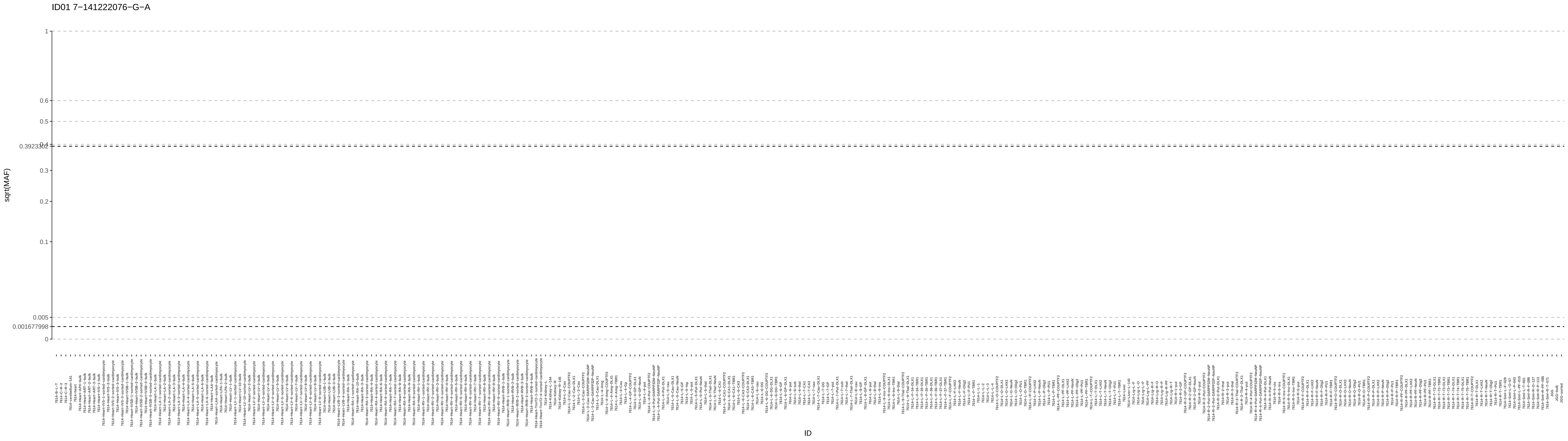
ID01 7−141222076−G−A
sqrt(MAF)
1
0.6
0.5
0.3923302
0.3
0.2
0.1
0.005
0.001677998
0
ID

ID01 7−150175576−A−C

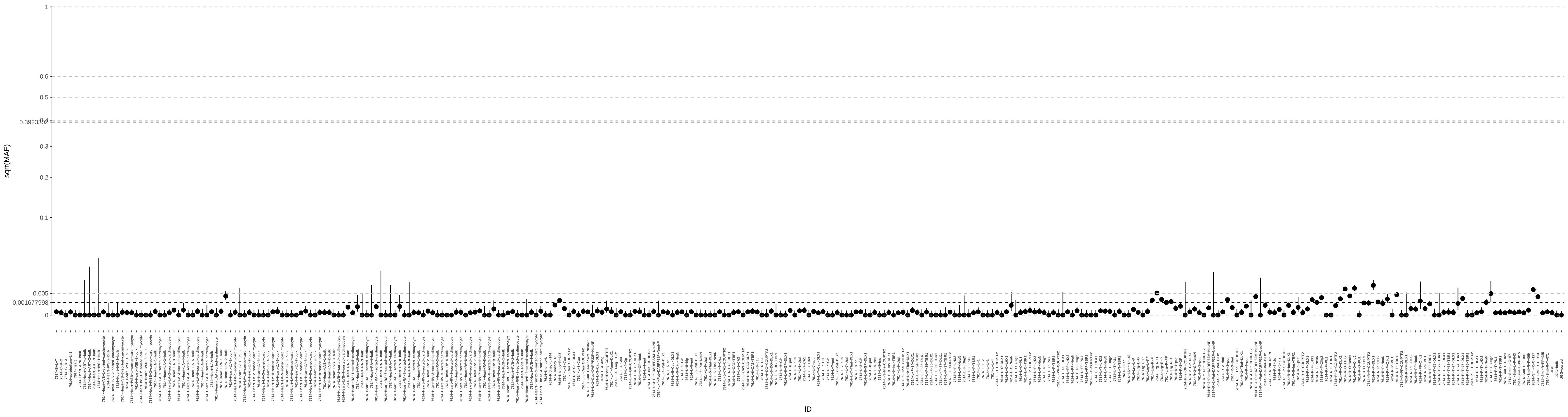

ID01 7−31796412−T−G

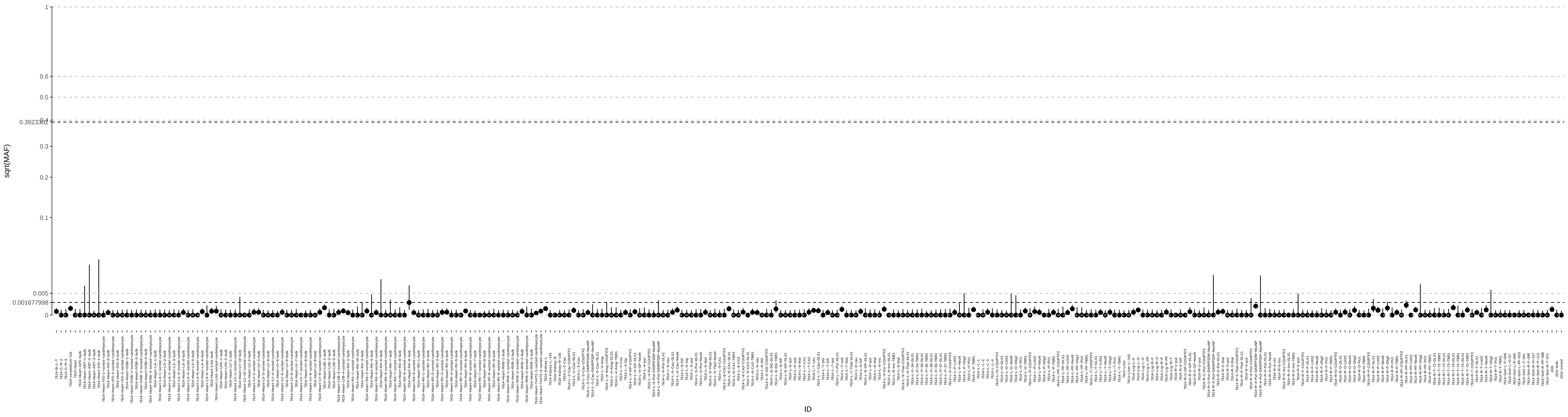

ID01 7−33765368−A−G

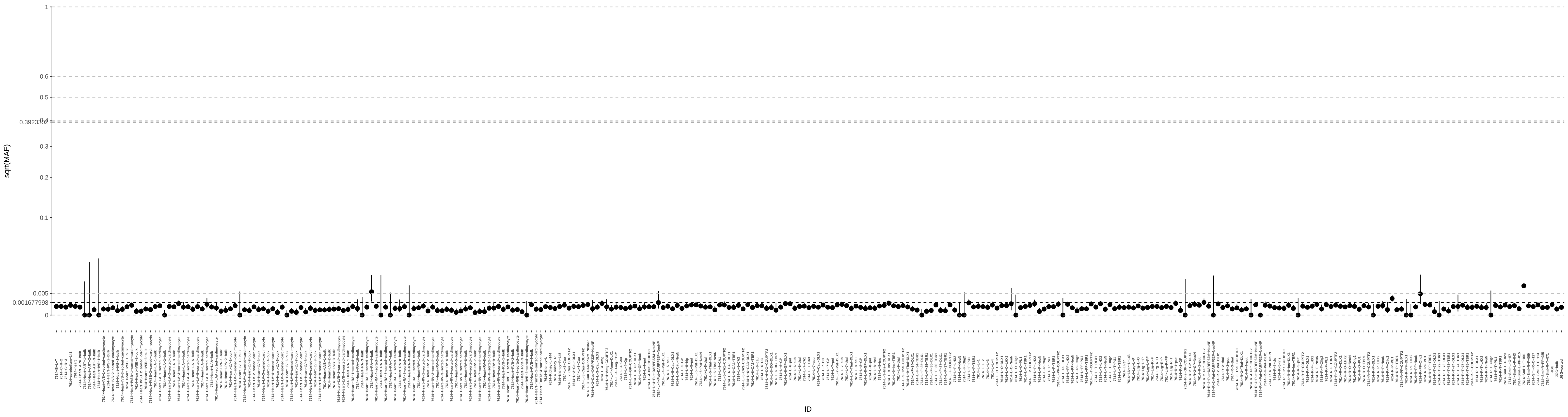

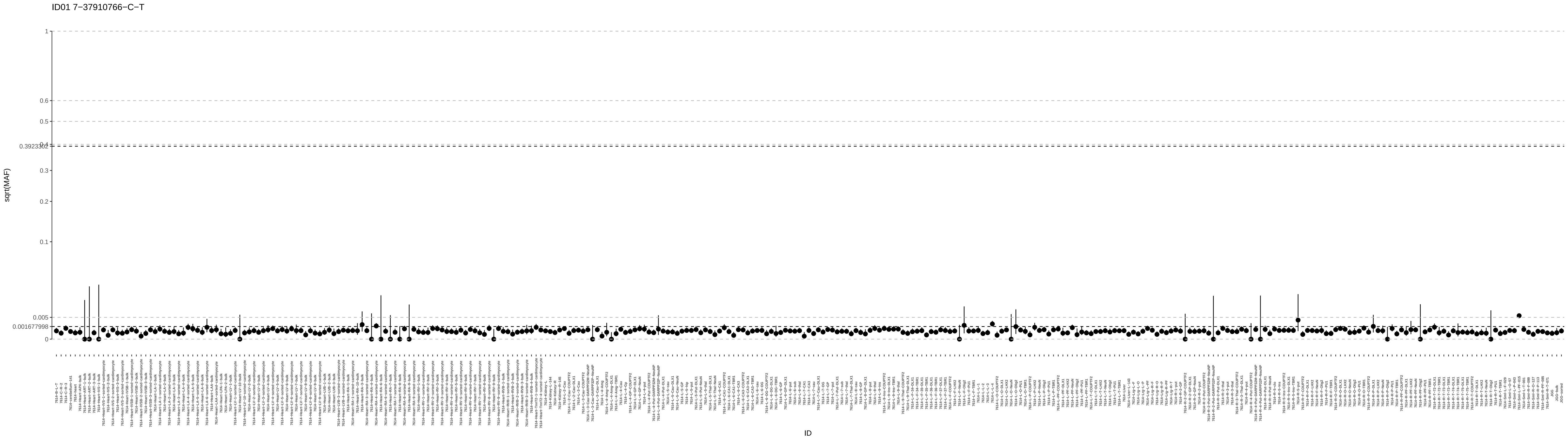
ID01 7−37910766−C−T
sqrt(MAF)
1
0.6
0.5
0.3923302
0.3
0.2
0.1
0.005
0.001677998
0
ID

ID01 7−43766364−T−G

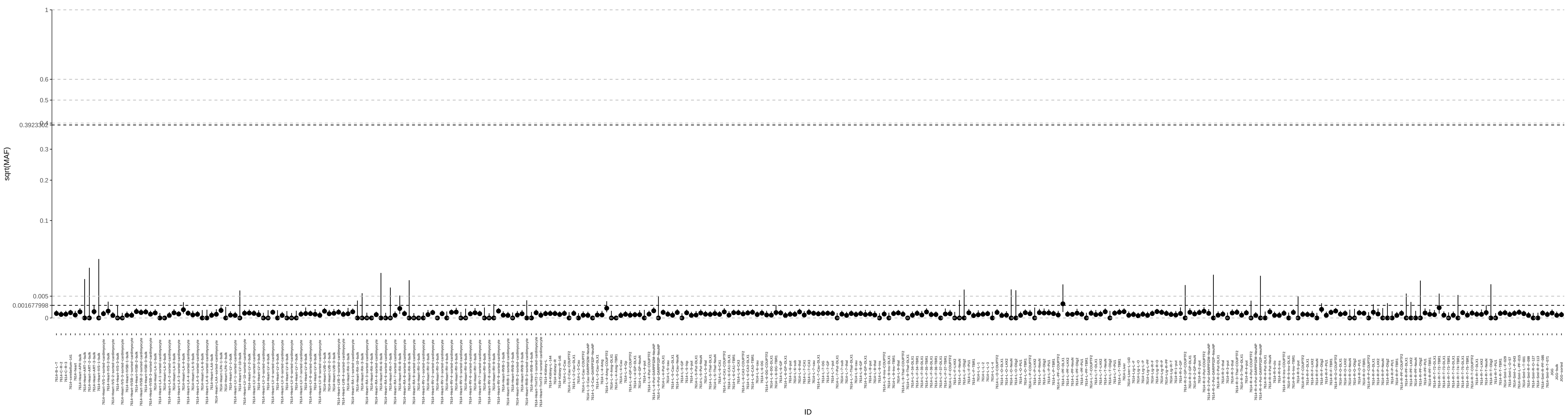

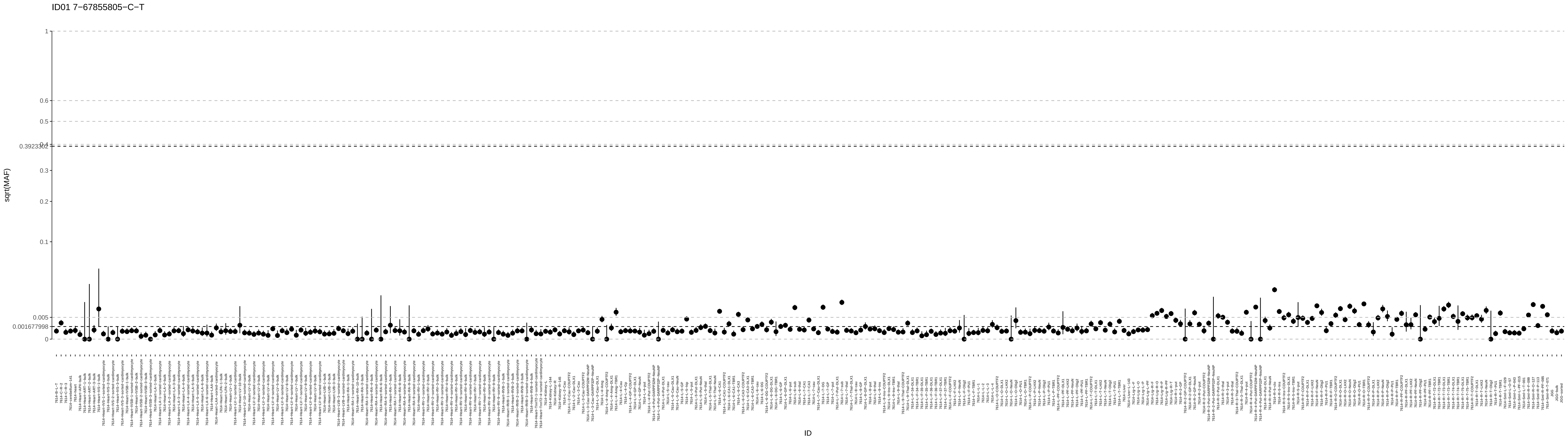
ID01 7−67855805−C−T
sqrt(MAF)
1
0.6
0.5
0.4
0.3923302
0.3
0.2
0.1
0.005
0.001677998
0
ID

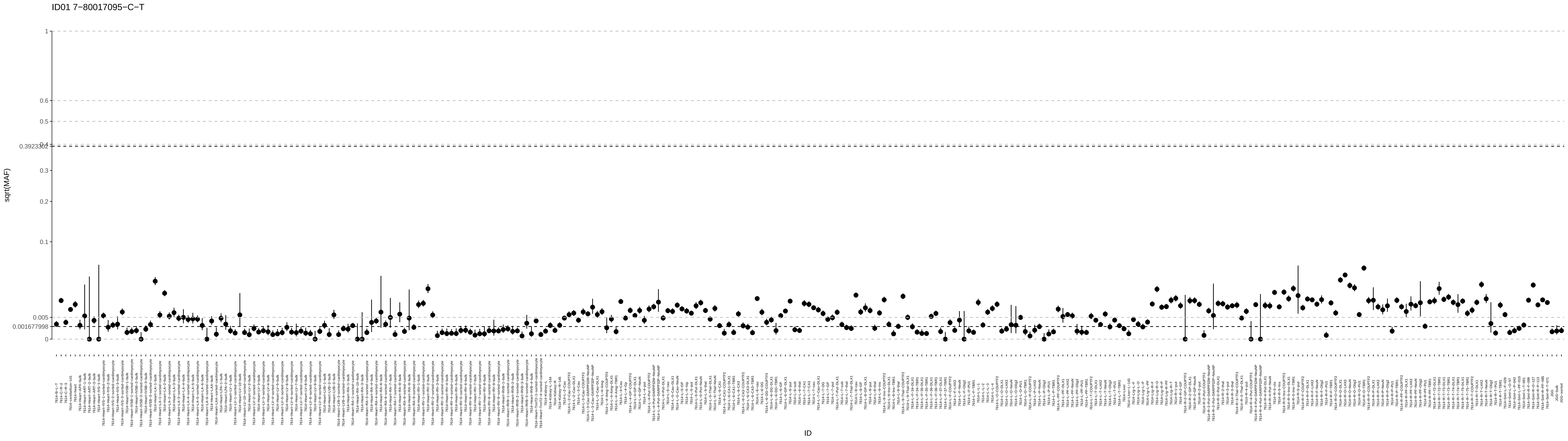

ID01 7−80017095−C−T
sqrt(MAF)
1
0.6
0.5
0.3923302
0.3
0.2
0.1
0.005
0.001677998
0
ID

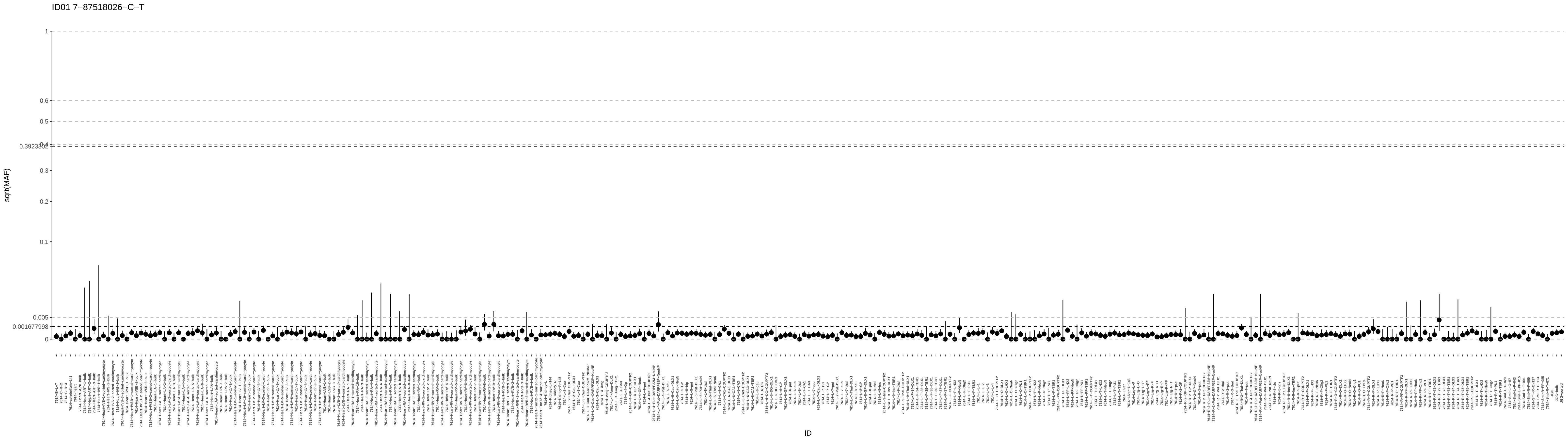
ID01 7−87518026−C−T
sqrt(MAF)
1
0.6
0.5
0.3923302
0.3
0.2
0.1
0.005
0.001677998
0
ID

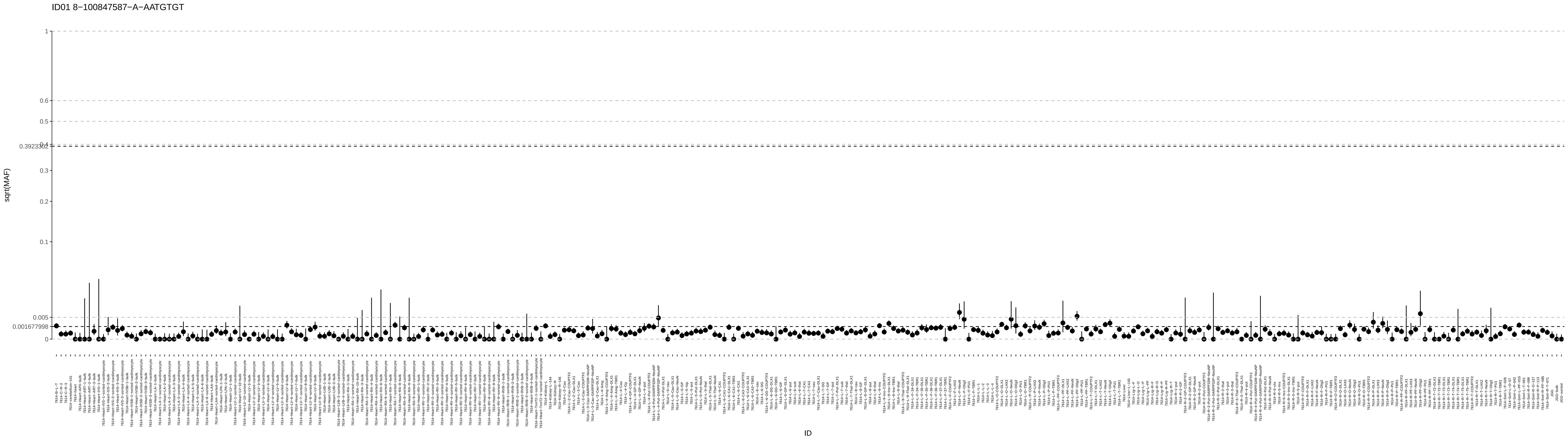
ID01 8−100847587−A−AATGTGT
sqrt(MAF)
1
0.6
0.5
0.4
0.3923302
0.3
0.2
0.1
0.005
0.001677998
0
ID

ID01 8−116848693−T−A

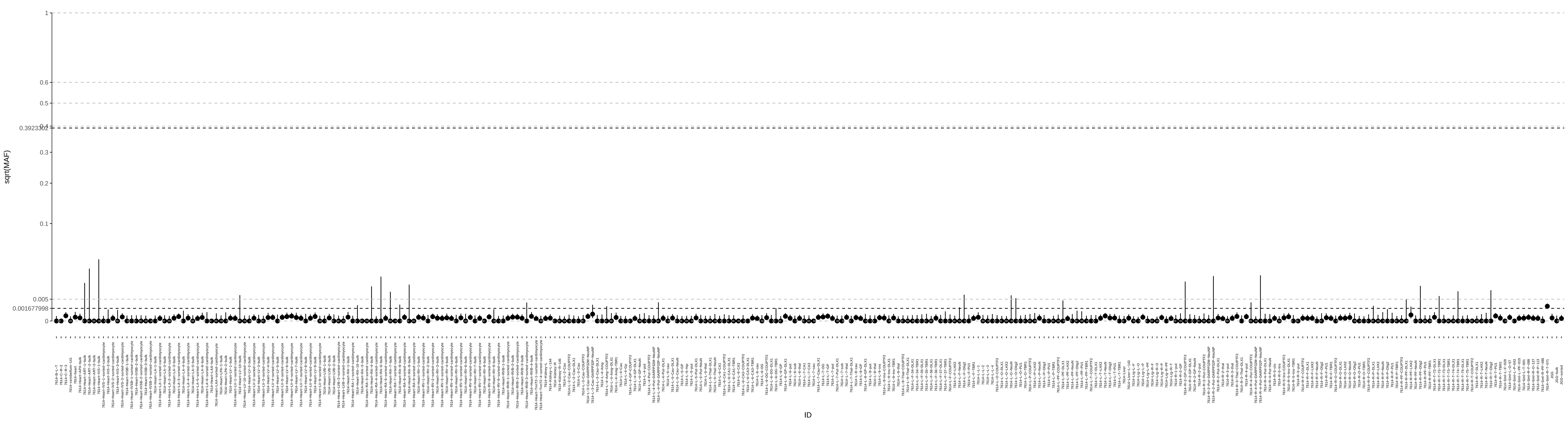

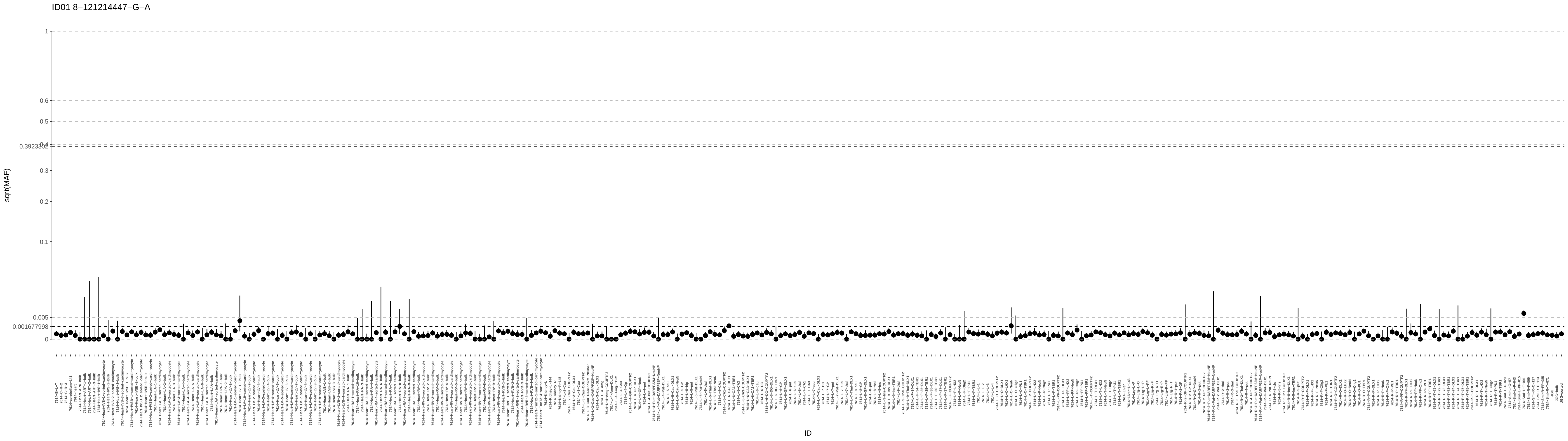
ID01 8−121214447−G−A
sqrt(MAF)
1
0.6
0.5
0.3923302
0.3
0.2
0.1
0.005
0.001677998
0
ID

ID01 8−122651370−C−A

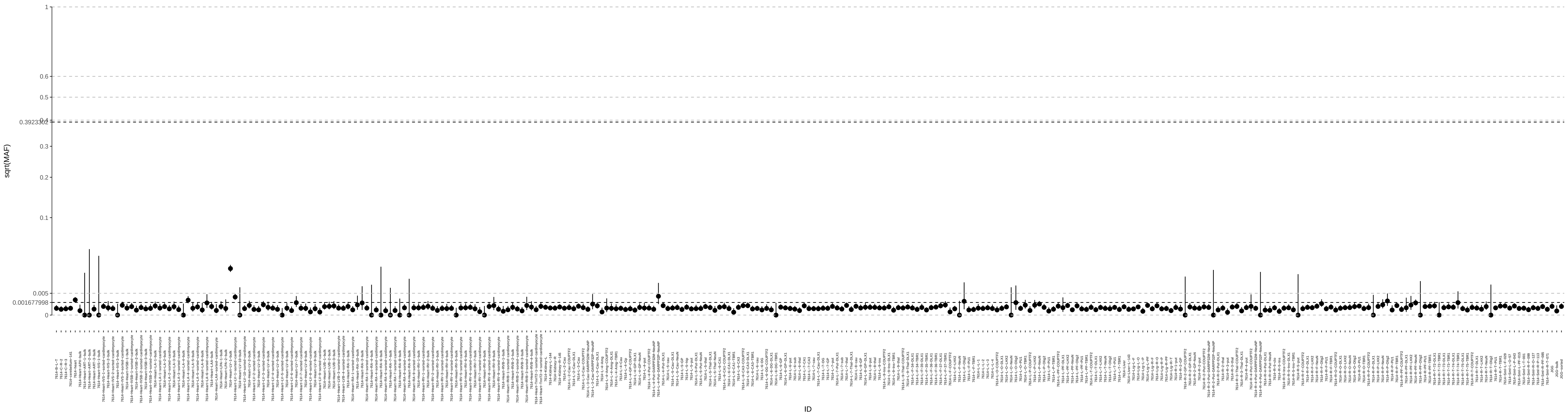

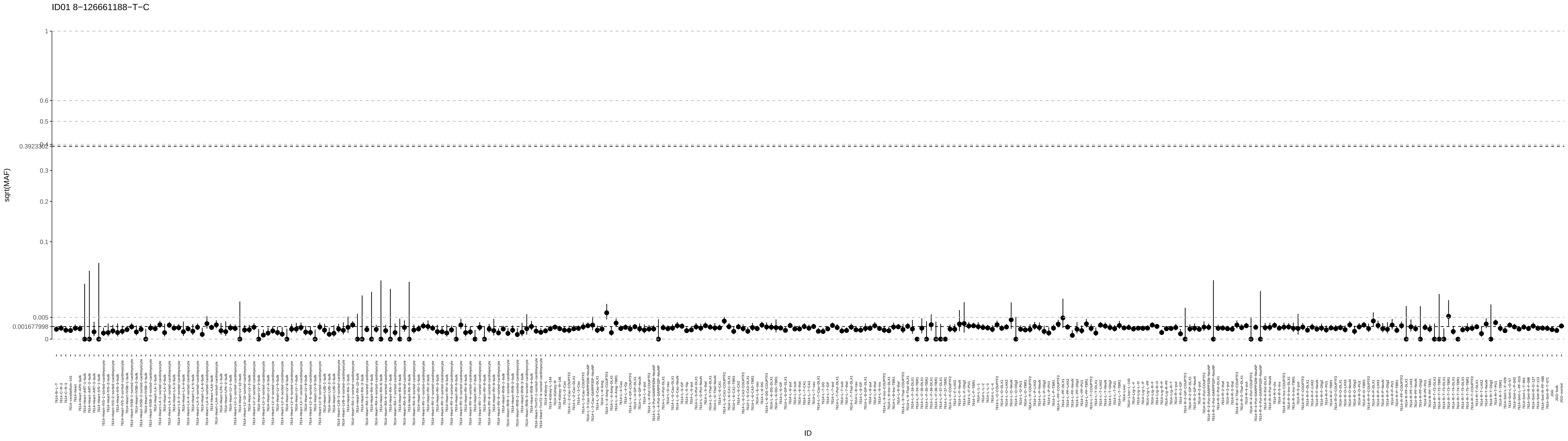
ID01 8−126661188−T−C
sqrt(MAF)
1
0.6
0.5
0.4
0.3923302
0.3
0.2
0.1
0.005
0.001677998
0
ID

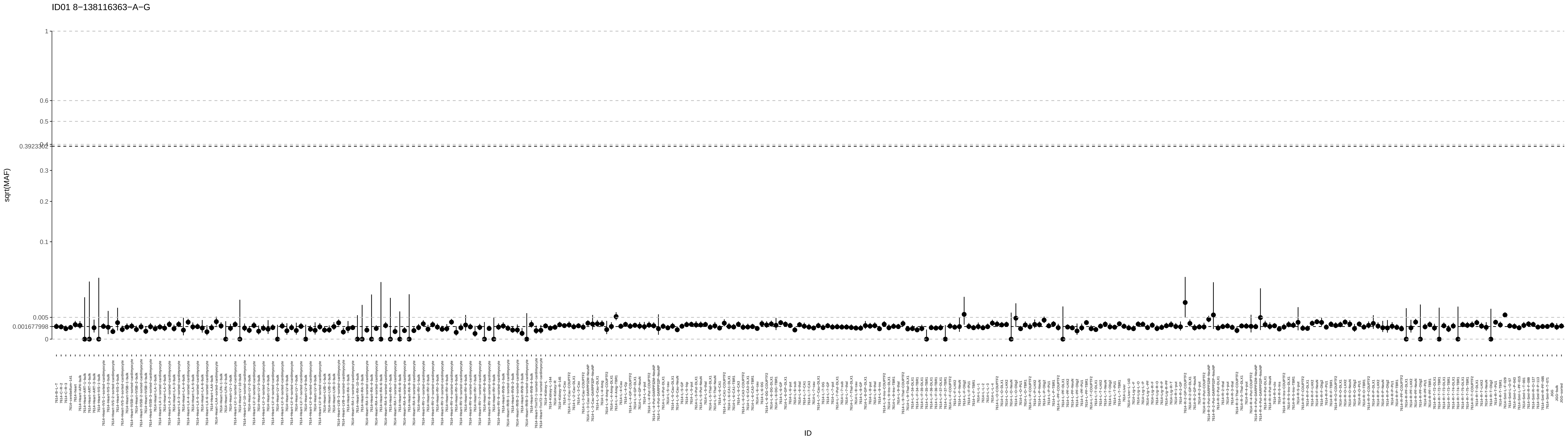
ID01 8−138116363−A−G
sqrt(MAF)
1
0.6
0.5
0.3923302
0.3
0.2
0.1
0.005
0.001677998
0
ID

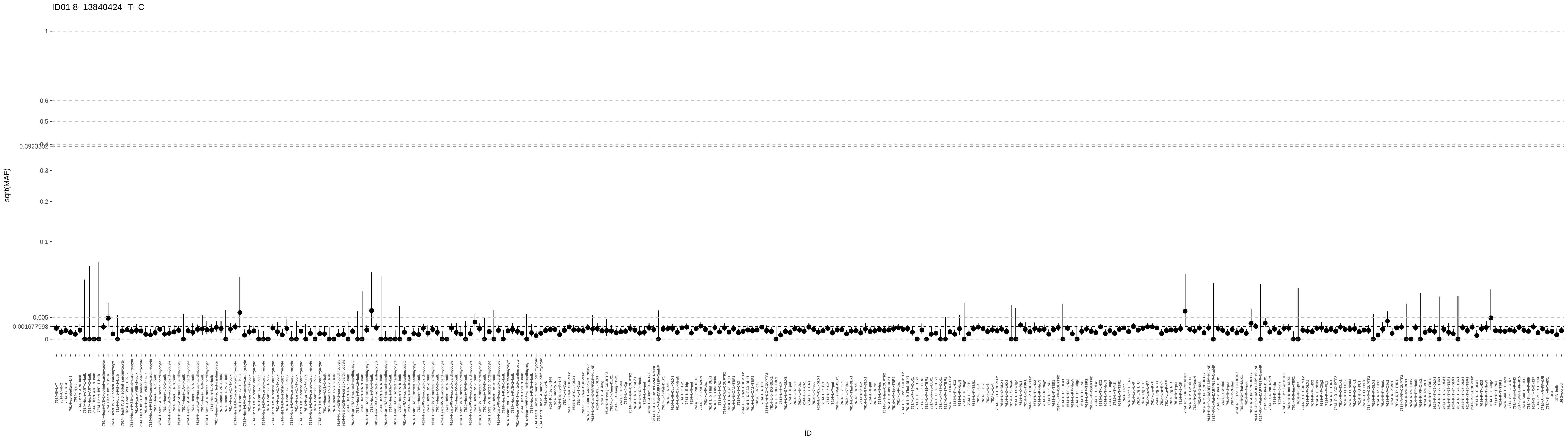
ID01 8−13840424−T−C
sqrt(MAF)
1
0.6
0.5
0.3923302
0.3
0.2
0.1
0.005
0.001677998
0
ID

ID01 8−14166252−C−T

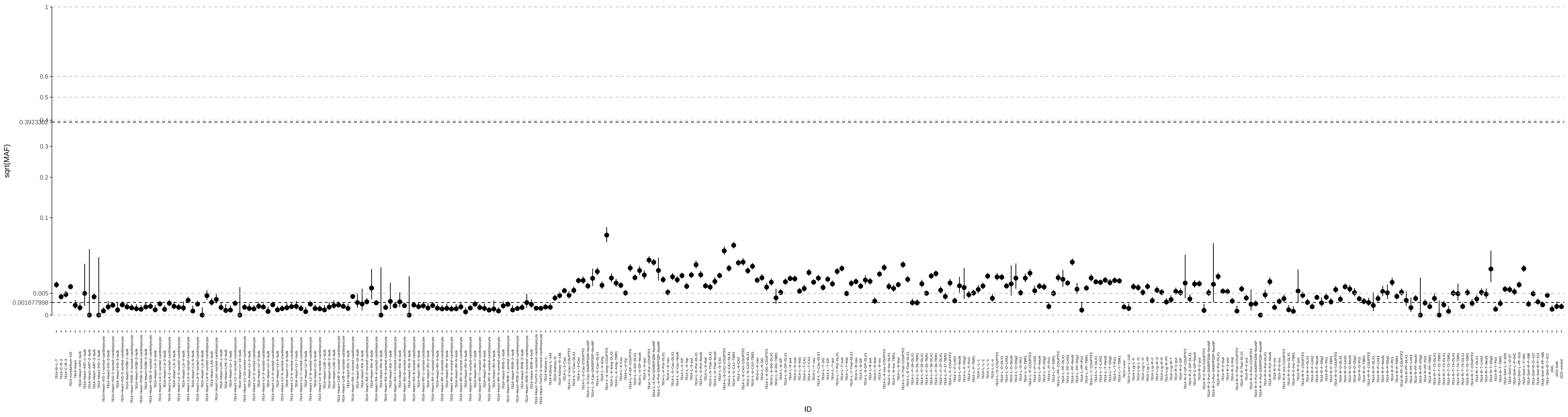

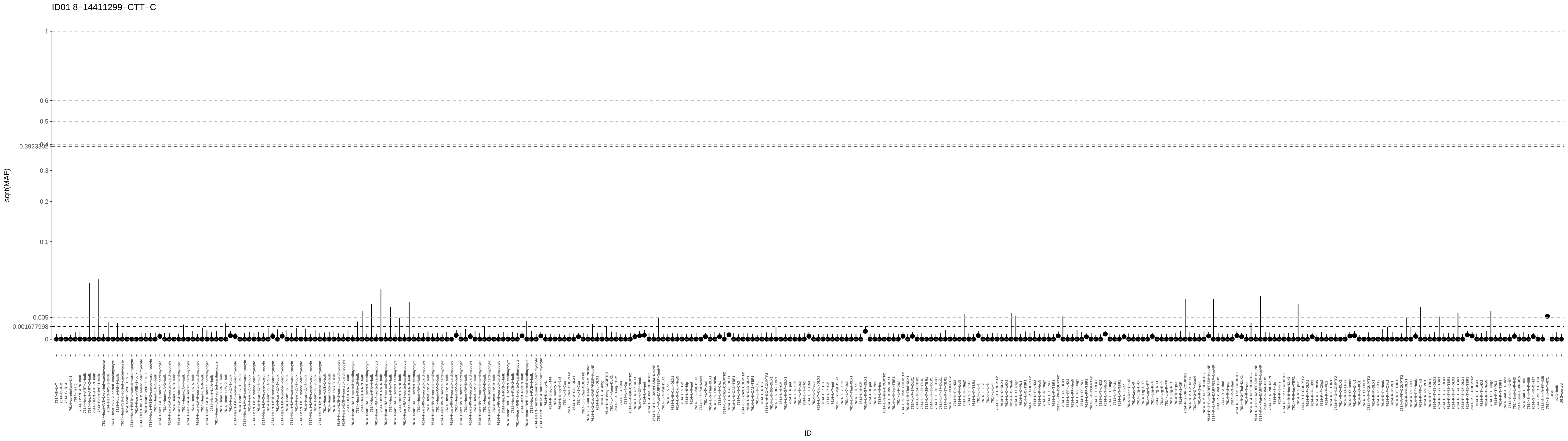
ID01 8−14411299−CTT−C
sqrt(MAF)
1
0.6
0.5
0.3923302
0.3
0.2
0.1
0.005
0.001677998
0
ID

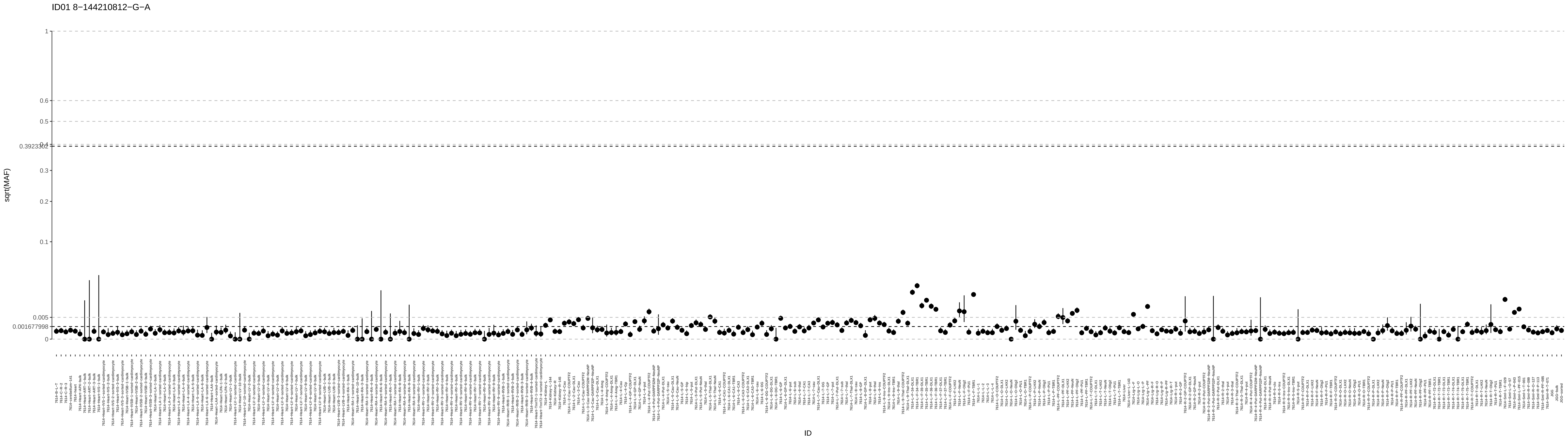

ID01 8−144210812−G−A
sqrt(MAF)
1
0.6
0.5
0.3923302
0.3
0.2
0.1
0.005
0.001677998
0
ID

ID01 8−2199684−G−A

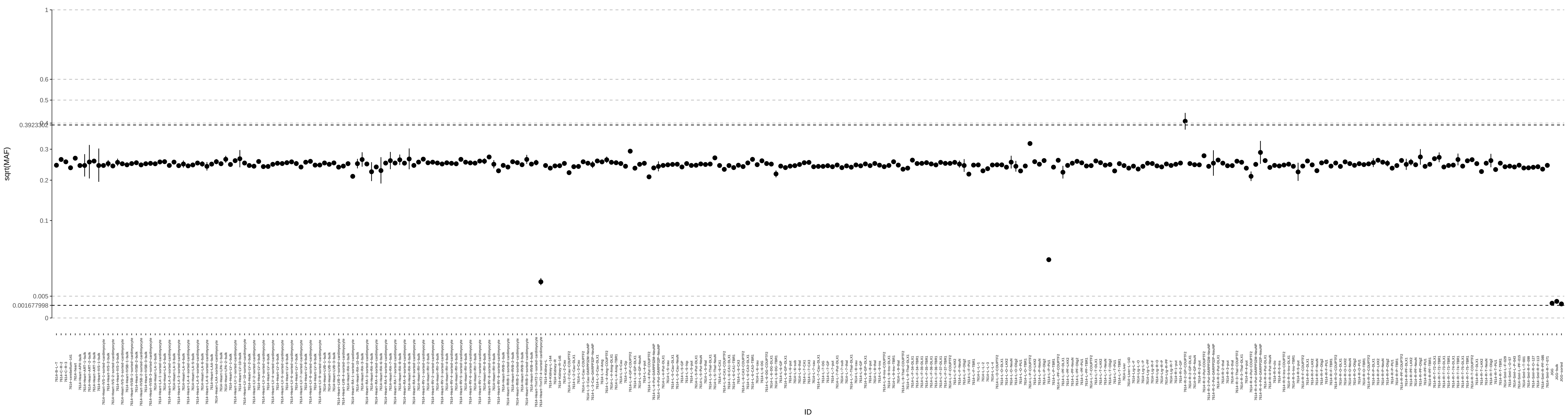

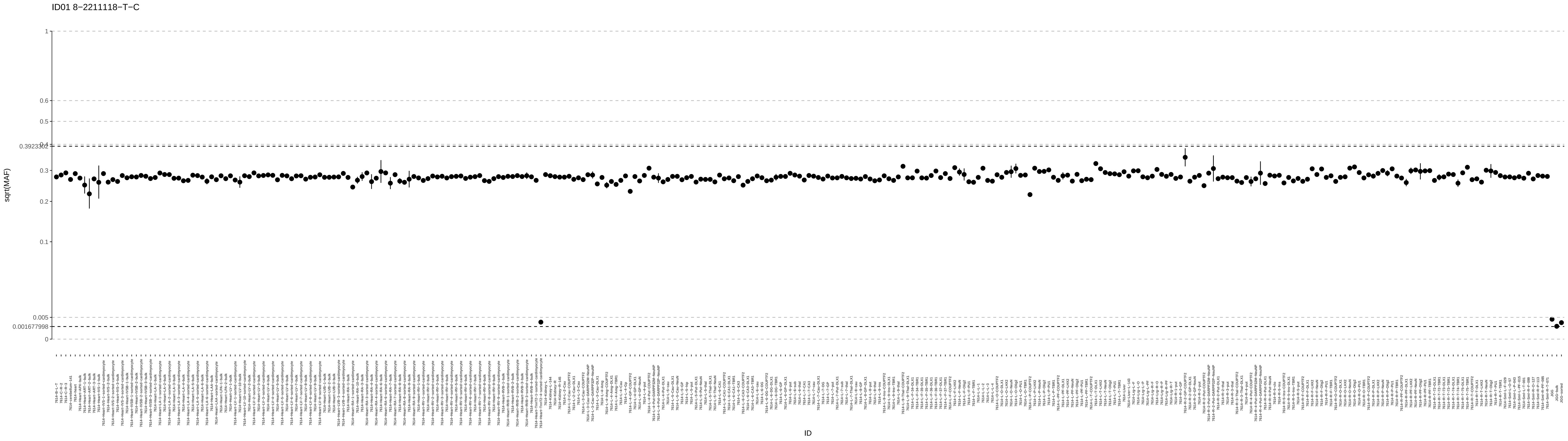
ID01 8−2211118−T−C
sqrt(MAF)
1
0.6
0.5
0.3923302
0.3
0.2
0.1
0.005
0.001677998
0
ID

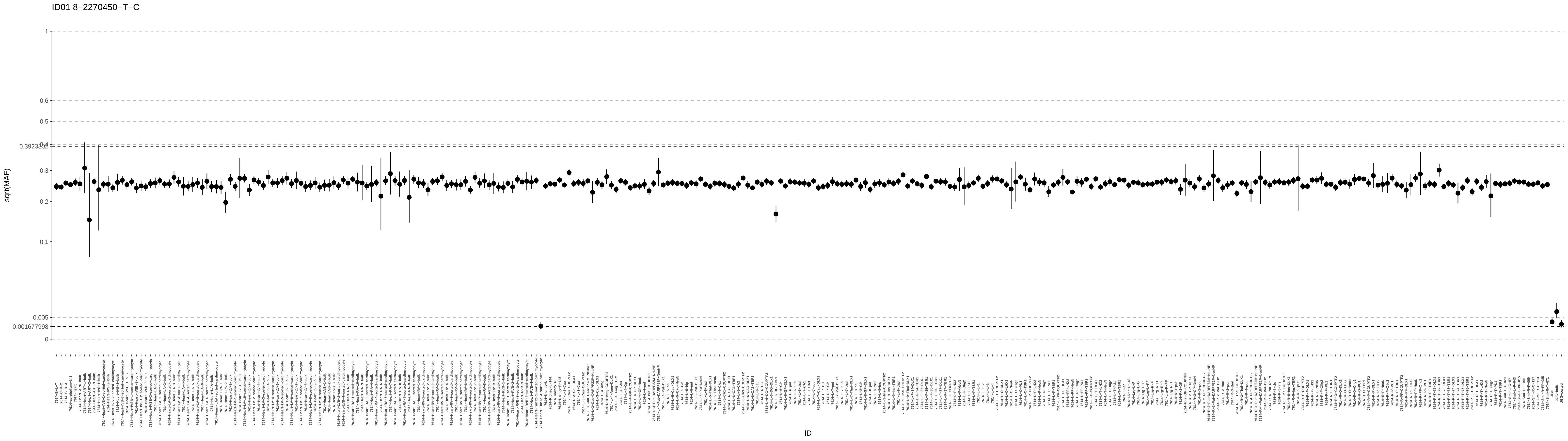
ID01 8−2270450−T−C
sqrt(MAF)
1
0.6
0.5
0.3923302
0.3
0.2
0.1
0.005
0.001677998
0
ID

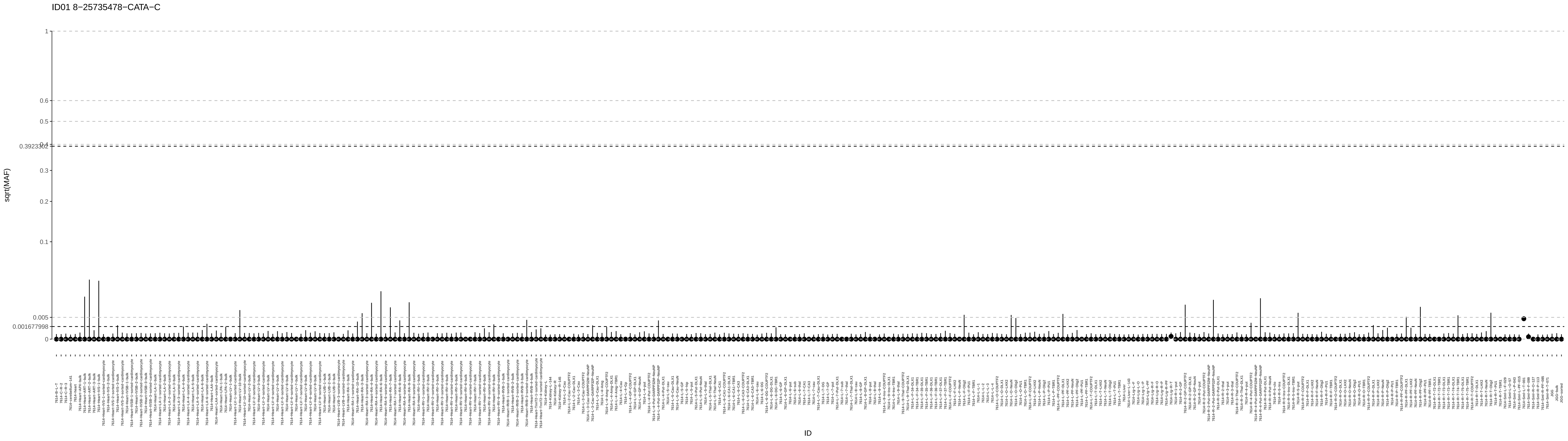
ID01 8−25735478−CATA−C
sqrt(MAF)
ID
0.3923302
0.001677998

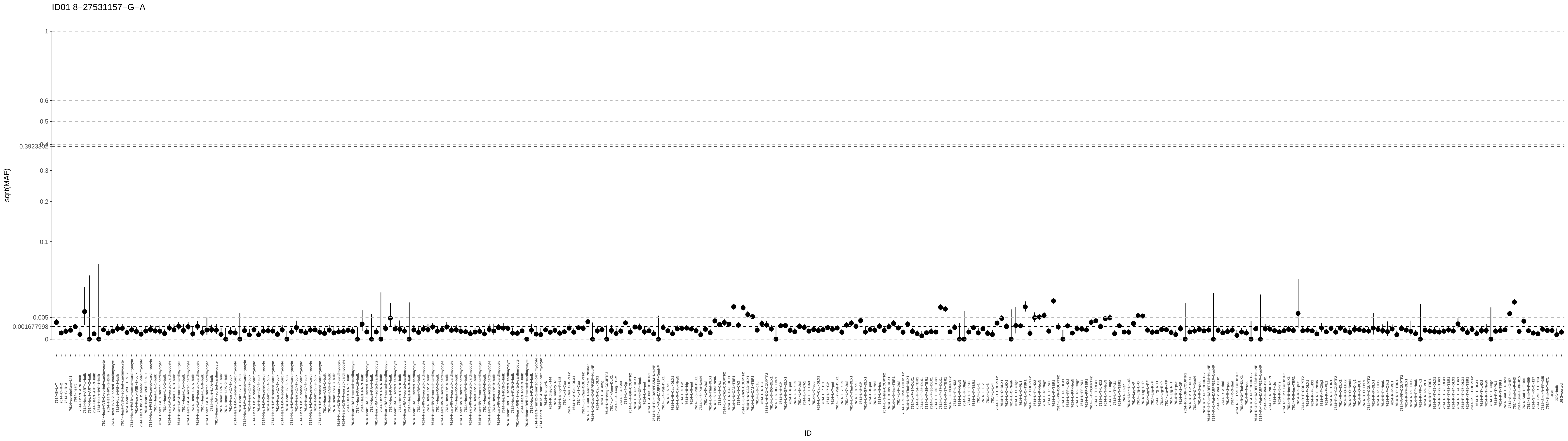

ID01 8−27531157−G−A
sqrt(MAF)
1
0.6
0.5
0.3923302
0.3
0.2
0.1
0.005
0.001677998
0
ID

ID01 8-28894088-T-G

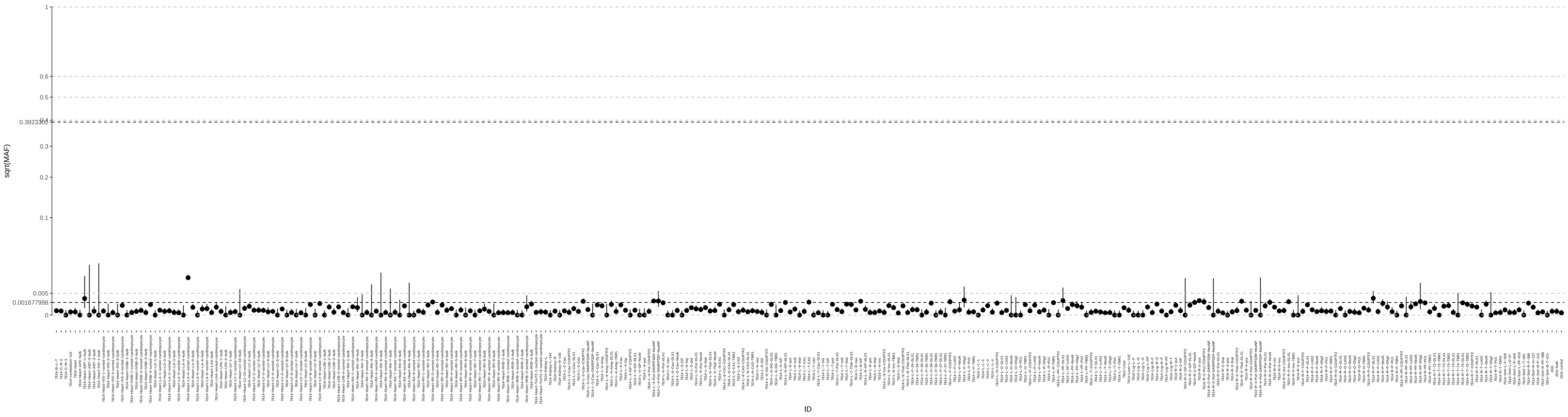

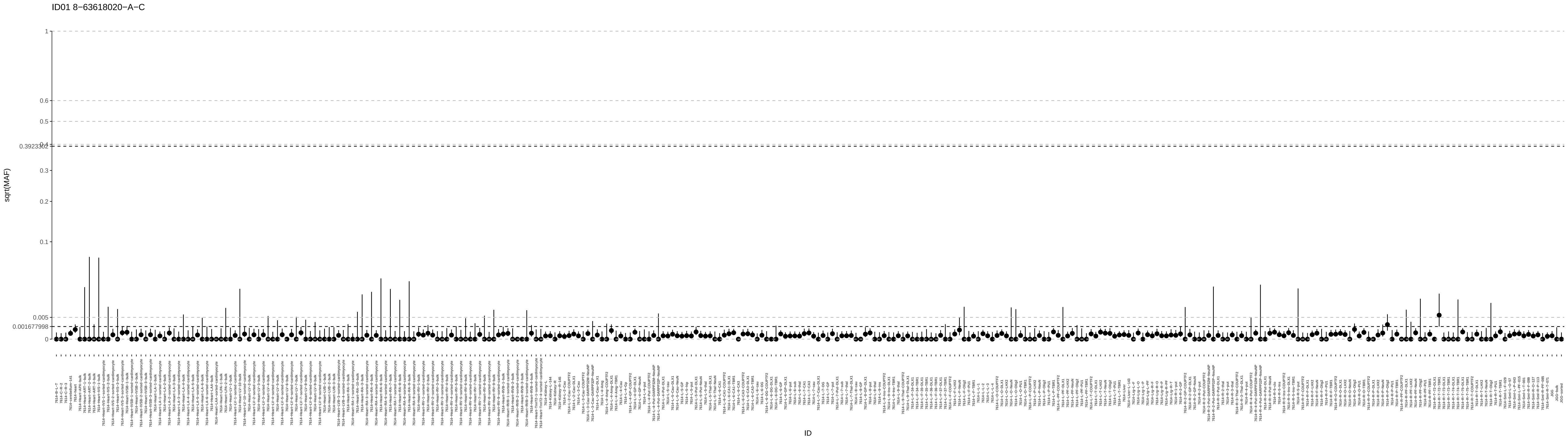
ID01 8−63618020−A−C
sqrt(MAF)
1
0.6
0.5
0.3923302
0.3
0.2
0.1
0.005
0.001677998
0
ID

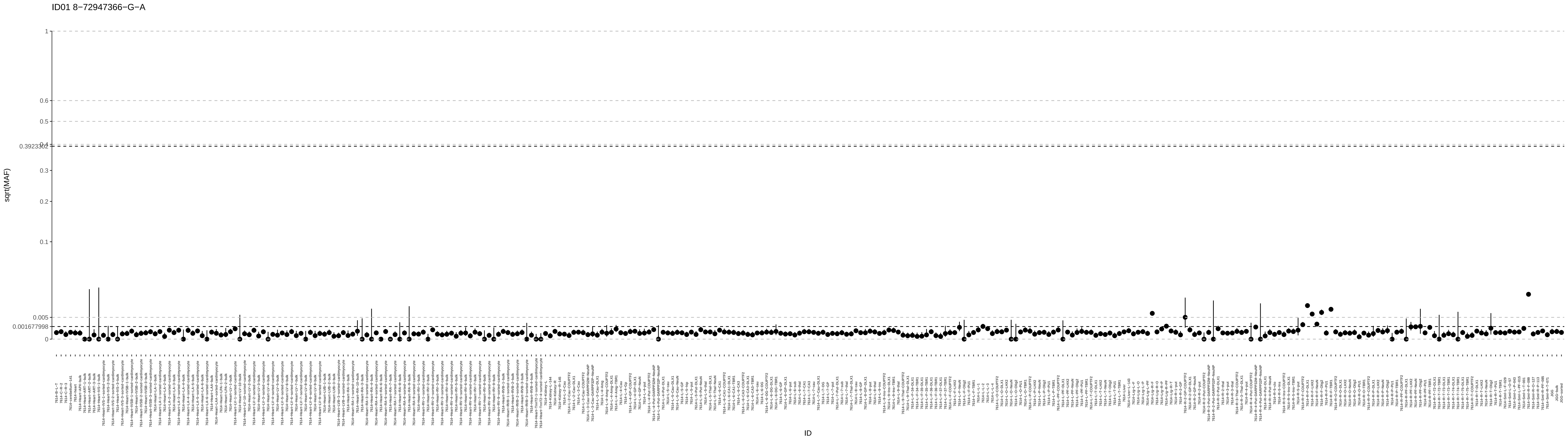
ID01 8−72947366−G−A
sqrt(MAF)
1
0.6
0.5
0.3923302
0.4
0.3
0.2
0.1
0.005
0.001677998
0
ID

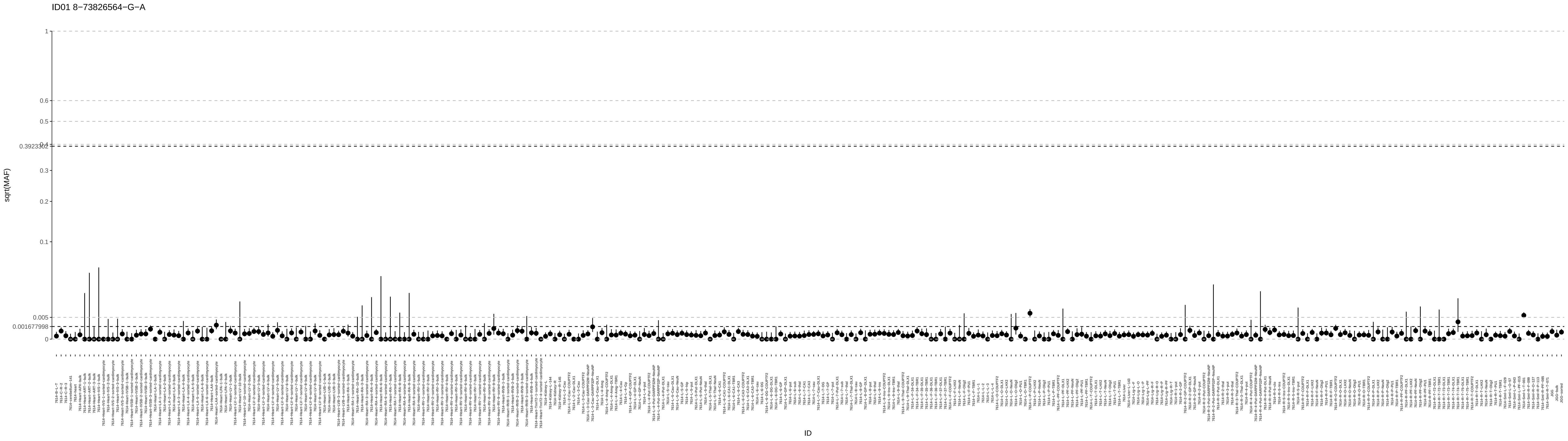
ID01 8−73826564−G−A
sqrt(MAF)
1
0.6
0.5
0.4
0.3923302
0.3
0.2
0.1
0.005
0.001677998
0
ID

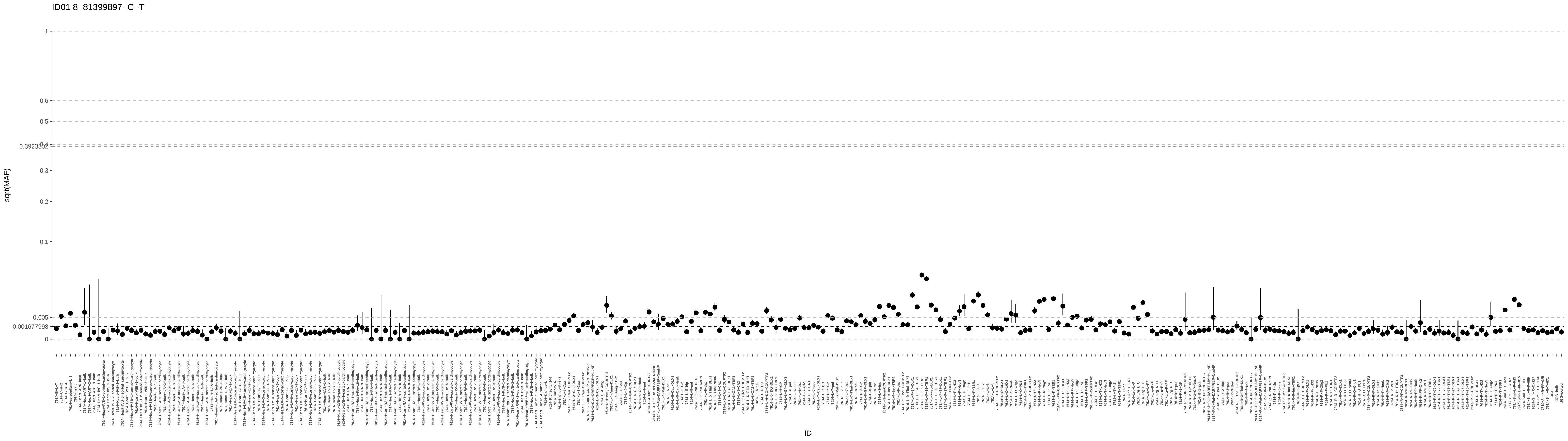
ID01 8−81399897−C−T
sqrt(MAF)
1
0.6
0.5
0.3923302
0.3
0.2
0.1
0.005
0.001677998
0
ID

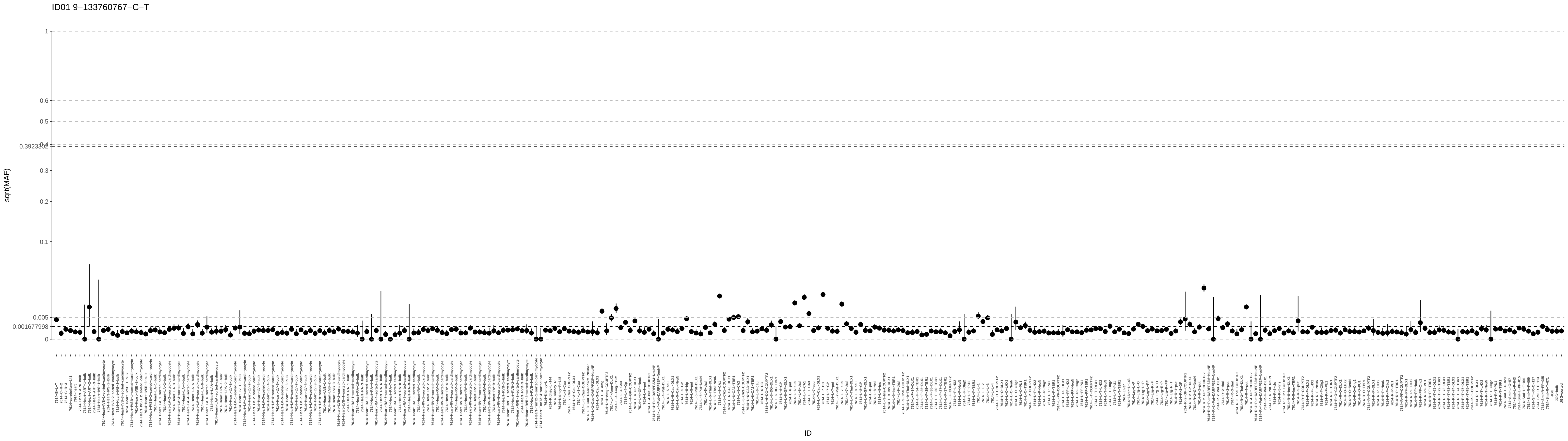
ID01 9-133760767-C-T
sqrt(MAF)
1
0.6
0.5
0.3923302
0.3
0.2
0.1
0.005
0.001677998
0
ID

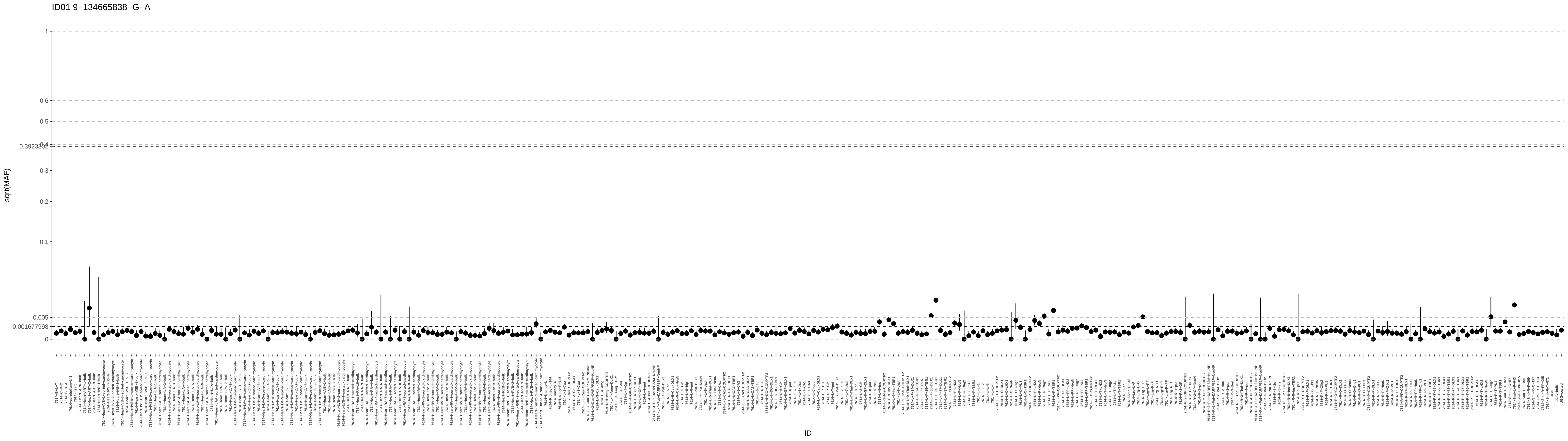
ID01 9−134665838−G−A
sqrt(MAF)
1
0.6
0.5
0.4
0.3923302
0.3
0.2
0.1
0.005
0.001677998
0
ID

ID01 9−134950488−G−T

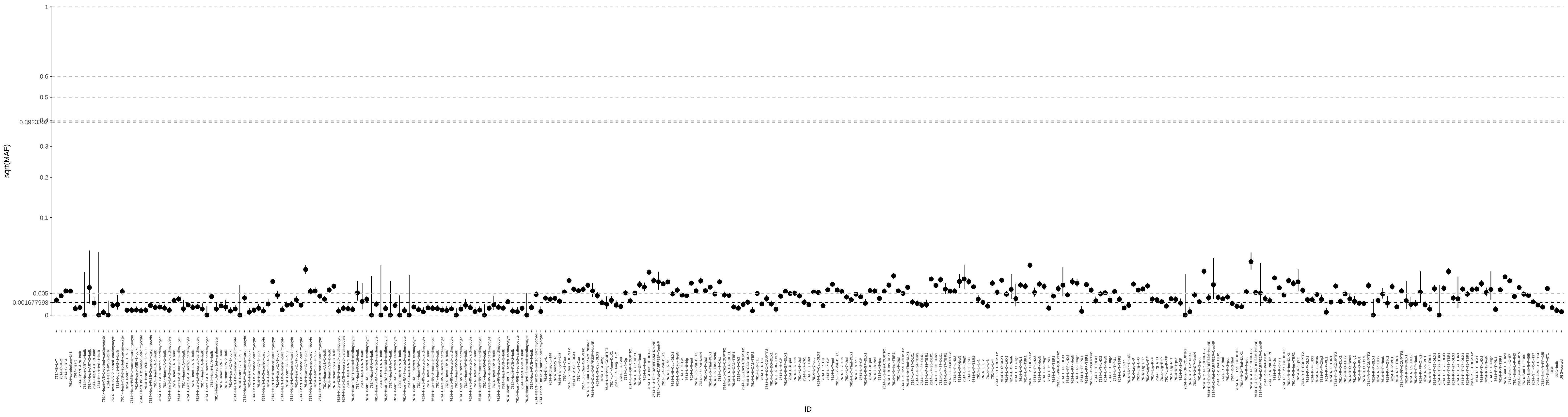

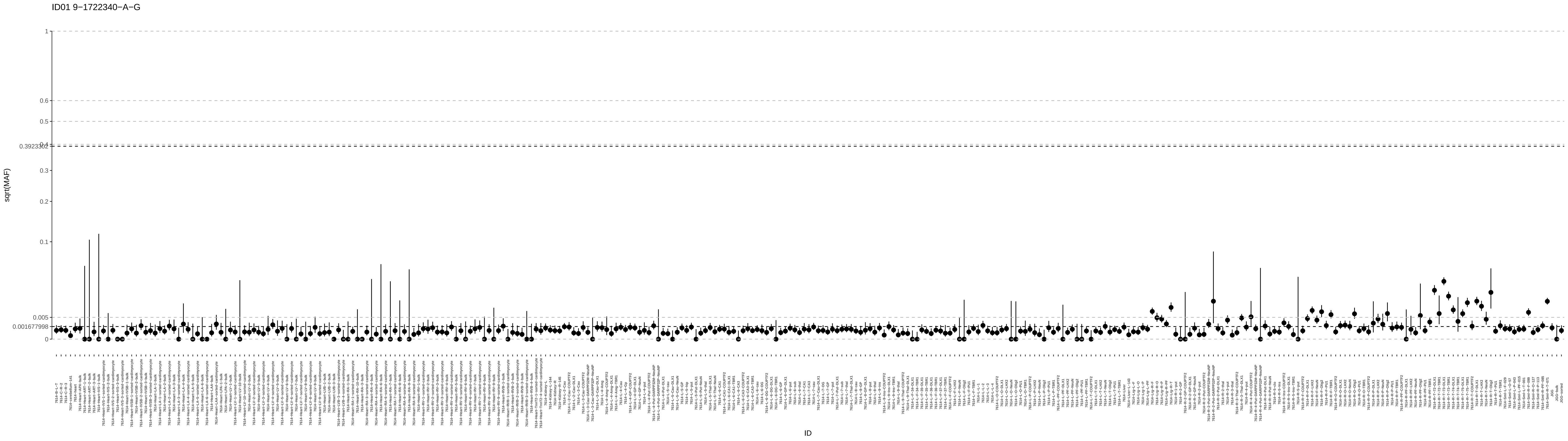
ID01 9-1722340-A-G
sqrt(MAF)
1
0.6
0.5
0.3923302
0.3
0.2
0.1
0.005
0.001677998
0
ID

ID01 9−38668272−C−T

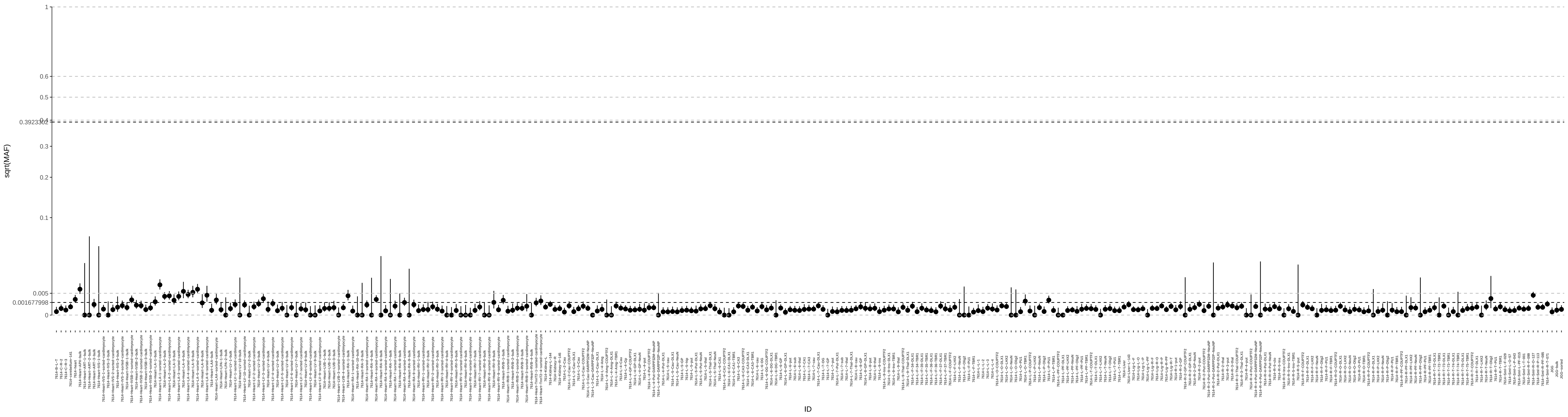

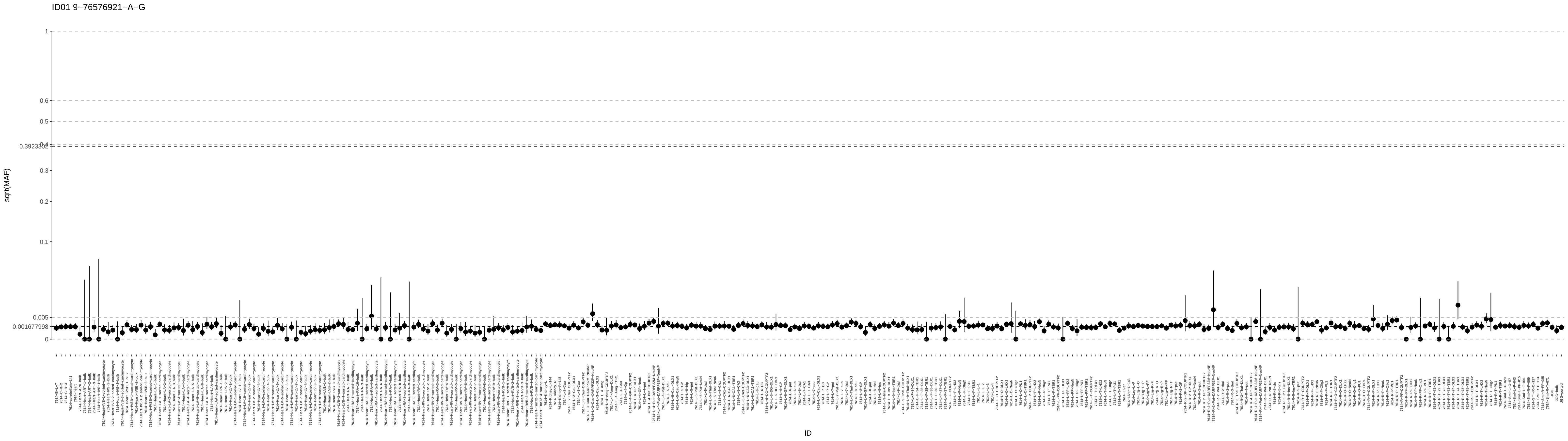
ID01 9−76576921−A−G
sqrt(MAF)
1
0.6
0.5
0.4
0.3923302
0.3
0.2
0.1
0.005
0.001677998
0
ID

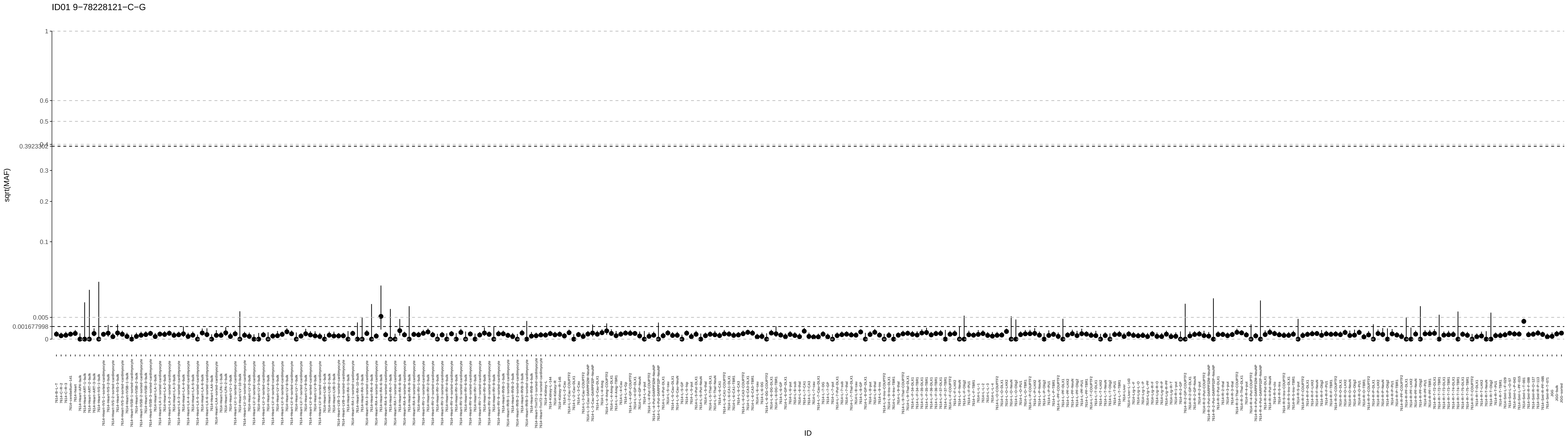
ID01 9−78228121−C−G
sqrt(MAF)
ID
1
0.6
0.5
0.3923302
0.3
0.2
0.1
0.005
0.001677998
0

ID01 9-7860755-C-T

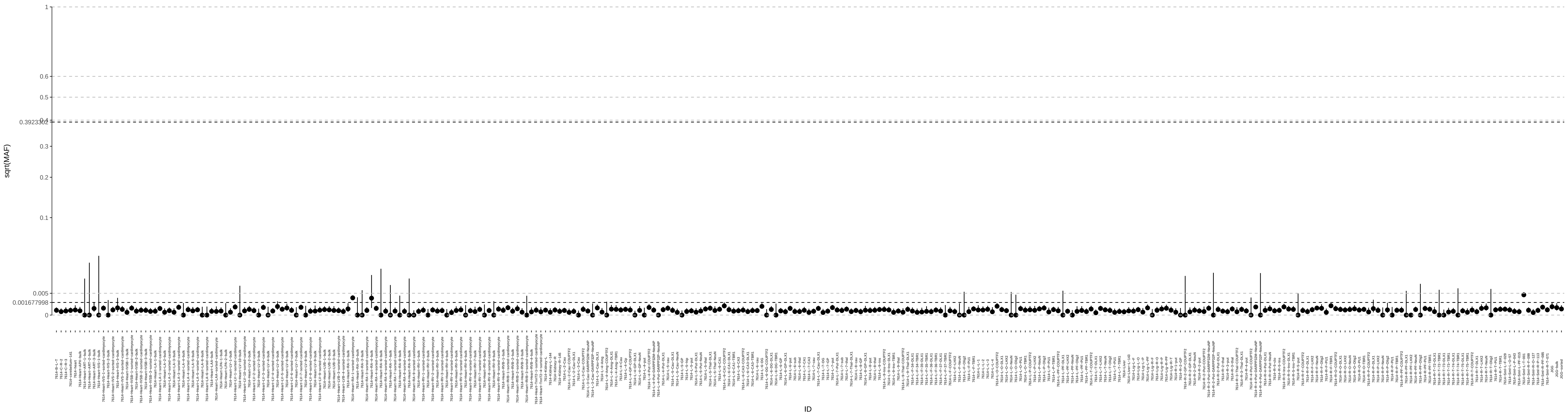

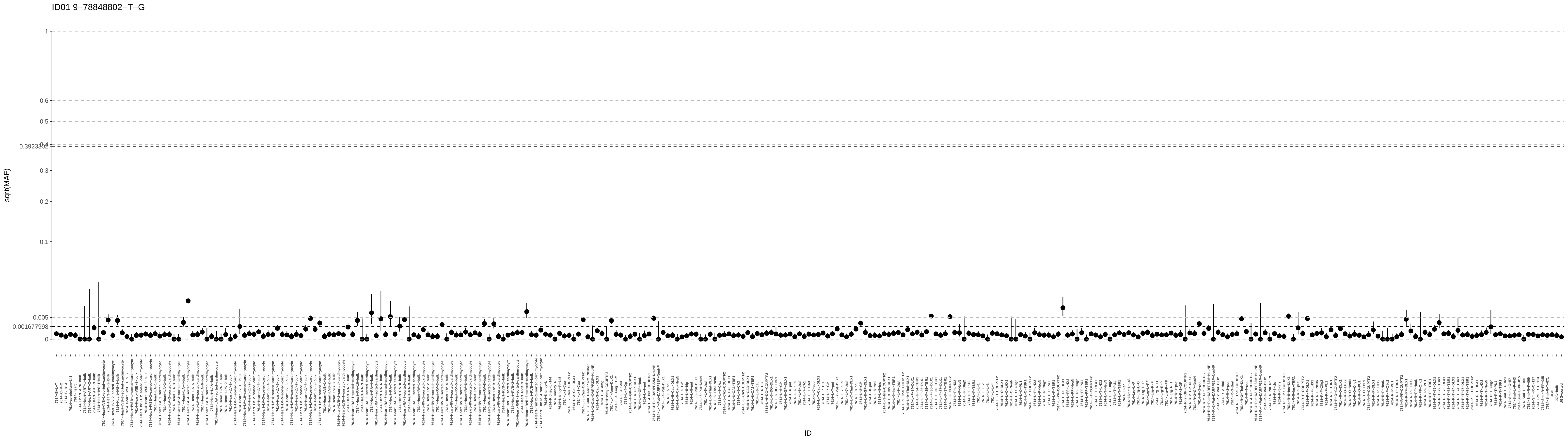
ID01 9−78848802−T−G
sqrt(MAF)
1
0.6
0.5
0.3923302
0.3
0.2
0.1
0.005
0.001677998
0
ID

ID01 9−88011667−A−C

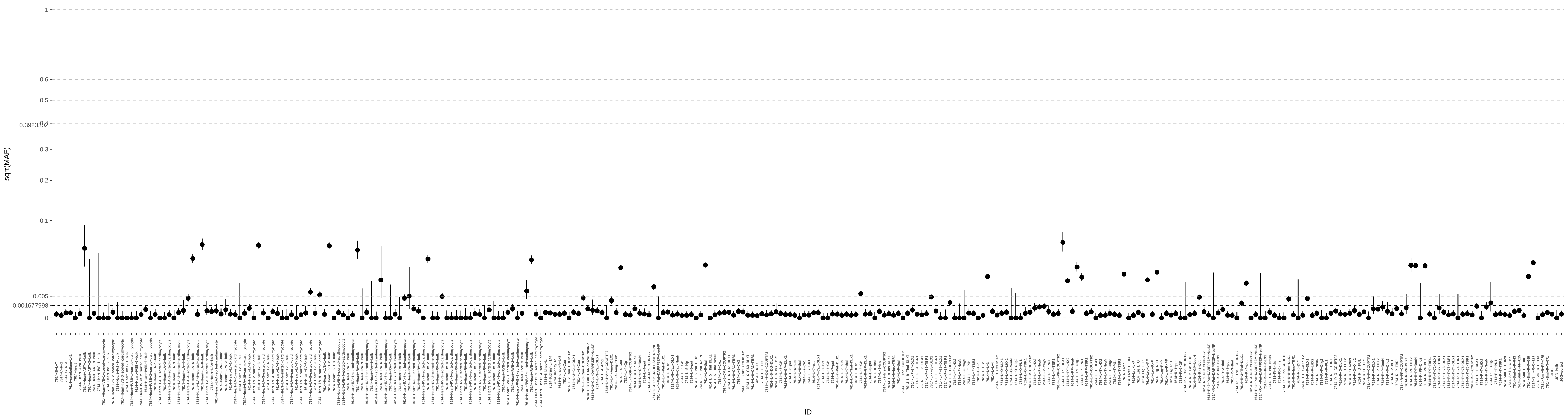

ID01 9–9457115–T–A

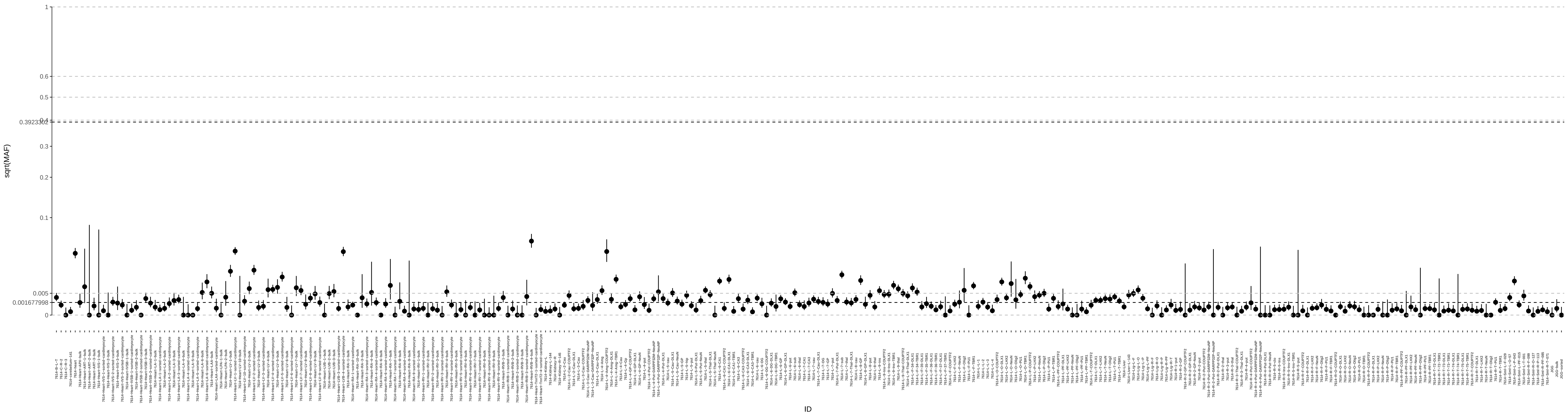

ID01 X−10480747−T−A

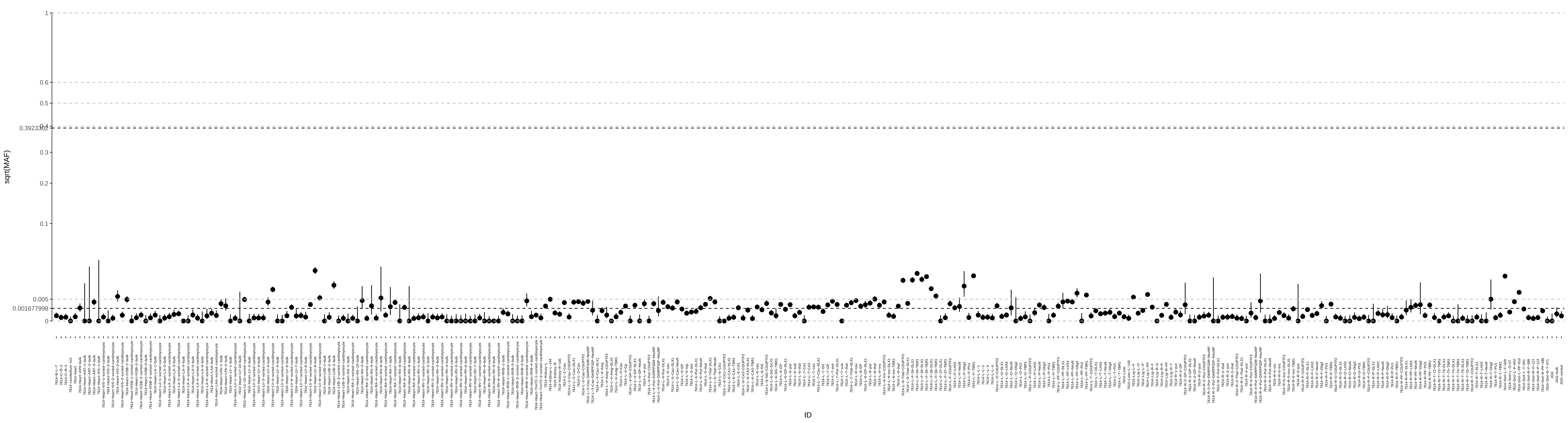

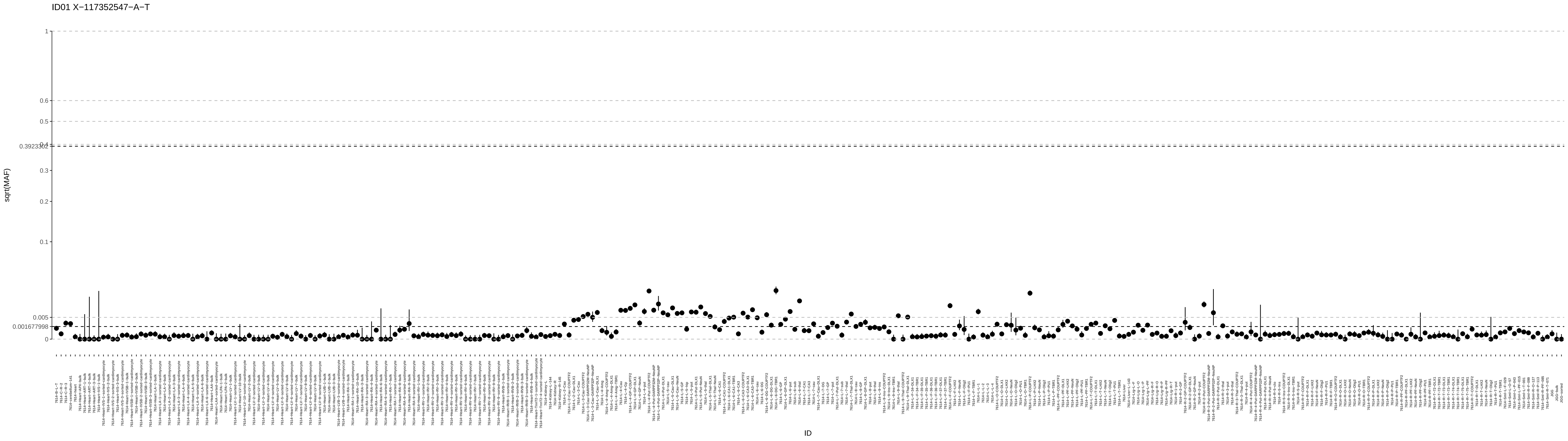
ID01 X−117352547−A−T
sqrt(MAF)
1
0.6
0.5
0.3923302
0.3
0.2
0.1
0.005
0.001677998
0
ID

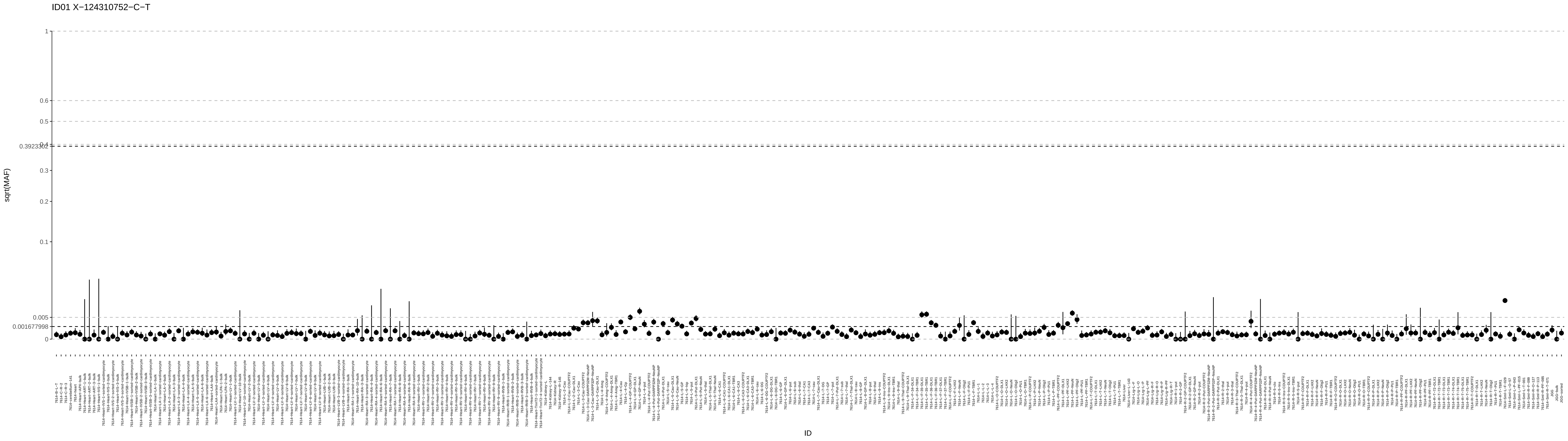
ID01 X−124310752−C−T
1
0.6
0.5
0.4
0.3923302
0.3
0.2
0.1
0.005
0.001677998
0
sqrt(MAF)
ID

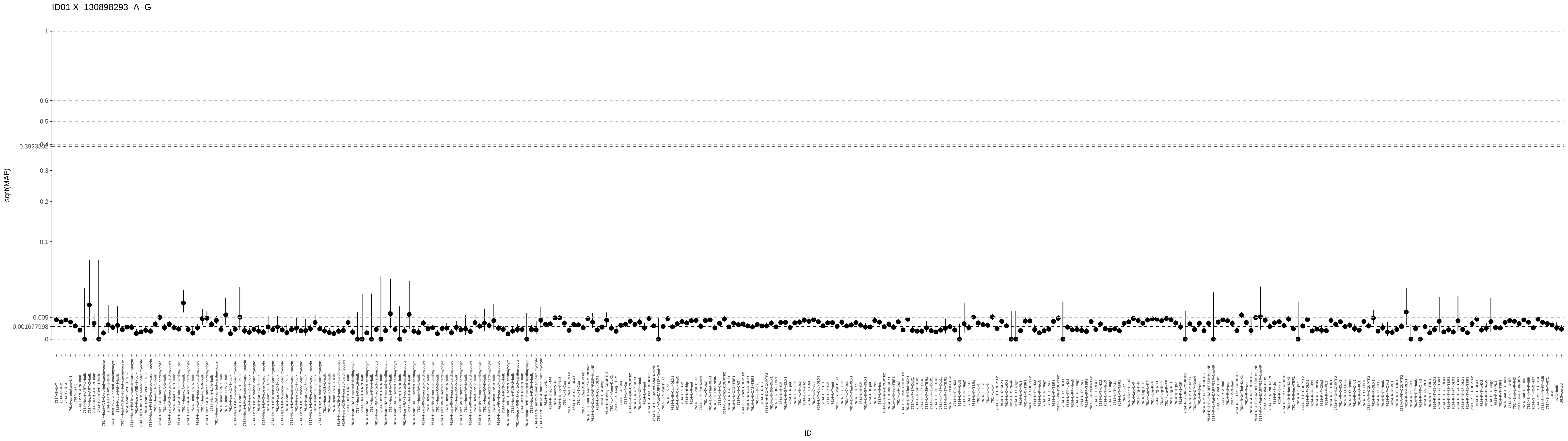
ID01 X−130898293−A−G
sqrt(MAF)
1
0.6
0.5
0.3923302
0.3
0.2
0.1
0.005
0.001677998
0
ID

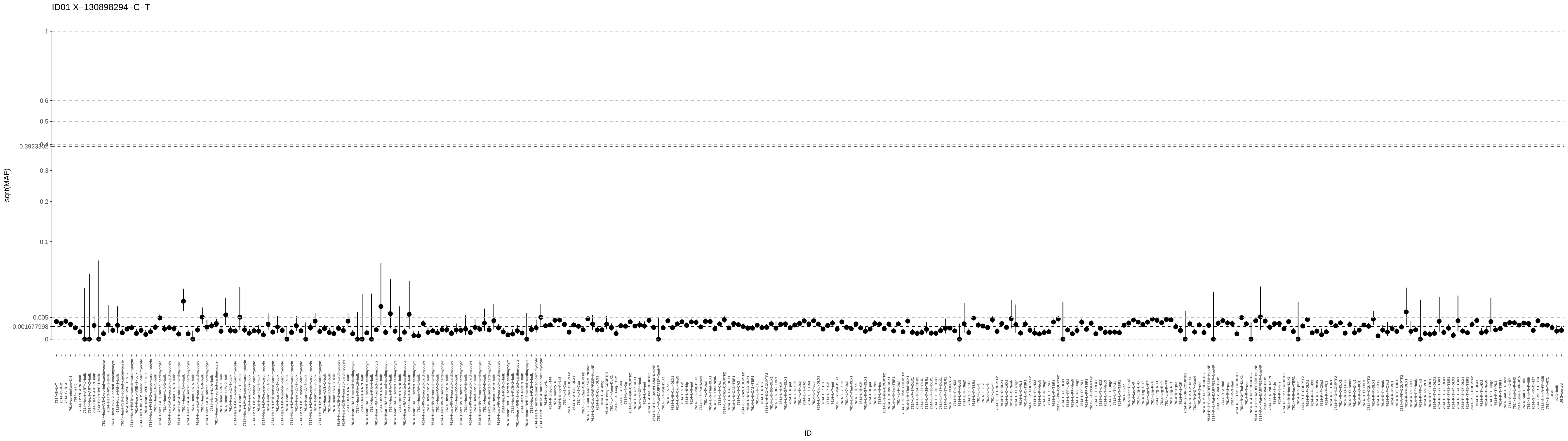
ID01 X−130898294−C−T
sqrt(MAF)
1
0.6
0.5
0.3923302
0.3
0.2
0.1
0.005
0.001677998
0
ID

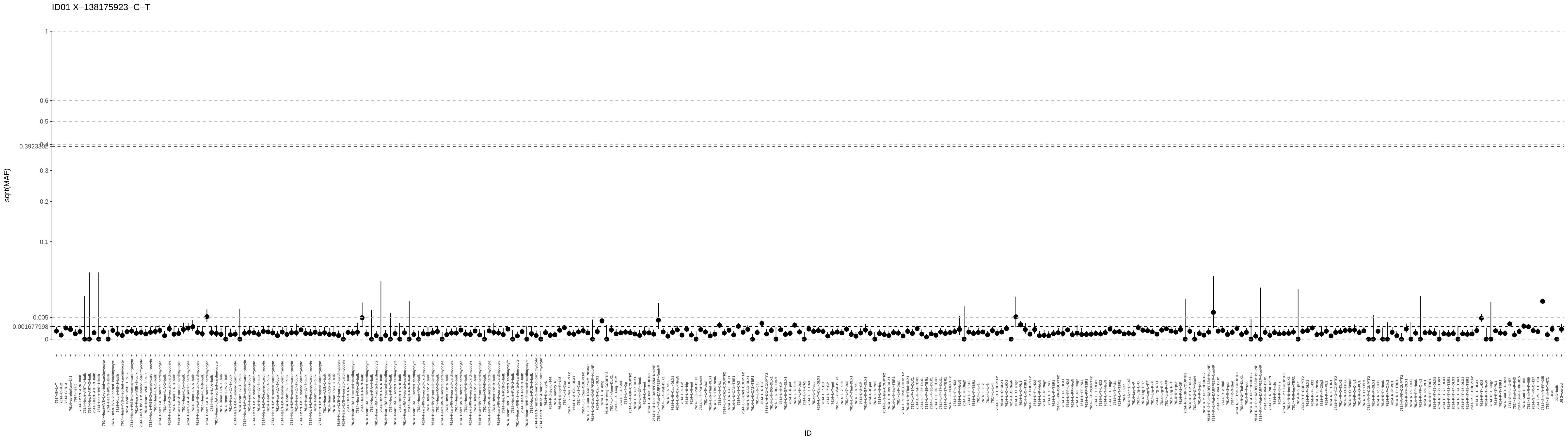
ID01 X−138175923−C−T
sqrt(MAF)
1
0.6
0.5
0.3923302
0.3
0.2
0.1
0.005
0.001677998
0
ID

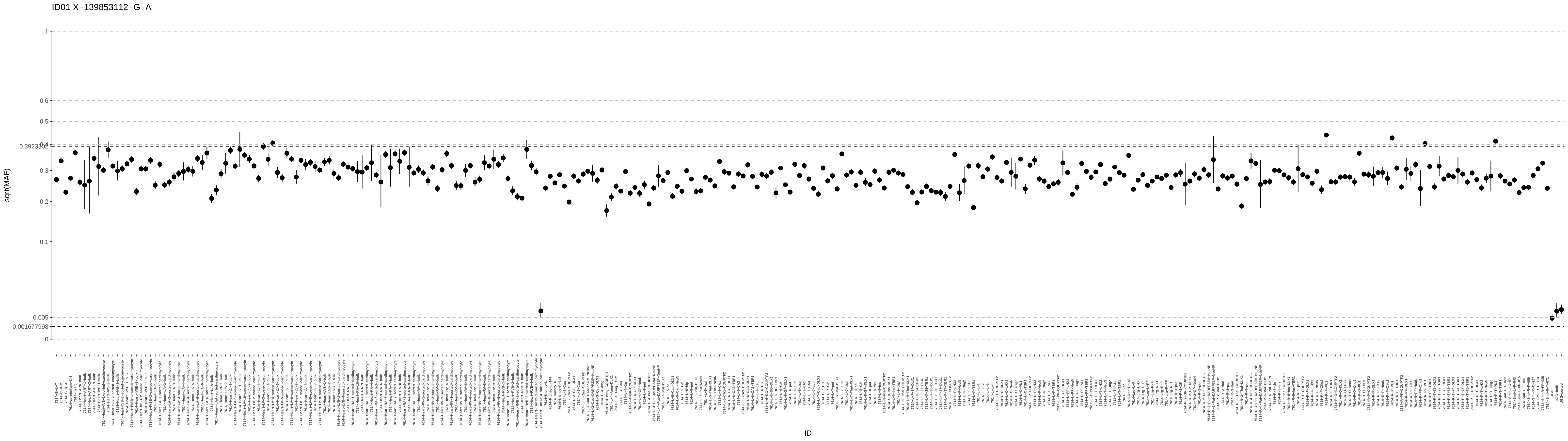
ID01 X−139853112−G−A
sqrt(MAF)
1
0.6
0.5
0.4
0.3923302
0.3
0.2
0.1
0.005
0.001677998
0
ID

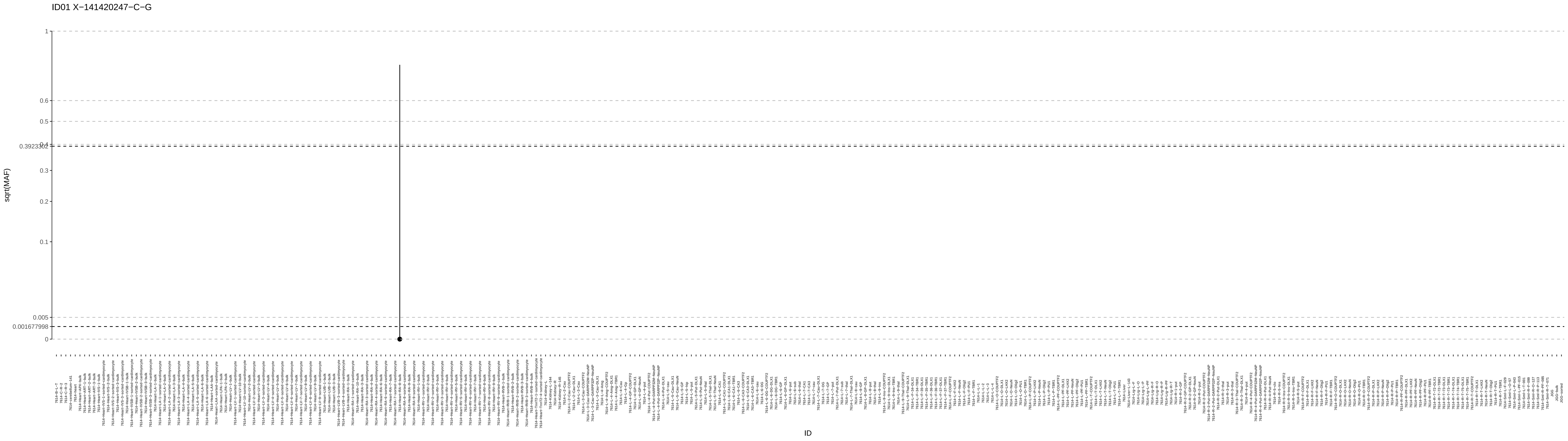
ID01 X−141420247−C−G
sqrt(MAF)
1
0.6
0.5
0.3923302
0.3
0.2
0.1
0.005
0.001677998
0
ID

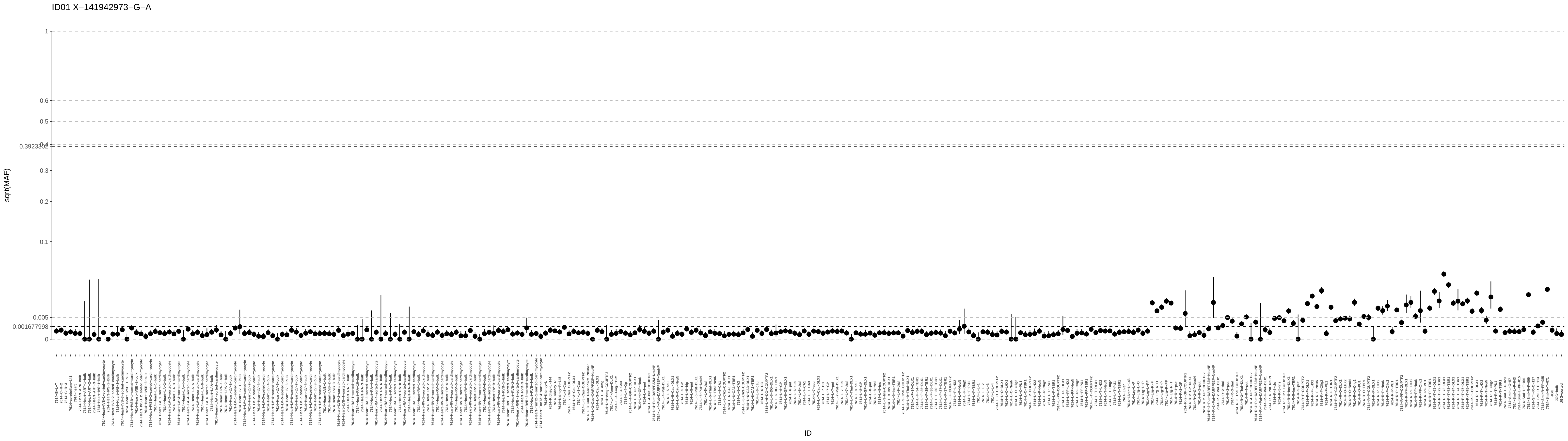

ID01 X−141942973−G−A
sqrt(MAF)
1
0.6
0.5
0.3923302
0.3
0.2
0.1
0.005
0.001677998
0
ID

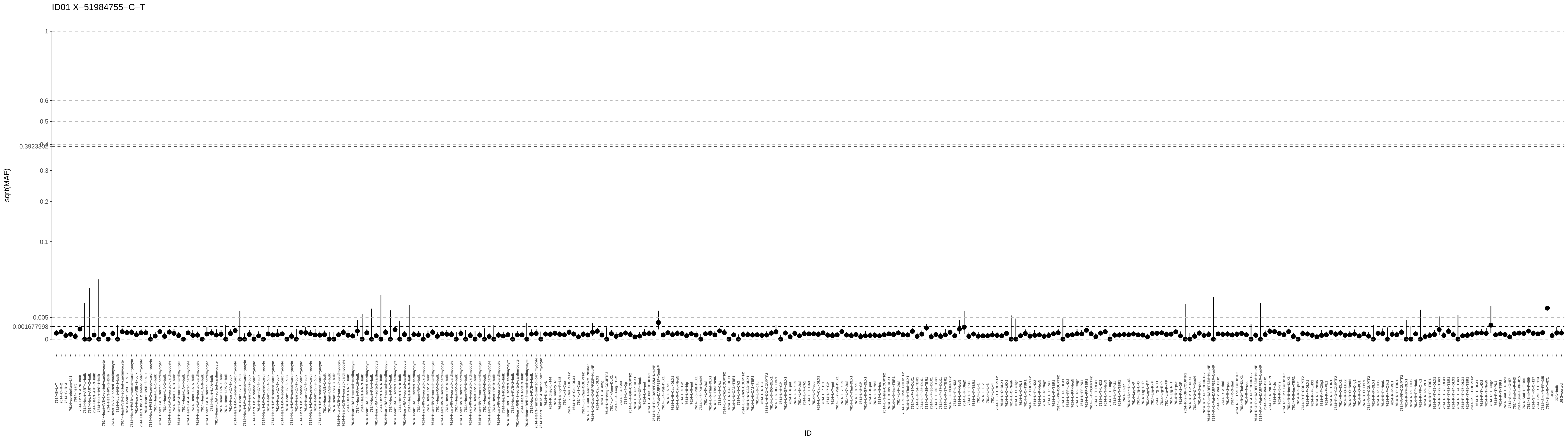
ID01 X−51984755−C−T
sqrt(MAF)
1
0.6
0.5
0.3923302
0.3
0.2
0.1
0.005
0.001677998
0
ID

ID01 X−54739596−C−G

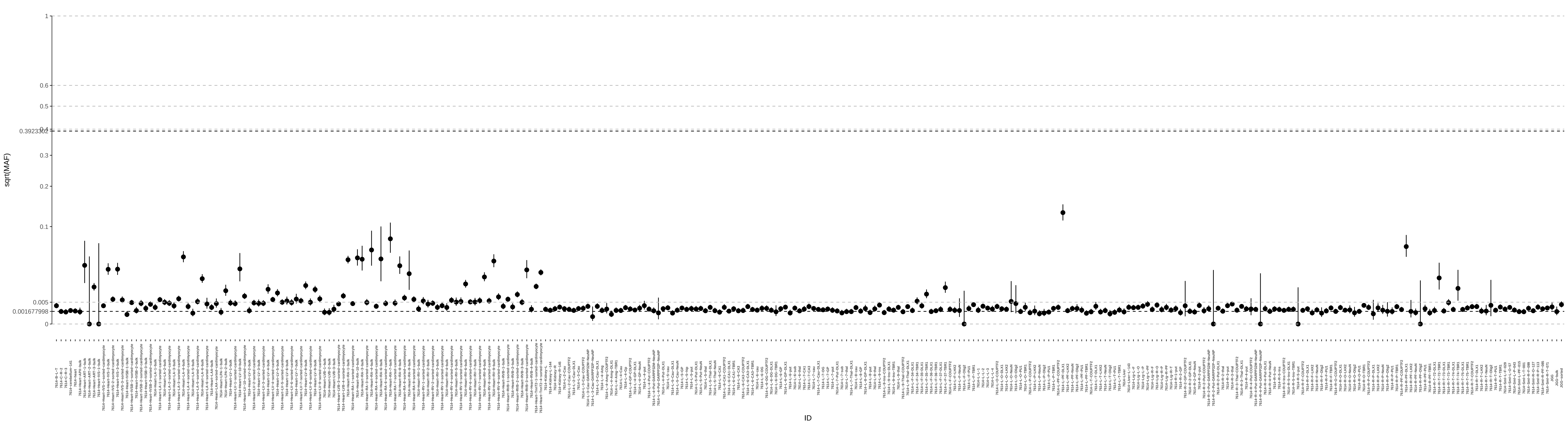

ID01 X−62980946−T−C

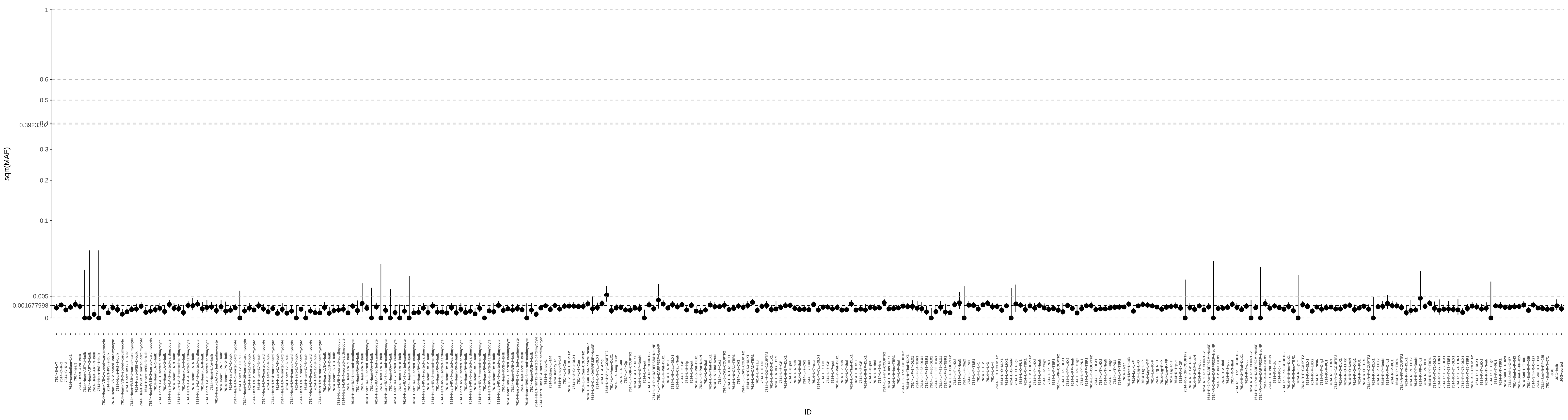

ID01 X-77453233-C-T

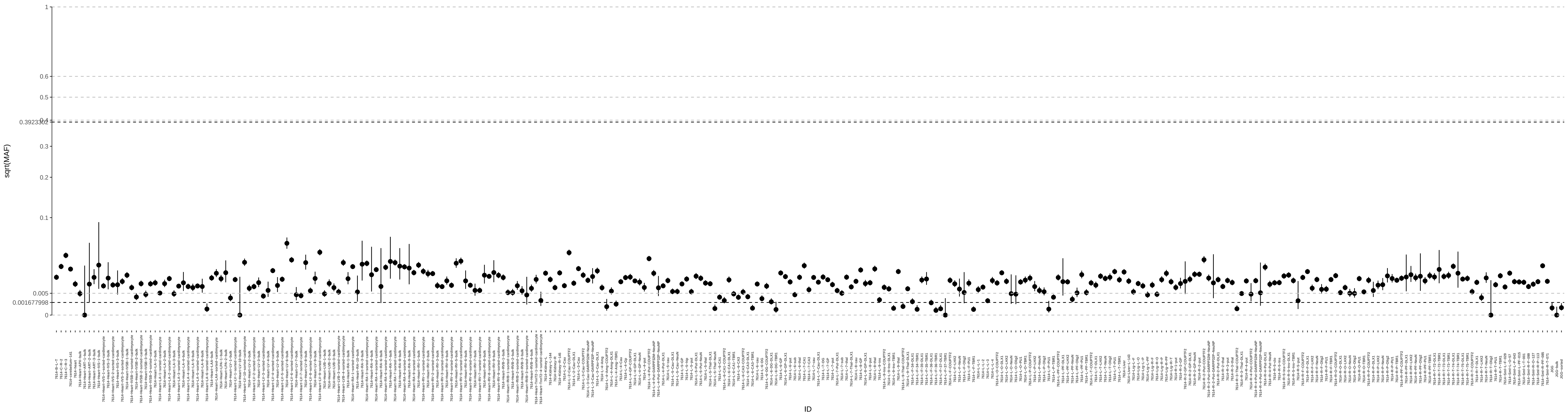

ID05 1−102152318−C−T

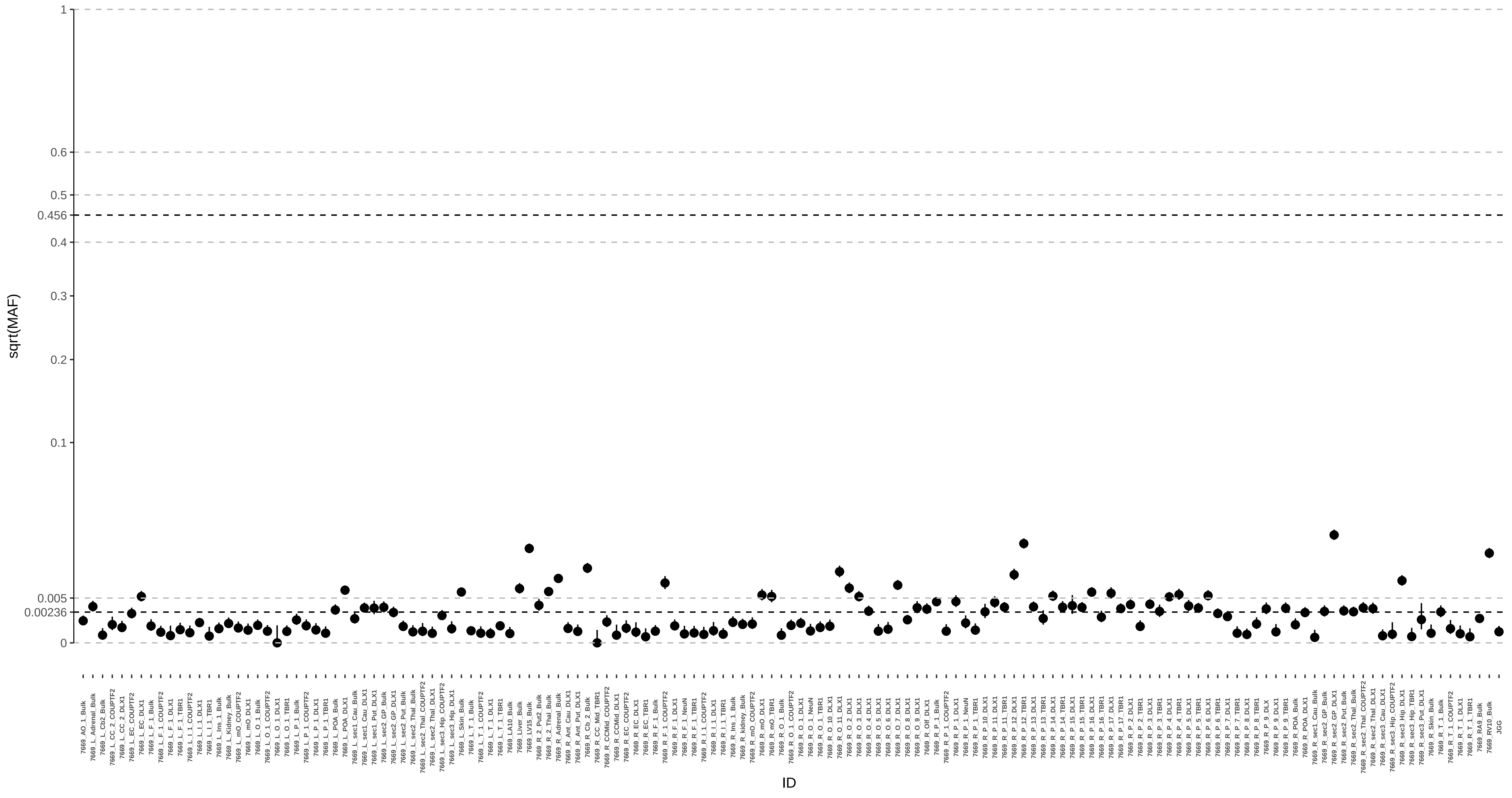

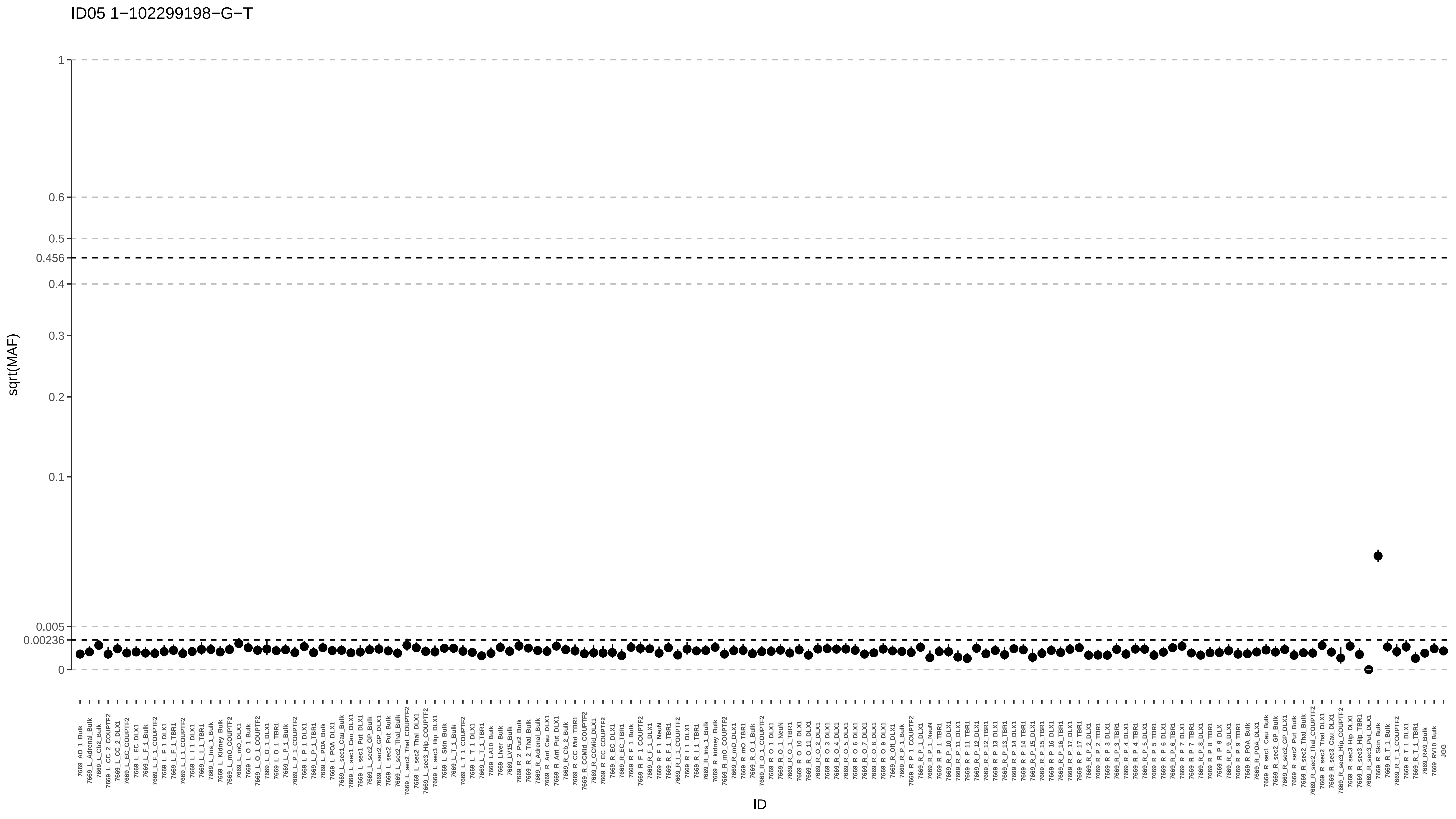
ID05 1−102299198−G−T
sqrt(MAF)
ID

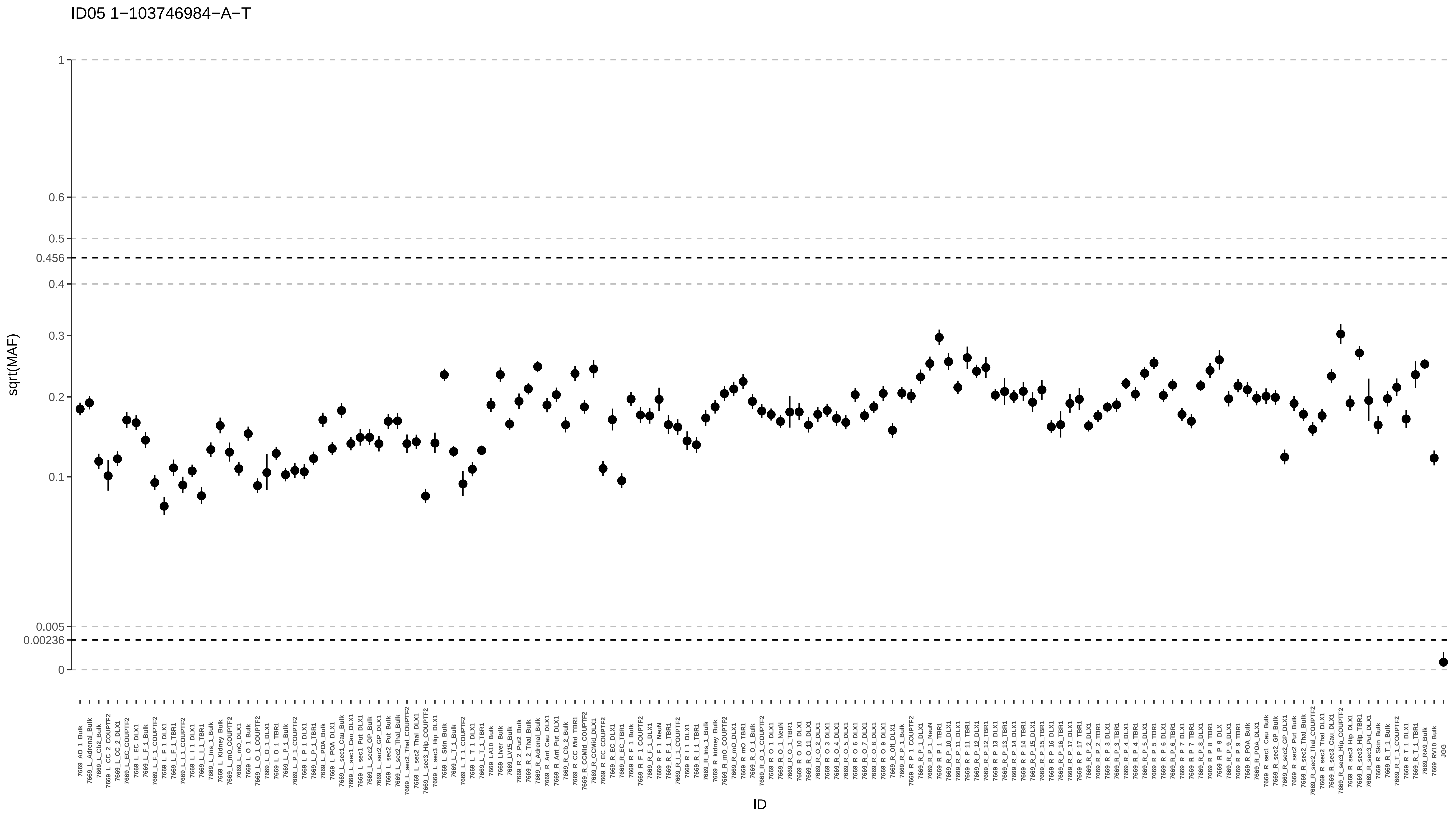
ID05 1−103746984−A−T
sqrt(MAF)
1
0.6
0.5
0.456
0.4
0.3
0.2
0.1
0.005
0.00236
0
ID

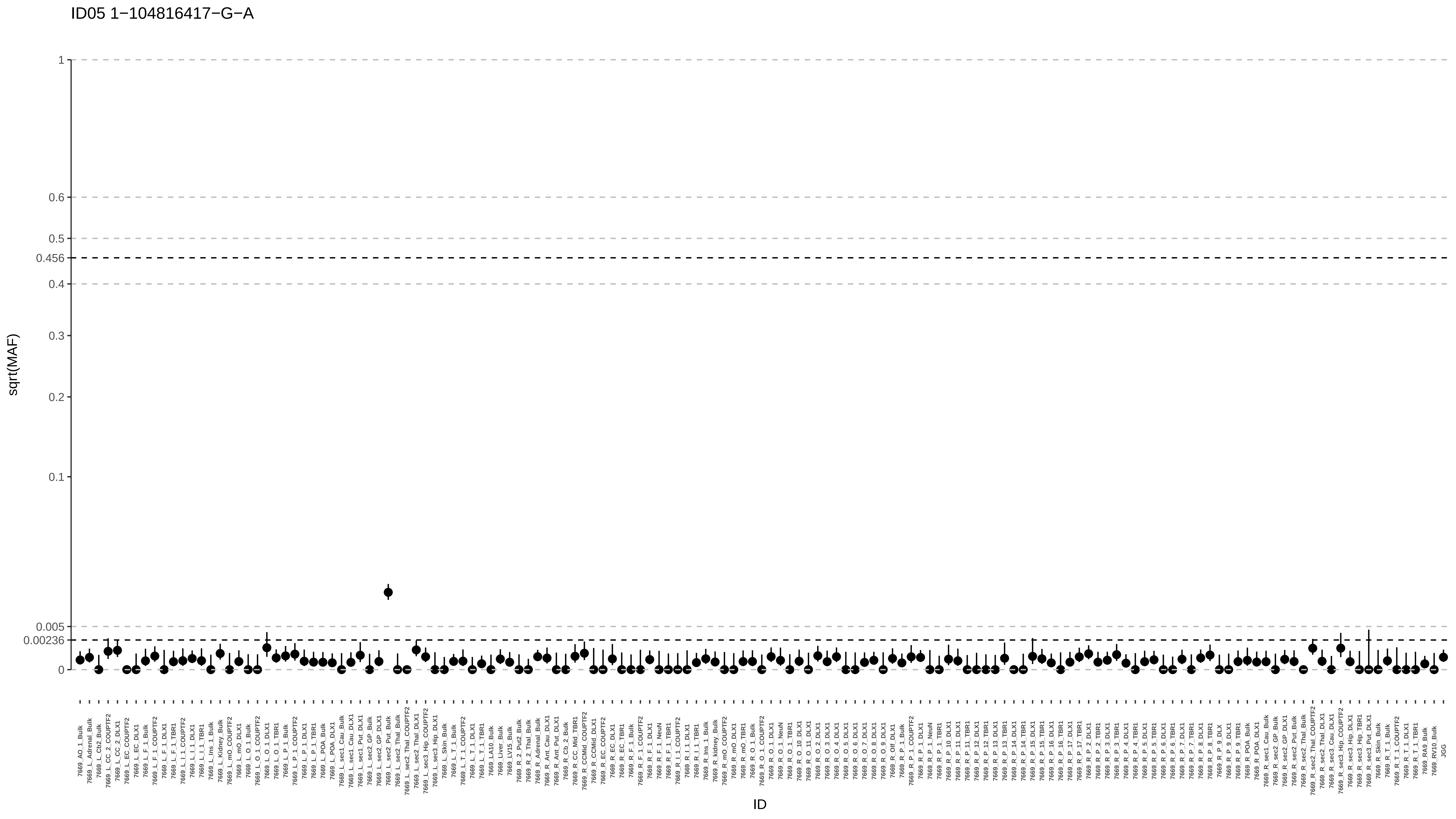
ID05 1−104816417−G−A
sqrt(MAF)
ID

ID05 1−105448120−C−T

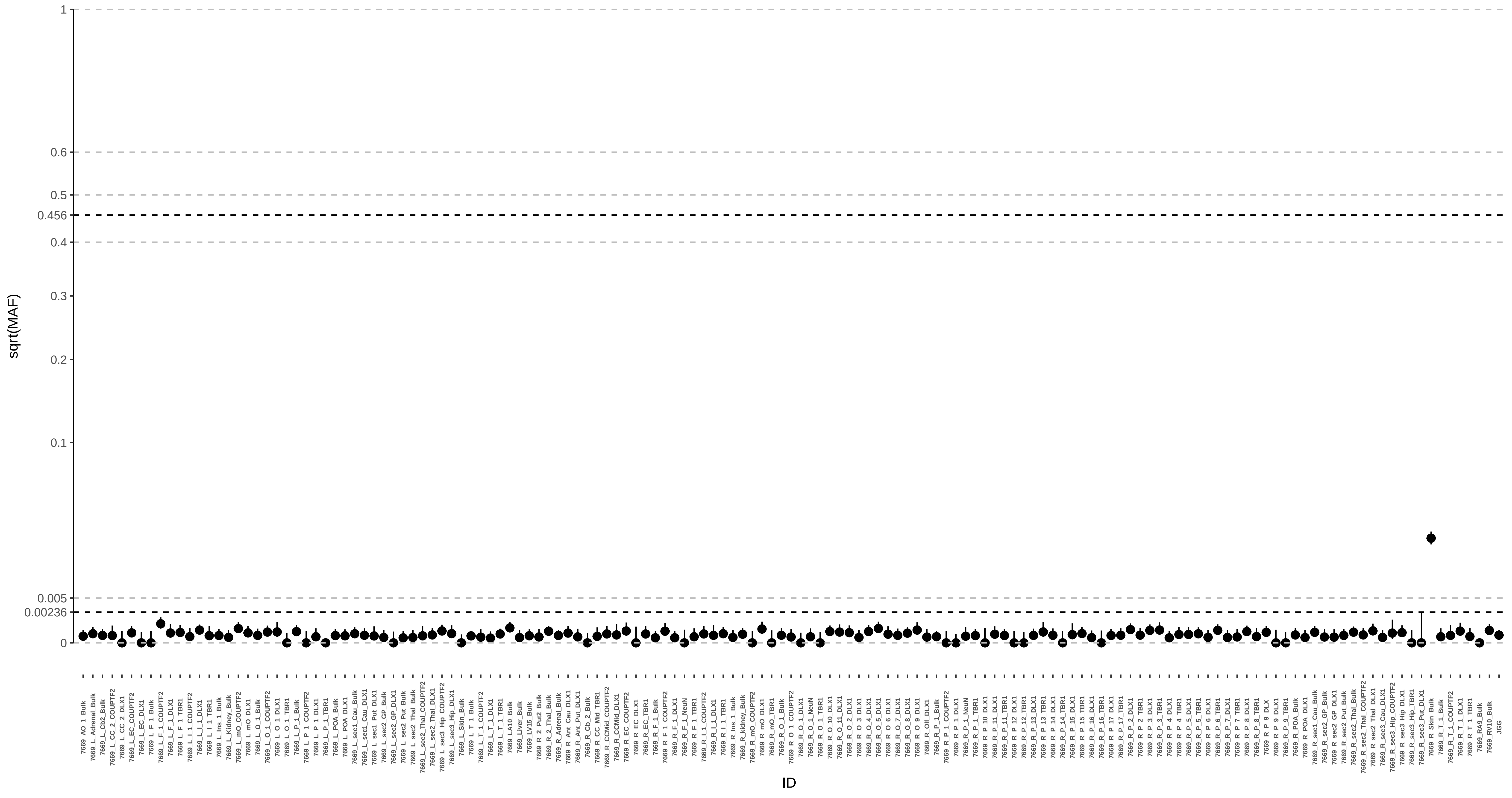

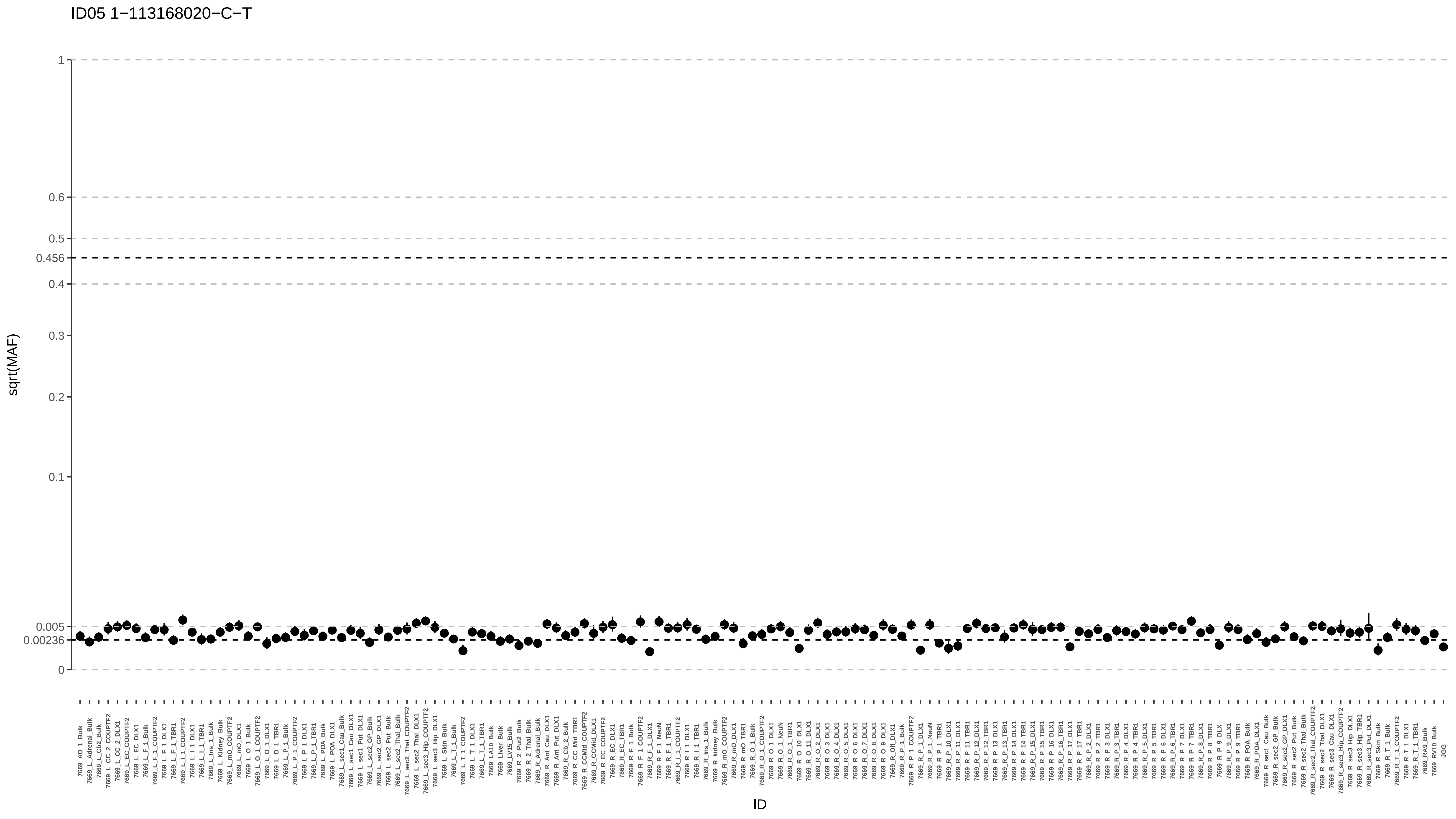
ID05 1−113168020−C−T
sqrt(MAF)
ID
1
0.6
0.5
0.456
0.4
0.3
0.2
0.1
0.005
0.00236
0

ID05 1−113168021−A−G

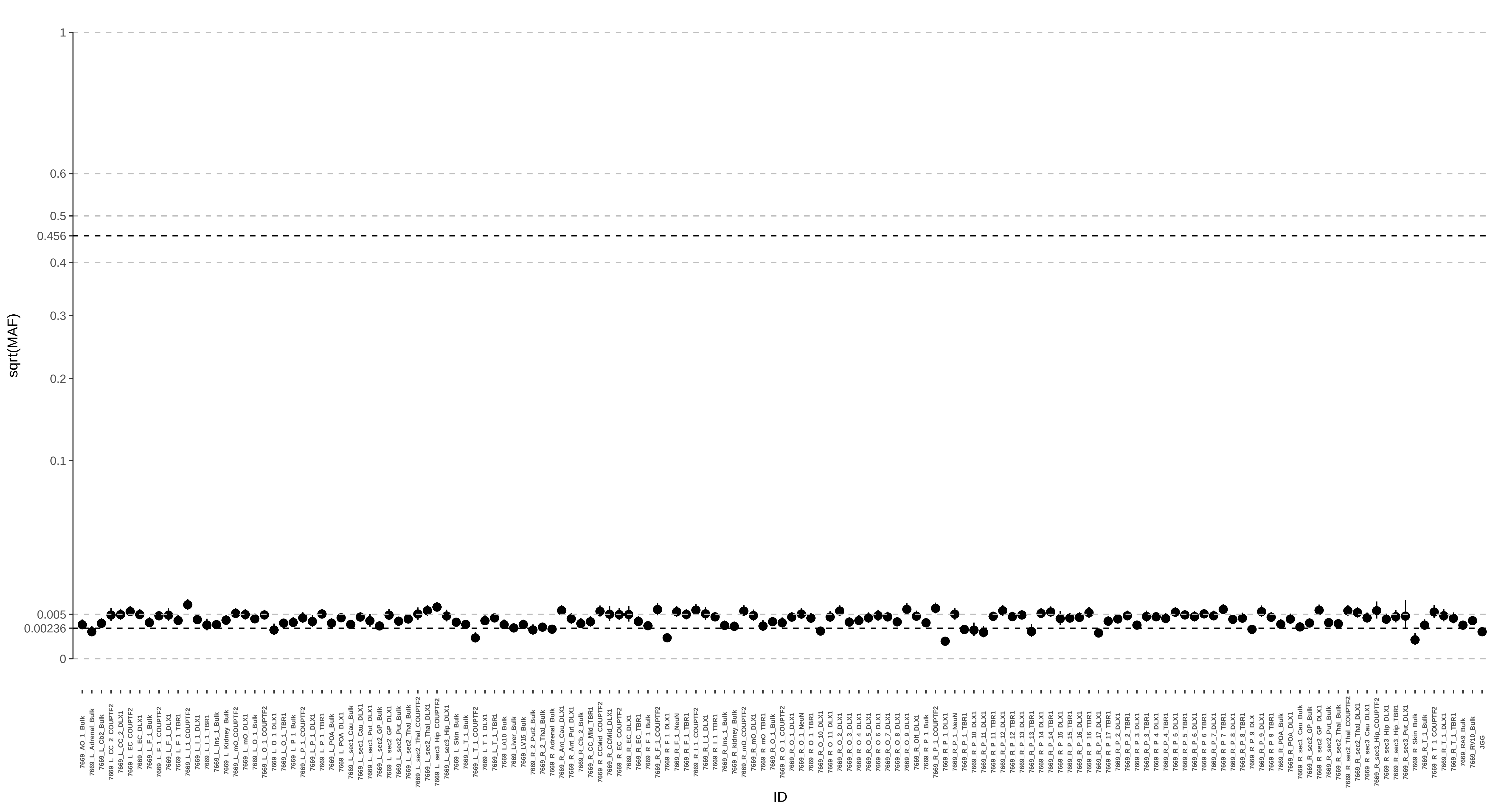

ID05 1−116835962−G−A

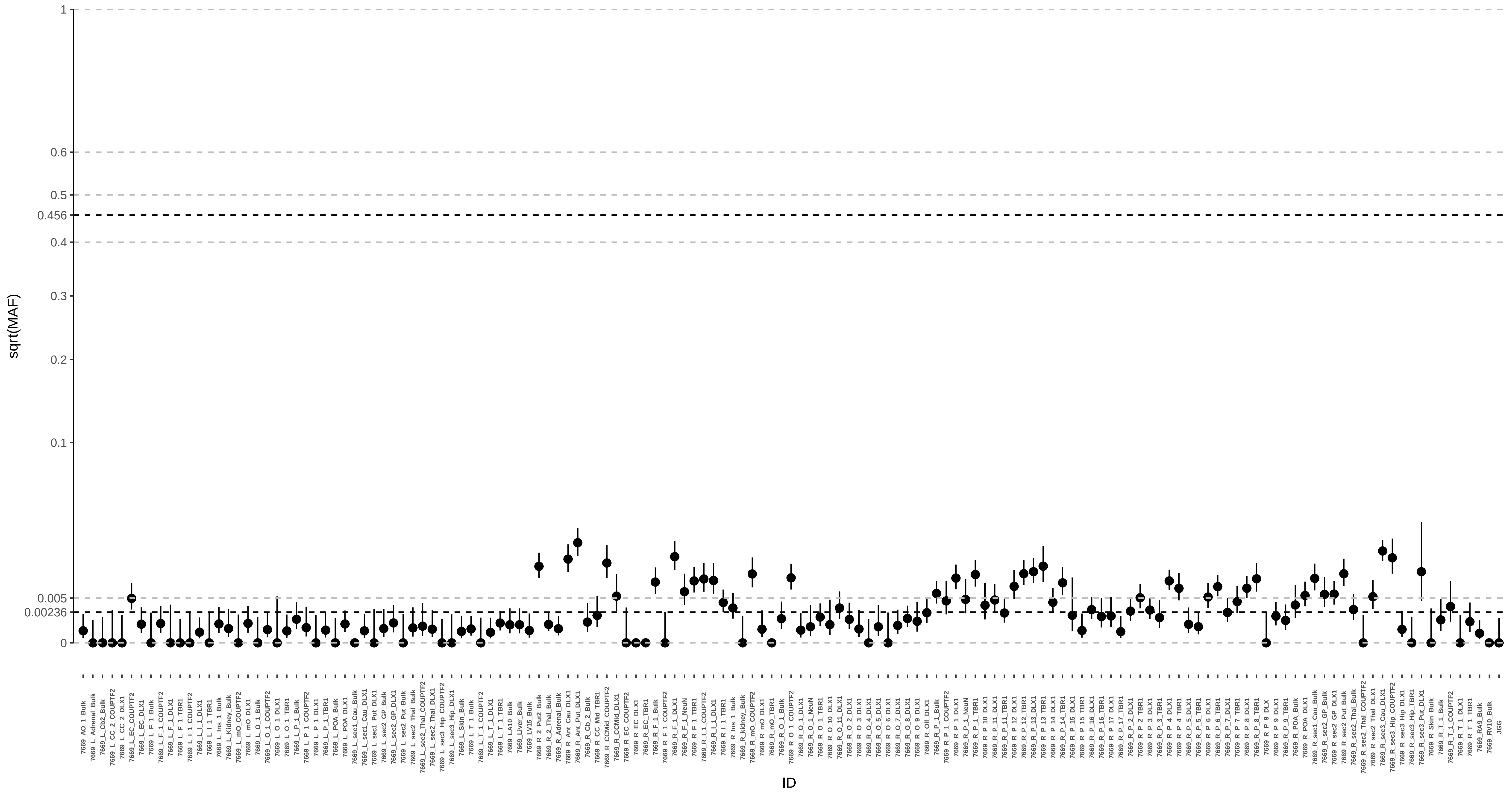

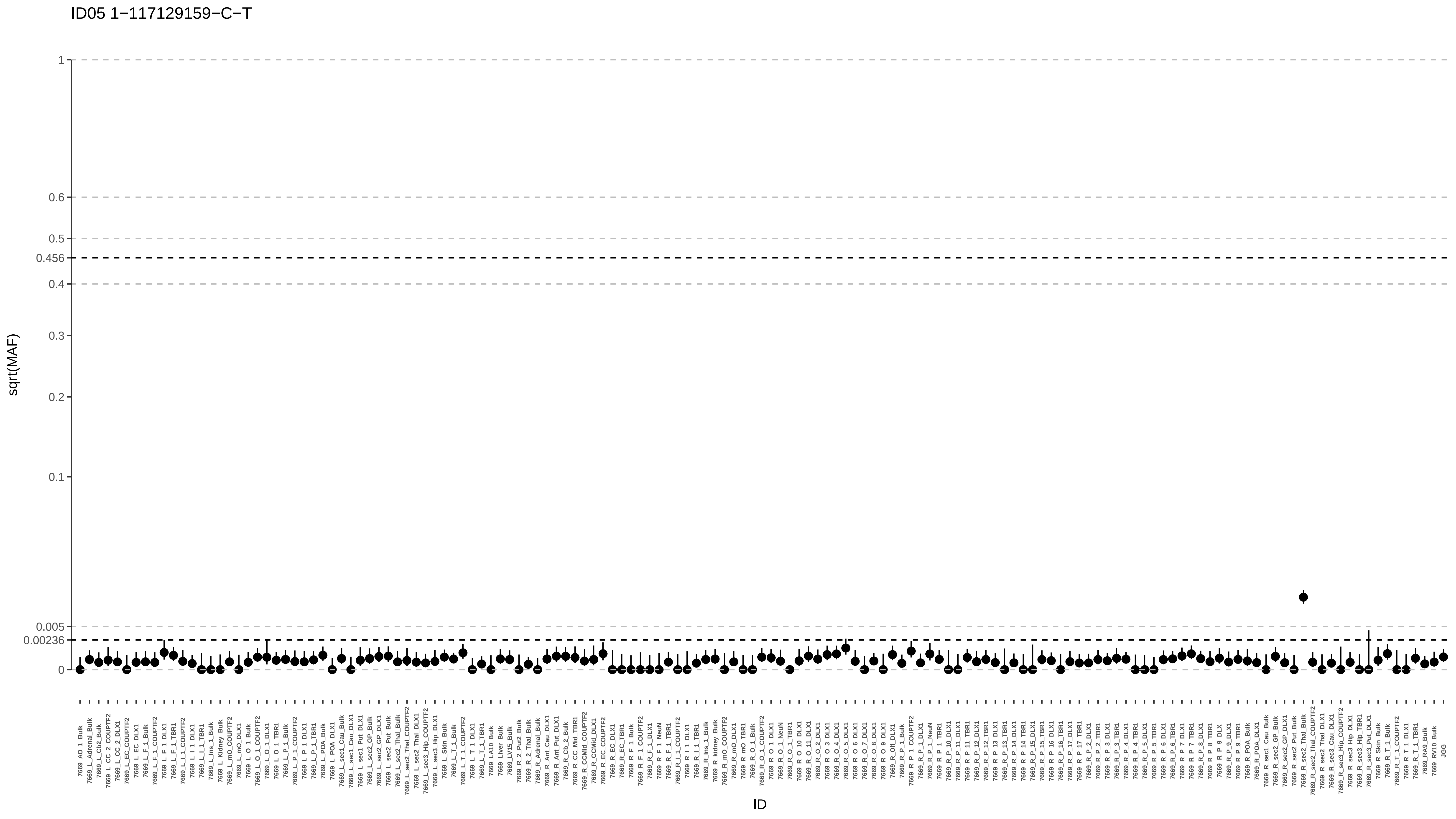
ID05 1−117129159−C−T
sqrt(MAF)
ID
0.456
0.00236

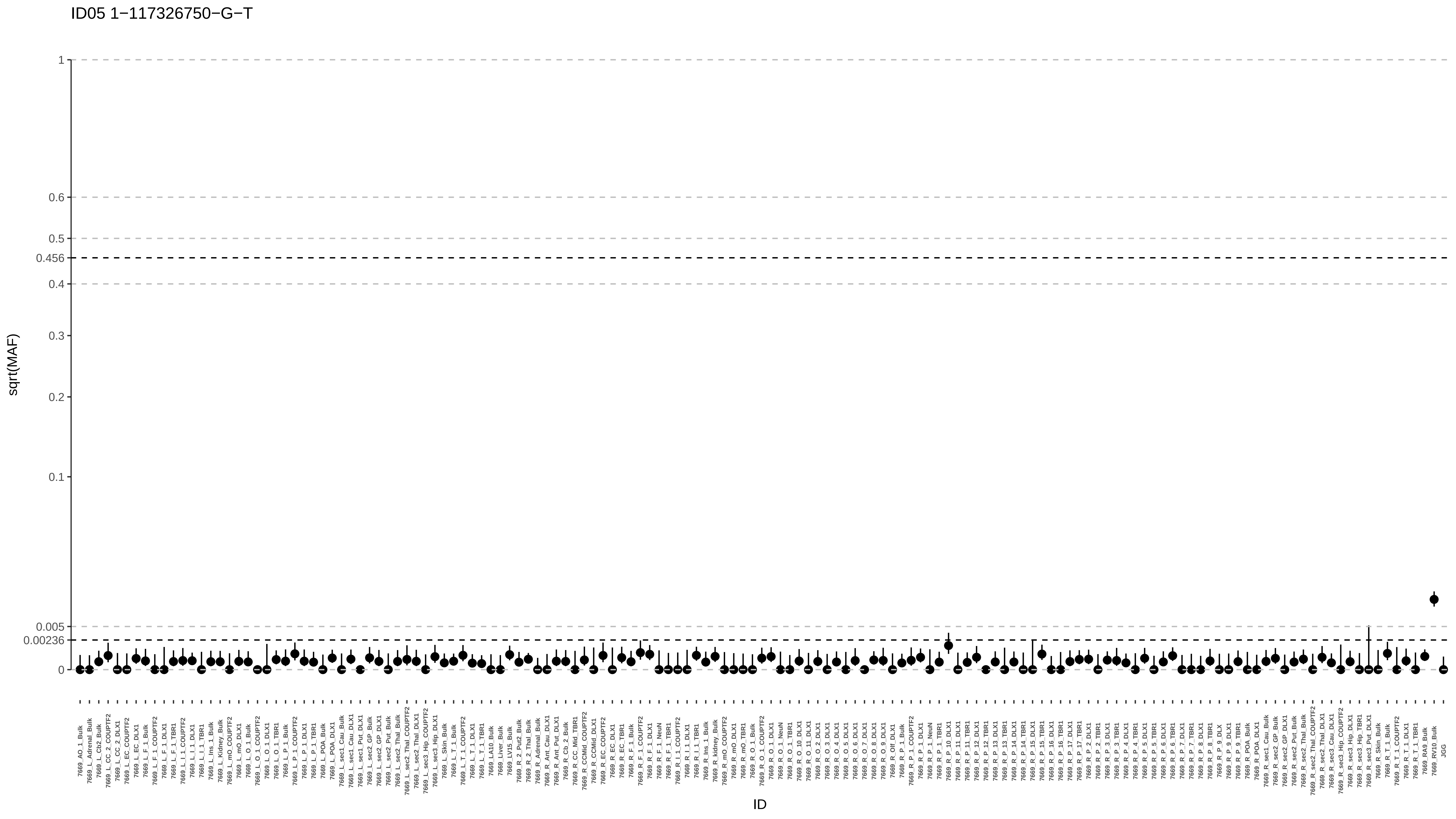

ID05 1−117326750−G−T
sqrt(MAF)
1
0.6
0.5
0.456
0.4
0.3
0.2
0.1
0.005
0.00236
0
ID
7669_AO_1_Bulk
7669_L_Adrenal_Bulk
7669_L_Cb2_Bulk
7669_L_CC_2_COUPTF2
7669_L_CC_2_DLX1
7669_L_EC_COUPTF2
7669_L_EC_DLX1
7669_L_F_1_Bulk
7669_L_F_1_COUPTF2
7669_L_F_1_DLX1
7669_L_F_1_TBR1
7669_L_I_1_COUPTF2
7669_L_I_1_DLX1
7669_L_I_1_TBR1
7669_L_Ins_1_Bulk
7669_L_Kidney_Bulk
7669_L_mO_COUPTF2
7669_L_mO_DLX1
7669_L_O_1_Bulk
7669_L_O_1_COUPTF2
7669_L_O_1_DLX1
7669_L_O_1_TBR1
7669_L_P_1_Bulk
7669_L_P_1_COUPTF2
7669_L_P_1_DLX1
7669_L_P_1_TBR1
7669_L_POA_Bulk
7669_L_POA_DLX1
7669_L_sec1_Cau_Bulk
7669_L_sec1_Cau_DLX1
7669_L_sec1_Put_DLX1
7669_L_sec2_GP_Bulk
7669_L_sec2_GP_DLX1
7669_L_sec2_Put_Bulk
7669_L_sec2_Thal_Bulk
7669_L_sec2_Thal_COUPTF2
7669_L_sec2_Thal_DLX1
7669_L_sec3_Hip_COUPTF2
7669_L_sec3_Hip_DLX1
7669_L_Skin_Bulk
7669_L_T_1_Bulk
7669_L_T_1_COUPTF2
7669_L_T_1_DLX1
7669_L_T_1_TBR1
7669_LA10_Bulk
7669_Liver_Bulk
7669_LV15_Bulk
7669_R_2_Put2_Bulk
7669_R_2_Thal_Bulk
7669_R_Adrenal_Bulk
7669_R_Ant_Cau_DLX1
7669_R_Ant_Put_DLX1
7669_R_Cb_2_Bulk
7669_R_CC_Mid_TBR1
7669_R_CCMid_COUPTF2
7669_R_CCMid_DLX1
7669_R_EC_COUPTF2
7669_R_EC_DLX1
7669_R_EC_TBR1
7669_R_F_1_Bulk
7669_R_F_1_COUPTF2
7669_R_F_1_DLX1
7669_R_F_1_NeuN
7669_R_F_1_TBR1
7669_R_I_1_COUPTF2
7669_R_I_1_DLX1
7669_R_I_1_TBR1
7669_R_Ins_1_Bulk
7669_R_kidney_Bulk
7669_R_mO_COUPTF2
7669_R_mO_DLX1
7669_R_mO_TBR1
7669_R_O_1_Bulk
7669_R_O_1_COUPTF2
7669_R_O_1_DLX1
7669_R_O_1_NeuN
7669_R_O_1_TBR1
7669_R_O_10_DLX1
7669_R_O_11_DLX1
7669_R_O_2_DLX1
7669_R_O_3_DLX1
7669_R_O_4_DLX1
7669_R_O_5_DLX1
7669_R_O_6_DLX1
7669_R_O_7_DLX1
7669_R_O_8_DLX1
7669_R_O_9_DLX1
7669_R_Olf_DLX1
7669_R_P_1_Bulk
7669_R_P_1_COUPTF2
7669_R_P_1_DLX1
7669_R_P_1_NeuN
7669_R_P_1_TBR1
7669_R_P_10_DLX1
7669_R_P_11_DLX1
7669_R_P_11_TBR1
7669_R_P_12_DLX1
7669_R_P_12_TBR1
7669_R_P_13_DLX1
7669_R_P_13_TBR1
7669_R_P_14_DLX1
7669_R_P_14_TBR1
7669_R_P_15_DLX1
7669_R_P_15_TBR1
7669_R_P_16_DLX1
7669_R_P_16_TBR1
7669_R_P_17_DLX1
7669_R_P_17_TBR1
7669_R_P_2_DLX1
7669_R_P_2_TBR1
7669_R_P_3_DLX1
7669_R_P_3_TBR1
7669_R_P_4_DLX1
7669_R_P_4_TBR1
7669_R_P_5_DLX1
7669_R_P_5_TBR1
7669_R_P_6_DLX1
7669_R_P_6_TBR1
7669_R_P_7_DLX1
7669_R_P_7_TBR1
7669_R_P_8_DLX1
7669_R_P_8_TBR1
7669_R_P_9_DLX
7669_R_P_9_DLX1
7669_R_P_9_TBR1
7669_R_POA_Bulk
7669_R_POA_DLX1
7669_R_sec1_Cau_Bulk
7669_R_sec2_GP_Bulk
7669_R_sec2_GP_DLX1
7669_R_sec2_Put_Bulk
7669_R_sec2_Thal_Bulk
7669_R_sec2_Thal_COUPTF2
7669_R_sec2_Thal_DLX1
7669_R_sec3_Cau_DLX1
7669_R_sec3_Hip_COUPTF2
7669_R_sec3_Hip_DLX1
7669_R_sec3_Hip_TBR1
7669_R_sec3_Put_DLX1
7669_R_Skin_Bulk
7669_R_T_1_Bulk
7669_R_T_1_COUPTF2
7669_R_T_1_DLX1
7669_R_T_1_TBR1
7669_RA9_Bulk
7669_RV10_Bulk
JGG

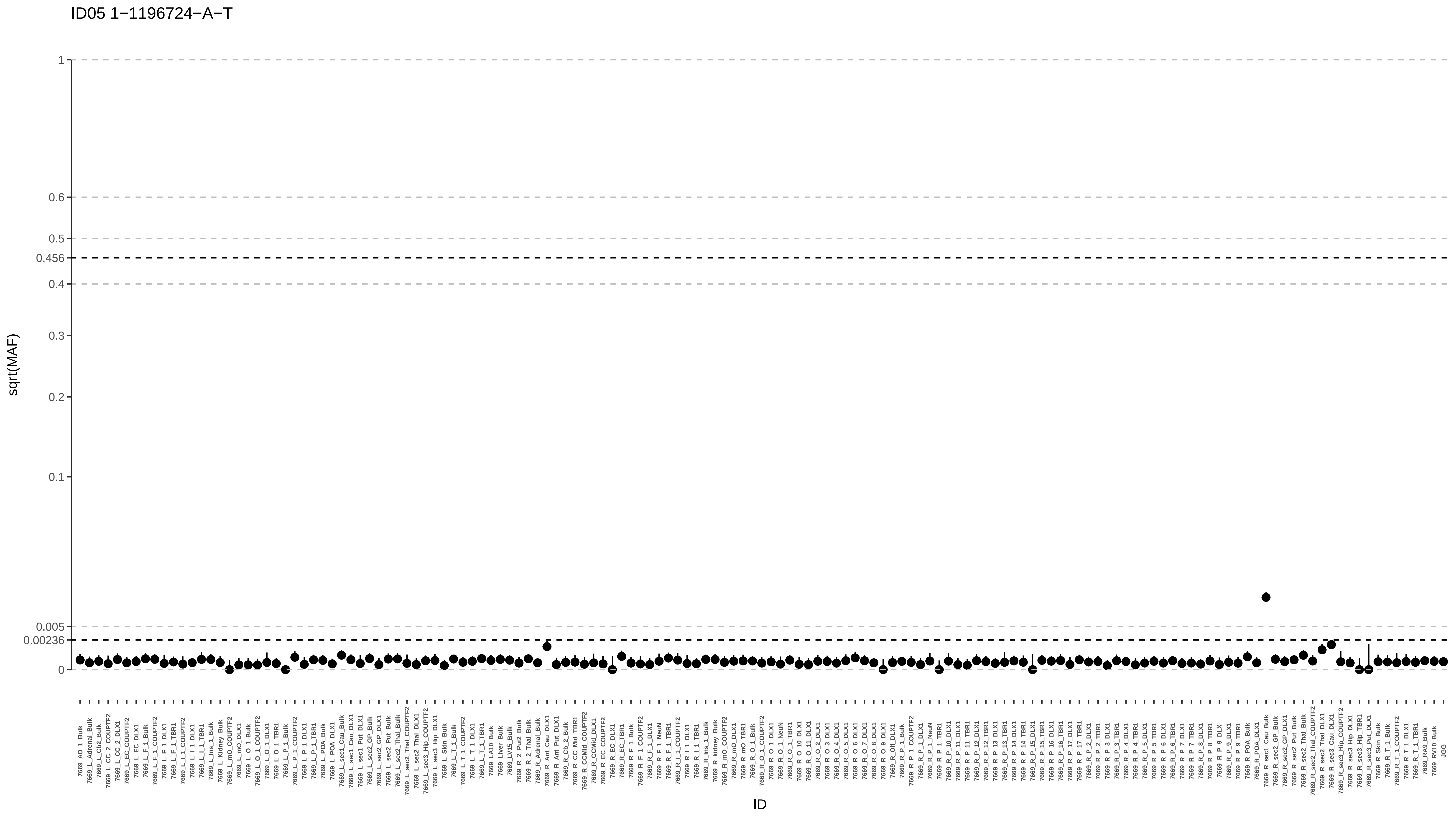
ID05 1−1196724−A−T
sqrt(MAF)
1
0.6
0.5
0.456
0.4
0.3
0.2
0.1
0.005
0.00236
0
ID

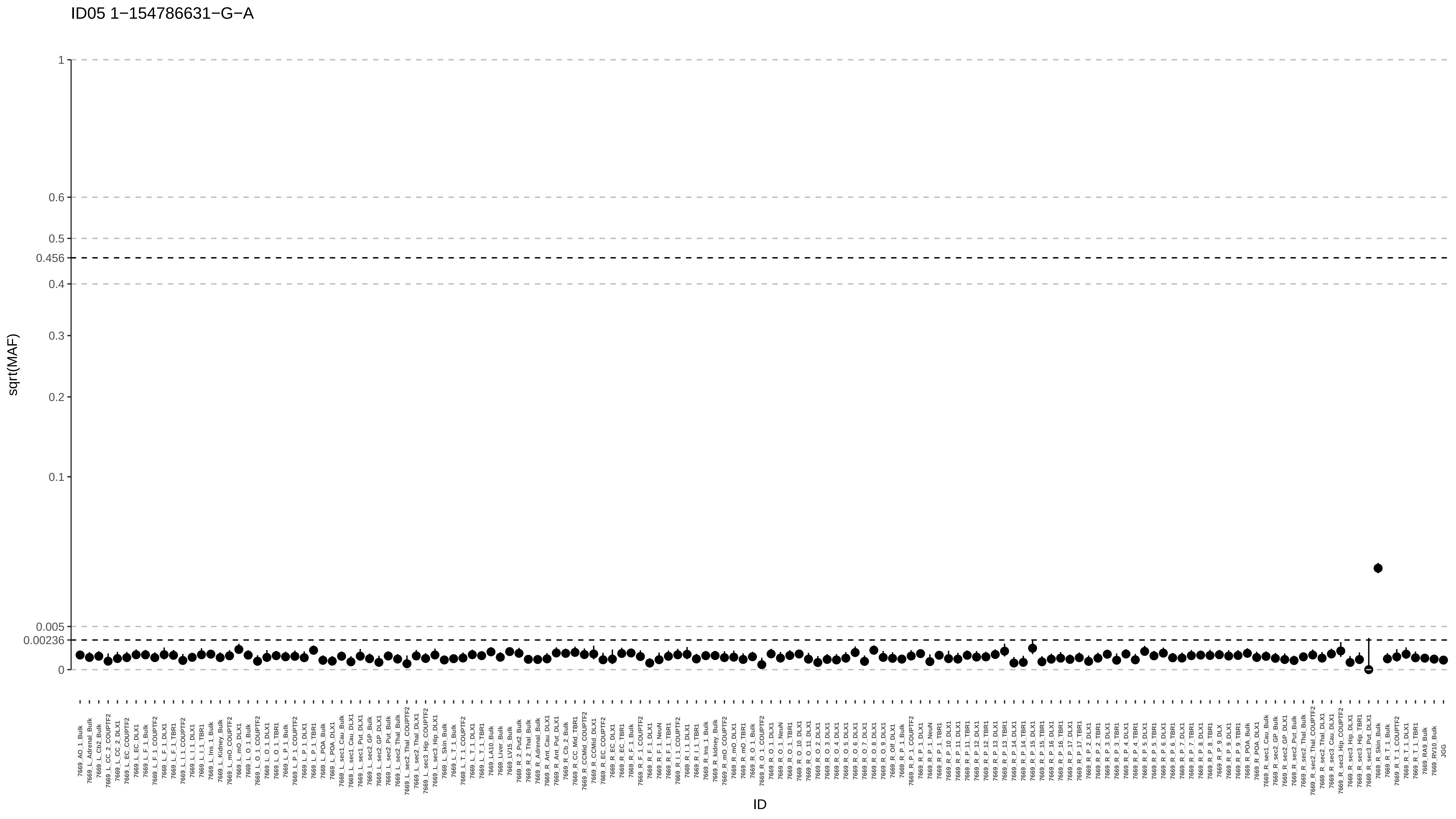

ID05 1−154786631−G−A
sqrt(MAF)
ID
1
0.6
0.5
0.456
0.4
0.3
0.2
0.1
0.005
0.00236
0

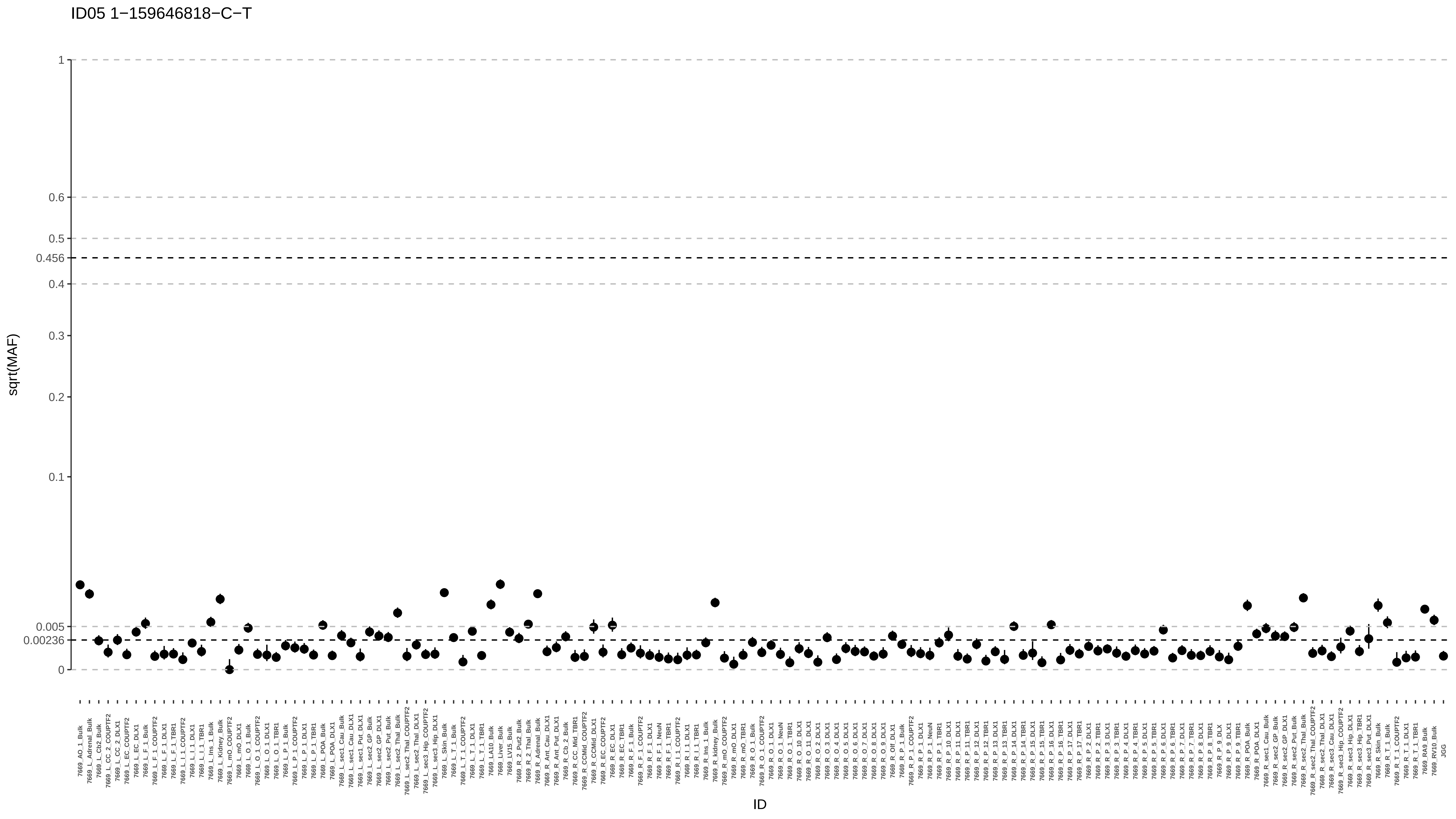
ID05 1−159646818−C−T
sqrt(MAF)
1
0.6
0.5
0.456
0.4
0.3
0.2
0.1
0.005
0.00236
0
ID

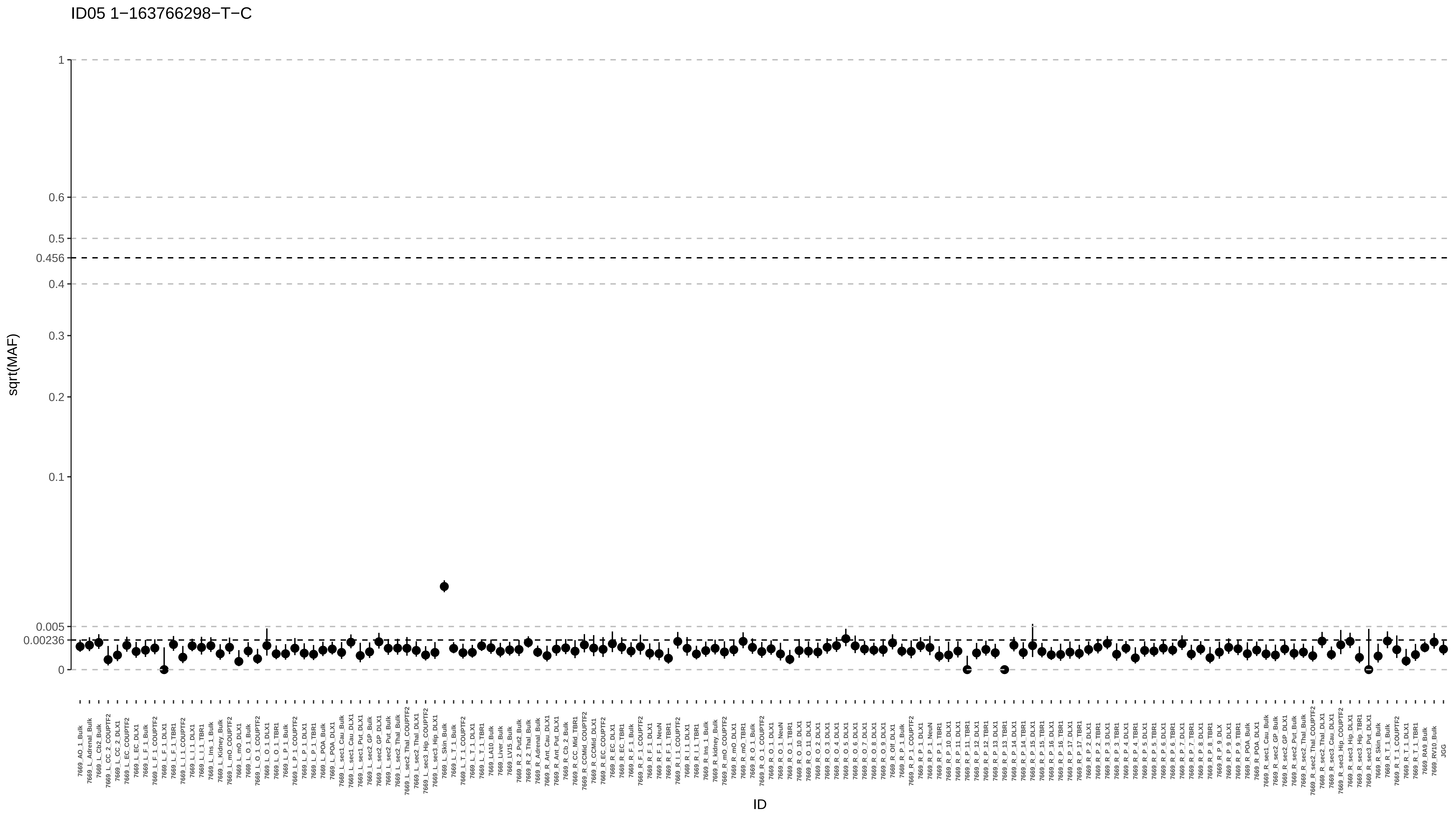
ID05 1−163766298−T−C
sqrt(MAF)
ID

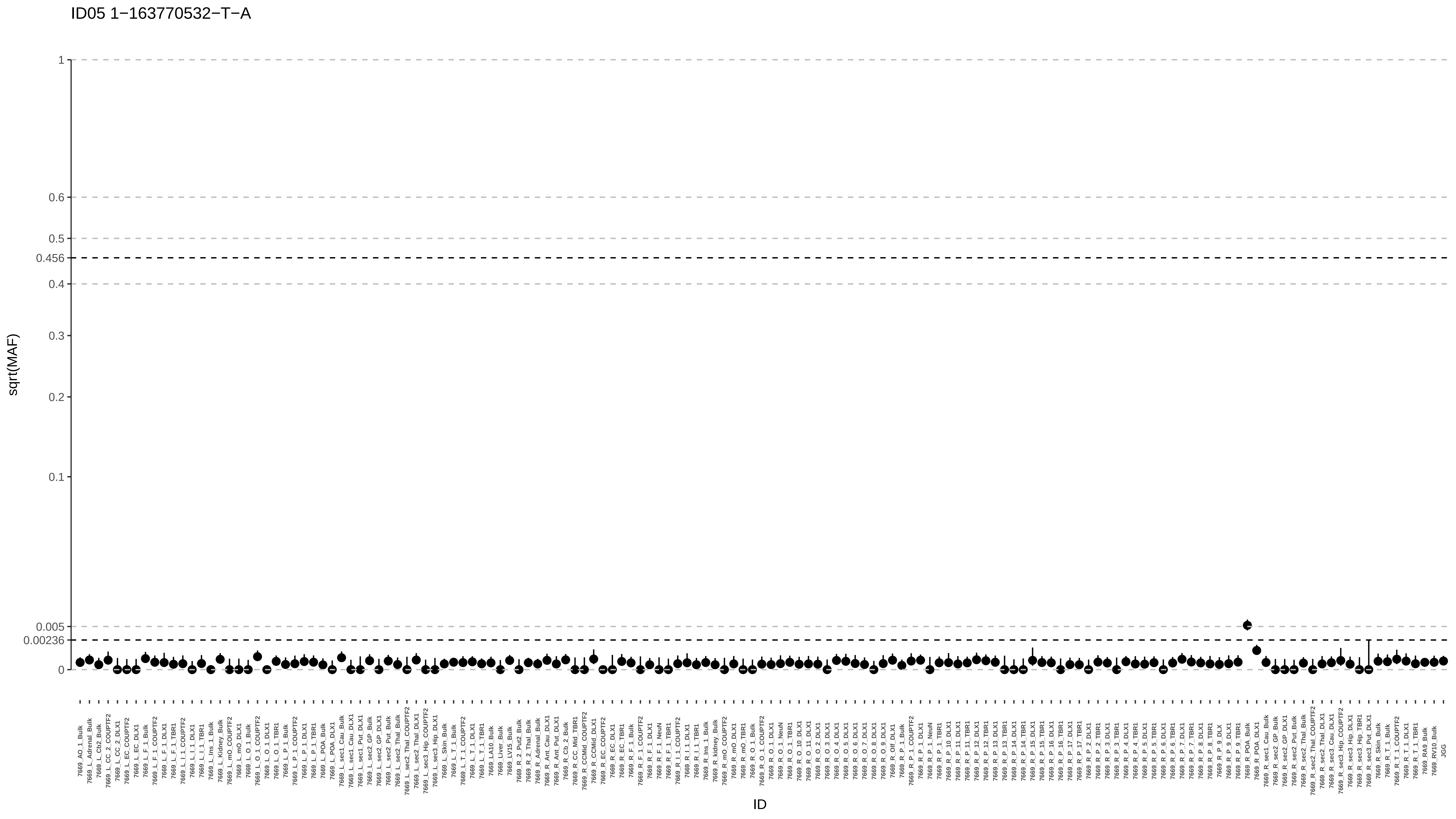
ID05 1−163770532−T−A
sqrt(MAF)
1
0.6
0.5
0.456
0.4
0.3
0.2
0.1
0.005
0.00236
0
ID
7669_AO_1_Bulk
7669_L_Adrenal_Bulk
7669_L_Cb2_Bulk
7669_L_CC_2_COUPTF2
7669_L_CC_2_DLX1
7669_L_EC_COUPTF2
7669_L_EC_DLX1
7669_L_F_1_Bulk
7669_L_F_1_COUPTF2
7669_L_F_1_DLX1
7669_L_F_1_TBR1
7669_L_I_1_COUPTF2
7669_L_I_1_DLX1
7669_L_I_1_TBR1
7669_L_Ins_1_Bulk
7669_L_Kidney_Bulk
7669_L_mO_COUPTF2
7669_L_mO_DLX1
7669_L_O_1_Bulk
7669_L_O_1_COUPTF2
7669_L_O_1_DLX1
7669_L_O_1_TBR1
7669_L_P_1_Bulk
7669_L_P_1_COUPTF2
7669_L_P_1_DLX1
7669_L_P_1_TBR1
7669_L_POA_Bulk
7669_L_POA_DLX1
7669_L_sec1_Cau_Bulk
7669_L_sec1_Cau_DLX1
7669_L_sec1_Put_DLX1
7669_L_sec2_GP_Bulk
7669_L_sec2_GP_DLX1
7669_L_sec2_Put_Bulk
7669_L_sec2_Thal_Bulk
7669_L_sec2_Thal_COUPTF2
7669_L_sec2_Thal_DLX1
7669_L_sec3_Hip_COUPTF2
7669_L_sec3_Hip_DLX1
7669_L_Skin_Bulk
7669_L_T_1_Bulk
7669_L_T_1_COUPTF2
7669_L_T_1_DLX1
7669_L_T_1_TBR1
7669_LA10_Bulk
7669_Liver_Bulk
7669_LV15_Bulk
7669_R_2_Put2_Bulk
7669_R_2_Thal_Bulk
7669_R_Adrenal_Bulk
7669_R_Ant_Cau_DLX1
7669_R_Ant_Put_DLX1
7669_R_Cb_2_Bulk
7669_R_CC_Mid_TBR1
7669_R_CCMid_COUPTF2
7669_R_CCMid_DLX1
7669_R_EC_COUPTF2
7669_R_EC_DLX1
7669_R_EC_TBR1
7669_R_F_1_Bulk
7669_R_F_1_COUPTF2
7669_R_F_1_DLX1
7669_R_F_1_NeuN
7669_R_F_1_TBR1
7669_R_I_1_COUPTF2
7669_R_I_1_DLX1
7669_R_I_1_TBR1
7669_R_Ins_1_Bulk
7669_R_kidney_Bulk
7669_R_mO_COUPTF2
7669_R_mO_DLX1
7669_R_mO_TBR1
7669_R_O_1_Bulk
7669_R_O_1_COUPTF2
7669_R_O_1_DLX1
7669_R_O_1_NeuN
7669_R_O_1_TBR1
7669_R_O_10_DLX1
7669_R_O_11_DLX1
7669_R_O_2_DLX1
7669_R_O_3_DLX1
7669_R_O_4_DLX1
7669_R_O_5_DLX1
7669_R_O_6_DLX1
7669_R_O_7_DLX1
7669_R_O_8_DLX1
7669_R_O_9_DLX1
7669_R_Olf_DLX1
7669_R_P_1_Bulk
7669_R_P_1_COUPTF2
7669_R_P_1_DLX1
7669_R_P_1_NeuN
7669_R_P_1_TBR1
7669_R_P_10_DLX1
7669_R_P_11_DLX1
7669_R_P_11_TBR1
7669_R_P_12_DLX1
7669_R_P_12_TBR1
7669_R_P_13_DLX1
7669_R_P_13_TBR1
7669_R_P_14_DLX1
7669_R_P_14_TBR1
7669_R_P_15_DLX1
7669_R_P_15_TBR1
7669_R_P_16_DLX1
7669_R_P_16_TBR1
7669_R_P_17_DLX1
7669_R_P_17_TBR1
7669_R_P_2_DLX1
7669_R_P_2_TBR1
7669_R_P_3_DLX1
7669_R_P_3_TBR1
7669_R_P_4_DLX1
7669_R_P_4_TBR1
7669_R_P_5_DLX1
7669_R_P_5_TBR1
7669_R_P_6_DLX1
7669_R_P_6_TBR1
7669_R_P_7_DLX1
7669_R_P_7_TBR1
7669_R_P_8_DLX1
7669_R_P_8_TBR1
7669_R_P_9_DLX
7669_R_P_9_DLX1
7669_R_P_9_TBR1
7669_R_POA_Bulk
7669_R_POA_DLX1
7669_R_sec1_Cau_Bulk
7669_R_sec2_GP_Bulk
7669_R_sec2_GP_DLX1
7669_R_sec2_Put_Bulk
7669_R_sec2_Thal_Bulk
7669_R_sec2_Thal_COUPTF2
7669_R_sec2_Thal_DLX1
7669_R_sec3_Cau_DLX1
7669_R_sec3_Hip_COUPTF2
7669_R_sec3_Hip_DLX1
7669_R_sec3_Hip_TBR1
7669_R_sec3_Put_DLX1
7669_R_Skin_Bulk
7669_R_T_1_Bulk
7669_R_T_1_COUPTF2
7669_R_T_1_DLX1
7669_R_T_1_TBR1
7669_RA9_Bulk
7669_RV10_Bulk
JGG

ID05 1−177430967−T−C

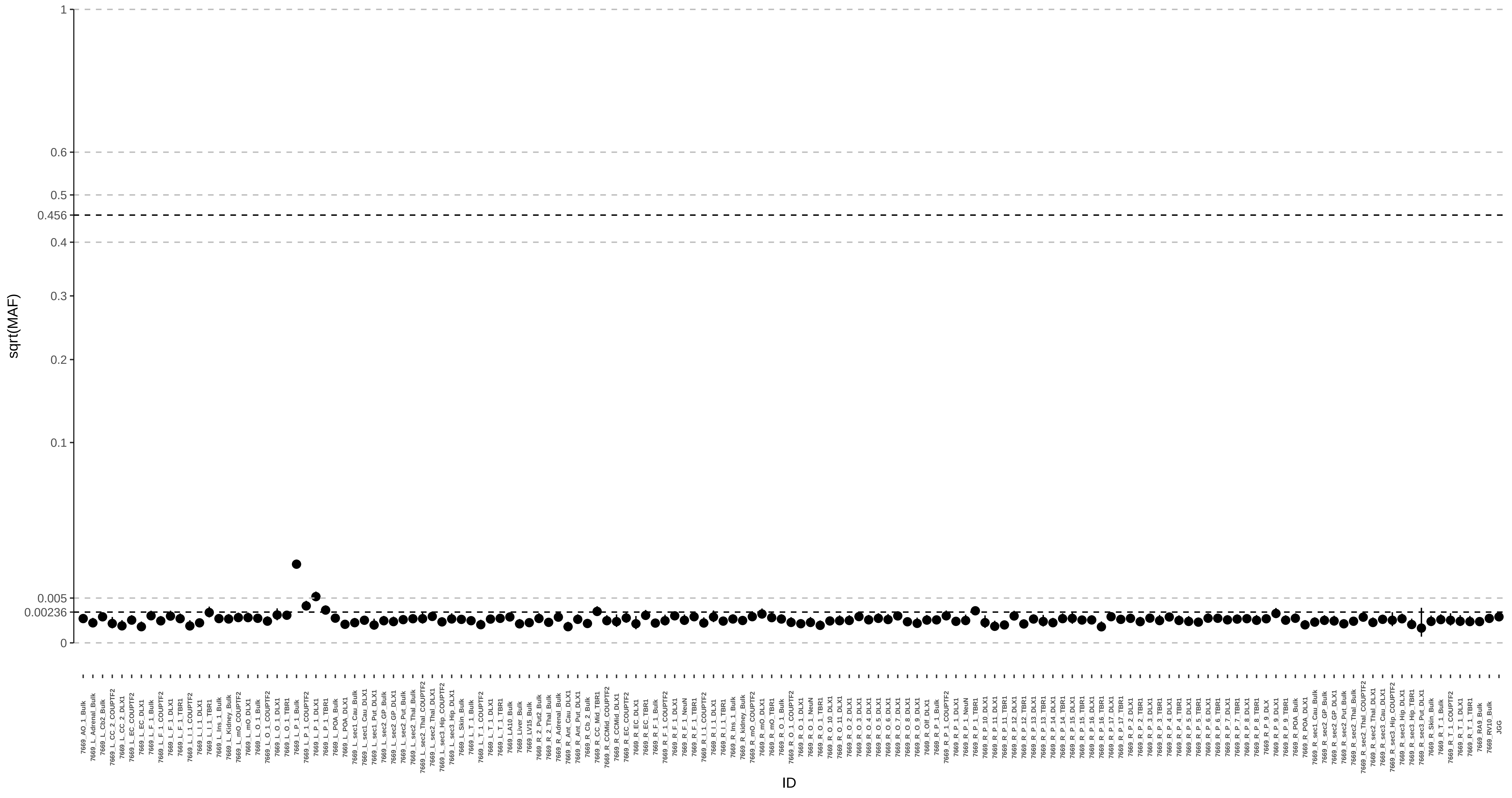

ID05 1−18387130−C−T

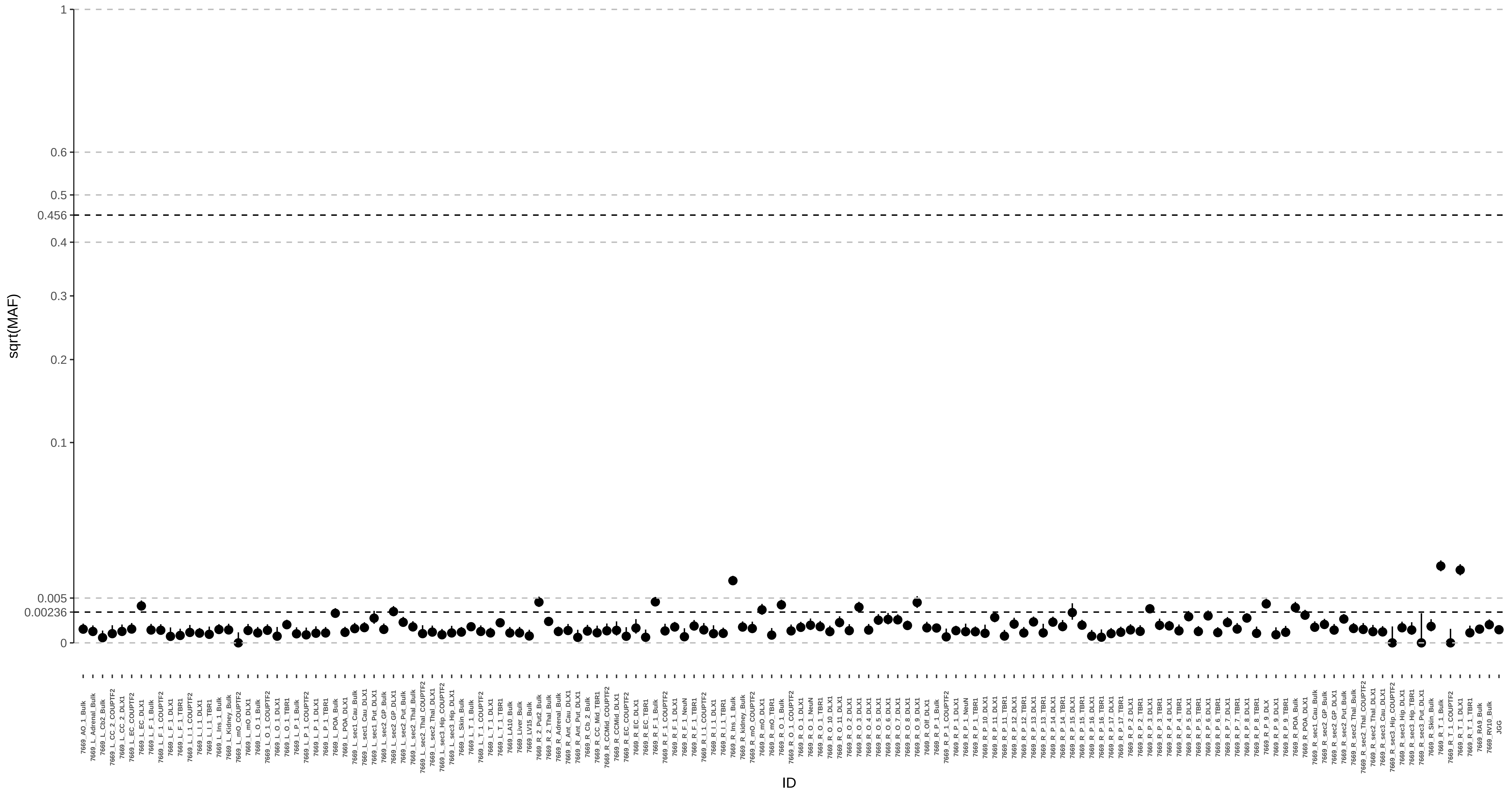

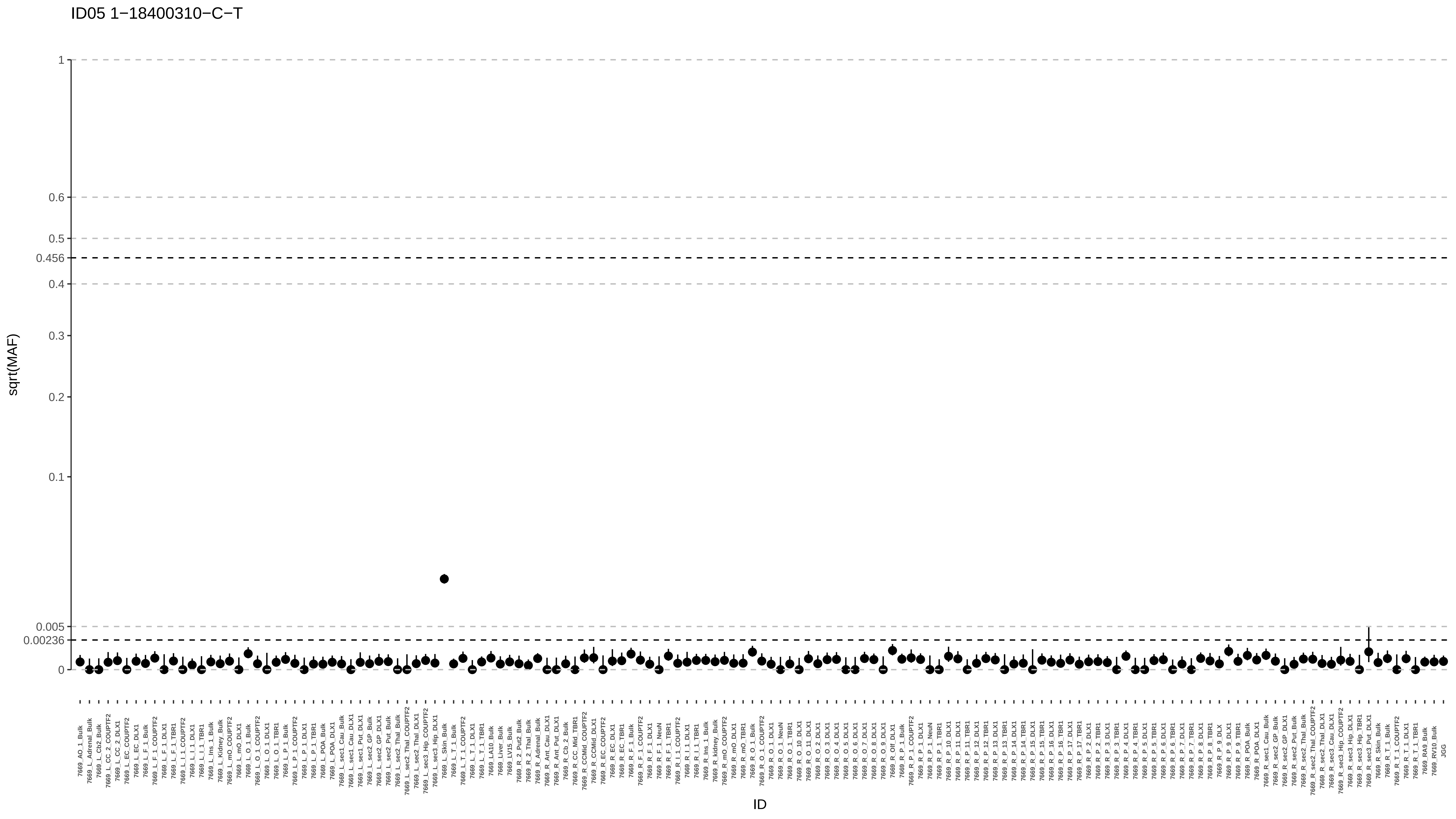

ID05 1−18400310−C−T
sqrt(MAF)
1
0.6
0.5
0.456
0.4
0.3
0.2
0.1
0.005
0.00236
0
ID

# ID05 1−188074317−T−C

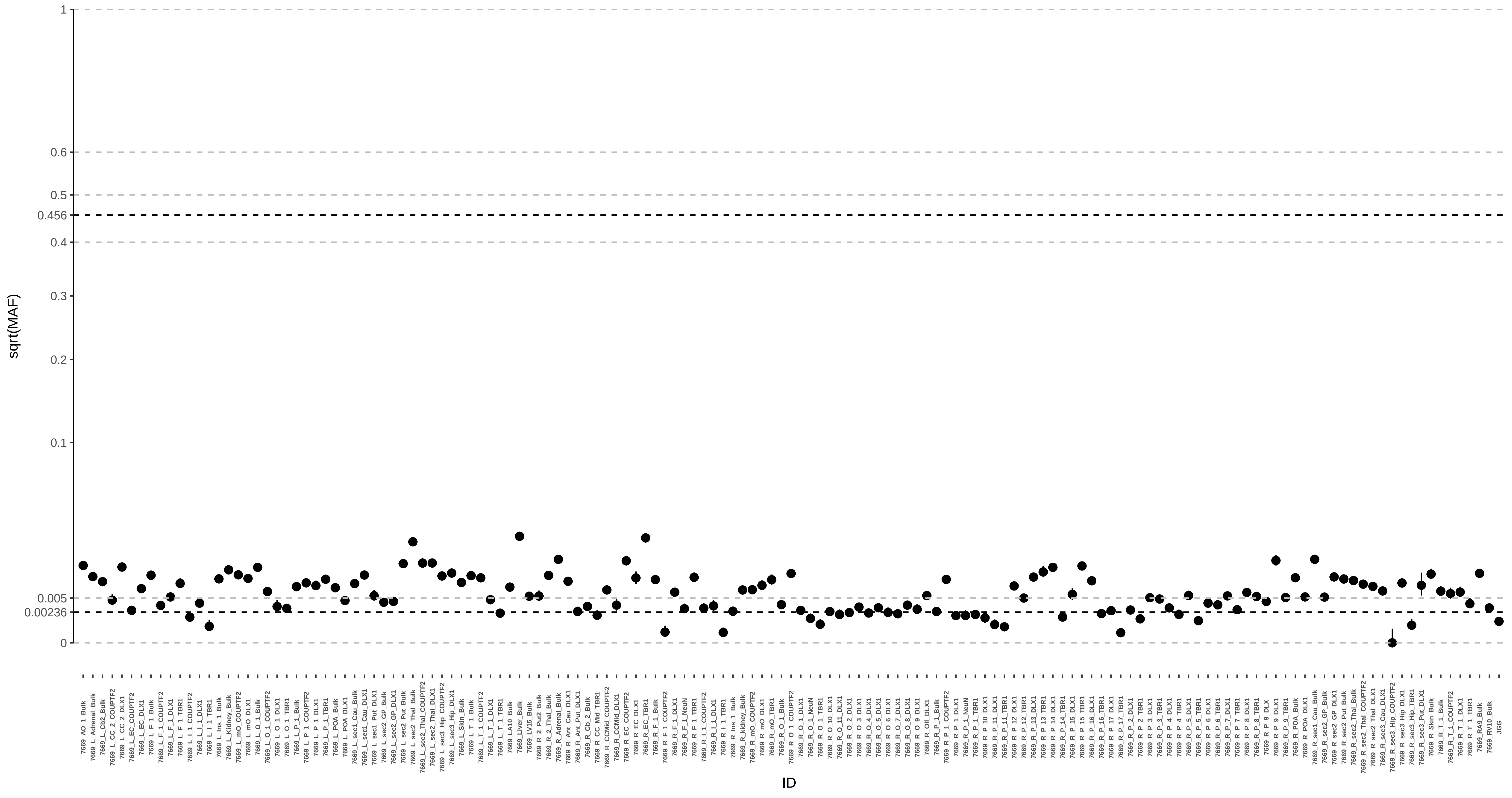

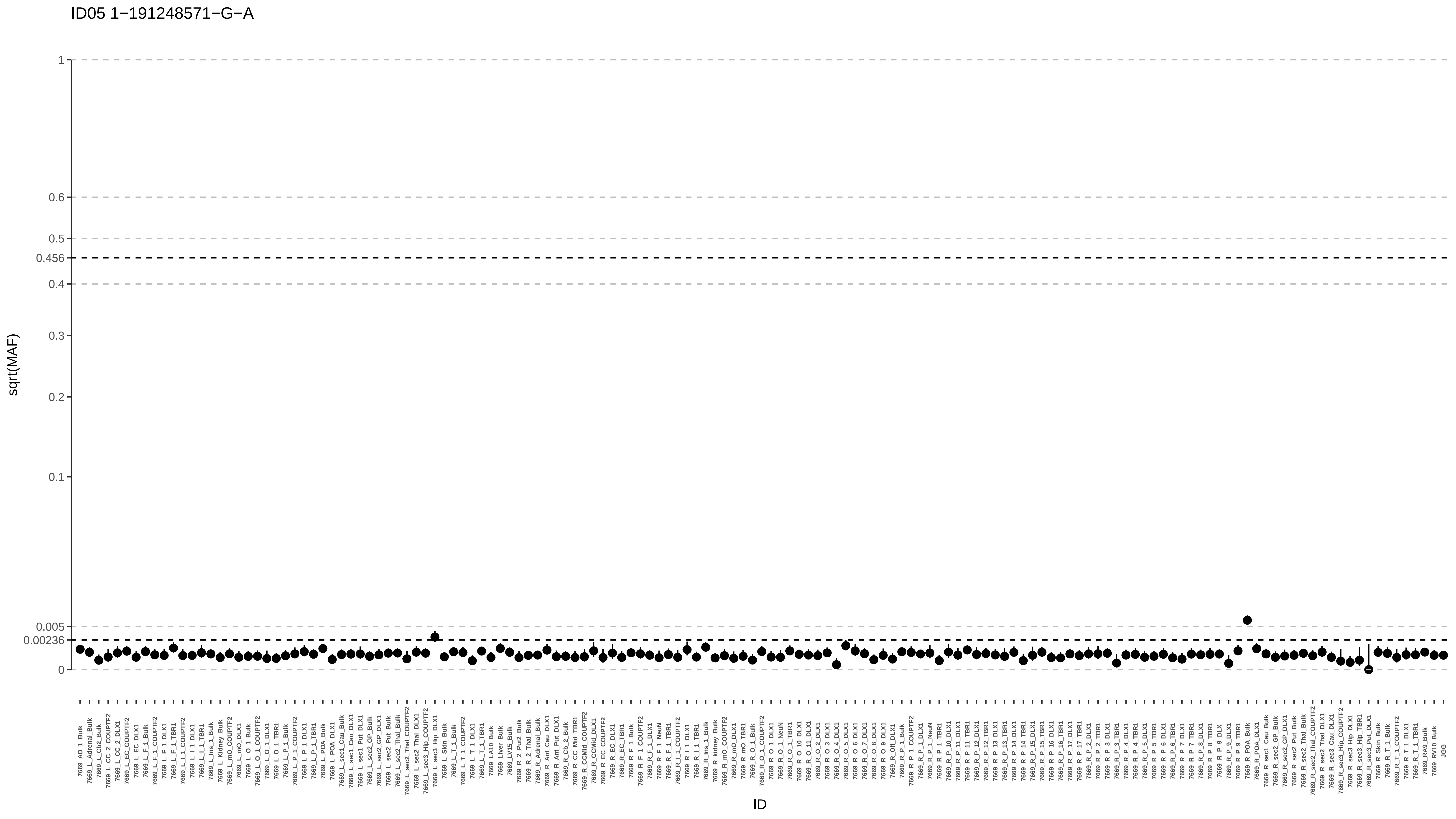
ID05 1−191248571−G−A
sqrt(MAF)
1
0.6
0.5
0.456
0.4
0.3
0.2
0.1
0.005
0.00236
0
ID
7669_AO_1_Bulk
7669_L_Adrenal_Bulk
7669_L_Cb2_Bulk
7669_L_CC_2_COUPTF2
7669_L_CC_2_DLX1
7669_L_EC_COUPTF2
7669_L_EC_DLX1
7669_L_F_1_Bulk
7669_L_F_1_COUPTF2
7669_L_F_1_DLX1
7669_L_F_1_TBR1
7669_L_I_1_COUPTF2
7669_L_I_1_DLX1
7669_L_I_1_TBR1
7669_L_Ins_1_Bulk
7669_L_Kidney_Bulk
7669_L_mO_COUPTF2
7669_L_mO_DLX1
7669_L_O_1_Bulk
7669_L_O_1_COUPTF2
7669_L_O_1_DLX1
7669_L_O_1_TBR1
7669_L_P_1_Bulk
7669_L_P_1_COUPTF2
7669_L_P_1_DLX1
7669_L_P_1_TBR1
7669_L_POA_Bulk
7669_L_POA_DLX1
7669_L_sec1_Cau_Bulk
7669_L_sec1_Cau_DLX1
7669_L_sec1_Put_DLX1
7669_L_sec2_GP_Bulk
7669_L_sec2_GP_DLX1
7669_L_sec2_Put_Bulk
7669_L_sec2_Thal_Bulk
7669_L_sec2_Thal_COUPTF2
7669_L_sec2_Thal_DLX1
7669_L_sec3_Hip_COUPTF2
7669_L_sec3_Hip_DLX1
7669_L_Skin_Bulk
7669_L_T_1_Bulk
7669_L_T_1_COUPTF2
7669_L_T_1_DLX1
7669_L_T_1_TBR1
7669_LA10_Bulk
7669_Liver_Bulk
7669_LV15_Bulk
7669_R_2_Put2_Bulk
7669_R_2_Thal_Bulk
7669_R_Adrenal_Bulk
7669_R_Ant_Cau_DLX1
7669_R_Ant_Put_DLX1
7669_R_Cb_2_Bulk
7669_R_CC_Mid_TBR1
7669_R_CCMid_COUPTF2
7669_R_CCMid_DLX1
7669_R_EC_COUPTF2
7669_R_EC_DLX1
7669_R_EC_TBR1
7669_R_F_1_Bulk
7669_R_F_1_COUPTF2
7669_R_F_1_DLX1
7669_R_F_1_NeuN
7669_R_F_1_TBR1
7669_R_I_1_COUPTF2
7669_R_I_1_DLX1
7669_R_I_1_TBR1
7669_R_Ins_1_Bulk
7669_R_kidney_Bulk
7669_R_mO_COUPTF2
7669_R_mO_DLX1
7669_R_mO_TBR1
7669_R_O_1_Bulk
7669_R_O_1_COUPTF2
7669_R_O_1_DLX1
7669_R_O_1_NeuN
7669_R_O_1_TBR1
7669_R_O_10_DLX1
7669_R_O_11_DLX1
7669_R_O_2_DLX1
7669_R_O_3_DLX1
7669_R_O_4_DLX1
7669_R_O_5_DLX1
7669_R_O_6_DLX1
7669_R_O_7_DLX1
7669_R_O_8_DLX1
7669_R_O_9_DLX1
7669_R_Olf_DLX1
7669_R_P_1_Bulk
7669_R_P_1_COUPTF2
7669_R_P_1_DLX1
7669_R_P_1_NeuN
7669_R_P_1_TBR1
7669_R_P_10_DLX1
7669_R_P_11_DLX1
7669_R_P_11_TBR1
7669_R_P_12_DLX1
7669_R_P_12_TBR1
7669_R_P_13_DLX1
7669_R_P_13_TBR1
7669_R_P_14_DLX1
7669_R_P_14_TBR1
7669_R_P_15_DLX1
7669_R_P_15_TBR1
7669_R_P_16_DLX1
7669_R_P_16_TBR1
7669_R_P_17_DLX1
7669_R_P_17_TBR1
7669_R_P_2_DLX1
7669_R_P_2_TBR1
7669_R_P_3_DLX1
7669_R_P_3_TBR1
7669_R_P_4_DLX1
7669_R_P_4_TBR1
7669_R_P_5_DLX1
7669_R_P_5_TBR1
7669_R_P_6_DLX1
7669_R_P_6_TBR1
7669_R_P_7_DLX1
7669_R_P_7_TBR1
7669_R_P_8_DLX1
7669_R_P_8_TBR1
7669_R_P_9_DLX
7669_R_P_9_DLX1
7669_R_P_9_TBR1
7669_R_POA_Bulk
7669_R_POA_DLX1
7669_R_sec1_Cau_Bulk
7669_R_sec2_GP_Bulk
7669_R_sec2_GP_DLX1
7669_R_sec2_Put_Bulk
7669_R_sec2_Thal_Bulk
7669_R_sec2_Thal_COUPTF2
7669_R_sec2_Thal_DLX1
7669_R_sec3_Cau_DLX1
7669_R_sec3_Hip_COUPTF2
7669_R_sec3_Hip_DLX1
7669_R_sec3_Hip_TBR1
7669_R_sec3_Put_DLX1
7669_R_Skin_Bulk
7669_R_T_1_Bulk
7669_R_T_1_COUPTF2
7669_R_T_1_DLX1
7669_R_T_1_TBR1
7669_RA9_Bulk
7669_RV10_Bulk
JGG

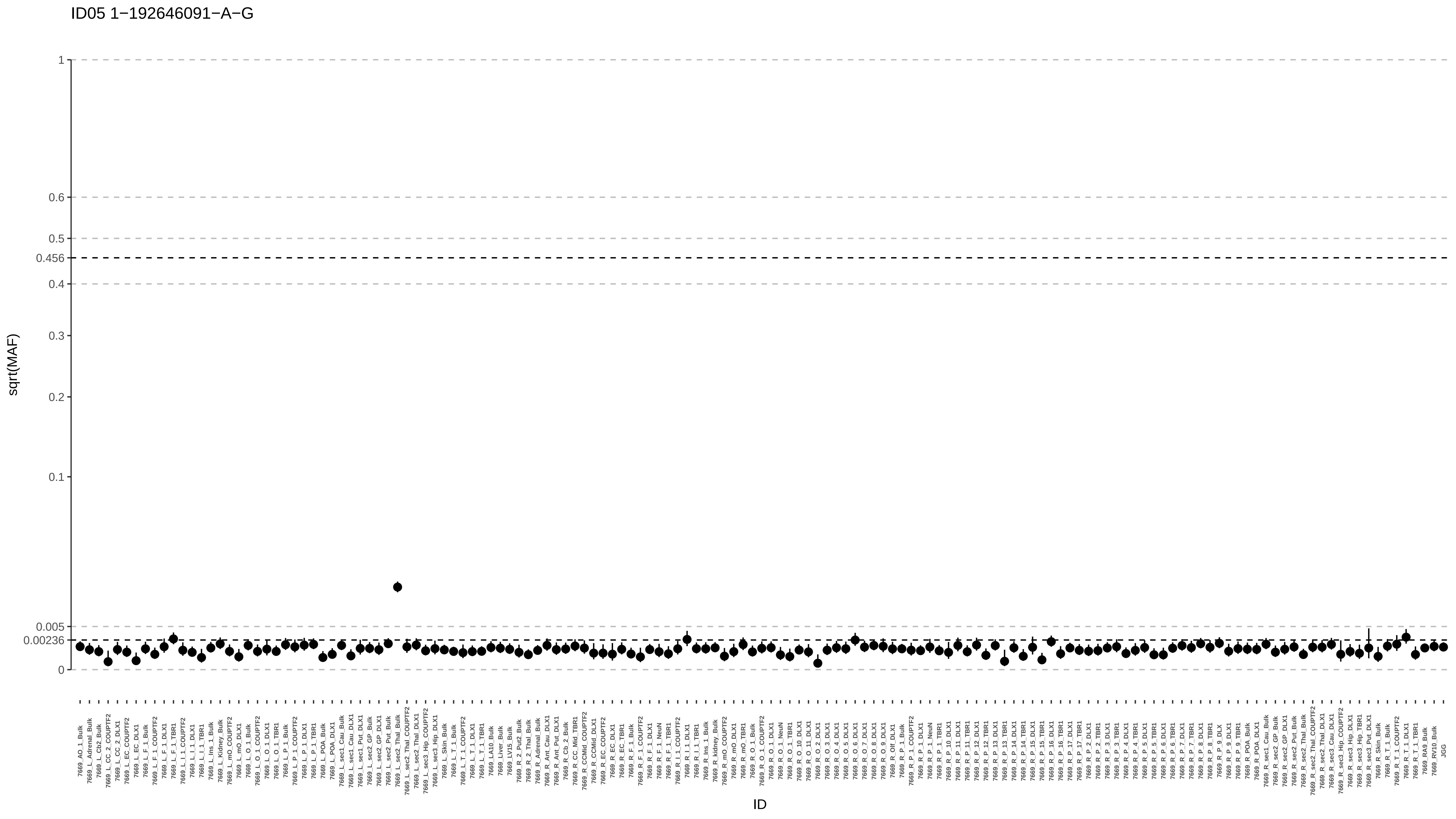
ID05 1−192646091−A−G
sqrt(MAF)
1
0.6
0.5
0.456
0.4
0.3
0.2
0.1
0.005
0.00236
0
ID

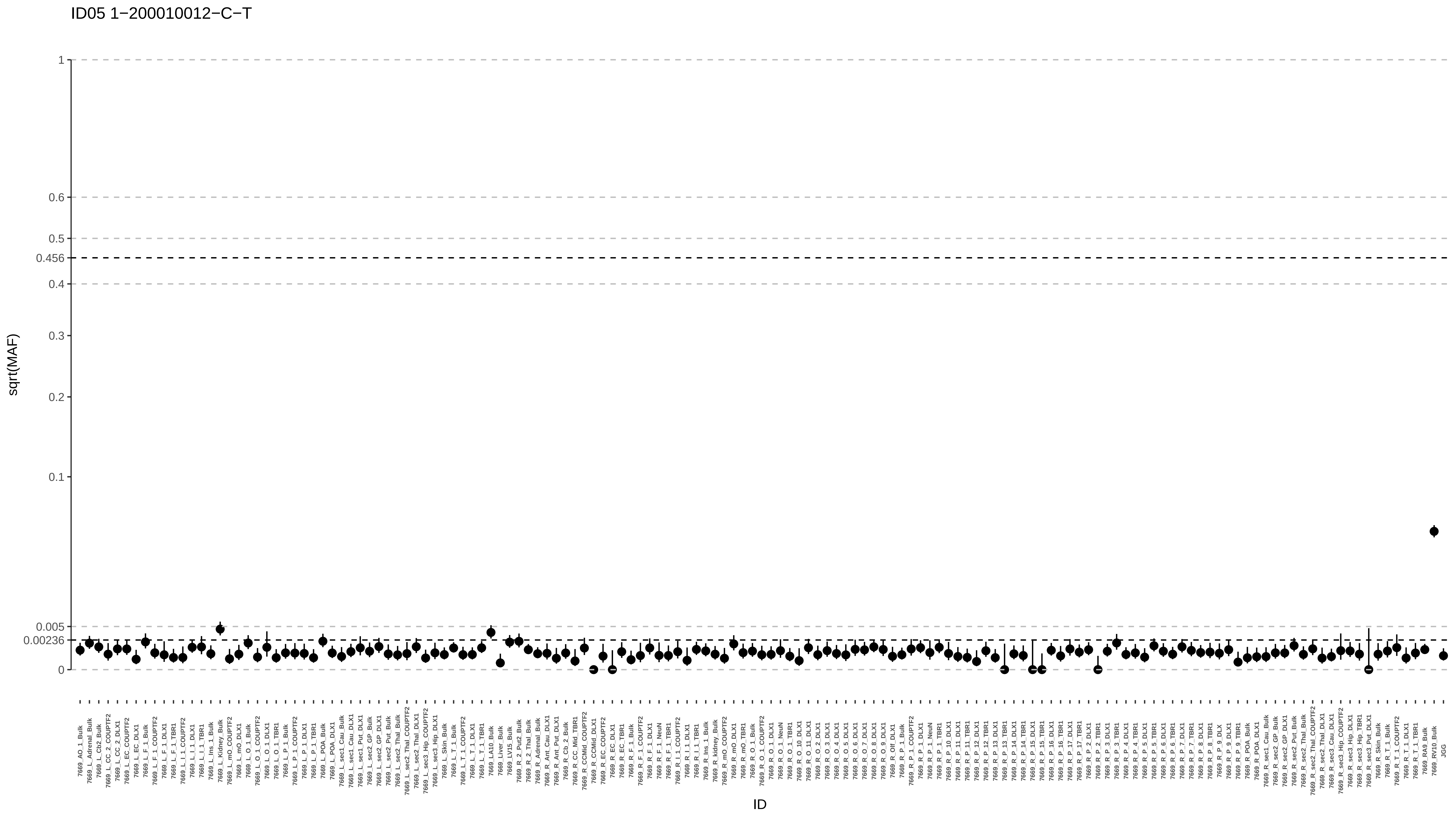
ID05 1−200010012−C−T
sqrt(MAF)
1
0.6
0.5
0.456
0.4
0.3
0.2
0.1
0.005
0.00236
0
ID
JGG

ID05 1−201266932−C−T

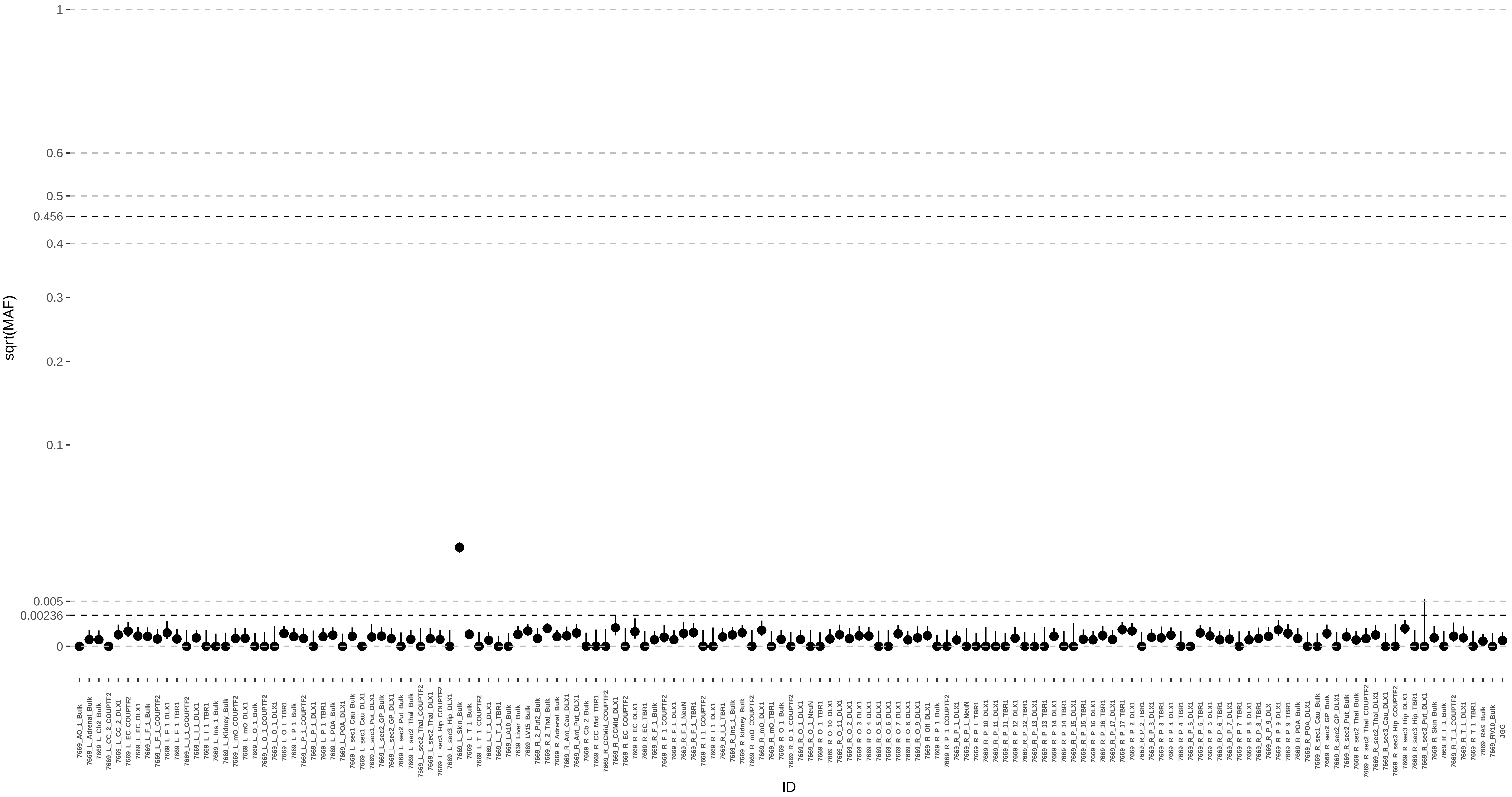

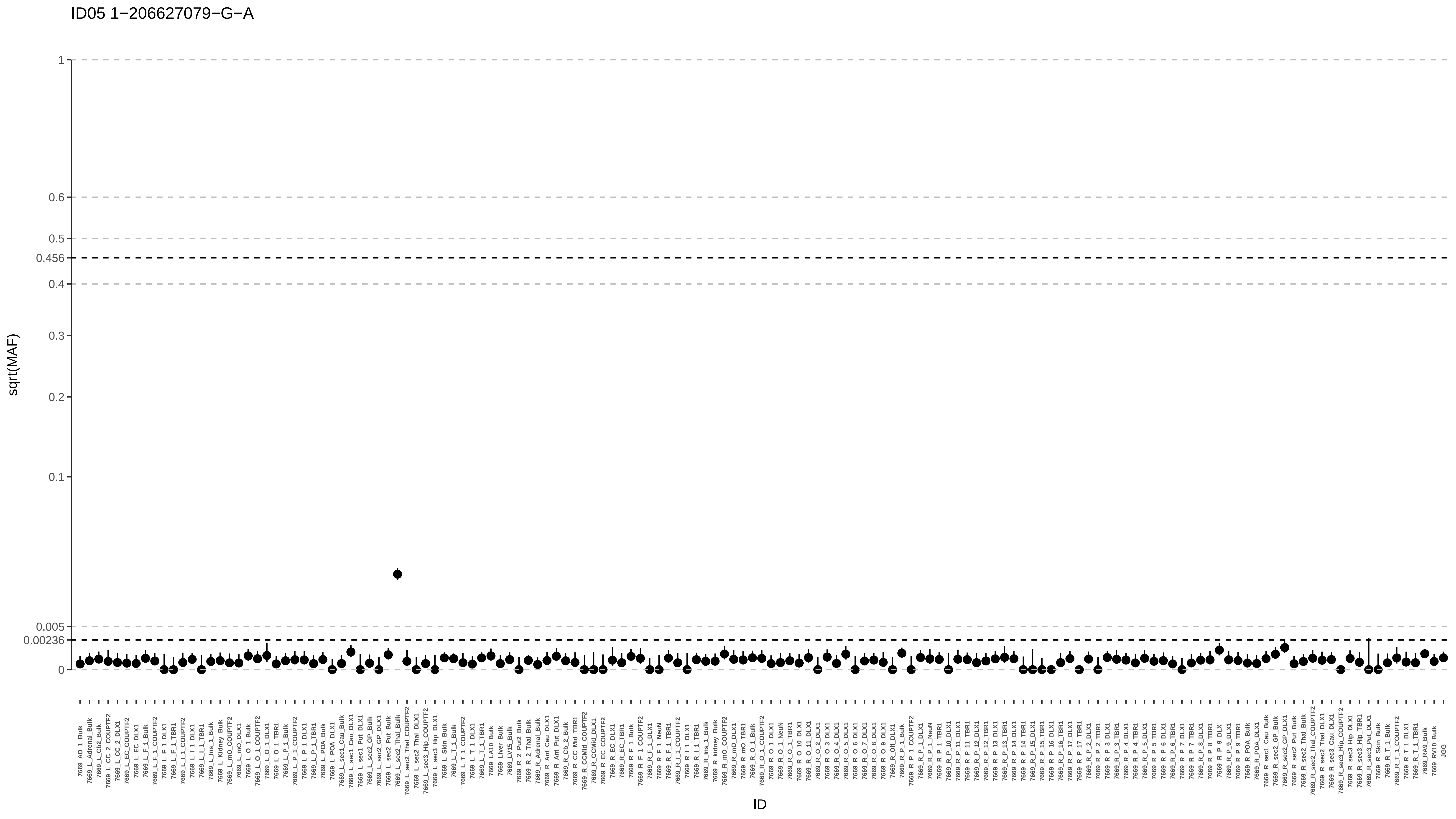
ID05 1−206627079−G−A
sqrt(MAF)
1
0.6
0.5
0.456
0.4
0.3
0.2
0.1
0.005
0.00236
0
ID

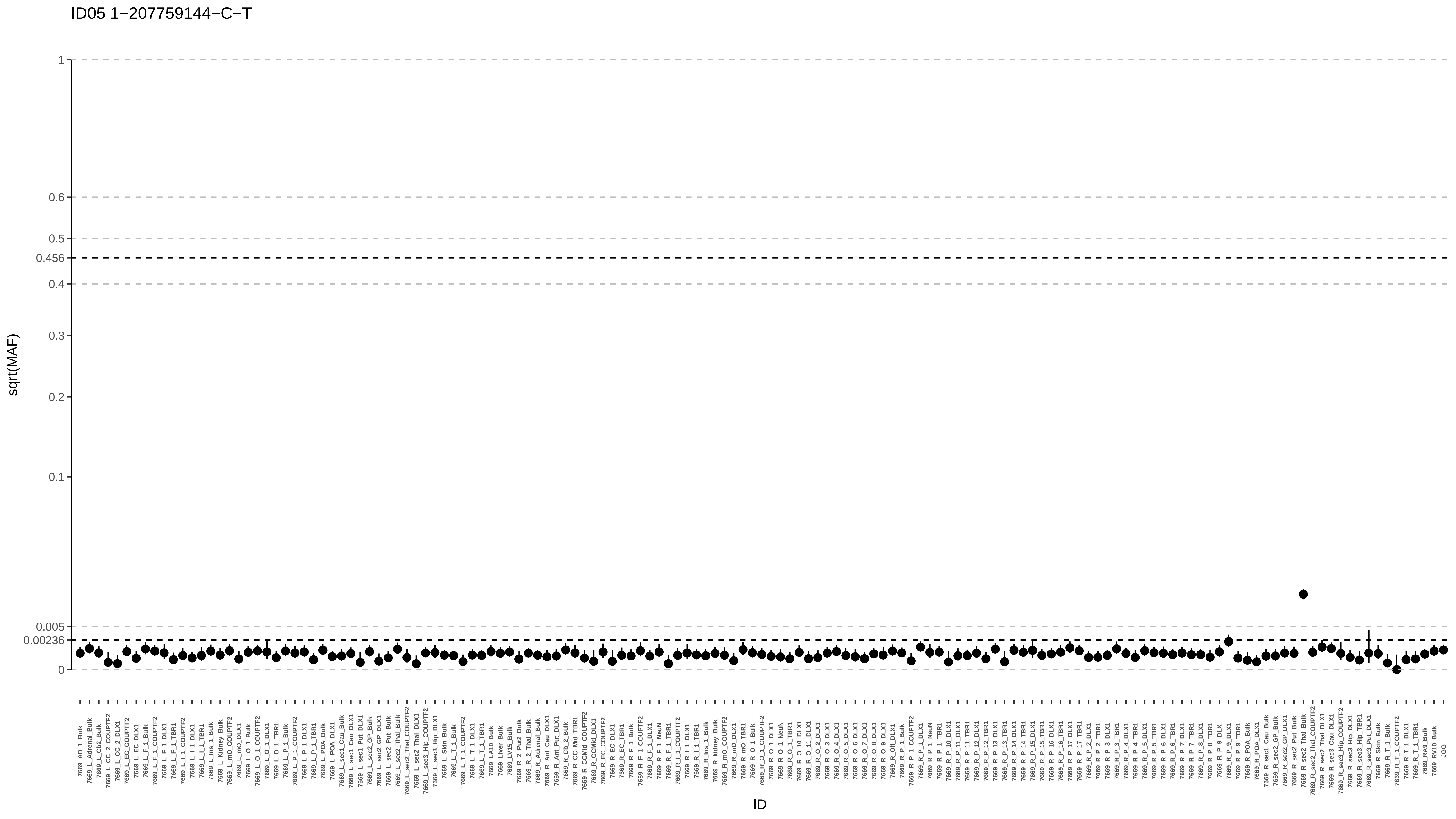
ID05 1−207759144−C−T
sqrt(MAF)
1
0.6
0.5
0.456
0.4
0.3
0.2
0.1
0.005
0.00236
0
ID

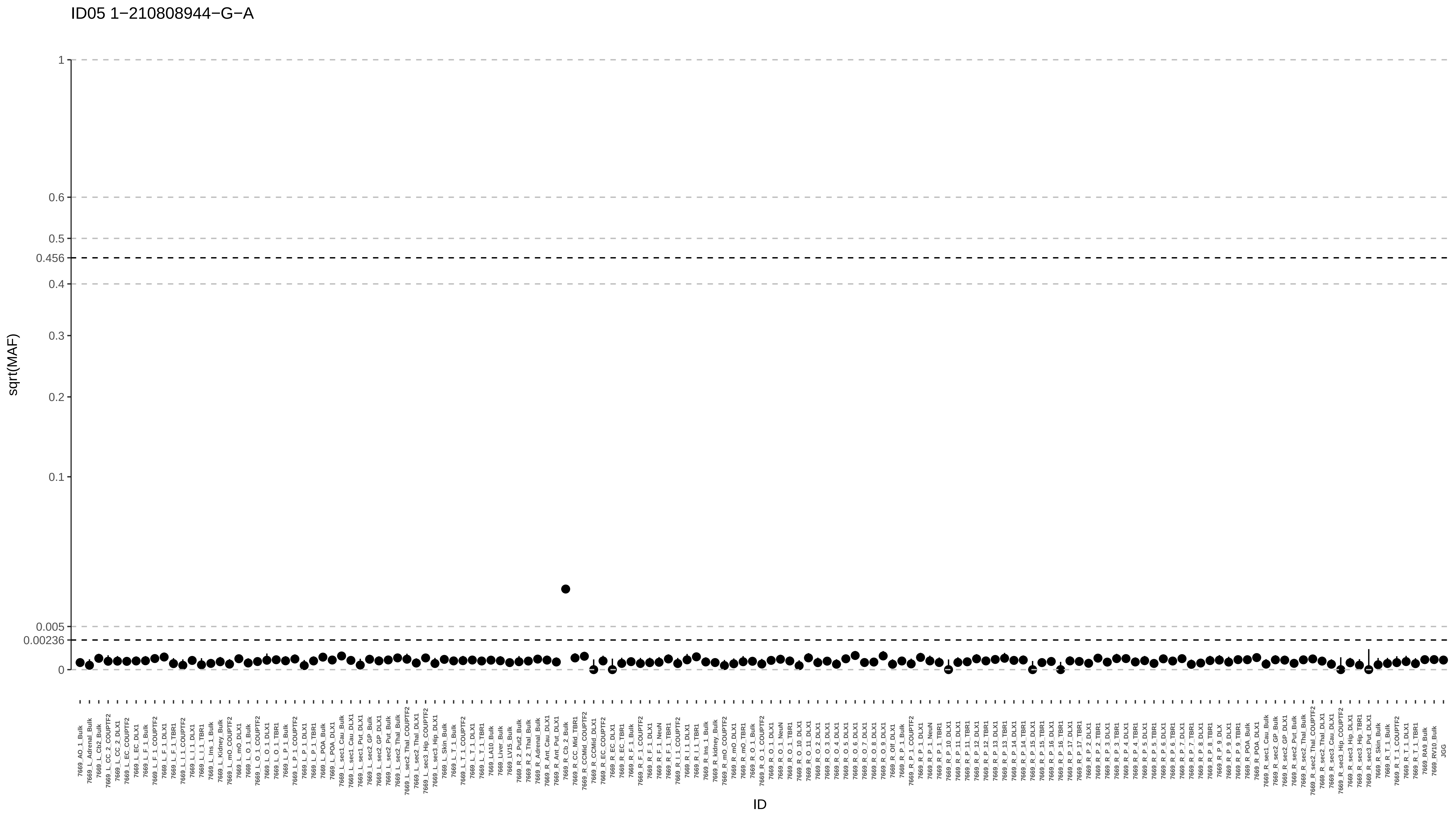
ID05 1−210808944−G−A
sqrt(MAF)
ID
1
0.6
0.5
0.456
0.4
0.3
0.2
0.1
0.005
0.00236
0

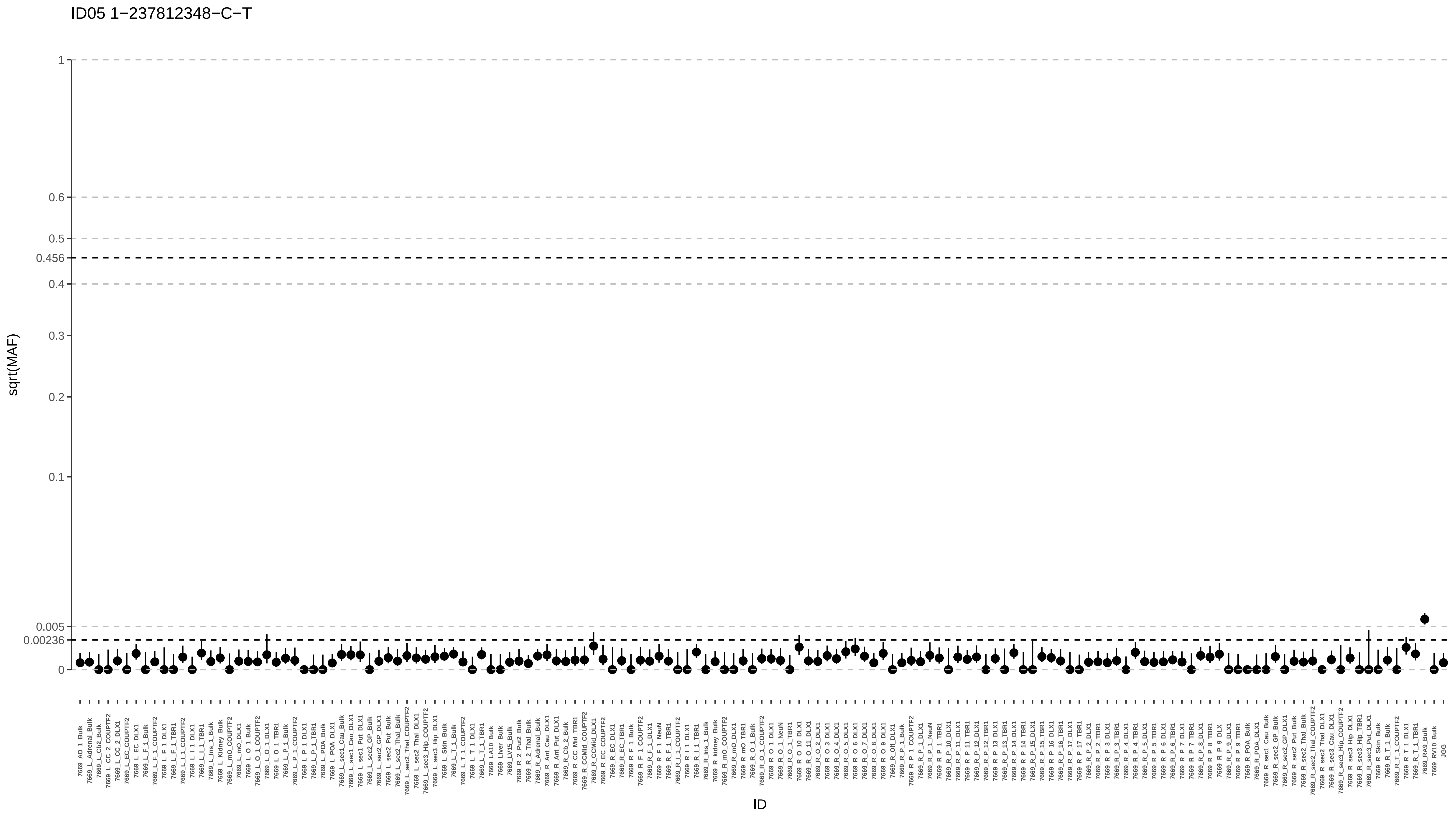
ID05 1−237812348−C−T
sqrt(MAF)
ID
1
0.6
0.5
0.456
0.4
0.3
0.2
0.1
0.005
0.00236
0

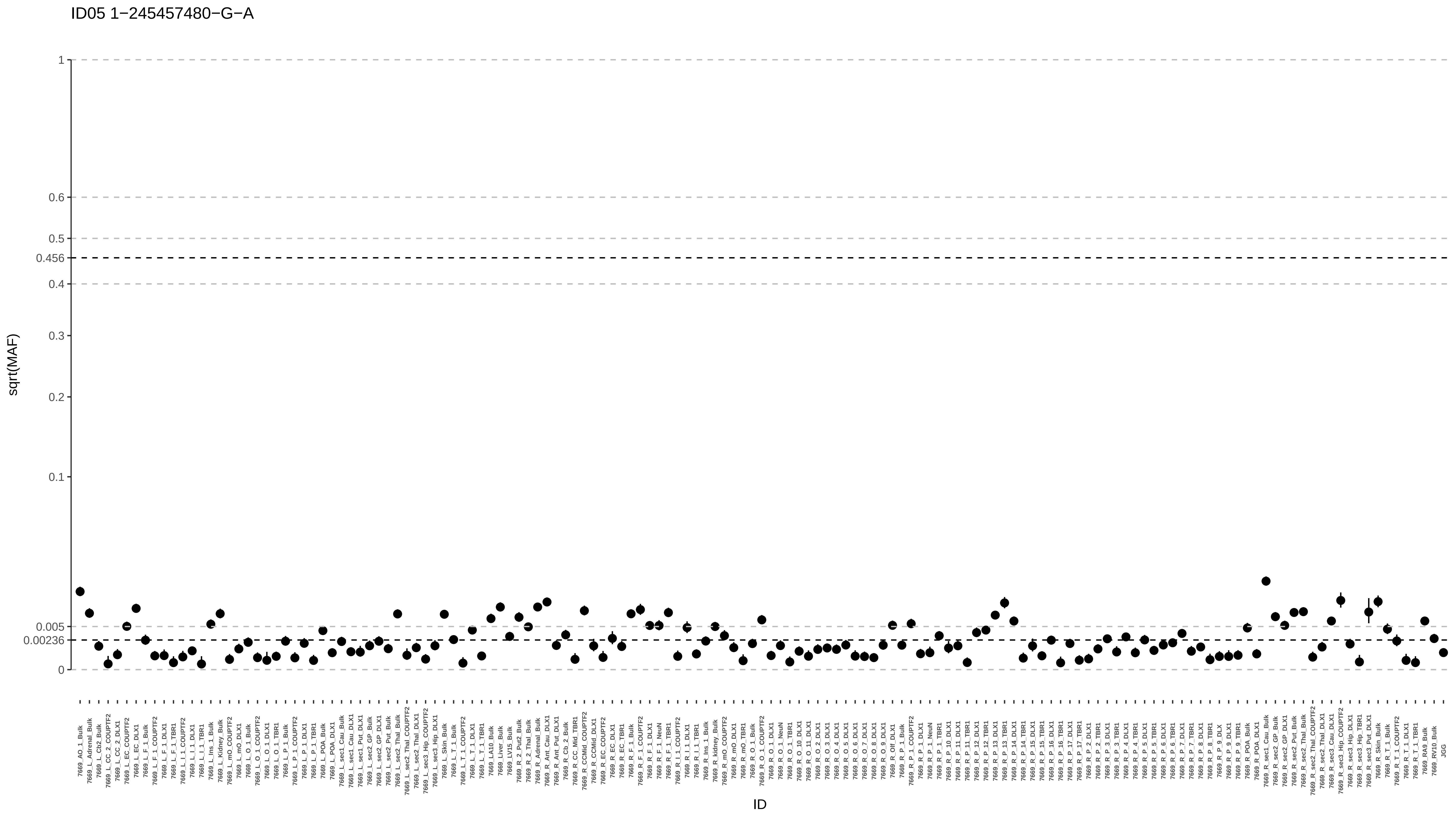
ID05 1−245457480−G−A
sqrt(MAF)
1
0.6
0.5
0.456
0.4
0.3
0.2
0.1
0.005
0.00236
0
ID

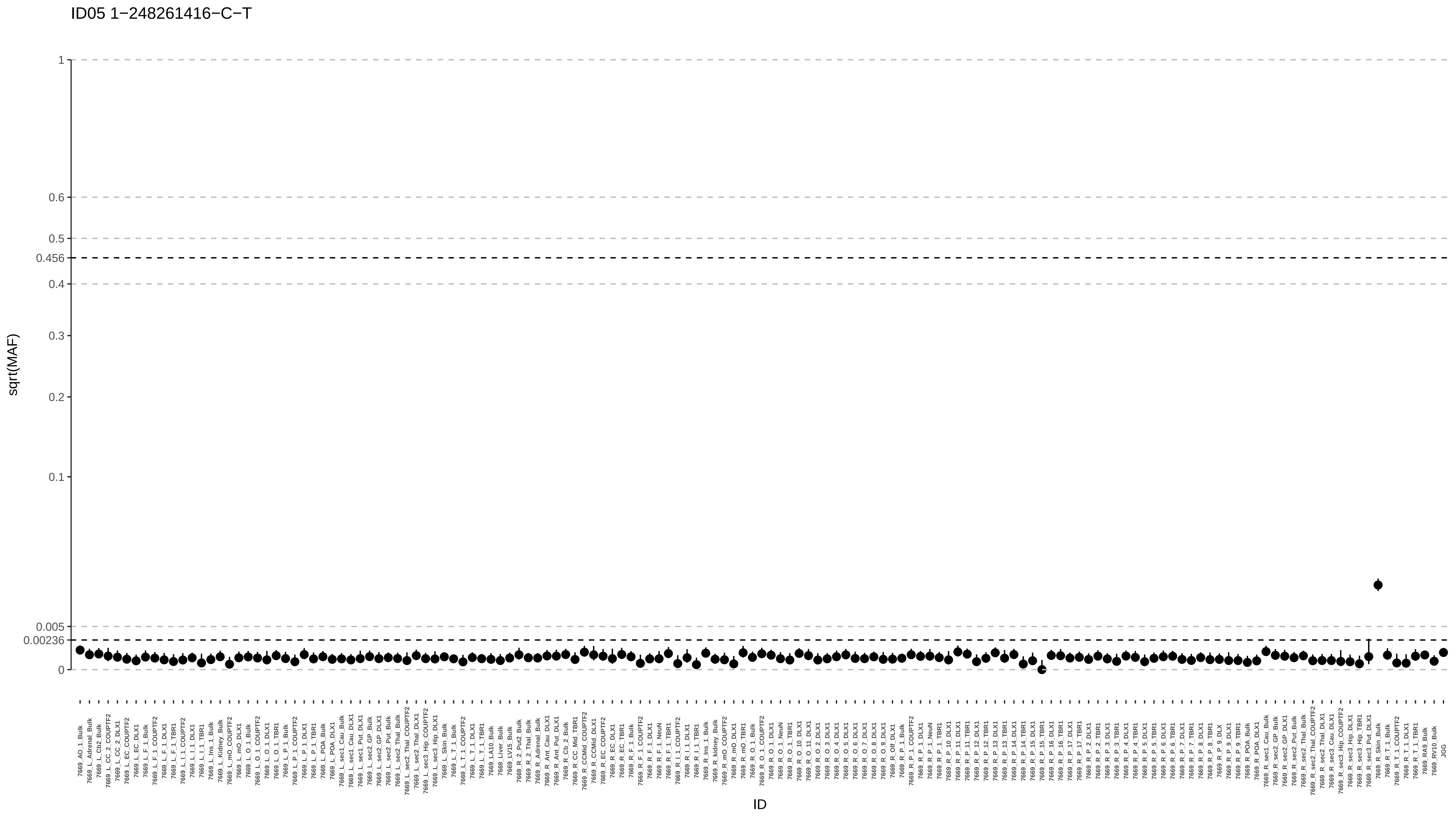
ID05 1−248261416−C−T
sqrt(MAF)
1
0.6
0.5
0.456
0.4
0.3
0.2
0.1
0.005
0.00236
0
ID

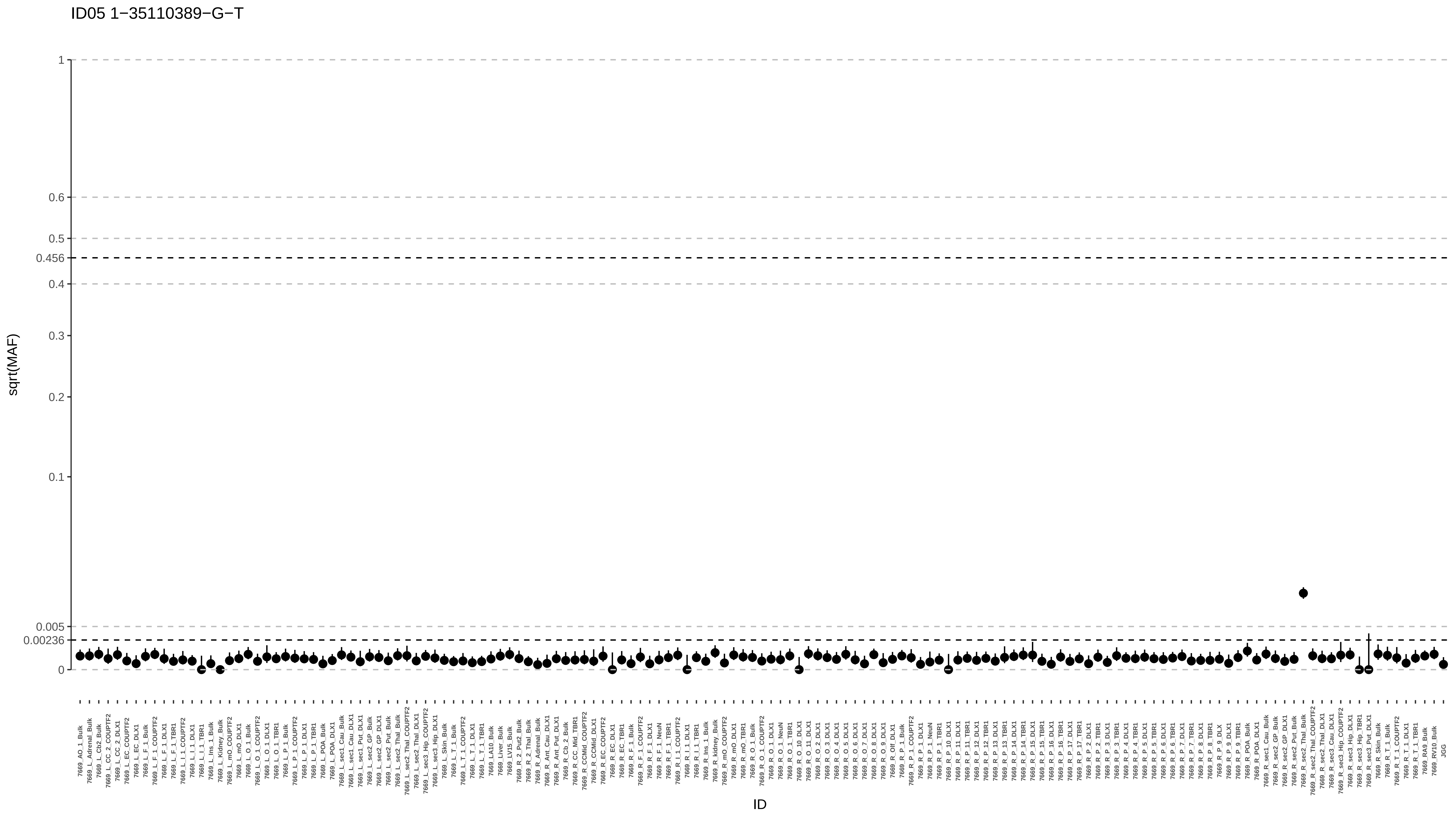
ID05 1−35110389−G−T
sqrt(MAF)
1
0.6
0.5
0.456
0.4
0.3
0.2
0.1
0.005
0.00236
0
ID
7669_AO_1_Bulk
7669_L_Adrenal_Bulk
7669_L_Cb2_Bulk
7669_L_CC_2_COUPTF2
7669_L_CC_2_DLX1
7669_L_EC_COUPTF2
7669_L_EC_DLX1
7669_L_F_1_Bulk
7669_L_F_1_COUPTF2
7669_L_F_1_DLX1
7669_L_F_1_TBR1
7669_L_I_1_COUPTF2
7669_L_I_1_DLX1
7669_L_I_1_TBR1
7669_L_Ins_1_Bulk
7669_L_Kidney_Bulk
7669_L_mO_COUPTF2
7669_L_mO_DLX1
7669_L_O_1_Bulk
7669_L_O_1_COUPTF2
7669_L_O_1_DLX1
7669_L_O_1_TBR1
7669_L_P_1_Bulk
7669_L_P_1_COUPTF2
7669_L_P_1_DLX1
7669_L_P_1_TBR1
7669_L_POA_Bulk
7669_L_POA_DLX1
7669_L_sec1_Cau_Bulk
7669_L_sec1_Cau_DLX1
7669_L_sec1_Put_DLX1
7669_L_sec2_GP_Bulk
7669_L_sec2_GP_DLX1
7669_L_sec2_Put_Bulk
7669_L_sec2_Thal_Bulk
7669_L_sec2_Thal_COUPTF2
7669_L_sec2_Thal_DLX1
7669_L_sec3_Hip_COUPTF2
7669_L_sec3_Hip_DLX1
7669_L_Skin_Bulk
7669_L_T_1_Bulk
7669_L_T_1_COUPTF2
7669_L_T_1_DLX1
7669_L_T_1_TBR1
7669_LA10_Bulk
7669_Liver_Bulk
7669_LV15_Bulk
7669_R_2_Put2_Bulk
7669_R_2_Thal_Bulk
7669_R_Adrenal_Bulk
7669_R_Ant_Cau_DLX1
7669_R_Ant_Put_DLX1
7669_R_Cb_2_Bulk
7669_R_CC_Mid_TBR1
7669_R_CCMid_COUPTF2
7669_R_CCMid_DLX1
7669_R_EC_COUPTF2
7669_R_EC_DLX1
7669_R_EC_TBR1
7669_R_F_1_Bulk
7669_R_F_1_COUPTF2
7669_R_F_1_DLX1
7669_R_F_1_NeuN
7669_R_F_1_TBR1
7669_R_I_1_COUPTF2
7669_R_I_1_DLX1
7669_R_I_1_TBR1
7669_R_Ins_1_Bulk
7669_R_kidney_Bulk
7669_R_mO_COUPTF2
7669_R_mO_DLX1
7669_R_mO_TBR1
7669_R_O_1_Bulk
7669_R_O_1_COUPTF2
7669_R_O_1_DLX1
7669_R_O_1_NeuN
7669_R_O_1_TBR1
7669_R_O_10_DLX1
7669_R_O_11_DLX1
7669_R_O_2_DLX1
7669_R_O_3_DLX1
7669_R_O_4_DLX1
7669_R_O_5_DLX1
7669_R_O_6_DLX1
7669_R_O_7_DLX1
7669_R_O_8_DLX1
7669_R_O_9_DLX1
7669_R_Olf_DLX1
7669_R_P_1_Bulk
7669_R_P_1_COUPTF2
7669_R_P_1_DLX1
7669_R_P_1_NeuN
7669_R_P_1_TBR1
7669_R_P_10_DLX1
7669_R_P_11_DLX1
7669_R_P_11_TBR1
7669_R_P_12_DLX1
7669_R_P_12_TBR1
7669_R_P_13_DLX1
7669_R_P_13_TBR1
7669_R_P_14_DLX1
7669_R_P_14_TBR1
7669_R_P_15_DLX1
7669_R_P_15_TBR1
7669_R_P_16_DLX1
7669_R_P_16_TBR1
7669_R_P_17_DLX1
7669_R_P_17_TBR1
7669_R_P_2_DLX1
7669_R_P_2_TBR1
7669_R_P_3_DLX1
7669_R_P_3_TBR1
7669_R_P_4_DLX1
7669_R_P_4_TBR1
7669_R_P_5_DLX1
7669_R_P_5_TBR1
7669_R_P_6_DLX1
7669_R_P_6_TBR1
7669_R_P_7_DLX1
7669_R_P_7_TBR1
7669_R_P_8_DLX1
7669_R_P_8_TBR1
7669_R_P_9_DLX
7669_R_P_9_DLX1
7669_R_P_9_TBR1
7669_R_POA_Bulk
7669_R_POA_DLX1
7669_R_sec1_Cau_Bulk
7669_R_sec2_GP_Bulk
7669_R_sec2_GP_DLX1
7669_R_sec2_Put_Bulk
7669_R_sec2_Thal_Bulk
7669_R_sec2_Thal_COUPTF2
7669_R_sec2_Thal_DLX1
7669_R_sec3_Cau_DLX1
7669_R_sec3_Hip_COUPTF2
7669_R_sec3_Hip_DLX1
7669_R_sec3_Hip_TBR1
7669_R_sec3_Put_DLX1
7669_R_Skin_Bulk
7669_R_T_1_Bulk
7669_R_T_1_COUPTF2
7669_R_T_1_DLX1
7669_R_T_1_TBR1
7669_RA9_Bulk
7669_RV10_Bulk
JGG

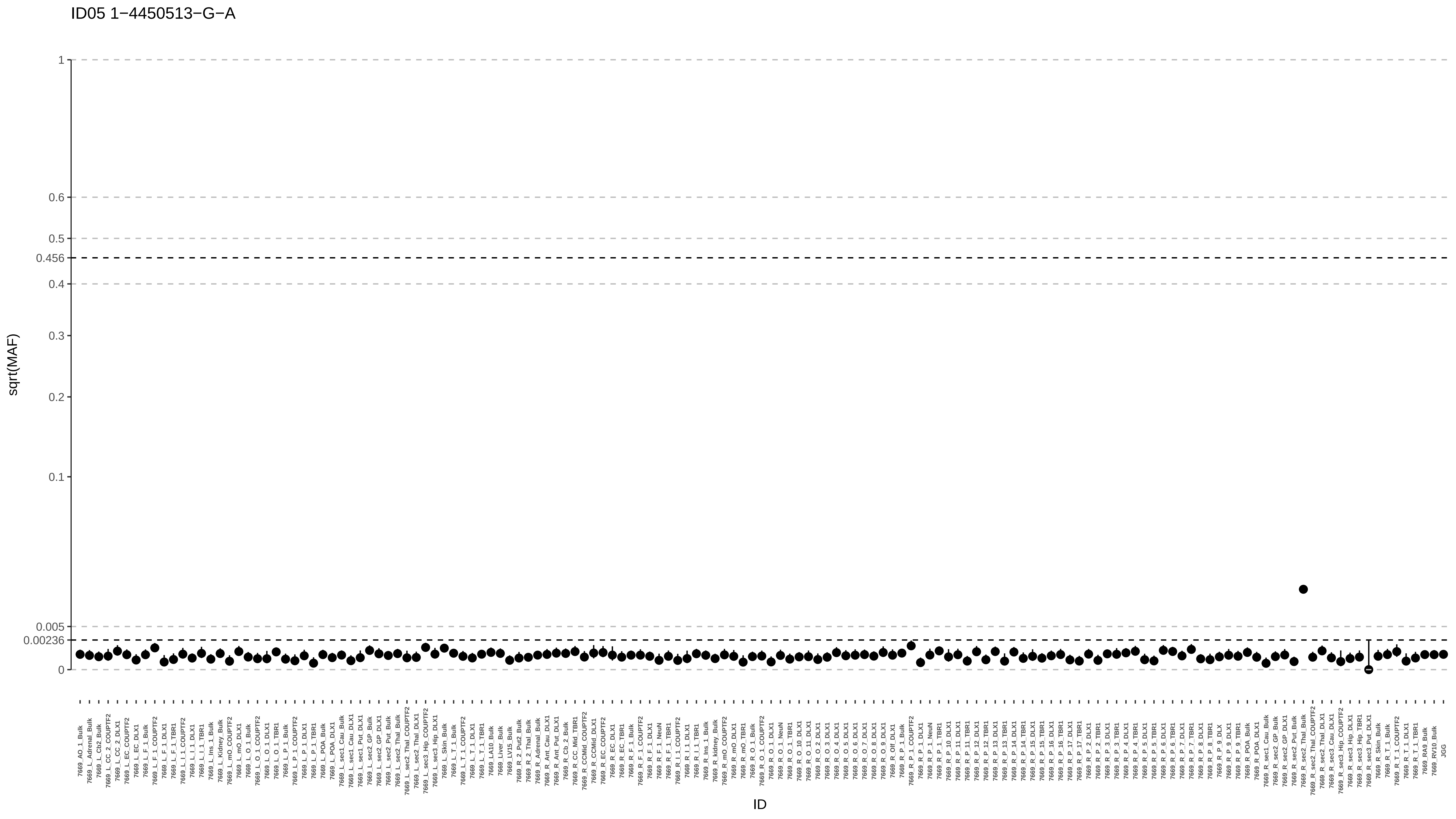

ID05 1−4450513−G−A
sqrt(MAF)
1
0.6
0.5
0.456
0.4
0.3
0.2
0.1
0.005
0.00236
0
ID

ID05 1−4465349−T−G

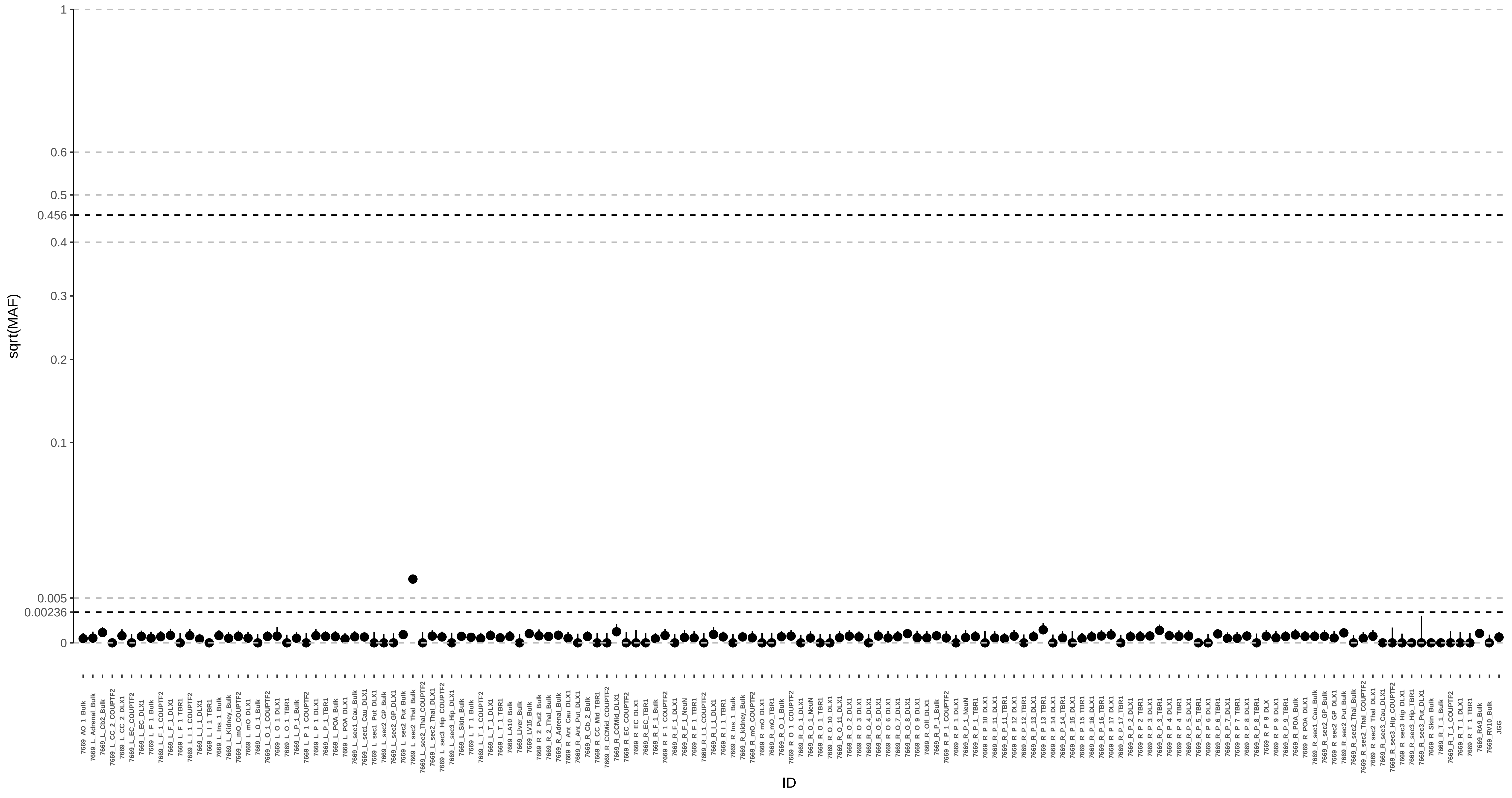

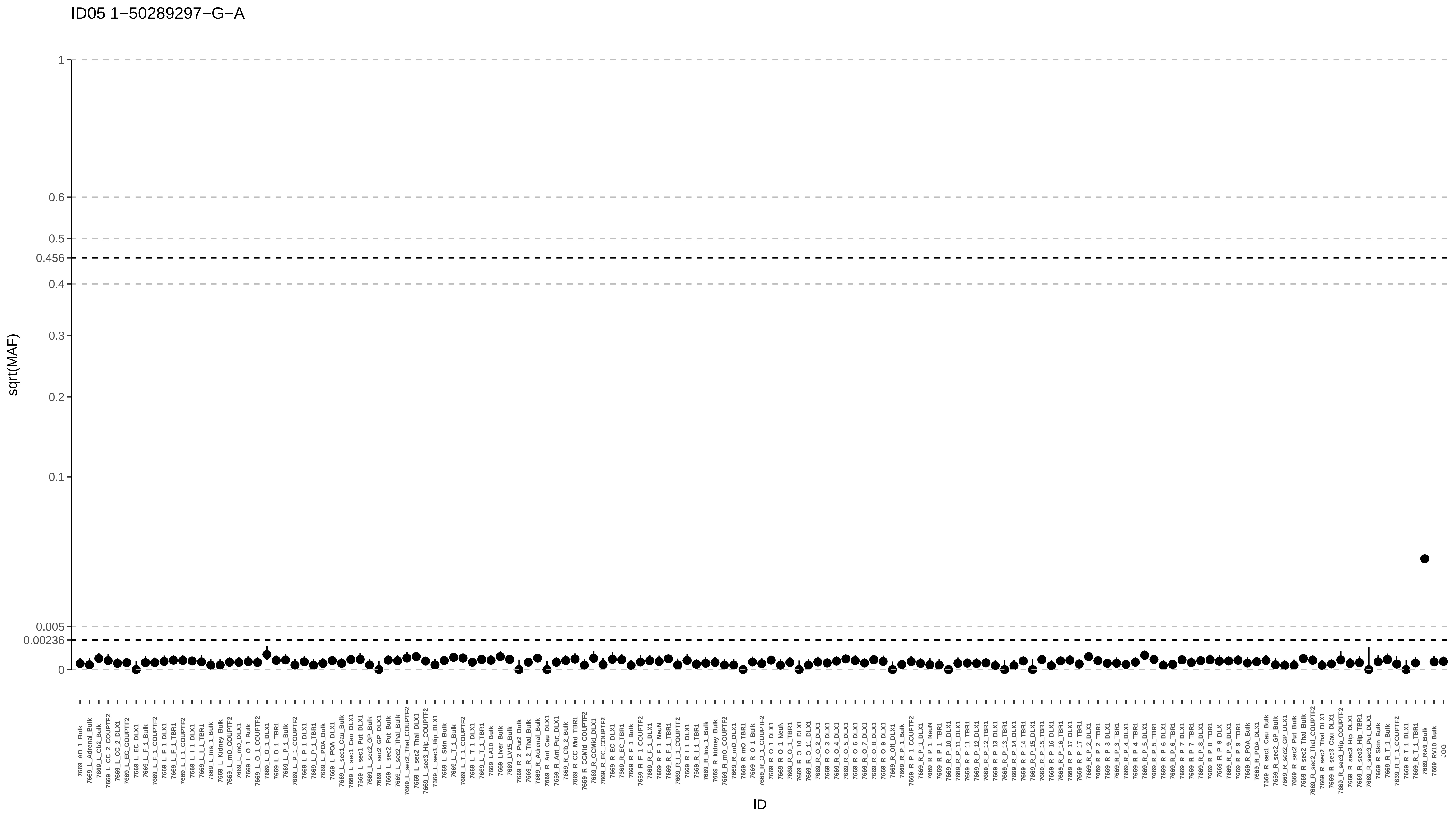

ID05 1−50289297−G−A
sqrt(MAF)
1
0.6
0.5
0.456
0.4
0.3
0.2
0.1
0.005
0.00236
0
ID

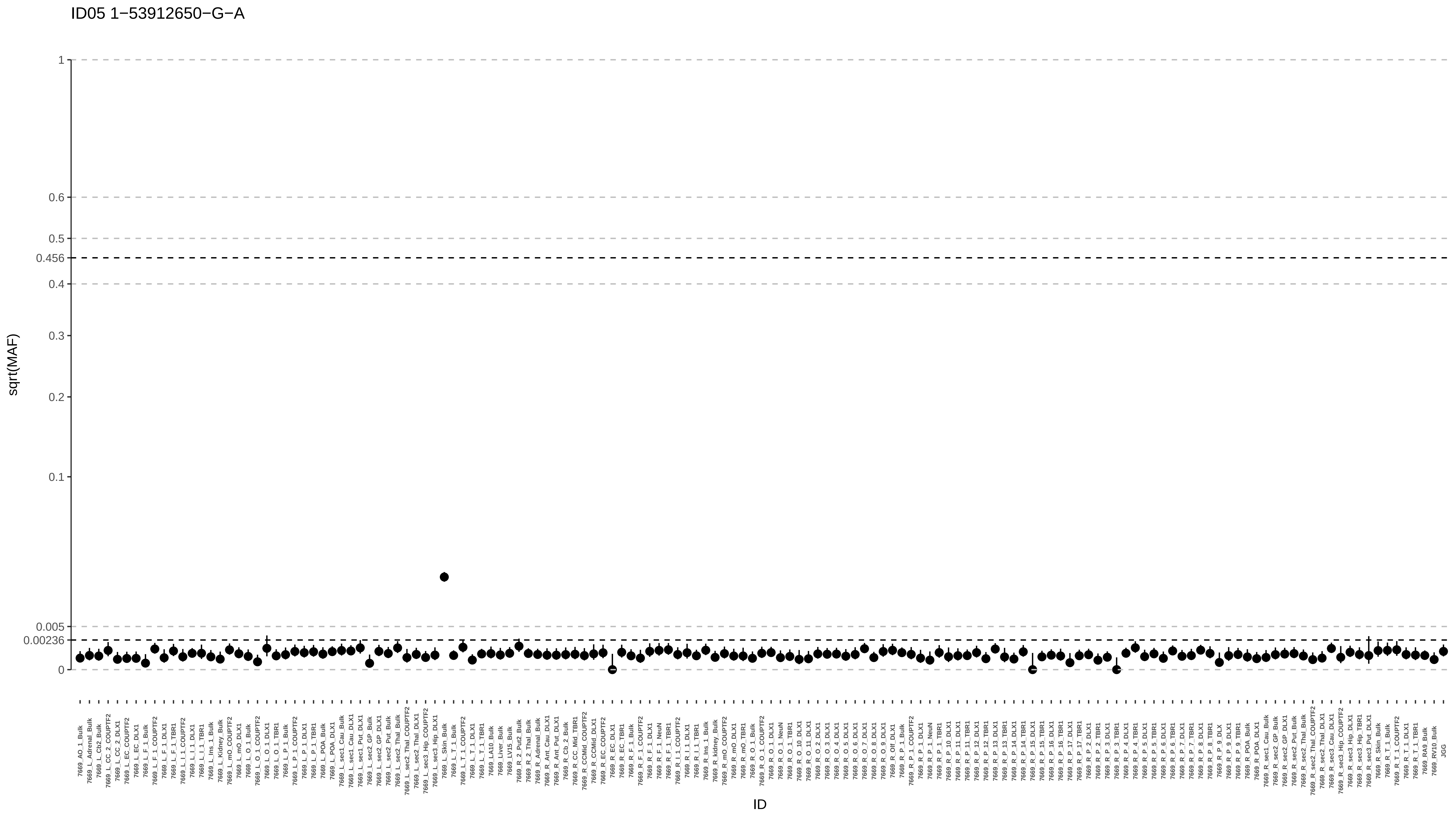

ID05 1−53912650−G−A
sqrt(MAF)
1
0.6
0.5
0.456
0.4
0.3
0.2
0.1
0.005
0.00236
0
ID

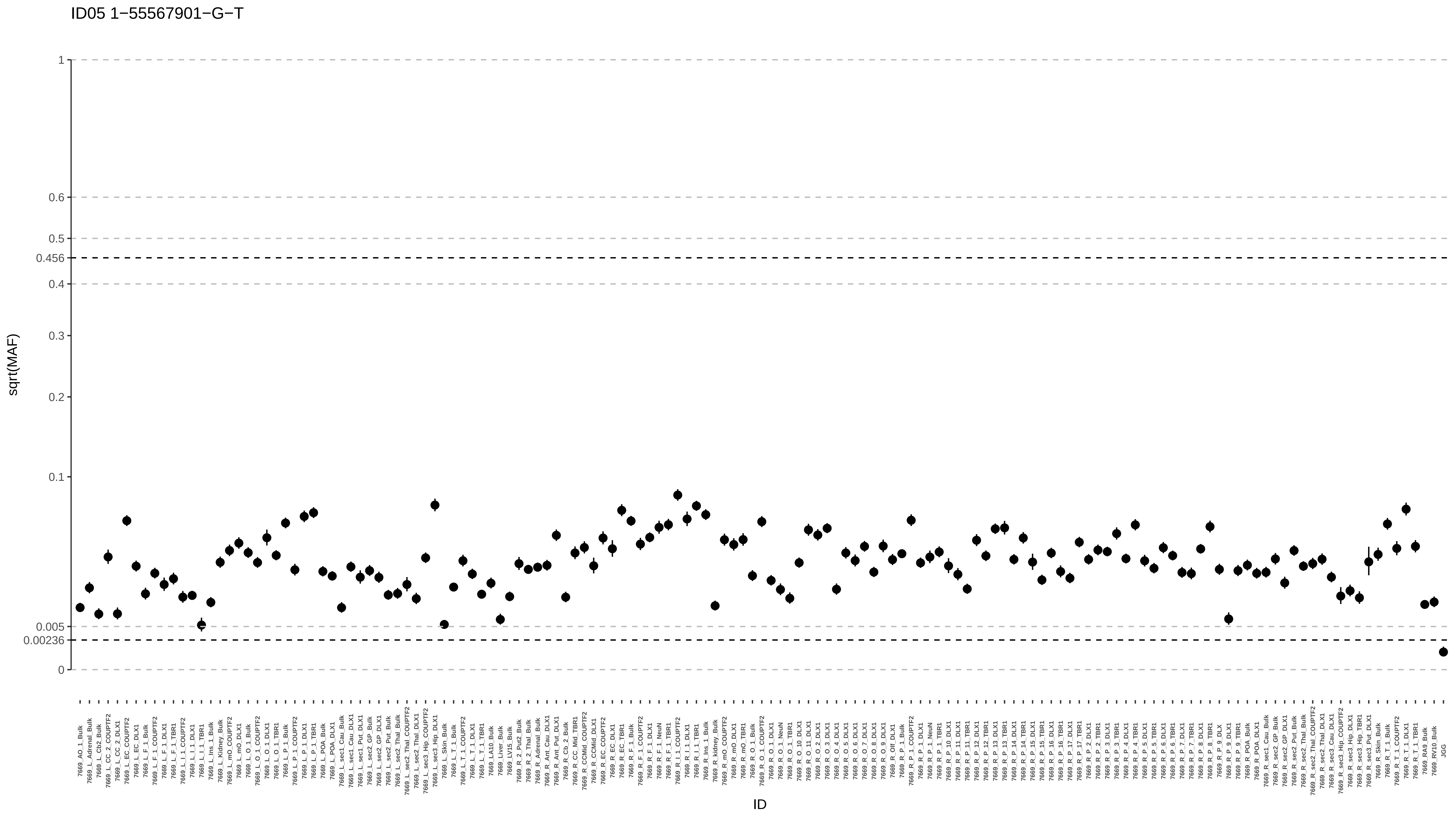
ID05 1−55567901−G−T
sqrt(MAF)
1
0.6
0.5
0.456
0.4
0.3
0.2
0.1
0.005
0.00236
0
ID

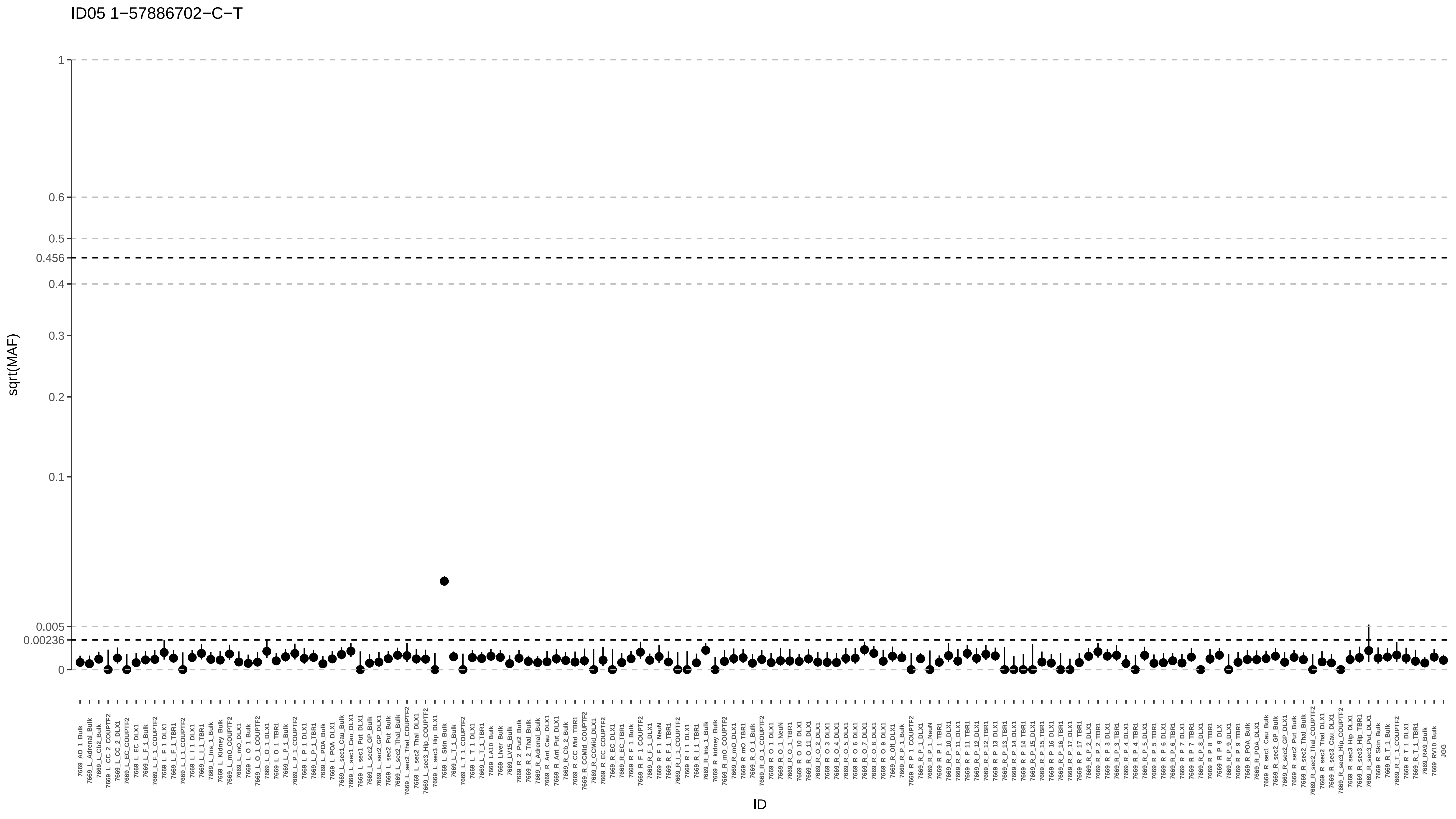
ID05 1−57886702−C−T
sqrt(MAF)
1
0.6
0.5
0.456
0.4
0.3
0.2
0.1
0.005
0.00236
0
ID
7669_AO_1_Bulk
7669_L_Adrenal_Bulk
7669_L_Cb2_Bulk
7669_L_CC_2_COUPTF2
7669_L_CC_2_DLX1
7669_L_EC_COUPTF2
7669_L_EC_DLX1
7669_L_F_1_Bulk
7669_L_F_1_COUPTF2
7669_L_F_1_DLX1
7669_L_F_1_TBR1
7669_L_I_1_COUPTF2
7669_L_I_1_DLX1
7669_L_I_1_TBR1
7669_L_Ins_1_Bulk
7669_L_Kidney_Bulk
7669_L_mO_COUPTF2
7669_L_mO_DLX1
7669_L_O_1_Bulk
7669_L_O_1_COUPTF2
7669_L_O_1_DLX1
7669_L_O_1_TBR1
7669_L_P_1_Bulk
7669_L_P_1_COUPTF2
7669_L_P_1_DLX1
7669_L_P_1_TBR1
7669_L_POA_Bulk
7669_L_POA_DLX1
7669_L_sec1_Cau_Bulk
7669_L_sec1_Cau_DLX1
7669_L_sec1_Put_DLX1
7669_L_sec2_GP_Bulk
7669_L_sec2_GP_DLX1
7669_L_sec2_Put_Bulk
7669_L_sec2_Thal_Bulk
7669_L_sec2_Thal_COUPTF2
7669_L_sec2_Thal_DLX1
7669_L_sec3_Hip_COUPTF2
7669_L_sec3_Hip_DLX1
7669_L_Skin_Bulk
7669_L_T_1_Bulk
7669_L_T_1_COUPTF2
7669_L_T_1_DLX1
7669_L_T_1_TBR1
7669_LA10_Bulk
7669_Liver_Bulk
7669_LV15_Bulk
7669_R_2_Put2_Bulk
7669_R_2_Thal_Bulk
7669_R_Adrenal_Bulk
7669_R_Ant_Cau_DLX1
7669_R_Ant_Put_DLX1
7669_R_Cb_2_Bulk
7669_R_CC_Mid_TBR1
7669_R_CCMid_COUPTF2
7669_R_CCMid_DLX1
7669_R_EC_COUPTF2
7669_R_EC_DLX1
7669_R_EC_TBR1
7669_R_F_1_Bulk
7669_R_F_1_COUPTF2
7669_R_F_1_DLX1
7669_R_F_1_NeuN
7669_R_F_1_TBR1
7669_R_I_1_COUPTF2
7669_R_I_1_DLX1
7669_R_I_1_TBR1
7669_R_Ins_1_Bulk
7669_R_kidney_Bulk
7669_R_mO_COUPTF2
7669_R_mO_DLX1
7669_R_mO_TBR1
7669_R_O_1_Bulk
7669_R_O_1_COUPTF2
7669_R_O_1_DLX1
7669_R_O_1_NeuN
7669_R_O_1_TBR1
7669_R_O_10_DLX1
7669_R_O_11_DLX1
7669_R_O_2_DLX1
7669_R_O_3_DLX1
7669_R_O_4_DLX1
7669_R_O_5_DLX1
7669_R_O_6_DLX1
7669_R_O_7_DLX1
7669_R_O_8_DLX1
7669_R_O_9_DLX1
7669_R_Olf_DLX1
7669_R_P_1_Bulk
7669_R_P_1_COUPTF2
7669_R_P_1_DLX1
7669_R_P_1_NeuN
7669_R_P_1_TBR1
7669_R_P_10_DLX1
7669_R_P_11_DLX1
7669_R_P_11_TBR1
7669_R_P_12_DLX1
7669_R_P_12_TBR1
7669_R_P_13_DLX1
7669_R_P_13_TBR1
7669_R_P_14_DLX1
7669_R_P_14_TBR1
7669_R_P_15_DLX1
7669_R_P_15_TBR1
7669_R_P_16_DLX1
7669_R_P_16_TBR1
7669_R_P_17_DLX1
7669_R_P_17_TBR1
7669_R_P_2_DLX1
7669_R_P_2_TBR1
7669_R_P_3_DLX1
7669_R_P_3_TBR1
7669_R_P_4_DLX1
7669_R_P_4_TBR1
7669_R_P_5_DLX1
7669_R_P_5_TBR1
7669_R_P_6_DLX1
7669_R_P_6_TBR1
7669_R_P_7_DLX1
7669_R_P_7_TBR1
7669_R_P_8_DLX1
7669_R_P_8_TBR1
7669_R_P_9_DLX
7669_R_P_9_DLX1
7669_R_P_9_TBR1
7669_R_POA_Bulk
7669_R_POA_DLX1
7669_R_sec1_Cau_Bulk
7669_R_sec2_GP_Bulk
7669_R_sec2_GP_DLX1
7669_R_sec2_Put_Bulk
7669_R_sec2_Thal_Bulk
7669_R_sec2_Thal_COUPTF2
7669_R_sec2_Thal_DLX1
7669_R_sec3_Cau_DLX1
7669_R_sec3_Hip_COUPTF2
7669_R_sec3_Hip_DLX1
7669_R_sec3_Hip_TBR1
7669_R_sec3_Put_DLX1
7669_R_Skin_Bulk
7669_R_T_1_Bulk
7669_R_T_1_COUPTF2
7669_R_T_1_DLX1
7669_R_T_1_TBR1
7669_RA9_Bulk
7669_RV10_Bulk
JGG

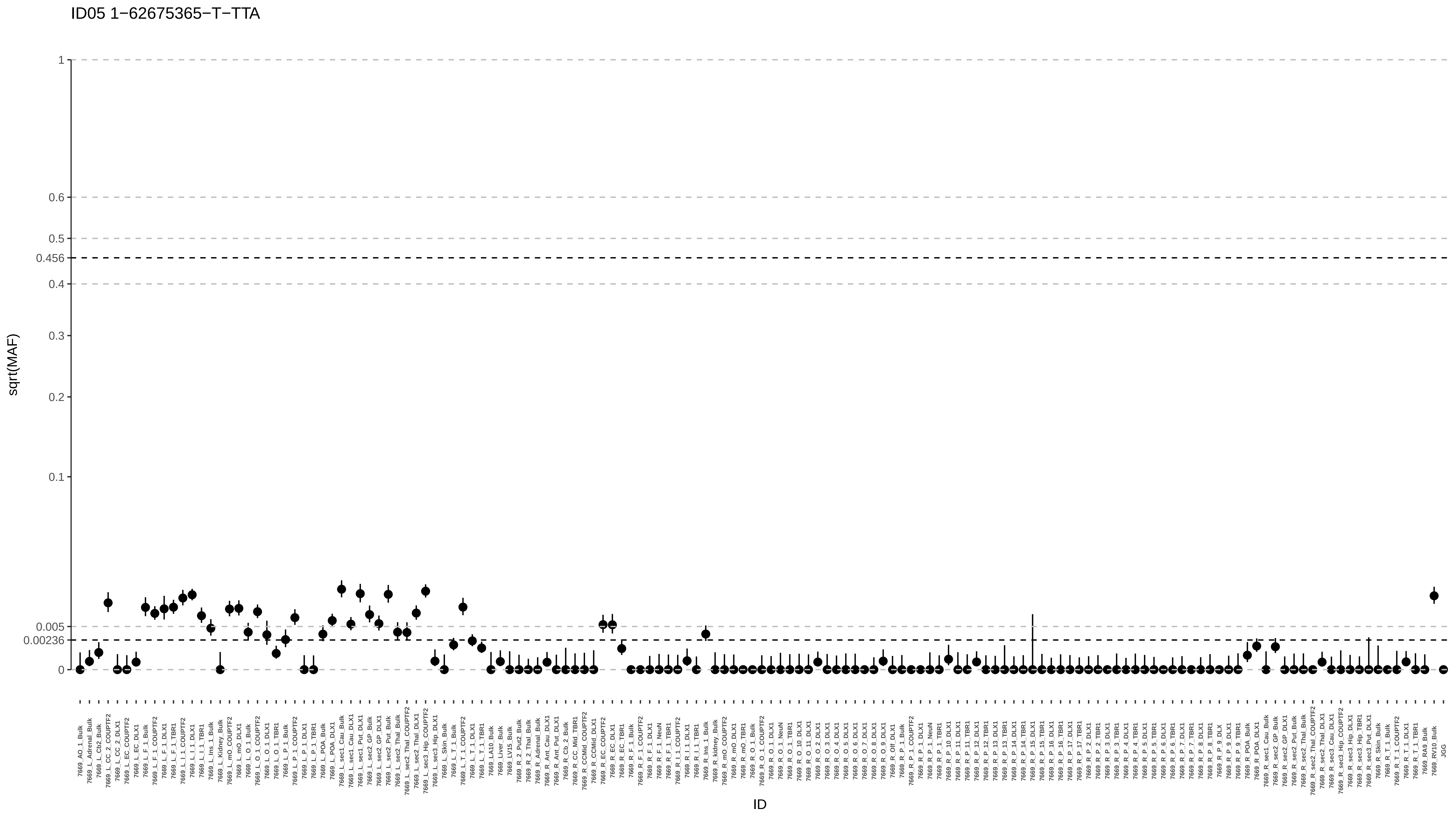
ID05 1−62675365−T−TTA
sqrt(MAF)
1
0.6
0.5
0.456
0.4
0.3
0.2
0.1
0.005
0.00236
0
ID

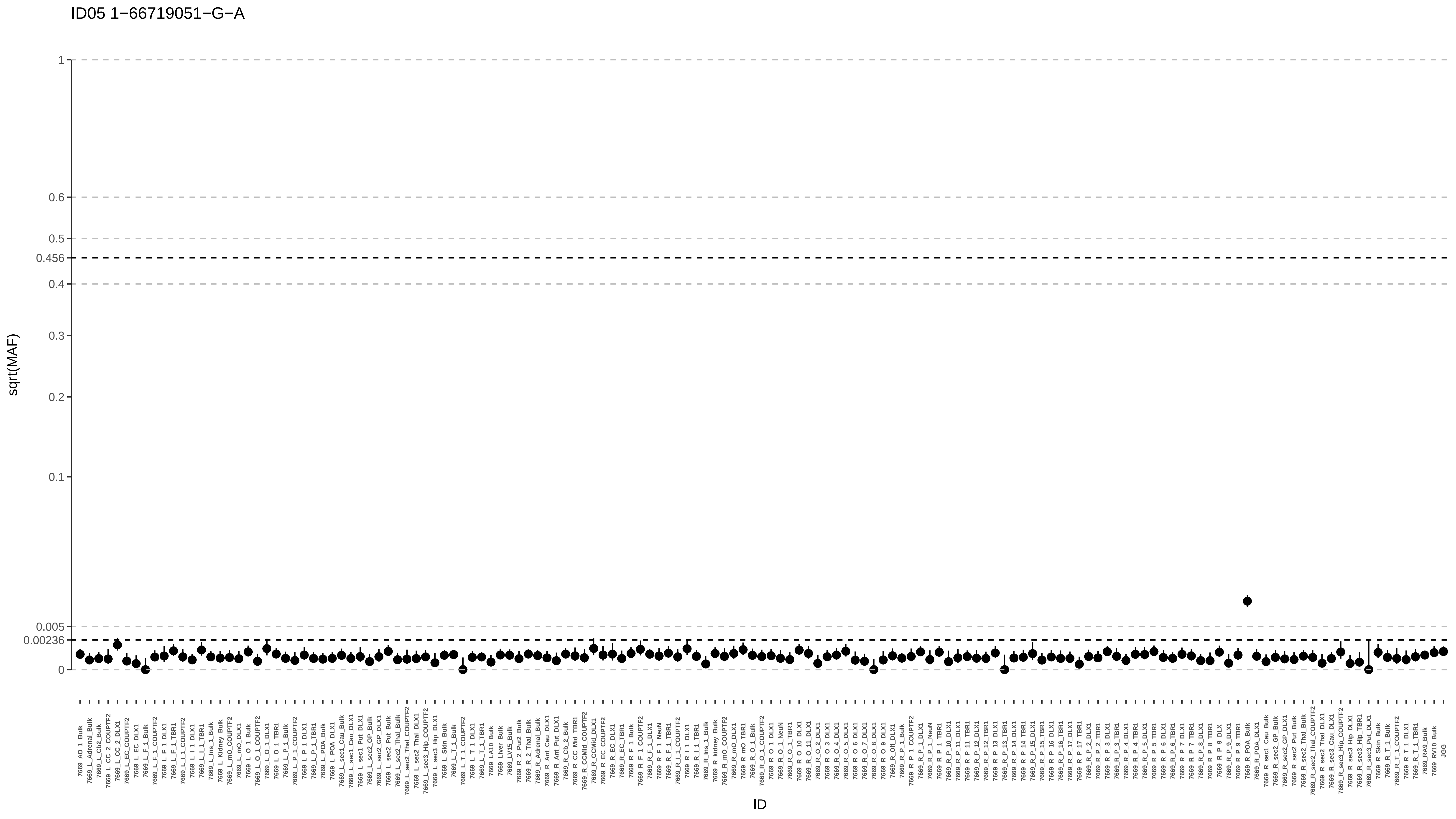
ID05 1−66719051−G−A
sqrt(MAF)
1
0.6
0.5
0.456
0.4
0.3
0.2
0.1
0.005
0.00236
0
ID

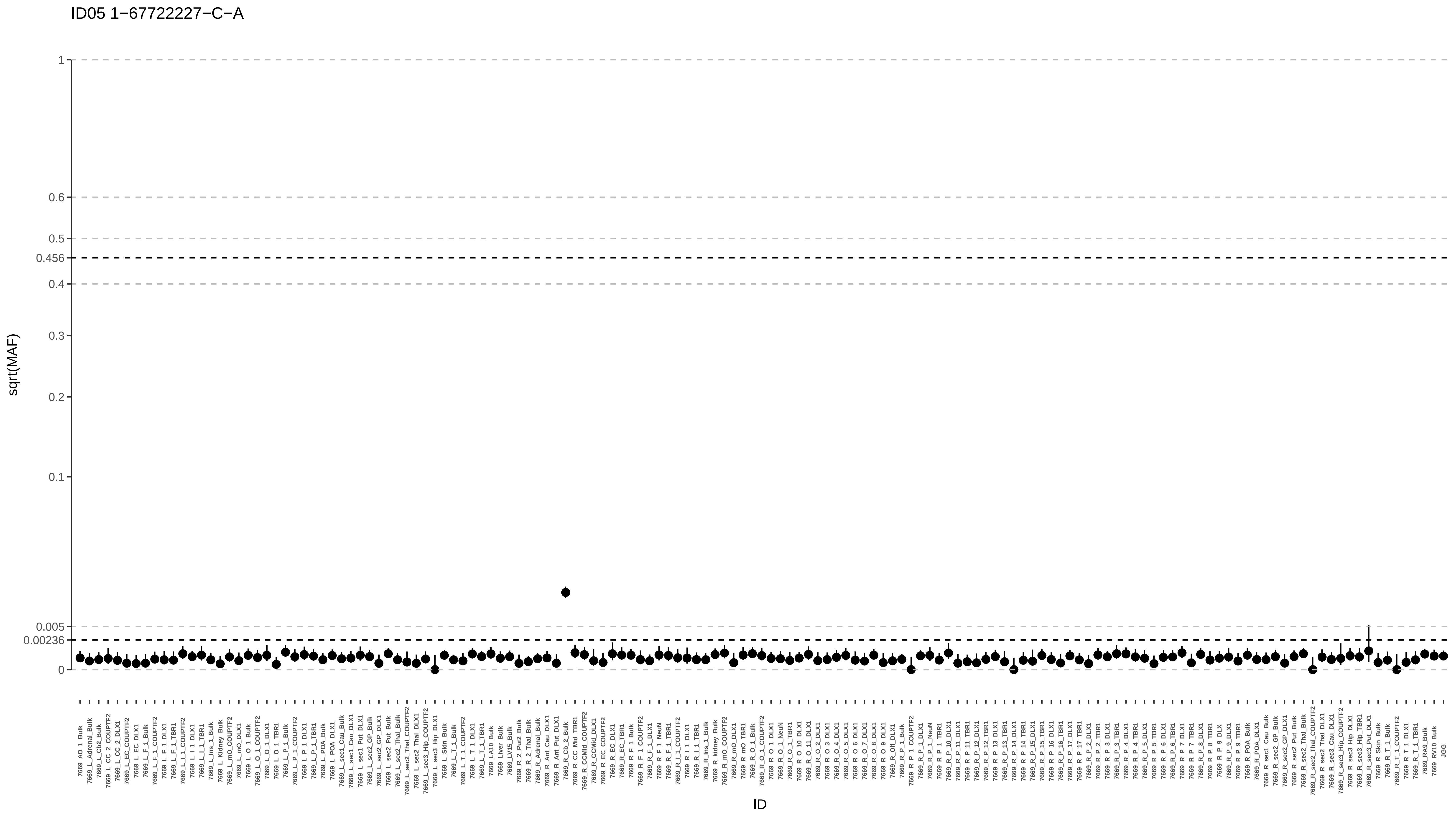
ID05 1−67722227−C−A
sqrt(MAF)
ID
1
0.6
0.5
0.456
0.4
0.3
0.2
0.1
0.005
0.00236
0

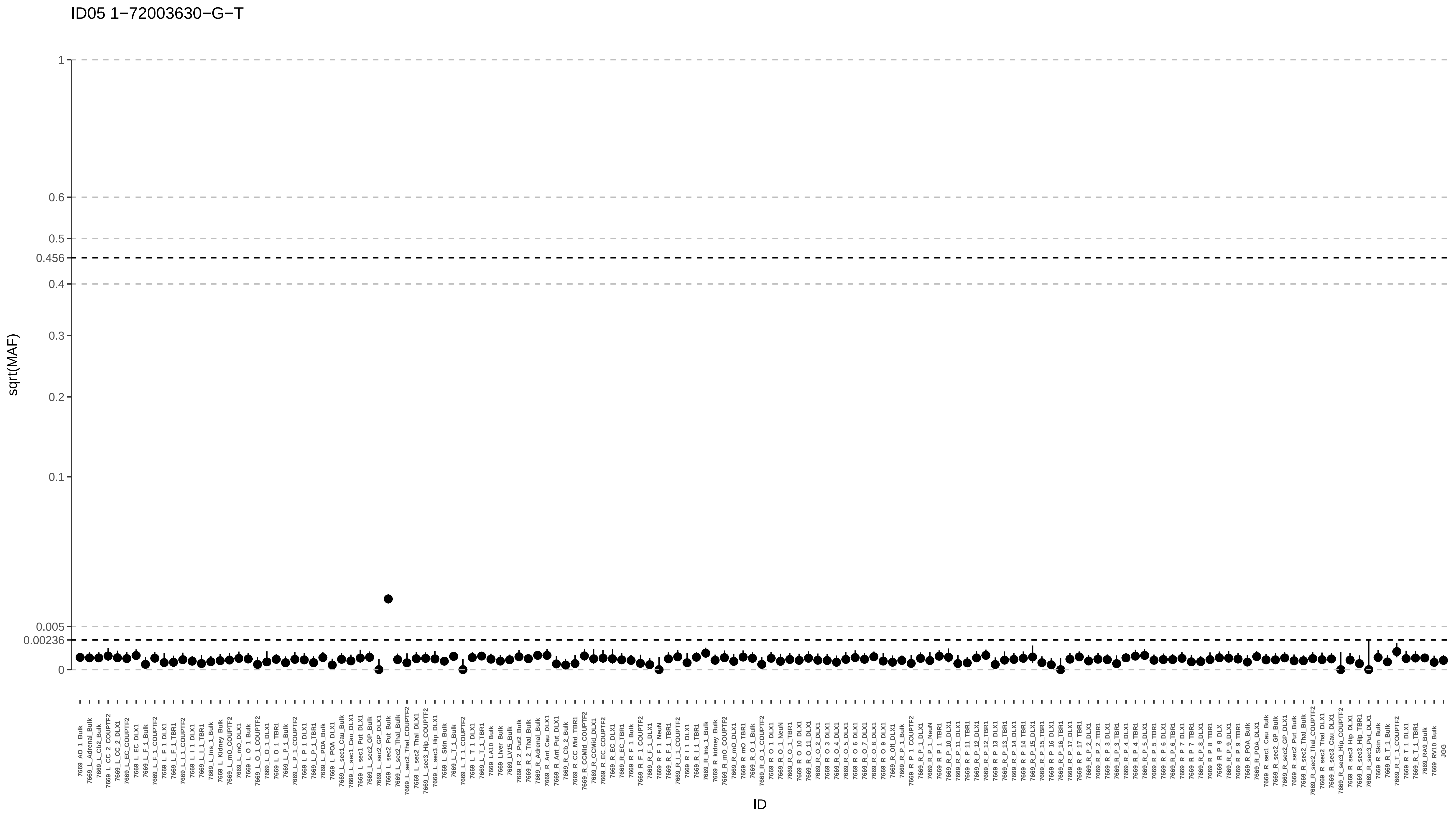
ID05 1−72003630−G−T
sqrt(MAF)
1
0.6
0.5
0.456
0.4
0.3
0.2
0.1
0.005
0.00236
0
ID

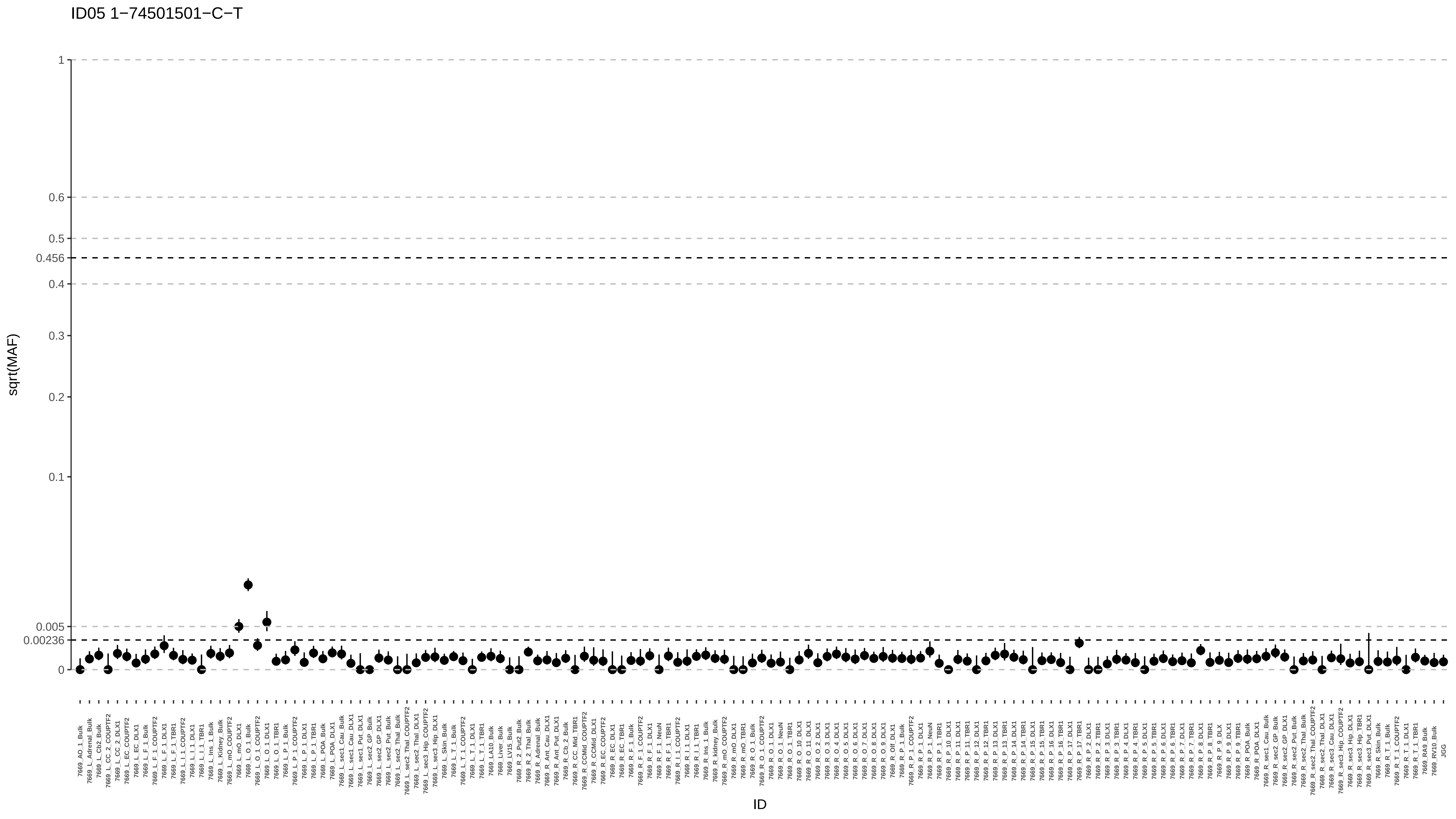
ID05 1−74501501−C−T
sqrt(MAF)
1
0.6
0.5
0.456
0.4
0.3
0.2
0.1
0.005
0.00236
0
ID
7669_AO_1_Bulk
7669_L_Adrenal_Bulk
7669_L_Cb2_Bulk
7669_L_CC_2_COUPTF2
7669_L_CC_2_DLX1
7669_L_EC_COUPTF2
7669_L_EC_DLX1
7669_L_F_1_Bulk
7669_L_F_1_COUPTF2
7669_L_F_1_DLX1
7669_L_F_1_TBR1
7669_L_I_1_COUPTF2
7669_L_I_1_DLX1
7669_L_I_1_TBR1
7669_L_Ins_1_Bulk
7669_L_Kidney_Bulk
7669_L_mO_COUPTF2
7669_L_mO_DLX1
7669_L_O_1_Bulk
7669_L_O_1_COUPTF2
7669_L_O_1_DLX1
7669_L_O_1_TBR1
7669_L_P_1_Bulk
7669_L_P_1_COUPTF2
7669_L_P_1_DLX1
7669_L_P_1_TBR1
7669_L_POA_Bulk
7669_L_POA_DLX1
7669_L_sec1_Cau_Bulk
7669_L_sec1_Cau_DLX1
7669_L_sec1_Put_DLX1
7669_L_sec2_GP_Bulk
7669_L_sec2_GP_DLX1
7669_L_sec2_Put_Bulk
7669_L_sec2_Thal_Bulk
7669_L_sec2_Thal_COUPTF2
7669_L_sec2_Thal_DLX1
7669_L_sec3_Hip_COUPTF2
7669_L_sec3_Hip_DLX1
7669_L_Skin_Bulk
7669_L_T_1_Bulk
7669_L_T_1_COUPTF2
7669_L_T_1_DLX1
7669_L_T_1_TBR1
7669_LA10_Bulk
7669_Liver_Bulk
7669_LV15_Bulk
7669_R_2_Put2_Bulk
7669_R_2_Thal_Bulk
7669_R_Adrenal_Bulk
7669_R_Ant_Cau_DLX1
7669_R_Ant_Put_DLX1
7669_R_Cb_2_Bulk
7669_R_CC_Mid_TBR1
7669_R_CCMid_COUPTF2
7669_R_CCMid_DLX1
7669_R_EC_COUPTF2
7669_R_EC_DLX1
7669_R_EC_TBR1
7669_R_F_1_Bulk
7669_R_F_1_COUPTF2
7669_R_F_1_DLX1
7669_R_F_1_NeuN
7669_R_F_1_TBR1
7669_R_I_1_COUPTF2
7669_R_I_1_DLX1
7669_R_I_1_TBR1
7669_R_Ins_1_Bulk
7669_R_kidney_Bulk
7669_R_mO_COUPTF2
7669_R_mO_DLX1
7669_R_mO_TBR1
7669_R_O_1_Bulk
7669_R_O_1_COUPTF2
7669_R_O_1_DLX1
7669_R_O_1_NeuN
7669_R_O_1_TBR1
7669_R_O_10_DLX1
7669_R_O_11_DLX1
7669_R_O_2_DLX1
7669_R_O_3_DLX1
7669_R_O_4_DLX1
7669_R_O_5_DLX1
7669_R_O_6_DLX1
7669_R_O_7_DLX1
7669_R_O_8_DLX1
7669_R_O_9_DLX1
7669_R_Olf_DLX1
7669_R_P_1_Bulk
7669_R_P_1_COUPTF2
7669_R_P_1_DLX1
7669_R_P_1_NeuN
7669_R_P_1_TBR1
7669_R_P_10_DLX1
7669_R_P_11_DLX1
7669_R_P_11_TBR1
7669_R_P_12_DLX1
7669_R_P_12_TBR1
7669_R_P_13_DLX1
7669_R_P_13_TBR1
7669_R_P_14_DLX1
7669_R_P_14_TBR1
7669_R_P_15_DLX1
7669_R_P_15_TBR1
7669_R_P_16_DLX1
7669_R_P_16_TBR1
7669_R_P_17_DLX1
7669_R_P_17_TBR1
7669_R_P_2_DLX1
7669_R_P_2_TBR1
7669_R_P_3_DLX1
7669_R_P_3_TBR1
7669_R_P_4_DLX1
7669_R_P_4_TBR1
7669_R_P_5_DLX1
7669_R_P_5_TBR1
7669_R_P_6_DLX1
7669_R_P_6_TBR1
7669_R_P_7_DLX1
7669_R_P_7_TBR1
7669_R_P_8_DLX1
7669_R_P_8_TBR1
7669_R_P_9_DLX
7669_R_P_9_DLX1
7669_R_P_9_TBR1
7669_R_POA_Bulk
7669_R_POA_DLX1
7669_R_sec1_Cau_Bulk
7669_R_sec2_GP_Bulk
7669_R_sec2_GP_DLX1
7669_R_sec2_Put_Bulk
7669_R_sec2_Thal_Bulk
7669_R_sec2_Thal_COUPTF2
7669_R_sec2_Thal_DLX1
7669_R_sec3_Cau_DLX1
7669_R_sec3_Hip_COUPTF2
7669_R_sec3_Hip_DLX1
7669_R_sec3_Hip_TBR1
7669_R_sec3_Put_DLX1
7669_R_Skin_Bulk
7669_R_T_1_Bulk
7669_R_T_1_COUPTF2
7669_R_T_1_DLX1
7669_R_T_1_TBR1
7669_RA9_Bulk
7669_RV10_Bulk
JGG

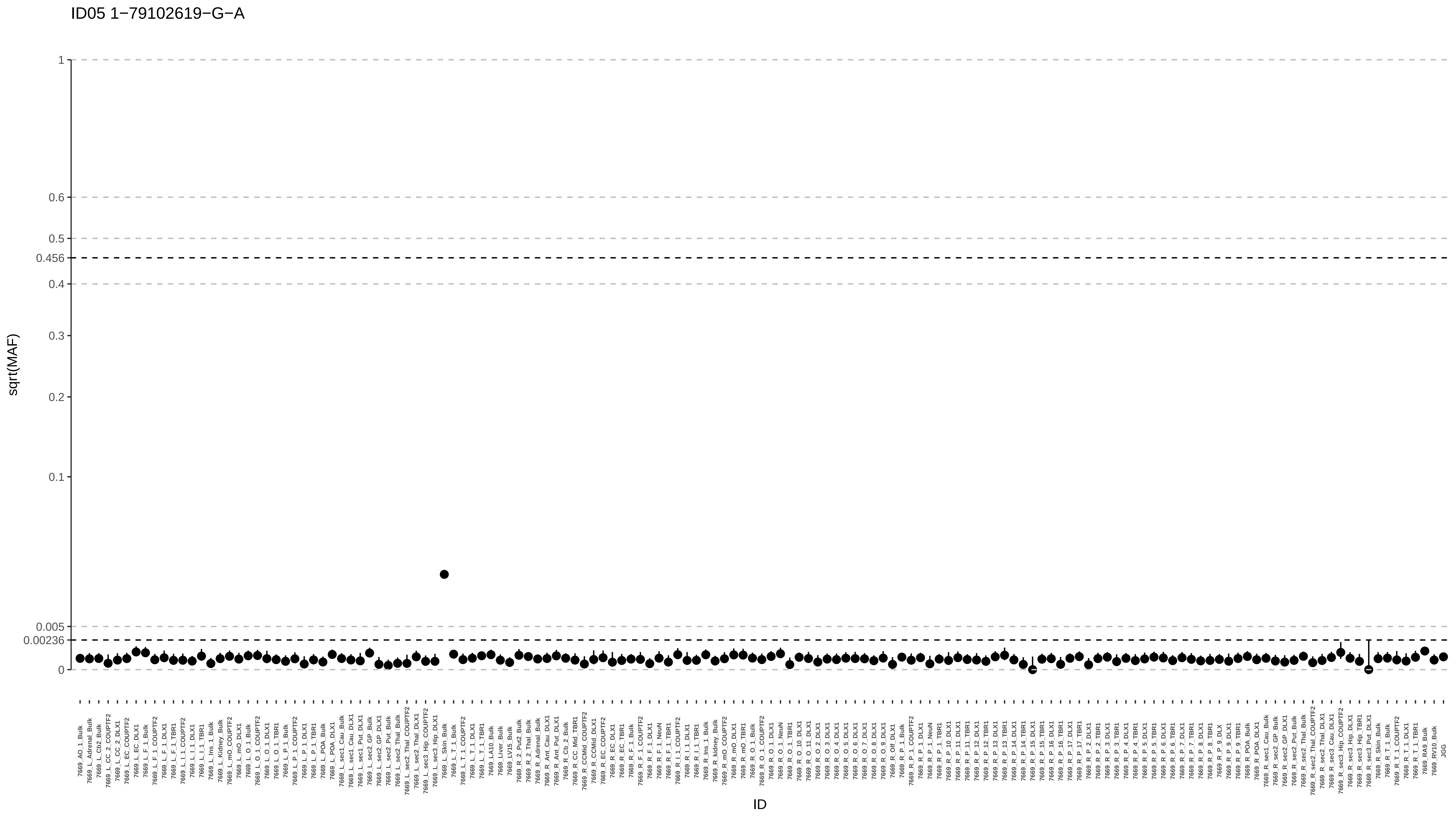
ID05 1−79102619−G−A
sqrt(MAF)
1
0.6
0.5
0.456
0.4
0.3
0.2
0.1
0.005
0.00236
0
ID

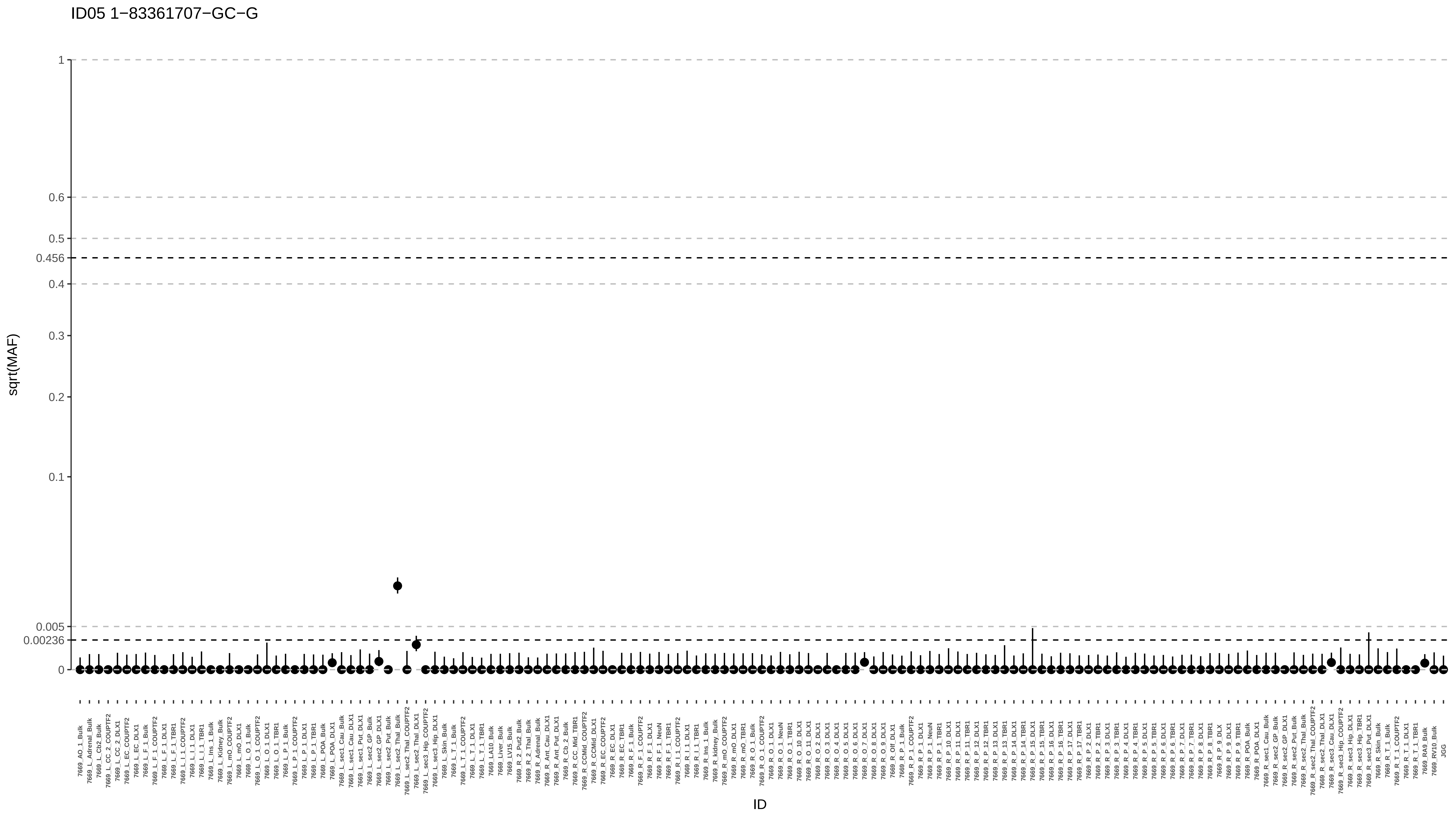
ID05 1−83361707−GC−G
sqrt(MAF)
1
0.6
0.5
0.456
0.4
0.3
0.2
0.1
0.005
0.00236
0
ID

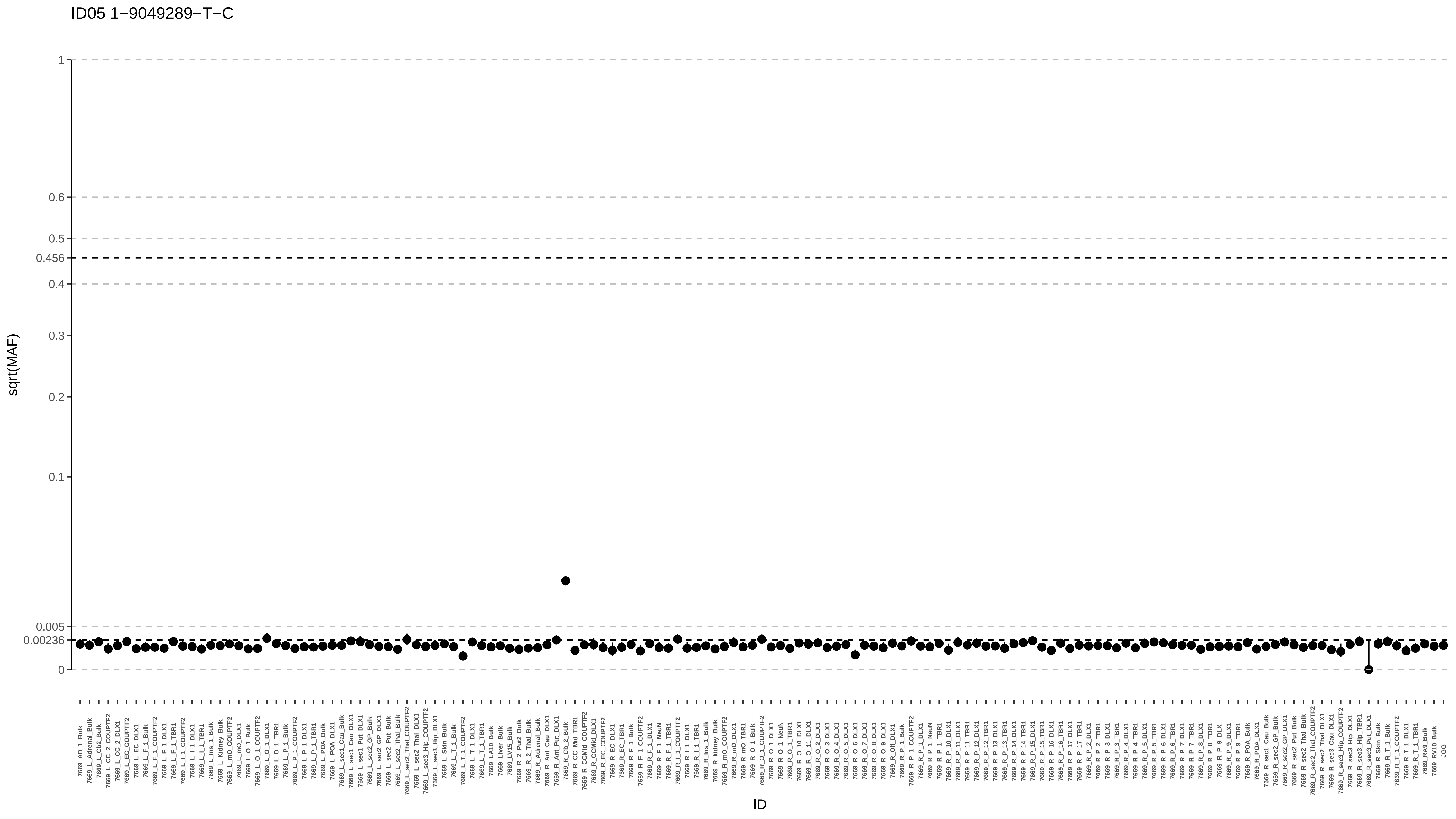
ID05 1−9049289−T−C
sqrt(MAF)
1
0.6
0.5
0.456
0.4
0.3
0.2
0.1
0.005
0.00236
0
ID

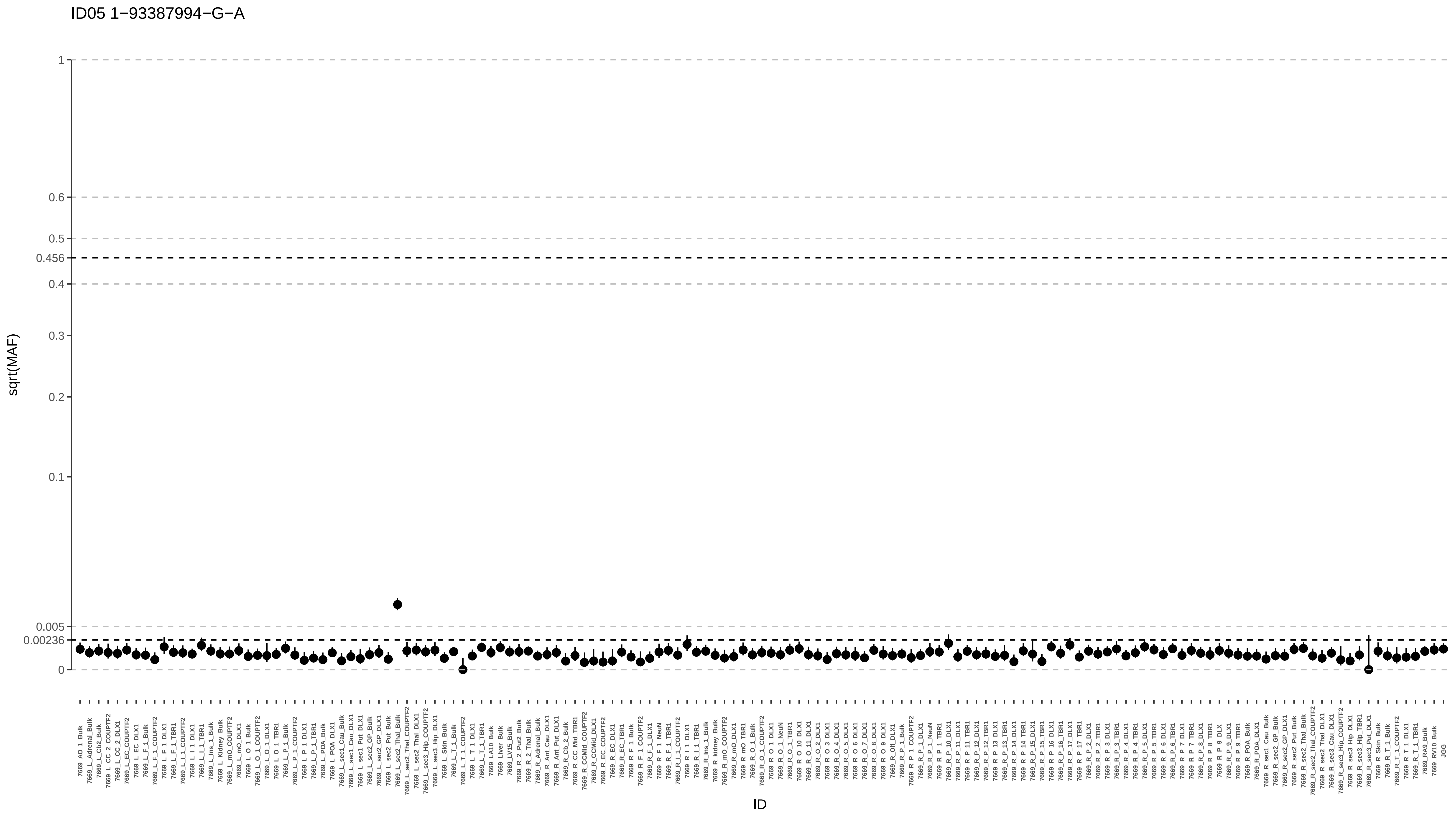
ID05 1−93387994−G−A
sqrt(MAF)
1
0.6
0.5
0.456
0.4
0.3
0.2
0.1
0.005
0.00236
0
ID

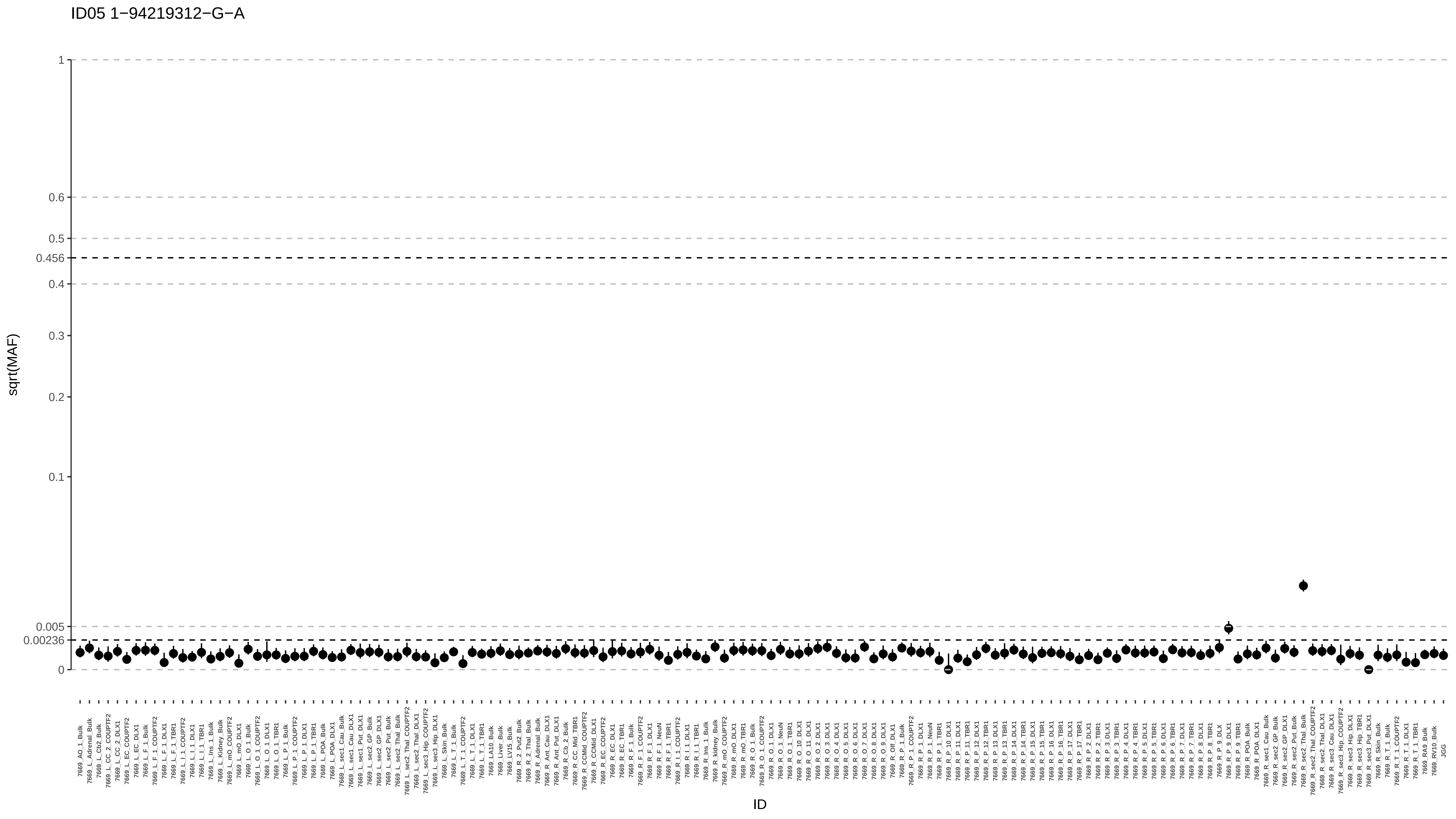
ID05 1−94219312−G−A
sqrt(MAF)
1
0.6
0.5
0.456
0.4
0.3
0.2
0.1
0.005
0.00236
0
ID

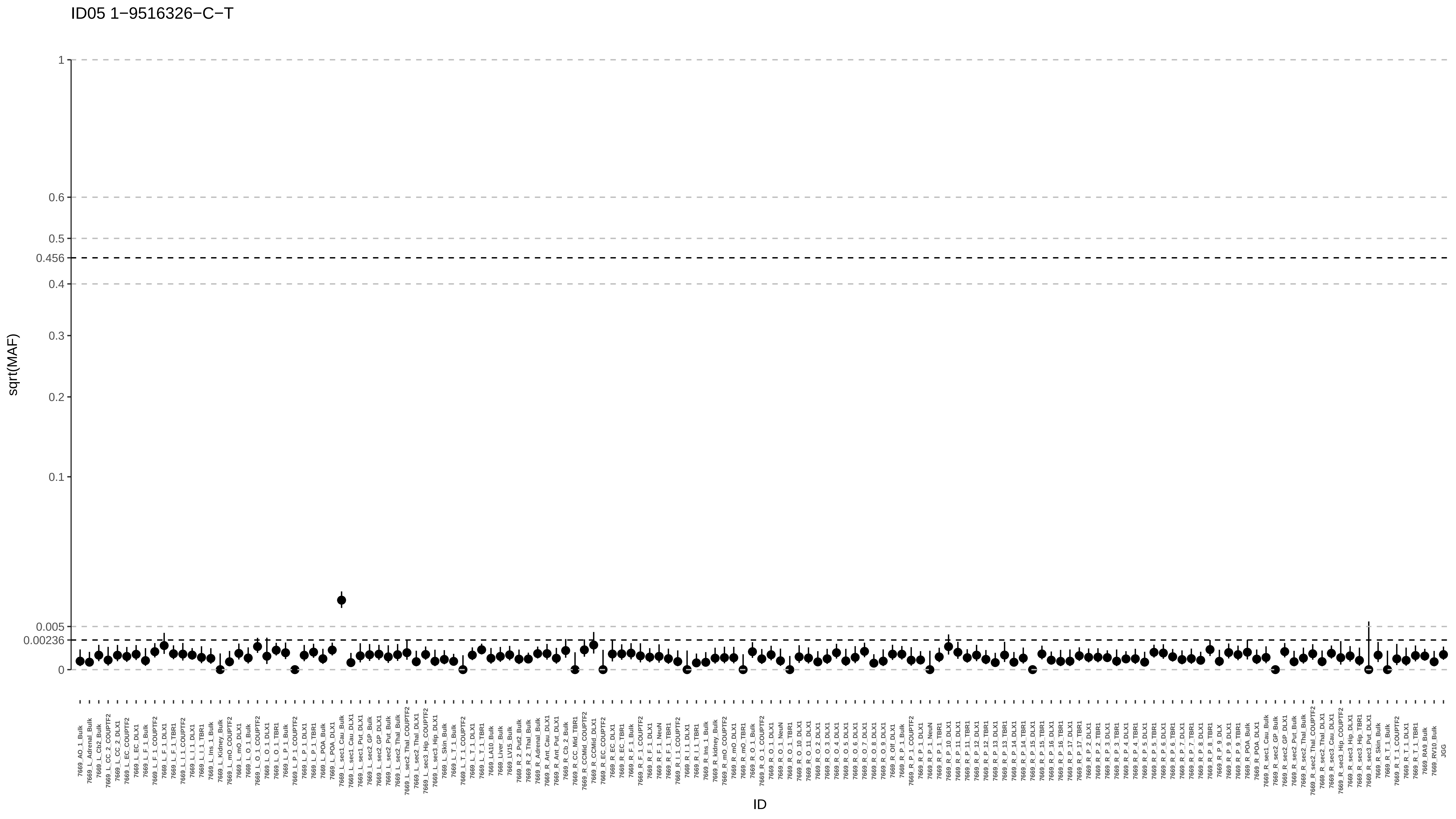

ID05 1−9516326−C−T
sqrt(MAF)
1
0.6
0.5
0.456
0.4
0.3
0.2
0.1
0.005
0.00236
0
ID

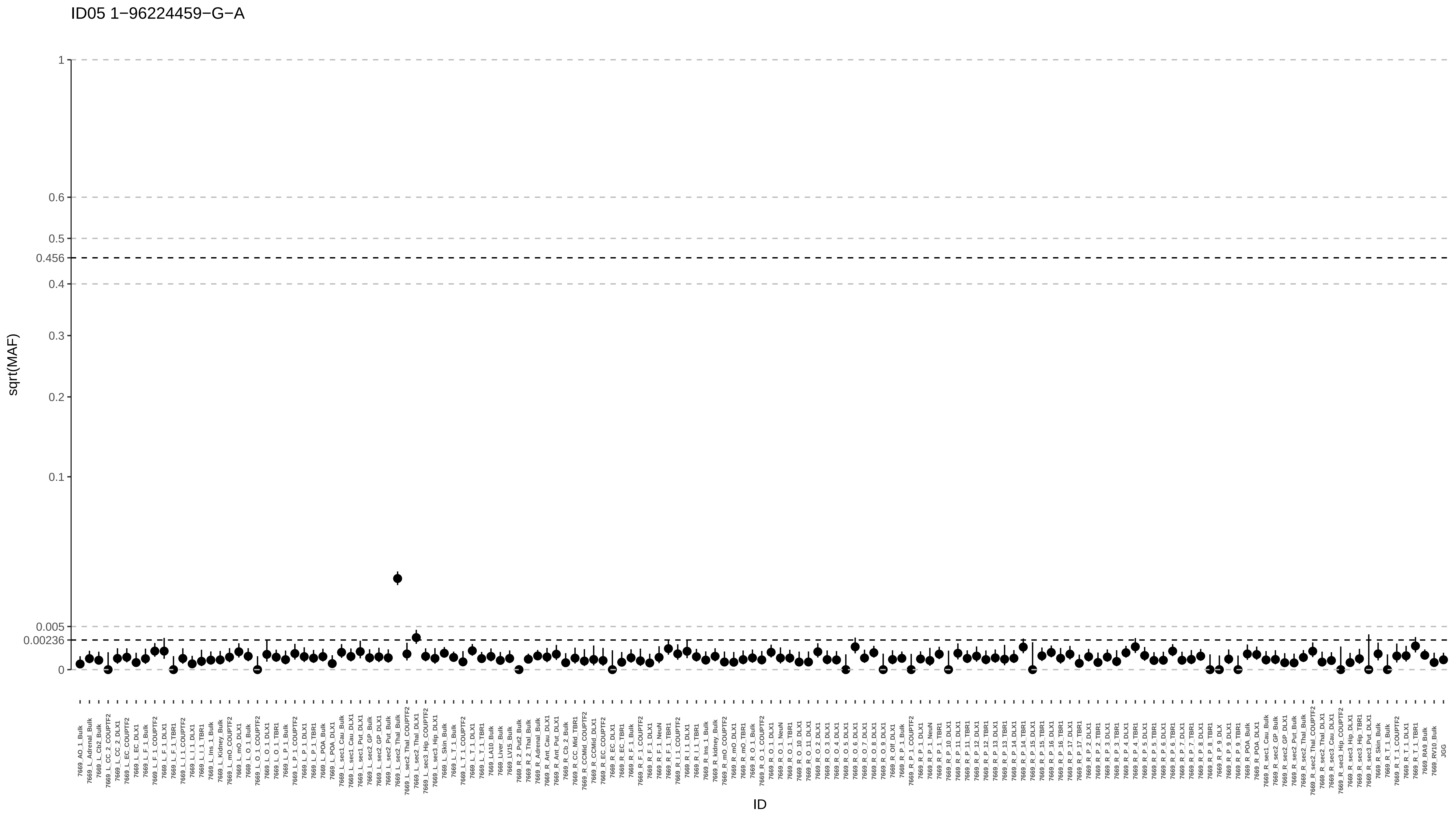
ID05 1−96224459−G−A
sqrt(MAF)
1
0.6
0.5
0.456
0.4
0.3
0.2
0.1
0.005
0.00236
0
ID

ID05 1−9672930−G−A

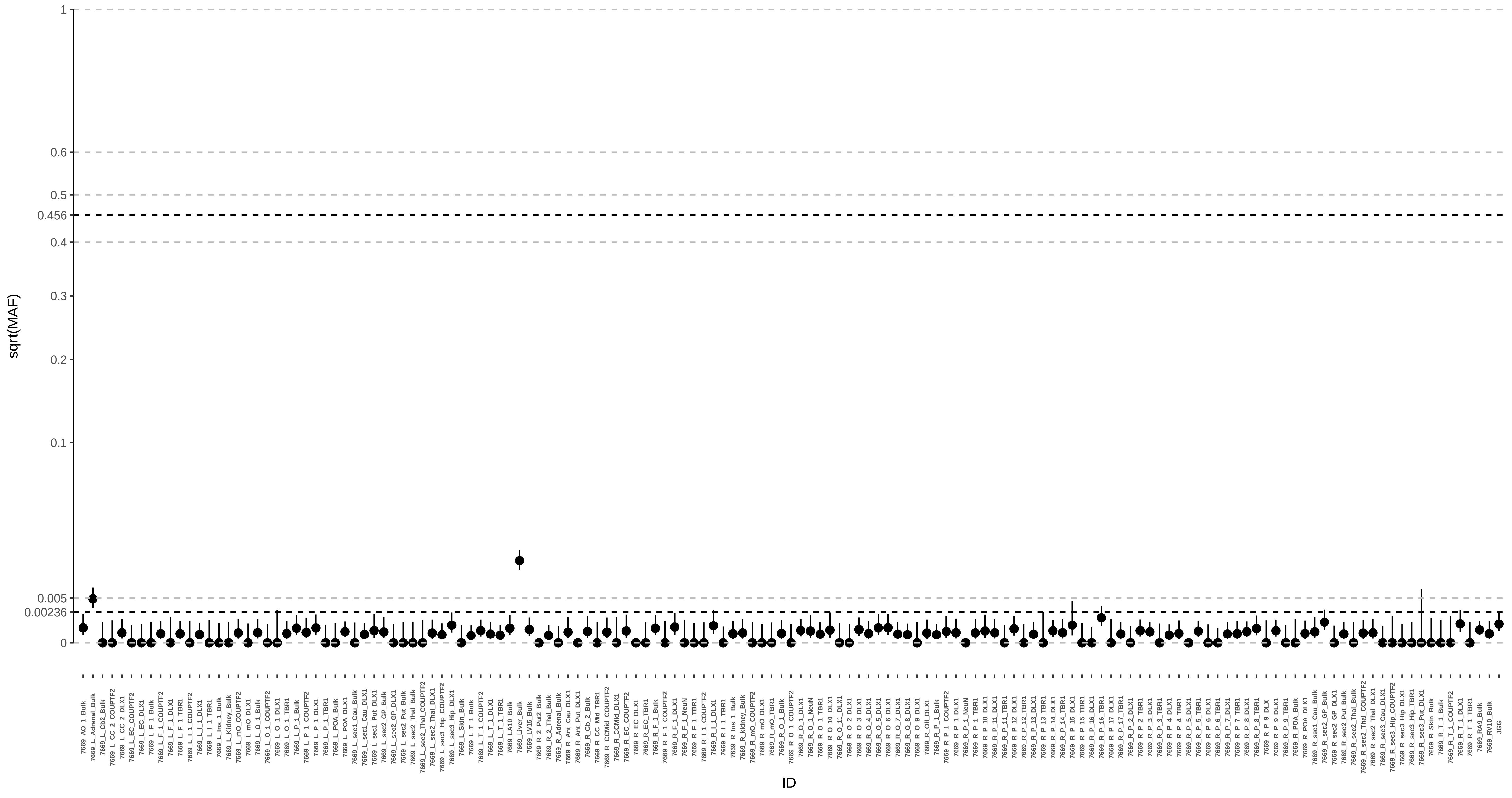

ID05 1−98301896−T−C

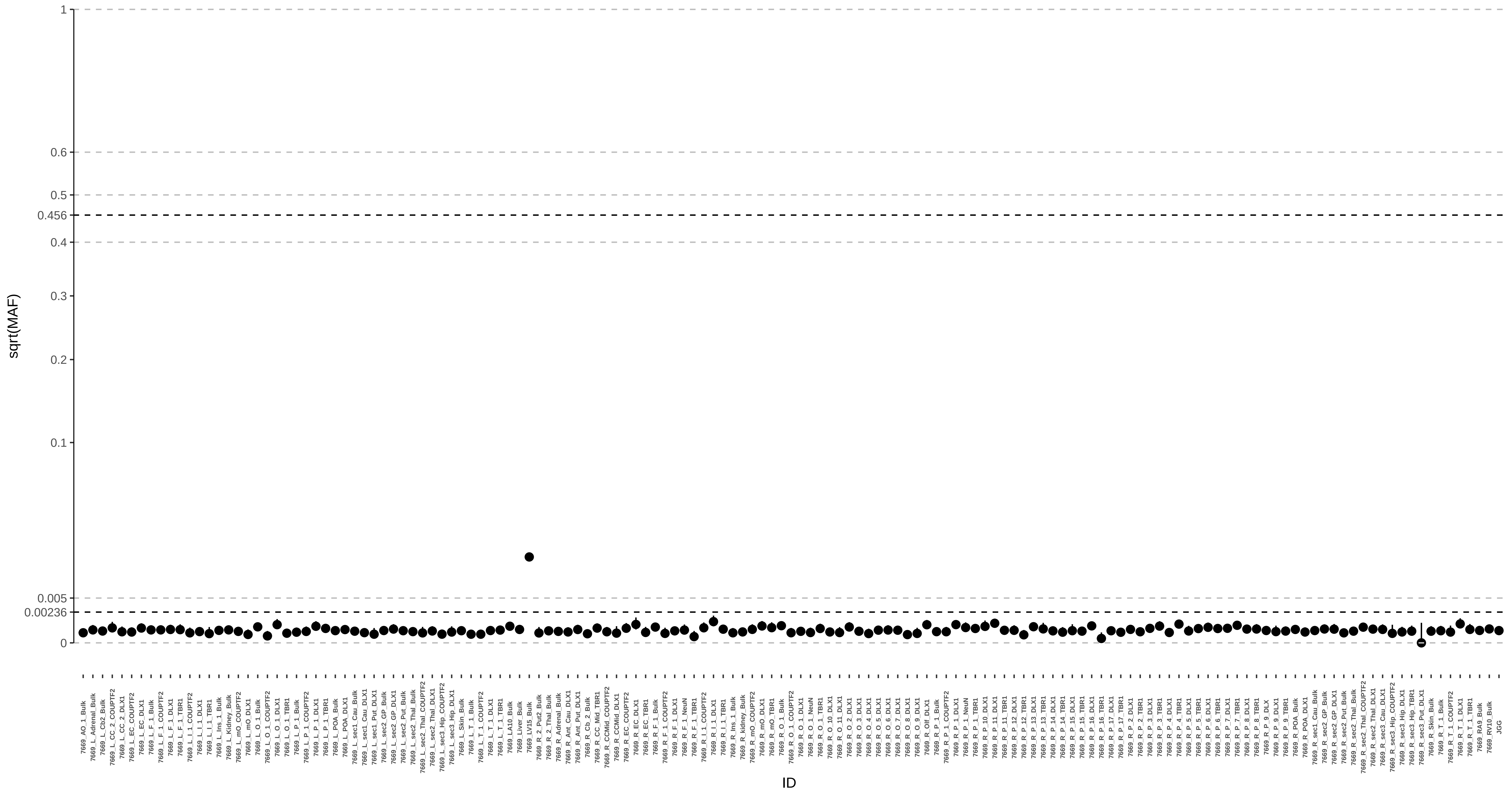

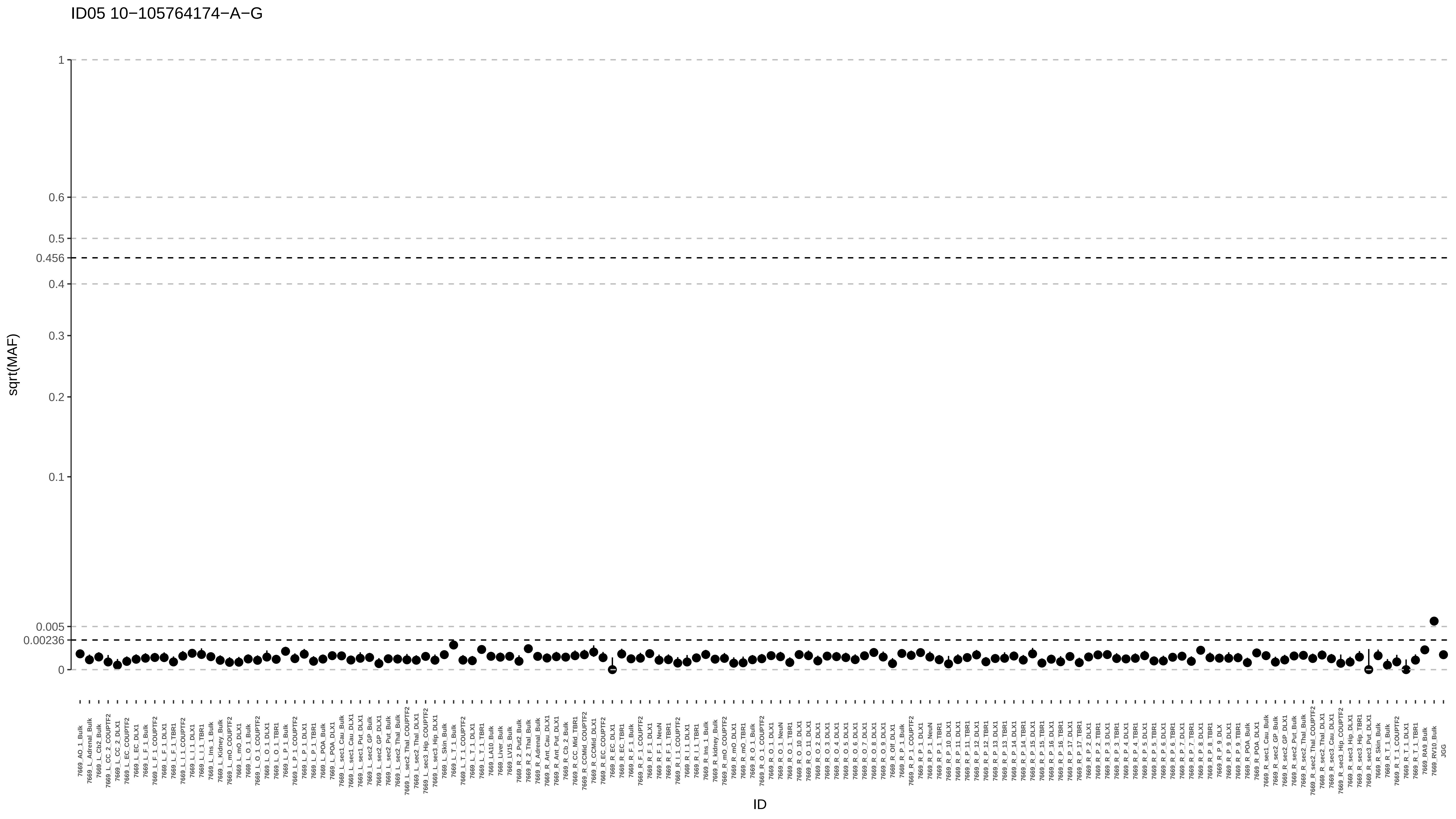
ID05 10−105764174−A−G
sqrt(MAF)
1
0.6
0.5
0.456
0.4
0.3
0.2
0.1
0.005
0.00236
0
ID
7669_AO_1_Bulk
7669_L_Adrenal_Bulk
7669_L_Cb2_Bulk
7669_L_CC_2_COUPTF2
7669_L_CC_2_DLX1
7669_L_EC_COUPTF2
7669_L_EC_DLX1
7669_L_F_1_Bulk
7669_L_F_1_COUPTF2
7669_L_F_1_DLX1
7669_L_F_1_TBR1
7669_L_I_1_COUPTF2
7669_L_I_1_DLX1
7669_L_I_1_TBR1
7669_L_Ins_1_Bulk
7669_L_Kidney_Bulk
7669_L_mO_COUPTF2
7669_L_mO_DLX1
7669_L_O_1_Bulk
7669_L_O_1_COUPTF2
7669_L_O_1_DLX1
7669_L_O_1_TBR1
7669_L_P_1_Bulk
7669_L_P_1_COUPTF2
7669_L_P_1_DLX1
7669_L_P_1_TBR1
7669_L_POA_Bulk
7669_L_POA_DLX1
7669_L_sec1_Cau_Bulk
7669_L_sec1_Cau_DLX1
7669_L_sec1_Put_DLX1
7669_L_sec2_GP_Bulk
7669_L_sec2_GP_DLX1
7669_L_sec2_Put_Bulk
7669_L_sec2_Thal_Bulk
7669_L_sec2_Thal_COUPTF2
7669_L_sec2_Thal_DLX1
7669_L_sec3_Hip_COUPTF2
7669_L_sec3_Hip_DLX1
7669_L_Skin_Bulk
7669_L_T_1_Bulk
7669_L_T_1_COUPTF2
7669_L_T_1_DLX1
7669_L_T_1_TBR1
7669_LA10_Bulk
7669_Liver_Bulk
7669_LV15_Bulk
7669_R_2_Put2_Bulk
7669_R_2_Thal_Bulk
7669_R_Adrenal_Bulk
7669_R_Ant_Cau_DLX1
7669_R_Ant_Put_DLX1
7669_R_Cb_2_Bulk
7669_R_CC_Mid_TBR1
7669_R_CCMid_COUPTF2
7669_R_CCMid_DLX1
7669_R_EC_COUPTF2
7669_R_EC_DLX1
7669_R_EC_TBR1
7669_R_F_1_Bulk
7669_R_F_1_COUPTF2
7669_R_F_1_DLX1
7669_R_F_1_NeuN
7669_R_F_1_TBR1
7669_R_I_1_COUPTF2
7669_R_I_1_DLX1
7669_R_I_1_TBR1
7669_R_Ins_1_Bulk
7669_R_kidney_Bulk
7669_R_mO_COUPTF2
7669_R_mO_DLX1
7669_R_mO_TBR1
7669_R_O_1_Bulk
7669_R_O_1_COUPTF2
7669_R_O_1_DLX1
7669_R_O_1_NeuN
7669_R_O_1_TBR1
7669_R_O_10_DLX1
7669_R_O_11_DLX1
7669_R_O_2_DLX1
7669_R_O_3_DLX1
7669_R_O_4_DLX1
7669_R_O_5_DLX1
7669_R_O_6_DLX1
7669_R_O_7_DLX1
7669_R_O_8_DLX1
7669_R_O_9_DLX1
7669_R_Olf_DLX1
7669_R_P_1_Bulk
7669_R_P_1_COUPTF2
7669_R_P_1_DLX1
7669_R_P_1_NeuN
7669_R_P_1_TBR1
7669_R_P_10_DLX1
7669_R_P_11_DLX1
7669_R_P_11_TBR1
7669_R_P_12_DLX1
7669_R_P_12_TBR1
7669_R_P_13_DLX1
7669_R_P_13_TBR1
7669_R_P_14_DLX1
7669_R_P_14_TBR1
7669_R_P_15_DLX1
7669_R_P_15_TBR1
7669_R_P_16_DLX1
7669_R_P_16_TBR1
7669_R_P_17_DLX1
7669_R_P_17_TBR1
7669_R_P_2_DLX1
7669_R_P_2_TBR1
7669_R_P_3_DLX1
7669_R_P_3_TBR1
7669_R_P_4_DLX1
7669_R_P_4_TBR1
7669_R_P_5_DLX1
7669_R_P_5_TBR1
7669_R_P_6_DLX1
7669_R_P_6_TBR1
7669_R_P_7_DLX1
7669_R_P_7_TBR1
7669_R_P_8_DLX1
7669_R_P_8_TBR1
7669_R_P_9_DLX
7669_R_P_9_DLX1
7669_R_P_9_TBR1
7669_R_POA_Bulk
7669_R_POA_DLX1
7669_R_sec1_Cau_Bulk
7669_R_sec2_GP_Bulk
7669_R_sec2_GP_DLX1
7669_R_sec2_Put_Bulk
7669_R_sec2_Thal_Bulk
7669_R_sec2_Thal_COUPTF2
7669_R_sec2_Thal_DLX1
7669_R_sec3_Cau_DLX1
7669_R_sec3_Hip_COUPTF2
7669_R_sec3_Hip_DLX1
7669_R_sec3_Hip_TBR1
7669_R_sec3_Put_DLX1
7669_R_Skin_Bulk
7669_R_T_1_Bulk
7669_R_T_1_COUPTF2
7669_R_T_1_DLX1
7669_R_T_1_TBR1
7669_RA9_Bulk
7669_RV10_Bulk
JGG

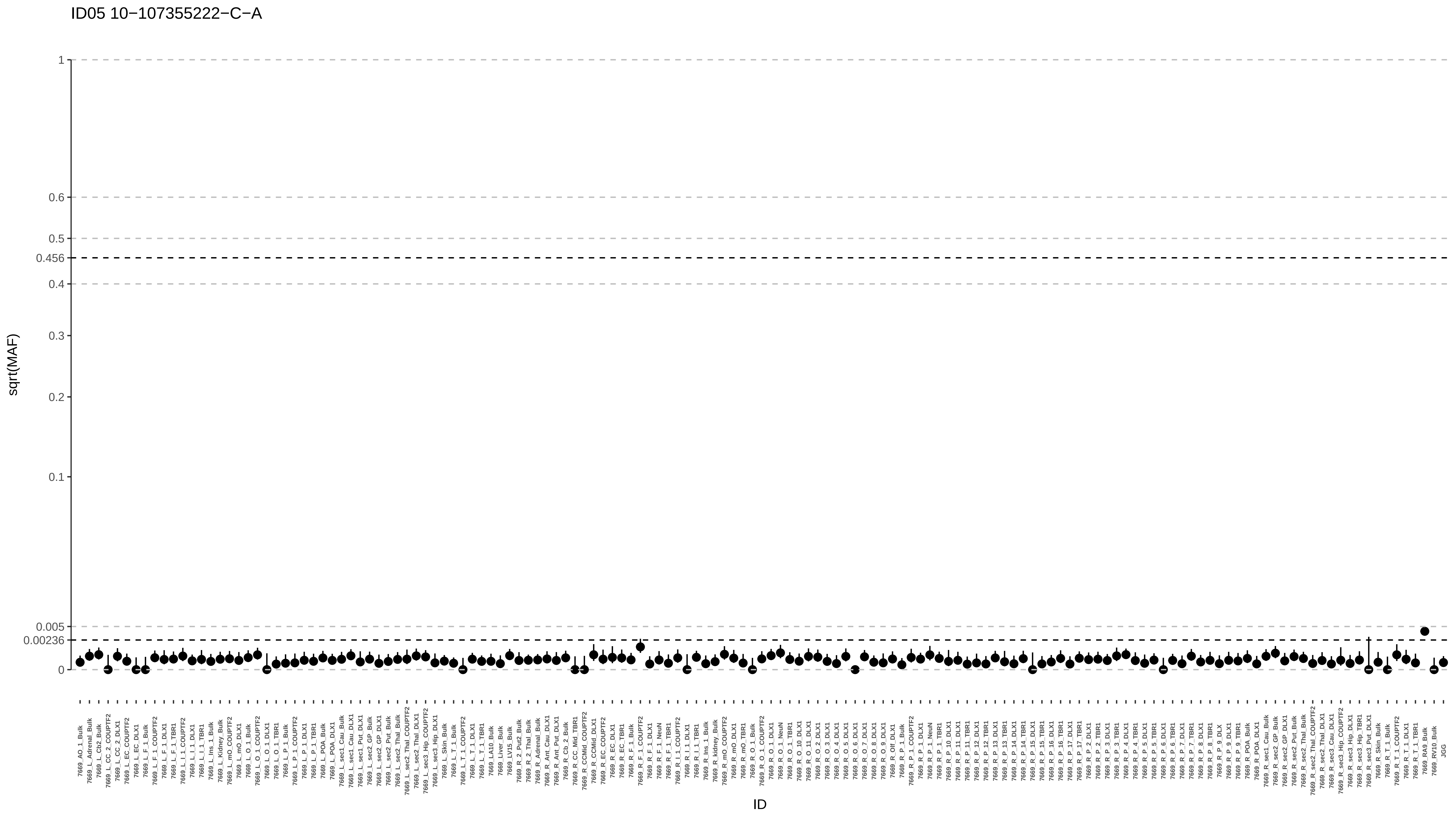
ID05 10−107355222−C−A
sqrt(MAF)
1
0.6
0.5
0.456
0.4
0.3
0.2
0.1
0.005
0.00236
0
ID

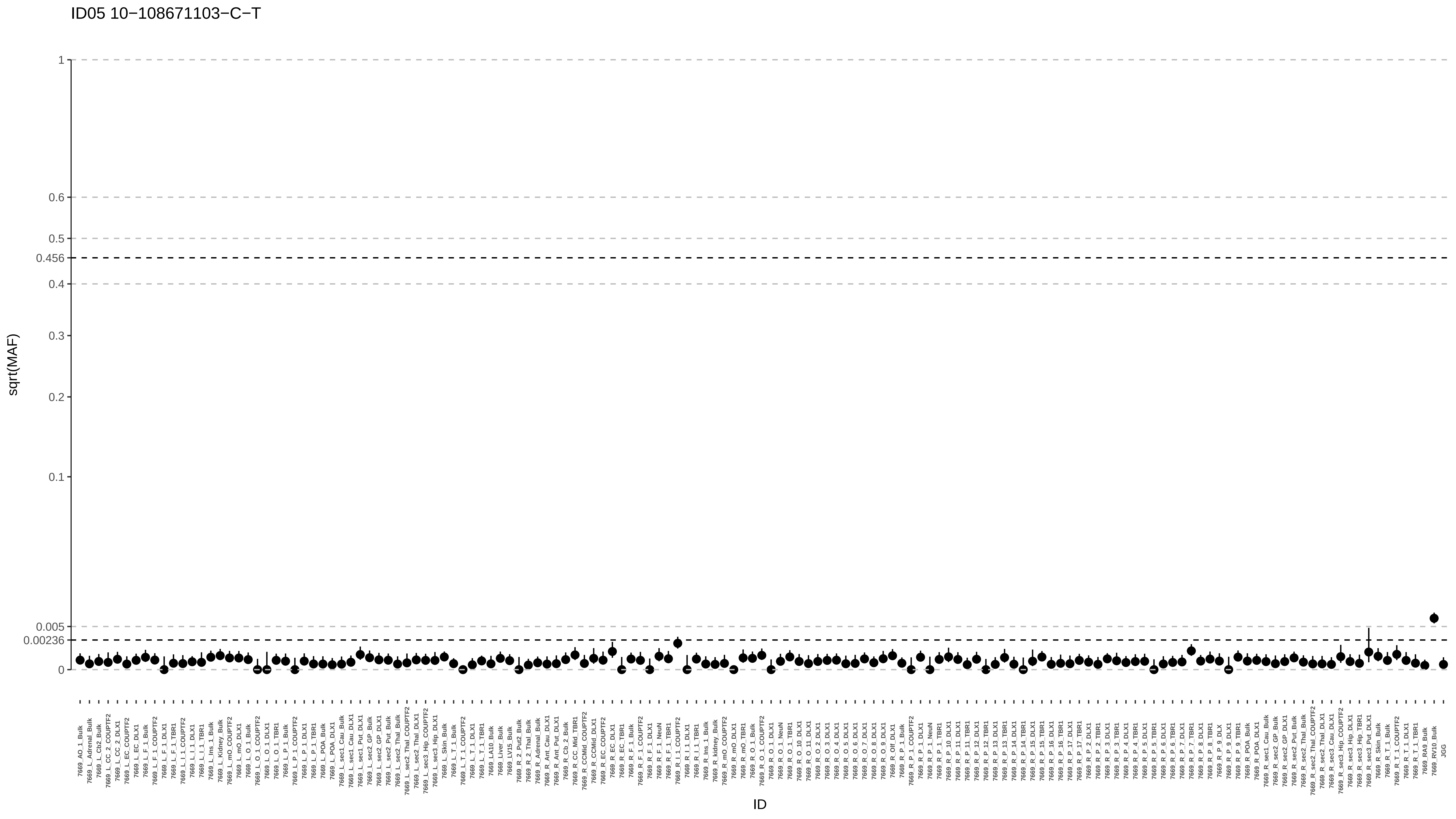
ID05 10−108671103−C−T
sqrt(MAF)
ID

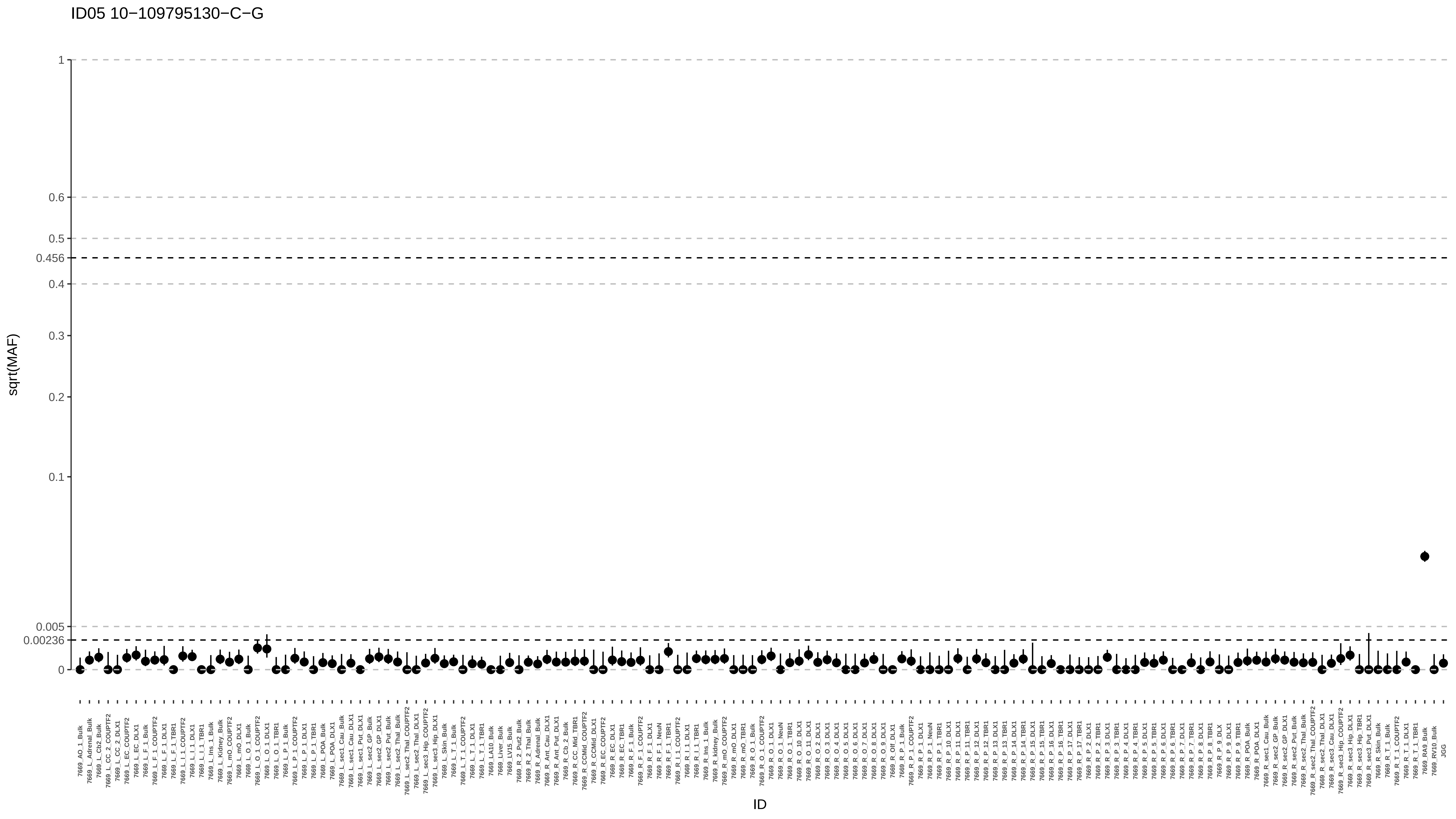

ID05 10−109795130−C−G
sqrt(MAF)
ID

ID05 10−113404344−G−A

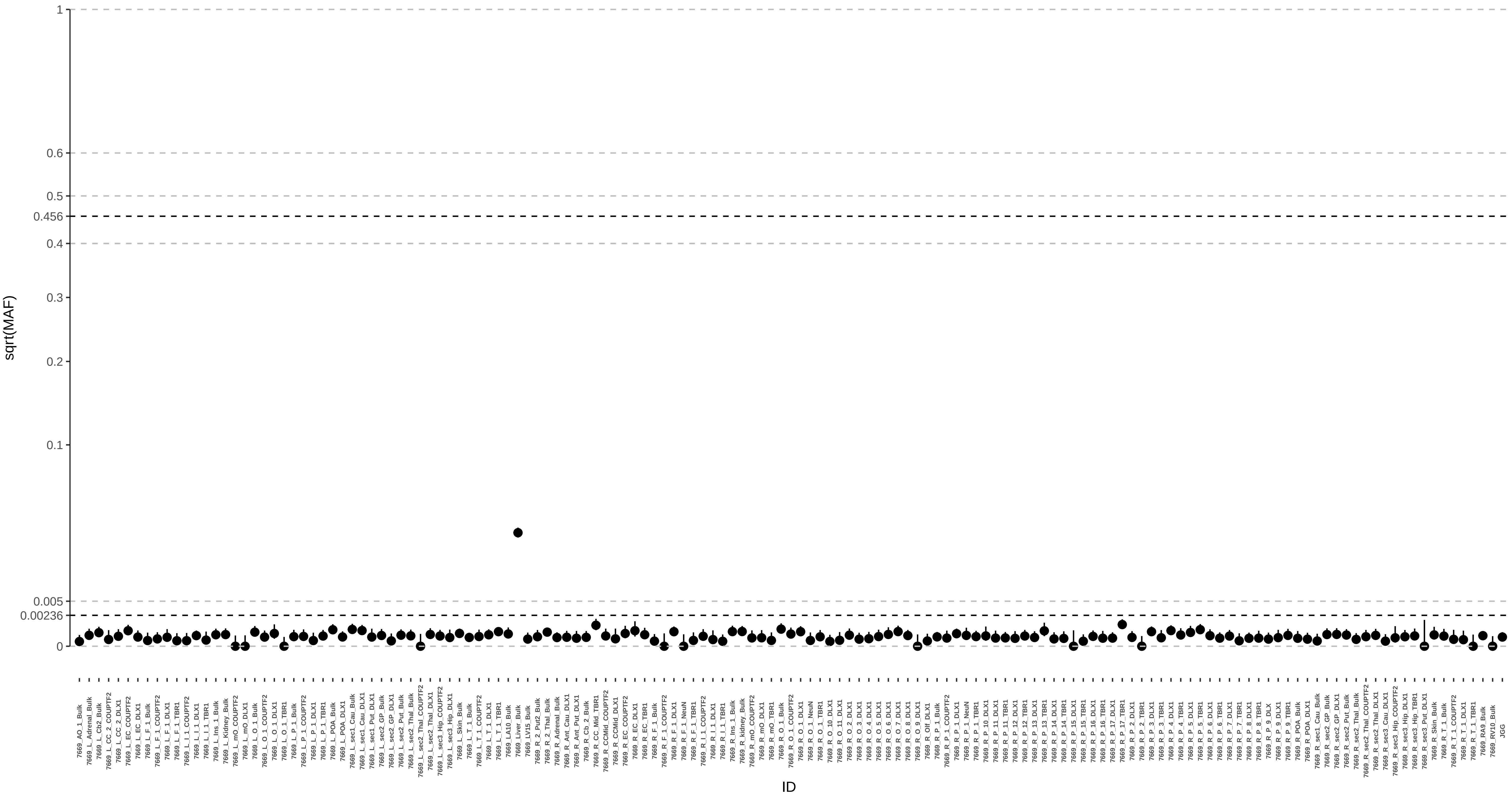

ID05 10−116677604−C−T

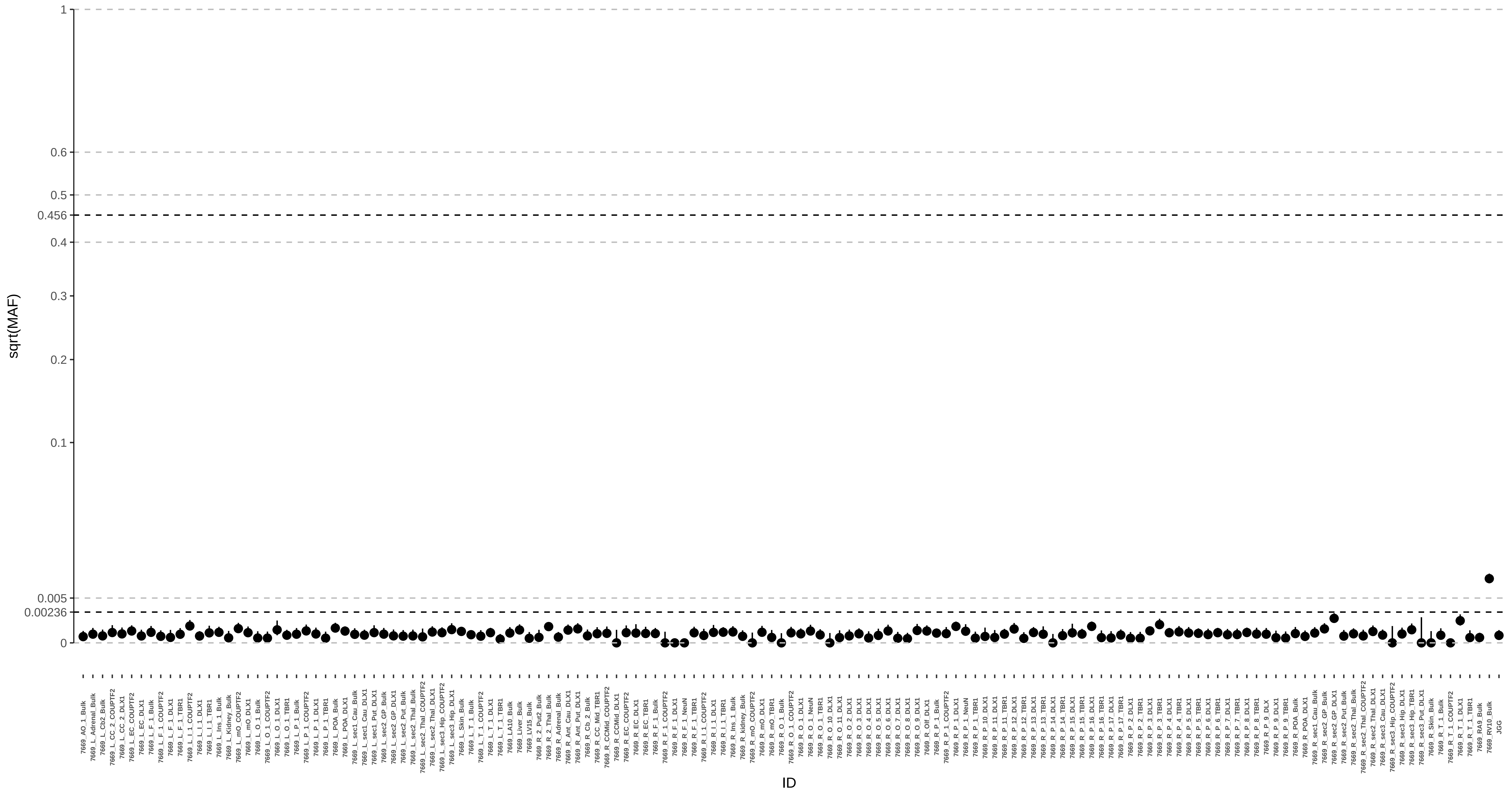

ID05 10−117540461−G−A

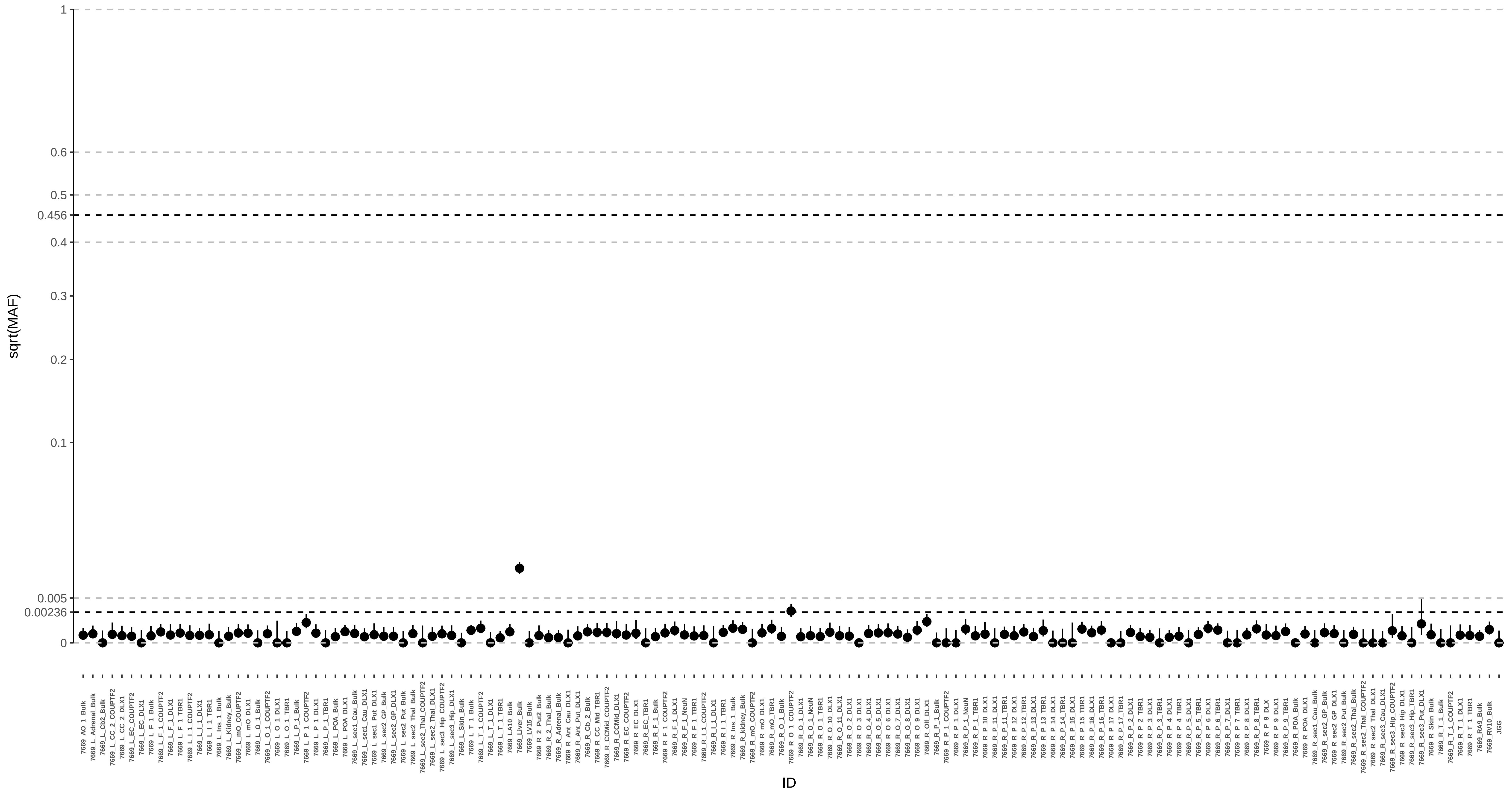

ID05 10−121073761−C−T

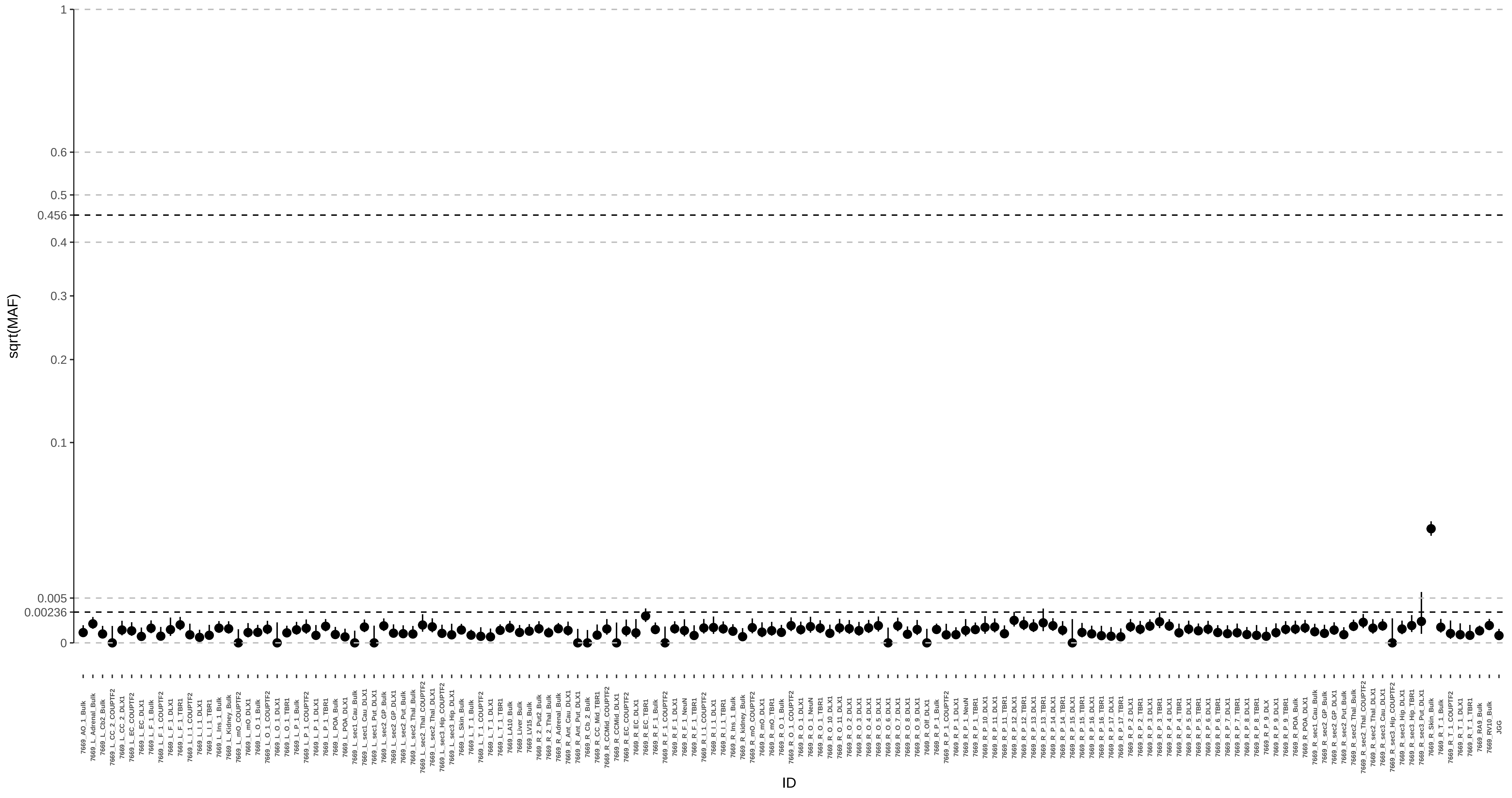

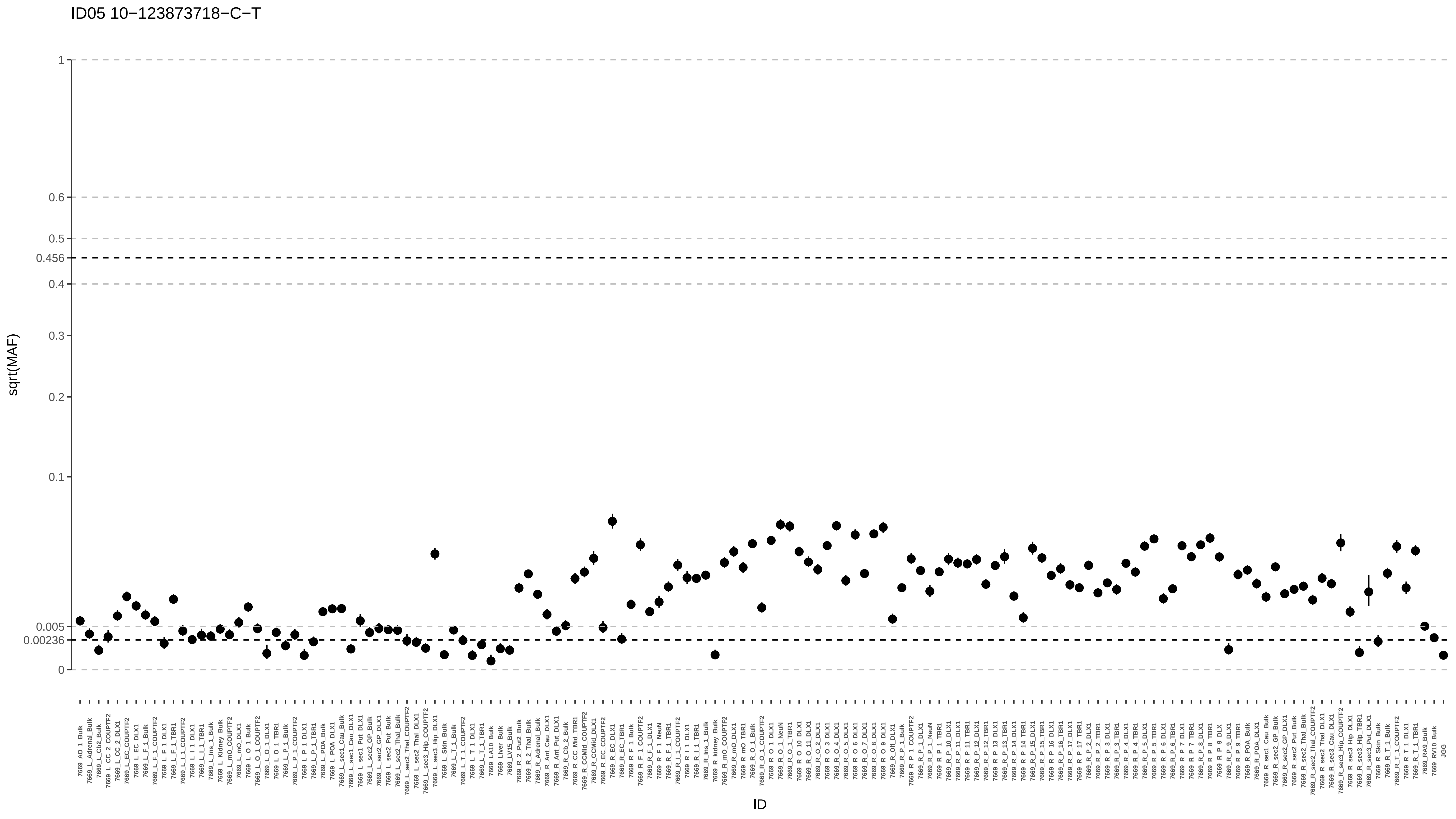
ID05 10−123873718−C−T
sqrt(MAF)
1
0.6
0.5
0.456
0.4
0.3
0.2
0.1
0.005
0.00236
0
ID
7669_AO_1_Bulk
7669_L_Adrenal_Bulk
7669_L_Cb2_Bulk
7669_L_CC_2_COUPTF2
7669_L_CC_2_DLX1
7669_L_EC_COUPTF2
7669_L_EC_DLX1
7669_L_F_1_Bulk
7669_L_F_1_COUPTF2
7669_L_F_1_DLX1
7669_L_F_1_TBR1
7669_L_I_1_COUPTF2
7669_L_I_1_DLX1
7669_L_I_1_TBR1
7669_L_Ins_1_Bulk
7669_L_Kidney_Bulk
7669_L_mO_COUPTF2
7669_L_mO_DLX1
7669_L_O_1_Bulk
7669_L_O_1_COUPTF2
7669_L_O_1_DLX1
7669_L_O_1_TBR1
7669_L_P_1_Bulk
7669_L_P_1_COUPTF2
7669_L_P_1_DLX1
7669_L_P_1_TBR1
7669_L_POA_Bulk
7669_L_POA_DLX1
7669_L_sec1_Cau_Bulk
7669_L_sec1_Cau_DLX1
7669_L_sec1_Put_DLX1
7669_L_sec2_GP_Bulk
7669_L_sec2_GP_DLX1
7669_L_sec2_Put_Bulk
7669_L_sec2_Thal_Bulk
7669_L_sec2_Thal_COUPTF2
7669_L_sec2_Thal_DLX1
7669_L_sec3_Hip_COUPTF2
7669_L_sec3_Hip_DLX1
7669_L_Skin_Bulk
7669_L_T_1_Bulk
7669_L_T_1_COUPTF2
7669_L_T_1_DLX1
7669_L_T_1_TBR1
7669_LA10_Bulk
7669_Liver_Bulk
7669_LV15_Bulk
7669_R_2_Put2_Bulk
7669_R_2_Thal_Bulk
7669_R_Adrenal_Bulk
7669_R_Ant_Cau_DLX1
7669_R_Ant_Put_DLX1
7669_R_Cb_2_Bulk
7669_R_CC_Mid_TBR1
7669_R_CCMid_COUPTF2
7669_R_CCMid_DLX1
7669_R_EC_COUPTF2
7669_R_EC_DLX1
7669_R_EC_TBR1
7669_R_F_1_Bulk
7669_R_F_1_COUPTF2
7669_R_F_1_DLX1
7669_R_F_1_NeuN
7669_R_F_1_TBR1
7669_R_I_1_COUPTF2
7669_R_I_1_DLX1
7669_R_I_1_TBR1
7669_R_Ins_1_Bulk
7669_R_kidney_Bulk
7669_R_mO_COUPTF2
7669_R_mO_DLX1
7669_R_mO_TBR1
7669_R_O_1_Bulk
7669_R_O_1_COUPTF2
7669_R_O_1_DLX1
7669_R_O_1_NeuN
7669_R_O_1_TBR1
7669_R_O_10_DLX1
7669_R_O_11_DLX1
7669_R_O_2_DLX1
7669_R_O_3_DLX1
7669_R_O_4_DLX1
7669_R_O_5_DLX1
7669_R_O_6_DLX1
7669_R_O_7_DLX1
7669_R_O_8_DLX1
7669_R_O_9_DLX1
7669_R_Olf_DLX1
7669_R_P_1_Bulk
7669_R_P_1_COUPTF2
7669_R_P_1_DLX1
7669_R_P_1_NeuN
7669_R_P_1_TBR1
7669_R_P_10_DLX1
7669_R_P_11_DLX1
7669_R_P_11_TBR1
7669_R_P_12_DLX1
7669_R_P_12_TBR1
7669_R_P_13_DLX1
7669_R_P_13_TBR1
7669_R_P_14_DLX1
7669_R_P_14_TBR1
7669_R_P_15_DLX1
7669_R_P_15_TBR1
7669_R_P_16_DLX1
7669_R_P_16_TBR1
7669_R_P_17_DLX1
7669_R_P_17_TBR1
7669_R_P_2_DLX1
7669_R_P_2_TBR1
7669_R_P_3_DLX1
7669_R_P_3_TBR1
7669_R_P_4_DLX1
7669_R_P_4_TBR1
7669_R_P_5_DLX1
7669_R_P_5_TBR1
7669_R_P_6_DLX1
7669_R_P_6_TBR1
7669_R_P_7_DLX1
7669_R_P_7_TBR1
7669_R_P_8_DLX1
7669_R_P_8_TBR1
7669_R_P_9_DLX
7669_R_P_9_DLX1
7669_R_P_9_TBR1
7669_R_POA_Bulk
7669_R_POA_DLX1
7669_R_sec1_Cau_Bulk
7669_R_sec2_GP_Bulk
7669_R_sec2_GP_DLX1
7669_R_sec2_Put_Bulk
7669_R_sec2_Thal_Bulk
7669_R_sec2_Thal_COUPTF2
7669_R_sec2_Thal_DLX1
7669_R_sec3_Cau_DLX1
7669_R_sec3_Hip_COUPTF2
7669_R_sec3_Hip_DLX1
7669_R_sec3_Hip_TBR1
7669_R_sec3_Put_DLX1
7669_R_Skin_Bulk
7669_R_T_1_Bulk
7669_R_T_1_COUPTF2
7669_R_T_1_DLX1
7669_R_T_1_TBR1
7669_RA9_Bulk
7669_RV10_Bulk
JGG

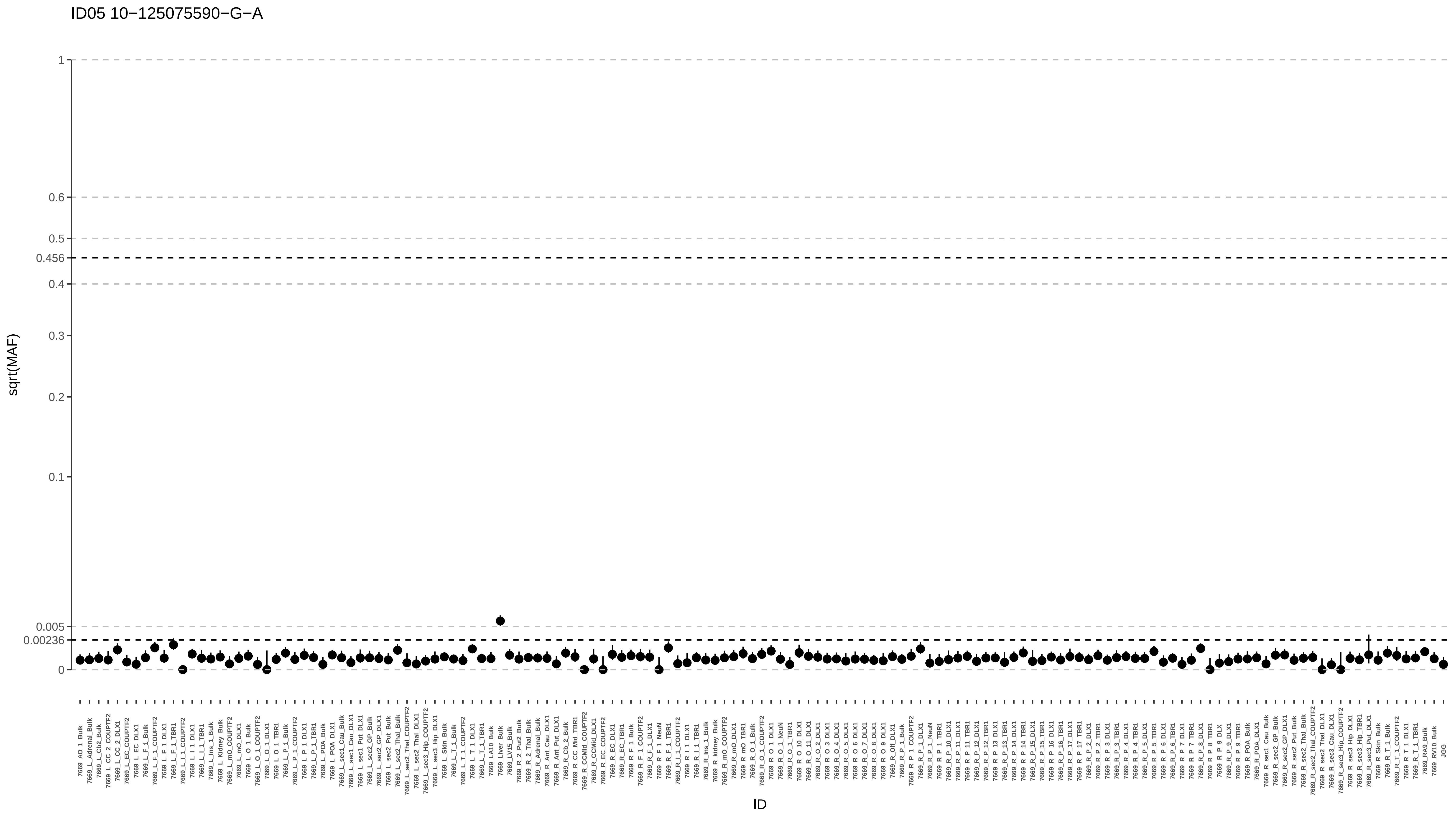
ID05 10−125075590−G−A
sqrt(MAF)
1
0.6
0.5
0.456
0.4
0.3
0.2
0.1
0.005
0.00236
0
ID

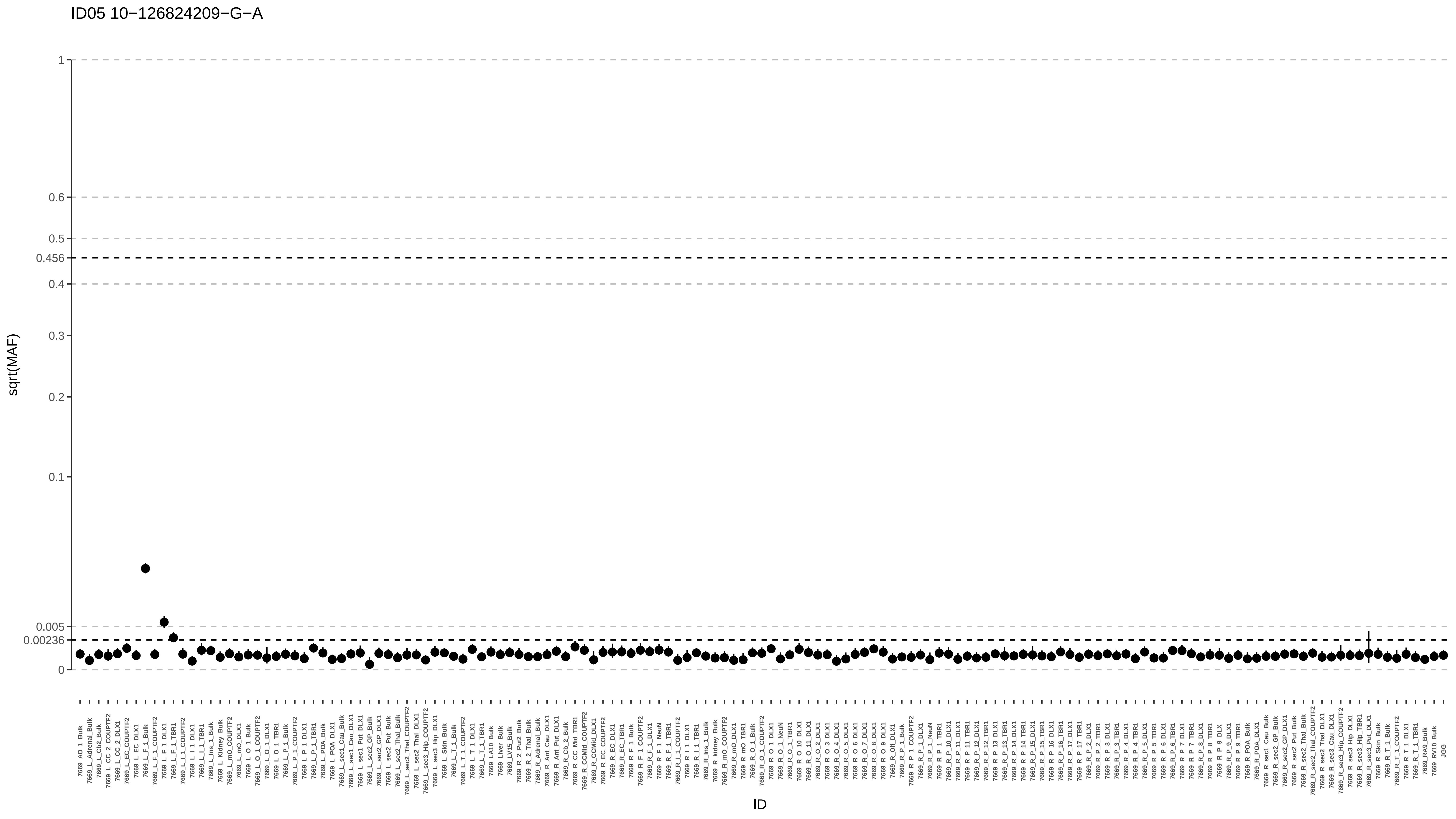

ID05 10−126824209−G−A
sqrt(MAF)
1
0.6
0.5
0.456
0.4
0.3
0.2
0.1
0.005
0.00236
0
ID
7669_AO_1_Bulk
7669_L_Adrenal_Bulk
7669_L_Cb2_Bulk
7669_L_CC_2_COUPTF2
7669_L_CC_2_DLX1
7669_L_EC_COUPTF2
7669_L_EC_DLX1
7669_L_F_1_Bulk
7669_L_F_1_COUPTF2
7669_L_F_1_DLX1
7669_L_F_1_TBR1
7669_L_I_1_COUPTF2
7669_L_I_1_DLX1
7669_L_I_1_TBR1
7669_L_Ins_1_Bulk
7669_L_Kidney_Bulk
7669_L_mO_COUPTF2
7669_L_mO_DLX1
7669_L_O_1_Bulk
7669_L_O_1_COUPTF2
7669_L_O_1_DLX1
7669_L_O_1_TBR1
7669_L_P_1_Bulk
7669_L_P_1_COUPTF2
7669_L_P_1_DLX1
7669_L_P_1_TBR1
7669_L_POA_Bulk
7669_L_POA_DLX1
7669_L_sec1_Cau_Bulk
7669_L_sec1_Cau_DLX1
7669_L_sec1_Put_DLX1
7669_L_sec2_GP_Bulk
7669_L_sec2_GP_DLX1
7669_L_sec2_Put_Bulk
7669_L_sec2_Thal_Bulk
7669_L_sec2_Thal_COUPTF2
7669_L_sec2_Thal_DLX1
7669_L_sec3_Hip_COUPTF2
7669_L_sec3_Hip_DLX1
7669_L_Skin_Bulk
7669_L_T_1_Bulk
7669_L_T_1_COUPTF2
7669_L_T_1_DLX1
7669_L_T_1_TBR1
7669_LA10_Bulk
7669_Liver_Bulk
7669_LV15_Bulk
7669_R_2_Put2_Bulk
7669_R_2_Thal_Bulk
7669_R_Adrenal_Bulk
7669_R_Ant_Cau_DLX1
7669_R_Ant_Put_DLX1
7669_R_Cb_2_Bulk
7669_R_CC_Mid_TBR1
7669_R_CCMid_COUPTF2
7669_R_CCMid_DLX1
7669_R_EC_COUPTF2
7669_R_EC_DLX1
7669_R_EC_TBR1
7669_R_F_1_Bulk
7669_R_F_1_COUPTF2
7669_R_F_1_DLX1
7669_R_F_1_NeuN
7669_R_F_1_TBR1
7669_R_I_1_COUPTF2
7669_R_I_1_DLX1
7669_R_I_1_TBR1
7669_R_Ins_1_Bulk
7669_R_kidney_Bulk
7669_R_mO_COUPTF2
7669_R_mO_DLX1
7669_R_mO_TBR1
7669_R_O_1_Bulk
7669_R_O_1_COUPTF2
7669_R_O_1_DLX1
7669_R_O_1_NeuN
7669_R_O_1_TBR1
7669_R_O_10_DLX1
7669_R_O_11_DLX1
7669_R_O_2_DLX1
7669_R_O_3_DLX1
7669_R_O_4_DLX1
7669_R_O_5_DLX1
7669_R_O_6_DLX1
7669_R_O_7_DLX1
7669_R_O_8_DLX1
7669_R_O_9_DLX1
7669_R_Olf_DLX1
7669_R_P_1_Bulk
7669_R_P_1_COUPTF2
7669_R_P_1_DLX1
7669_R_P_1_NeuN
7669_R_P_1_TBR1
7669_R_P_10_DLX1
7669_R_P_11_DLX1
7669_R_P_11_TBR1
7669_R_P_12_DLX1
7669_R_P_12_TBR1
7669_R_P_13_DLX1
7669_R_P_13_TBR1
7669_R_P_14_DLX1
7669_R_P_14_TBR1
7669_R_P_15_DLX1
7669_R_P_15_TBR1
7669_R_P_16_DLX1
7669_R_P_16_TBR1
7669_R_P_17_DLX1
7669_R_P_17_TBR1
7669_R_P_2_DLX1
7669_R_P_2_TBR1
7669_R_P_3_DLX1
7669_R_P_3_TBR1
7669_R_P_4_DLX1
7669_R_P_4_TBR1
7669_R_P_5_DLX1
7669_R_P_5_TBR1
7669_R_P_6_DLX1
7669_R_P_6_TBR1
7669_R_P_7_DLX1
7669_R_P_7_TBR1
7669_R_P_8_DLX1
7669_R_P_8_TBR1
7669_R_P_9_DLX
7669_R_P_9_DLX1
7669_R_P_9_TBR1
7669_R_POA_Bulk
7669_R_POA_DLX1
7669_R_sec1_Cau_Bulk
7669_R_sec2_GP_Bulk
7669_R_sec2_GP_DLX1
7669_R_sec2_Put_Bulk
7669_R_sec2_Thal_Bulk
7669_R_sec2_Thal_COUPTF2
7669_R_sec2_Thal_DLX1
7669_R_sec3_Cau_DLX1
7669_R_sec3_Hip_COUPTF2
7669_R_sec3_Hip_DLX1
7669_R_sec3_Hip_TBR1
7669_R_sec3_Put_DLX1
7669_R_Skin_Bulk
7669_R_T_1_Bulk
7669_R_T_1_COUPTF2
7669_R_T_1_DLX1
7669_R_T_1_TBR1
7669_RA9_Bulk
7669_RV10_Bulk
JGG

ID05 10−12696593−A−G

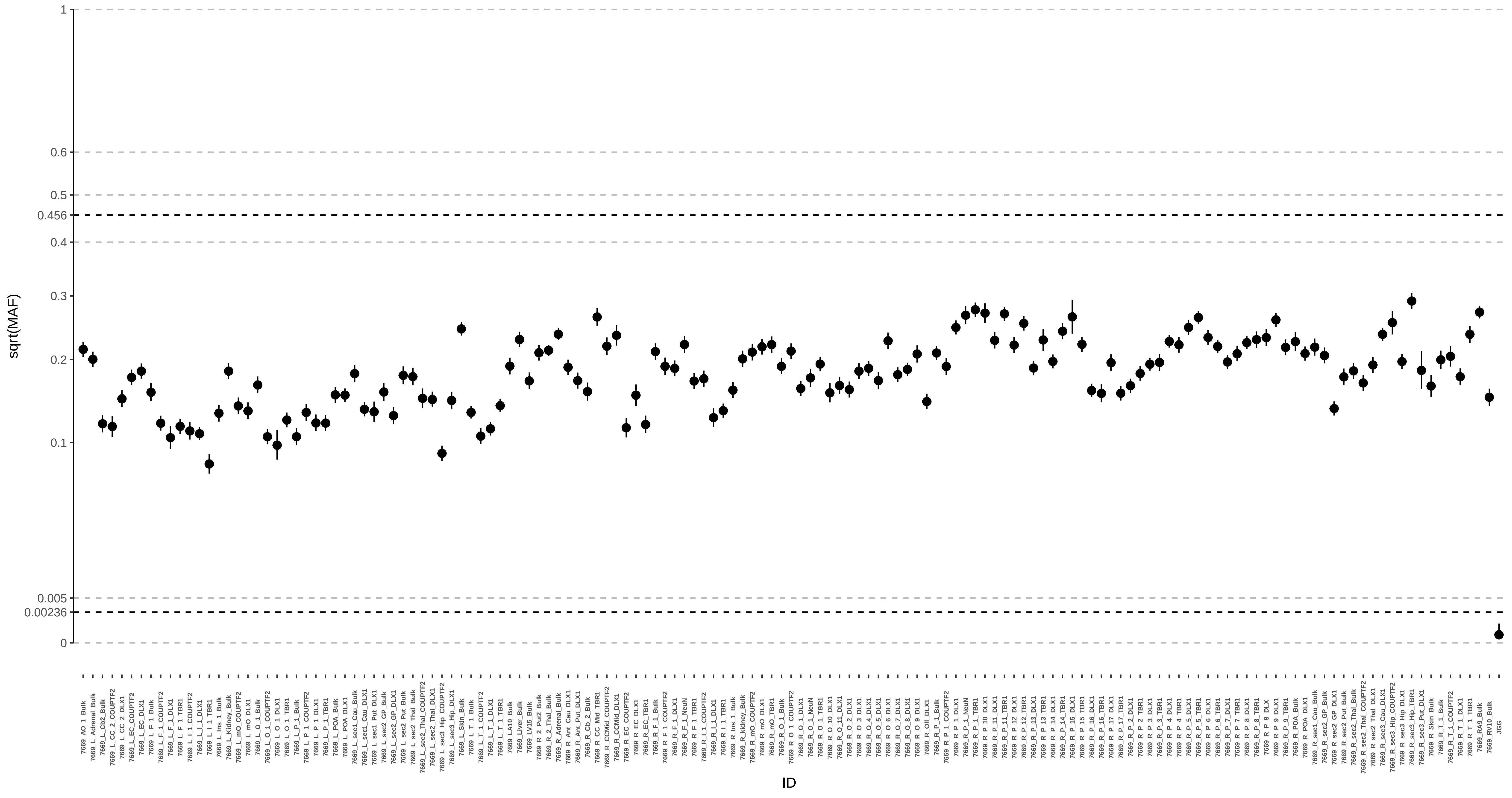

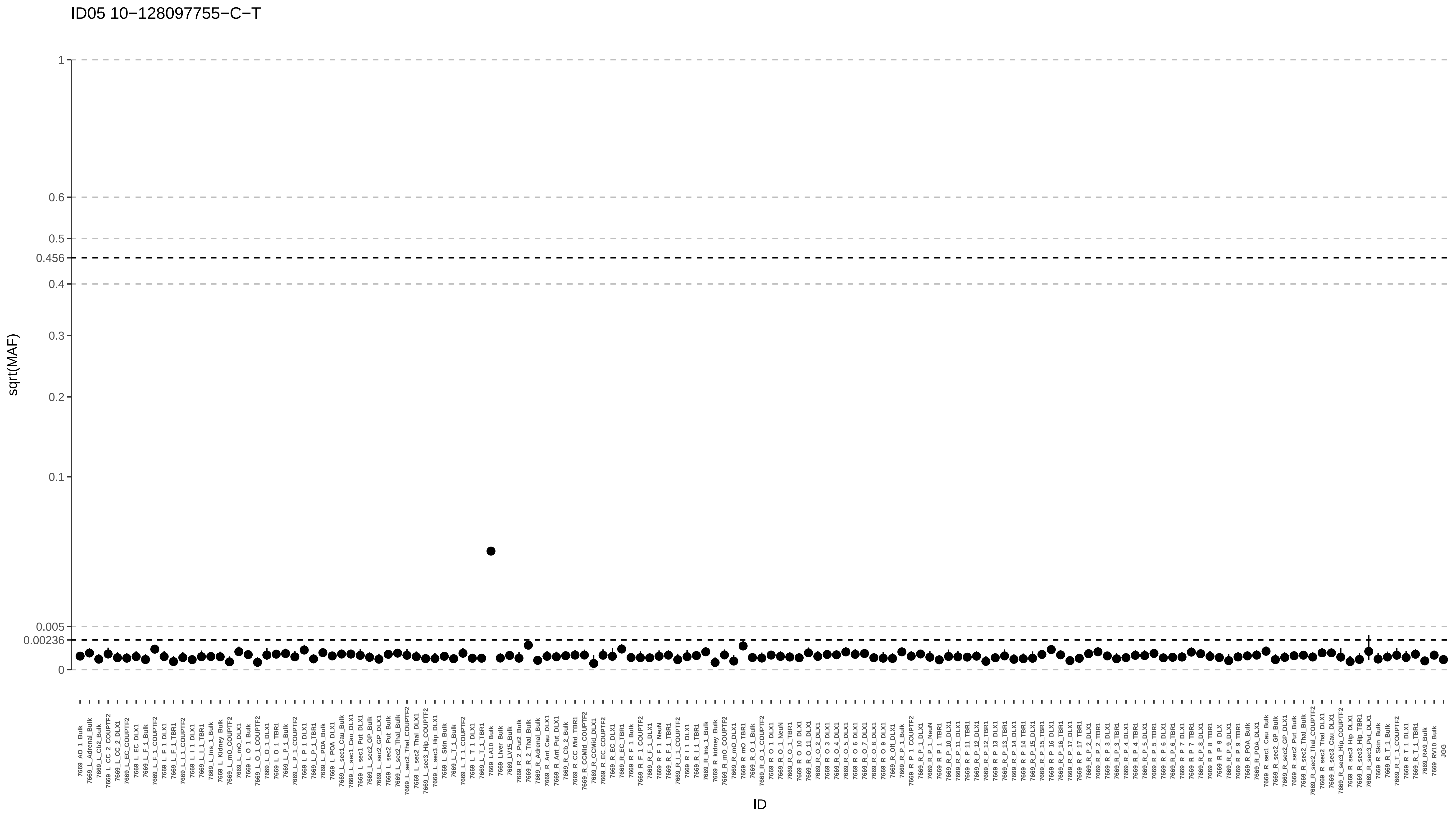
ID05 10−128097755−C−T
sqrt(MAF)
1
0.6
0.5
0.456
0.4
0.3
0.2
0.1
0.005
0.00236
0
ID

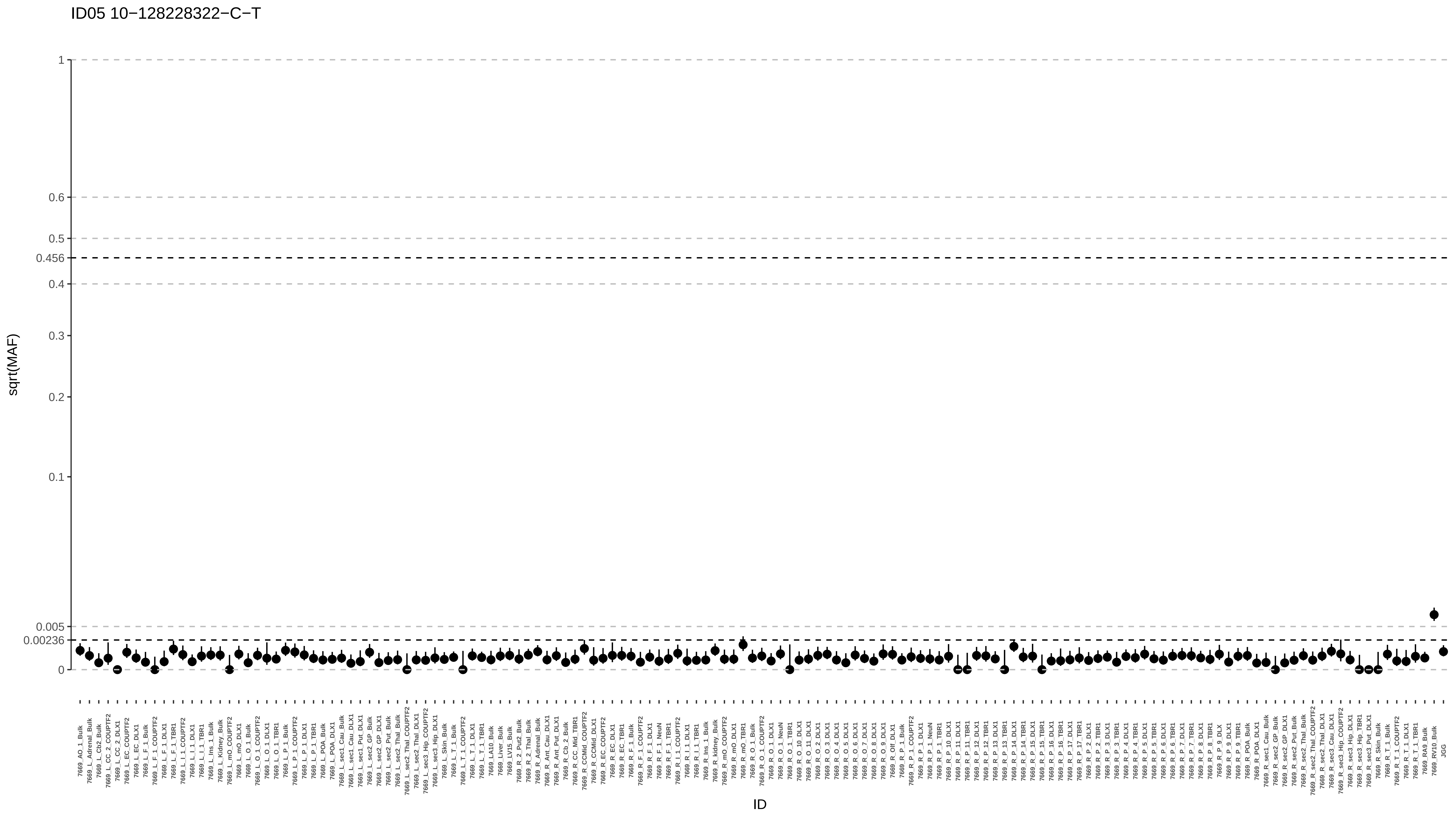
ID05 10−128228322−C−T
sqrt(MAF)
1
0.6
0.5
0.456
0.4
0.3
0.2
0.1
0.005
0.00236
0
ID

ID05 10−129376060−T−G

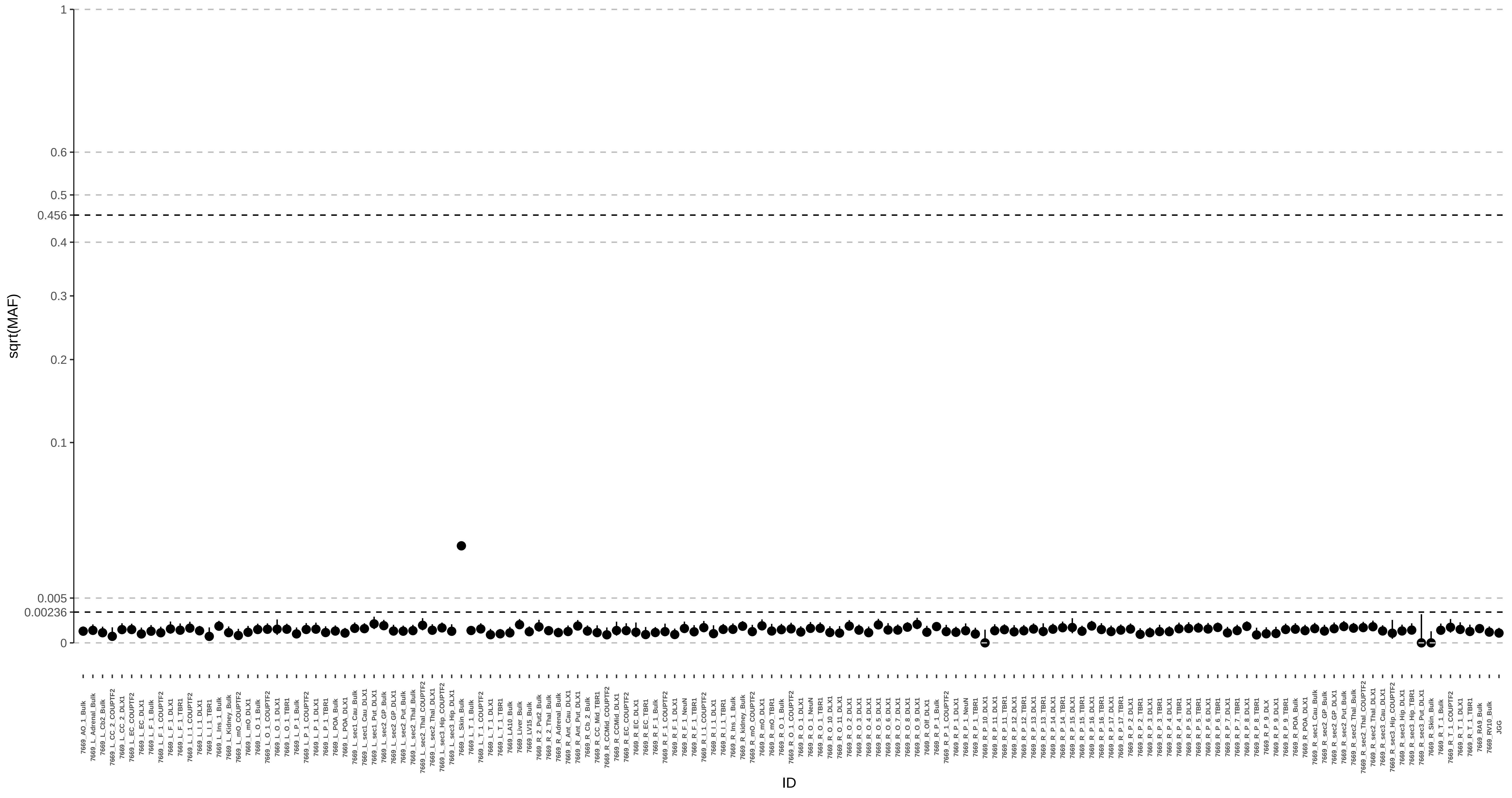

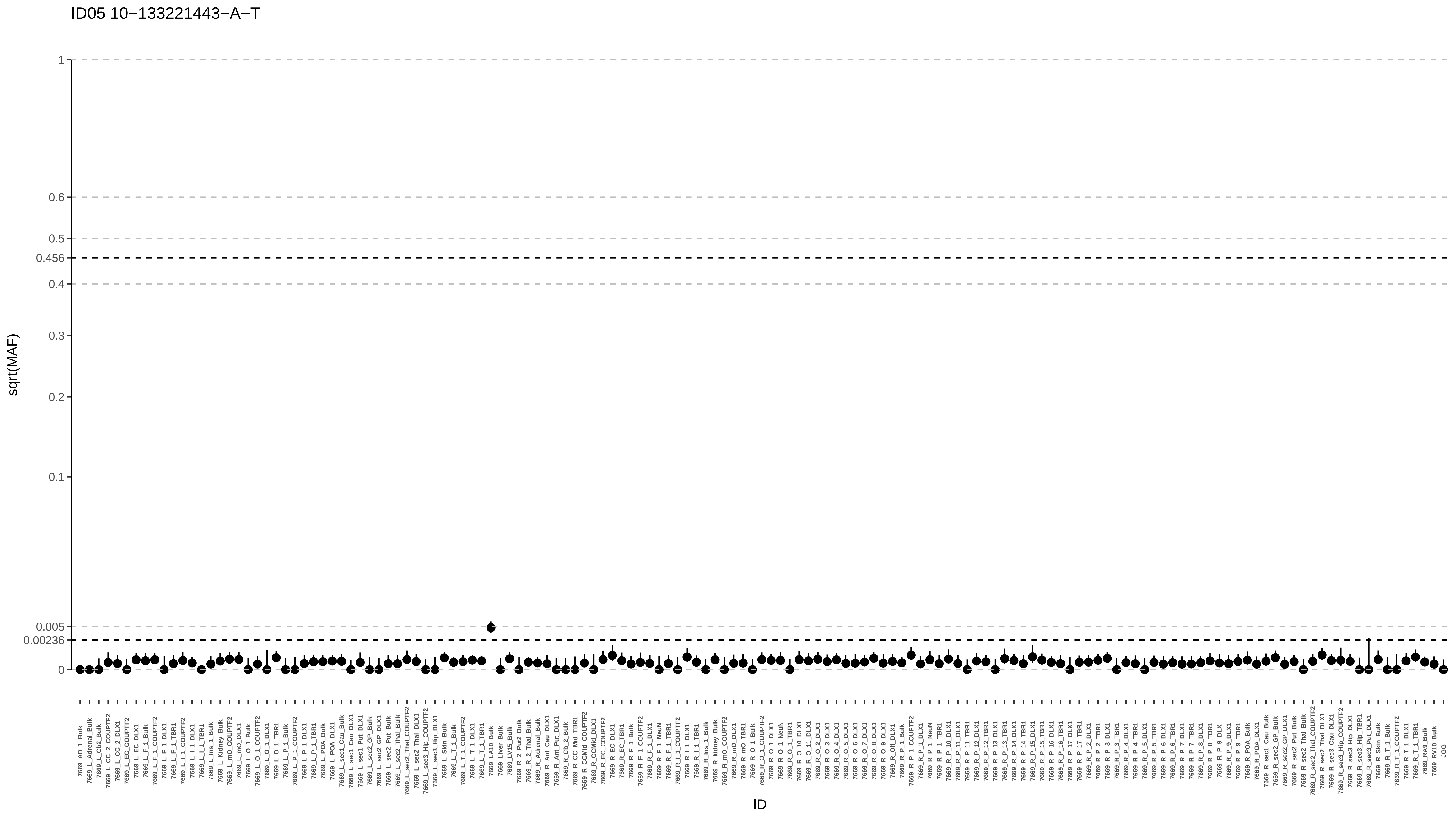
ID05 10−133221443−A−T
sqrt(MAF)
ID

ID05 10−134211625−C−T

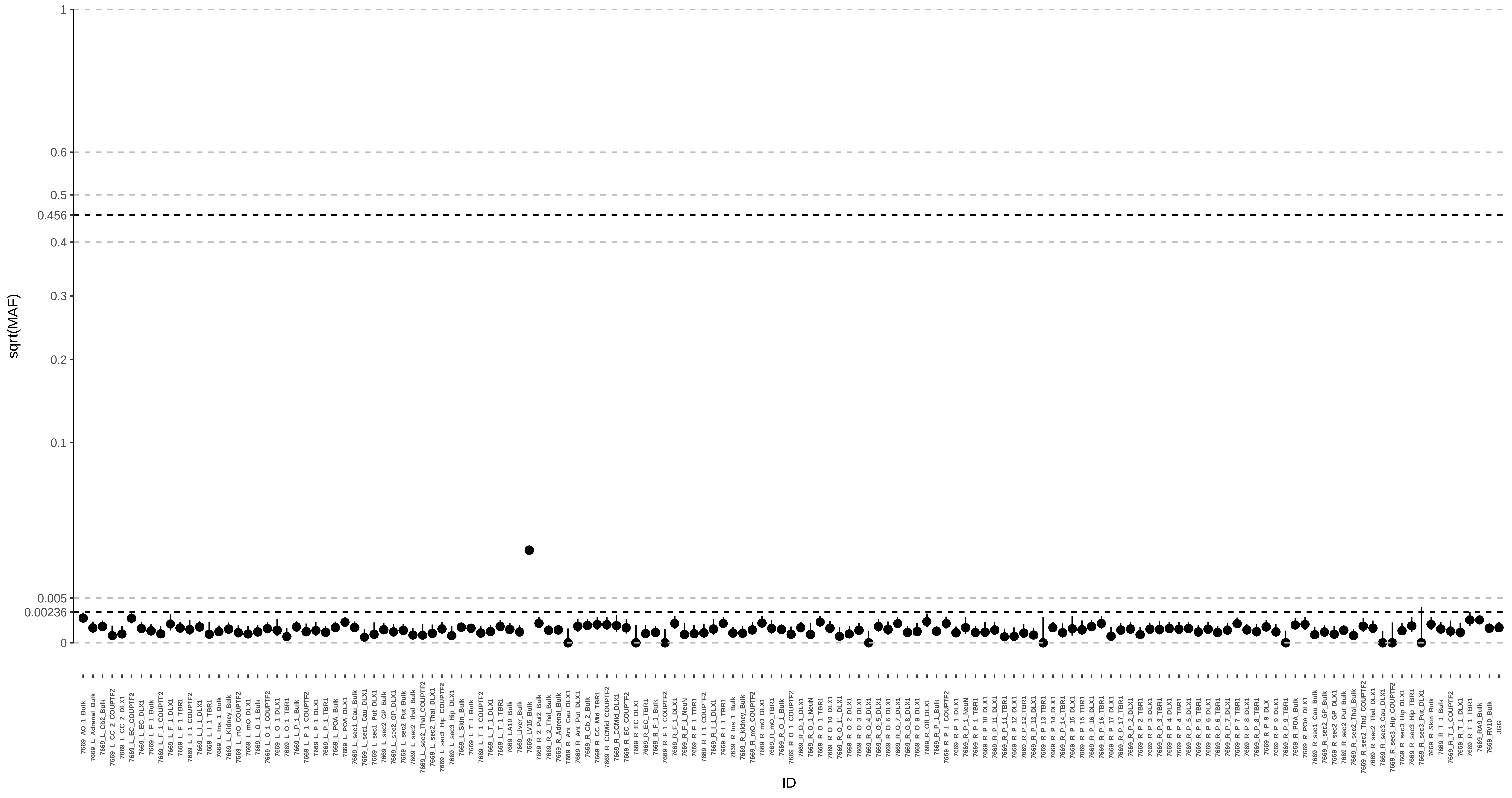

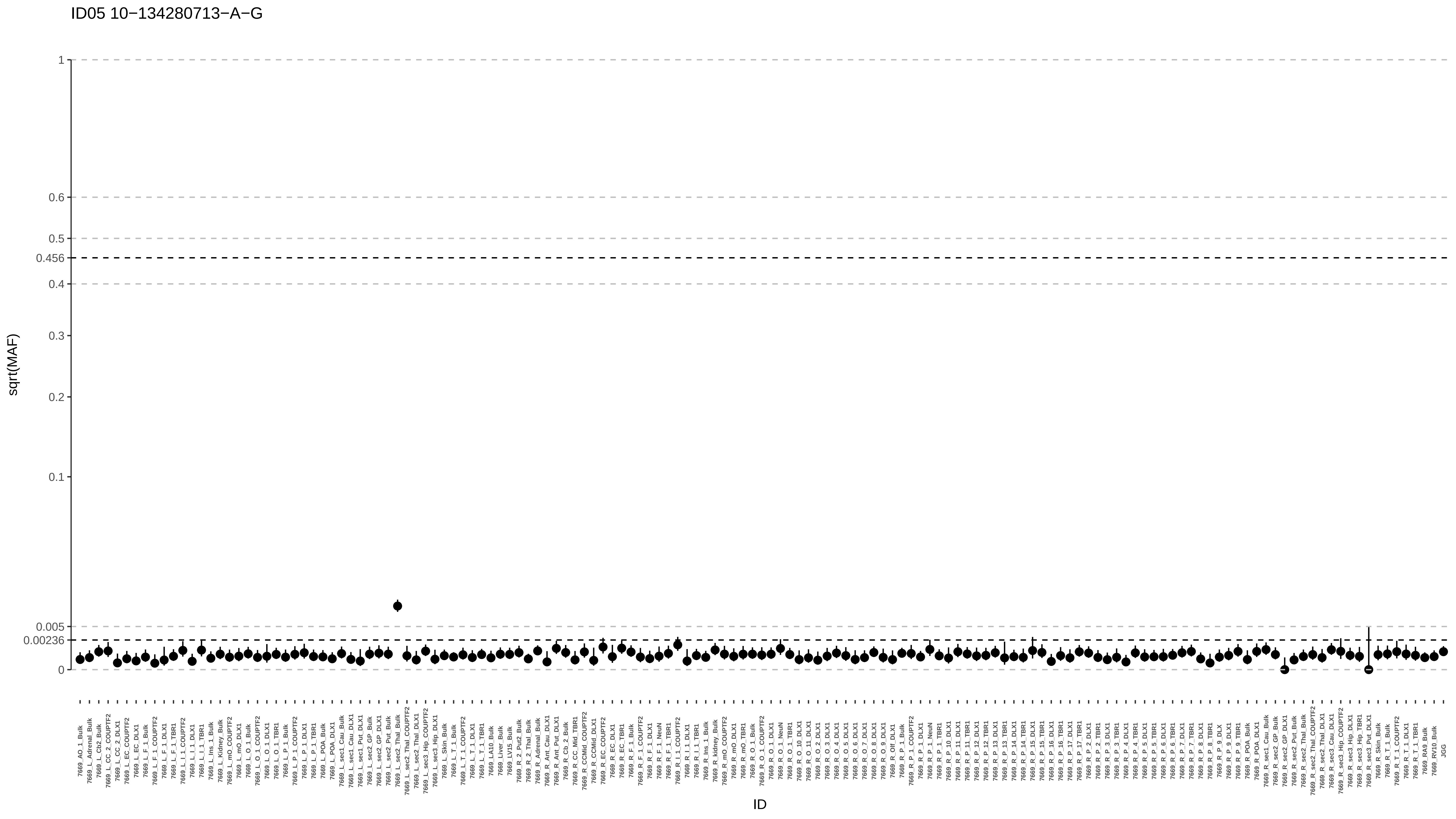
ID05 10−134280713−A−G
sqrt(MAF)
1
0.6
0.5
0.456
0.4
0.3
0.2
0.1
0.005
0.00236
0
ID

ID05 10−13796054−C−T

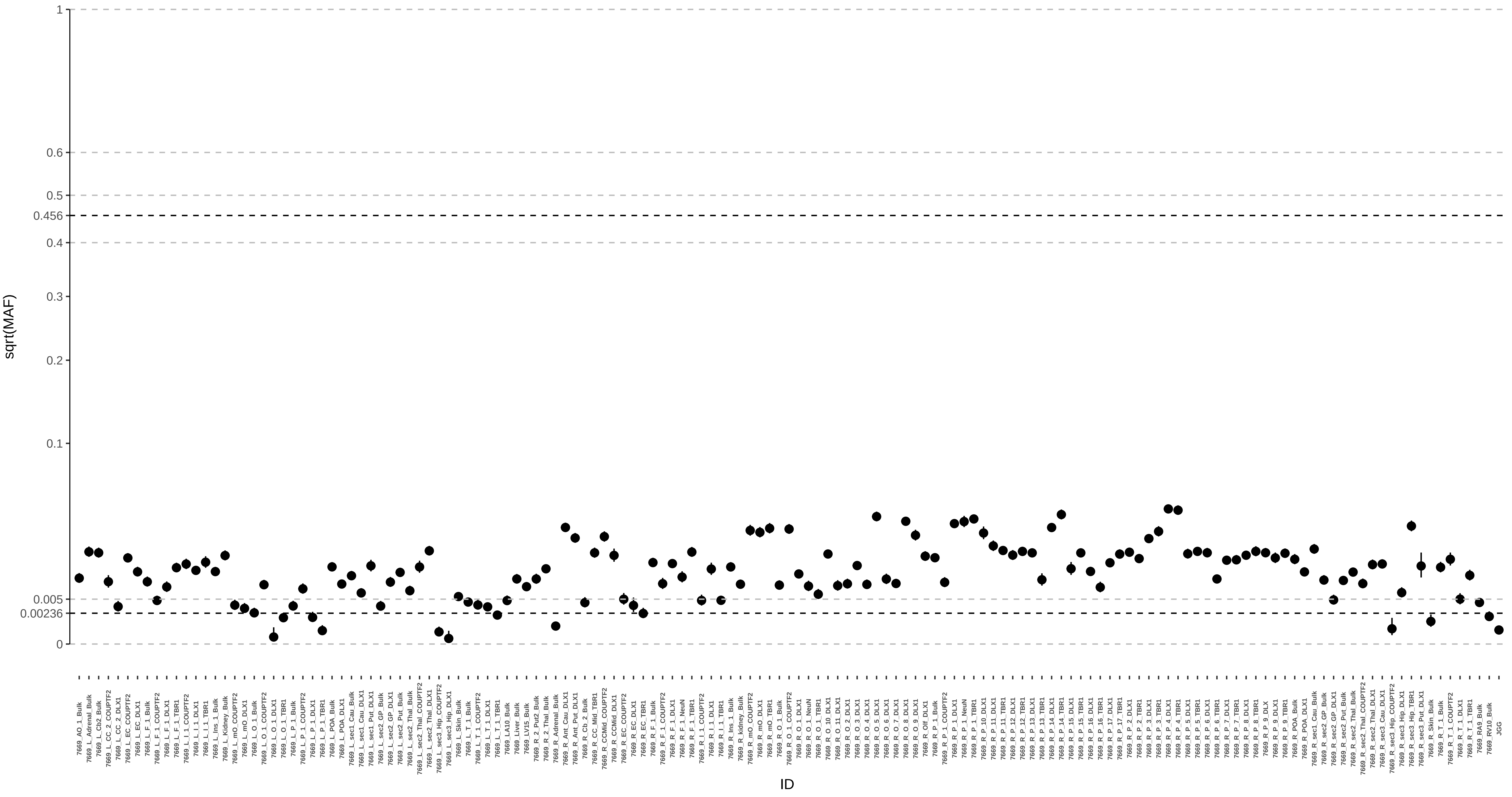

ID05 10−14490737−G−A

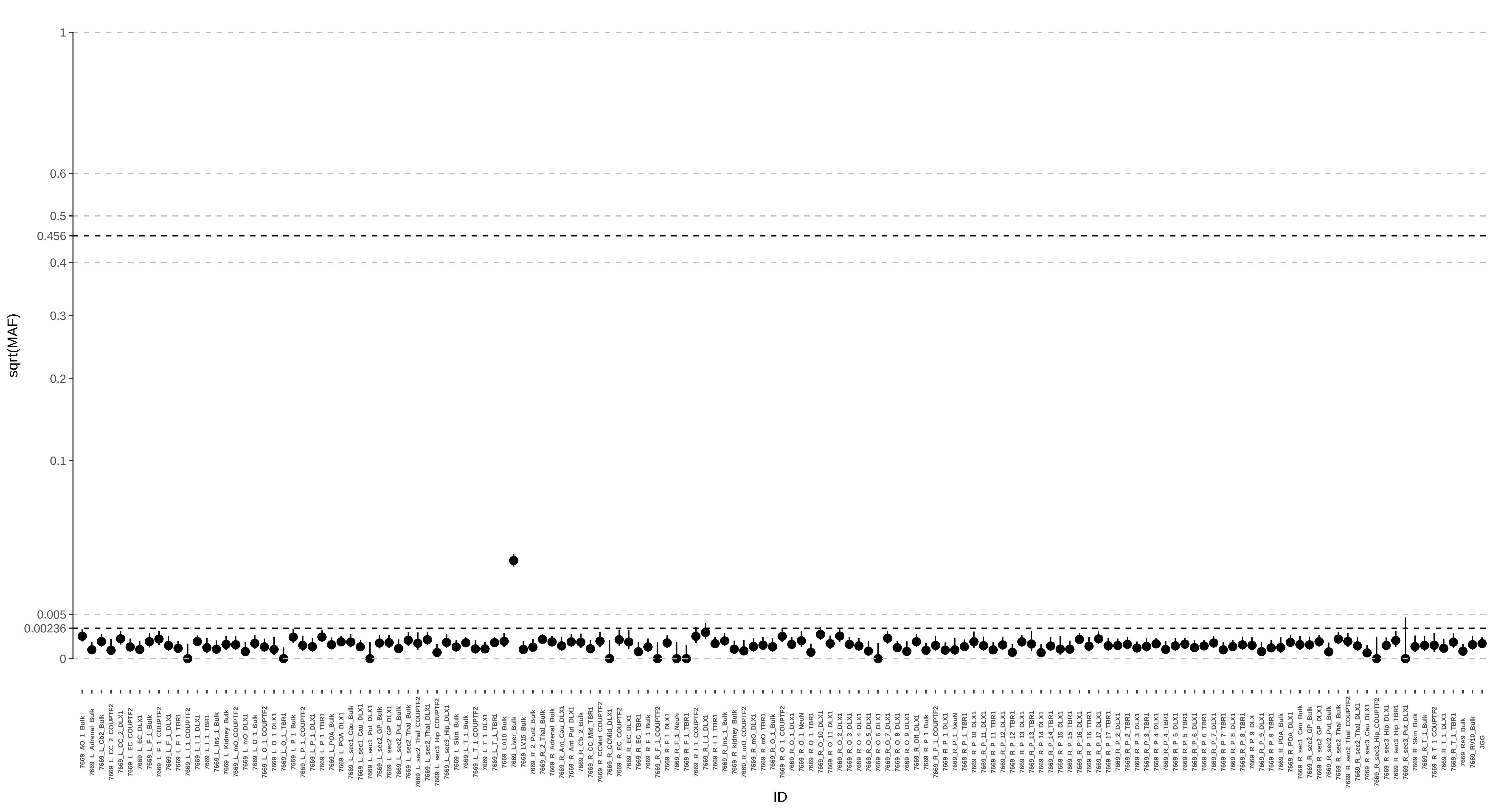

ID05 10−18693563−A−G

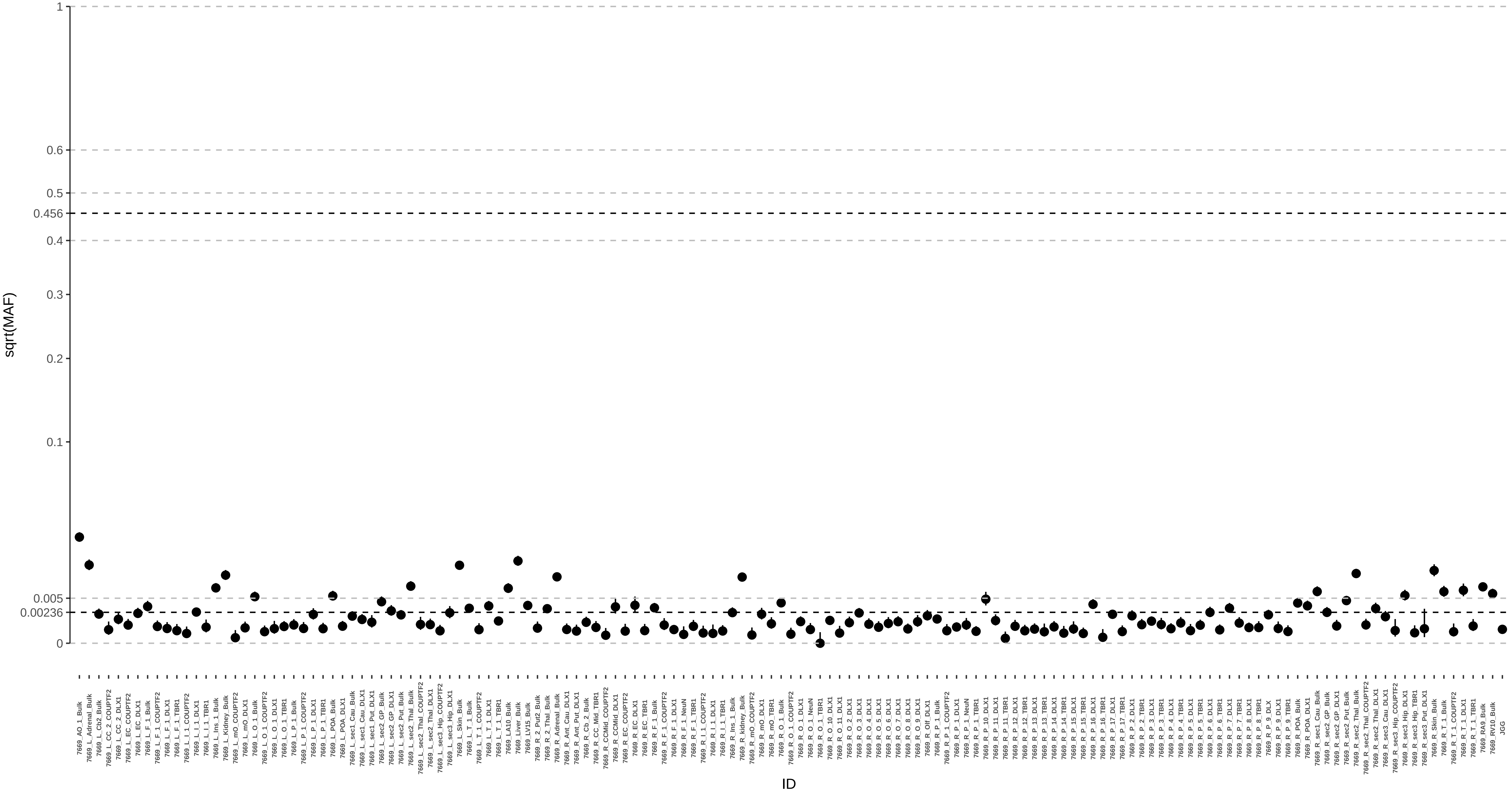

ID05 10−18760480−C−T

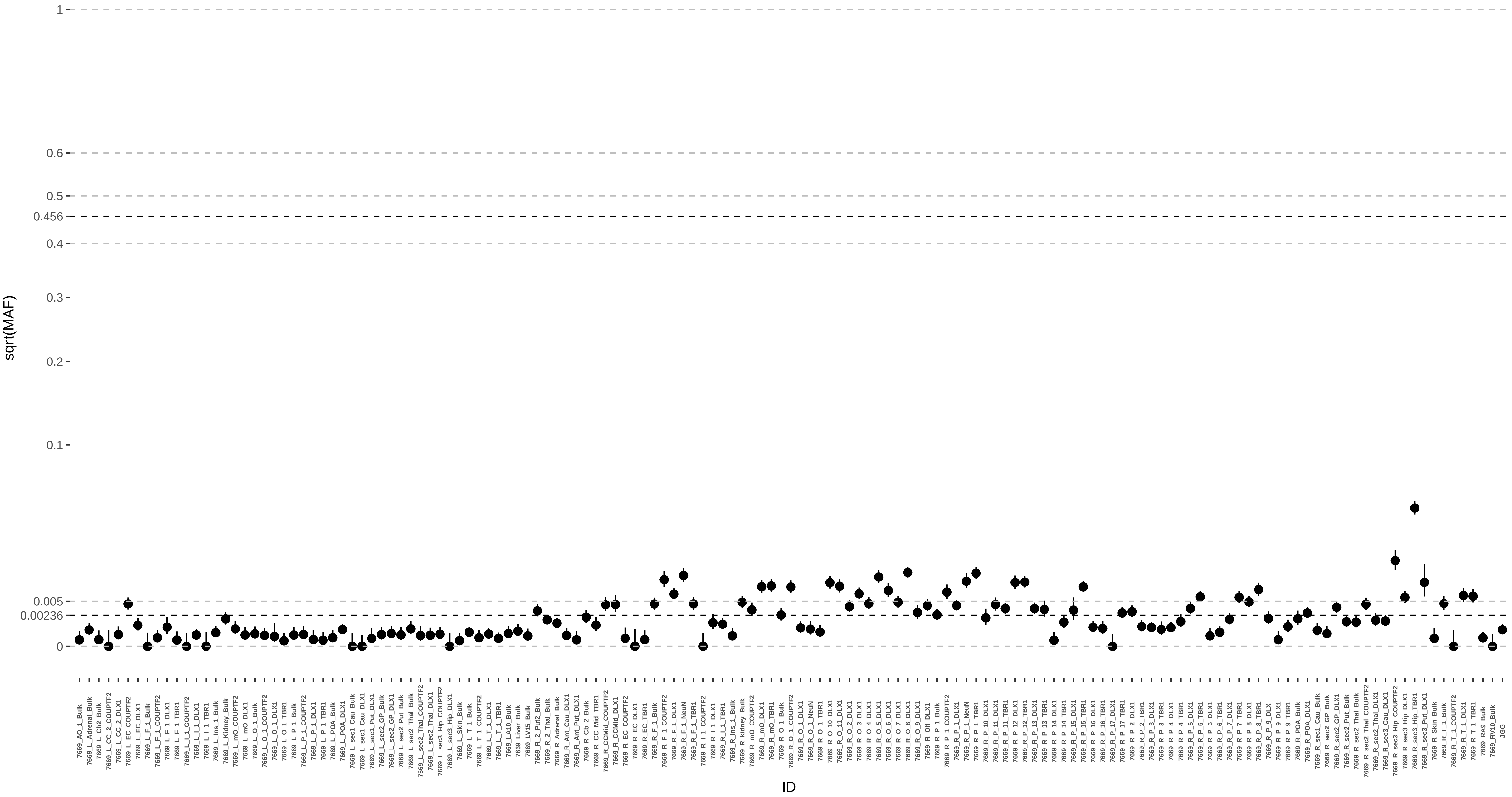

ID05 10−2261697−A−C

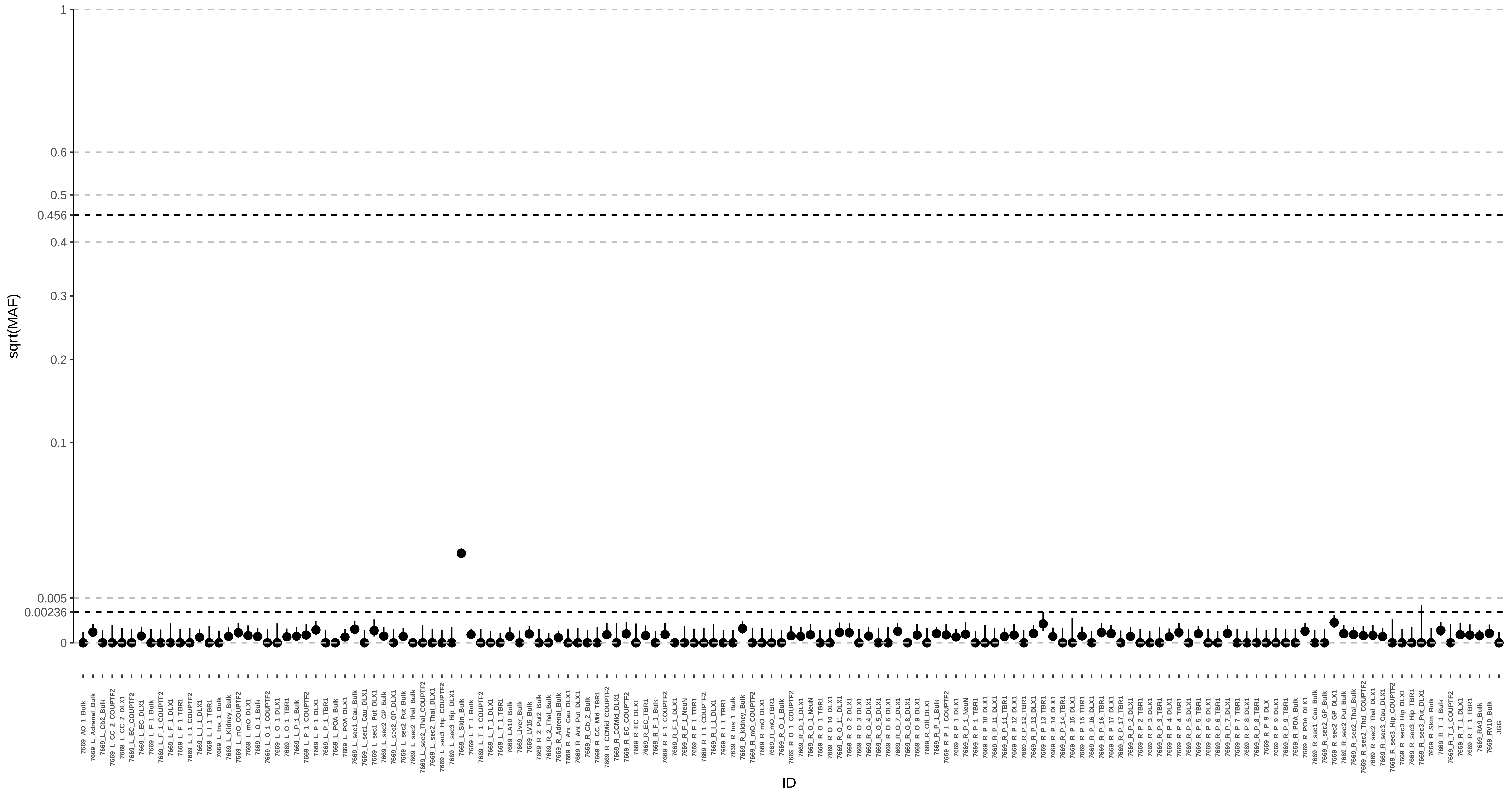

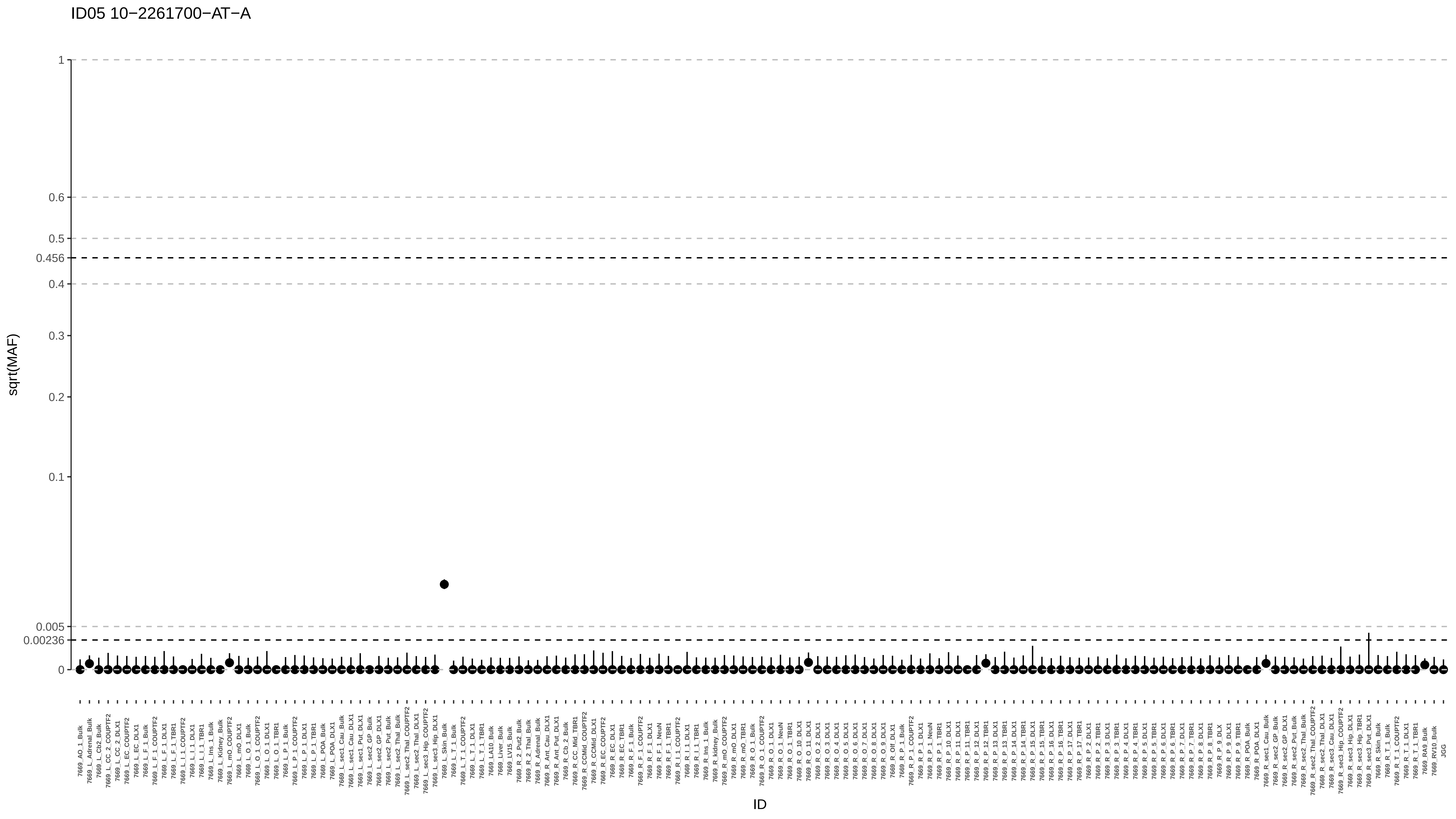

ID05 10−2261700−AT−A
sqrt(MAF)
1
0.6
0.5
0.456
0.4
0.3
0.2
0.1
0.005
0.00236
0
ID

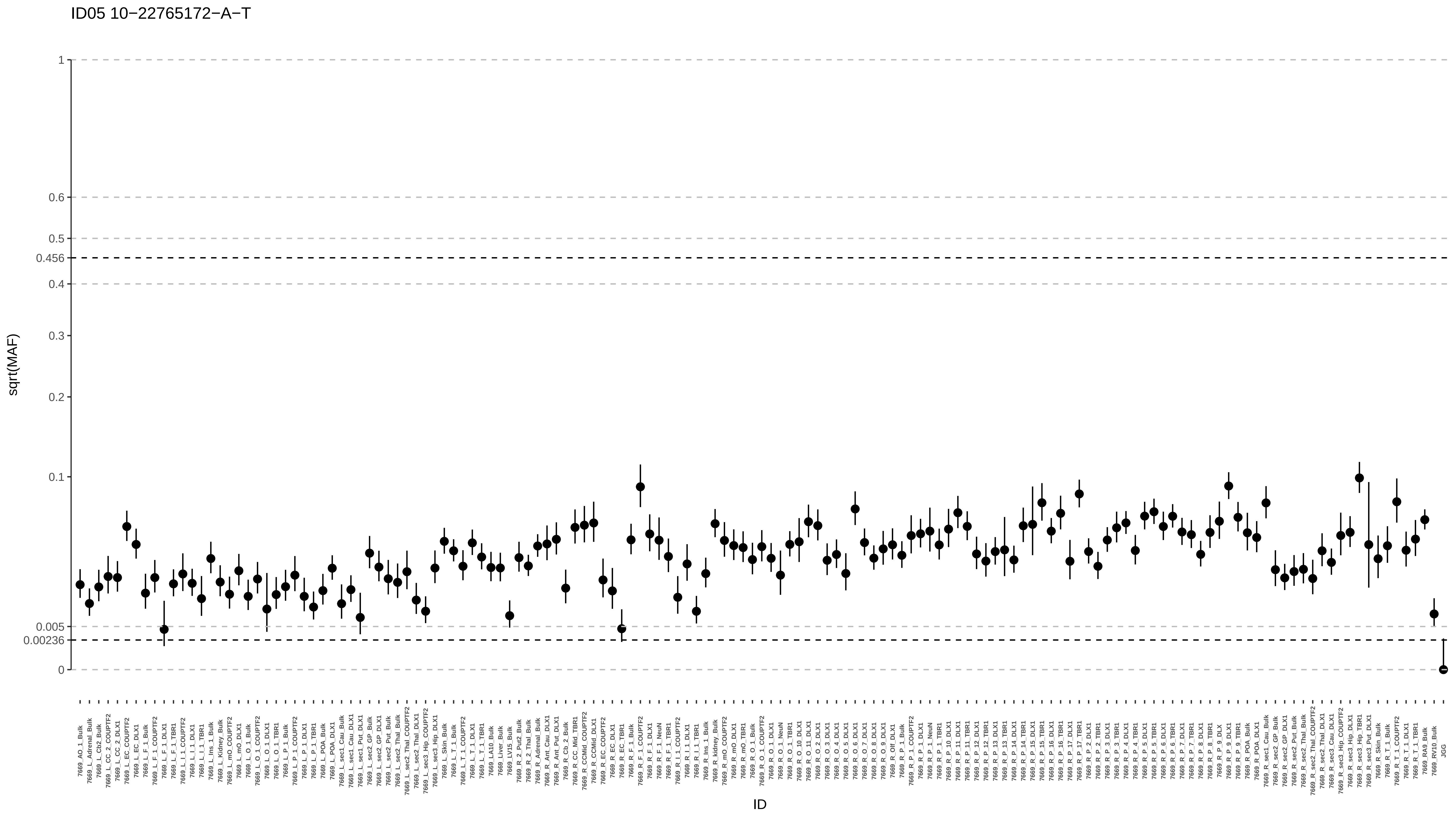
ID05 10−22765172−A−T
sqrt(MAF)
1
0.6
0.5
0.456
0.4
0.3
0.2
0.1
0.005
0.00236
0
ID

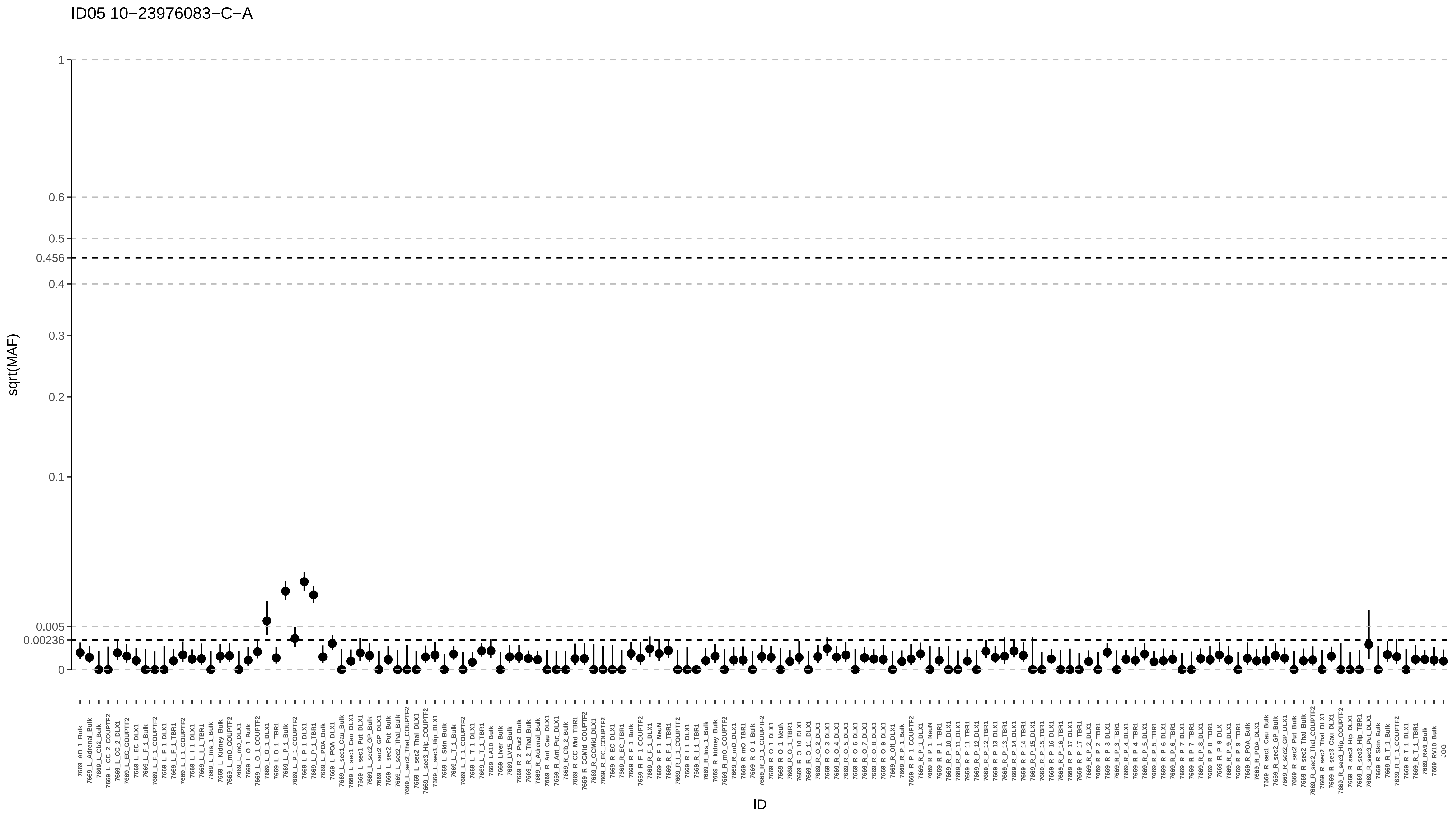
ID05 10−23976083−C−A
sqrt(MAF)
1
0.6
0.5
0.456
0.4
0.3
0.2
0.1
0.005
0.00236
0
ID

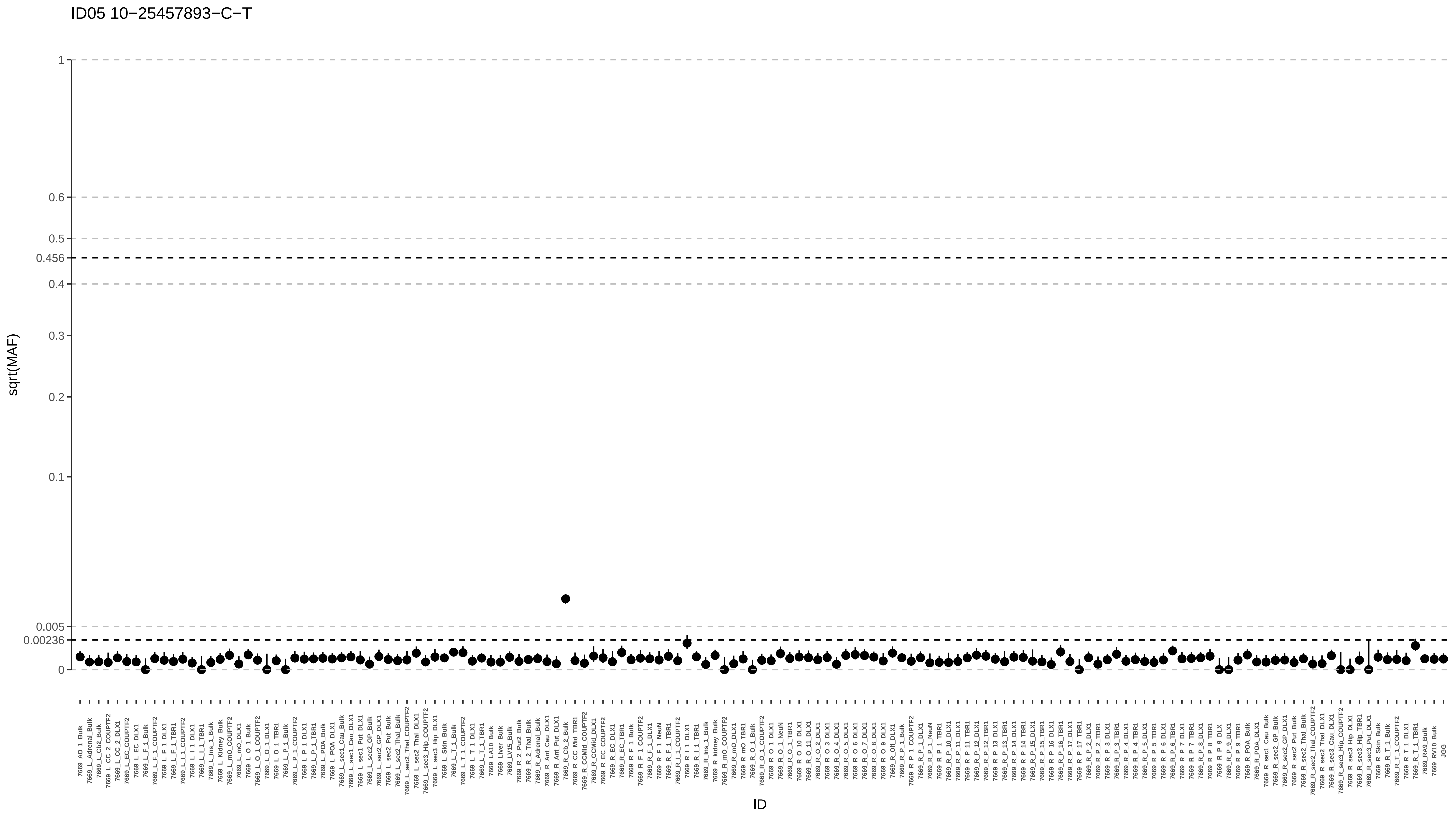
ID05 10−25457893−C−T
sqrt(MAF)
1
0.6
0.5
0.456
0.4
0.3
0.2
0.1
0.005
0.00236
0
ID

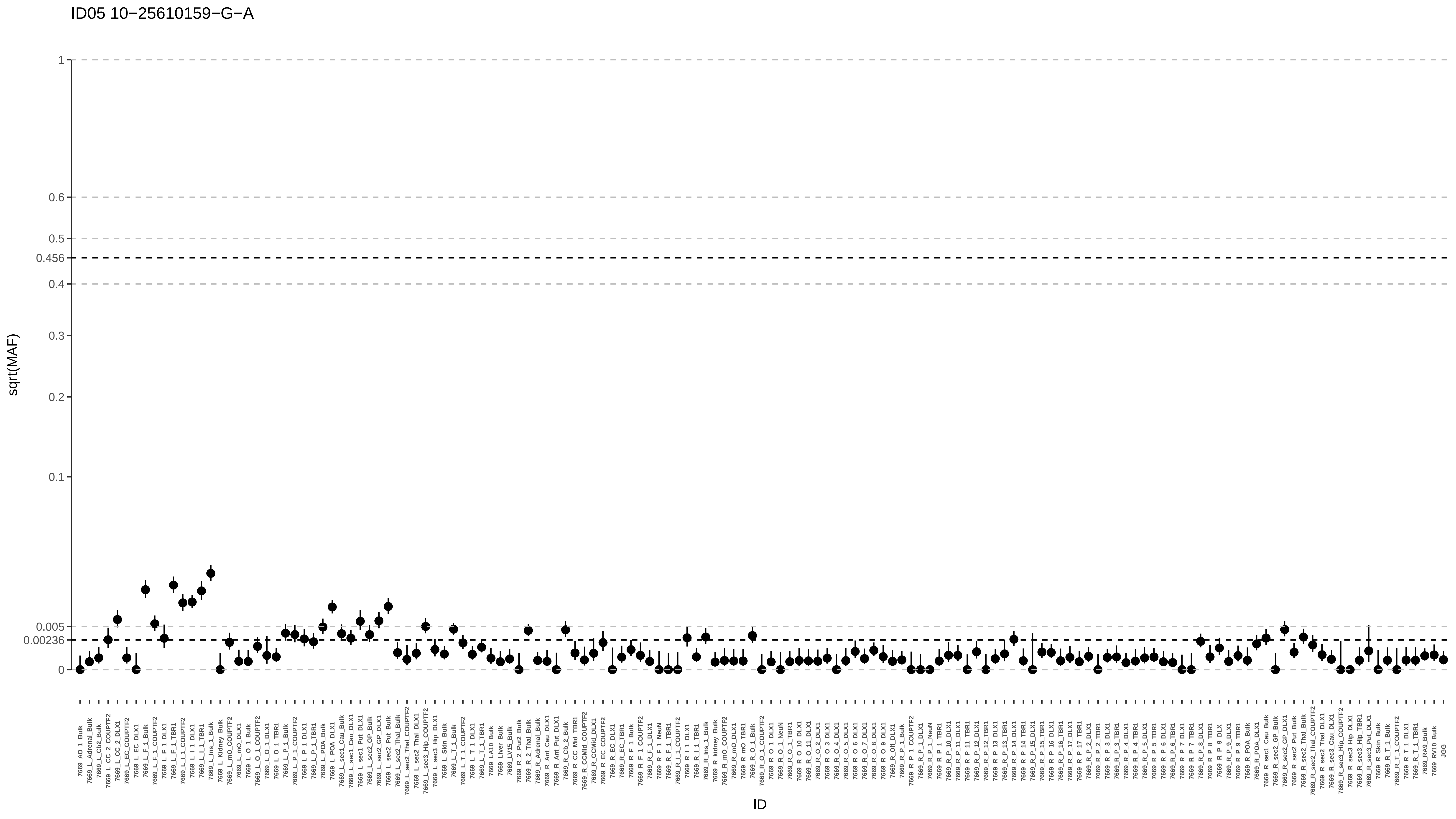
ID05 10−25610159−G−A
sqrt(MAF)
1
0.6
0.5
0.456
0.4
0.3
0.2
0.1
0.005
0.00236
0
ID

ID05 10−34121088−A−G

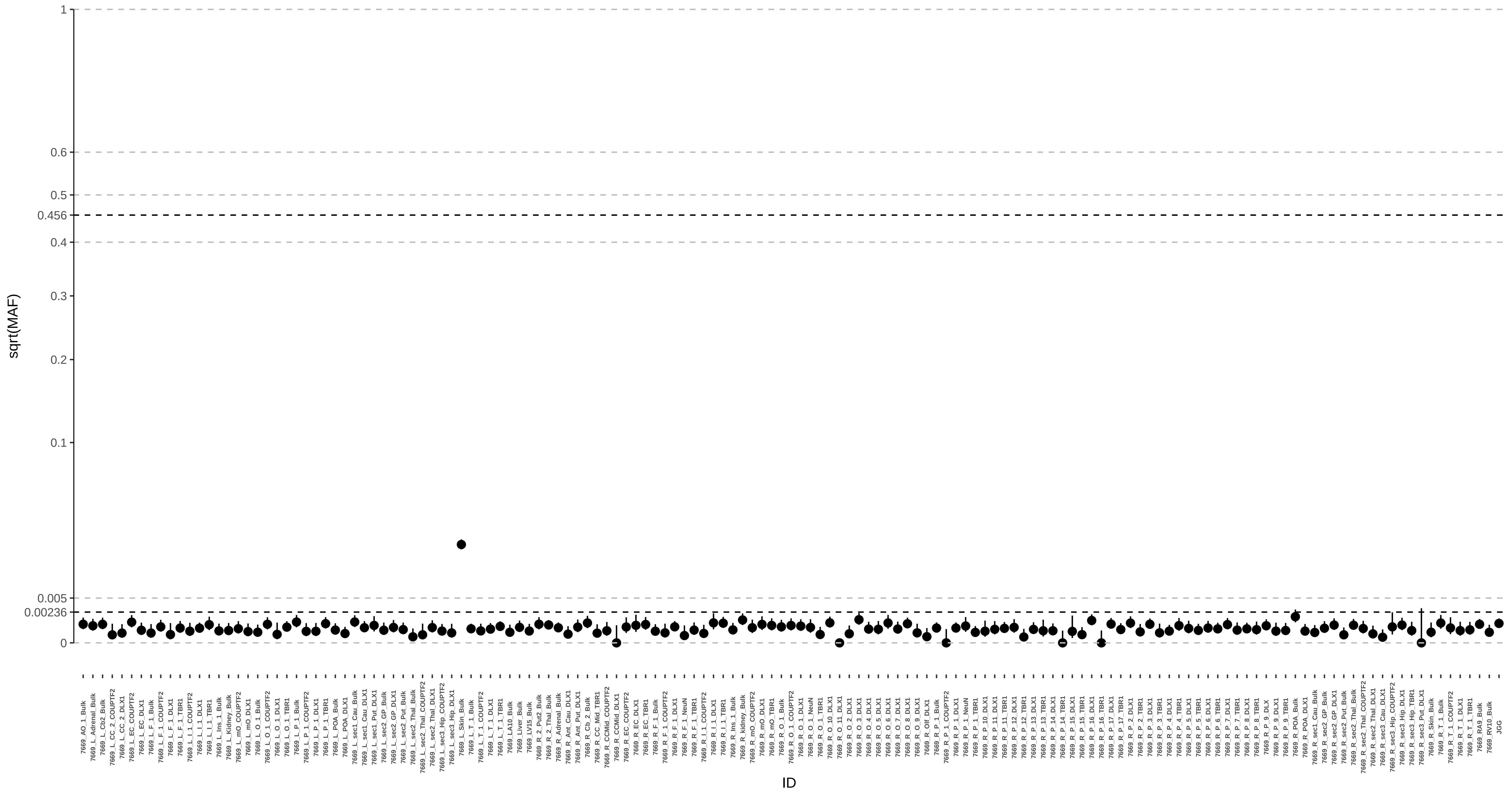

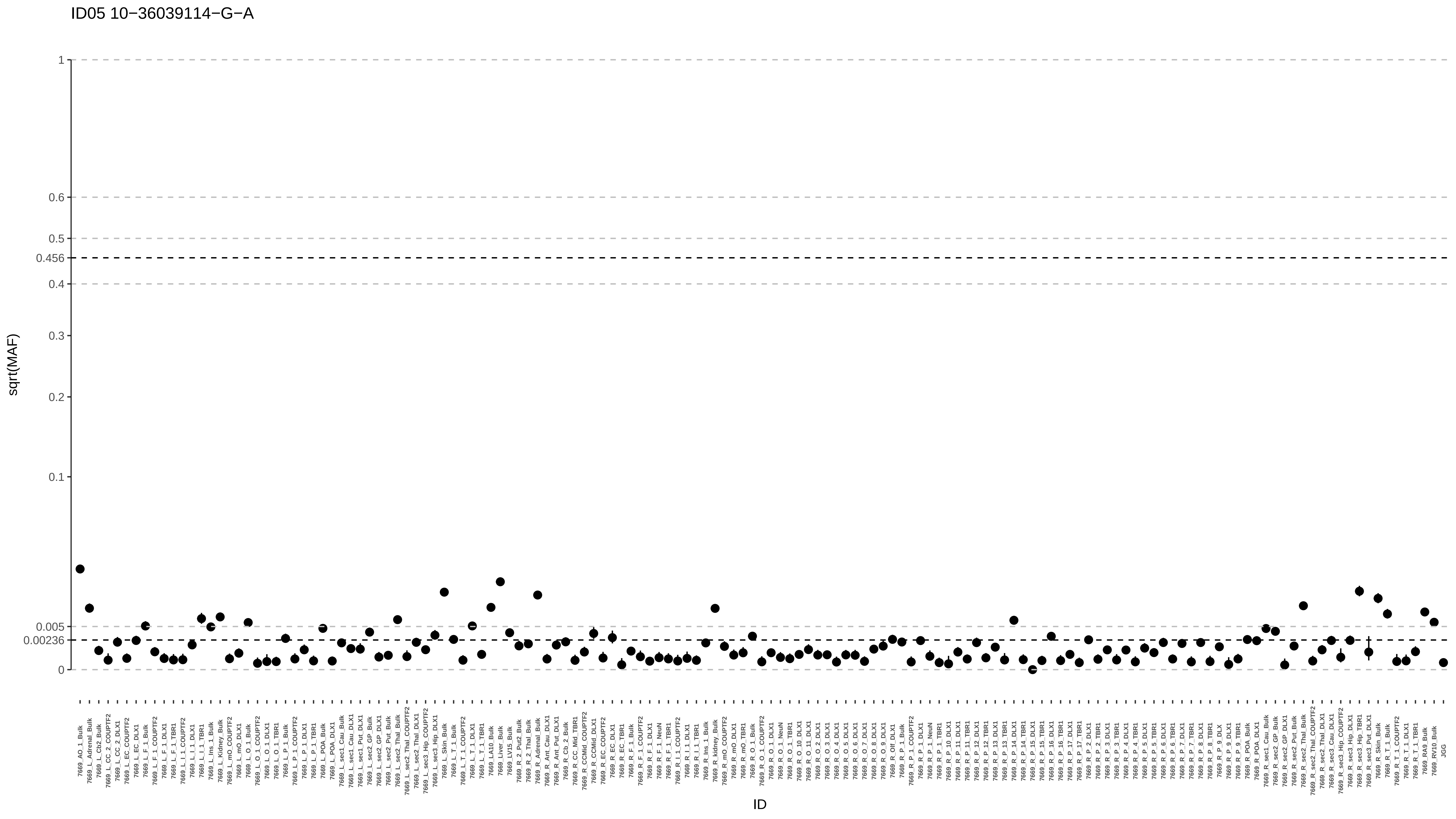
ID05 10−36039114−G−A
sqrt(MAF)
1
0.6
0.5
0.456
0.4
0.3
0.2
0.1
0.005
0.00236
0
ID

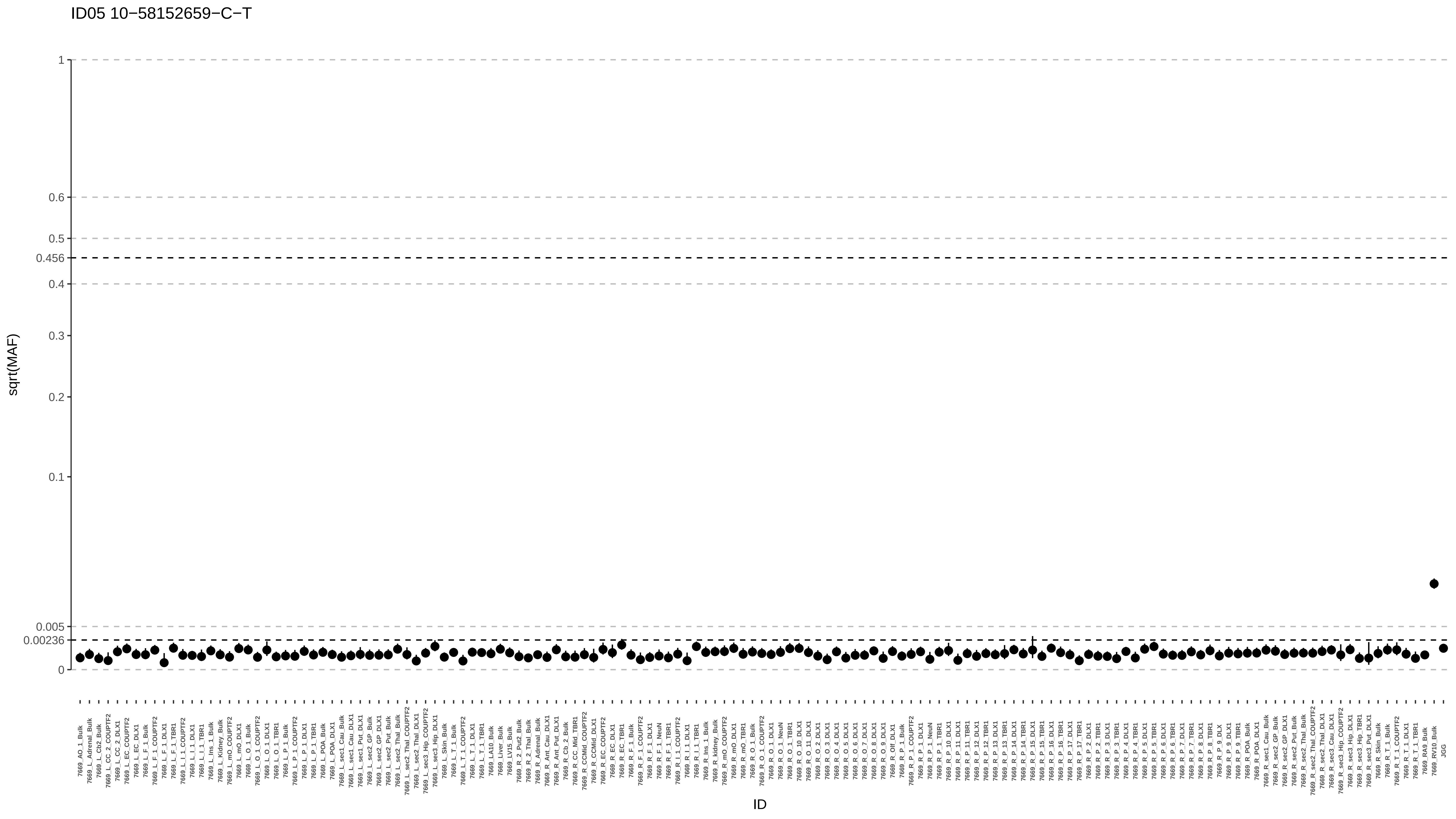
ID05 10−58152659−C−T
sqrt(MAF)
1
0.6
0.5
0.456
0.4
0.3
0.2
0.1
0.005
0.00236
0
ID

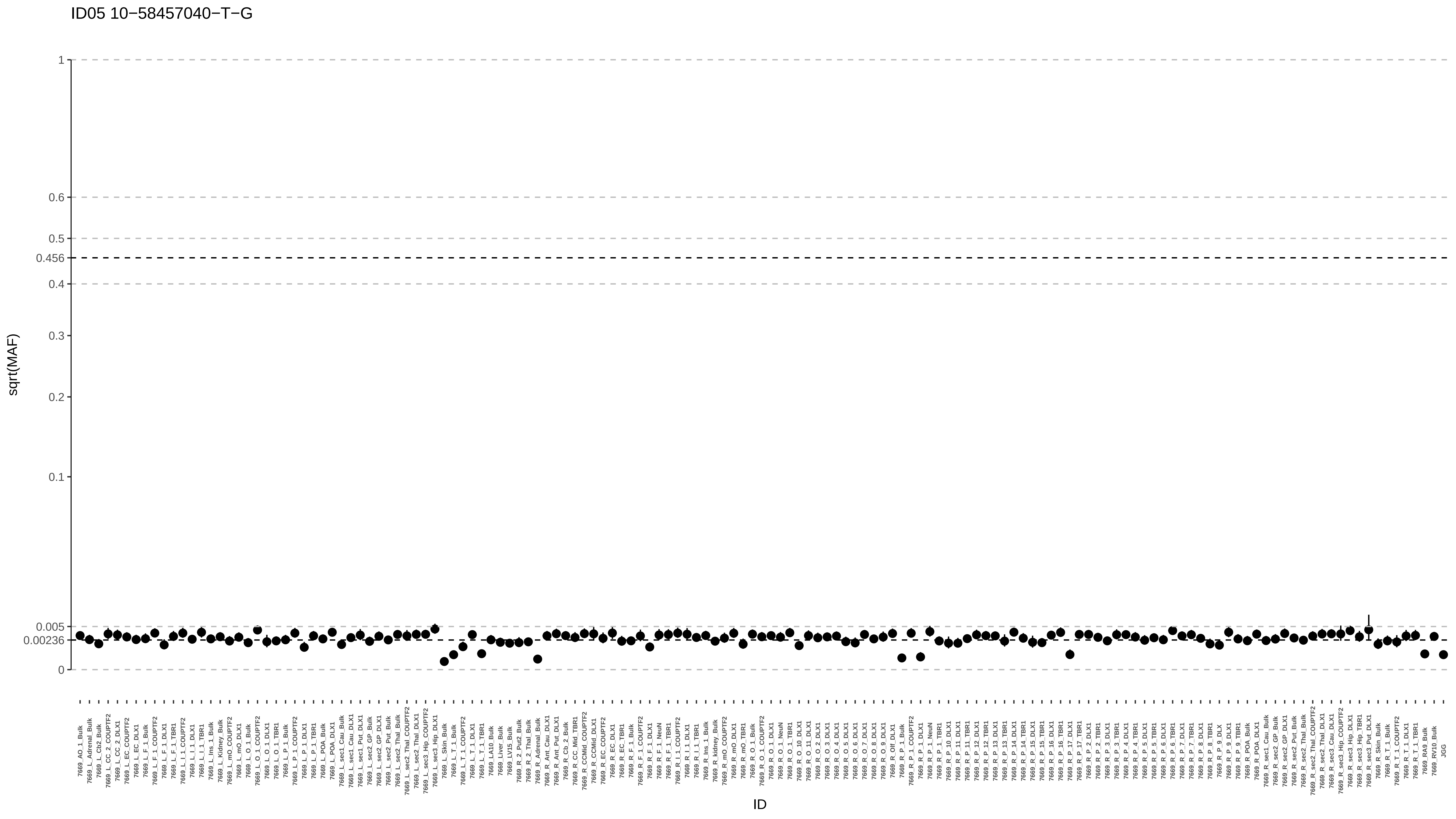
ID05 10−58457040−T−G
sqrt(MAF)
1
0.6
0.5
0.456
0.4
0.3
0.2
0.1
0.005
0.00236
0
ID

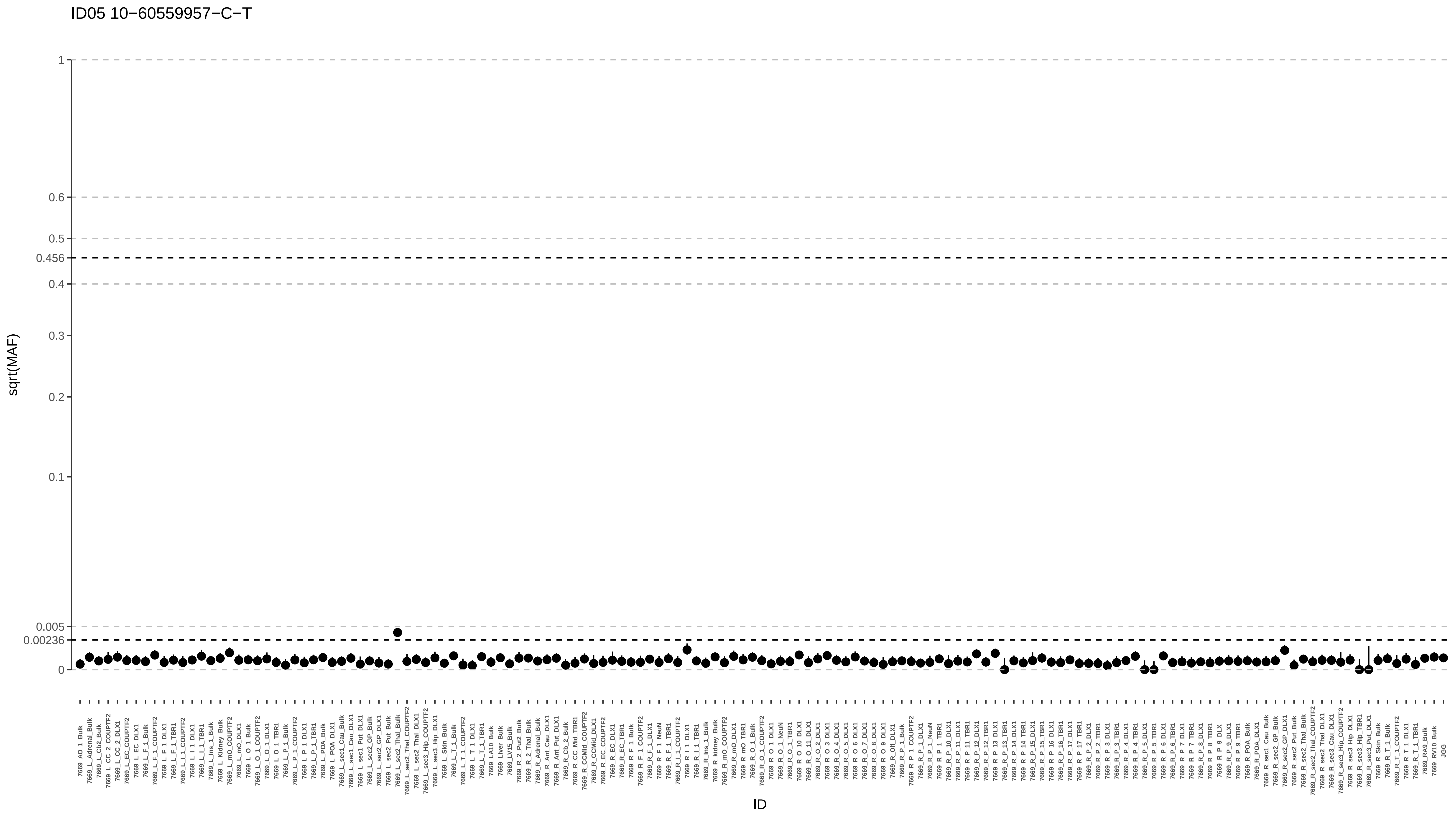
ID05 10−60559957−C−T
sqrt(MAF)
ID

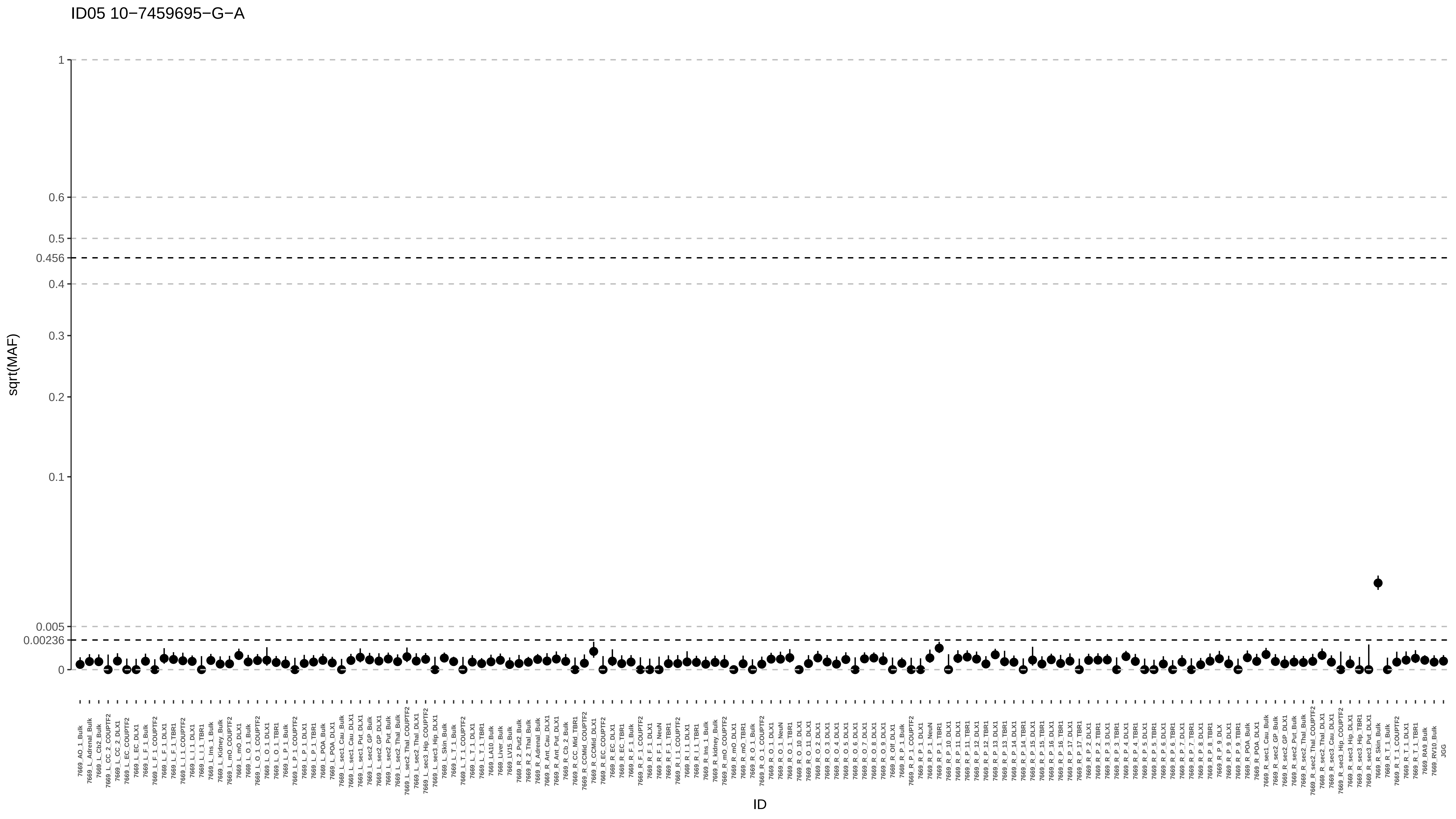
ID05 10−7459695−G−A
sqrt(MAF)
1
0.6
0.5
0.456
0.4
0.3
0.2
0.1
0.005
0.00236
0
ID

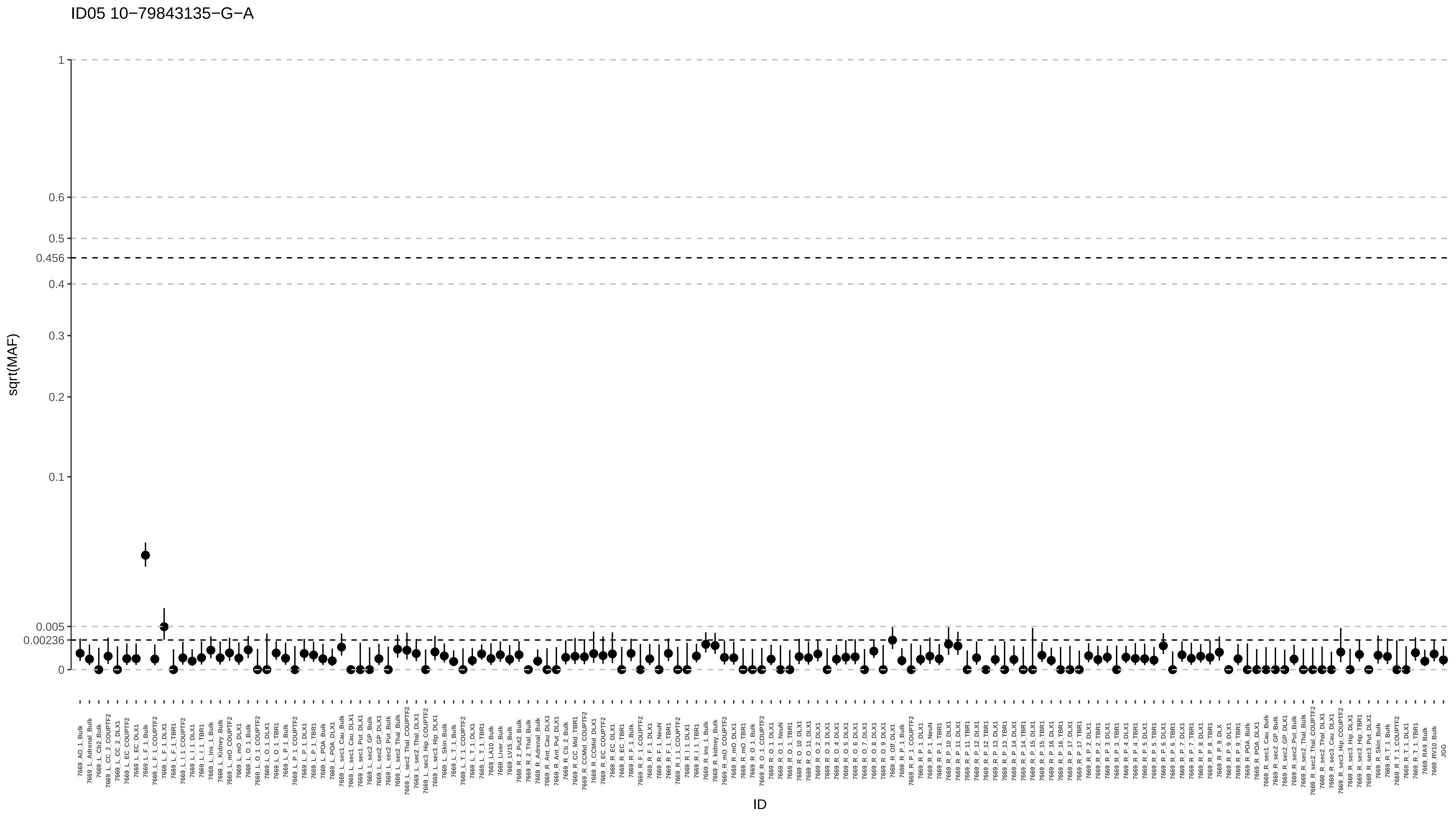
ID05 10−79843135−G−A
sqrt(MAF)
1
0.6
0.5
0.456
0.4
0.3
0.2
0.1
0.005
0.00236
0
ID

ID05 10−86000116−G−A

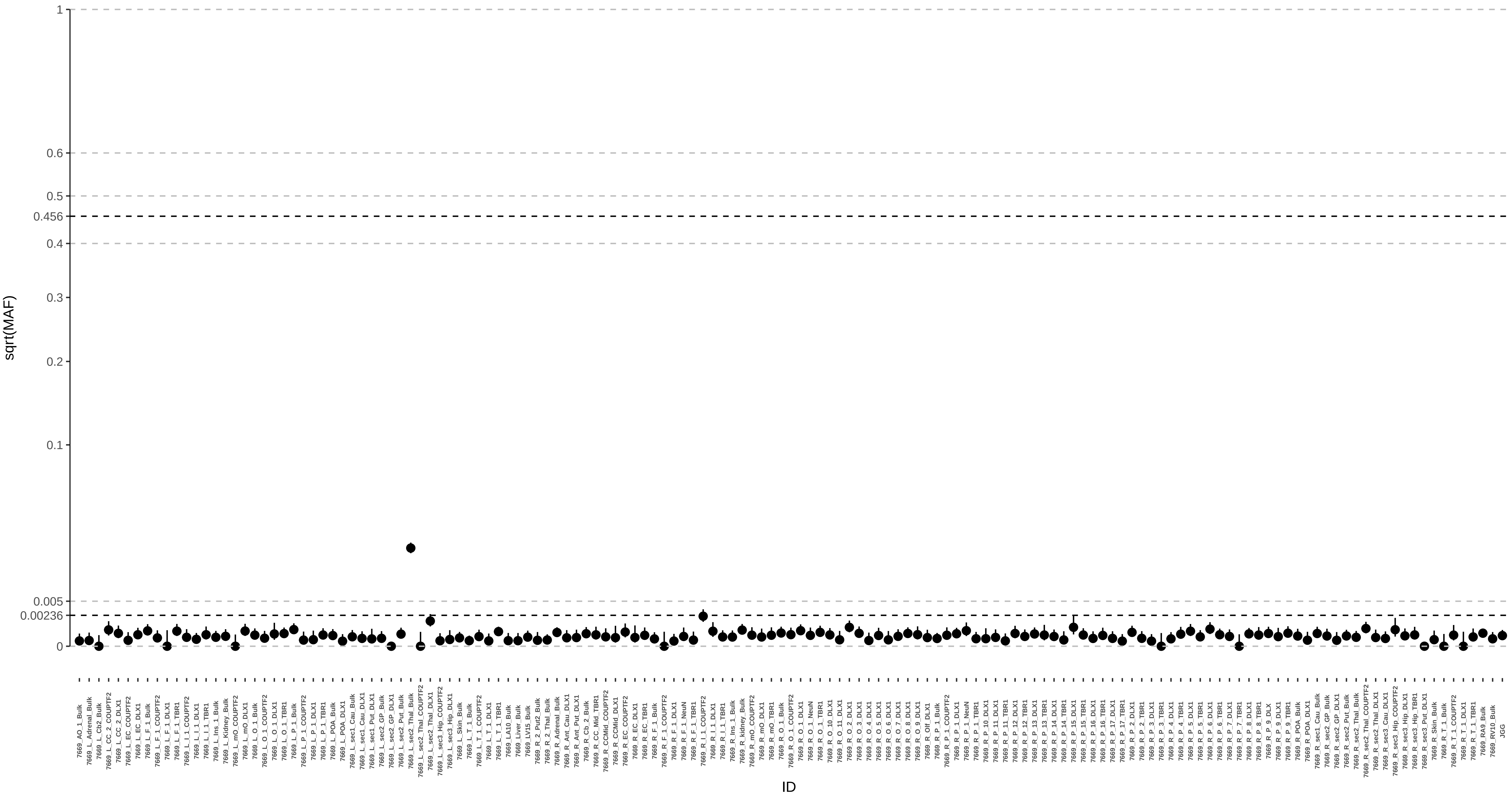

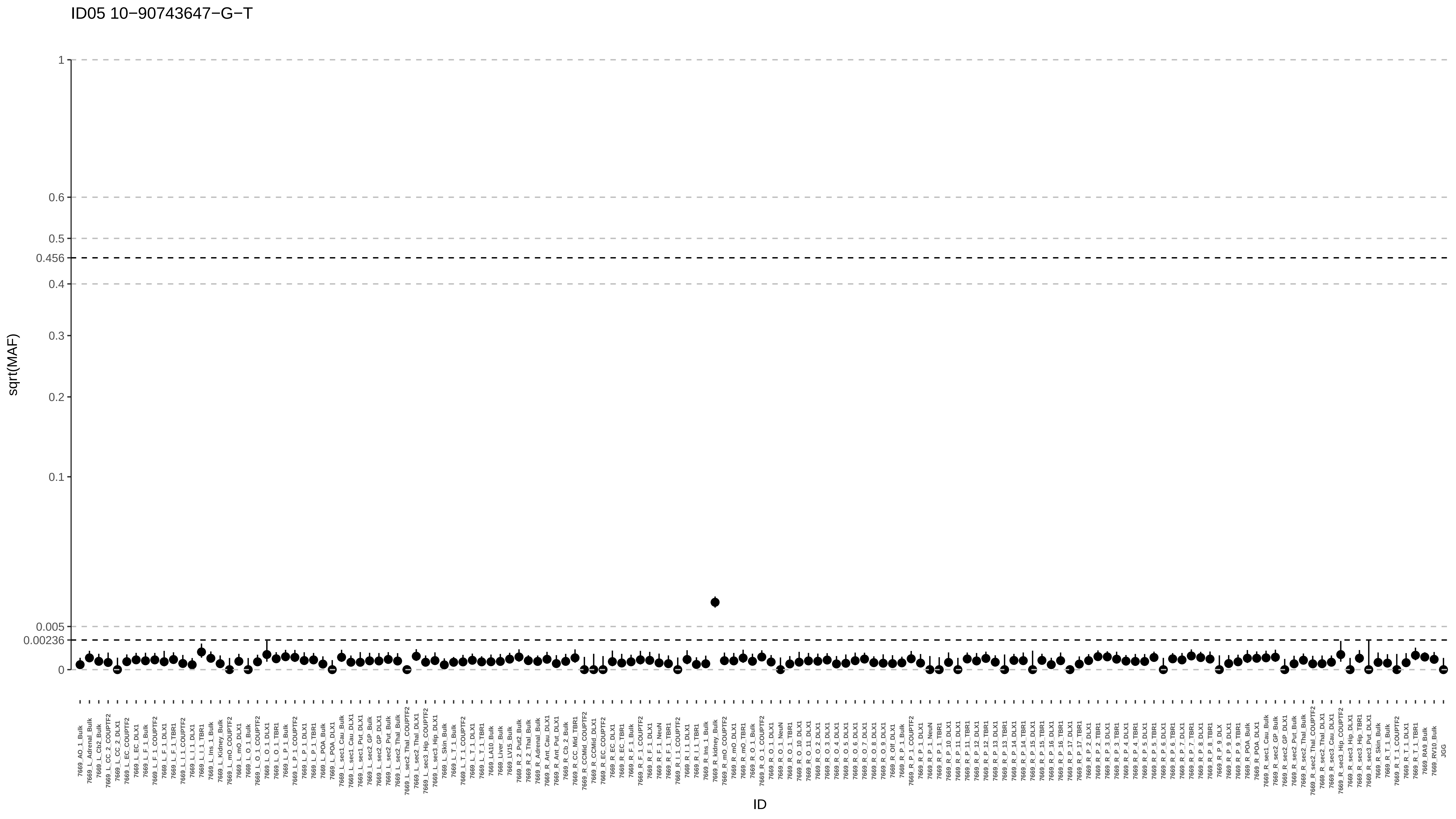

ID05 10−90743647−G−T
sqrt(MAF)
1
0.6
0.5
0.456
0.4
0.3
0.2
0.1
0.005
0.00236
0
ID

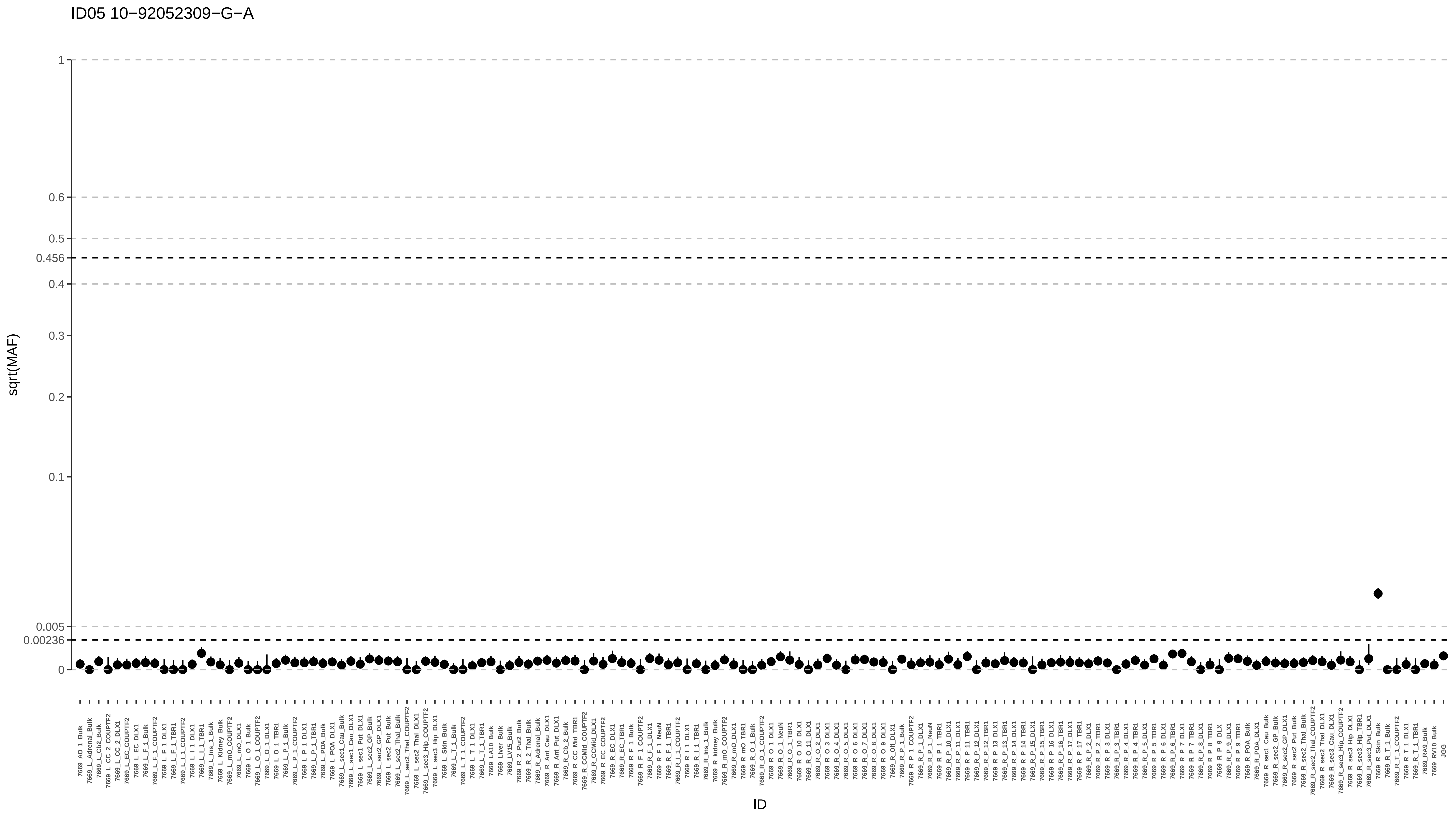
ID05 10−92052309−G−A
sqrt(MAF)
1
0.6
0.5
0.456
0.4
0.3
0.2
0.1
0.005
0.00236
0
ID

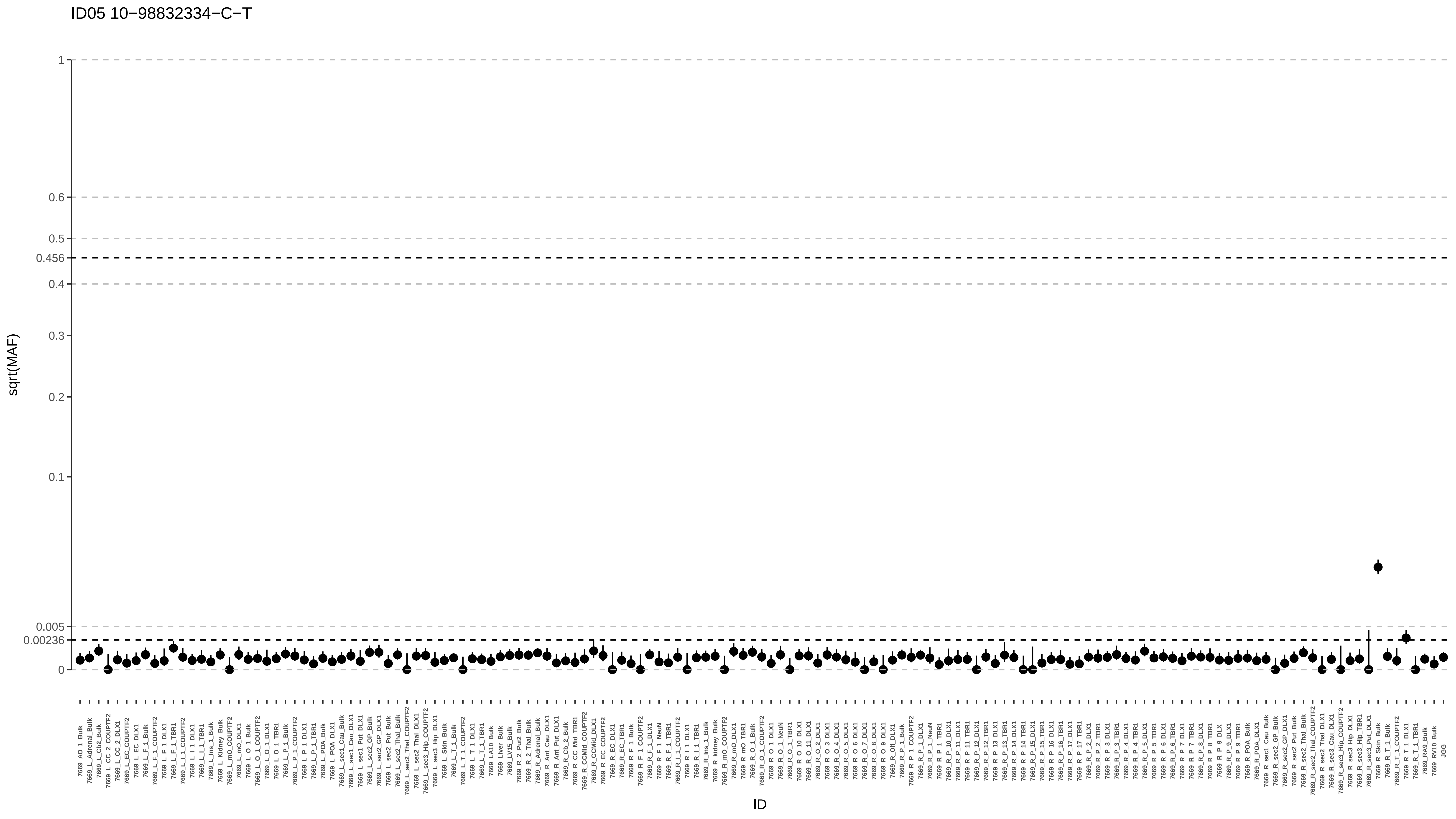

ID05 10−98832334−C−T
sqrt(MAF)
1
0.6
0.5
0.456
0.4
0.3
0.2
0.1
0.005
0.00236
0
ID

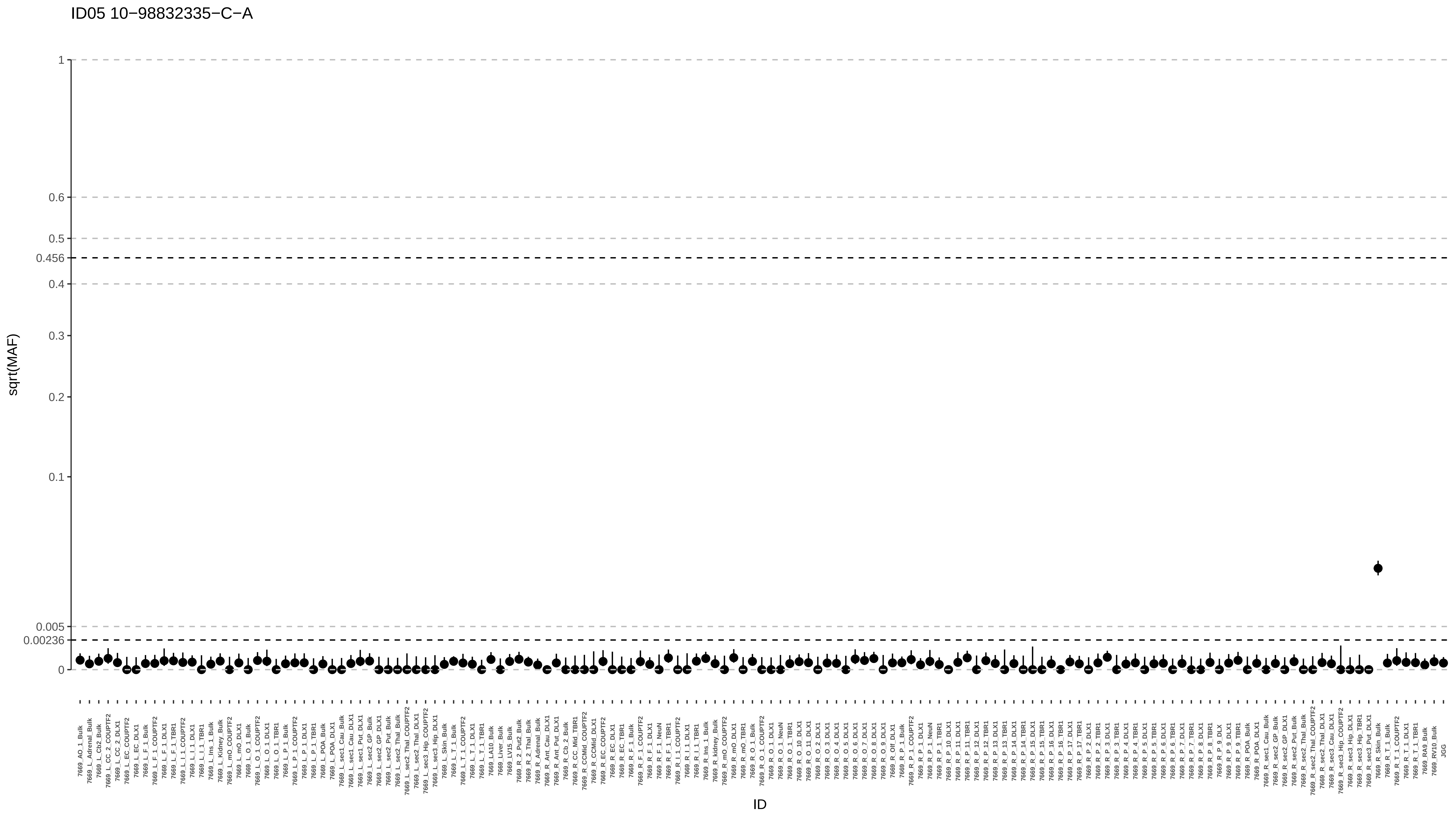
ID05 10−98832335−C−A
sqrt(MAF)
1
0.6
0.5
0.456
0.4
0.3
0.2
0.1
0.005
0.00236
0
ID

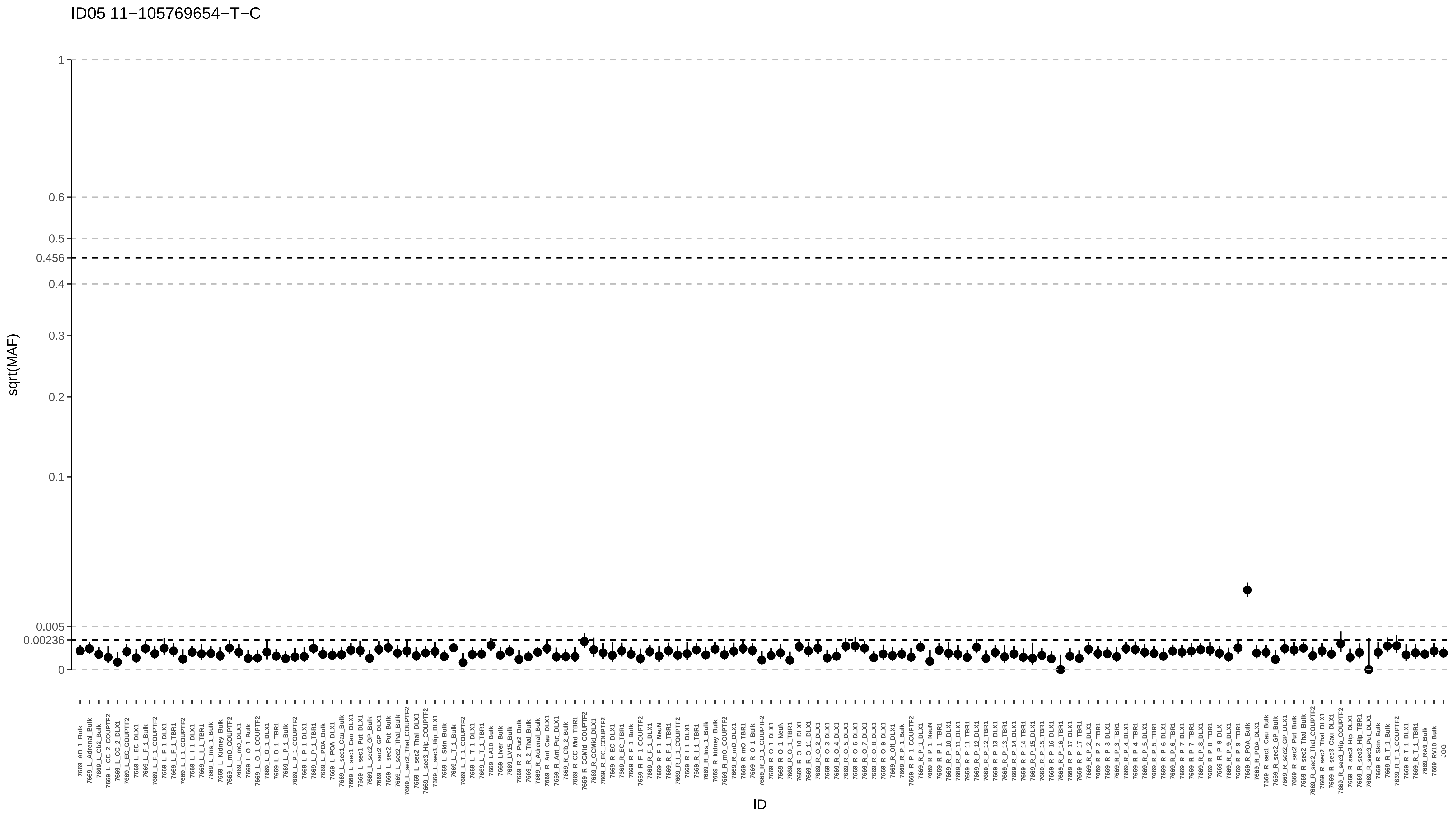

ID05 11−105769654−T−C
sqrt(MAF)
1
0.6
0.5
0.456
0.4
0.3
0.2
0.1
0.005
0.00236
0
ID

ID05 11−109546001−G−C

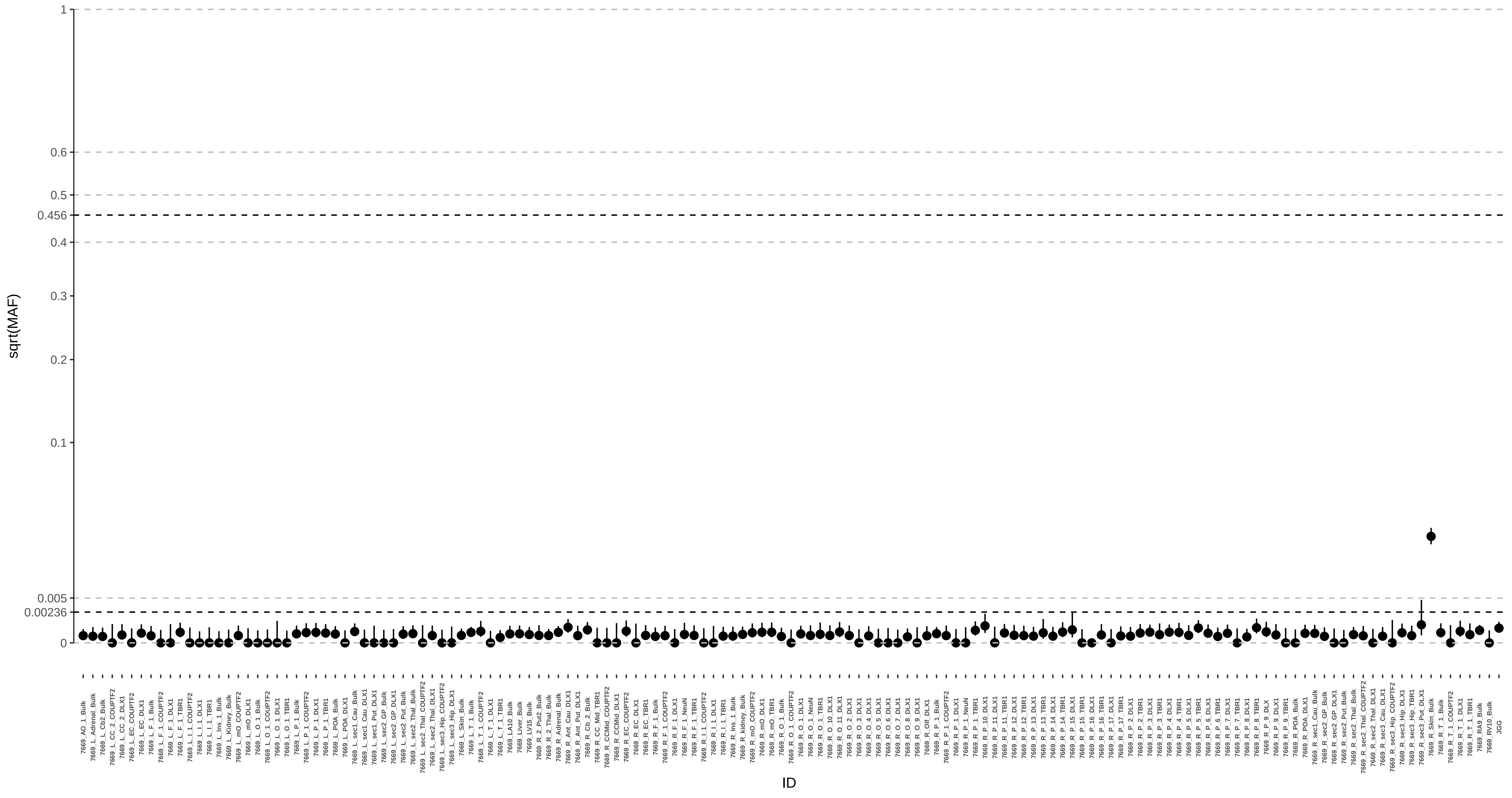

ID05 11−111580997−G−A

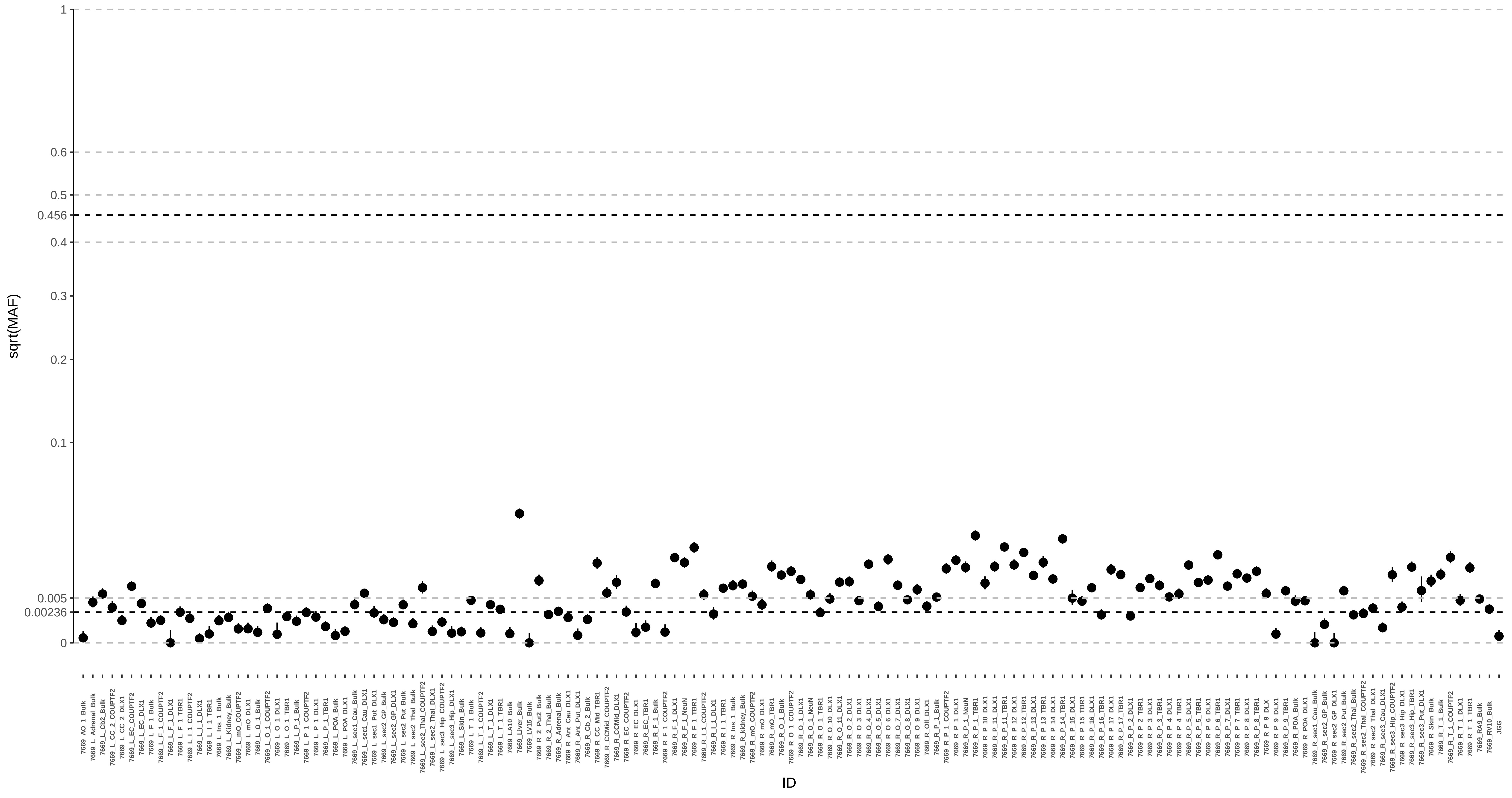

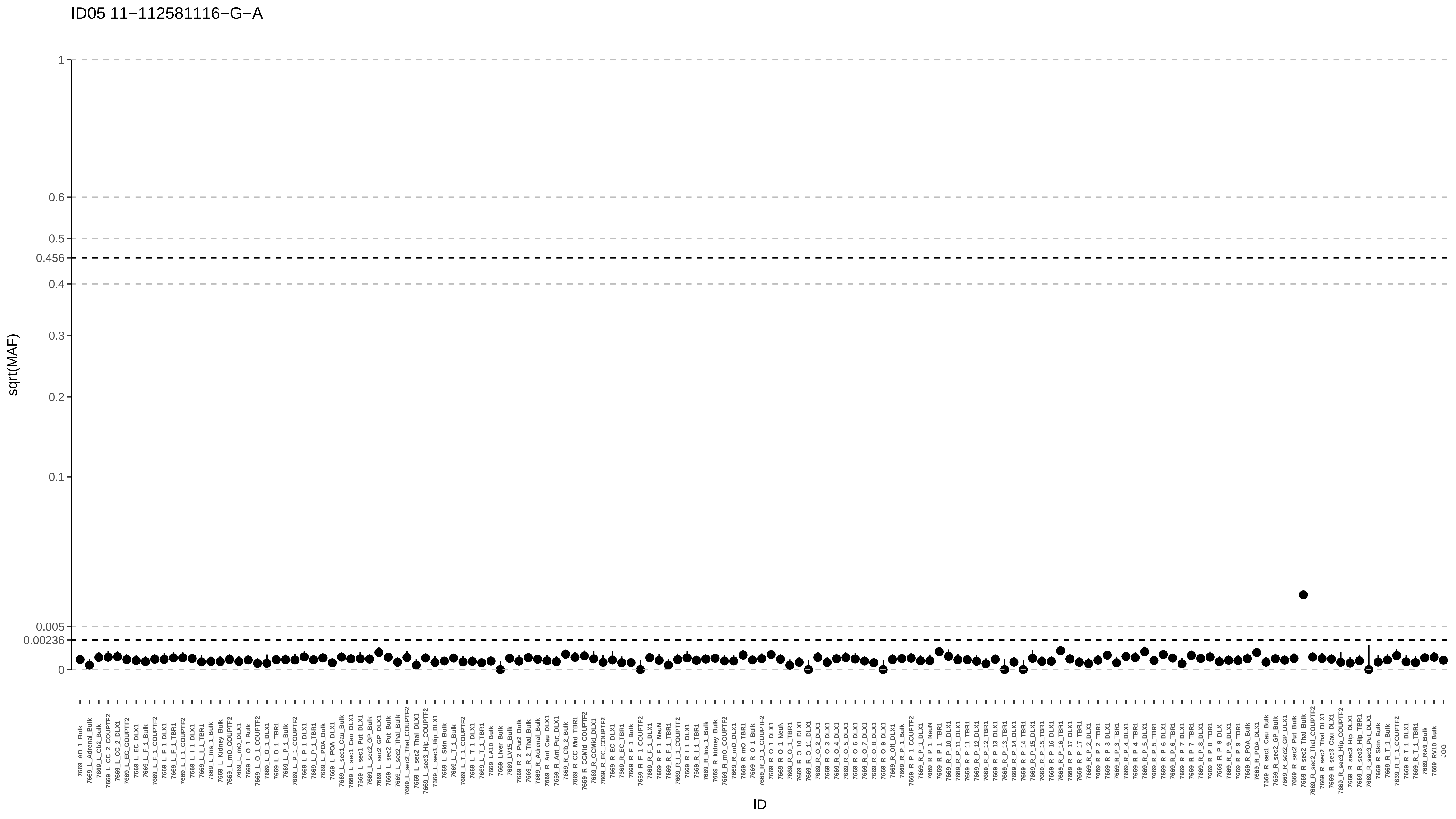
ID05 11−112581116−G−A
sqrt(MAF)
1
0.6
0.5
0.456
0.4
0.3
0.2
0.1
0.005
0.00236
0
ID

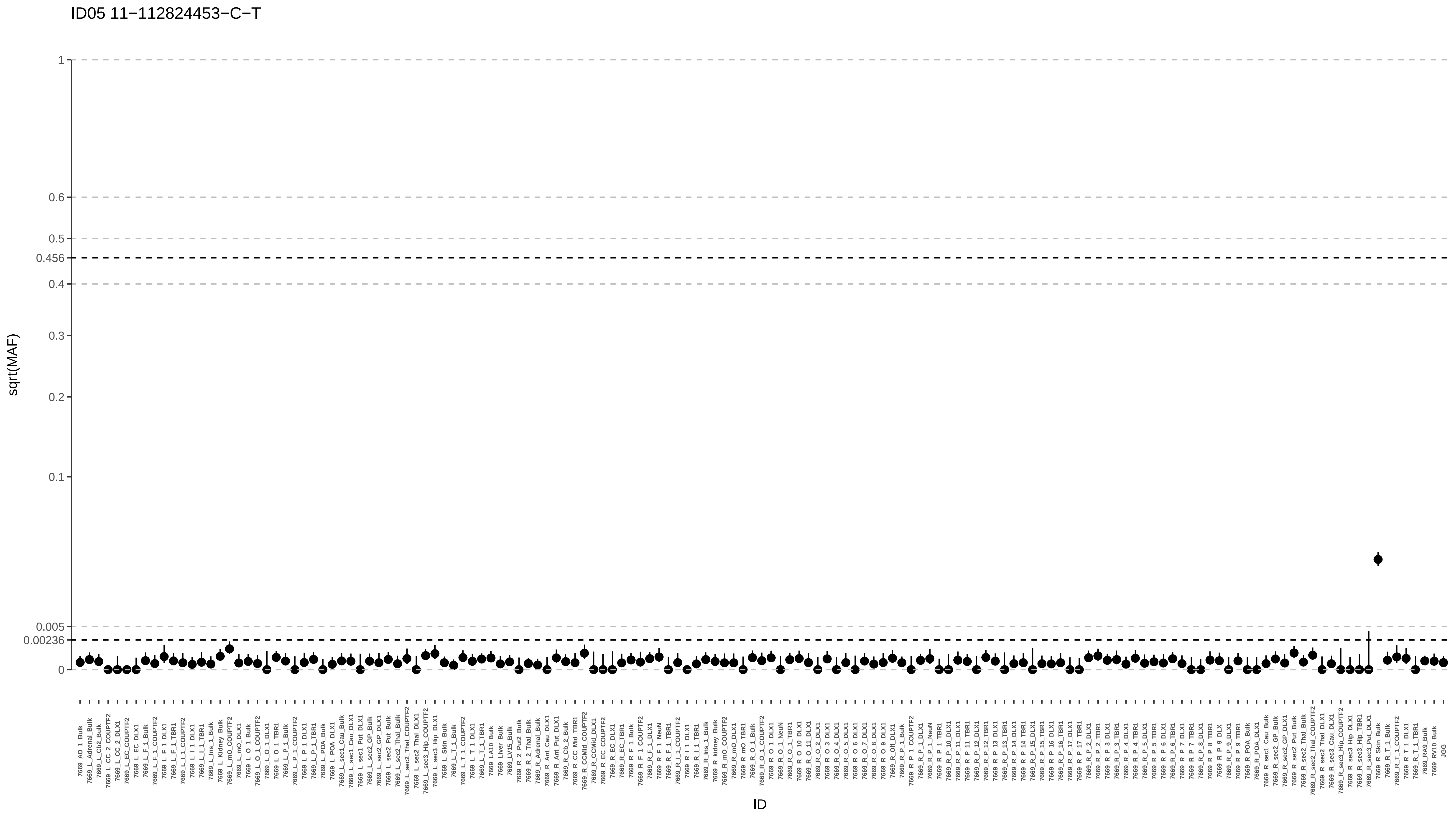

ID05 11−112824453−C−T
sqrt(MAF)
1
0.6
0.5
0.456
0.4
0.3
0.2
0.1
0.005
0.00236
0
ID

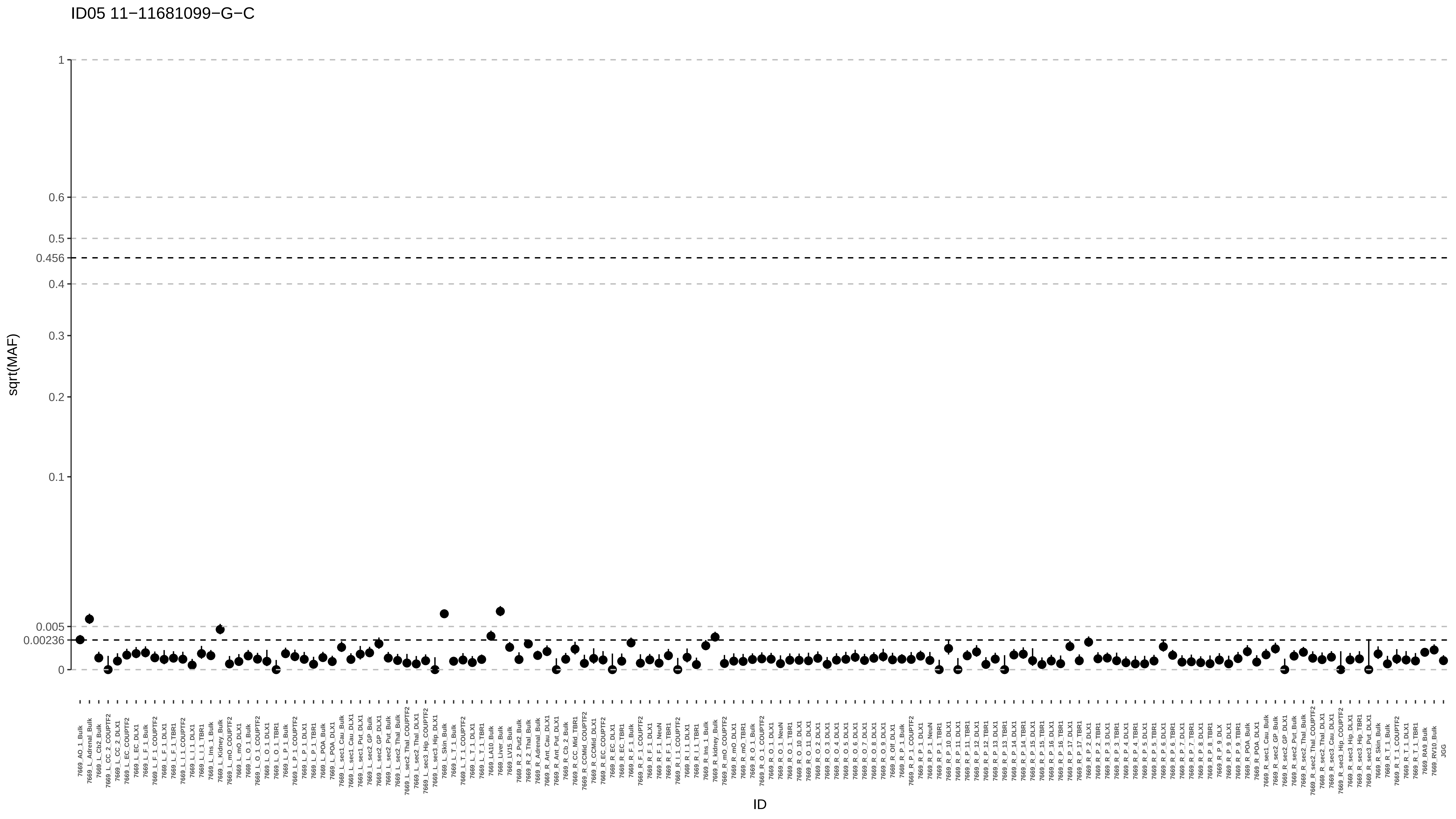

ID05 11−11681099−G−C
sqrt(MAF)
1
0.6
0.5
0.456
0.4
0.3
0.2
0.1
0.005
0.00236
0
ID

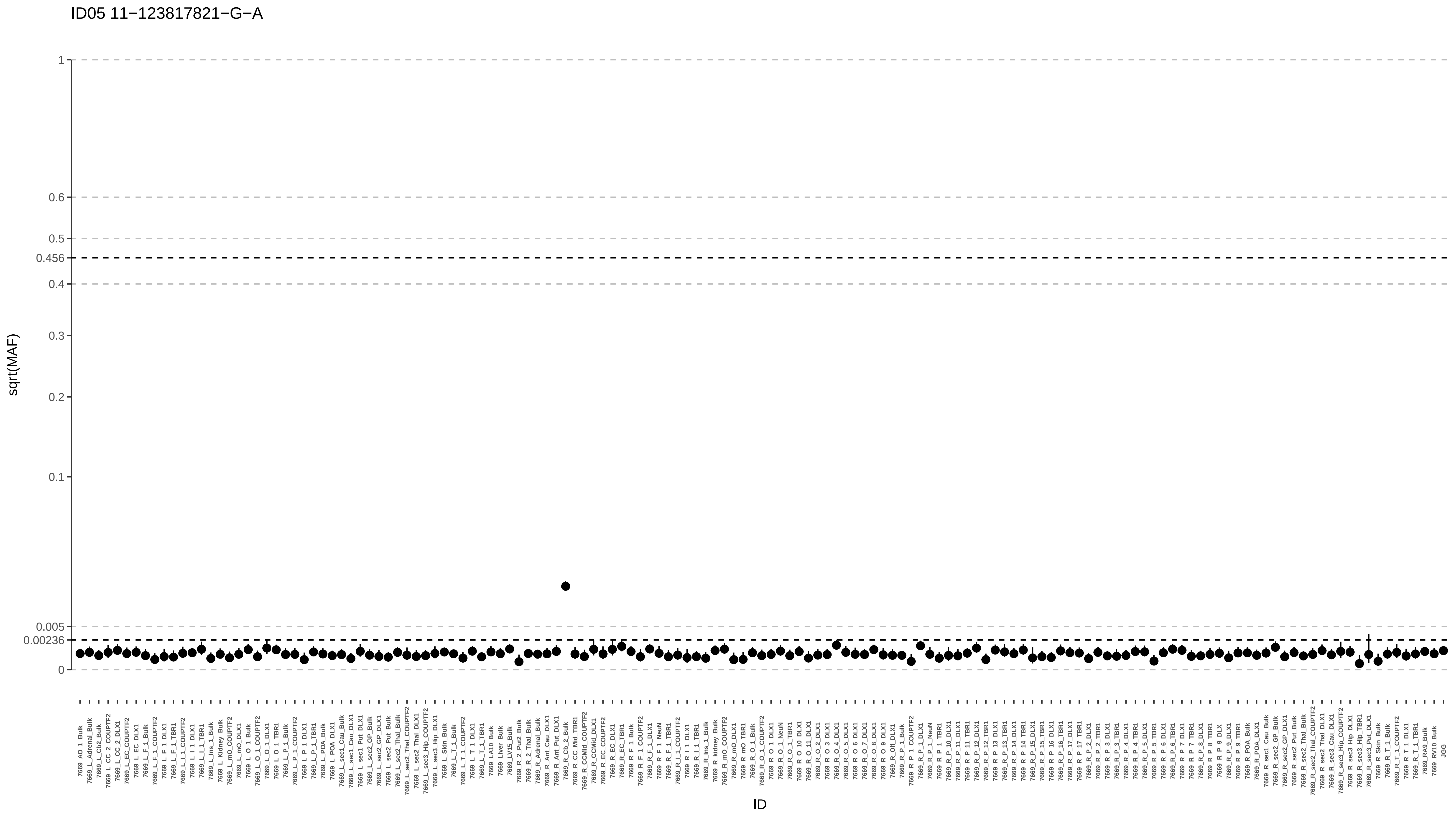
ID05 11−123817821−G−A
sqrt(MAF)
1
0.6
0.5
0.456
0.4
0.3
0.2
0.1
0.005
0.00236
0
ID
7669_AO_1_Bulk
7669_L_Adrenal_Bulk
7669_L_Cb2_Bulk
7669_L_CC_2_COUPTF2
7669_L_CC_2_DLX1
7669_L_EC_COUPTF2
7669_L_EC_DLX1
7669_L_F_1_Bulk
7669_L_F_1_COUPTF2
7669_L_F_1_DLX1
7669_L_F_1_TBR1
7669_L_I_1_COUPTF2
7669_L_I_1_DLX1
7669_L_I_1_TBR1
7669_L_Ins_1_Bulk
7669_L_Kidney_Bulk
7669_L_mO_COUPTF2
7669_L_mO_DLX1
7669_L_O_1_Bulk
7669_L_O_1_COUPTF2
7669_L_O_1_DLX1
7669_L_O_1_TBR1
7669_L_P_1_Bulk
7669_L_P_1_COUPTF2
7669_L_P_1_DLX1
7669_L_P_1_TBR1
7669_L_POA_Bulk
7669_L_POA_DLX1
7669_L_sec1_Cau_Bulk
7669_L_sec1_Cau_DLX1
7669_L_sec1_Put_DLX1
7669_L_sec2_GP_Bulk
7669_L_sec2_GP_DLX1
7669_L_sec2_Put_Bulk
7669_L_sec2_Thal_Bulk
7669_L_sec2_Thal_COUPTF2
7669_L_sec2_Thal_DLX1
7669_L_sec3_Hip_COUPTF2
7669_L_sec3_Hip_DLX1
7669_L_Skin_Bulk
7669_L_T_1_Bulk
7669_L_T_1_COUPTF2
7669_L_T_1_DLX1
7669_L_T_1_TBR1
7669_LA10_Bulk
7669_Liver_Bulk
7669_LV15_Bulk
7669_R_2_Put2_Bulk
7669_R_2_Thal_Bulk
7669_R_Adrenal_Bulk
7669_R_Ant_Cau_DLX1
7669_R_Ant_Put_DLX1
7669_R_Cb_2_Bulk
7669_R_CC_Mid_TBR1
7669_R_CCMid_COUPTF2
7669_R_CCMid_DLX1
7669_R_EC_COUPTF2
7669_R_EC_DLX1
7669_R_EC_TBR1
7669_R_F_1_Bulk
7669_R_F_1_COUPTF2
7669_R_F_1_DLX1
7669_R_F_1_NeuN
7669_R_F_1_TBR1
7669_R_I_1_COUPTF2
7669_R_I_1_DLX1
7669_R_I_1_TBR1
7669_R_Ins_1_Bulk
7669_R_kidney_Bulk
7669_R_mO_COUPTF2
7669_R_mO_DLX1
7669_R_mO_TBR1
7669_R_O_1_Bulk
7669_R_O_1_COUPTF2
7669_R_O_1_DLX1
7669_R_O_1_NeuN
7669_R_O_1_TBR1
7669_R_O_10_DLX1
7669_R_O_11_DLX1
7669_R_O_2_DLX1
7669_R_O_3_DLX1
7669_R_O_4_DLX1
7669_R_O_5_DLX1
7669_R_O_6_DLX1
7669_R_O_7_DLX1
7669_R_O_8_DLX1
7669_R_O_9_DLX1
7669_R_Olf_DLX1
7669_R_P_1_Bulk
7669_R_P_1_COUPTF2
7669_R_P_1_DLX1
7669_R_P_1_NeuN
7669_R_P_1_TBR1
7669_R_P_10_DLX1
7669_R_P_11_DLX1
7669_R_P_11_TBR1
7669_R_P_12_DLX1
7669_R_P_12_TBR1
7669_R_P_13_DLX1
7669_R_P_13_TBR1
7669_R_P_14_DLX1
7669_R_P_14_TBR1
7669_R_P_15_DLX1
7669_R_P_15_TBR1
7669_R_P_16_DLX1
7669_R_P_16_TBR1
7669_R_P_17_DLX1
7669_R_P_17_TBR1
7669_R_P_2_DLX1
7669_R_P_2_TBR1
7669_R_P_3_DLX1
7669_R_P_3_TBR1
7669_R_P_4_DLX1
7669_R_P_4_TBR1
7669_R_P_5_DLX1
7669_R_P_5_TBR1
7669_R_P_6_DLX1
7669_R_P_6_TBR1
7669_R_P_7_DLX1
7669_R_P_7_TBR1
7669_R_P_8_DLX1
7669_R_P_8_TBR1
7669_R_P_9_DLX
7669_R_P_9_DLX1
7669_R_P_9_TBR1
7669_R_POA_Bulk
7669_R_POA_DLX1
7669_R_sec1_Cau_Bulk
7669_R_sec2_GP_Bulk
7669_R_sec2_GP_DLX1
7669_R_sec2_Put_Bulk
7669_R_sec2_Thal_Bulk
7669_R_sec2_Thal_COUPTF2
7669_R_sec2_Thal_DLX1
7669_R_sec3_Cau_DLX1
7669_R_sec3_Hip_COUPTF2
7669_R_sec3_Hip_DLX1
7669_R_sec3_Hip_TBR1
7669_R_sec3_Put_DLX1
7669_R_Skin_Bulk
7669_R_T_1_Bulk
7669_R_T_1_COUPTF2
7669_R_T_1_DLX1
7669_R_T_1_TBR1
7669_RA9_Bulk
7669_RV10_Bulk
JGG

ID05 11−125271170−G−C

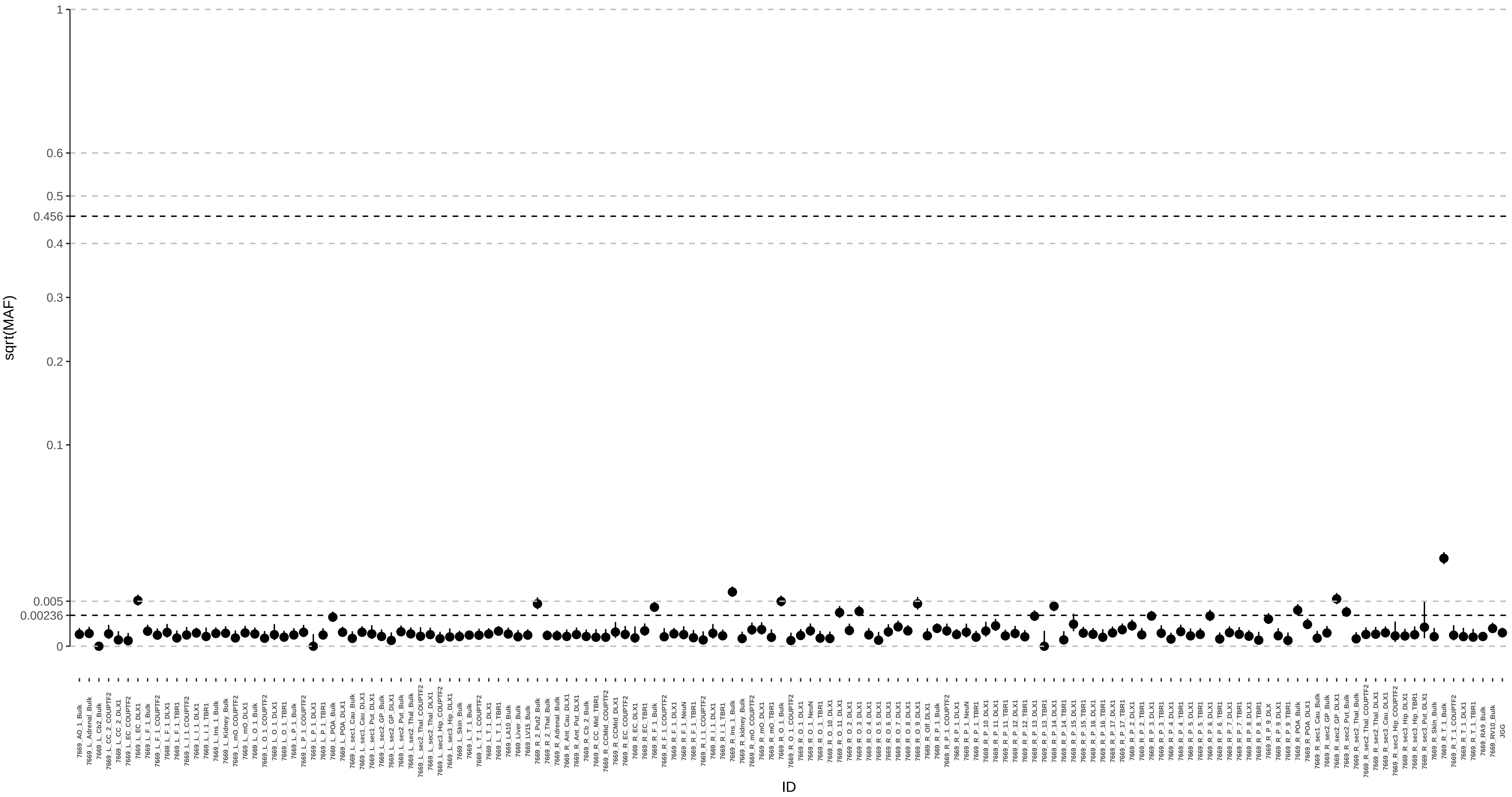

ID05 11−127350815−C−T

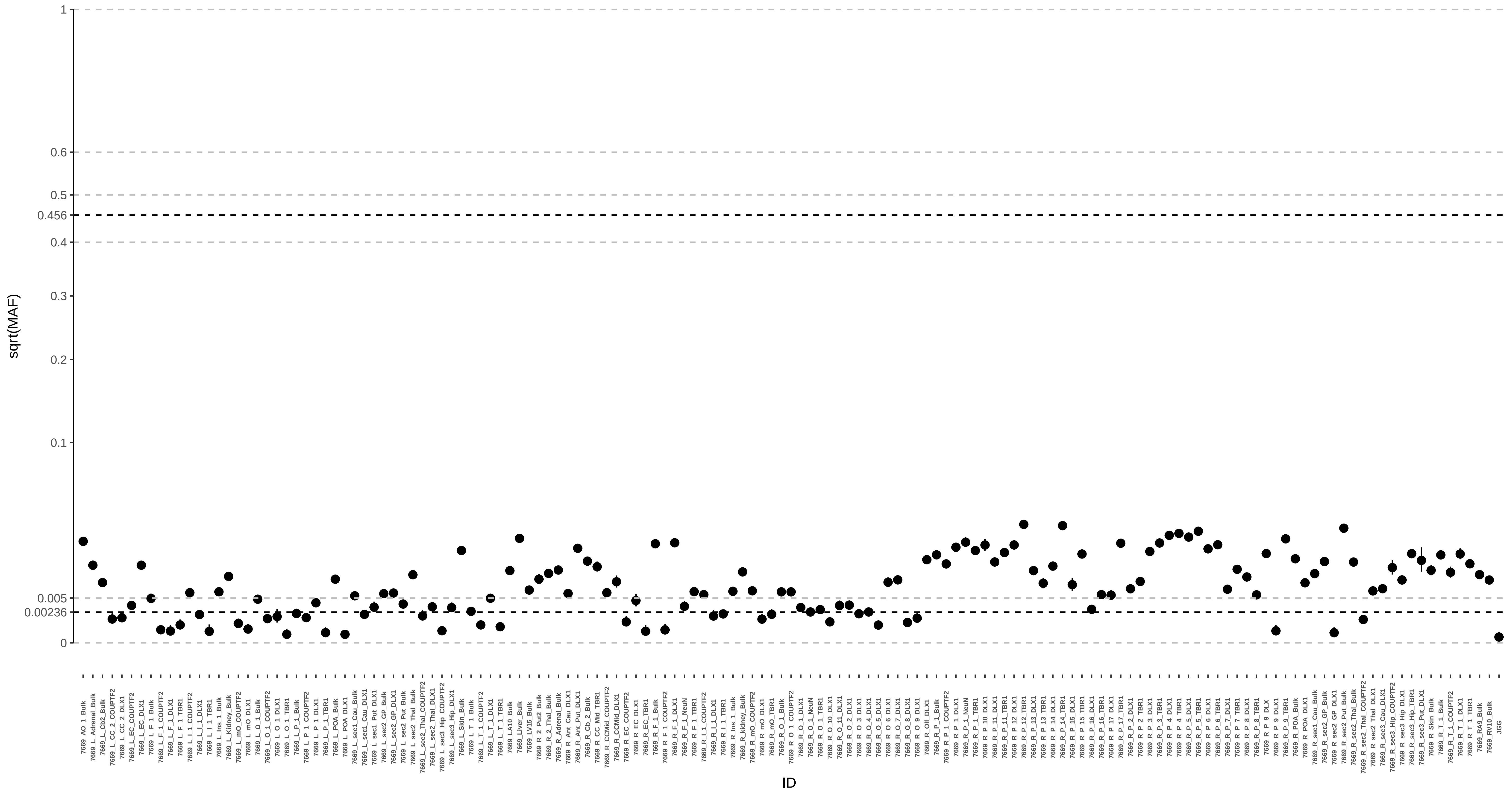

ID05 11−133014880−G−C

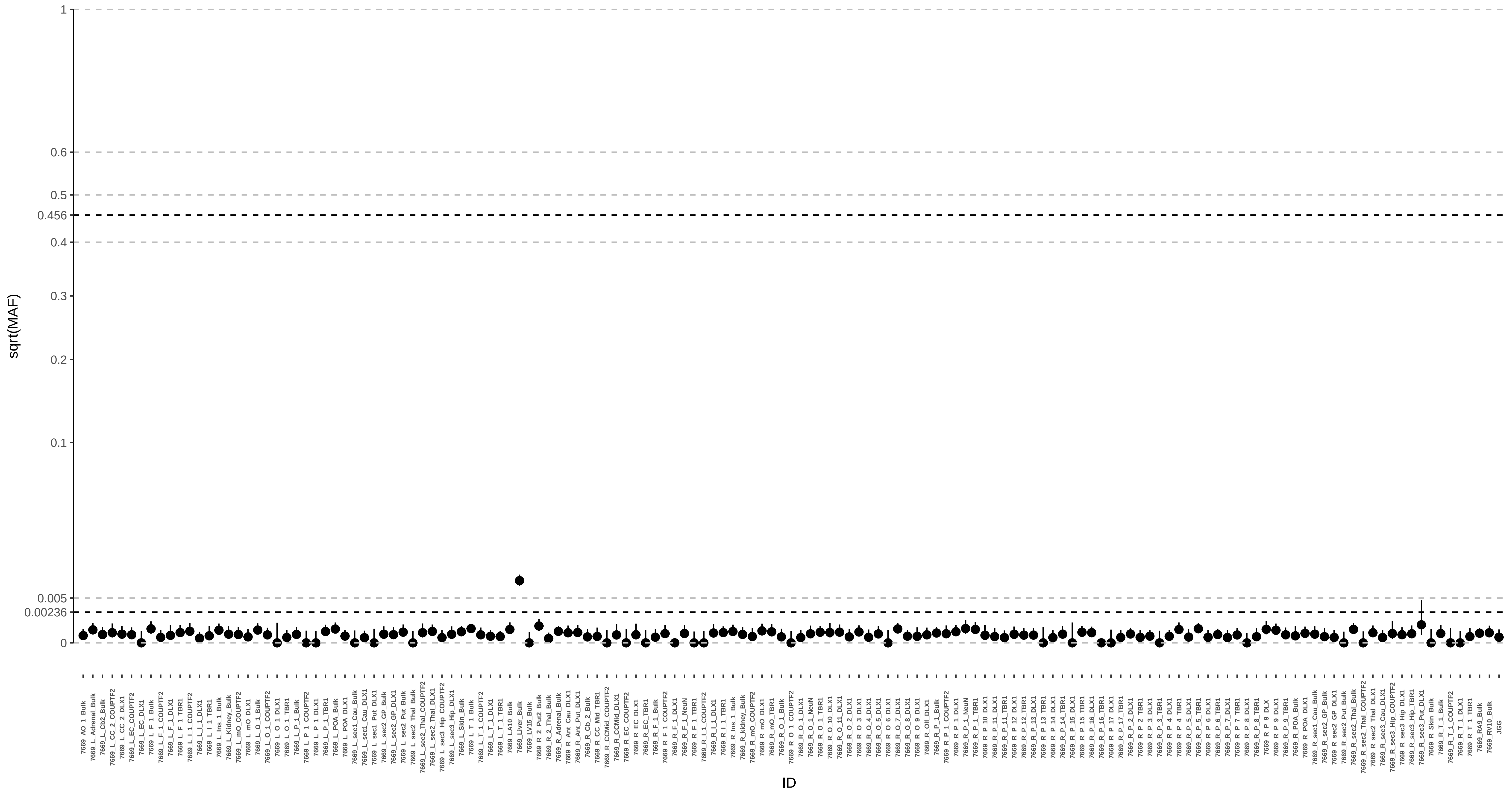

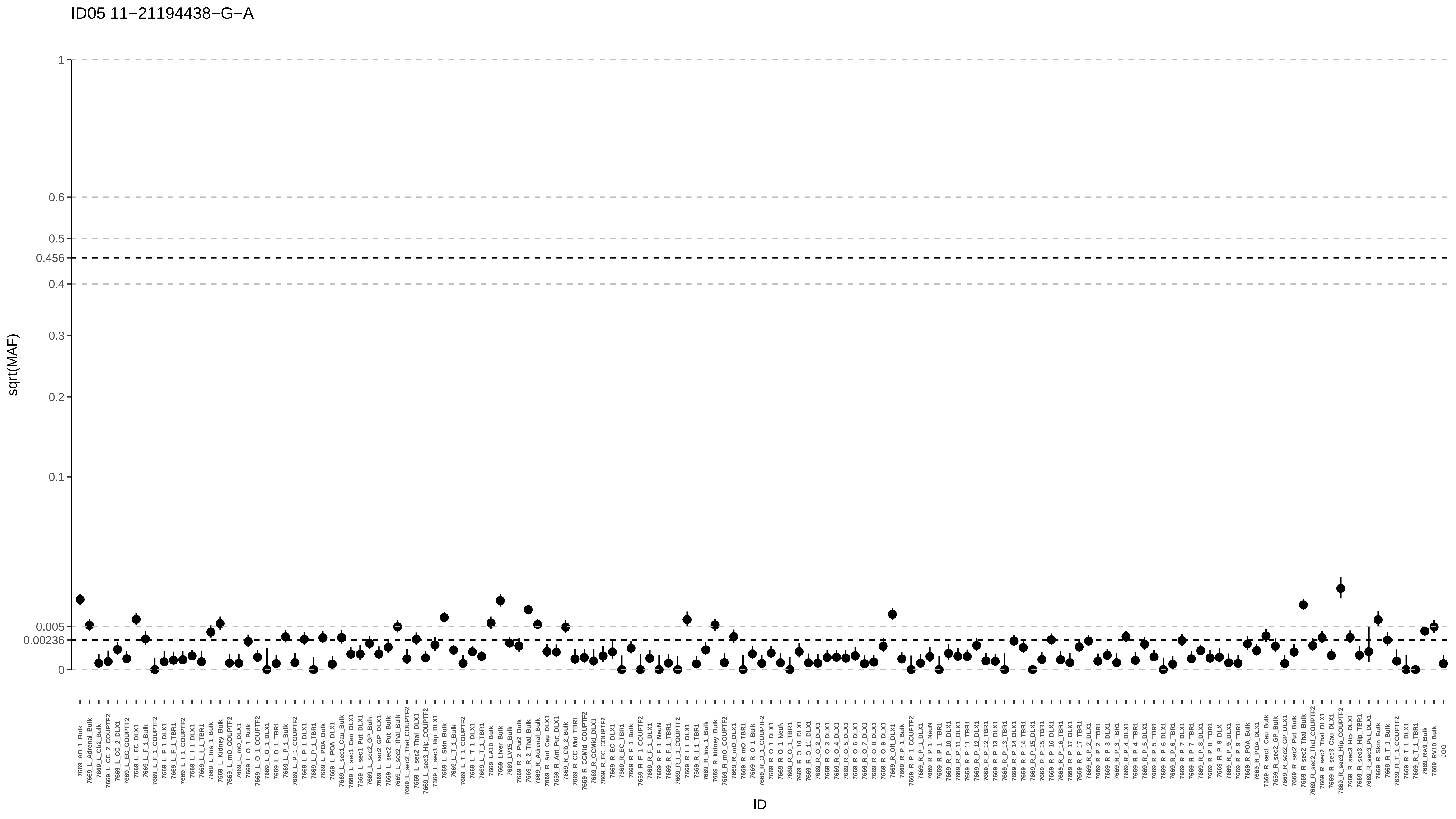
ID05 11−21194438−G−A
sqrt(MAF)
1
0.6
0.5
0.456
0.4
0.3
0.2
0.1
0.005
0.00236
0
ID

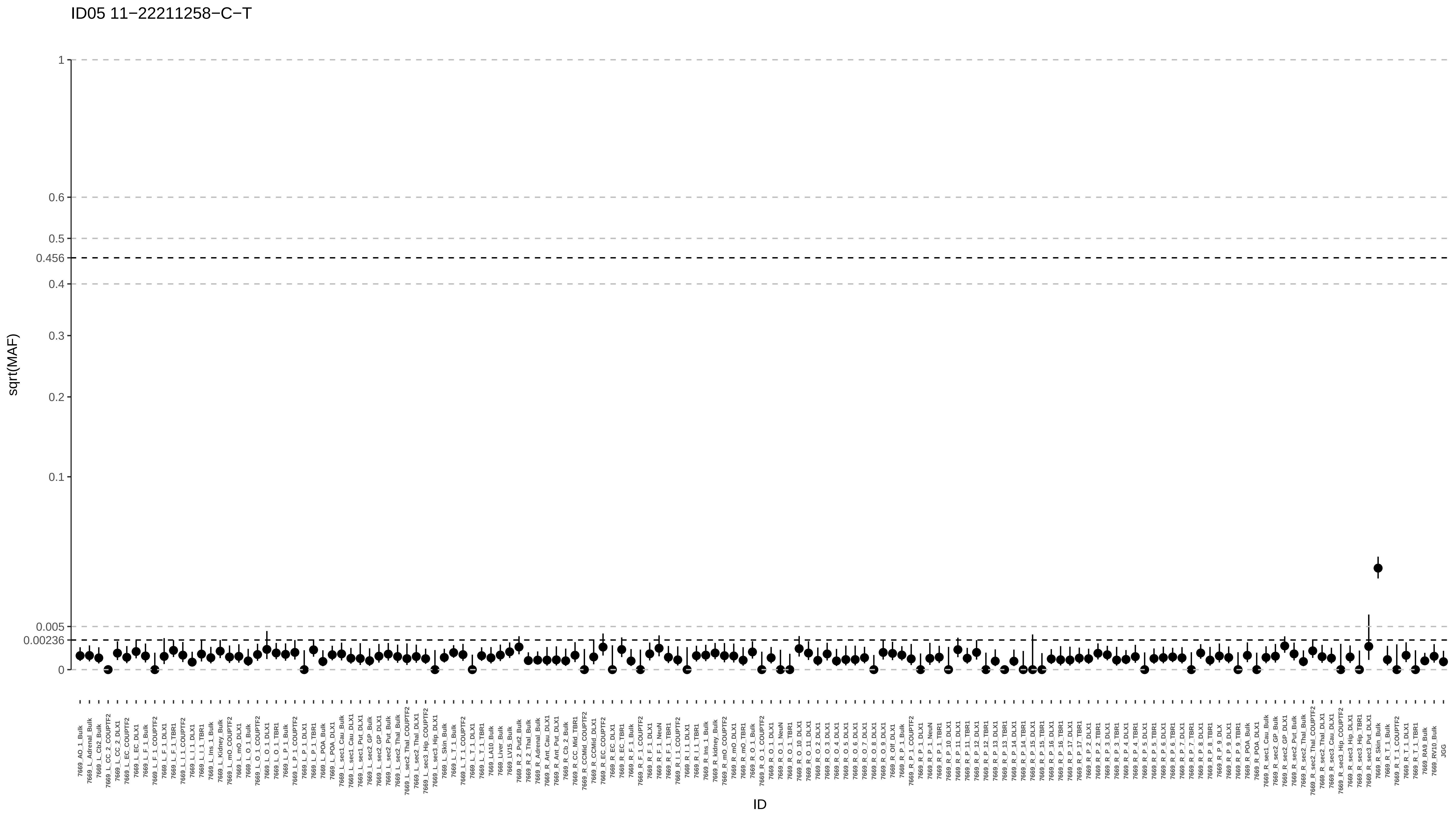

ID05 11−22211258−C−T
sqrt(MAF)
ID

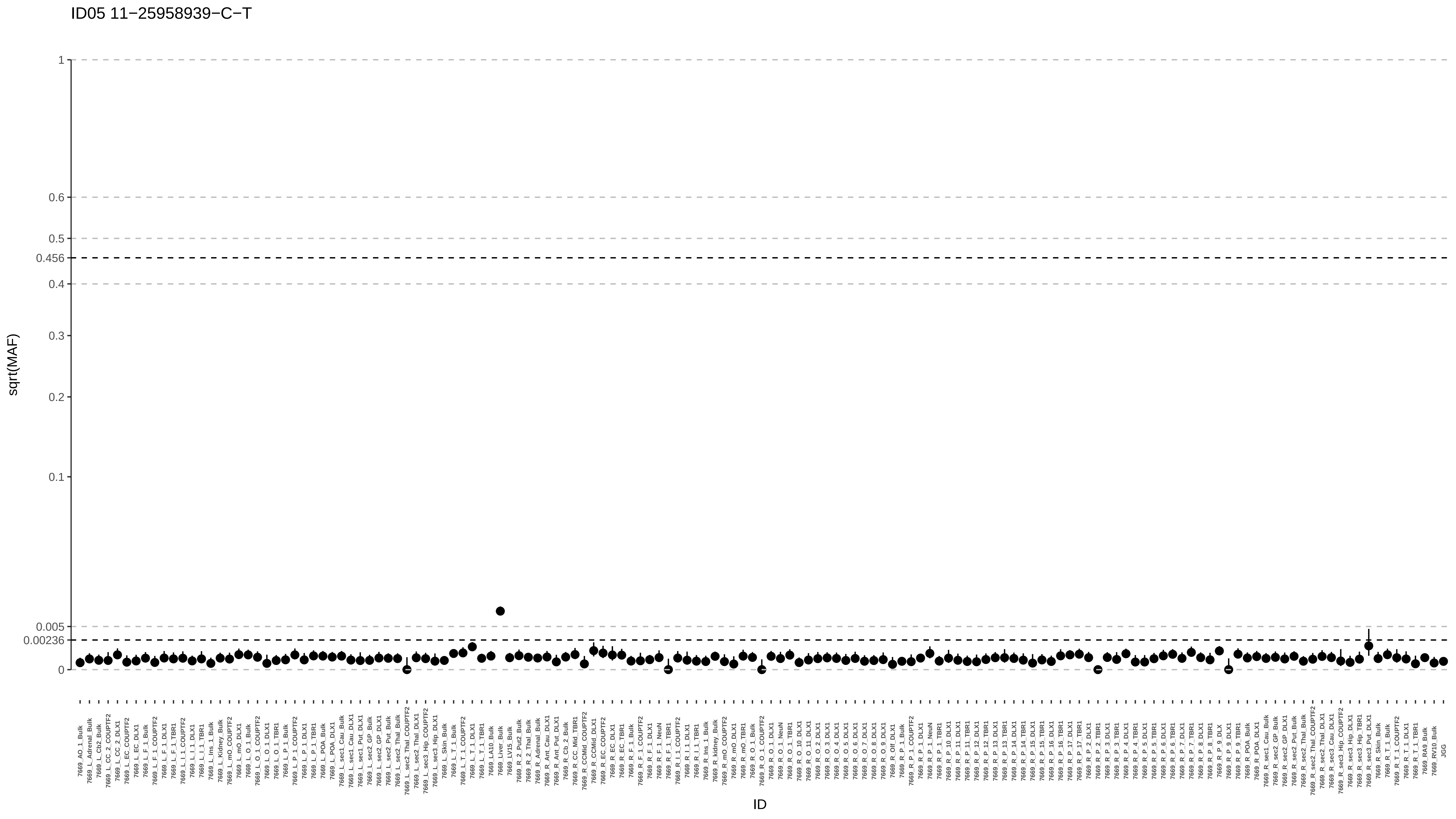
ID05 11−25958939−C−T
sqrt(MAF)
ID

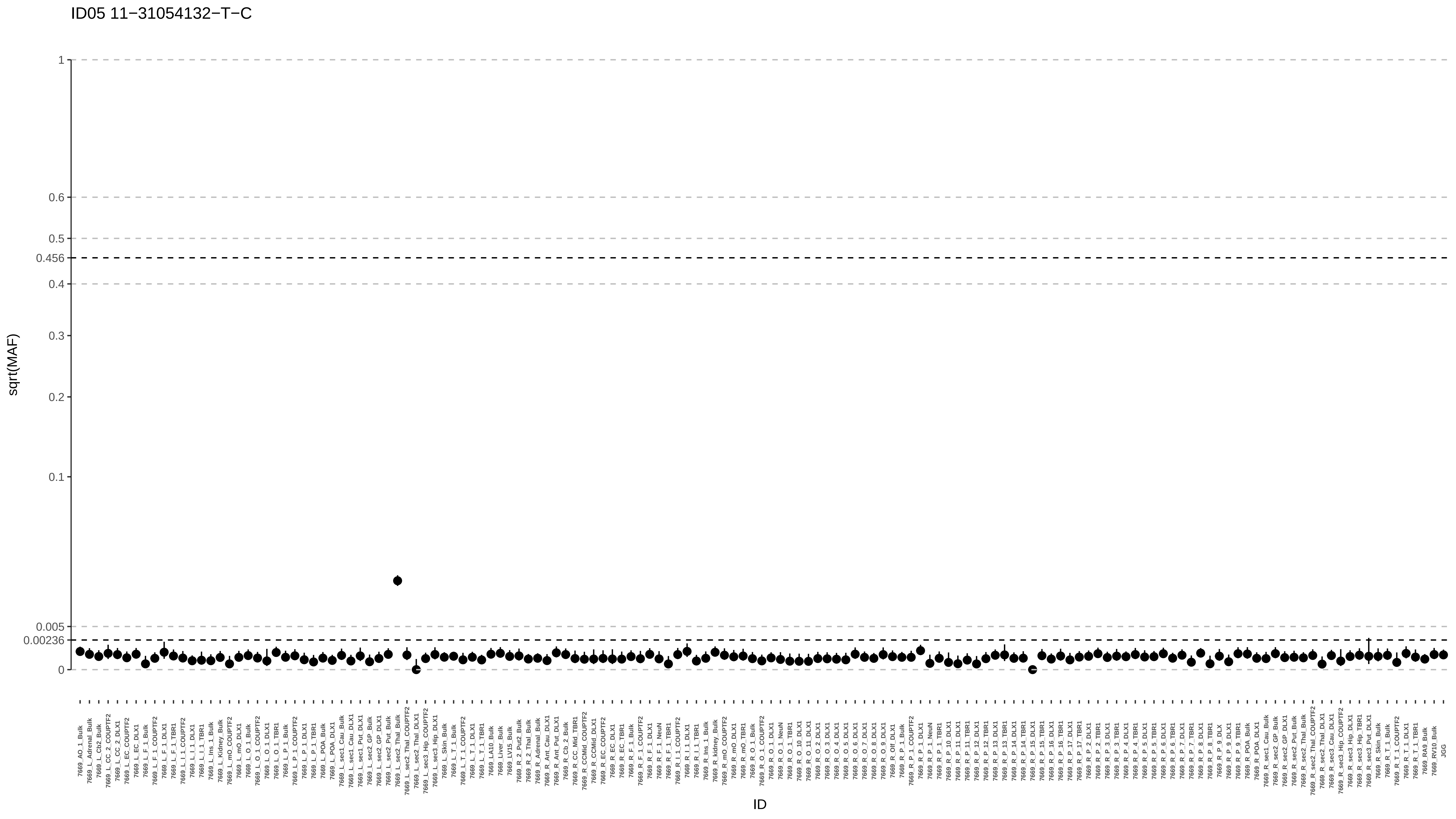
ID05 11−31054132−T−C
sqrt(MAF)
ID
1
0.6
0.5
0.456
0.4
0.3
0.2
0.1
0.005
0.00236
0

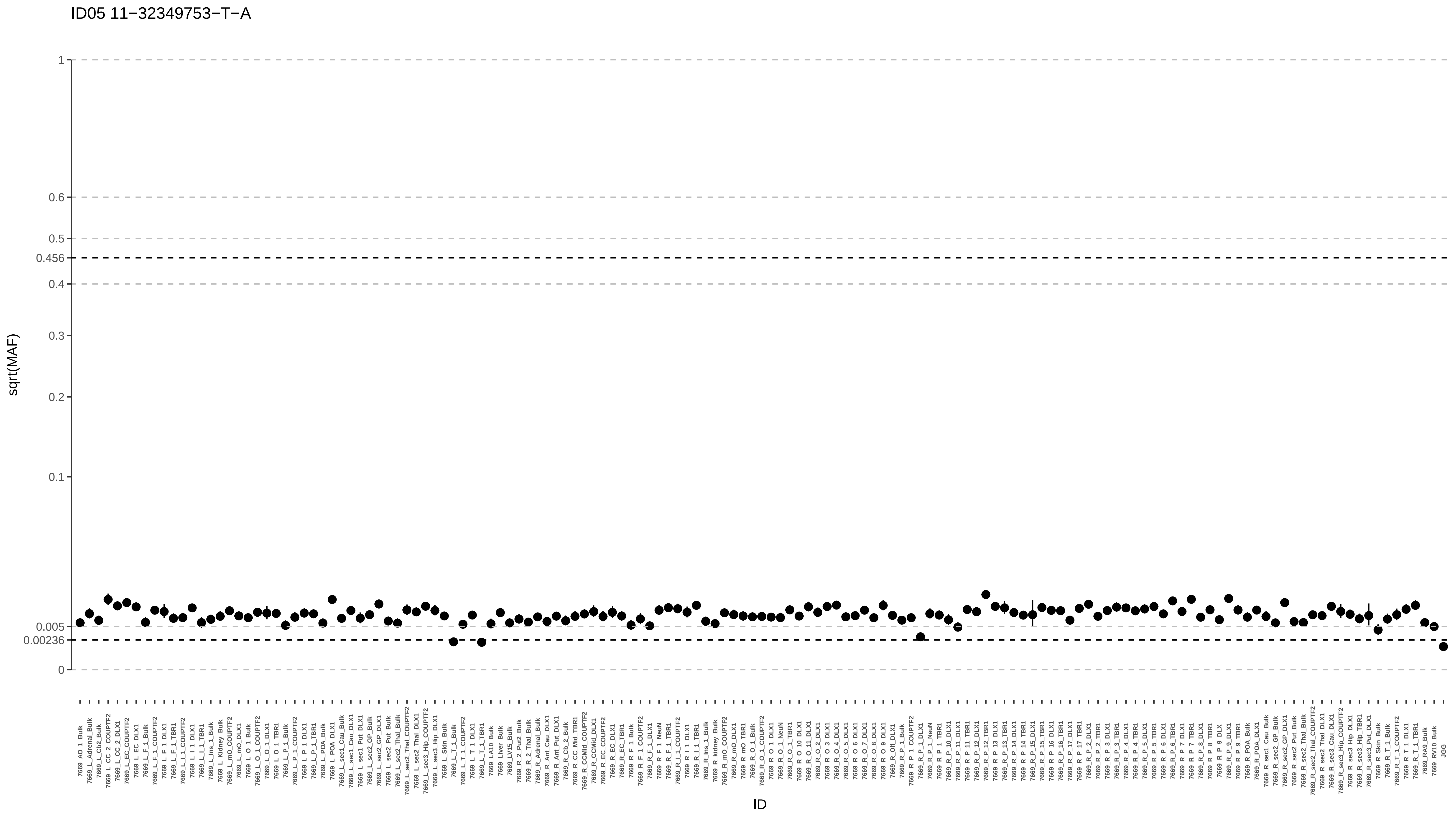
ID05 11−32349753−T−A
sqrt(MAF)
1
0.6
0.5
0.456
0.4
0.3
0.2
0.1
0.005
0.00236
0
ID
7669_AO_1_Bulk
7669_L_Adrenal_Bulk
7669_L_Cb2_Bulk
7669_L_CC_2_COUPTF2
7669_L_CC_2_DLX1
7669_L_EC_COUPTF2
7669_L_EC_DLX1
7669_L_F_1_Bulk
7669_L_F_1_COUPTF2
7669_L_F_1_DLX1
7669_L_F_1_TBR1
7669_L_I_1_COUPTF2
7669_L_I_1_DLX1
7669_L_I_1_TBR1
7669_L_Ins_1_Bulk
7669_L_Kidney_Bulk
7669_L_mO_COUPTF2
7669_L_mO_DLX1
7669_L_O_1_Bulk
7669_L_O_1_COUPTF2
7669_L_O_1_DLX1
7669_L_O_1_TBR1
7669_L_P_1_Bulk
7669_L_P_1_COUPTF2
7669_L_P_1_DLX1
7669_L_P_1_TBR1
7669_L_POA_Bulk
7669_L_POA_DLX1
7669_L_sec1_Cau_Bulk
7669_L_sec1_Cau_DLX1
7669_L_sec1_Put_DLX1
7669_L_sec2_GP_Bulk
7669_L_sec2_GP_DLX1
7669_L_sec2_Put_Bulk
7669_L_sec2_Thal_Bulk
7669_L_sec2_Thal_COUPTF2
7669_L_sec2_Thal_DLX1
7669_L_sec3_Hip_COUPTF2
7669_L_sec3_Hip_DLX1
7669_L_Skin_Bulk
7669_L_T_1_Bulk
7669_L_T_1_COUPTF2
7669_L_T_1_DLX1
7669_L_T_1_TBR1
7669_LA10_Bulk
7669_Liver_Bulk
7669_LV15_Bulk
7669_R_2_Put2_Bulk
7669_R_2_Thal_Bulk
7669_R_Adrenal_Bulk
7669_R_Ant_Cau_DLX1
7669_R_Ant_Put_DLX1
7669_R_Cb_2_Bulk
7669_R_CC_Mid_TBR1
7669_R_CCMid_COUPTF2
7669_R_CCMid_DLX1
7669_R_EC_COUPTF2
7669_R_EC_DLX1
7669_R_EC_TBR1
7669_R_F_1_Bulk
7669_R_F_1_COUPTF2
7669_R_F_1_DLX1
7669_R_F_1_NeuN
7669_R_F_1_TBR1
7669_R_I_1_COUPTF2
7669_R_I_1_DLX1
7669_R_I_1_TBR1
7669_R_Ins_1_Bulk
7669_R_kidney_Bulk
7669_R_mO_COUPTF2
7669_R_mO_DLX1
7669_R_mO_TBR1
7669_R_O_1_Bulk
7669_R_O_1_COUPTF2
7669_R_O_1_DLX1
7669_R_O_1_NeuN
7669_R_O_1_TBR1
7669_R_O_10_DLX1
7669_R_O_11_DLX1
7669_R_O_2_DLX1
7669_R_O_3_DLX1
7669_R_O_4_DLX1
7669_R_O_5_DLX1
7669_R_O_6_DLX1
7669_R_O_7_DLX1
7669_R_O_8_DLX1
7669_R_O_9_DLX1
7669_R_Olf_DLX1
7669_R_P_1_Bulk
7669_R_P_1_COUPTF2
7669_R_P_1_DLX1
7669_R_P_1_NeuN
7669_R_P_1_TBR1
7669_R_P_10_DLX1
7669_R_P_11_DLX1
7669_R_P_11_TBR1
7669_R_P_12_DLX1
7669_R_P_12_TBR1
7669_R_P_13_DLX1
7669_R_P_13_TBR1
7669_R_P_14_DLX1
7669_R_P_14_TBR1
7669_R_P_15_DLX1
7669_R_P_15_TBR1
7669_R_P_16_DLX1
7669_R_P_16_TBR1
7669_R_P_17_DLX1
7669_R_P_17_TBR1
7669_R_P_2_DLX1
7669_R_P_2_TBR1
7669_R_P_3_DLX1
7669_R_P_3_TBR1
7669_R_P_4_DLX1
7669_R_P_4_TBR1
7669_R_P_5_DLX1
7669_R_P_5_TBR1
7669_R_P_6_DLX1
7669_R_P_6_TBR1
7669_R_P_7_DLX1
7669_R_P_7_TBR1
7669_R_P_8_DLX1
7669_R_P_8_TBR1
7669_R_P_9_DLX
7669_R_P_9_DLX1
7669_R_P_9_TBR1
7669_R_POA_Bulk
7669_R_POA_DLX1
7669_R_sec1_Cau_Bulk
7669_R_sec2_GP_Bulk
7669_R_sec2_GP_DLX1
7669_R_sec2_Put_Bulk
7669_R_sec2_Thal_Bulk
7669_R_sec2_Thal_COUPTF2
7669_R_sec2_Thal_DLX1
7669_R_sec3_Cau_DLX1
7669_R_sec3_Hip_COUPTF2
7669_R_sec3_Hip_DLX1
7669_R_sec3_Hip_TBR1
7669_R_sec3_Put_DLX1
7669_R_Skin_Bulk
7669_R_T_1_Bulk
7669_R_T_1_COUPTF2
7669_R_T_1_DLX1
7669_R_T_1_TBR1
7669_RA9_Bulk
7669_RV10_Bulk
JGG

ID05 11−32349754−T−A

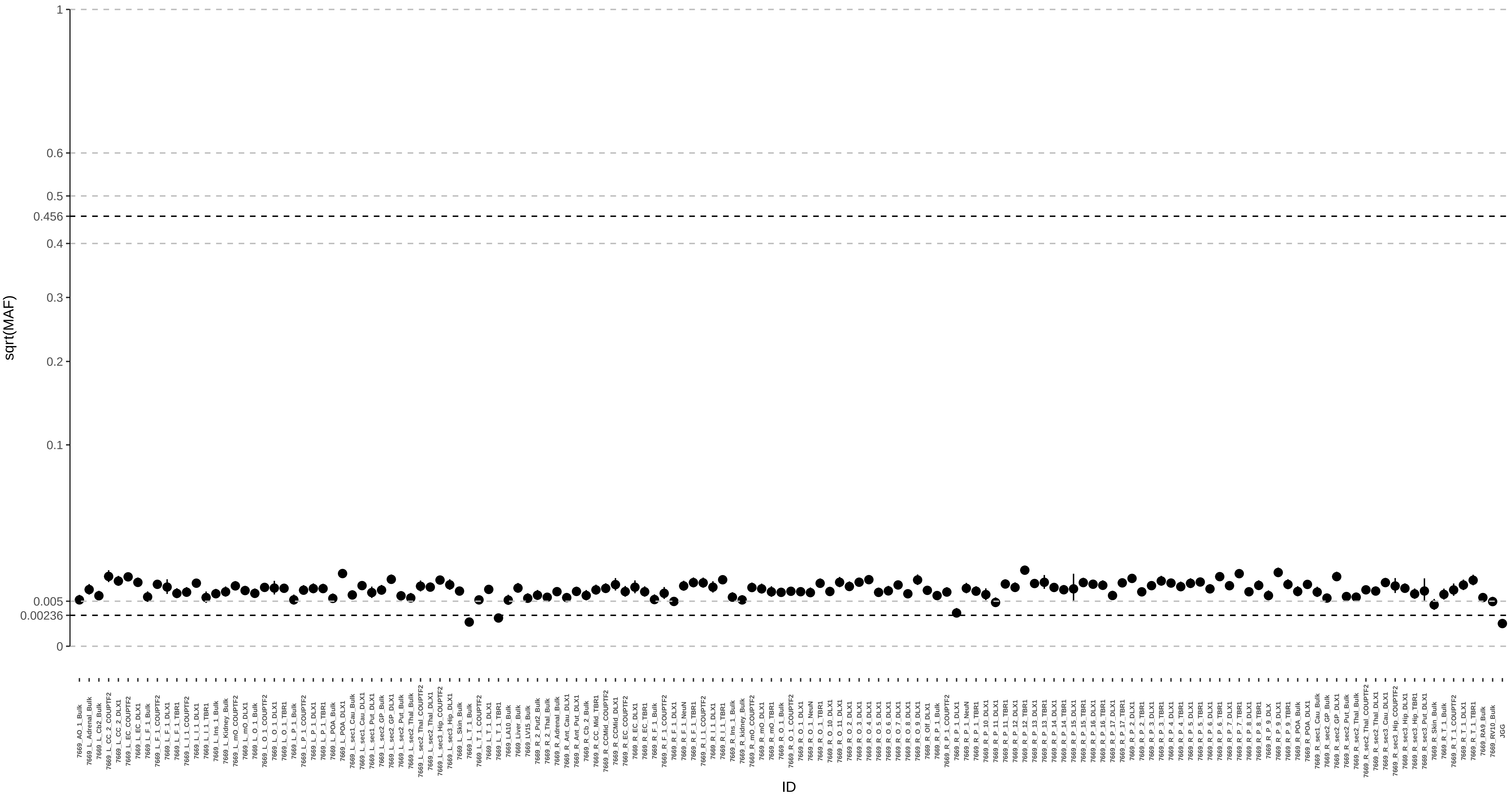

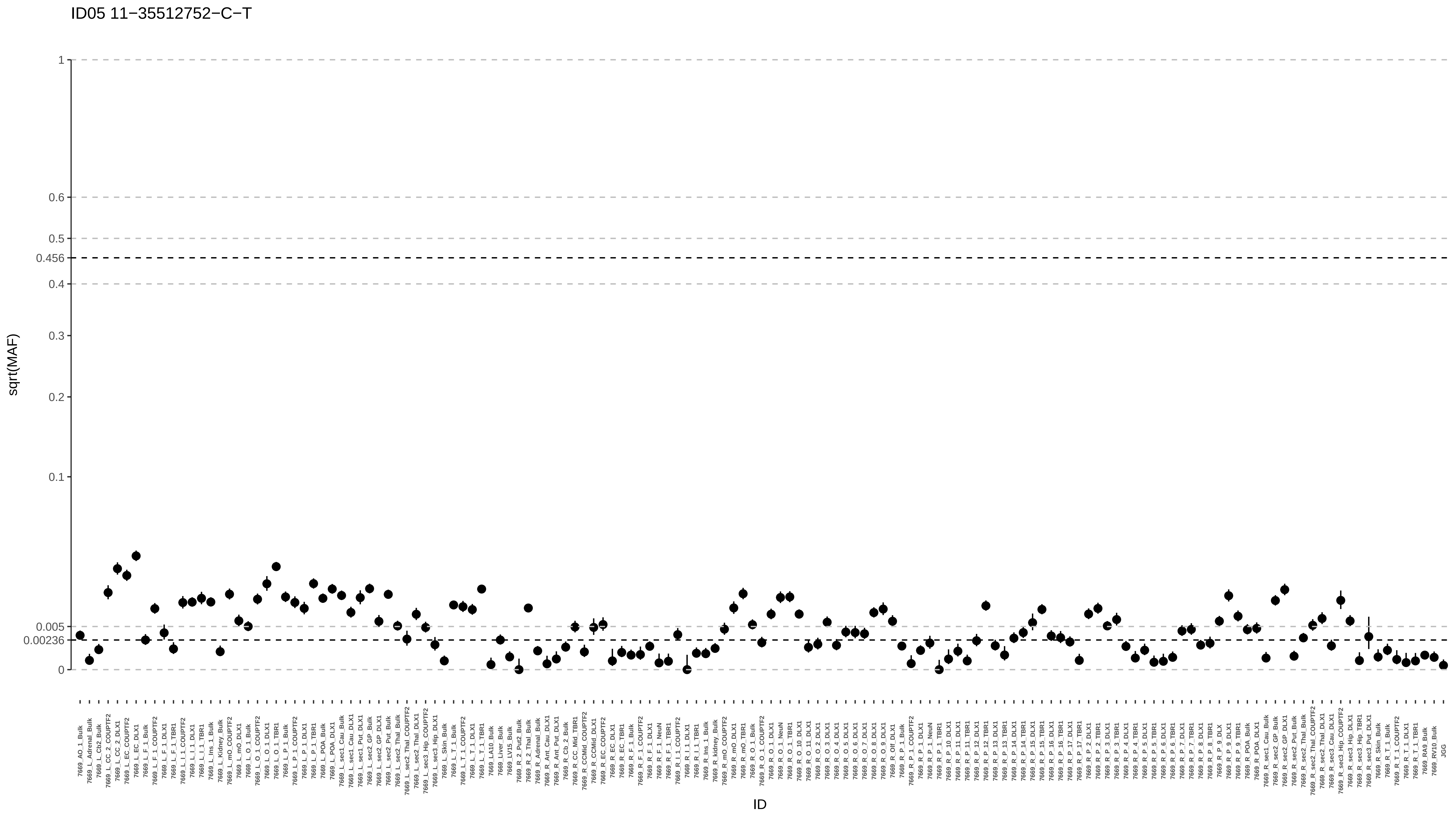
ID05 11−35512752−C−T
sqrt(MAF)
ID
1
0.6
0.5
0.456
0.4
0.3
0.2
0.1
0.005
0.00236
0

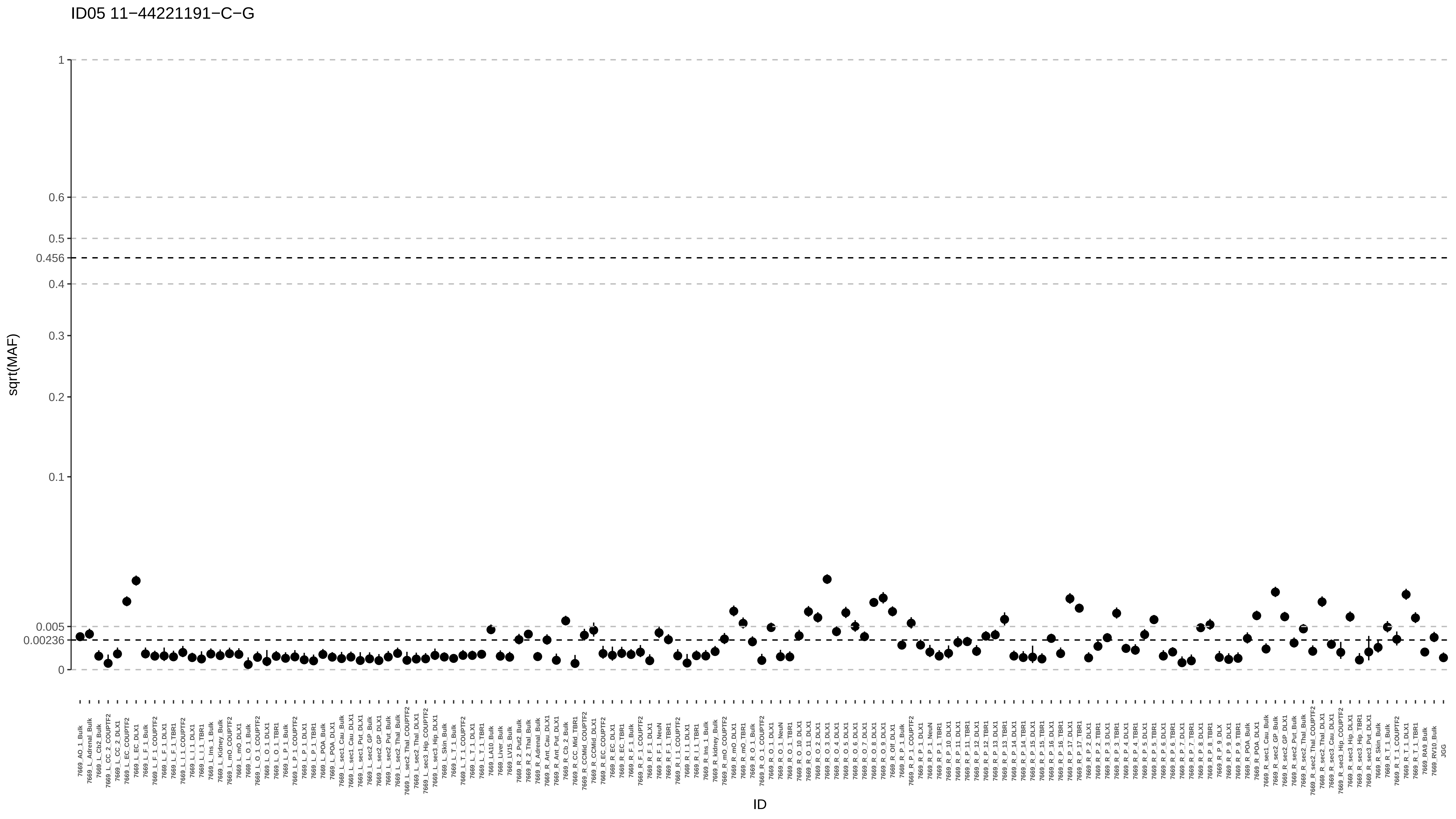
ID05 11−44221191−C−G
sqrt(MAF)
1
0.6
0.5
0.456
0.4
0.3
0.2
0.1
0.005
0.00236
0
ID

ID05 11−45873419−G−A

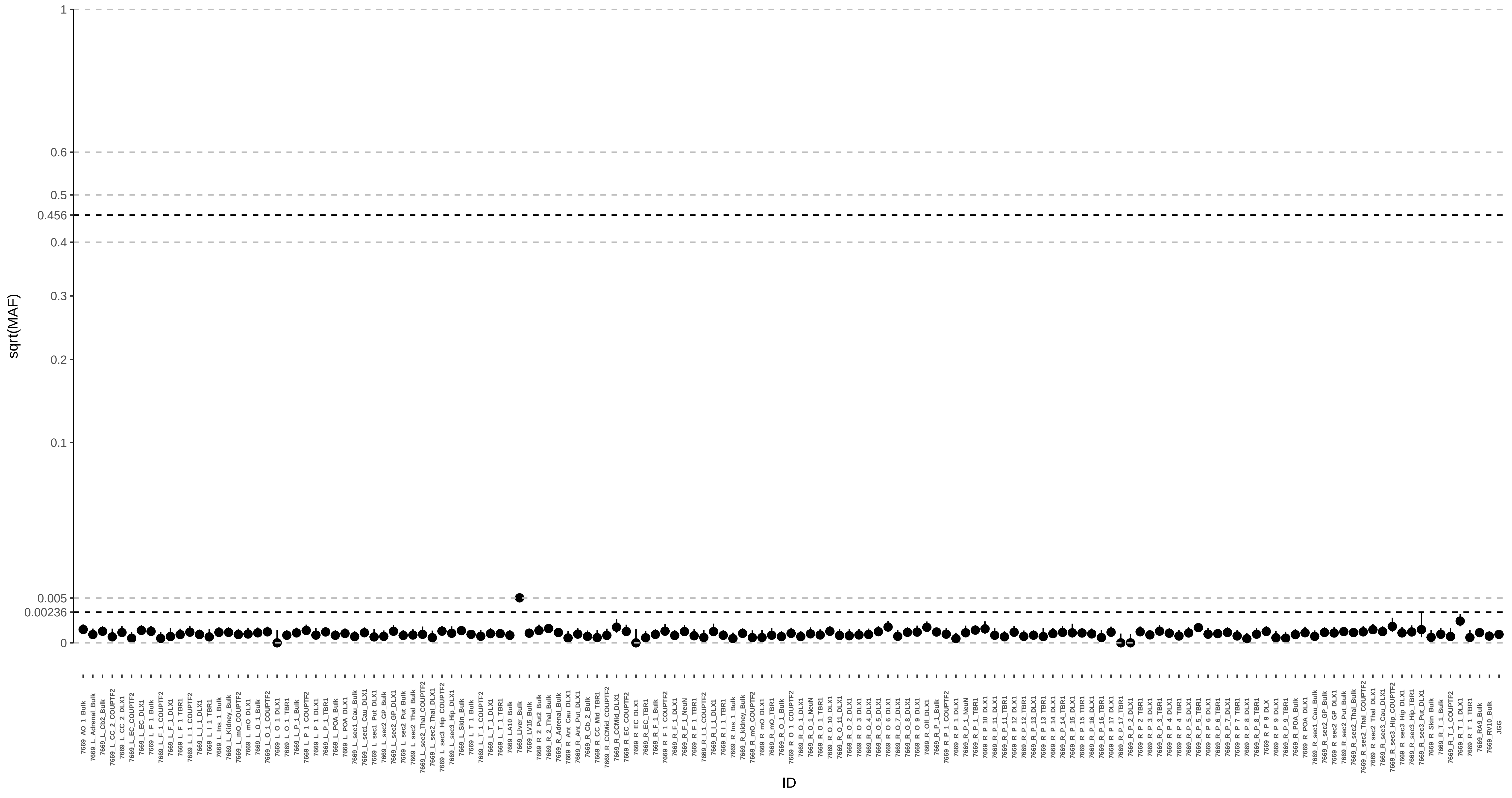

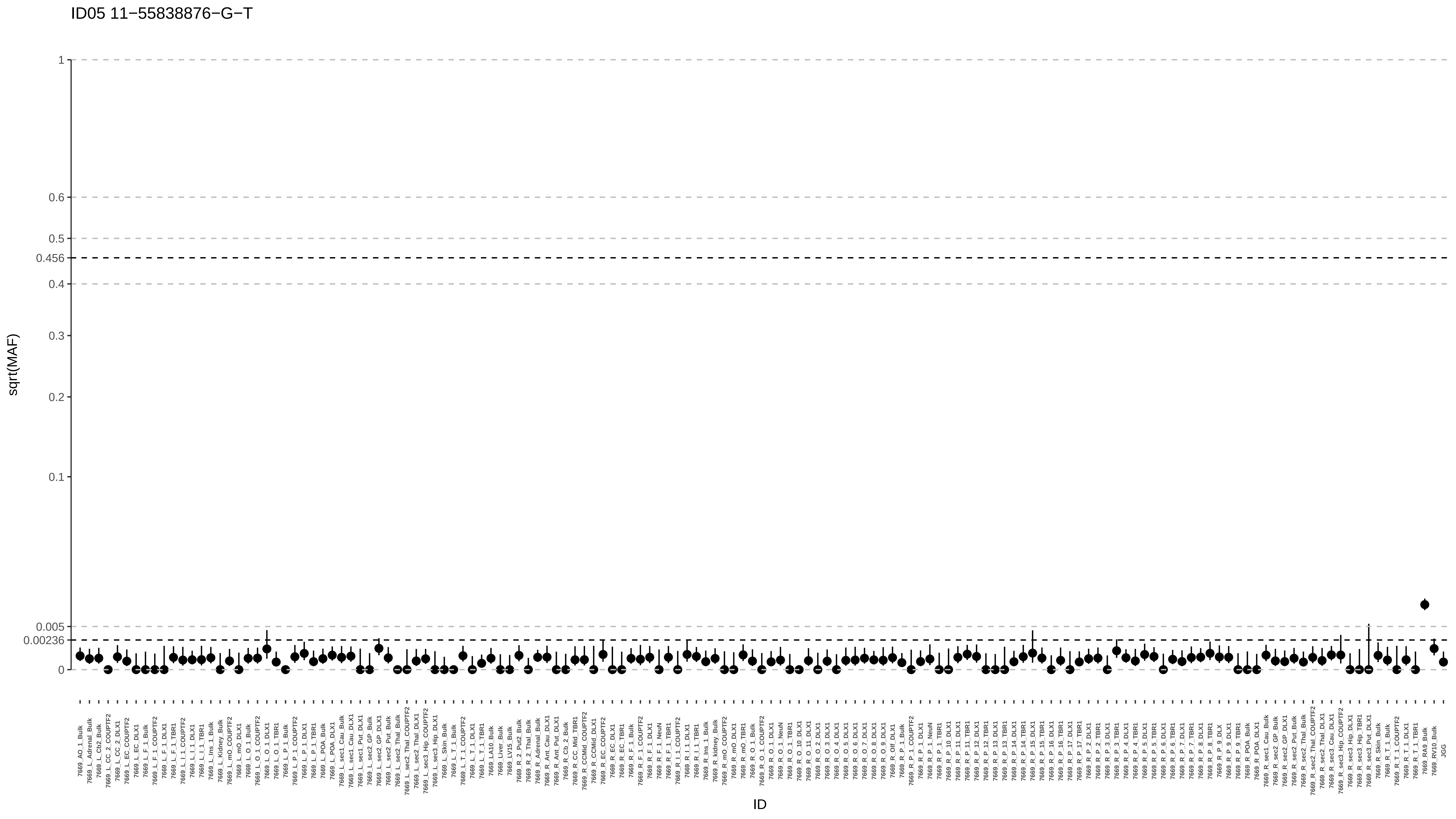

ID05 11−55838876−G−T
sqrt(MAF)
1
0.6
0.5
0.456
0.4
0.3
0.2
0.1
0.005
0.00236
0
ID

ID05 11−55997990−C−T

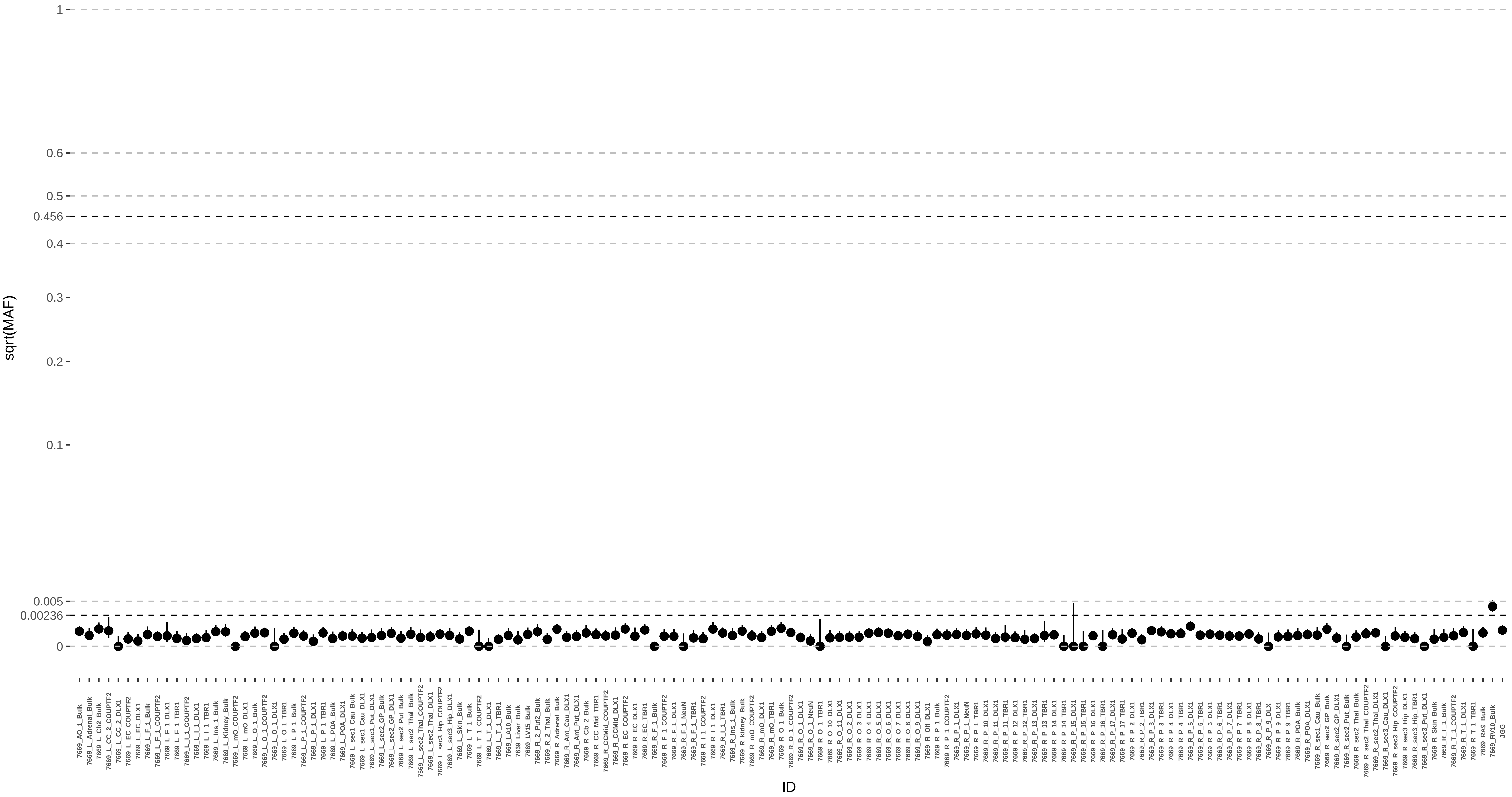

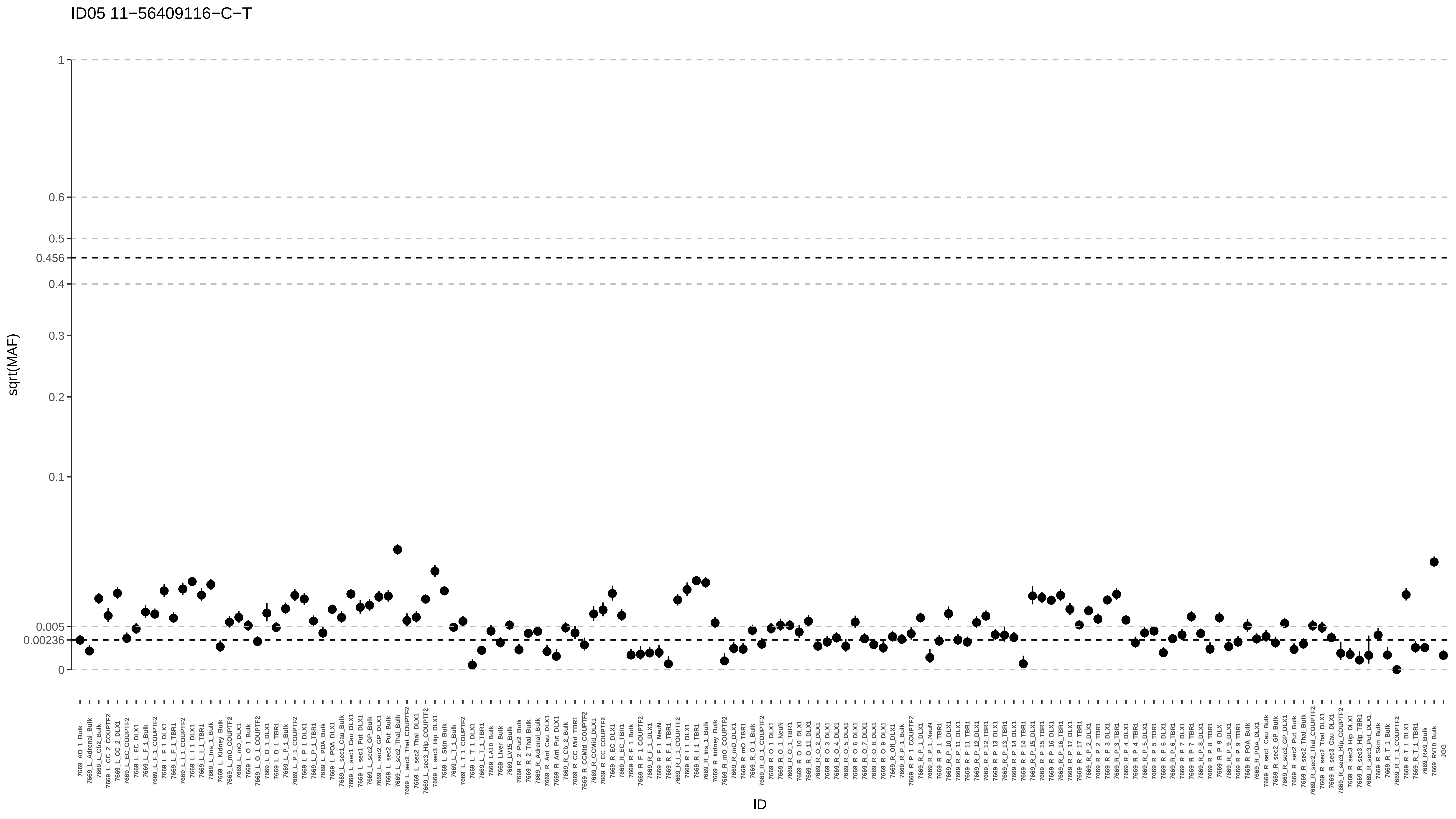
ID05 11−56409116−C−T
sqrt(MAF)
1
0.6
0.5
0.456
0.4
0.3
0.2
0.1
0.005
0.00236
0
ID

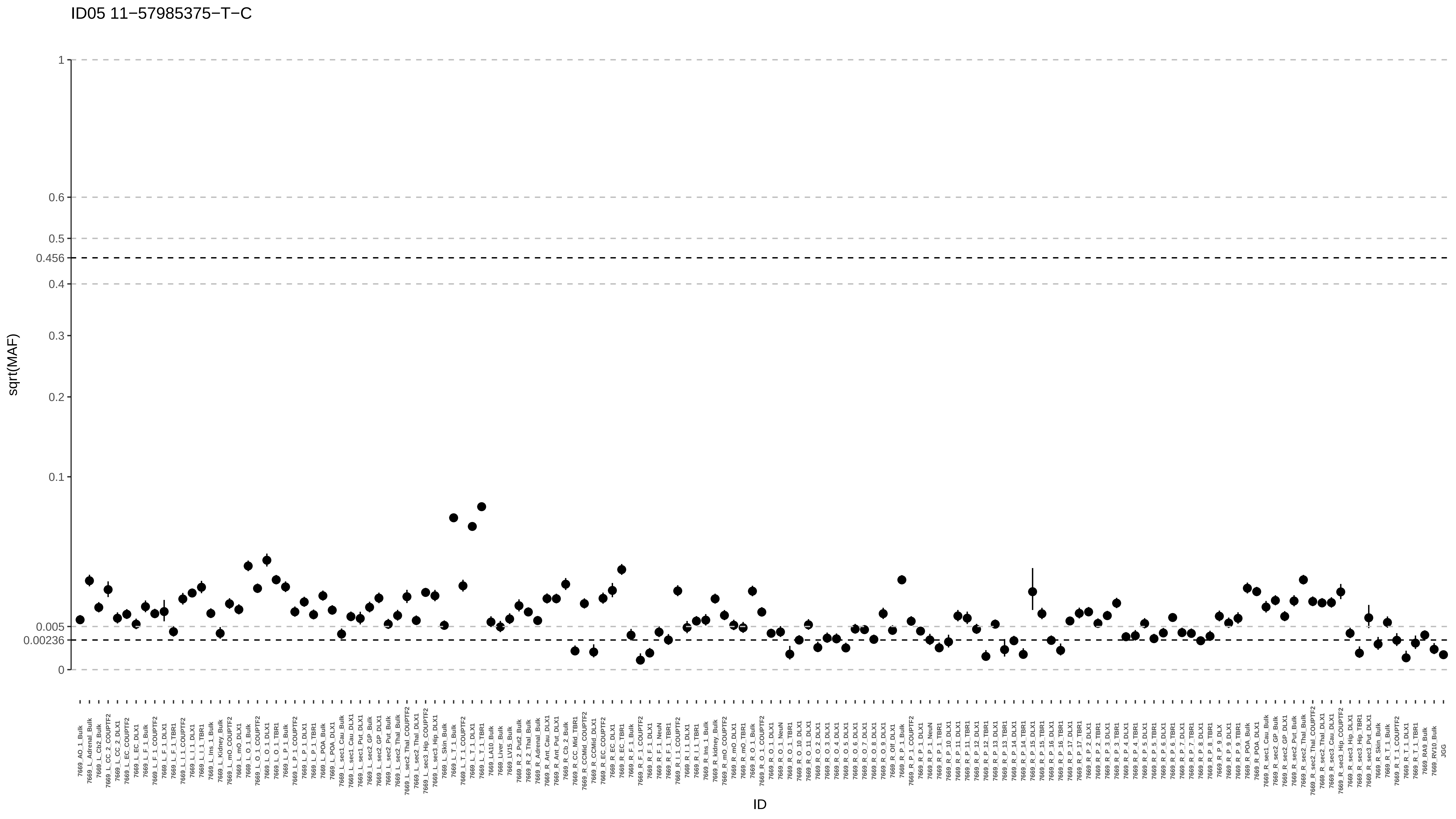
ID05 11−57985375−T−C
sqrt(MAF)
1
0.6
0.5
0.456
0.4
0.3
0.2
0.1
0.005
0.00236
0
ID

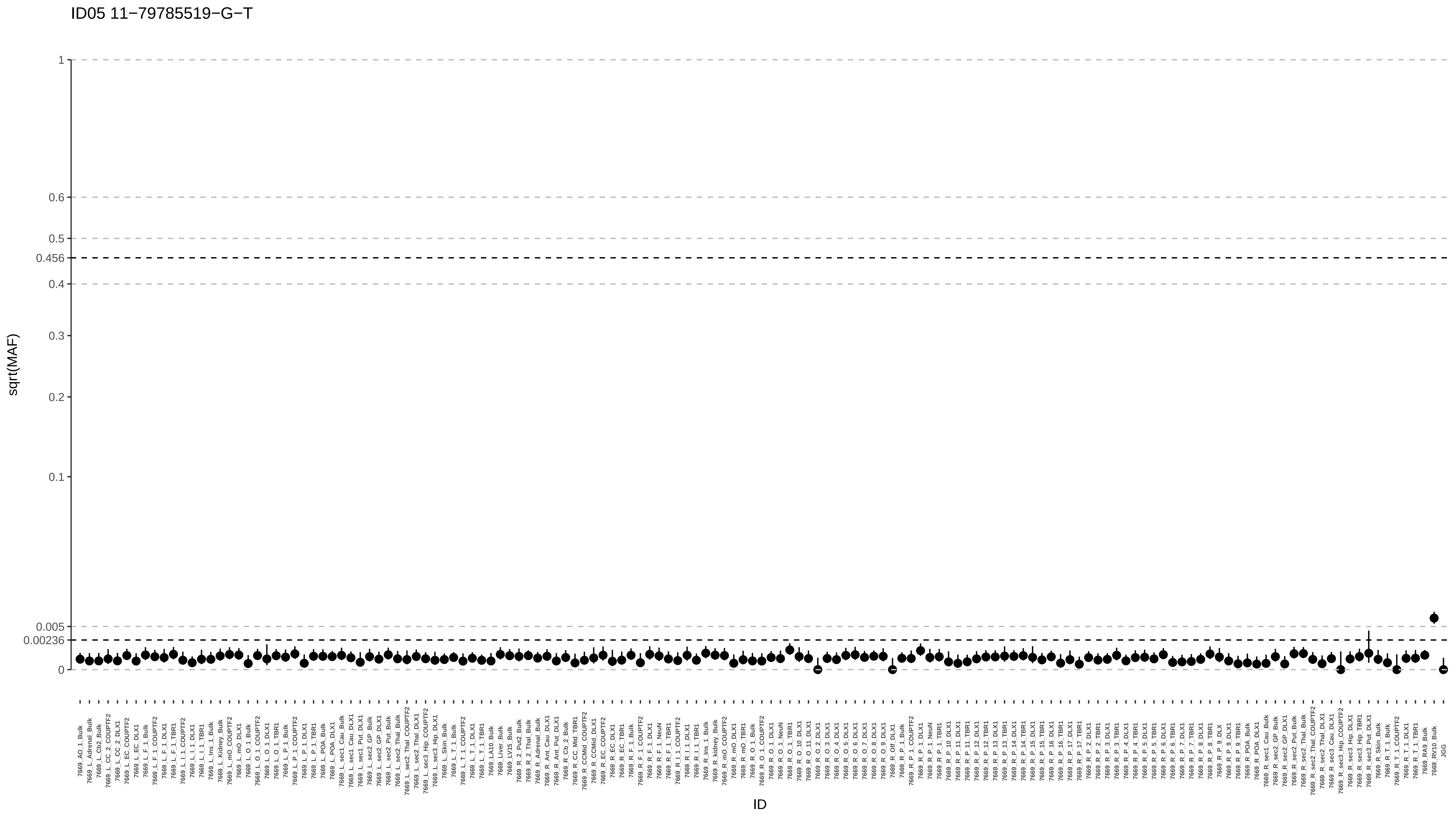

ID05 11−79785519−G−T
sqrt(MAF)
1
0.6
0.5
0.456
0.4
0.3
0.2
0.1
0.005
0.00236
0
ID

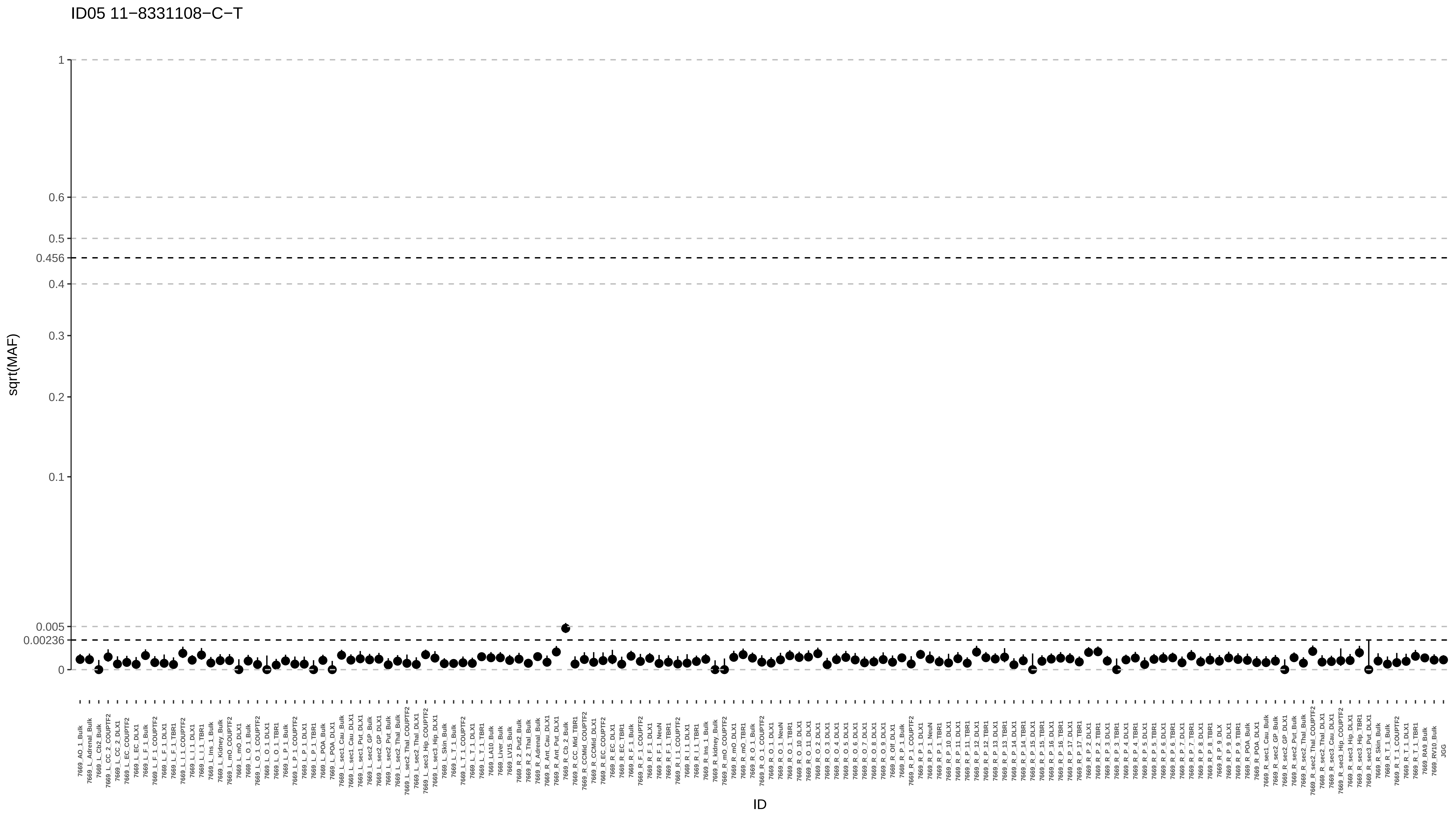
ID05 11−8331108−C−T
sqrt(MAF)
1
0.6
0.5
0.456
0.4
0.3
0.2
0.1
0.005
0.00236
0
ID

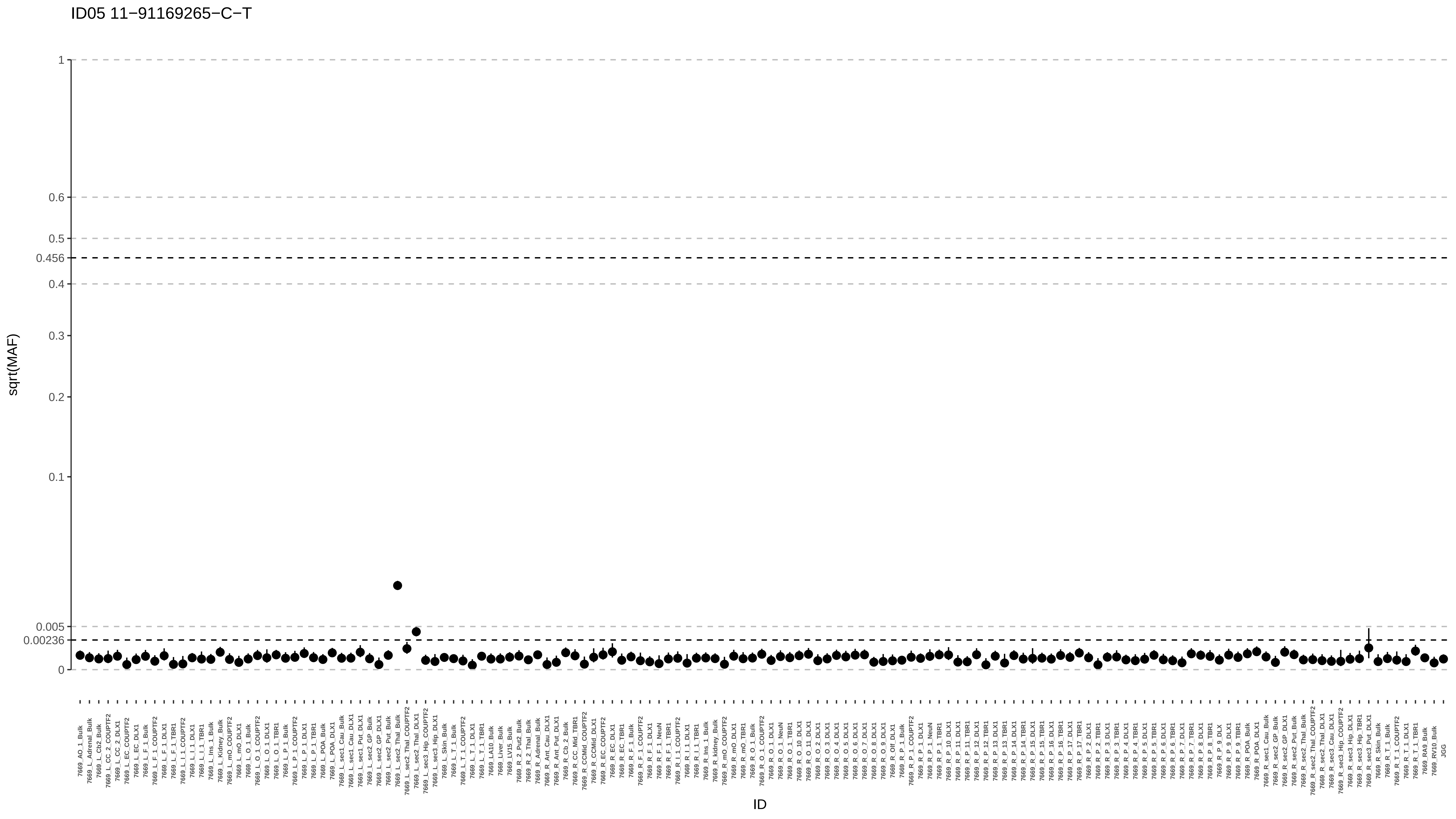
ID05 11−91169265−C−T
sqrt(MAF)
1
0.6
0.5
0.456
0.4
0.3
0.2
0.1
0.005
0.00236
0
ID

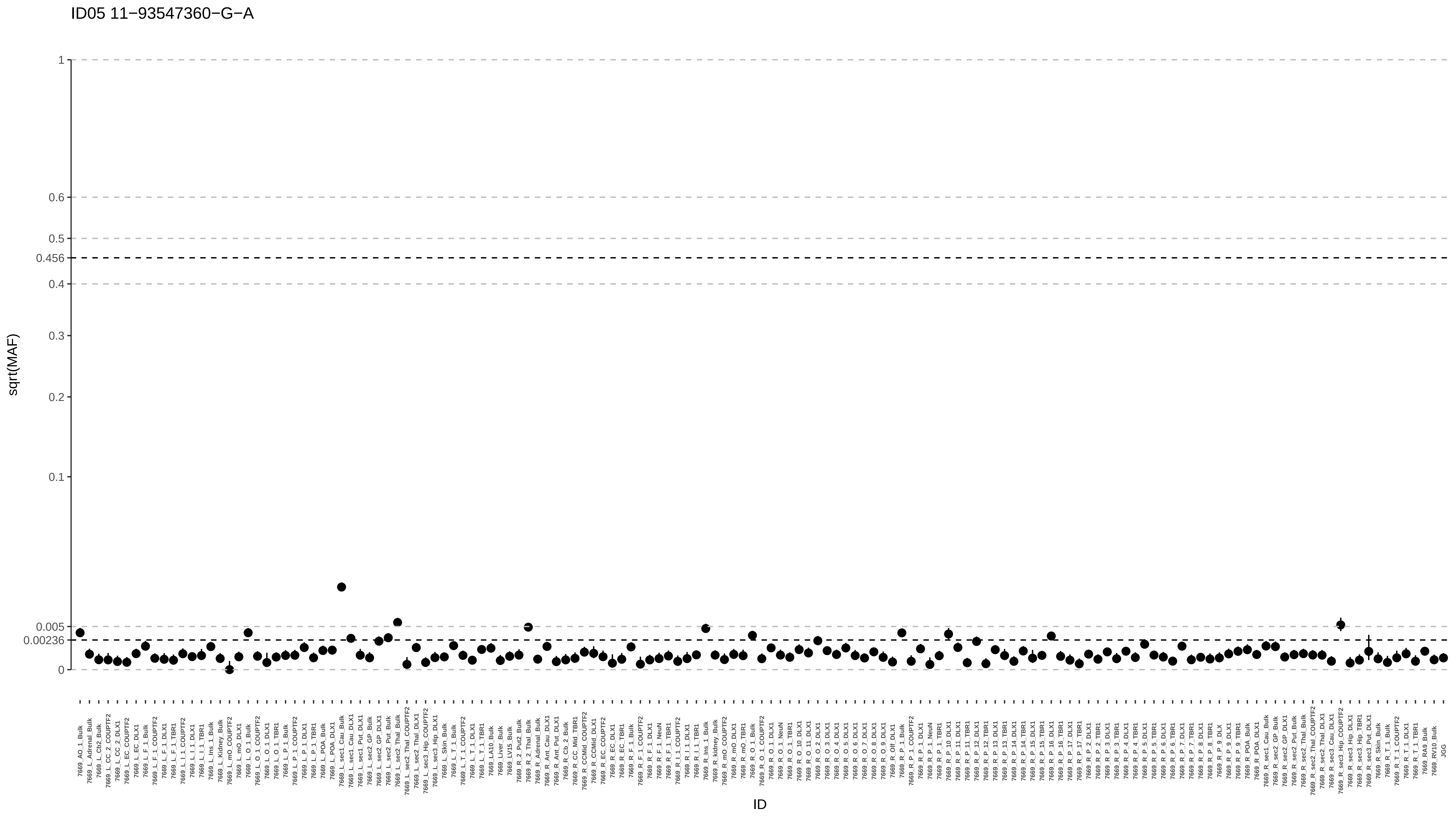
ID05 11−93547360−G−A
sqrt(MAF)
1
0.6
0.5
0.456
0.4
0.3
0.2
0.1
0.005
0.00236
0
ID

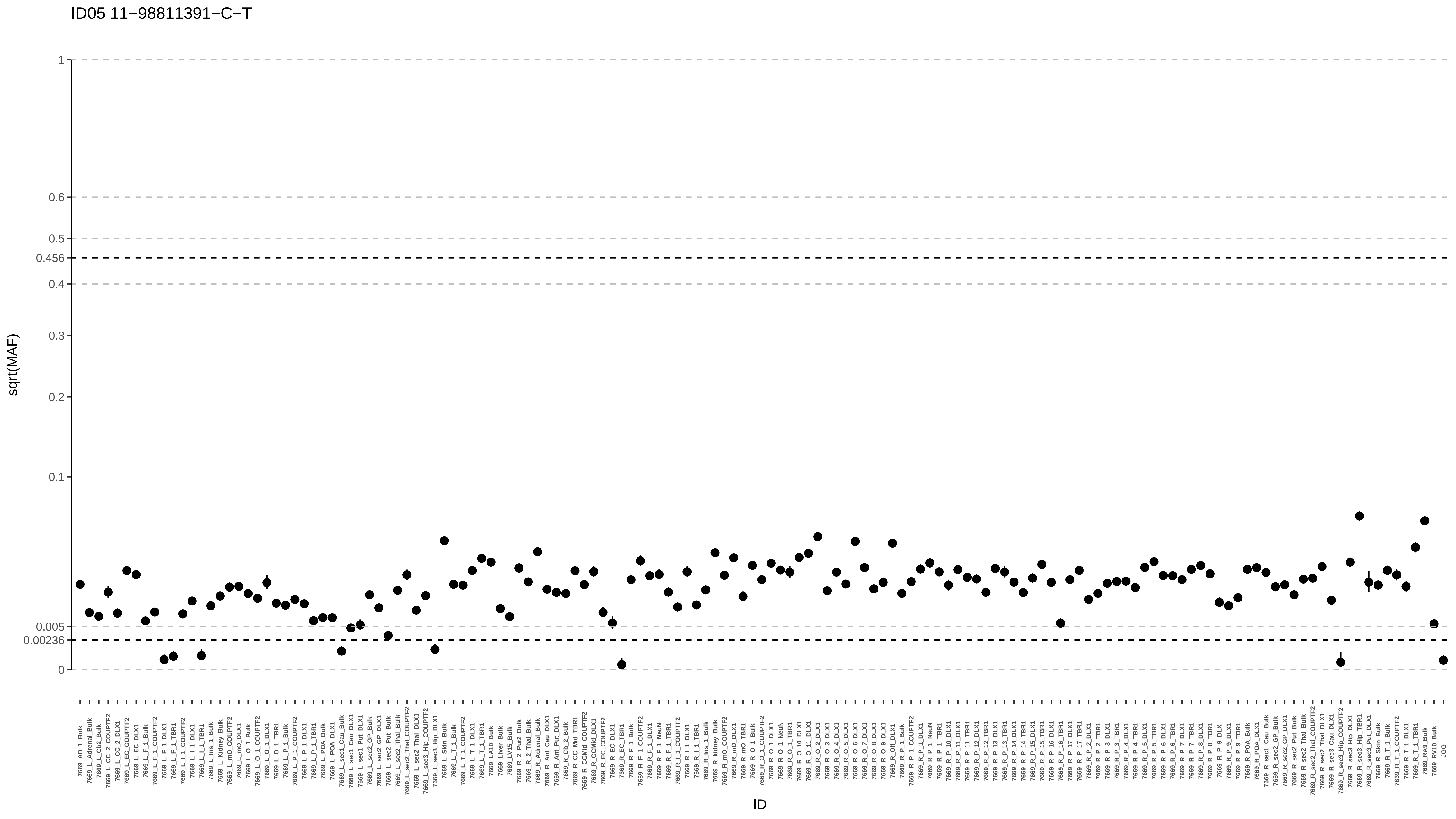
ID05 11−98811391−C−T
sqrt(MAF)
1
0.6
0.5
0.456
0.4
0.3
0.2
0.1
0.005
0.00236
0
7669_AO_1_Bulk
7669_L_Adrenal_Bulk
7669_L_Cb2_Bulk
7669_L_CC_2_COUPTF2
7669_L_CC_2_DLX1
7669_L_EC_COUPTF2
7669_L_EC_DLX1
7669_L_F_1_Bulk
7669_L_F_1_COUPTF2
7669_L_F_1_DLX1
7669_L_F_1_TBR1
7669_L_I_1_COUPTF2
7669_L_I_1_DLX1
7669_L_I_1_TBR1
7669_L_Ins_1_Bulk
7669_L_Kidney_Bulk
7669_L_mO_COUPTF2
7669_L_mO_DLX1
7669_L_O_1_Bulk
7669_L_O_1_COUPTF2
7669_L_O_1_DLX1
7669_L_O_1_TBR1
7669_L_P_1_Bulk
7669_L_P_1_COUPTF2
7669_L_P_1_DLX1
7669_L_P_1_TBR1
7669_L_POA_Bulk
7669_L_POA_DLX1
7669_L_sec1_Cau_Bulk
7669_L_sec1_Cau_DLX1
7669_L_sec1_Put_DLX1
7669_L_sec2_GP_Bulk
7669_L_sec2_GP_DLX1
7669_L_sec2_Put_Bulk
7669_L_sec2_Thal_Bulk
7669_L_sec2_Thal_COUPTF2
7669_L_sec2_Thal_DLX1
7669_L_sec3_Hip_COUPTF2
7669_L_sec3_Hip_DLX1
7669_L_Skin_Bulk
7669_L_T_1_Bulk
7669_L_T_1_COUPTF2
7669_L_T_1_DLX1
7669_L_T_1_TBR1
7669_LA10_Bulk
7669_Liver_Bulk
7669_LV15_Bulk
7669_R_2_Put2_Bulk
7669_R_2_Thal_Bulk
7669_R_Adrenal_Bulk
7669_R_Ant_Cau_DLX1
7669_R_Ant_Put_DLX1
7669_R_Cb_2_Bulk
7669_R_CC_Mid_TBR1
7669_R_CCMid_COUPTF2
7669_R_CCMid_DLX1
7669_R_EC_COUPTF2
7669_R_EC_DLX1
7669_R_EC_TBR1
7669_R_F_1_Bulk
7669_R_F_1_COUPTF2
7669_R_F_1_DLX1
7669_R_F_1_NeuN
7669_R_F_1_TBR1
7669_R_I_1_COUPTF2
7669_R_I_1_DLX1
7669_R_I_1_TBR1
7669_R_Ins_1_Bulk
7669_R_kidney_Bulk
7669_R_mO_COUPTF2
7669_R_mO_DLX1
7669_R_mO_TBR1
7669_R_O_1_Bulk
7669_R_O_1_COUPTF2
7669_R_O_1_DLX1
7669_R_O_1_NeuN
7669_R_O_1_TBR1
7669_R_O_10_DLX1
7669_R_O_11_DLX1
7669_R_O_2_DLX1
7669_R_O_3_DLX1
7669_R_O_4_DLX1
7669_R_O_5_DLX1
7669_R_O_6_DLX1
7669_R_O_7_DLX1
7669_R_O_8_DLX1
7669_R_O_9_DLX1
7669_R_Olf_DLX1
7669_R_P_1_Bulk
7669_R_P_1_COUPTF2
7669_R_P_1_DLX1
7669_R_P_1_NeuN
7669_R_P_1_TBR1
7669_R_P_10_DLX1
7669_R_P_11_DLX1
7669_R_P_11_TBR1
7669_R_P_12_DLX1
7669_R_P_12_TBR1
7669_R_P_13_DLX1
7669_R_P_13_TBR1
7669_R_P_14_DLX1
7669_R_P_14_TBR1
7669_R_P_15_DLX1
7669_R_P_15_TBR1
7669_R_P_16_DLX1
7669_R_P_16_TBR1
7669_R_P_17_DLX1
7669_R_P_17_TBR1
7669_R_P_2_DLX1
7669_R_P_2_TBR1
7669_R_P_3_DLX1
7669_R_P_3_TBR1
7669_R_P_4_DLX1
7669_R_P_4_TBR1
7669_R_P_5_DLX1
7669_R_P_5_TBR1
7669_R_P_6_DLX1
7669_R_P_6_TBR1
7669_R_P_7_DLX1
7669_R_P_7_TBR1
7669_R_P_8_DLX1
7669_R_P_8_TBR1
7669_R_P_9_DLX
7669_R_P_9_DLX1
7669_R_P_9_TBR1
7669_R_POA_Bulk
7669_R_POA_DLX1
7669_R_sec1_Cau_Bulk
7669_R_sec2_GP_Bulk
7669_R_sec2_GP_DLX1
7669_R_sec2_Put_Bulk
7669_R_sec2_Thal_Bulk
7669_R_sec2_Thal_COUPTF2
7669_R_sec2_Thal_DLX1
7669_R_sec3_Cau_DLX1
7669_R_sec3_Hip_COUPTF2
7669_R_sec3_Hip_DLX1
7669_R_sec3_Hip_TBR1
7669_R_sec3_Put_DLX1
7669_R_Skin_Bulk
7669_R_T_1_Bulk
7669_R_T_1_COUPTF2
7669_R_T_1_DLX1
7669_R_T_1_TBR1
7669_RA9_Bulk
7669_RV10_Bulk
JGG
ID

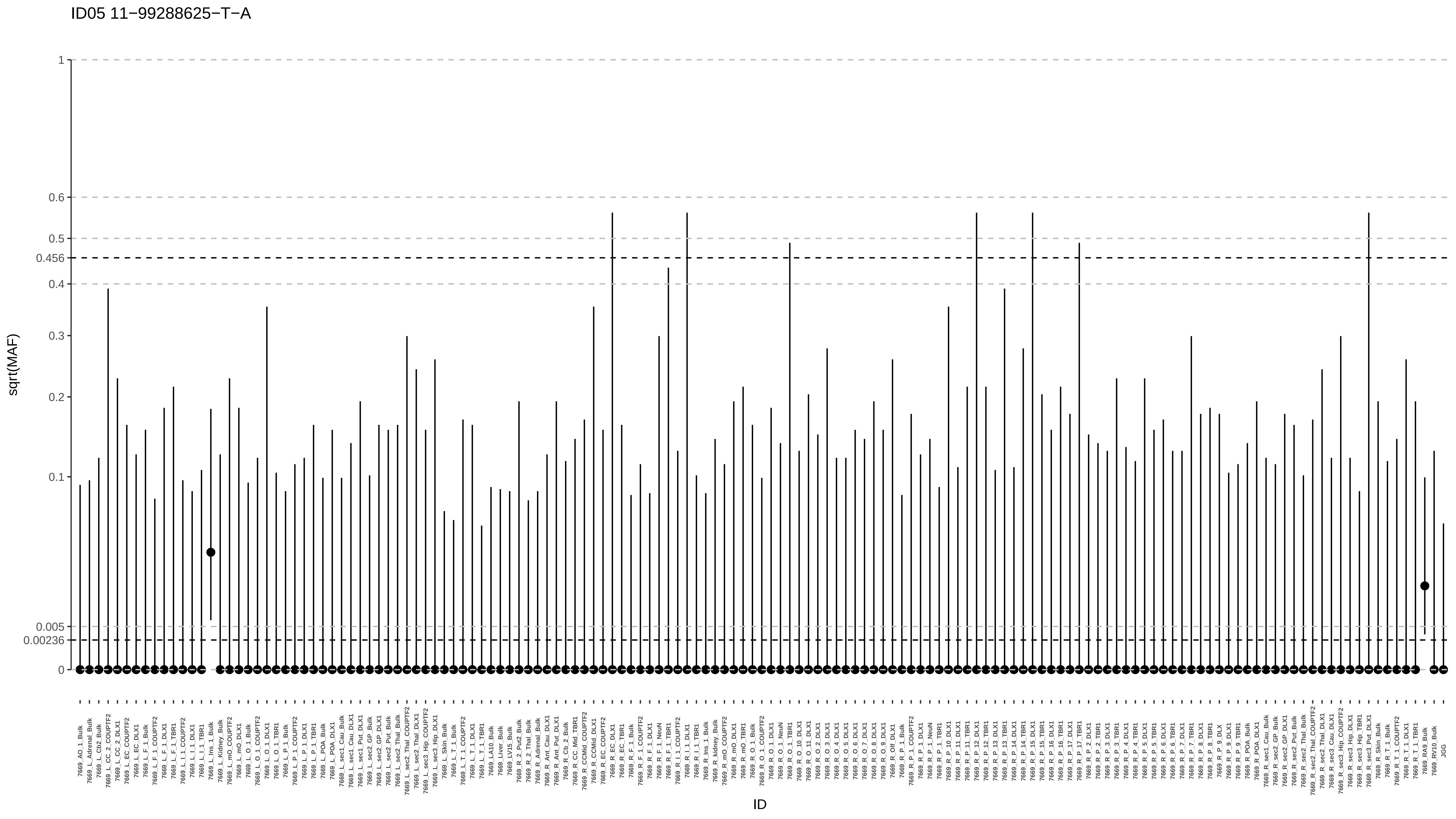
ID05 11−99288625−T−A
sqrt(MAF)
1
0.6
0.5
0.456
0.4
0.3
0.2
0.1
0.005
0.00236
0
ID

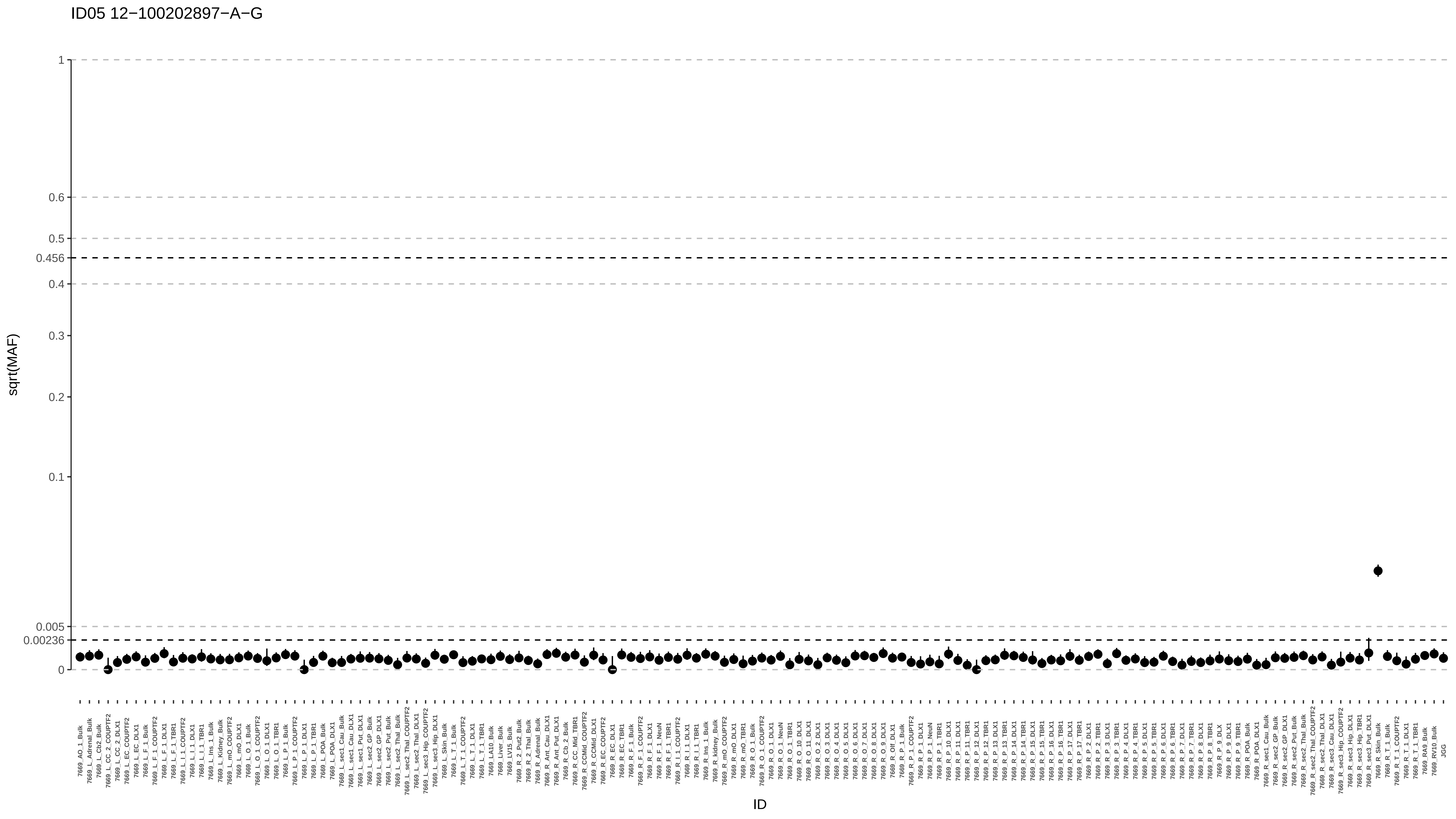

ID05 12−100202897−A−G
sqrt(MAF)
1
0.6
0.5
0.456
0.4
0.3
0.2
0.1
0.005
0.00236
0
ID

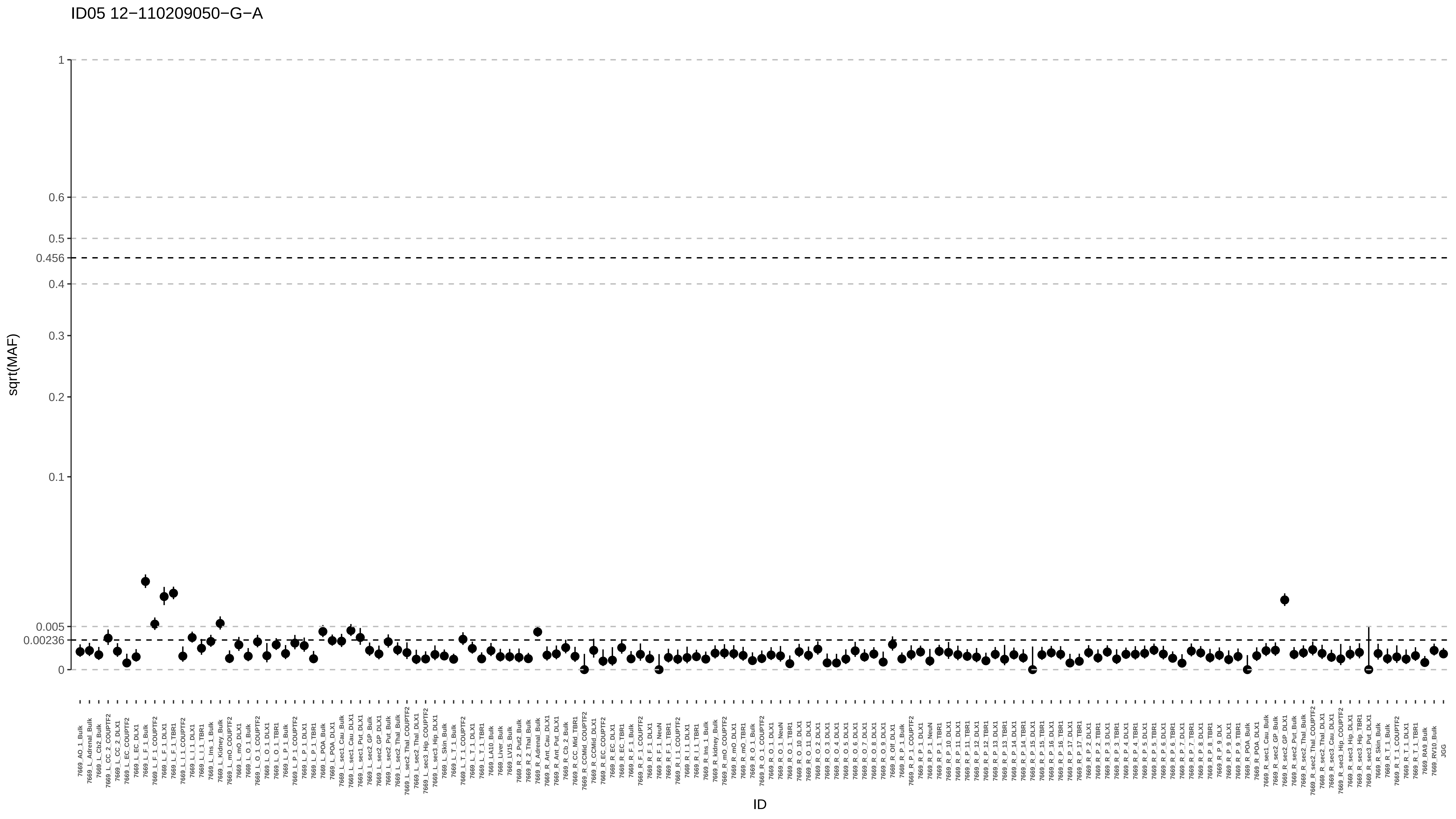
ID05 12−110209050−G−A
sqrt(MAF)
1
0.6
0.5
0.456
0.4
0.3
0.2
0.1
0.005
0.00236
0
ID

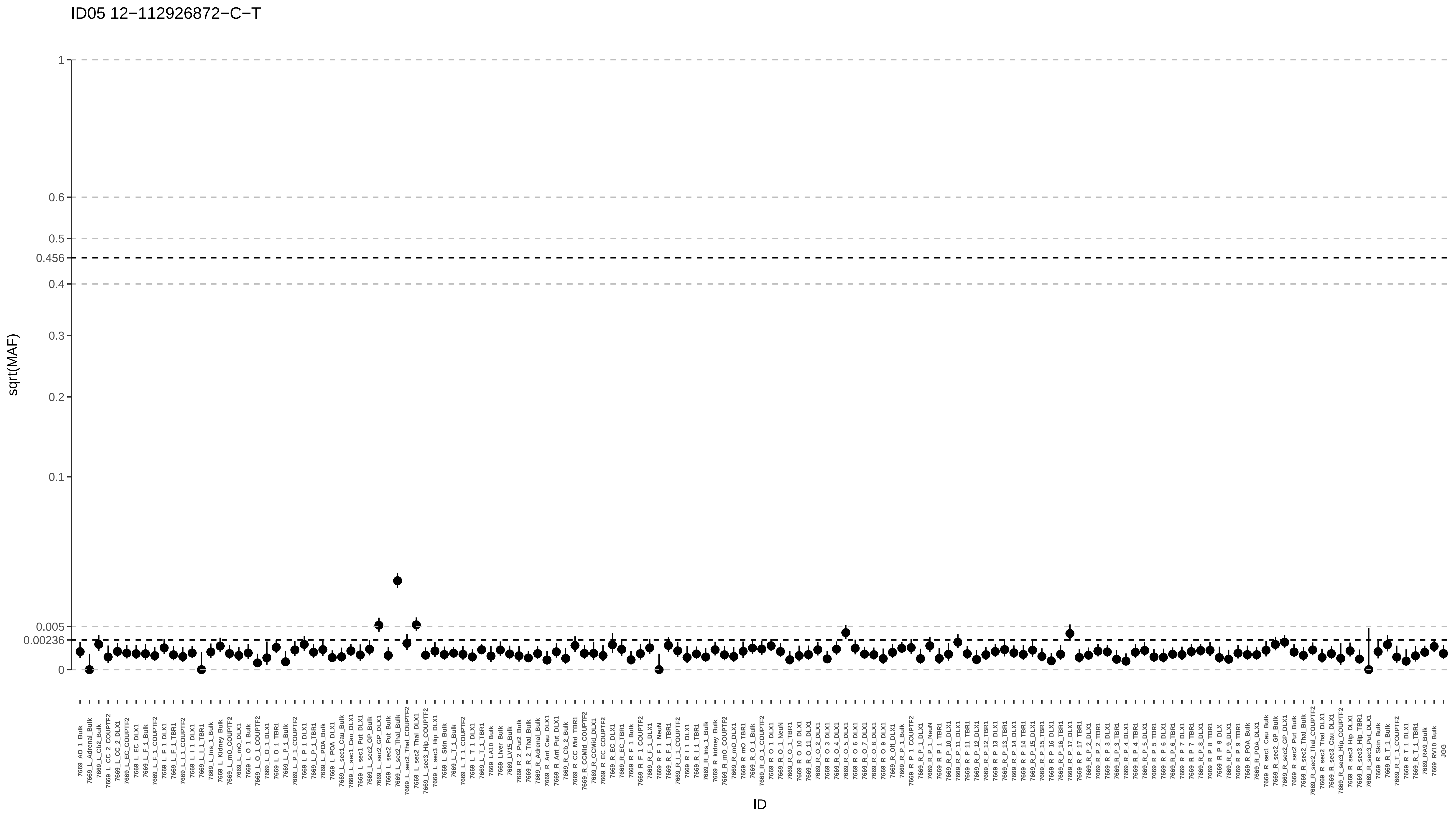
ID05 12−112926872−C−T
sqrt(MAF)
ID

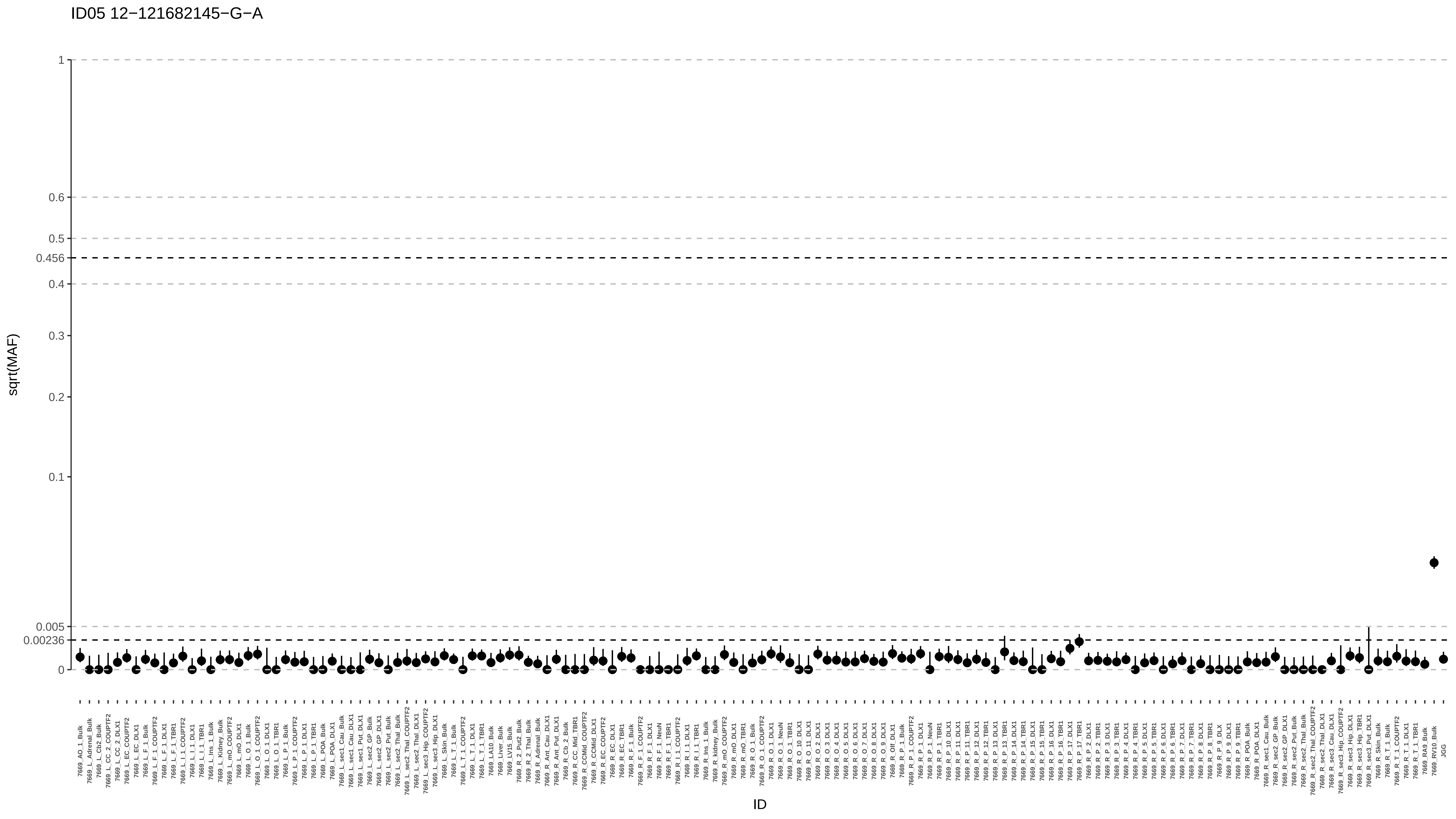
ID05 12−121682145−G−A
sqrt(MAF)
1
0.6
0.5
0.456
0.4
0.3
0.2
0.1
0.005
0.00236
0
ID

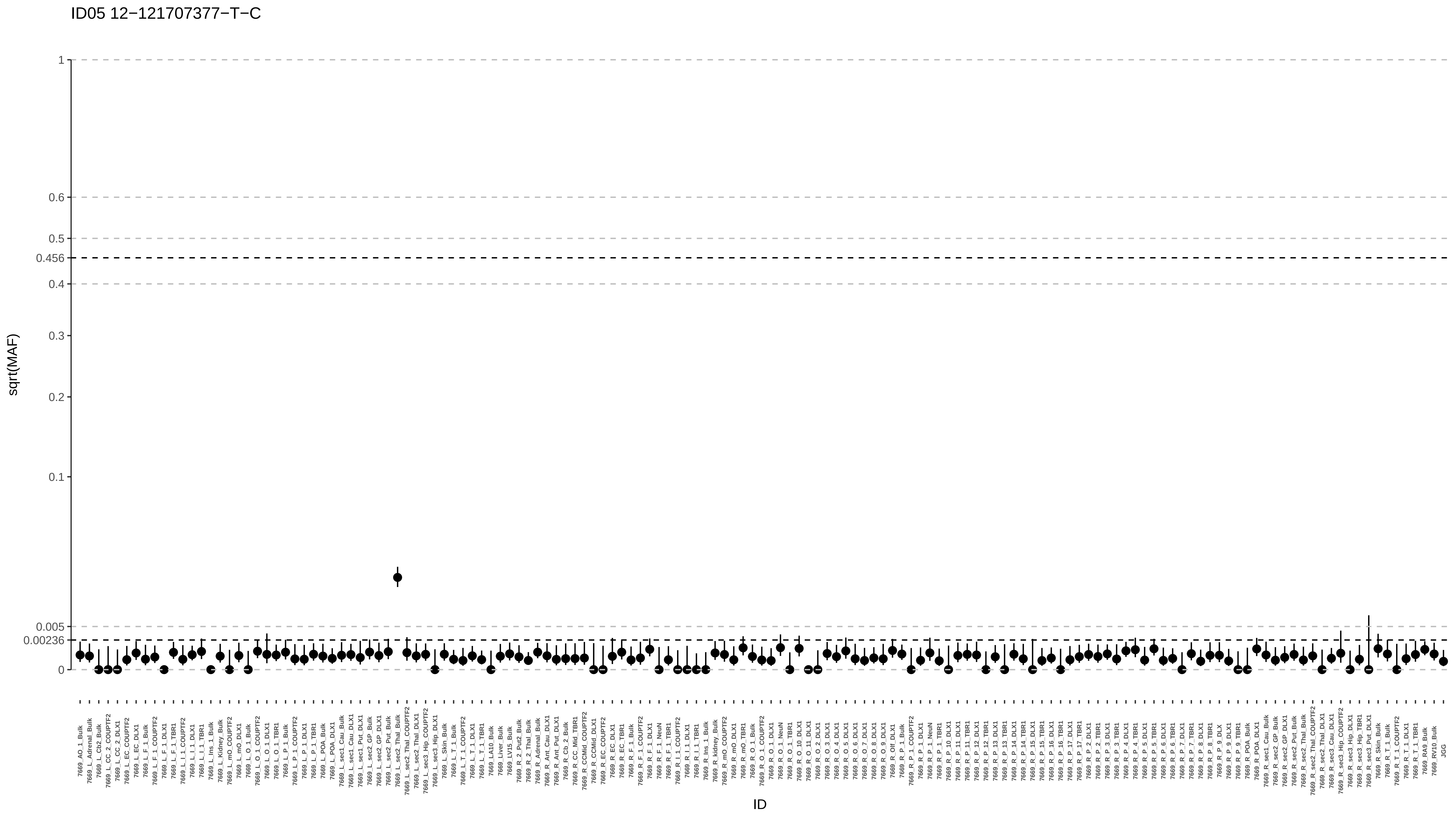

ID05 12−121707377−T−C
sqrt(MAF)
1
0.6
0.5
0.456
0.4
0.3
0.2
0.1
0.005
0.00236
0
ID

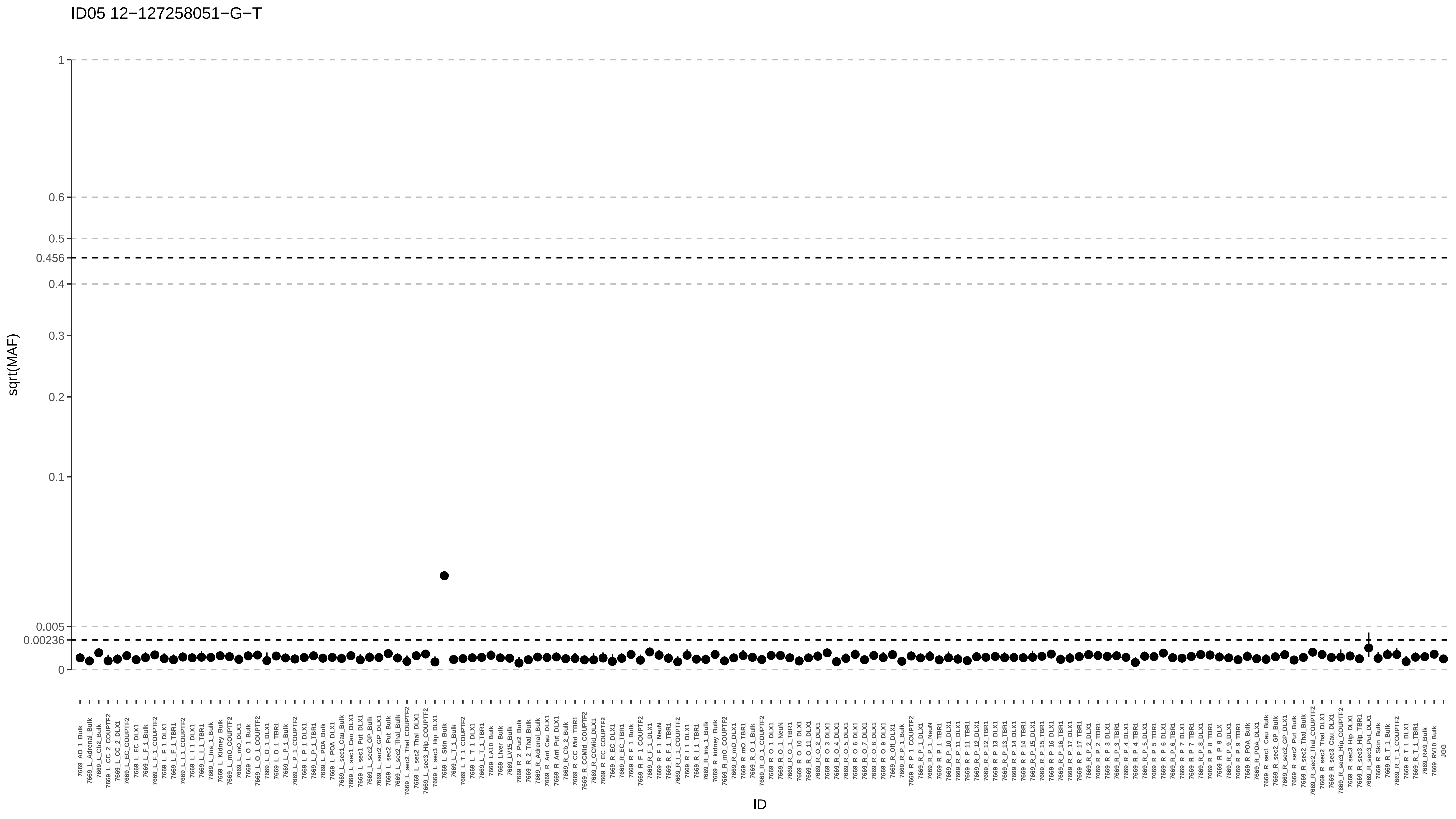
ID05 12−127258051−G−T
sqrt(MAF)
1
0.6
0.5
0.456
0.4
0.3
0.2
0.1
0.005
0.00236
0
ID

ID05 12−128703875−C−T

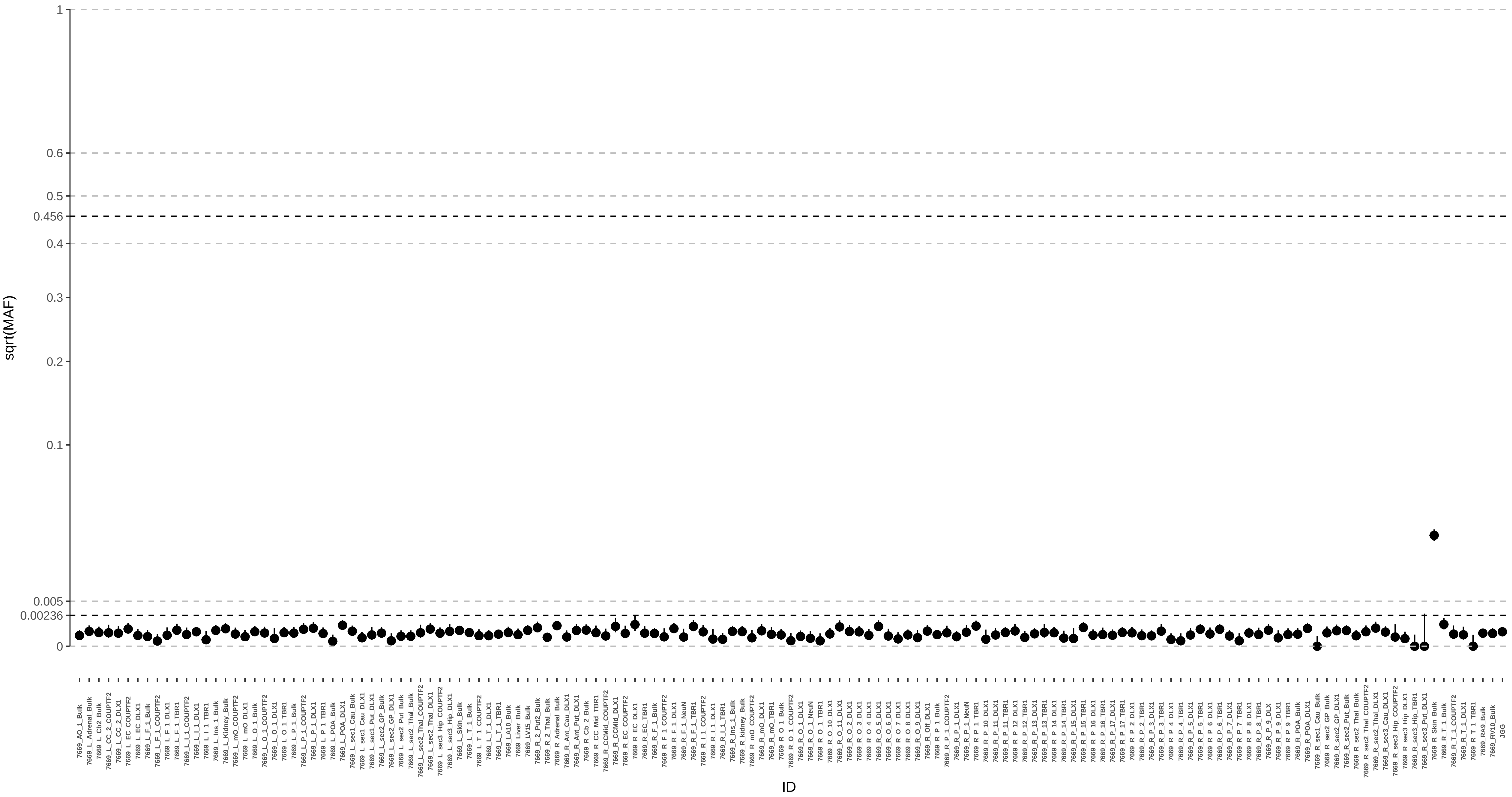

ID05 12−130892713−C−T

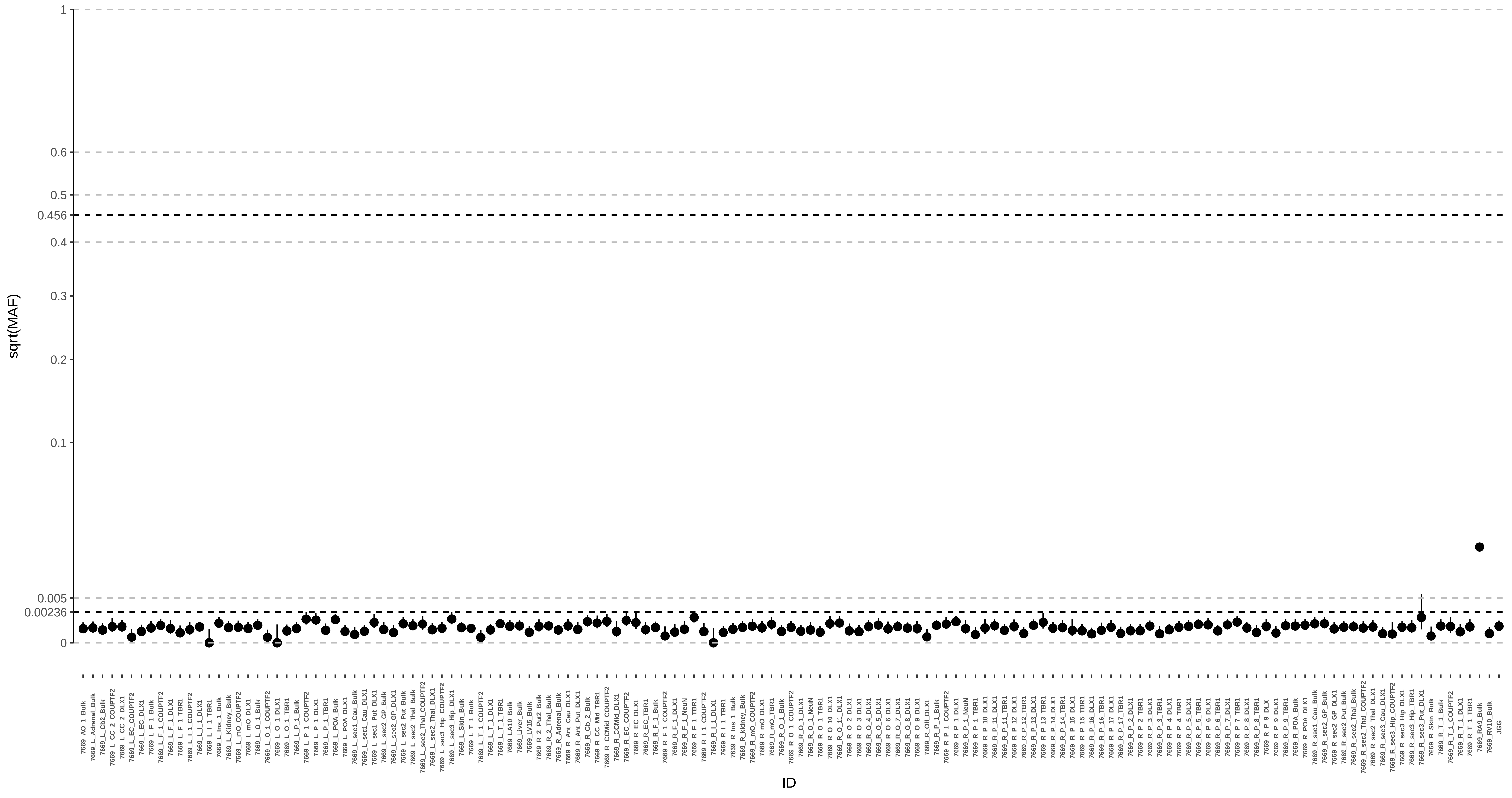

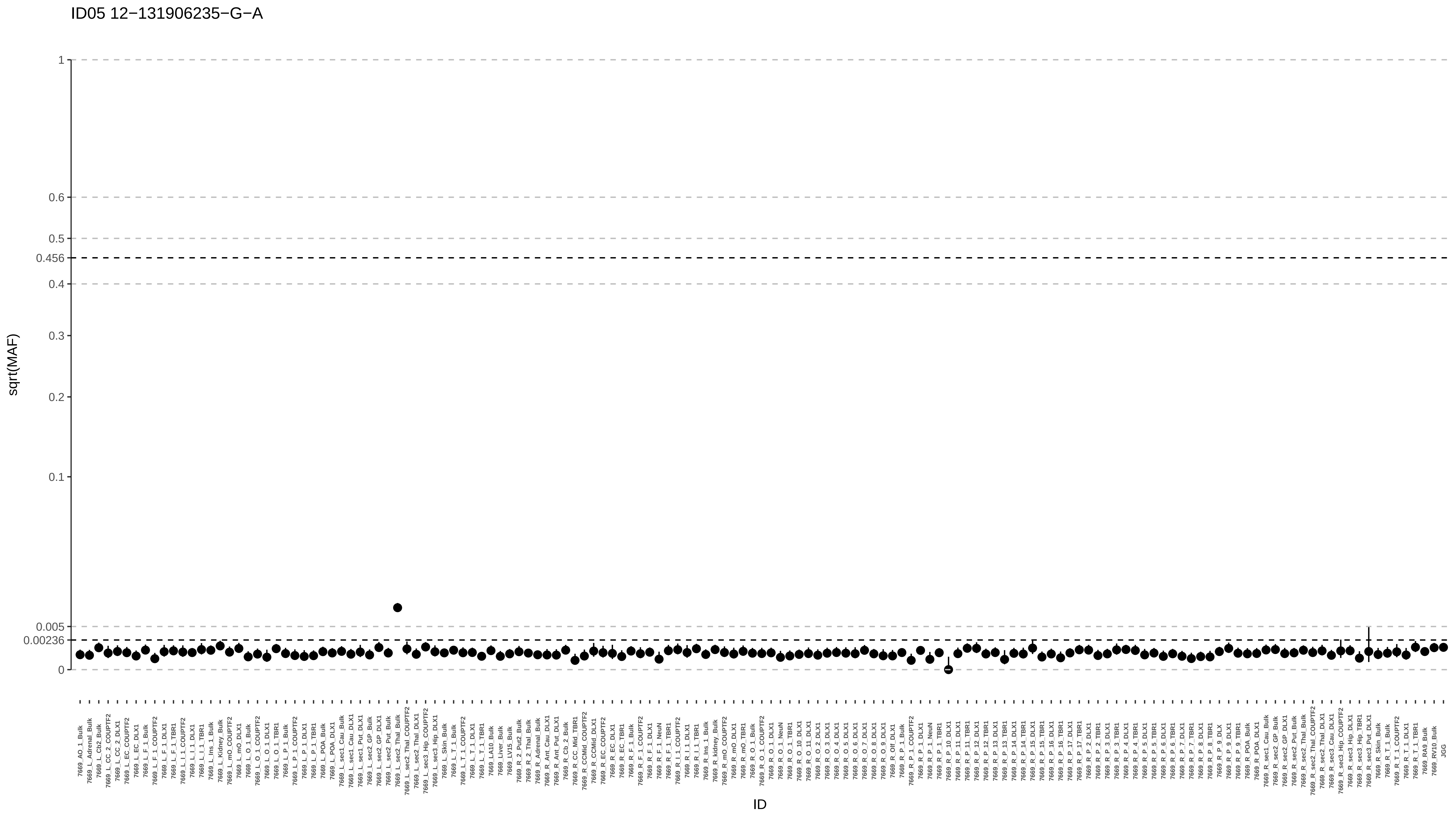

ID05 12−131906235−G−A
sqrt(MAF)
1
0.6
0.5
0.456
0.4
0.3
0.2
0.1
0.005
0.00236
0
ID

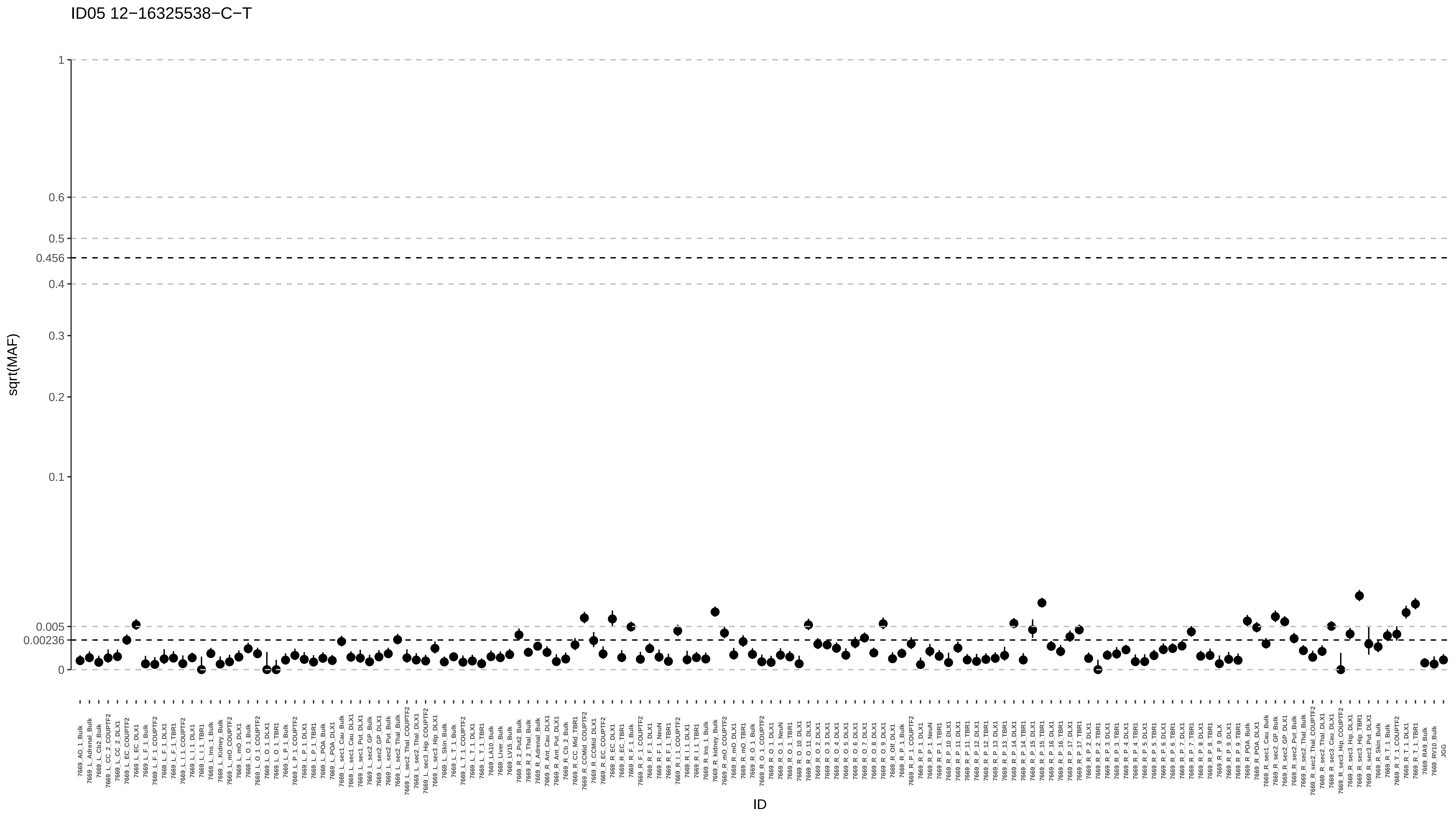
ID05 12−16325538−C−T
sqrt(MAF)
1
0.6
0.5
0.456
0.4
0.3
0.2
0.1
0.005
0.00236
0
ID

ID05 12−17467014−G−T

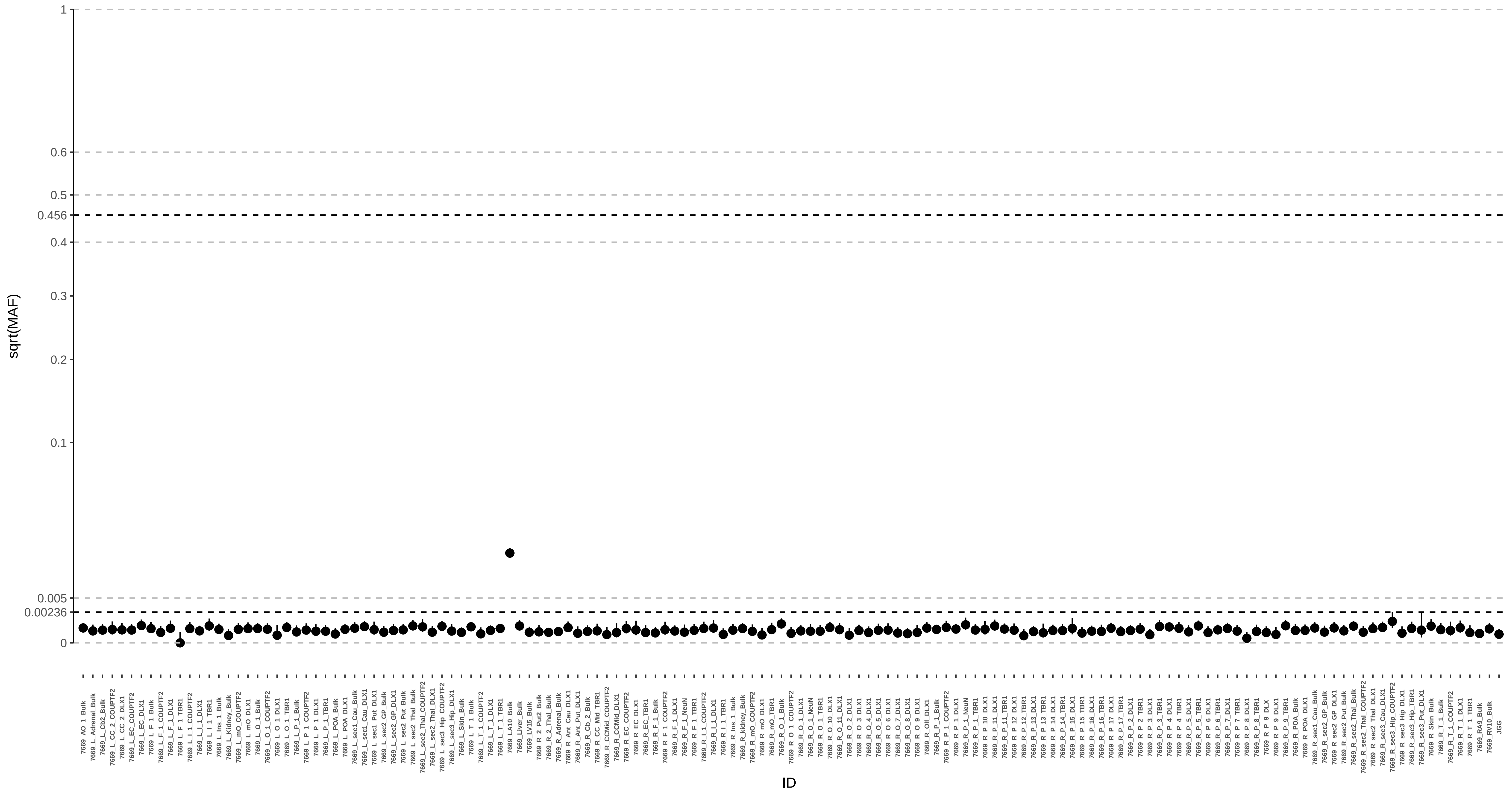

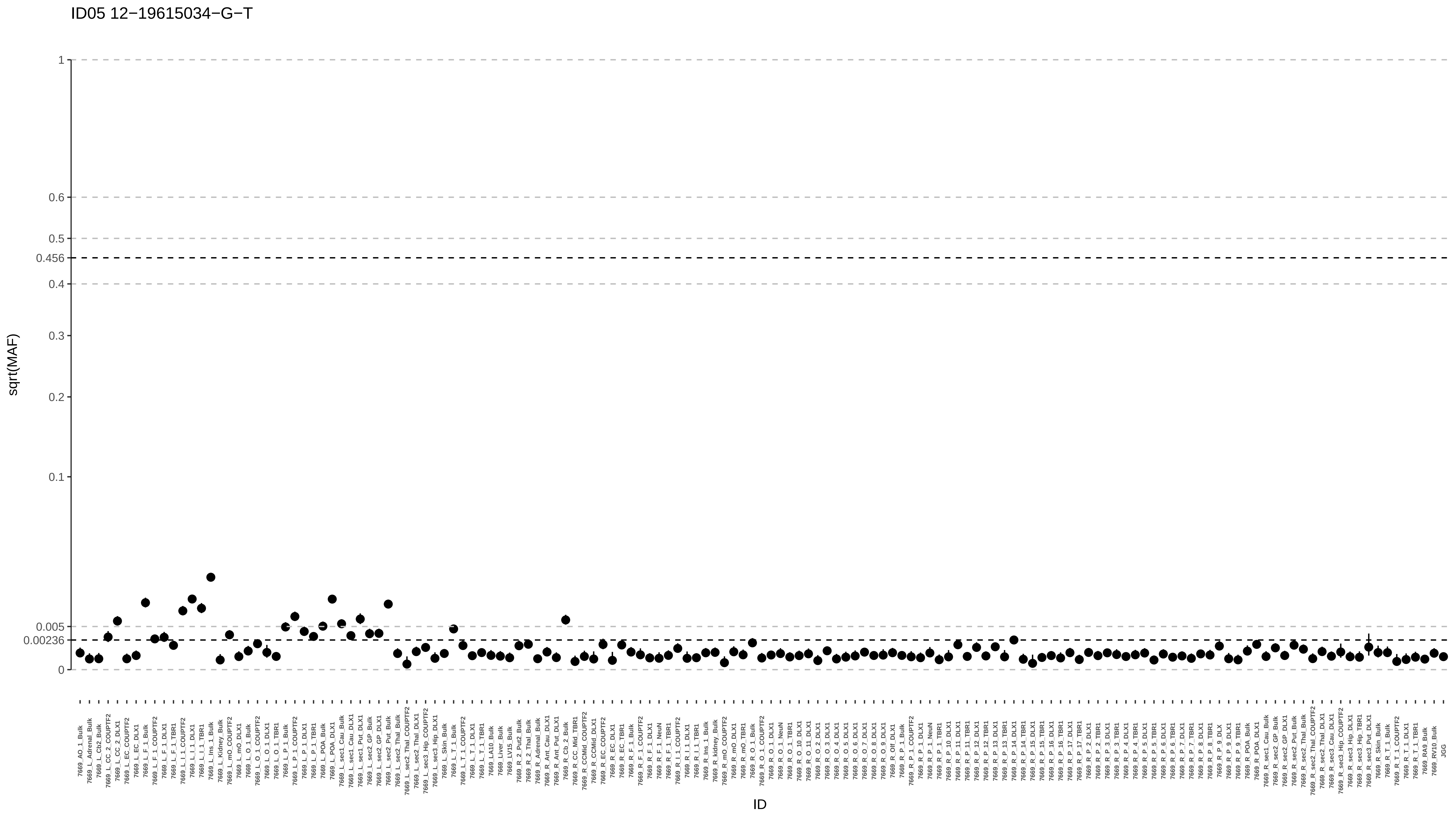
ID05 12−19615034−G−T
sqrt(MAF)
1
0.6
0.5
0.456
0.4
0.3
0.2
0.1
0.005
0.00236
0
ID

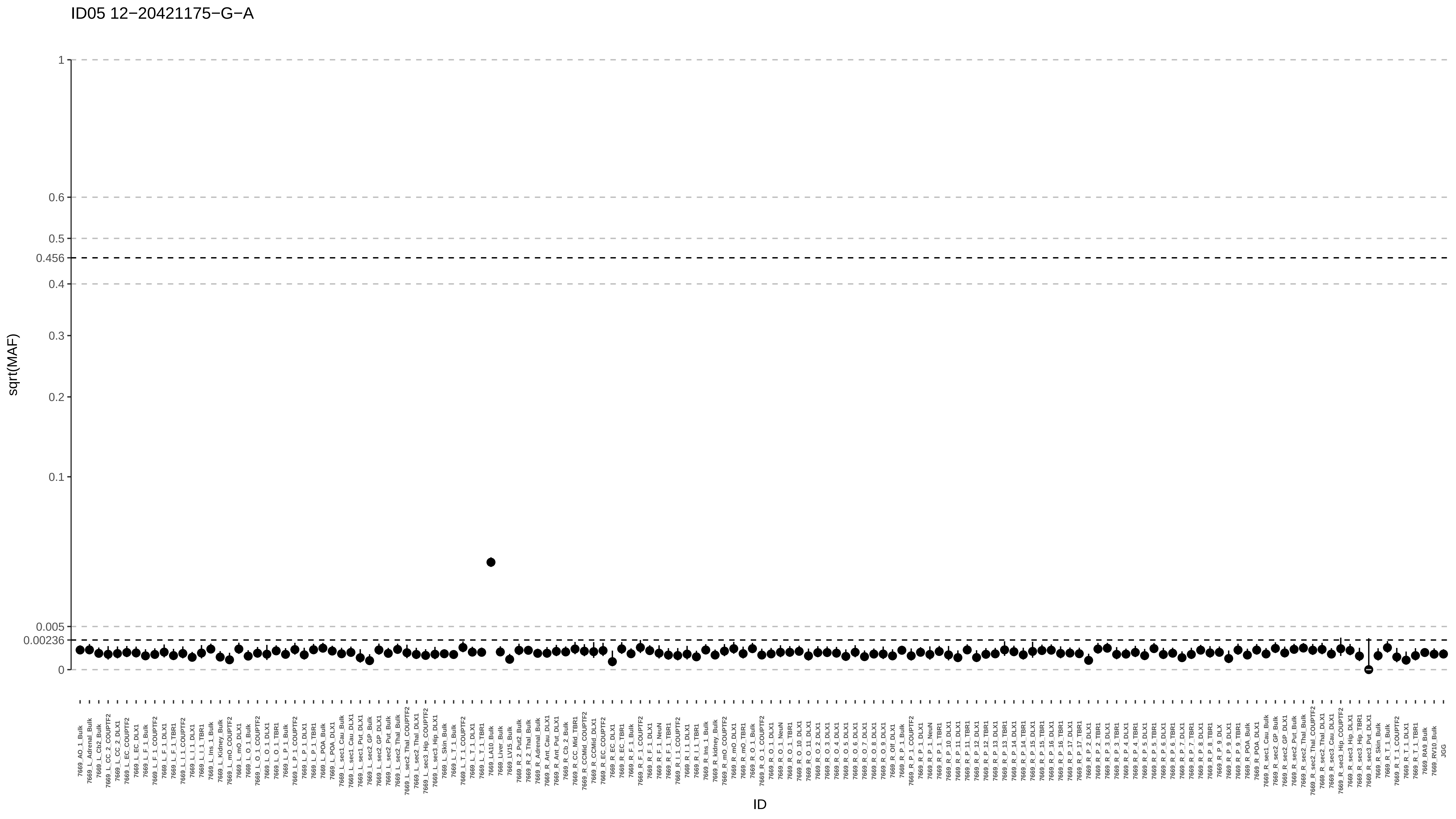
ID05 12−20421175−G−A
sqrt(MAF)
1
0.6
0.5
0.456
0.4
0.3
0.2
0.1
0.005
0.00236
0
ID

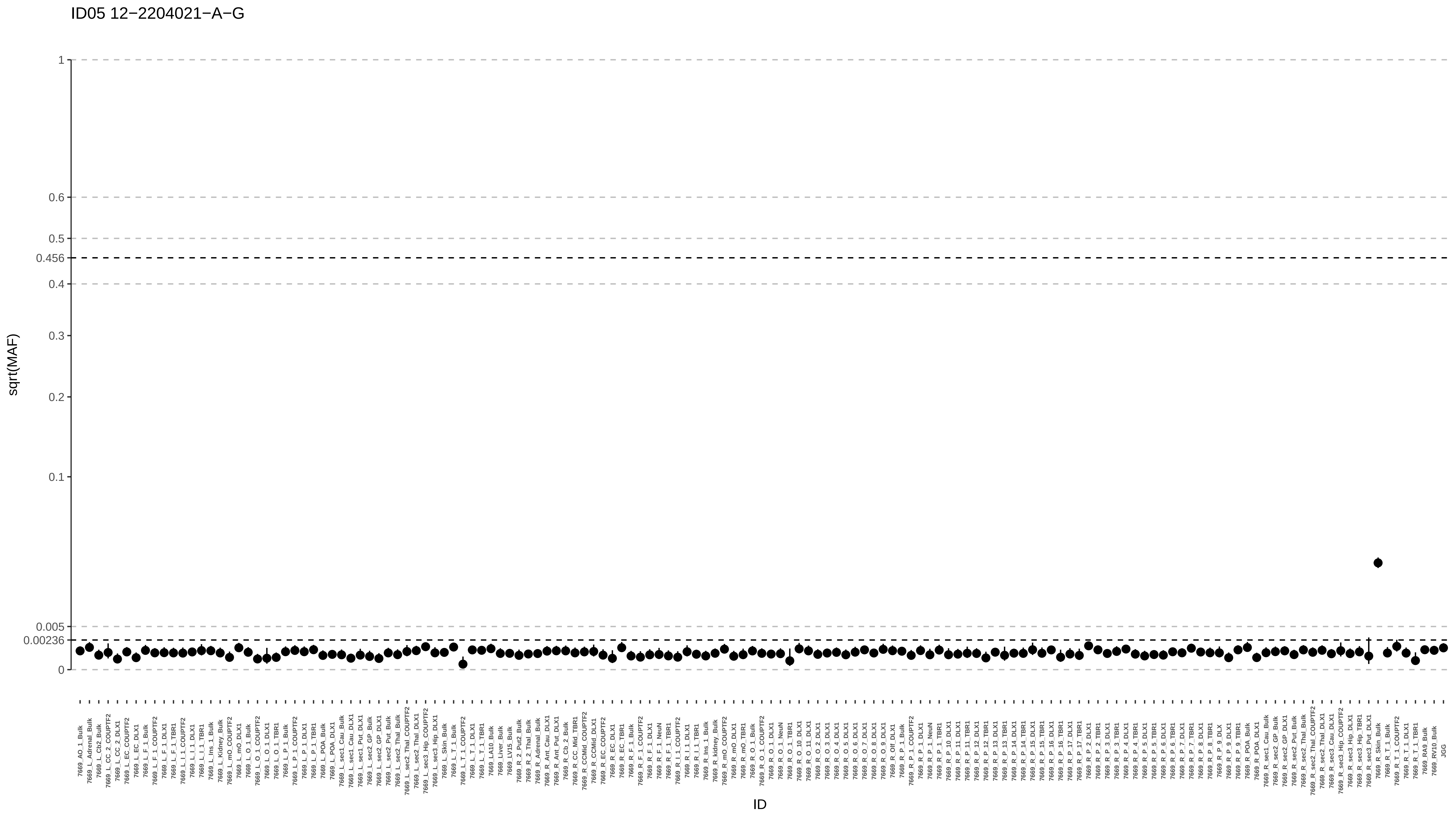
ID05 12−2204021−A−G
sqrt(MAF)
1
0.6
0.5
0.456
0.4
0.3
0.2
0.1
0.005
0.00236
0
ID

ID05 12−24436884−C−T

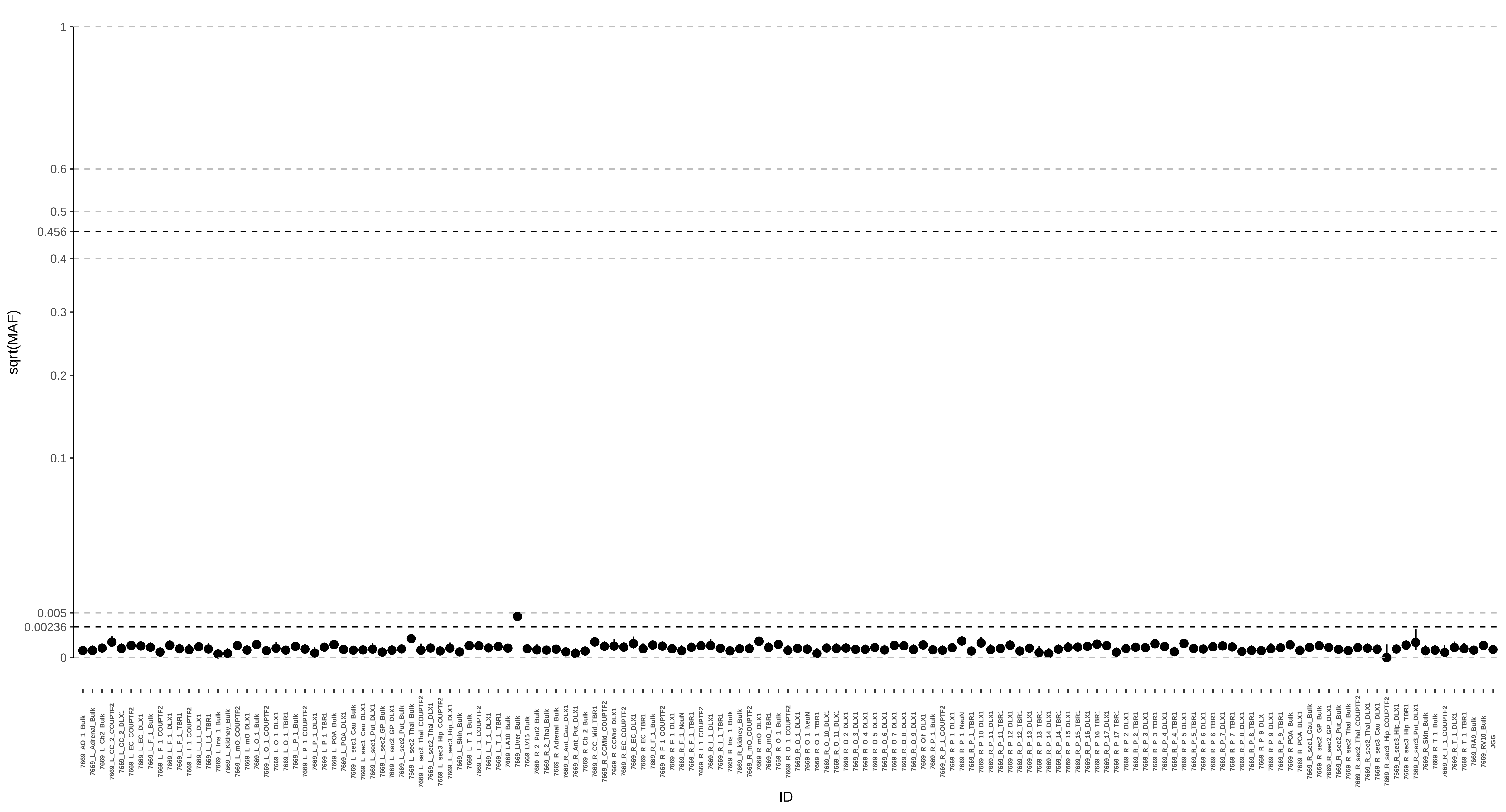

ID05 12−55677268−C−A

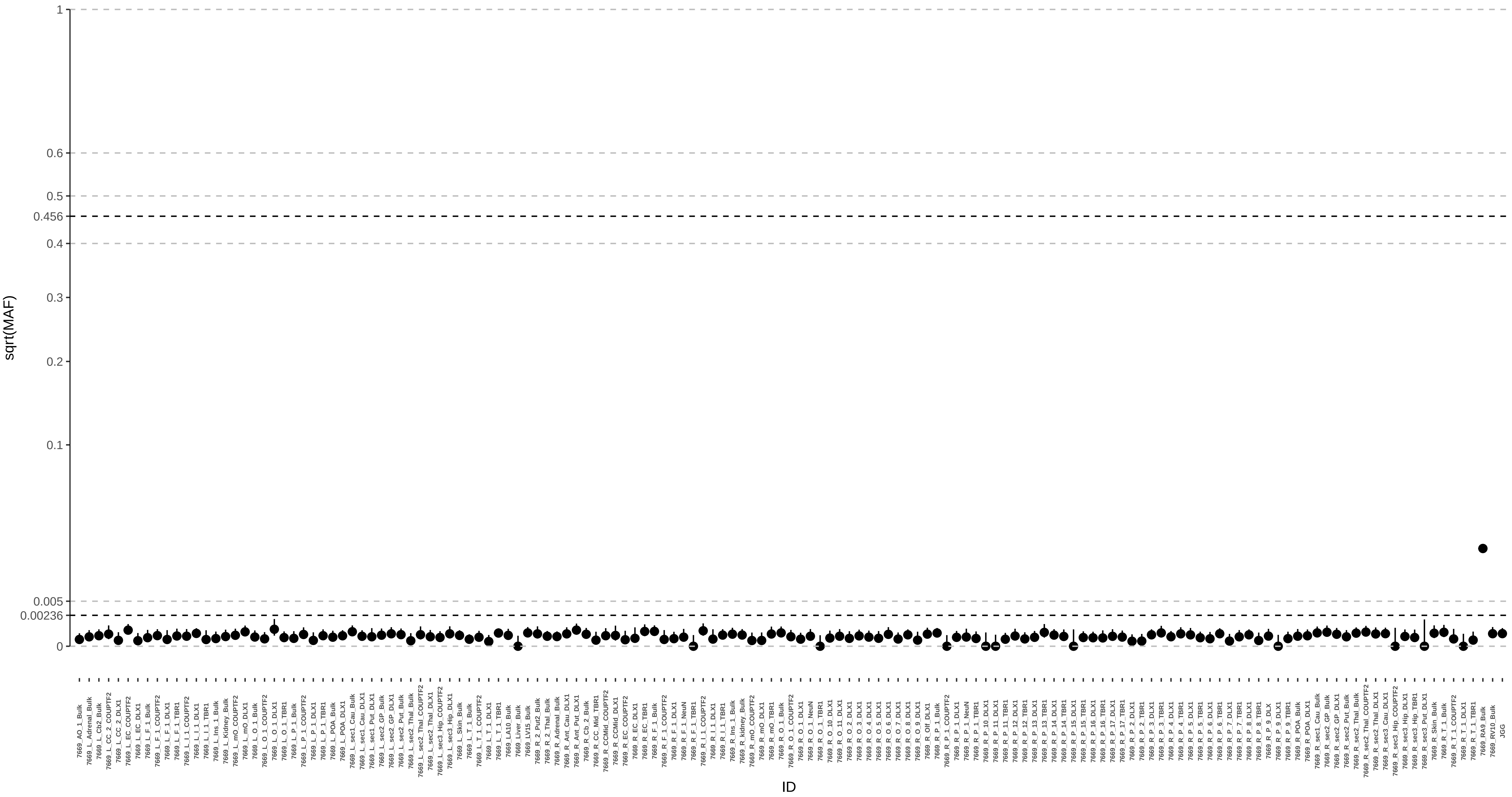

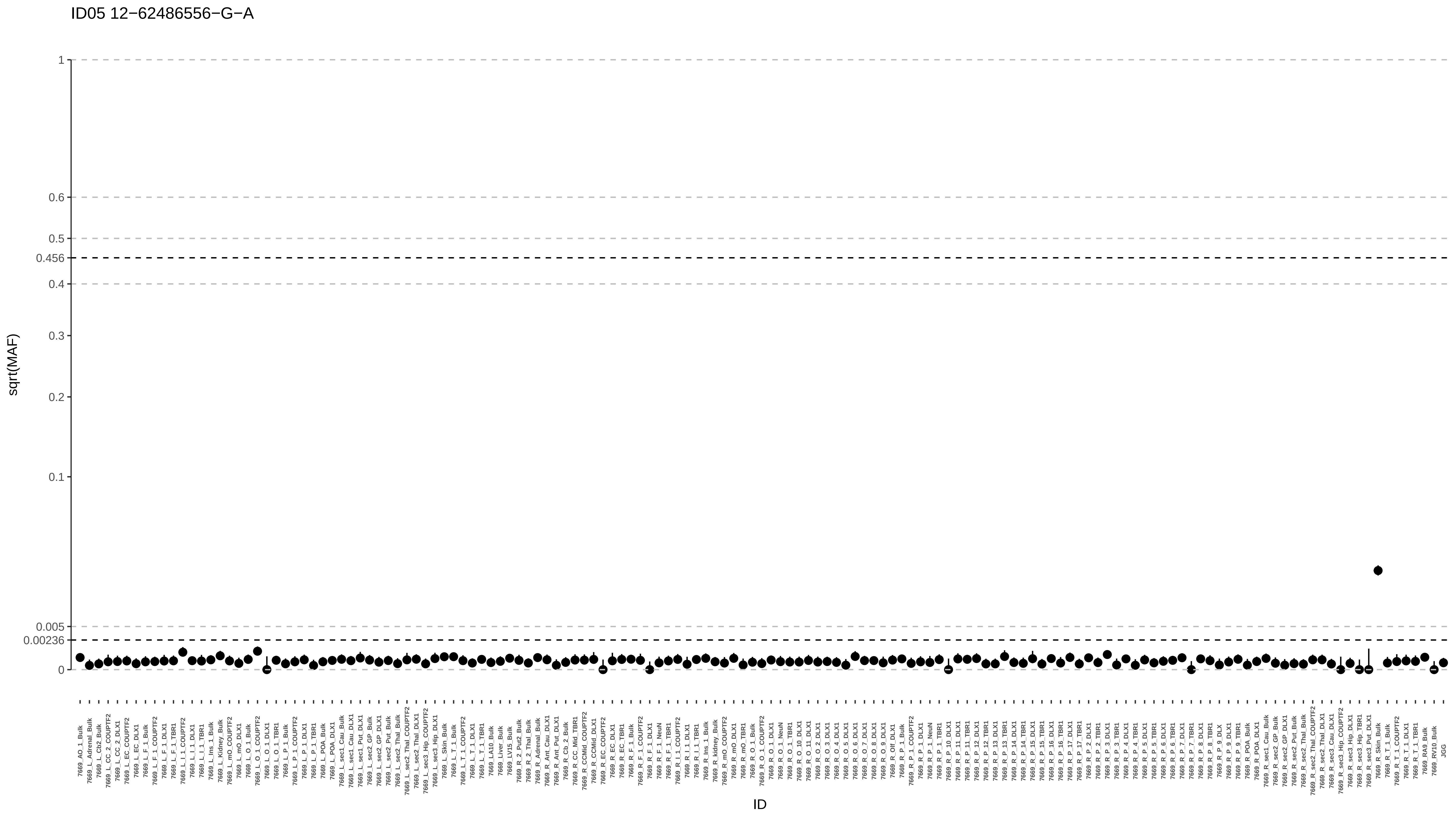
ID05 12−62486556−G−A
sqrt(MAF)
1
0.6
0.5
0.456
0.4
0.3
0.2
0.1
0.005
0.00236
0
ID

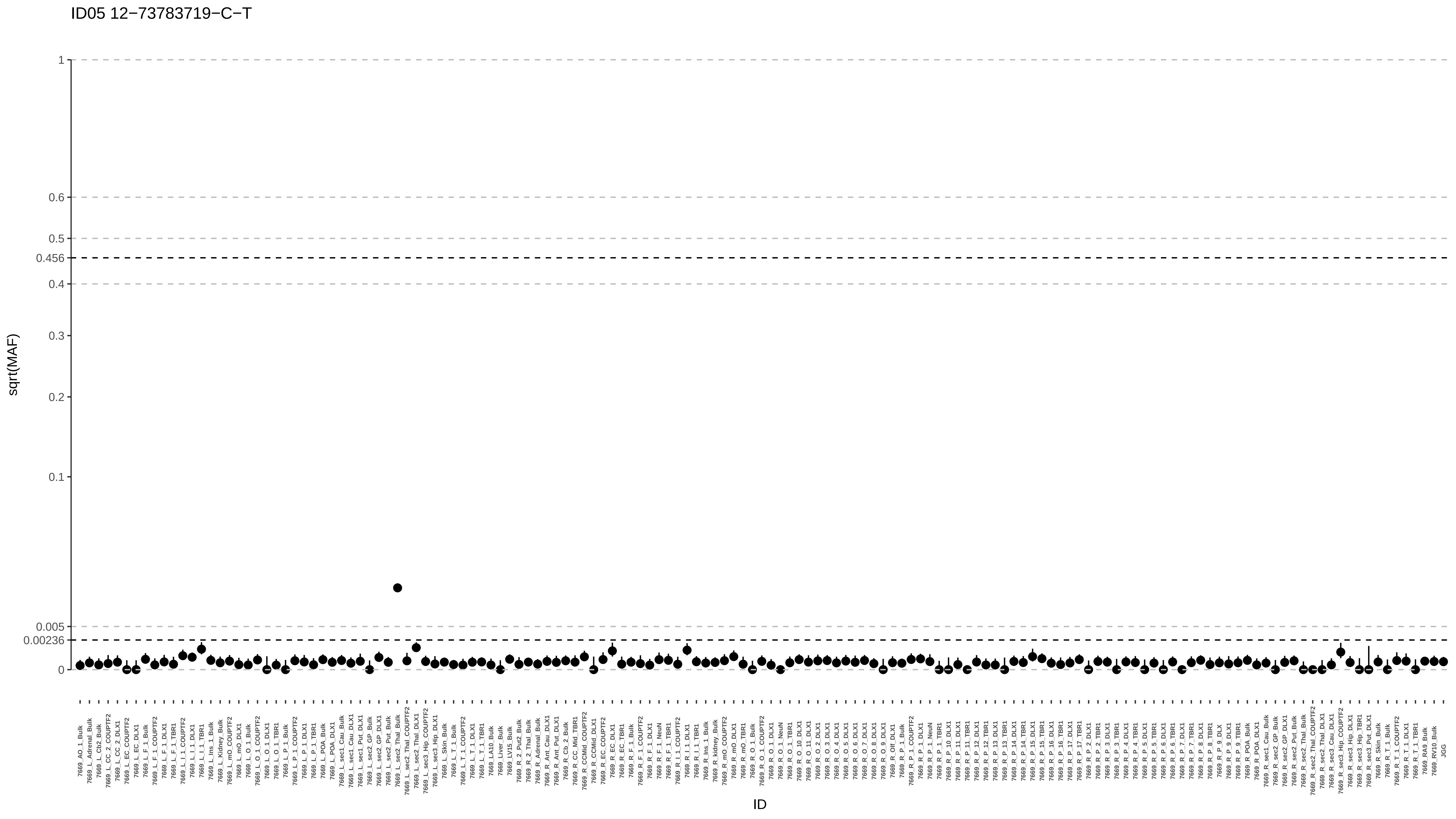
ID05 12−73783719−C−T
sqrt(MAF)
ID
1
0.6
0.5
0.456
0.4
0.3
0.2
0.1
0.005
0.00236
0

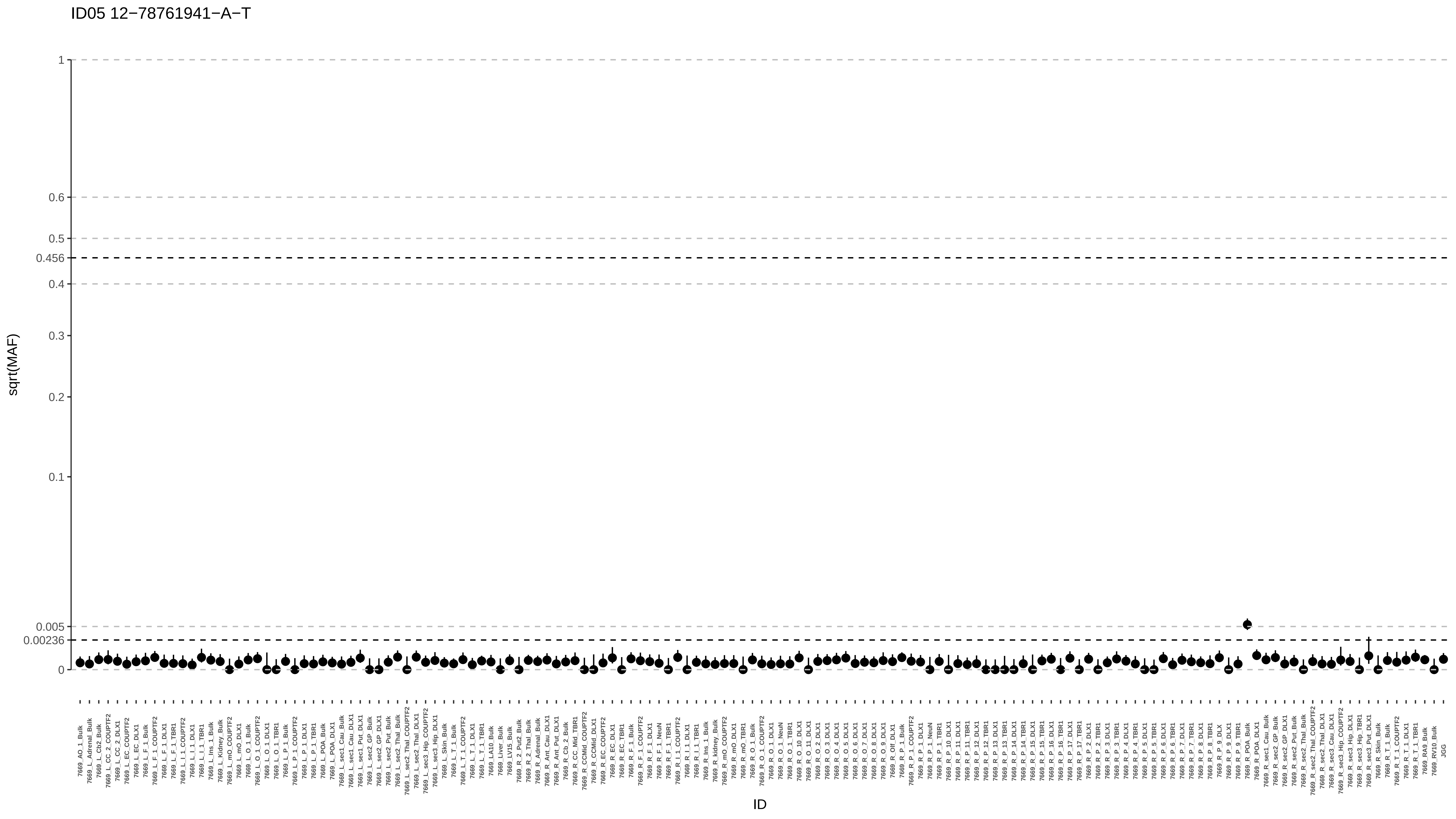
ID05 12−78761941−A−T
sqrt(MAF)
1
0.6
0.5
0.456
0.4
0.3
0.2
0.1
0.005
0.00236
0
ID
7669_AO_1_Bulk
7669_L_Adrenal_Bulk
7669_L_Cb2_Bulk
7669_L_CC_2_COUPTF2
7669_L_CC_2_DLX1
7669_L_EC_COUPTF2
7669_L_EC_DLX1
7669_L_F_1_Bulk
7669_L_F_1_COUPTF2
7669_L_F_1_DLX1
7669_L_F_1_TBR1
7669_L_I_1_COUPTF2
7669_L_I_1_DLX1
7669_L_I_1_TBR1
7669_L_Ins_1_Bulk
7669_L_Kidney_Bulk
7669_L_mO_COUPTF2
7669_L_mO_DLX1
7669_L_O_1_Bulk
7669_L_O_1_COUPTF2
7669_L_O_1_DLX1
7669_L_O_1_TBR1
7669_L_P_1_Bulk
7669_L_P_1_COUPTF2
7669_L_P_1_DLX1
7669_L_P_1_TBR1
7669_L_POA_Bulk
7669_L_POA_DLX1
7669_L_sec1_Cau_Bulk
7669_L_sec1_Cau_DLX1
7669_L_sec1_Put_DLX1
7669_L_sec2_GP_Bulk
7669_L_sec2_GP_DLX1
7669_L_sec2_Put_Bulk
7669_L_sec2_Thal_Bulk
7669_L_sec2_Thal_COUPTF2
7669_L_sec2_Thal_DLX1
7669_L_sec3_Hip_COUPTF2
7669_L_sec3_Hip_DLX1
7669_L_Skin_Bulk
7669_L_T_1_Bulk
7669_L_T_1_COUPTF2
7669_L_T_1_DLX1
7669_L_T_1_TBR1
7669_LA10_Bulk
7669_Liver_Bulk
7669_LV15_Bulk
7669_R_2_Put2_Bulk
7669_R_2_Thal_Bulk
7669_R_Adrenal_Bulk
7669_R_Ant_Cau_DLX1
7669_R_Ant_Put_DLX1
7669_R_Cb_2_Bulk
7669_R_CC_Mid_TBR1
7669_R_CCMid_COUPTF2
7669_R_CCMid_DLX1
7669_R_EC_COUPTF2
7669_R_EC_DLX1
7669_R_EC_TBR1
7669_R_F_1_Bulk
7669_R_F_1_COUPTF2
7669_R_F_1_DLX1
7669_R_F_1_NeuN
7669_R_F_1_TBR1
7669_R_I_1_COUPTF2
7669_R_I_1_DLX1
7669_R_I_1_TBR1
7669_R_Ins_1_Bulk
7669_R_kidney_Bulk
7669_R_mO_COUPTF2
7669_R_mO_DLX1
7669_R_mO_TBR1
7669_R_O_1_Bulk
7669_R_O_1_COUPTF2
7669_R_O_1_DLX1
7669_R_O_1_NeuN
7669_R_O_1_TBR1
7669_R_O_10_DLX1
7669_R_O_11_DLX1
7669_R_O_2_DLX1
7669_R_O_3_DLX1
7669_R_O_4_DLX1
7669_R_O_5_DLX1
7669_R_O_6_DLX1
7669_R_O_7_DLX1
7669_R_O_8_DLX1
7669_R_O_9_DLX1
7669_R_Olf_DLX1
7669_R_P_1_Bulk
7669_R_P_1_COUPTF2
7669_R_P_1_DLX1
7669_R_P_1_NeuN
7669_R_P_1_TBR1
7669_R_P_10_DLX1
7669_R_P_11_DLX1
7669_R_P_11_TBR1
7669_R_P_12_DLX1
7669_R_P_12_TBR1
7669_R_P_13_DLX1
7669_R_P_13_TBR1
7669_R_P_14_DLX1
7669_R_P_14_TBR1
7669_R_P_15_DLX1
7669_R_P_15_TBR1
7669_R_P_16_DLX1
7669_R_P_16_TBR1
7669_R_P_17_DLX1
7669_R_P_17_TBR1
7669_R_P_2_DLX1
7669_R_P_2_TBR1
7669_R_P_3_DLX1
7669_R_P_3_TBR1
7669_R_P_4_DLX1
7669_R_P_4_TBR1
7669_R_P_5_DLX1
7669_R_P_5_TBR1
7669_R_P_6_DLX1
7669_R_P_6_TBR1
7669_R_P_7_DLX1
7669_R_P_7_TBR1
7669_R_P_8_DLX1
7669_R_P_8_TBR1
7669_R_P_9_DLX
7669_R_P_9_DLX1
7669_R_P_9_TBR1
7669_R_POA_Bulk
7669_R_POA_DLX1
7669_R_sec1_Cau_Bulk
7669_R_sec2_GP_Bulk
7669_R_sec2_GP_DLX1
7669_R_sec2_Put_Bulk
7669_R_sec2_Thal_Bulk
7669_R_sec2_Thal_COUPTF2
7669_R_sec2_Thal_DLX1
7669_R_sec3_Cau_DLX1
7669_R_sec3_Hip_COUPTF2
7669_R_sec3_Hip_DLX1
7669_R_sec3_Hip_TBR1
7669_R_sec3_Put_DLX1
7669_R_Skin_Bulk
7669_R_T_1_Bulk
7669_R_T_1_COUPTF2
7669_R_T_1_DLX1
7669_R_T_1_TBR1
7669_RA9_Bulk
7669_RV10_Bulk
JGG

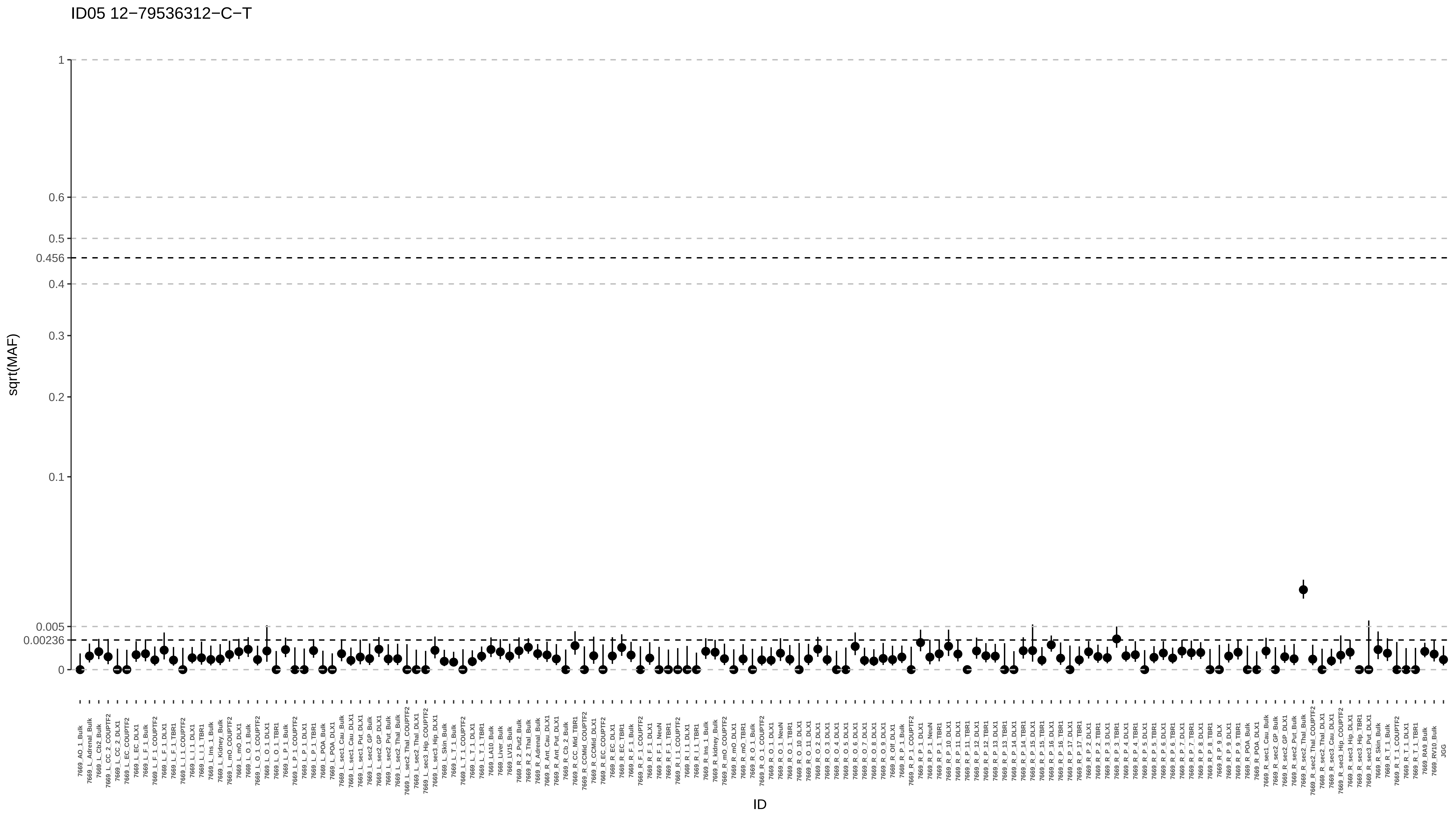
ID05 12−79536312−C−T
sqrt(MAF)
1
0.6
0.5
0.456
0.4
0.3
0.2
0.1
0.005
0.00236
0
ID
7669_AO_1_Bulk
7669_L_Adrenal_Bulk
7669_L_Cb2_Bulk
7669_L_CC_2_COUPTF2
7669_L_CC_2_DLX1
7669_L_EC_COUPTF2
7669_L_EC_DLX1
7669_L_F_1_Bulk
7669_L_F_1_COUPTF2
7669_L_F_1_DLX1
7669_L_F_1_TBR1
7669_L_I_1_COUPTF2
7669_L_I_1_DLX1
7669_L_I_1_TBR1
7669_L_Ins_1_Bulk
7669_L_Kidney_Bulk
7669_L_mO_COUPTF2
7669_L_mO_DLX1
7669_L_O_1_Bulk
7669_L_O_1_COUPTF2
7669_L_O_1_DLX1
7669_L_O_1_TBR1
7669_L_P_1_Bulk
7669_L_P_1_COUPTF2
7669_L_P_1_DLX1
7669_L_P_1_TBR1
7669_L_POA_Bulk
7669_L_POA_DLX1
7669_L_sec1_Cau_Bulk
7669_L_sec1_Cau_DLX1
7669_L_sec1_Put_DLX1
7669_L_sec2_GP_Bulk
7669_L_sec2_GP_DLX1
7669_L_sec2_Put_Bulk
7669_L_sec2_Thal_Bulk
7669_L_sec2_Thal_COUPTF2
7669_L_sec2_Thal_DLX1
7669_L_sec3_Hip_COUPTF2
7669_L_sec3_Hip_DLX1
7669_L_Skin_Bulk
7669_L_T_1_Bulk
7669_L_T_1_COUPTF2
7669_L_T_1_DLX1
7669_L_T_1_TBR1
7669_LA10_Bulk
7669_Liver_Bulk
7669_LV15_Bulk
7669_R_2_Put2_Bulk
7669_R_2_Thal_Bulk
7669_R_Adrenal_Bulk
7669_R_Ant_Cau_DLX1
7669_R_Ant_Put_DLX1
7669_R_Cb_2_Bulk
7669_R_CC_Mid_TBR1
7669_R_CCMid_COUPTF2
7669_R_CCMid_DLX1
7669_R_EC_COUPTF2
7669_R_EC_DLX1
7669_R_EC_TBR1
7669_R_F_1_Bulk
7669_R_F_1_COUPTF2
7669_R_F_1_DLX1
7669_R_F_1_NeuN
7669_R_F_1_TBR1
7669_R_I_1_COUPTF2
7669_R_I_1_DLX1
7669_R_I_1_TBR1
7669_R_Ins_1_Bulk
7669_R_kidney_Bulk
7669_R_mO_COUPTF2
7669_R_mO_DLX1
7669_R_mO_TBR1
7669_R_O_1_Bulk
7669_R_O_1_COUPTF2
7669_R_O_1_DLX1
7669_R_O_1_NeuN
7669_R_O_1_TBR1
7669_R_O_10_DLX1
7669_R_O_11_DLX1
7669_R_O_2_DLX1
7669_R_O_3_DLX1
7669_R_O_4_DLX1
7669_R_O_5_DLX1
7669_R_O_6_DLX1
7669_R_O_7_DLX1
7669_R_O_8_DLX1
7669_R_O_9_DLX1
7669_R_Olf_DLX1
7669_R_P_1_Bulk
7669_R_P_1_COUPTF2
7669_R_P_1_DLX1
7669_R_P_1_NeuN
7669_R_P_1_TBR1
7669_R_P_10_DLX1
7669_R_P_11_DLX1
7669_R_P_11_TBR1
7669_R_P_12_DLX1
7669_R_P_12_TBR1
7669_R_P_13_DLX1
7669_R_P_13_TBR1
7669_R_P_14_DLX1
7669_R_P_14_TBR1
7669_R_P_15_DLX1
7669_R_P_15_TBR1
7669_R_P_16_DLX1
7669_R_P_16_TBR1
7669_R_P_17_DLX1
7669_R_P_17_TBR1
7669_R_P_2_DLX1
7669_R_P_2_TBR1
7669_R_P_3_DLX1
7669_R_P_3_TBR1
7669_R_P_4_DLX1
7669_R_P_4_TBR1
7669_R_P_5_DLX1
7669_R_P_5_TBR1
7669_R_P_6_DLX1
7669_R_P_6_TBR1
7669_R_P_7_DLX1
7669_R_P_7_TBR1
7669_R_P_8_DLX1
7669_R_P_8_TBR1
7669_R_P_9_DLX
7669_R_P_9_DLX1
7669_R_P_9_TBR1
7669_R_POA_Bulk
7669_R_POA_DLX1
7669_R_sec1_Cau_Bulk
7669_R_sec2_GP_Bulk
7669_R_sec2_GP_DLX1
7669_R_sec2_Put_Bulk
7669_R_sec2_Thal_Bulk
7669_R_sec2_Thal_COUPTF2
7669_R_sec2_Thal_DLX1
7669_R_sec3_Cau_DLX1
7669_R_sec3_Hip_COUPTF2
7669_R_sec3_Hip_DLX1
7669_R_sec3_Hip_TBR1
7669_R_sec3_Put_DLX1
7669_R_Skin_Bulk
7669_R_T_1_Bulk
7669_R_T_1_COUPTF2
7669_R_T_1_DLX1
7669_R_T_1_TBR1
7669_RA9_Bulk
7669_RV10_Bulk
JGG

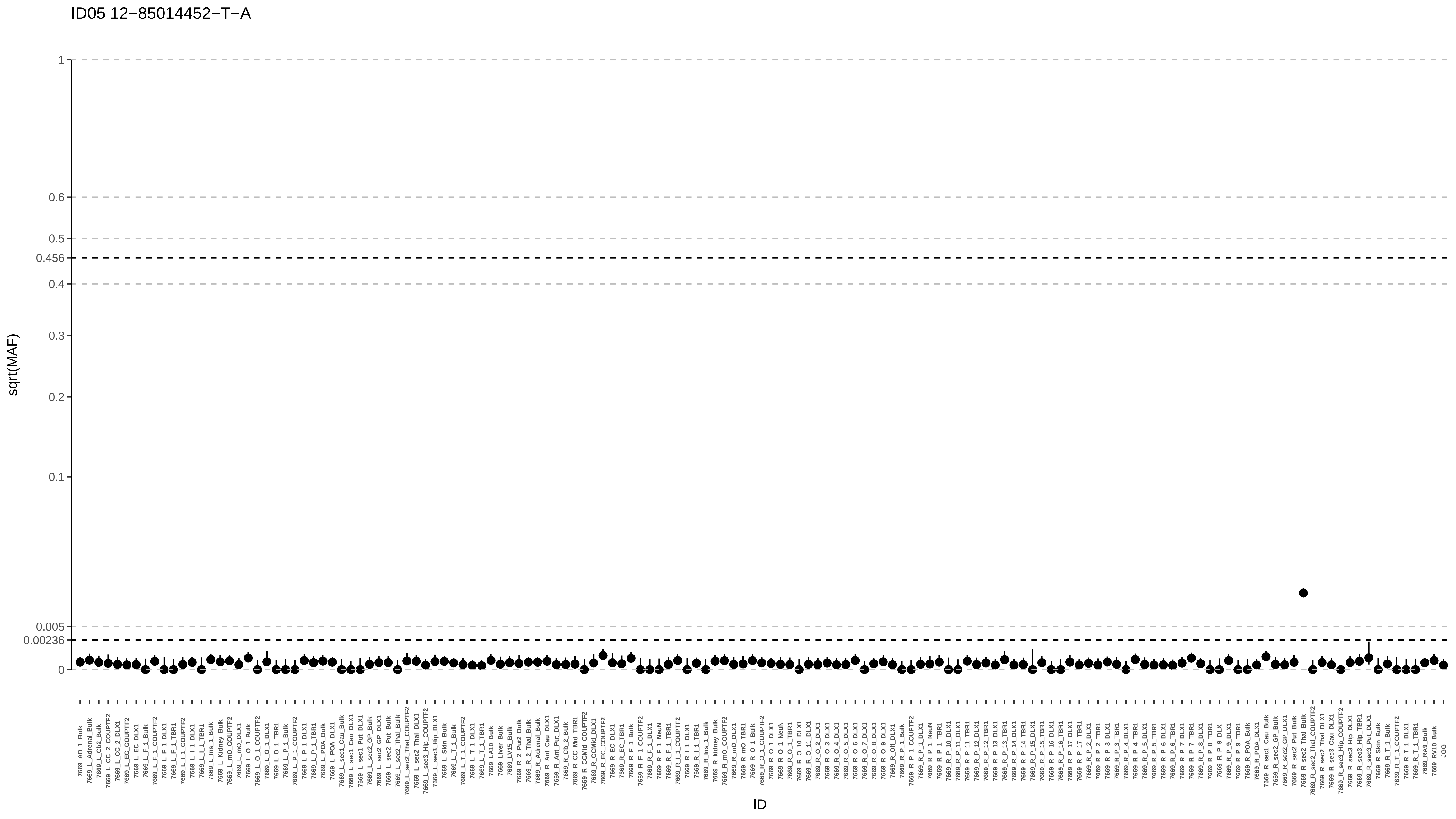
ID05 12−85014452−T−A
sqrt(MAF)
1
0.6
0.5
0.456
0.4
0.3
0.2
0.1
0.005
0.00236
0
ID

ID05 12−85787527−A−T

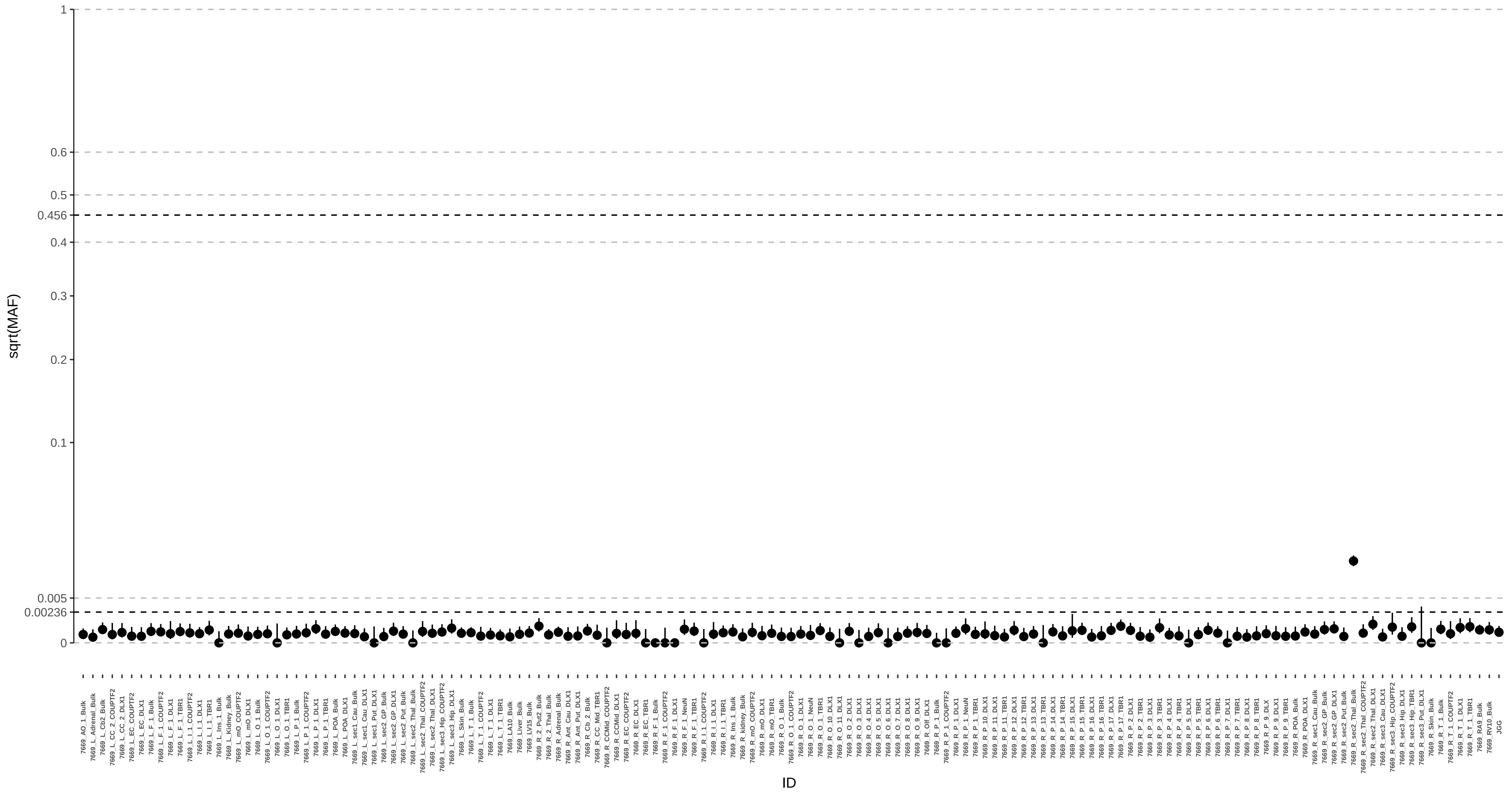

ID05 12−90774790−C−T

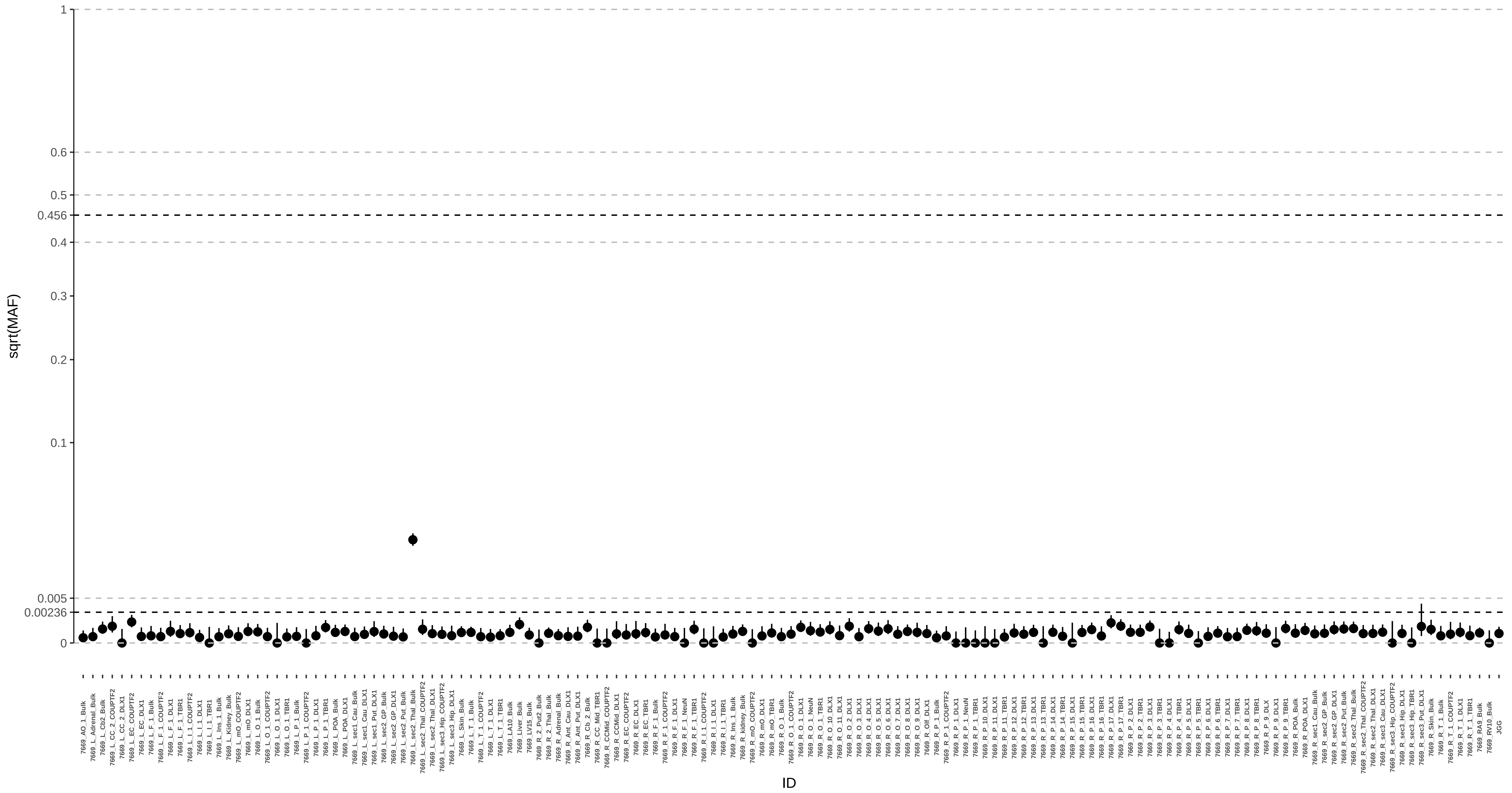

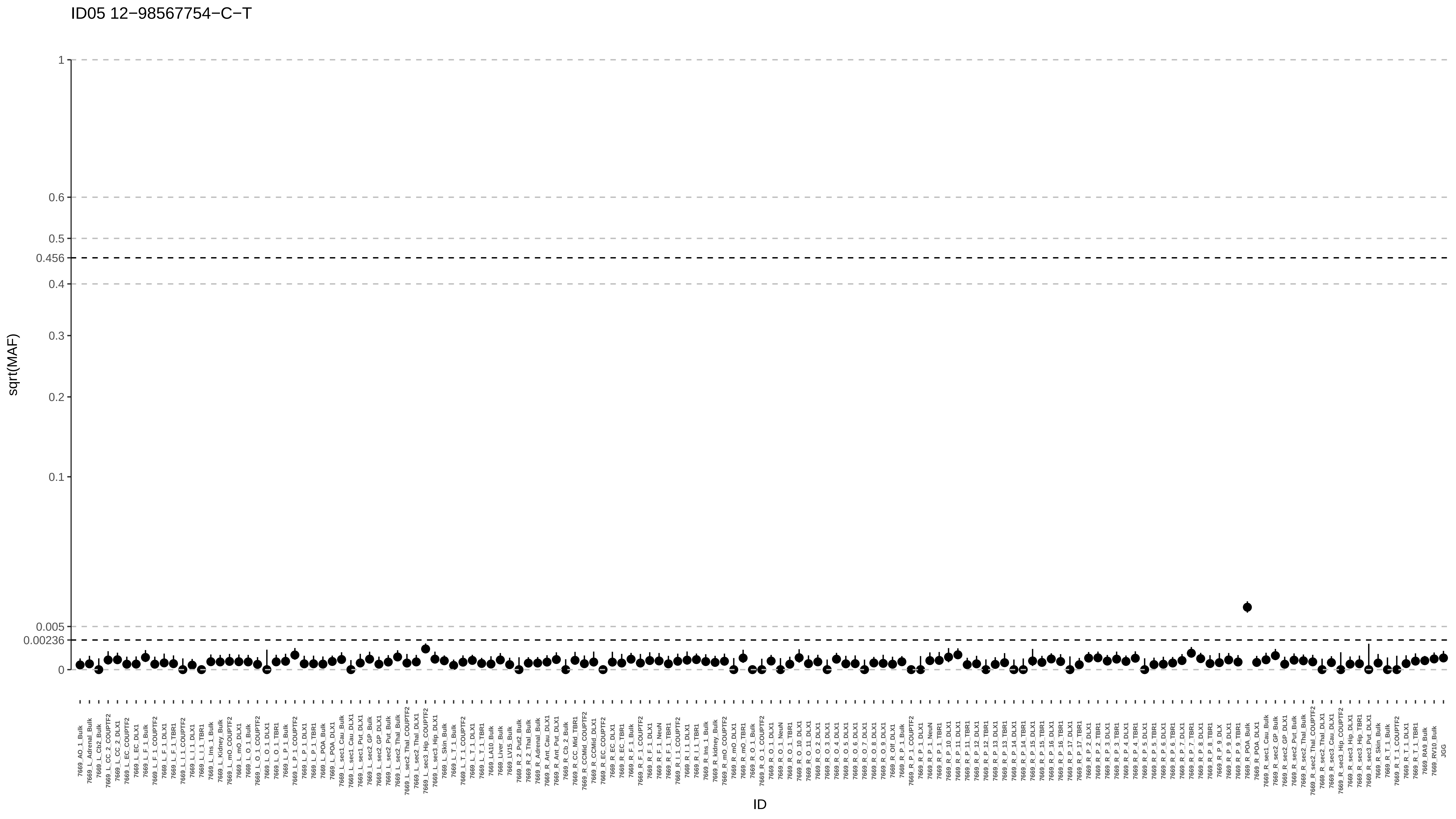
ID05 12−98567754−C−T
sqrt(MAF)
1
0.6
0.5
0.456
0.4
0.3
0.2
0.1
0.005
0.00236
0
ID
7669_AO_1_Bulk
7669_L_Adrenal_Bulk
7669_L_Cb2_Bulk
7669_L_CC_2_COUPTF2
7669_L_CC_2_DLX1
7669_L_EC_COUPTF2
7669_L_EC_DLX1
7669_L_F_1_Bulk
7669_L_F_1_COUPTF2
7669_L_F_1_DLX1
7669_L_F_1_TBR1
7669_L_I_1_COUPTF2
7669_L_I_1_DLX1
7669_L_I_1_TBR1
7669_L_Ins_1_Bulk
7669_L_Kidney_Bulk
7669_L_mO_COUPTF2
7669_L_mO_DLX1
7669_L_O_1_Bulk
7669_L_O_1_COUPTF2
7669_L_O_1_DLX1
7669_L_O_1_TBR1
7669_L_P_1_Bulk
7669_L_P_1_COUPTF2
7669_L_P_1_DLX1
7669_L_P_1_TBR1
7669_L_POA_Bulk
7669_L_POA_DLX1
7669_L_sec1_Cau_Bulk
7669_L_sec1_Cau_DLX1
7669_L_sec1_Put_DLX1
7669_L_sec2_GP_Bulk
7669_L_sec2_GP_DLX1
7669_L_sec2_Put_Bulk
7669_L_sec2_Thal_Bulk
7669_L_sec2_Thal_COUPTF2
7669_L_sec2_Thal_DLX1
7669_L_sec3_Hip_COUPTF2
7669_L_sec3_Hip_DLX1
7669_L_Skin_Bulk
7669_L_T_1_Bulk
7669_L_T_1_COUPTF2
7669_L_T_1_DLX1
7669_L_T_1_TBR1
7669_LA10_Bulk
7669_Liver_Bulk
7669_LV15_Bulk
7669_R_2_Put2_Bulk
7669_R_2_Thal_Bulk
7669_R_Adrenal_Bulk
7669_R_Ant_Cau_DLX1
7669_R_Ant_Put_DLX1
7669_R_Cb_2_Bulk
7669_R_CC_Mid_TBR1
7669_R_CCMid_COUPTF2
7669_R_CCMid_DLX1
7669_R_EC_COUPTF2
7669_R_EC_DLX1
7669_R_EC_TBR1
7669_R_F_1_Bulk
7669_R_F_1_COUPTF2
7669_R_F_1_DLX1
7669_R_F_1_NeuN
7669_R_F_1_TBR1
7669_R_I_1_COUPTF2
7669_R_I_1_DLX1
7669_R_I_1_TBR1
7669_R_Ins_1_Bulk
7669_R_kidney_Bulk
7669_R_mO_COUPTF2
7669_R_mO_DLX1
7669_R_mO_TBR1
7669_R_O_1_Bulk
7669_R_O_1_COUPTF2
7669_R_O_1_DLX1
7669_R_O_1_NeuN
7669_R_O_1_TBR1
7669_R_O_10_DLX1
7669_R_O_11_DLX1
7669_R_O_2_DLX1
7669_R_O_3_DLX1
7669_R_O_4_DLX1
7669_R_O_5_DLX1
7669_R_O_6_DLX1
7669_R_O_7_DLX1
7669_R_O_8_DLX1
7669_R_O_9_DLX1
7669_R_Olf_DLX1
7669_R_P_1_Bulk
7669_R_P_1_COUPTF2
7669_R_P_1_DLX1
7669_R_P_1_NeuN
7669_R_P_1_TBR1
7669_R_P_10_DLX1
7669_R_P_11_DLX1
7669_R_P_11_TBR1
7669_R_P_12_DLX1
7669_R_P_12_TBR1
7669_R_P_13_DLX1
7669_R_P_13_TBR1
7669_R_P_14_DLX1
7669_R_P_14_TBR1
7669_R_P_15_DLX1
7669_R_P_15_TBR1
7669_R_P_16_DLX1
7669_R_P_16_TBR1
7669_R_P_17_DLX1
7669_R_P_17_TBR1
7669_R_P_2_DLX1
7669_R_P_2_TBR1
7669_R_P_3_DLX1
7669_R_P_3_TBR1
7669_R_P_4_DLX1
7669_R_P_4_TBR1
7669_R_P_5_DLX1
7669_R_P_5_TBR1
7669_R_P_6_DLX1
7669_R_P_6_TBR1
7669_R_P_7_DLX1
7669_R_P_7_TBR1
7669_R_P_8_DLX1
7669_R_P_8_TBR1
7669_R_P_9_DLX
7669_R_P_9_DLX1
7669_R_P_9_TBR1
7669_R_POA_Bulk
7669_R_POA_DLX1
7669_R_sec1_Cau_Bulk
7669_R_sec2_GP_Bulk
7669_R_sec2_GP_DLX1
7669_R_sec2_Put_Bulk
7669_R_sec2_Thal_Bulk
7669_R_sec2_Thal_COUPTF2
7669_R_sec2_Thal_DLX1
7669_R_sec3_Cau_DLX1
7669_R_sec3_Hip_COUPTF2
7669_R_sec3_Hip_DLX1
7669_R_sec3_Hip_TBR1
7669_R_sec3_Put_DLX1
7669_R_Skin_Bulk
7669_R_T_1_Bulk
7669_R_T_1_COUPTF2
7669_R_T_1_DLX1
7669_R_T_1_TBR1
7669_RA9_Bulk
7669_RV10_Bulk
JGG

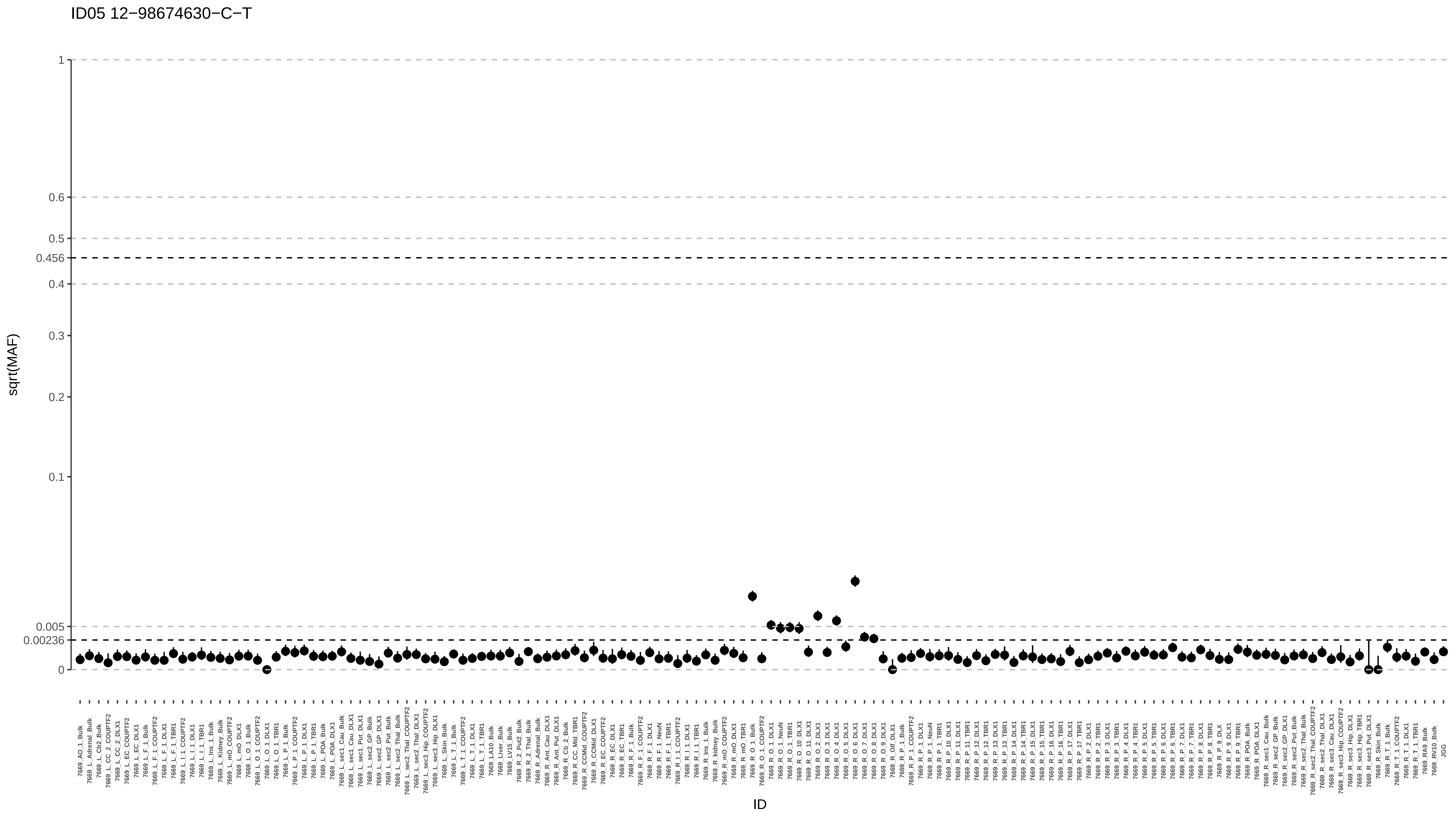

ID05 12−98674630−C−T
sqrt(MAF)
1
0.6
0.5
0.456
0.4
0.3
0.2
0.1
0.005
0.00236
0
ID

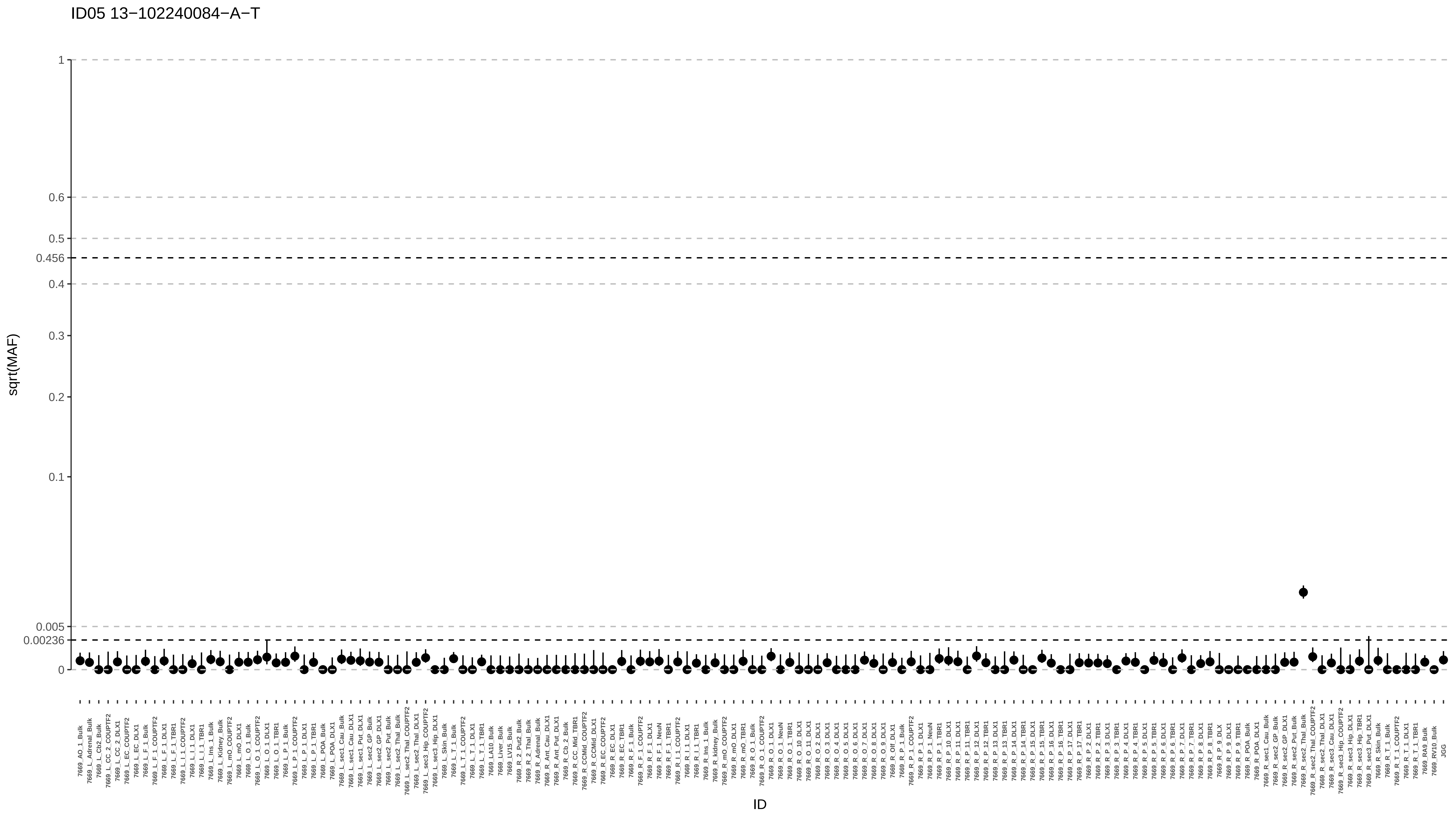
ID05 13−102240084−A−T
sqrt(MAF)
1
0.6
0.5
0.456
0.4
0.3
0.2
0.1
0.005
0.00236
0
ID

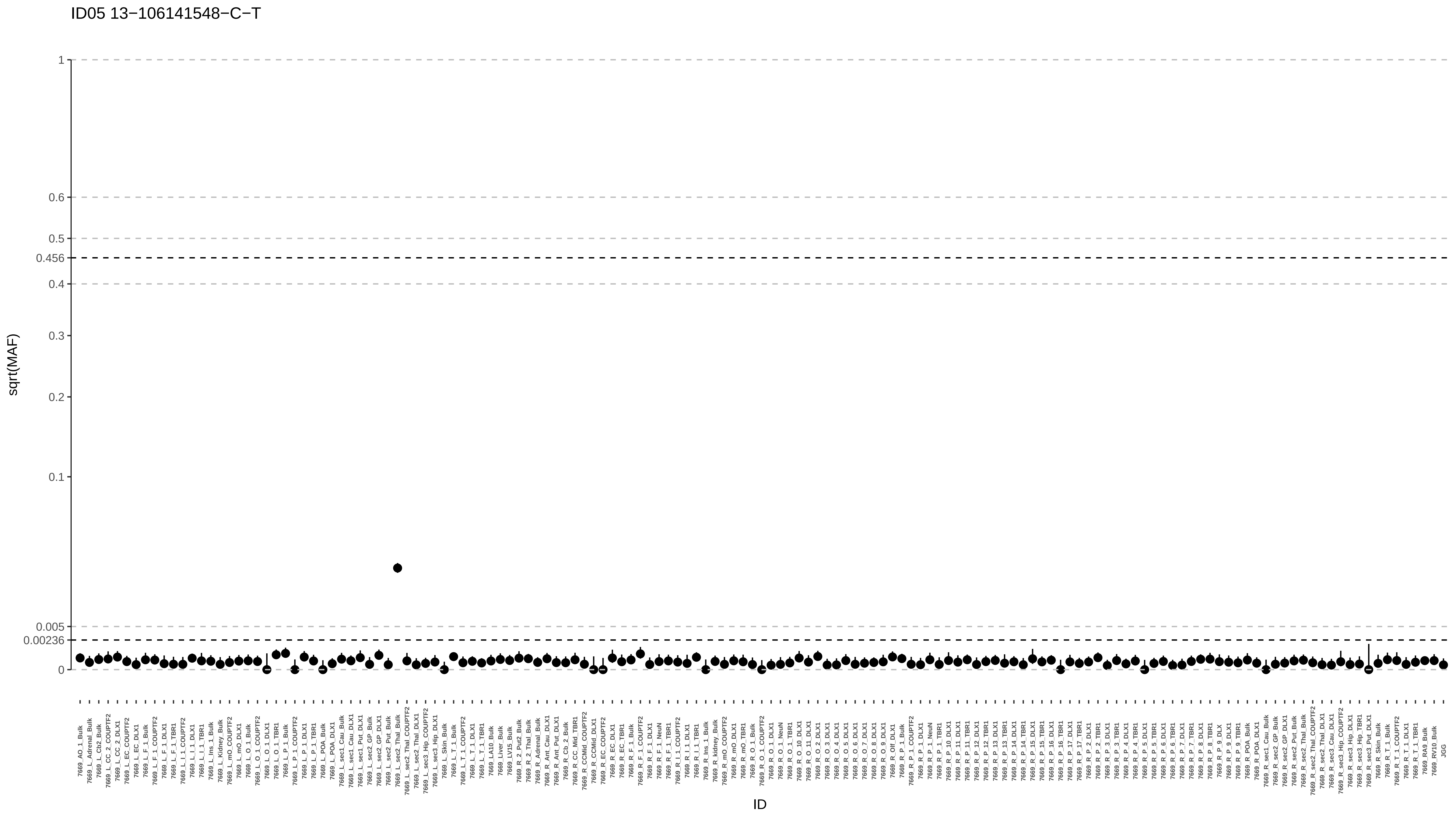

ID05 13−106141548−C−T
sqrt(MAF)
1
0.6
0.5
0.456
0.4
0.3
0.2
0.1
0.005
0.00236
0
ID
7669_AO_1_Bulk
7669_L_Adrenal_Bulk
7669_L_Cb2_Bulk
7669_L_CC_2_COUPTF2
7669_L_CC_2_DLX1
7669_L_EC_COUPTF2
7669_L_EC_DLX1
7669_L_F_1_Bulk
7669_L_F_1_COUPTF2
7669_L_F_1_DLX1
7669_L_F_1_TBR1
7669_L_I_1_COUPTF2
7669_L_I_1_DLX1
7669_L_I_1_TBR1
7669_L_Ins_1_Bulk
7669_L_Kidney_Bulk
7669_L_mO_COUPTF2
7669_L_mO_DLX1
7669_L_O_1_Bulk
7669_L_O_1_COUPTF2
7669_L_O_1_DLX1
7669_L_O_1_TBR1
7669_L_P_1_Bulk
7669_L_P_1_COUPTF2
7669_L_P_1_DLX1
7669_L_P_1_TBR1
7669_L_POA_Bulk
7669_L_POA_DLX1
7669_L_sec1_Cau_Bulk
7669_L_sec1_Cau_DLX1
7669_L_sec1_Put_DLX1
7669_L_sec2_GP_Bulk
7669_L_sec2_GP_DLX1
7669_L_sec2_Put_Bulk
7669_L_sec2_Thal_Bulk
7669_L_sec2_Thal_COUPTF2
7669_L_sec2_Thal_DLX1
7669_L_sec3_Hip_COUPTF2
7669_L_sec3_Hip_DLX1
7669_L_Skin_Bulk
7669_L_T_1_Bulk
7669_L_T_1_COUPTF2
7669_L_T_1_DLX1
7669_L_T_1_TBR1
7669_LA10_Bulk
7669_Liver_Bulk
7669_LV15_Bulk
7669_R_2_Put2_Bulk
7669_R_2_Thal_Bulk
7669_R_Adrenal_Bulk
7669_R_Ant_Cau_DLX1
7669_R_Ant_Put_DLX1
7669_R_Cb_2_Bulk
7669_R_CC_Mid_TBR1
7669_R_CCMid_COUPTF2
7669_R_CCMid_DLX1
7669_R_EC_COUPTF2
7669_R_EC_DLX1
7669_R_EC_TBR1
7669_R_F_1_Bulk
7669_R_F_1_COUPTF2
7669_R_F_1_DLX1
7669_R_F_1_NeuN
7669_R_F_1_TBR1
7669_R_I_1_COUPTF2
7669_R_I_1_DLX1
7669_R_I_1_TBR1
7669_R_Ins_1_Bulk
7669_R_kidney_Bulk
7669_R_mO_COUPTF2
7669_R_mO_DLX1
7669_R_mO_TBR1
7669_R_O_1_Bulk
7669_R_O_1_COUPTF2
7669_R_O_1_DLX1
7669_R_O_1_NeuN
7669_R_O_1_TBR1
7669_R_O_10_DLX1
7669_R_O_11_DLX1
7669_R_O_2_DLX1
7669_R_O_3_DLX1
7669_R_O_4_DLX1
7669_R_O_5_DLX1
7669_R_O_6_DLX1
7669_R_O_7_DLX1
7669_R_O_8_DLX1
7669_R_O_9_DLX1
7669_R_Olf_DLX1
7669_R_P_1_Bulk
7669_R_P_1_COUPTF2
7669_R_P_1_DLX1
7669_R_P_1_NeuN
7669_R_P_1_TBR1
7669_R_P_10_DLX1
7669_R_P_11_DLX1
7669_R_P_11_TBR1
7669_R_P_12_DLX1
7669_R_P_12_TBR1
7669_R_P_13_DLX1
7669_R_P_13_TBR1
7669_R_P_14_DLX1
7669_R_P_14_TBR1
7669_R_P_15_DLX1
7669_R_P_15_TBR1
7669_R_P_16_DLX1
7669_R_P_16_TBR1
7669_R_P_17_DLX1
7669_R_P_17_TBR1
7669_R_P_2_DLX1
7669_R_P_2_TBR1
7669_R_P_3_DLX1
7669_R_P_3_TBR1
7669_R_P_4_DLX1
7669_R_P_4_TBR1
7669_R_P_5_DLX1
7669_R_P_5_TBR1
7669_R_P_6_DLX1
7669_R_P_6_TBR1
7669_R_P_7_DLX1
7669_R_P_7_TBR1
7669_R_P_8_DLX1
7669_R_P_8_TBR1
7669_R_P_9_DLX
7669_R_P_9_DLX1
7669_R_P_9_TBR1
7669_R_POA_Bulk
7669_R_POA_DLX1
7669_R_sec1_Cau_Bulk
7669_R_sec2_GP_Bulk
7669_R_sec2_GP_DLX1
7669_R_sec2_Put_Bulk
7669_R_sec2_Thal_Bulk
7669_R_sec2_Thal_COUPTF2
7669_R_sec2_Thal_DLX1
7669_R_sec3_Cau_DLX1
7669_R_sec3_Hip_COUPTF2
7669_R_sec3_Hip_DLX1
7669_R_sec3_Hip_TBR1
7669_R_sec3_Put_DLX1
7669_R_Skin_Bulk
7669_R_T_1_Bulk
7669_R_T_1_COUPTF2
7669_R_T_1_DLX1
7669_R_T_1_TBR1
7669_RA9_Bulk
7669_RV10_Bulk
JGG

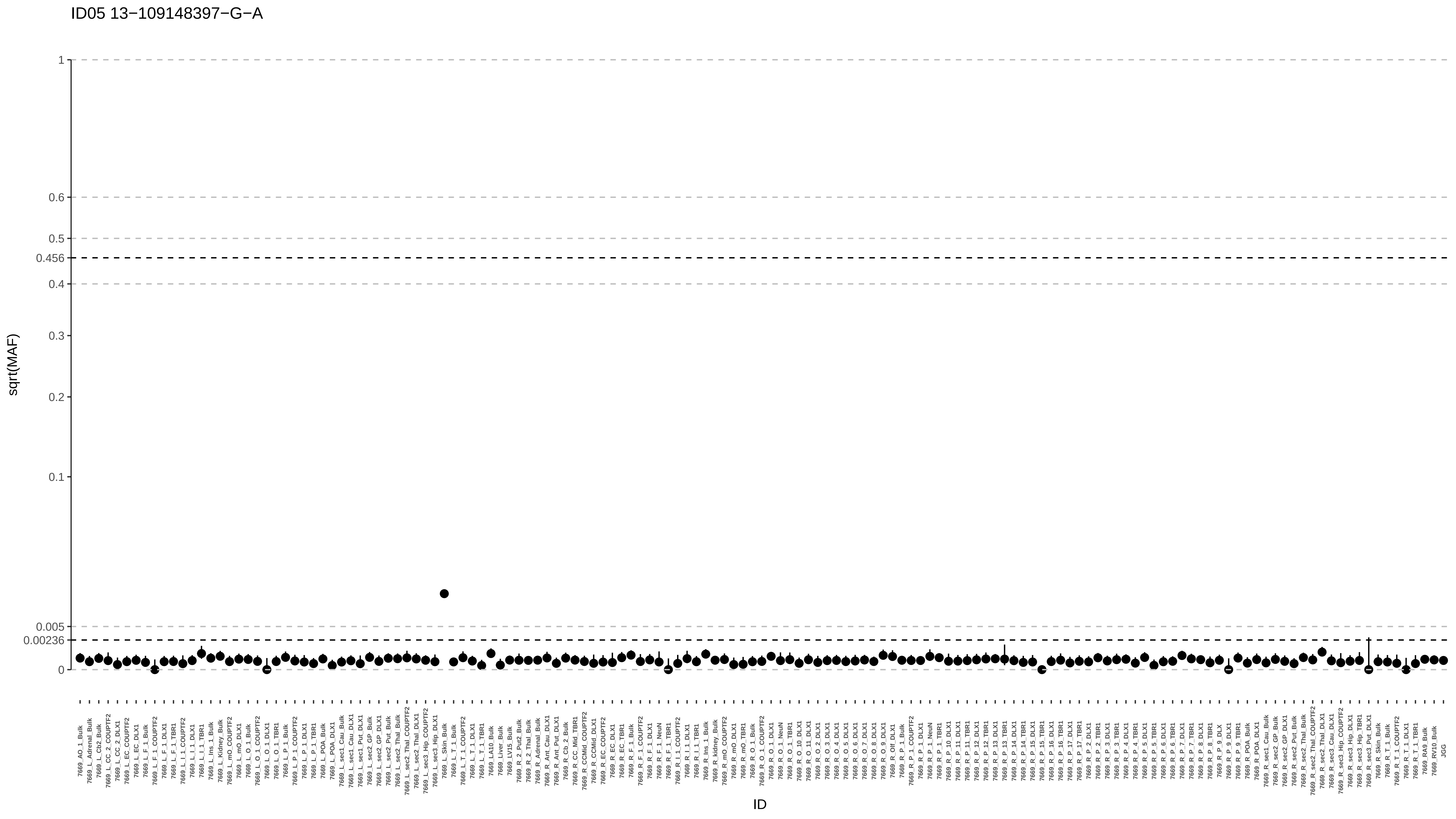
ID05 13−109148397−G−A
sqrt(MAF)
1
0.6
0.5
0.456
0.4
0.3
0.2
0.1
0.005
0.00236
0
ID

ID05 13−109292761−G−A

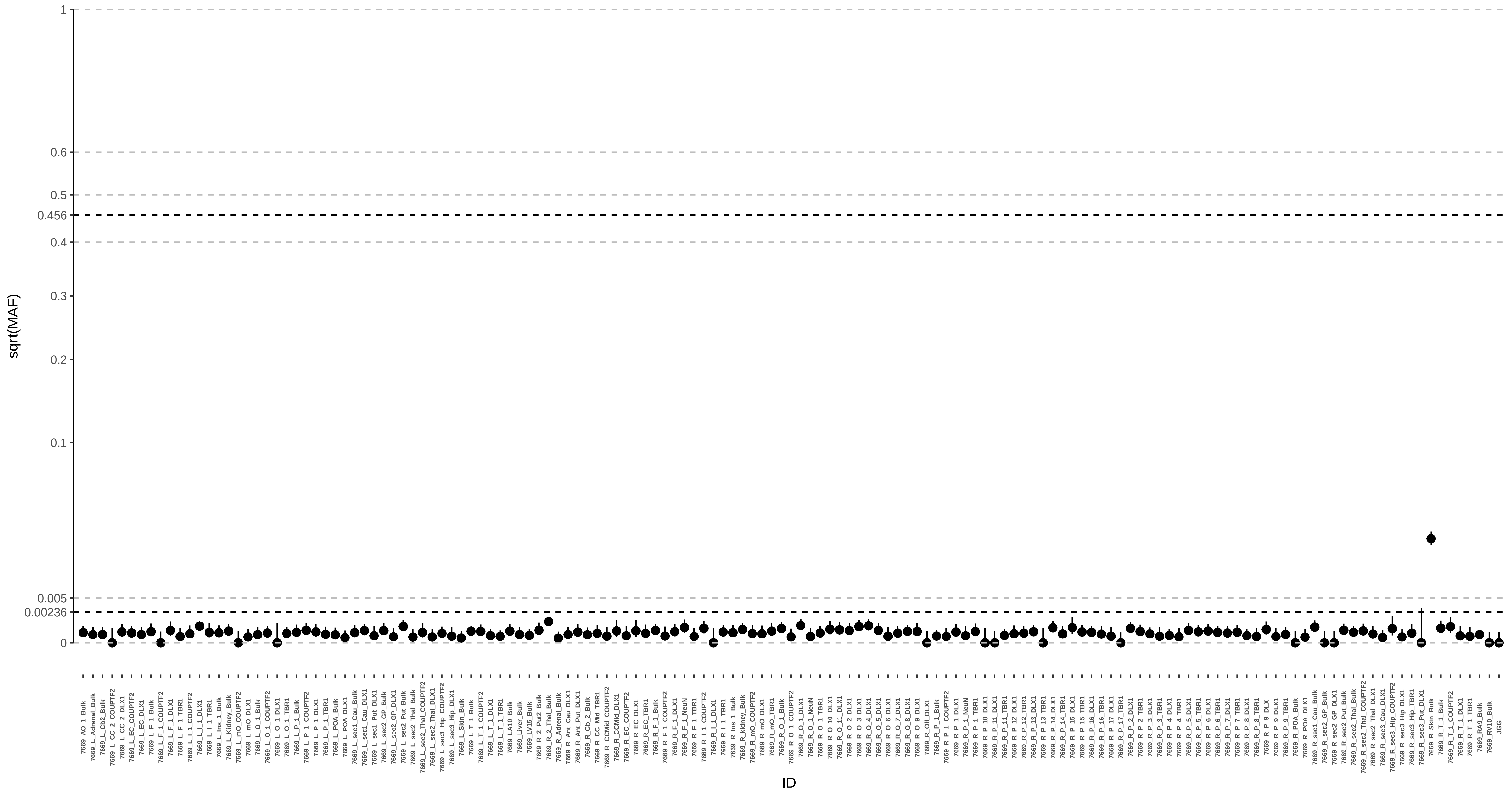

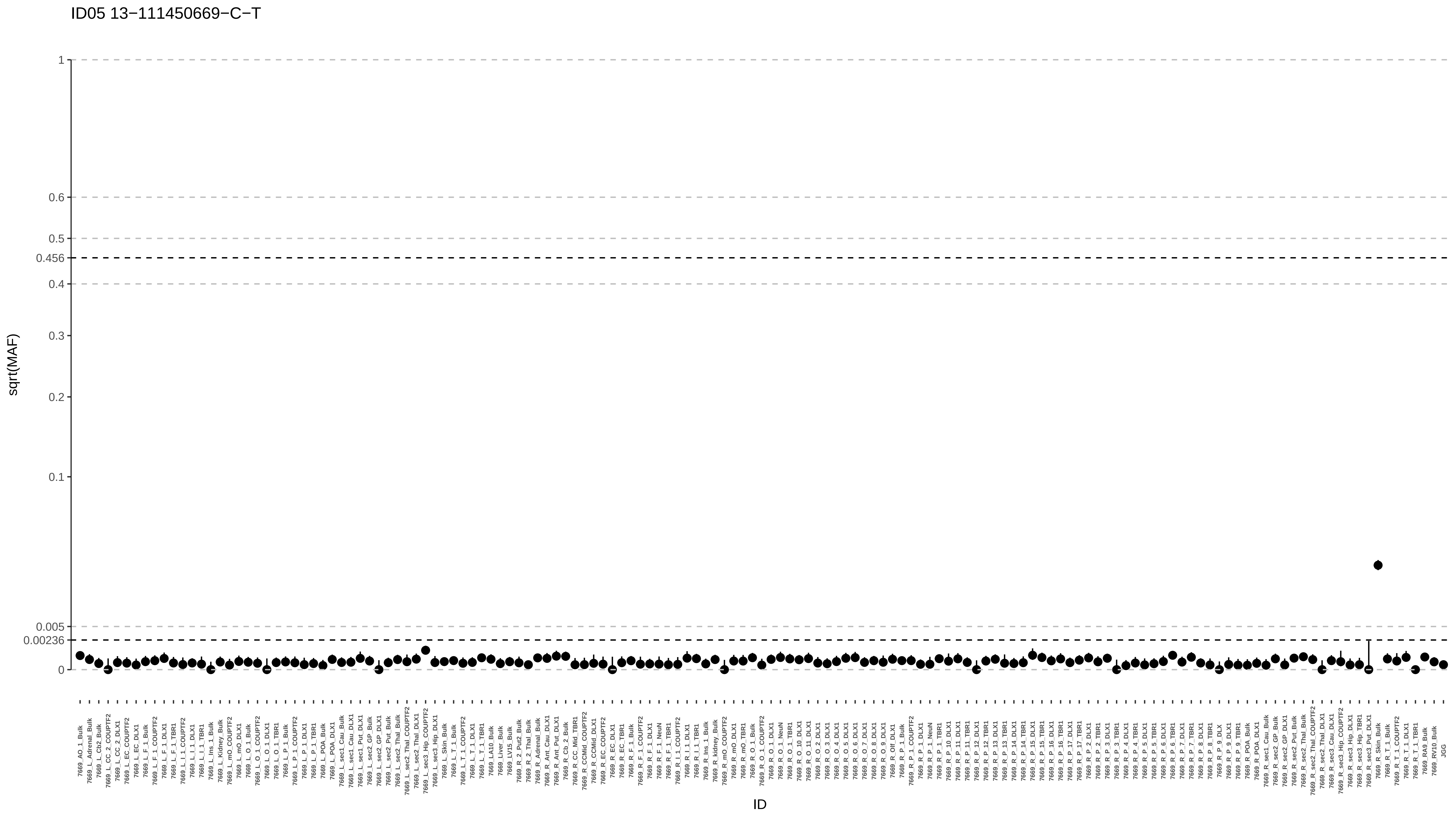

ID05 13−111450669−C−T
sqrt(MAF)
ID

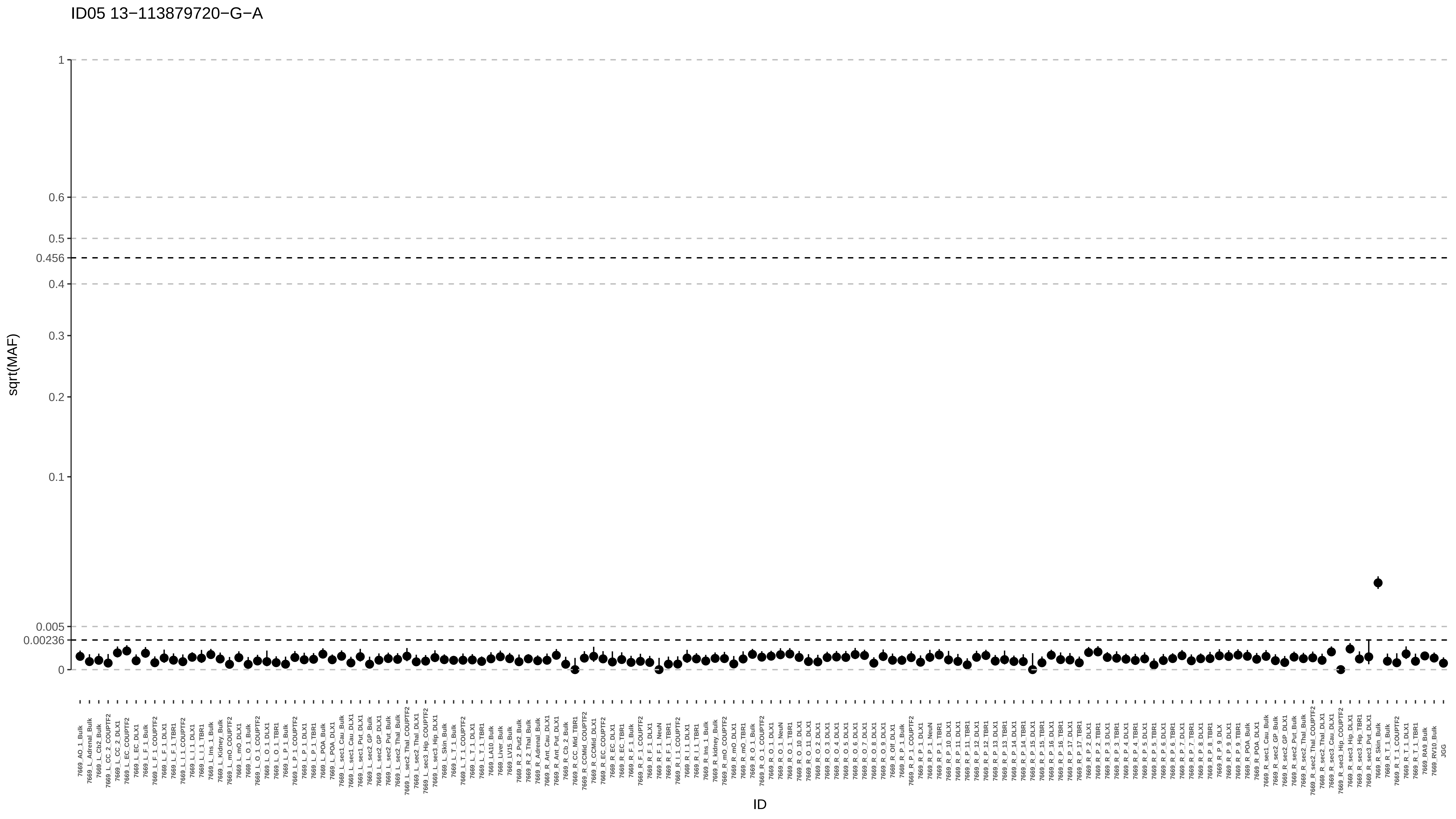
ID05 13−113879720−G−A
sqrt(MAF)
1
0.6
0.5
0.456
0.4
0.3
0.2
0.1
0.005
0.00236
0
ID

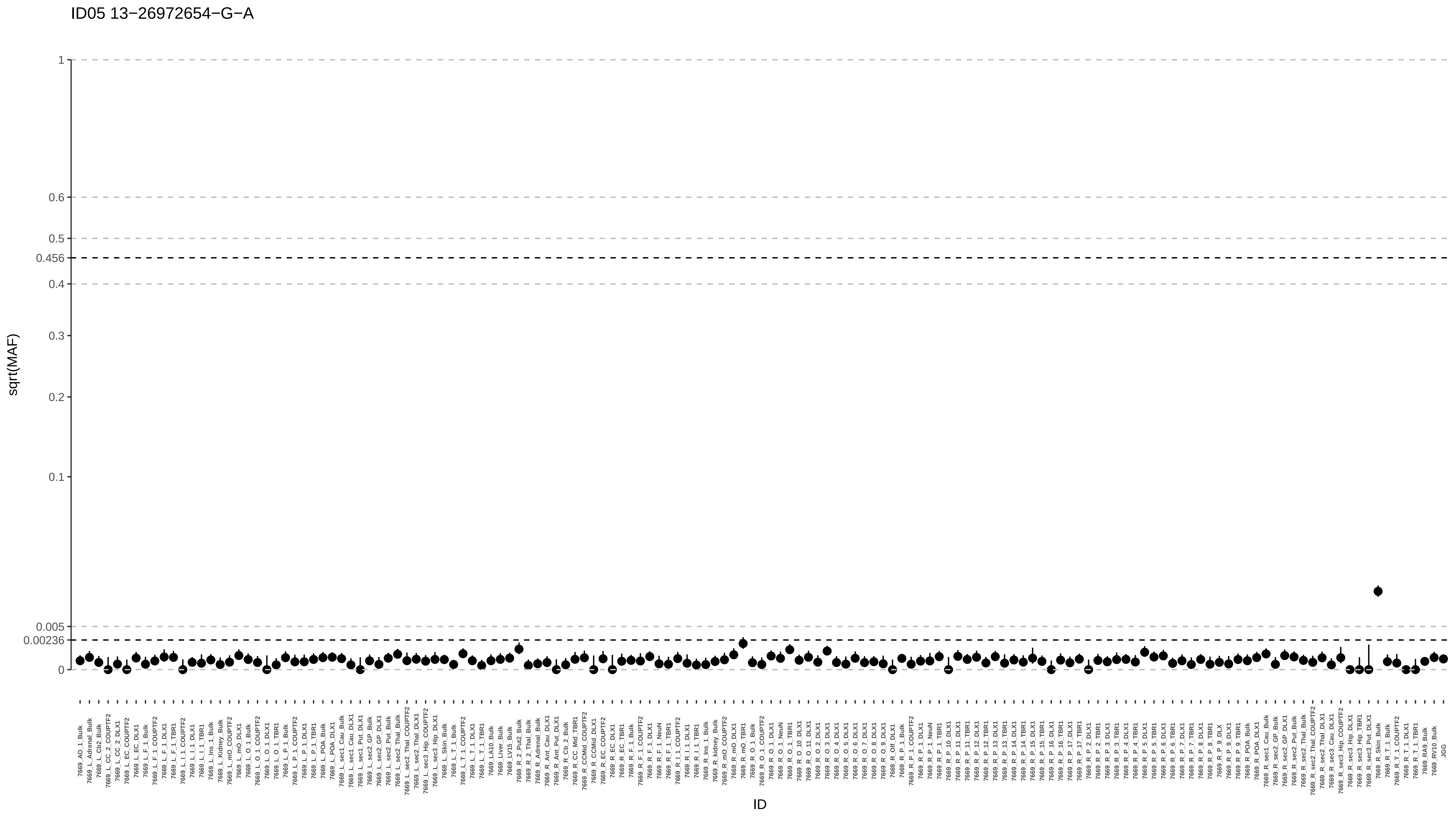
ID05 13−26972654−G−A
sqrt(MAF)
1
0.6
0.5
0.456
0.4
0.3
0.2
0.1
0.005
0.00236
0
ID

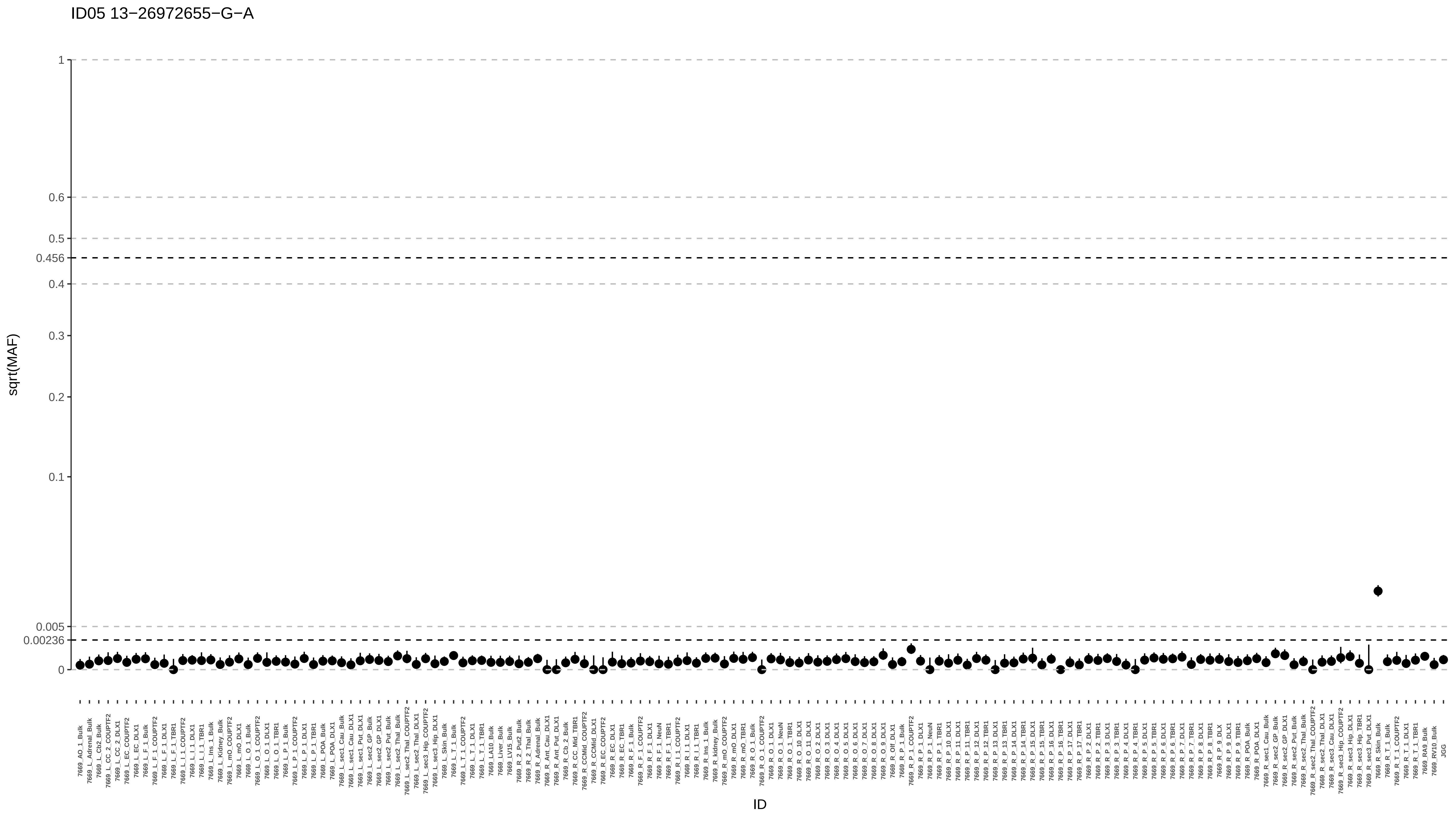

ID05 13−26972655−G−A
sqrt(MAF)
1
0.6
0.5
0.456
0.4
0.3
0.2
0.1
0.005
0.00236
0
ID

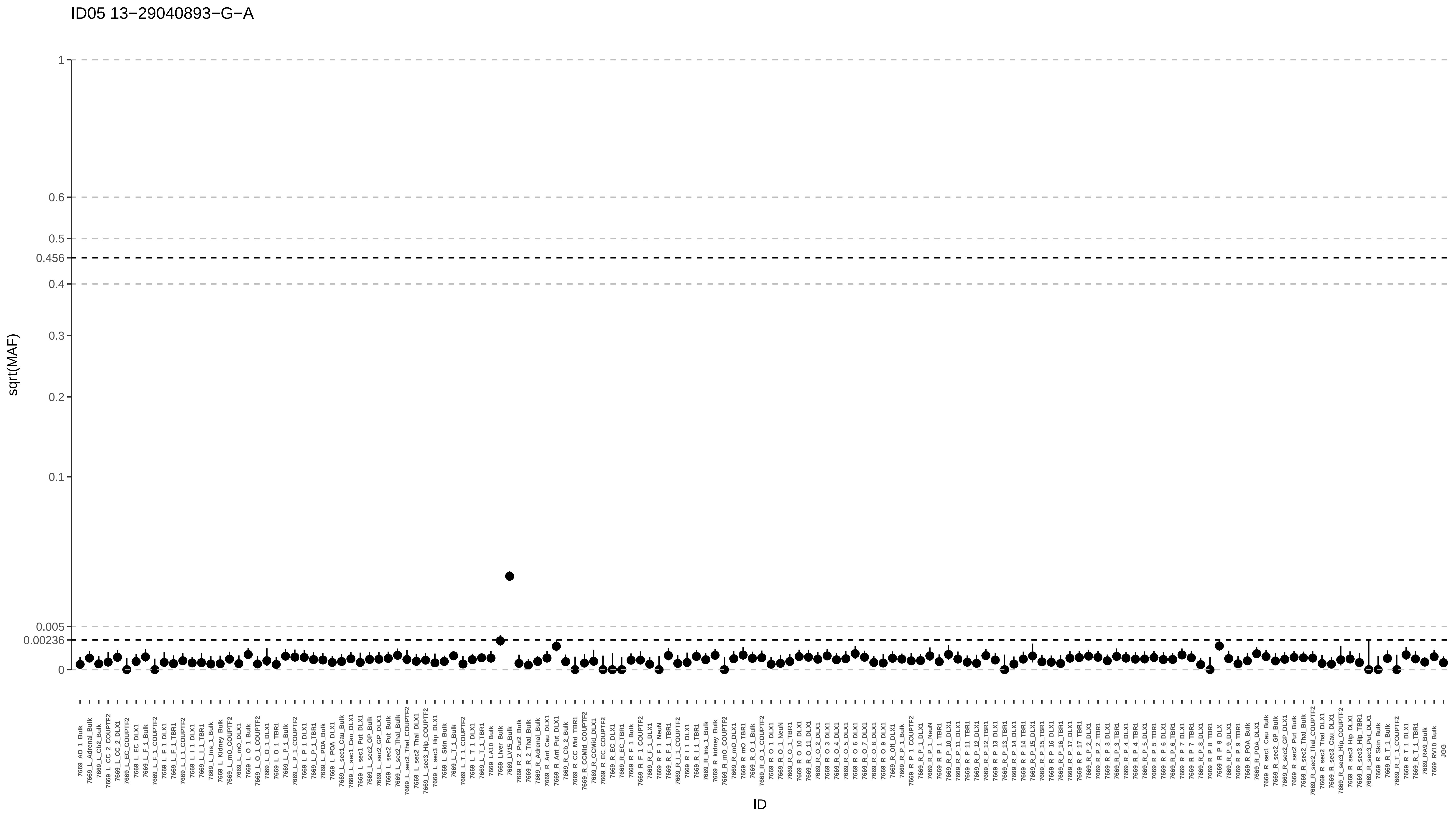
ID05 13−29040893−G−A
sqrt(MAF)
ID
1
0.6
0.5
0.456
0.4
0.3
0.2
0.1
0.005
0.00236
0

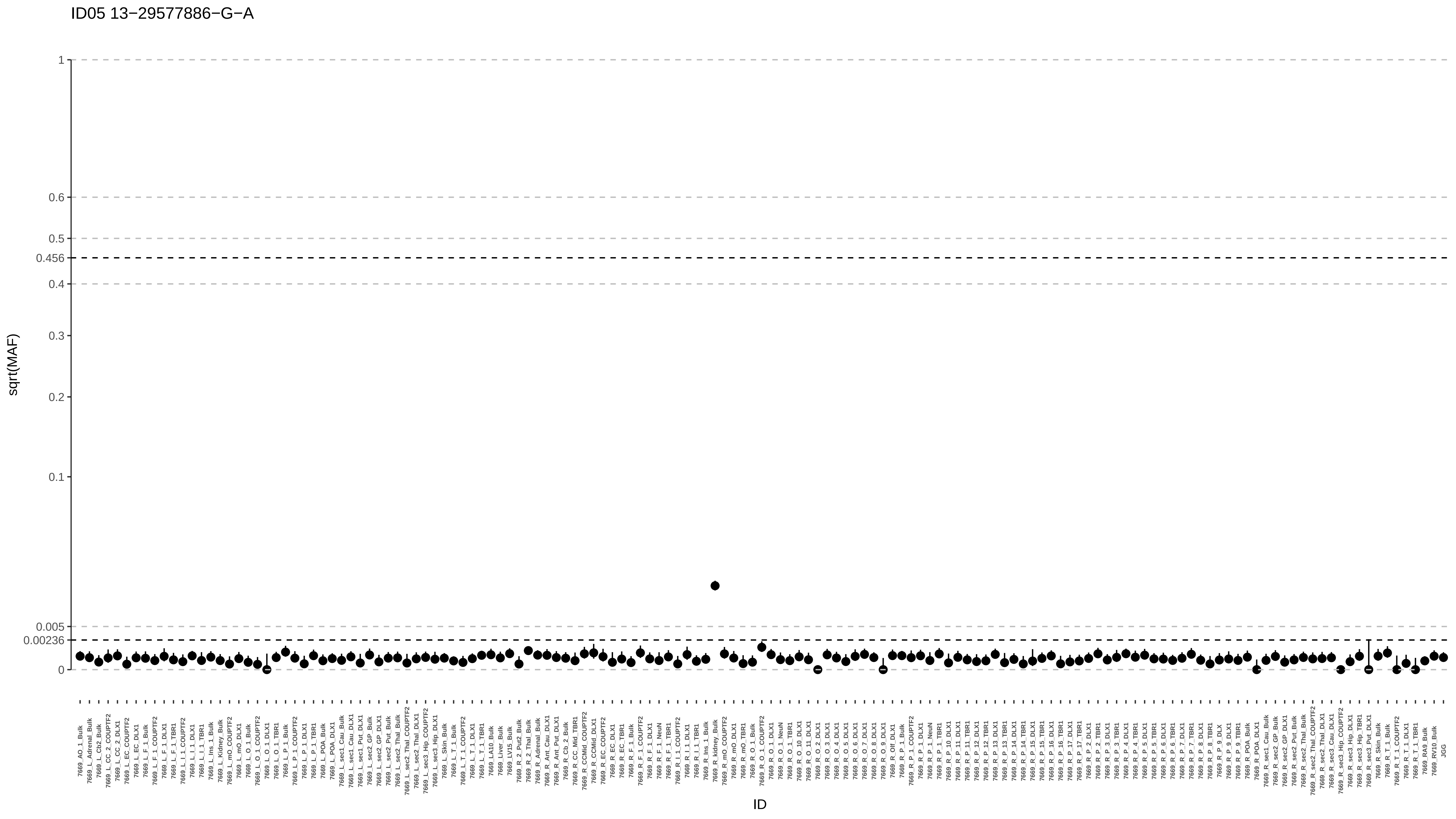
ID05 13−29577886−G−A
sqrt(MAF)
1
0.6
0.5
0.456
0.4
0.3
0.2
0.1
0.005
0.00236
0
ID

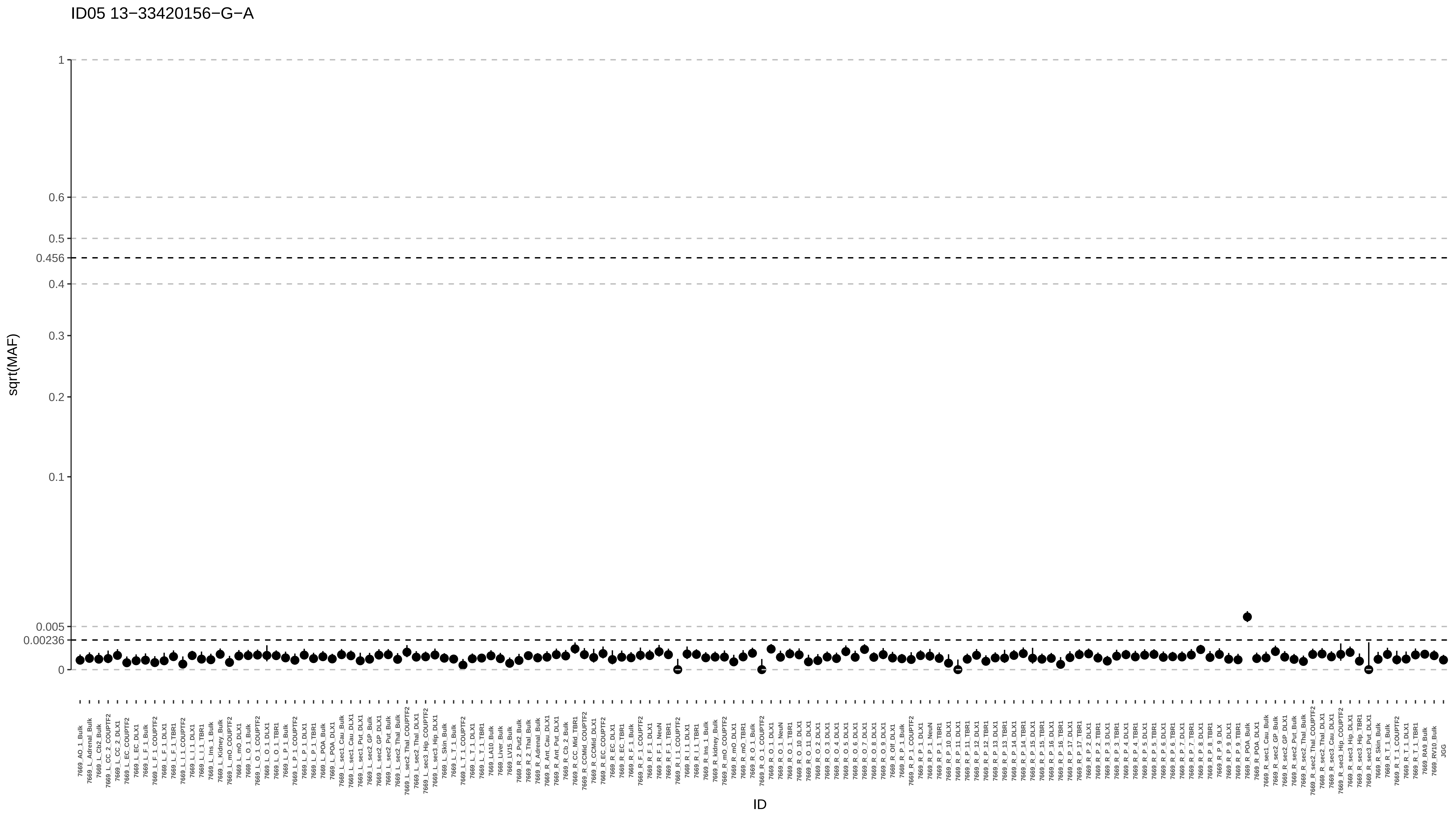

ID05 13−33420156−G−A
sqrt(MAF)
1
0.6
0.5
0.456
0.4
0.3
0.2
0.1
0.005
0.00236
0
ID

ID05 13−41556737−G−C

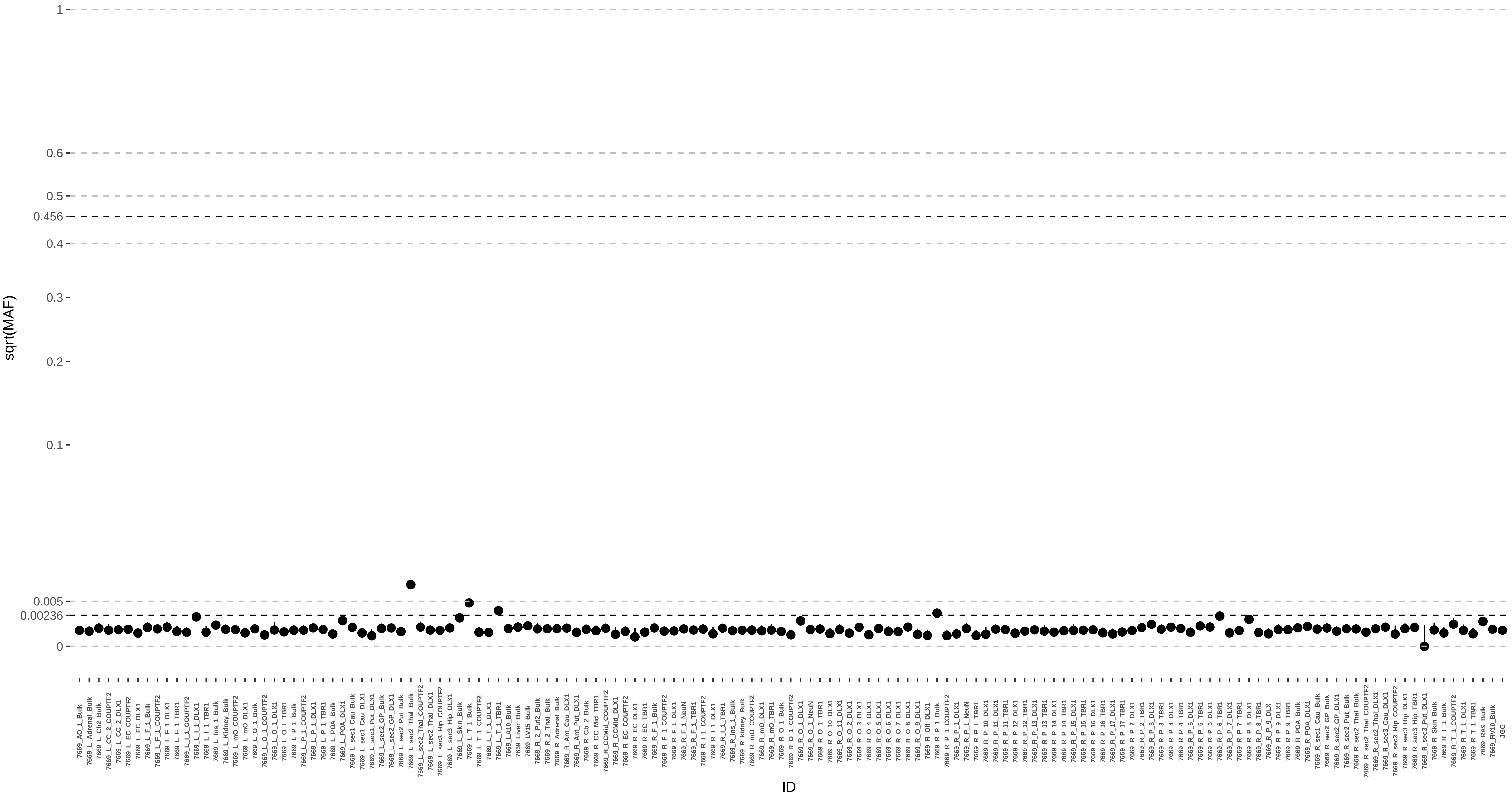

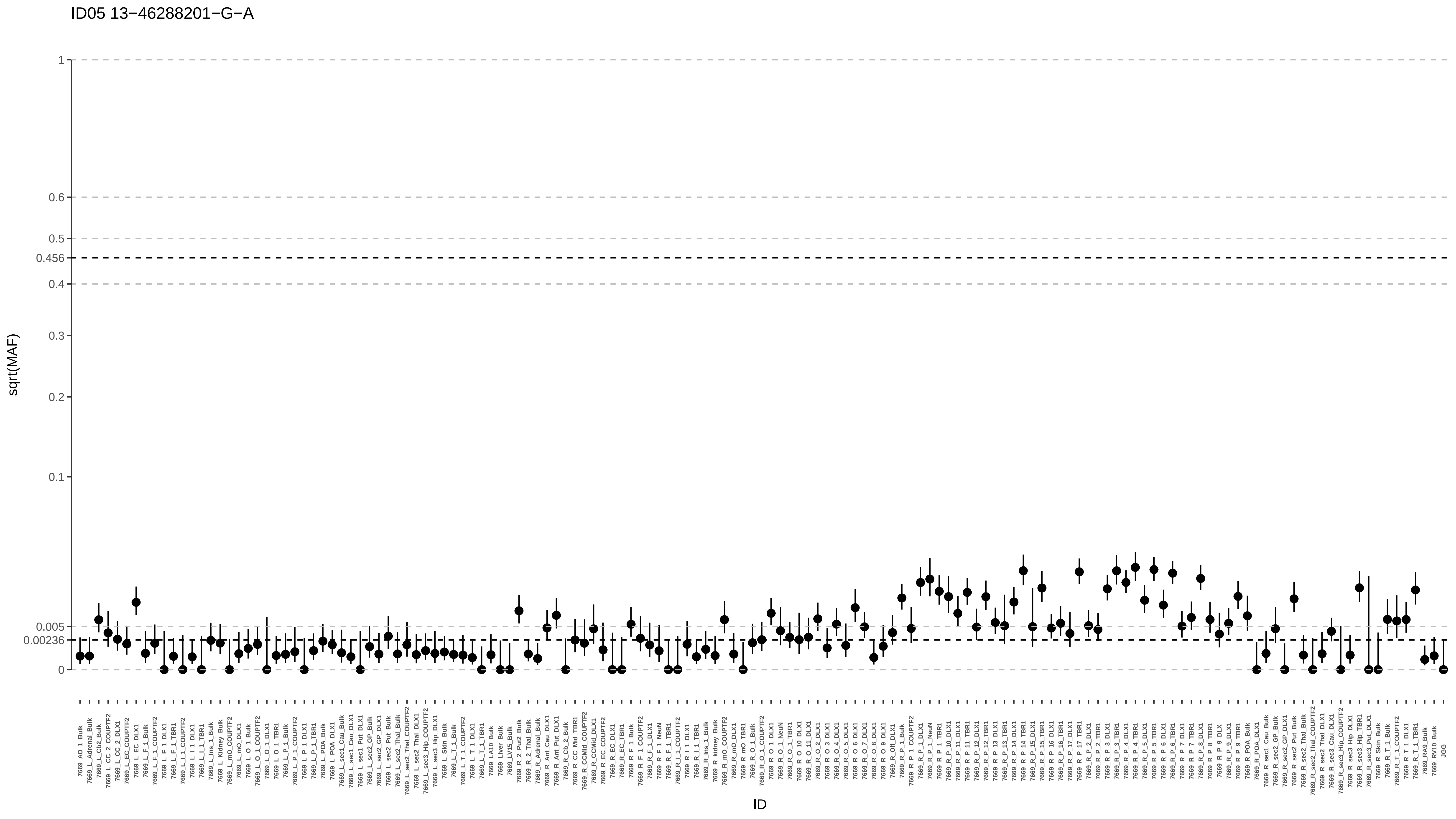

ID05 13−46288201−G−A
sqrt(MAF)
1
0.6
0.5
0.456
0.4
0.3
0.2
0.1
0.005
0.00236
0
ID
7669_AO_1_Bulk
7669_L_Adrenal_Bulk
7669_L_Cb2_Bulk
7669_L_CC_2_COUPTF2
7669_L_CC_2_DLX1
7669_L_EC_COUPTF2
7669_L_EC_DLX1
7669_L_F_1_Bulk
7669_L_F_1_COUPTF2
7669_L_F_1_DLX1
7669_L_F_1_TBR1
7669_L_I_1_COUPTF2
7669_L_I_1_DLX1
7669_L_I_1_TBR1
7669_L_Ins_1_Bulk
7669_L_Kidney_Bulk
7669_L_mO_COUPTF2
7669_L_mO_DLX1
7669_L_O_1_Bulk
7669_L_O_1_COUPTF2
7669_L_O_1_DLX1
7669_L_O_1_TBR1
7669_L_P_1_Bulk
7669_L_P_1_COUPTF2
7669_L_P_1_DLX1
7669_L_P_1_TBR1
7669_L_POA_Bulk
7669_L_POA_DLX1
7669_L_sec1_Cau_Bulk
7669_L_sec1_Cau_DLX1
7669_L_sec1_Put_DLX1
7669_L_sec2_GP_Bulk
7669_L_sec2_GP_DLX1
7669_L_sec2_Put_Bulk
7669_L_sec2_Thal_Bulk
7669_L_sec2_Thal_COUPTF2
7669_L_sec2_Thal_DLX1
7669_L_sec3_Hip_COUPTF2
7669_L_sec3_Hip_DLX1
7669_L_Skin_Bulk
7669_L_T_1_Bulk
7669_L_T_1_COUPTF2
7669_L_T_1_DLX1
7669_L_T_1_TBR1
7669_LA10_Bulk
7669_Liver_Bulk
7669_LV15_Bulk
7669_R_2_Put2_Bulk
7669_R_2_Thal_Bulk
7669_R_Adrenal_Bulk
7669_R_Ant_Cau_DLX1
7669_R_Ant_Put_DLX1
7669_R_Cb_2_Bulk
7669_R_CC_Mid_TBR1
7669_R_CCMid_COUPTF2
7669_R_CCMid_DLX1
7669_R_EC_COUPTF2
7669_R_EC_DLX1
7669_R_EC_TBR1
7669_R_F_1_Bulk
7669_R_F_1_COUPTF2
7669_R_F_1_DLX1
7669_R_F_1_NeuN
7669_R_F_1_TBR1
7669_R_I_1_COUPTF2
7669_R_I_1_DLX1
7669_R_I_1_TBR1
7669_R_Ins_1_Bulk
7669_R_kidney_Bulk
7669_R_mO_COUPTF2
7669_R_mO_DLX1
7669_R_mO_TBR1
7669_R_O_1_Bulk
7669_R_O_1_COUPTF2
7669_R_O_1_DLX1
7669_R_O_1_NeuN
7669_R_O_1_TBR1
7669_R_O_10_DLX1
7669_R_O_11_DLX1
7669_R_O_2_DLX1
7669_R_O_3_DLX1
7669_R_O_4_DLX1
7669_R_O_5_DLX1
7669_R_O_6_DLX1
7669_R_O_7_DLX1
7669_R_O_8_DLX1
7669_R_O_9_DLX1
7669_R_Olf_DLX1
7669_R_P_1_Bulk
7669_R_P_1_COUPTF2
7669_R_P_1_DLX1
7669_R_P_1_NeuN
7669_R_P_1_TBR1
7669_R_P_10_DLX1
7669_R_P_11_DLX1
7669_R_P_11_TBR1
7669_R_P_12_DLX1
7669_R_P_12_TBR1
7669_R_P_13_DLX1
7669_R_P_13_TBR1
7669_R_P_14_DLX1
7669_R_P_14_TBR1
7669_R_P_15_DLX1
7669_R_P_15_TBR1
7669_R_P_16_DLX1
7669_R_P_16_TBR1
7669_R_P_17_DLX1
7669_R_P_17_TBR1
7669_R_P_2_DLX1
7669_R_P_2_TBR1
7669_R_P_3_DLX1
7669_R_P_3_TBR1
7669_R_P_4_DLX1
7669_R_P_4_TBR1
7669_R_P_5_DLX1
7669_R_P_5_TBR1
7669_R_P_6_DLX1
7669_R_P_6_TBR1
7669_R_P_7_DLX1
7669_R_P_7_TBR1
7669_R_P_8_DLX1
7669_R_P_8_TBR1
7669_R_P_9_DLX
7669_R_P_9_DLX1
7669_R_P_9_TBR1
7669_R_POA_Bulk
7669_R_POA_DLX1
7669_R_sec1_Cau_Bulk
7669_R_sec2_GP_Bulk
7669_R_sec2_GP_DLX1
7669_R_sec2_Put_Bulk
7669_R_sec2_Thal_Bulk
7669_R_sec2_Thal_COUPTF2
7669_R_sec2_Thal_DLX1
7669_R_sec3_Cau_DLX1
7669_R_sec3_Hip_COUPTF2
7669_R_sec3_Hip_DLX1
7669_R_sec3_Hip_TBR1
7669_R_sec3_Put_DLX1
7669_R_Skin_Bulk
7669_R_T_1_Bulk
7669_R_T_1_COUPTF2
7669_R_T_1_DLX1
7669_R_T_1_TBR1
7669_RA9_Bulk
7669_RV10_Bulk
JGG

ID05 13−55140512−C−T

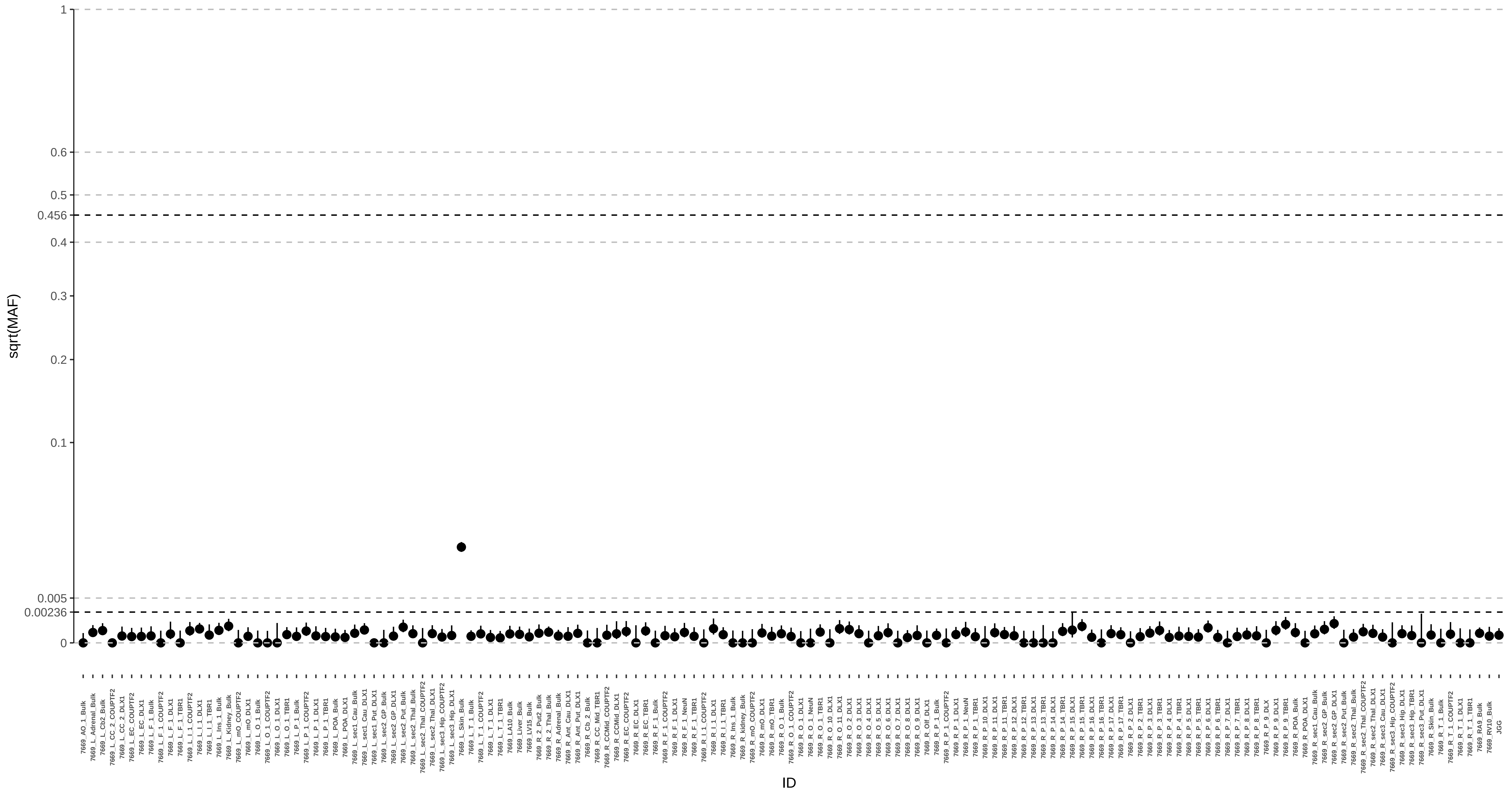

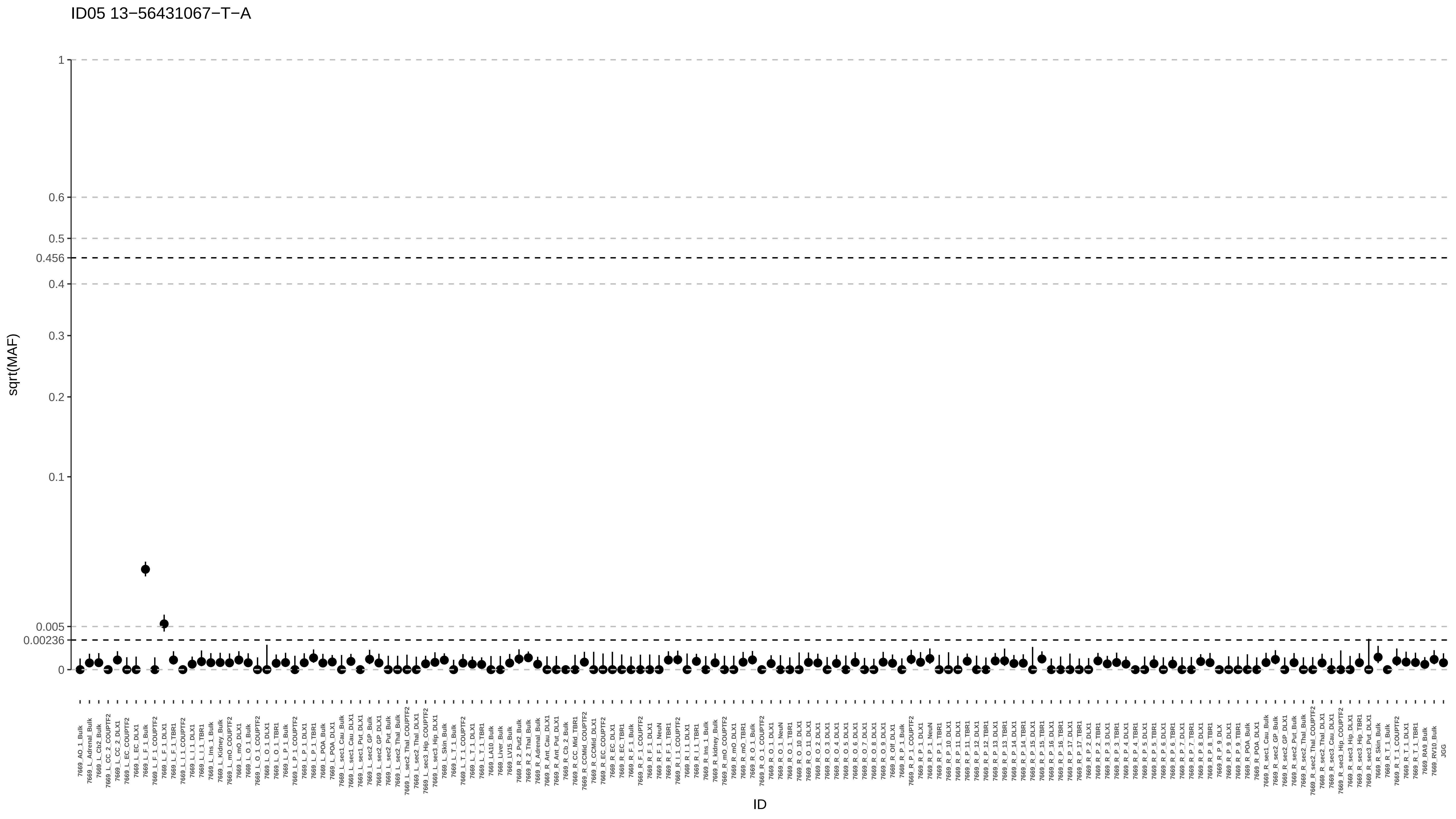
ID05 13−56431067−T−A
sqrt(MAF)
1
0.6
0.5
0.456
0.4
0.3
0.2
0.1
0.005
0.00236
0
ID

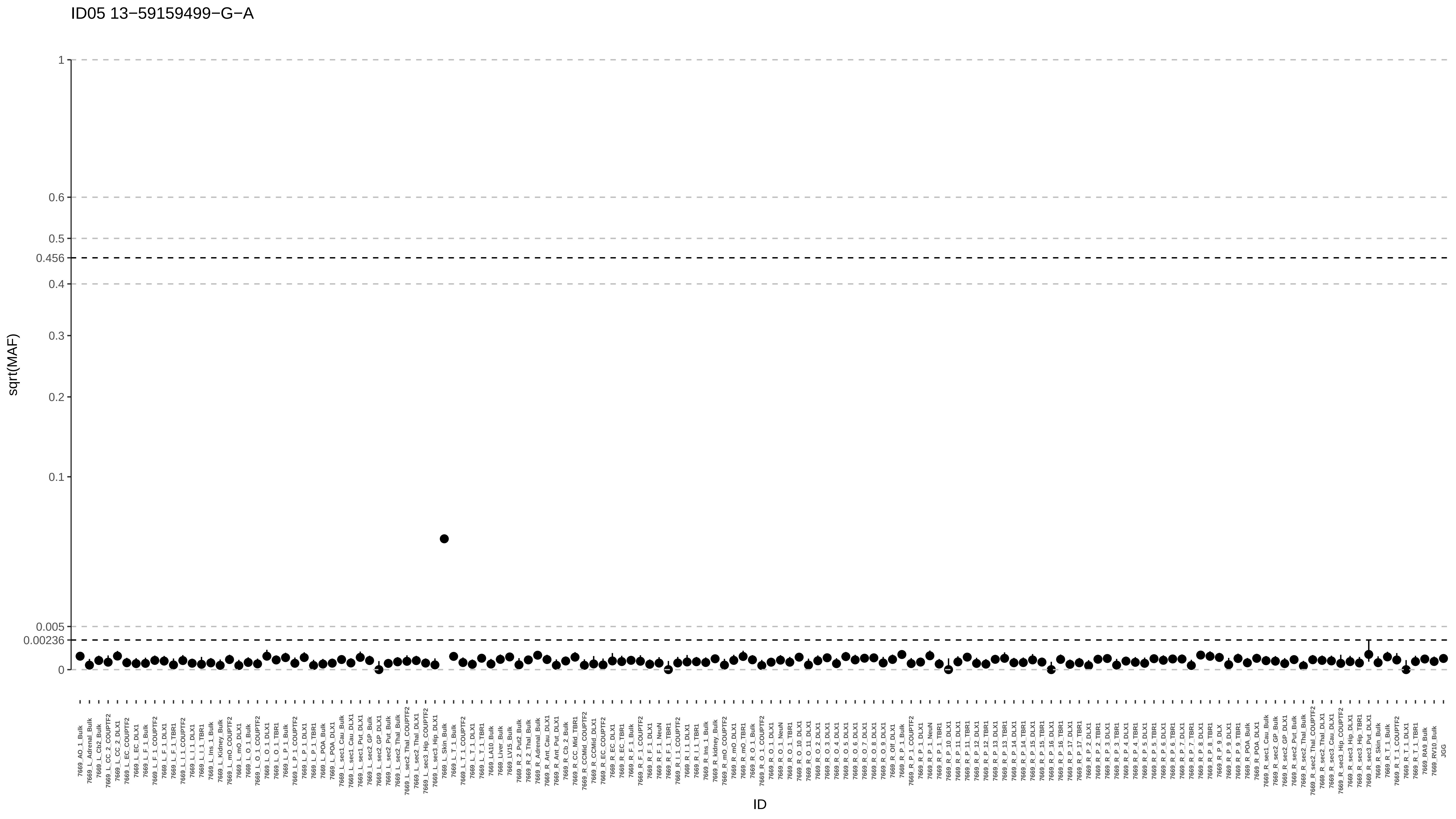
ID05 13−59159499−G−A
sqrt(MAF)
1
0.6
0.5
0.456
0.4
0.3
0.2
0.1
0.005
0.00236
0
ID

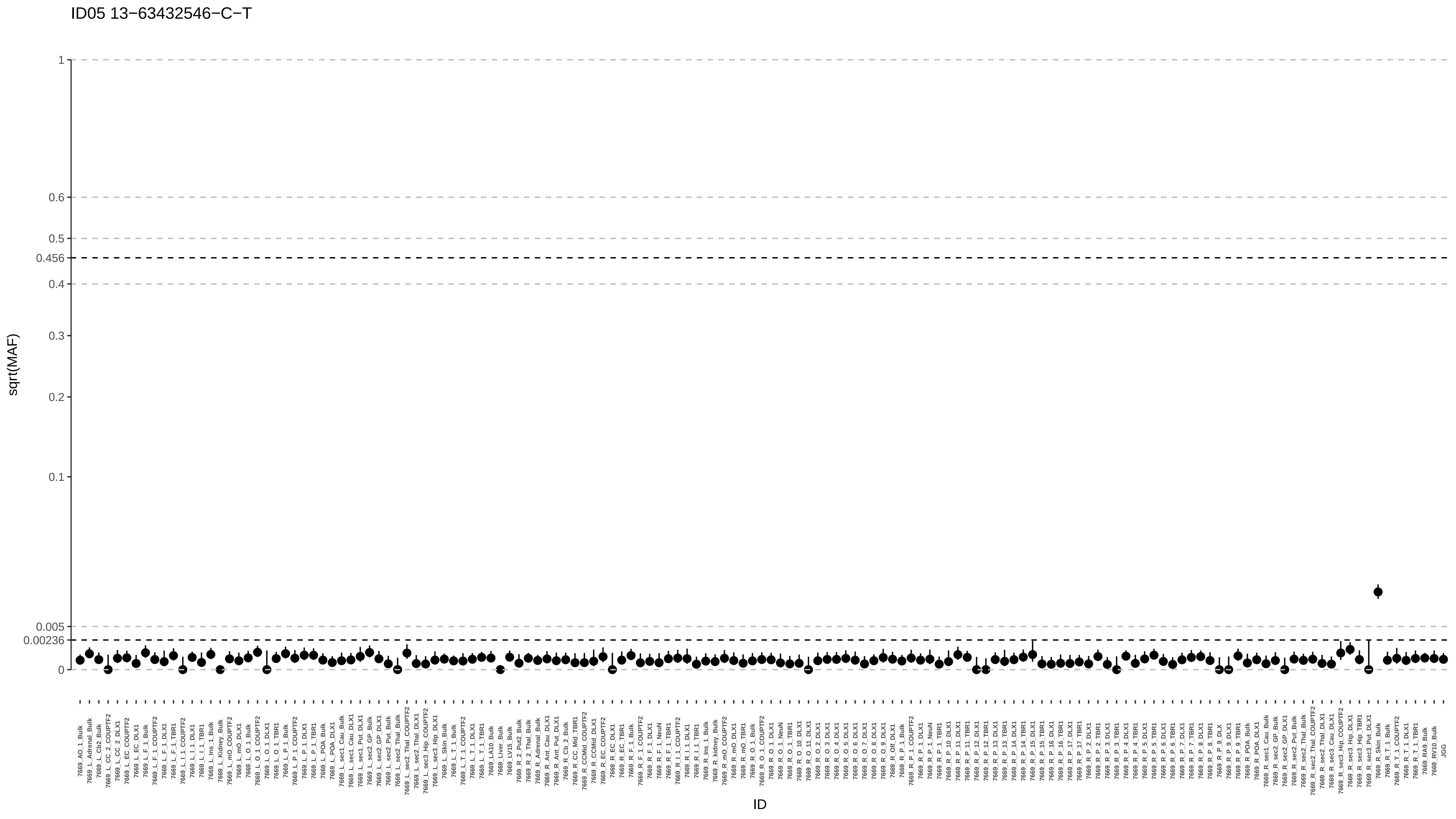
ID05 13−63432546−C−T
sqrt(MAF)
1
0.6
0.5
0.456
0.4
0.3
0.2
0.1
0.005
0.00236
0
ID
7669_AO_1_Bulk
7669_L_Adrenal_Bulk
7669_L_Cb2_Bulk
7669_L_CC_2_COUPTF2
7669_L_CC_2_DLX1
7669_L_EC_COUPTF2
7669_L_EC_DLX1
7669_L_F_1_Bulk
7669_L_F_1_COUPTF2
7669_L_F_1_DLX1
7669_L_F_1_TBR1
7669_L_I_1_COUPTF2
7669_L_I_1_DLX1
7669_L_I_1_TBR1
7669_L_Ins_1_Bulk
7669_L_Kidney_Bulk
7669_L_mO_COUPTF2
7669_L_mO_DLX1
7669_L_O_1_Bulk
7669_L_O_1_COUPTF2
7669_L_O_1_DLX1
7669_L_O_1_TBR1
7669_L_P_1_Bulk
7669_L_P_1_COUPTF2
7669_L_P_1_DLX1
7669_L_P_1_TBR1
7669_L_POA_Bulk
7669_L_POA_DLX1
7669_L_sec1_Cau_Bulk
7669_L_sec1_Cau_DLX1
7669_L_sec1_Put_DLX1
7669_L_sec2_GP_Bulk
7669_L_sec2_GP_DLX1
7669_L_sec2_Put_Bulk
7669_L_sec2_Thal_Bulk
7669_L_sec2_Thal_COUPTF2
7669_L_sec2_Thal_DLX1
7669_L_sec3_Hip_COUPTF2
7669_L_sec3_Hip_DLX1
7669_L_Skin_Bulk
7669_L_T_1_Bulk
7669_L_T_1_COUPTF2
7669_L_T_1_DLX1
7669_L_T_1_TBR1
7669_LA10_Bulk
7669_Liver_Bulk
7669_LV15_Bulk
7669_R_2_Put2_Bulk
7669_R_2_Thal_Bulk
7669_R_Adrenal_Bulk
7669_R_Ant_Cau_DLX1
7669_R_Ant_Put_DLX1
7669_R_Cb_2_Bulk
7669_R_CC_Mid_TBR1
7669_R_CCMid_COUPTF2
7669_R_CCMid_DLX1
7669_R_EC_COUPTF2
7669_R_EC_DLX1
7669_R_EC_TBR1
7669_R_F_1_Bulk
7669_R_F_1_COUPTF2
7669_R_F_1_DLX1
7669_R_F_1_NeuN
7669_R_F_1_TBR1
7669_R_I_1_COUPTF2
7669_R_I_1_DLX1
7669_R_I_1_TBR1
7669_R_Ins_1_Bulk
7669_R_kidney_Bulk
7669_R_mO_COUPTF2
7669_R_mO_DLX1
7669_R_mO_TBR1
7669_R_O_1_Bulk
7669_R_O_1_COUPTF2
7669_R_O_1_DLX1
7669_R_O_1_NeuN
7669_R_O_1_TBR1
7669_R_O_10_DLX1
7669_R_O_11_DLX1
7669_R_O_2_DLX1
7669_R_O_3_DLX1
7669_R_O_4_DLX1
7669_R_O_5_DLX1
7669_R_O_6_DLX1
7669_R_O_7_DLX1
7669_R_O_8_DLX1
7669_R_O_9_DLX1
7669_R_Olf_DLX1
7669_R_P_1_Bulk
7669_R_P_1_COUPTF2
7669_R_P_1_DLX1
7669_R_P_1_NeuN
7669_R_P_1_TBR1
7669_R_P_10_DLX1
7669_R_P_11_DLX1
7669_R_P_11_TBR1
7669_R_P_12_DLX1
7669_R_P_12_TBR1
7669_R_P_13_DLX1
7669_R_P_13_TBR1
7669_R_P_14_DLX1
7669_R_P_14_TBR1
7669_R_P_15_DLX1
7669_R_P_15_TBR1
7669_R_P_16_DLX1
7669_R_P_16_TBR1
7669_R_P_17_DLX1
7669_R_P_17_TBR1
7669_R_P_2_DLX1
7669_R_P_2_TBR1
7669_R_P_3_DLX1
7669_R_P_3_TBR1
7669_R_P_4_DLX1
7669_R_P_4_TBR1
7669_R_P_5_DLX1
7669_R_P_5_TBR1
7669_R_P_6_DLX1
7669_R_P_6_TBR1
7669_R_P_7_DLX1
7669_R_P_7_TBR1
7669_R_P_8_DLX1
7669_R_P_8_TBR1
7669_R_P_9_DLX
7669_R_P_9_DLX1
7669_R_P_9_TBR1
7669_R_POA_Bulk
7669_R_POA_DLX1
7669_R_sec1_Cau_Bulk
7669_R_sec2_GP_Bulk
7669_R_sec2_GP_DLX1
7669_R_sec2_Put_Bulk
7669_R_sec2_Thal_Bulk
7669_R_sec2_Thal_COUPTF2
7669_R_sec2_Thal_DLX1
7669_R_sec3_Cau_DLX1
7669_R_sec3_Hip_COUPTF2
7669_R_sec3_Hip_DLX1
7669_R_sec3_Hip_TBR1
7669_R_sec3_Put_DLX1
7669_R_Skin_Bulk
7669_R_T_1_Bulk
7669_R_T_1_COUPTF2
7669_R_T_1_DLX1
7669_R_T_1_TBR1
7669_RA9_Bulk
7669_RV10_Bulk
JGG

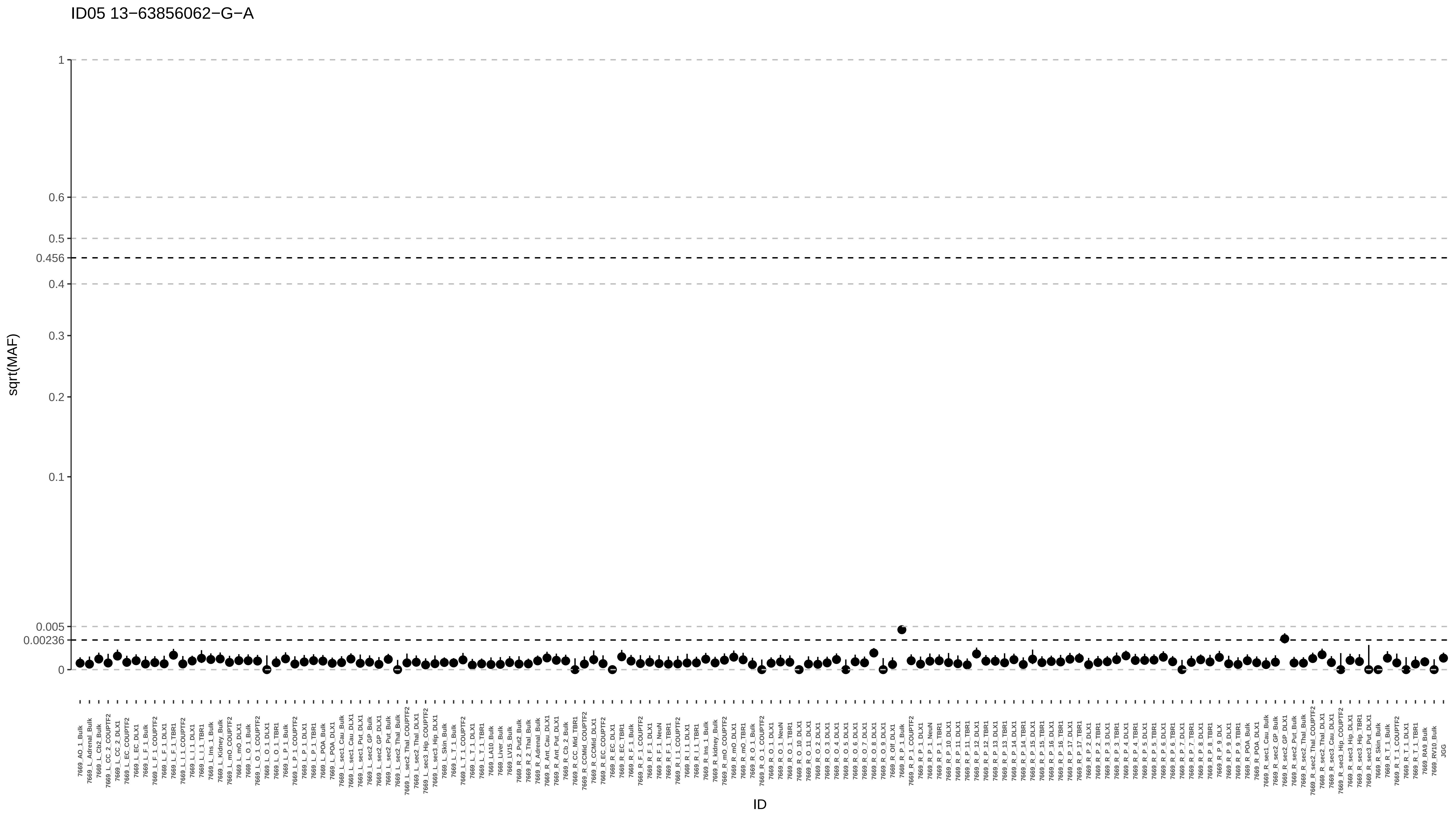
ID05 13−63856062−G−A
sqrt(MAF)
ID
1
0.6
0.5
0.456
0.4
0.3
0.2
0.1
0.005
0.00236
0

ID05 13−65586962−C−A

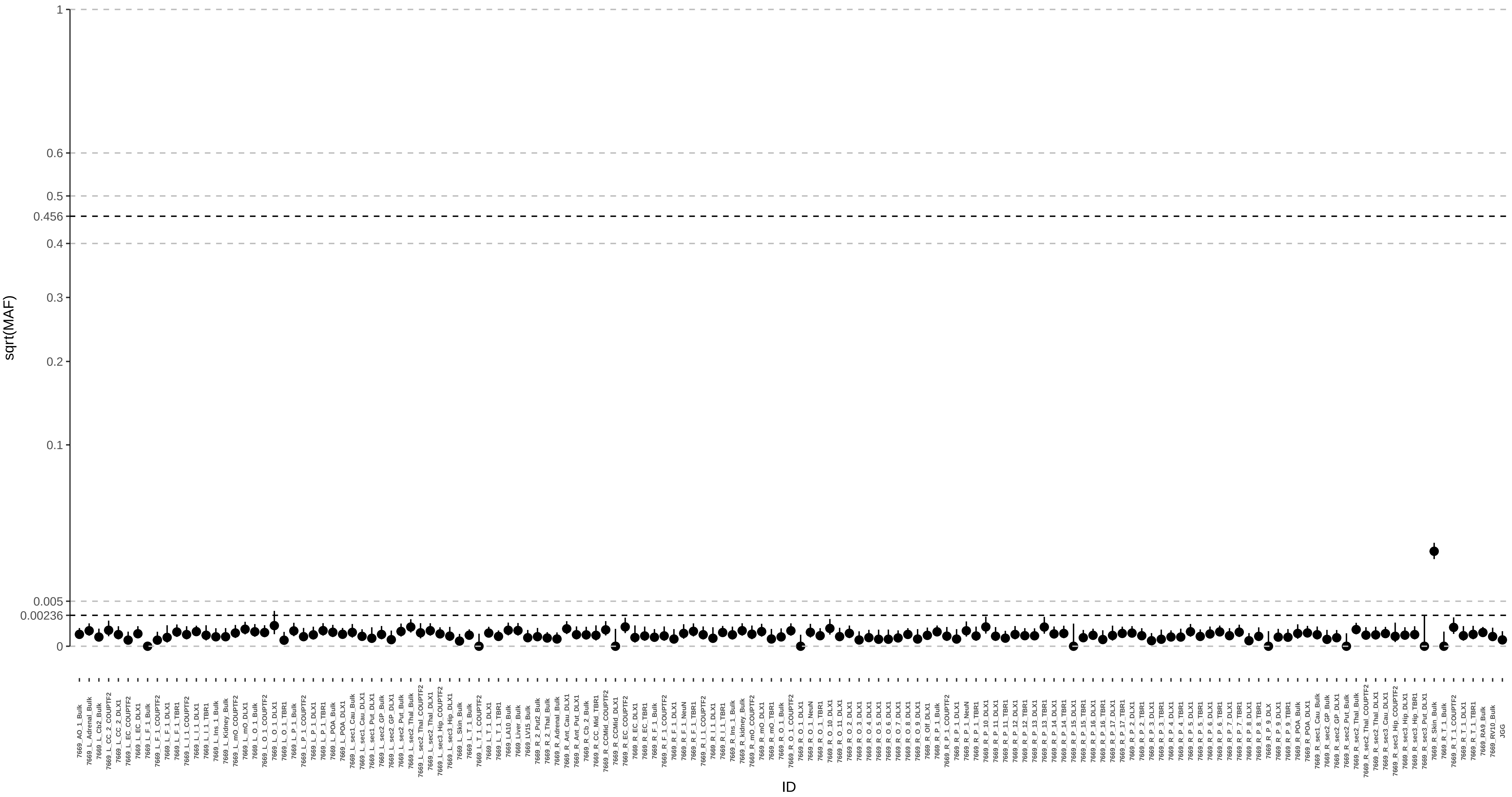

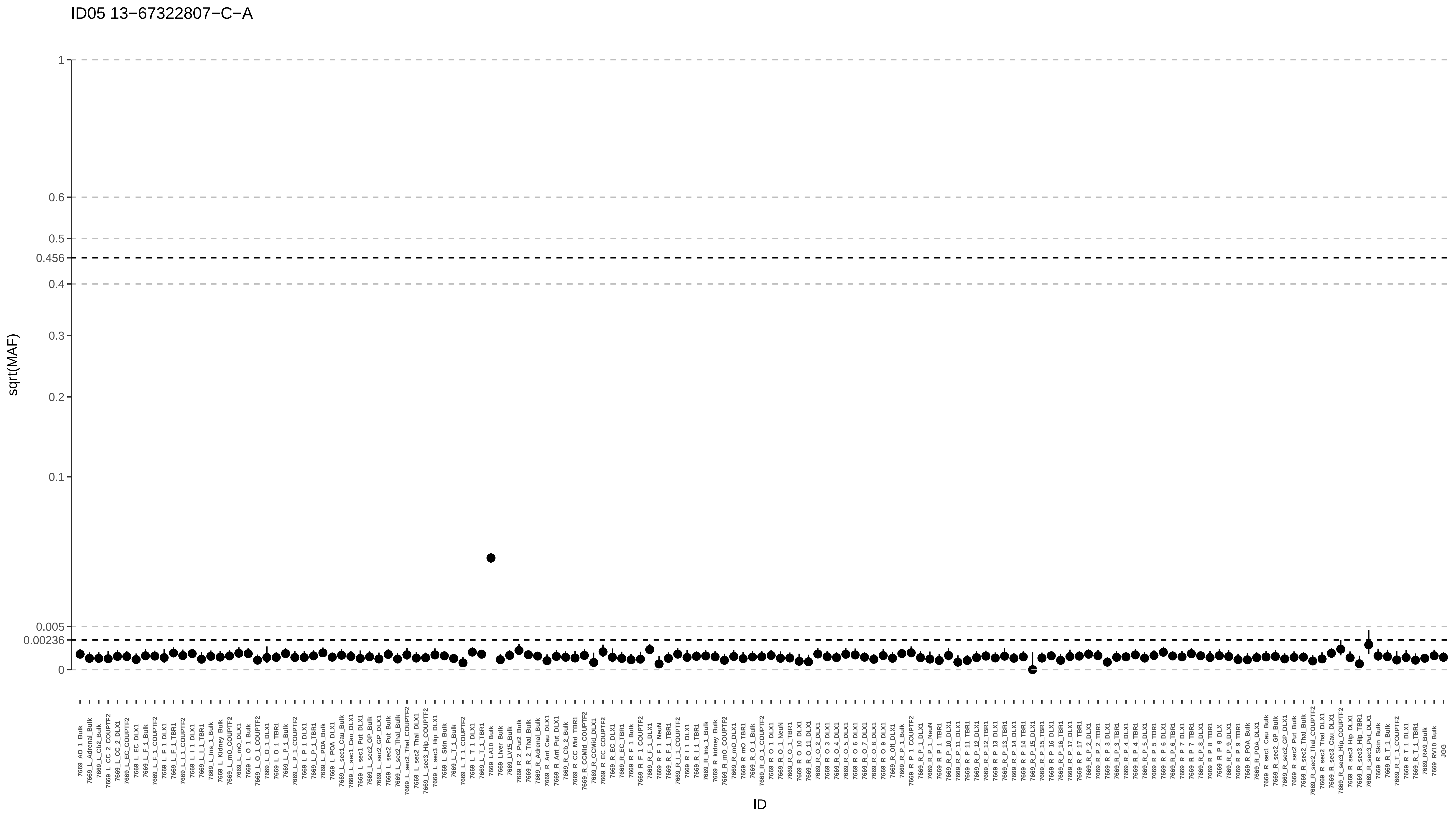
ID05 13−67322807−C−A
sqrt(MAF)
ID

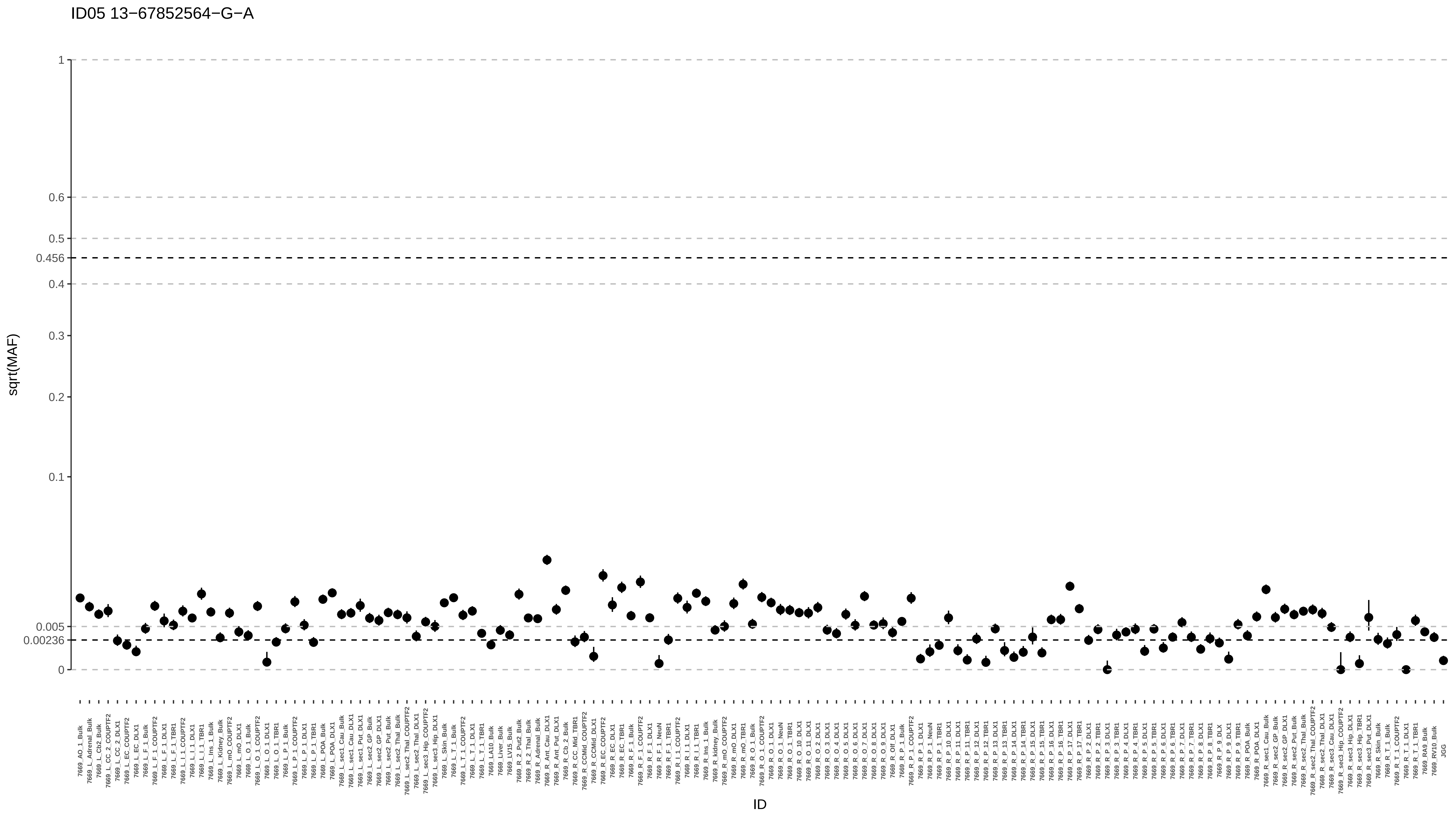
ID05 13−67852564−G−A
sqrt(MAF)
1
0.6
0.5
0.456
0.4
0.3
0.2
0.1
0.005
0.00236
0
ID

ID05 13−69093869−G−A

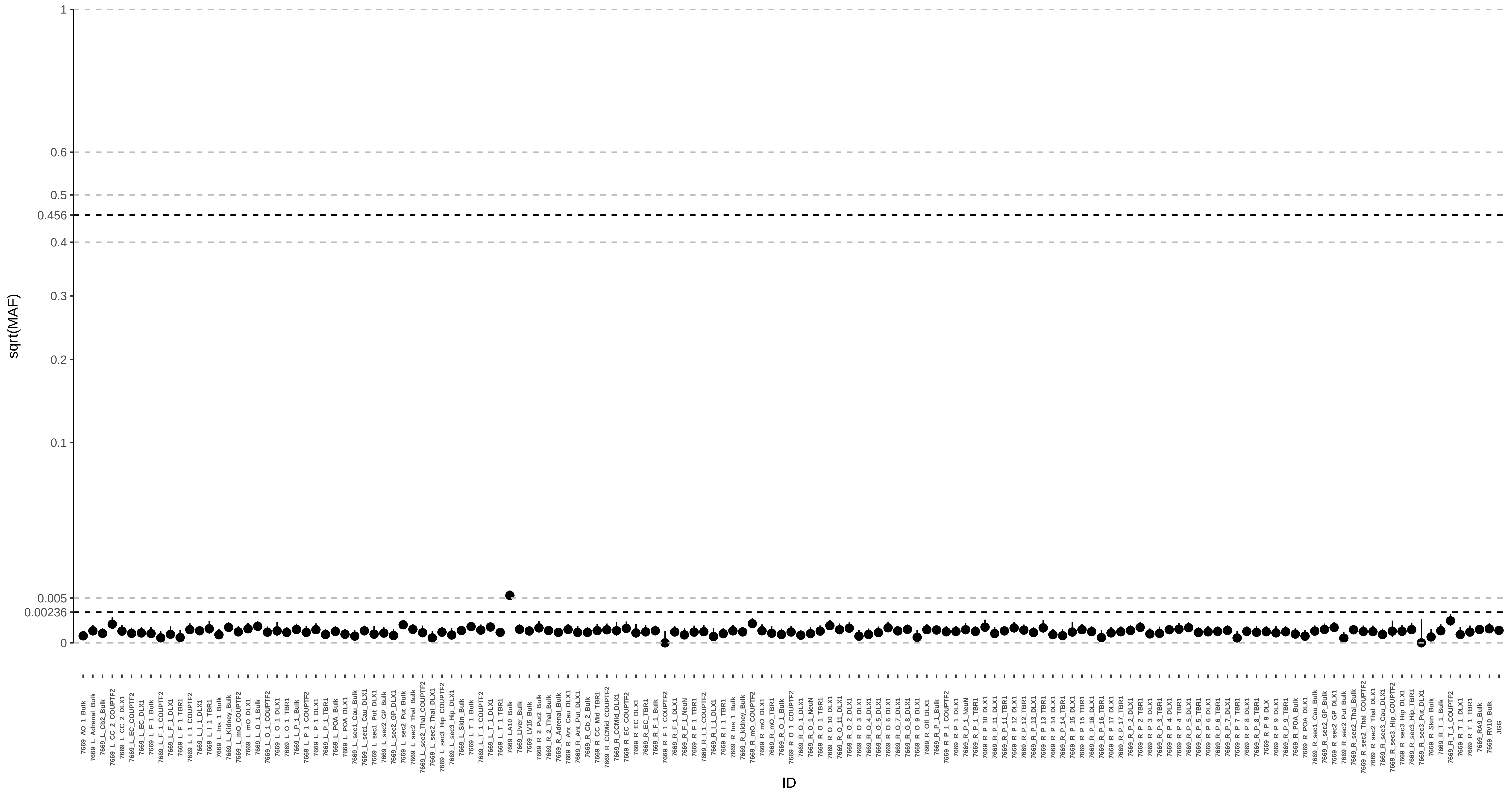

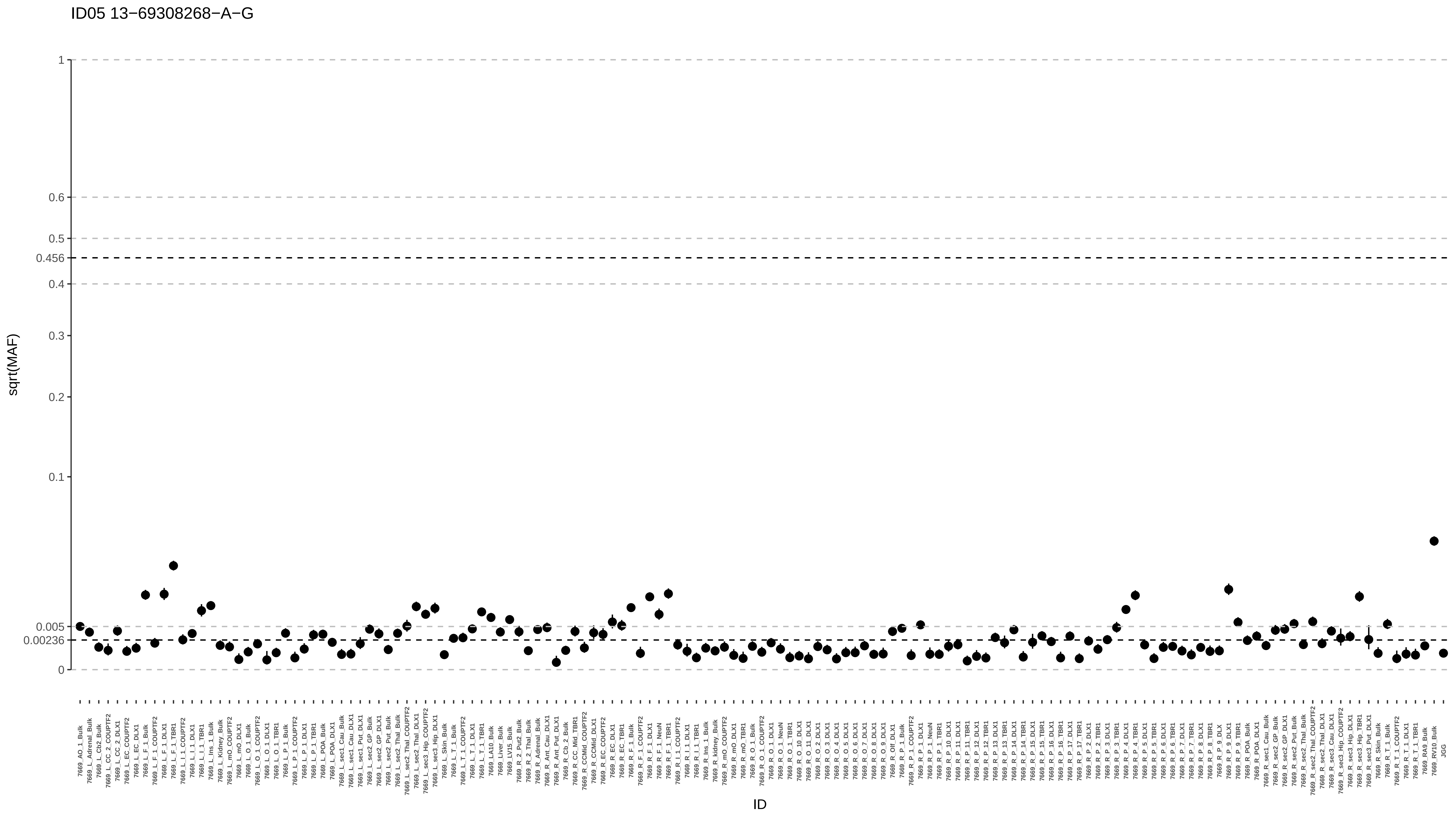

ID05 13−69308268−A−G
sqrt(MAF)
ID

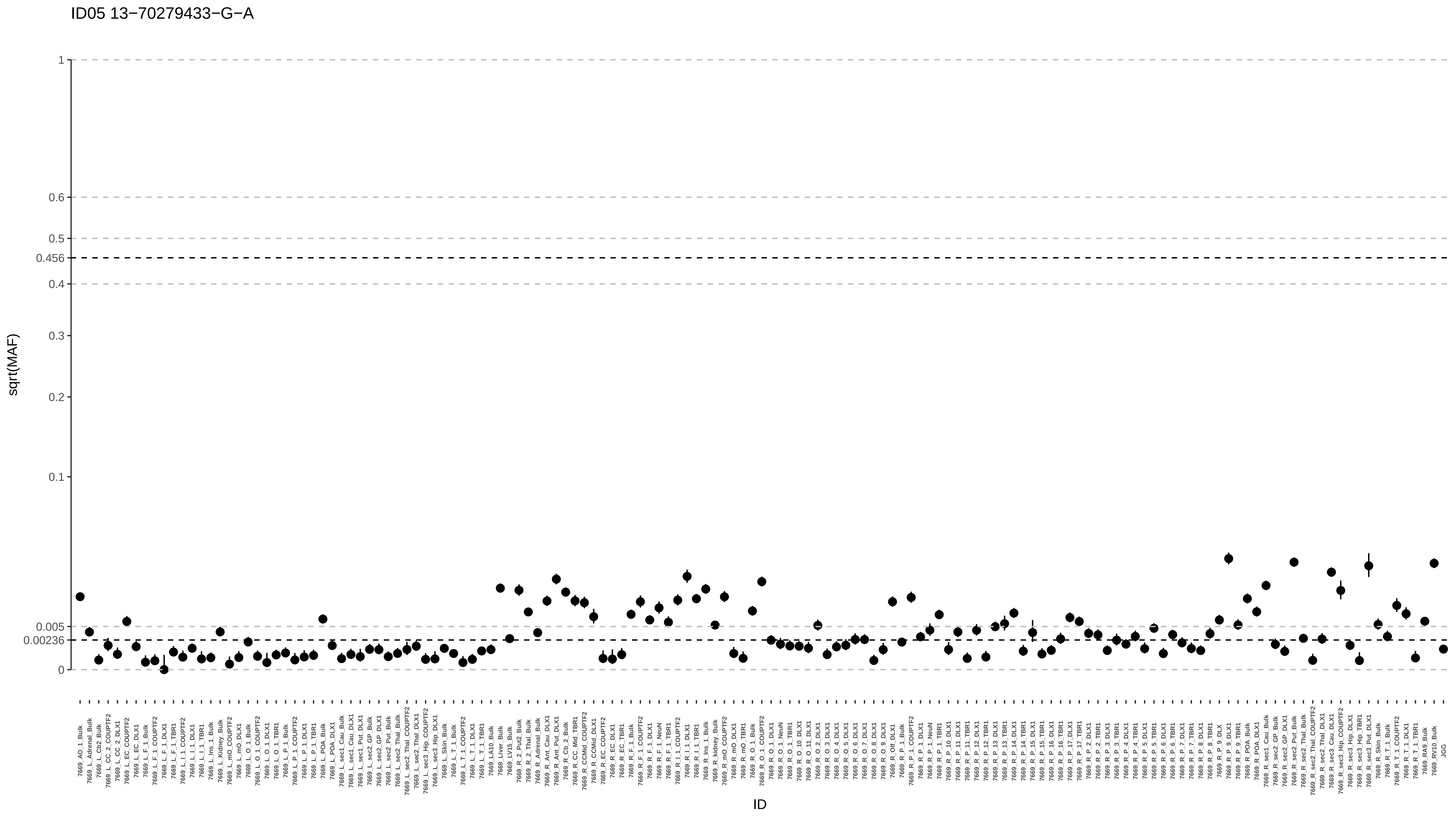
ID05 13−70279433−G−A
sqrt(MAF)
1
0.6
0.5
0.456
0.4
0.3
0.2
0.1
0.005
0.00236
0
ID

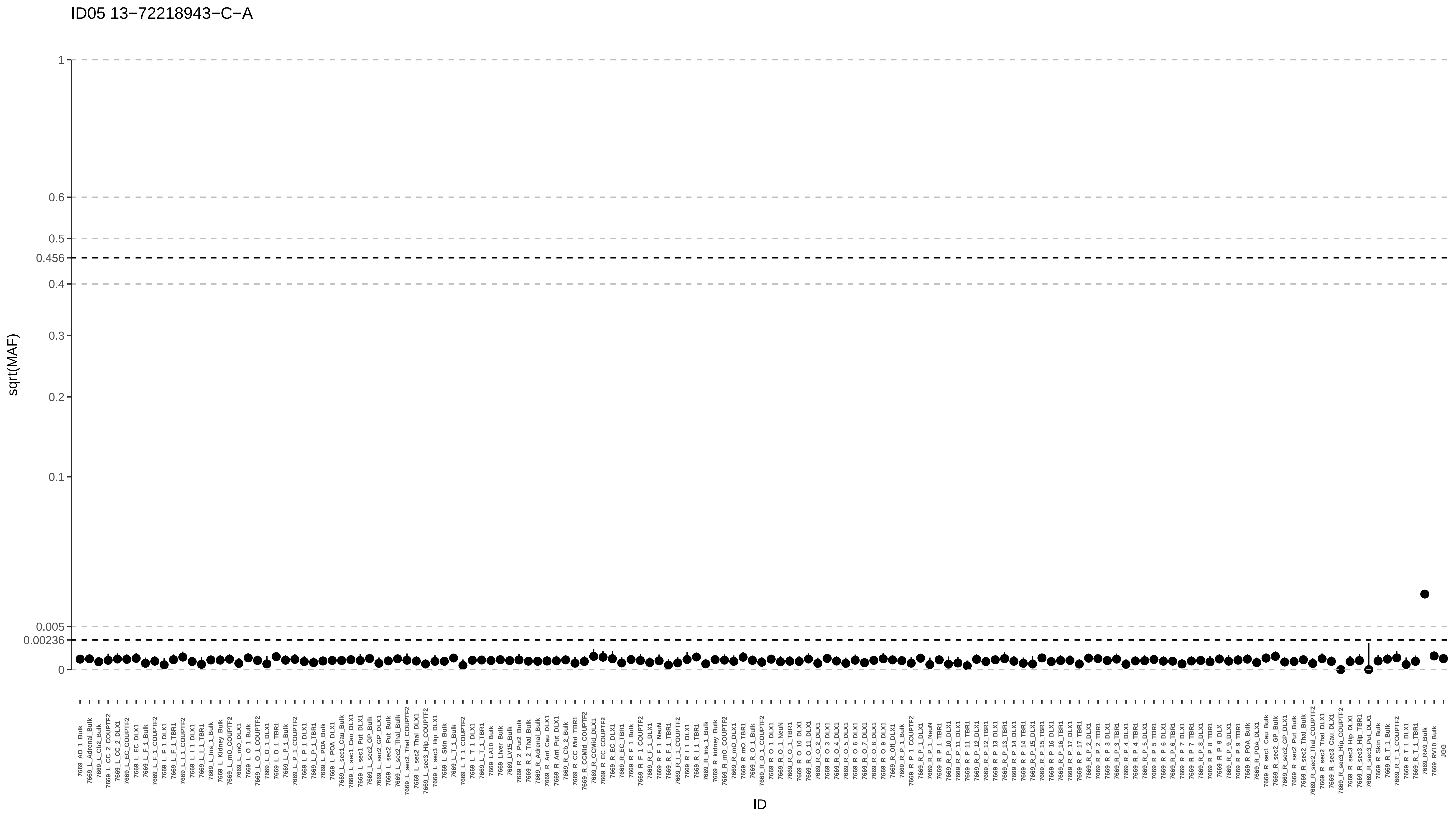
ID05 13−72218943−C−A
sqrt(MAF)
ID

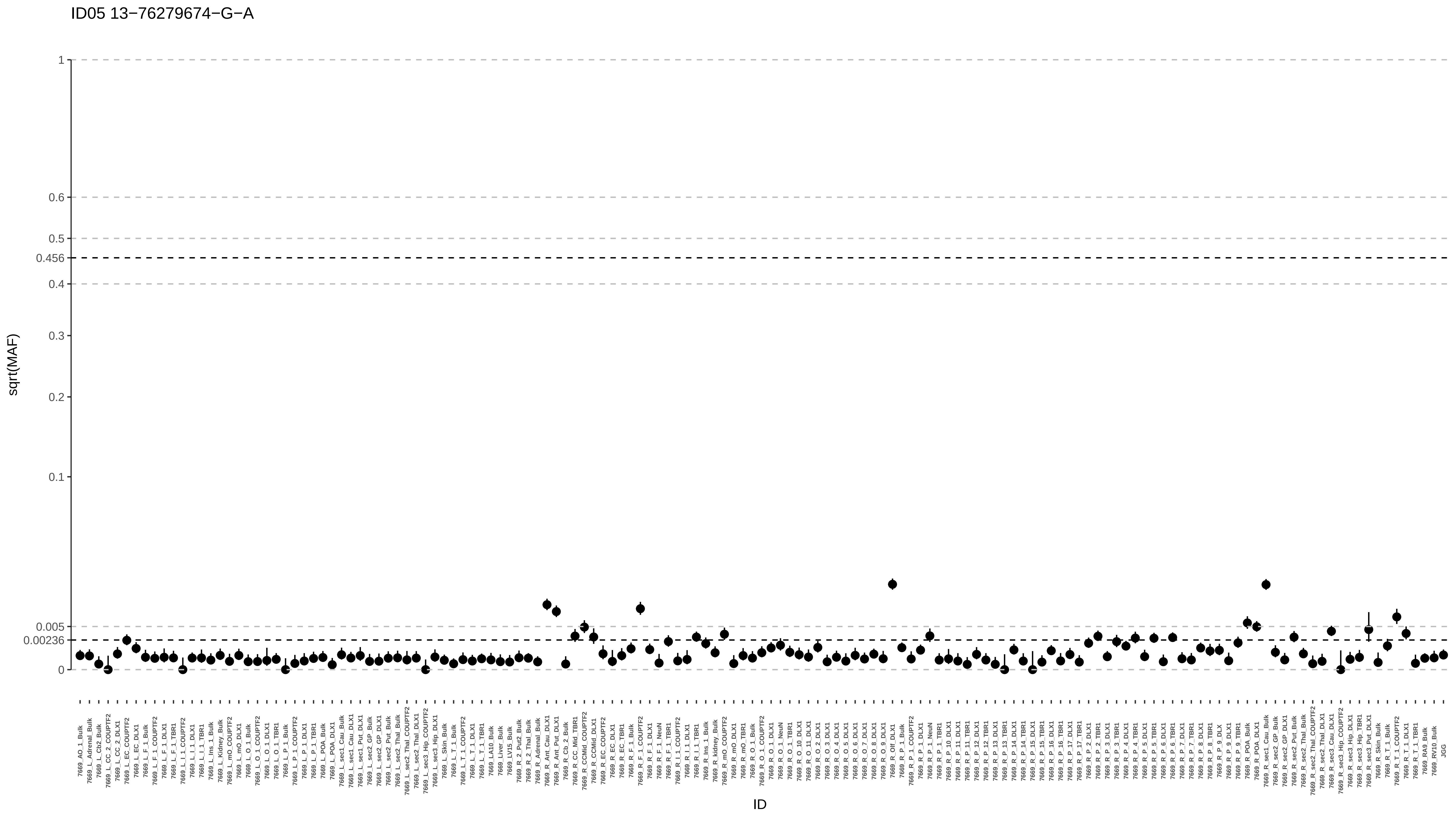
ID05 13−76279674−G−A
sqrt(MAF)
1
0.6
0.5
0.456
0.4
0.3
0.2
0.1
0.005
0.00236
0
ID
7669_AO_1_Bulk
7669_L_Adrenal_Bulk
7669_L_Cb2_Bulk
7669_L_CC_2_COUPTF2
7669_L_CC_2_DLX1
7669_L_EC_COUPTF2
7669_L_EC_DLX1
7669_L_F_1_Bulk
7669_L_F_1_COUPTF2
7669_L_F_1_DLX1
7669_L_F_1_TBR1
7669_L_I_1_COUPTF2
7669_L_I_1_DLX1
7669_L_I_1_TBR1
7669_L_Ins_1_Bulk
7669_L_Kidney_Bulk
7669_L_mO_COUPTF2
7669_L_mO_DLX1
7669_L_O_1_Bulk
7669_L_O_1_COUPTF2
7669_L_O_1_DLX1
7669_L_O_1_TBR1
7669_L_P_1_Bulk
7669_L_P_1_COUPTF2
7669_L_P_1_DLX1
7669_L_P_1_TBR1
7669_L_POA_Bulk
7669_L_POA_DLX1
7669_L_sec1_Cau_Bulk
7669_L_sec1_Cau_DLX1
7669_L_sec1_Put_DLX1
7669_L_sec2_GP_Bulk
7669_L_sec2_GP_DLX1
7669_L_sec2_Put_Bulk
7669_L_sec2_Thal_Bulk
7669_L_sec2_Thal_COUPTF2
7669_L_sec2_Thal_DLX1
7669_L_sec3_Hip_COUPTF2
7669_L_sec3_Hip_DLX1
7669_L_Skin_Bulk
7669_L_T_1_Bulk
7669_L_T_1_COUPTF2
7669_L_T_1_DLX1
7669_L_T_1_TBR1
7669_LA10_Bulk
7669_Liver_Bulk
7669_LV15_Bulk
7669_R_2_Put2_Bulk
7669_R_2_Thal_Bulk
7669_R_Adrenal_Bulk
7669_R_Ant_Cau_DLX1
7669_R_Ant_Put_DLX1
7669_R_Cb_2_Bulk
7669_R_CC_Mid_TBR1
7669_R_CCMid_COUPTF2
7669_R_CCMid_DLX1
7669_R_EC_COUPTF2
7669_R_EC_DLX1
7669_R_EC_TBR1
7669_R_F_1_Bulk
7669_R_F_1_COUPTF2
7669_R_F_1_DLX1
7669_R_F_1_NeuN
7669_R_F_1_TBR1
7669_R_I_1_COUPTF2
7669_R_I_1_DLX1
7669_R_I_1_TBR1
7669_R_Ins_1_Bulk
7669_R_kidney_Bulk
7669_R_mO_COUPTF2
7669_R_mO_DLX1
7669_R_mO_TBR1
7669_R_O_1_Bulk
7669_R_O_1_COUPTF2
7669_R_O_1_DLX1
7669_R_O_1_NeuN
7669_R_O_1_TBR1
7669_R_O_10_DLX1
7669_R_O_11_DLX1
7669_R_O_2_DLX1
7669_R_O_3_DLX1
7669_R_O_4_DLX1
7669_R_O_5_DLX1
7669_R_O_6_DLX1
7669_R_O_7_DLX1
7669_R_O_8_DLX1
7669_R_O_9_DLX1
7669_R_Olf_DLX1
7669_R_P_1_Bulk
7669_R_P_1_COUPTF2
7669_R_P_1_DLX1
7669_R_P_1_NeuN
7669_R_P_1_TBR1
7669_R_P_10_DLX1
7669_R_P_11_DLX1
7669_R_P_11_TBR1
7669_R_P_12_DLX1
7669_R_P_12_TBR1
7669_R_P_13_DLX1
7669_R_P_13_TBR1
7669_R_P_14_DLX1
7669_R_P_14_TBR1
7669_R_P_15_DLX1
7669_R_P_15_TBR1
7669_R_P_16_DLX1
7669_R_P_16_TBR1
7669_R_P_17_DLX1
7669_R_P_17_TBR1
7669_R_P_2_DLX1
7669_R_P_2_TBR1
7669_R_P_3_DLX1
7669_R_P_3_TBR1
7669_R_P_4_DLX1
7669_R_P_4_TBR1
7669_R_P_5_DLX1
7669_R_P_5_TBR1
7669_R_P_6_DLX1
7669_R_P_6_TBR1
7669_R_P_7_DLX1
7669_R_P_7_TBR1
7669_R_P_8_DLX1
7669_R_P_8_TBR1
7669_R_P_9_DLX
7669_R_P_9_DLX1
7669_R_P_9_TBR1
7669_R_POA_Bulk
7669_R_POA_DLX1
7669_R_sec1_Cau_Bulk
7669_R_sec2_GP_Bulk
7669_R_sec2_GP_DLX1
7669_R_sec2_Put_Bulk
7669_R_sec2_Thal_Bulk
7669_R_sec2_Thal_COUPTF2
7669_R_sec2_Thal_DLX1
7669_R_sec3_Cau_DLX1
7669_R_sec3_Hip_COUPTF2
7669_R_sec3_Hip_DLX1
7669_R_sec3_Hip_TBR1
7669_R_sec3_Put_DLX1
7669_R_Skin_Bulk
7669_R_T_1_Bulk
7669_R_T_1_COUPTF2
7669_R_T_1_DLX1
7669_R_T_1_TBR1
7669_RA9_Bulk
7669_RV10_Bulk
JGG

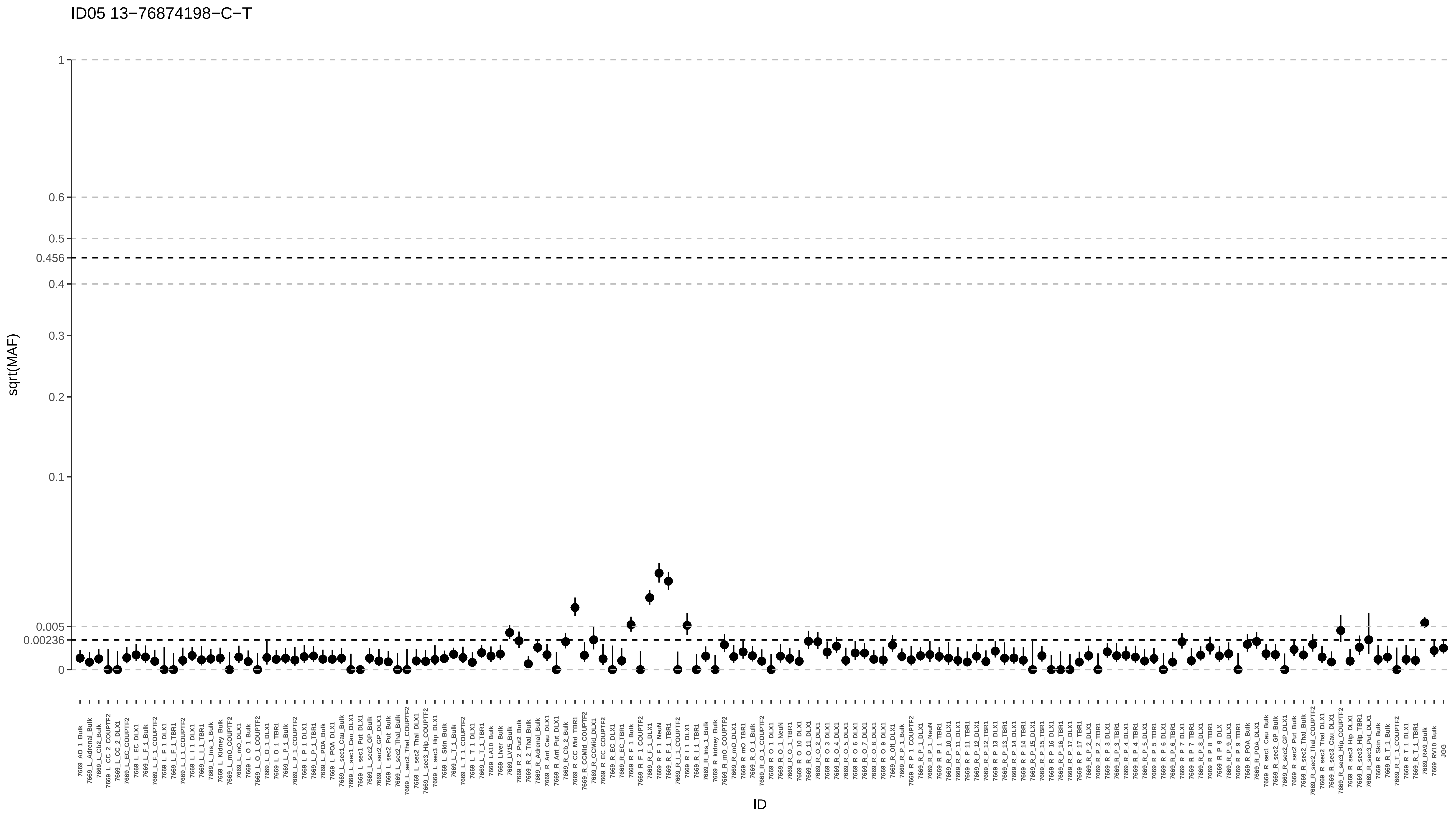
ID05 13−76874198−C−T
sqrt(MAF)
1
0.6
0.5
0.456
0.4
0.3
0.2
0.1
0.005
0.00236
0
ID

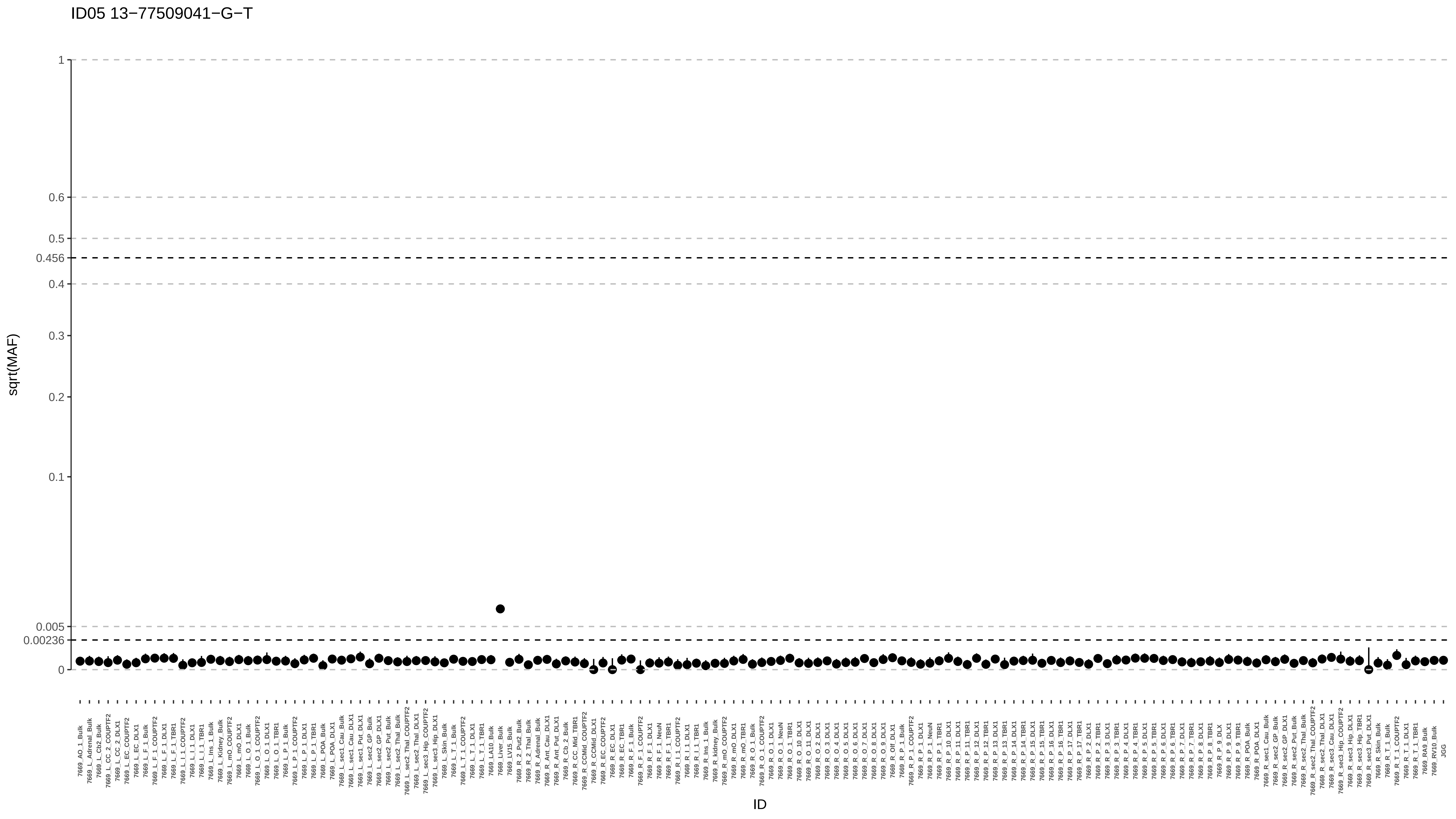
ID05 13−77509041−G−T
sqrt(MAF)
1
0.6
0.5
0.456
0.4
0.3
0.2
0.1
0.005
0.00236
0
ID

ID05 13−83875706−C−T

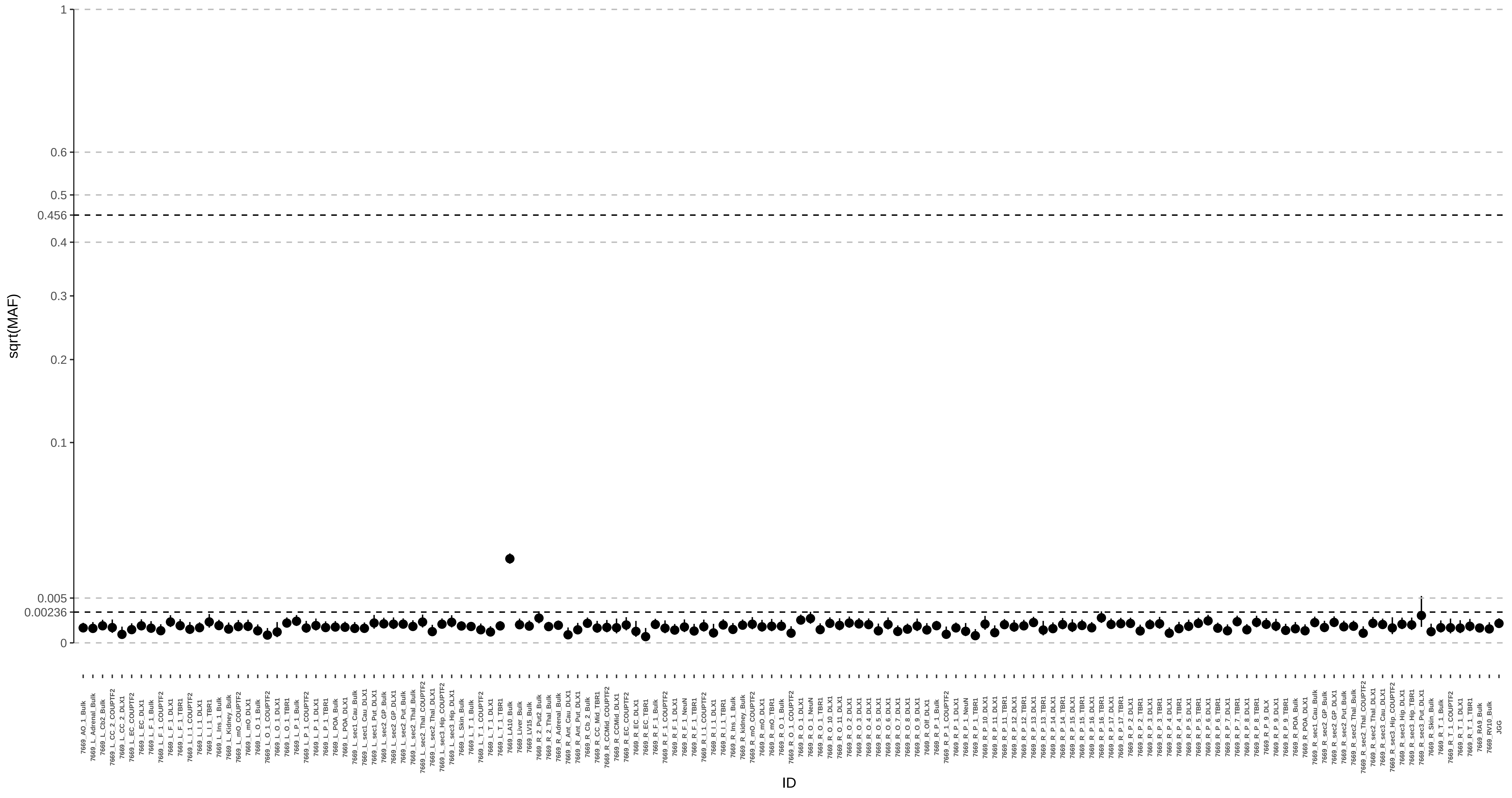

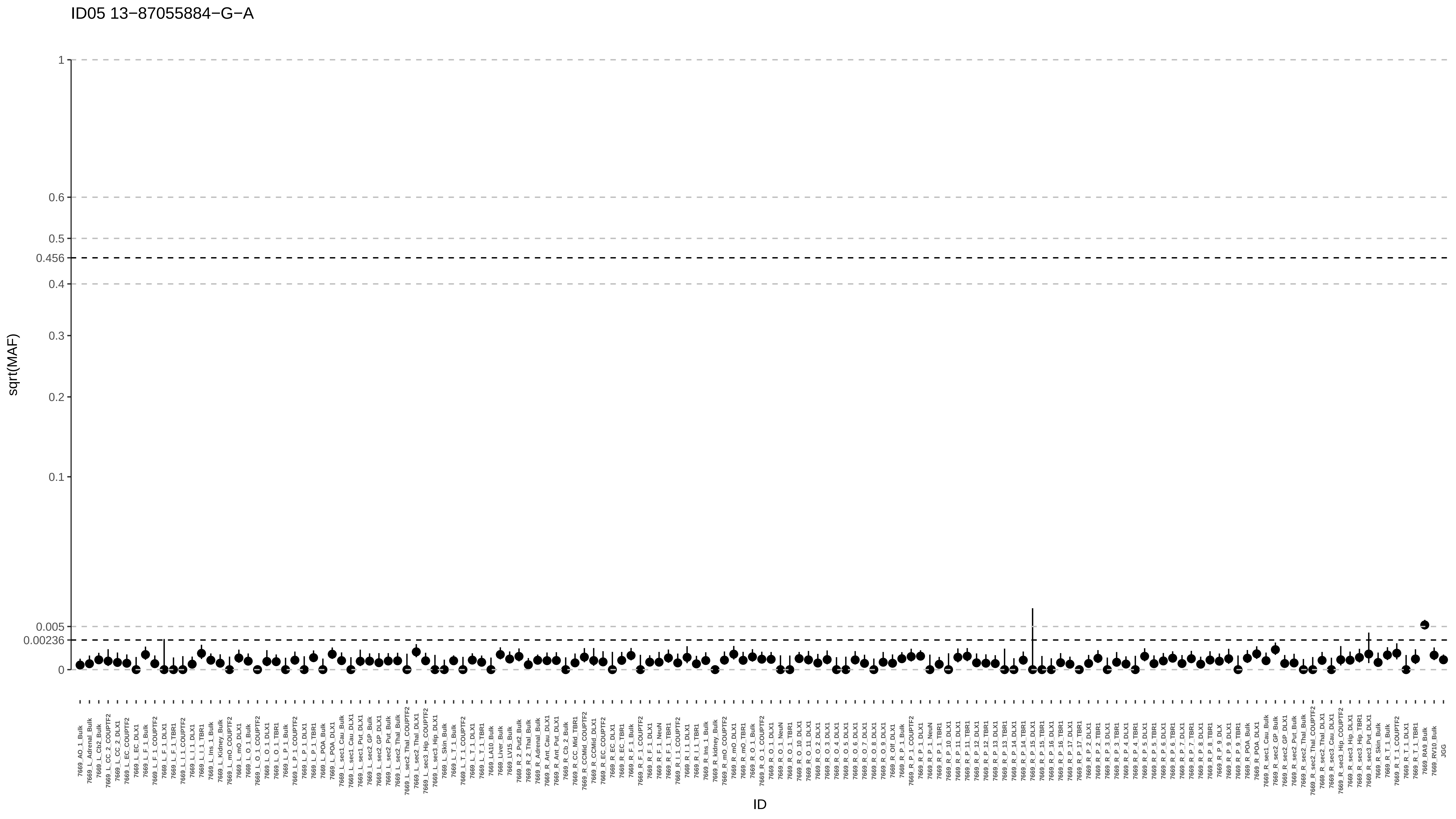
ID05 13−87055884−G−A
sqrt(MAF)
1
0.6
0.5
0.456
0.4
0.3
0.2
0.1
0.005
0.00236
0
ID
7669_AO_1_Bulk
7669_L_Adrenal_Bulk
7669_L_Cb2_Bulk
7669_L_CC_2_COUPTF2
7669_L_CC_2_DLX1
7669_L_EC_COUPTF2
7669_L_EC_DLX1
7669_L_F_1_Bulk
7669_L_F_1_COUPTF2
7669_L_F_1_DLX1
7669_L_F_1_TBR1
7669_L_I_1_COUPTF2
7669_L_I_1_DLX1
7669_L_I_1_TBR1
7669_L_Ins_1_Bulk
7669_L_Kidney_Bulk
7669_L_mO_COUPTF2
7669_L_mO_DLX1
7669_L_O_1_Bulk
7669_L_O_1_COUPTF2
7669_L_O_1_DLX1
7669_L_O_1_TBR1
7669_L_P_1_Bulk
7669_L_P_1_COUPTF2
7669_L_P_1_DLX1
7669_L_P_1_TBR1
7669_L_POA_Bulk
7669_L_POA_DLX1
7669_L_sec1_Cau_Bulk
7669_L_sec1_Cau_DLX1
7669_L_sec1_Put_DLX1
7669_L_sec2_GP_Bulk
7669_L_sec2_GP_DLX1
7669_L_sec2_Put_Bulk
7669_L_sec2_Thal_Bulk
7669_L_sec2_Thal_COUPTF2
7669_L_sec2_Thal_DLX1
7669_L_sec3_Hip_COUPTF2
7669_L_sec3_Hip_DLX1
7669_L_Skin_Bulk
7669_L_T_1_Bulk
7669_L_T_1_COUPTF2
7669_L_T_1_DLX1
7669_L_T_1_TBR1
7669_LA10_Bulk
7669_Liver_Bulk
7669_LV15_Bulk
7669_R_2_Put2_Bulk
7669_R_2_Thal_Bulk
7669_R_Adrenal_Bulk
7669_R_Ant_Cau_DLX1
7669_R_Ant_Put_DLX1
7669_R_Cb_2_Bulk
7669_R_CC_Mid_TBR1
7669_R_CCMid_COUPTF2
7669_R_CCMid_DLX1
7669_R_EC_COUPTF2
7669_R_EC_DLX1
7669_R_EC_TBR1
7669_R_F_1_Bulk
7669_R_F_1_COUPTF2
7669_R_F_1_DLX1
7669_R_F_1_NeuN
7669_R_F_1_TBR1
7669_R_I_1_COUPTF2
7669_R_I_1_DLX1
7669_R_I_1_TBR1
7669_R_Ins_1_Bulk
7669_R_kidney_Bulk
7669_R_mO_COUPTF2
7669_R_mO_DLX1
7669_R_mO_TBR1
7669_R_O_1_Bulk
7669_R_O_1_COUPTF2
7669_R_O_1_DLX1
7669_R_O_1_NeuN
7669_R_O_1_TBR1
7669_R_O_10_DLX1
7669_R_O_11_DLX1
7669_R_O_2_DLX1
7669_R_O_3_DLX1
7669_R_O_4_DLX1
7669_R_O_5_DLX1
7669_R_O_6_DLX1
7669_R_O_7_DLX1
7669_R_O_8_DLX1
7669_R_O_9_DLX1
7669_R_Olf_DLX1
7669_R_P_1_Bulk
7669_R_P_1_COUPTF2
7669_R_P_1_DLX1
7669_R_P_1_NeuN
7669_R_P_1_TBR1
7669_R_P_10_DLX1
7669_R_P_11_DLX1
7669_R_P_11_TBR1
7669_R_P_12_DLX1
7669_R_P_12_TBR1
7669_R_P_13_DLX1
7669_R_P_13_TBR1
7669_R_P_14_DLX1
7669_R_P_14_TBR1
7669_R_P_15_DLX1
7669_R_P_15_TBR1
7669_R_P_16_DLX1
7669_R_P_16_TBR1
7669_R_P_17_DLX1
7669_R_P_17_TBR1
7669_R_P_2_DLX1
7669_R_P_2_TBR1
7669_R_P_3_DLX1
7669_R_P_3_TBR1
7669_R_P_4_DLX1
7669_R_P_4_TBR1
7669_R_P_5_DLX1
7669_R_P_5_TBR1
7669_R_P_6_DLX1
7669_R_P_6_TBR1
7669_R_P_7_DLX1
7669_R_P_7_TBR1
7669_R_P_8_DLX1
7669_R_P_8_TBR1
7669_R_P_9_DLX
7669_R_P_9_DLX1
7669_R_P_9_TBR1
7669_R_POA_Bulk
7669_R_POA_DLX1
7669_R_sec1_Cau_Bulk
7669_R_sec2_GP_Bulk
7669_R_sec2_GP_DLX1
7669_R_sec2_Put_Bulk
7669_R_sec2_Thal_Bulk
7669_R_sec2_Thal_COUPTF2
7669_R_sec2_Thal_DLX1
7669_R_sec3_Cau_DLX1
7669_R_sec3_Hip_COUPTF2
7669_R_sec3_Hip_DLX1
7669_R_sec3_Hip_TBR1
7669_R_sec3_Put_DLX1
7669_R_Skin_Bulk
7669_R_T_1_Bulk
7669_R_T_1_COUPTF2
7669_R_T_1_DLX1
7669_R_T_1_TBR1
7669_RA9_Bulk
7669_RV10_Bulk
JGG

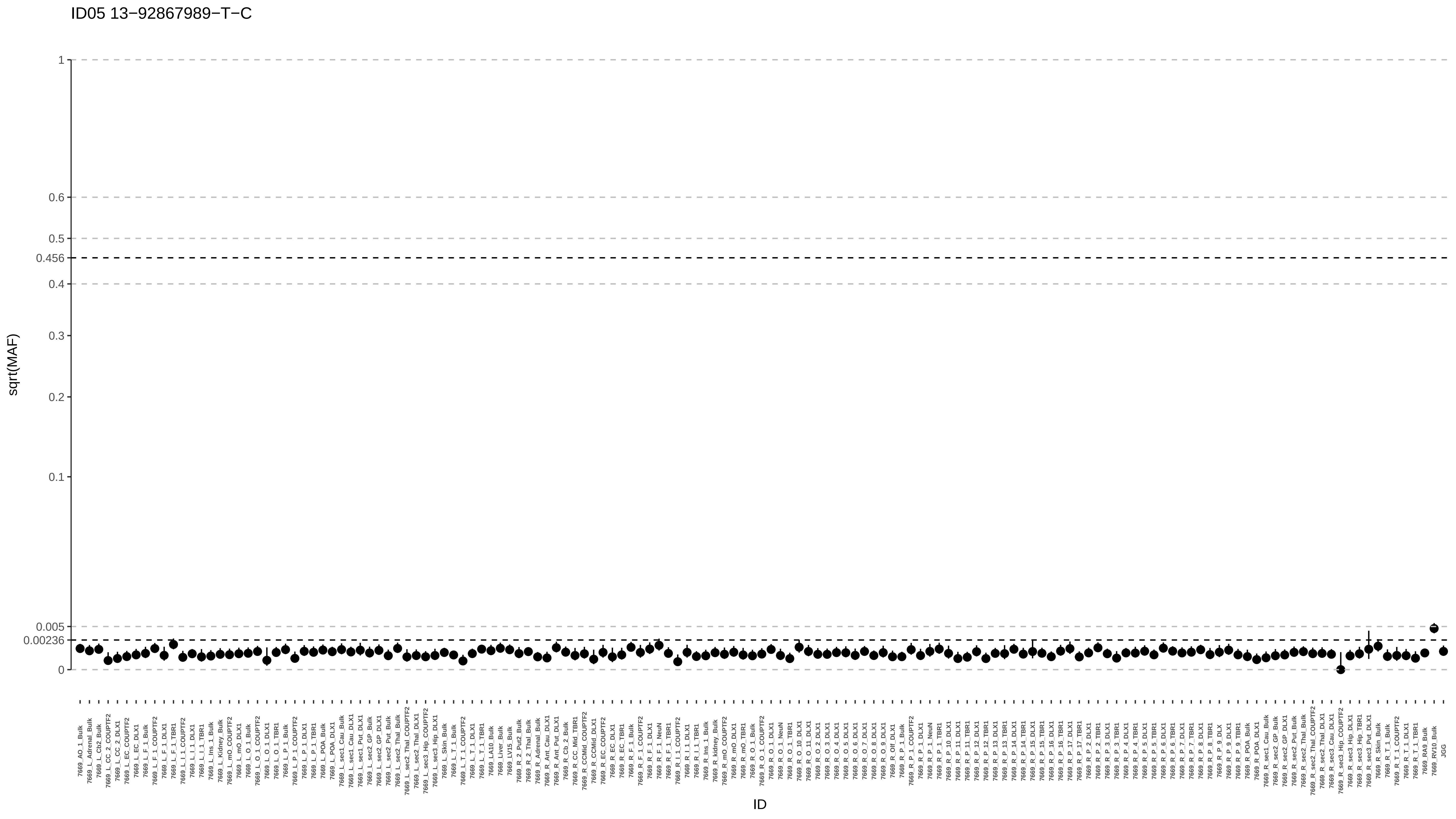
ID05 13−92867989−T−C
sqrt(MAF)
ID

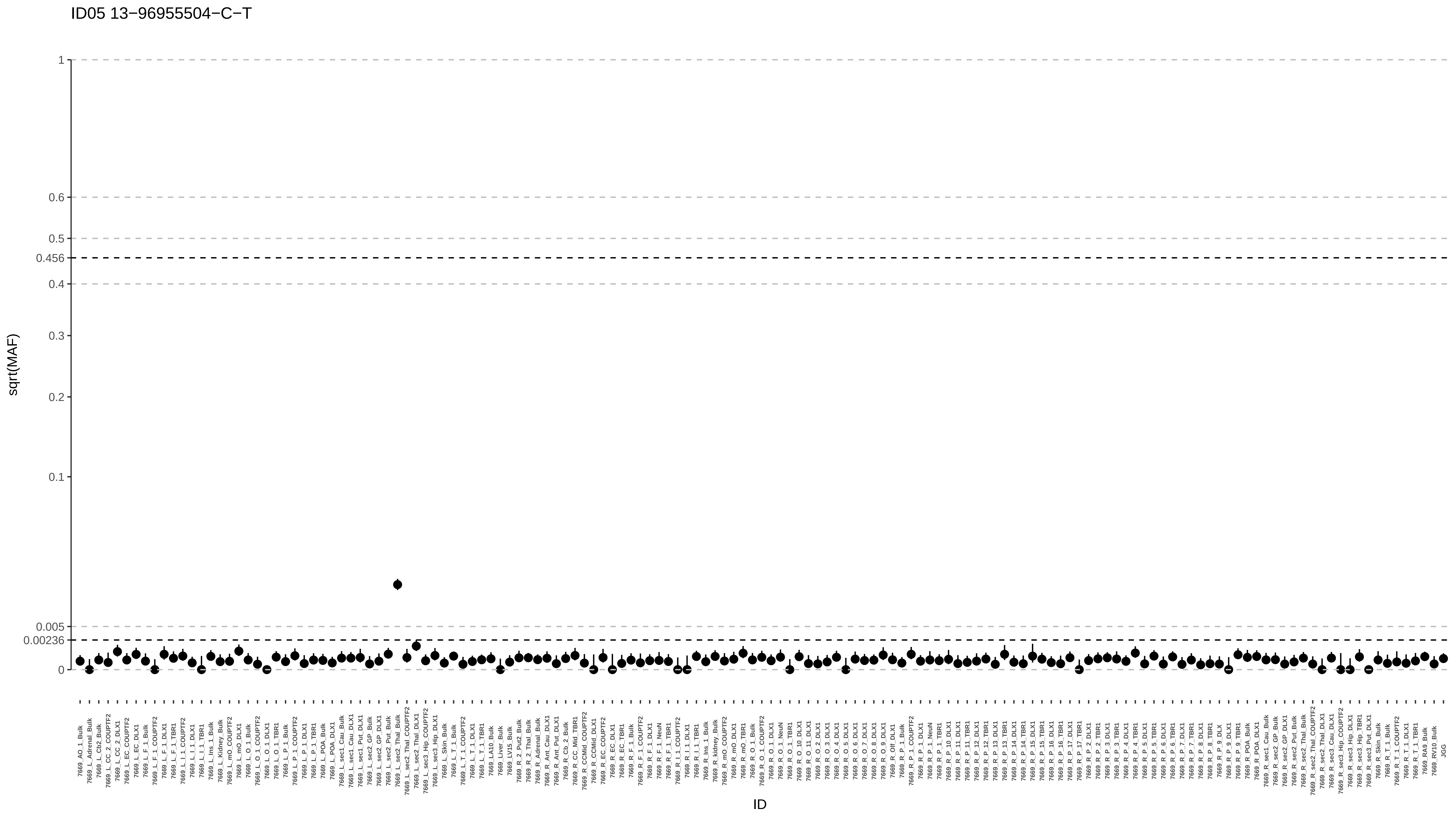
ID05 13−96955504−C−T
sqrt(MAF)
1
0.6
0.5
0.456
0.4
0.3
0.2
0.1
0.005
0.00236
0
ID

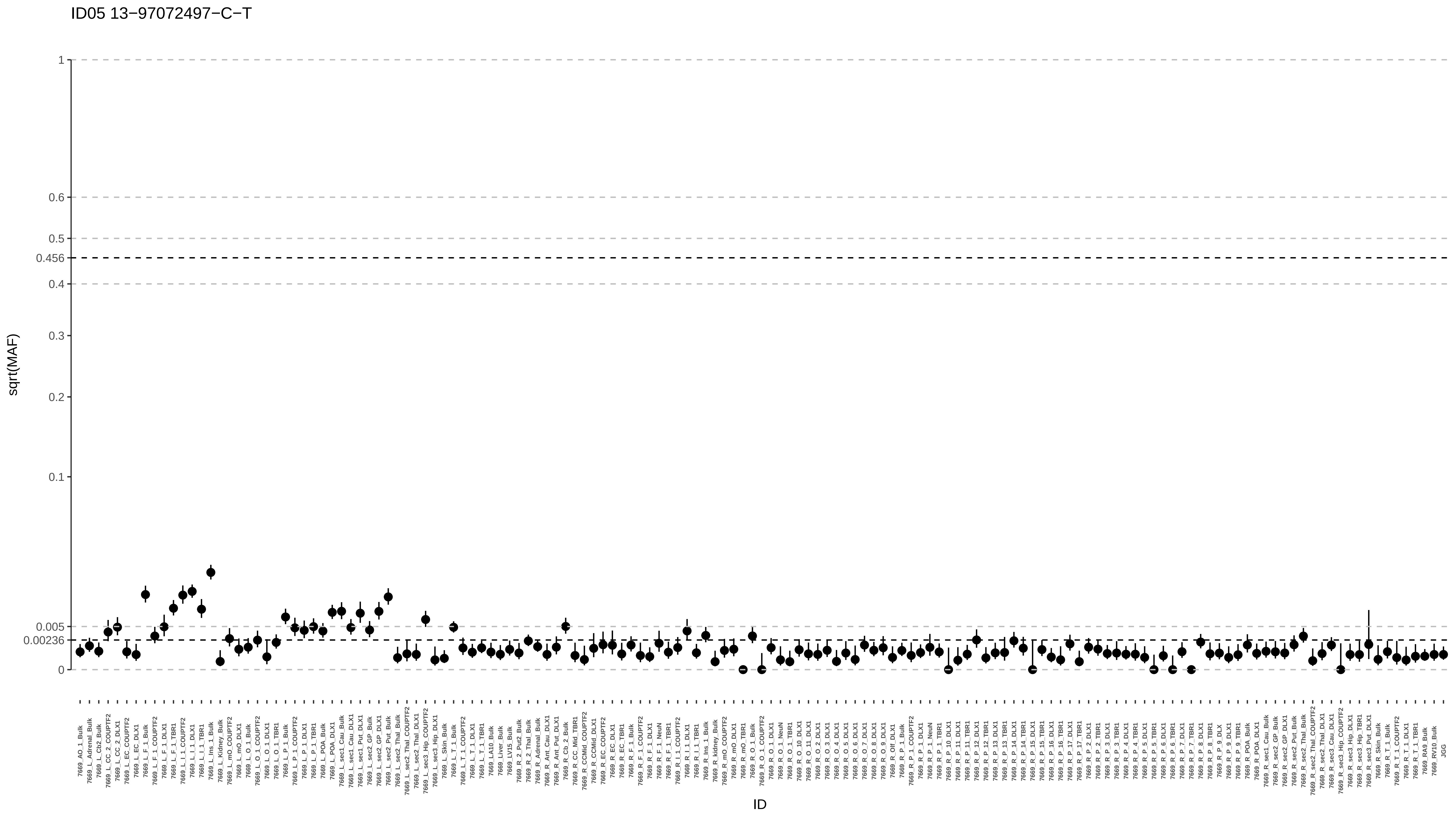
ID05 13−97072497−C−T
sqrt(MAF)
1
0.6
0.5
0.456
0.4
0.3
0.2
0.1
0.005
0.00236
0
ID
7669_AO_1_Bulk
7669_L_Adrenal_Bulk
7669_L_Cb2_Bulk
7669_L_CC_2_COUPTF2
7669_L_CC_2_DLX1
7669_L_EC_COUPTF2
7669_L_EC_DLX1
7669_L_F_1_Bulk
7669_L_F_1_COUPTF2
7669_L_F_1_DLX1
7669_L_F_1_TBR1
7669_L_I_1_COUPTF2
7669_L_I_1_DLX1
7669_L_I_1_TBR1
7669_L_Ins_1_Bulk
7669_L_Kidney_Bulk
7669_L_mO_COUPTF2
7669_L_mO_DLX1
7669_L_O_1_Bulk
7669_L_O_1_COUPTF2
7669_L_O_1_DLX1
7669_L_O_1_TBR1
7669_L_P_1_Bulk
7669_L_P_1_COUPTF2
7669_L_P_1_DLX1
7669_L_P_1_TBR1
7669_L_POA_Bulk
7669_L_POA_DLX1
7669_L_sec1_Cau_Bulk
7669_L_sec1_Cau_DLX1
7669_L_sec1_Put_DLX1
7669_L_sec2_GP_Bulk
7669_L_sec2_GP_DLX1
7669_L_sec2_Put_Bulk
7669_L_sec2_Thal_Bulk
7669_L_sec2_Thal_COUPTF2
7669_L_sec2_Thal_DLX1
7669_L_sec3_Hip_COUPTF2
7669_L_sec3_Hip_DLX1
7669_L_Skin_Bulk
7669_L_T_1_Bulk
7669_L_T_1_COUPTF2
7669_L_T_1_DLX1
7669_L_T_1_TBR1
7669_LA10_Bulk
7669_Liver_Bulk
7669_LV15_Bulk
7669_R_2_Put2_Bulk
7669_R_2_Thal_Bulk
7669_R_Adrenal_Bulk
7669_R_Ant_Cau_DLX1
7669_R_Ant_Put_DLX1
7669_R_Cb_2_Bulk
7669_R_CC_Mid_TBR1
7669_R_CCMid_COUPTF2
7669_R_CCMid_DLX1
7669_R_EC_COUPTF2
7669_R_EC_DLX1
7669_R_EC_TBR1
7669_R_F_1_Bulk
7669_R_F_1_COUPTF2
7669_R_F_1_DLX1
7669_R_F_1_NeuN
7669_R_F_1_TBR1
7669_R_I_1_COUPTF2
7669_R_I_1_DLX1
7669_R_I_1_TBR1
7669_R_Ins_1_Bulk
7669_R_kidney_Bulk
7669_R_mO_COUPTF2
7669_R_mO_DLX1
7669_R_mO_TBR1
7669_R_O_1_Bulk
7669_R_O_1_COUPTF2
7669_R_O_1_DLX1
7669_R_O_1_NeuN
7669_R_O_1_TBR1
7669_R_O_10_DLX1
7669_R_O_11_DLX1
7669_R_O_2_DLX1
7669_R_O_3_DLX1
7669_R_O_4_DLX1
7669_R_O_5_DLX1
7669_R_O_6_DLX1
7669_R_O_7_DLX1
7669_R_O_8_DLX1
7669_R_O_9_DLX1
7669_R_Olf_DLX1
7669_R_P_1_Bulk
7669_R_P_1_COUPTF2
7669_R_P_1_DLX1
7669_R_P_1_NeuN
7669_R_P_1_TBR1
7669_R_P_10_DLX1
7669_R_P_11_DLX1
7669_R_P_11_TBR1
7669_R_P_12_DLX1
7669_R_P_12_TBR1
7669_R_P_13_DLX1
7669_R_P_13_TBR1
7669_R_P_14_DLX1
7669_R_P_14_TBR1
7669_R_P_15_DLX1
7669_R_P_15_TBR1
7669_R_P_16_DLX1
7669_R_P_16_TBR1
7669_R_P_17_DLX1
7669_R_P_17_TBR1
7669_R_P_2_DLX1
7669_R_P_2_TBR1
7669_R_P_3_DLX1
7669_R_P_3_TBR1
7669_R_P_4_DLX1
7669_R_P_4_TBR1
7669_R_P_5_DLX1
7669_R_P_5_TBR1
7669_R_P_6_DLX1
7669_R_P_6_TBR1
7669_R_P_7_DLX1
7669_R_P_7_TBR1
7669_R_P_8_DLX1
7669_R_P_8_TBR1
7669_R_P_9_DLX
7669_R_P_9_DLX1
7669_R_P_9_TBR1
7669_R_POA_Bulk
7669_R_POA_DLX1
7669_R_sec1_Cau_Bulk
7669_R_sec2_GP_Bulk
7669_R_sec2_GP_DLX1
7669_R_sec2_Put_Bulk
7669_R_sec2_Thal_Bulk
7669_R_sec2_Thal_COUPTF2
7669_R_sec2_Thal_DLX1
7669_R_sec3_Cau_DLX1
7669_R_sec3_Hip_COUPTF2
7669_R_sec3_Hip_DLX1
7669_R_sec3_Hip_TBR1
7669_R_sec3_Put_DLX1
7669_R_Skin_Bulk
7669_R_T_1_Bulk
7669_R_T_1_COUPTF2
7669_R_T_1_DLX1
7669_R_T_1_TBR1
7669_RA9_Bulk
7669_RV10_Bulk
JGG

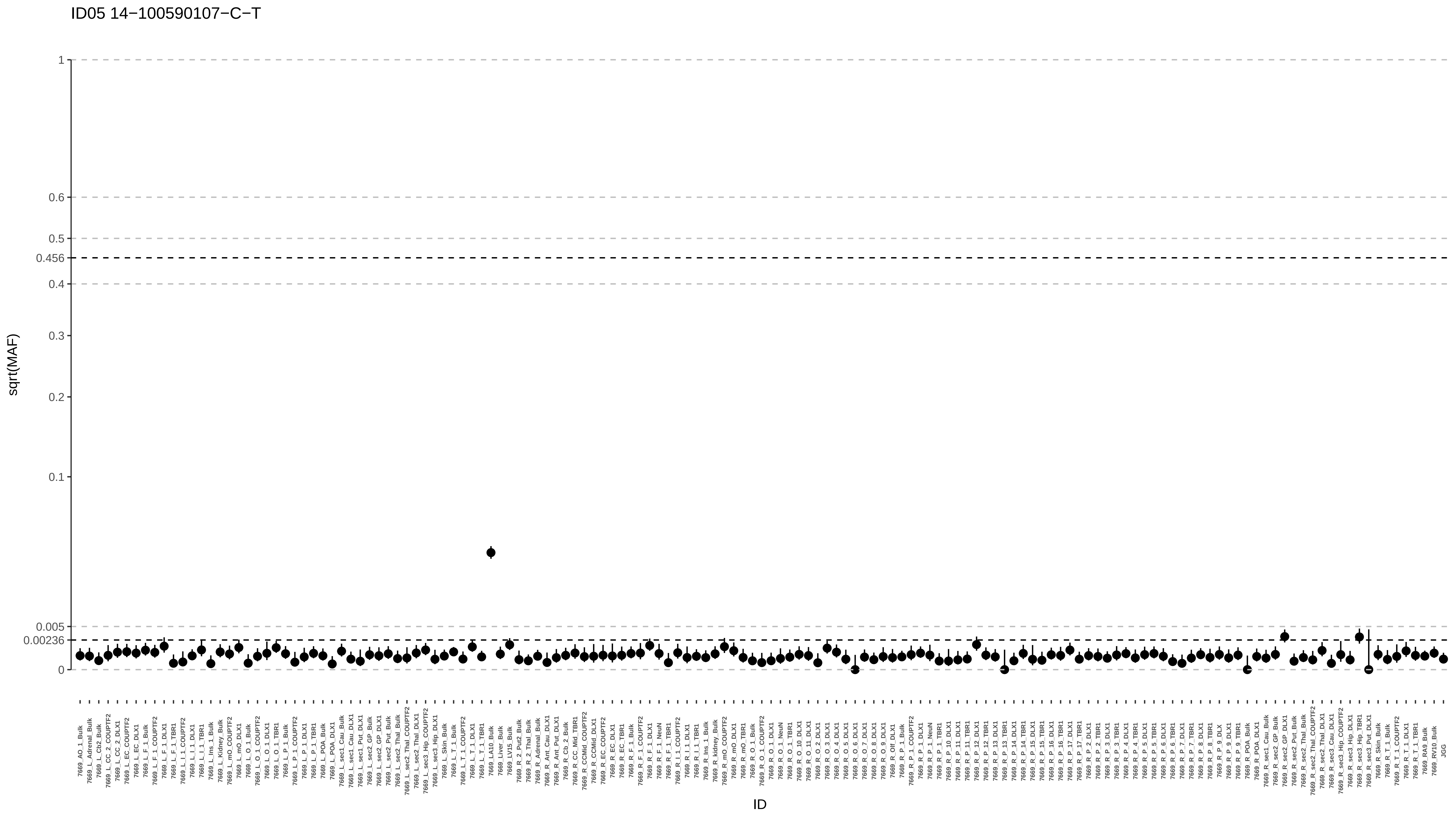
ID05 14−100590107−C−T
sqrt(MAF)
1
0.6
0.5
0.456
0.4
0.3
0.2
0.1
0.005
0.00236
0
ID

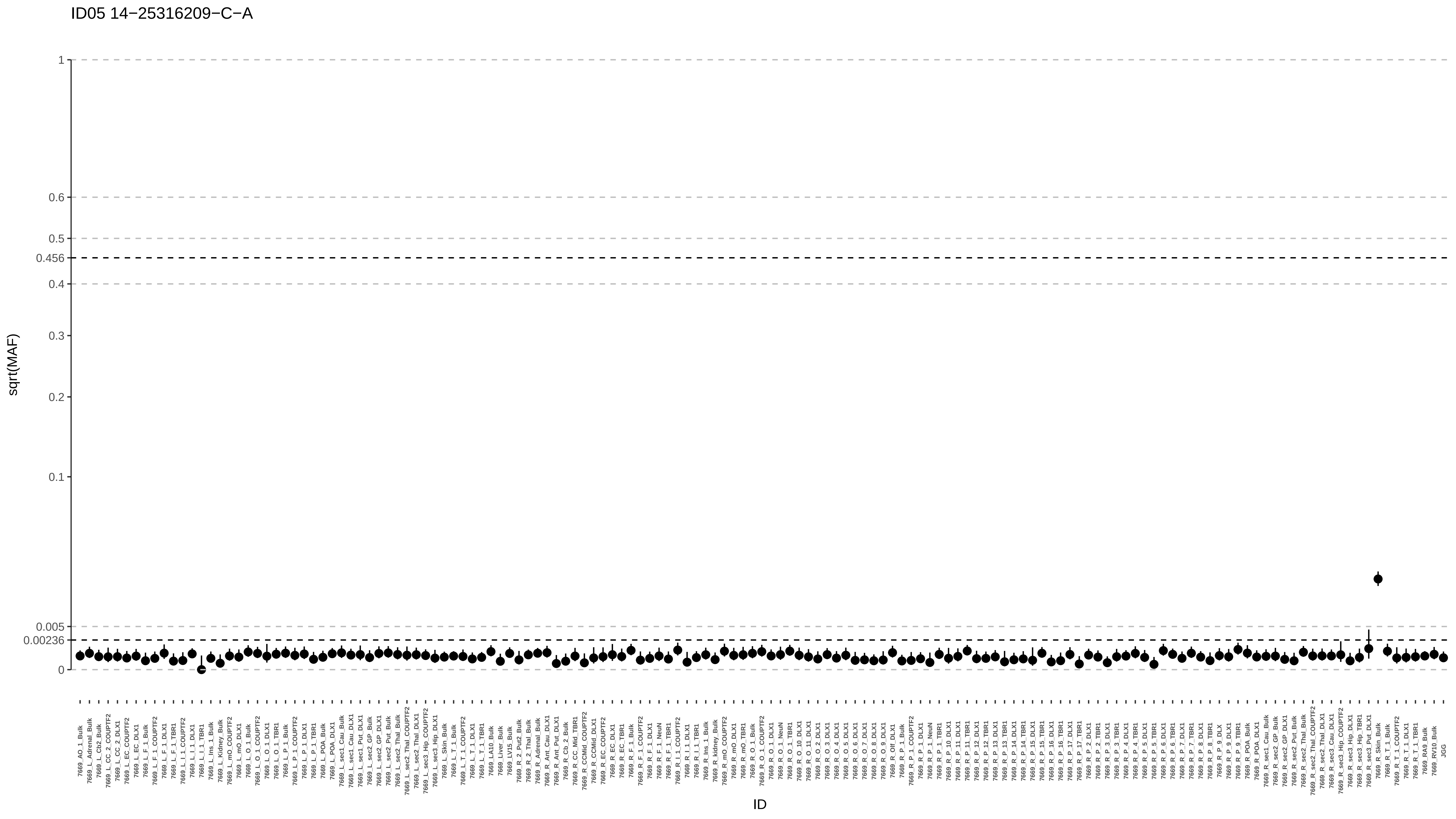
ID05 14−25316209−C−A
sqrt(MAF)
1
0.6
0.5
0.456
0.4
0.3
0.2
0.1
0.005
0.00236
0
ID

ID05 14−27412390−G−A

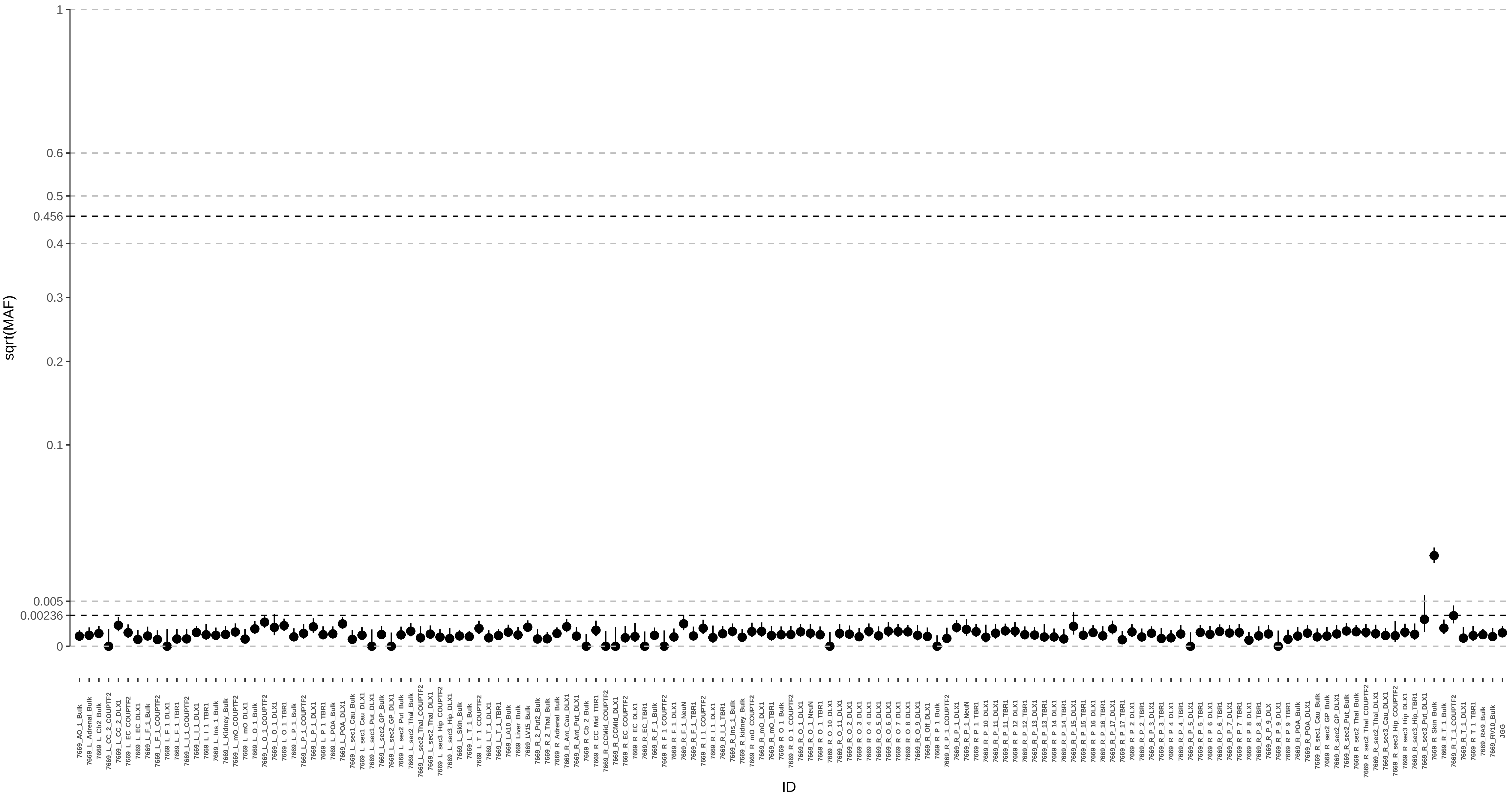

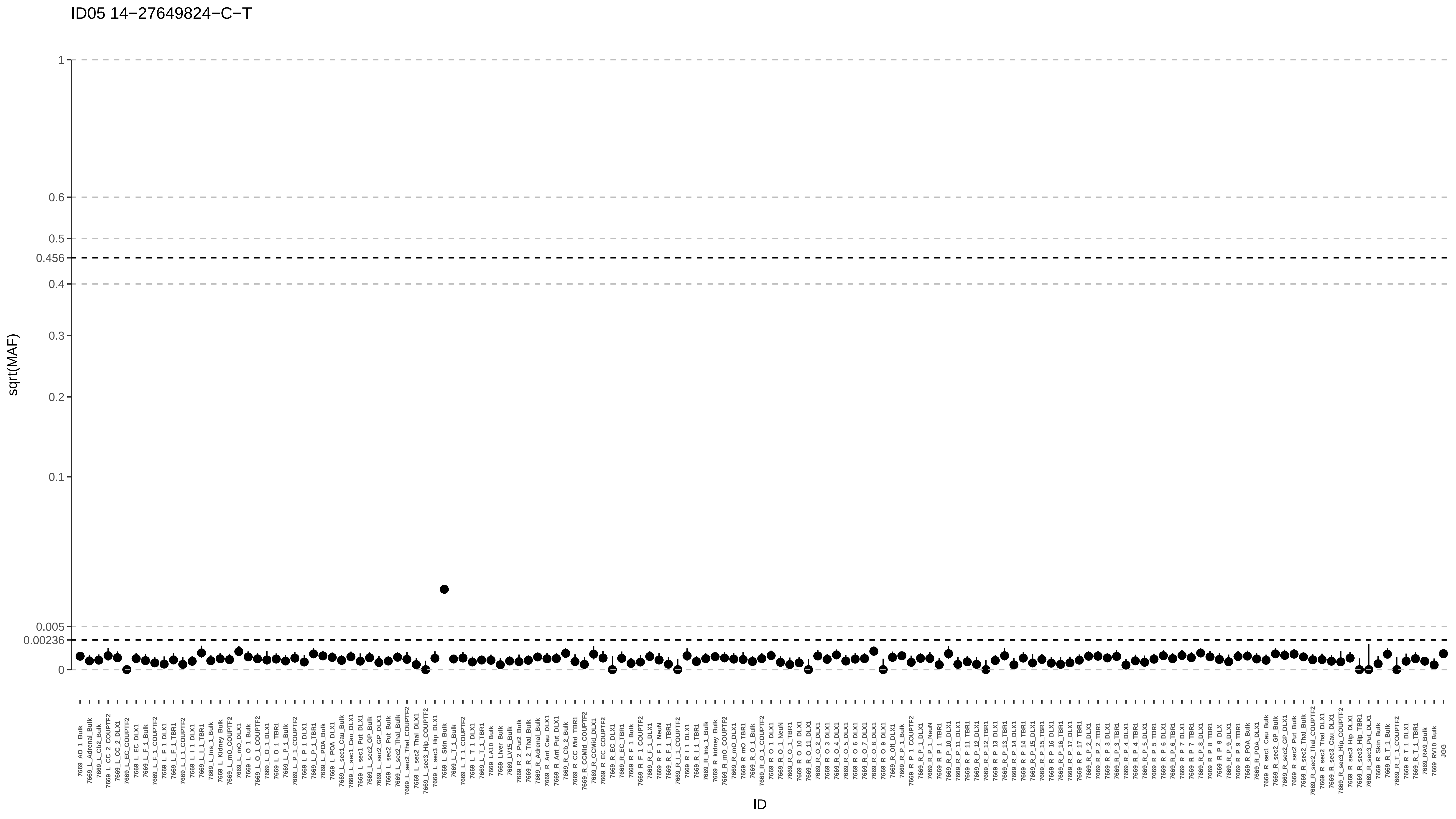
ID05 14−27649824−C−T
sqrt(MAF)
1
0.6
0.5
0.456
0.4
0.3
0.2
0.1
0.005
0.00236
0
ID

ID05 14−36924791−G−C

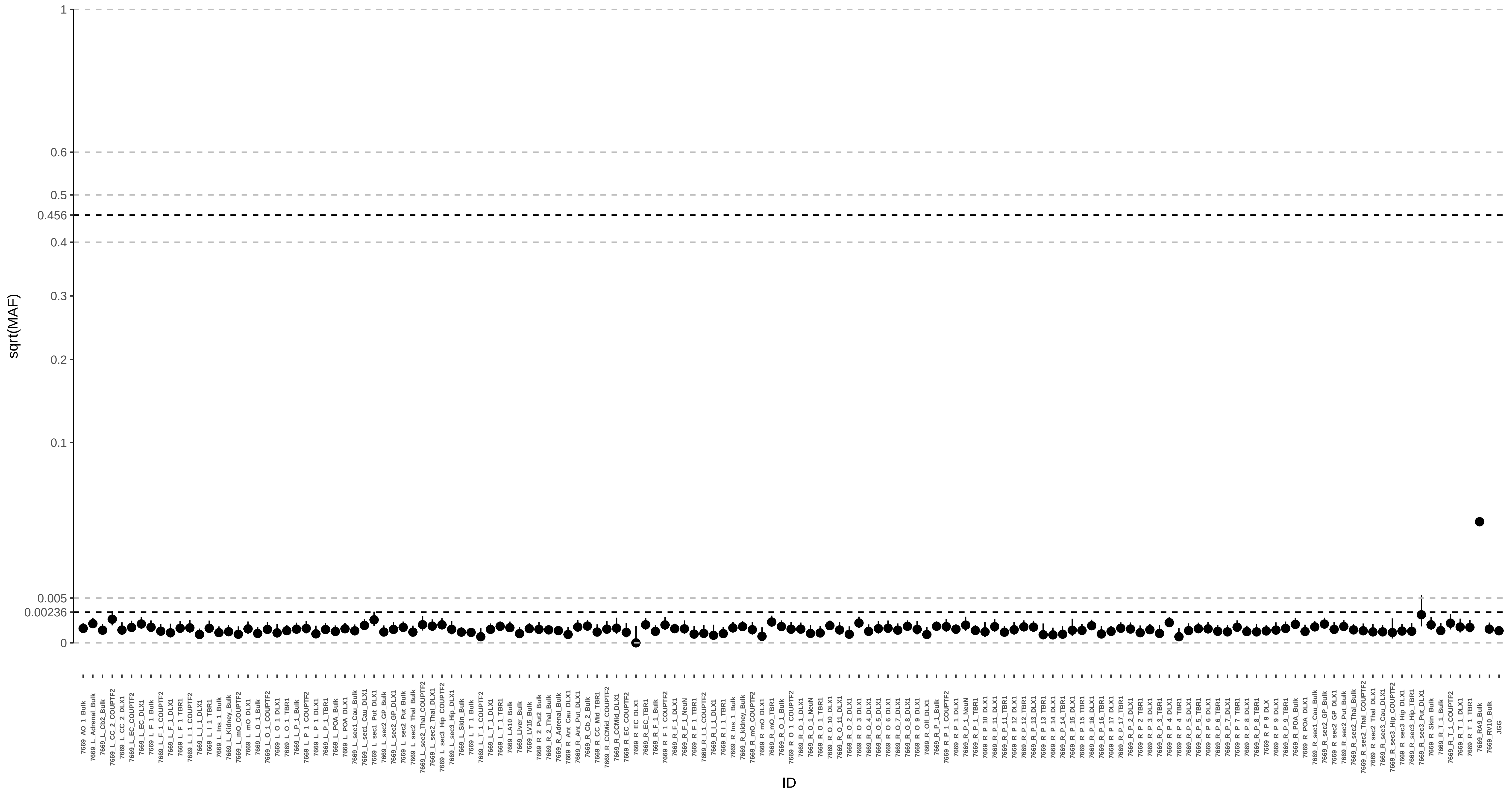

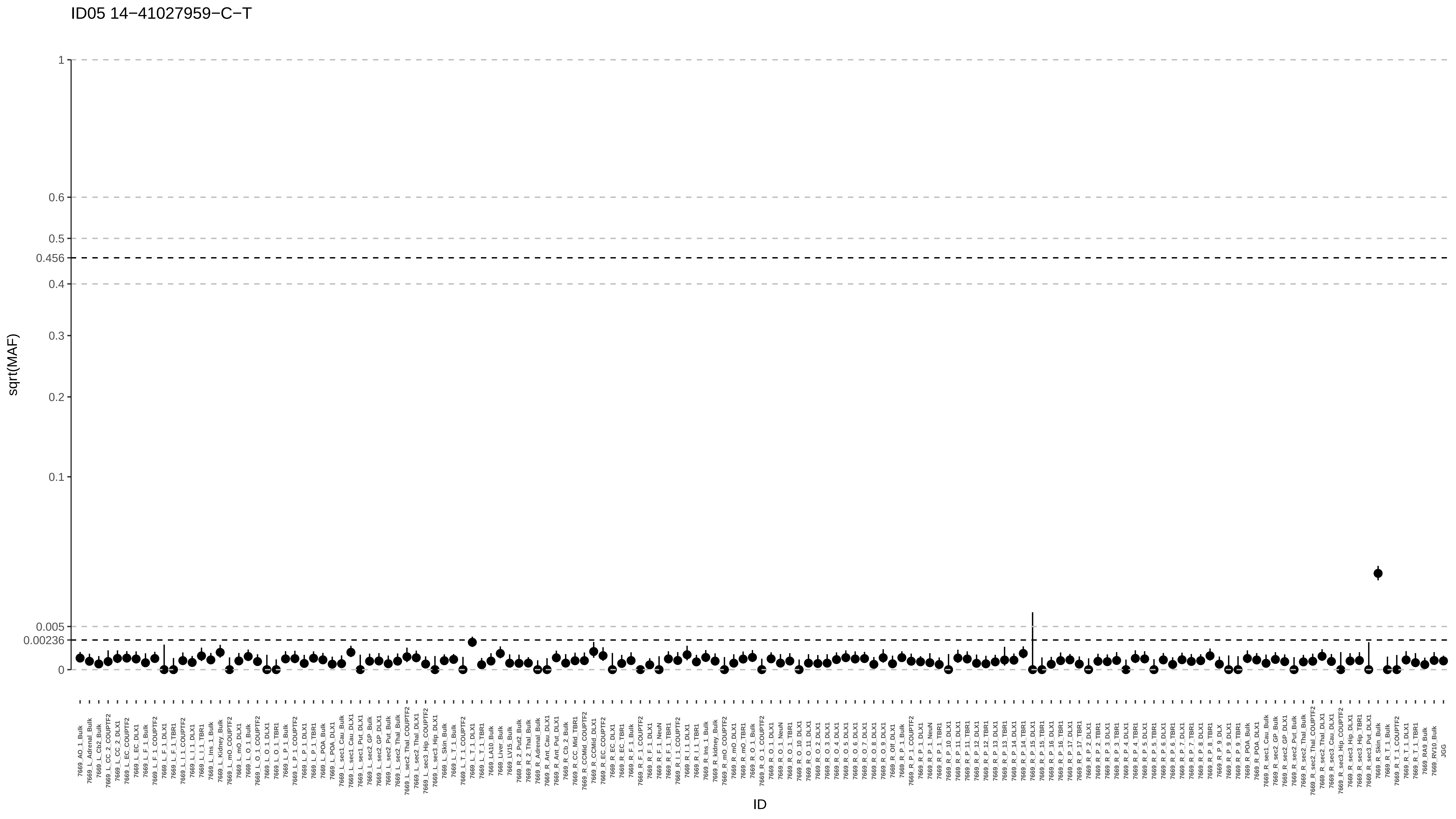
ID05 14−41027959−C−T
sqrt(MAF)
1
0.6
0.5
0.456
0.4
0.3
0.2
0.1
0.005
0.00236
0
ID

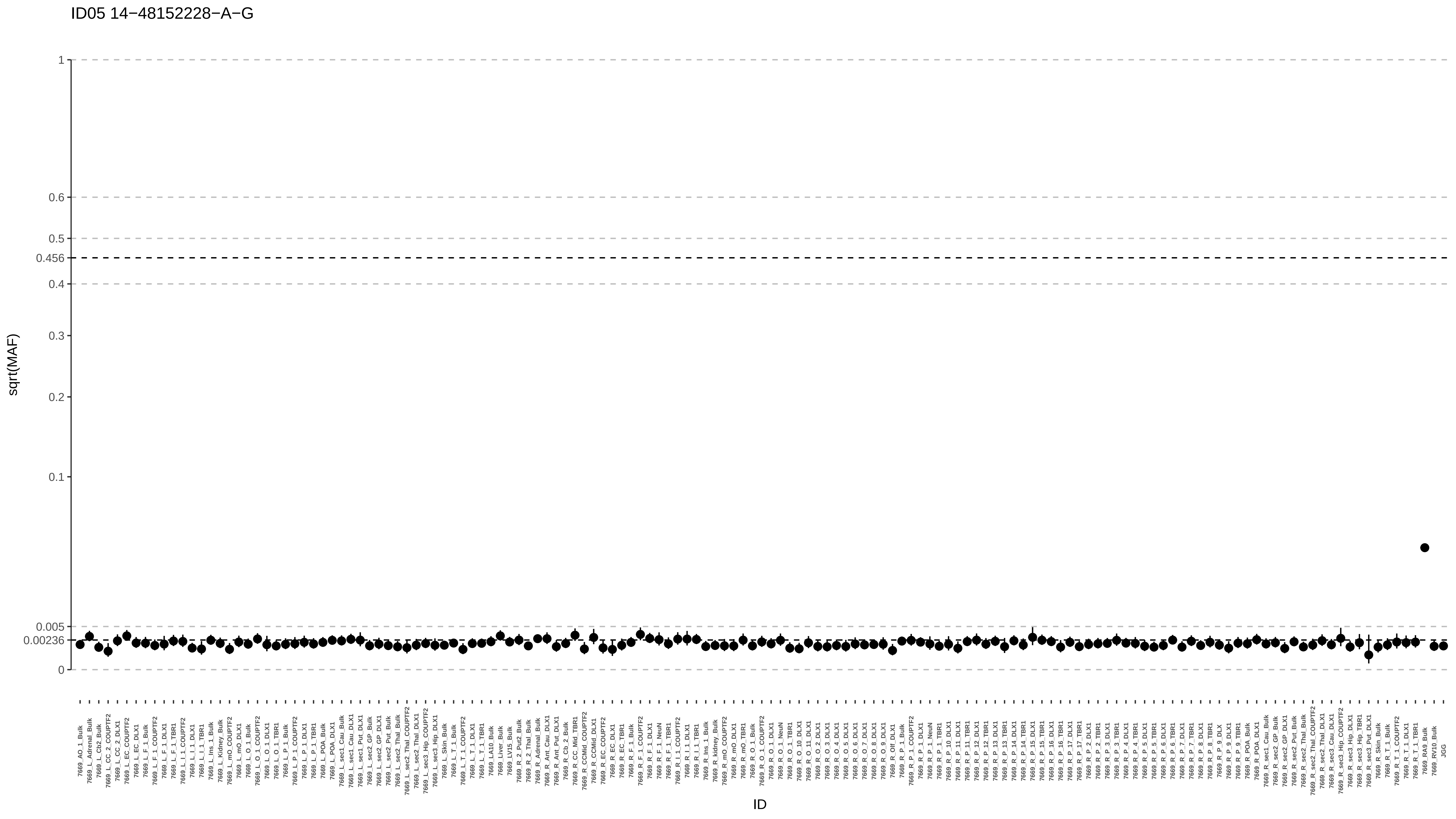
ID05 14−48152228−A−G
sqrt(MAF)
1
0.6
0.5
0.456
0.4
0.3
0.2
0.1
0.005
0.00236
0
ID

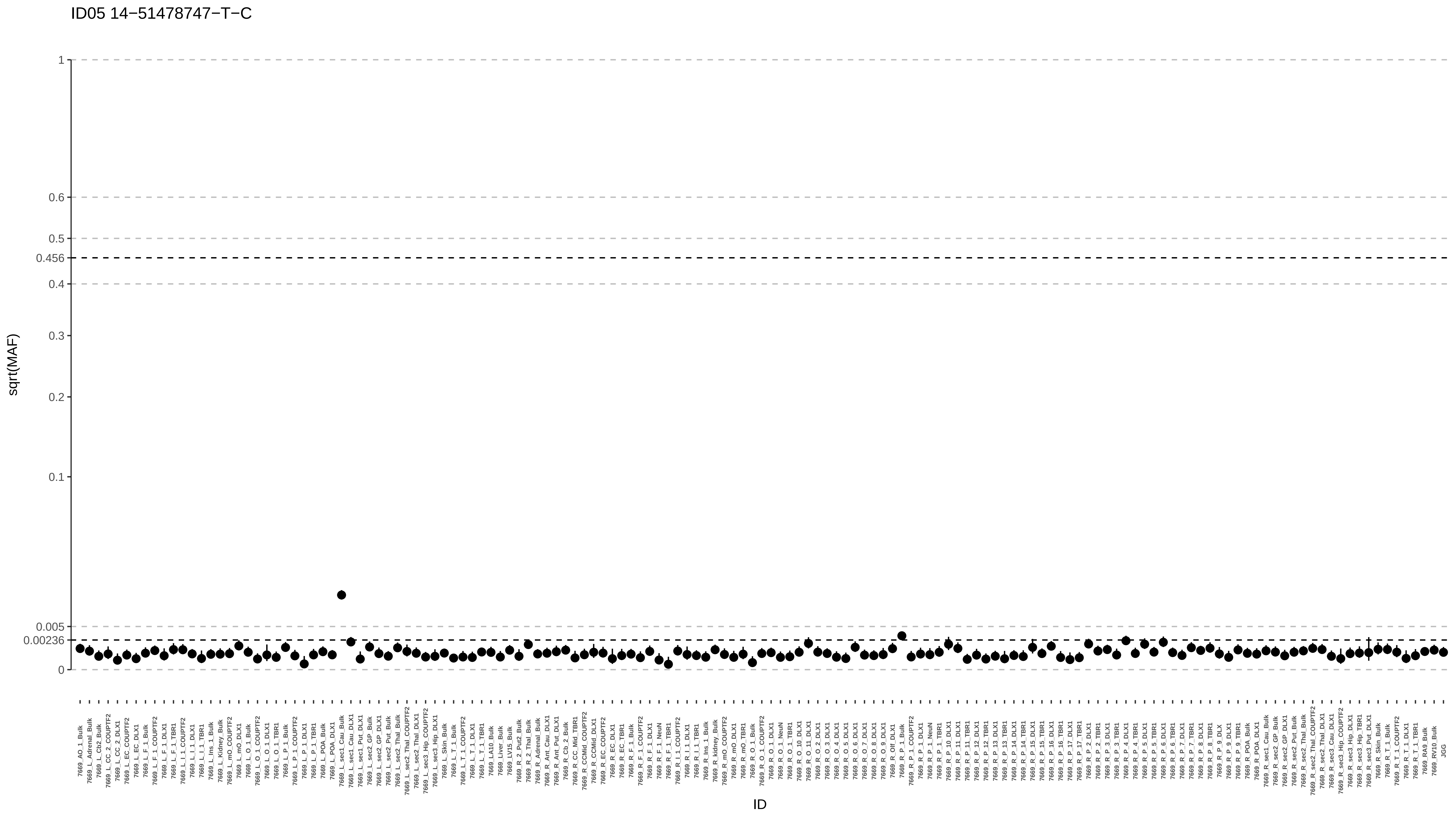
ID05 14−51478747−T−C
sqrt(MAF)
1
0.6
0.5
0.456
0.4
0.3
0.2
0.1
0.005
0.00236
0
ID

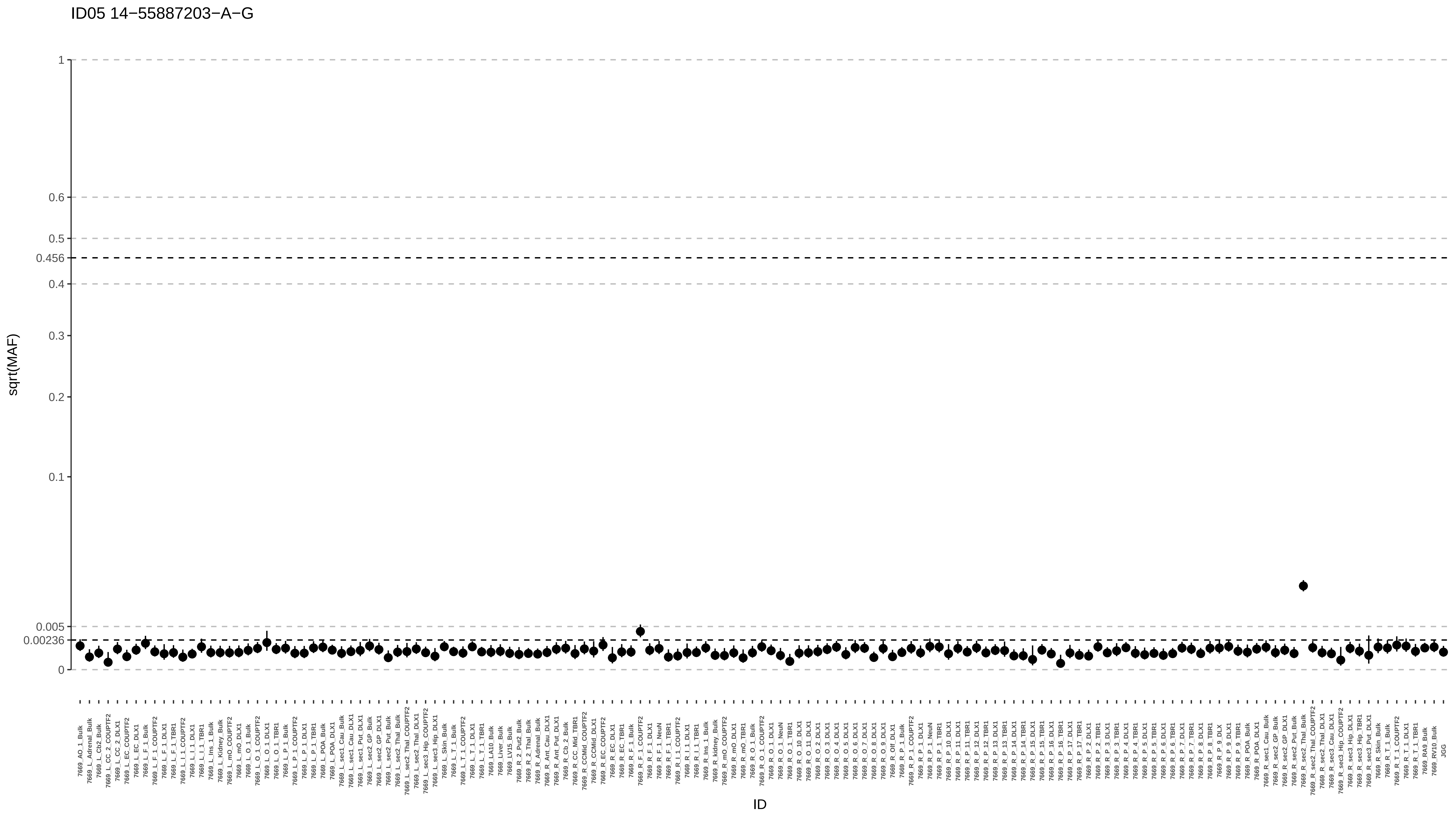
ID05 14−55887203−A−G
sqrt(MAF)
1
0.6
0.5
0.456
0.4
0.3
0.2
0.1
0.005
0.00236
0
ID

ID05 14−59827441−C−T

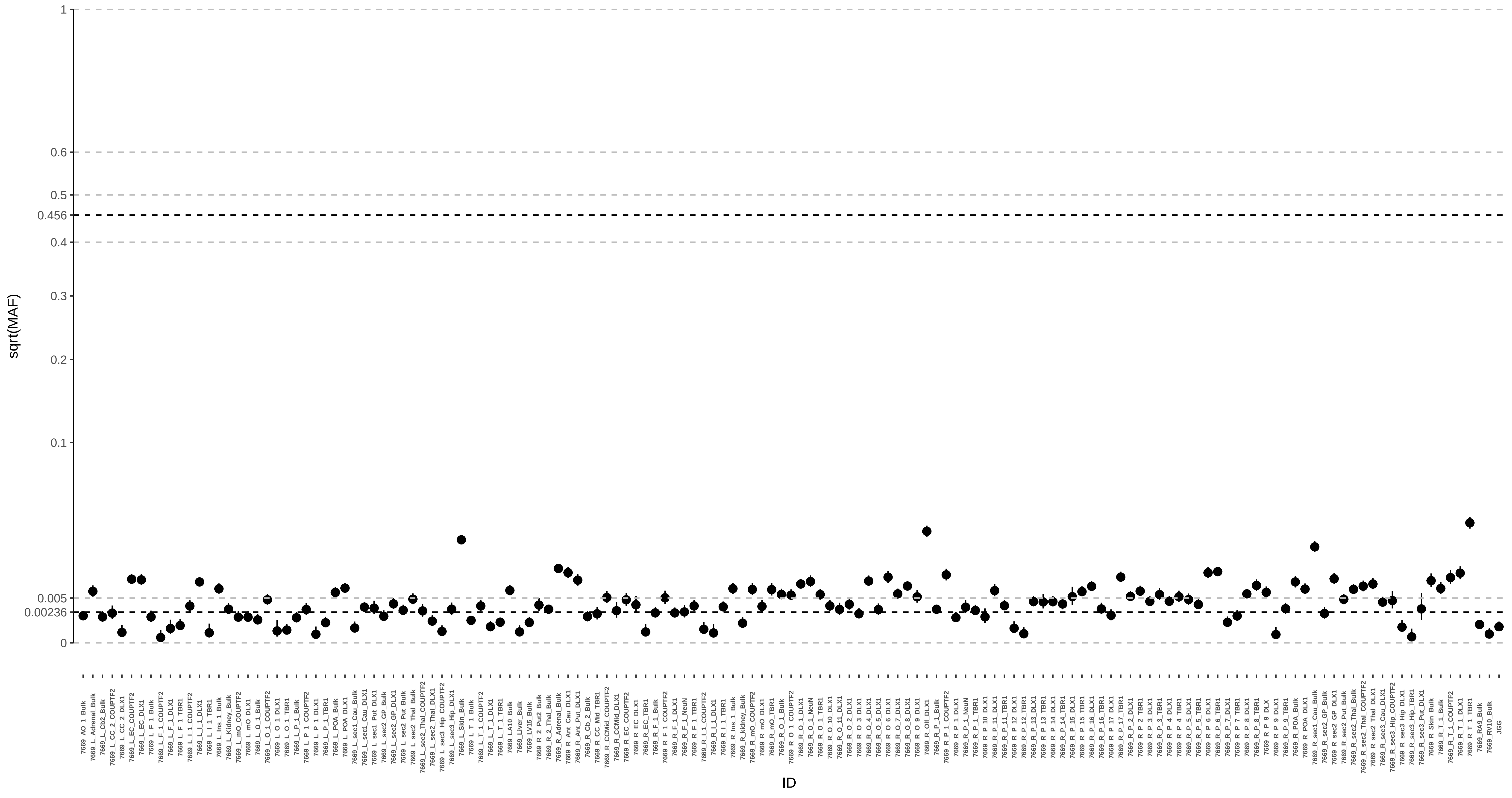

ID05 14−65264866−C−T

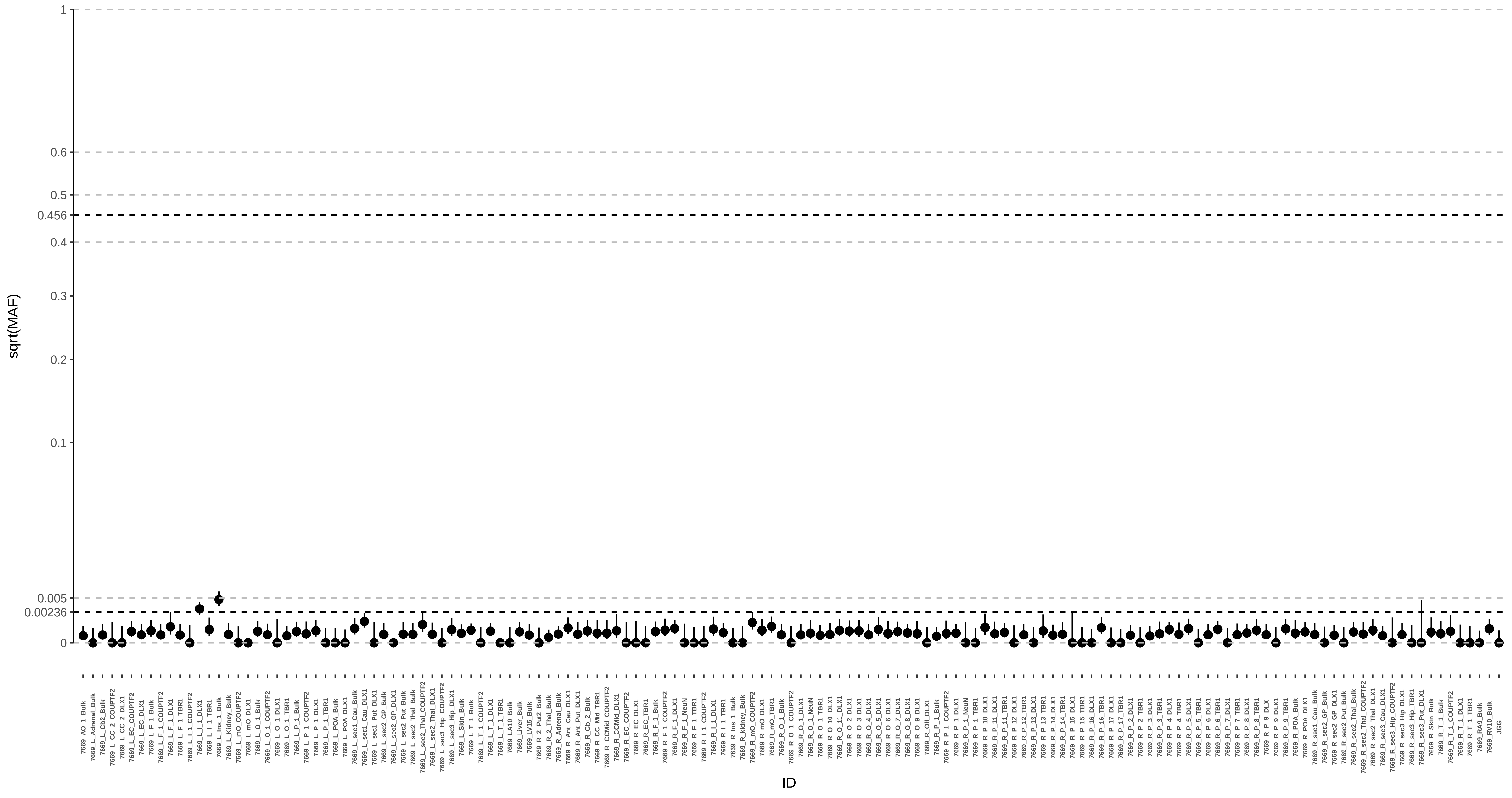

ID05 14−70538632−G−A

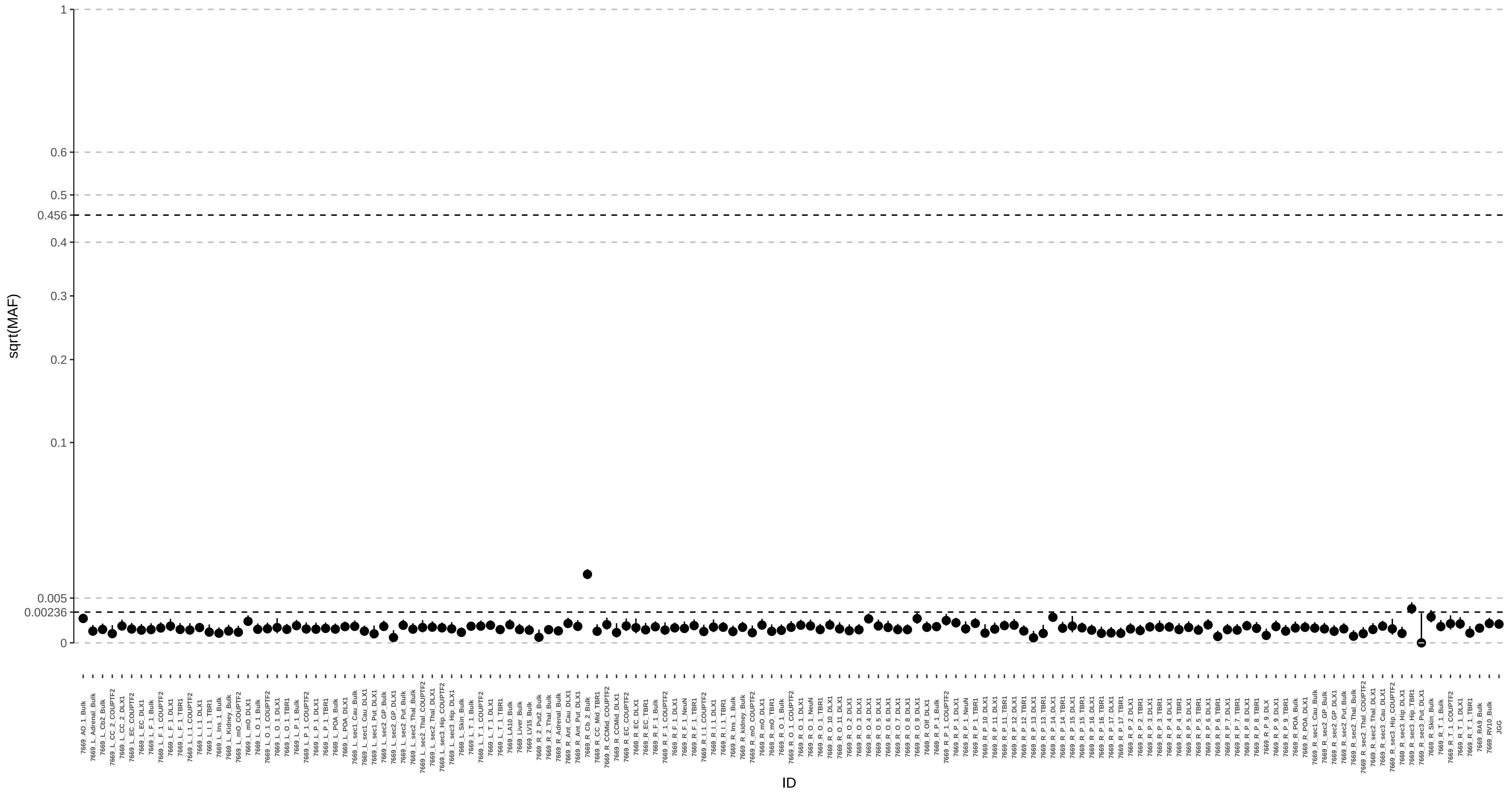

ID05 14−77126705−G−A

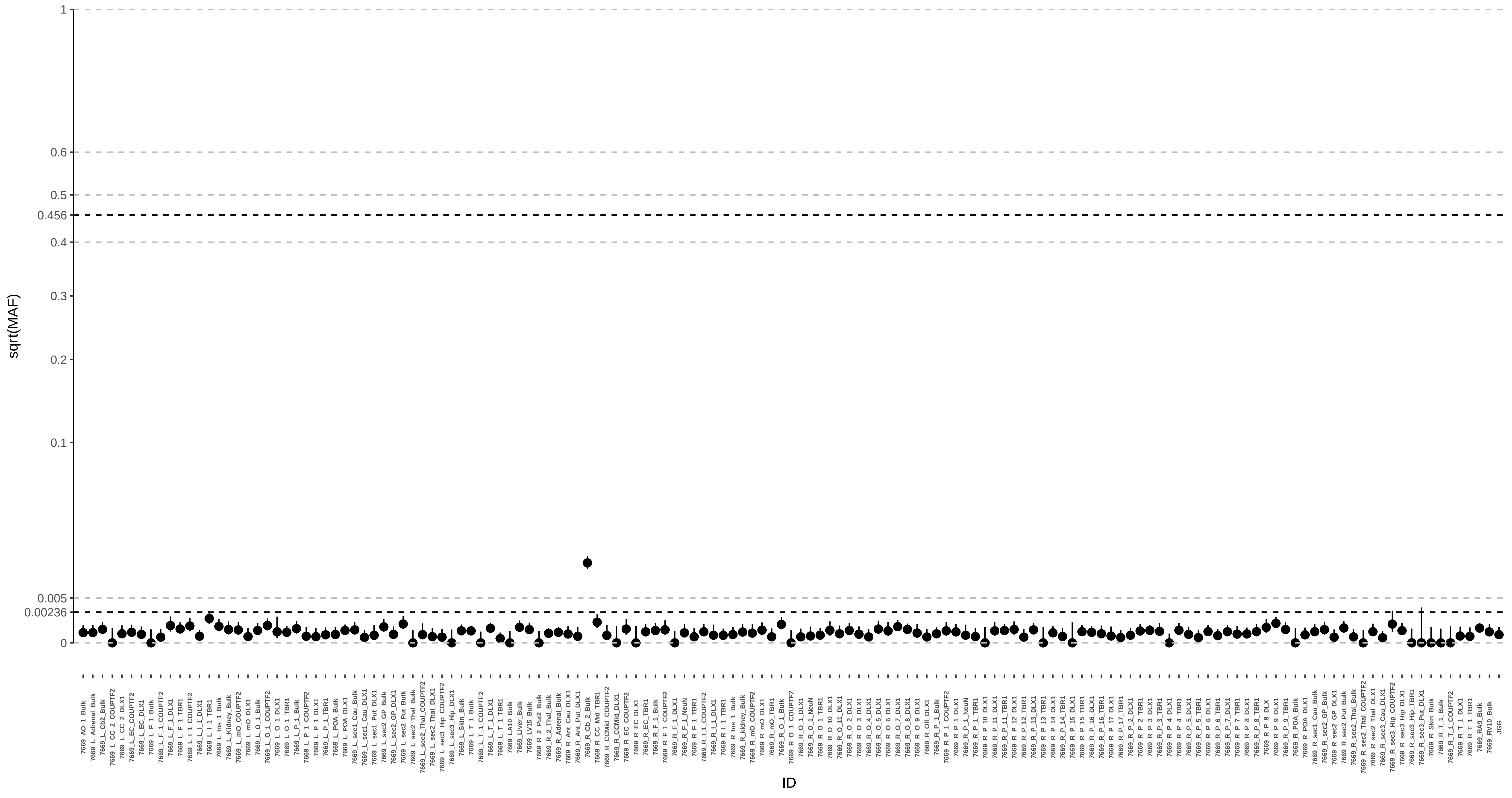

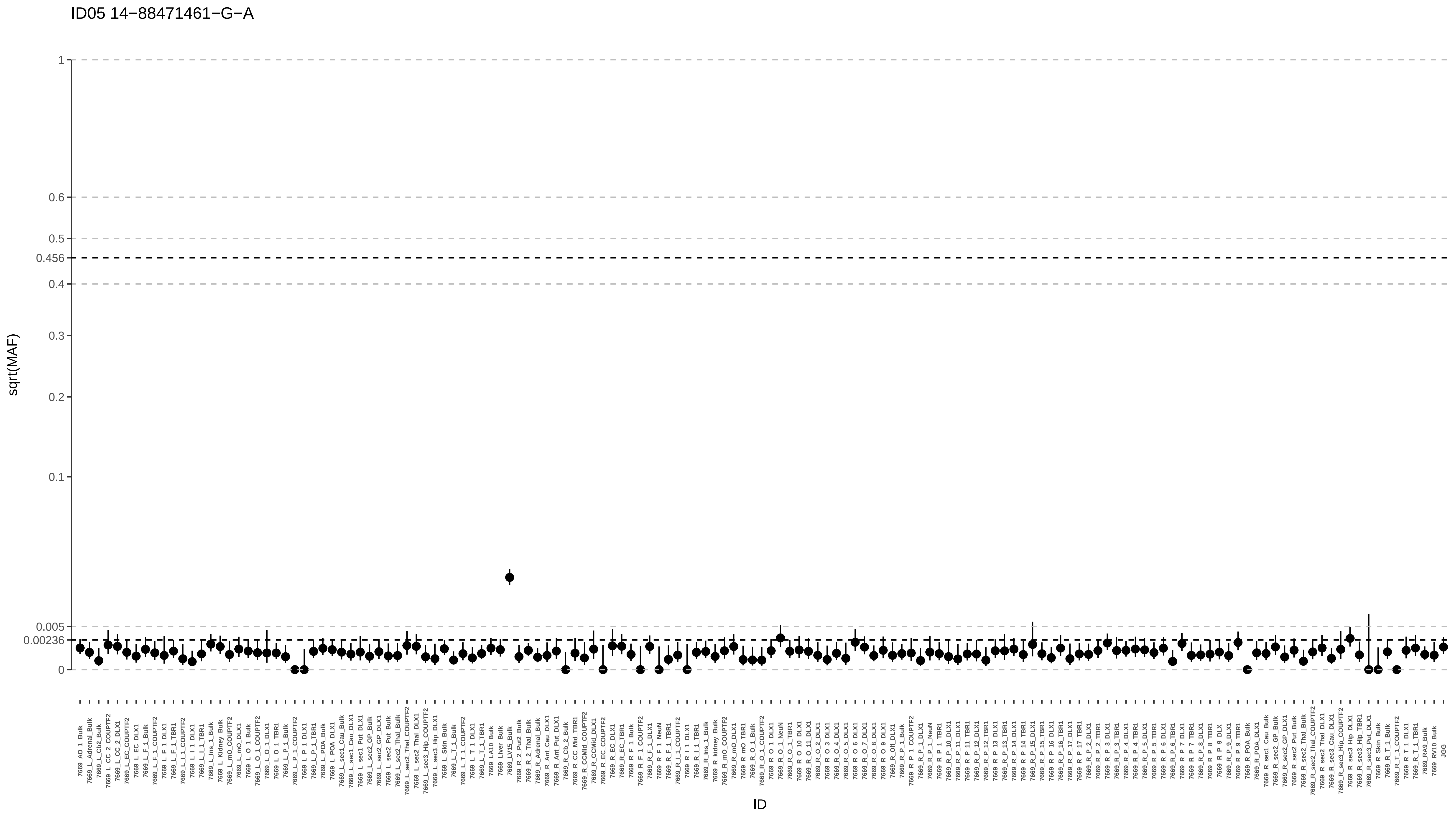
ID05 14−88471461−G−A
sqrt(MAF)
ID
1
0.6
0.5
0.456
0.4
0.3
0.2
0.1
0.005
0.00236
0

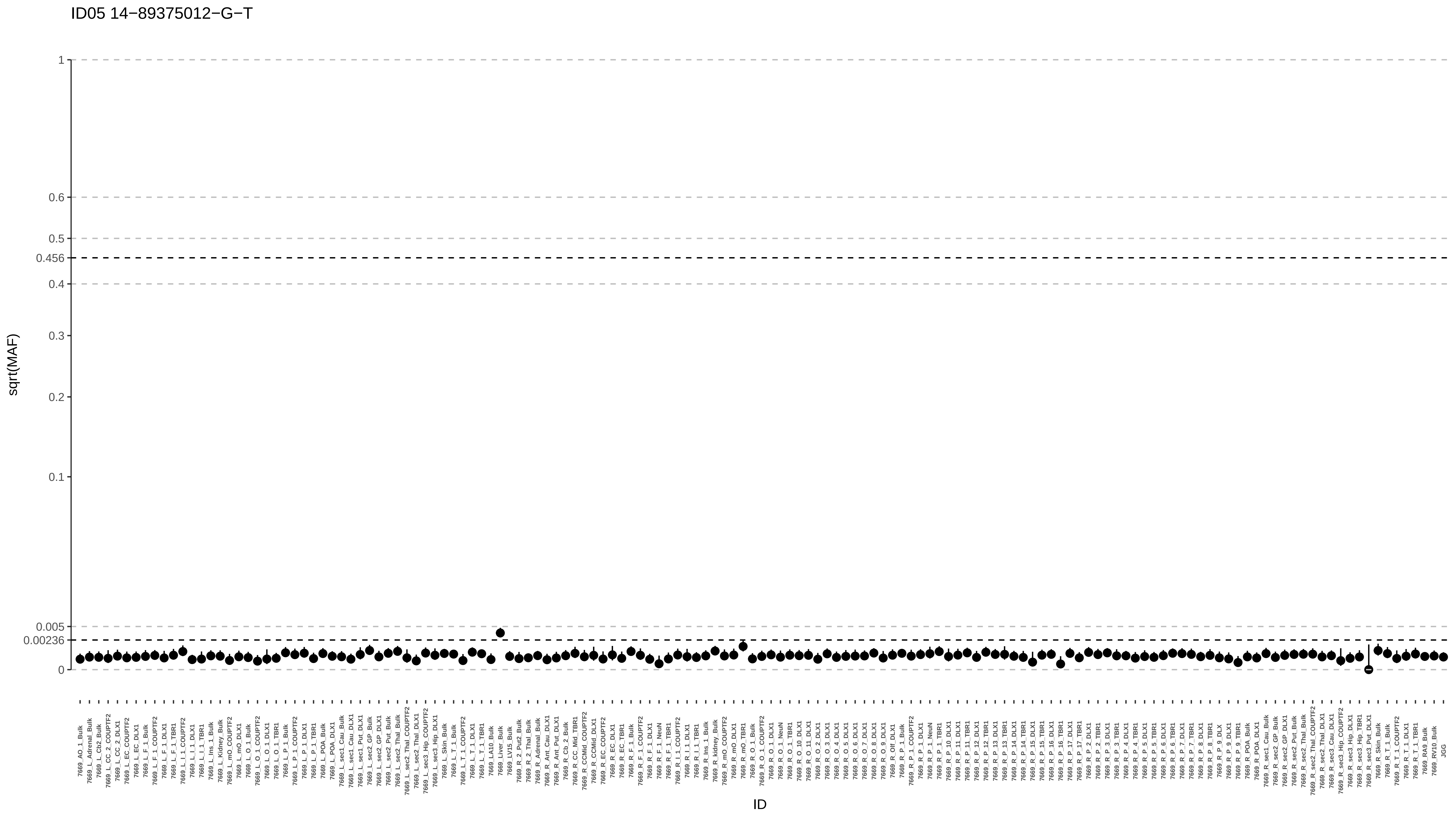
ID05 14−89375012−G−T
sqrt(MAF)
1
0.6
0.5
0.456
0.4
0.3
0.2
0.1
0.005
0.00236
0
ID

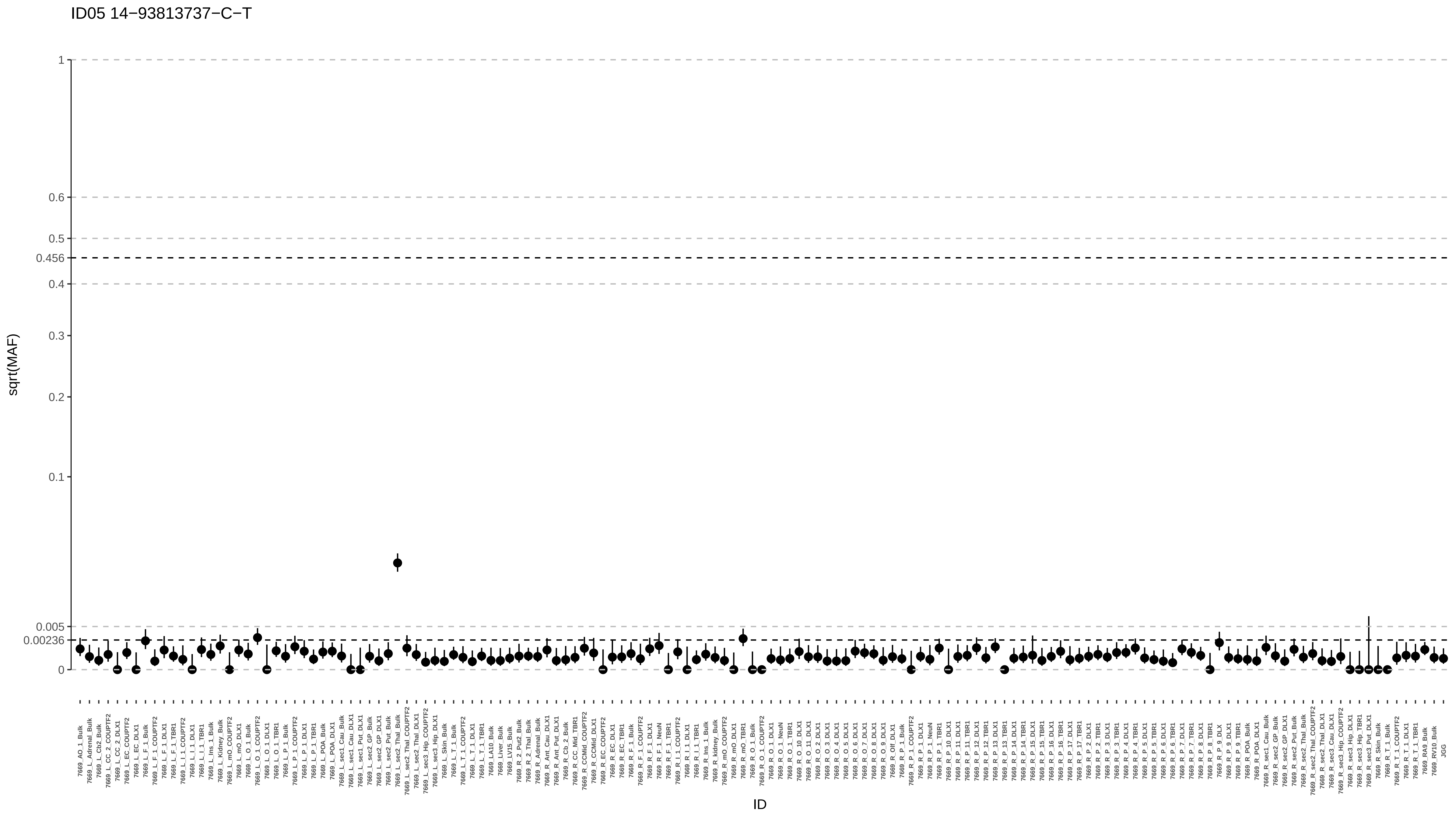
ID05 14−93813737−C−T
sqrt(MAF)
1
0.6
0.5
0.456
0.4
0.3
0.2
0.1
0.005
0.00236
0
ID

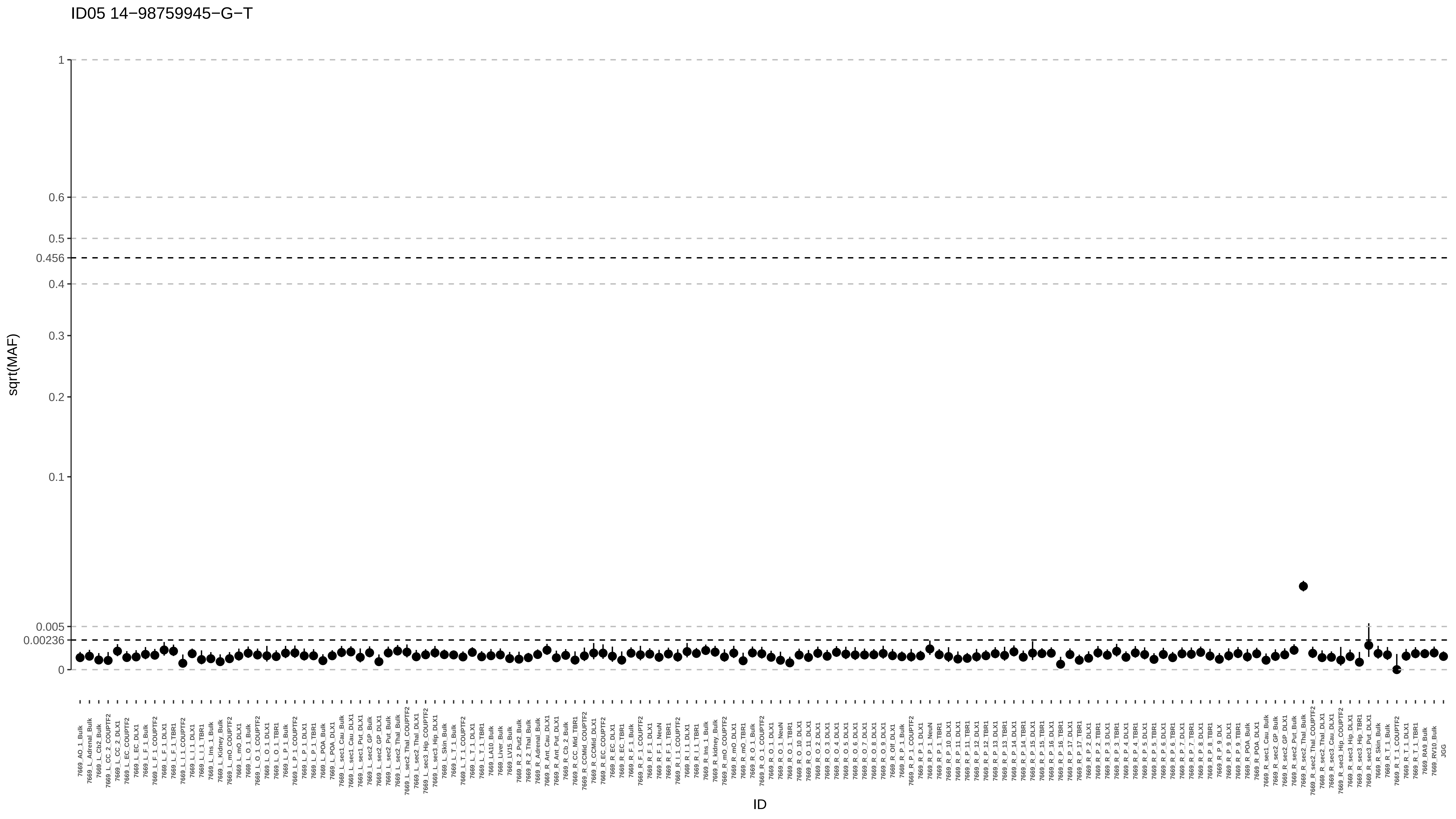
ID05 14−98759945−G−T
sqrt(MAF)
1
0.6
0.5
0.456
0.4
0.3
0.2
0.1
0.005
0.00236
0
ID

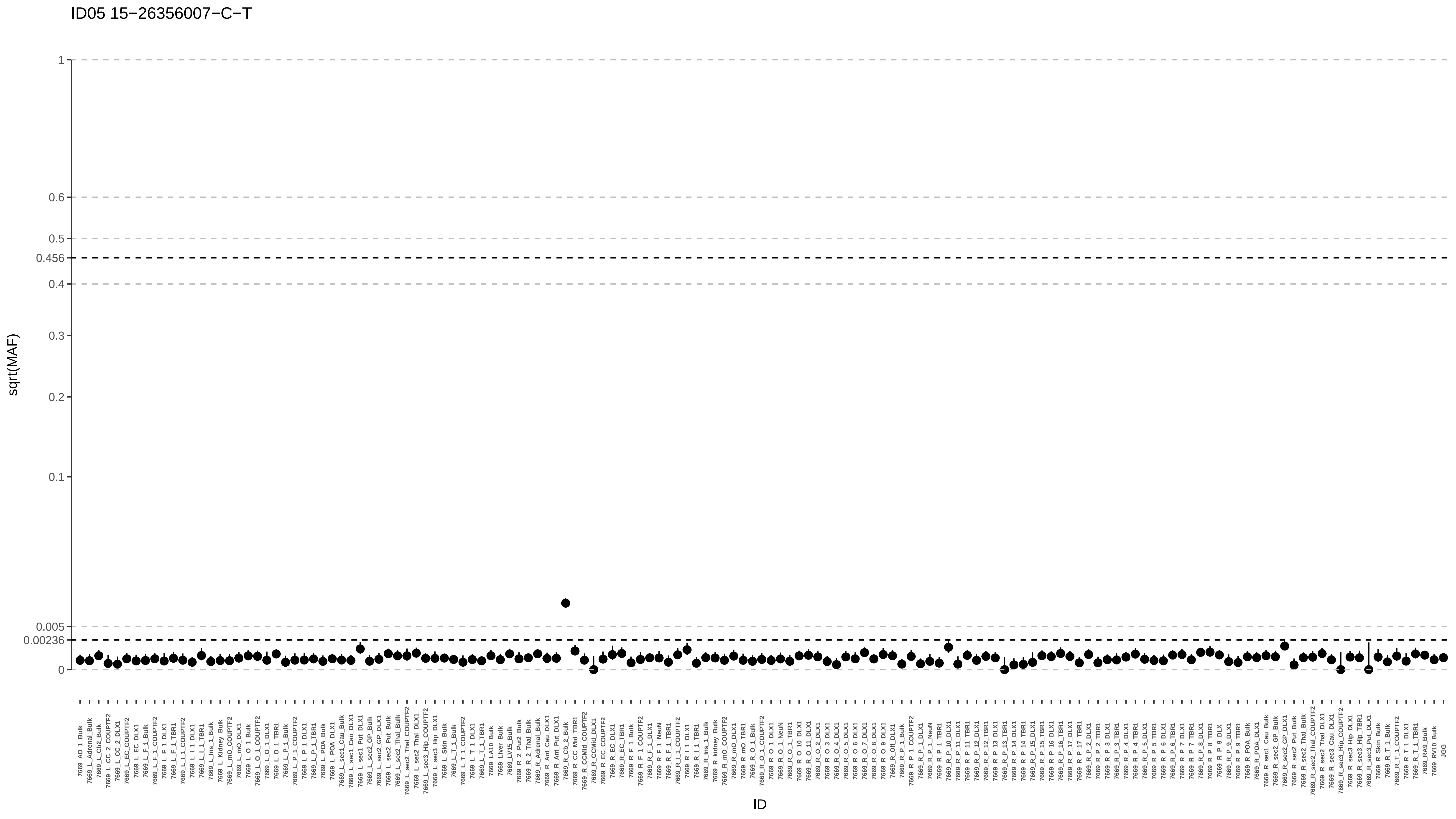
ID05 15−26356007−C−T
sqrt(MAF)
1
0.6
0.5
0.456
0.4
0.3
0.2
0.1
0.005
0.00236
0
ID

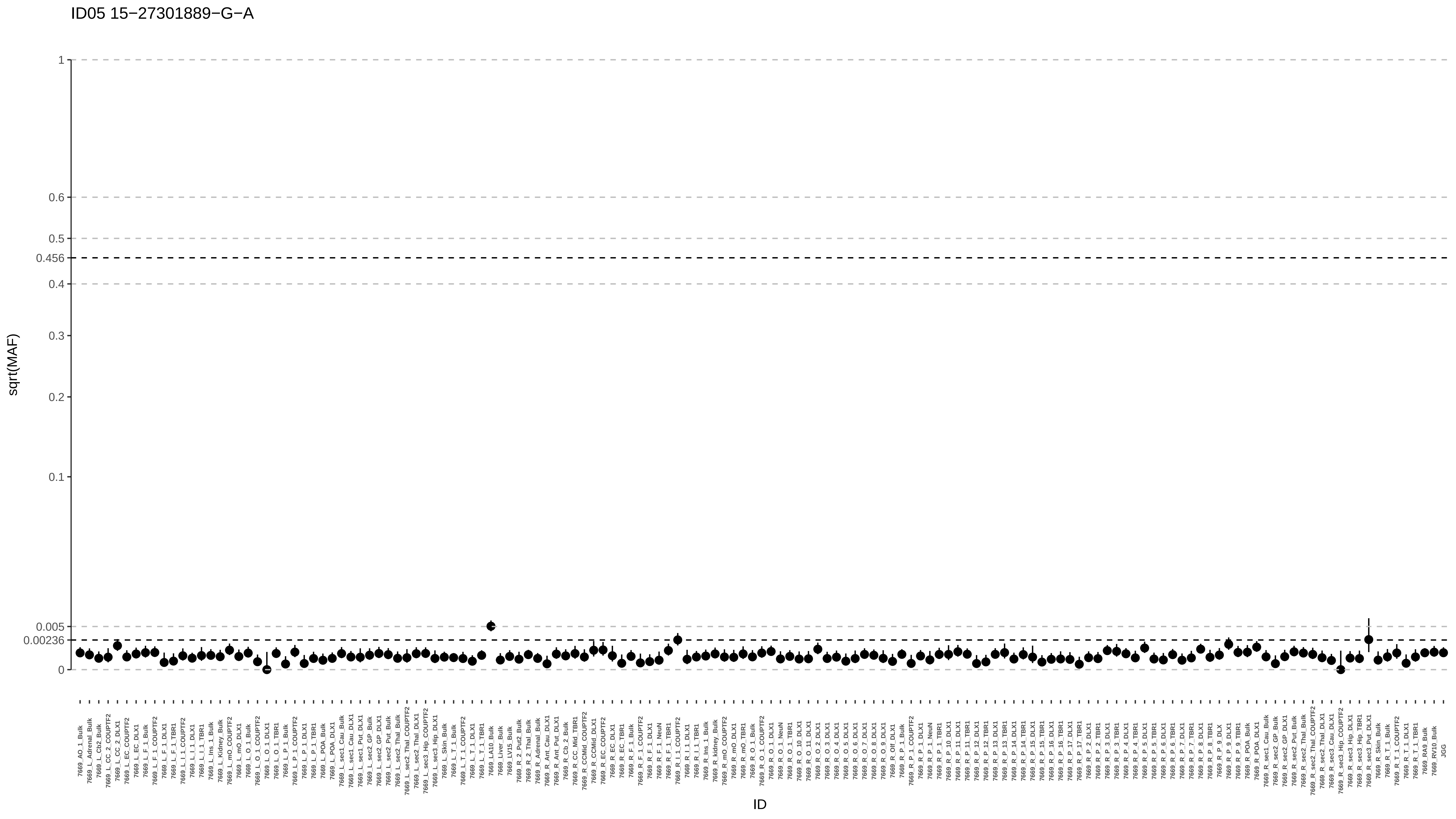

ID05 15−27301889−G−A
sqrt(MAF)
1
0.6
0.5
0.456
0.4
0.3
0.2
0.1
0.005
0.00236
0
ID

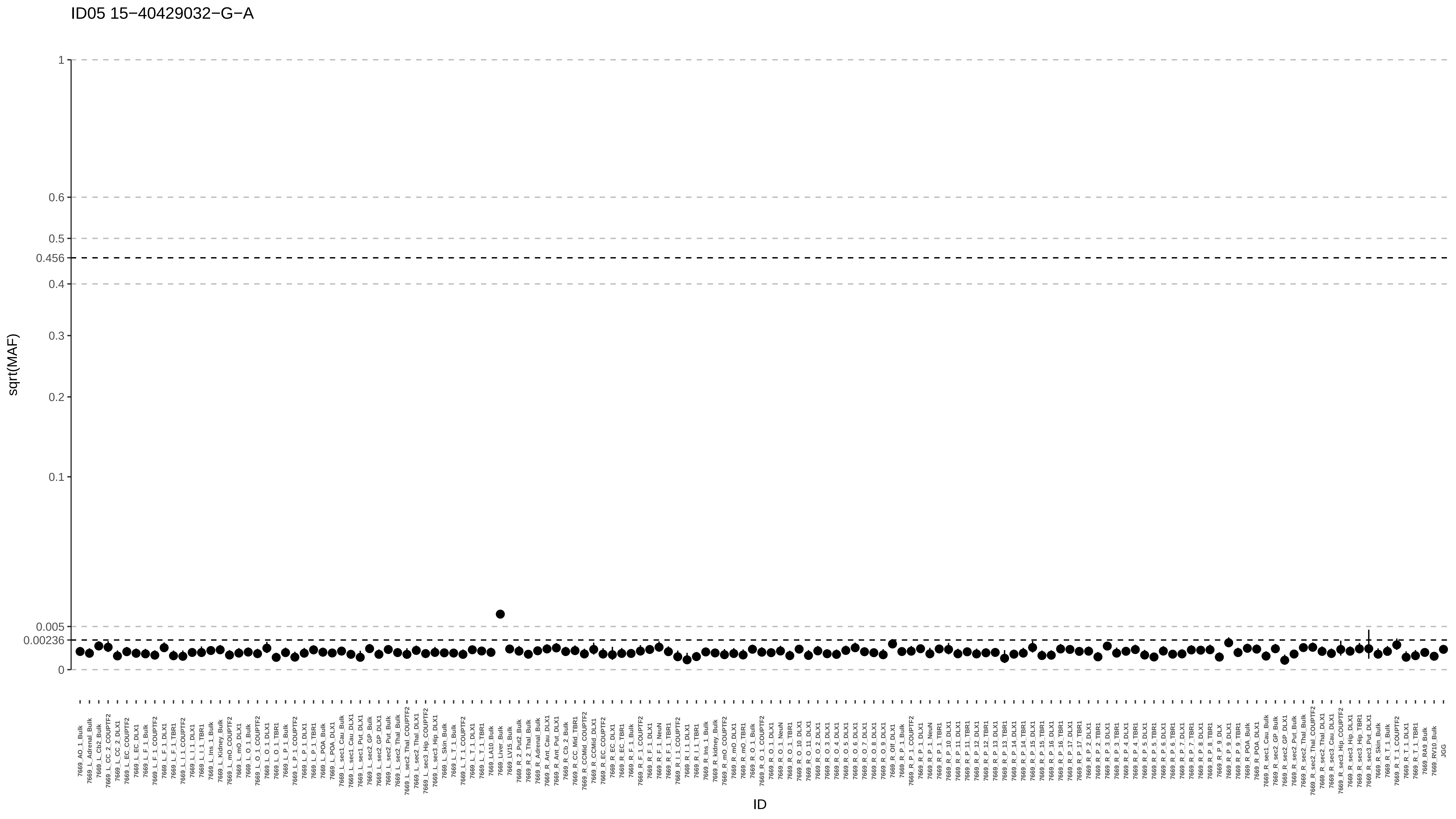
ID05 15−40429032−G−A
sqrt(MAF)
1
0.6
0.5
0.456
0.4
0.3
0.2
0.1
0.005
0.00236
0
ID

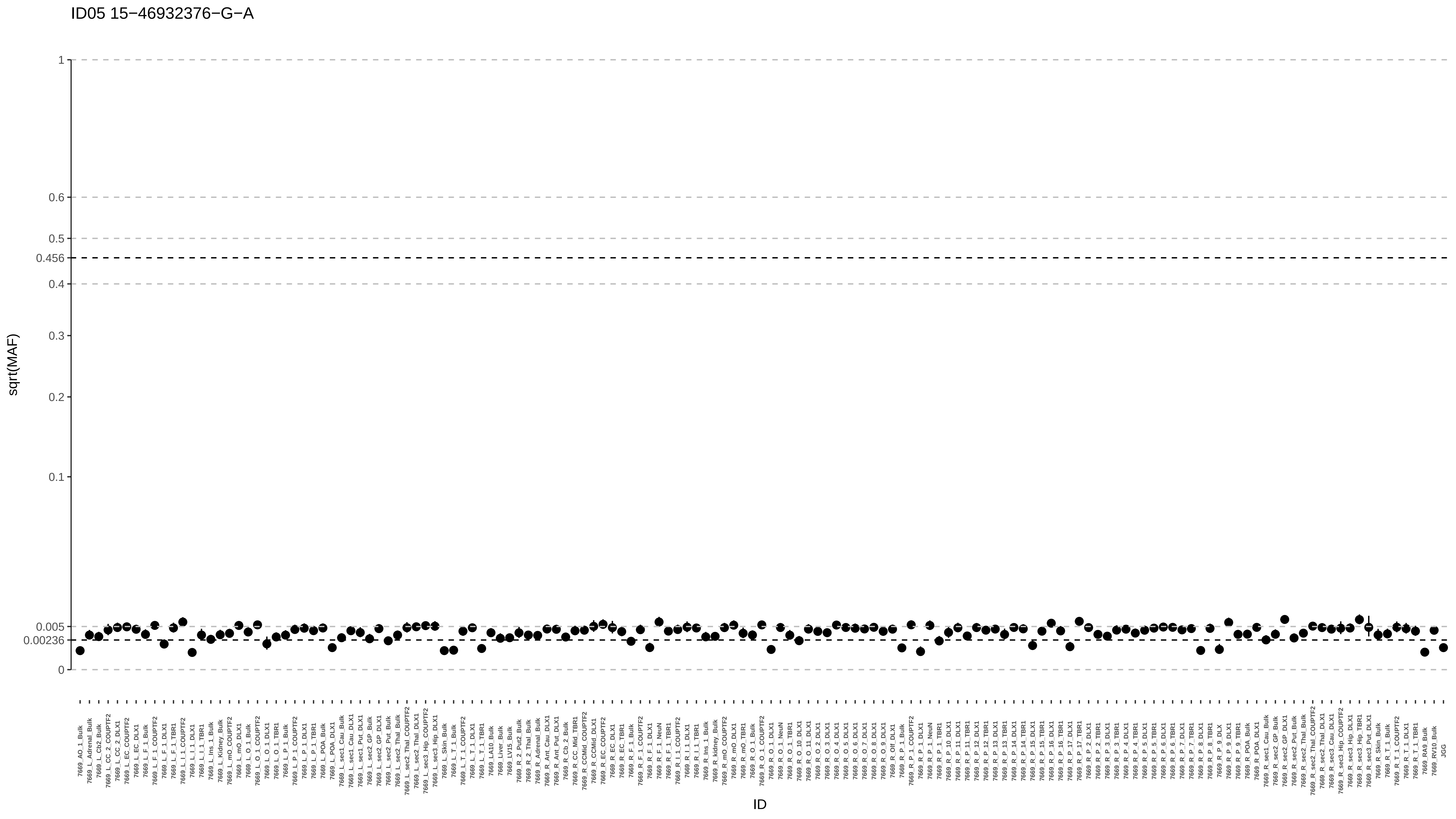
ID05 15−46932376−G−A
sqrt(MAF)
ID
1
0.6
0.5
0.456
0.4
0.3
0.2
0.1
0.005
0.00236
0

ID05 15−46962188−T−A

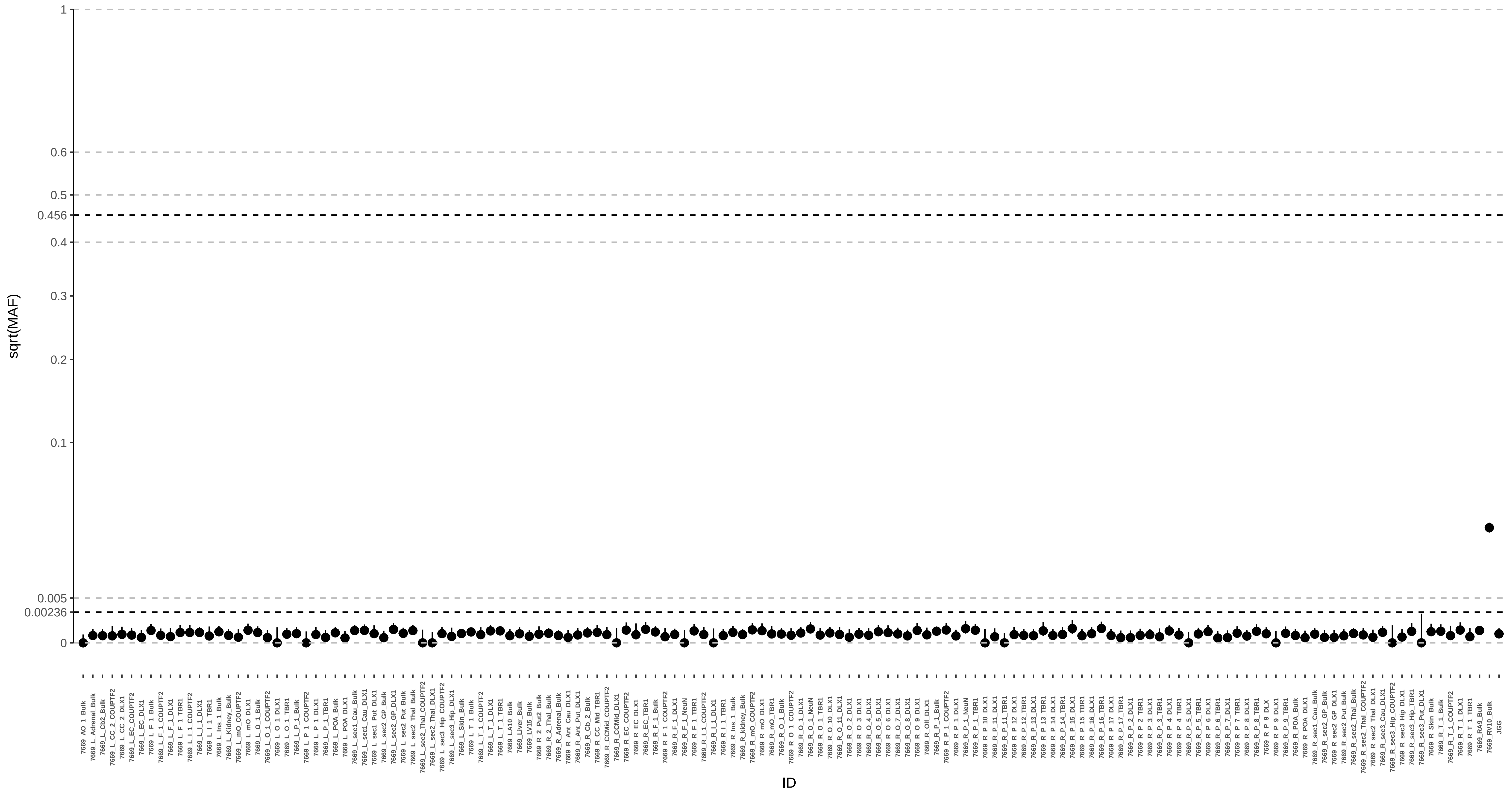

ID05 15−51489592−C−T

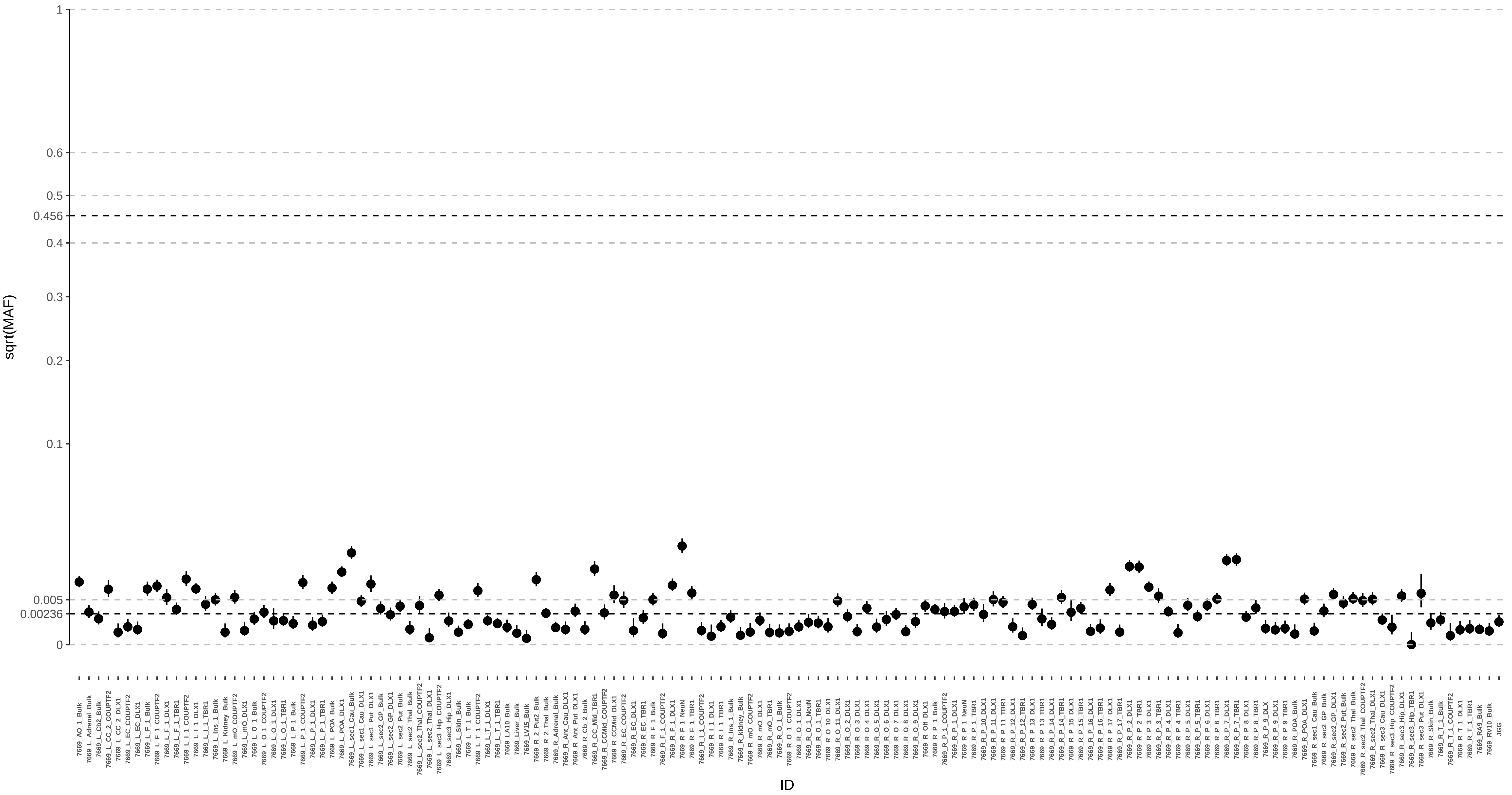

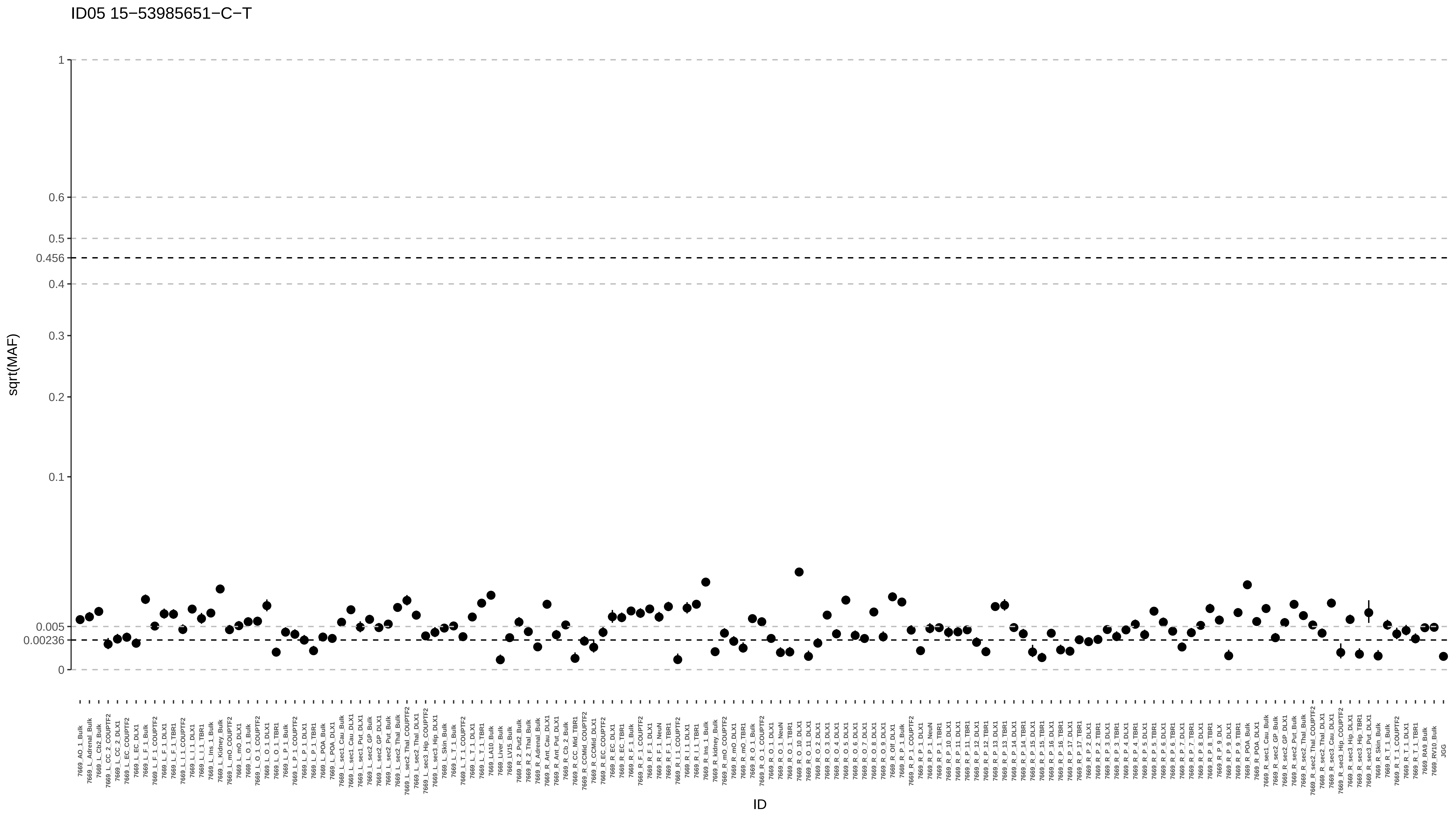
ID05 15−53985651−C−T
sqrt(MAF)
1
0.6
0.5
0.456
0.4
0.3
0.2
0.1
0.005
0.00236
0
ID

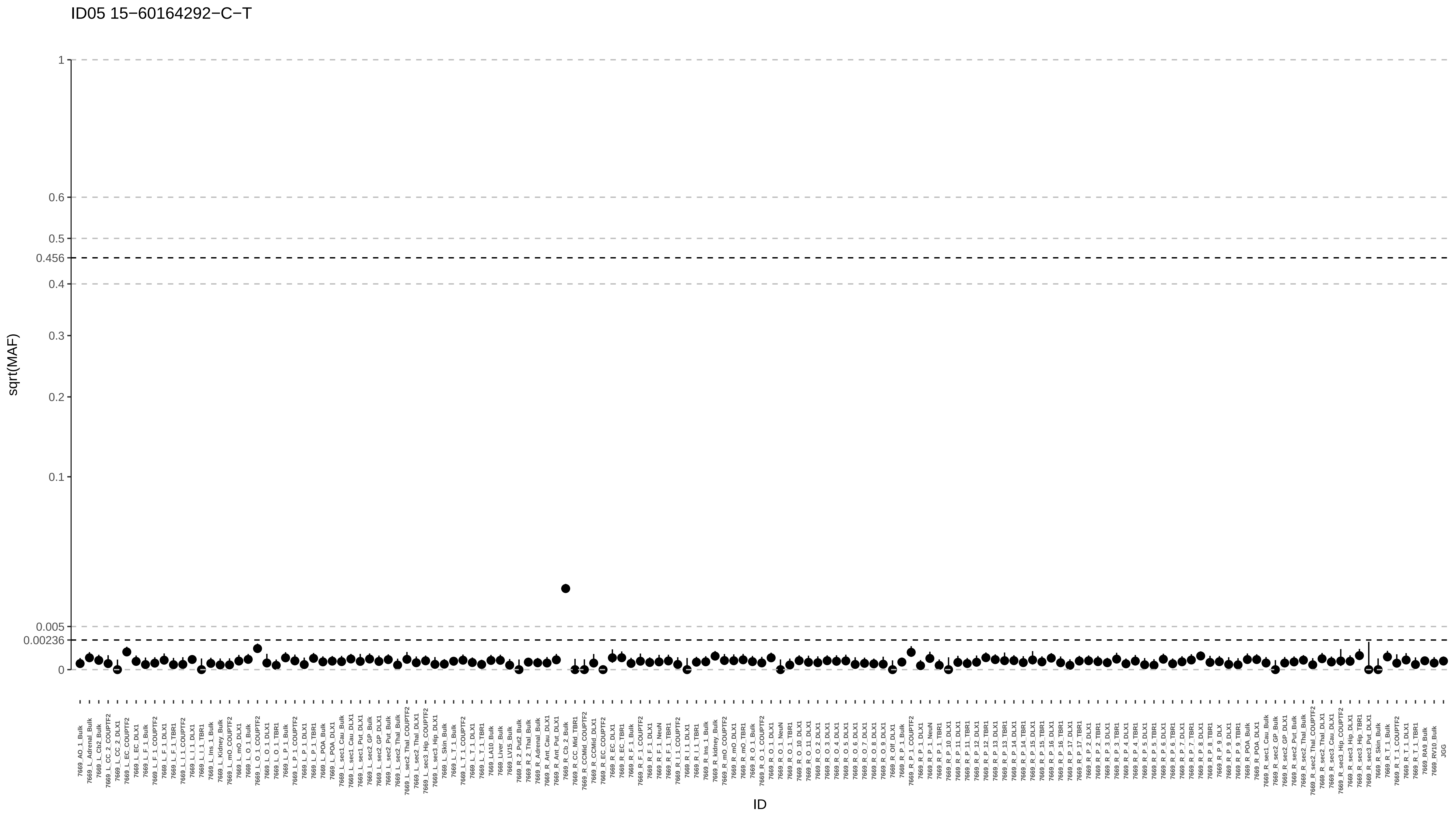
ID05 15−60164292−C−T
sqrt(MAF)
1
0.6
0.5
0.456
0.4
0.3
0.2
0.1
0.005
0.00236
0
ID

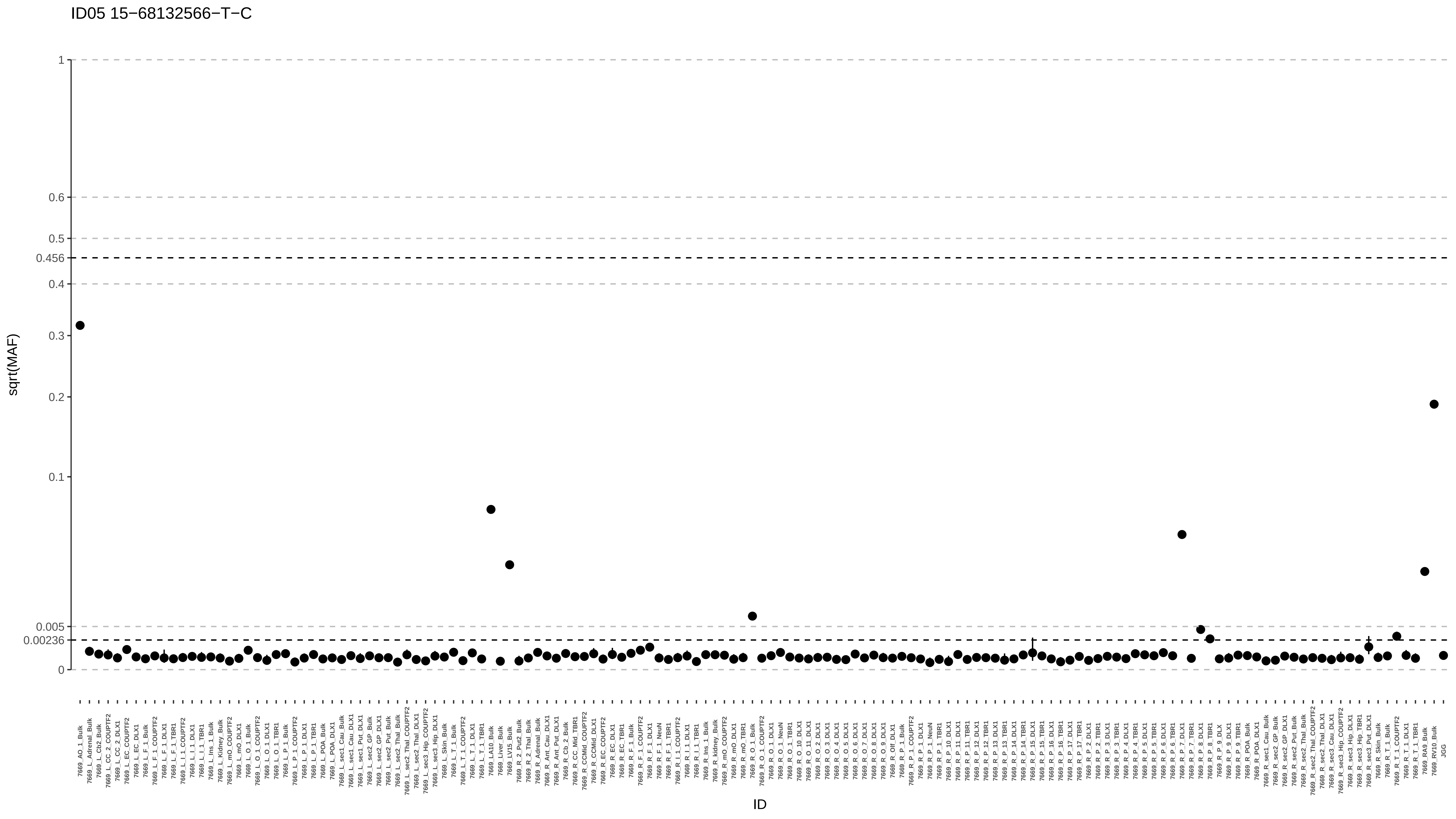
ID05 15−68132566−T−C
sqrt(MAF)
1
0.6
0.5
0.456
0.4
0.3
0.2
0.1
0.005
0.00236
0
ID

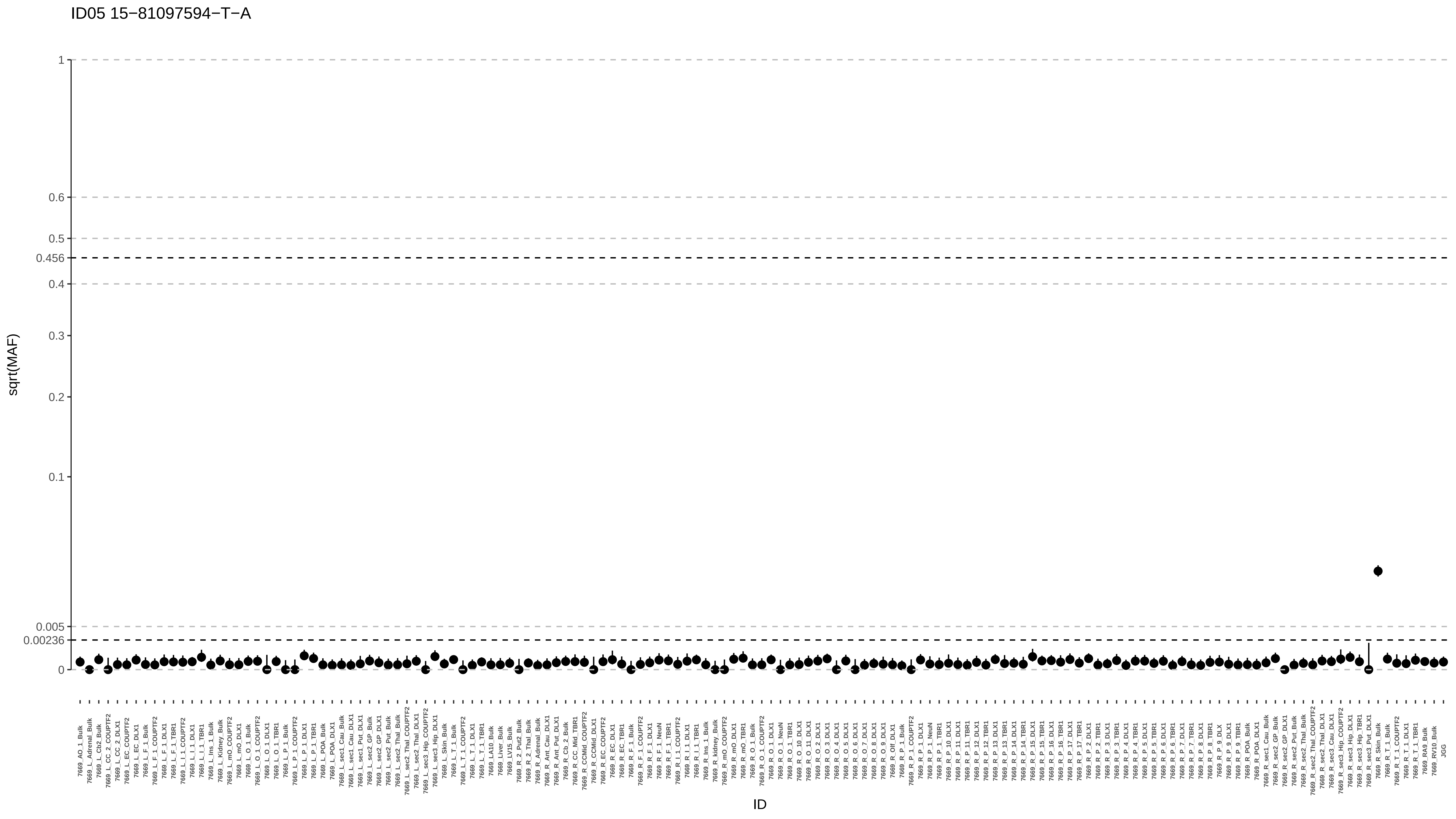
ID05 15−81097594−T−A
sqrt(MAF)
1
0.6
0.5
0.456
0.4
0.3
0.2
0.1
0.005
0.00236
0
ID
7669_AO_1_Bulk
7669_L_Adrenal_Bulk
7669_L_Cb2_Bulk
7669_L_CC_2_COUPTF2
7669_L_CC_2_DLX1
7669_L_EC_COUPTF2
7669_L_EC_DLX1
7669_L_F_1_Bulk
7669_L_F_1_COUPTF2
7669_L_F_1_DLX1
7669_L_F_1_TBR1
7669_L_I_1_COUPTF2
7669_L_I_1_DLX1
7669_L_I_1_TBR1
7669_L_Ins_1_Bulk
7669_L_Kidney_Bulk
7669_L_mO_COUPTF2
7669_L_mO_DLX1
7669_L_O_1_Bulk
7669_L_O_1_COUPTF2
7669_L_O_1_DLX1
7669_L_O_1_TBR1
7669_L_P_1_Bulk
7669_L_P_1_COUPTF2
7669_L_P_1_DLX1
7669_L_P_1_TBR1
7669_L_POA_Bulk
7669_L_POA_DLX1
7669_L_sec1_Cau_Bulk
7669_L_sec1_Cau_DLX1
7669_L_sec1_Put_DLX1
7669_L_sec2_GP_Bulk
7669_L_sec2_GP_DLX1
7669_L_sec2_Put_Bulk
7669_L_sec2_Thal_Bulk
7669_L_sec2_Thal_COUPTF2
7669_L_sec2_Thal_DLX1
7669_L_sec3_Hip_COUPTF2
7669_L_sec3_Hip_DLX1
7669_L_Skin_Bulk
7669_L_T_1_Bulk
7669_L_T_1_COUPTF2
7669_L_T_1_DLX1
7669_L_T_1_TBR1
7669_LA10_Bulk
7669_Liver_Bulk
7669_LV15_Bulk
7669_R_2_Put2_Bulk
7669_R_2_Thal_Bulk
7669_R_Adrenal_Bulk
7669_R_Ant_Cau_DLX1
7669_R_Ant_Put_DLX1
7669_R_Cb_2_Bulk
7669_R_CC_Mid_TBR1
7669_R_CCMid_COUPTF2
7669_R_CCMid_DLX1
7669_R_EC_COUPTF2
7669_R_EC_DLX1
7669_R_EC_TBR1
7669_R_F_1_Bulk
7669_R_F_1_COUPTF2
7669_R_F_1_DLX1
7669_R_F_1_NeuN
7669_R_F_1_TBR1
7669_R_I_1_COUPTF2
7669_R_I_1_DLX1
7669_R_I_1_TBR1
7669_R_Ins_1_Bulk
7669_R_kidney_Bulk
7669_R_mO_COUPTF2
7669_R_mO_DLX1
7669_R_mO_TBR1
7669_R_O_1_Bulk
7669_R_O_1_COUPTF2
7669_R_O_1_DLX1
7669_R_O_1_NeuN
7669_R_O_1_TBR1
7669_R_O_10_DLX1
7669_R_O_11_DLX1
7669_R_O_2_DLX1
7669_R_O_3_DLX1
7669_R_O_4_DLX1
7669_R_O_5_DLX1
7669_R_O_6_DLX1
7669_R_O_7_DLX1
7669_R_O_8_DLX1
7669_R_O_9_DLX1
7669_R_Olf_DLX1
7669_R_P_1_Bulk
7669_R_P_1_COUPTF2
7669_R_P_1_DLX1
7669_R_P_1_NeuN
7669_R_P_1_TBR1
7669_R_P_10_DLX1
7669_R_P_11_DLX1
7669_R_P_11_TBR1
7669_R_P_12_DLX1
7669_R_P_12_TBR1
7669_R_P_13_DLX1
7669_R_P_13_TBR1
7669_R_P_14_DLX1
7669_R_P_14_TBR1
7669_R_P_15_DLX1
7669_R_P_15_TBR1
7669_R_P_16_DLX1
7669_R_P_16_TBR1
7669_R_P_17_DLX1
7669_R_P_17_TBR1
7669_R_P_2_DLX1
7669_R_P_2_TBR1
7669_R_P_3_DLX1
7669_R_P_3_TBR1
7669_R_P_4_DLX1
7669_R_P_4_TBR1
7669_R_P_5_DLX1
7669_R_P_5_TBR1
7669_R_P_6_DLX1
7669_R_P_6_TBR1
7669_R_P_7_DLX1
7669_R_P_7_TBR1
7669_R_P_8_DLX1
7669_R_P_8_TBR1
7669_R_P_9_DLX
7669_R_P_9_DLX1
7669_R_P_9_TBR1
7669_R_POA_Bulk
7669_R_POA_DLX1
7669_R_sec1_Cau_Bulk
7669_R_sec2_GP_Bulk
7669_R_sec2_GP_DLX1
7669_R_sec2_Put_Bulk
7669_R_sec2_Thal_Bulk
7669_R_sec2_Thal_COUPTF2
7669_R_sec2_Thal_DLX1
7669_R_sec3_Cau_DLX1
7669_R_sec3_Hip_COUPTF2
7669_R_sec3_Hip_DLX1
7669_R_sec3_Hip_TBR1
7669_R_sec3_Put_DLX1
7669_R_Skin_Bulk
7669_R_T_1_Bulk
7669_R_T_1_COUPTF2
7669_R_T_1_DLX1
7669_R_T_1_TBR1
7669_RA9_Bulk
7669_RV10_Bulk
JGG

ID05 15−82103153−G−A

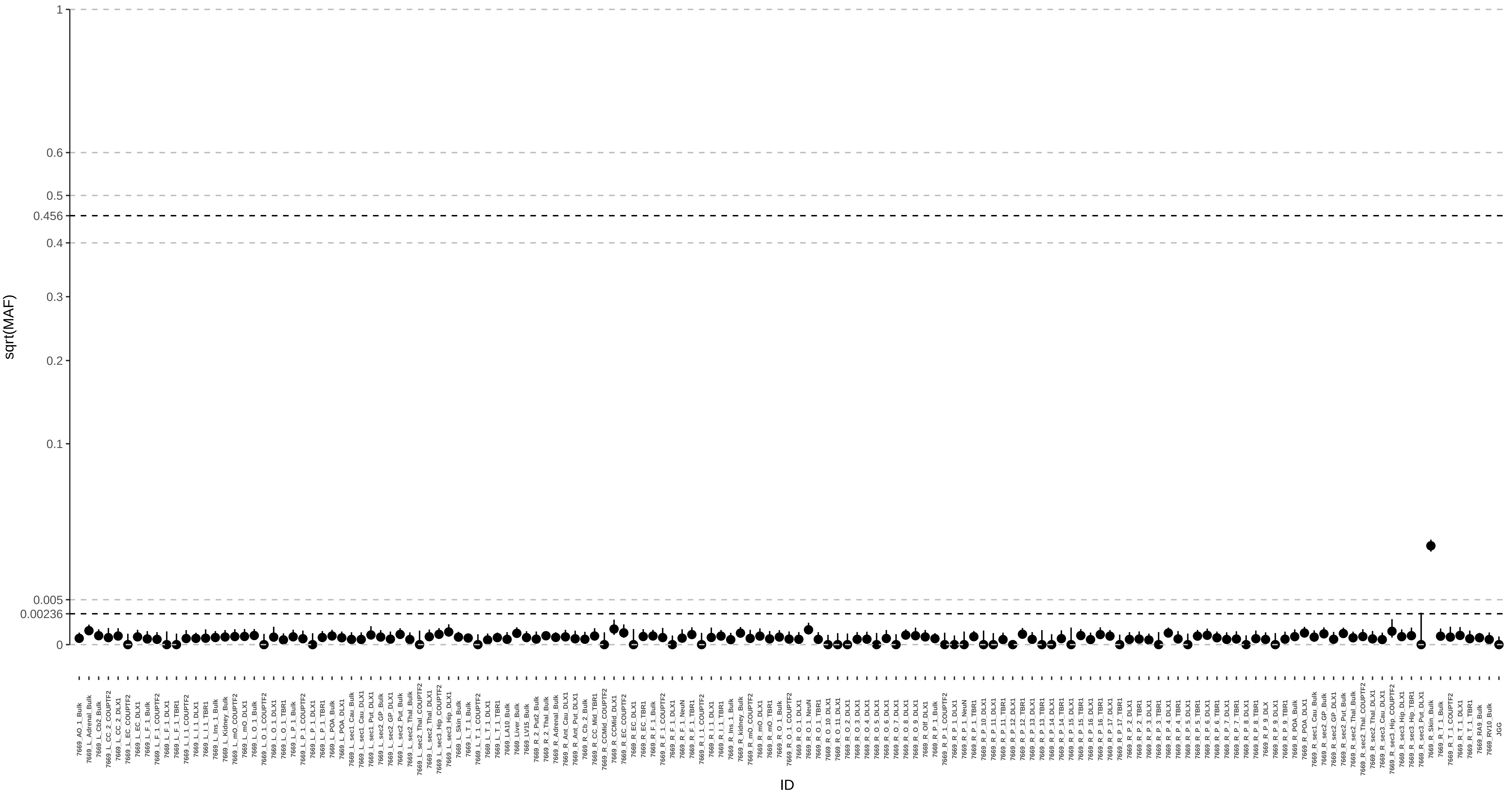

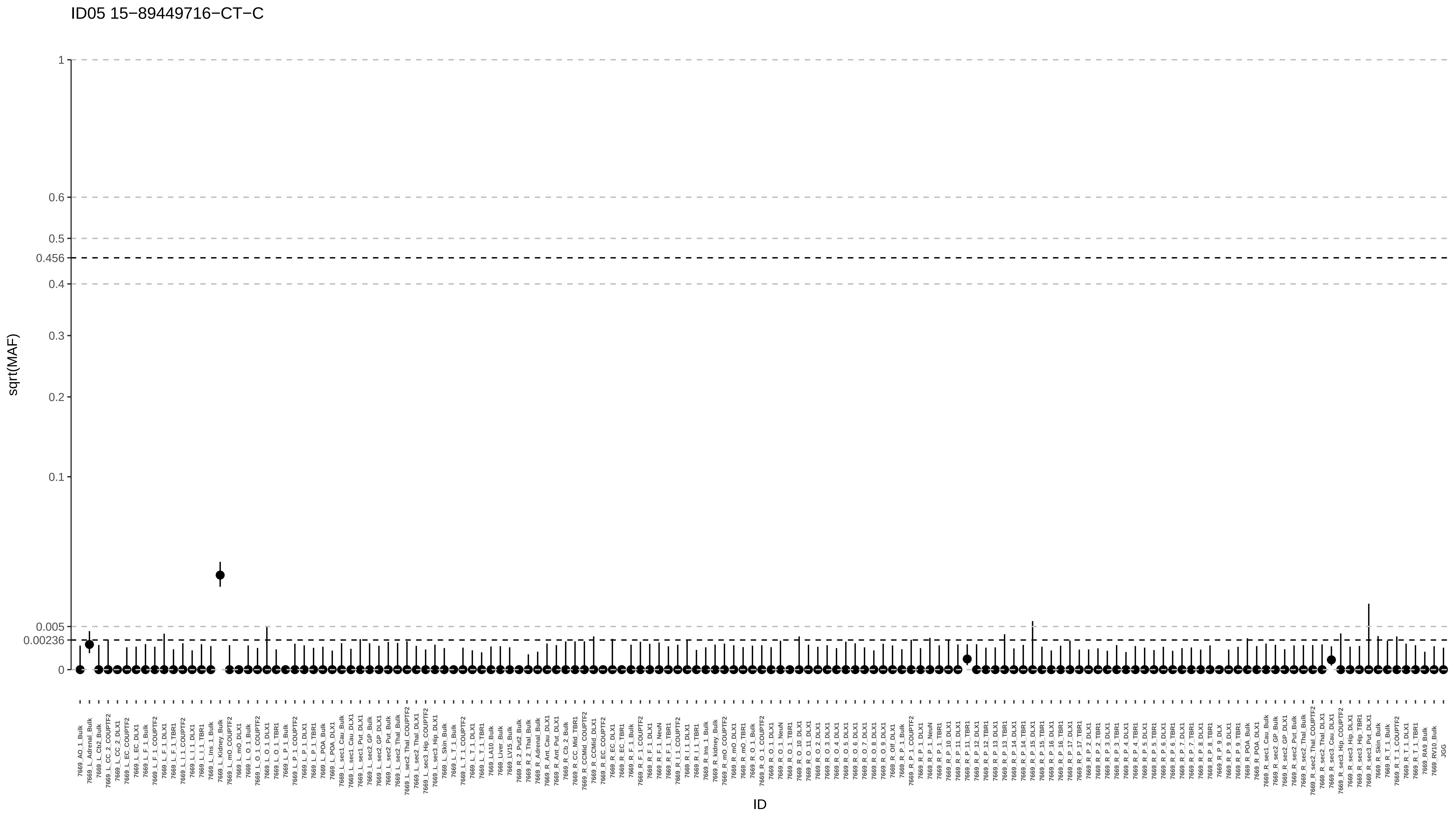
ID05 15−89449716−CT−C
sqrt(MAF)
1
0.6
0.5
0.456
0.4
0.3
0.2
0.1
0.005
0.00236
0
ID

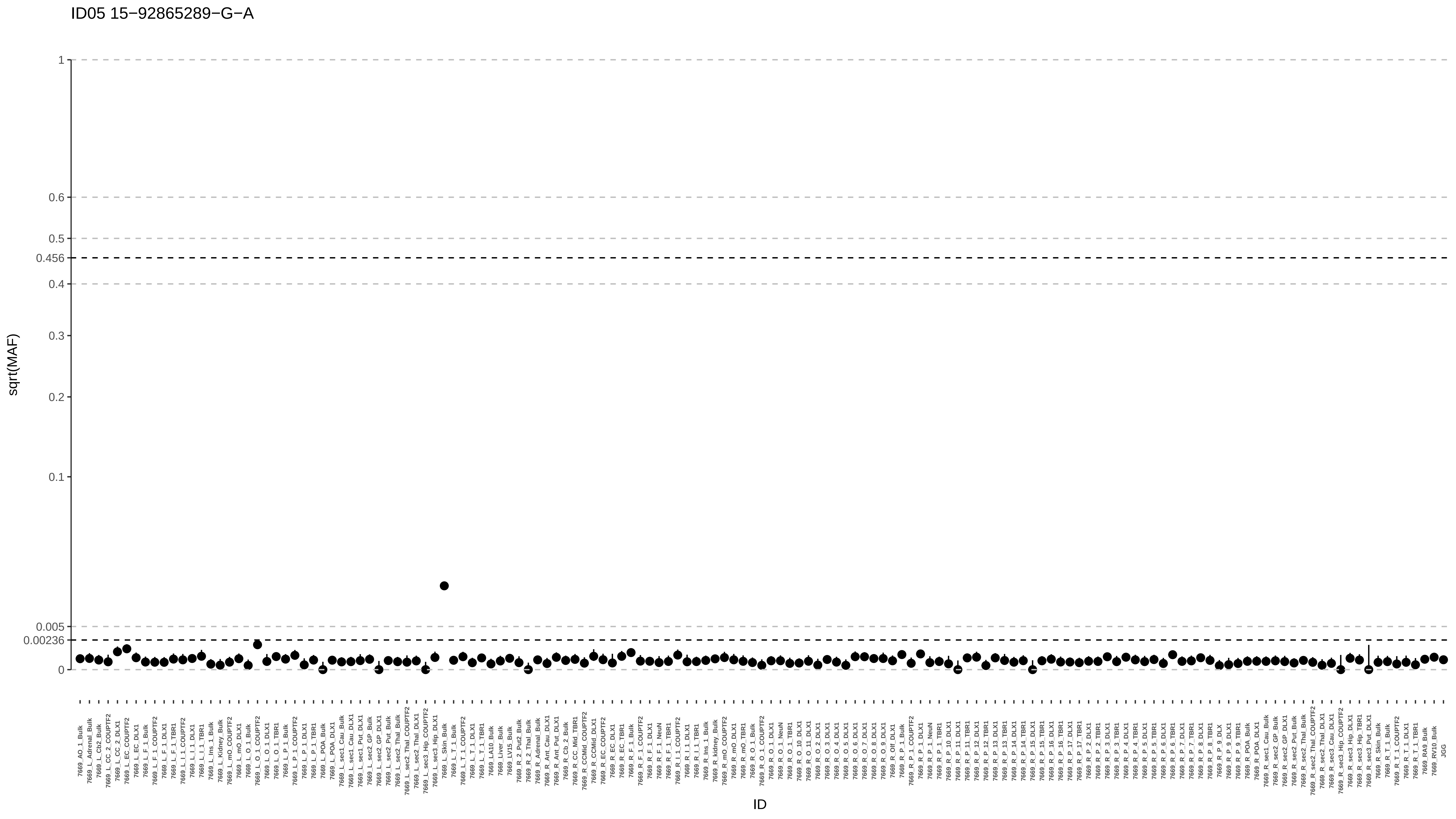
ID05 15−92865289−G−A
sqrt(MAF)
1
0.6
0.5
0.456
0.4
0.3
0.2
0.1
0.005
0.00236
0
ID

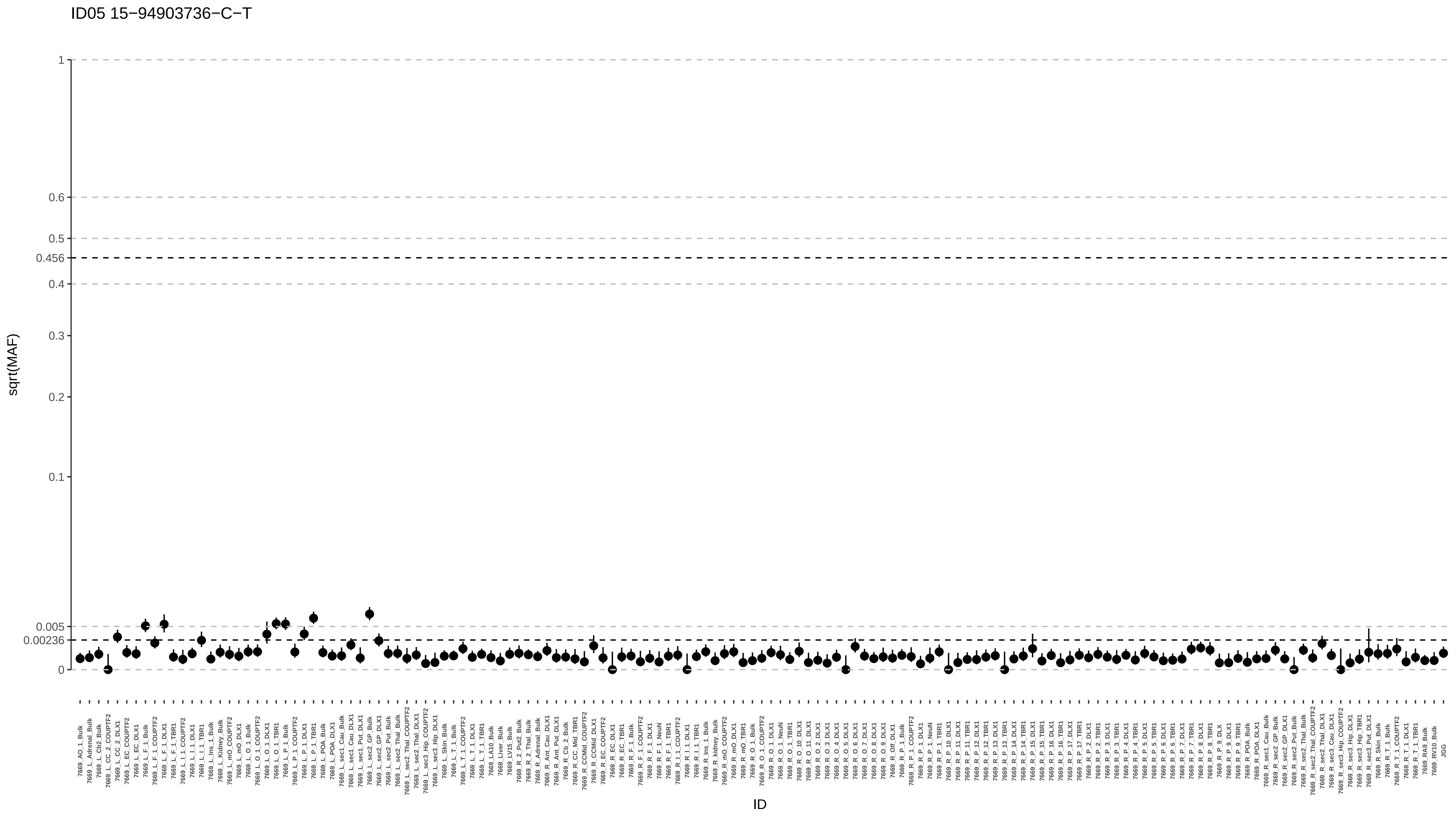
ID05 15−94903736−C−T
sqrt(MAF)
1
0.6
0.5
0.456
0.4
0.3
0.2
0.1
0.005
0.00236
0
ID

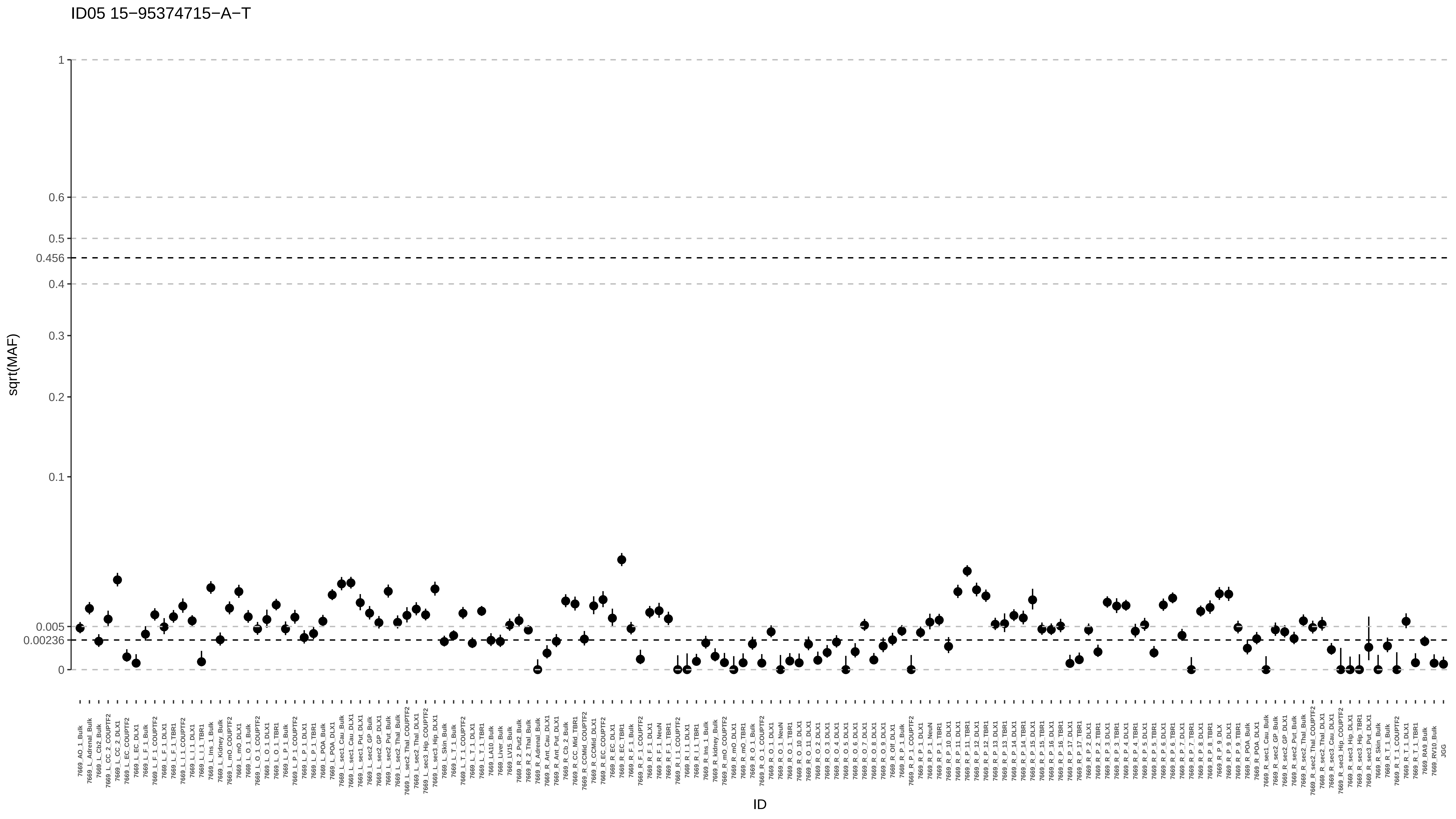
ID05 15−95374715−A−T
sqrt(MAF)
1
0.6
0.5
0.456
0.4
0.3
0.2
0.1
0.005
0.00236
0
ID

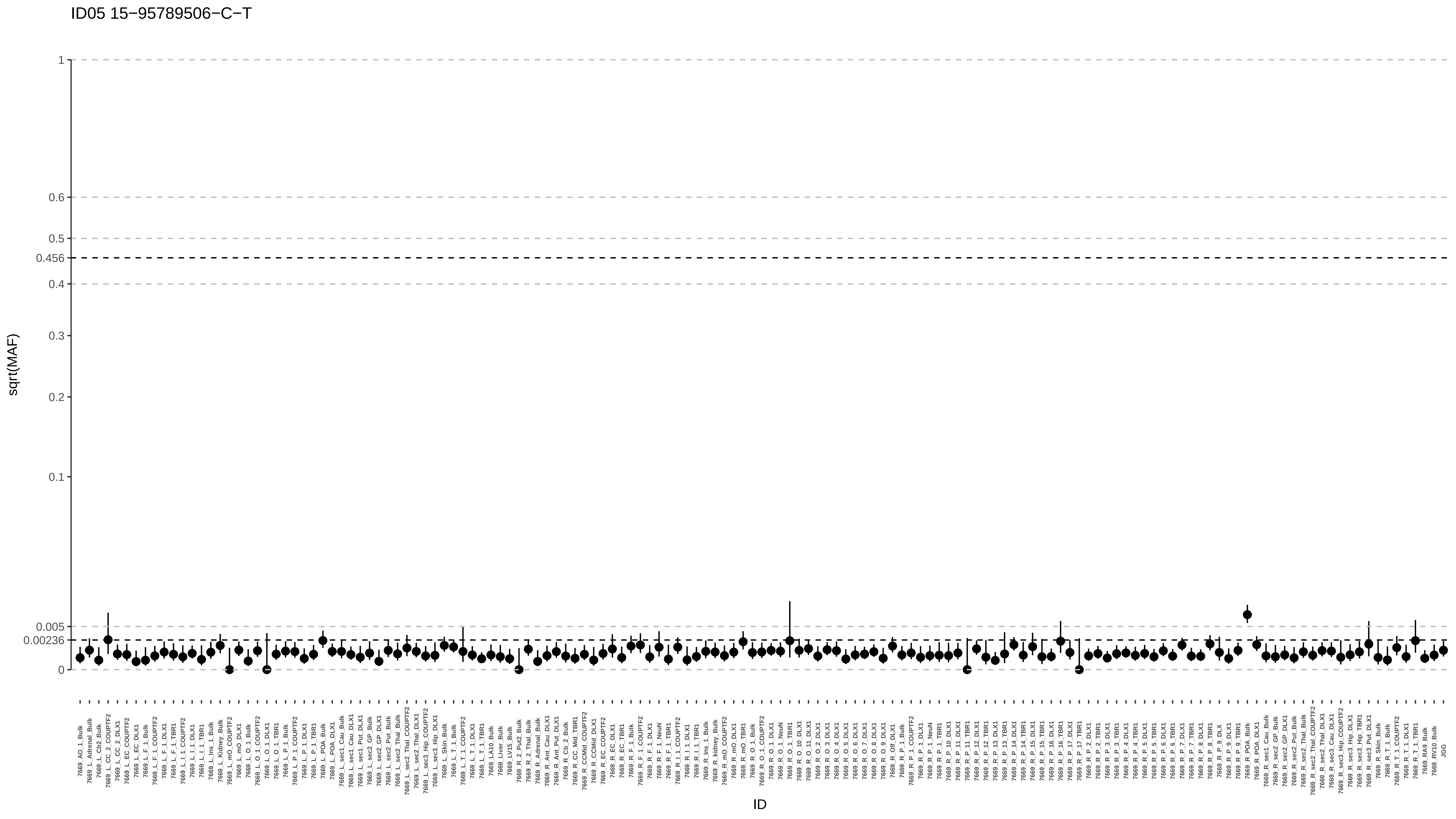
ID05 15−95789506−C−T
1
0.6
0.5
0.456
0.4
0.3
0.2
0.1
0.005
0.00236
0
sqrt(MAF)
ID

ID05 15−95947987−C−G

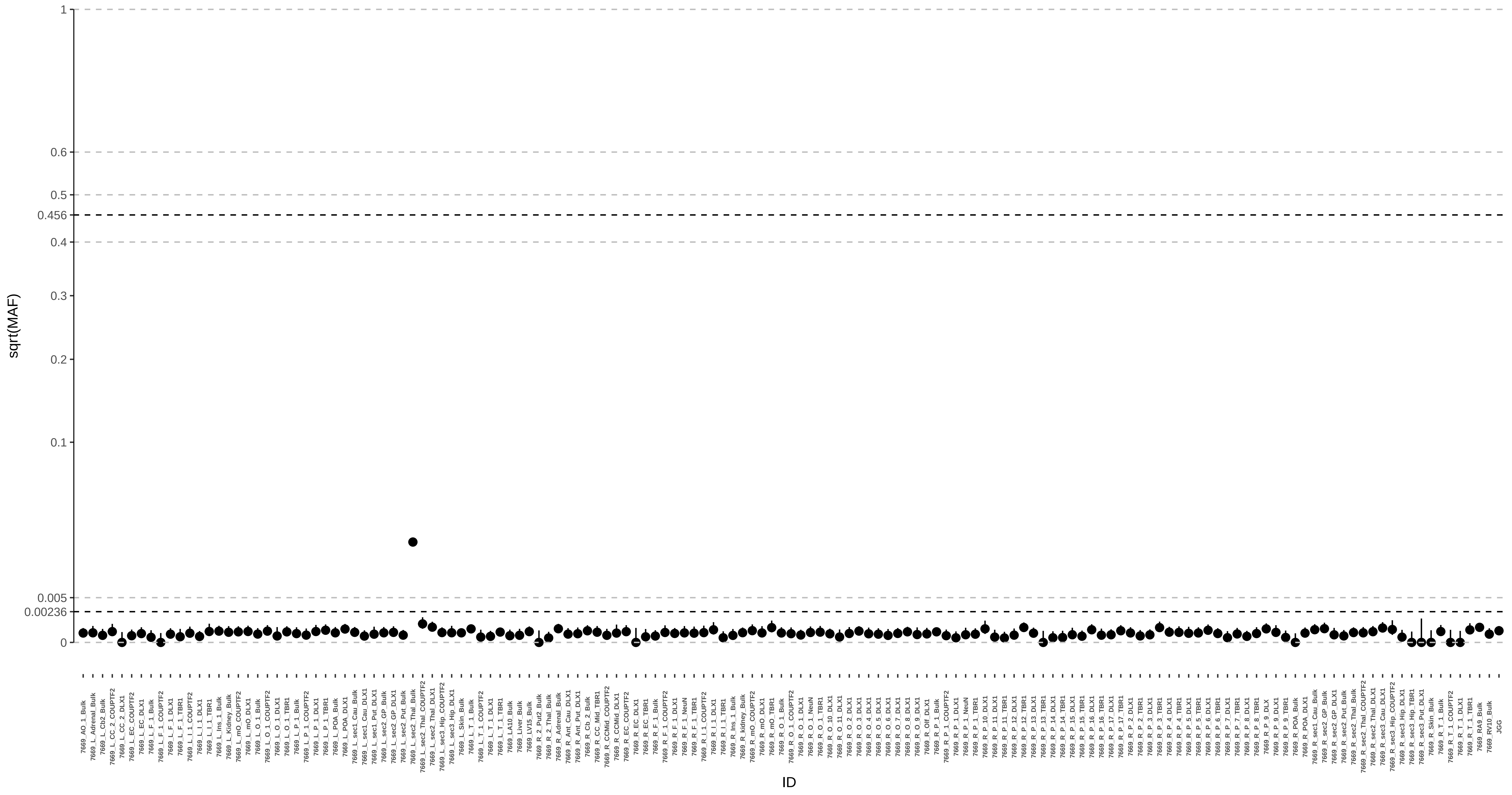

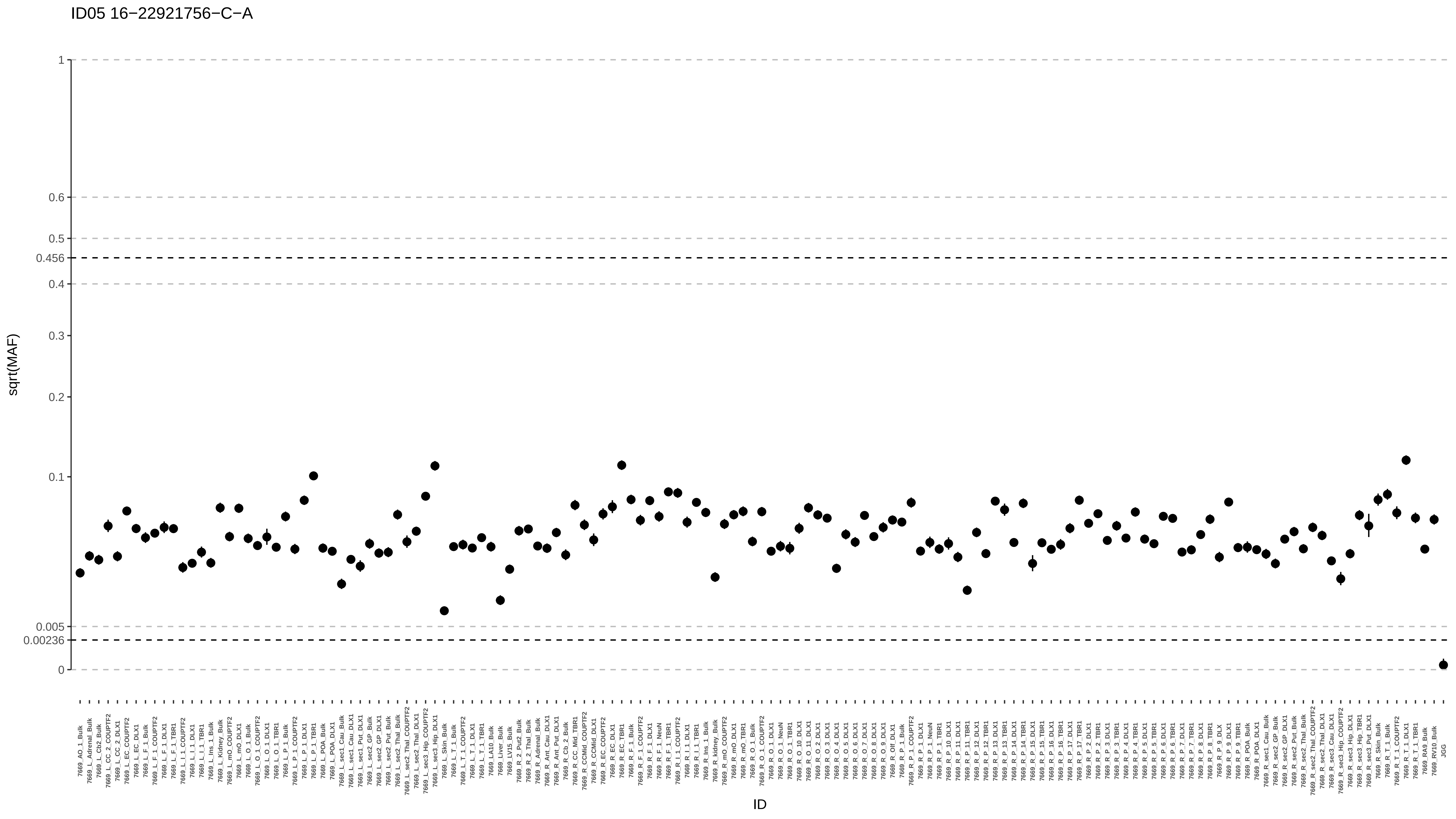
ID05 16−22921756−C−A
sqrt(MAF)
ID
1
0.6
0.5
0.456
0.4
0.3
0.2
0.1
0.005
0.00236
0

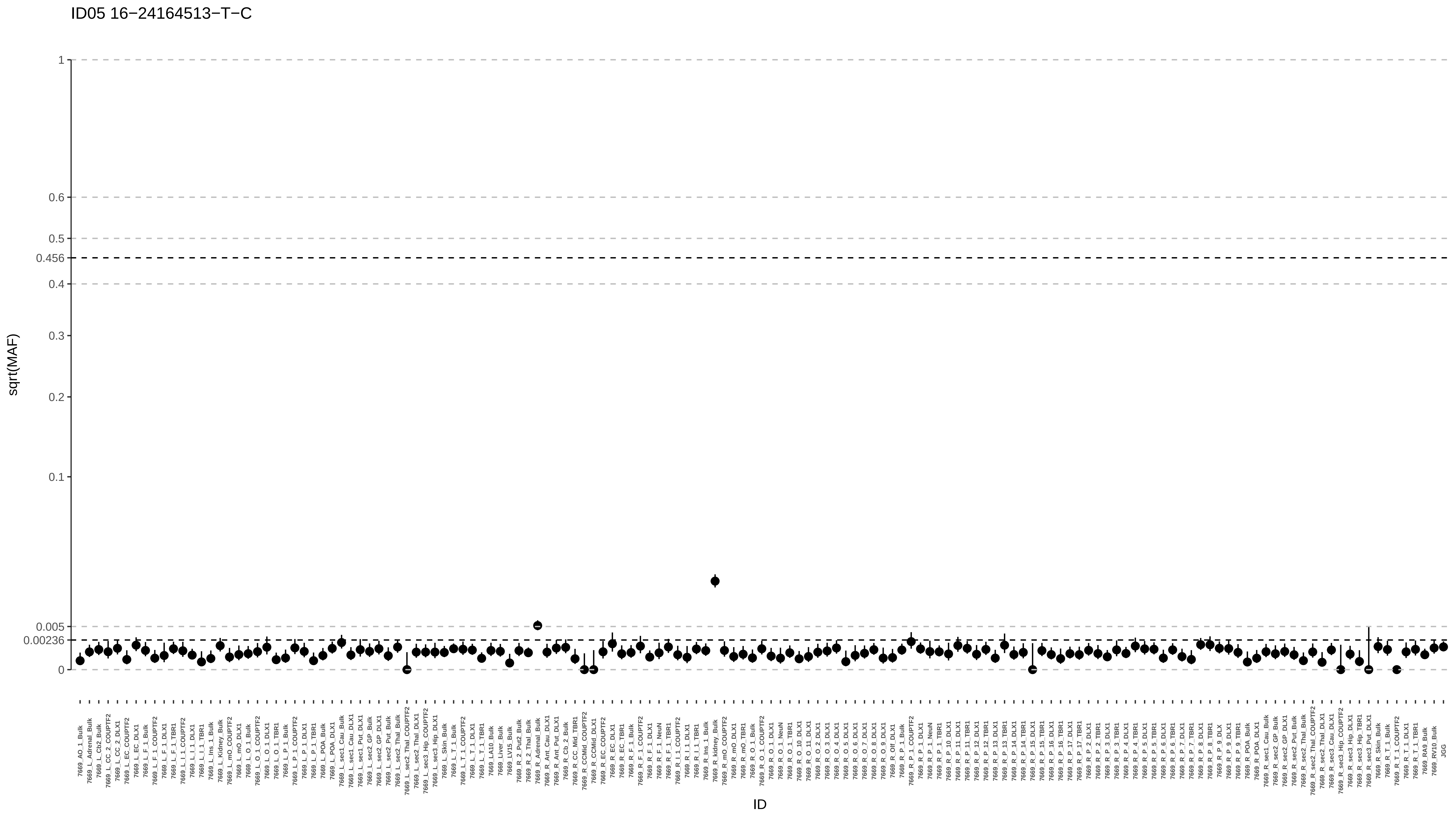
ID05 16−24164513−T−C
sqrt(MAF)
ID
1
0.6
0.5
0.456
0.4
0.3
0.2
0.1
0.005
0.00236
0

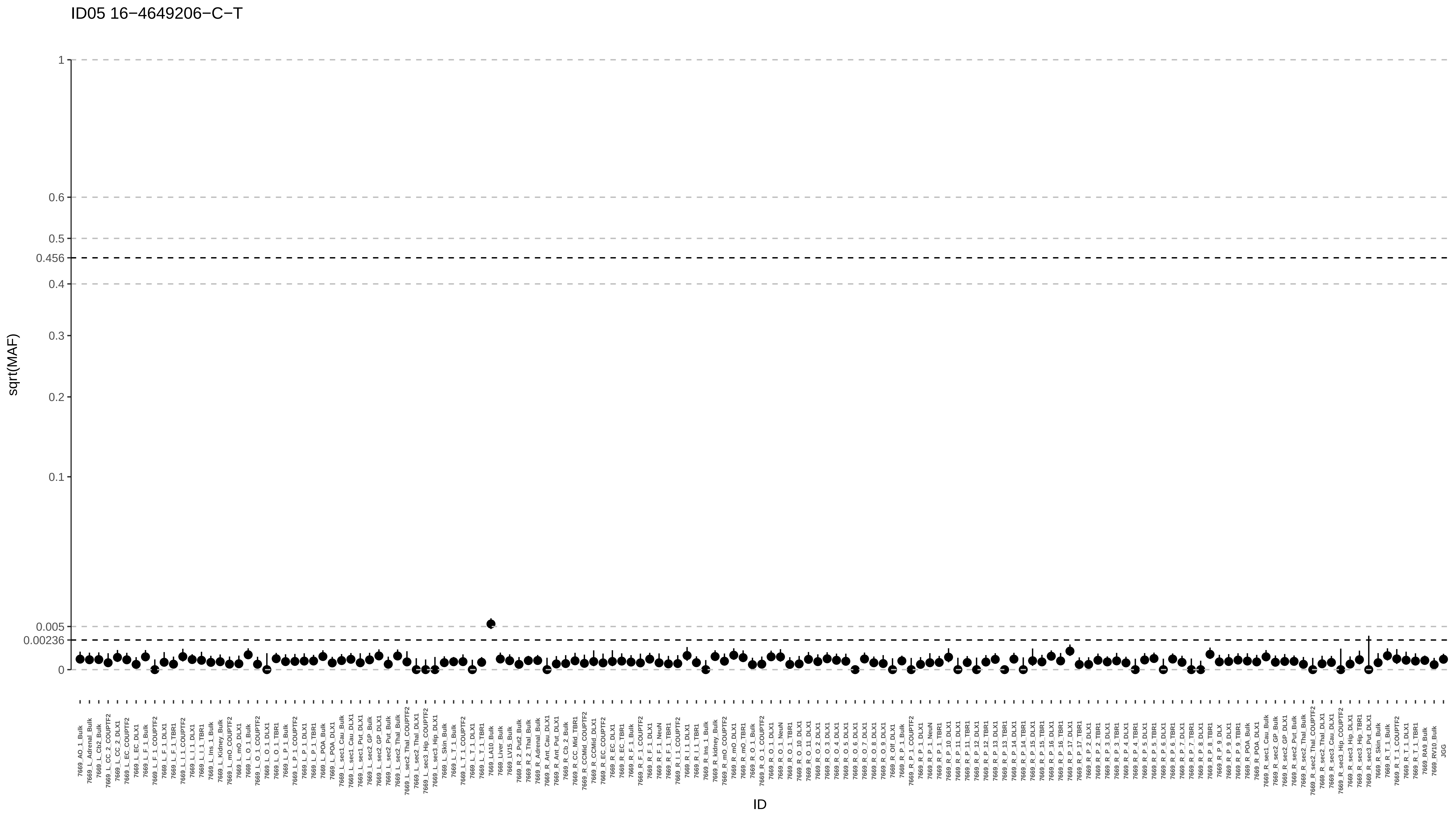
ID05 16−4649206−C−T
sqrt(MAF)
1
0.6
0.5
0.456
0.4
0.3
0.2
0.1
0.005
0.00236
0
ID

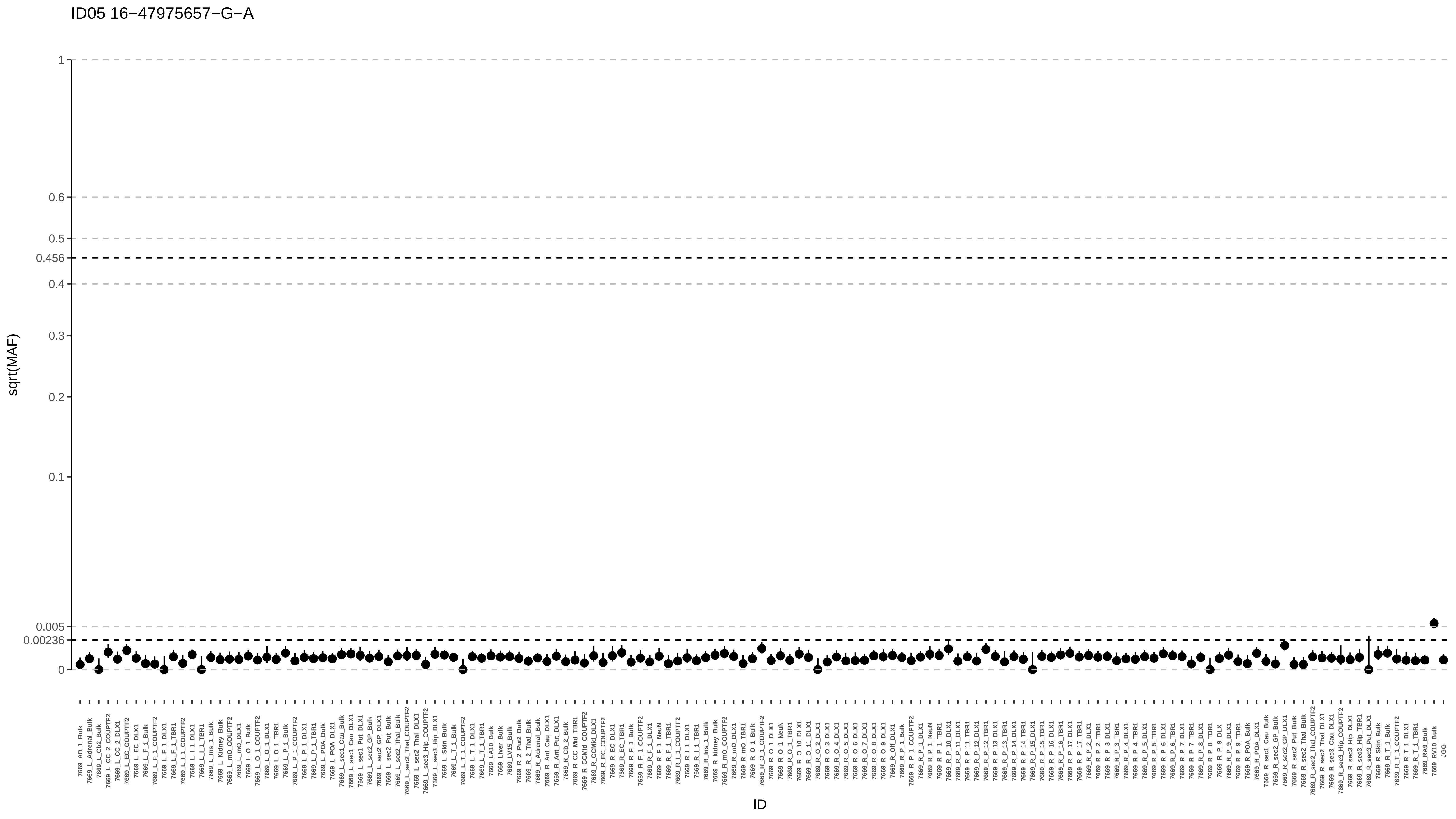

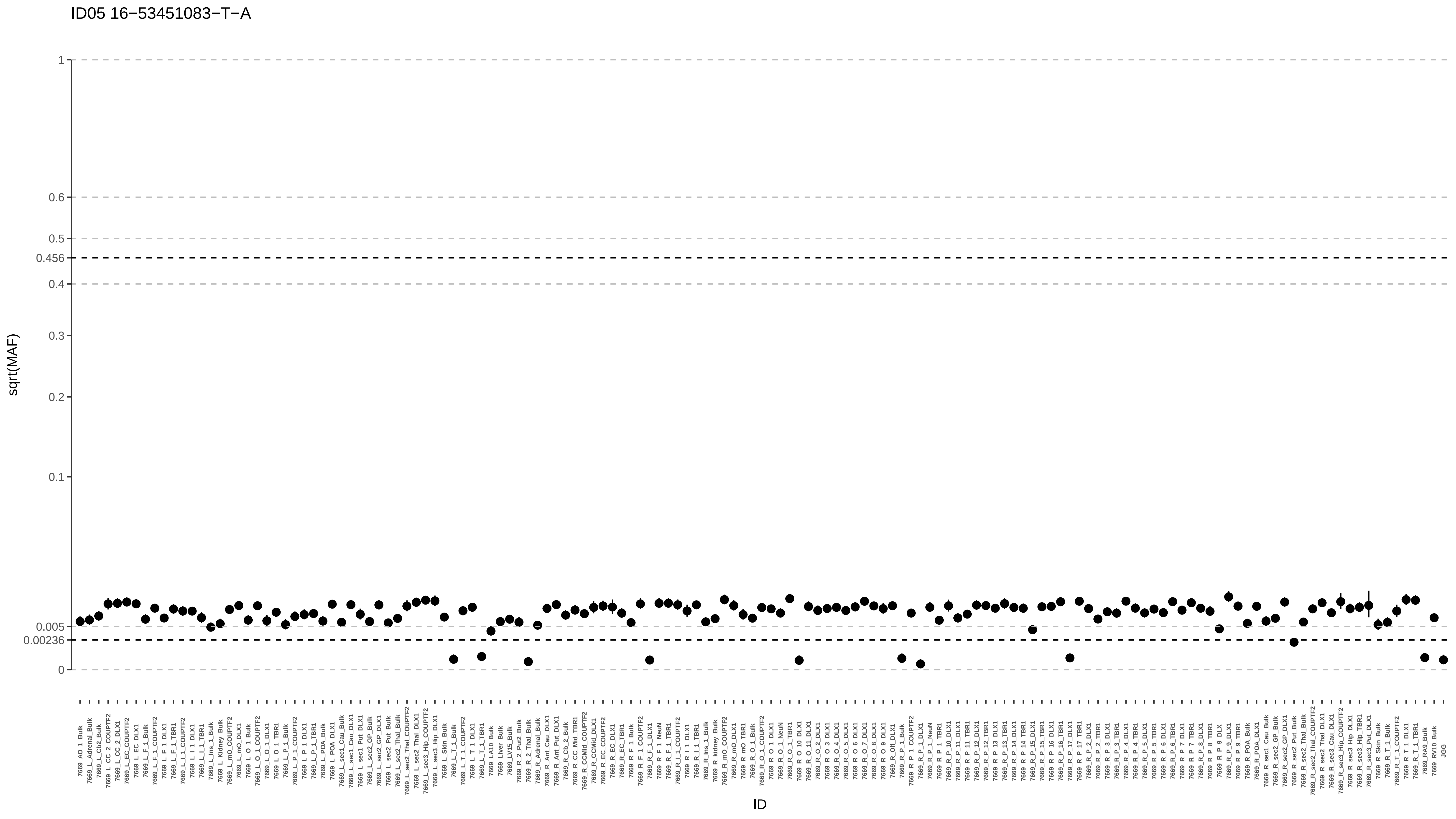
ID05 16−53451083−T−A
sqrt(MAF)
1
0.6
0.5
0.456
0.4
0.3
0.2
0.1
0.005
0.00236
0
ID

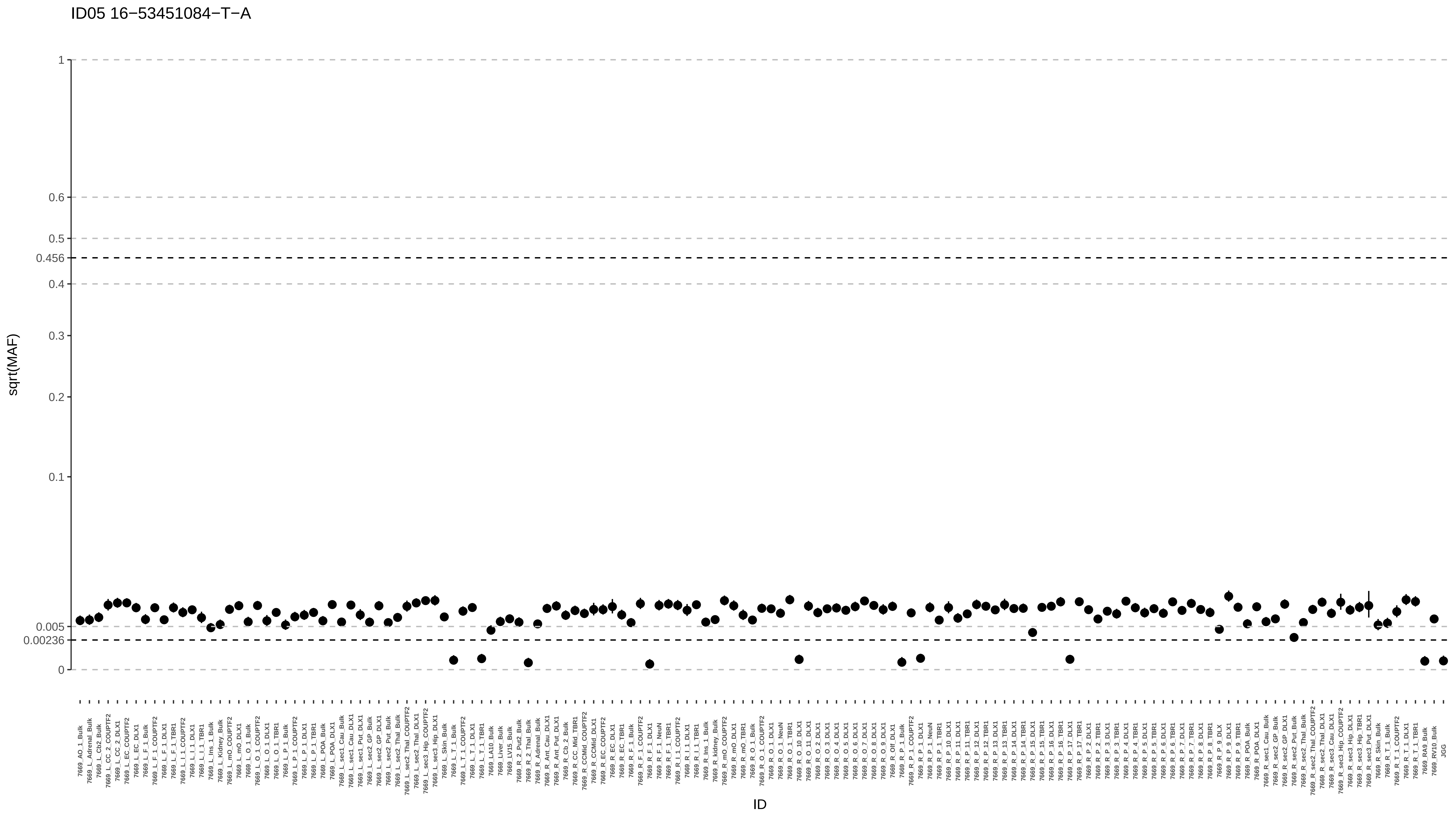
ID05 16−53451084−T−A
sqrt(MAF)
ID
1
0.6
0.5
0.456
0.4
0.3
0.2
0.1
0.005
0.00236
0

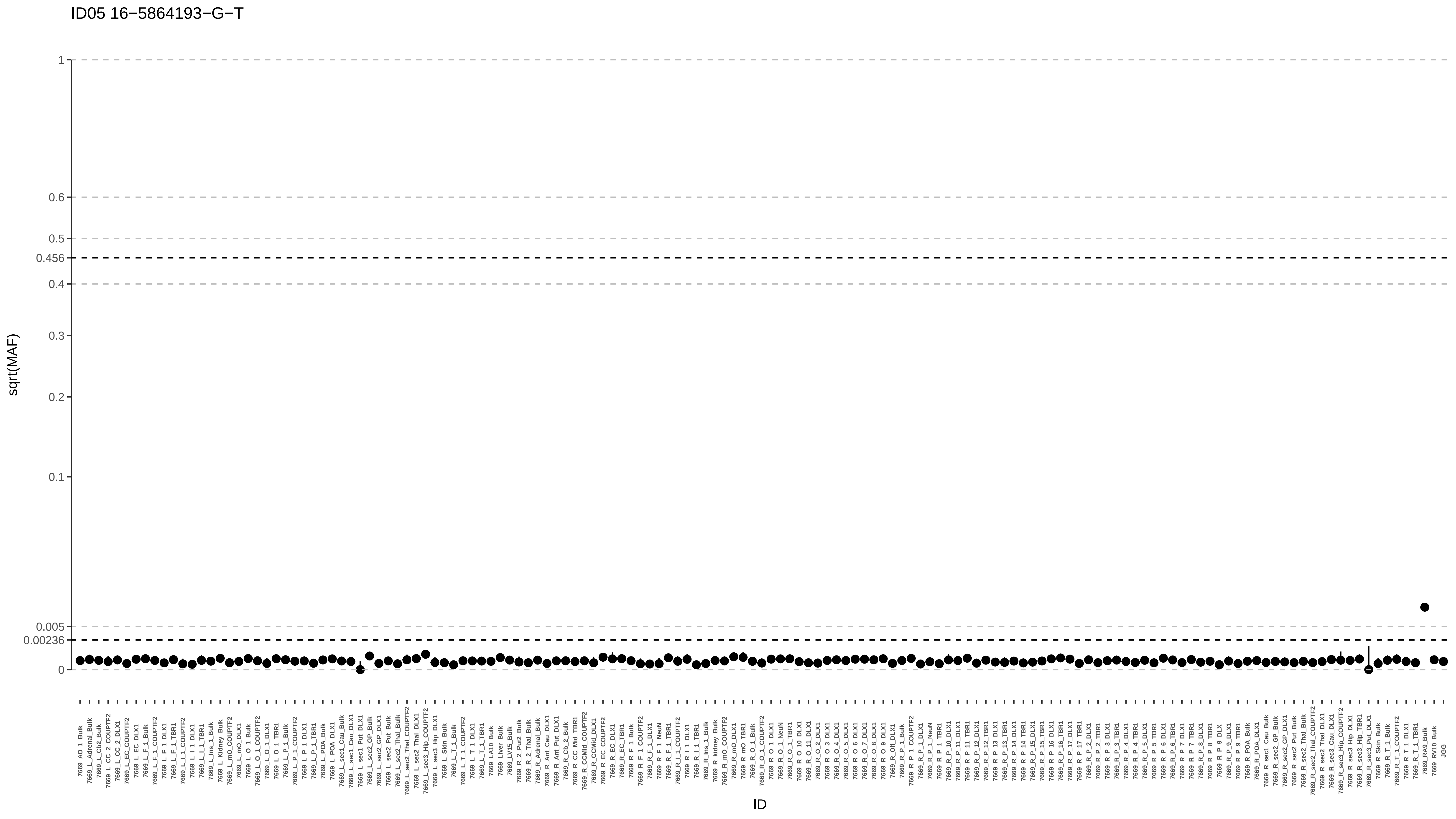
ID05 16−5864193−G−T
sqrt(MAF)
1
0.6
0.5
0.456
0.4
0.3
0.2
0.1
0.005
0.00236
0
ID

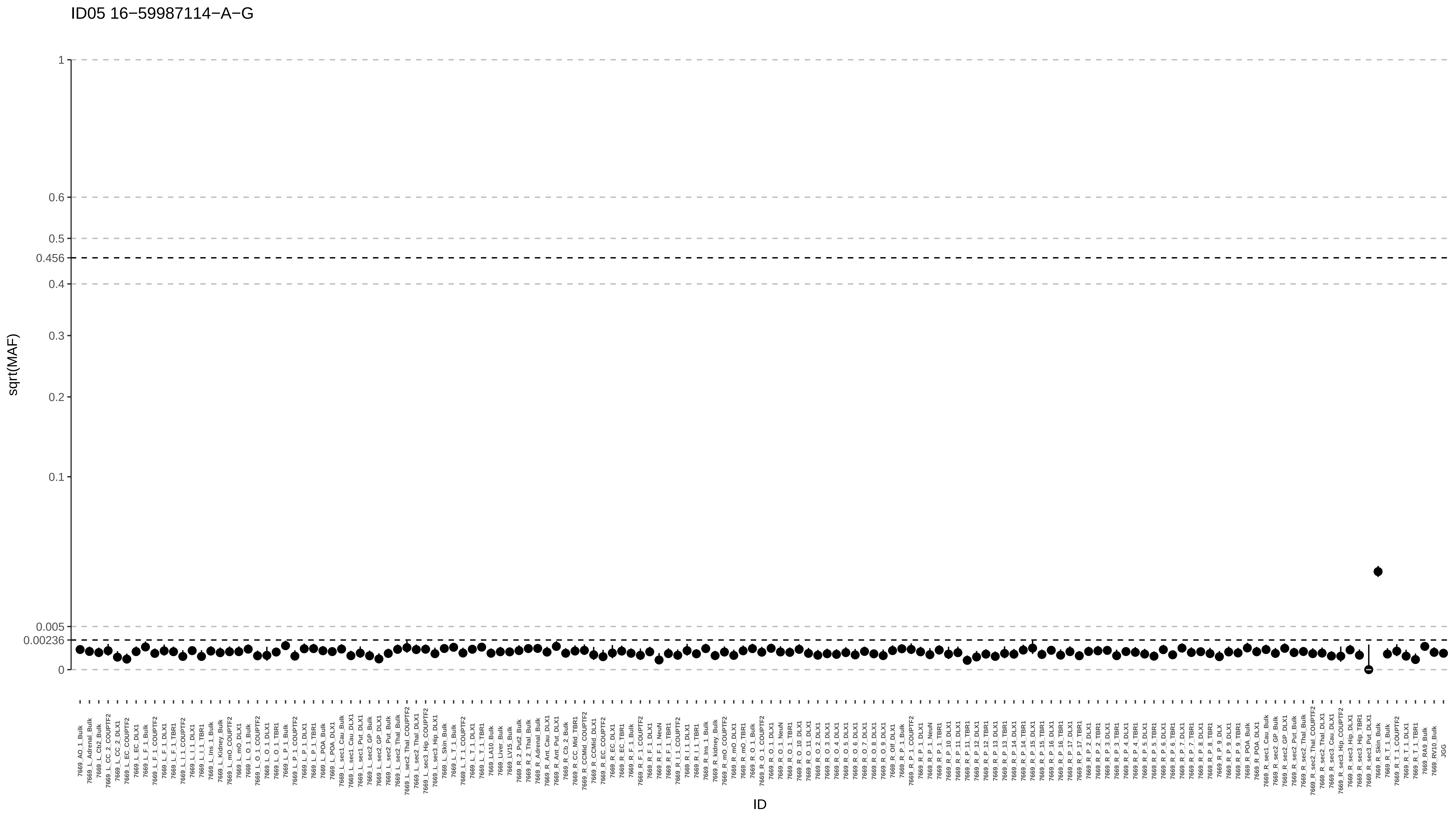

ID05 16−59987114−A−G
sqrt(MAF)
1
0.6
0.5
0.456
0.4
0.3
0.2
0.1
0.005
0.00236
0
ID

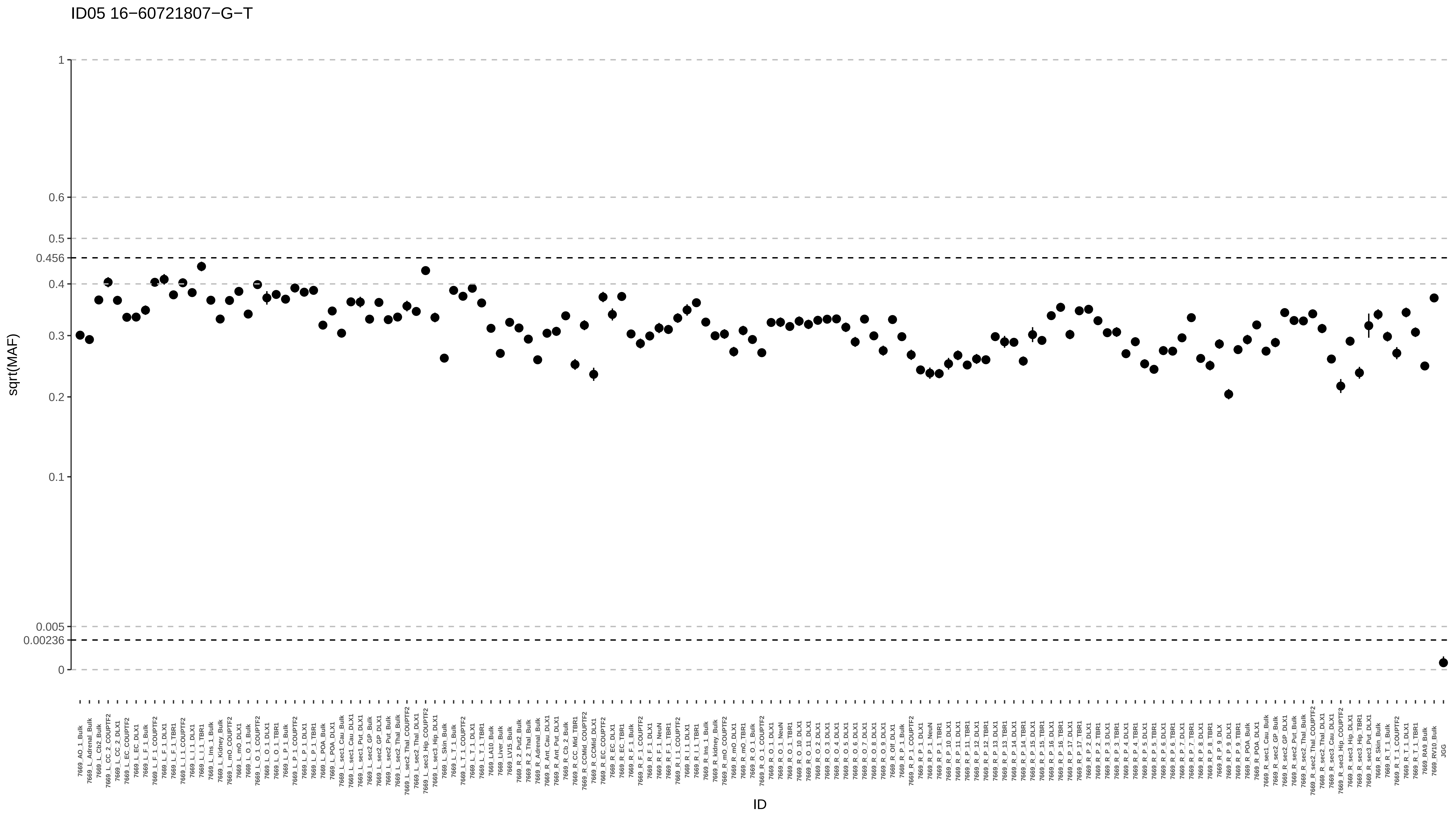
ID05 16−60721807−G−T
sqrt(MAF)
1
0.6
0.5
0.456
0.4
0.3
0.2
0.1
0.005
0.00236
0
ID
7669_AO_1_Bulk
7669_L_Adrenal_Bulk
7669_L_Cb2_Bulk
7669_L_CC_2_COUPTF2
7669_L_CC_2_DLX1
7669_L_EC_COUPTF2
7669_L_EC_DLX1
7669_L_F_1_Bulk
7669_L_F_1_COUPTF2
7669_L_F_1_DLX1
7669_L_F_1_TBR1
7669_L_I_1_COUPTF2
7669_L_I_1_DLX1
7669_L_I_1_TBR1
7669_L_Ins_1_Bulk
7669_L_Kidney_Bulk
7669_L_mO_COUPTF2
7669_L_mO_DLX1
7669_L_O_1_Bulk
7669_L_O_1_COUPTF2
7669_L_O_1_DLX1
7669_L_O_1_TBR1
7669_L_P_1_Bulk
7669_L_P_1_COUPTF2
7669_L_P_1_DLX1
7669_L_P_1_TBR1
7669_L_POA_Bulk
7669_L_POA_DLX1
7669_L_sec1_Cau_Bulk
7669_L_sec1_Cau_DLX1
7669_L_sec1_Put_DLX1
7669_L_sec2_GP_Bulk
7669_L_sec2_GP_DLX1
7669_L_sec2_Put_Bulk
7669_L_sec2_Thal_Bulk
7669_L_sec2_Thal_COUPTF2
7669_L_sec2_Thal_DLX1
7669_L_sec3_Hip_COUPTF2
7669_L_sec3_Hip_DLX1
7669_L_Skin_Bulk
7669_L_T_1_Bulk
7669_L_T_1_COUPTF2
7669_L_T_1_DLX1
7669_L_T_1_TBR1
7669_LA10_Bulk
7669_Liver_Bulk
7669_LV15_Bulk
7669_R_2_Put2_Bulk
7669_R_2_Thal_Bulk
7669_R_Adrenal_Bulk
7669_R_Ant_Cau_DLX1
7669_R_Ant_Put_DLX1
7669_R_Cb_2_Bulk
7669_R_CC_Mid_TBR1
7669_R_CCMid_COUPTF2
7669_R_CCMid_DLX1
7669_R_EC_COUPTF2
7669_R_EC_DLX1
7669_R_EC_TBR1
7669_R_F_1_Bulk
7669_R_F_1_COUPTF2
7669_R_F_1_DLX1
7669_R_F_1_NeuN
7669_R_F_1_TBR1
7669_R_I_1_COUPTF2
7669_R_I_1_DLX1
7669_R_I_1_TBR1
7669_R_Ins_1_Bulk
7669_R_kidney_Bulk
7669_R_mO_COUPTF2
7669_R_mO_DLX1
7669_R_mO_TBR1
7669_R_O_1_Bulk
7669_R_O_1_COUPTF2
7669_R_O_1_DLX1
7669_R_O_1_NeuN
7669_R_O_1_TBR1
7669_R_O_10_DLX1
7669_R_O_11_DLX1
7669_R_O_2_DLX1
7669_R_O_3_DLX1
7669_R_O_4_DLX1
7669_R_O_5_DLX1
7669_R_O_6_DLX1
7669_R_O_7_DLX1
7669_R_O_8_DLX1
7669_R_O_9_DLX1
7669_R_Olf_DLX1
7669_R_P_1_Bulk
7669_R_P_1_COUPTF2
7669_R_P_1_DLX1
7669_R_P_1_NeuN
7669_R_P_1_TBR1
7669_R_P_10_DLX1
7669_R_P_11_DLX1
7669_R_P_11_TBR1
7669_R_P_12_DLX1
7669_R_P_12_TBR1
7669_R_P_13_DLX1
7669_R_P_13_TBR1
7669_R_P_14_DLX1
7669_R_P_14_TBR1
7669_R_P_15_DLX1
7669_R_P_15_TBR1
7669_R_P_16_DLX1
7669_R_P_16_TBR1
7669_R_P_17_DLX1
7669_R_P_17_TBR1
7669_R_P_2_DLX1
7669_R_P_2_TBR1
7669_R_P_3_DLX1
7669_R_P_3_TBR1
7669_R_P_4_DLX1
7669_R_P_4_TBR1
7669_R_P_5_DLX1
7669_R_P_5_TBR1
7669_R_P_6_DLX1
7669_R_P_6_TBR1
7669_R_P_7_DLX1
7669_R_P_7_TBR1
7669_R_P_8_DLX1
7669_R_P_8_TBR1
7669_R_P_9_DLX
7669_R_P_9_DLX1
7669_R_P_9_TBR1
7669_R_POA_Bulk
7669_R_POA_DLX1
7669_R_sec1_Cau_Bulk
7669_R_sec2_GP_Bulk
7669_R_sec2_GP_DLX1
7669_R_sec2_Put_Bulk
7669_R_sec2_Thal_Bulk
7669_R_sec2_Thal_COUPTF2
7669_R_sec2_Thal_DLX1
7669_R_sec3_Cau_DLX1
7669_R_sec3_Hip_COUPTF2
7669_R_sec3_Hip_DLX1
7669_R_sec3_Hip_TBR1
7669_R_sec3_Put_DLX1
7669_R_Skin_Bulk
7669_R_T_1_Bulk
7669_R_T_1_COUPTF2
7669_R_T_1_DLX1
7669_R_T_1_TBR1
7669_RA9_Bulk
7669_RV10_Bulk
JGG

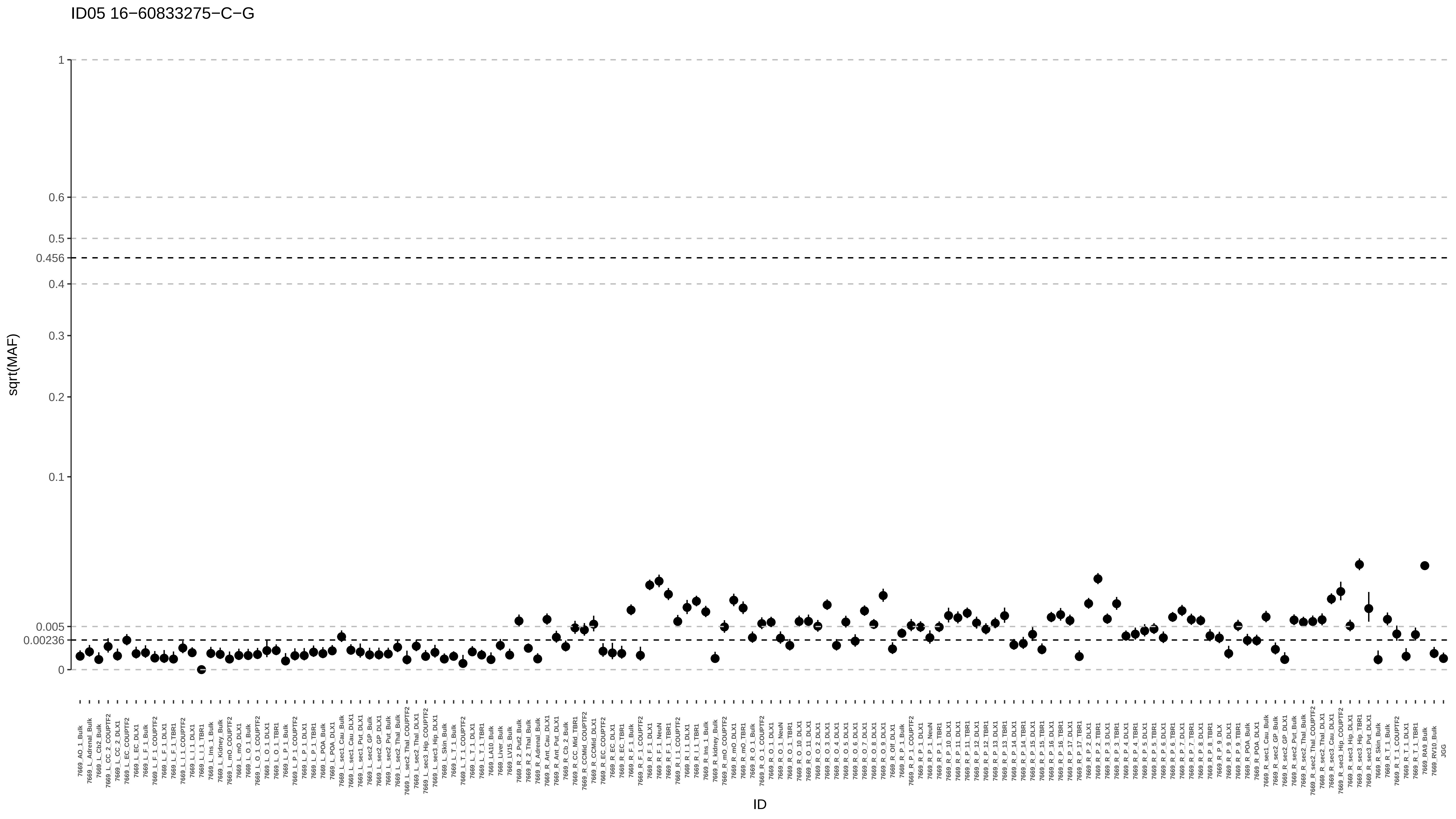
ID05 16−60833275−C−G
sqrt(MAF)
1
0.6
0.5
0.456
0.4
0.3
0.2
0.1
0.005
0.00236
0
ID
7669_AO_1_Bulk
7669_L_Adrenal_Bulk
7669_L_Cb2_Bulk
7669_L_CC_2_COUPTF2
7669_L_CC_2_DLX1
7669_L_EC_COUPTF2
7669_L_EC_DLX1
7669_L_F_1_Bulk
7669_L_F_1_COUPTF2
7669_L_F_1_DLX1
7669_L_F_1_TBR1
7669_L_I_1_COUPTF2
7669_L_I_1_DLX1
7669_L_I_1_TBR1
7669_L_Ins_1_Bulk
7669_L_Kidney_Bulk
7669_L_mO_COUPTF2
7669_L_mO_DLX1
7669_L_O_1_Bulk
7669_L_O_1_COUPTF2
7669_L_O_1_DLX1
7669_L_O_1_TBR1
7669_L_P_1_Bulk
7669_L_P_1_COUPTF2
7669_L_P_1_DLX1
7669_L_P_1_TBR1
7669_L_POA_Bulk
7669_L_POA_DLX1
7669_L_sec1_Cau_Bulk
7669_L_sec1_Cau_DLX1
7669_L_sec1_Put_DLX1
7669_L_sec2_GP_Bulk
7669_L_sec2_Put_Bulk
7669_L_sec2_Thal_Bulk
7669_L_sec2_Thal_COUPTF2
7669_L_sec2_Thal_DLX1
7669_L_sec3_Hip_COUPTF2
7669_L_sec3_Hip_DLX1
7669_L_Skin_Bulk
7669_L_T_1_Bulk
7669_L_T_1_COUPTF2
7669_L_T_1_DLX1
7669_L_T_1_TBR1
7669_LA10_Bulk
7669_Liver_Bulk
7669_LV15_Bulk
7669_R_2_Put2_Bulk
7669_R_2_Thal_Bulk
7669_R_Adrenal_Bulk
7669_R_Ant_Cau_DLX1
7669_R_Ant_Put_DLX1
7669_R_Cb_2_Bulk
7669_R_CC_Mid_TBR1
7669_R_CCMid_COUPTF2
7669_R_CCMid_DLX1
7669_R_EC_COUPTF2
7669_R_EC_DLX1
7669_R_EC_TBR1
7669_R_F_1_Bulk
7669_R_F_1_COUPTF2
7669_R_F_1_DLX1
7669_R_F_1_NeuN
7669_R_F_1_TBR1
7669_R_I_1_COUPTF2
7669_R_I_1_DLX1
7669_R_I_1_TBR1
7669_R_Ins_1_Bulk
7669_R_kidney_Bulk
7669_R_mO_COUPTF2
7669_R_mO_DLX1
7669_R_mO_TBR1
7669_R_O_1_Bulk
7669_R_O_1_COUPTF2
7669_R_O_1_DLX1
7669_R_O_1_NeuN
7669_R_O_1_TBR1
7669_R_O_10_DLX1
7669_R_O_11_DLX1
7669_R_O_2_DLX1
7669_R_O_3_DLX1
7669_R_O_4_DLX1
7669_R_O_5_DLX1
7669_R_O_6_DLX1
7669_R_O_7_DLX1
7669_R_O_8_DLX1
7669_R_O_9_DLX1
7669_R_Olf_DLX1
7669_R_P_1_Bulk
7669_R_P_1_COUPTF2
7669_R_P_1_DLX1
7669_R_P_1_NeuN
7669_R_P_1_TBR1
7669_R_P_10_DLX1
7669_R_P_11_DLX1
7669_R_P_11_TBR1
7669_R_P_12_DLX1
7669_R_P_12_TBR1
7669_R_P_13_DLX1
7669_R_P_13_TBR1
7669_R_P_14_DLX1
7669_R_P_14_TBR1
7669_R_P_15_DLX1
7669_R_P_15_TBR1
7669_R_P_16_DLX1
7669_R_P_16_TBR1
7669_R_P_17_DLX1
7669_R_P_17_TBR1
7669_R_P_2_DLX1
7669_R_P_2_TBR1
7669_R_P_3_DLX1
7669_R_P_3_TBR1
7669_R_P_4_DLX1
7669_R_P_4_TBR1
7669_R_P_5_DLX1
7669_R_P_5_TBR1
7669_R_P_6_DLX1
7669_R_P_6_TBR1
7669_R_P_7_DLX1
7669_R_P_7_TBR1
7669_R_P_8_DLX1
7669_R_P_8_TBR1
7669_R_P_9_DLX
7669_R_P_9_DLX1
7669_R_P_9_TBR1
7669_R_POA_Bulk
7669_R_POA_DLX1
7669_R_sec1_Cau_Bulk
7669_R_sec2_GP_Bulk
7669_R_sec2_GP_DLX1
7669_R_sec2_Put_Bulk
7669_R_sec2_Thal_Bulk
7669_R_sec2_Thal_COUPTF2
7669_R_sec2_Thal_DLX1
7669_R_sec3_Cau_DLX1
7669_R_sec3_Hip_COUPTF2
7669_R_sec3_Hip_DLX1
7669_R_sec3_Hip_TBR1
7669_R_sec3_Put_DLX1
7669_R_Skin_Bulk
7669_R_T_1_Bulk
7669_R_T_1_COUPTF2
7669_R_T_1_DLX1
7669_R_T_1_TBR1
7669_RA9_Bulk
7669_RV10_Bulk
JGG

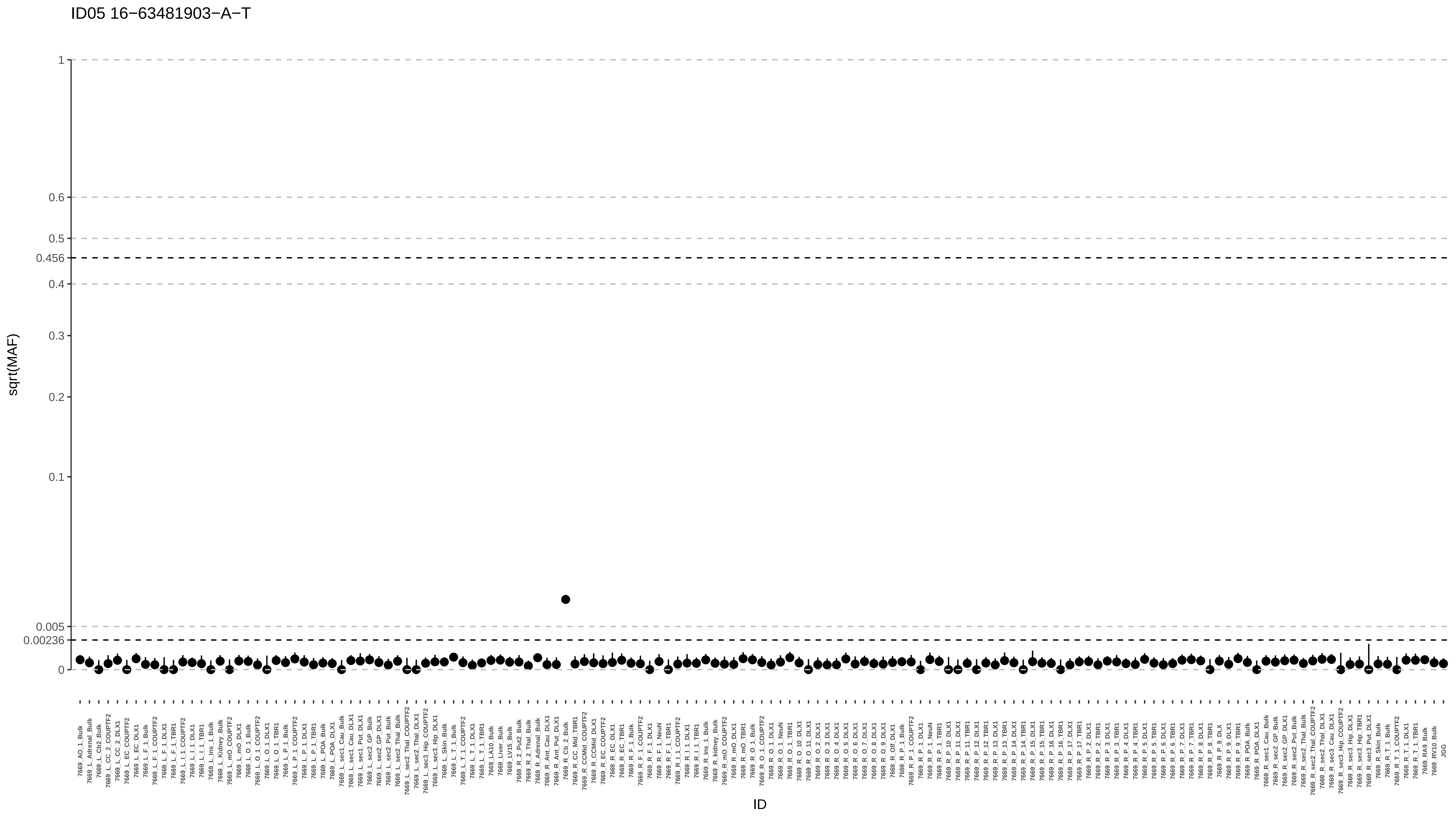
ID05 16−63481903−A−T
sqrt(MAF)
1
0.6
0.5
0.456
0.4
0.3
0.2
0.1
0.005
0.00236
0
ID

ID05 16−64667272−C−T

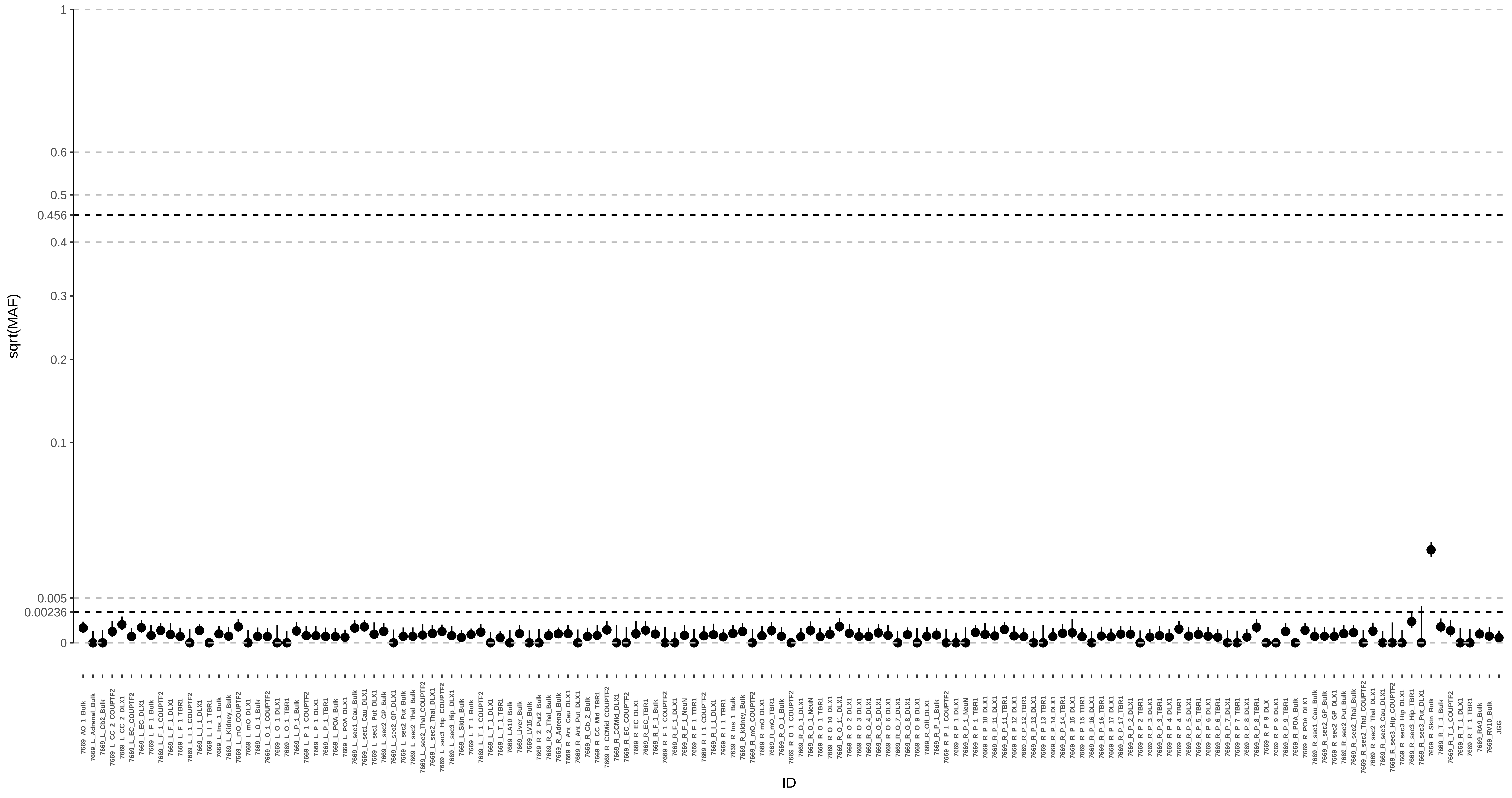

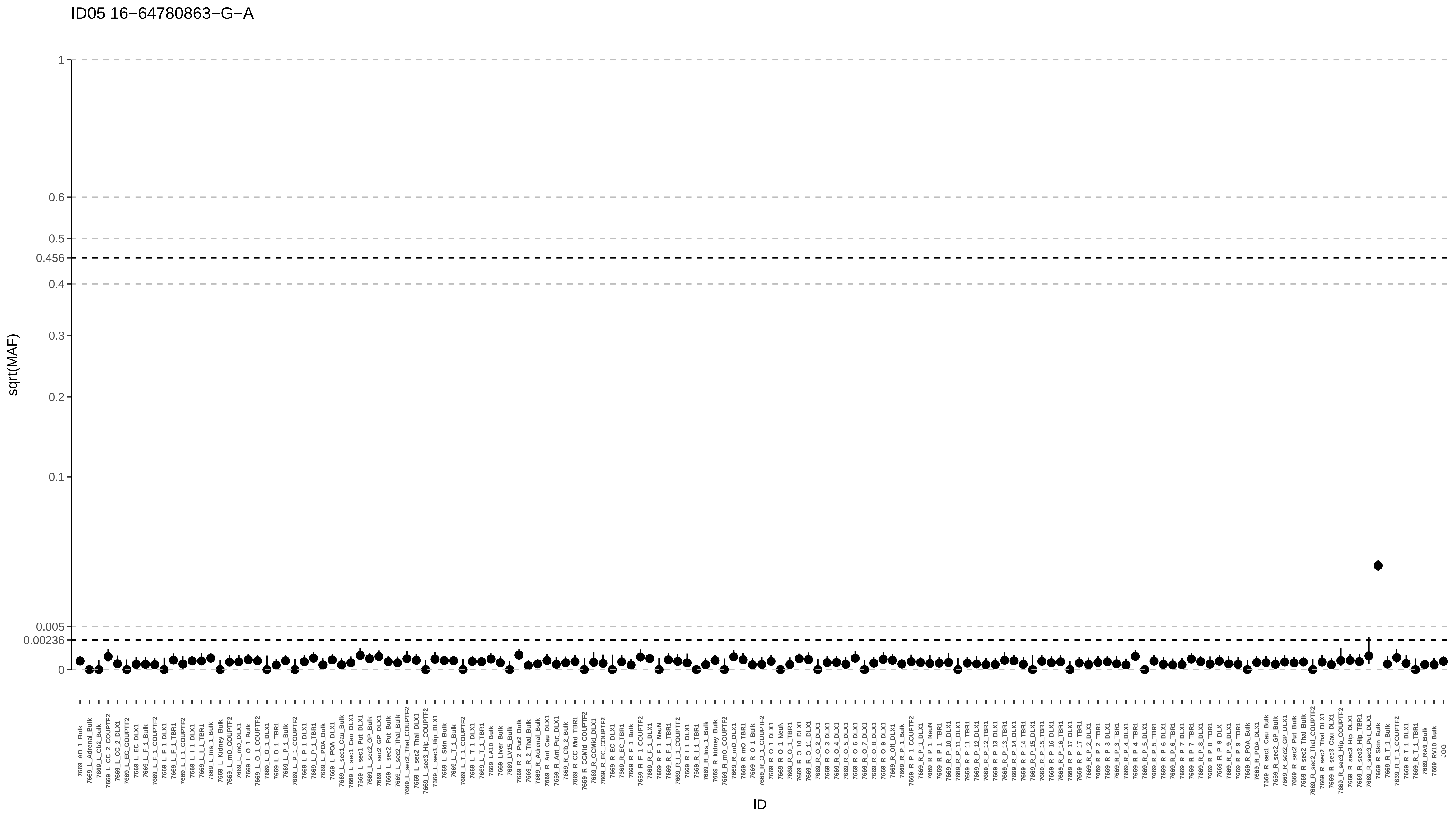
ID05 16−64780863−G−A
sqrt(MAF)
1
0.6
0.5
0.456
0.4
0.3
0.2
0.1
0.005
0.00236
0
ID
7669_AO_1_Bulk
7669_L_Adrenal_Bulk
7669_L_Cb2_Bulk
7669_L_CC_2_COUPTF2
7669_L_CC_2_DLX1
7669_L_EC_COUPTF2
7669_L_EC_DLX1
7669_L_F_1_Bulk
7669_L_F_1_COUPTF2
7669_L_F_1_DLX1
7669_L_F_1_TBR1
7669_L_I_1_COUPTF2
7669_L_I_1_DLX1
7669_L_I_1_TBR1
7669_L_Ins_1_Bulk
7669_L_Kidney_Bulk
7669_L_mO_COUPTF2
7669_L_mO_DLX1
7669_L_O_1_Bulk
7669_L_O_1_COUPTF2
7669_L_O_1_DLX1
7669_L_O_1_TBR1
7669_L_P_1_Bulk
7669_L_P_1_COUPTF2
7669_L_P_1_DLX1
7669_L_P_1_TBR1
7669_L_POA_Bulk
7669_L_POA_DLX1
7669_L_sec1_Cau_Bulk
7669_L_sec1_Cau_DLX1
7669_L_sec1_Put_DLX1
7669_L_sec2_GP_Bulk
7669_L_sec2_GP_DLX1
7669_L_sec2_Put_Bulk
7669_L_sec2_Thal_Bulk
7669_L_sec2_Thal_COUPTF2
7669_L_sec2_Thal_DLX1
7669_L_sec3_Hip_COUPTF2
7669_L_sec3_Hip_DLX1
7669_L_Skin_Bulk
7669_L_T_1_Bulk
7669_L_T_1_COUPTF2
7669_L_T_1_DLX1
7669_L_T_1_TBR1
7669_LA10_Bulk
7669_Liver_Bulk
7669_LV15_Bulk
7669_R_2_Put2_Bulk
7669_R_2_Thal_Bulk
7669_R_Adrenal_Bulk
7669_R_Ant_Cau_DLX1
7669_R_Ant_Put_DLX1
7669_R_Cb_2_Bulk
7669_R_CC_Mid_TBR1
7669_R_CCMid_COUPTF2
7669_R_CCMid_DLX1
7669_R_EC_COUPTF2
7669_R_EC_DLX1
7669_R_EC_TBR1
7669_R_F_1_Bulk
7669_R_F_1_COUPTF2
7669_R_F_1_DLX1
7669_R_F_1_NeuN
7669_R_F_1_TBR1
7669_R_I_1_COUPTF2
7669_R_I_1_DLX1
7669_R_I_1_TBR1
7669_R_Ins_1_Bulk
7669_R_kidney_Bulk
7669_R_mO_COUPTF2
7669_R_mO_DLX1
7669_R_mO_TBR1
7669_R_O_1_Bulk
7669_R_O_1_COUPTF2
7669_R_O_1_DLX1
7669_R_O_1_NeuN
7669_R_O_1_TBR1
7669_R_O_10_DLX1
7669_R_O_11_DLX1
7669_R_O_2_DLX1
7669_R_O_3_DLX1
7669_R_O_4_DLX1
7669_R_O_5_DLX1
7669_R_O_6_DLX1
7669_R_O_7_DLX1
7669_R_O_8_DLX1
7669_R_O_9_DLX1
7669_R_Olf_DLX1
7669_R_P_1_Bulk
7669_R_P_1_COUPTF2
7669_R_P_1_DLX1
7669_R_P_1_NeuN
7669_R_P_1_TBR1
7669_R_P_10_DLX1
7669_R_P_11_DLX1
7669_R_P_11_TBR1
7669_R_P_12_DLX1
7669_R_P_12_TBR1
7669_R_P_13_DLX1
7669_R_P_13_TBR1
7669_R_P_14_DLX1
7669_R_P_14_TBR1
7669_R_P_15_DLX1
7669_R_P_15_TBR1
7669_R_P_16_DLX1
7669_R_P_16_TBR1
7669_R_P_17_DLX1
7669_R_P_17_TBR1
7669_R_P_2_DLX1
7669_R_P_2_TBR1
7669_R_P_3_DLX1
7669_R_P_3_TBR1
7669_R_P_4_DLX1
7669_R_P_4_TBR1
7669_R_P_5_DLX1
7669_R_P_5_TBR1
7669_R_P_6_DLX1
7669_R_P_6_TBR1
7669_R_P_7_DLX1
7669_R_P_7_TBR1
7669_R_P_8_DLX1
7669_R_P_8_TBR1
7669_R_P_9_DLX
7669_R_P_9_DLX1
7669_R_P_9_TBR1
7669_R_POA_Bulk
7669_R_POA_DLX1
7669_R_sec1_Cau_Bulk
7669_R_sec2_GP_Bulk
7669_R_sec2_GP_DLX1
7669_R_sec2_Put_Bulk
7669_R_sec2_Thal_Bulk
7669_R_sec2_Thal_COUPTF2
7669_R_sec2_Thal_DLX1
7669_R_sec3_Cau_DLX1
7669_R_sec3_Hip_COUPTF2
7669_R_sec3_Hip_DLX1
7669_R_sec3_Hip_TBR1
7669_R_sec3_Put_DLX1
7669_R_Skin_Bulk
7669_R_T_1_Bulk
7669_R_T_1_COUPTF2
7669_R_T_1_DLX1
7669_R_T_1_TBR1
7669_RA9_Bulk
7669_RV10_Bulk
JGG

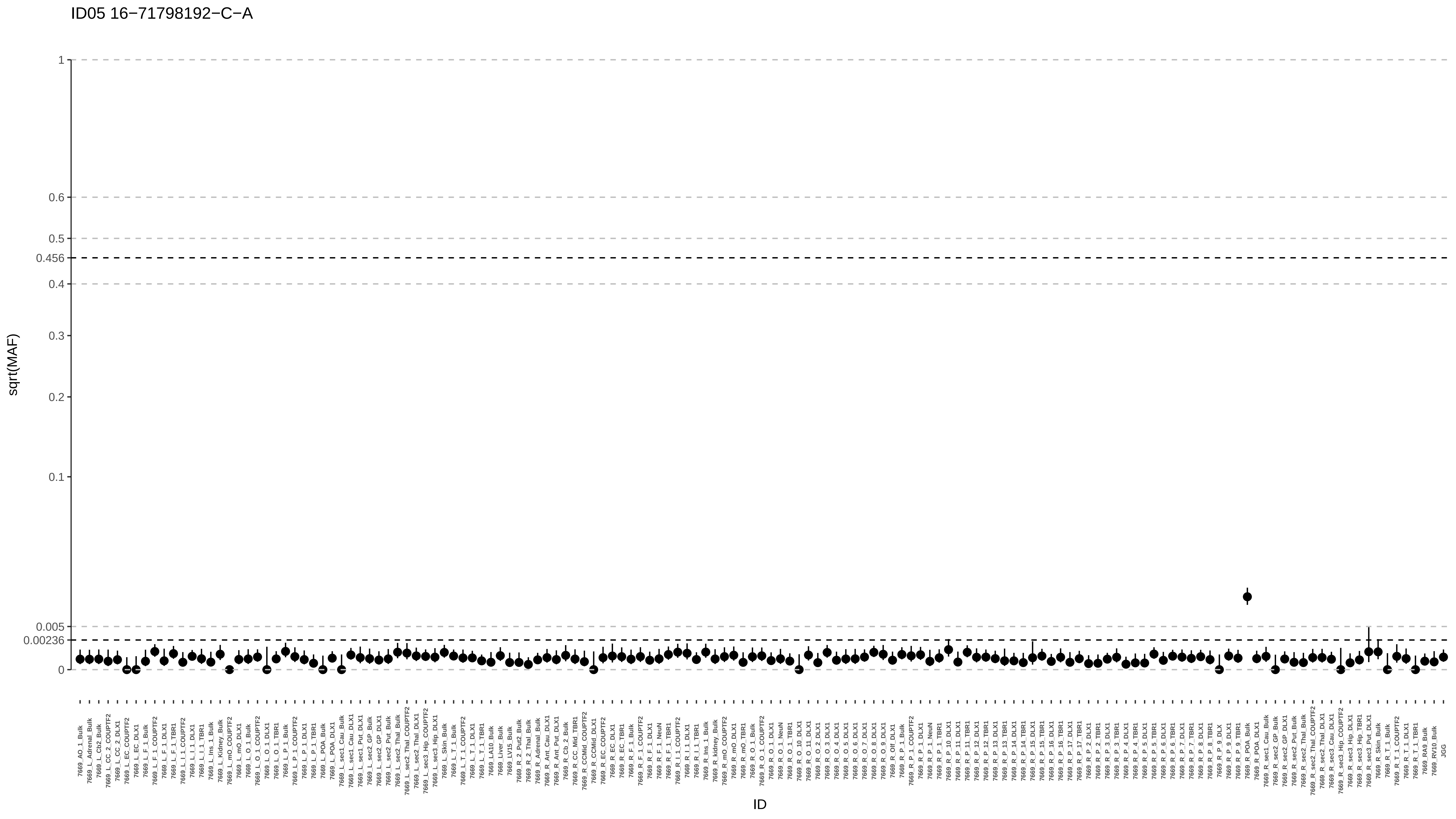
ID05 16−71798192−C−A
sqrt(MAF)
1
0.6
0.5
0.456
0.4
0.3
0.2
0.1
0.005
0.00236
0
ID
7669_AO_1_Bulk
7669_L_Adrenal_Bulk
7669_L_Cb2_Bulk
7669_L_CC_2_COUPTF2
7669_L_CC_2_DLX1
7669_L_EC_COUPTF2
7669_L_EC_DLX1
7669_L_F_1_Bulk
7669_L_F_1_COUPTF2
7669_L_F_1_DLX1
7669_L_F_1_TBR1
7669_L_I_1_COUPTF2
7669_L_I_1_DLX1
7669_L_I_1_TBR1
7669_L_Ins_1_Bulk
7669_L_Kidney_Bulk
7669_L_mO_COUPTF2
7669_L_mO_DLX1
7669_L_O_1_Bulk
7669_L_O_1_COUPTF2
7669_L_O_1_DLX1
7669_L_O_1_TBR1
7669_L_P_1_Bulk
7669_L_P_1_COUPTF2
7669_L_P_1_DLX1
7669_L_P_1_TBR1
7669_L_POA_Bulk
7669_L_POA_DLX1
7669_L_sec1_Cau_Bulk
7669_L_sec1_Cau_DLX1
7669_L_sec1_Put_DLX1
7669_L_sec2_GP_Bulk
7669_L_sec2_GP_DLX1
7669_L_sec2_Put_Bulk
7669_L_sec2_Thal_Bulk
7669_L_sec2_Thal_COUPTF2
7669_L_sec2_Thal_DLX1
7669_L_sec3_Hip_COUPTF2
7669_L_sec3_Hip_DLX1
7669_L_Skin_Bulk
7669_L_T_1_Bulk
7669_L_T_1_COUPTF2
7669_L_T_1_DLX1
7669_L_T_1_TBR1
7669_LA10_Bulk
7669_Liver_Bulk
7669_LV15_Bulk
7669_R_2_Put2_Bulk
7669_R_2_Thal_Bulk
7669_R_Adrenal_Bulk
7669_R_Ant_Cau_DLX1
7669_R_Ant_Put_DLX1
7669_R_Cb_2_Bulk
7669_R_CC_Mid_TBR1
7669_R_CCMid_COUPTF2
7669_R_CCMid_DLX1
7669_R_EC_COUPTF2
7669_R_EC_DLX1
7669_R_EC_TBR1
7669_R_F_1_Bulk
7669_R_F_1_COUPTF2
7669_R_F_1_DLX1
7669_R_F_1_NeuN
7669_R_F_1_TBR1
7669_R_I_1_COUPTF2
7669_R_I_1_DLX1
7669_R_I_1_TBR1
7669_R_Ins_1_Bulk
7669_R_kidney_Bulk
7669_R_mO_COUPTF2
7669_R_mO_DLX1
7669_R_mO_TBR1
7669_R_O_1_Bulk
7669_R_O_1_COUPTF2
7669_R_O_1_DLX1
7669_R_O_1_NeuN
7669_R_O_1_TBR1
7669_R_O_10_DLX1
7669_R_O_11_DLX1
7669_R_O_2_DLX1
7669_R_O_3_DLX1
7669_R_O_4_DLX1
7669_R_O_5_DLX1
7669_R_O_6_DLX1
7669_R_O_7_DLX1
7669_R_O_8_DLX1
7669_R_O_9_DLX1
7669_R_Olf_DLX1
7669_R_P_1_Bulk
7669_R_P_1_COUPTF2
7669_R_P_1_DLX1
7669_R_P_1_NeuN
7669_R_P_1_TBR1
7669_R_P_10_DLX1
7669_R_P_11_DLX1
7669_R_P_11_TBR1
7669_R_P_12_DLX1
7669_R_P_12_TBR1
7669_R_P_13_DLX1
7669_R_P_13_TBR1
7669_R_P_14_DLX1
7669_R_P_14_TBR1
7669_R_P_15_DLX1
7669_R_P_15_TBR1
7669_R_P_16_DLX1
7669_R_P_16_TBR1
7669_R_P_17_DLX1
7669_R_P_17_TBR1
7669_R_P_2_DLX1
7669_R_P_2_TBR1
7669_R_P_3_DLX1
7669_R_P_3_TBR1
7669_R_P_4_DLX1
7669_R_P_4_TBR1
7669_R_P_5_DLX1
7669_R_P_5_TBR1
7669_R_P_6_DLX1
7669_R_P_6_TBR1
7669_R_P_7_DLX1
7669_R_P_7_TBR1
7669_R_P_8_DLX1
7669_R_P_8_TBR1
7669_R_P_9_DLX
7669_R_P_9_DLX1
7669_R_P_9_TBR1
7669_R_POA_Bulk
7669_R_POA_DLX1
7669_R_sec1_Cau_Bulk
7669_R_sec2_GP_Bulk
7669_R_sec2_GP_DLX1
7669_R_sec2_Put_Bulk
7669_R_sec2_Thal_Bulk
7669_R_sec2_Thal_COUPTF2
7669_R_sec2_Thal_DLX1
7669_R_sec3_Cau_DLX1
7669_R_sec3_Hip_COUPTF2
7669_R_sec3_Hip_DLX1
7669_R_sec3_Hip_TBR1
7669_R_sec3_Put_DLX1
7669_R_Skin_Bulk
7669_R_T_1_Bulk
7669_R_T_1_COUPTF2
7669_R_T_1_DLX1
7669_R_T_1_TBR1
7669_RA9_Bulk
7669_RV10_Bulk
JGG

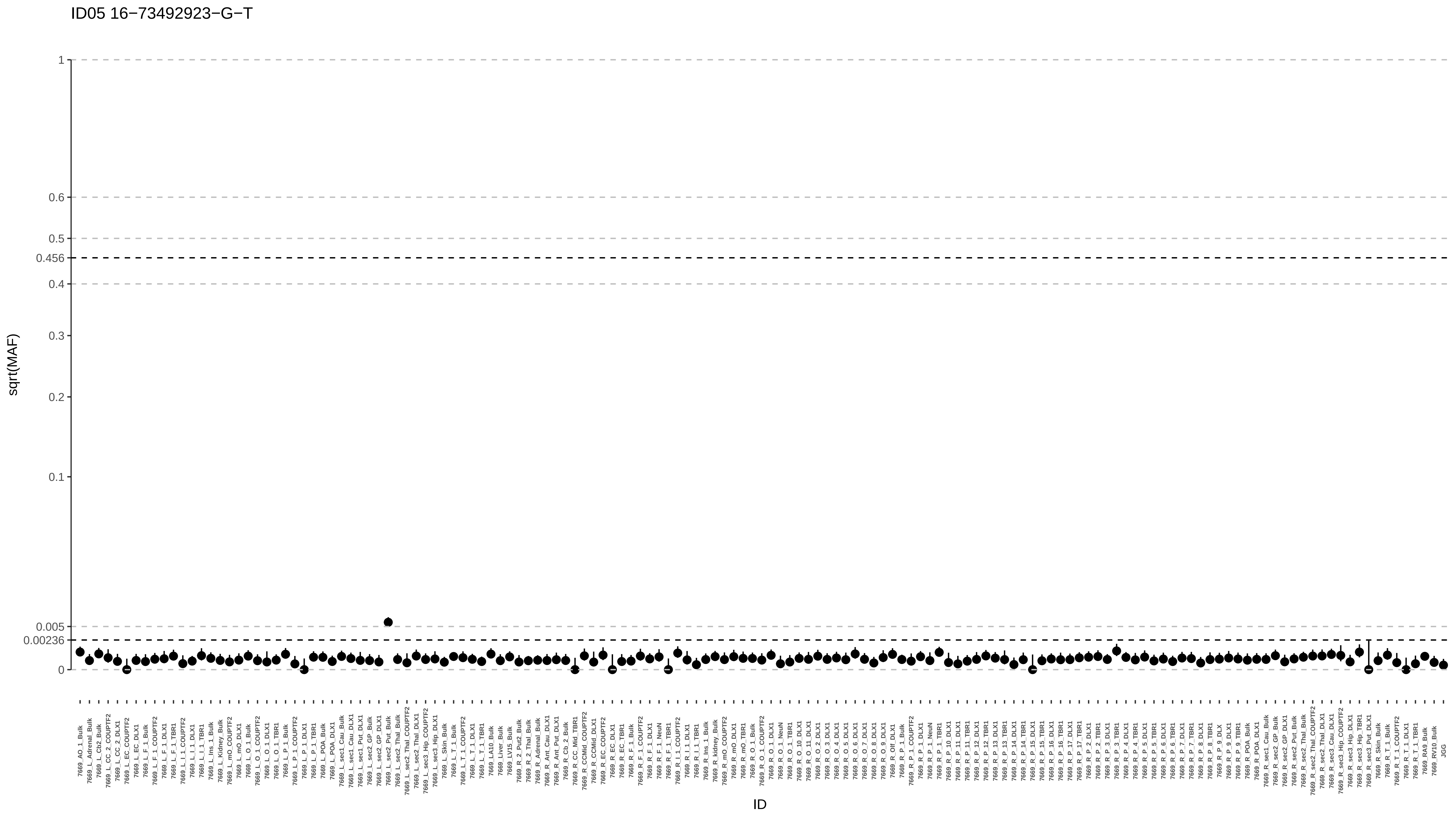
ID05 16−73492923−G−T
sqrt(MAF)
1
0.6
0.5
0.456
0.4
0.3
0.2
0.1
0.005
0.00236
0
ID

ID05 16−77437520−G−T

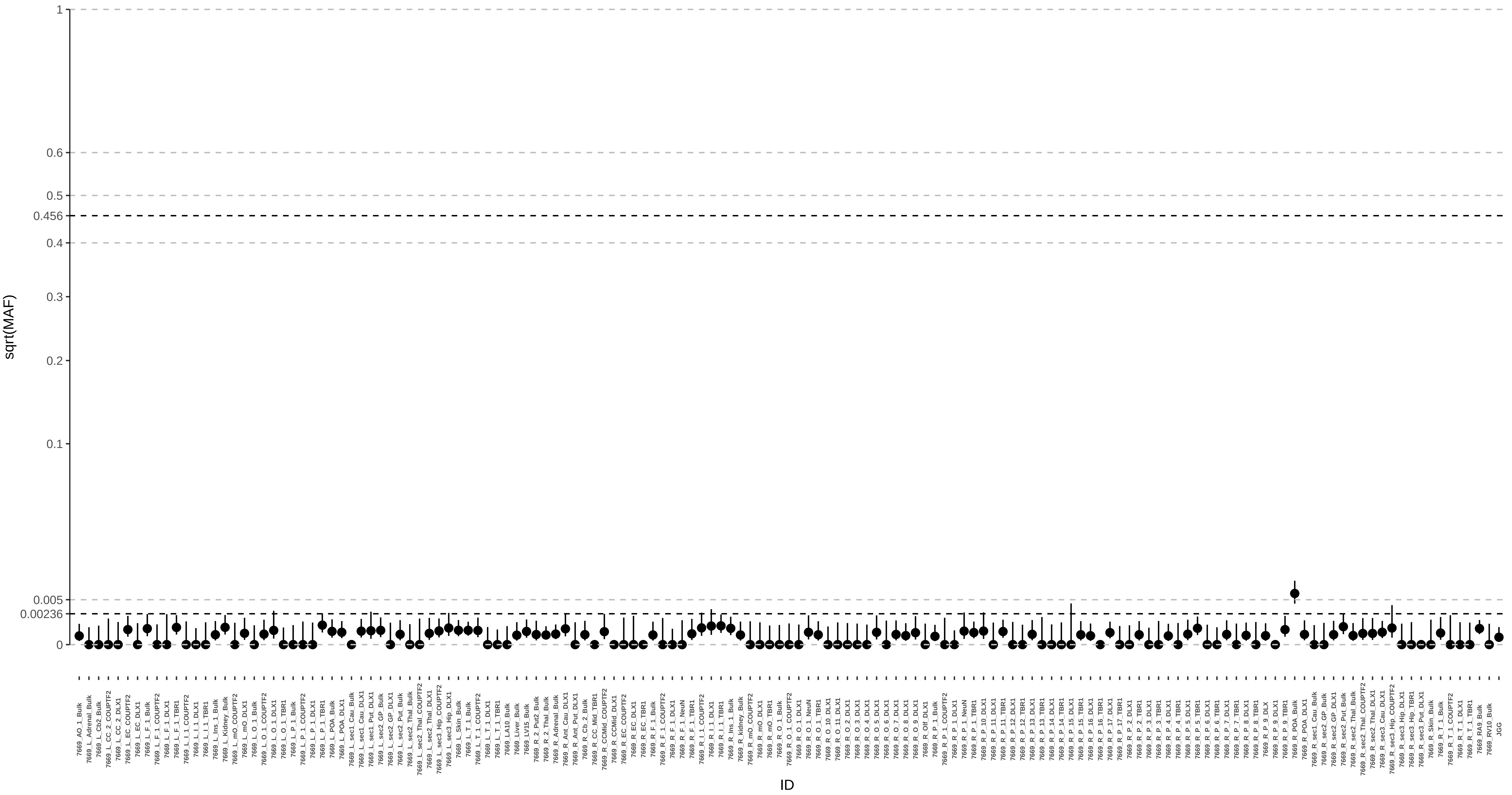

ID05 16−83153418−C−T

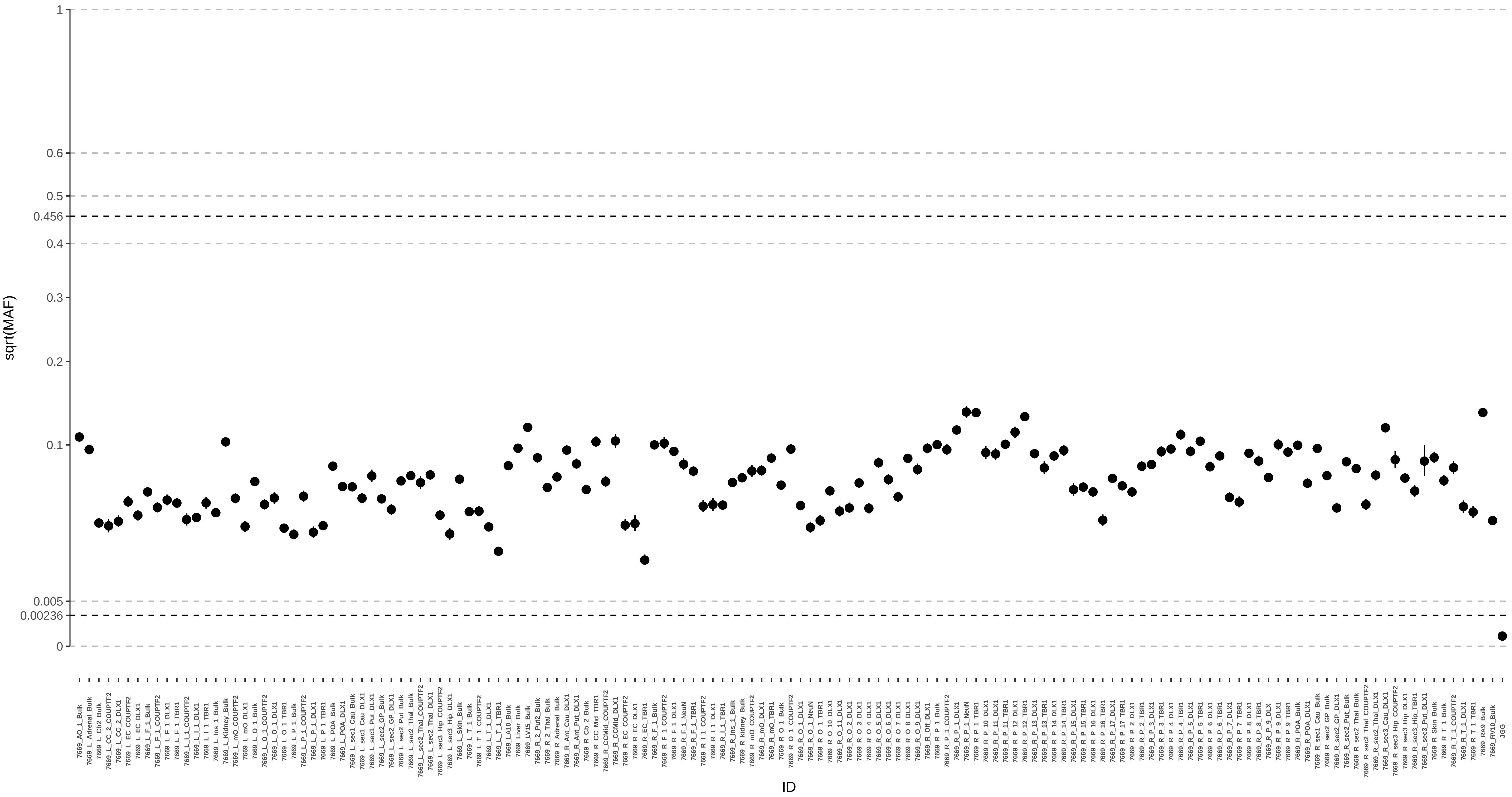

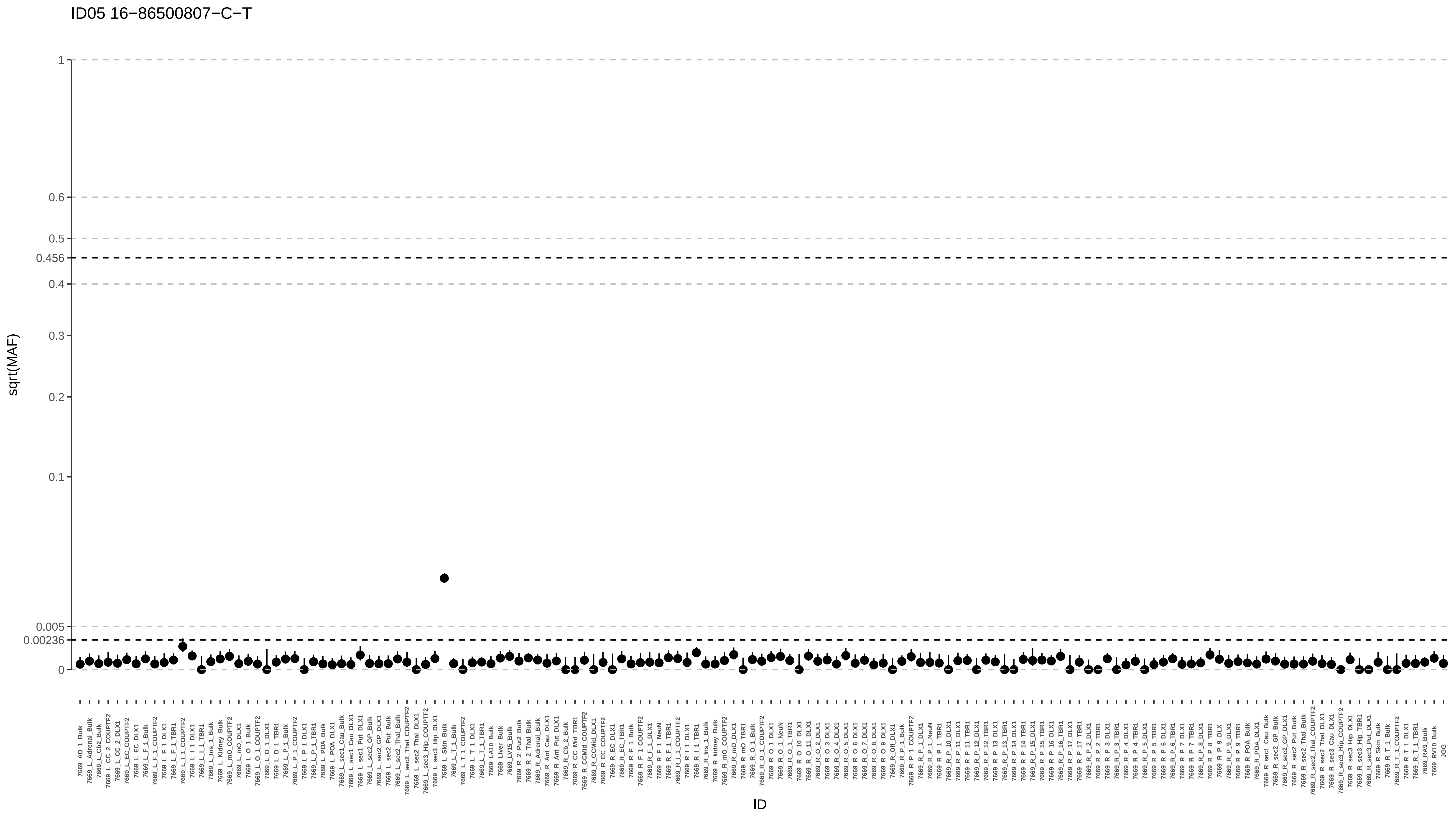
ID05 16−86500807−C−T
sqrt(MAF)
1
0.6
0.5
0.456
0.4
0.3
0.2
0.1
0.005
0.00236
0
ID

ID05 16−8802550−C−T

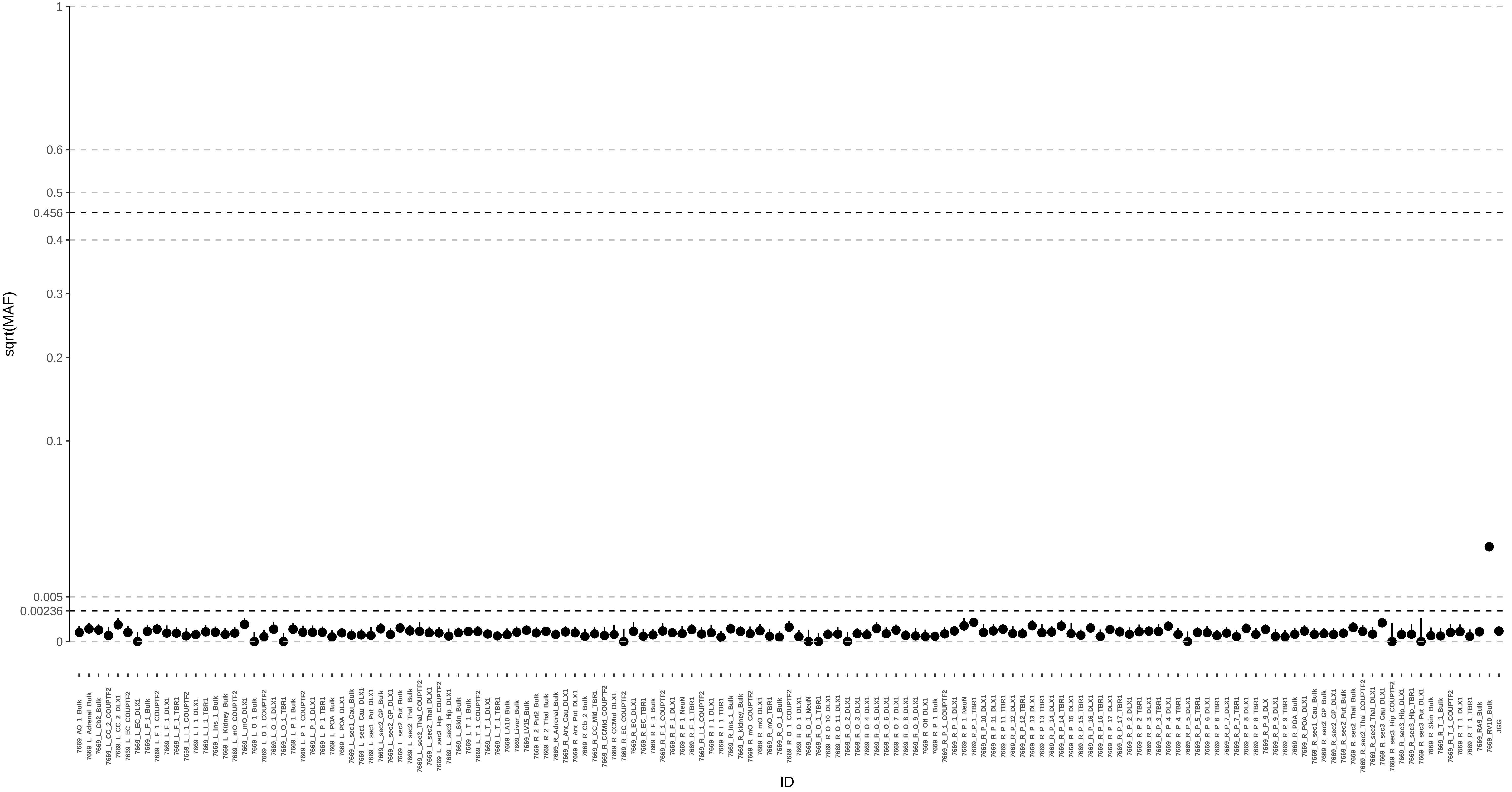

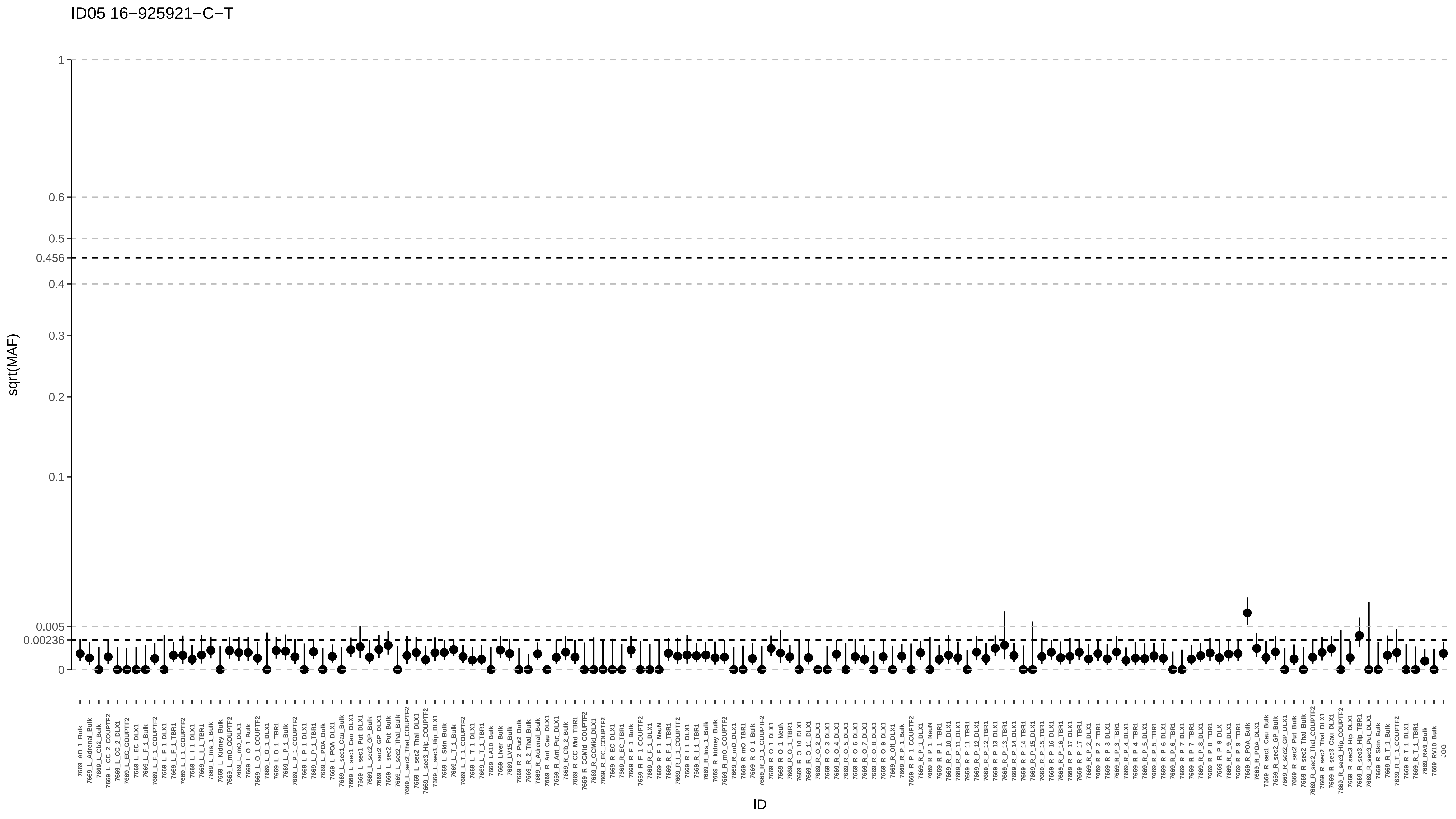

ID05 16−925921−C−T
sqrt(MAF)
1
0.6
0.5
0.456
0.4
0.3
0.2
0.1
0.005
0.00236
0
ID

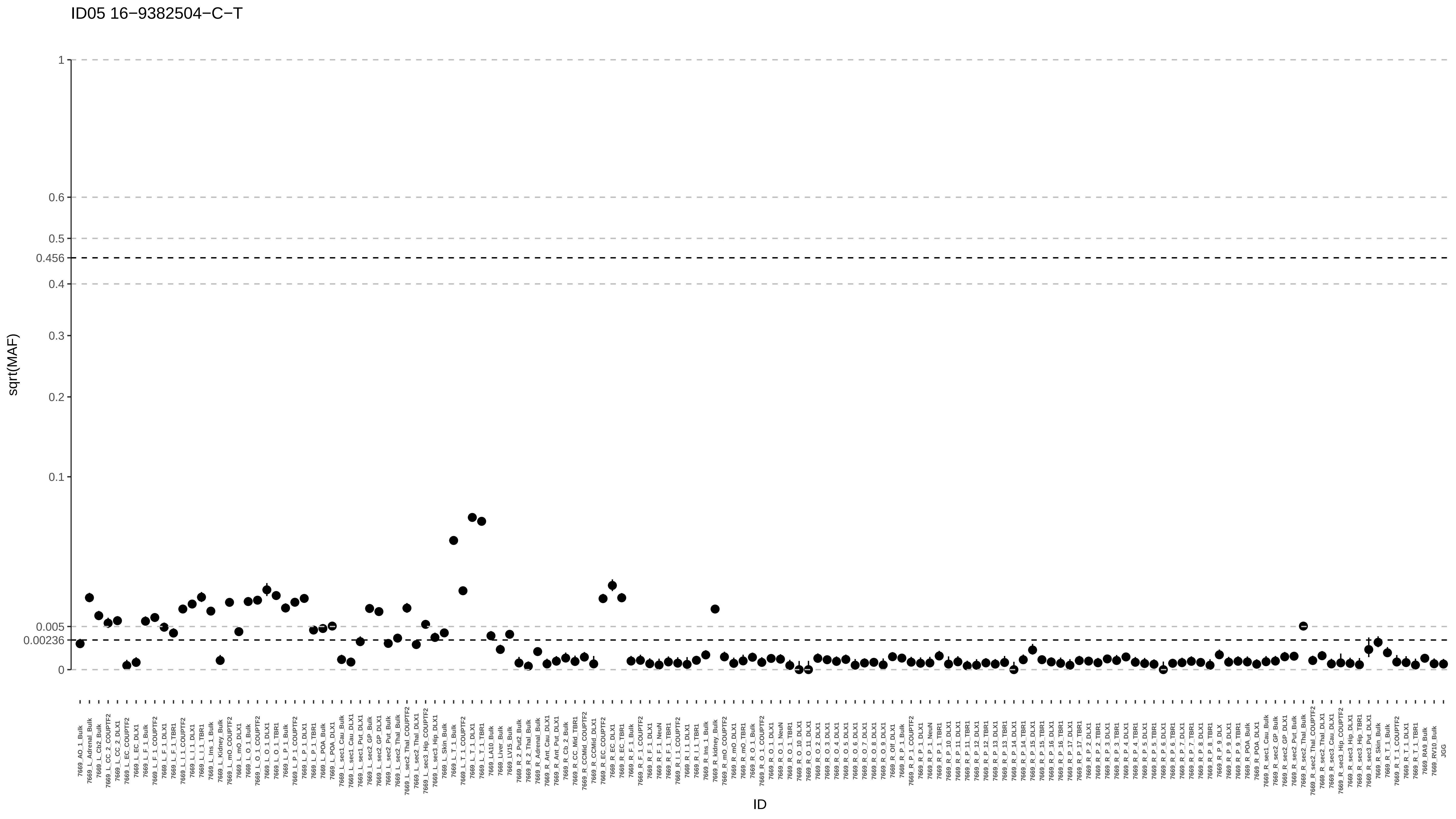
ID05 16−9382504−C−T
sqrt(MAF)
ID
1
0.6
0.5
0.456
0.4
0.3
0.2
0.1
0.005
0.00236
0

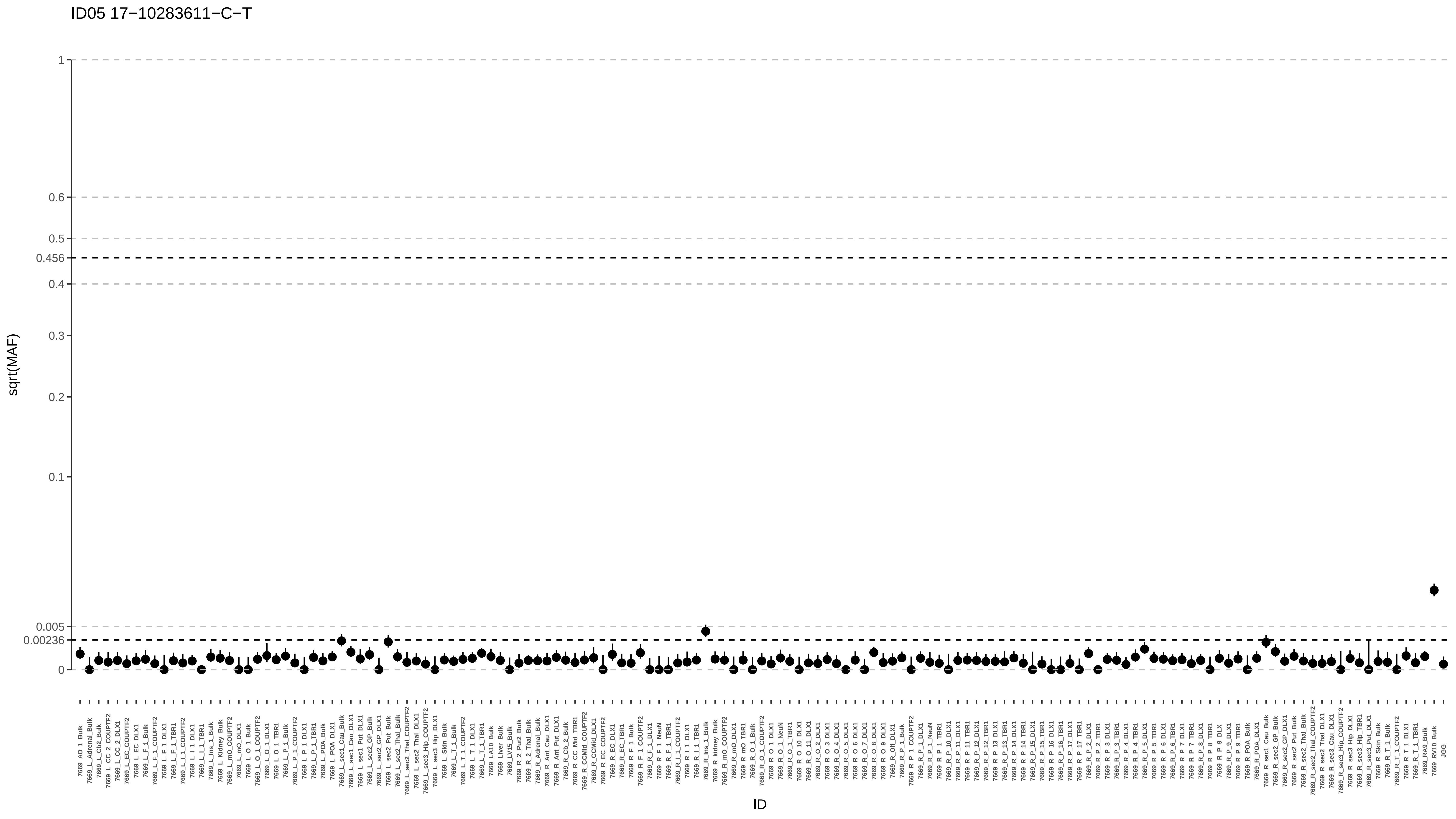
ID05 17−10283611−C−T
sqrt(MAF)
ID

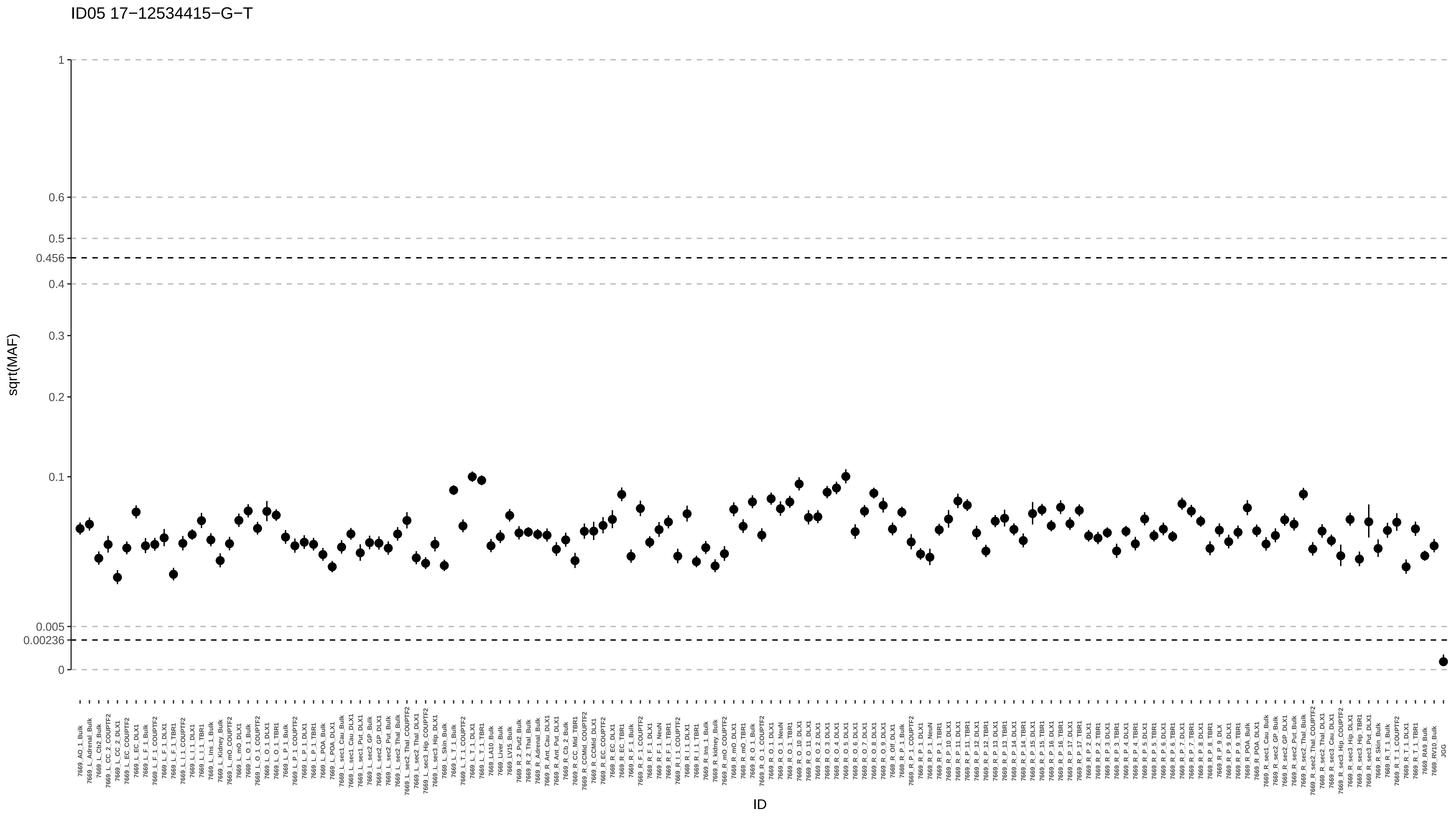
ID05 17−12534415−G−T
sqrt(MAF)
ID
1
0.6
0.5
0.456
0.4
0.3
0.2
0.1
0.005
0.00236
0

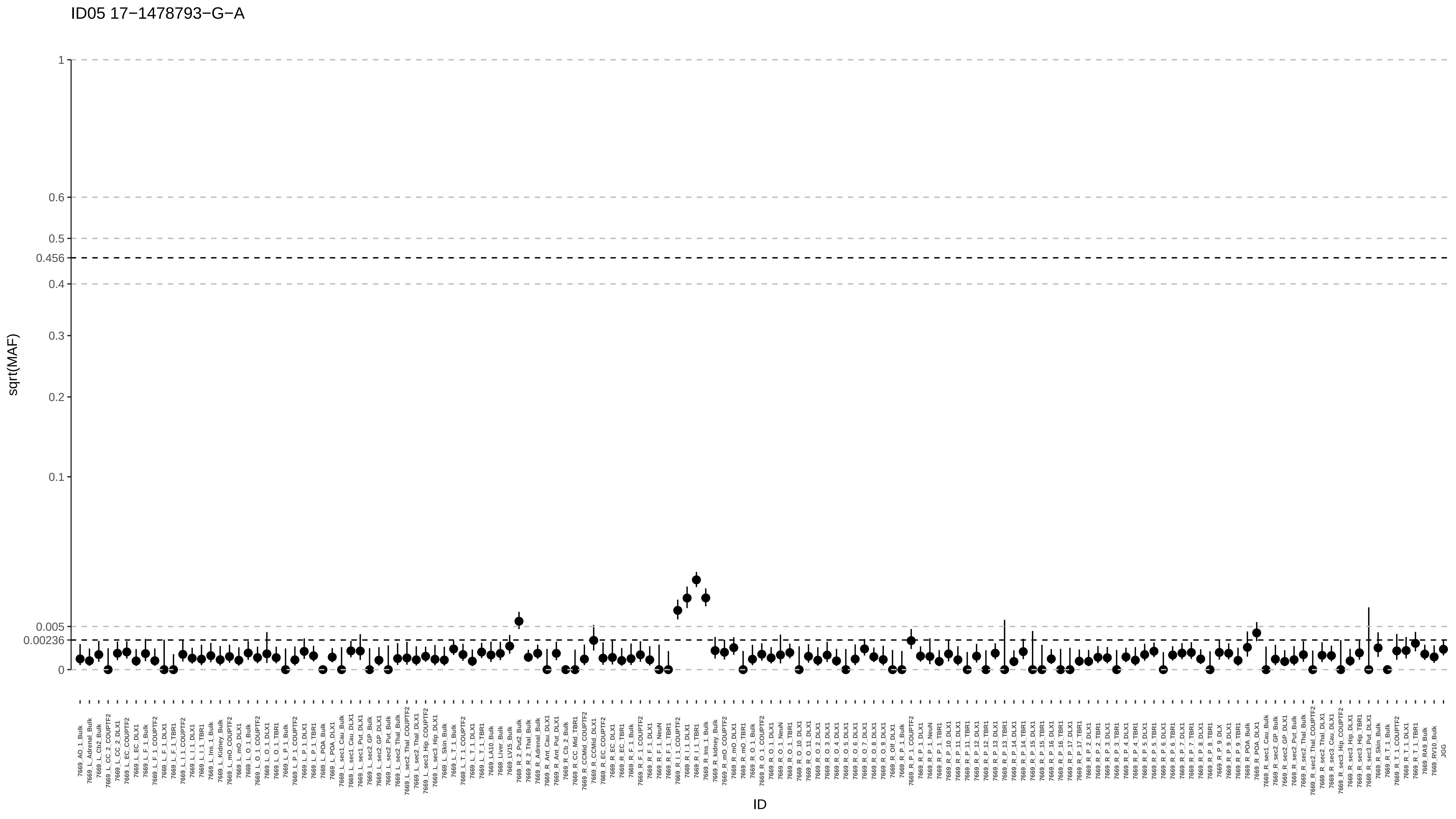
ID05 17−1478793−G−A
sqrt(MAF)
1
0.6
0.5
0.456
0.4
0.3
0.2
0.1
0.005
0.00236
0
ID
7669_AO_1_Bulk
7669_L_Adrenal_Bulk
7669_L_Cb2_Bulk
7669_L_CC_2_COUPTF2
7669_L_CC_2_DLX1
7669_L_EC_COUPTF2
7669_L_EC_DLX1
7669_L_F_1_Bulk
7669_L_F_1_COUPTF2
7669_L_F_1_DLX1
7669_L_F_1_TBR1
7669_L_I_1_COUPTF2
7669_L_I_1_DLX1
7669_L_I_1_TBR1
7669_L_Ins_1_Bulk
7669_L_Kidney_Bulk
7669_L_mO_COUPTF2
7669_L_mO_DLX1
7669_L_O_1_Bulk
7669_L_O_1_COUPTF2
7669_L_O_1_DLX1
7669_L_O_1_TBR1
7669_L_P_1_Bulk
7669_L_P_1_COUPTF2
7669_L_P_1_DLX1
7669_L_P_1_TBR1
7669_L_POA_Bulk
7669_L_POA_DLX1
7669_L_sec1_Cau_Bulk
7669_L_sec1_Cau_DLX1
7669_L_sec1_Put_DLX1
7669_L_sec2_GP_Bulk
7669_L_sec2_GP_DLX1
7669_L_sec2_Put_Bulk
7669_L_sec2_Thal_Bulk
7669_L_sec2_Thal_COUPTF2
7669_L_sec2_Thal_DLX1
7669_L_sec3_Hip_COUPTF2
7669_L_sec3_Hip_DLX1
7669_L_Skin_Bulk
7669_L_T_1_Bulk
7669_L_T_1_COUPTF2
7669_L_T_1_DLX1
7669_L_T_1_TBR1
7669_LA10_Bulk
7669_Liver_Bulk
7669_LV15_Bulk
7669_R_2_Put2_Bulk
7669_R_2_Thal_Bulk
7669_R_Adrenal_Bulk
7669_R_Ant_Cau_DLX1
7669_R_Ant_Put_DLX1
7669_R_Cb_2_Bulk
7669_R_CC_Mid_TBR1
7669_R_CCMid_COUPTF2
7669_R_CCMid_DLX1
7669_R_EC_COUPTF2
7669_R_EC_DLX1
7669_R_EC_TBR1
7669_R_F_1_Bulk
7669_R_F_1_COUPTF2
7669_R_F_1_DLX1
7669_R_F_1_NeuN
7669_R_F_1_TBR1
7669_R_I_1_COUPTF2
7669_R_I_1_DLX1
7669_R_I_1_TBR1
7669_R_Ins_1_Bulk
7669_R_kidney_Bulk
7669_R_mO_COUPTF2
7669_R_mO_DLX1
7669_R_mO_TBR1
7669_R_O_1_Bulk
7669_R_O_1_COUPTF2
7669_R_O_1_DLX1
7669_R_O_1_NeuN
7669_R_O_1_TBR1
7669_R_O_10_DLX1
7669_R_O_11_DLX1
7669_R_O_2_DLX1
7669_R_O_3_DLX1
7669_R_O_4_DLX1
7669_R_O_5_DLX1
7669_R_O_6_DLX1
7669_R_O_7_DLX1
7669_R_O_8_DLX1
7669_R_O_9_DLX1
7669_R_Olf_DLX1
7669_R_P_1_Bulk
7669_R_P_1_COUPTF2
7669_R_P_1_DLX1
7669_R_P_1_NeuN
7669_R_P_1_TBR1
7669_R_P_10_DLX1
7669_R_P_11_DLX1
7669_R_P_11_TBR1
7669_R_P_12_DLX1
7669_R_P_12_TBR1
7669_R_P_13_DLX1
7669_R_P_13_TBR1
7669_R_P_14_DLX1
7669_R_P_14_TBR1
7669_R_P_15_DLX1
7669_R_P_15_TBR1
7669_R_P_16_DLX1
7669_R_P_16_TBR1
7669_R_P_17_DLX1
7669_R_P_17_TBR1
7669_R_P_2_DLX1
7669_R_P_2_TBR1
7669_R_P_3_DLX1
7669_R_P_3_TBR1
7669_R_P_4_DLX1
7669_R_P_4_TBR1
7669_R_P_5_DLX1
7669_R_P_5_TBR1
7669_R_P_6_DLX1
7669_R_P_6_TBR1
7669_R_P_7_DLX1
7669_R_P_7_TBR1
7669_R_P_8_DLX1
7669_R_P_8_TBR1
7669_R_P_9_DLX
7669_R_P_9_DLX1
7669_R_P_9_TBR1
7669_R_POA_Bulk
7669_R_POA_DLX1
7669_R_sec1_Cau_Bulk
7669_R_sec2_GP_Bulk
7669_R_sec2_GP_DLX1
7669_R_sec2_Put_Bulk
7669_R_sec2_Thal_Bulk
7669_R_sec2_Thal_COUPTF2
7669_R_sec2_Thal_DLX1
7669_R_sec3_Cau_DLX1
7669_R_sec3_Hip_COUPTF2
7669_R_sec3_Hip_DLX1
7669_R_sec3_Hip_TBR1
7669_R_sec3_Put_DLX1
7669_R_Skin_Bulk
7669_R_T_1_Bulk
7669_R_T_1_COUPTF2
7669_R_T_1_DLX1
7669_R_T_1_TBR1
7669_RA9_Bulk
7669_RV10_Bulk
JGG

ID05 17−1620800−G−T

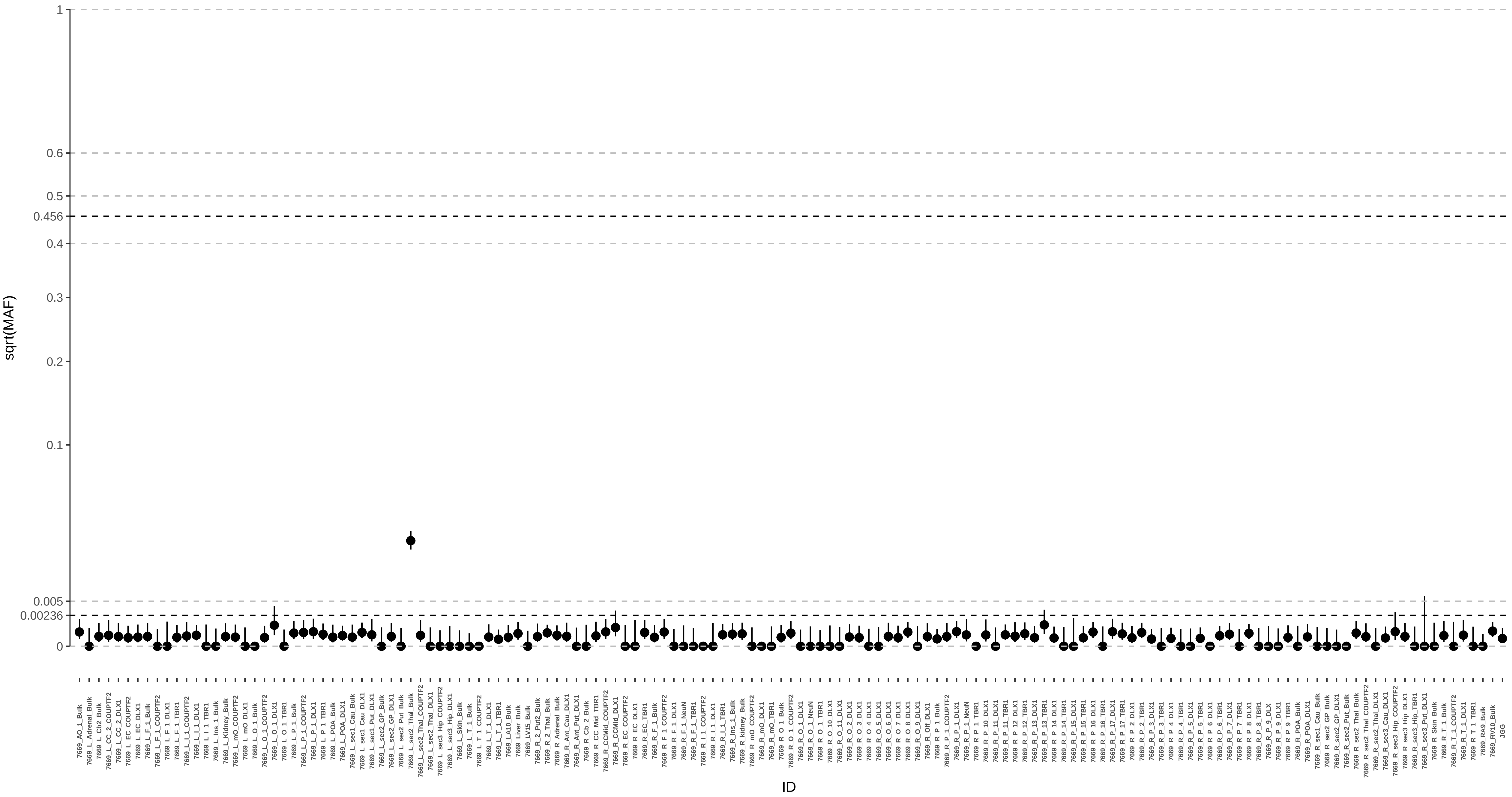

ID05 17−17411221−G−A

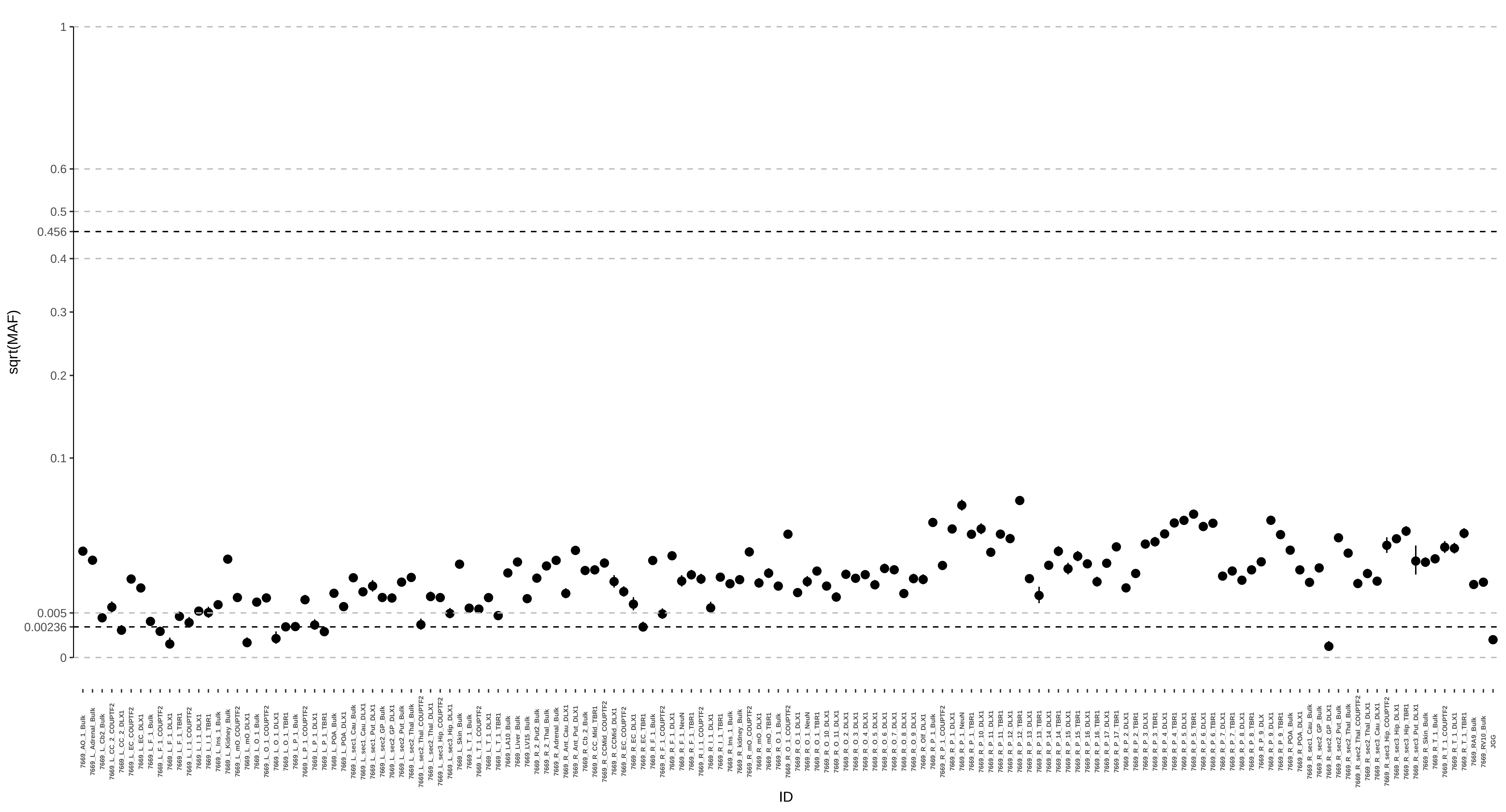

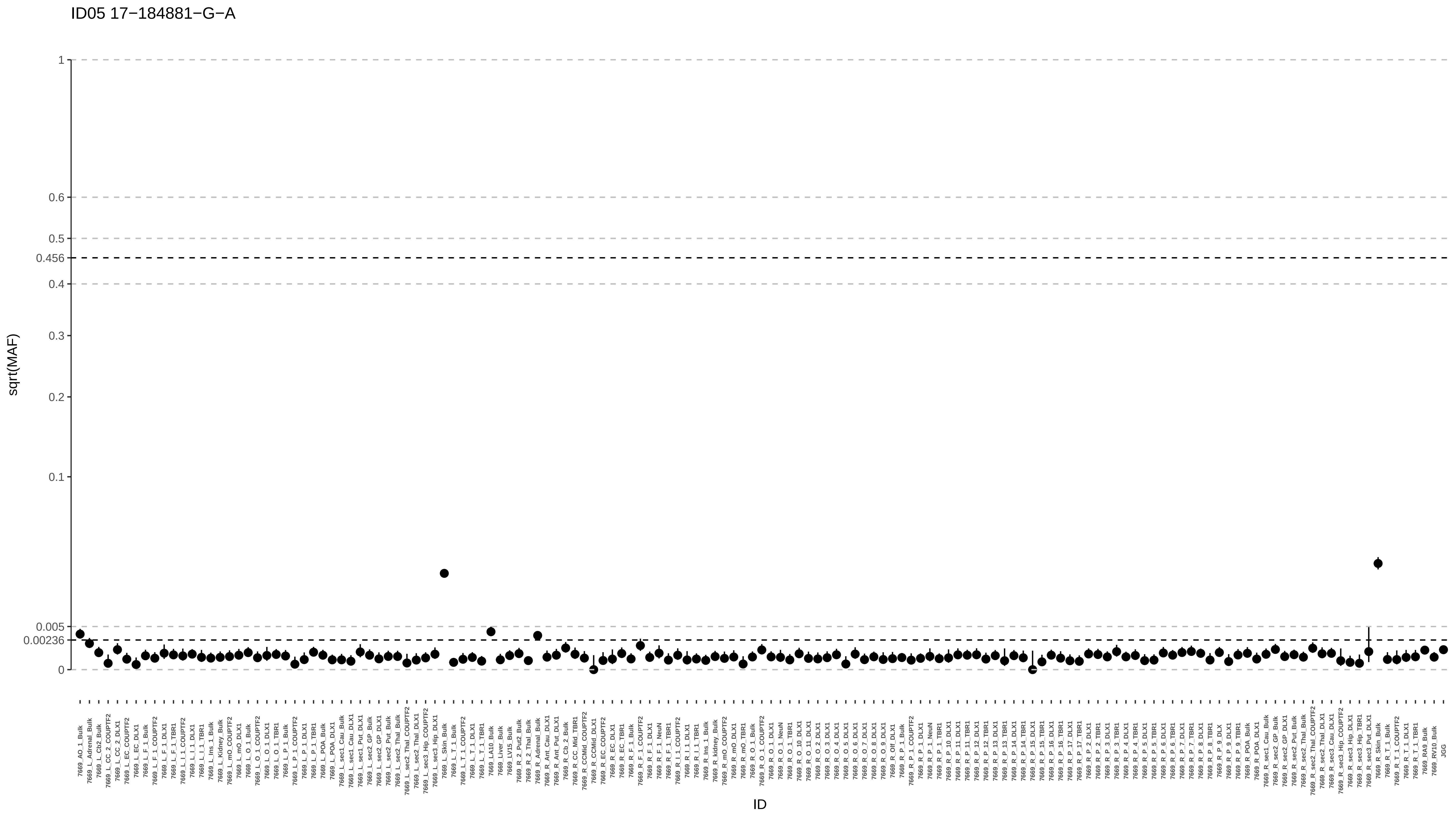
ID05 17−184881−G−A
sqrt(MAF)
1
0.6
0.5
0.456
0.4
0.3
0.2
0.1
0.005
0.00236
0
ID

ID05 17−20005504−G−A

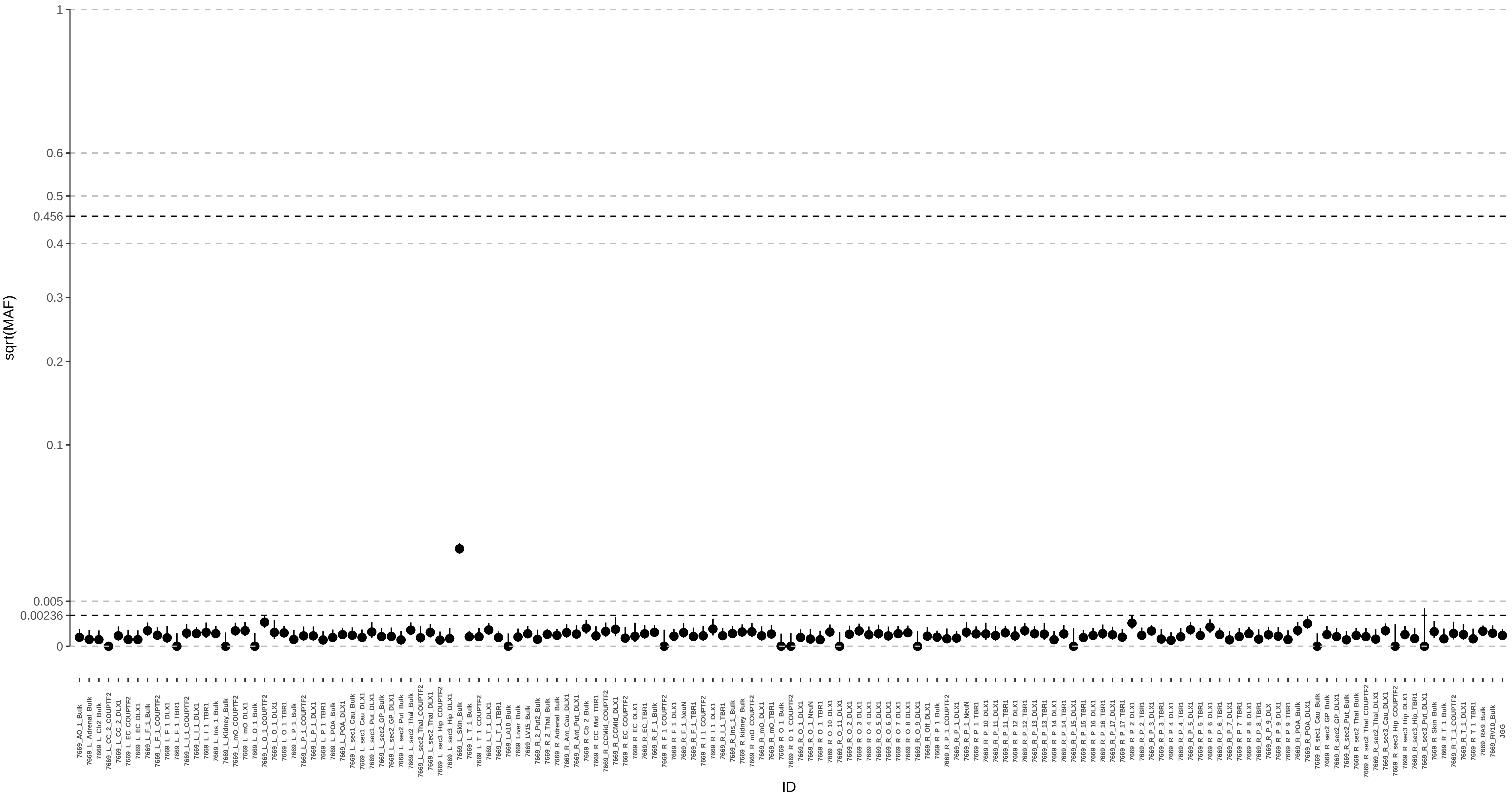

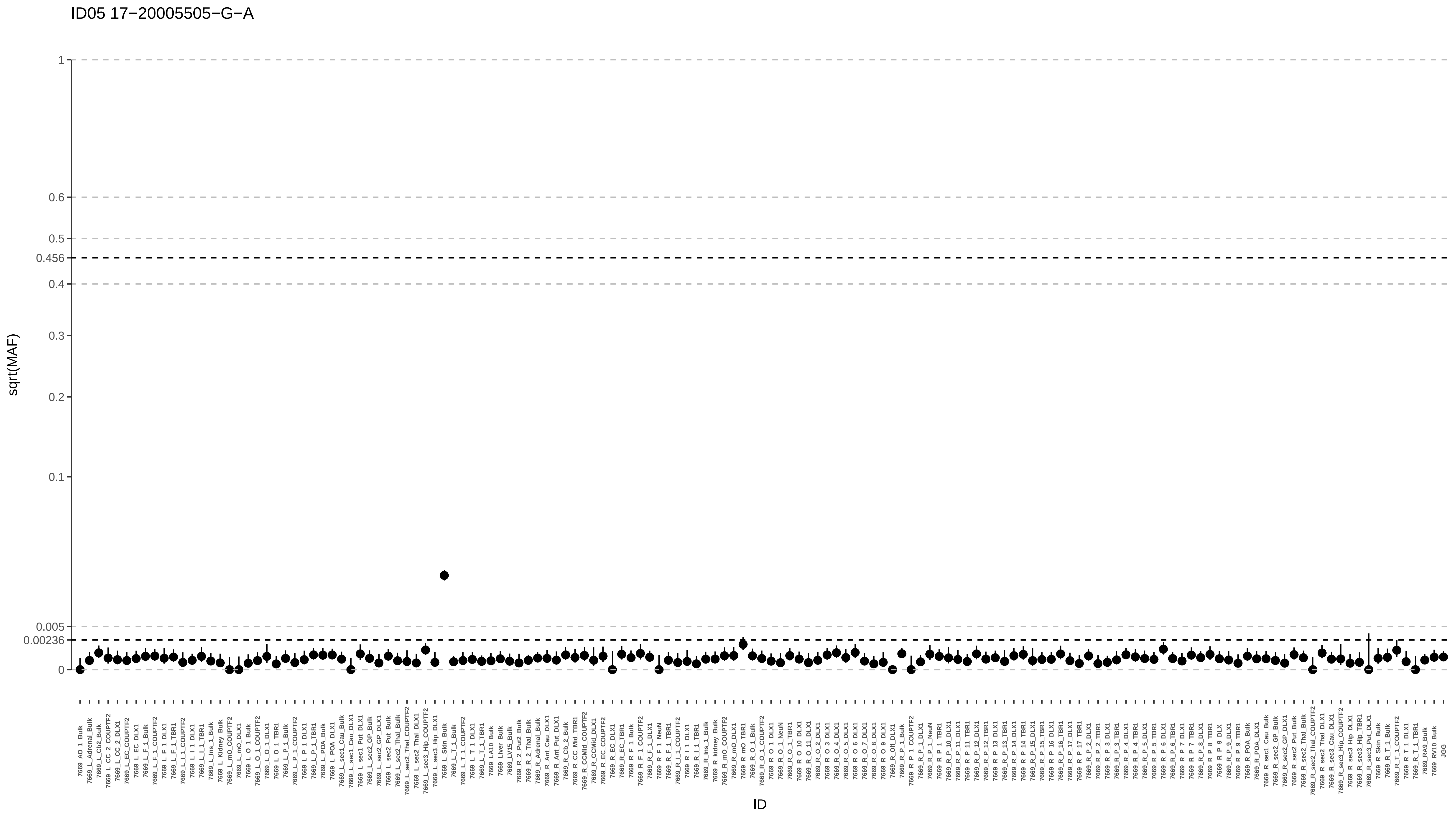
ID05 17−20005505−G−A
sqrt(MAF)
1
0.6
0.5
0.456
0.4
0.3
0.2
0.1
0.005
0.00236
0
ID

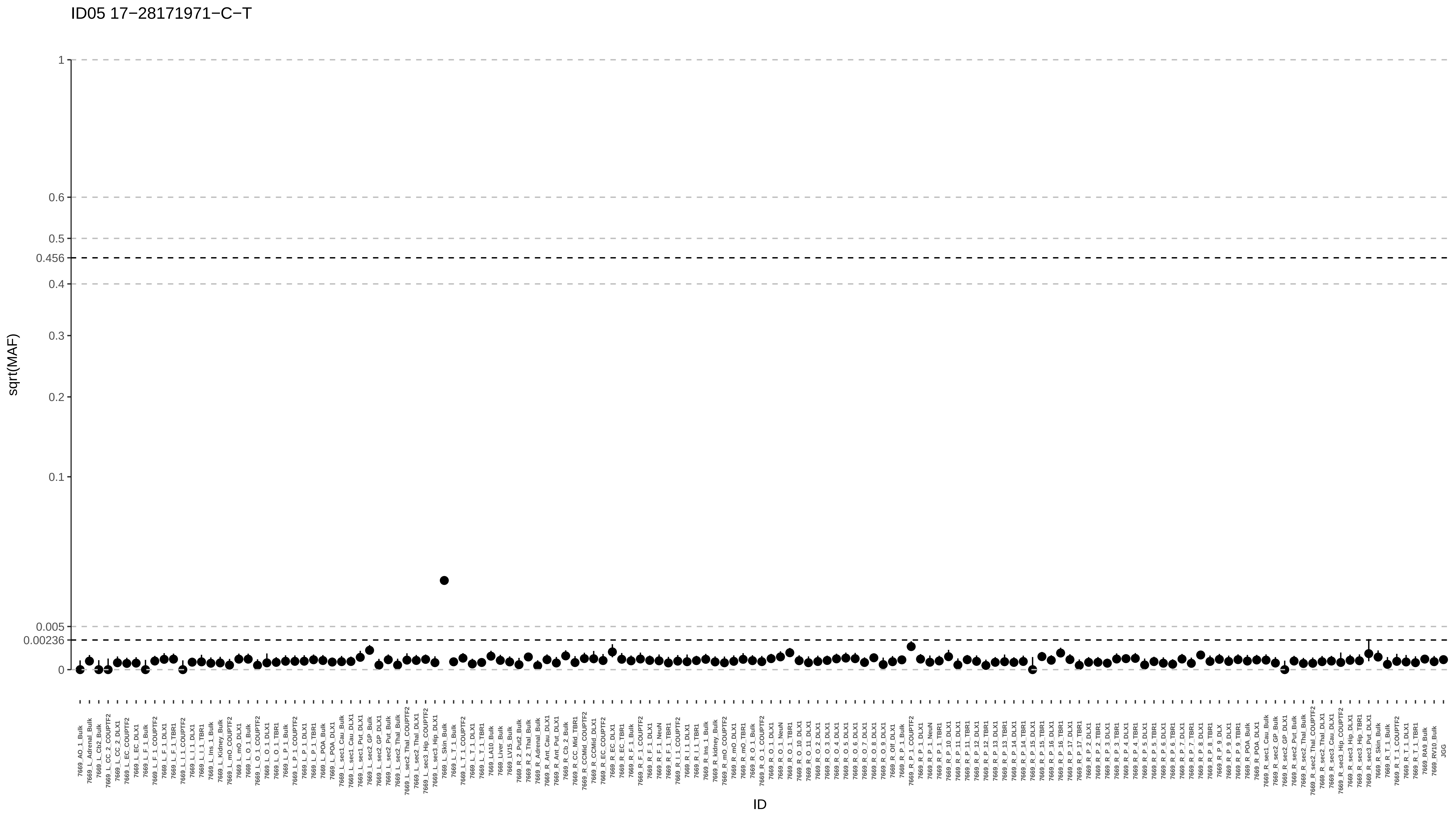
ID05 17−28171971−C−T
sqrt(MAF)
1
0.6
0.5
0.456
0.4
0.3
0.2
0.1
0.005
0.00236
0
ID

ID05 17−40192535−C−G

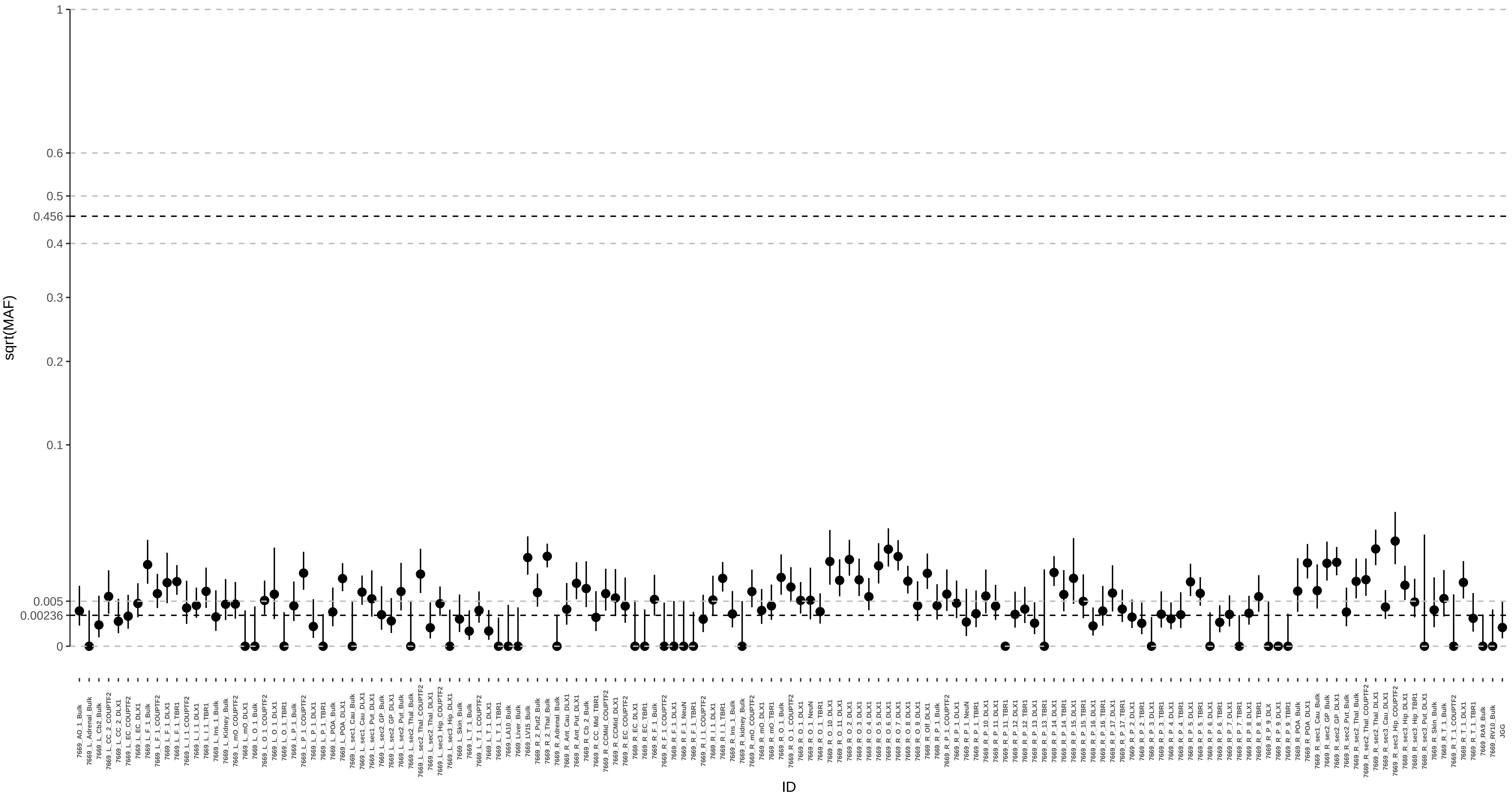

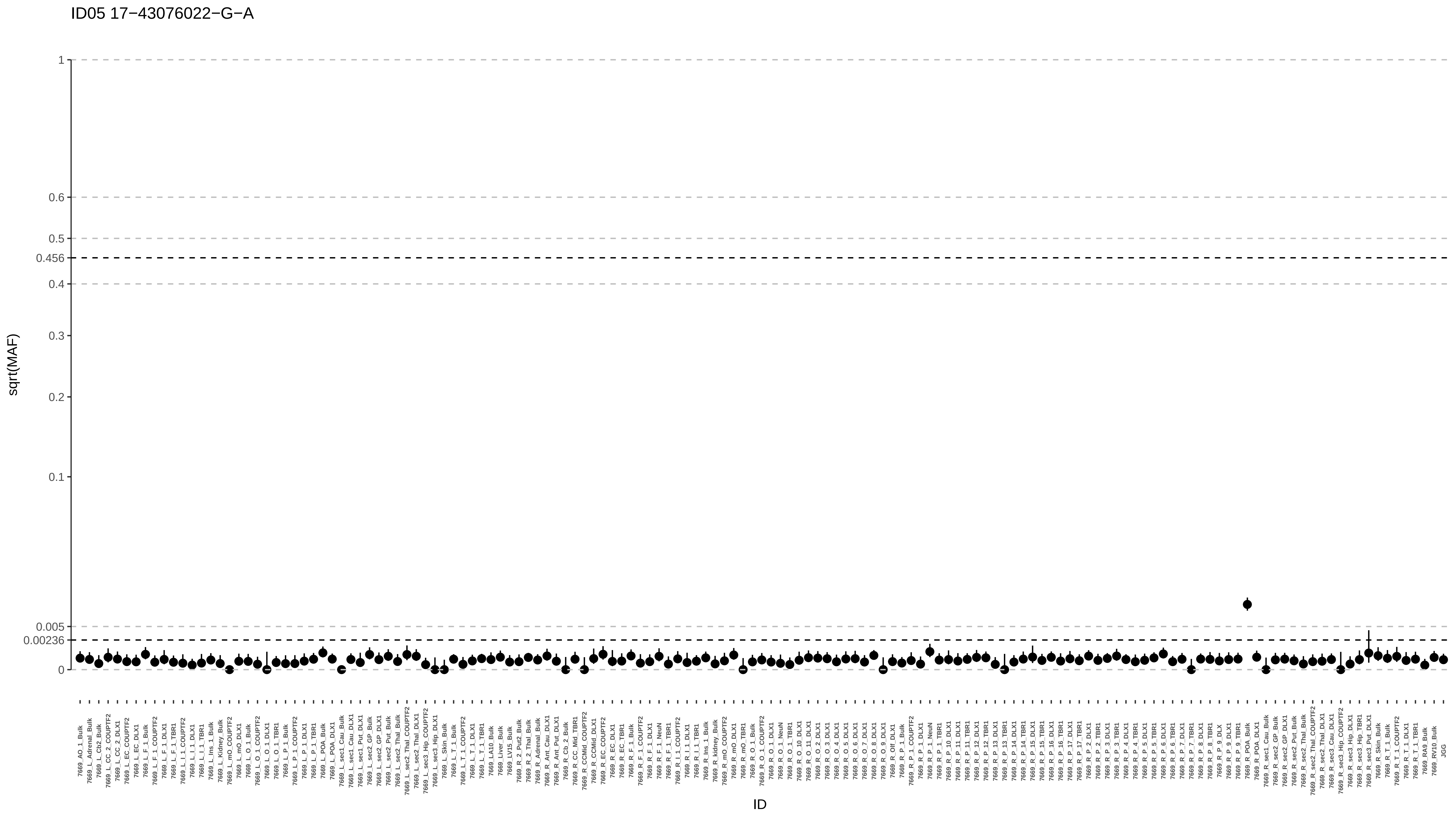
ID05 17−43076022−G−A
sqrt(MAF)
1
0.6
0.5
0.456
0.4
0.3
0.2
0.1
0.005
0.00236
0
ID
7669_AO_1_Bulk
7669_L_Adrenal_Bulk
7669_L_Cb2_Bulk
7669_L_CC_2_COUPTF2
7669_L_CC_2_DLX1
7669_L_EC_COUPTF2
7669_L_EC_DLX1
7669_L_F_1_Bulk
7669_L_F_1_COUPTF2
7669_L_F_1_DLX1
7669_L_F_1_TBR1
7669_L_I_1_COUPTF2
7669_L_I_1_DLX1
7669_L_I_1_TBR1
7669_L_Ins_1_Bulk
7669_L_Kidney_Bulk
7669_L_mO_COUPTF2
7669_L_mO_DLX1
7669_L_O_1_Bulk
7669_L_O_1_COUPTF2
7669_L_O_1_DLX1
7669_L_O_1_TBR1
7669_L_P_1_Bulk
7669_L_P_1_COUPTF2
7669_L_P_1_DLX1
7669_L_P_1_TBR1
7669_L_POA_Bulk
7669_L_POA_DLX1
7669_L_sec1_Cau_Bulk
7669_L_sec1_Cau_DLX1
7669_L_sec1_Put_DLX1
7669_L_sec2_GP_Bulk
7669_L_sec2_GP_DLX1
7669_L_sec2_Put_Bulk
7669_L_sec2_Thal_Bulk
7669_L_sec2_Thal_COUPTF2
7669_L_sec2_Thal_DLX1
7669_L_sec3_Hip_COUPTF2
7669_L_sec3_Hip_DLX1
7669_L_Skin_Bulk
7669_L_T_1_Bulk
7669_L_T_1_COUPTF2
7669_L_T_1_DLX1
7669_L_T_1_TBR1
7669_LA10_Bulk
7669_Liver_Bulk
7669_LV15_Bulk
7669_R_2_Put2_Bulk
7669_R_2_Thal_Bulk
7669_R_Adrenal_Bulk
7669_R_Ant_Cau_DLX1
7669_R_Ant_Put_DLX1
7669_R_Cb_2_Bulk
7669_R_CC_Mid_TBR1
7669_R_CCMid_COUPTF2
7669_R_CCMid_DLX1
7669_R_EC_COUPTF2
7669_R_EC_DLX1
7669_R_EC_TBR1
7669_R_F_1_Bulk
7669_R_F_1_COUPTF2
7669_R_F_1_DLX1
7669_R_F_1_NeuN
7669_R_F_1_TBR1
7669_R_I_1_COUPTF2
7669_R_I_1_DLX1
7669_R_I_1_TBR1
7669_R_Ins_1_Bulk
7669_R_kidney_Bulk
7669_R_mO_COUPTF2
7669_R_mO_DLX1
7669_R_mO_TBR1
7669_R_O_1_Bulk
7669_R_O_1_COUPTF2
7669_R_O_1_DLX1
7669_R_O_1_NeuN
7669_R_O_1_TBR1
7669_R_O_10_DLX1
7669_R_O_11_DLX1
7669_R_O_2_DLX1
7669_R_O_3_DLX1
7669_R_O_4_DLX1
7669_R_O_5_DLX1
7669_R_O_6_DLX1
7669_R_O_7_DLX1
7669_R_O_8_DLX1
7669_R_O_9_DLX1
7669_R_Olf_DLX1
7669_R_P_1_Bulk
7669_R_P_1_COUPTF2
7669_R_P_1_DLX1
7669_R_P_1_NeuN
7669_R_P_1_TBR1
7669_R_P_10_DLX1
7669_R_P_11_DLX1
7669_R_P_11_TBR1
7669_R_P_12_DLX1
7669_R_P_12_TBR1
7669_R_P_13_DLX1
7669_R_P_13_TBR1
7669_R_P_14_DLX1
7669_R_P_14_TBR1
7669_R_P_15_DLX1
7669_R_P_15_TBR1
7669_R_P_16_DLX1
7669_R_P_16_TBR1
7669_R_P_17_DLX1
7669_R_P_17_TBR1
7669_R_P_2_DLX1
7669_R_P_2_TBR1
7669_R_P_3_DLX1
7669_R_P_3_TBR1
7669_R_P_4_DLX1
7669_R_P_4_TBR1
7669_R_P_5_DLX1
7669_R_P_5_TBR1
7669_R_P_6_DLX1
7669_R_P_6_TBR1
7669_R_P_7_DLX1
7669_R_P_7_TBR1
7669_R_P_8_DLX1
7669_R_P_8_TBR1
7669_R_P_9_DLX
7669_R_P_9_DLX1
7669_R_P_9_TBR1
7669_R_POA_Bulk
7669_R_POA_DLX1
7669_R_sec1_Cau_Bulk
7669_R_sec2_GP_Bulk
7669_R_sec2_GP_DLX1
7669_R_sec2_Put_Bulk
7669_R_sec2_Thal_Bulk
7669_R_sec2_Thal_COUPTF2
7669_R_sec2_Thal_DLX1
7669_R_sec3_Cau_DLX1
7669_R_sec3_Hip_COUPTF2
7669_R_sec3_Hip_DLX1
7669_R_sec3_Hip_TBR1
7669_R_sec3_Put_DLX1
7669_R_Skin_Bulk
7669_R_T_1_Bulk
7669_R_T_1_COUPTF2
7669_R_T_1_DLX1
7669_R_T_1_TBR1
7669_RA9_Bulk
7669_RV10_Bulk
JGG

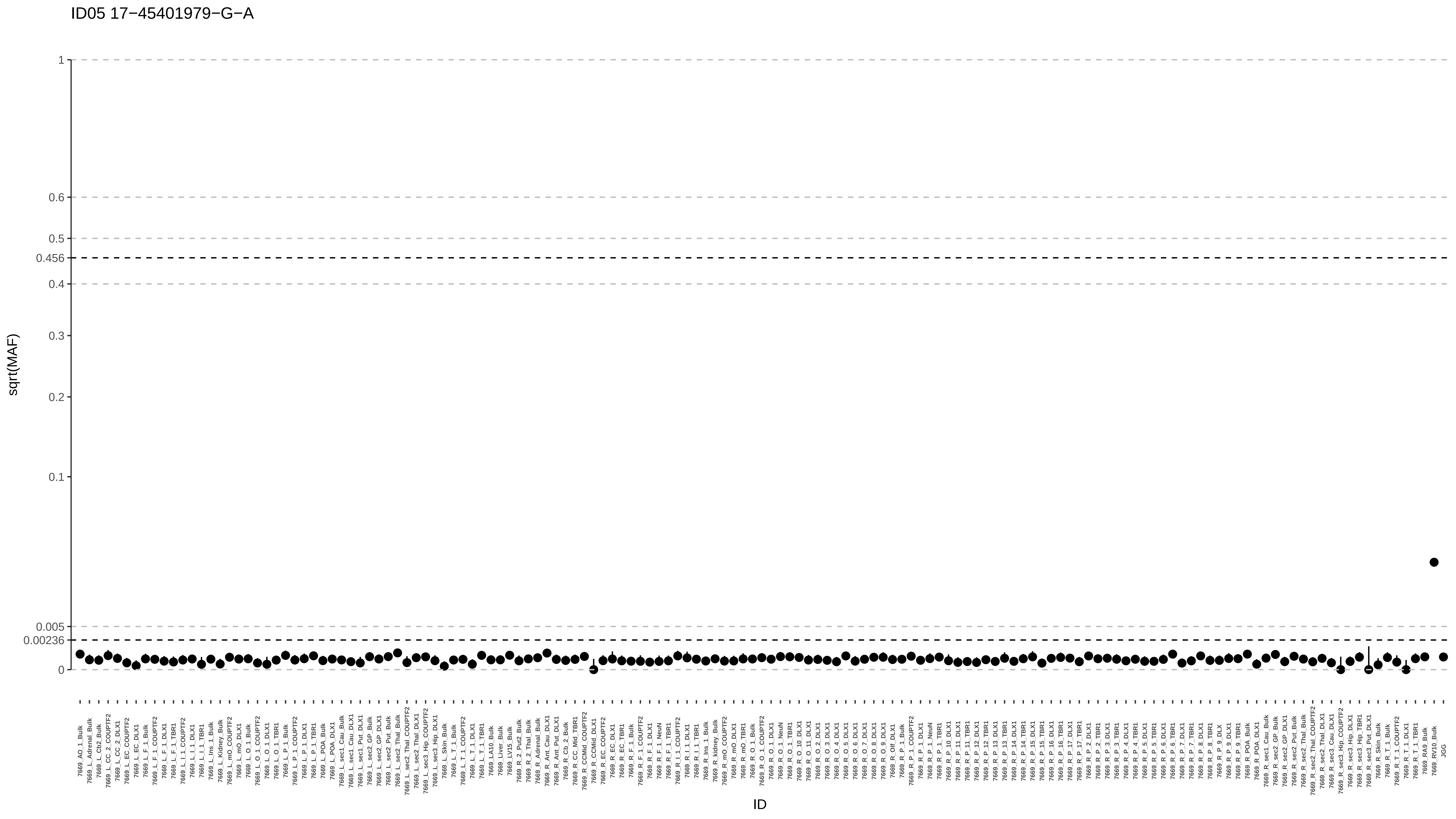

ID05 17−45401979−G−A
sqrt(MAF)
1
0.6
0.5
0.456
0.4
0.3
0.2
0.1
0.005
0.00236
0
ID
7669_AO_1_Bulk
7669_L_Adrenal_Bulk
7669_L_Cb2_Bulk
7669_L_CC_2_COUPTF2
7669_L_CC_2_DLX1
7669_L_EC_COUPTF2
7669_L_EC_DLX1
7669_L_F_1_Bulk
7669_L_F_1_COUPTF2
7669_L_F_1_DLX1
7669_L_F_1_TBR1
7669_L_I_1_COUPTF2
7669_L_I_1_DLX1
7669_L_I_1_TBR1
7669_L_Ins_1_Bulk
7669_L_Kidney_Bulk
7669_L_mO_COUPTF2
7669_L_mO_DLX1
7669_L_O_1_Bulk
7669_L_O_1_COUPTF2
7669_L_O_1_DLX1
7669_L_O_1_TBR1
7669_L_P_1_Bulk
7669_L_P_1_COUPTF2
7669_L_P_1_DLX1
7669_L_P_1_TBR1
7669_L_POA_Bulk
7669_L_POA_DLX1
7669_L_sec1_Cau_Bulk
7669_L_sec1_Cau_DLX1
7669_L_sec1_Put_DLX1
7669_L_sec2_GP_Bulk
7669_L_sec2_GP_DLX1
7669_L_sec2_Put_Bulk
7669_L_sec2_Thal_Bulk
7669_L_sec2_Thal_COUPTF2
7669_L_sec2_Thal_DLX1
7669_L_sec3_Hip_COUPTF2
7669_L_sec3_Hip_DLX1
7669_L_Skin_Bulk
7669_L_T_1_Bulk
7669_L_T_1_COUPTF2
7669_L_T_1_DLX1
7669_L_T_1_TBR1
7669_LA10_Bulk
7669_Liver_Bulk
7669_LV15_Bulk
7669_R_2_Put2_Bulk
7669_R_2_Thal_Bulk
7669_R_Adrenal_Bulk
7669_R_Ant_Cau_DLX1
7669_R_Ant_Put_DLX1
7669_R_Cb_2_Bulk
7669_R_CC_Mid_TBR1
7669_R_CCMid_COUPTF2
7669_R_CCMid_DLX1
7669_R_EC_COUPTF2
7669_R_EC_DLX1
7669_R_EC_TBR1
7669_R_F_1_Bulk
7669_R_F_1_COUPTF2
7669_R_F_1_DLX1
7669_R_F_1_NeuN
7669_R_F_1_TBR1
7669_R_I_1_COUPTF2
7669_R_I_1_DLX1
7669_R_I_1_TBR1
7669_R_Ins_1_Bulk
7669_R_kidney_Bulk
7669_R_mO_COUPTF2
7669_R_mO_DLX1
7669_R_mO_TBR1
7669_R_O_1_Bulk
7669_R_O_1_COUPTF2
7669_R_O_1_DLX1
7669_R_O_1_NeuN
7669_R_O_1_TBR1
7669_R_O_10_DLX1
7669_R_O_11_DLX1
7669_R_O_2_DLX1
7669_R_O_3_DLX1
7669_R_O_4_DLX1
7669_R_O_5_DLX1
7669_R_O_6_DLX1
7669_R_O_7_DLX1
7669_R_O_8_DLX1
7669_R_O_9_DLX1
7669_R_Olf_DLX1
7669_R_P_1_Bulk
7669_R_P_1_COUPTF2
7669_R_P_1_DLX1
7669_R_P_1_NeuN
7669_R_P_1_TBR1
7669_R_P_10_DLX1
7669_R_P_11_DLX1
7669_R_P_11_TBR1
7669_R_P_12_DLX1
7669_R_P_12_TBR1
7669_R_P_13_DLX1
7669_R_P_13_TBR1
7669_R_P_14_DLX1
7669_R_P_14_TBR1
7669_R_P_15_DLX1
7669_R_P_15_TBR1
7669_R_P_16_DLX1
7669_R_P_16_TBR1
7669_R_P_17_DLX1
7669_R_P_17_TBR1
7669_R_P_2_DLX1
7669_R_P_2_TBR1
7669_R_P_3_DLX1
7669_R_P_3_TBR1
7669_R_P_4_DLX1
7669_R_P_4_TBR1
7669_R_P_5_DLX1
7669_R_P_5_TBR1
7669_R_P_6_DLX1
7669_R_P_6_TBR1
7669_R_P_7_DLX1
7669_R_P_7_TBR1
7669_R_P_8_DLX1
7669_R_P_8_TBR1
7669_R_P_9_DLX
7669_R_P_9_DLX1
7669_R_P_9_TBR1
7669_R_POA_Bulk
7669_R_POA_DLX1
7669_R_sec1_Cau_Bulk
7669_R_sec2_GP_Bulk
7669_R_sec2_GP_DLX1
7669_R_sec2_Put_Bulk
7669_R_sec2_Thal_Bulk
7669_R_sec2_Thal_COUPTF2
7669_R_sec2_Thal_DLX1
7669_R_sec3_Cau_DLX1
7669_R_sec3_Hip_COUPTF2
7669_R_sec3_Hip_DLX1
7669_R_sec3_Hip_TBR1
7669_R_sec3_Put_DLX1
7669_R_Skin_Bulk
7669_R_T_1_Bulk
7669_R_T_1_COUPTF2
7669_R_T_1_DLX1
7669_R_T_1_TBR1
7669_RA9_Bulk
7669_RV10_Bulk
JGG

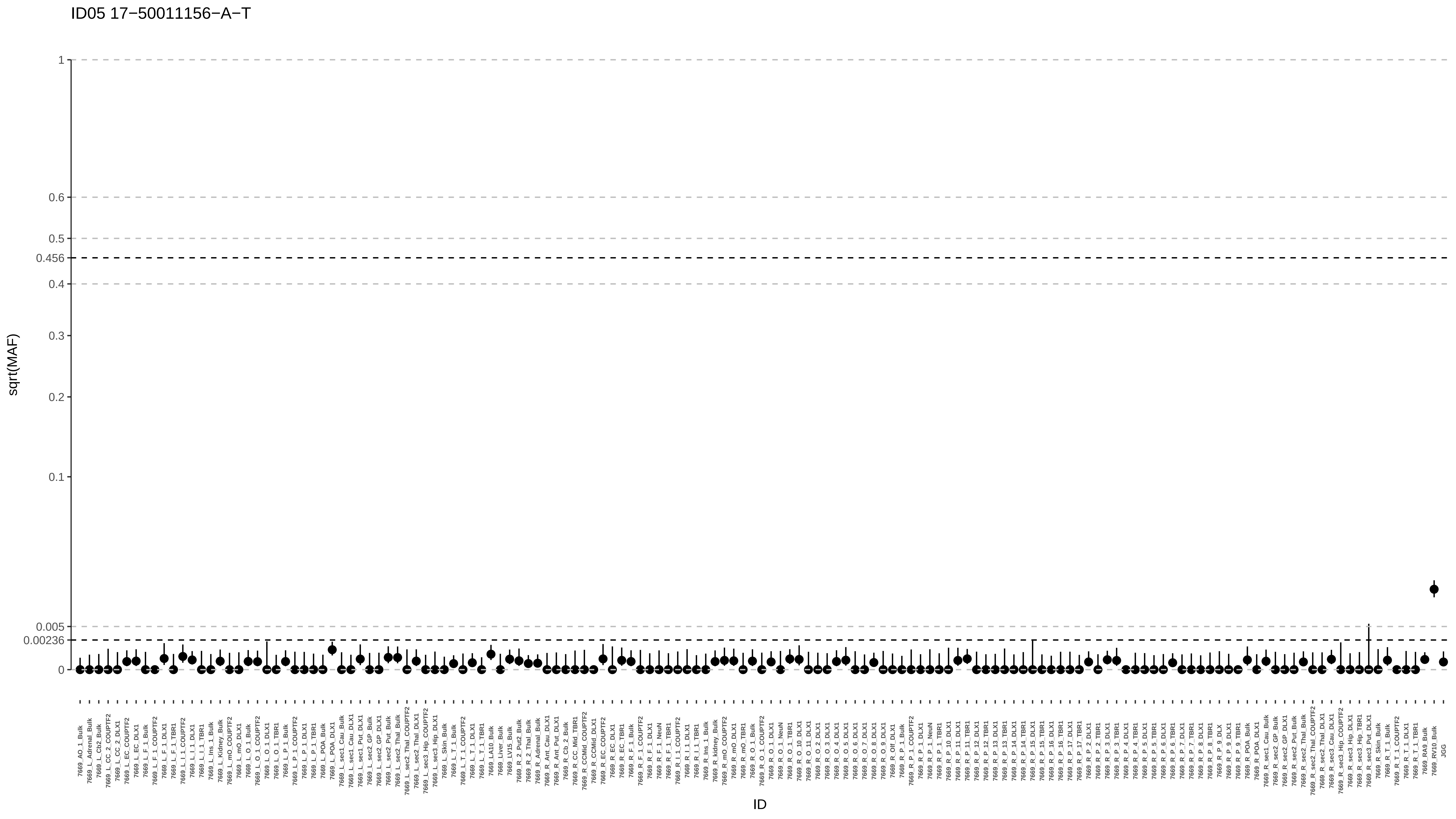
ID05 17−50011156−A−T
sqrt(MAF)
1
0.6
0.5
0.456
0.4
0.3
0.2
0.1
0.005
0.00236
0
ID

ID05 17−59031927−C−T

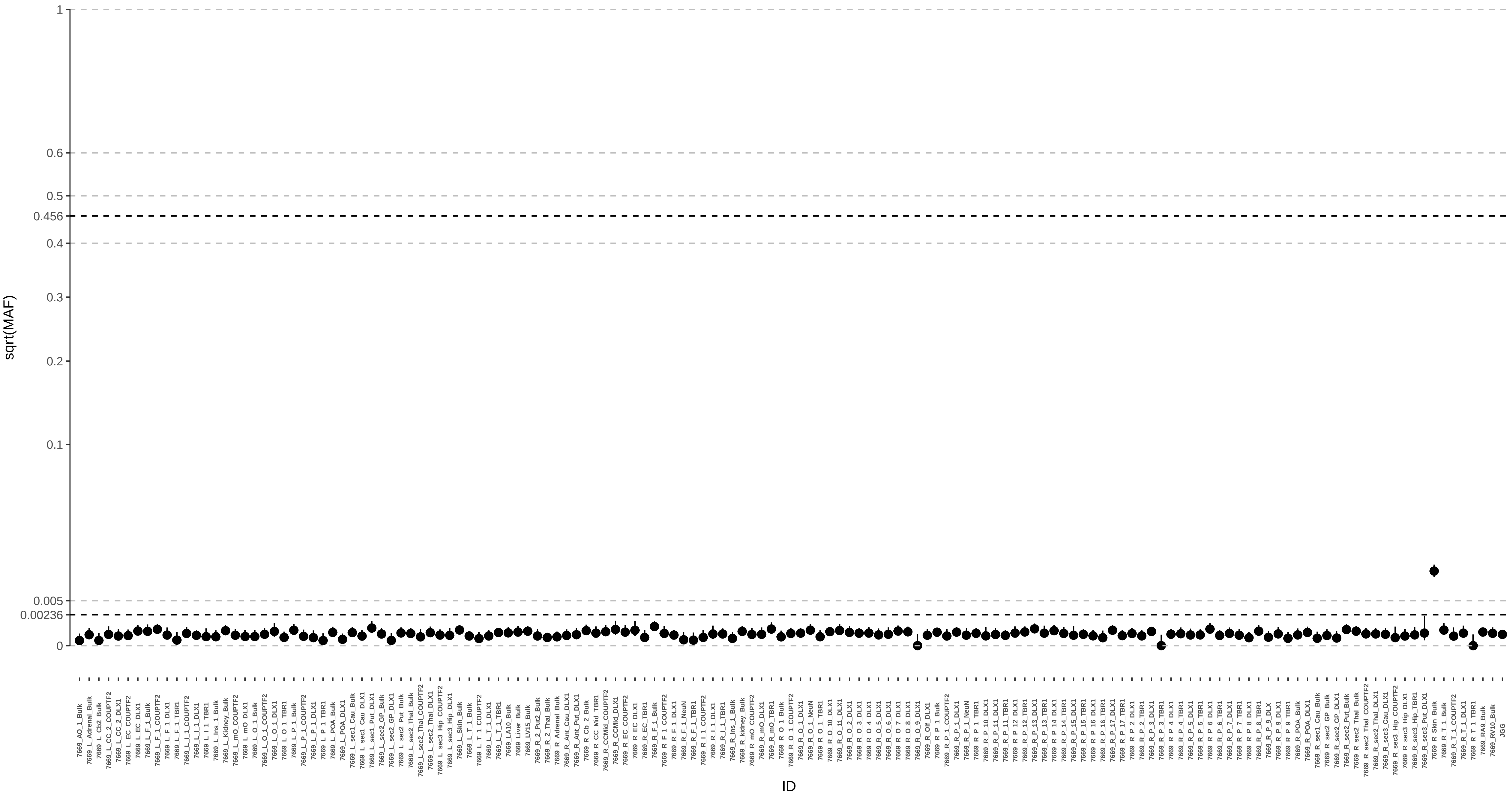

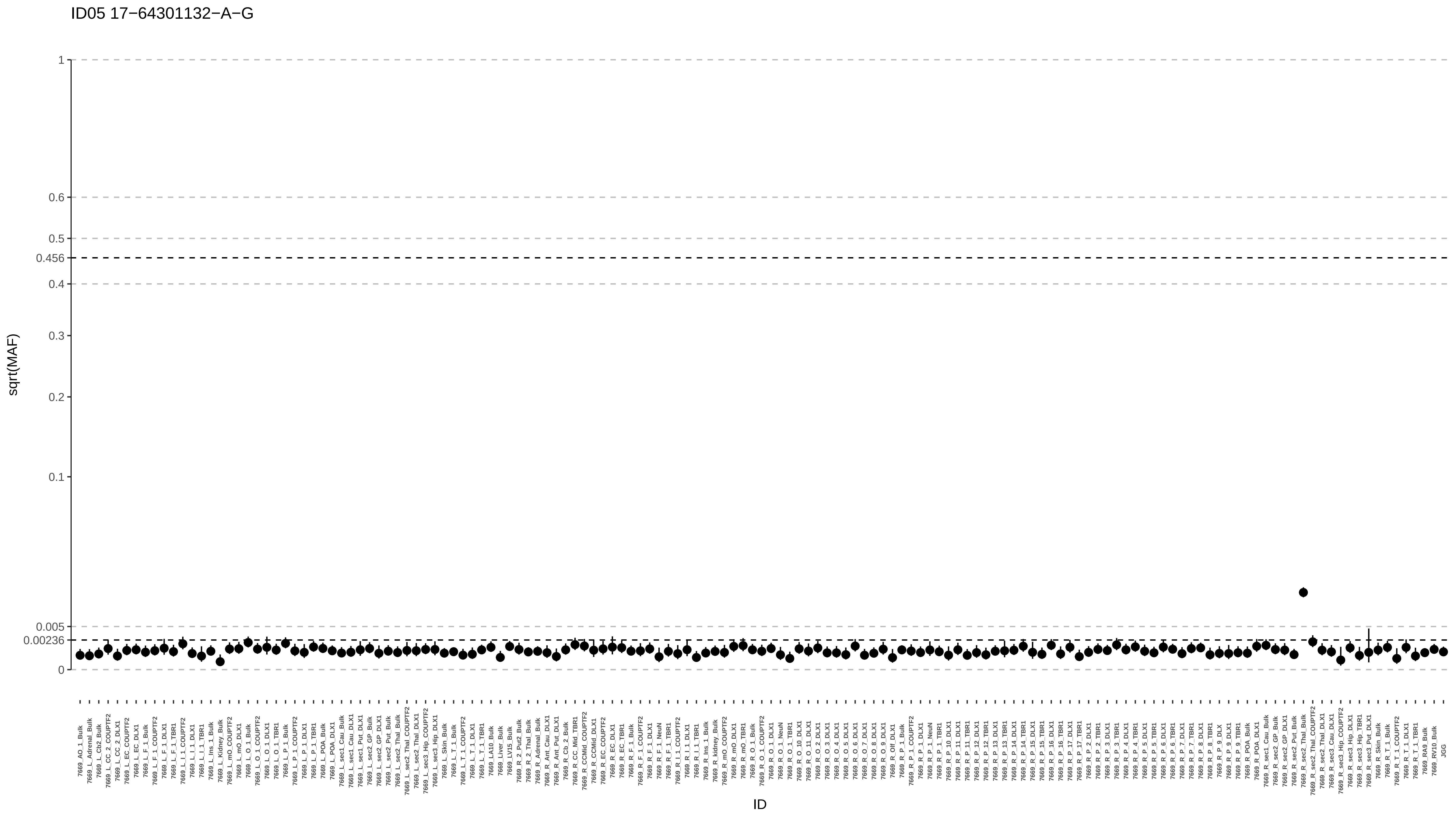
ID05 17−64301132−A−G
sqrt(MAF)
1
0.6
0.5
0.456
0.4
0.3
0.2
0.1
0.005
0.00236
0
ID

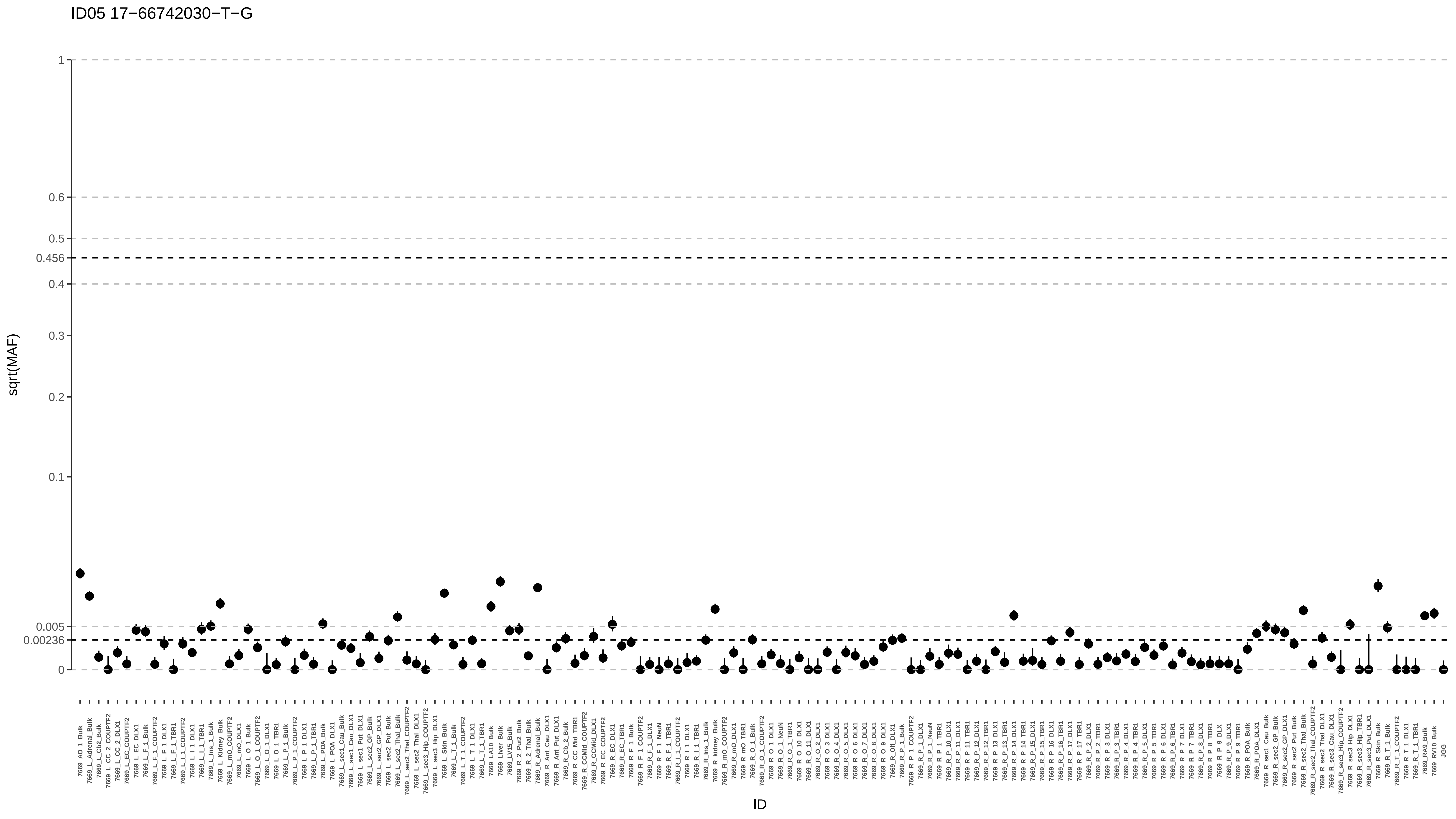
ID05 17−66742030−T−G
sqrt(MAF)
1
0.6
0.5
0.456
0.4
0.3
0.2
0.1
0.005
0.00236
0
ID
7669_AO_1_Bulk
7669_L_Adrenal_Bulk
7669_L_Cb2_Bulk
7669_L_CC_2_COUPTF2
7669_L_CC_2_DLX1
7669_L_EC_COUPTF2
7669_L_EC_DLX1
7669_L_F_1_Bulk
7669_L_F_1_COUPTF2
7669_L_F_1_DLX1
7669_L_F_1_TBR1
7669_L_I_1_COUPTF2
7669_L_I_1_DLX1
7669_L_I_1_TBR1
7669_L_Ins_1_Bulk
7669_L_Kidney_Bulk
7669_L_mO_COUPTF2
7669_L_mO_DLX1
7669_L_O_1_Bulk
7669_L_O_1_COUPTF2
7669_L_O_1_DLX1
7669_L_O_1_TBR1
7669_L_P_1_Bulk
7669_L_P_1_COUPTF2
7669_L_P_1_DLX1
7669_L_P_1_TBR1
7669_L_POA_Bulk
7669_L_POA_DLX1
7669_L_sec1_Cau_Bulk
7669_L_sec1_Cau_DLX1
7669_L_sec1_Put_DLX1
7669_L_sec2_GP_Bulk
7669_L_sec2_GP_DLX1
7669_L_sec2_Put_Bulk
7669_L_sec2_Thal_Bulk
7669_L_sec2_Thal_COUPTF2
7669_L_sec2_Thal_DLX1
7669_L_sec3_Hip_COUPTF2
7669_L_sec3_Hip_DLX1
7669_L_Skin_Bulk
7669_L_T_1_Bulk
7669_L_T_1_COUPTF2
7669_L_T_1_DLX1
7669_L_T_1_TBR1
7669_LA10_Bulk
7669_Liver_Bulk
7669_LV15_Bulk
7669_R_2_Put2_Bulk
7669_R_2_Thal_Bulk
7669_R_Adrenal_Bulk
7669_R_Ant_Cau_DLX1
7669_R_Ant_Put_DLX1
7669_R_Cb_2_Bulk
7669_R_CC_Mid_TBR1
7669_R_CCMid_COUPTF2
7669_R_CCMid_DLX1
7669_R_EC_COUPTF2
7669_R_EC_DLX1
7669_R_EC_TBR1
7669_R_F_1_Bulk
7669_R_F_1_COUPTF2
7669_R_F_1_DLX1
7669_R_F_1_NeuN
7669_R_F_1_TBR1
7669_R_I_1_COUPTF2
7669_R_I_1_DLX1
7669_R_I_1_TBR1
7669_R_Ins_1_Bulk
7669_R_kidney_Bulk
7669_R_mO_COUPTF2
7669_R_mO_DLX1
7669_R_mO_TBR1
7669_R_O_1_Bulk
7669_R_O_1_COUPTF2
7669_R_O_1_DLX1
7669_R_O_1_NeuN
7669_R_O_1_TBR1
7669_R_O_10_DLX1
7669_R_O_11_DLX1
7669_R_O_2_DLX1
7669_R_O_3_DLX1
7669_R_O_4_DLX1
7669_R_O_5_DLX1
7669_R_O_6_DLX1
7669_R_O_7_DLX1
7669_R_O_8_DLX1
7669_R_O_9_DLX1
7669_R_Olf_DLX1
7669_R_P_1_Bulk
7669_R_P_1_COUPTF2
7669_R_P_1_DLX1
7669_R_P_1_NeuN
7669_R_P_1_TBR1
7669_R_P_10_DLX1
7669_R_P_11_DLX1
7669_R_P_11_TBR1
7669_R_P_12_DLX1
7669_R_P_12_TBR1
7669_R_P_13_DLX1
7669_R_P_13_TBR1
7669_R_P_14_DLX1
7669_R_P_14_TBR1
7669_R_P_15_DLX1
7669_R_P_15_TBR1
7669_R_P_16_DLX1
7669_R_P_16_TBR1
7669_R_P_17_DLX1
7669_R_P_17_TBR1
7669_R_P_2_DLX1
7669_R_P_2_TBR1
7669_R_P_3_DLX1
7669_R_P_3_TBR1
7669_R_P_4_DLX1
7669_R_P_4_TBR1
7669_R_P_5_DLX1
7669_R_P_5_TBR1
7669_R_P_6_DLX1
7669_R_P_6_TBR1
7669_R_P_7_DLX1
7669_R_P_7_TBR1
7669_R_P_8_DLX1
7669_R_P_8_TBR1
7669_R_P_9_DLX
7669_R_P_9_DLX1
7669_R_P_9_TBR1
7669_R_POA_Bulk
7669_R_POA_DLX1
7669_R_sec1_Cau_Bulk
7669_R_sec2_GP_Bulk
7669_R_sec2_GP_DLX1
7669_R_sec2_Put_Bulk
7669_R_sec2_Thal_Bulk
7669_R_sec2_Thal_COUPTF2
7669_R_sec2_Thal_DLX1
7669_R_sec3_Cau_DLX1
7669_R_sec3_Hip_COUPTF2
7669_R_sec3_Hip_DLX1
7669_R_sec3_Hip_TBR1
7669_R_sec3_Put_DLX1
7669_R_Skin_Bulk
7669_R_T_1_Bulk
7669_R_T_1_COUPTF2
7669_R_T_1_DLX1
7669_R_T_1_TBR1
7669_RA9_Bulk
7669_RV10_Bulk
JGG

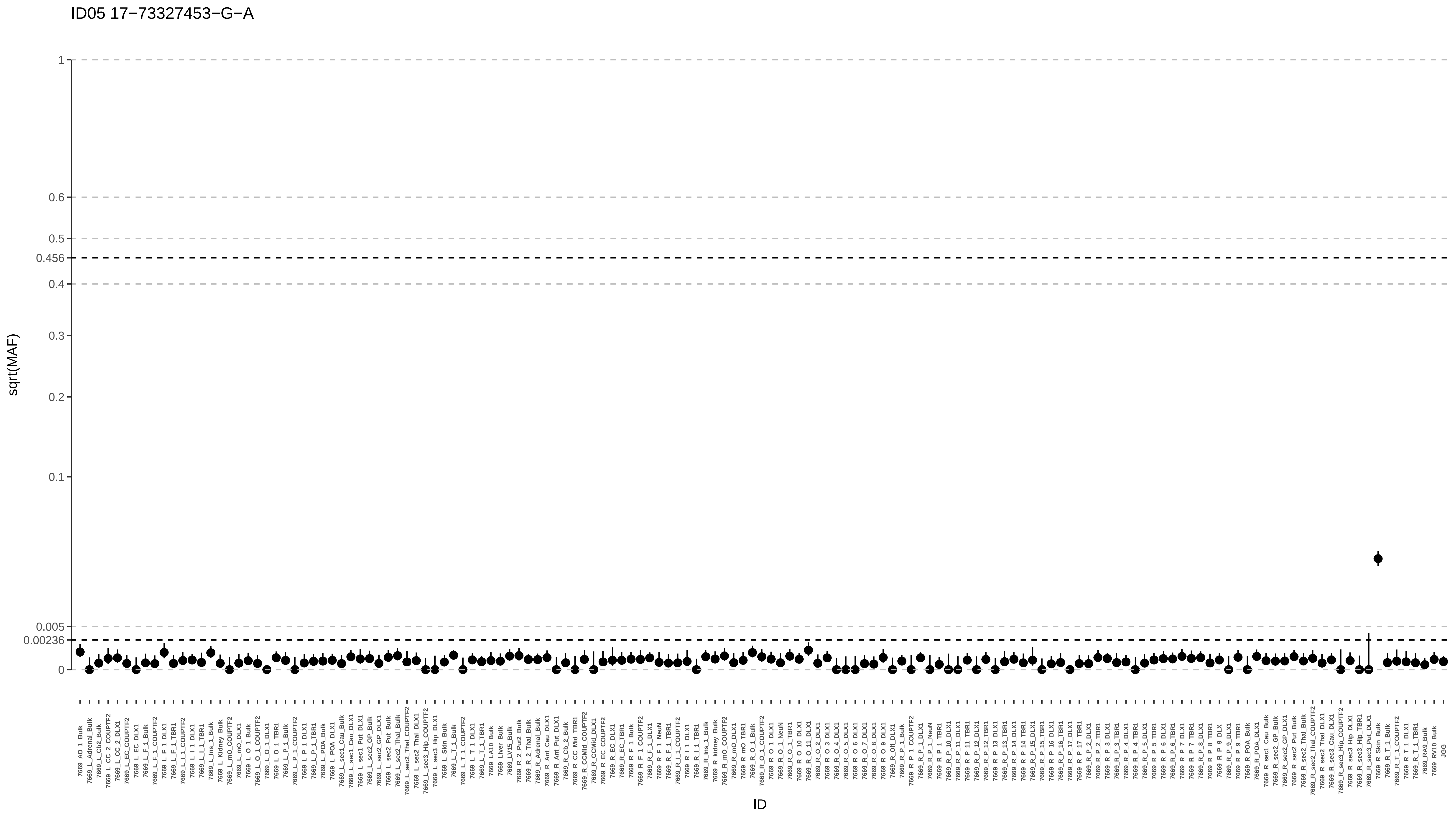
ID05 17−73327453−G−A
sqrt(MAF)
1
0.6
0.5
0.456
0.4
0.3
0.2
0.1
0.005
0.00236
0
ID

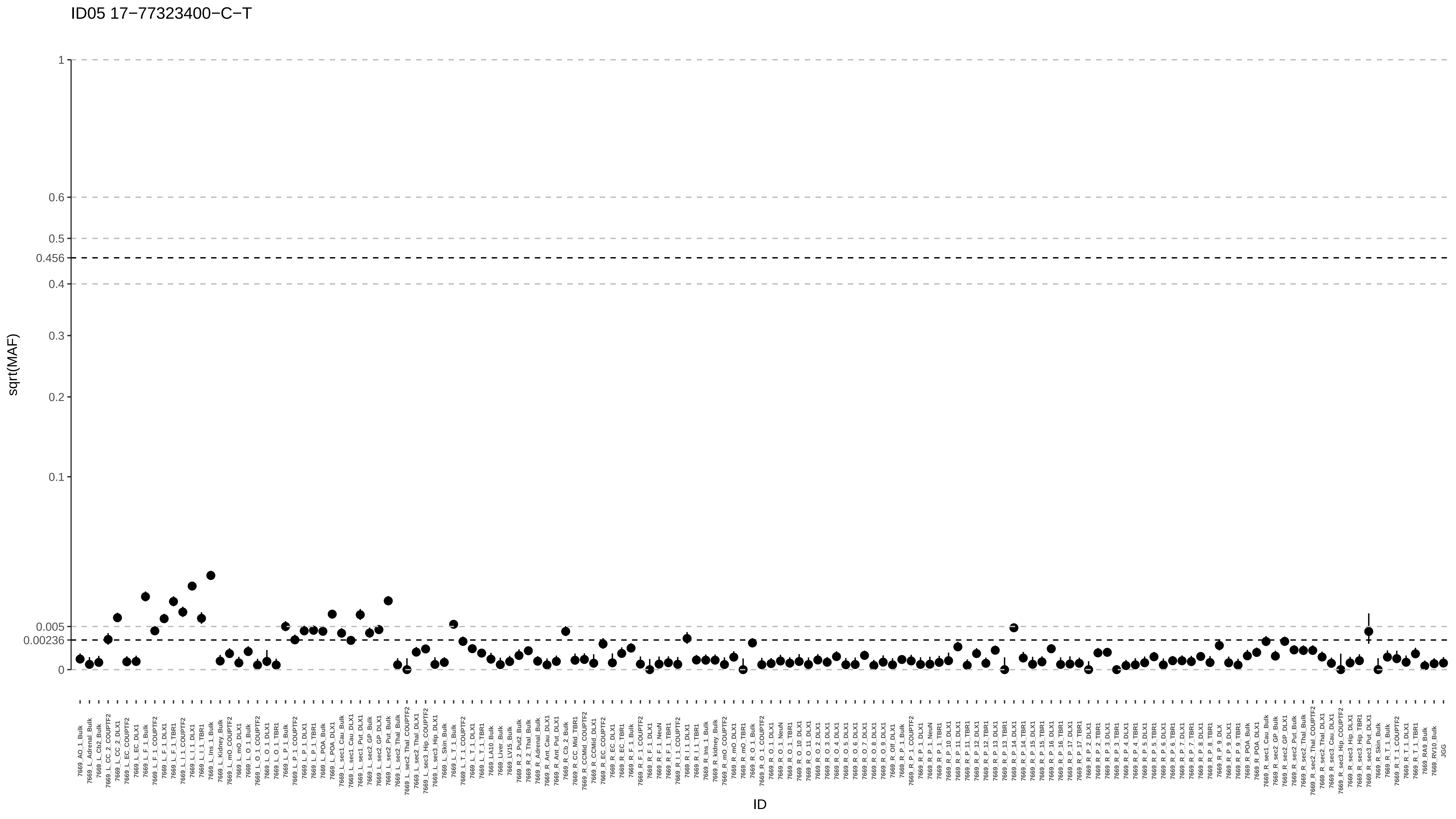

ID05 17−77323400−C−T
sqrt(MAF)
1
0.6
0.5
0.456
0.4
0.3
0.2
0.1
0.005
0.00236
0
ID

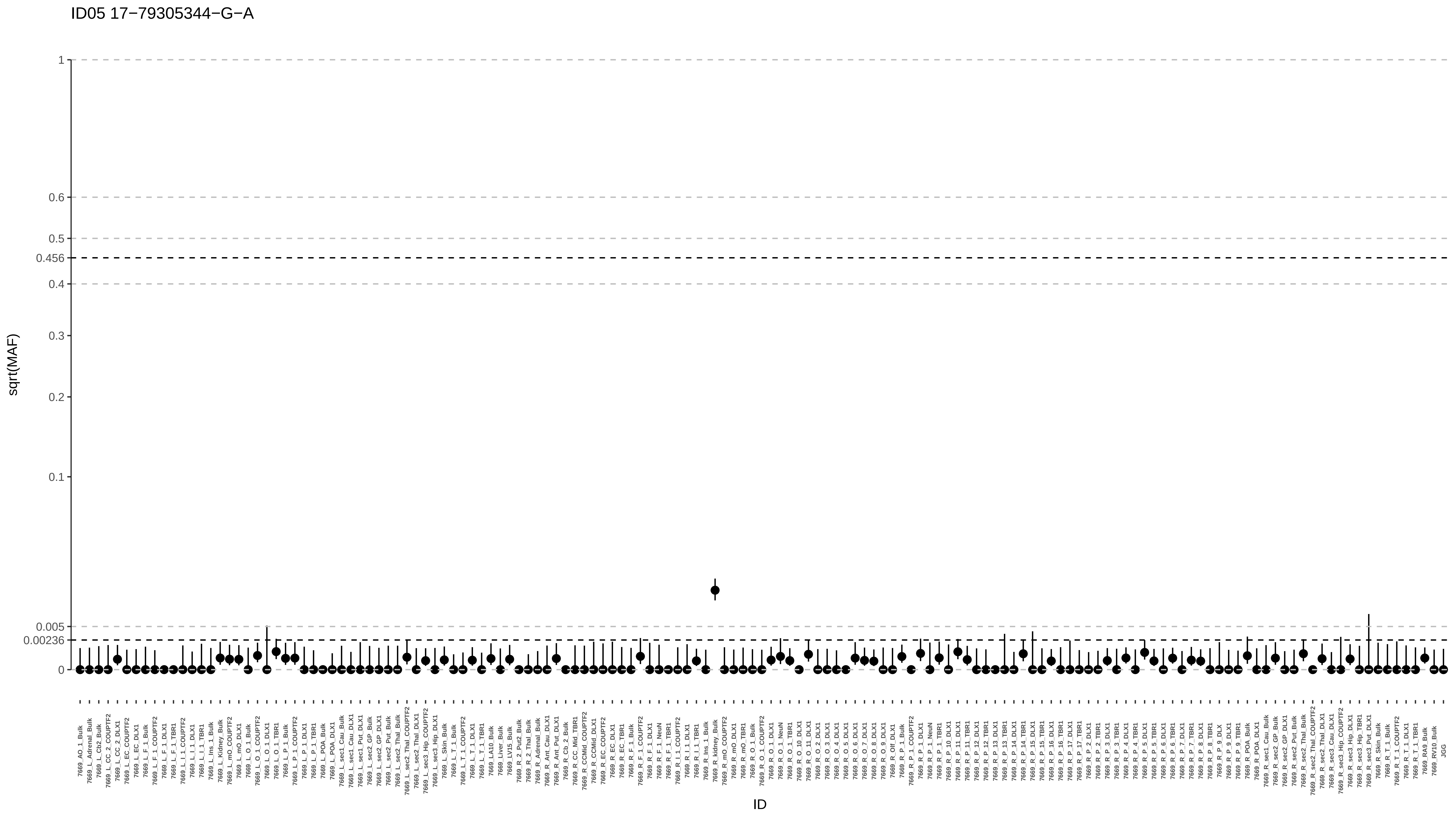
ID05 17−79305344−G−A
sqrt(MAF)
1
0.6
0.5
0.456
0.4
0.3
0.2
0.1
0.005
0.00236
0
ID
7669_AO_1_Bulk
7669_L_Adrenal_Bulk
7669_L_Cb2_Bulk
7669_L_CC_2_COUPTF2
7669_L_CC_2_DLX1
7669_L_EC_COUPTF2
7669_L_EC_DLX1
7669_L_F_1_Bulk
7669_L_F_1_COUPTF2
7669_L_F_1_DLX1
7669_L_F_1_TBR1
7669_L_I_1_COUPTF2
7669_L_I_1_DLX1
7669_L_I_1_TBR1
7669_L_Ins_1_Bulk
7669_L_Kidney_Bulk
7669_L_mO_COUPTF2
7669_L_mO_DLX1
7669_L_O_1_Bulk
7669_L_O_1_COUPTF2
7669_L_O_1_DLX1
7669_L_O_1_TBR1
7669_L_P_1_Bulk
7669_L_P_1_COUPTF2
7669_L_P_1_DLX1
7669_L_P_1_TBR1
7669_L_POA_Bulk
7669_L_POA_DLX1
7669_L_sec1_Cau_Bulk
7669_L_sec1_Cau_DLX1
7669_L_sec1_Put_DLX1
7669_L_sec2_GP_Bulk
7669_L_sec2_GP_DLX1
7669_L_sec2_Put_Bulk
7669_L_sec2_Thal_Bulk
7669_L_sec2_Thal_COUPTF2
7669_L_sec2_Thal_DLX1
7669_L_sec3_Hip_COUPTF2
7669_L_sec3_Hip_DLX1
7669_L_Skin_Bulk
7669_L_T_1_Bulk
7669_L_T_1_COUPTF2
7669_L_T_1_DLX1
7669_L_T_1_TBR1
7669_LA10_Bulk
7669_Liver_Bulk
7669_LV15_Bulk
7669_R_2_Put2_Bulk
7669_R_2_Thal_Bulk
7669_R_Adrenal_Bulk
7669_R_Ant_Cau_DLX1
7669_R_Ant_Put_DLX1
7669_R_Cb_2_Bulk
7669_R_CC_Mid_TBR1
7669_R_CCMid_COUPTF2
7669_R_CCMid_DLX1
7669_R_EC_COUPTF2
7669_R_EC_DLX1
7669_R_EC_TBR1
7669_R_F_1_Bulk
7669_R_F_1_COUPTF2
7669_R_F_1_DLX1
7669_R_F_1_NeuN
7669_R_F_1_TBR1
7669_R_I_1_COUPTF2
7669_R_I_1_DLX1
7669_R_I_1_TBR1
7669_R_Ins_1_Bulk
7669_R_kidney_Bulk
7669_R_mO_COUPTF2
7669_R_mO_DLX1
7669_R_mO_TBR1
7669_R_O_1_Bulk
7669_R_O_1_COUPTF2
7669_R_O_1_DLX1
7669_R_O_1_NeuN
7669_R_O_1_TBR1
7669_R_O_10_DLX1
7669_R_O_11_DLX1
7669_R_O_2_DLX1
7669_R_O_3_DLX1
7669_R_O_4_DLX1
7669_R_O_5_DLX1
7669_R_O_6_DLX1
7669_R_O_7_DLX1
7669_R_O_8_DLX1
7669_R_O_9_DLX1
7669_R_Olf_DLX1
7669_R_P_1_Bulk
7669_R_P_1_COUPTF2
7669_R_P_1_DLX1
7669_R_P_1_NeuN
7669_R_P_1_TBR1
7669_R_P_10_DLX1
7669_R_P_11_DLX1
7669_R_P_11_TBR1
7669_R_P_12_DLX1
7669_R_P_12_TBR1
7669_R_P_13_DLX1
7669_R_P_13_TBR1
7669_R_P_14_DLX1
7669_R_P_14_TBR1
7669_R_P_15_DLX1
7669_R_P_15_TBR1
7669_R_P_16_DLX1
7669_R_P_16_TBR1
7669_R_P_17_DLX1
7669_R_P_17_TBR1
7669_R_P_2_DLX1
7669_R_P_2_TBR1
7669_R_P_3_DLX1
7669_R_P_3_TBR1
7669_R_P_4_DLX1
7669_R_P_4_TBR1
7669_R_P_5_DLX1
7669_R_P_5_TBR1
7669_R_P_6_DLX1
7669_R_P_6_TBR1
7669_R_P_7_DLX1
7669_R_P_7_TBR1
7669_R_P_8_DLX1
7669_R_P_8_TBR1
7669_R_P_9_DLX
7669_R_P_9_DLX1
7669_R_P_9_TBR1
7669_R_POA_Bulk
7669_R_POA_DLX1
7669_R_sec1_Cau_Bulk
7669_R_sec2_GP_Bulk
7669_R_sec2_GP_DLX1
7669_R_sec2_Put_Bulk
7669_R_sec2_Thal_Bulk
7669_R_sec2_Thal_COUPTF2
7669_R_sec2_Thal_DLX1
7669_R_sec3_Cau_DLX1
7669_R_sec3_Hip_COUPTF2
7669_R_sec3_Hip_DLX1
7669_R_sec3_Hip_TBR1
7669_R_sec3_Put_DLX1
7669_R_Skin_Bulk
7669_R_T_1_Bulk
7669_R_T_1_COUPTF2
7669_R_T_1_DLX1
7669_R_T_1_TBR1
7669_RA9_Bulk
7669_RV10_Bulk
JGG

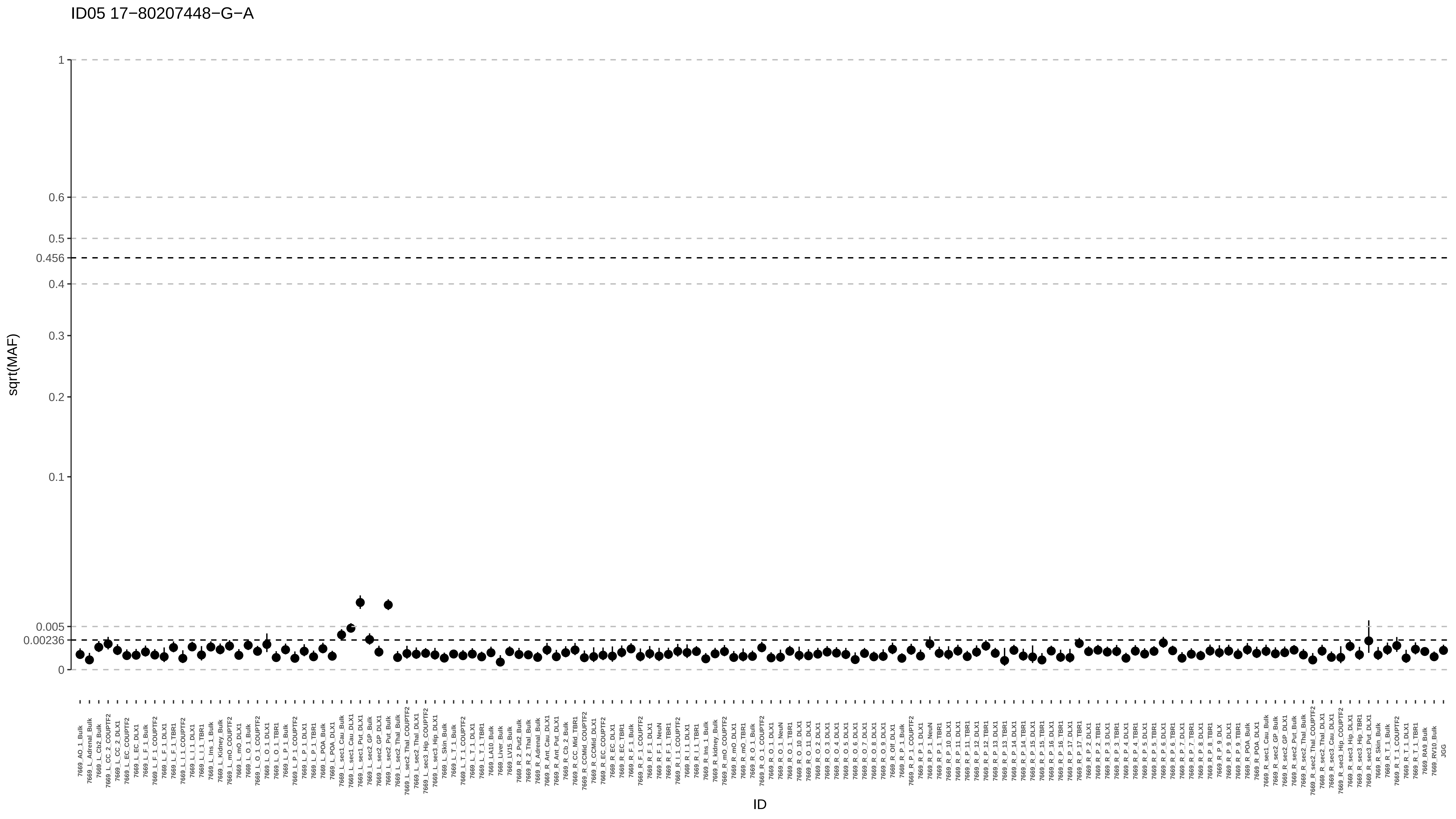
ID05 17−80207448−G−A
sqrt(MAF)
ID
1
0.6
0.5
0.456
0.4
0.3
0.2
0.1
0.005
0.00236
0

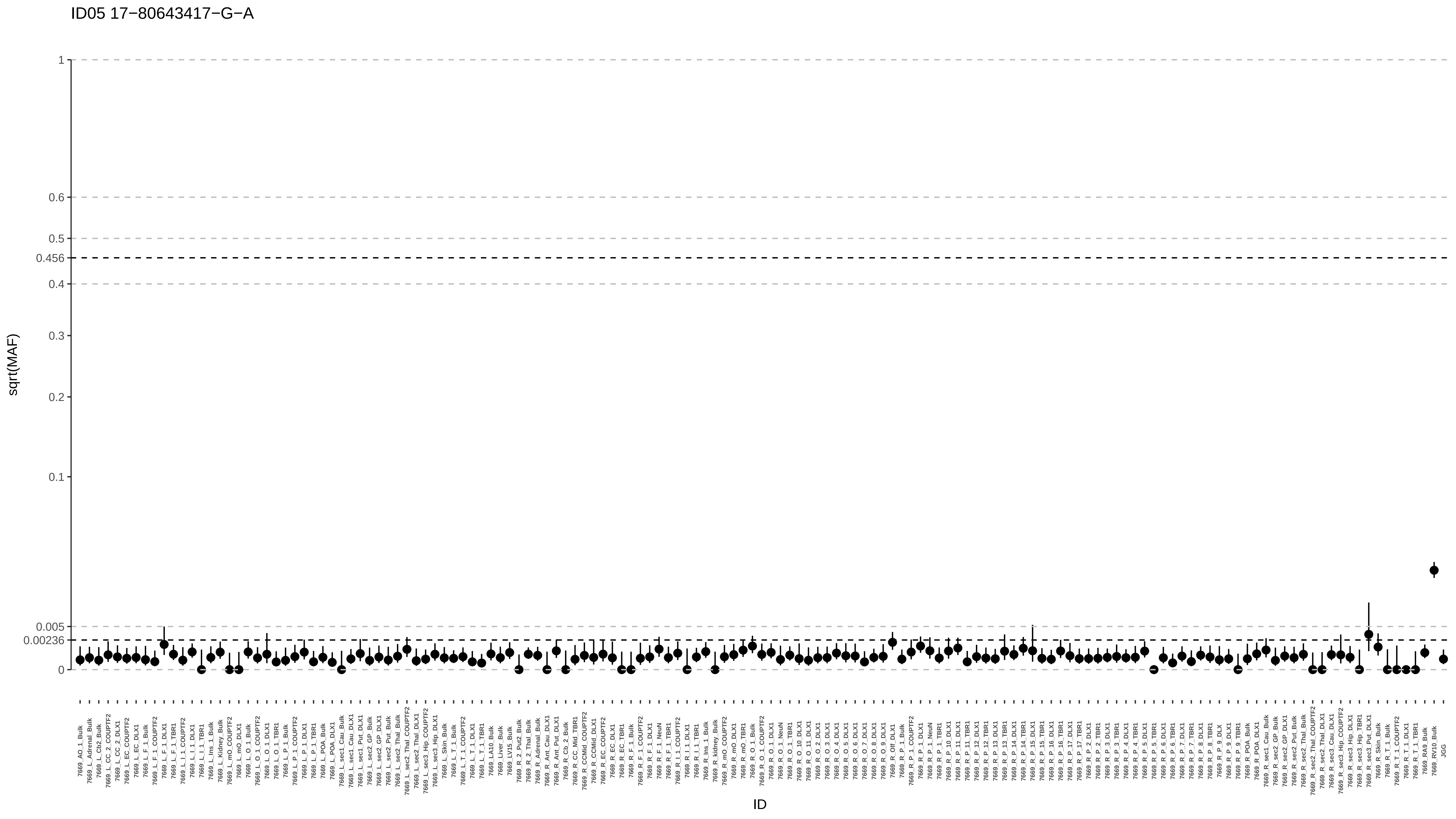
ID05 17−80643417−G−A
sqrt(MAF)
1
0.6
0.5
0.456
0.4
0.3
0.2
0.1
0.005
0.00236
0
ID
7669_AO_1_Bulk
7669_L_Adrenal_Bulk
7669_L_Cb2_Bulk
7669_L_CC_2_COUPTF2
7669_L_CC_2_DLX1
7669_L_EC_COUPTF2
7669_L_EC_DLX1
7669_L_F_1_Bulk
7669_L_F_1_COUPTF2
7669_L_F_1_DLX1
7669_L_F_1_TBR1
7669_L_I_1_COUPTF2
7669_L_I_1_DLX1
7669_L_I_1_TBR1
7669_L_Ins_1_Bulk
7669_L_Kidney_Bulk
7669_L_mO_COUPTF2
7669_L_mO_DLX1
7669_L_O_1_Bulk
7669_L_O_1_COUPTF2
7669_L_O_1_DLX1
7669_L_O_1_TBR1
7669_L_P_1_Bulk
7669_L_P_1_COUPTF2
7669_L_P_1_DLX1
7669_L_P_1_TBR1
7669_L_POA_Bulk
7669_L_POA_DLX1
7669_L_sec1_Cau_Bulk
7669_L_sec1_Cau_DLX1
7669_L_sec1_Put_DLX1
7669_L_sec2_GP_Bulk
7669_L_sec2_GP_DLX1
7669_L_sec2_Put_Bulk
7669_L_sec2_Thal_Bulk
7669_L_sec2_Thal_COUPTF2
7669_L_sec2_Thal_DLX1
7669_L_sec3_Hip_COUPTF2
7669_L_sec3_Hip_DLX1
7669_L_Skin_Bulk
7669_L_T_1_Bulk
7669_L_T_1_COUPTF2
7669_L_T_1_DLX1
7669_L_T_1_TBR1
7669_LA10_Bulk
7669_Liver_Bulk
7669_LV15_Bulk
7669_R_2_Put2_Bulk
7669_R_2_Thal_Bulk
7669_R_Adrenal_Bulk
7669_R_Ant_Cau_DLX1
7669_R_Ant_Put_DLX1
7669_R_Cb_2_Bulk
7669_R_CC_Mid_TBR1
7669_R_CCMid_COUPTF2
7669_R_CCMid_DLX1
7669_R_EC_COUPTF2
7669_R_EC_DLX1
7669_R_EC_TBR1
7669_R_F_1_Bulk
7669_R_F_1_COUPTF2
7669_R_F_1_DLX1
7669_R_F_1_NeuN
7669_R_F_1_TBR1
7669_R_I_1_COUPTF2
7669_R_I_1_DLX1
7669_R_I_1_TBR1
7669_R_Ins_1_Bulk
7669_R_kidney_Bulk
7669_R_mO_COUPTF2
7669_R_mO_DLX1
7669_R_mO_TBR1
7669_R_O_1_Bulk
7669_R_O_1_COUPTF2
7669_R_O_1_DLX1
7669_R_O_1_NeuN
7669_R_O_1_TBR1
7669_R_O_10_DLX1
7669_R_O_11_DLX1
7669_R_O_2_DLX1
7669_R_O_3_DLX1
7669_R_O_4_DLX1
7669_R_O_5_DLX1
7669_R_O_6_DLX1
7669_R_O_7_DLX1
7669_R_O_8_DLX1
7669_R_O_9_DLX1
7669_R_Olf_DLX1
7669_R_P_1_Bulk
7669_R_P_1_COUPTF2
7669_R_P_1_DLX1
7669_R_P_1_NeuN
7669_R_P_1_TBR1
7669_R_P_10_DLX1
7669_R_P_11_DLX1
7669_R_P_11_TBR1
7669_R_P_12_DLX1
7669_R_P_12_TBR1
7669_R_P_13_DLX1
7669_R_P_13_TBR1
7669_R_P_14_DLX1
7669_R_P_14_TBR1
7669_R_P_15_DLX1
7669_R_P_15_TBR1
7669_R_P_16_DLX1
7669_R_P_16_TBR1
7669_R_P_17_DLX1
7669_R_P_17_TBR1
7669_R_P_2_DLX1
7669_R_P_2_TBR1
7669_R_P_3_DLX1
7669_R_P_3_TBR1
7669_R_P_4_DLX1
7669_R_P_4_TBR1
7669_R_P_5_DLX1
7669_R_P_5_TBR1
7669_R_P_6_DLX1
7669_R_P_6_TBR1
7669_R_P_7_DLX1
7669_R_P_7_TBR1
7669_R_P_8_DLX1
7669_R_P_8_TBR1
7669_R_P_9_DLX
7669_R_P_9_DLX1
7669_R_P_9_TBR1
7669_R_POA_Bulk
7669_R_POA_DLX1
7669_R_sec1_Cau_Bulk
7669_R_sec2_GP_Bulk
7669_R_sec2_GP_DLX1
7669_R_sec2_Put_Bulk
7669_R_sec2_Thal_Bulk
7669_R_sec2_Thal_COUPTF2
7669_R_sec2_Thal_DLX1
7669_R_sec3_Cau_DLX1
7669_R_sec3_Hip_COUPTF2
7669_R_sec3_Hip_DLX1
7669_R_sec3_Hip_TBR1
7669_R_sec3_Put_DLX1
7669_R_Skin_Bulk
7669_R_T_1_Bulk
7669_R_T_1_COUPTF2
7669_R_T_1_DLX1
7669_R_T_1_TBR1
7669_RA9_Bulk
7669_RV10_Bulk
JGG

ID05 17−8815877−G−A

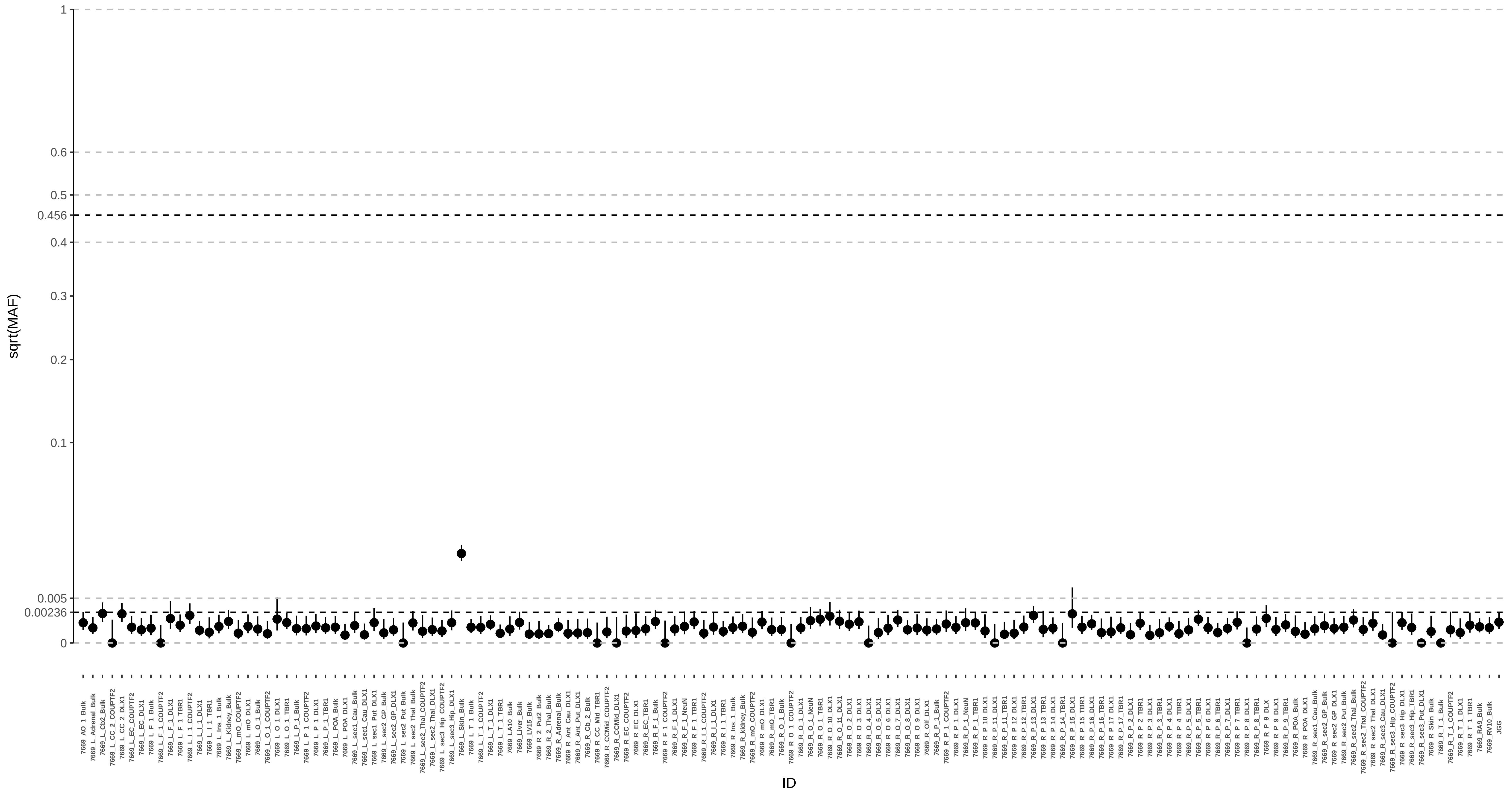

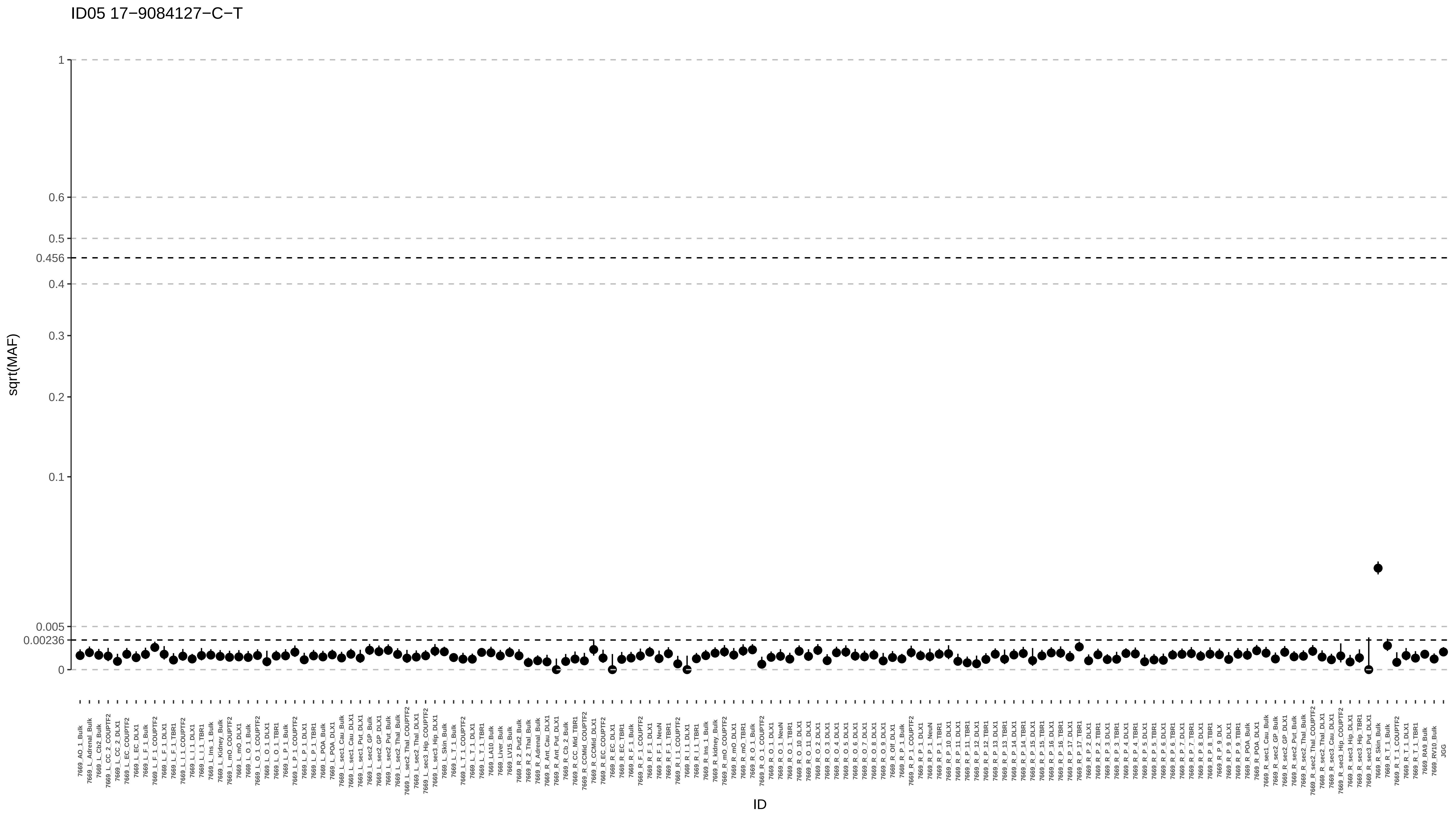
ID05 17−9084127−C−T
sqrt(MAF)
1
0.6
0.5
0.456
0.4
0.3
0.2
0.1
0.005
0.00236
0
ID

ID05 18−1735890−A−C

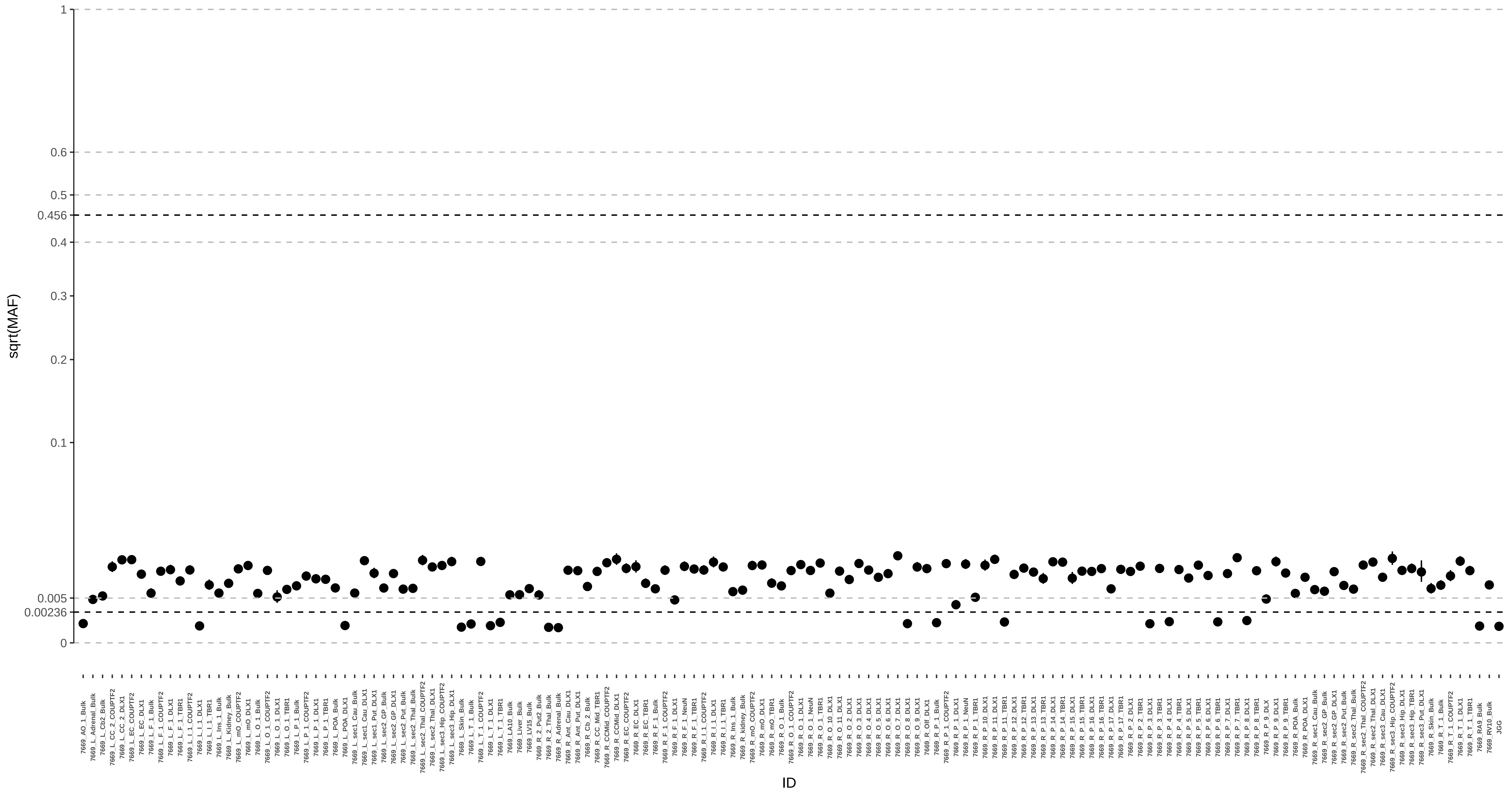

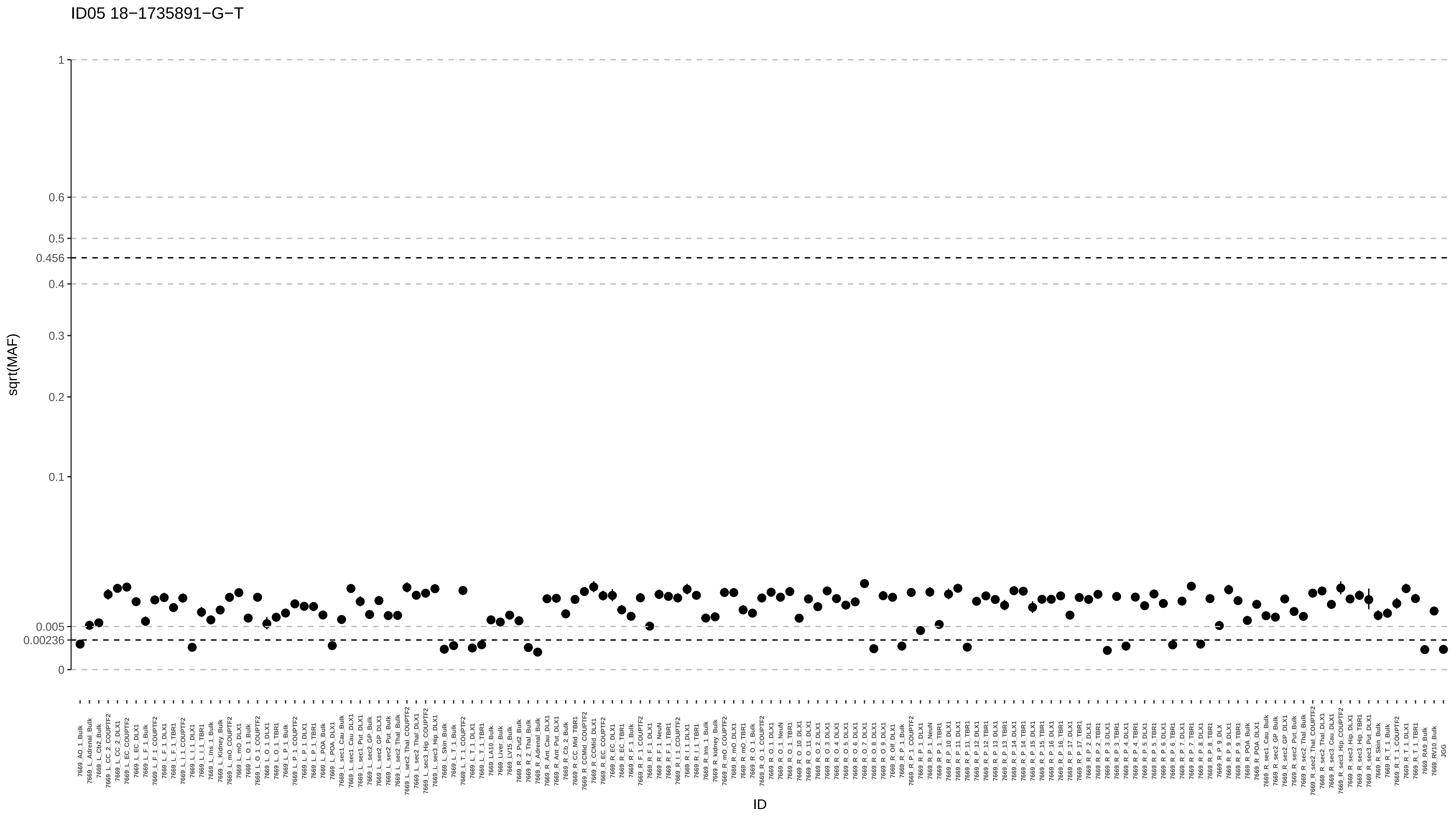
ID05 18−1735891−G−T
sqrt(MAF)
1
0.6
0.5
0.456
0.4
0.3
0.2
0.1
0.005
0.00236
0
ID

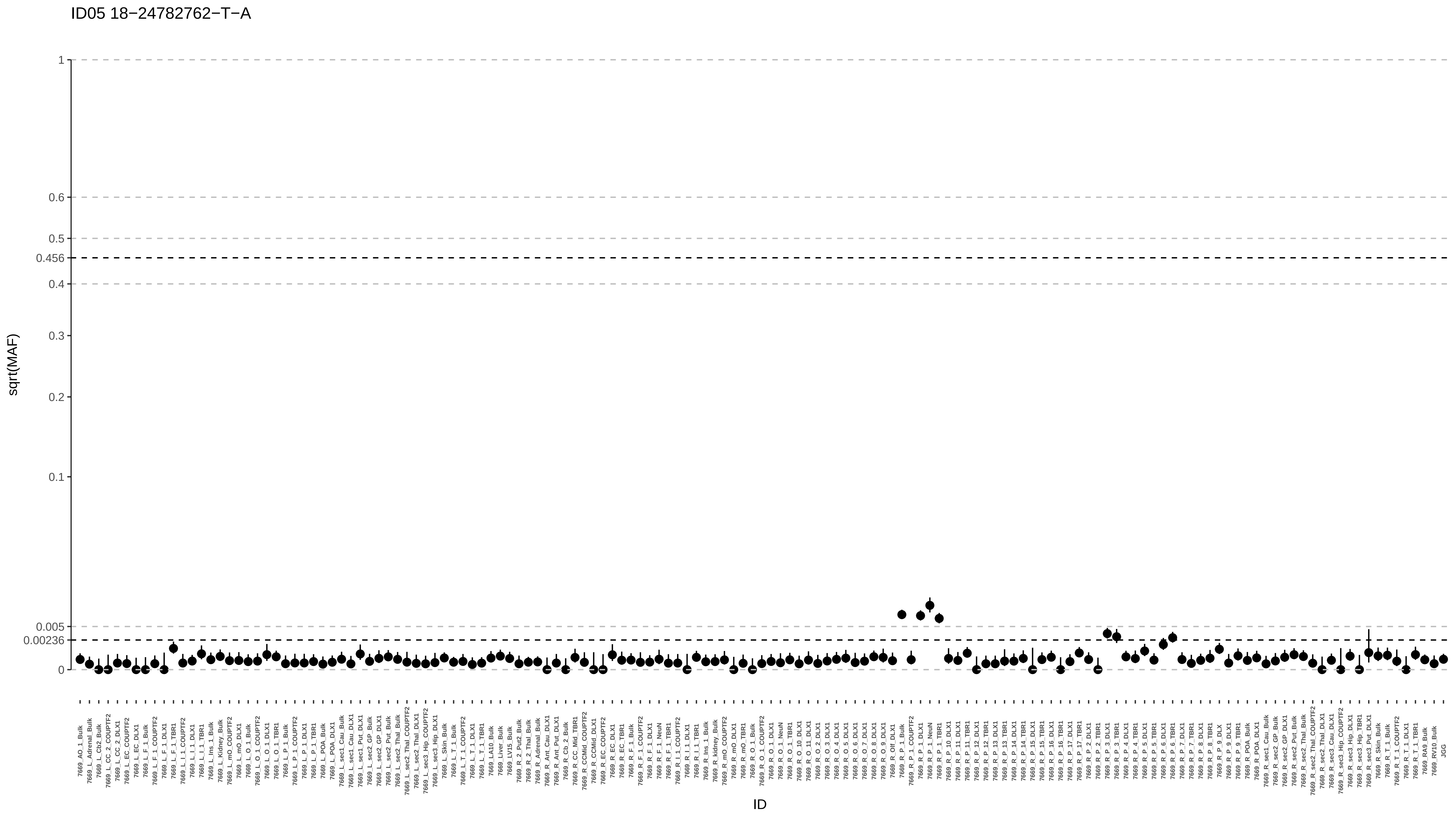
ID05 18−24782762−T−A
sqrt(MAF)
1
0.6
0.5
0.456
0.4
0.3
0.2
0.1
0.005
0.00236
0
ID

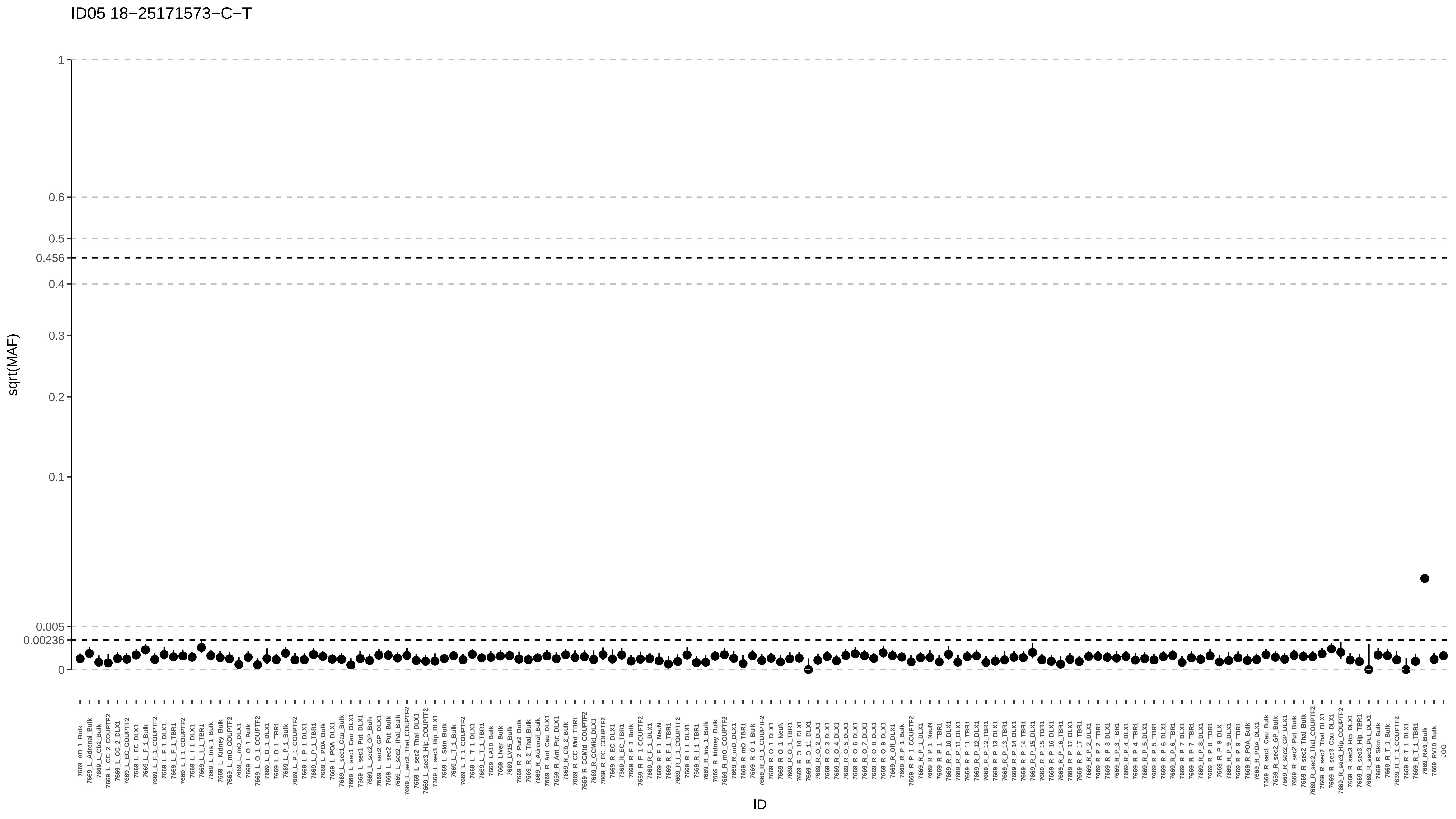

ID05 18−25171573−C−T
1
0.6
0.5
0.456
0.4
0.3
0.2
0.1
0.005
0.00236
0
sqrt(MAF)
ID

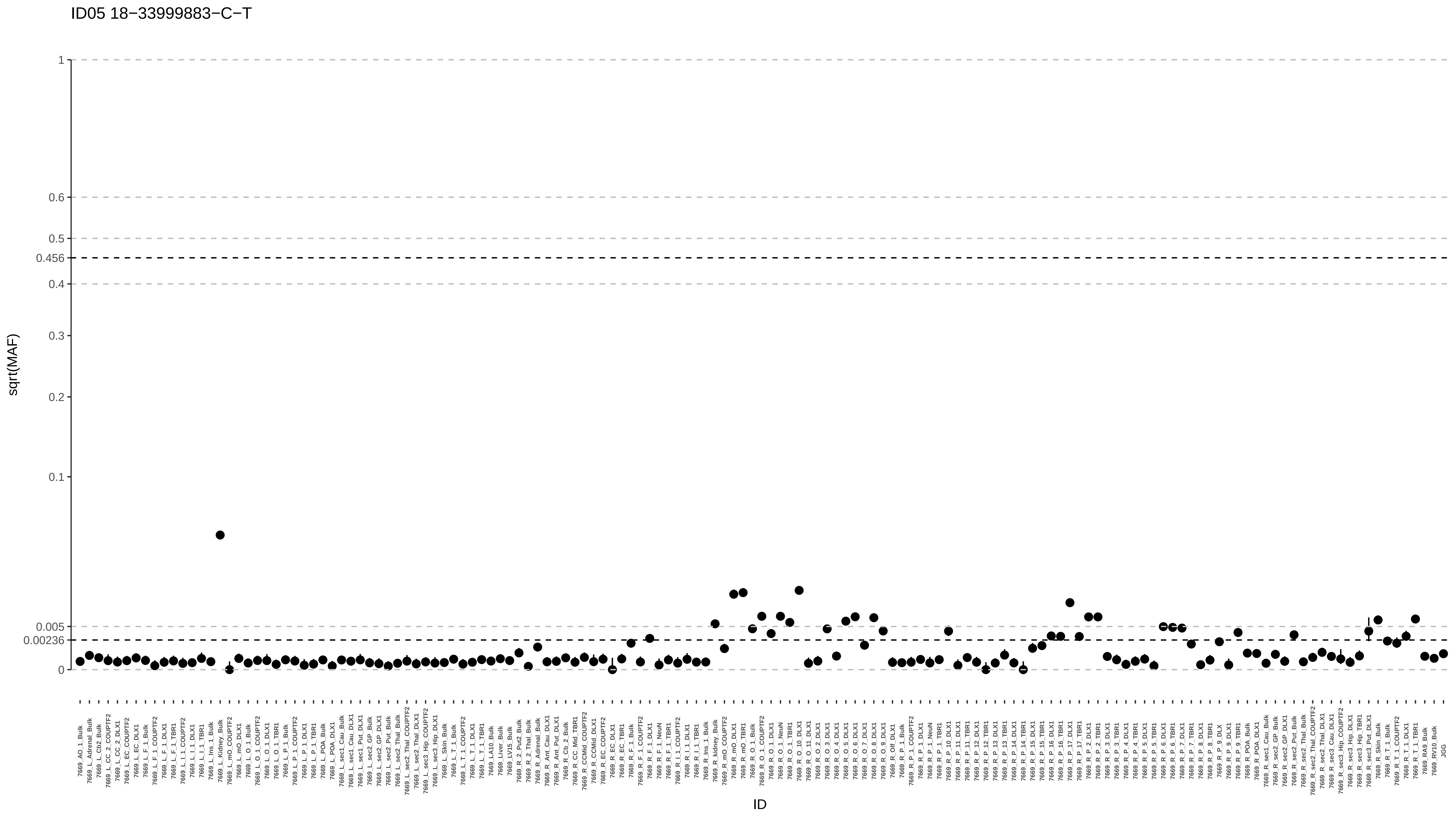
ID05 18−33999883−C−T
sqrt(MAF)
1
0.6
0.5
0.456
0.4
0.3
0.2
0.1
0.005
0.00236
0
ID
7669_AO_1_Bulk
7669_L_Adrenal_Bulk
7669_L_Cb2_Bulk
7669_L_CC_2_COUPTF2
7669_L_CC_2_DLX1
7669_L_EC_COUPTF2
7669_L_EC_DLX1
7669_L_F_1_Bulk
7669_L_F_1_COUPTF2
7669_L_F_1_DLX1
7669_L_F_1_TBR1
7669_L_I_1_COUPTF2
7669_L_I_1_DLX1
7669_L_I_1_TBR1
7669_L_Ins_1_Bulk
7669_L_Kidney_Bulk
7669_L_mO_COUPTF2
7669_L_mO_DLX1
7669_L_O_1_Bulk
7669_L_O_1_COUPTF2
7669_L_O_1_DLX1
7669_L_O_1_TBR1
7669_L_P_1_Bulk
7669_L_P_1_COUPTF2
7669_L_P_1_DLX1
7669_L_P_1_TBR1
7669_L_POA_Bulk
7669_L_POA_DLX1
7669_L_sec1_Cau_Bulk
7669_L_sec1_Cau_DLX1
7669_L_sec1_Put_DLX1
7669_L_sec2_GP_Bulk
7669_L_sec2_GP_DLX1
7669_L_sec2_Put_Bulk
7669_L_sec2_Thal_Bulk
7669_L_sec2_Thal_COUPTF2
7669_L_sec2_Thal_DLX1
7669_L_sec3_Hip_COUPTF2
7669_L_sec3_Hip_DLX1
7669_L_Skin_Bulk
7669_L_T_1_Bulk
7669_L_T_1_COUPTF2
7669_L_T_1_DLX1
7669_L_T_1_TBR1
7669_LA10_Bulk
7669_Liver_Bulk
7669_LV15_Bulk
7669_R_2_Put2_Bulk
7669_R_2_Thal_Bulk
7669_R_Adrenal_Bulk
7669_R_Ant_Cau_DLX1
7669_R_Ant_Put_DLX1
7669_R_Cb_2_Bulk
7669_R_CC_Mid_TBR1
7669_R_CCMid_COUPTF2
7669_R_CCMid_DLX1
7669_R_EC_COUPTF2
7669_R_EC_DLX1
7669_R_EC_TBR1
7669_R_F_1_Bulk
7669_R_F_1_COUPTF2
7669_R_F_1_DLX1
7669_R_F_1_NeuN
7669_R_F_1_TBR1
7669_R_I_1_COUPTF2
7669_R_I_1_DLX1
7669_R_I_1_TBR1
7669_R_Ins_1_Bulk
7669_R_kidney_Bulk
7669_R_mO_COUPTF2
7669_R_mO_DLX1
7669_R_mO_TBR1
7669_R_O_1_Bulk
7669_R_O_1_COUPTF2
7669_R_O_1_DLX1
7669_R_O_1_NeuN
7669_R_O_1_TBR1
7669_R_O_10_DLX1
7669_R_O_11_DLX1
7669_R_O_2_DLX1
7669_R_O_3_DLX1
7669_R_O_4_DLX1
7669_R_O_5_DLX1
7669_R_O_6_DLX1
7669_R_O_7_DLX1
7669_R_O_8_DLX1
7669_R_O_9_DLX1
7669_R_Olf_DLX1
7669_R_P_1_Bulk
7669_R_P_1_COUPTF2
7669_R_P_1_DLX1
7669_R_P_1_NeuN
7669_R_P_1_TBR1
7669_R_P_10_DLX1
7669_R_P_11_DLX1
7669_R_P_11_TBR1
7669_R_P_12_DLX1
7669_R_P_12_TBR1
7669_R_P_13_DLX1
7669_R_P_13_TBR1
7669_R_P_14_DLX1
7669_R_P_14_TBR1
7669_R_P_15_DLX1
7669_R_P_15_TBR1
7669_R_P_16_DLX1
7669_R_P_16_TBR1
7669_R_P_17_DLX1
7669_R_P_17_TBR1
7669_R_P_2_DLX1
7669_R_P_2_TBR1
7669_R_P_3_DLX1
7669_R_P_3_TBR1
7669_R_P_4_DLX1
7669_R_P_4_TBR1
7669_R_P_5_DLX1
7669_R_P_5_TBR1
7669_R_P_6_DLX1
7669_R_P_6_TBR1
7669_R_P_7_DLX1
7669_R_P_7_TBR1
7669_R_P_8_DLX1
7669_R_P_8_TBR1
7669_R_P_9_DLX
7669_R_P_9_DLX1
7669_R_P_9_TBR1
7669_R_POA_Bulk
7669_R_POA_DLX1
7669_R_sec1_Cau_Bulk
7669_R_sec2_GP_Bulk
7669_R_sec2_GP_DLX1
7669_R_sec2_Put_Bulk
7669_R_sec2_Thal_Bulk
7669_R_sec2_Thal_COUPTF2
7669_R_sec2_Thal_DLX1
7669_R_sec3_Cau_DLX1
7669_R_sec3_Hip_COUPTF2
7669_R_sec3_Hip_DLX1
7669_R_sec3_Hip_TBR1
7669_R_sec3_Put_DLX1
7669_R_Skin_Bulk
7669_R_T_1_Bulk
7669_R_T_1_COUPTF2
7669_R_T_1_DLX1
7669_R_T_1_TBR1
7669_RA9_Bulk
7669_RV10_Bulk
JGG

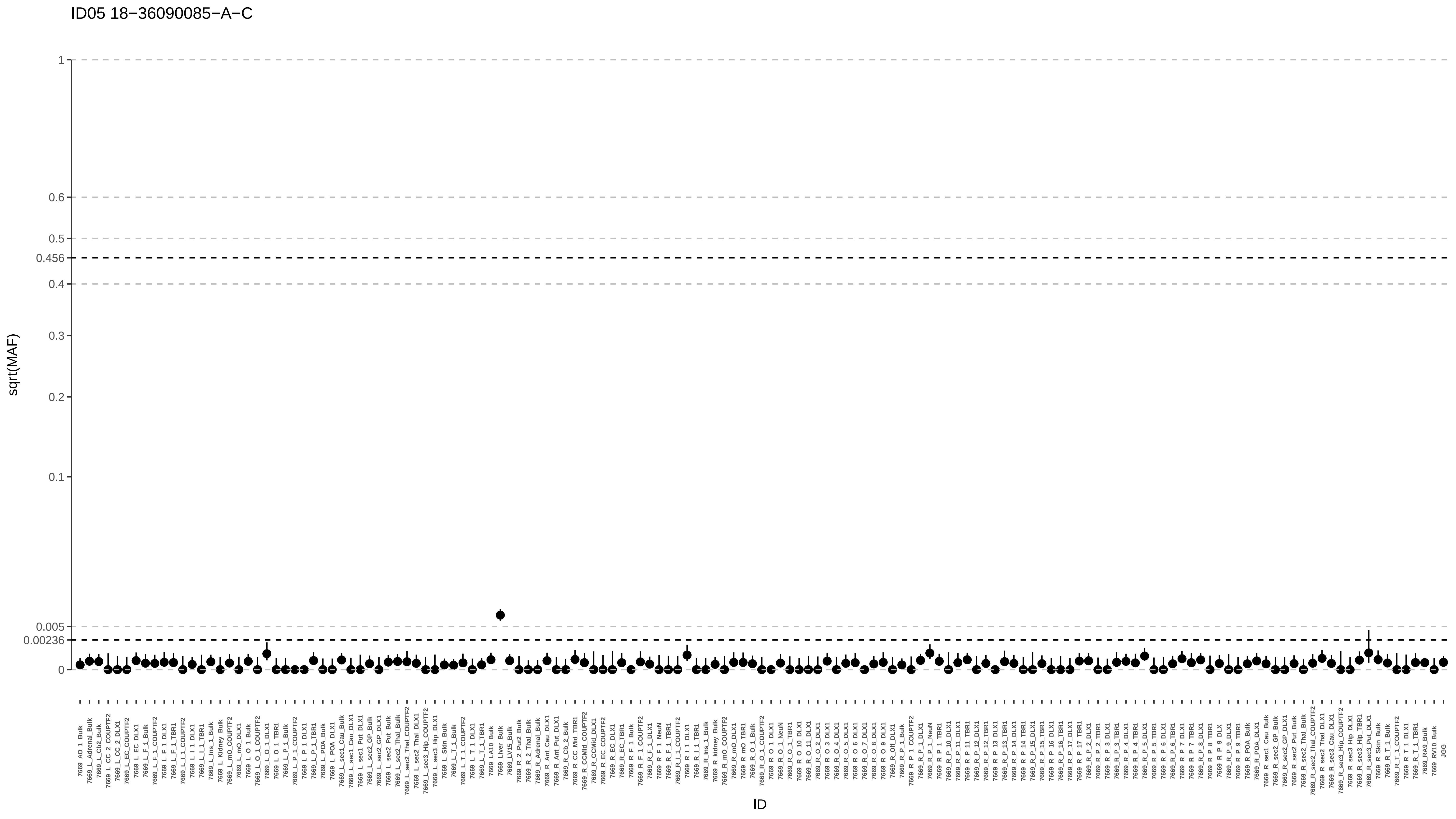
ID05 18−36090085−A−C
sqrt(MAF)
1
0.6
0.5
0.456
0.4
0.3
0.2
0.1
0.005
0.00236
0
ID
7669_AO_1_Bulk
7669_L_Adrenal_Bulk
7669_L_Cb2_Bulk
7669_L_CC_2_COUPTF2
7669_L_CC_2_DLX1
7669_L_EC_COUPTF2
7669_L_EC_DLX1
7669_L_F_1_Bulk
7669_L_F_1_COUPTF2
7669_L_F_1_DLX1
7669_L_F_1_TBR1
7669_L_I_1_COUPTF2
7669_L_I_1_DLX1
7669_L_I_1_TBR1
7669_L_Ins_1_Bulk
7669_L_Kidney_Bulk
7669_L_mO_COUPTF2
7669_L_mO_DLX1
7669_L_O_1_Bulk
7669_L_O_1_COUPTF2
7669_L_O_1_DLX1
7669_L_O_1_TBR1
7669_L_P_1_Bulk
7669_L_P_1_COUPTF2
7669_L_P_1_DLX1
7669_L_P_1_TBR1
7669_L_POA_Bulk
7669_L_POA_DLX1
7669_L_sec1_Cau_Bulk
7669_L_sec1_Cau_DLX1
7669_L_sec1_Put_DLX1
7669_L_sec2_GP_Bulk
7669_L_sec2_GP_DLX1
7669_L_sec2_Put_Bulk
7669_L_sec2_Thal_Bulk
7669_L_sec2_Thal_COUPTF2
7669_L_sec2_Thal_DLX1
7669_L_sec3_Hip_COUPTF2
7669_L_sec3_Hip_DLX1
7669_L_Skin_Bulk
7669_L_T_1_Bulk
7669_L_T_1_COUPTF2
7669_L_T_1_DLX1
7669_L_T_1_TBR1
7669_LA10_Bulk
7669_Liver_Bulk
7669_LV15_Bulk
7669_R_2_Put2_Bulk
7669_R_2_Thal_Bulk
7669_R_Adrenal_Bulk
7669_R_Ant_Cau_DLX1
7669_R_Ant_Put_DLX1
7669_R_Cb_2_Bulk
7669_R_CC_Mid_TBR1
7669_R_CCMid_COUPTF2
7669_R_CCMid_DLX1
7669_R_EC_COUPTF2
7669_R_EC_DLX1
7669_R_EC_TBR1
7669_R_F_1_Bulk
7669_R_F_1_COUPTF2
7669_R_F_1_DLX1
7669_R_F_1_NeuN
7669_R_F_1_TBR1
7669_R_I_1_COUPTF2
7669_R_I_1_DLX1
7669_R_I_1_TBR1
7669_R_Ins_1_Bulk
7669_R_kidney_Bulk
7669_R_mO_COUPTF2
7669_R_mO_DLX1
7669_R_mO_TBR1
7669_R_O_1_Bulk
7669_R_O_1_COUPTF2
7669_R_O_1_DLX1
7669_R_O_1_NeuN
7669_R_O_1_TBR1
7669_R_O_10_DLX1
7669_R_O_11_DLX1
7669_R_O_2_DLX1
7669_R_O_3_DLX1
7669_R_O_4_DLX1
7669_R_O_5_DLX1
7669_R_O_6_DLX1
7669_R_O_7_DLX1
7669_R_O_8_DLX1
7669_R_O_9_DLX1
7669_R_Olf_DLX1
7669_R_P_1_Bulk
7669_R_P_1_COUPTF2
7669_R_P_1_DLX1
7669_R_P_1_NeuN
7669_R_P_1_TBR1
7669_R_P_10_DLX1
7669_R_P_11_DLX1
7669_R_P_11_TBR1
7669_R_P_12_DLX1
7669_R_P_12_TBR1
7669_R_P_13_DLX1
7669_R_P_13_TBR1
7669_R_P_14_DLX1
7669_R_P_14_TBR1
7669_R_P_15_DLX1
7669_R_P_15_TBR1
7669_R_P_16_DLX1
7669_R_P_16_TBR1
7669_R_P_17_DLX1
7669_R_P_17_TBR1
7669_R_P_2_DLX1
7669_R_P_2_TBR1
7669_R_P_3_DLX1
7669_R_P_3_TBR1
7669_R_P_4_DLX1
7669_R_P_4_TBR1
7669_R_P_5_DLX1
7669_R_P_5_TBR1
7669_R_P_6_DLX1
7669_R_P_6_TBR1
7669_R_P_7_DLX1
7669_R_P_7_TBR1
7669_R_P_8_DLX1
7669_R_P_8_TBR1
7669_R_P_9_DLX
7669_R_P_9_DLX1
7669_R_P_9_TBR1
7669_R_POA_Bulk
7669_R_POA_DLX1
7669_R_sec1_Cau_Bulk
7669_R_sec2_GP_Bulk
7669_R_sec2_GP_DLX1
7669_R_sec2_Put_Bulk
7669_R_sec2_Thal_Bulk
7669_R_sec2_Thal_COUPTF2
7669_R_sec2_Thal_DLX1
7669_R_sec3_Cau_DLX1
7669_R_sec3_Hip_COUPTF2
7669_R_sec3_Hip_DLX1
7669_R_sec3_Hip_TBR1
7669_R_sec3_Put_DLX1
7669_R_Skin_Bulk
7669_R_T_1_Bulk
7669_R_T_1_COUPTF2
7669_R_T_1_DLX1
7669_R_T_1_TBR1
7669_RA9_Bulk
7669_RV10_Bulk
JGG

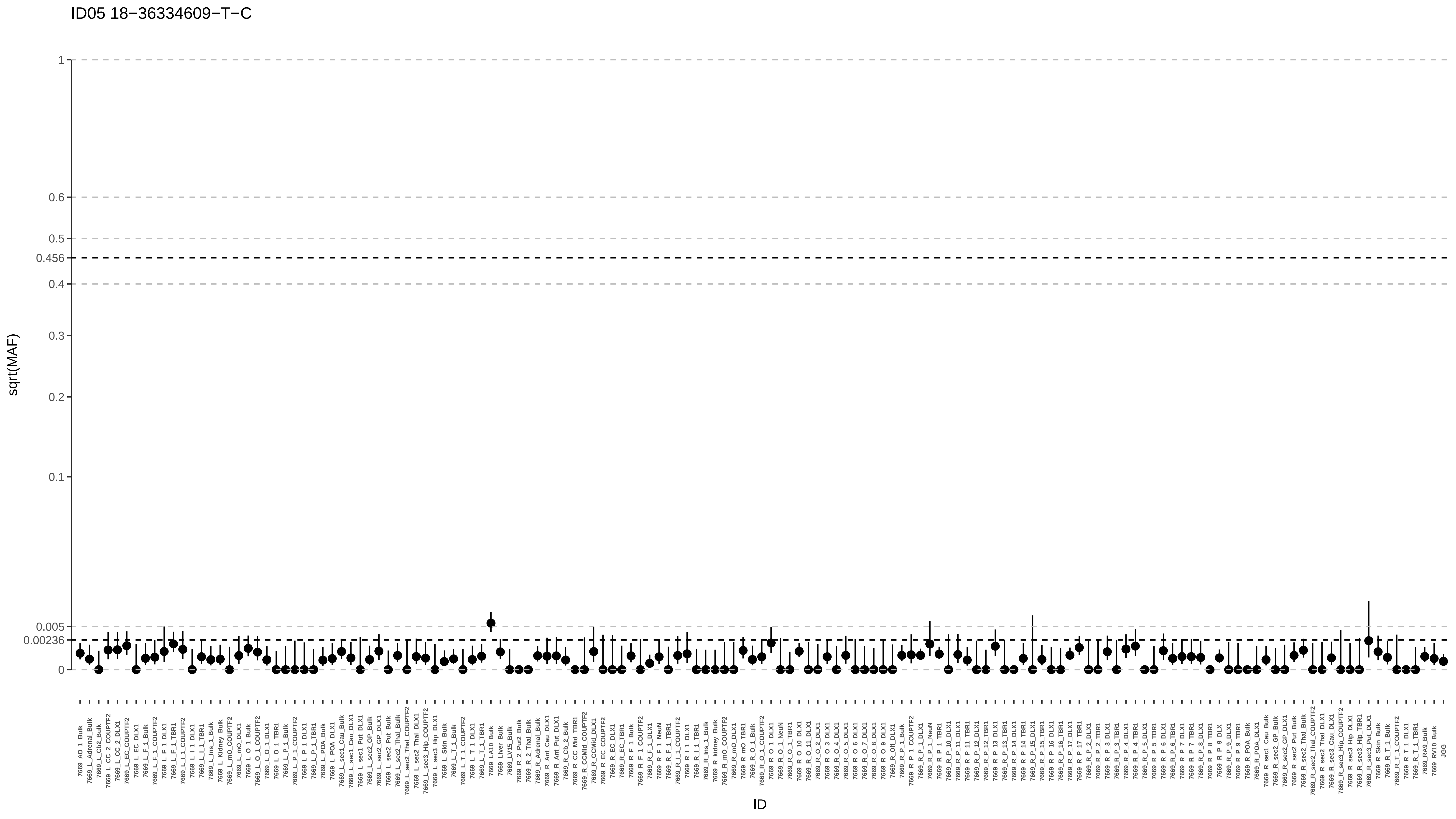

ID05 18−36334609−T−C
sqrt(MAF)
1
0.6
0.5
0.456
0.4
0.3
0.2
0.1
0.005
0.00236
0
ID

ID05 18−37021461−T−C

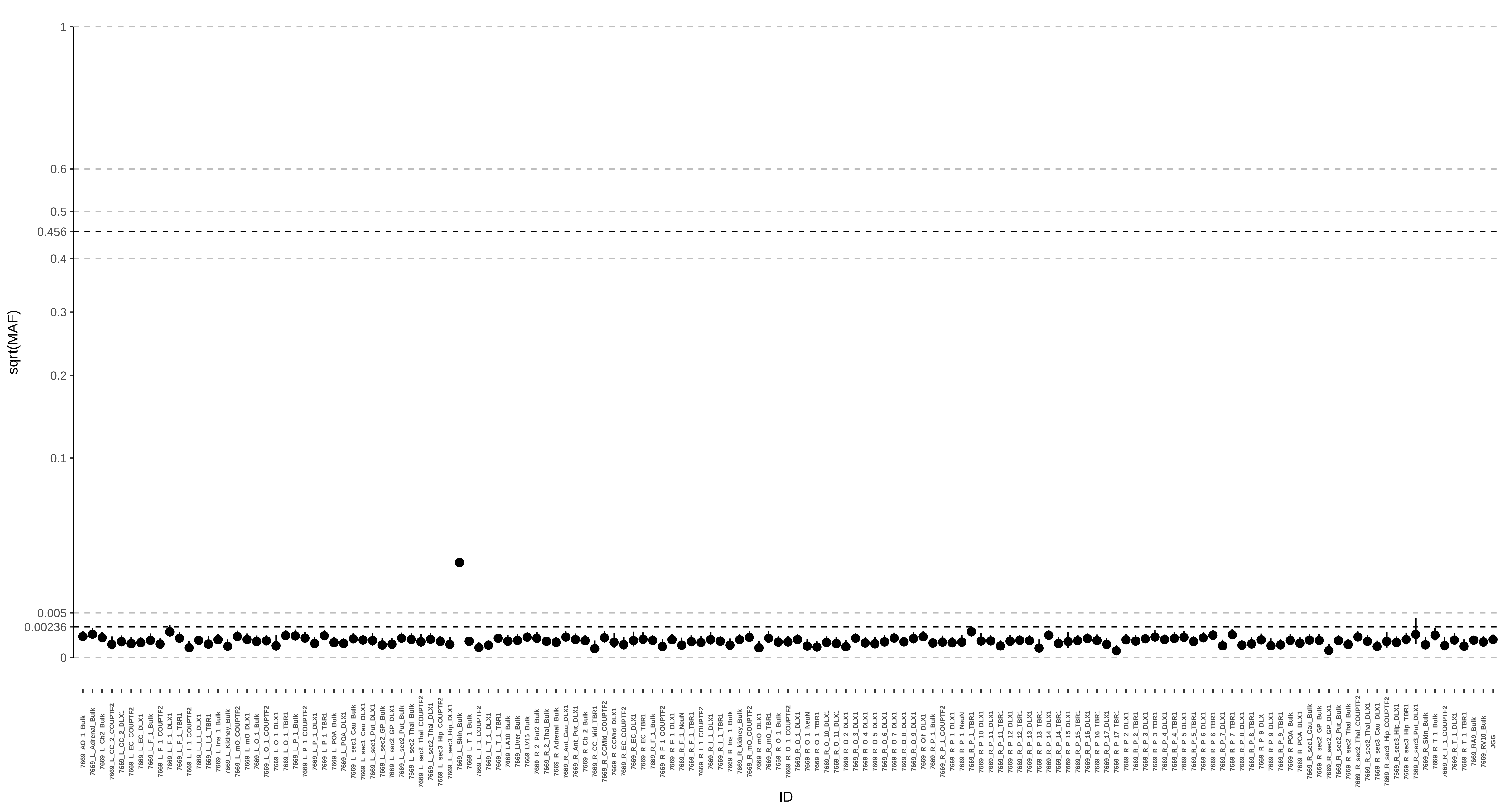

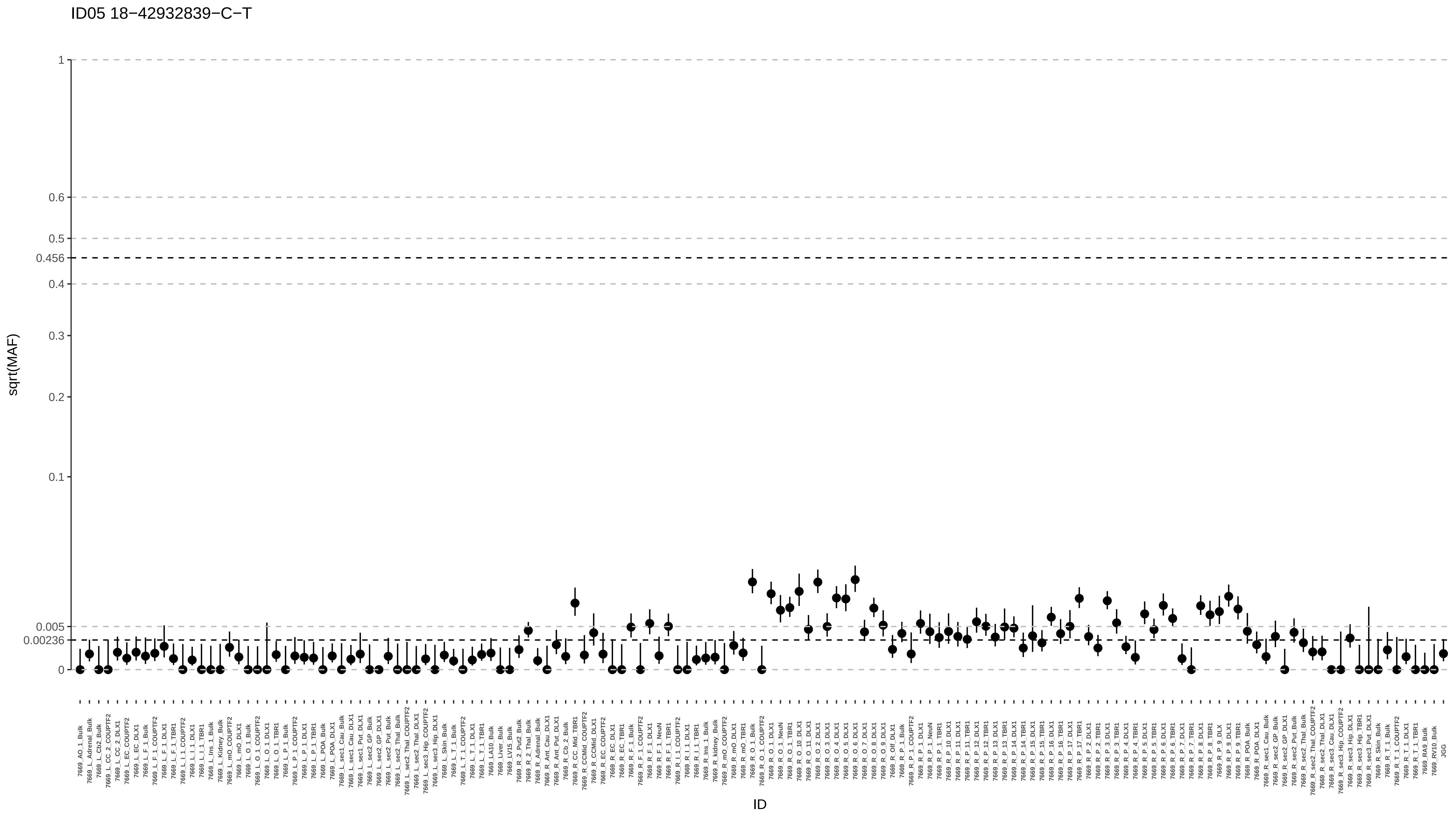
ID05 18−42932839−C−T
sqrt(MAF)
1
0.6
0.5
0.456
0.4
0.3
0.2
0.1
0.005
0.00236
0
ID

ID05 18−49194474−G−A

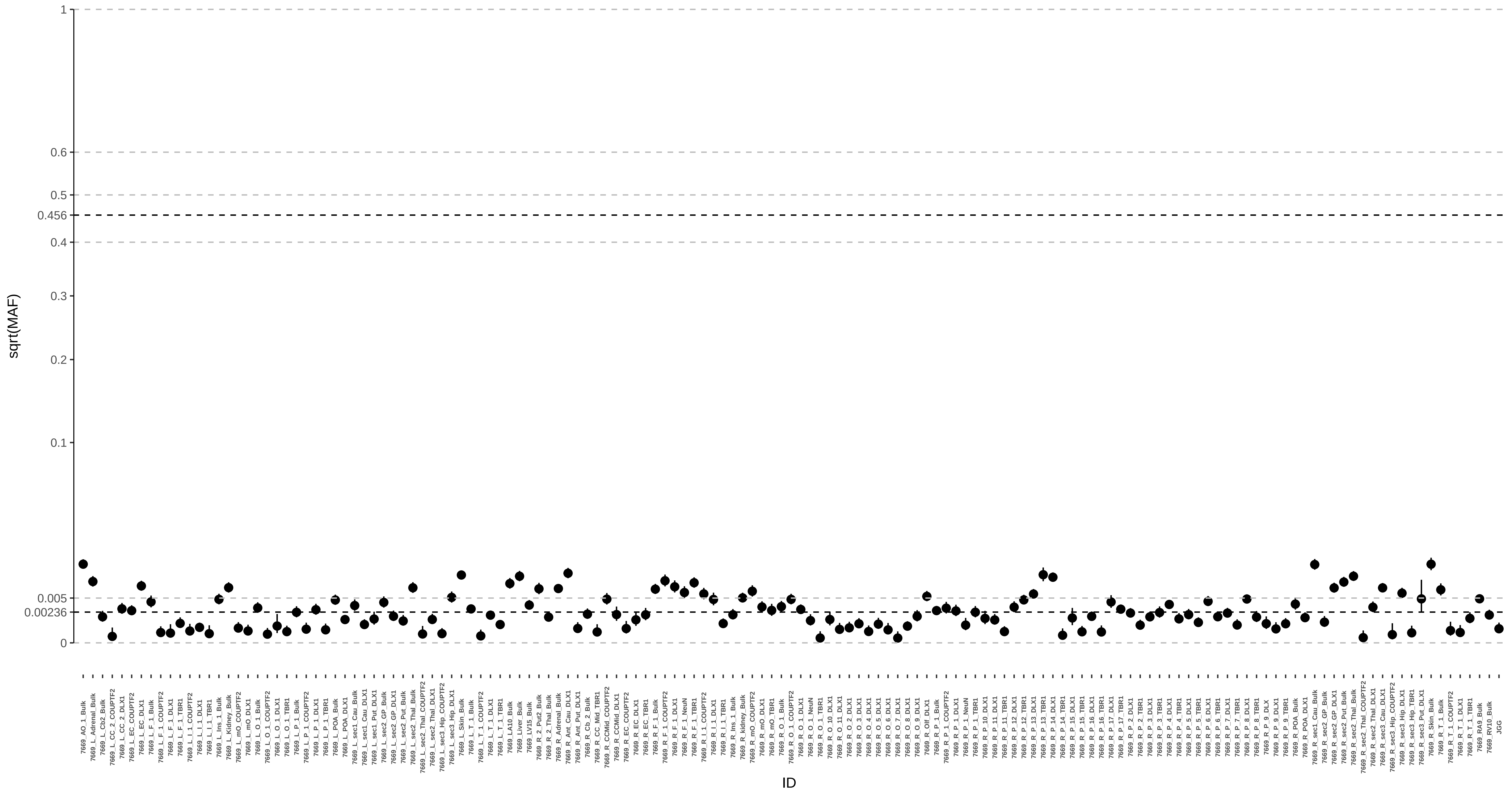

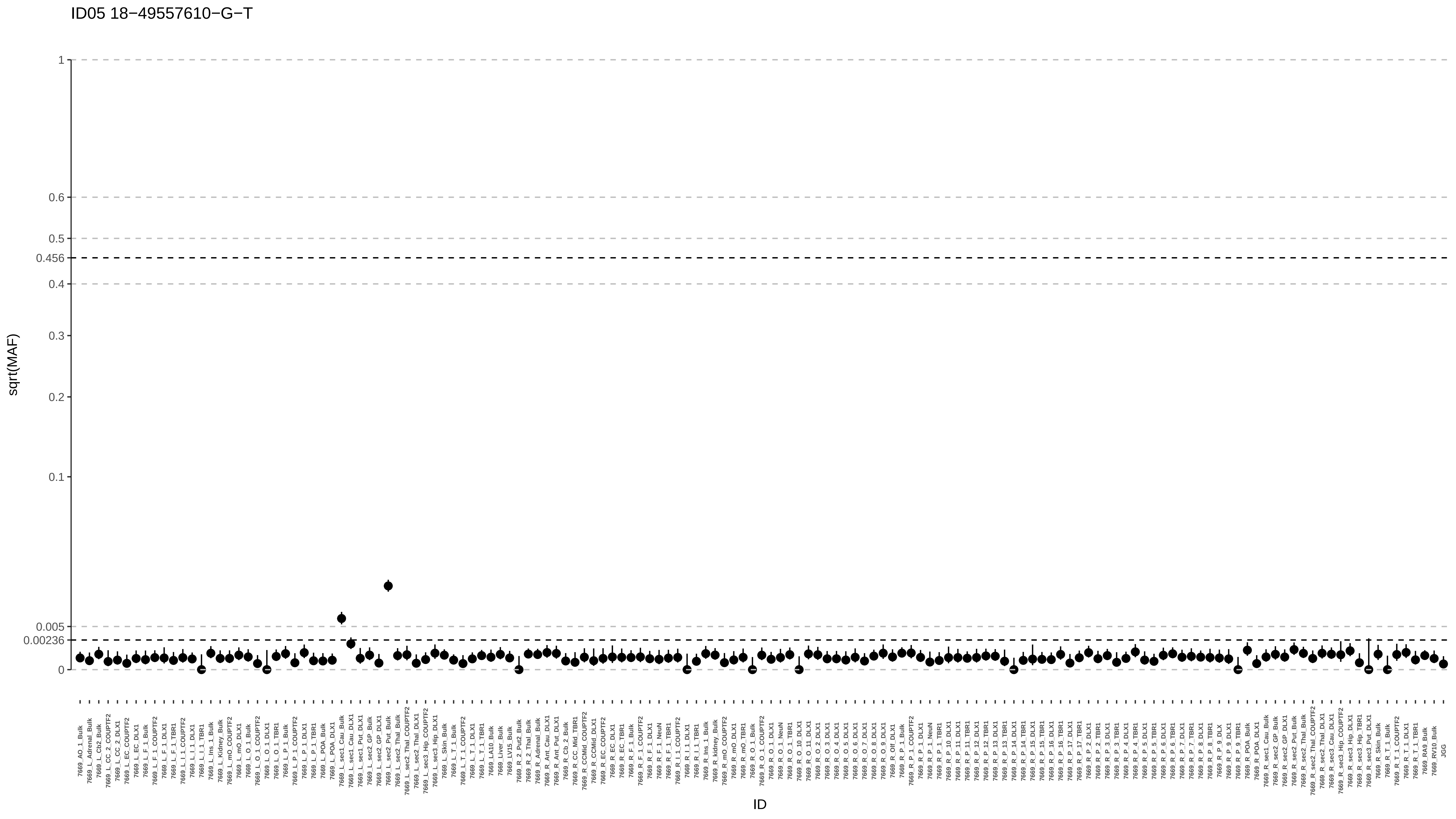
ID05 18−49557610−G−T
sqrt(MAF)
1
0.6
0.5
0.456
0.4
0.3
0.2
0.1
0.005
0.00236
0
ID

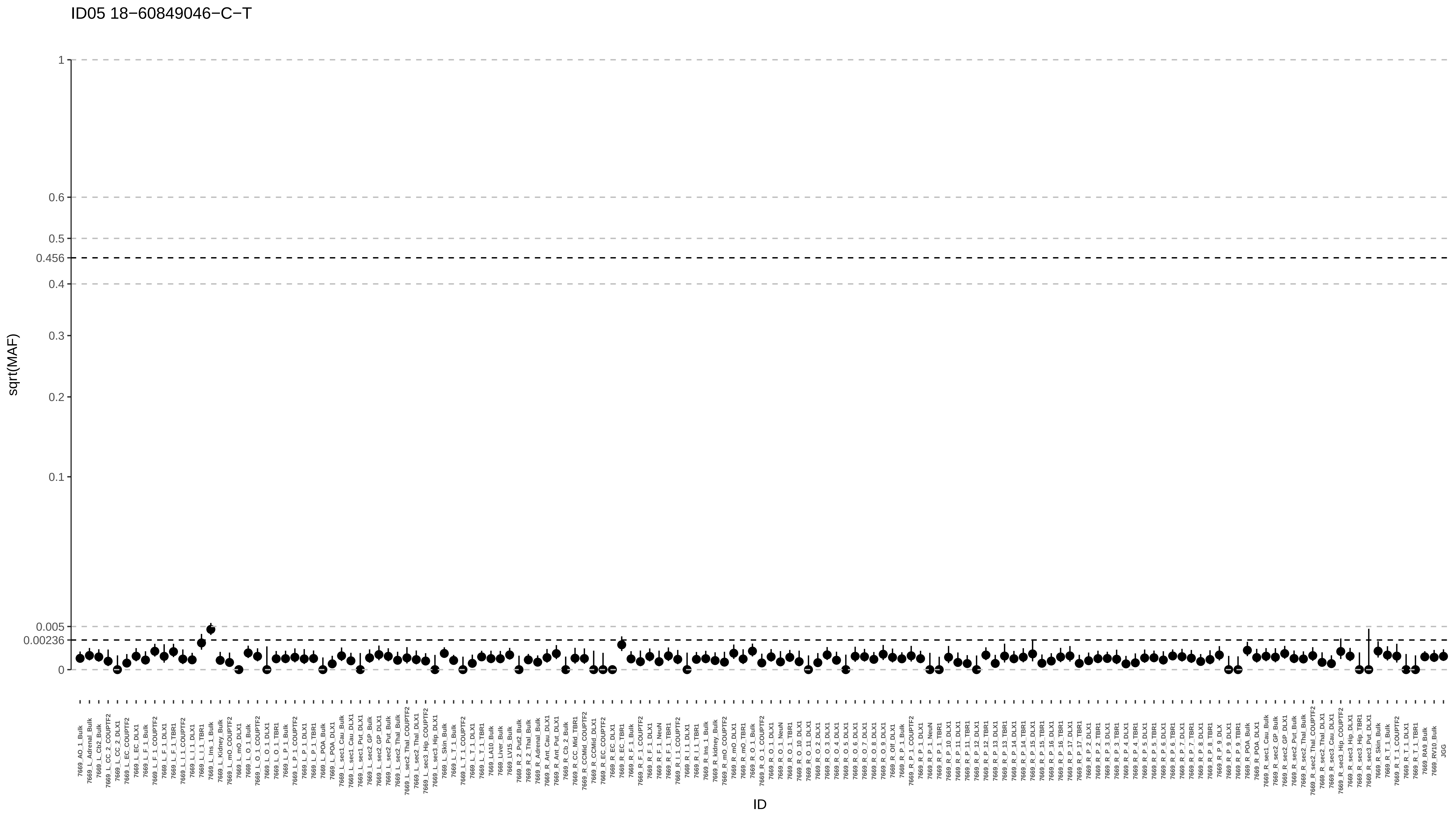

ID05 18−60849046−C−T
sqrt(MAF)
1
0.6
0.5
0.456
0.4
0.3
0.2
0.1
0.005
0.00236
0
ID

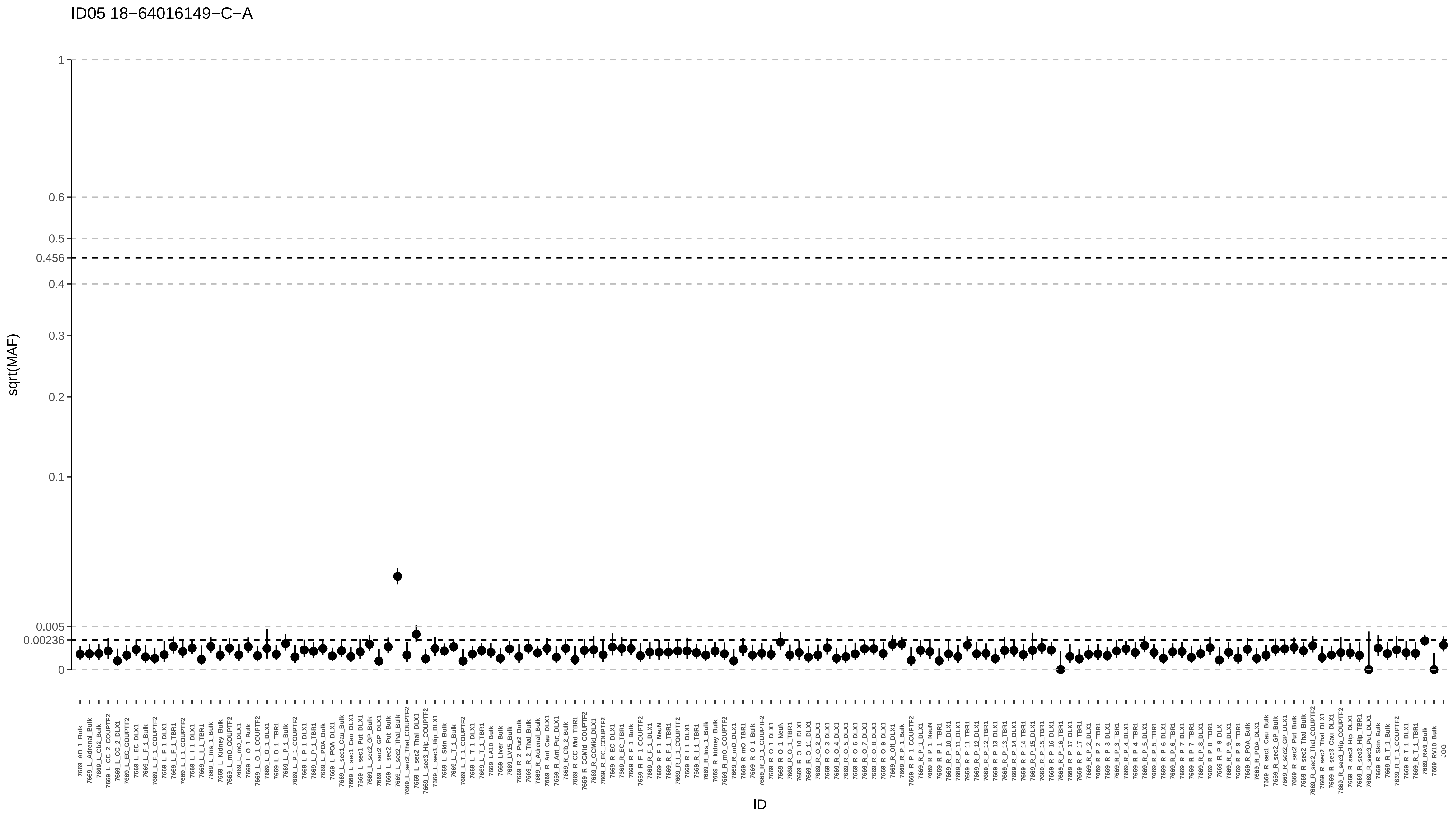

ID05 18−64016149−C−A
sqrt(MAF)
ID

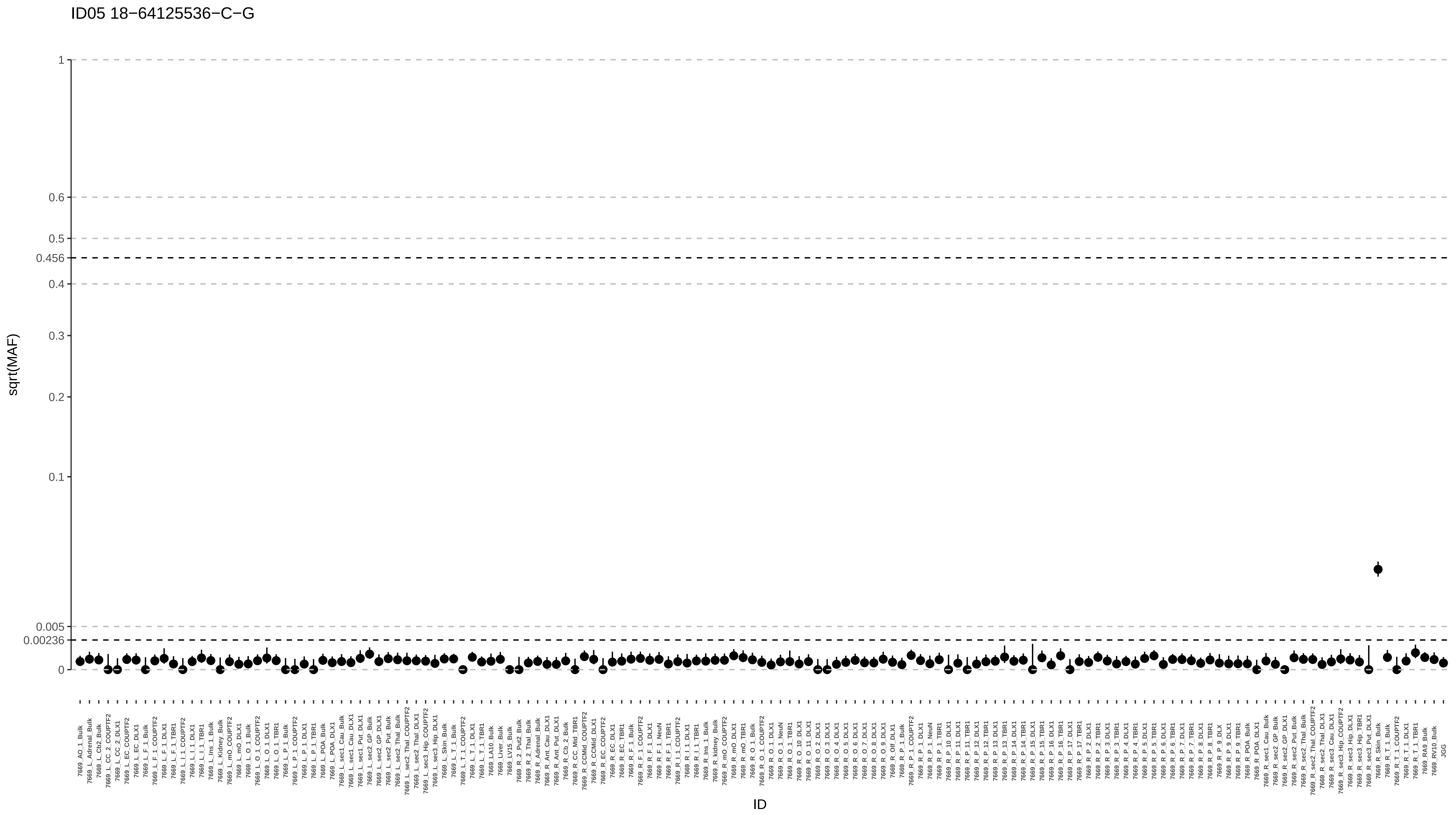

ID05 18−64125536−C−G
sqrt(MAF)
1
0.6
0.5
0.456
0.4
0.3
0.2
0.1
0.005
0.00236
0
ID

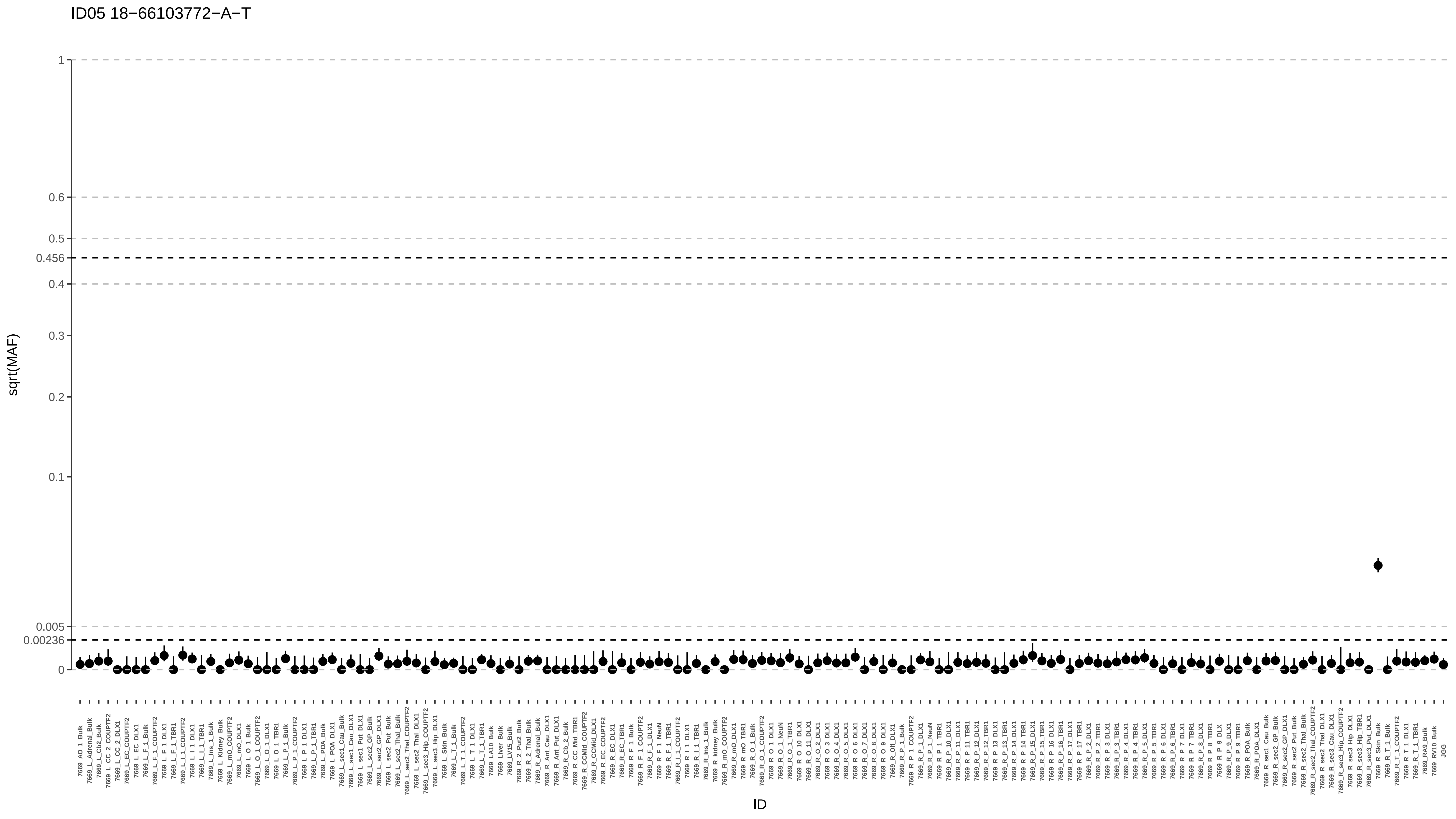
ID05 18−66103772−A−T
sqrt(MAF)
1
0.6
0.5
0.456
0.4
0.3
0.2
0.1
0.005
0.00236
0
7669_AO_1_Bulk
7669_L_Adrenal_Bulk
7669_L_Cb2_Bulk
7669_L_CC_2_COUPTF2
7669_L_CC_2_DLX1
7669_L_EC_COUPTF2
7669_L_EC_DLX1
7669_L_F_1_Bulk
7669_L_F_1_COUPTF2
7669_L_F_1_DLX1
7669_L_F_1_TBR1
7669_L_I_1_COUPTF2
7669_L_I_1_DLX1
7669_L_I_1_TBR1
7669_L_Ins_1_Bulk
7669_L_Kidney_Bulk
7669_L_mO_COUPTF2
7669_L_mO_DLX1
7669_L_O_1_Bulk
7669_L_O_1_COUPTF2
7669_L_O_1_DLX1
7669_L_O_1_TBR1
7669_L_P_1_Bulk
7669_L_P_1_COUPTF2
7669_L_P_1_DLX1
7669_L_P_1_TBR1
7669_L_POA_Bulk
7669_L_POA_DLX1
7669_L_sec1_Cau_Bulk
7669_L_sec1_Cau_DLX1
7669_L_sec1_Put_DLX1
7669_L_sec2_GP_Bulk
7669_L_sec2_GP_DLX1
7669_L_sec2_Put_Bulk
7669_L_sec2_Thal_Bulk
7669_L_sec2_Thal_COUPTF2
7669_L_sec2_Thal_DLX1
7669_L_sec3_Hip_COUPTF2
7669_L_sec3_Hip_DLX1
7669_L_Skin_Bulk
7669_L_T_1_Bulk
7669_L_T_1_COUPTF2
7669_L_T_1_DLX1
7669_L_T_1_TBR1
7669_LA10_Bulk
7669_Liver_Bulk
7669_LV15_Bulk
7669_R_2_Put2_Bulk
7669_R_2_Thal_Bulk
7669_R_Adrenal_Bulk
7669_R_Ant_Cau_DLX1
7669_R_Ant_Put_DLX1
7669_R_Cb_2_Bulk
7669_R_CC_Mid_TBR1
7669_R_CCMid_COUPTF2
7669_R_CCMid_DLX1
7669_R_EC_COUPTF2
7669_R_EC_DLX1
7669_R_EC_TBR1
7669_R_F_1_Bulk
7669_R_F_1_COUPTF2
7669_R_F_1_DLX1
7669_R_F_1_NeuN
7669_R_F_1_TBR1
7669_R_I_1_COUPTF2
7669_R_I_1_DLX1
7669_R_I_1_TBR1
7669_R_Ins_1_Bulk
7669_R_kidney_Bulk
7669_R_mO_COUPTF2
7669_R_mO_DLX1
7669_R_mO_TBR1
7669_R_O_1_Bulk
7669_R_O_1_COUPTF2
7669_R_O_1_DLX1
7669_R_O_1_NeuN
7669_R_O_1_TBR1
7669_R_O_10_DLX1
7669_R_O_11_DLX1
7669_R_O_2_DLX1
7669_R_O_3_DLX1
7669_R_O_4_DLX1
7669_R_O_5_DLX1
7669_R_O_6_DLX1
7669_R_O_7_DLX1
7669_R_O_8_DLX1
7669_R_O_9_DLX1
7669_R_Olf_DLX1
7669_R_P_1_Bulk
7669_R_P_1_COUPTF2
7669_R_P_1_DLX1
7669_R_P_1_NeuN
7669_R_P_1_TBR1
7669_R_P_10_DLX1
7669_R_P_11_DLX1
7669_R_P_11_TBR1
7669_R_P_12_DLX1
7669_R_P_12_TBR1
7669_R_P_13_DLX1
7669_R_P_13_TBR1
7669_R_P_14_DLX1
7669_R_P_14_TBR1
7669_R_P_15_DLX1
7669_R_P_15_TBR1
7669_R_P_16_DLX1
7669_R_P_16_TBR1
7669_R_P_17_DLX1
7669_R_P_17_TBR1
7669_R_P_2_DLX1
7669_R_P_2_TBR1
7669_R_P_3_DLX1
7669_R_P_3_TBR1
7669_R_P_4_DLX1
7669_R_P_4_TBR1
7669_R_P_5_DLX1
7669_R_P_5_TBR1
7669_R_P_6_DLX1
7669_R_P_6_TBR1
7669_R_P_7_DLX1
7669_R_P_7_TBR1
7669_R_P_8_DLX1
7669_R_P_8_TBR1
7669_R_P_9_DLX
7669_R_P_9_DLX1
7669_R_P_9_TBR1
7669_R_POA_Bulk
7669_R_POA_DLX1
7669_R_sec1_Cau_Bulk
7669_R_sec2_GP_Bulk
7669_R_sec2_GP_DLX1
7669_R_sec2_Put_Bulk
7669_R_sec2_Thal_Bulk
7669_R_sec2_Thal_COUPTF2
7669_R_sec2_Thal_DLX1
7669_R_sec3_Cau_DLX1
7669_R_sec3_Hip_COUPTF2
7669_R_sec3_Hip_DLX1
7669_R_sec3_Hip_TBR1
7669_R_sec3_Put_DLX1
7669_R_Skin_Bulk
7669_R_T_1_Bulk
7669_R_T_1_COUPTF2
7669_R_T_1_DLX1
7669_R_T_1_TBR1
7669_RA9_Bulk
7669_RV10_Bulk
JGG
ID

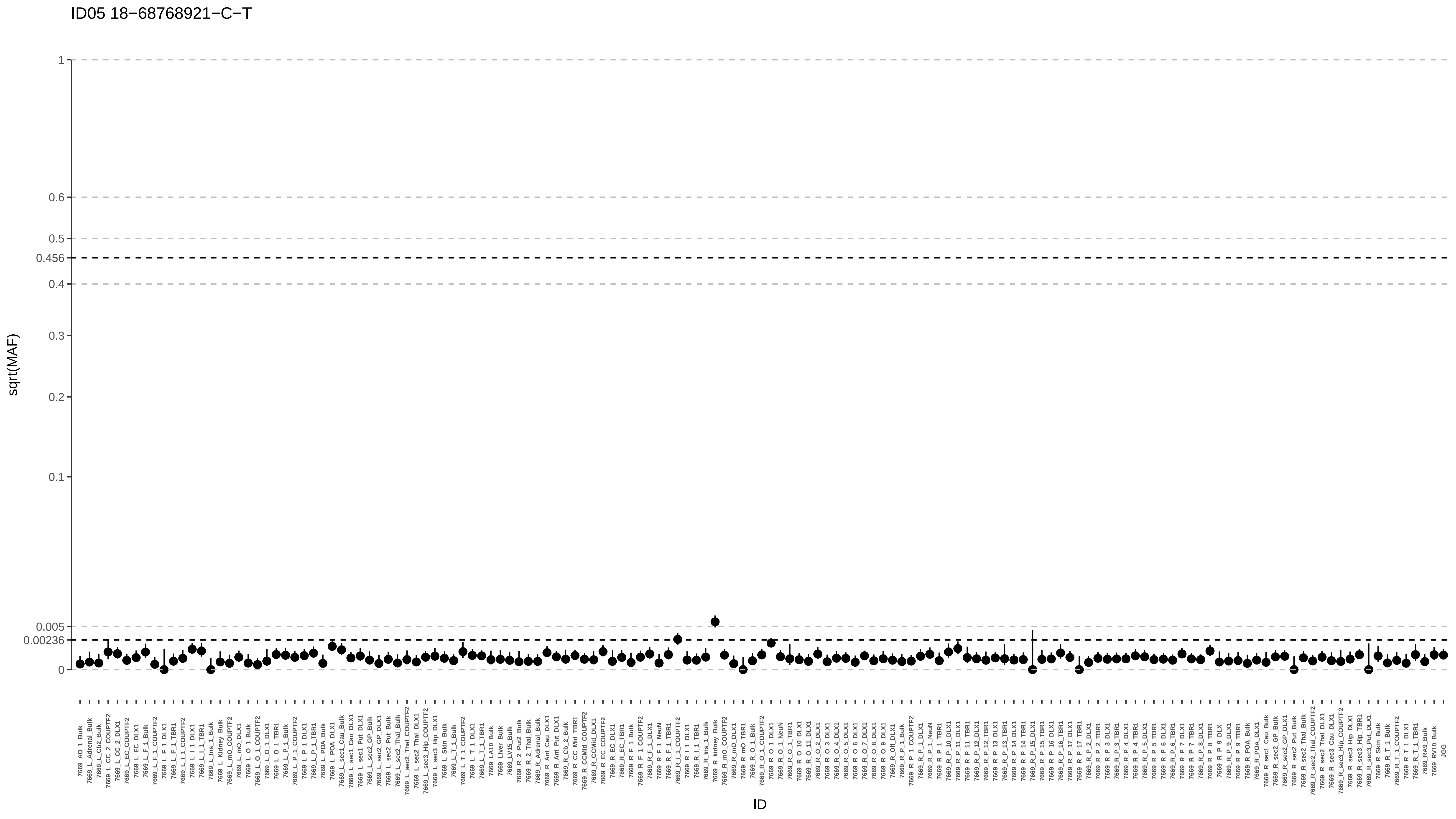

ID05 18−68768921−C−T
sqrt(MAF)
1
0.6
0.5
0.456
0.4
0.3
0.2
0.1
0.005
0.00236
0
ID

ID05 19−1124954−C−T

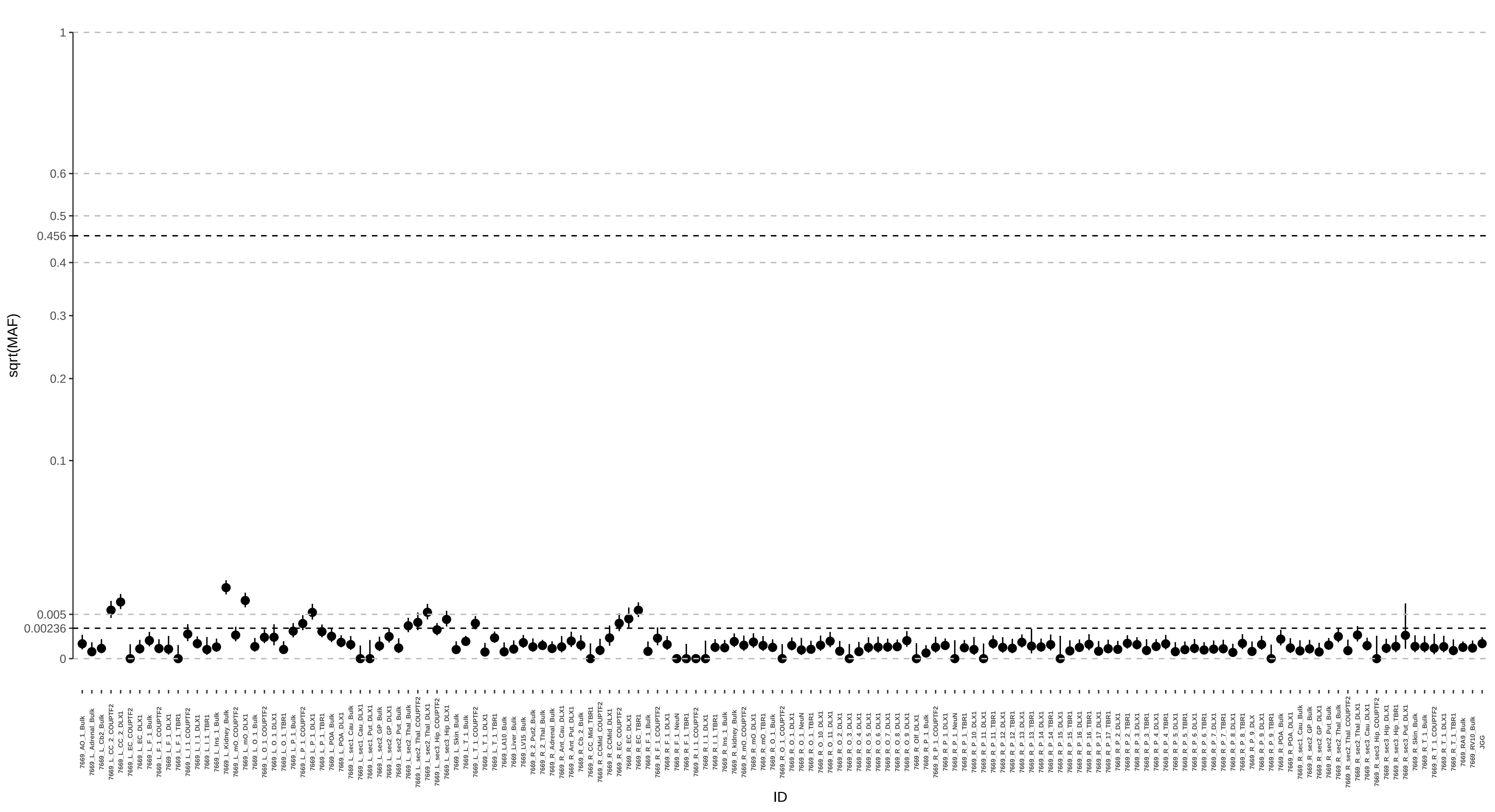

ID05 19−13941229−G−A

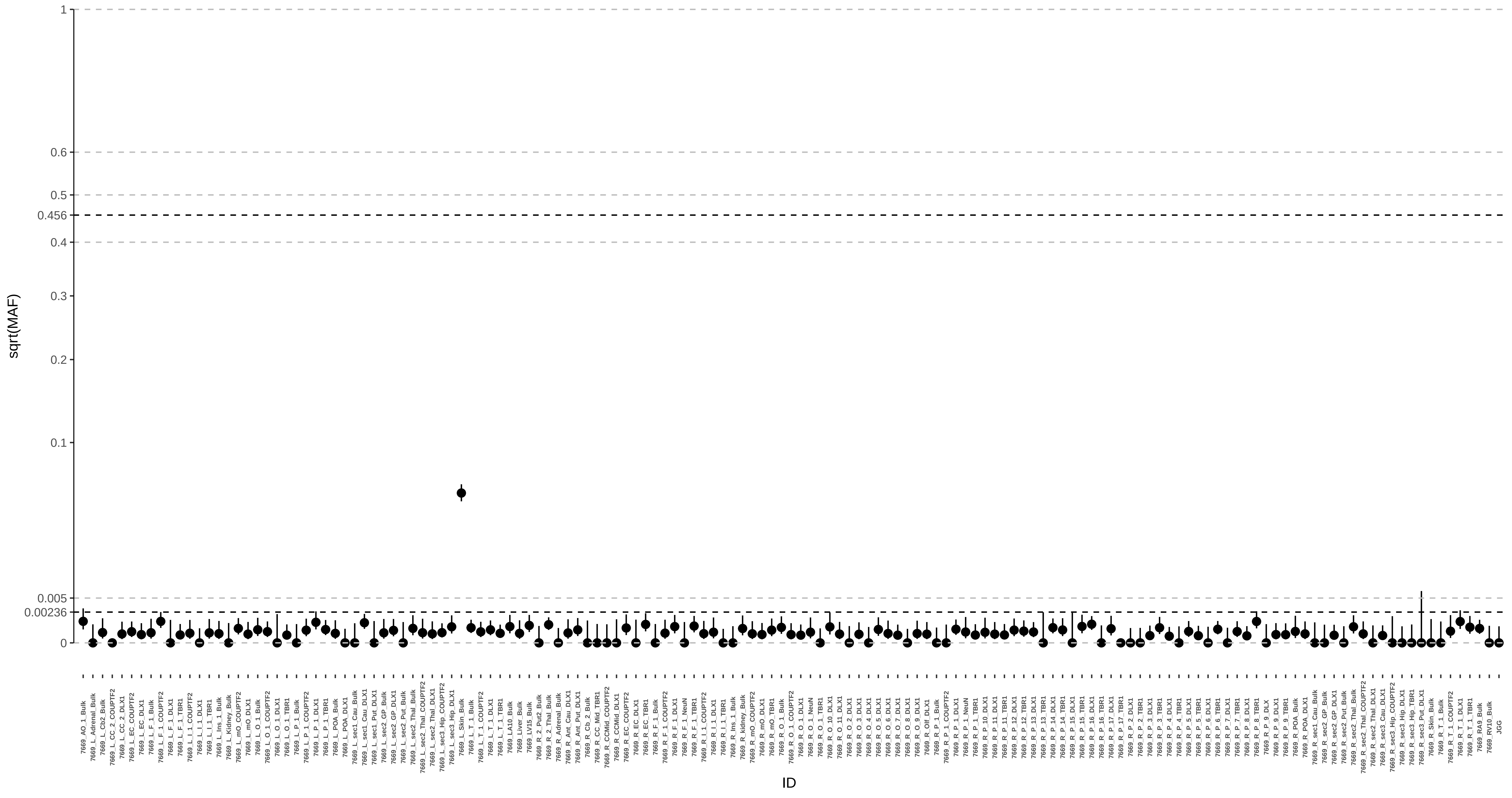

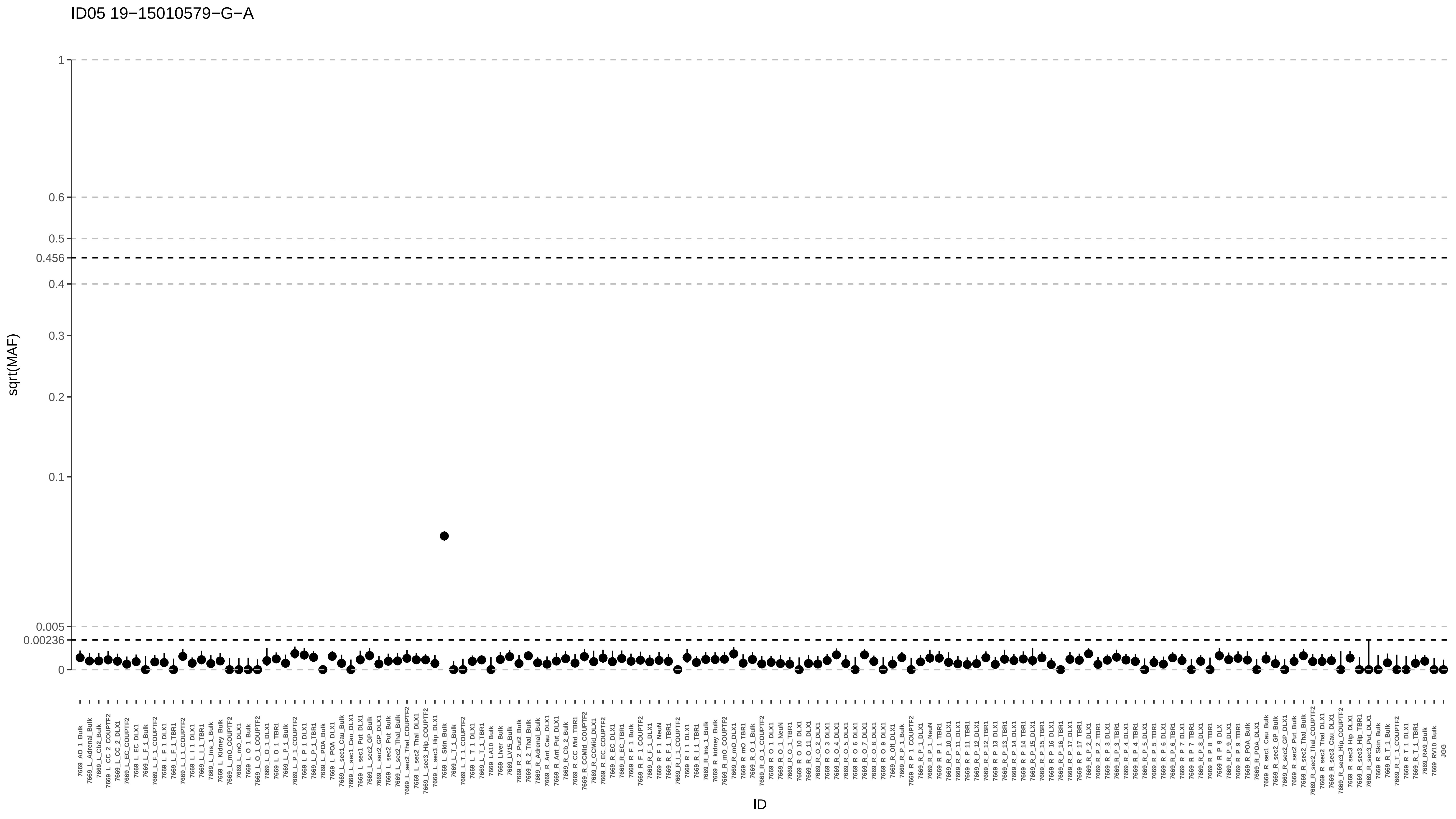
ID05 19−15010579−G−A
sqrt(MAF)
1
0.6
0.5
0.456
0.4
0.3
0.2
0.1
0.005
0.00236
0
ID

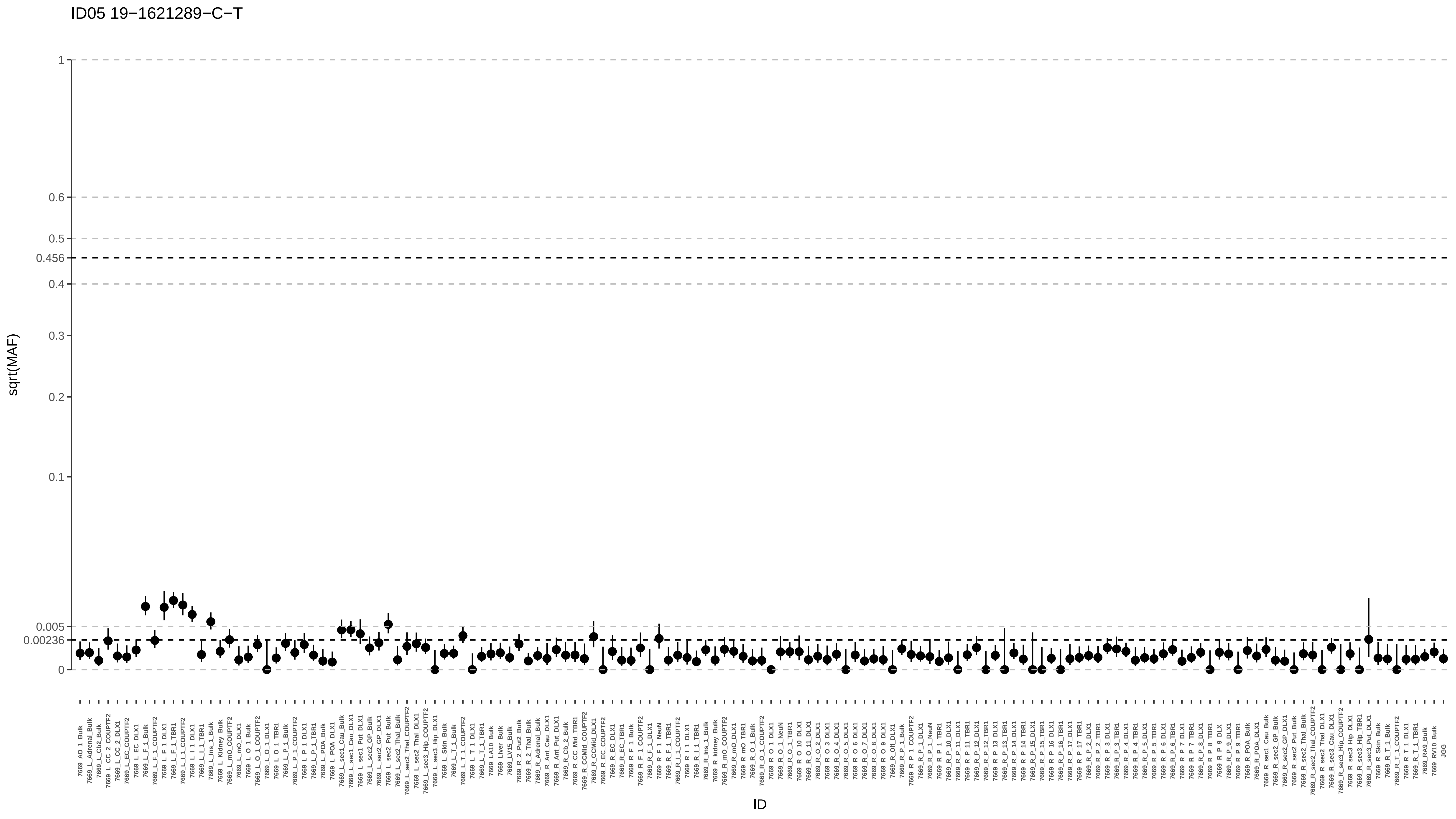
ID05 19−1621289−C−T
sqrt(MAF)
1
0.6
0.5
0.456
0.4
0.3
0.2
0.1
0.005
0.00236
0
ID

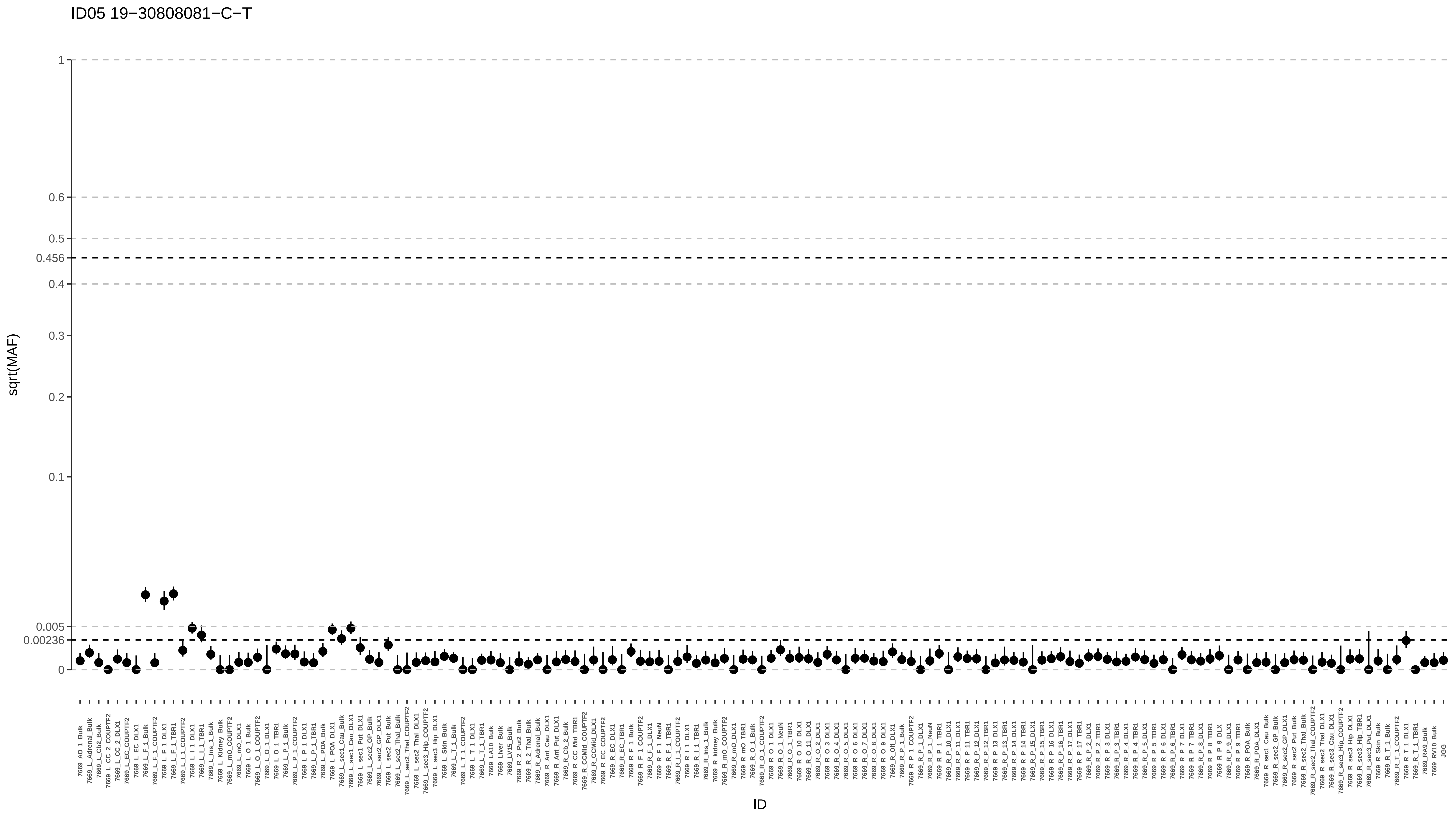
ID05 19−30808081−C−T
sqrt(MAF)
1
0.6
0.5
0.456
0.4
0.3
0.2
0.1
0.005
0.00236
0
ID

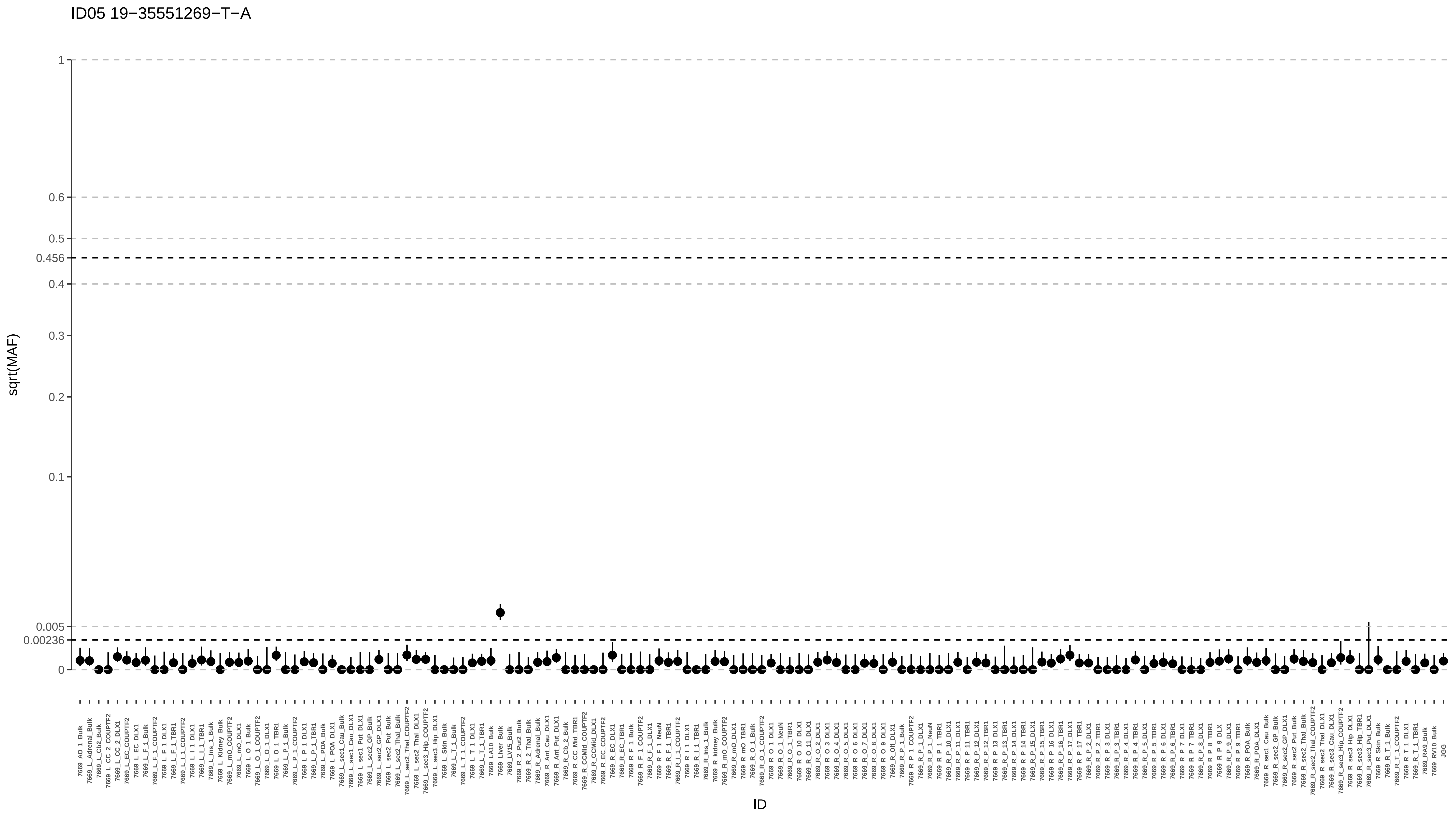
ID05 19−35551269−T−A
sqrt(MAF)
1
0.6
0.5
0.456
0.4
0.3
0.2
0.1
0.005
0.00236
0
ID
JGG

ID05 19−36516973−C−T

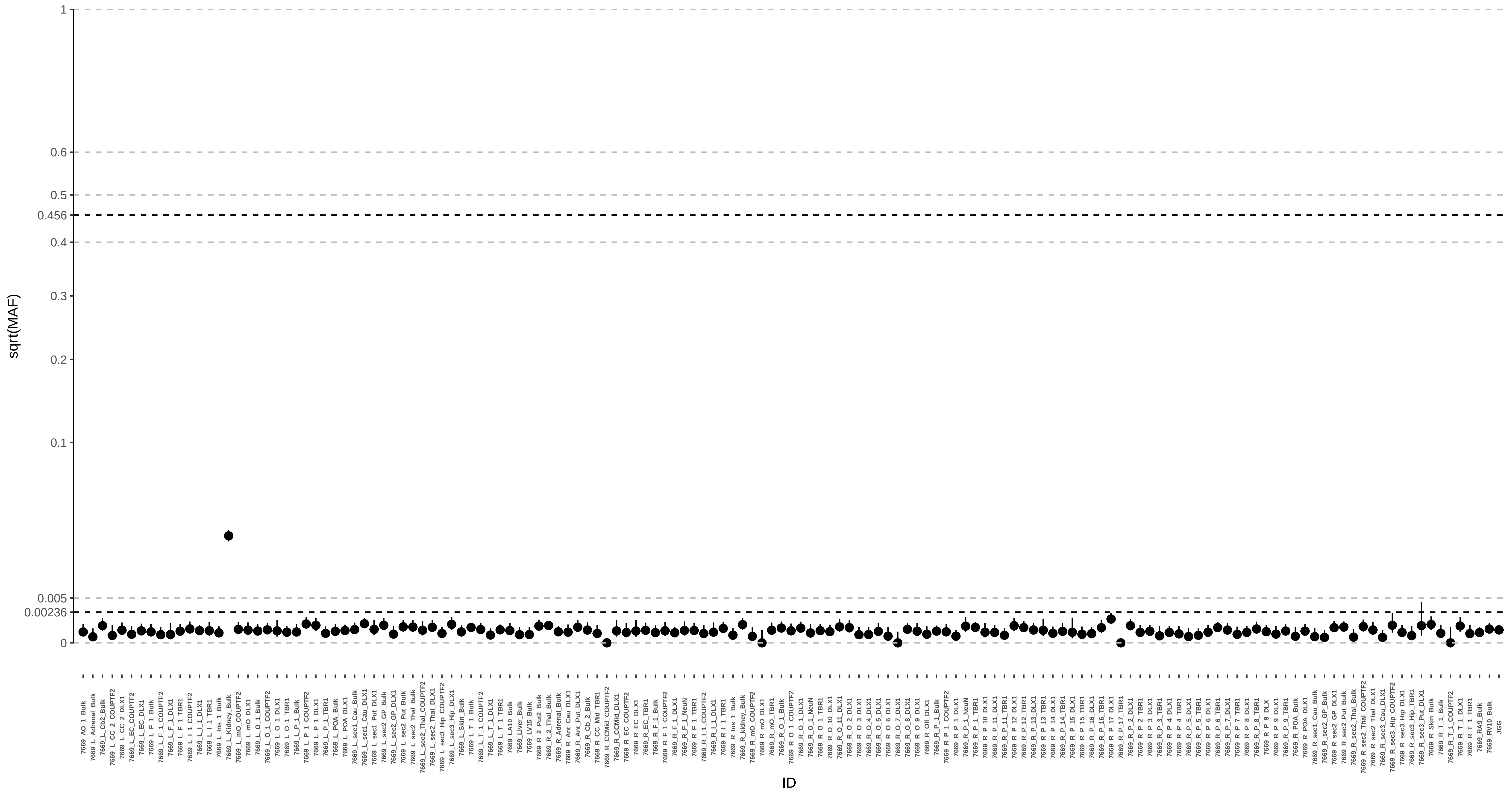

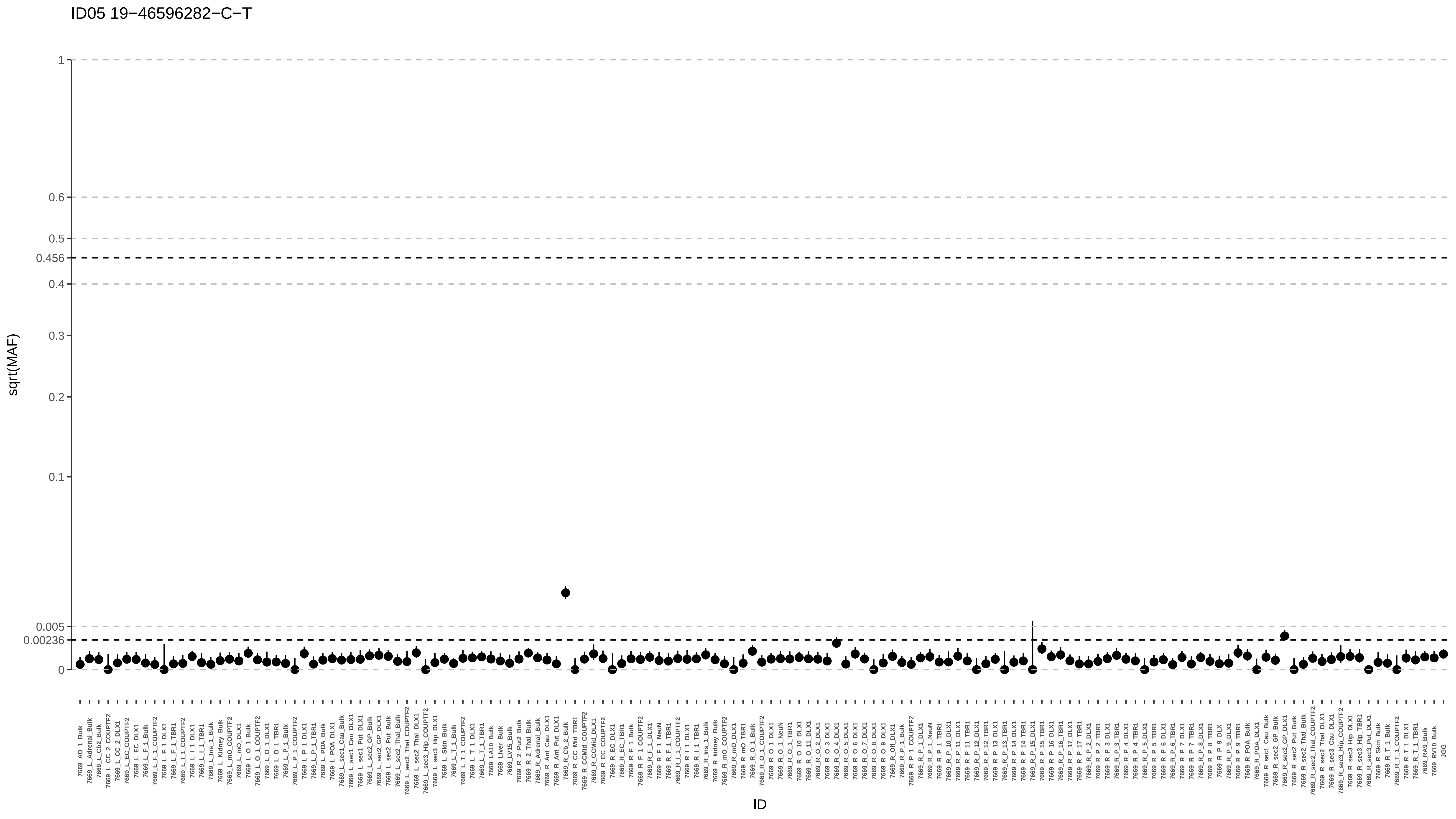
ID05 19−46596282−C−T
sqrt(MAF)
1
0.6
0.5
0.456
0.4
0.3
0.2
0.1
0.005
0.00236
0
ID

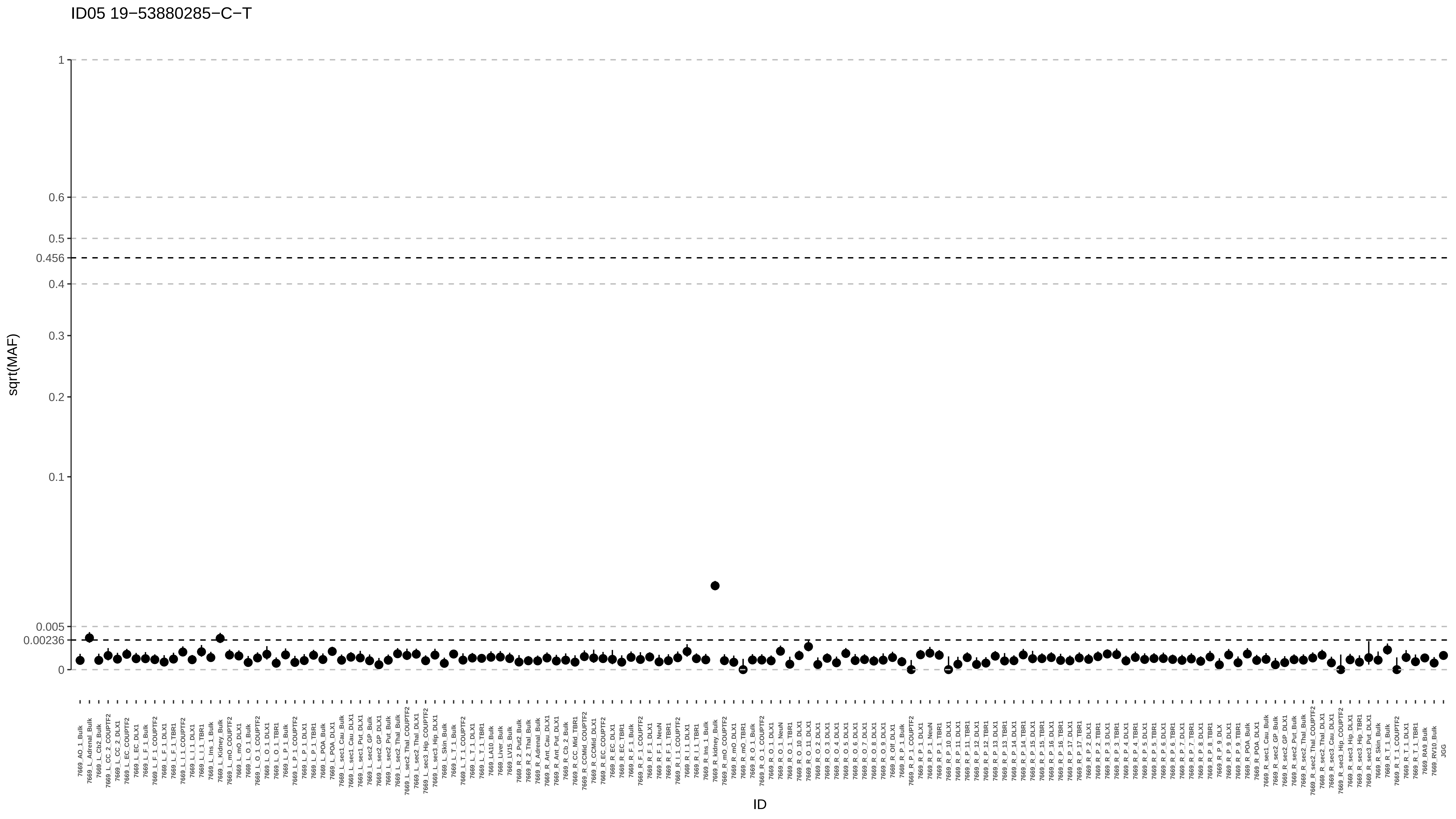
ID05 19−53880285−C−T
sqrt(MAF)
ID

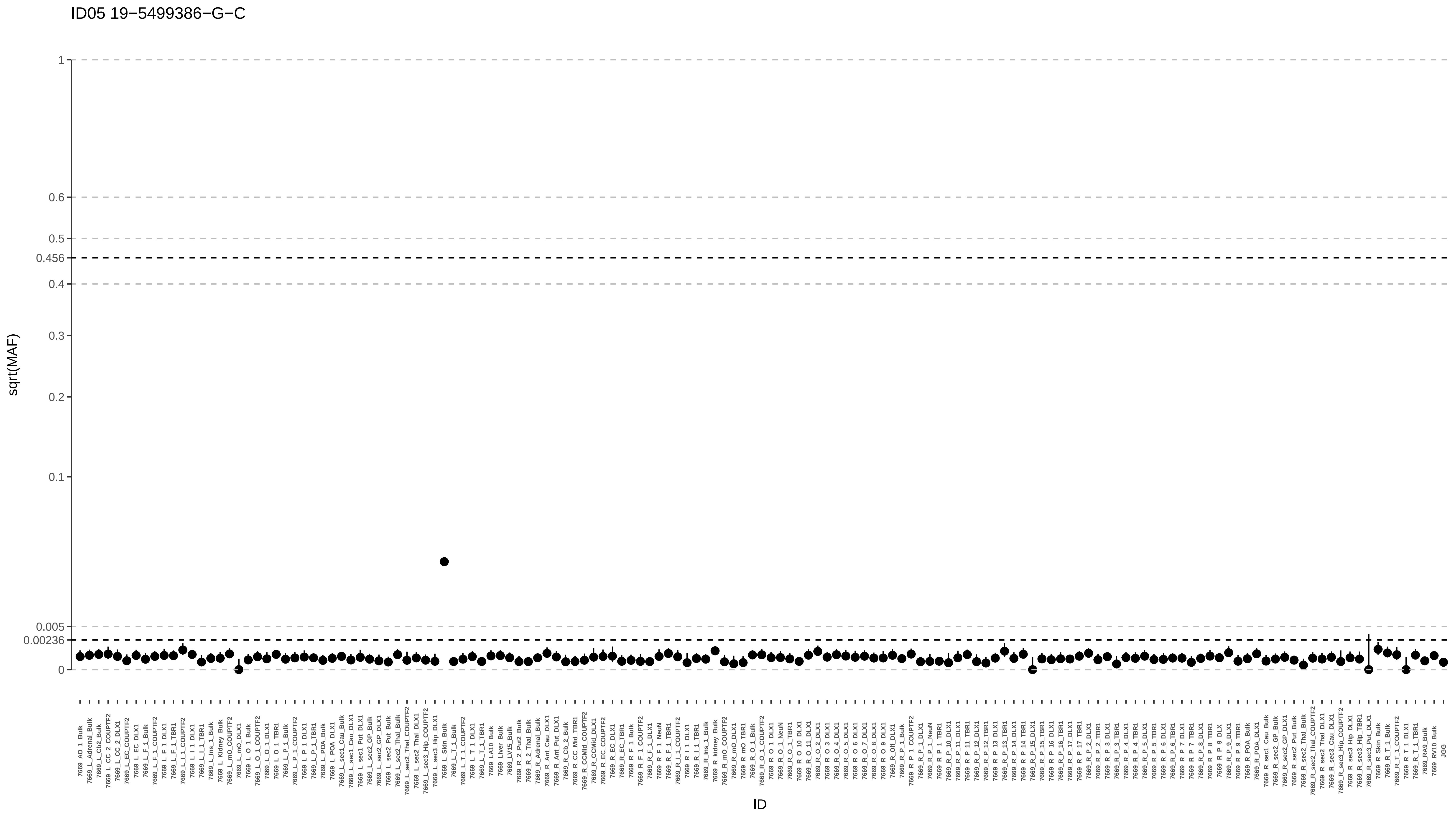

ID05 19−5499386−G−C
sqrt(MAF)
1
0.6
0.5
0.456
0.4
0.3
0.2
0.1
0.005
0.00236
0
ID

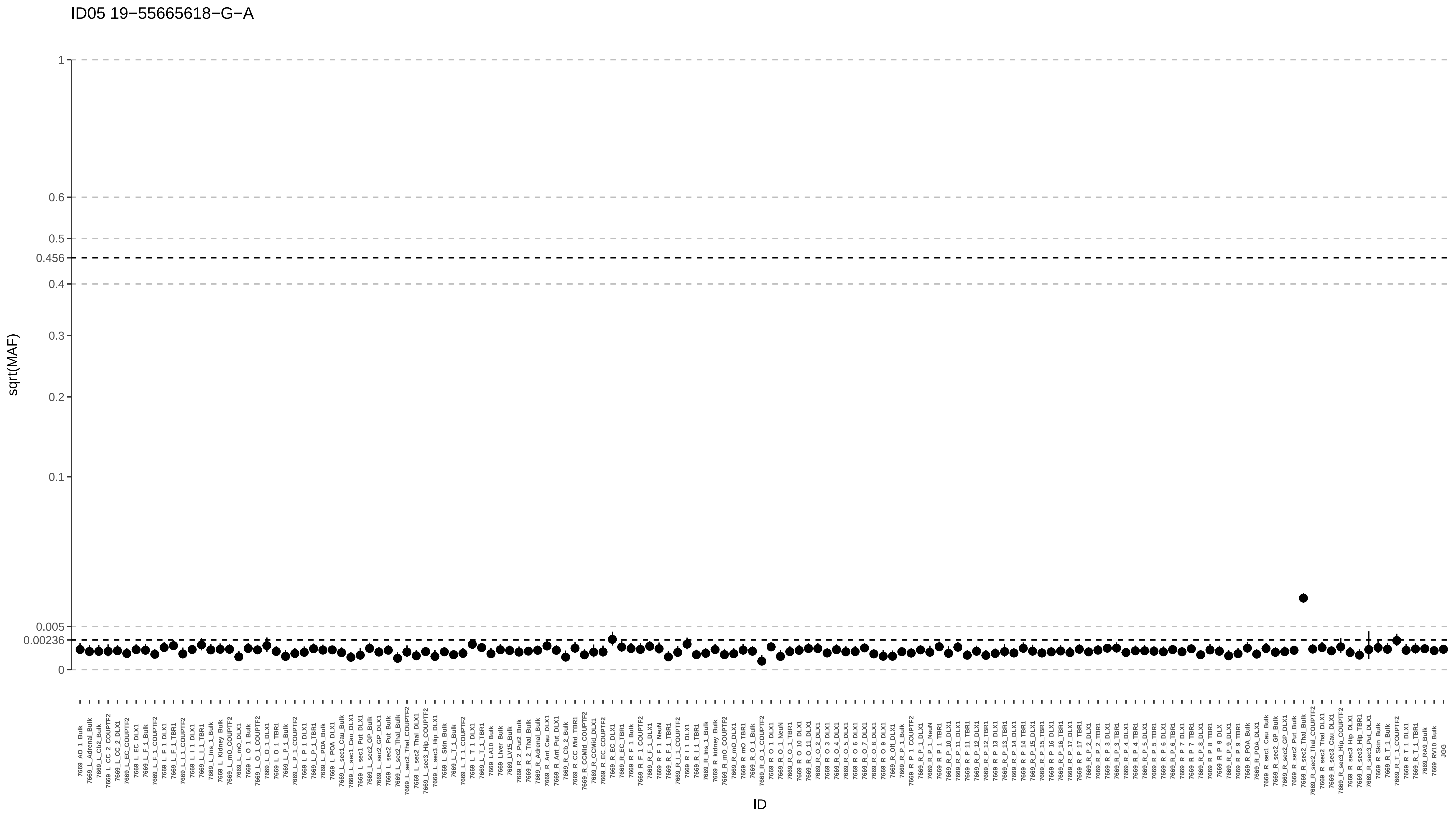
ID05 19−55665618−G−A
sqrt(MAF)
1
0.6
0.5
0.456
0.4
0.3
0.2
0.1
0.005
0.00236
0
ID

ID05 19−57342142−G−A

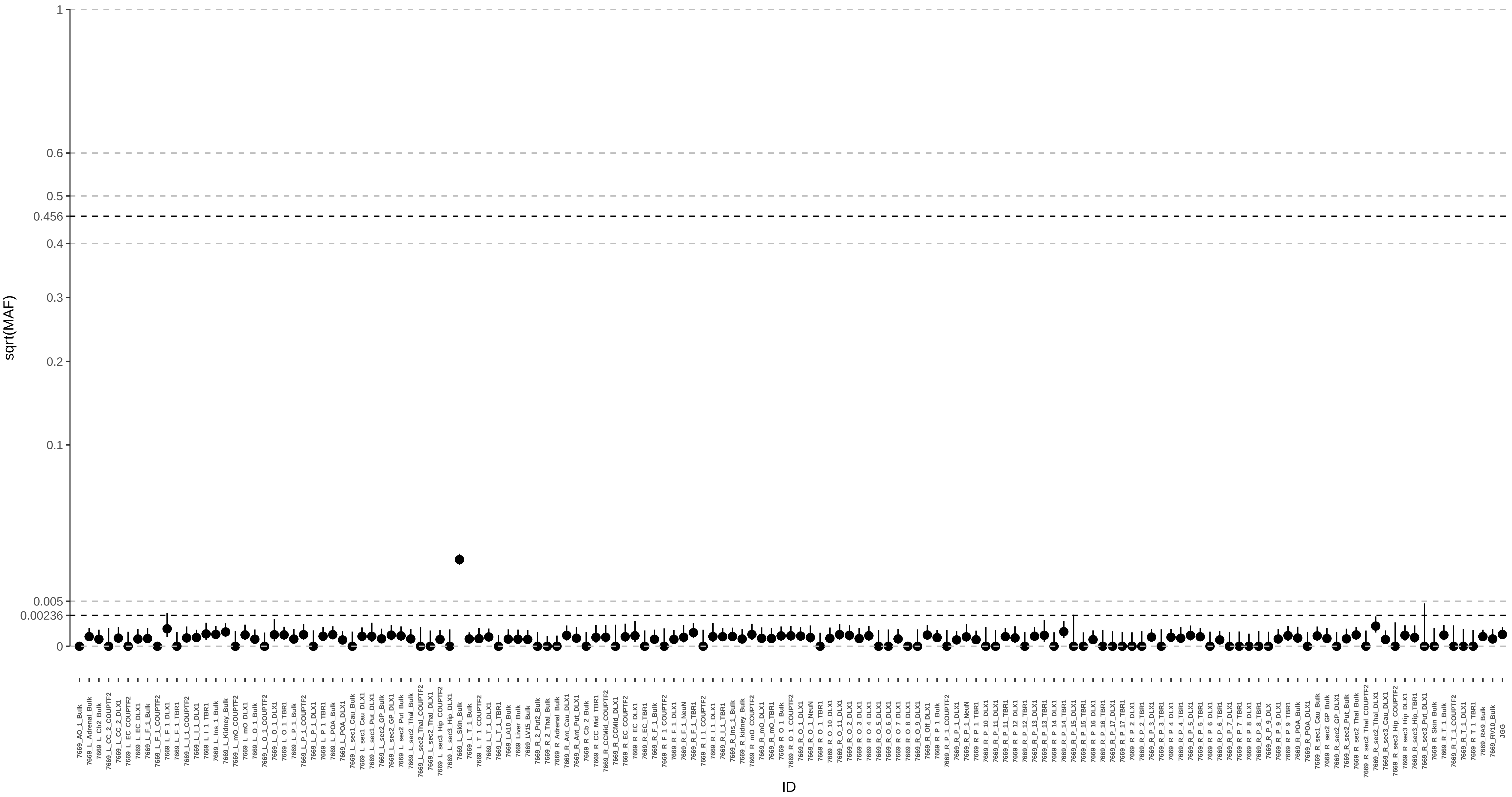

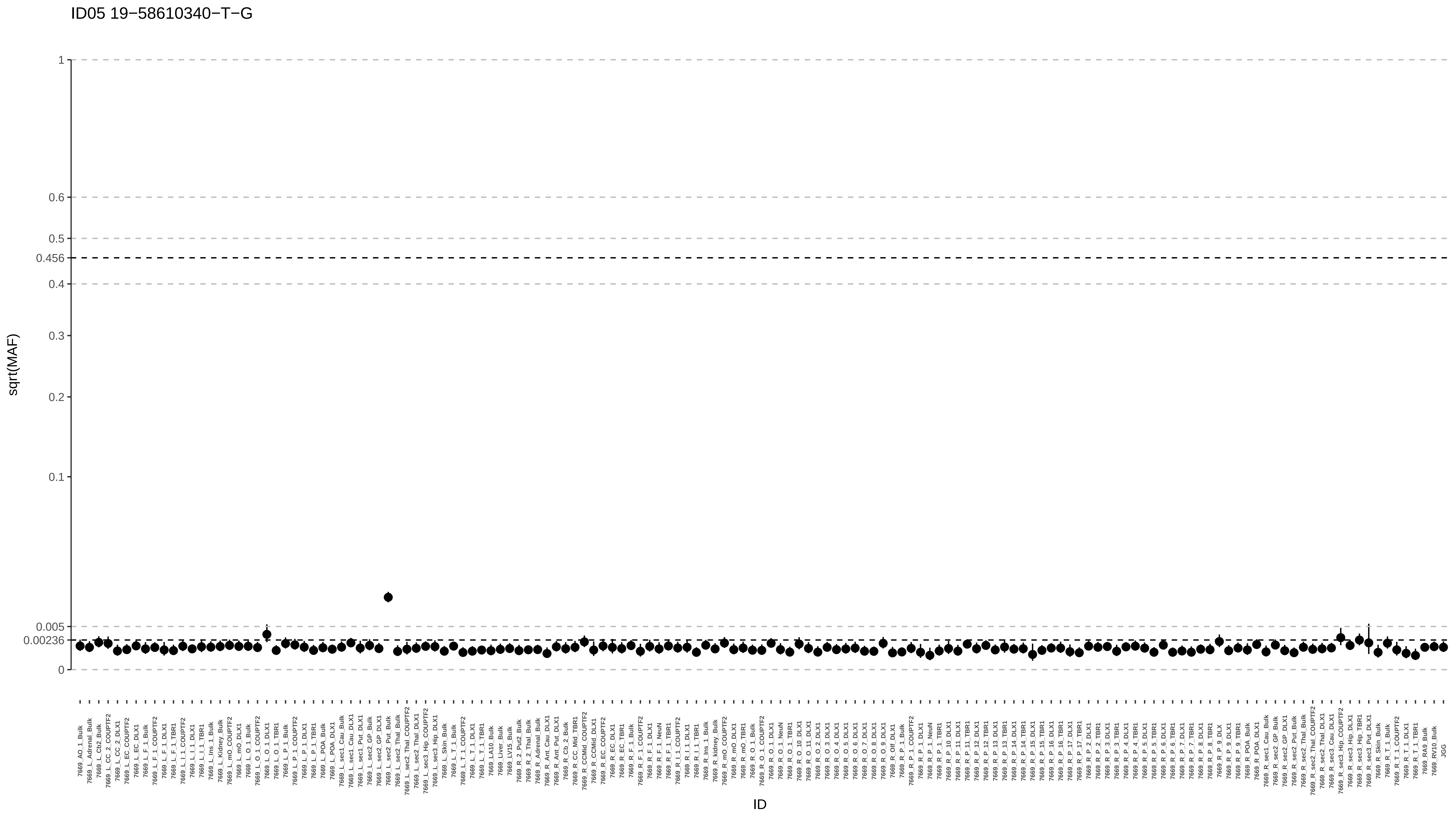
ID05 19−58610340−T−G
sqrt(MAF)
1
0.6
0.5
0.456
0.4
0.3
0.2
0.1
0.005
0.00236
0
ID

ID05 19−5879037−C−T

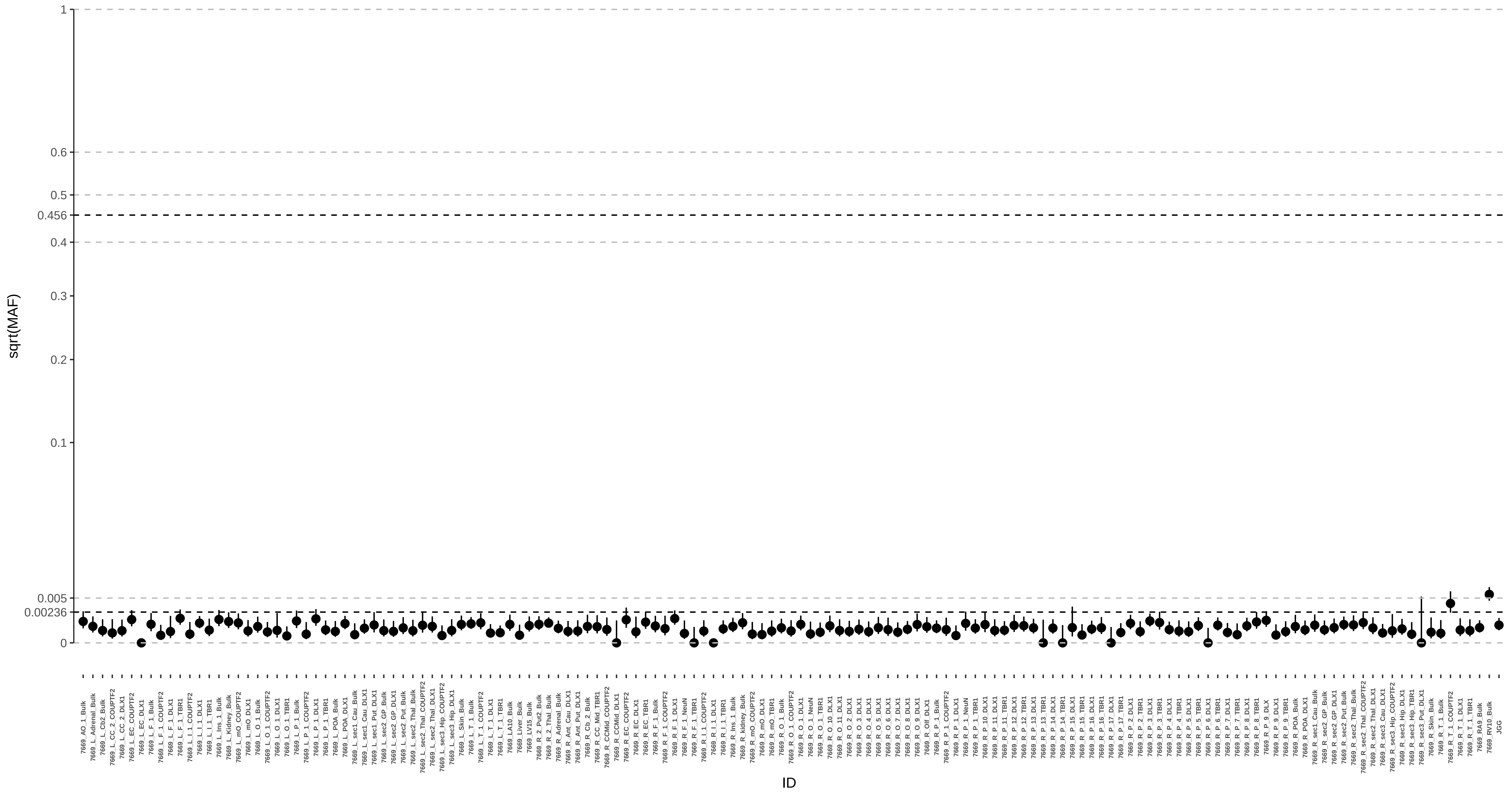

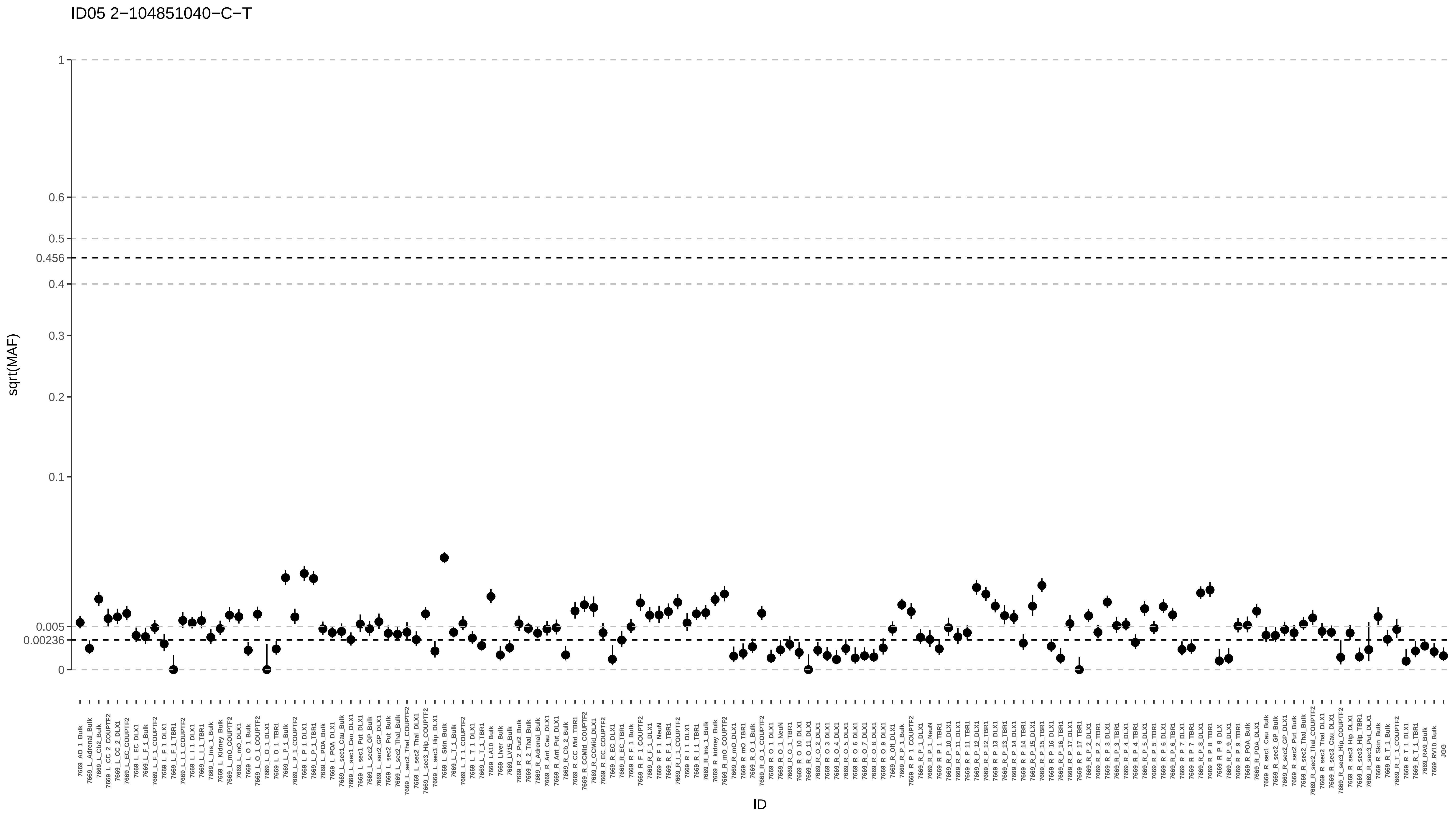
ID05 2−104851040−C−T
sqrt(MAF)
1
0.6
0.5
0.456
0.4
0.3
0.2
0.1
0.005
0.00236
0
ID

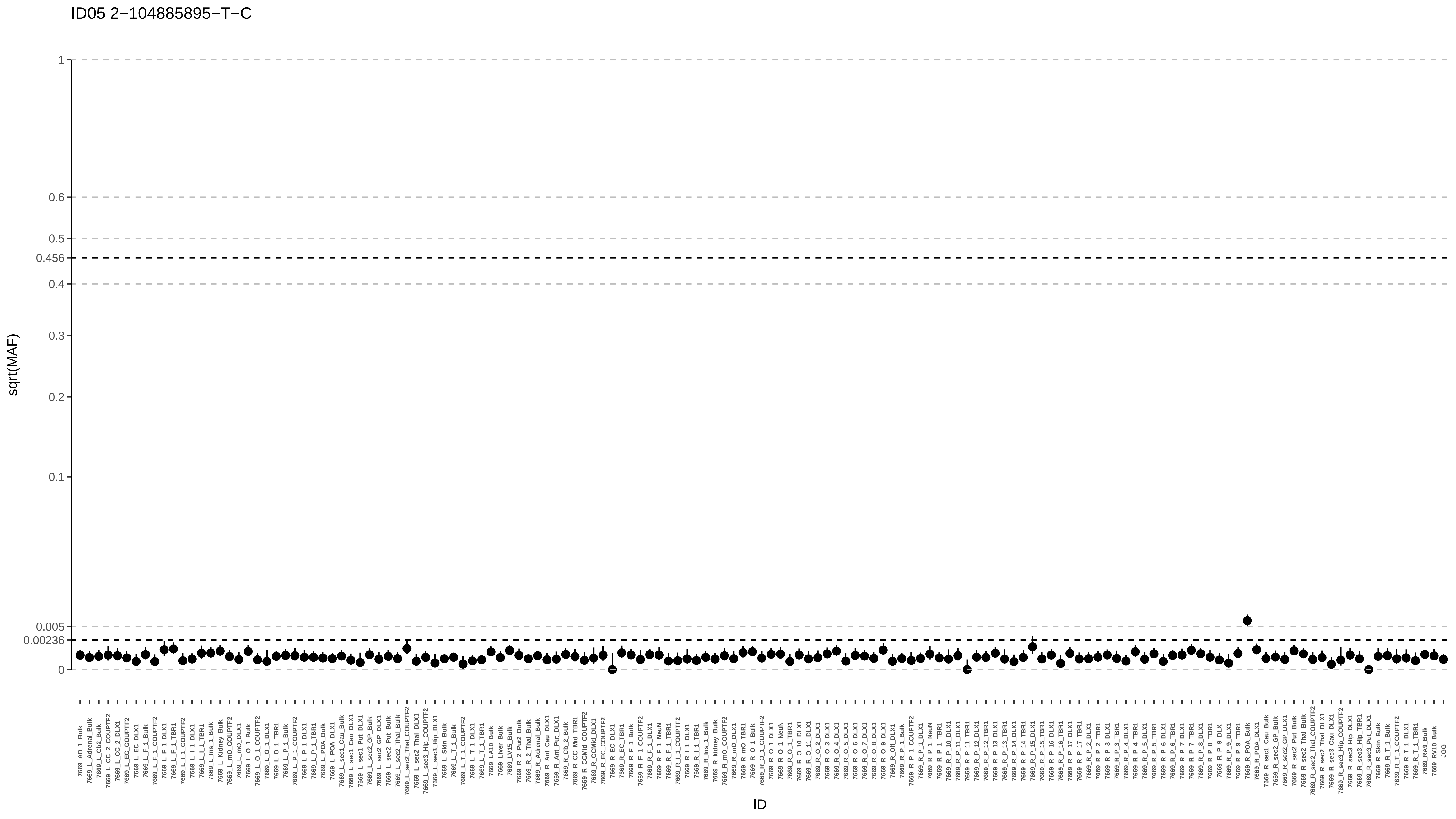
ID05 2−104885895−T−C
sqrt(MAF)
1
0.6
0.5
0.456
0.4
0.3
0.2
0.1
0.005
0.00236
0
ID

ID05 2−105083647−G−A

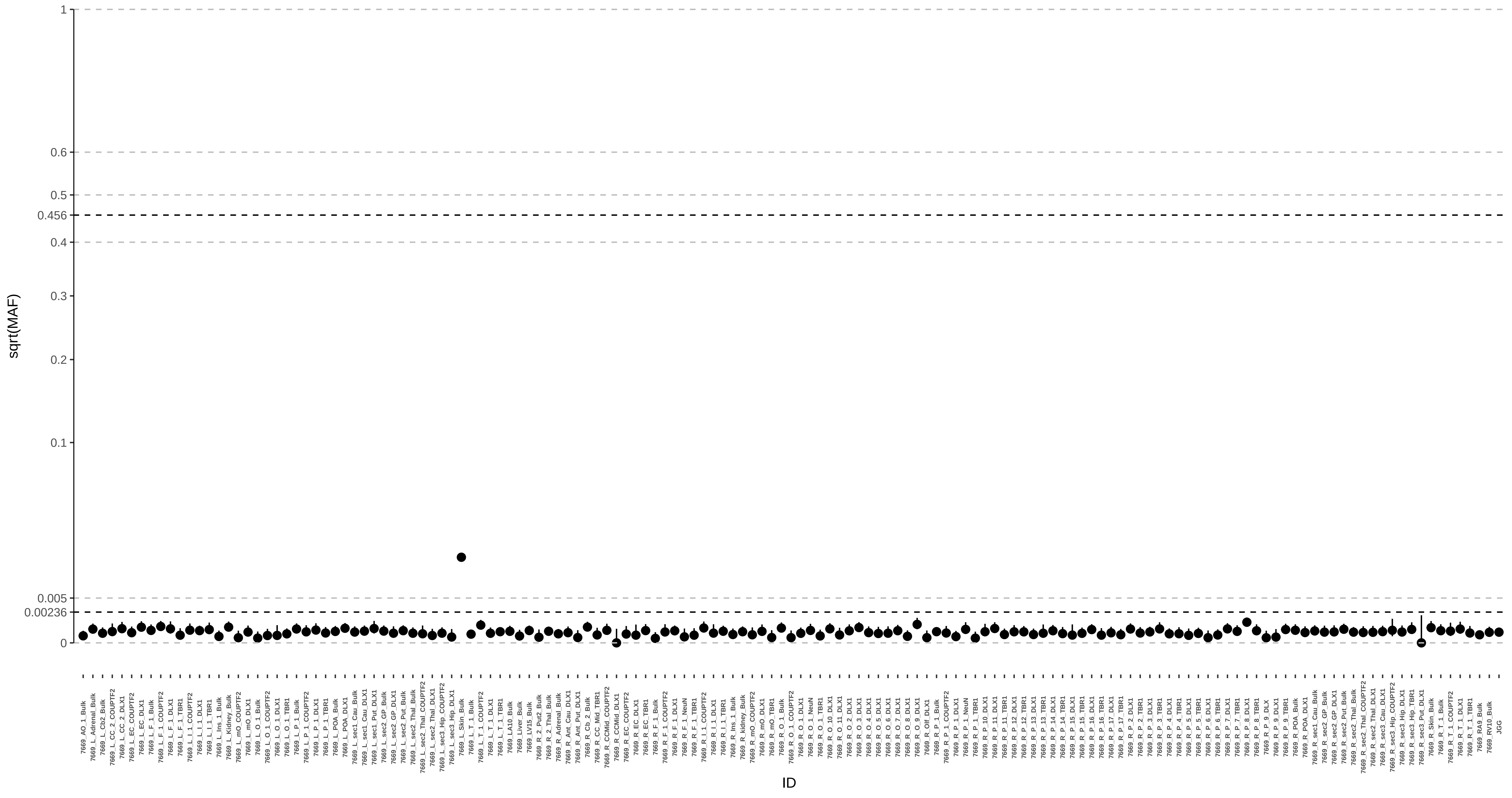

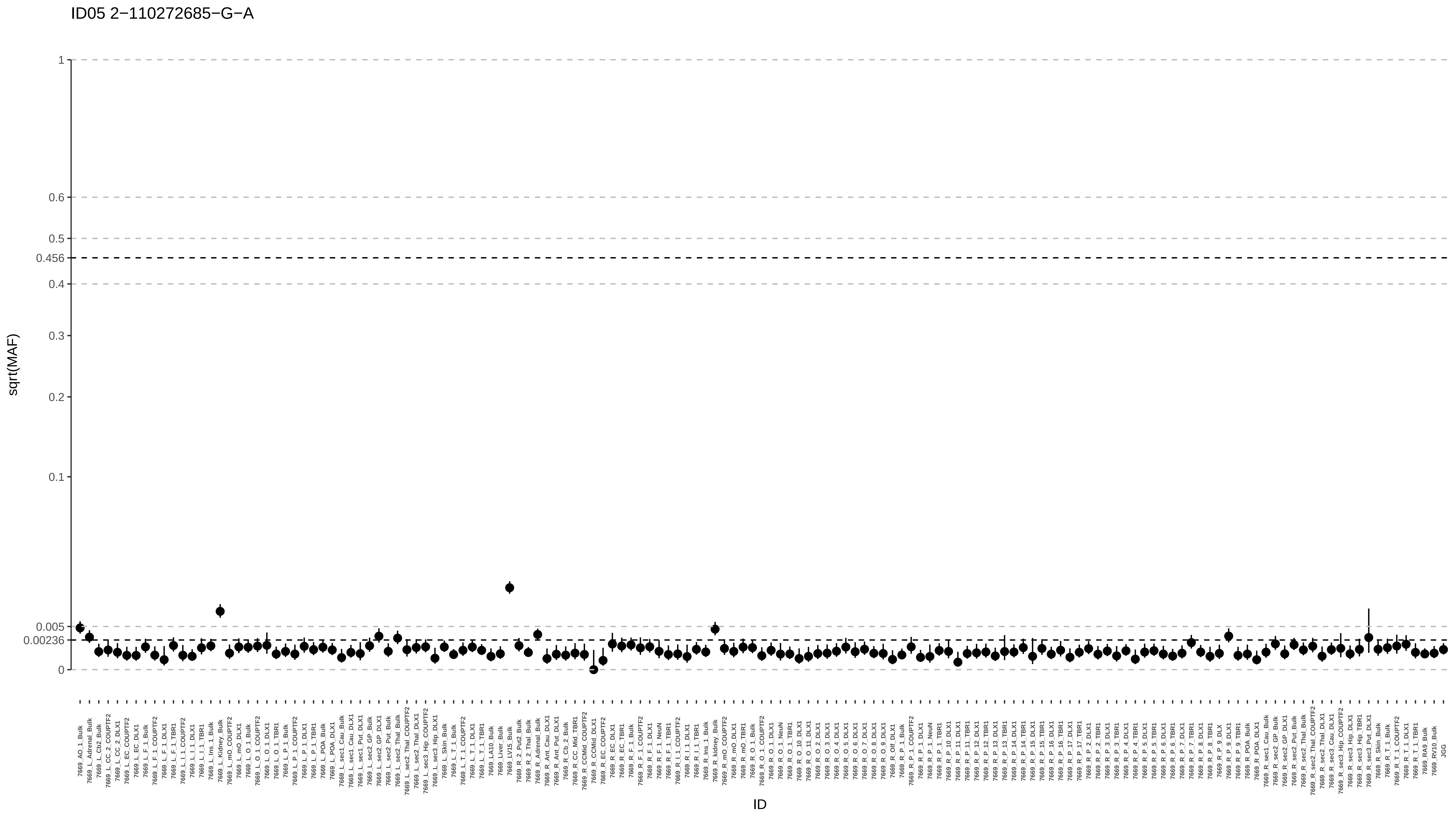
ID05 2−110272685−G−A
sqrt(MAF)
ID

ID05 2−115000633−A−G

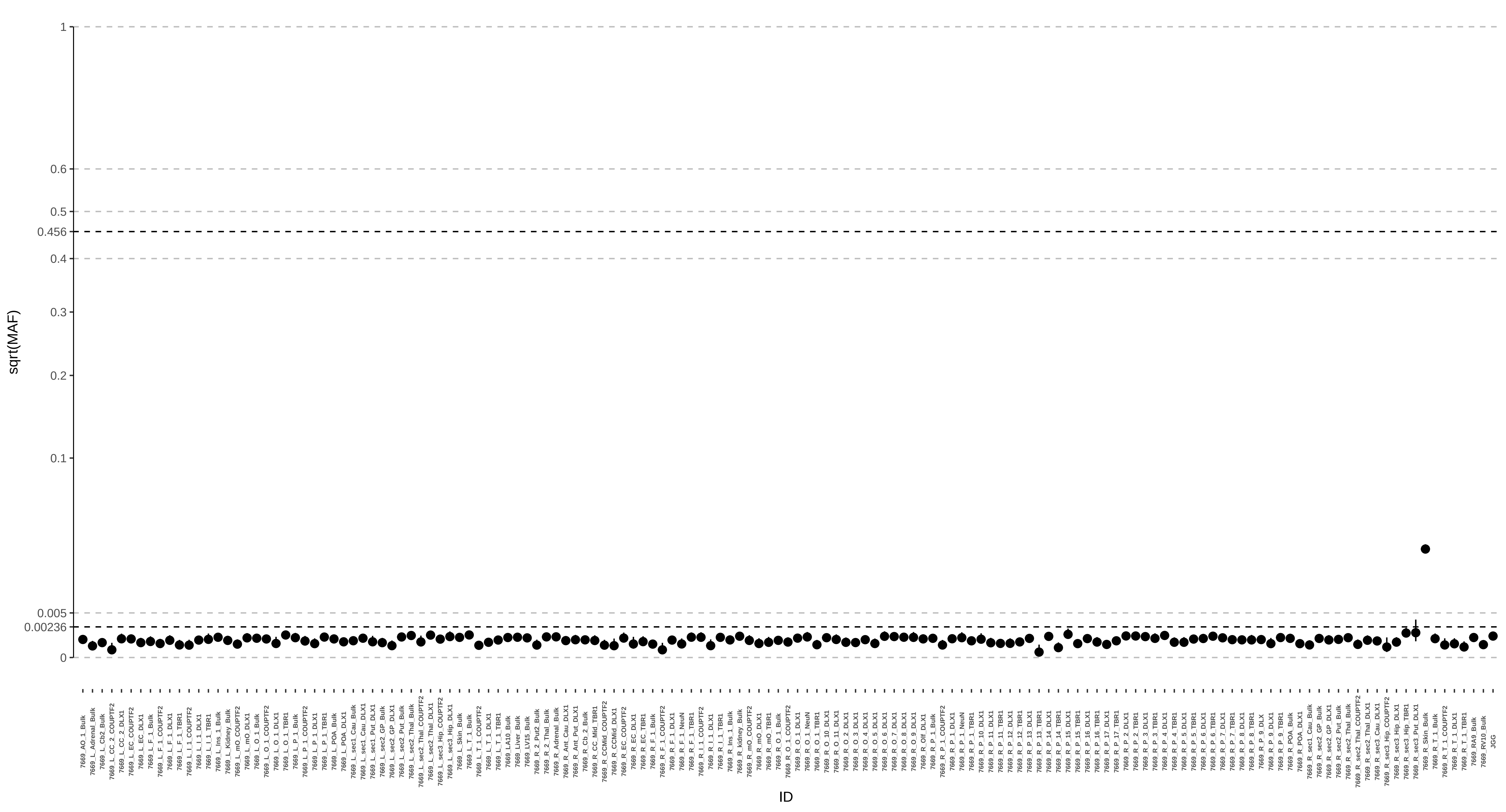

ID05 2−115027303−G−A

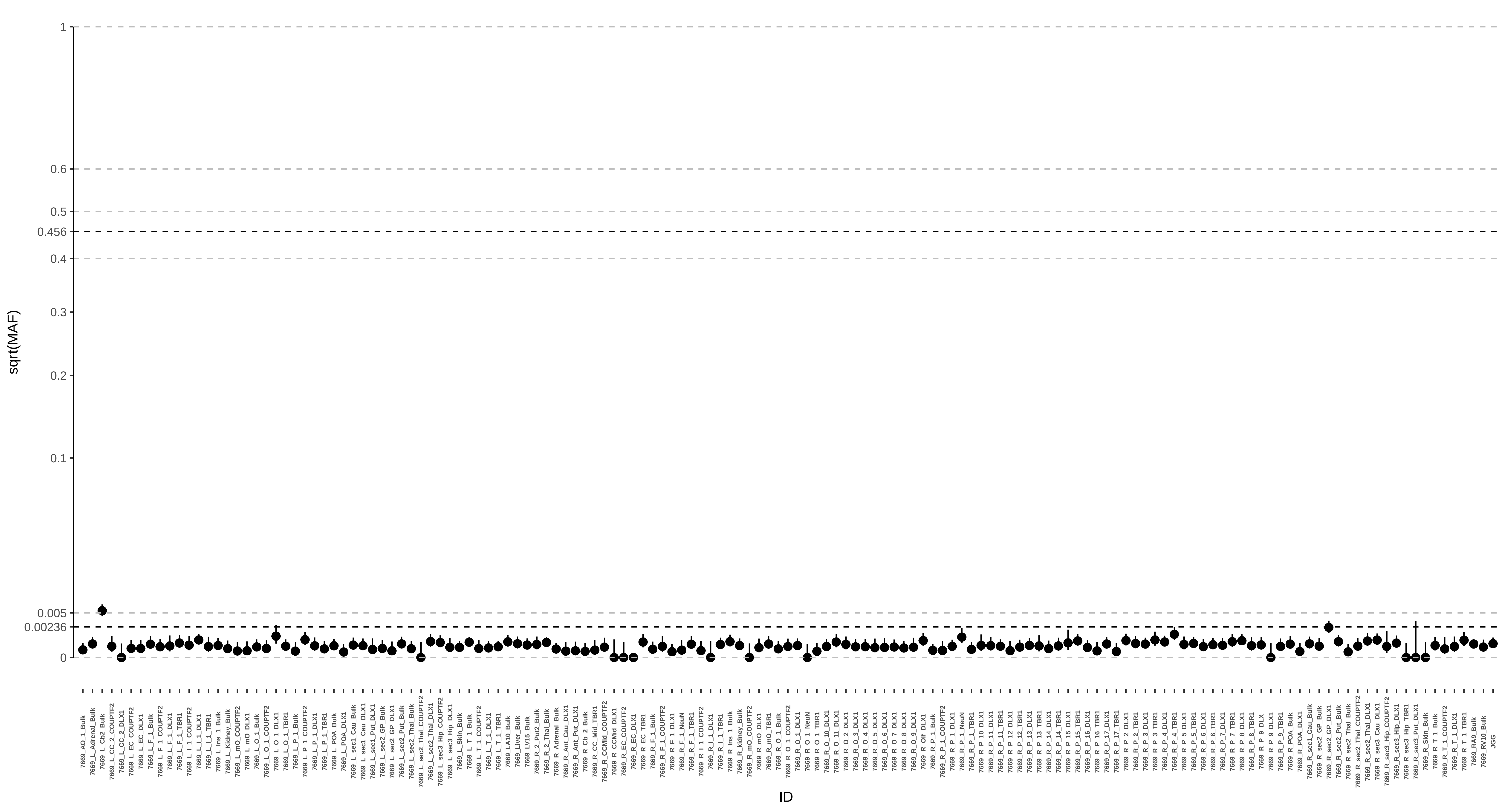

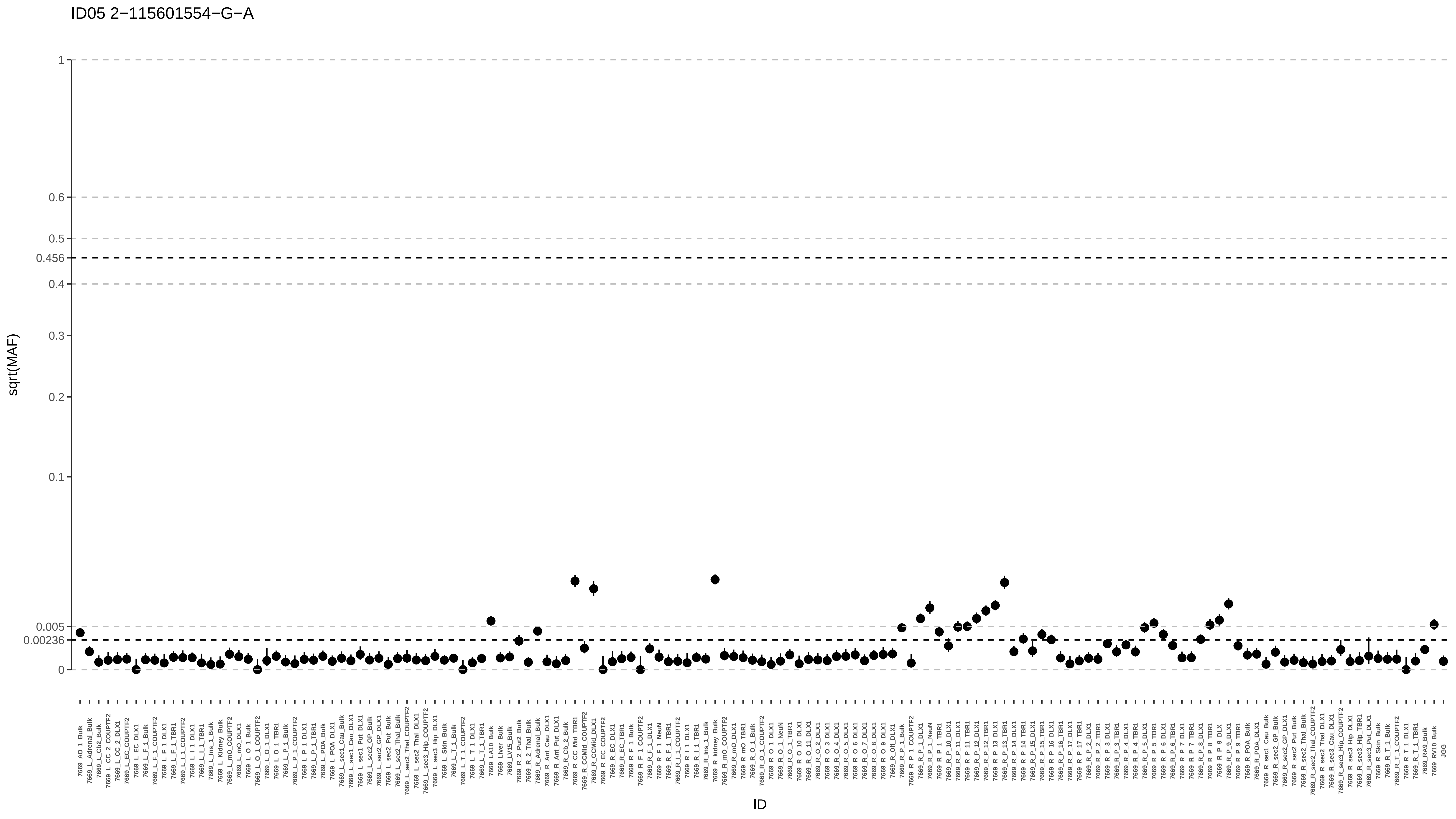
ID05 2−115601554−G−A
sqrt(MAF)
1
0.6
0.5
0.456
0.4
0.3
0.2
0.1
0.005
0.00236
0
ID

ID05 2−115708162−C−T

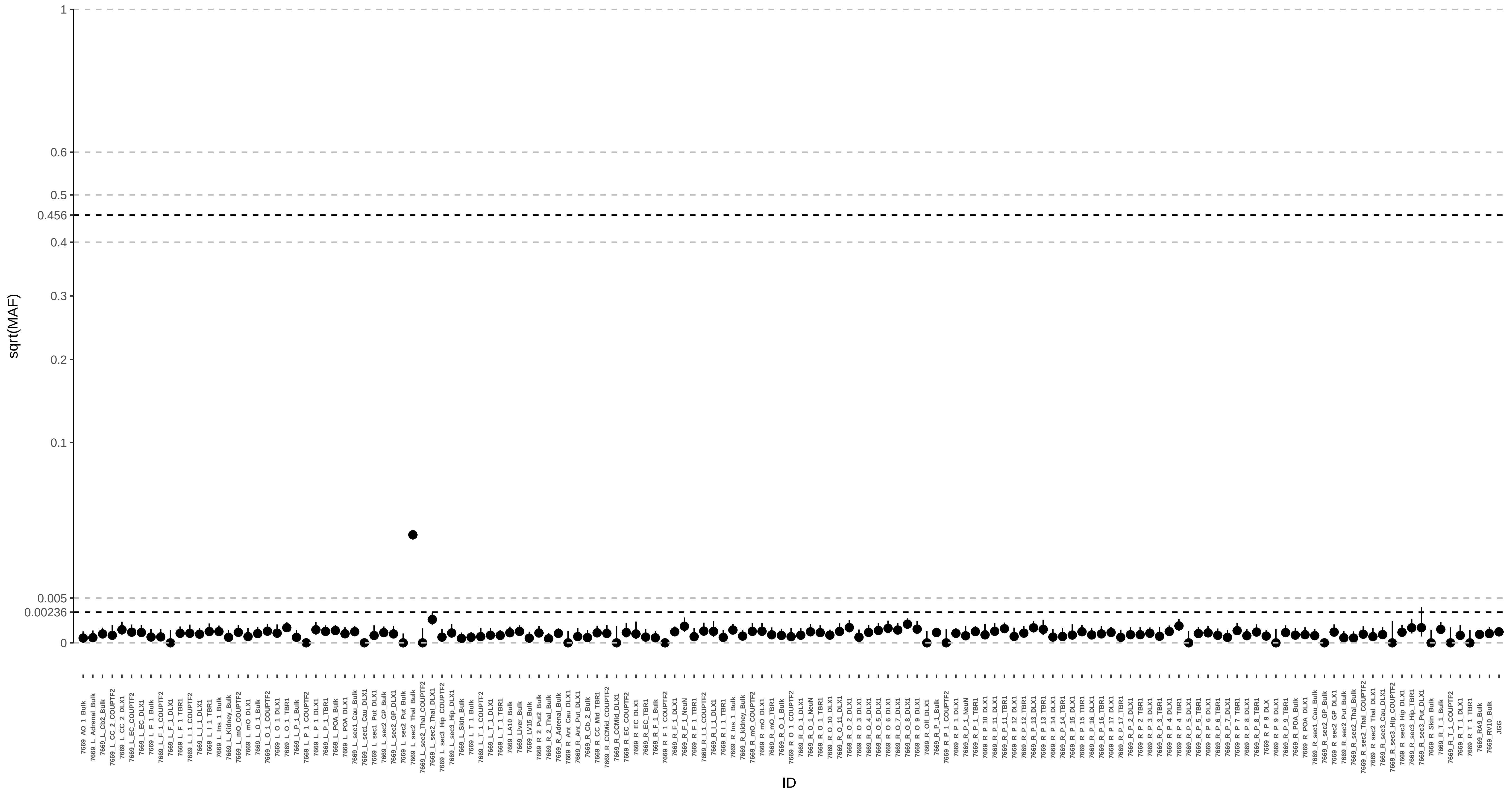

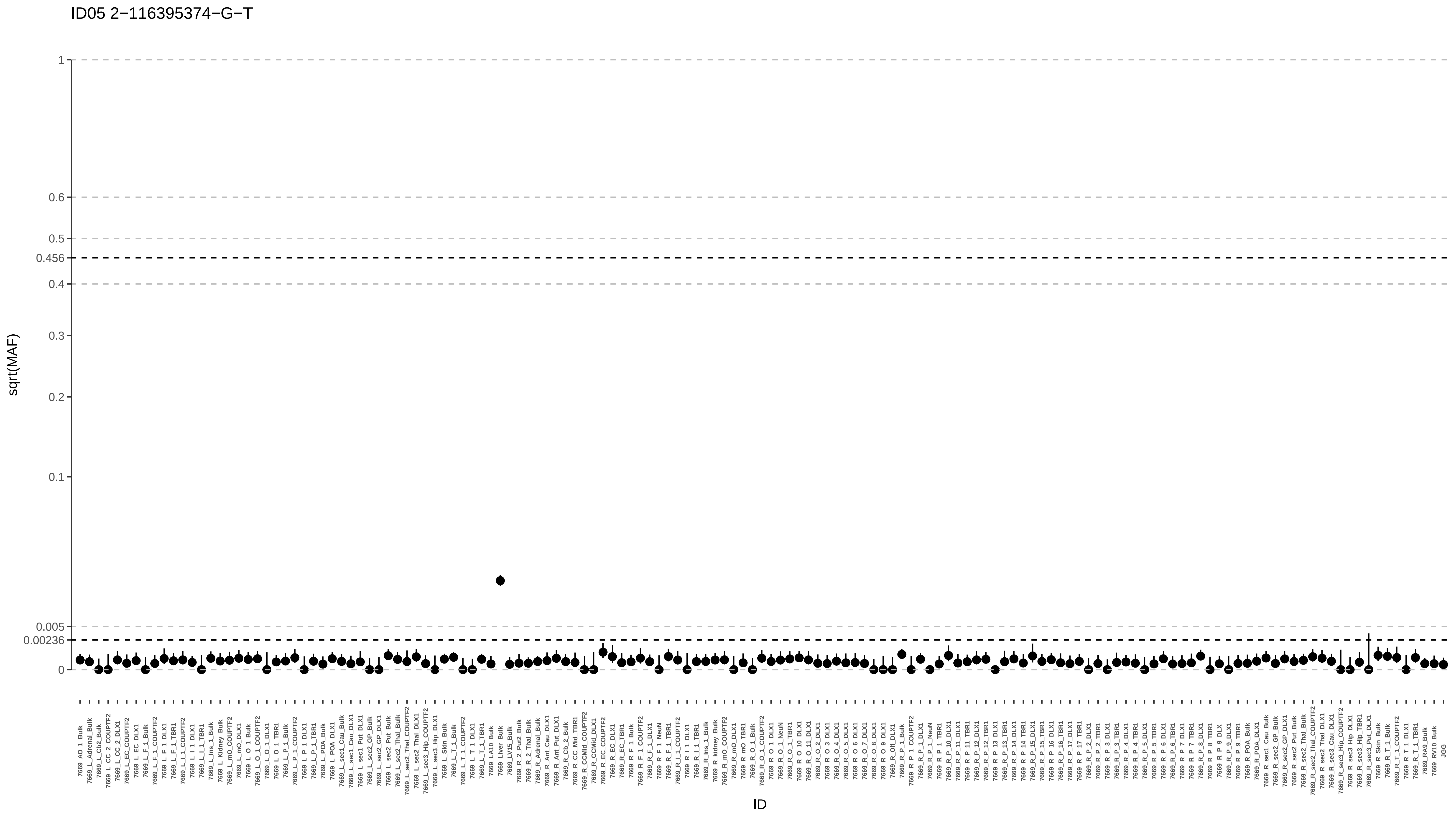
ID05 2−116395374−G−T
sqrt(MAF)
1
0.6
0.5
0.456
0.4
0.3
0.2
0.1
0.005
0.00236
0
ID

ID05 2−118425584−C−T

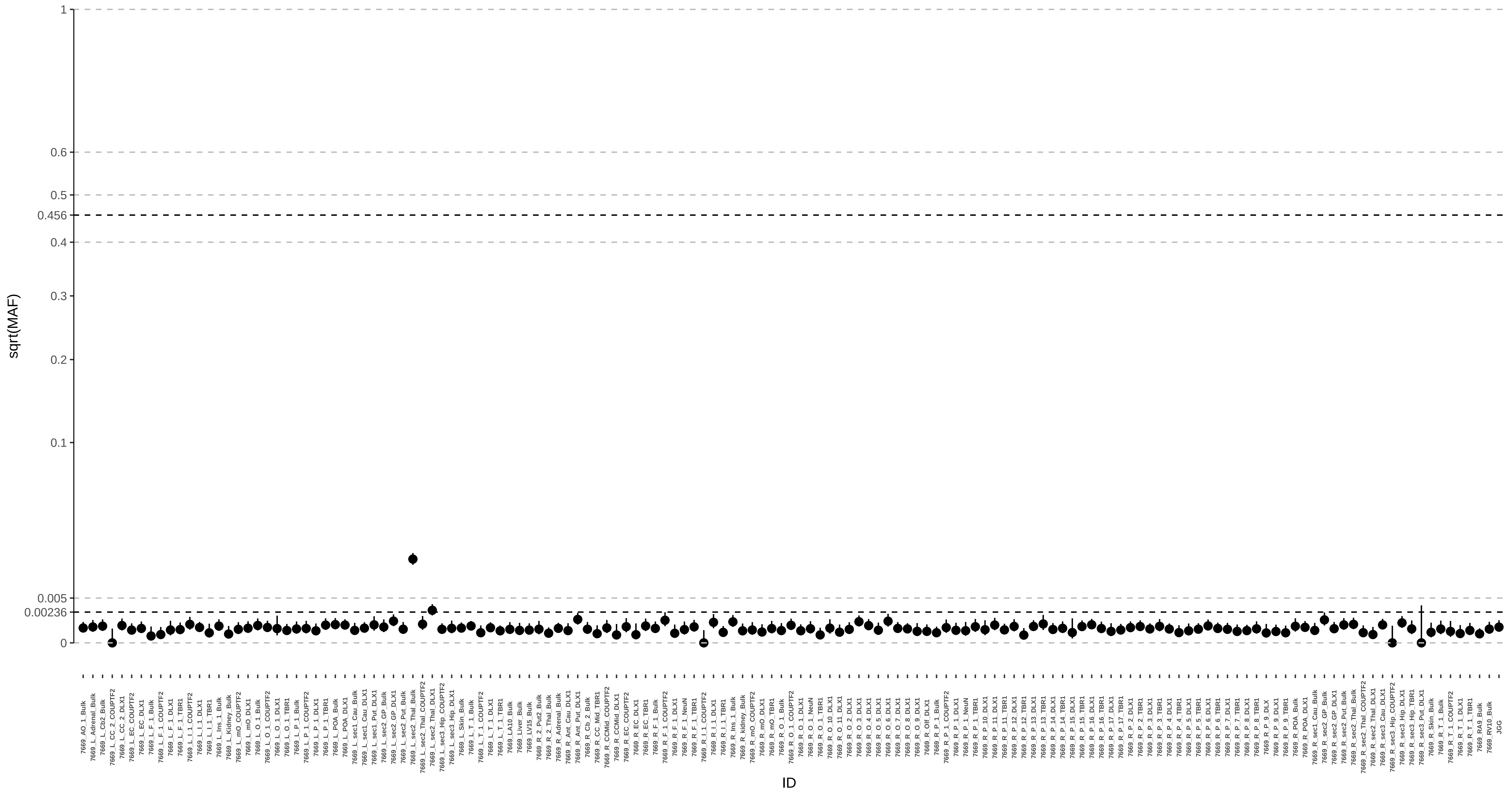

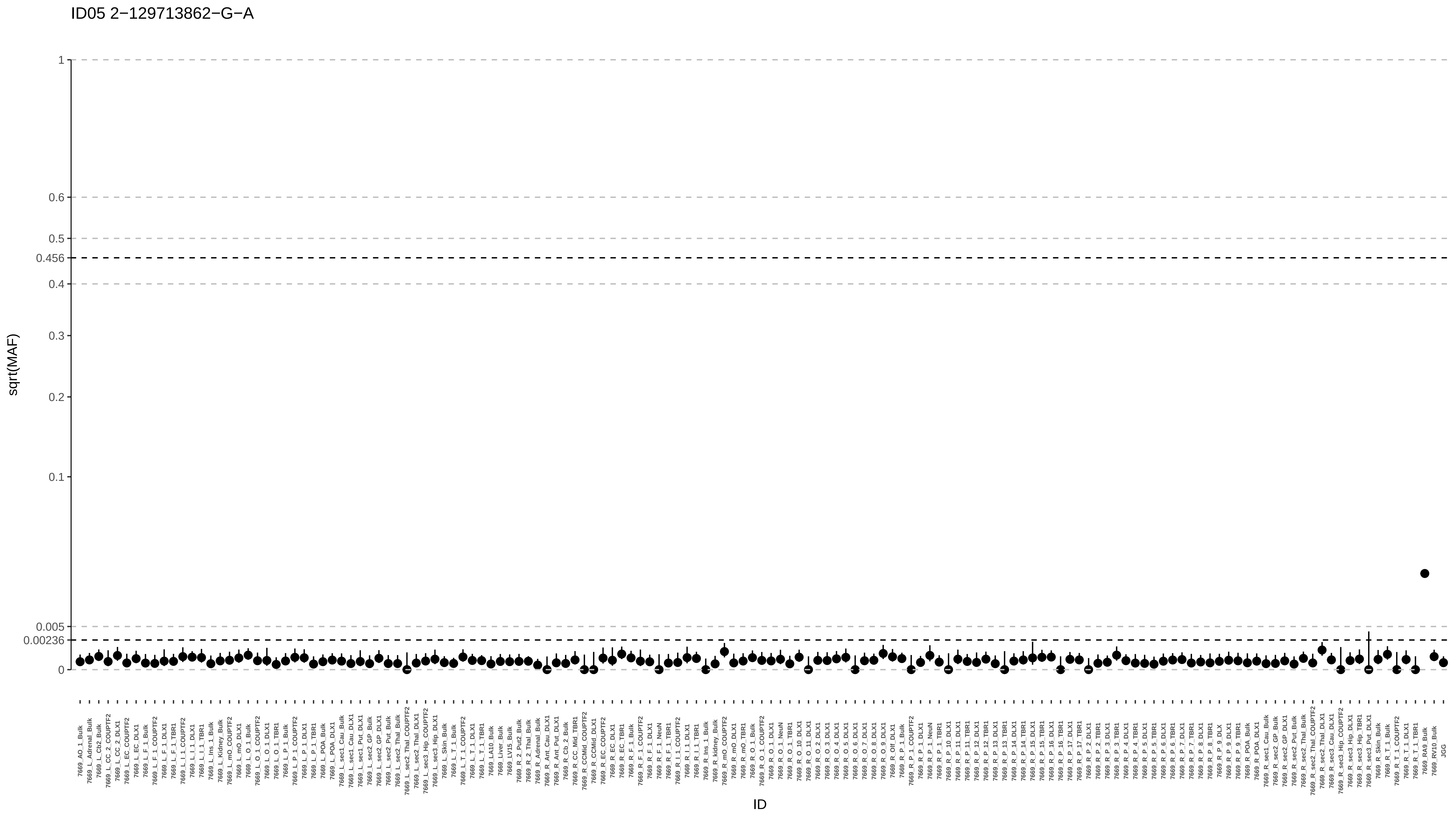
ID05 2−129713862−G−A
sqrt(MAF)
1
0.6
0.5
0.456
0.4
0.3
0.2
0.1
0.005
0.00236
0
ID

ID05 2−130483876−C−T

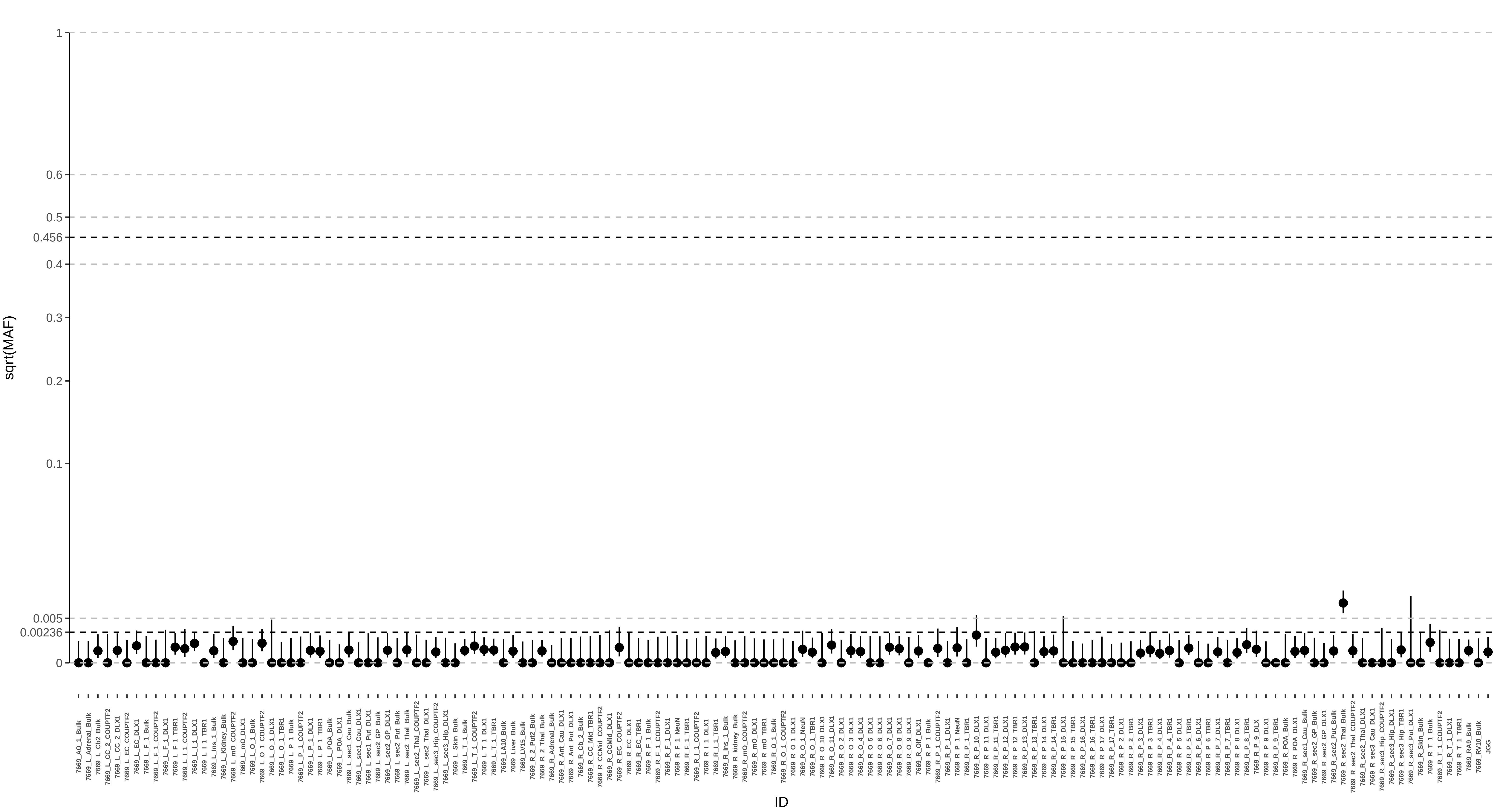

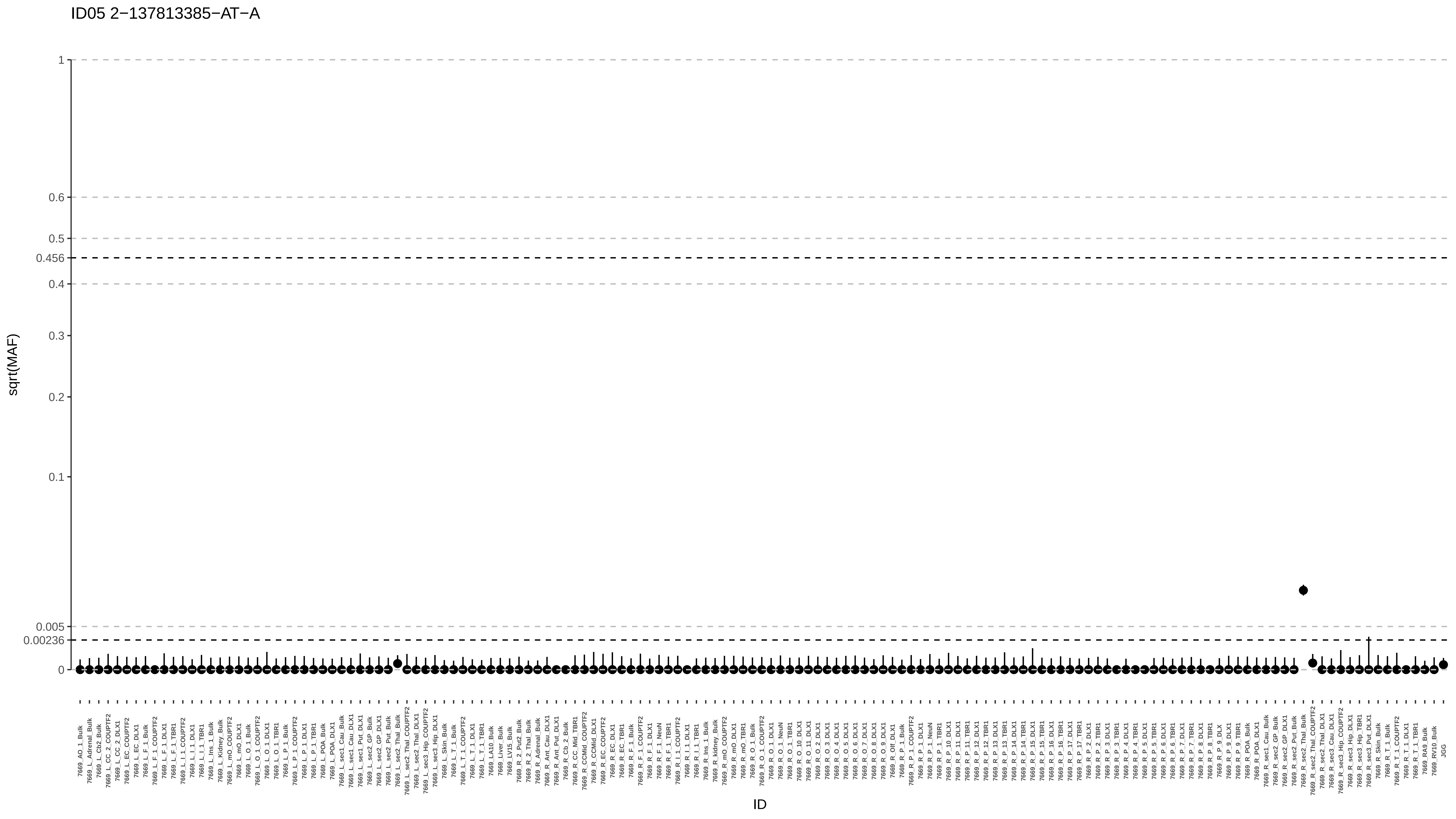
ID05 2−137813385−AT−A
sqrt(MAF)
1
0.6
0.5
0.456
0.4
0.3
0.2
0.1
0.005
0.00236
0
ID
7669_AO_1_Bulk
7669_L_Adrenal_Bulk
7669_L_Cb2_Bulk
7669_L_CC_2_COUPTF2
7669_L_CC_2_DLX1
7669_L_EC_COUPTF2
7669_L_EC_DLX1
7669_L_F_1_Bulk
7669_L_F_1_COUPTF2
7669_L_F_1_DLX1
7669_L_F_1_TBR1
7669_L_I_1_COUPTF2
7669_L_I_1_DLX1
7669_L_I_1_TBR1
7669_L_Ins_1_Bulk
7669_L_Kidney_Bulk
7669_L_mO_COUPTF2
7669_L_mO_DLX1
7669_L_O_1_Bulk
7669_L_O_1_COUPTF2
7669_L_O_1_DLX1
7669_L_O_1_TBR1
7669_L_P_1_Bulk
7669_L_P_1_COUPTF2
7669_L_P_1_DLX1
7669_L_P_1_TBR1
7669_L_POA_Bulk
7669_L_POA_DLX1
7669_L_sec1_Cau_Bulk
7669_L_sec1_Cau_DLX1
7669_L_sec1_Put_DLX1
7669_L_sec2_GP_Bulk
7669_L_sec2_GP_DLX1
7669_L_sec2_Put_Bulk
7669_L_sec2_Thal_Bulk
7669_L_sec2_Thal_COUPTF2
7669_L_sec2_Thal_DLX1
7669_L_sec3_Hip_COUPTF2
7669_L_sec3_Hip_DLX1
7669_L_Skin_Bulk
7669_L_T_1_Bulk
7669_L_T_1_COUPTF2
7669_L_T_1_DLX1
7669_L_T_1_TBR1
7669_LA10_Bulk
7669_Liver_Bulk
7669_LV15_Bulk
7669_R_2_Put2_Bulk
7669_R_2_Thal_Bulk
7669_R_Adrenal_Bulk
7669_R_Ant_Cau_DLX1
7669_R_Ant_Put_DLX1
7669_R_Cb_2_Bulk
7669_R_CC_Mid_TBR1
7669_R_CCMid_COUPTF2
7669_R_CCMid_DLX1
7669_R_EC_COUPTF2
7669_R_EC_DLX1
7669_R_EC_TBR1
7669_R_F_1_Bulk
7669_R_F_1_COUPTF2
7669_R_F_1_DLX1
7669_R_F_1_NeuN
7669_R_F_1_TBR1
7669_R_I_1_COUPTF2
7669_R_I_1_DLX1
7669_R_I_1_TBR1
7669_R_Ins_1_Bulk
7669_R_kidney_Bulk
7669_R_mO_COUPTF2
7669_R_mO_DLX1
7669_R_mO_TBR1
7669_R_O_1_Bulk
7669_R_O_1_COUPTF2
7669_R_O_1_DLX1
7669_R_O_1_NeuN
7669_R_O_1_TBR1
7669_R_O_10_DLX1
7669_R_O_11_DLX1
7669_R_O_2_DLX1
7669_R_O_3_DLX1
7669_R_O_4_DLX1
7669_R_O_5_DLX1
7669_R_O_6_DLX1
7669_R_O_7_DLX1
7669_R_O_8_DLX1
7669_R_O_9_DLX1
7669_R_Olf_DLX1
7669_R_P_1_Bulk
7669_R_P_1_COUPTF2
7669_R_P_1_DLX1
7669_R_P_1_NeuN
7669_R_P_1_TBR1
7669_R_P_10_DLX1
7669_R_P_11_DLX1
7669_R_P_11_TBR1
7669_R_P_12_DLX1
7669_R_P_12_TBR1
7669_R_P_13_DLX1
7669_R_P_13_TBR1
7669_R_P_14_DLX1
7669_R_P_14_TBR1
7669_R_P_15_DLX1
7669_R_P_15_TBR1
7669_R_P_16_DLX1
7669_R_P_16_TBR1
7669_R_P_17_DLX1
7669_R_P_17_TBR1
7669_R_P_2_DLX1
7669_R_P_2_TBR1
7669_R_P_3_DLX1
7669_R_P_3_TBR1
7669_R_P_4_DLX1
7669_R_P_4_TBR1
7669_R_P_5_DLX1
7669_R_P_5_TBR1
7669_R_P_6_DLX1
7669_R_P_6_TBR1
7669_R_P_7_DLX1
7669_R_P_7_TBR1
7669_R_P_8_DLX1
7669_R_P_8_TBR1
7669_R_P_9_DLX
7669_R_P_9_DLX1
7669_R_P_9_TBR1
7669_R_POA_Bulk
7669_R_POA_DLX1
7669_R_sec1_Cau_Bulk
7669_R_sec2_GP_Bulk
7669_R_sec2_GP_DLX1
7669_R_sec2_Put_Bulk
7669_R_sec2_Thal_Bulk
7669_R_sec2_Thal_COUPTF2
7669_R_sec2_Thal_DLX1
7669_R_sec3_Cau_DLX1
7669_R_sec3_Hip_COUPTF2
7669_R_sec3_Hip_DLX1
7669_R_sec3_Hip_TBR1
7669_R_sec3_Put_DLX1
7669_R_Skin_Bulk
7669_R_T_1_Bulk
7669_R_T_1_COUPTF2
7669_R_T_1_DLX1
7669_R_T_1_TBR1
7669_RA9_Bulk
7669_RV10_Bulk
JGG

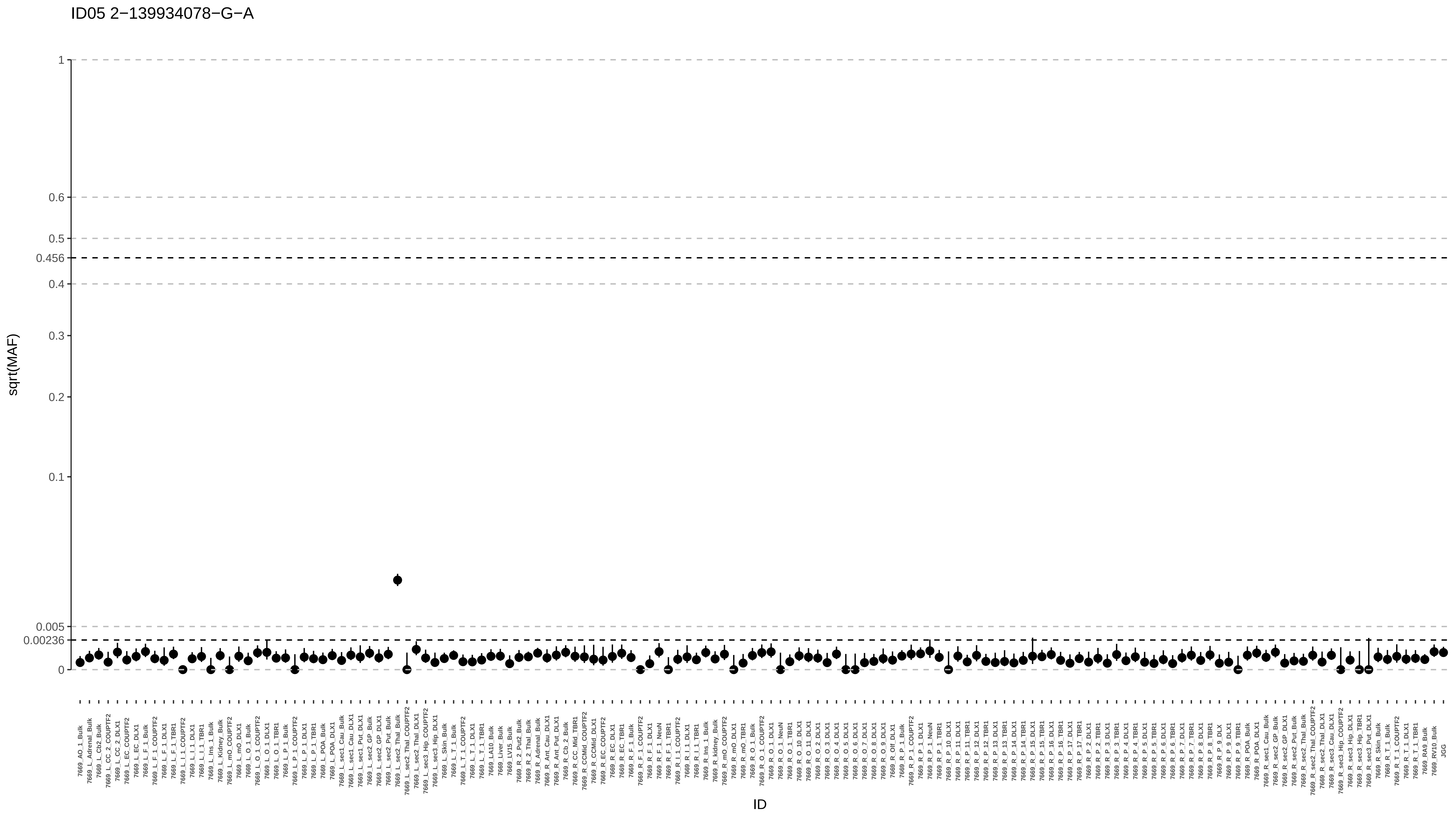
ID05 2−139934078−G−A
sqrt(MAF)
ID

ID05 2−141914766−A−T

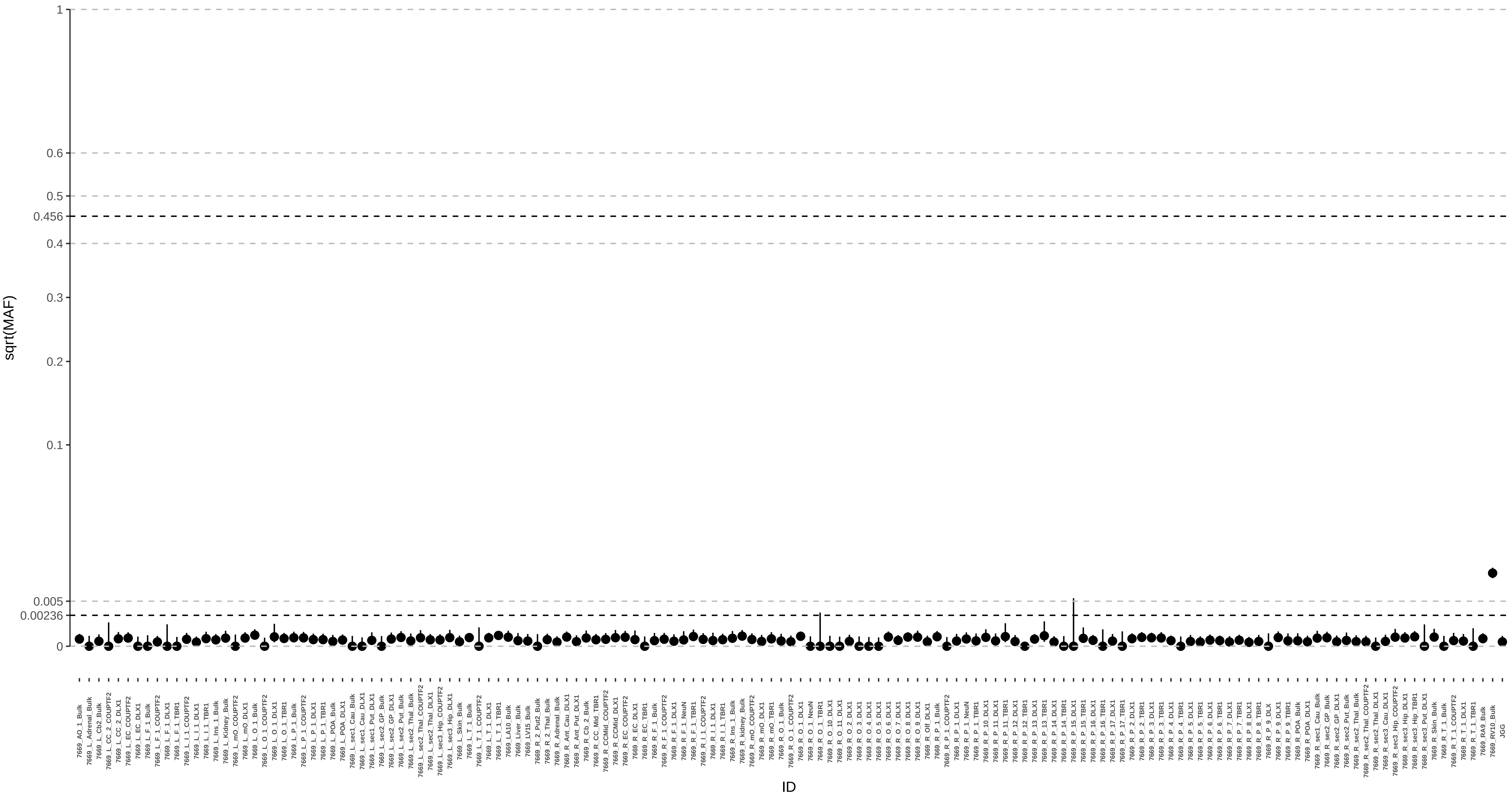

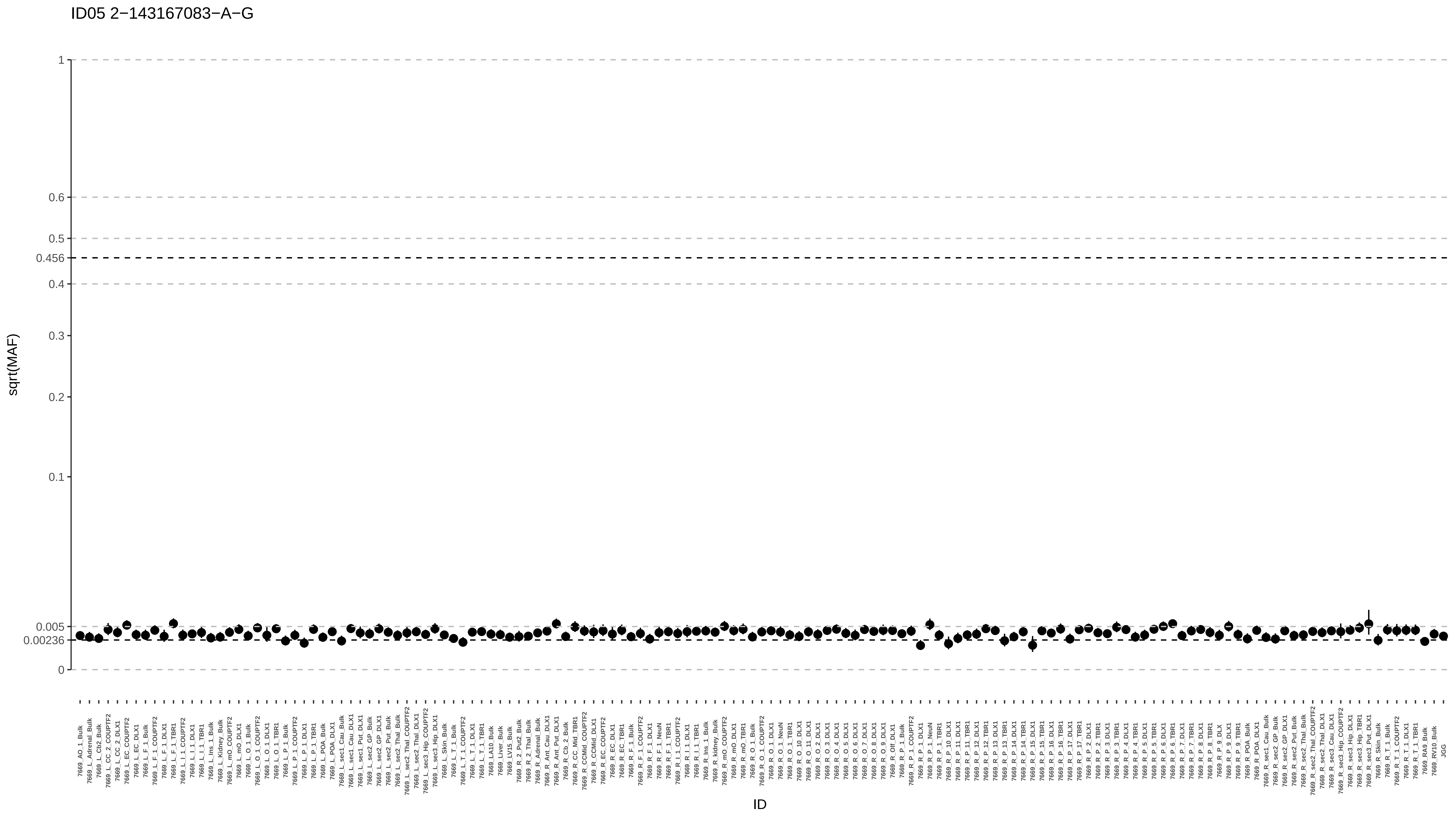
ID05 2−143167083−A−G
sqrt(MAF)
1
0.6
0.5
0.456
0.4
0.3
0.2
0.1
0.005
0.00236
0
ID
7669_AO_1_Bulk
7669_L_Adrenal_Bulk
7669_L_Cb2_Bulk
7669_L_CC_2_COUPTF2
7669_L_CC_2_DLX1
7669_L_EC_COUPTF2
7669_L_EC_DLX1
7669_L_F_1_Bulk
7669_L_F_1_COUPTF2
7669_L_F_1_DLX1
7669_L_F_1_TBR1
7669_L_I_1_COUPTF2
7669_L_I_1_DLX1
7669_L_I_1_TBR1
7669_L_Ins_1_Bulk
7669_L_Kidney_Bulk
7669_L_mO_COUPTF2
7669_L_mO_DLX1
7669_L_O_1_Bulk
7669_L_O_1_COUPTF2
7669_L_O_1_DLX1
7669_L_O_1_TBR1
7669_L_P_1_Bulk
7669_L_P_1_COUPTF2
7669_L_P_1_DLX1
7669_L_P_1_TBR1
7669_L_POA_Bulk
7669_L_POA_DLX1
7669_L_sec1_Cau_Bulk
7669_L_sec1_Cau_DLX1
7669_L_sec1_Put_DLX1
7669_L_sec2_GP_Bulk
7669_L_sec2_GP_DLX1
7669_L_sec2_Put_Bulk
7669_L_sec2_Thal_Bulk
7669_L_sec2_Thal_COUPTF2
7669_L_sec2_Thal_DLX1
7669_L_sec3_Hip_COUPTF2
7669_L_sec3_Hip_DLX1
7669_L_Skin_Bulk
7669_L_T_1_Bulk
7669_L_T_1_COUPTF2
7669_L_T_1_DLX1
7669_L_T_1_TBR1
7669_LA10_Bulk
7669_Liver_Bulk
7669_LV15_Bulk
7669_R_2_Put2_Bulk
7669_R_2_Thal_Bulk
7669_R_Adrenal_Bulk
7669_R_Ant_Cau_DLX1
7669_R_Ant_Put_DLX1
7669_R_Cb_2_Bulk
7669_R_CC_Mid_TBR1
7669_R_CCMid_COUPTF2
7669_R_CCMid_DLX1
7669_R_EC_COUPTF2
7669_R_EC_DLX1
7669_R_EC_TBR1
7669_R_F_1_Bulk
7669_R_F_1_COUPTF2
7669_R_F_1_DLX1
7669_R_F_1_NeuN
7669_R_F_1_TBR1
7669_R_I_1_COUPTF2
7669_R_I_1_DLX1
7669_R_I_1_TBR1
7669_R_Ins_1_Bulk
7669_R_kidney_Bulk
7669_R_mO_COUPTF2
7669_R_mO_DLX1
7669_R_mO_TBR1
7669_R_O_1_Bulk
7669_R_O_1_COUPTF2
7669_R_O_1_DLX1
7669_R_O_1_NeuN
7669_R_O_1_TBR1
7669_R_O_10_DLX1
7669_R_O_11_DLX1
7669_R_O_2_DLX1
7669_R_O_3_DLX1
7669_R_O_4_DLX1
7669_R_O_5_DLX1
7669_R_O_6_DLX1
7669_R_O_7_DLX1
7669_R_O_8_DLX1
7669_R_O_9_DLX1
7669_R_Olf_DLX1
7669_R_P_1_Bulk
7669_R_P_1_COUPTF2
7669_R_P_1_DLX1
7669_R_P_1_NeuN
7669_R_P_1_TBR1
7669_R_P_10_DLX1
7669_R_P_11_DLX1
7669_R_P_11_TBR1
7669_R_P_12_DLX1
7669_R_P_12_TBR1
7669_R_P_13_DLX1
7669_R_P_13_TBR1
7669_R_P_14_DLX1
7669_R_P_14_TBR1
7669_R_P_15_DLX1
7669_R_P_15_TBR1
7669_R_P_16_DLX1
7669_R_P_16_TBR1
7669_R_P_17_DLX1
7669_R_P_17_TBR1
7669_R_P_2_DLX1
7669_R_P_2_TBR1
7669_R_P_3_DLX1
7669_R_P_3_TBR1
7669_R_P_4_DLX1
7669_R_P_4_TBR1
7669_R_P_5_DLX1
7669_R_P_5_TBR1
7669_R_P_6_DLX1
7669_R_P_6_TBR1
7669_R_P_7_DLX1
7669_R_P_7_TBR1
7669_R_P_8_DLX1
7669_R_P_8_TBR1
7669_R_P_9_DLX
7669_R_P_9_DLX1
7669_R_P_9_TBR1
7669_R_POA_Bulk
7669_R_POA_DLX1
7669_R_sec1_Cau_Bulk
7669_R_sec2_GP_Bulk
7669_R_sec2_GP_DLX1
7669_R_sec2_Put_Bulk
7669_R_sec2_Thal_Bulk
7669_R_sec2_Thal_COUPTF2
7669_R_sec2_Thal_DLX1
7669_R_sec3_Cau_DLX1
7669_R_sec3_Hip_COUPTF2
7669_R_sec3_Hip_DLX1
7669_R_sec3_Hip_TBR1
7669_R_sec3_Put_DLX1
7669_R_Skin_Bulk
7669_R_T_1_Bulk
7669_R_T_1_COUPTF2
7669_R_T_1_DLX1
7669_R_T_1_TBR1
7669_RA9_Bulk
7669_RV10_Bulk
JGG

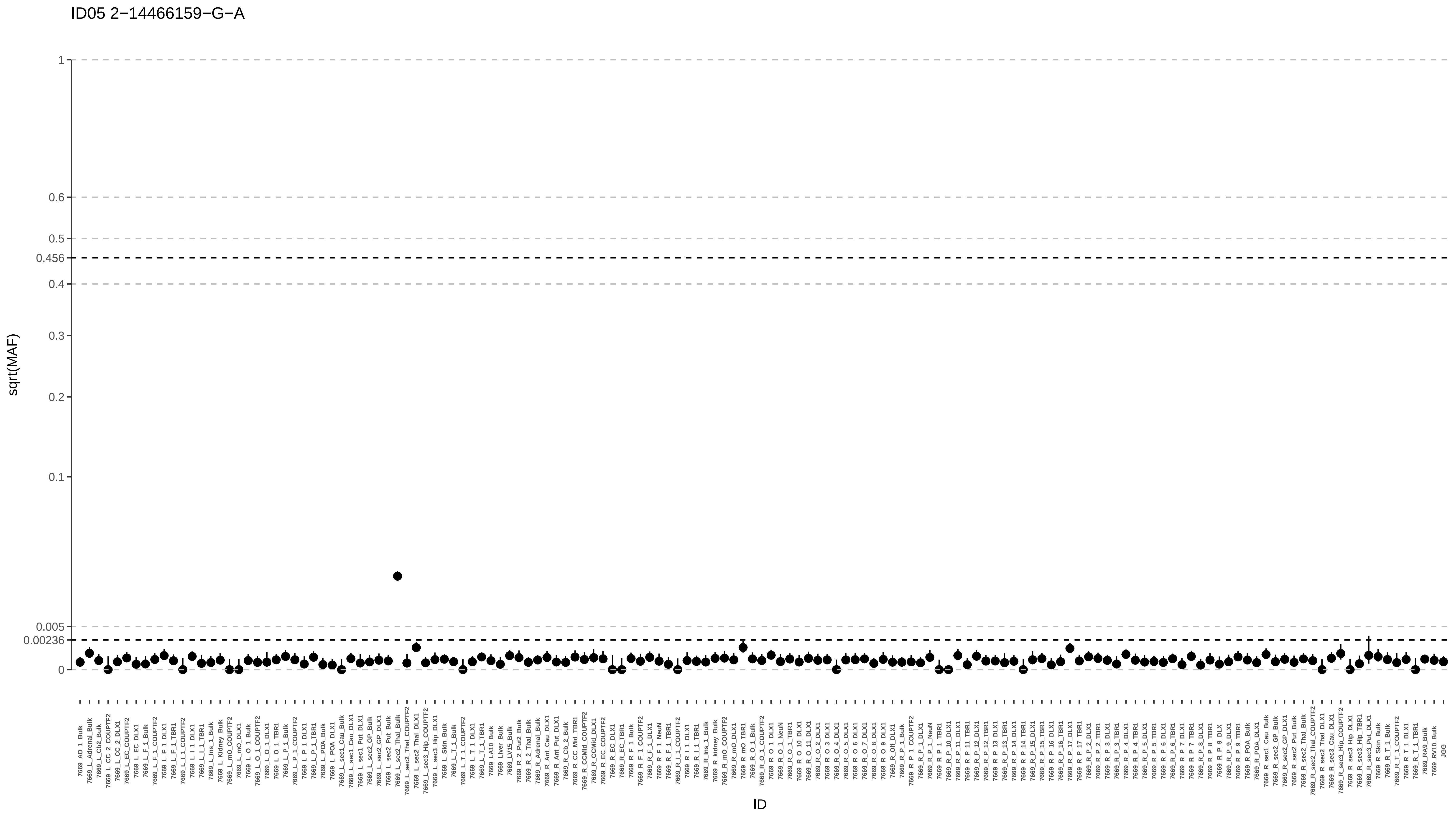
ID05 2−14466159−G−A
1
0.6
0.5
0.456
0.4
0.3
0.2
0.1
0.005
0.00236
0
sqrt(MAF)
ID

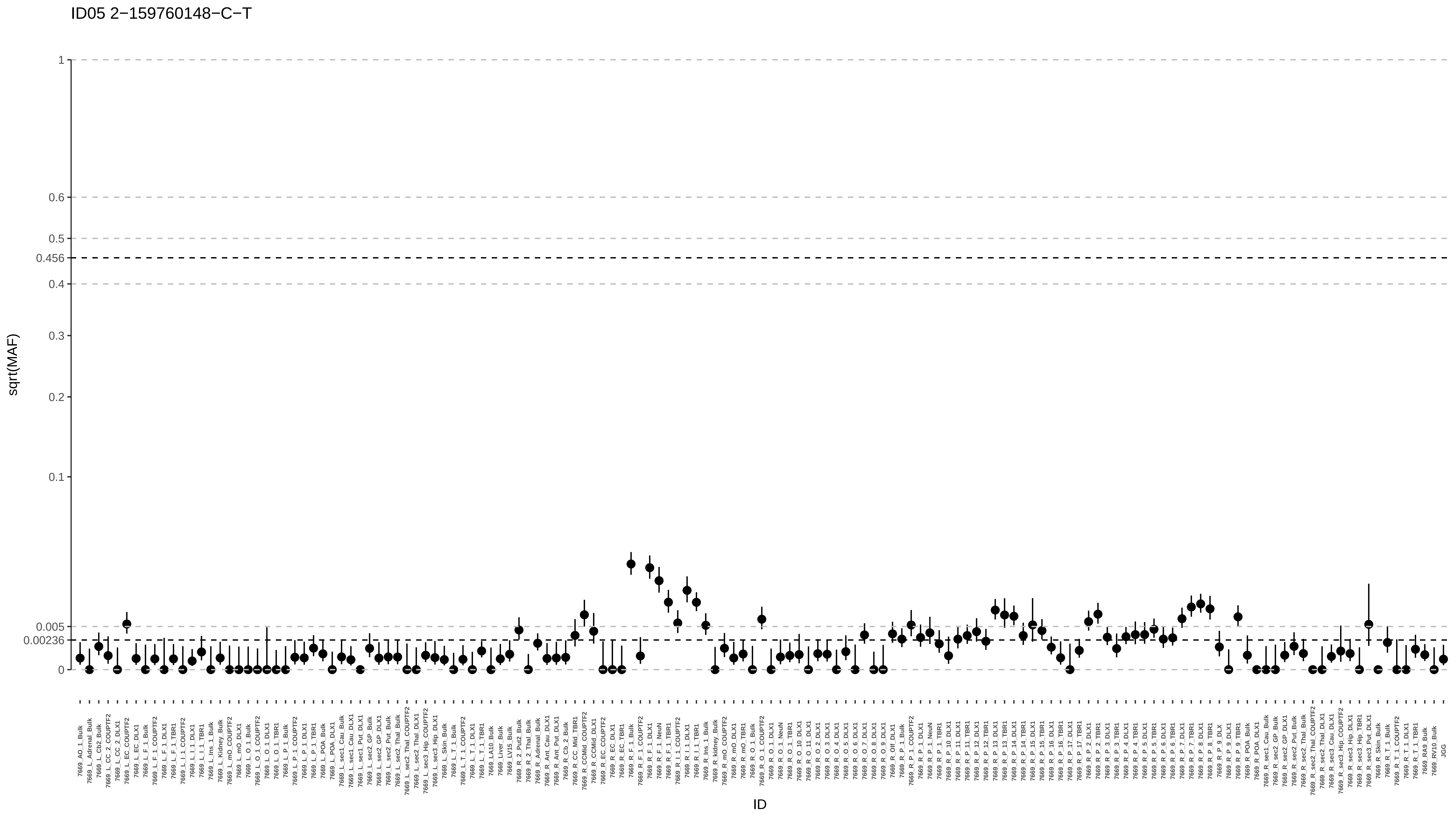
ID05 2−159760148−C−T
sqrt(MAF)
1
0.6
0.5
0.456
0.4
0.3
0.2
0.1
0.005
0.00236
0
ID
7669_AO_1_Bulk
7669_L_Adrenal_Bulk
7669_L_Cb2_Bulk
7669_L_CC_2_COUPTF2
7669_L_CC_2_DLX1
7669_L_EC_COUPTF2
7669_L_EC_DLX1
7669_L_F_1_Bulk
7669_L_F_1_COUPTF2
7669_L_F_1_DLX1
7669_L_F_1_TBR1
7669_L_I_1_COUPTF2
7669_L_I_1_DLX1
7669_L_I_1_TBR1
7669_L_Ins_1_Bulk
7669_L_Kidney_Bulk
7669_L_mO_COUPTF2
7669_L_mO_DLX1
7669_L_O_1_Bulk
7669_L_O_1_COUPTF2
7669_L_O_1_DLX1
7669_L_O_1_TBR1
7669_L_P_1_Bulk
7669_L_P_1_COUPTF2
7669_L_P_1_DLX1
7669_L_P_1_TBR1
7669_L_POA_Bulk
7669_L_POA_DLX1
7669_L_sec1_Cau_Bulk
7669_L_sec1_Cau_DLX1
7669_L_sec1_Put_DLX1
7669_L_sec2_GP_Bulk
7669_L_sec2_GP_DLX1
7669_L_sec2_Put_Bulk
7669_L_sec2_Thal_Bulk
7669_L_sec2_Thal_COUPTF2
7669_L_sec2_Thal_DLX1
7669_L_sec3_Hip_COUPTF2
7669_L_sec3_Hip_DLX1
7669_L_Skin_Bulk
7669_L_T_1_Bulk
7669_L_T_1_COUPTF2
7669_L_T_1_DLX1
7669_L_T_1_TBR1
7669_LA10_Bulk
7669_Liver_Bulk
7669_LV15_Bulk
7669_R_2_Put2_Bulk
7669_R_2_Thal_Bulk
7669_R_Adrenal_Bulk
7669_R_Ant_Cau_DLX1
7669_R_Ant_Put_DLX1
7669_R_Cb_2_Bulk
7669_R_CC_Mid_TBR1
7669_R_CCMid_COUPTF2
7669_R_CCMid_DLX1
7669_R_EC_COUPTF2
7669_R_EC_DLX1
7669_R_EC_TBR1
7669_R_F_1_Bulk
7669_R_F_1_COUPTF2
7669_R_F_1_DLX1
7669_R_F_1_NeuN
7669_R_F_1_TBR1
7669_R_I_1_COUPTF2
7669_R_I_1_DLX1
7669_R_I_1_TBR1
7669_R_Ins_1_Bulk
7669_R_kidney_Bulk
7669_R_mO_COUPTF2
7669_R_mO_DLX1
7669_R_mO_TBR1
7669_R_O_1_Bulk
7669_R_O_1_COUPTF2
7669_R_O_1_DLX1
7669_R_O_1_NeuN
7669_R_O_1_TBR1
7669_R_O_10_DLX1
7669_R_O_11_DLX1
7669_R_O_2_DLX1
7669_R_O_3_DLX1
7669_R_O_4_DLX1
7669_R_O_5_DLX1
7669_R_O_6_DLX1
7669_R_O_7_DLX1
7669_R_O_8_DLX1
7669_R_O_9_DLX1
7669_R_Olf_DLX1
7669_R_P_1_Bulk
7669_R_P_1_COUPTF2
7669_R_P_1_DLX1
7669_R_P_1_NeuN
7669_R_P_1_TBR1
7669_R_P_10_DLX1
7669_R_P_11_DLX1
7669_R_P_11_TBR1
7669_R_P_12_DLX1
7669_R_P_12_TBR1
7669_R_P_13_DLX1
7669_R_P_13_TBR1
7669_R_P_14_DLX1
7669_R_P_14_TBR1
7669_R_P_15_DLX1
7669_R_P_15_TBR1
7669_R_P_16_DLX1
7669_R_P_16_TBR1
7669_R_P_17_DLX1
7669_R_P_17_TBR1
7669_R_P_2_DLX1
7669_R_P_2_TBR1
7669_R_P_3_DLX1
7669_R_P_3_TBR1
7669_R_P_4_DLX1
7669_R_P_4_TBR1
7669_R_P_5_DLX1
7669_R_P_5_TBR1
7669_R_P_6_DLX1
7669_R_P_6_TBR1
7669_R_P_7_DLX1
7669_R_P_7_TBR1
7669_R_P_8_DLX1
7669_R_P_8_TBR1
7669_R_P_9_DLX
7669_R_P_9_DLX1
7669_R_P_9_TBR1
7669_R_POA_Bulk
7669_R_POA_DLX1
7669_R_sec1_Cau_Bulk
7669_R_sec2_GP_Bulk
7669_R_sec2_GP_DLX1
7669_R_sec2_Put_Bulk
7669_R_sec2_Thal_Bulk
7669_R_sec2_Thal_COUPTF2
7669_R_sec2_Thal_DLX1
7669_R_sec3_Cau_DLX1
7669_R_sec3_Hip_COUPTF2
7669_R_sec3_Hip_DLX1
7669_R_sec3_Hip_TBR1
7669_R_sec3_Put_DLX1
7669_R_Skin_Bulk
7669_R_T_1_Bulk
7669_R_T_1_COUPTF2
7669_R_T_1_DLX1
7669_R_T_1_TBR1
7669_RA9_Bulk
7669_RV10_Bulk
JGG

ID05 2−163073318−G−A

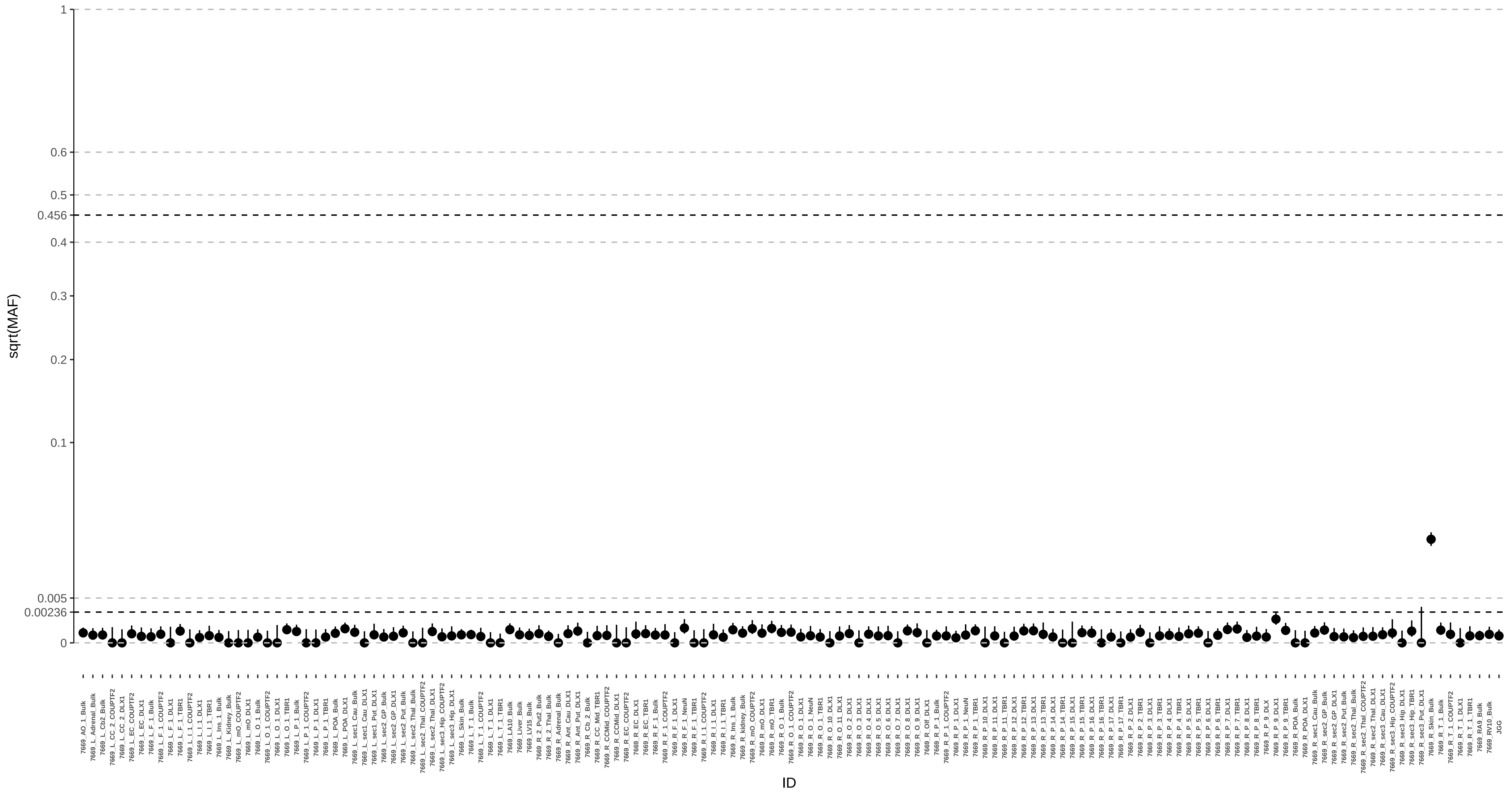

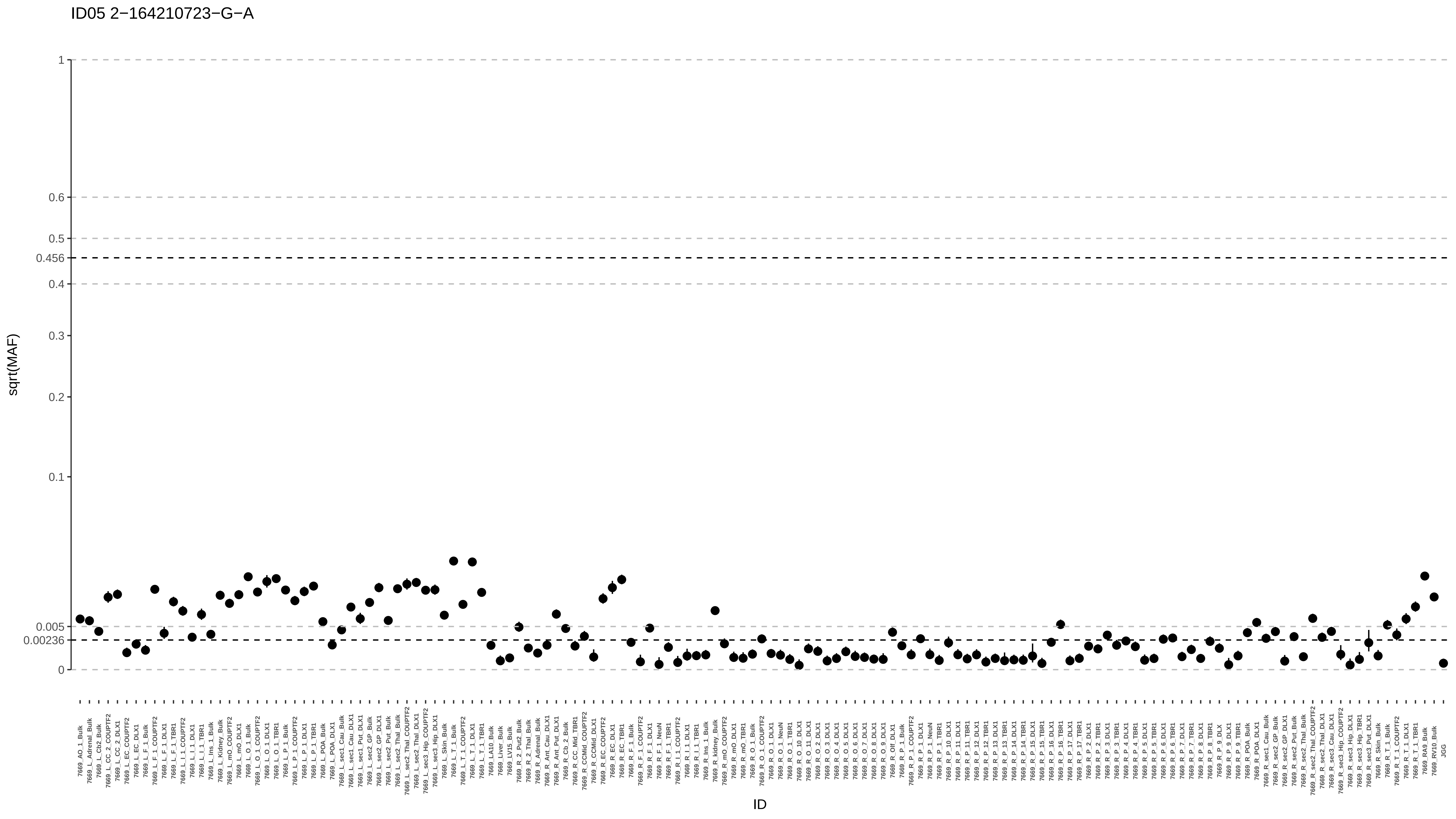
ID05 2−164210723−G−A
sqrt(MAF)
1
0.6
0.5
0.456
0.4
0.3
0.2
0.1
0.005
0.00236
0
ID

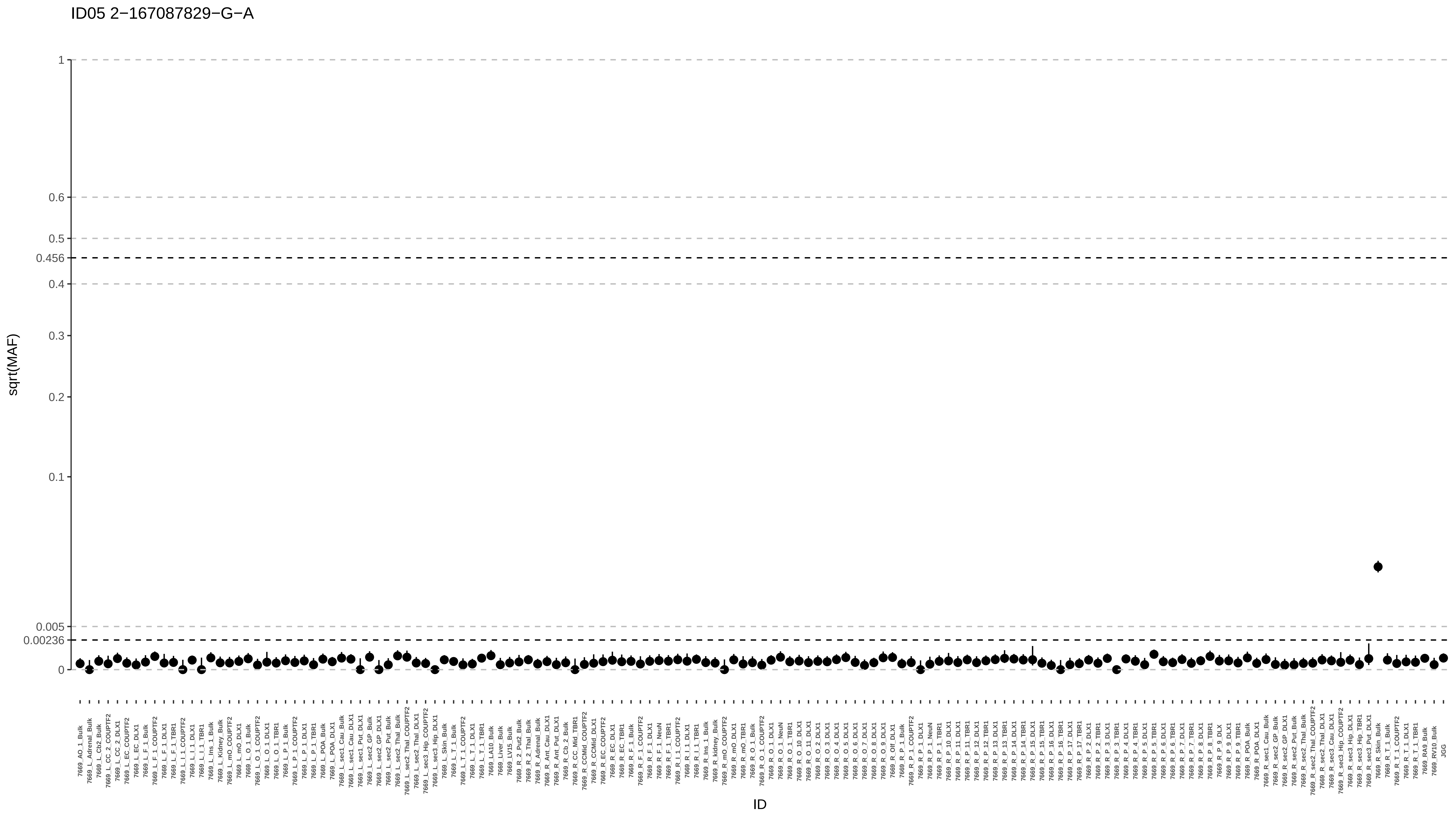
ID05 2−167087829−G−A
sqrt(MAF)
ID
1
0.6
0.5
0.456
0.4
0.3
0.2
0.1
0.005
0.00236
0

ID05 2−167342653−C−T

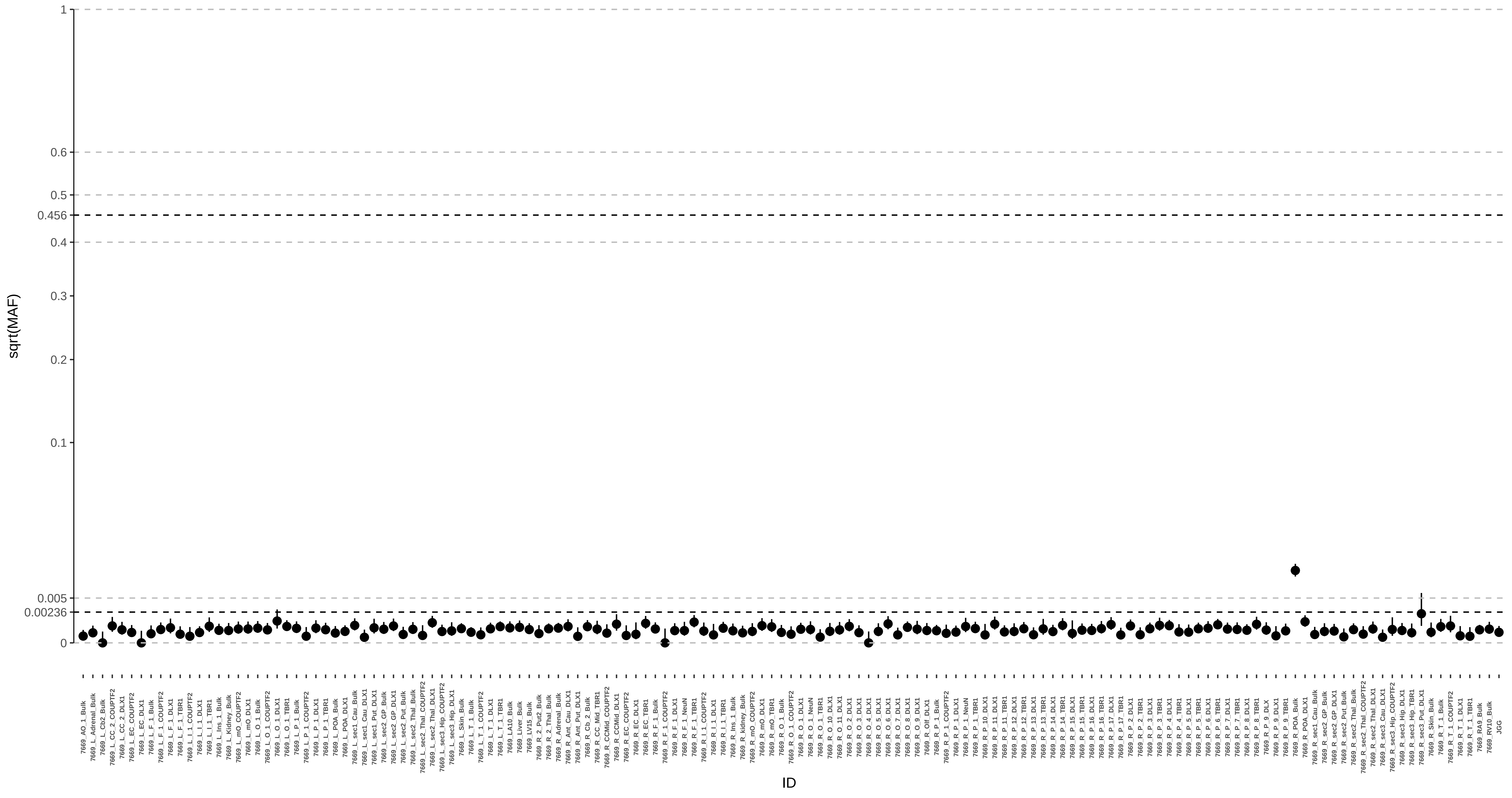

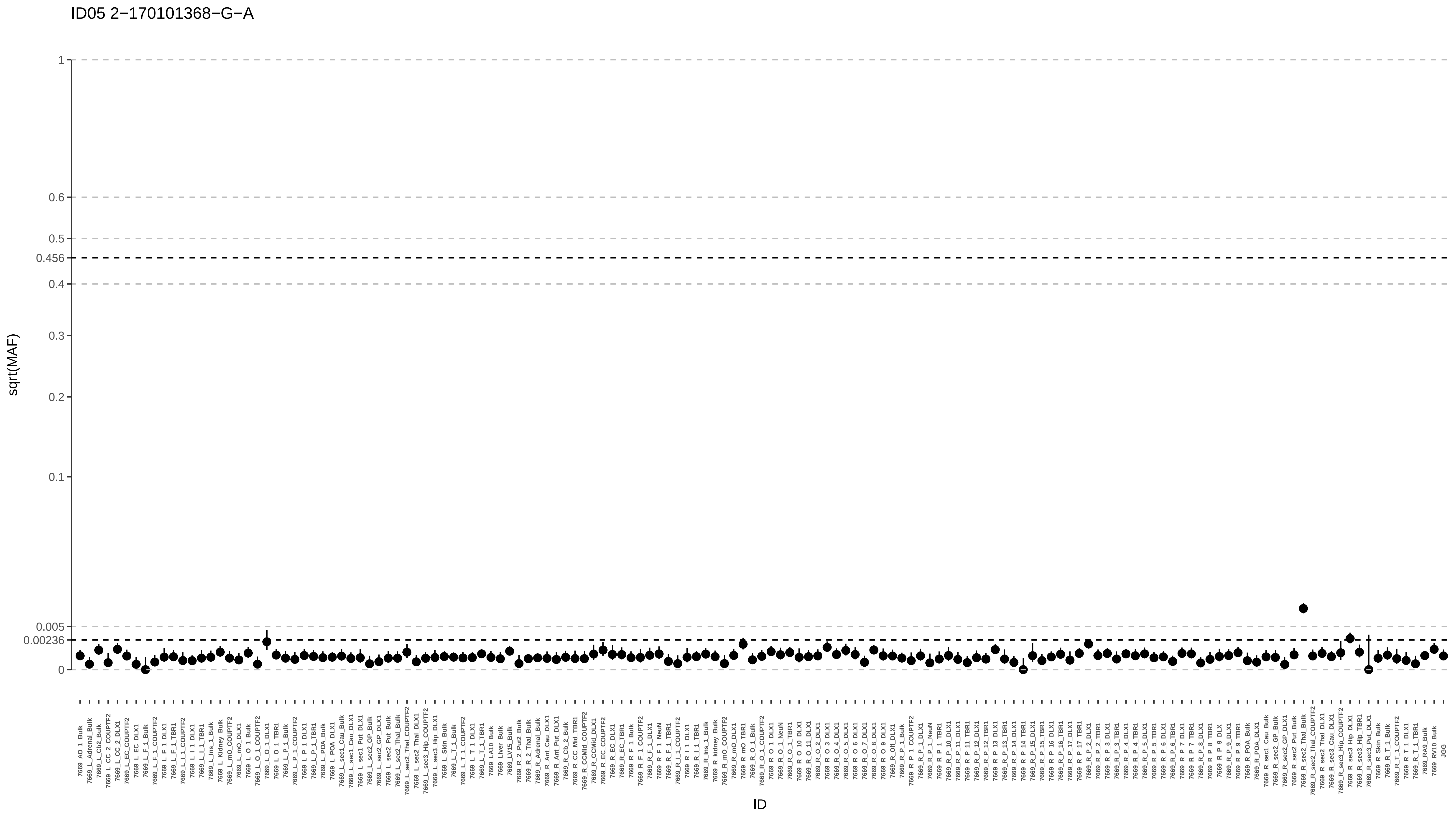
ID05 2−170101368−G−A
sqrt(MAF)
ID

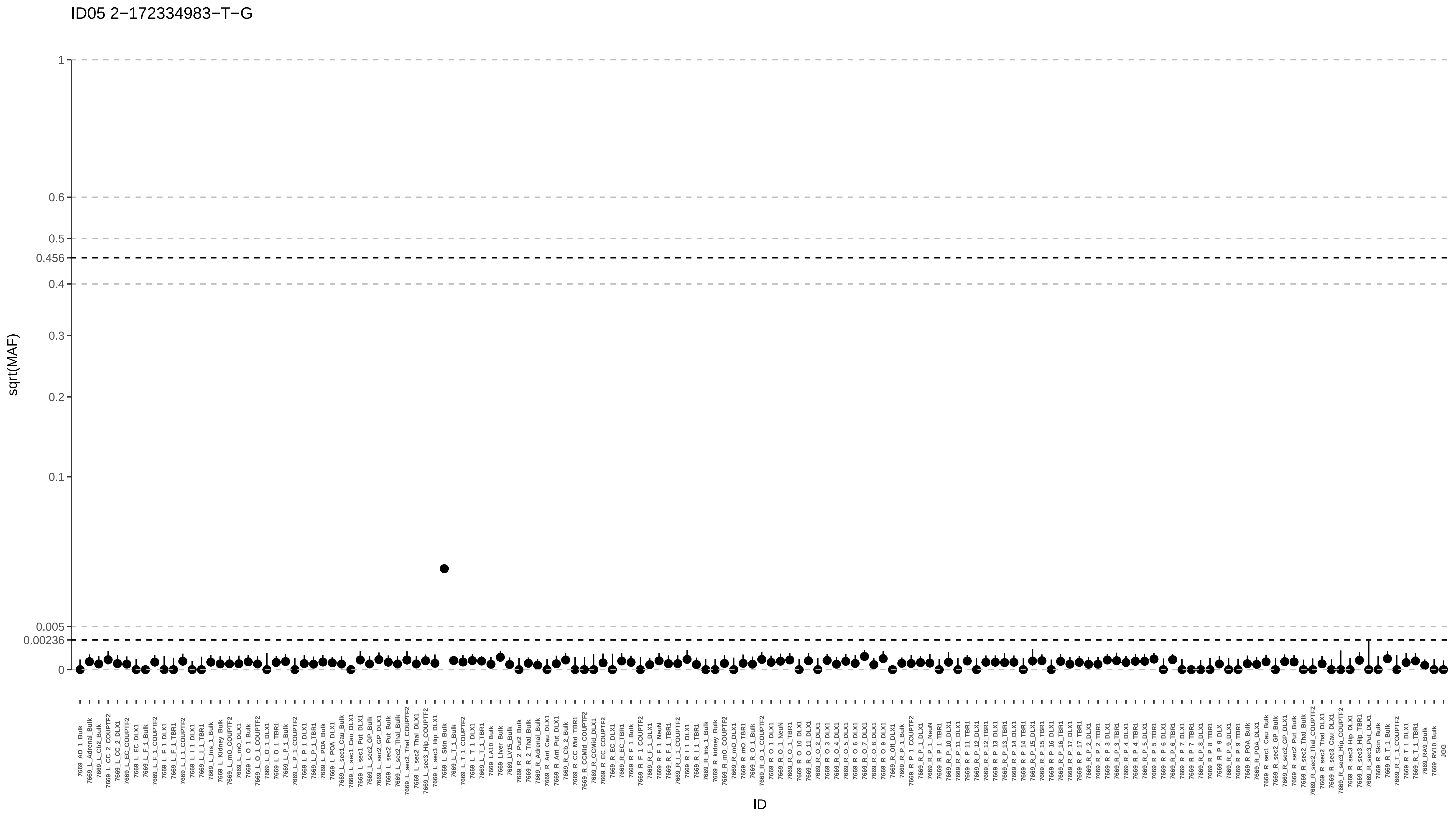

ID05 2−172334983−T−G
sqrt(MAF)
1
0.6
0.5
0.456
0.4
0.3
0.2
0.1
0.005
0.00236
0
ID
7669_AO_1_Bulk
7669_L_Adrenal_Bulk
7669_L_Cb2_Bulk
7669_L_CC_2_COUPTF2
7669_L_CC_2_DLX1
7669_L_EC_COUPTF2
7669_L_EC_DLX1
7669_L_F_1_Bulk
7669_L_F_1_COUPTF2
7669_L_F_1_DLX1
7669_L_F_1_TBR1
7669_L_I_1_COUPTF2
7669_L_I_1_DLX1
7669_L_I_1_TBR1
7669_L_Ins_1_Bulk
7669_L_Kidney_Bulk
7669_L_mO_COUPTF2
7669_L_mO_DLX1
7669_L_O_1_Bulk
7669_L_O_1_COUPTF2
7669_L_O_1_DLX1
7669_L_O_1_TBR1
7669_L_P_1_Bulk
7669_L_P_1_COUPTF2
7669_L_P_1_DLX1
7669_L_P_1_TBR1
7669_L_POA_Bulk
7669_L_POA_DLX1
7669_L_sec1_Cau_Bulk
7669_L_sec1_Cau_DLX1
7669_L_sec1_Put_DLX1
7669_L_sec2_GP_Bulk
7669_L_sec2_GP_DLX1
7669_L_sec2_Put_Bulk
7669_L_sec2_Thal_Bulk
7669_L_sec2_Thal_COUPTF2
7669_L_sec2_Thal_DLX1
7669_L_sec3_Hip_COUPTF2
7669_L_sec3_Hip_DLX1
7669_L_Skin_Bulk
7669_L_T_1_Bulk
7669_L_T_1_COUPTF2
7669_L_T_1_DLX1
7669_L_T_1_TBR1
7669_LA10_Bulk
7669_Liver_Bulk
7669_LV15_Bulk
7669_R_2_Put2_Bulk
7669_R_2_Thal_Bulk
7669_R_Adrenal_Bulk
7669_R_Ant_Cau_DLX1
7669_R_Ant_Put_DLX1
7669_R_Cb_2_Bulk
7669_R_CC_Mid_TBR1
7669_R_CCMid_COUPTF2
7669_R_CCMid_DLX1
7669_R_EC_COUPTF2
7669_R_EC_DLX1
7669_R_EC_TBR1
7669_R_F_1_Bulk
7669_R_F_1_COUPTF2
7669_R_F_1_DLX1
7669_R_F_1_NeuN
7669_R_F_1_TBR1
7669_R_I_1_COUPTF2
7669_R_I_1_DLX1
7669_R_I_1_TBR1
7669_R_Ins_1_Bulk
7669_R_kidney_Bulk
7669_R_mO_COUPTF2
7669_R_mO_DLX1
7669_R_mO_TBR1
7669_R_O_1_Bulk
7669_R_O_1_COUPTF2
7669_R_O_1_DLX1
7669_R_O_1_NeuN
7669_R_O_1_TBR1
7669_R_O_10_DLX1
7669_R_O_11_DLX1
7669_R_O_2_DLX1
7669_R_O_3_DLX1
7669_R_O_4_DLX1
7669_R_O_5_DLX1
7669_R_O_6_DLX1
7669_R_O_7_DLX1
7669_R_O_8_DLX1
7669_R_O_9_DLX1
7669_R_Olf_DLX1
7669_R_P_1_Bulk
7669_R_P_1_COUPTF2
7669_R_P_1_DLX1
7669_R_P_1_NeuN
7669_R_P_1_TBR1
7669_R_P_10_DLX1
7669_R_P_11_DLX1
7669_R_P_11_TBR1
7669_R_P_12_DLX1
7669_R_P_12_TBR1
7669_R_P_13_DLX1
7669_R_P_13_TBR1
7669_R_P_14_DLX1
7669_R_P_14_TBR1
7669_R_P_15_DLX1
7669_R_P_15_TBR1
7669_R_P_16_DLX1
7669_R_P_16_TBR1
7669_R_P_17_DLX1
7669_R_P_17_TBR1
7669_R_P_2_DLX1
7669_R_P_2_TBR1
7669_R_P_3_DLX1
7669_R_P_3_TBR1
7669_R_P_4_DLX1
7669_R_P_4_TBR1
7669_R_P_5_DLX1
7669_R_P_5_TBR1
7669_R_P_6_DLX1
7669_R_P_6_TBR1
7669_R_P_7_DLX1
7669_R_P_7_TBR1
7669_R_P_8_DLX1
7669_R_P_8_TBR1
7669_R_P_9_DLX
7669_R_P_9_DLX1
7669_R_P_9_TBR1
7669_R_POA_Bulk
7669_R_POA_DLX1
7669_R_sec1_Cau_Bulk
7669_R_sec2_GP_Bulk
7669_R_sec2_GP_DLX1
7669_R_sec2_Put_Bulk
7669_R_sec2_Thal_Bulk
7669_R_sec2_Thal_COUPTF2
7669_R_sec2_Thal_DLX1
7669_R_sec3_Cau_DLX1
7669_R_sec3_Hip_COUPTF2
7669_R_sec3_Hip_DLX1
7669_R_sec3_Hip_TBR1
7669_R_sec3_Put_DLX1
7669_R_Skin_Bulk
7669_R_T_1_Bulk
7669_R_T_1_COUPTF2
7669_R_T_1_DLX1
7669_R_T_1_TBR1
7669_RA9_Bulk
7669_RV10_Bulk
JGG

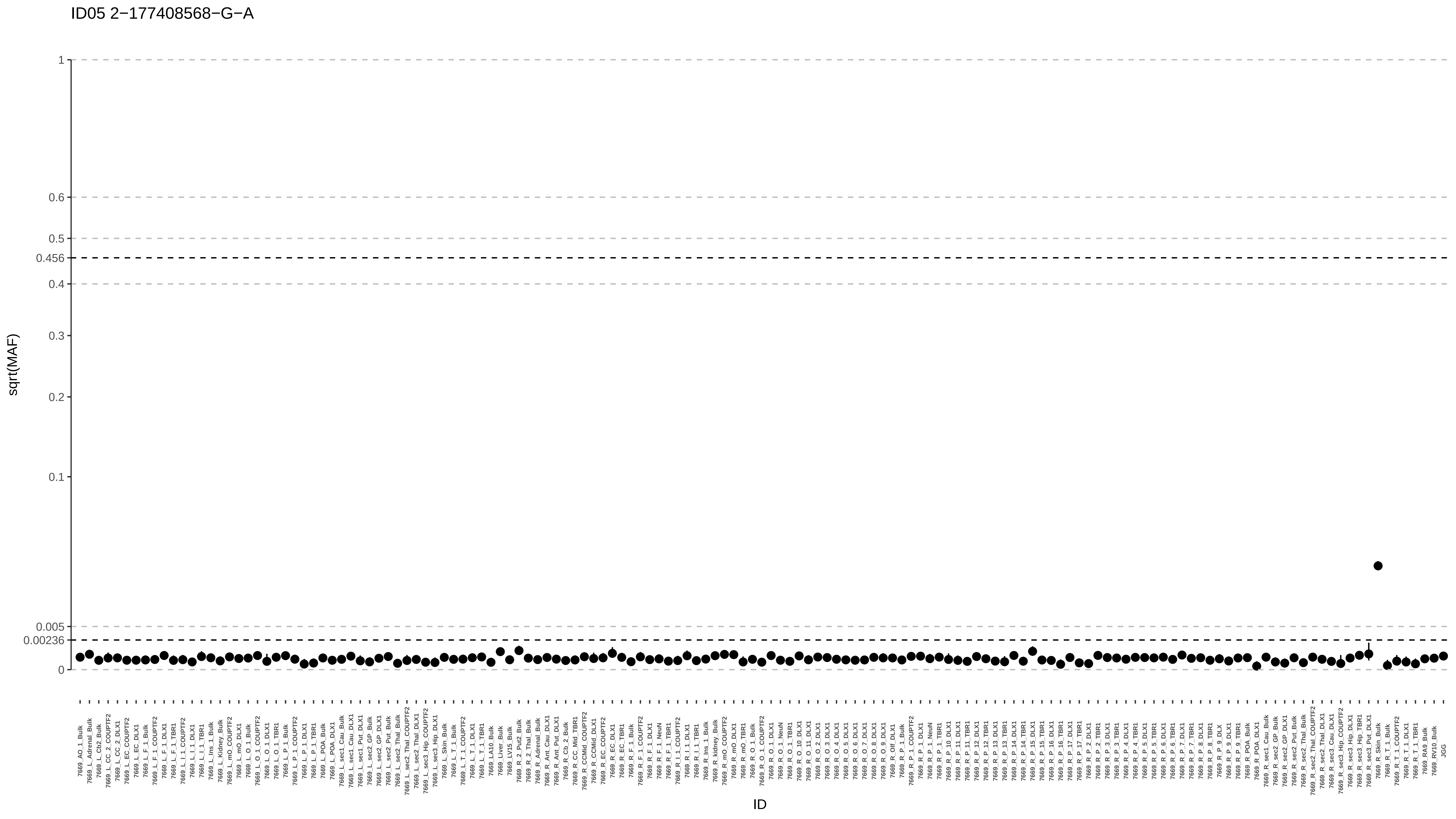
ID05 2−177408568−G−A
sqrt(MAF)
ID
1
0.6
0.5
0.456
0.4
0.3
0.2
0.1
0.005
0.00236
0

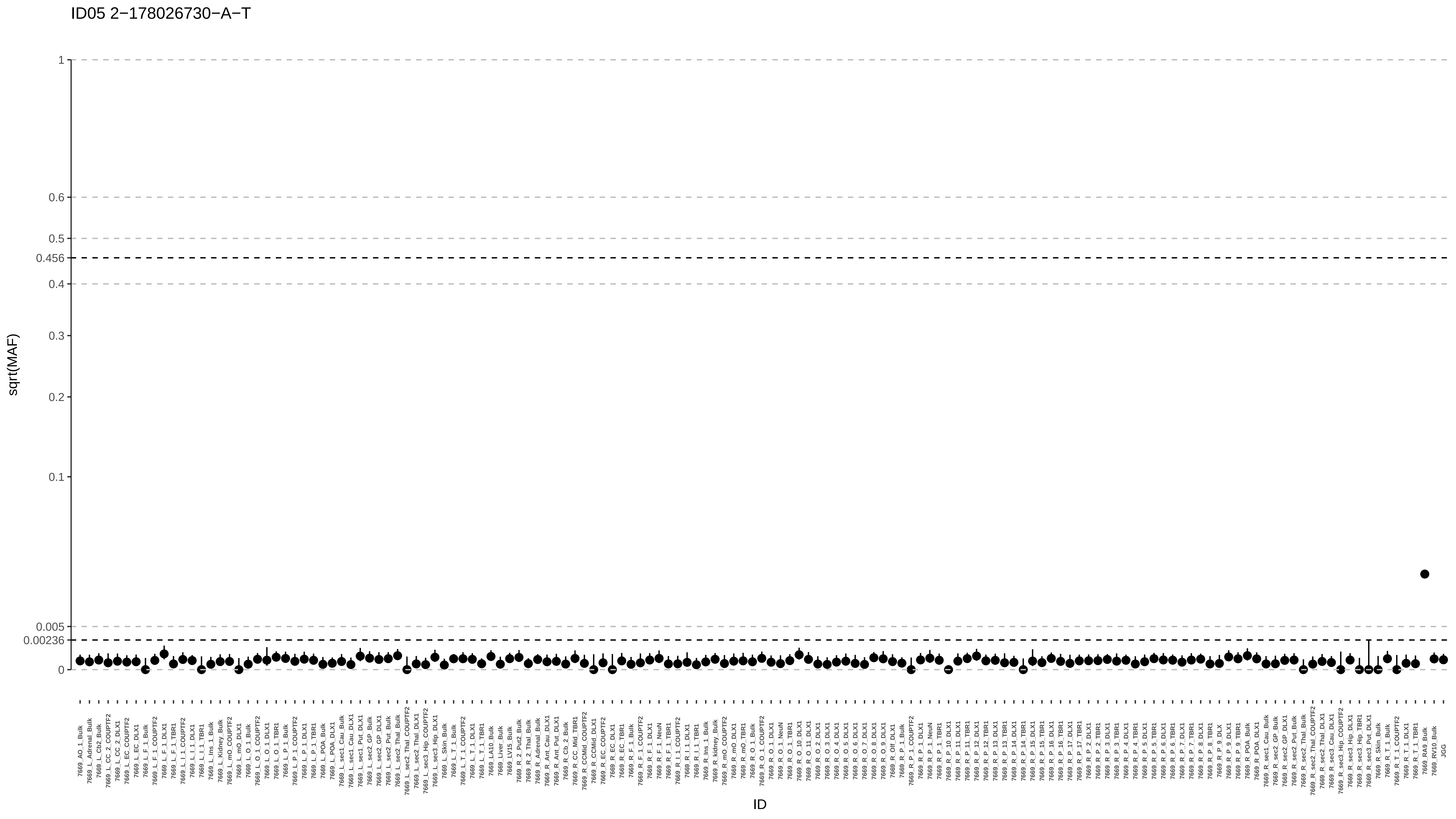
ID05 2−178026730−A−T
sqrt(MAF)
1
0.6
0.5
0.456
0.4
0.3
0.2
0.1
0.005
0.00236
0
ID

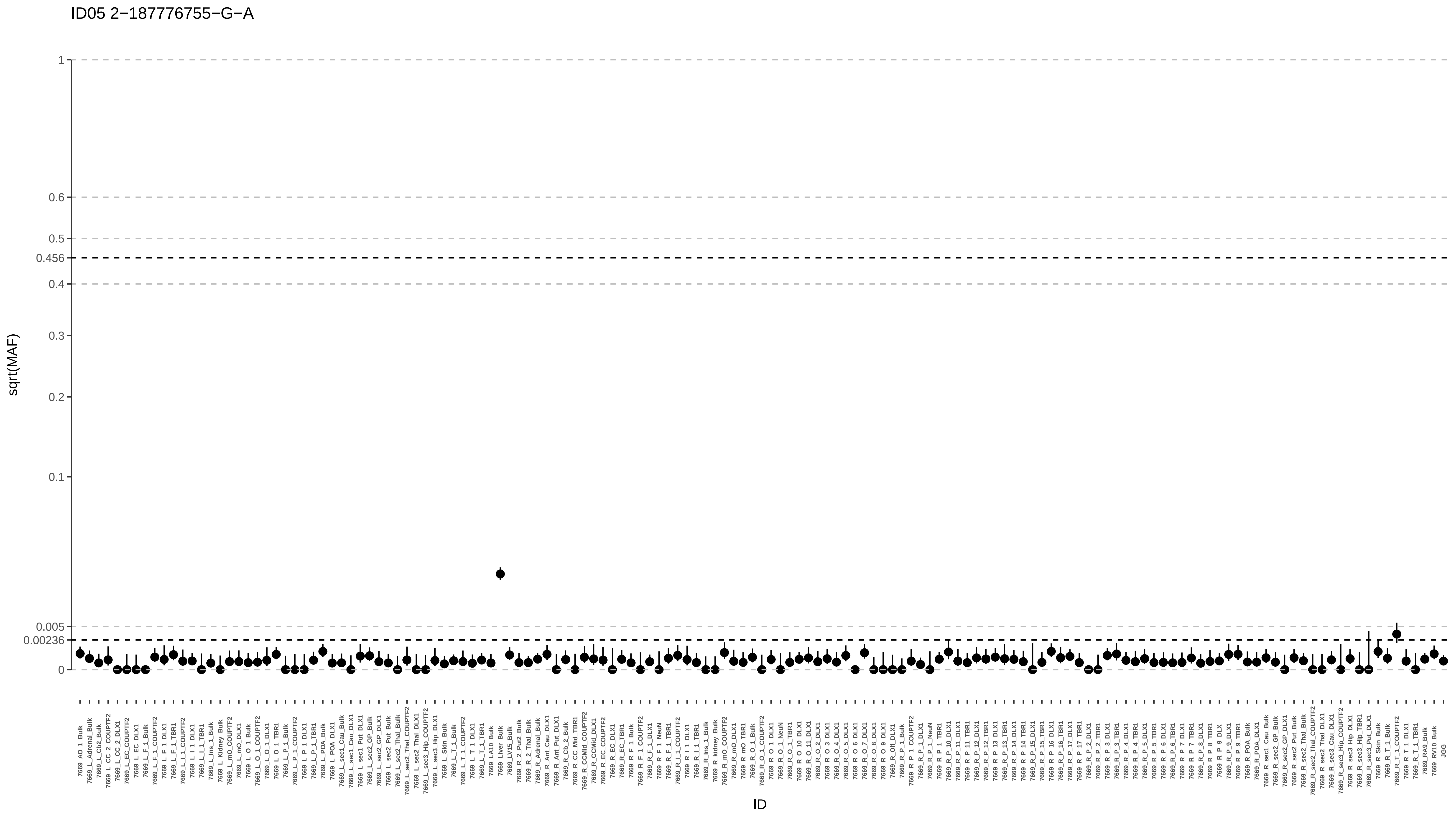
ID05 2−187776755−G−A
sqrt(MAF)
ID
1
0.6
0.5
0.456
0.4
0.3
0.2
0.1
0.005
0.00236
0

ID05 2−190501942−A−G

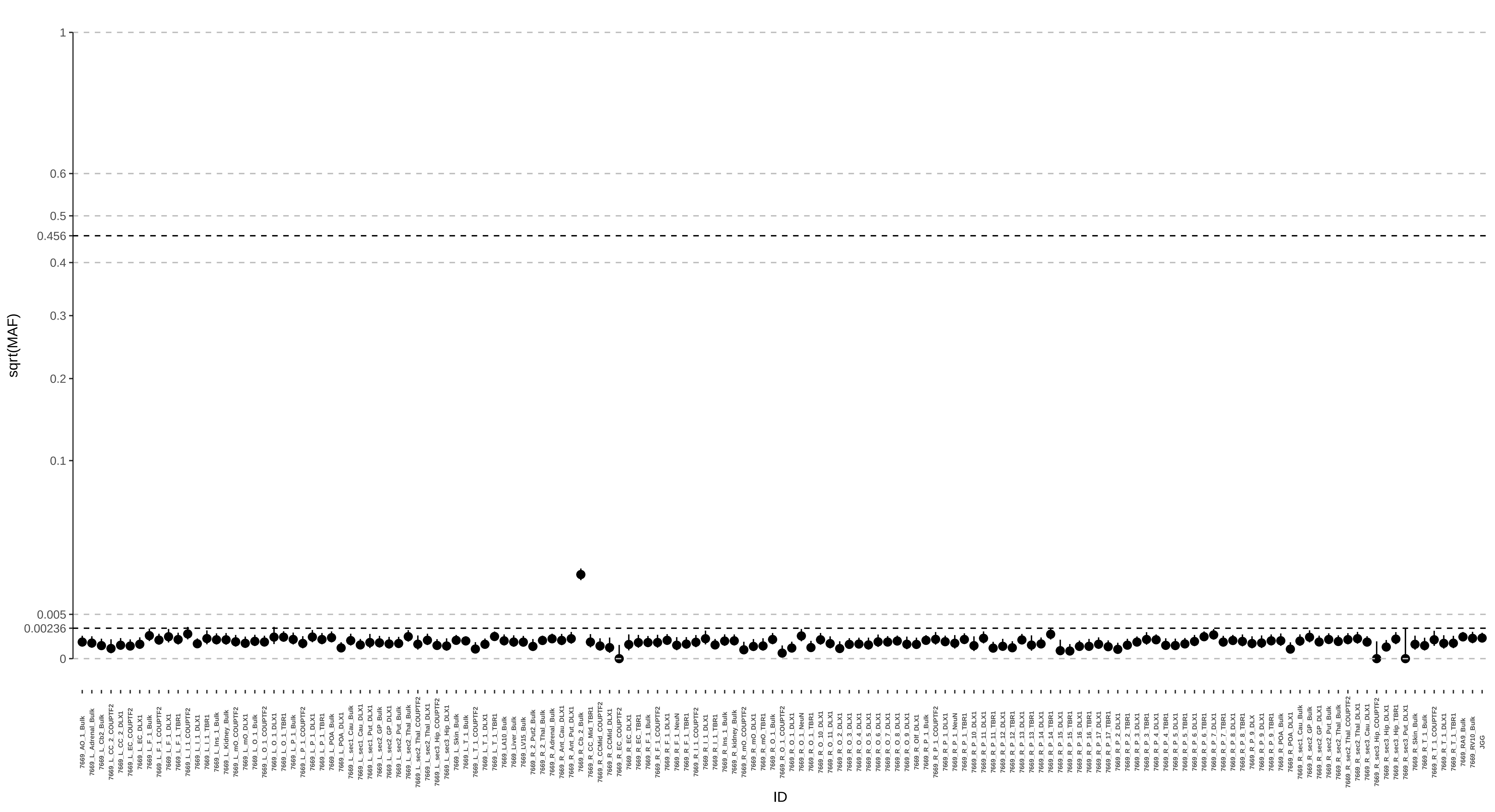

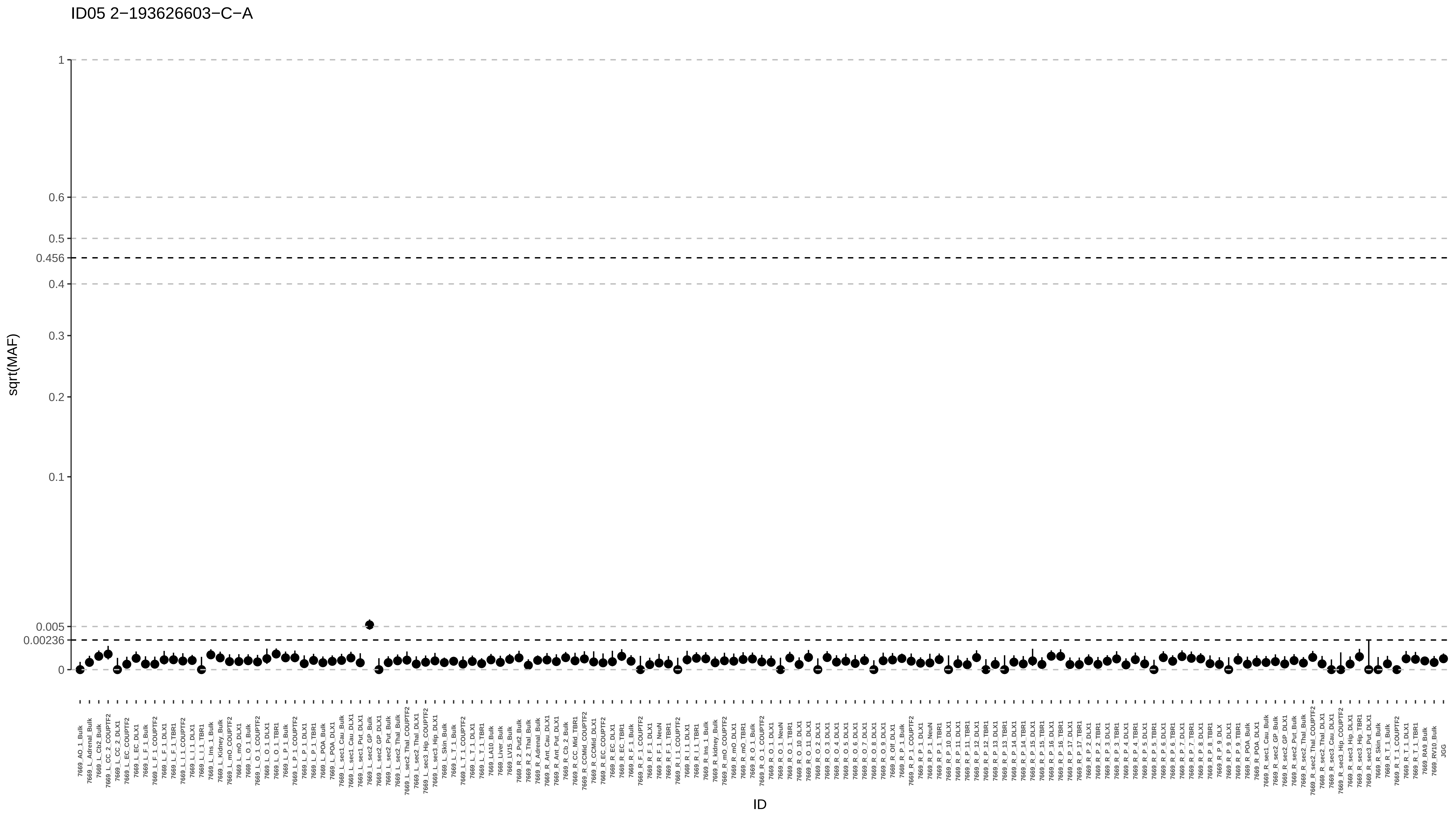
ID05 2−193626603−C−A
sqrt(MAF)
ID

ID05 2−193831190−T−A

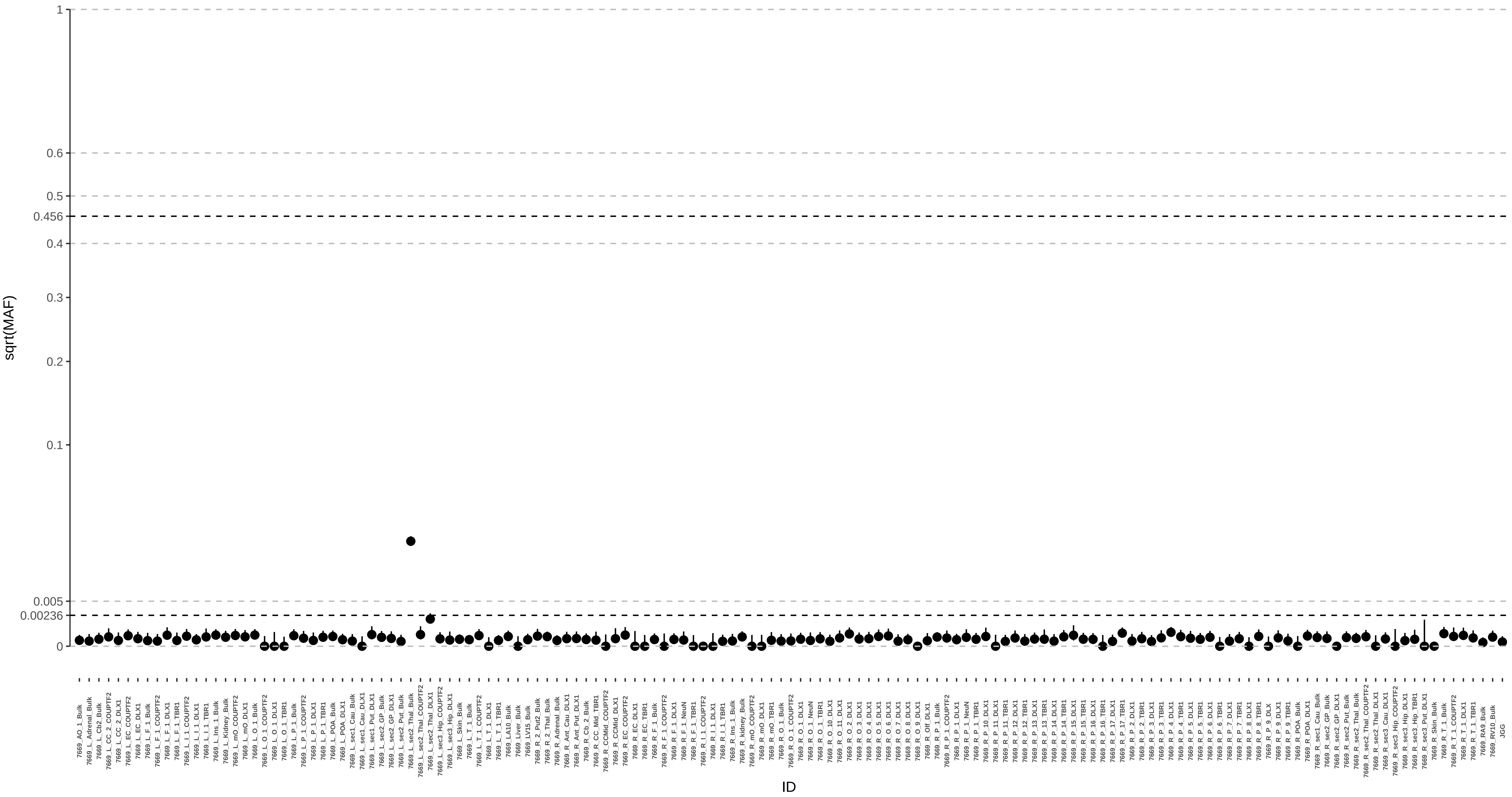

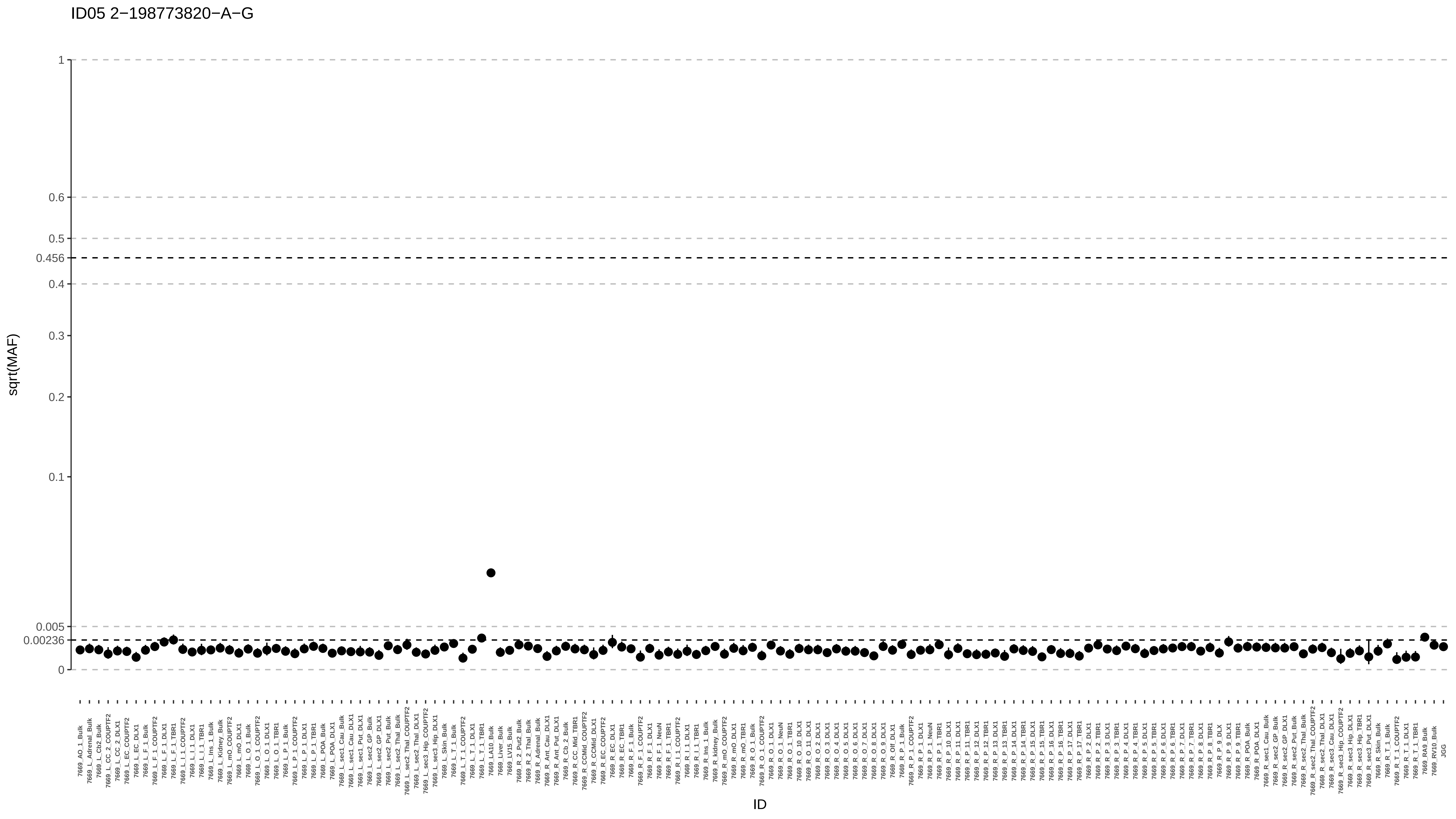

ID05 2−198773820−A−G
sqrt(MAF)
ID

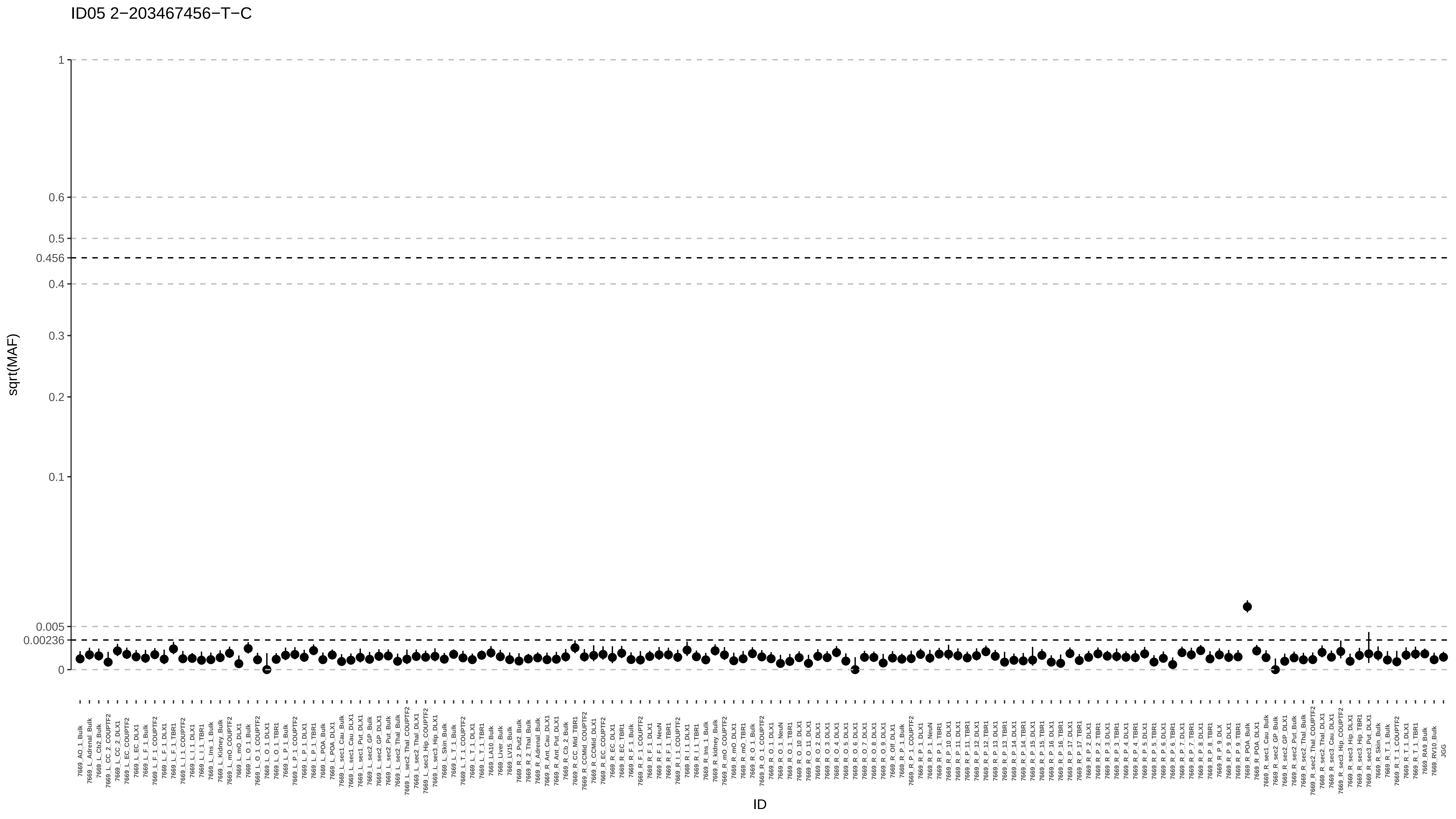
ID05 2−203467456−T−C
sqrt(MAF)
1
0.6
0.5
0.456
0.4
0.3
0.2
0.1
0.005
0.00236
0
ID

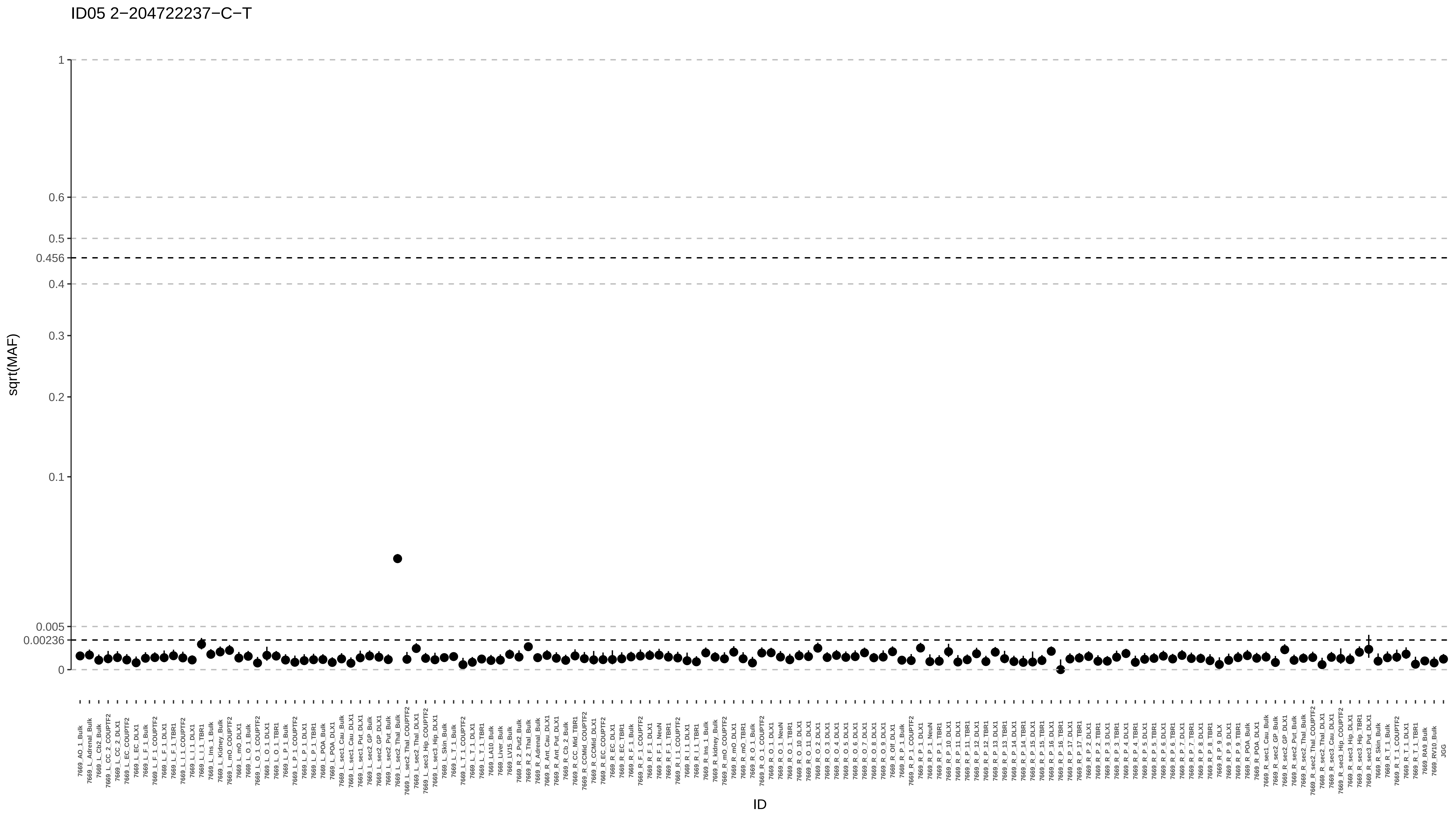
ID05 2−204722237−C−T
sqrt(MAF)
1
0.6
0.5
0.456
0.4
0.3
0.2
0.1
0.005
0.00236
0
ID

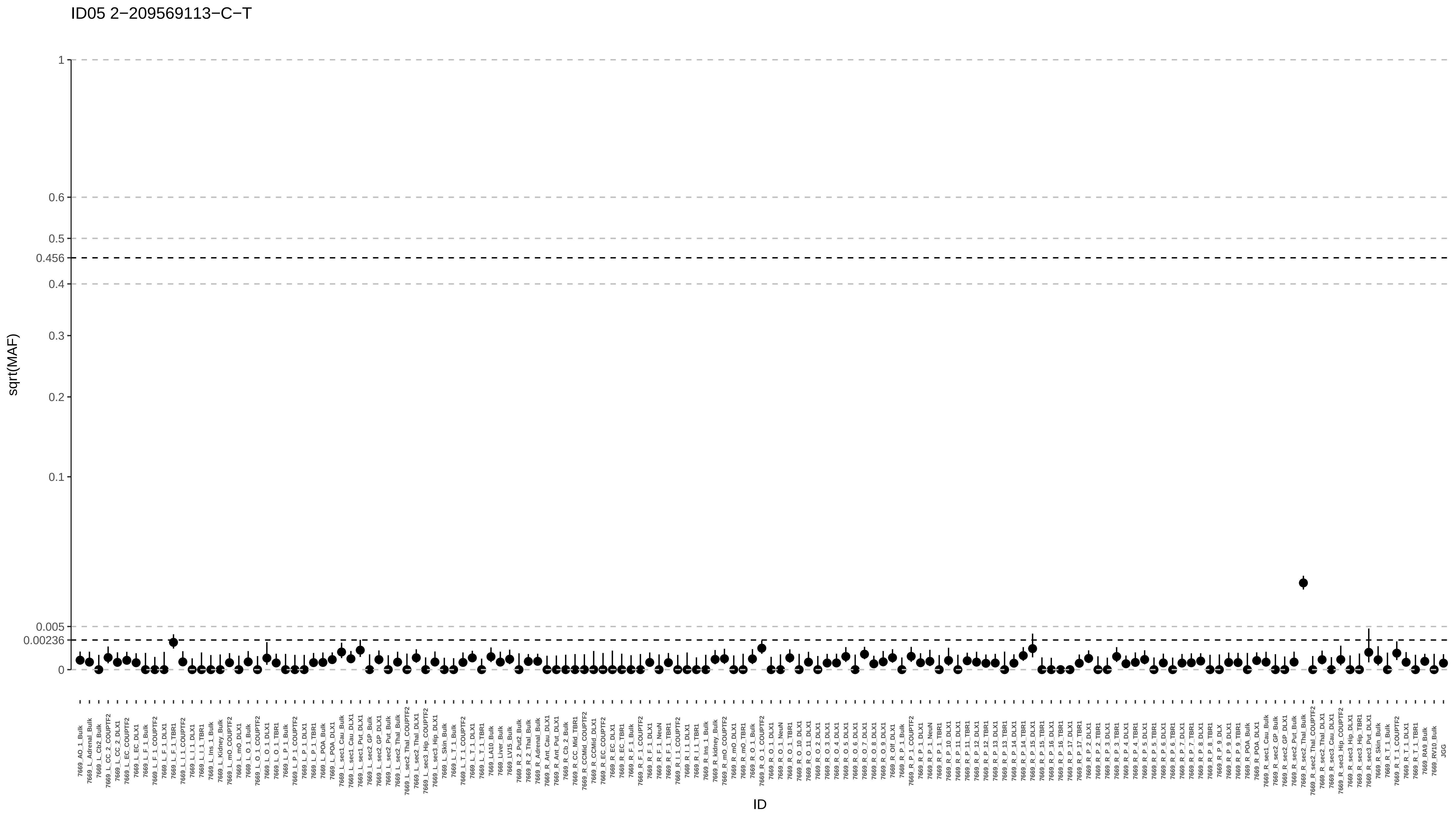
ID05 2−209569113−C−T
sqrt(MAF)
1
0.6
0.5
0.456
0.4
0.3
0.2
0.1
0.005
0.00236
0
ID

ID05 2−213063972−C−T

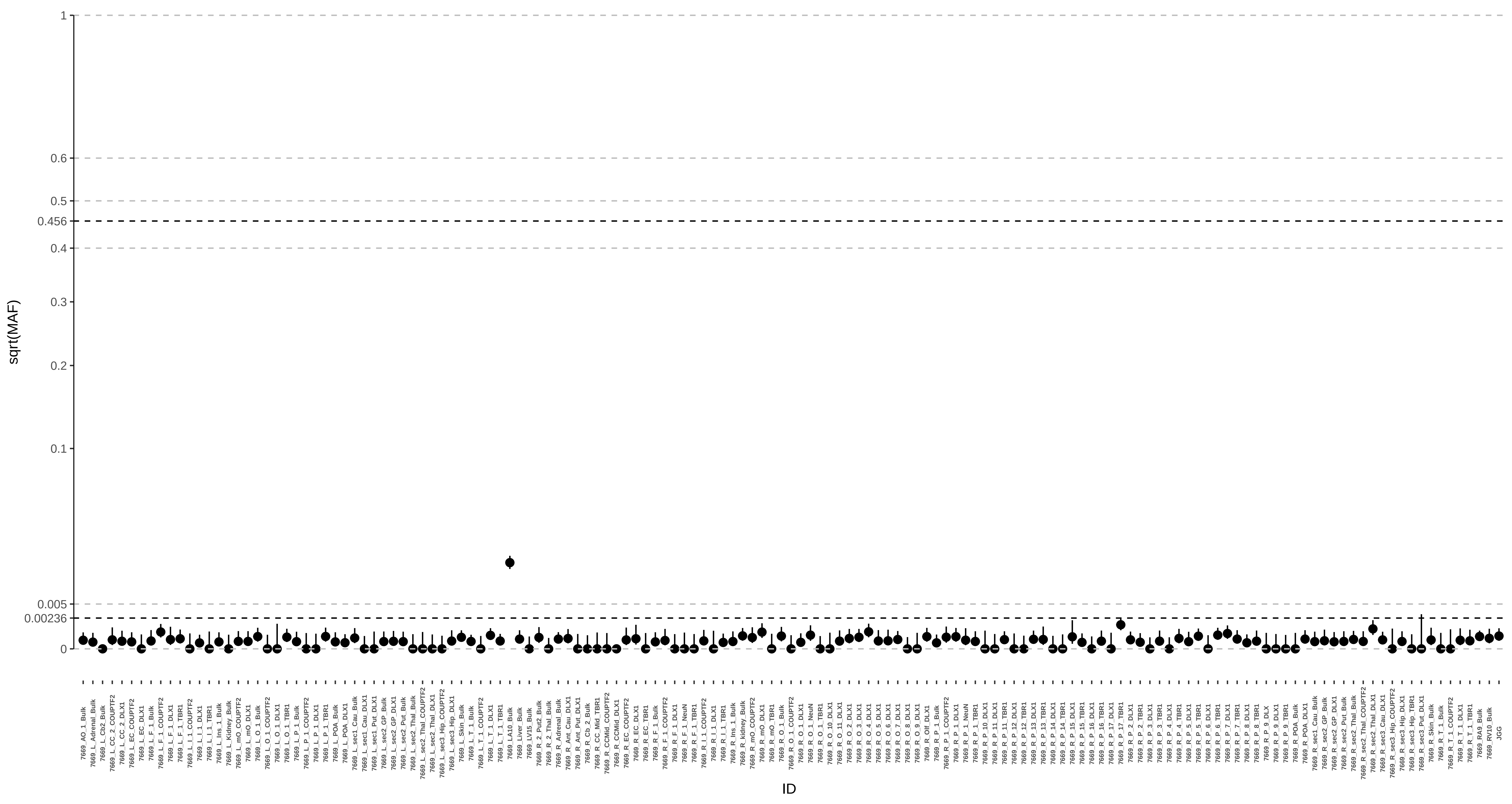

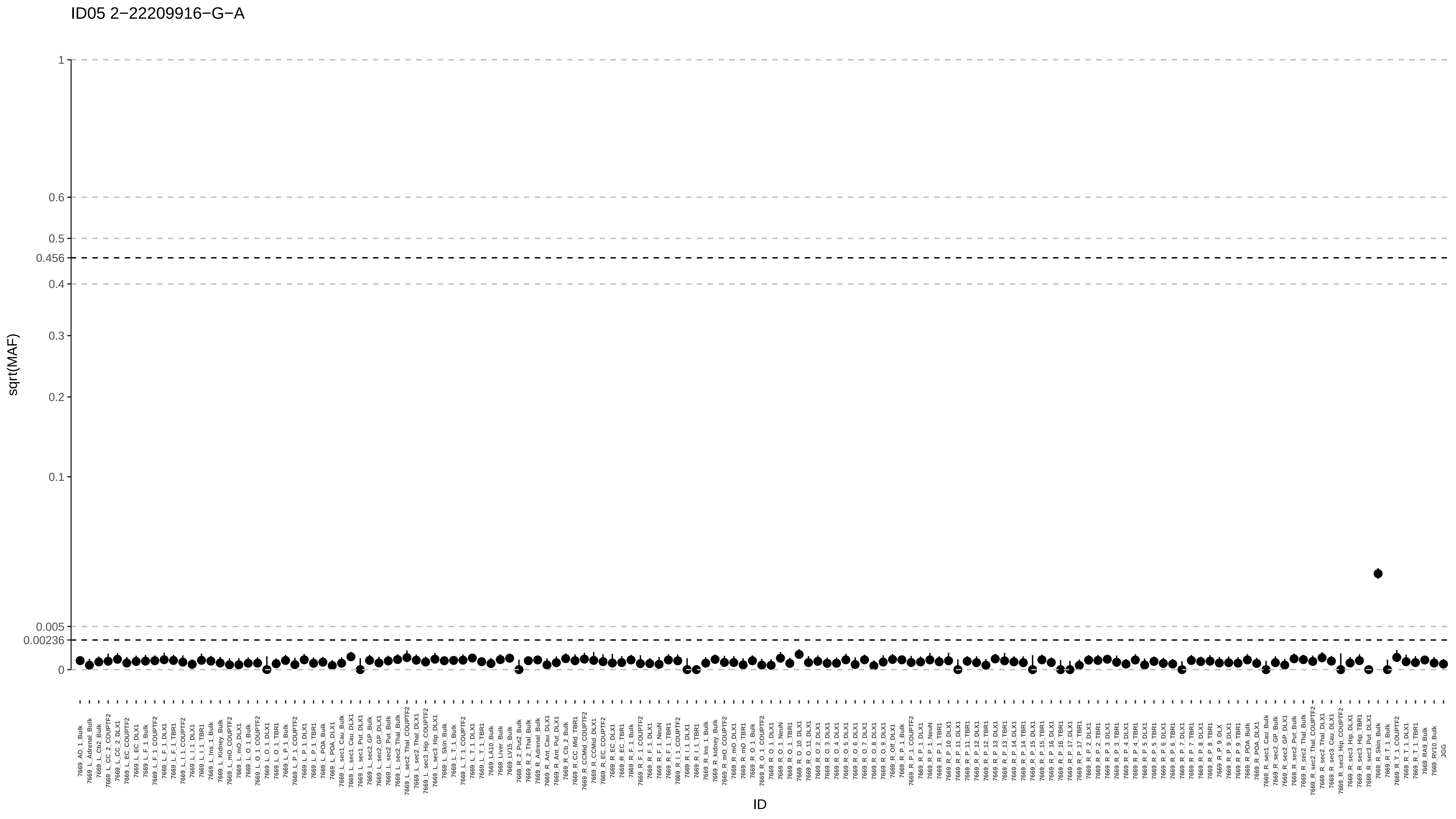

ID05 2−22209916−G−A
sqrt(MAF)
ID
1
0.6
0.5
0.456
0.4
0.3
0.2
0.1
0.005
0.00236
0

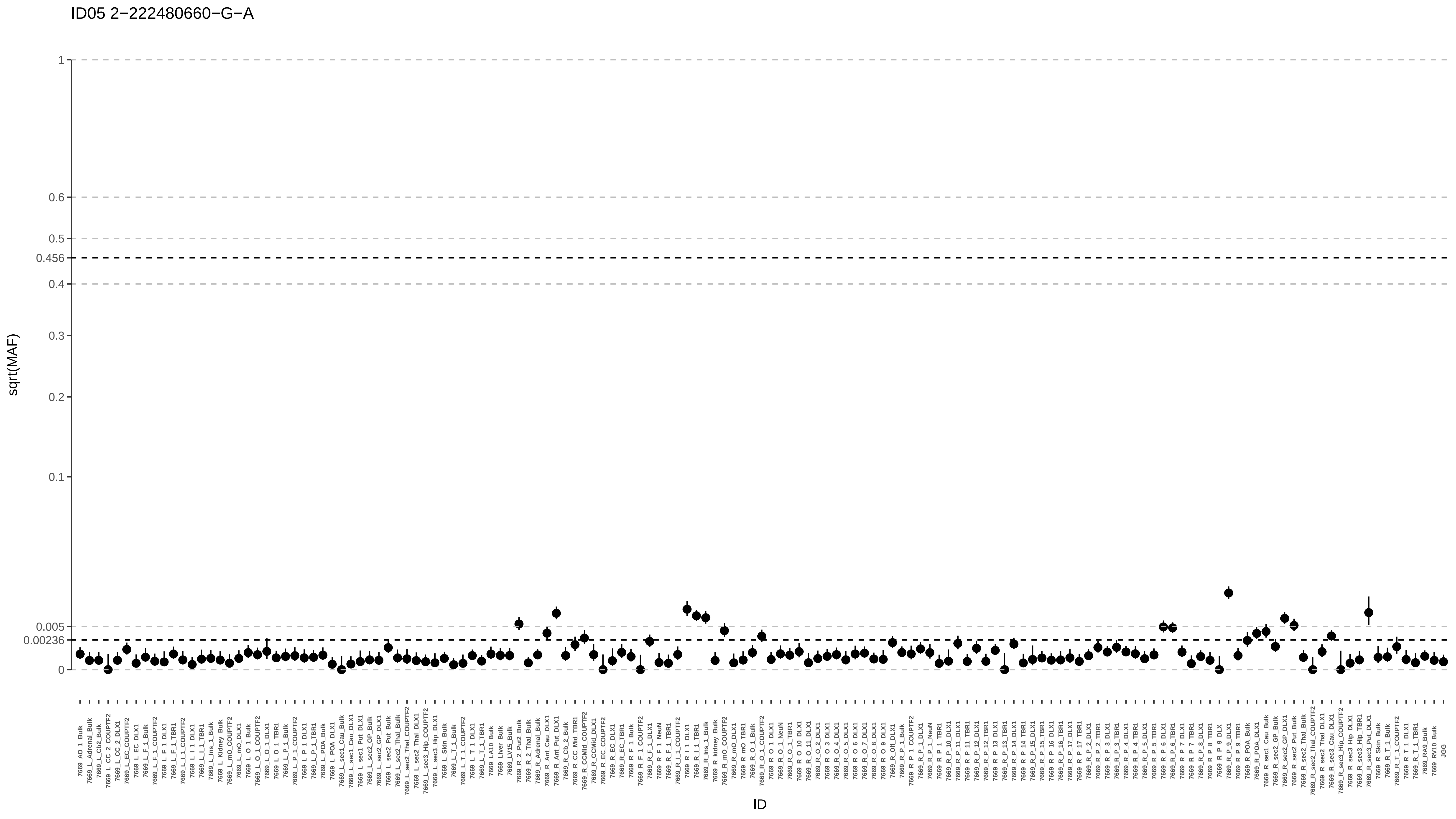
ID05 2−222480660−G−A
1
0.6
0.5
0.456
0.4
0.3
0.2
0.1
0.005
0.00236
0
sqrt(MAF)
ID

ID05 2−225875953−A−G

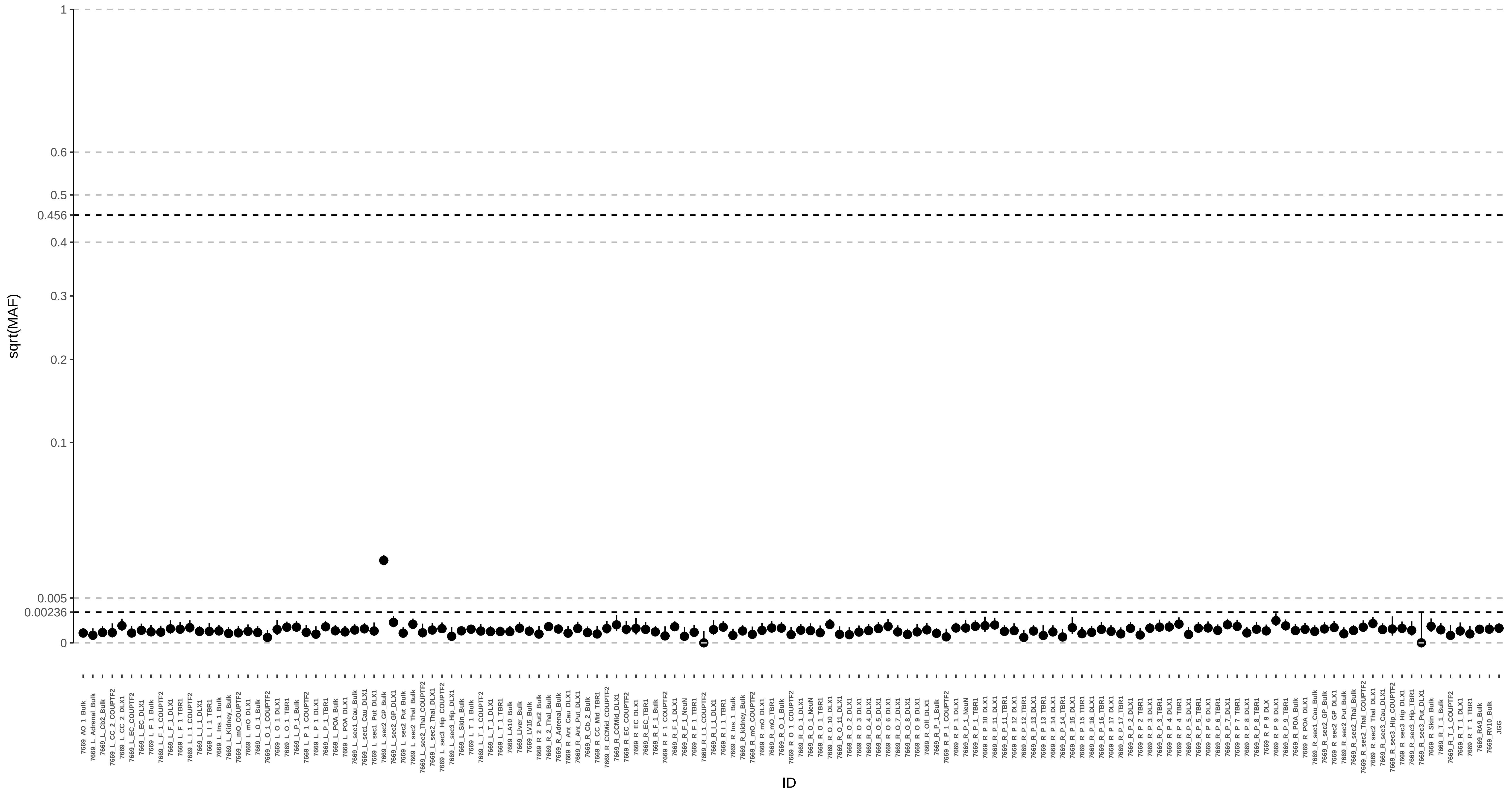

ID05 2−226074050−C−A

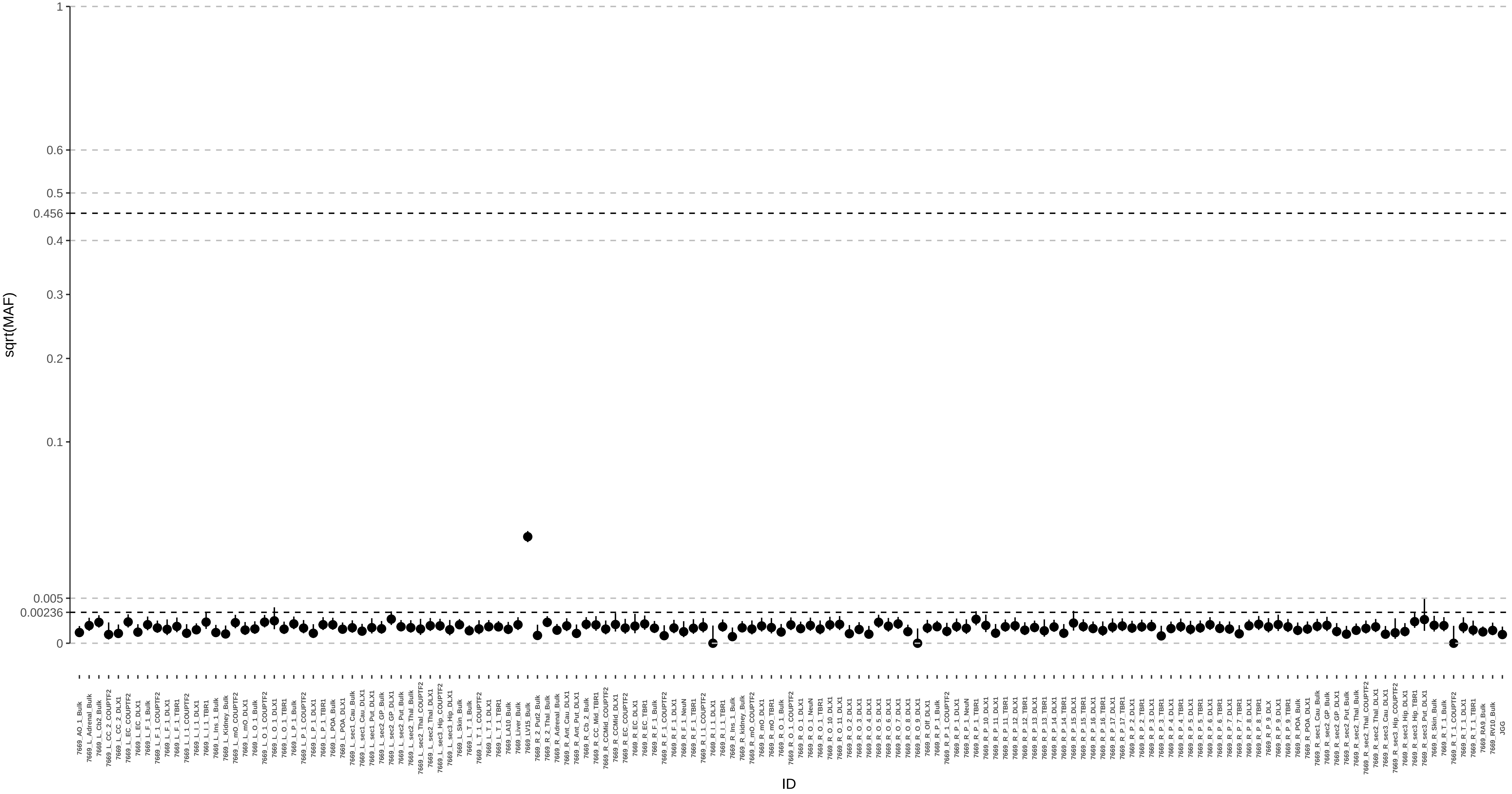

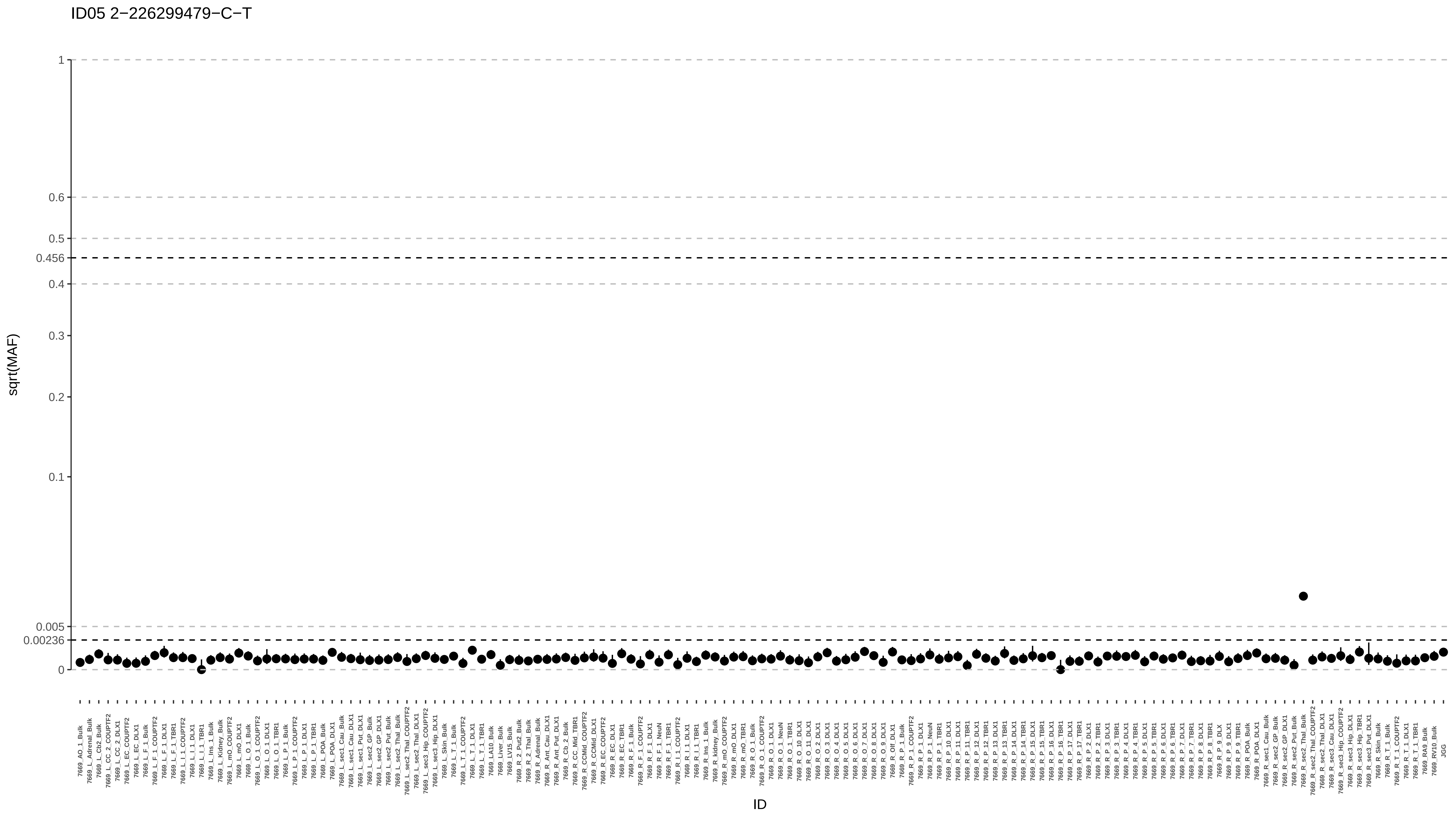
ID05 2−226299479−C−T
sqrt(MAF)
1
0.6
0.5
0.456
0.4
0.3
0.2
0.1
0.005
0.00236
0
ID
7669_AO_1_Bulk
7669_L_Adrenal_Bulk
7669_L_Cb2_Bulk
7669_L_CC_2_COUPTF2
7669_L_CC_2_DLX1
7669_L_EC_COUPTF2
7669_L_EC_DLX1
7669_L_F_1_Bulk
7669_L_F_1_COUPTF2
7669_L_F_1_DLX1
7669_L_F_1_TBR1
7669_L_I_1_COUPTF2
7669_L_I_1_DLX1
7669_L_I_1_TBR1
7669_L_Ins_1_Bulk
7669_L_Kidney_Bulk
7669_L_mO_COUPTF2
7669_L_mO_DLX1
7669_L_O_1_Bulk
7669_L_O_1_COUPTF2
7669_L_O_1_DLX1
7669_L_O_1_TBR1
7669_L_P_1_Bulk
7669_L_P_1_COUPTF2
7669_L_P_1_DLX1
7669_L_P_1_TBR1
7669_L_POA_Bulk
7669_L_POA_DLX1
7669_L_sec1_Cau_Bulk
7669_L_sec1_Cau_DLX1
7669_L_sec1_Put_DLX1
7669_L_sec2_GP_Bulk
7669_L_sec2_GP_DLX1
7669_L_sec2_Put_Bulk
7669_L_sec2_Thal_Bulk
7669_L_sec2_Thal_COUPTF2
7669_L_sec2_Thal_DLX1
7669_L_sec3_Hip_COUPTF2
7669_L_sec3_Hip_DLX1
7669_L_Skin_Bulk
7669_L_T_1_Bulk
7669_L_T_1_COUPTF2
7669_L_T_1_DLX1
7669_L_T_1_TBR1
7669_LA10_Bulk
7669_Liver_Bulk
7669_LV15_Bulk
7669_R_2_Put2_Bulk
7669_R_2_Thal_Bulk
7669_R_Adrenal_Bulk
7669_R_Ant_Cau_DLX1
7669_R_Ant_Put_DLX1
7669_R_Cb_2_Bulk
7669_R_CC_Mid_TBR1
7669_R_CCMid_COUPTF2
7669_R_CCMid_DLX1
7669_R_EC_COUPTF2
7669_R_EC_DLX1
7669_R_EC_TBR1
7669_R_F_1_Bulk
7669_R_F_1_COUPTF2
7669_R_F_1_DLX1
7669_R_F_1_NeuN
7669_R_F_1_TBR1
7669_R_I_1_COUPTF2
7669_R_I_1_DLX1
7669_R_I_1_TBR1
7669_R_Ins_1_Bulk
7669_R_kidney_Bulk
7669_R_mO_COUPTF2
7669_R_mO_DLX1
7669_R_mO_TBR1
7669_R_O_1_Bulk
7669_R_O_1_COUPTF2
7669_R_O_1_DLX1
7669_R_O_1_NeuN
7669_R_O_1_TBR1
7669_R_O_10_DLX1
7669_R_O_11_DLX1
7669_R_O_2_DLX1
7669_R_O_3_DLX1
7669_R_O_4_DLX1
7669_R_O_5_DLX1
7669_R_O_6_DLX1
7669_R_O_7_DLX1
7669_R_O_8_DLX1
7669_R_O_9_DLX1
7669_R_Olf_DLX1
7669_R_P_1_Bulk
7669_R_P_1_COUPTF2
7669_R_P_1_DLX1
7669_R_P_1_NeuN
7669_R_P_1_TBR1
7669_R_P_10_DLX1
7669_R_P_11_DLX1
7669_R_P_11_TBR1
7669_R_P_12_DLX1
7669_R_P_12_TBR1
7669_R_P_13_DLX1
7669_R_P_13_TBR1
7669_R_P_14_DLX1
7669_R_P_14_TBR1
7669_R_P_15_DLX1
7669_R_P_15_TBR1
7669_R_P_16_DLX1
7669_R_P_16_TBR1
7669_R_P_17_DLX1
7669_R_P_17_TBR1
7669_R_P_2_DLX1
7669_R_P_2_TBR1
7669_R_P_3_DLX1
7669_R_P_3_TBR1
7669_R_P_4_DLX1
7669_R_P_4_TBR1
7669_R_P_5_DLX1
7669_R_P_5_TBR1
7669_R_P_6_DLX1
7669_R_P_6_TBR1
7669_R_P_7_DLX1
7669_R_P_7_TBR1
7669_R_P_8_DLX1
7669_R_P_8_TBR1
7669_R_P_9_DLX
7669_R_P_9_DLX1
7669_R_P_9_TBR1
7669_R_POA_Bulk
7669_R_POA_DLX1
7669_R_sec1_Cau_Bulk
7669_R_sec2_GP_Bulk
7669_R_sec2_GP_DLX1
7669_R_sec2_Put_Bulk
7669_R_sec2_Thal_Bulk
7669_R_sec2_Thal_COUPTF2
7669_R_sec2_Thal_DLX1
7669_R_sec3_Cau_DLX1
7669_R_sec3_Hip_COUPTF2
7669_R_sec3_Hip_DLX1
7669_R_sec3_Hip_TBR1
7669_R_sec3_Put_DLX1
7669_R_Skin_Bulk
7669_R_T_1_Bulk
7669_R_T_1_COUPTF2
7669_R_T_1_DLX1
7669_R_T_1_TBR1
7669_RA9_Bulk
7669_RV10_Bulk
JGG

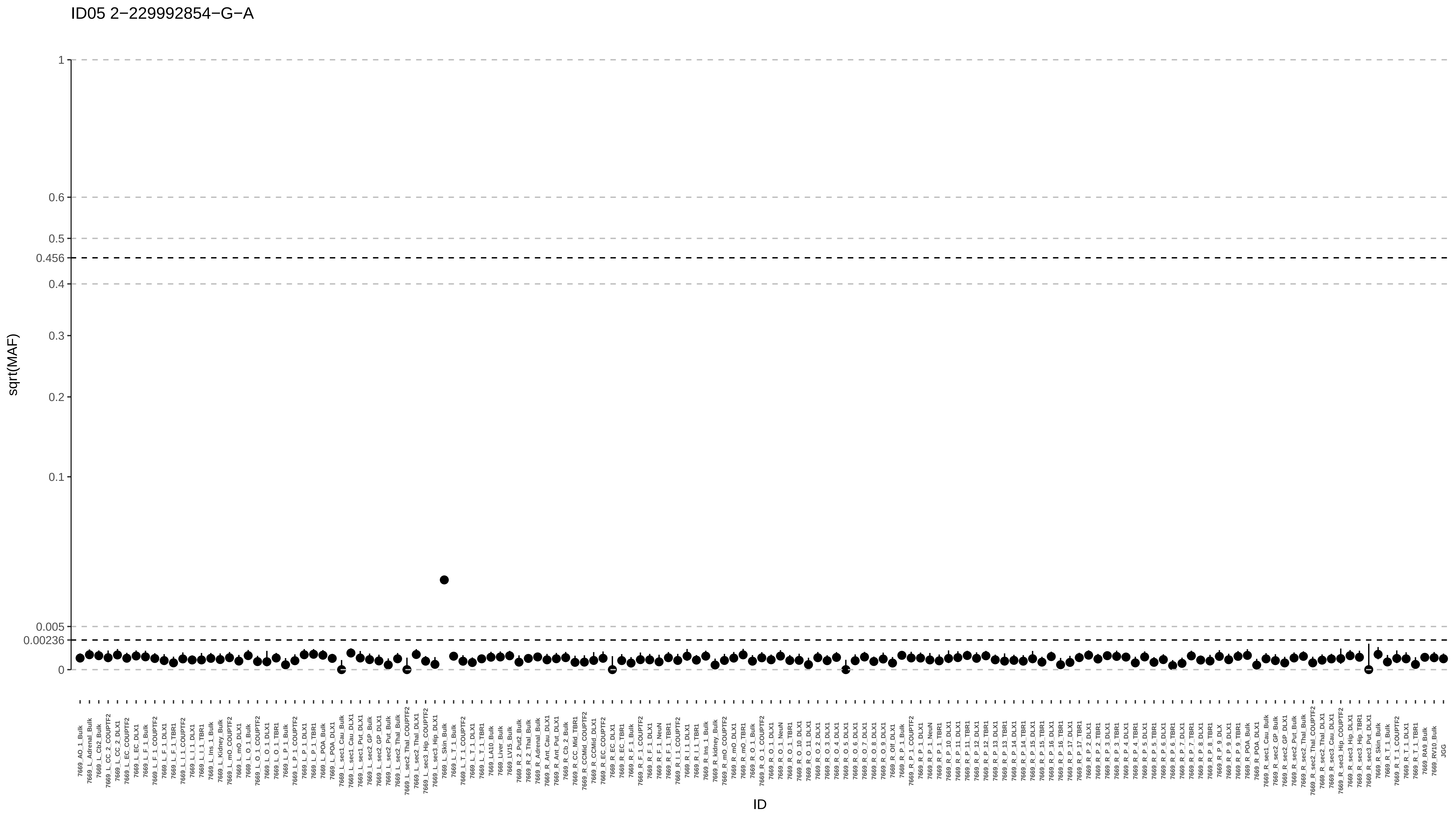
ID05 2−229992854−G−A
sqrt(MAF)
ID
1
0.6
0.5
0.456
0.4
0.3
0.2
0.1
0.005
0.00236
0

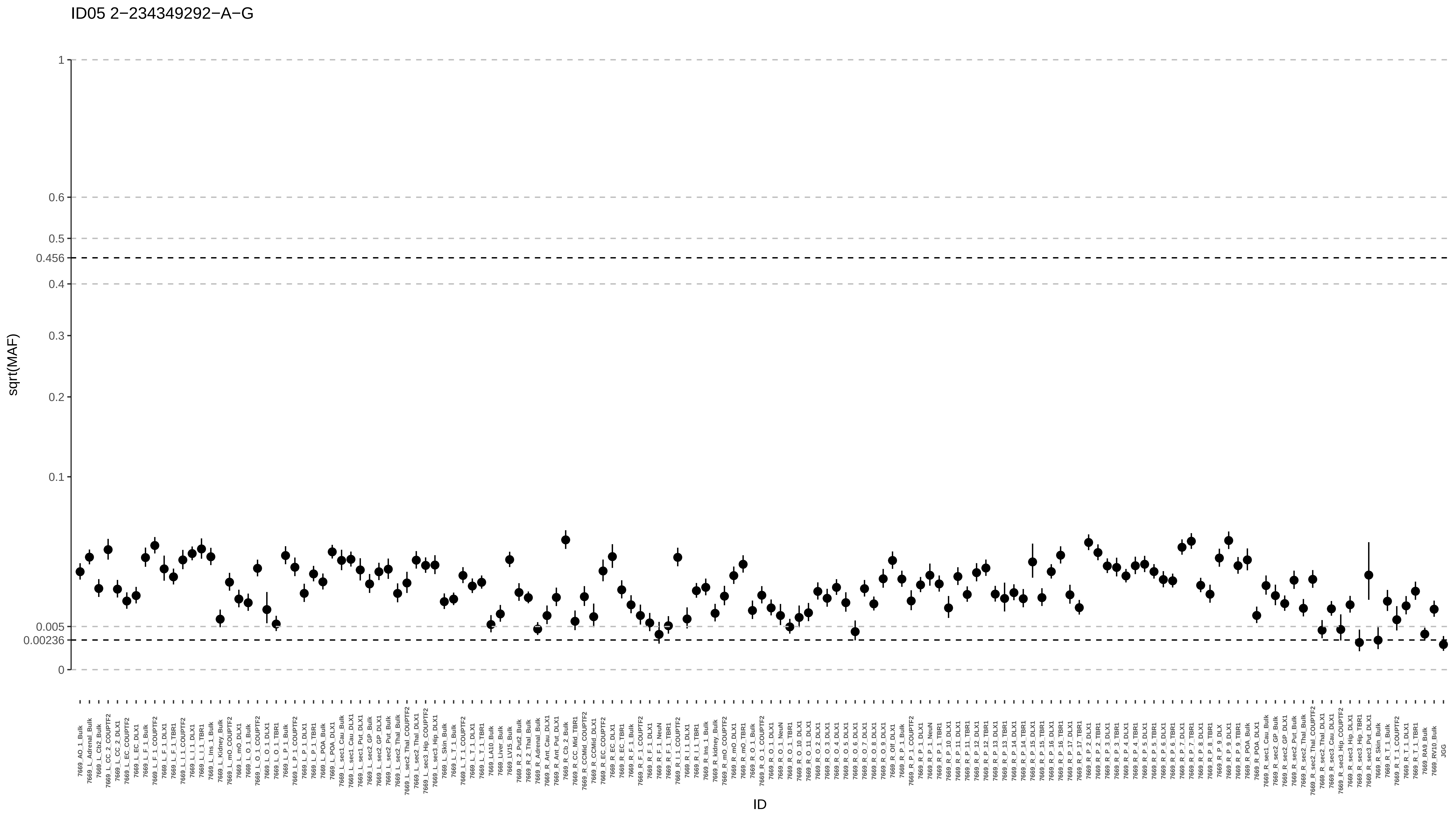

ID05 2−234349292−A−G
sqrt(MAF)
1
0.6
0.5
0.456
0.4
0.3
0.2
0.1
0.005
0.00236
0
ID

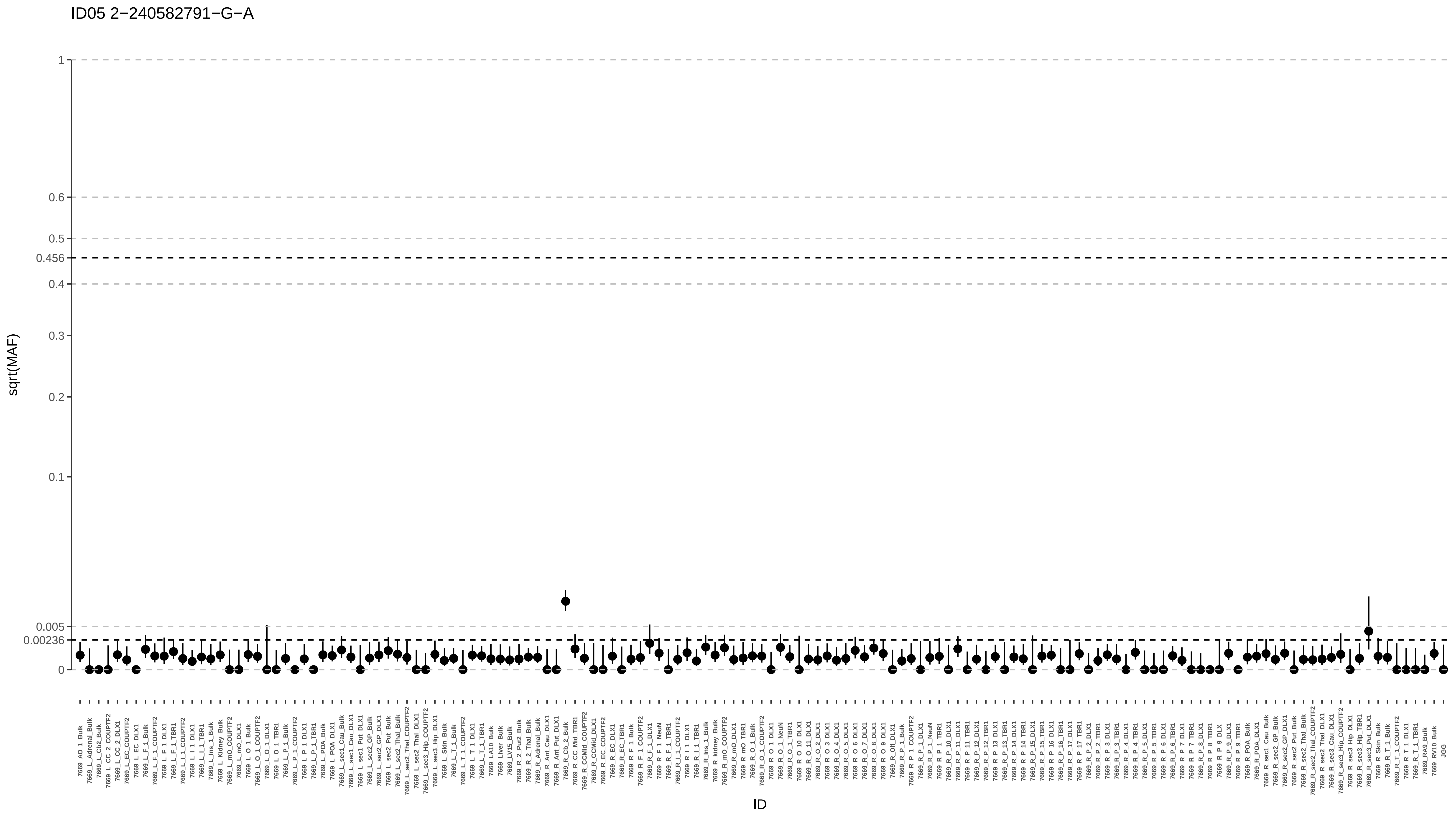

ID05 2−240582791−G−A
sqrt(MAF)
1
0.6
0.5
0.456
0.4
0.3
0.2
0.1
0.005
0.00236
0
ID

ID05 2−241015446−G−A

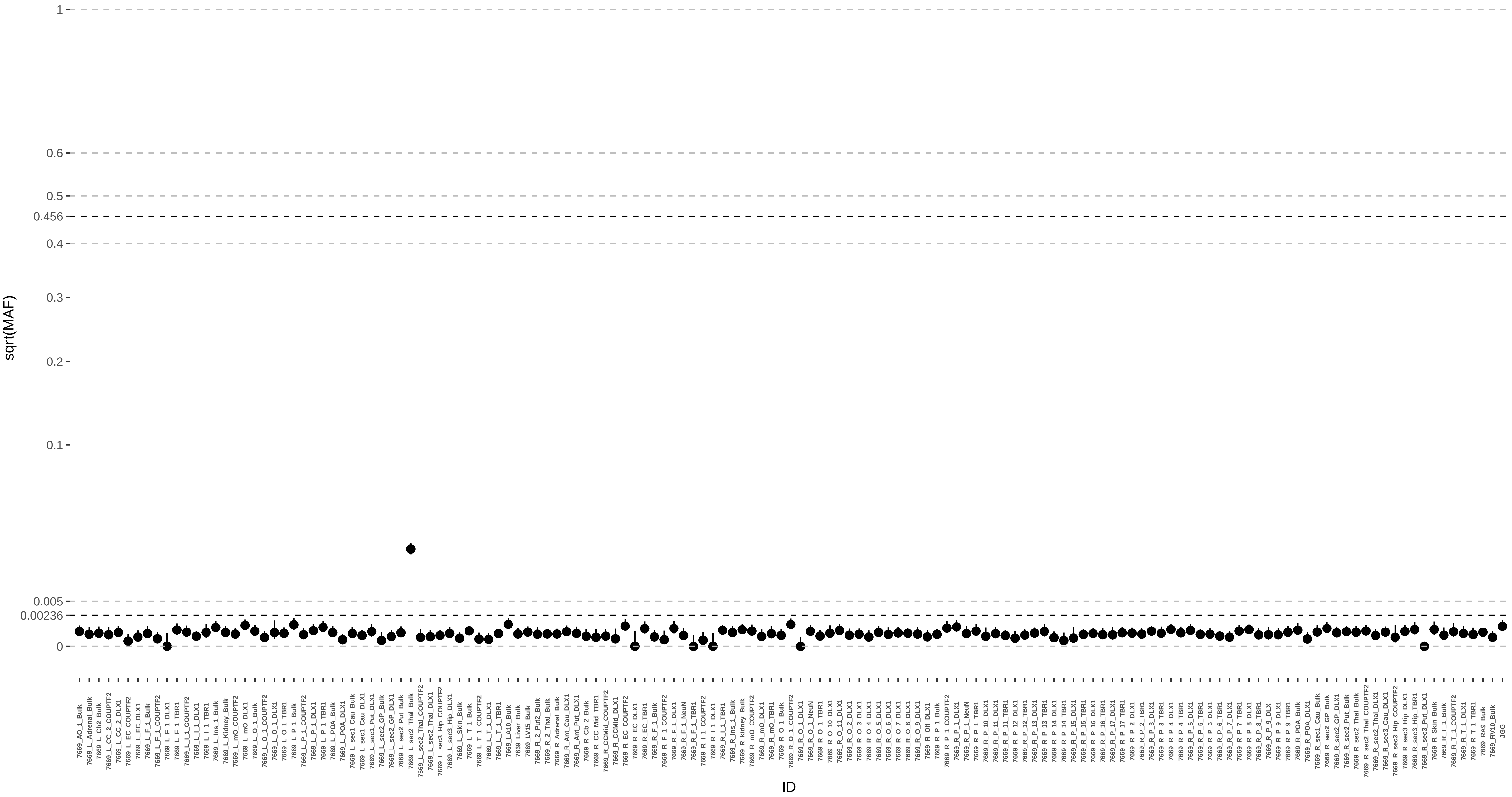

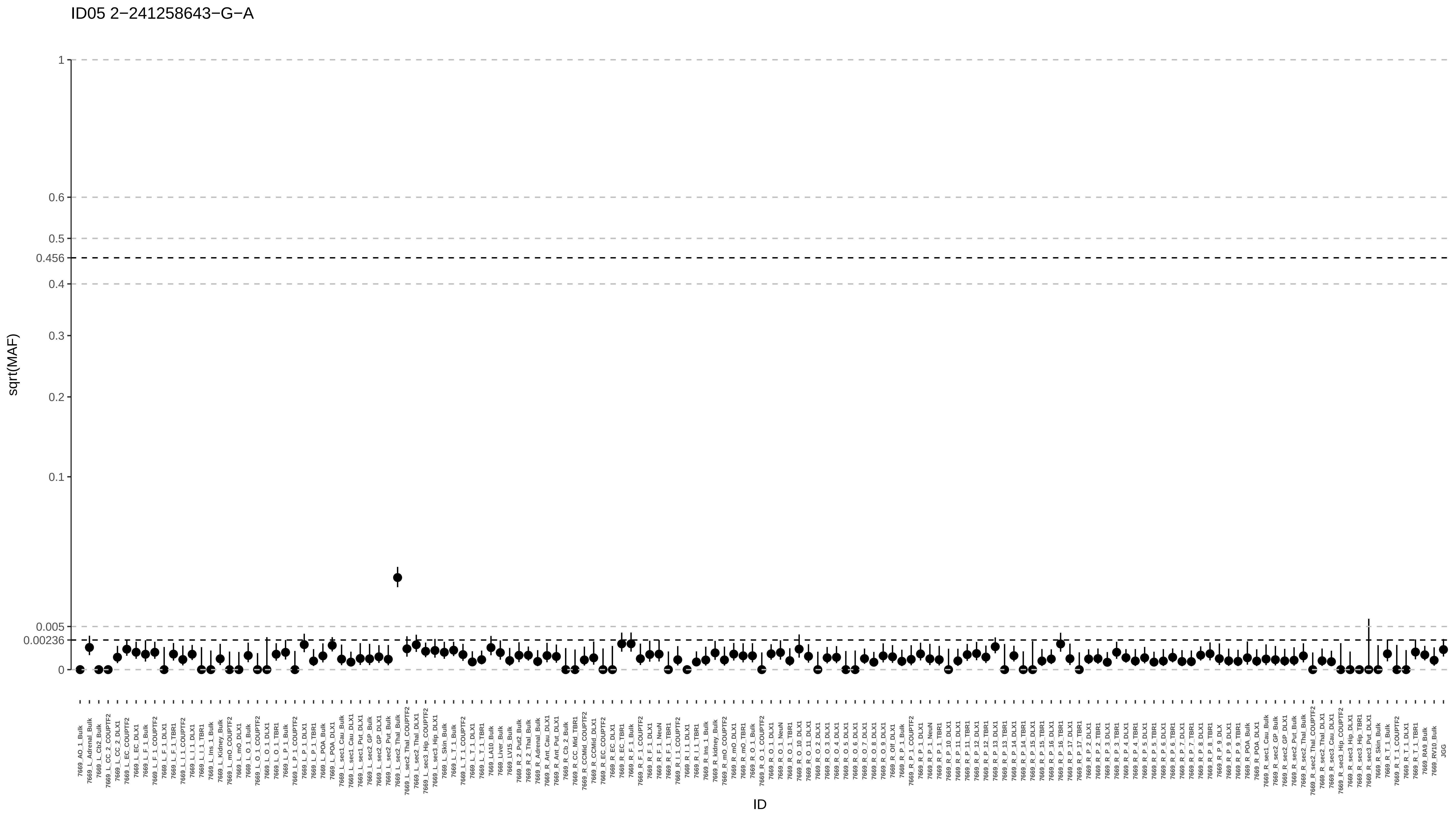
ID05 2−241258643−G−A
sqrt(MAF)
1
0.6
0.5
0.456
0.4
0.3
0.2
0.1
0.005
0.00236
0
ID

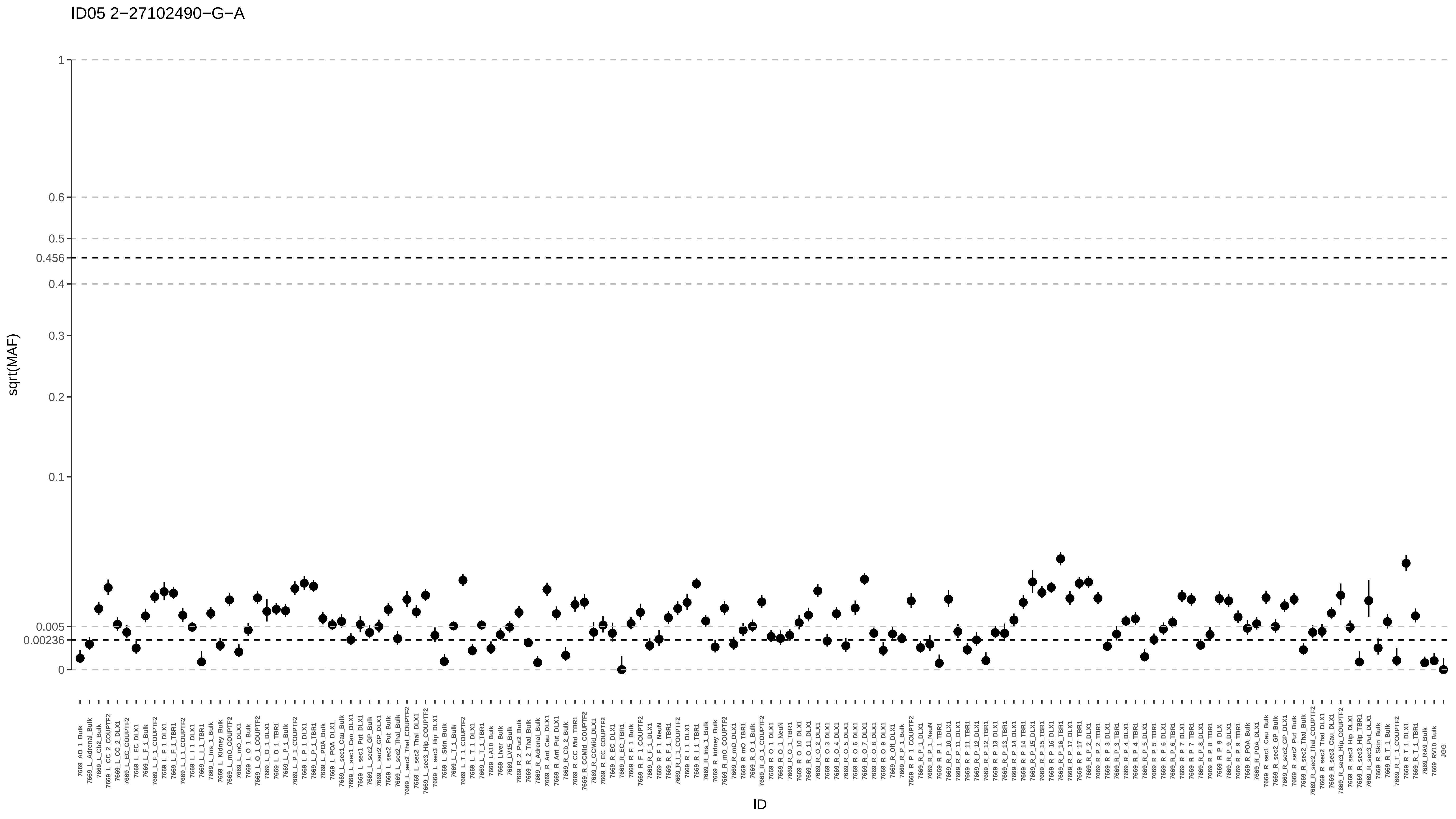
ID05 2−27102490−G−A
sqrt(MAF)
1
0.6
0.5
0.456
0.4
0.3
0.2
0.1
0.005
0.00236
0
ID
7669_AO_1_Bulk
7669_L_Adrenal_Bulk
7669_L_Cb2_Bulk
7669_L_CC_2_COUPTF2
7669_L_CC_2_DLX1
7669_L_EC_COUPTF2
7669_L_EC_DLX1
7669_L_F_1_Bulk
7669_L_F_1_COUPTF2
7669_L_F_1_DLX1
7669_L_F_1_TBR1
7669_L_I_1_COUPTF2
7669_L_I_1_DLX1
7669_L_I_1_TBR1
7669_L_Ins_1_Bulk
7669_L_Kidney_Bulk
7669_L_mO_COUPTF2
7669_L_mO_DLX1
7669_L_O_1_Bulk
7669_L_O_1_COUPTF2
7669_L_O_1_DLX1
7669_L_O_1_TBR1
7669_L_P_1_Bulk
7669_L_P_1_COUPTF2
7669_L_P_1_DLX1
7669_L_P_1_TBR1
7669_L_POA_Bulk
7669_L_POA_DLX1
7669_L_sec1_Cau_Bulk
7669_L_sec1_Cau_DLX1
7669_L_sec1_Put_DLX1
7669_L_sec2_GP_Bulk
7669_L_sec2_GP_DLX1
7669_L_sec2_Put_Bulk
7669_L_sec2_Thal_Bulk
7669_L_sec2_Thal_COUPTF2
7669_L_sec2_Thal_DLX1
7669_L_sec3_Hip_COUPTF2
7669_L_sec3_Hip_DLX1
7669_L_Skin_Bulk
7669_L_T_1_Bulk
7669_L_T_1_COUPTF2
7669_L_T_1_DLX1
7669_L_T_1_TBR1
7669_LA10_Bulk
7669_Liver_Bulk
7669_LV15_Bulk
7669_R_2_Put2_Bulk
7669_R_2_Thal_Bulk
7669_R_Adrenal_Bulk
7669_R_Ant_Cau_DLX1
7669_R_Ant_Put_DLX1
7669_R_Cb_2_Bulk
7669_R_CC_Mid_TBR1
7669_R_CCMid_COUPTF2
7669_R_CCMid_DLX1
7669_R_EC_COUPTF2
7669_R_EC_DLX1
7669_R_EC_TBR1
7669_R_F_1_Bulk
7669_R_F_1_COUPTF2
7669_R_F_1_DLX1
7669_R_F_1_NeuN
7669_R_F_1_TBR1
7669_R_I_1_COUPTF2
7669_R_I_1_DLX1
7669_R_I_1_TBR1
7669_R_Ins_1_Bulk
7669_R_kidney_Bulk
7669_R_mO_COUPTF2
7669_R_mO_DLX1
7669_R_mO_TBR1
7669_R_O_1_Bulk
7669_R_O_1_COUPTF2
7669_R_O_1_DLX1
7669_R_O_1_NeuN
7669_R_O_1_TBR1
7669_R_O_10_DLX1
7669_R_O_11_DLX1
7669_R_O_2_DLX1
7669_R_O_3_DLX1
7669_R_O_4_DLX1
7669_R_O_5_DLX1
7669_R_O_6_DLX1
7669_R_O_7_DLX1
7669_R_O_8_DLX1
7669_R_O_9_DLX1
7669_R_Olf_DLX1
7669_R_P_1_Bulk
7669_R_P_1_COUPTF2
7669_R_P_1_DLX1
7669_R_P_1_NeuN
7669_R_P_1_TBR1
7669_R_P_10_DLX1
7669_R_P_11_DLX1
7669_R_P_11_TBR1
7669_R_P_12_DLX1
7669_R_P_12_TBR1
7669_R_P_13_DLX1
7669_R_P_13_TBR1
7669_R_P_14_DLX1
7669_R_P_14_TBR1
7669_R_P_15_DLX1
7669_R_P_15_TBR1
7669_R_P_16_DLX1
7669_R_P_16_TBR1
7669_R_P_17_DLX1
7669_R_P_17_TBR1
7669_R_P_2_DLX1
7669_R_P_2_TBR1
7669_R_P_3_DLX1
7669_R_P_3_TBR1
7669_R_P_4_DLX1
7669_R_P_4_TBR1
7669_R_P_5_DLX1
7669_R_P_5_TBR1
7669_R_P_6_DLX1
7669_R_P_6_TBR1
7669_R_P_7_DLX1
7669_R_P_7_TBR1
7669_R_P_8_DLX1
7669_R_P_8_TBR1
7669_R_P_9_DLX
7669_R_P_9_DLX1
7669_R_P_9_TBR1
7669_R_POA_Bulk
7669_R_POA_DLX1
7669_R_sec1_Cau_Bulk
7669_R_sec2_GP_Bulk
7669_R_sec2_GP_DLX1
7669_R_sec2_Put_Bulk
7669_R_sec2_Thal_Bulk
7669_R_sec2_Thal_COUPTF2
7669_R_sec2_Thal_DLX1
7669_R_sec3_Cau_DLX1
7669_R_sec3_Hip_COUPTF2
7669_R_sec3_Hip_DLX1
7669_R_sec3_Hip_TBR1
7669_R_sec3_Put_DLX1
7669_R_Skin_Bulk
7669_R_T_1_Bulk
7669_R_T_1_COUPTF2
7669_R_T_1_DLX1
7669_R_T_1_TBR1
7669_RA9_Bulk
7669_RV10_Bulk
JGG

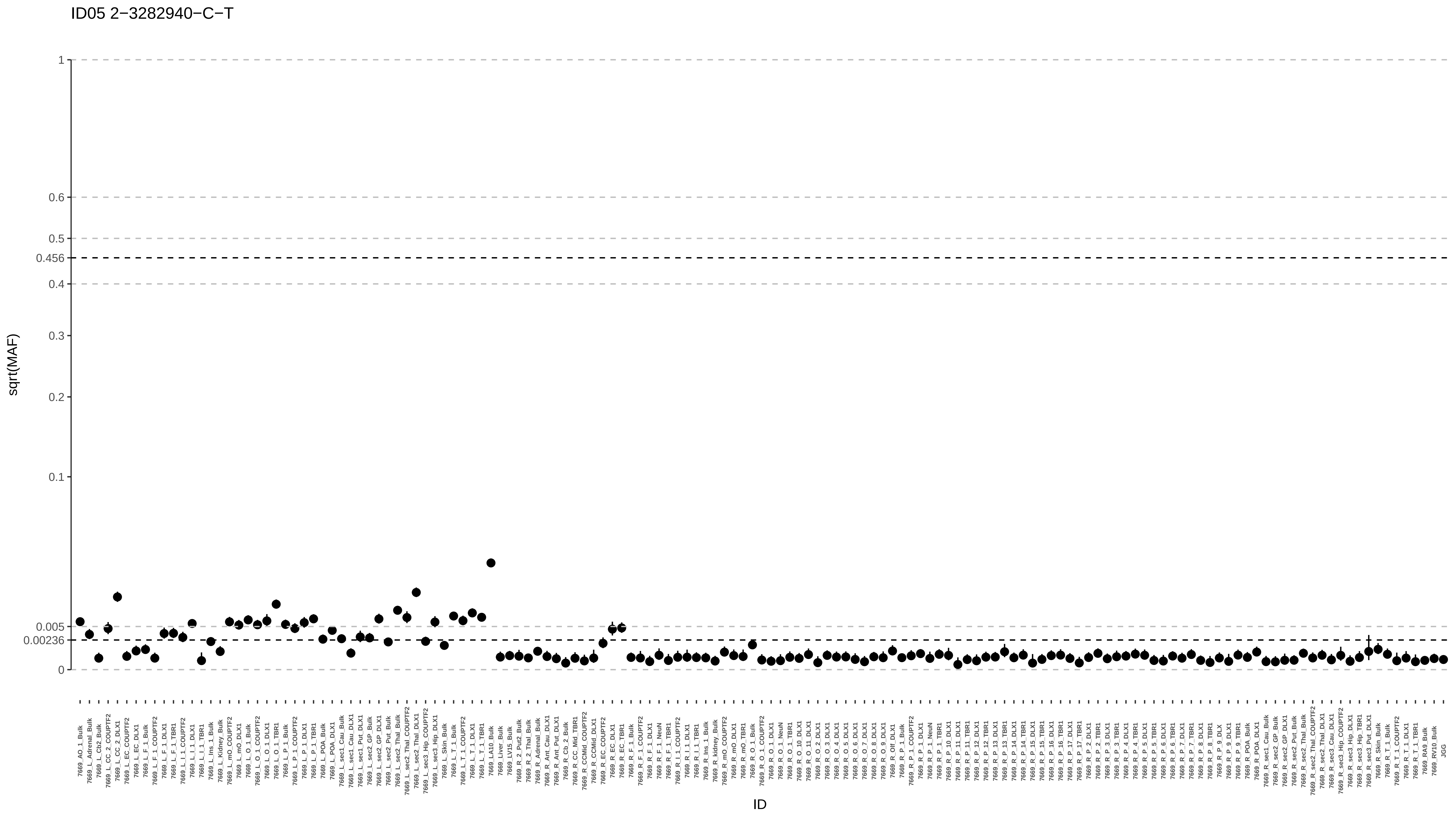
ID05 2−3282940−C−T
sqrt(MAF)
1
0.6
0.5
0.456
0.4
0.3
0.2
0.1
0.005
0.00236
0
ID

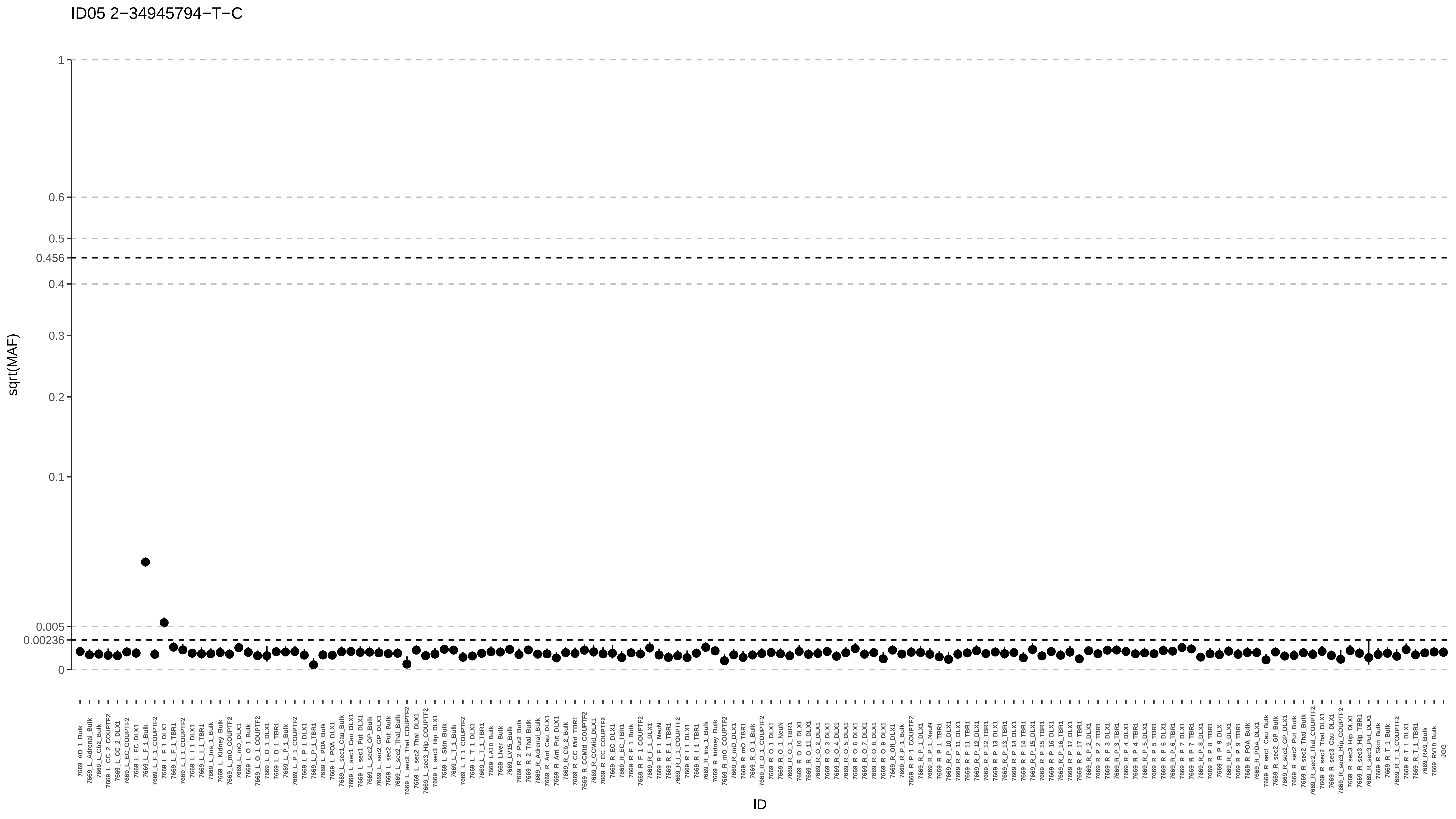
ID05 2−34945794−T−C
sqrt(MAF)
1
0.6
0.5
0.456
0.4
0.3
0.2
0.1
0.005
0.00236
0
ID

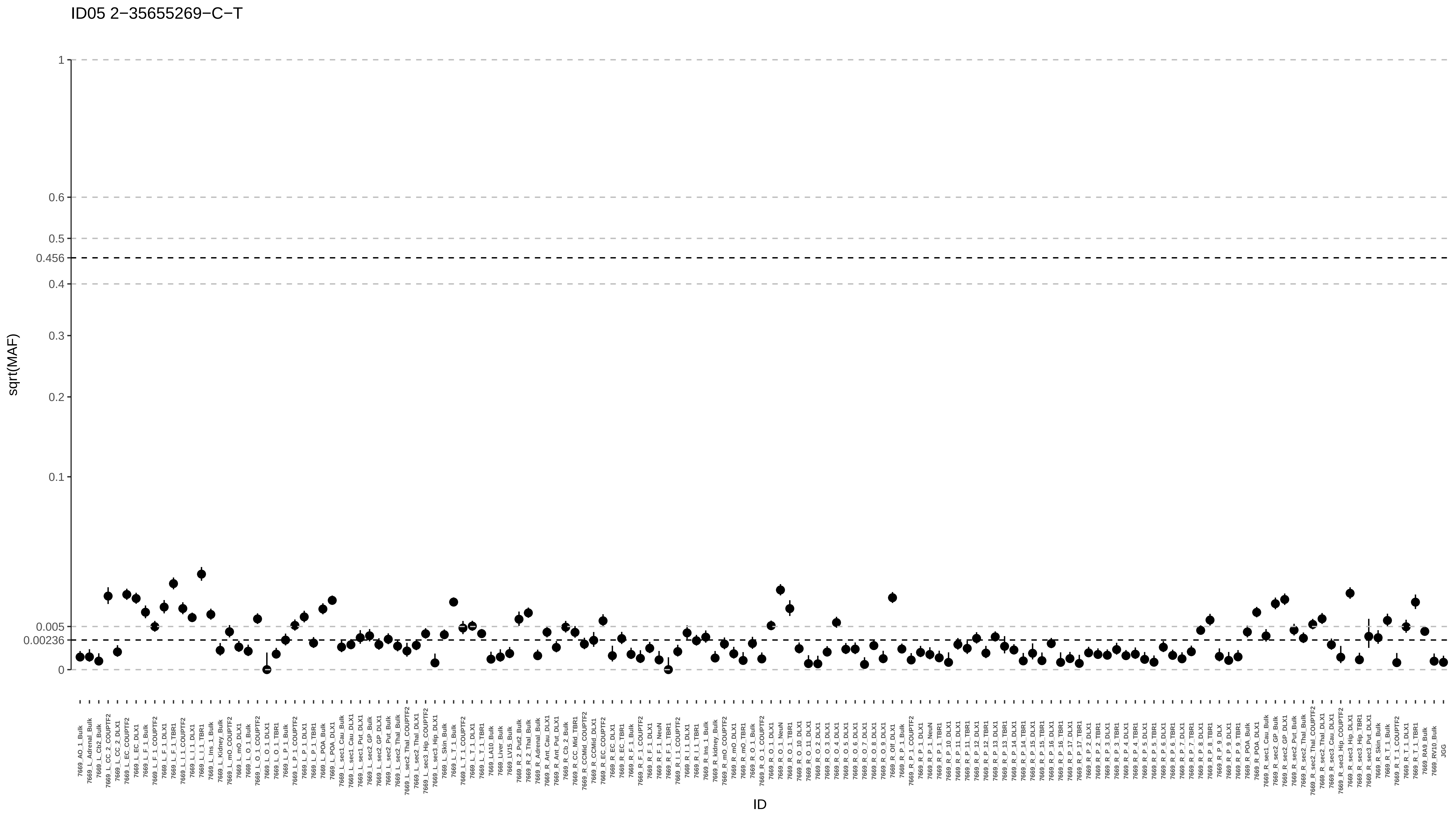
ID05 2−35655269−C−T
sqrt(MAF)
ID
1
0.6
0.5
0.456
0.4
0.3
0.2
0.1
0.005
0.00236
0

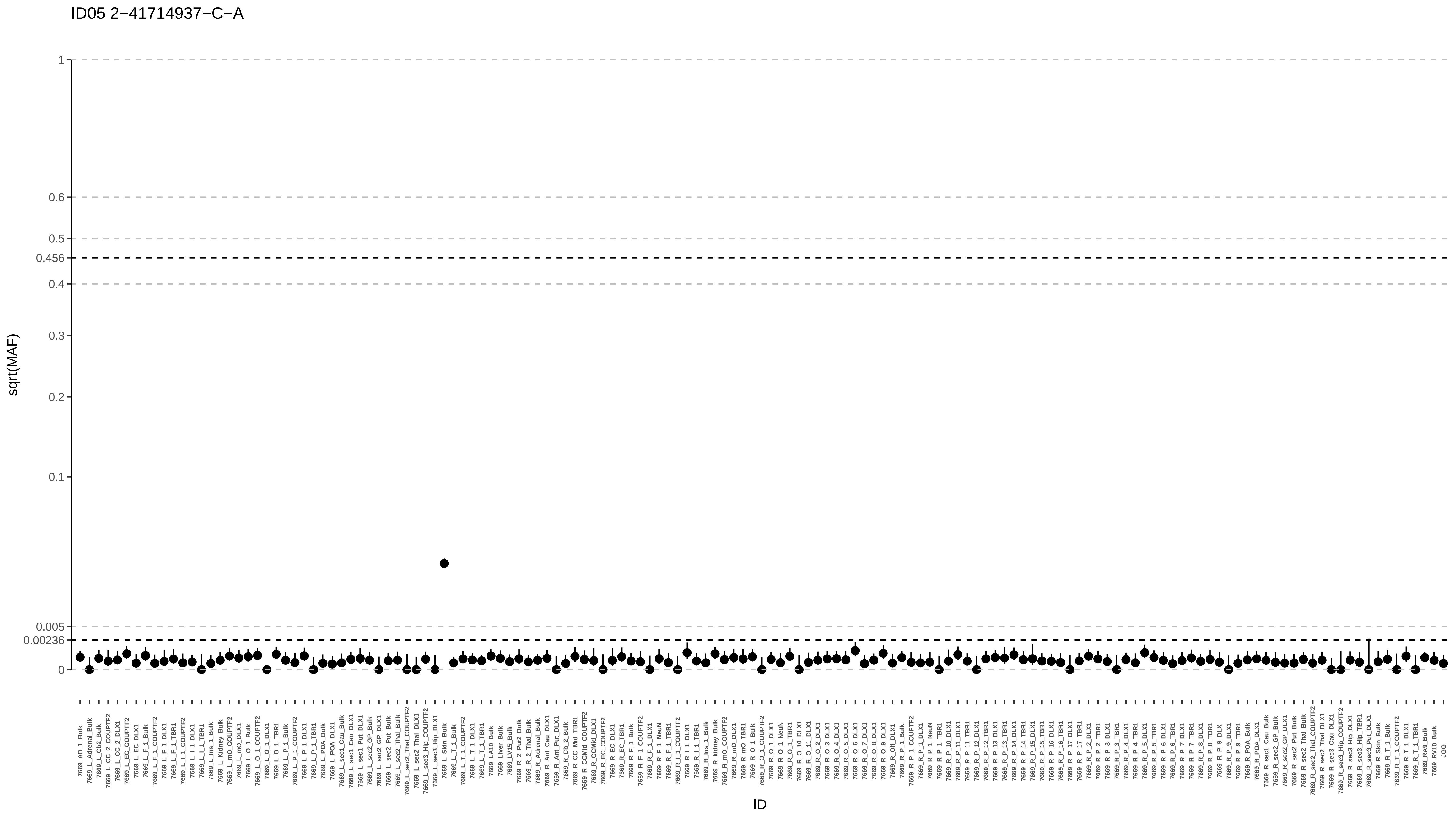
ID05 2−41714937−C−A
sqrt(MAF)
1
0.6
0.5
0.456
0.4
0.3
0.2
0.1
0.005
0.00236
0
ID

ID05 2−424135−C−T

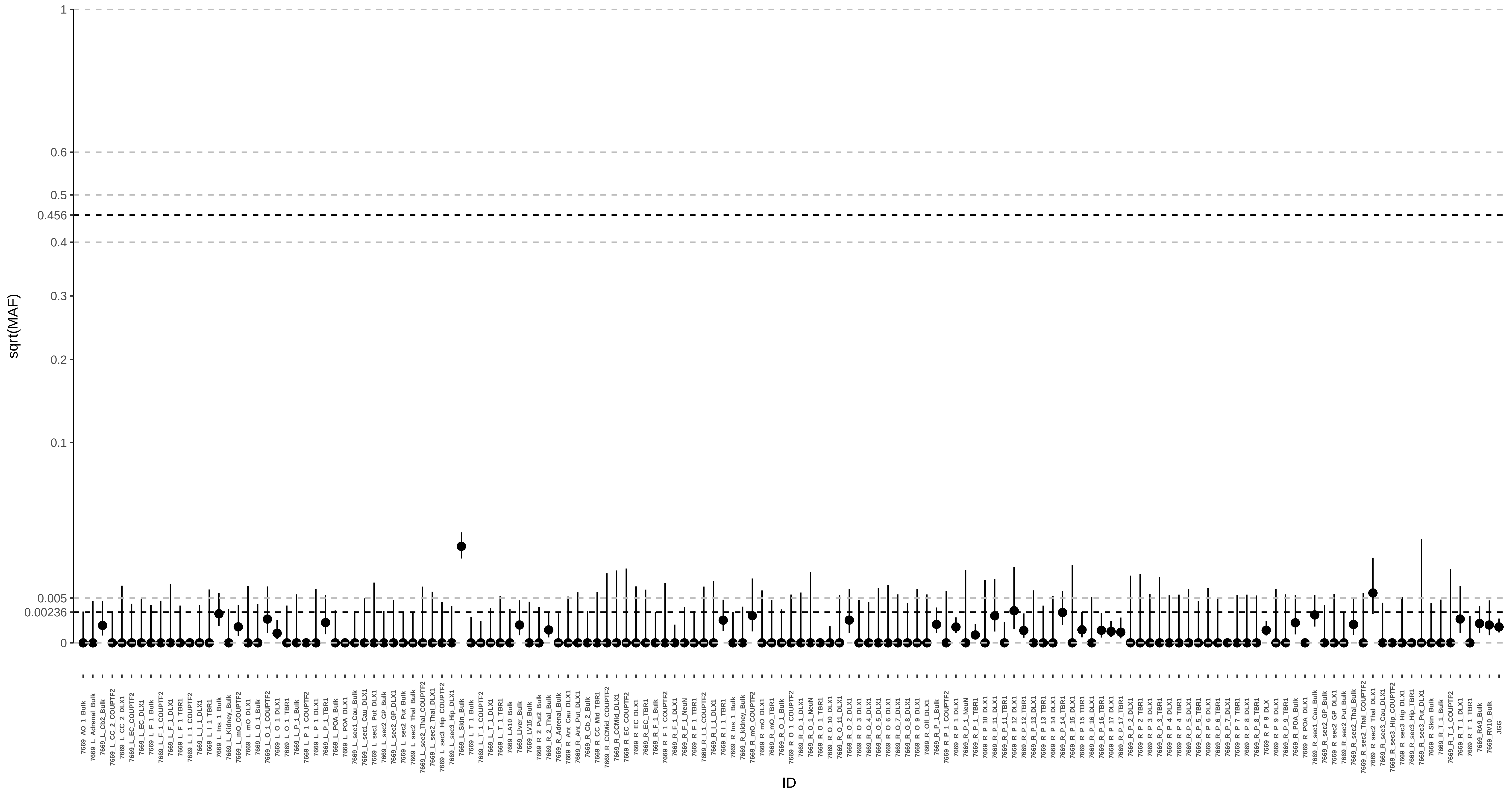

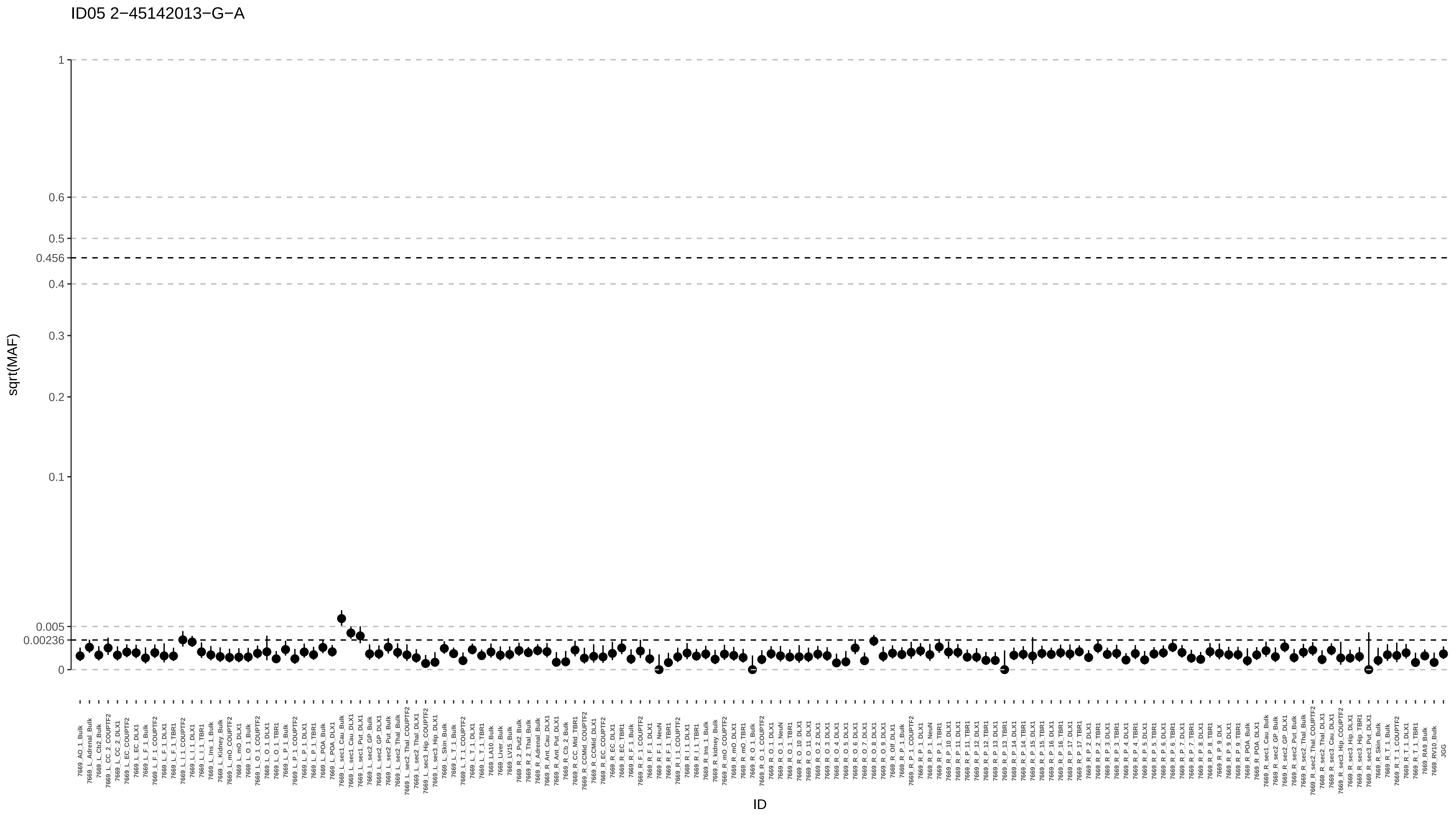
ID05 2−45142013−G−A
sqrt(MAF)
1
0.6
0.5
0.456
0.4
0.3
0.2
0.1
0.005
0.00236
0
ID
7669_AO_1_Bulk
7669_L_Adrenal_Bulk
7669_L_Cb2_Bulk
7669_L_CC_2_COUPTF2
7669_L_CC_2_DLX1
7669_L_EC_COUPTF2
7669_L_EC_DLX1
7669_L_F_1_Bulk
7669_L_F_1_COUPTF2
7669_L_F_1_DLX1
7669_L_F_1_TBR1
7669_L_I_1_COUPTF2
7669_L_I_1_DLX1
7669_L_I_1_TBR1
7669_L_Ins_1_Bulk
7669_L_Kidney_Bulk
7669_L_mO_COUPTF2
7669_L_mO_DLX1
7669_L_O_1_Bulk
7669_L_O_1_COUPTF2
7669_L_O_1_DLX1
7669_L_O_1_TBR1
7669_L_P_1_Bulk
7669_L_P_1_COUPTF2
7669_L_P_1_DLX1
7669_L_P_1_TBR1
7669_L_POA_Bulk
7669_L_POA_DLX1
7669_L_sec1_Cau_Bulk
7669_L_sec1_Cau_DLX1
7669_L_sec1_Put_DLX1
7669_L_sec2_GP_Bulk
7669_L_sec2_GP_DLX1
7669_L_sec2_Put_Bulk
7669_L_sec2_Thal_Bulk
7669_L_sec2_Thal_COUPTF2
7669_L_sec2_Thal_DLX1
7669_L_sec3_Hip_COUPTF2
7669_L_sec3_Hip_DLX1
7669_L_Skin_Bulk
7669_L_T_1_Bulk
7669_L_T_1_COUPTF2
7669_L_T_1_DLX1
7669_L_T_1_TBR1
7669_LA10_Bulk
7669_Liver_Bulk
7669_LV15_Bulk
7669_R_2_Put2_Bulk
7669_R_2_Thal_Bulk
7669_R_Adrenal_Bulk
7669_R_Ant_Cau_DLX1
7669_R_Ant_Put_DLX1
7669_R_Cb_2_Bulk
7669_R_CC_Mid_TBR1
7669_R_CCMid_COUPTF2
7669_R_CCMid_DLX1
7669_R_EC_COUPTF2
7669_R_EC_DLX1
7669_R_EC_TBR1
7669_R_F_1_Bulk
7669_R_F_1_COUPTF2
7669_R_F_1_DLX1
7669_R_F_1_NeuN
7669_R_F_1_TBR1
7669_R_I_1_COUPTF2
7669_R_I_1_DLX1
7669_R_I_1_TBR1
7669_R_Ins_1_Bulk
7669_R_kidney_Bulk
7669_R_mO_COUPTF2
7669_R_mO_DLX1
7669_R_mO_TBR1
7669_R_O_1_Bulk
7669_R_O_1_COUPTF2
7669_R_O_1_DLX1
7669_R_O_1_NeuN
7669_R_O_1_TBR1
7669_R_O_10_DLX1
7669_R_O_11_DLX1
7669_R_O_2_DLX1
7669_R_O_3_DLX1
7669_R_O_4_DLX1
7669_R_O_5_DLX1
7669_R_O_6_DLX1
7669_R_O_7_DLX1
7669_R_O_8_DLX1
7669_R_O_9_DLX1
7669_R_Olf_DLX1
7669_R_P_1_Bulk
7669_R_P_1_COUPTF2
7669_R_P_1_DLX1
7669_R_P_1_NeuN
7669_R_P_1_TBR1
7669_R_P_10_DLX1
7669_R_P_11_DLX1
7669_R_P_11_TBR1
7669_R_P_12_DLX1
7669_R_P_12_TBR1
7669_R_P_13_DLX1
7669_R_P_13_TBR1
7669_R_P_14_DLX1
7669_R_P_14_TBR1
7669_R_P_15_DLX1
7669_R_P_15_TBR1
7669_R_P_16_DLX1
7669_R_P_16_TBR1
7669_R_P_17_DLX1
7669_R_P_17_TBR1
7669_R_P_2_DLX1
7669_R_P_2_TBR1
7669_R_P_3_DLX1
7669_R_P_3_TBR1
7669_R_P_4_DLX1
7669_R_P_4_TBR1
7669_R_P_5_DLX1
7669_R_P_5_TBR1
7669_R_P_6_DLX1
7669_R_P_6_TBR1
7669_R_P_7_DLX1
7669_R_P_7_TBR1
7669_R_P_8_DLX1
7669_R_P_8_TBR1
7669_R_P_9_DLX
7669_R_P_9_DLX1
7669_R_P_9_TBR1
7669_R_POA_Bulk
7669_R_POA_DLX1
7669_R_sec1_Cau_Bulk
7669_R_sec2_GP_Bulk
7669_R_sec2_GP_DLX1
7669_R_sec2_Put_Bulk
7669_R_sec2_Thal_Bulk
7669_R_sec2_Thal_COUPTF2
7669_R_sec2_Thal_DLX1
7669_R_sec3_Cau_DLX1
7669_R_sec3_Hip_COUPTF2
7669_R_sec3_Hip_DLX1
7669_R_sec3_Hip_TBR1
7669_R_sec3_Put_DLX1
7669_R_Skin_Bulk
7669_R_T_1_Bulk
7669_R_T_1_COUPTF2
7669_R_T_1_DLX1
7669_R_T_1_TBR1
7669_RA9_Bulk
7669_RV10_Bulk
JGG

ID05 2−48643619−C−T

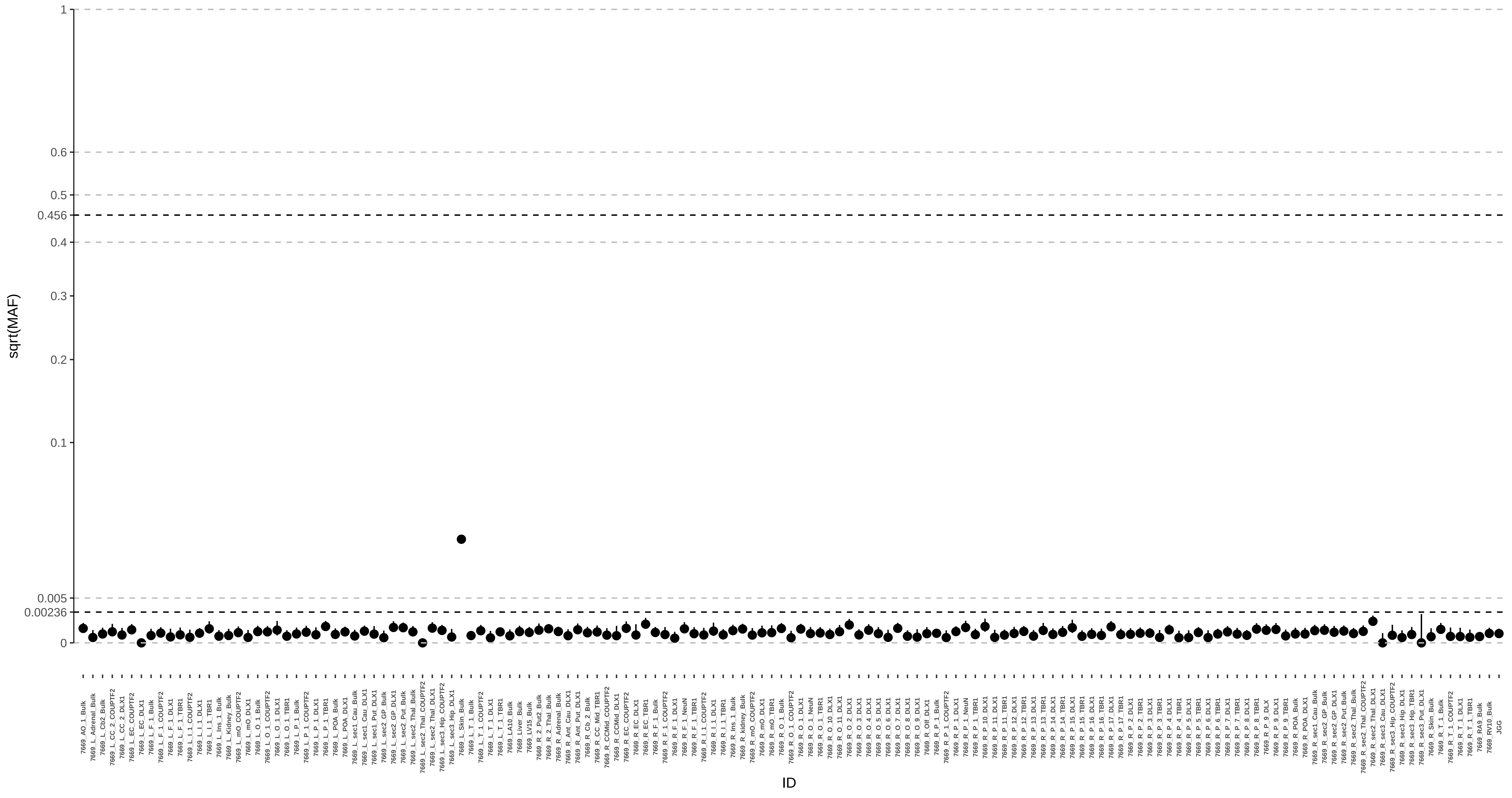

ID05 2−52452769−G−A

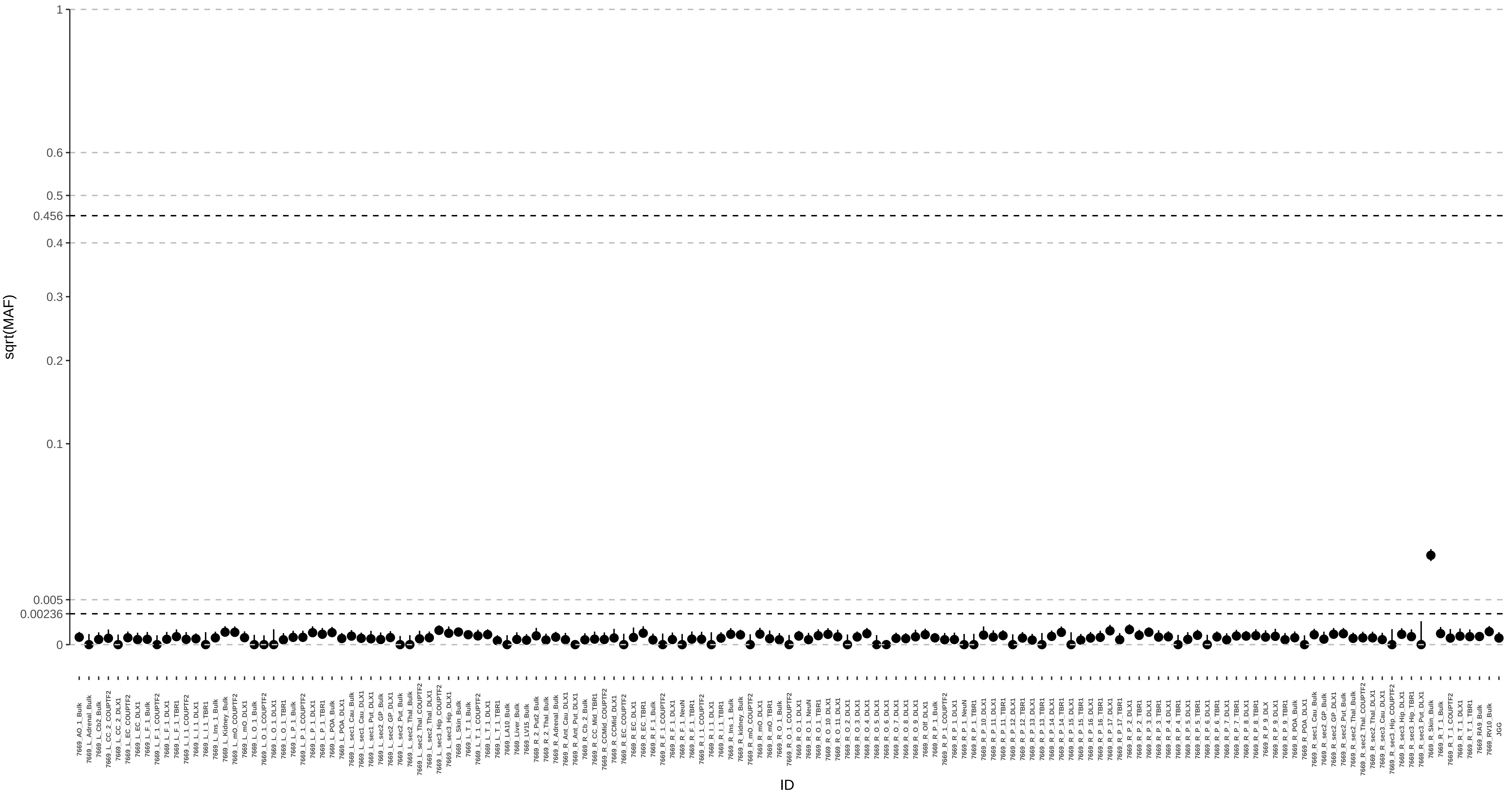

ID05 2−53839419−G−A

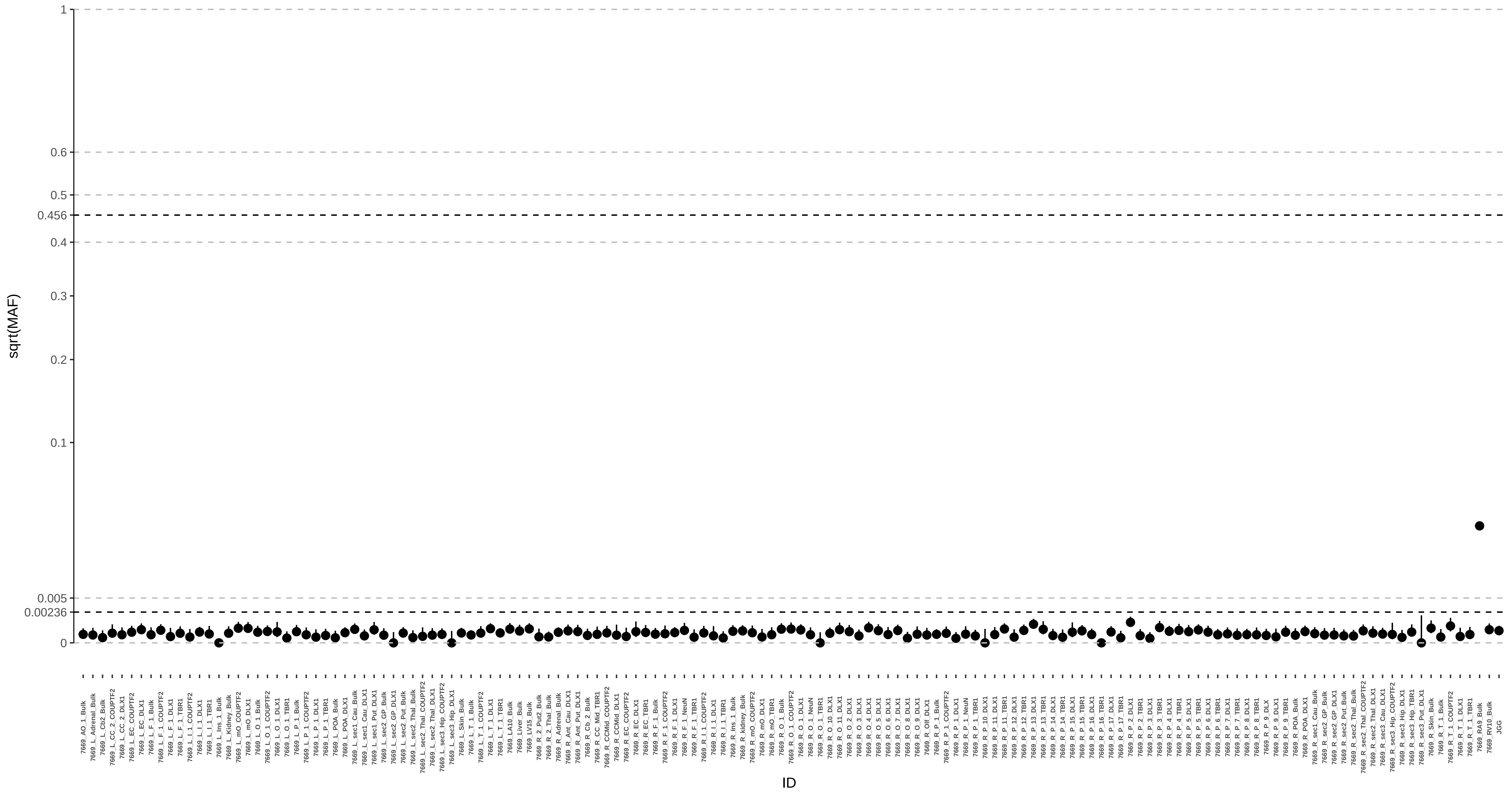

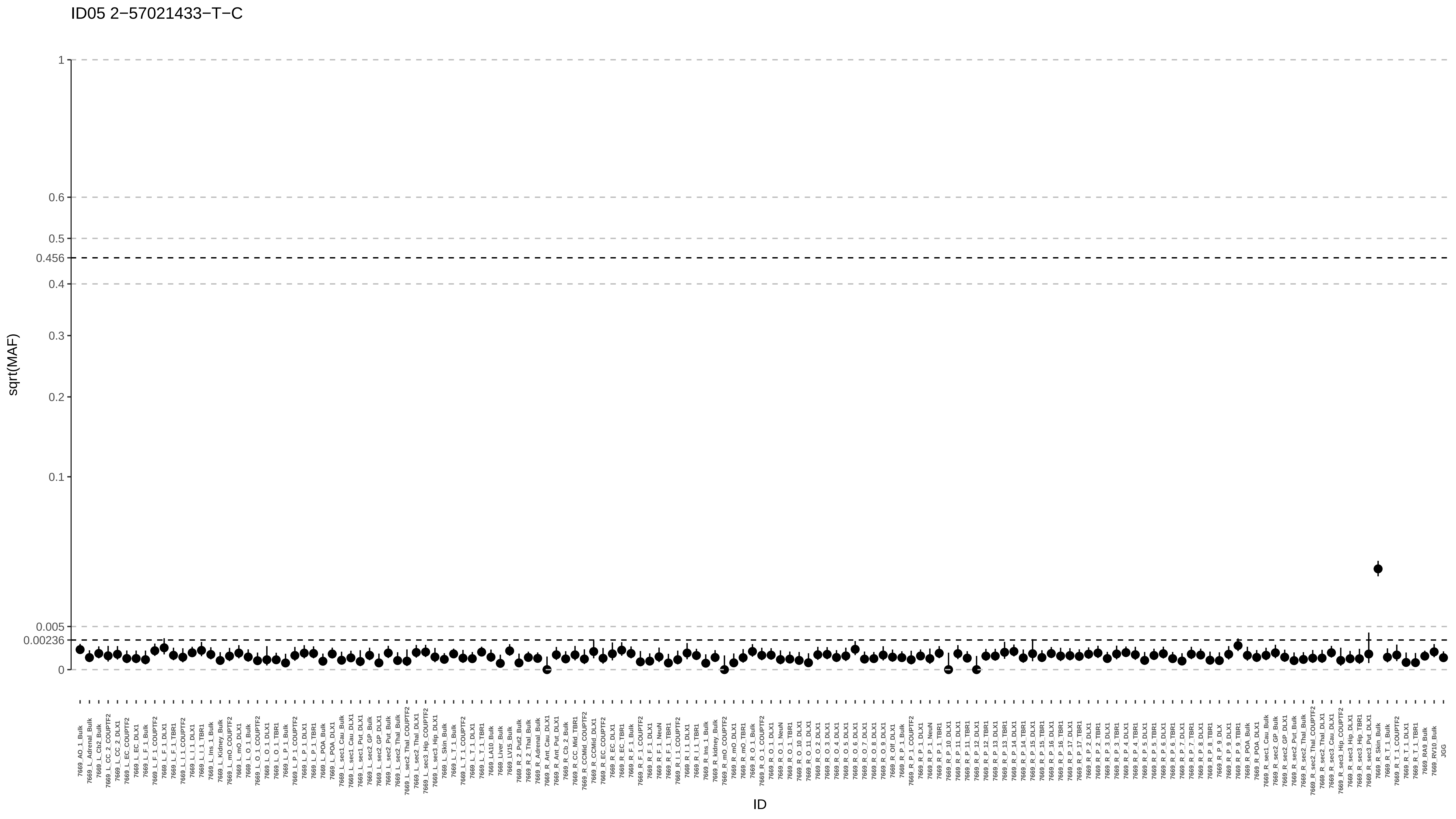

ID05 2−57021433−T−C
sqrt(MAF)
1
0.6
0.5
0.456
0.4
0.3
0.2
0.1
0.005
0.00236
0
ID

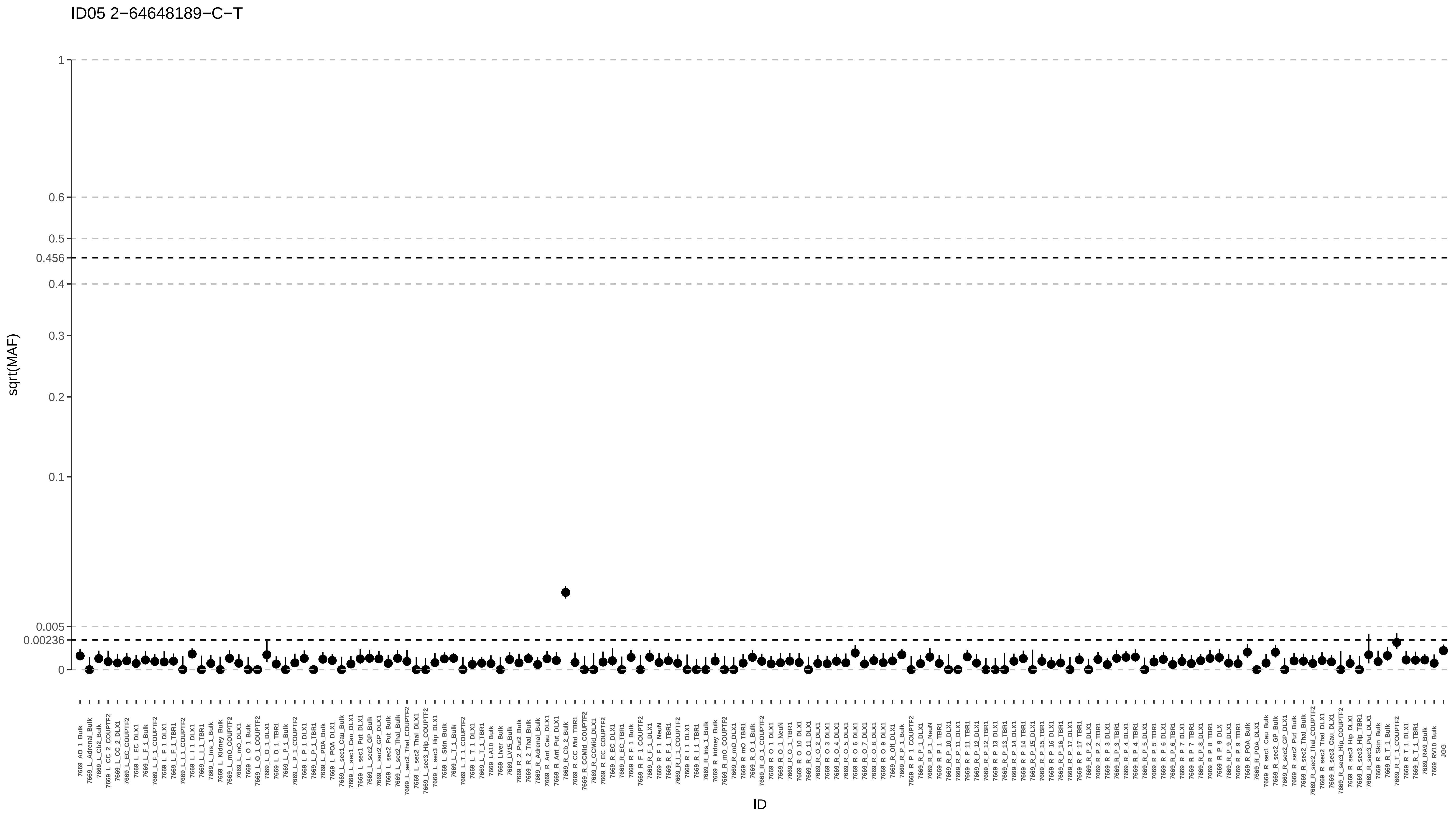
ID05 2−64648189−C−T
sqrt(MAF)
1
0.6
0.5
0.456
0.4
0.3
0.2
0.1
0.005
0.00236
0
ID

ID05 2−72858074−C−A

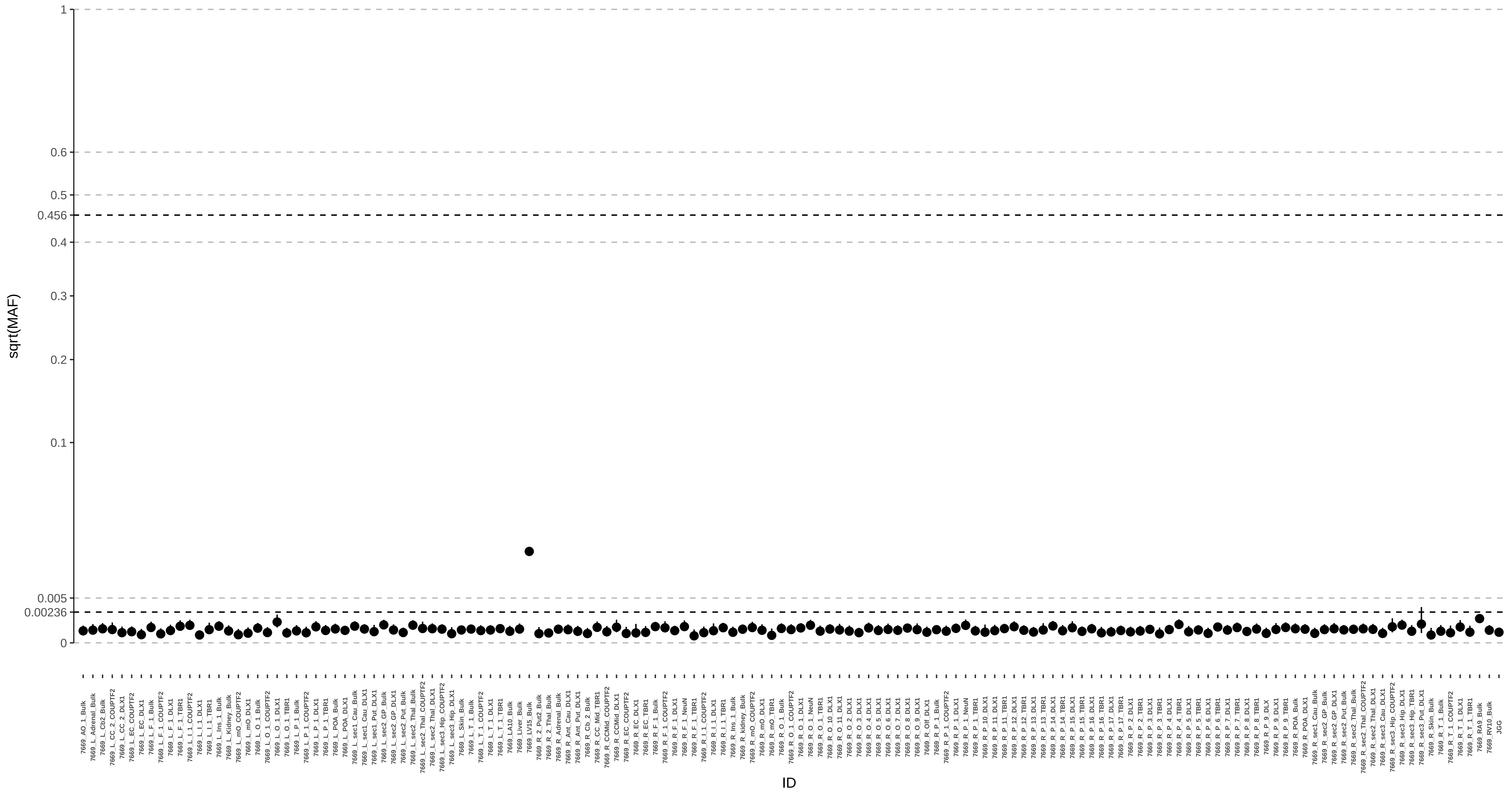

ID05 2−78747329−C−A

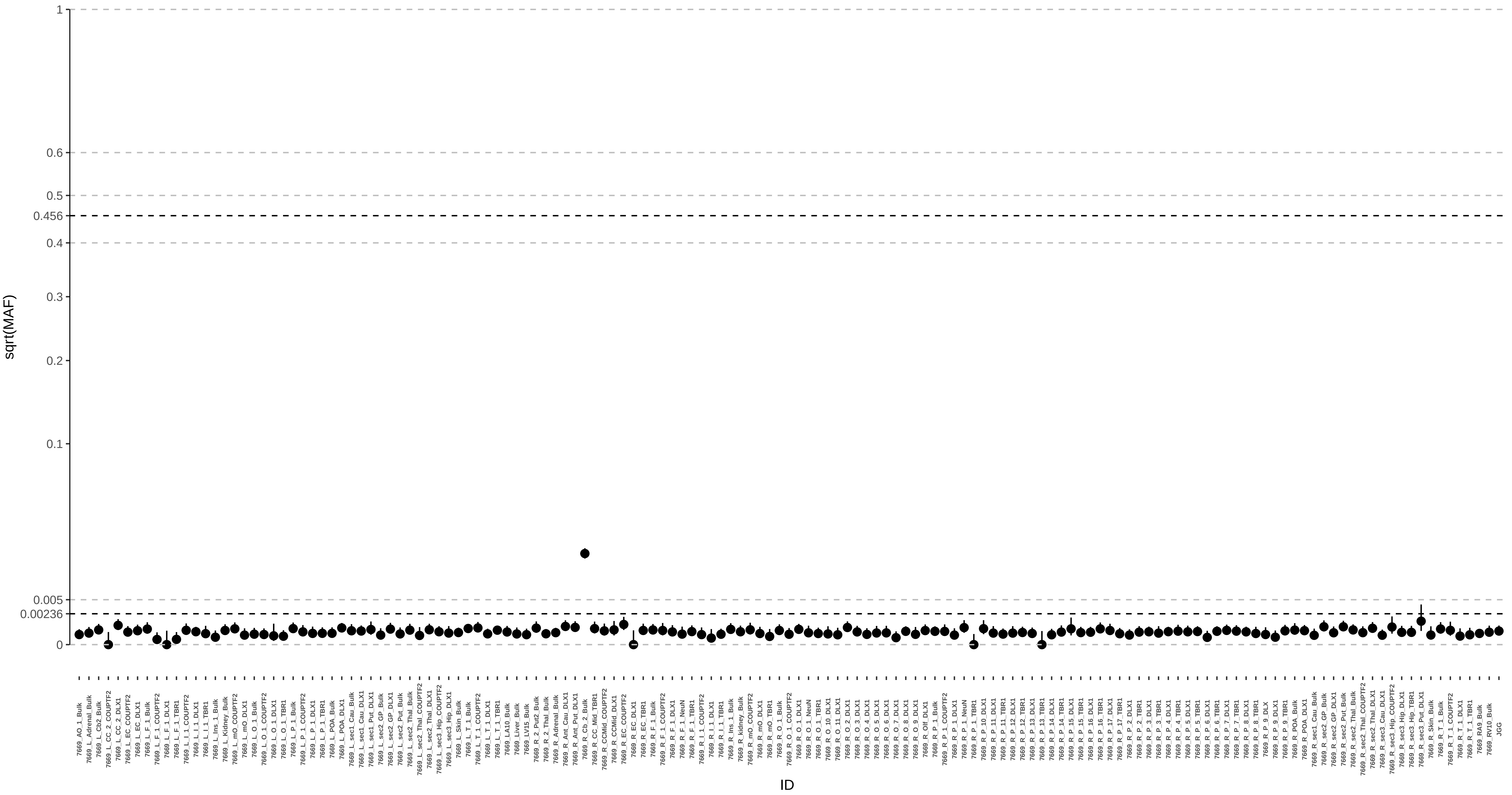

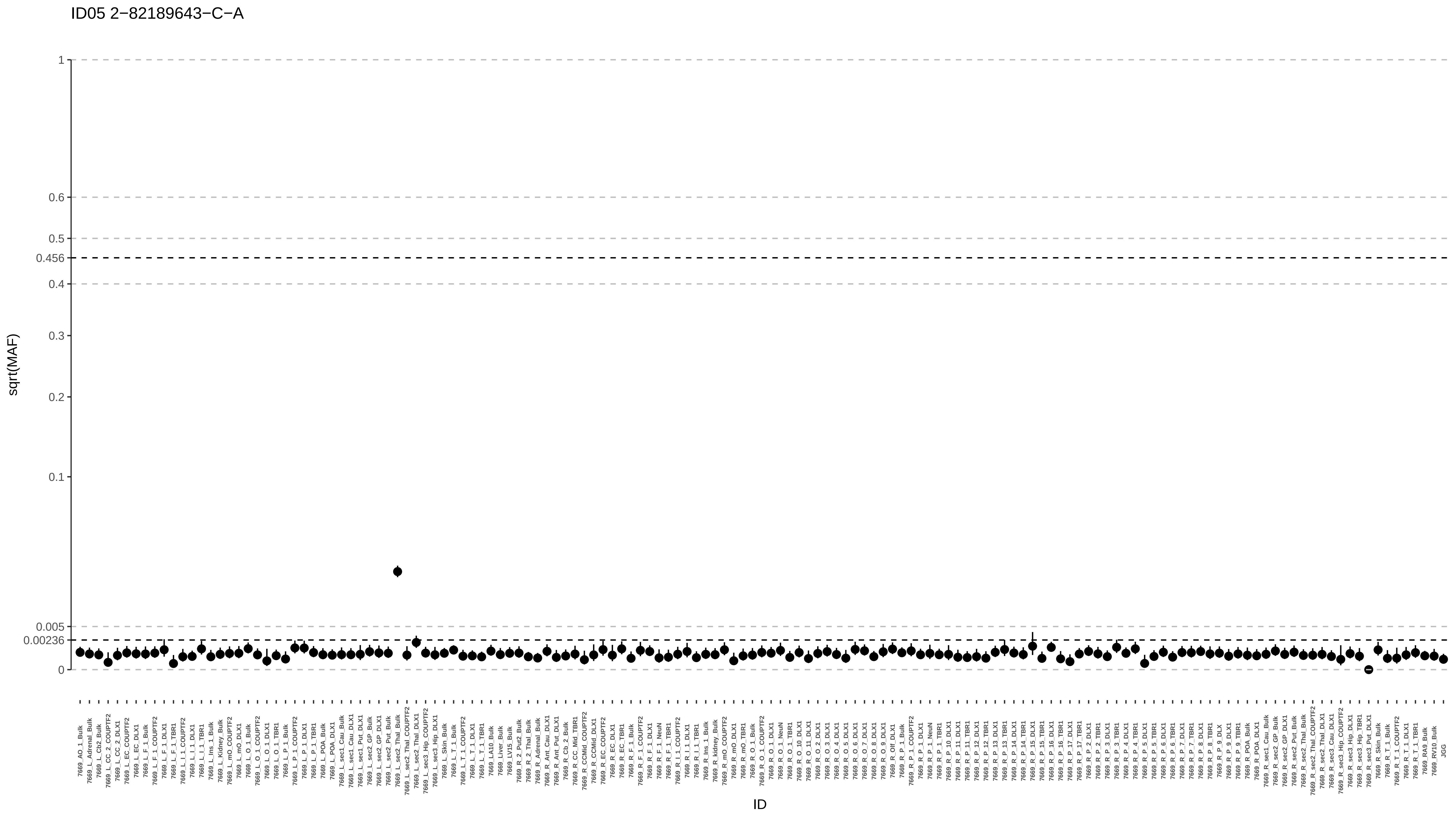
ID05 2−82189643−C−A
sqrt(MAF)
1
0.6
0.5
0.456
0.4
0.3
0.2
0.1
0.005
0.00236
0
ID

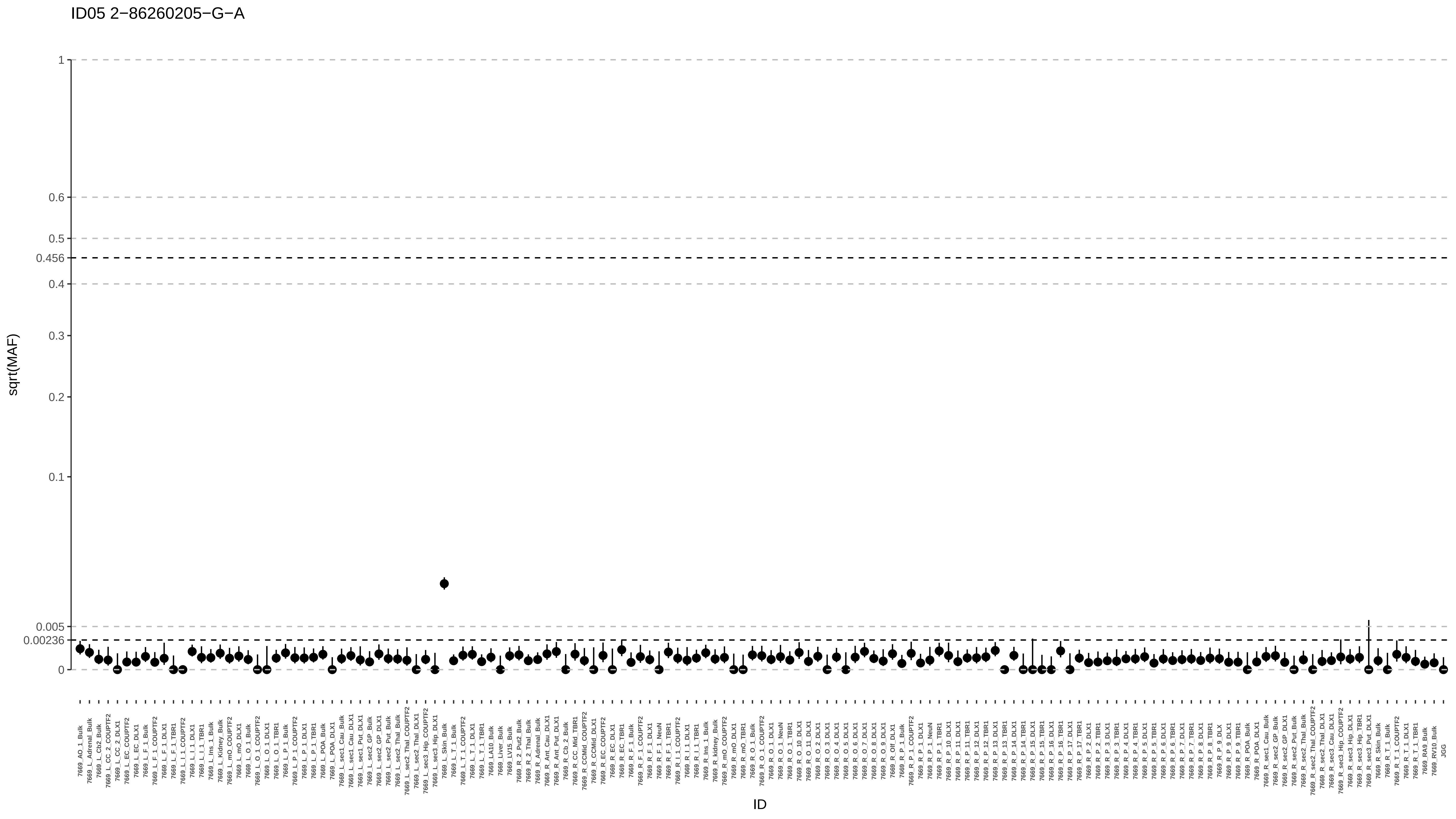
ID05 2−86260205−G−A
sqrt(MAF)
1
0.6
0.5
0.456
0.4
0.3
0.2
0.1
0.005
0.00236
0
ID
7669_AO_1_Bulk
7669_L_Adrenal_Bulk
7669_L_Cb2_Bulk
7669_L_CC_2_COUPTF2
7669_L_CC_2_DLX1
7669_L_EC_COUPTF2
7669_L_EC_DLX1
7669_L_F_1_Bulk
7669_L_F_1_COUPTF2
7669_L_F_1_DLX1
7669_L_F_1_TBR1
7669_L_I_1_COUPTF2
7669_L_I_1_DLX1
7669_L_I_1_TBR1
7669_L_Ins_1_Bulk
7669_L_Kidney_Bulk
7669_L_mO_COUPTF2
7669_L_mO_DLX1
7669_L_O_1_Bulk
7669_L_O_1_COUPTF2
7669_L_O_1_DLX1
7669_L_O_1_TBR1
7669_L_P_1_Bulk
7669_L_P_1_COUPTF2
7669_L_P_1_DLX1
7669_L_P_1_TBR1
7669_L_POA_Bulk
7669_L_POA_DLX1
7669_L_sec1_Cau_Bulk
7669_L_sec1_Cau_DLX1
7669_L_sec1_Put_DLX1
7669_L_sec2_GP_Bulk
7669_L_sec2_GP_DLX1
7669_L_sec2_Put_Bulk
7669_L_sec2_Thal_Bulk
7669_L_sec2_Thal_COUPTF2
7669_L_sec2_Thal_DLX1
7669_L_sec3_Hip_COUPTF2
7669_L_sec3_Hip_DLX1
7669_L_Skin_Bulk
7669_L_T_1_Bulk
7669_L_T_1_COUPTF2
7669_L_T_1_DLX1
7669_L_T_1_TBR1
7669_LA10_Bulk
7669_Liver_Bulk
7669_LV15_Bulk
7669_R_2_Put2_Bulk
7669_R_2_Thal_Bulk
7669_R_Adrenal_Bulk
7669_R_Ant_Cau_DLX1
7669_R_Ant_Put_DLX1
7669_R_Cb_2_Bulk
7669_R_CC_Mid_TBR1
7669_R_CCMid_COUPTF2
7669_R_CCMid_DLX1
7669_R_EC_COUPTF2
7669_R_EC_DLX1
7669_R_EC_TBR1
7669_R_F_1_Bulk
7669_R_F_1_COUPTF2
7669_R_F_1_DLX1
7669_R_F_1_NeuN
7669_R_F_1_TBR1
7669_R_I_1_COUPTF2
7669_R_I_1_DLX1
7669_R_I_1_TBR1
7669_R_Ins_1_Bulk
7669_R_kidney_Bulk
7669_R_mO_COUPTF2
7669_R_mO_DLX1
7669_R_mO_TBR1
7669_R_O_1_Bulk
7669_R_O_1_COUPTF2
7669_R_O_1_DLX1
7669_R_O_1_NeuN
7669_R_O_1_TBR1
7669_R_O_10_DLX1
7669_R_O_11_DLX1
7669_R_O_2_DLX1
7669_R_O_3_DLX1
7669_R_O_4_DLX1
7669_R_O_5_DLX1
7669_R_O_6_DLX1
7669_R_O_7_DLX1
7669_R_O_8_DLX1
7669_R_O_9_DLX1
7669_R_Olf_DLX1
7669_R_P_1_Bulk
7669_R_P_1_COUPTF2
7669_R_P_1_DLX1
7669_R_P_1_NeuN
7669_R_P_1_TBR1
7669_R_P_10_DLX1
7669_R_P_11_DLX1
7669_R_P_11_TBR1
7669_R_P_12_DLX1
7669_R_P_12_TBR1
7669_R_P_13_DLX1
7669_R_P_13_TBR1
7669_R_P_14_DLX1
7669_R_P_14_TBR1
7669_R_P_15_DLX1
7669_R_P_15_TBR1
7669_R_P_16_DLX1
7669_R_P_16_TBR1
7669_R_P_17_DLX1
7669_R_P_17_TBR1
7669_R_P_2_DLX1
7669_R_P_2_TBR1
7669_R_P_3_DLX1
7669_R_P_3_TBR1
7669_R_P_4_DLX1
7669_R_P_4_TBR1
7669_R_P_5_DLX1
7669_R_P_5_TBR1
7669_R_P_6_DLX1
7669_R_P_6_TBR1
7669_R_P_7_DLX1
7669_R_P_7_TBR1
7669_R_P_8_DLX1
7669_R_P_8_TBR1
7669_R_P_9_DLX
7669_R_P_9_DLX1
7669_R_P_9_TBR1
7669_R_POA_Bulk
7669_R_POA_DLX1
7669_R_sec1_Cau_Bulk
7669_R_sec2_GP_Bulk
7669_R_sec2_GP_DLX1
7669_R_sec2_Put_Bulk
7669_R_sec2_Thal_Bulk
7669_R_sec2_Thal_COUPTF2
7669_R_sec2_Thal_DLX1
7669_R_sec3_Cau_DLX1
7669_R_sec3_Hip_COUPTF2
7669_R_sec3_Hip_DLX1
7669_R_sec3_Hip_TBR1
7669_R_sec3_Put_DLX1
7669_R_Skin_Bulk
7669_R_T_1_Bulk
7669_R_T_1_COUPTF2
7669_R_T_1_DLX1
7669_R_T_1_TBR1
7669_RA9_Bulk
7669_RV10_Bulk
JGG

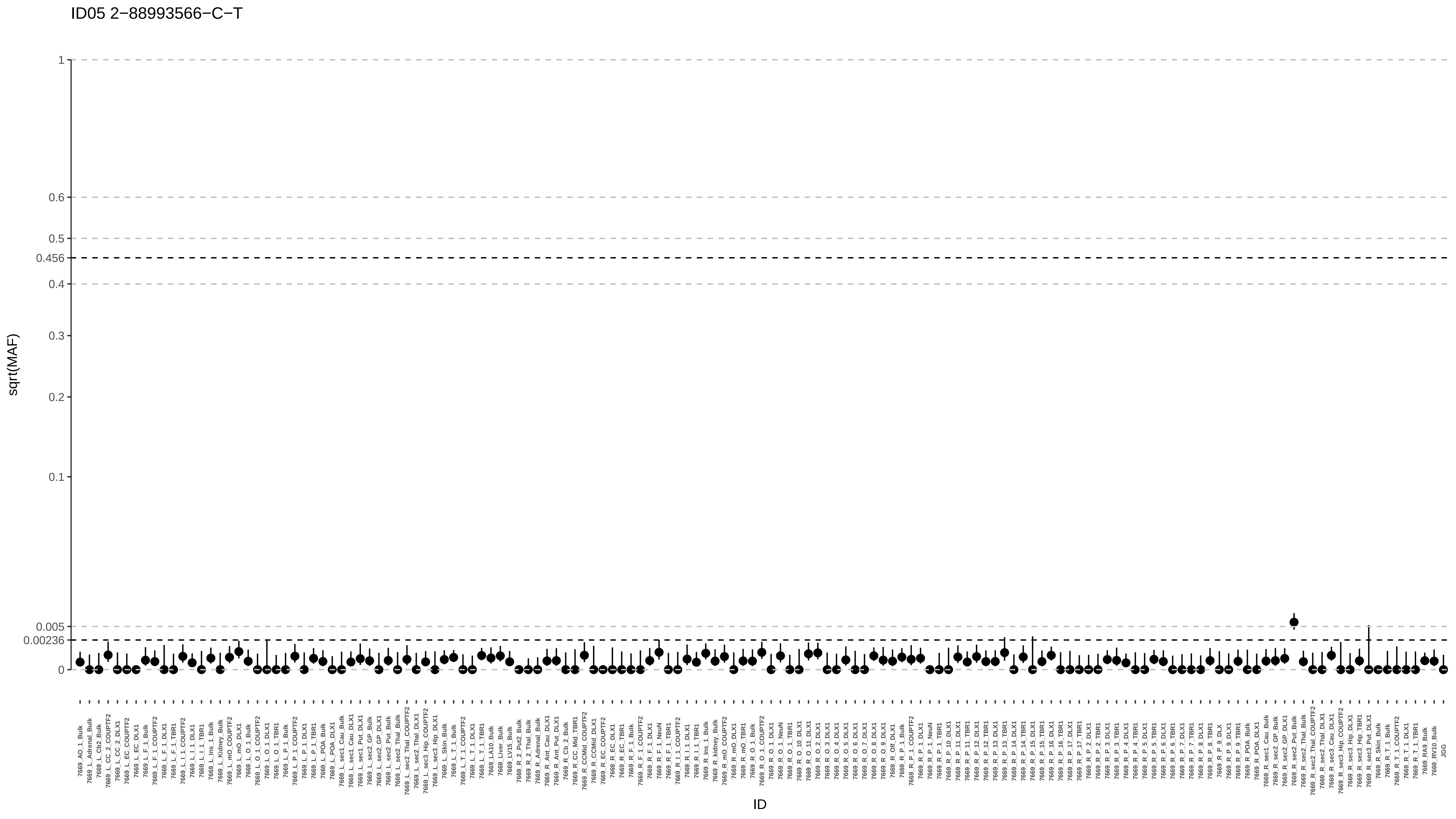
ID05 2−88993566−C−T
sqrt(MAF)
ID
1
0.6
0.5
0.456
0.4
0.3
0.2
0.1
0.005
0.00236
0

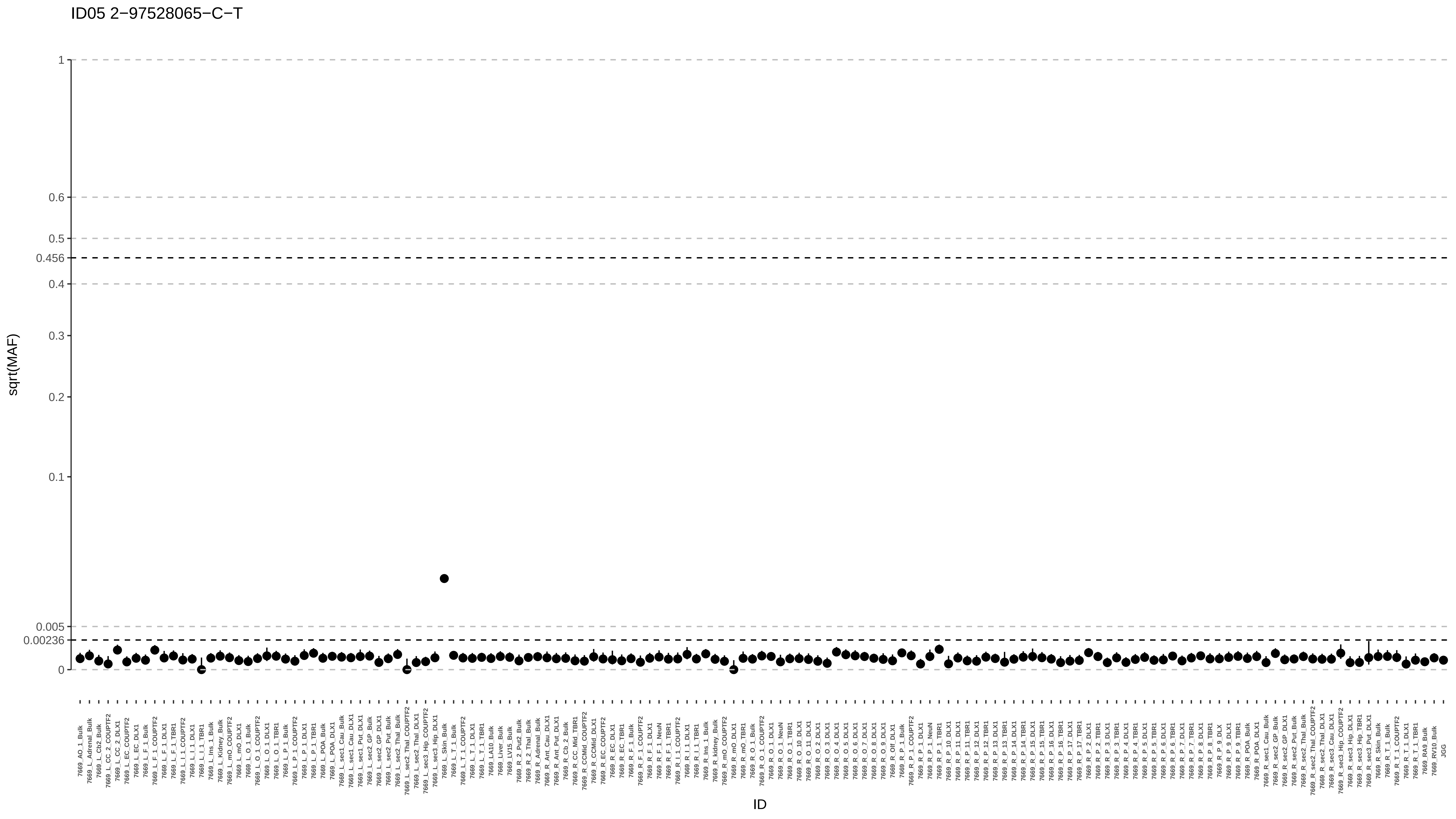
ID05 2−97528065−C−T
sqrt(MAF)
1
0.6
0.5
0.456
0.4
0.3
0.2
0.1
0.005
0.00236
0
ID

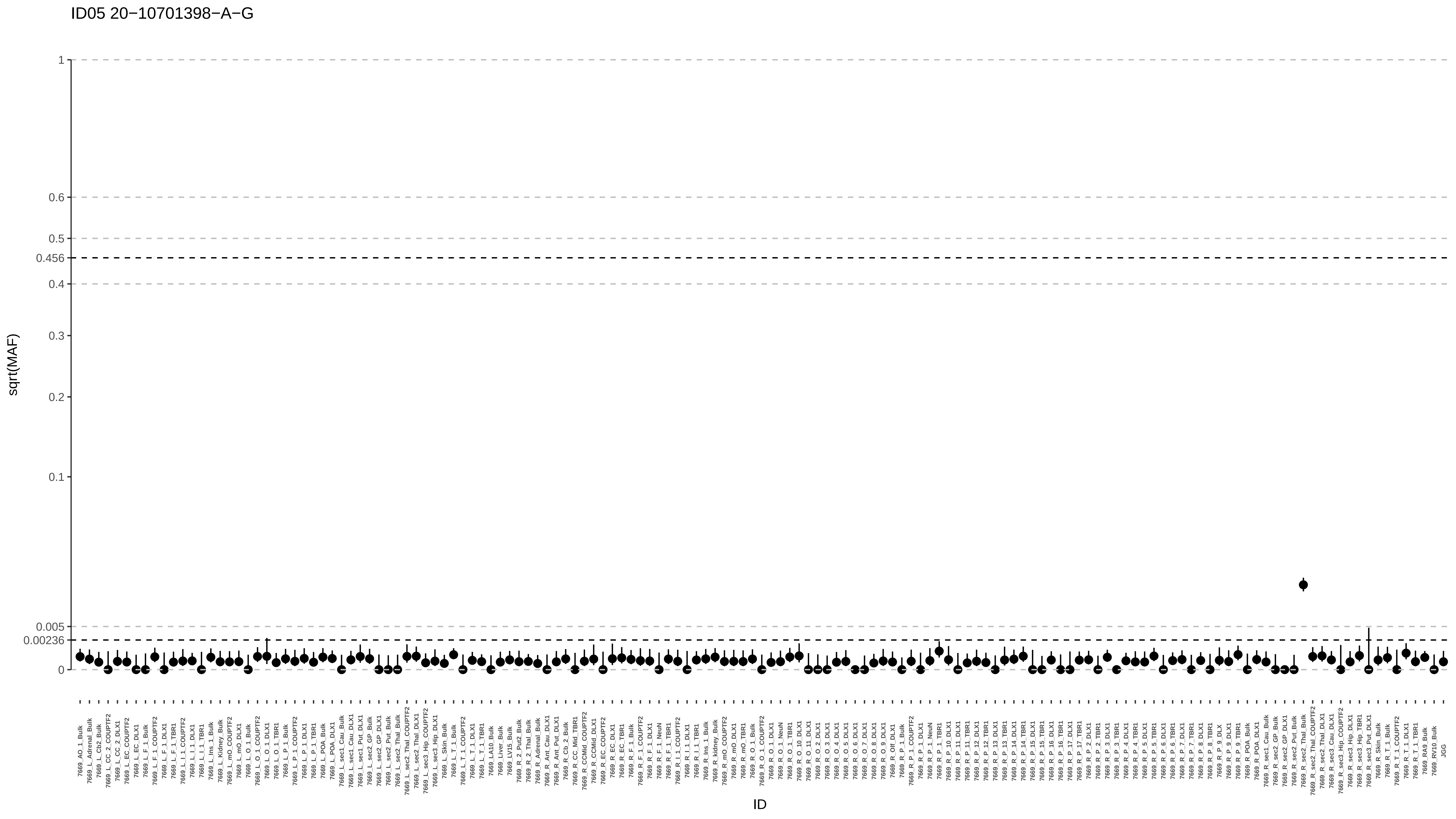
ID05 20−10701398−A−G
sqrt(MAF)
1
0.6
0.5
0.456
0.4
0.3
0.2
0.1
0.005
0.00236
0
ID
7669_AO_1_Bulk
7669_L_Adrenal_Bulk
7669_L_Cb2_Bulk
7669_L_CC_2_COUPTF2
7669_L_CC_2_DLX1
7669_L_EC_COUPTF2
7669_L_EC_DLX1
7669_L_F_1_Bulk
7669_L_F_1_COUPTF2
7669_L_F_1_DLX1
7669_L_F_1_TBR1
7669_L_I_1_COUPTF2
7669_L_I_1_DLX1
7669_L_I_1_TBR1
7669_L_Ins_1_Bulk
7669_L_Kidney_Bulk
7669_L_mO_COUPTF2
7669_L_mO_DLX1
7669_L_O_1_Bulk
7669_L_O_1_COUPTF2
7669_L_O_1_DLX1
7669_L_O_1_TBR1
7669_L_P_1_Bulk
7669_L_P_1_COUPTF2
7669_L_P_1_DLX1
7669_L_P_1_TBR1
7669_L_POA_Bulk
7669_L_POA_DLX1
7669_L_sec1_Cau_Bulk
7669_L_sec1_Cau_DLX1
7669_L_sec1_Put_DLX1
7669_L_sec2_GP_Bulk
7669_L_sec2_GP_DLX1
7669_L_sec2_Put_Bulk
7669_L_sec2_Thal_Bulk
7669_L_sec2_Thal_COUPTF2
7669_L_sec2_Thal_DLX1
7669_L_sec3_Hip_COUPTF2
7669_L_sec3_Hip_DLX1
7669_L_Skin_Bulk
7669_L_T_1_Bulk
7669_L_T_1_COUPTF2
7669_L_T_1_DLX1
7669_L_T_1_TBR1
7669_LA10_Bulk
7669_Liver_Bulk
7669_LV15_Bulk
7669_R_2_Put2_Bulk
7669_R_2_Thal_Bulk
7669_R_Adrenal_Bulk
7669_R_Ant_Cau_DLX1
7669_R_Ant_Put_DLX1
7669_R_Cb_2_Bulk
7669_R_CC_Mid_TBR1
7669_R_CCMid_COUPTF2
7669_R_CCMid_DLX1
7669_R_EC_COUPTF2
7669_R_EC_DLX1
7669_R_EC_TBR1
7669_R_F_1_Bulk
7669_R_F_1_COUPTF2
7669_R_F_1_DLX1
7669_R_F_1_NeuN
7669_R_F_1_TBR1
7669_R_I_1_COUPTF2
7669_R_I_1_DLX1
7669_R_I_1_TBR1
7669_R_Ins_1_Bulk
7669_R_kidney_Bulk
7669_R_mO_COUPTF2
7669_R_mO_DLX1
7669_R_mO_TBR1
7669_R_O_1_Bulk
7669_R_O_1_COUPTF2
7669_R_O_1_DLX1
7669_R_O_1_NeuN
7669_R_O_1_TBR1
7669_R_O_10_DLX1
7669_R_O_11_DLX1
7669_R_O_2_DLX1
7669_R_O_3_DLX1
7669_R_O_4_DLX1
7669_R_O_5_DLX1
7669_R_O_6_DLX1
7669_R_O_7_DLX1
7669_R_O_8_DLX1
7669_R_O_9_DLX1
7669_R_Olf_DLX1
7669_R_P_1_Bulk
7669_R_P_1_COUPTF2
7669_R_P_1_DLX1
7669_R_P_1_NeuN
7669_R_P_1_TBR1
7669_R_P_10_DLX1
7669_R_P_11_DLX1
7669_R_P_11_TBR1
7669_R_P_12_DLX1
7669_R_P_12_TBR1
7669_R_P_13_DLX1
7669_R_P_13_TBR1
7669_R_P_14_DLX1
7669_R_P_14_TBR1
7669_R_P_15_DLX1
7669_R_P_15_TBR1
7669_R_P_16_DLX1
7669_R_P_16_TBR1
7669_R_P_17_DLX1
7669_R_P_17_TBR1
7669_R_P_2_DLX1
7669_R_P_2_TBR1
7669_R_P_3_DLX1
7669_R_P_3_TBR1
7669_R_P_4_DLX1
7669_R_P_4_TBR1
7669_R_P_5_DLX1
7669_R_P_5_TBR1
7669_R_P_6_DLX1
7669_R_P_6_TBR1
7669_R_P_7_DLX1
7669_R_P_7_TBR1
7669_R_P_8_DLX1
7669_R_P_8_TBR1
7669_R_P_9_DLX
7669_R_P_9_DLX1
7669_R_P_9_TBR1
7669_R_POA_Bulk
7669_R_POA_DLX1
7669_R_sec1_Cau_Bulk
7669_R_sec2_GP_Bulk
7669_R_sec2_GP_DLX1
7669_R_sec2_Put_Bulk
7669_R_sec2_Thal_Bulk
7669_R_sec2_Thal_COUPTF2
7669_R_sec2_Thal_DLX1
7669_R_sec3_Cau_DLX1
7669_R_sec3_Hip_COUPTF2
7669_R_sec3_Hip_DLX1
7669_R_sec3_Hip_TBR1
7669_R_sec3_Put_DLX1
7669_R_Skin_Bulk
7669_R_T_1_Bulk
7669_R_T_1_COUPTF2
7669_R_T_1_DLX1
7669_R_T_1_TBR1
7669_RA9_Bulk
7669_RV10_Bulk
JGG

ID05 20−10742088−G−A

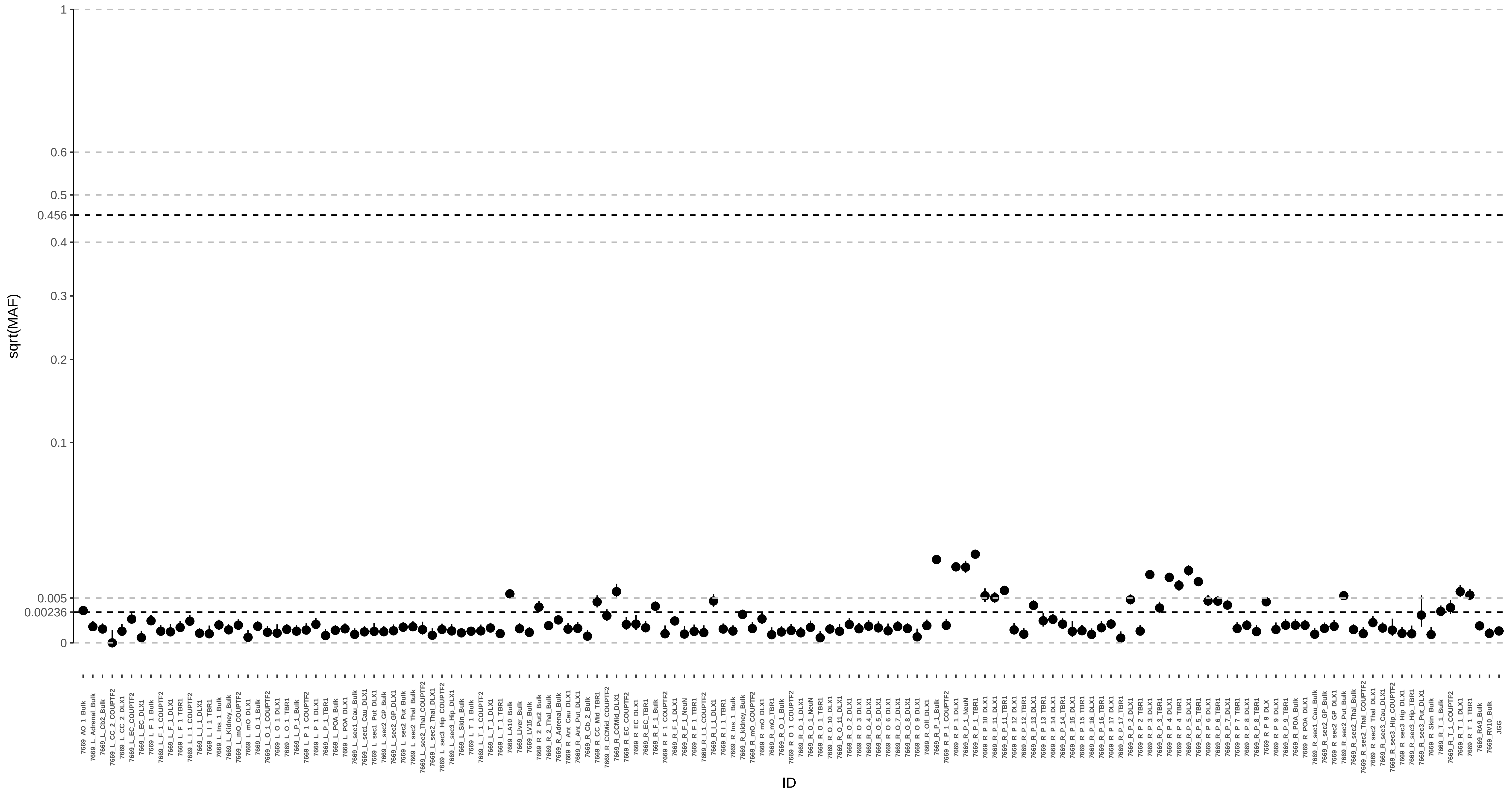

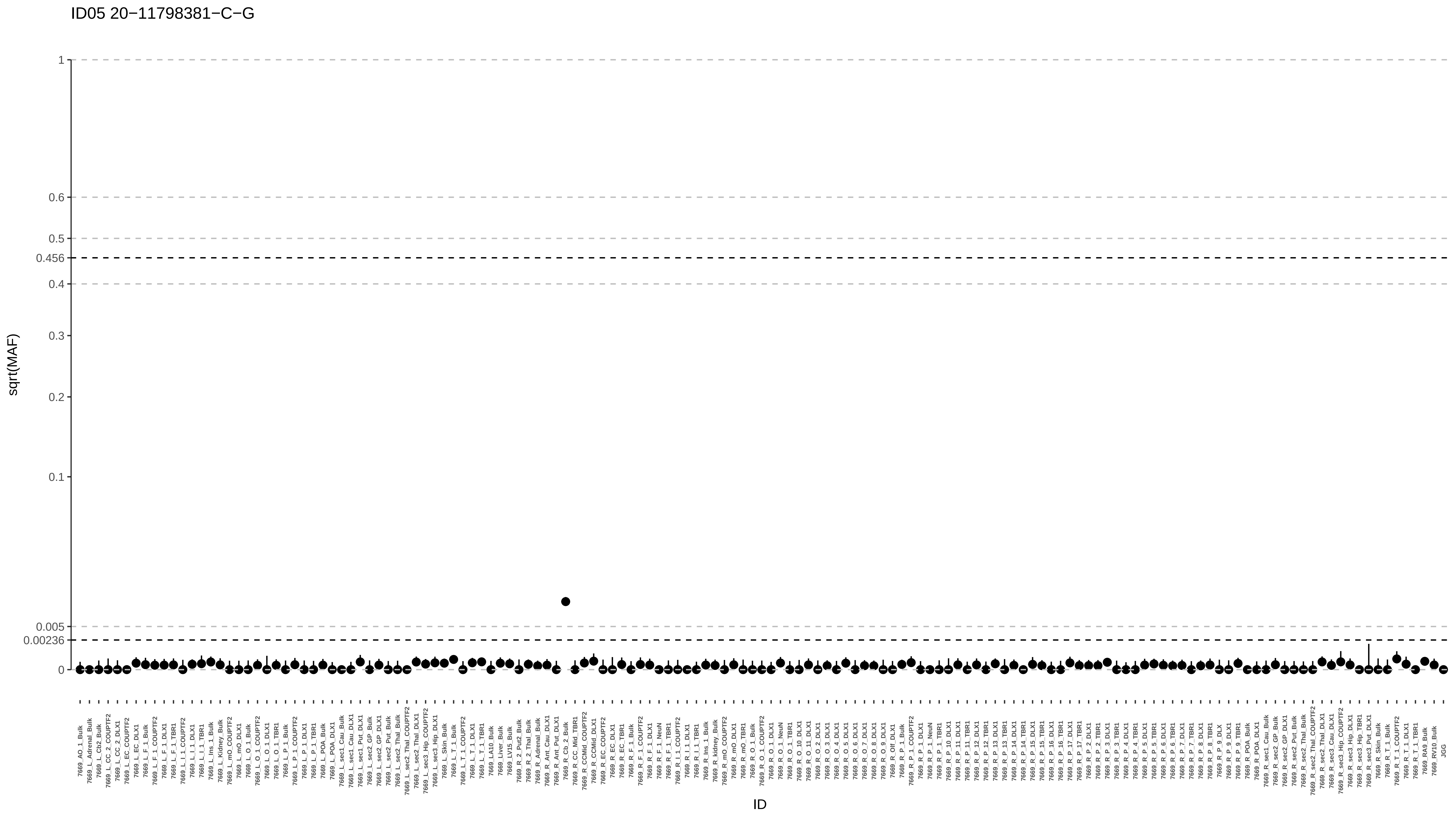
ID05 20−11798381−C−G
sqrt(MAF)
1
0.6
0.5
0.456
0.4
0.3
0.2
0.1
0.005
0.00236
0
ID

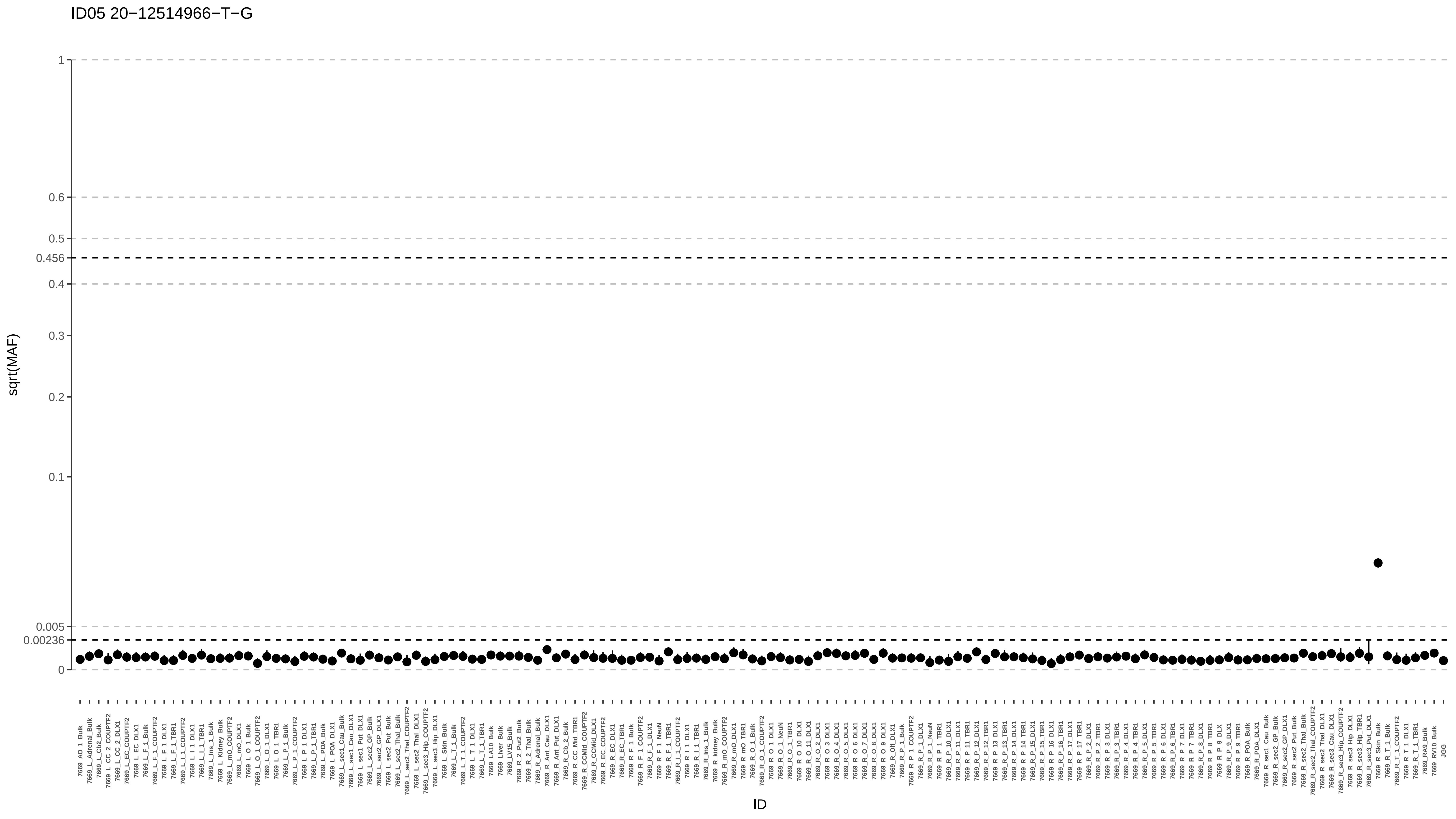
ID05 20−12514966−T−G
sqrt(MAF)
1
0.6
0.5
0.456
0.4
0.3
0.2
0.1
0.005
0.00236
0
ID
7669_AO_1_Bulk
7669_L_Adrenal_Bulk
7669_L_Cb2_Bulk
7669_L_CC_2_COUPTF2
7669_L_CC_2_DLX1
7669_L_EC_COUPTF2
7669_L_EC_DLX1
7669_L_F_1_Bulk
7669_L_F_1_COUPTF2
7669_L_F_1_DLX1
7669_L_F_1_TBR1
7669_L_I_1_COUPTF2
7669_L_I_1_DLX1
7669_L_I_1_TBR1
7669_L_Ins_1_Bulk
7669_L_Kidney_Bulk
7669_L_mO_COUPTF2
7669_L_mO_DLX1
7669_L_O_1_Bulk
7669_L_O_1_COUPTF2
7669_L_O_1_DLX1
7669_L_O_1_TBR1
7669_L_P_1_Bulk
7669_L_P_1_COUPTF2
7669_L_P_1_DLX1
7669_L_P_1_TBR1
7669_L_POA_Bulk
7669_L_POA_DLX1
7669_L_sec1_Cau_Bulk
7669_L_sec1_Cau_DLX1
7669_L_sec1_Put_DLX1
7669_L_sec2_GP_Bulk
7669_L_sec2_GP_DLX1
7669_L_sec2_Put_Bulk
7669_L_sec2_Thal_Bulk
7669_L_sec2_Thal_COUPTF2
7669_L_sec2_Thal_DLX1
7669_L_sec3_Hip_COUPTF2
7669_L_sec3_Hip_DLX1
7669_L_Skin_Bulk
7669_L_T_1_Bulk
7669_L_T_1_COUPTF2
7669_L_T_1_DLX1
7669_L_T_1_TBR1
7669_LA10_Bulk
7669_Liver_Bulk
7669_LV15_Bulk
7669_R_2_Put2_Bulk
7669_R_2_Thal_Bulk
7669_R_Adrenal_Bulk
7669_R_Ant_Cau_DLX1
7669_R_Ant_Put_DLX1
7669_R_Cb_2_Bulk
7669_R_CC_Mid_TBR1
7669_R_CCMid_COUPTF2
7669_R_CCMid_DLX1
7669_R_EC_COUPTF2
7669_R_EC_DLX1
7669_R_EC_TBR1
7669_R_F_1_Bulk
7669_R_F_1_COUPTF2
7669_R_F_1_DLX1
7669_R_F_1_NeuN
7669_R_F_1_TBR1
7669_R_I_1_COUPTF2
7669_R_I_1_DLX1
7669_R_I_1_TBR1
7669_R_Ins_1_Bulk
7669_R_kidney_Bulk
7669_R_mO_COUPTF2
7669_R_mO_DLX1
7669_R_mO_TBR1
7669_R_O_1_Bulk
7669_R_O_1_COUPTF2
7669_R_O_1_DLX1
7669_R_O_1_NeuN
7669_R_O_1_TBR1
7669_R_O_10_DLX1
7669_R_O_11_DLX1
7669_R_O_2_DLX1
7669_R_O_3_DLX1
7669_R_O_4_DLX1
7669_R_O_5_DLX1
7669_R_O_6_DLX1
7669_R_O_7_DLX1
7669_R_O_8_DLX1
7669_R_O_9_DLX1
7669_R_Olf_DLX1
7669_R_P_1_Bulk
7669_R_P_1_COUPTF2
7669_R_P_1_DLX1
7669_R_P_1_NeuN
7669_R_P_1_TBR1
7669_R_P_10_DLX1
7669_R_P_11_DLX1
7669_R_P_11_TBR1
7669_R_P_12_DLX1
7669_R_P_12_TBR1
7669_R_P_13_DLX1
7669_R_P_13_TBR1
7669_R_P_14_DLX1
7669_R_P_14_TBR1
7669_R_P_15_DLX1
7669_R_P_15_TBR1
7669_R_P_16_DLX1
7669_R_P_16_TBR1
7669_R_P_17_DLX1
7669_R_P_17_TBR1
7669_R_P_2_DLX1
7669_R_P_2_TBR1
7669_R_P_3_DLX1
7669_R_P_3_TBR1
7669_R_P_4_DLX1
7669_R_P_4_TBR1
7669_R_P_5_DLX1
7669_R_P_5_TBR1
7669_R_P_6_DLX1
7669_R_P_6_TBR1
7669_R_P_7_DLX1
7669_R_P_7_TBR1
7669_R_P_8_DLX1
7669_R_P_8_TBR1
7669_R_P_9_DLX
7669_R_P_9_DLX1
7669_R_P_9_TBR1
7669_R_POA_Bulk
7669_R_POA_DLX1
7669_R_sec1_Cau_Bulk
7669_R_sec2_GP_Bulk
7669_R_sec2_GP_DLX1
7669_R_sec2_Put_Bulk
7669_R_sec2_Thal_Bulk
7669_R_sec2_Thal_COUPTF2
7669_R_sec2_Thal_DLX1
7669_R_sec3_Cau_DLX1
7669_R_sec3_Hip_COUPTF2
7669_R_sec3_Hip_DLX1
7669_R_sec3_Hip_TBR1
7669_R_sec3_Put_DLX1
7669_R_Skin_Bulk
7669_R_T_1_Bulk
7669_R_T_1_COUPTF2
7669_R_T_1_DLX1
7669_R_T_1_TBR1
7669_RA9_Bulk
7669_RV10_Bulk
JGG

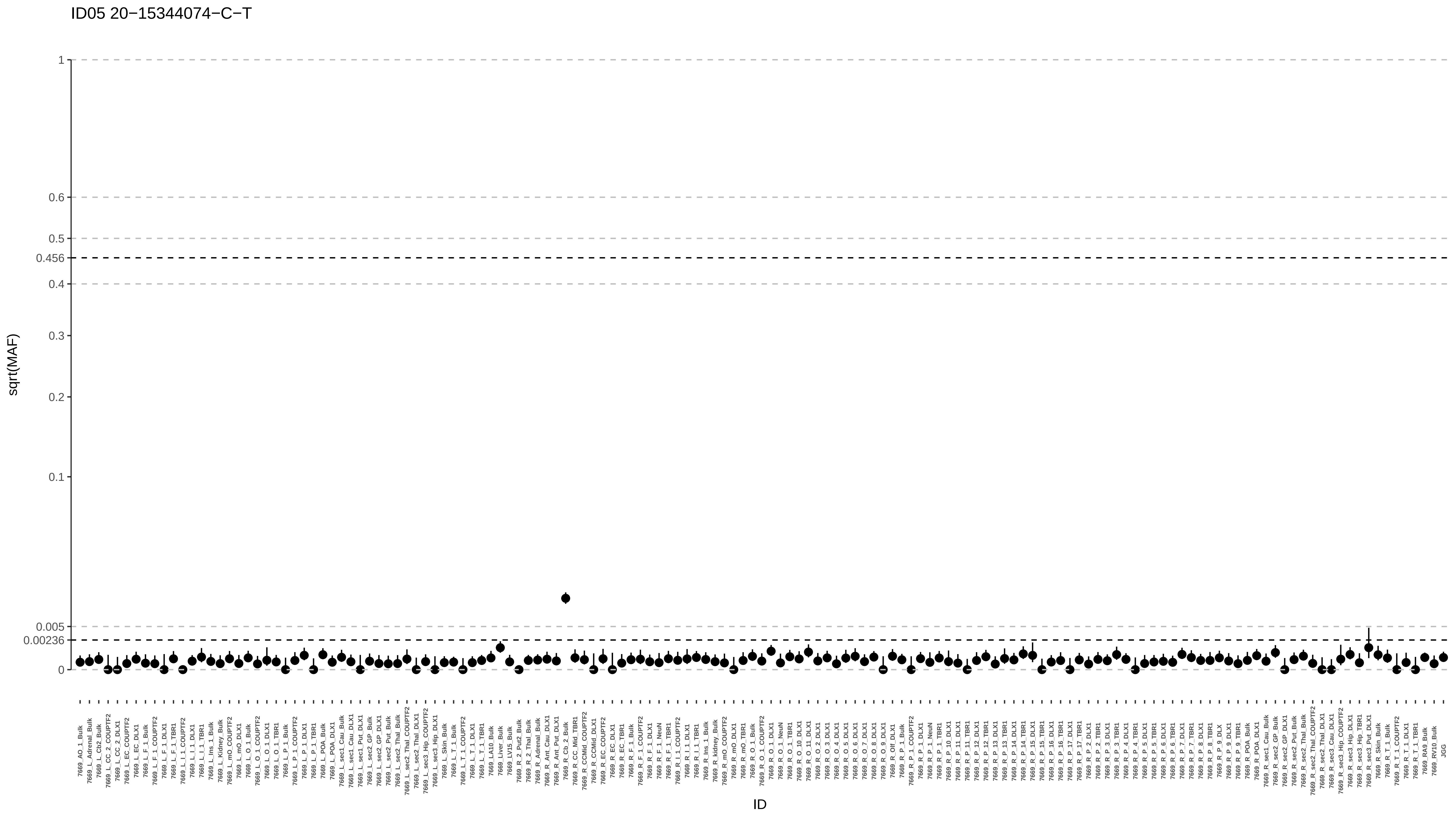

ID05 20−15344074−C−T
sqrt(MAF)
1
0.6
0.5
0.456
0.4
0.3
0.2
0.1
0.005
0.00236
0
ID

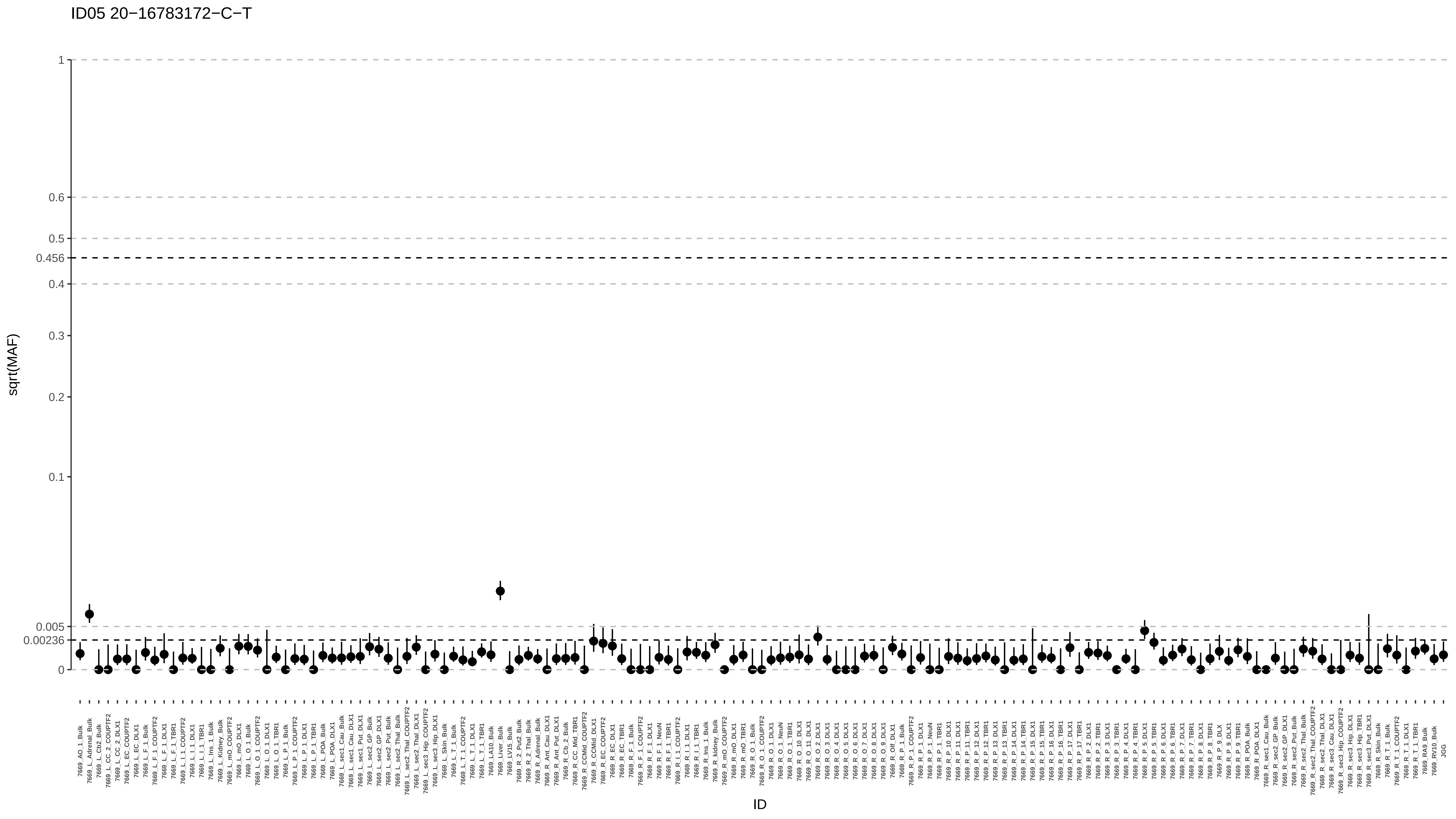
ID05 20−16783172−C−T
sqrt(MAF)
1
0.6
0.5
0.456
0.4
0.3
0.2
0.1
0.005
0.00236
0
ID
7669_AO_1_Bulk
7669_L_Adrenal_Bulk
7669_L_Cb2_Bulk
7669_L_CC_2_COUPTF2
7669_L_CC_2_DLX1
7669_L_EC_COUPTF2
7669_L_EC_DLX1
7669_L_F_1_Bulk
7669_L_F_1_COUPTF2
7669_L_F_1_DLX1
7669_L_F_1_TBR1
7669_L_I_1_COUPTF2
7669_L_I_1_DLX1
7669_L_I_1_TBR1
7669_L_Ins_1_Bulk
7669_L_Kidney_Bulk
7669_L_mO_COUPTF2
7669_L_mO_DLX1
7669_L_O_1_Bulk
7669_L_O_1_COUPTF2
7669_L_O_1_DLX1
7669_L_O_1_TBR1
7669_L_P_1_Bulk
7669_L_P_1_COUPTF2
7669_L_P_1_DLX1
7669_L_P_1_TBR1
7669_L_POA_Bulk
7669_L_POA_DLX1
7669_L_sec1_Cau_Bulk
7669_L_sec1_Cau_DLX1
7669_L_sec1_Put_DLX1
7669_L_sec2_GP_Bulk
7669_L_sec2_GP_DLX1
7669_L_sec2_Put_Bulk
7669_L_sec2_Thal_Bulk
7669_L_sec2_Thal_COUPTF2
7669_L_sec2_Thal_DLX1
7669_L_sec3_Hip_COUPTF2
7669_L_sec3_Hip_DLX1
7669_L_Skin_Bulk
7669_L_T_1_Bulk
7669_L_T_1_COUPTF2
7669_L_T_1_DLX1
7669_L_T_1_TBR1
7669_LA10_Bulk
7669_Liver_Bulk
7669_LV15_Bulk
7669_R_2_Put2_Bulk
7669_R_2_Thal_Bulk
7669_R_Adrenal_Bulk
7669_R_Ant_Cau_DLX1
7669_R_Ant_Put_DLX1
7669_R_Cb_2_Bulk
7669_R_CC_Mid_TBR1
7669_R_CCMid_COUPTF2
7669_R_CCMid_DLX1
7669_R_EC_COUPTF2
7669_R_EC_DLX1
7669_R_EC_TBR1
7669_R_F_1_Bulk
7669_R_F_1_COUPTF2
7669_R_F_1_DLX1
7669_R_F_1_NeuN
7669_R_F_1_TBR1
7669_R_I_1_COUPTF2
7669_R_I_1_DLX1
7669_R_I_1_TBR1
7669_R_Ins_1_Bulk
7669_R_kidney_Bulk
7669_R_mO_COUPTF2
7669_R_mO_DLX1
7669_R_mO_TBR1
7669_R_O_1_Bulk
7669_R_O_1_COUPTF2
7669_R_O_1_DLX1
7669_R_O_1_NeuN
7669_R_O_1_TBR1
7669_R_O_10_DLX1
7669_R_O_11_DLX1
7669_R_O_2_DLX1
7669_R_O_3_DLX1
7669_R_O_4_DLX1
7669_R_O_5_DLX1
7669_R_O_6_DLX1
7669_R_O_7_DLX1
7669_R_O_8_DLX1
7669_R_O_9_DLX1
7669_R_Olf_DLX1
7669_R_P_1_Bulk
7669_R_P_1_COUPTF2
7669_R_P_1_DLX1
7669_R_P_1_NeuN
7669_R_P_1_TBR1
7669_R_P_10_DLX1
7669_R_P_11_DLX1
7669_R_P_11_TBR1
7669_R_P_12_DLX1
7669_R_P_12_TBR1
7669_R_P_13_DLX1
7669_R_P_13_TBR1
7669_R_P_14_DLX1
7669_R_P_14_TBR1
7669_R_P_15_DLX1
7669_R_P_15_TBR1
7669_R_P_16_DLX1
7669_R_P_16_TBR1
7669_R_P_17_DLX1
7669_R_P_17_TBR1
7669_R_P_2_DLX1
7669_R_P_2_TBR1
7669_R_P_3_DLX1
7669_R_P_3_TBR1
7669_R_P_4_DLX1
7669_R_P_4_TBR1
7669_R_P_5_DLX1
7669_R_P_5_TBR1
7669_R_P_6_DLX1
7669_R_P_6_TBR1
7669_R_P_7_DLX1
7669_R_P_7_TBR1
7669_R_P_8_DLX1
7669_R_P_8_TBR1
7669_R_P_9_DLX
7669_R_P_9_DLX1
7669_R_P_9_TBR1
7669_R_POA_Bulk
7669_R_POA_DLX1
7669_R_sec1_Cau_Bulk
7669_R_sec2_GP_Bulk
7669_R_sec2_GP_DLX1
7669_R_sec2_Put_Bulk
7669_R_sec2_Thal_Bulk
7669_R_sec2_Thal_COUPTF2
7669_R_sec2_Thal_DLX1
7669_R_sec3_Cau_DLX1
7669_R_sec3_Hip_COUPTF2
7669_R_sec3_Hip_DLX1
7669_R_sec3_Hip_TBR1
7669_R_sec3_Put_DLX1
7669_R_Skin_Bulk
7669_R_T_1_Bulk
7669_R_T_1_COUPTF2
7669_R_T_1_DLX1
7669_R_T_1_TBR1
7669_RA9_Bulk
7669_RV10_Bulk
JGG

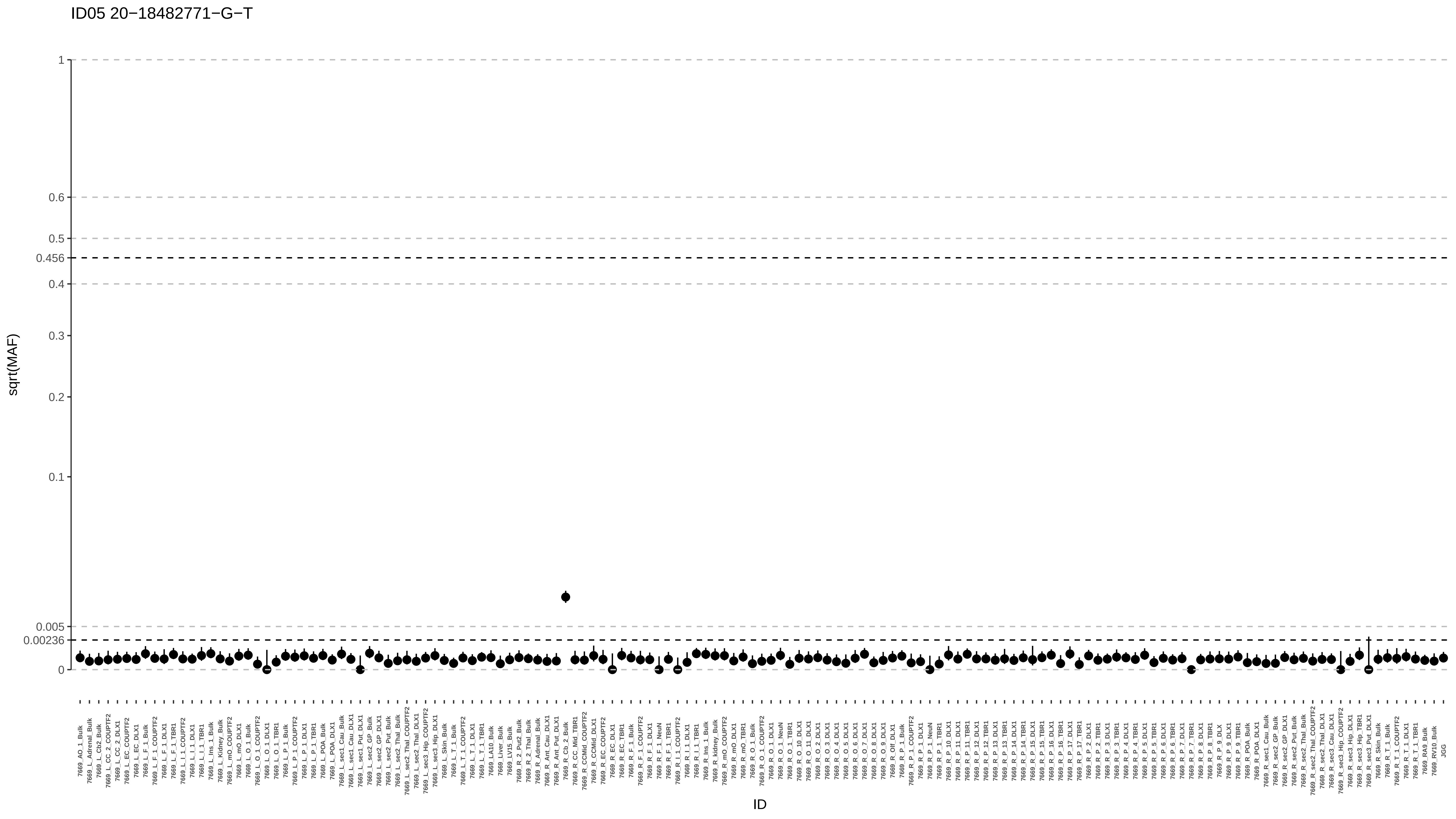
ID05 20−18482771−G−T
sqrt(MAF)
ID

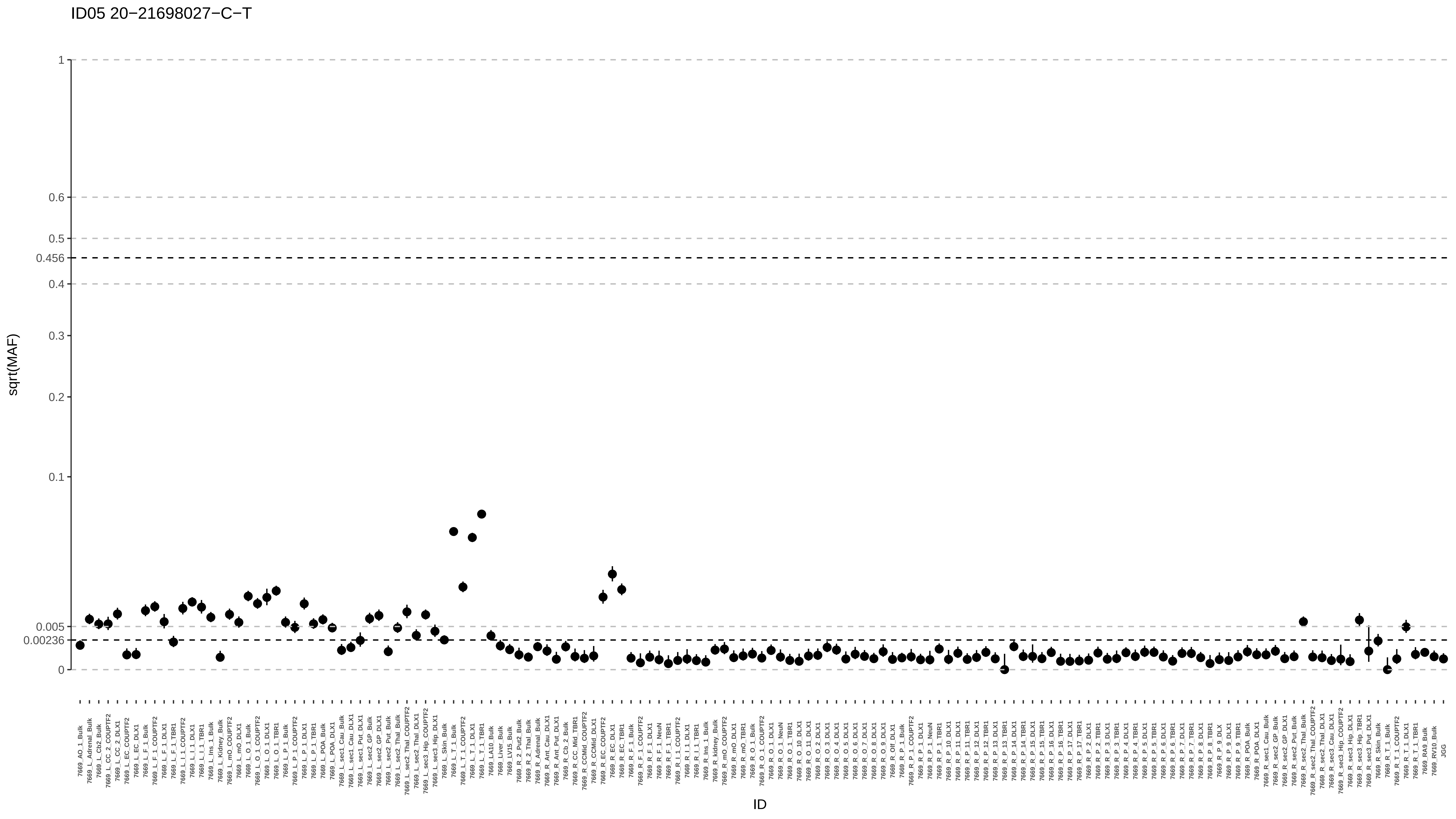
ID05 20−21698027−C−T
sqrt(MAF)
1
0.6
0.5
0.456
0.4
0.3
0.2
0.1
0.005
0.00236
0
ID

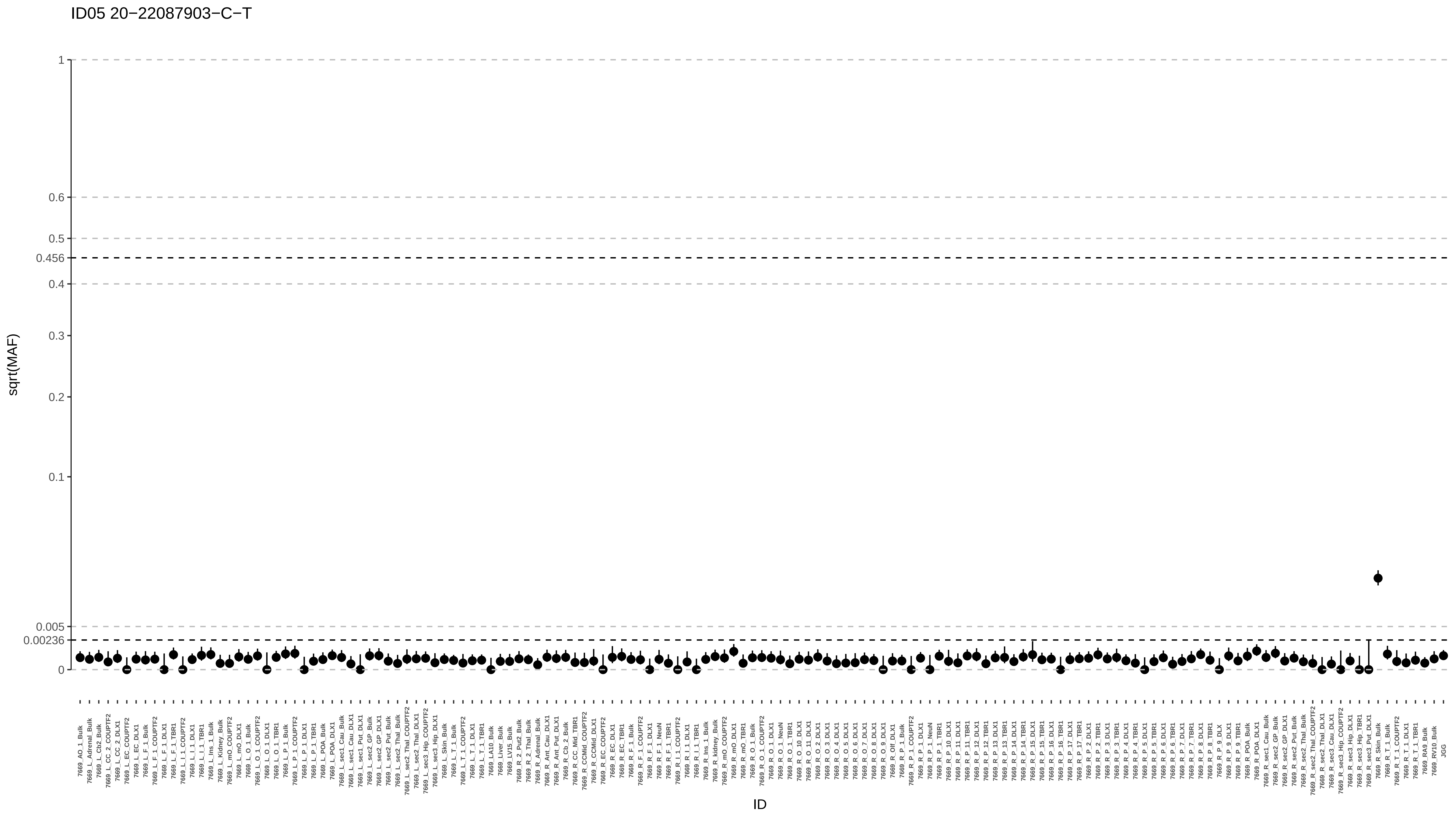

ID05 20−22087903−C−T
sqrt(MAF)
1
0.6
0.5
0.456
0.4
0.3
0.2
0.1
0.005
0.00236
0
ID

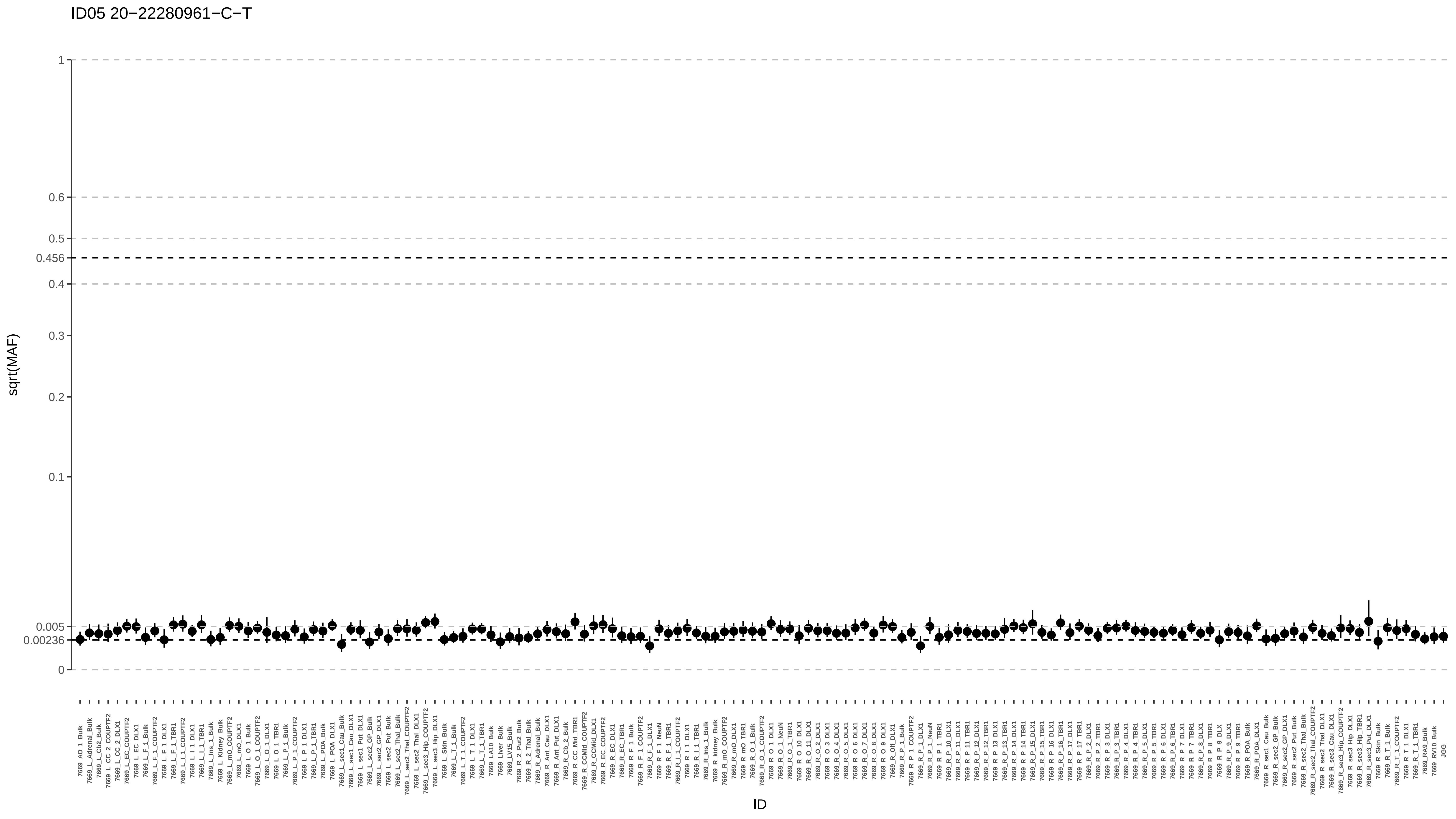
ID05 20−22280961−C−T
sqrt(MAF)
1
0.6
0.5
0.456
0.4
0.3
0.2
0.1
0.005
0.00236
0
ID
7669_AO_1_Bulk
7669_L_Adrenal_Bulk
7669_L_Cb2_Bulk
7669_L_CC_2_COUPTF2
7669_L_CC_2_DLX1
7669_L_EC_COUPTF2
7669_L_EC_DLX1
7669_L_F_1_Bulk
7669_L_F_1_COUPTF2
7669_L_F_1_DLX1
7669_L_F_1_TBR1
7669_L_I_1_COUPTF2
7669_L_I_1_DLX1
7669_L_I_1_TBR1
7669_L_Ins_1_Bulk
7669_L_Kidney_Bulk
7669_L_mO_COUPTF2
7669_L_mO_DLX1
7669_L_O_1_Bulk
7669_L_O_1_COUPTF2
7669_L_O_1_DLX1
7669_L_O_1_TBR1
7669_L_P_1_Bulk
7669_L_P_1_COUPTF2
7669_L_P_1_DLX1
7669_L_P_1_TBR1
7669_L_POA_Bulk
7669_L_POA_DLX1
7669_L_sec1_Cau_Bulk
7669_L_sec1_Cau_DLX1
7669_L_sec1_Put_DLX1
7669_L_sec2_GP_Bulk
7669_L_sec2_GP_DLX1
7669_L_sec2_Put_Bulk
7669_L_sec2_Thal_Bulk
7669_L_sec2_Thal_COUPTF2
7669_L_sec2_Thal_DLX1
7669_L_sec3_Hip_COUPTF2
7669_L_sec3_Hip_DLX1
7669_L_Skin_Bulk
7669_L_T_1_Bulk
7669_L_T_1_COUPTF2
7669_L_T_1_DLX1
7669_L_T_1_TBR1
7669_LA10_Bulk
7669_Liver_Bulk
7669_LV15_Bulk
7669_R_2_Put2_Bulk
7669_R_2_Thal_Bulk
7669_R_Adrenal_Bulk
7669_R_Ant_Cau_DLX1
7669_R_Ant_Put_DLX1
7669_R_Cb_2_Bulk
7669_R_CC_Mid_TBR1
7669_R_CCMid_COUPTF2
7669_R_CCMid_DLX1
7669_R_EC_COUPTF2
7669_R_EC_DLX1
7669_R_EC_TBR1
7669_R_F_1_Bulk
7669_R_F_1_COUPTF2
7669_R_F_1_DLX1
7669_R_F_1_NeuN
7669_R_F_1_TBR1
7669_R_I_1_COUPTF2
7669_R_I_1_DLX1
7669_R_I_1_TBR1
7669_R_Ins_1_Bulk
7669_R_kidney_Bulk
7669_R_mO_COUPTF2
7669_R_mO_DLX1
7669_R_mO_TBR1
7669_R_O_1_Bulk
7669_R_O_1_COUPTF2
7669_R_O_1_DLX1
7669_R_O_1_NeuN
7669_R_O_1_TBR1
7669_R_O_10_DLX1
7669_R_O_11_DLX1
7669_R_O_2_DLX1
7669_R_O_3_DLX1
7669_R_O_4_DLX1
7669_R_O_5_DLX1
7669_R_O_6_DLX1
7669_R_O_7_DLX1
7669_R_O_8_DLX1
7669_R_O_9_DLX1
7669_R_Olf_DLX1
7669_R_P_1_Bulk
7669_R_P_1_COUPTF2
7669_R_P_1_DLX1
7669_R_P_1_NeuN
7669_R_P_1_TBR1
7669_R_P_10_DLX1
7669_R_P_11_DLX1
7669_R_P_11_TBR1
7669_R_P_12_DLX1
7669_R_P_12_TBR1
7669_R_P_13_DLX1
7669_R_P_13_TBR1
7669_R_P_14_DLX1
7669_R_P_14_TBR1
7669_R_P_15_DLX1
7669_R_P_15_TBR1
7669_R_P_16_DLX1
7669_R_P_16_TBR1
7669_R_P_17_DLX1
7669_R_P_17_TBR1
7669_R_P_2_DLX1
7669_R_P_2_TBR1
7669_R_P_3_DLX1
7669_R_P_3_TBR1
7669_R_P_4_DLX1
7669_R_P_4_TBR1
7669_R_P_5_DLX1
7669_R_P_5_TBR1
7669_R_P_6_DLX1
7669_R_P_6_TBR1
7669_R_P_7_DLX1
7669_R_P_7_TBR1
7669_R_P_8_DLX1
7669_R_P_8_TBR1
7669_R_P_9_DLX
7669_R_P_9_DLX1
7669_R_P_9_TBR1
7669_R_POA_Bulk
7669_R_POA_DLX1
7669_R_sec1_Cau_Bulk
7669_R_sec2_GP_Bulk
7669_R_sec2_GP_DLX1
7669_R_sec2_Put_Bulk
7669_R_sec2_Thal_Bulk
7669_R_sec2_Thal_COUPTF2
7669_R_sec2_Thal_DLX1
7669_R_sec3_Cau_DLX1
7669_R_sec3_Hip_COUPTF2
7669_R_sec3_Hip_DLX1
7669_R_sec3_Hip_TBR1
7669_R_sec3_Put_DLX1
7669_R_Skin_Bulk
7669_R_T_1_Bulk
7669_R_T_1_COUPTF2
7669_R_T_1_DLX1
7669_R_T_1_TBR1
7669_RA9_Bulk
7669_RV10_Bulk
JGG

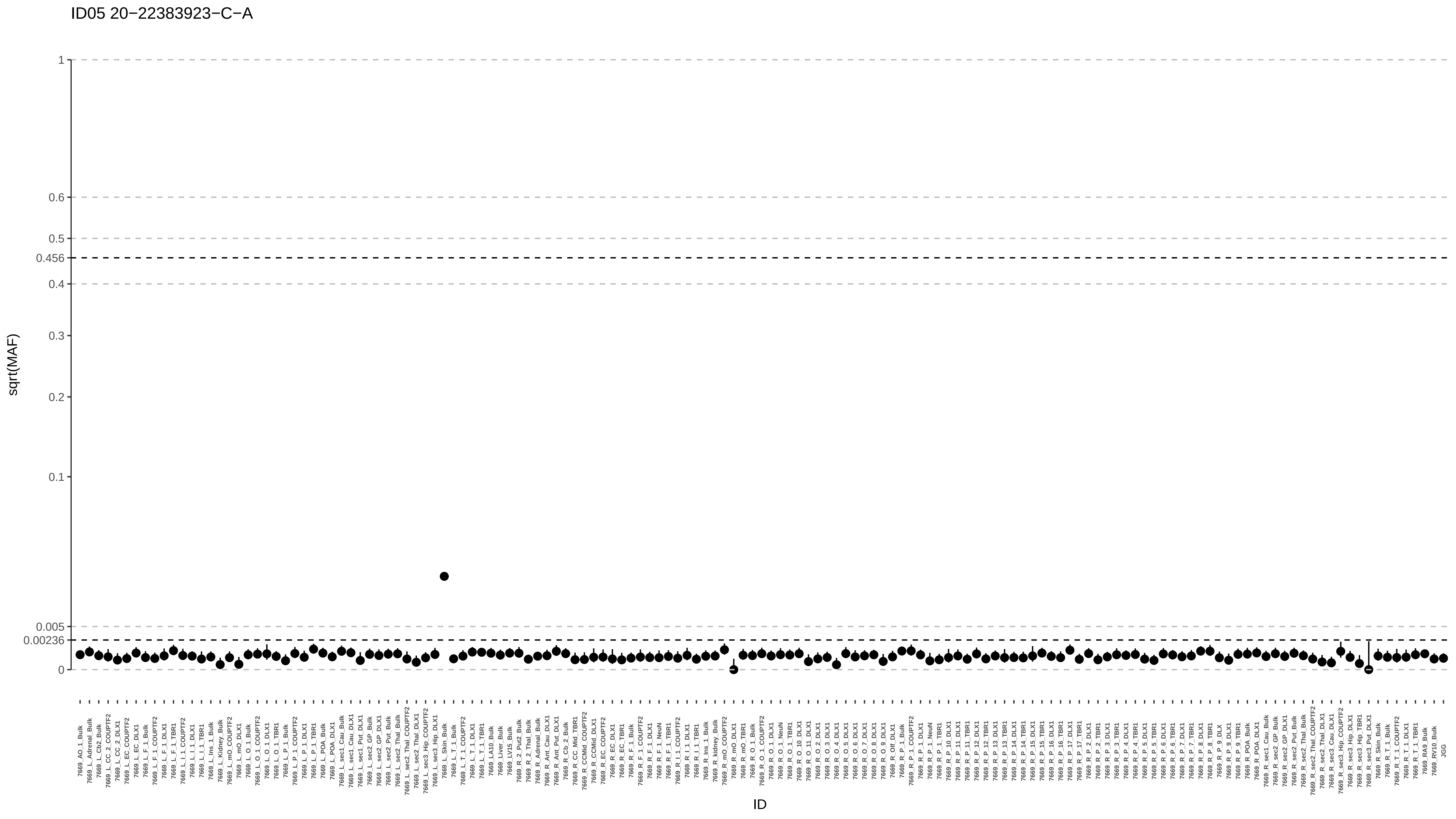
ID05 20−22383923−C−A
sqrt(MAF)
1
0.6
0.5
0.456
0.4
0.3
0.2
0.1
0.005
0.00236
0
ID

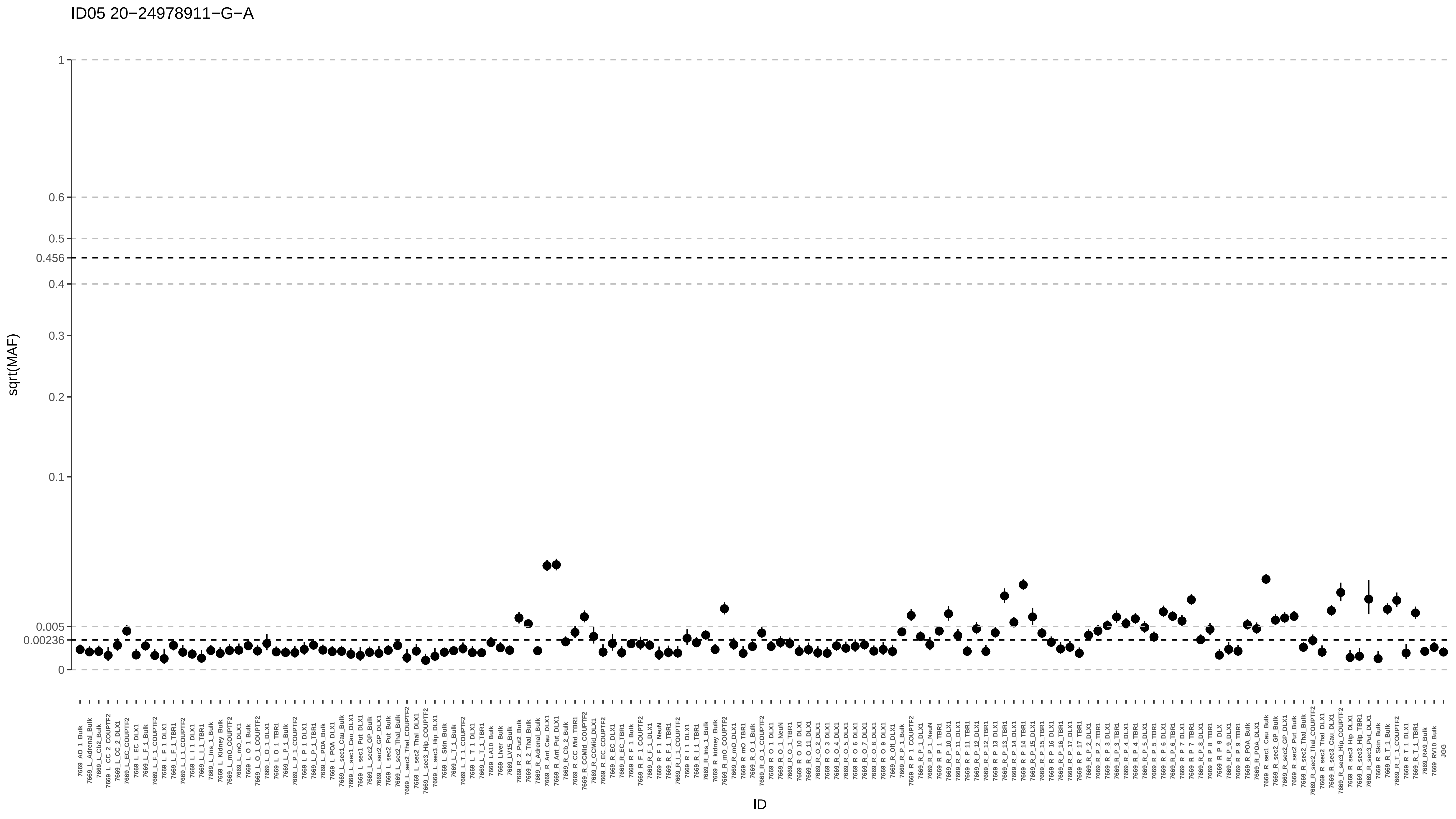
ID05 20−24978911−G−A
sqrt(MAF)
1
0.6
0.5
0.456
0.4
0.3
0.2
0.1
0.005
0.00236
0
ID

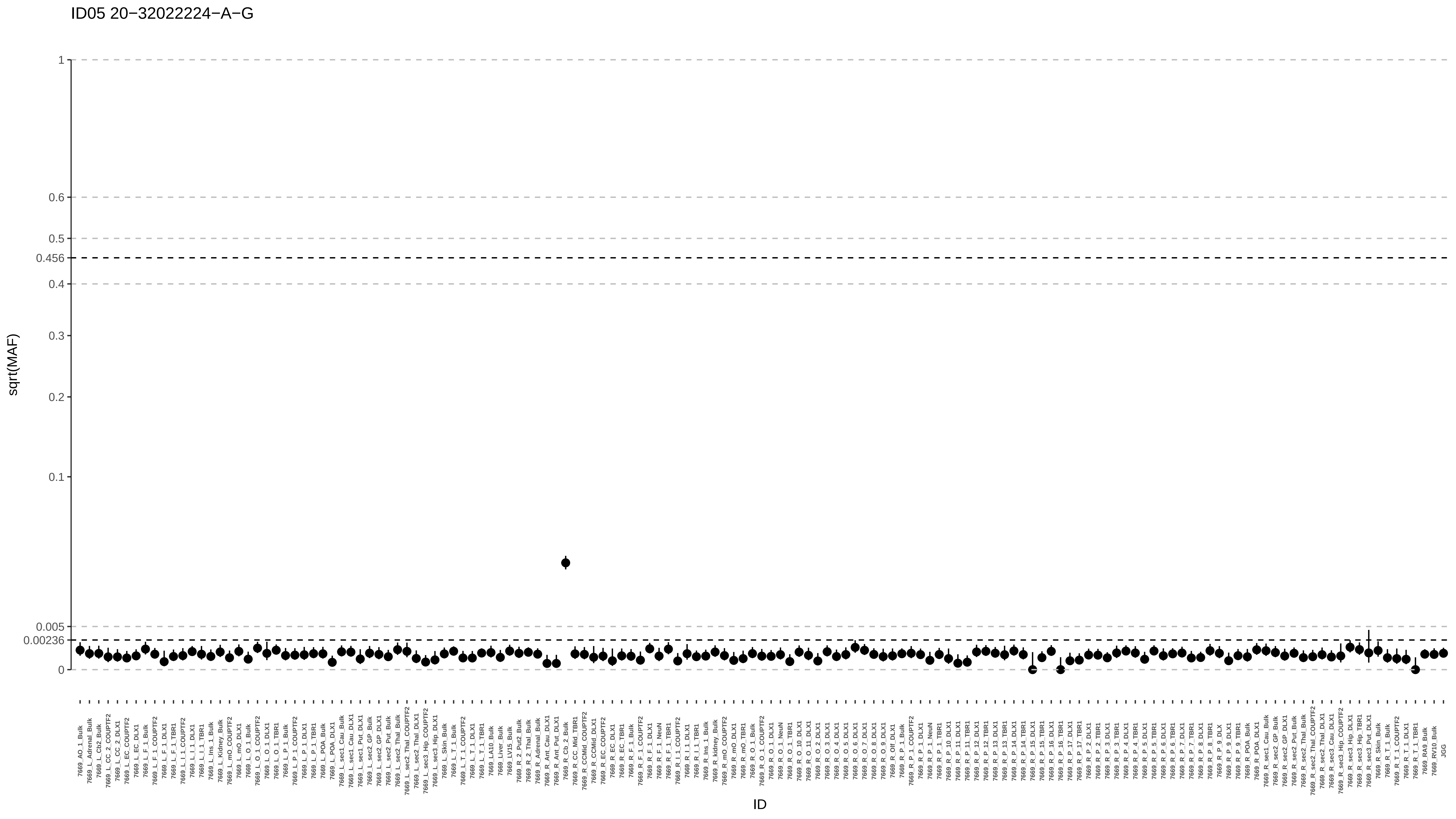
ID05 20−32022224−A−G
sqrt(MAF)
1
0.6
0.5
0.456
0.4
0.3
0.2
0.1
0.005
0.00236
0
ID

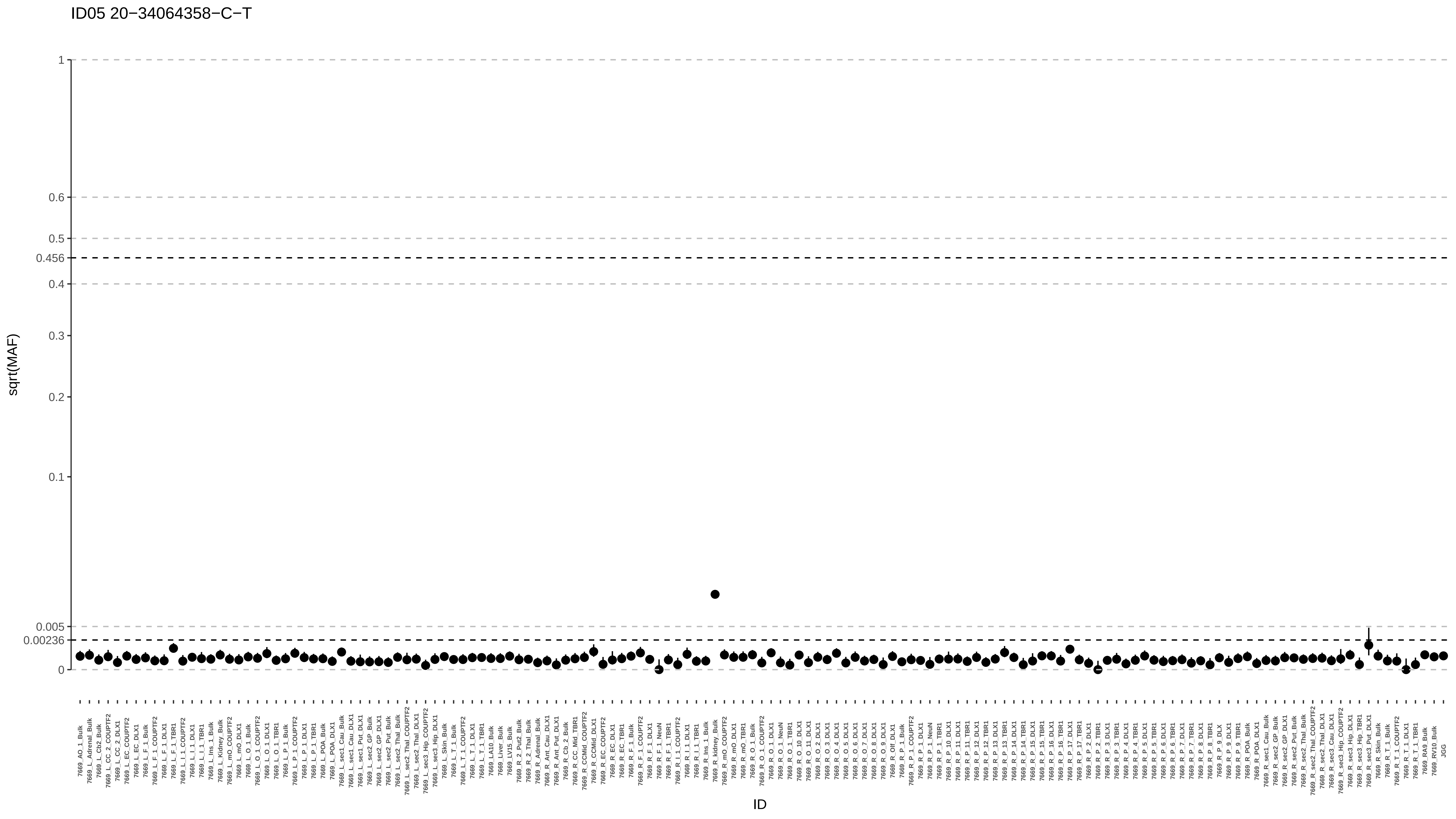

ID05 20−34064358−C−T
sqrt(MAF)
1
0.6
0.5
0.456
0.4
0.3
0.2
0.1
0.005
0.00236
0
ID

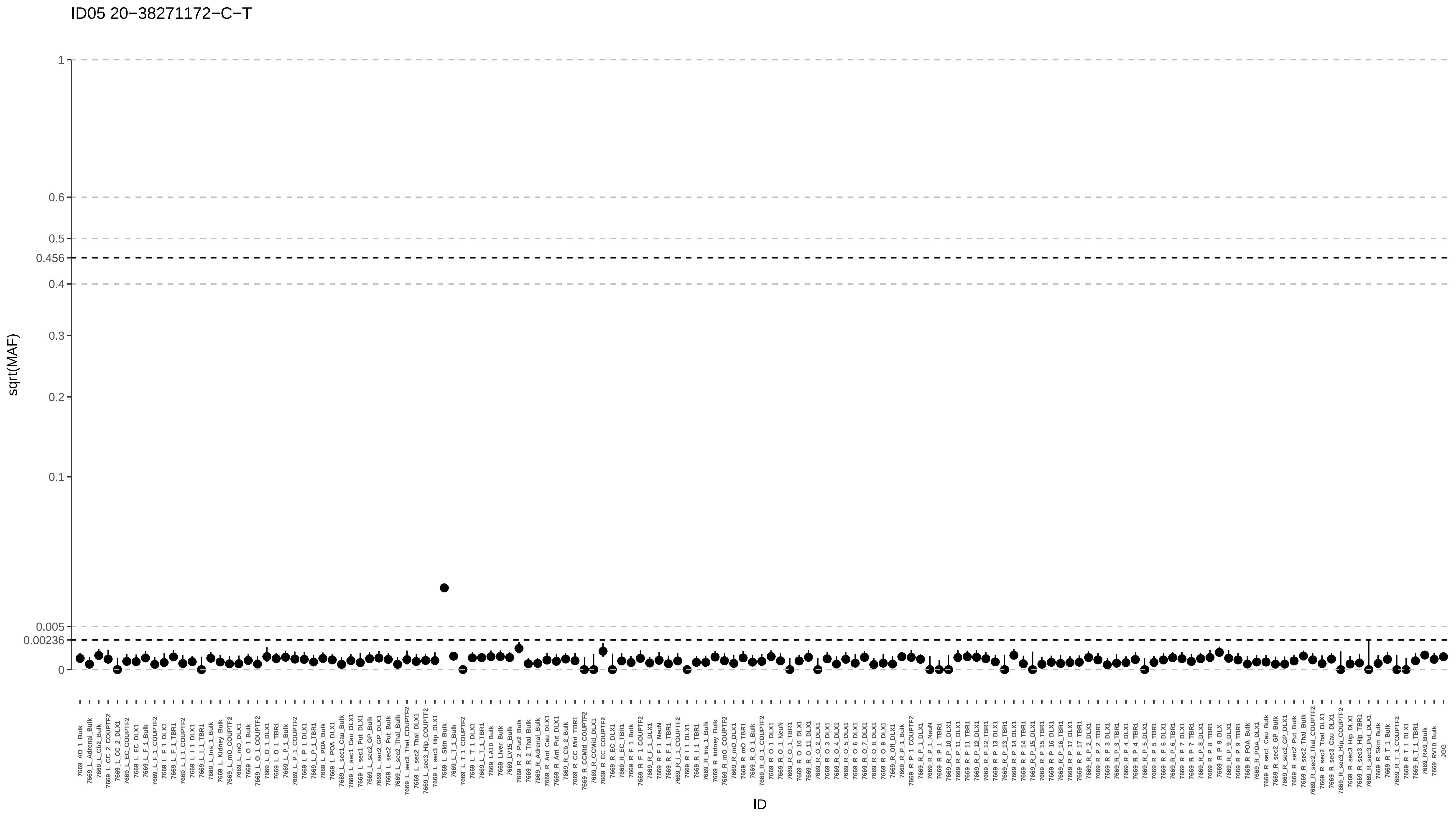
ID05 20−38271172−C−T
sqrt(MAF)
1
0.6
0.5
0.456
0.4
0.3
0.2
0.1
0.005
0.00236
0
ID

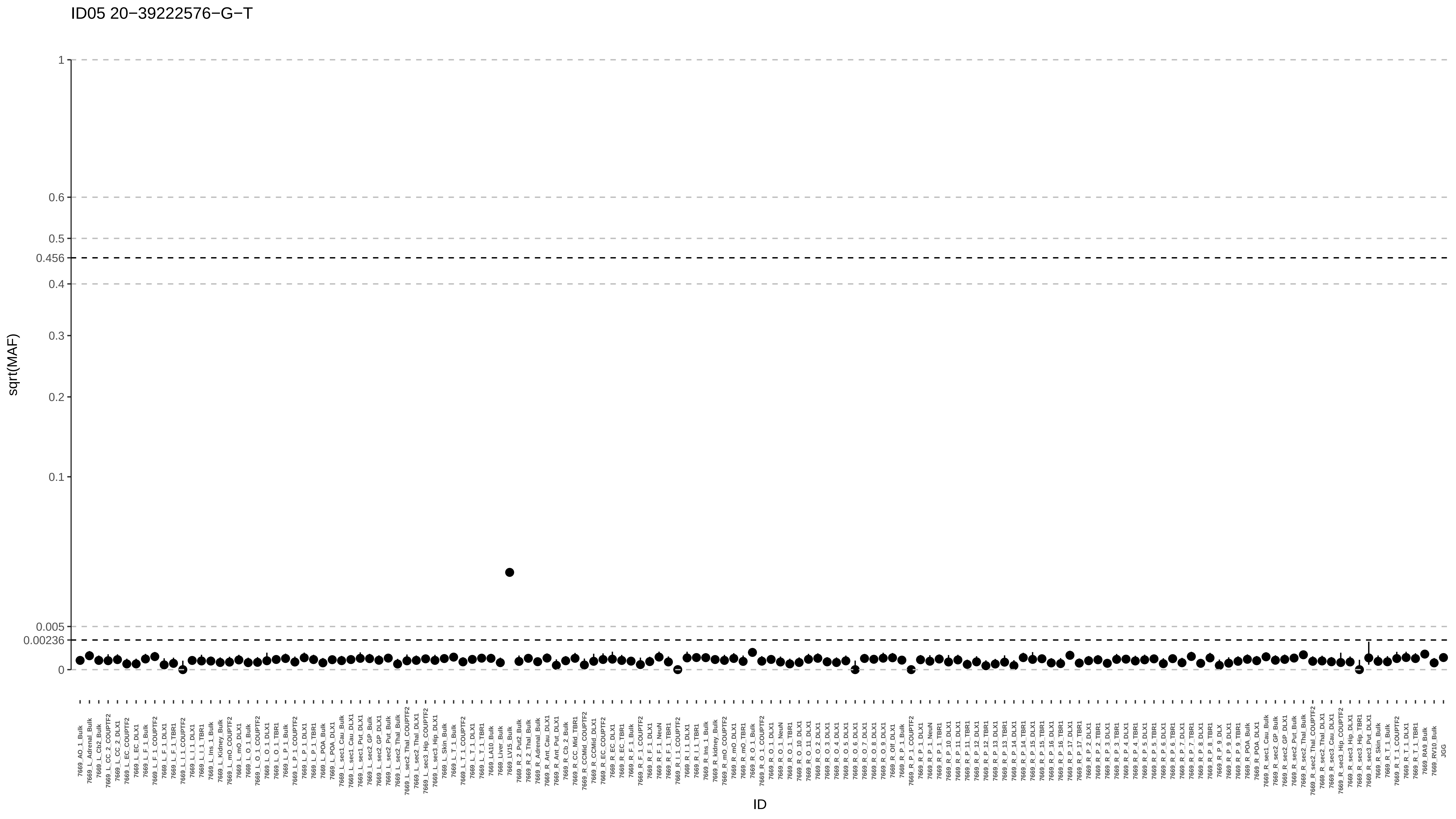
ID05 20−39222576−G−T
sqrt(MAF)
1
0.6
0.5
0.456
0.4
0.3
0.2
0.1
0.005
0.00236
0
ID

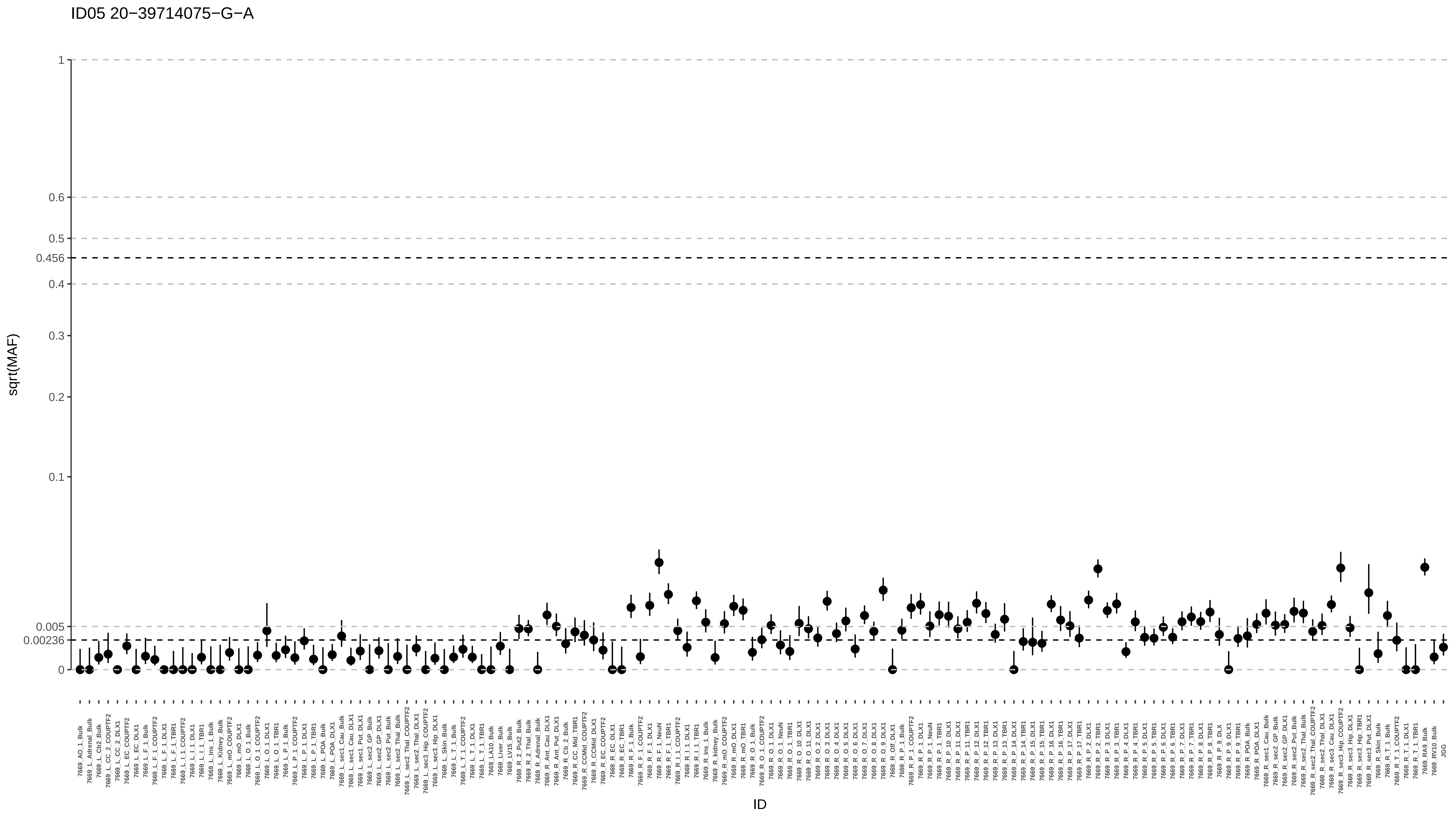

ID05 20−39714075−G−A
sqrt(MAF)
1
0.6
0.5
0.456
0.4
0.3
0.2
0.1
0.005
0.00236
0
ID

ID05 20−41427297−T−C

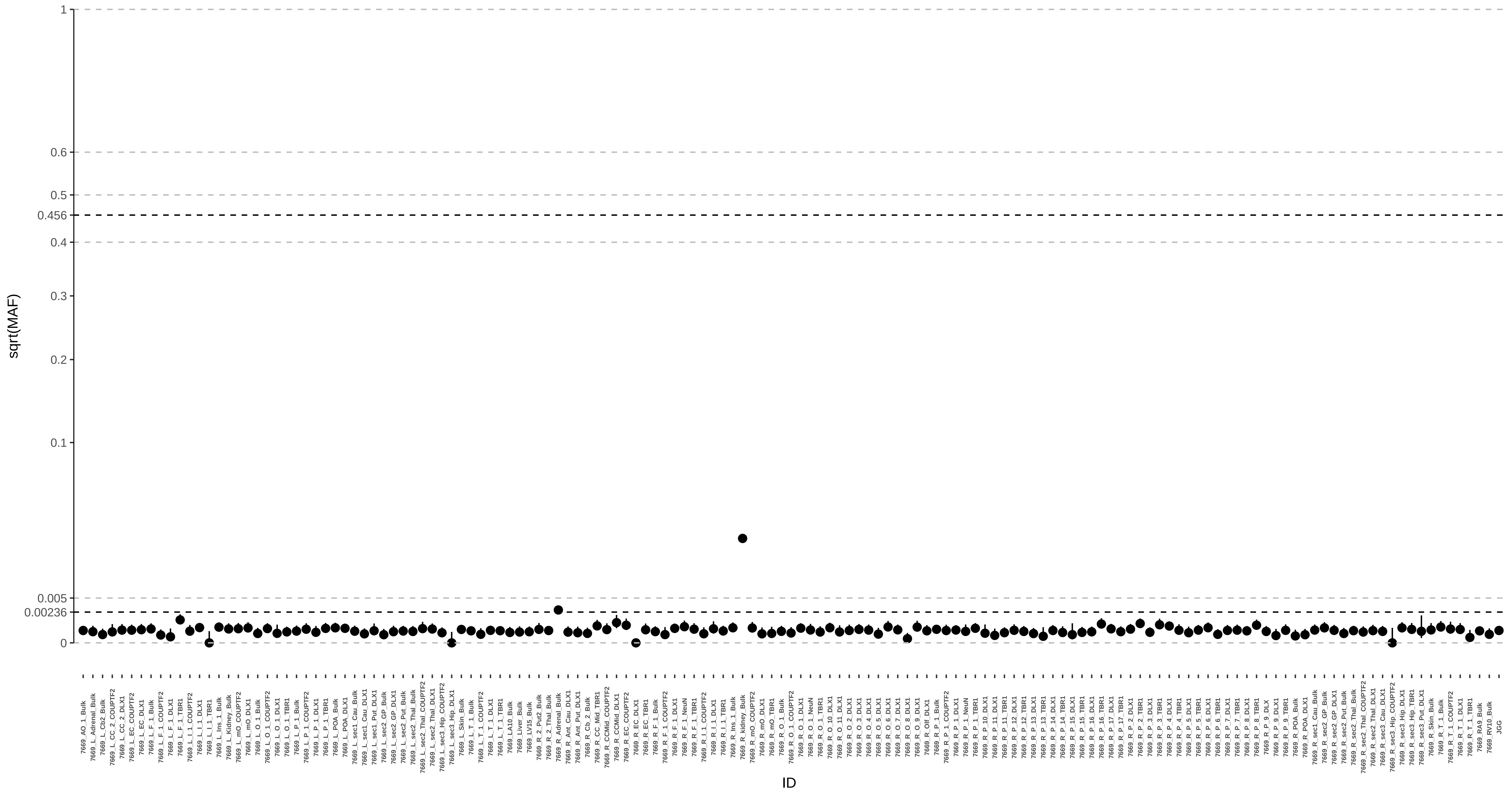

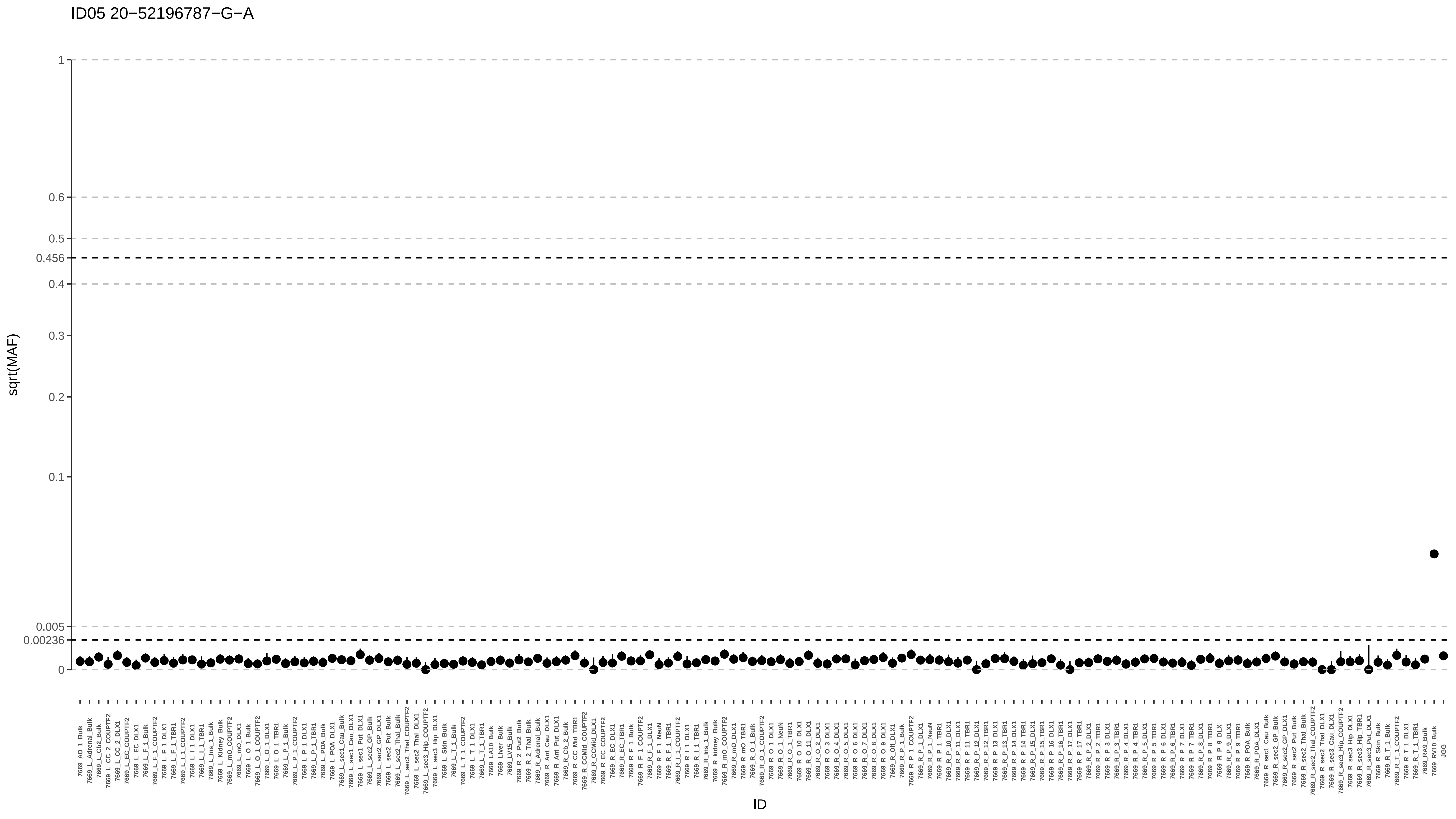
ID05 20−52196787−G−A
sqrt(MAF)
1
0.6
0.5
0.456
0.4
0.3
0.2
0.1
0.005
0.00236
0
ID
7669_AO_1_Bulk
7669_L_Adrenal_Bulk
7669_L_Cb2_Bulk
7669_L_CC_2_COUPTF2
7669_L_CC_2_DLX1
7669_L_EC_COUPTF2
7669_L_EC_DLX1
7669_L_F_1_Bulk
7669_L_F_1_COUPTF2
7669_L_F_1_DLX1
7669_L_F_1_TBR1
7669_L_I_1_COUPTF2
7669_L_I_1_DLX1
7669_L_I_1_TBR1
7669_L_Ins_1_Bulk
7669_L_Kidney_Bulk
7669_L_mO_COUPTF2
7669_L_mO_DLX1
7669_L_O_1_Bulk
7669_L_O_1_COUPTF2
7669_L_O_1_DLX1
7669_L_O_1_TBR1
7669_L_P_1_Bulk
7669_L_P_1_COUPTF2
7669_L_P_1_DLX1
7669_L_P_1_TBR1
7669_L_POA_Bulk
7669_L_POA_DLX1
7669_L_sec1_Cau_Bulk
7669_L_sec1_Cau_DLX1
7669_L_sec1_Put_DLX1
7669_L_sec2_GP_Bulk
7669_L_sec2_GP_DLX1
7669_L_sec2_Put_Bulk
7669_L_sec2_Thal_Bulk
7669_L_sec2_Thal_COUPTF2
7669_L_sec2_Thal_DLX1
7669_L_sec3_Hip_COUPTF2
7669_L_sec3_Hip_DLX1
7669_L_Skin_Bulk
7669_L_T_1_Bulk
7669_L_T_1_COUPTF2
7669_L_T_1_DLX1
7669_L_T_1_TBR1
7669_LA10_Bulk
7669_Liver_Bulk
7669_LV15_Bulk
7669_R_2_Put2_Bulk
7669_R_2_Thal_Bulk
7669_R_Adrenal_Bulk
7669_R_Ant_Cau_DLX1
7669_R_Ant_Put_DLX1
7669_R_Cb_2_Bulk
7669_R_CC_Mid_TBR1
7669_R_CCMid_COUPTF2
7669_R_CCMid_DLX1
7669_R_EC_COUPTF2
7669_R_EC_DLX1
7669_R_EC_TBR1
7669_R_F_1_Bulk
7669_R_F_1_COUPTF2
7669_R_F_1_DLX1
7669_R_F_1_NeuN
7669_R_F_1_TBR1
7669_R_I_1_COUPTF2
7669_R_I_1_DLX1
7669_R_I_1_TBR1
7669_R_Ins_1_Bulk
7669_R_kidney_Bulk
7669_R_mO_COUPTF2
7669_R_mO_DLX1
7669_R_mO_TBR1
7669_R_O_1_Bulk
7669_R_O_1_COUPTF2
7669_R_O_1_DLX1
7669_R_O_1_NeuN
7669_R_O_1_TBR1
7669_R_O_10_DLX1
7669_R_O_11_DLX1
7669_R_O_2_DLX1
7669_R_O_3_DLX1
7669_R_O_4_DLX1
7669_R_O_5_DLX1
7669_R_O_6_DLX1
7669_R_O_7_DLX1
7669_R_O_8_DLX1
7669_R_O_9_DLX1
7669_R_Olf_DLX1
7669_R_P_1_Bulk
7669_R_P_1_COUPTF2
7669_R_P_1_DLX1
7669_R_P_1_NeuN
7669_R_P_1_TBR1
7669_R_P_10_DLX1
7669_R_P_11_DLX1
7669_R_P_11_TBR1
7669_R_P_12_DLX1
7669_R_P_12_TBR1
7669_R_P_13_DLX1
7669_R_P_13_TBR1
7669_R_P_14_DLX1
7669_R_P_14_TBR1
7669_R_P_15_DLX1
7669_R_P_15_TBR1
7669_R_P_16_DLX1
7669_R_P_16_TBR1
7669_R_P_17_DLX1
7669_R_P_17_TBR1
7669_R_P_2_DLX1
7669_R_P_2_TBR1
7669_R_P_3_DLX1
7669_R_P_3_TBR1
7669_R_P_4_DLX1
7669_R_P_4_TBR1
7669_R_P_5_DLX1
7669_R_P_5_TBR1
7669_R_P_6_DLX1
7669_R_P_6_TBR1
7669_R_P_7_DLX1
7669_R_P_7_TBR1
7669_R_P_8_DLX1
7669_R_P_8_TBR1
7669_R_P_9_DLX
7669_R_P_9_DLX1
7669_R_P_9_TBR1
7669_R_POA_Bulk
7669_R_POA_DLX1
7669_R_sec1_Cau_Bulk
7669_R_sec2_GP_Bulk
7669_R_sec2_GP_DLX1
7669_R_sec2_Put_Bulk
7669_R_sec2_Thal_Bulk
7669_R_sec2_Thal_COUPTF2
7669_R_sec2_Thal_DLX1
7669_R_sec3_Cau_DLX1
7669_R_sec3_Hip_COUPTF2
7669_R_sec3_Hip_DLX1
7669_R_sec3_Hip_TBR1
7669_R_sec3_Put_DLX1
7669_R_Skin_Bulk
7669_R_T_1_Bulk
7669_R_T_1_COUPTF2
7669_R_T_1_DLX1
7669_R_T_1_TBR1
7669_RA9_Bulk
7669_RV10_Bulk
JGG

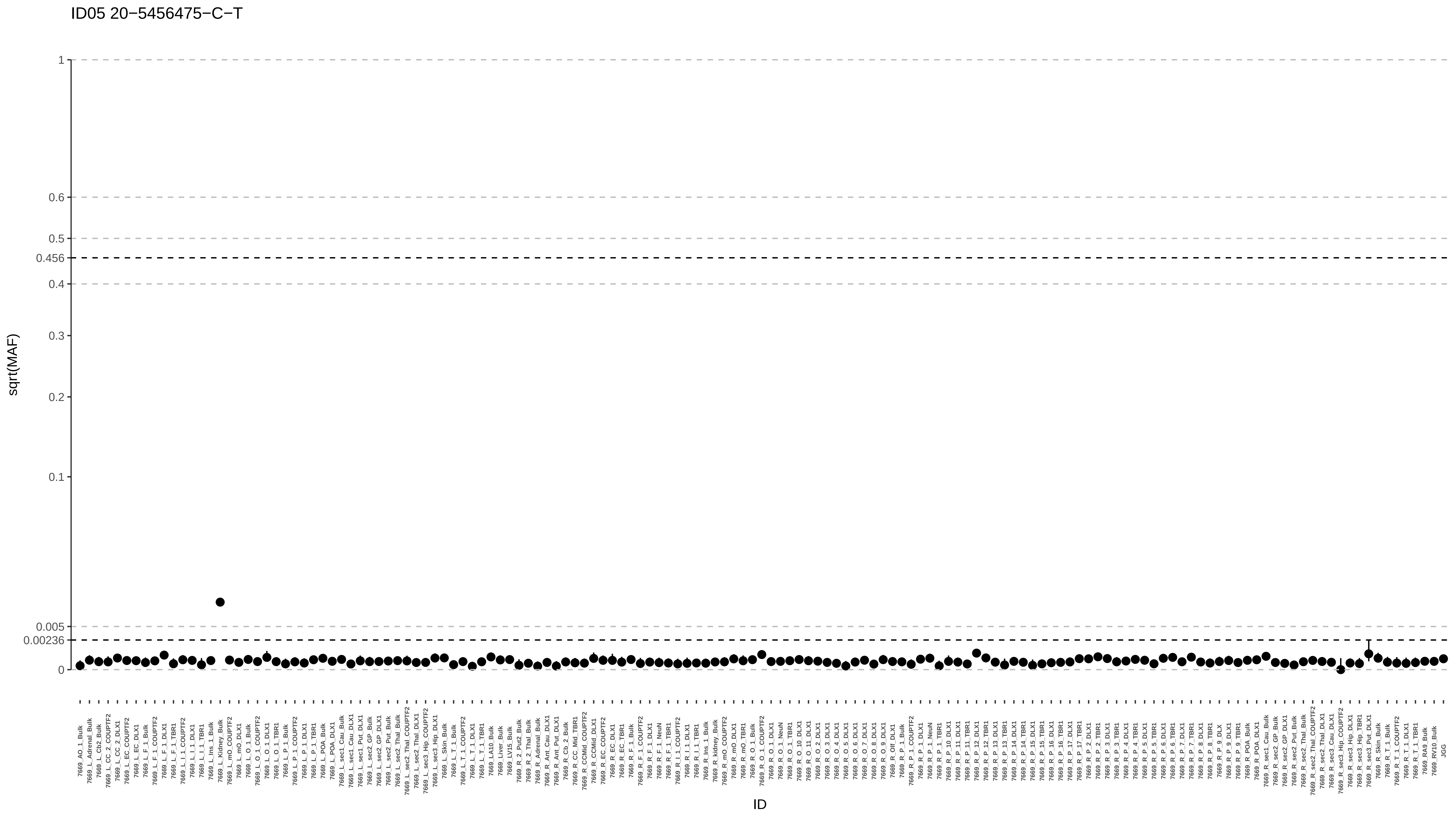

ID05 20−5456475−C−T
sqrt(MAF)
1
0.6
0.5
0.456
0.4
0.3
0.2
0.1
0.005
0.00236
0
ID

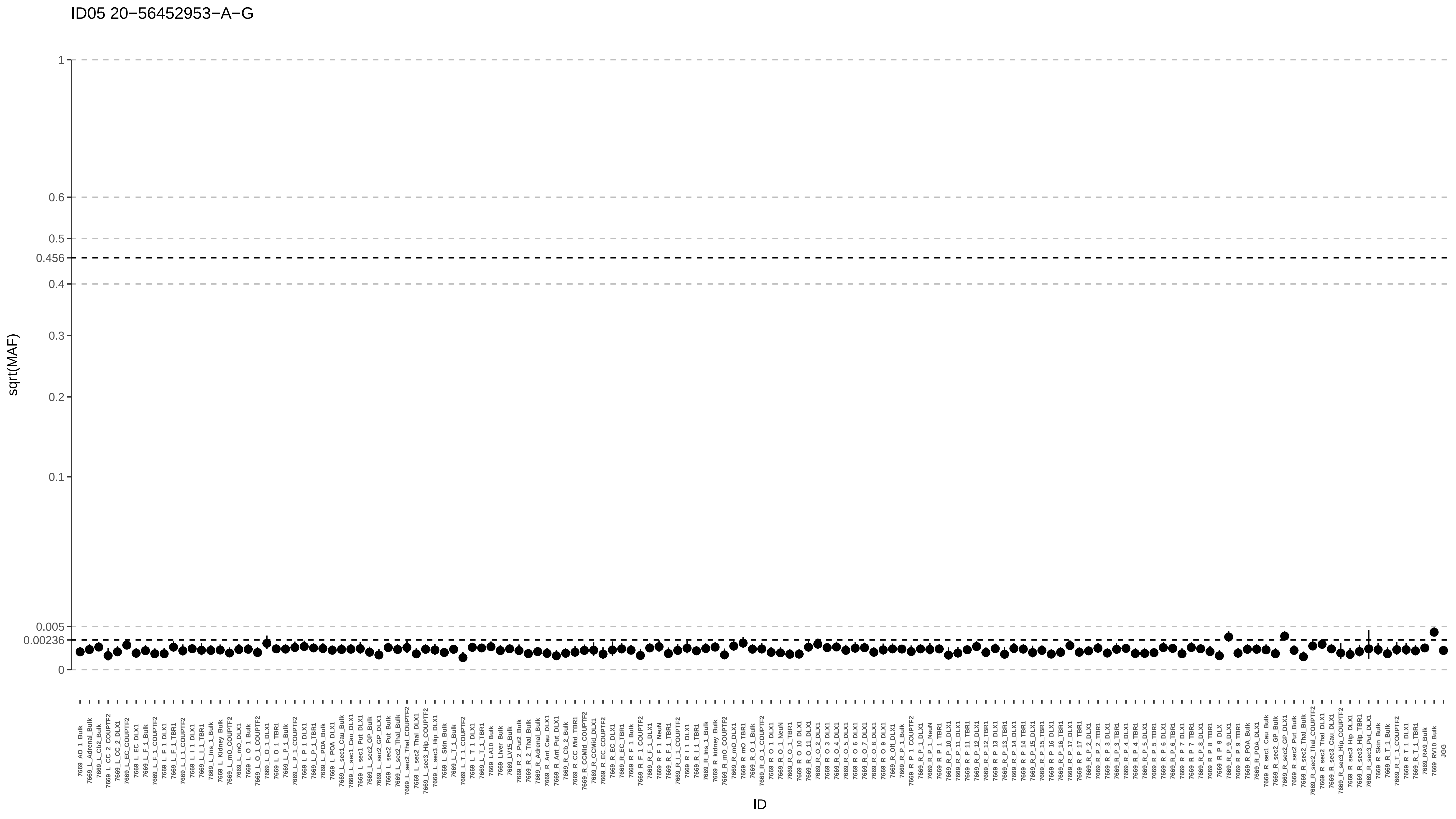
ID05 20−56452953−A−G
sqrt(MAF)
1
0.6
0.5
0.456
0.4
0.3
0.2
0.1
0.005
0.00236
0
ID

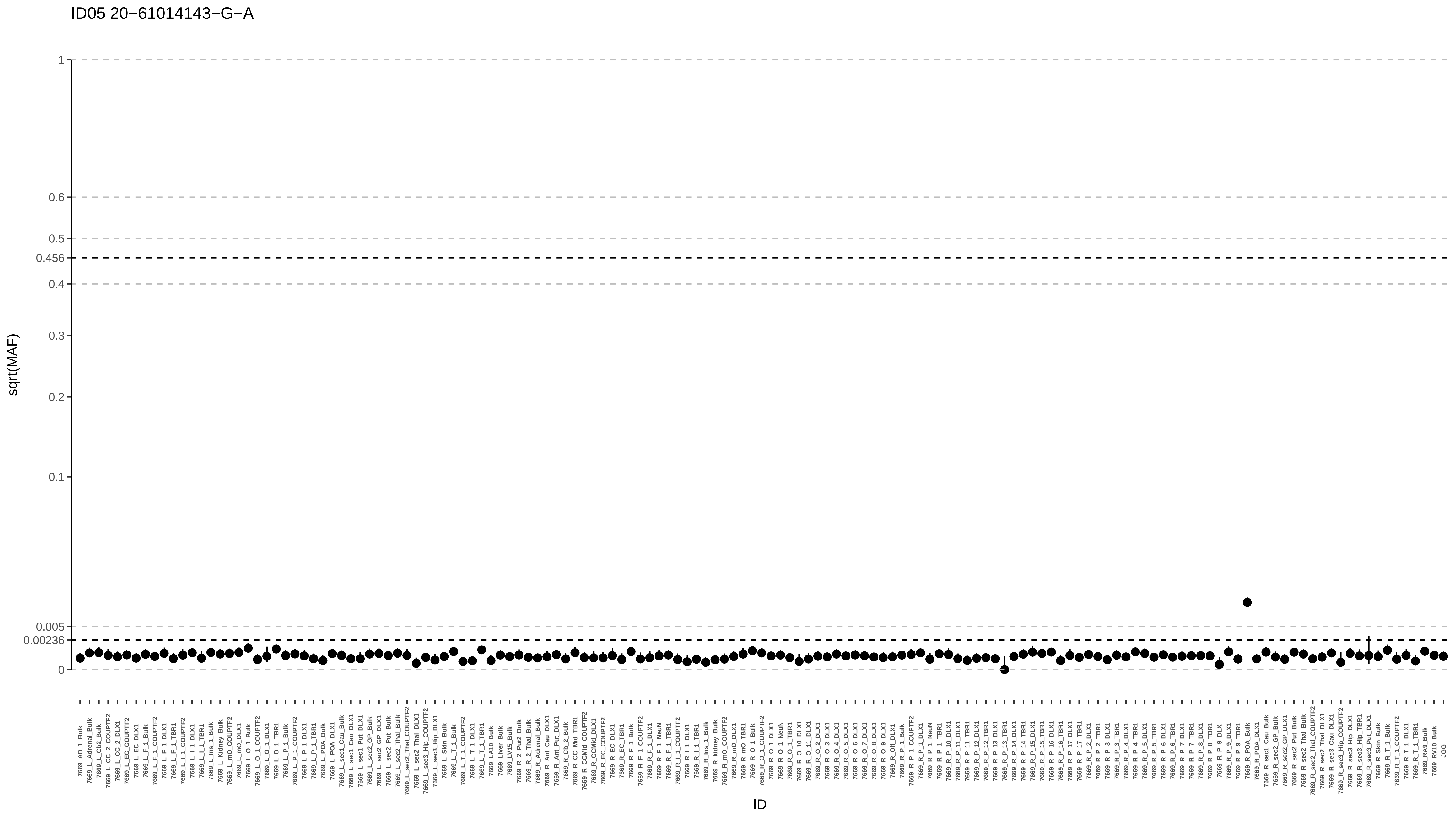

ID05 20−61014143−G−A
sqrt(MAF)
1
0.6
0.5
0.456
0.4
0.3
0.2
0.1
0.005
0.00236
0
ID

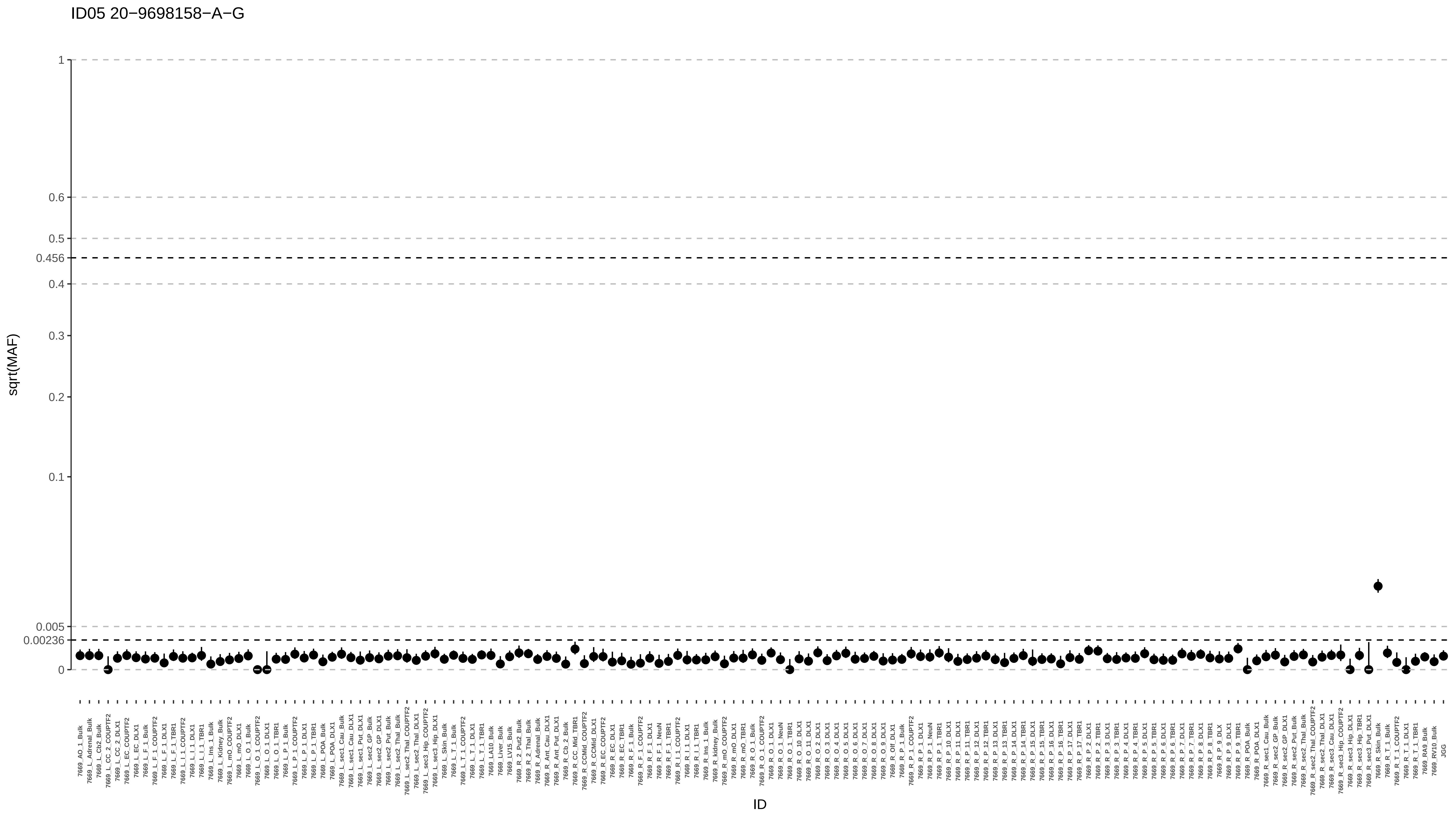

ID05 20−9698158−A−G
sqrt(MAF)
1
0.6
0.5
0.456
0.4
0.3
0.2
0.1
0.005
0.00236
0
ID

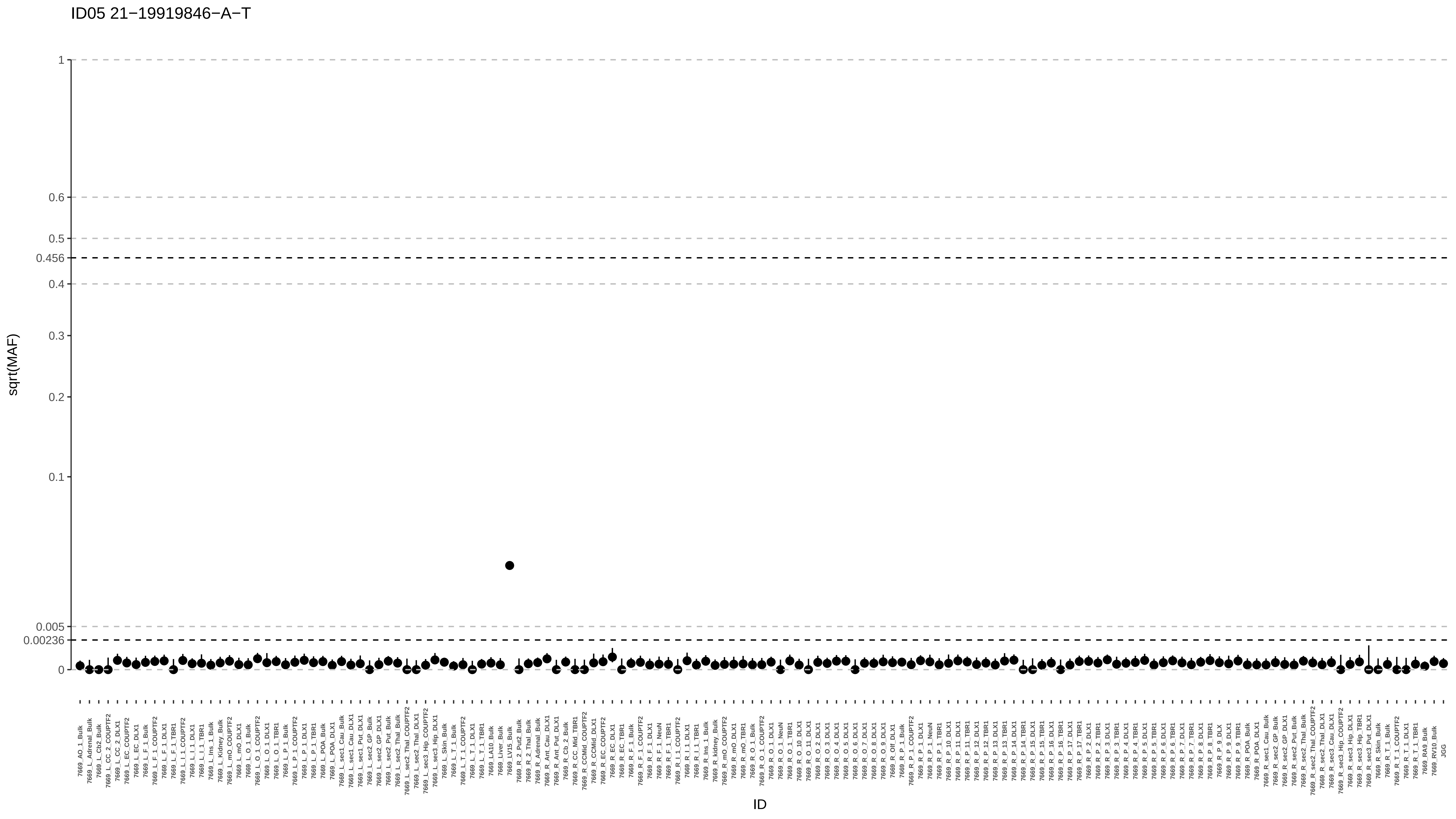
ID05 21−19919846−A−T
sqrt(MAF)
1
0.6
0.5
0.456
0.4
0.3
0.2
0.1
0.005
0.00236
0
ID

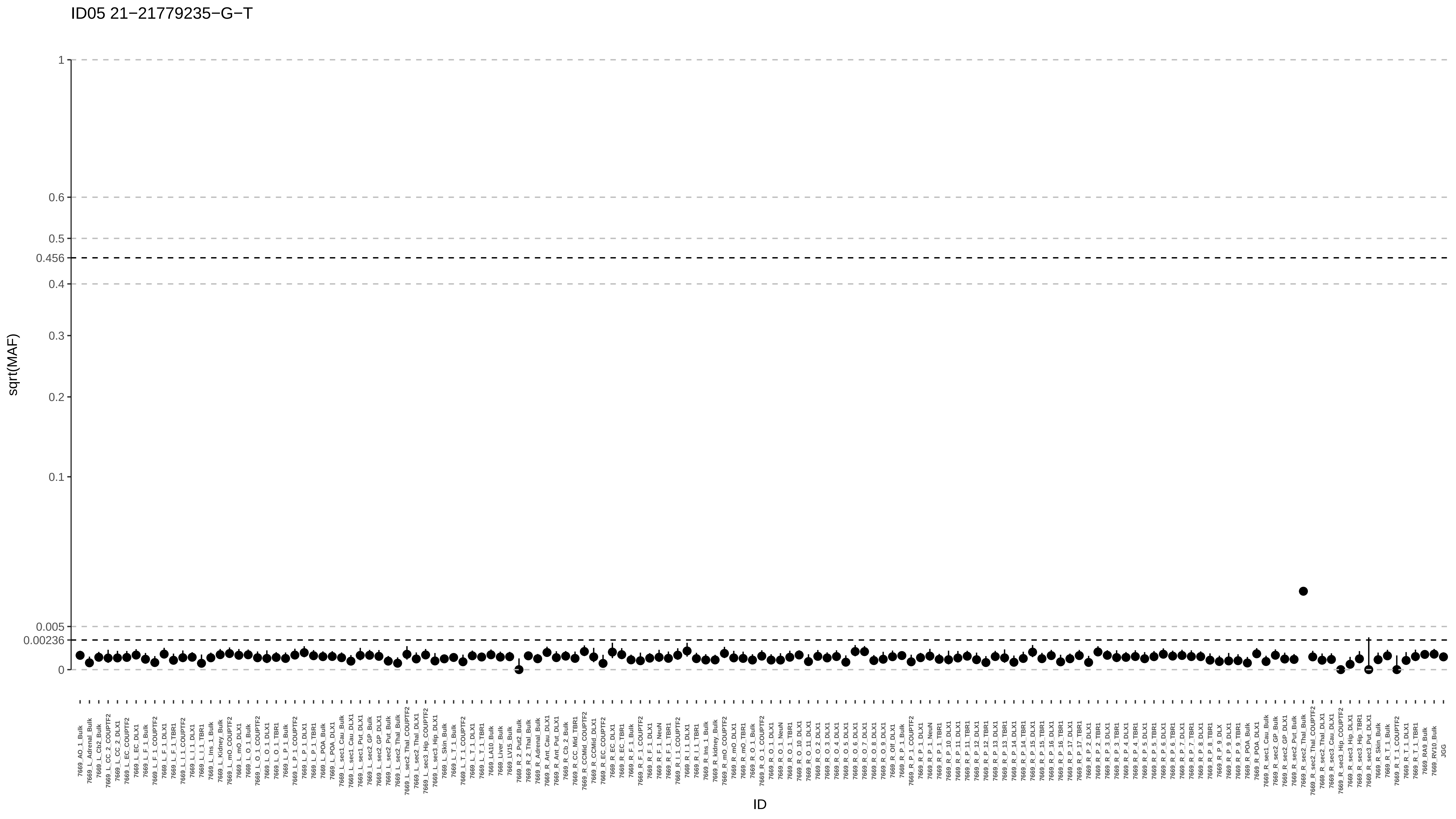

ID05 21−21779235−G−T
sqrt(MAF)
1
0.6
0.5
0.456
0.4
0.3
0.2
0.1
0.005
0.00236
0
ID
7669_AO_1_Bulk
7669_L_Adrenal_Bulk
7669_L_Cb2_Bulk
7669_L_CC_2_COUPTF2
7669_L_CC_2_DLX1
7669_L_EC_COUPTF2
7669_L_EC_DLX1
7669_L_F_1_Bulk
7669_L_F_1_COUPTF2
7669_L_F_1_DLX1
7669_L_F_1_TBR1
7669_L_I_1_COUPTF2
7669_L_I_1_DLX1
7669_L_I_1_TBR1
7669_L_Ins_1_Bulk
7669_L_Kidney_Bulk
7669_L_mO_COUPTF2
7669_L_mO_DLX1
7669_L_O_1_Bulk
7669_L_O_1_COUPTF2
7669_L_O_1_DLX1
7669_L_O_1_TBR1
7669_L_P_1_Bulk
7669_L_P_1_COUPTF2
7669_L_P_1_DLX1
7669_L_P_1_TBR1
7669_L_POA_Bulk
7669_L_POA_DLX1
7669_L_sec1_Cau_Bulk
7669_L_sec1_Cau_DLX1
7669_L_sec1_Put_DLX1
7669_L_sec2_GP_Bulk
7669_L_sec2_GP_DLX1
7669_L_sec2_Put_Bulk
7669_L_sec2_Thal_Bulk
7669_L_sec2_Thal_COUPTF2
7669_L_sec2_Thal_DLX1
7669_L_sec3_Hip_COUPTF2
7669_L_sec3_Hip_DLX1
7669_L_Skin_Bulk
7669_L_T_1_Bulk
7669_L_T_1_COUPTF2
7669_L_T_1_DLX1
7669_L_T_1_TBR1
7669_LA10_Bulk
7669_Liver_Bulk
7669_LV15_Bulk
7669_R_2_Put2_Bulk
7669_R_2_Thal_Bulk
7669_R_Adrenal_Bulk
7669_R_Ant_Cau_DLX1
7669_R_Ant_Put_DLX1
7669_R_Cb_2_Bulk
7669_R_CC_Mid_TBR1
7669_R_CCMid_COUPTF2
7669_R_CCMid_DLX1
7669_R_EC_COUPTF2
7669_R_EC_DLX1
7669_R_EC_TBR1
7669_R_F_1_Bulk
7669_R_F_1_COUPTF2
7669_R_F_1_DLX1
7669_R_F_1_NeuN
7669_R_F_1_TBR1
7669_R_I_1_COUPTF2
7669_R_I_1_DLX1
7669_R_I_1_TBR1
7669_R_Ins_1_Bulk
7669_R_kidney_Bulk
7669_R_mO_COUPTF2
7669_R_mO_DLX1
7669_R_mO_TBR1
7669_R_O_1_Bulk
7669_R_O_1_COUPTF2
7669_R_O_1_DLX1
7669_R_O_1_NeuN
7669_R_O_1_TBR1
7669_R_O_10_DLX1
7669_R_O_11_DLX1
7669_R_O_2_DLX1
7669_R_O_3_DLX1
7669_R_O_4_DLX1
7669_R_O_5_DLX1
7669_R_O_6_DLX1
7669_R_O_7_DLX1
7669_R_O_8_DLX1
7669_R_O_9_DLX1
7669_R_Olf_DLX1
7669_R_P_1_Bulk
7669_R_P_1_COUPTF2
7669_R_P_1_DLX1
7669_R_P_1_NeuN
7669_R_P_1_TBR1
7669_R_P_10_DLX1
7669_R_P_11_DLX1
7669_R_P_11_TBR1
7669_R_P_12_DLX1
7669_R_P_12_TBR1
7669_R_P_13_DLX1
7669_R_P_13_TBR1
7669_R_P_14_DLX1
7669_R_P_14_TBR1
7669_R_P_15_DLX1
7669_R_P_15_TBR1
7669_R_P_16_DLX1
7669_R_P_16_TBR1
7669_R_P_17_DLX1
7669_R_P_17_TBR1
7669_R_P_2_DLX1
7669_R_P_2_TBR1
7669_R_P_3_DLX1
7669_R_P_3_TBR1
7669_R_P_4_DLX1
7669_R_P_4_TBR1
7669_R_P_5_DLX1
7669_R_P_5_TBR1
7669_R_P_6_DLX1
7669_R_P_6_TBR1
7669_R_P_7_DLX1
7669_R_P_7_TBR1
7669_R_P_8_DLX1
7669_R_P_8_TBR1
7669_R_P_9_DLX
7669_R_P_9_DLX1
7669_R_P_9_TBR1
7669_R_POA_Bulk
7669_R_POA_DLX1
7669_R_sec1_Cau_Bulk
7669_R_sec2_GP_Bulk
7669_R_sec2_GP_DLX1
7669_R_sec2_Put_Bulk
7669_R_sec2_Thal_Bulk
7669_R_sec2_Thal_COUPTF2
7669_R_sec2_Thal_DLX1
7669_R_sec3_Cau_DLX1
7669_R_sec3_Hip_COUPTF2
7669_R_sec3_Hip_DLX1
7669_R_sec3_Hip_TBR1
7669_R_sec3_Put_DLX1
7669_R_Skin_Bulk
7669_R_T_1_Bulk
7669_R_T_1_COUPTF2
7669_R_T_1_DLX1
7669_R_T_1_TBR1
7669_RA9_Bulk
7669_RV10_Bulk
JGG

ID05 21−24508933−T−C

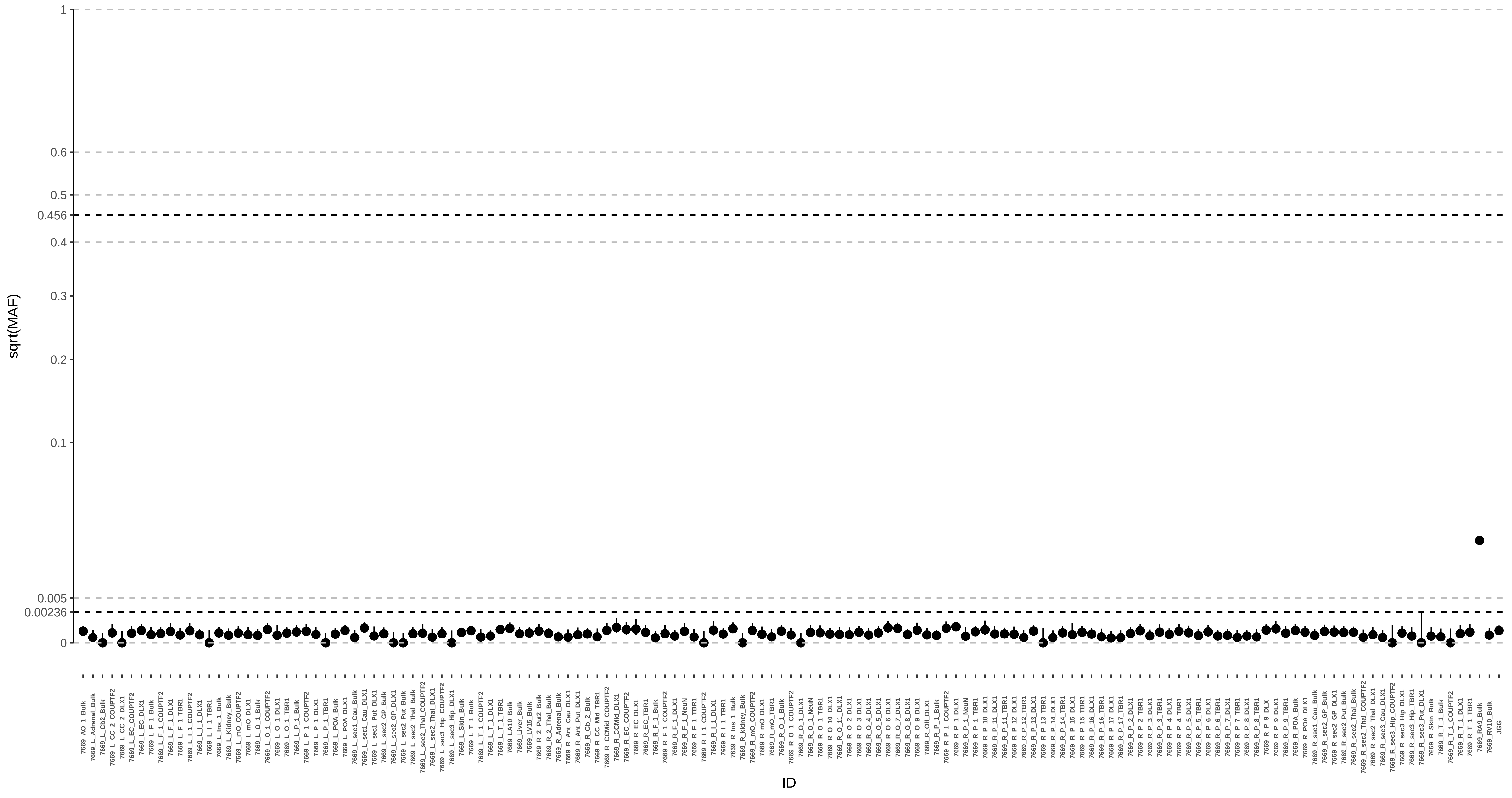

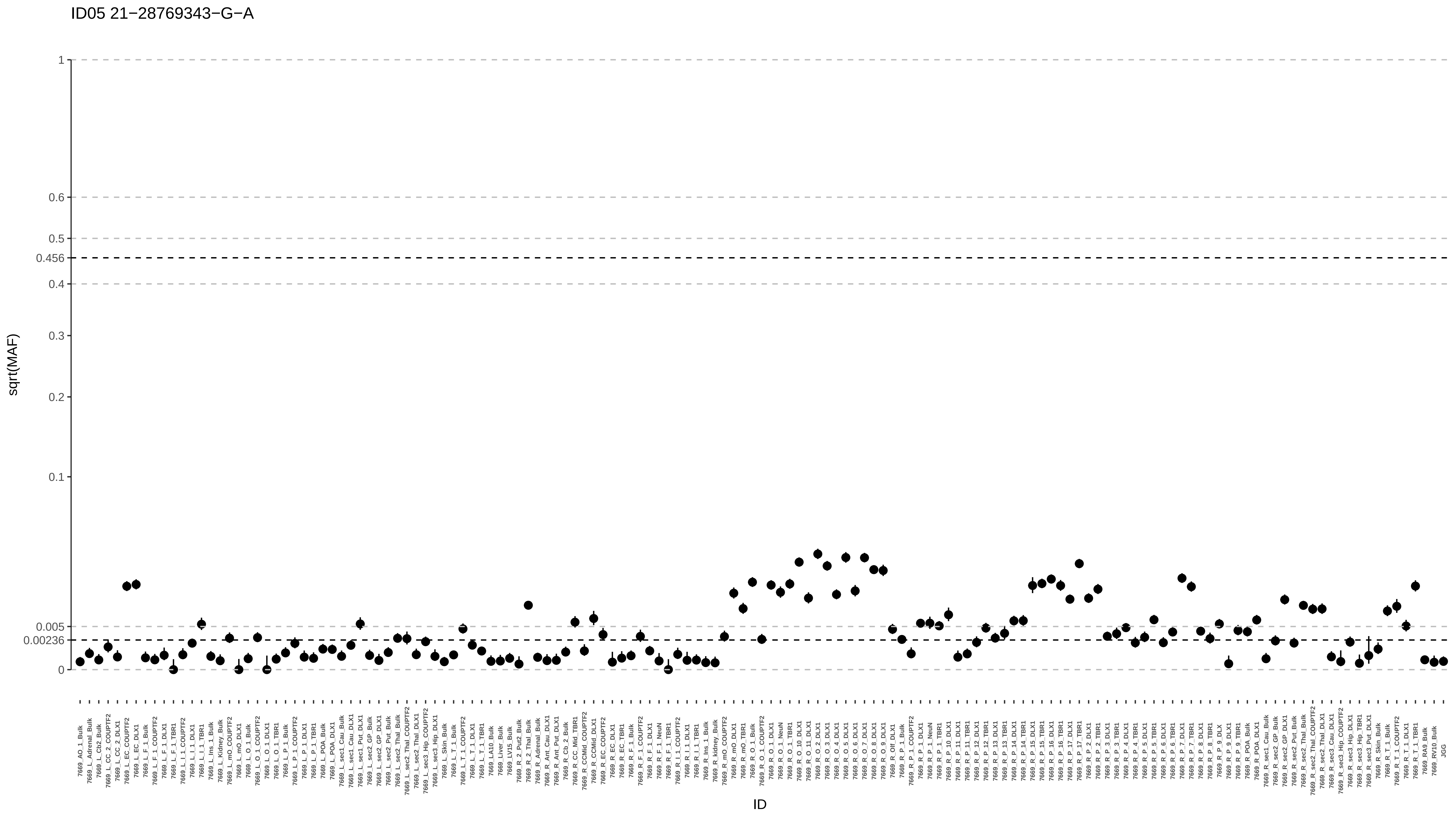
ID05 21−28769343−G−A
sqrt(MAF)
1
0.6
0.5
0.456
0.4
0.3
0.2
0.1
0.005
0.00236
0
ID

ID05 21−32316236−C−T

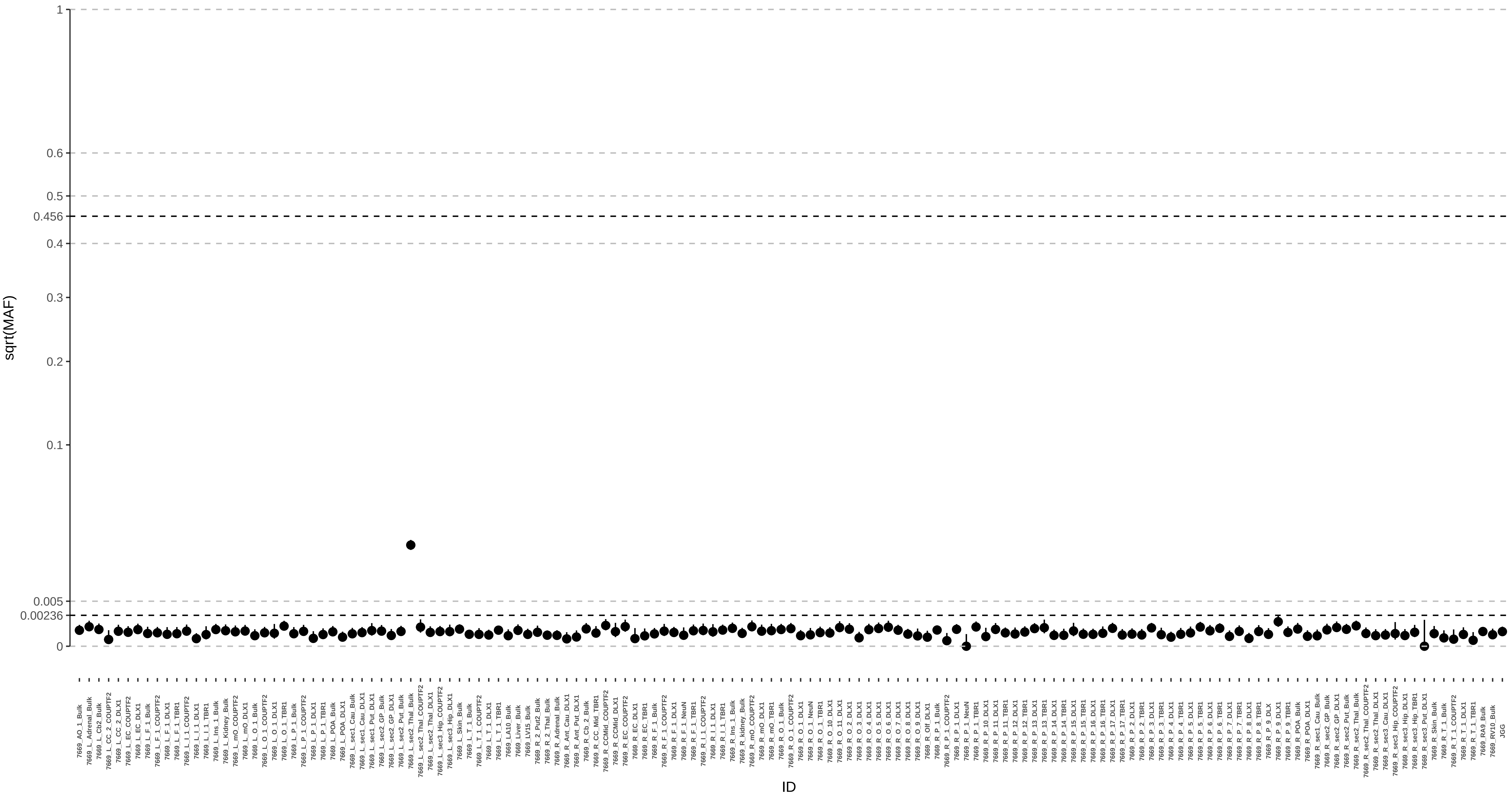

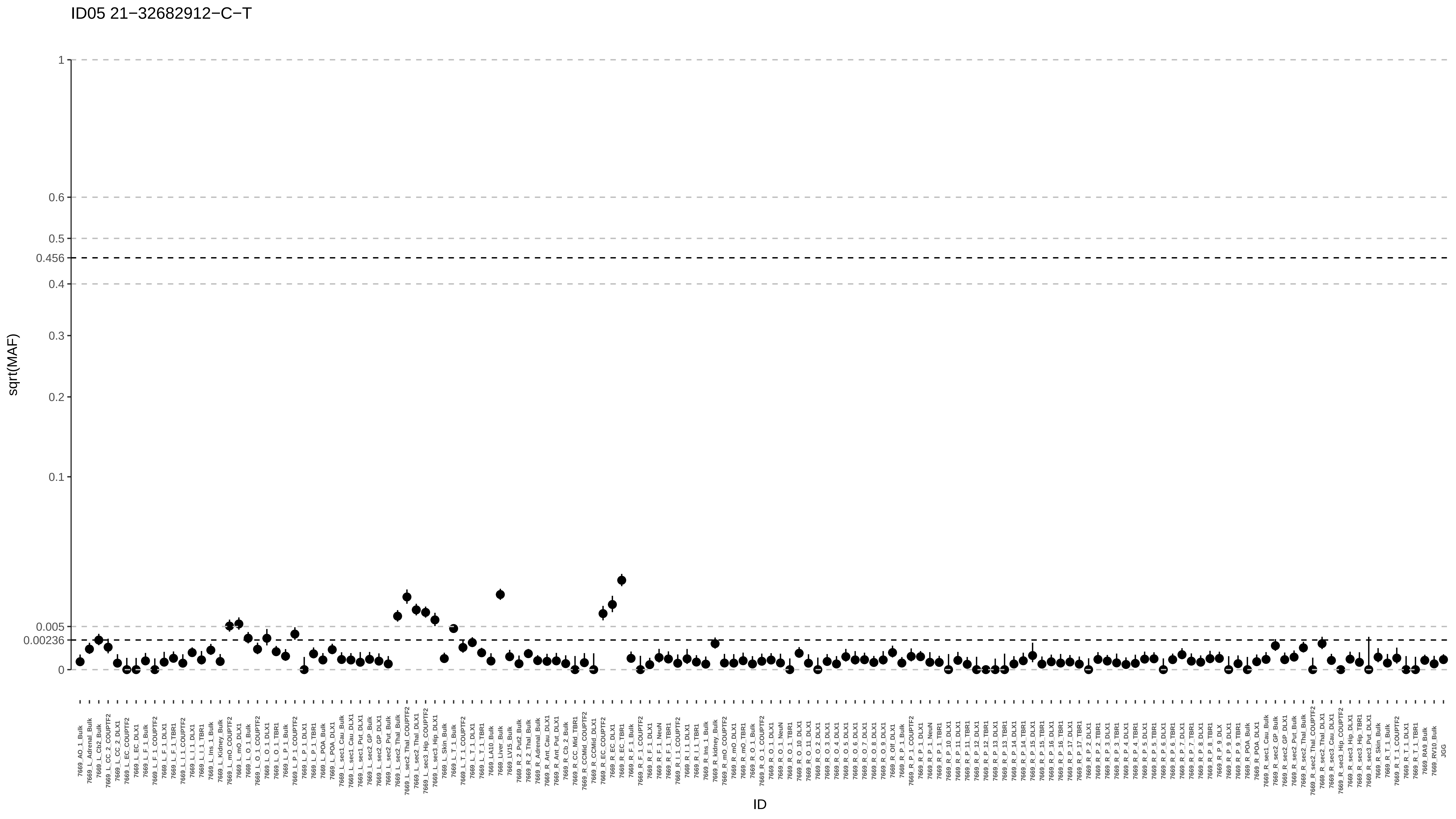
ID05 21−32682912−C−T
sqrt(MAF)
ID

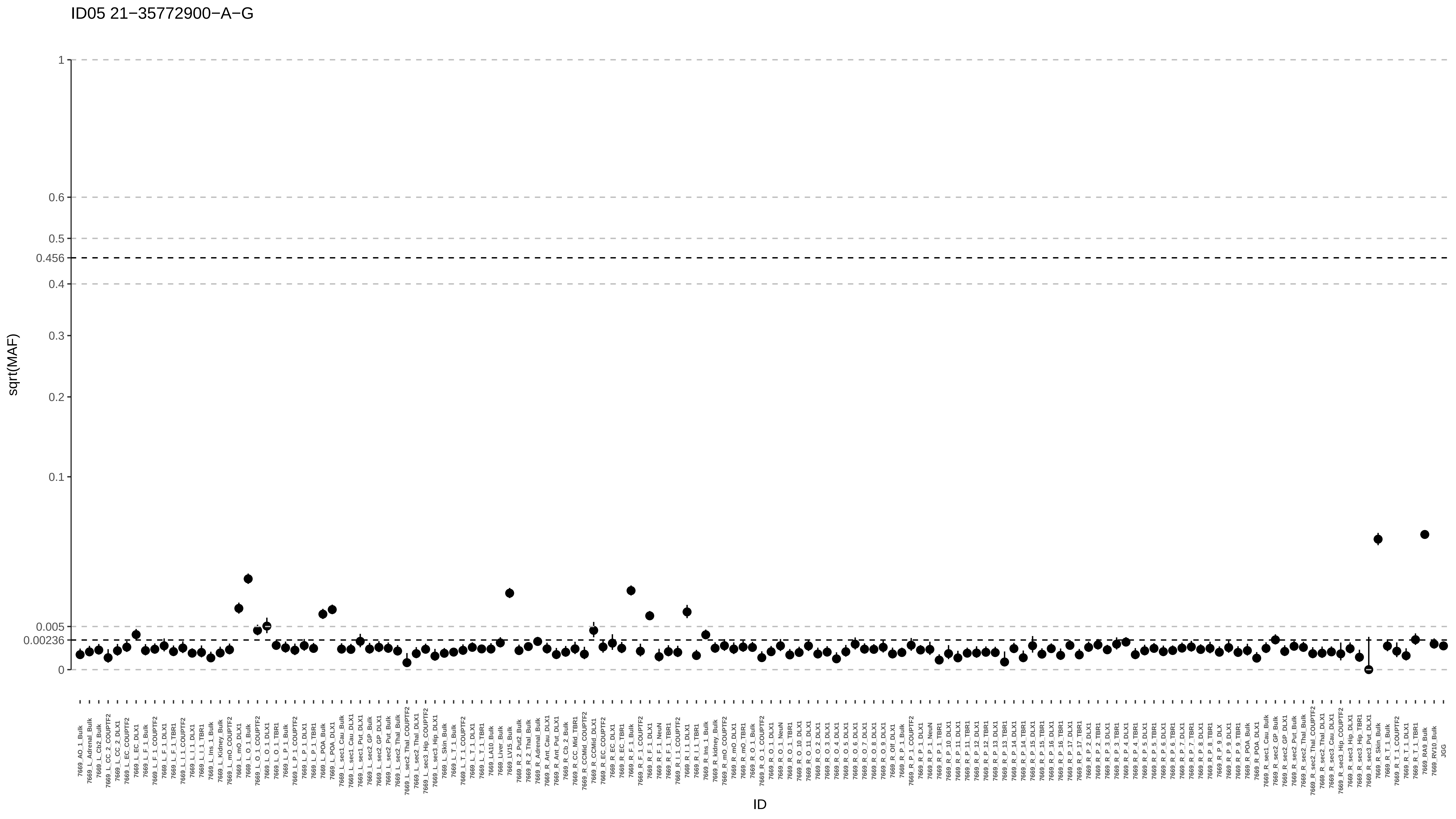
ID05 21−35772900−A−G
sqrt(MAF)
1
0.6
0.5
0.456
0.4
0.3
0.2
0.1
0.005
0.00236
0
ID

ID05 21−41023573−C−T

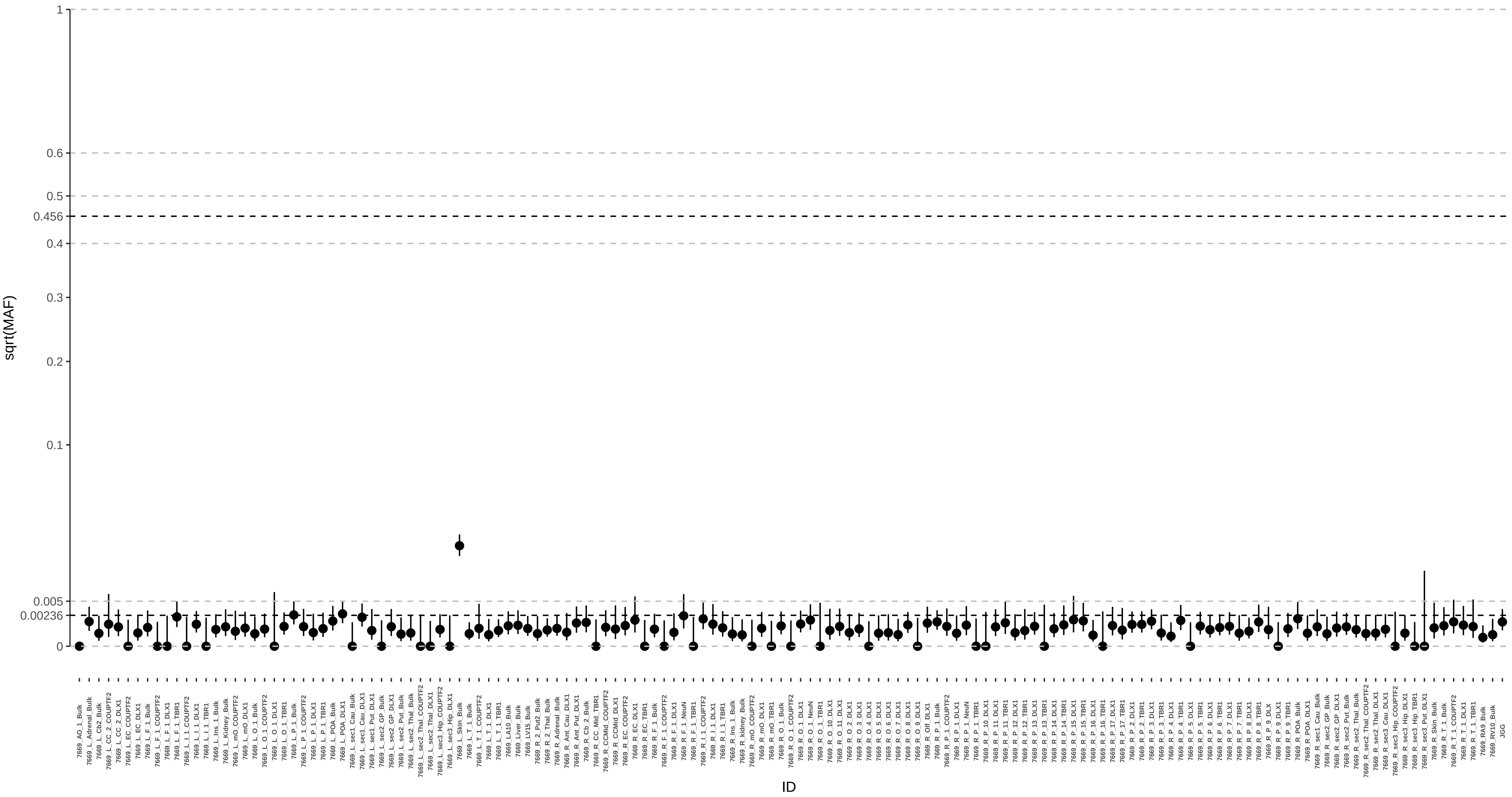

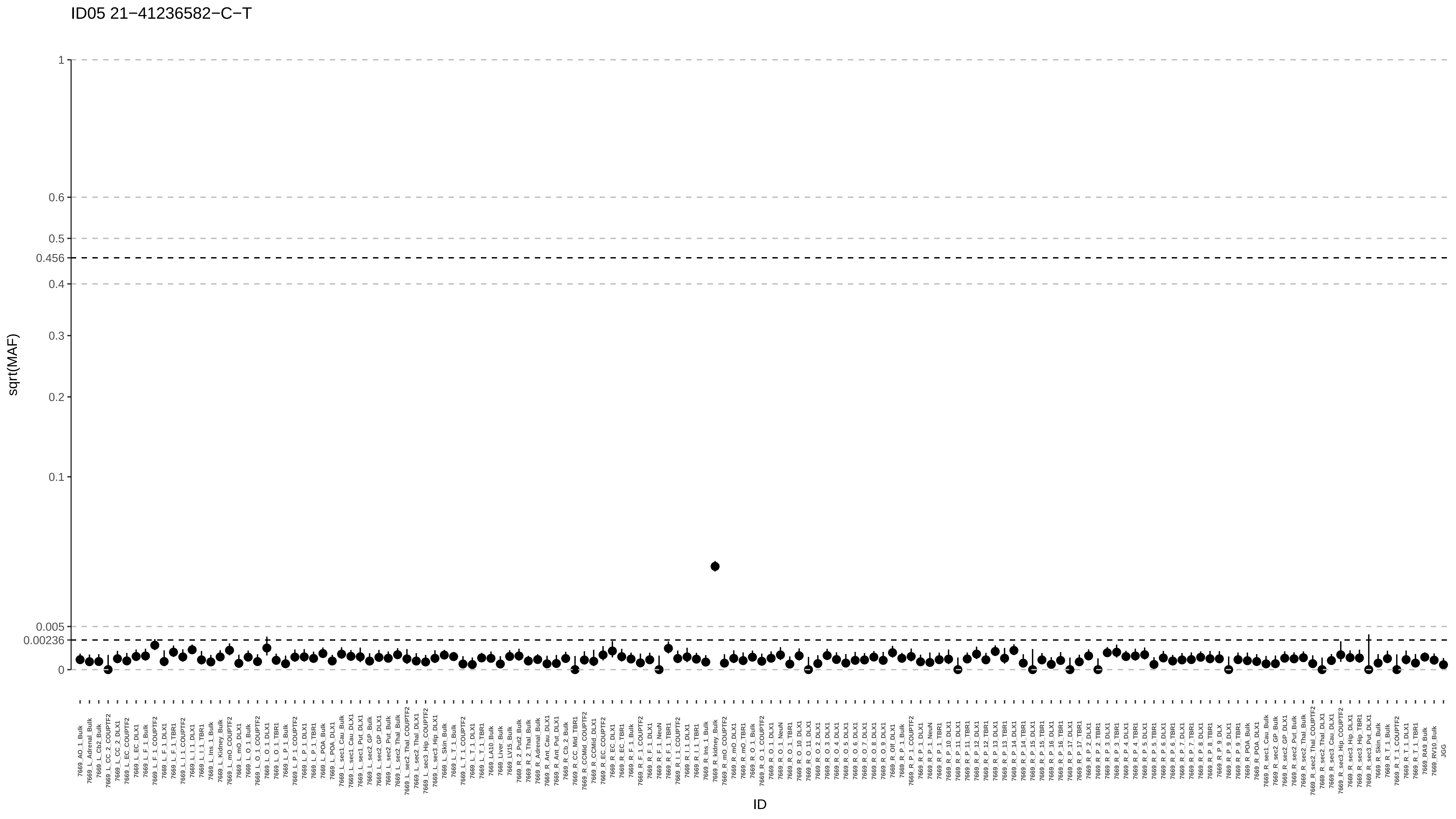
ID05 21−41236582−C−T
sqrt(MAF)
1
0.6
0.5
0.456
0.4
0.3
0.2
0.1
0.005
0.00236
0
ID

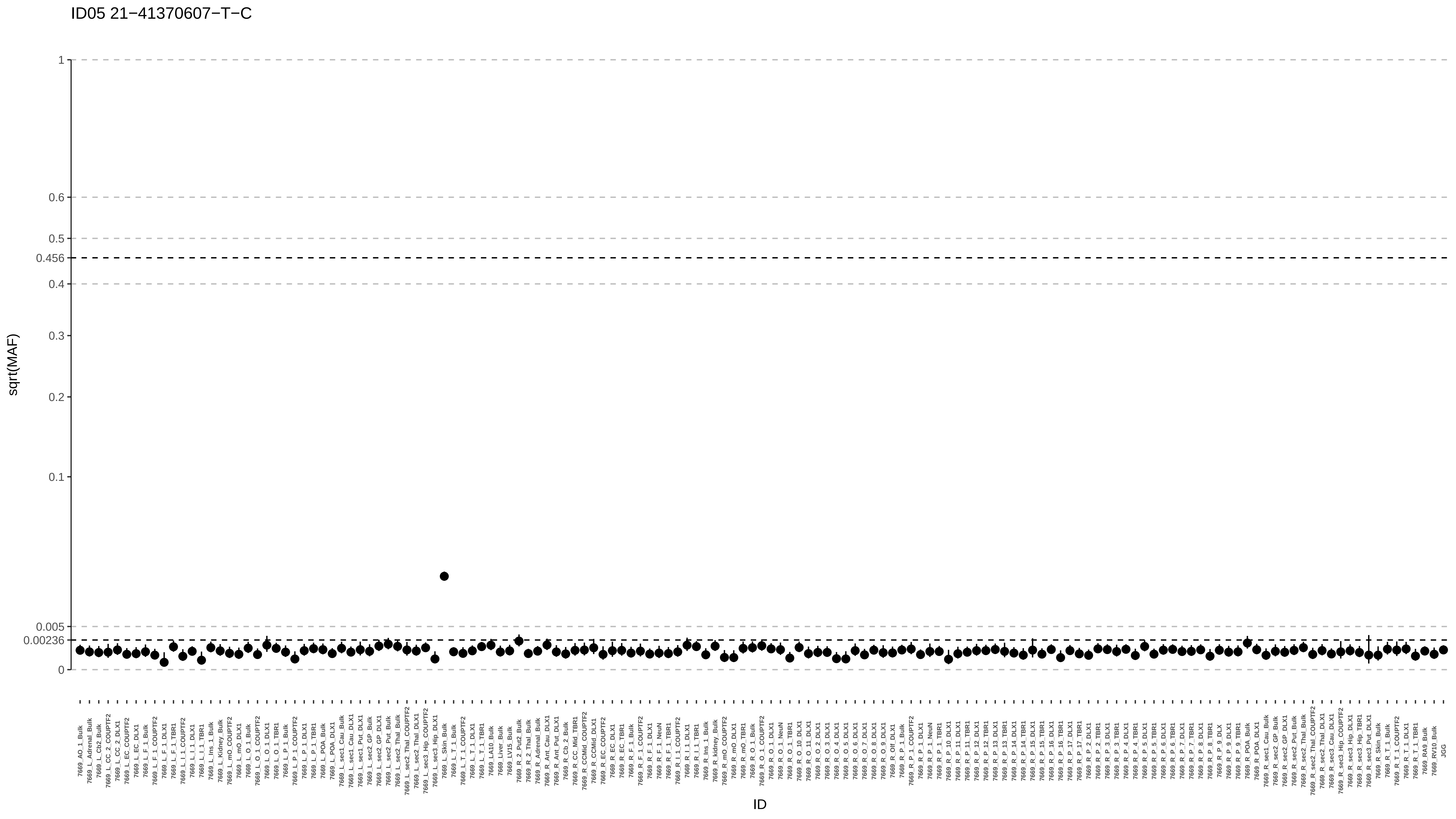
ID05 21−41370607−T−C
sqrt(MAF)
1
0.6
0.5
0.456
0.4
0.3
0.2
0.1
0.005
0.00236
0
ID

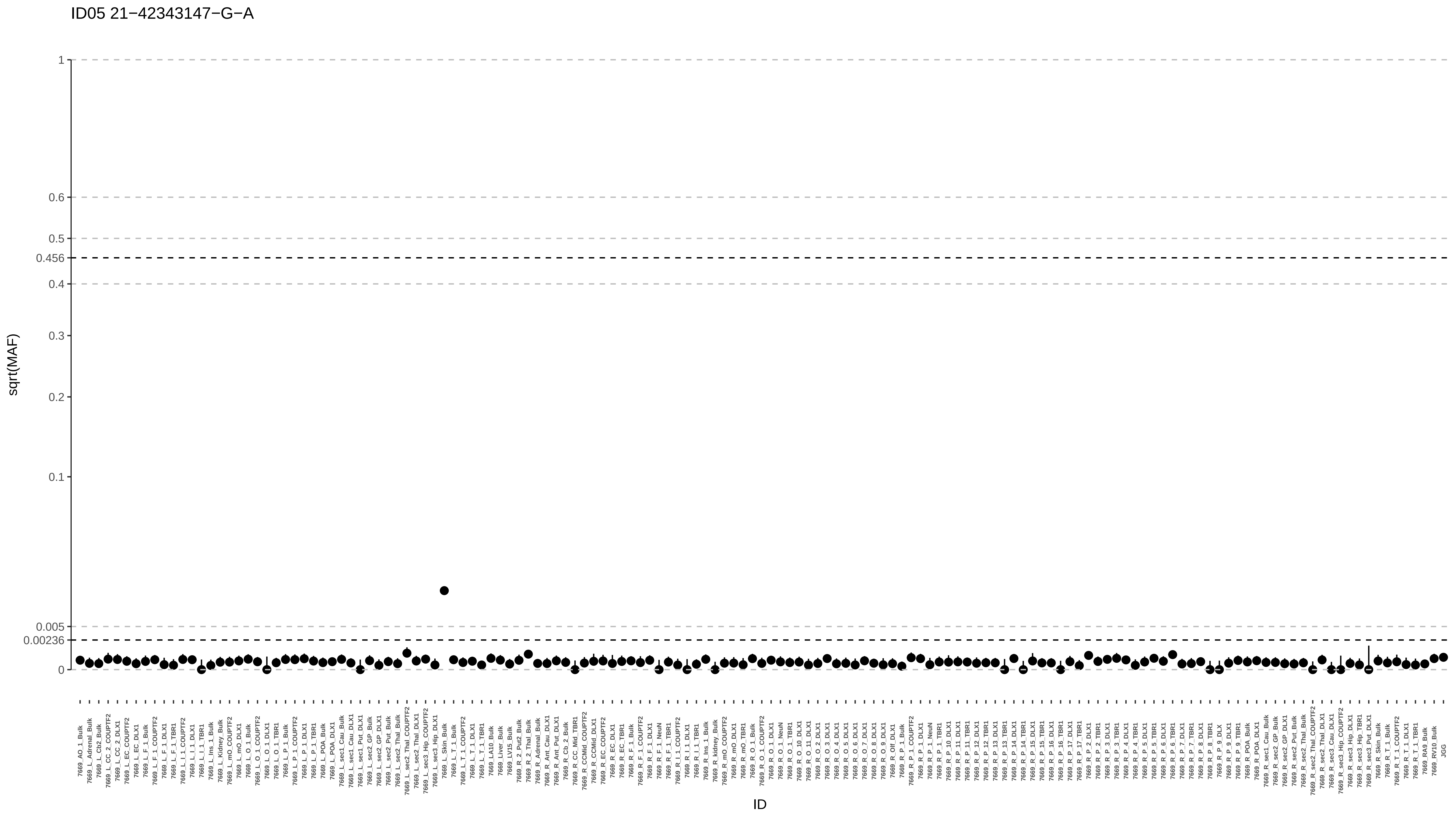

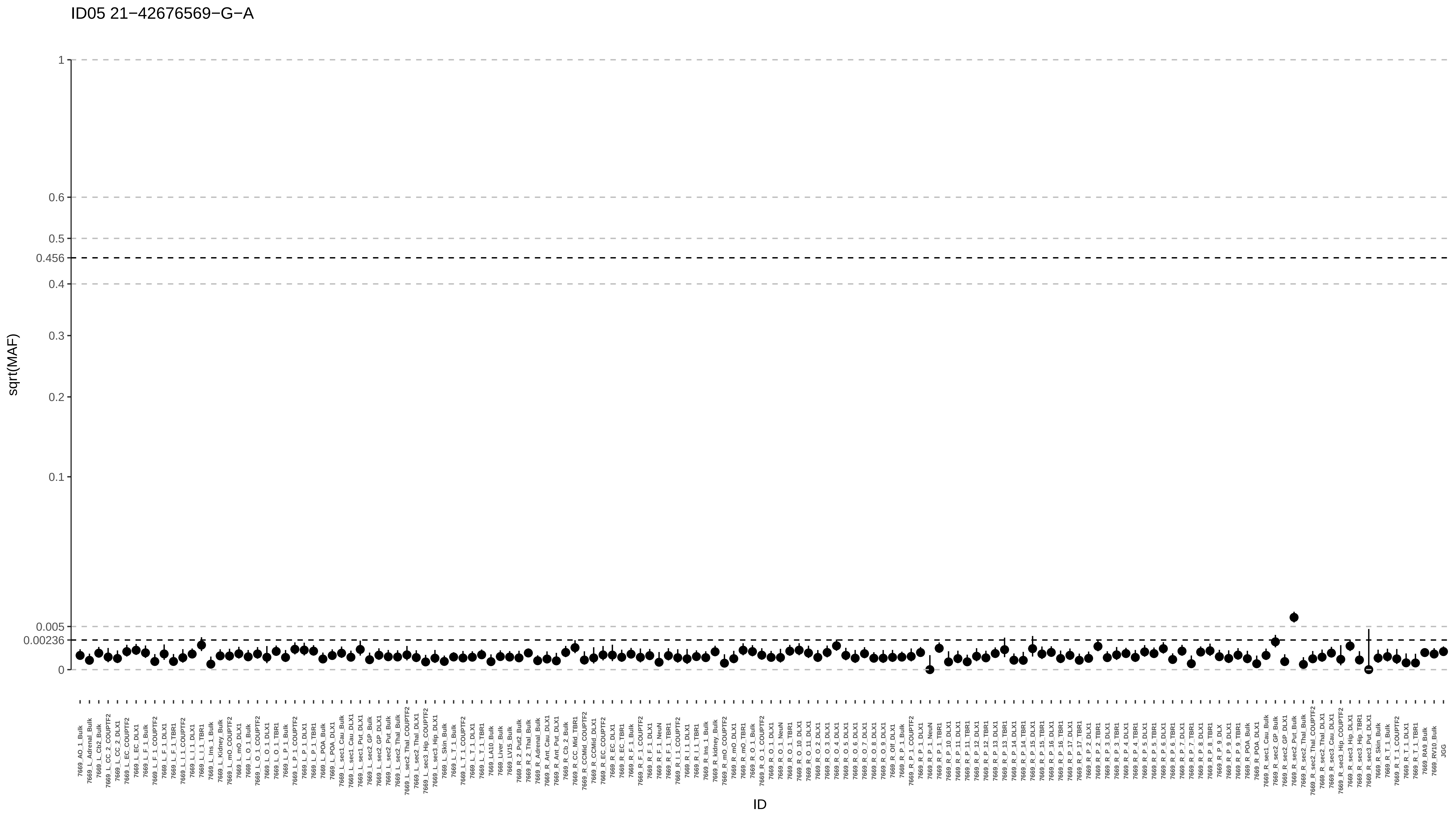
ID05 21−42676569−G−A
sqrt(MAF)
ID
1
0.6
0.5
0.456
0.4
0.3
0.2
0.1
0.005
0.00236
0

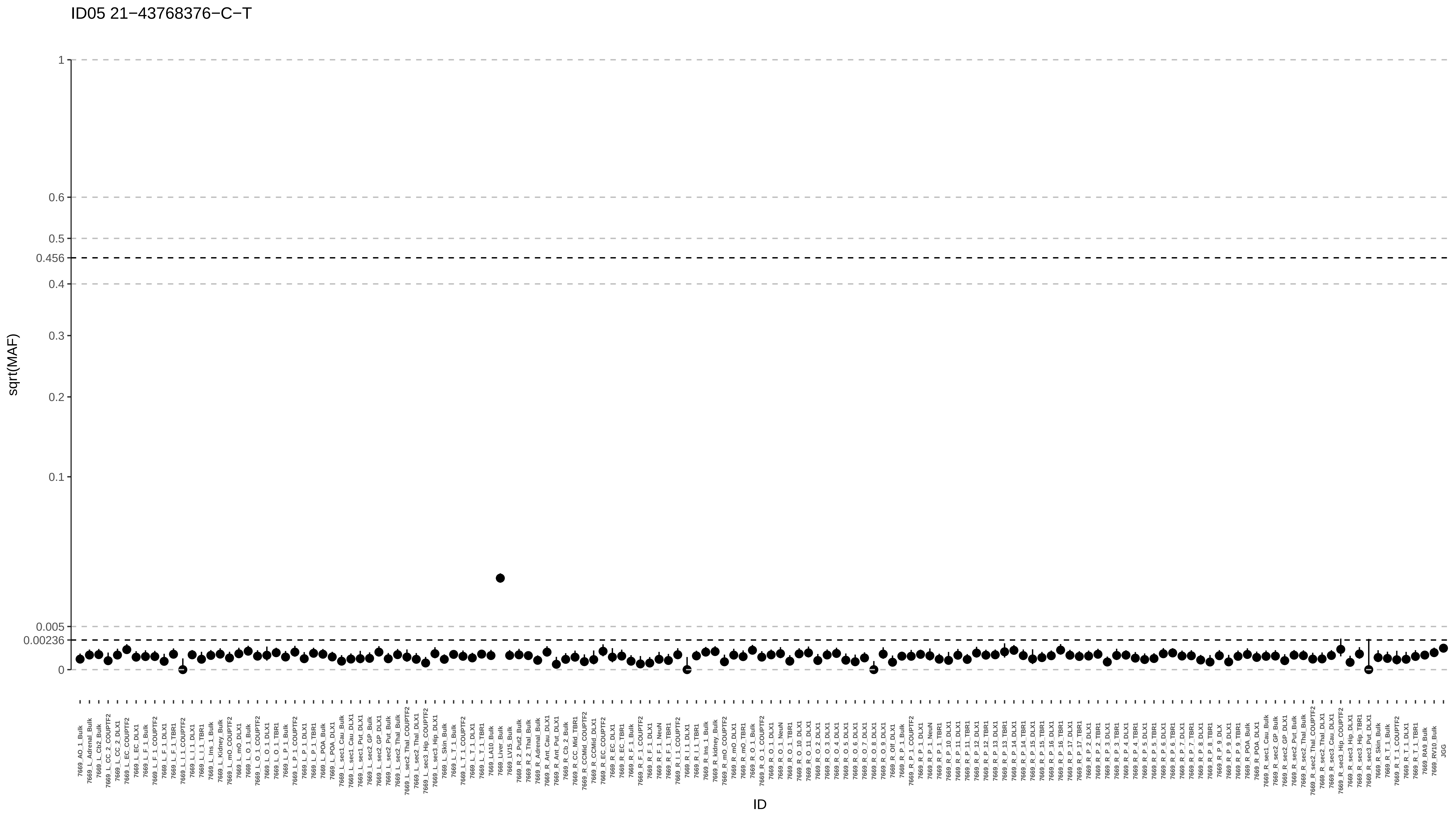
ID05 21−43768376−C−T
sqrt(MAF)
1
0.6
0.5
0.456
0.4
0.3
0.2
0.1
0.005
0.00236
0
ID

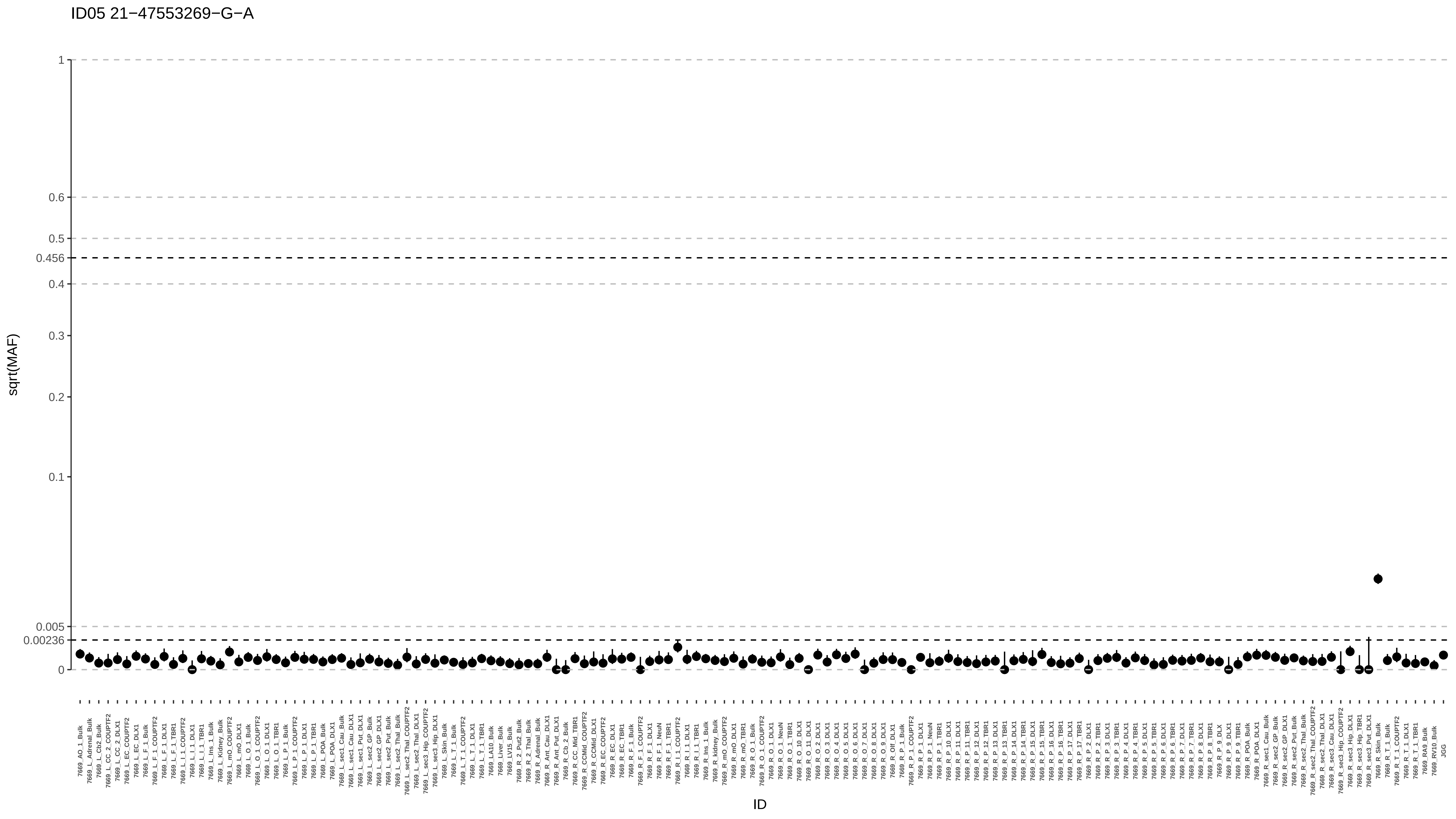
ID05 21−47553269−G−A
sqrt(MAF)
1
0.6
0.5
0.456
0.4
0.3
0.2
0.1
0.005
0.00236
0
ID

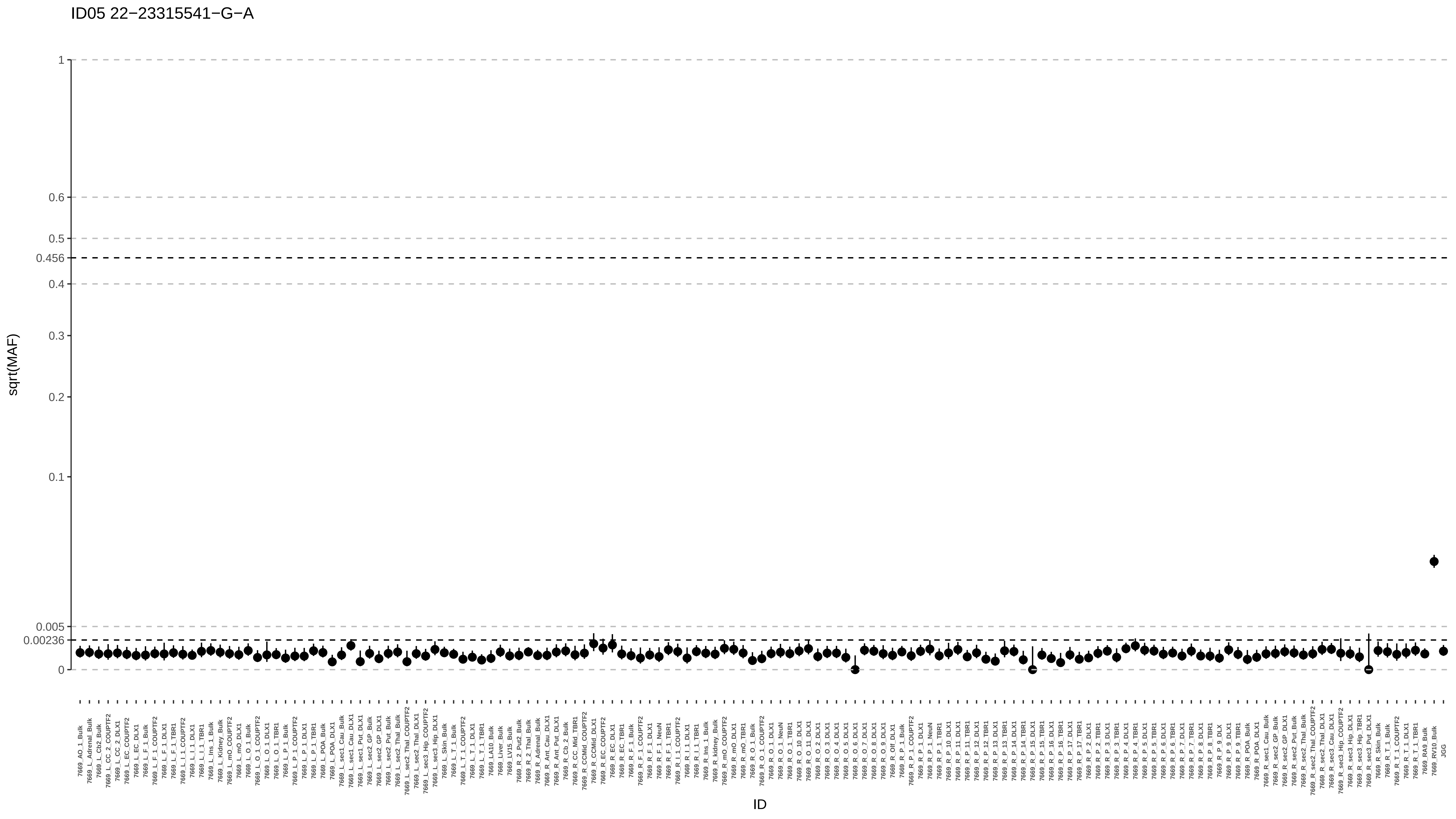
ID05 22−23315541−G−A
sqrt(MAF)
1
0.6
0.5
0.456
0.4
0.3
0.2
0.1
0.005
0.00236
0
ID

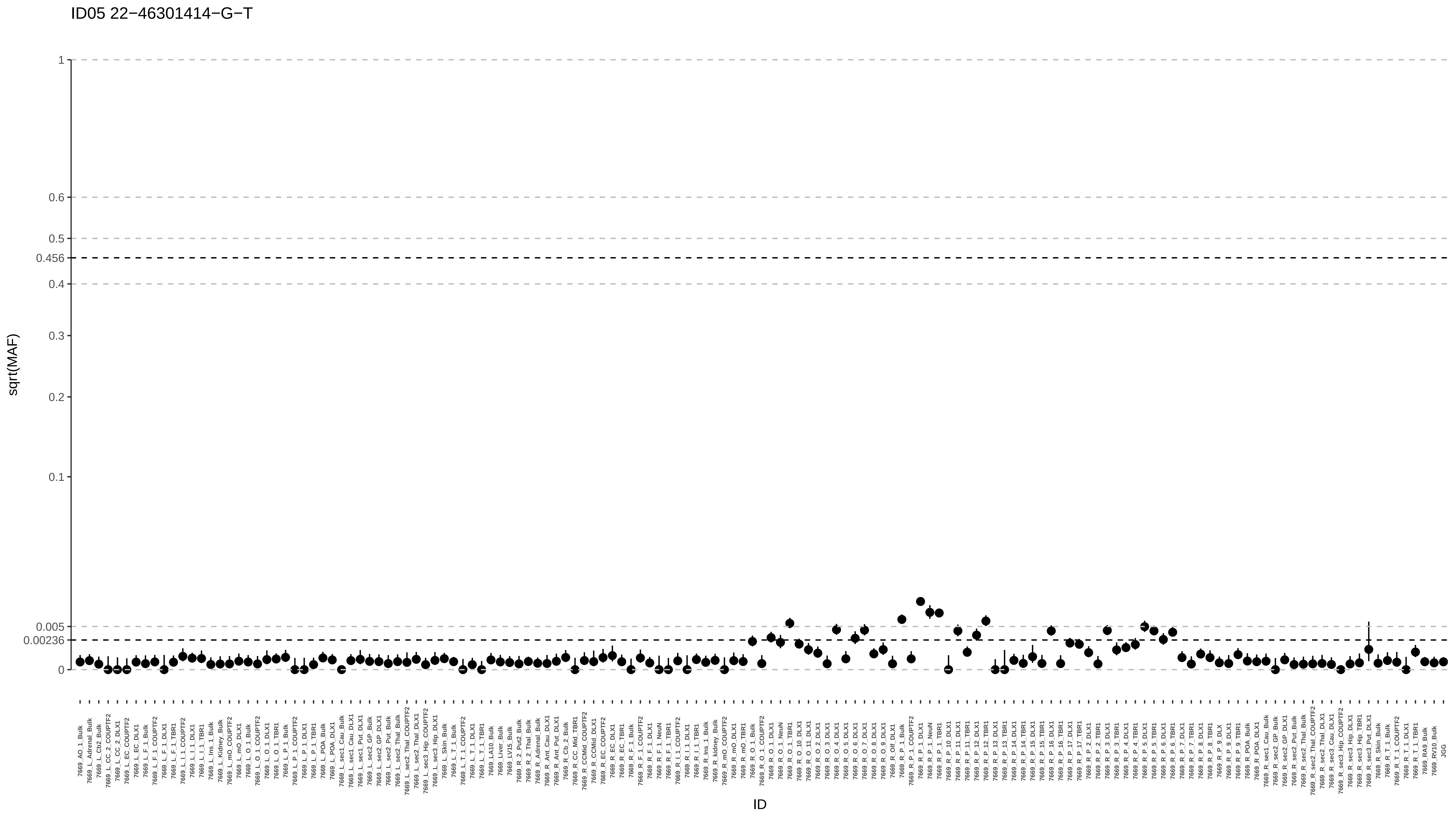
ID05 22−46301414−G−T
sqrt(MAF)
1
0.6
0.5
0.456
0.4
0.3
0.2
0.1
0.005
0.00236
0
ID

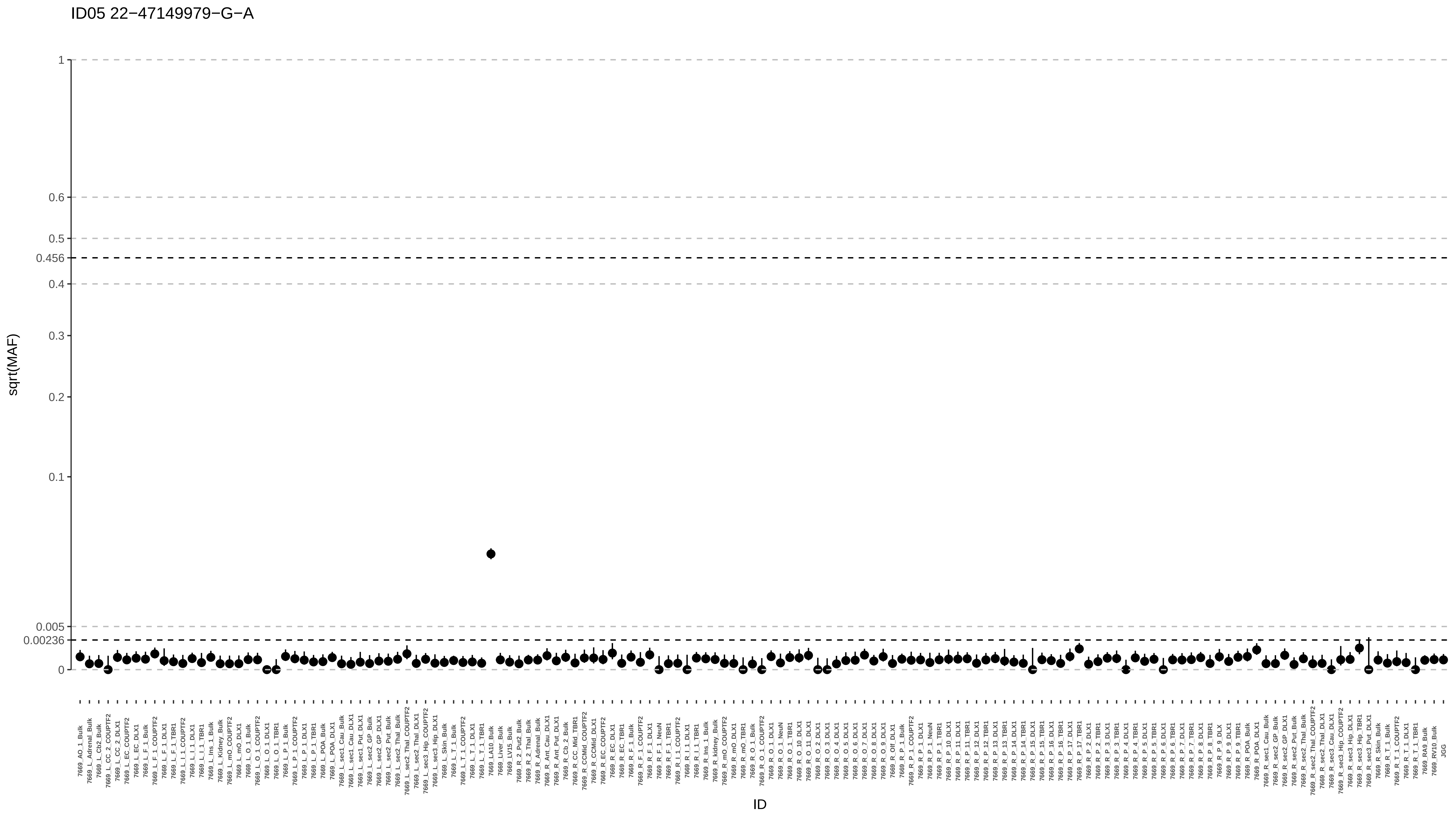
ID05 22−47149979−G−A
sqrt(MAF)
1
0.6
0.5
0.456
0.4
0.3
0.2
0.1
0.005
0.00236
0
ID
7669_AO_1_Bulk
7669_L_Adrenal_Bulk
7669_L_Cb2_Bulk
7669_L_CC_2_COUPTF2
7669_L_CC_2_DLX1
7669_L_EC_COUPTF2
7669_L_EC_DLX1
7669_L_F_1_Bulk
7669_L_F_1_COUPTF2
7669_L_F_1_DLX1
7669_L_F_1_TBR1
7669_L_I_1_COUPTF2
7669_L_I_1_DLX1
7669_L_I_1_TBR1
7669_L_Ins_1_Bulk
7669_L_Kidney_Bulk
7669_L_mO_COUPTF2
7669_L_mO_DLX1
7669_L_O_1_Bulk
7669_L_O_1_COUPTF2
7669_L_O_1_DLX1
7669_L_O_1_TBR1
7669_L_P_1_Bulk
7669_L_P_1_COUPTF2
7669_L_P_1_DLX1
7669_L_P_1_TBR1
7669_L_POA_Bulk
7669_L_POA_DLX1
7669_L_sec1_Cau_Bulk
7669_L_sec1_Cau_DLX1
7669_L_sec1_Put_DLX1
7669_L_sec2_GP_Bulk
7669_L_sec2_GP_DLX1
7669_L_sec2_Put_Bulk
7669_L_sec2_Thal_Bulk
7669_L_sec2_Thal_COUPTF2
7669_L_sec2_Thal_DLX1
7669_L_sec3_Hip_COUPTF2
7669_L_sec3_Hip_DLX1
7669_L_Skin_Bulk
7669_L_T_1_Bulk
7669_L_T_1_COUPTF2
7669_L_T_1_DLX1
7669_L_T_1_TBR1
7669_LA10_Bulk
7669_Liver_Bulk
7669_LV15_Bulk
7669_R_2_Put2_Bulk
7669_R_2_Thal_Bulk
7669_R_Adrenal_Bulk
7669_R_Ant_Cau_DLX1
7669_R_Ant_Put_DLX1
7669_R_Cb_2_Bulk
7669_R_CC_Mid_TBR1
7669_R_CCMid_COUPTF2
7669_R_CCMid_DLX1
7669_R_EC_COUPTF2
7669_R_EC_DLX1
7669_R_EC_TBR1
7669_R_F_1_Bulk
7669_R_F_1_COUPTF2
7669_R_F_1_DLX1
7669_R_F_1_NeuN
7669_R_F_1_TBR1
7669_R_I_1_COUPTF2
7669_R_I_1_DLX1
7669_R_I_1_TBR1
7669_R_Ins_1_Bulk
7669_R_kidney_Bulk
7669_R_mO_COUPTF2
7669_R_mO_DLX1
7669_R_mO_TBR1
7669_R_O_1_Bulk
7669_R_O_1_COUPTF2
7669_R_O_1_DLX1
7669_R_O_1_NeuN
7669_R_O_1_TBR1
7669_R_O_10_DLX1
7669_R_O_11_DLX1
7669_R_O_2_DLX1
7669_R_O_3_DLX1
7669_R_O_4_DLX1
7669_R_O_5_DLX1
7669_R_O_6_DLX1
7669_R_O_7_DLX1
7669_R_O_8_DLX1
7669_R_O_9_DLX1
7669_R_Olf_DLX1
7669_R_P_1_Bulk
7669_R_P_1_COUPTF2
7669_R_P_1_DLX1
7669_R_P_1_NeuN
7669_R_P_1_TBR1
7669_R_P_10_DLX1
7669_R_P_11_DLX1
7669_R_P_11_TBR1
7669_R_P_12_DLX1
7669_R_P_12_TBR1
7669_R_P_13_DLX1
7669_R_P_13_TBR1
7669_R_P_14_DLX1
7669_R_P_14_TBR1
7669_R_P_15_DLX1
7669_R_P_15_TBR1
7669_R_P_16_DLX1
7669_R_P_16_TBR1
7669_R_P_17_DLX1
7669_R_P_17_TBR1
7669_R_P_2_DLX1
7669_R_P_2_TBR1
7669_R_P_3_DLX1
7669_R_P_3_TBR1
7669_R_P_4_DLX1
7669_R_P_4_TBR1
7669_R_P_5_DLX1
7669_R_P_5_TBR1
7669_R_P_6_DLX1
7669_R_P_6_TBR1
7669_R_P_7_DLX1
7669_R_P_7_TBR1
7669_R_P_8_DLX1
7669_R_P_8_TBR1
7669_R_P_9_DLX
7669_R_P_9_DLX1
7669_R_P_9_TBR1
7669_R_POA_Bulk
7669_R_POA_DLX1
7669_R_sec1_Cau_Bulk
7669_R_sec2_GP_Bulk
7669_R_sec2_GP_DLX1
7669_R_sec2_Put_Bulk
7669_R_sec2_Thal_Bulk
7669_R_sec2_Thal_COUPTF2
7669_R_sec2_Thal_DLX1
7669_R_sec3_Cau_DLX1
7669_R_sec3_Hip_COUPTF2
7669_R_sec3_Hip_DLX1
7669_R_sec3_Hip_TBR1
7669_R_sec3_Put_DLX1
7669_R_Skin_Bulk
7669_R_T_1_Bulk
7669_R_T_1_COUPTF2
7669_R_T_1_DLX1
7669_R_T_1_TBR1
7669_RA9_Bulk
7669_RV10_Bulk
JGG

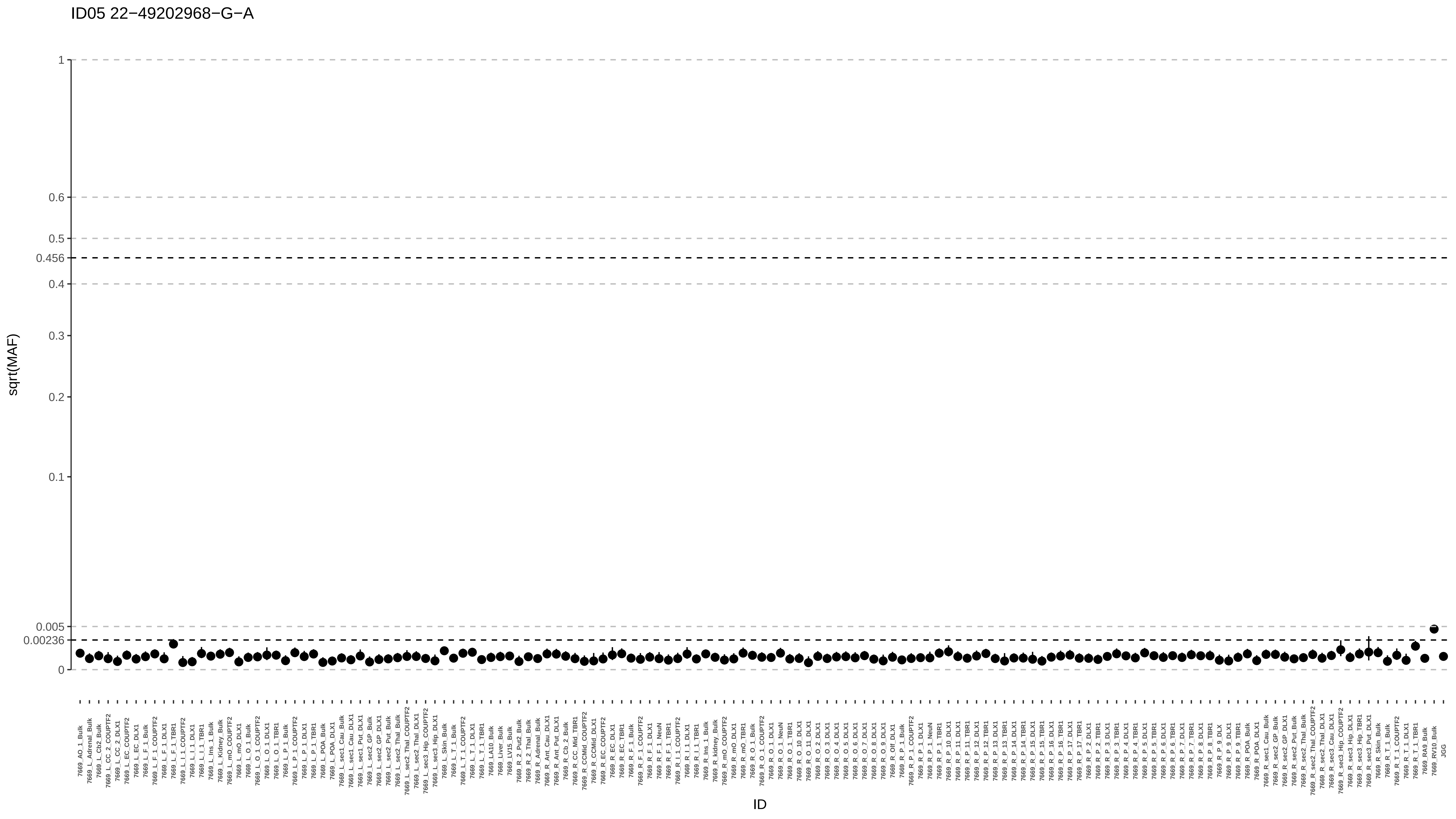
ID05 22−49202968−G−A
sqrt(MAF)
1
0.6
0.5
0.456
0.4
0.3
0.2
0.1
0.005
0.00236
0
ID

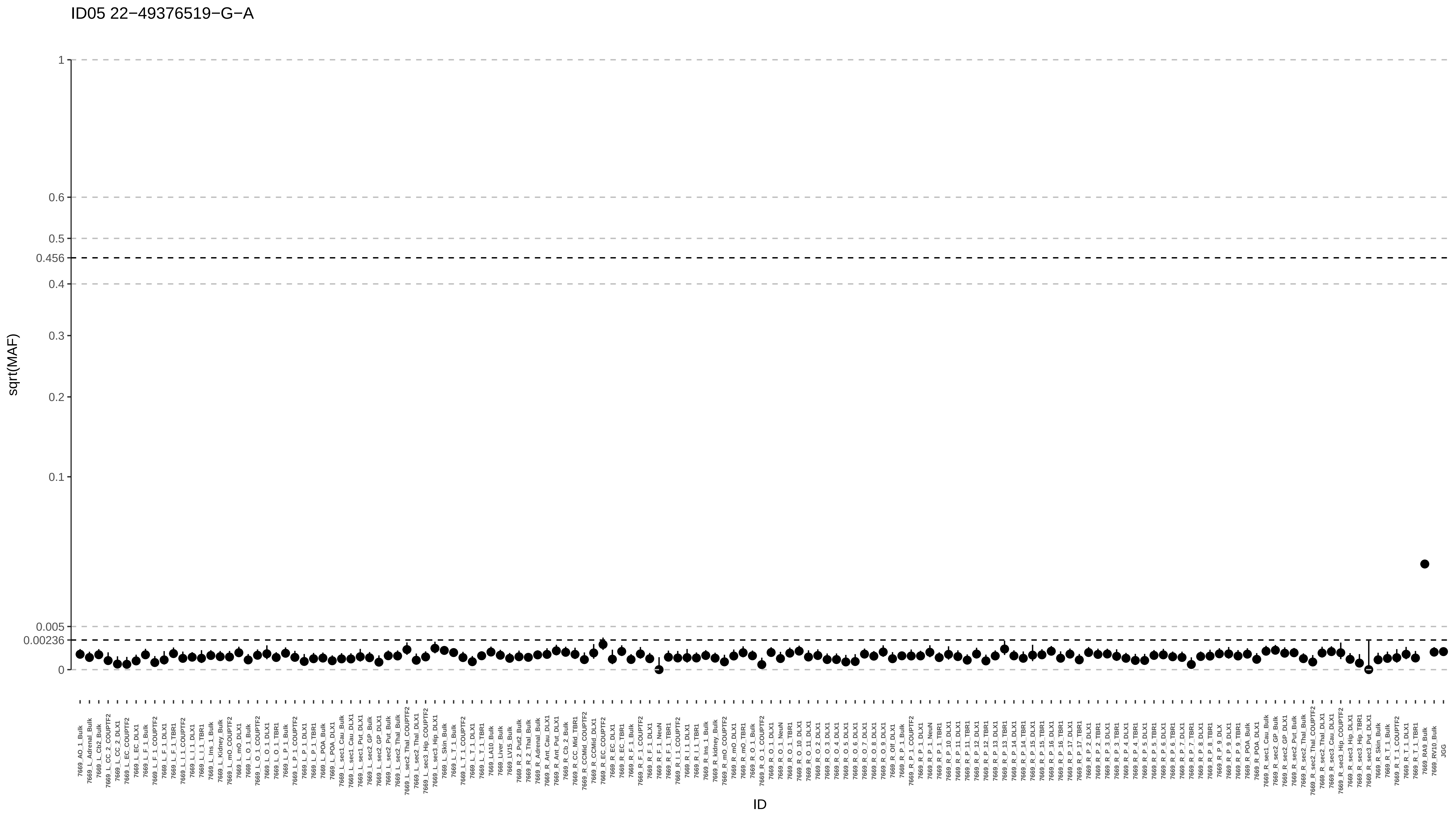
ID05 22−49376519−G−A
sqrt(MAF)
1
0.6
0.5
0.456
0.4
0.3
0.2
0.1
0.005
0.00236
0
ID

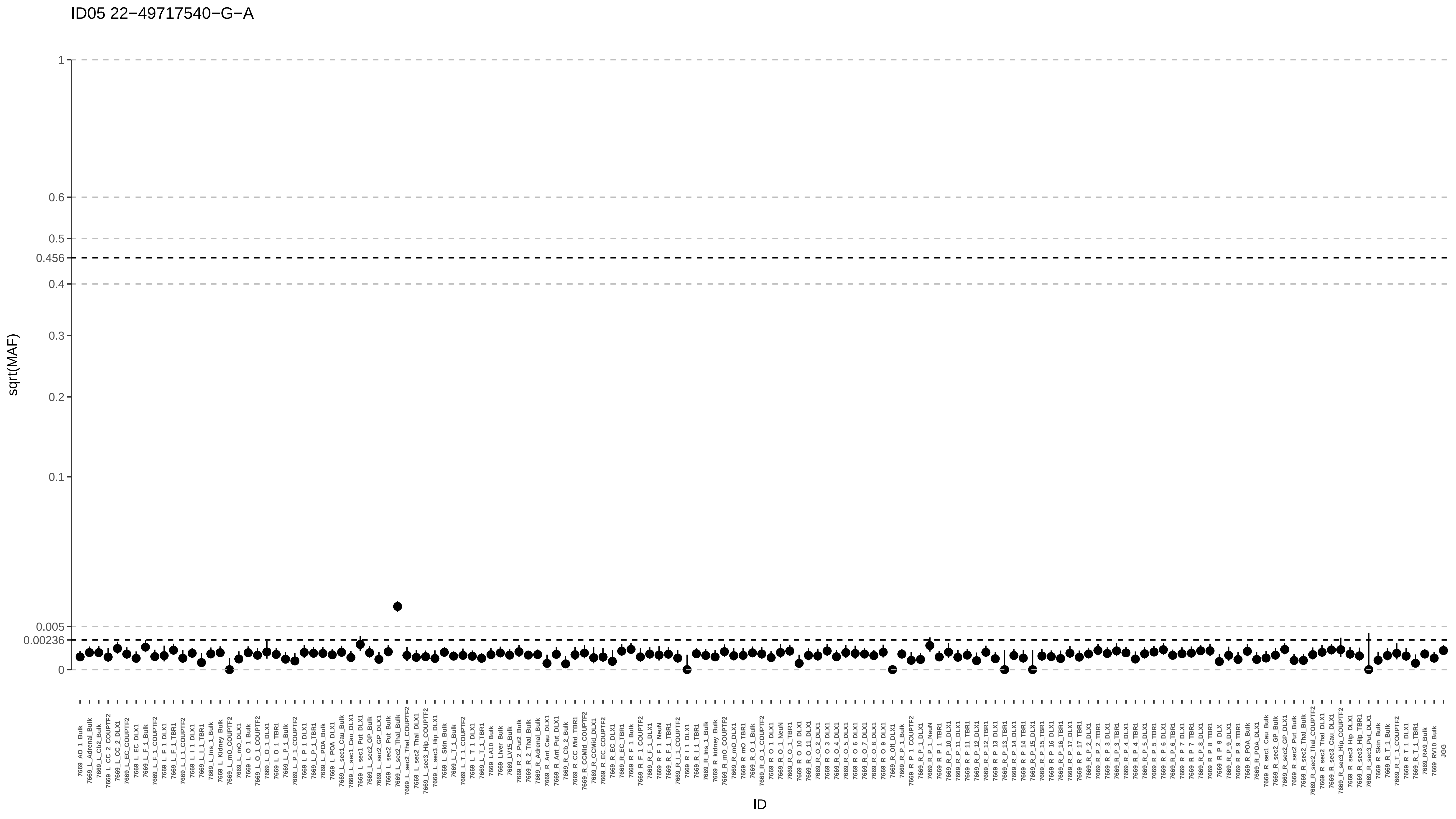

ID05 22−49717540−G−A
sqrt(MAF)
ID

ID05 22−49847904−G−A

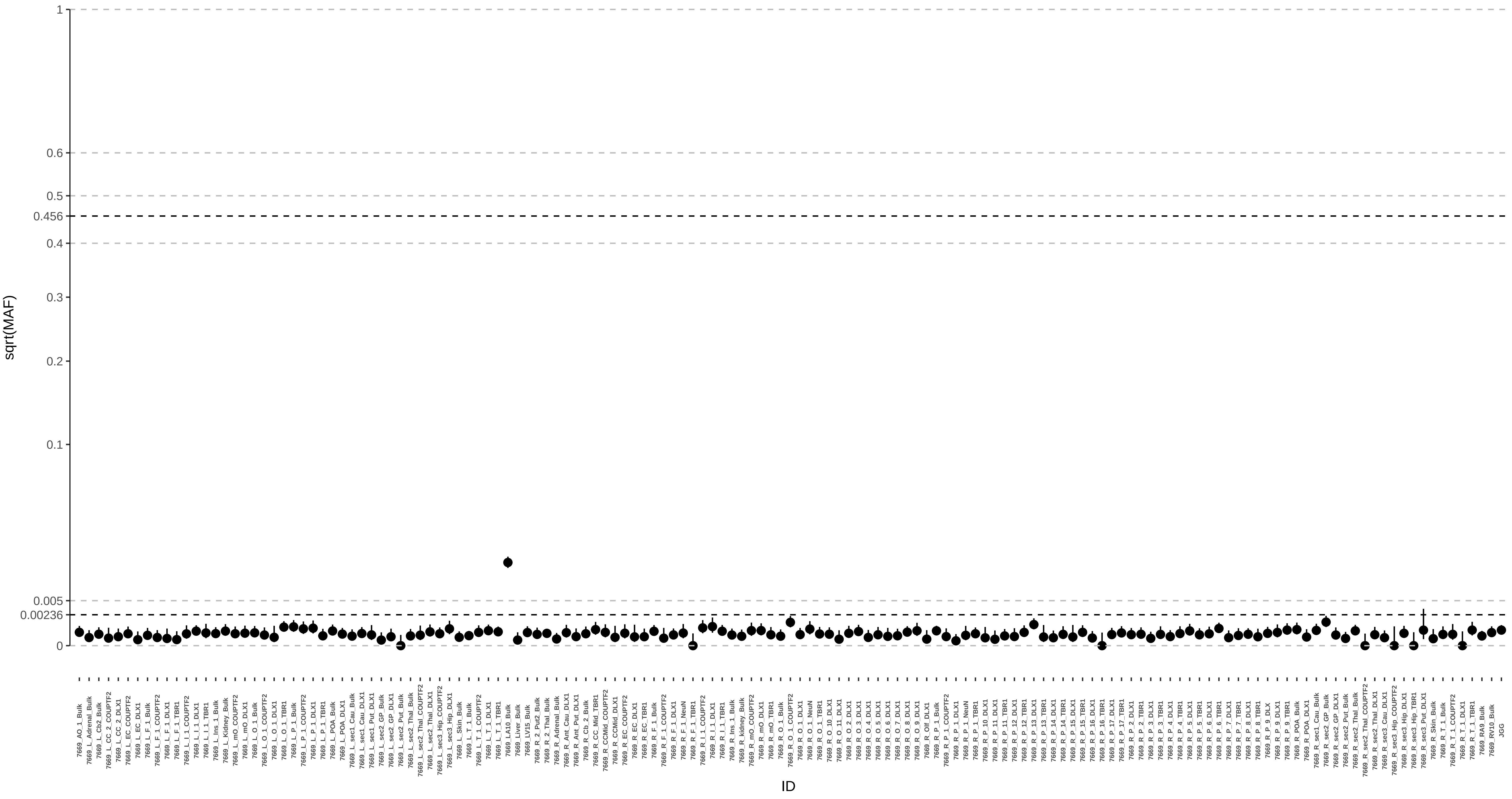

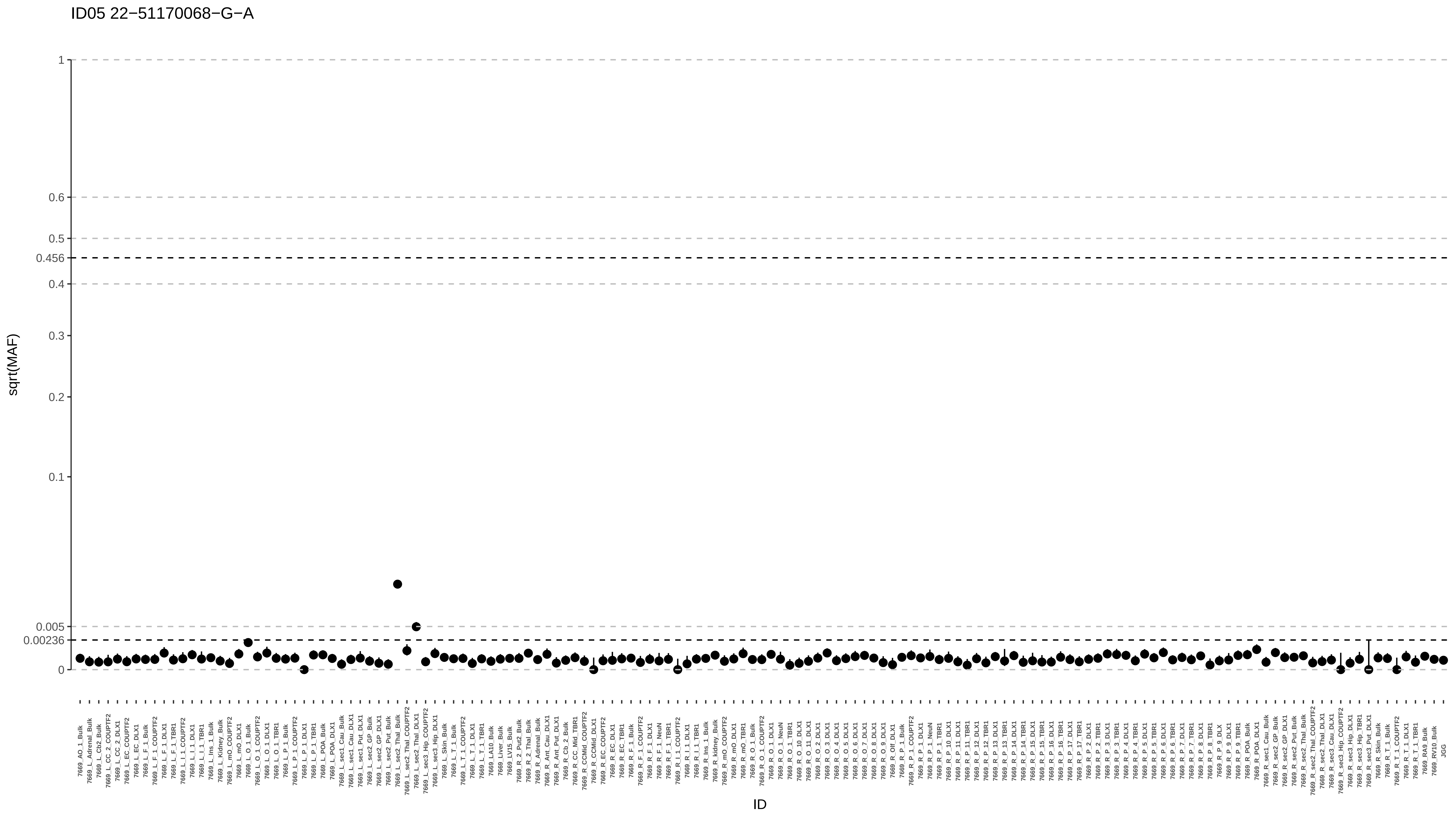
ID05 22−51170068−G−A
1
0.6
0.5
0.456
0.4
0.3
0.2
0.1
0.005
0.00236
0
sqrt(MAF)
7669_AO_1_Bulk
7669_L_Adrenal_Bulk
7669_L_Cb2_Bulk
7669_L_CC_2_COUPTF2
7669_L_CC_2_DLX1
7669_L_EC_COUPTF2
7669_L_EC_DLX1
7669_L_F_1_Bulk
7669_L_F_1_COUPTF2
7669_L_F_1_DLX1
7669_L_F_1_TBR1
7669_L_I_1_COUPTF2
7669_L_I_1_DLX1
7669_L_I_1_TBR1
7669_L_Ins_1_Bulk
7669_L_Kidney_Bulk
7669_L_mO_COUPTF2
7669_L_mO_DLX1
7669_L_O_1_Bulk
7669_L_O_1_COUPTF2
7669_L_O_1_DLX1
7669_L_O_1_TBR1
7669_L_P_1_Bulk
7669_L_P_1_COUPTF2
7669_L_P_1_DLX1
7669_L_P_1_TBR1
7669_L_POA_Bulk
7669_L_POA_DLX1
7669_L_sec1_Cau_Bulk
7669_L_sec1_Cau_DLX1
7669_L_sec1_Put_DLX1
7669_L_sec2_GP_Bulk
7669_L_sec2_GP_DLX1
7669_L_sec2_Put_Bulk
7669_L_sec2_Thal_Bulk
7669_L_sec2_Thal_COUPTF2
7669_L_sec2_Thal_DLX1
7669_L_sec3_Hip_COUPTF2
7669_L_sec3_Hip_DLX1
7669_L_Skin_Bulk
7669_L_T_1_Bulk
7669_L_T_1_COUPTF2
7669_L_T_1_DLX1
7669_L_T_1_TBR1
7669_LA10_Bulk
7669_Liver_Bulk
7669_LV15_Bulk
7669_R_2_Put2_Bulk
7669_R_2_Thal_Bulk
7669_R_Adrenal_Bulk
7669_R_Ant_Cau_DLX1
7669_R_Ant_Put_DLX1
7669_R_Cb_2_Bulk
7669_R_CC_Mid_TBR1
7669_R_CCMid_COUPTF2
7669_R_CCMid_DLX1
7669_R_EC_COUPTF2
7669_R_EC_DLX1
7669_R_EC_TBR1
7669_R_F_1_Bulk
7669_R_F_1_COUPTF2
7669_R_F_1_DLX1
7669_R_F_1_NeuN
7669_R_F_1_TBR1
7669_R_I_1_COUPTF2
7669_R_I_1_DLX1
7669_R_I_1_TBR1
7669_R_Ins_1_Bulk
7669_R_kidney_Bulk
7669_R_mO_COUPTF2
7669_R_mO_DLX1
7669_R_mO_TBR1
7669_R_O_1_Bulk
7669_R_O_1_COUPTF2
7669_R_O_1_DLX1
7669_R_O_1_NeuN
7669_R_O_1_TBR1
7669_R_O_10_DLX1
7669_R_O_11_DLX1
7669_R_O_2_DLX1
7669_R_O_3_DLX1
7669_R_O_4_DLX1
7669_R_O_5_DLX1
7669_R_O_6_DLX1
7669_R_O_7_DLX1
7669_R_O_8_DLX1
7669_R_O_9_DLX1
7669_R_Olf_DLX1
7669_R_P_1_Bulk
7669_R_P_1_COUPTF2
7669_R_P_1_DLX1
7669_R_P_1_NeuN
7669_R_P_1_TBR1
7669_R_P_10_DLX1
7669_R_P_11_DLX1
7669_R_P_11_TBR1
7669_R_P_12_DLX1
7669_R_P_12_TBR1
7669_R_P_13_DLX1
7669_R_P_13_TBR1
7669_R_P_14_DLX1
7669_R_P_14_TBR1
7669_R_P_15_DLX1
7669_R_P_15_TBR1
7669_R_P_16_DLX1
7669_R_P_16_TBR1
7669_R_P_17_DLX1
7669_R_P_17_TBR1
7669_R_P_2_DLX1
7669_R_P_2_TBR1
7669_R_P_3_DLX1
7669_R_P_3_TBR1
7669_R_P_4_DLX1
7669_R_P_4_TBR1
7669_R_P_5_DLX1
7669_R_P_5_TBR1
7669_R_P_6_DLX1
7669_R_P_6_TBR1
7669_R_P_7_DLX1
7669_R_P_7_TBR1
7669_R_P_8_DLX1
7669_R_P_8_TBR1
7669_R_P_9_DLX
7669_R_P_9_DLX1
7669_R_P_9_TBR1
7669_R_POA_Bulk
7669_R_POA_DLX1
7669_R_sec1_Cau_Bulk
7669_R_sec2_GP_Bulk
7669_R_sec2_GP_DLX1
7669_R_sec2_Put_Bulk
7669_R_sec2_Thal_Bulk
7669_R_sec2_Thal_COUPTF2
7669_R_sec2_Thal_DLX1
7669_R_sec3_Cau_DLX1
7669_R_sec3_Hip_COUPTF2
7669_R_sec3_Hip_DLX1
7669_R_sec3_Hip_TBR1
7669_R_sec3_Put_DLX1
7669_R_Skin_Bulk
7669_R_T_1_Bulk
7669_R_T_1_COUPTF2
7669_R_T_1_DLX1
7669_R_T_1_TBR1
7669_RA9_Bulk
7669_RV10_Bulk
JGG
ID

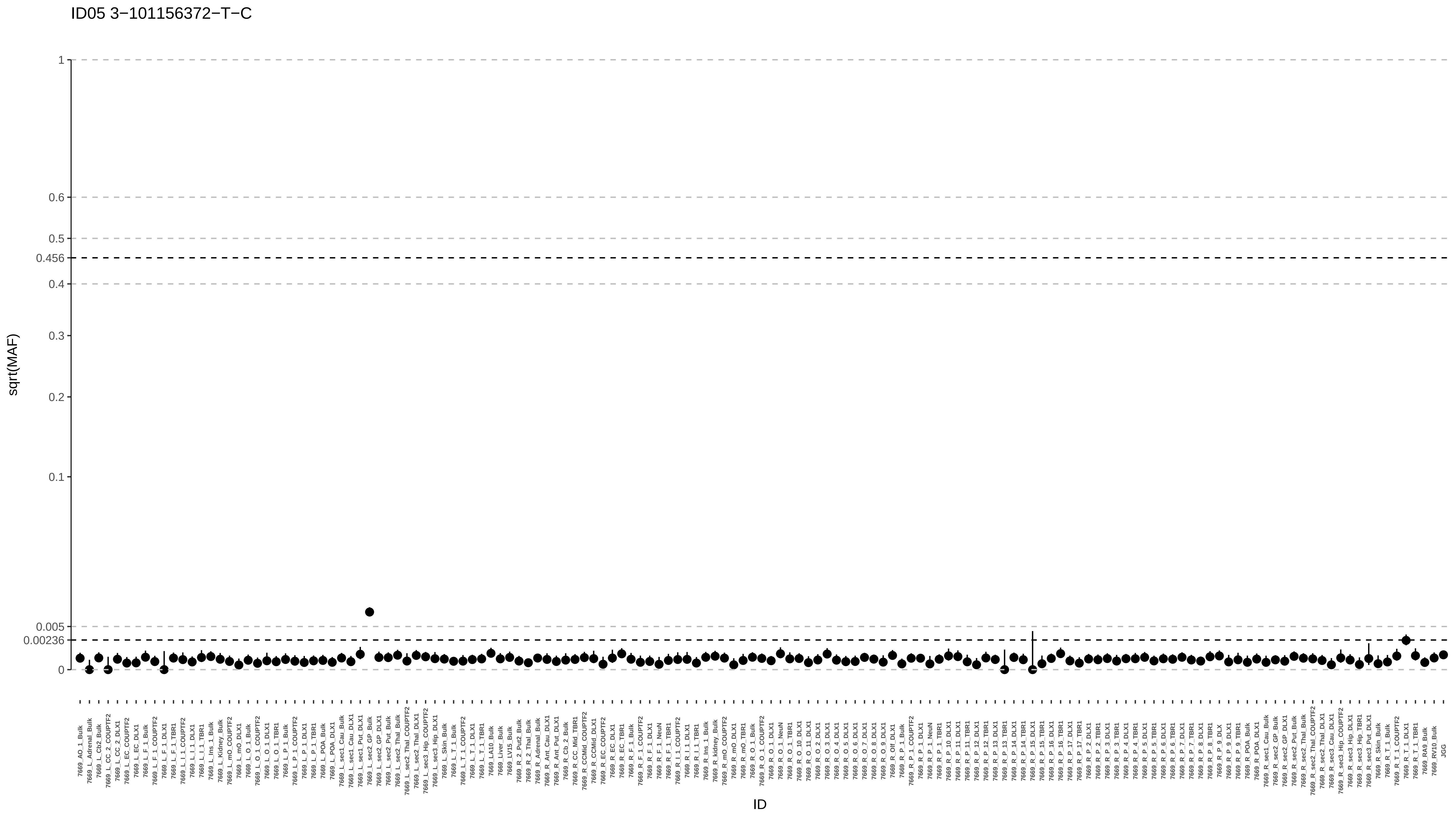

ID05 3−101156372−T−C
sqrt(MAF)
1
0.6
0.5
0.456
0.4
0.3
0.2
0.1
0.005
0.00236
0
ID

## ID05 3−103294509−T−G

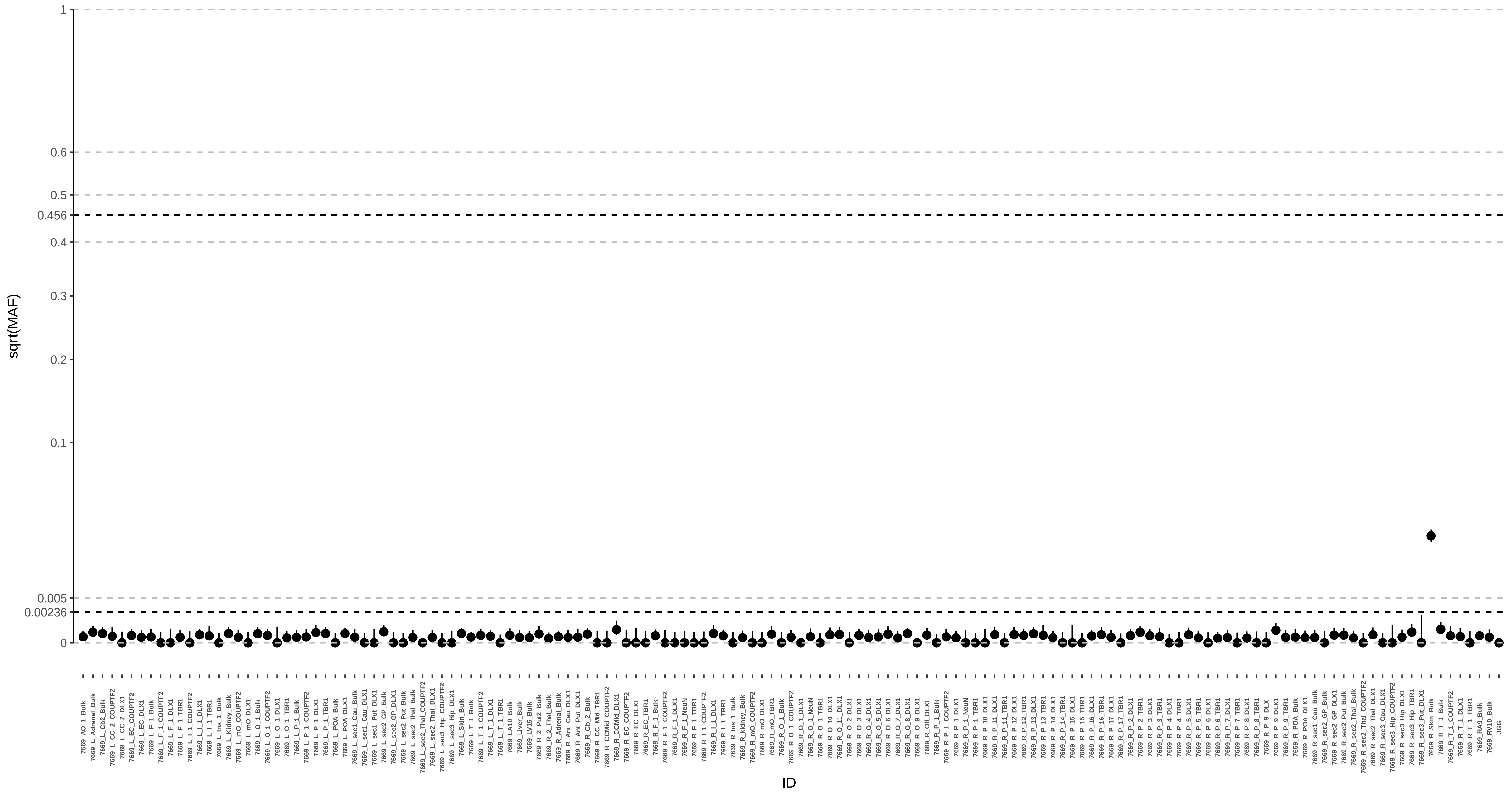

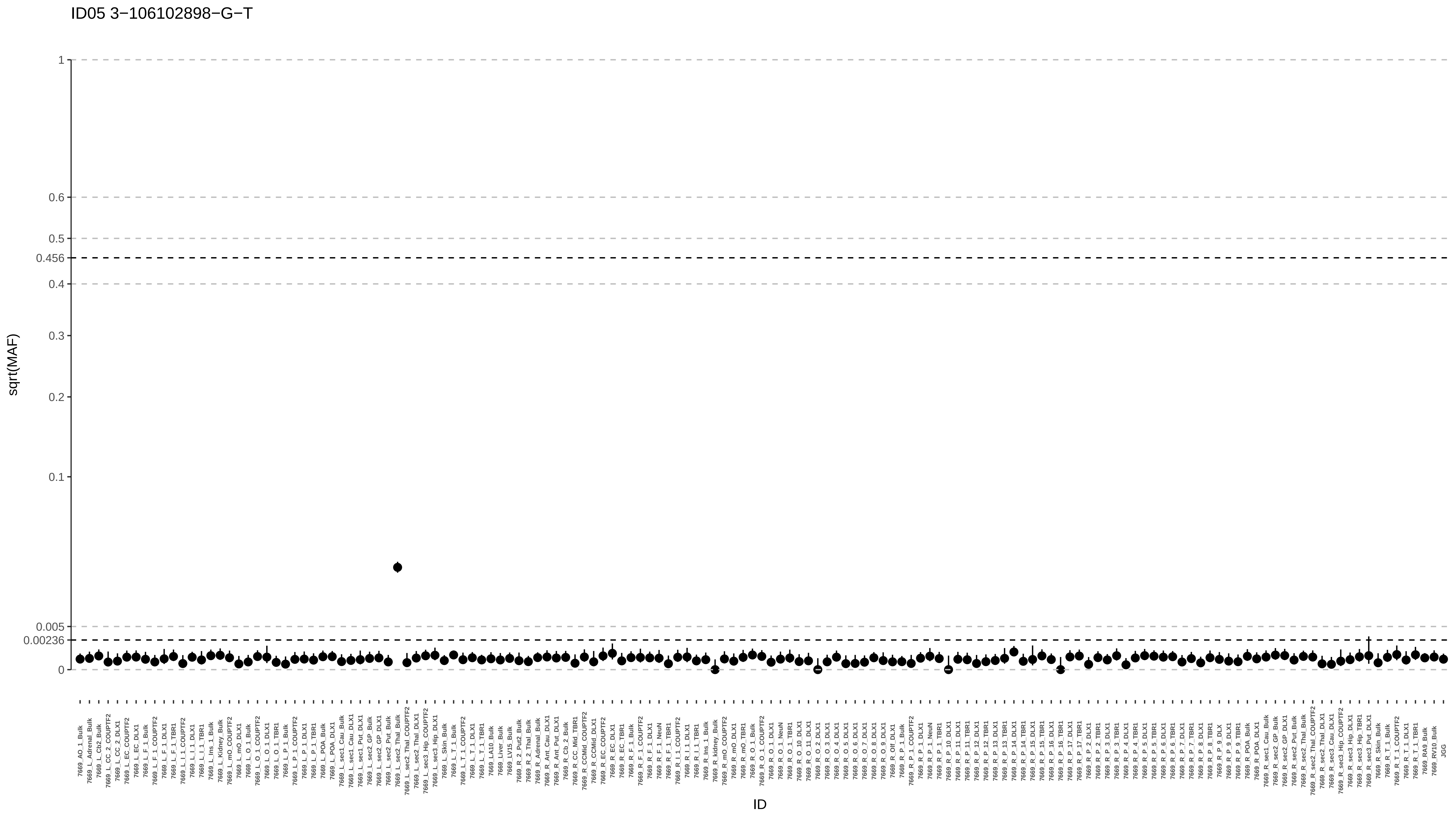

ID05 3−106102898−G−T
sqrt(MAF)
1
0.6
0.5
0.456
0.4
0.3
0.2
0.1
0.005
0.00236
0
ID

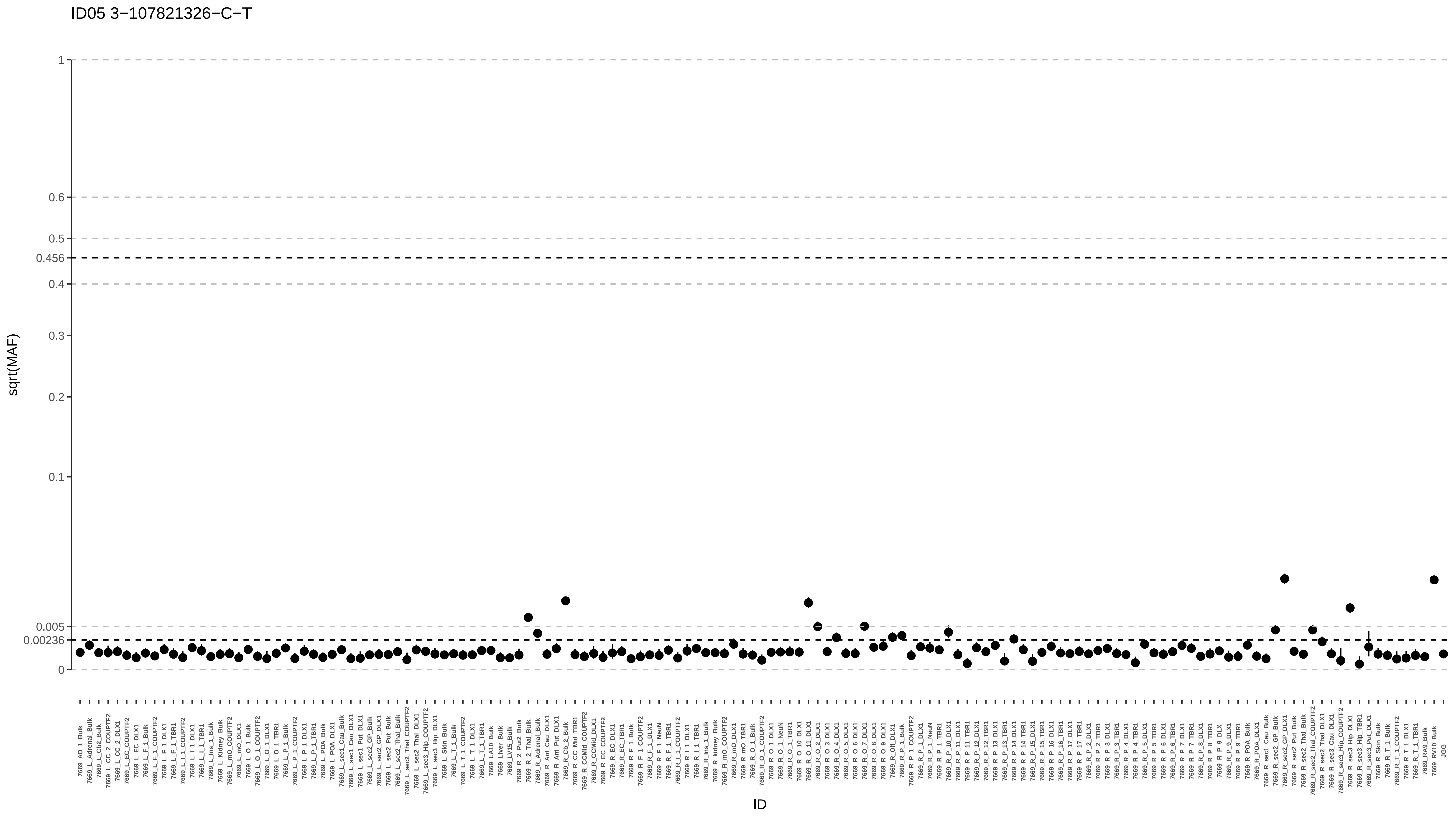
ID05 3−107821326−C−T
sqrt(MAF)
1
0.6
0.5
0.456
0.4
0.3
0.2
0.1
0.005
0.00236
0
ID

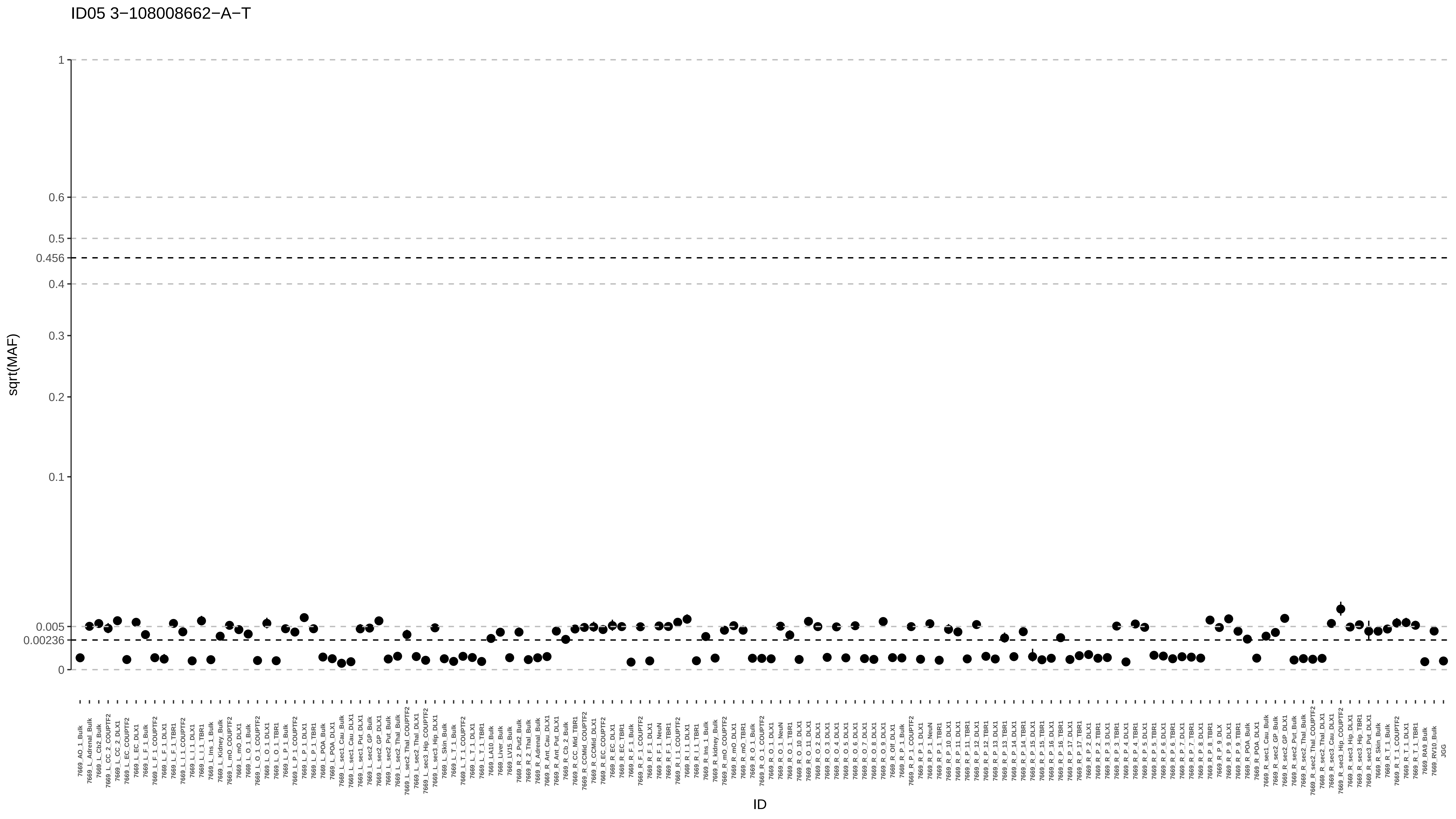
ID05 3−108008662−A−T
sqrt(MAF)
1
0.6
0.5
0.456
0.4
0.3
0.2
0.1
0.005
0.00236
0
ID

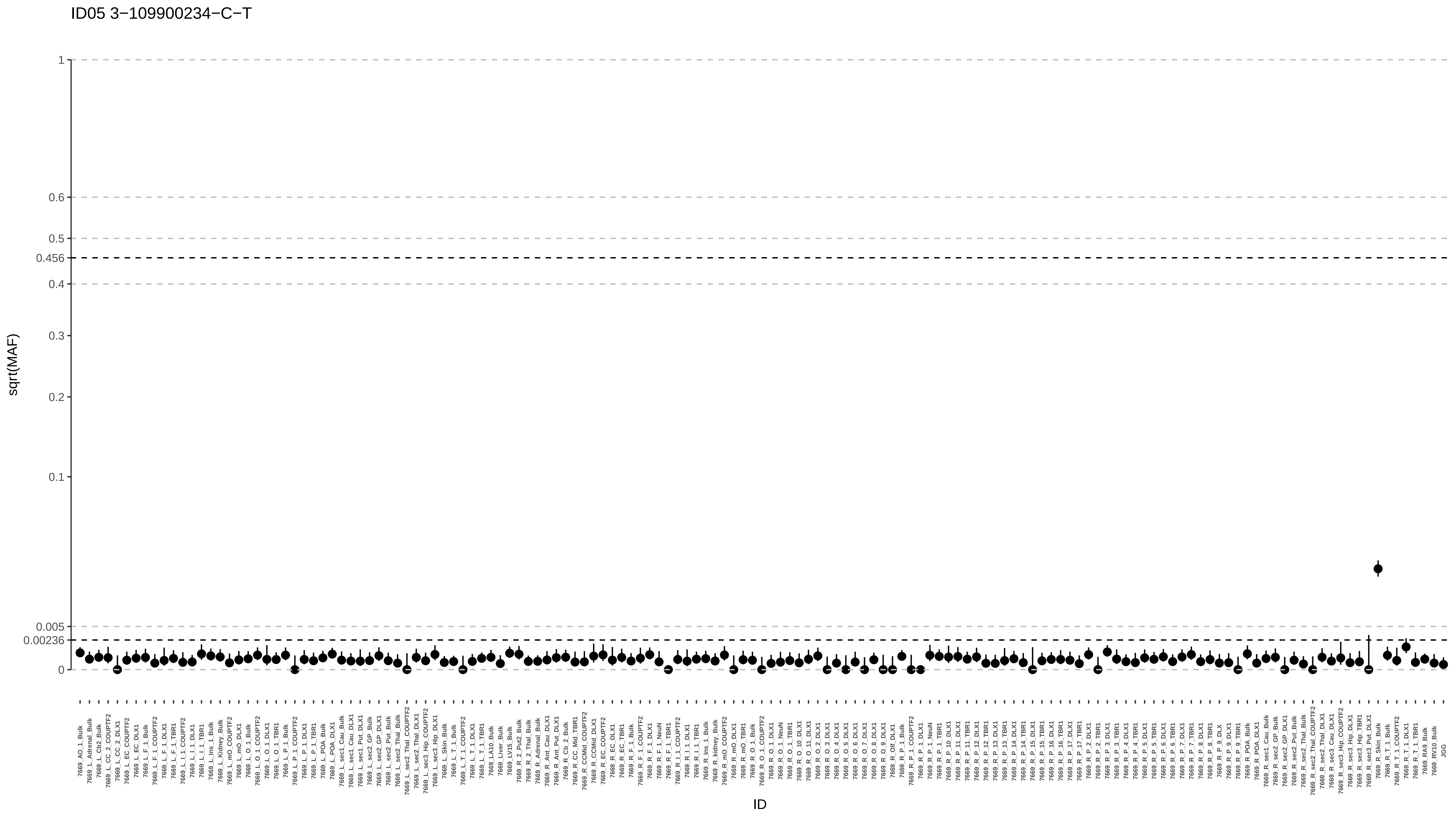
ID05 3−109900234−C−T
sqrt(MAF)
1
0.6
0.5
0.456
0.4
0.3
0.2
0.1
0.005
0.00236
0
ID

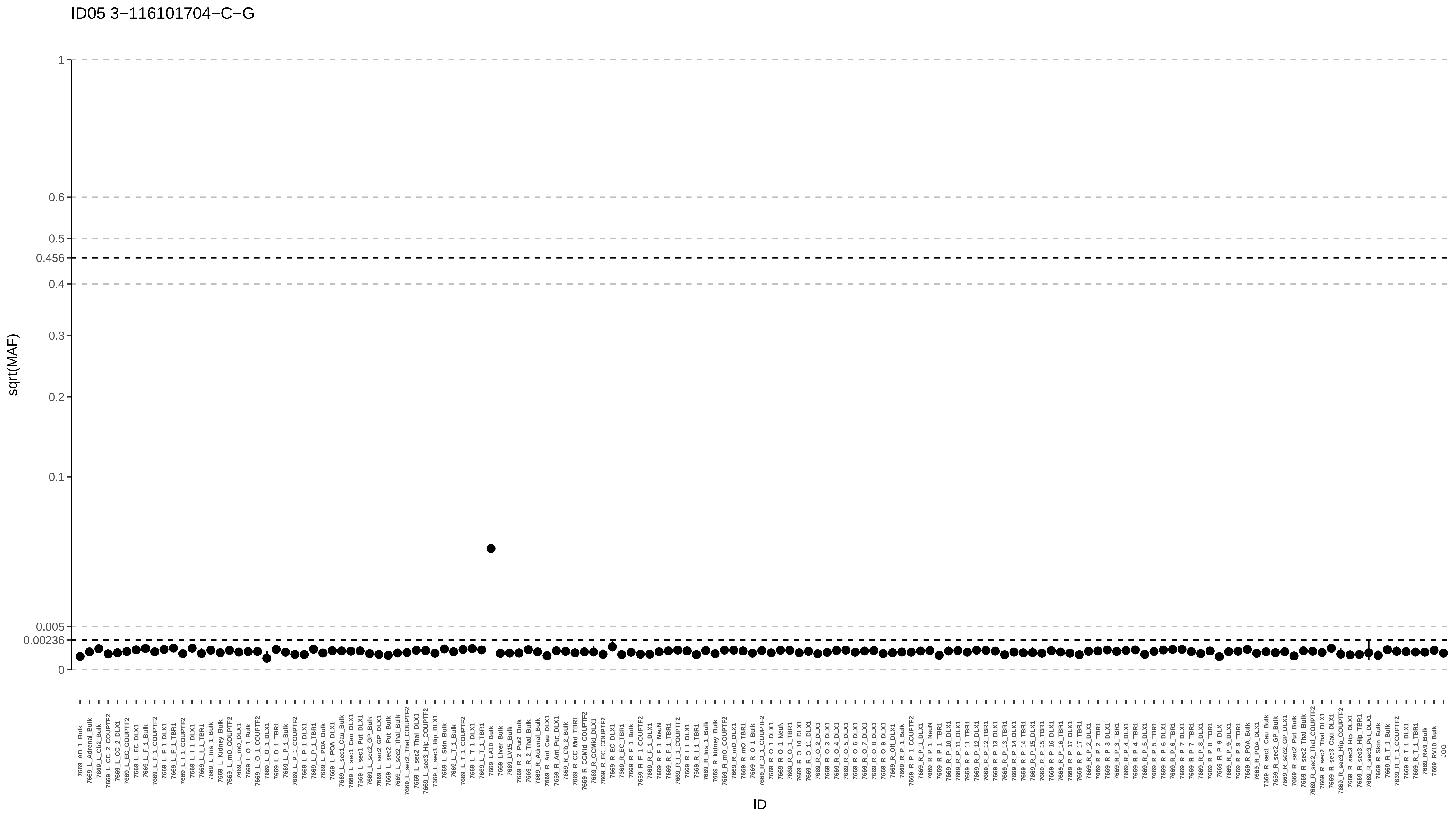
ID05 3−116101704−C−G
sqrt(MAF)
1
0.6
0.5
0.456
0.4
0.3
0.2
0.1
0.005
0.00236
0
ID

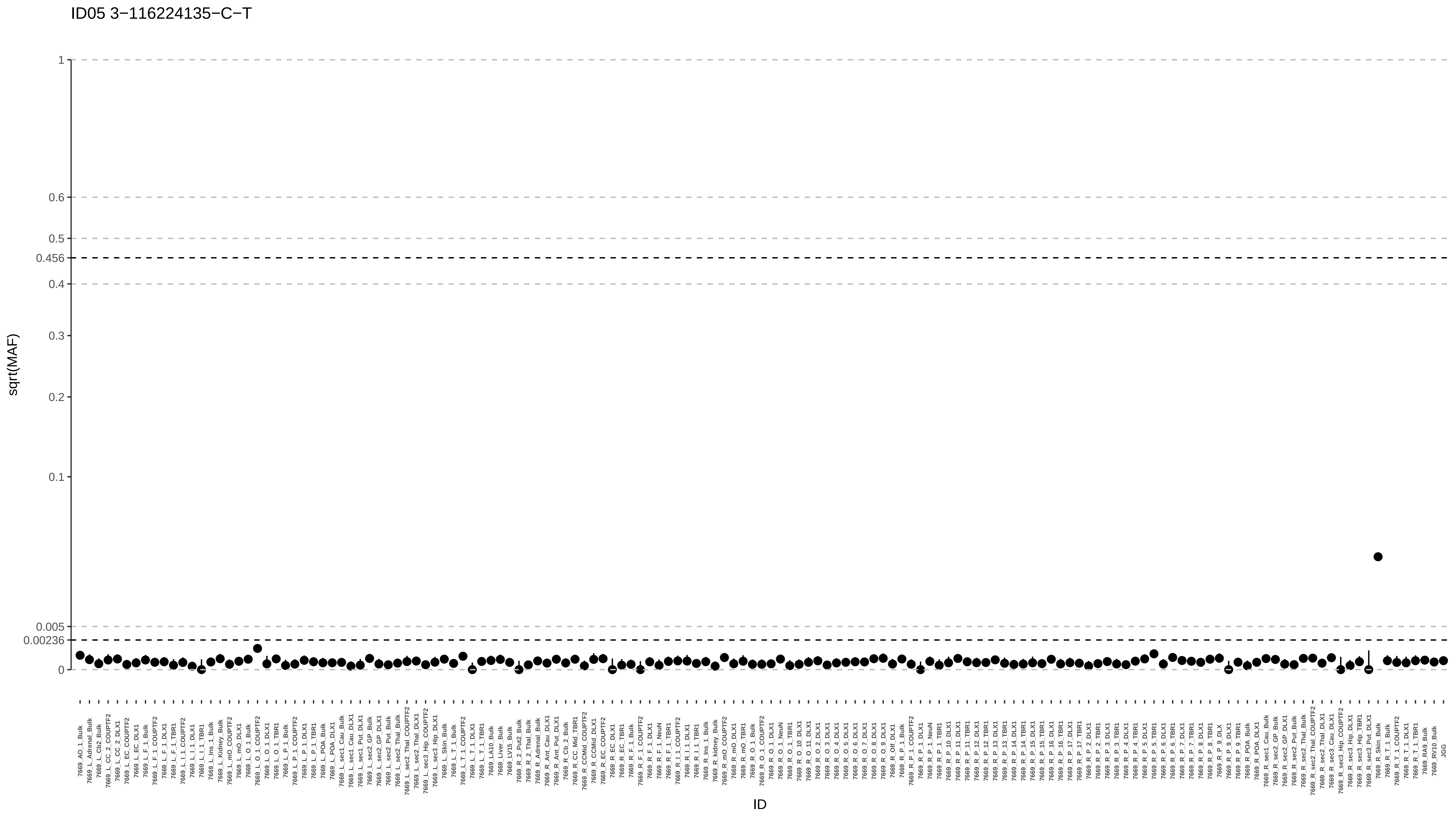
ID05 3−116224135−C−T
sqrt(MAF)
1
0.6
0.5
0.456
0.4
0.3
0.2
0.1
0.005
0.00236
0
ID

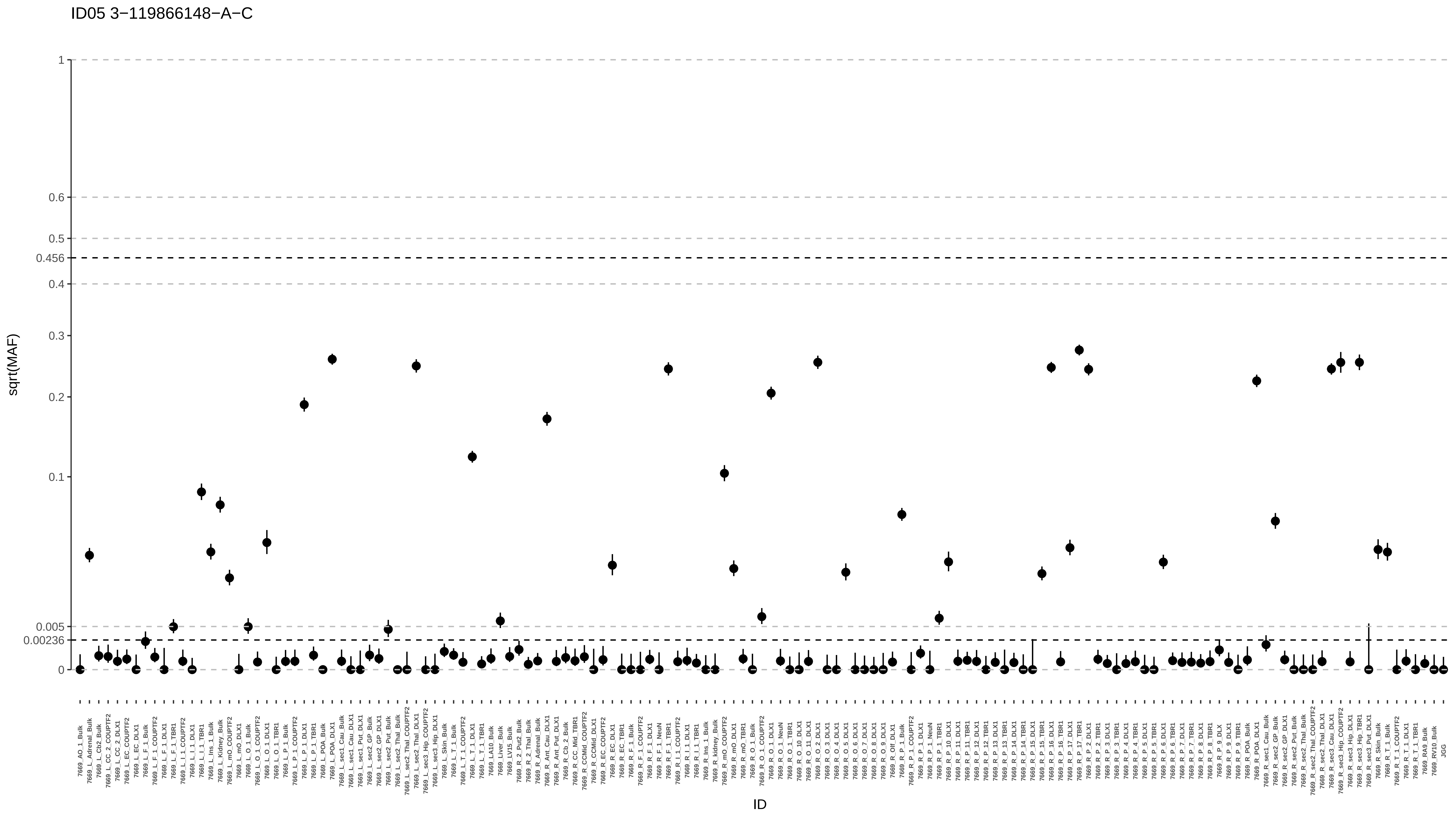

ID05 3−119866148−A−C
sqrt(MAF)
1
0.6
0.5
0.456
0.4
0.3
0.2
0.1
0.005
0.00236
0
ID
7669_AO_1_Bulk
7669_L_Adrenal_Bulk
7669_L_Cb2_Bulk
7669_L_CC_2_COUPTF2
7669_L_CC_2_DLX1
7669_L_EC_COUPTF2
7669_L_EC_DLX1
7669_L_F_1_Bulk
7669_L_F_1_COUPTF2
7669_L_F_1_DLX1
7669_L_F_1_TBR1
7669_L_I_1_COUPTF2
7669_L_I_1_DLX1
7669_L_I_1_TBR1
7669_L_Ins_1_Bulk
7669_L_Kidney_Bulk
7669_L_mO_COUPTF2
7669_L_mO_DLX1
7669_L_O_1_Bulk
7669_L_O_1_COUPTF2
7669_L_O_1_DLX1
7669_L_O_1_TBR1
7669_L_P_1_Bulk
7669_L_P_1_COUPTF2
7669_L_P_1_DLX1
7669_L_P_1_TBR1
7669_L_POA_Bulk
7669_L_POA_DLX1
7669_L_sec1_Cau_Bulk
7669_L_sec1_Cau_DLX1
7669_L_sec1_Put_DLX1
7669_L_sec2_GP_Bulk
7669_L_sec2_GP_DLX1
7669_L_sec2_Put_Bulk
7669_L_sec2_Thal_Bulk
7669_L_sec2_Thal_COUPTF2
7669_L_sec2_Thal_DLX1
7669_L_sec3_Hip_COUPTF2
7669_L_sec3_Hip_DLX1
7669_L_Skin_Bulk
7669_L_T_1_Bulk
7669_L_T_1_COUPTF2
7669_L_T_1_DLX1
7669_L_T_1_TBR1
7669_LA10_Bulk
7669_Liver_Bulk
7669_LV15_Bulk
7669_R_2_Put2_Bulk
7669_R_2_Thal_Bulk
7669_R_Adrenal_Bulk
7669_R_Ant_Cau_DLX1
7669_R_Ant_Put_DLX1
7669_R_Cb_2_Bulk
7669_R_CC_Mid_TBR1
7669_R_CCMid_COUPTF2
7669_R_CCMid_DLX1
7669_R_EC_COUPTF2
7669_R_EC_DLX1
7669_R_EC_TBR1
7669_R_F_1_Bulk
7669_R_F_1_COUPTF2
7669_R_F_1_DLX1
7669_R_F_1_NeuN
7669_R_F_1_TBR1
7669_R_I_1_COUPTF2
7669_R_I_1_DLX1
7669_R_I_1_TBR1
7669_R_Ins_1_Bulk
7669_R_kidney_Bulk
7669_R_mO_COUPTF2
7669_R_mO_DLX1
7669_R_mO_TBR1
7669_R_O_1_Bulk
7669_R_O_1_COUPTF2
7669_R_O_1_DLX1
7669_R_O_1_NeuN
7669_R_O_1_TBR1
7669_R_O_10_DLX1
7669_R_O_11_DLX1
7669_R_O_2_DLX1
7669_R_O_3_DLX1
7669_R_O_4_DLX1
7669_R_O_5_DLX1
7669_R_O_6_DLX1
7669_R_O_7_DLX1
7669_R_O_8_DLX1
7669_R_O_9_DLX1
7669_R_Olf_DLX1
7669_R_P_1_Bulk
7669_R_P_1_COUPTF2
7669_R_P_1_DLX1
7669_R_P_1_NeuN
7669_R_P_1_TBR1
7669_R_P_10_DLX1
7669_R_P_11_DLX1
7669_R_P_11_TBR1
7669_R_P_12_DLX1
7669_R_P_12_TBR1
7669_R_P_13_DLX1
7669_R_P_13_TBR1
7669_R_P_14_DLX1
7669_R_P_14_TBR1
7669_R_P_15_DLX1
7669_R_P_15_TBR1
7669_R_P_16_DLX1
7669_R_P_16_TBR1
7669_R_P_17_DLX1
7669_R_P_17_TBR1
7669_R_P_2_DLX1
7669_R_P_2_TBR1
7669_R_P_3_DLX1
7669_R_P_3_TBR1
7669_R_P_4_DLX1
7669_R_P_4_TBR1
7669_R_P_5_DLX1
7669_R_P_5_TBR1
7669_R_P_6_DLX1
7669_R_P_6_TBR1
7669_R_P_7_DLX1
7669_R_P_7_TBR1
7669_R_P_8_DLX1
7669_R_P_8_TBR1
7669_R_P_9_DLX
7669_R_P_9_DLX1
7669_R_P_9_TBR1
7669_R_POA_Bulk
7669_R_POA_DLX1
7669_R_sec1_Cau_Bulk
7669_R_sec2_GP_Bulk
7669_R_sec2_GP_DLX1
7669_R_sec2_Put_Bulk
7669_R_sec2_Thal_Bulk
7669_R_sec2_Thal_COUPTF2
7669_R_sec2_Thal_DLX1
7669_R_sec3_Cau_DLX1
7669_R_sec3_Hip_COUPTF2
7669_R_sec3_Hip_DLX1
7669_R_sec3_Hip_TBR1
7669_R_sec3_Put_DLX1
7669_R_Skin_Bulk
7669_R_T_1_Bulk
7669_R_T_1_COUPTF2
7669_R_T_1_DLX1
7669_R_T_1_TBR1
7669_RA9_Bulk
7669_RV10_Bulk
JGG

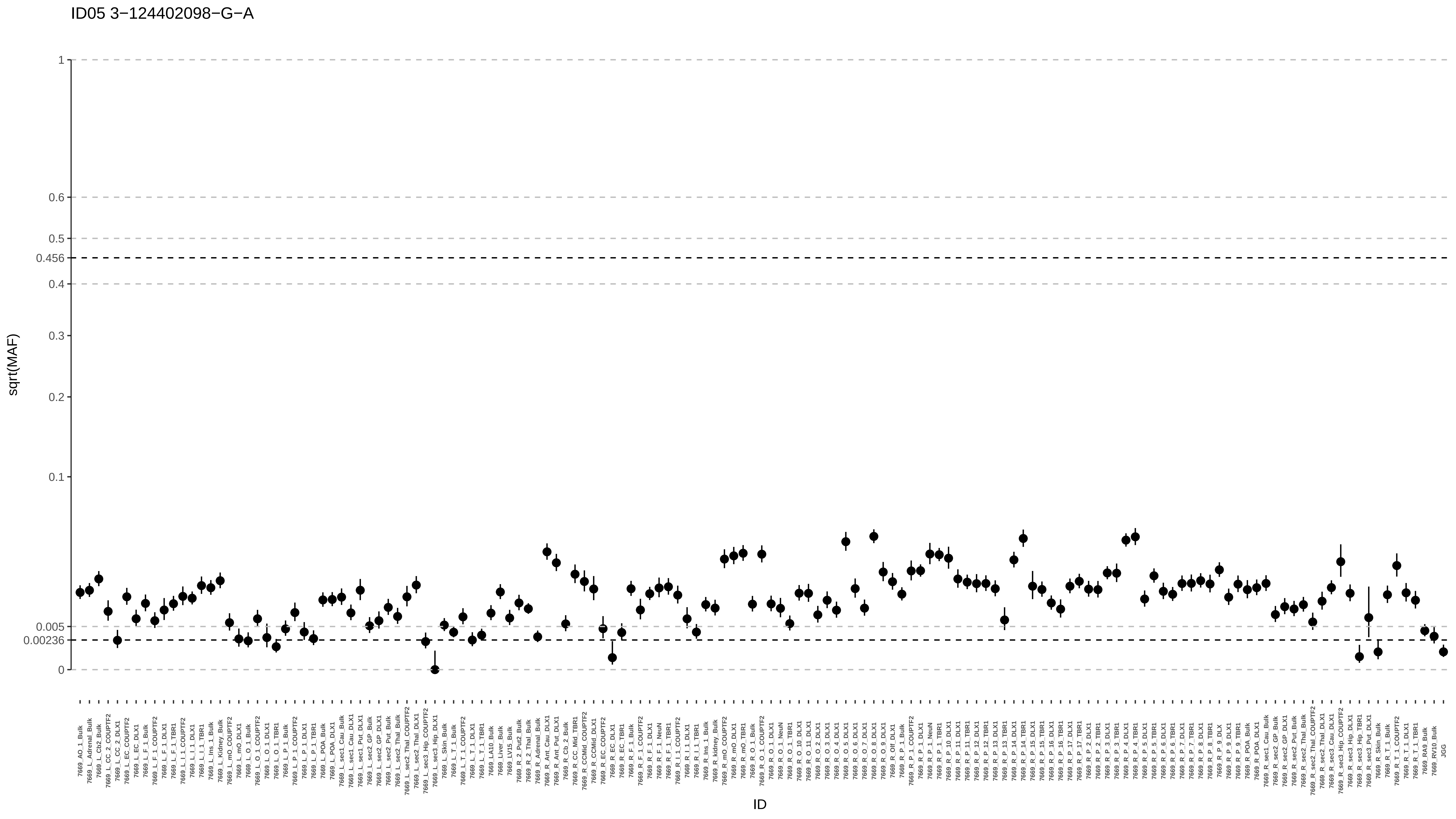
ID05 3−124402098−G−A
sqrt(MAF)
1
0.6
0.5
0.456
0.4
0.3
0.2
0.1
0.005
0.00236
0
ID
7669_AO_1_Bulk
7669_L_Adrenal_Bulk
7669_L_Cb2_Bulk
7669_L_CC_2_COUPTF2
7669_L_CC_2_DLX1
7669_L_EC_COUPTF2
7669_L_EC_DLX1
7669_L_F_1_Bulk
7669_L_F_1_COUPTF2
7669_L_F_1_DLX1
7669_L_F_1_TBR1
7669_L_I_1_COUPTF2
7669_L_I_1_DLX1
7669_L_I_1_TBR1
7669_L_Ins_1_Bulk
7669_L_Kidney_Bulk
7669_L_mO_COUPTF2
7669_L_mO_DLX1
7669_L_O_1_Bulk
7669_L_O_1_COUPTF2
7669_L_O_1_DLX1
7669_L_O_1_TBR1
7669_L_P_1_Bulk
7669_L_P_1_COUPTF2
7669_L_P_1_DLX1
7669_L_P_1_TBR1
7669_L_POA_Bulk
7669_L_POA_DLX1
7669_L_sec1_Cau_Bulk
7669_L_sec1_Cau_DLX1
7669_L_sec1_Put_DLX1
7669_L_sec2_GP_Bulk
7669_L_sec2_GP_DLX1
7669_L_sec2_Put_Bulk
7669_L_sec2_Thal_Bulk
7669_L_sec2_Thal_COUPTF2
7669_L_sec2_Thal_DLX1
7669_L_sec3_Hip_COUPTF2
7669_L_sec3_Hip_DLX1
7669_L_Skin_Bulk
7669_L_T_1_Bulk
7669_L_T_1_COUPTF2
7669_L_T_1_DLX1
7669_L_T_1_TBR1
7669_LA10_Bulk
7669_Liver_Bulk
7669_LV15_Bulk
7669_R_2_Put2_Bulk
7669_R_2_Thal_Bulk
7669_R_Adrenal_Bulk
7669_R_Ant_Cau_DLX1
7669_R_Ant_Put_DLX1
7669_R_Cb_2_Bulk
7669_R_CC_Mid_TBR1
7669_R_CCMid_COUPTF2
7669_R_CCMid_DLX1
7669_R_EC_COUPTF2
7669_R_EC_DLX1
7669_R_EC_TBR1
7669_R_F_1_Bulk
7669_R_F_1_COUPTF2
7669_R_F_1_DLX1
7669_R_F_1_NeuN
7669_R_F_1_TBR1
7669_R_I_1_COUPTF2
7669_R_I_1_DLX1
7669_R_I_1_TBR1
7669_R_Ins_1_Bulk
7669_R_kidney_Bulk
7669_R_mO_COUPTF2
7669_R_mO_DLX1
7669_R_mO_TBR1
7669_R_O_1_Bulk
7669_R_O_1_COUPTF2
7669_R_O_1_DLX1
7669_R_O_1_NeuN
7669_R_O_1_TBR1
7669_R_O_10_DLX1
7669_R_O_11_DLX1
7669_R_O_2_DLX1
7669_R_O_3_DLX1
7669_R_O_4_DLX1
7669_R_O_5_DLX1
7669_R_O_6_DLX1
7669_R_O_7_DLX1
7669_R_O_8_DLX1
7669_R_O_9_DLX1
7669_R_Olf_DLX1
7669_R_P_1_Bulk
7669_R_P_1_COUPTF2
7669_R_P_1_DLX1
7669_R_P_1_NeuN
7669_R_P_1_TBR1
7669_R_P_10_DLX1
7669_R_P_11_DLX1
7669_R_P_11_TBR1
7669_R_P_12_DLX1
7669_R_P_12_TBR1
7669_R_P_13_DLX1
7669_R_P_13_TBR1
7669_R_P_14_DLX1
7669_R_P_14_TBR1
7669_R_P_15_DLX1
7669_R_P_15_TBR1
7669_R_P_16_DLX1
7669_R_P_16_TBR1
7669_R_P_17_DLX1
7669_R_P_17_TBR1
7669_R_P_2_DLX1
7669_R_P_2_TBR1
7669_R_P_3_DLX1
7669_R_P_3_TBR1
7669_R_P_4_DLX1
7669_R_P_4_TBR1
7669_R_P_5_DLX1
7669_R_P_5_TBR1
7669_R_P_6_DLX1
7669_R_P_6_TBR1
7669_R_P_7_DLX1
7669_R_P_7_TBR1
7669_R_P_8_DLX1
7669_R_P_8_TBR1
7669_R_P_9_DLX
7669_R_P_9_DLX1
7669_R_P_9_TBR1
7669_R_POA_Bulk
7669_R_POA_DLX1
7669_R_sec1_Cau_Bulk
7669_R_sec2_GP_Bulk
7669_R_sec2_GP_DLX1
7669_R_sec2_Put_Bulk
7669_R_sec2_Thal_Bulk
7669_R_sec2_Thal_COUPTF2
7669_R_sec2_Thal_DLX1
7669_R_sec3_Cau_DLX1
7669_R_sec3_Hip_COUPTF2
7669_R_sec3_Hip_DLX1
7669_R_sec3_Hip_TBR1
7669_R_sec3_Put_DLX1
7669_R_Skin_Bulk
7669_R_T_1_Bulk
7669_R_T_1_COUPTF2
7669_R_T_1_DLX1
7669_R_T_1_TBR1
7669_RA9_Bulk
7669_RV10_Bulk
JGG

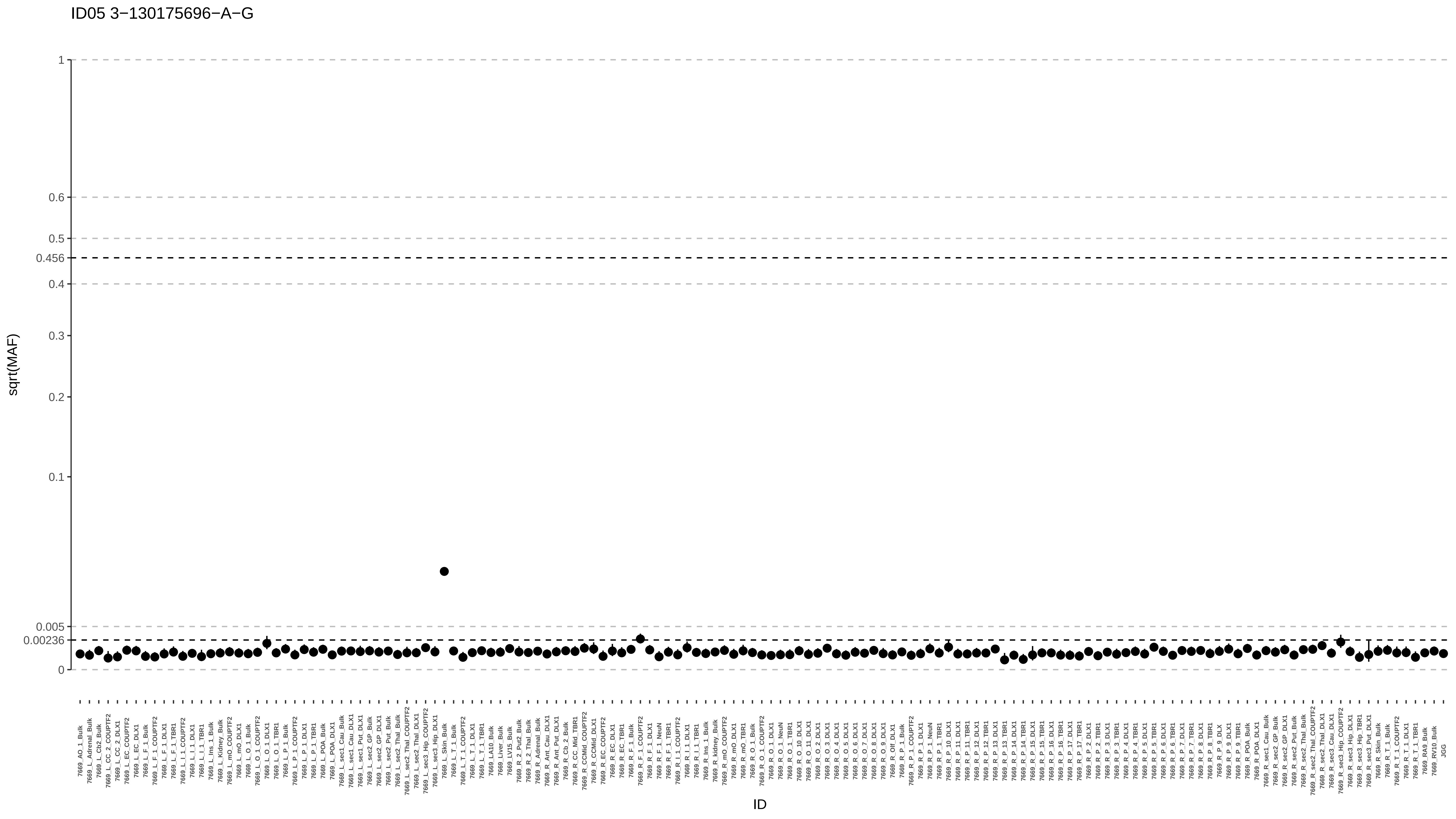
ID05 3−130175696−A−G
sqrt(MAF)
ID
1
0.6
0.5
0.456
0.4
0.3
0.2
0.1
0.005
0.00236
0

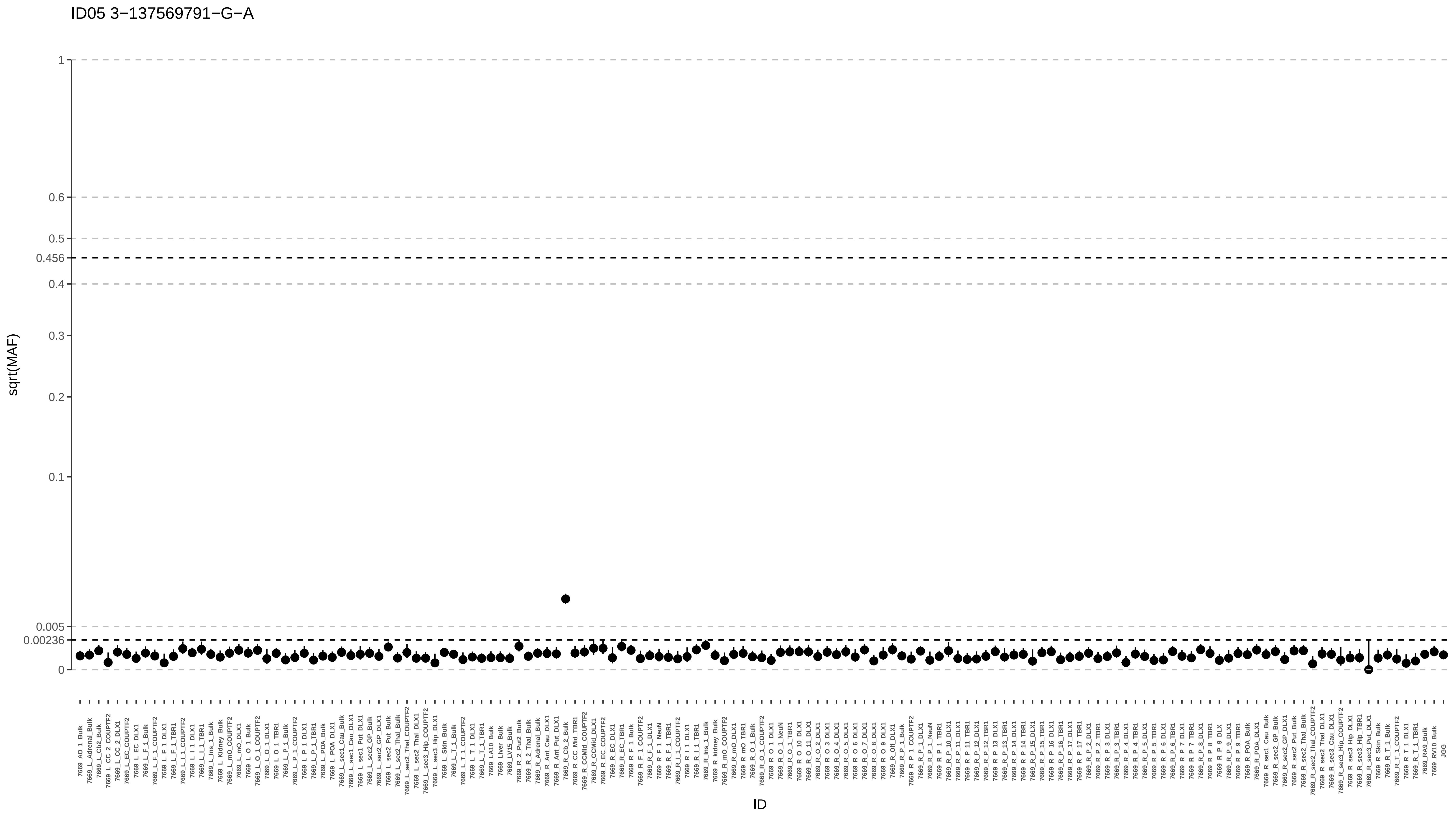

ID05 3−137569791−G−A
sqrt(MAF)
1
0.6
0.5
0.456
0.4
0.3
0.2
0.1
0.005
0.00236
0
ID

ID05 3−145391171−A−G

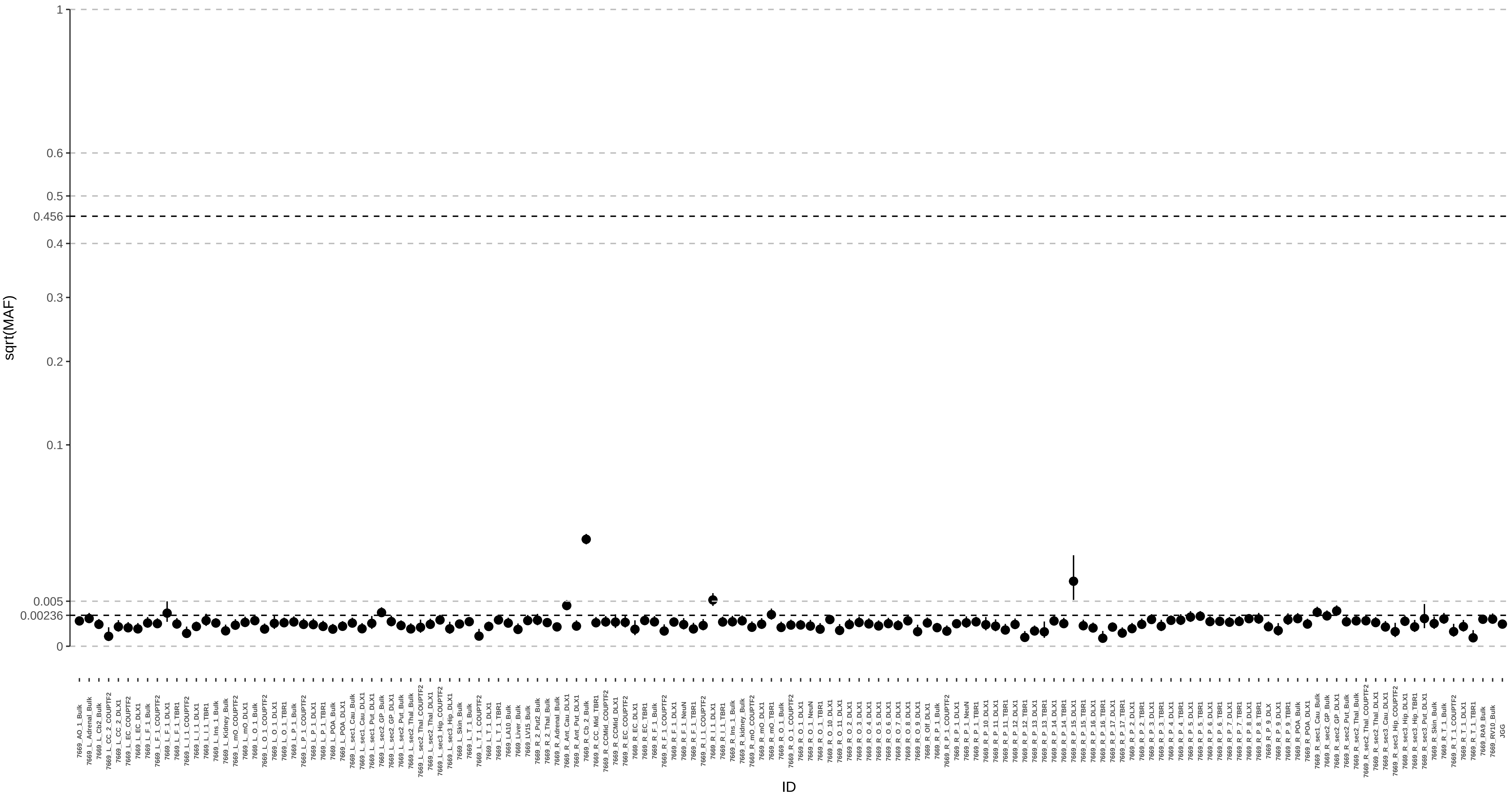

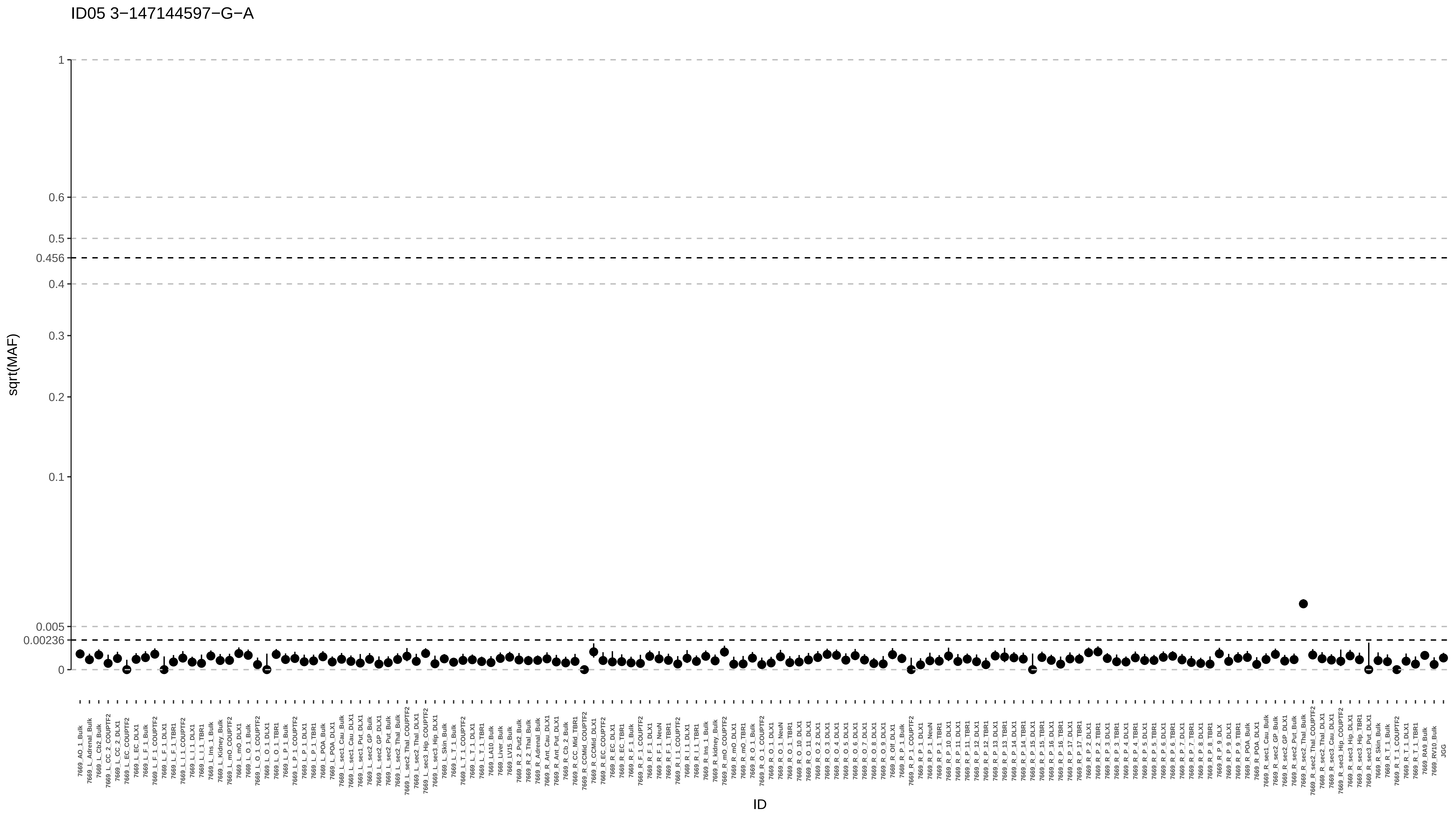
ID05 3−147144597−G−A
sqrt(MAF)
1
0.6
0.5
0.456
0.4
0.3
0.2
0.1
0.005
0.00236
0
ID

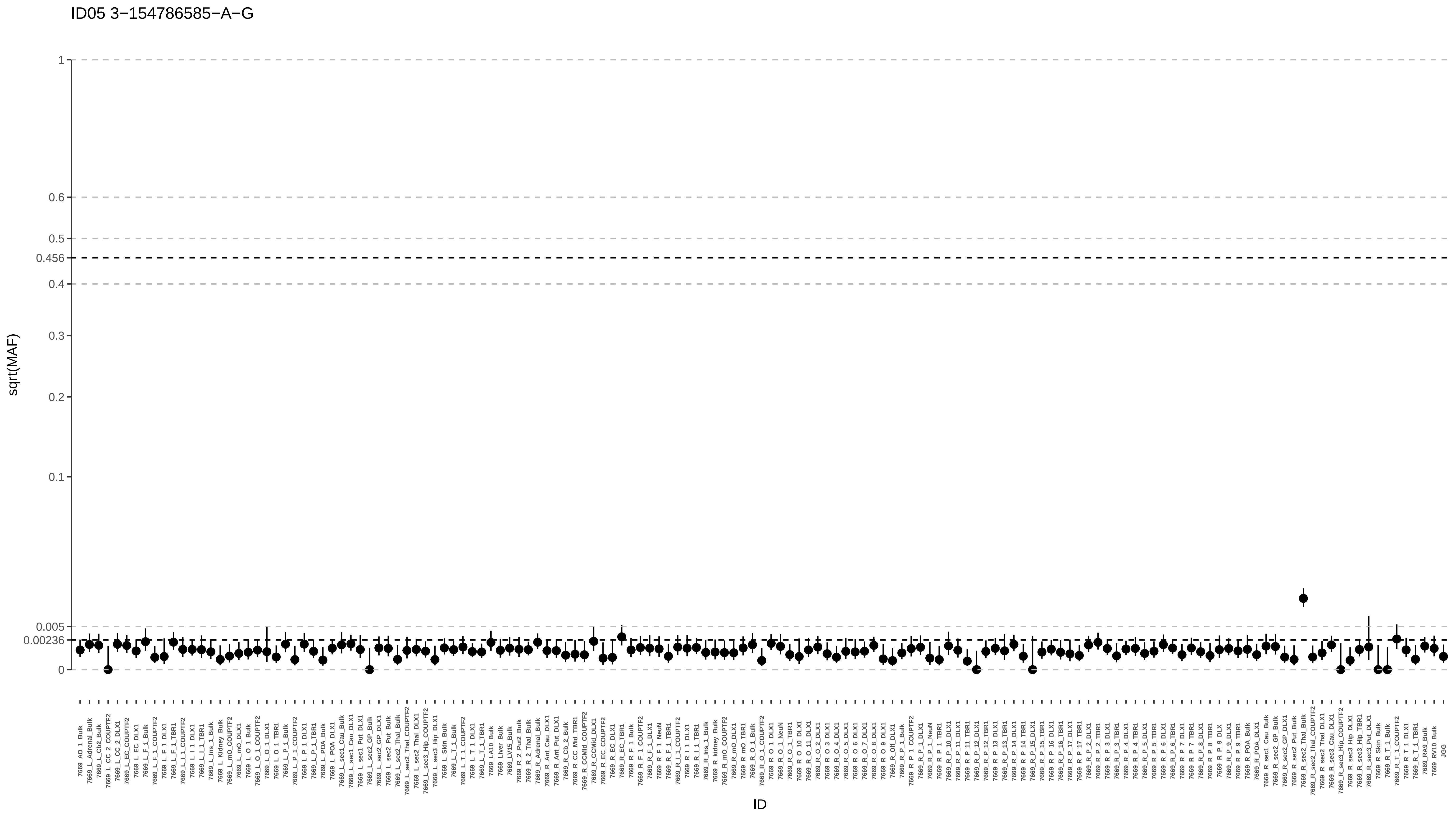
ID05 3−154786585−A−G
sqrt(MAF)
1
0.6
0.5
0.456
0.4
0.3
0.2
0.1
0.005
0.00236
0
ID
JGG

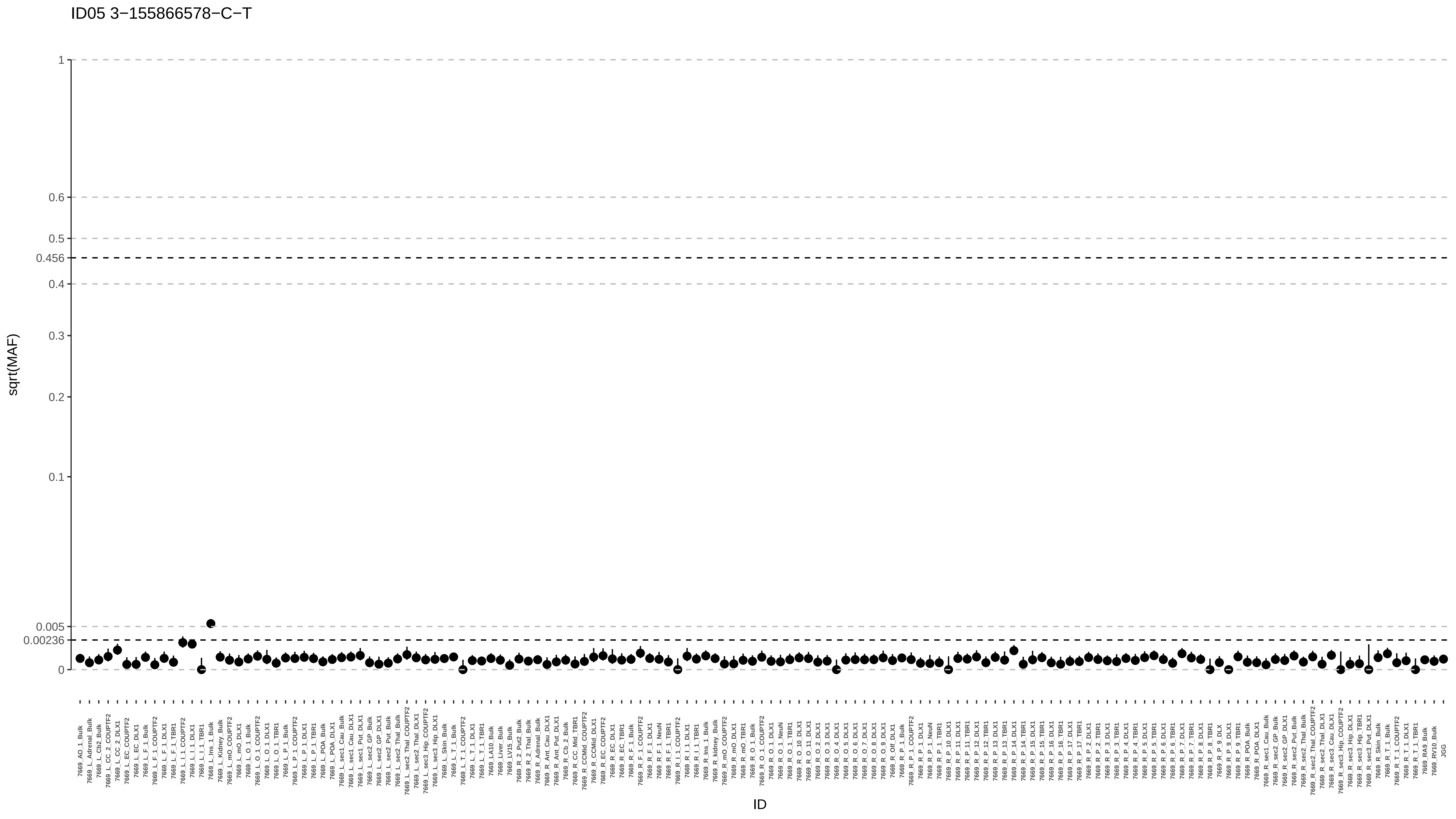
ID05 3−155866578−C−T
sqrt(MAF)
ID

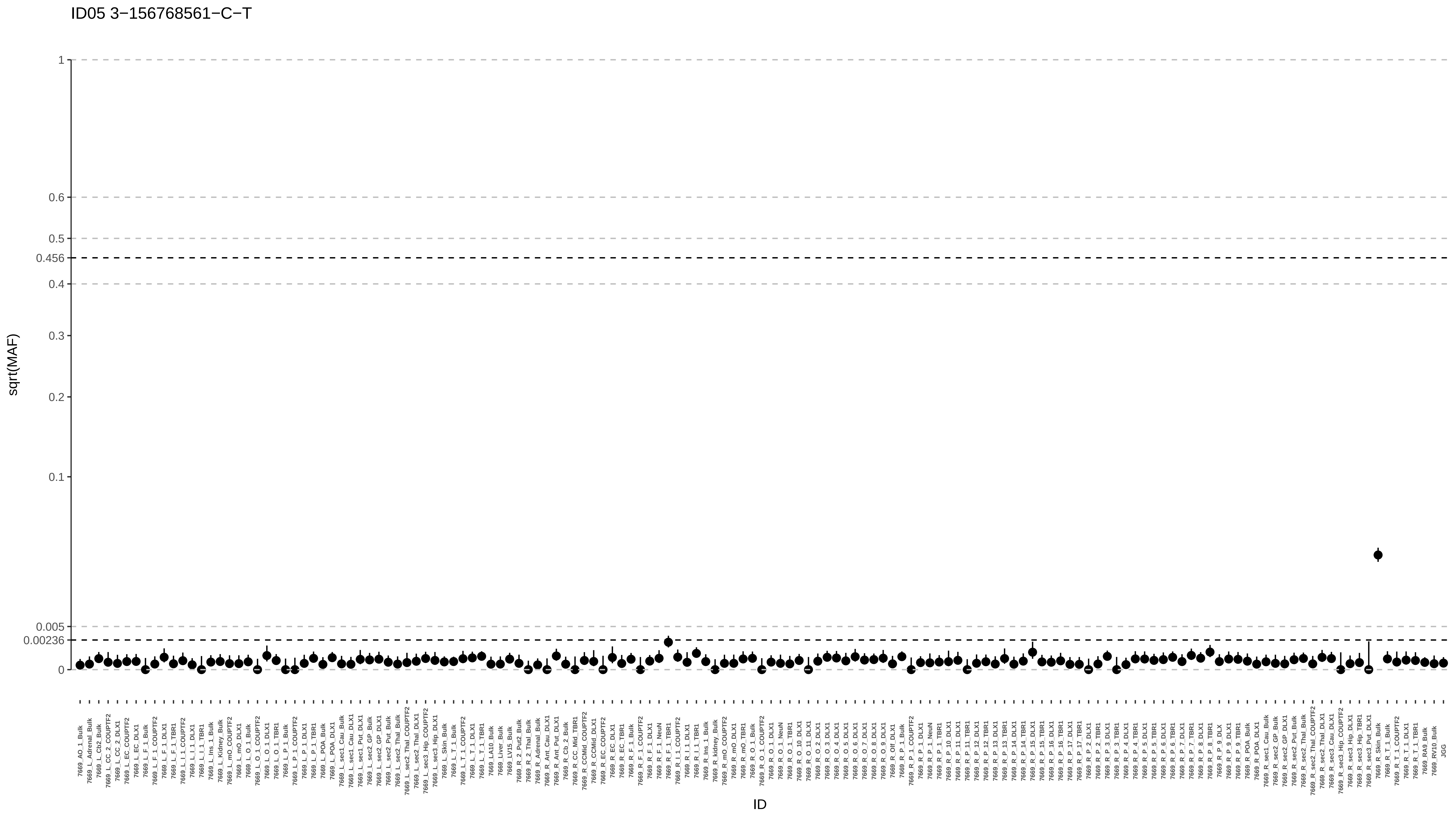
ID05 3−156768561−C−T
sqrt(MAF)
1
0.6
0.5
0.456
0.4
0.3
0.2
0.1
0.005
0.00236
0
ID

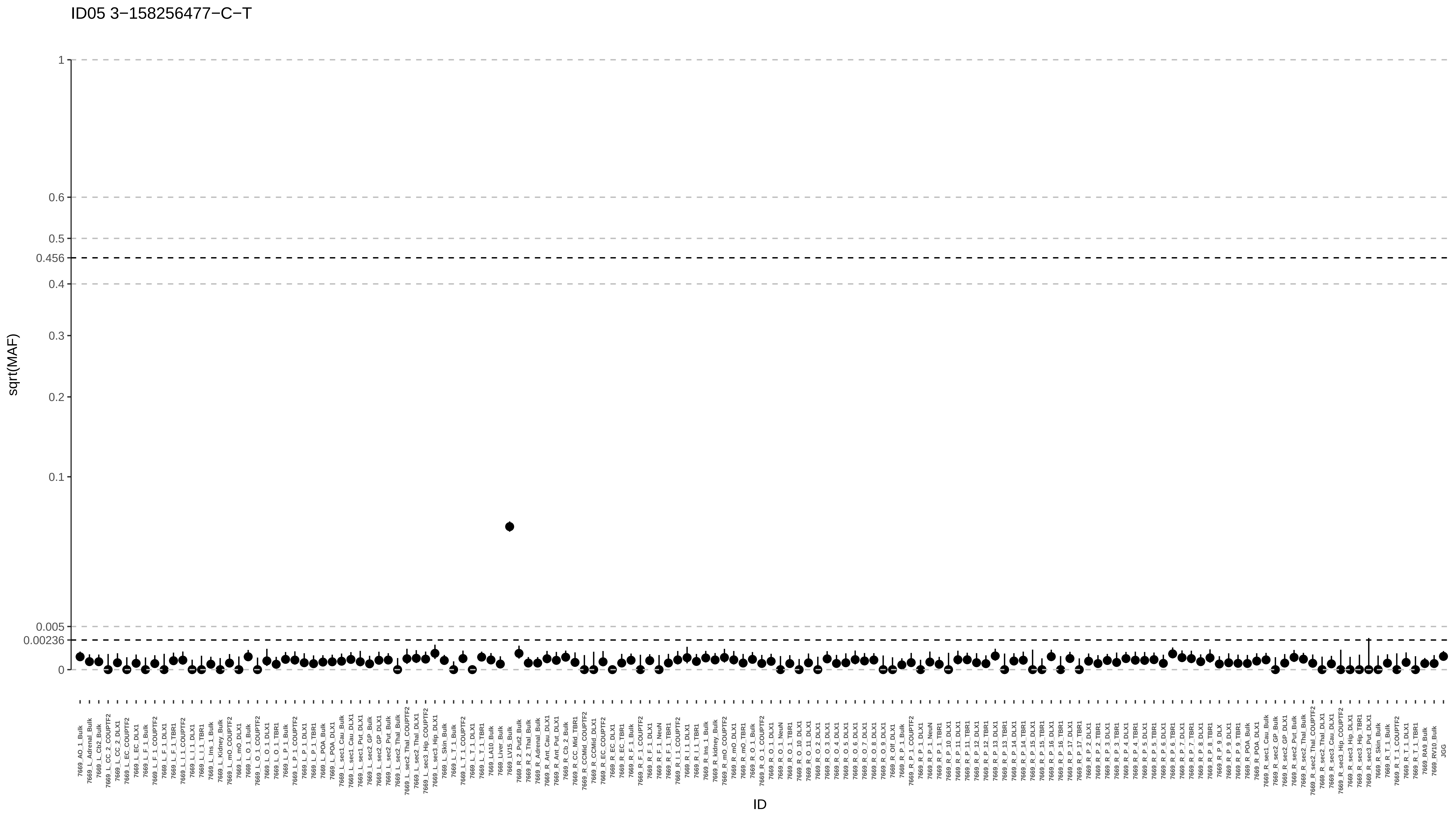
ID05 3−158256477−C−T
sqrt(MAF)
1
0.6
0.5
0.456
0.4
0.3
0.2
0.1
0.005
0.00236
0
ID

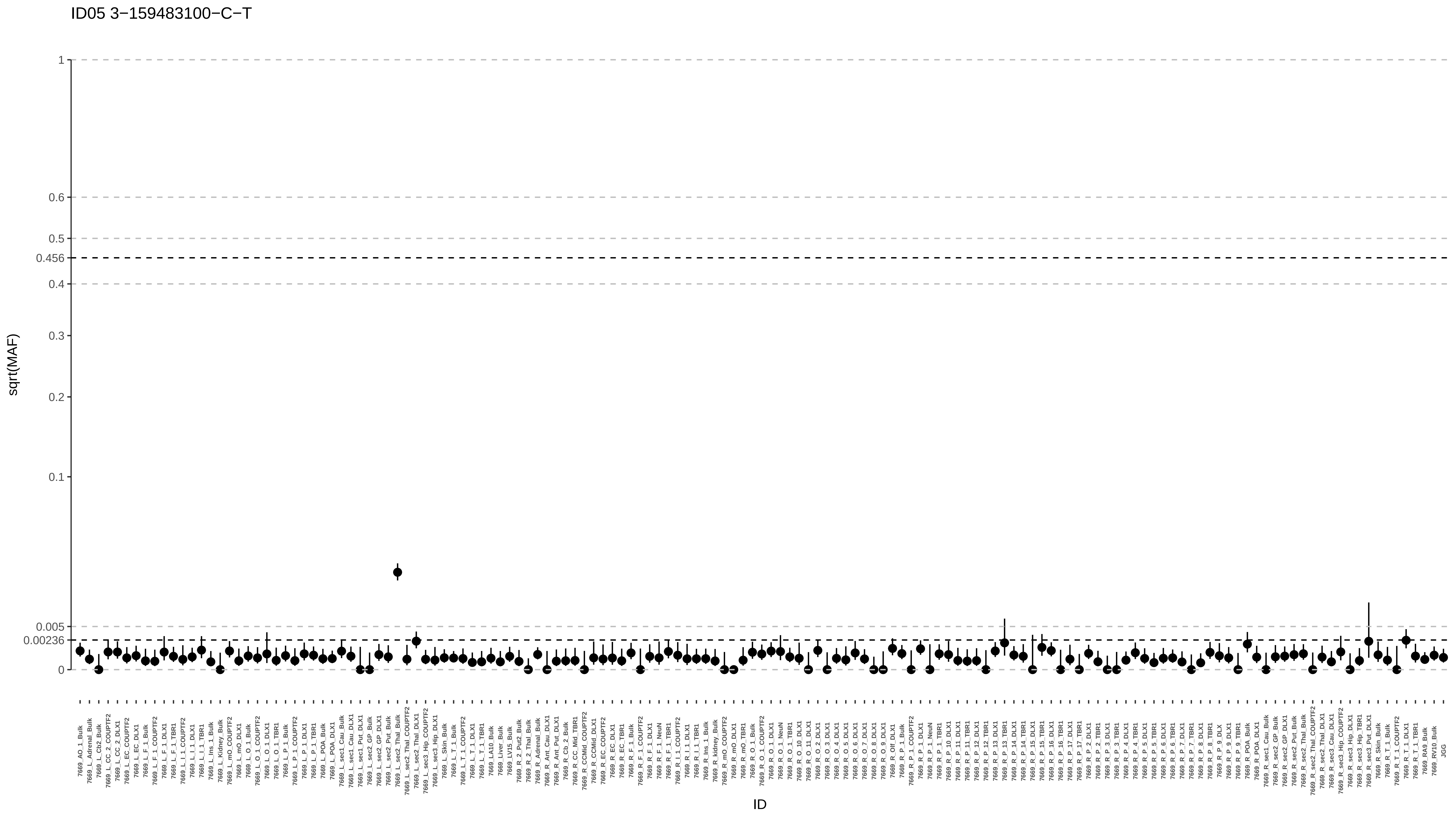
ID05 3−159483100−C−T
sqrt(MAF)
1
0.6
0.5
0.456
0.4
0.3
0.2
0.1
0.005
0.00236
0
ID

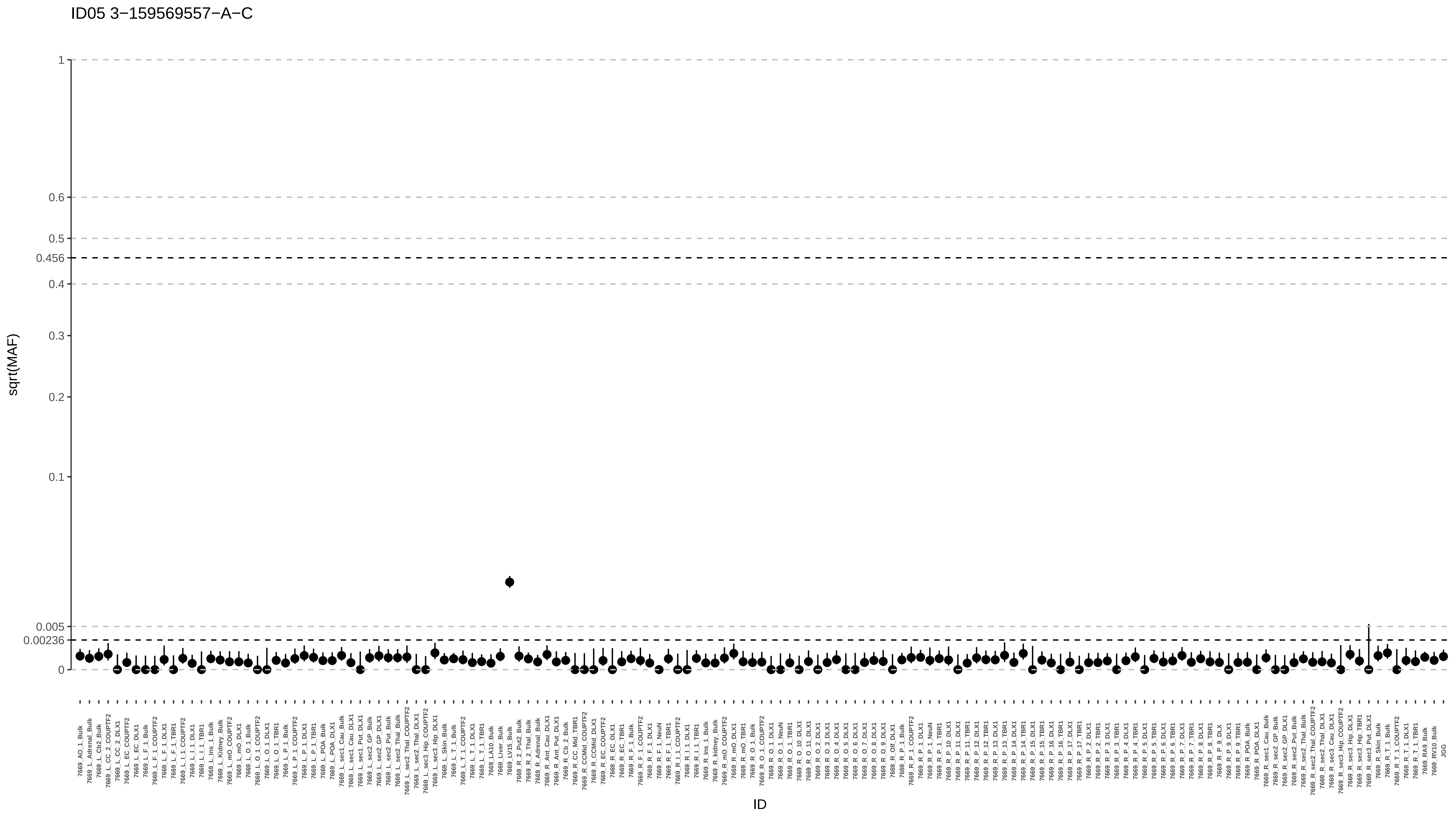
ID05 3−159569557−A−C
sqrt(MAF)
ID
1
0.6
0.5
0.456
0.4
0.3
0.2
0.1
0.005
0.00236
0

ID05 3−164879470−C−T

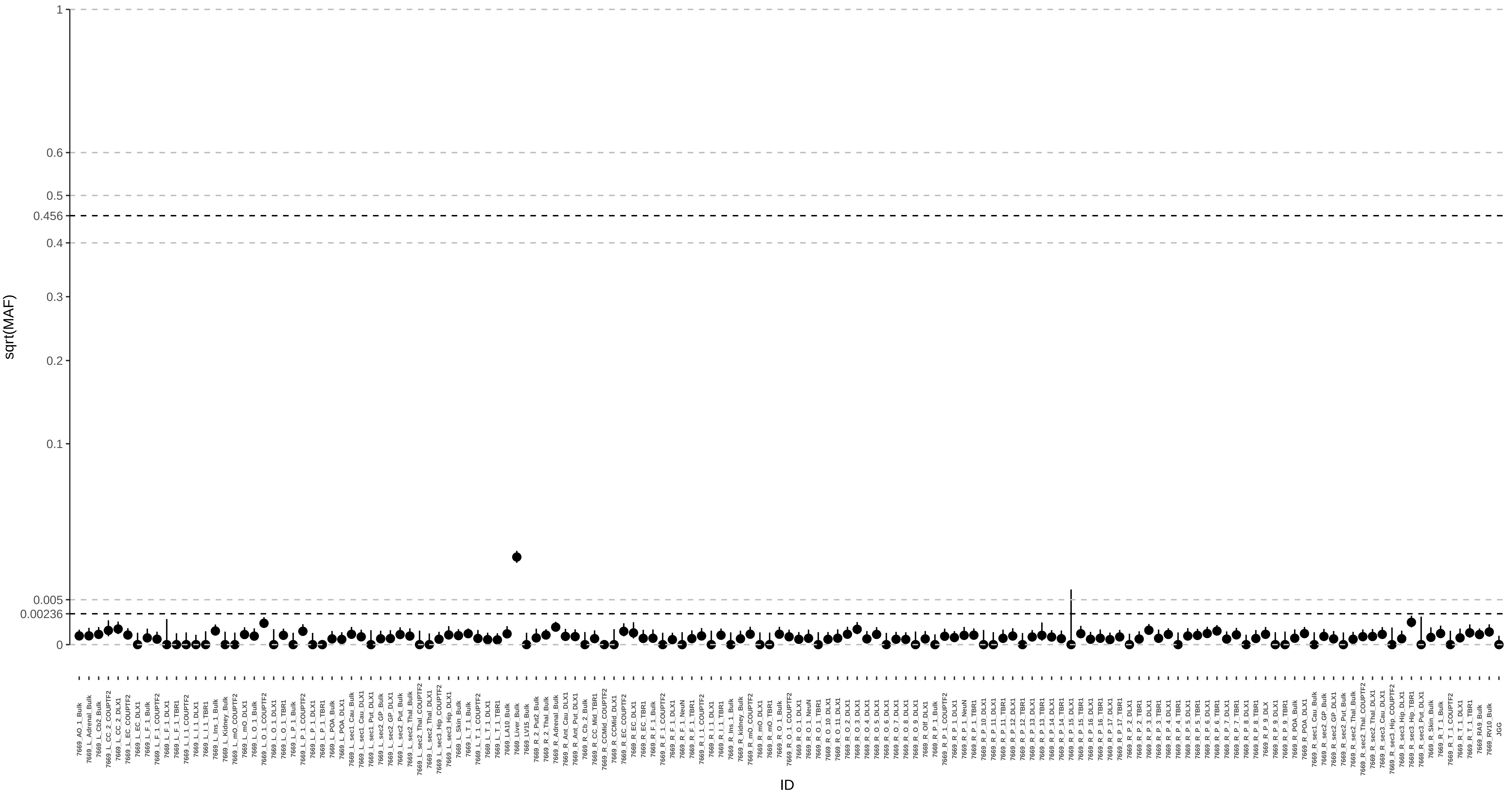

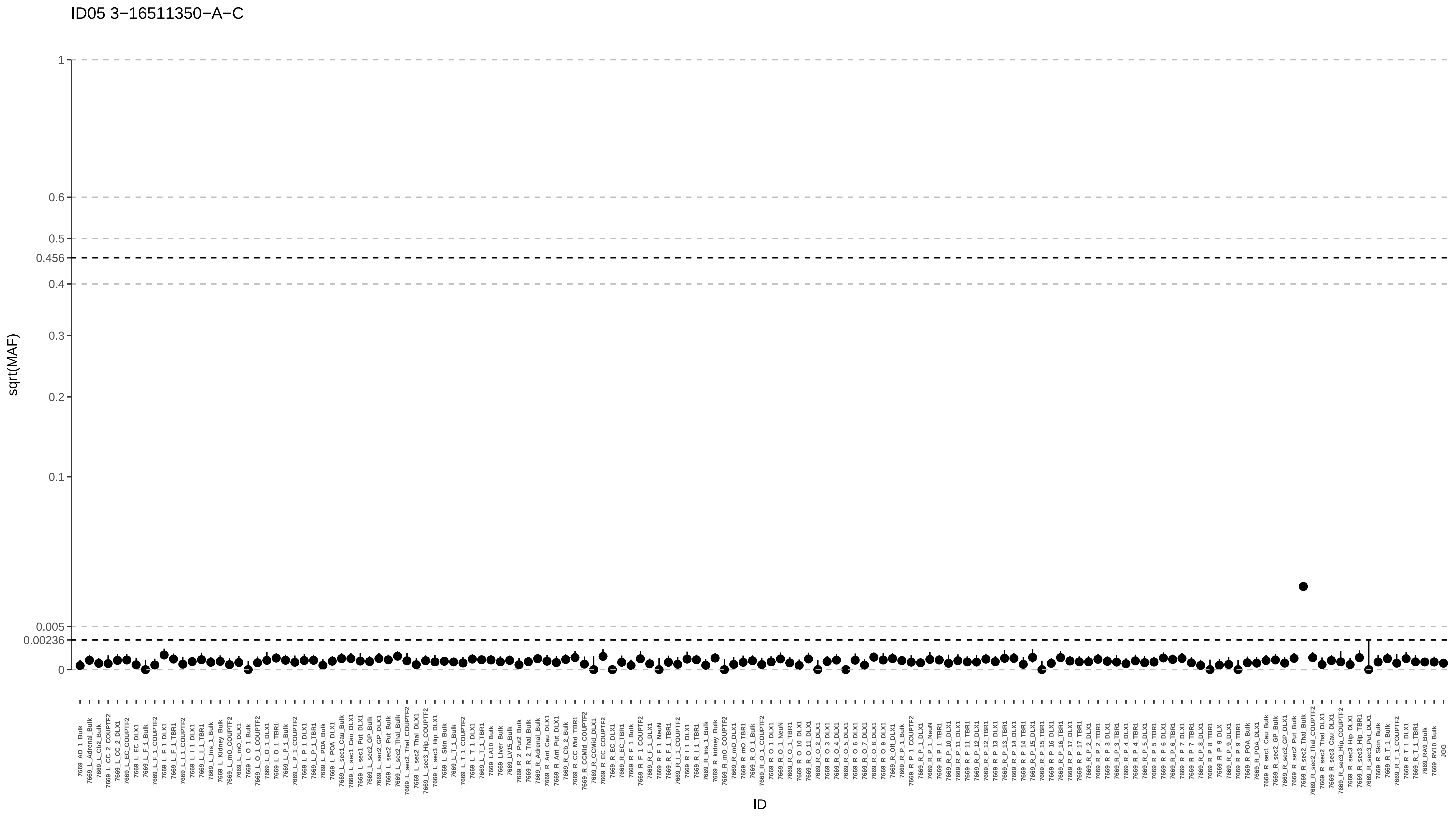
ID05 3−16511350−A−C
sqrt(MAF)
1
0.6
0.5
0.456
0.4
0.3
0.2
0.1
0.005
0.00236
0
ID

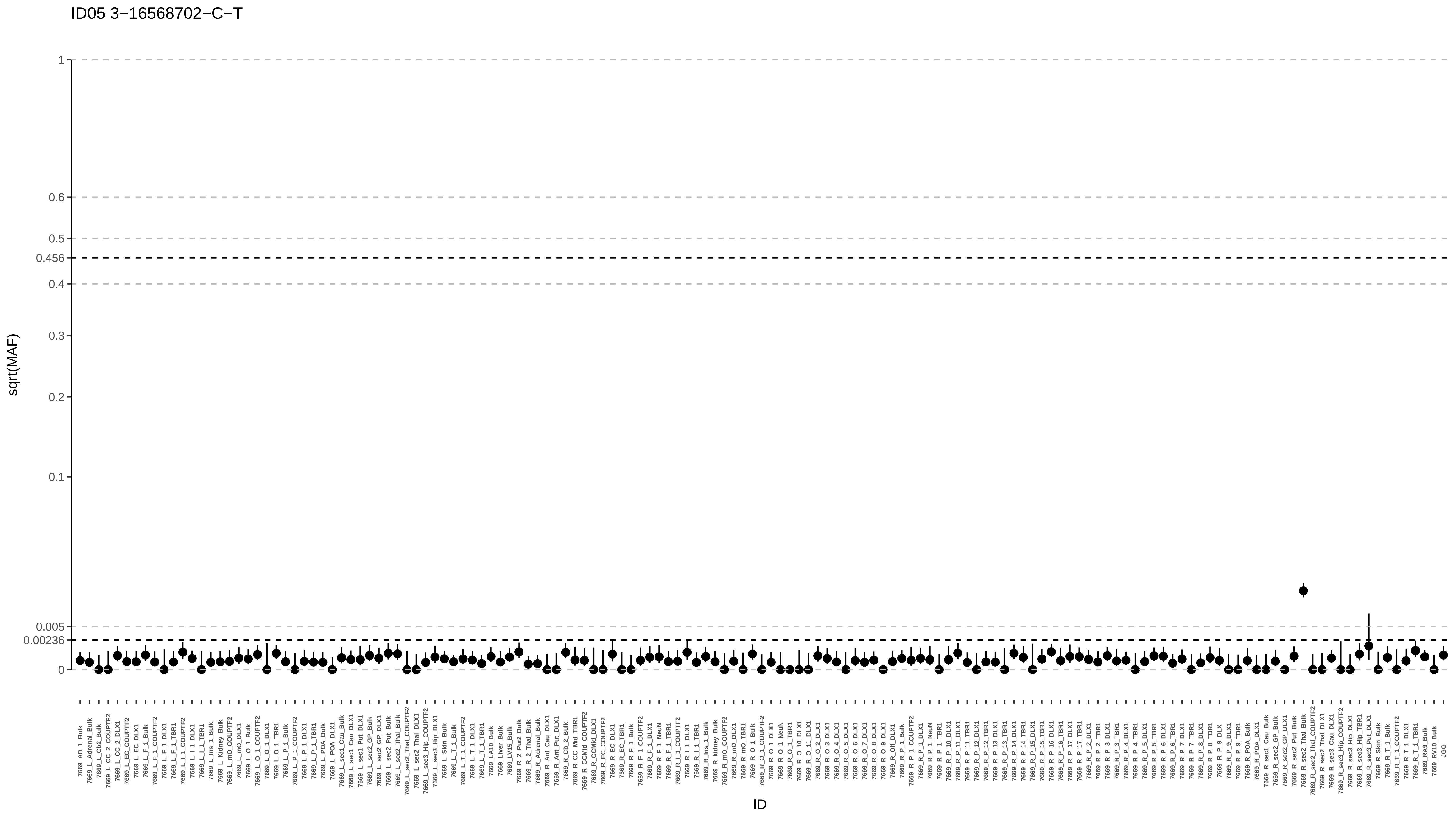
ID05 3−16568702−C−T
sqrt(MAF)
1
0.6
0.5
0.456
0.4
0.3
0.2
0.1
0.005
0.00236
0
ID

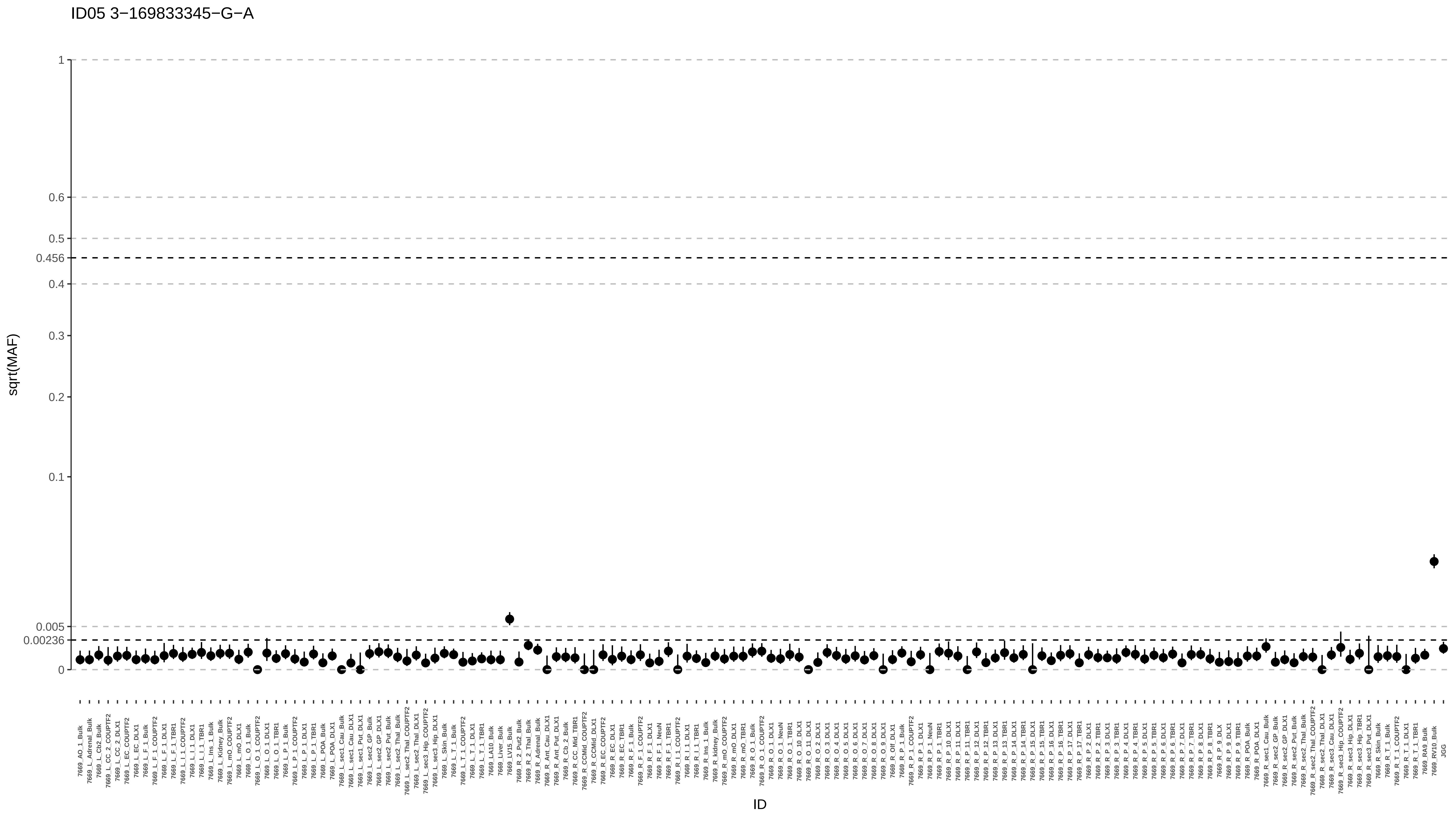
ID05 3−169833345−G−A
sqrt(MAF)
1
0.6
0.5
0.456
0.4
0.3
0.2
0.1
0.005
0.00236
0
ID

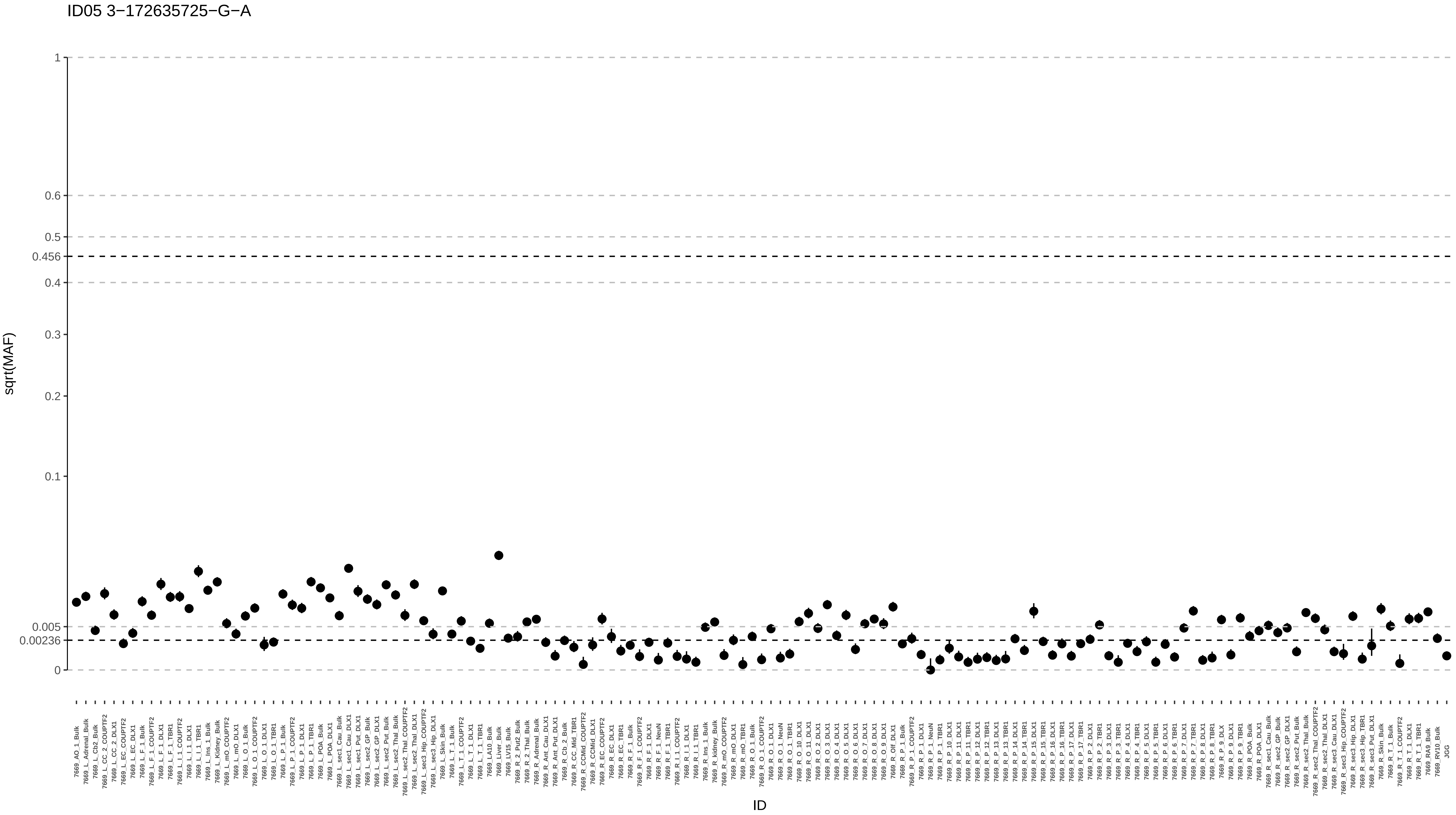
ID05 3−172635725−G−A
sqrt(MAF)
1
0.6
0.5
0.456
0.4
0.3
0.2
0.1
0.005
0.00236
0
ID
7669_AO_1_Bulk
7669_L_Adrenal_Bulk
7669_L_Cb2_Bulk
7669_L_CC_2_COUPTF2
7669_L_CC_2_DLX1
7669_L_EC_COUPTF2
7669_L_EC_DLX1
7669_L_F_1_Bulk
7669_L_F_1_COUPTF2
7669_L_F_1_DLX1
7669_L_F_1_TBR1
7669_L_I_1_COUPTF2
7669_L_I_1_DLX1
7669_L_I_1_TBR1
7669_L_Ins_1_Bulk
7669_L_Kidney_Bulk
7669_L_mO_COUPTF2
7669_L_mO_DLX1
7669_L_O_1_Bulk
7669_L_O_1_COUPTF2
7669_L_O_1_DLX1
7669_L_O_1_TBR1
7669_L_P_1_Bulk
7669_L_P_1_COUPTF2
7669_L_P_1_DLX1
7669_L_P_1_TBR1
7669_L_POA_Bulk
7669_L_POA_DLX1
7669_L_sec1_Cau_Bulk
7669_L_sec1_Cau_DLX1
7669_L_sec1_Put_DLX1
7669_L_sec2_GP_Bulk
7669_L_sec2_GP_DLX1
7669_L_sec2_Put_Bulk
7669_L_sec2_Thal_Bulk
7669_L_sec2_Thal_COUPTF2
7669_L_sec2_Thal_DLX1
7669_L_sec3_Hip_COUPTF2
7669_L_sec3_Hip_DLX1
7669_L_Skin_Bulk
7669_L_T_1_Bulk
7669_L_T_1_COUPTF2
7669_L_T_1_DLX1
7669_L_T_1_TBR1
7669_LA10_Bulk
7669_Liver_Bulk
7669_LV15_Bulk
7669_R_2_Put2_Bulk
7669_R_2_Thal_Bulk
7669_R_Adrenal_Bulk
7669_R_Ant_Cau_DLX1
7669_R_Ant_Put_DLX1
7669_R_Cb_2_Bulk
7669_R_CC_Mid_TBR1
7669_R_CCMid_COUPTF2
7669_R_CCMid_DLX1
7669_R_EC_COUPTF2
7669_R_EC_DLX1
7669_R_EC_TBR1
7669_R_F_1_Bulk
7669_R_F_1_COUPTF2
7669_R_F_1_DLX1
7669_R_F_1_NeuN
7669_R_F_1_TBR1
7669_R_I_1_COUPTF2
7669_R_I_1_DLX1
7669_R_I_1_TBR1
7669_R_Ins_1_Bulk
7669_R_kidney_Bulk
7669_R_mO_COUPTF2
7669_R_mO_DLX1
7669_R_mO_TBR1
7669_R_O_1_Bulk
7669_R_O_1_COUPTF2
7669_R_O_1_DLX1
7669_R_O_1_NeuN
7669_R_O_1_TBR1
7669_R_O_10_DLX1
7669_R_O_11_DLX1
7669_R_O_2_DLX1
7669_R_O_3_DLX1
7669_R_O_4_DLX1
7669_R_O_5_DLX1
7669_R_O_6_DLX1
7669_R_O_7_DLX1
7669_R_O_8_DLX1
7669_R_O_9_DLX1
7669_R_Olf_DLX1
7669_R_P_1_Bulk
7669_R_P_1_COUPTF2
7669_R_P_1_DLX1
7669_R_P_1_NeuN
7669_R_P_1_TBR1
7669_R_P_10_DLX1
7669_R_P_11_DLX1
7669_R_P_11_TBR1
7669_R_P_12_DLX1
7669_R_P_12_TBR1
7669_R_P_13_DLX1
7669_R_P_13_TBR1
7669_R_P_14_DLX1
7669_R_P_14_TBR1
7669_R_P_15_DLX1
7669_R_P_15_TBR1
7669_R_P_16_DLX1
7669_R_P_16_TBR1
7669_R_P_17_DLX1
7669_R_P_17_TBR1
7669_R_P_2_DLX1
7669_R_P_2_TBR1
7669_R_P_3_DLX1
7669_R_P_3_TBR1
7669_R_P_4_DLX1
7669_R_P_4_TBR1
7669_R_P_5_DLX1
7669_R_P_5_TBR1
7669_R_P_6_DLX1
7669_R_P_6_TBR1
7669_R_P_7_DLX1
7669_R_P_7_TBR1
7669_R_P_8_DLX1
7669_R_P_8_TBR1
7669_R_P_9_DLX
7669_R_P_9_DLX1
7669_R_P_9_TBR1
7669_R_POA_Bulk
7669_R_POA_DLX1
7669_R_sec1_Cau_Bulk
7669_R_sec2_GP_Bulk
7669_R_sec2_GP_DLX1
7669_R_sec2_Put_Bulk
7669_R_sec2_Thal_Bulk
7669_R_sec2_Thal_COUPTF2
7669_R_sec2_Thal_DLX1
7669_R_sec3_Cau_DLX1
7669_R_sec3_Hip_COUPTF2
7669_R_sec3_Hip_DLX1
7669_R_sec3_Hip_TBR1
7669_R_sec3_Put_DLX1
7669_R_Skin_Bulk
7669_R_T_1_Bulk
7669_R_T_1_COUPTF2
7669_R_T_1_DLX1
7669_R_T_1_TBR1
7669_RA9_Bulk
7669_RV10_Bulk
JGG

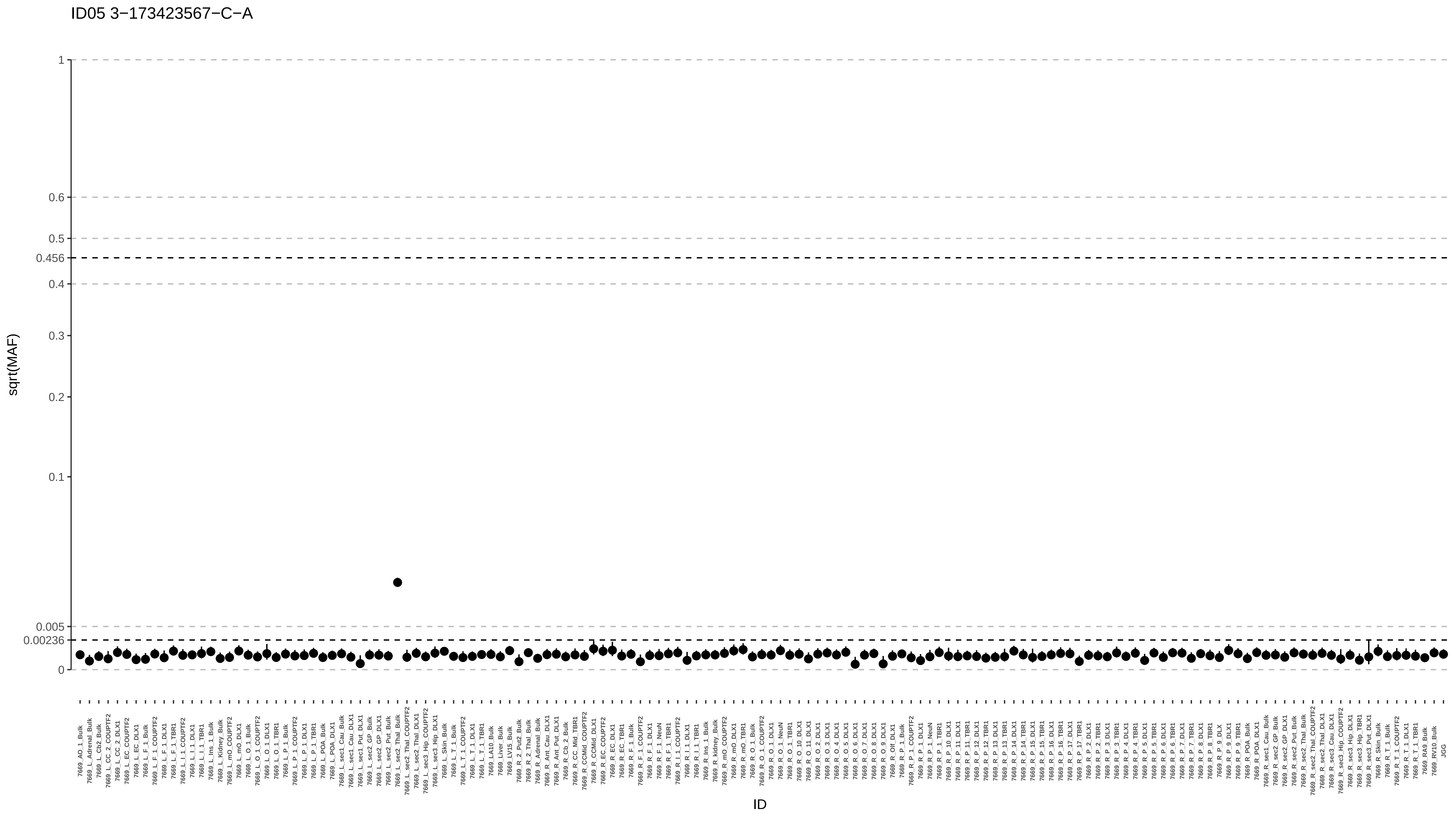
ID05 3−173423567−C−A
sqrt(MAF)
1
0.6
0.5
0.456
0.4
0.3
0.2
0.1
0.005
0.00236
0
ID
7669_AO_1_Bulk
7669_L_Adrenal_Bulk
7669_L_Cb2_Bulk
7669_L_CC_2_COUPTF2
7669_L_CC_2_DLX1
7669_L_EC_COUPTF2
7669_L_EC_DLX1
7669_L_F_1_Bulk
7669_L_F_1_COUPTF2
7669_L_F_1_DLX1
7669_L_F_1_TBR1
7669_L_I_1_COUPTF2
7669_L_I_1_DLX1
7669_L_I_1_TBR1
7669_L_Ins_1_Bulk
7669_L_Kidney_Bulk
7669_L_mO_COUPTF2
7669_L_mO_DLX1
7669_L_O_1_Bulk
7669_L_O_1_COUPTF2
7669_L_O_1_DLX1
7669_L_O_1_TBR1
7669_L_P_1_Bulk
7669_L_P_1_COUPTF2
7669_L_P_1_DLX1
7669_L_P_1_TBR1
7669_L_POA_Bulk
7669_L_POA_DLX1
7669_L_sec1_Cau_Bulk
7669_L_sec1_Cau_DLX1
7669_L_sec1_Put_DLX1
7669_L_sec2_GP_Bulk
7669_L_sec2_GP_DLX1
7669_L_sec2_Put_Bulk
7669_L_sec2_Thal_Bulk
7669_L_sec2_Thal_COUPTF2
7669_L_sec2_Thal_DLX1
7669_L_sec3_Hip_COUPTF2
7669_L_sec3_Hip_DLX1
7669_L_Skin_Bulk
7669_L_T_1_Bulk
7669_L_T_1_COUPTF2
7669_L_T_1_DLX1
7669_L_T_1_TBR1
7669_LA10_Bulk
7669_Liver_Bulk
7669_LV15_Bulk
7669_R_2_Put2_Bulk
7669_R_2_Thal_Bulk
7669_R_Adrenal_Bulk
7669_R_Ant_Cau_DLX1
7669_R_Ant_Put_DLX1
7669_R_Cb_2_Bulk
7669_R_CC_Mid_TBR1
7669_R_CCMid_COUPTF2
7669_R_CCMid_DLX1
7669_R_EC_COUPTF2
7669_R_EC_DLX1
7669_R_EC_TBR1
7669_R_F_1_Bulk
7669_R_F_1_COUPTF2
7669_R_F_1_DLX1
7669_R_F_1_NeuN
7669_R_F_1_TBR1
7669_R_I_1_COUPTF2
7669_R_I_1_DLX1
7669_R_I_1_TBR1
7669_R_Ins_1_Bulk
7669_R_kidney_Bulk
7669_R_mO_COUPTF2
7669_R_mO_DLX1
7669_R_mO_TBR1
7669_R_O_1_Bulk
7669_R_O_1_COUPTF2
7669_R_O_1_DLX1
7669_R_O_1_NeuN
7669_R_O_1_TBR1
7669_R_O_10_DLX1
7669_R_O_11_DLX1
7669_R_O_2_DLX1
7669_R_O_3_DLX1
7669_R_O_4_DLX1
7669_R_O_5_DLX1
7669_R_O_6_DLX1
7669_R_O_7_DLX1
7669_R_O_8_DLX1
7669_R_O_9_DLX1
7669_R_Olf_DLX1
7669_R_P_1_Bulk
7669_R_P_1_COUPTF2
7669_R_P_1_DLX1
7669_R_P_1_NeuN
7669_R_P_1_TBR1
7669_R_P_10_DLX1
7669_R_P_11_DLX1
7669_R_P_11_TBR1
7669_R_P_12_DLX1
7669_R_P_12_TBR1
7669_R_P_13_DLX1
7669_R_P_13_TBR1
7669_R_P_14_DLX1
7669_R_P_14_TBR1
7669_R_P_15_DLX1
7669_R_P_15_TBR1
7669_R_P_16_DLX1
7669_R_P_16_TBR1
7669_R_P_17_DLX1
7669_R_P_17_TBR1
7669_R_P_2_DLX1
7669_R_P_2_TBR1
7669_R_P_3_DLX1
7669_R_P_3_TBR1
7669_R_P_4_DLX1
7669_R_P_4_TBR1
7669_R_P_5_DLX1
7669_R_P_5_TBR1
7669_R_P_6_DLX1
7669_R_P_6_TBR1
7669_R_P_7_DLX1
7669_R_P_7_TBR1
7669_R_P_8_DLX1
7669_R_P_8_TBR1
7669_R_P_9_DLX
7669_R_P_9_DLX1
7669_R_P_9_TBR1
7669_R_POA_Bulk
7669_R_POA_DLX1
7669_R_sec1_Cau_Bulk
7669_R_sec2_GP_Bulk
7669_R_sec2_GP_DLX1
7669_R_sec2_Put_Bulk
7669_R_sec2_Thal_Bulk
7669_R_sec2_Thal_COUPTF2
7669_R_sec2_Thal_DLX1
7669_R_sec3_Cau_DLX1
7669_R_sec3_Hip_COUPTF2
7669_R_sec3_Hip_DLX1
7669_R_sec3_Hip_TBR1
7669_R_sec3_Put_DLX1
7669_R_Skin_Bulk
7669_R_T_1_Bulk
7669_R_T_1_COUPTF2
7669_R_T_1_DLX1
7669_R_T_1_TBR1
7669_RA9_Bulk
7669_RV10_Bulk
JGG

ID05 3−181259475−C−T

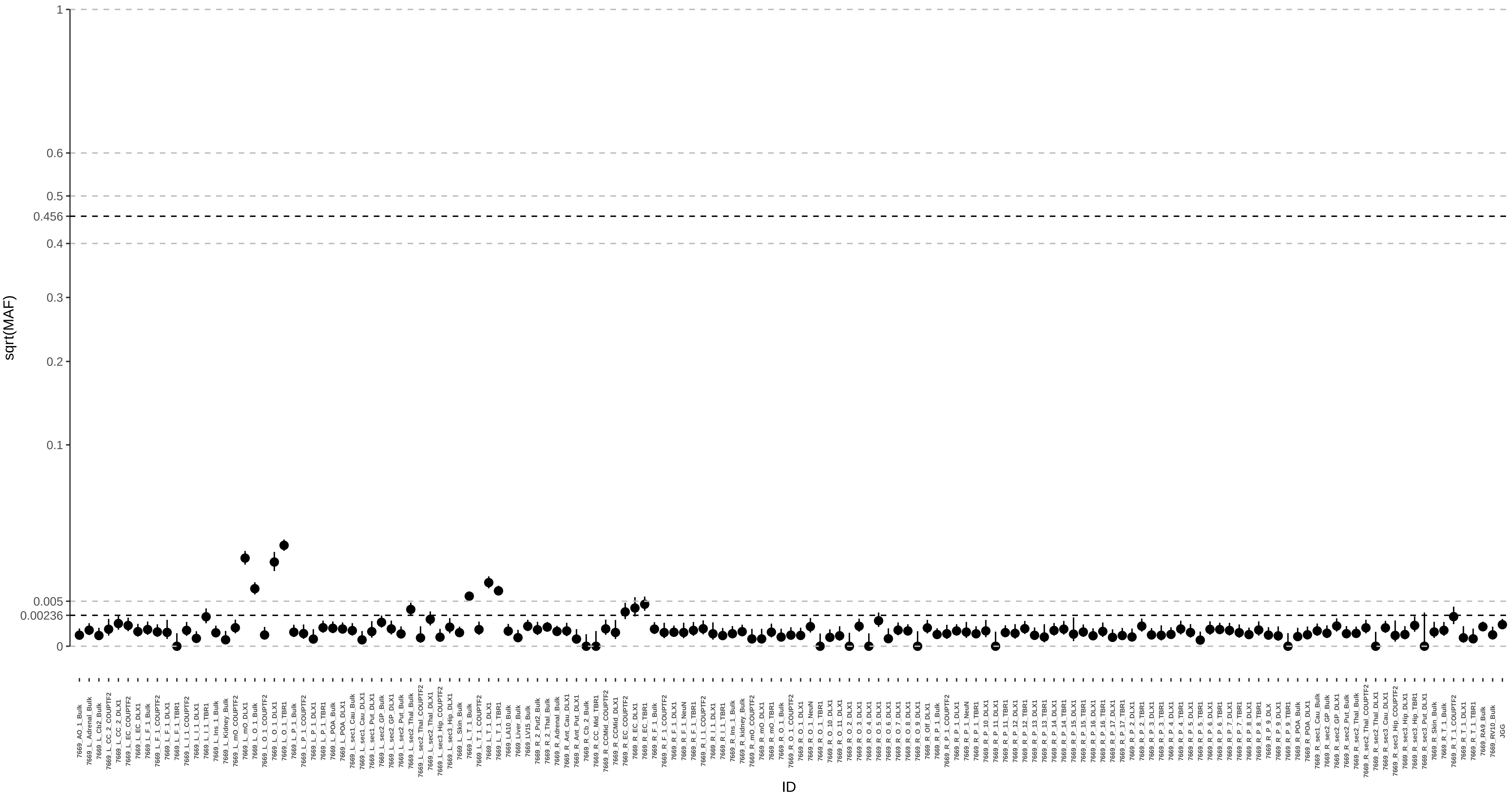

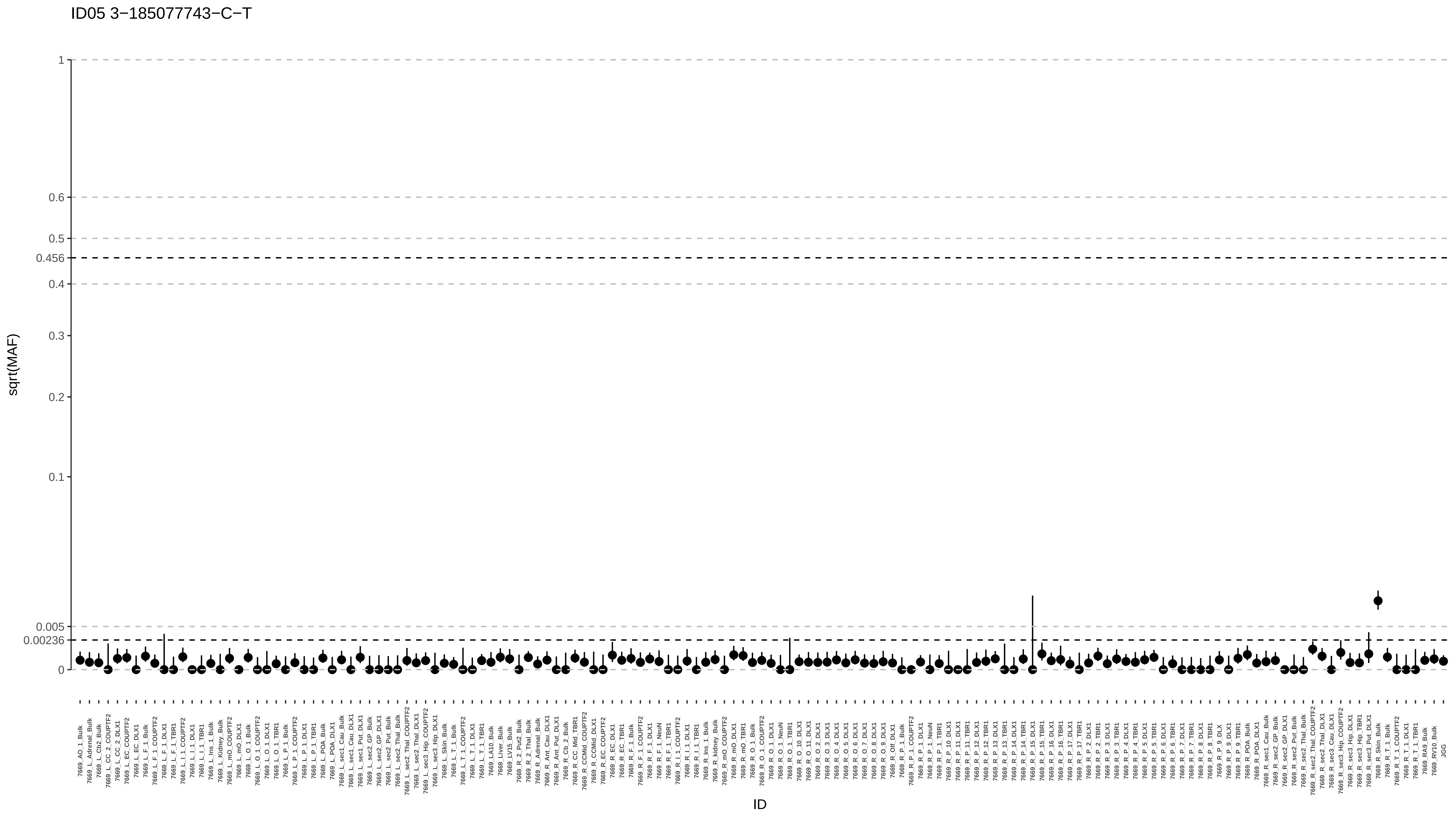
ID05 3−185077743−C−T
sqrt(MAF)
ID
1
0.6
0.5
0.456
0.4
0.3
0.2
0.1
0.005
0.00236
0

ID05 3−189834708−G−T

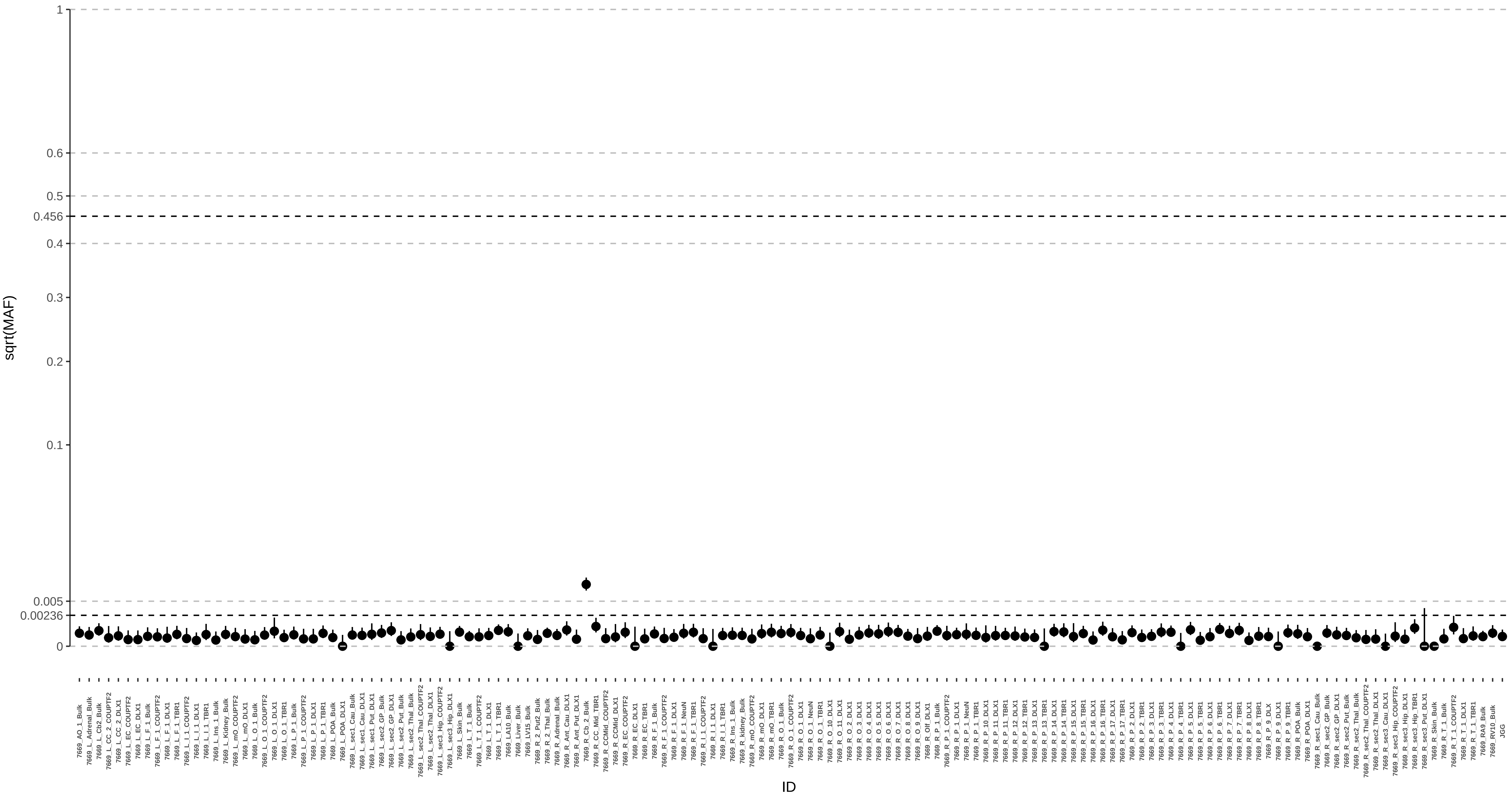

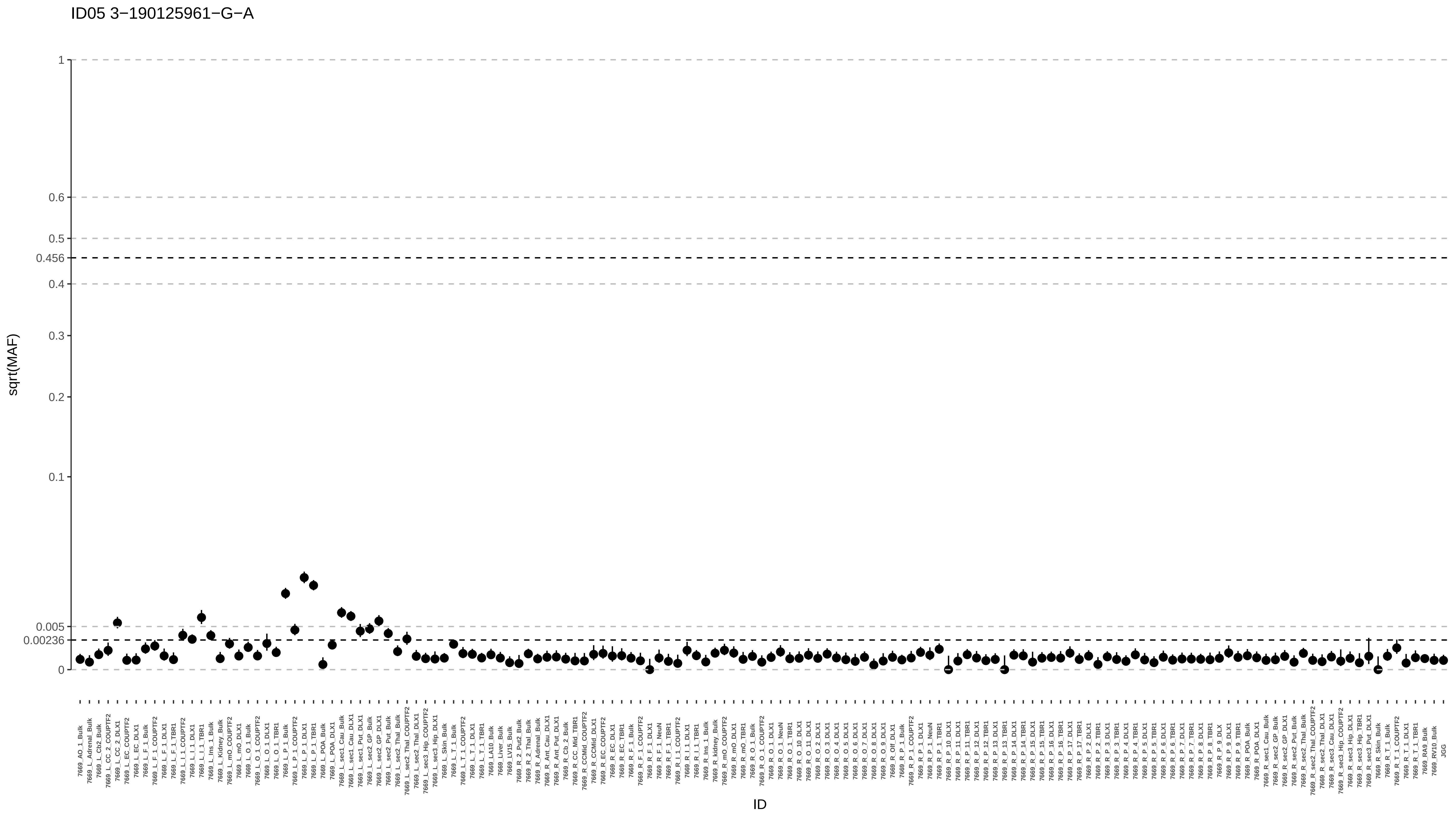
ID05 3−190125961−G−A
sqrt(MAF)
1
0.6
0.5
0.456
0.4
0.3
0.2
0.1
0.005
0.00236
0
ID

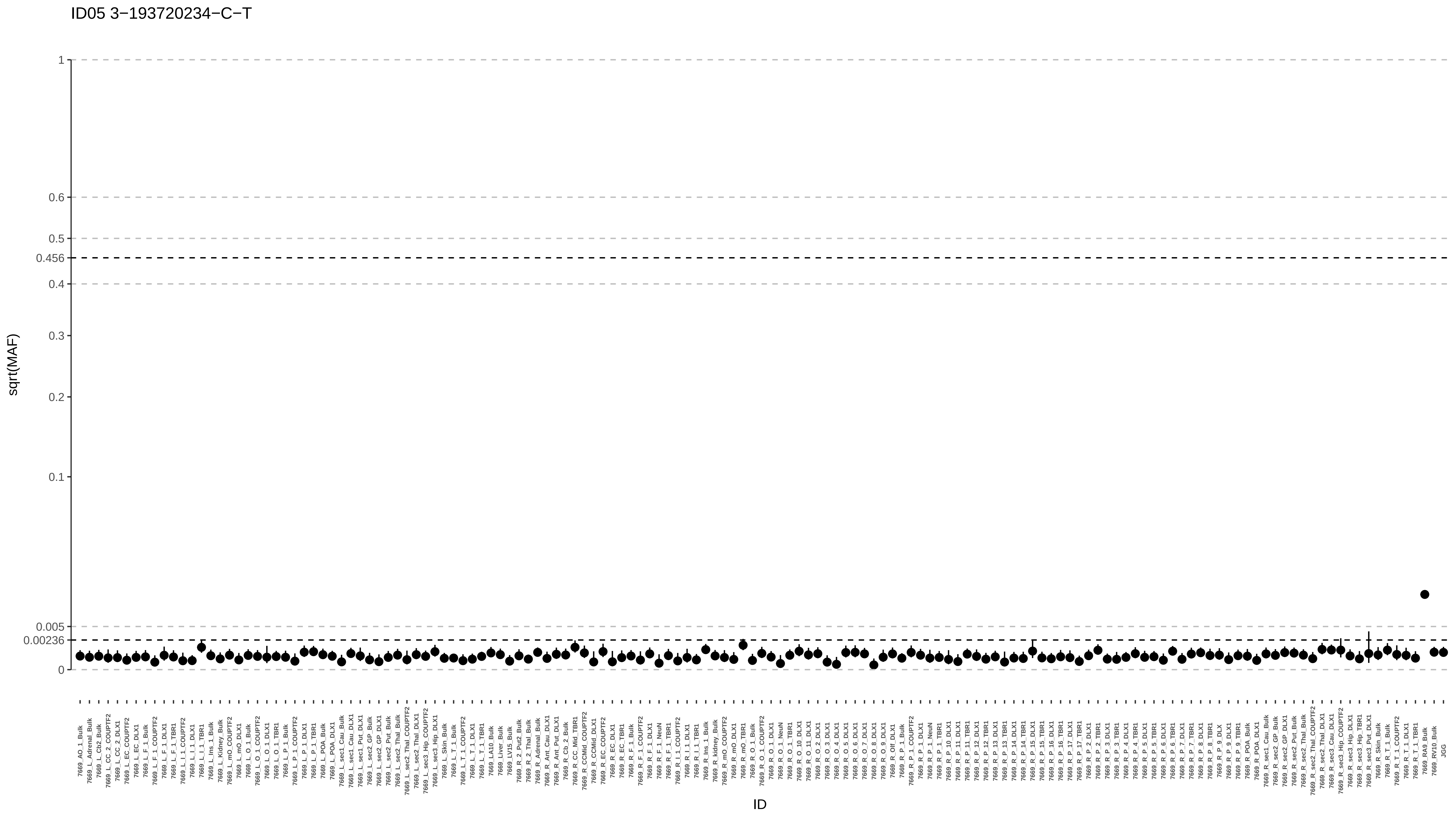
ID05 3−193720234−C−T
sqrt(MAF)
ID
1
0.6
0.5
0.456
0.4
0.3
0.2
0.1
0.005
0.00236
0

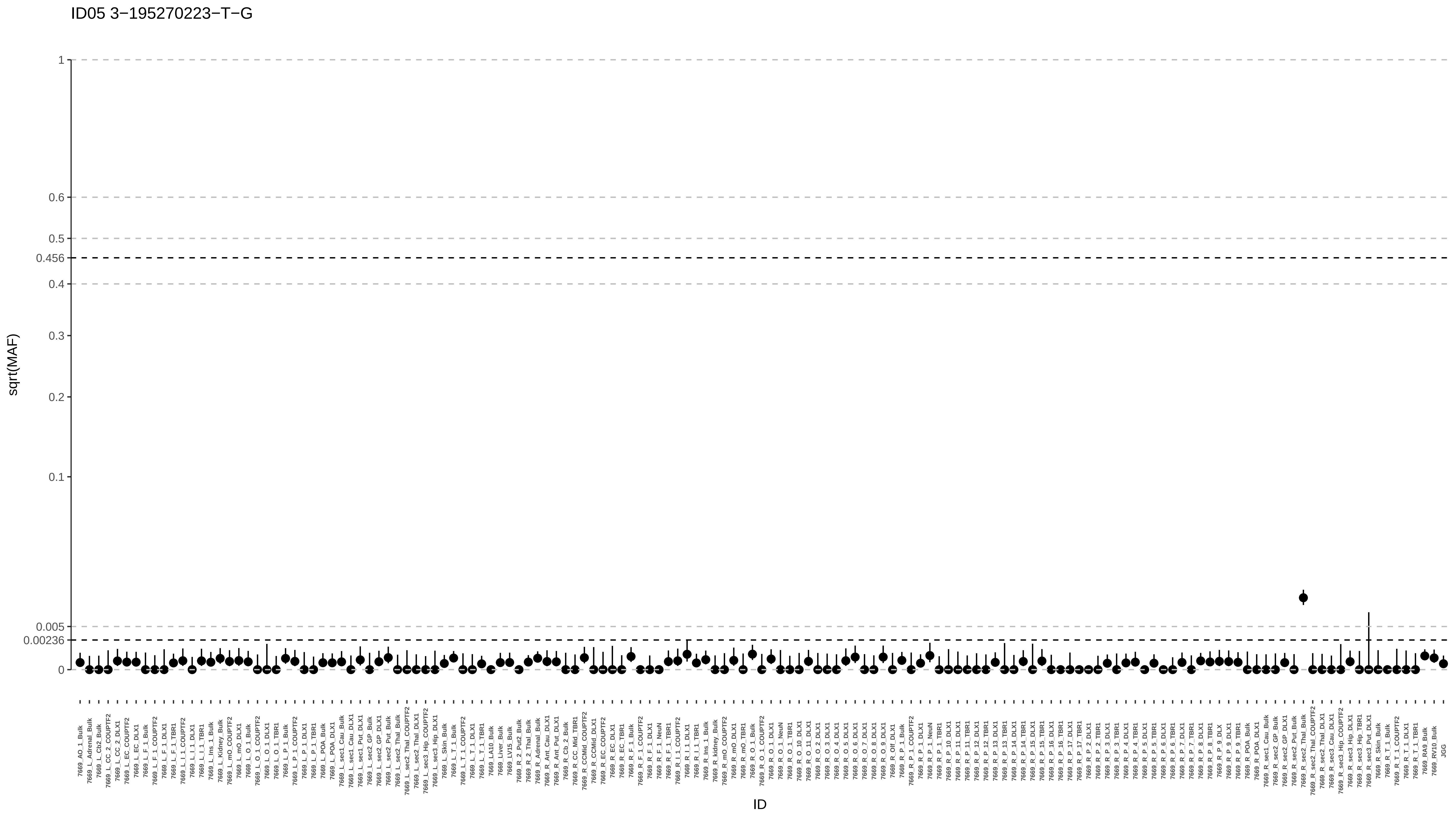
ID05 3−195270223−T−G
sqrt(MAF)
1
0.6
0.5
0.456
0.4
0.3
0.2
0.1
0.005
0.00236
0
ID

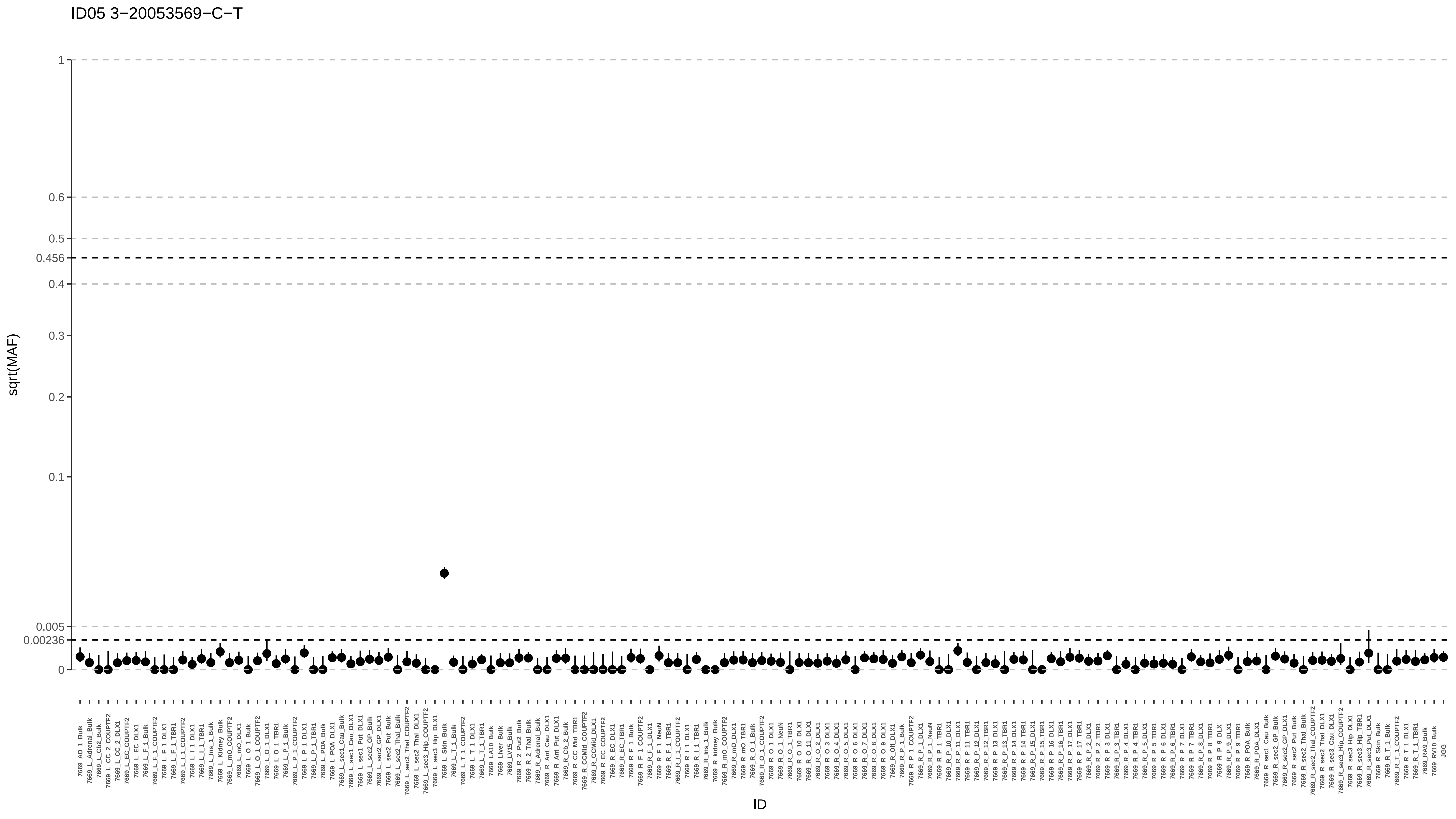

ID05 3−20053569−C−T
sqrt(MAF)
1
0.6
0.5
0.456
0.4
0.3
0.2
0.1
0.005
0.00236
0
ID

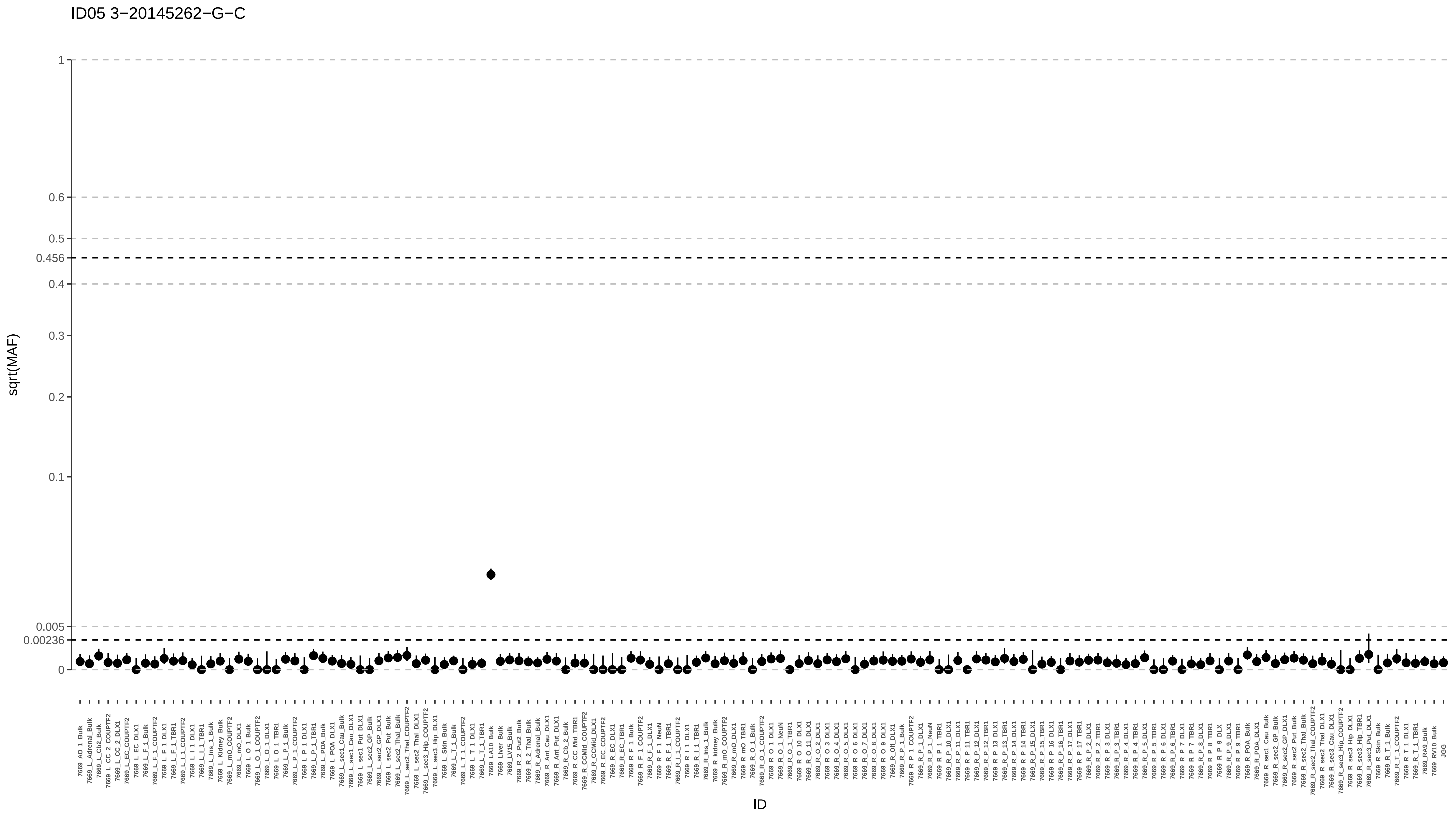

ID05 3−20145262−G−C
sqrt(MAF)
1
0.6
0.5
0.456
0.4
0.3
0.2
0.1
0.005
0.00236
0
ID

ID05 3−20222654−C−T

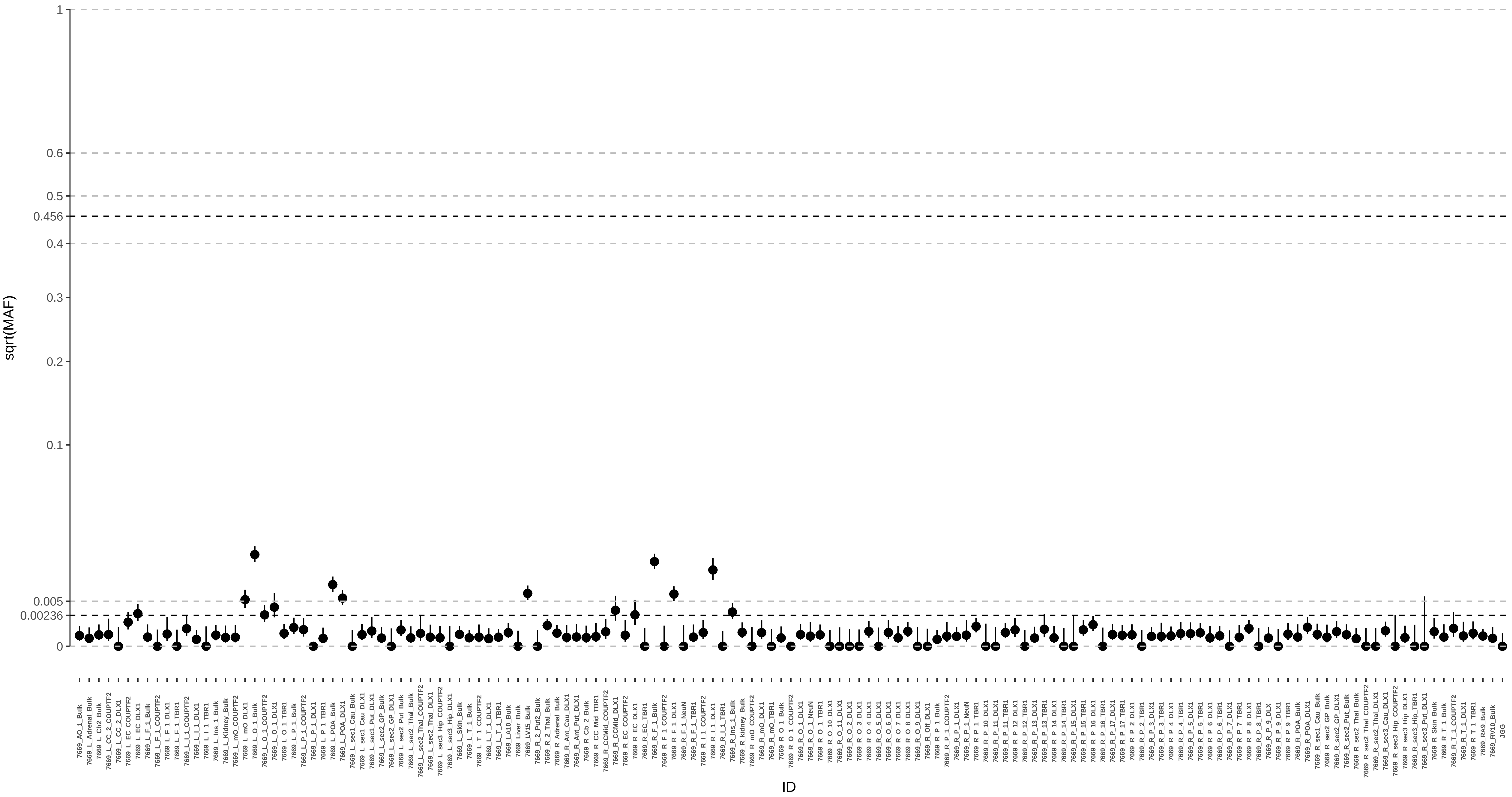

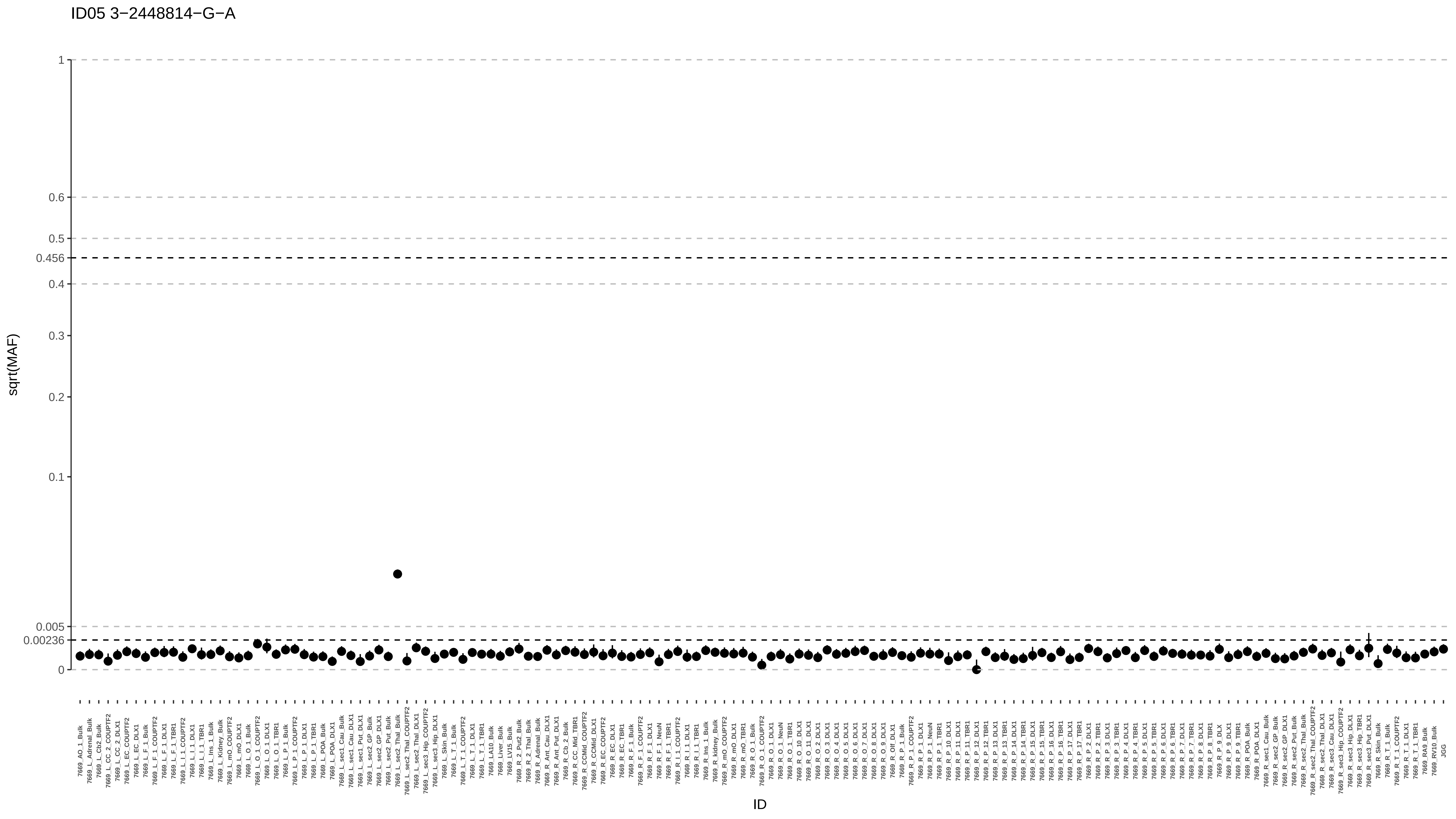
ID05 3−2448814−G−A
sqrt(MAF)
1
0.6
0.5
0.456
0.4
0.3
0.2
0.1
0.005
0.00236
0
ID

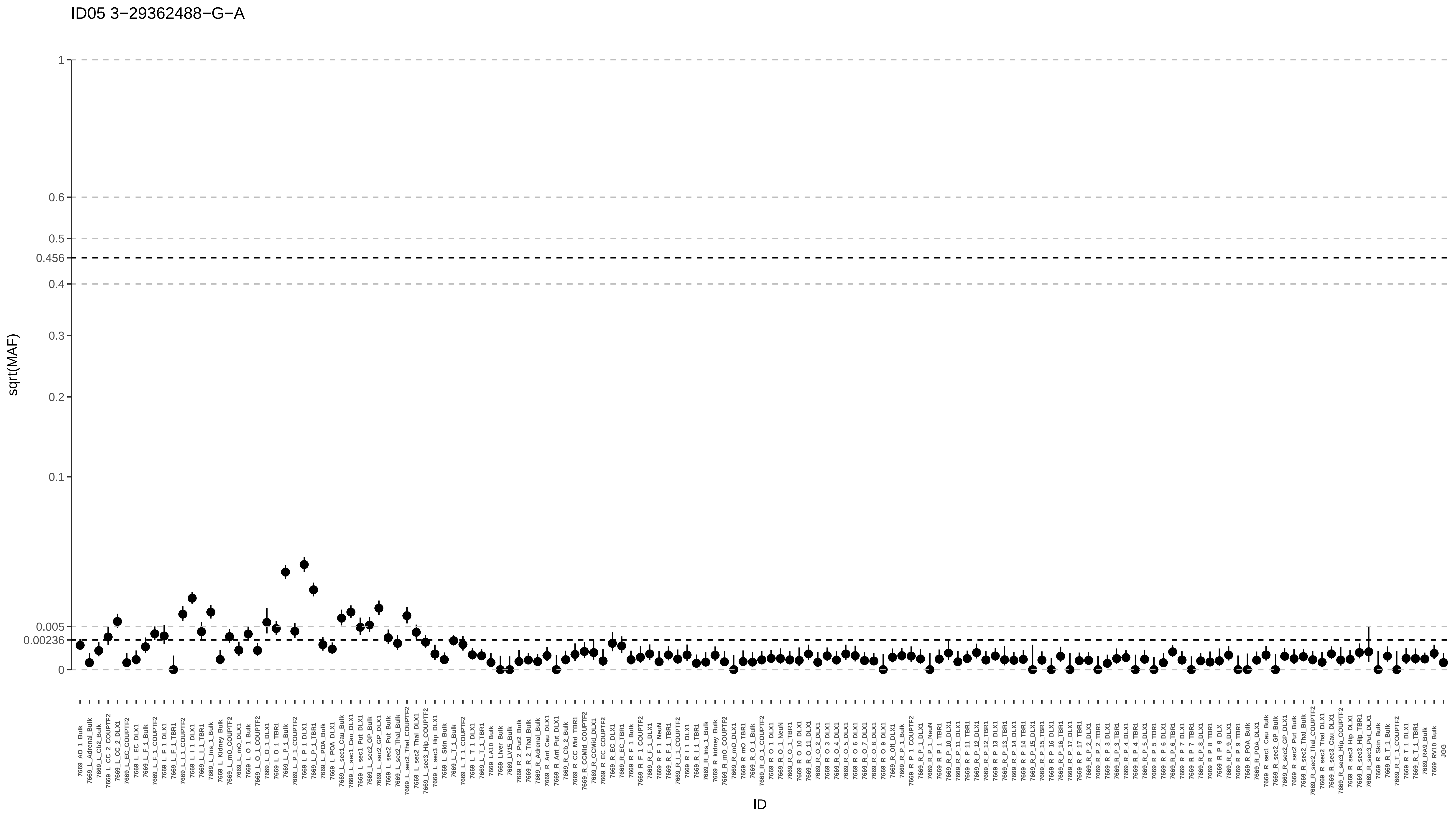
ID05 3−29362488−G−A
sqrt(MAF)
1
0.6
0.5
0.456
0.4
0.3
0.2
0.1
0.005
0.00236
0
ID

ID05 3−29763350−T−C

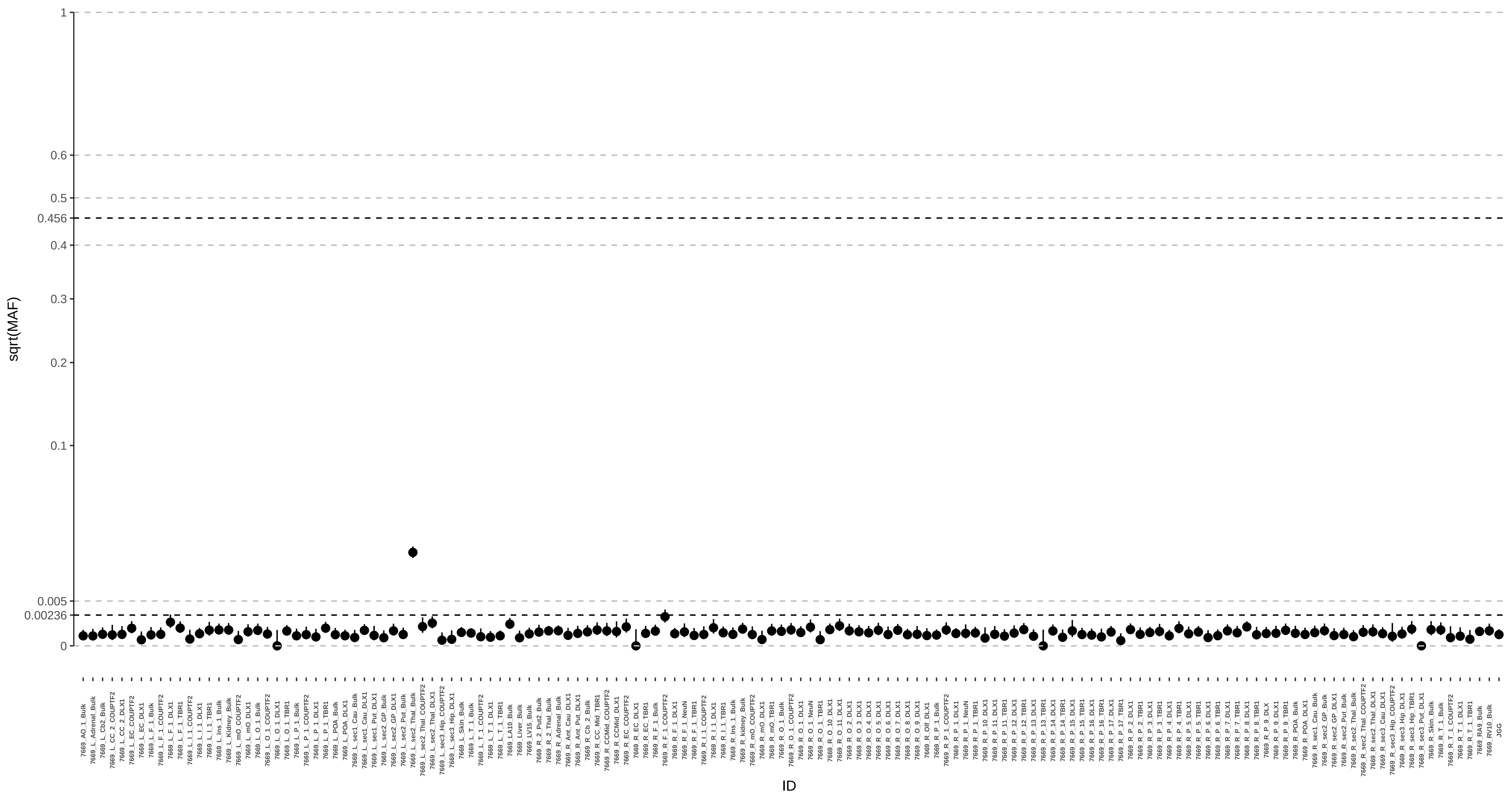

ID05 3−32863852−G−A

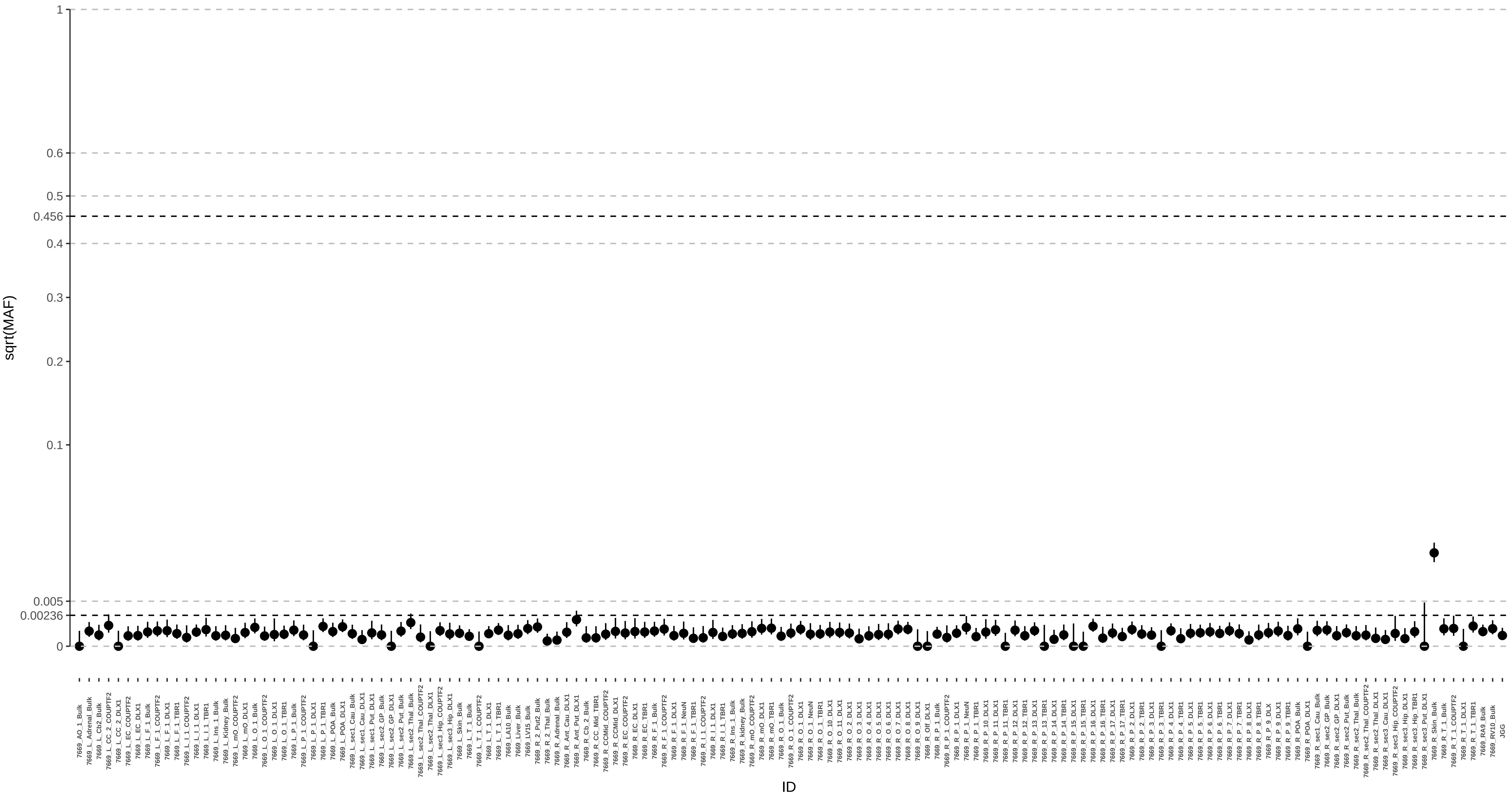

ID05 3−3458468−TG−T

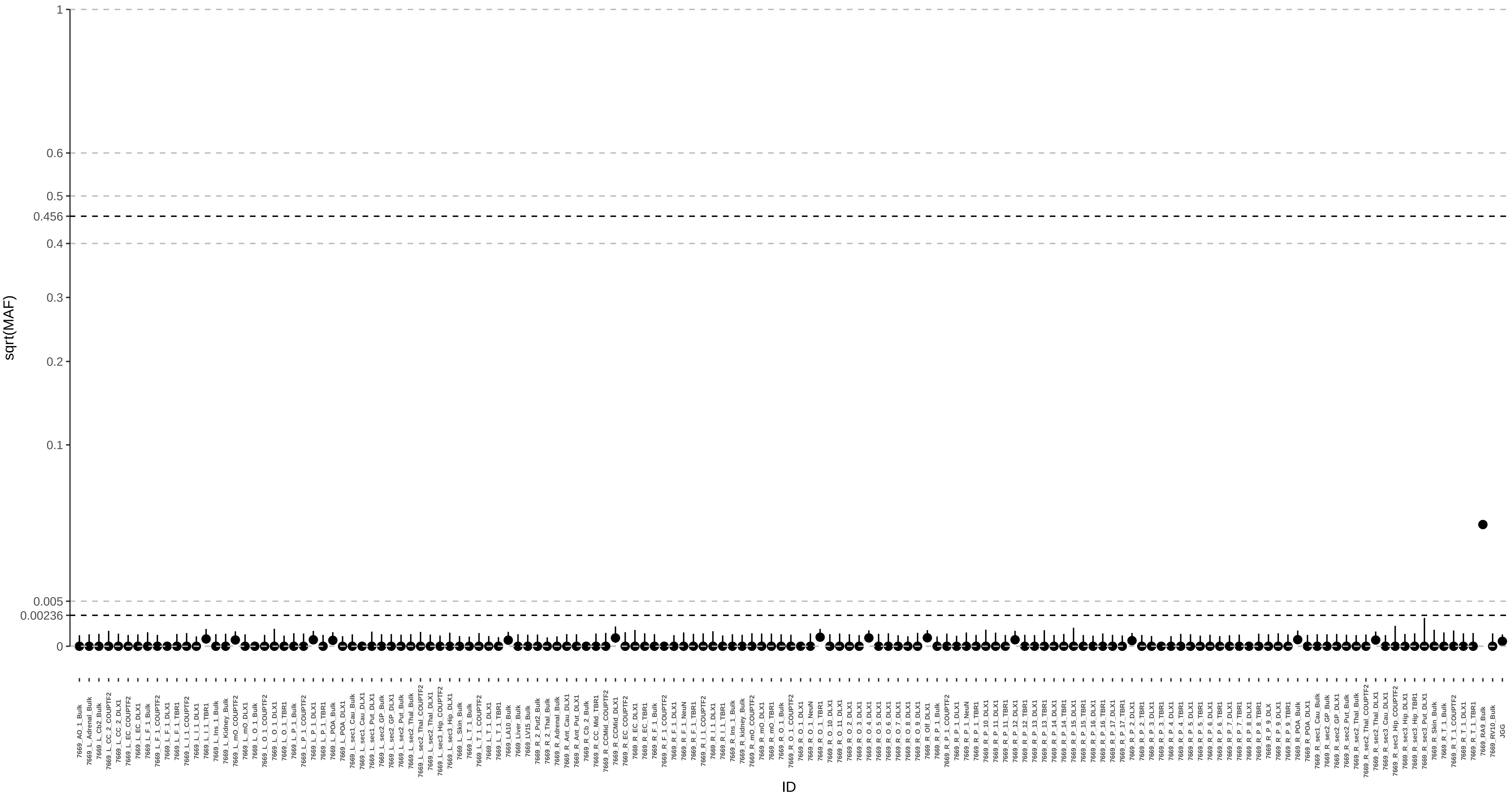

ID05 3−36389918−C−T

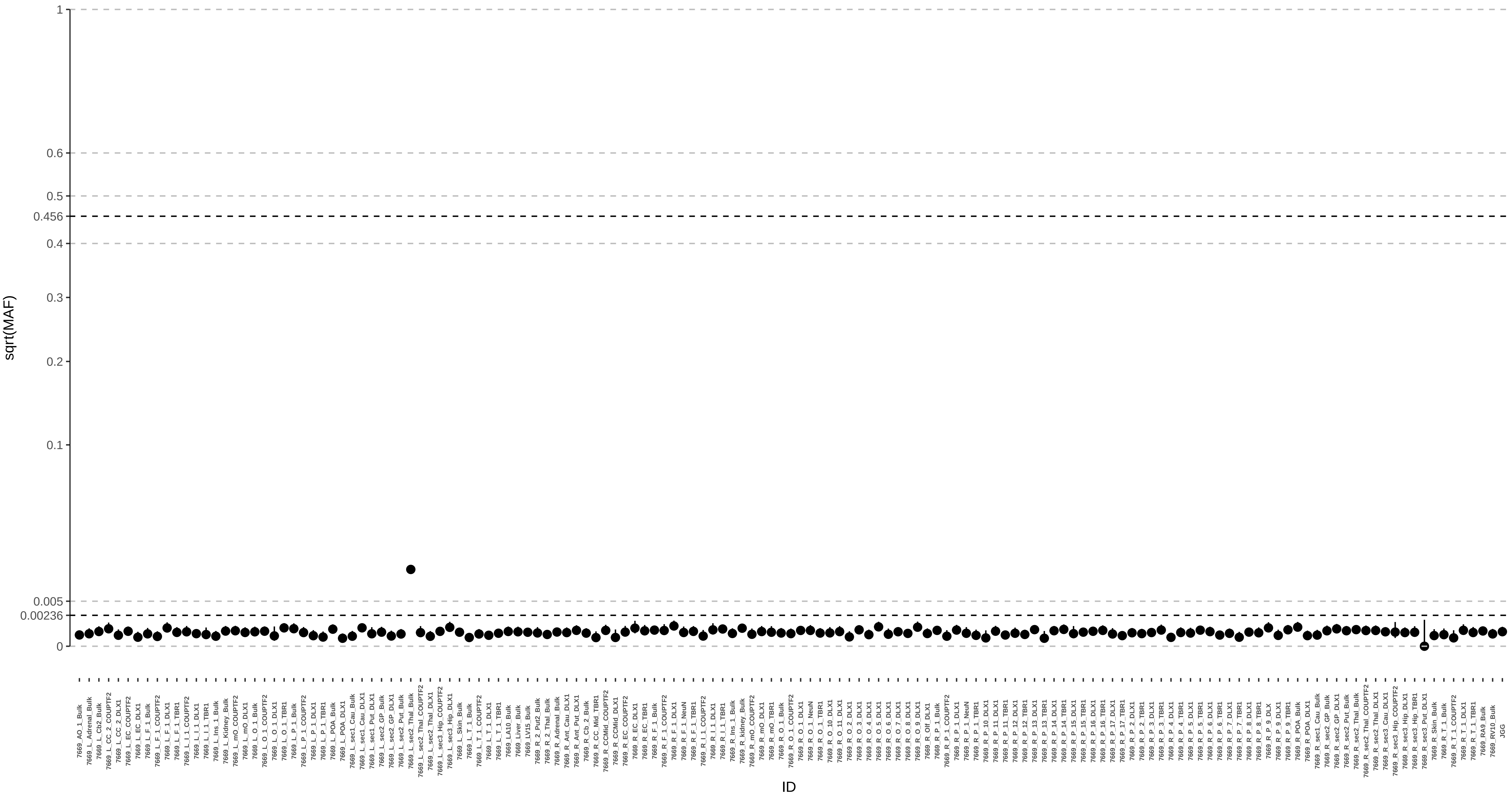

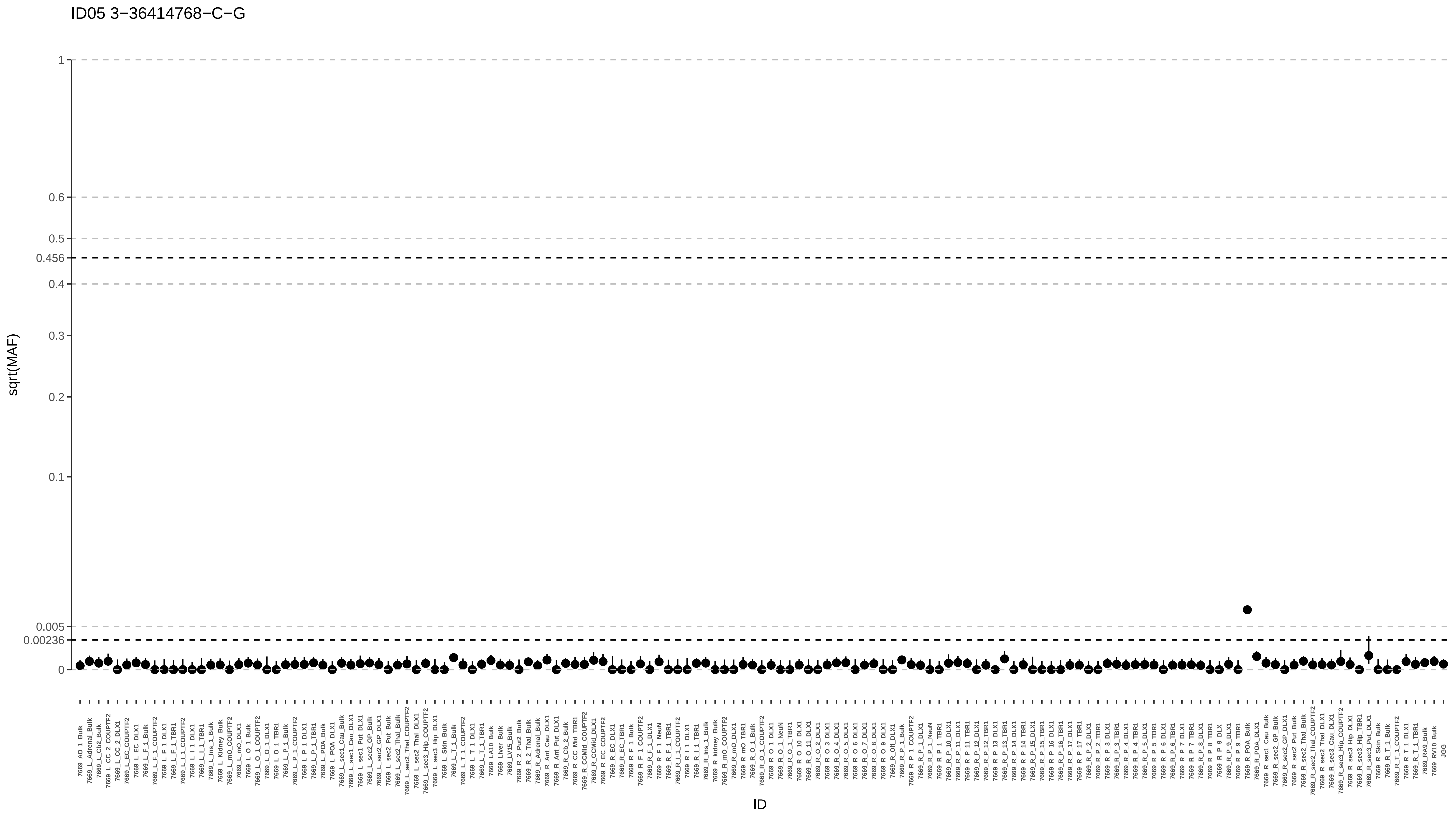
ID05 3−36414768−C−G
sqrt(MAF)
1
0.6
0.5
0.456
0.4
0.3
0.2
0.1
0.005
0.00236
0
ID

ID05 3−40086182−G−T

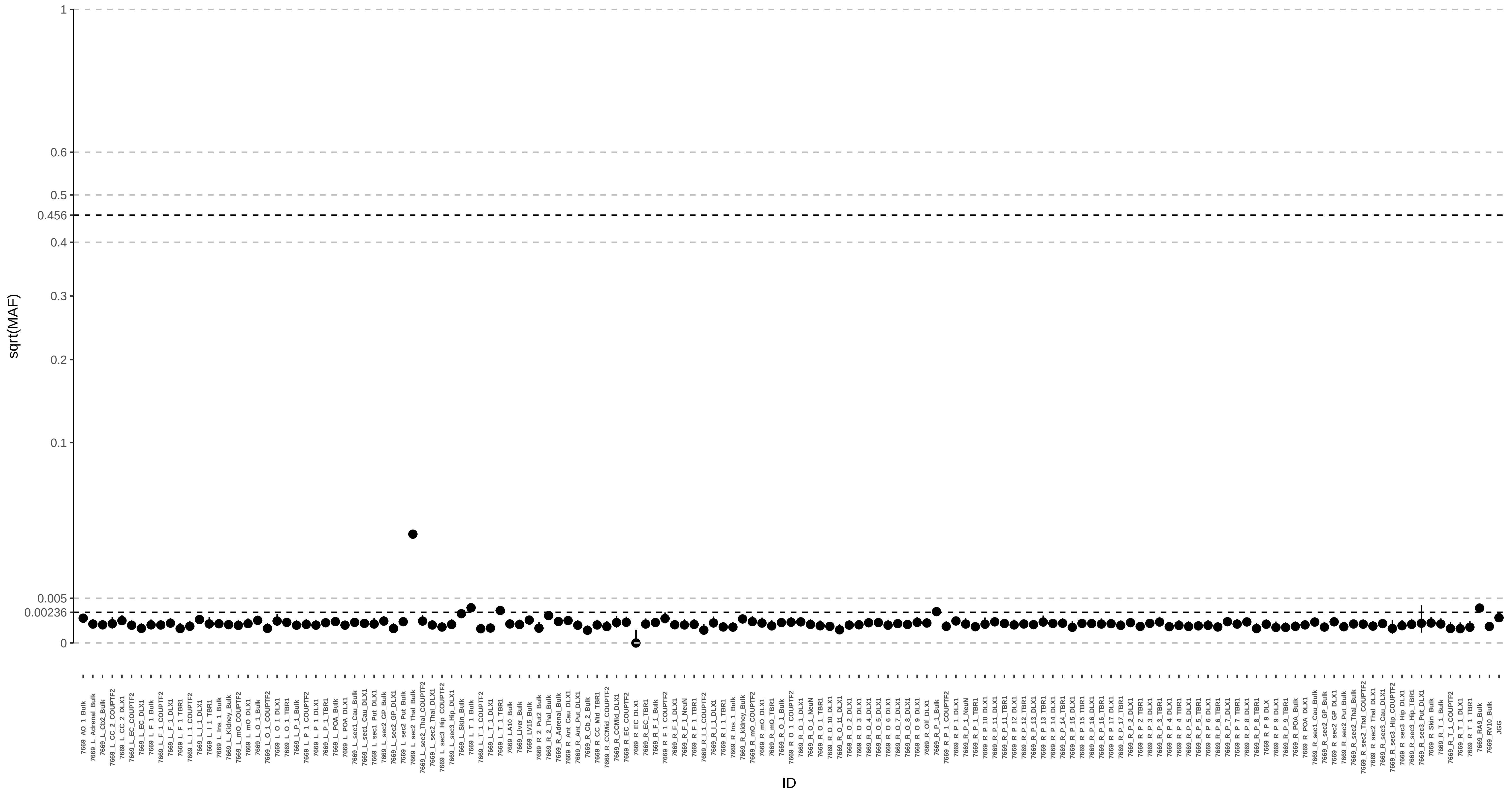

ID05 3−44078495−G−A

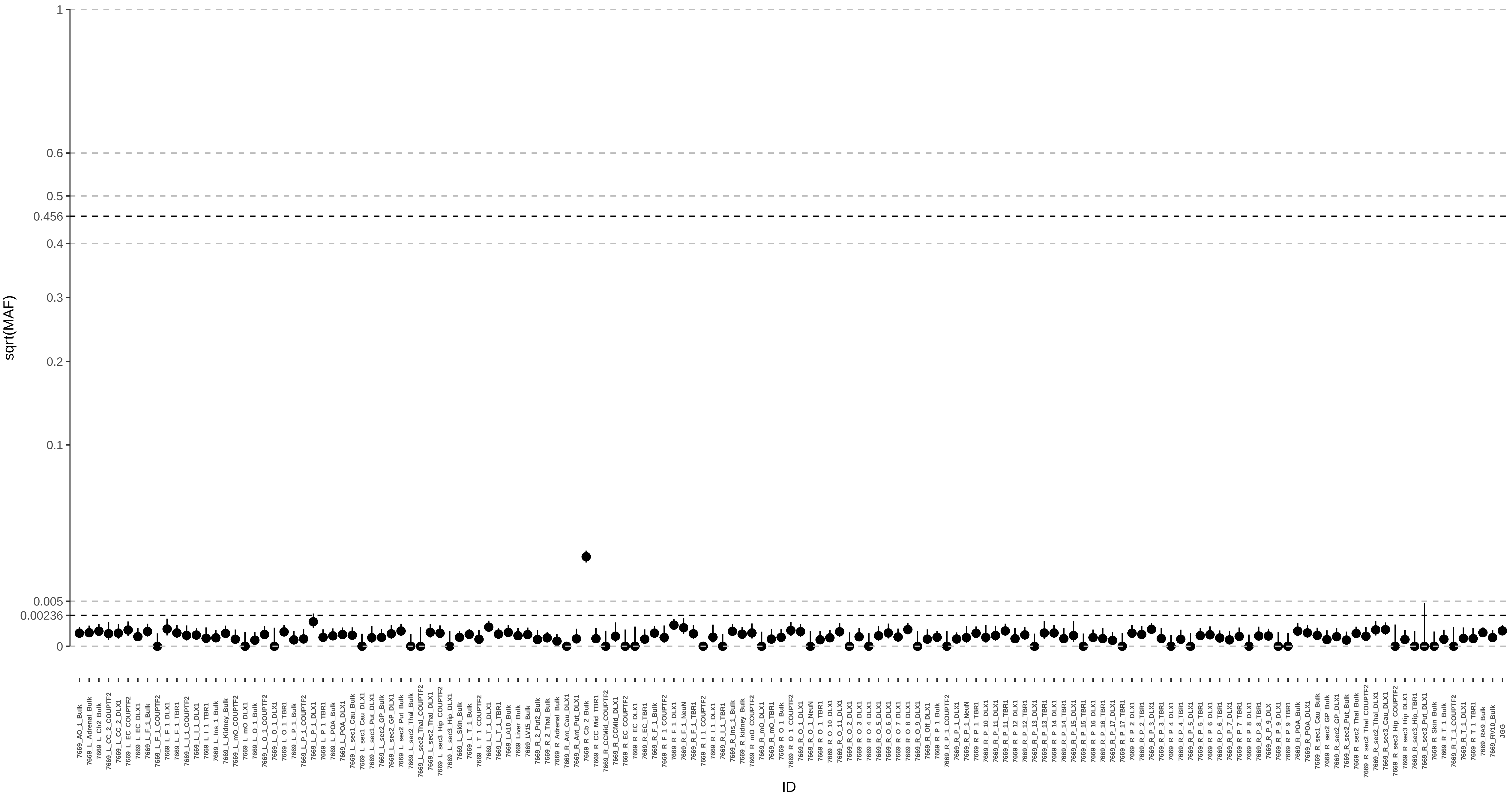

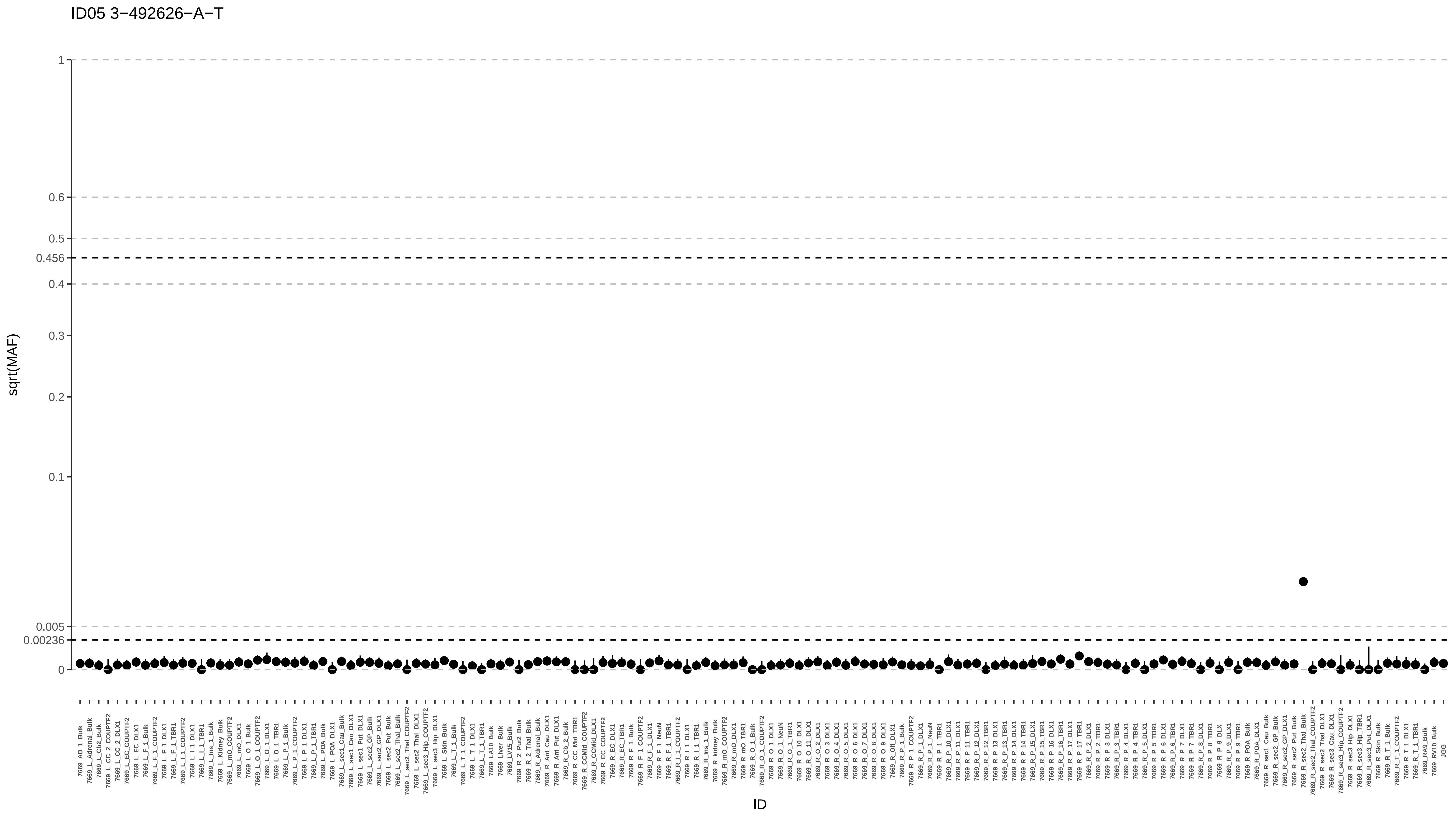
ID05 3−492626−A−T
sqrt(MAF)
1
0.6
0.5
0.456
0.4
0.3
0.2
0.1
0.005
0.00236
0
ID

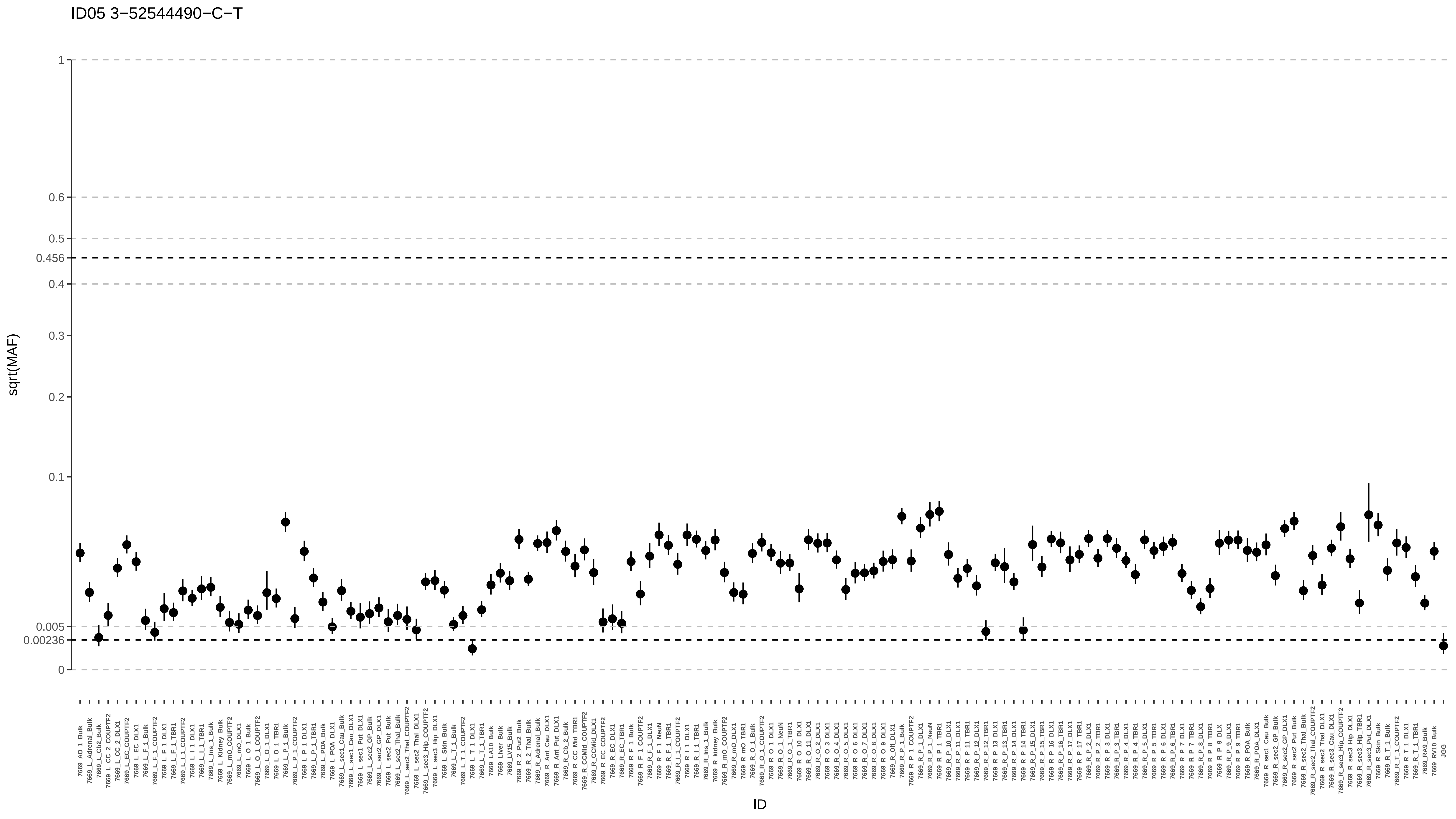
ID05 3−52544490−C−T
sqrt(MAF)
1
0.6
0.5
0.456
0.4
0.3
0.2
0.1
0.005
0.00236
0
ID

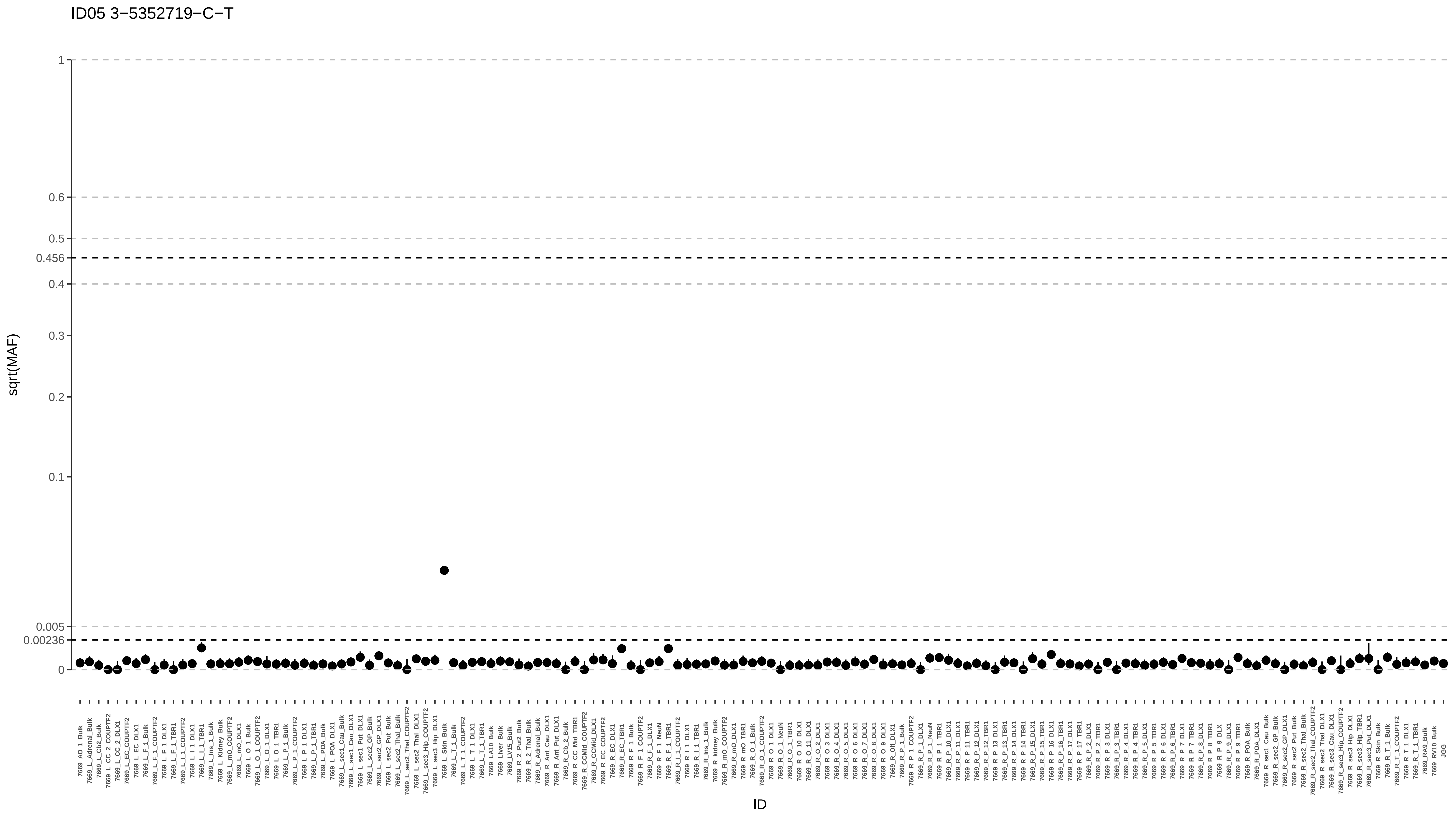
ID05 3−5352719−C−T
sqrt(MAF)
1
0.6
0.5
0.456
0.4
0.3
0.2
0.1
0.005
0.00236
0
ID

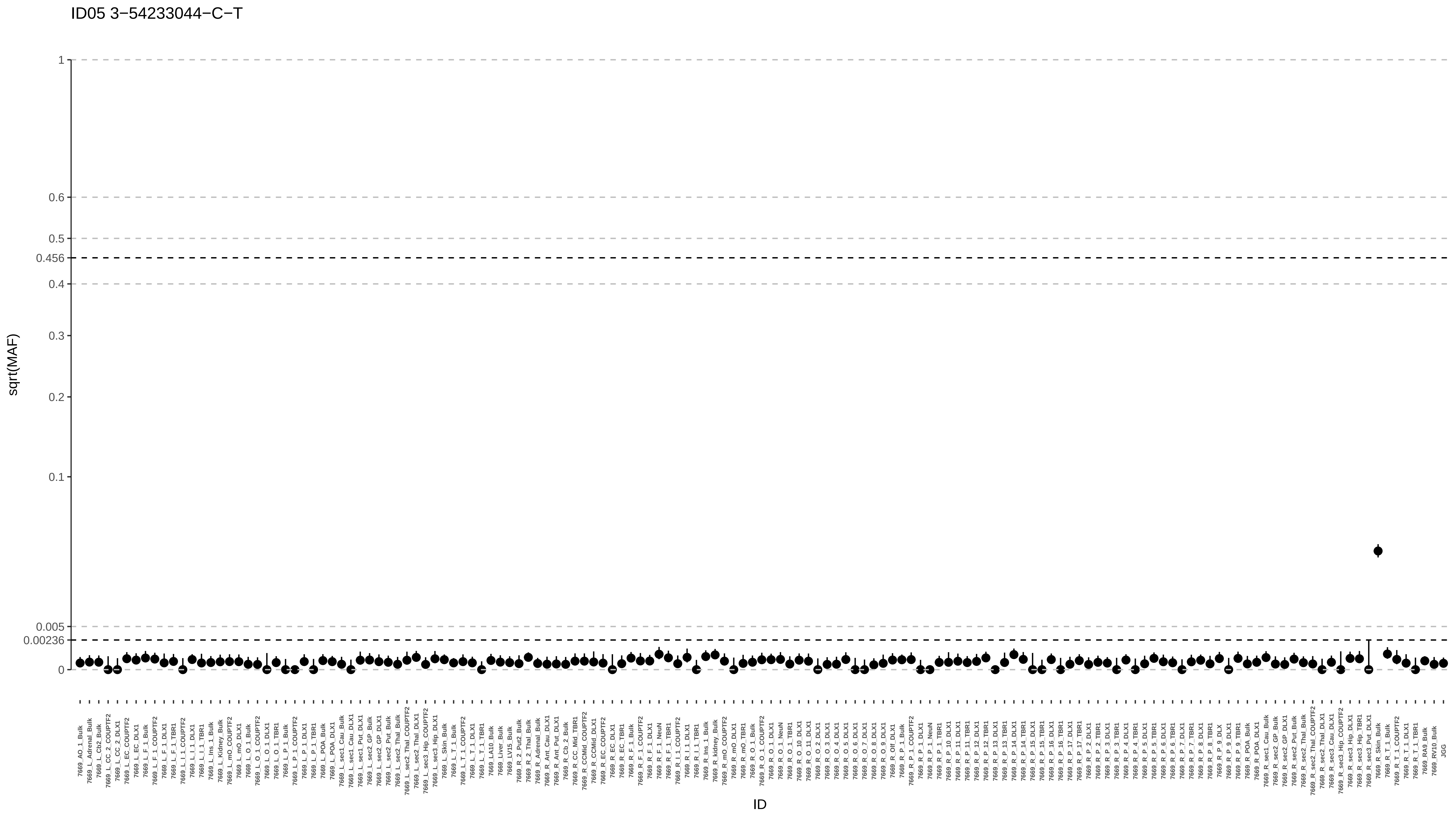
ID05 3−54233044−C−T
sqrt(MAF)
1
0.6
0.5
0.456
0.4
0.3
0.2
0.1
0.005
0.00236
0
ID

ID05 3−57851309−G−A

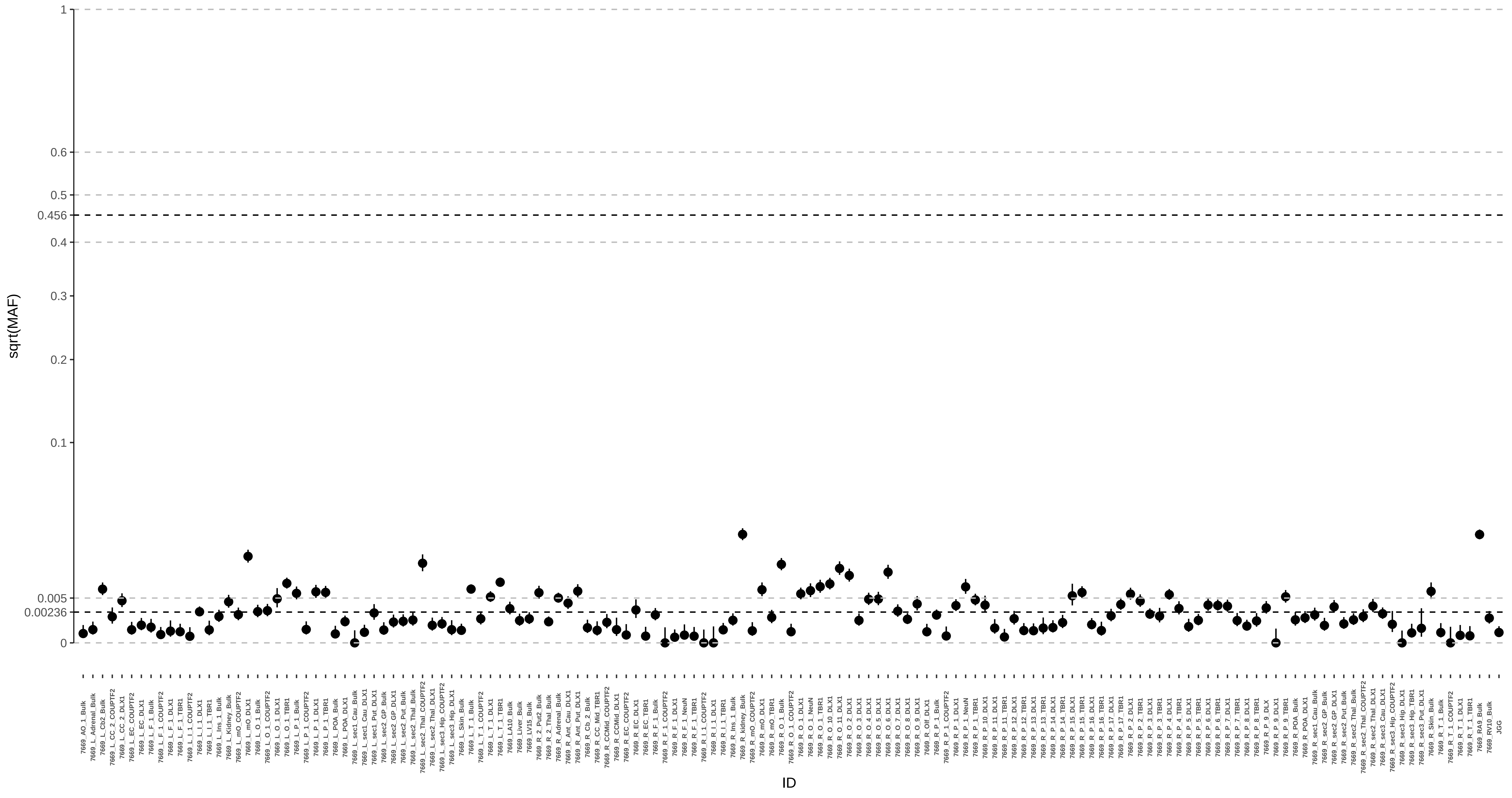

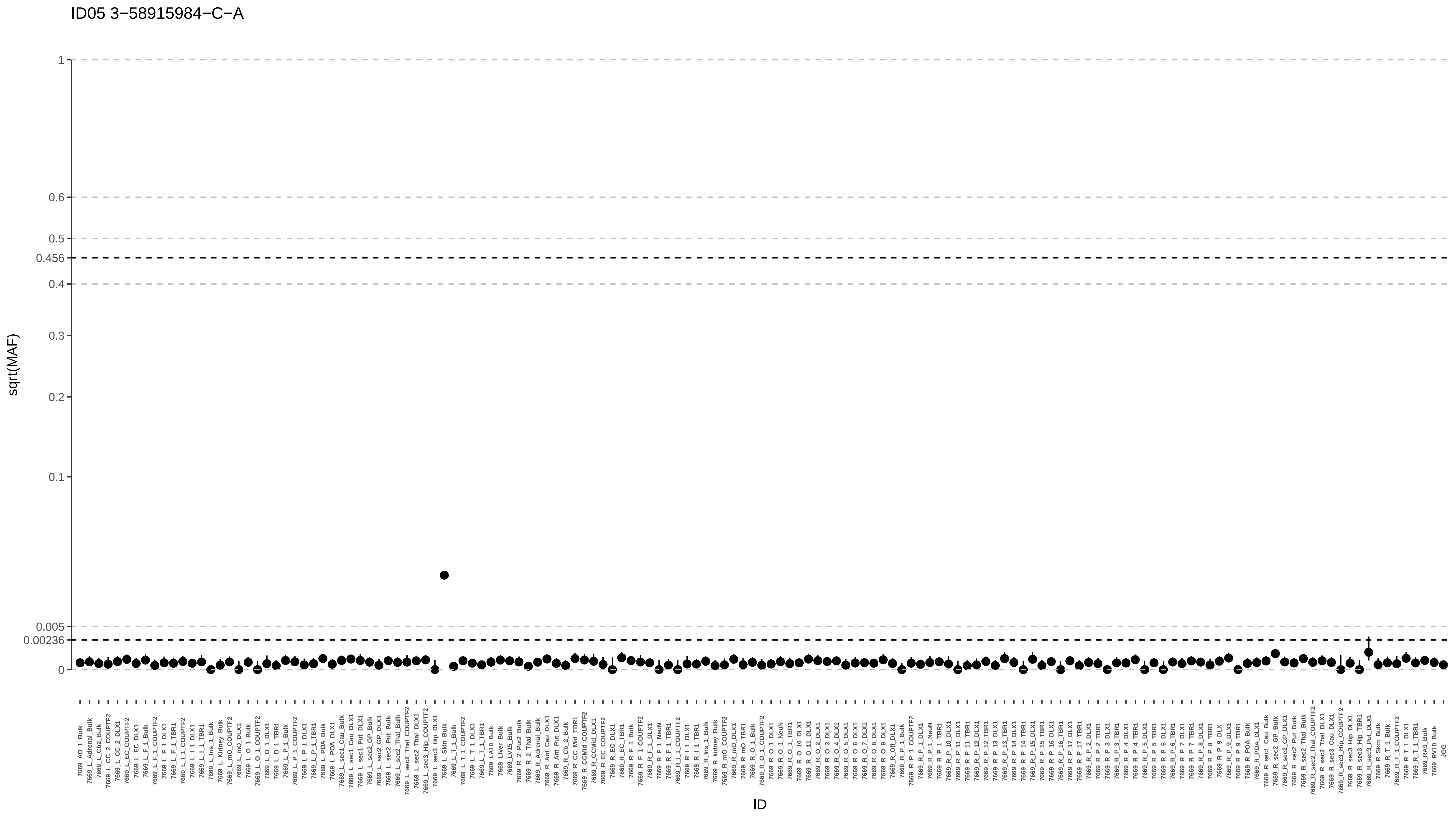

ID05 3−58915984−C−A
sqrt(MAF)
ID

ID05 3−61853127−C−T

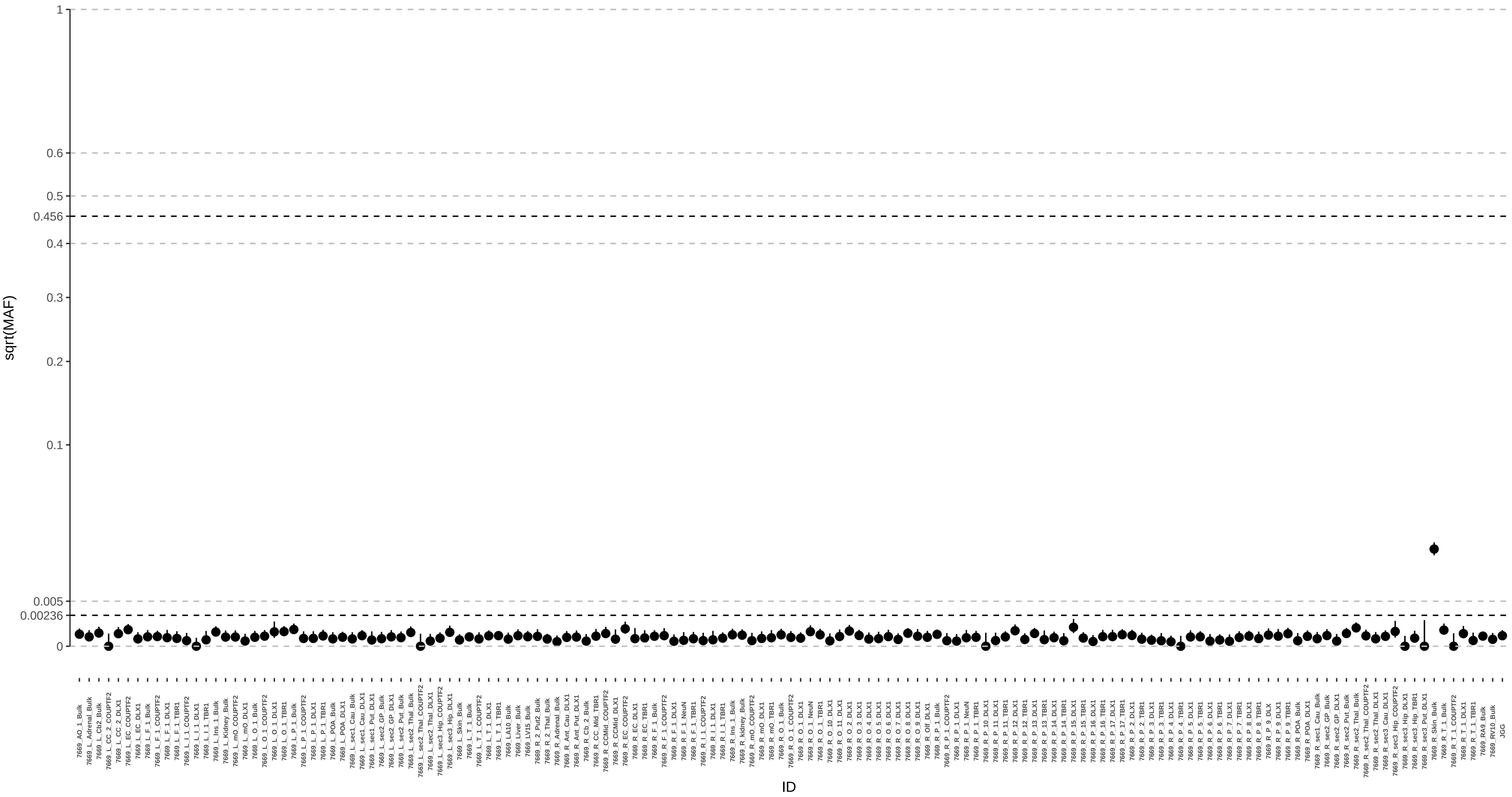

ID05 3−69001163−C−T

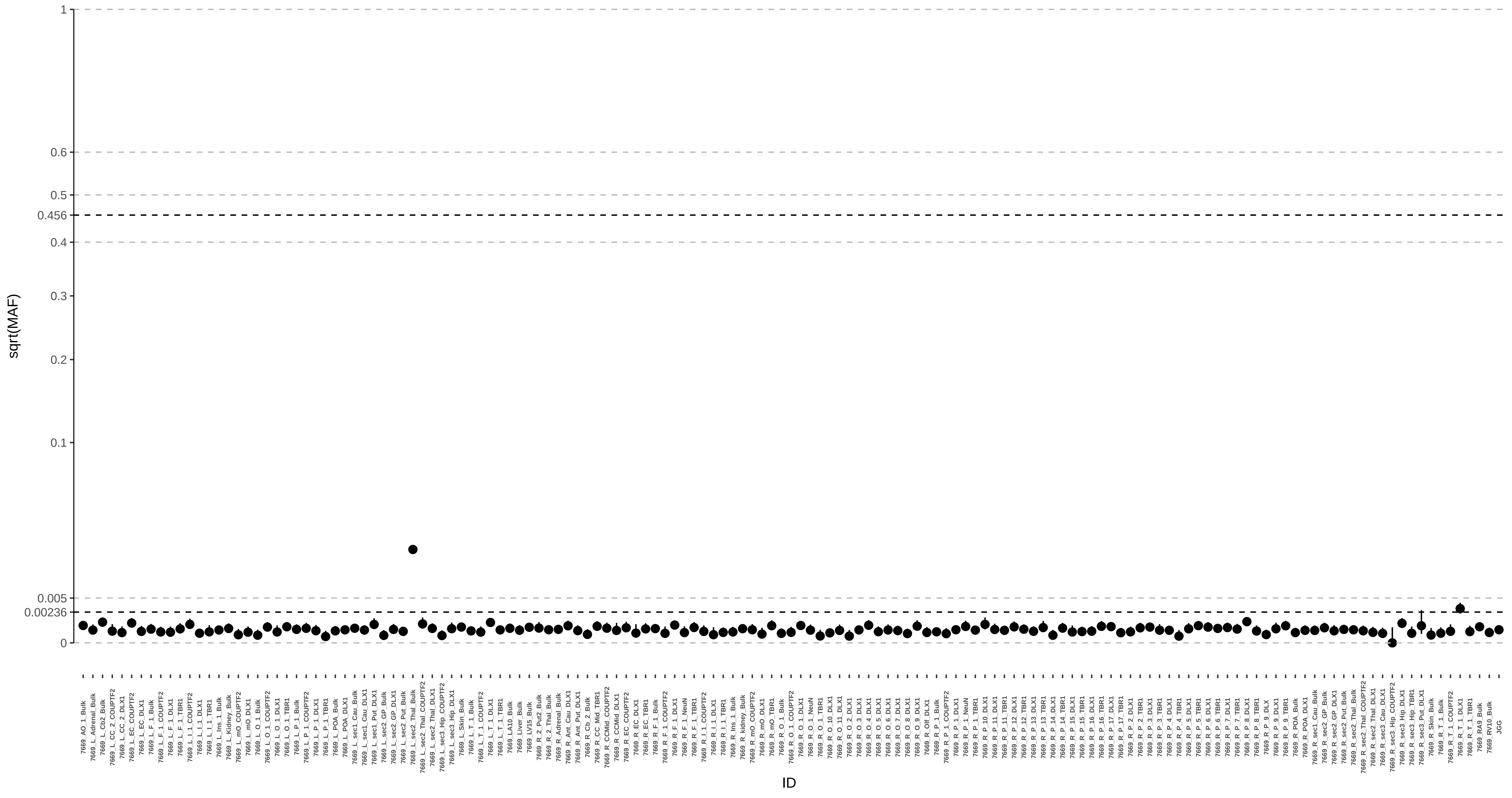

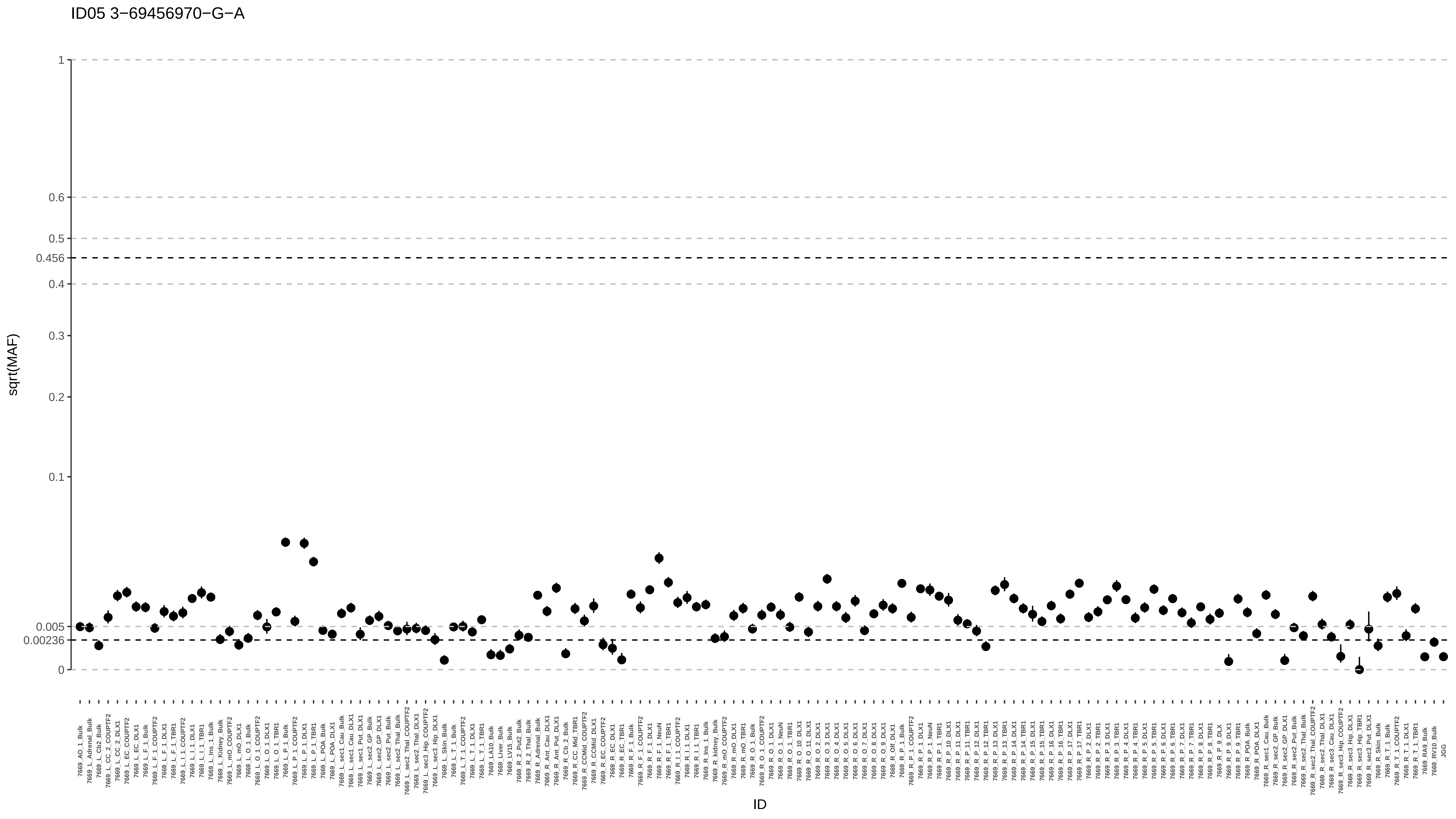

ID05 3−69456970−G−A
sqrt(MAF)
1
0.6
0.5
0.456
0.4
0.3
0.2
0.1
0.005
0.00236
0
ID

ID05 3−78738978−T−C

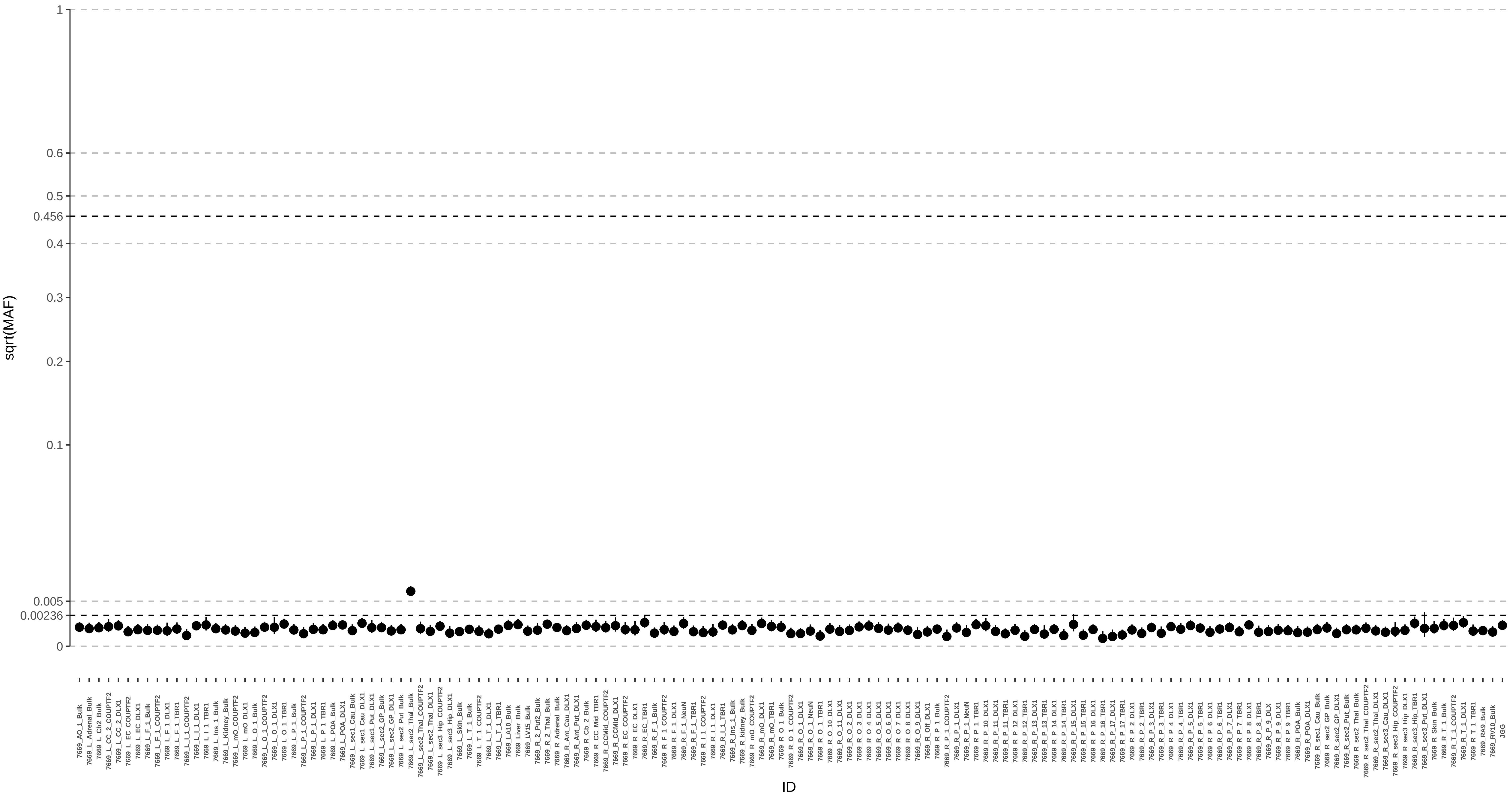

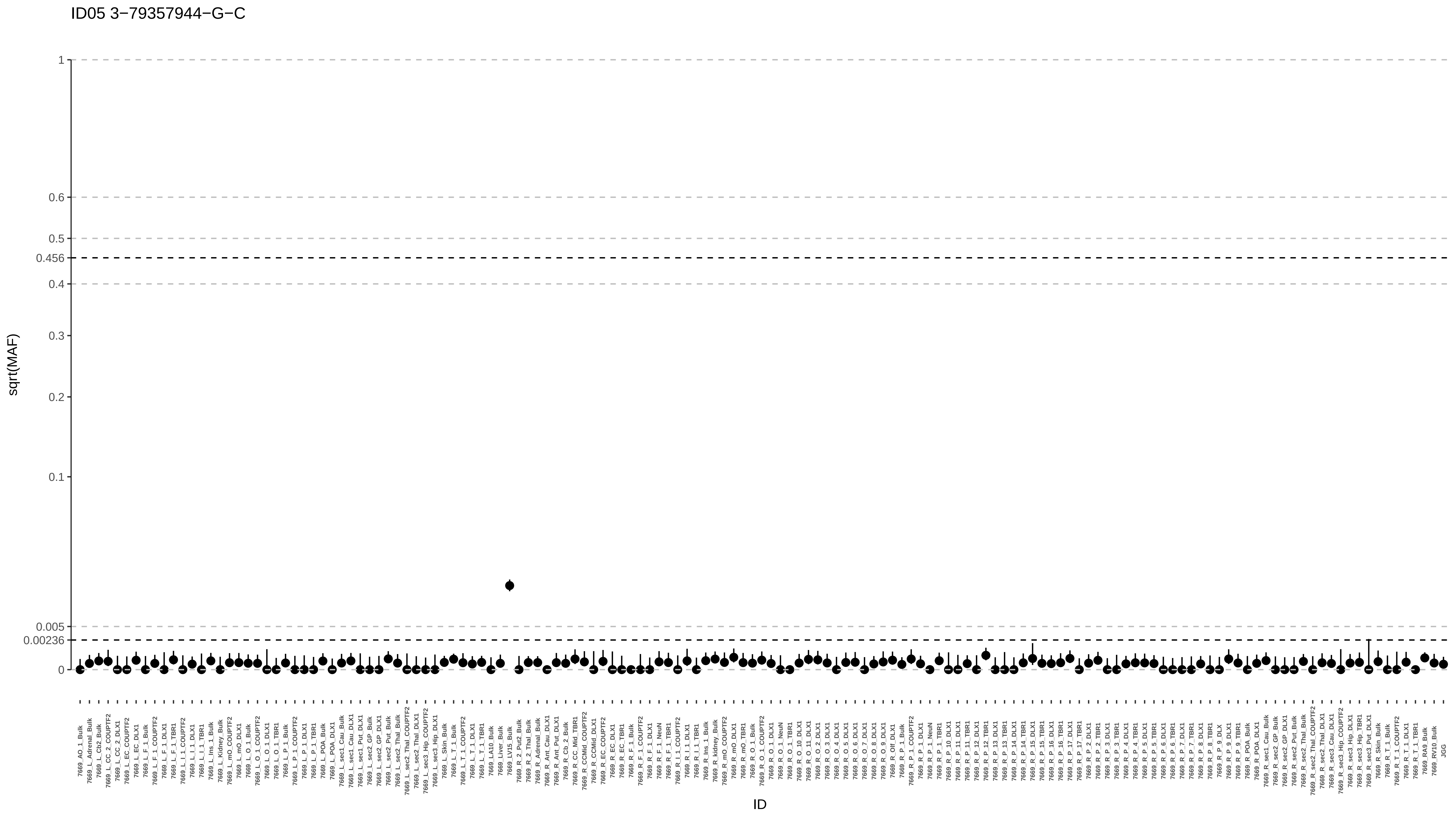
ID05 3−79357944−G−C
sqrt(MAF)
1
0.6
0.5
0.456
0.4
0.3
0.2
0.1
0.005
0.00236
0
ID

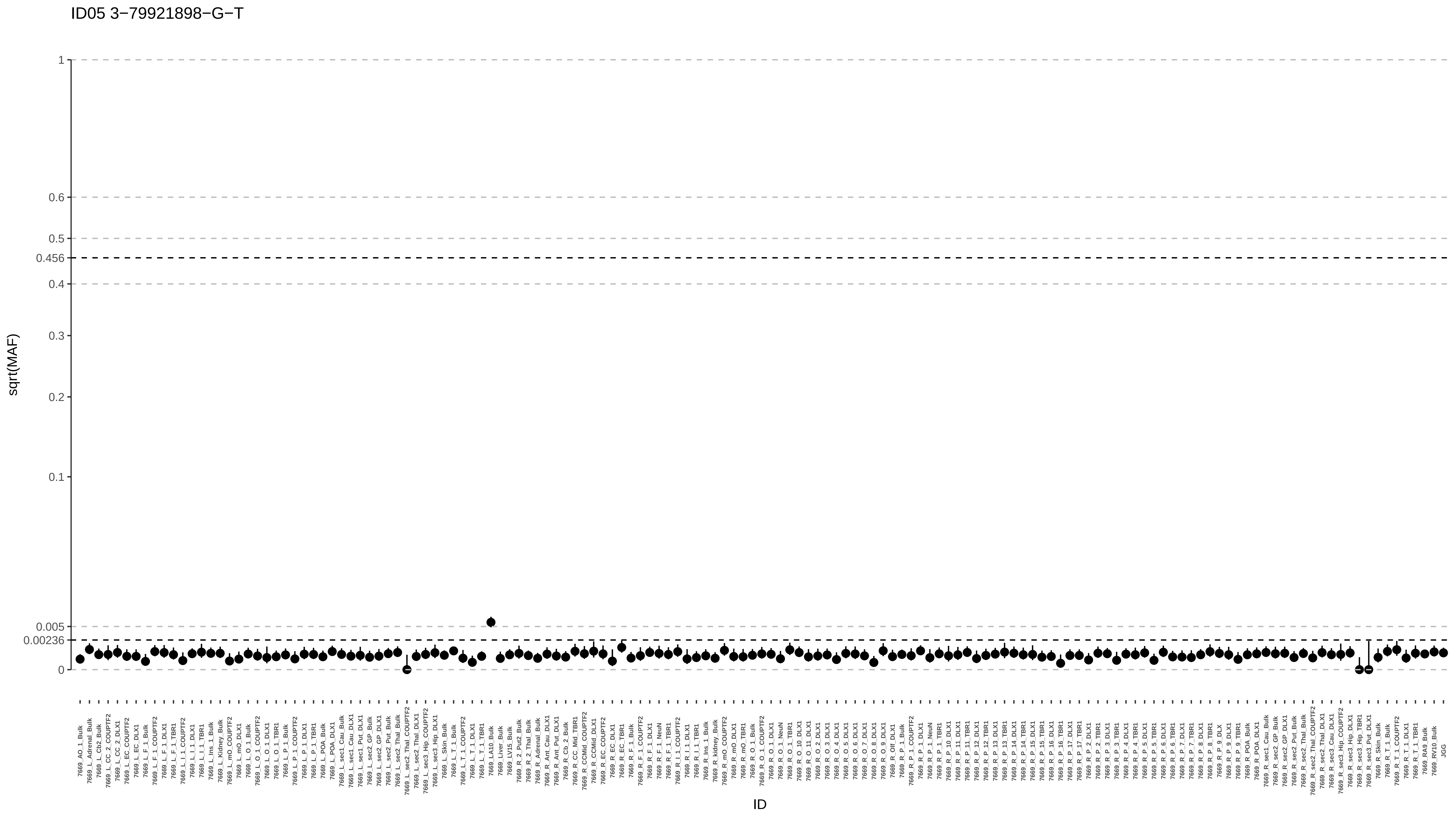
ID05 3−79921898−G−T
sqrt(MAF)
ID
1
0.6
0.5
0.456
0.4
0.3
0.2
0.1
0.005
0.00236
0

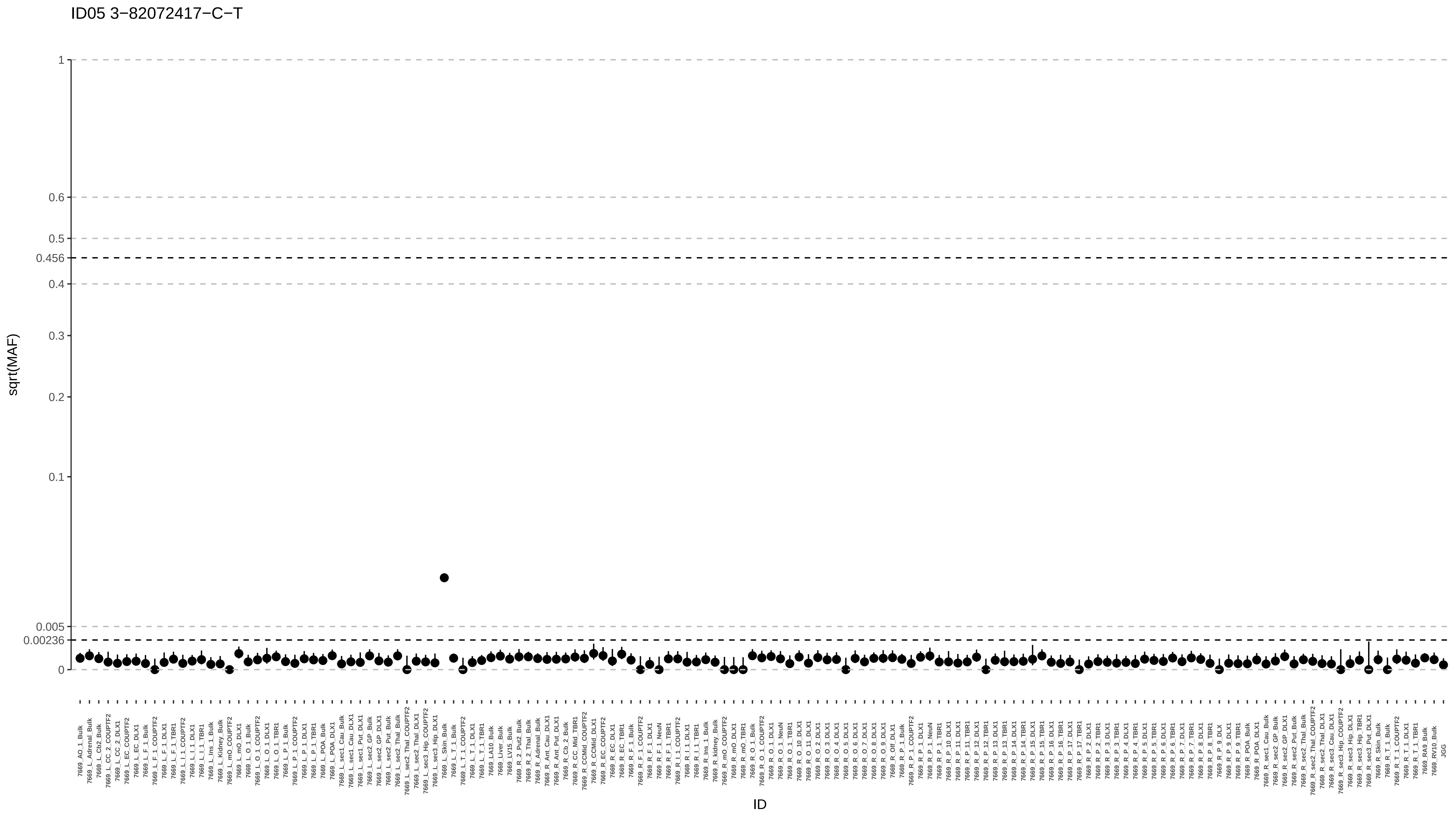

ID05 3−82072417−C−T
sqrt(MAF)
1
0.6
0.5
0.456
0.4
0.3
0.2
0.1
0.005
0.00236
0
ID

ID05 3−82901114−C−T

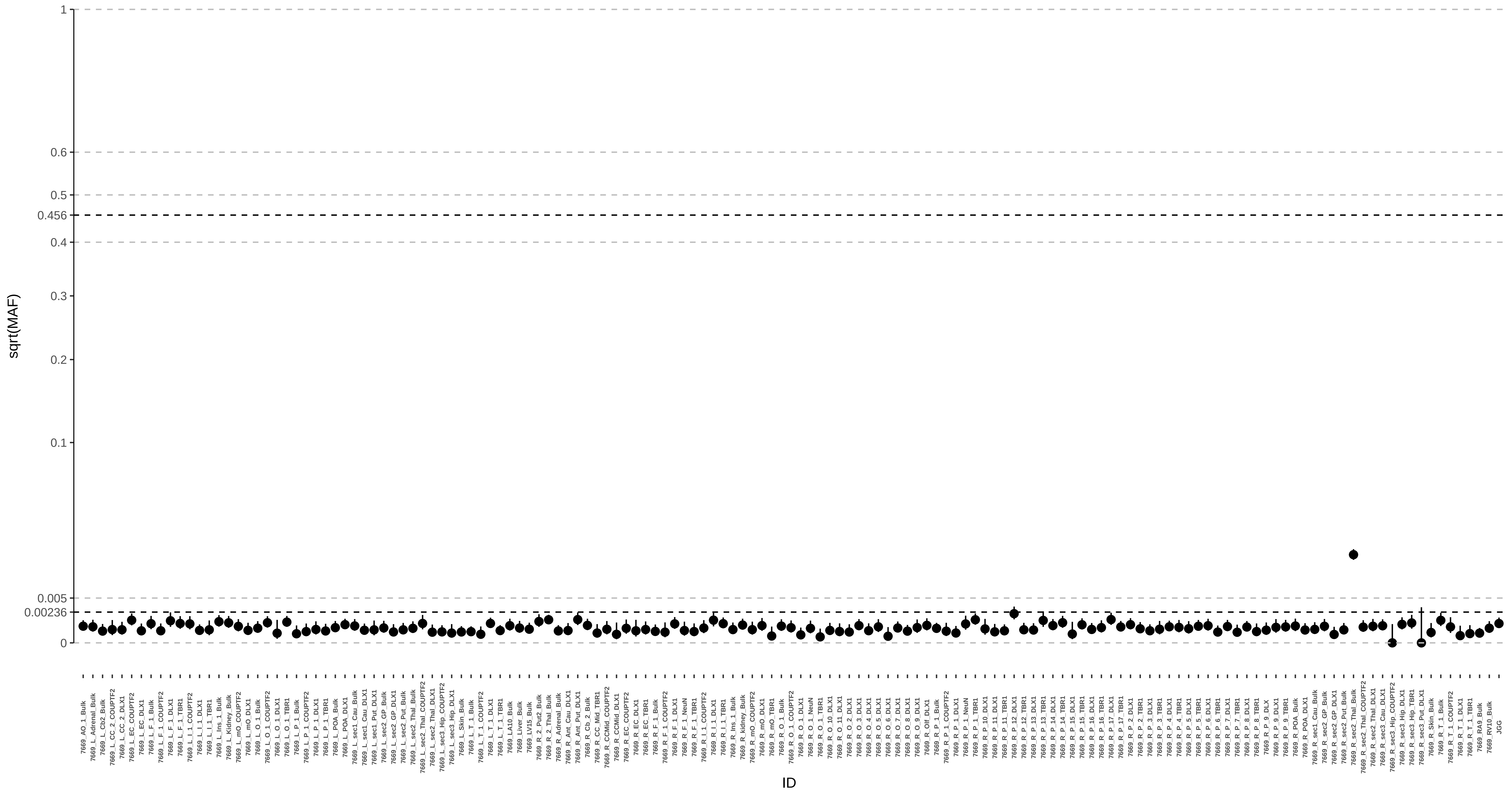

ID05 3−83516288−A−C

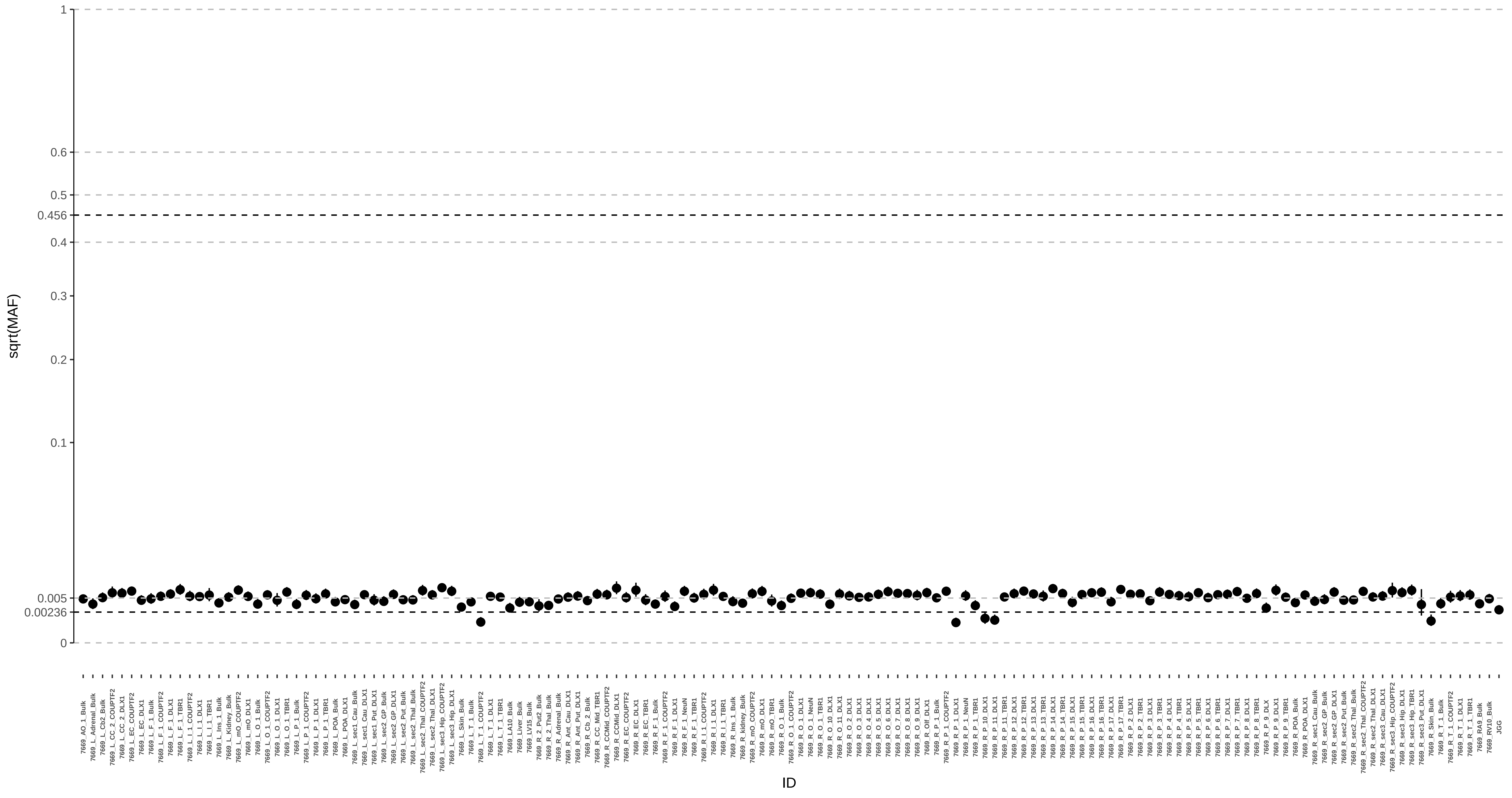

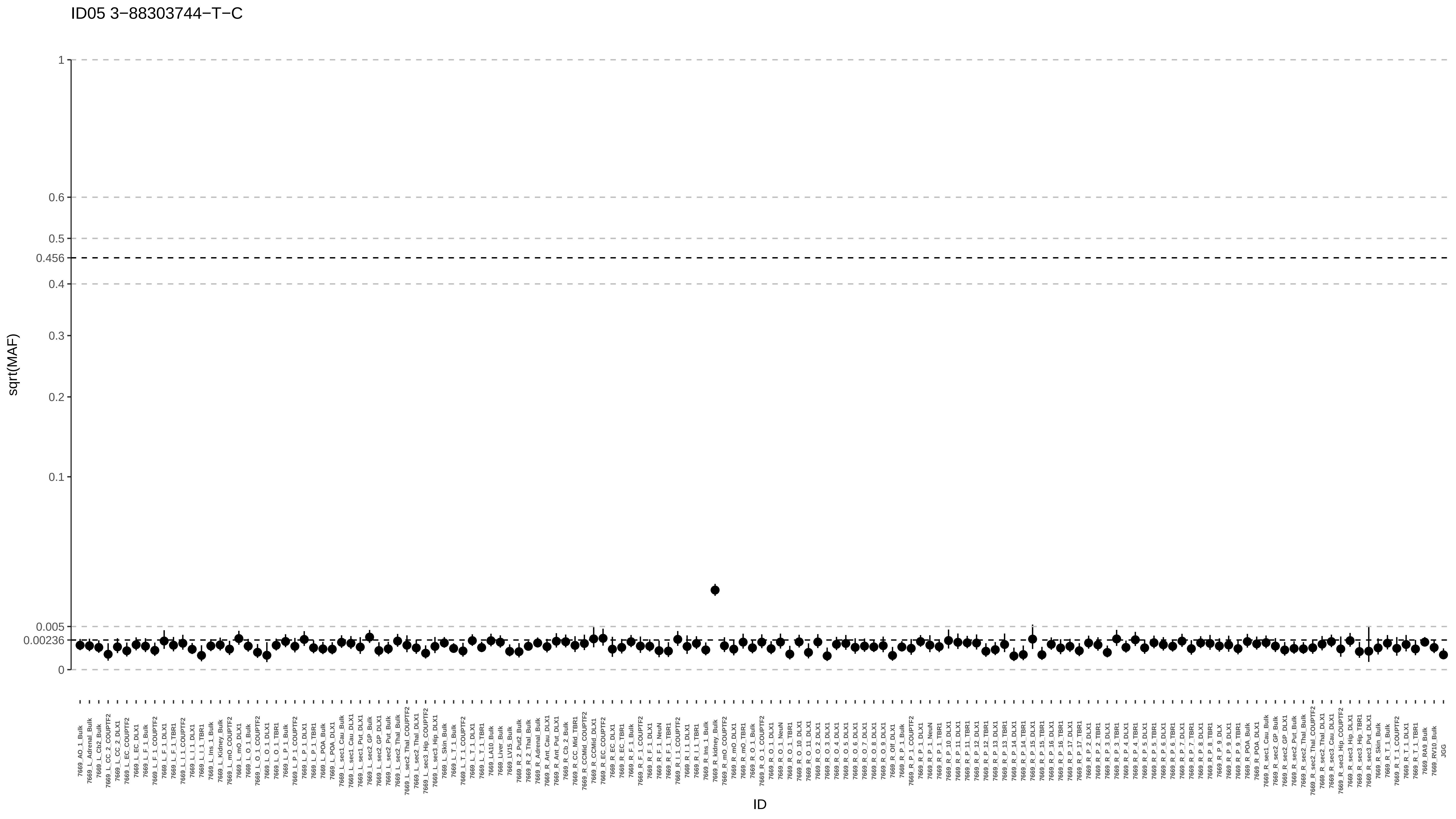
ID05 3−88303744−T−C
sqrt(MAF)
1
0.6
0.5
0.456
0.4
0.3
0.2
0.1
0.005
0.00236
0
ID

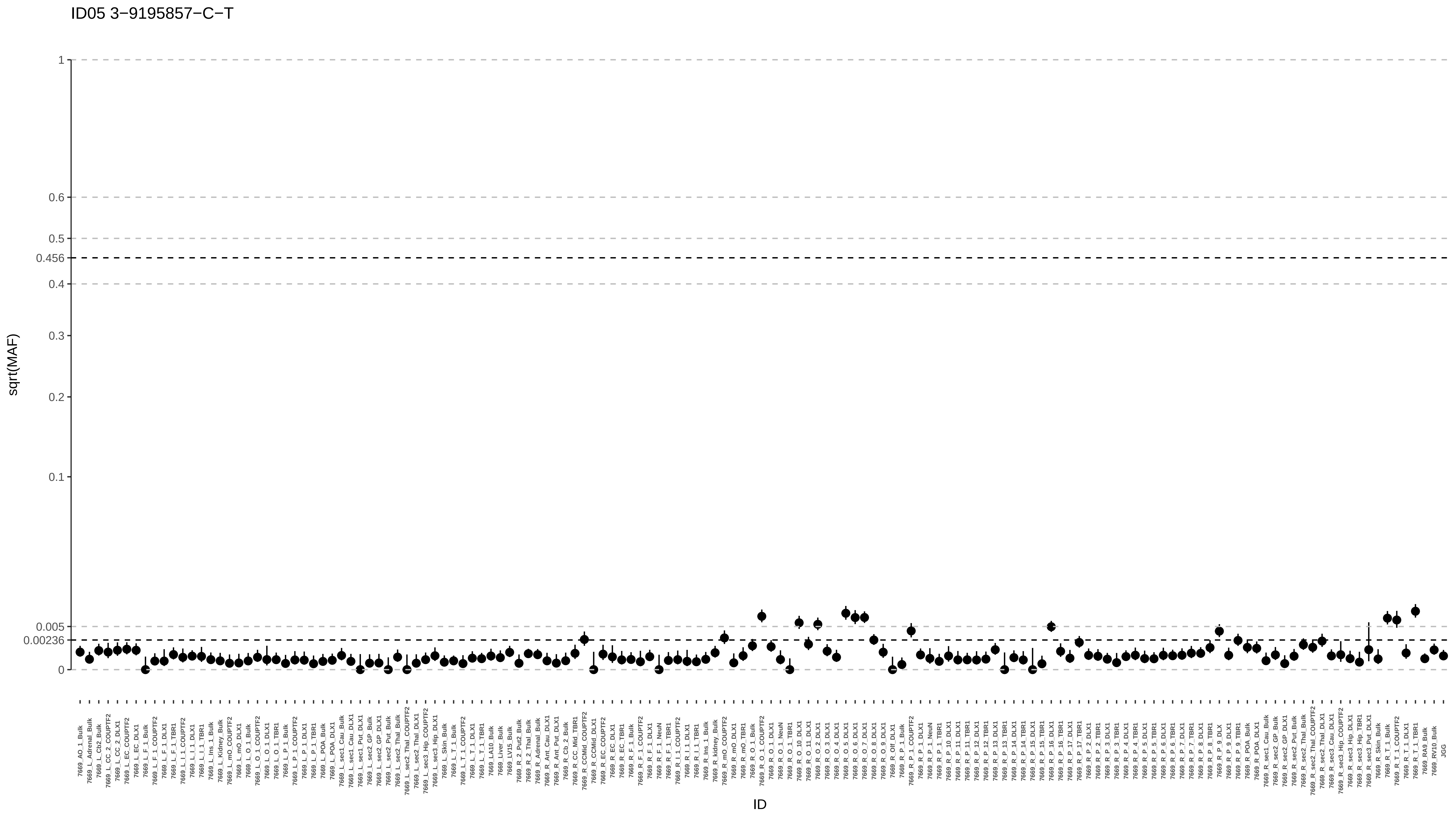
ID05 3−9195857−C−T
sqrt(MAF)
1
0.6
0.5
0.456
0.4
0.3
0.2
0.1
0.005
0.00236
0
ID

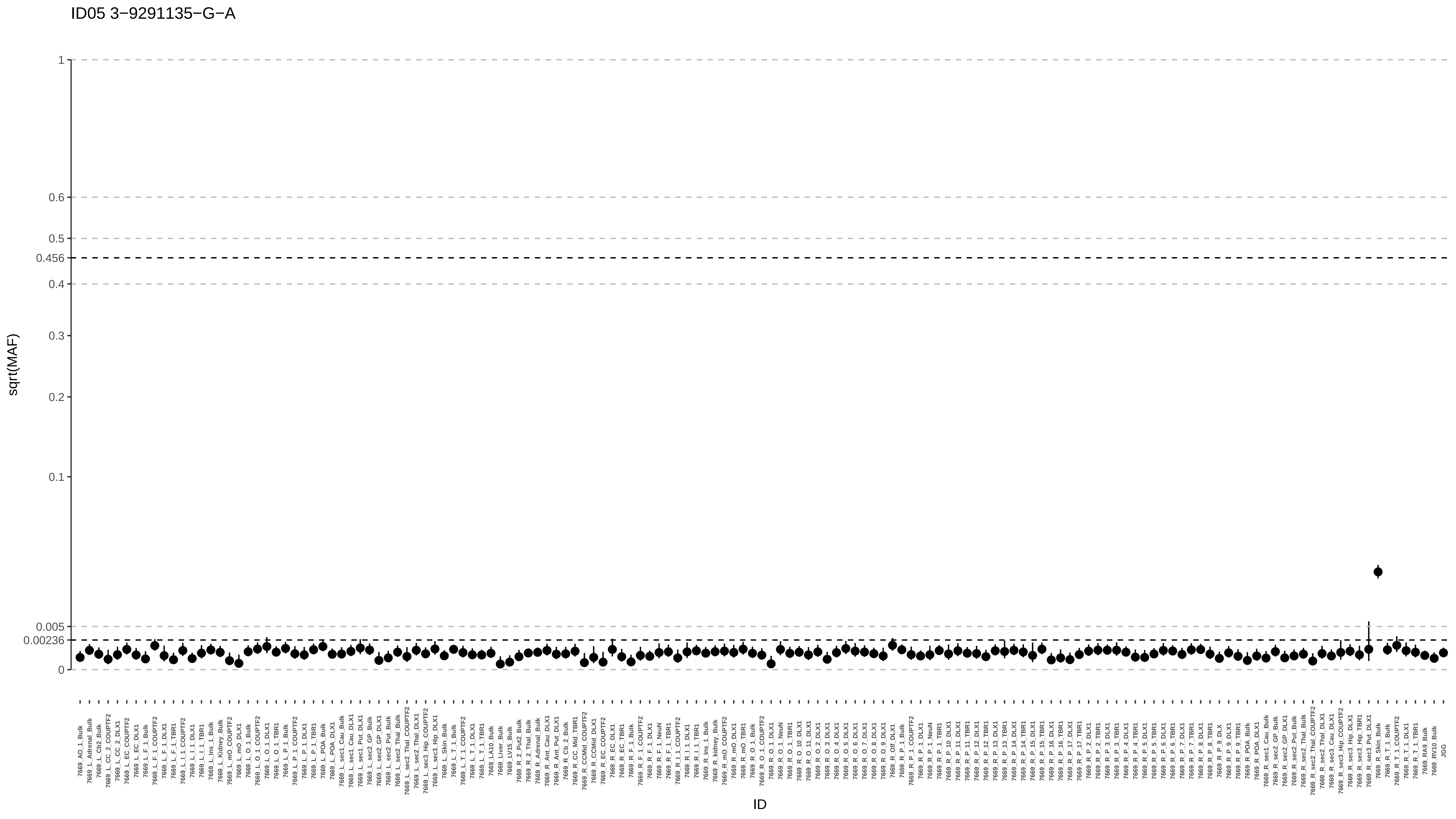

ID05 3−9291135−G−A
sqrt(MAF)
1
0.6
0.5
0.456
0.4
0.3
0.2
0.1
0.005
0.00236
0
ID

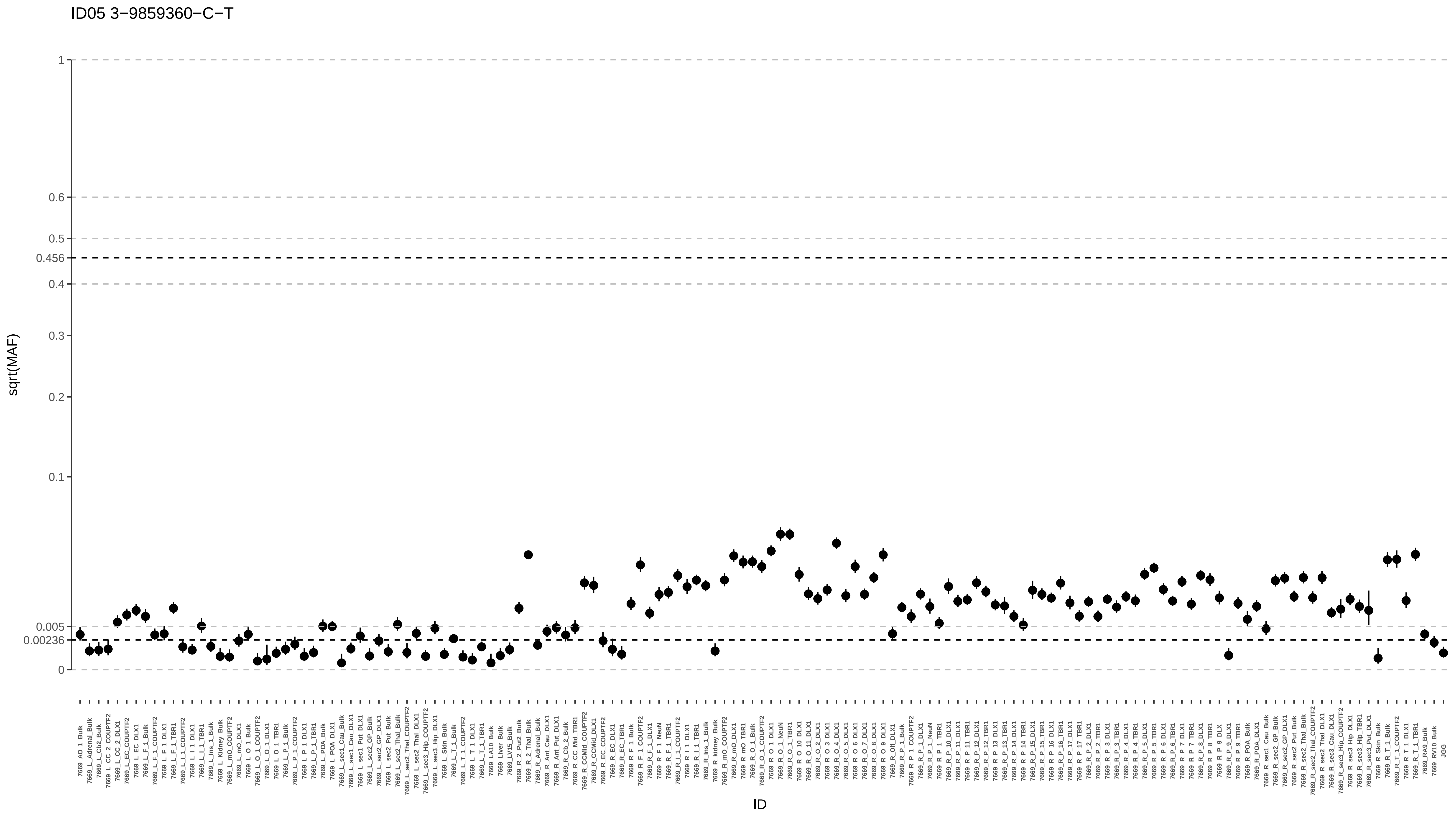
ID05 3−9859360−C−T
sqrt(MAF)
1
0.6
0.5
0.456
0.4
0.3
0.2
0.1
0.005
0.00236
0
ID
7669_AO_1_Bulk
7669_L_Adrenal_Bulk
7669_L_Cb2_Bulk
7669_L_CC_2_COUPTF2
7669_L_CC_2_DLX1
7669_L_EC_COUPTF2
7669_L_EC_DLX1
7669_L_F_1_Bulk
7669_L_F_1_COUPTF2
7669_L_F_1_DLX1
7669_L_F_1_TBR1
7669_L_I_1_COUPTF2
7669_L_I_1_DLX1
7669_L_I_1_TBR1
7669_L_Ins_1_Bulk
7669_L_Kidney_Bulk
7669_L_mO_COUPTF2
7669_L_mO_DLX1
7669_L_O_1_Bulk
7669_L_O_1_COUPTF2
7669_L_O_1_DLX1
7669_L_O_1_TBR1
7669_L_P_1_Bulk
7669_L_P_1_COUPTF2
7669_L_P_1_DLX1
7669_L_P_1_TBR1
7669_L_POA_Bulk
7669_L_POA_DLX1
7669_L_sec1_Cau_Bulk
7669_L_sec1_Cau_DLX1
7669_L_sec1_Put_DLX1
7669_L_sec2_GP_Bulk
7669_L_sec2_GP_DLX1
7669_L_sec2_Put_Bulk
7669_L_sec2_Thal_Bulk
7669_L_sec2_Thal_COUPTF2
7669_L_sec2_Thal_DLX1
7669_L_sec3_Hip_COUPTF2
7669_L_sec3_Hip_DLX1
7669_L_Skin_Bulk
7669_L_T_1_Bulk
7669_L_T_1_COUPTF2
7669_L_T_1_DLX1
7669_L_T_1_TBR1
7669_LA10_Bulk
7669_Liver_Bulk
7669_LV15_Bulk
7669_R_2_Put2_Bulk
7669_R_2_Thal_Bulk
7669_R_Adrenal_Bulk
7669_R_Ant_Cau_DLX1
7669_R_Ant_Put_DLX1
7669_R_Cb_2_Bulk
7669_R_CC_Mid_TBR1
7669_R_CCMid_COUPTF2
7669_R_CCMid_DLX1
7669_R_EC_COUPTF2
7669_R_EC_DLX1
7669_R_EC_TBR1
7669_R_F_1_Bulk
7669_R_F_1_COUPTF2
7669_R_F_1_DLX1
7669_R_F_1_NeuN
7669_R_F_1_TBR1
7669_R_I_1_COUPTF2
7669_R_I_1_DLX1
7669_R_I_1_TBR1
7669_R_Ins_1_Bulk
7669_R_kidney_Bulk
7669_R_mO_COUPTF2
7669_R_mO_DLX1
7669_R_mO_TBR1
7669_R_O_1_Bulk
7669_R_O_1_COUPTF2
7669_R_O_1_DLX1
7669_R_O_1_NeuN
7669_R_O_1_TBR1
7669_R_O_10_DLX1
7669_R_O_11_DLX1
7669_R_O_2_DLX1
7669_R_O_3_DLX1
7669_R_O_4_DLX1
7669_R_O_5_DLX1
7669_R_O_6_DLX1
7669_R_O_7_DLX1
7669_R_O_8_DLX1
7669_R_O_9_DLX1
7669_R_Olf_DLX1
7669_R_P_1_Bulk
7669_R_P_1_COUPTF2
7669_R_P_1_DLX1
7669_R_P_1_NeuN
7669_R_P_1_TBR1
7669_R_P_10_DLX1
7669_R_P_11_DLX1
7669_R_P_11_TBR1
7669_R_P_12_DLX1
7669_R_P_12_TBR1
7669_R_P_13_DLX1
7669_R_P_13_TBR1
7669_R_P_14_DLX1
7669_R_P_14_TBR1
7669_R_P_15_DLX1
7669_R_P_15_TBR1
7669_R_P_16_DLX1
7669_R_P_16_TBR1
7669_R_P_17_DLX1
7669_R_P_17_TBR1
7669_R_P_2_DLX1
7669_R_P_2_TBR1
7669_R_P_3_DLX1
7669_R_P_3_TBR1
7669_R_P_4_DLX1
7669_R_P_4_TBR1
7669_R_P_5_DLX1
7669_R_P_5_TBR1
7669_R_P_6_DLX1
7669_R_P_6_TBR1
7669_R_P_7_DLX1
7669_R_P_7_TBR1
7669_R_P_8_DLX1
7669_R_P_8_TBR1
7669_R_P_9_DLX
7669_R_P_9_DLX1
7669_R_P_9_TBR1
7669_R_POA_Bulk
7669_R_POA_DLX1
7669_R_sec1_Cau_Bulk
7669_R_sec2_GP_Bulk
7669_R_sec2_GP_DLX1
7669_R_sec2_Put_Bulk
7669_R_sec2_Thal_Bulk
7669_R_sec2_Thal_COUPTF2
7669_R_sec2_Thal_DLX1
7669_R_sec3_Cau_DLX1
7669_R_sec3_Hip_COUPTF2
7669_R_sec3_Hip_DLX1
7669_R_sec3_Hip_TBR1
7669_R_sec3_Put_DLX1
7669_R_Skin_Bulk
7669_R_T_1_Bulk
7669_R_T_1_COUPTF2
7669_R_T_1_DLX1
7669_R_T_1_TBR1
7669_RA9_Bulk
7669_RV10_Bulk
JGG

ID05 3−99869280−C−T

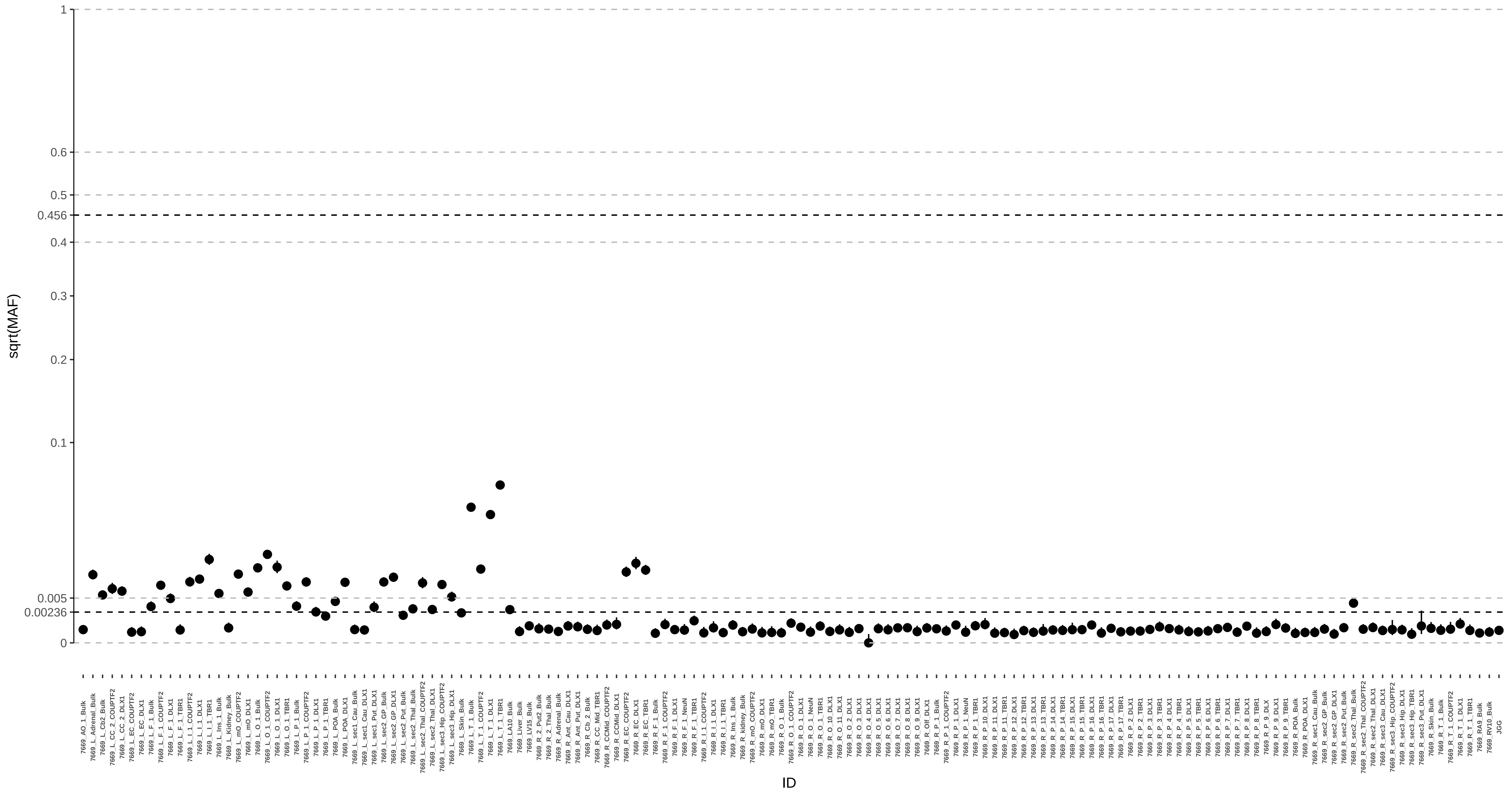

ID05 4−100236559−G−A

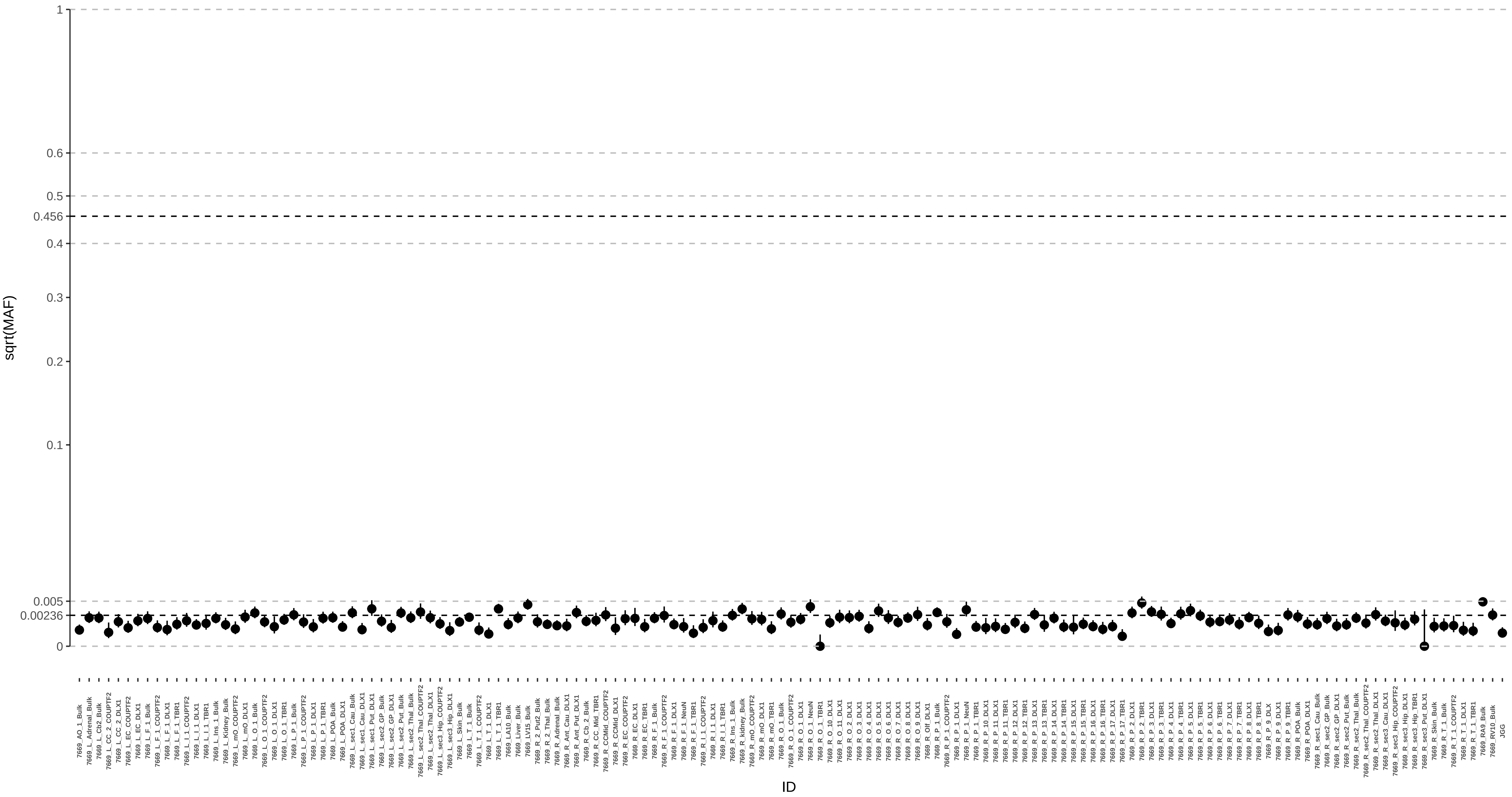

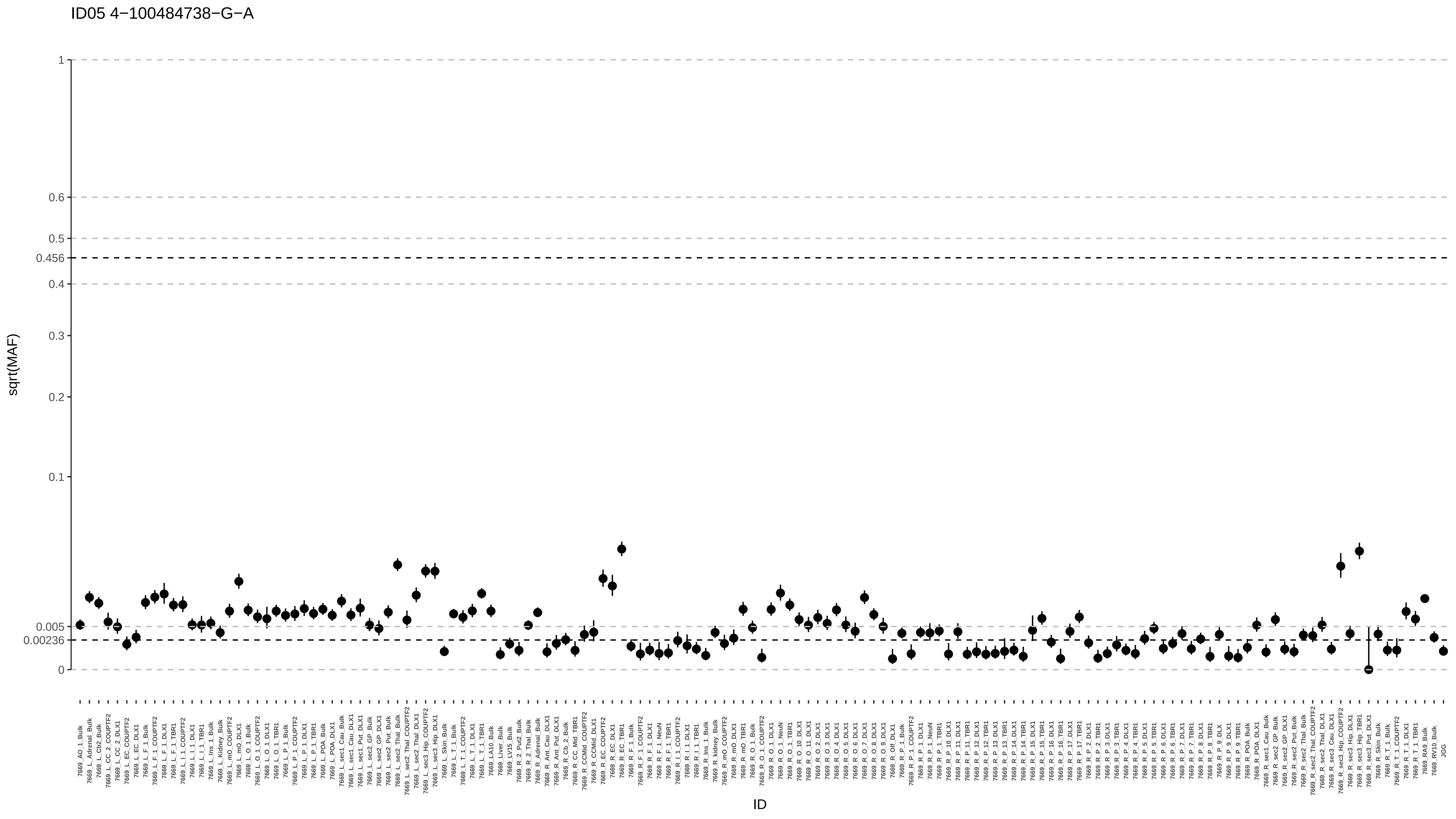
ID05 4−100484738−G−A
sqrt(MAF)
1
0.6
0.5
0.456
0.4
0.3
0.2
0.1
0.005
0.00236
0
ID

ID05 4−1028303−G−A

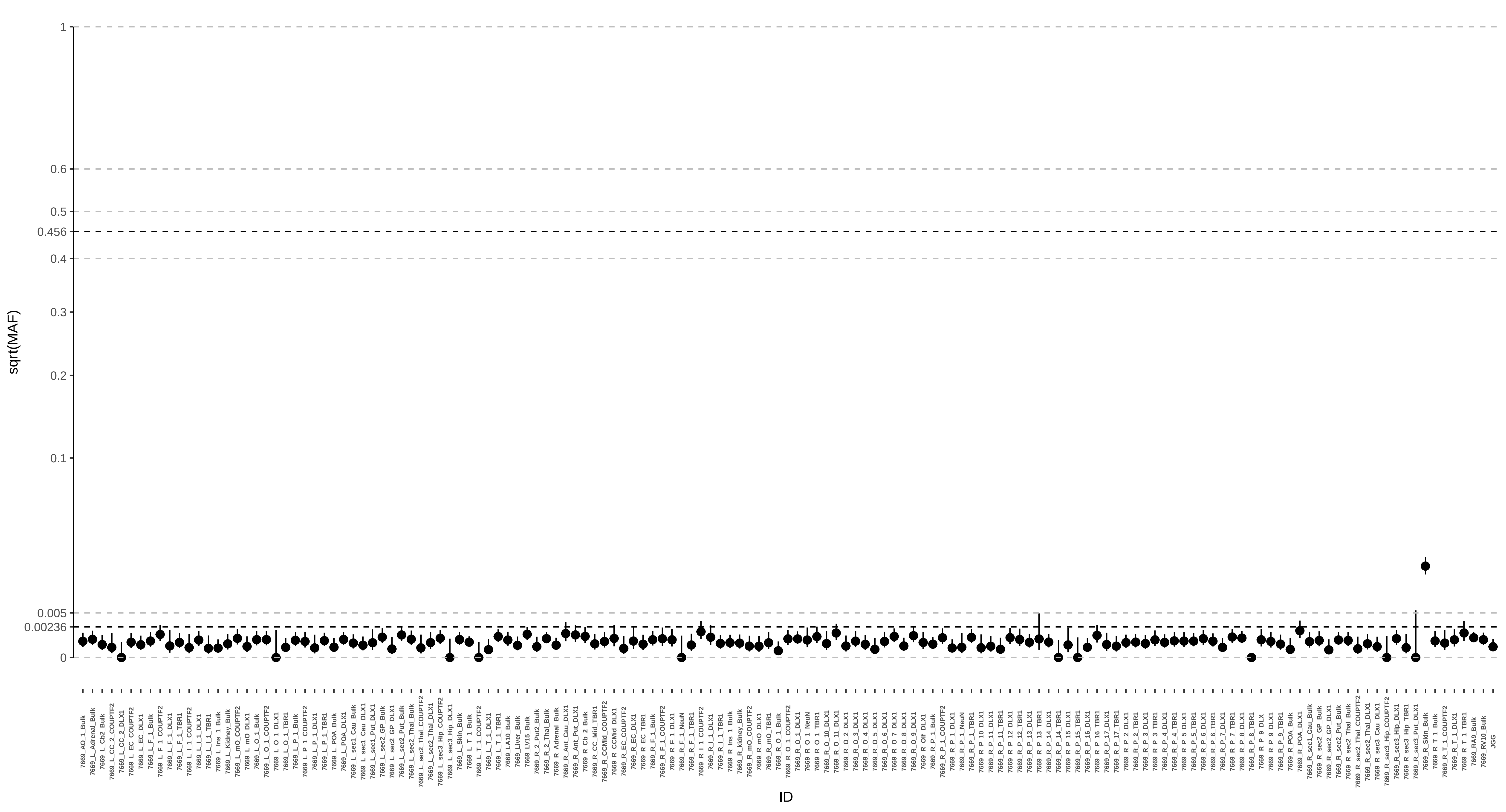

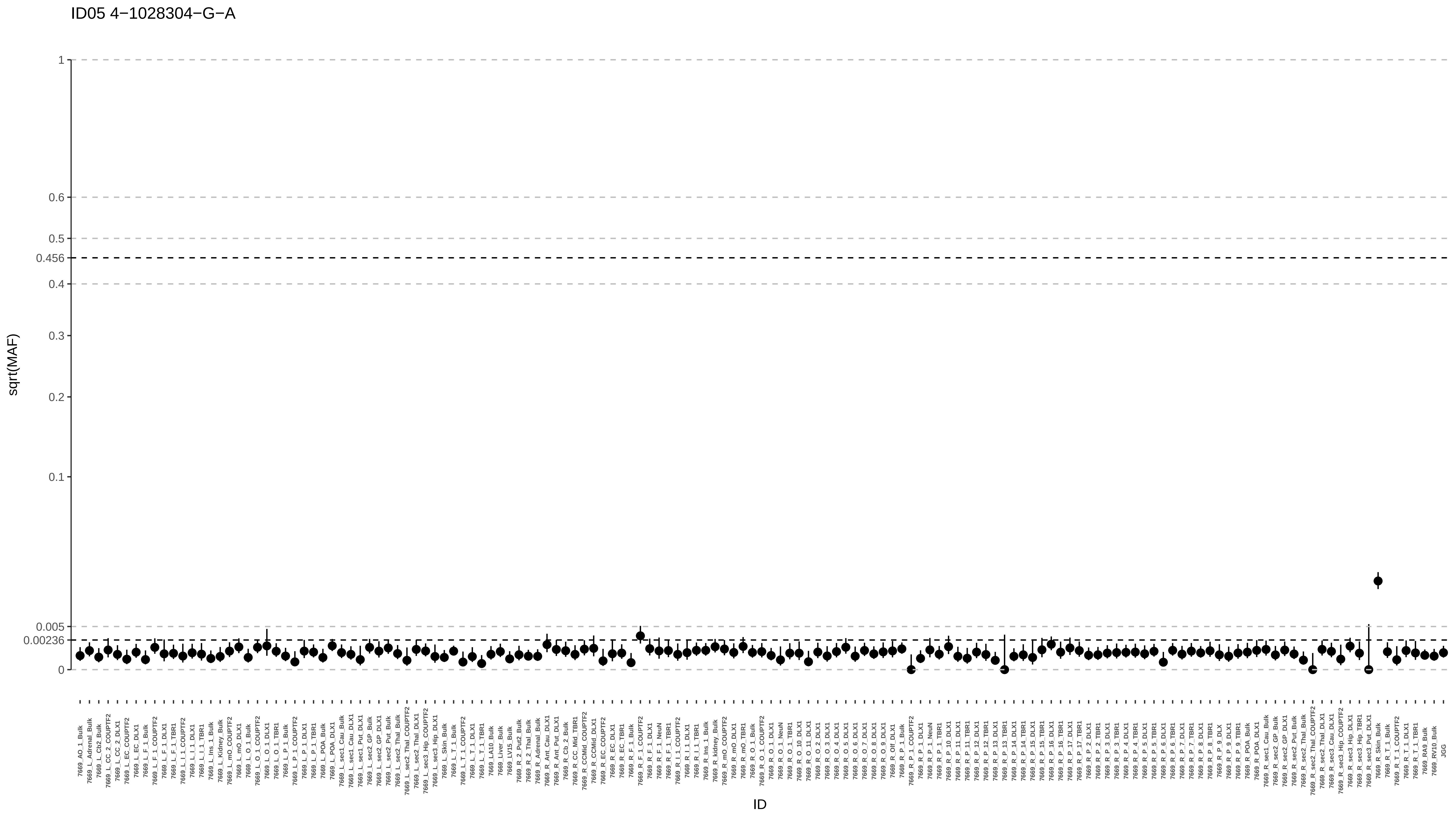
ID05 4−1028304−G−A
sqrt(MAF)
ID
1
0.6
0.5
0.456
0.4
0.3
0.2
0.1
0.005
0.00236
0

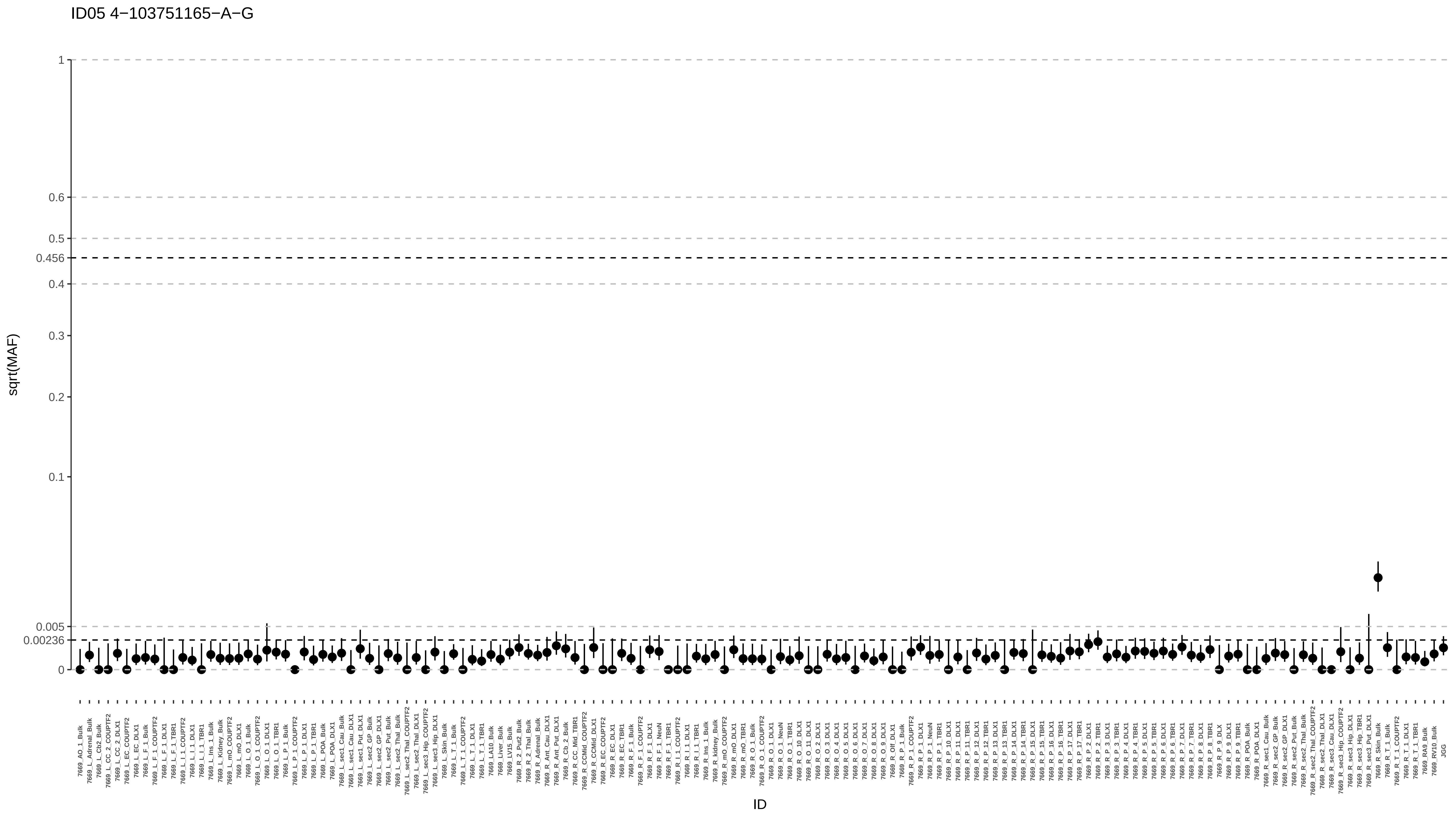
ID05 4−103751165−A−G
sqrt(MAF)
1
0.6
0.5
0.456
0.4
0.3
0.2
0.1
0.005
0.00236
0
ID

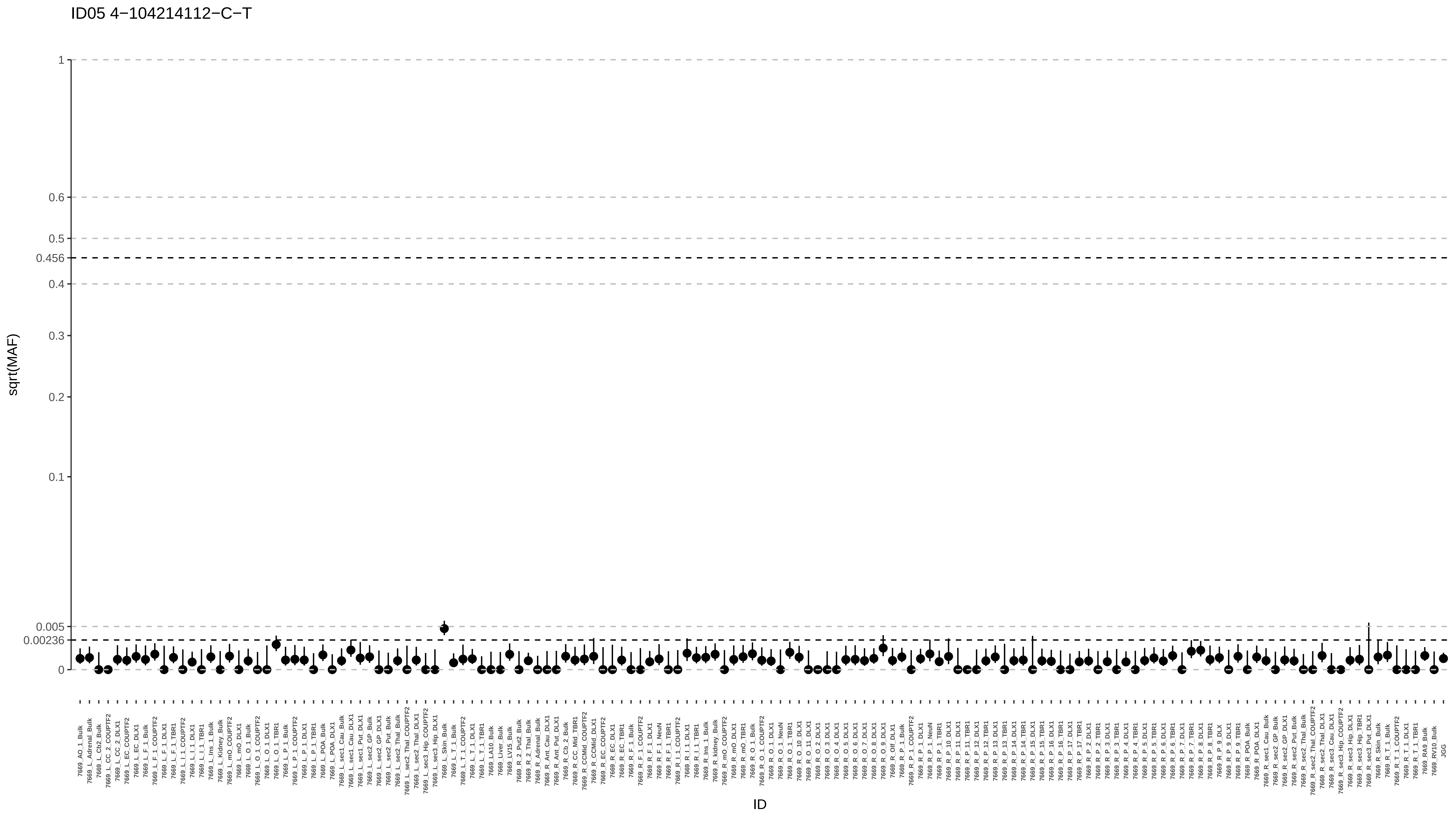
ID05 4−104214112−C−T
sqrt(MAF)
1
0.6
0.5
0.456
0.4
0.3
0.2
0.1
0.005
0.00236
0
ID

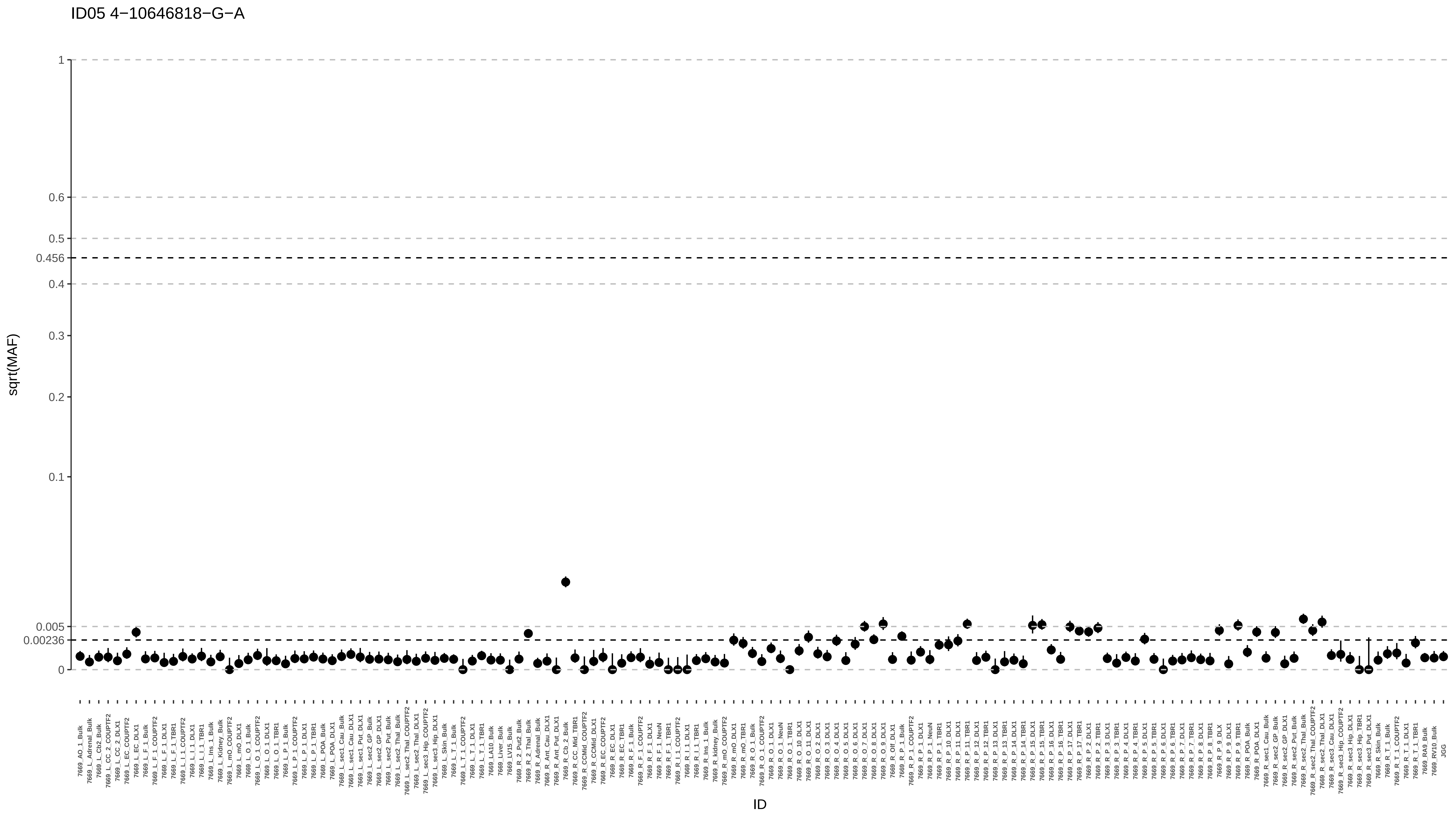

ID05 4−10646818−G−A
sqrt(MAF)
1
0.6
0.5
0.456
0.4
0.3
0.2
0.1
0.005
0.00236
0
ID

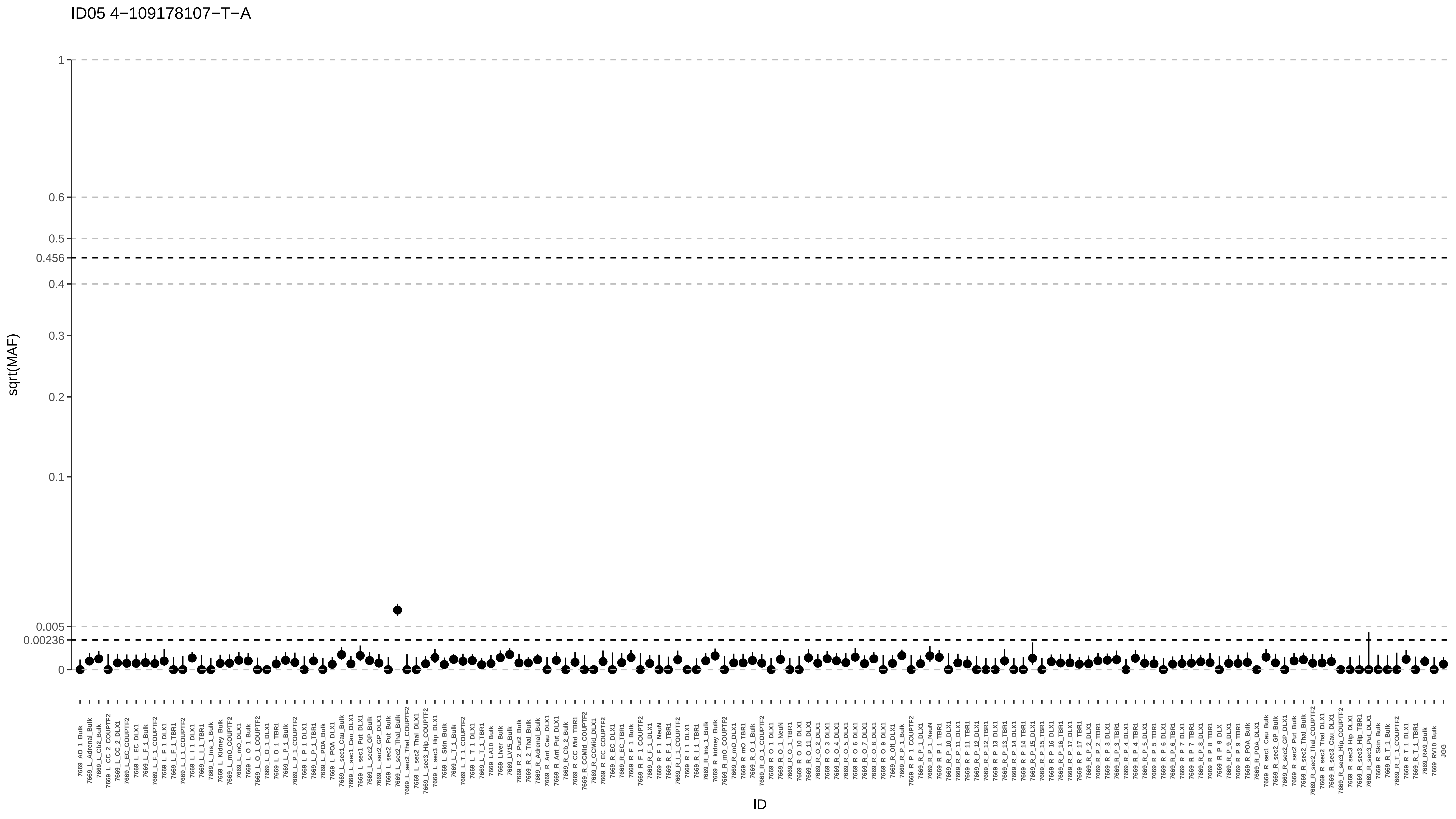
ID05 4−109178107−T−A
sqrt(MAF)
1
0.6
0.5
0.456
0.4
0.3
0.2
0.1
0.005
0.00236
0
ID

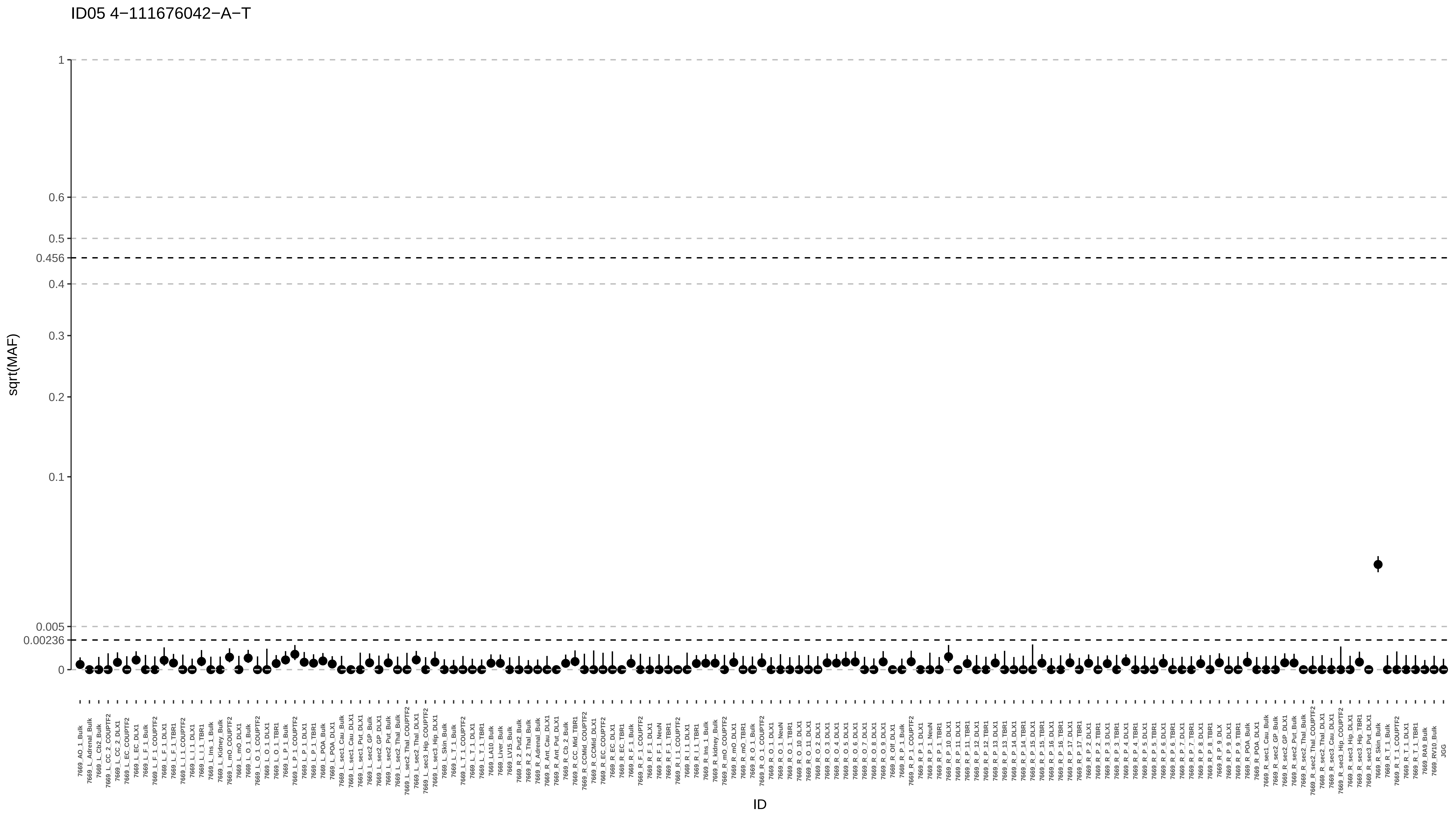
ID05 4−111676042−A−T
sqrt(MAF)
1
0.6
0.5
0.456
0.4
0.3
0.2
0.1
0.005
0.00236
0
ID

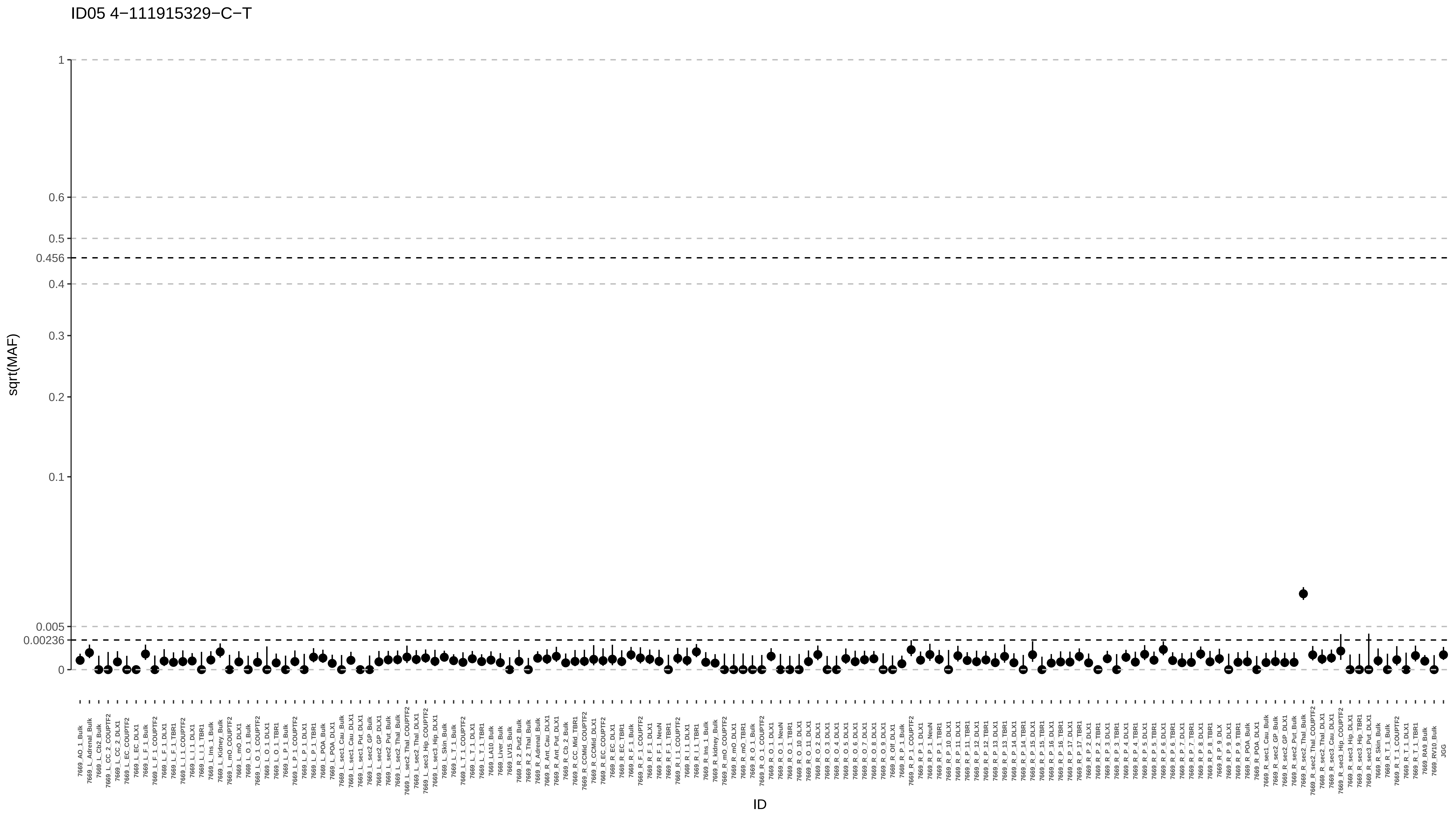
ID05 4−111915329−C−T
sqrt(MAF)
1
0.6
0.5
0.456
0.4
0.3
0.2
0.1
0.005
0.00236
0
ID

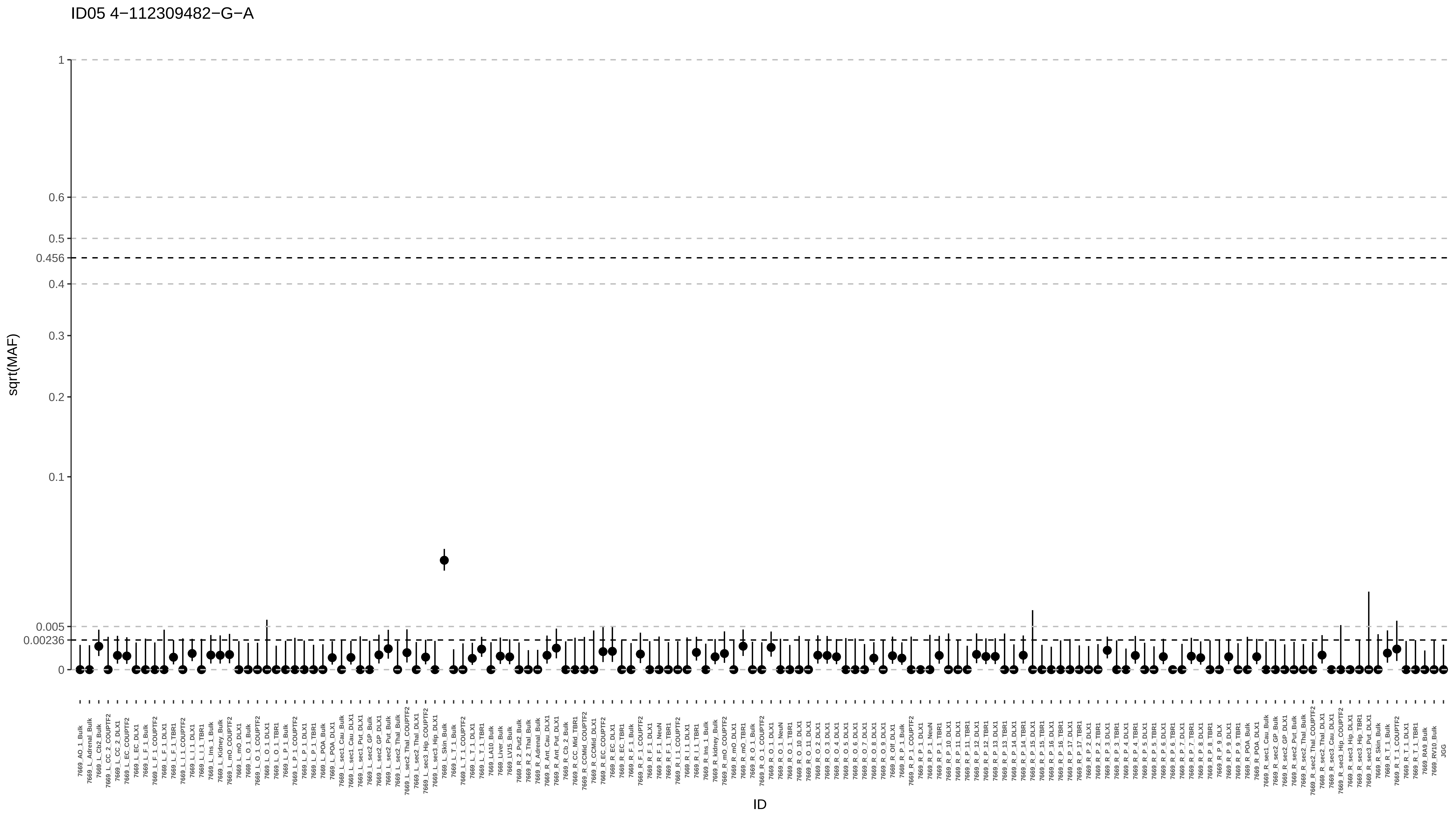
ID05 4−112309482−G−A
sqrt(MAF)
1
0.6
0.5
0.456
0.4
0.3
0.2
0.1
0.005
0.00236
0
ID

ID05 4−116228416−A−G

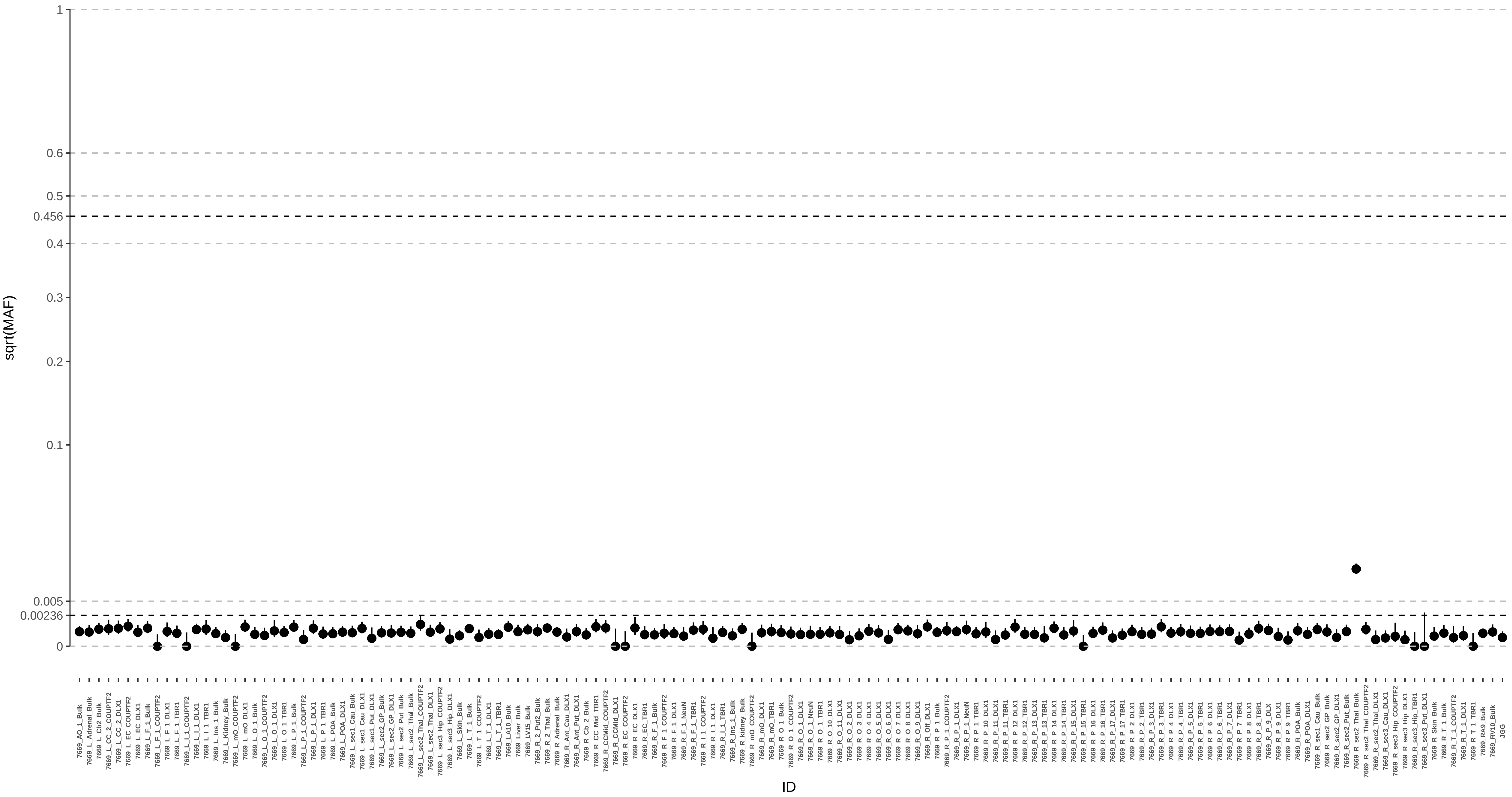

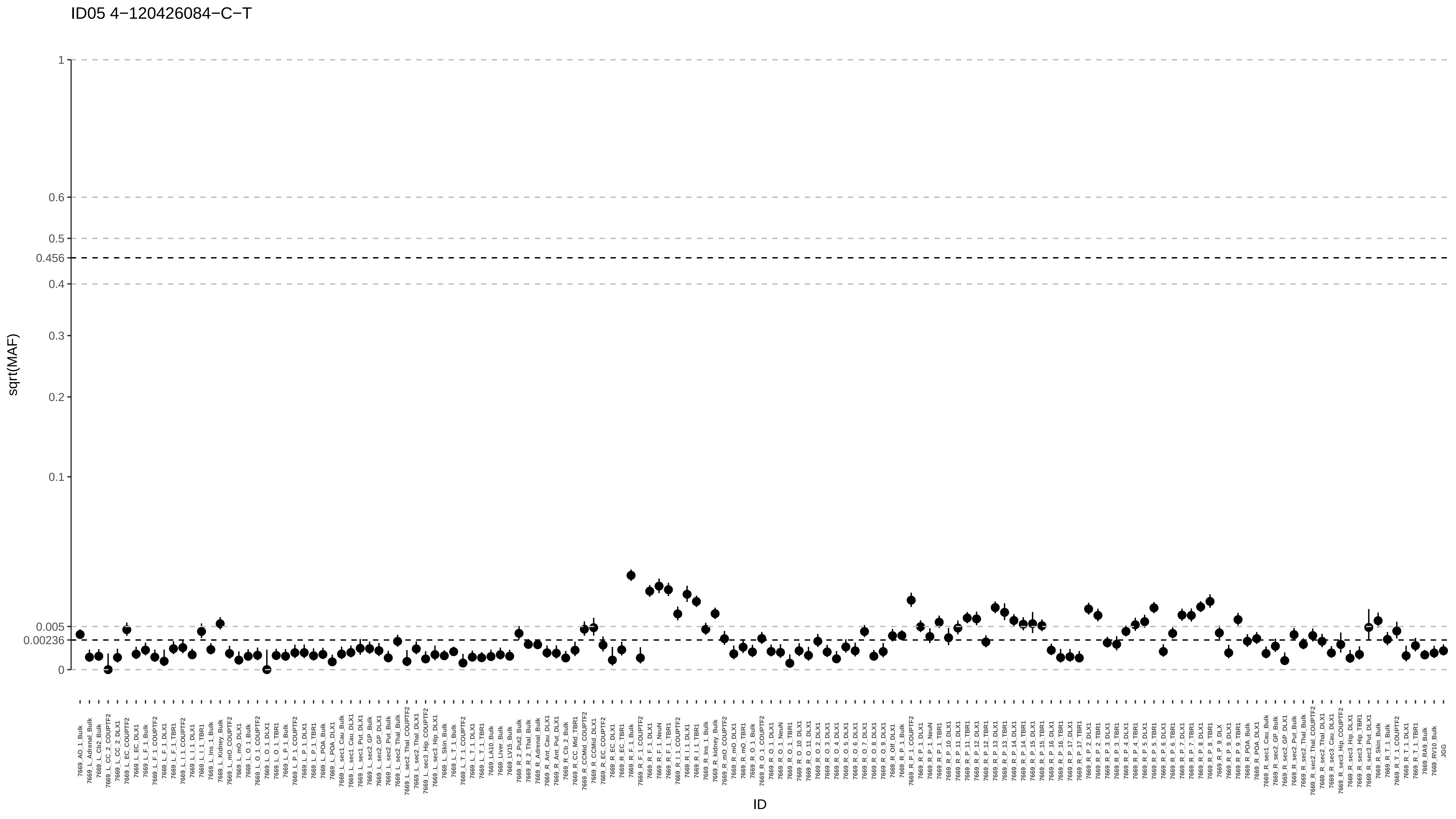
ID05 4−120426084−C−T
sqrt(MAF)
1
0.6
0.5
0.456
0.4
0.3
0.2
0.1
0.005
0.00236
0
ID

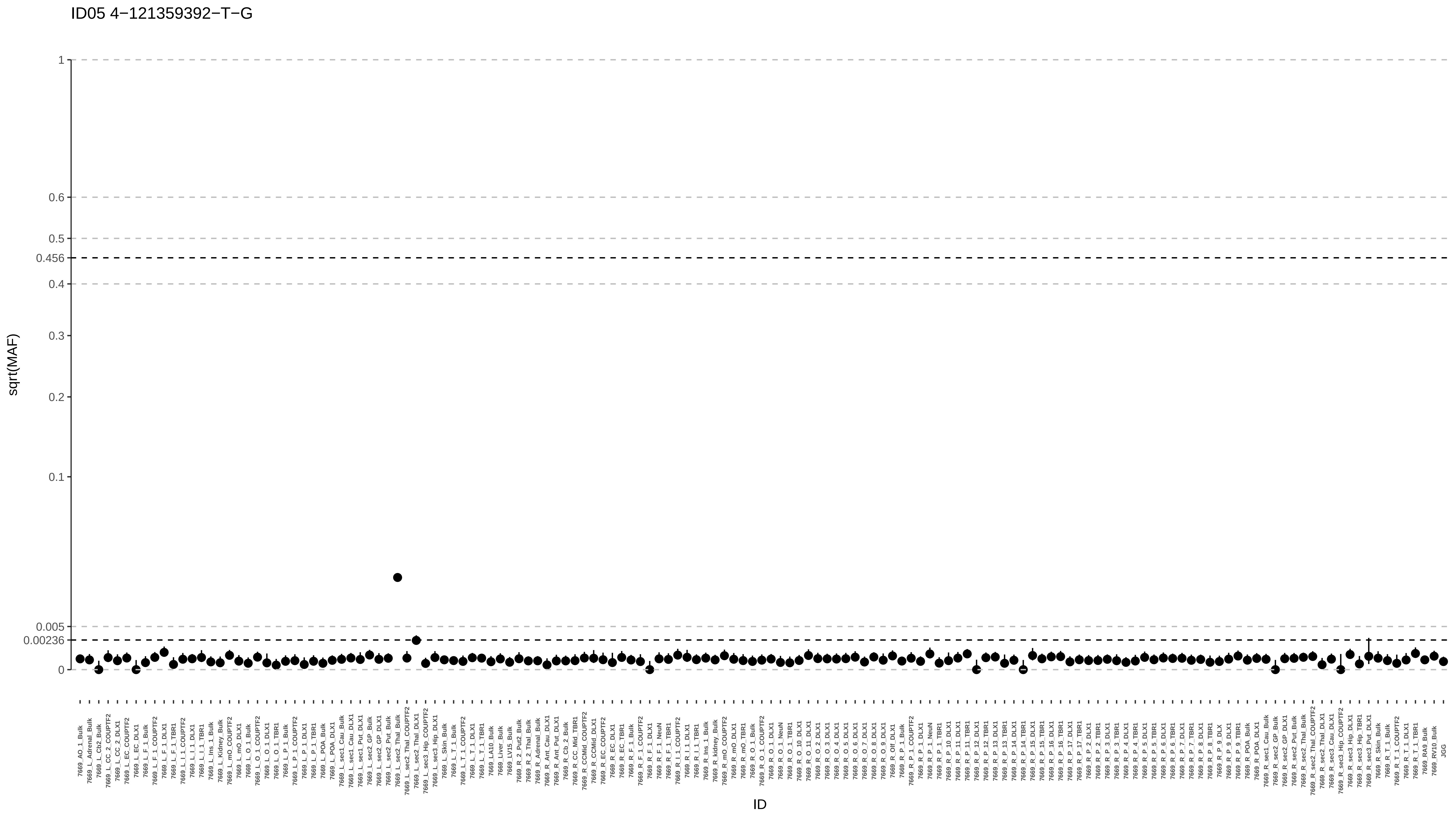

ID05 4−121359392−T−G
sqrt(MAF)
1
0.6
0.5
0.456
0.4
0.3
0.2
0.1
0.005
0.00236
0
ID

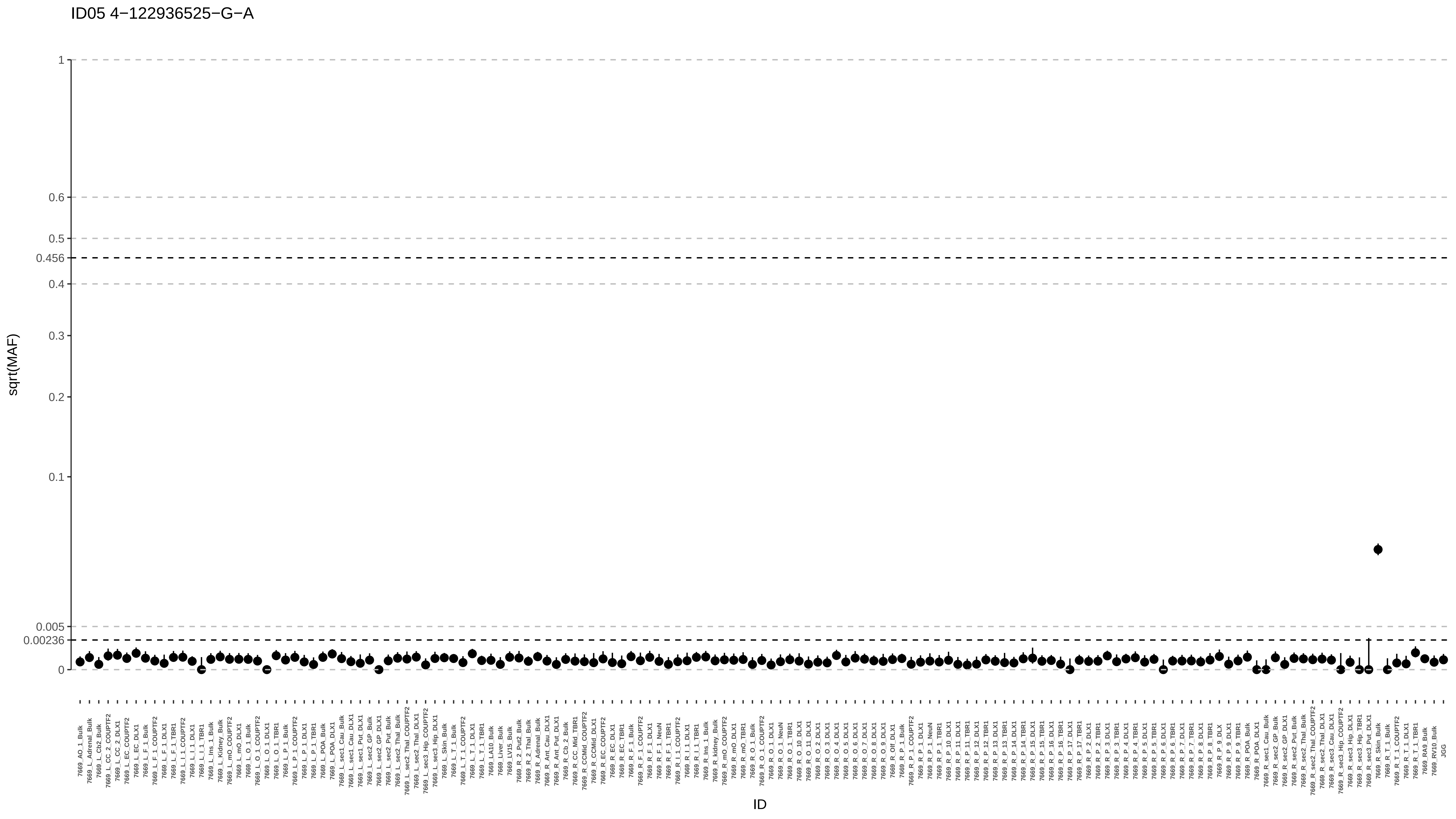
ID05 4−122936525−G−A
sqrt(MAF)
ID
1
0.6
0.5
0.456
0.4
0.3
0.2
0.1
0.005
0.00236
0

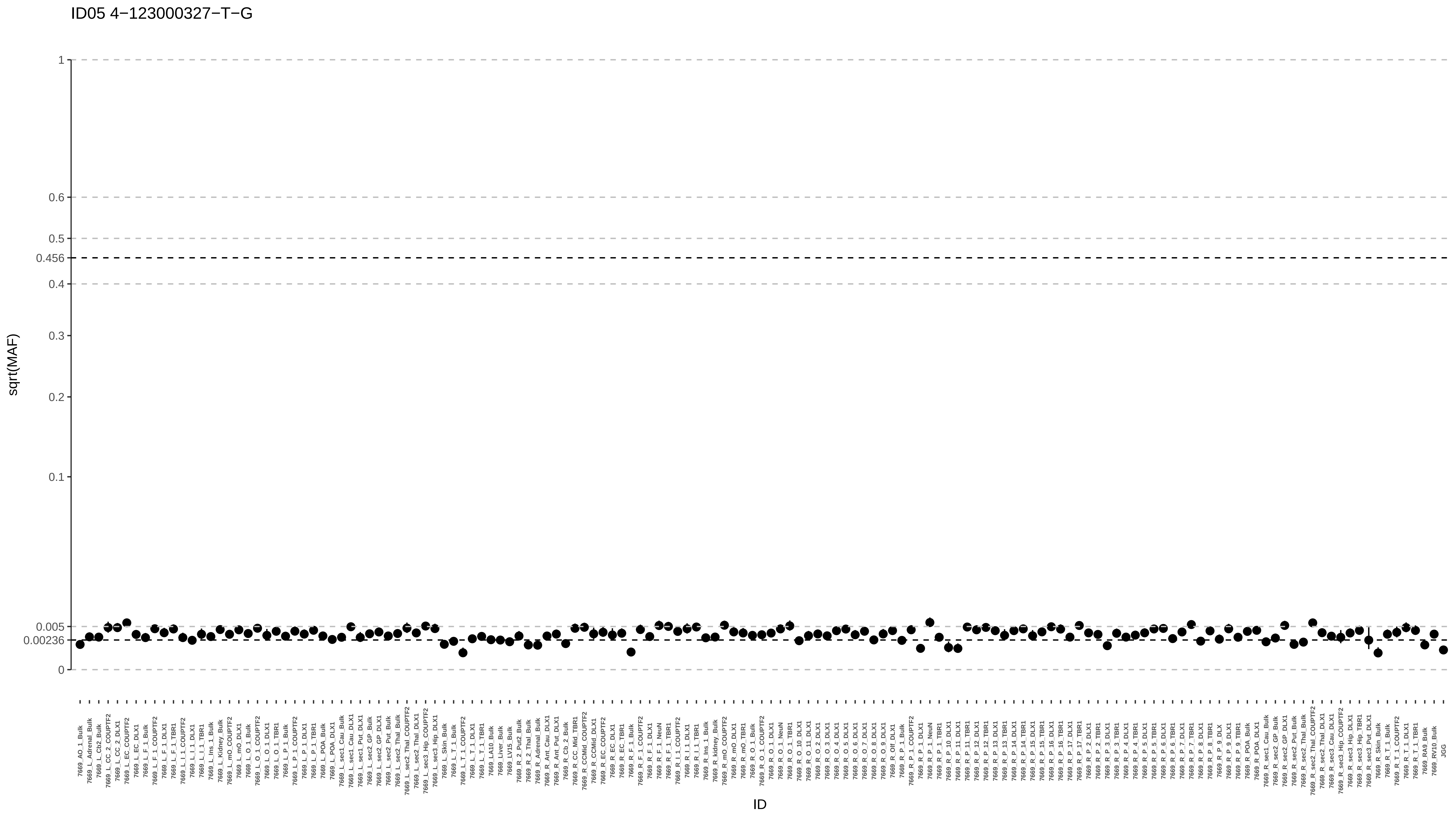
ID05 4−123000327−T−G
sqrt(MAF)
1
0.6
0.5
0.456
0.4
0.3
0.2
0.1
0.005
0.00236
0
ID

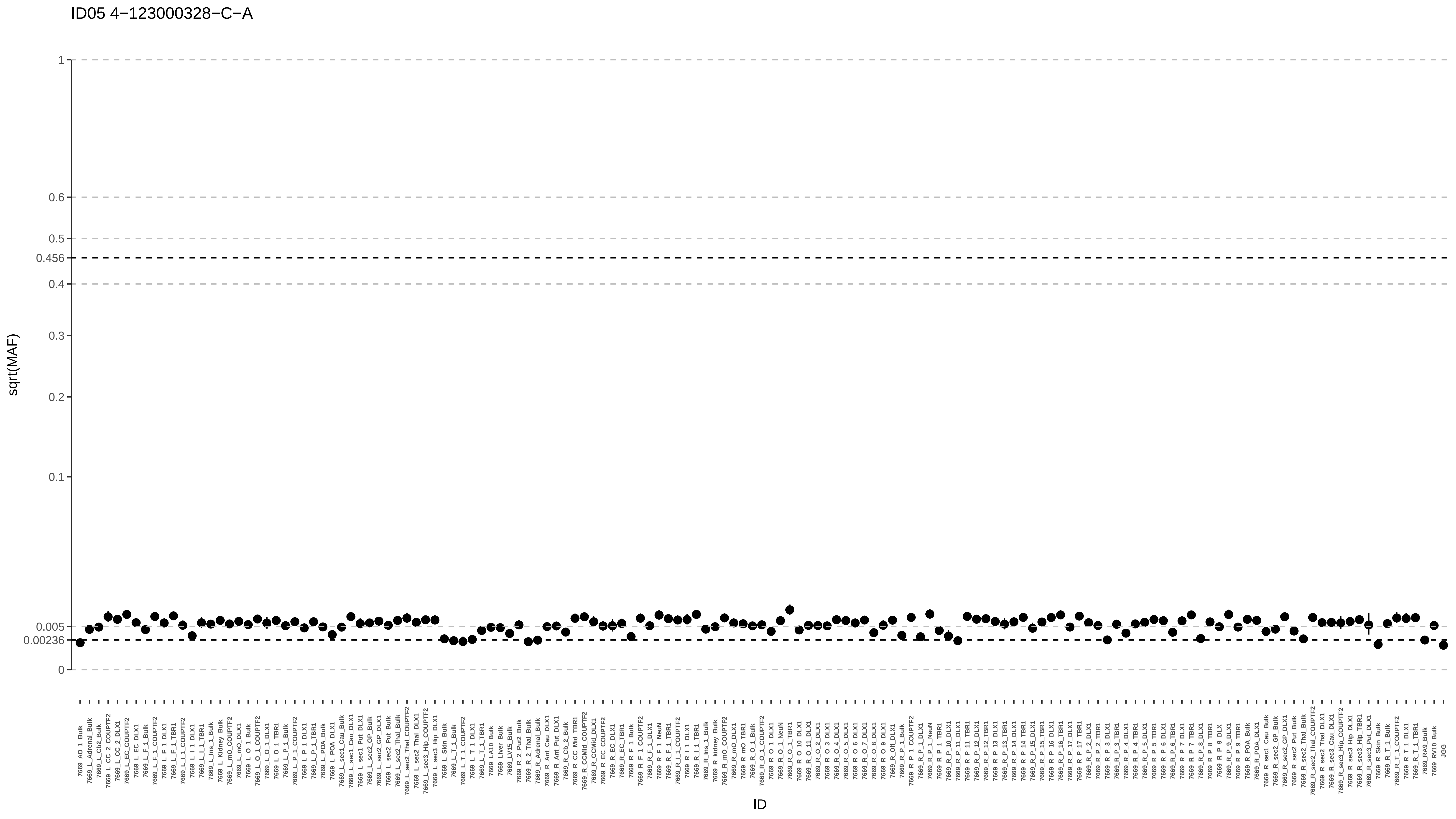
ID05 4−123000328−C−A
sqrt(MAF)
1
0.6
0.5
0.456
0.4
0.3
0.2
0.1
0.005
0.00236
0
ID

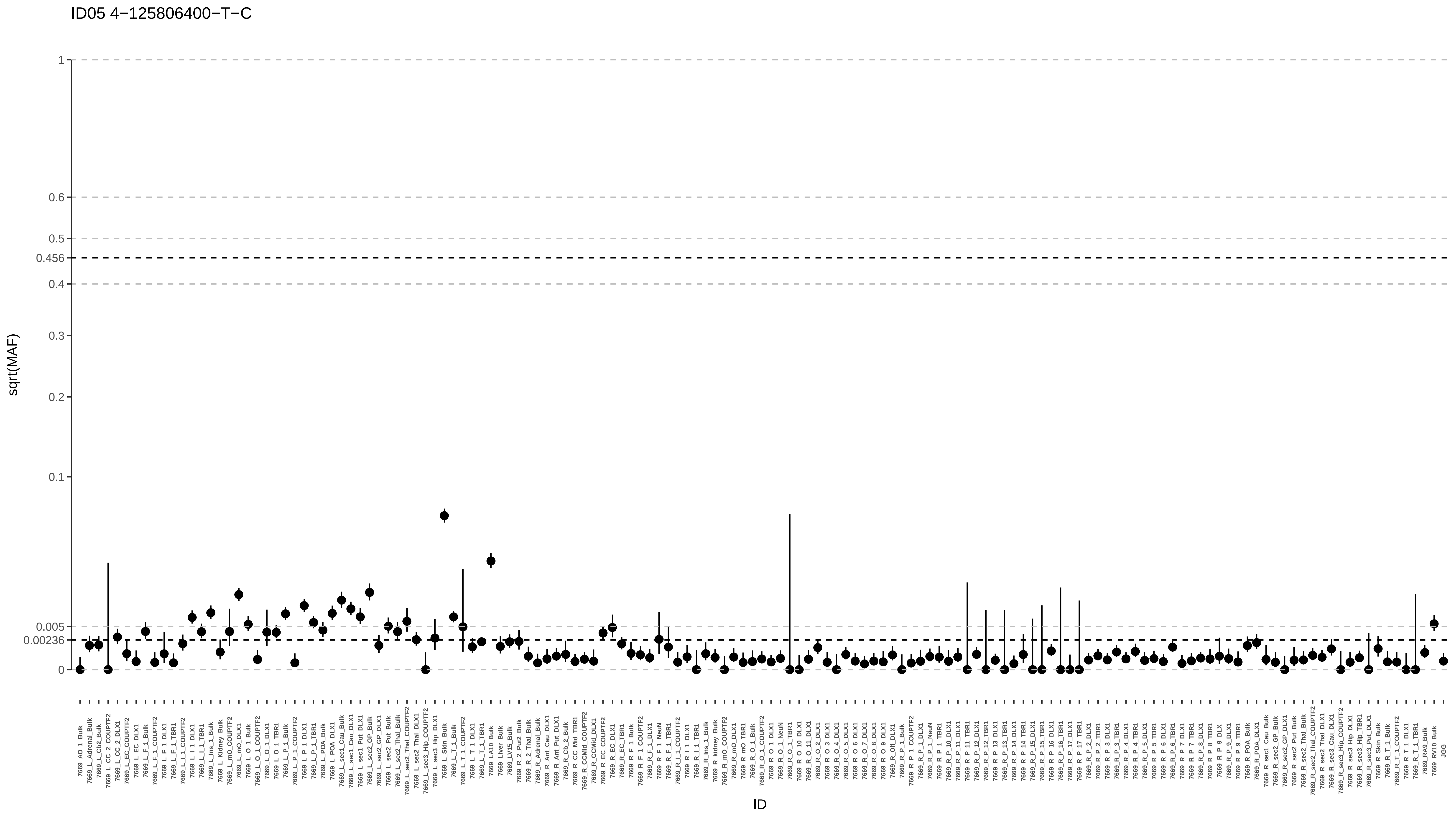

ID05 4−125806400−T−C
1
0.6
0.5
0.456
0.4
0.3
0.2
0.1
0.005
0.00236
0
sqrt(MAF)
ID

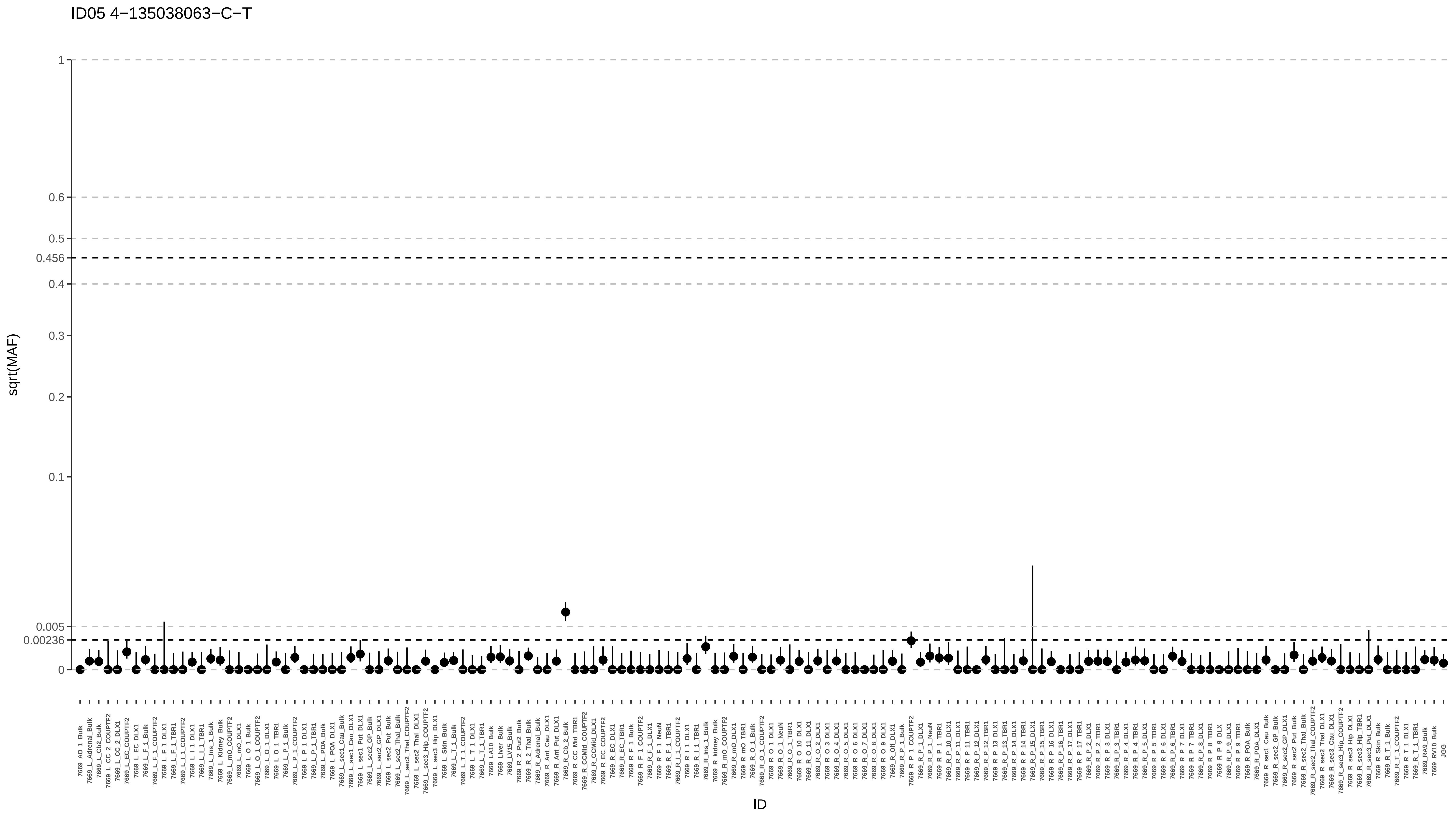

ID05 4−135038063−C−T
sqrt(MAF)
1
0.6
0.5
0.456
0.4
0.3
0.2
0.1
0.005
0.00236
0
ID

ID05 4−137937295−G−T

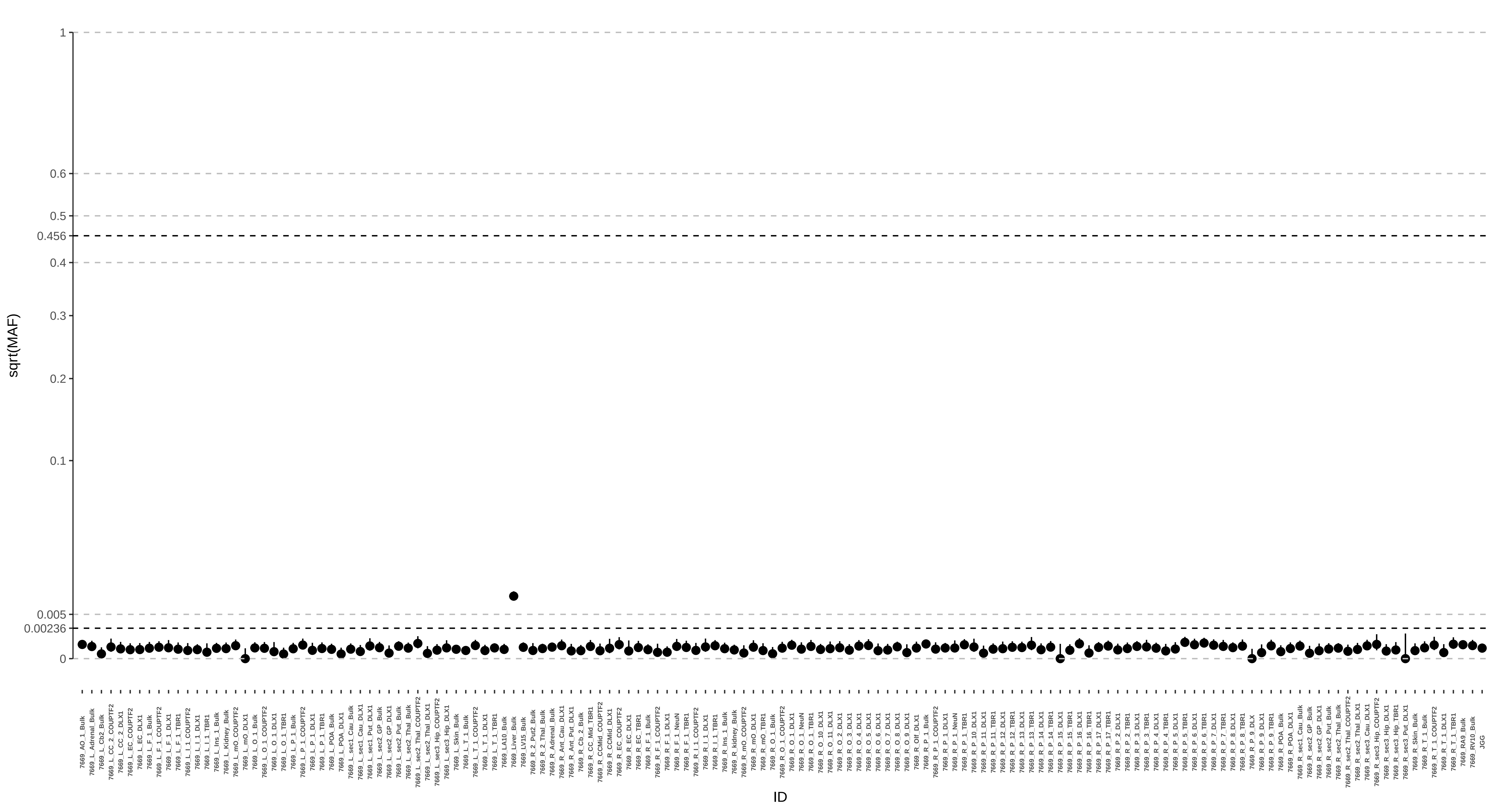

ID05 4−139482903−G−A

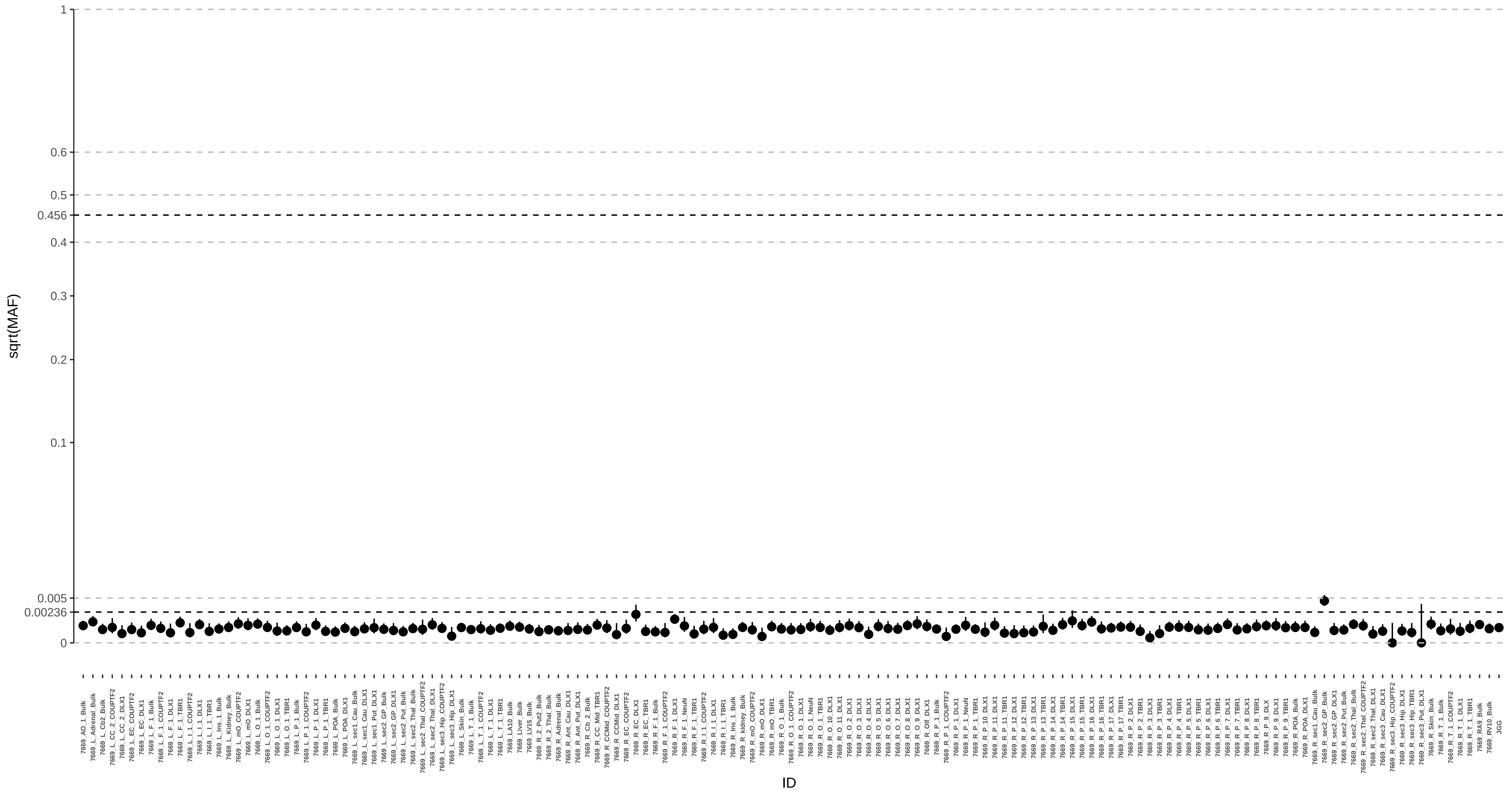

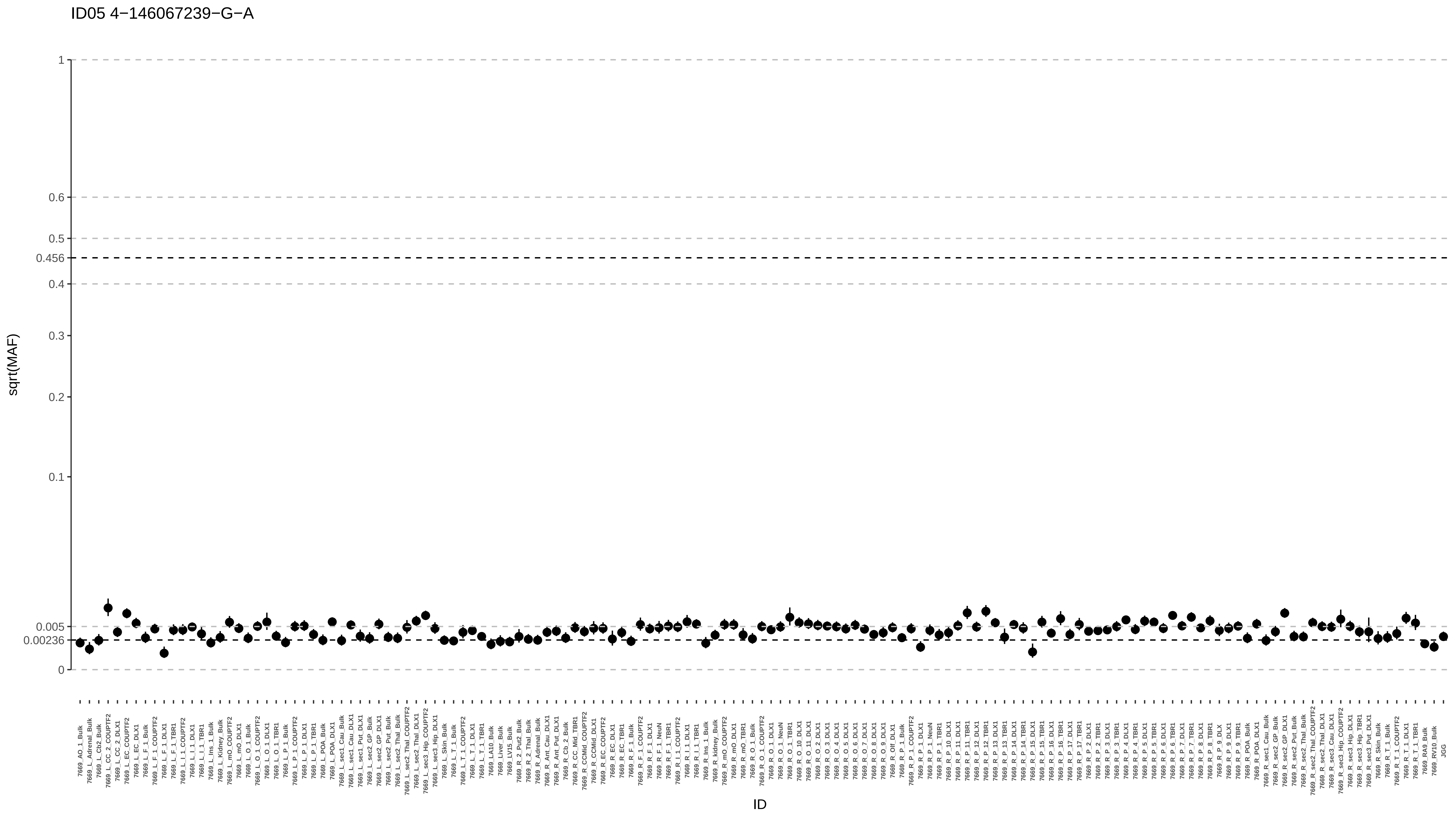
ID05 4−146067239−G−A
sqrt(MAF)
ID
1
0.6
0.5
0.456
0.4
0.3
0.2
0.1
0.005
0.00236
0

ID05 4−152509643−C−A

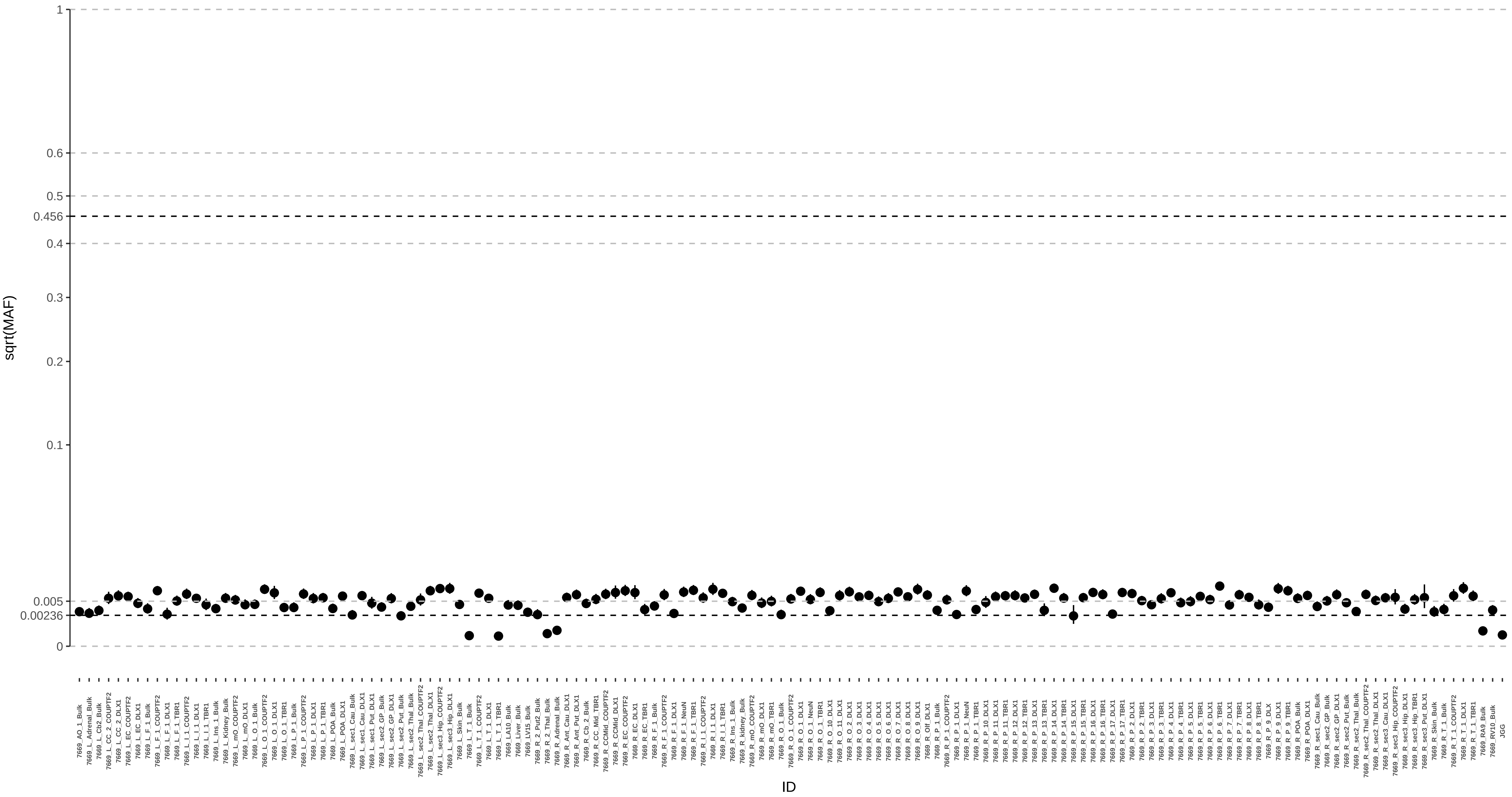

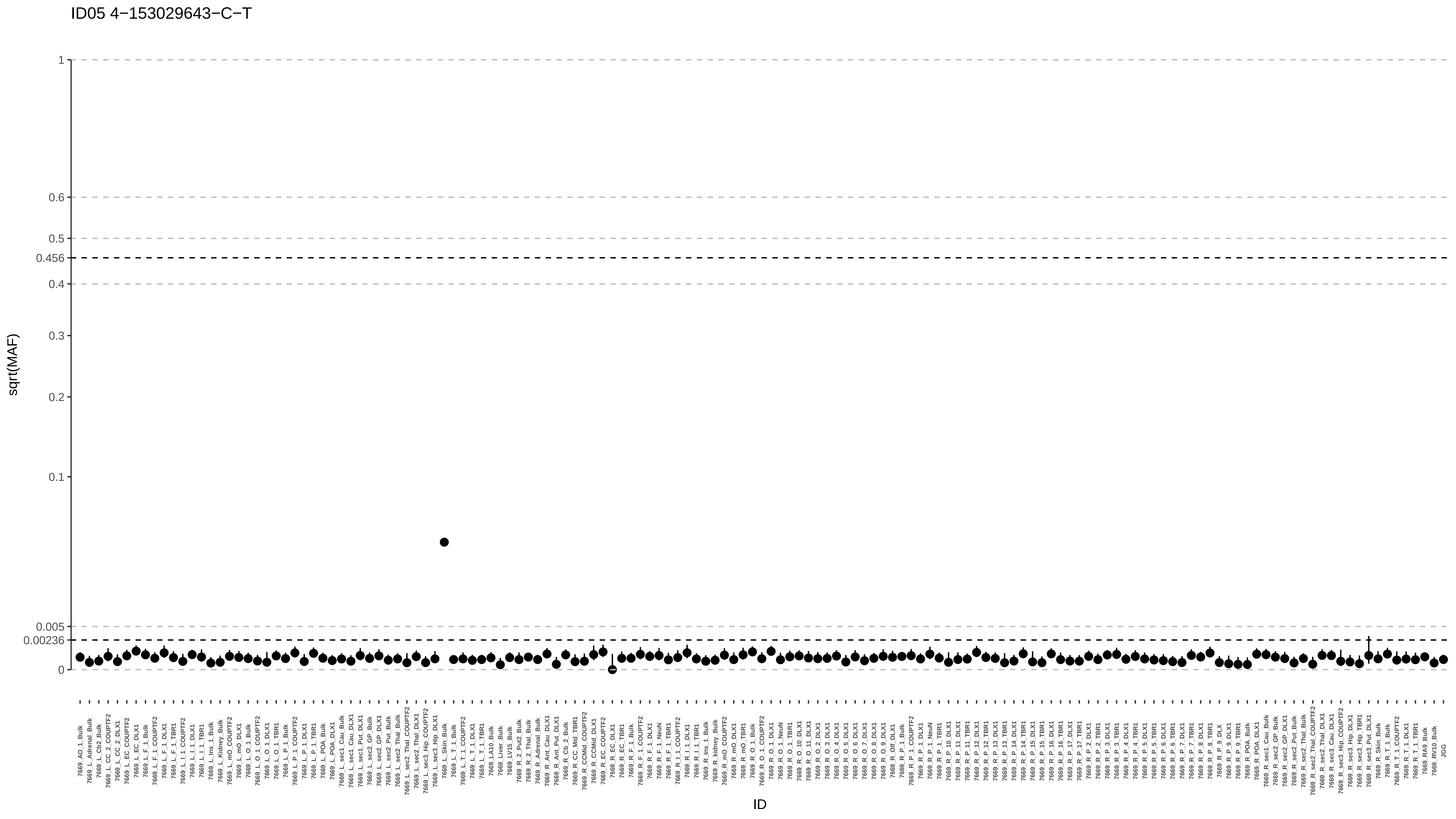

ID05 4−153029643−C−T
sqrt(MAF)
1
0.6
0.5
0.456
0.4
0.3
0.2
0.1
0.005
0.00236
0
ID

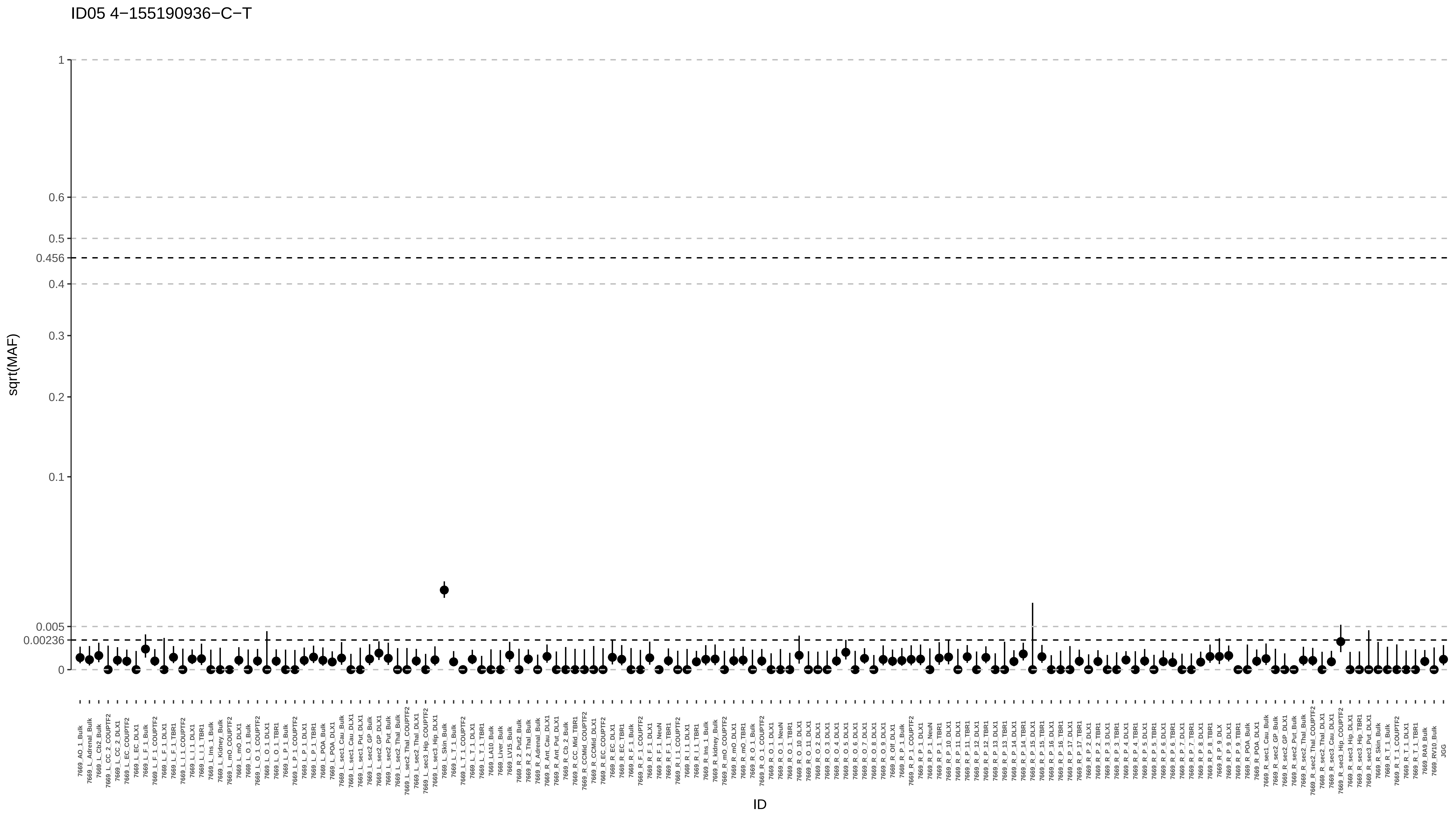

ID05 4−155190936−C−T
sqrt(MAF)
ID

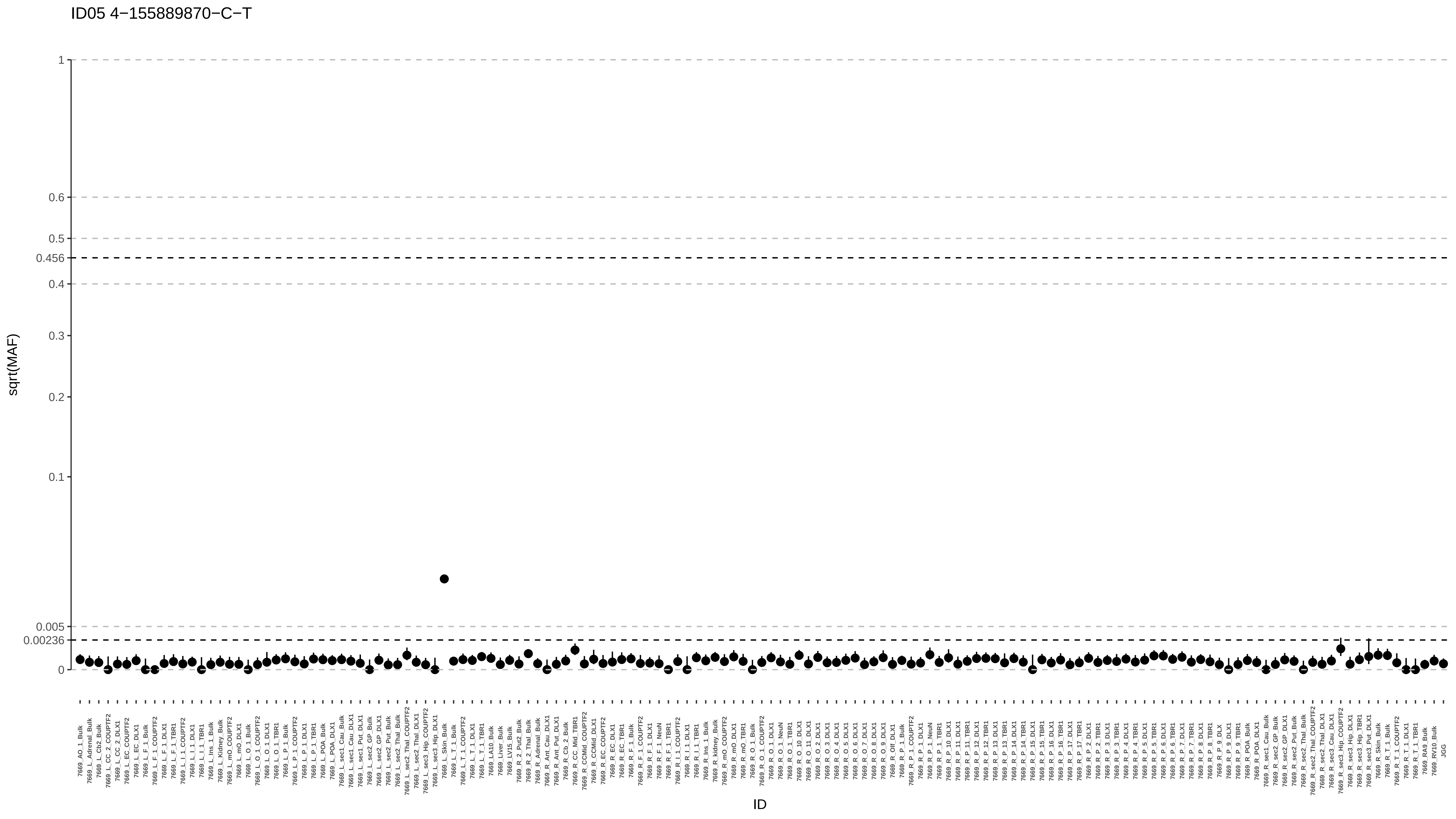
ID05 4−155889870−C−T
sqrt(MAF)
1
0.6
0.5
0.456
0.4
0.3
0.2
0.1
0.005
0.00236
0
ID

ID05 4−155904014−T−G

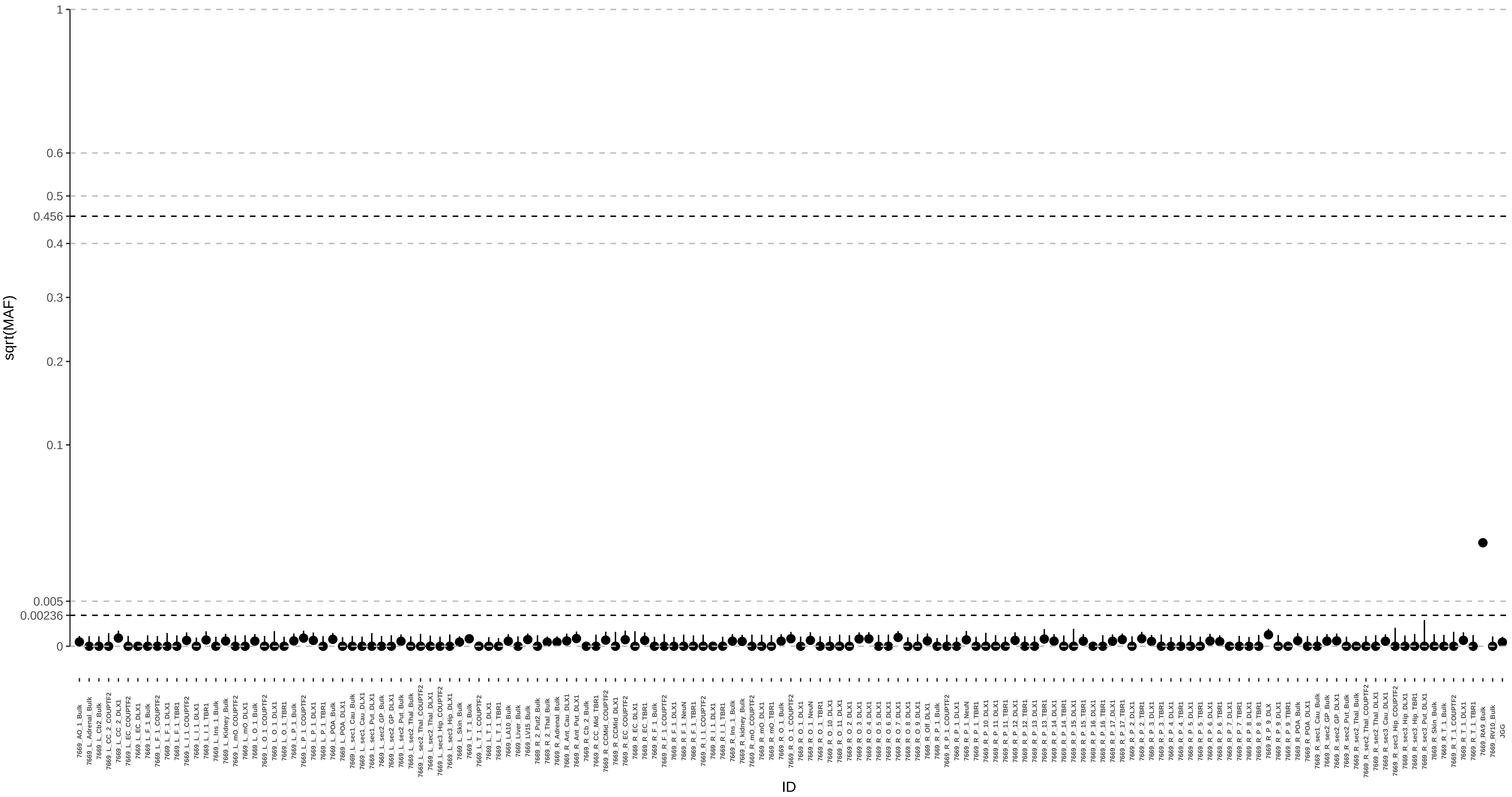

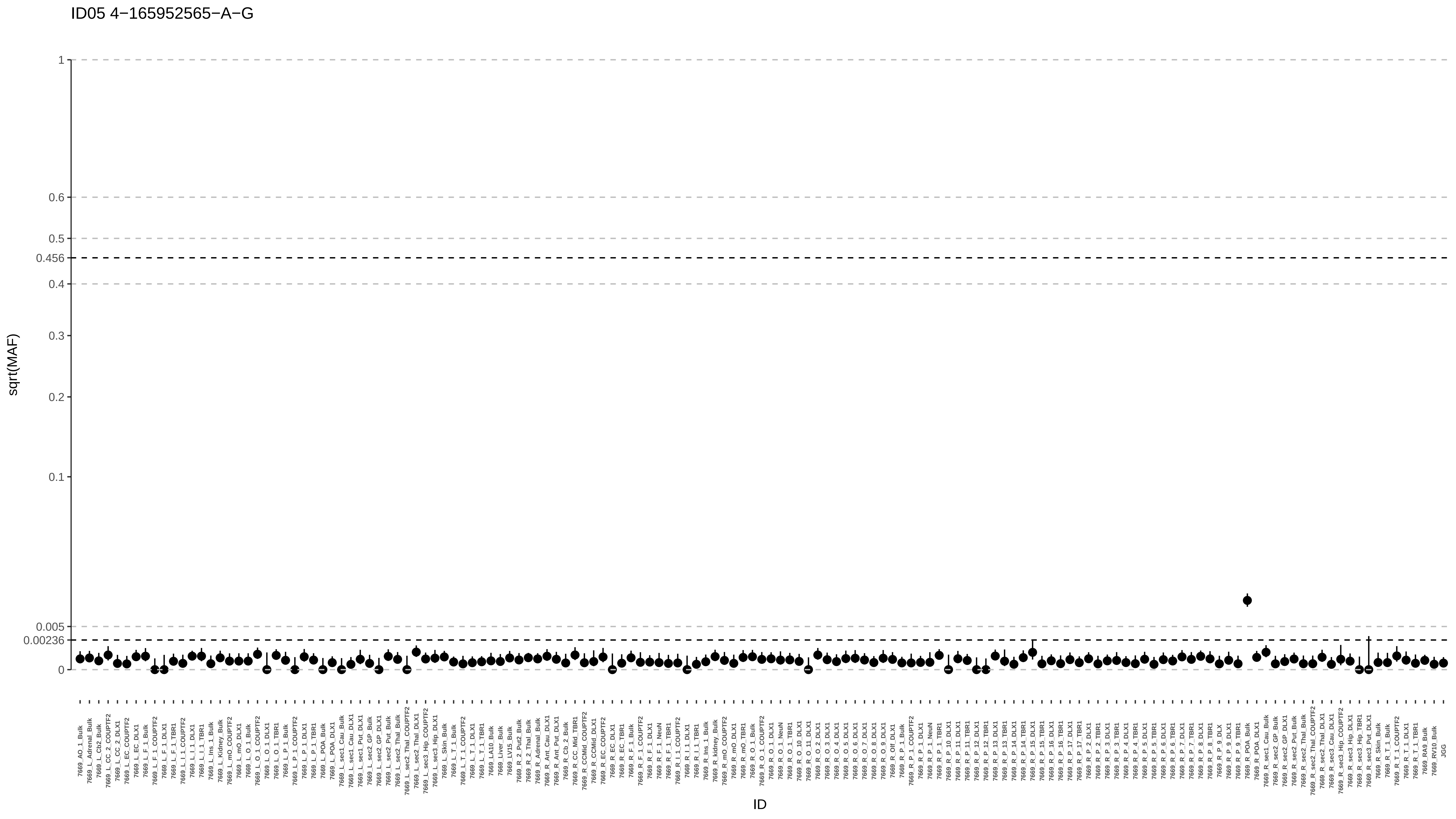

ID05 4−165952565−A−G
sqrt(MAF)
1
0.6
0.5
0.456
0.4
0.3
0.2
0.1
0.005
0.00236
0
ID

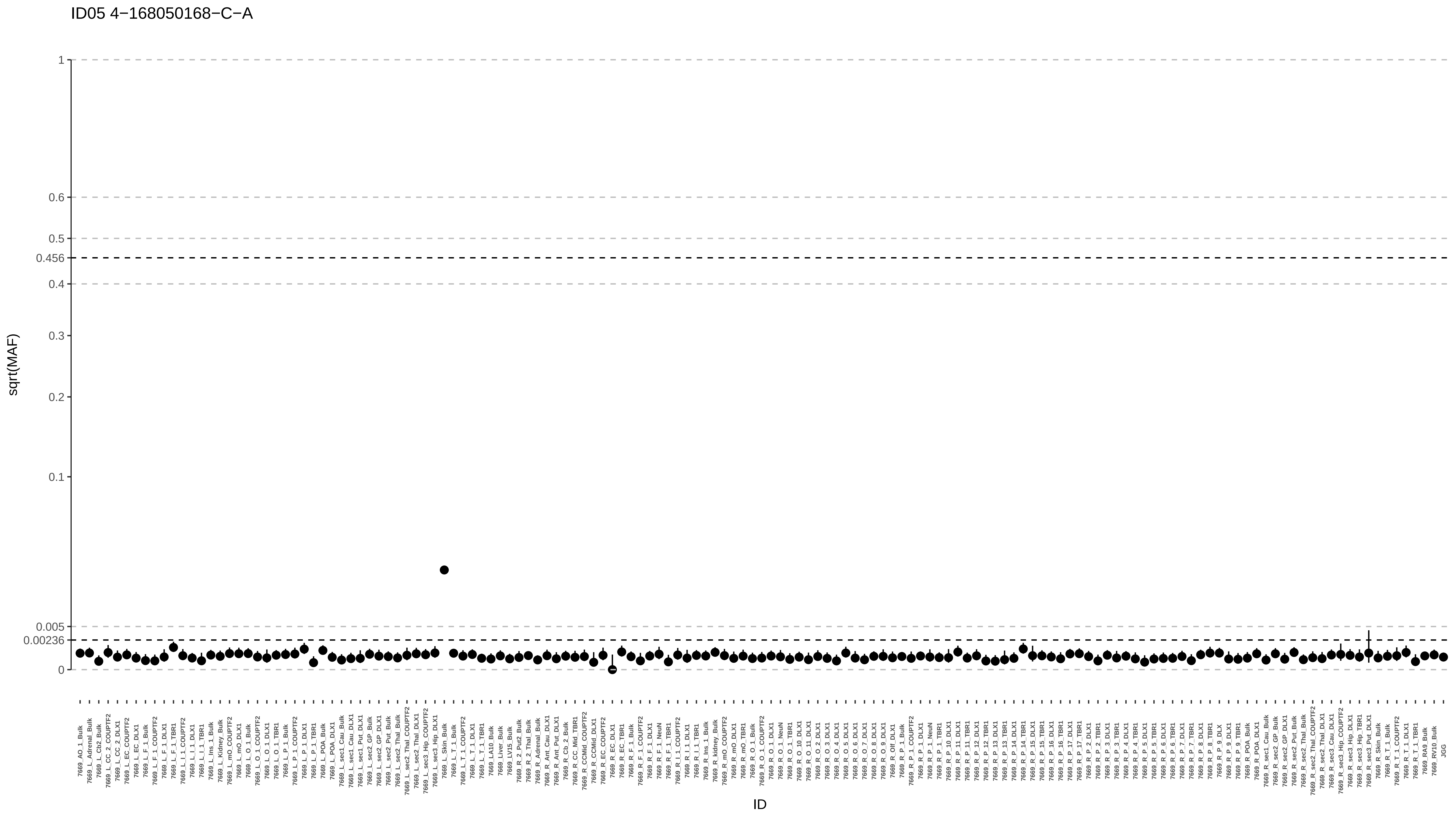
ID05 4−168050168−C−A
sqrt(MAF)
1
0.6
0.5
0.456
0.4
0.3
0.2
0.1
0.005
0.00236
0
ID

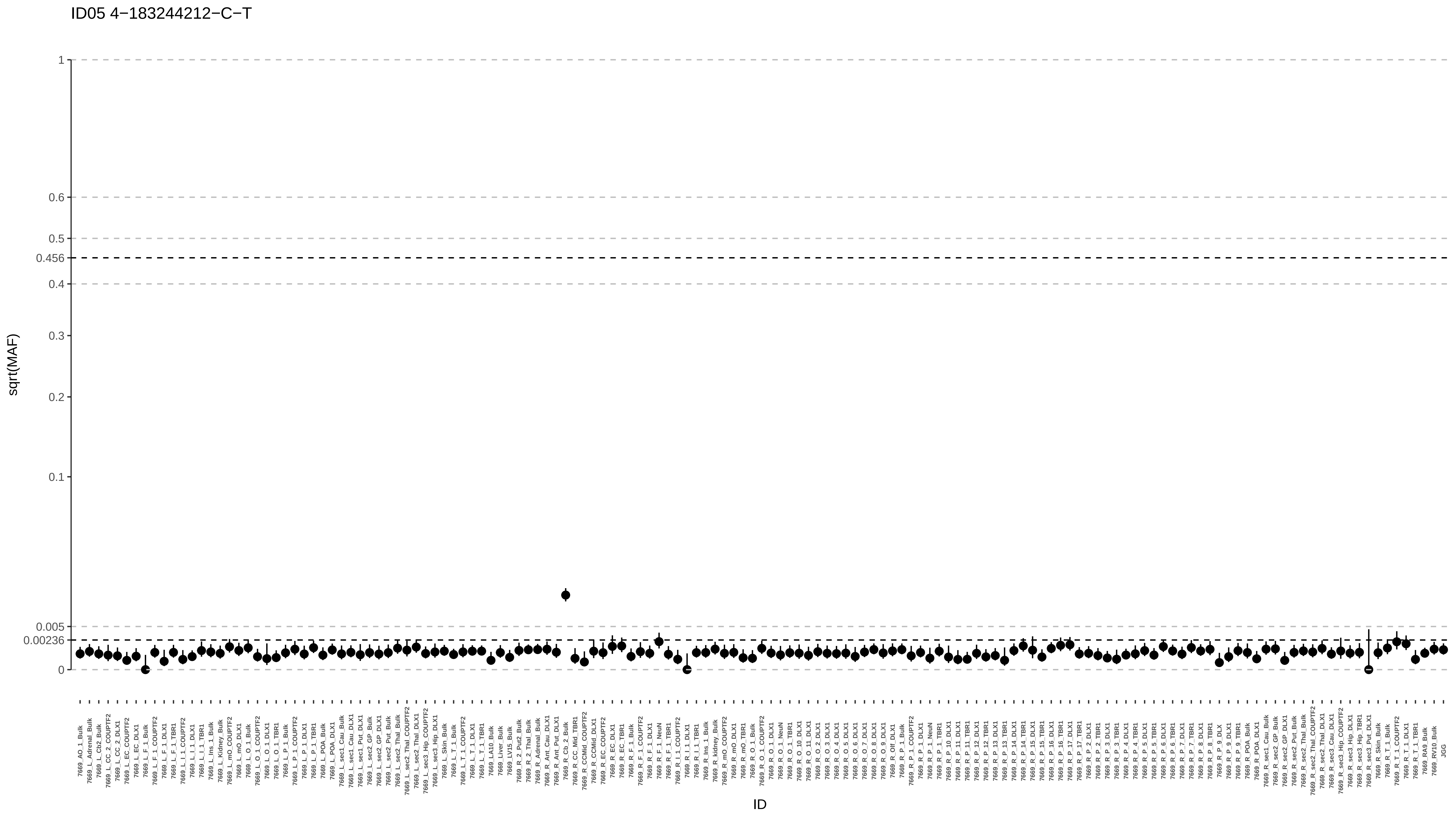

ID05 4−183244212−C−T
sqrt(MAF)
1
0.6
0.5
0.456
0.4
0.3
0.2
0.1
0.005
0.00236
0
ID

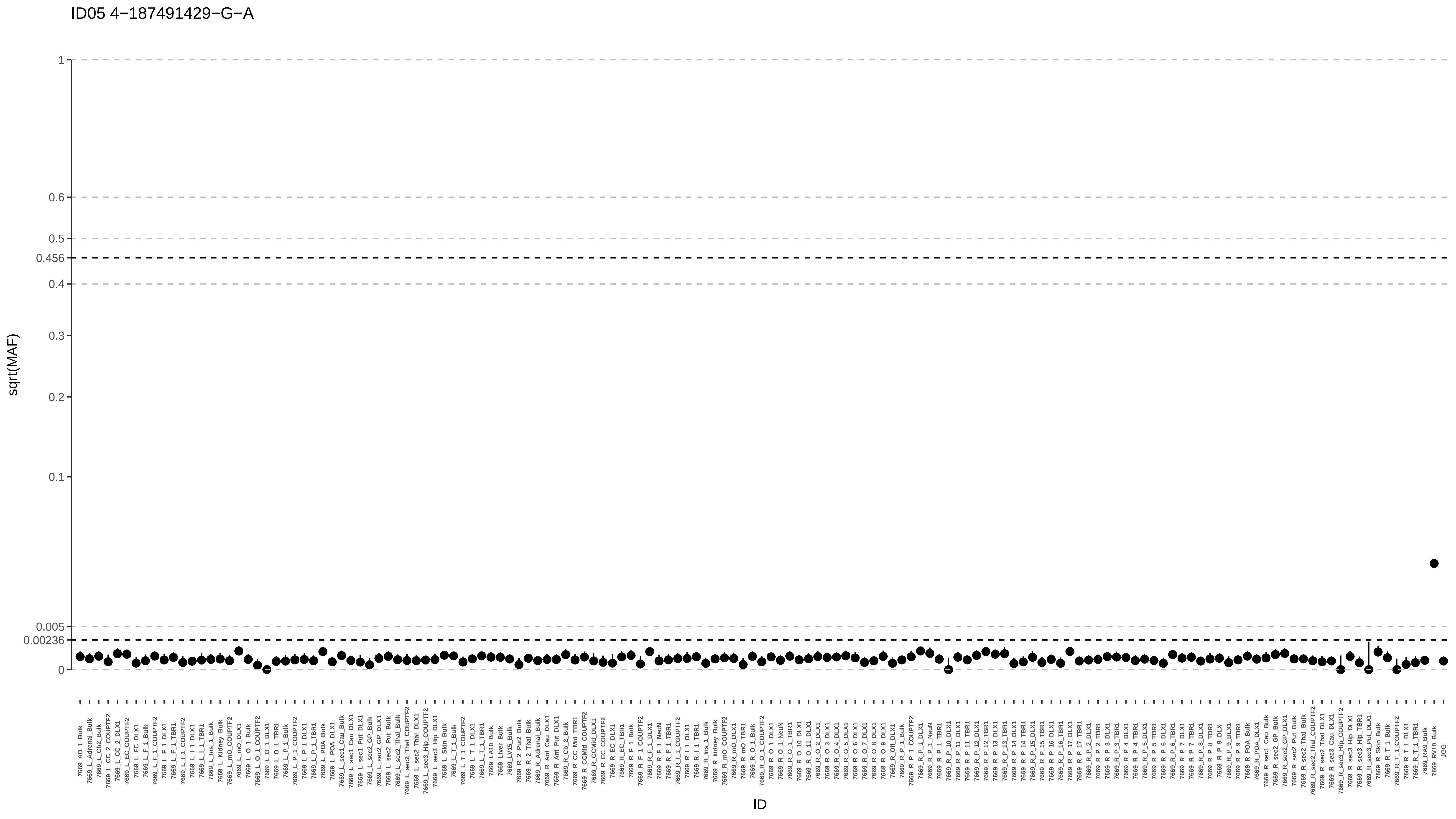
ID05 4−187491429−G−A
sqrt(MAF)
ID
1
0.6
0.5
0.456
0.4
0.3
0.2
0.1
0.005
0.00236
0

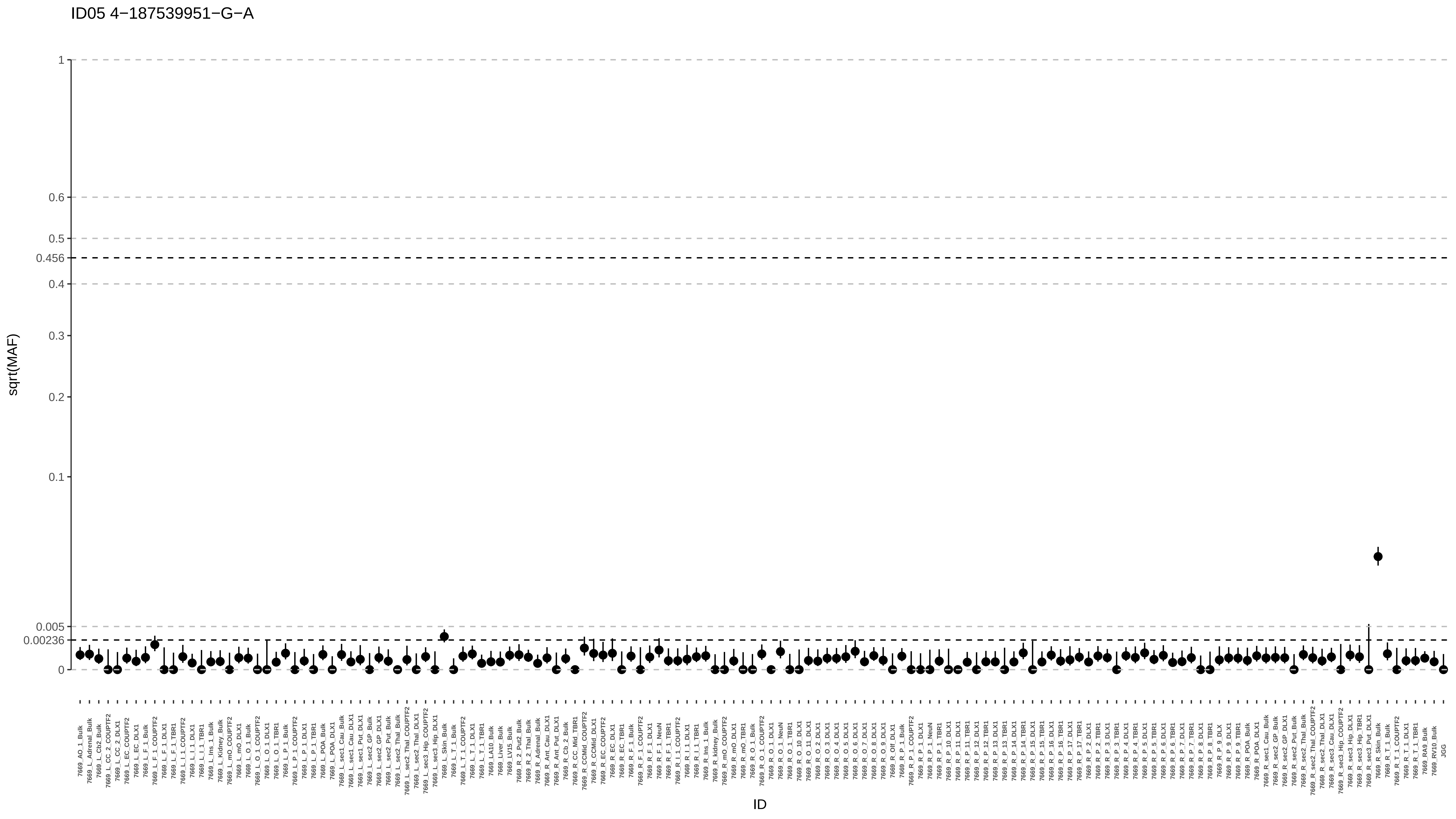
ID05 4−187539951−G−A
sqrt(MAF)
1
0.6
0.5
0.456
0.4
0.3
0.2
0.1
0.005
0.00236
0
ID

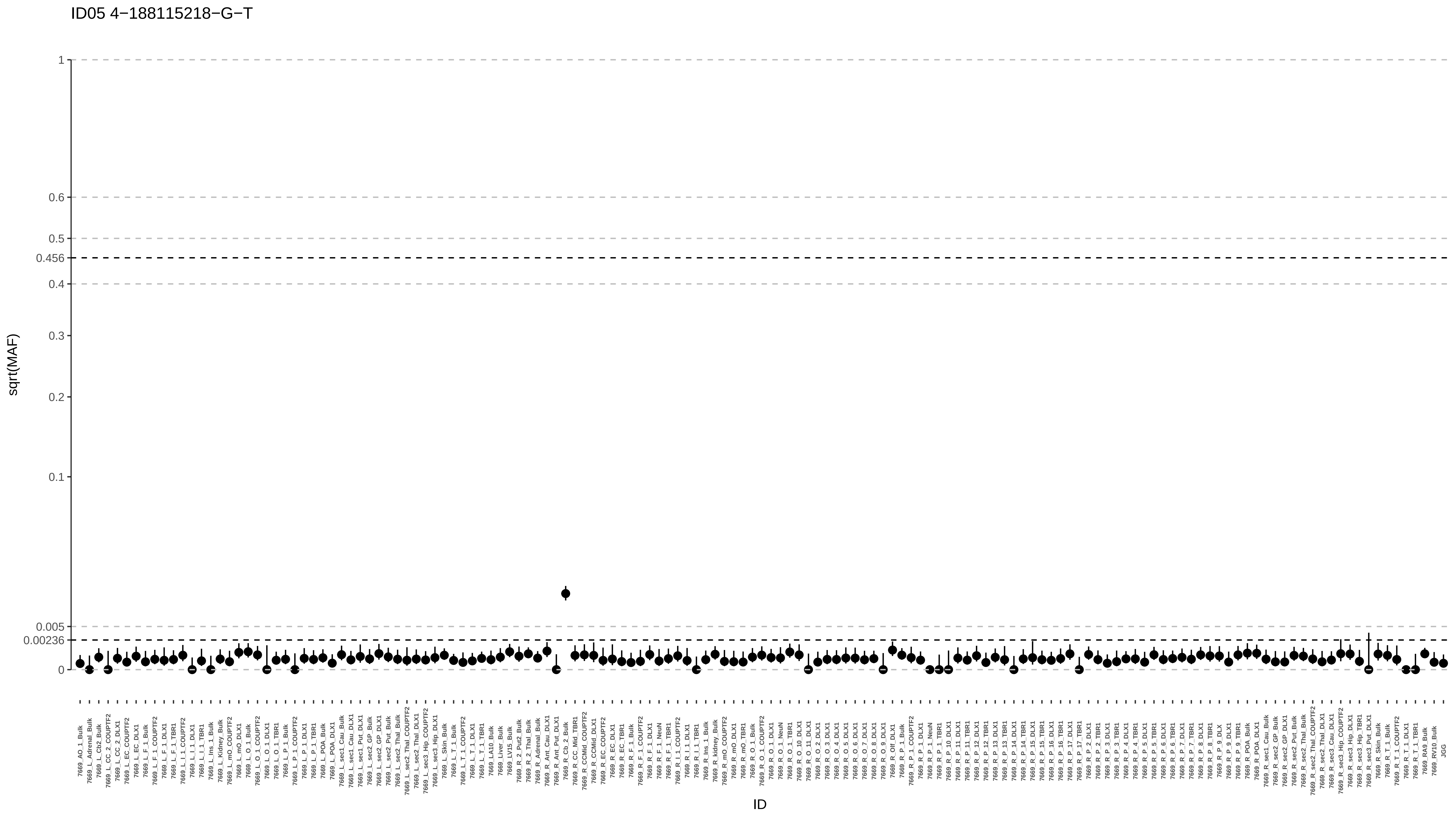
ID05 4−188115218−G−T
sqrt(MAF)
1
0.6
0.5
0.456
0.4
0.3
0.2
0.1
0.005
0.00236
0
ID

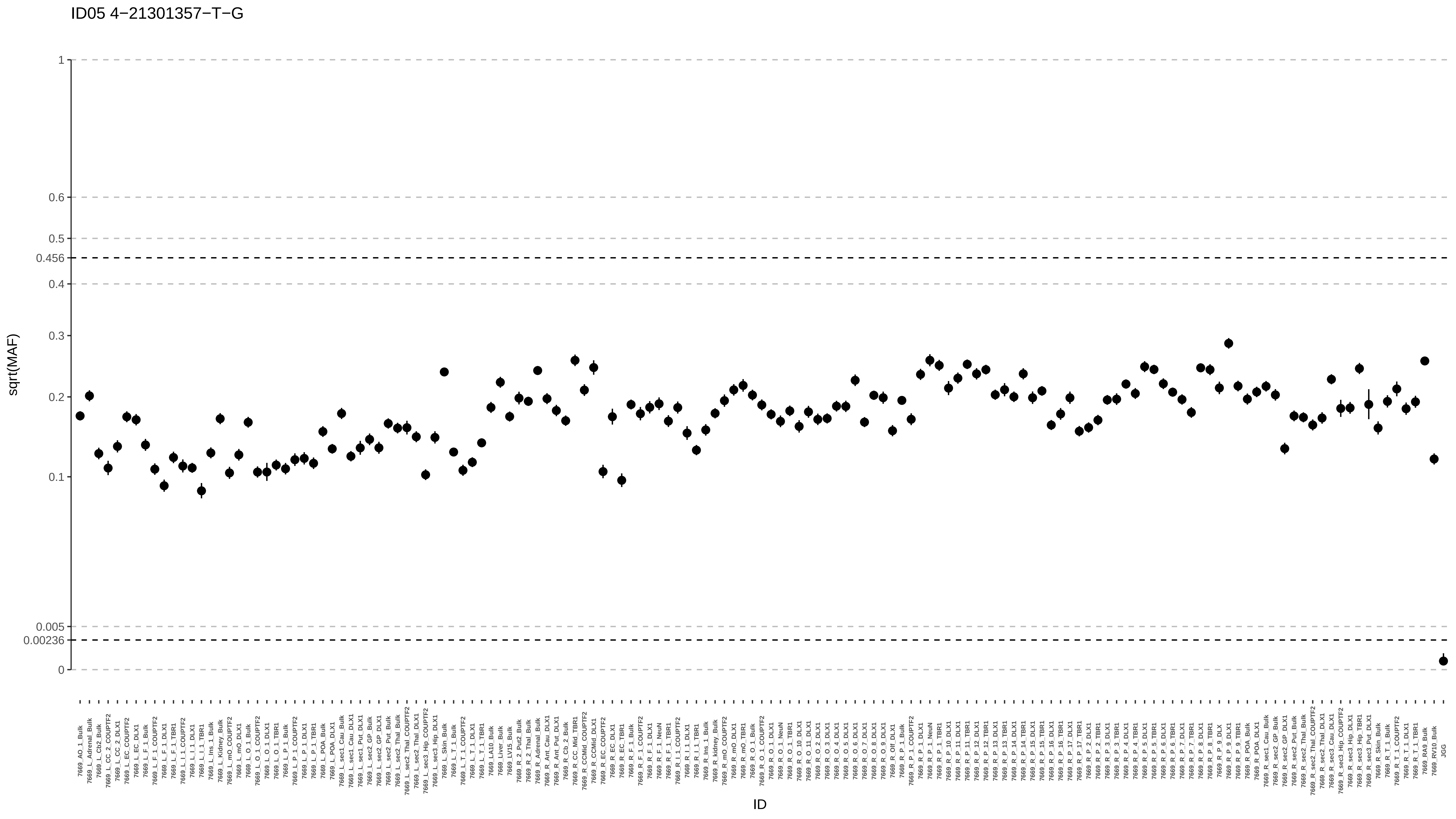
ID05 4−21301357−T−G
sqrt(MAF)
1
0.6
0.5
0.456
0.4
0.3
0.2
0.1
0.005
0.00236
0
ID
7669_AO_1_Bulk
7669_L_Adrenal_Bulk
7669_L_Cb2_Bulk
7669_L_CC_2_COUPTF2
7669_L_CC_2_DLX1
7669_L_EC_COUPTF2
7669_L_EC_DLX1
7669_L_F_1_Bulk
7669_L_F_1_COUPTF2
7669_L_F_1_DLX1
7669_L_F_1_TBR1
7669_L_I_1_COUPTF2
7669_L_I_1_DLX1
7669_L_I_1_TBR1
7669_L_Ins_1_Bulk
7669_L_Kidney_Bulk
7669_L_mO_COUPTF2
7669_L_mO_DLX1
7669_L_O_1_Bulk
7669_L_O_1_COUPTF2
7669_L_O_1_DLX1
7669_L_O_1_TBR1
7669_L_P_1_Bulk
7669_L_P_1_COUPTF2
7669_L_P_1_DLX1
7669_L_P_1_TBR1
7669_L_POA_Bulk
7669_L_POA_DLX1
7669_L_sec1_Cau_Bulk
7669_L_sec1_Cau_DLX1
7669_L_sec1_Put_DLX1
7669_L_sec2_GP_Bulk
7669_L_sec2_GP_DLX1
7669_L_sec2_Put_Bulk
7669_L_sec2_Thal_Bulk
7669_L_sec2_Thal_COUPTF2
7669_L_sec2_Thal_DLX1
7669_L_sec3_Hip_COUPTF2
7669_L_sec3_Hip_DLX1
7669_L_Skin_Bulk
7669_L_T_1_Bulk
7669_L_T_1_COUPTF2
7669_L_T_1_DLX1
7669_L_T_1_TBR1
7669_LA10_Bulk
7669_Liver_Bulk
7669_LV15_Bulk
7669_R_2_Put2_Bulk
7669_R_2_Thal_Bulk
7669_R_Adrenal_Bulk
7669_R_Ant_Cau_DLX1
7669_R_Ant_Put_DLX1
7669_R_Cb_2_Bulk
7669_R_CC_Mid_TBR1
7669_R_CCMid_COUPTF2
7669_R_CCMid_DLX1
7669_R_EC_COUPTF2
7669_R_EC_DLX1
7669_R_EC_TBR1
7669_R_F_1_Bulk
7669_R_F_1_COUPTF2
7669_R_F_1_DLX1
7669_R_F_1_NeuN
7669_R_F_1_TBR1
7669_R_I_1_COUPTF2
7669_R_I_1_DLX1
7669_R_I_1_TBR1
7669_R_Ins_1_Bulk
7669_R_kidney_Bulk
7669_R_mO_COUPTF2
7669_R_mO_DLX1
7669_R_mO_TBR1
7669_R_O_1_Bulk
7669_R_O_1_COUPTF2
7669_R_O_1_DLX1
7669_R_O_1_NeuN
7669_R_O_1_TBR1
7669_R_O_10_DLX1
7669_R_O_11_DLX1
7669_R_O_2_DLX1
7669_R_O_3_DLX1
7669_R_O_4_DLX1
7669_R_O_5_DLX1
7669_R_O_6_DLX1
7669_R_O_7_DLX1
7669_R_O_8_DLX1
7669_R_O_9_DLX1
7669_R_Olf_DLX1
7669_R_P_1_Bulk
7669_R_P_1_COUPTF2
7669_R_P_1_DLX1
7669_R_P_1_NeuN
7669_R_P_1_TBR1
7669_R_P_10_DLX1
7669_R_P_11_DLX1
7669_R_P_11_TBR1
7669_R_P_12_DLX1
7669_R_P_12_TBR1
7669_R_P_13_DLX1
7669_R_P_13_TBR1
7669_R_P_14_DLX1
7669_R_P_14_TBR1
7669_R_P_15_DLX1
7669_R_P_15_TBR1
7669_R_P_16_DLX1
7669_R_P_16_TBR1
7669_R_P_17_DLX1
7669_R_P_17_TBR1
7669_R_P_2_DLX1
7669_R_P_2_TBR1
7669_R_P_3_DLX1
7669_R_P_3_TBR1
7669_R_P_4_DLX1
7669_R_P_4_TBR1
7669_R_P_5_DLX1
7669_R_P_5_TBR1
7669_R_P_6_DLX1
7669_R_P_6_TBR1
7669_R_P_7_DLX1
7669_R_P_7_TBR1
7669_R_P_8_DLX1
7669_R_P_8_TBR1
7669_R_P_9_DLX
7669_R_P_9_DLX1
7669_R_P_9_TBR1
7669_R_POA_Bulk
7669_R_POA_DLX1
7669_R_sec1_Cau_Bulk
7669_R_sec2_GP_Bulk
7669_R_sec2_GP_DLX1
7669_R_sec2_Put_Bulk
7669_R_sec2_Thal_Bulk
7669_R_sec2_Thal_COUPTF2
7669_R_sec2_Thal_DLX1
7669_R_sec3_Cau_DLX1
7669_R_sec3_Hip_COUPTF2
7669_R_sec3_Hip_DLX1
7669_R_sec3_Hip_TBR1
7669_R_sec3_Put_DLX1
7669_R_Skin_Bulk
7669_R_T_1_Bulk
7669_R_T_1_COUPTF2
7669_R_T_1_DLX1
7669_R_T_1_TBR1
7669_RA9_Bulk
7669_RV10_Bulk
JGG

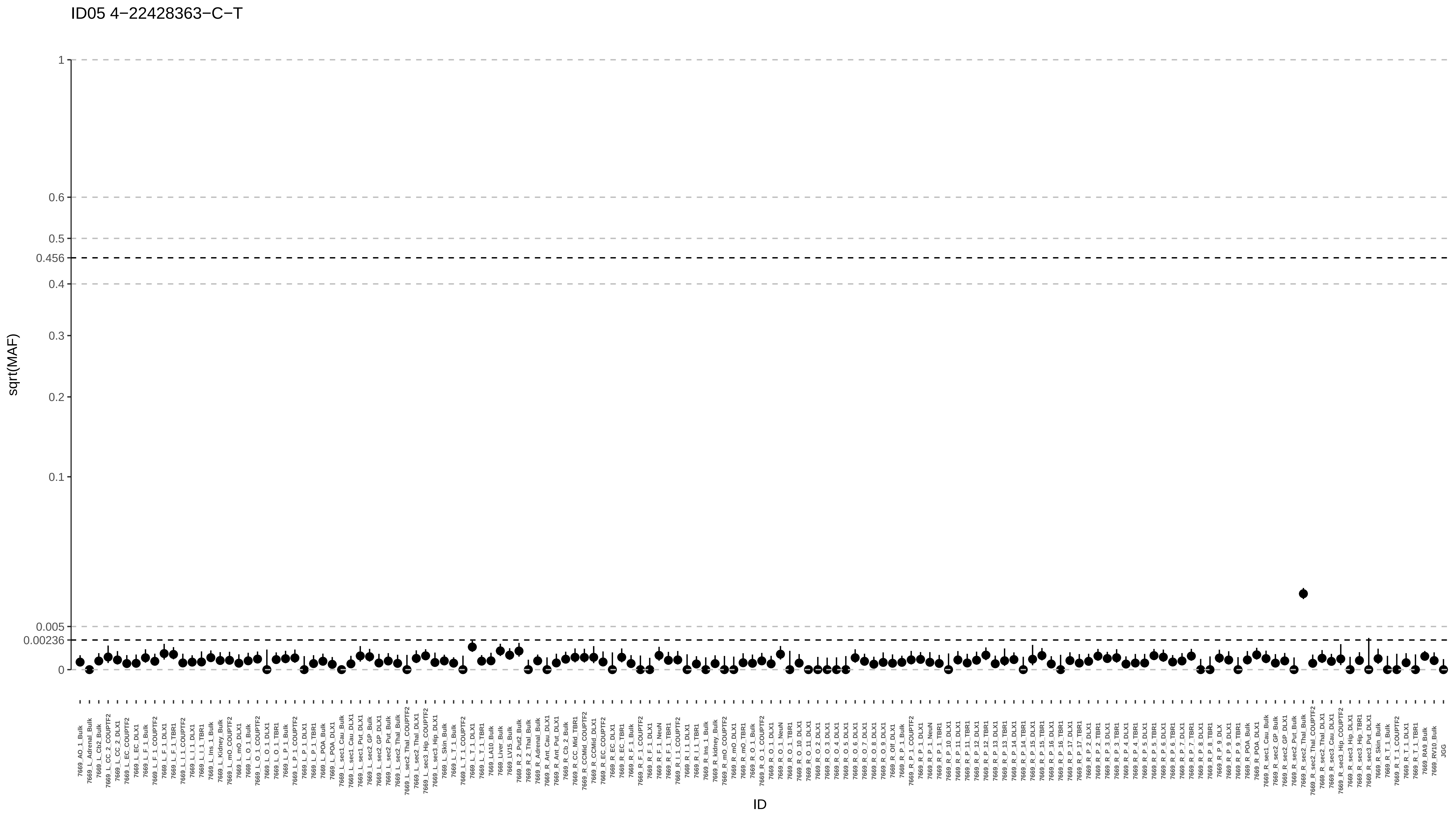
ID05 4−22428363−C−T
sqrt(MAF)
1
0.6
0.5
0.456
0.4
0.3
0.2
0.1
0.005
0.00236
0
ID

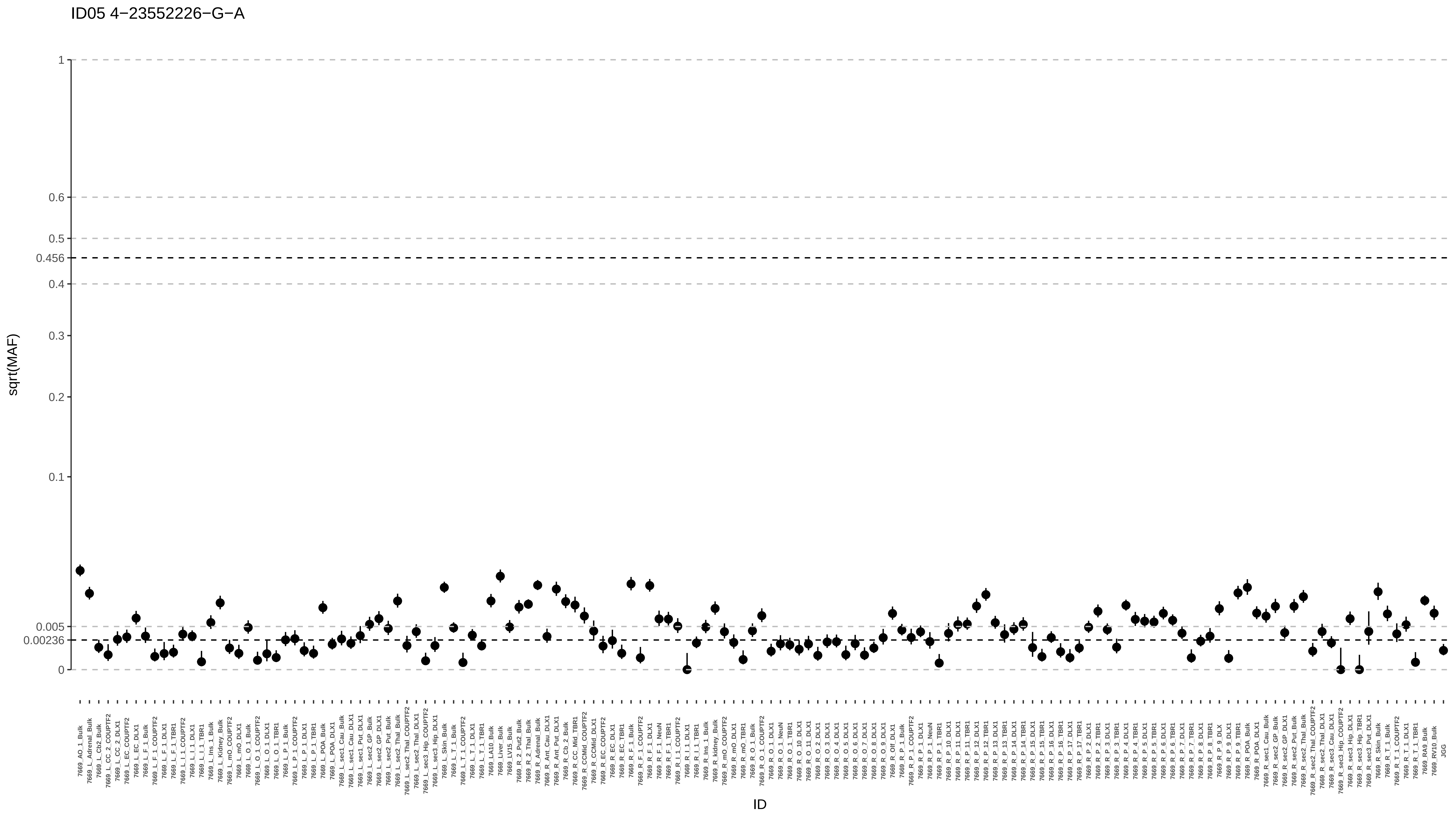

ID05 4−23552226−G−A
sqrt(MAF)
1
0.6
0.5
0.456
0.4
0.3
0.2
0.1
0.005
0.00236
0
ID

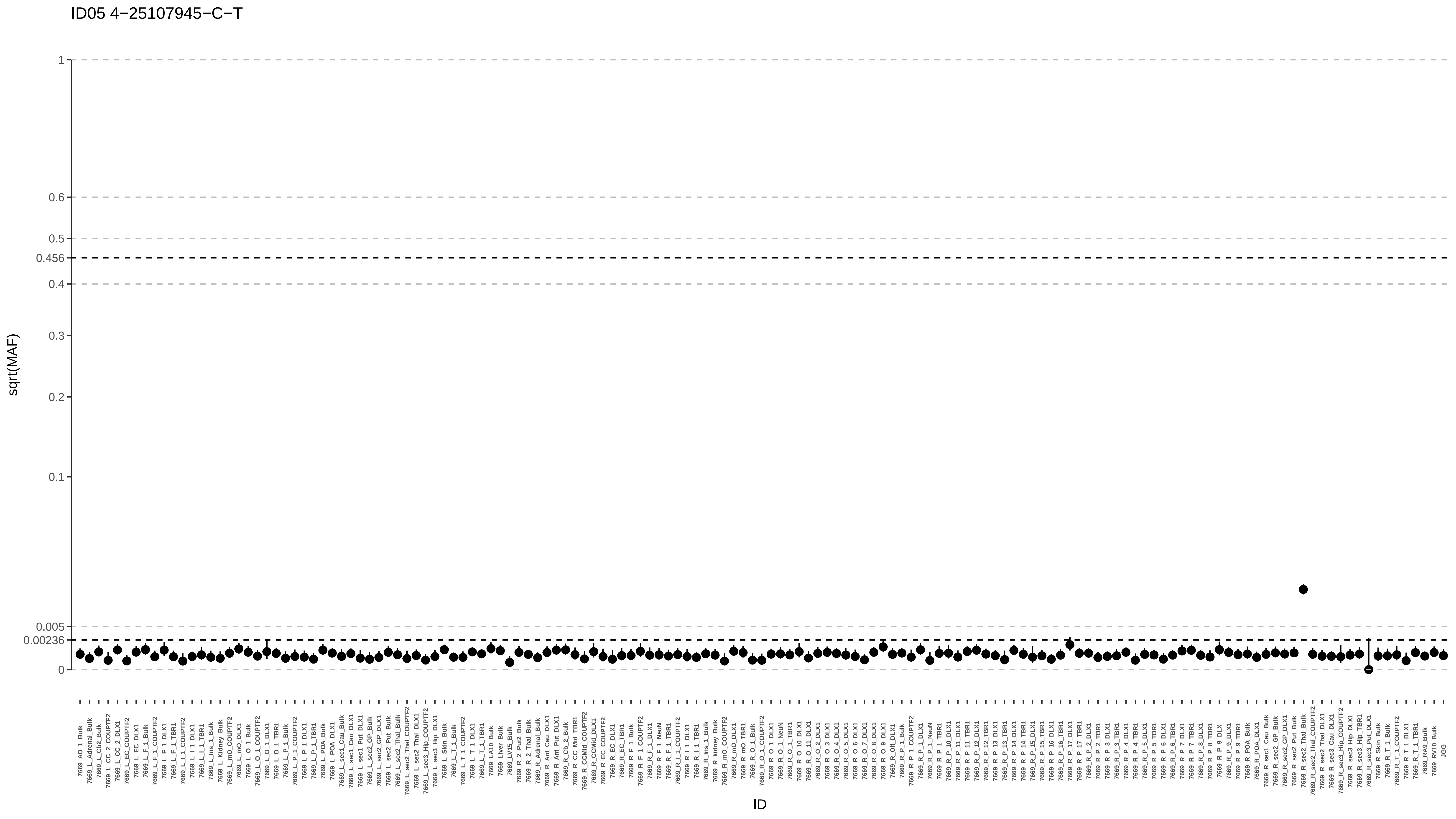

ID05 4−25107945−C−T
sqrt(MAF)
ID

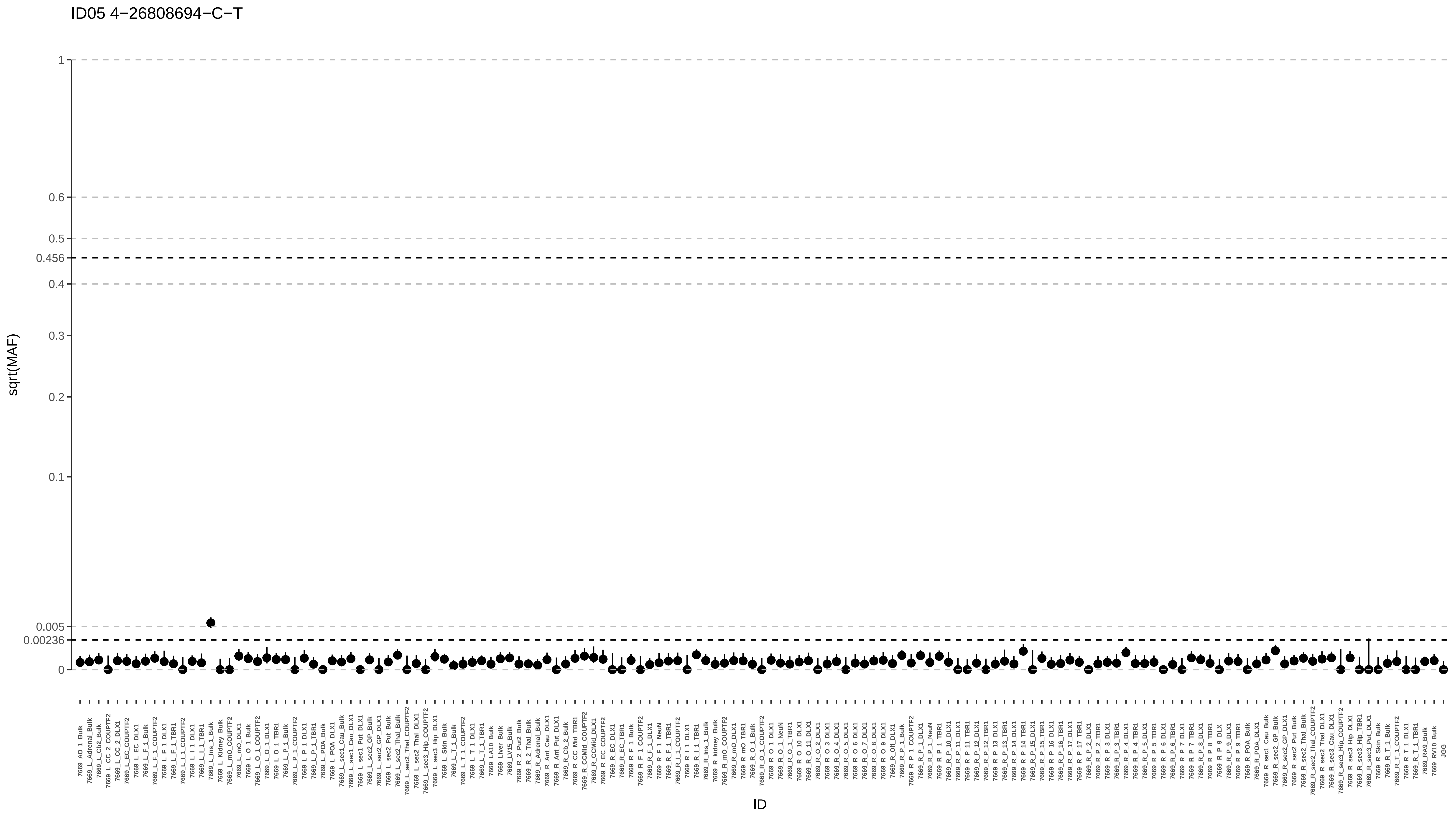

ID05 4−26808694−C−T
sqrt(MAF)
1
0.6
0.5
0.456
0.4
0.3
0.2
0.1
0.005
0.00236
0
ID

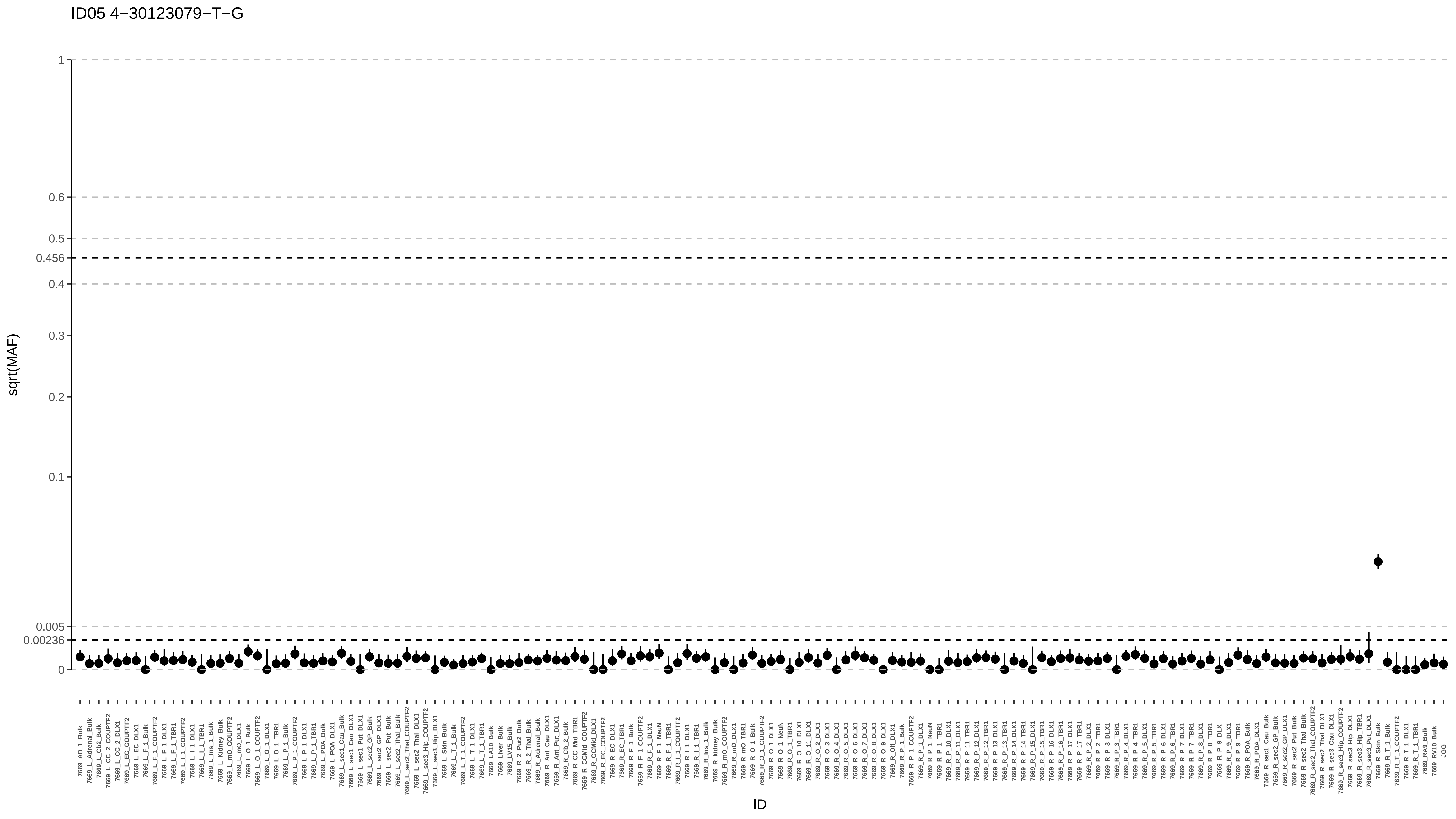
ID05 4−30123079−T−G
sqrt(MAF)
1
0.6
0.5
0.456
0.4
0.3
0.2
0.1
0.005
0.00236
0
ID

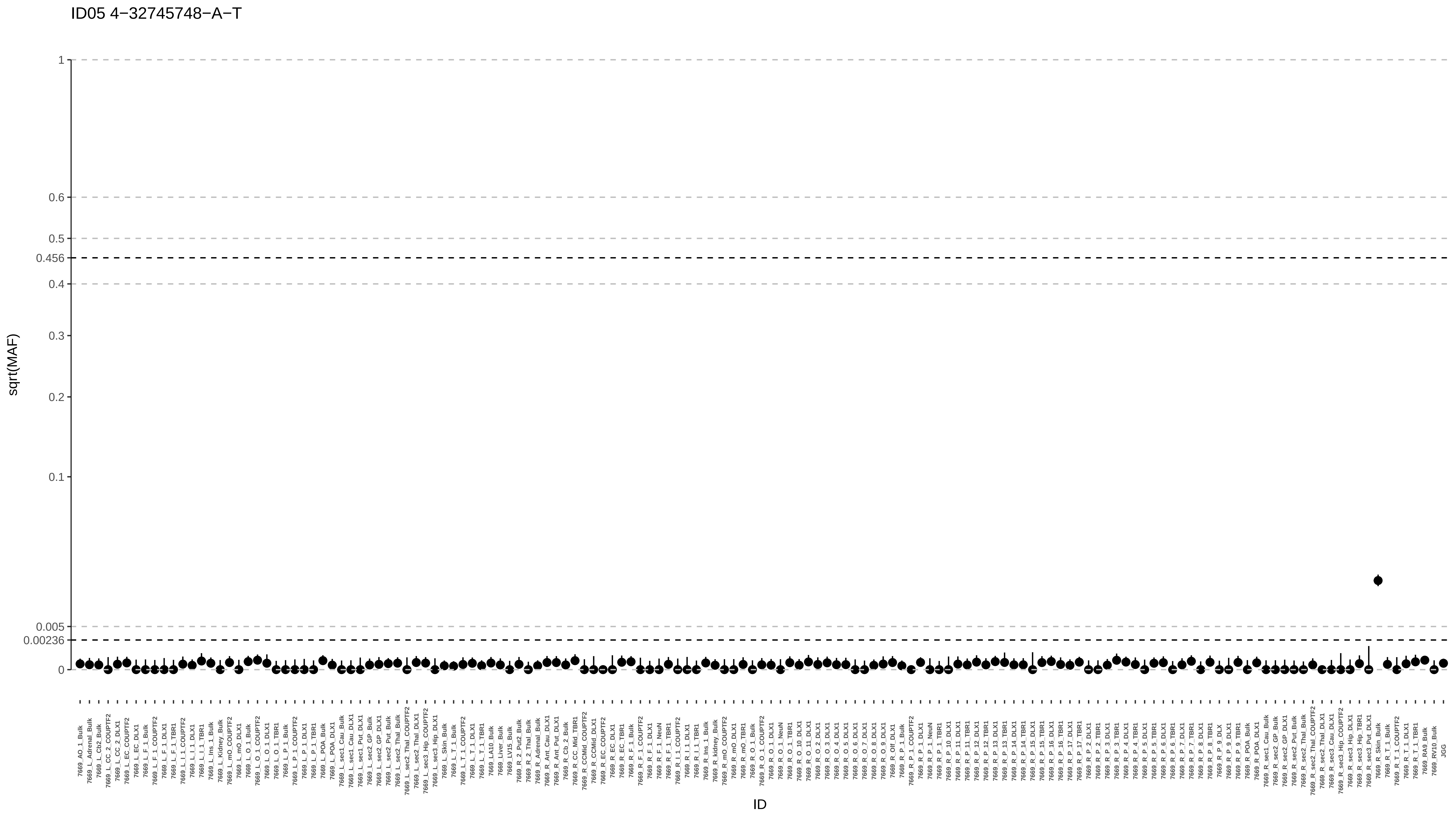
ID05 4−32745748−A−T
sqrt(MAF)
1
0.6
0.5
0.456
0.4
0.3
0.2
0.1
0.005
0.00236
0
ID

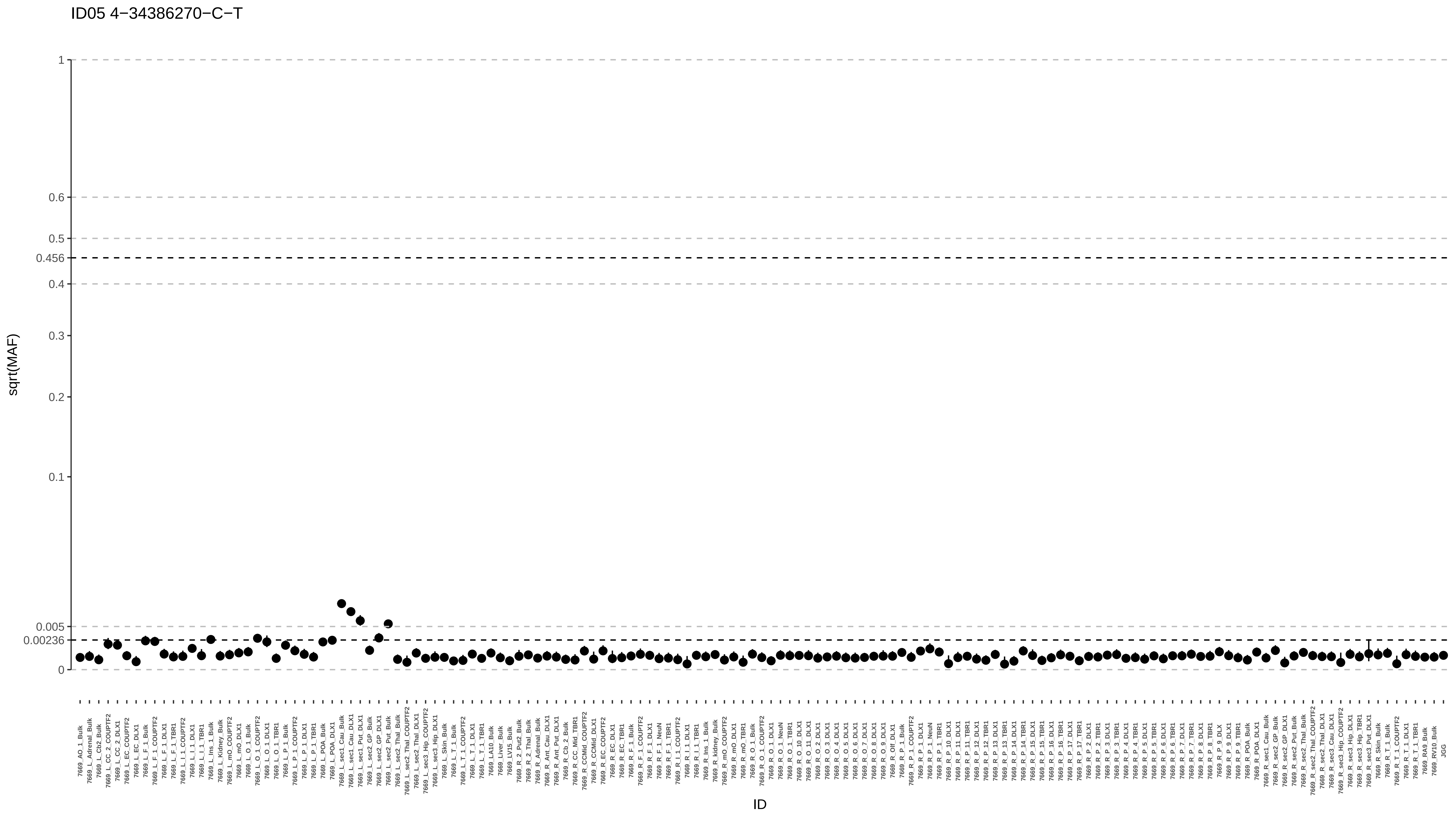
ID05 4−34386270−C−T
sqrt(MAF)
1
0.6
0.5
0.456
0.4
0.3
0.2
0.1
0.005
0.00236
0
ID

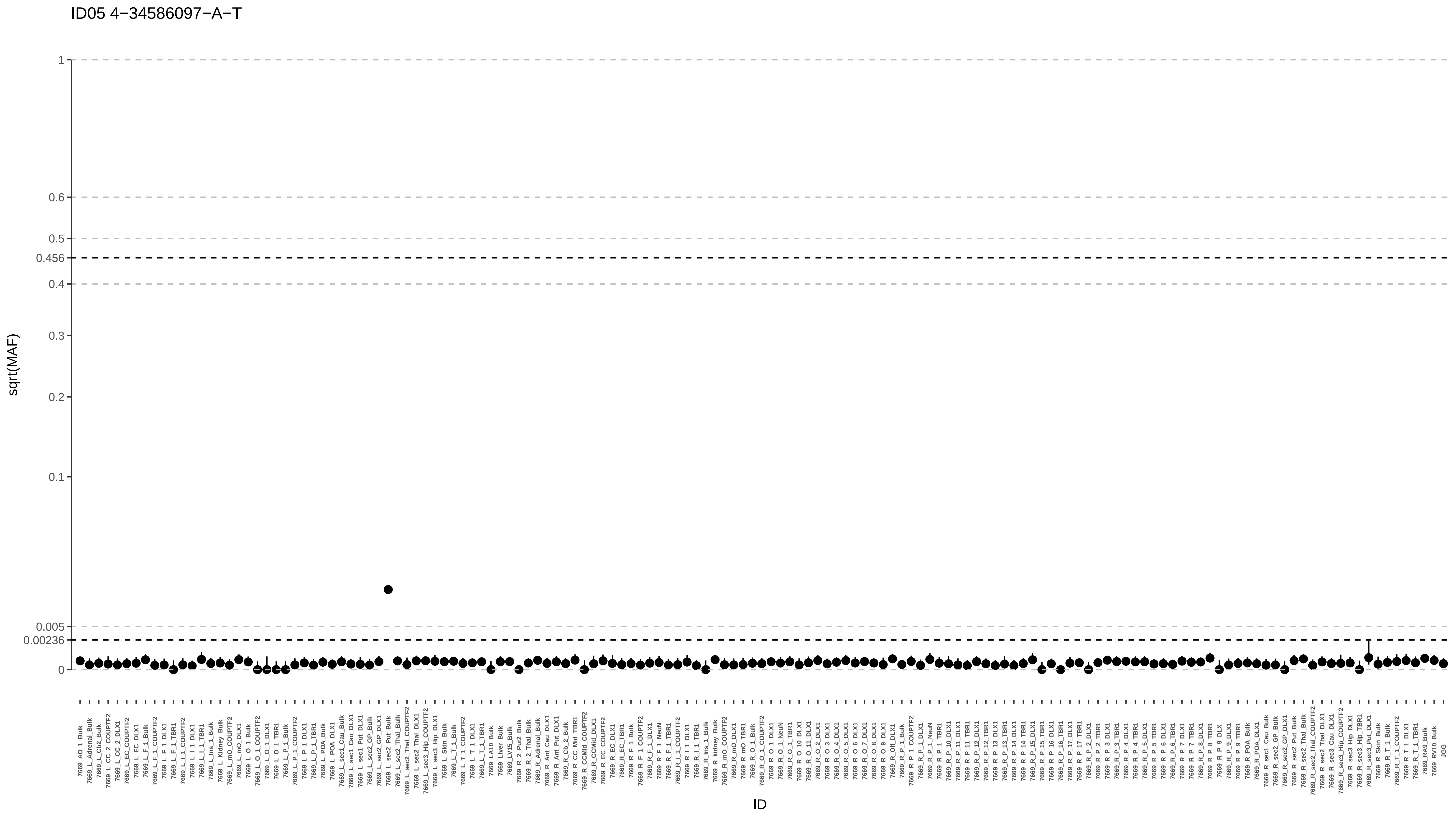
ID05 4−34586097−A−T
sqrt(MAF)
1
0.6
0.5
0.456
0.4
0.3
0.2
0.1
0.005
0.00236
0
ID

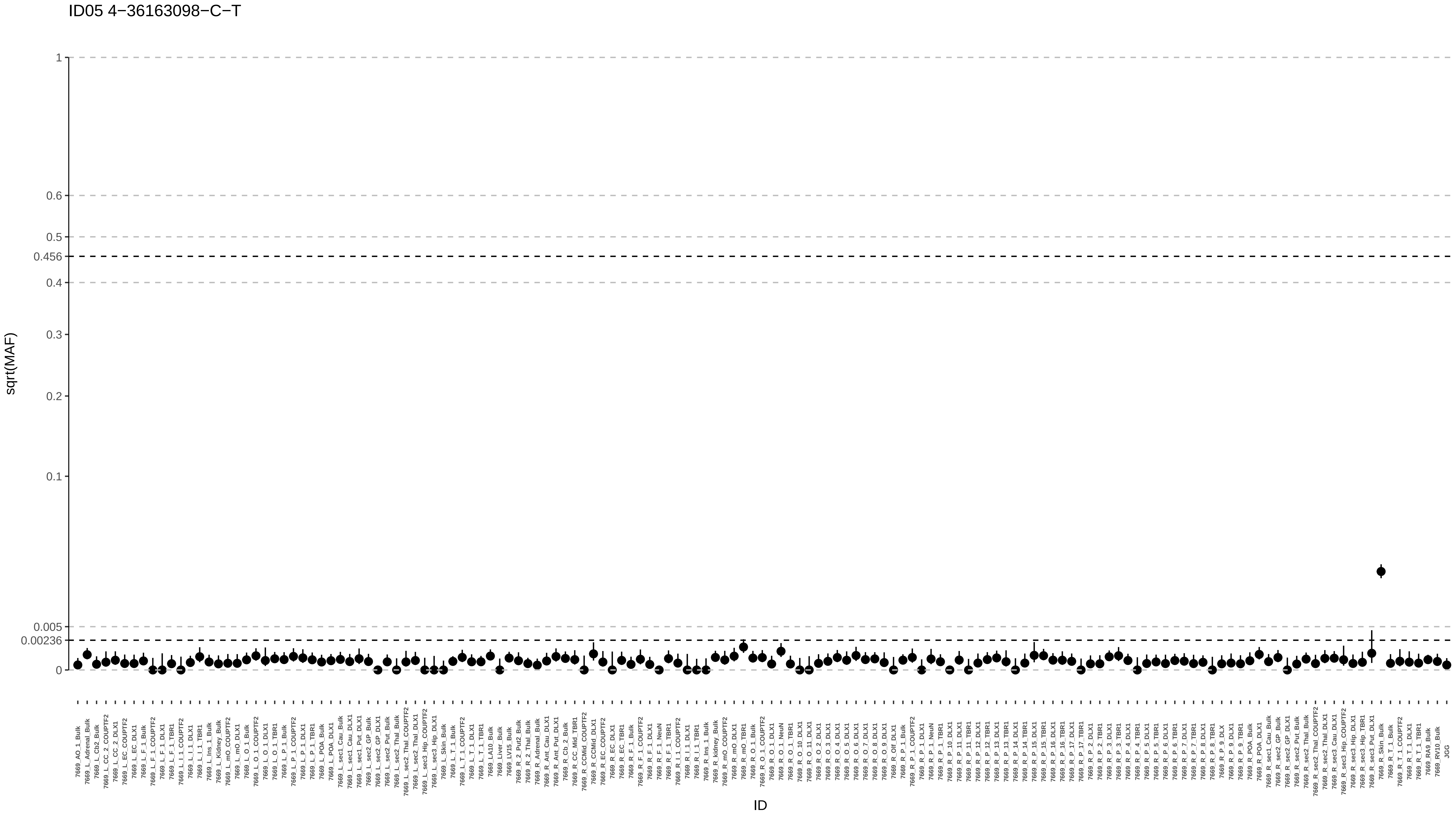
ID05 4−36163098−C−T
sqrt(MAF)
1
0.6
0.5
0.456
0.4
0.3
0.2
0.1
0.005
0.00236
0
ID
7669_AO_1_Bulk
7669_L_Adrenal_Bulk
7669_L_Cb2_Bulk
7669_L_CC_2_COUPTF2
7669_L_CC_2_DLX1
7669_L_EC_COUPTF2
7669_L_EC_DLX1
7669_L_F_1_Bulk
7669_L_F_1_COUPTF2
7669_L_F_1_DLX1
7669_L_F_1_TBR1
7669_L_I_1_COUPTF2
7669_L_I_1_DLX1
7669_L_I_1_TBR1
7669_L_Ins_1_Bulk
7669_L_Kidney_Bulk
7669_L_mO_COUPTF2
7669_L_mO_DLX1
7669_L_O_1_Bulk
7669_L_O_1_COUPTF2
7669_L_O_1_DLX1
7669_L_O_1_TBR1
7669_L_P_1_Bulk
7669_L_P_1_COUPTF2
7669_L_P_1_DLX1
7669_L_P_1_TBR1
7669_L_POA_Bulk
7669_L_POA_DLX1
7669_L_sec1_Cau_Bulk
7669_L_sec1_Cau_DLX1
7669_L_sec1_Put_DLX1
7669_L_sec2_GP_Bulk
7669_L_sec2_GP_DLX1
7669_L_sec2_Put_Bulk
7669_L_sec2_Thal_Bulk
7669_L_sec2_Thal_COUPTF2
7669_L_sec2_Thal_DLX1
7669_L_sec3_Hip_COUPTF2
7669_L_sec3_Hip_DLX1
7669_L_Skin_Bulk
7669_L_T_1_Bulk
7669_L_T_1_COUPTF2
7669_L_T_1_DLX1
7669_L_T_1_TBR1
7669_LA10_Bulk
7669_Liver_Bulk
7669_LV15_Bulk
7669_R_2_Put2_Bulk
7669_R_2_Thal_Bulk
7669_R_Adrenal_Bulk
7669_R_Ant_Cau_DLX1
7669_R_Ant_Put_DLX1
7669_R_Cb_2_Bulk
7669_R_CC_Mid_TBR1
7669_R_CCMid_COUPTF2
7669_R_CCMid_DLX1
7669_R_EC_COUPTF2
7669_R_EC_DLX1
7669_R_EC_TBR1
7669_R_F_1_Bulk
7669_R_F_1_COUPTF2
7669_R_F_1_DLX1
7669_R_F_1_NeuN
7669_R_F_1_TBR1
7669_R_I_1_COUPTF2
7669_R_I_1_DLX1
7669_R_I_1_TBR1
7669_R_Ins_1_Bulk
7669_R_kidney_Bulk
7669_R_mO_COUPTF2
7669_R_mO_DLX1
7669_R_mO_TBR1
7669_R_O_1_Bulk
7669_R_O_1_COUPTF2
7669_R_O_1_DLX1
7669_R_O_1_NeuN
7669_R_O_1_TBR1
7669_R_O_10_DLX1
7669_R_O_11_DLX1
7669_R_O_2_DLX1
7669_R_O_3_DLX1
7669_R_O_4_DLX1
7669_R_O_5_DLX1
7669_R_O_6_DLX1
7669_R_O_7_DLX1
7669_R_O_8_DLX1
7669_R_O_9_DLX1
7669_R_Olf_DLX1
7669_R_P_1_Bulk
7669_R_P_1_COUPTF2
7669_R_P_1_DLX1
7669_R_P_1_NeuN
7669_R_P_1_TBR1
7669_R_P_10_DLX1
7669_R_P_11_DLX1
7669_R_P_11_TBR1
7669_R_P_12_DLX1
7669_R_P_12_TBR1
7669_R_P_13_DLX1
7669_R_P_13_TBR1
7669_R_P_14_DLX1
7669_R_P_14_TBR1
7669_R_P_15_DLX1
7669_R_P_15_TBR1
7669_R_P_16_DLX1
7669_R_P_16_TBR1
7669_R_P_17_DLX1
7669_R_P_17_TBR1
7669_R_P_2_DLX1
7669_R_P_2_TBR1
7669_R_P_3_DLX1
7669_R_P_3_TBR1
7669_R_P_4_DLX1
7669_R_P_4_TBR1
7669_R_P_5_DLX1
7669_R_P_5_TBR1
7669_R_P_6_DLX1
7669_R_P_6_TBR1
7669_R_P_7_DLX1
7669_R_P_7_TBR1
7669_R_P_8_DLX1
7669_R_P_8_TBR1
7669_R_P_9_DLX
7669_R_P_9_DLX1
7669_R_P_9_TBR1
7669_R_POA_Bulk
7669_R_POA_DLX1
7669_R_sec1_Cau_Bulk
7669_R_sec2_GP_Bulk
7669_R_sec2_GP_DLX1
7669_R_sec2_Put_Bulk
7669_R_sec2_Thal_Bulk
7669_R_sec2_Thal_COUPTF2
7669_R_sec2_Thal_DLX1
7669_R_sec3_Cau_DLX1
7669_R_sec3_Hip_COUPTF2
7669_R_sec3_Hip_DLX1
7669_R_sec3_Hip_TBR1
7669_R_sec3_Put_DLX1
7669_R_Skin_Bulk
7669_R_T_1_Bulk
7669_R_T_1_COUPTF2
7669_R_T_1_DLX1
7669_R_T_1_TBR1
7669_RA9_Bulk
7669_RV10_Bulk
JGG

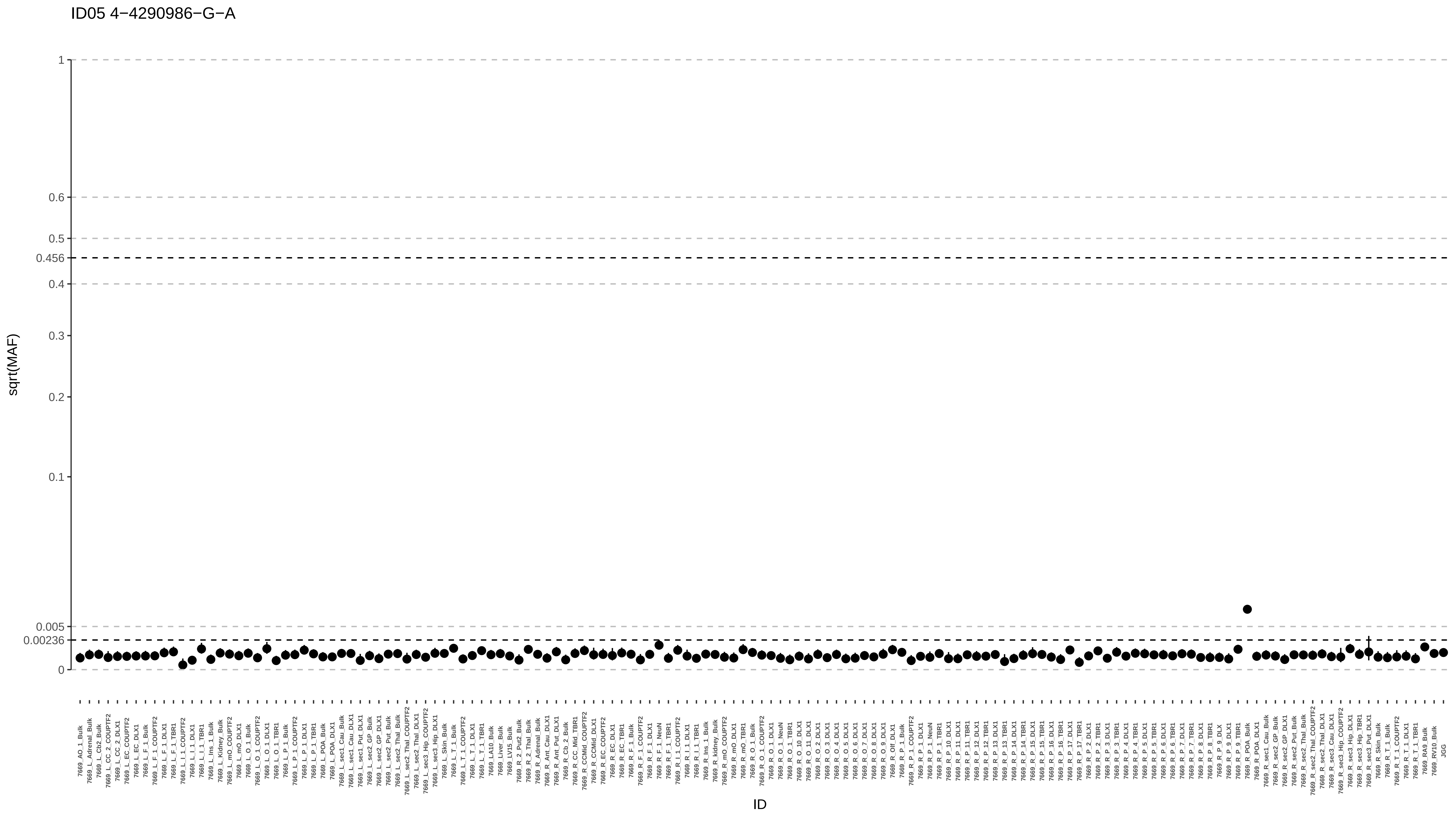

ID05 4−4290986−G−A
sqrt(MAF)
1
0.6
0.5
0.456
0.4
0.3
0.2
0.1
0.005
0.00236
0
ID

ID05 4−44006646−G−A

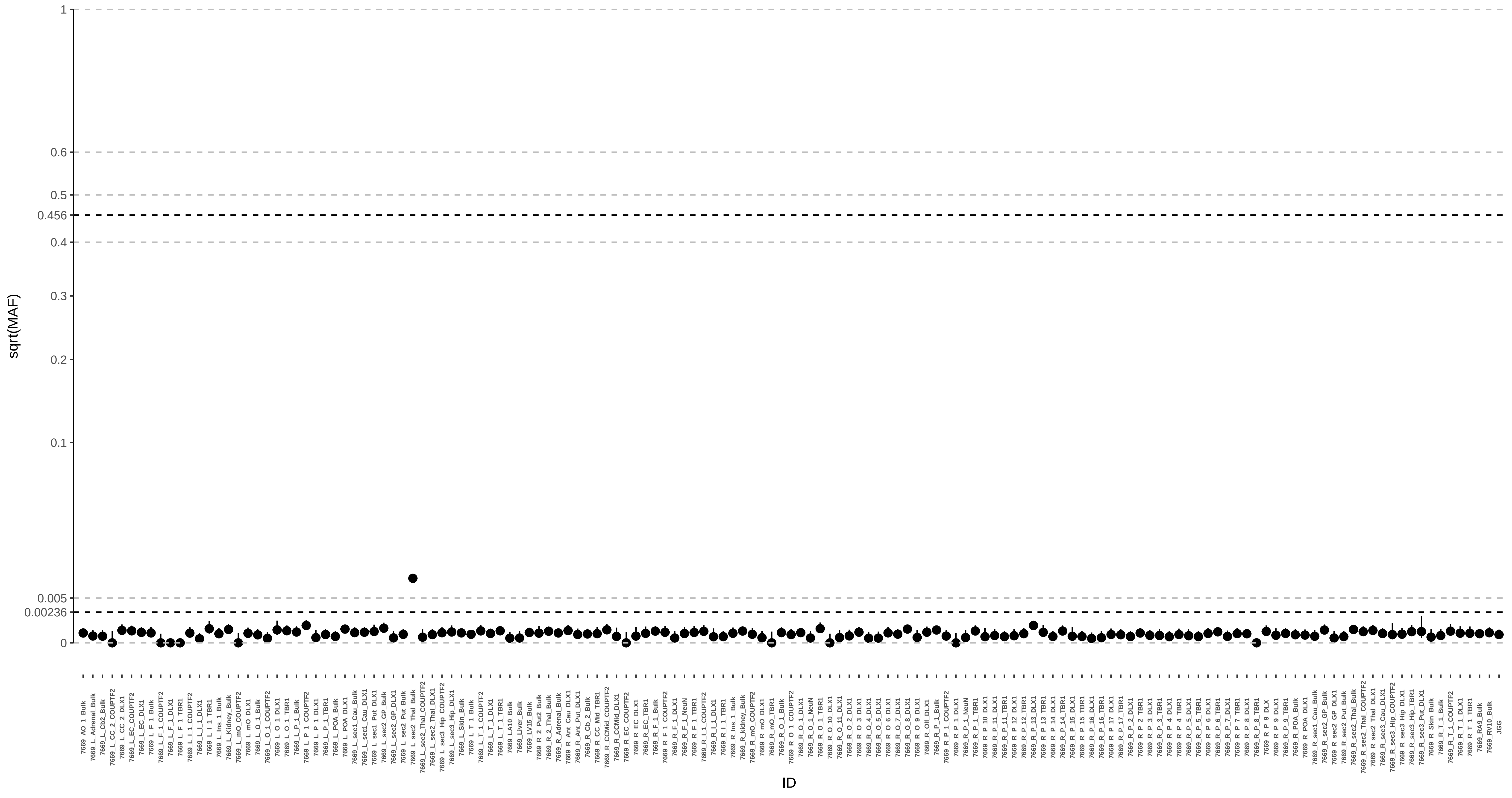

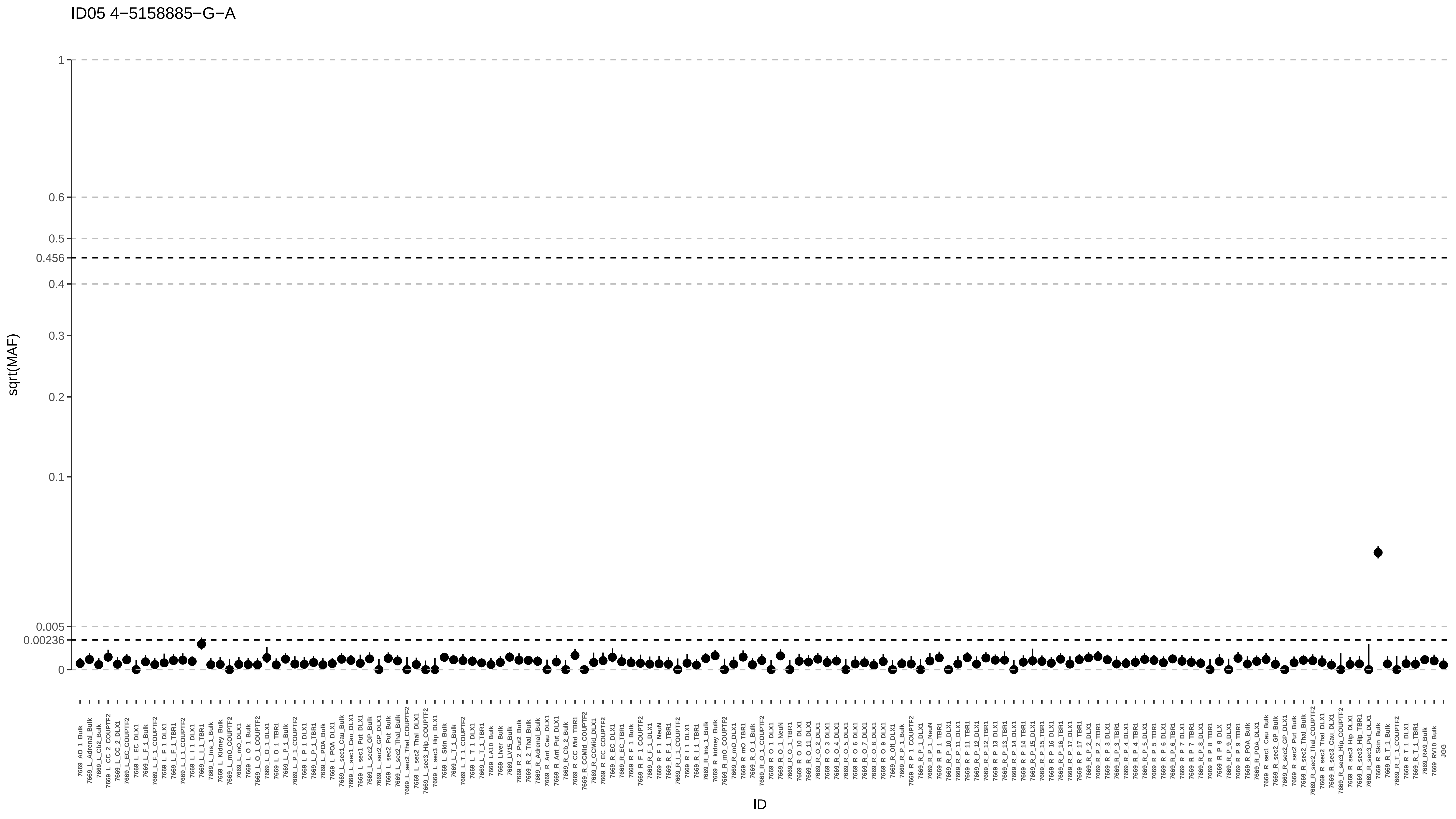
ID05 4−5158885−G−A
sqrt(MAF)
1
0.6
0.5
0.456
0.4
0.3
0.2
0.1
0.005
0.00236
0
ID

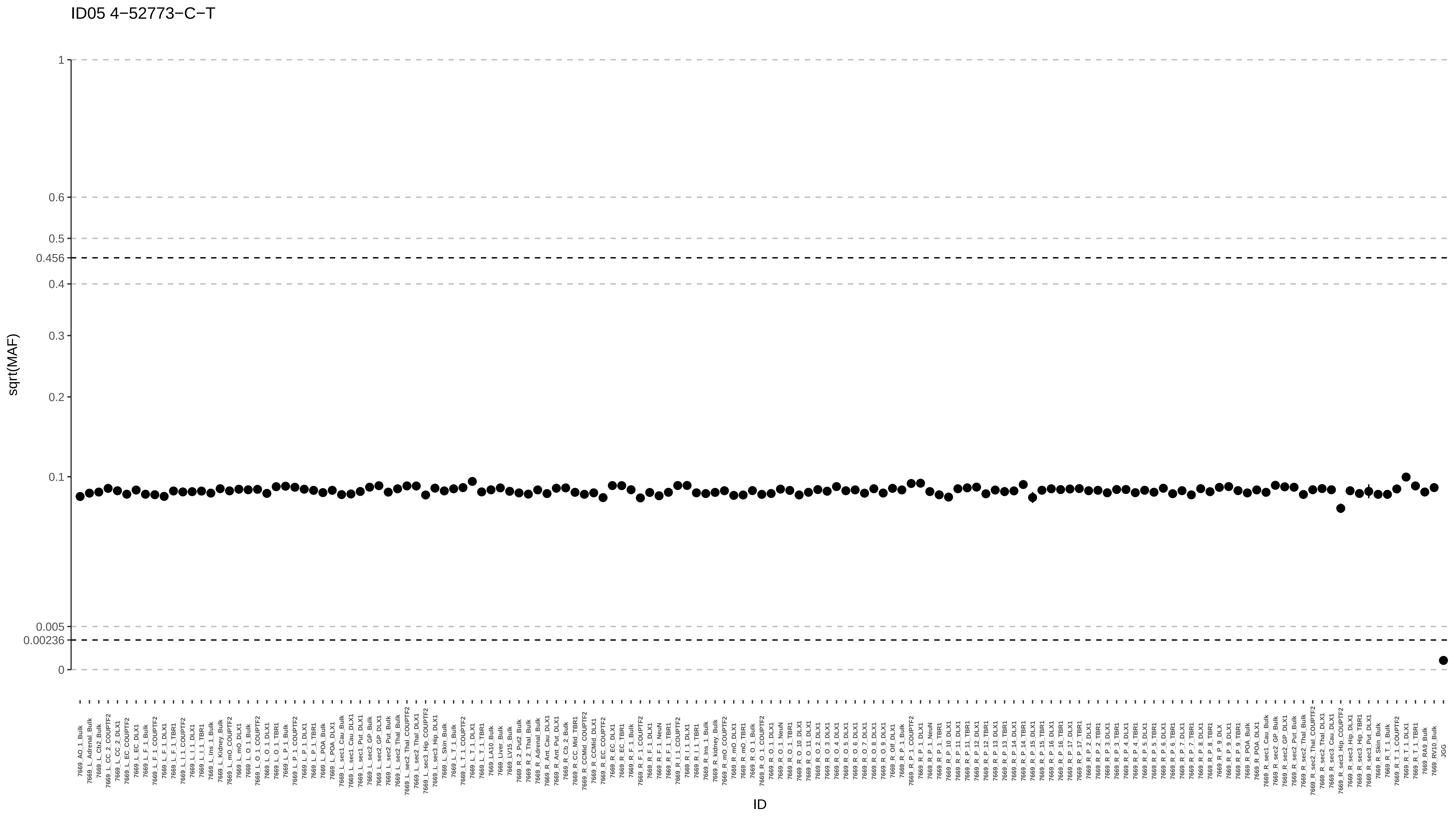
ID05 4−52773−C−T
sqrt(MAF)
1
0.6
0.5
0.456
0.4
0.3
0.2
0.1
0.005
0.00236
0
ID

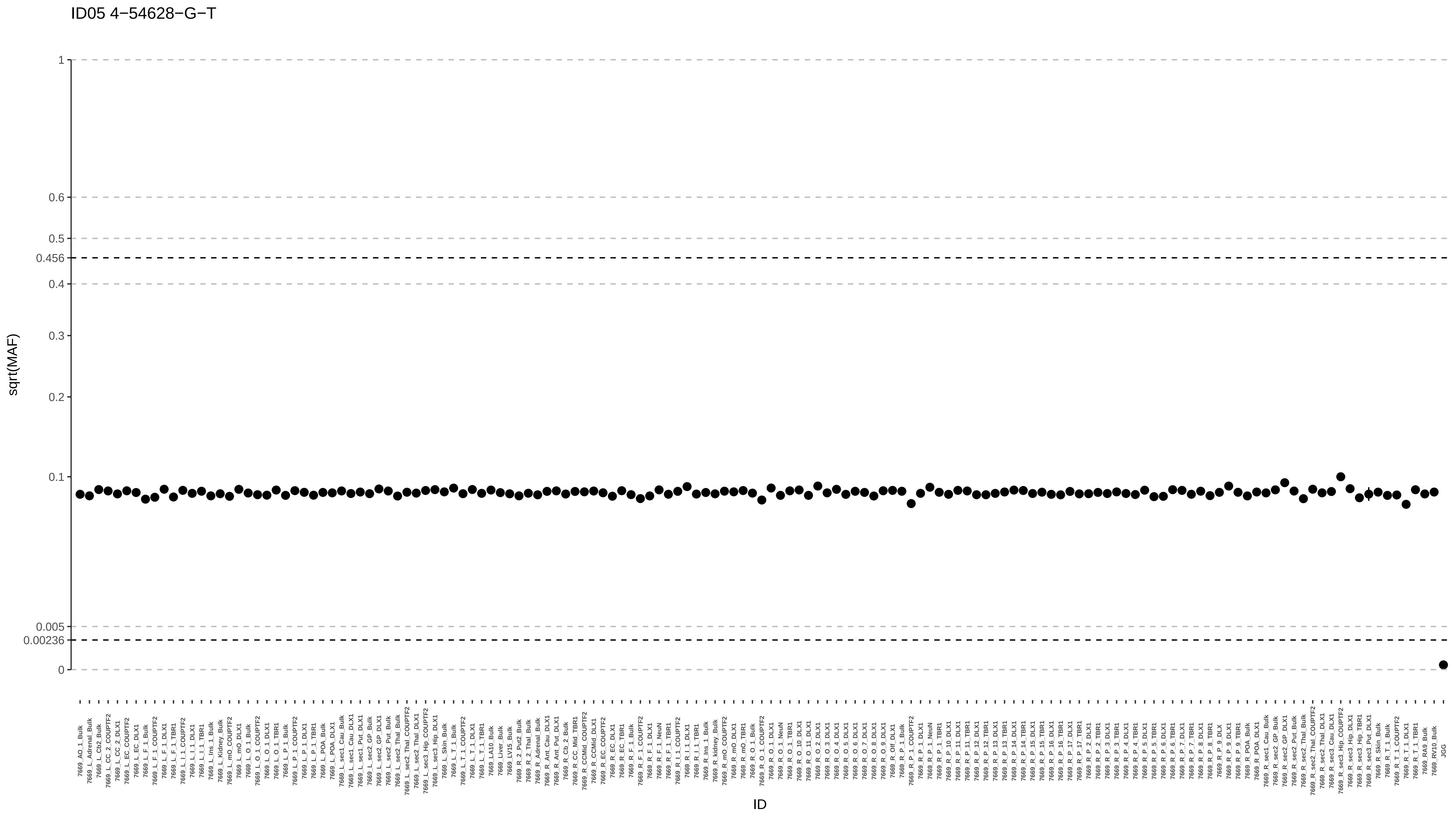

ID05 4−54628−G−T
sqrt(MAF)
1
0.6
0.5
0.456
0.4
0.3
0.2
0.1
0.005
0.00236
0
ID
7669_AO_1_Bulk
7669_L_Adrenal_Bulk
7669_L_Cb2_Bulk
7669_L_CC_2_COUPTF2
7669_L_CC_2_DLX1
7669_L_EC_COUPTF2
7669_L_EC_DLX1
7669_L_F_1_Bulk
7669_L_F_1_COUPTF2
7669_L_F_1_DLX1
7669_L_F_1_TBR1
7669_L_I_1_COUPTF2
7669_L_I_1_DLX1
7669_L_I_1_TBR1
7669_L_Ins_1_Bulk
7669_L_Kidney_Bulk
7669_L_mO_COUPTF2
7669_L_mO_DLX1
7669_L_O_1_Bulk
7669_L_O_1_COUPTF2
7669_L_O_1_DLX1
7669_L_O_1_TBR1
7669_L_P_1_Bulk
7669_L_P_1_COUPTF2
7669_L_P_1_DLX1
7669_L_P_1_TBR1
7669_L_POA_Bulk
7669_L_POA_DLX1
7669_L_sec1_Cau_Bulk
7669_L_sec1_Cau_DLX1
7669_L_sec1_Put_DLX1
7669_L_sec2_GP_Bulk
7669_L_sec2_GP_DLX1
7669_L_sec2_Put_Bulk
7669_L_sec2_Thal_Bulk
7669_L_sec2_Thal_COUPTF2
7669_L_sec2_Thal_DLX1
7669_L_sec3_Hip_COUPTF2
7669_L_sec3_Hip_DLX1
7669_L_Skin_Bulk
7669_L_T_1_Bulk
7669_L_T_1_COUPTF2
7669_L_T_1_DLX1
7669_L_T_1_TBR1
7669_LA10_Bulk
7669_Liver_Bulk
7669_LV15_Bulk
7669_R_2_Put2_Bulk
7669_R_2_Thal_Bulk
7669_R_Adrenal_Bulk
7669_R_Ant_Cau_DLX1
7669_R_Ant_Put_DLX1
7669_R_Cb_2_Bulk
7669_R_CC_Mid_TBR1
7669_R_CCMid_COUPTF2
7669_R_CCMid_DLX1
7669_R_EC_COUPTF2
7669_R_EC_DLX1
7669_R_EC_TBR1
7669_R_F_1_Bulk
7669_R_F_1_COUPTF2
7669_R_F_1_DLX1
7669_R_F_1_NeuN
7669_R_F_1_TBR1
7669_R_I_1_COUPTF2
7669_R_I_1_DLX1
7669_R_I_1_TBR1
7669_R_Ins_1_Bulk
7669_R_kidney_Bulk
7669_R_mO_COUPTF2
7669_R_mO_DLX1
7669_R_mO_TBR1
7669_R_O_1_Bulk
7669_R_O_1_COUPTF2
7669_R_O_1_DLX1
7669_R_O_1_NeuN
7669_R_O_1_TBR1
7669_R_O_10_DLX1
7669_R_O_11_DLX1
7669_R_O_2_DLX1
7669_R_O_3_DLX1
7669_R_O_4_DLX1
7669_R_O_5_DLX1
7669_R_O_6_DLX1
7669_R_O_7_DLX1
7669_R_O_8_DLX1
7669_R_O_9_DLX1
7669_R_Olf_DLX1
7669_R_P_1_Bulk
7669_R_P_1_COUPTF2
7669_R_P_1_DLX1
7669_R_P_1_NeuN
7669_R_P_1_TBR1
7669_R_P_10_DLX1
7669_R_P_11_DLX1
7669_R_P_11_TBR1
7669_R_P_12_DLX1
7669_R_P_12_TBR1
7669_R_P_13_DLX1
7669_R_P_13_TBR1
7669_R_P_14_DLX1
7669_R_P_14_TBR1
7669_R_P_15_DLX1
7669_R_P_15_TBR1
7669_R_P_16_DLX1
7669_R_P_16_TBR1
7669_R_P_17_DLX1
7669_R_P_17_TBR1
7669_R_P_2_DLX1
7669_R_P_2_TBR1
7669_R_P_3_DLX1
7669_R_P_3_TBR1
7669_R_P_4_DLX1
7669_R_P_4_TBR1
7669_R_P_5_DLX1
7669_R_P_5_TBR1
7669_R_P_6_DLX1
7669_R_P_6_TBR1
7669_R_P_7_DLX1
7669_R_P_7_TBR1
7669_R_P_8_DLX1
7669_R_P_8_TBR1
7669_R_P_9_DLX
7669_R_P_9_DLX1
7669_R_P_9_TBR1
7669_R_POA_Bulk
7669_R_POA_DLX1
7669_R_sec1_Cau_Bulk
7669_R_sec2_GP_Bulk
7669_R_sec2_GP_DLX1
7669_R_sec2_Put_Bulk
7669_R_sec2_Thal_Bulk
7669_R_sec2_Thal_COUPTF2
7669_R_sec2_Thal_DLX1
7669_R_sec3_Cau_DLX1
7669_R_sec3_Hip_COUPTF2
7669_R_sec3_Hip_DLX1
7669_R_sec3_Hip_TBR1
7669_R_sec3_Put_DLX1
7669_R_Skin_Bulk
7669_R_T_1_Bulk
7669_R_T_1_COUPTF2
7669_R_T_1_DLX1
7669_R_T_1_TBR1
7669_RA9_Bulk
7669_RV10_Bulk
JGG

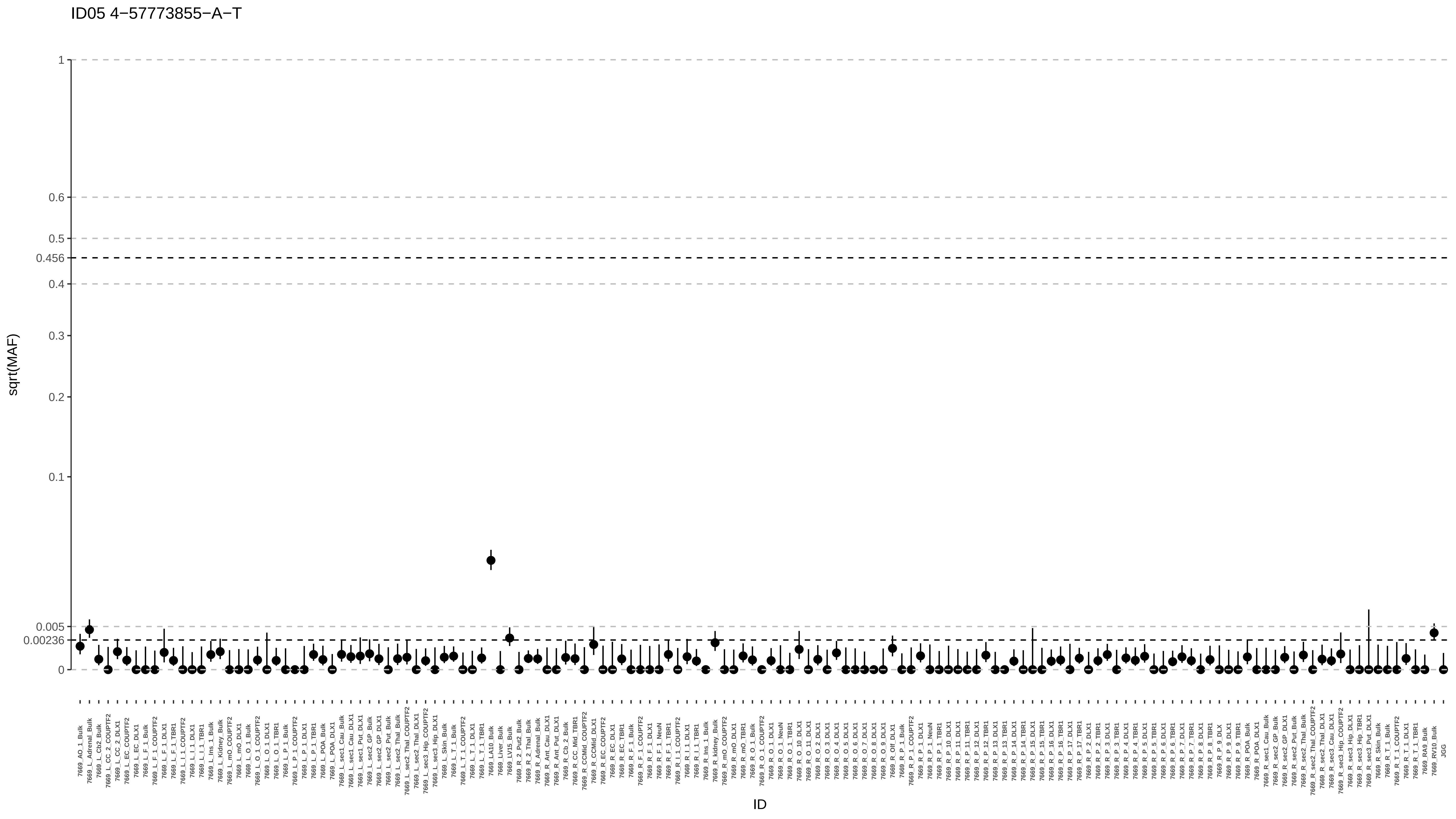
ID05 4−57773855−A−T
sqrt(MAF)
1
0.6
0.5
0.456
0.4
0.3
0.2
0.1
0.005
0.00236
0
ID

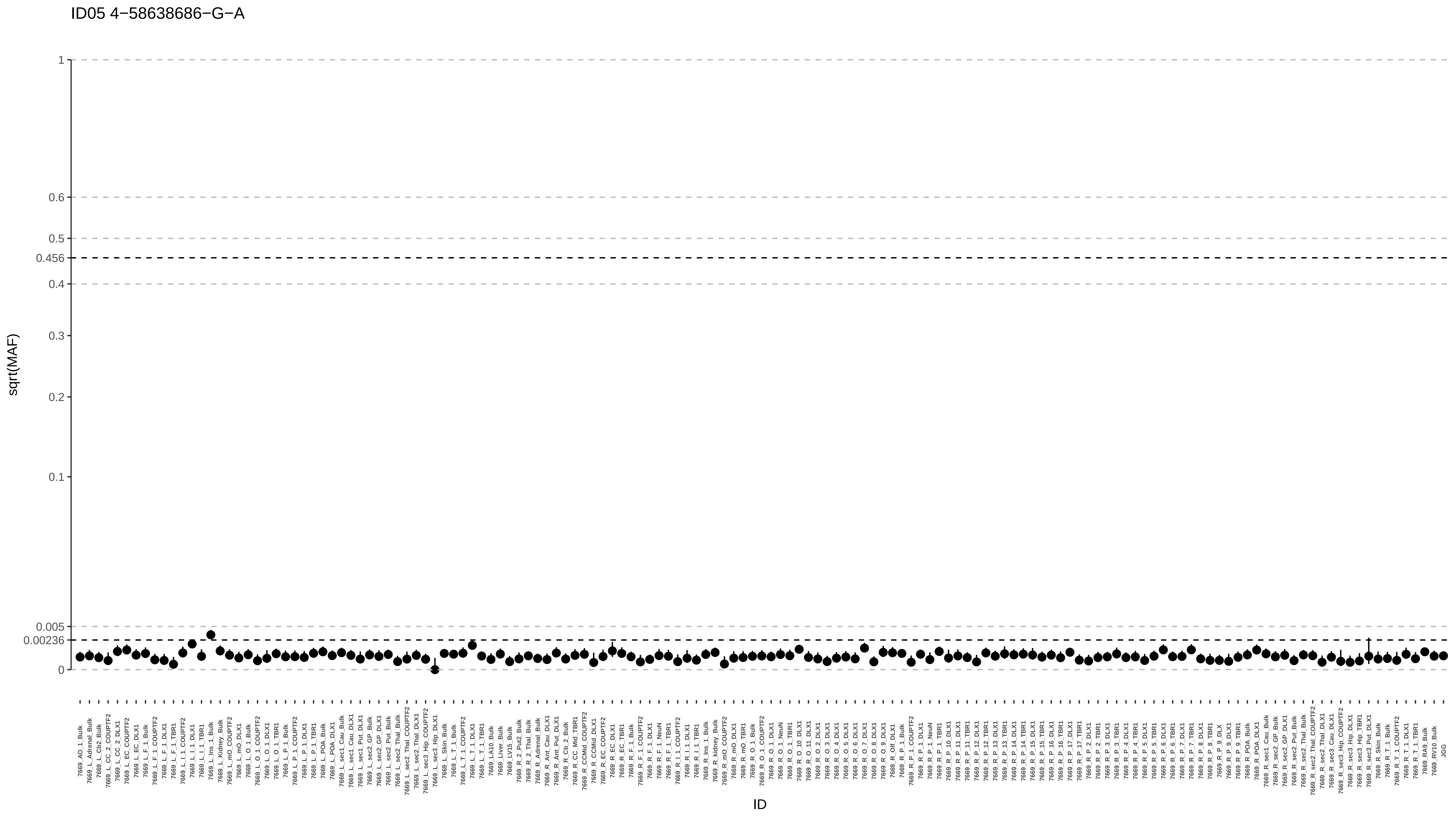
ID05 4−58638686−G−A
sqrt(MAF)
1
0.6
0.5
0.456
0.4
0.3
0.2
0.1
0.005
0.00236
0
ID

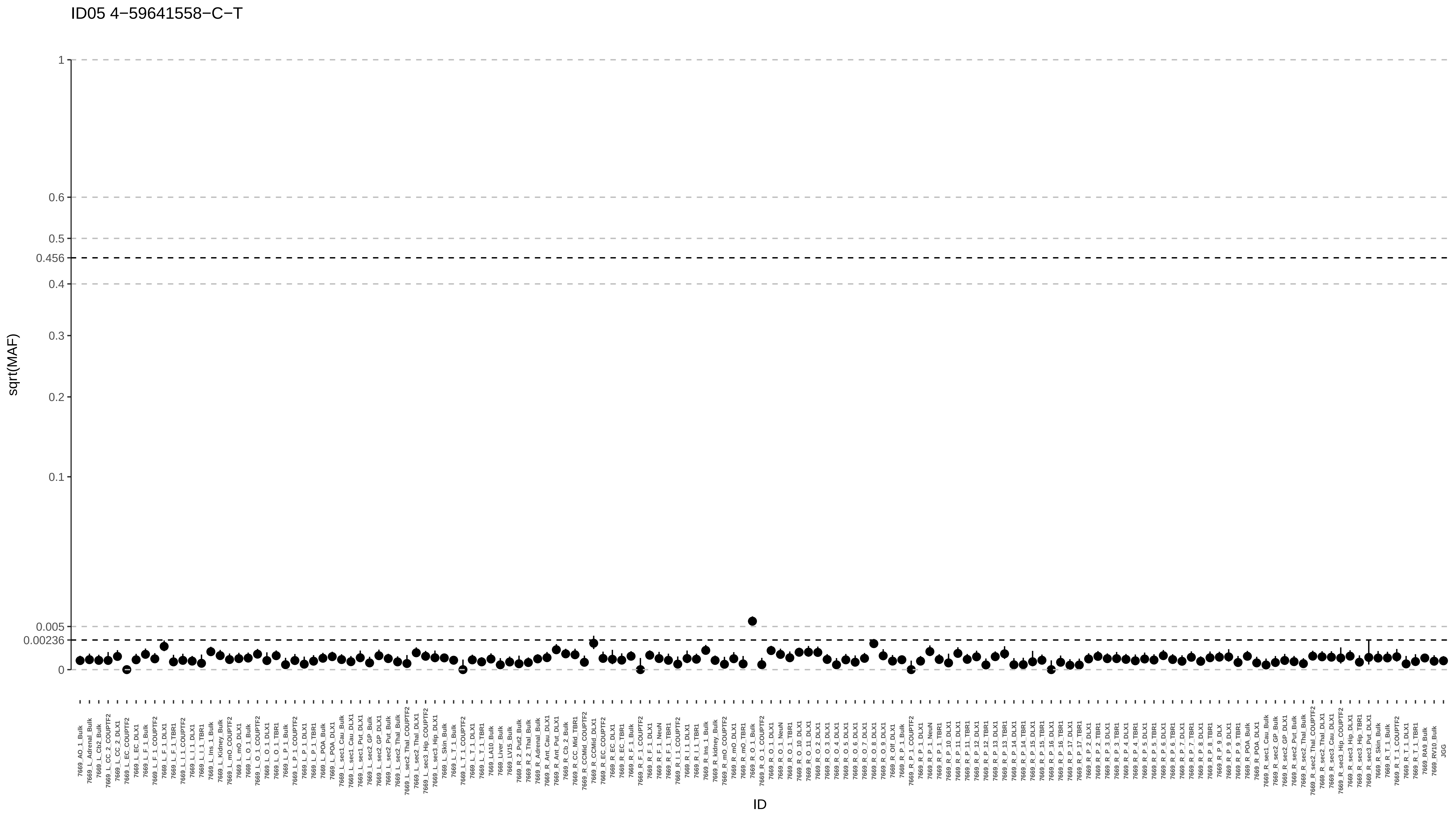

ID05 4−59641558−C−T
sqrt(MAF)
ID
1
0.6
0.5
0.456
0.4
0.3
0.2
0.1
0.005
0.00236
0

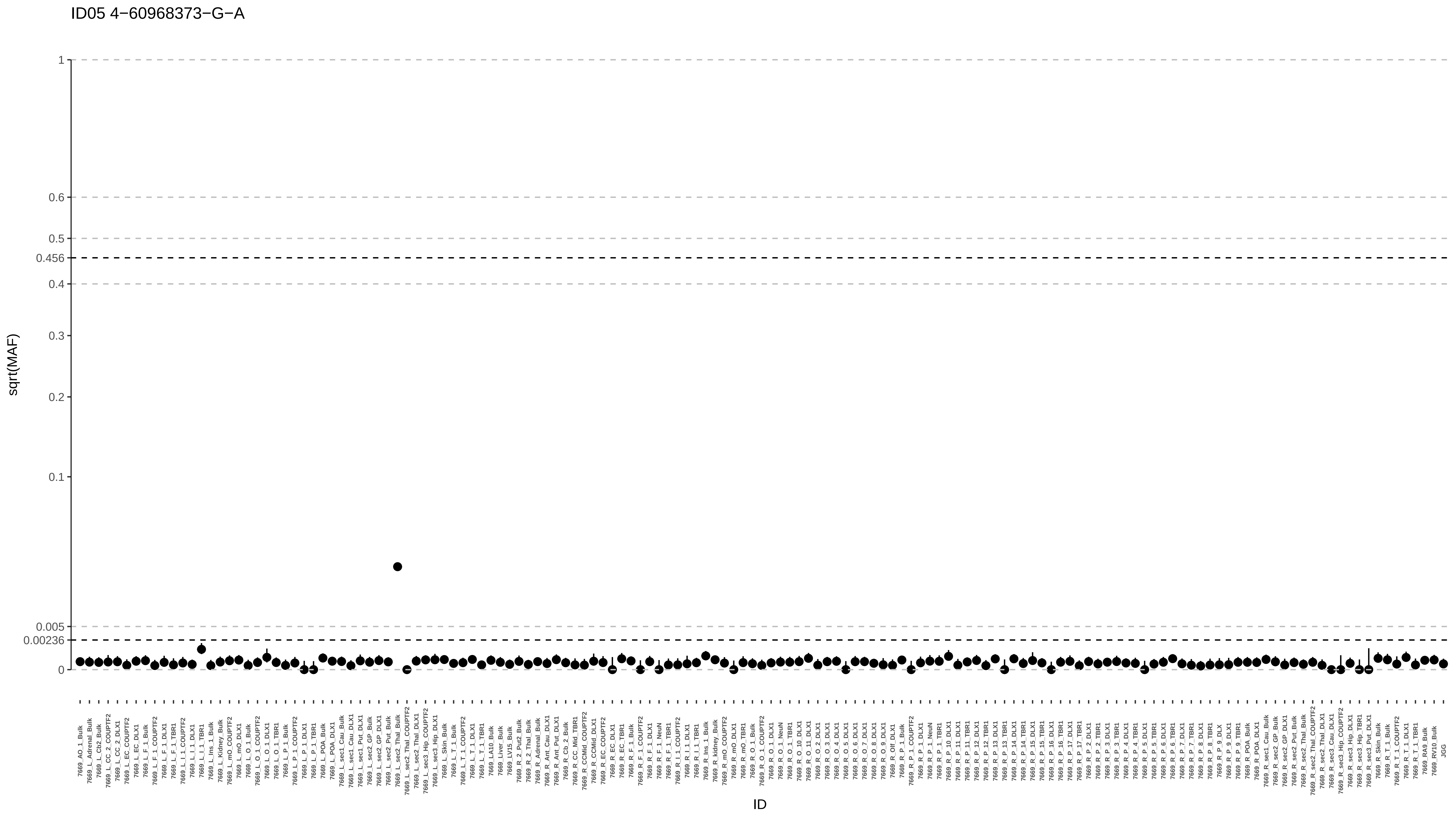
ID05 4−60968373−G−A
sqrt(MAF)
1
0.6
0.5
0.456
0.4
0.3
0.2
0.1
0.005
0.00236
0
ID

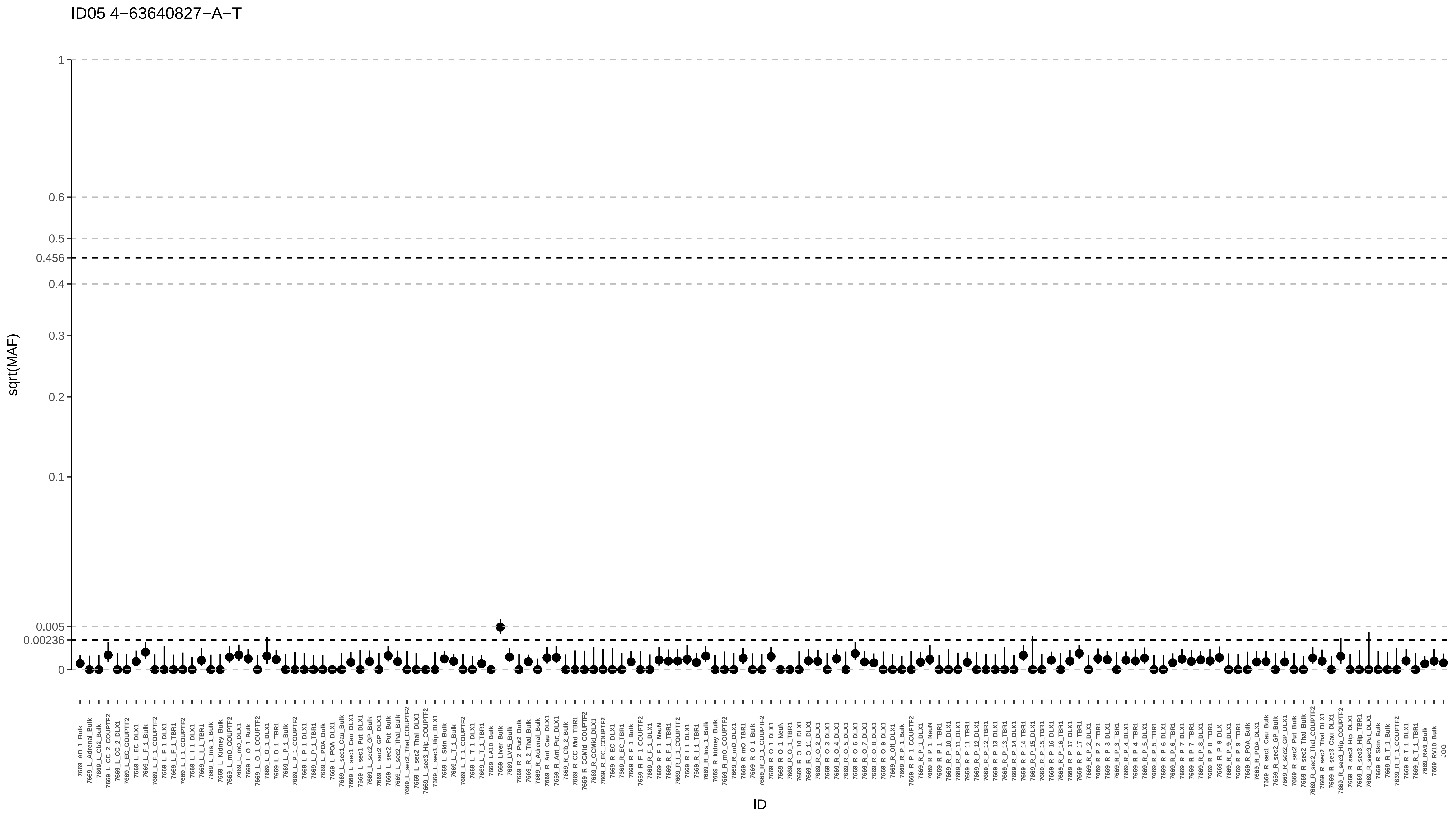
ID05 4−63640827−A−T
sqrt(MAF)
1
0.6
0.5
0.456
0.4
0.3
0.2
0.1
0.005
0.00236
0
ID

ID05 4−64742736−G−A

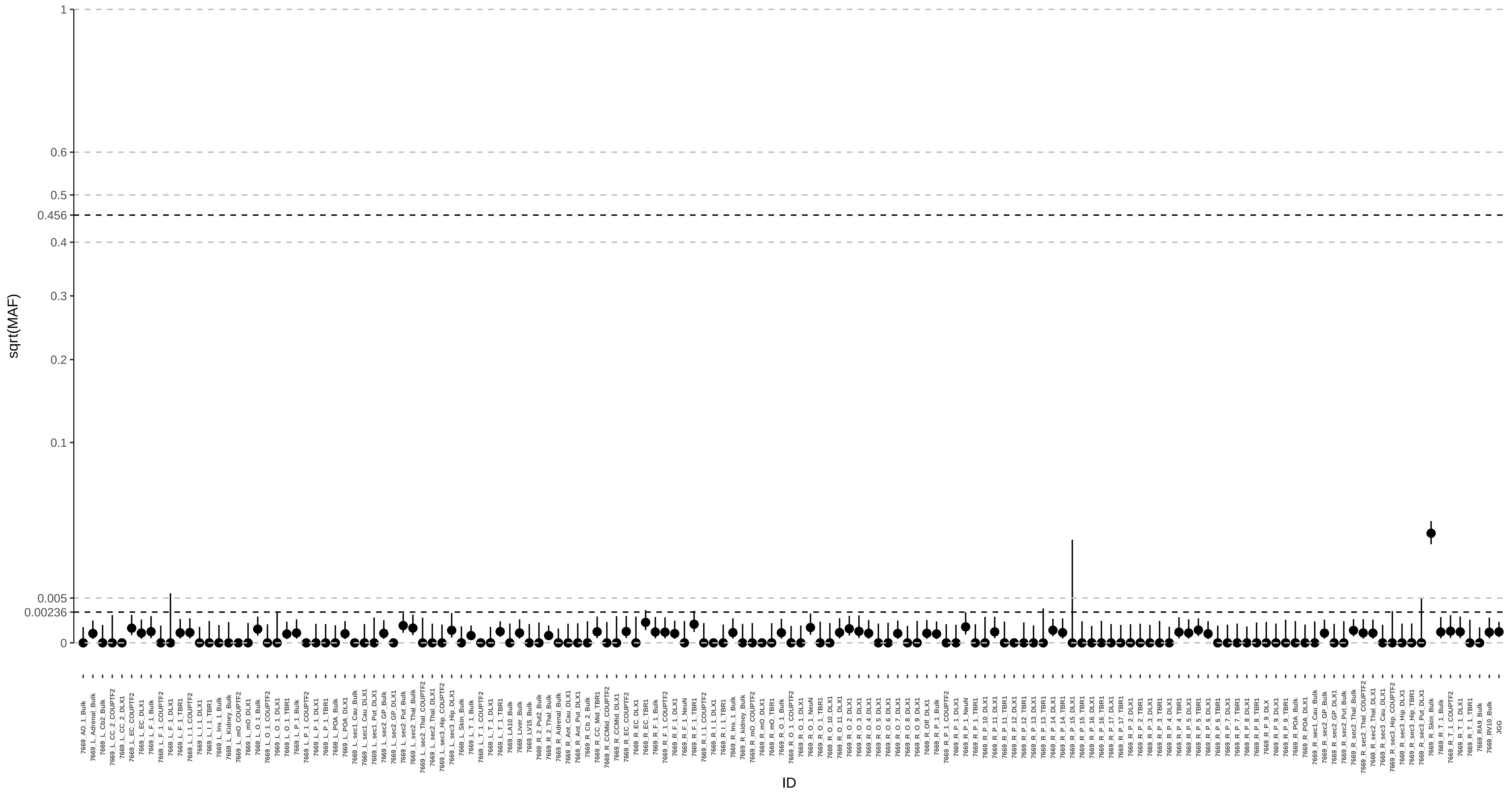

ID05 4−66798015−A−T

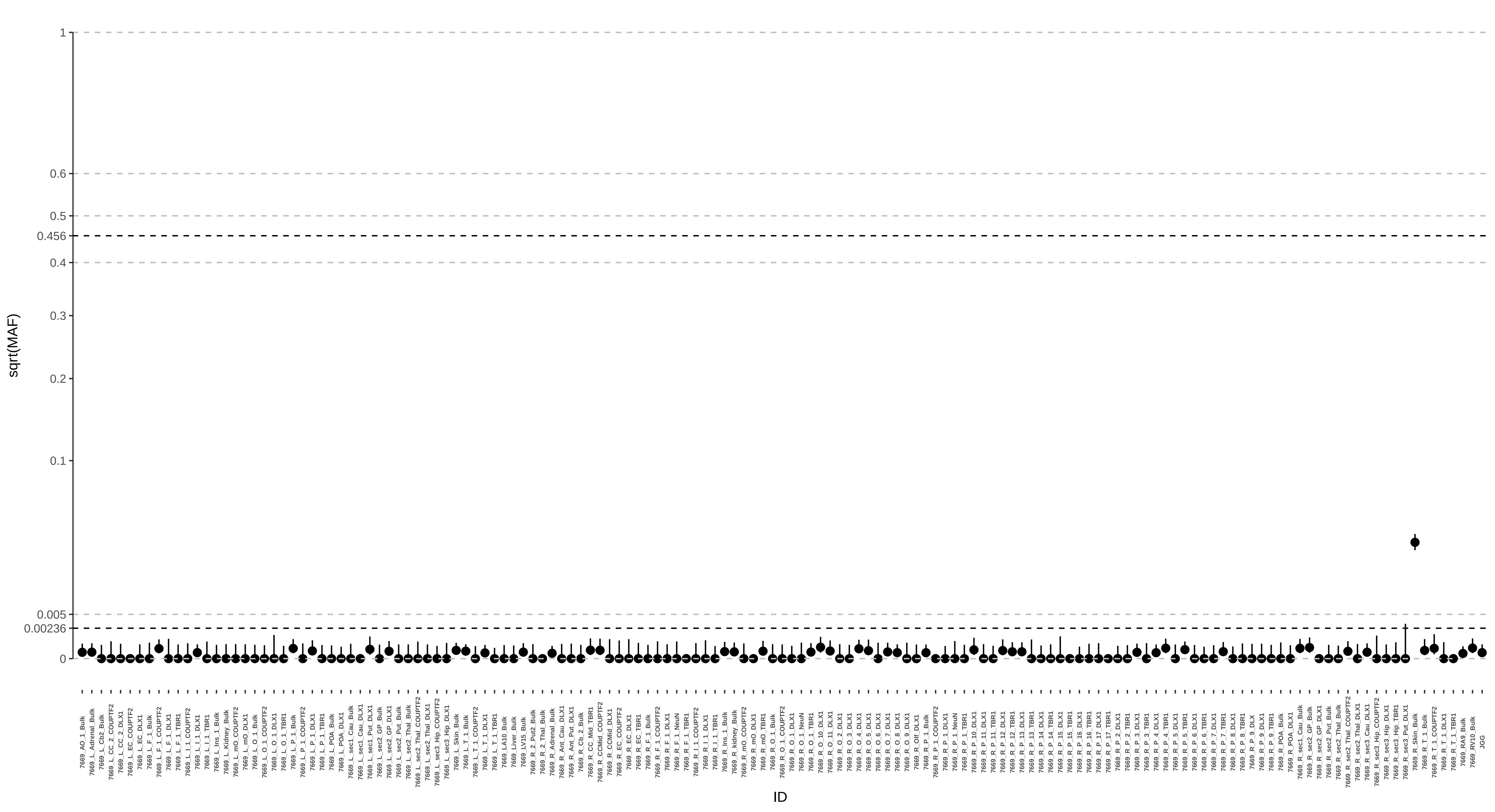

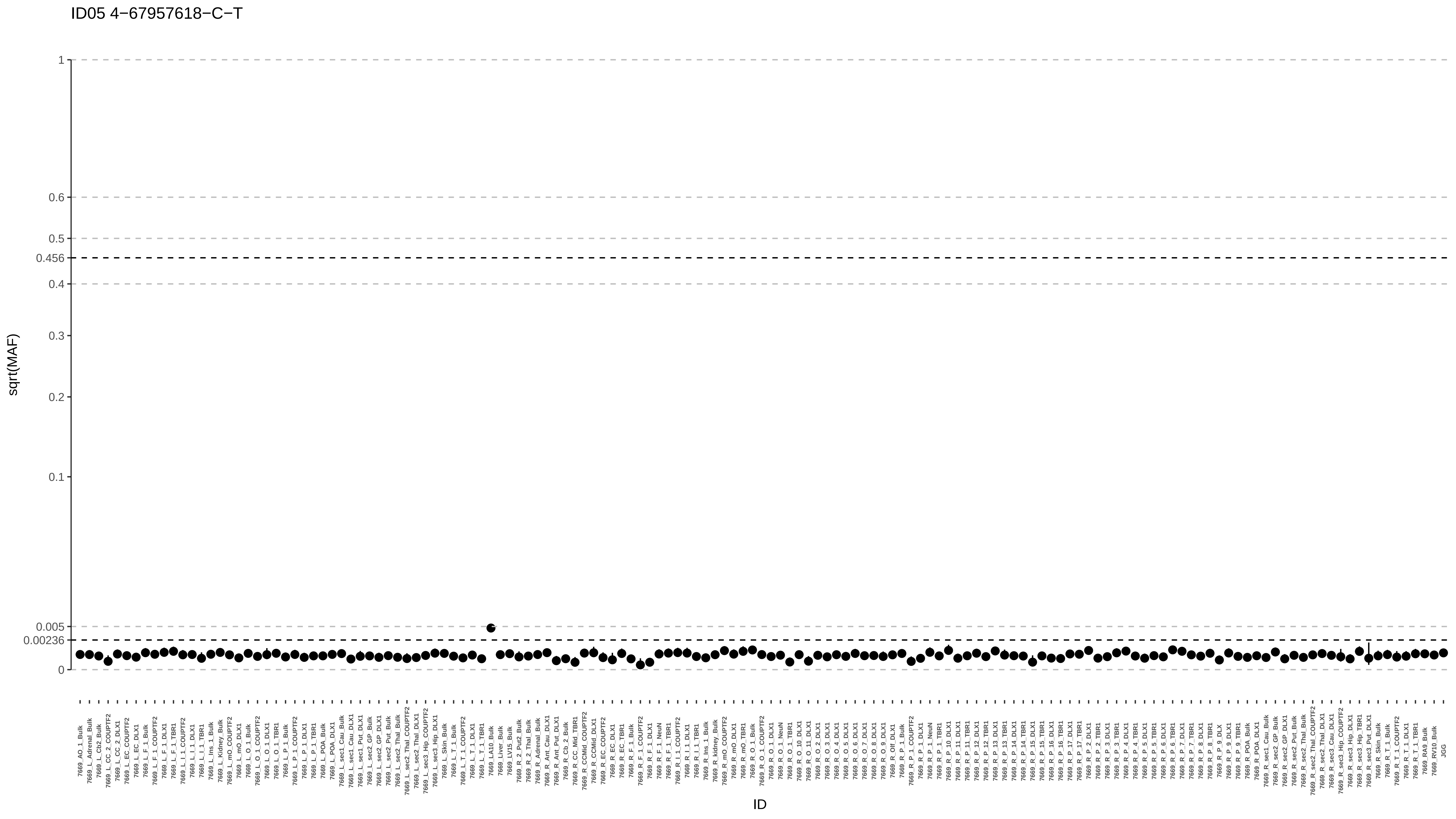
ID05 4−67957618−C−T
sqrt(MAF)
1
0.6
0.5
0.456
0.4
0.3
0.2
0.1
0.005
0.00236
0
ID
7669_AO_1_Bulk
7669_L_Adrenal_Bulk
7669_L_Cb2_Bulk
7669_L_CC_2_COUPTF2
7669_L_CC_2_DLX1
7669_L_EC_COUPTF2
7669_L_EC_DLX1
7669_L_F_1_Bulk
7669_L_F_1_COUPTF2
7669_L_F_1_DLX1
7669_L_F_1_TBR1
7669_L_I_1_COUPTF2
7669_L_I_1_DLX1
7669_L_I_1_TBR1
7669_L_Ins_1_Bulk
7669_L_Kidney_Bulk
7669_L_mO_COUPTF2
7669_L_mO_DLX1
7669_L_O_1_Bulk
7669_L_O_1_COUPTF2
7669_L_O_1_DLX1
7669_L_O_1_TBR1
7669_L_P_1_Bulk
7669_L_P_1_COUPTF2
7669_L_P_1_DLX1
7669_L_P_1_TBR1
7669_L_POA_Bulk
7669_L_POA_DLX1
7669_L_sec1_Cau_Bulk
7669_L_sec1_Cau_DLX1
7669_L_sec1_Put_DLX1
7669_L_sec2_GP_Bulk
7669_L_sec2_GP_DLX1
7669_L_sec2_Put_Bulk
7669_L_sec2_Thal_Bulk
7669_L_sec2_Thal_COUPTF2
7669_L_sec2_Thal_DLX1
7669_L_sec3_Hip_COUPTF2
7669_L_sec3_Hip_DLX1
7669_L_Skin_Bulk
7669_L_T_1_Bulk
7669_L_T_1_COUPTF2
7669_L_T_1_DLX1
7669_L_T_1_TBR1
7669_LA10_Bulk
7669_Liver_Bulk
7669_LV15_Bulk
7669_R_2_Put2_Bulk
7669_R_2_Thal_Bulk
7669_R_Adrenal_Bulk
7669_R_Ant_Cau_DLX1
7669_R_Ant_Put_DLX1
7669_R_Cb_2_Bulk
7669_R_CC_Mid_TBR1
7669_R_CCMid_COUPTF2
7669_R_CCMid_DLX1
7669_R_EC_COUPTF2
7669_R_EC_DLX1
7669_R_EC_TBR1
7669_R_F_1_Bulk
7669_R_F_1_COUPTF2
7669_R_F_1_DLX1
7669_R_F_1_NeuN
7669_R_F_1_TBR1
7669_R_I_1_COUPTF2
7669_R_I_1_DLX1
7669_R_I_1_TBR1
7669_R_Ins_1_Bulk
7669_R_kidney_Bulk
7669_R_mO_COUPTF2
7669_R_mO_DLX1
7669_R_mO_TBR1
7669_R_O_1_Bulk
7669_R_O_1_COUPTF2
7669_R_O_1_DLX1
7669_R_O_1_NeuN
7669_R_O_1_TBR1
7669_R_O_10_DLX1
7669_R_O_11_DLX1
7669_R_O_2_DLX1
7669_R_O_3_DLX1
7669_R_O_4_DLX1
7669_R_O_5_DLX1
7669_R_O_6_DLX1
7669_R_O_7_DLX1
7669_R_O_8_DLX1
7669_R_O_9_DLX1
7669_R_Olf_DLX1
7669_R_P_1_Bulk
7669_R_P_1_COUPTF2
7669_R_P_1_DLX1
7669_R_P_1_NeuN
7669_R_P_1_TBR1
7669_R_P_10_DLX1
7669_R_P_11_DLX1
7669_R_P_11_TBR1
7669_R_P_12_DLX1
7669_R_P_12_TBR1
7669_R_P_13_DLX1
7669_R_P_13_TBR1
7669_R_P_14_DLX1
7669_R_P_14_TBR1
7669_R_P_15_DLX1
7669_R_P_15_TBR1
7669_R_P_16_DLX1
7669_R_P_16_TBR1
7669_R_P_17_DLX1
7669_R_P_17_TBR1
7669_R_P_2_DLX1
7669_R_P_2_TBR1
7669_R_P_3_DLX1
7669_R_P_3_TBR1
7669_R_P_4_DLX1
7669_R_P_4_TBR1
7669_R_P_5_DLX1
7669_R_P_5_TBR1
7669_R_P_6_DLX1
7669_R_P_6_TBR1
7669_R_P_7_DLX1
7669_R_P_7_TBR1
7669_R_P_8_DLX1
7669_R_P_8_TBR1
7669_R_P_9_DLX
7669_R_P_9_DLX1
7669_R_P_9_TBR1
7669_R_POA_Bulk
7669_R_POA_DLX1
7669_R_sec1_Cau_Bulk
7669_R_sec2_GP_Bulk
7669_R_sec2_GP_DLX1
7669_R_sec2_Put_Bulk
7669_R_sec2_Thal_Bulk
7669_R_sec2_Thal_COUPTF2
7669_R_sec2_Thal_DLX1
7669_R_sec3_Cau_DLX1
7669_R_sec3_Hip_COUPTF2
7669_R_sec3_Hip_DLX1
7669_R_sec3_Hip_TBR1
7669_R_sec3_Put_DLX1
7669_R_Skin_Bulk
7669_R_T_1_Bulk
7669_R_T_1_COUPTF2
7669_R_T_1_DLX1
7669_R_T_1_TBR1
7669_RA9_Bulk
7669_RV10_Bulk
JGG

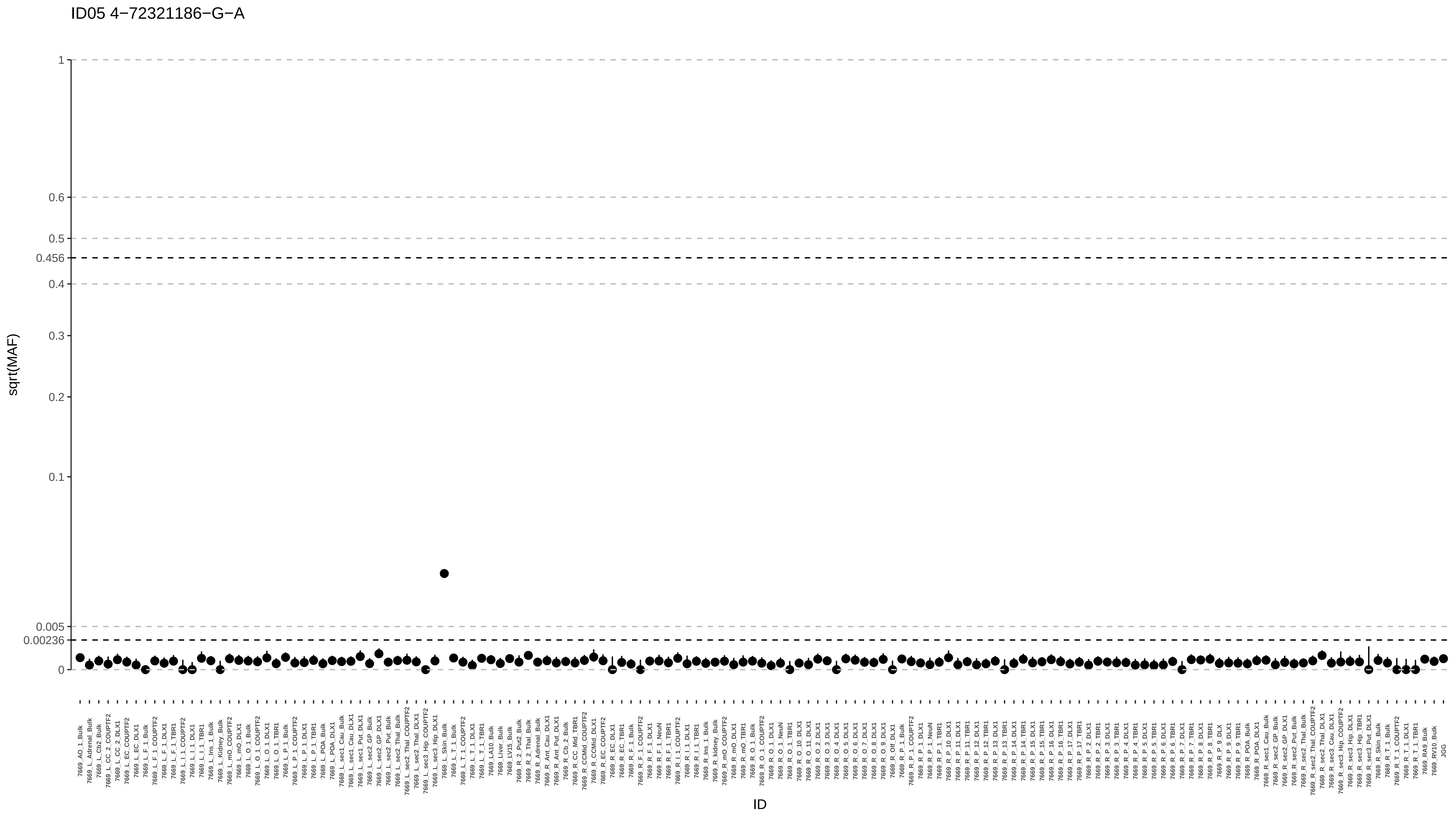
ID05 4−72321186−G−A
sqrt(MAF)
1
0.6
0.5
0.456
0.4
0.3
0.2
0.1
0.005
0.00236
0
ID

ID05 4−72321187−G−A

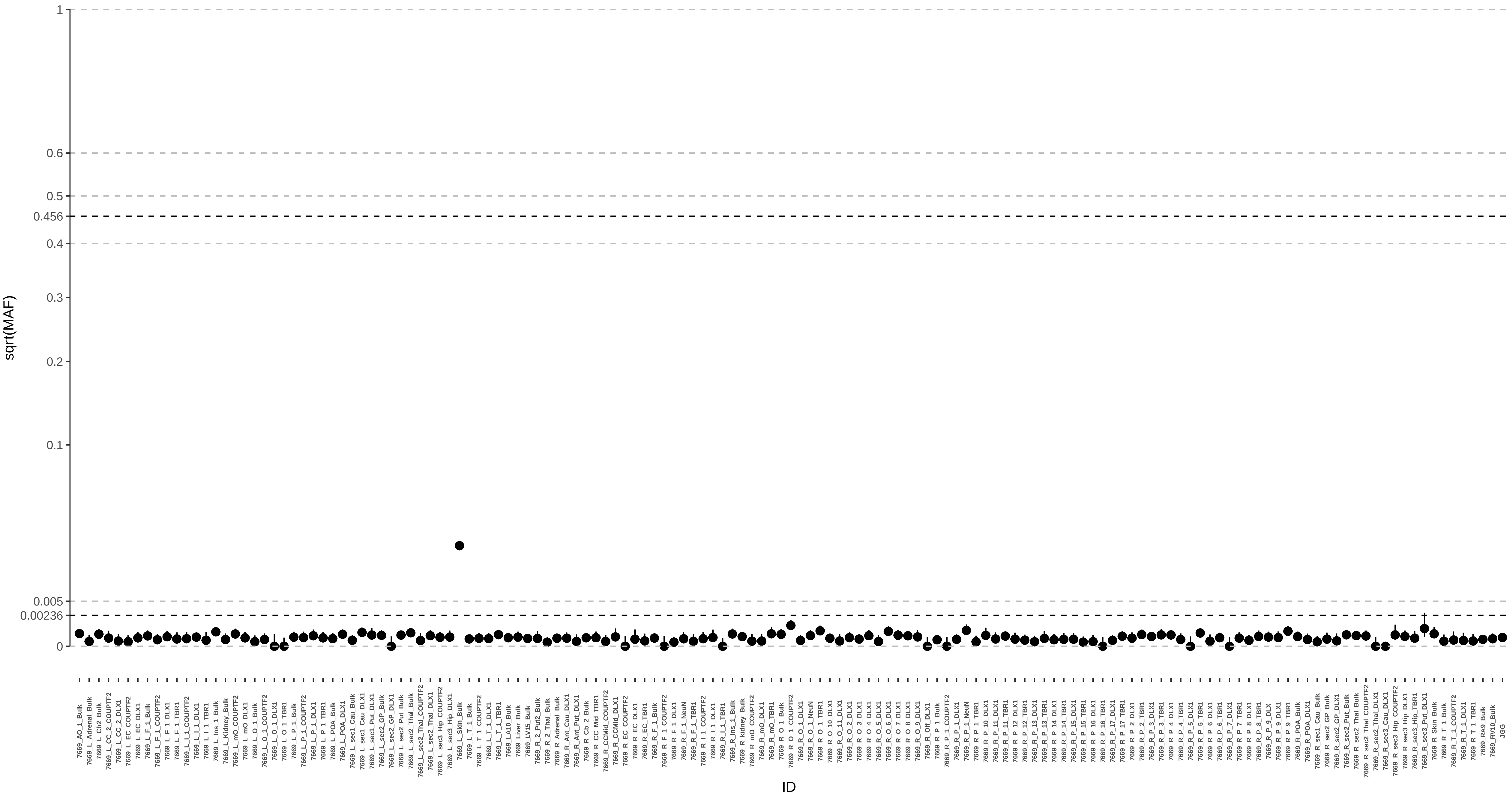

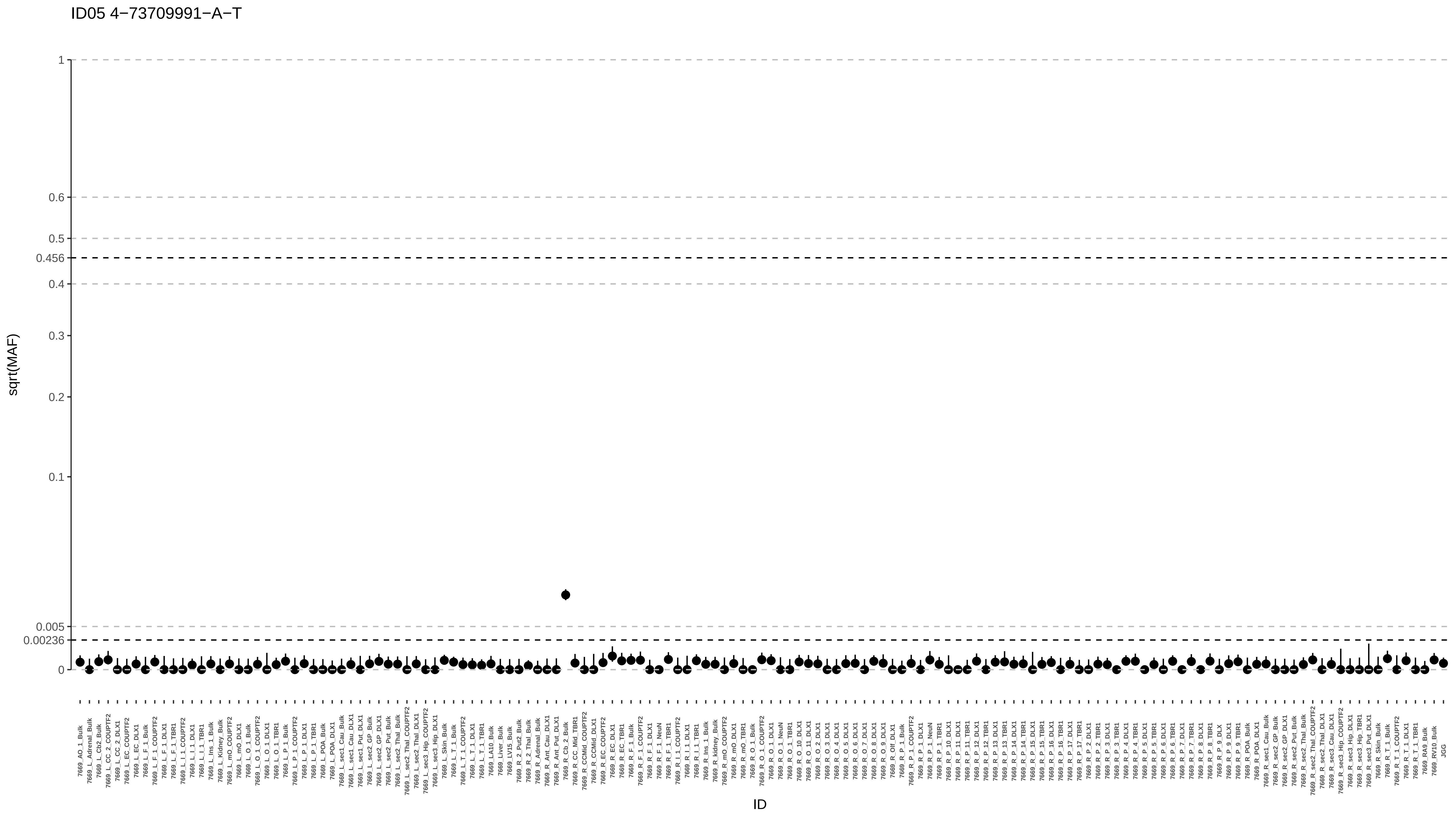
ID05 4−73709991−A−T
sqrt(MAF)
ID
1
0.6
0.5
0.456
0.4
0.3
0.2
0.1
0.005
0.00236
0

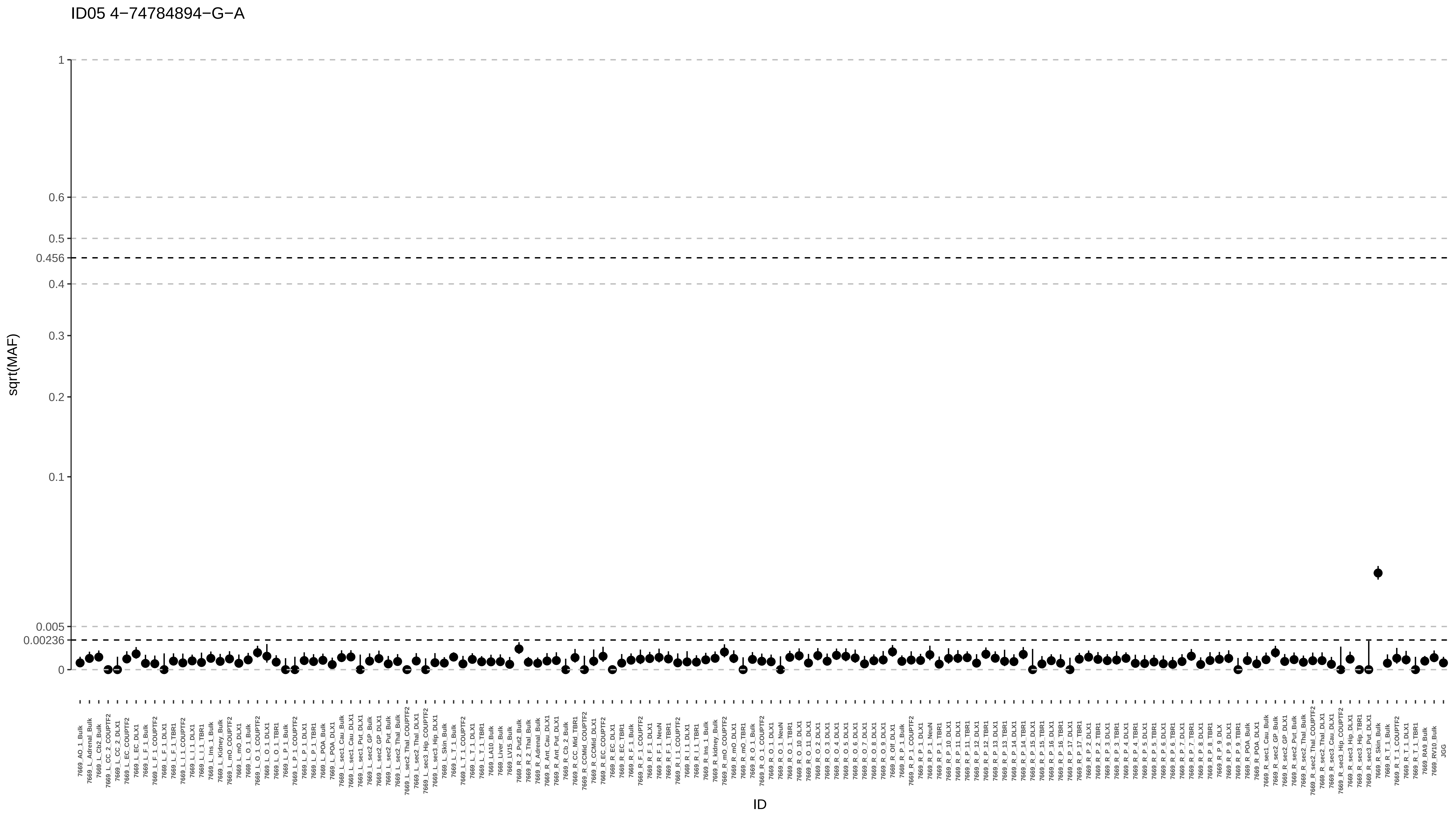
ID05 4−74784894−G−A
sqrt(MAF)
1
0.6
0.5
0.456
0.4
0.3
0.2
0.1
0.005
0.00236
0
ID
7669_AO_1_Bulk
7669_L_Adrenal_Bulk
7669_L_Cb2_Bulk
7669_L_CC_2_COUPTF2
7669_L_CC_2_DLX1
7669_L_EC_COUPTF2
7669_L_EC_DLX1
7669_L_F_1_Bulk
7669_L_F_1_COUPTF2
7669_L_F_1_DLX1
7669_L_F_1_TBR1
7669_L_I_1_COUPTF2
7669_L_I_1_DLX1
7669_L_I_1_TBR1
7669_L_Ins_1_Bulk
7669_L_Kidney_Bulk
7669_L_mO_COUPTF2
7669_L_mO_DLX1
7669_L_O_1_Bulk
7669_L_O_1_COUPTF2
7669_L_O_1_DLX1
7669_L_O_1_TBR1
7669_L_P_1_Bulk
7669_L_P_1_COUPTF2
7669_L_P_1_DLX1
7669_L_P_1_TBR1
7669_L_POA_Bulk
7669_L_POA_DLX1
7669_L_sec1_Cau_Bulk
7669_L_sec1_Cau_DLX1
7669_L_sec1_Put_DLX1
7669_L_sec2_GP_Bulk
7669_L_sec2_GP_DLX1
7669_L_sec2_Put_Bulk
7669_L_sec2_Thal_Bulk
7669_L_sec2_Thal_COUPTF2
7669_L_sec2_Thal_DLX1
7669_L_sec3_Hip_COUPTF2
7669_L_sec3_Hip_DLX1
7669_L_Skin_Bulk
7669_L_T_1_Bulk
7669_L_T_1_COUPTF2
7669_L_T_1_DLX1
7669_L_T_1_TBR1
7669_LA10_Bulk
7669_Liver_Bulk
7669_LV15_Bulk
7669_R_2_Put2_Bulk
7669_R_2_Thal_Bulk
7669_R_Adrenal_Bulk
7669_R_Ant_Cau_DLX1
7669_R_Ant_Put_DLX1
7669_R_Cb_2_Bulk
7669_R_CC_Mid_TBR1
7669_R_CCMid_COUPTF2
7669_R_CCMid_DLX1
7669_R_EC_COUPTF2
7669_R_EC_DLX1
7669_R_EC_TBR1
7669_R_F_1_Bulk
7669_R_F_1_COUPTF2
7669_R_F_1_DLX1
7669_R_F_1_NeuN
7669_R_F_1_TBR1
7669_R_I_1_COUPTF2
7669_R_I_1_DLX1
7669_R_I_1_TBR1
7669_R_Ins_1_Bulk
7669_R_kidney_Bulk
7669_R_mO_COUPTF2
7669_R_mO_DLX1
7669_R_mO_TBR1
7669_R_O_1_Bulk
7669_R_O_1_COUPTF2
7669_R_O_1_DLX1
7669_R_O_1_NeuN
7669_R_O_1_TBR1
7669_R_O_10_DLX1
7669_R_O_11_DLX1
7669_R_O_2_DLX1
7669_R_O_3_DLX1
7669_R_O_4_DLX1
7669_R_O_5_DLX1
7669_R_O_6_DLX1
7669_R_O_7_DLX1
7669_R_O_8_DLX1
7669_R_O_9_DLX1
7669_R_Olf_DLX1
7669_R_P_1_Bulk
7669_R_P_1_COUPTF2
7669_R_P_1_DLX1
7669_R_P_1_NeuN
7669_R_P_1_TBR1
7669_R_P_10_DLX1
7669_R_P_11_DLX1
7669_R_P_11_TBR1
7669_R_P_12_DLX1
7669_R_P_12_TBR1
7669_R_P_13_DLX1
7669_R_P_13_TBR1
7669_R_P_14_DLX1
7669_R_P_14_TBR1
7669_R_P_15_DLX1
7669_R_P_15_TBR1
7669_R_P_16_DLX1
7669_R_P_16_TBR1
7669_R_P_17_DLX1
7669_R_P_17_TBR1
7669_R_P_2_DLX1
7669_R_P_2_TBR1
7669_R_P_3_DLX1
7669_R_P_3_TBR1
7669_R_P_4_DLX1
7669_R_P_4_TBR1
7669_R_P_5_DLX1
7669_R_P_5_TBR1
7669_R_P_6_DLX1
7669_R_P_6_TBR1
7669_R_P_7_DLX1
7669_R_P_7_TBR1
7669_R_P_8_DLX1
7669_R_P_8_TBR1
7669_R_P_9_DLX
7669_R_P_9_DLX1
7669_R_P_9_TBR1
7669_R_POA_Bulk
7669_R_POA_DLX1
7669_R_sec1_Cau_Bulk
7669_R_sec2_GP_Bulk
7669_R_sec2_GP_DLX1
7669_R_sec2_Put_Bulk
7669_R_sec2_Thal_Bulk
7669_R_sec2_Thal_COUPTF2
7669_R_sec2_Thal_DLX1
7669_R_sec3_Cau_DLX1
7669_R_sec3_Hip_COUPTF2
7669_R_sec3_Hip_DLX1
7669_R_sec3_Hip_TBR1
7669_R_sec3_Put_DLX1
7669_R_Skin_Bulk
7669_R_T_1_Bulk
7669_R_T_1_COUPTF2
7669_R_T_1_DLX1
7669_R_T_1_TBR1
7669_RA9_Bulk
7669_RV10_Bulk
JGG

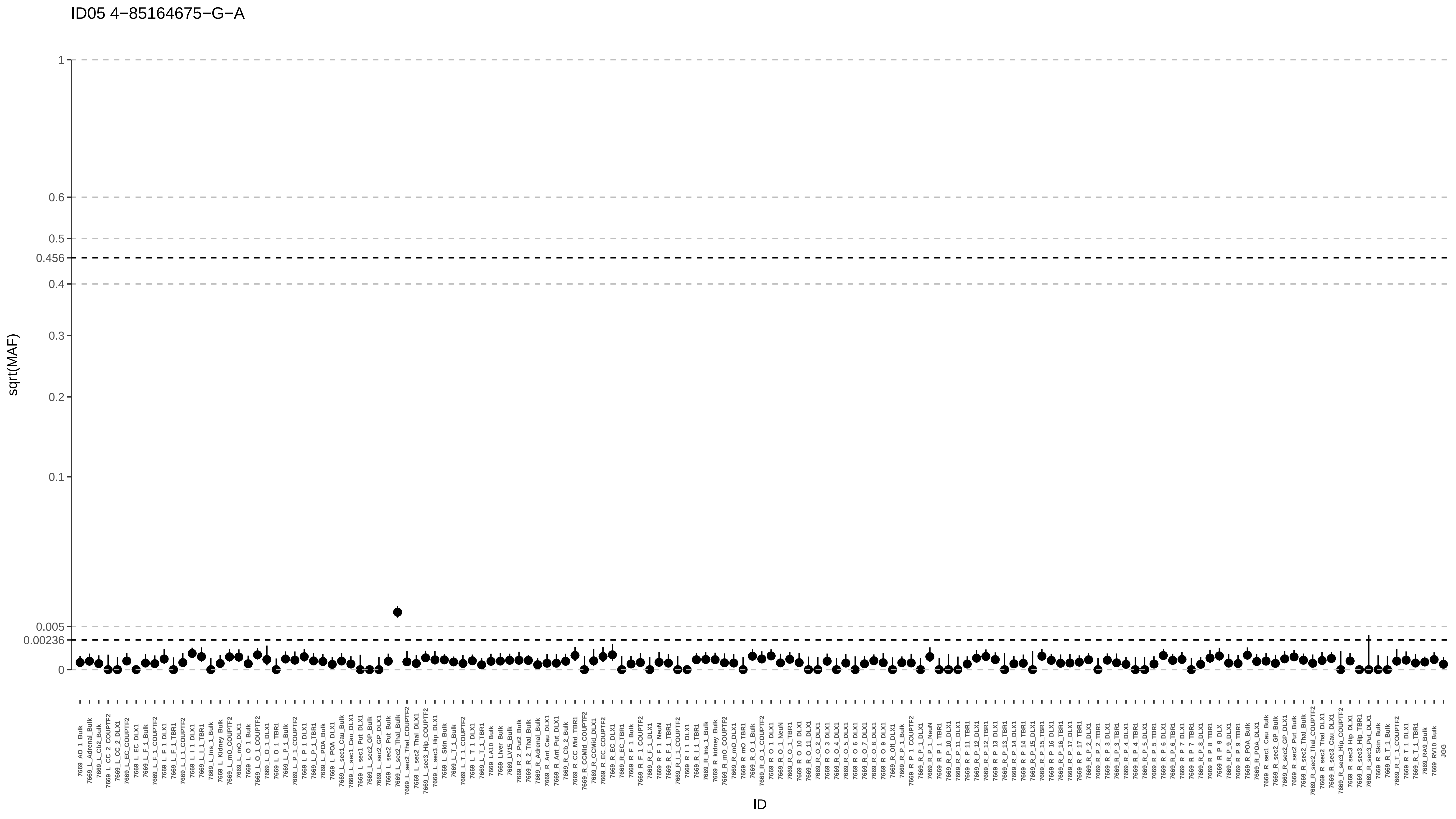
ID05 4−85164675−G−A
sqrt(MAF)
1
0.6
0.5
0.456
0.4
0.3
0.2
0.1
0.005
0.00236
0
ID

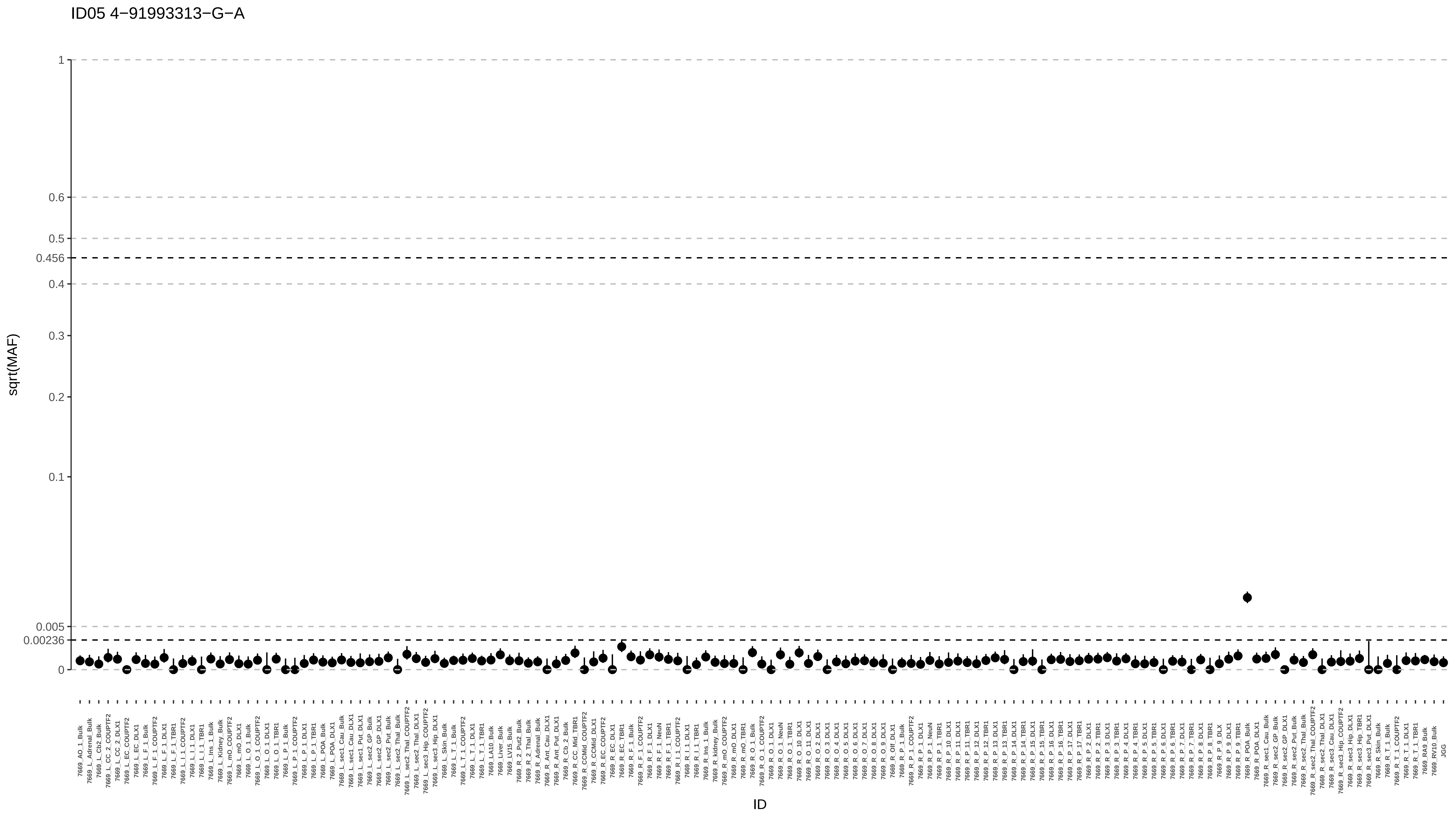
ID05 4−91993313−G−A
sqrt(MAF)
1
0.6
0.5
0.456
0.4
0.3
0.2
0.1
0.005
0.00236
0
ID
7669_AO_1_Bulk
7669_L_Adrenal_Bulk
7669_L_Cb2_Bulk
7669_L_CC_2_COUPTF2
7669_L_CC_2_DLX1
7669_L_EC_COUPTF2
7669_L_EC_DLX1
7669_L_F_1_Bulk
7669_L_F_1_COUPTF2
7669_L_F_1_DLX1
7669_L_F_1_TBR1
7669_L_I_1_COUPTF2
7669_L_I_1_DLX1
7669_L_I_1_TBR1
7669_L_Ins_1_Bulk
7669_L_Kidney_Bulk
7669_L_mO_COUPTF2
7669_L_mO_DLX1
7669_L_O_1_Bulk
7669_L_O_1_COUPTF2
7669_L_O_1_DLX1
7669_L_O_1_TBR1
7669_L_P_1_Bulk
7669_L_P_1_COUPTF2
7669_L_P_1_DLX1
7669_L_P_1_TBR1
7669_L_POA_Bulk
7669_L_POA_DLX1
7669_L_sec1_Cau_Bulk
7669_L_sec1_Cau_DLX1
7669_L_sec1_Put_DLX1
7669_L_sec2_GP_Bulk
7669_L_sec2_GP_DLX1
7669_L_sec2_Put_Bulk
7669_L_sec2_Thal_Bulk
7669_L_sec2_Thal_COUPTF2
7669_L_sec2_Thal_DLX1
7669_L_sec3_Hip_COUPTF2
7669_L_sec3_Hip_DLX1
7669_L_Skin_Bulk
7669_L_T_1_Bulk
7669_L_T_1_COUPTF2
7669_L_T_1_DLX1
7669_L_T_1_TBR1
7669_LA10_Bulk
7669_Liver_Bulk
7669_LV15_Bulk
7669_R_2_Put2_Bulk
7669_R_2_Thal_Bulk
7669_R_Adrenal_Bulk
7669_R_Ant_Cau_DLX1
7669_R_Ant_Put_DLX1
7669_R_Cb_2_Bulk
7669_R_CC_Mid_TBR1
7669_R_CCMid_COUPTF2
7669_R_CCMid_DLX1
7669_R_EC_COUPTF2
7669_R_EC_DLX1
7669_R_EC_TBR1
7669_R_F_1_Bulk
7669_R_F_1_COUPTF2
7669_R_F_1_DLX1
7669_R_F_1_NeuN
7669_R_F_1_TBR1
7669_R_I_1_COUPTF2
7669_R_I_1_DLX1
7669_R_I_1_TBR1
7669_R_Ins_1_Bulk
7669_R_kidney_Bulk
7669_R_mO_COUPTF2
7669_R_mO_DLX1
7669_R_mO_TBR1
7669_R_O_1_Bulk
7669_R_O_1_COUPTF2
7669_R_O_1_DLX1
7669_R_O_1_NeuN
7669_R_O_1_TBR1
7669_R_O_10_DLX1
7669_R_O_11_DLX1
7669_R_O_2_DLX1
7669_R_O_3_DLX1
7669_R_O_4_DLX1
7669_R_O_5_DLX1
7669_R_O_6_DLX1
7669_R_O_7_DLX1
7669_R_O_8_DLX1
7669_R_O_9_DLX1
7669_R_Olf_DLX1
7669_R_P_1_Bulk
7669_R_P_1_COUPTF2
7669_R_P_1_DLX1
7669_R_P_1_NeuN
7669_R_P_1_TBR1
7669_R_P_10_DLX1
7669_R_P_11_DLX1
7669_R_P_11_TBR1
7669_R_P_12_DLX1
7669_R_P_12_TBR1
7669_R_P_13_DLX1
7669_R_P_13_TBR1
7669_R_P_14_DLX1
7669_R_P_14_TBR1
7669_R_P_15_DLX1
7669_R_P_15_TBR1
7669_R_P_16_DLX1
7669_R_P_16_TBR1
7669_R_P_17_DLX1
7669_R_P_17_TBR1
7669_R_P_2_DLX1
7669_R_P_2_TBR1
7669_R_P_3_DLX1
7669_R_P_3_TBR1
7669_R_P_4_DLX1
7669_R_P_4_TBR1
7669_R_P_5_DLX1
7669_R_P_5_TBR1
7669_R_P_6_DLX1
7669_R_P_6_TBR1
7669_R_P_7_DLX1
7669_R_P_7_TBR1
7669_R_P_8_DLX1
7669_R_P_8_TBR1
7669_R_P_9_DLX
7669_R_P_9_DLX1
7669_R_P_9_TBR1
7669_R_POA_Bulk
7669_R_POA_DLX1
7669_R_sec1_Cau_Bulk
7669_R_sec2_GP_Bulk
7669_R_sec2_GP_DLX1
7669_R_sec2_Put_Bulk
7669_R_sec2_Thal_Bulk
7669_R_sec2_Thal_COUPTF2
7669_R_sec2_Thal_DLX1
7669_R_sec3_Cau_DLX1
7669_R_sec3_Hip_COUPTF2
7669_R_sec3_Hip_DLX1
7669_R_sec3_Hip_TBR1
7669_R_sec3_Put_DLX1
7669_R_Skin_Bulk
7669_R_T_1_Bulk
7669_R_T_1_COUPTF2
7669_R_T_1_DLX1
7669_R_T_1_TBR1
7669_RA9_Bulk
7669_RV10_Bulk
JGG

ID05 4−92330313−C−G

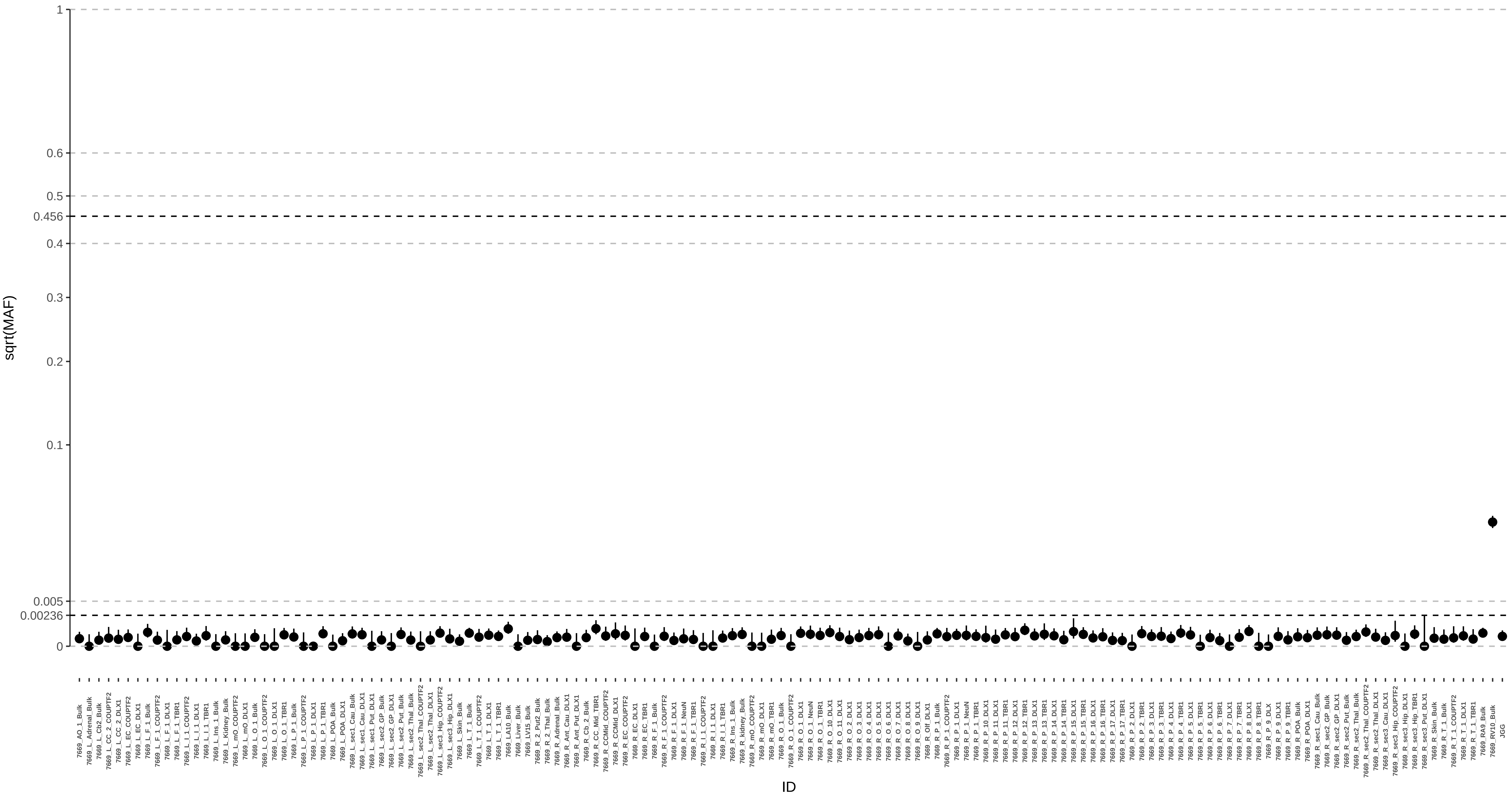

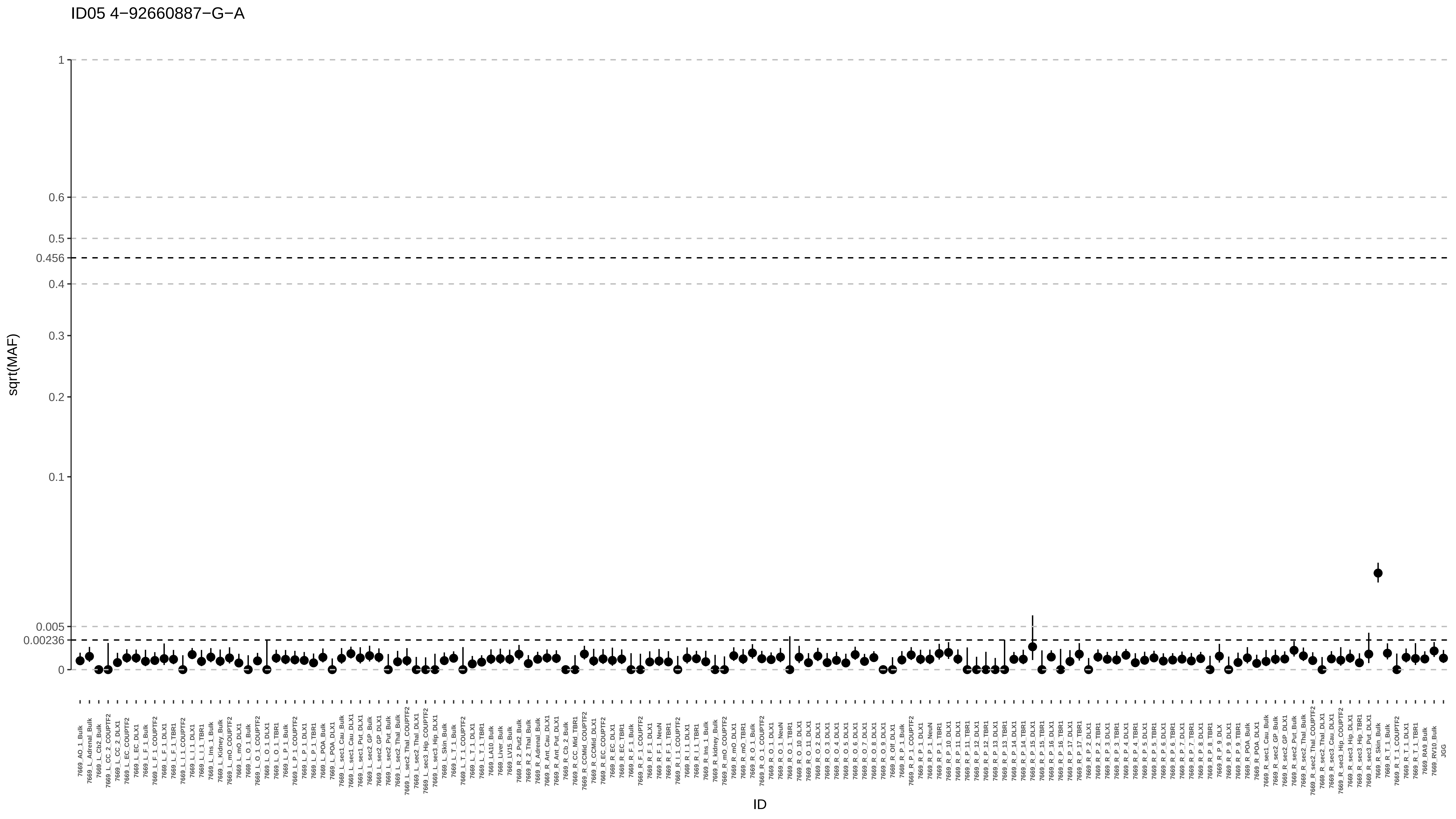

ID05 4−92660887−G−A
sqrt(MAF)
ID
1
0.6
0.5
0.456
0.4
0.3
0.2
0.1
0.005
0.00236
0

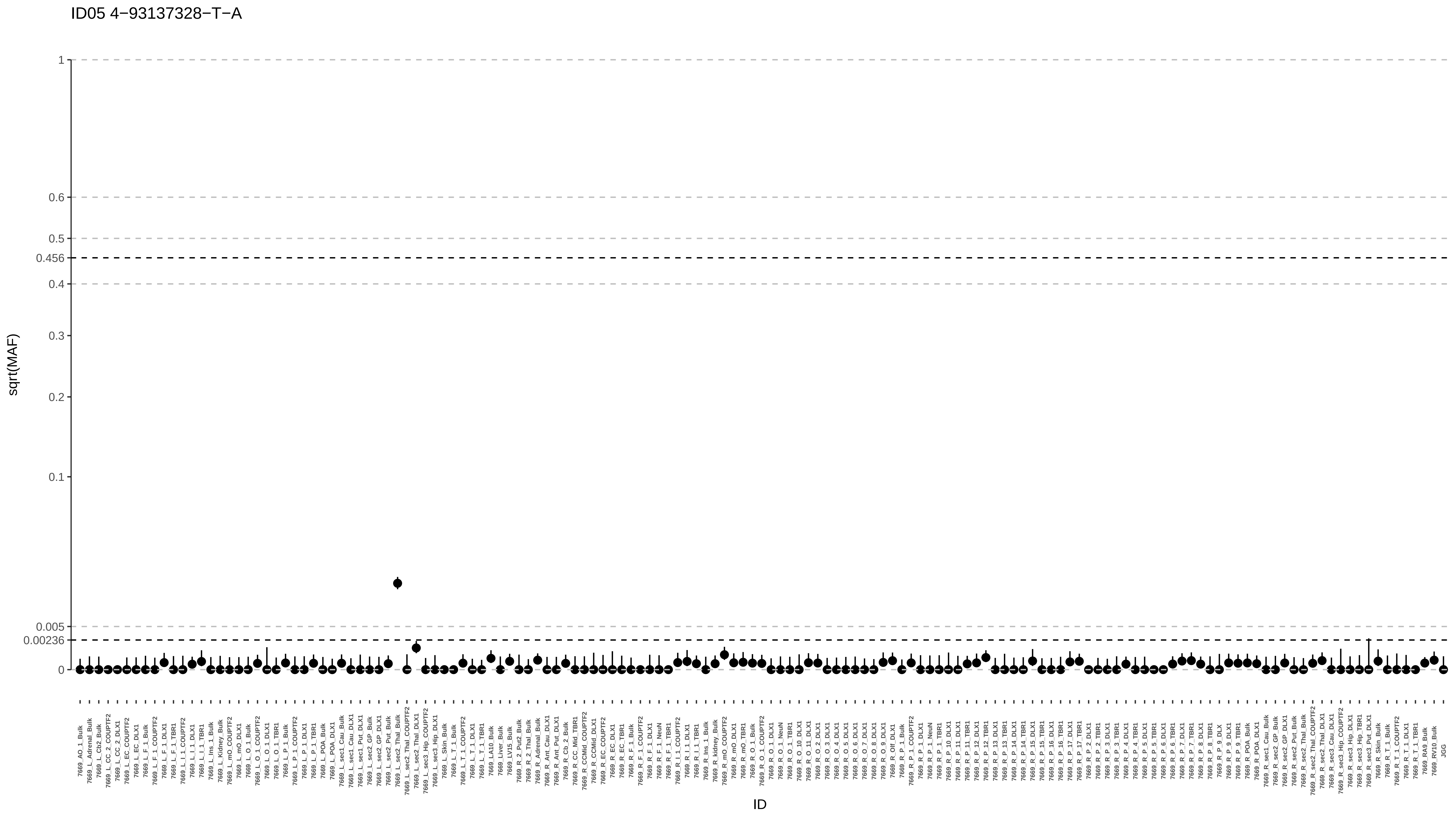
ID05 4−93137328−T−A
sqrt(MAF)
ID
1
0.6
0.5
0.456
0.4
0.3
0.2
0.1
0.005
0.00236
0

ID05 4−94277150−C−T

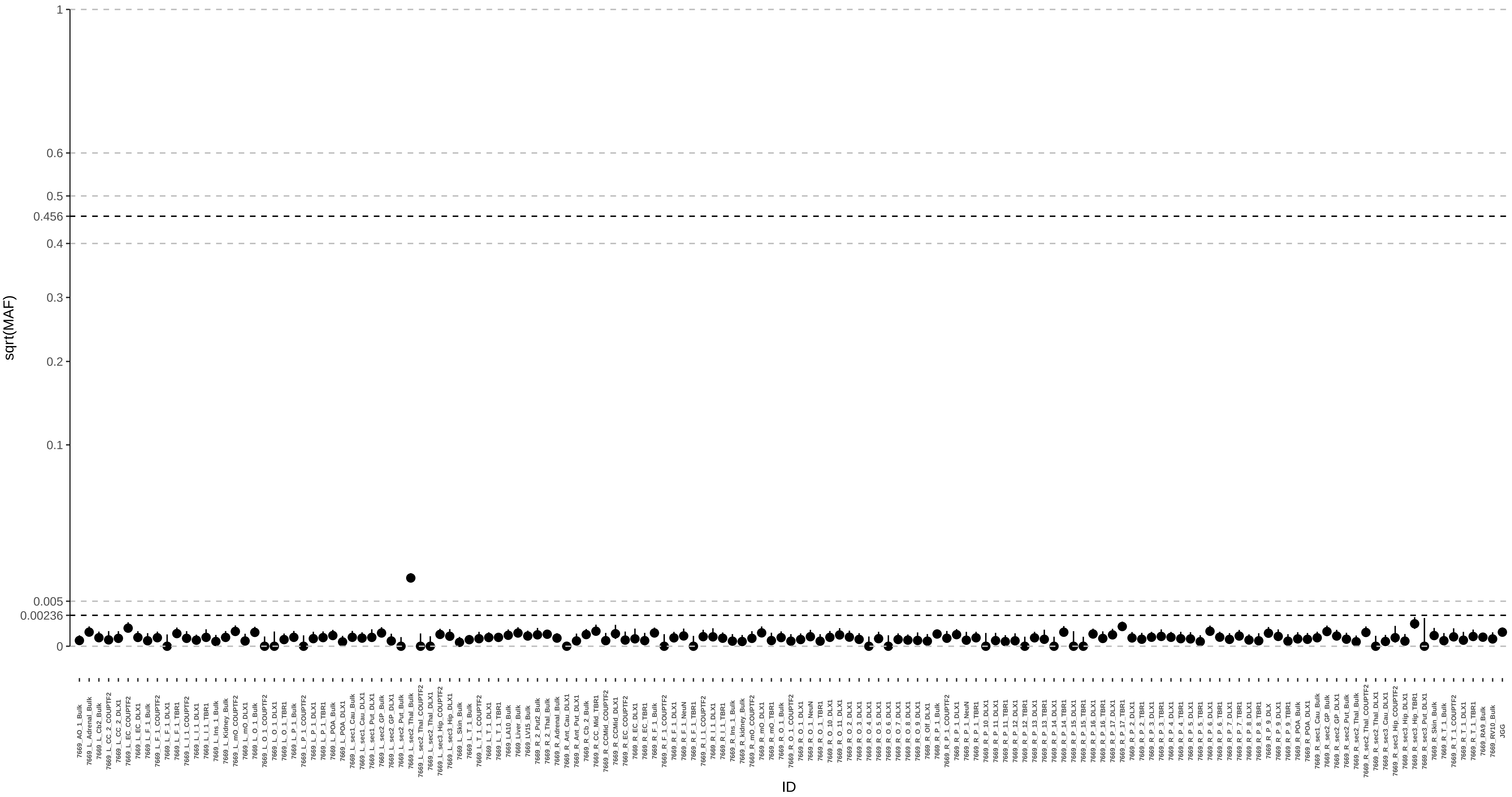

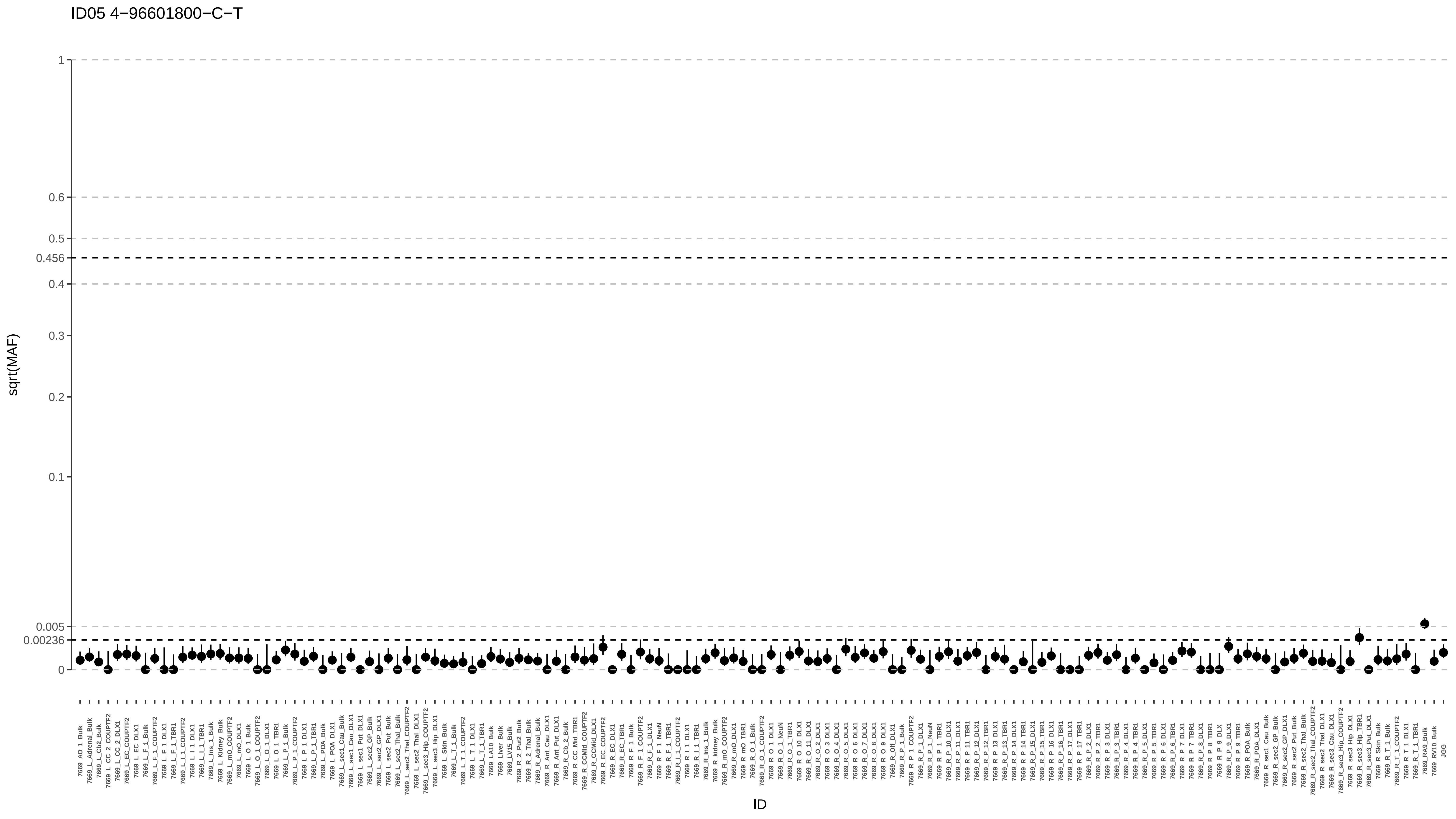
ID05 4−96601800−C−T
sqrt(MAF)
1
0.6
0.5
0.456
0.4
0.3
0.2
0.1
0.005
0.00236
0
ID

ID05 5−10808399−G−A

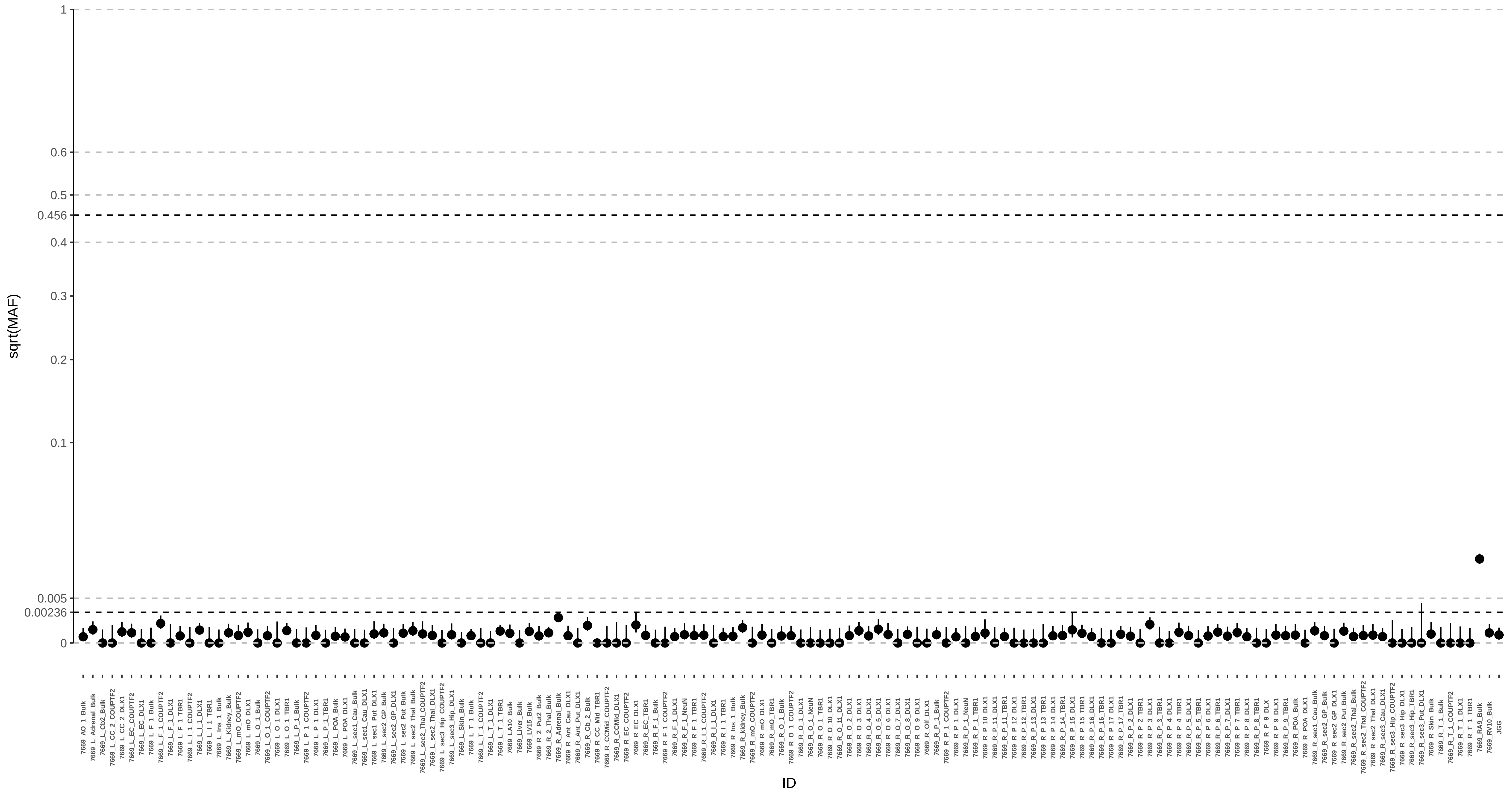

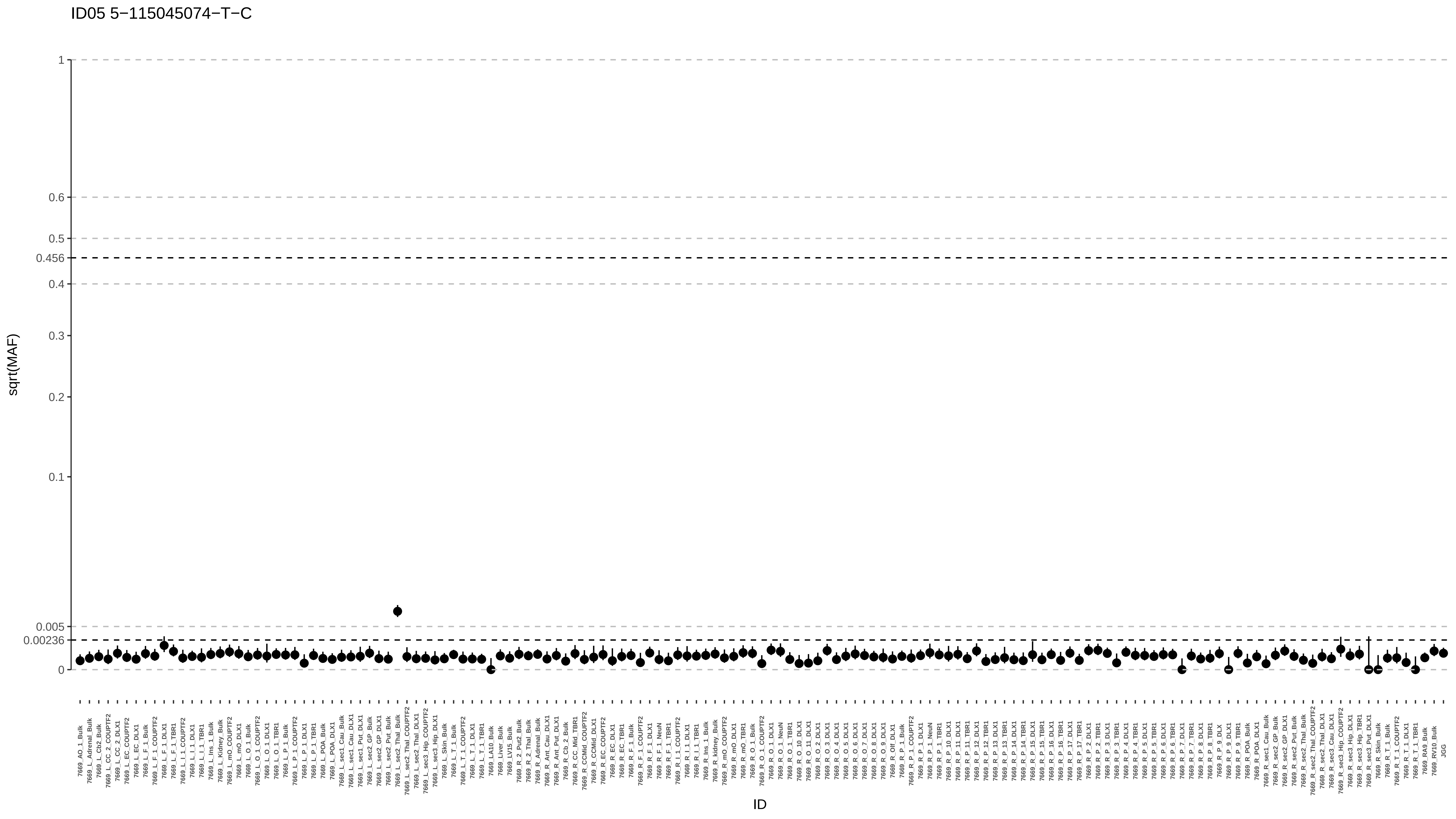
ID05 5−115045074−T−C
sqrt(MAF)
1
0.6
0.5
0.456
0.4
0.3
0.2
0.1
0.005
0.00236
0
ID

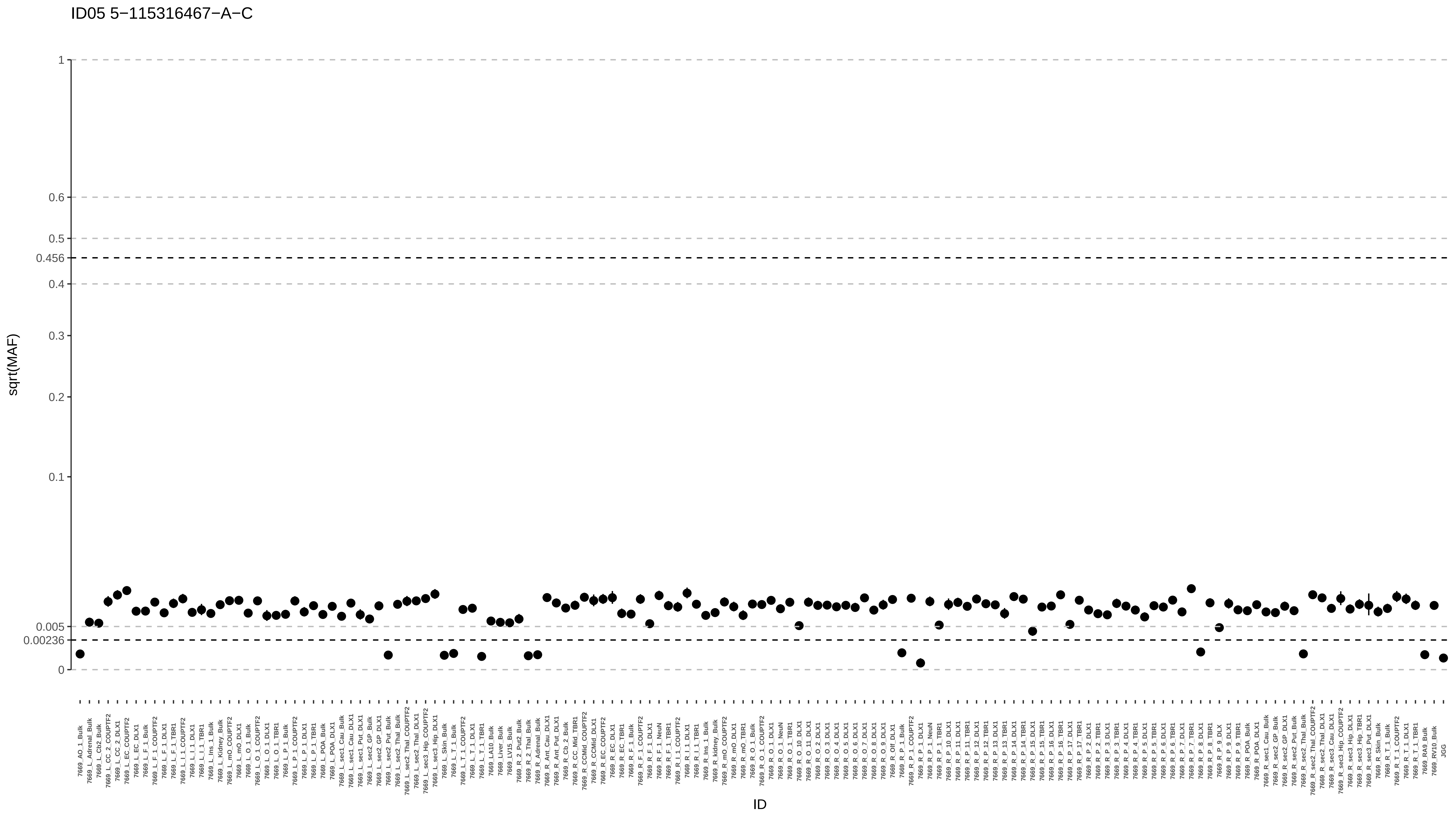
ID05 5−115316467−A−C
sqrt(MAF)
1
0.6
0.5
0.456
0.4
0.3
0.2
0.1
0.005
0.00236
0
ID

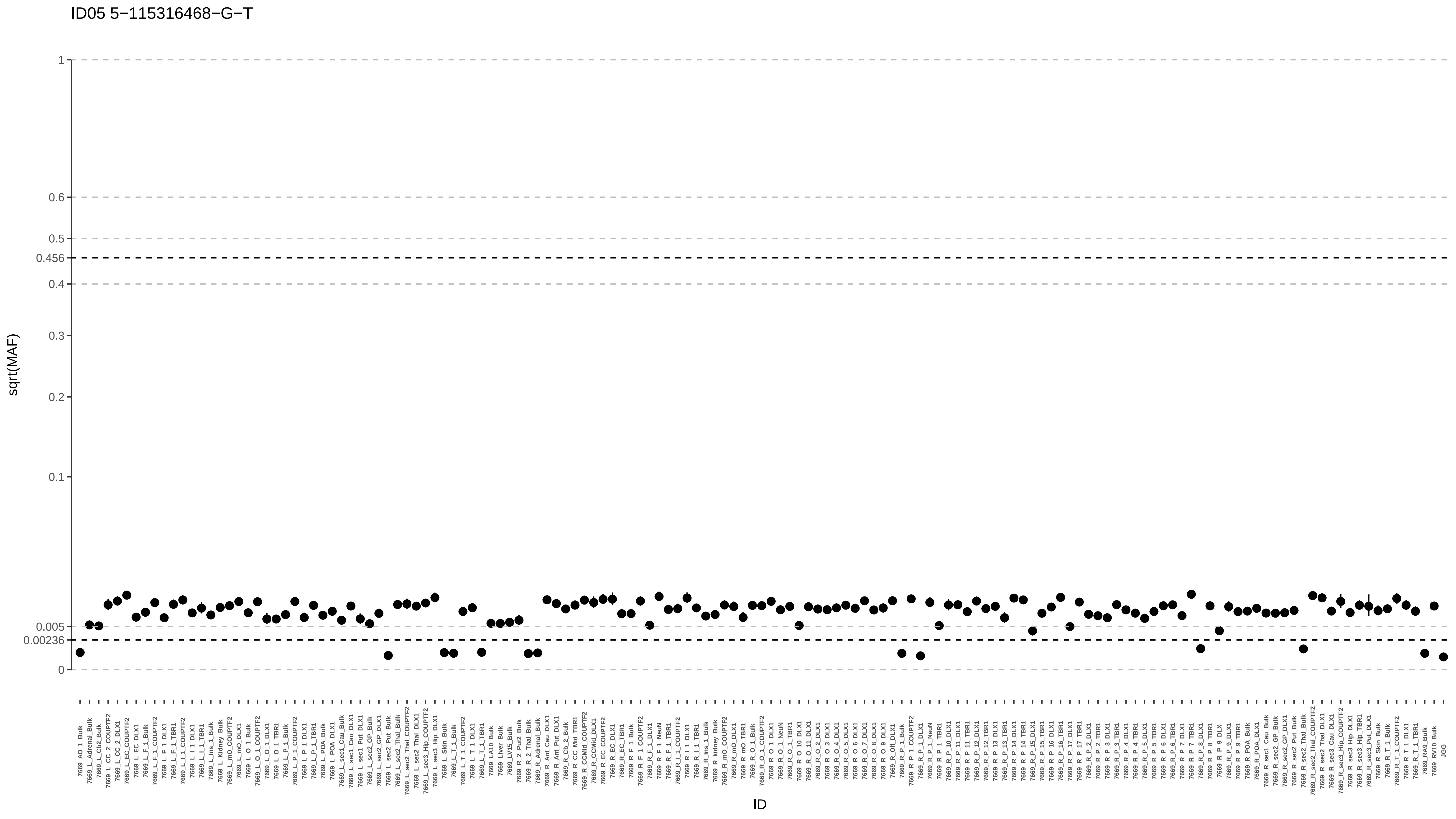
ID05 5−115316468−G−T
sqrt(MAF)
ID
1
0.6
0.5
0.456
0.4
0.3
0.2
0.1
0.005
0.00236
0

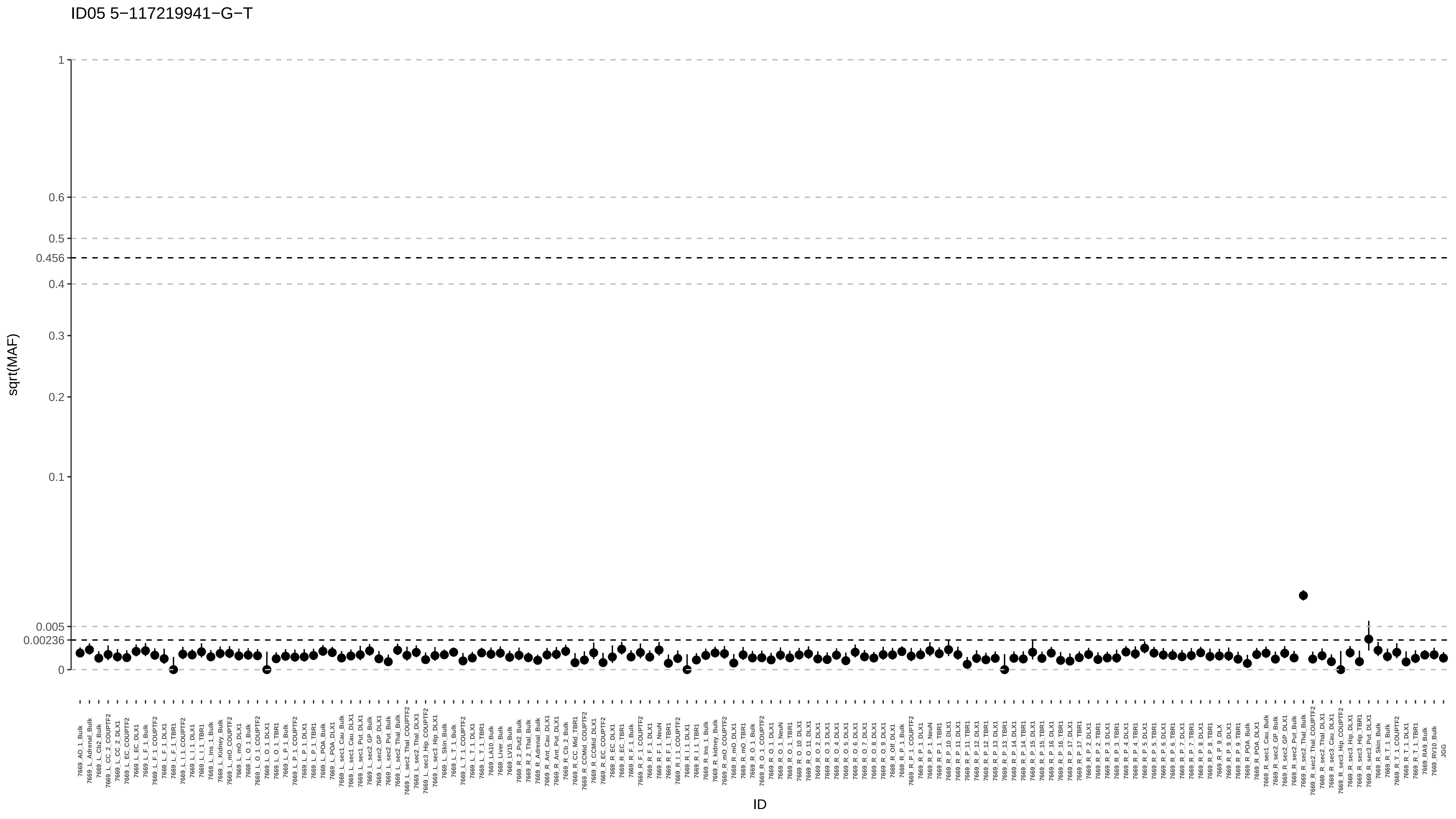
ID05 5−117219941−G−T
sqrt(MAF)
1
0.6
0.5
0.456
0.4
0.3
0.2
0.1
0.005
0.00236
0
ID

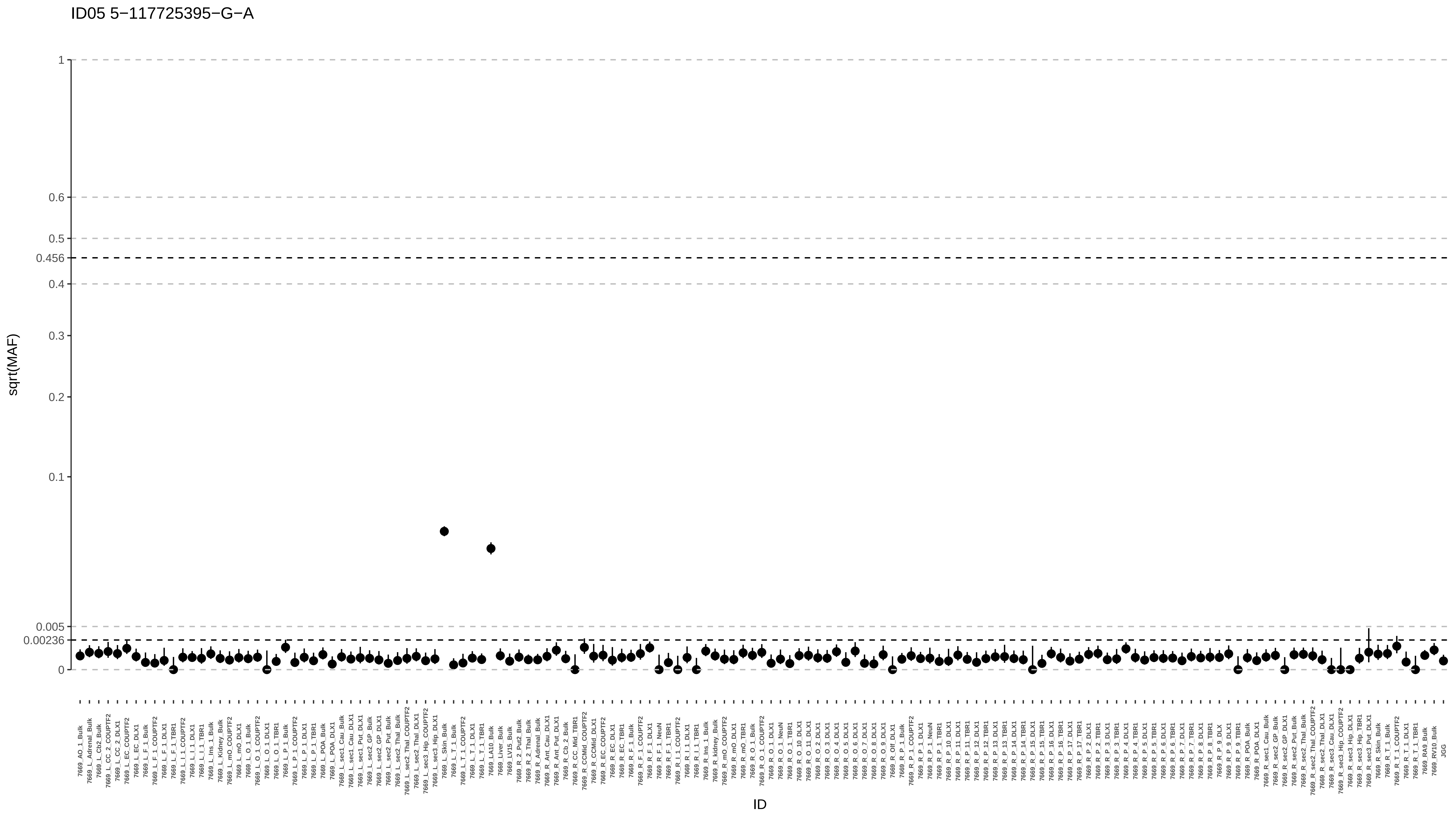
ID05 5−117725395−G−A
sqrt(MAF)
ID
1
0.6
0.5
0.456
0.4
0.3
0.2
0.1
0.005
0.00236
0

ID05 5−121446225−C−T

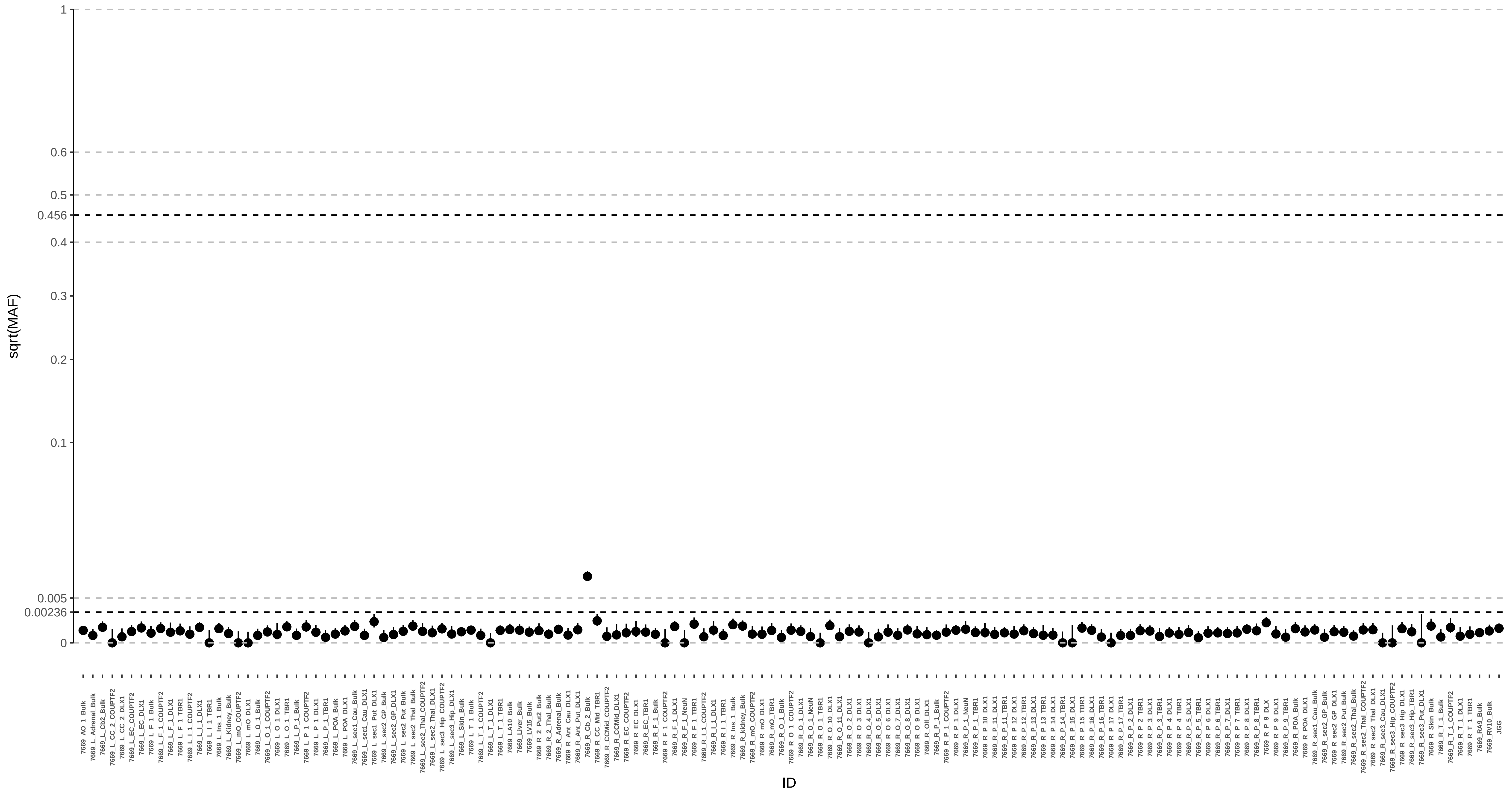

ID05 5−121518814−G−A

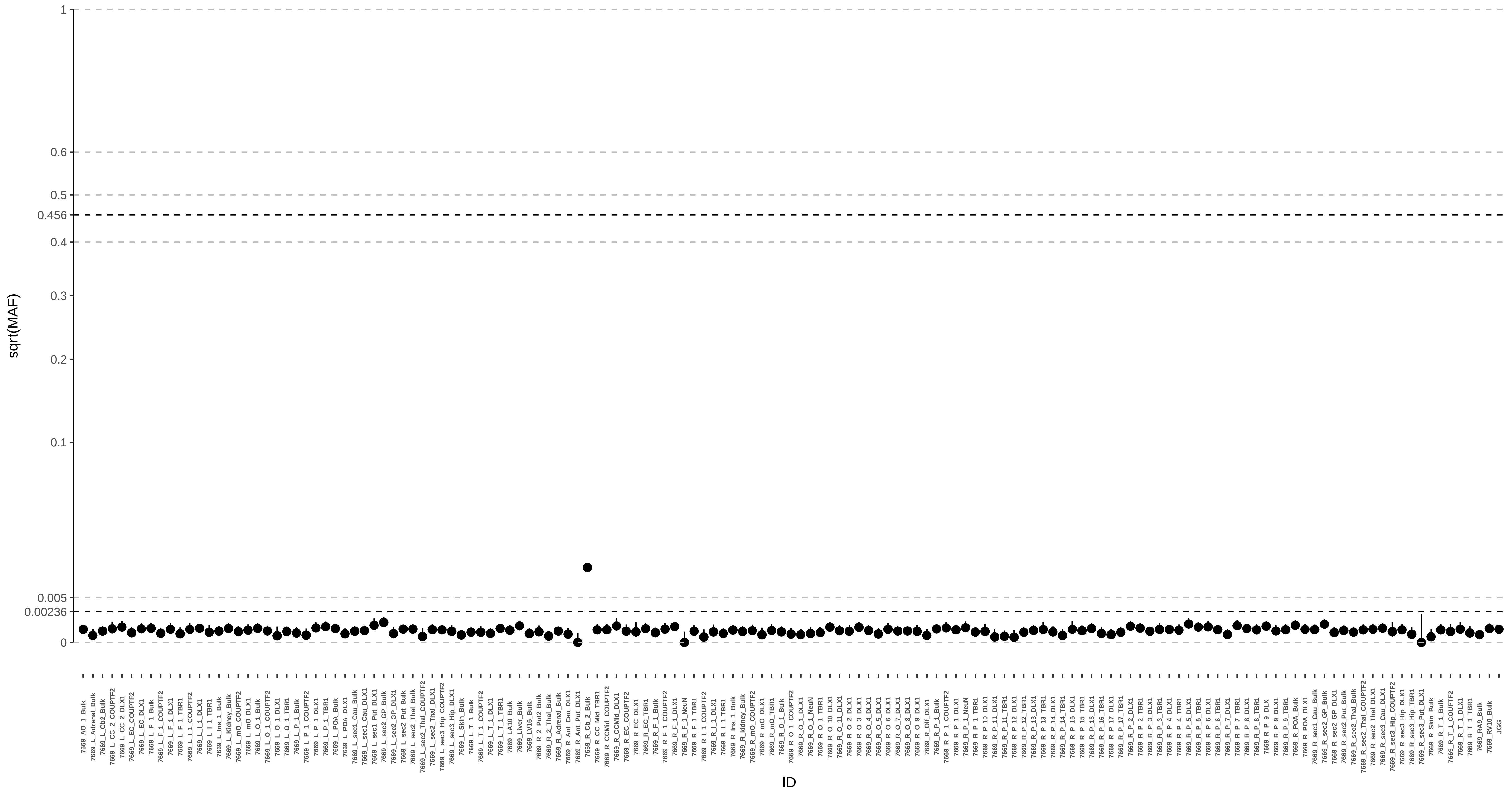

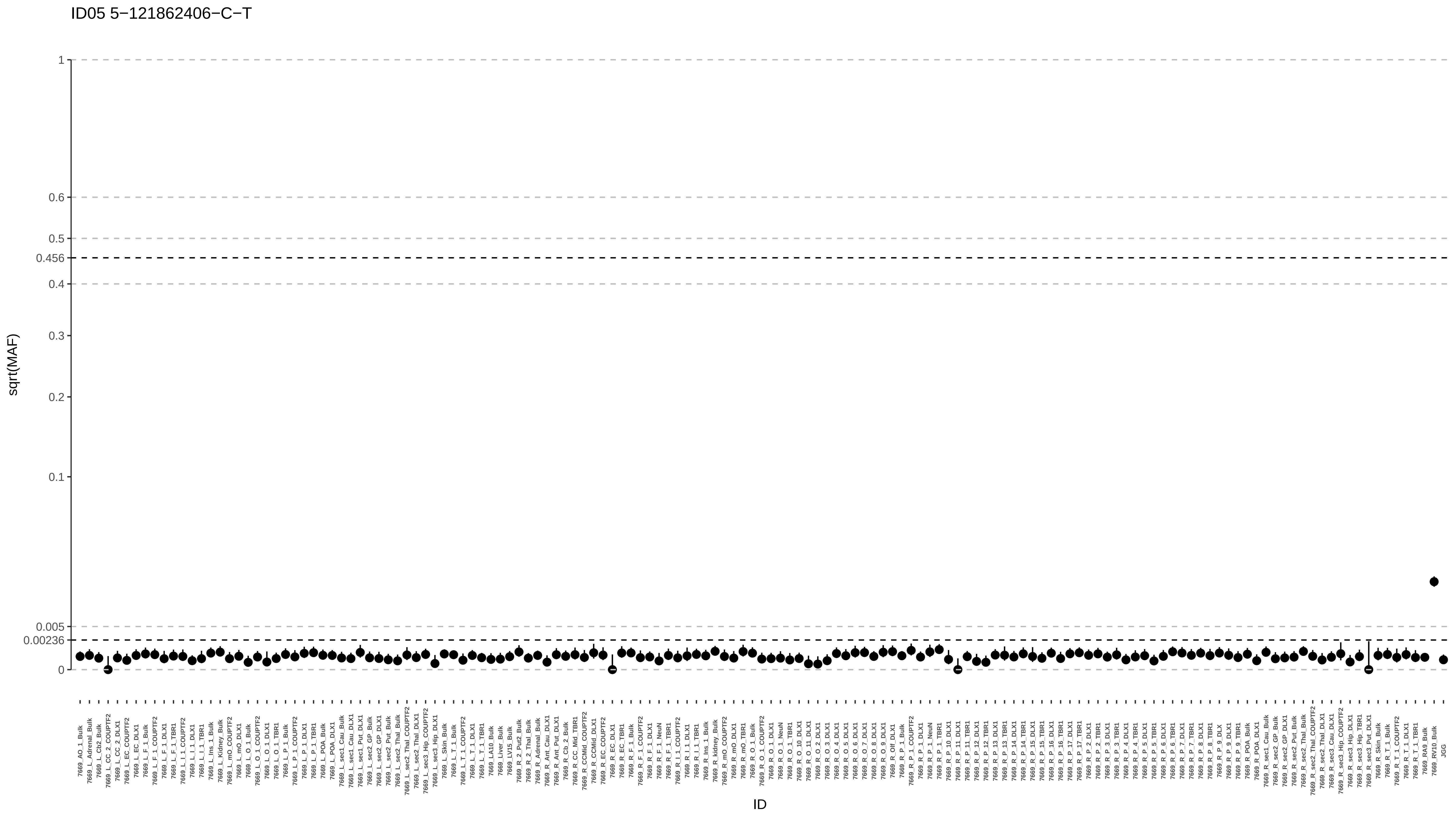
ID05 5−121862406−C−T
sqrt(MAF)
1
0.6
0.5
0.456
0.4
0.3
0.2
0.1
0.005
0.00236
0
ID
7669_AO_1_Bulk
7669_L_Adrenal_Bulk
7669_L_Cb2_Bulk
7669_L_CC_2_COUPTF2
7669_L_CC_2_DLX1
7669_L_EC_COUPTF2
7669_L_EC_DLX1
7669_L_F_1_Bulk
7669_L_F_1_COUPTF2
7669_L_F_1_DLX1
7669_L_F_1_TBR1
7669_L_I_1_COUPTF2
7669_L_I_1_DLX1
7669_L_I_1_TBR1
7669_L_Ins_1_Bulk
7669_L_Kidney_Bulk
7669_L_mO_COUPTF2
7669_L_mO_DLX1
7669_L_O_1_Bulk
7669_L_O_1_COUPTF2
7669_L_O_1_DLX1
7669_L_O_1_TBR1
7669_L_P_1_Bulk
7669_L_P_1_COUPTF2
7669_L_P_1_DLX1
7669_L_P_1_TBR1
7669_L_POA_Bulk
7669_L_POA_DLX1
7669_L_sec1_Cau_Bulk
7669_L_sec1_Cau_DLX1
7669_L_sec1_Put_DLX1
7669_L_sec2_GP_Bulk
7669_L_sec2_GP_DLX1
7669_L_sec2_Put_Bulk
7669_L_sec2_Thal_Bulk
7669_L_sec2_Thal_COUPTF2
7669_L_sec2_Thal_DLX1
7669_L_sec3_Hip_COUPTF2
7669_L_sec3_Hip_DLX1
7669_L_Skin_Bulk
7669_L_T_1_Bulk
7669_L_T_1_COUPTF2
7669_L_T_1_DLX1
7669_L_T_1_TBR1
7669_LA10_Bulk
7669_Liver_Bulk
7669_LV15_Bulk
7669_R_2_Put2_Bulk
7669_R_2_Thal_Bulk
7669_R_Adrenal_Bulk
7669_R_Ant_Cau_DLX1
7669_R_Ant_Put_DLX1
7669_R_Cb_2_Bulk
7669_R_CC_Mid_TBR1
7669_R_CCMid_COUPTF2
7669_R_CCMid_DLX1
7669_R_EC_COUPTF2
7669_R_EC_DLX1
7669_R_EC_TBR1
7669_R_F_1_Bulk
7669_R_F_1_COUPTF2
7669_R_F_1_DLX1
7669_R_F_1_NeuN
7669_R_F_1_TBR1
7669_R_I_1_COUPTF2
7669_R_I_1_DLX1
7669_R_I_1_TBR1
7669_R_Ins_1_Bulk
7669_R_kidney_Bulk
7669_R_mO_COUPTF2
7669_R_mO_DLX1
7669_R_mO_TBR1
7669_R_O_1_Bulk
7669_R_O_1_COUPTF2
7669_R_O_1_DLX1
7669_R_O_1_NeuN
7669_R_O_1_TBR1
7669_R_O_10_DLX1
7669_R_O_11_DLX1
7669_R_O_2_DLX1
7669_R_O_3_DLX1
7669_R_O_4_DLX1
7669_R_O_5_DLX1
7669_R_O_6_DLX1
7669_R_O_7_DLX1
7669_R_O_8_DLX1
7669_R_O_9_DLX1
7669_R_Olf_DLX1
7669_R_P_1_Bulk
7669_R_P_1_COUPTF2
7669_R_P_1_DLX1
7669_R_P_1_NeuN
7669_R_P_1_TBR1
7669_R_P_10_DLX1
7669_R_P_11_DLX1
7669_R_P_11_TBR1
7669_R_P_12_DLX1
7669_R_P_12_TBR1
7669_R_P_13_DLX1
7669_R_P_13_TBR1
7669_R_P_14_DLX1
7669_R_P_14_TBR1
7669_R_P_15_DLX1
7669_R_P_15_TBR1
7669_R_P_16_DLX1
7669_R_P_16_TBR1
7669_R_P_17_DLX1
7669_R_P_17_TBR1
7669_R_P_2_DLX1
7669_R_P_2_TBR1
7669_R_P_3_DLX1
7669_R_P_3_TBR1
7669_R_P_4_DLX1
7669_R_P_4_TBR1
7669_R_P_5_DLX1
7669_R_P_5_TBR1
7669_R_P_6_DLX1
7669_R_P_6_TBR1
7669_R_P_7_DLX1
7669_R_P_7_TBR1
7669_R_P_8_DLX1
7669_R_P_8_TBR1
7669_R_P_9_DLX
7669_R_P_9_DLX1
7669_R_P_9_TBR1
7669_R_POA_Bulk
7669_R_POA_DLX1
7669_R_sec1_Cau_Bulk
7669_R_sec2_GP_Bulk
7669_R_sec2_GP_DLX1
7669_R_sec2_Put_Bulk
7669_R_sec2_Thal_Bulk
7669_R_sec2_Thal_COUPTF2
7669_R_sec2_Thal_DLX1
7669_R_sec3_Cau_DLX1
7669_R_sec3_Hip_COUPTF2
7669_R_sec3_Hip_DLX1
7669_R_sec3_Hip_TBR1
7669_R_sec3_Put_DLX1
7669_R_Skin_Bulk
7669_R_T_1_Bulk
7669_R_T_1_COUPTF2
7669_R_T_1_DLX1
7669_R_T_1_TBR1
7669_RA9_Bulk
7669_RV10_Bulk
JGG

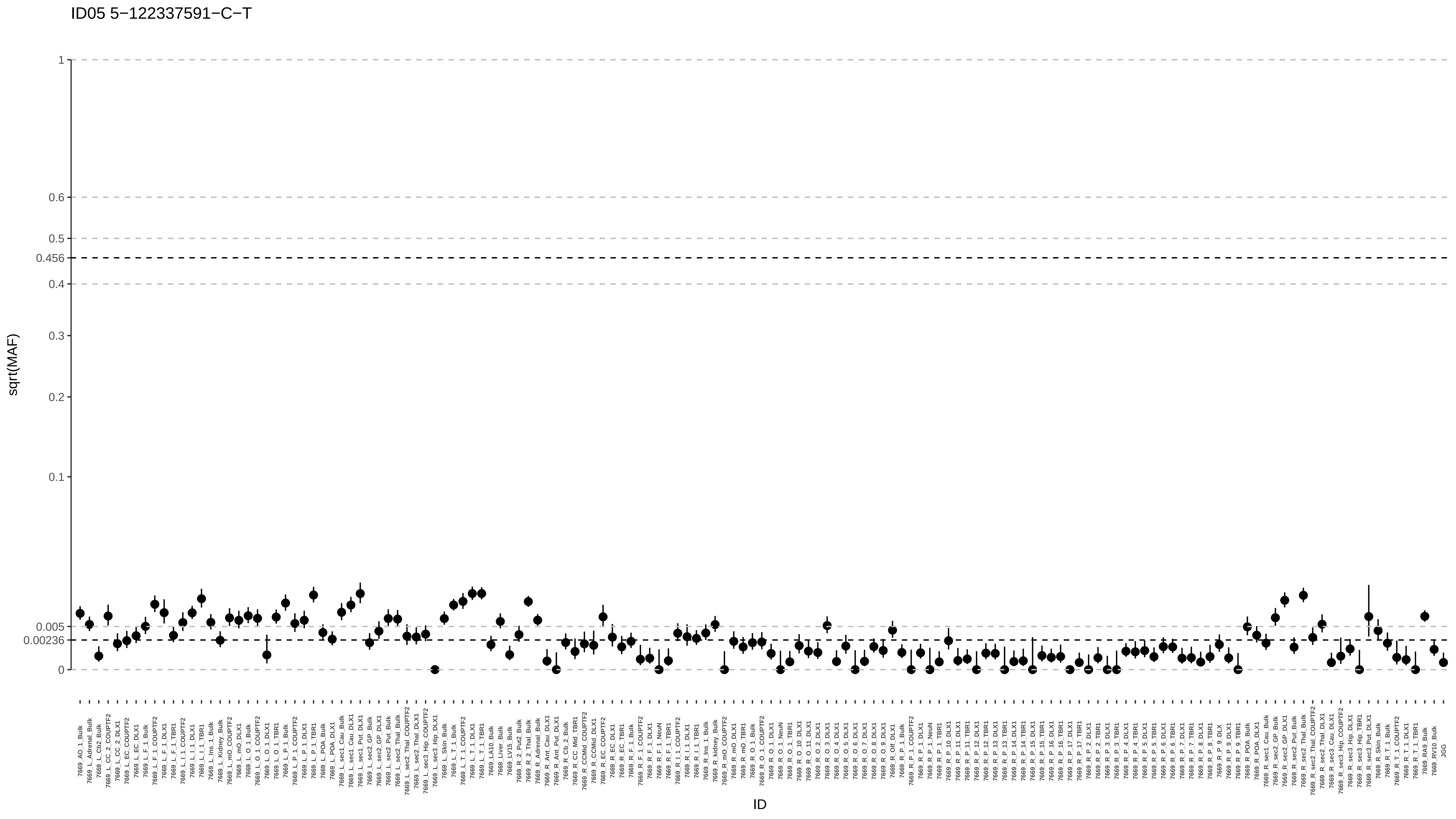
ID05 5−122337591−C−T
sqrt(MAF)
ID
1
0.6
0.5
0.456
0.4
0.3
0.2
0.1
0.005
0.00236
0

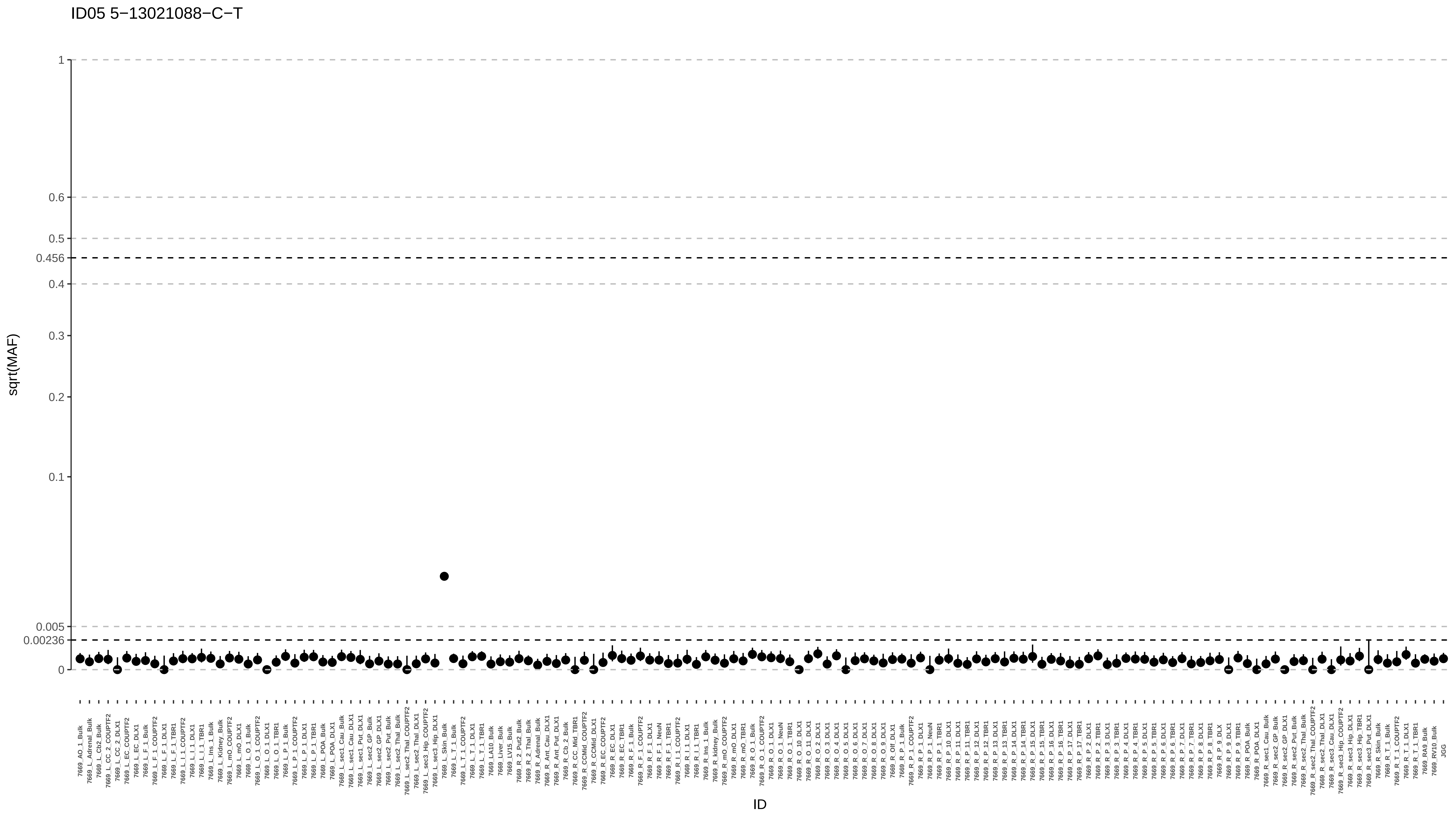
ID05 5−13021088−C−T
sqrt(MAF)
1
0.6
0.5
0.456
0.4
0.3
0.2
0.1
0.005
0.00236
0
ID

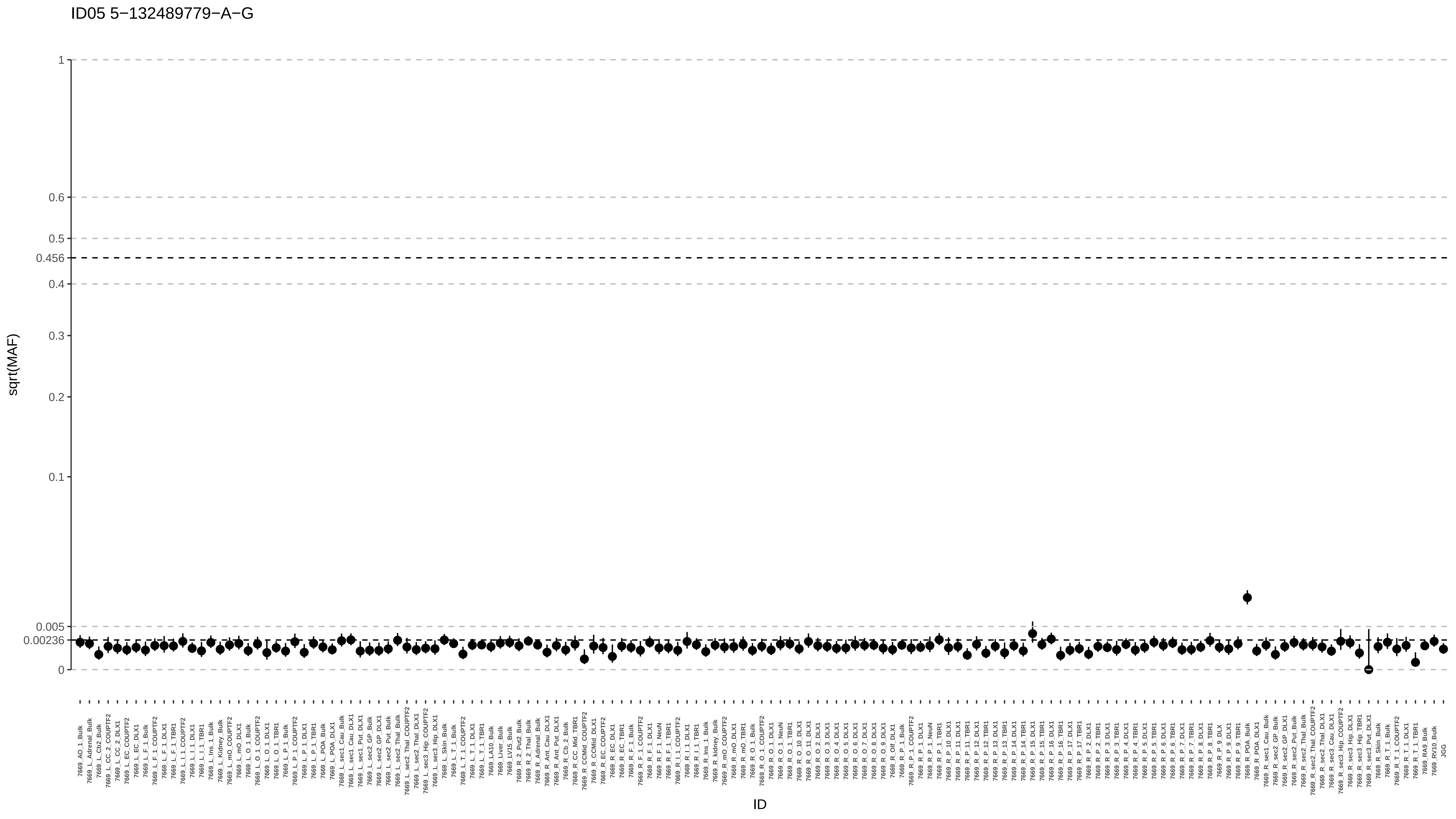
ID05 5−132489779−A−G
sqrt(MAF)
ID
1
0.6
0.5
0.456
0.4
0.3
0.2
0.1
0.005
0.00236
0

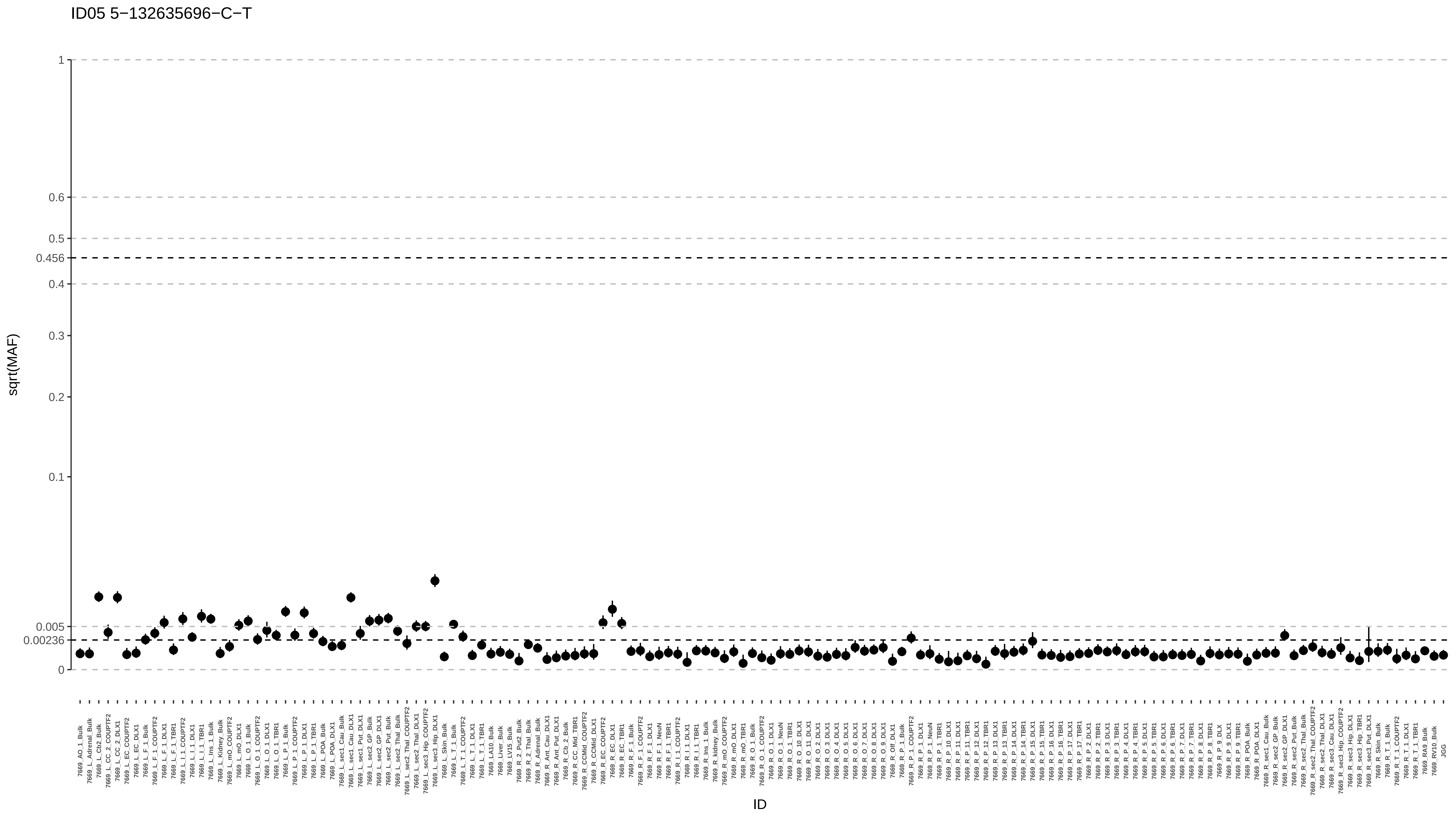
ID05 5−132635696−C−T
sqrt(MAF)
1
0.6
0.5
0.456
0.4
0.3
0.2
0.1
0.005
0.00236
0
ID

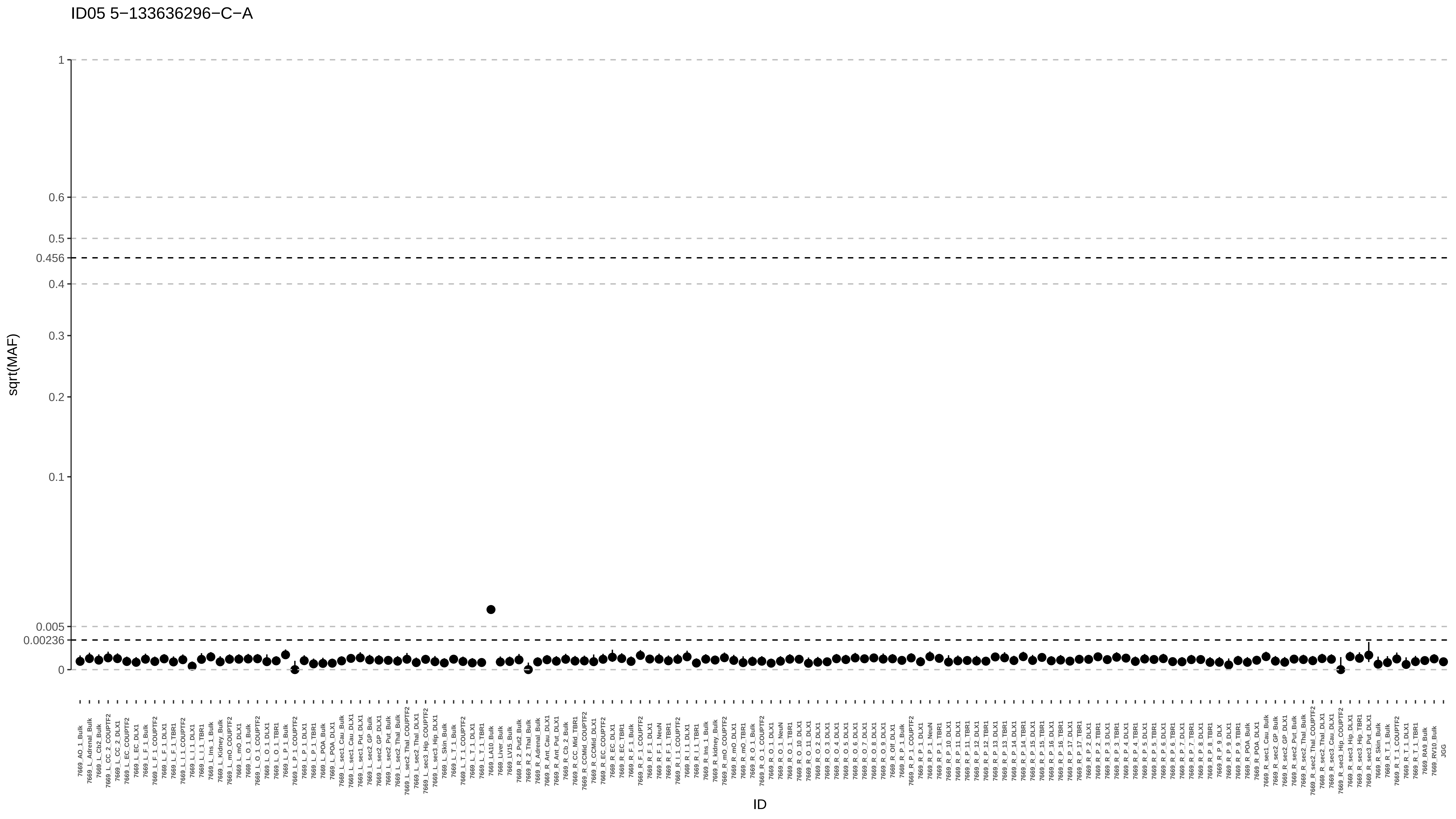
ID05 5−133636296−C−A
sqrt(MAF)
1
0.6
0.5
0.456
0.4
0.3
0.2
0.1
0.005
0.00236
0
ID

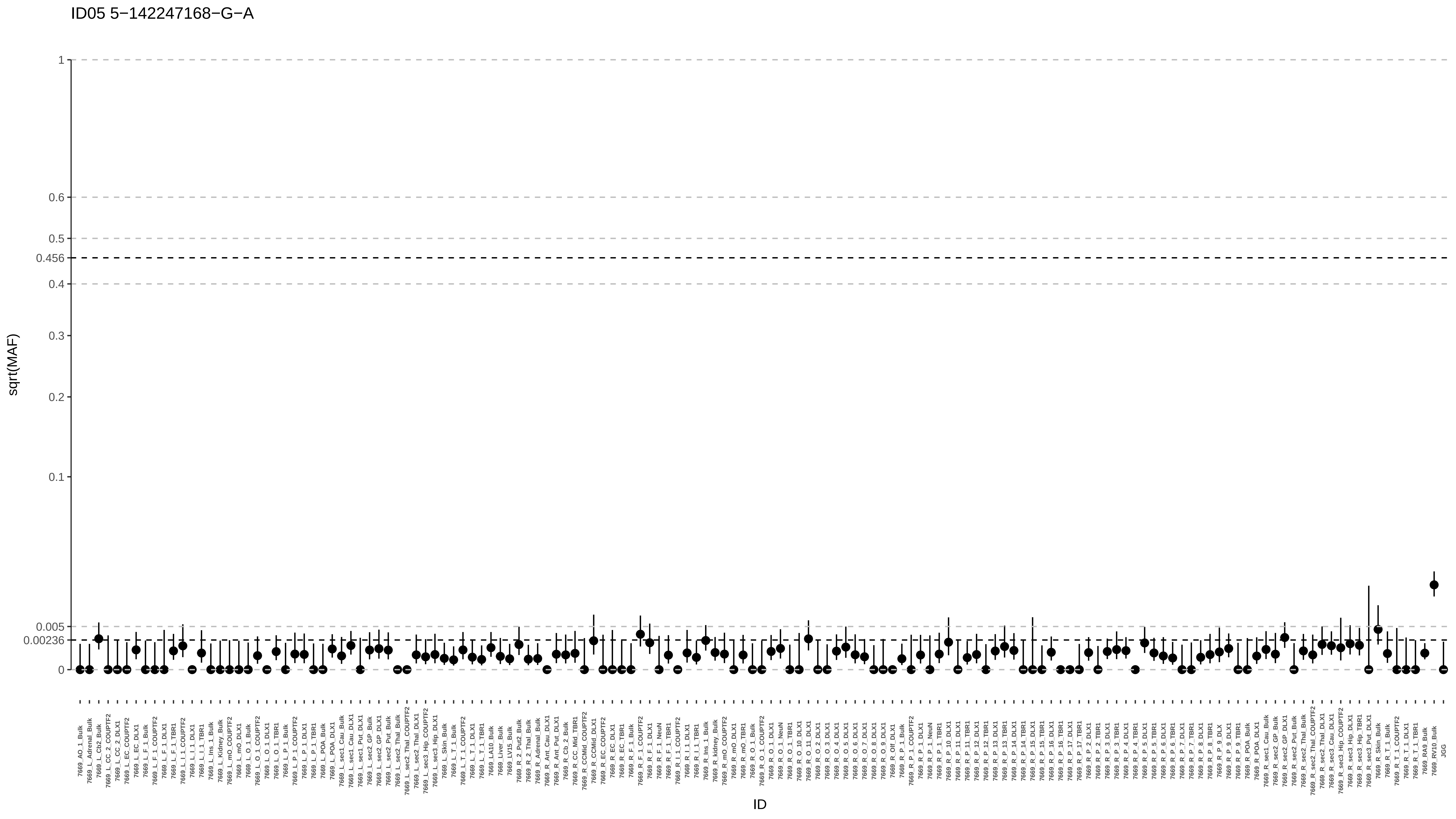

ID05 5−142247168−G−A
sqrt(MAF)
1
0.6
0.5
0.456
0.4
0.3
0.2
0.1
0.005
0.00236
0
ID

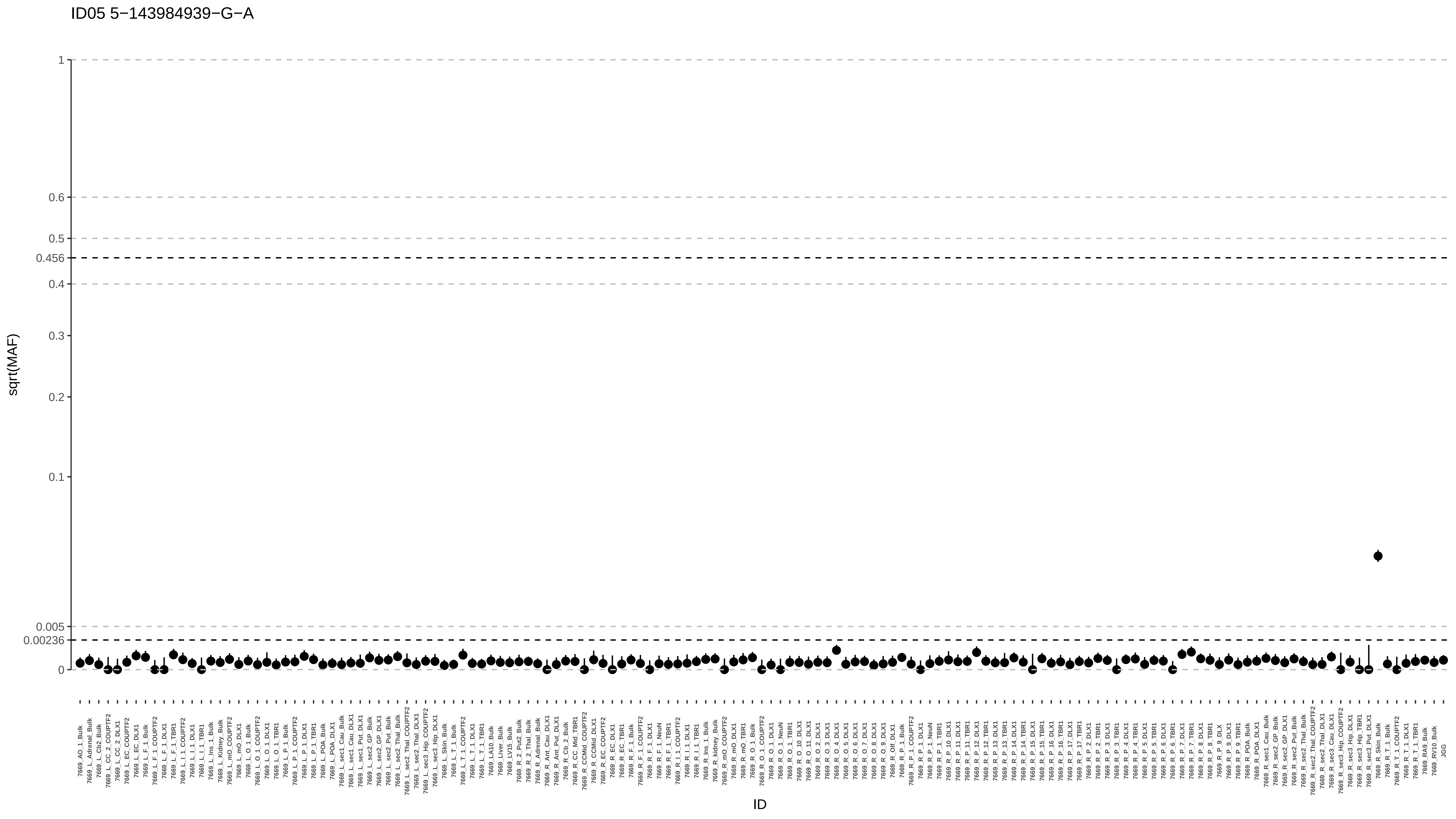
ID05 5−143984939−G−A
sqrt(MAF)
1
0.6
0.5
0.456
0.4
0.3
0.2
0.1
0.005
0.00236
0
ID

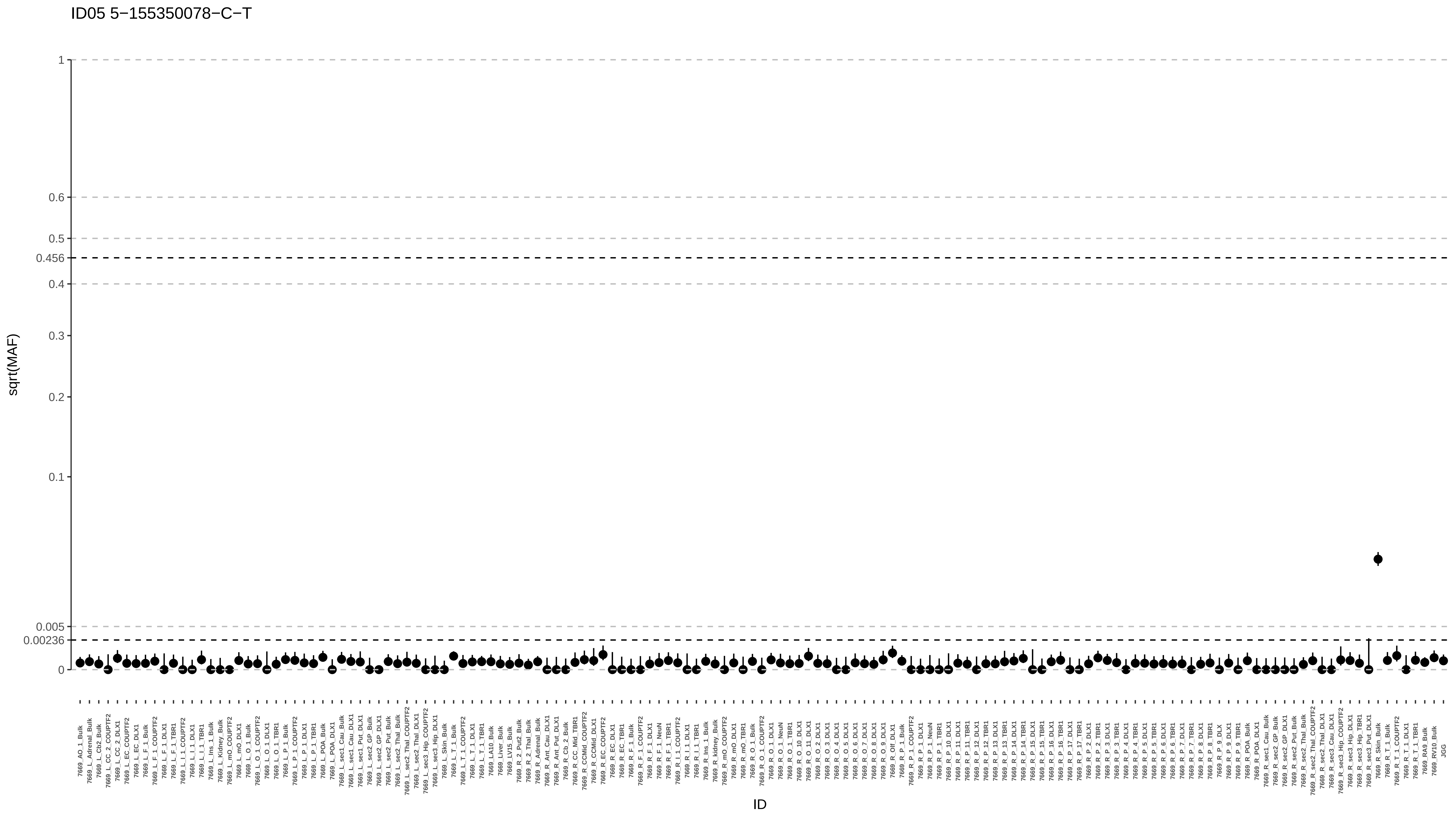

ID05 5−155350078−C−T
sqrt(MAF)
1
0.6
0.5
0.456
0.4
0.3
0.2
0.1
0.005
0.00236
0
ID

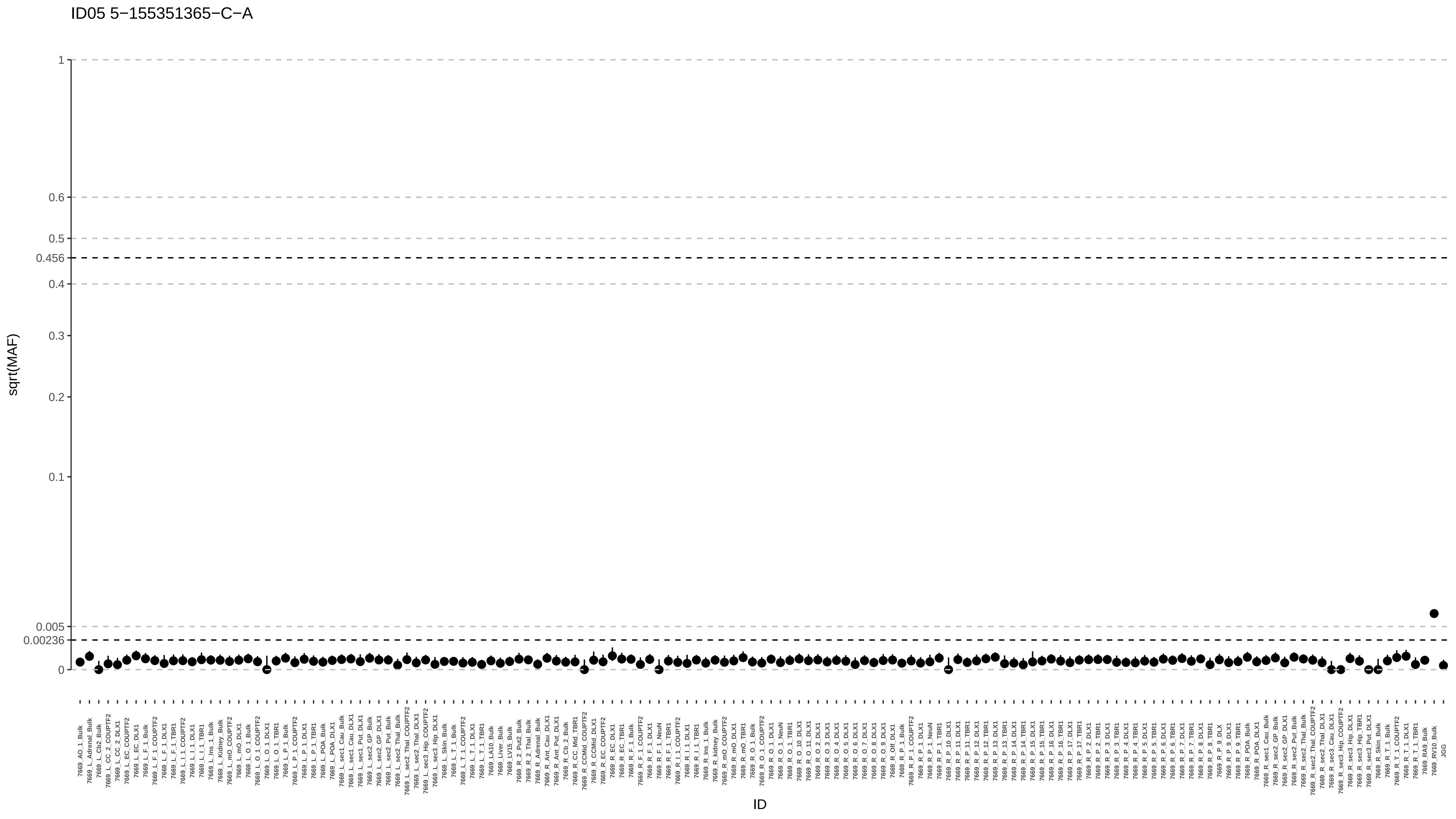
ID05 5−155351365−C−A
sqrt(MAF)
1
0.6
0.5
0.456
0.4
0.3
0.2
0.1
0.005
0.00236
0
ID

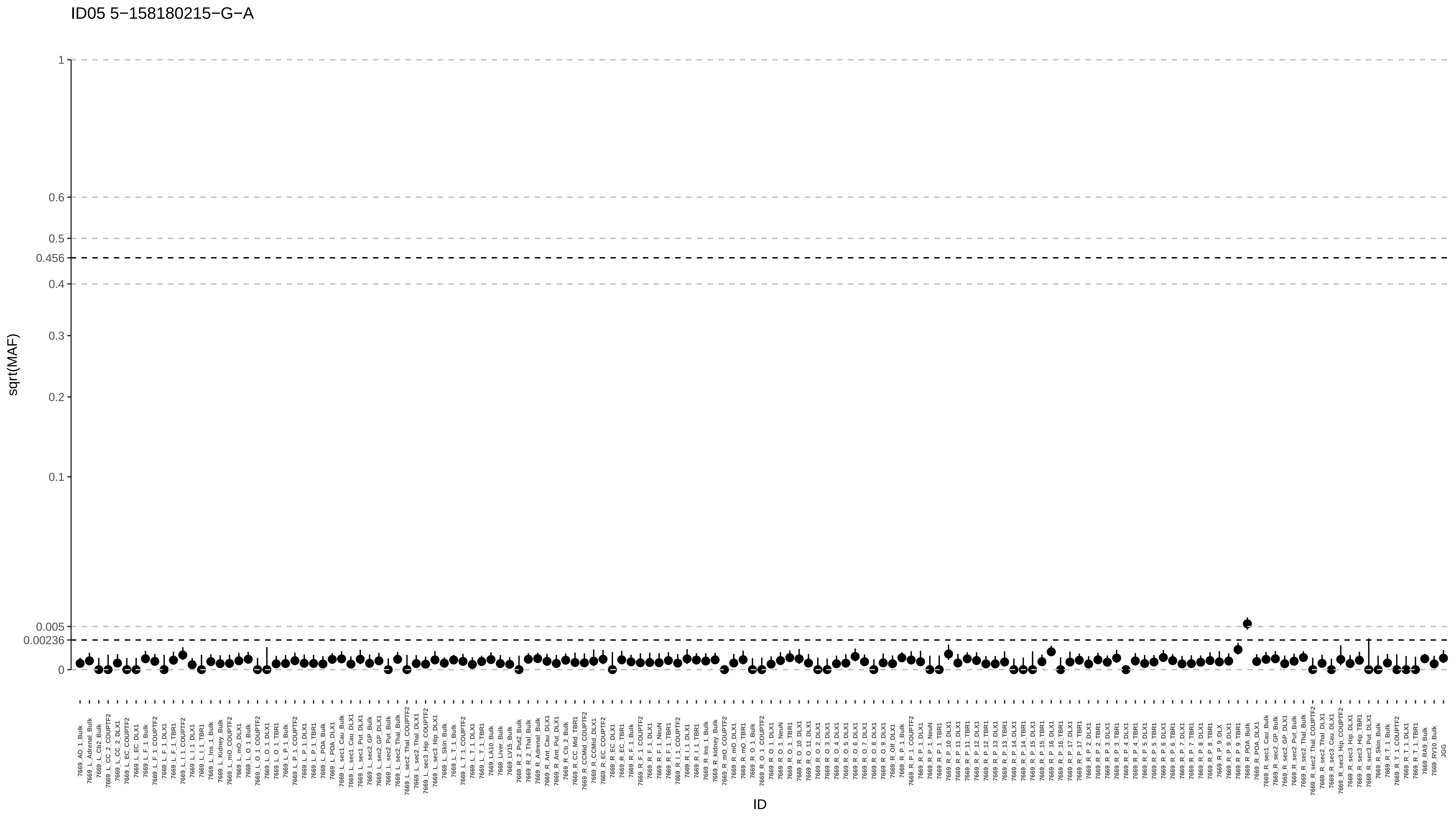
ID05 5−158180215−G−A
sqrt(MAF)
1
0.6
0.5
0.456
0.4
0.3
0.2
0.1
0.005
0.00236
0
ID

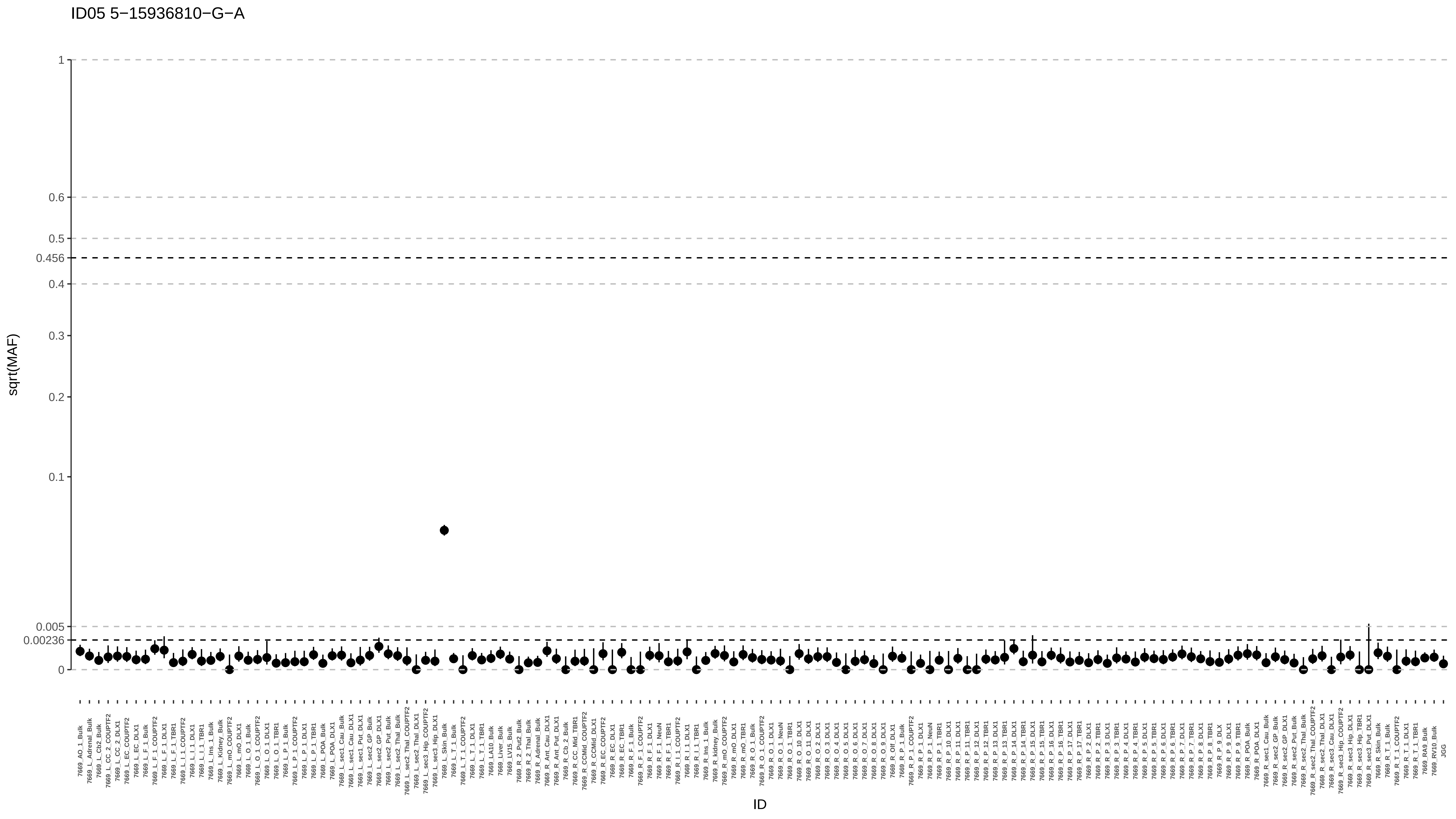
ID05 5−15936810−G−A
sqrt(MAF)
1
0.6
0.5
0.456
0.4
0.3
0.2
0.1
0.005
0.00236
0
ID

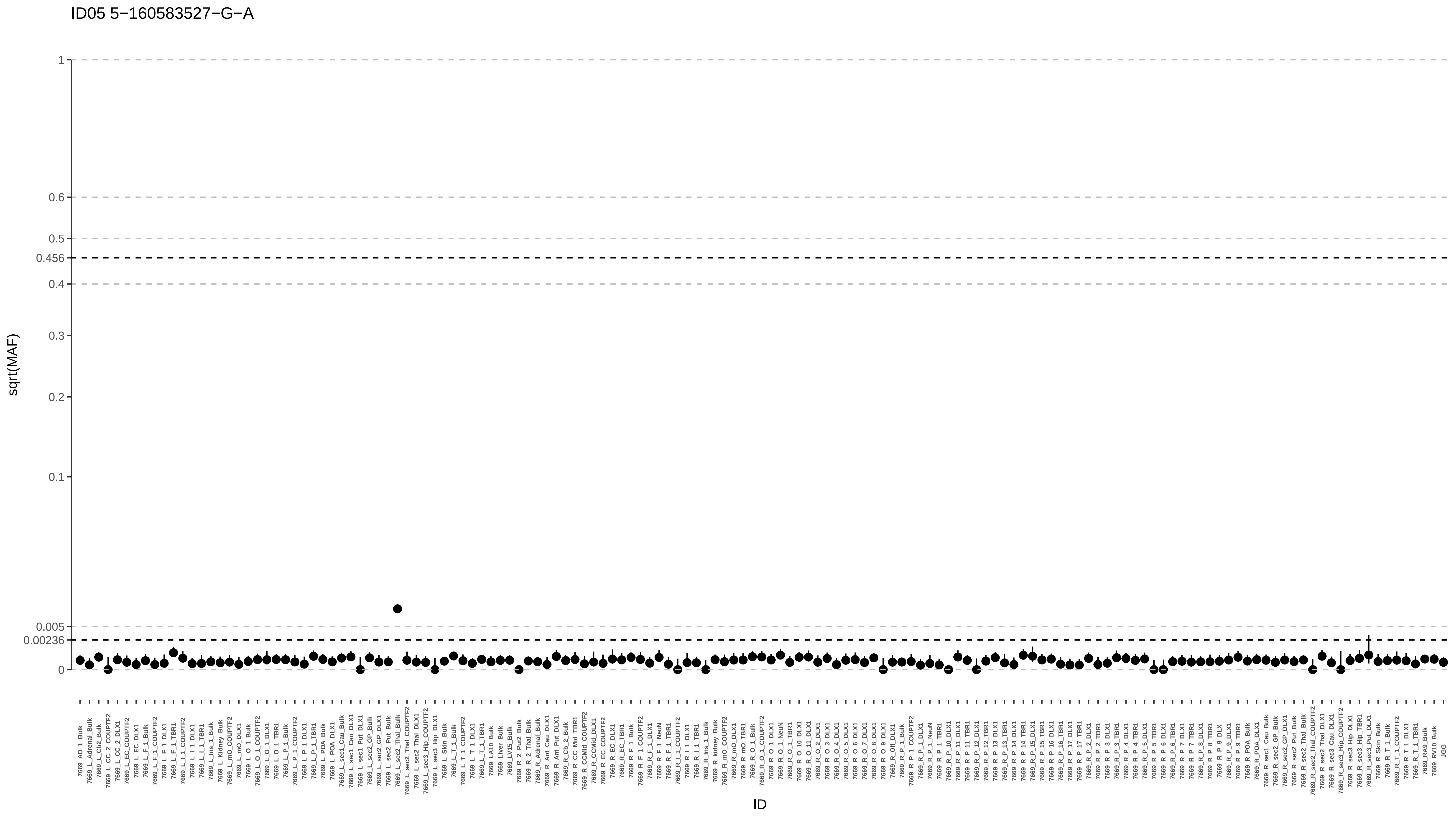
ID05 5−160583527−G−A
sqrt(MAF)
1
0.6
0.5
0.456
0.4
0.3
0.2
0.1
0.005
0.00236
0
ID

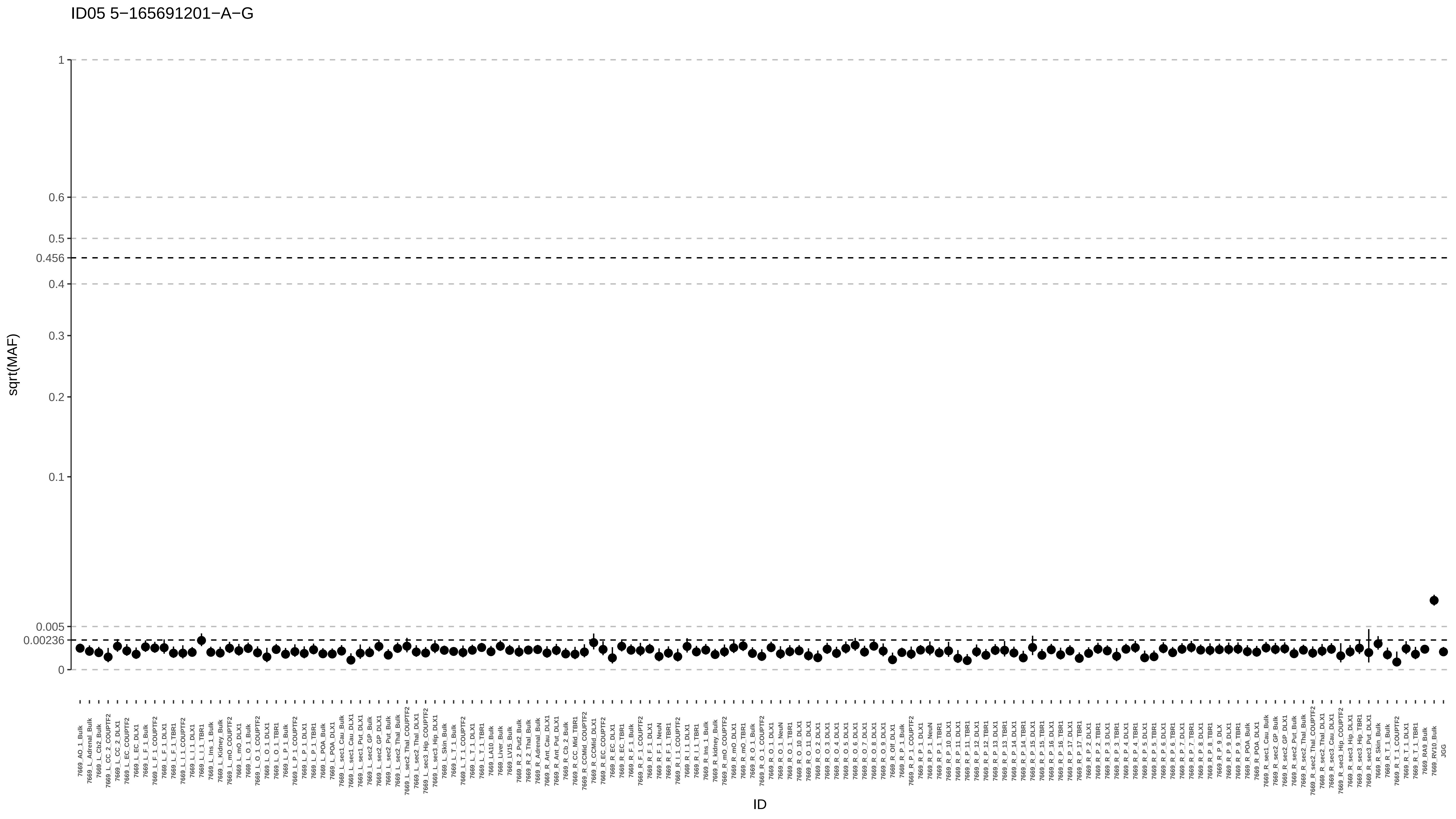
ID05 5−165691201−A−G
sqrt(MAF)
1
0.6
0.5
0.456
0.4
0.3
0.2
0.1
0.005
0.00236
0
ID

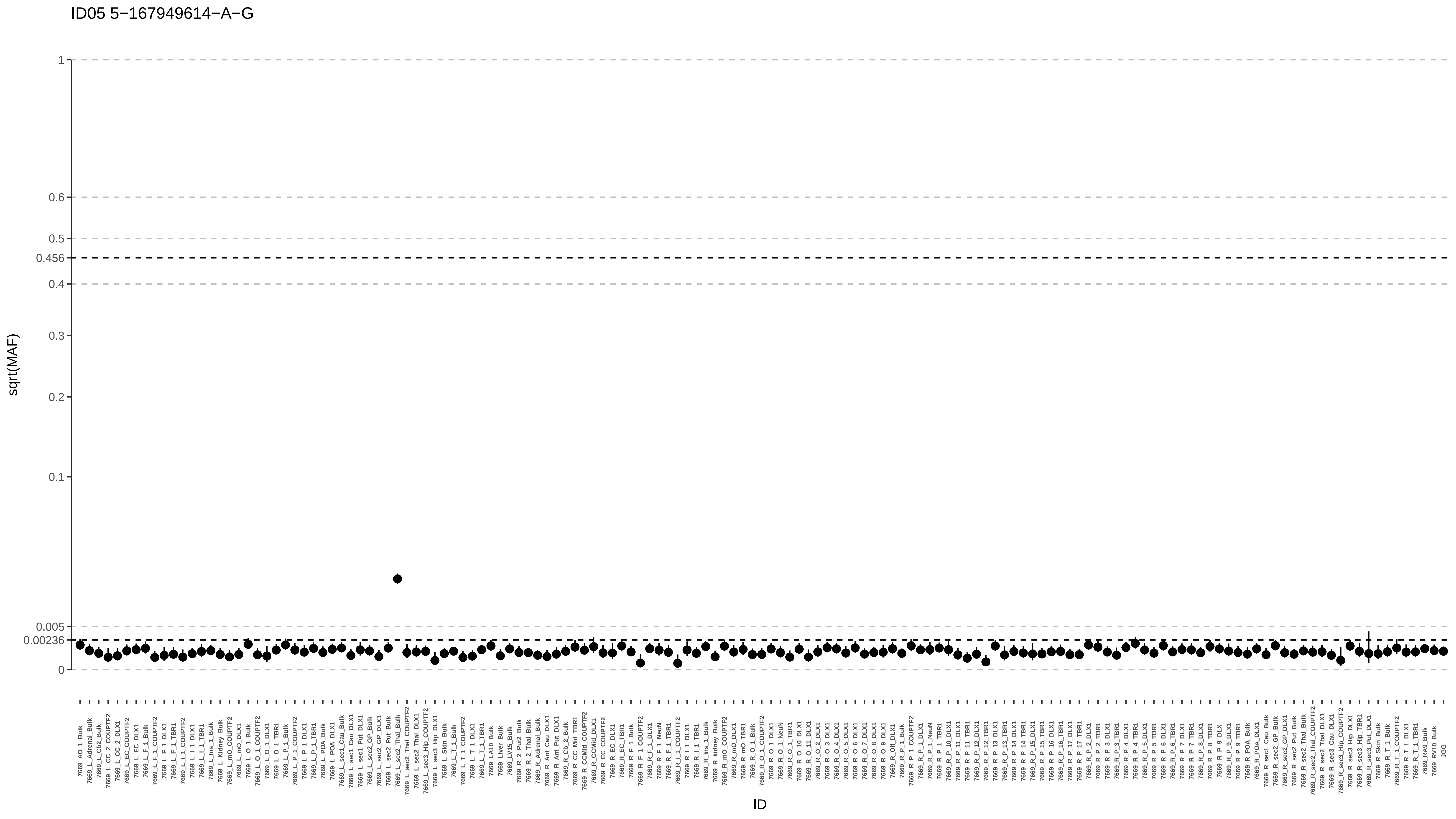
ID05 5−167949614−A−G
sqrt(MAF)
1
0.6
0.5
0.456
0.4
0.3
0.2
0.1
0.005
0.00236
0
ID

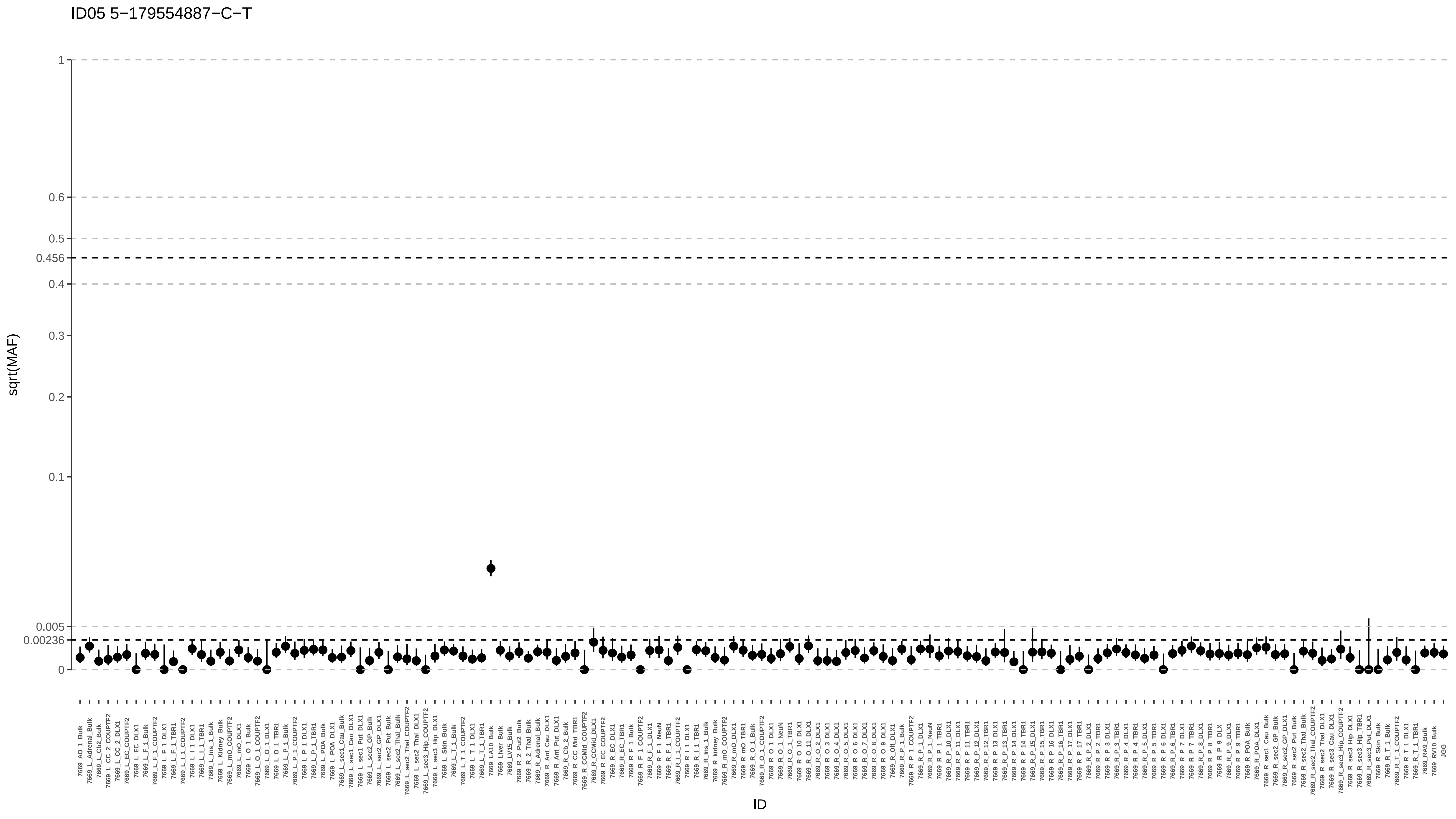
ID05 5−179554887−C−T
sqrt(MAF)
1
0.6
0.5
0.456
0.4
0.3
0.2
0.1
0.005
0.00236
0
ID

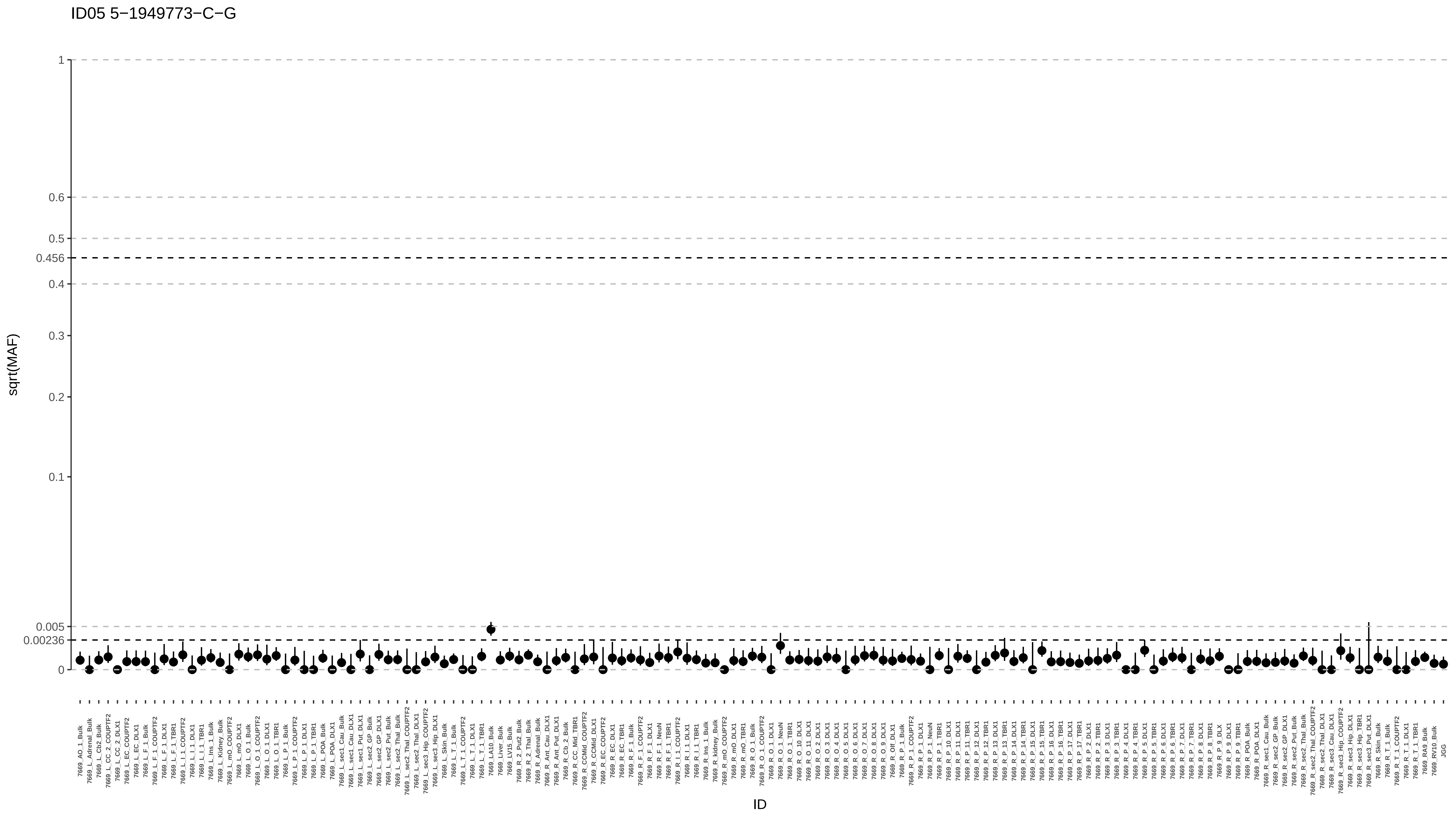
ID05 5−1949773−C−G
sqrt(MAF)
1
0.6
0.5
0.456
0.4
0.3
0.2
0.1
0.005
0.00236
0
ID
7669_AO_1_Bulk
7669_L_Adrenal_Bulk
7669_L_Cb2_Bulk
7669_L_CC_2_COUPTF2
7669_L_CC_2_DLX1
7669_L_EC_COUPTF2
7669_L_EC_DLX1
7669_L_F_1_Bulk
7669_L_F_1_COUPTF2
7669_L_F_1_DLX1
7669_L_F_1_TBR1
7669_L_I_1_COUPTF2
7669_L_I_1_DLX1
7669_L_I_1_TBR1
7669_L_Ins_1_Bulk
7669_L_Kidney_Bulk
7669_L_mO_COUPTF2
7669_L_mO_DLX1
7669_L_O_1_Bulk
7669_L_O_1_COUPTF2
7669_L_O_1_DLX1
7669_L_O_1_TBR1
7669_L_P_1_Bulk
7669_L_P_1_COUPTF2
7669_L_P_1_DLX1
7669_L_P_1_TBR1
7669_L_POA_Bulk
7669_L_POA_DLX1
7669_L_sec1_Cau_Bulk
7669_L_sec1_Cau_DLX1
7669_L_sec1_Put_DLX1
7669_L_sec2_GP_Bulk
7669_L_sec2_GP_DLX1
7669_L_sec2_Put_Bulk
7669_L_sec2_Thal_Bulk
7669_L_sec2_Thal_COUPTF2
7669_L_sec2_Thal_DLX1
7669_L_sec3_Hip_COUPTF2
7669_L_sec3_Hip_DLX1
7669_L_Skin_Bulk
7669_L_T_1_Bulk
7669_L_T_1_COUPTF2
7669_L_T_1_DLX1
7669_L_T_1_TBR1
7669_LA10_Bulk
7669_Liver_Bulk
7669_LV15_Bulk
7669_R_2_Put2_Bulk
7669_R_2_Thal_Bulk
7669_R_Adrenal_Bulk
7669_R_Ant_Cau_DLX1
7669_R_Ant_Put_DLX1
7669_R_Cb_2_Bulk
7669_R_CC_Mid_TBR1
7669_R_CCMid_COUPTF2
7669_R_CCMid_DLX1
7669_R_EC_COUPTF2
7669_R_EC_DLX1
7669_R_EC_TBR1
7669_R_F_1_Bulk
7669_R_F_1_COUPTF2
7669_R_F_1_DLX1
7669_R_F_1_NeuN
7669_R_F_1_TBR1
7669_R_I_1_COUPTF2
7669_R_I_1_DLX1
7669_R_I_1_TBR1
7669_R_Ins_1_Bulk
7669_R_kidney_Bulk
7669_R_mO_COUPTF2
7669_R_mO_DLX1
7669_R_mO_TBR1
7669_R_O_1_Bulk
7669_R_O_1_COUPTF2
7669_R_O_1_DLX1
7669_R_O_1_NeuN
7669_R_O_1_TBR1
7669_R_O_10_DLX1
7669_R_O_11_DLX1
7669_R_O_2_DLX1
7669_R_O_3_DLX1
7669_R_O_4_DLX1
7669_R_O_5_DLX1
7669_R_O_6_DLX1
7669_R_O_7_DLX1
7669_R_O_8_DLX1
7669_R_O_9_DLX1
7669_R_Olf_DLX1
7669_R_P_1_Bulk
7669_R_P_1_COUPTF2
7669_R_P_1_DLX1
7669_R_P_1_NeuN
7669_R_P_1_TBR1
7669_R_P_10_DLX1
7669_R_P_11_DLX1
7669_R_P_11_TBR1
7669_R_P_12_DLX1
7669_R_P_12_TBR1
7669_R_P_13_DLX1
7669_R_P_13_TBR1
7669_R_P_14_DLX1
7669_R_P_14_TBR1
7669_R_P_15_DLX1
7669_R_P_15_TBR1
7669_R_P_16_DLX1
7669_R_P_16_TBR1
7669_R_P_17_DLX1
7669_R_P_17_TBR1
7669_R_P_2_DLX1
7669_R_P_2_TBR1
7669_R_P_3_DLX1
7669_R_P_3_TBR1
7669_R_P_4_DLX1
7669_R_P_4_TBR1
7669_R_P_5_DLX1
7669_R_P_5_TBR1
7669_R_P_6_DLX1
7669_R_P_6_TBR1
7669_R_P_7_DLX1
7669_R_P_7_TBR1
7669_R_P_8_DLX1
7669_R_P_8_TBR1
7669_R_P_9_DLX
7669_R_P_9_DLX1
7669_R_P_9_TBR1
7669_R_POA_Bulk
7669_R_POA_DLX1
7669_R_sec1_Cau_Bulk
7669_R_sec2_GP_Bulk
7669_R_sec2_GP_DLX1
7669_R_sec2_Put_Bulk
7669_R_sec2_Thal_Bulk
7669_R_sec2_Thal_COUPTF2
7669_R_sec2_Thal_DLX1
7669_R_sec3_Cau_DLX1
7669_R_sec3_Hip_COUPTF2
7669_R_sec3_Hip_DLX1
7669_R_sec3_Hip_TBR1
7669_R_sec3_Put_DLX1
7669_R_Skin_Bulk
7669_R_T_1_Bulk
7669_R_T_1_COUPTF2
7669_R_T_1_DLX1
7669_R_T_1_TBR1
7669_RA9_Bulk
7669_RV10_Bulk
JGG

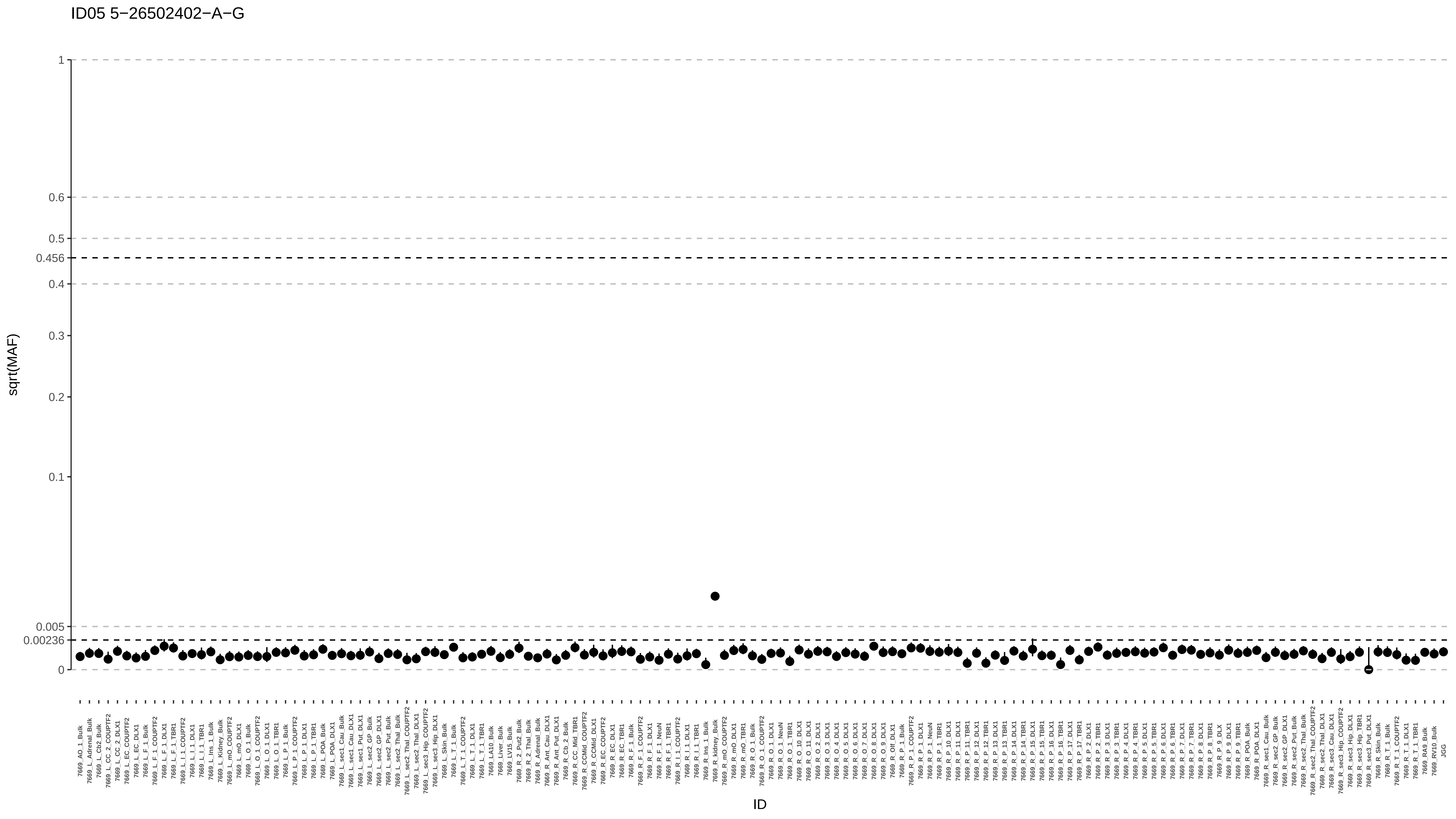

ID05 5−26502402−A−G
sqrt(MAF)
ID

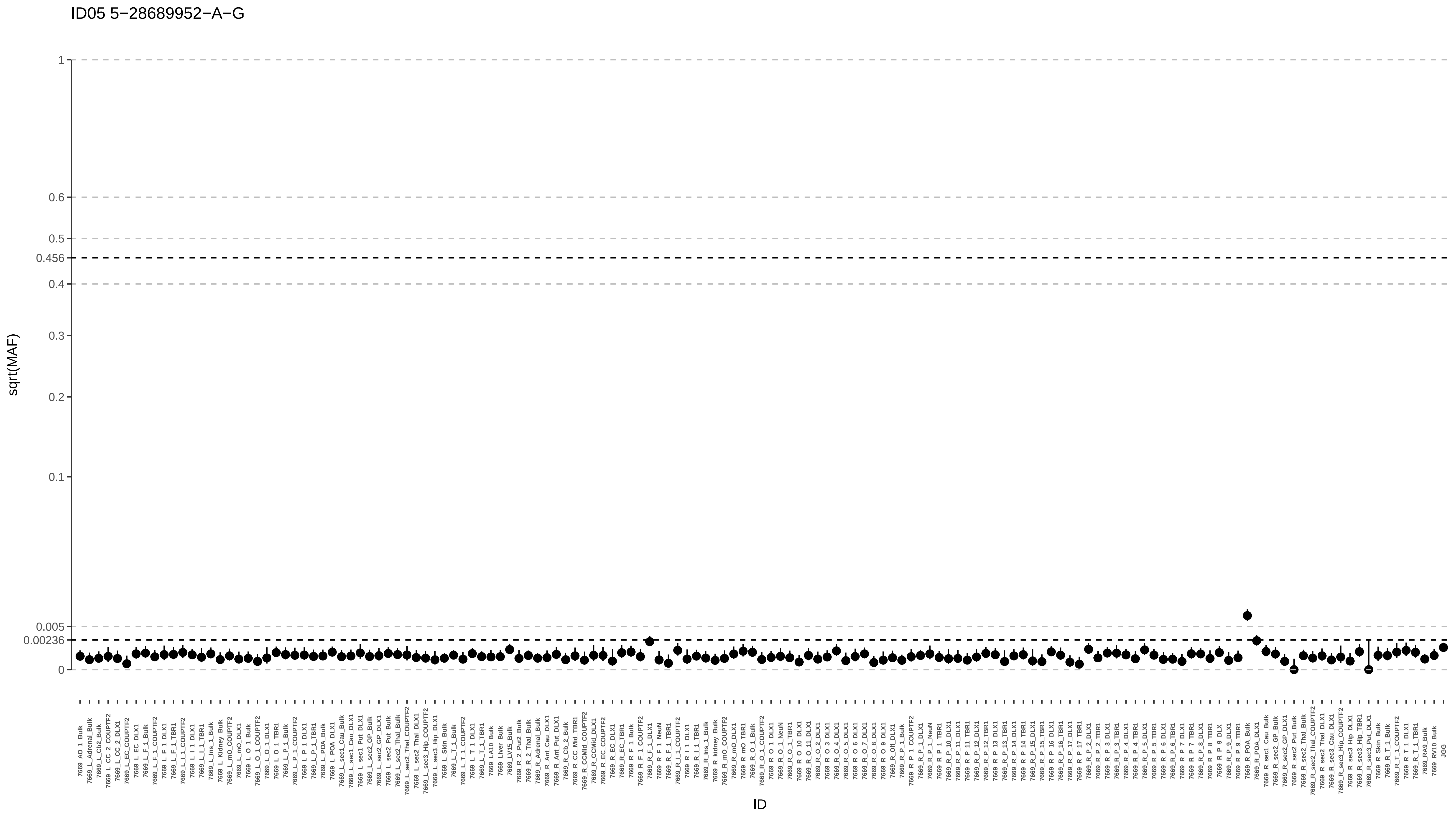
ID05 5−28689952−A−G
sqrt(MAF)
1
0.6
0.5
0.456
0.4
0.3
0.2
0.1
0.005
0.00236
0
ID

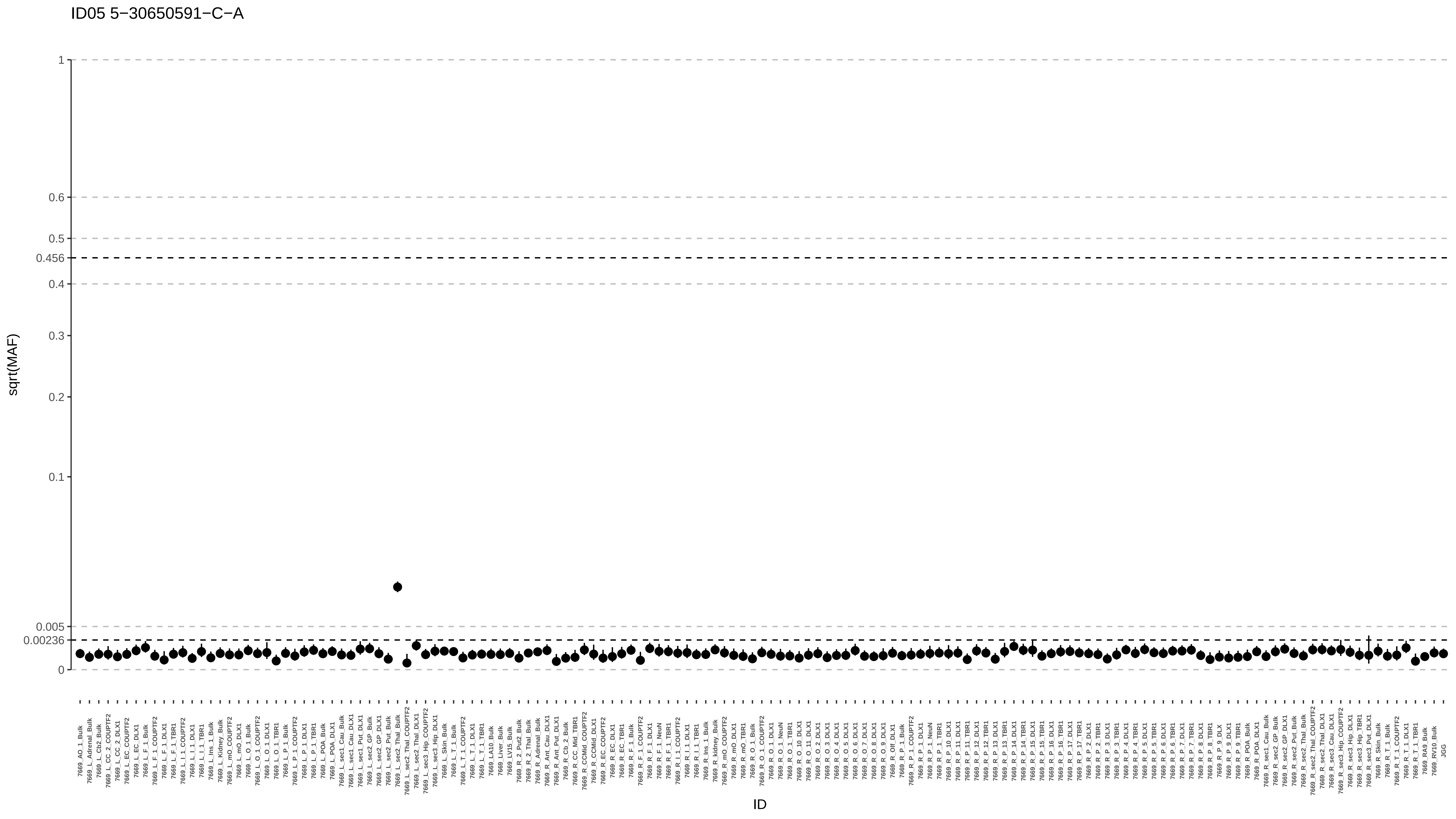

ID05 5−30650591−C−A
sqrt(MAF)
1
0.6
0.5
0.456
0.4
0.3
0.2
0.1
0.005
0.00236
0
ID

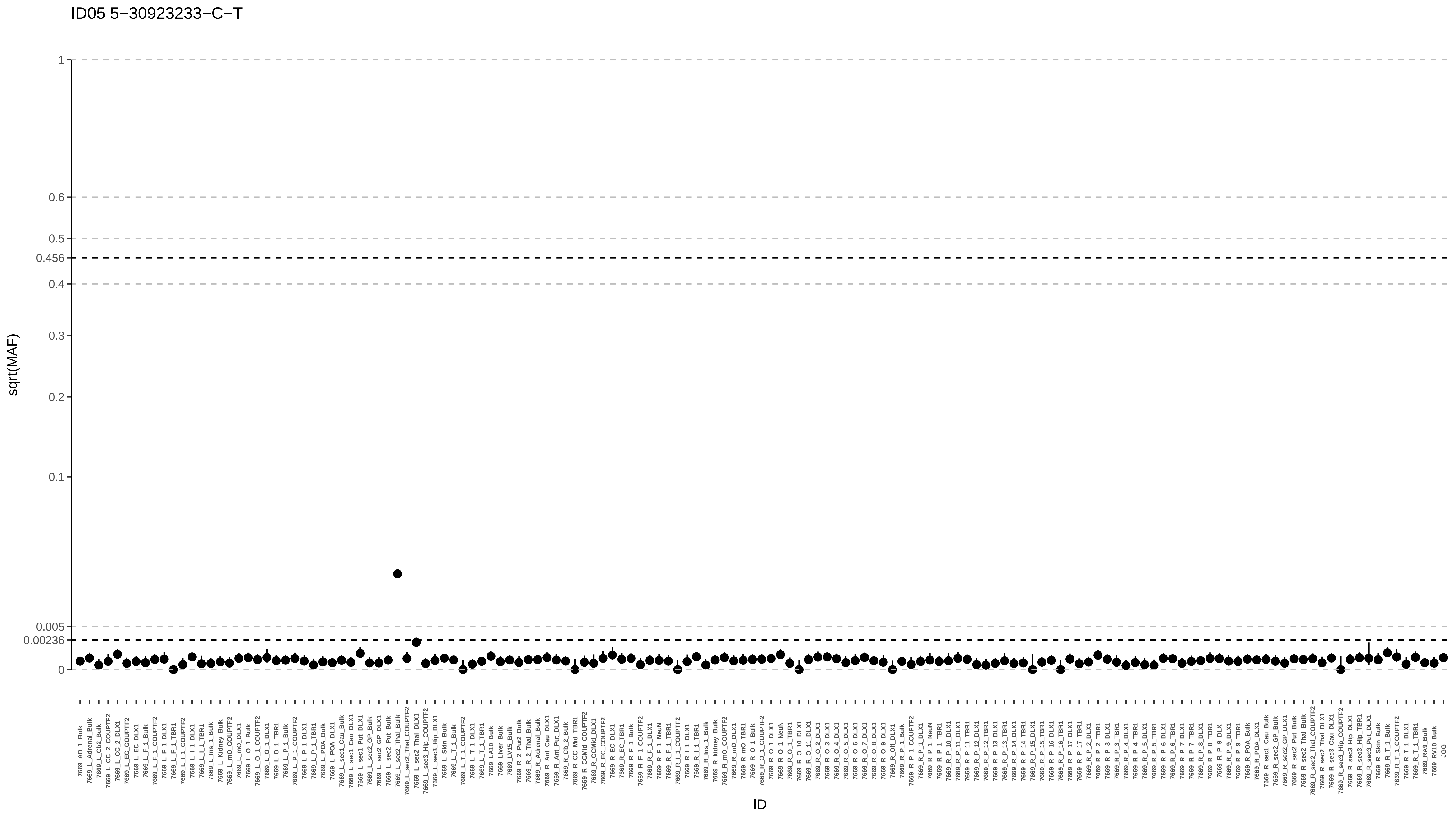
ID05 5−30923233−C−T
sqrt(MAF)
1
0.6
0.5
0.456
0.4
0.3
0.2
0.1
0.005
0.00236
0
ID

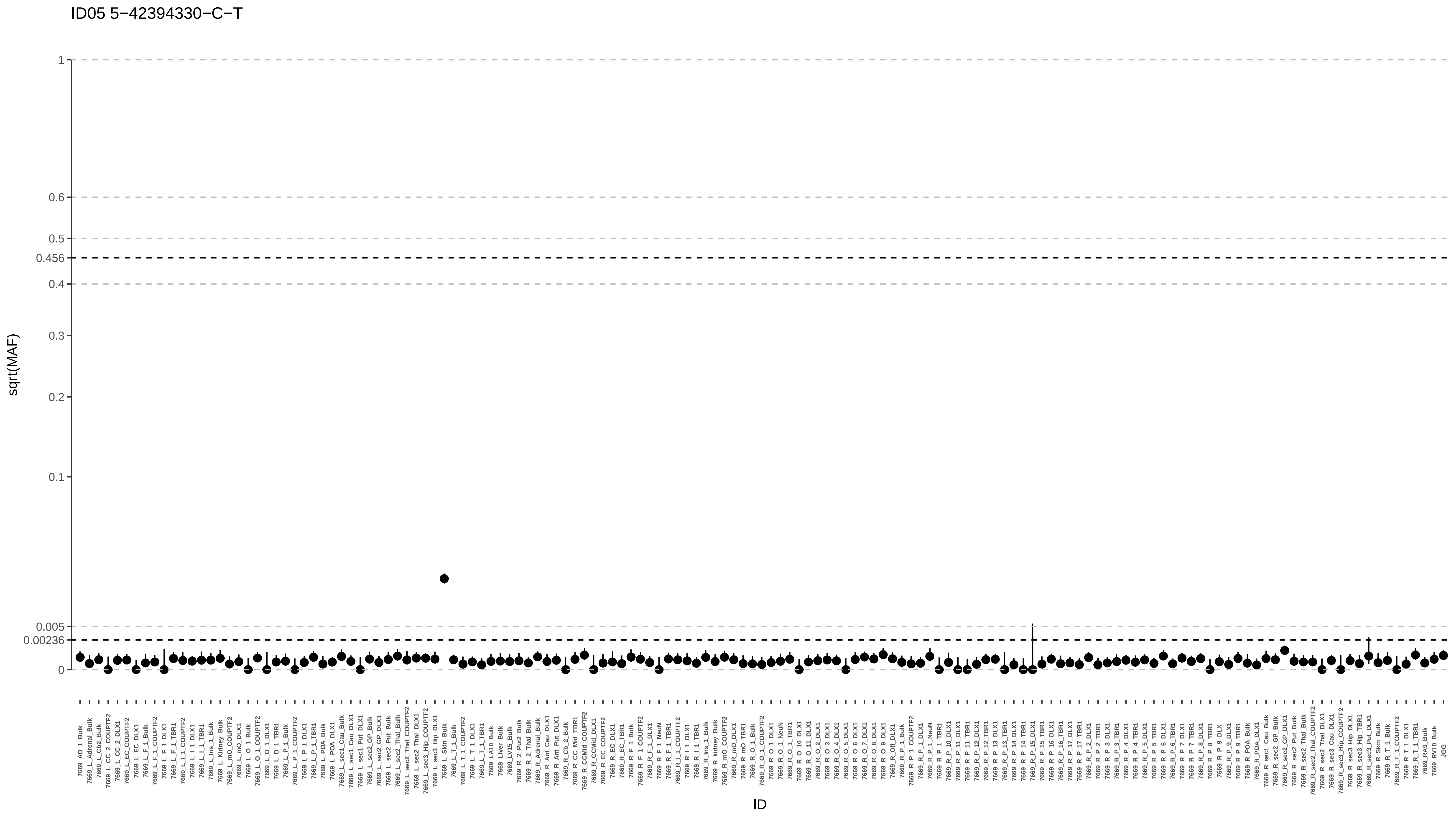
ID05 5−42394330−C−T
sqrt(MAF)
1
0.6
0.5
0.456
0.4
0.3
0.2
0.1
0.005
0.00236
0
ID
7669_AO_1_Bulk
7669_L_Adrenal_Bulk
7669_L_Cb2_Bulk
7669_L_CC_2_COUPTF2
7669_L_CC_2_DLX1
7669_L_EC_COUPTF2
7669_L_EC_DLX1
7669_L_F_1_Bulk
7669_L_F_1_COUPTF2
7669_L_F_1_DLX1
7669_L_F_1_TBR1
7669_L_I_1_COUPTF2
7669_L_I_1_DLX1
7669_L_I_1_TBR1
7669_L_Ins_1_Bulk
7669_L_Kidney_Bulk
7669_L_mO_COUPTF2
7669_L_mO_DLX1
7669_L_O_1_Bulk
7669_L_O_1_COUPTF2
7669_L_O_1_DLX1
7669_L_O_1_TBR1
7669_L_P_1_Bulk
7669_L_P_1_COUPTF2
7669_L_P_1_DLX1
7669_L_P_1_TBR1
7669_L_POA_Bulk
7669_L_POA_DLX1
7669_L_sec1_Cau_Bulk
7669_L_sec1_Cau_DLX1
7669_L_sec1_Put_DLX1
7669_L_sec2_GP_Bulk
7669_L_sec2_GP_DLX1
7669_L_sec2_Put_Bulk
7669_L_sec2_Thal_Bulk
7669_L_sec2_Thal_COUPTF2
7669_L_sec2_Thal_DLX1
7669_L_sec3_Hip_COUPTF2
7669_L_sec3_Hip_DLX1
7669_L_Skin_Bulk
7669_L_T_1_Bulk
7669_L_T_1_COUPTF2
7669_L_T_1_DLX1
7669_L_T_1_TBR1
7669_LA10_Bulk
7669_Liver_Bulk
7669_LV15_Bulk
7669_R_2_Put2_Bulk
7669_R_2_Thal_Bulk
7669_R_Adrenal_Bulk
7669_R_Ant_Cau_DLX1
7669_R_Ant_Put_DLX1
7669_R_Cb_2_Bulk
7669_R_CC_Mid_TBR1
7669_R_CCMid_COUPTF2
7669_R_CCMid_DLX1
7669_R_EC_COUPTF2
7669_R_EC_DLX1
7669_R_EC_TBR1
7669_R_F_1_Bulk
7669_R_F_1_COUPTF2
7669_R_F_1_DLX1
7669_R_F_1_NeuN
7669_R_F_1_TBR1
7669_R_I_1_COUPTF2
7669_R_I_1_DLX1
7669_R_I_1_TBR1
7669_R_Ins_1_Bulk
7669_R_kidney_Bulk
7669_R_mO_COUPTF2
7669_R_mO_DLX1
7669_R_mO_TBR1
7669_R_O_1_Bulk
7669_R_O_1_COUPTF2
7669_R_O_1_DLX1
7669_R_O_1_NeuN
7669_R_O_1_TBR1
7669_R_O_10_DLX1
7669_R_O_11_DLX1
7669_R_O_2_DLX1
7669_R_O_3_DLX1
7669_R_O_4_DLX1
7669_R_O_5_DLX1
7669_R_O_6_DLX1
7669_R_O_7_DLX1
7669_R_O_8_DLX1
7669_R_O_9_DLX1
7669_R_Olf_DLX1
7669_R_P_1_Bulk
7669_R_P_1_COUPTF2
7669_R_P_1_DLX1
7669_R_P_1_NeuN
7669_R_P_1_TBR1
7669_R_P_10_DLX1
7669_R_P_11_DLX1
7669_R_P_11_TBR1
7669_R_P_12_DLX1
7669_R_P_12_TBR1
7669_R_P_13_DLX1
7669_R_P_13_TBR1
7669_R_P_14_DLX1
7669_R_P_14_TBR1
7669_R_P_15_DLX1
7669_R_P_15_TBR1
7669_R_P_16_DLX1
7669_R_P_16_TBR1
7669_R_P_17_DLX1
7669_R_P_17_TBR1
7669_R_P_2_DLX1
7669_R_P_2_TBR1
7669_R_P_3_DLX1
7669_R_P_3_TBR1
7669_R_P_4_DLX1
7669_R_P_4_TBR1
7669_R_P_5_DLX1
7669_R_P_5_TBR1
7669_R_P_6_DLX1
7669_R_P_6_TBR1
7669_R_P_7_DLX1
7669_R_P_7_TBR1
7669_R_P_8_DLX1
7669_R_P_8_TBR1
7669_R_P_9_DLX
7669_R_P_9_DLX1
7669_R_P_9_TBR1
7669_R_POA_Bulk
7669_R_POA_DLX1
7669_R_sec1_Cau_Bulk
7669_R_sec2_GP_Bulk
7669_R_sec2_GP_DLX1
7669_R_sec2_Put_Bulk
7669_R_sec2_Thal_Bulk
7669_R_sec2_Thal_COUPTF2
7669_R_sec2_Thal_DLX1
7669_R_sec3_Cau_DLX1
7669_R_sec3_Hip_COUPTF2
7669_R_sec3_Hip_DLX1
7669_R_sec3_Hip_TBR1
7669_R_sec3_Put_DLX1
7669_R_Skin_Bulk
7669_R_T_1_Bulk
7669_R_T_1_COUPTF2
7669_R_T_1_DLX1
7669_R_T_1_TBR1
7669_RA9_Bulk
7669_RV10_Bulk
JGG

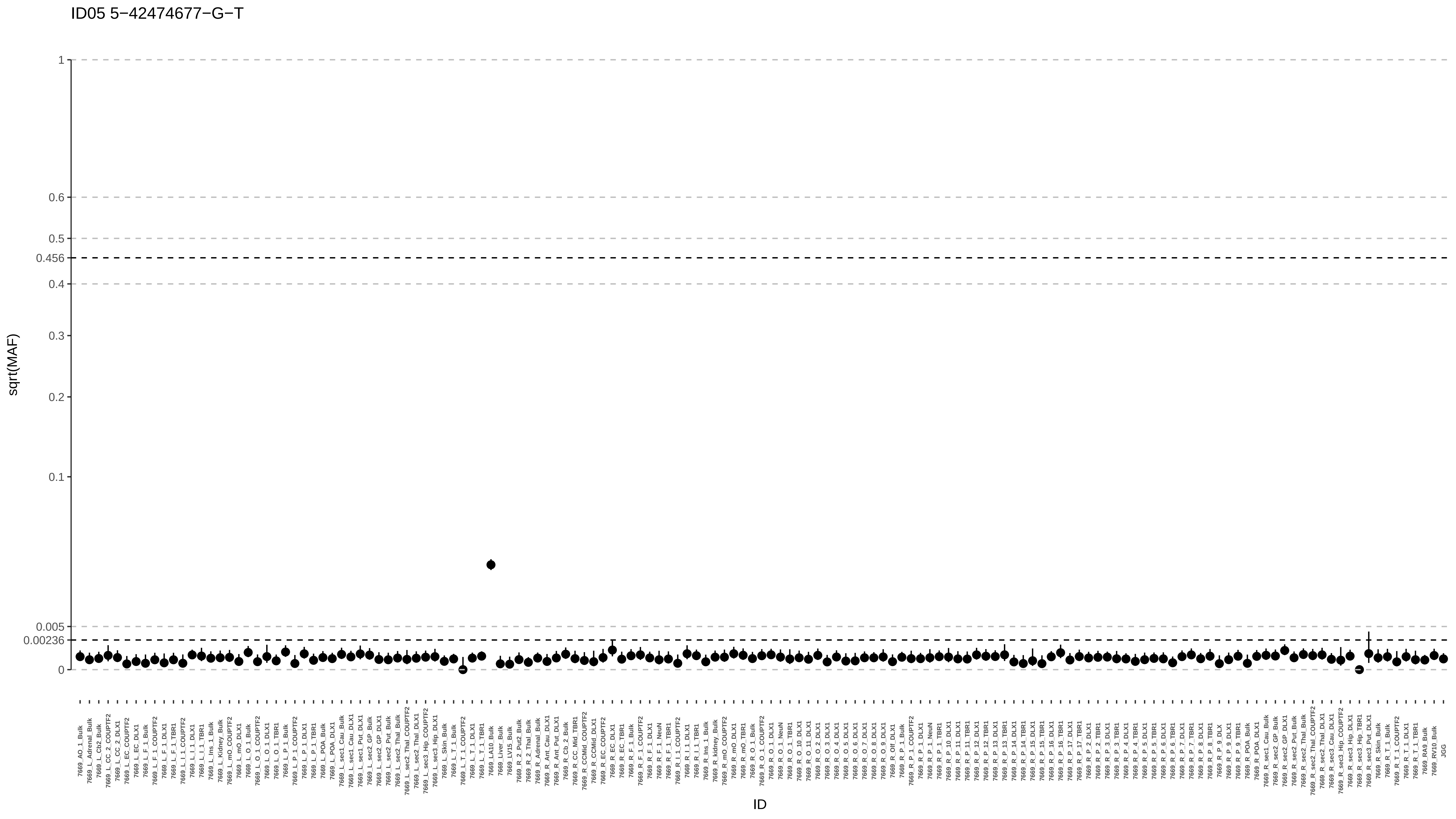
ID05 5−42474677−G−T
sqrt(MAF)
1
0.6
0.5
0.456
0.4
0.3
0.2
0.1
0.005
0.00236
0
ID
7669_AO_1_Bulk
7669_L_Adrenal_Bulk
7669_L_Cb2_Bulk
7669_L_CC_2_COUPTF2
7669_L_CC_2_DLX1
7669_L_EC_COUPTF2
7669_L_EC_DLX1
7669_L_F_1_Bulk
7669_L_F_1_COUPTF2
7669_L_F_1_DLX1
7669_L_F_1_TBR1
7669_L_I_1_COUPTF2
7669_L_I_1_DLX1
7669_L_I_1_TBR1
7669_L_Ins_1_Bulk
7669_L_Kidney_Bulk
7669_L_mO_COUPTF2
7669_L_mO_DLX1
7669_L_O_1_Bulk
7669_L_O_1_COUPTF2
7669_L_O_1_DLX1
7669_L_O_1_TBR1
7669_L_P_1_Bulk
7669_L_P_1_COUPTF2
7669_L_P_1_DLX1
7669_L_P_1_TBR1
7669_L_POA_Bulk
7669_L_POA_DLX1
7669_L_sec1_Cau_Bulk
7669_L_sec1_Cau_DLX1
7669_L_sec1_Put_DLX1
7669_L_sec2_GP_Bulk
7669_L_sec2_GP_DLX1
7669_L_sec2_Put_Bulk
7669_L_sec2_Thal_Bulk
7669_L_sec2_Thal_COUPTF2
7669_L_sec2_Thal_DLX1
7669_L_sec3_Hip_COUPTF2
7669_L_sec3_Hip_DLX1
7669_L_Skin_Bulk
7669_L_T_1_Bulk
7669_L_T_1_COUPTF2
7669_L_T_1_DLX1
7669_L_T_1_TBR1
7669_LA10_Bulk
7669_Liver_Bulk
7669_LV15_Bulk
7669_R_2_Put2_Bulk
7669_R_2_Thal_Bulk
7669_R_Adrenal_Bulk
7669_R_Ant_Cau_DLX1
7669_R_Ant_Put_DLX1
7669_R_Cb_2_Bulk
7669_R_CC_Mid_TBR1
7669_R_CCMid_COUPTF2
7669_R_CCMid_DLX1
7669_R_EC_COUPTF2
7669_R_EC_DLX1
7669_R_EC_TBR1
7669_R_F_1_Bulk
7669_R_F_1_COUPTF2
7669_R_F_1_DLX1
7669_R_F_1_NeuN
7669_R_F_1_TBR1
7669_R_I_1_COUPTF2
7669_R_I_1_DLX1
7669_R_I_1_TBR1
7669_R_Ins_1_Bulk
7669_R_kidney_Bulk
7669_R_mO_COUPTF2
7669_R_mO_DLX1
7669_R_mO_TBR1
7669_R_O_1_Bulk
7669_R_O_1_COUPTF2
7669_R_O_1_DLX1
7669_R_O_1_NeuN
7669_R_O_1_TBR1
7669_R_O_10_DLX1
7669_R_O_11_DLX1
7669_R_O_2_DLX1
7669_R_O_3_DLX1
7669_R_O_4_DLX1
7669_R_O_5_DLX1
7669_R_O_6_DLX1
7669_R_O_7_DLX1
7669_R_O_8_DLX1
7669_R_O_9_DLX1
7669_R_Olf_DLX1
7669_R_P_1_Bulk
7669_R_P_1_COUPTF2
7669_R_P_1_DLX1
7669_R_P_1_NeuN
7669_R_P_1_TBR1
7669_R_P_10_DLX1
7669_R_P_11_DLX1
7669_R_P_11_TBR1
7669_R_P_12_DLX1
7669_R_P_12_TBR1
7669_R_P_13_DLX1
7669_R_P_13_TBR1
7669_R_P_14_DLX1
7669_R_P_14_TBR1
7669_R_P_15_DLX1
7669_R_P_15_TBR1
7669_R_P_16_DLX1
7669_R_P_16_TBR1
7669_R_P_17_DLX1
7669_R_P_17_TBR1
7669_R_P_2_DLX1
7669_R_P_2_TBR1
7669_R_P_3_DLX1
7669_R_P_3_TBR1
7669_R_P_4_DLX1
7669_R_P_4_TBR1
7669_R_P_5_DLX1
7669_R_P_5_TBR1
7669_R_P_6_DLX1
7669_R_P_6_TBR1
7669_R_P_7_DLX1
7669_R_P_7_TBR1
7669_R_P_8_DLX1
7669_R_P_8_TBR1
7669_R_P_9_DLX
7669_R_P_9_DLX1
7669_R_P_9_TBR1
7669_R_POA_Bulk
7669_R_POA_DLX1
7669_R_sec1_Cau_Bulk
7669_R_sec2_GP_Bulk
7669_R_sec2_GP_DLX1
7669_R_sec2_Put_Bulk
7669_R_sec2_Thal_Bulk
7669_R_sec2_Thal_COUPTF2
7669_R_sec2_Thal_DLX1
7669_R_sec3_Cau_DLX1
7669_R_sec3_Hip_COUPTF2
7669_R_sec3_Hip_DLX1
7669_R_sec3_Hip_TBR1
7669_R_sec3_Put_DLX1
7669_R_Skin_Bulk
7669_R_T_1_Bulk
7669_R_T_1_COUPTF2
7669_R_T_1_DLX1
7669_R_T_1_TBR1
7669_RA9_Bulk
7669_RV10_Bulk
JGG

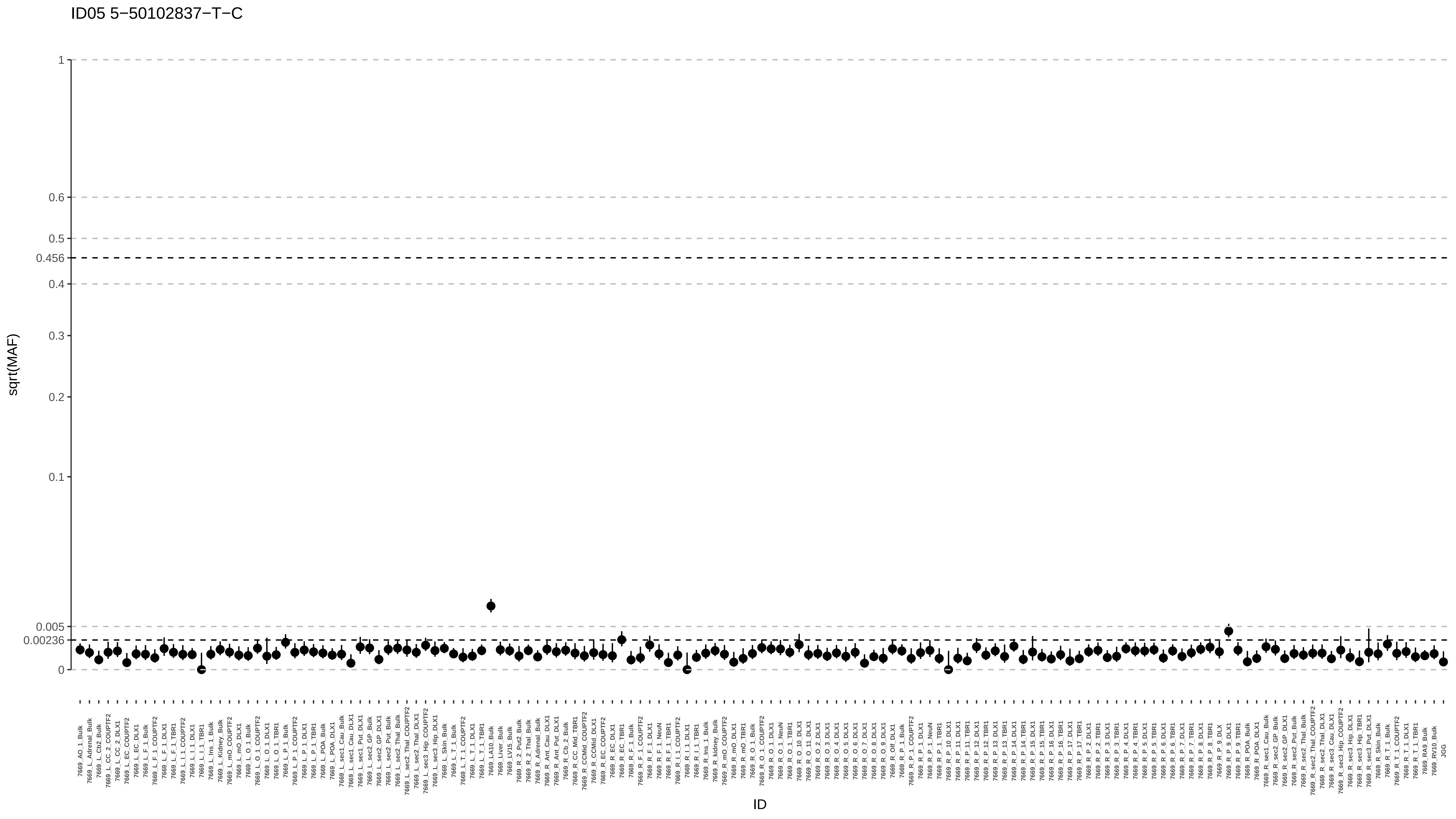
ID05 5−50102837−T−C
sqrt(MAF)
1
0.6
0.5
0.456
0.4
0.3
0.2
0.1
0.005
0.00236
0
ID

ID05 5−53535260−A−G

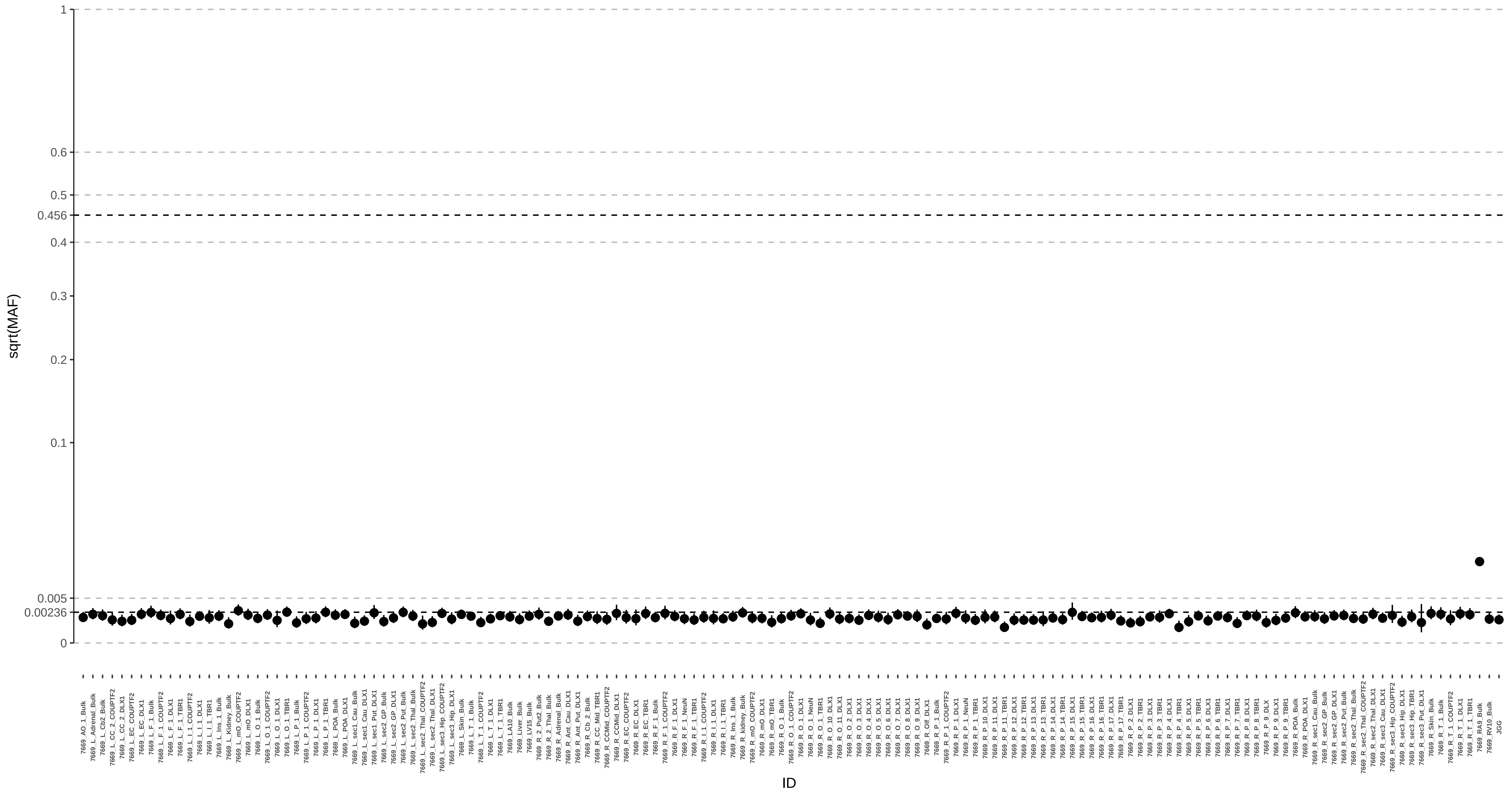

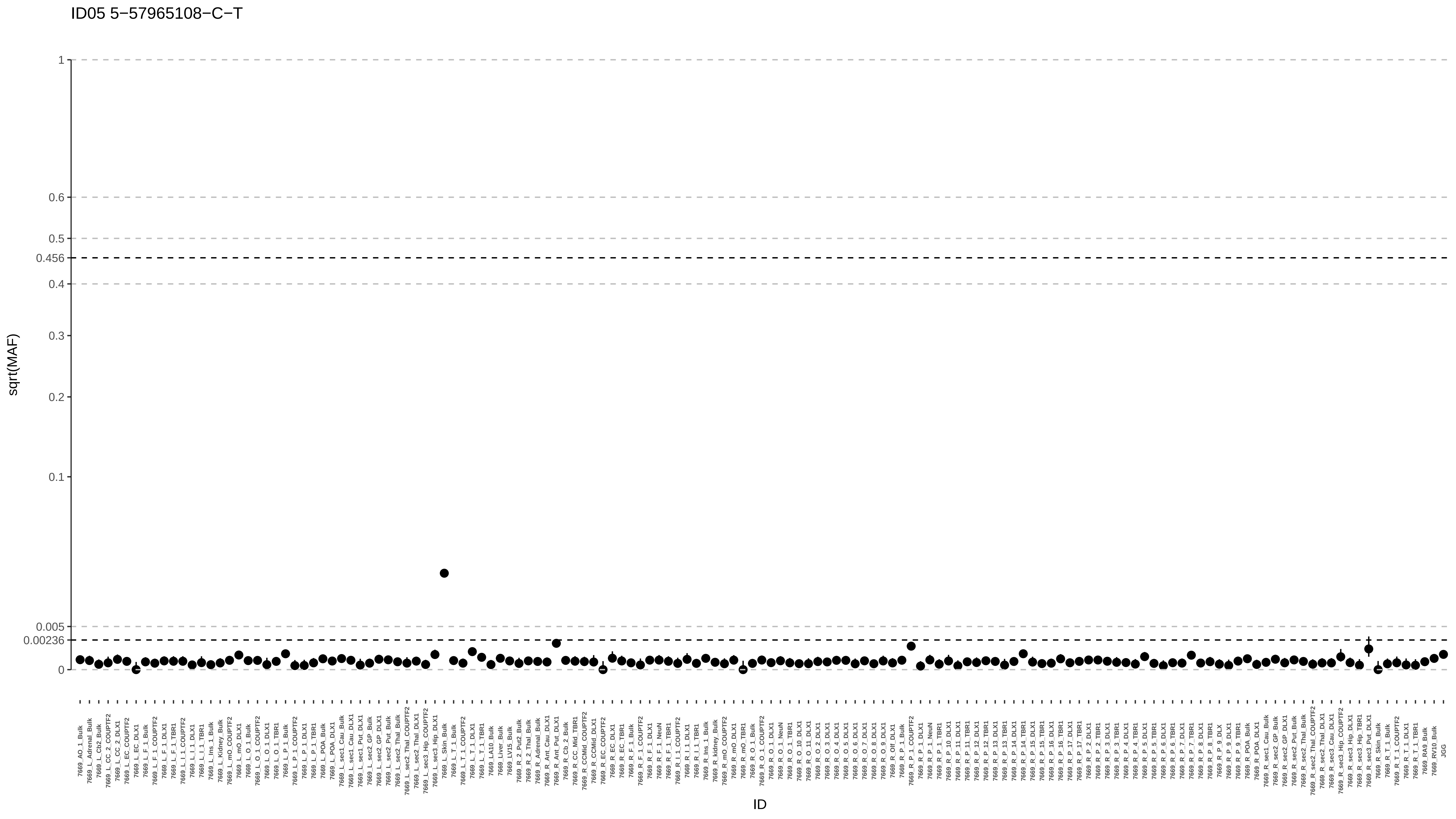
ID05 5−57965108−C−T
sqrt(MAF)
1
0.6
0.5
0.456
0.4
0.3
0.2
0.1
0.005
0.00236
0
ID

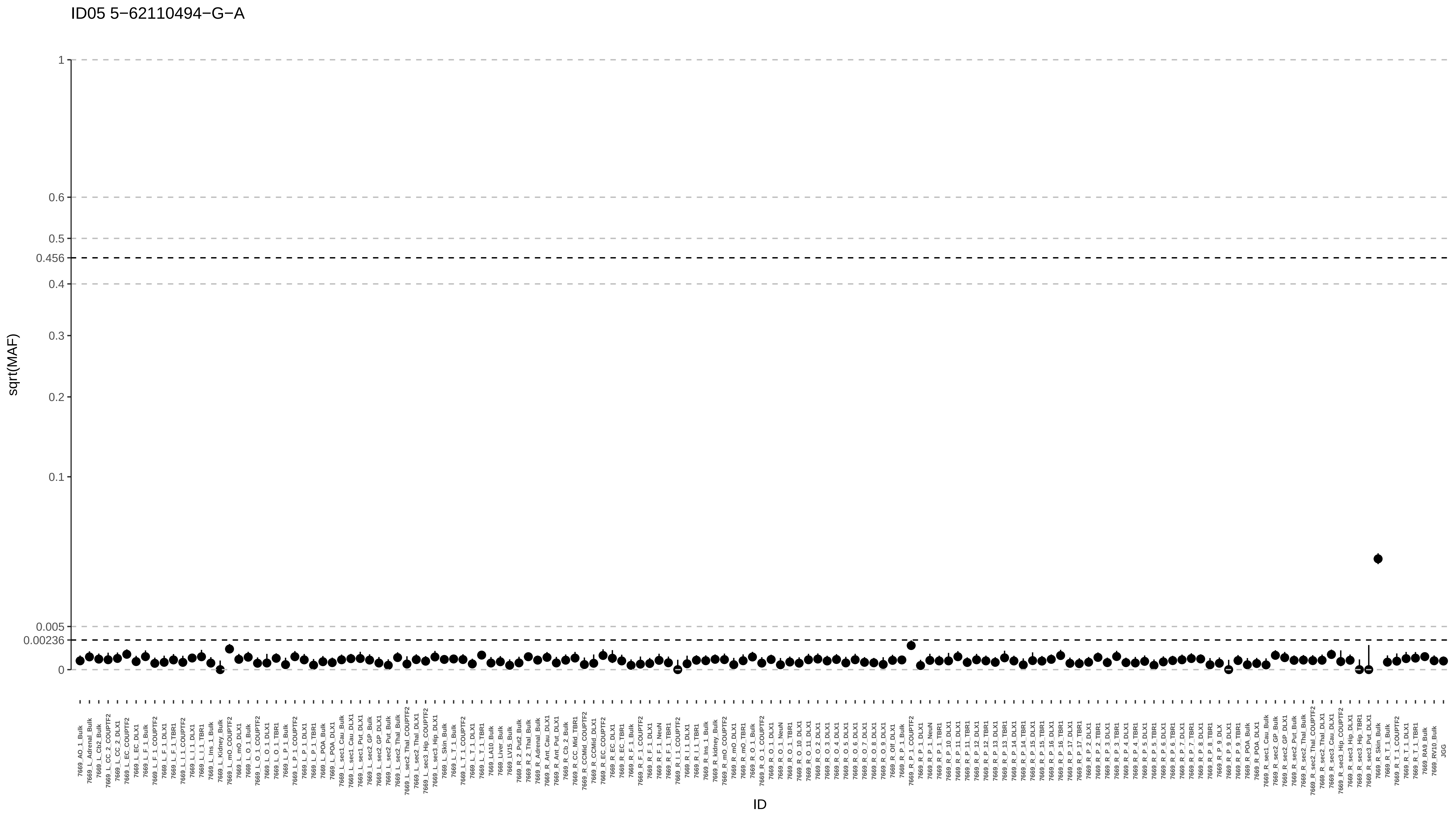
ID05 5−62110494−G−A
sqrt(MAF)
ID
1
0.6
0.5
0.456
0.4
0.3
0.2
0.1
0.005
0.00236
0

ID05 5−64328747−C−T

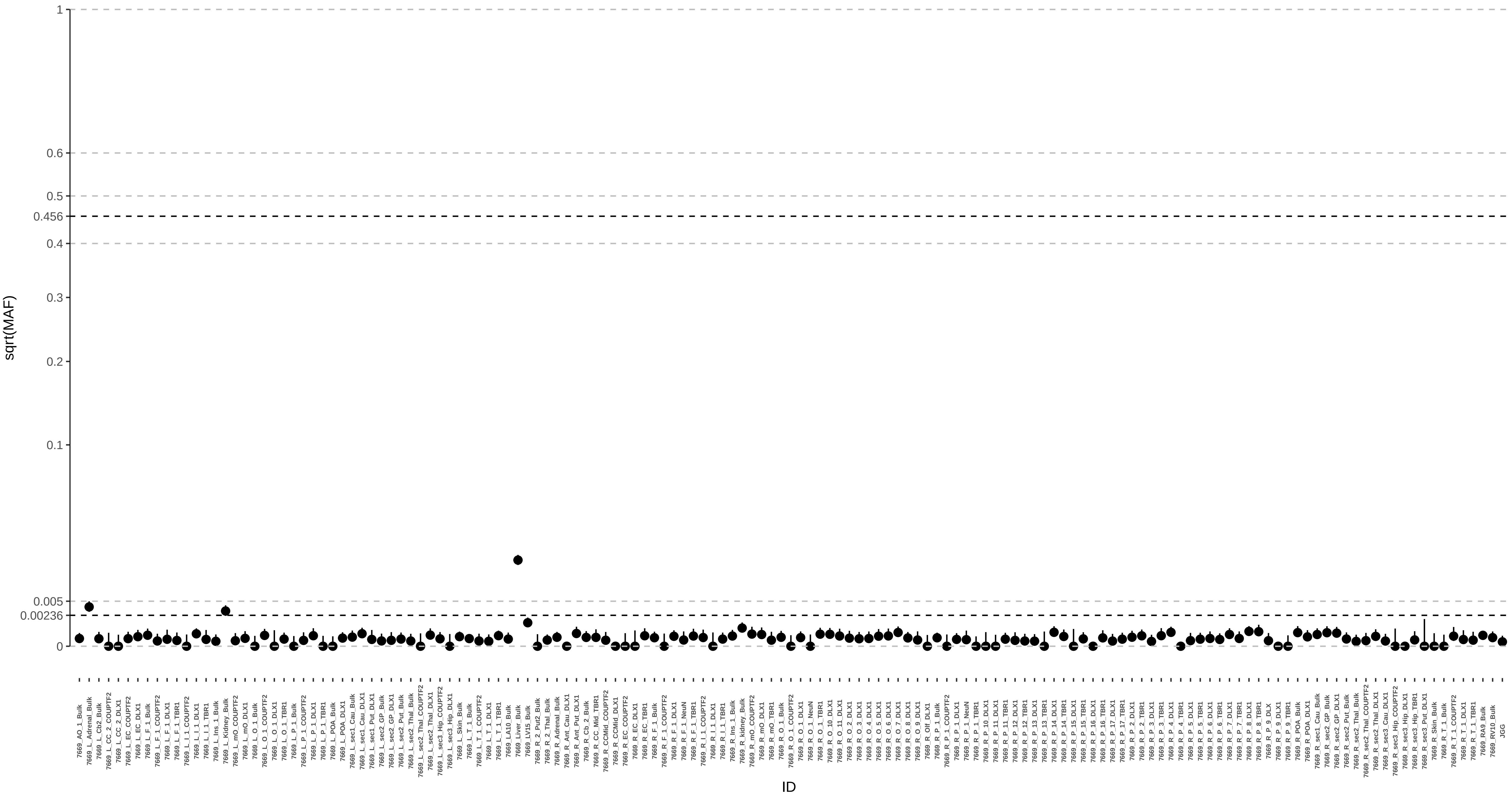

ID05 5−66922121−G−T

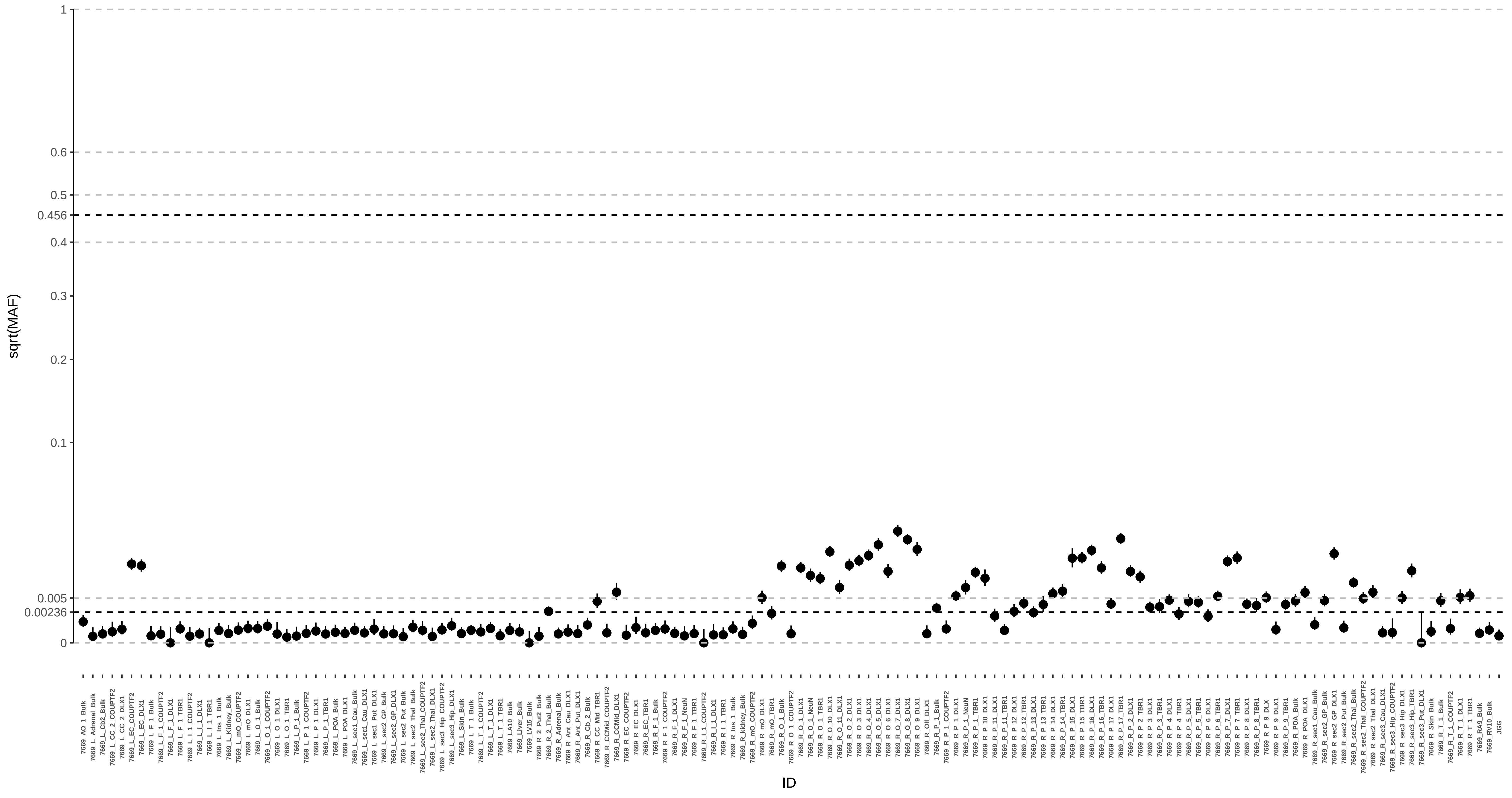

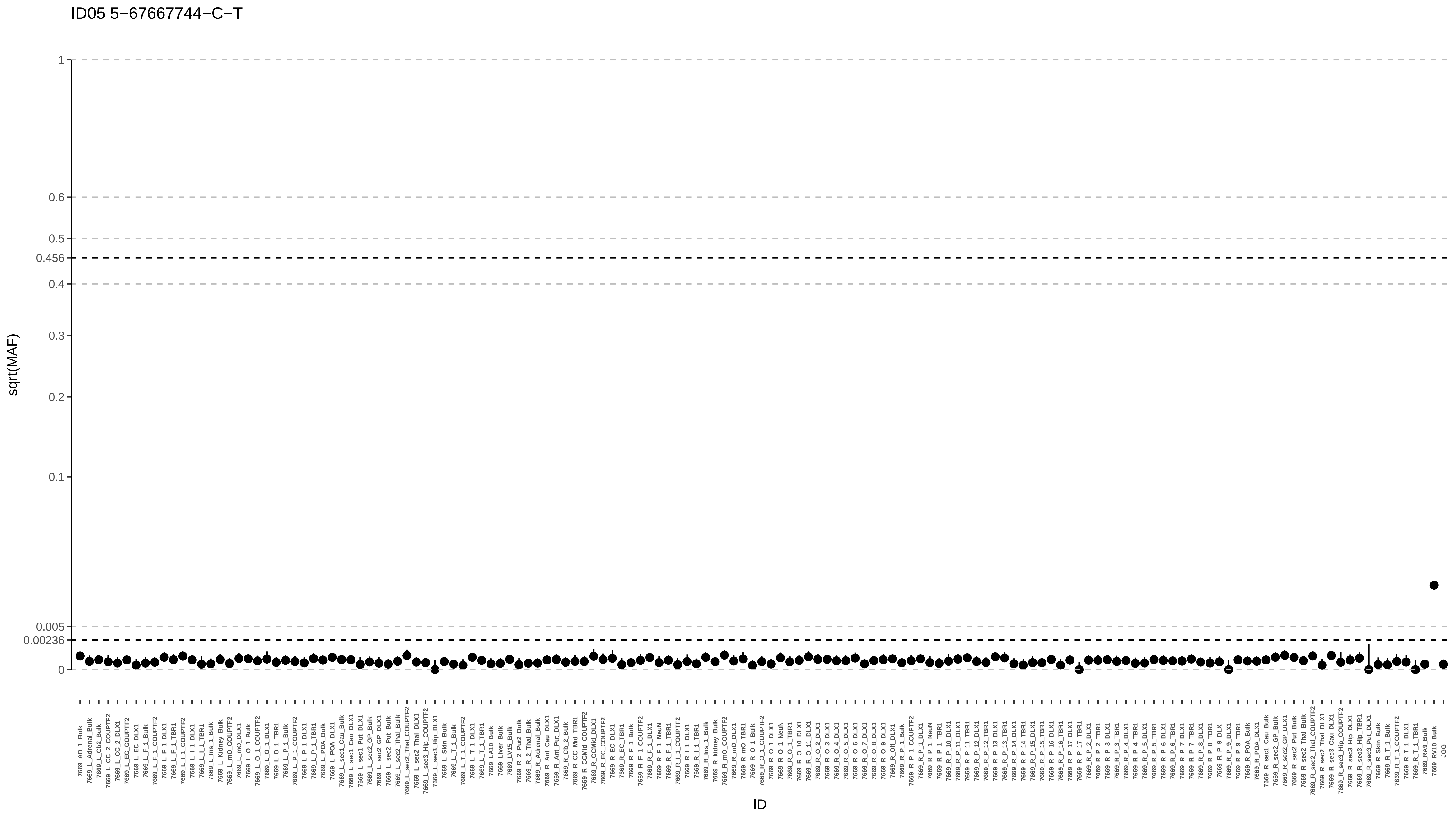
ID05 5−67667744−C−T
sqrt(MAF)
1
0.6
0.5
0.456
0.4
0.3
0.2
0.1
0.005
0.00236
0
ID

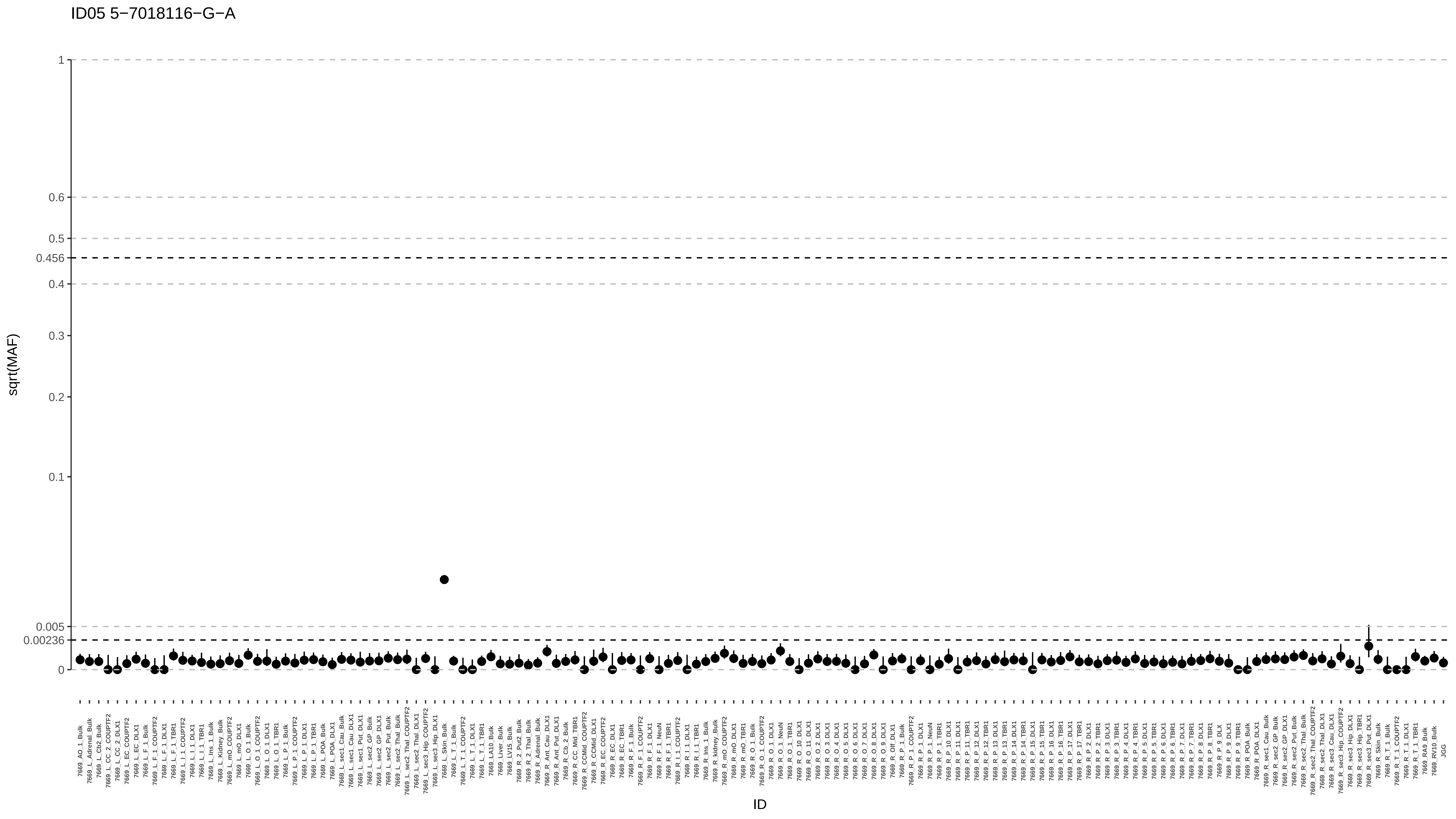
ID05 5−7018116−G−A
sqrt(MAF)
1
0.6
0.5
0.456
0.4
0.3
0.2
0.1
0.005
0.00236
0
ID

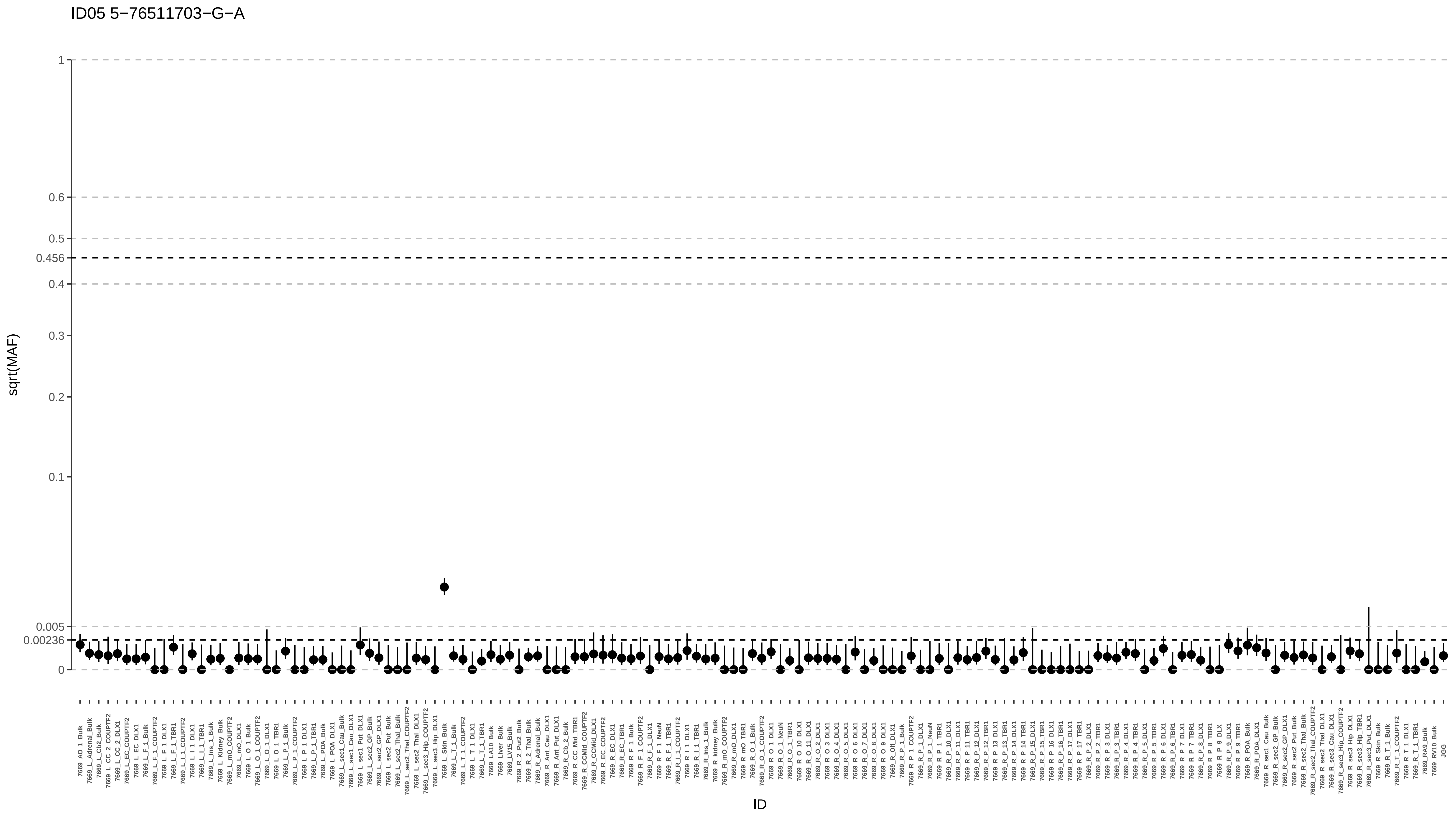
ID05 5−76511703−G−A
sqrt(MAF)
ID
1
0.6
0.5
0.456
0.4
0.3
0.2
0.1
0.005
0.00236
0

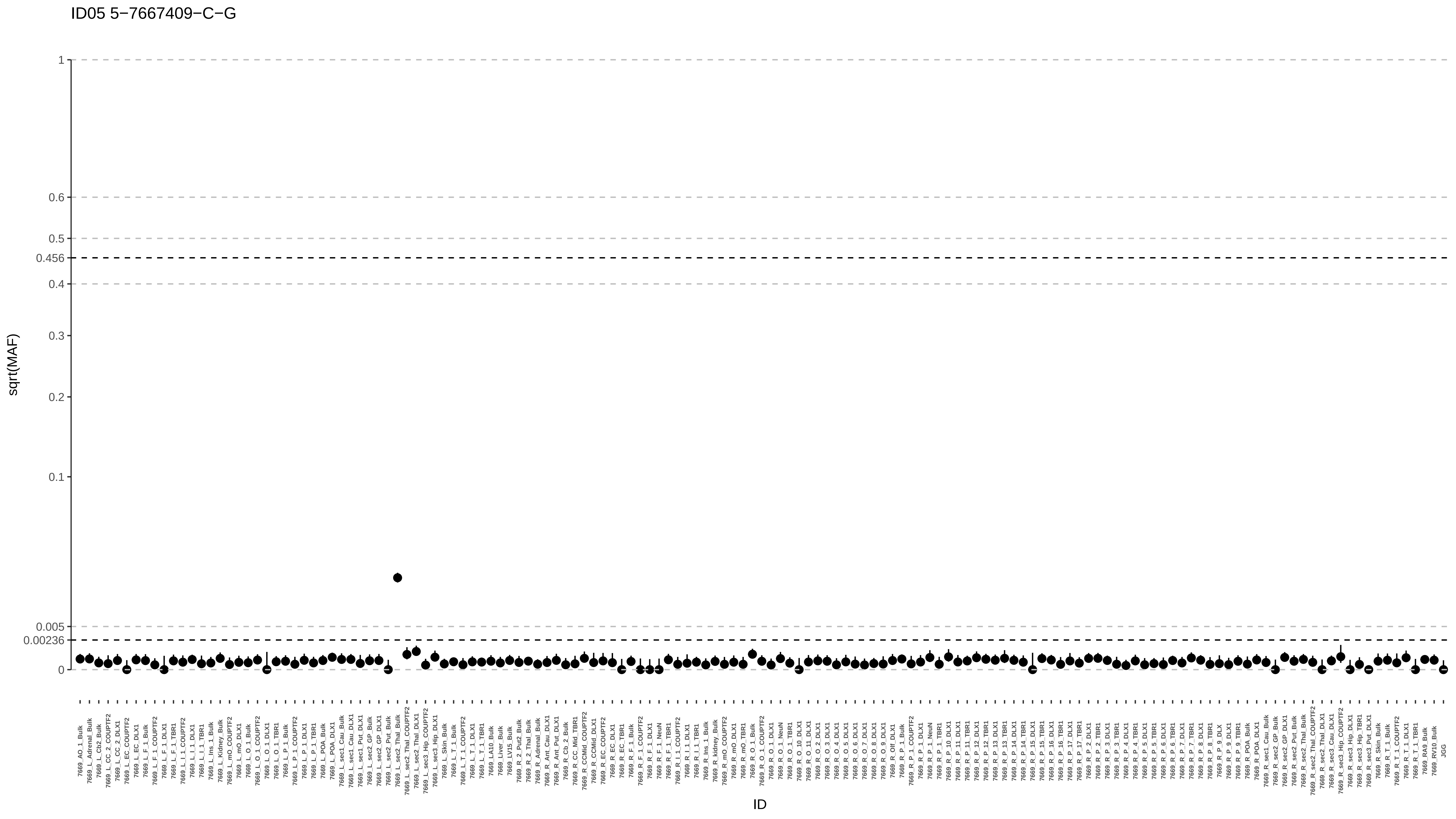

ID05 5−7667409−C−G
sqrt(MAF)
ID
1
0.6
0.5
0.456
0.4
0.3
0.2
0.1
0.005
0.00236
0

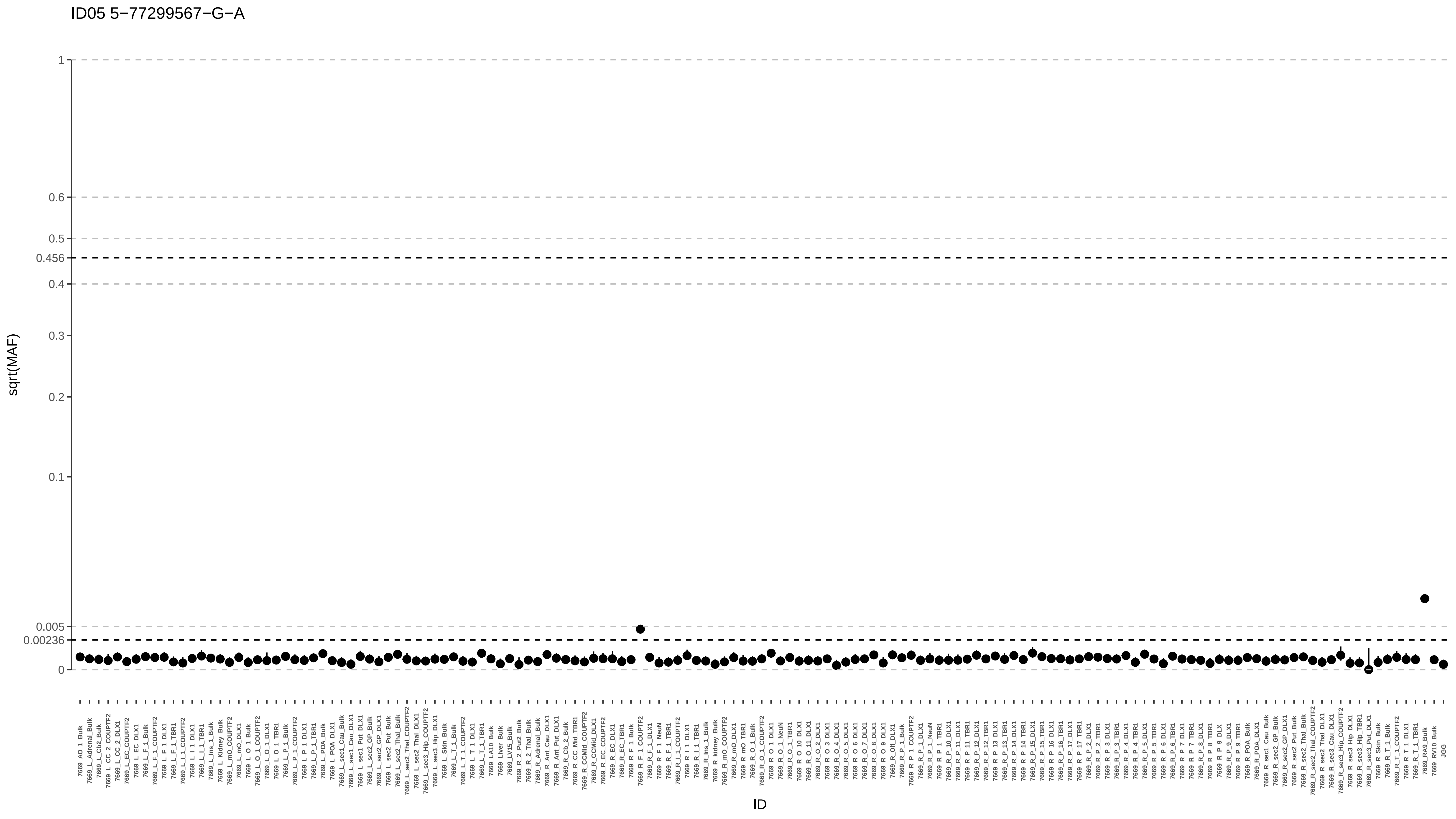

ID05 5−77299567−G−A
sqrt(MAF)
1
0.6
0.5
0.456
0.4
0.3
0.2
0.1
0.005
0.00236
0
ID

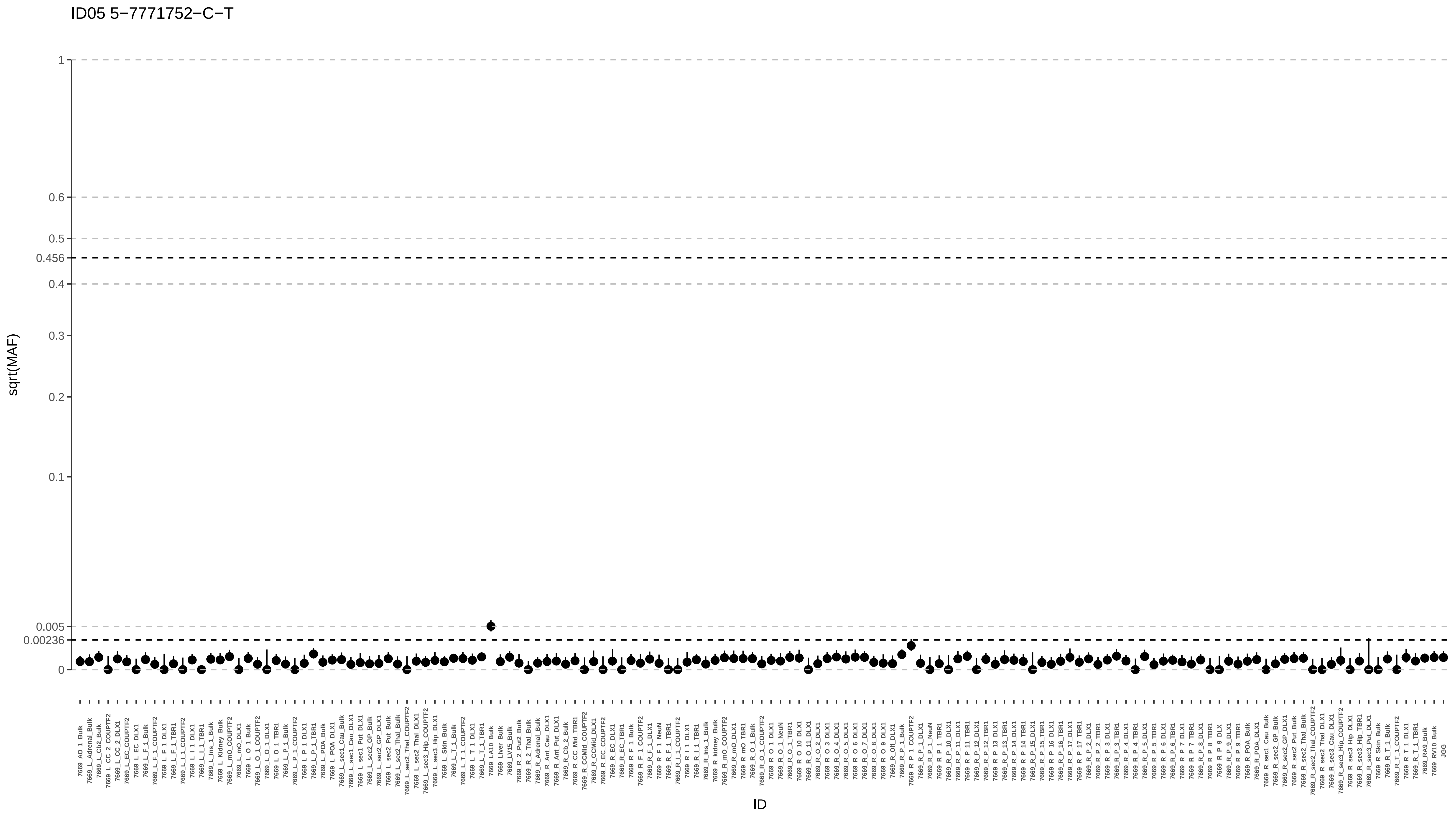
ID05 5−7771752−C−T
sqrt(MAF)
1
0.6
0.5
0.456
0.4
0.3
0.2
0.1
0.005
0.00236
0
ID

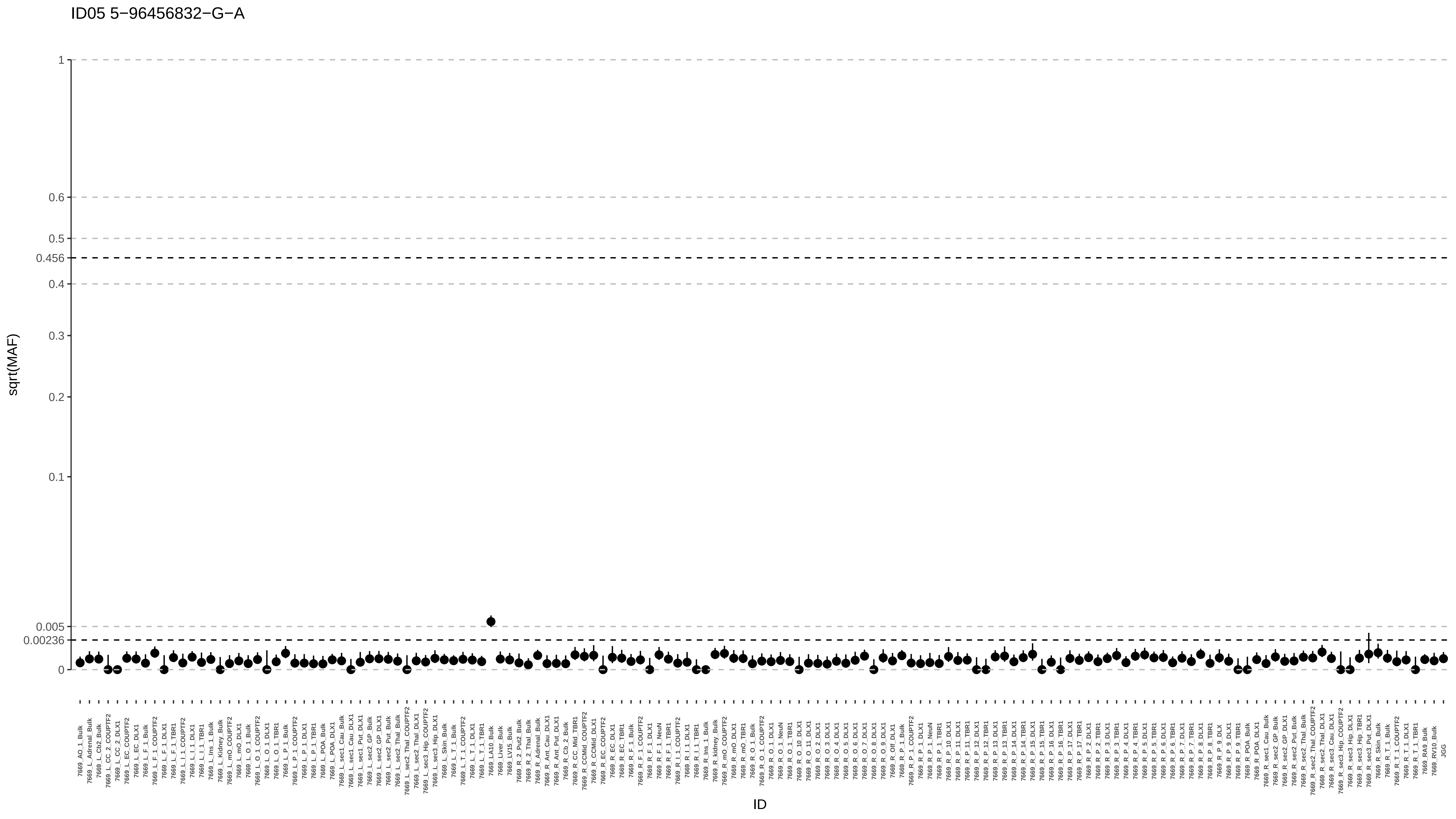
ID05 5−96456832−G−A
sqrt(MAF)
ID
1
0.6
0.5
0.456
0.4
0.3
0.2
0.1
0.005
0.00236
0

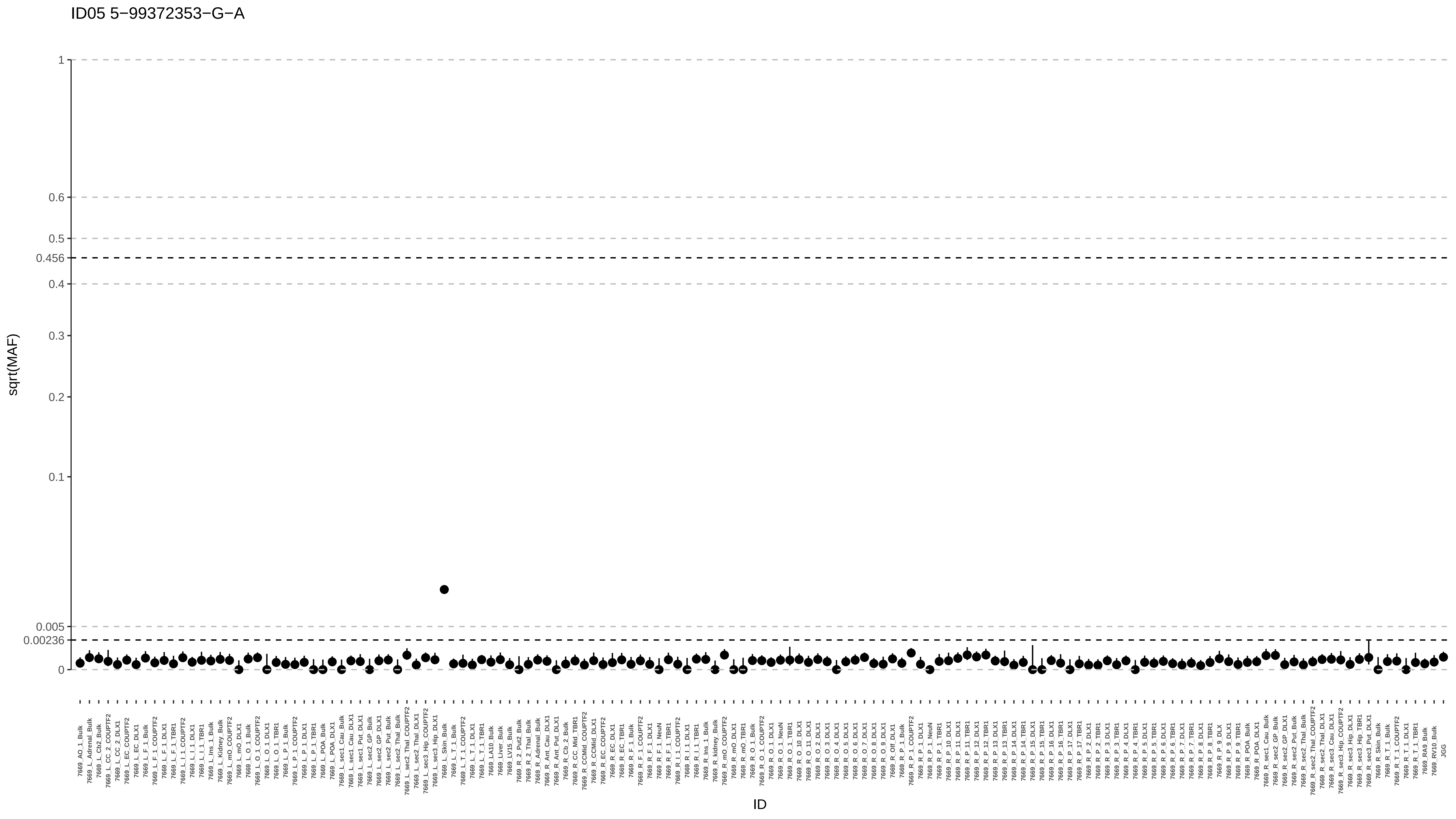
ID05 5−99372353−G−A
sqrt(MAF)
ID

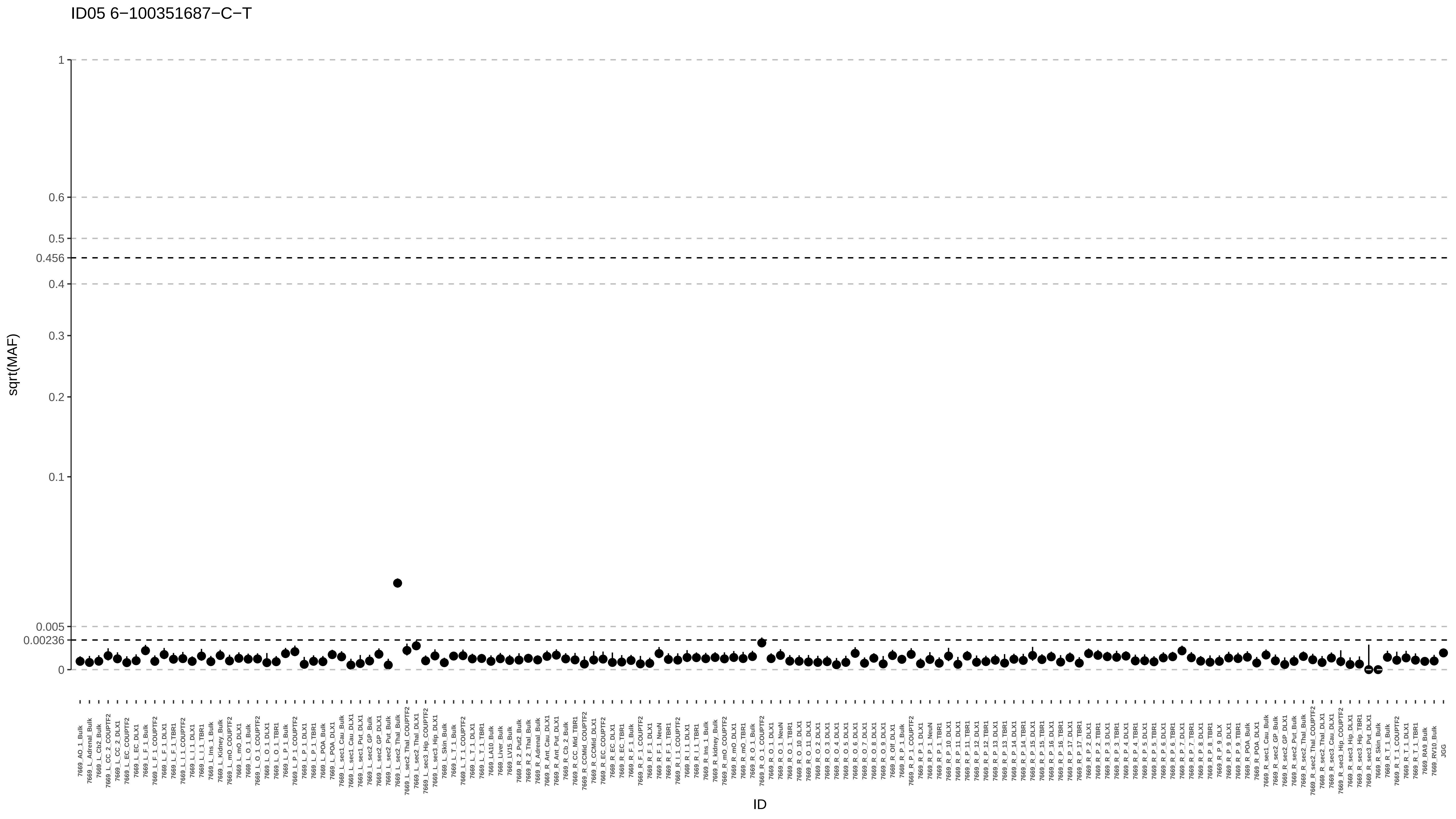
ID05 6−100351687−C−T
sqrt(MAF)
1
0.6
0.5
0.456
0.4
0.3
0.2
0.1
0.005
0.00236
0
ID

ID05 6−102878754−T−A

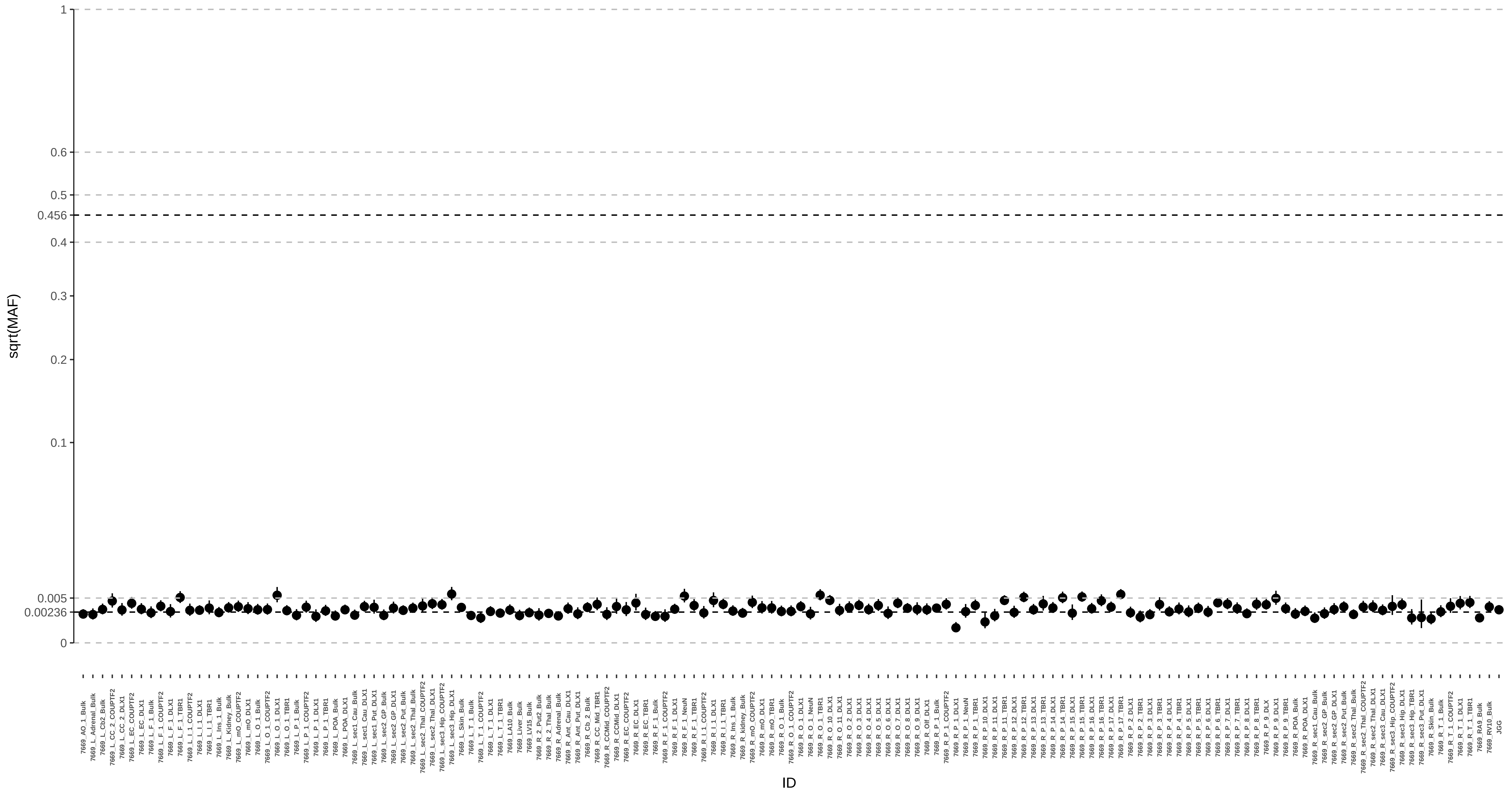

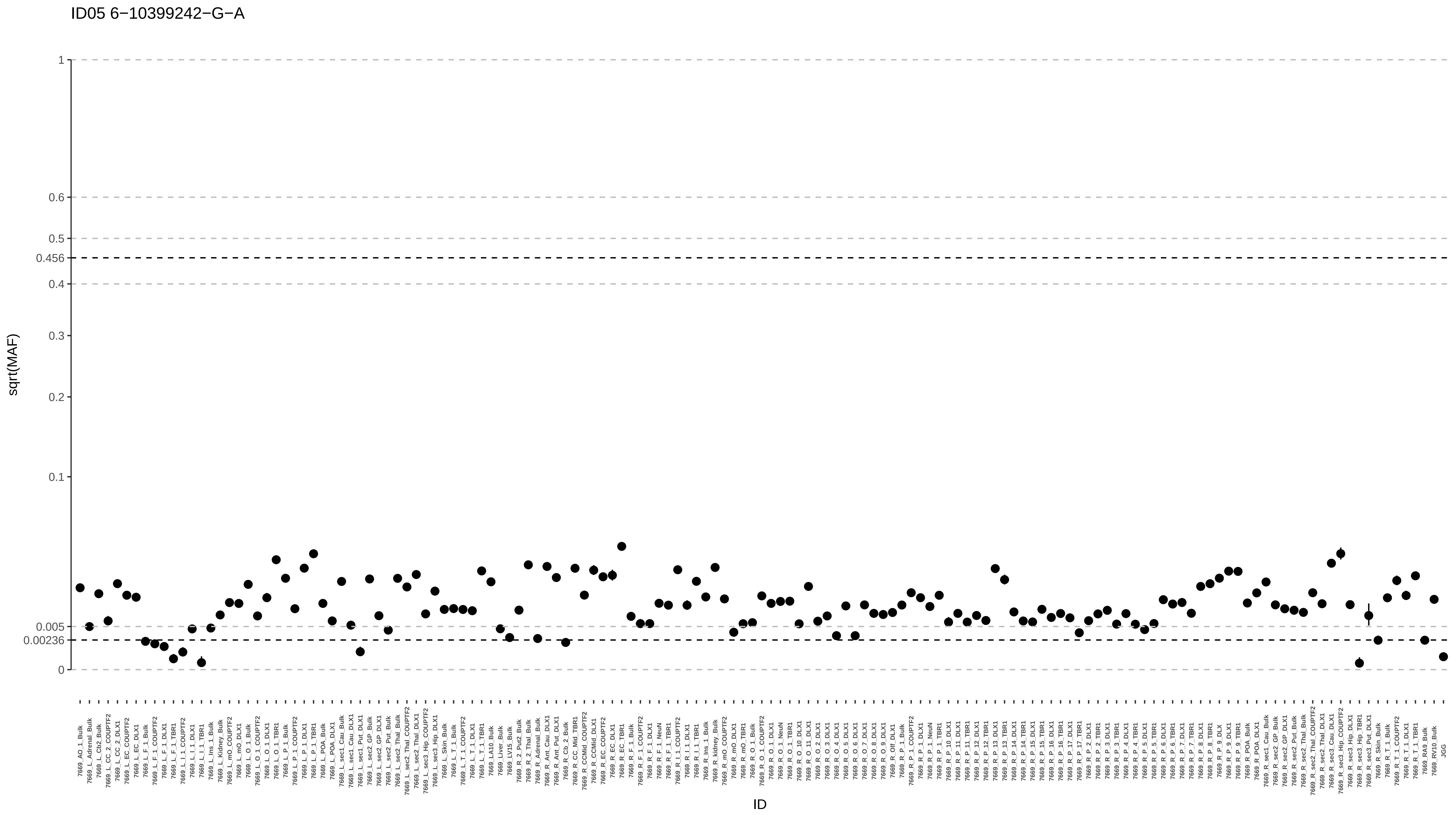
ID05 6−10399242−G−A
sqrt(MAF)
1
0.6
0.5
0.456
0.4
0.3
0.2
0.1
0.005
0.00236
0
ID

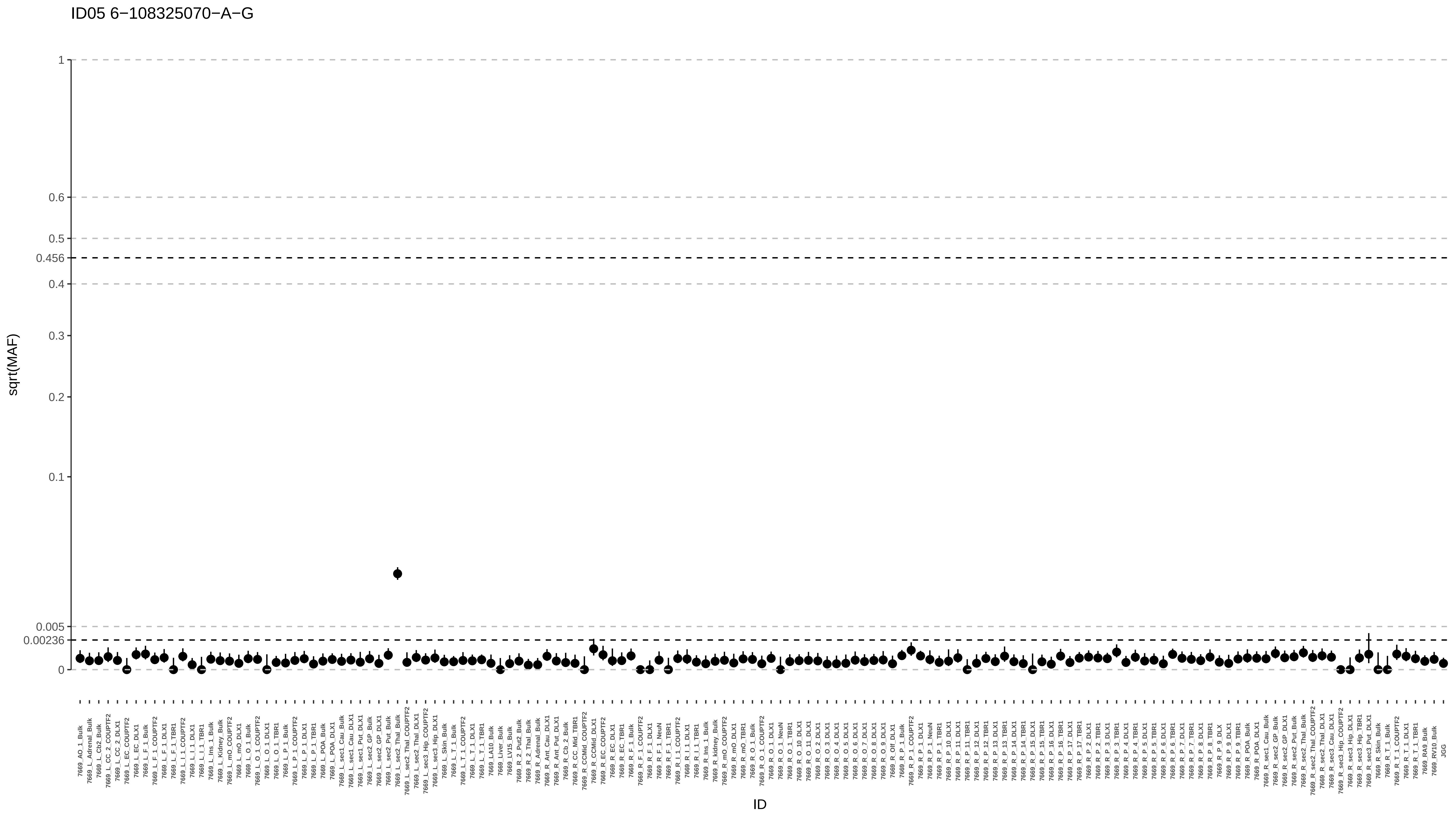

ID05 6−108325070−A−G
sqrt(MAF)
1
0.6
0.5
0.456
0.4
0.3
0.2
0.1
0.005
0.00236
0
7669_AO_1_Bulk
7669_L_Adrenal_Bulk
7669_L_Cb2_Bulk
7669_L_CC_2_COUPTF2
7669_L_CC_2_DLX1
7669_L_EC_COUPTF2
7669_L_EC_DLX1
7669_L_F_1_Bulk
7669_L_F_1_COUPTF2
7669_L_F_1_DLX1
7669_L_F_1_TBR1
7669_L_I_1_COUPTF2
7669_L_I_1_DLX1
7669_L_I_1_TBR1
7669_L_Ins_1_Bulk
7669_L_Kidney_Bulk
7669_L_mO_COUPTF2
7669_L_mO_DLX1
7669_L_O_1_Bulk
7669_L_O_1_COUPTF2
7669_L_O_1_DLX1
7669_L_O_1_TBR1
7669_L_P_1_Bulk
7669_L_P_1_COUPTF2
7669_L_P_1_DLX1
7669_L_P_1_TBR1
7669_L_POA_Bulk
7669_L_POA_DLX1
7669_L_sec1_Cau_Bulk
7669_L_sec1_Cau_DLX1
7669_L_sec1_Put_DLX1
7669_L_sec2_GP_Bulk
7669_L_sec2_GP_DLX1
7669_L_sec2_Put_Bulk
7669_L_sec2_Thal_Bulk
7669_L_sec2_Thal_COUPTF2
7669_L_sec2_Thal_DLX1
7669_L_sec3_Hip_COUPTF2
7669_L_sec3_Hip_DLX1
7669_L_Skin_Bulk
7669_L_T_1_Bulk
7669_L_T_1_COUPTF2
7669_L_T_1_DLX1
7669_L_T_1_TBR1
7669_LA10_Bulk
7669_Liver_Bulk
7669_LV15_Bulk
7669_R_2_Put2_Bulk
7669_R_2_Thal_Bulk
7669_R_Adrenal_Bulk
7669_R_Ant_Cau_DLX1
7669_R_Ant_Put_DLX1
7669_R_Cb_2_Bulk
7669_R_CC_Mid_TBR1
7669_R_CCMid_COUPTF2
7669_R_CCMid_DLX1
7669_R_EC_COUPTF2
7669_R_EC_DLX1
7669_R_EC_TBR1
7669_R_F_1_Bulk
7669_R_F_1_COUPTF2
7669_R_F_1_DLX1
7669_R_F_1_NeuN
7669_R_F_1_TBR1
7669_R_I_1_COUPTF2
7669_R_I_1_DLX1
7669_R_I_1_TBR1
7669_R_Ins_1_Bulk
7669_R_kidney_Bulk
7669_R_mO_COUPTF2
7669_R_mO_DLX1
7669_R_mO_TBR1
7669_R_O_1_Bulk
7669_R_O_1_COUPTF2
7669_R_O_1_DLX1
7669_R_O_1_NeuN
7669_R_O_1_TBR1
7669_R_O_10_DLX1
7669_R_O_11_DLX1
7669_R_O_2_DLX1
7669_R_O_3_DLX1
7669_R_O_4_DLX1
7669_R_O_5_DLX1
7669_R_O_6_DLX1
7669_R_O_7_DLX1
7669_R_O_8_DLX1
7669_R_O_9_DLX1
7669_R_Olf_DLX1
7669_R_P_1_Bulk
7669_R_P_1_COUPTF2
7669_R_P_1_DLX1
7669_R_P_1_NeuN
7669_R_P_1_TBR1
7669_R_P_10_DLX1
7669_R_P_11_DLX1
7669_R_P_11_TBR1
7669_R_P_12_DLX1
7669_R_P_12_TBR1
7669_R_P_13_DLX1
7669_R_P_13_TBR1
7669_R_P_14_DLX1
7669_R_P_14_TBR1
7669_R_P_15_DLX1
7669_R_P_15_TBR1
7669_R_P_16_DLX1
7669_R_P_16_TBR1
7669_R_P_17_DLX1
7669_R_P_17_TBR1
7669_R_P_2_DLX1
7669_R_P_2_TBR1
7669_R_P_3_DLX1
7669_R_P_3_TBR1
7669_R_P_4_DLX1
7669_R_P_4_TBR1
7669_R_P_5_DLX1
7669_R_P_5_TBR1
7669_R_P_6_DLX1
7669_R_P_6_TBR1
7669_R_P_7_DLX1
7669_R_P_7_TBR1
7669_R_P_8_DLX1
7669_R_P_8_TBR1
7669_R_P_9_DLX
7669_R_P_9_DLX1
7669_R_P_9_TBR1
7669_R_POA_Bulk
7669_R_POA_DLX1
7669_R_sec1_Cau_Bulk
7669_R_sec2_GP_Bulk
7669_R_sec2_GP_DLX1
7669_R_sec2_Put_Bulk
7669_R_sec2_Thal_Bulk
7669_R_sec2_Thal_COUPTF2
7669_R_sec2_Thal_DLX1
7669_R_sec3_Cau_DLX1
7669_R_sec3_Hip_COUPTF2
7669_R_sec3_Hip_DLX1
7669_R_sec3_Hip_TBR1
7669_R_sec3_Put_DLX1
7669_R_Skin_Bulk
7669_R_T_1_Bulk
7669_R_T_1_COUPTF2
7669_R_T_1_DLX1
7669_R_T_1_TBR1
7669_RA9_Bulk
7669_RV10_Bulk
JGG
ID

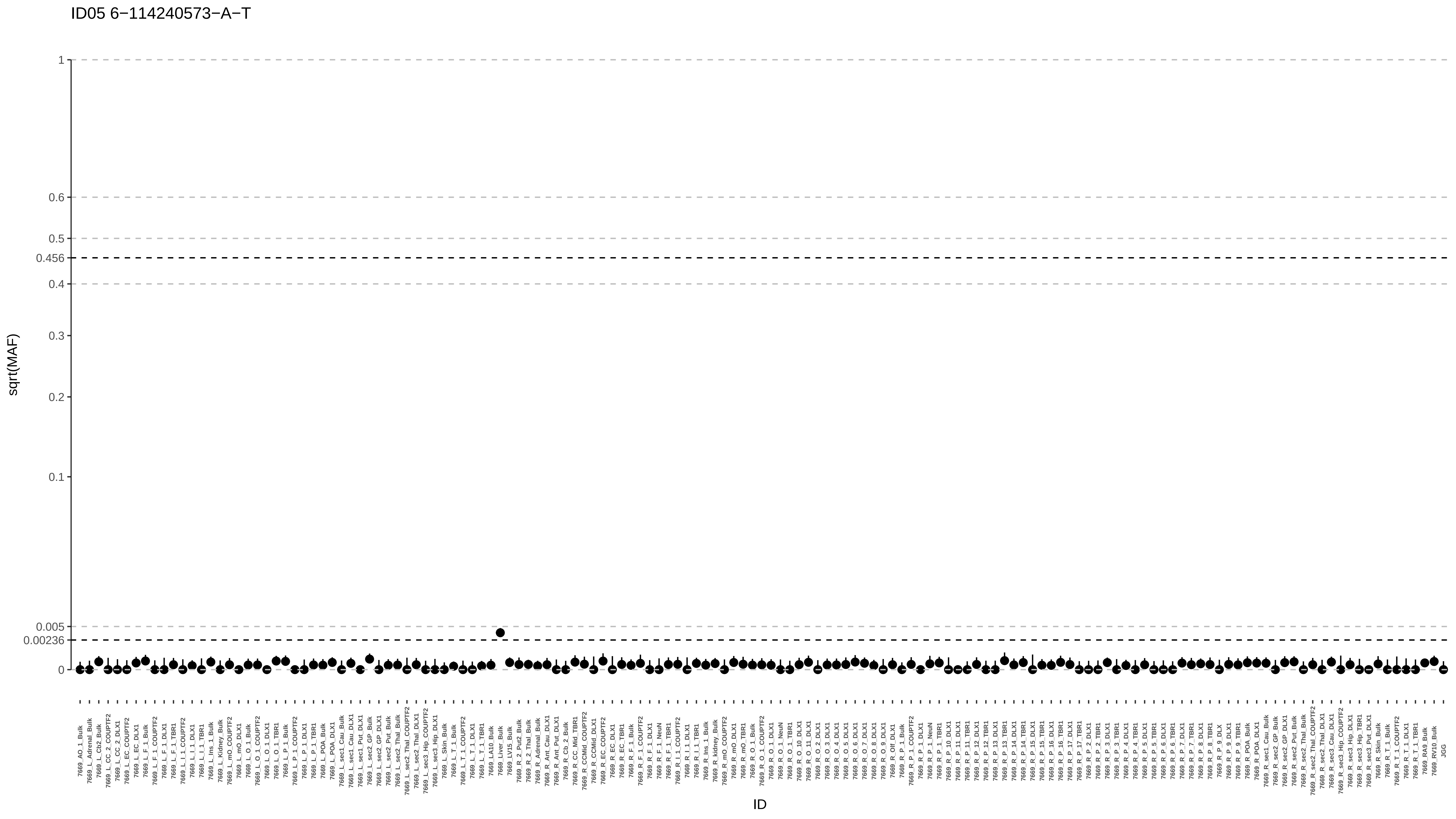
ID05 6−114240573−A−T
sqrt(MAF)
1
0.6
0.5
0.456
0.4
0.3
0.2
0.1
0.005
0.00236
0
ID

ID05 6−115212381−G−A

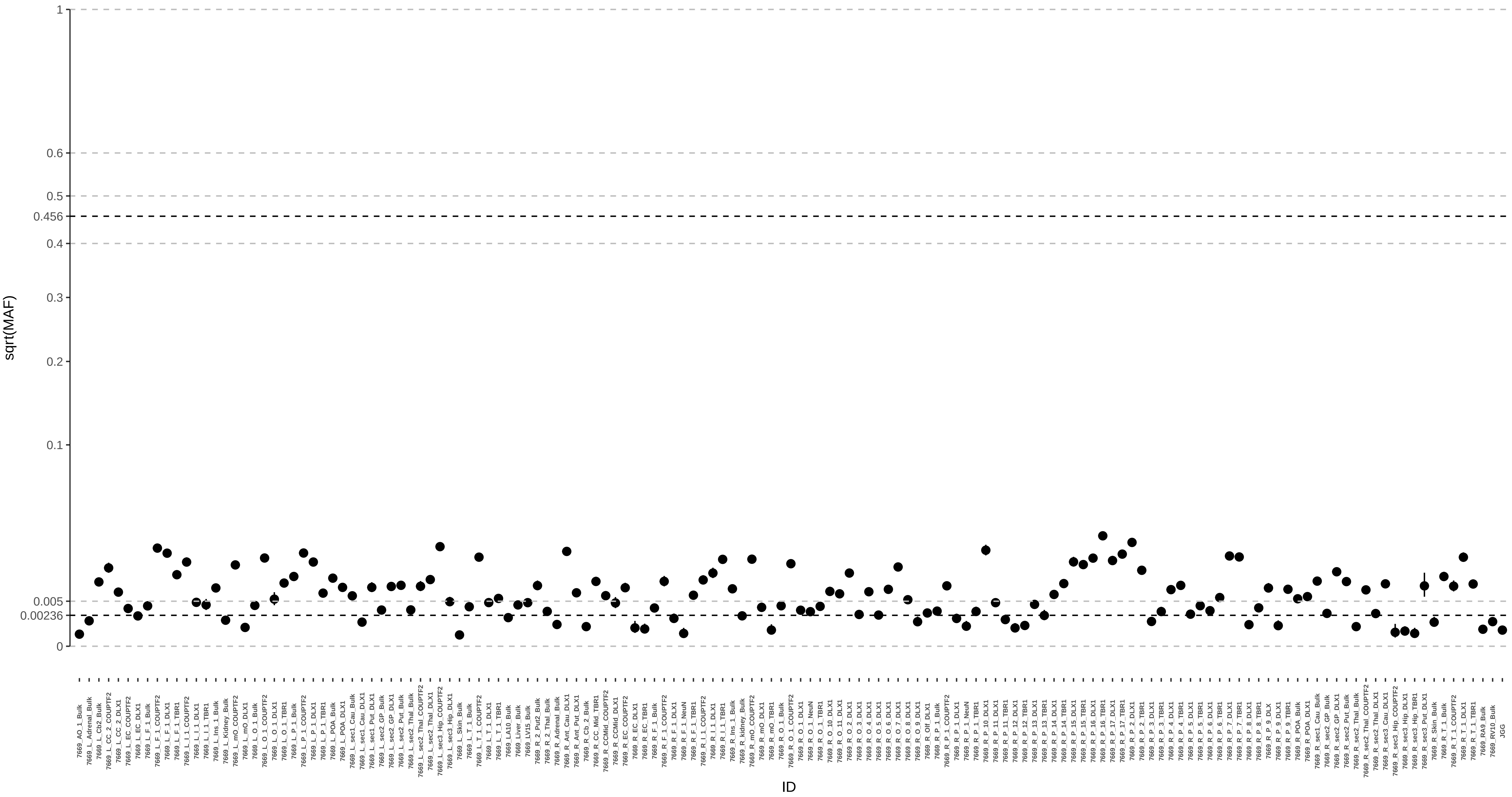

ID05 6−115259744−C−T

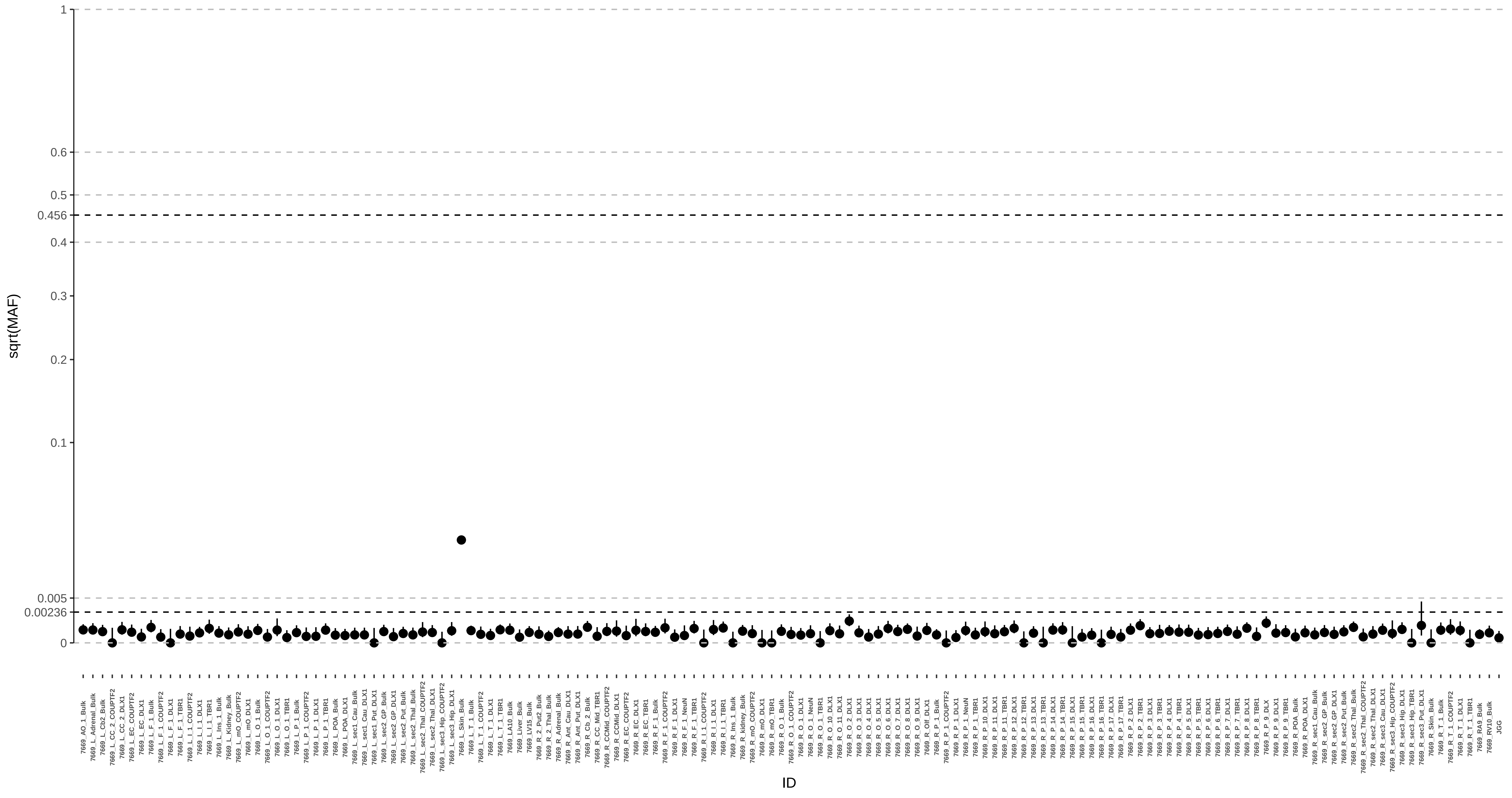

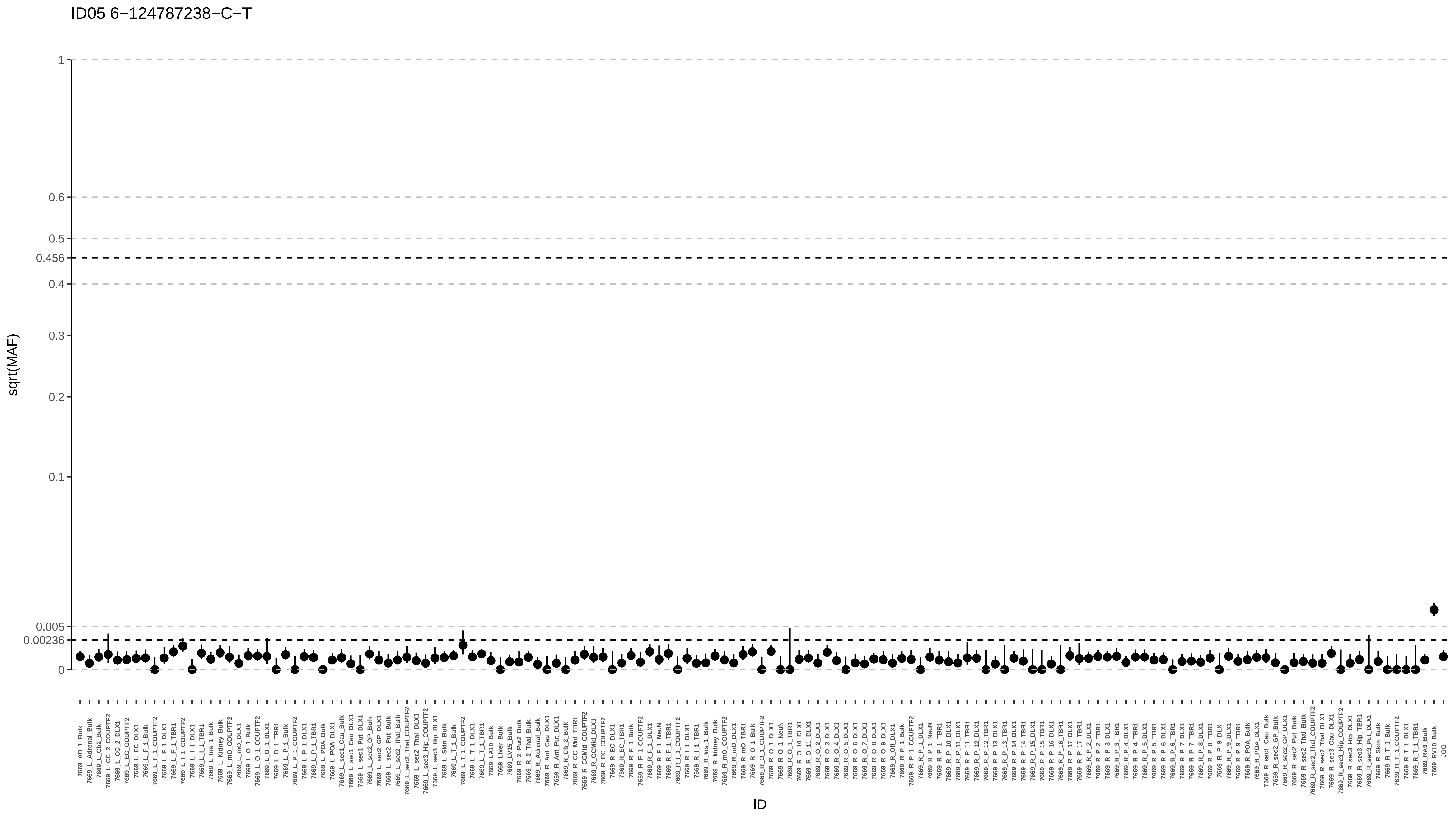

ID05 6−124787238−C−T
sqrt(MAF)
1
0.6
0.5
0.456
0.4
0.3
0.2
0.1
0.005
0.00236
0
ID

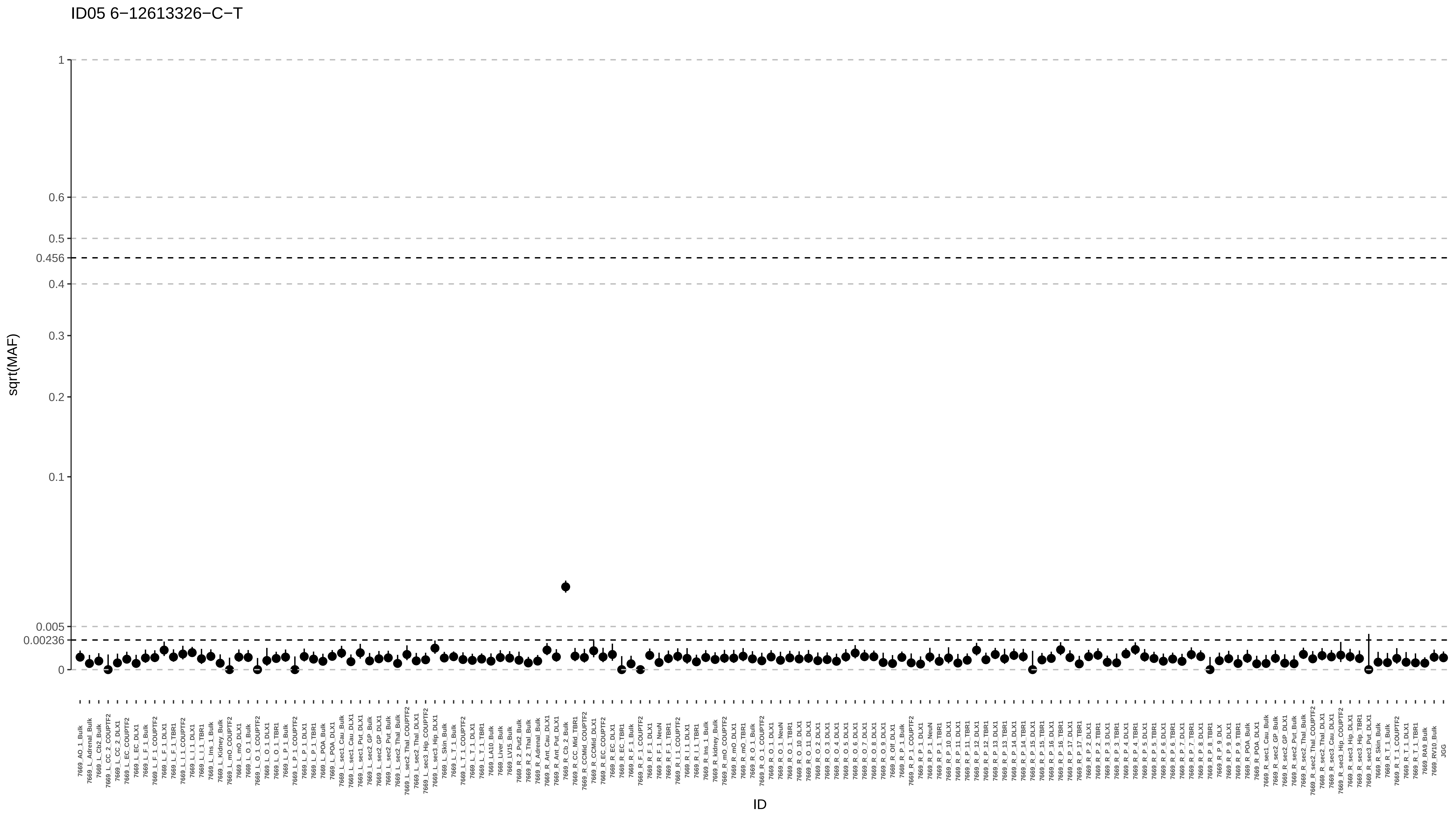
ID05 6−12613326−C−T
sqrt(MAF)
1
0.6
0.5
0.456
0.4
0.3
0.2
0.1
0.005
0.00236
0
ID

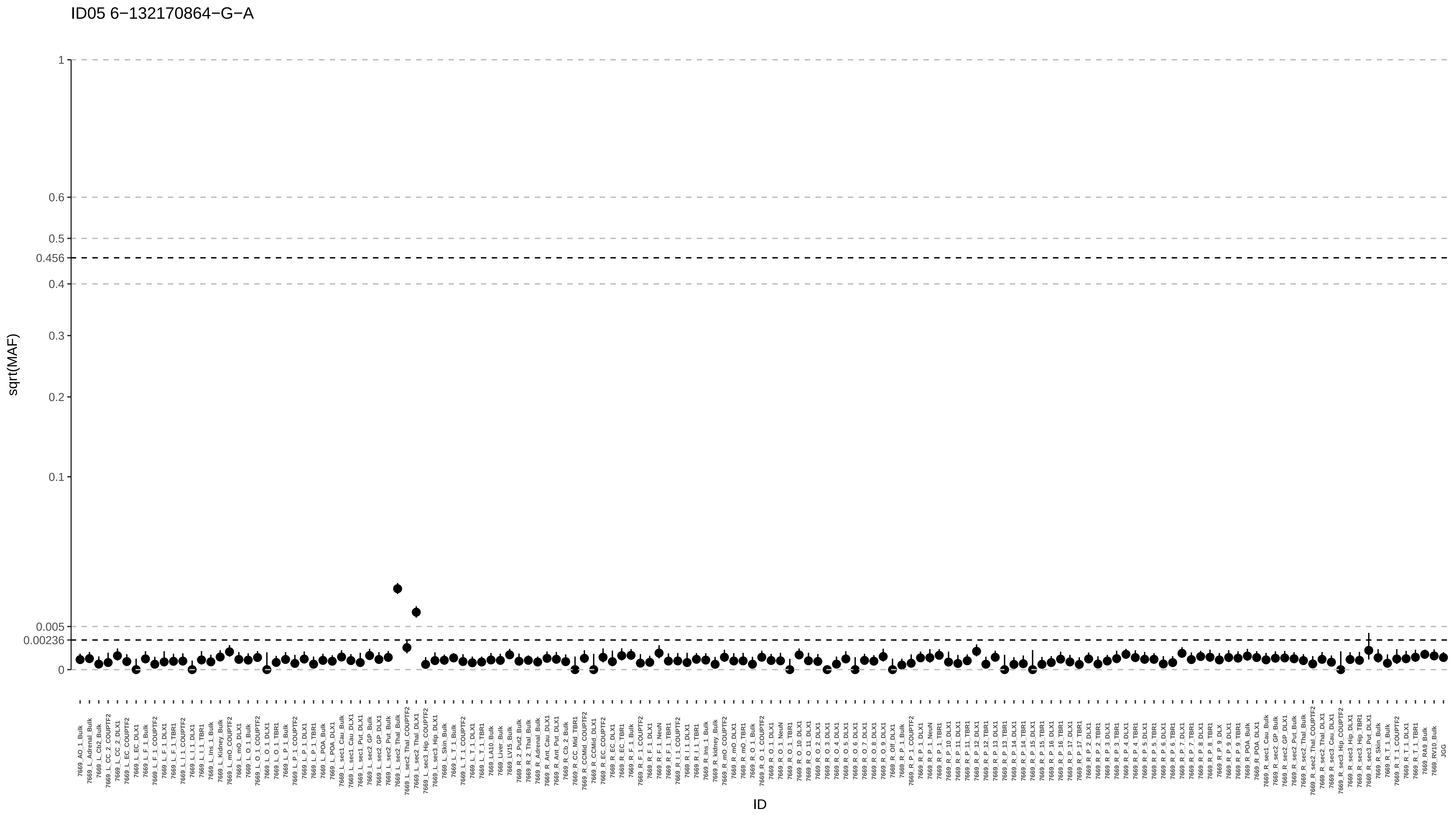
ID05 6−132170864−G−A
sqrt(MAF)
1
0.6
0.5
0.456
0.4
0.3
0.2
0.1
0.005
0.00236
0
ID

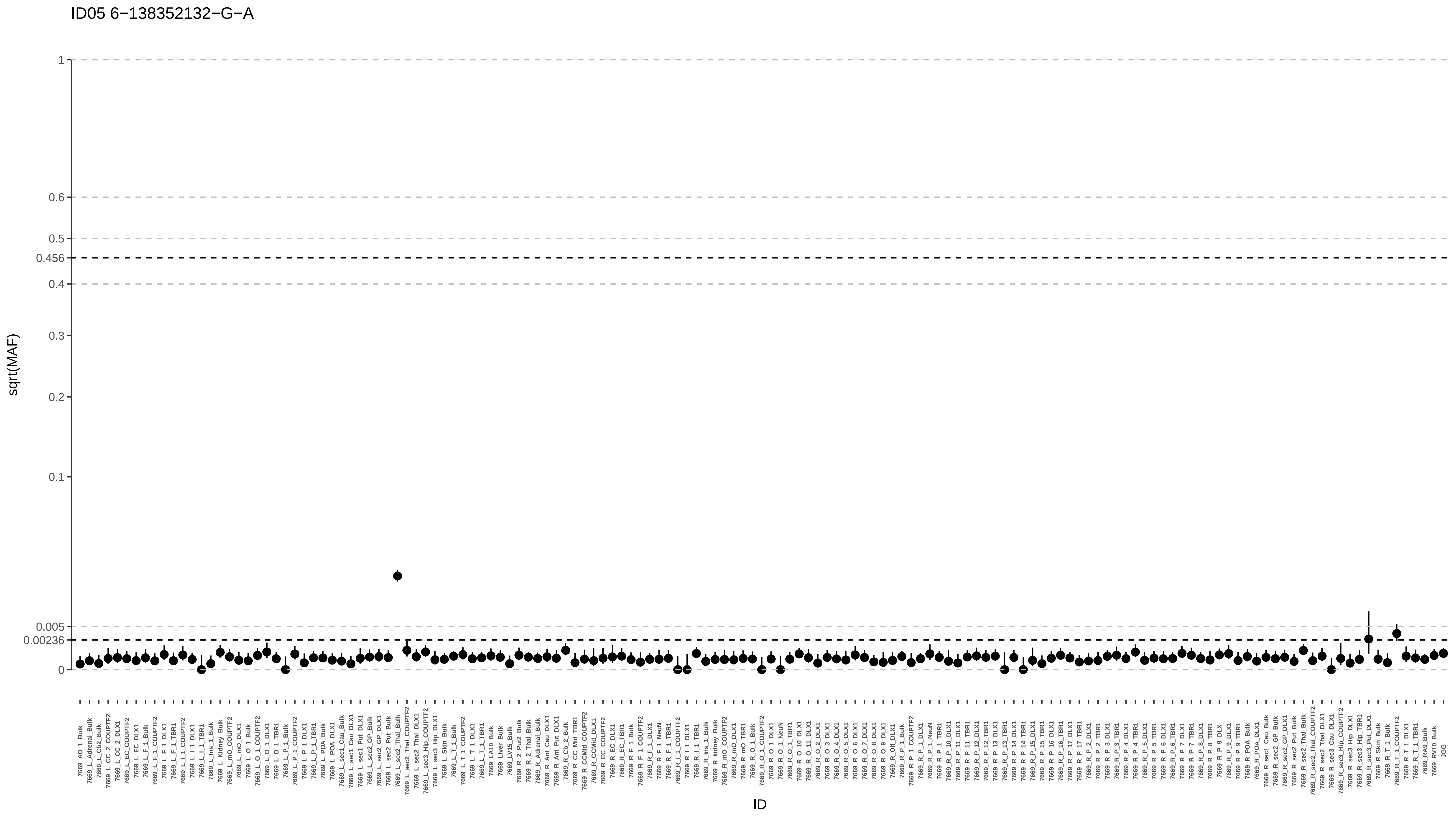
ID05 6−138352132−G−A
sqrt(MAF)
1
0.6
0.5
0.456
0.4
0.3
0.2
0.1
0.005
0.00236
0
ID

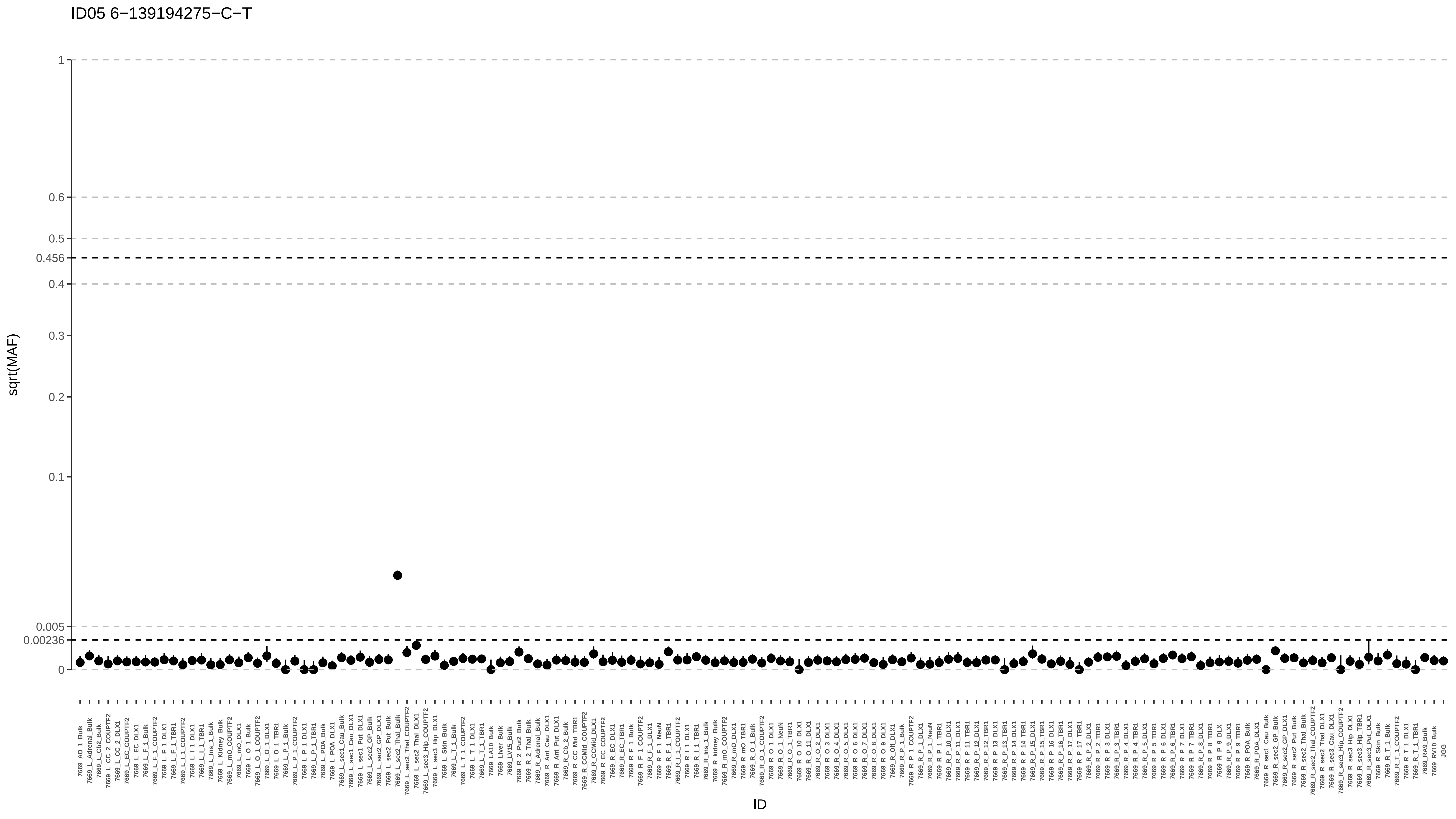
ID05 6−139194275−C−T
sqrt(MAF)
1
0.6
0.5
0.456
0.4
0.3
0.2
0.1
0.005
0.00236
0
ID

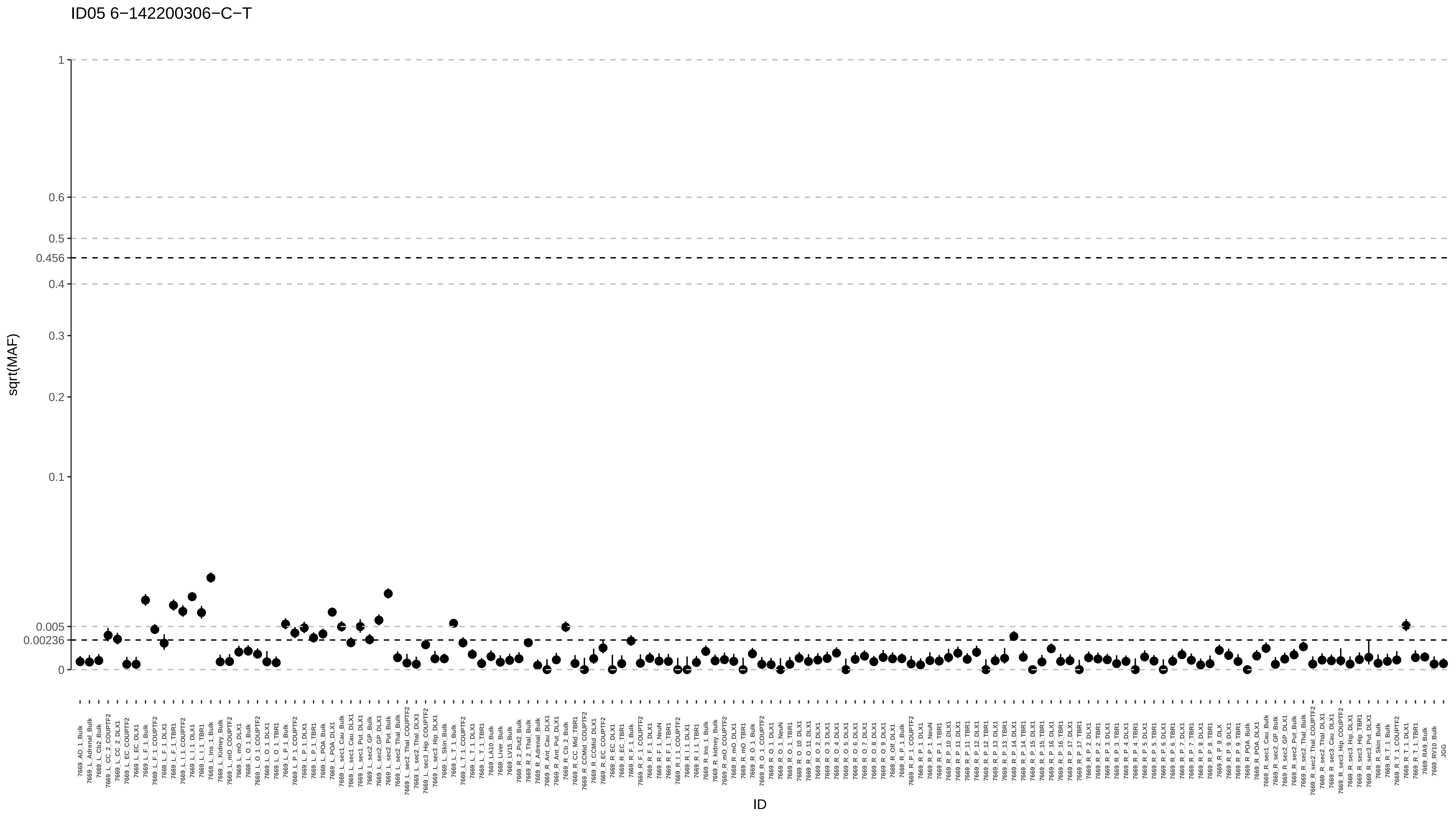

ID05 6−142200306−C−T
sqrt(MAF)
1
0.6
0.5
0.456
0.4
0.3
0.2
0.1
0.005
0.00236
0
ID

ID05 6−144775505−C−T

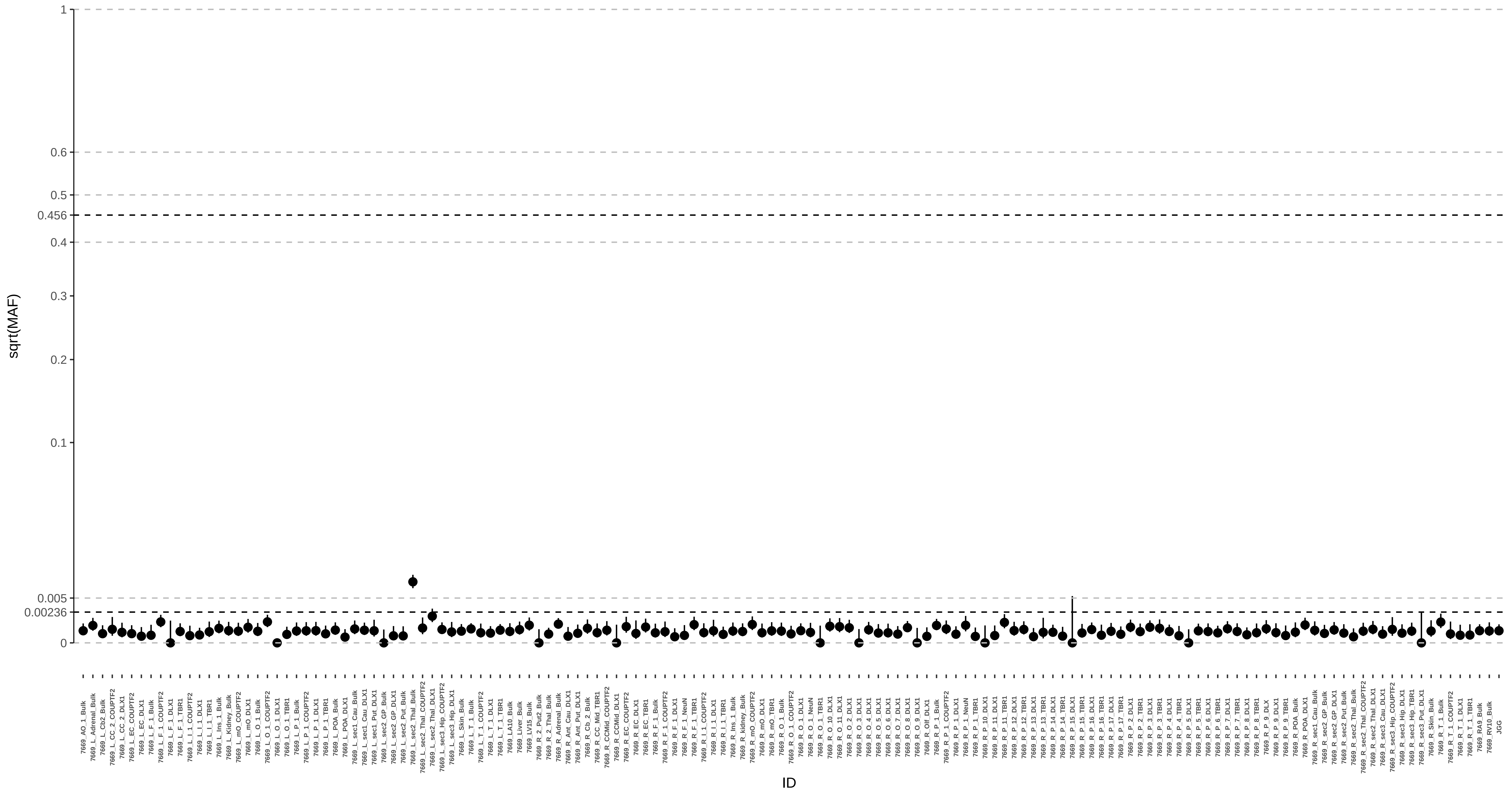

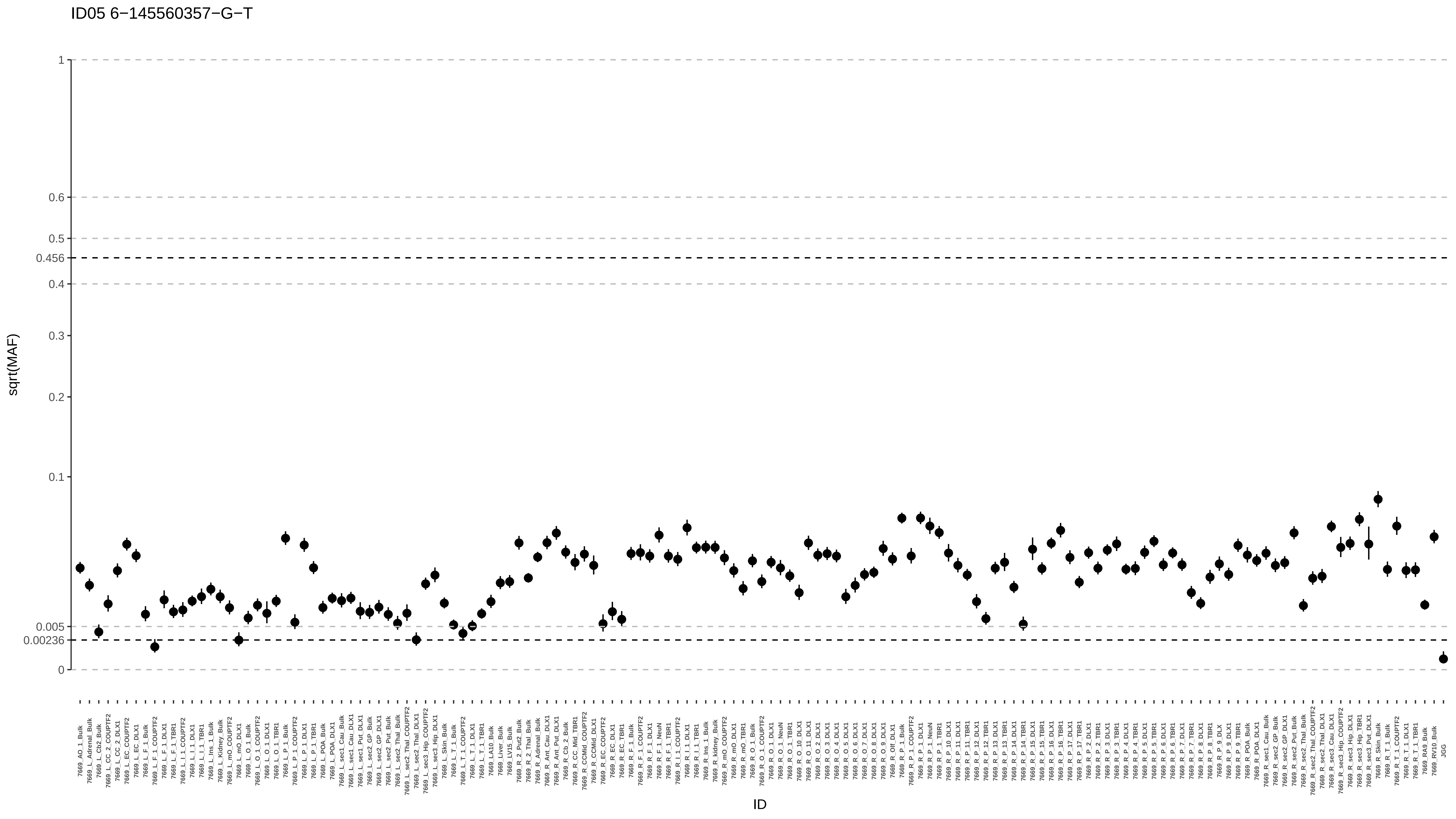

ID05 6−145560357−G−T
sqrt(MAF)
1
0.6
0.5
0.456
0.4
0.3
0.2
0.1
0.005
0.00236
0
ID
7669_AO_1_Bulk
7669_L_Adrenal_Bulk
7669_L_Cb2_Bulk
7669_L_CC_2_COUPTF2
7669_L_CC_2_DLX1
7669_L_EC_COUPTF2
7669_L_EC_DLX1
7669_L_F_1_Bulk
7669_L_F_1_COUPTF2
7669_L_F_1_DLX1
7669_L_F_1_TBR1
7669_L_I_1_COUPTF2
7669_L_I_1_DLX1
7669_L_I_1_TBR1
7669_L_Ins_1_Bulk
7669_L_Kidney_Bulk
7669_L_mO_COUPTF2
7669_L_mO_DLX1
7669_L_O_1_Bulk
7669_L_O_1_COUPTF2
7669_L_O_1_DLX1
7669_L_O_1_TBR1
7669_L_P_1_Bulk
7669_L_P_1_COUPTF2
7669_L_P_1_DLX1
7669_L_P_1_TBR1
7669_L_POA_Bulk
7669_L_POA_DLX1
7669_L_sec1_Cau_Bulk
7669_L_sec1_Cau_DLX1
7669_L_sec1_Put_DLX1
7669_L_sec2_GP_Bulk
7669_L_sec2_GP_DLX1
7669_L_sec2_Put_Bulk
7669_L_sec2_Thal_Bulk
7669_L_sec2_Thal_COUPTF2
7669_L_sec2_Thal_DLX1
7669_L_sec3_Hip_COUPTF2
7669_L_sec3_Hip_DLX1
7669_L_Skin_Bulk
7669_L_T_1_Bulk
7669_L_T_1_COUPTF2
7669_L_T_1_DLX1
7669_L_T_1_TBR1
7669_LA10_Bulk
7669_Liver_Bulk
7669_LV15_Bulk
7669_R_2_Put2_Bulk
7669_R_2_Thal_Bulk
7669_R_Adrenal_Bulk
7669_R_Ant_Cau_DLX1
7669_R_Ant_Put_DLX1
7669_R_Cb_2_Bulk
7669_R_CC_Mid_TBR1
7669_R_CCMid_COUPTF2
7669_R_CCMid_DLX1
7669_R_EC_COUPTF2
7669_R_EC_DLX1
7669_R_EC_TBR1
7669_R_F_1_Bulk
7669_R_F_1_COUPTF2
7669_R_F_1_DLX1
7669_R_F_1_NeuN
7669_R_F_1_TBR1
7669_R_I_1_COUPTF2
7669_R_I_1_DLX1
7669_R_I_1_TBR1
7669_R_Ins_1_Bulk
7669_R_kidney_Bulk
7669_R_mO_COUPTF2
7669_R_mO_DLX1
7669_R_mO_TBR1
7669_R_O_1_Bulk
7669_R_O_1_COUPTF2
7669_R_O_1_DLX1
7669_R_O_1_NeuN
7669_R_O_1_TBR1
7669_R_O_10_DLX1
7669_R_O_11_DLX1
7669_R_O_2_DLX1
7669_R_O_3_DLX1
7669_R_O_4_DLX1
7669_R_O_5_DLX1
7669_R_O_6_DLX1
7669_R_O_7_DLX1
7669_R_O_8_DLX1
7669_R_O_9_DLX1
7669_R_Olf_DLX1
7669_R_P_1_Bulk
7669_R_P_1_COUPTF2
7669_R_P_1_DLX1
7669_R_P_1_NeuN
7669_R_P_1_TBR1
7669_R_P_10_DLX1
7669_R_P_11_DLX1
7669_R_P_11_TBR1
7669_R_P_12_DLX1
7669_R_P_12_TBR1
7669_R_P_13_DLX1
7669_R_P_13_TBR1
7669_R_P_14_DLX1
7669_R_P_14_TBR1
7669_R_P_15_DLX1
7669_R_P_15_TBR1
7669_R_P_16_DLX1
7669_R_P_16_TBR1
7669_R_P_17_DLX1
7669_R_P_17_TBR1
7669_R_P_2_DLX1
7669_R_P_2_TBR1
7669_R_P_3_DLX1
7669_R_P_3_TBR1
7669_R_P_4_DLX1
7669_R_P_4_TBR1
7669_R_P_5_DLX1
7669_R_P_5_TBR1
7669_R_P_6_DLX1
7669_R_P_6_TBR1
7669_R_P_7_DLX1
7669_R_P_7_TBR1
7669_R_P_8_DLX1
7669_R_P_8_TBR1
7669_R_P_9_DLX
7669_R_P_9_DLX1
7669_R_P_9_TBR1
7669_R_POA_Bulk
7669_R_POA_DLX1
7669_R_sec1_Cau_Bulk
7669_R_sec2_GP_Bulk
7669_R_sec2_GP_DLX1
7669_R_sec2_Put_Bulk
7669_R_sec2_Thal_Bulk
7669_R_sec2_Thal_COUPTF2
7669_R_sec2_Thal_DLX1
7669_R_sec3_Cau_DLX1
7669_R_sec3_Hip_COUPTF2
7669_R_sec3_Hip_DLX1
7669_R_sec3_Hip_TBR1
7669_R_sec3_Put_DLX1
7669_R_Skin_Bulk
7669_R_T_1_Bulk
7669_R_T_1_COUPTF2
7669_R_T_1_DLX1
7669_R_T_1_TBR1
7669_RA9_Bulk
7669_RV10_Bulk
JGG

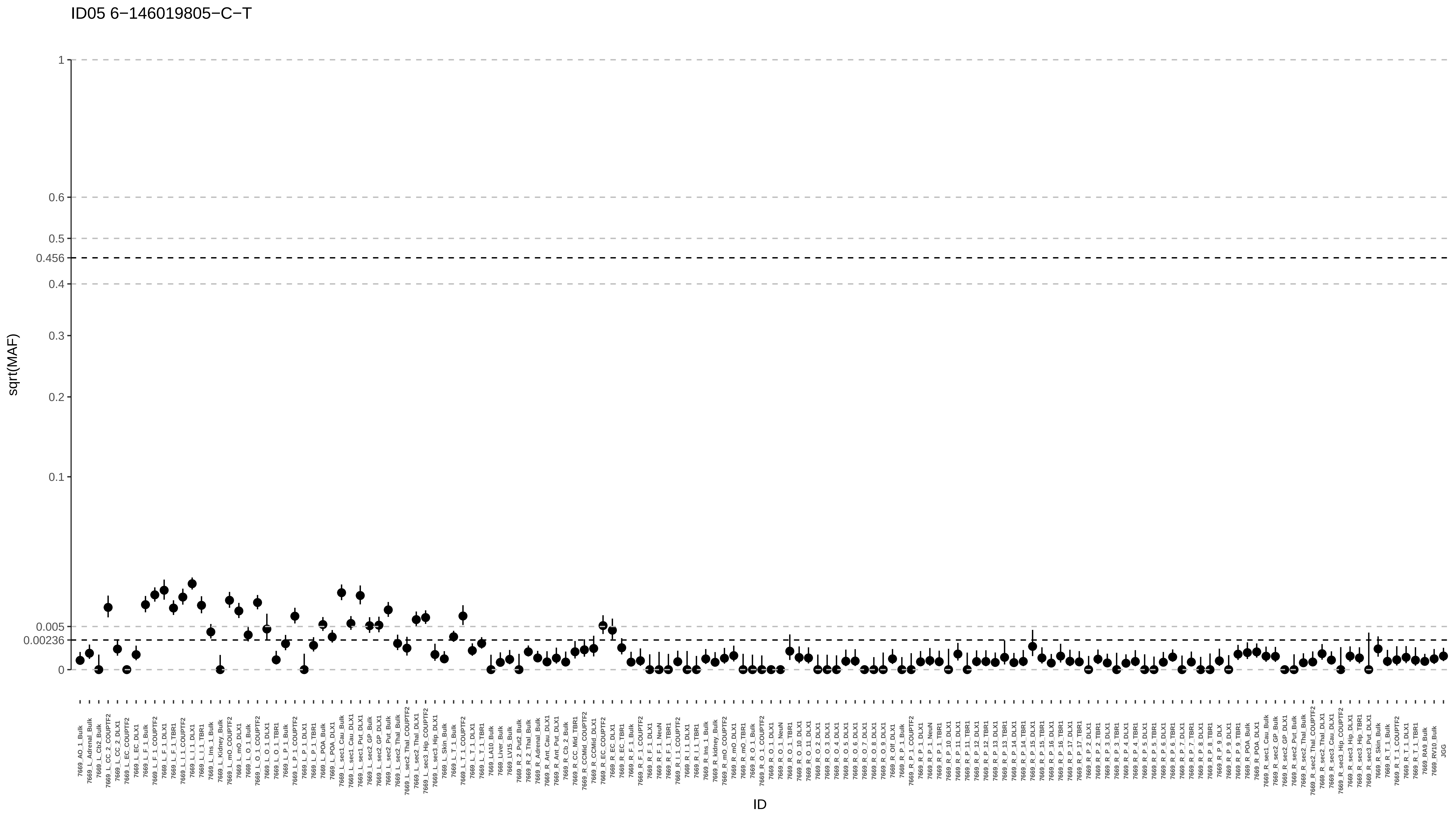
ID05 6−146019805−C−T
sqrt(MAF)
ID
1
0.6
0.5
0.456
0.4
0.3
0.2
0.1
0.005
0.00236
0

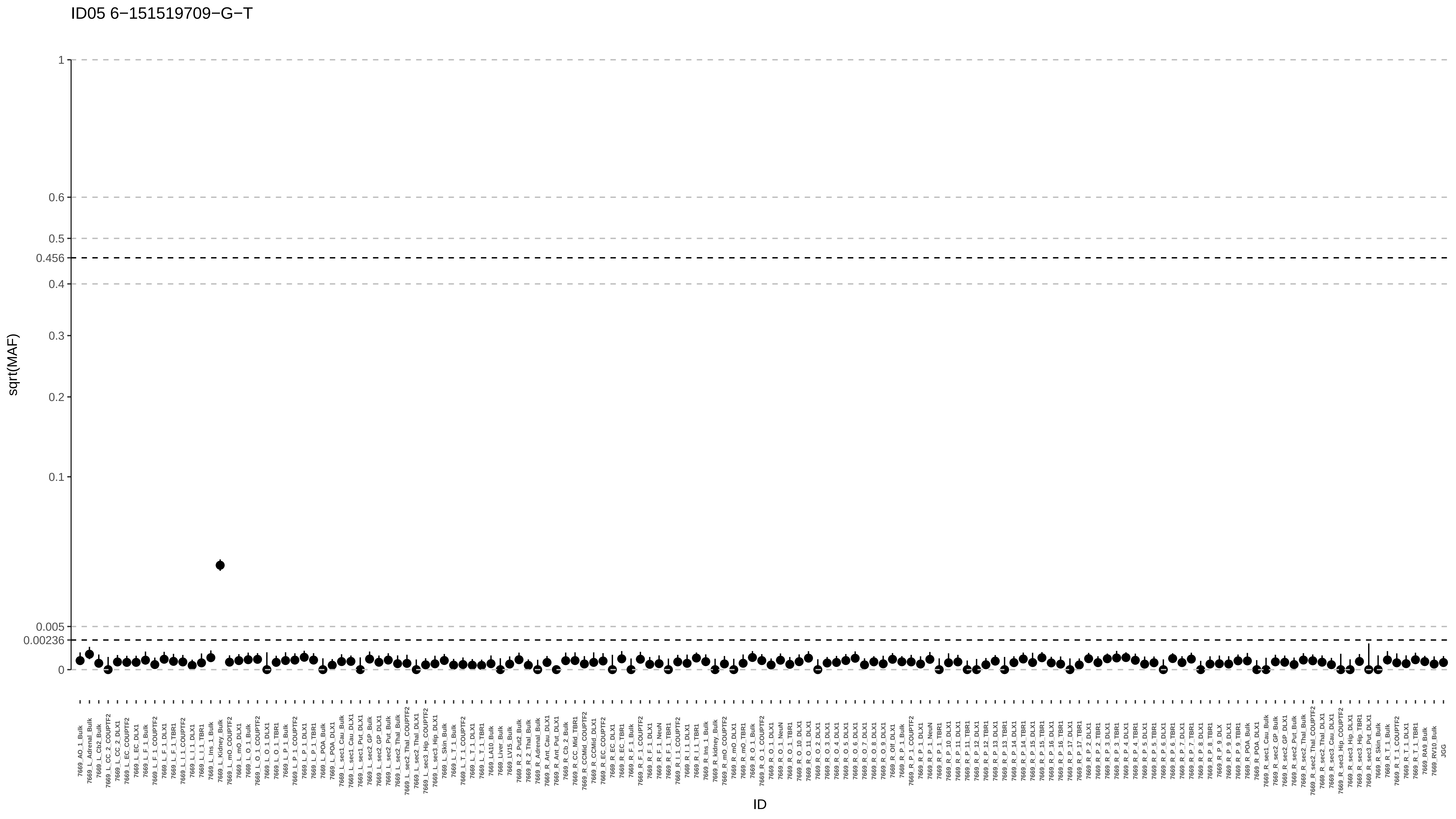
ID05 6−151519709−G−T
sqrt(MAF)
1
0.6
0.5
0.456
0.4
0.3
0.2
0.1
0.005
0.00236
0
ID

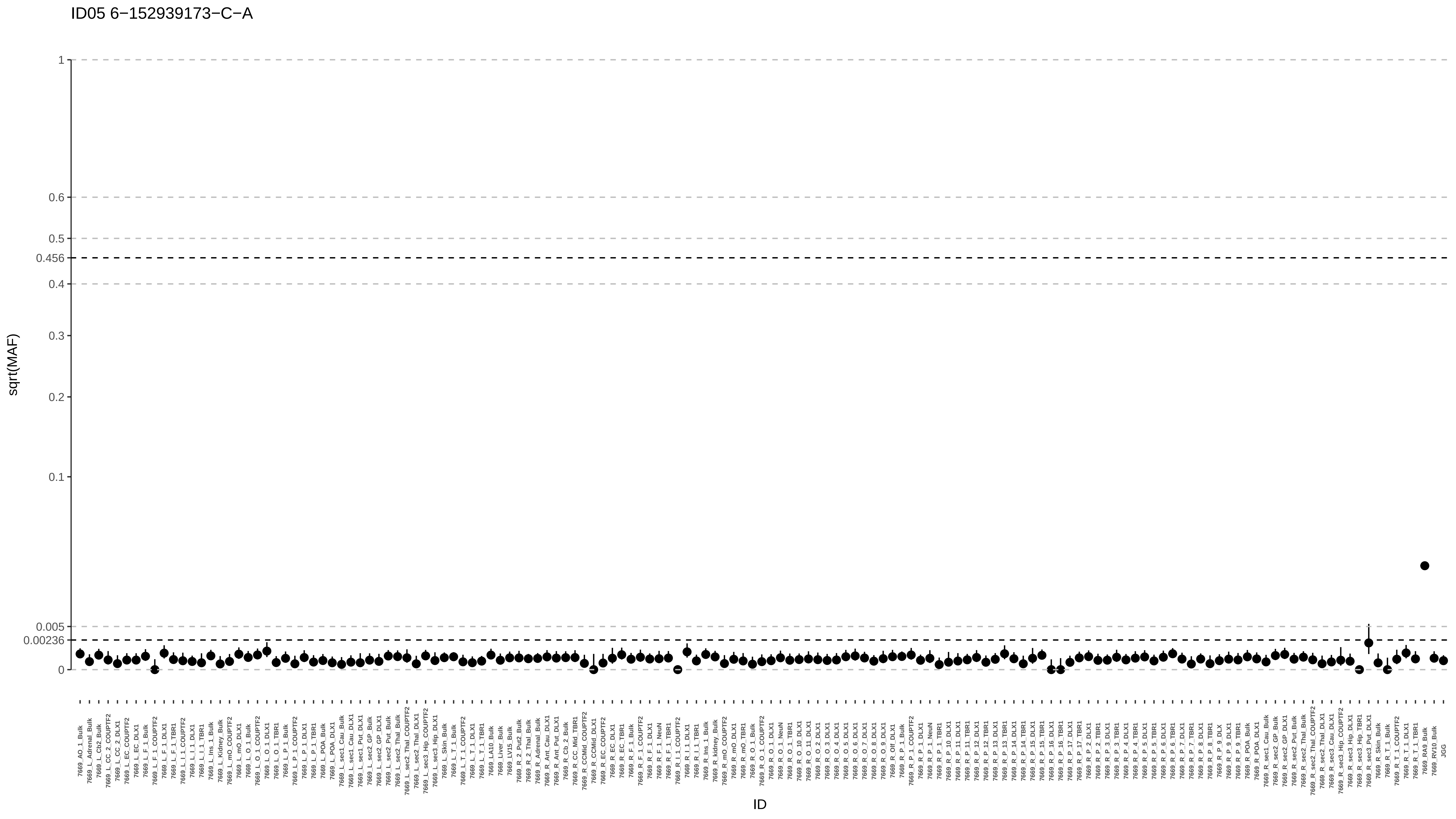
ID05 6−152939173−C−A
sqrt(MAF)
1
0.6
0.5
0.456
0.4
0.3
0.2
0.1
0.005
0.00236
0
ID

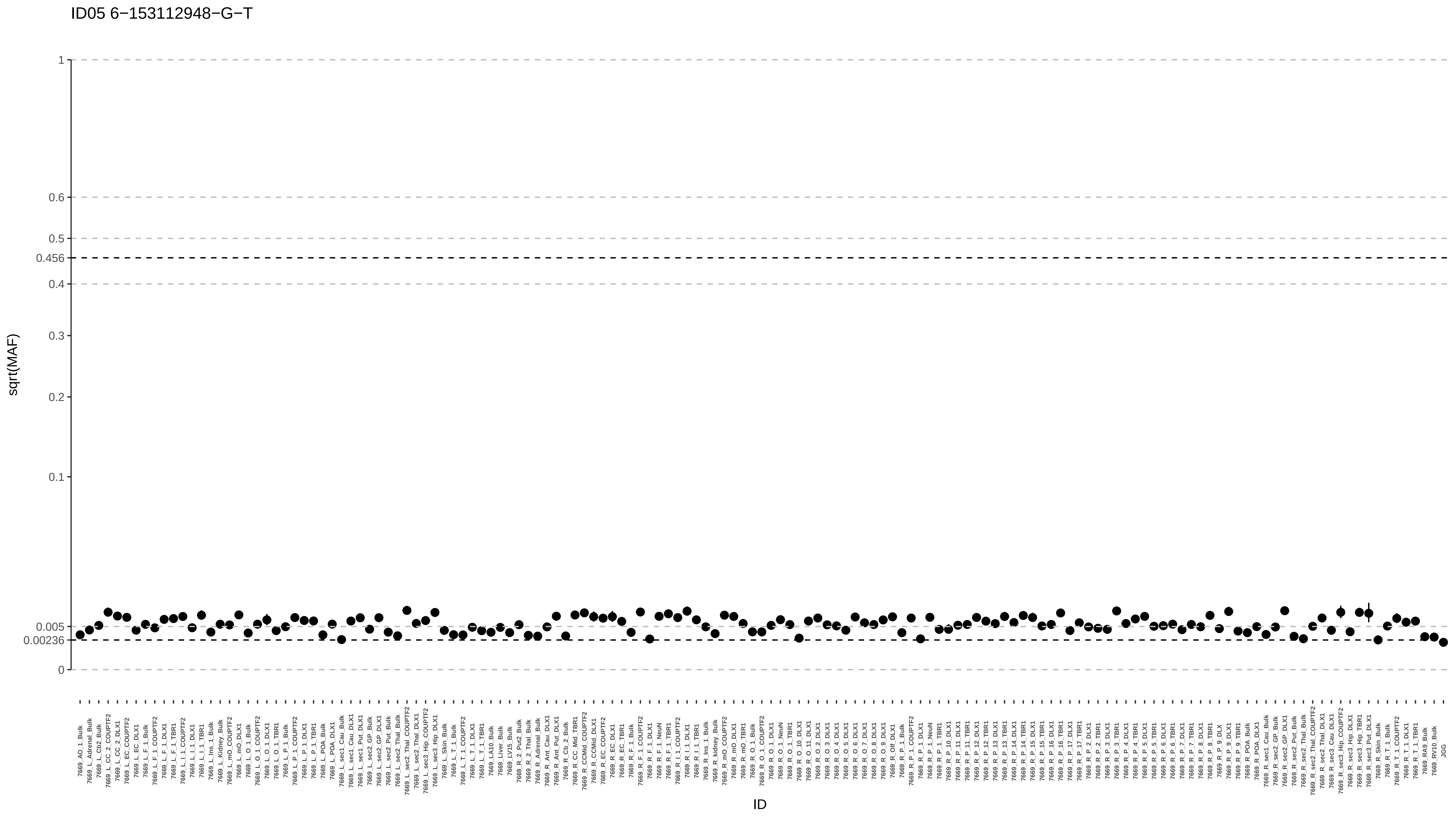
ID05 6−153112948−G−T
sqrt(MAF)
1
0.6
0.5
0.456
0.4
0.3
0.2
0.1
0.005
0.00236
0
ID

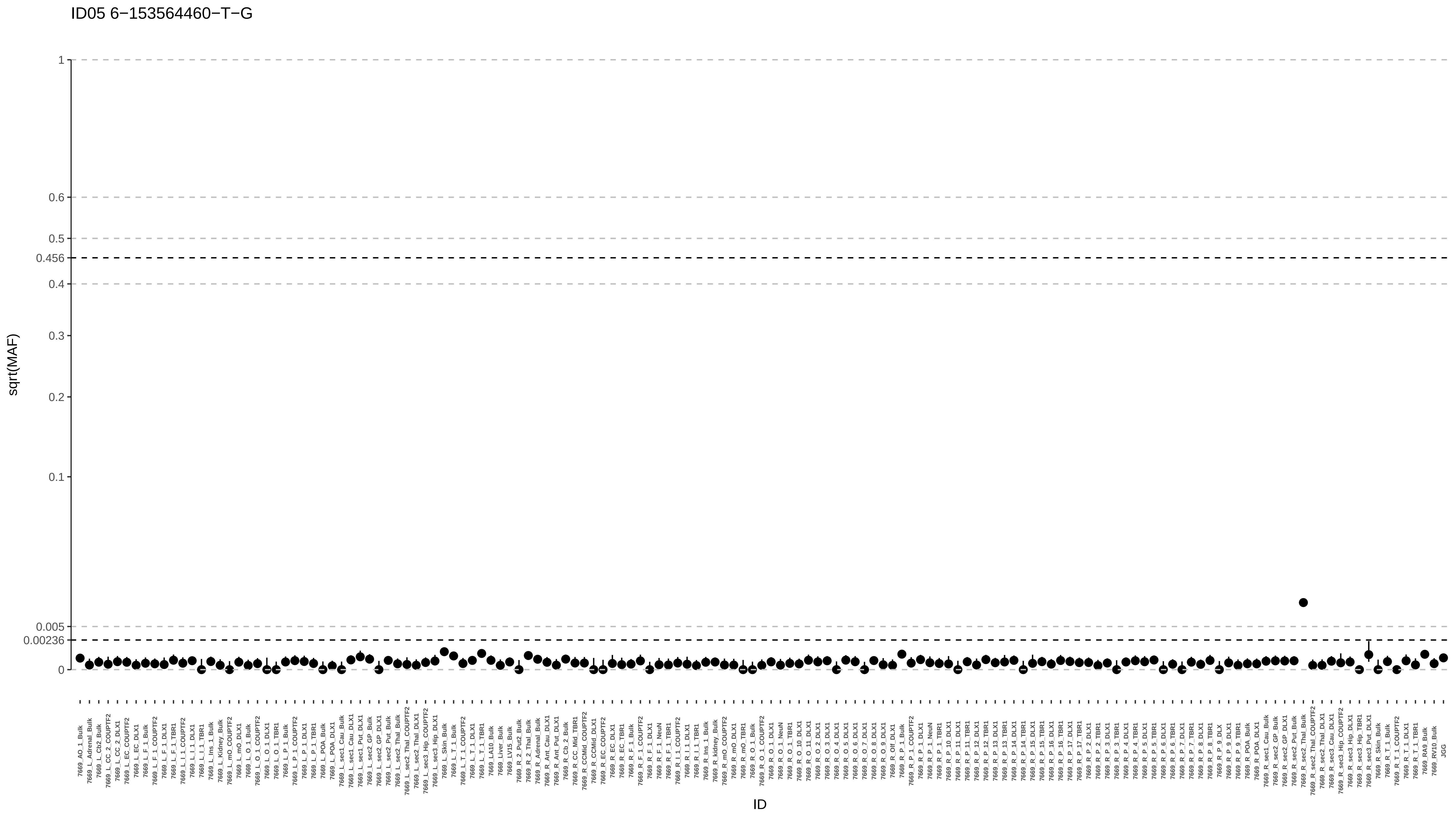
ID05 6−153564460−T−G
sqrt(MAF)
ID

ID05 6−155986031−C−T

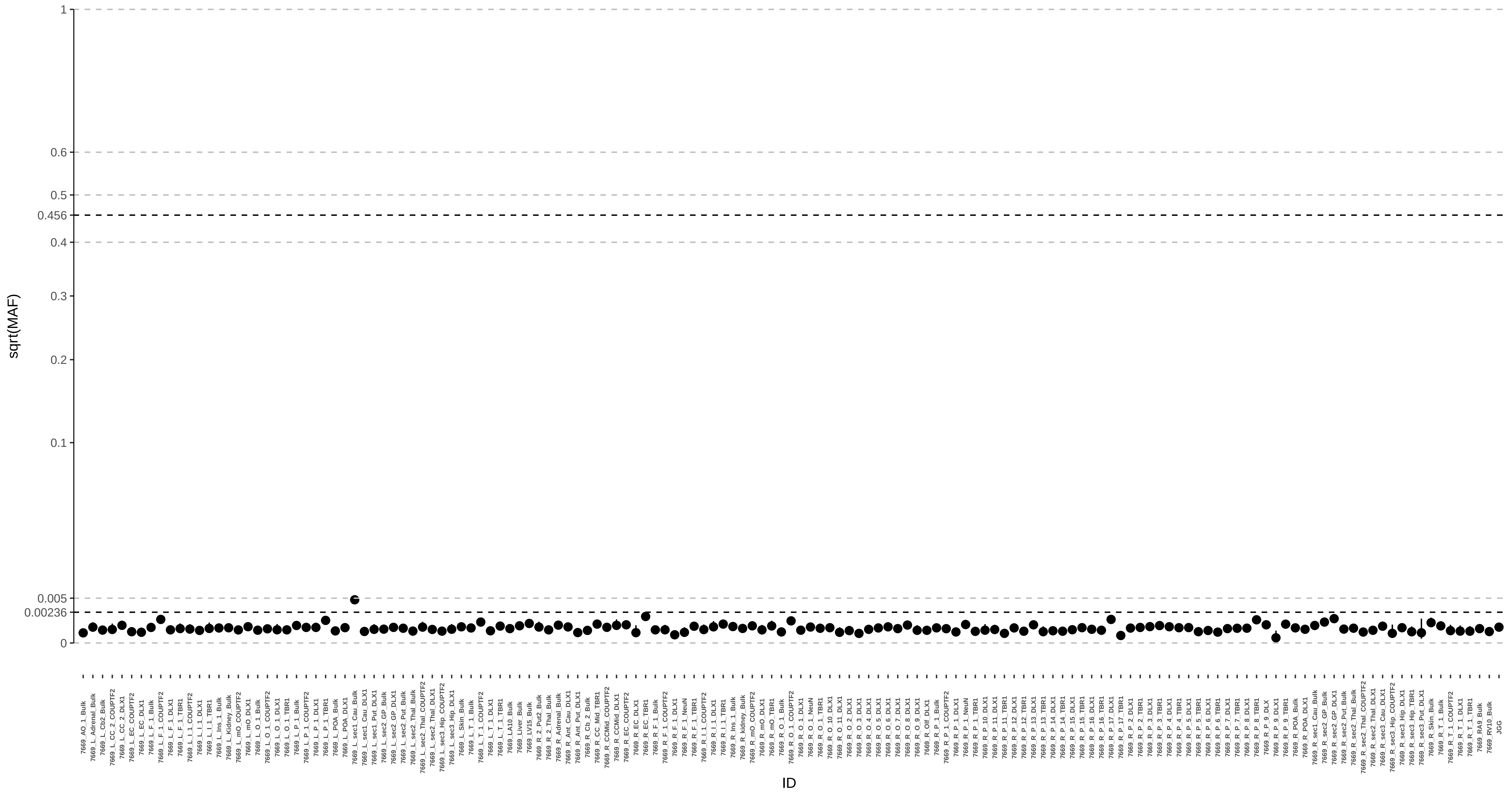

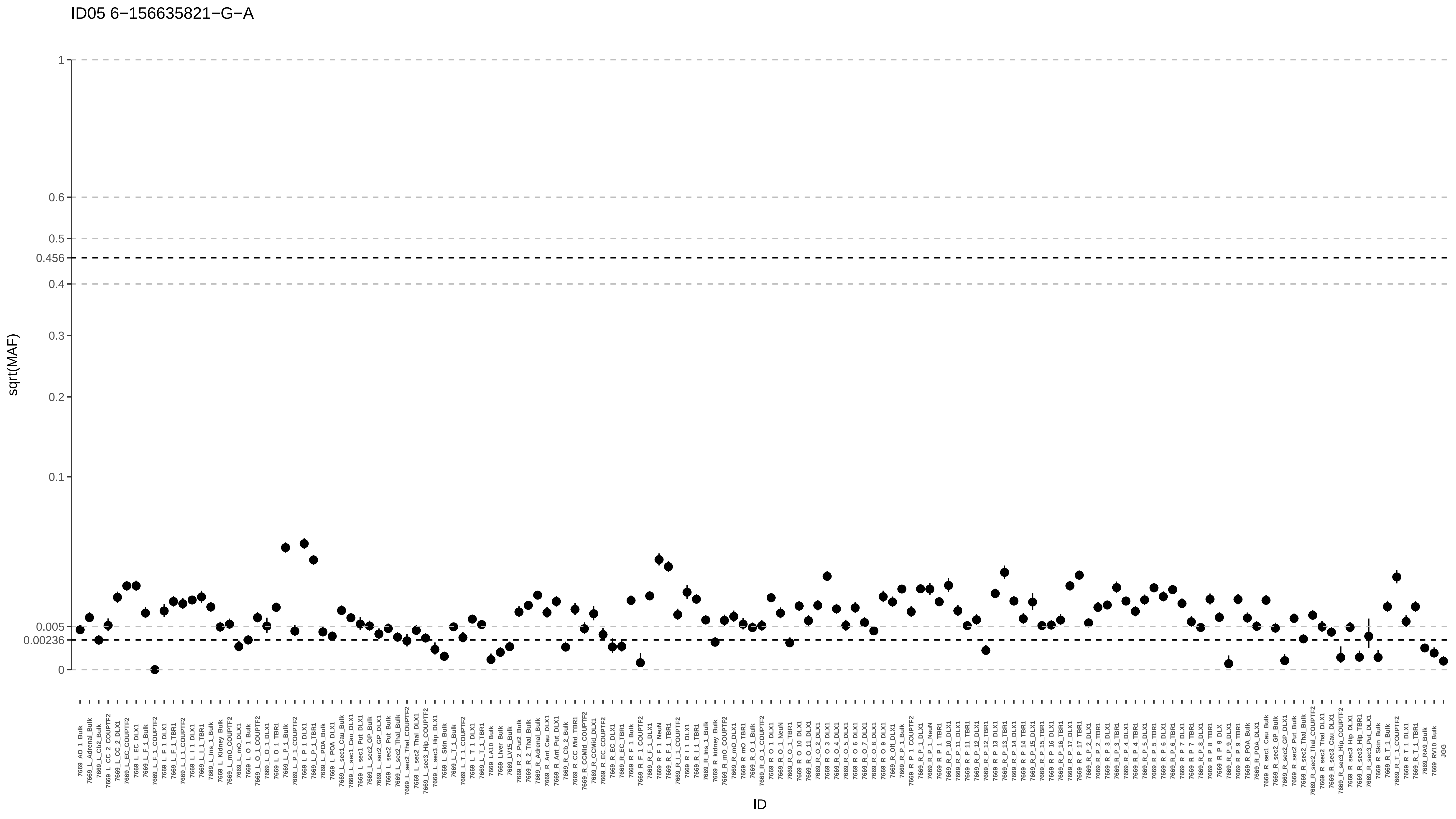
ID05 6−156635821−G−A
sqrt(MAF)
1
0.6
0.5
0.456
0.4
0.3
0.2
0.1
0.005
0.00236
0
ID

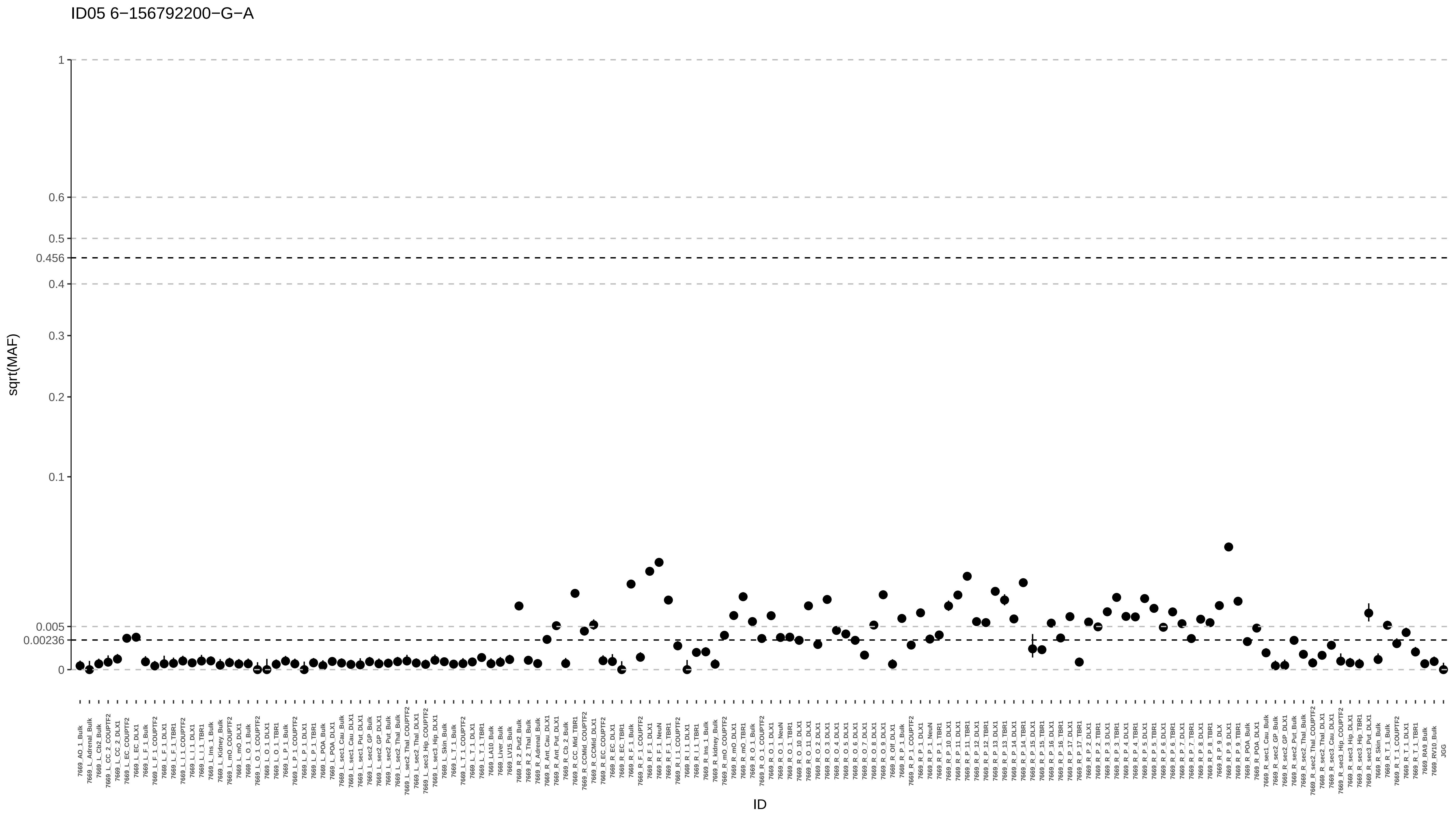
ID05 6−156792200−G−A
sqrt(MAF)
ID
1
0.6
0.5
0.456
0.4
0.3
0.2
0.1
0.005
0.00236
0

ID05 6−159935200−C−T

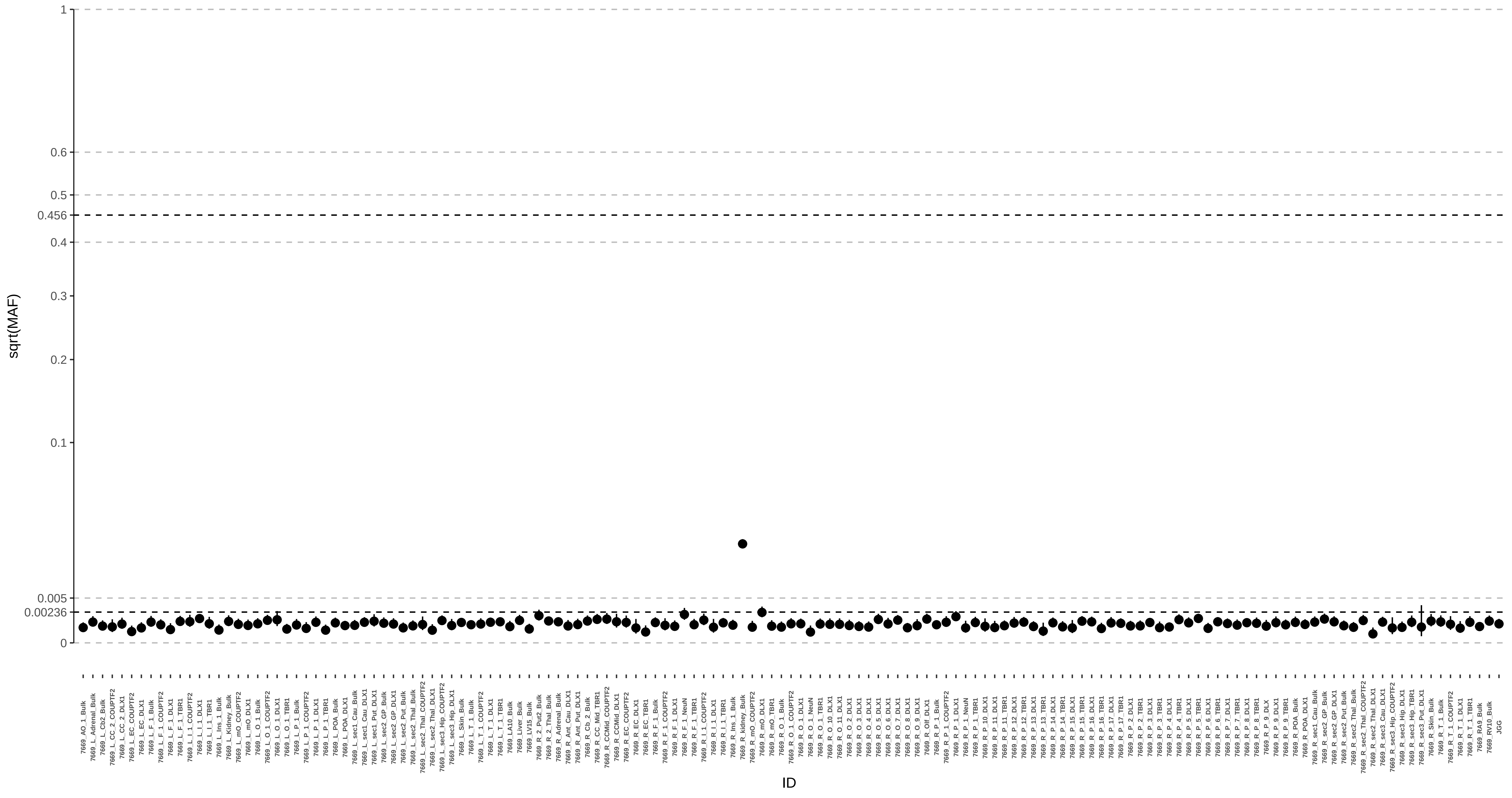

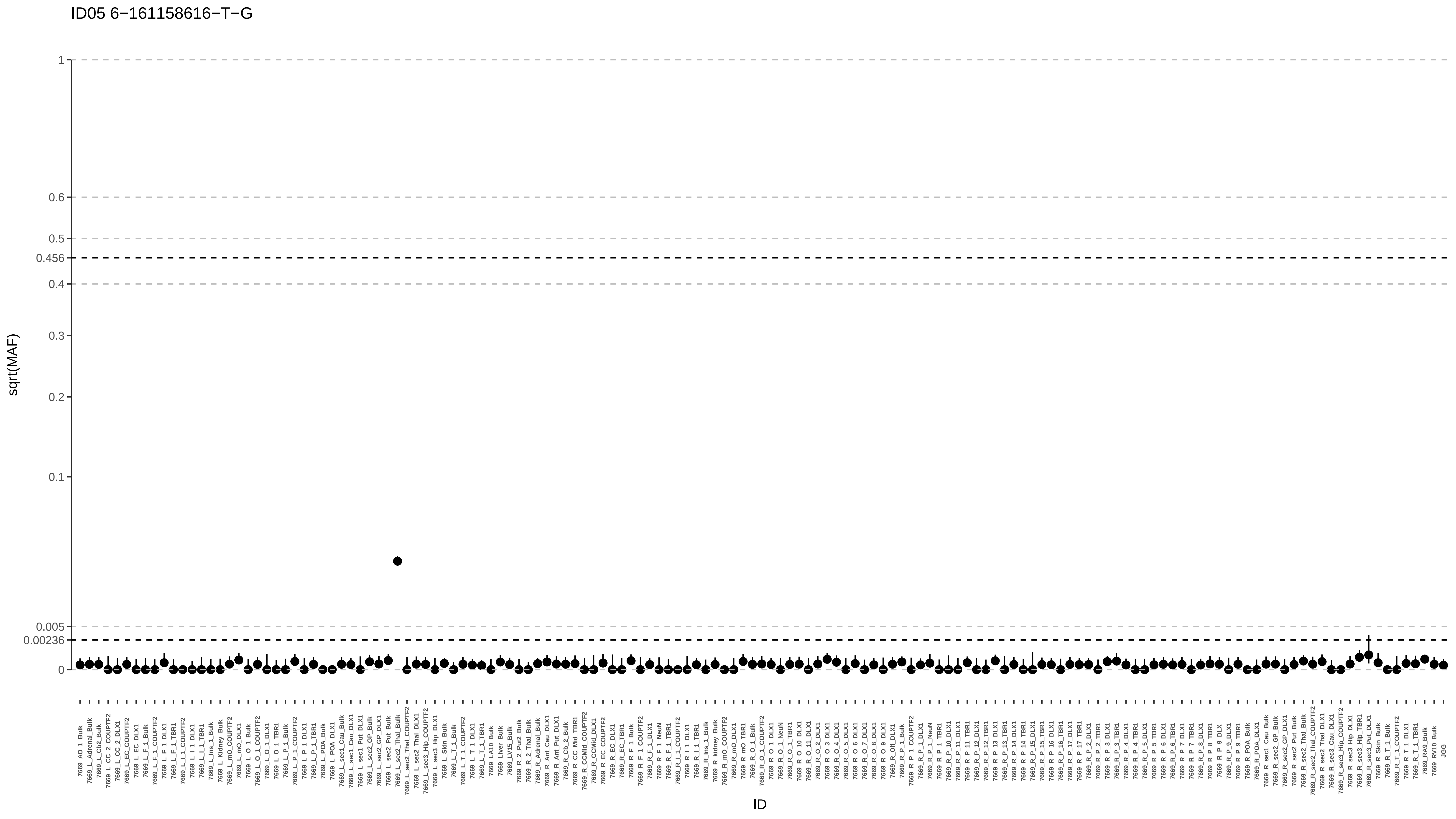
ID05 6−161158616−T−G
sqrt(MAF)
ID
1
0.6
0.5
0.456
0.4
0.3
0.2
0.1
0.005
0.00236
0

ID05 6−162806617−C−T

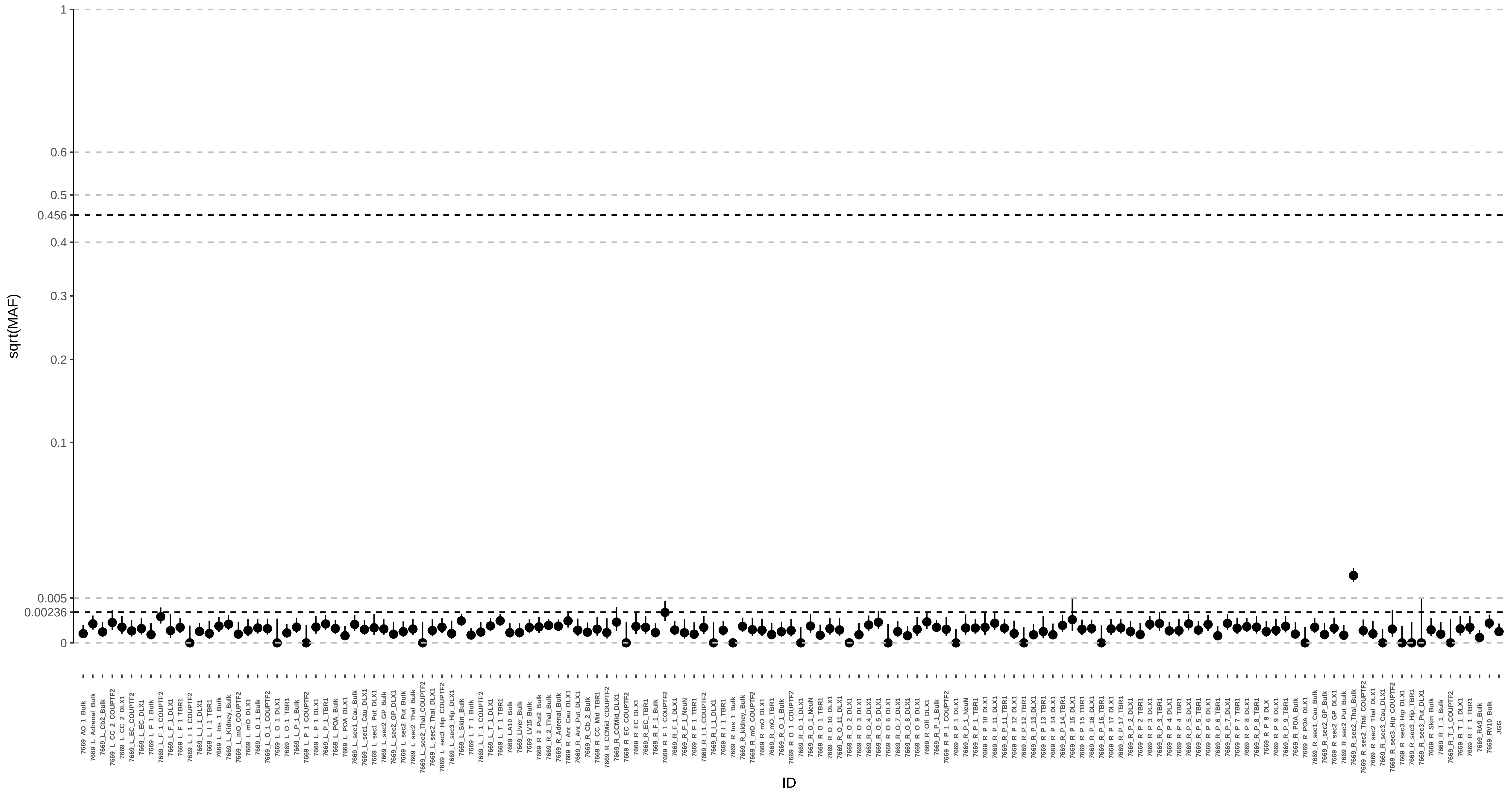

# ID05 6−1661827−C−T

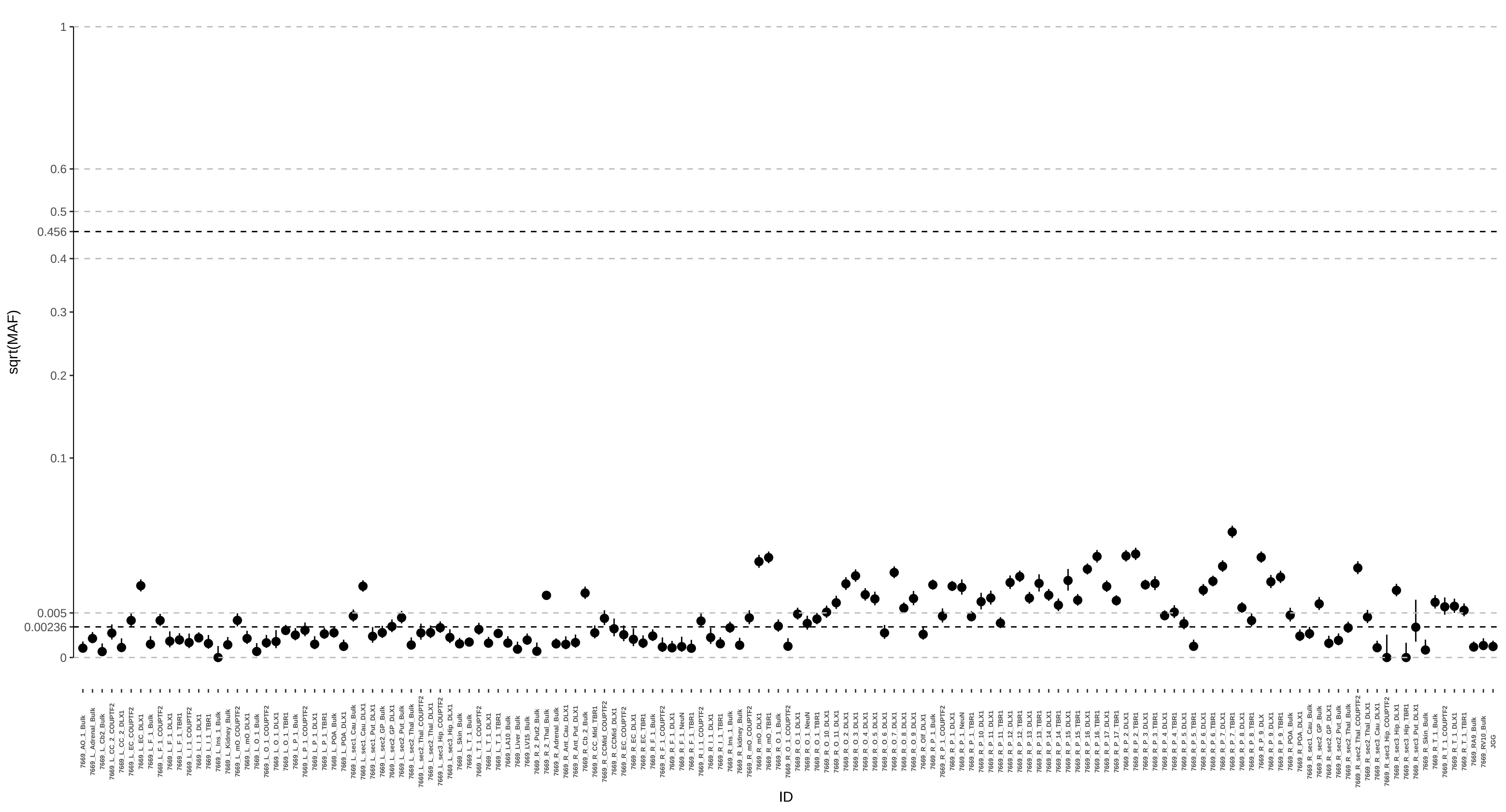

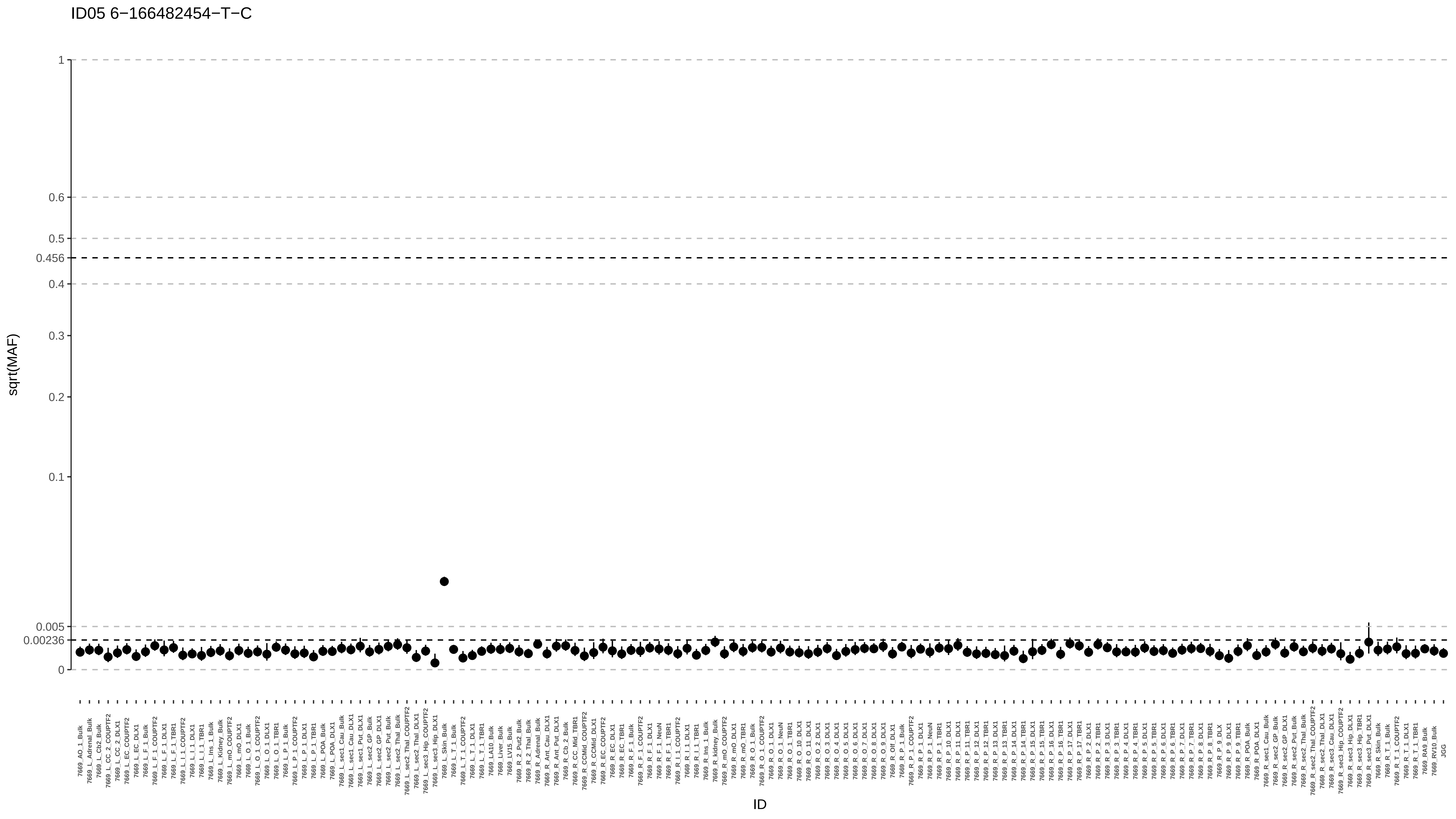
ID05 6−166482454−T−C
sqrt(MAF)
1
0.6
0.5
0.456
0.4
0.3
0.2
0.1
0.005
0.00236
0
ID

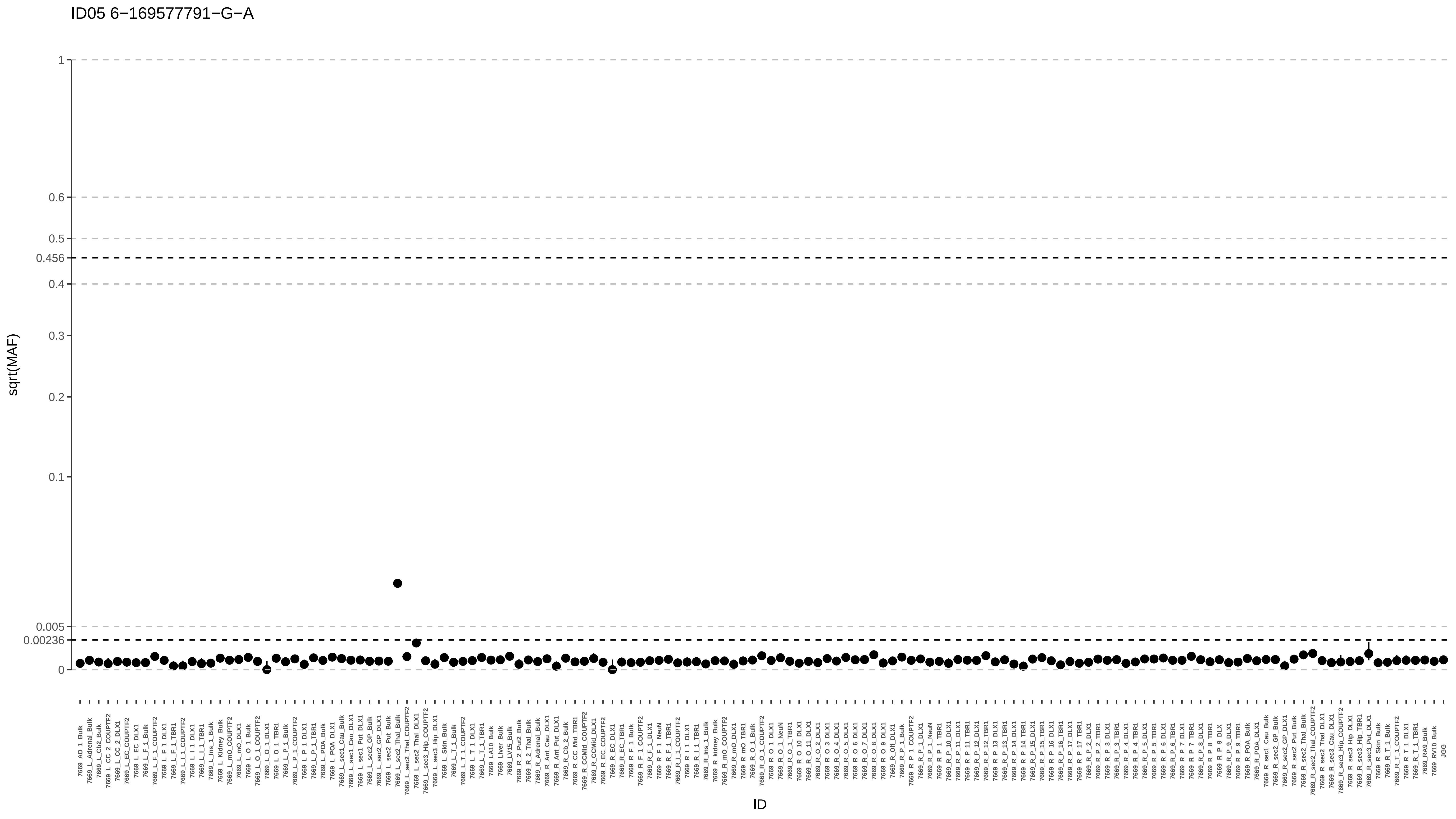
ID05 6−169577791−G−A
sqrt(MAF)
1
0.6
0.5
0.456
0.4
0.3
0.2
0.1
0.005
0.00236
0
ID
7669_AO_1_Bulk
7669_L_Adrenal_Bulk
7669_L_Cb2_Bulk
7669_L_CC_2_COUPTF2
7669_L_CC_2_DLX1
7669_L_EC_COUPTF2
7669_L_EC_DLX1
7669_L_F_1_Bulk
7669_L_F_1_COUPTF2
7669_L_F_1_DLX1
7669_L_F_1_TBR1
7669_L_I_1_COUPTF2
7669_L_I_1_DLX1
7669_L_I_1_TBR1
7669_L_Ins_1_Bulk
7669_L_Kidney_Bulk
7669_L_mO_COUPTF2
7669_L_mO_DLX1
7669_L_O_1_Bulk
7669_L_O_1_COUPTF2
7669_L_O_1_DLX1
7669_L_O_1_TBR1
7669_L_P_1_Bulk
7669_L_P_1_COUPTF2
7669_L_P_1_DLX1
7669_L_P_1_TBR1
7669_L_POA_Bulk
7669_L_POA_DLX1
7669_L_sec1_Cau_Bulk
7669_L_sec1_Cau_DLX1
7669_L_sec1_Put_DLX1
7669_L_sec2_GP_Bulk
7669_L_sec2_GP_DLX1
7669_L_sec2_Put_Bulk
7669_L_sec2_Thal_Bulk
7669_L_sec2_Thal_COUPTF2
7669_L_sec2_Thal_DLX1
7669_L_sec3_Hip_COUPTF2
7669_L_sec3_Hip_DLX1
7669_L_Skin_Bulk
7669_L_T_1_Bulk
7669_L_T_1_COUPTF2
7669_L_T_1_DLX1
7669_L_T_1_TBR1
7669_LA10_Bulk
7669_Liver_Bulk
7669_LV15_Bulk
7669_R_2_Put2_Bulk
7669_R_2_Thal_Bulk
7669_R_Adrenal_Bulk
7669_R_Ant_Cau_DLX1
7669_R_Ant_Put_DLX1
7669_R_Cb_2_Bulk
7669_R_CC_Mid_TBR1
7669_R_CCMid_COUPTF2
7669_R_CCMid_DLX1
7669_R_EC_COUPTF2
7669_R_EC_DLX1
7669_R_EC_TBR1
7669_R_F_1_Bulk
7669_R_F_1_COUPTF2
7669_R_F_1_DLX1
7669_R_F_1_NeuN
7669_R_F_1_TBR1
7669_R_I_1_COUPTF2
7669_R_I_1_DLX1
7669_R_I_1_TBR1
7669_R_Ins_1_Bulk
7669_R_kidney_Bulk
7669_R_mO_COUPTF2
7669_R_mO_DLX1
7669_R_mO_TBR1
7669_R_O_1_Bulk
7669_R_O_1_COUPTF2
7669_R_O_1_DLX1
7669_R_O_1_NeuN
7669_R_O_1_TBR1
7669_R_O_10_DLX1
7669_R_O_11_DLX1
7669_R_O_2_DLX1
7669_R_O_3_DLX1
7669_R_O_4_DLX1
7669_R_O_5_DLX1
7669_R_O_6_DLX1
7669_R_O_7_DLX1
7669_R_O_8_DLX1
7669_R_O_9_DLX1
7669_R_Olf_DLX1
7669_R_P_1_Bulk
7669_R_P_1_COUPTF2
7669_R_P_1_DLX1
7669_R_P_1_NeuN
7669_R_P_1_TBR1
7669_R_P_10_DLX1
7669_R_P_11_DLX1
7669_R_P_11_TBR1
7669_R_P_12_DLX1
7669_R_P_12_TBR1
7669_R_P_13_DLX1
7669_R_P_13_TBR1
7669_R_P_14_DLX1
7669_R_P_14_TBR1
7669_R_P_15_DLX1
7669_R_P_15_TBR1
7669_R_P_16_DLX1
7669_R_P_16_TBR1
7669_R_P_17_DLX1
7669_R_P_17_TBR1
7669_R_P_2_DLX1
7669_R_P_2_TBR1
7669_R_P_3_DLX1
7669_R_P_3_TBR1
7669_R_P_4_DLX1
7669_R_P_4_TBR1
7669_R_P_5_DLX1
7669_R_P_5_TBR1
7669_R_P_6_DLX1
7669_R_P_6_TBR1
7669_R_P_7_DLX1
7669_R_P_7_TBR1
7669_R_P_8_DLX1
7669_R_P_8_TBR1
7669_R_P_9_DLX
7669_R_P_9_DLX1
7669_R_P_9_TBR1
7669_R_POA_Bulk
7669_R_POA_DLX1
7669_R_sec1_Cau_Bulk
7669_R_sec2_GP_Bulk
7669_R_sec2_GP_DLX1
7669_R_sec2_Put_Bulk
7669_R_sec2_Thal_Bulk
7669_R_sec2_Thal_COUPTF2
7669_R_sec2_Thal_DLX1
7669_R_sec3_Cau_DLX1
7669_R_sec3_Hip_COUPTF2
7669_R_sec3_Hip_DLX1
7669_R_sec3_Hip_TBR1
7669_R_sec3_Put_DLX1
7669_R_Skin_Bulk
7669_R_T_1_Bulk
7669_R_T_1_COUPTF2
7669_R_T_1_DLX1
7669_R_T_1_TBR1
7669_RA9_Bulk
7669_RV10_Bulk
JGG

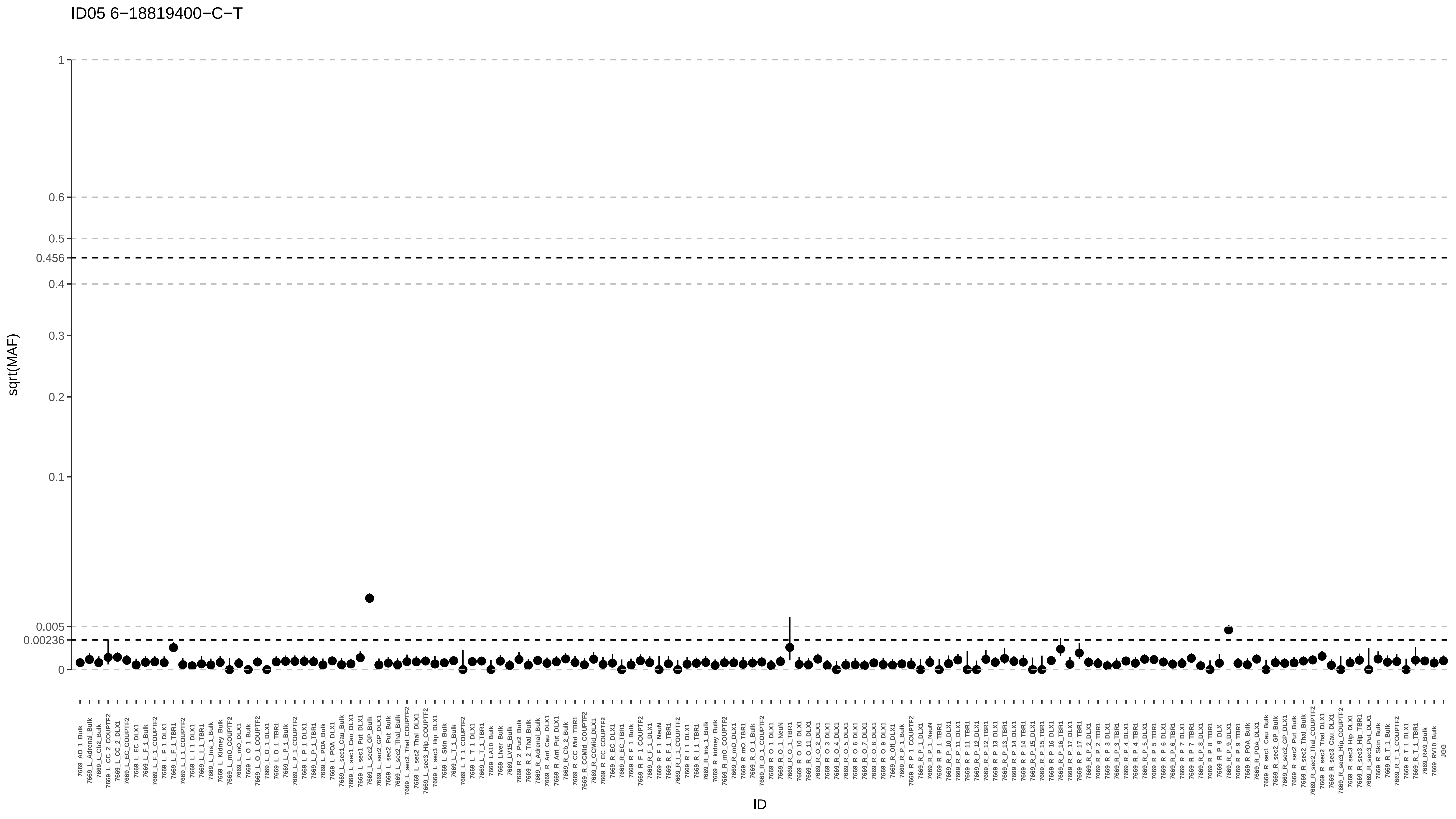
ID05 6−18819400−C−T
sqrt(MAF)
1
0.6
0.5
0.456
0.4
0.3
0.2
0.1
0.005
0.00236
0
ID
7669_AO_1_Bulk
7669_L_Adrenal_Bulk
7669_L_Cb2_Bulk
7669_L_CC_2_COUPTF2
7669_L_CC_2_DLX1
7669_L_EC_COUPTF2
7669_L_EC_DLX1
7669_L_F_1_Bulk
7669_L_F_1_COUPTF2
7669_L_F_1_DLX1
7669_L_F_1_TBR1
7669_L_I_1_COUPTF2
7669_L_I_1_DLX1
7669_L_I_1_TBR1
7669_L_Ins_1_Bulk
7669_L_Kidney_Bulk
7669_L_mO_COUPTF2
7669_L_mO_DLX1
7669_L_O_1_Bulk
7669_L_O_1_COUPTF2
7669_L_O_1_DLX1
7669_L_O_1_TBR1
7669_L_P_1_Bulk
7669_L_P_1_COUPTF2
7669_L_P_1_DLX1
7669_L_P_1_TBR1
7669_L_POA_Bulk
7669_L_POA_DLX1
7669_L_sec1_Cau_Bulk
7669_L_sec1_Cau_DLX1
7669_L_sec1_Put_DLX1
7669_L_sec2_GP_Bulk
7669_L_sec2_GP_DLX1
7669_L_sec2_Put_Bulk
7669_L_sec2_Thal_Bulk
7669_L_sec2_Thal_COUPTF2
7669_L_sec2_Thal_DLX1
7669_L_sec3_Hip_COUPTF2
7669_L_sec3_Hip_DLX1
7669_L_Skin_Bulk
7669_L_T_1_Bulk
7669_L_T_1_COUPTF2
7669_L_T_1_DLX1
7669_L_T_1_TBR1
7669_LA10_Bulk
7669_Liver_Bulk
7669_LV15_Bulk
7669_R_2_Put2_Bulk
7669_R_2_Thal_Bulk
7669_R_Adrenal_Bulk
7669_R_Ant_Cau_DLX1
7669_R_Ant_Put_DLX1
7669_R_Cb_2_Bulk
7669_R_CC_Mid_TBR1
7669_R_CCMid_COUPTF2
7669_R_CCMid_DLX1
7669_R_EC_COUPTF2
7669_R_EC_DLX1
7669_R_EC_TBR1
7669_R_F_1_Bulk
7669_R_F_1_COUPTF2
7669_R_F_1_DLX1
7669_R_F_1_NeuN
7669_R_F_1_TBR1
7669_R_I_1_COUPTF2
7669_R_I_1_DLX1
7669_R_I_1_TBR1
7669_R_Ins_1_Bulk
7669_R_kidney_Bulk
7669_R_mO_COUPTF2
7669_R_mO_DLX1
7669_R_mO_TBR1
7669_R_O_1_Bulk
7669_R_O_1_COUPTF2
7669_R_O_1_DLX1
7669_R_O_1_NeuN
7669_R_O_1_TBR1
7669_R_O_10_DLX1
7669_R_O_11_DLX1
7669_R_O_2_DLX1
7669_R_O_3_DLX1
7669_R_O_4_DLX1
7669_R_O_5_DLX1
7669_R_O_6_DLX1
7669_R_O_7_DLX1
7669_R_O_8_DLX1
7669_R_O_9_DLX1
7669_R_Olf_DLX1
7669_R_P_1_Bulk
7669_R_P_1_COUPTF2
7669_R_P_1_DLX1
7669_R_P_1_NeuN
7669_R_P_1_TBR1
7669_R_P_10_DLX1
7669_R_P_11_DLX1
7669_R_P_11_TBR1
7669_R_P_12_DLX1
7669_R_P_12_TBR1
7669_R_P_13_DLX1
7669_R_P_13_TBR1
7669_R_P_14_DLX1
7669_R_P_14_TBR1
7669_R_P_15_DLX1
7669_R_P_15_TBR1
7669_R_P_16_DLX1
7669_R_P_16_TBR1
7669_R_P_17_DLX1
7669_R_P_17_TBR1
7669_R_P_2_DLX1
7669_R_P_2_TBR1
7669_R_P_3_DLX1
7669_R_P_3_TBR1
7669_R_P_4_DLX1
7669_R_P_4_TBR1
7669_R_P_5_DLX1
7669_R_P_5_TBR1
7669_R_P_6_DLX1
7669_R_P_6_TBR1
7669_R_P_7_DLX1
7669_R_P_7_TBR1
7669_R_P_8_DLX1
7669_R_P_8_TBR1
7669_R_P_9_DLX
7669_R_P_9_DLX1
7669_R_P_9_TBR1
7669_R_POA_Bulk
7669_R_POA_DLX1
7669_R_sec1_Cau_Bulk
7669_R_sec2_GP_Bulk
7669_R_sec2_GP_DLX1
7669_R_sec2_Put_Bulk
7669_R_sec2_Thal_Bulk
7669_R_sec2_Thal_COUPTF2
7669_R_sec2_Thal_DLX1
7669_R_sec3_Cau_DLX1
7669_R_sec3_Hip_COUPTF2
7669_R_sec3_Hip_DLX1
7669_R_sec3_Hip_TBR1
7669_R_sec3_Put_DLX1
7669_R_Skin_Bulk
7669_R_T_1_Bulk
7669_R_T_1_COUPTF2
7669_R_T_1_DLX1
7669_R_T_1_TBR1
7669_RA9_Bulk
7669_RV10_Bulk
JGG

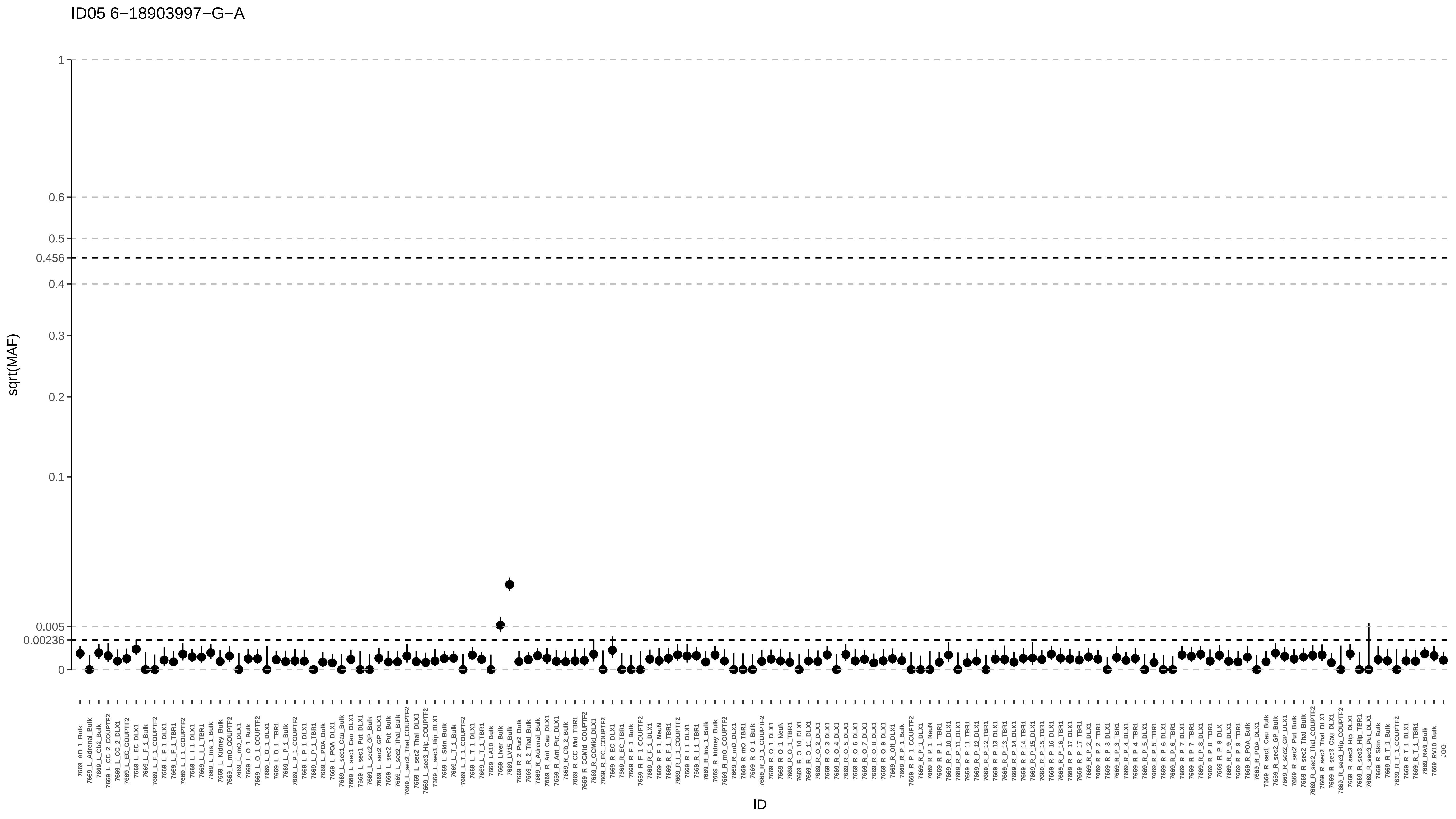
ID05 6−18903997−G−A
sqrt(MAF)
1
0.6
0.5
0.456
0.4
0.3
0.2
0.1
0.005
0.00236
0
ID

ID05 6−24537268−G−A

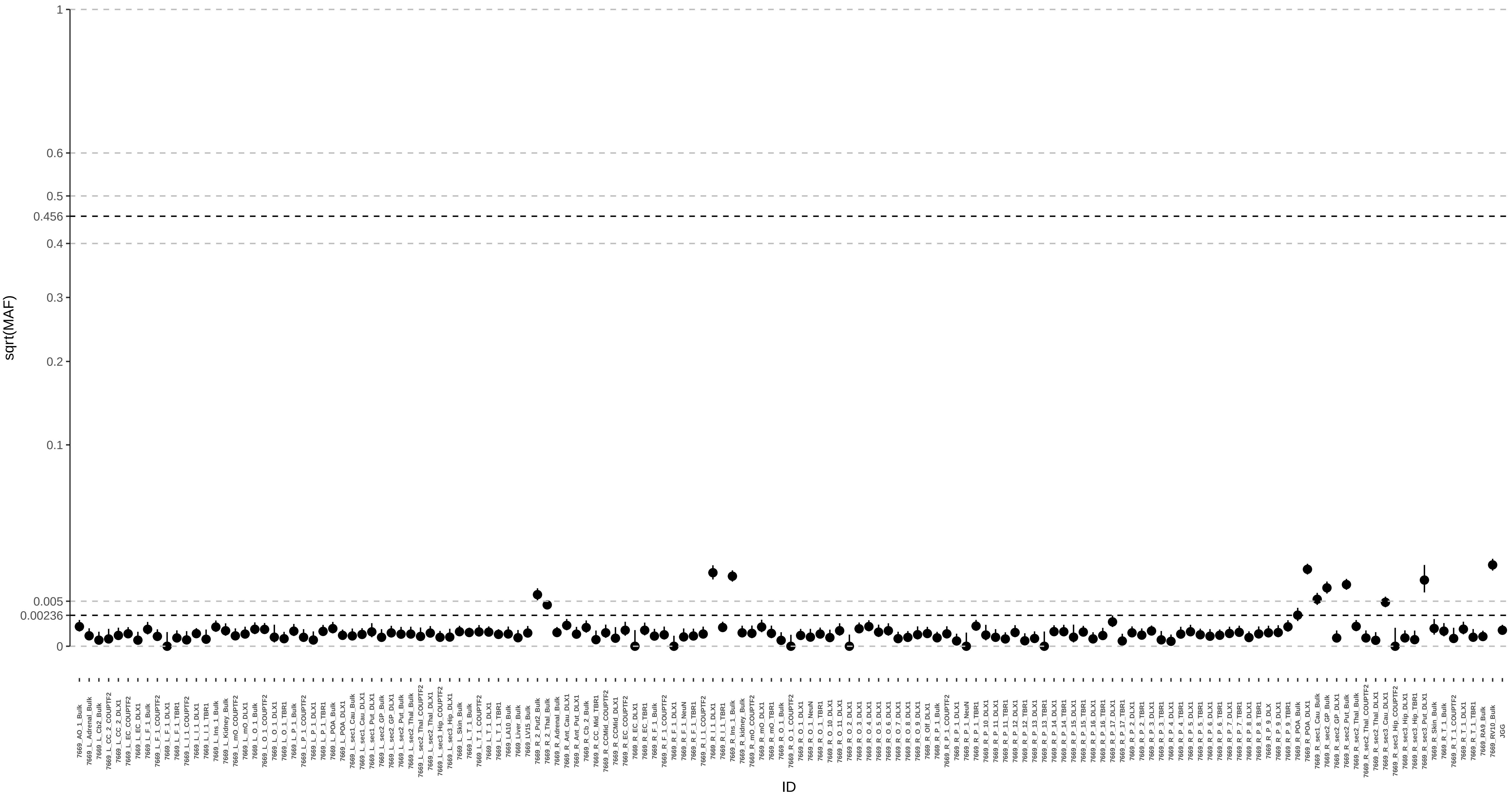

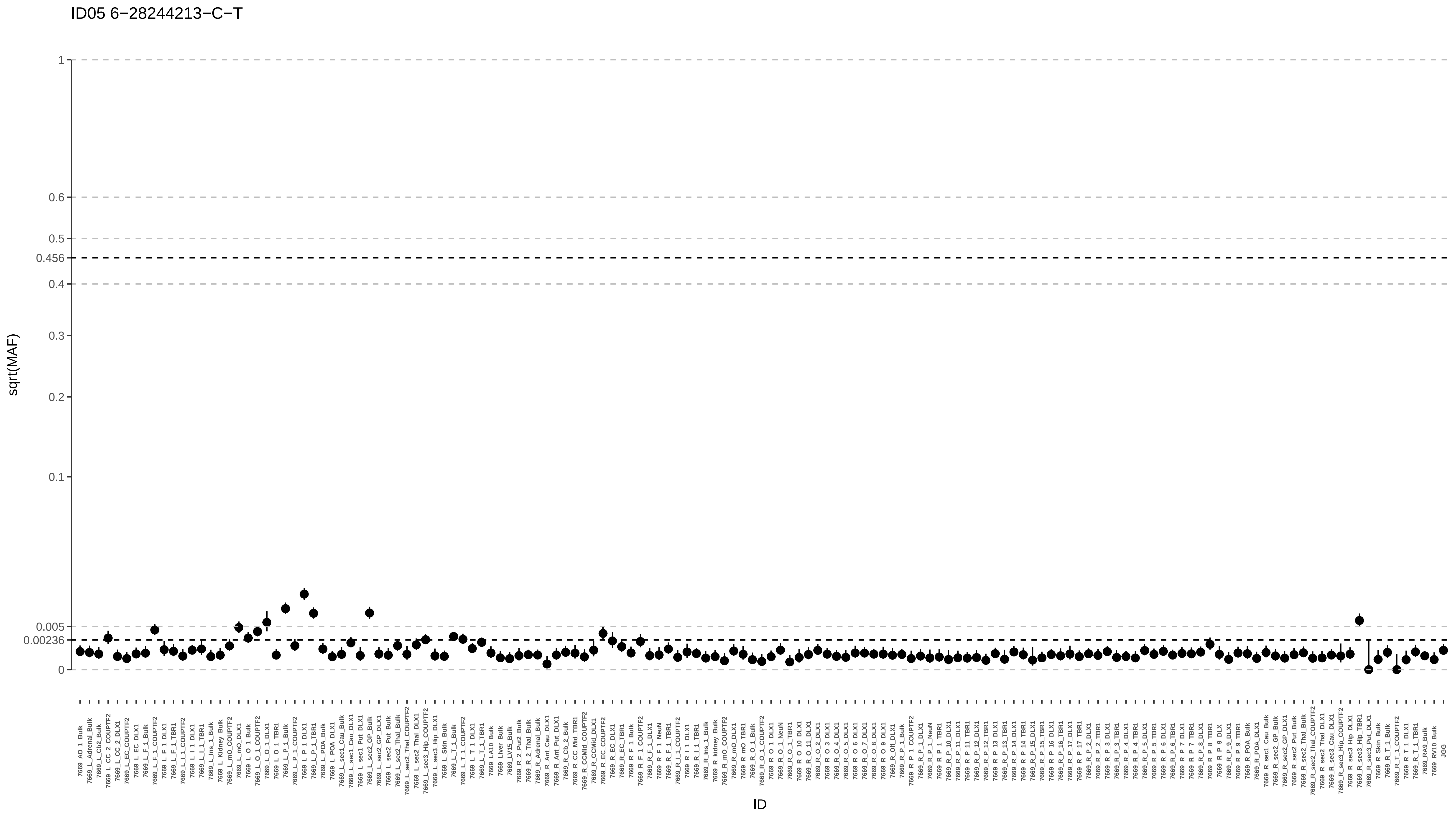

ID05 6−28244213−C−T
sqrt(MAF)
1
0.6
0.5
0.456
0.4
0.3
0.2
0.1
0.005
0.00236
0
ID

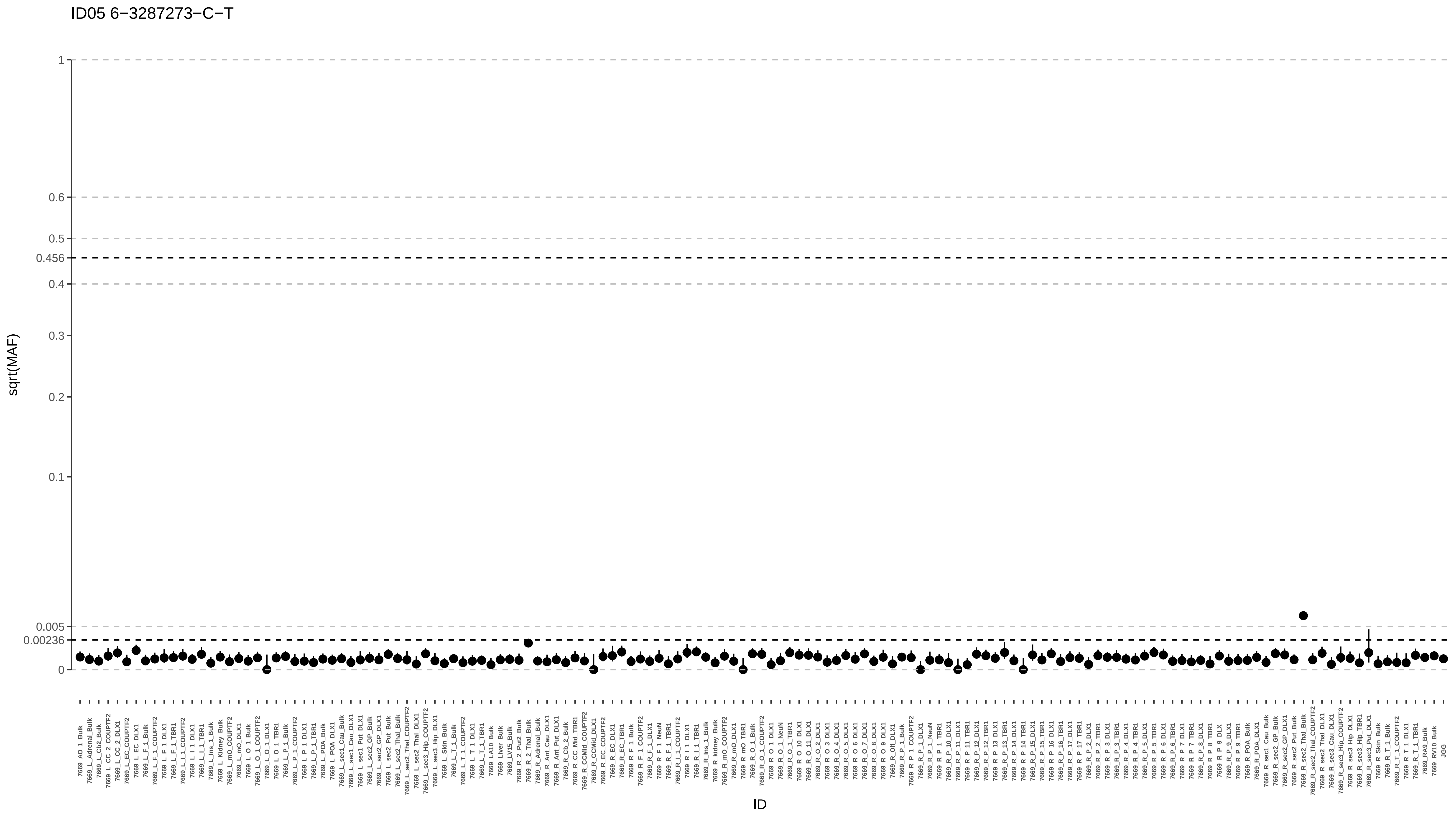
ID05 6−3287273−C−T
sqrt(MAF)
1
0.6
0.5
0.456
0.4
0.3
0.2
0.1
0.005
0.00236
0
ID

ID05 6−36931639−G−C

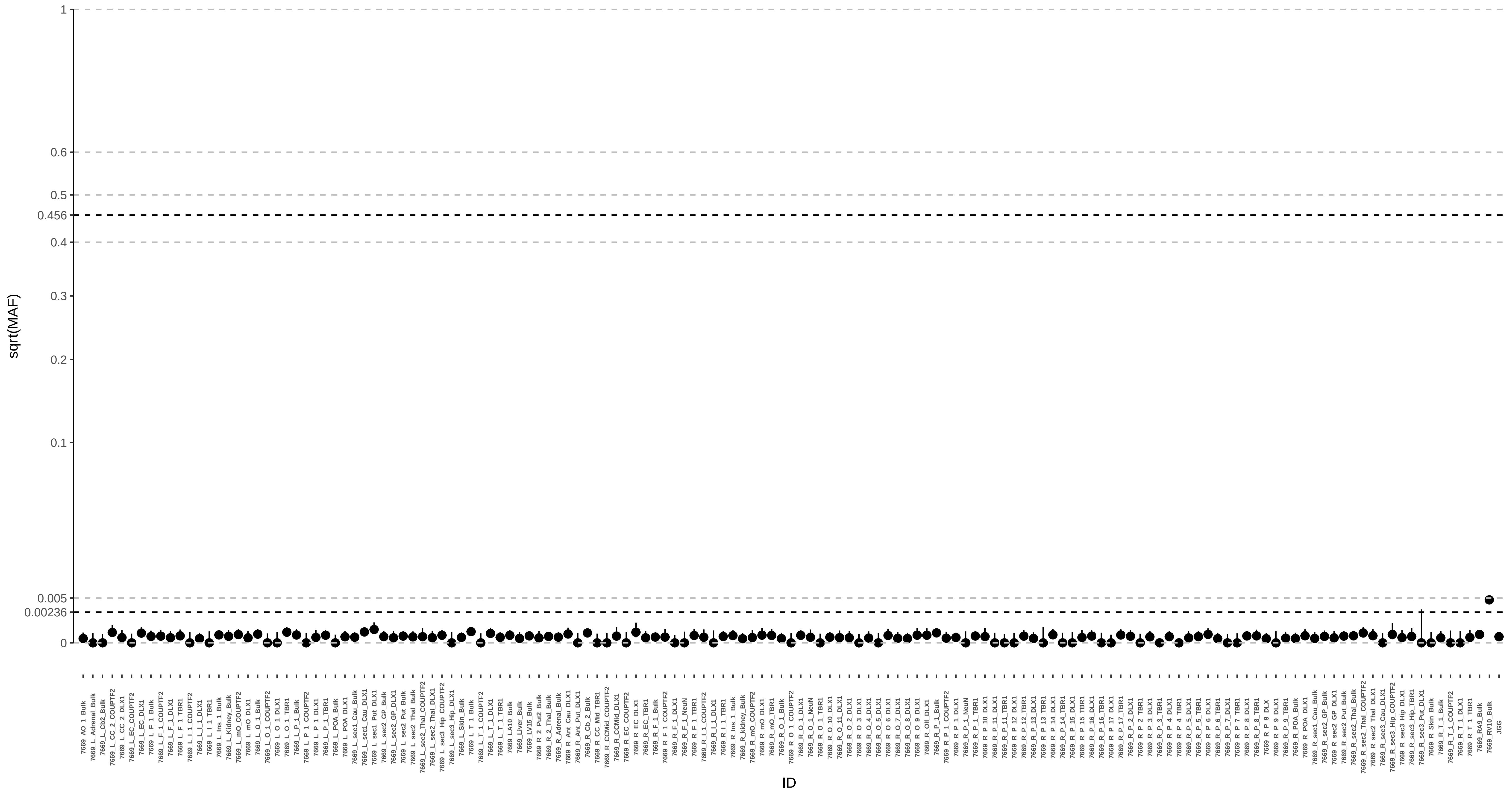

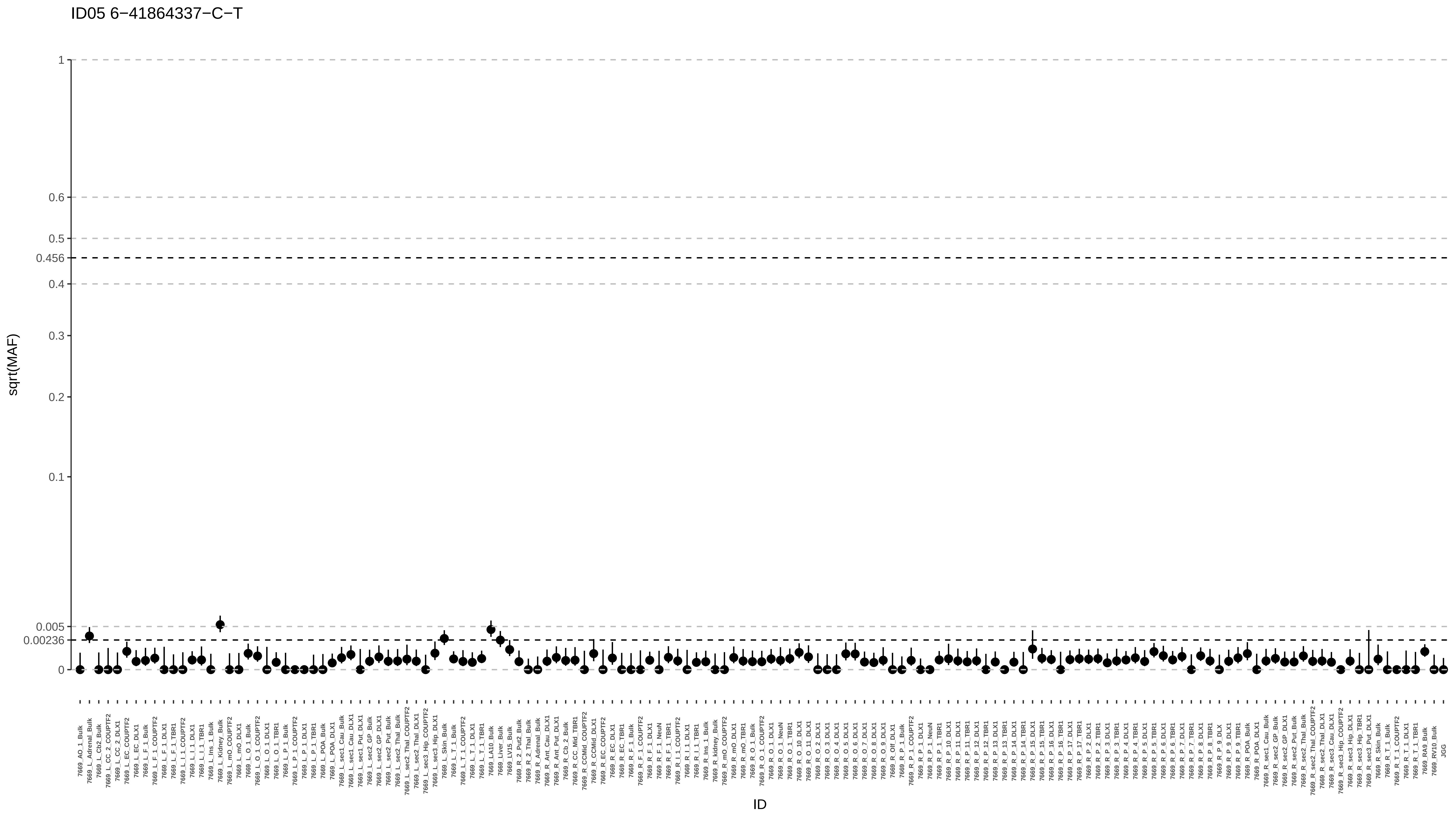
ID05 6−41864337−C−T
sqrt(MAF)
1
0.6
0.5
0.456
0.4
0.3
0.2
0.1
0.005
0.00236
0
ID

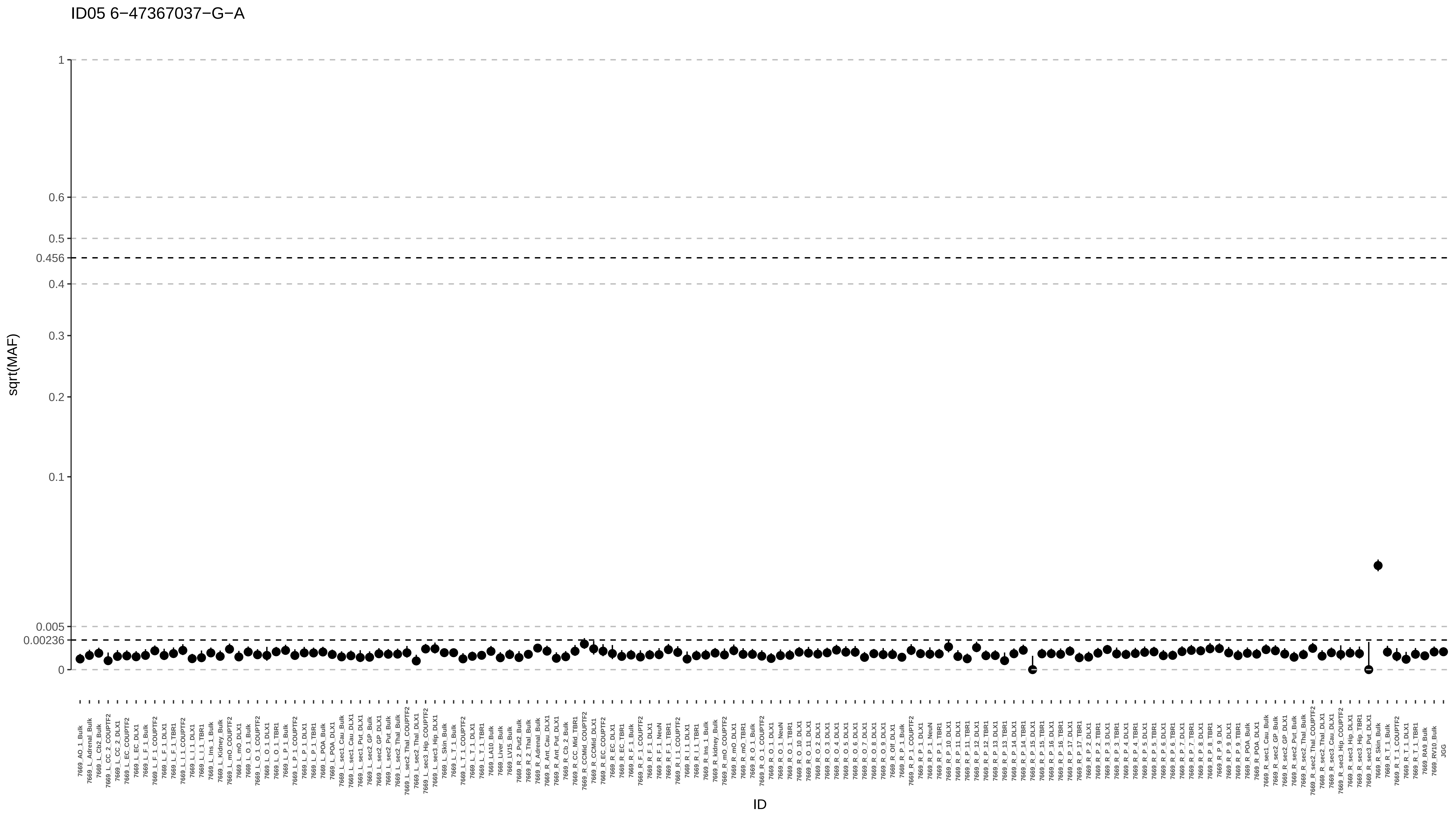
ID05 6−47367037−G−A
sqrt(MAF)
1
0.6
0.5
0.456
0.4
0.3
0.2
0.1
0.005
0.00236
0
ID

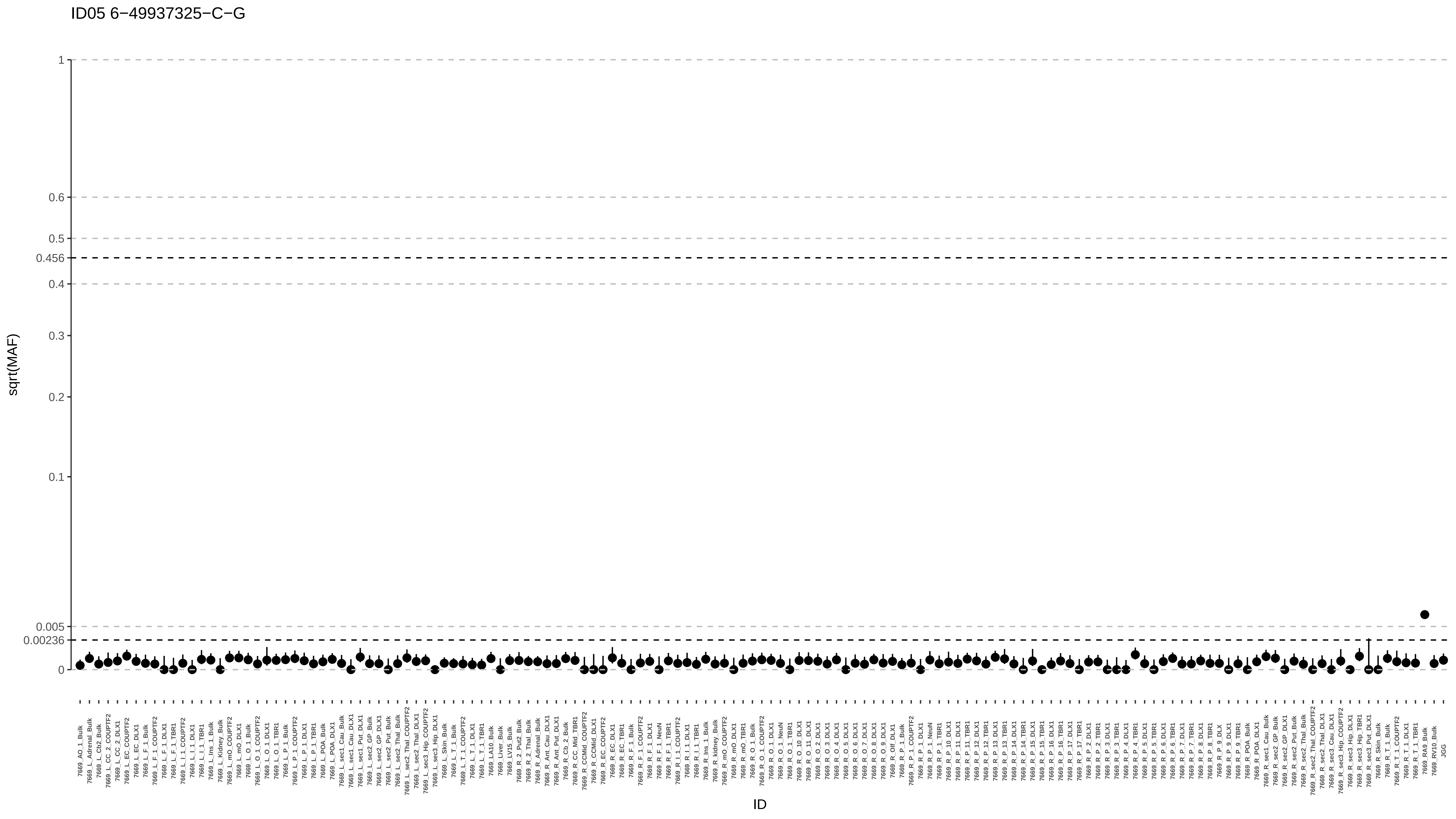
ID05 6−49937325−C−G
sqrt(MAF)
ID
1
0.6
0.5
0.456
0.4
0.3
0.2
0.1
0.005
0.00236
0

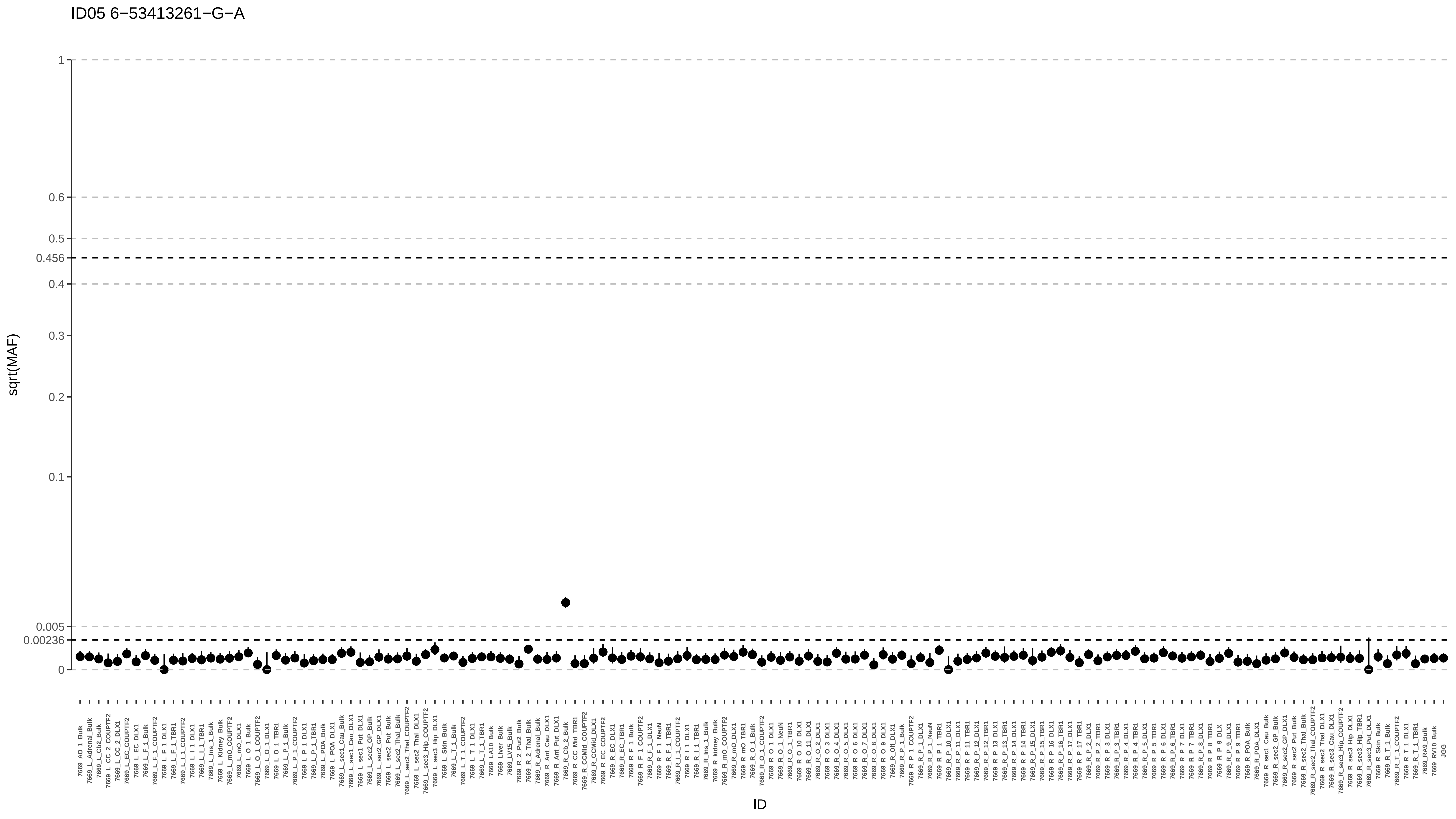
ID05 6−53413261−G−A
sqrt(MAF)
1
0.6
0.5
0.456
0.4
0.3
0.2
0.1
0.005
0.00236
0
ID

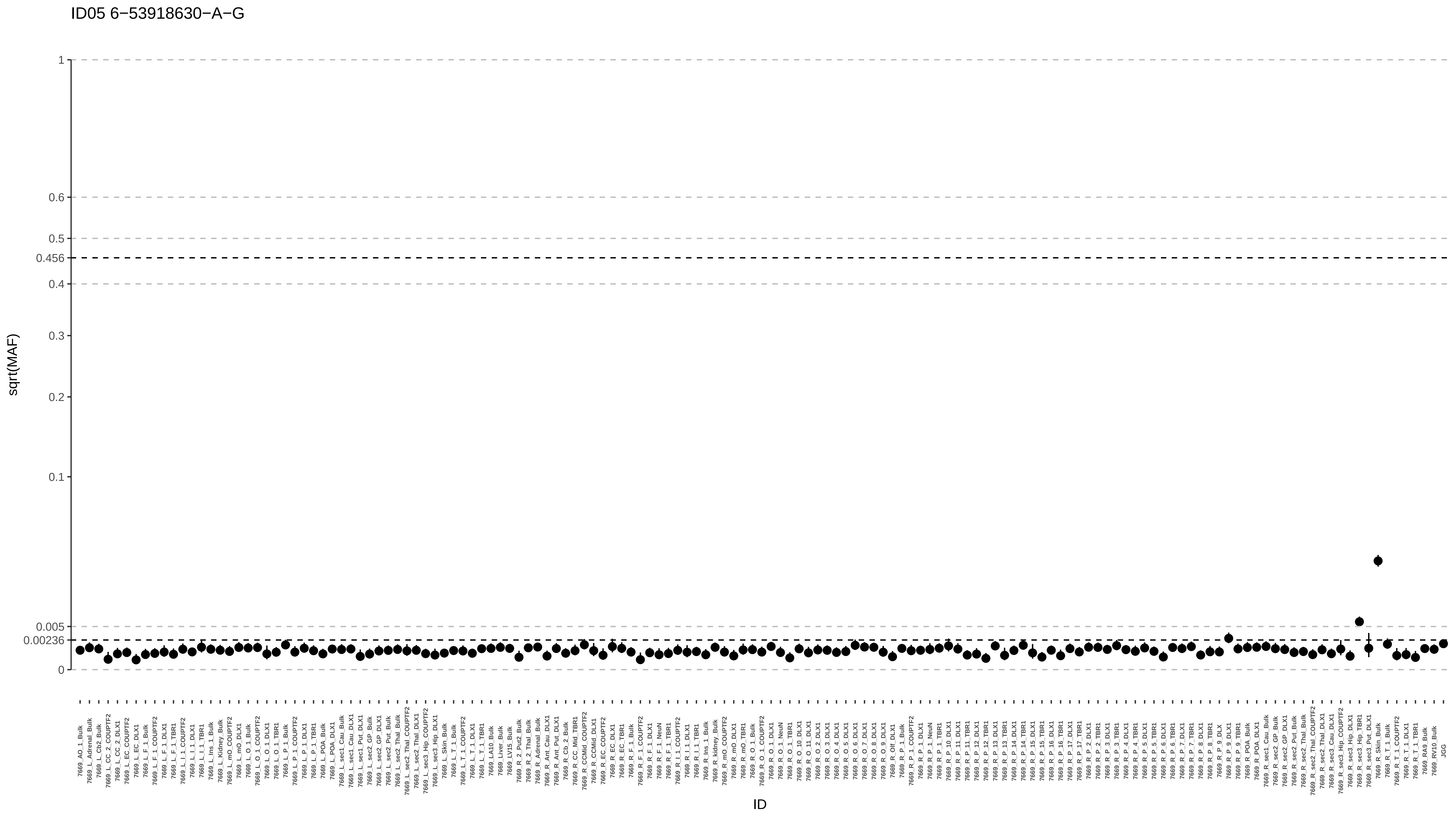

ID05 6−53918630−A−G
sqrt(MAF)
1
0.6
0.5
0.456
0.4
0.3
0.2
0.1
0.005
0.00236
0
ID

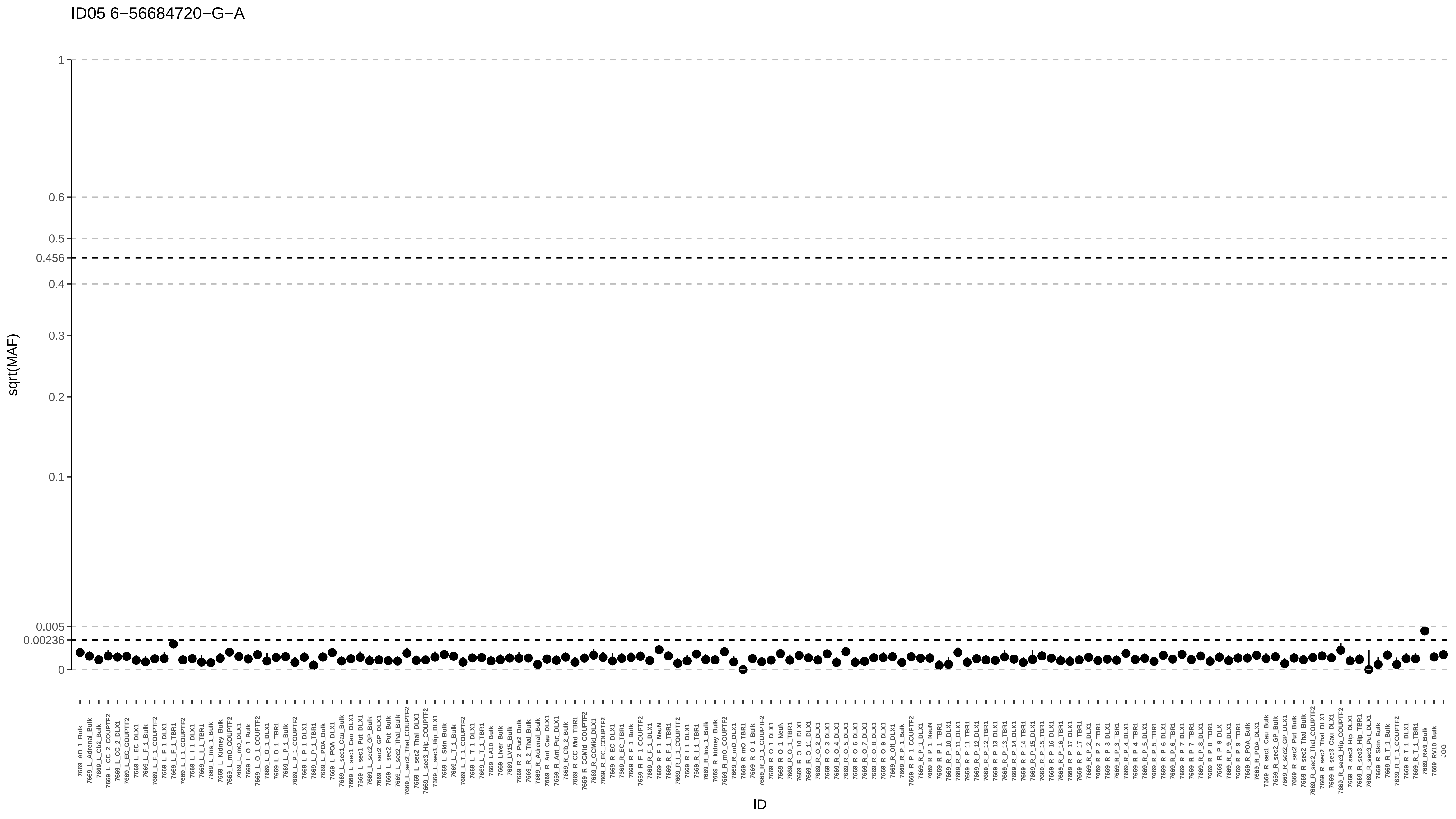
ID05 6−56684720−G−A
sqrt(MAF)
ID
1
0.6
0.5
0.456
0.4
0.3
0.2
0.1
0.005
0.00236
0

ID05 6−57165516−G−A

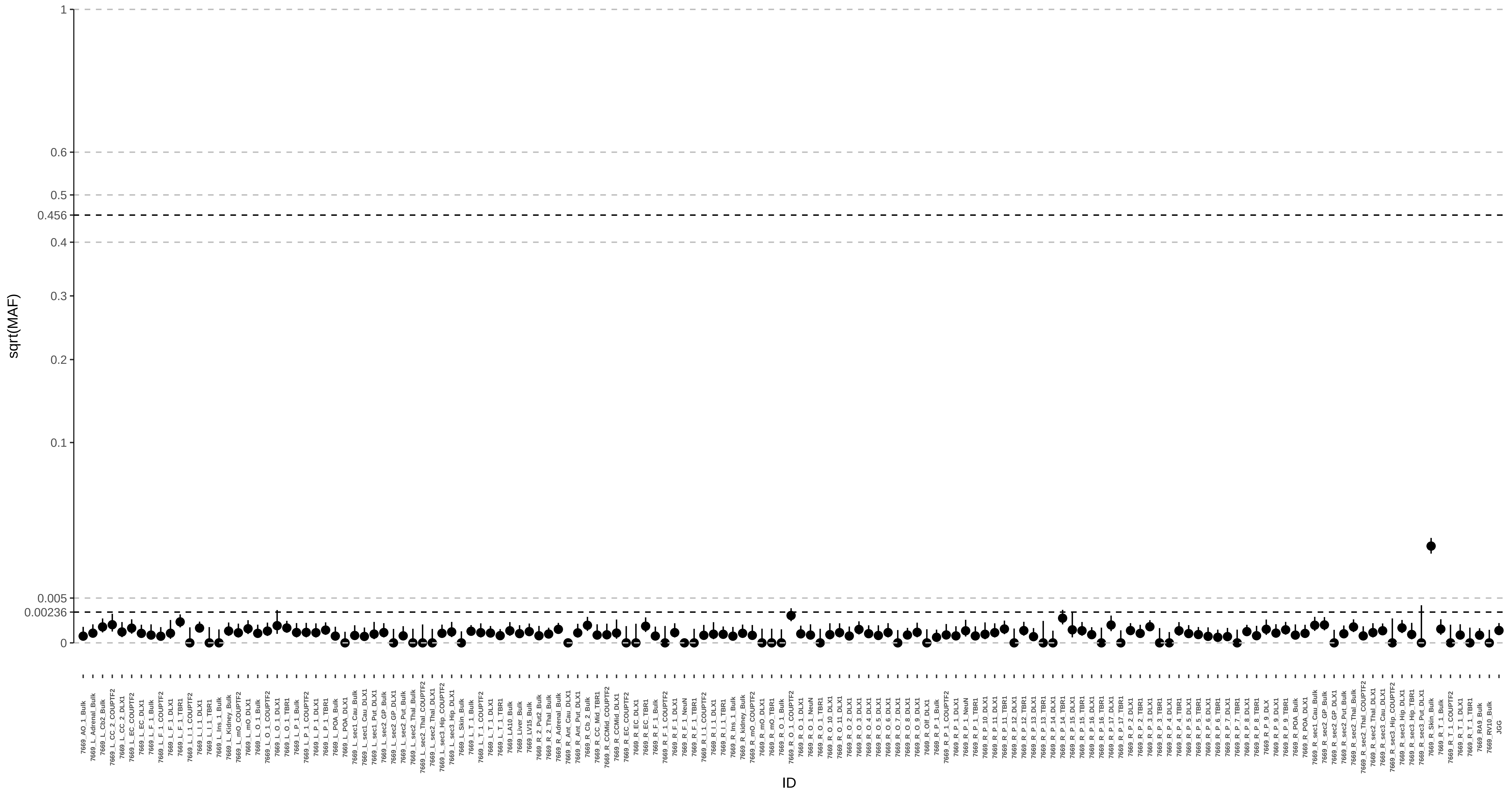

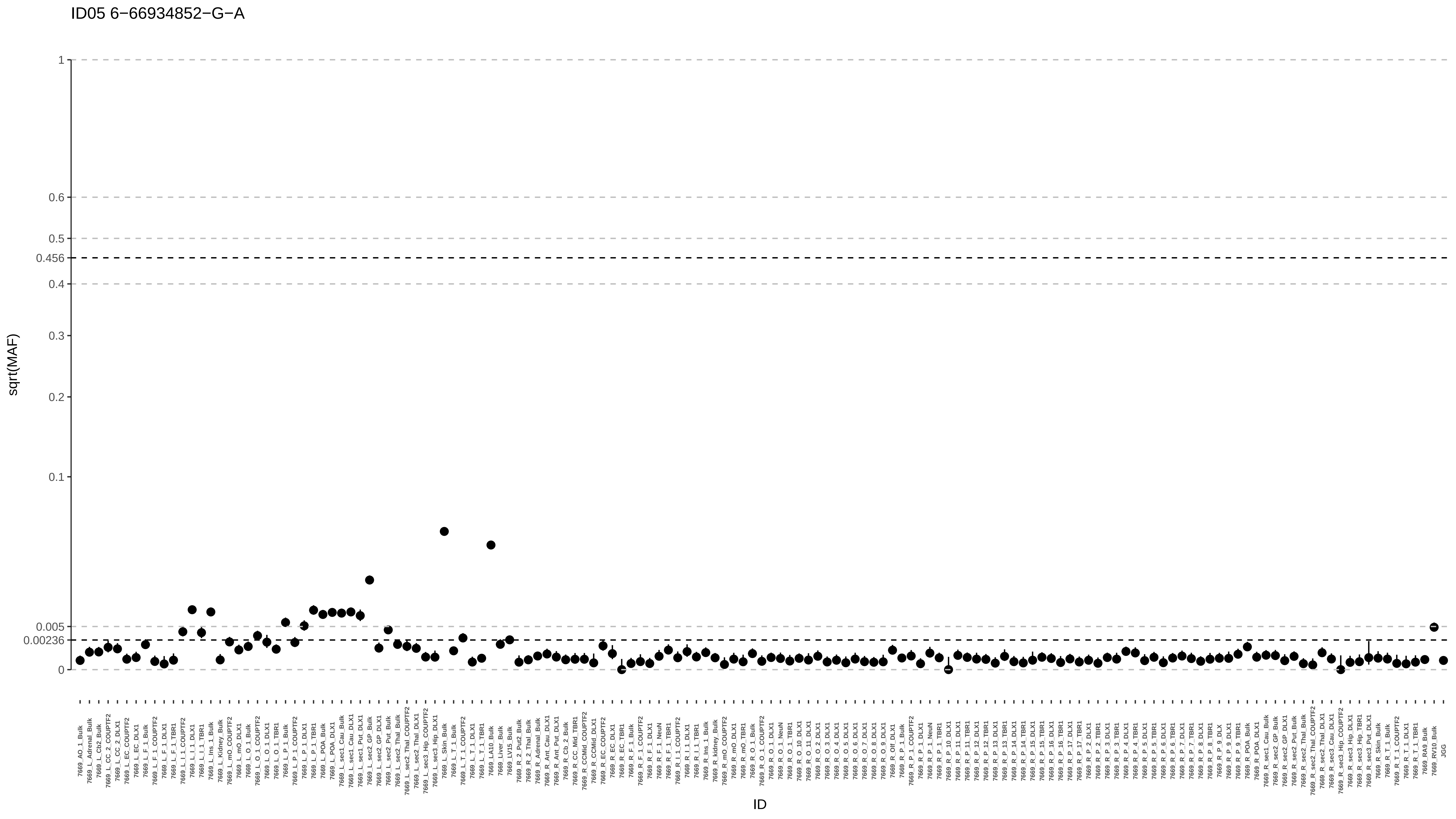
ID05 6−66934852−G−A
sqrt(MAF)
ID
1
0.6
0.5
0.456
0.4
0.3
0.2
0.1
0.005
0.00236
0

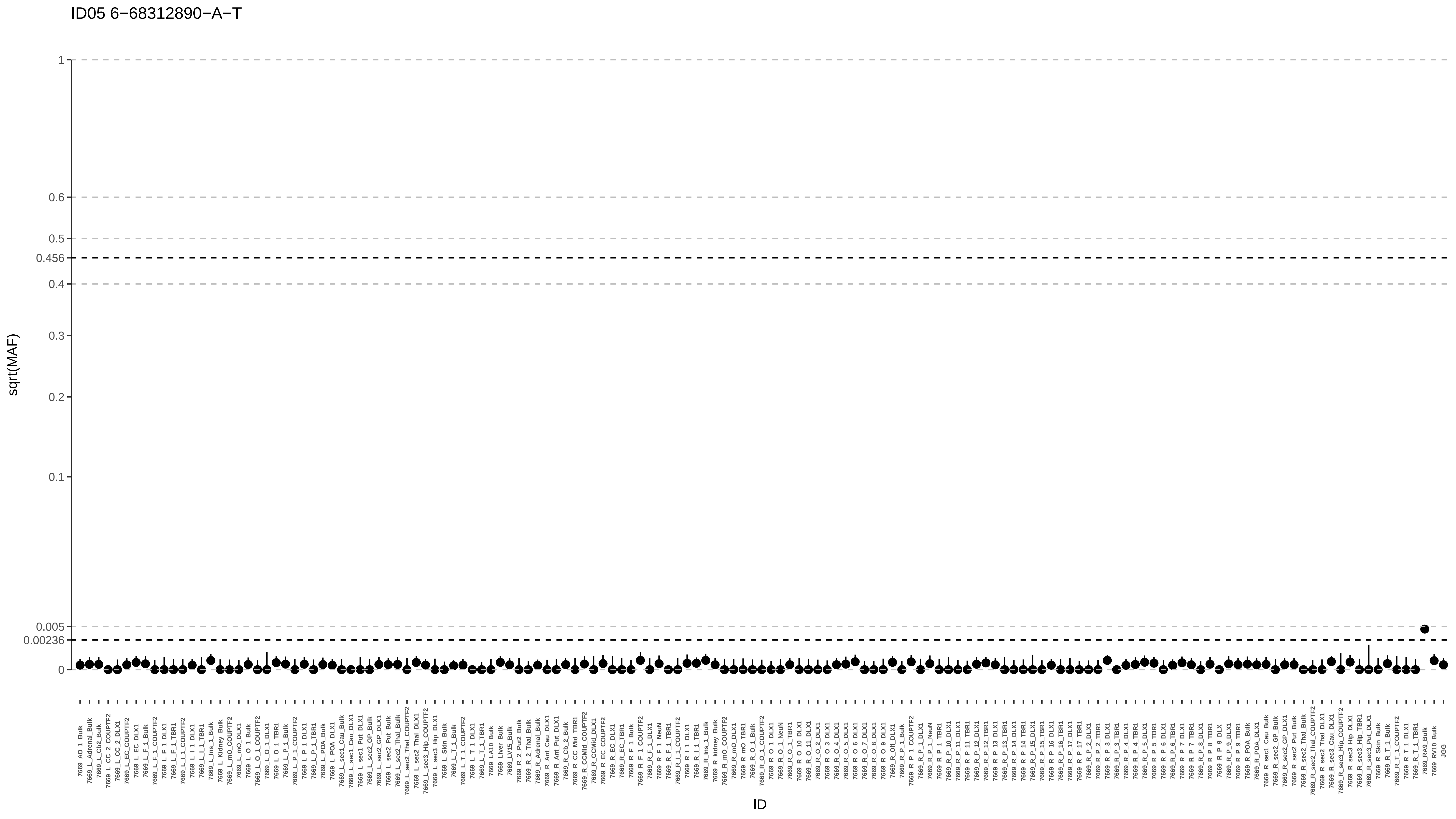

ID05 6−68312890−A−T
sqrt(MAF)
1
0.6
0.5
0.456
0.4
0.3
0.2
0.1
0.005
0.00236
0
ID
7669_AO_1_Bulk
7669_L_Adrenal_Bulk
7669_L_Cb2_Bulk
7669_L_CC_2_COUPTF2
7669_L_CC_2_DLX1
7669_L_EC_COUPTF2
7669_L_EC_DLX1
7669_L_F_1_Bulk
7669_L_F_1_COUPTF2
7669_L_F_1_DLX1
7669_L_F_1_TBR1
7669_L_I_1_COUPTF2
7669_L_I_1_DLX1
7669_L_I_1_TBR1
7669_L_Ins_1_Bulk
7669_L_Kidney_Bulk
7669_L_mO_COUPTF2
7669_L_mO_DLX1
7669_L_O_1_Bulk
7669_L_O_1_COUPTF2
7669_L_O_1_DLX1
7669_L_O_1_TBR1
7669_L_P_1_Bulk
7669_L_P_1_COUPTF2
7669_L_P_1_DLX1
7669_L_P_1_TBR1
7669_L_POA_Bulk
7669_L_POA_DLX1
7669_L_sec1_Cau_Bulk
7669_L_sec1_Cau_DLX1
7669_L_sec1_Put_DLX1
7669_L_sec2_GP_Bulk
7669_L_sec2_GP_DLX1
7669_L_sec2_Put_Bulk
7669_L_sec2_Thal_Bulk
7669_L_sec2_Thal_COUPTF2
7669_L_sec2_Thal_DLX1
7669_L_sec3_Hip_COUPTF2
7669_L_sec3_Hip_DLX1
7669_L_Skin_Bulk
7669_L_T_1_Bulk
7669_L_T_1_COUPTF2
7669_L_T_1_DLX1
7669_L_T_1_TBR1
7669_LA10_Bulk
7669_Liver_Bulk
7669_LV15_Bulk
7669_R_2_Put2_Bulk
7669_R_2_Thal_Bulk
7669_R_Adrenal_Bulk
7669_R_Ant_Cau_DLX1
7669_R_Ant_Put_DLX1
7669_R_Cb_2_Bulk
7669_R_CC_Mid_TBR1
7669_R_CCMid_COUPTF2
7669_R_CCMid_DLX1
7669_R_EC_COUPTF2
7669_R_EC_DLX1
7669_R_EC_TBR1
7669_R_F_1_Bulk
7669_R_F_1_COUPTF2
7669_R_F_1_DLX1
7669_R_F_1_NeuN
7669_R_F_1_TBR1
7669_R_I_1_COUPTF2
7669_R_I_1_DLX1
7669_R_I_1_TBR1
7669_R_Ins_1_Bulk
7669_R_kidney_Bulk
7669_R_mO_COUPTF2
7669_R_mO_DLX1
7669_R_mO_TBR1
7669_R_O_1_Bulk
7669_R_O_1_COUPTF2
7669_R_O_1_DLX1
7669_R_O_1_NeuN
7669_R_O_1_TBR1
7669_R_O_10_DLX1
7669_R_O_11_DLX1
7669_R_O_2_DLX1
7669_R_O_3_DLX1
7669_R_O_4_DLX1
7669_R_O_5_DLX1
7669_R_O_6_DLX1
7669_R_O_7_DLX1
7669_R_O_8_DLX1
7669_R_O_9_DLX1
7669_R_Olf_DLX1
7669_R_P_1_Bulk
7669_R_P_1_COUPTF2
7669_R_P_1_DLX1
7669_R_P_1_NeuN
7669_R_P_1_TBR1
7669_R_P_10_DLX1
7669_R_P_11_DLX1
7669_R_P_11_TBR1
7669_R_P_12_DLX1
7669_R_P_12_TBR1
7669_R_P_13_DLX1
7669_R_P_13_TBR1
7669_R_P_14_DLX1
7669_R_P_14_TBR1
7669_R_P_15_DLX1
7669_R_P_15_TBR1
7669_R_P_16_DLX1
7669_R_P_16_TBR1
7669_R_P_17_DLX1
7669_R_P_17_TBR1
7669_R_P_2_DLX1
7669_R_P_2_TBR1
7669_R_P_3_DLX1
7669_R_P_3_TBR1
7669_R_P_4_DLX1
7669_R_P_4_TBR1
7669_R_P_5_DLX1
7669_R_P_5_TBR1
7669_R_P_6_DLX1
7669_R_P_6_TBR1
7669_R_P_7_DLX1
7669_R_P_7_TBR1
7669_R_P_8_DLX1
7669_R_P_8_TBR1
7669_R_P_9_DLX
7669_R_P_9_DLX1
7669_R_P_9_TBR1
7669_R_POA_Bulk
7669_R_POA_DLX1
7669_R_sec1_Cau_Bulk
7669_R_sec2_GP_Bulk
7669_R_sec2_GP_DLX1
7669_R_sec2_Put_Bulk
7669_R_sec2_Thal_Bulk
7669_R_sec2_Thal_COUPTF2
7669_R_sec2_Thal_DLX1
7669_R_sec3_Cau_DLX1
7669_R_sec3_Hip_COUPTF2
7669_R_sec3_Hip_DLX1
7669_R_sec3_Hip_TBR1
7669_R_sec3_Put_DLX1
7669_R_Skin_Bulk
7669_R_T_1_Bulk
7669_R_T_1_COUPTF2
7669_R_T_1_DLX1
7669_R_T_1_TBR1
7669_RA9_Bulk
7669_RV10_Bulk
JGG

ID05 6−91839541−C−T

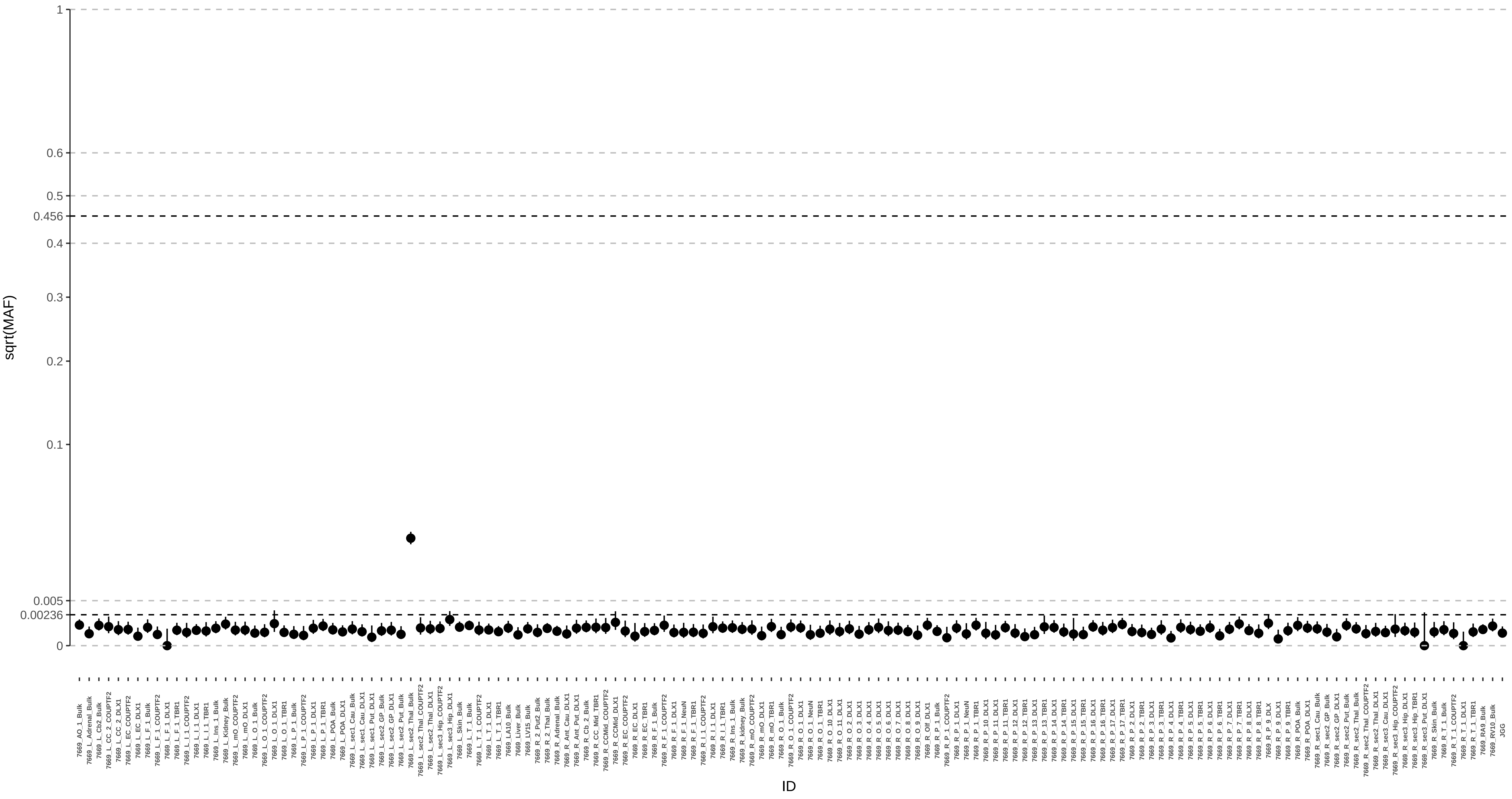

ID05 6−92211371−C−T

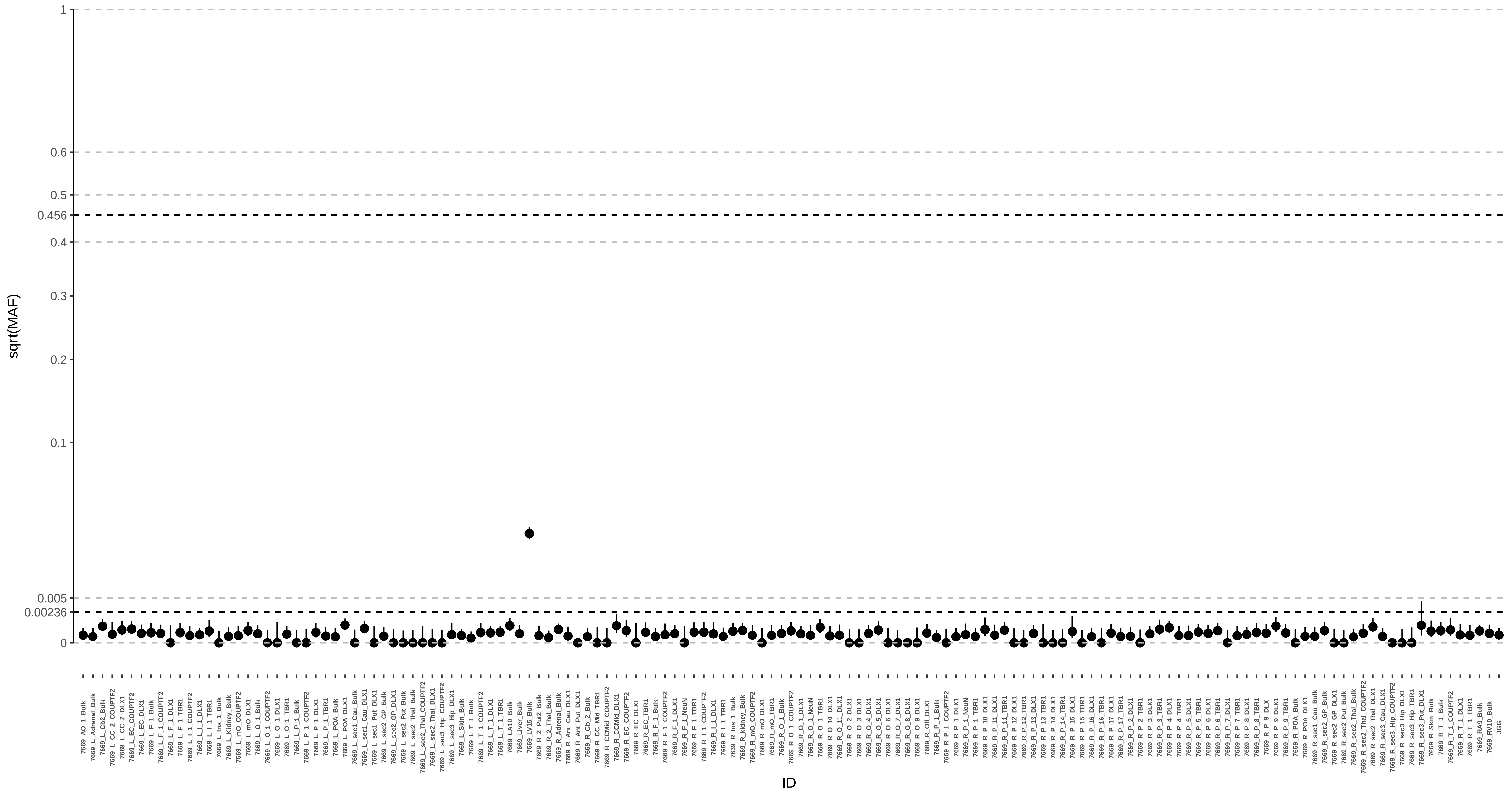

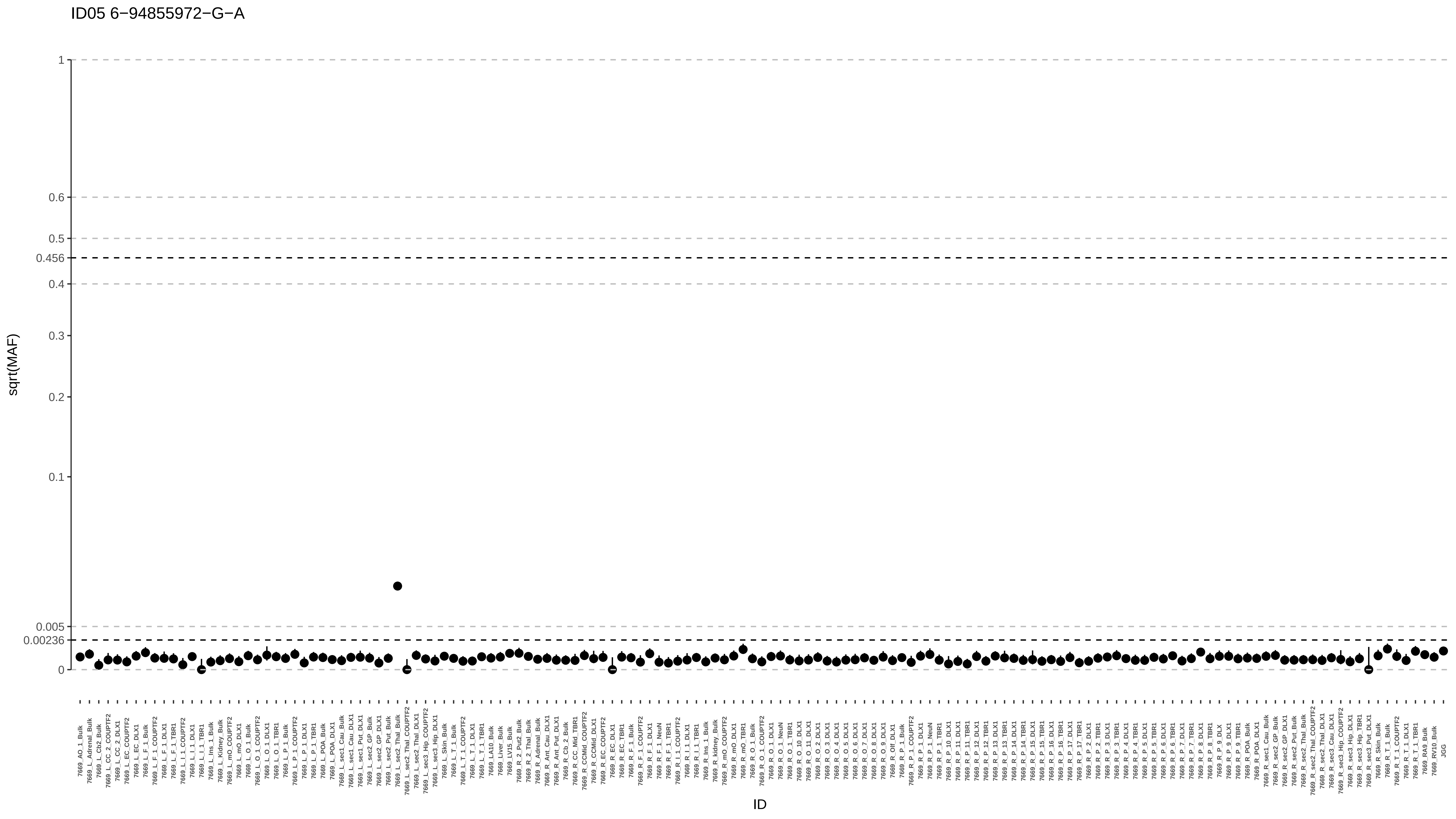
ID05 6−94855972−G−A
sqrt(MAF)
1
0.6
0.5
0.456
0.4
0.3
0.2
0.1
0.005
0.00236
0
ID

ID05 6−96176402−T−C

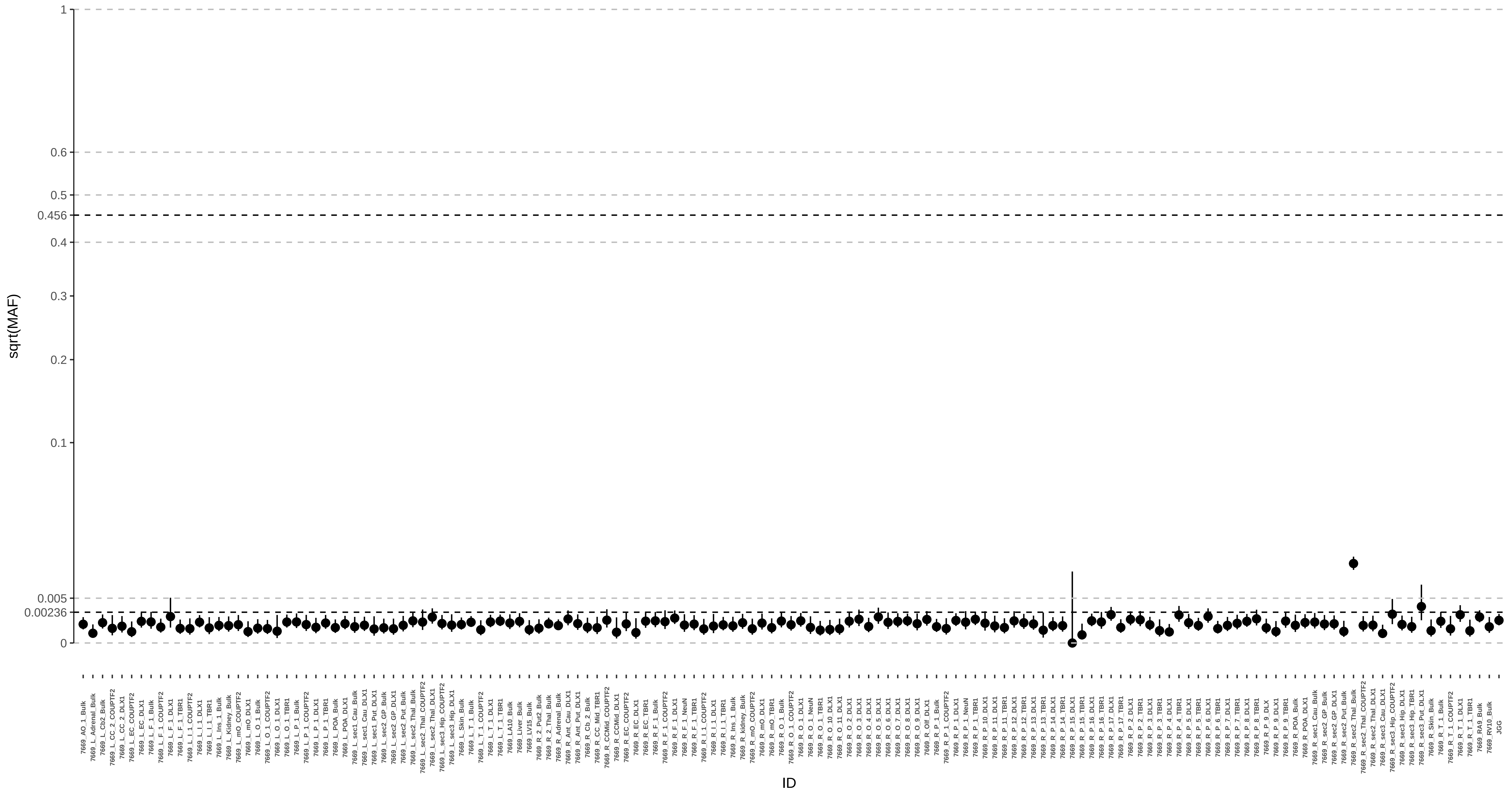

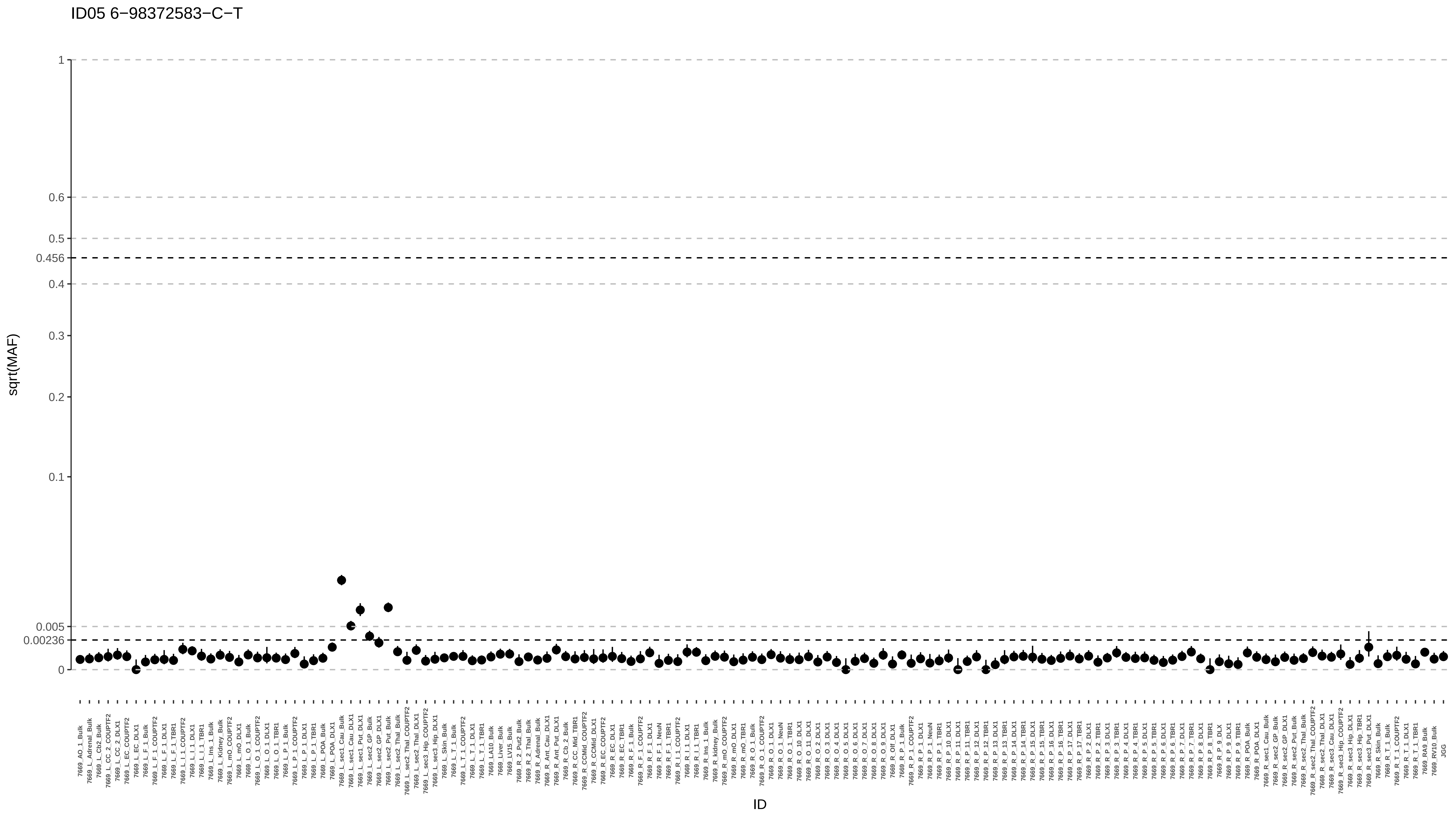
ID05 6−98372583−C−T
sqrt(MAF)
1
0.6
0.5
0.456
0.4
0.3
0.2
0.1
0.005
0.00236
0
ID

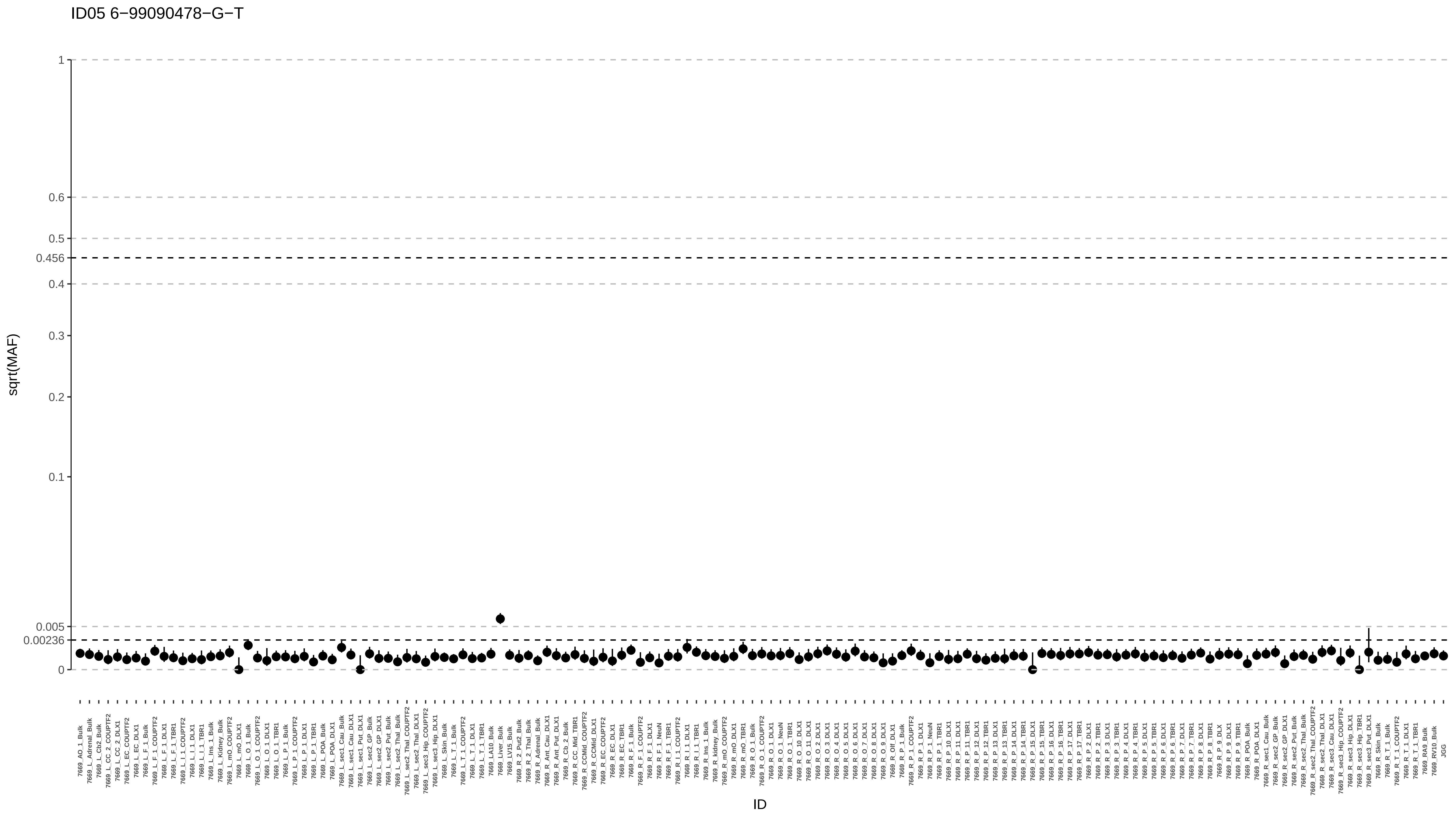
ID05 6−99090478−G−T
sqrt(MAF)
1
0.6
0.5
0.456
0.4
0.3
0.2
0.1
0.005
0.00236
0
ID

ID05 6−99303058−CCT−C

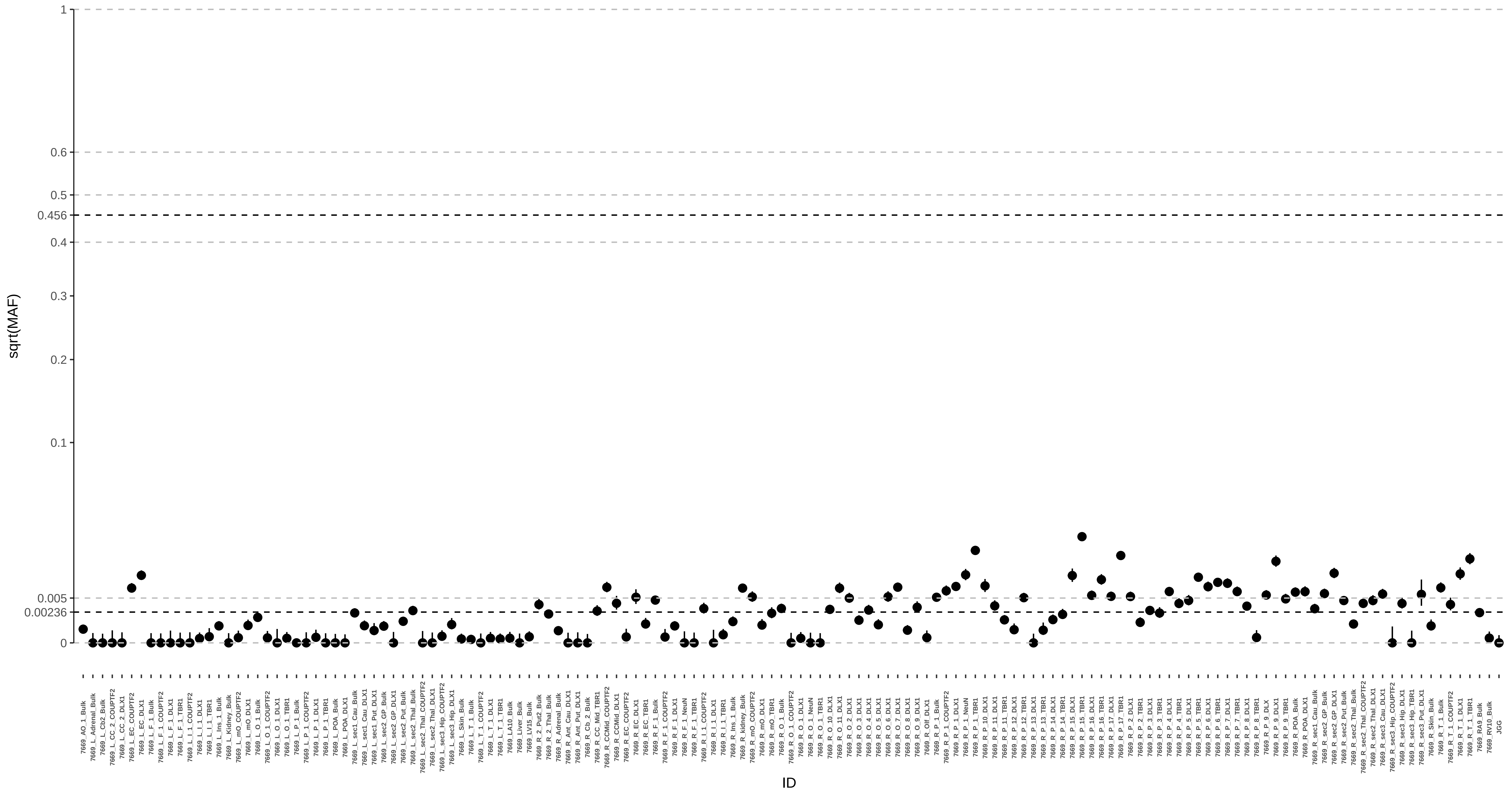

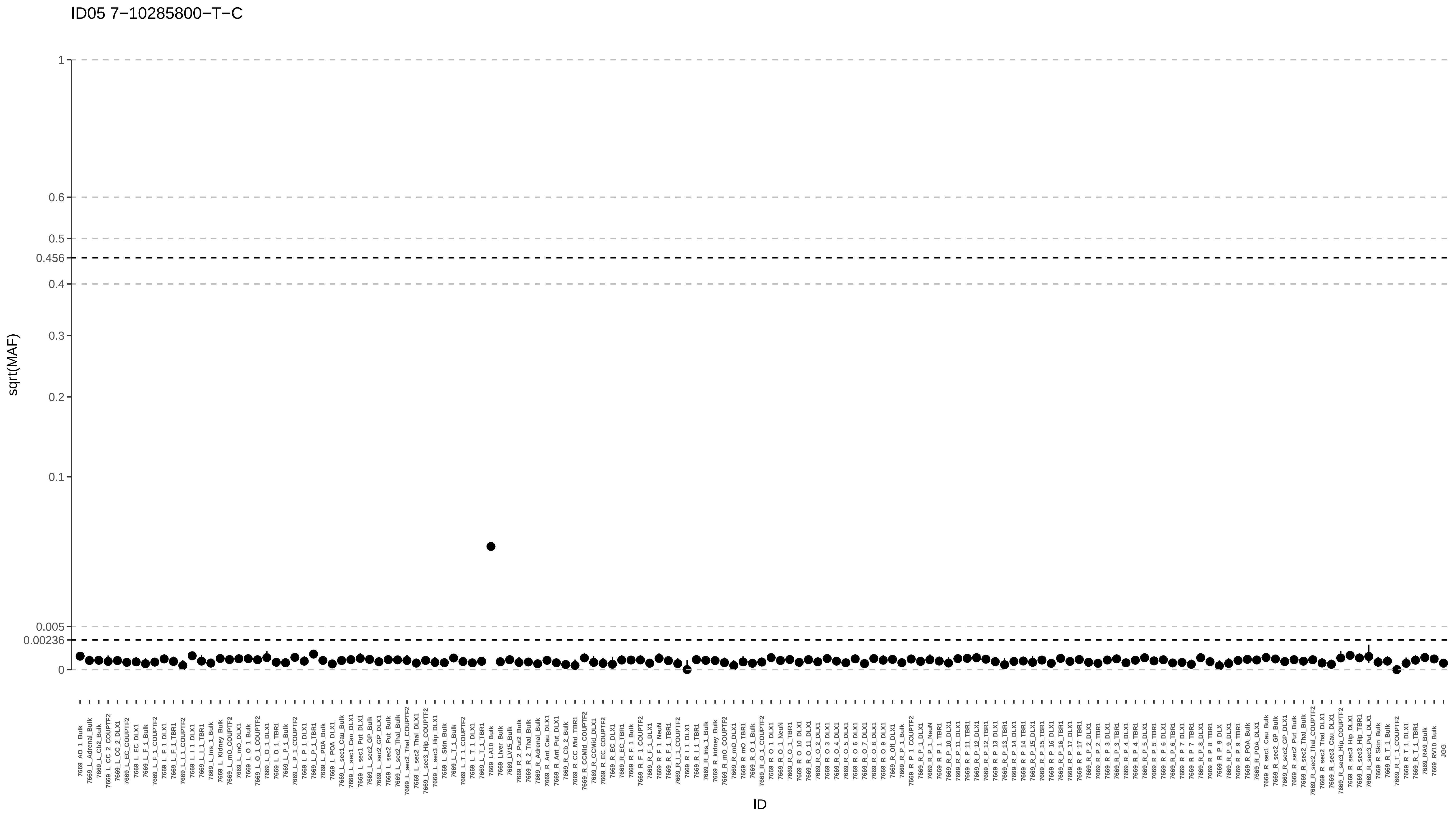
ID05 7−10285800−T−C
sqrt(MAF)
1
0.6
0.5
0.456
0.4
0.3
0.2
0.1
0.005
0.00236
0
ID
7669_AO_1_Bulk
7669_L_Adrenal_Bulk
7669_L_Cb2_Bulk
7669_L_CC_2_COUPTF2
7669_L_CC_2_DLX1
7669_L_EC_COUPTF2
7669_L_EC_DLX1
7669_L_F_1_Bulk
7669_L_F_1_COUPTF2
7669_L_F_1_DLX1
7669_L_F_1_TBR1
7669_L_I_1_COUPTF2
7669_L_I_1_DLX1
7669_L_I_1_TBR1
7669_L_Ins_1_Bulk
7669_L_Kidney_Bulk
7669_L_mO_COUPTF2
7669_L_mO_DLX1
7669_L_O_1_Bulk
7669_L_O_1_COUPTF2
7669_L_O_1_DLX1
7669_L_O_1_TBR1
7669_L_P_1_Bulk
7669_L_P_1_COUPTF2
7669_L_P_1_DLX1
7669_L_P_1_TBR1
7669_L_POA_Bulk
7669_L_POA_DLX1
7669_L_sec1_Cau_Bulk
7669_L_sec1_Cau_DLX1
7669_L_sec1_Put_DLX1
7669_L_sec2_GP_Bulk
7669_L_sec2_GP_DLX1
7669_L_sec2_Put_Bulk
7669_L_sec2_Thal_Bulk
7669_L_sec2_Thal_COUPTF2
7669_L_sec2_Thal_DLX1
7669_L_sec3_Hip_COUPTF2
7669_L_sec3_Hip_DLX1
7669_L_Skin_Bulk
7669_L_T_1_Bulk
7669_L_T_1_COUPTF2
7669_L_T_1_DLX1
7669_L_T_1_TBR1
7669_LA10_Bulk
7669_Liver_Bulk
7669_LV15_Bulk
7669_R_2_Put2_Bulk
7669_R_2_Thal_Bulk
7669_R_Adrenal_Bulk
7669_R_Ant_Cau_DLX1
7669_R_Ant_Put_DLX1
7669_R_Cb_2_Bulk
7669_R_CC_Mid_TBR1
7669_R_CCMid_COUPTF2
7669_R_CCMid_DLX1
7669_R_EC_COUPTF2
7669_R_EC_DLX1
7669_R_EC_TBR1
7669_R_F_1_Bulk
7669_R_F_1_COUPTF2
7669_R_F_1_DLX1
7669_R_F_1_NeuN
7669_R_F_1_TBR1
7669_R_I_1_COUPTF2
7669_R_I_1_DLX1
7669_R_I_1_TBR1
7669_R_Ins_1_Bulk
7669_R_kidney_Bulk
7669_R_mO_COUPTF2
7669_R_mO_DLX1
7669_R_mO_TBR1
7669_R_O_1_Bulk
7669_R_O_1_COUPTF2
7669_R_O_1_DLX1
7669_R_O_1_NeuN
7669_R_O_1_TBR1
7669_R_O_10_DLX1
7669_R_O_11_DLX1
7669_R_O_2_DLX1
7669_R_O_3_DLX1
7669_R_O_4_DLX1
7669_R_O_5_DLX1
7669_R_O_6_DLX1
7669_R_O_7_DLX1
7669_R_O_8_DLX1
7669_R_O_9_DLX1
7669_R_Olf_DLX1
7669_R_P_1_Bulk
7669_R_P_1_COUPTF2
7669_R_P_1_DLX1
7669_R_P_1_NeuN
7669_R_P_1_TBR1
7669_R_P_10_DLX1
7669_R_P_11_DLX1
7669_R_P_11_TBR1
7669_R_P_12_DLX1
7669_R_P_12_TBR1
7669_R_P_13_DLX1
7669_R_P_13_TBR1
7669_R_P_14_DLX1
7669_R_P_14_TBR1
7669_R_P_15_DLX1
7669_R_P_15_TBR1
7669_R_P_16_DLX1
7669_R_P_16_TBR1
7669_R_P_17_DLX1
7669_R_P_17_TBR1
7669_R_P_2_DLX1
7669_R_P_2_TBR1
7669_R_P_3_DLX1
7669_R_P_3_TBR1
7669_R_P_4_DLX1
7669_R_P_4_TBR1
7669_R_P_5_DLX1
7669_R_P_5_TBR1
7669_R_P_6_DLX1
7669_R_P_6_TBR1
7669_R_P_7_DLX1
7669_R_P_7_TBR1
7669_R_P_8_DLX1
7669_R_P_8_TBR1
7669_R_P_9_DLX
7669_R_P_9_DLX1
7669_R_P_9_TBR1
7669_R_POA_Bulk
7669_R_POA_DLX1
7669_R_sec1_Cau_Bulk
7669_R_sec2_GP_Bulk
7669_R_sec2_GP_DLX1
7669_R_sec2_Put_Bulk
7669_R_sec2_Thal_Bulk
7669_R_sec2_Thal_COUPTF2
7669_R_sec2_Thal_DLX1
7669_R_sec3_Cau_DLX1
7669_R_sec3_Hip_COUPTF2
7669_R_sec3_Hip_DLX1
7669_R_sec3_Hip_TBR1
7669_R_sec3_Put_DLX1
7669_R_Skin_Bulk
7669_R_T_1_Bulk
7669_R_T_1_COUPTF2
7669_R_T_1_DLX1
7669_R_T_1_TBR1
7669_RA9_Bulk
7669_RV10_Bulk
JGG

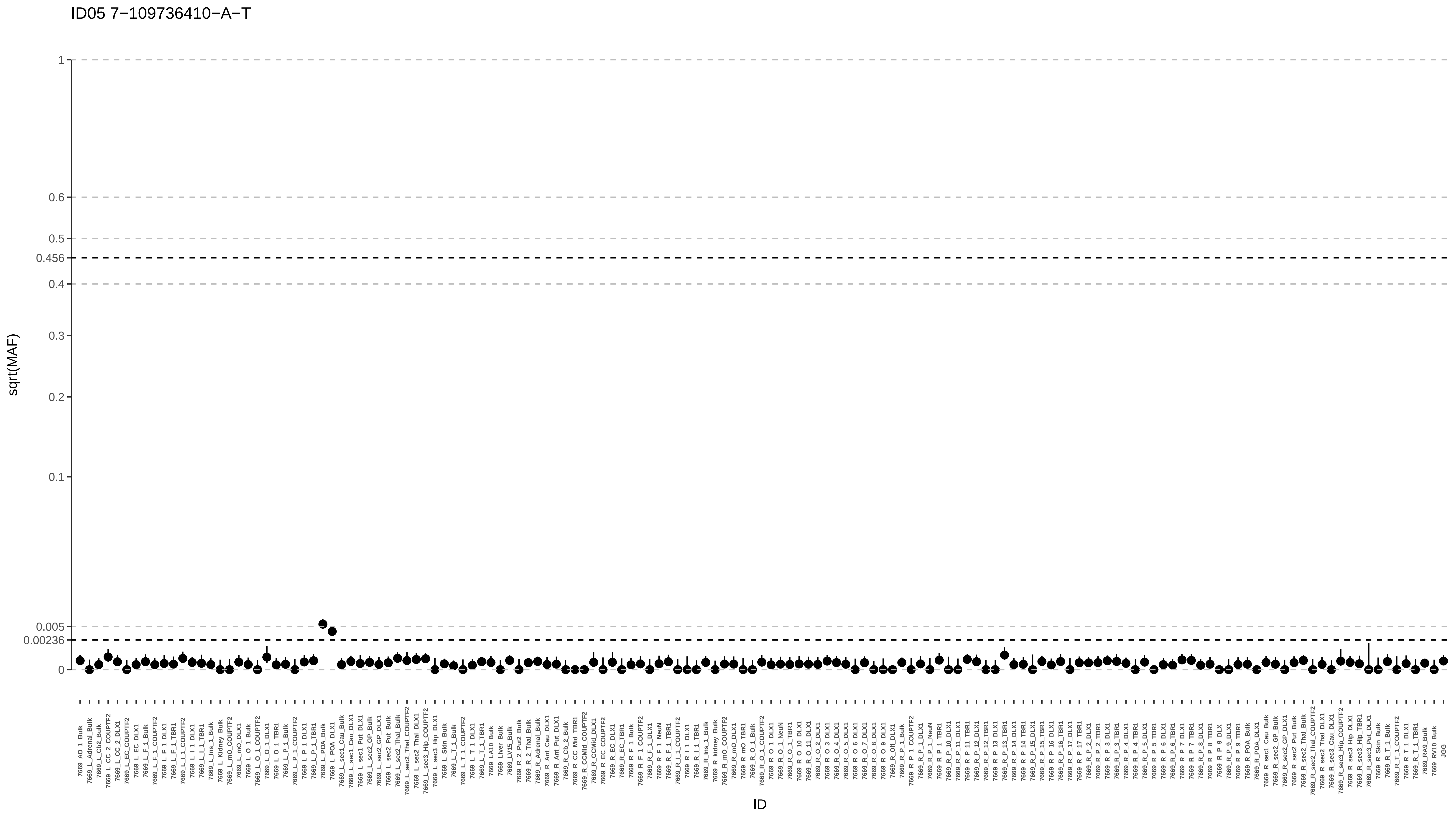
ID05 7−109736410−A−T
sqrt(MAF)
1
0.6
0.5
0.456
0.4
0.3
0.2
0.1
0.005
0.00236
0
ID

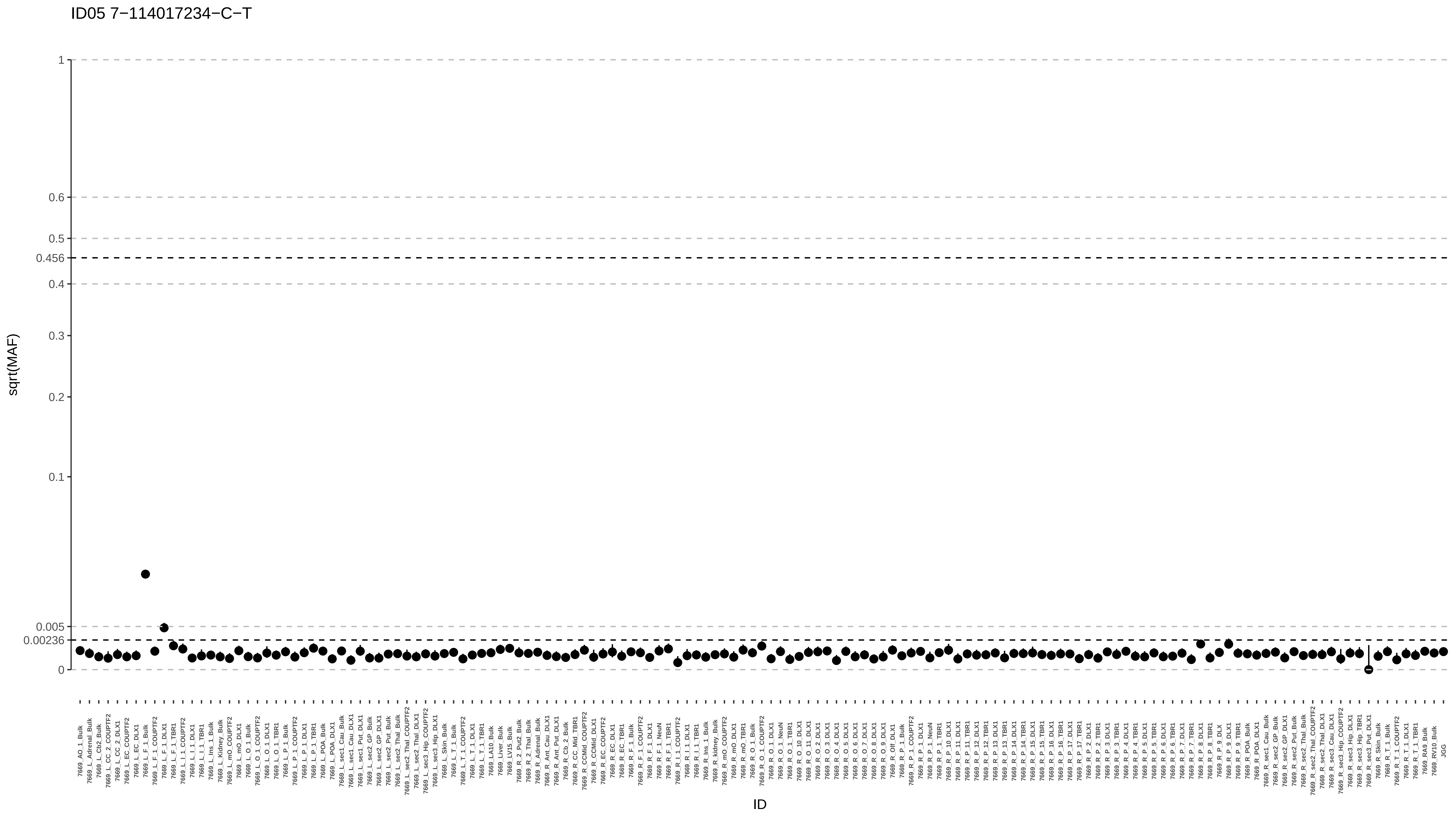
ID05 7−114017234−C−T
sqrt(MAF)
1
0.6
0.5
0.456
0.4
0.3
0.2
0.1
0.005
0.00236
0
ID

ID05 7−114658860−A−G

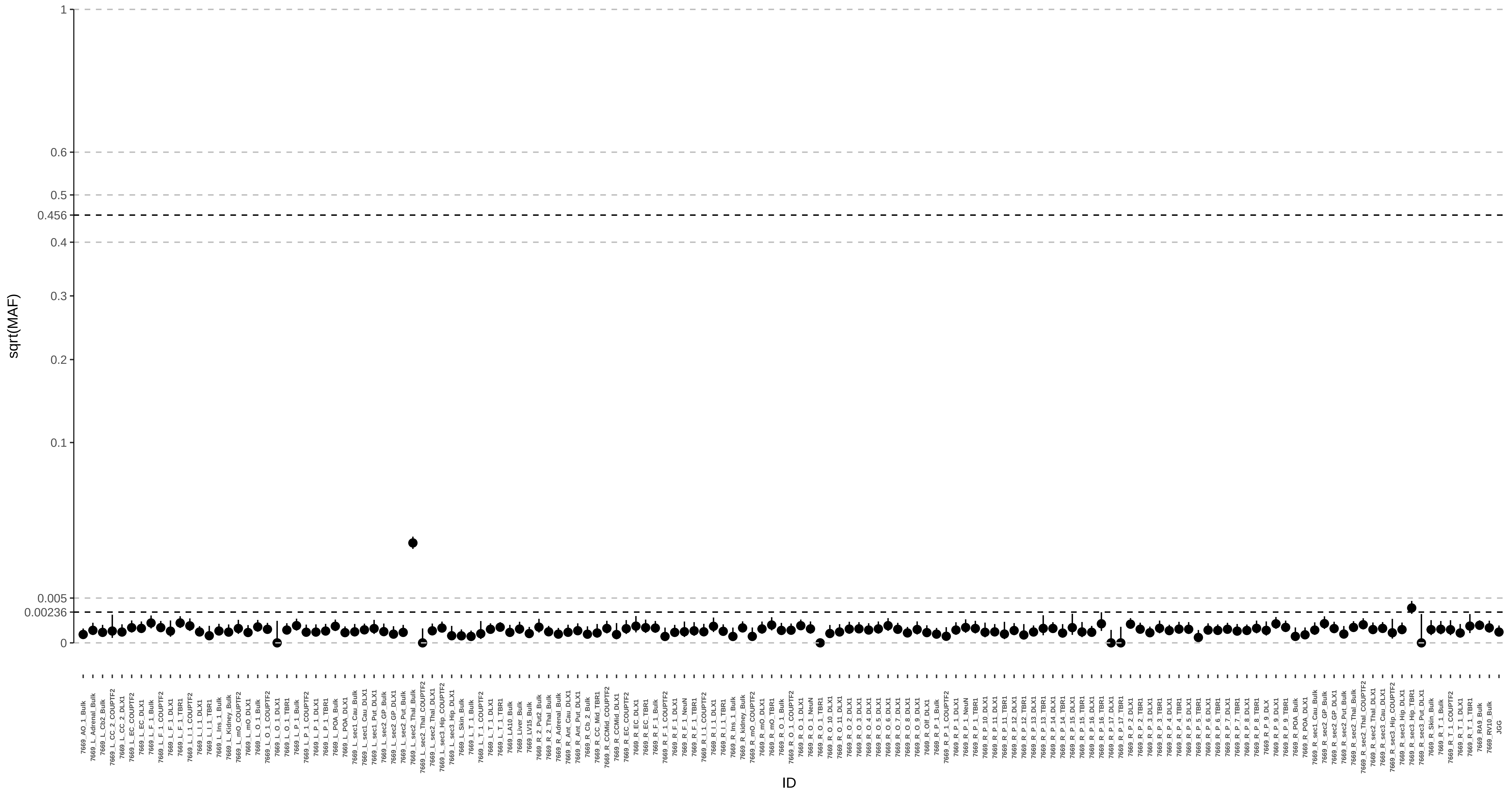

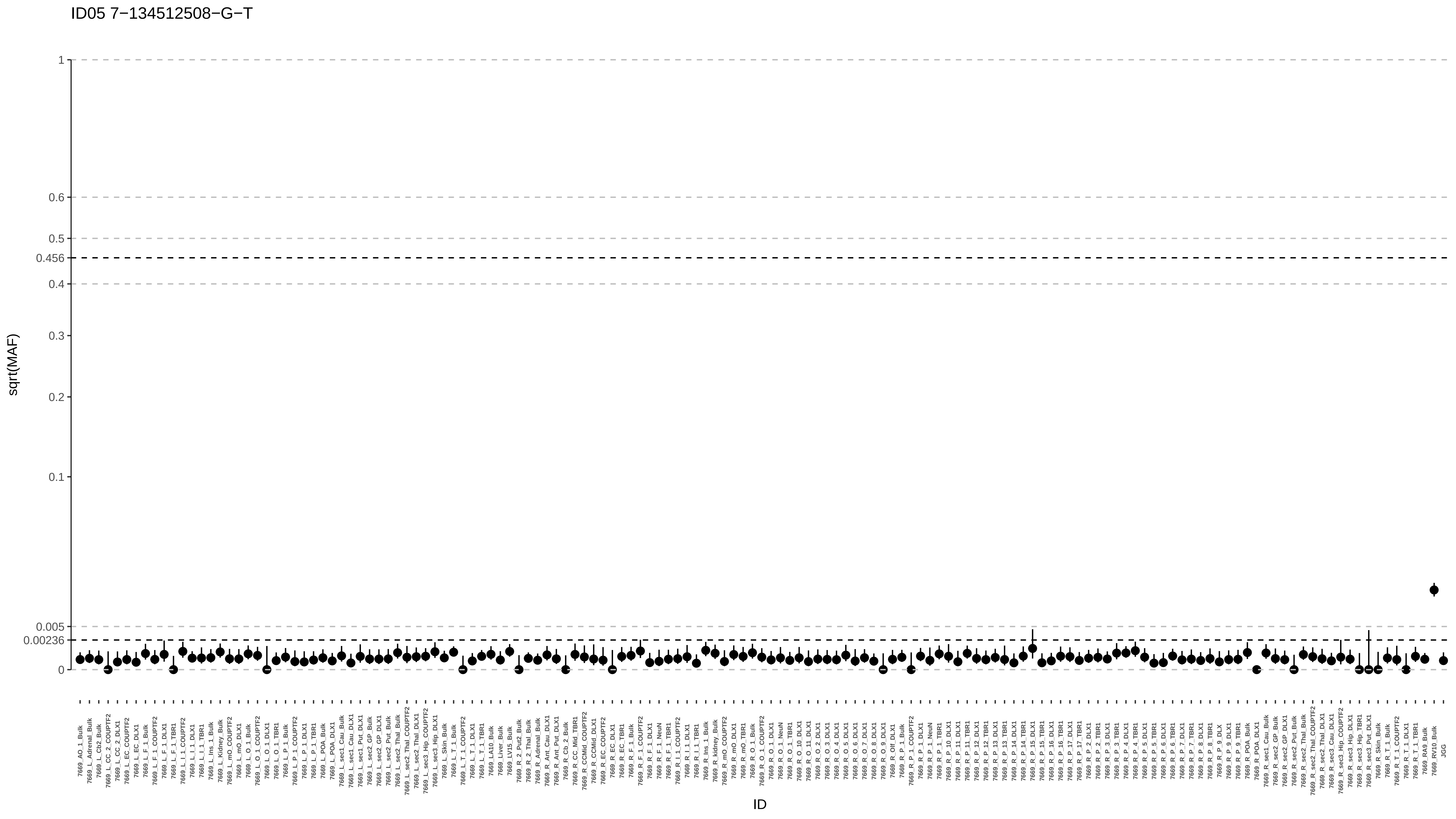
ID05 7−134512508−G−T
1
0.6
0.5
0.456
0.4
0.3
0.2
0.1
0.005
0.00236
0
sqrt(MAF)
ID

ID05 7−137220703−G−A

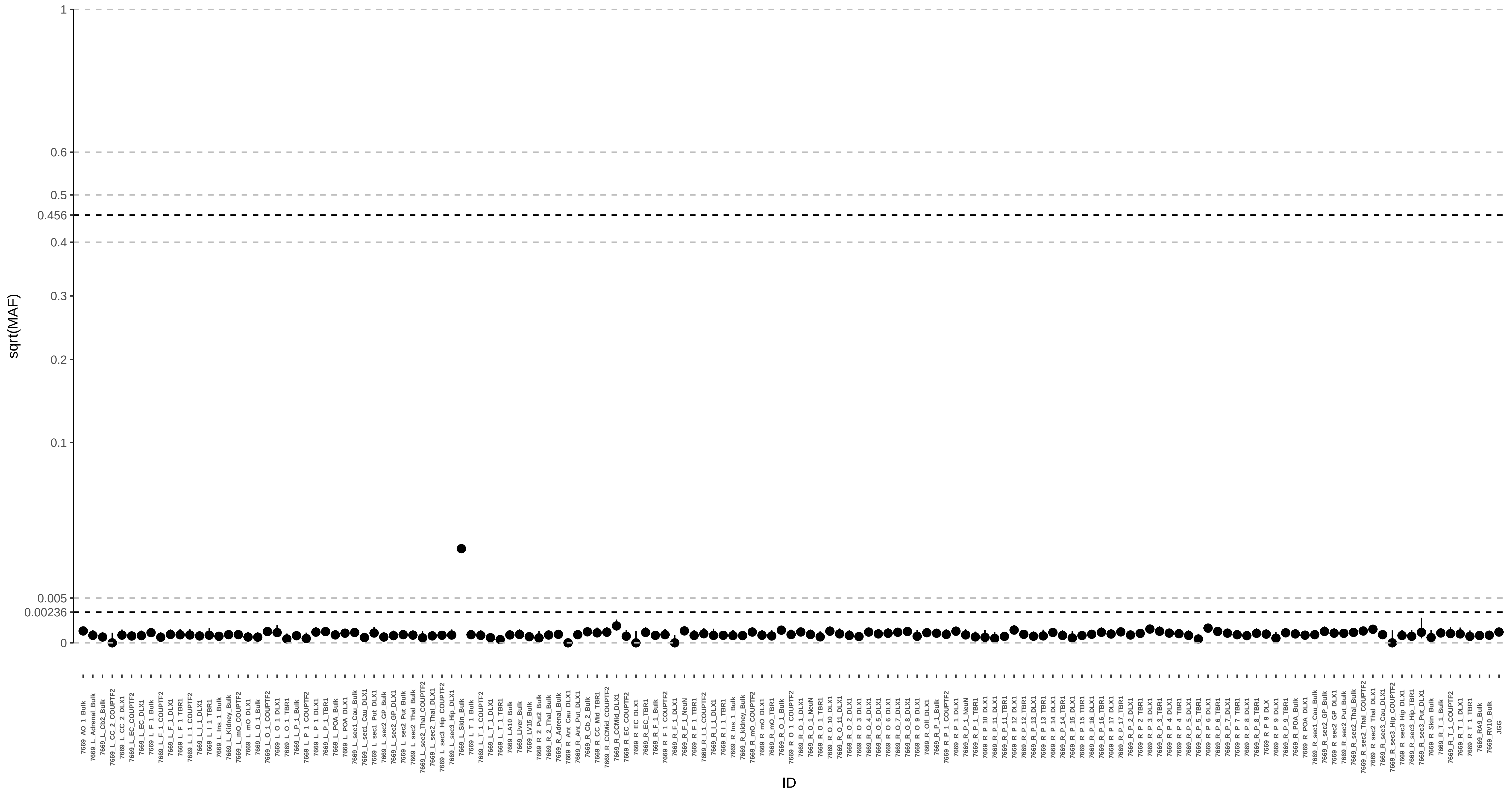

ID05 7−139192862−C−T

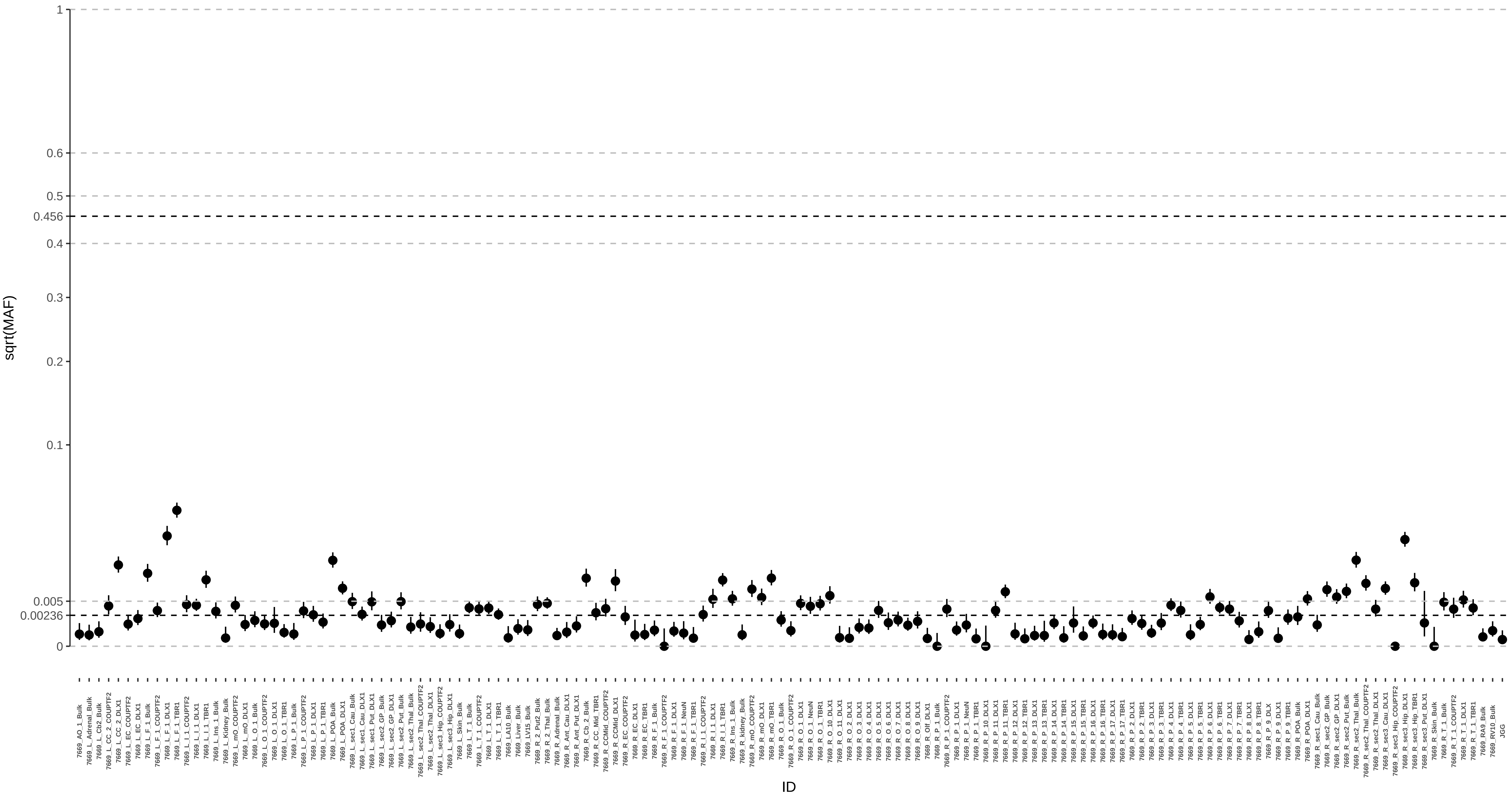

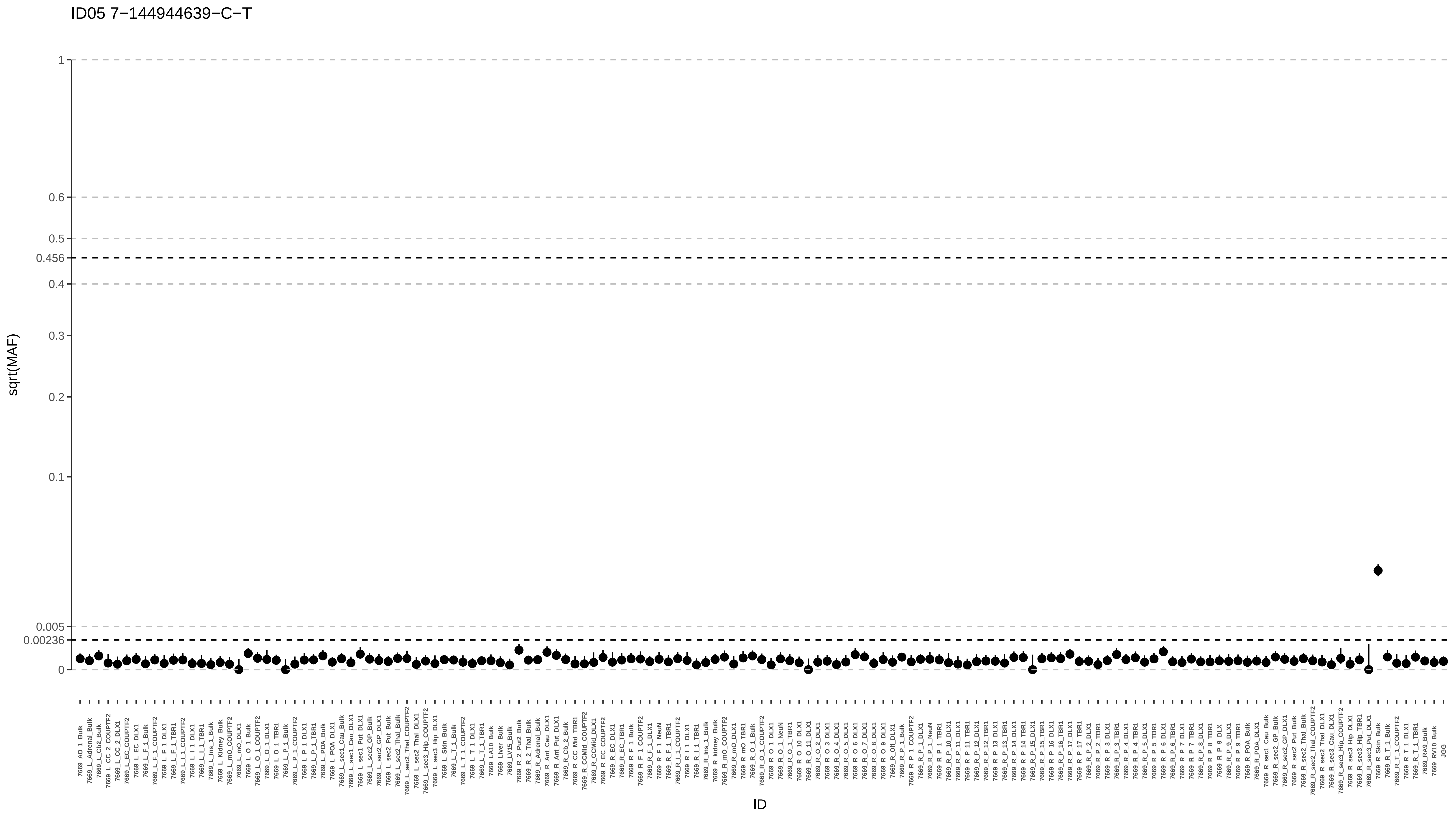
ID05 7−144944639−C−T
sqrt(MAF)
1
0.6
0.5
0.456
0.4
0.3
0.2
0.1
0.005
0.00236
0
ID

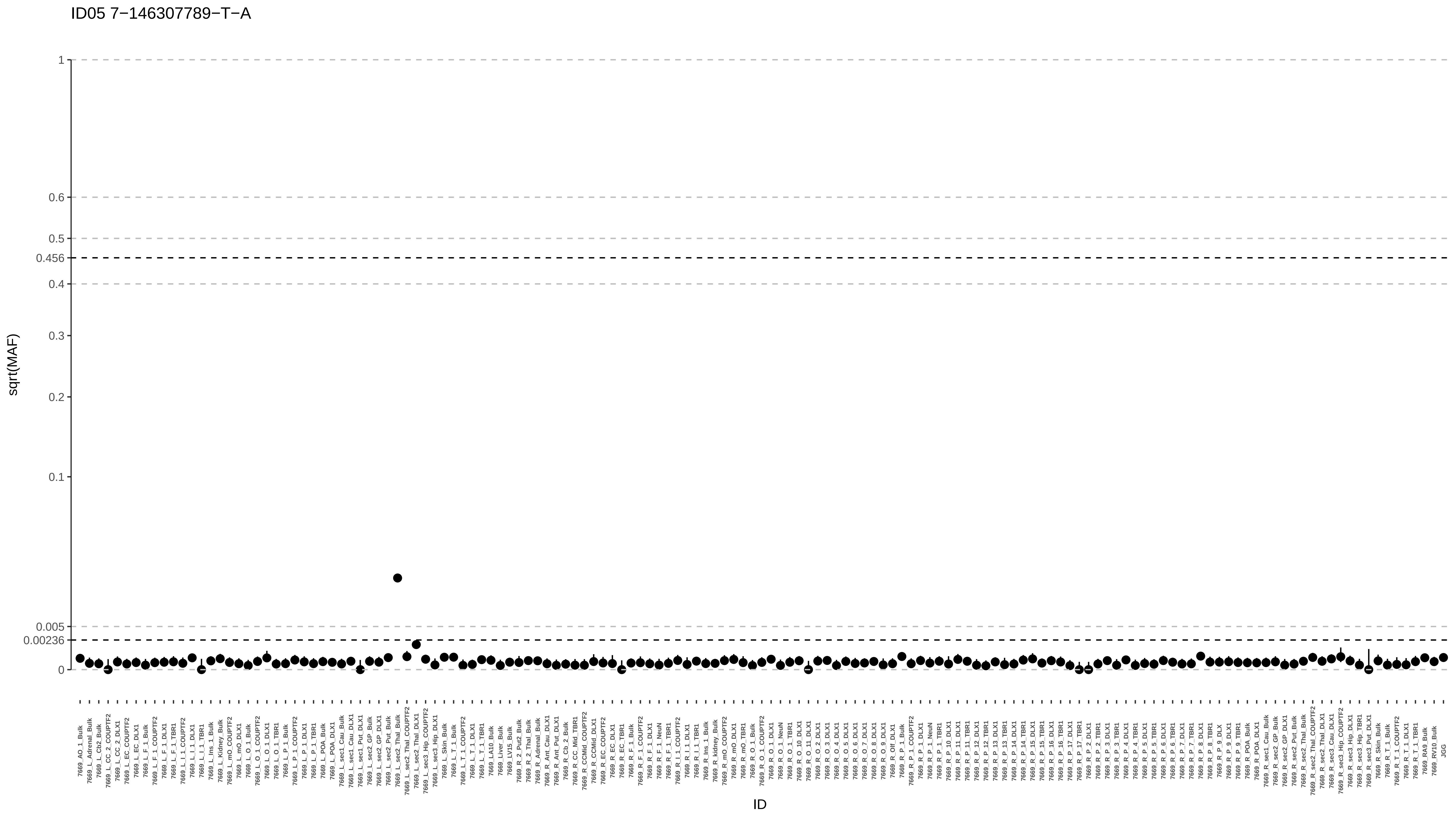
ID05 7−146307789−T−A
sqrt(MAF)
ID
1
0.6
0.5
0.456
0.4
0.3
0.2
0.1
0.005
0.00236
0

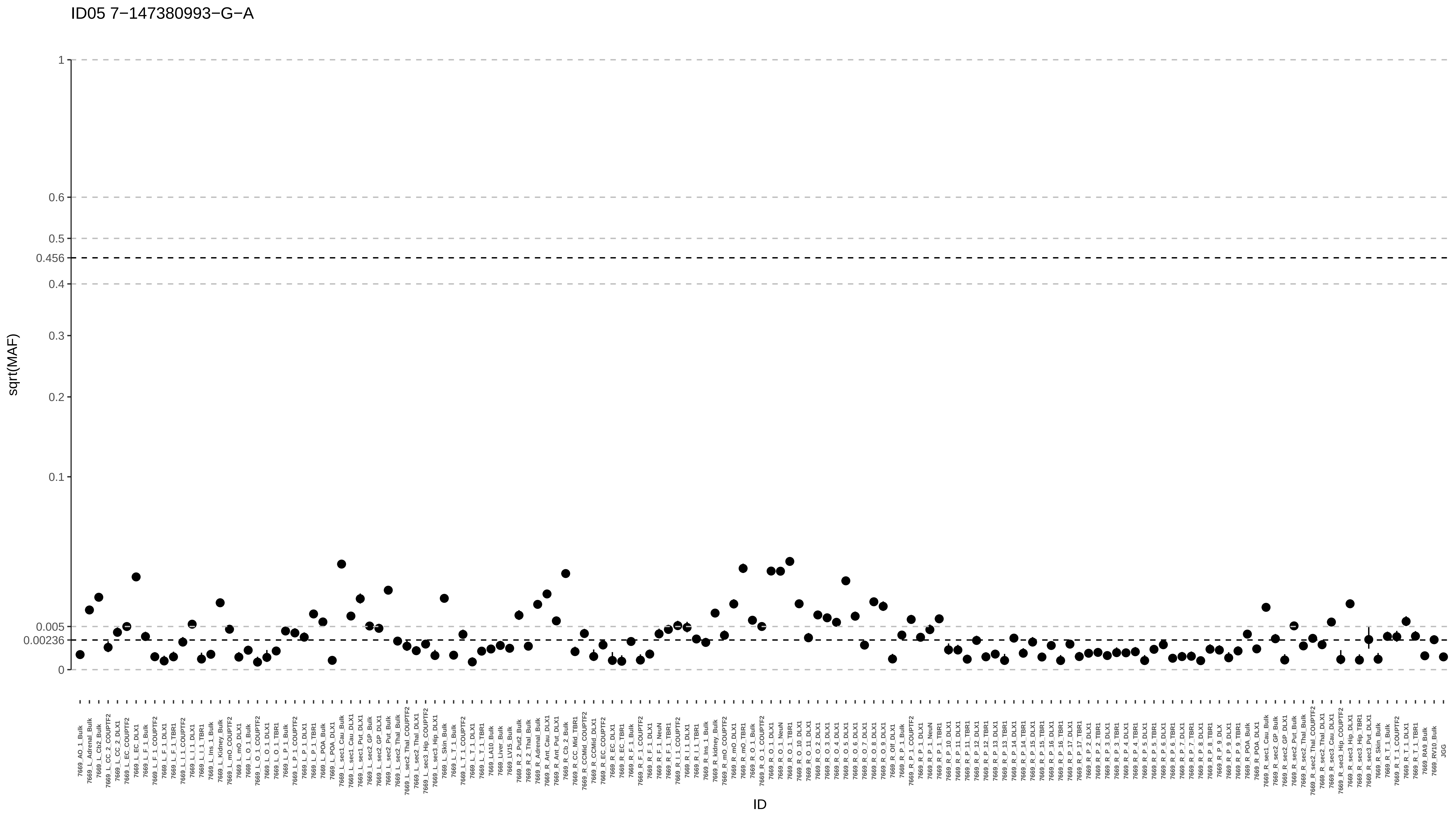

ID05 7−147380993−G−A
sqrt(MAF)
1
0.6
0.5
0.456
0.4
0.3
0.2
0.1
0.005
0.00236
0
ID

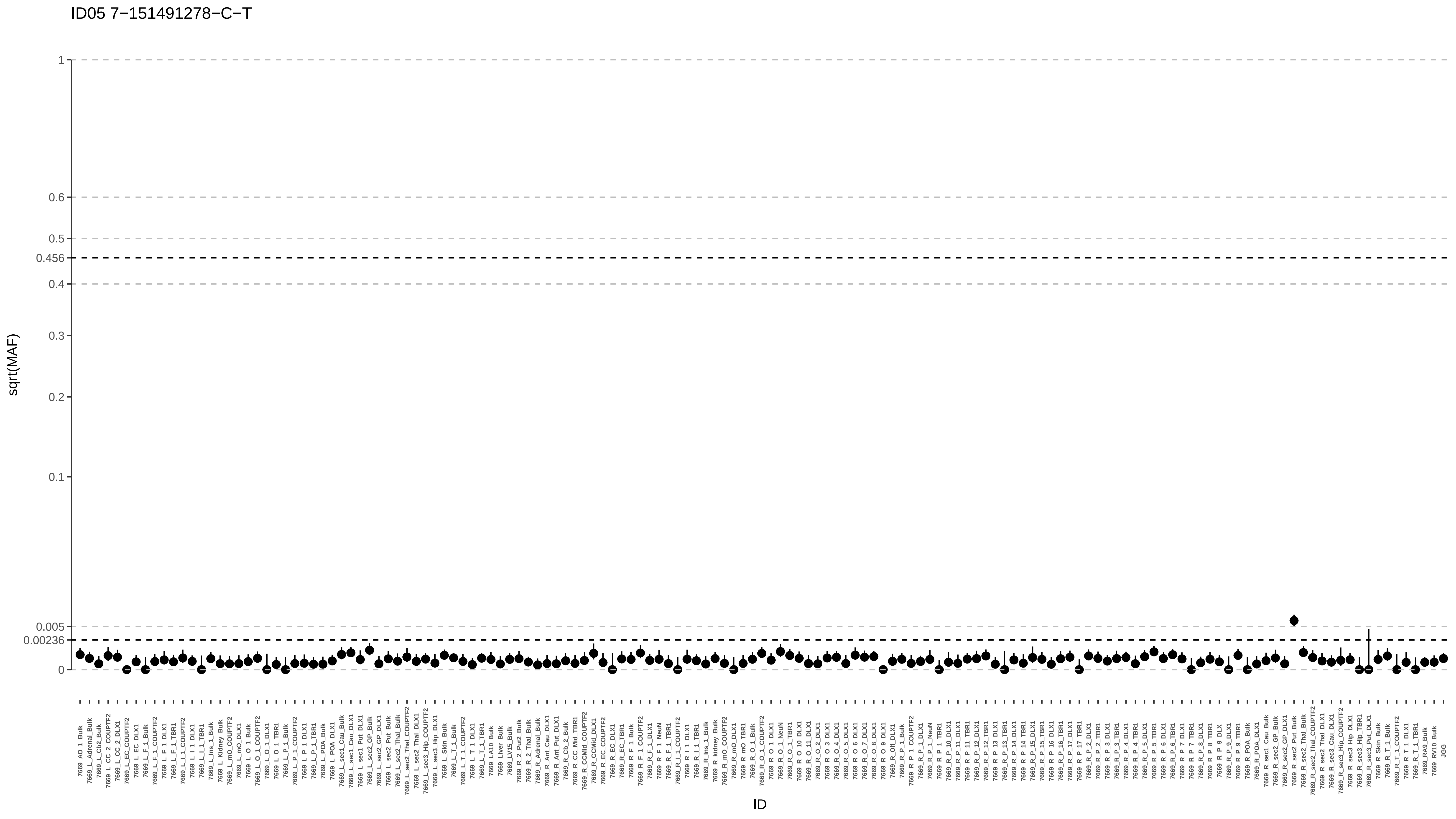
ID05 7−151491278−C−T
sqrt(MAF)
ID
1
0.6
0.5
0.456
0.4
0.3
0.2
0.1
0.005
0.00236
0

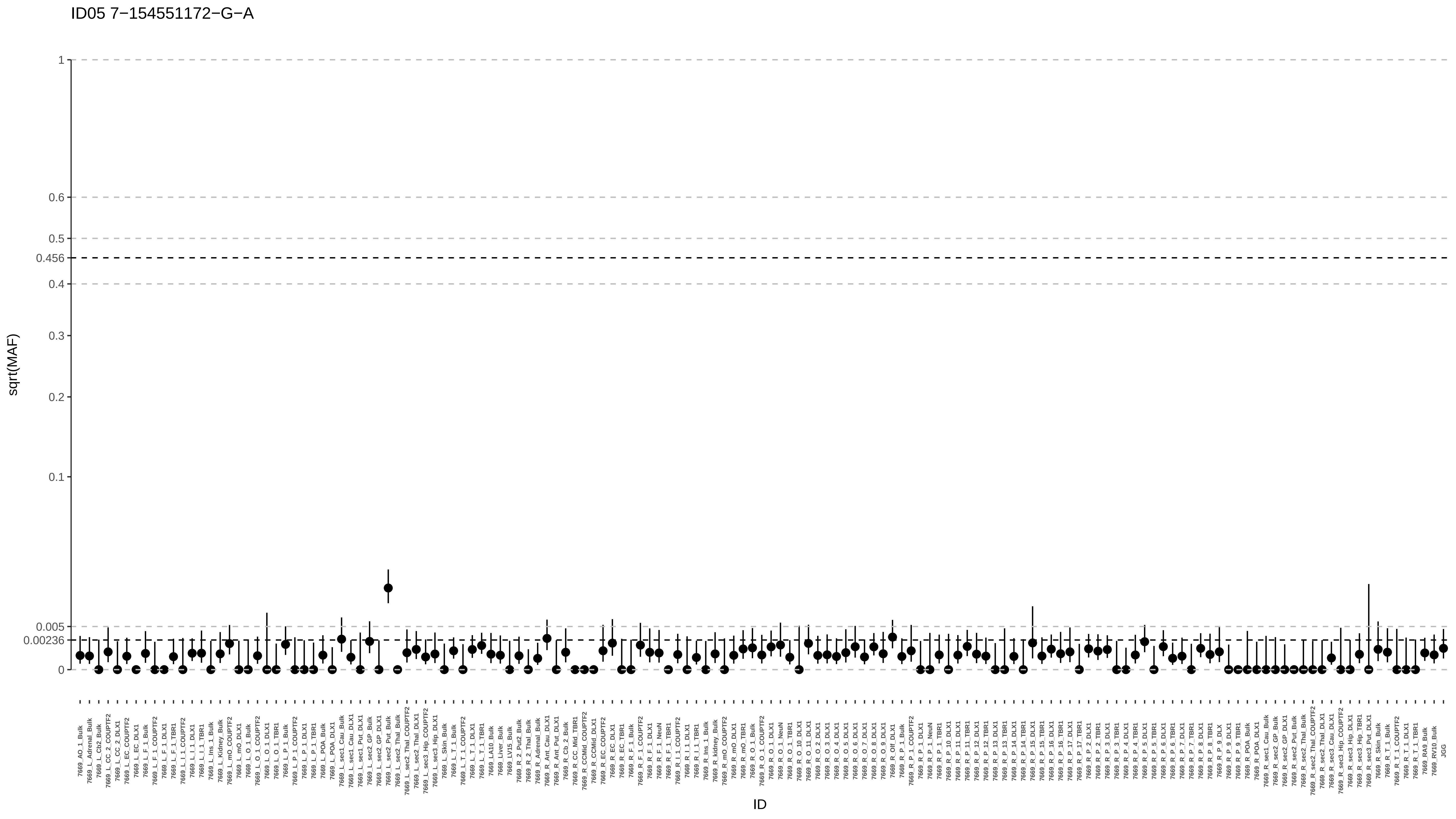

ID05 7−154551172−G−A
sqrt(MAF)
1
0.6
0.5
0.456
0.4
0.3
0.2
0.1
0.005
0.00236
0
ID

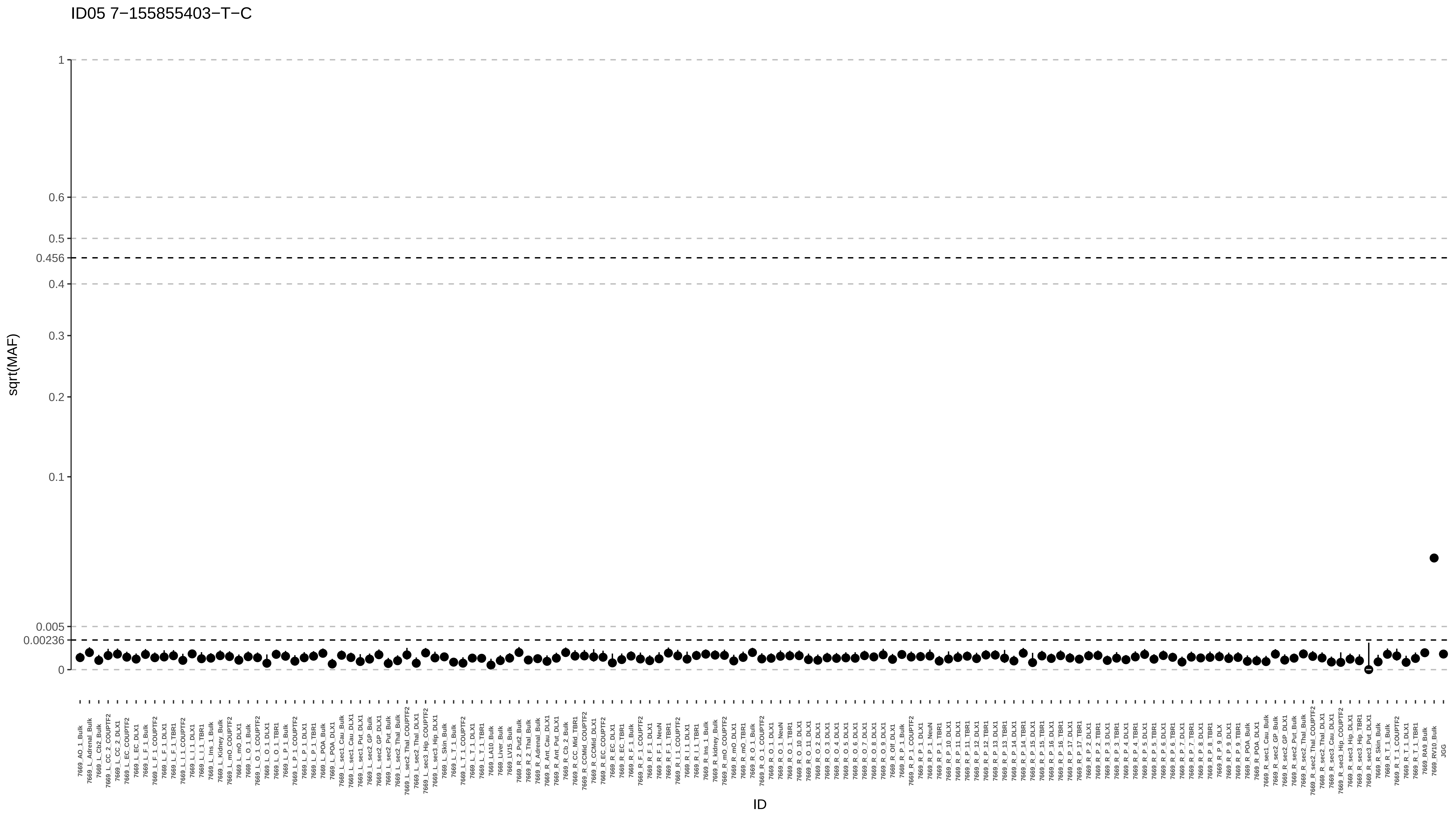
ID05 7−155855403−T−C
sqrt(MAF)
ID

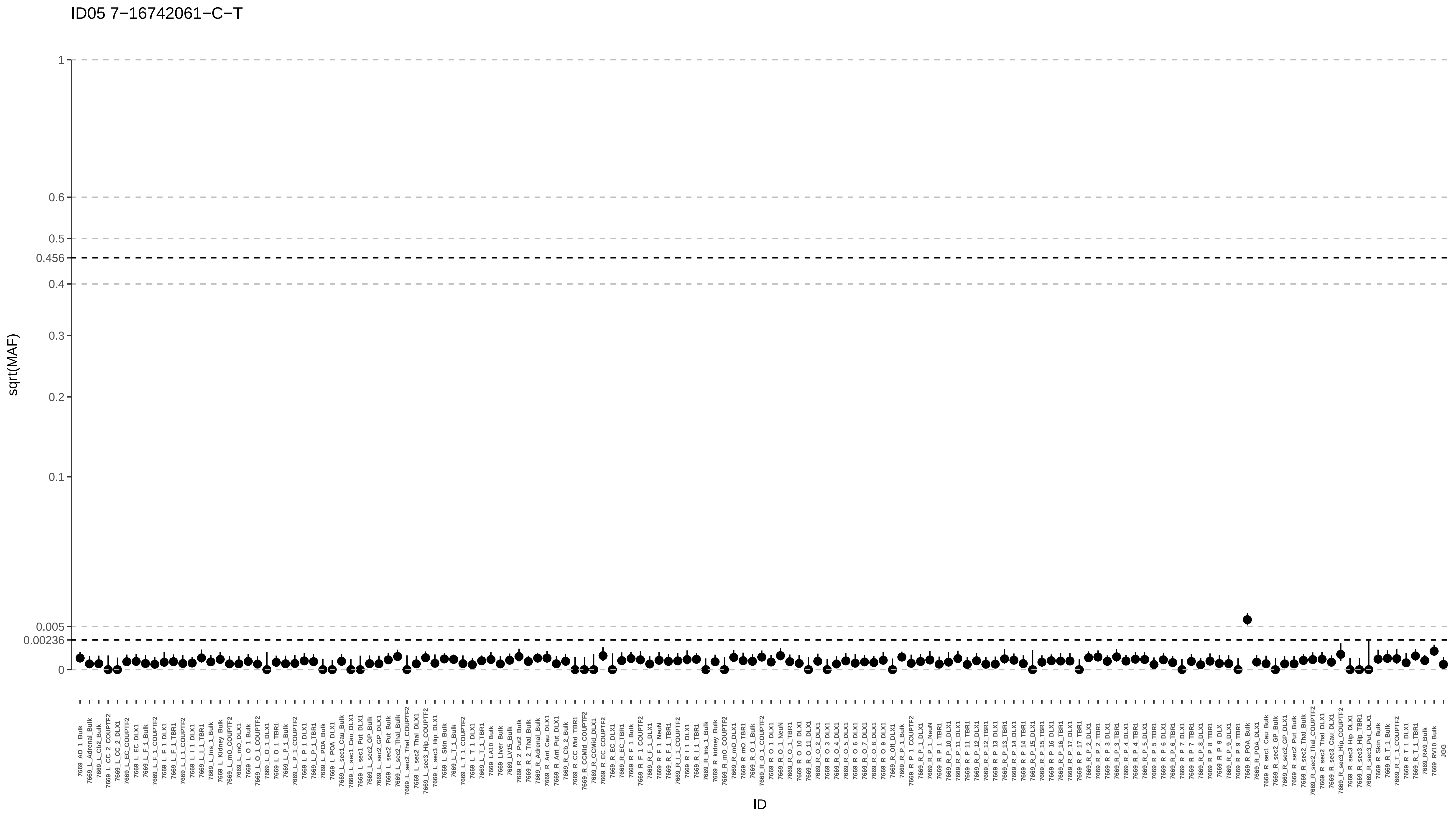

ID05 7−16742061−C−T
sqrt(MAF)
1
0.6
0.5
0.456
0.4
0.3
0.2
0.1
0.005
0.00236
0
ID

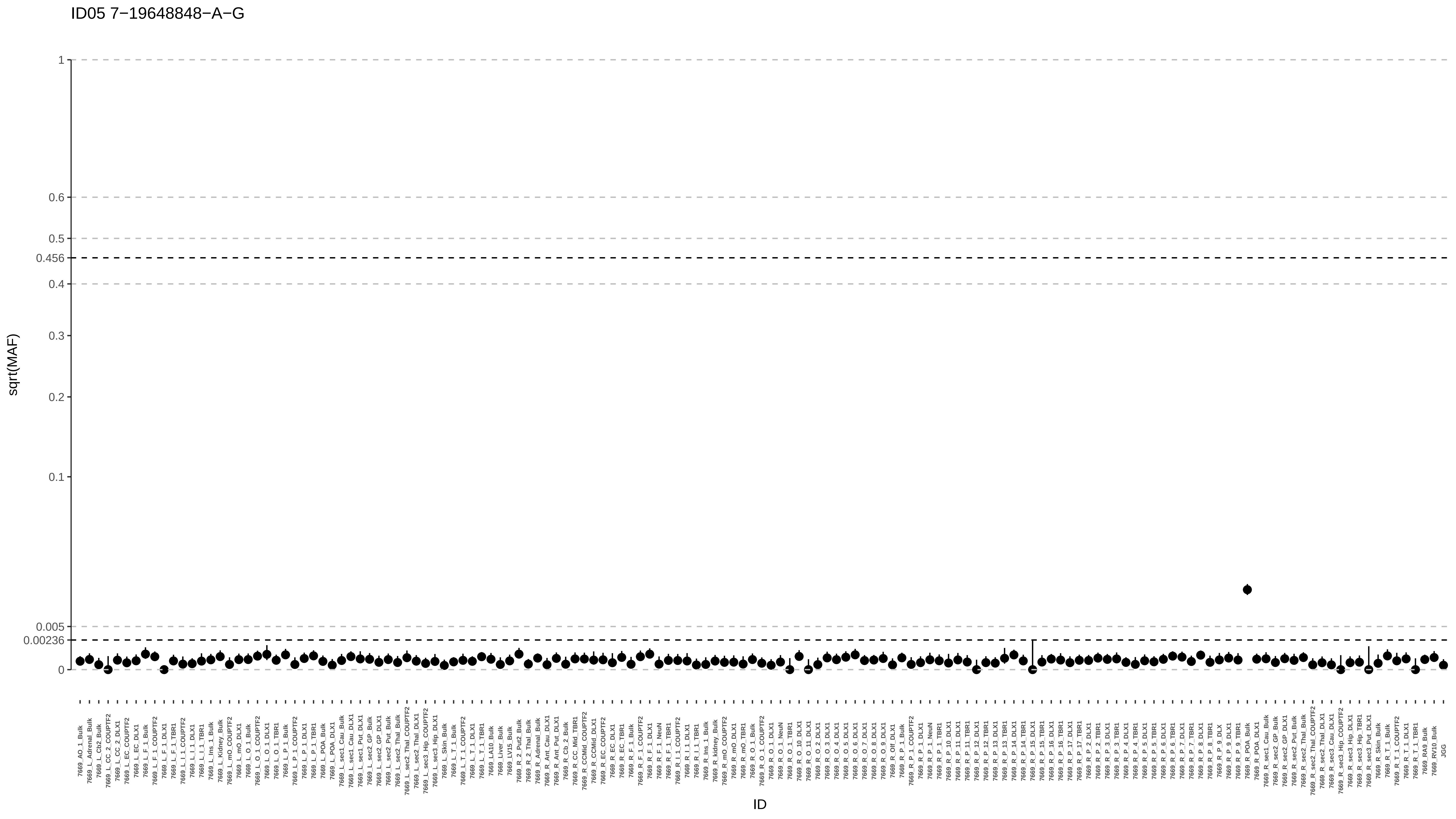
ID05 7−19648848−A−G
sqrt(MAF)
1
0.6
0.5
0.456
0.4
0.3
0.2
0.1
0.005
0.00236
0
ID

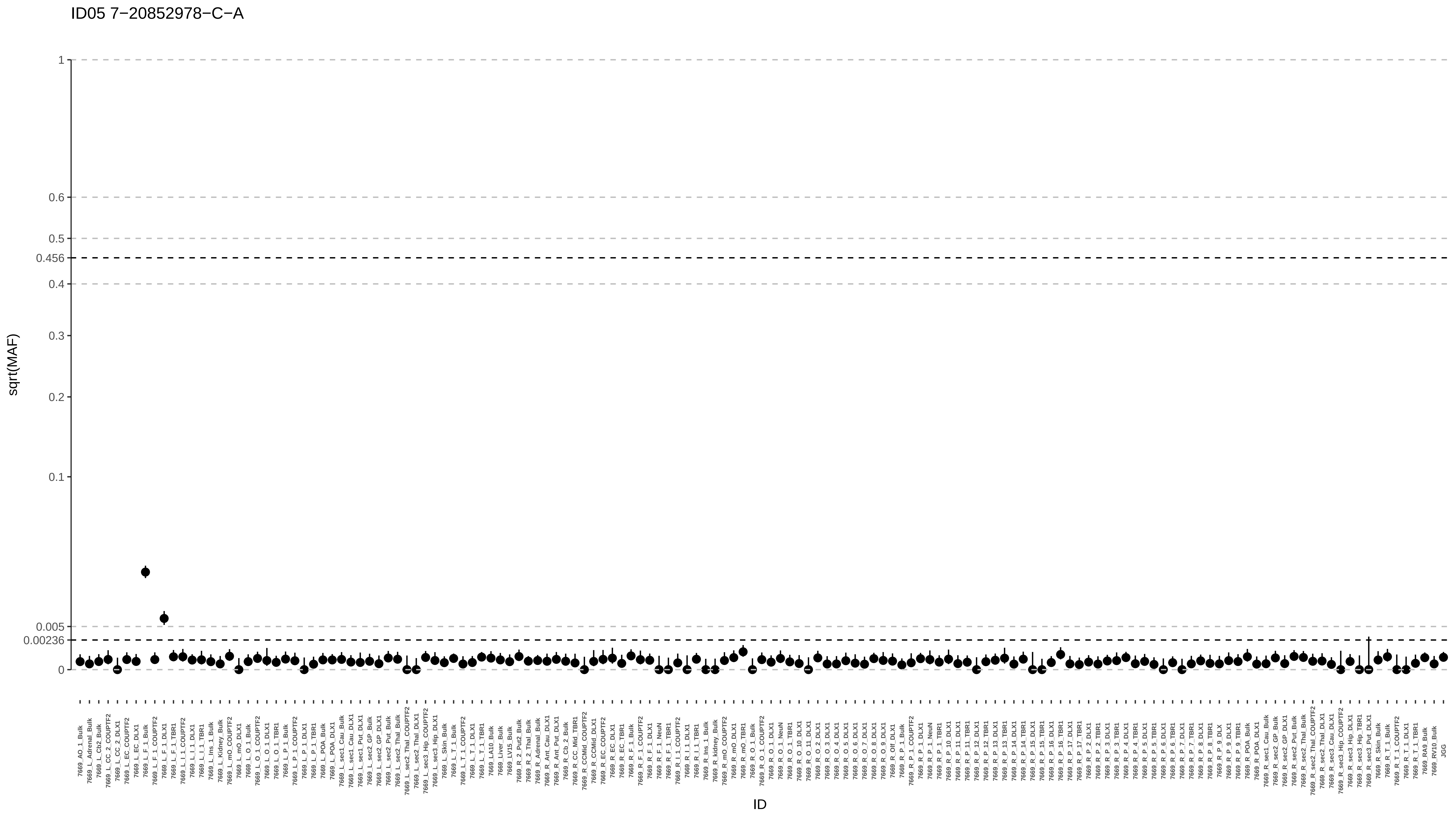

ID05 7−20852978−C−A
sqrt(MAF)
1
0.6
0.5
0.456
0.4
0.3
0.2
0.1
0.005
0.00236
0
ID

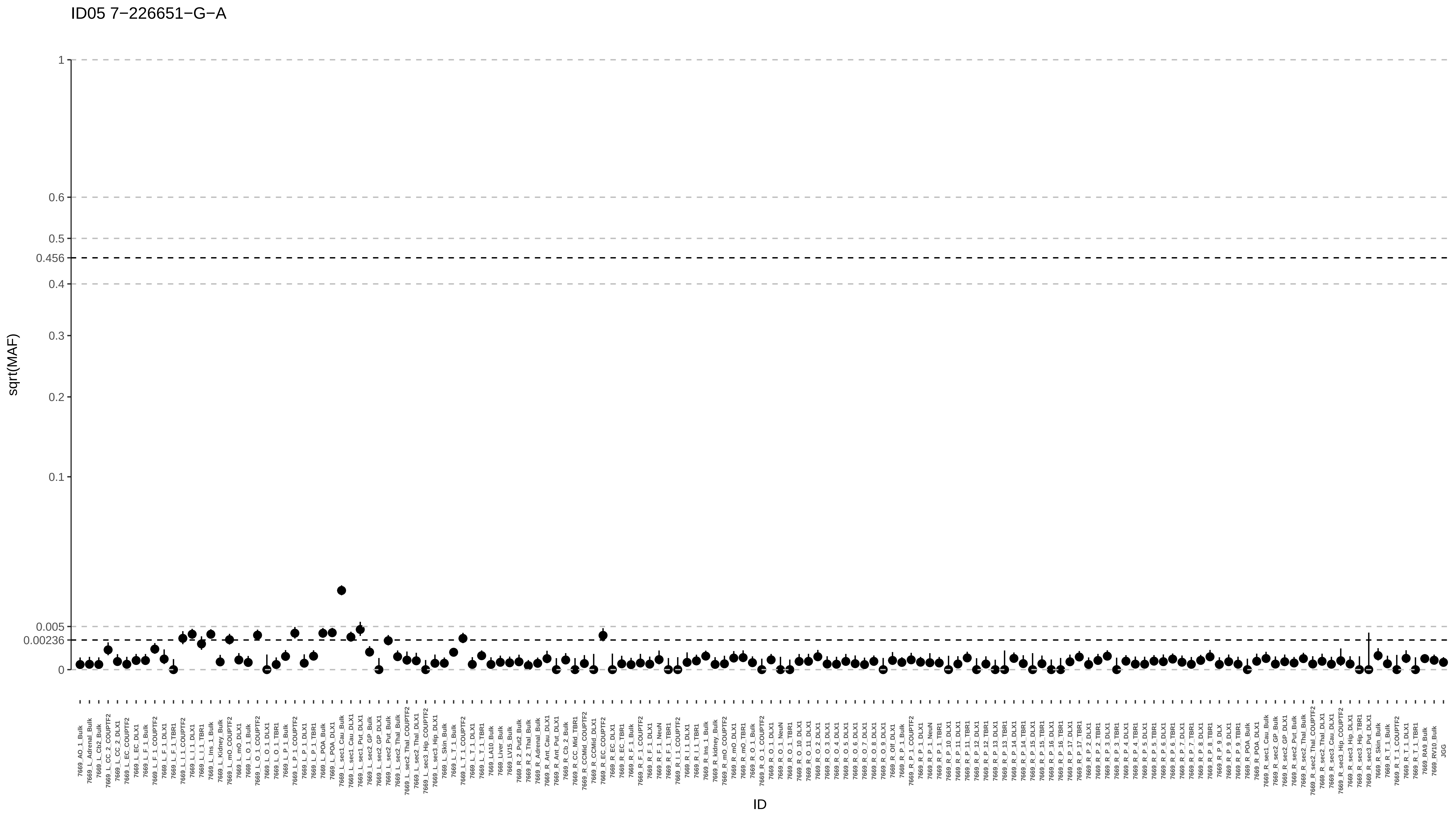
ID05 7−226651−G−A
sqrt(MAF)
1
0.6
0.5
0.456
0.4
0.3
0.2
0.1
0.005
0.00236
0
ID

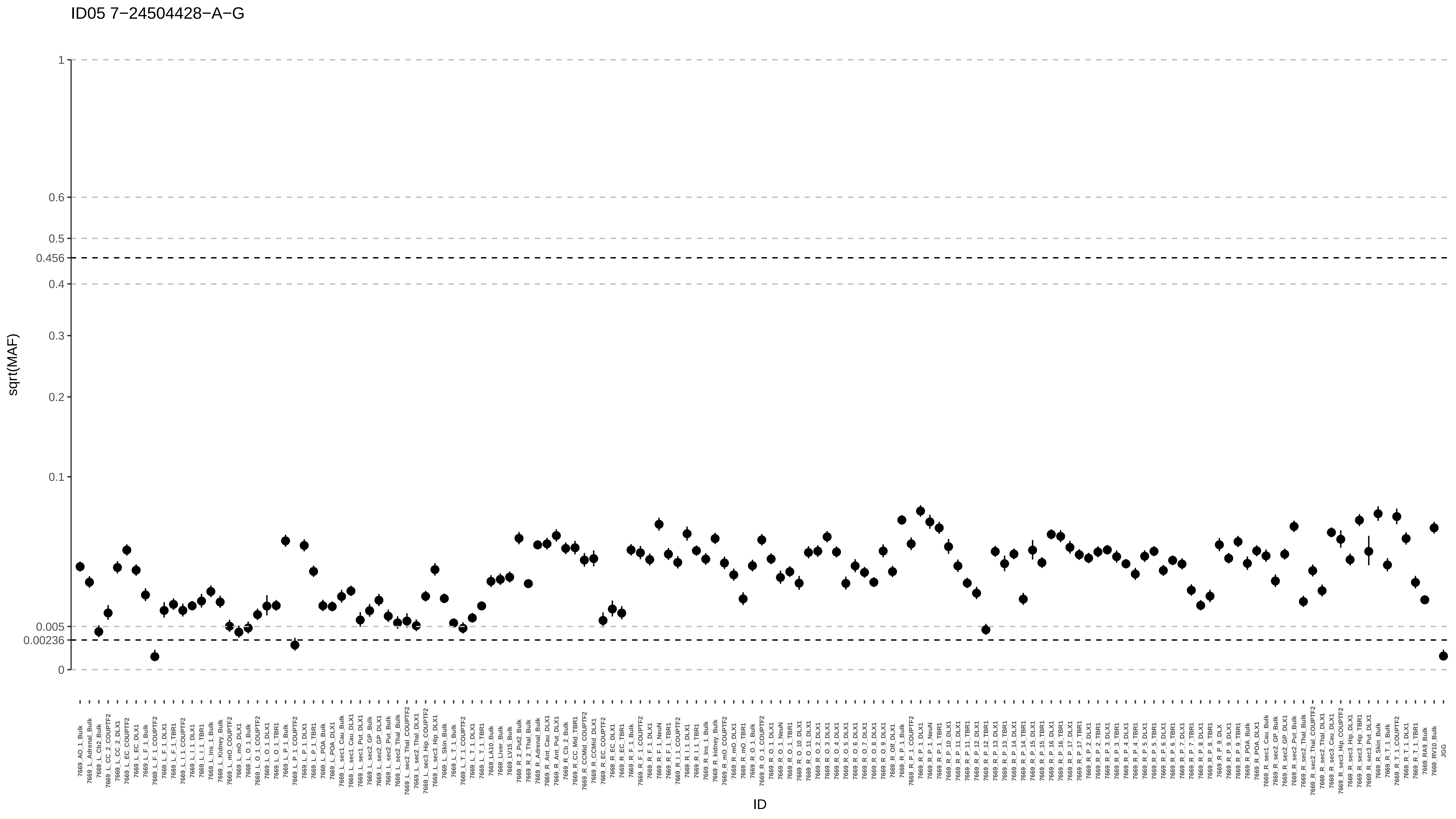
ID05 7−24504428−A−G
sqrt(MAF)
1
0.6
0.5
0.456
0.4
0.3
0.2
0.1
0.005
0.00236
0
ID

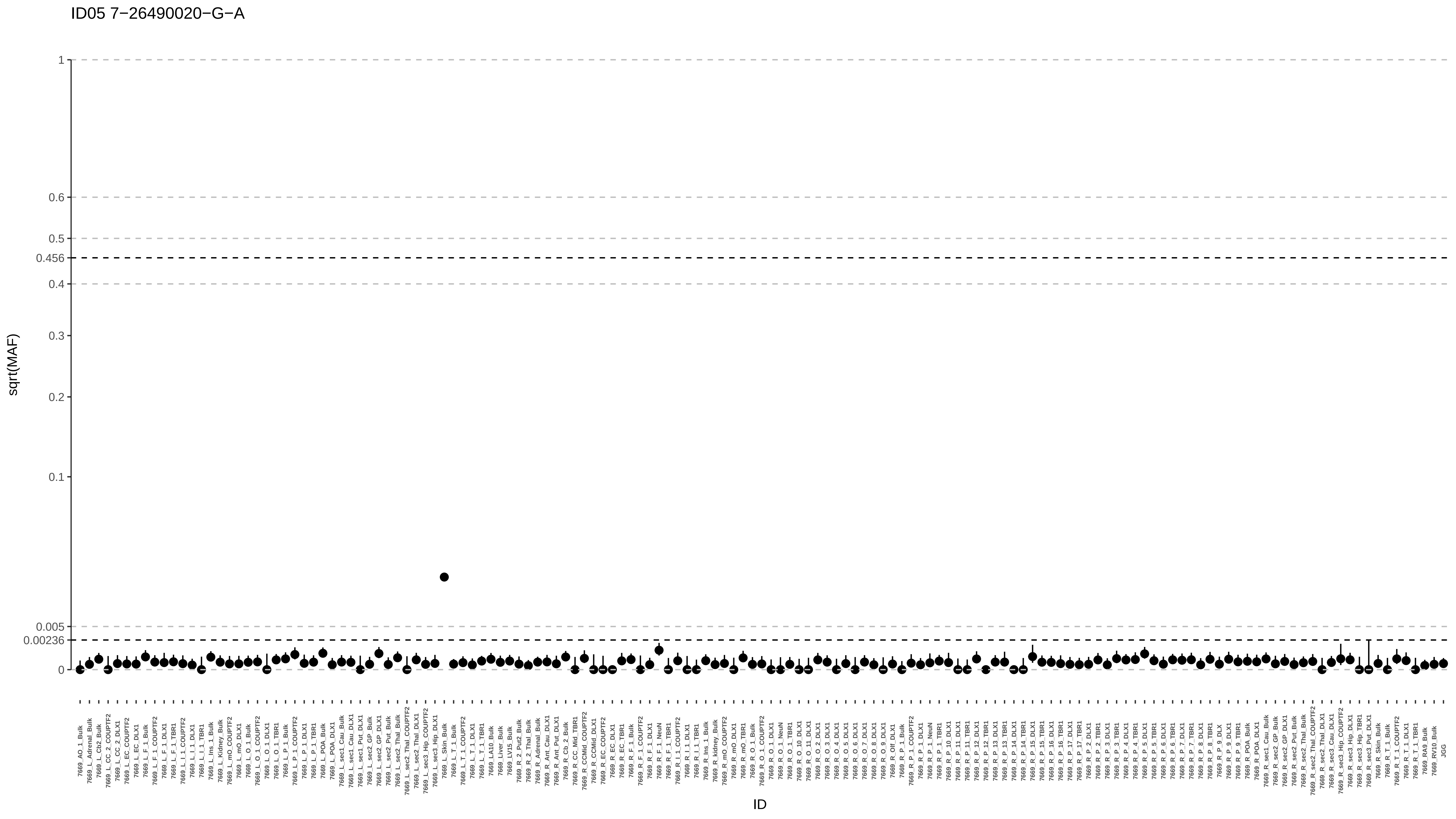

ID05 7−26490020−G−A
sqrt(MAF)
ID

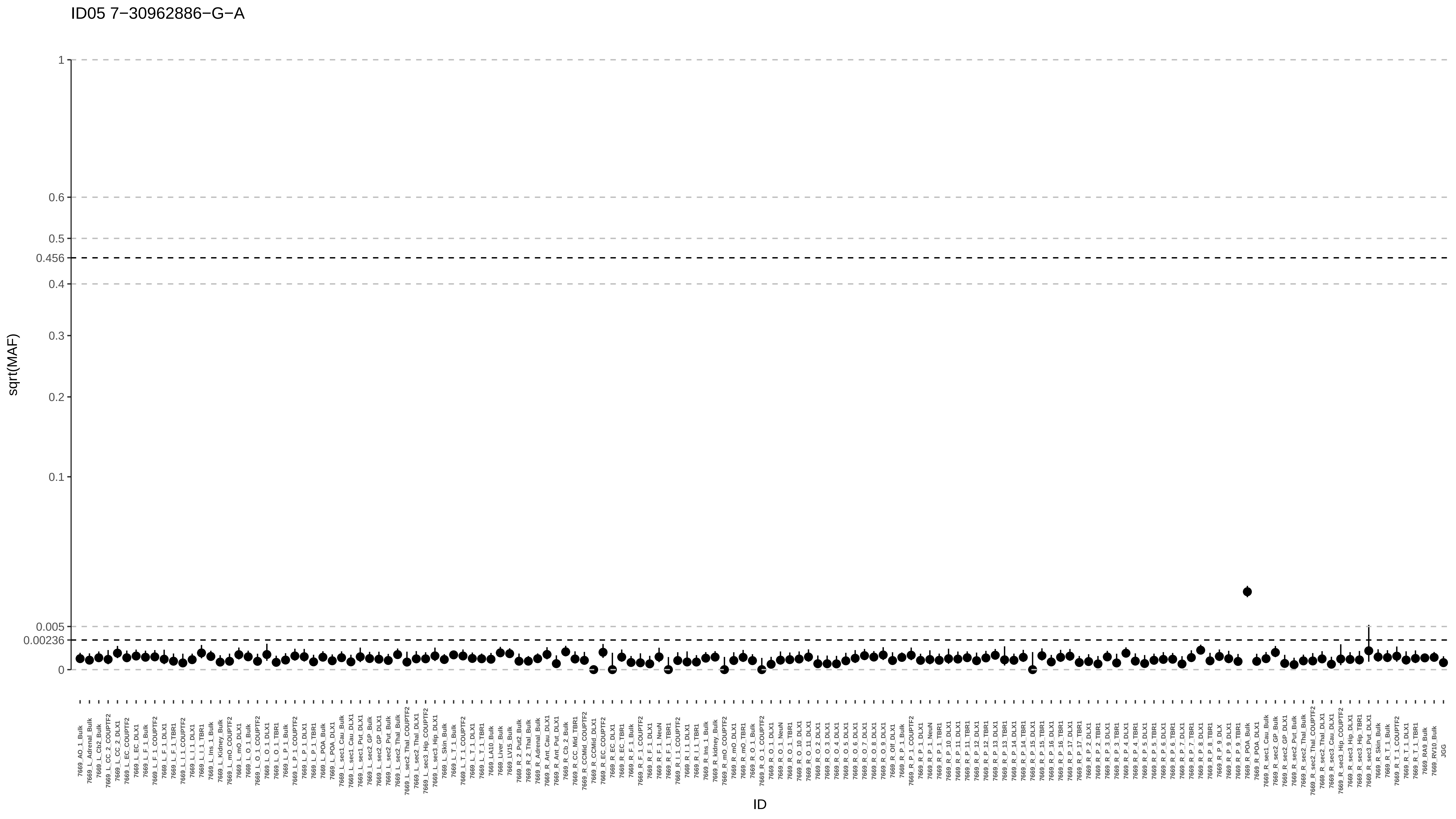
ID05 7−30962886−G−A
sqrt(MAF)
1
0.6
0.5
0.456
0.4
0.3
0.2
0.1
0.005
0.00236
0
ID
7669_AO_1_Bulk
7669_L_Adrenal_Bulk
7669_L_Cb2_Bulk
7669_L_CC_2_COUPTF2
7669_L_CC_2_DLX1
7669_L_EC_COUPTF2
7669_L_EC_DLX1
7669_L_F_1_Bulk
7669_L_F_1_COUPTF2
7669_L_F_1_DLX1
7669_L_F_1_TBR1
7669_L_I_1_COUPTF2
7669_L_I_1_DLX1
7669_L_I_1_TBR1
7669_L_Ins_1_Bulk
7669_L_Kidney_Bulk
7669_L_mO_COUPTF2
7669_L_mO_DLX1
7669_L_O_1_Bulk
7669_L_O_1_COUPTF2
7669_L_O_1_DLX1
7669_L_O_1_TBR1
7669_L_P_1_Bulk
7669_L_P_1_COUPTF2
7669_L_P_1_DLX1
7669_L_P_1_TBR1
7669_L_POA_Bulk
7669_L_POA_DLX1
7669_L_sec1_Cau_Bulk
7669_L_sec1_Cau_DLX1
7669_L_sec1_Put_DLX1
7669_L_sec2_GP_Bulk
7669_L_sec2_GP_DLX1
7669_L_sec2_Put_Bulk
7669_L_sec2_Thal_Bulk
7669_L_sec2_Thal_COUPTF2
7669_L_sec2_Thal_DLX1
7669_L_sec3_Hip_COUPTF2
7669_L_sec3_Hip_DLX1
7669_L_Skin_Bulk
7669_L_T_1_Bulk
7669_L_T_1_COUPTF2
7669_L_T_1_DLX1
7669_L_T_1_TBR1
7669_LA10_Bulk
7669_Liver_Bulk
7669_LV15_Bulk
7669_R_2_Put2_Bulk
7669_R_2_Thal_Bulk
7669_R_Adrenal_Bulk
7669_R_Ant_Cau_DLX1
7669_R_Ant_Put_DLX1
7669_R_Cb_2_Bulk
7669_R_CC_Mid_TBR1
7669_R_CCMid_COUPTF2
7669_R_CCMid_DLX1
7669_R_EC_COUPTF2
7669_R_EC_DLX1
7669_R_EC_TBR1
7669_R_F_1_Bulk
7669_R_F_1_COUPTF2
7669_R_F_1_DLX1
7669_R_F_1_NeuN
7669_R_F_1_TBR1
7669_R_I_1_COUPTF2
7669_R_I_1_DLX1
7669_R_I_1_TBR1
7669_R_Ins_1_Bulk
7669_R_kidney_Bulk
7669_R_mO_COUPTF2
7669_R_mO_DLX1
7669_R_mO_TBR1
7669_R_O_1_Bulk
7669_R_O_1_COUPTF2
7669_R_O_1_DLX1
7669_R_O_1_NeuN
7669_R_O_1_TBR1
7669_R_O_10_DLX1
7669_R_O_11_DLX1
7669_R_O_2_DLX1
7669_R_O_3_DLX1
7669_R_O_4_DLX1
7669_R_O_5_DLX1
7669_R_O_6_DLX1
7669_R_O_7_DLX1
7669_R_O_8_DLX1
7669_R_O_9_DLX1
7669_R_Olf_DLX1
7669_R_P_1_Bulk
7669_R_P_1_COUPTF2
7669_R_P_1_DLX1
7669_R_P_1_NeuN
7669_R_P_1_TBR1
7669_R_P_10_DLX1
7669_R_P_11_DLX1
7669_R_P_11_TBR1
7669_R_P_12_DLX1
7669_R_P_12_TBR1
7669_R_P_13_DLX1
7669_R_P_13_TBR1
7669_R_P_14_DLX1
7669_R_P_14_TBR1
7669_R_P_15_DLX1
7669_R_P_15_TBR1
7669_R_P_16_DLX1
7669_R_P_16_TBR1
7669_R_P_17_DLX1
7669_R_P_17_TBR1
7669_R_P_2_DLX1
7669_R_P_2_TBR1
7669_R_P_3_DLX1
7669_R_P_3_TBR1
7669_R_P_4_DLX1
7669_R_P_4_TBR1
7669_R_P_5_DLX1
7669_R_P_5_TBR1
7669_R_P_6_DLX1
7669_R_P_6_TBR1
7669_R_P_7_DLX1
7669_R_P_7_TBR1
7669_R_P_8_DLX1
7669_R_P_8_TBR1
7669_R_P_9_DLX
7669_R_P_9_DLX1
7669_R_P_9_TBR1
7669_R_POA_Bulk
7669_R_POA_DLX1
7669_R_sec1_Cau_Bulk
7669_R_sec2_GP_Bulk
7669_R_sec2_GP_DLX1
7669_R_sec2_Put_Bulk
7669_R_sec2_Thal_Bulk
7669_R_sec2_Thal_COUPTF2
7669_R_sec2_Thal_DLX1
7669_R_sec3_Cau_DLX1
7669_R_sec3_Hip_COUPTF2
7669_R_sec3_Hip_DLX1
7669_R_sec3_Hip_TBR1
7669_R_sec3_Put_DLX1
7669_R_Skin_Bulk
7669_R_T_1_Bulk
7669_R_T_1_COUPTF2
7669_R_T_1_DLX1
7669_R_T_1_TBR1
7669_RA9_Bulk
7669_RV10_Bulk
JGG

ID05 7−31462228−G−A

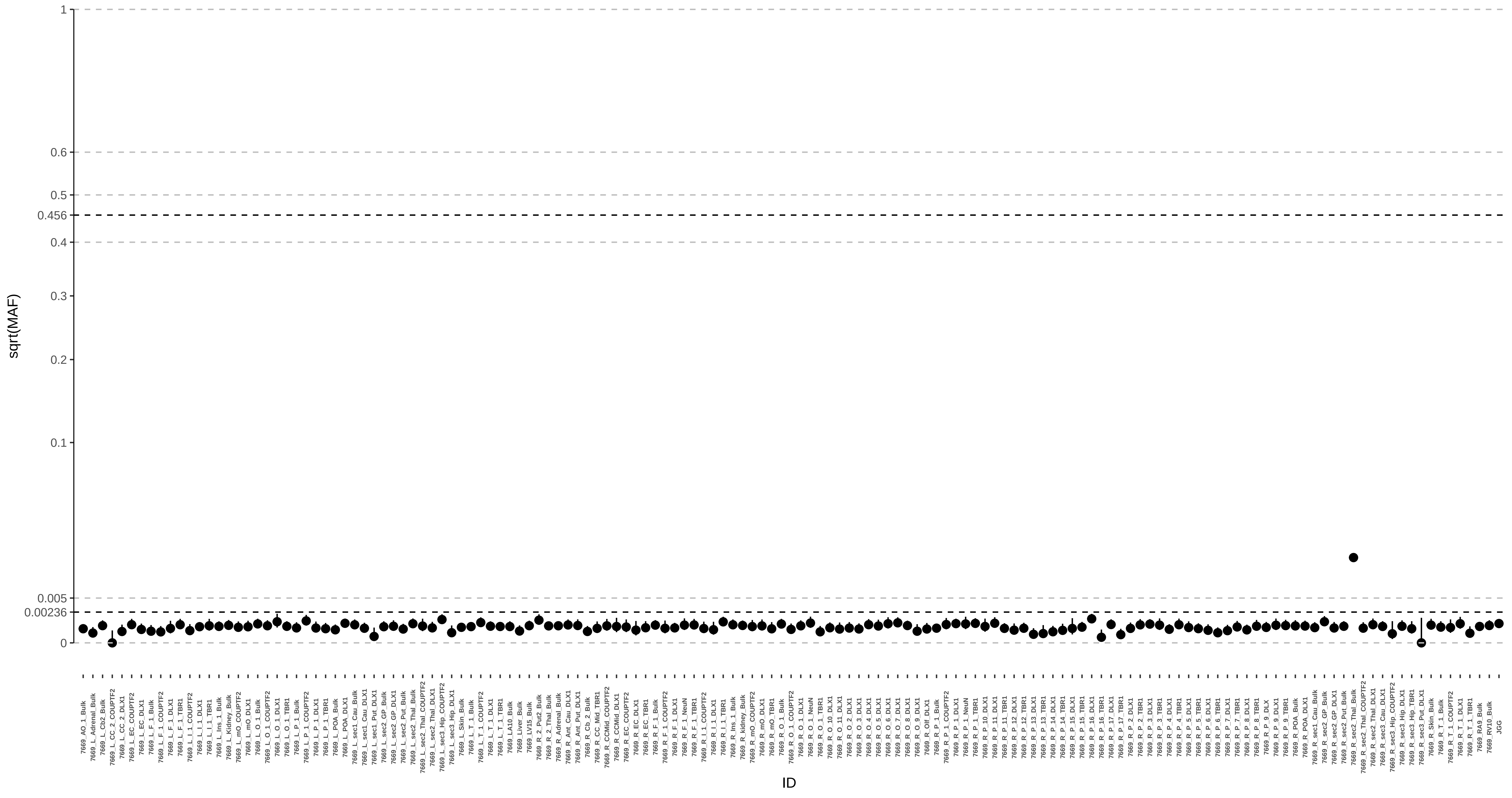

ID05 7−35583026−G−C

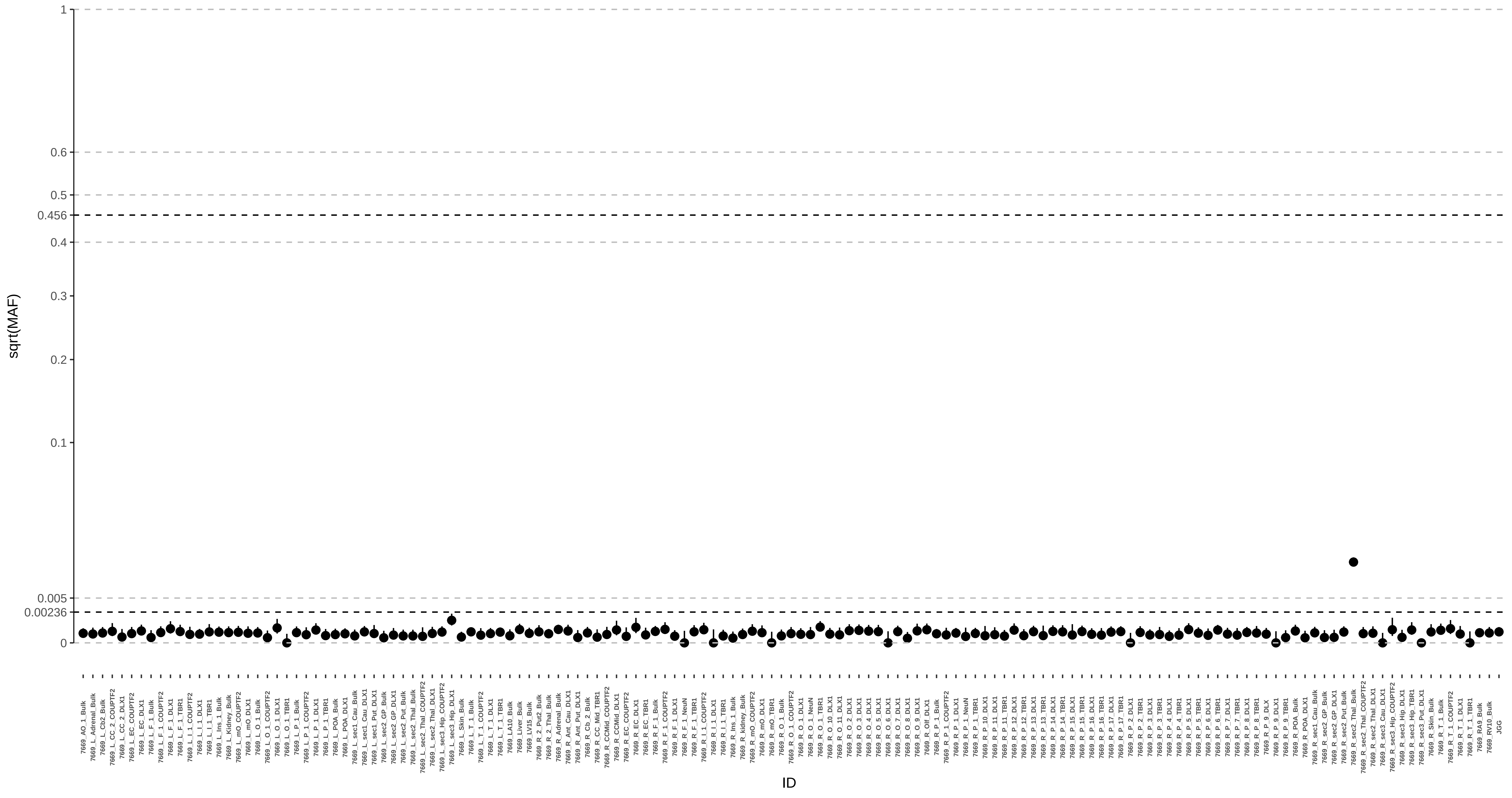

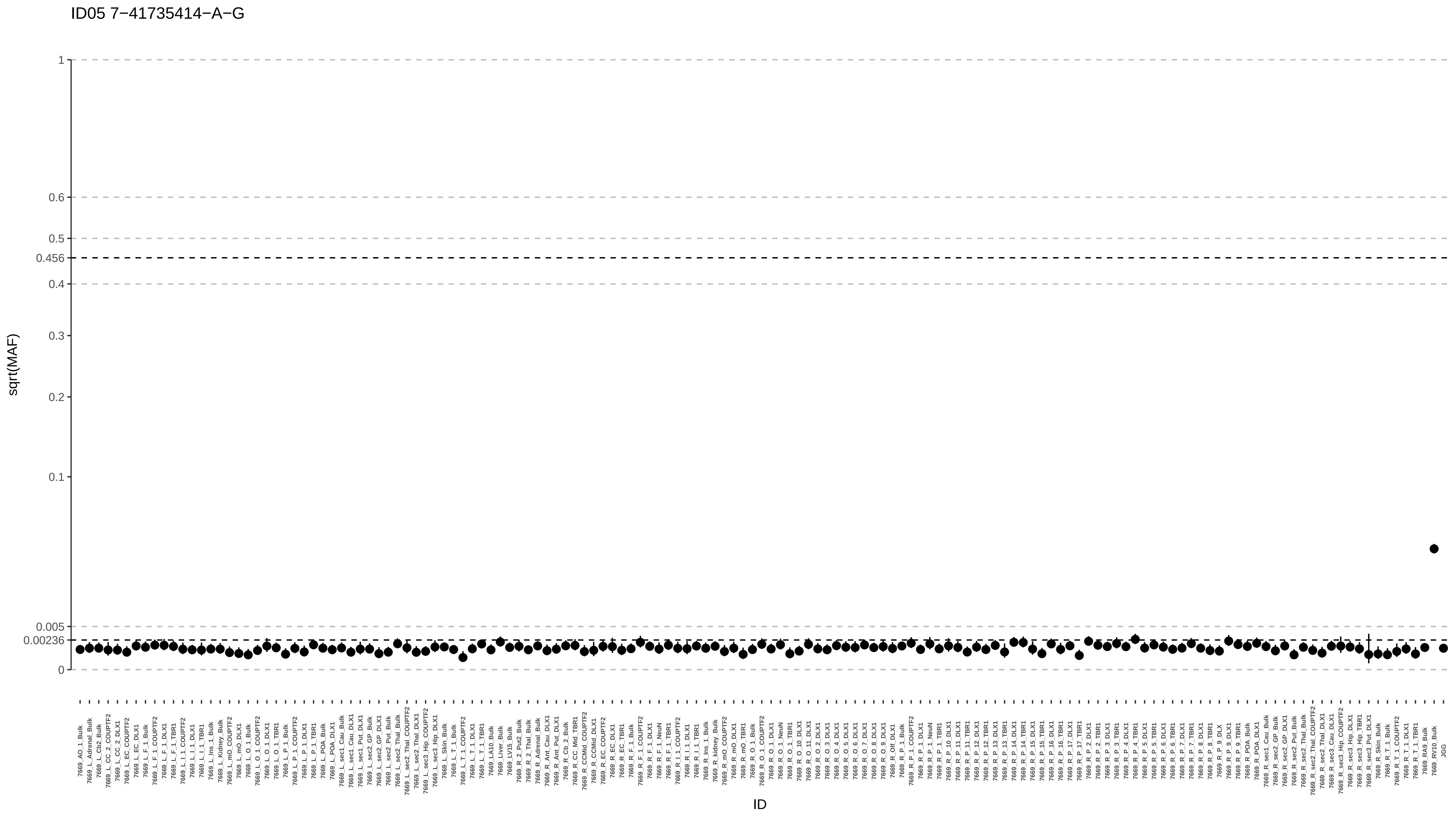
ID05 7−41735414−A−G
sqrt(MAF)
1
0.6
0.5
0.456
0.4
0.3
0.2
0.1
0.005
0.00236
0
ID
7669_AO_1_Bulk
7669_L_Adrenal_Bulk
7669_L_Cb2_Bulk
7669_L_CC_2_COUPTF2
7669_L_CC_2_DLX1
7669_L_EC_COUPTF2
7669_L_EC_DLX1
7669_L_F_1_Bulk
7669_L_F_1_COUPTF2
7669_L_F_1_DLX1
7669_L_F_1_TBR1
7669_L_I_1_COUPTF2
7669_L_I_1_DLX1
7669_L_I_1_TBR1
7669_L_Ins_1_Bulk
7669_L_Kidney_Bulk
7669_L_mO_COUPTF2
7669_L_mO_DLX1
7669_L_O_1_Bulk
7669_L_O_1_COUPTF2
7669_L_O_1_DLX1
7669_L_O_1_TBR1
7669_L_P_1_Bulk
7669_L_P_1_COUPTF2
7669_L_P_1_DLX1
7669_L_P_1_TBR1
7669_L_POA_Bulk
7669_L_POA_DLX1
7669_L_sec1_Cau_Bulk
7669_L_sec1_Cau_DLX1
7669_L_sec1_Put_DLX1
7669_L_sec2_GP_Bulk
7669_L_sec2_GP_DLX1
7669_L_sec2_Put_Bulk
7669_L_sec2_Thal_Bulk
7669_L_sec2_Thal_COUPTF2
7669_L_sec2_Thal_DLX1
7669_L_sec3_Hip_COUPTF2
7669_L_sec3_Hip_DLX1
7669_L_Skin_Bulk
7669_L_T_1_Bulk
7669_L_T_1_COUPTF2
7669_L_T_1_DLX1
7669_L_T_1_TBR1
7669_LA10_Bulk
7669_Liver_Bulk
7669_LV15_Bulk
7669_R_2_Put2_Bulk
7669_R_2_Thal_Bulk
7669_R_Adrenal_Bulk
7669_R_Ant_Cau_DLX1
7669_R_Ant_Put_DLX1
7669_R_Cb_2_Bulk
7669_R_CC_Mid_TBR1
7669_R_CCMid_COUPTF2
7669_R_CCMid_DLX1
7669_R_EC_COUPTF2
7669_R_EC_DLX1
7669_R_EC_TBR1
7669_R_F_1_Bulk
7669_R_F_1_COUPTF2
7669_R_F_1_DLX1
7669_R_F_1_NeuN
7669_R_F_1_TBR1
7669_R_I_1_COUPTF2
7669_R_I_1_DLX1
7669_R_I_1_TBR1
7669_R_Ins_1_Bulk
7669_R_kidney_Bulk
7669_R_mO_COUPTF2
7669_R_mO_DLX1
7669_R_mO_TBR1
7669_R_O_1_Bulk
7669_R_O_1_COUPTF2
7669_R_O_1_DLX1
7669_R_O_1_NeuN
7669_R_O_1_TBR1
7669_R_O_10_DLX1
7669_R_O_11_DLX1
7669_R_O_2_DLX1
7669_R_O_3_DLX1
7669_R_O_4_DLX1
7669_R_O_5_DLX1
7669_R_O_6_DLX1
7669_R_O_7_DLX1
7669_R_O_8_DLX1
7669_R_O_9_DLX1
7669_R_Olf_DLX1
7669_R_P_1_Bulk
7669_R_P_1_COUPTF2
7669_R_P_1_DLX1
7669_R_P_1_NeuN
7669_R_P_1_TBR1
7669_R_P_10_DLX1
7669_R_P_11_DLX1
7669_R_P_11_TBR1
7669_R_P_12_DLX1
7669_R_P_12_TBR1
7669_R_P_13_DLX1
7669_R_P_13_TBR1
7669_R_P_14_DLX1
7669_R_P_14_TBR1
7669_R_P_15_DLX1
7669_R_P_15_TBR1
7669_R_P_16_DLX1
7669_R_P_16_TBR1
7669_R_P_17_DLX1
7669_R_P_17_TBR1
7669_R_P_2_DLX1
7669_R_P_2_TBR1
7669_R_P_3_DLX1
7669_R_P_3_TBR1
7669_R_P_4_DLX1
7669_R_P_4_TBR1
7669_R_P_5_DLX1
7669_R_P_5_TBR1
7669_R_P_6_DLX1
7669_R_P_6_TBR1
7669_R_P_7_DLX1
7669_R_P_7_TBR1
7669_R_P_8_DLX1
7669_R_P_8_TBR1
7669_R_P_9_DLX
7669_R_P_9_DLX1
7669_R_P_9_TBR1
7669_R_POA_Bulk
7669_R_POA_DLX1
7669_R_sec1_Cau_Bulk
7669_R_sec2_GP_Bulk
7669_R_sec2_GP_DLX1
7669_R_sec2_Put_Bulk
7669_R_sec2_Thal_Bulk
7669_R_sec2_Thal_COUPTF2
7669_R_sec2_Thal_DLX1
7669_R_sec3_Cau_DLX1
7669_R_sec3_Hip_COUPTF2
7669_R_sec3_Hip_DLX1
7669_R_sec3_Hip_TBR1
7669_R_sec3_Put_DLX1
7669_R_Skin_Bulk
7669_R_T_1_Bulk
7669_R_T_1_COUPTF2
7669_R_T_1_DLX1
7669_R_T_1_TBR1
7669_RA9_Bulk
7669_RV10_Bulk
JGG

ID05 7−46623362−G−C

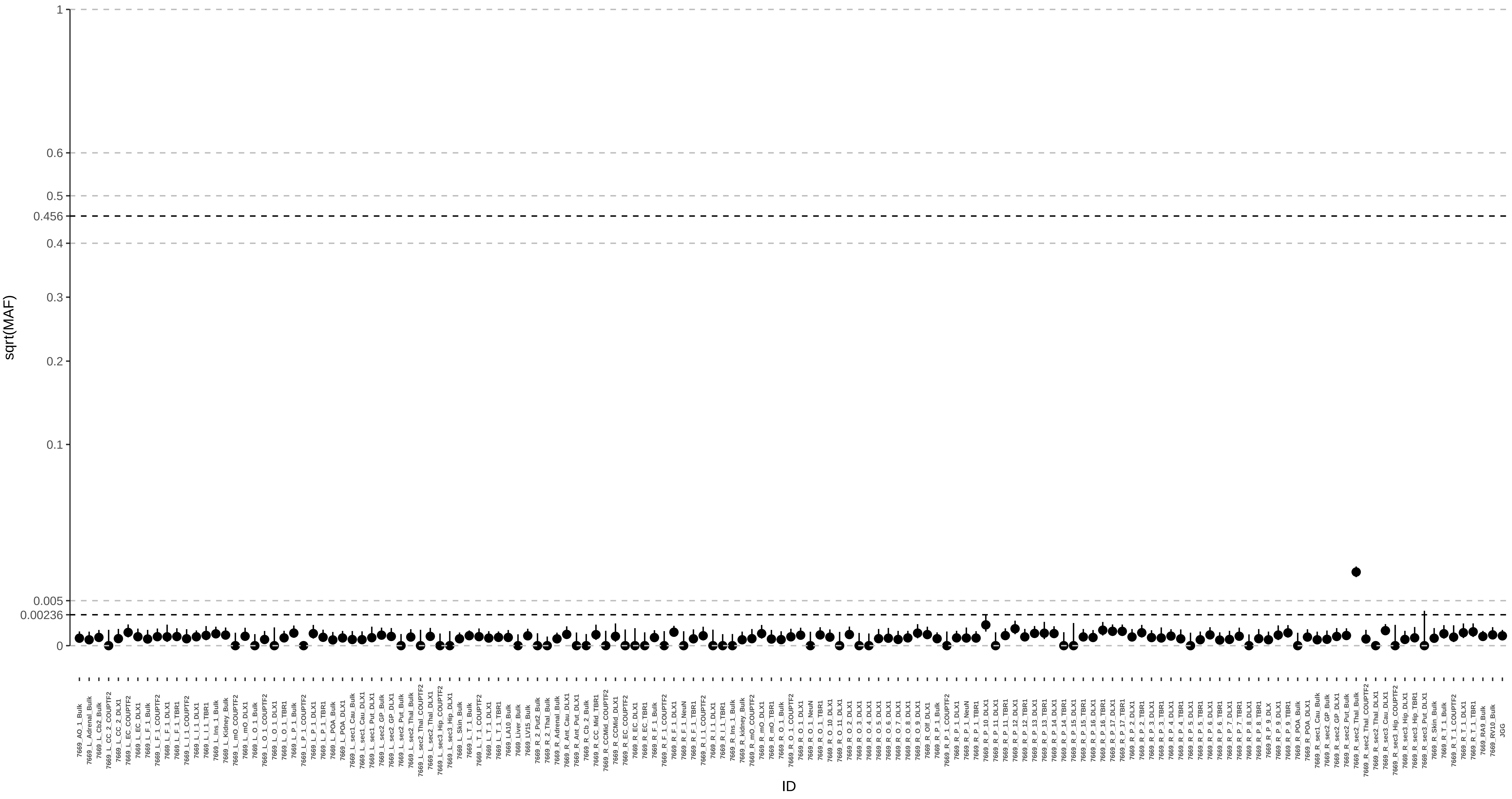

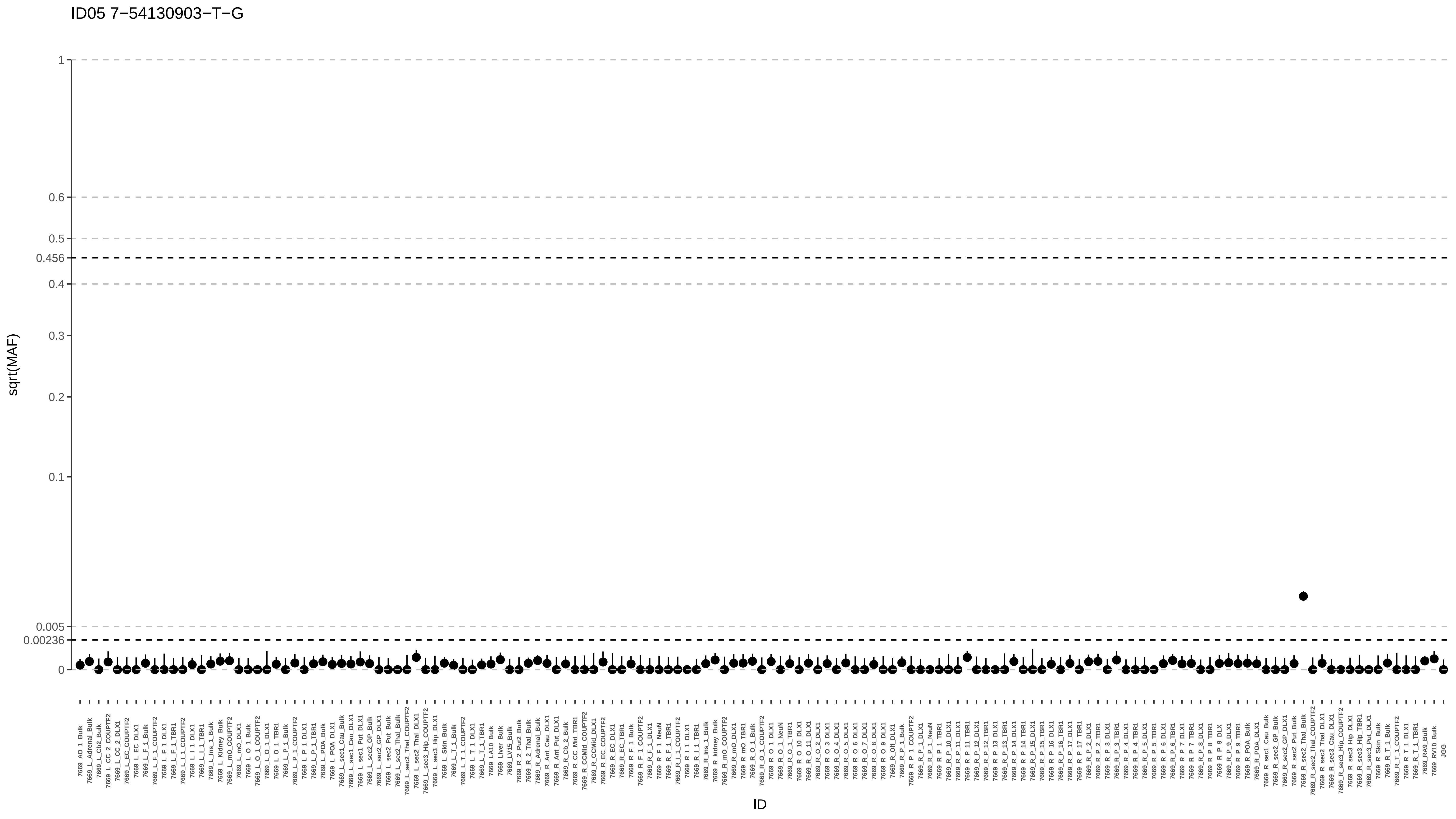
ID05 7−54130903−T−G
sqrt(MAF)
1
0.6
0.5
0.456
0.4
0.3
0.2
0.1
0.005
0.00236
0
ID

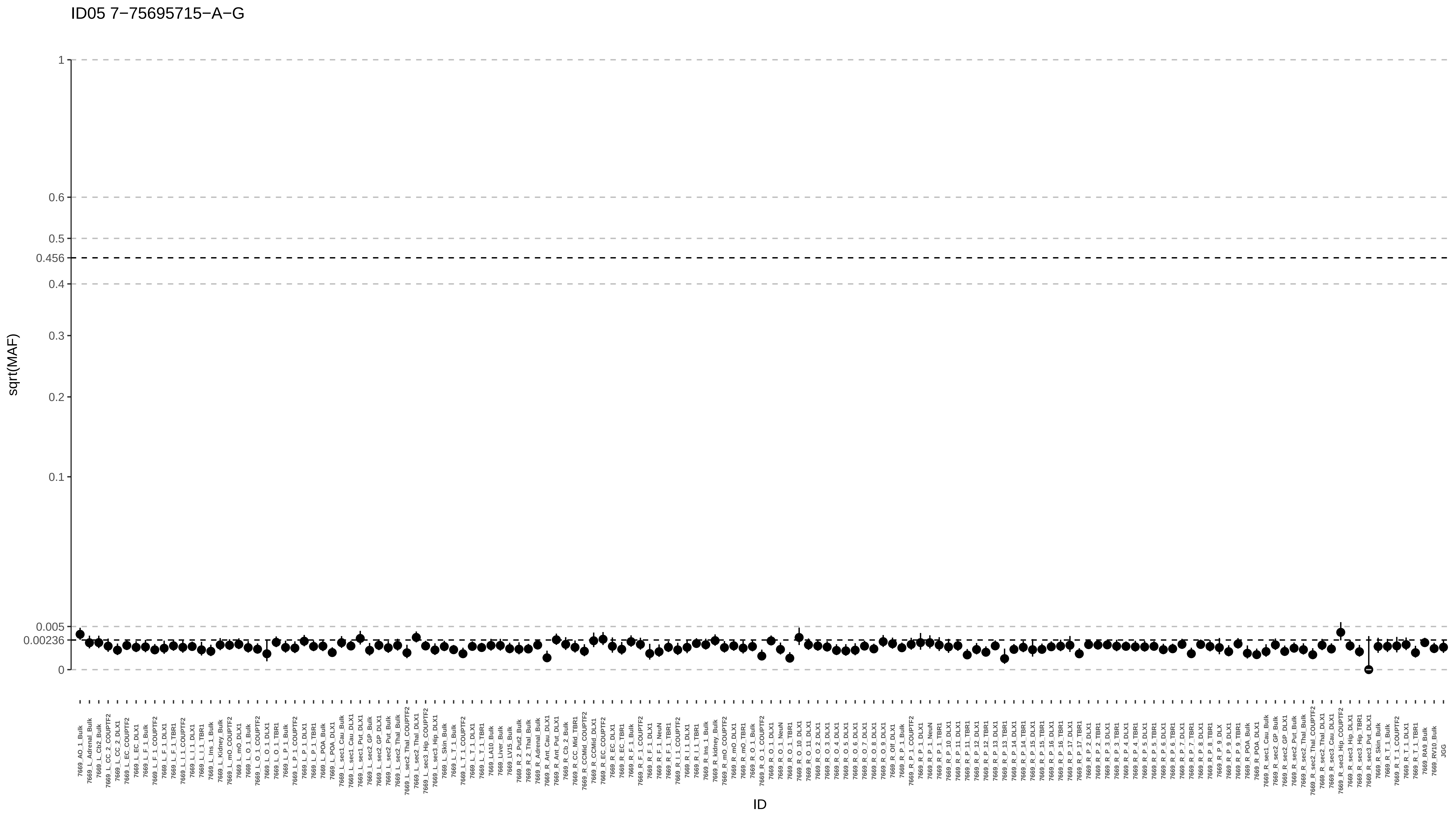
ID05 7−75695715−A−G
sqrt(MAF)
1
0.6
0.5
0.456
0.4
0.3
0.2
0.1
0.005
0.00236
0
ID

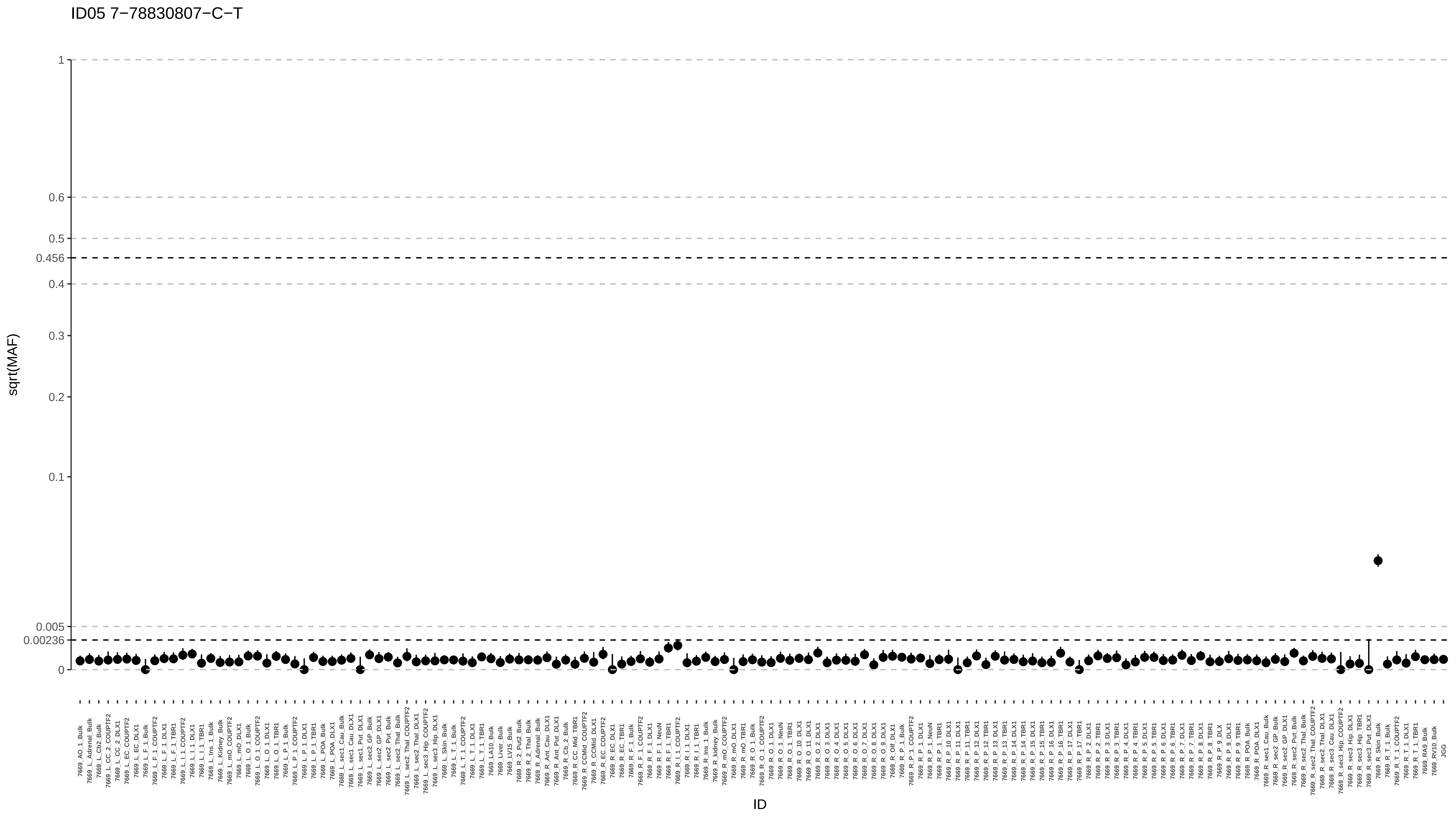
ID05 7−78830807−C−T
sqrt(MAF)
1
0.6
0.5
0.456
0.4
0.3
0.2
0.1
0.005
0.00236
0
ID

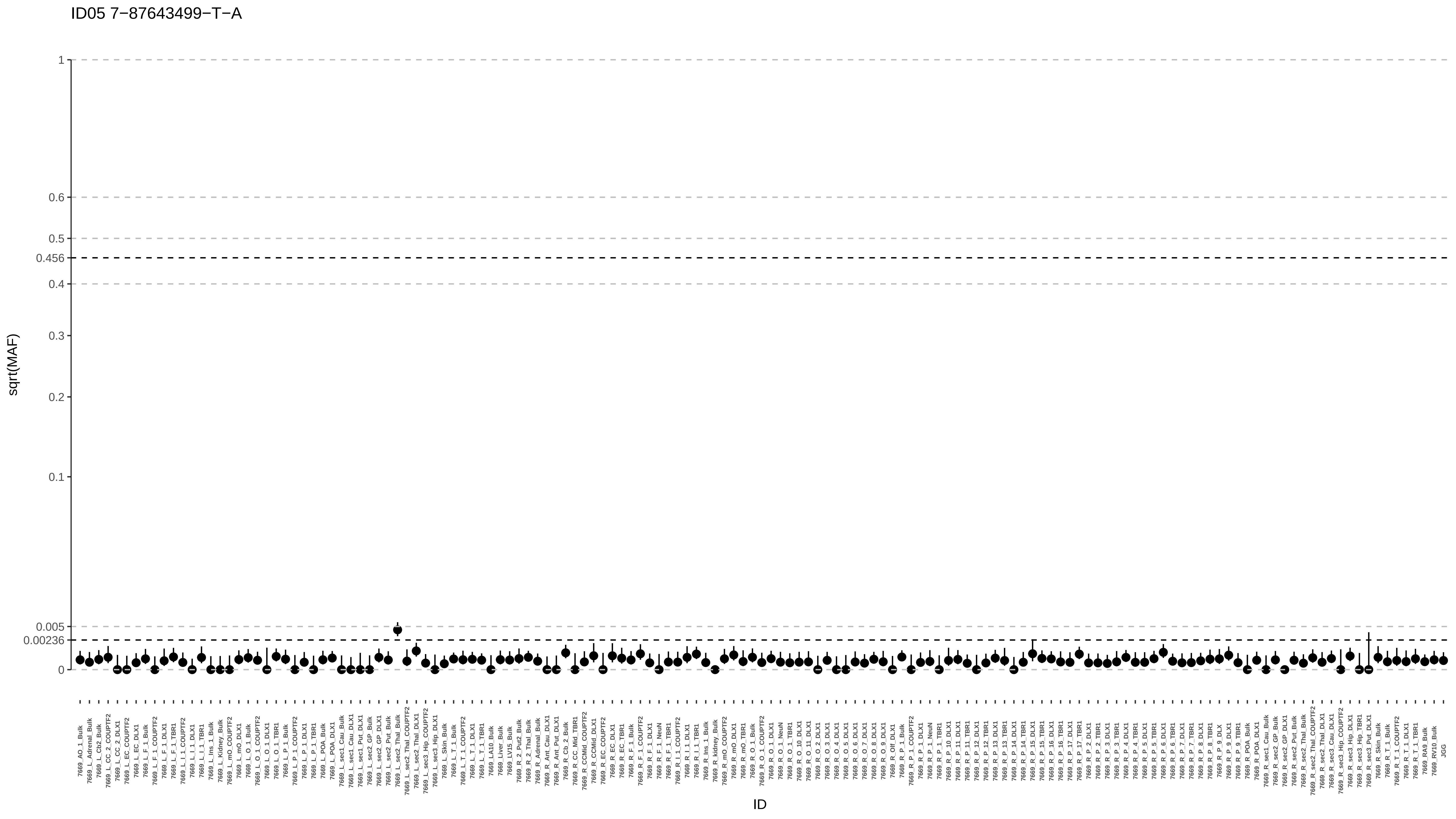
ID05 7−87643499−T−A
sqrt(MAF)
1
0.6
0.5
0.456
0.4
0.3
0.2
0.1
0.005
0.00236
0
ID
7669_AO_1_Bulk
7669_L_Adrenal_Bulk
7669_L_Cb2_Bulk
7669_L_CC_2_COUPTF2
7669_L_CC_2_DLX1
7669_L_EC_COUPTF2
7669_L_EC_DLX1
7669_L_F_1_Bulk
7669_L_F_1_COUPTF2
7669_L_F_1_DLX1
7669_L_F_1_TBR1
7669_L_I_1_COUPTF2
7669_L_I_1_DLX1
7669_L_I_1_TBR1
7669_L_Ins_1_Bulk
7669_L_Kidney_Bulk
7669_L_mO_COUPTF2
7669_L_mO_DLX1
7669_L_O_1_Bulk
7669_L_O_1_COUPTF2
7669_L_O_1_DLX1
7669_L_O_1_TBR1
7669_L_P_1_Bulk
7669_L_P_1_COUPTF2
7669_L_P_1_DLX1
7669_L_P_1_TBR1
7669_L_POA_Bulk
7669_L_POA_DLX1
7669_L_sec1_Cau_Bulk
7669_L_sec1_Cau_DLX1
7669_L_sec1_Put_DLX1
7669_L_sec2_GP_Bulk
7669_L_sec2_GP_DLX1
7669_L_sec2_Put_Bulk
7669_L_sec2_Thal_Bulk
7669_L_sec2_Thal_COUPTF2
7669_L_sec2_Thal_DLX1
7669_L_sec3_Hip_COUPTF2
7669_L_sec3_Hip_DLX1
7669_L_Skin_Bulk
7669_L_T_1_Bulk
7669_L_T_1_COUPTF2
7669_L_T_1_DLX1
7669_L_T_1_TBR1
7669_LA10_Bulk
7669_Liver_Bulk
7669_LV15_Bulk
7669_R_2_Put2_Bulk
7669_R_2_Thal_Bulk
7669_R_Adrenal_Bulk
7669_R_Ant_Cau_DLX1
7669_R_Ant_Put_DLX1
7669_R_Cb_2_Bulk
7669_R_CC_Mid_TBR1
7669_R_CCMid_COUPTF2
7669_R_CCMid_DLX1
7669_R_EC_COUPTF2
7669_R_EC_DLX1
7669_R_EC_TBR1
7669_R_F_1_Bulk
7669_R_F_1_COUPTF2
7669_R_F_1_DLX1
7669_R_F_1_NeuN
7669_R_F_1_TBR1
7669_R_I_1_COUPTF2
7669_R_I_1_DLX1
7669_R_I_1_TBR1
7669_R_Ins_1_Bulk
7669_R_kidney_Bulk
7669_R_mO_COUPTF2
7669_R_mO_DLX1
7669_R_mO_TBR1
7669_R_O_1_Bulk
7669_R_O_1_COUPTF2
7669_R_O_1_DLX1
7669_R_O_1_NeuN
7669_R_O_1_TBR1
7669_R_O_10_DLX1
7669_R_O_11_DLX1
7669_R_O_2_DLX1
7669_R_O_3_DLX1
7669_R_O_4_DLX1
7669_R_O_5_DLX1
7669_R_O_6_DLX1
7669_R_O_7_DLX1
7669_R_O_8_DLX1
7669_R_O_9_DLX1
7669_R_Olf_DLX1
7669_R_P_1_Bulk
7669_R_P_1_COUPTF2
7669_R_P_1_DLX1
7669_R_P_1_NeuN
7669_R_P_1_TBR1
7669_R_P_10_DLX1
7669_R_P_11_DLX1
7669_R_P_11_TBR1
7669_R_P_12_DLX1
7669_R_P_12_TBR1
7669_R_P_13_DLX1
7669_R_P_13_TBR1
7669_R_P_14_DLX1
7669_R_P_14_TBR1
7669_R_P_15_DLX1
7669_R_P_15_TBR1
7669_R_P_16_DLX1
7669_R_P_16_TBR1
7669_R_P_17_DLX1
7669_R_P_17_TBR1
7669_R_P_2_DLX1
7669_R_P_2_TBR1
7669_R_P_3_DLX1
7669_R_P_3_TBR1
7669_R_P_4_DLX1
7669_R_P_4_TBR1
7669_R_P_5_DLX1
7669_R_P_5_TBR1
7669_R_P_6_DLX1
7669_R_P_6_TBR1
7669_R_P_7_DLX1
7669_R_P_7_TBR1
7669_R_P_8_DLX1
7669_R_P_8_TBR1
7669_R_P_9_DLX
7669_R_P_9_DLX1
7669_R_P_9_TBR1
7669_R_POA_Bulk
7669_R_POA_DLX1
7669_R_sec1_Cau_Bulk
7669_R_sec2_GP_Bulk
7669_R_sec2_GP_DLX1
7669_R_sec2_Put_Bulk
7669_R_sec2_Thal_Bulk
7669_R_sec2_Thal_COUPTF2
7669_R_sec2_Thal_DLX1
7669_R_sec3_Cau_DLX1
7669_R_sec3_Hip_COUPTF2
7669_R_sec3_Hip_DLX1
7669_R_sec3_Hip_TBR1
7669_R_sec3_Put_DLX1
7669_R_Skin_Bulk
7669_R_T_1_Bulk
7669_R_T_1_COUPTF2
7669_R_T_1_DLX1
7669_R_T_1_TBR1
7669_RA9_Bulk
7669_RV10_Bulk
JGG

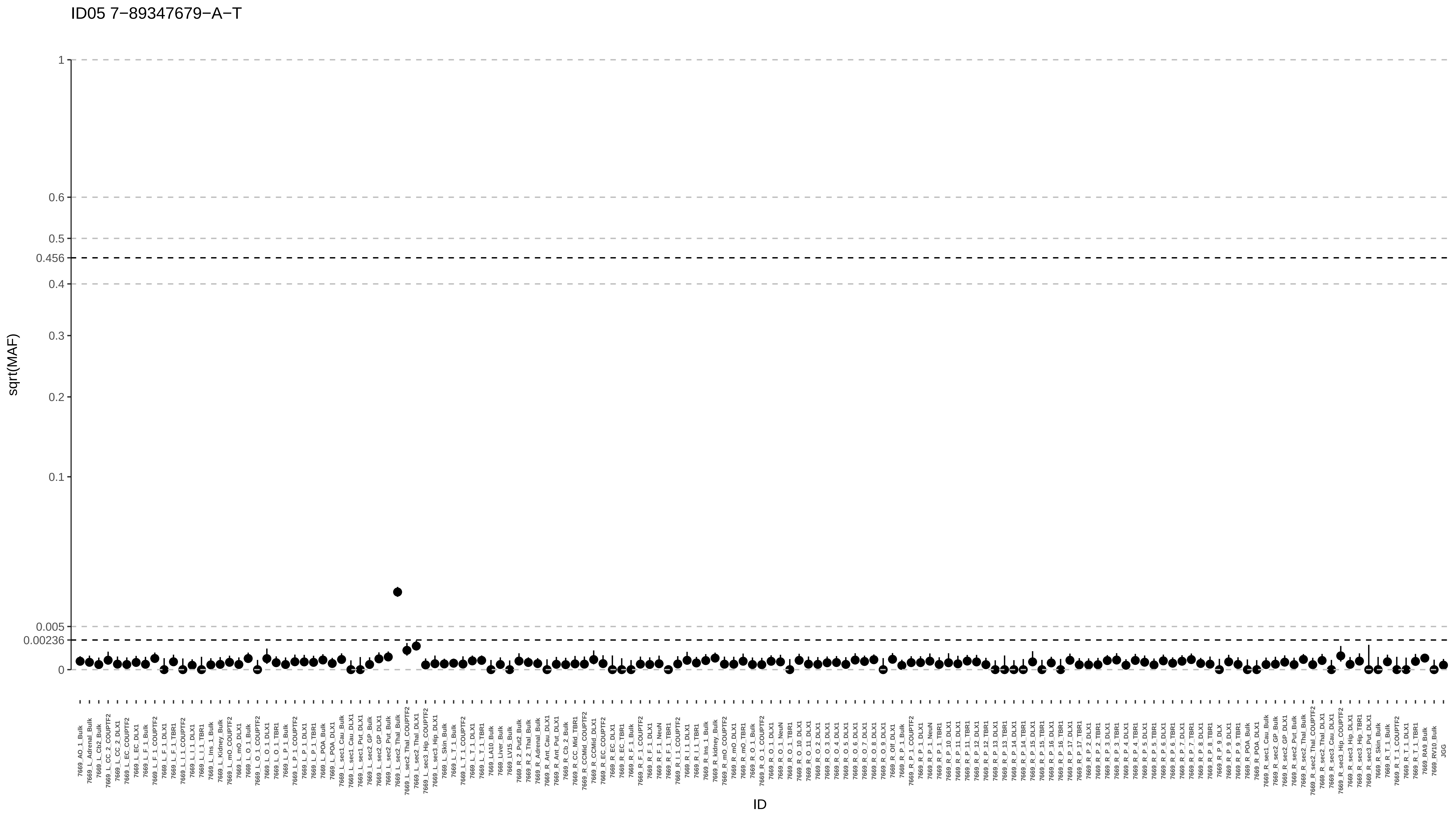

ID05 7−89347679−A−T
sqrt(MAF)
1
0.6
0.5
0.456
0.4
0.3
0.2
0.1
0.005
0.00236
0
ID

ID05 7−9192631−C−T

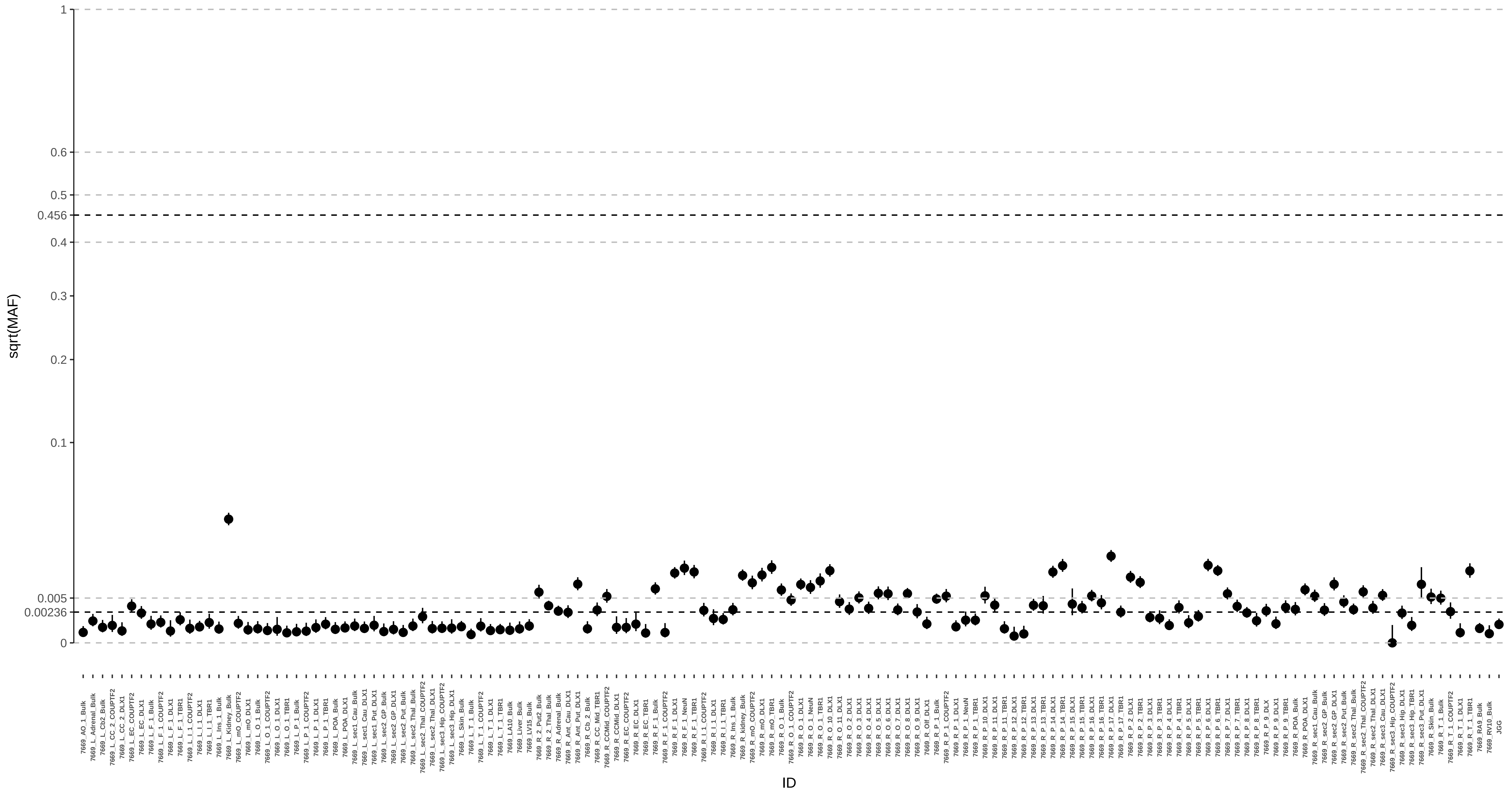

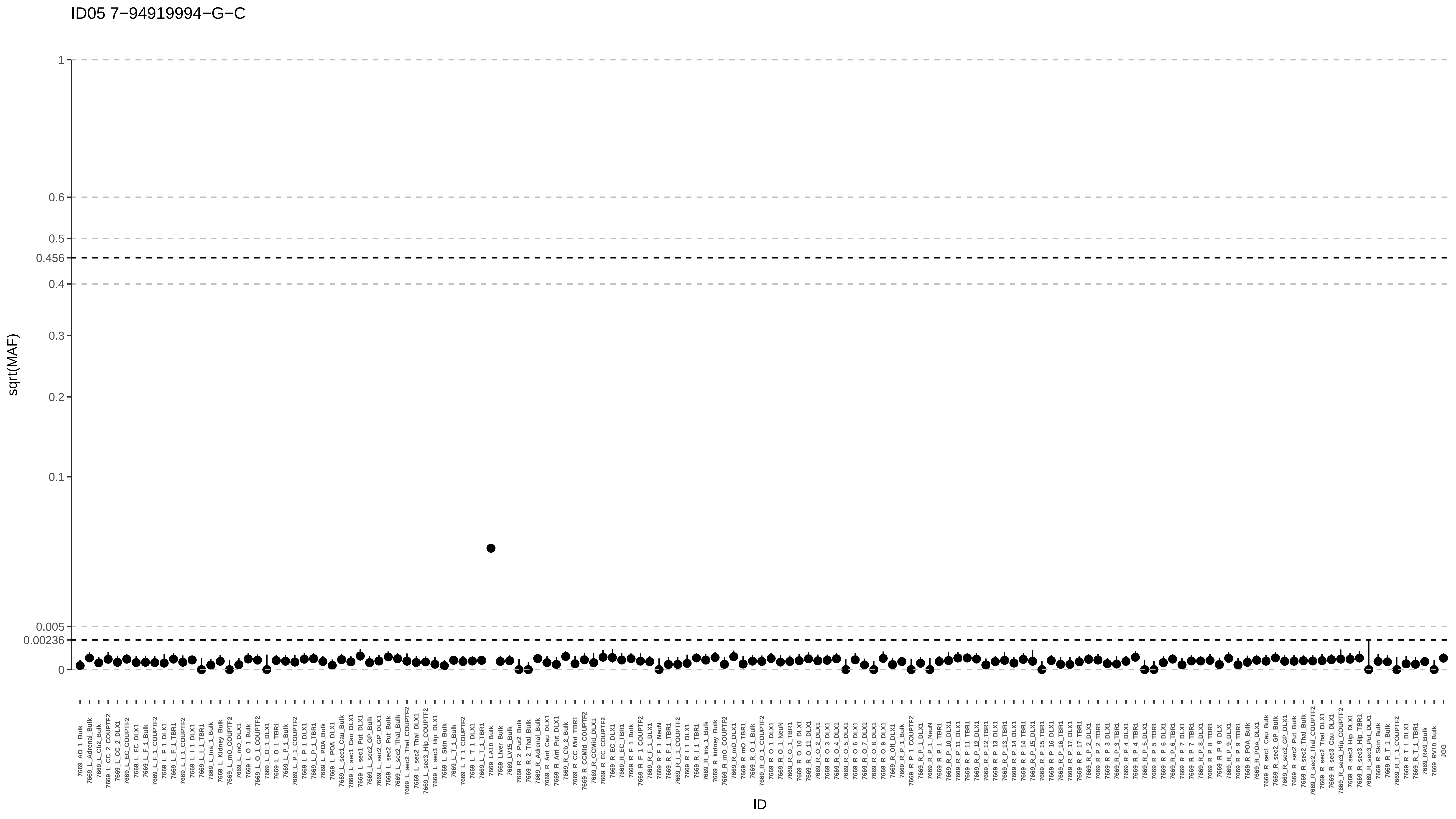
ID05 7−94919994−G−C
sqrt(MAF)
1
0.6
0.5
0.456
0.4
0.3
0.2
0.1
0.005
0.00236
0
ID

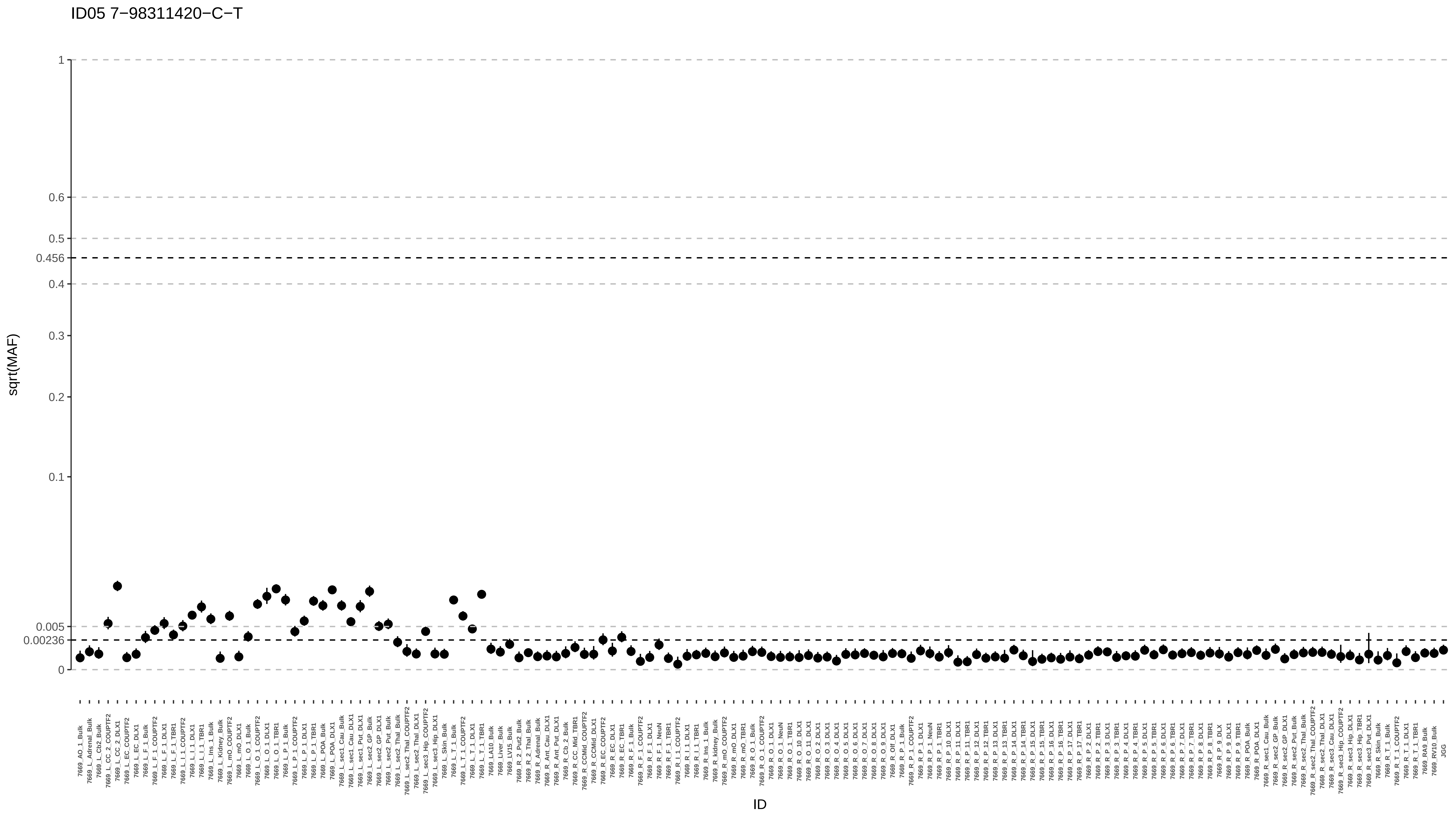
ID05 7−98311420−C−T
sqrt(MAF)
1
0.6
0.5
0.456
0.4
0.3
0.2
0.1
0.005
0.00236
0
7669_AO_1_Bulk
7669_L_Adrenal_Bulk
7669_L_Cb2_Bulk
7669_L_CC_2_COUPTF2
7669_L_CC_2_DLX1
7669_L_EC_COUPTF2
7669_L_EC_DLX1
7669_L_F_1_Bulk
7669_L_F_1_COUPTF2
7669_L_F_1_DLX1
7669_L_F_1_TBR1
7669_L_I_1_COUPTF2
7669_L_I_1_DLX1
7669_L_I_1_TBR1
7669_L_Ins_1_Bulk
7669_L_Kidney_Bulk
7669_L_mO_COUPTF2
7669_L_mO_DLX1
7669_L_O_1_Bulk
7669_L_O_1_COUPTF2
7669_L_O_1_DLX1
7669_L_O_1_TBR1
7669_L_P_1_Bulk
7669_L_P_1_COUPTF2
7669_L_P_1_DLX1
7669_L_P_1_TBR1
7669_L_POA_Bulk
7669_L_POA_DLX1
7669_L_sec1_Cau_Bulk
7669_L_sec1_Cau_DLX1
7669_L_sec1_Put_DLX1
7669_L_sec2_GP_Bulk
7669_L_sec2_GP_DLX1
7669_L_sec2_Put_Bulk
7669_L_sec2_Thal_Bulk
7669_L_sec2_Thal_COUPTF2
7669_L_sec2_Thal_DLX1
7669_L_sec3_Hip_COUPTF2
7669_L_sec3_Hip_DLX1
7669_L_Skin_Bulk
7669_L_T_1_Bulk
7669_L_T_1_COUPTF2
7669_L_T_1_DLX1
7669_L_T_1_TBR1
7669_LA10_Bulk
7669_Liver_Bulk
7669_LV15_Bulk
7669_R_2_Put2_Bulk
7669_R_2_Thal_Bulk
7669_R_Adrenal_Bulk
7669_R_Ant_Cau_DLX1
7669_R_Ant_Put_DLX1
7669_R_Cb_2_Bulk
7669_R_CC_Mid_TBR1
7669_R_CCMid_COUPTF2
7669_R_CCMid_DLX1
7669_R_EC_COUPTF2
7669_R_EC_DLX1
7669_R_EC_TBR1
7669_R_F_1_Bulk
7669_R_F_1_COUPTF2
7669_R_F_1_DLX1
7669_R_F_1_NeuN
7669_R_F_1_TBR1
7669_R_I_1_COUPTF2
7669_R_I_1_DLX1
7669_R_I_1_TBR1
7669_R_Ins_1_Bulk
7669_R_kidney_Bulk
7669_R_mO_COUPTF2
7669_R_mO_DLX1
7669_R_mO_TBR1
7669_R_O_1_Bulk
7669_R_O_1_COUPTF2
7669_R_O_1_DLX1
7669_R_O_1_NeuN
7669_R_O_1_TBR1
7669_R_O_10_DLX1
7669_R_O_11_DLX1
7669_R_O_2_DLX1
7669_R_O_3_DLX1
7669_R_O_4_DLX1
7669_R_O_5_DLX1
7669_R_O_6_DLX1
7669_R_O_7_DLX1
7669_R_O_8_DLX1
7669_R_O_9_DLX1
7669_R_Olf_DLX1
7669_R_P_1_Bulk
7669_R_P_1_COUPTF2
7669_R_P_1_DLX1
7669_R_P_1_NeuN
7669_R_P_1_TBR1
7669_R_P_10_DLX1
7669_R_P_11_DLX1
7669_R_P_11_TBR1
7669_R_P_12_DLX1
7669_R_P_12_TBR1
7669_R_P_13_DLX1
7669_R_P_13_TBR1
7669_R_P_14_DLX1
7669_R_P_14_TBR1
7669_R_P_15_DLX1
7669_R_P_15_TBR1
7669_R_P_16_DLX1
7669_R_P_16_TBR1
7669_R_P_17_DLX1
7669_R_P_17_TBR1
7669_R_P_2_DLX1
7669_R_P_2_TBR1
7669_R_P_3_DLX1
7669_R_P_3_TBR1
7669_R_P_4_DLX1
7669_R_P_4_TBR1
7669_R_P_5_DLX1
7669_R_P_5_TBR1
7669_R_P_6_DLX1
7669_R_P_6_TBR1
7669_R_P_7_DLX1
7669_R_P_7_TBR1
7669_R_P_8_DLX1
7669_R_P_8_TBR1
7669_R_P_9_DLX
7669_R_P_9_DLX1
7669_R_P_9_TBR1
7669_R_POA_Bulk
7669_R_POA_DLX1
7669_R_sec1_Cau_Bulk
7669_R_sec2_GP_Bulk
7669_R_sec2_GP_DLX1
7669_R_sec2_Put_Bulk
7669_R_sec2_Thal_Bulk
7669_R_sec2_Thal_COUPTF2
7669_R_sec2_Thal_DLX1
7669_R_sec3_Cau_DLX1
7669_R_sec3_Hip_COUPTF2
7669_R_sec3_Hip_DLX1
7669_R_sec3_Hip_TBR1
7669_R_sec3_Put_DLX1
7669_R_Skin_Bulk
7669_R_T_1_Bulk
7669_R_T_1_COUPTF2
7669_R_T_1_DLX1
7669_R_T_1_TBR1
7669_RA9_Bulk
7669_RV10_Bulk
JGG
ID

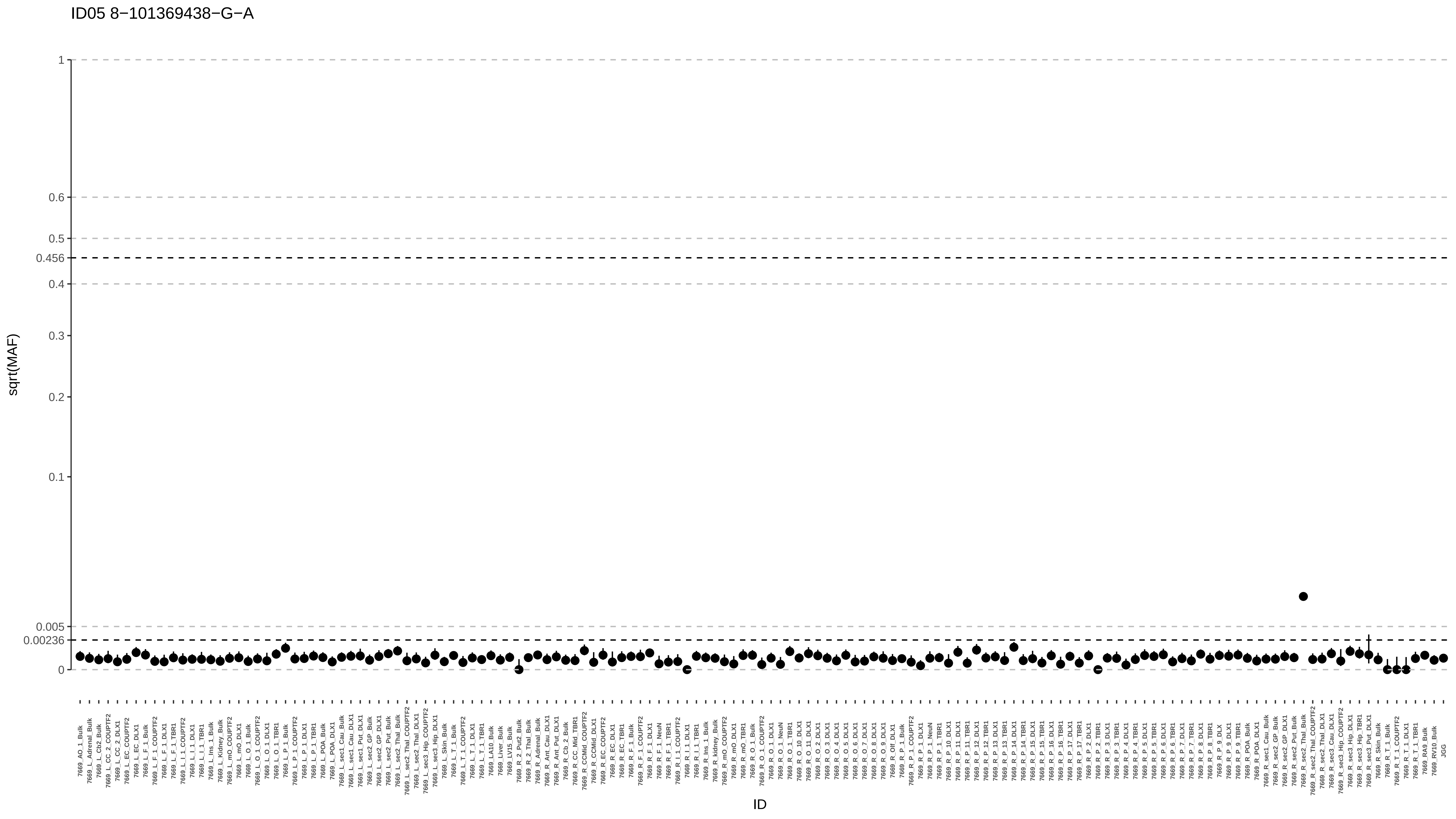
ID05 8−101369438−G−A
sqrt(MAF)
ID

ID05 8−101774660−C−A

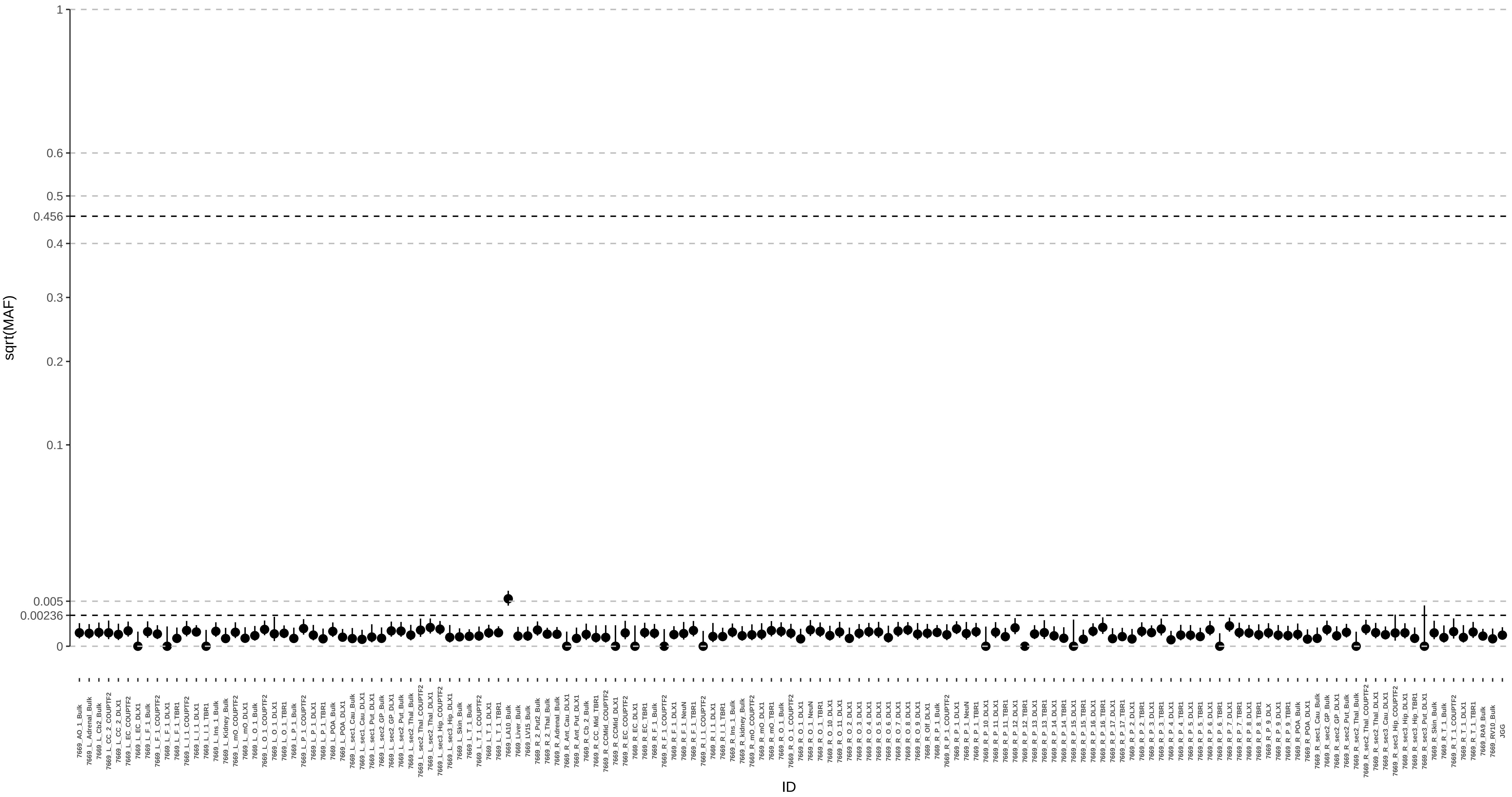

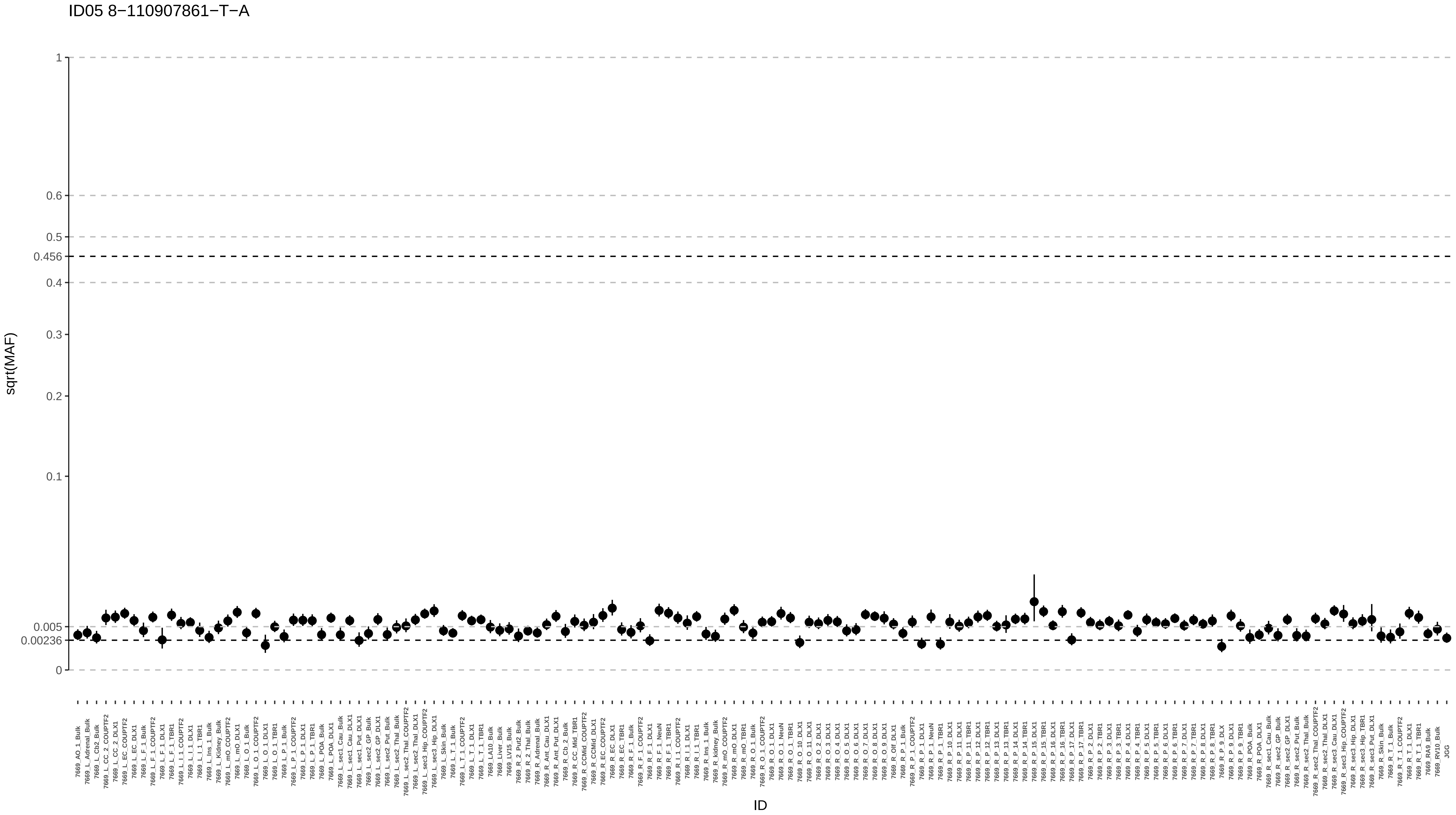
ID05 8−110907861−T−A
sqrt(MAF)
1
0.6
0.5
0.456
0.4
0.3
0.2
0.1
0.005
0.00236
0
ID
7669_AO_1_Bulk
7669_L_Adrenal_Bulk
7669_L_Cb2_Bulk
7669_L_CC_2_COUPTF2
7669_L_CC_2_DLX1
7669_L_EC_COUPTF2
7669_L_EC_DLX1
7669_L_F_1_Bulk
7669_L_F_1_COUPTF2
7669_L_F_1_DLX1
7669_L_F_1_TBR1
7669_L_I_1_COUPTF2
7669_L_I_1_DLX1
7669_L_I_1_TBR1
7669_L_Ins_1_Bulk
7669_L_Kidney_Bulk
7669_L_mO_COUPTF2
7669_L_mO_DLX1
7669_L_O_1_Bulk
7669_L_O_1_COUPTF2
7669_L_O_1_DLX1
7669_L_O_1_TBR1
7669_L_P_1_Bulk
7669_L_P_1_COUPTF2
7669_L_P_1_DLX1
7669_L_P_1_TBR1
7669_L_POA_Bulk
7669_L_POA_DLX1
7669_L_sec1_Cau_Bulk
7669_L_sec1_Cau_DLX1
7669_L_sec1_Put_DLX1
7669_L_sec2_GP_Bulk
7669_L_sec2_GP_DLX1
7669_L_sec2_Put_Bulk
7669_L_sec2_Thal_Bulk
7669_L_sec2_Thal_COUPTF2
7669_L_sec2_Thal_DLX1
7669_L_sec3_Hip_COUPTF2
7669_L_sec3_Hip_DLX1
7669_L_Skin_Bulk
7669_L_T_1_Bulk
7669_L_T_1_COUPTF2
7669_L_T_1_DLX1
7669_L_T_1_TBR1
7669_LA10_Bulk
7669_Liver_Bulk
7669_LV15_Bulk
7669_R_2_Put2_Bulk
7669_R_2_Thal_Bulk
7669_R_Adrenal_Bulk
7669_R_Ant_Cau_DLX1
7669_R_Ant_Put_DLX1
7669_R_Cb_2_Bulk
7669_R_CC_Mid_TBR1
7669_R_CCMid_COUPTF2
7669_R_CCMid_DLX1
7669_R_EC_COUPTF2
7669_R_EC_DLX1
7669_R_EC_TBR1
7669_R_F_1_Bulk
7669_R_F_1_COUPTF2
7669_R_F_1_DLX1
7669_R_F_1_NeuN
7669_R_F_1_TBR1
7669_R_I_1_COUPTF2
7669_R_I_1_DLX1
7669_R_I_1_TBR1
7669_R_Ins_1_Bulk
7669_R_kidney_Bulk
7669_R_mO_COUPTF2
7669_R_mO_DLX1
7669_R_mO_TBR1
7669_R_O_1_Bulk
7669_R_O_1_COUPTF2
7669_R_O_1_DLX1
7669_R_O_1_NeuN
7669_R_O_1_TBR1
7669_R_O_10_DLX1
7669_R_O_11_DLX1
7669_R_O_2_DLX1
7669_R_O_3_DLX1
7669_R_O_4_DLX1
7669_R_O_5_DLX1
7669_R_O_6_DLX1
7669_R_O_7_DLX1
7669_R_O_8_DLX1
7669_R_O_9_DLX1
7669_R_Olf_DLX1
7669_R_P_1_Bulk
7669_R_P_1_COUPTF2
7669_R_P_1_DLX1
7669_R_P_1_NeuN
7669_R_P_1_TBR1
7669_R_P_10_DLX1
7669_R_P_11_DLX1
7669_R_P_11_TBR1
7669_R_P_12_DLX1
7669_R_P_12_TBR1
7669_R_P_13_DLX1
7669_R_P_13_TBR1
7669_R_P_14_DLX1
7669_R_P_14_TBR1
7669_R_P_15_DLX1
7669_R_P_15_TBR1
7669_R_P_16_DLX1
7669_R_P_16_TBR1
7669_R_P_17_DLX1
7669_R_P_17_TBR1
7669_R_P_2_DLX1
7669_R_P_2_TBR1
7669_R_P_3_DLX1
7669_R_P_3_TBR1
7669_R_P_4_DLX1
7669_R_P_4_TBR1
7669_R_P_5_DLX1
7669_R_P_5_TBR1
7669_R_P_6_DLX1
7669_R_P_6_TBR1
7669_R_P_7_DLX1
7669_R_P_7_TBR1
7669_R_P_8_DLX1
7669_R_P_8_TBR1
7669_R_P_9_DLX
7669_R_P_9_DLX1
7669_R_P_9_TBR1
7669_R_POA_Bulk
7669_R_POA_DLX1
7669_R_sec1_Cau_Bulk
7669_R_sec2_GP_Bulk
7669_R_sec2_GP_DLX1
7669_R_sec2_Put_Bulk
7669_R_sec2_Thal_Bulk
7669_R_sec2_Thal_COUPTF2
7669_R_sec2_Thal_DLX1
7669_R_sec3_Cau_DLX1
7669_R_sec3_Hip_COUPTF2
7669_R_sec3_Hip_DLX1
7669_R_sec3_Hip_TBR1
7669_R_sec3_Put_DLX1
7669_R_Skin_Bulk
7669_R_T_1_Bulk
7669_R_T_1_COUPTF2
7669_R_T_1_DLX1
7669_R_T_1_TBR1
7669_RA9_Bulk
7669_RV10_Bulk
JGG

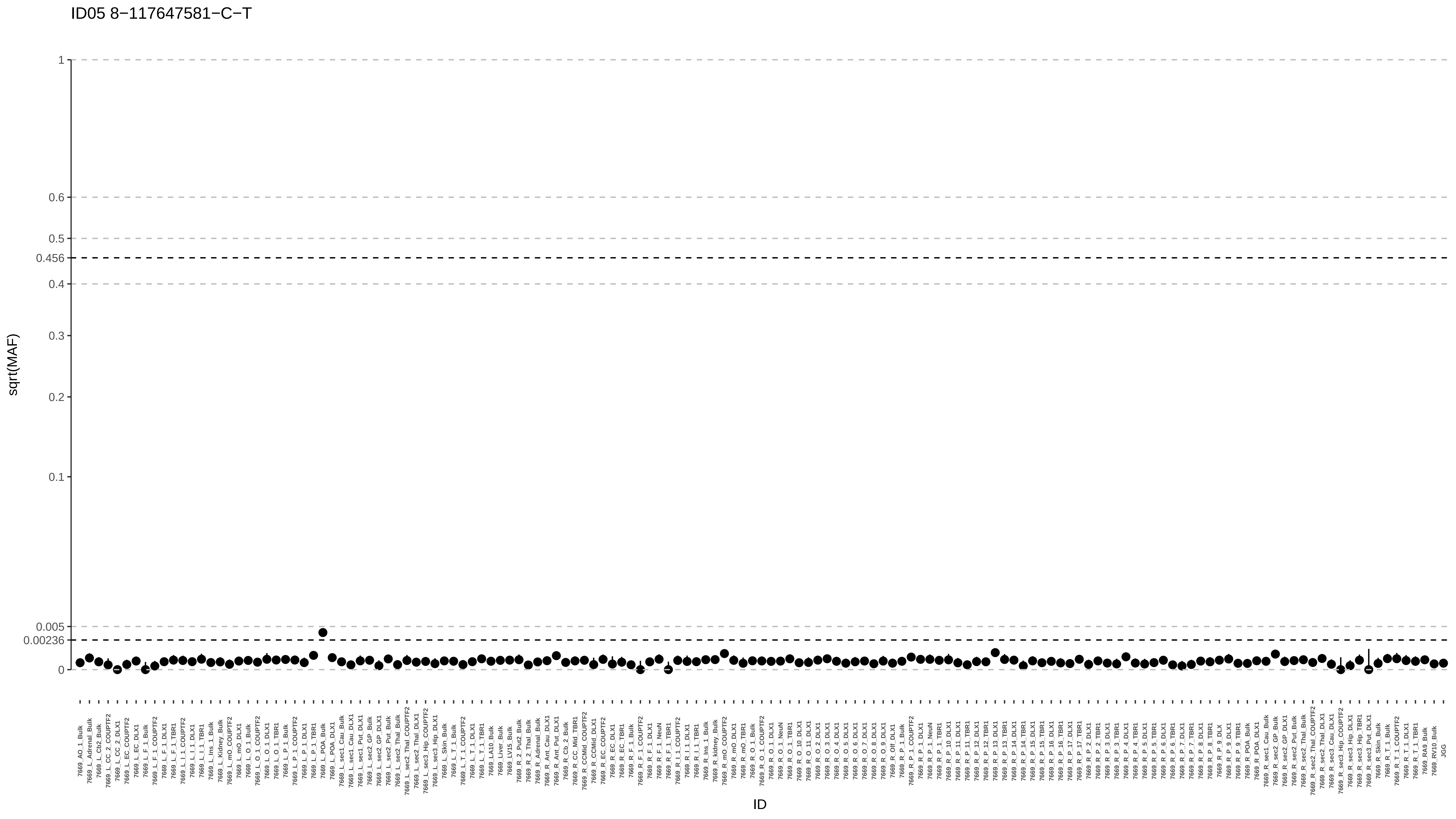

ID05 8−117647581−C−T
sqrt(MAF)
ID

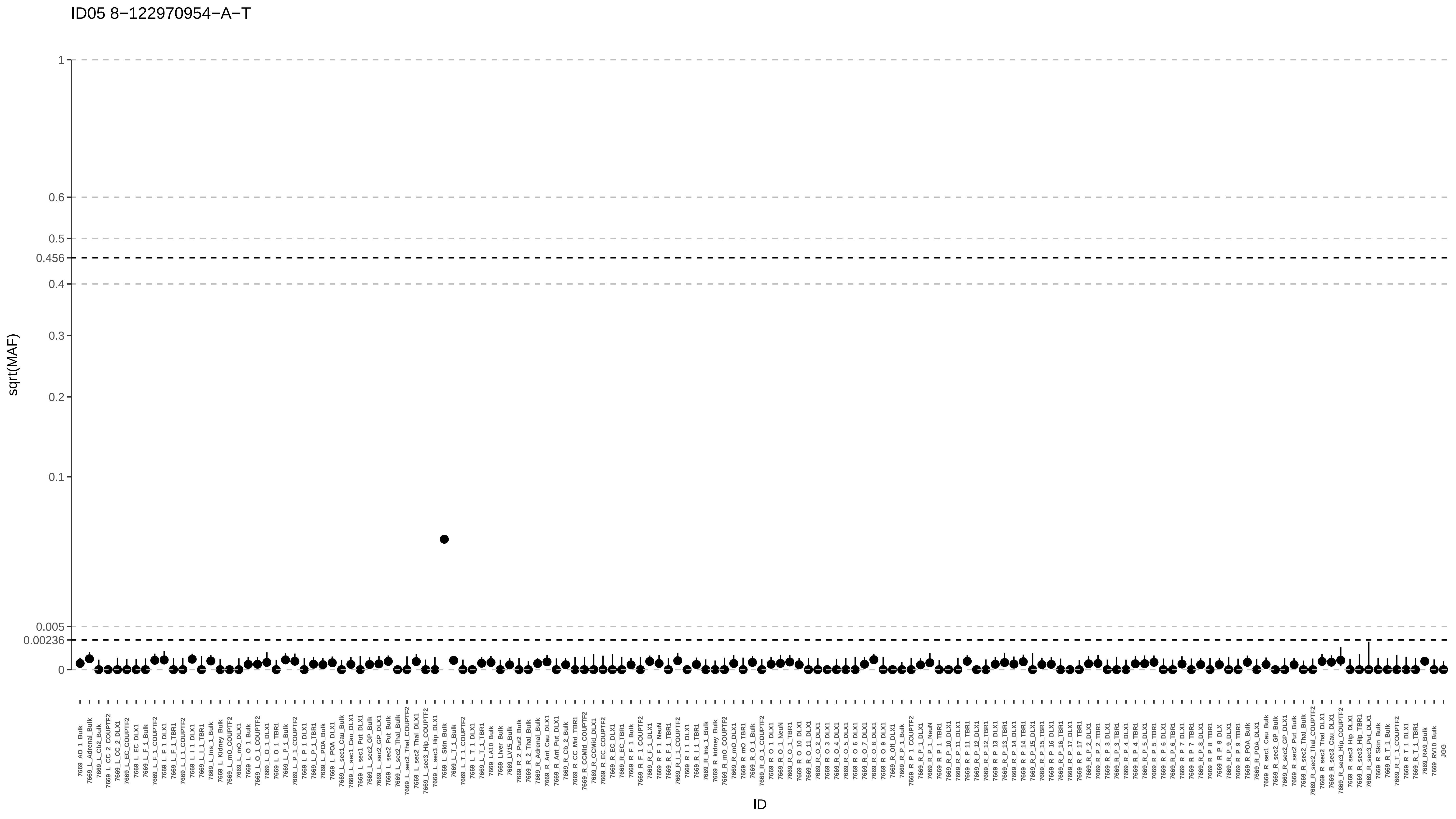
ID05 8−122970954−A−T
sqrt(MAF)
1
0.6
0.5
0.456
0.4
0.3
0.2
0.1
0.005
0.00236
0
ID

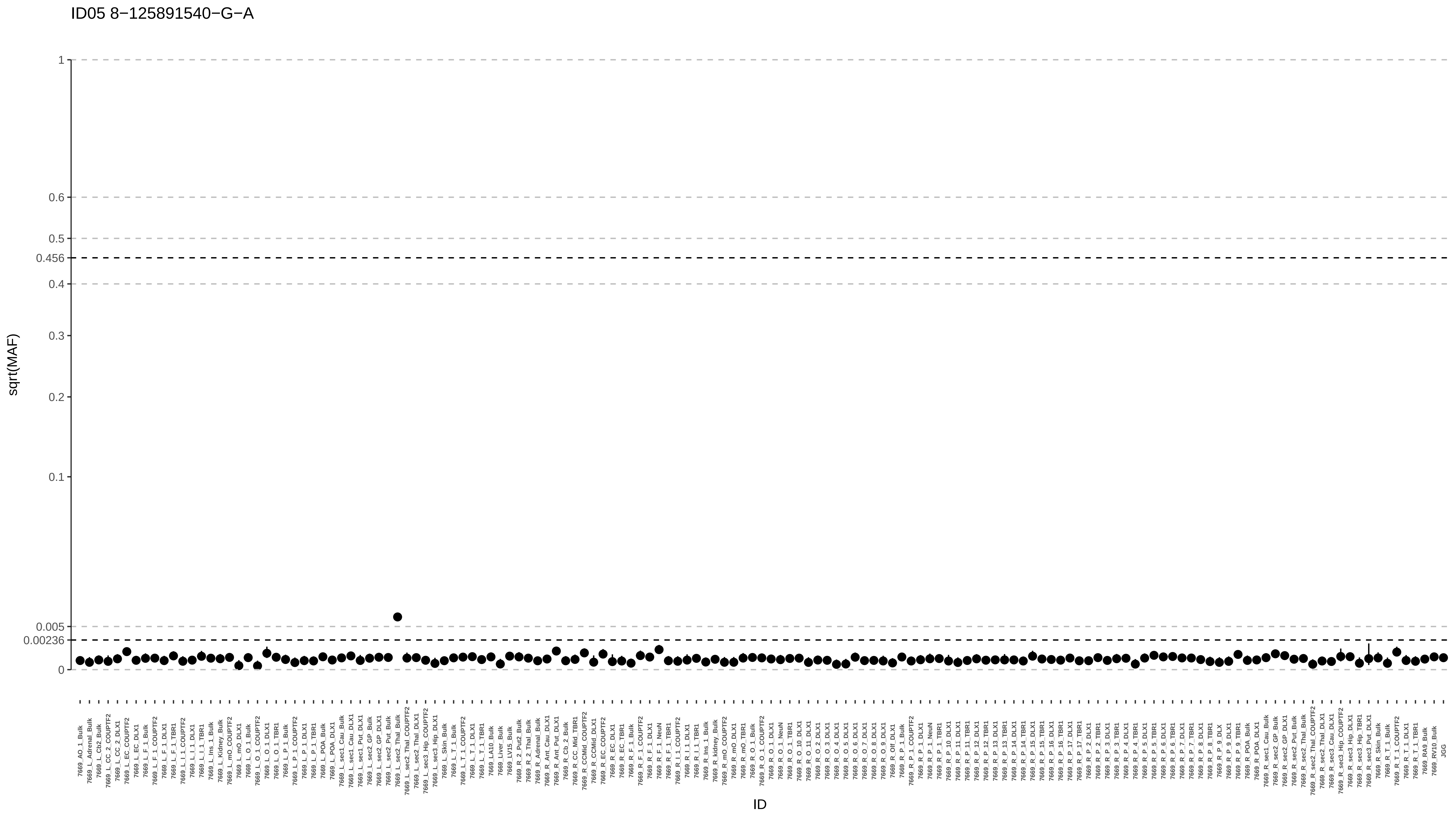
ID05 8−125891540−G−A
sqrt(MAF)
1
0.6
0.5
0.456
0.4
0.3
0.2
0.1
0.005
0.00236
0
ID

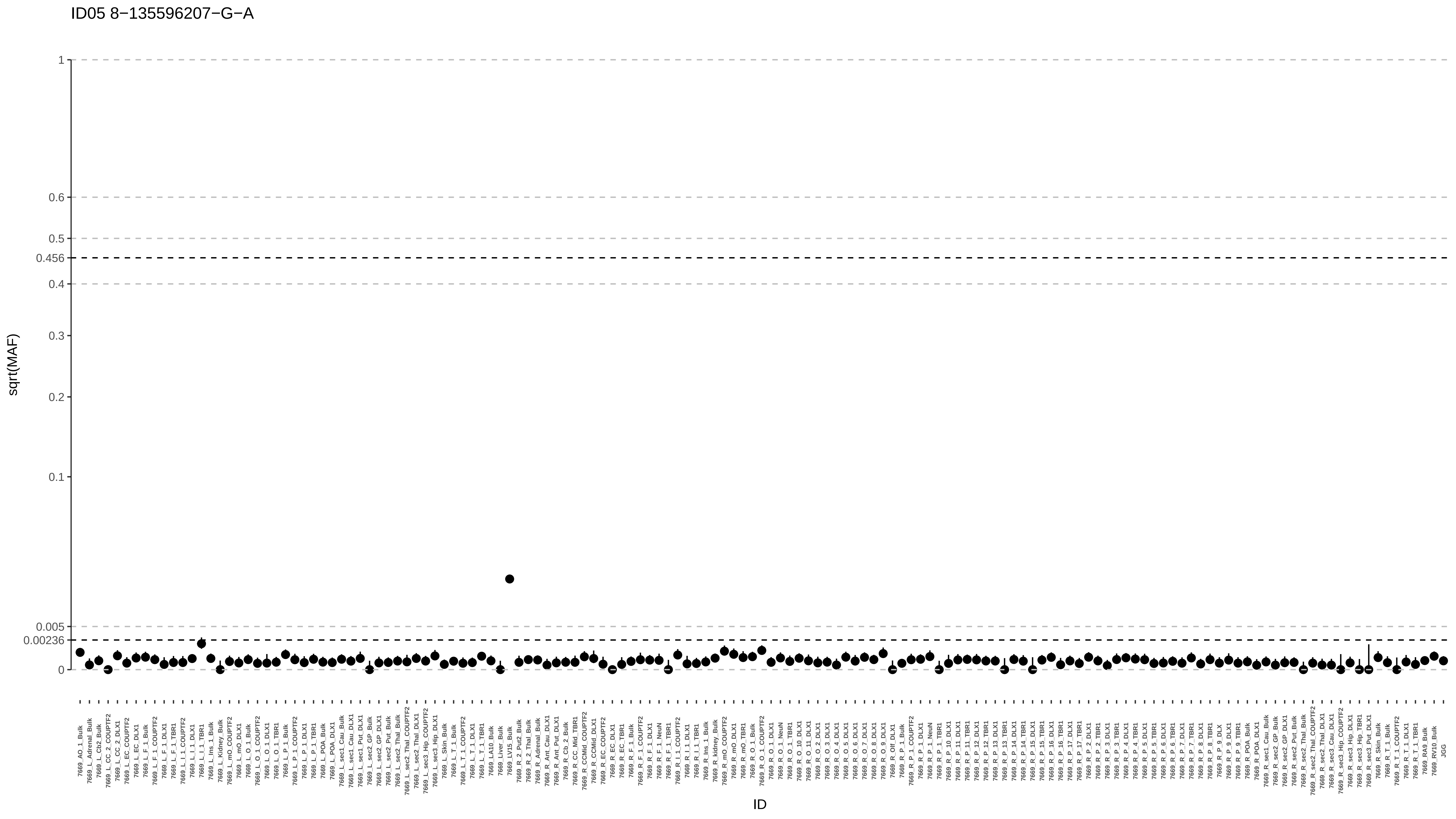
ID05 8−135596207−G−A
sqrt(MAF)
ID

ID05 8−140637848−C−A

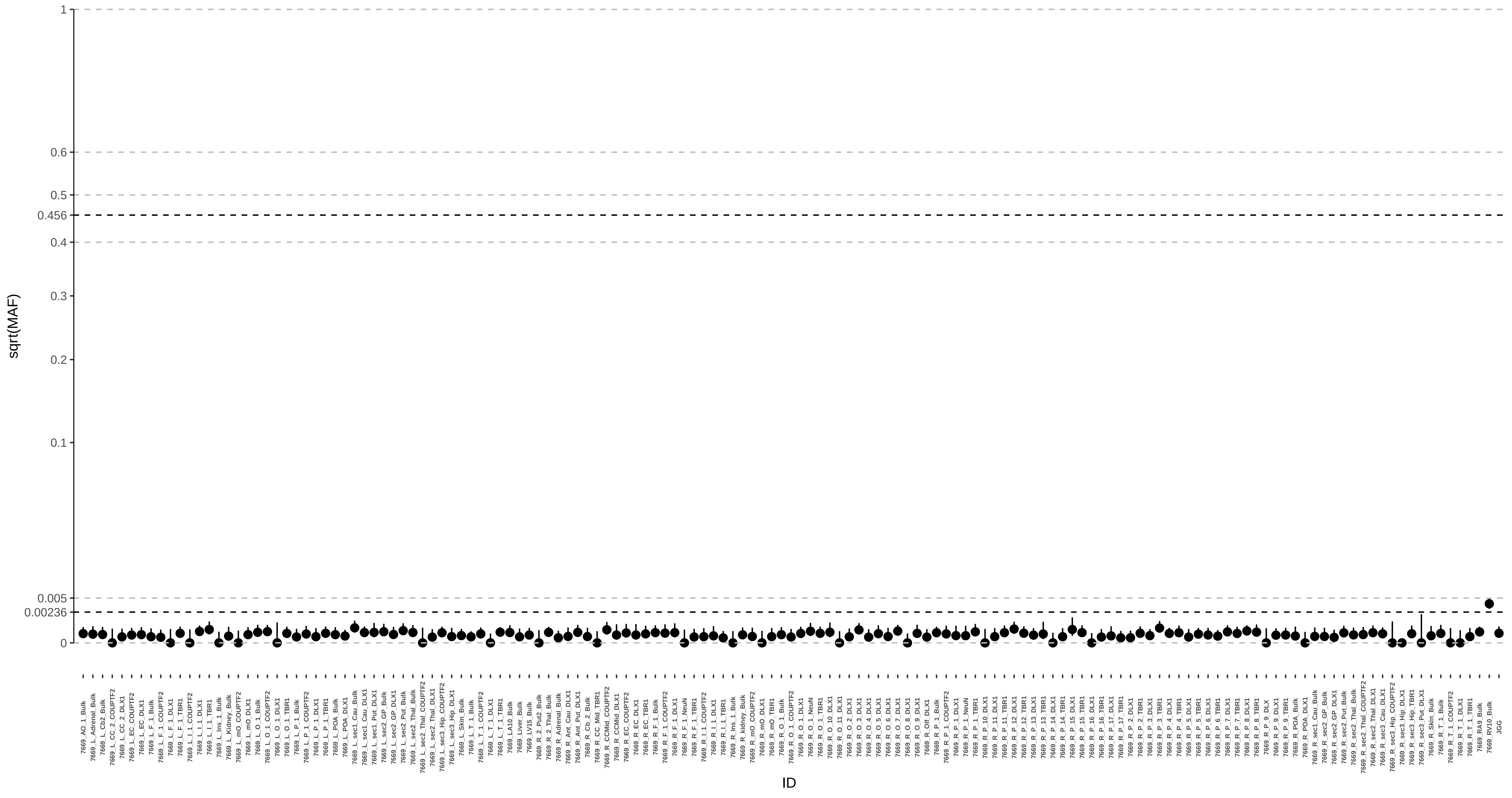

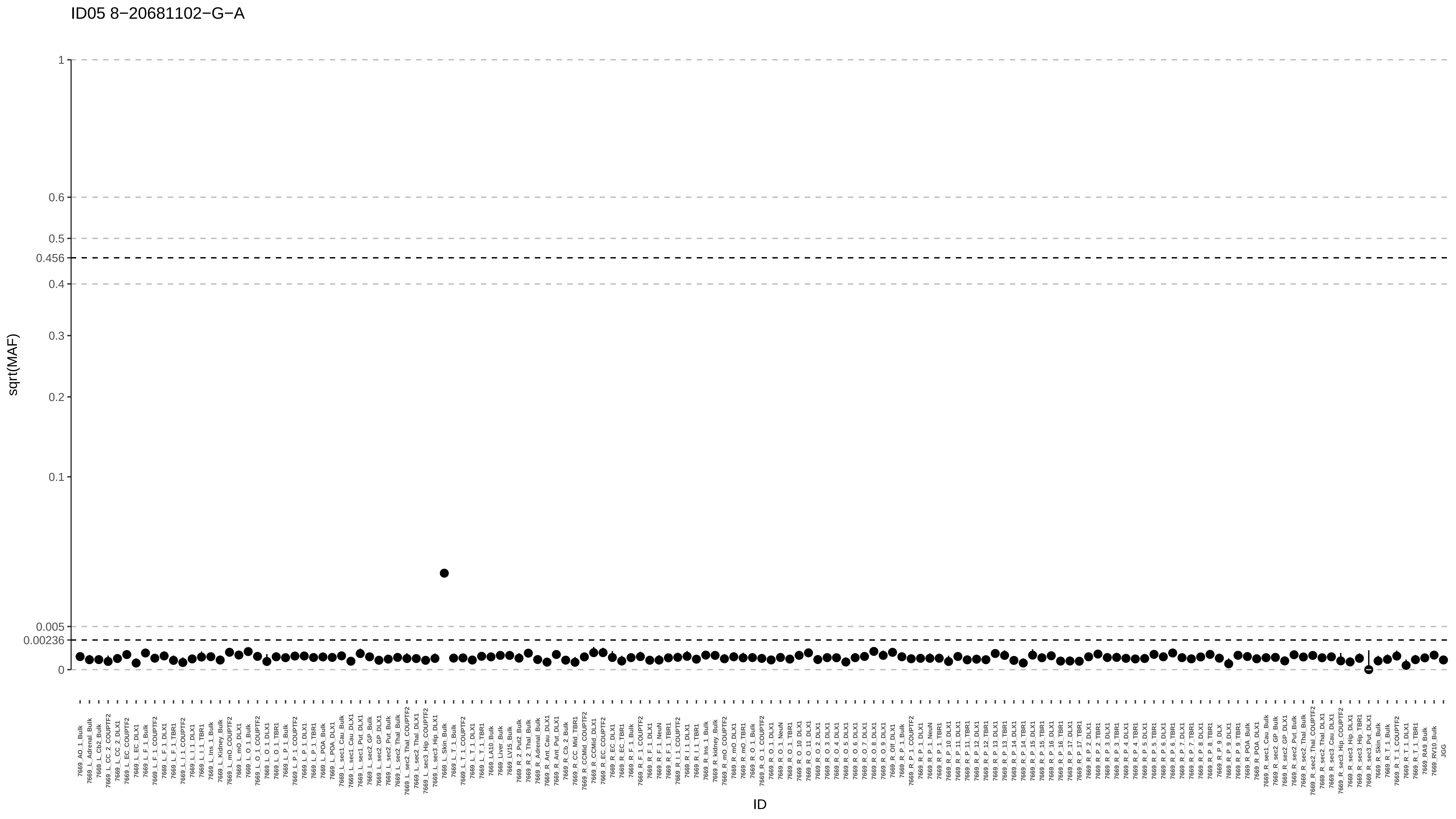

ID05 8−20681102−G−A
sqrt(MAF)
1
0.6
0.5
0.456
0.4
0.3
0.2
0.1
0.005
0.00236
0
ID

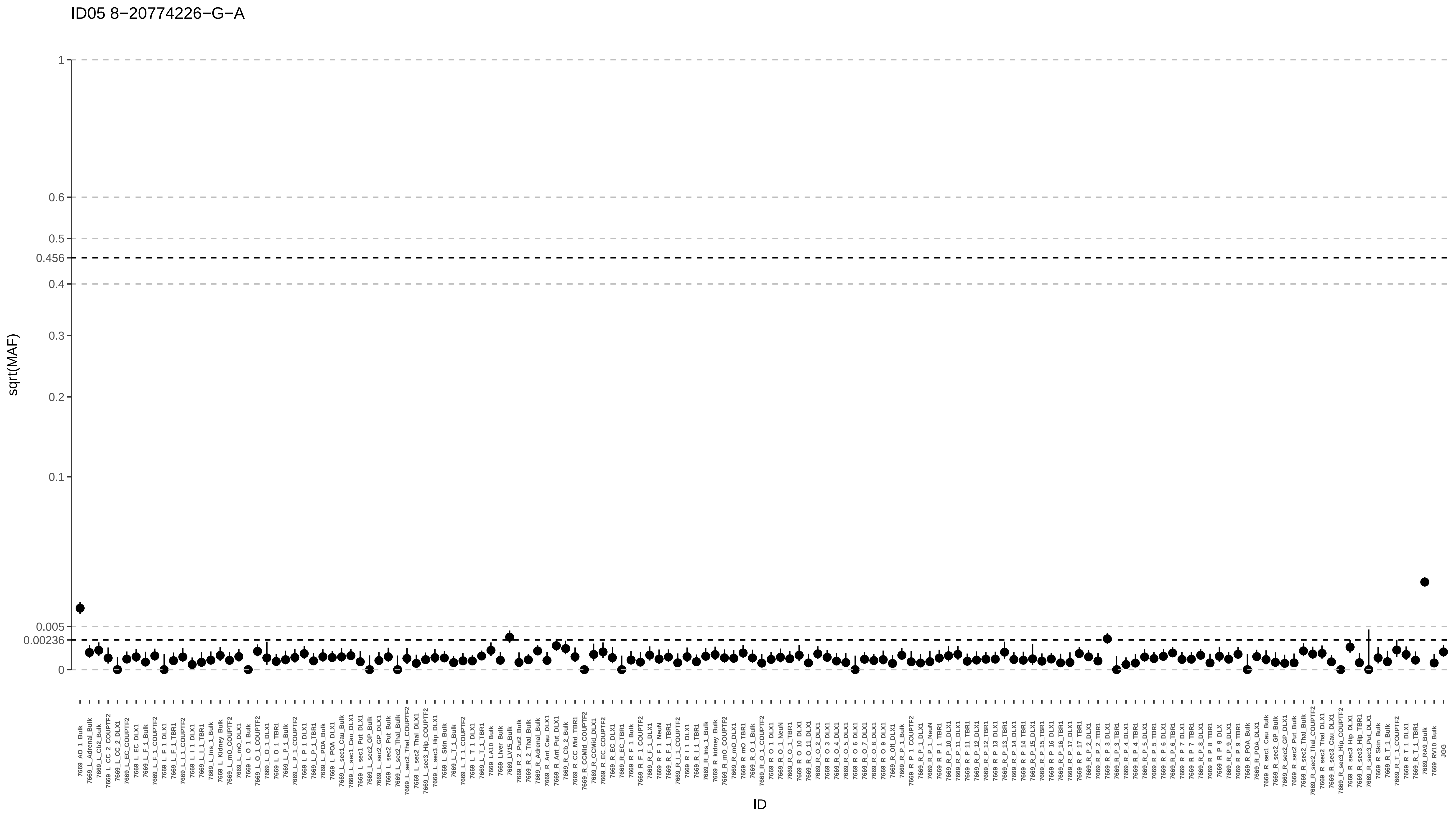
ID05 8−20774226−G−A
sqrt(MAF)
1
0.6
0.5
0.456
0.4
0.3
0.2
0.1
0.005
0.00236
0
ID

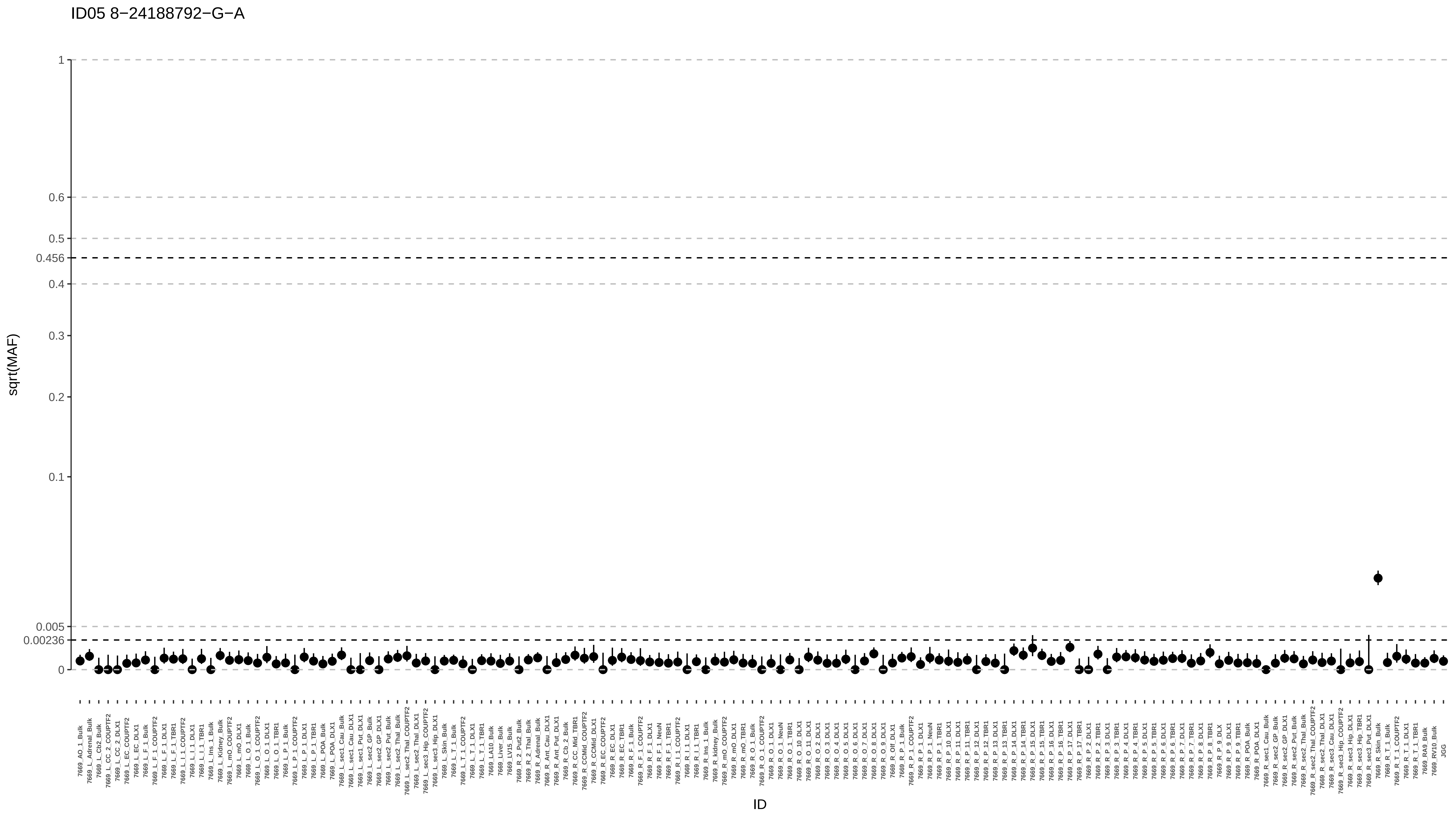
ID05 8−24188792−G−A
sqrt(MAF)
ID

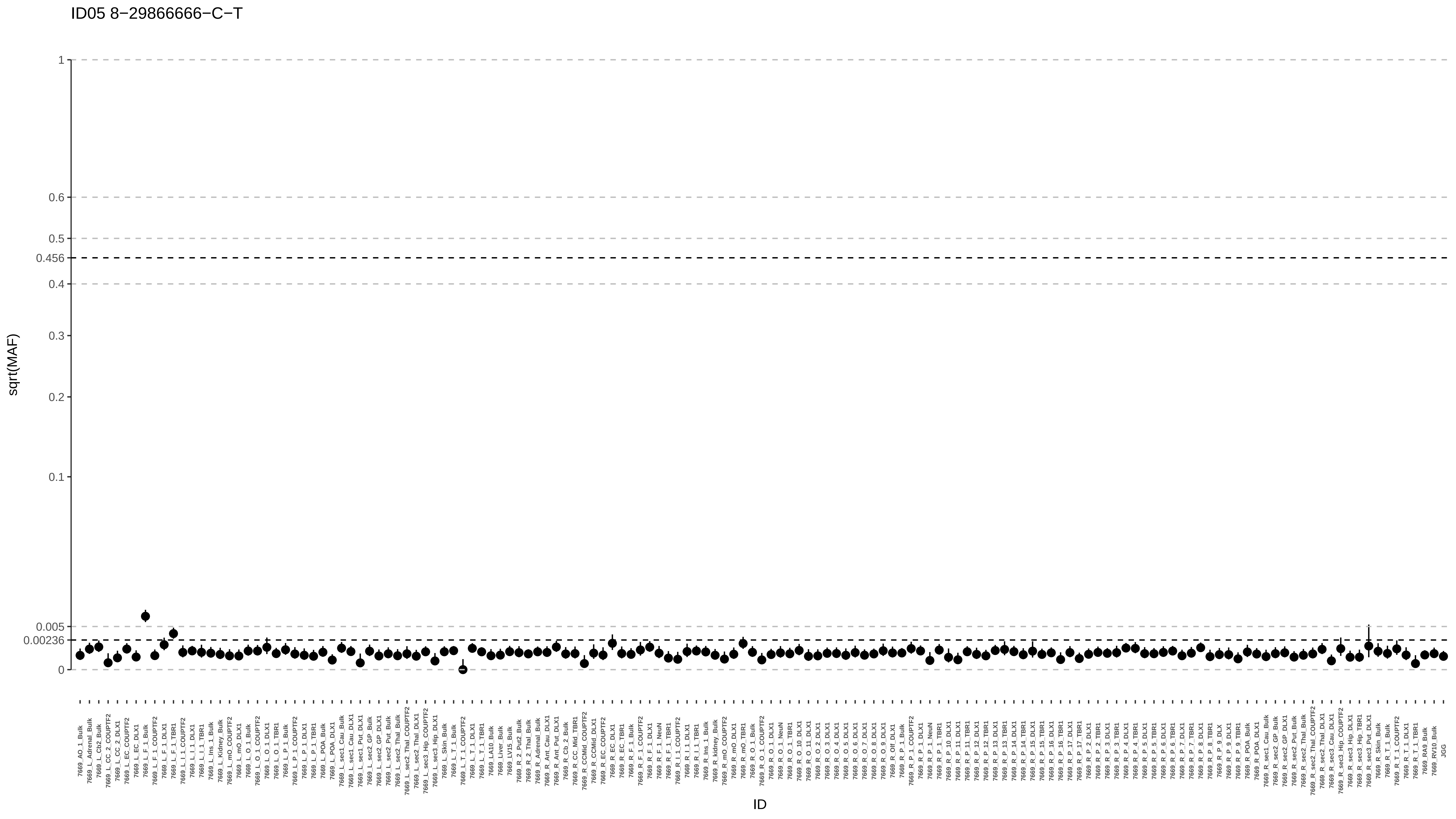

ID05 8−29866666−C−T
sqrt(MAF)
ID

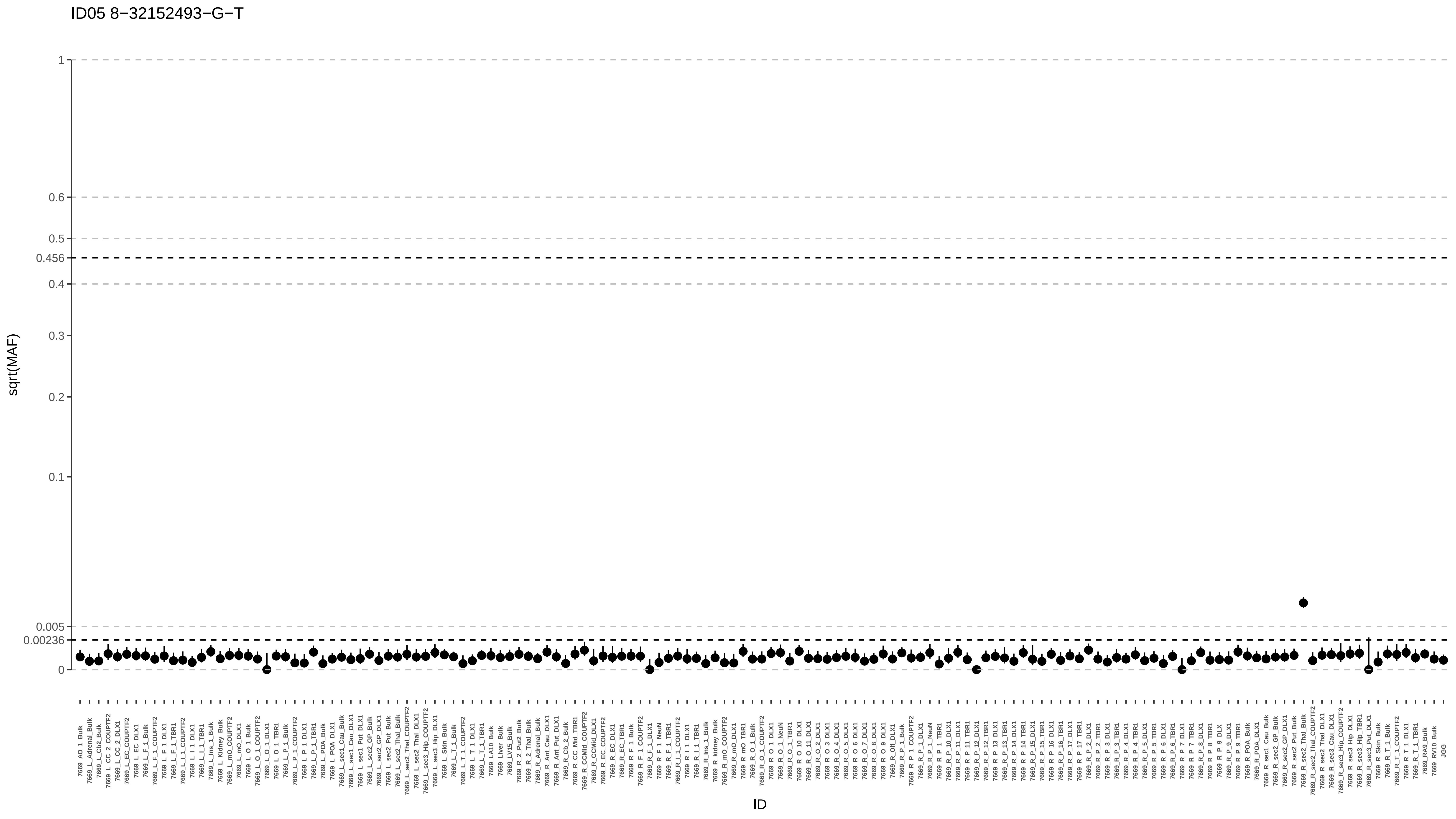
ID05 8−32152493−G−T
sqrt(MAF)
1
0.6
0.5
0.456
0.4
0.3
0.2
0.1
0.005
0.00236
0
ID

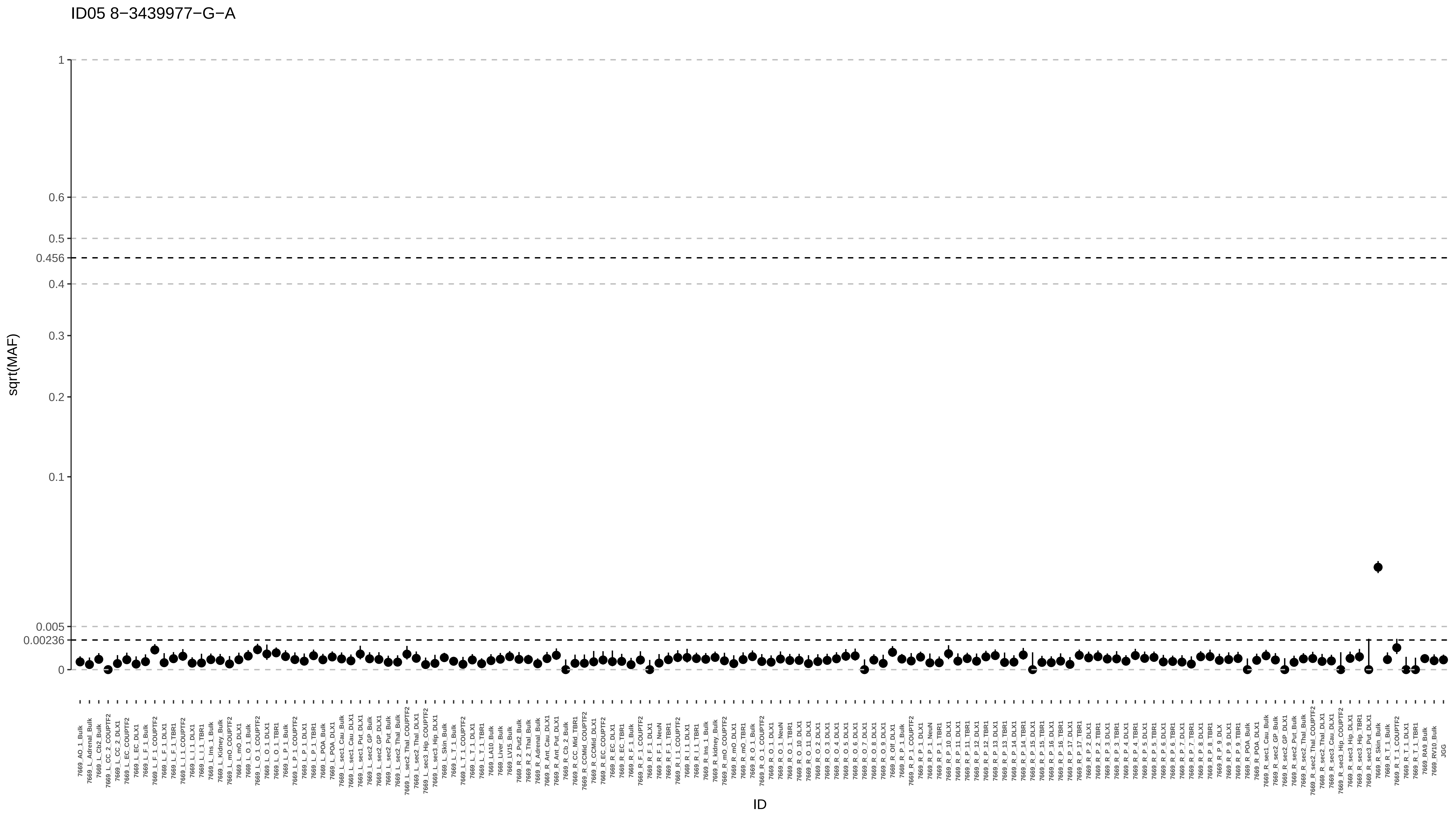
ID05 8−3439977−G−A
sqrt(MAF)
1
0.6
0.5
0.456
0.4
0.3
0.2
0.1
0.005
0.00236
0
ID

ID05 8−35923762−C−T

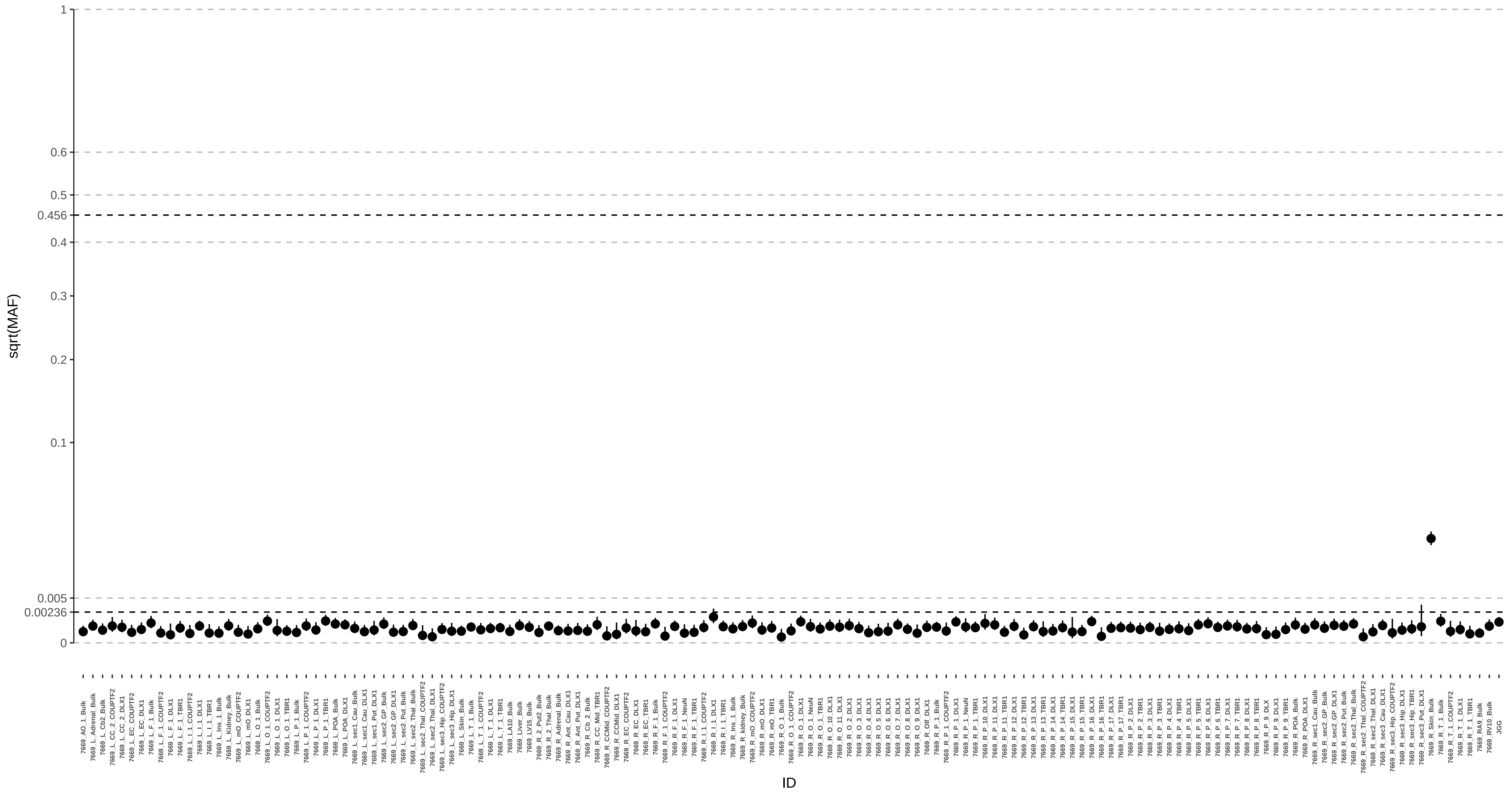

ID05 8−36571735−C−T

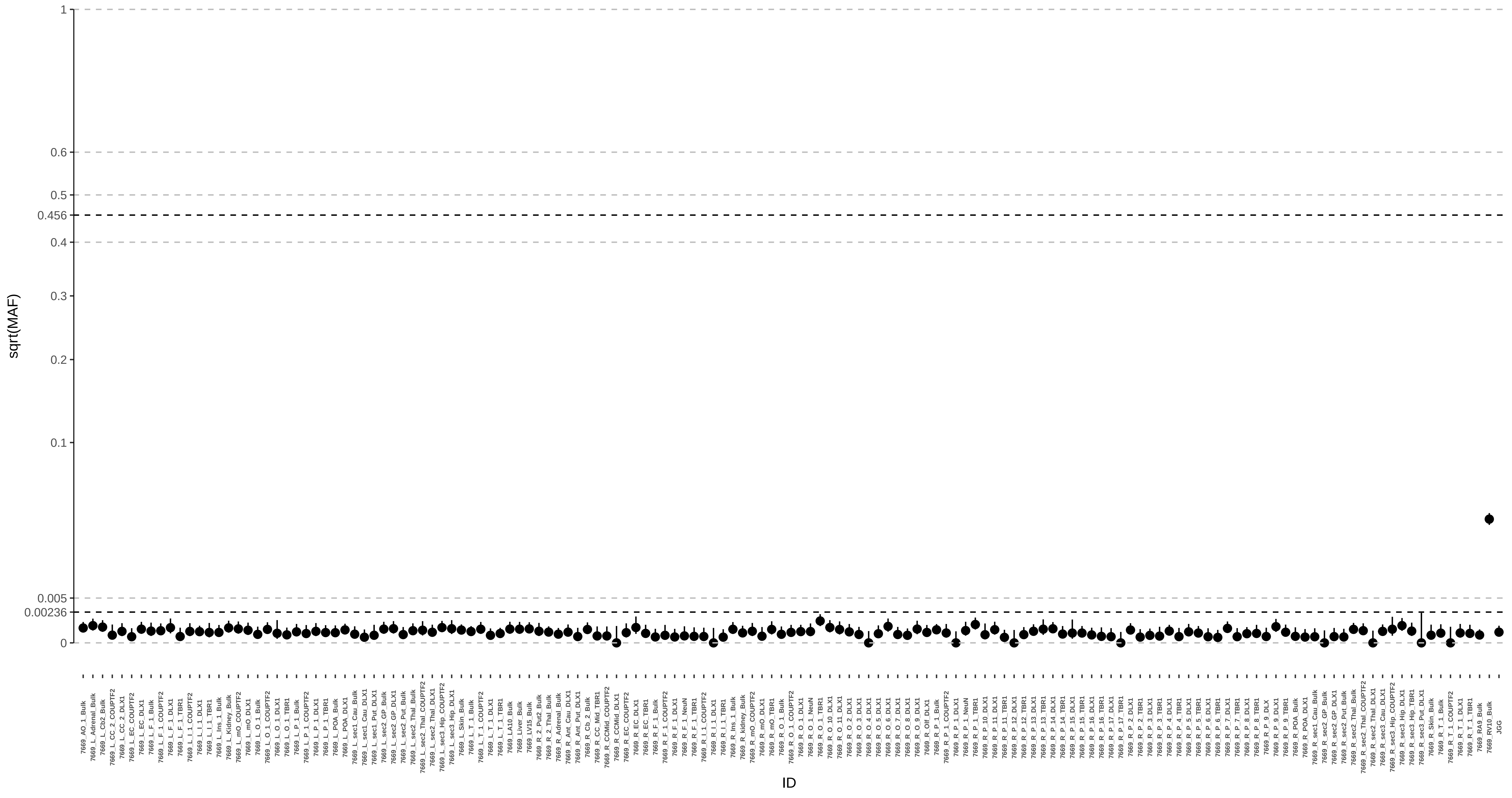

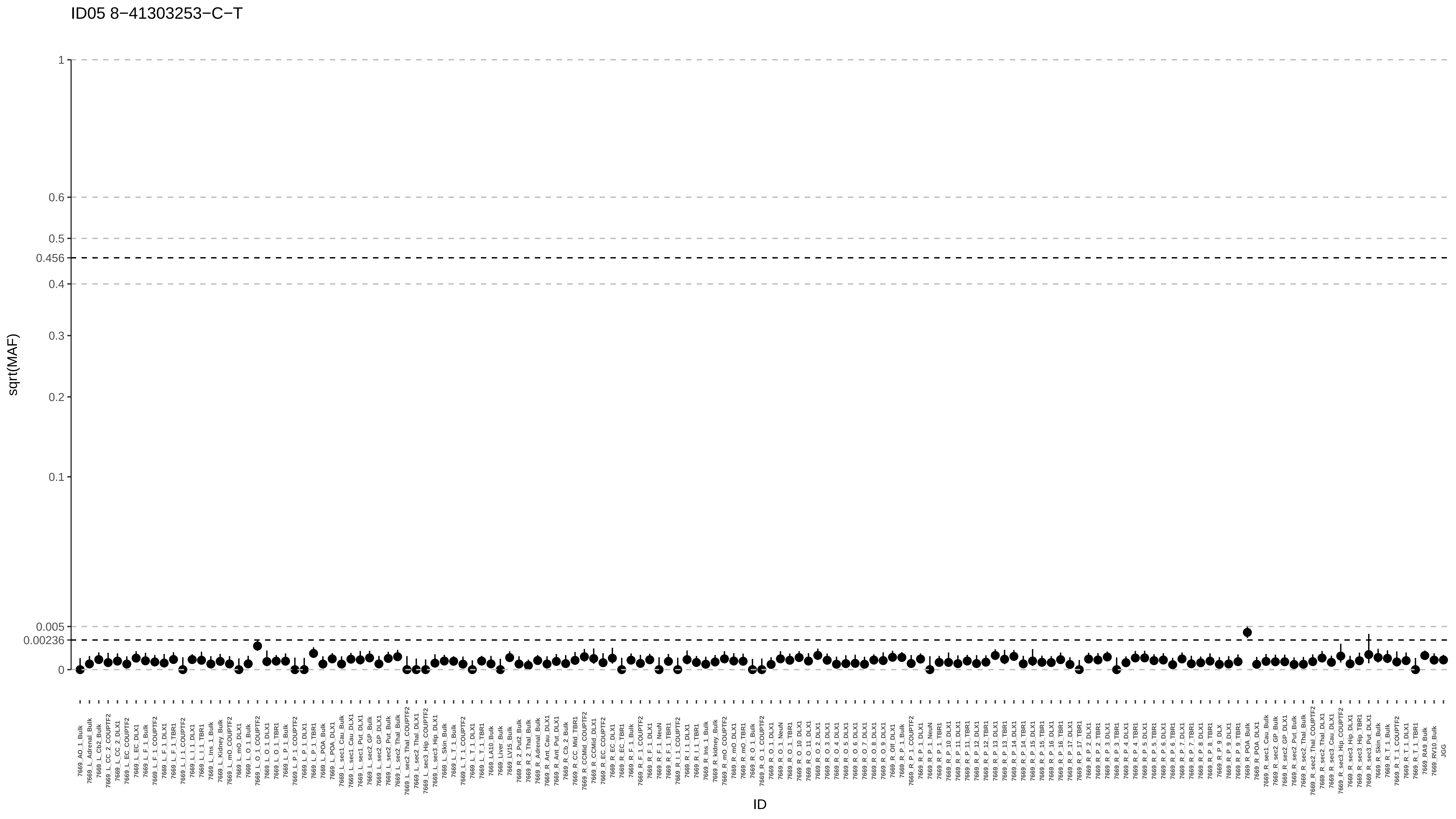
ID05 8−41303253−C−T
sqrt(MAF)
1
0.6
0.5
0.456
0.4
0.3
0.2
0.1
0.005
0.00236
0
ID

ID05 8−4601722−C−G

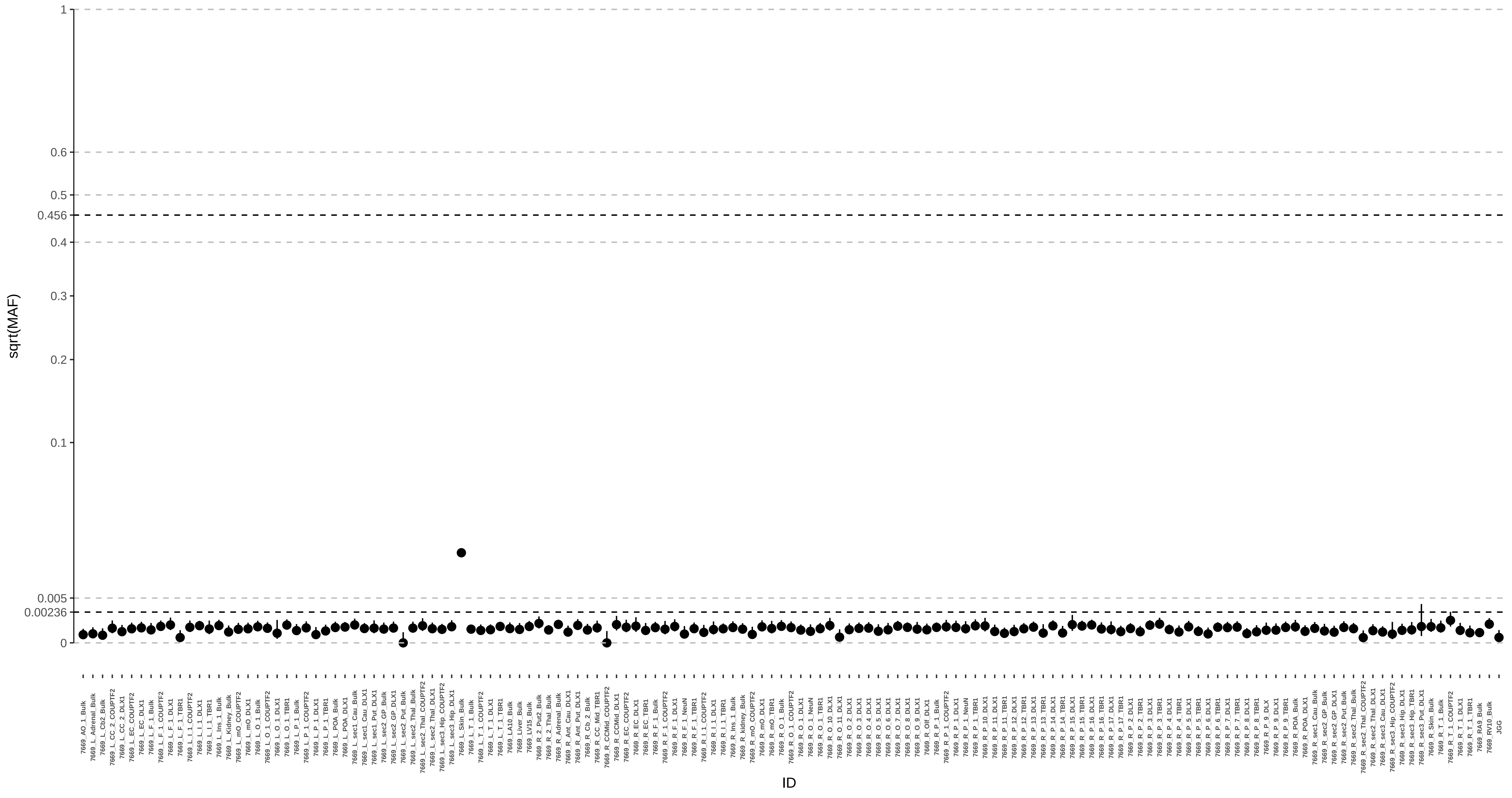

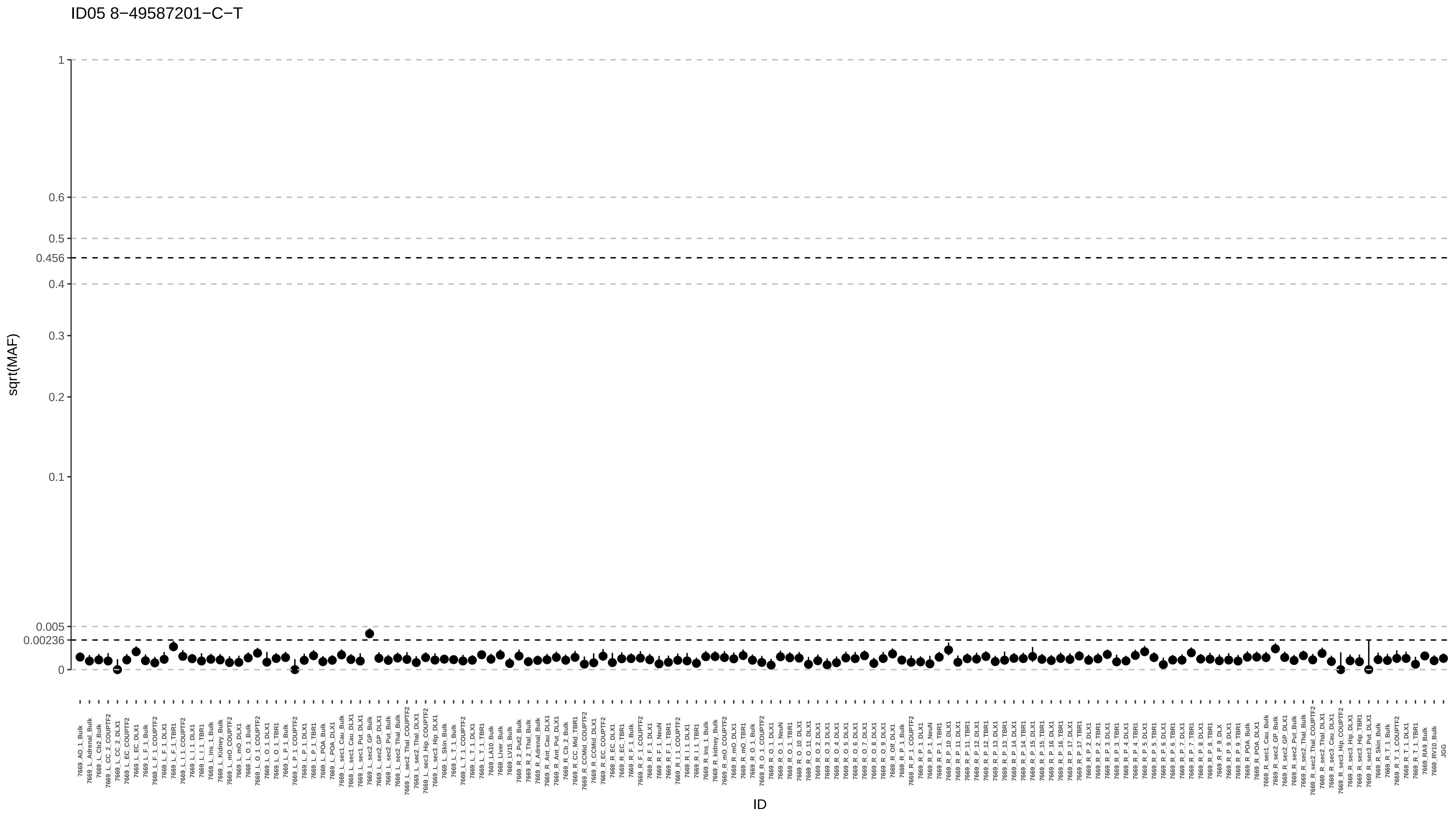
ID05 8−49587201−C−T
sqrt(MAF)
ID
1
0.6
0.5
0.456
0.4
0.3
0.2
0.1
0.005
0.00236
0

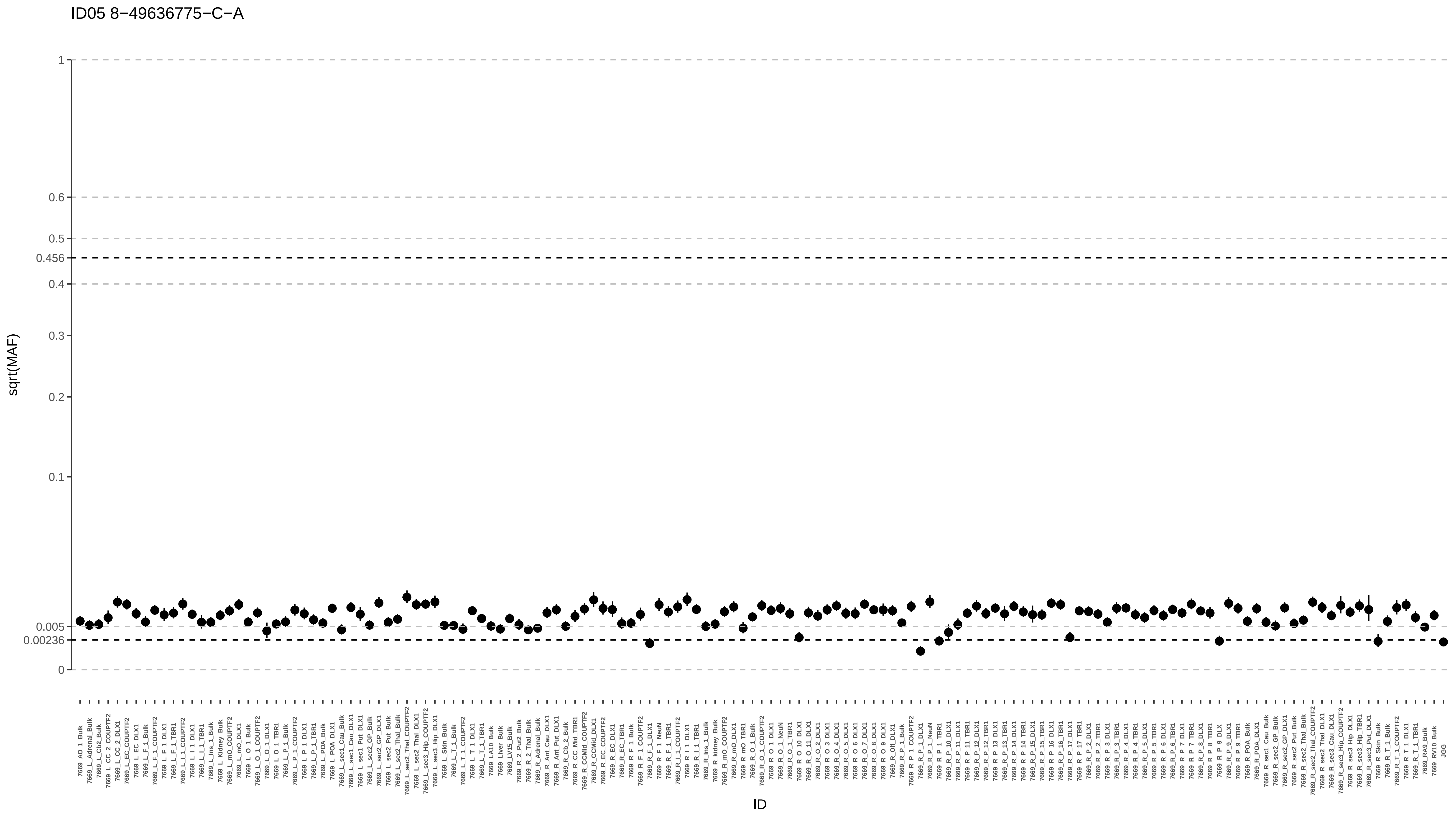
ID05 8−49636775−C−A
sqrt(MAF)
1
0.6
0.5
0.456
0.4
0.3
0.2
0.1
0.005
0.00236
0
ID
7669_AO_1_Bulk
7669_L_Adrenal_Bulk
7669_L_Cb2_Bulk
7669_L_CC_2_COUPTF2
7669_L_CC_2_DLX1
7669_L_EC_COUPTF2
7669_L_EC_DLX1
7669_L_F_1_Bulk
7669_L_F_1_COUPTF2
7669_L_F_1_DLX1
7669_L_F_1_TBR1
7669_L_I_1_COUPTF2
7669_L_I_1_DLX1
7669_L_I_1_TBR1
7669_L_Ins_1_Bulk
7669_L_Kidney_Bulk
7669_L_mO_COUPTF2
7669_L_mO_DLX1
7669_L_O_1_Bulk
7669_L_O_1_COUPTF2
7669_L_O_1_DLX1
7669_L_O_1_TBR1
7669_L_P_1_Bulk
7669_L_P_1_COUPTF2
7669_L_P_1_DLX1
7669_L_P_1_TBR1
7669_L_POA_Bulk
7669_L_POA_DLX1
7669_L_sec1_Cau_Bulk
7669_L_sec1_Cau_DLX1
7669_L_sec1_Put_DLX1
7669_L_sec2_GP_Bulk
7669_L_sec2_GP_DLX1
7669_L_sec2_Put_Bulk
7669_L_sec2_Thal_Bulk
7669_L_sec2_Thal_COUPTF2
7669_L_sec2_Thal_DLX1
7669_L_sec3_Hip_COUPTF2
7669_L_sec3_Hip_DLX1
7669_L_Skin_Bulk
7669_L_T_1_Bulk
7669_L_T_1_COUPTF2
7669_L_T_1_DLX1
7669_L_T_1_TBR1
7669_LA10_Bulk
7669_Liver_Bulk
7669_LV15_Bulk
7669_R_2_Put2_Bulk
7669_R_2_Thal_Bulk
7669_R_Adrenal_Bulk
7669_R_Ant_Cau_DLX1
7669_R_Ant_Put_DLX1
7669_R_Cb_2_Bulk
7669_R_CC_Mid_TBR1
7669_R_CCMid_COUPTF2
7669_R_CCMid_DLX1
7669_R_EC_COUPTF2
7669_R_EC_DLX1
7669_R_EC_TBR1
7669_R_F_1_Bulk
7669_R_F_1_COUPTF2
7669_R_F_1_DLX1
7669_R_F_1_NeuN
7669_R_F_1_TBR1
7669_R_I_1_COUPTF2
7669_R_I_1_DLX1
7669_R_I_1_TBR1
7669_R_Ins_1_Bulk
7669_R_kidney_Bulk
7669_R_mO_COUPTF2
7669_R_mO_DLX1
7669_R_mO_TBR1
7669_R_O_1_Bulk
7669_R_O_1_COUPTF2
7669_R_O_1_DLX1
7669_R_O_1_NeuN
7669_R_O_1_TBR1
7669_R_O_10_DLX1
7669_R_O_11_DLX1
7669_R_O_2_DLX1
7669_R_O_3_DLX1
7669_R_O_4_DLX1
7669_R_O_5_DLX1
7669_R_O_6_DLX1
7669_R_O_7_DLX1
7669_R_O_8_DLX1
7669_R_O_9_DLX1
7669_R_Olf_DLX1
7669_R_P_1_Bulk
7669_R_P_1_COUPTF2
7669_R_P_1_DLX1
7669_R_P_1_NeuN
7669_R_P_1_TBR1
7669_R_P_10_DLX1
7669_R_P_11_DLX1
7669_R_P_11_TBR1
7669_R_P_12_DLX1
7669_R_P_12_TBR1
7669_R_P_13_DLX1
7669_R_P_13_TBR1
7669_R_P_14_DLX1
7669_R_P_14_TBR1
7669_R_P_15_DLX1
7669_R_P_15_TBR1
7669_R_P_16_DLX1
7669_R_P_16_TBR1
7669_R_P_17_DLX1
7669_R_P_17_TBR1
7669_R_P_2_DLX1
7669_R_P_2_TBR1
7669_R_P_3_DLX1
7669_R_P_3_TBR1
7669_R_P_4_DLX1
7669_R_P_4_TBR1
7669_R_P_5_DLX1
7669_R_P_5_TBR1
7669_R_P_6_DLX1
7669_R_P_6_TBR1
7669_R_P_7_DLX1
7669_R_P_7_TBR1
7669_R_P_8_DLX1
7669_R_P_8_TBR1
7669_R_P_9_DLX
7669_R_P_9_DLX1
7669_R_P_9_TBR1
7669_R_POA_Bulk
7669_R_POA_DLX1
7669_R_sec1_Cau_Bulk
7669_R_sec2_GP_Bulk
7669_R_sec2_GP_DLX1
7669_R_sec2_Put_Bulk
7669_R_sec2_Thal_Bulk
7669_R_sec2_Thal_COUPTF2
7669_R_sec2_Thal_DLX1
7669_R_sec3_Cau_DLX1
7669_R_sec3_Hip_COUPTF2
7669_R_sec3_Hip_DLX1
7669_R_sec3_Hip_TBR1
7669_R_sec3_Put_DLX1
7669_R_Skin_Bulk
7669_R_T_1_Bulk
7669_R_T_1_COUPTF2
7669_R_T_1_DLX1
7669_R_T_1_TBR1
7669_RA9_Bulk
7669_RV10_Bulk
JGG

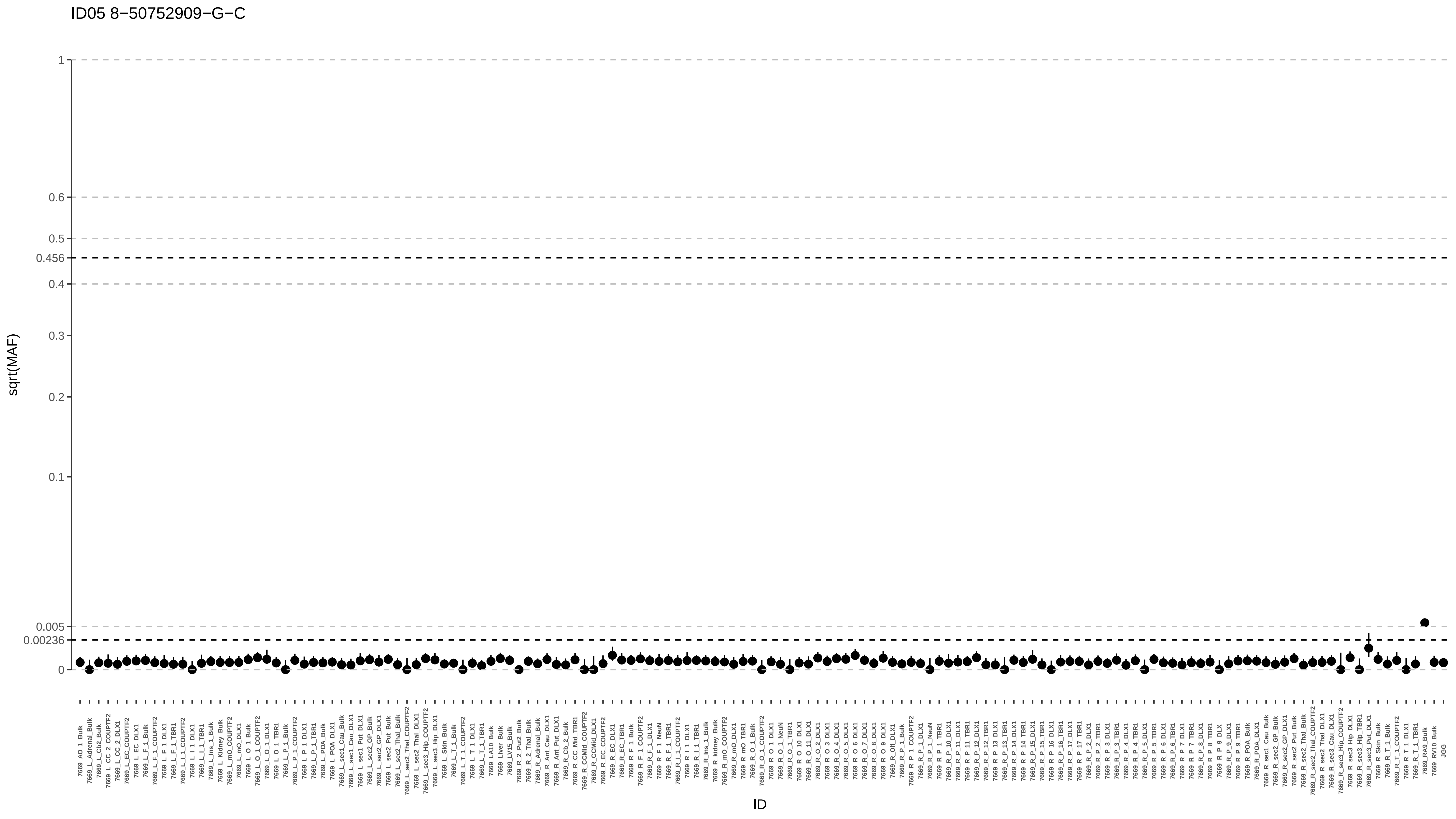
ID05 8−50752909−G−C
sqrt(MAF)
1
0.6
0.5
0.456
0.4
0.3
0.2
0.1
0.005
0.00236
0
ID

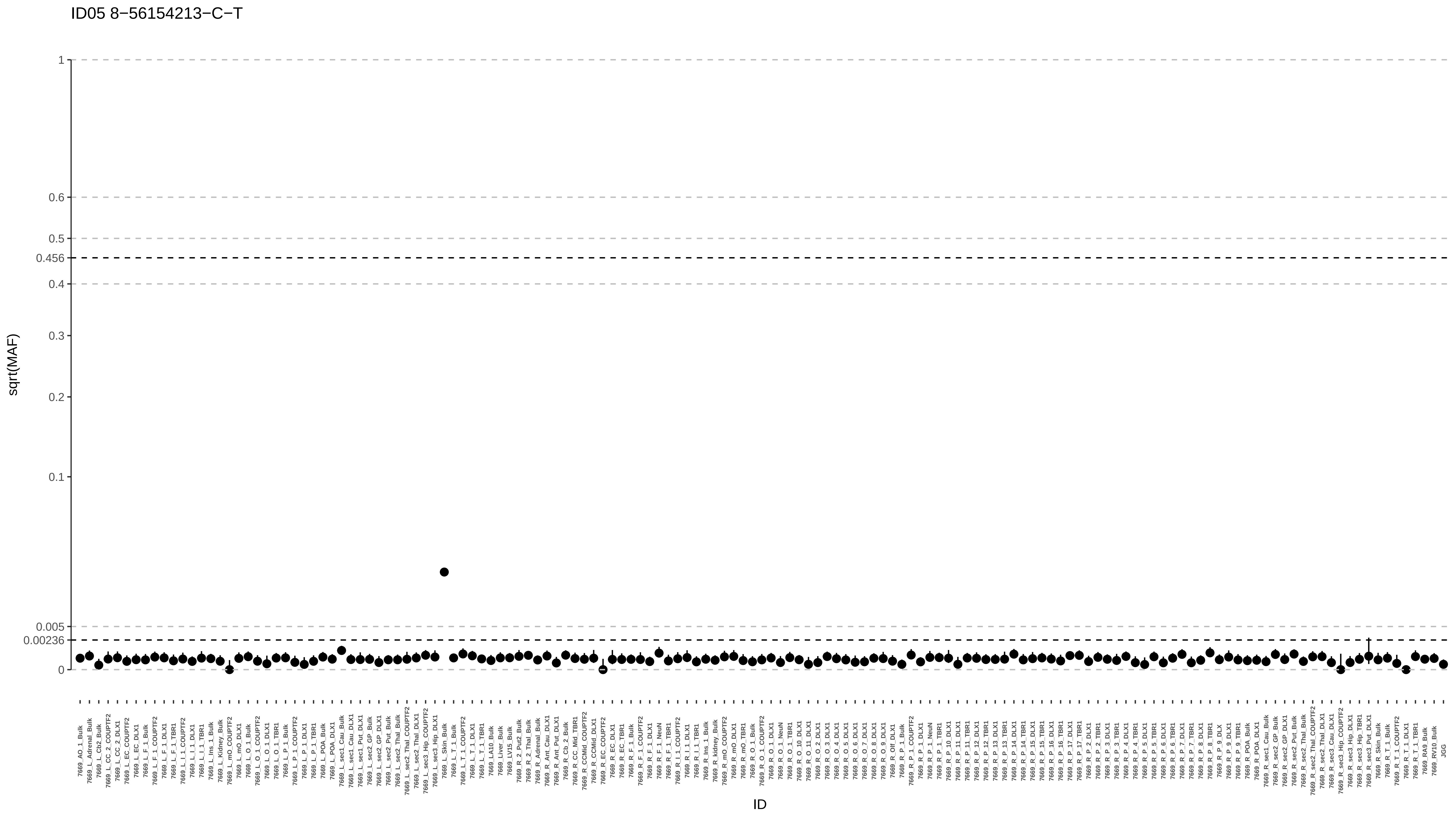
ID05 8−56154213−C−T
sqrt(MAF)
1
0.6
0.5
0.456
0.4
0.3
0.2
0.1
0.005
0.00236
0
ID
7669_AO_1_Bulk
7669_L_Adrenal_Bulk
7669_L_Cb2_Bulk
7669_L_CC_2_COUPTF2
7669_L_CC_2_DLX1
7669_L_EC_COUPTF2
7669_L_EC_DLX1
7669_L_F_1_Bulk
7669_L_F_1_COUPTF2
7669_L_F_1_DLX1
7669_L_F_1_TBR1
7669_L_I_1_COUPTF2
7669_L_I_1_DLX1
7669_L_I_1_TBR1
7669_L_Ins_1_Bulk
7669_L_Kidney_Bulk
7669_L_mO_COUPTF2
7669_L_mO_DLX1
7669_L_O_1_Bulk
7669_L_O_1_COUPTF2
7669_L_O_1_DLX1
7669_L_O_1_TBR1
7669_L_P_1_Bulk
7669_L_P_1_COUPTF2
7669_L_P_1_DLX1
7669_L_P_1_TBR1
7669_L_POA_Bulk
7669_L_POA_DLX1
7669_L_sec1_Cau_Bulk
7669_L_sec1_Cau_DLX1
7669_L_sec1_Put_DLX1
7669_L_sec2_GP_Bulk
7669_L_sec2_GP_DLX1
7669_L_sec2_Put_Bulk
7669_L_sec2_Thal_Bulk
7669_L_sec2_Thal_COUPTF2
7669_L_sec2_Thal_DLX1
7669_L_sec3_Hip_COUPTF2
7669_L_sec3_Hip_DLX1
7669_L_Skin_Bulk
7669_L_T_1_Bulk
7669_L_T_1_COUPTF2
7669_L_T_1_DLX1
7669_L_T_1_TBR1
7669_LA10_Bulk
7669_Liver_Bulk
7669_LV15_Bulk
7669_R_2_Put2_Bulk
7669_R_2_Thal_Bulk
7669_R_Adrenal_Bulk
7669_R_Ant_Cau_DLX1
7669_R_Ant_Put_DLX1
7669_R_Cb_2_Bulk
7669_R_CC_Mid_TBR1
7669_R_CCMid_COUPTF2
7669_R_CCMid_DLX1
7669_R_EC_COUPTF2
7669_R_EC_DLX1
7669_R_EC_TBR1
7669_R_F_1_Bulk
7669_R_F_1_COUPTF2
7669_R_F_1_DLX1
7669_R_F_1_NeuN
7669_R_F_1_TBR1
7669_R_I_1_COUPTF2
7669_R_I_1_DLX1
7669_R_I_1_TBR1
7669_R_Ins_1_Bulk
7669_R_kidney_Bulk
7669_R_mO_COUPTF2
7669_R_mO_DLX1
7669_R_mO_TBR1
7669_R_O_1_Bulk
7669_R_O_1_COUPTF2
7669_R_O_1_DLX1
7669_R_O_1_NeuN
7669_R_O_1_TBR1
7669_R_O_10_DLX1
7669_R_O_11_DLX1
7669_R_O_2_DLX1
7669_R_O_3_DLX1
7669_R_O_4_DLX1
7669_R_O_5_DLX1
7669_R_O_6_DLX1
7669_R_O_7_DLX1
7669_R_O_8_DLX1
7669_R_O_9_DLX1
7669_R_Olf_DLX1
7669_R_P_1_Bulk
7669_R_P_1_COUPTF2
7669_R_P_1_DLX1
7669_R_P_1_NeuN
7669_R_P_1_TBR1
7669_R_P_10_DLX1
7669_R_P_11_DLX1
7669_R_P_11_TBR1
7669_R_P_12_DLX1
7669_R_P_12_TBR1
7669_R_P_13_DLX1
7669_R_P_13_TBR1
7669_R_P_14_DLX1
7669_R_P_14_TBR1
7669_R_P_15_DLX1
7669_R_P_15_TBR1
7669_R_P_16_DLX1
7669_R_P_16_TBR1
7669_R_P_17_DLX1
7669_R_P_17_TBR1
7669_R_P_2_DLX1
7669_R_P_2_TBR1
7669_R_P_3_DLX1
7669_R_P_3_TBR1
7669_R_P_4_DLX1
7669_R_P_4_TBR1
7669_R_P_5_DLX1
7669_R_P_5_TBR1
7669_R_P_6_DLX1
7669_R_P_6_TBR1
7669_R_P_7_DLX1
7669_R_P_7_TBR1
7669_R_P_8_DLX1
7669_R_P_8_TBR1
7669_R_P_9_DLX
7669_R_P_9_DLX1
7669_R_P_9_TBR1
7669_R_POA_Bulk
7669_R_POA_DLX1
7669_R_sec1_Cau_Bulk
7669_R_sec2_GP_Bulk
7669_R_sec2_GP_DLX1
7669_R_sec2_Put_Bulk
7669_R_sec2_Thal_Bulk
7669_R_sec2_Thal_COUPTF2
7669_R_sec2_Thal_DLX1
7669_R_sec3_Cau_DLX1
7669_R_sec3_Hip_COUPTF2
7669_R_sec3_Hip_DLX1
7669_R_sec3_Hip_TBR1
7669_R_sec3_Put_DLX1
7669_R_Skin_Bulk
7669_R_T_1_Bulk
7669_R_T_1_COUPTF2
7669_R_T_1_DLX1
7669_R_T_1_TBR1
7669_RA9_Bulk
7669_RV10_Bulk
JGG

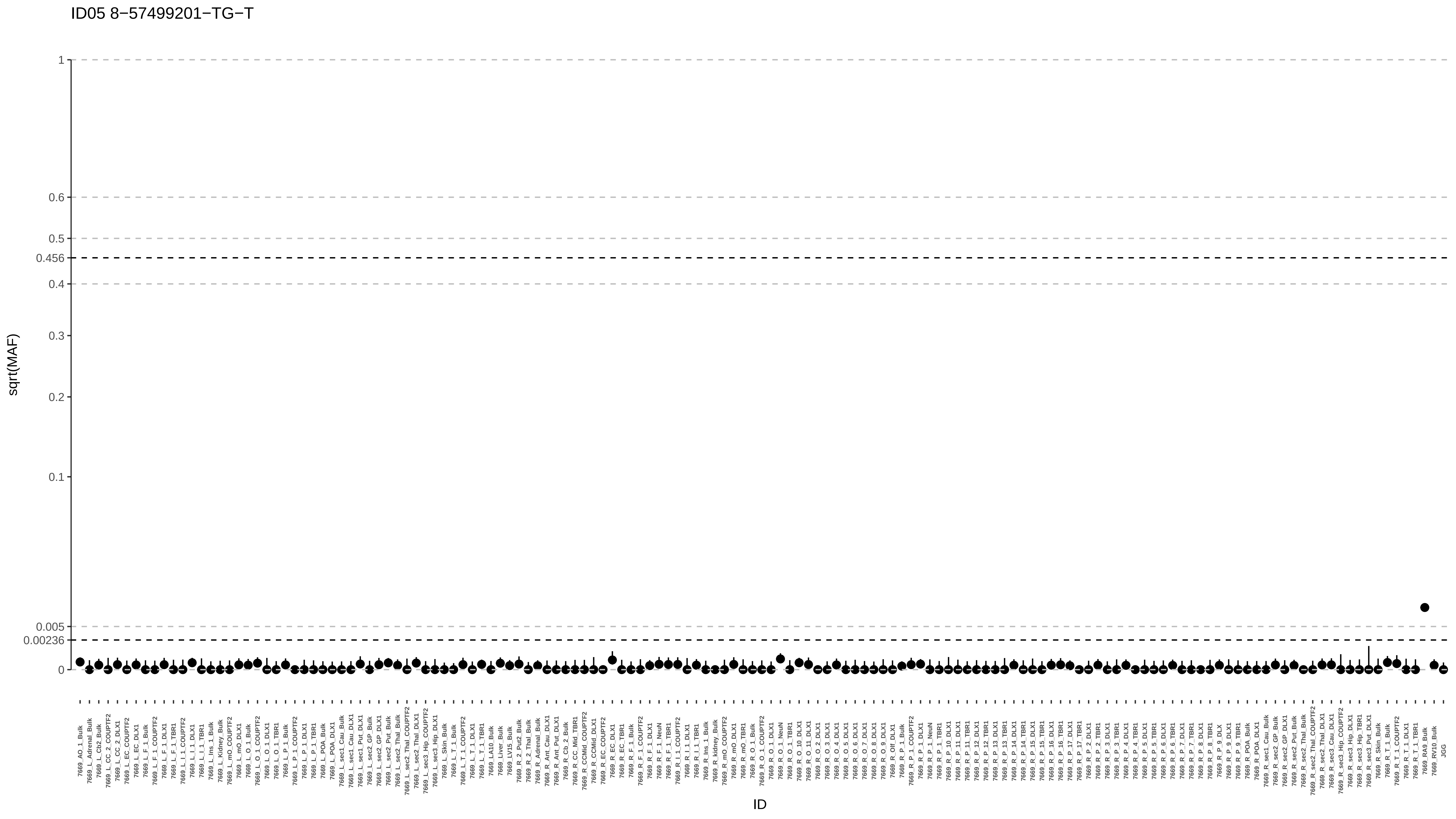
ID05 8−57499201−TG−T
sqrt(MAF)
ID
1
0.6
0.5
0.456
0.4
0.3
0.2
0.1
0.005
0.00236
0

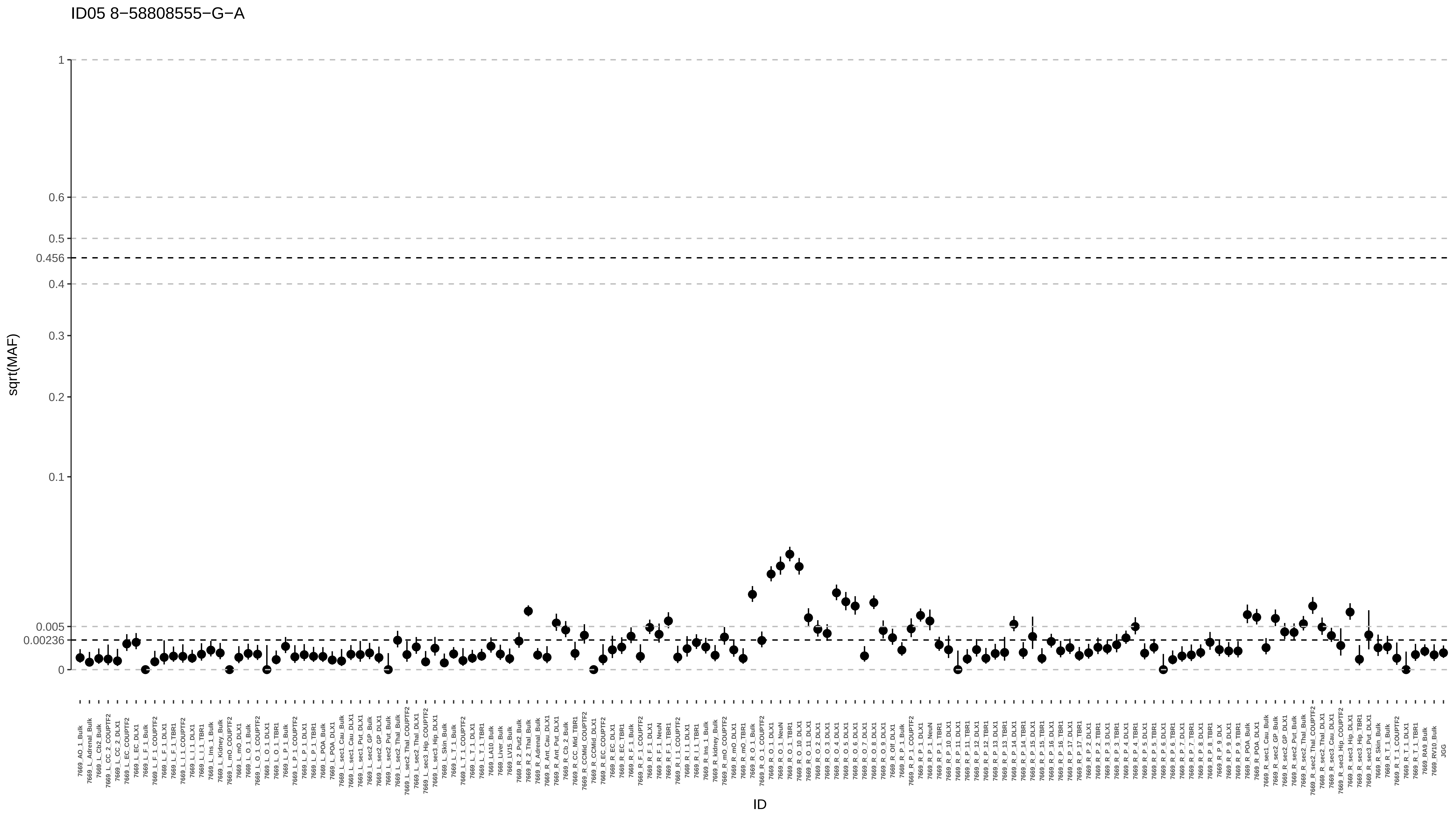
ID05 8−58808555−G−A
sqrt(MAF)
1
0.6
0.5
0.456
0.4
0.3
0.2
0.1
0.005
0.00236
0
ID
7669_AO_1_Bulk
7669_L_Adrenal_Bulk
7669_L_Cb2_Bulk
7669_L_CC_2_COUPTF2
7669_L_CC_2_DLX1
7669_L_EC_COUPTF2
7669_L_EC_DLX1
7669_L_F_1_Bulk
7669_L_F_1_COUPTF2
7669_L_F_1_DLX1
7669_L_F_1_TBR1
7669_L_I_1_COUPTF2
7669_L_I_1_DLX1
7669_L_I_1_TBR1
7669_L_Ins_1_Bulk
7669_L_Kidney_Bulk
7669_L_mO_COUPTF2
7669_L_mO_DLX1
7669_L_O_1_Bulk
7669_L_O_1_COUPTF2
7669_L_O_1_DLX1
7669_L_O_1_TBR1
7669_L_P_1_Bulk
7669_L_P_1_COUPTF2
7669_L_P_1_DLX1
7669_L_P_1_TBR1
7669_L_POA_Bulk
7669_L_POA_DLX1
7669_L_sec1_Cau_Bulk
7669_L_sec1_Cau_DLX1
7669_L_sec1_Put_DLX1
7669_L_sec2_GP_Bulk
7669_L_sec2_GP_DLX1
7669_L_sec2_Put_Bulk
7669_L_sec2_Thal_Bulk
7669_L_sec2_Thal_COUPTF2
7669_L_sec2_Thal_DLX1
7669_L_sec3_Hip_COUPTF2
7669_L_sec3_Hip_DLX1
7669_L_Skin_Bulk
7669_L_T_1_Bulk
7669_L_T_1_COUPTF2
7669_L_T_1_DLX1
7669_L_T_1_TBR1
7669_LA10_Bulk
7669_Liver_Bulk
7669_LV15_Bulk
7669_R_2_Put2_Bulk
7669_R_2_Thal_Bulk
7669_R_Adrenal_Bulk
7669_R_Ant_Cau_DLX1
7669_R_Ant_Put_DLX1
7669_R_Cb_2_Bulk
7669_R_CC_Mid_TBR1
7669_R_CCMid_COUPTF2
7669_R_CCMid_DLX1
7669_R_EC_COUPTF2
7669_R_EC_DLX1
7669_R_EC_TBR1
7669_R_F_1_Bulk
7669_R_F_1_COUPTF2
7669_R_F_1_DLX1
7669_R_F_1_NeuN
7669_R_F_1_TBR1
7669_R_I_1_COUPTF2
7669_R_I_1_DLX1
7669_R_I_1_TBR1
7669_R_Ins_1_Bulk
7669_R_kidney_Bulk
7669_R_mO_COUPTF2
7669_R_mO_DLX1
7669_R_mO_TBR1
7669_R_O_1_Bulk
7669_R_O_1_COUPTF2
7669_R_O_1_DLX1
7669_R_O_1_NeuN
7669_R_O_1_TBR1
7669_R_O_10_DLX1
7669_R_O_11_DLX1
7669_R_O_2_DLX1
7669_R_O_3_DLX1
7669_R_O_4_DLX1
7669_R_O_5_DLX1
7669_R_O_6_DLX1
7669_R_O_7_DLX1
7669_R_O_8_DLX1
7669_R_O_9_DLX1
7669_R_Olf_DLX1
7669_R_P_1_Bulk
7669_R_P_1_COUPTF2
7669_R_P_1_DLX1
7669_R_P_1_NeuN
7669_R_P_1_TBR1
7669_R_P_10_DLX1
7669_R_P_11_DLX1
7669_R_P_11_TBR1
7669_R_P_12_DLX1
7669_R_P_12_TBR1
7669_R_P_13_DLX1
7669_R_P_13_TBR1
7669_R_P_14_DLX1
7669_R_P_14_TBR1
7669_R_P_15_DLX1
7669_R_P_15_TBR1
7669_R_P_16_DLX1
7669_R_P_16_TBR1
7669_R_P_17_DLX1
7669_R_P_17_TBR1
7669_R_P_2_DLX1
7669_R_P_2_TBR1
7669_R_P_3_DLX1
7669_R_P_3_TBR1
7669_R_P_4_DLX1
7669_R_P_4_TBR1
7669_R_P_5_DLX1
7669_R_P_5_TBR1
7669_R_P_6_DLX1
7669_R_P_6_TBR1
7669_R_P_7_DLX1
7669_R_P_7_TBR1
7669_R_P_8_DLX1
7669_R_P_8_TBR1
7669_R_P_9_DLX
7669_R_P_9_DLX1
7669_R_P_9_TBR1
7669_R_POA_Bulk
7669_R_POA_DLX1
7669_R_sec1_Cau_Bulk
7669_R_sec2_GP_Bulk
7669_R_sec2_GP_DLX1
7669_R_sec2_Put_Bulk
7669_R_sec2_Thal_Bulk
7669_R_sec2_Thal_COUPTF2
7669_R_sec2_Thal_DLX1
7669_R_sec3_Cau_DLX1
7669_R_sec3_Hip_COUPTF2
7669_R_sec3_Hip_DLX1
7669_R_sec3_Hip_TBR1
7669_R_sec3_Put_DLX1
7669_R_Skin_Bulk
7669_R_T_1_Bulk
7669_R_T_1_COUPTF2
7669_R_T_1_DLX1
7669_R_T_1_TBR1
7669_RA9_Bulk
7669_RV10_Bulk
JGG

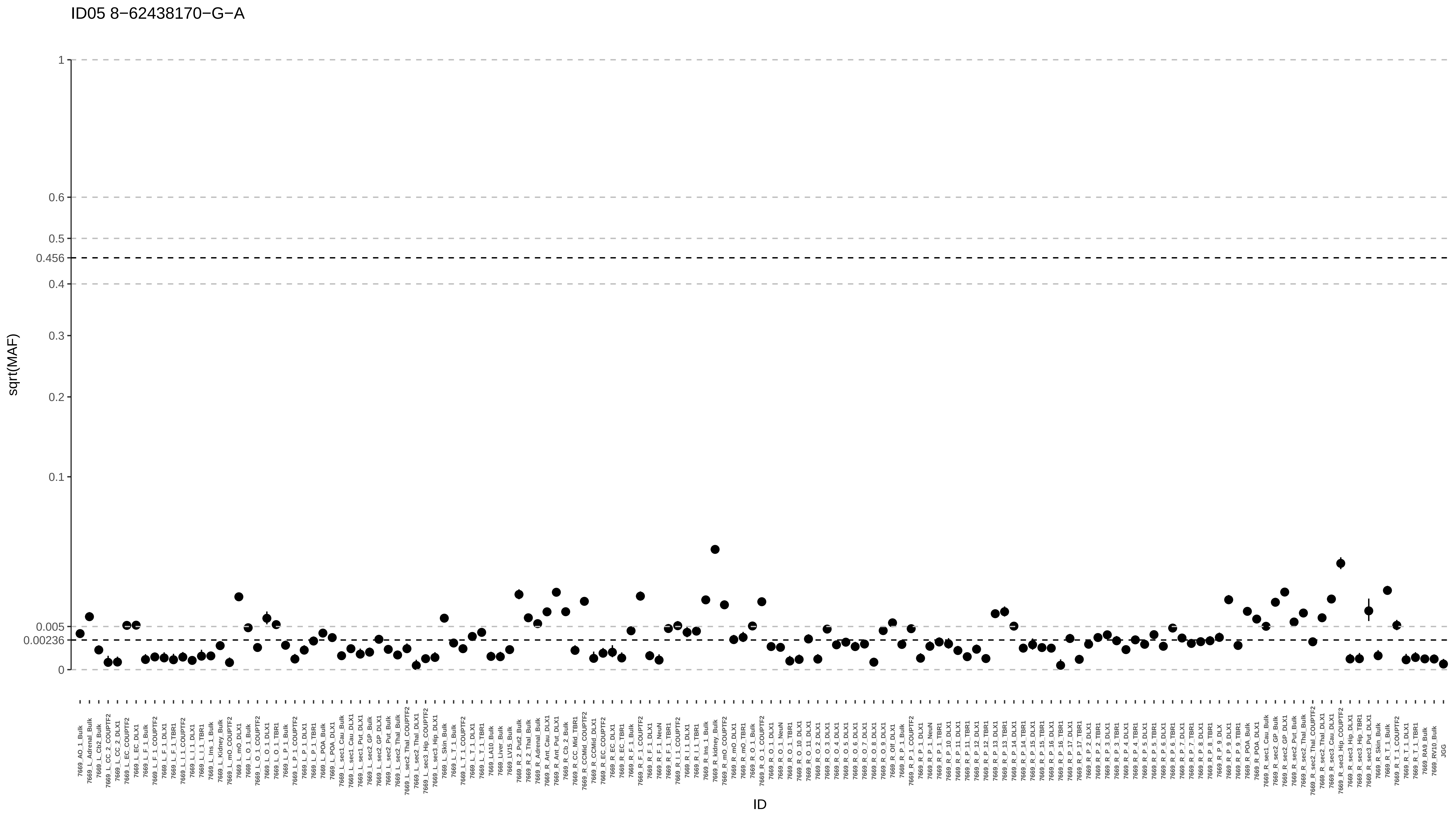
ID05 8−62438170−G−A
sqrt(MAF)
1
0.6
0.5
0.456
0.4
0.3
0.2
0.1
0.005
0.00236
0
ID

ID05 8−65149924−C−T

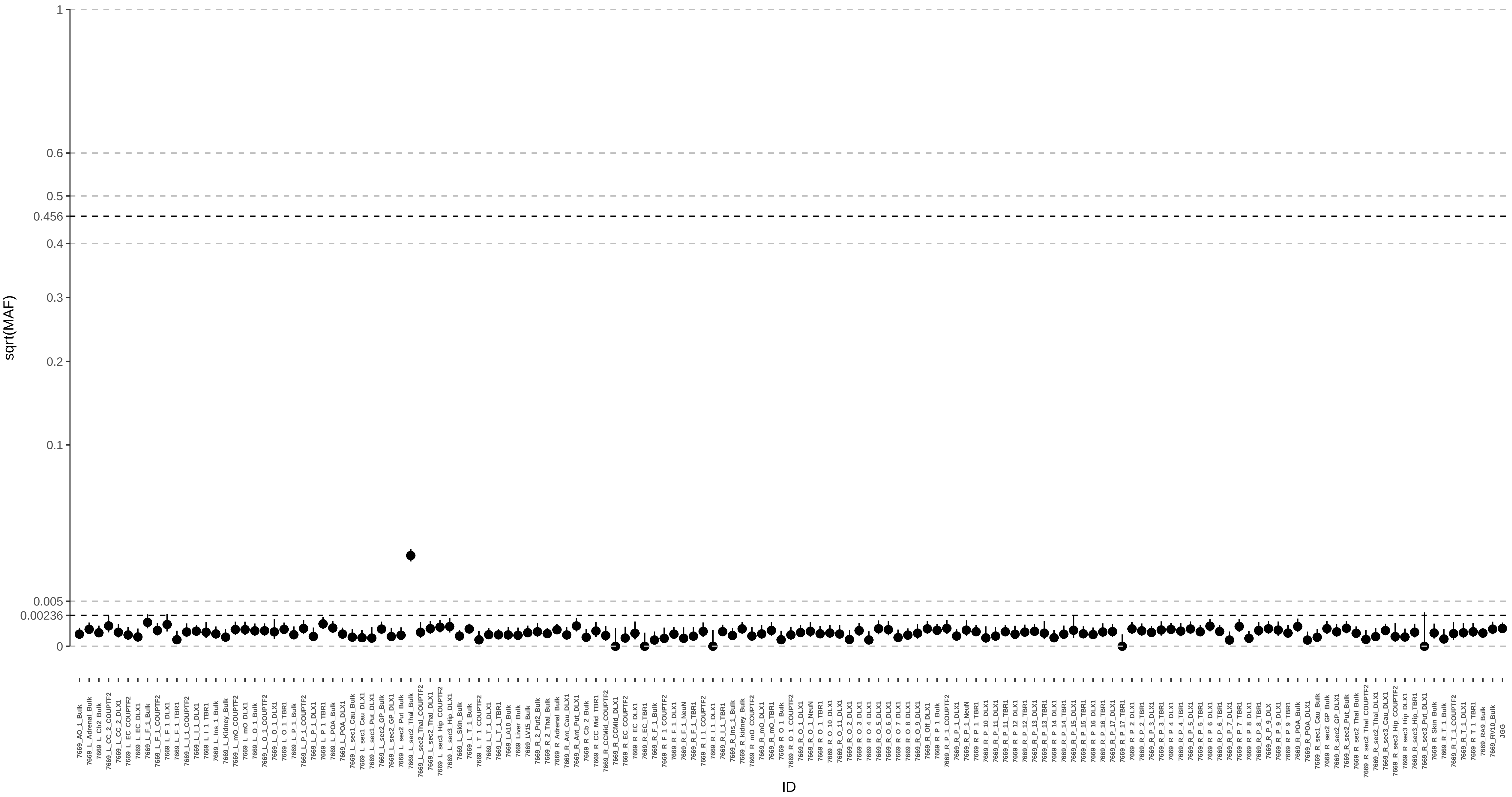

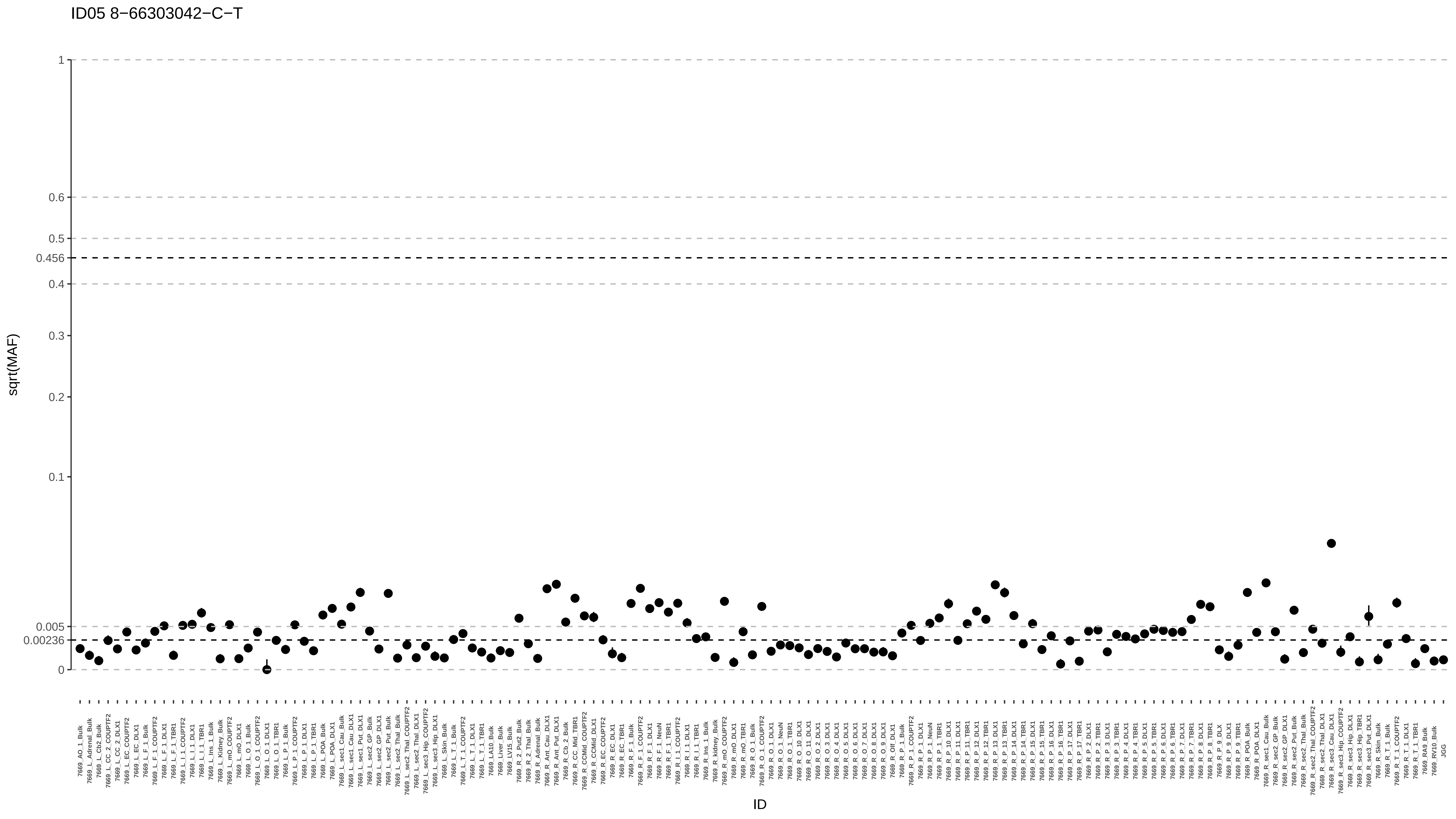
ID05 8−66303042−C−T
sqrt(MAF)
1
0.6
0.5
0.456
0.4
0.3
0.2
0.1
0.005
0.00236
0
ID

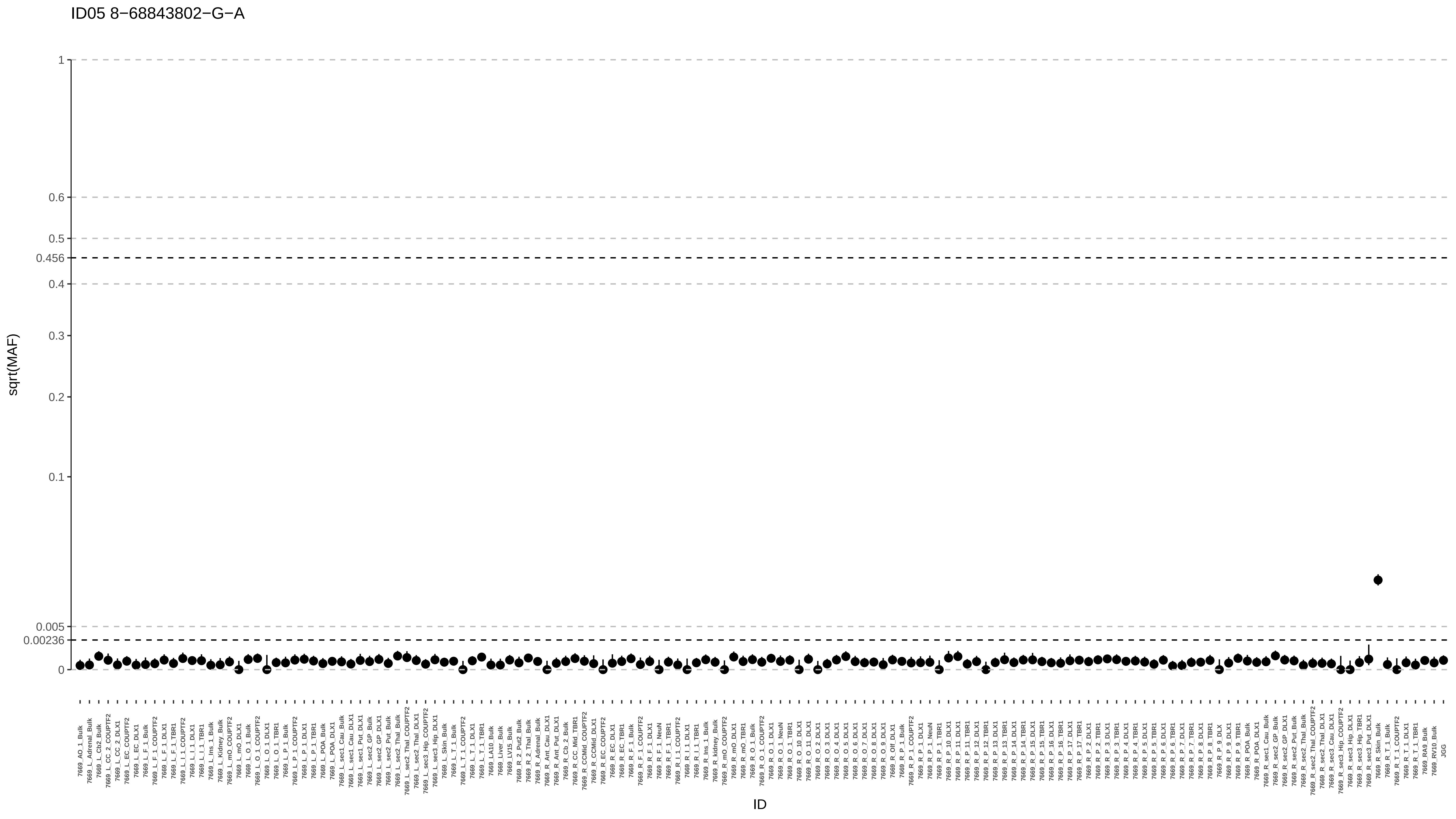
ID05 8−68843802−G−A
sqrt(MAF)
1
0.6
0.5
0.456
0.4
0.3
0.2
0.1
0.005
0.00236
0
ID

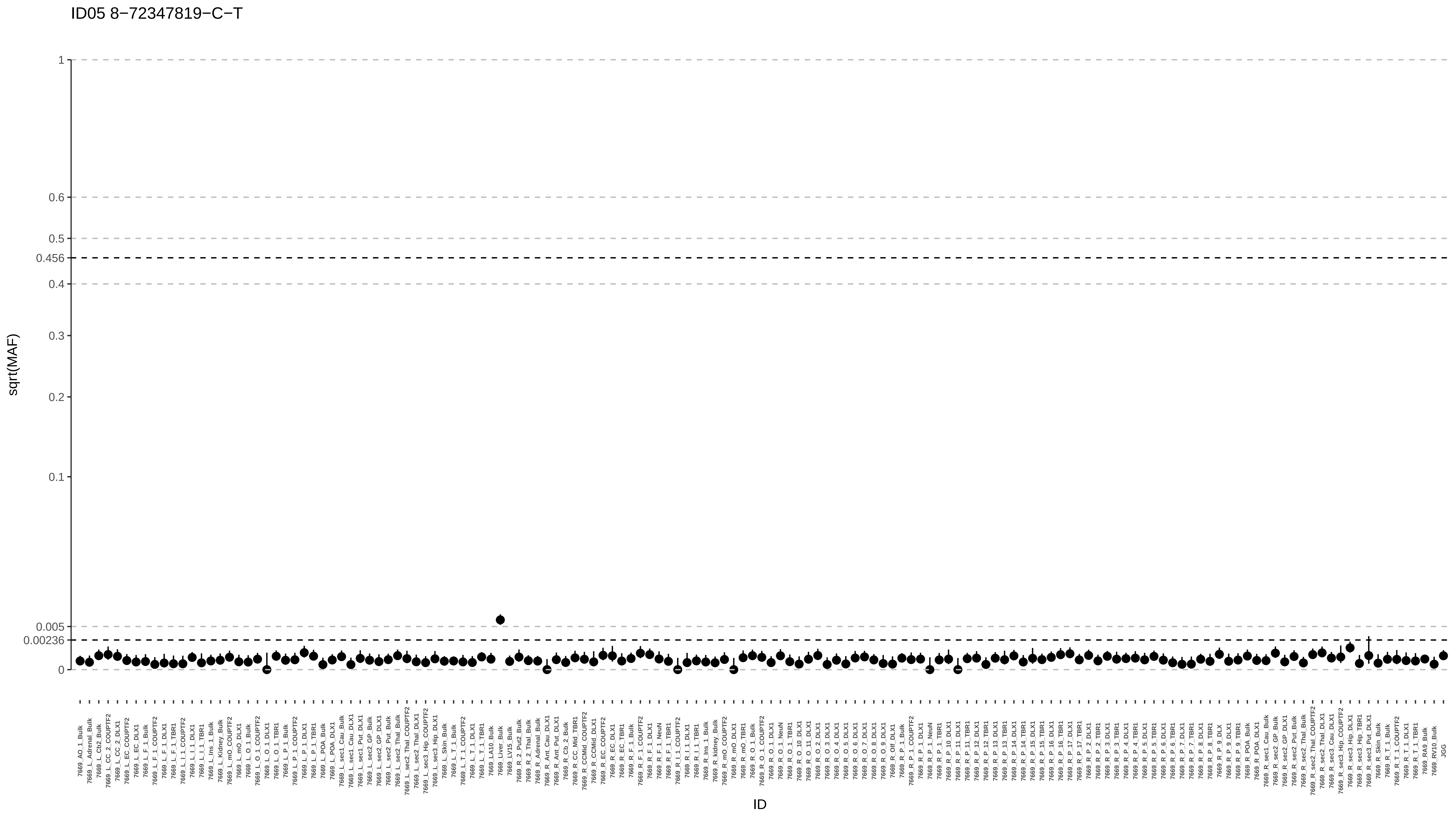

ID05 8−72347819−C−T
sqrt(MAF)
1
0.6
0.5
0.456
0.4
0.3
0.2
0.1
0.005
0.00236
0
ID

ID05 8−73048428−C−T

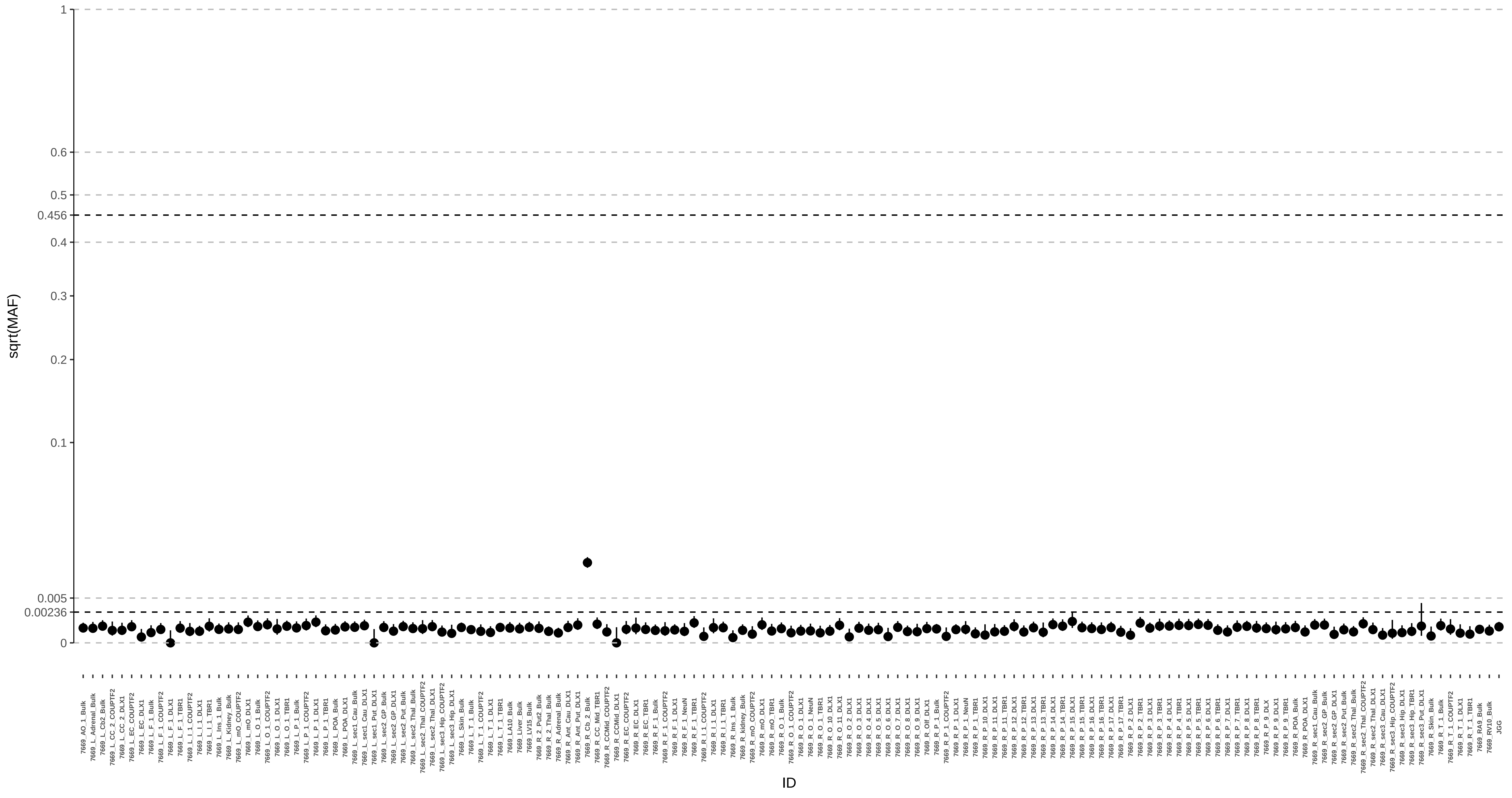

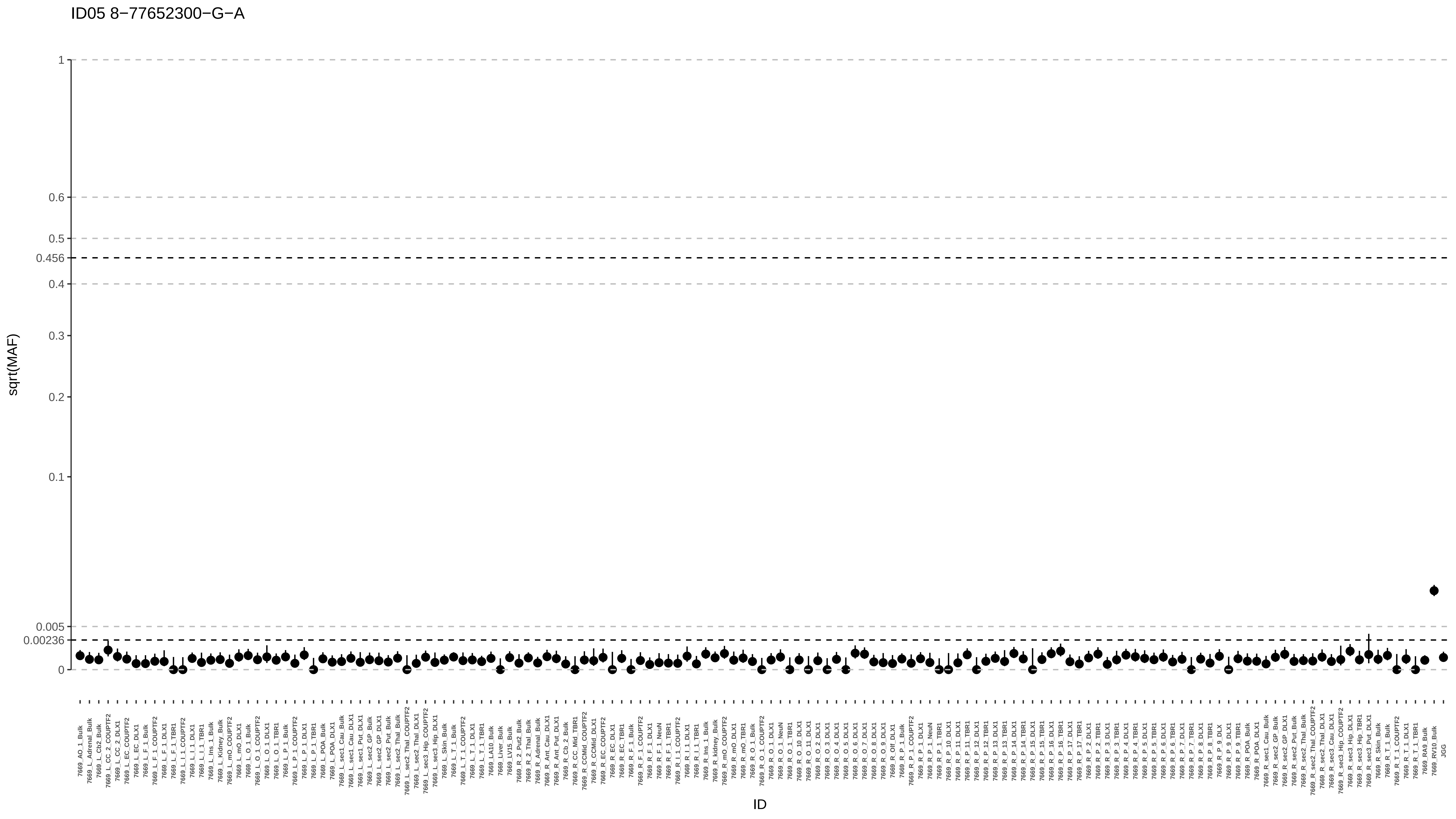
ID05 8−77652300−G−A
sqrt(MAF)
1
0.6
0.5
0.456
0.4
0.3
0.2
0.1
0.005
0.00236
0
ID
JGG

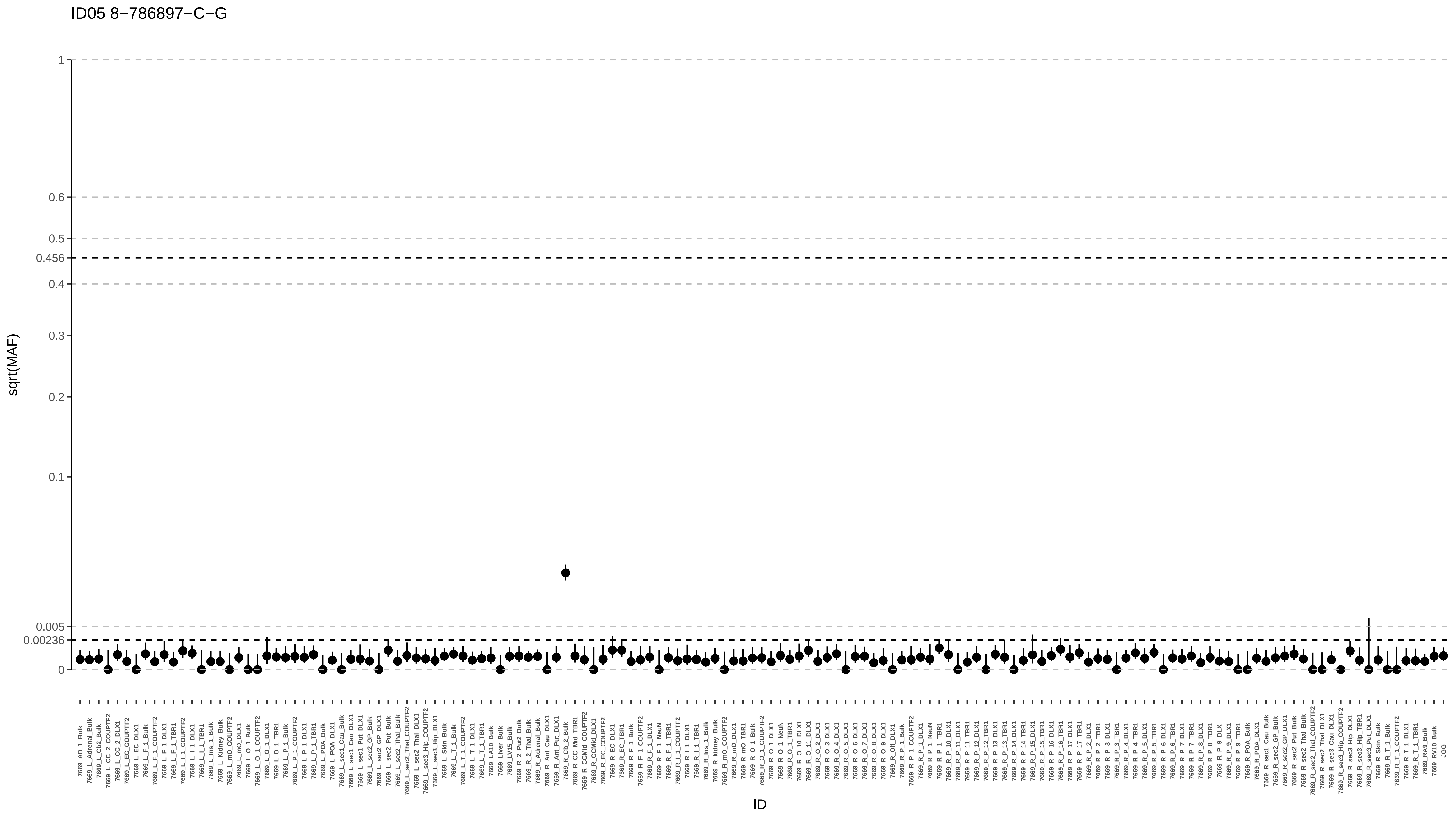

ID05 8−786897−C−G
sqrt(MAF)
ID

ID05 8−80121537−G−A

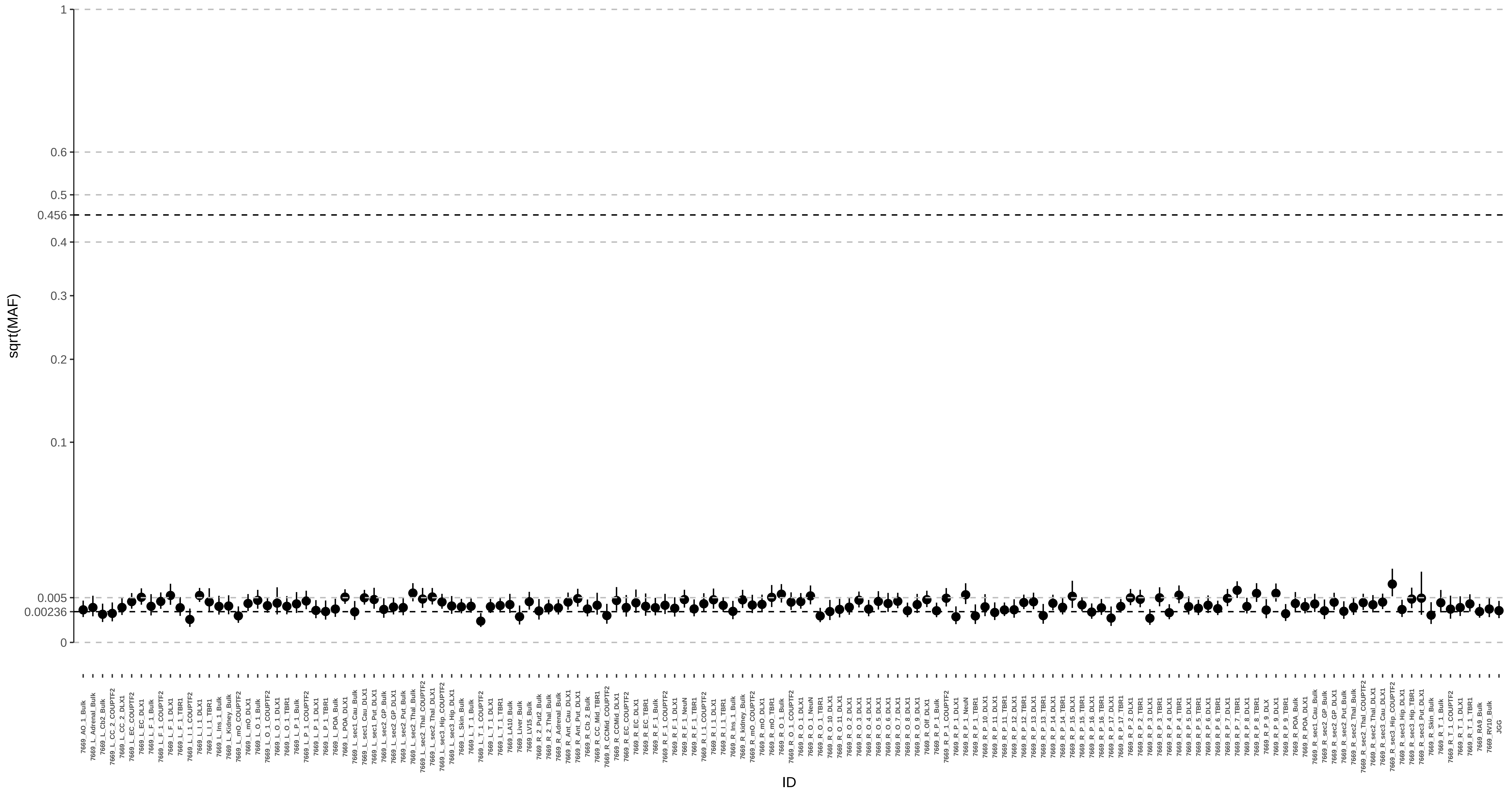

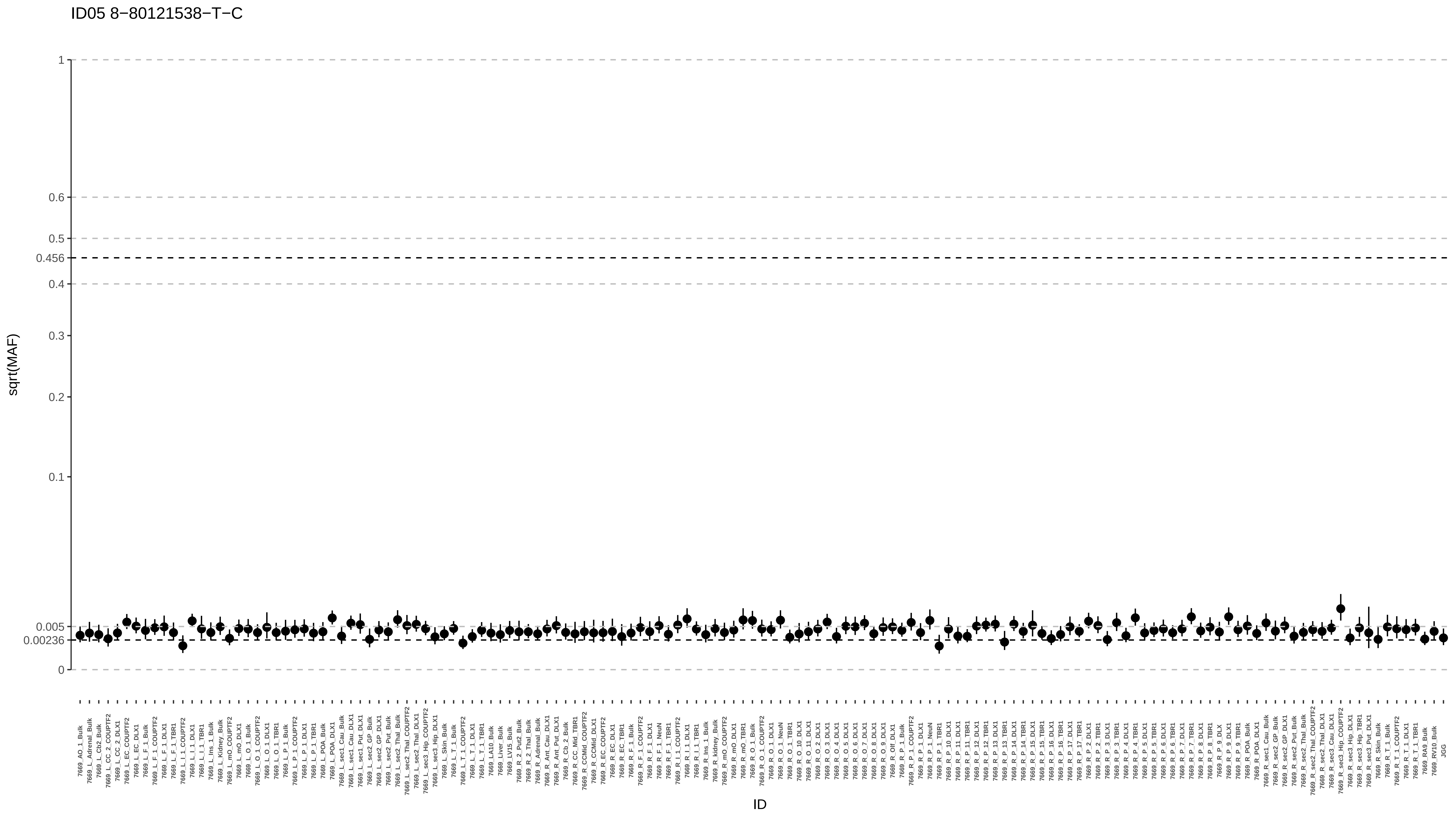
ID05 8−80121538−T−C
sqrt(MAF)
ID
1
0.6
0.5
0.456
0.4
0.3
0.2
0.1
0.005
0.00236
0

ID05 8−83768131−G−A

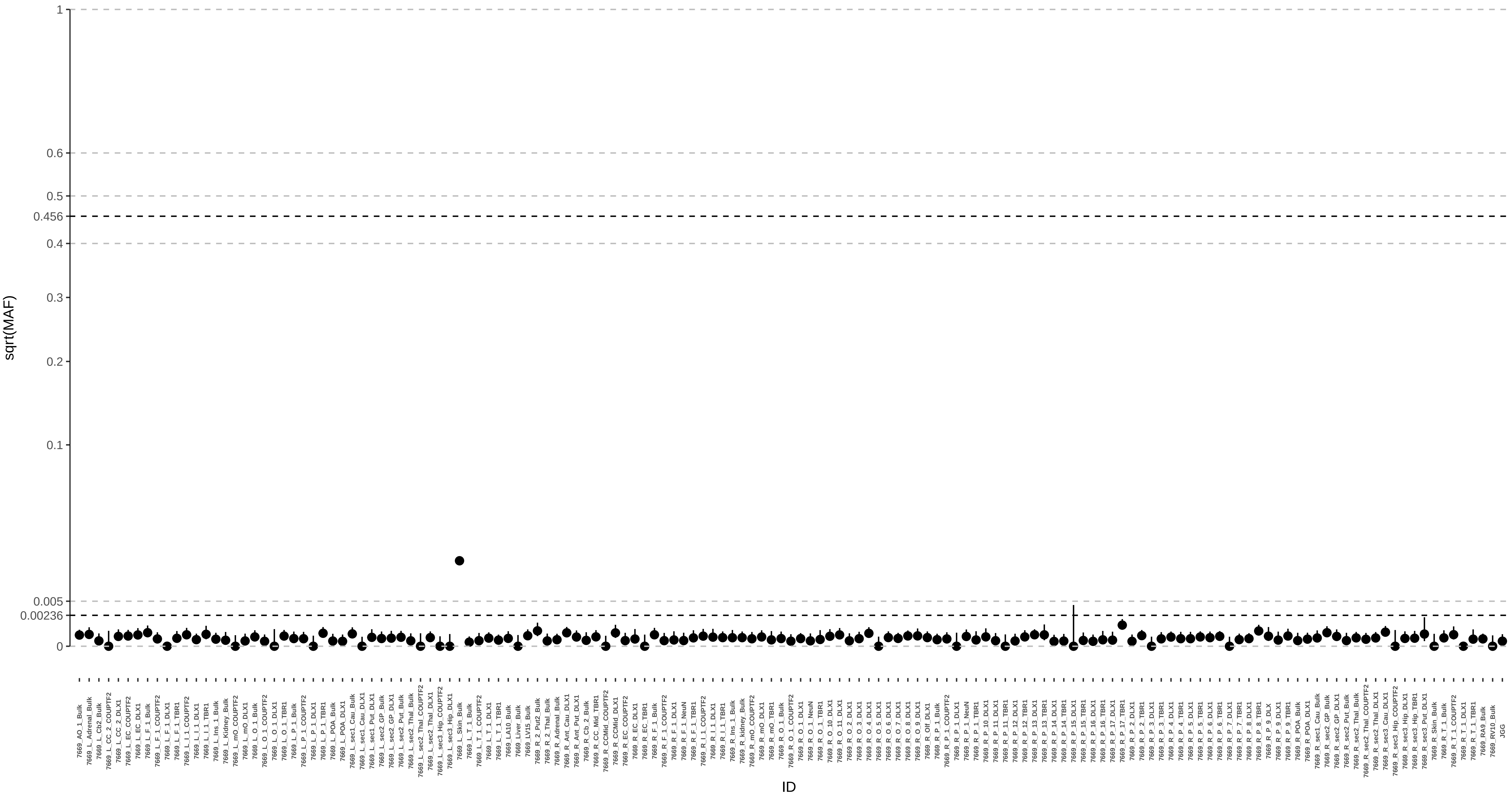

ID05 8−89142864−G−A

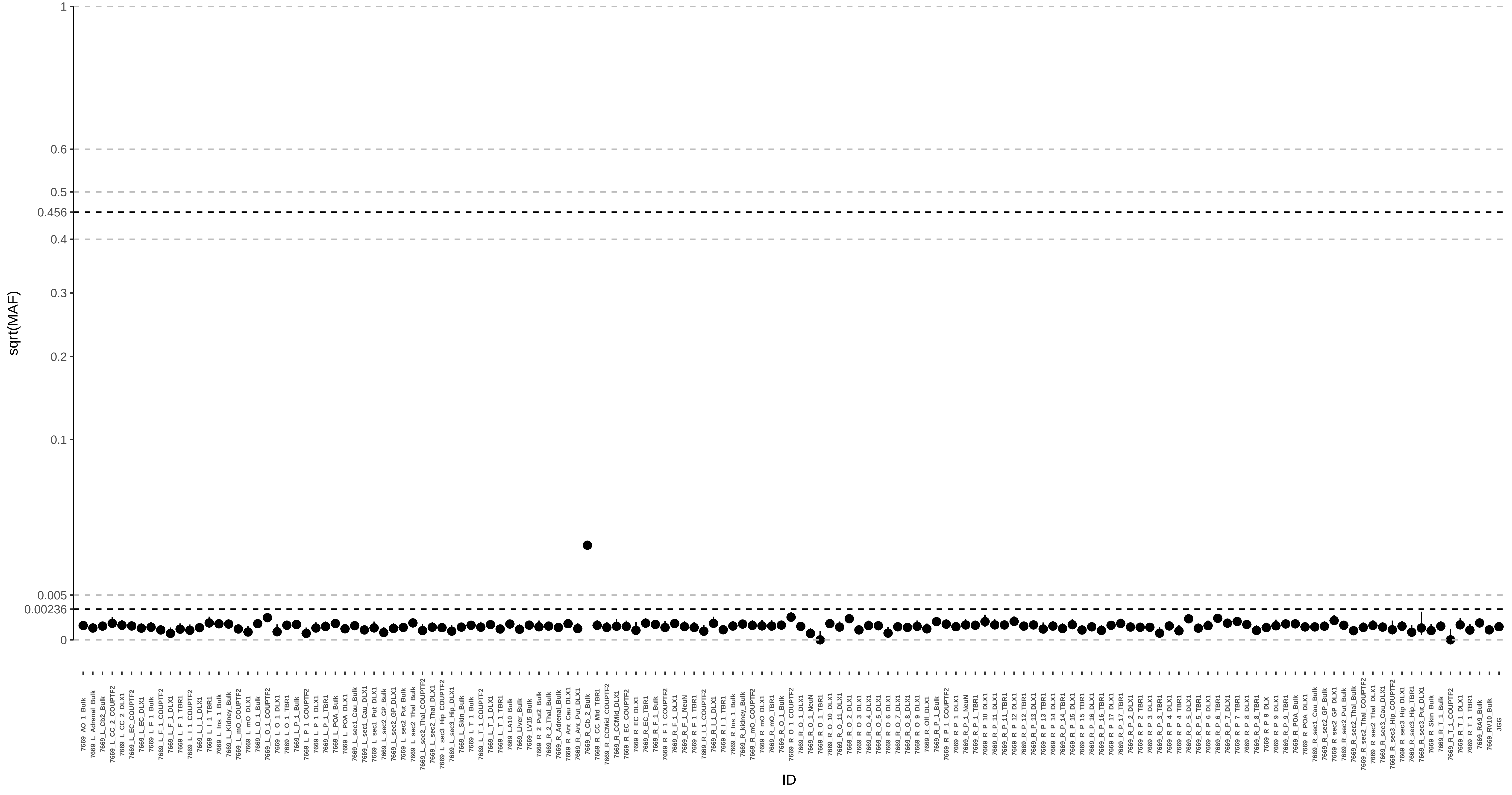

ID05 8−89612305−G−A

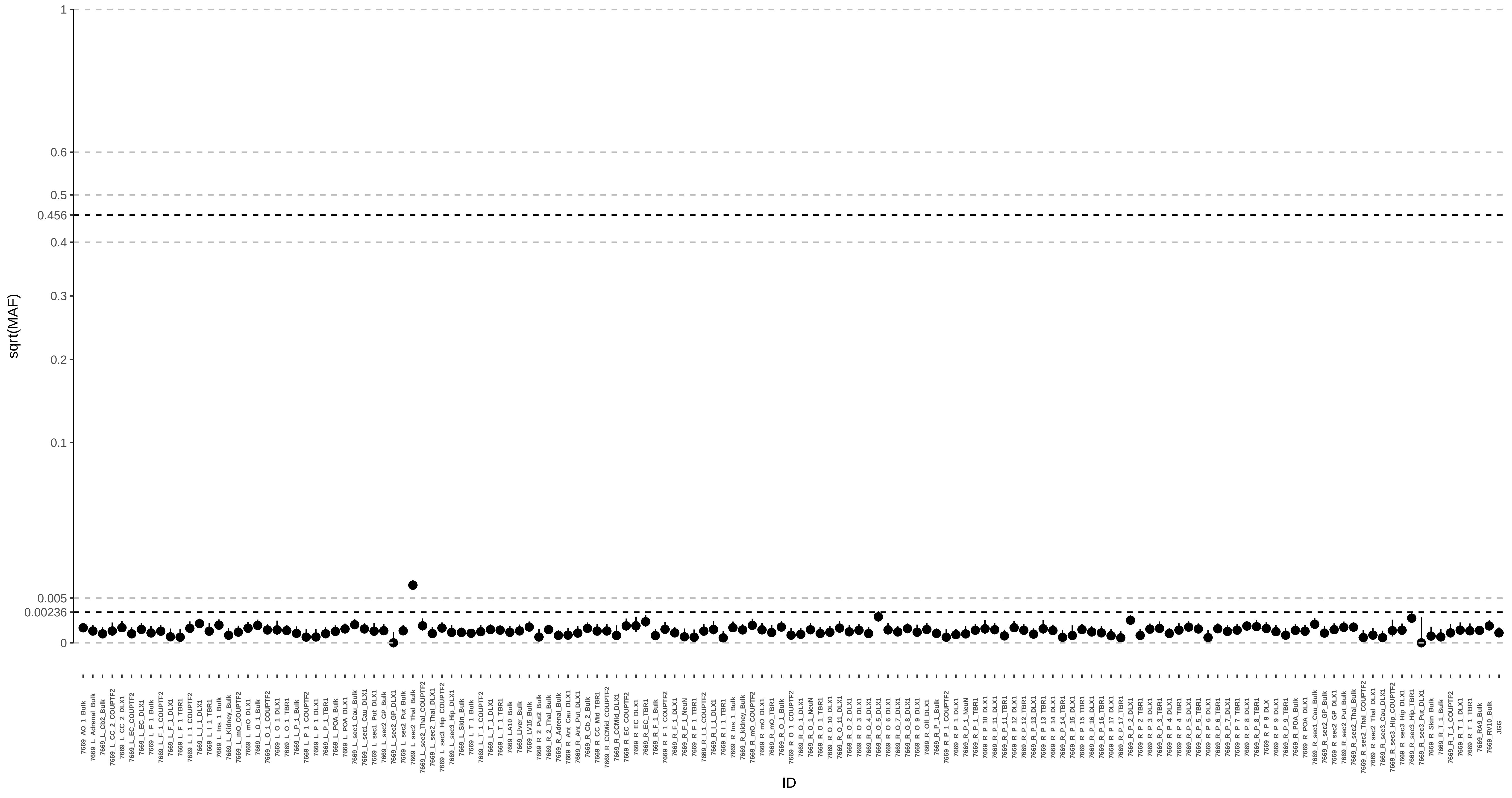

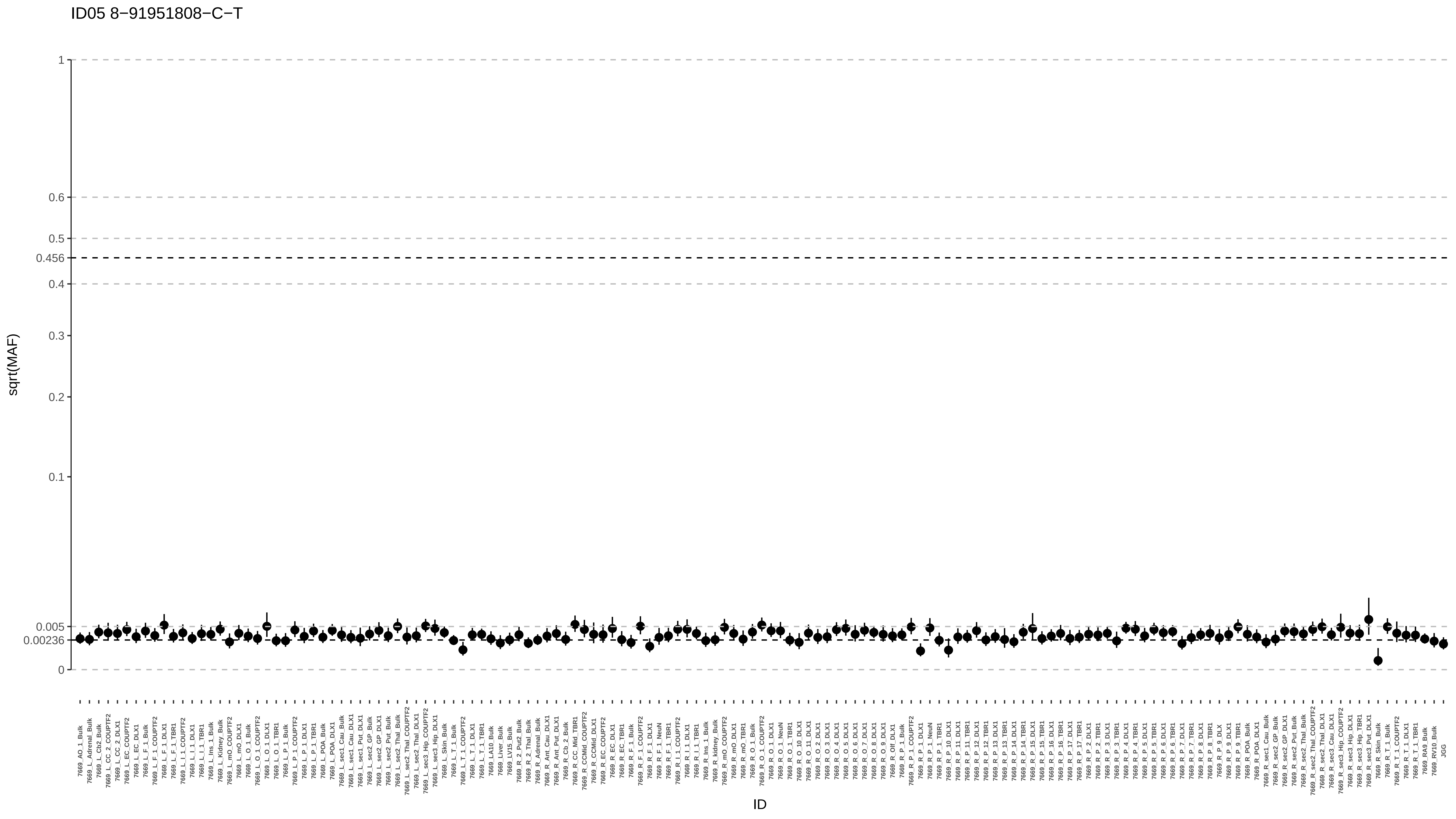

ID05 8−91951808−C−T
sqrt(MAF)
1
0.6
0.5
0.456
0.4
0.3
0.2
0.1
0.005
0.00236
0
ID

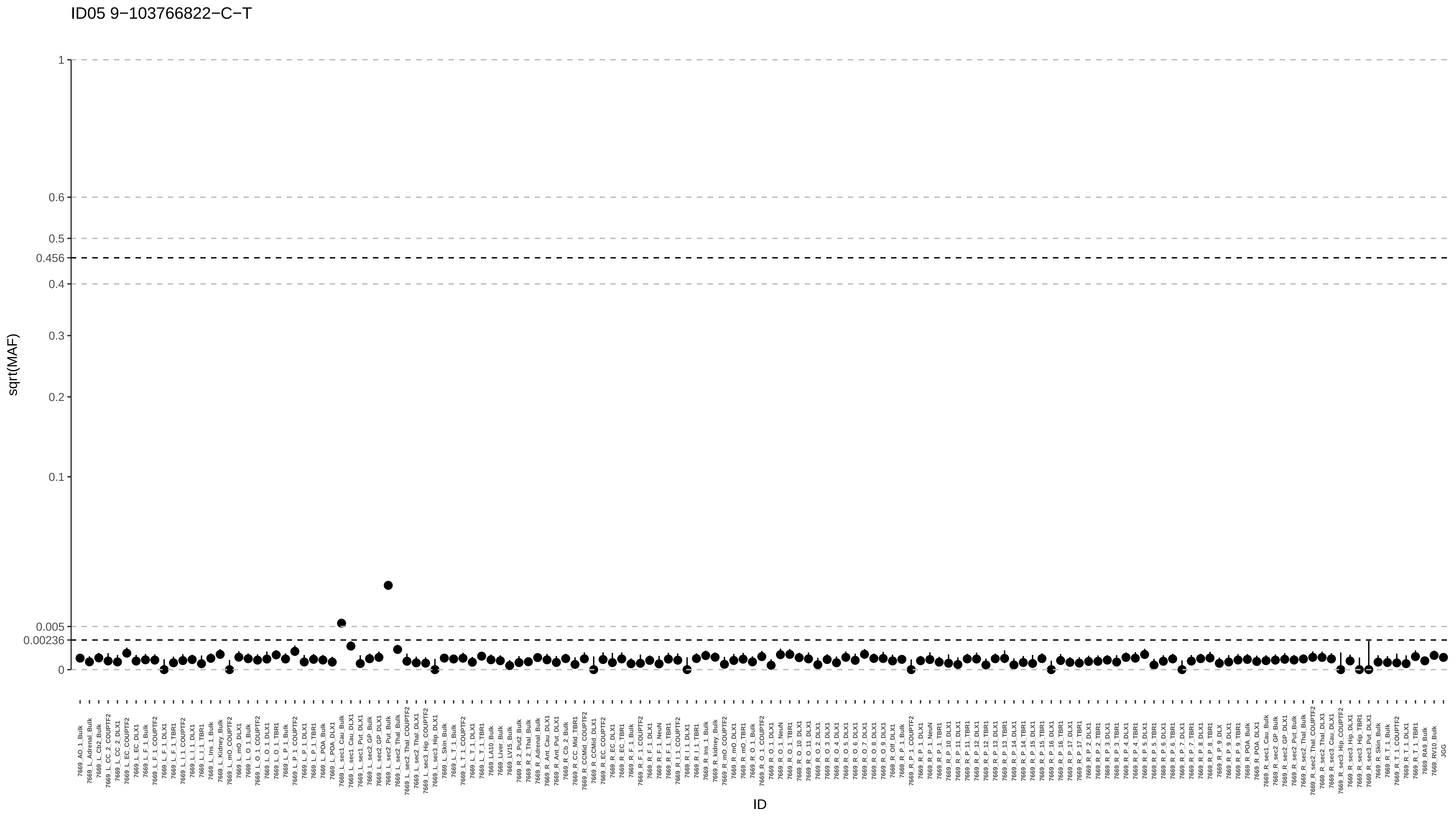
ID05 9−103766822−C−T
sqrt(MAF)
ID
1
0.6
0.5
0.456
0.4
0.3
0.2
0.1
0.005
0.00236
0

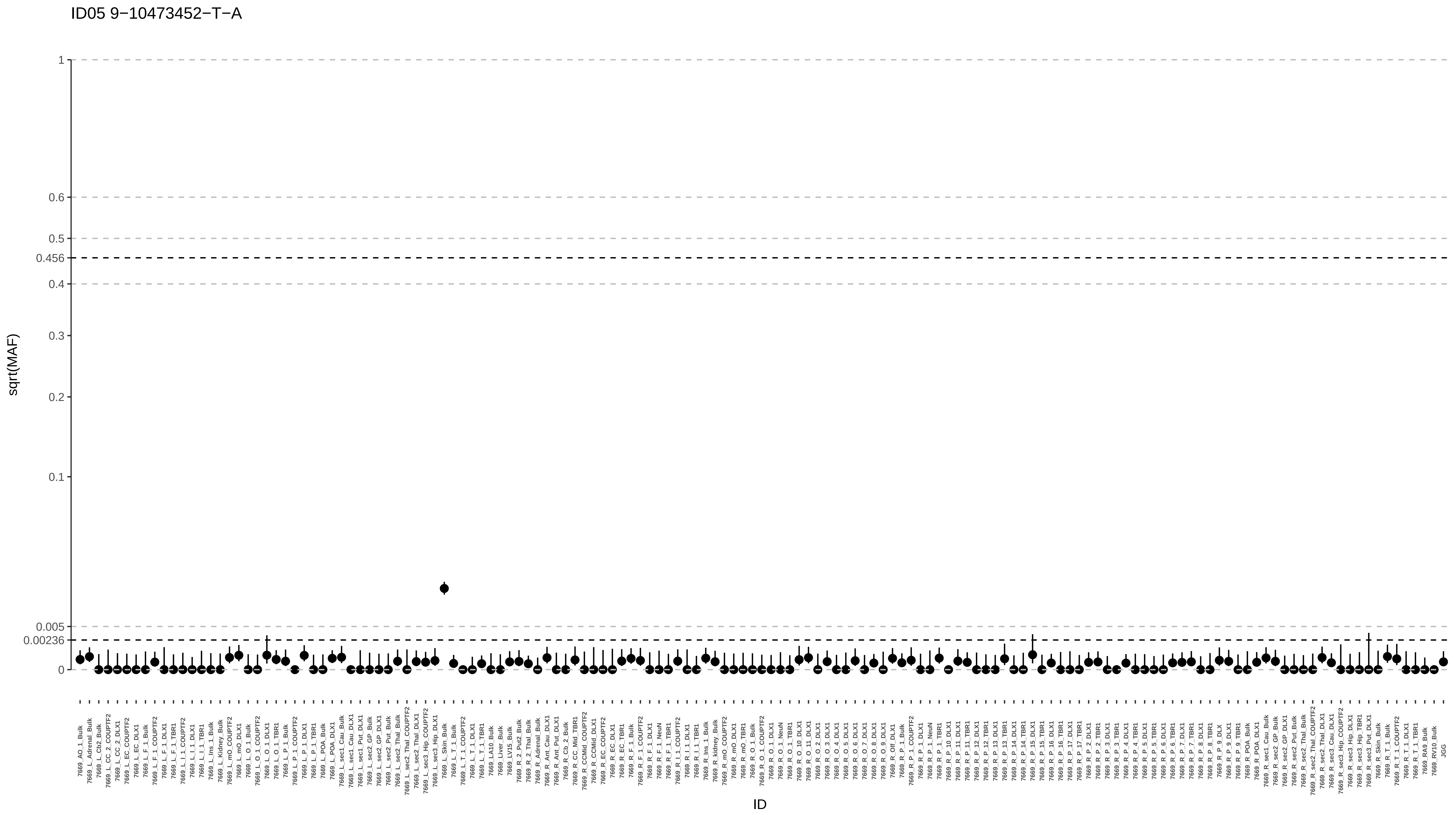
ID05 9−10473452−T−A
sqrt(MAF)
1
0.6
0.5
0.456
0.4
0.3
0.2
0.1
0.005
0.00236
0
ID

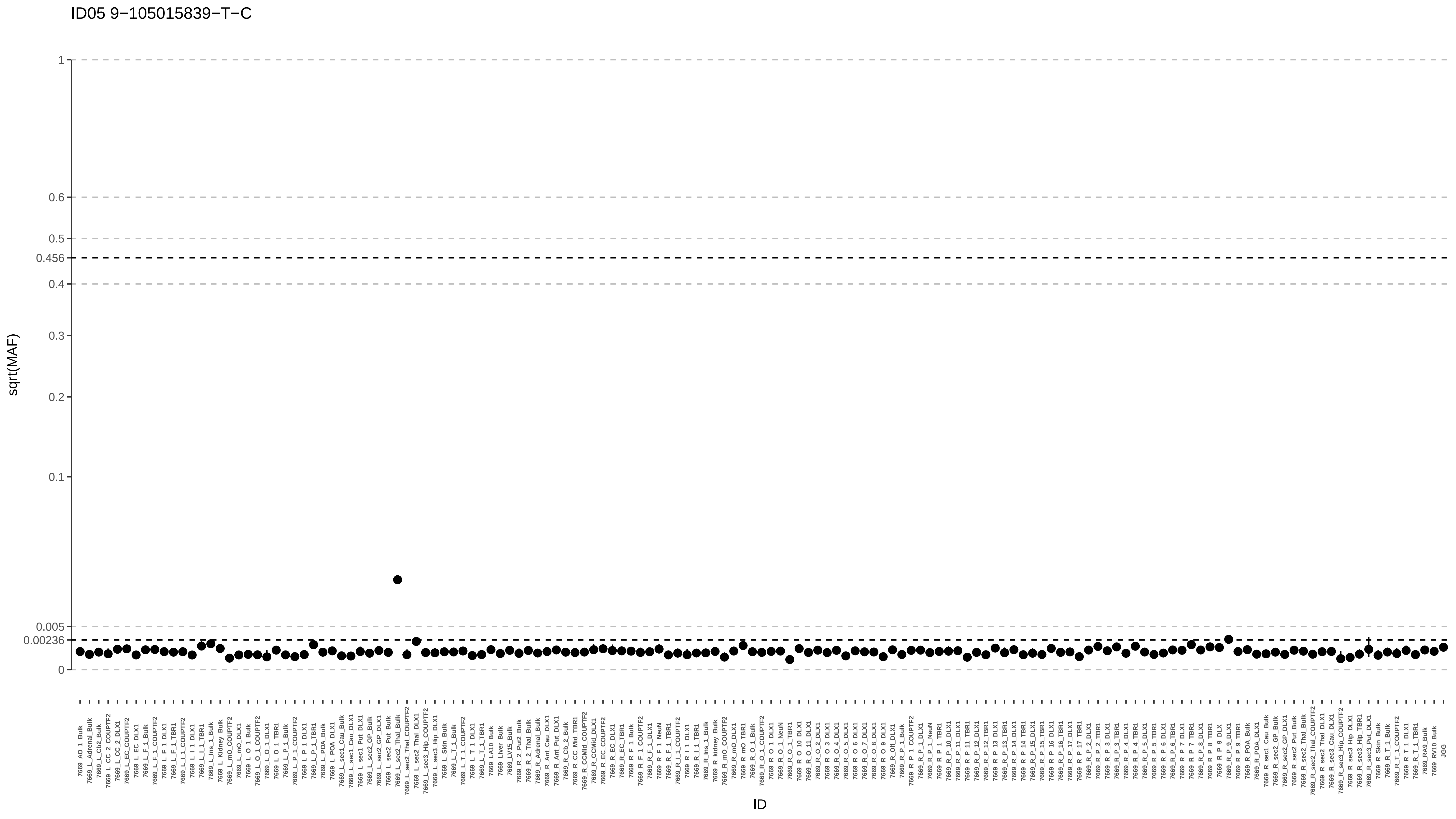
ID05 9−105015839−T−C
sqrt(MAF)
ID
1
0.6
0.5
0.456
0.4
0.3
0.2
0.1
0.005
0.00236
0

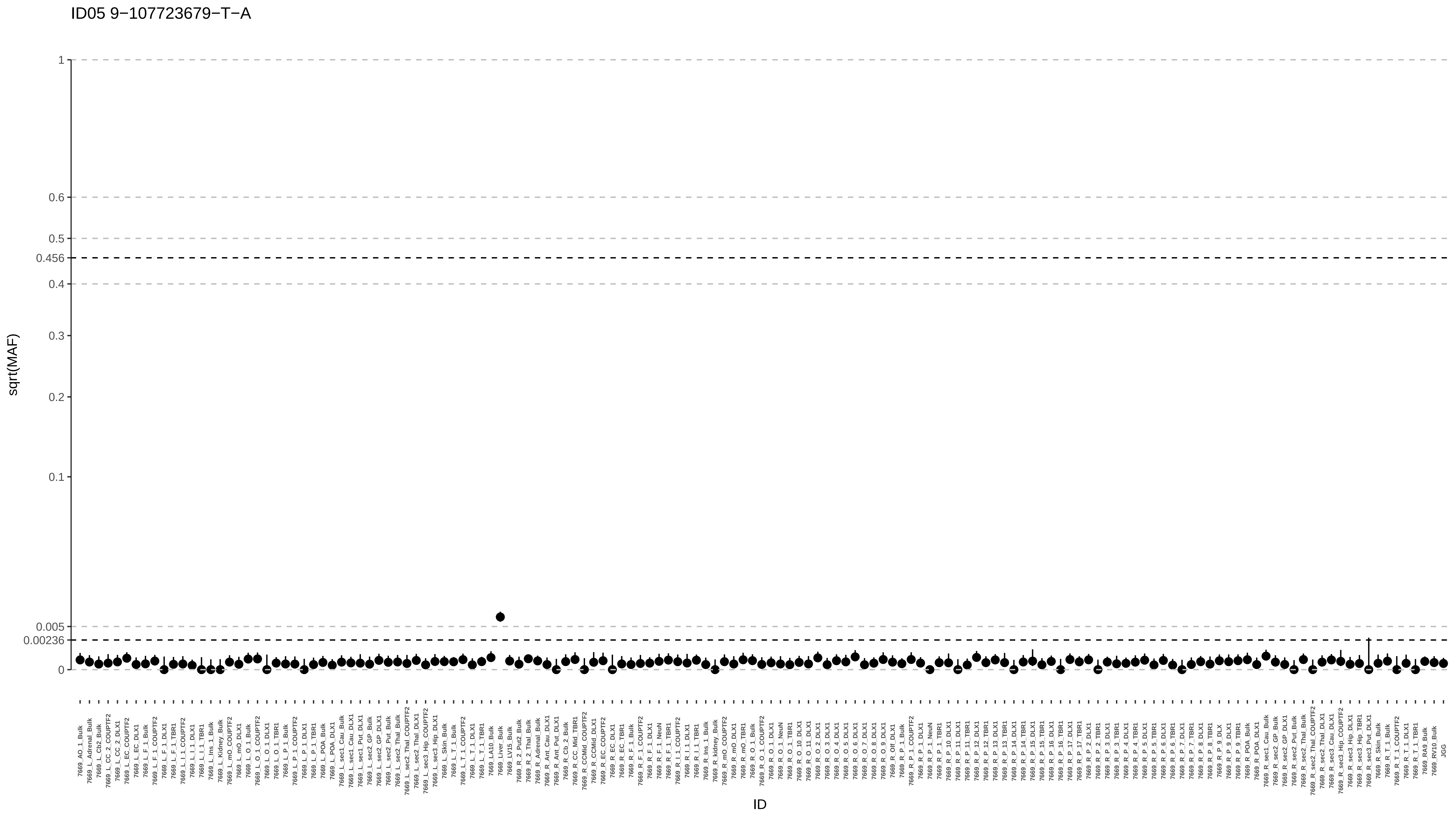
ID05 9−107723679−T−A
sqrt(MAF)
ID
1
0.6
0.5
0.456
0.4
0.3
0.2
0.1
0.005
0.00236
0

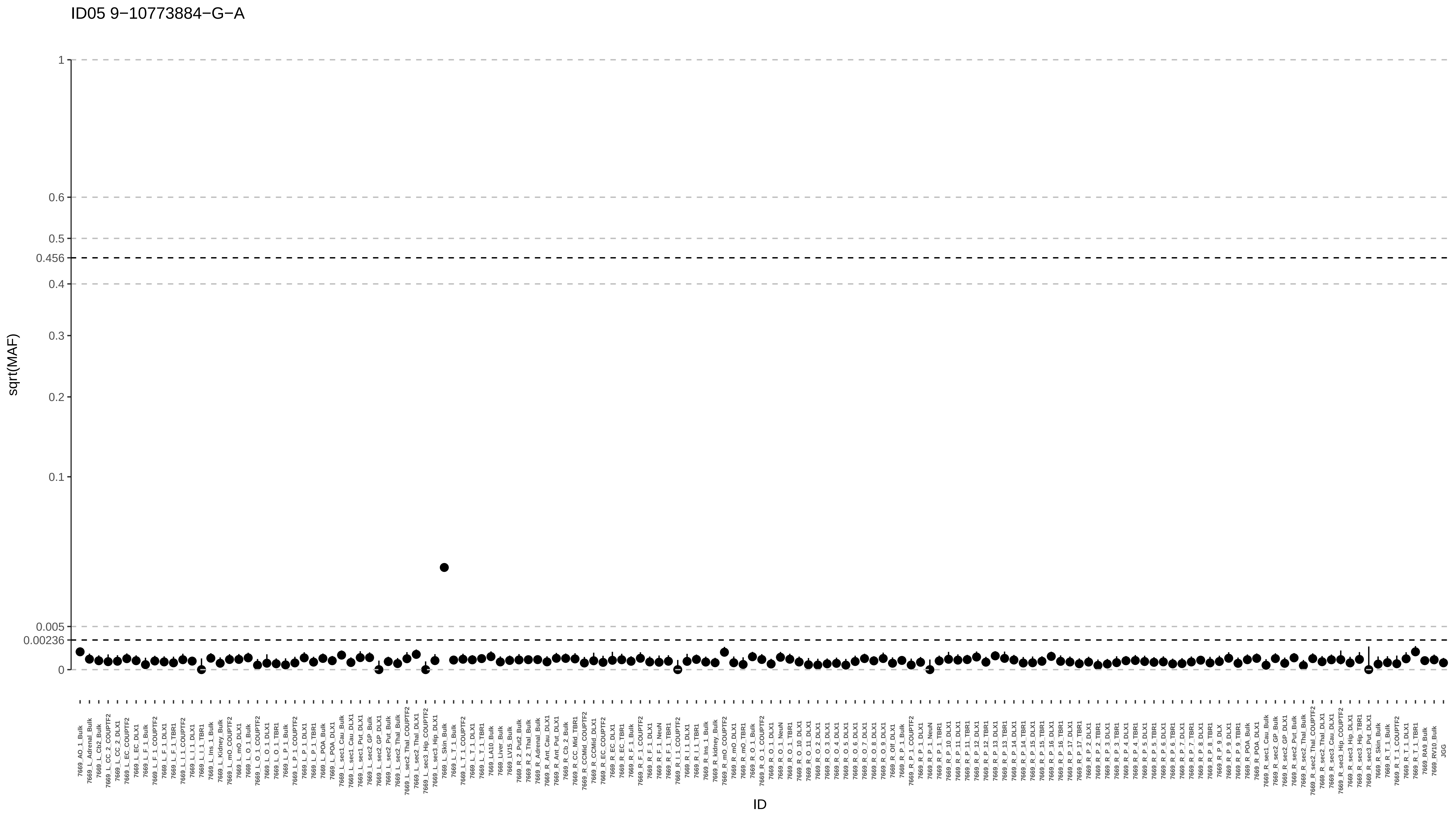
ID05 9−10773884−G−A
sqrt(MAF)
1
0.6
0.5
0.456
0.4
0.3
0.2
0.1
0.005
0.00236
0
ID

ID05 9−119721731−C−T

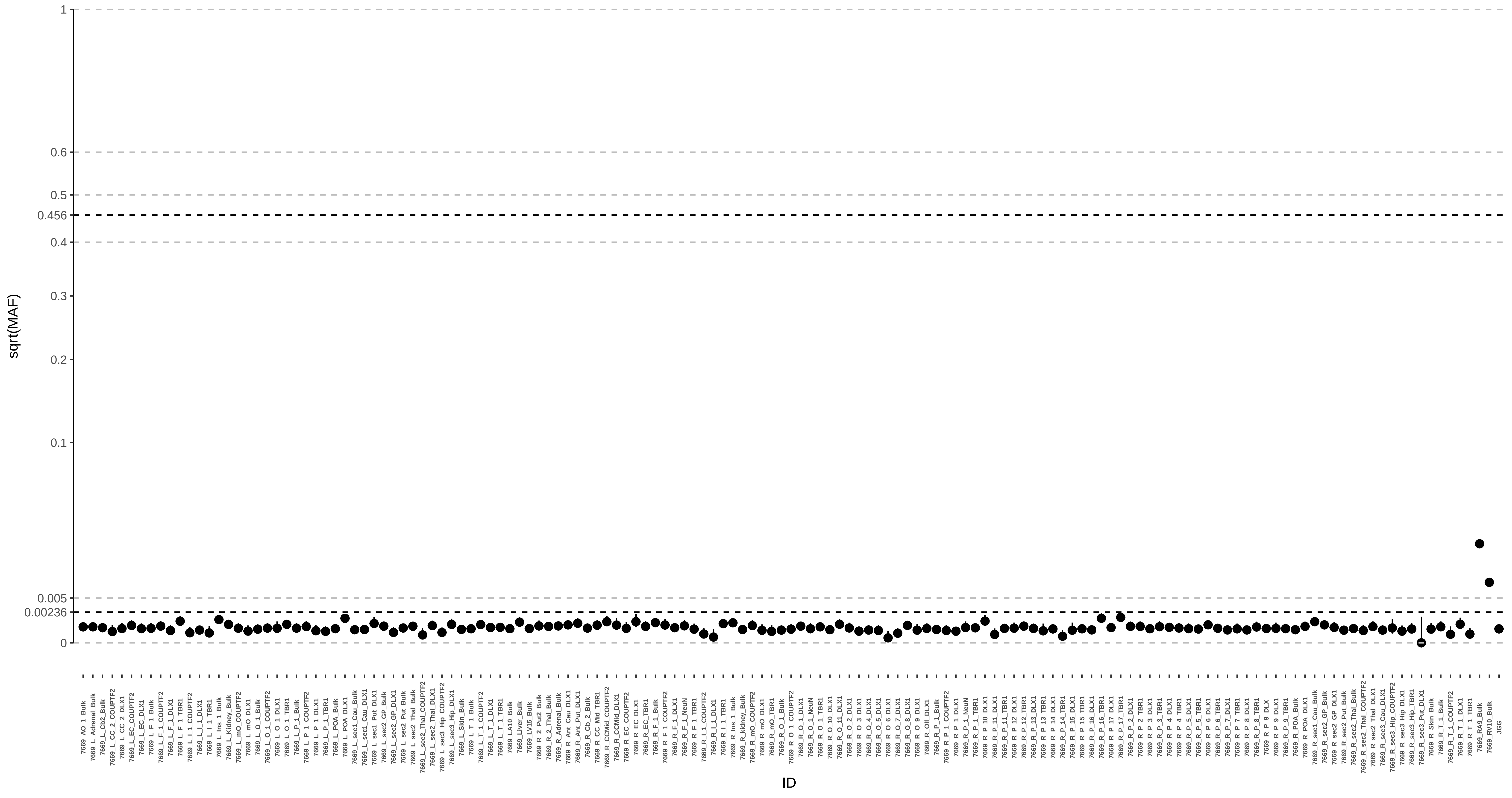

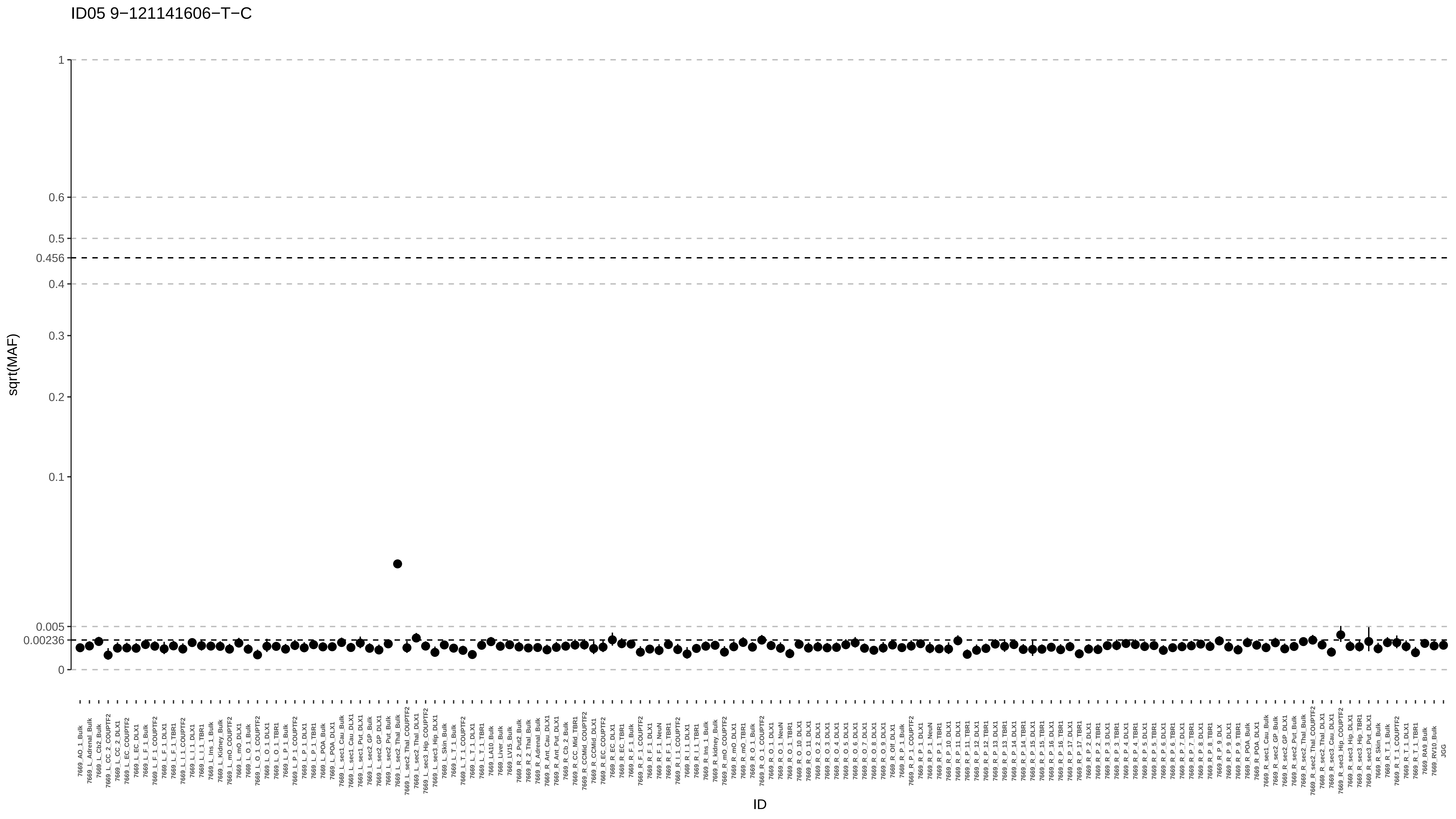
ID05 9−121141606−T−C
sqrt(MAF)
1
0.6
0.5
0.456
0.4
0.3
0.2
0.1
0.005
0.00236
0
ID
7669_AO_1_Bulk
7669_L_Adrenal_Bulk
7669_L_Cb2_Bulk
7669_L_CC_2_COUPTF2
7669_L_CC_2_DLX1
7669_L_EC_COUPTF2
7669_L_EC_DLX1
7669_L_F_1_Bulk
7669_L_F_1_COUPTF2
7669_L_F_1_DLX1
7669_L_F_1_TBR1
7669_L_I_1_COUPTF2
7669_L_I_1_DLX1
7669_L_I_1_TBR1
7669_L_Ins_1_Bulk
7669_L_Kidney_Bulk
7669_L_mO_COUPTF2
7669_L_mO_DLX1
7669_L_O_1_Bulk
7669_L_O_1_COUPTF2
7669_L_O_1_DLX1
7669_L_O_1_TBR1
7669_L_P_1_Bulk
7669_L_P_1_COUPTF2
7669_L_P_1_DLX1
7669_L_P_1_TBR1
7669_L_POA_Bulk
7669_L_POA_DLX1
7669_L_sec1_Cau_Bulk
7669_L_sec1_Cau_DLX1
7669_L_sec1_Put_DLX1
7669_L_sec2_GP_Bulk
7669_L_sec2_GP_DLX1
7669_L_sec2_Put_Bulk
7669_L_sec2_Thal_Bulk
7669_L_sec2_Thal_COUPTF2
7669_L_sec2_Thal_DLX1
7669_L_sec3_Hip_COUPTF2
7669_L_sec3_Hip_DLX1
7669_L_Skin_Bulk
7669_L_T_1_Bulk
7669_L_T_1_COUPTF2
7669_L_T_1_DLX1
7669_L_T_1_TBR1
7669_LA10_Bulk
7669_Liver_Bulk
7669_LV15_Bulk
7669_R_2_Put2_Bulk
7669_R_2_Thal_Bulk
7669_R_Adrenal_Bulk
7669_R_Ant_Cau_DLX1
7669_R_Ant_Put_DLX1
7669_R_Cb_2_Bulk
7669_R_CC_Mid_TBR1
7669_R_CCMid_COUPTF2
7669_R_CCMid_DLX1
7669_R_EC_COUPTF2
7669_R_EC_DLX1
7669_R_EC_TBR1
7669_R_F_1_Bulk
7669_R_F_1_COUPTF2
7669_R_F_1_DLX1
7669_R_F_1_NeuN
7669_R_F_1_TBR1
7669_R_I_1_COUPTF2
7669_R_I_1_DLX1
7669_R_I_1_TBR1
7669_R_Ins_1_Bulk
7669_R_kidney_Bulk
7669_R_mO_COUPTF2
7669_R_mO_DLX1
7669_R_mO_TBR1
7669_R_O_1_Bulk
7669_R_O_1_COUPTF2
7669_R_O_1_DLX1
7669_R_O_1_NeuN
7669_R_O_1_TBR1
7669_R_O_10_DLX1
7669_R_O_11_DLX1
7669_R_O_2_DLX1
7669_R_O_3_DLX1
7669_R_O_4_DLX1
7669_R_O_5_DLX1
7669_R_O_6_DLX1
7669_R_O_7_DLX1
7669_R_O_8_DLX1
7669_R_O_9_DLX1
7669_R_Olf_DLX1
7669_R_P_1_Bulk
7669_R_P_1_COUPTF2
7669_R_P_1_DLX1
7669_R_P_1_NeuN
7669_R_P_1_TBR1
7669_R_P_10_DLX1
7669_R_P_11_DLX1
7669_R_P_11_TBR1
7669_R_P_12_DLX1
7669_R_P_12_TBR1
7669_R_P_13_DLX1
7669_R_P_13_TBR1
7669_R_P_14_DLX1
7669_R_P_14_TBR1
7669_R_P_15_DLX1
7669_R_P_15_TBR1
7669_R_P_16_DLX1
7669_R_P_16_TBR1
7669_R_P_17_DLX1
7669_R_P_17_TBR1
7669_R_P_2_DLX1
7669_R_P_2_TBR1
7669_R_P_3_DLX1
7669_R_P_3_TBR1
7669_R_P_4_DLX1
7669_R_P_4_TBR1
7669_R_P_5_DLX1
7669_R_P_5_TBR1
7669_R_P_6_DLX1
7669_R_P_6_TBR1
7669_R_P_7_DLX1
7669_R_P_7_TBR1
7669_R_P_8_DLX1
7669_R_P_8_TBR1
7669_R_P_9_DLX
7669_R_P_9_DLX1
7669_R_P_9_TBR1
7669_R_POA_Bulk
7669_R_POA_DLX1
7669_R_sec1_Cau_Bulk
7669_R_sec2_GP_Bulk
7669_R_sec2_GP_DLX1
7669_R_sec2_Put_Bulk
7669_R_sec2_Thal_Bulk
7669_R_sec2_Thal_COUPTF2
7669_R_sec2_Thal_DLX1
7669_R_sec3_Cau_DLX1
7669_R_sec3_Hip_COUPTF2
7669_R_sec3_Hip_DLX1
7669_R_sec3_Hip_TBR1
7669_R_sec3_Put_DLX1
7669_R_Skin_Bulk
7669_R_T_1_Bulk
7669_R_T_1_COUPTF2
7669_R_T_1_DLX1
7669_R_T_1_TBR1
7669_RA9_Bulk
7669_RV10_Bulk
JGG

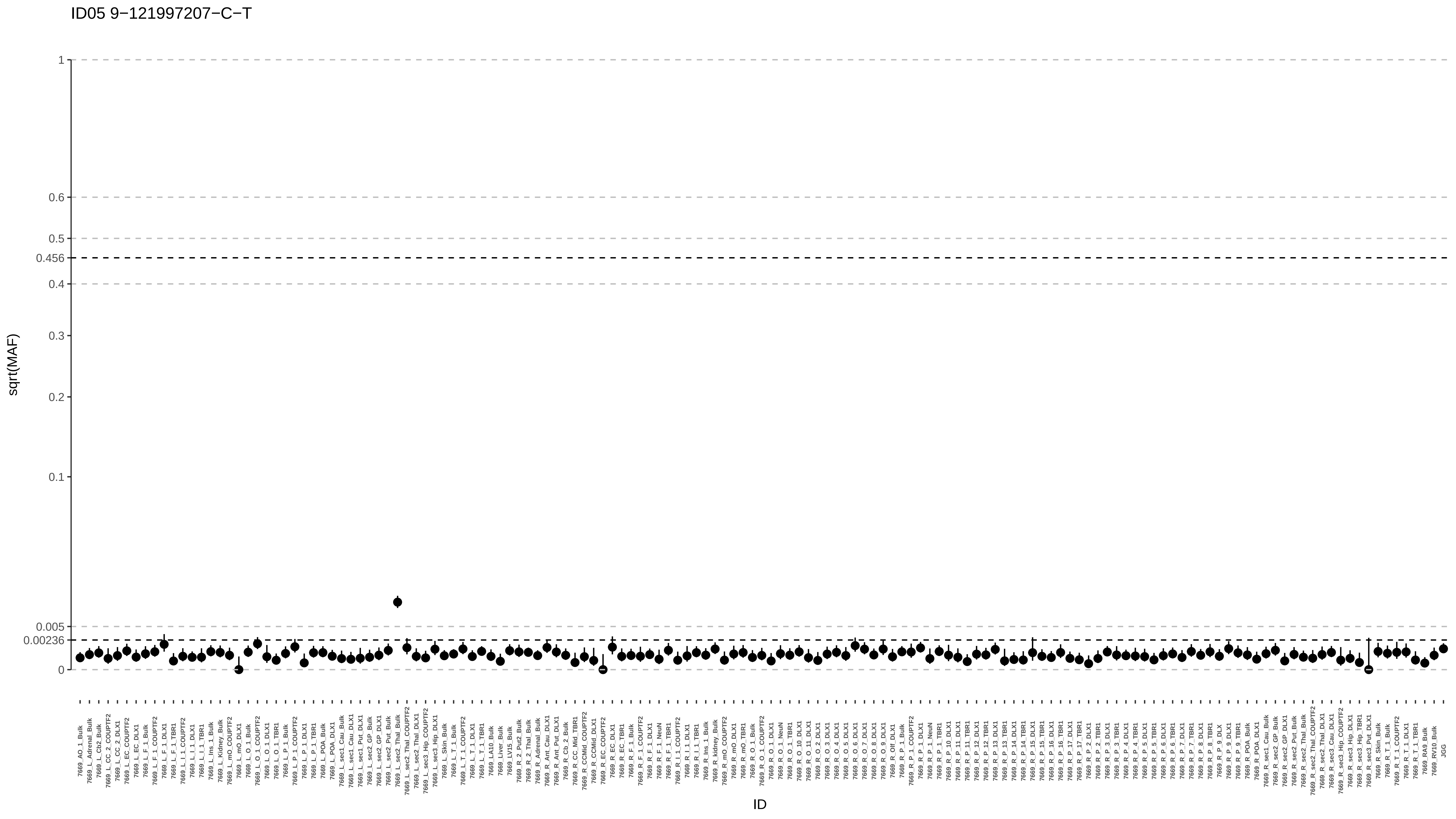
ID05 9−121997207−C−T
sqrt(MAF)
ID
1
0.6
0.5
0.456
0.4
0.3
0.2
0.1
0.005
0.00236
0

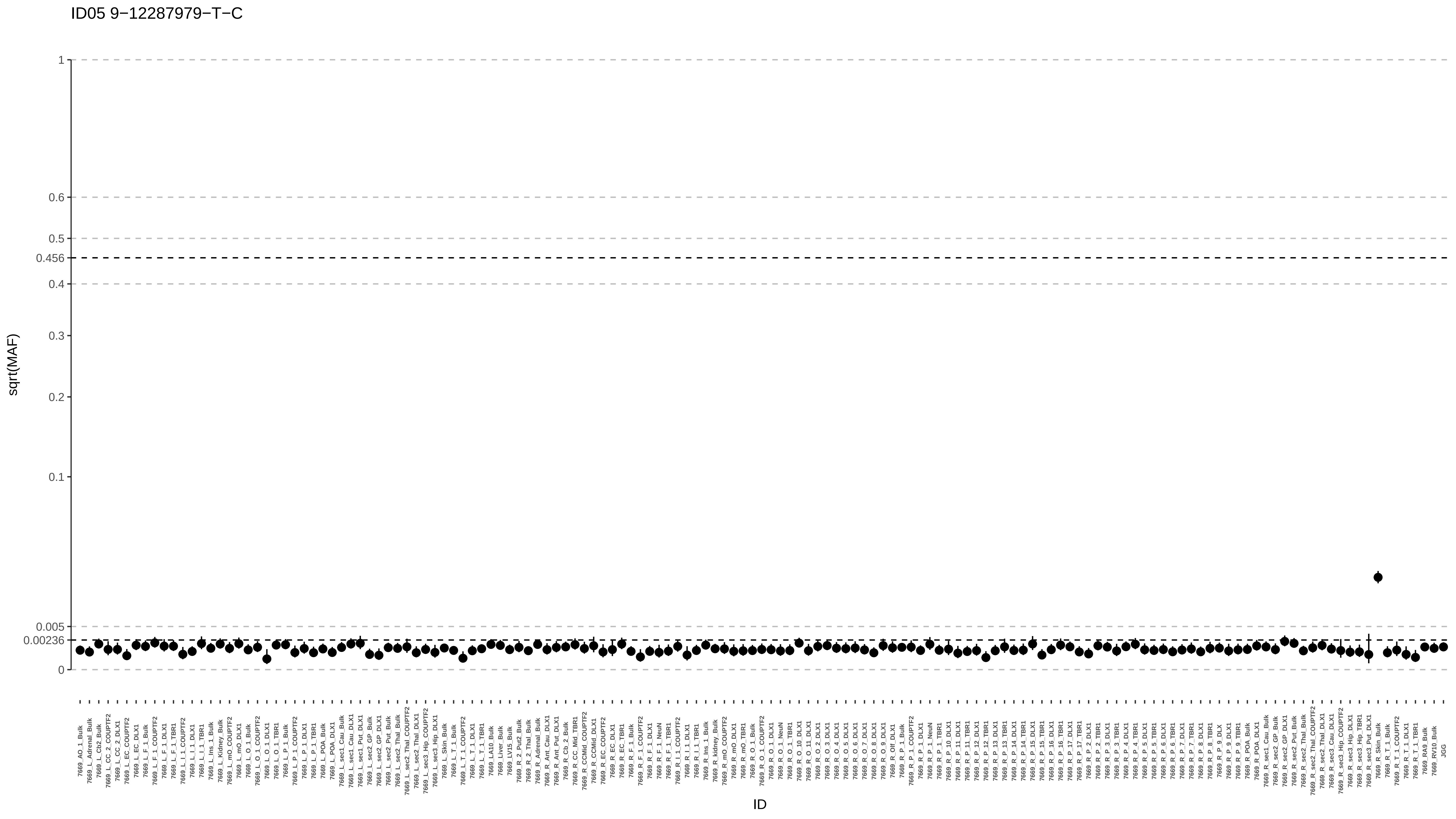

ID05 9−12287979−T−C
sqrt(MAF)
1
0.6
0.5
0.456
0.4
0.3
0.2
0.1
0.005
0.00236
0
ID

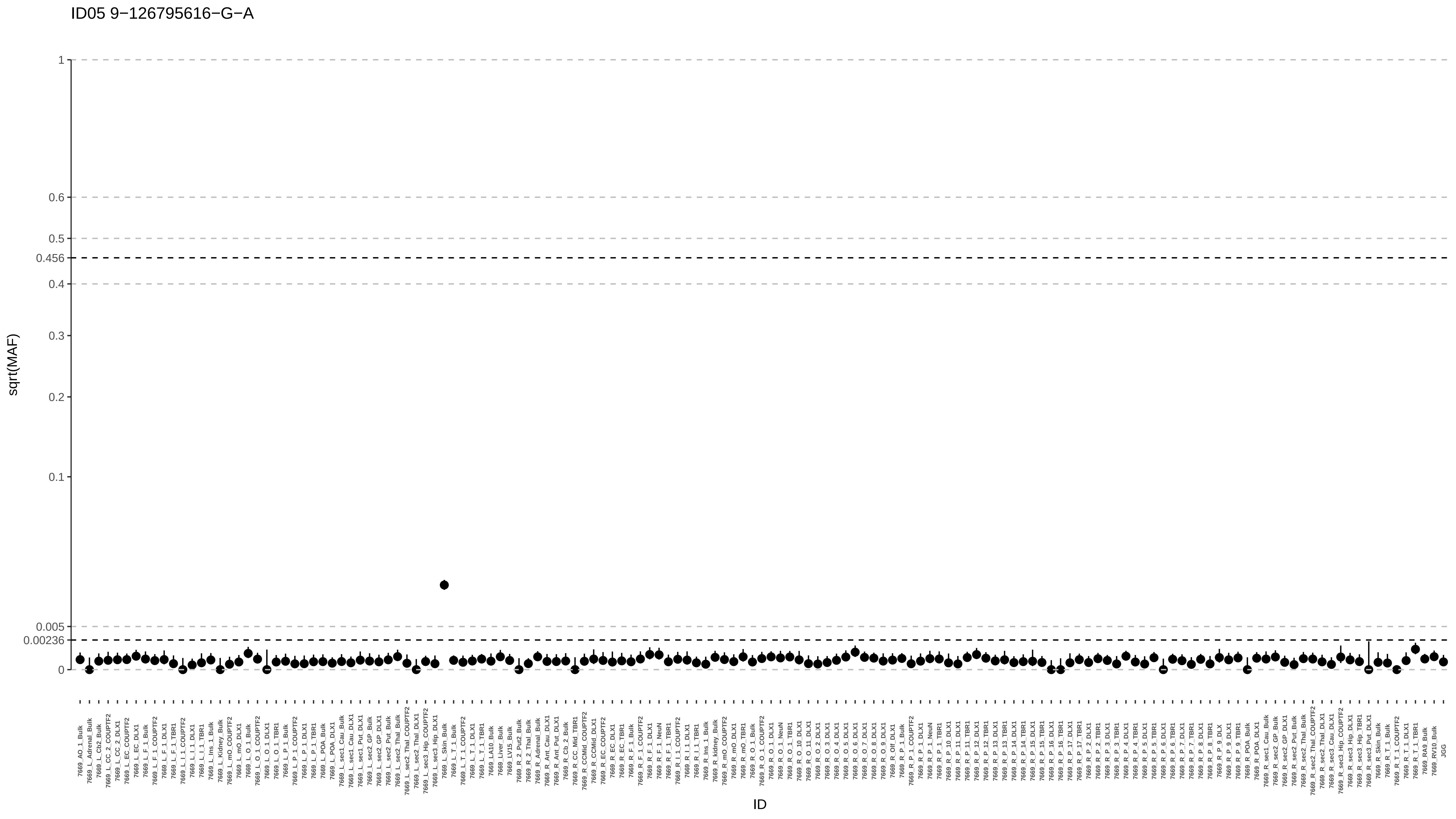
ID05 9−126795616−G−A
sqrt(MAF)
1
0.6
0.5
0.456
0.4
0.3
0.2
0.1
0.005
0.00236
0
ID

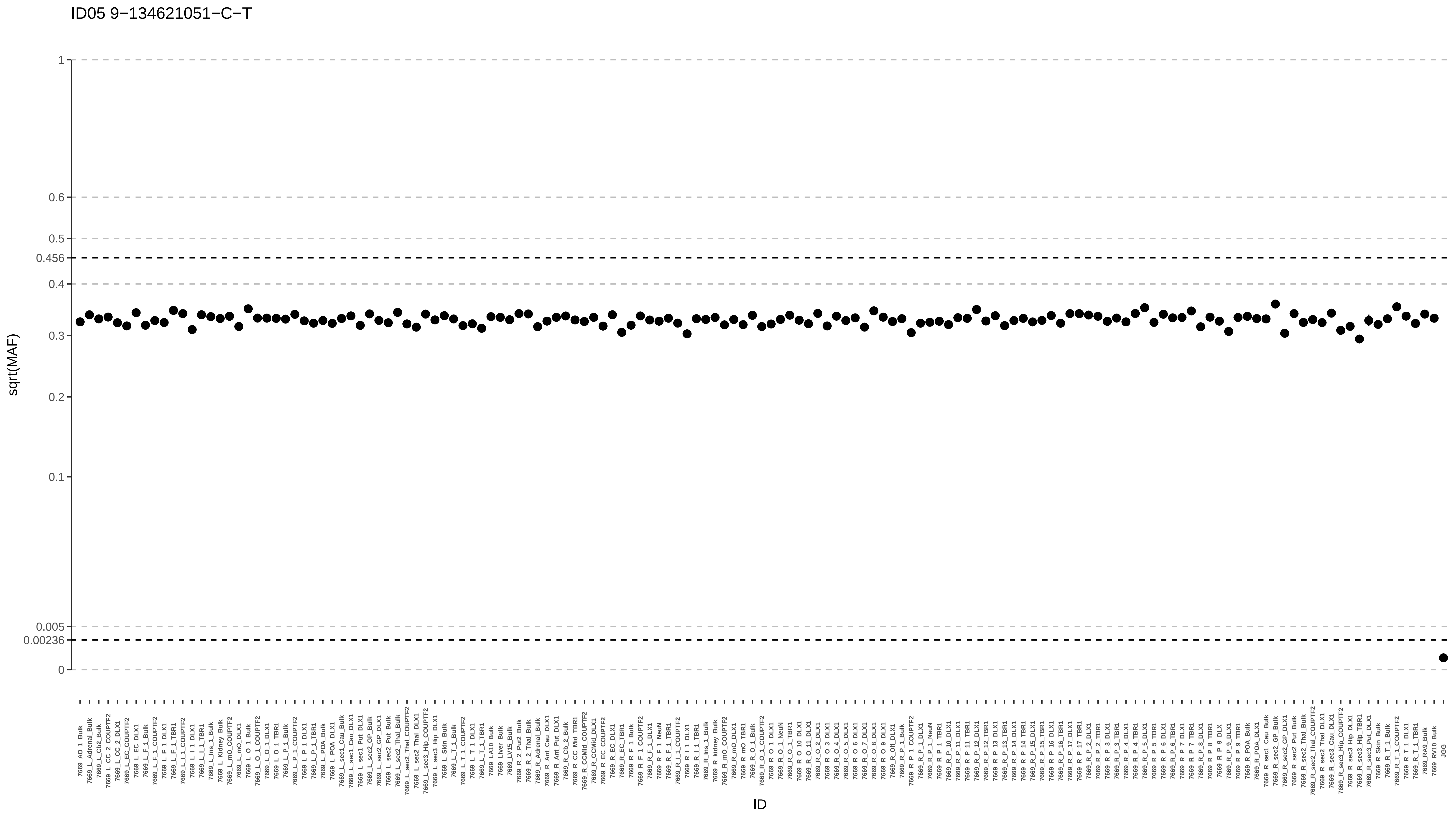
ID05 9−134621051−C−T
1
0.6
0.5
0.456
0.4
0.3
0.2
0.1
0.005
0.00236
0
sqrt(MAF)
ID
7669_AO_1_Bulk
7669_L_Adrenal_Bulk
7669_L_Cb2_Bulk
7669_L_CC_2_COUPTF2
7669_L_CC_2_DLX1
7669_L_EC_COUPTF2
7669_L_EC_DLX1
7669_L_F_1_Bulk
7669_L_F_1_COUPTF2
7669_L_F_1_DLX1
7669_L_F_1_TBR1
7669_L_I_1_COUPTF2
7669_L_I_1_DLX1
7669_L_I_1_TBR1
7669_L_Ins_1_Bulk
7669_L_Kidney_Bulk
7669_L_mO_COUPTF2
7669_L_mO_DLX1
7669_L_O_1_Bulk
7669_L_O_1_COUPTF2
7669_L_O_1_DLX1
7669_L_O_1_TBR1
7669_L_P_1_Bulk
7669_L_P_1_COUPTF2
7669_L_P_1_DLX1
7669_L_P_1_TBR1
7669_L_POA_Bulk
7669_L_POA_DLX1
7669_L_sec1_Cau_Bulk
7669_L_sec1_Cau_DLX1
7669_L_sec1_Put_DLX1
7669_L_sec2_GP_Bulk
7669_L_sec2_GP_DLX1
7669_L_sec2_Put_Bulk
7669_L_sec2_Thal_Bulk
7669_L_sec2_Thal_COUPTF2
7669_L_sec2_Thal_DLX1
7669_L_sec3_Hip_COUPTF2
7669_L_sec3_Hip_DLX1
7669_L_Skin_Bulk
7669_L_T_1_Bulk
7669_L_T_1_COUPTF2
7669_L_T_1_DLX1
7669_L_T_1_TBR1
7669_LA10_Bulk
7669_Liver_Bulk
7669_LV15_Bulk
7669_R_2_Put2_Bulk
7669_R_2_Thal_Bulk
7669_R_Adrenal_Bulk
7669_R_Ant_Cau_DLX1
7669_R_Ant_Put_DLX1
7669_R_Cb_2_Bulk
7669_R_CC_Mid_TBR1
7669_R_CCMid_COUPTF2
7669_R_CCMid_DLX1
7669_R_EC_COUPTF2
7669_R_EC_DLX1
7669_R_EC_TBR1
7669_R_F_1_Bulk
7669_R_F_1_COUPTF2
7669_R_F_1_DLX1
7669_R_F_1_NeuN
7669_R_F_1_TBR1
7669_R_I_1_COUPTF2
7669_R_I_1_DLX1
7669_R_I_1_TBR1
7669_R_Ins_1_Bulk
7669_R_kidney_Bulk
7669_R_mO_COUPTF2
7669_R_mO_DLX1
7669_R_mO_TBR1
7669_R_O_1_Bulk
7669_R_O_1_COUPTF2
7669_R_O_1_DLX1
7669_R_O_1_NeuN
7669_R_O_1_TBR1
7669_R_O_10_DLX1
7669_R_O_11_DLX1
7669_R_O_2_DLX1
7669_R_O_3_DLX1
7669_R_O_4_DLX1
7669_R_O_5_DLX1
7669_R_O_6_DLX1
7669_R_O_7_DLX1
7669_R_O_8_DLX1
7669_R_O_9_DLX1
7669_R_Olf_DLX1
7669_R_P_1_Bulk
7669_R_P_1_COUPTF2
7669_R_P_1_DLX1
7669_R_P_1_NeuN
7669_R_P_1_TBR1
7669_R_P_10_DLX1
7669_R_P_11_DLX1
7669_R_P_11_TBR1
7669_R_P_12_DLX1
7669_R_P_12_TBR1
7669_R_P_13_DLX1
7669_R_P_13_TBR1
7669_R_P_14_DLX1
7669_R_P_14_TBR1
7669_R_P_15_DLX1
7669_R_P_15_TBR1
7669_R_P_16_DLX1
7669_R_P_16_TBR1
7669_R_P_17_DLX1
7669_R_P_17_TBR1
7669_R_P_2_DLX1
7669_R_P_2_TBR1
7669_R_P_3_DLX1
7669_R_P_3_TBR1
7669_R_P_4_DLX1
7669_R_P_4_TBR1
7669_R_P_5_DLX1
7669_R_P_5_TBR1
7669_R_P_6_DLX1
7669_R_P_6_TBR1
7669_R_P_7_DLX1
7669_R_P_7_TBR1
7669_R_P_8_DLX1
7669_R_P_8_TBR1
7669_R_P_9_DLX
7669_R_P_9_DLX1
7669_R_P_9_TBR1
7669_R_POA_Bulk
7669_R_POA_DLX1
7669_R_sec1_Cau_Bulk
7669_R_sec2_GP_Bulk
7669_R_sec2_GP_DLX1
7669_R_sec2_Put_Bulk
7669_R_sec2_Thal_Bulk
7669_R_sec2_Thal_COUPTF2
7669_R_sec2_Thal_DLX1
7669_R_sec3_Cau_DLX1
7669_R_sec3_Hip_COUPTF2
7669_R_sec3_Hip_DLX1
7669_R_sec3_Hip_TBR1
7669_R_sec3_Put_DLX1
7669_R_Skin_Bulk
7669_R_T_1_Bulk
7669_R_T_1_COUPTF2
7669_R_T_1_DLX1
7669_R_T_1_TBR1
7669_RA9_Bulk
7669_RV10_Bulk
JGG

ID05 9−13509007−G−A

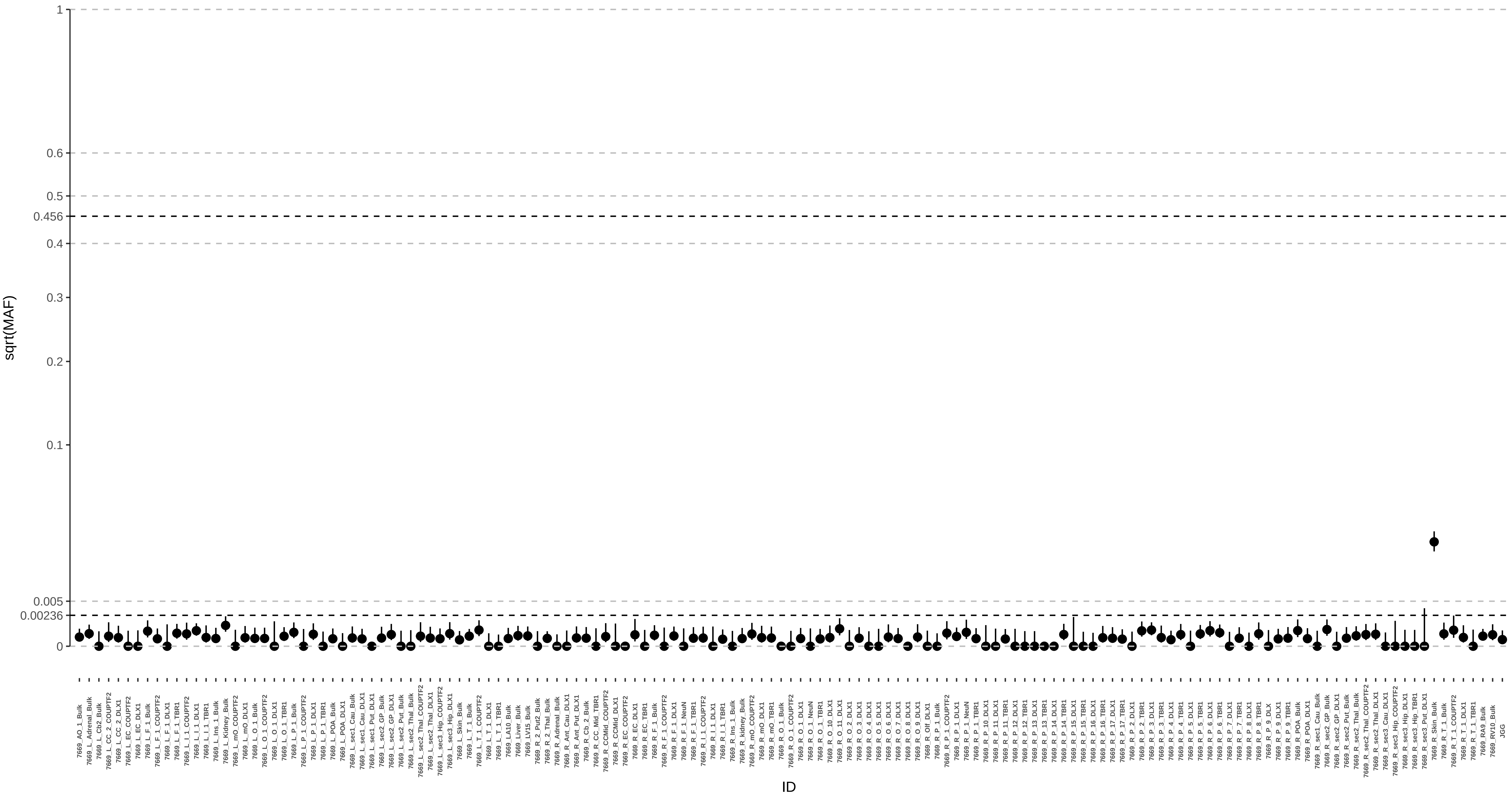

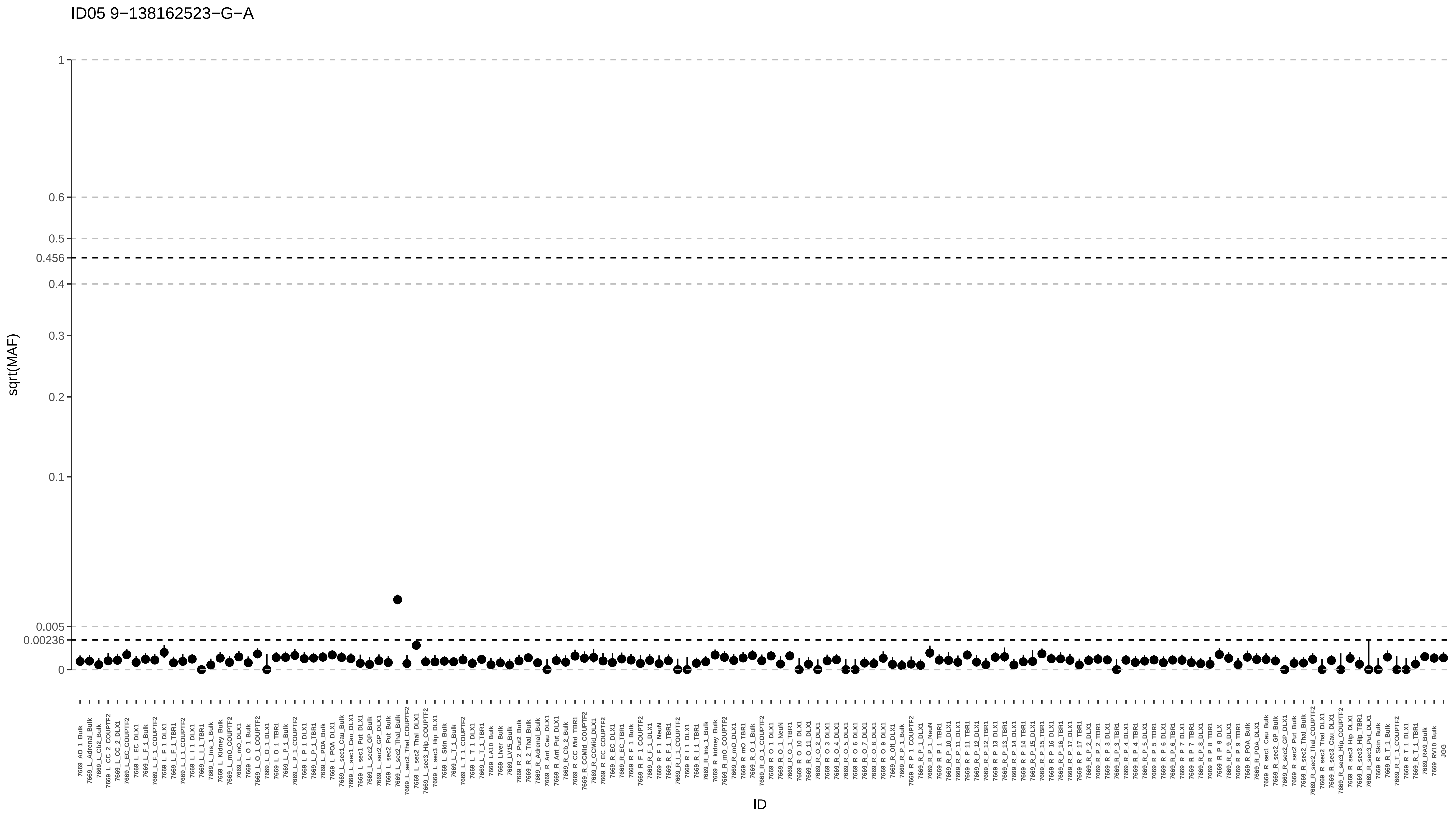
ID05 9−138162523−G−A
sqrt(MAF)
ID

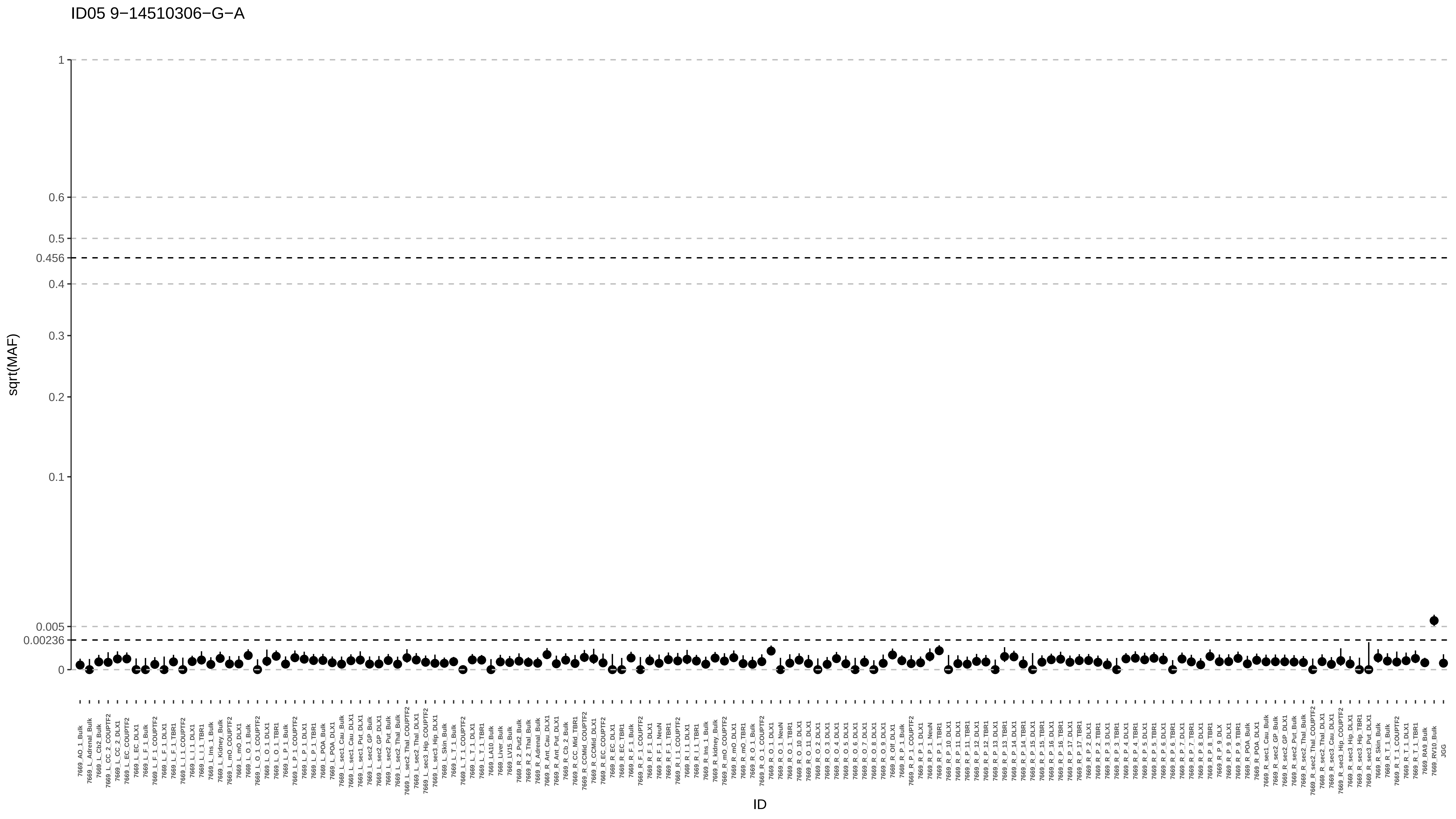
ID05 9−14510306−G−A
sqrt(MAF)
ID

ID05 9−19957121−C−T

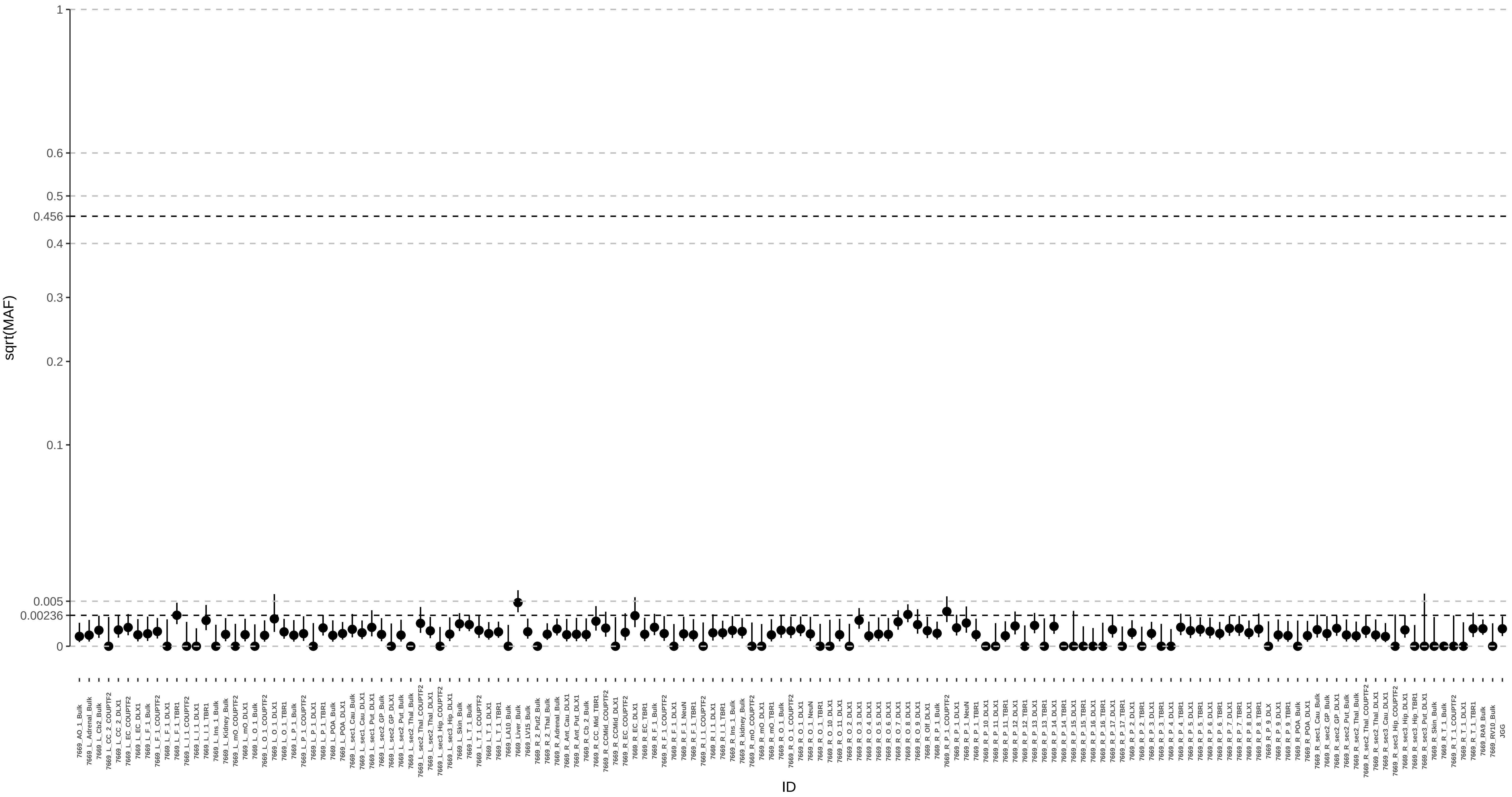

ID05 9−21154273−T−C

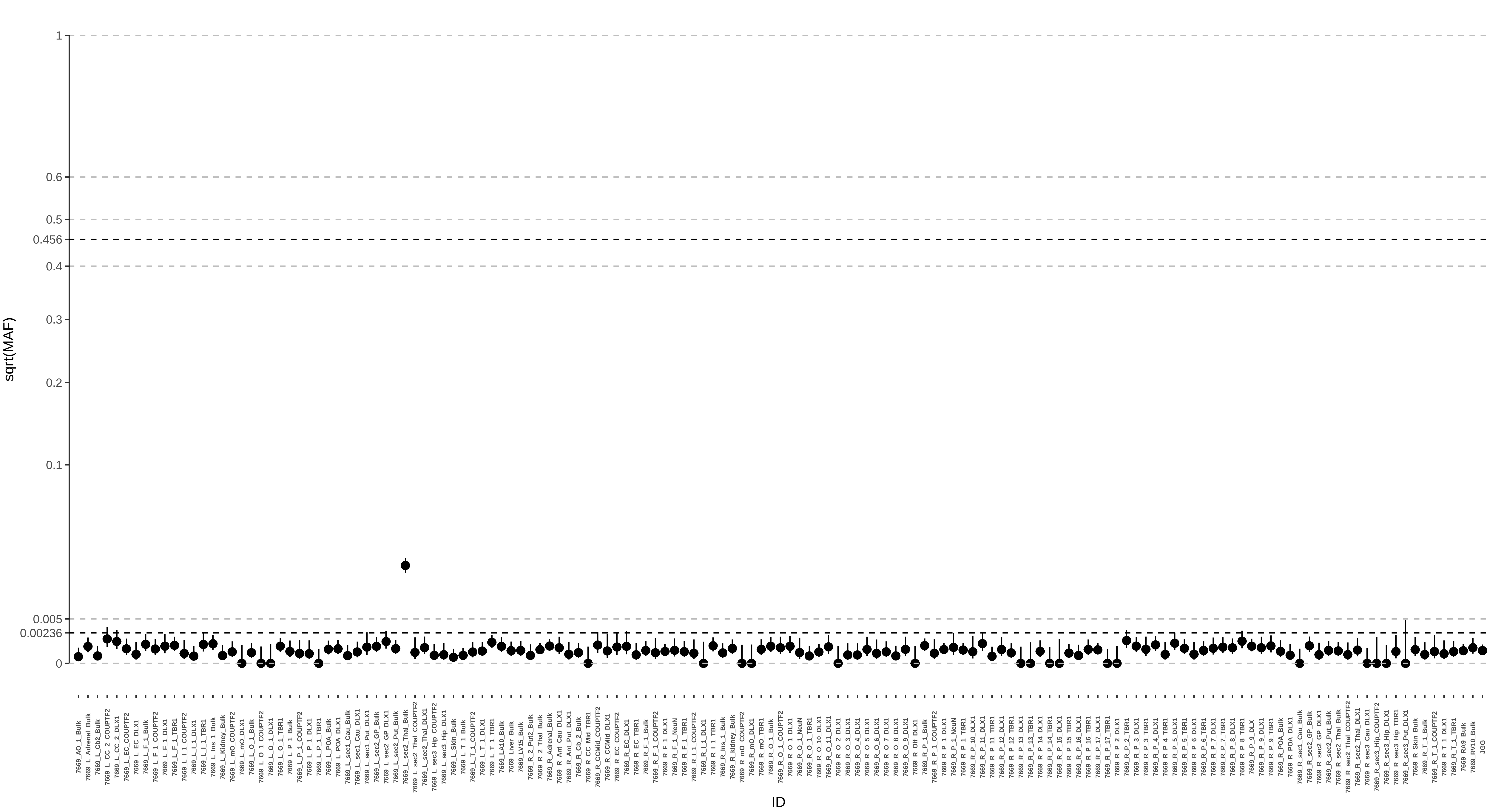

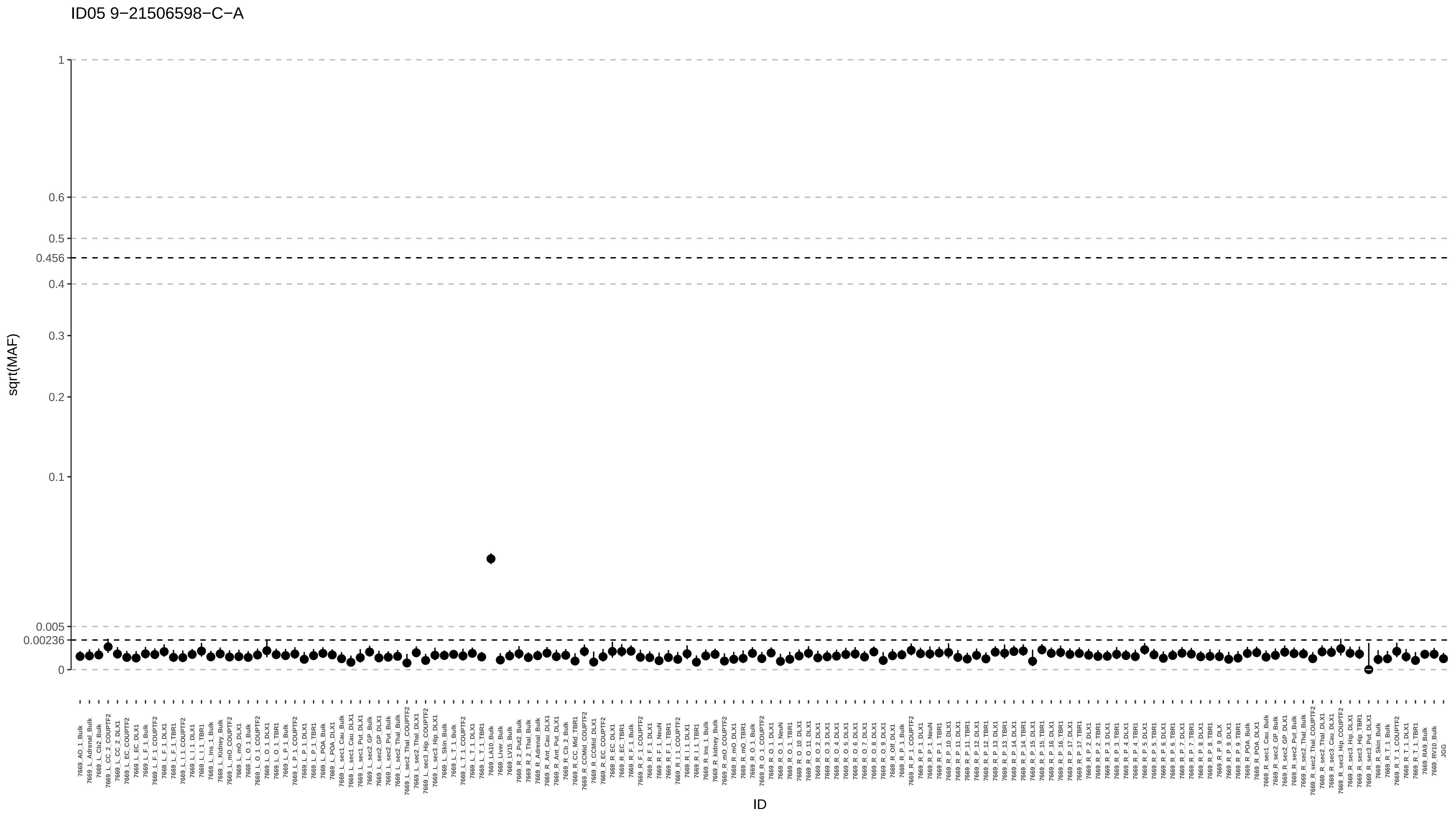
ID05 9−21506598−C−A
sqrt(MAF)
1
0.6
0.5
0.456
0.4
0.3
0.2
0.1
0.005
0.00236
0
ID

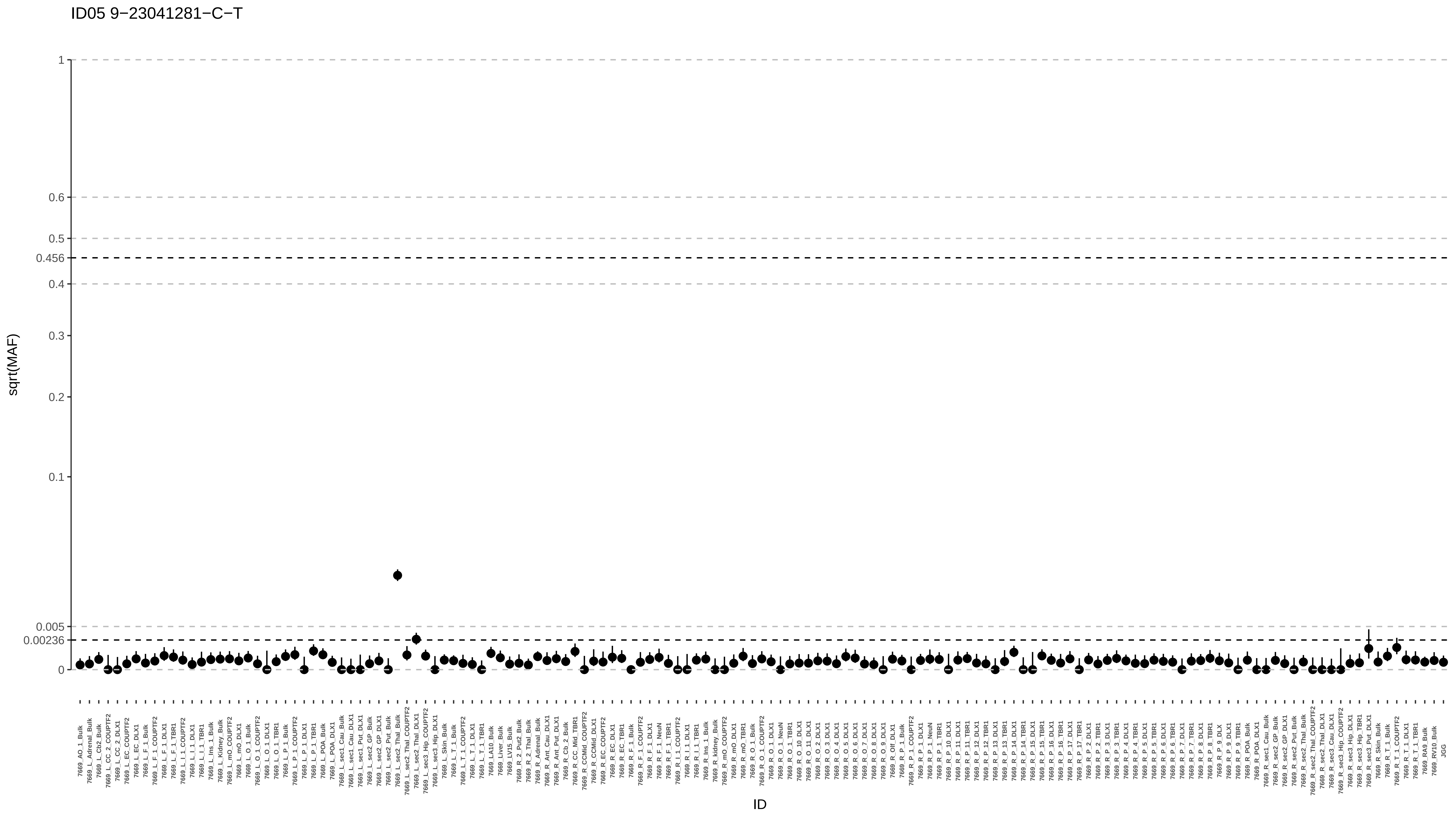
ID05 9−23041281−C−T
sqrt(MAF)
ID

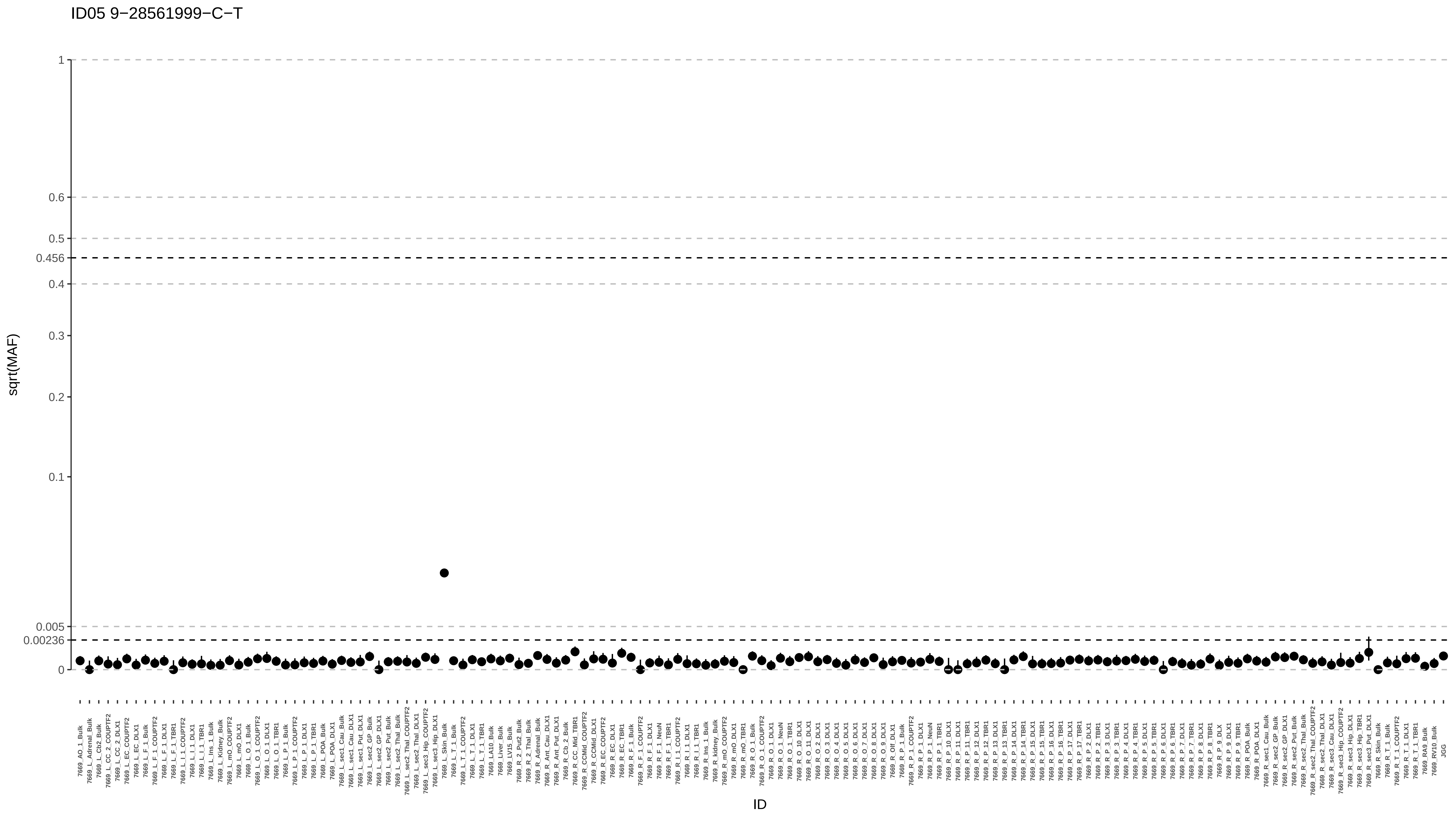
ID05 9−28561999−C−T
sqrt(MAF)
1
0.6
0.5
0.456
0.4
0.3
0.2
0.1
0.005
0.00236
0
ID

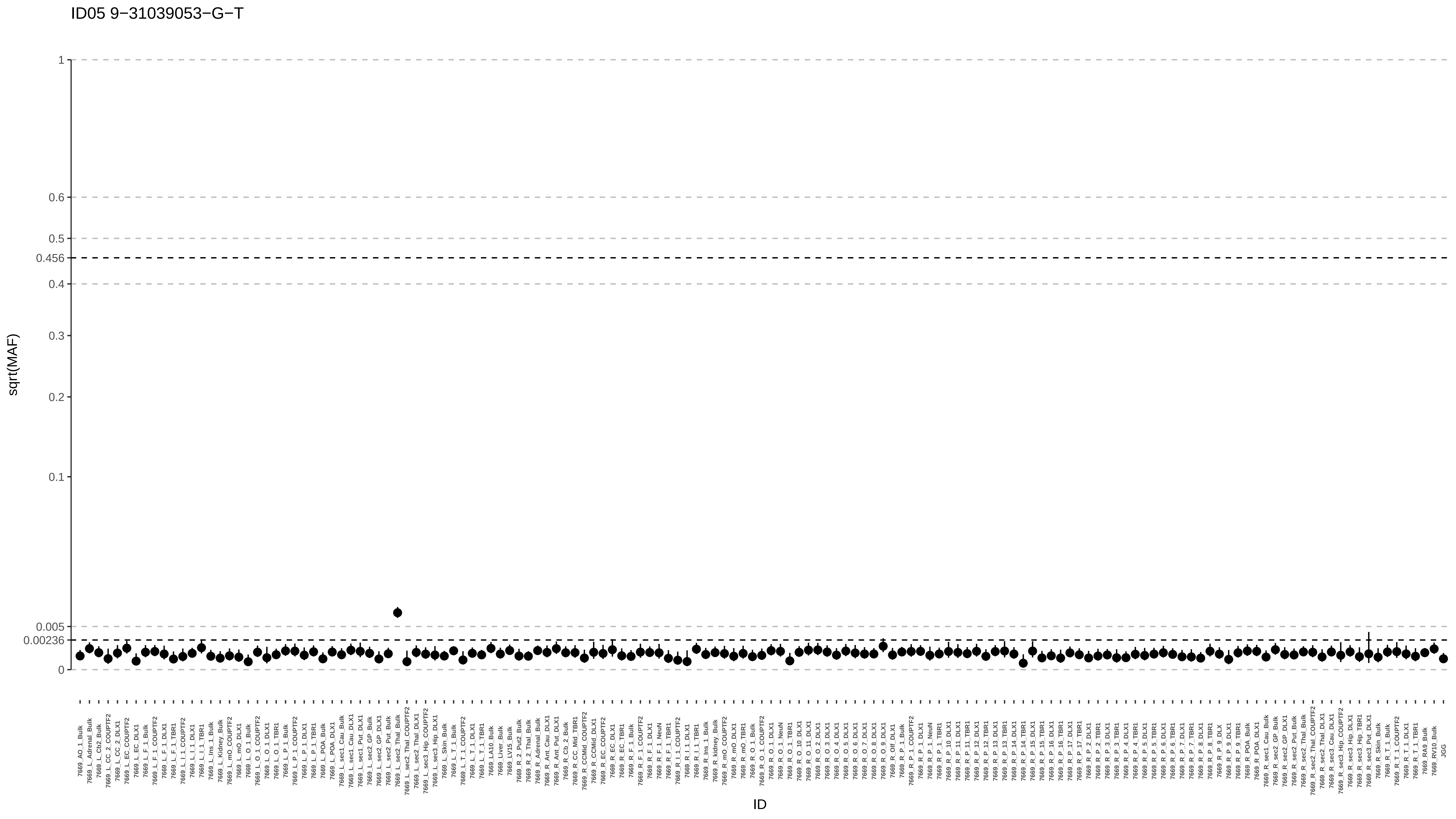
ID05 9−31039053−G−T
sqrt(MAF)
ID
1
0.6
0.5
0.456
0.4
0.3
0.2
0.1
0.005
0.00236
0

ID05 9−36967923−G−A

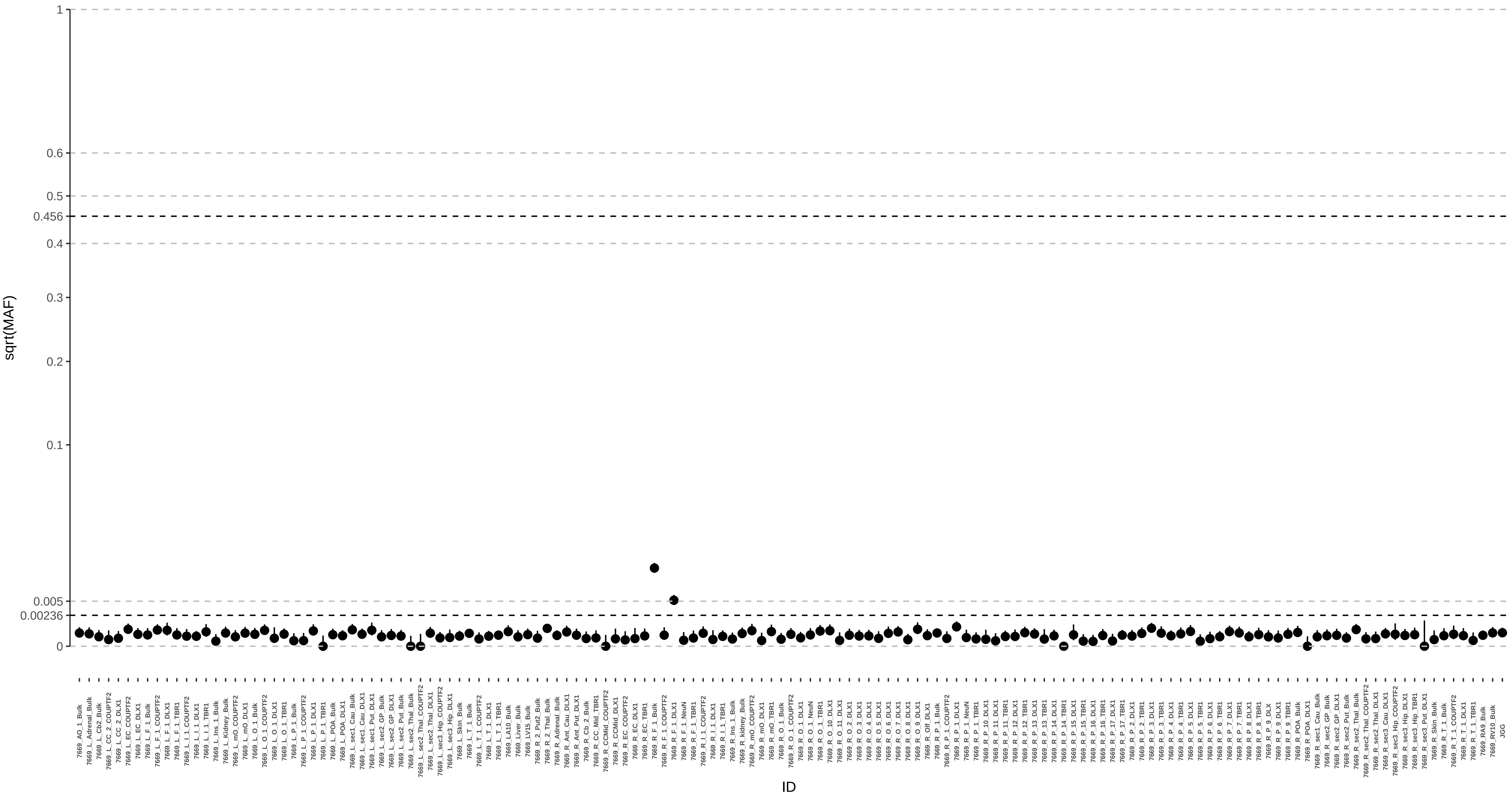

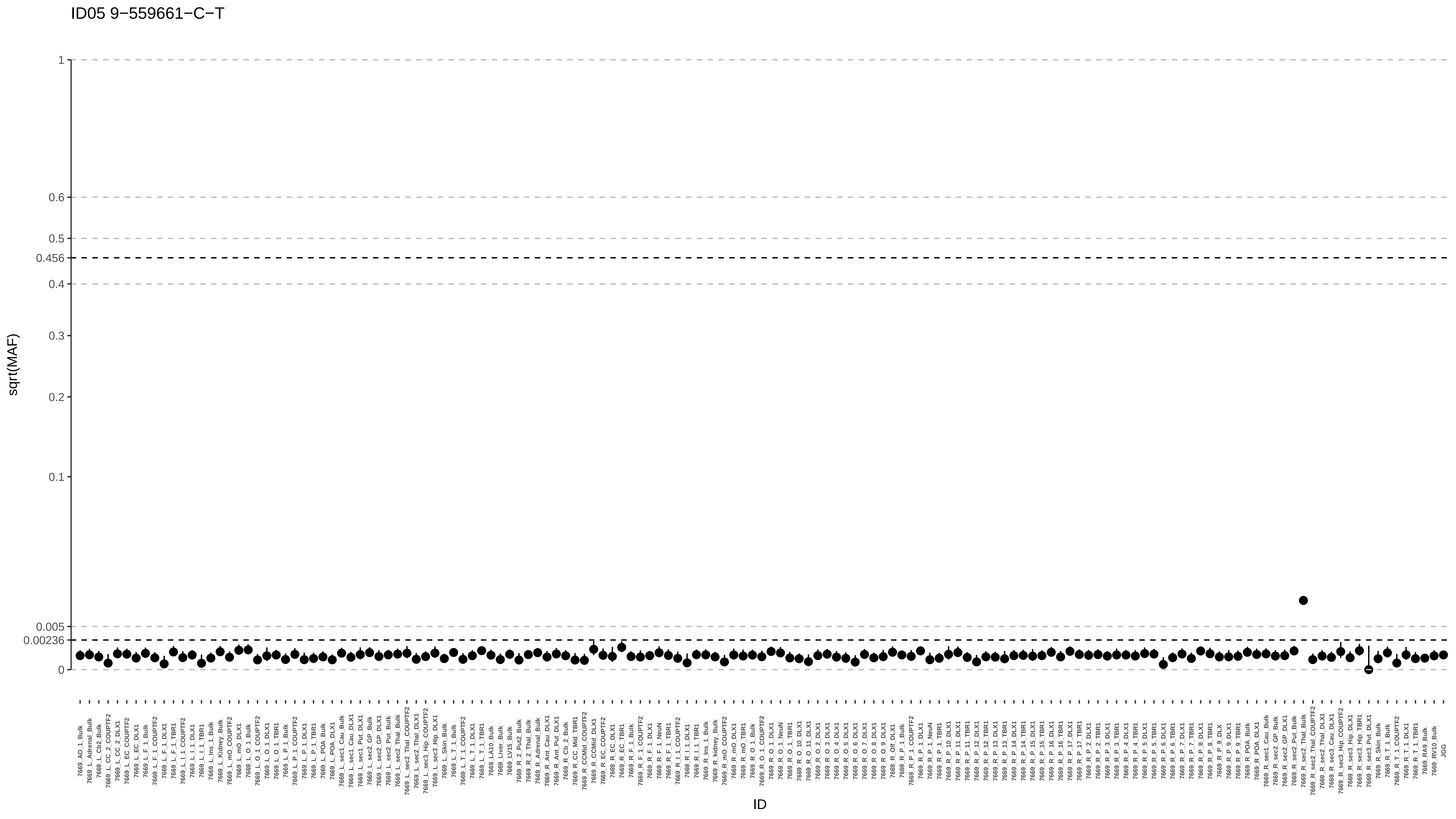
ID05 9−559661−C−T
sqrt(MAF)
1
0.6
0.5
0.456
0.4
0.3
0.2
0.1
0.005
0.00236
0
ID

ID05 9−74407663−G−A

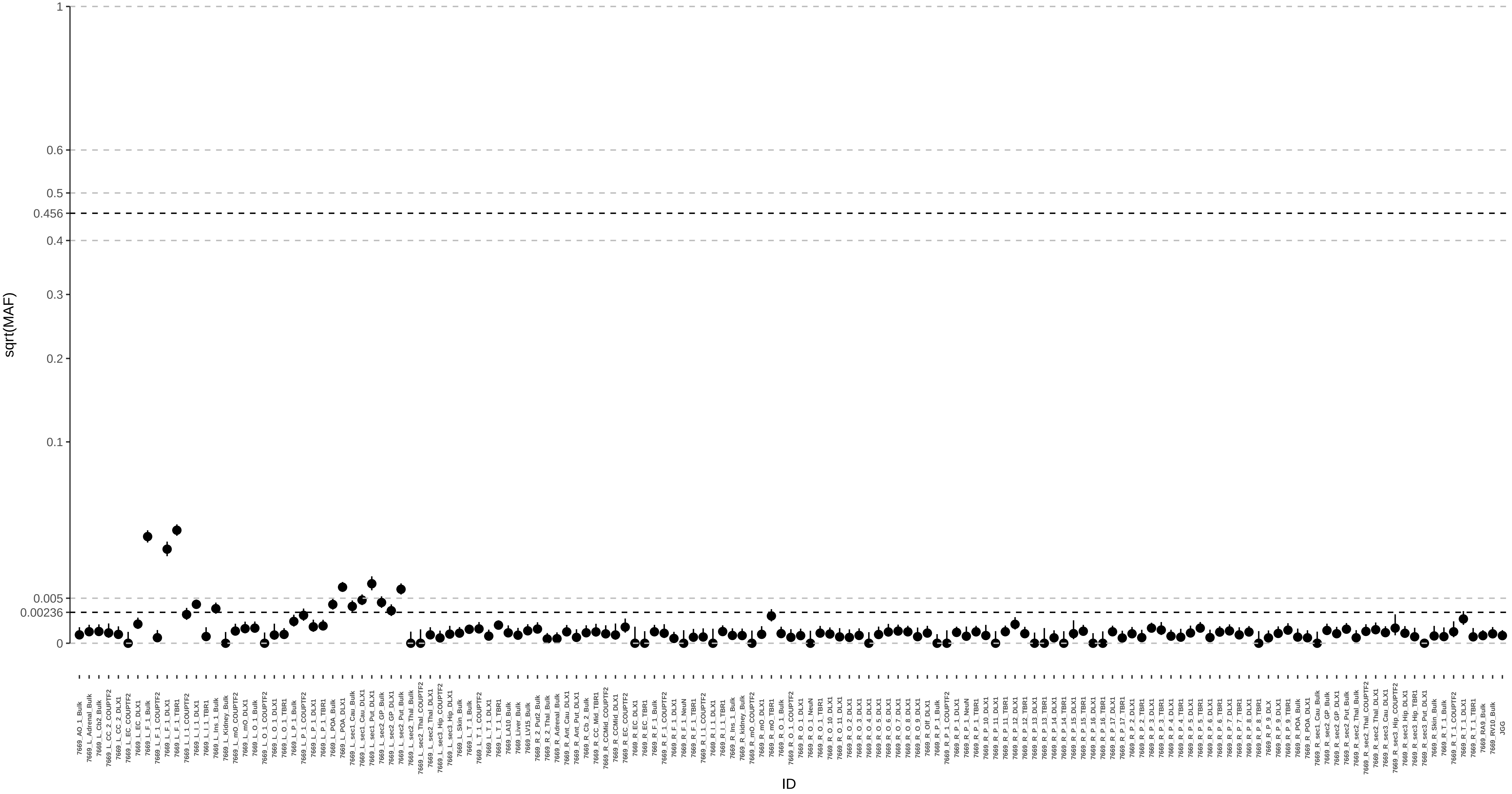

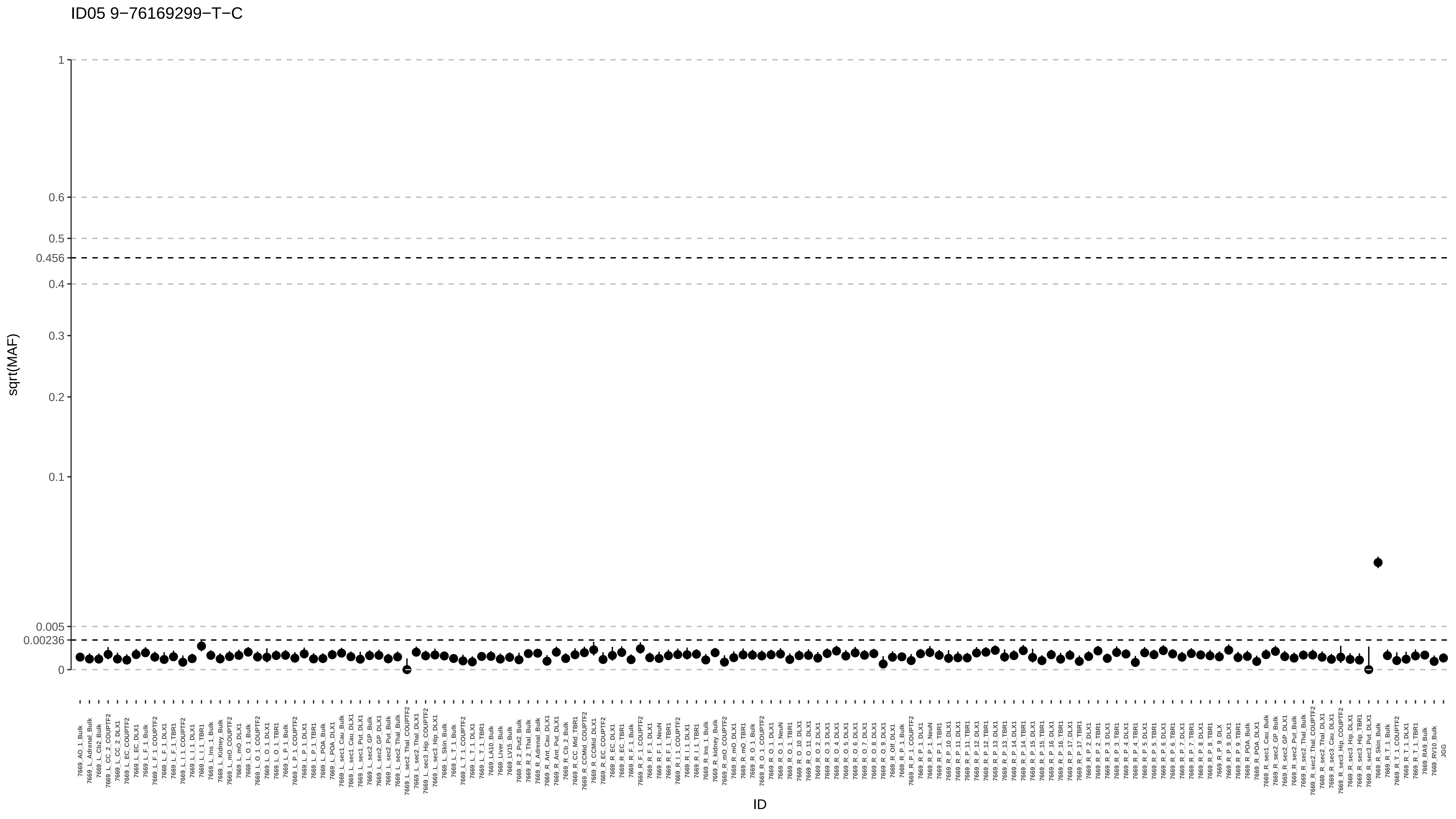

ID05 9−76169299−T−C
sqrt(MAF)
1
0.6
0.5
0.456
0.4
0.3
0.2
0.1
0.005
0.00236
0
ID
7669_AO_1_Bulk
7669_L_Adrenal_Bulk
7669_L_Cb2_Bulk
7669_L_CC_2_COUPTF2
7669_L_CC_2_DLX1
7669_L_EC_COUPTF2
7669_L_EC_DLX1
7669_L_F_1_Bulk
7669_L_F_1_COUPTF2
7669_L_F_1_DLX1
7669_L_F_1_TBR1
7669_L_I_1_COUPTF2
7669_L_I_1_DLX1
7669_L_I_1_TBR1
7669_L_Ins_1_Bulk
7669_L_Kidney_Bulk
7669_L_mO_COUPTF2
7669_L_mO_DLX1
7669_L_O_1_Bulk
7669_L_O_1_COUPTF2
7669_L_O_1_DLX1
7669_L_O_1_TBR1
7669_L_P_1_Bulk
7669_L_P_1_COUPTF2
7669_L_P_1_DLX1
7669_L_P_1_TBR1
7669_L_POA_Bulk
7669_L_POA_DLX1
7669_L_sec1_Cau_Bulk
7669_L_sec1_Cau_DLX1
7669_L_sec1_Put_DLX1
7669_L_sec2_GP_Bulk
7669_L_sec2_GP_DLX1
7669_L_sec2_Put_Bulk
7669_L_sec2_Thal_Bulk
7669_L_sec2_Thal_COUPTF2
7669_L_sec2_Thal_DLX1
7669_L_sec3_Hip_COUPTF2
7669_L_sec3_Hip_DLX1
7669_L_Skin_Bulk
7669_L_T_1_Bulk
7669_L_T_1_COUPTF2
7669_L_T_1_DLX1
7669_L_T_1_TBR1
7669_LA10_Bulk
7669_Liver_Bulk
7669_LV15_Bulk
7669_R_2_Put2_Bulk
7669_R_2_Thal_Bulk
7669_R_Adrenal_Bulk
7669_R_Ant_Cau_DLX1
7669_R_Ant_Put_DLX1
7669_R_Cb_2_Bulk
7669_R_CC_Mid_TBR1
7669_R_CCMid_COUPTF2
7669_R_CCMid_DLX1
7669_R_EC_COUPTF2
7669_R_EC_DLX1
7669_R_EC_TBR1
7669_R_F_1_Bulk
7669_R_F_1_COUPTF2
7669_R_F_1_DLX1
7669_R_F_1_NeuN
7669_R_F_1_TBR1
7669_R_I_1_COUPTF2
7669_R_I_1_DLX1
7669_R_I_1_TBR1
7669_R_Ins_1_Bulk
7669_R_kidney_Bulk
7669_R_mO_COUPTF2
7669_R_mO_DLX1
7669_R_mO_TBR1
7669_R_O_1_Bulk
7669_R_O_1_COUPTF2
7669_R_O_1_DLX1
7669_R_O_1_NeuN
7669_R_O_1_TBR1
7669_R_O_10_DLX1
7669_R_O_11_DLX1
7669_R_O_2_DLX1
7669_R_O_3_DLX1
7669_R_O_4_DLX1
7669_R_O_5_DLX1
7669_R_O_6_DLX1
7669_R_O_7_DLX1
7669_R_O_8_DLX1
7669_R_O_9_DLX1
7669_R_Olf_DLX1
7669_R_P_1_Bulk
7669_R_P_1_COUPTF2
7669_R_P_1_DLX1
7669_R_P_1_NeuN
7669_R_P_1_TBR1
7669_R_P_10_DLX1
7669_R_P_11_DLX1
7669_R_P_11_TBR1
7669_R_P_12_DLX1
7669_R_P_12_TBR1
7669_R_P_13_DLX1
7669_R_P_13_TBR1
7669_R_P_14_DLX1
7669_R_P_14_TBR1
7669_R_P_15_DLX1
7669_R_P_15_TBR1
7669_R_P_16_DLX1
7669_R_P_16_TBR1
7669_R_P_17_DLX1
7669_R_P_17_TBR1
7669_R_P_2_DLX1
7669_R_P_2_TBR1
7669_R_P_3_DLX1
7669_R_P_3_TBR1
7669_R_P_4_DLX1
7669_R_P_4_TBR1
7669_R_P_5_DLX1
7669_R_P_5_TBR1
7669_R_P_6_DLX1
7669_R_P_6_TBR1
7669_R_P_7_DLX1
7669_R_P_7_TBR1
7669_R_P_8_DLX1
7669_R_P_8_TBR1
7669_R_P_9_DLX
7669_R_P_9_DLX1
7669_R_P_9_TBR1
7669_R_POA_Bulk
7669_R_POA_DLX1
7669_R_sec1_Cau_Bulk
7669_R_sec2_GP_Bulk
7669_R_sec2_GP_DLX1
7669_R_sec2_Put_Bulk
7669_R_sec2_Thal_Bulk
7669_R_sec2_Thal_COUPTF2
7669_R_sec2_Thal_DLX1
7669_R_sec3_Cau_DLX1
7669_R_sec3_Hip_COUPTF2
7669_R_sec3_Hip_DLX1
7669_R_sec3_Hip_TBR1
7669_R_sec3_Put_DLX1
7669_R_Skin_Bulk
7669_R_T_1_Bulk
7669_R_T_1_COUPTF2
7669_R_T_1_DLX1
7669_R_T_1_TBR1
7669_RA9_Bulk
7669_RV10_Bulk
JGG

ID05 9−77140705−C−T

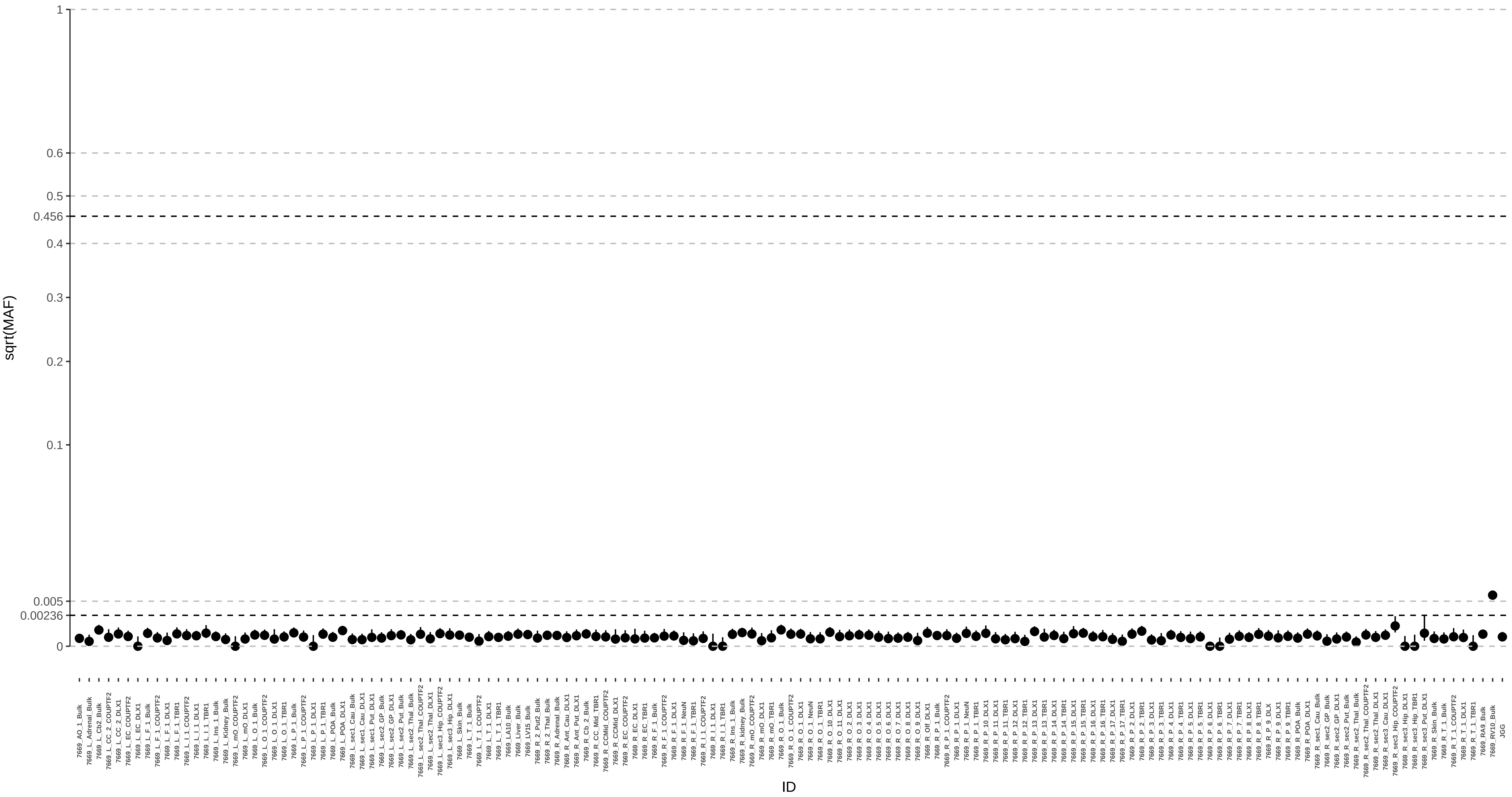

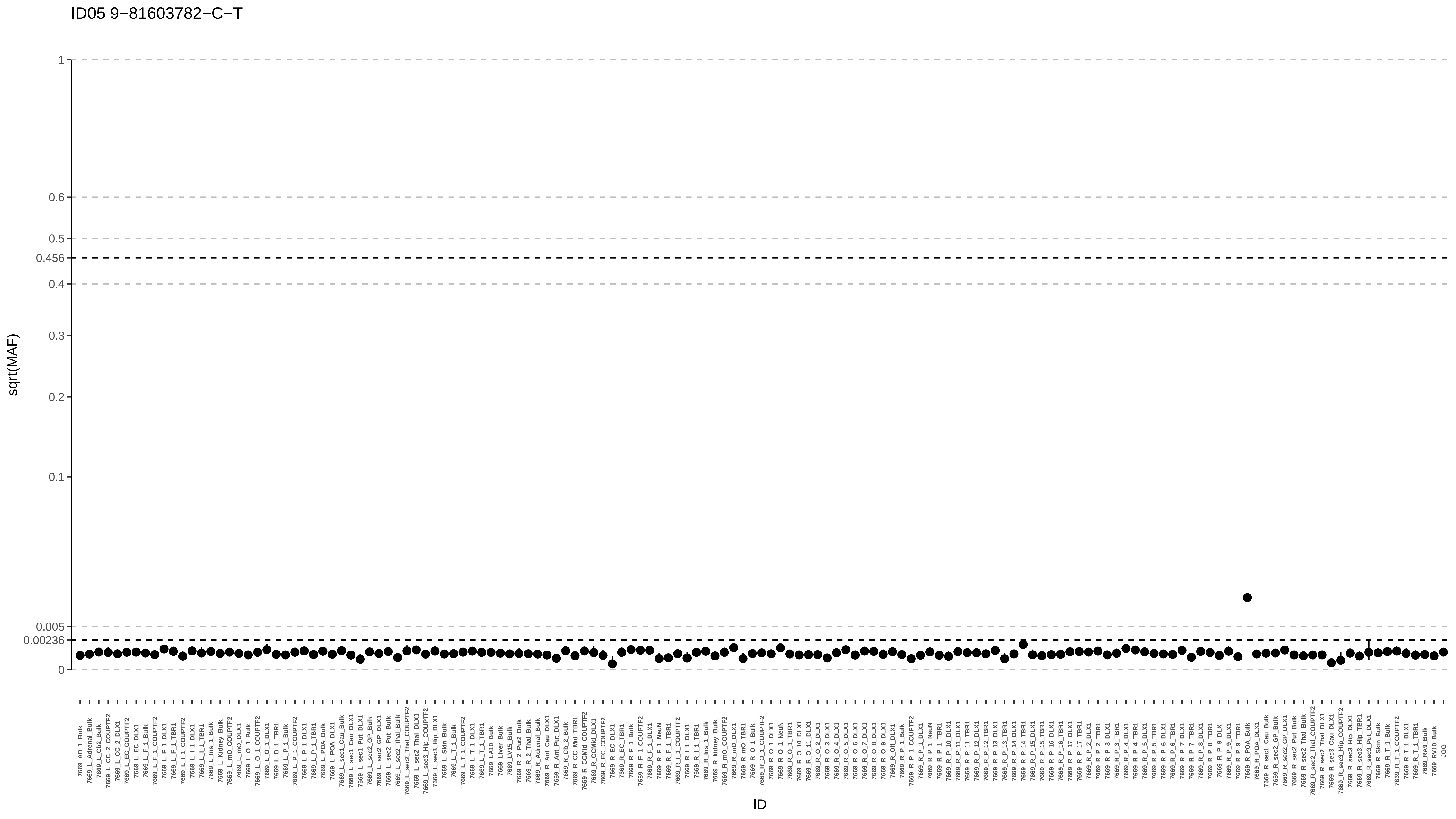
ID05 9−81603782−C−T
sqrt(MAF)
1
0.6
0.5
0.456
0.4
0.3
0.2
0.1
0.005
0.00236
0
ID
7669_AO_1_Bulk
7669_L_Adrenal_Bulk
7669_L_Cb2_Bulk
7669_L_CC_2_COUPTF2
7669_L_CC_2_DLX1
7669_L_EC_COUPTF2
7669_L_EC_DLX1
7669_L_F_1_Bulk
7669_L_F_1_COUPTF2
7669_L_F_1_DLX1
7669_L_F_1_TBR1
7669_L_I_1_COUPTF2
7669_L_I_1_DLX1
7669_L_I_1_TBR1
7669_L_Ins_1_Bulk
7669_L_Kidney_Bulk
7669_L_mO_COUPTF2
7669_L_mO_DLX1
7669_L_O_1_Bulk
7669_L_O_1_COUPTF2
7669_L_O_1_DLX1
7669_L_O_1_TBR1
7669_L_P_1_Bulk
7669_L_P_1_COUPTF2
7669_L_P_1_DLX1
7669_L_P_1_TBR1
7669_L_POA_Bulk
7669_L_POA_DLX1
7669_L_sec1_Cau_Bulk
7669_L_sec1_Cau_DLX1
7669_L_sec1_Put_DLX1
7669_L_sec2_GP_Bulk
7669_L_sec2_GP_DLX1
7669_L_sec2_Put_Bulk
7669_L_sec2_Thal_Bulk
7669_L_sec2_Thal_COUPTF2
7669_L_sec2_Thal_DLX1
7669_L_sec3_Hip_COUPTF2
7669_L_sec3_Hip_DLX1
7669_L_Skin_Bulk
7669_L_T_1_Bulk
7669_L_T_1_COUPTF2
7669_L_T_1_DLX1
7669_L_T_1_TBR1
7669_LA10_Bulk
7669_Liver_Bulk
7669_LV15_Bulk
7669_R_2_Put2_Bulk
7669_R_2_Thal_Bulk
7669_R_Adrenal_Bulk
7669_R_Ant_Cau_DLX1
7669_R_Ant_Put_DLX1
7669_R_Cb_2_Bulk
7669_R_CC_Mid_TBR1
7669_R_CCMid_COUPTF2
7669_R_CCMid_DLX1
7669_R_EC_COUPTF2
7669_R_EC_DLX1
7669_R_EC_TBR1
7669_R_F_1_Bulk
7669_R_F_1_COUPTF2
7669_R_F_1_DLX1
7669_R_F_1_NeuN
7669_R_F_1_TBR1
7669_R_I_1_COUPTF2
7669_R_I_1_DLX1
7669_R_I_1_TBR1
7669_R_Ins_1_Bulk
7669_R_kidney_Bulk
7669_R_mO_COUPTF2
7669_R_mO_DLX1
7669_R_mO_TBR1
7669_R_O_1_Bulk
7669_R_O_1_COUPTF2
7669_R_O_1_DLX1
7669_R_O_1_NeuN
7669_R_O_1_TBR1
7669_R_O_10_DLX1
7669_R_O_11_DLX1
7669_R_O_2_DLX1
7669_R_O_3_DLX1
7669_R_O_4_DLX1
7669_R_O_5_DLX1
7669_R_O_6_DLX1
7669_R_O_7_DLX1
7669_R_O_8_DLX1
7669_R_O_9_DLX1
7669_R_Olf_DLX1
7669_R_P_1_Bulk
7669_R_P_1_COUPTF2
7669_R_P_1_DLX1
7669_R_P_1_NeuN
7669_R_P_1_TBR1
7669_R_P_10_DLX1
7669_R_P_11_DLX1
7669_R_P_11_TBR1
7669_R_P_12_DLX1
7669_R_P_12_TBR1
7669_R_P_13_DLX1
7669_R_P_13_TBR1
7669_R_P_14_DLX1
7669_R_P_14_TBR1
7669_R_P_15_DLX1
7669_R_P_15_TBR1
7669_R_P_16_DLX1
7669_R_P_16_TBR1
7669_R_P_17_DLX1
7669_R_P_17_TBR1
7669_R_P_2_DLX1
7669_R_P_2_TBR1
7669_R_P_3_DLX1
7669_R_P_3_TBR1
7669_R_P_4_DLX1
7669_R_P_4_TBR1
7669_R_P_5_DLX1
7669_R_P_5_TBR1
7669_R_P_6_DLX1
7669_R_P_6_TBR1
7669_R_P_7_DLX1
7669_R_P_7_TBR1
7669_R_P_8_DLX1
7669_R_P_8_TBR1
7669_R_P_9_DLX
7669_R_P_9_DLX1
7669_R_P_9_TBR1
7669_R_POA_Bulk
7669_R_POA_DLX1
7669_R_sec1_Cau_Bulk
7669_R_sec2_GP_Bulk
7669_R_sec2_GP_DLX1
7669_R_sec2_Put_Bulk
7669_R_sec2_Thal_Bulk
7669_R_sec2_Thal_COUPTF2
7669_R_sec2_Thal_DLX1
7669_R_sec3_Cau_DLX1
7669_R_sec3_Hip_COUPTF2
7669_R_sec3_Hip_DLX1
7669_R_sec3_Hip_TBR1
7669_R_sec3_Put_DLX1
7669_R_Skin_Bulk
7669_R_T_1_Bulk
7669_R_T_1_COUPTF2
7669_R_T_1_DLX1
7669_R_T_1_TBR1
7669_RA9_Bulk
7669_RV10_Bulk
JGG

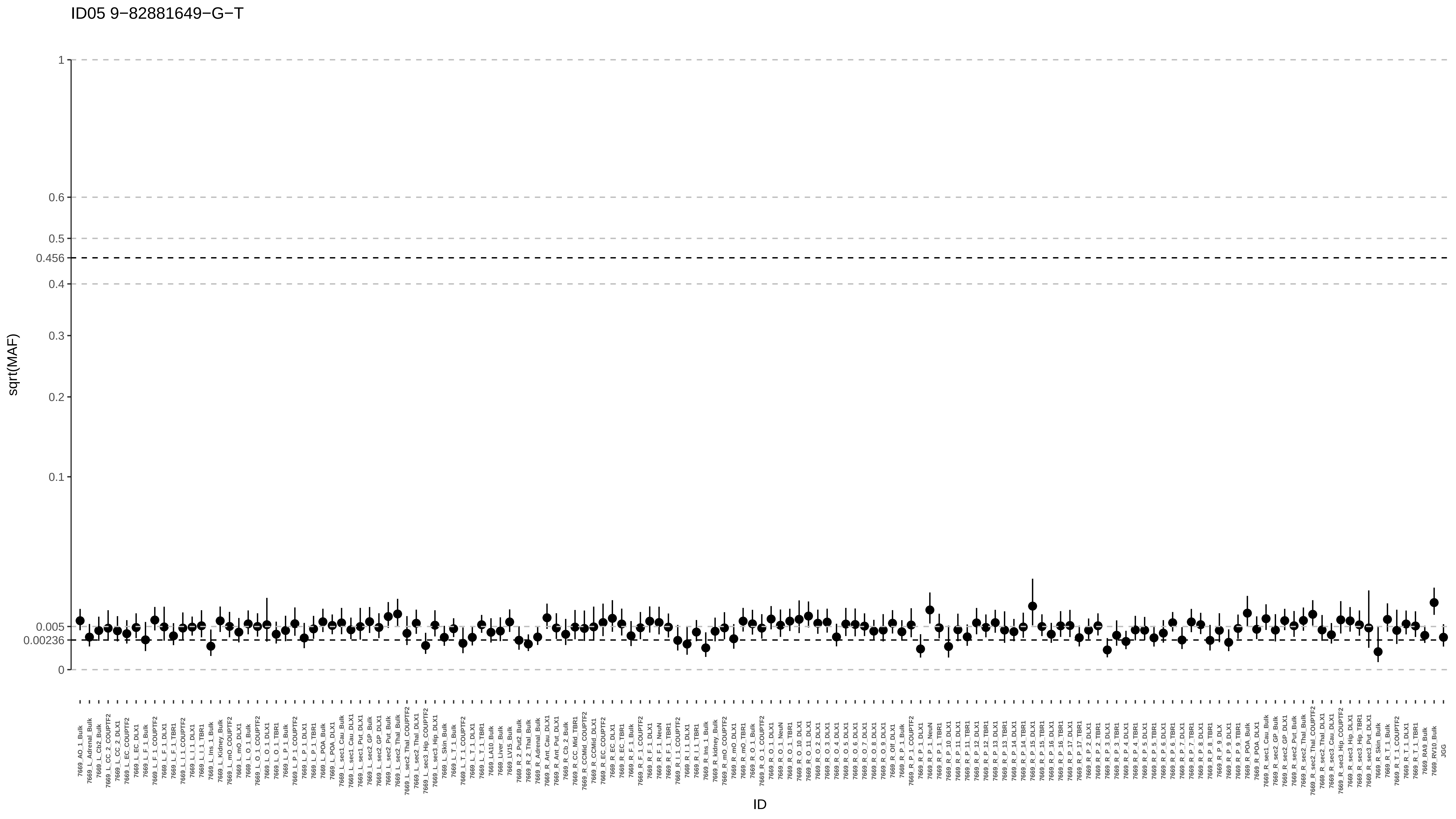
ID05 9−82881649−G−T
sqrt(MAF)
1
0.6
0.5
0.456
0.4
0.3
0.2
0.1
0.005
0.00236
0
ID

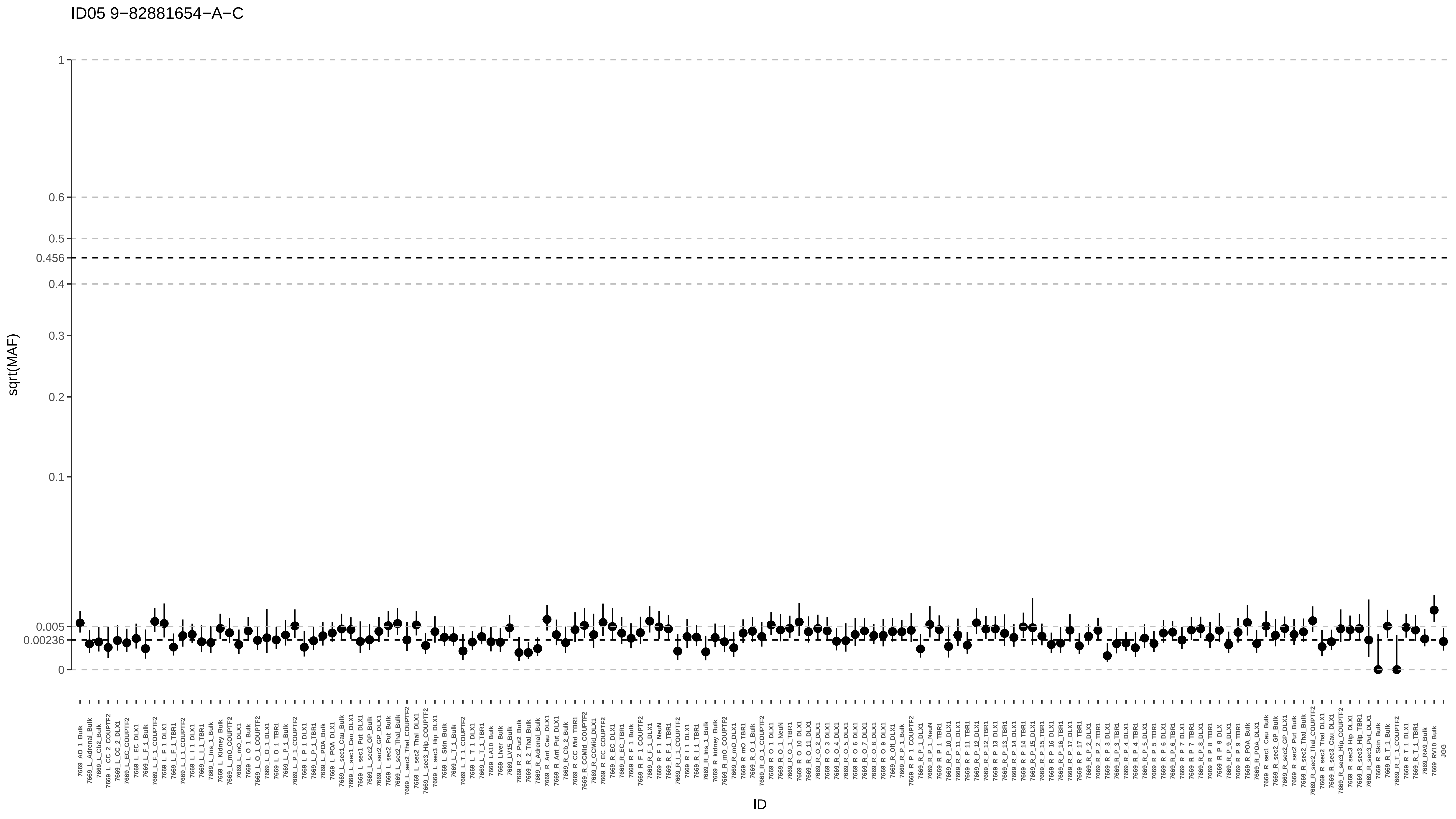

ID05 9−82881654−A−C
sqrt(MAF)
1
0.6
0.5
0.456
0.4
0.3
0.2
0.1
0.005
0.00236
0
ID
7669_AO_1_Bulk
7669_L_Adrenal_Bulk
7669_L_Cb2_Bulk
7669_L_CC_2_COUPTF2
7669_L_CC_2_DLX1
7669_L_EC_COUPTF2
7669_L_EC_DLX1
7669_L_F_1_Bulk
7669_L_F_1_COUPTF2
7669_L_F_1_DLX1
7669_L_F_1_TBR1
7669_L_I_1_COUPTF2
7669_L_I_1_DLX1
7669_L_I_1_TBR1
7669_L_Ins_1_Bulk
7669_L_Kidney_Bulk
7669_L_mO_COUPTF2
7669_L_mO_DLX1
7669_L_O_1_Bulk
7669_L_O_1_COUPTF2
7669_L_O_1_DLX1
7669_L_O_1_TBR1
7669_L_P_1_Bulk
7669_L_P_1_COUPTF2
7669_L_P_1_DLX1
7669_L_P_1_TBR1
7669_L_POA_Bulk
7669_L_POA_DLX1
7669_L_sec1_Cau_Bulk
7669_L_sec1_Cau_DLX1
7669_L_sec1_Put_DLX1
7669_L_sec2_GP_Bulk
7669_L_sec2_GP_DLX1
7669_L_sec2_Put_Bulk
7669_L_sec2_Thal_Bulk
7669_L_sec2_Thal_COUPTF2
7669_L_sec2_Thal_DLX1
7669_L_sec3_Hip_COUPTF2
7669_L_sec3_Hip_DLX1
7669_L_Skin_Bulk
7669_L_T_1_Bulk
7669_L_T_1_COUPTF2
7669_L_T_1_DLX1
7669_L_T_1_TBR1
7669_LA10_Bulk
7669_Liver_Bulk
7669_LV15_Bulk
7669_R_2_Put2_Bulk
7669_R_2_Thal_Bulk
7669_R_Adrenal_Bulk
7669_R_Ant_Cau_DLX1
7669_R_Ant_Put_DLX1
7669_R_Cb_2_Bulk
7669_R_CC_Mid_TBR1
7669_R_CCMid_COUPTF2
7669_R_CCMid_DLX1
7669_R_EC_COUPTF2
7669_R_EC_DLX1
7669_R_EC_TBR1
7669_R_F_1_Bulk
7669_R_F_1_COUPTF2
7669_R_F_1_DLX1
7669_R_F_1_NeuN
7669_R_F_1_TBR1
7669_R_I_1_COUPTF2
7669_R_I_1_DLX1
7669_R_I_1_TBR1
7669_R_Ins_1_Bulk
7669_R_kidney_Bulk
7669_R_mO_COUPTF2
7669_R_mO_DLX1
7669_R_mO_TBR1
7669_R_O_1_Bulk
7669_R_O_1_COUPTF2
7669_R_O_1_DLX1
7669_R_O_1_NeuN
7669_R_O_1_TBR1
7669_R_O_10_DLX1
7669_R_O_11_DLX1
7669_R_O_2_DLX1
7669_R_O_3_DLX1
7669_R_O_4_DLX1
7669_R_O_5_DLX1
7669_R_O_6_DLX1
7669_R_O_7_DLX1
7669_R_O_8_DLX1
7669_R_O_9_DLX1
7669_R_Olf_DLX1
7669_R_P_1_Bulk
7669_R_P_1_COUPTF2
7669_R_P_1_DLX1
7669_R_P_1_NeuN
7669_R_P_1_TBR1
7669_R_P_10_DLX1
7669_R_P_11_DLX1
7669_R_P_11_TBR1
7669_R_P_12_DLX1
7669_R_P_12_TBR1
7669_R_P_13_DLX1
7669_R_P_13_TBR1
7669_R_P_14_DLX1
7669_R_P_14_TBR1
7669_R_P_15_DLX1
7669_R_P_15_TBR1
7669_R_P_16_DLX1
7669_R_P_16_TBR1
7669_R_P_17_DLX1
7669_R_P_17_TBR1
7669_R_P_2_DLX1
7669_R_P_2_TBR1
7669_R_P_3_DLX1
7669_R_P_3_TBR1
7669_R_P_4_DLX1
7669_R_P_4_TBR1
7669_R_P_5_DLX1
7669_R_P_5_TBR1
7669_R_P_6_DLX1
7669_R_P_6_TBR1
7669_R_P_7_DLX1
7669_R_P_7_TBR1
7669_R_P_8_DLX1
7669_R_P_8_TBR1
7669_R_P_9_DLX
7669_R_P_9_DLX1
7669_R_P_9_TBR1
7669_R_POA_Bulk
7669_R_POA_DLX1
7669_R_sec1_Cau_Bulk
7669_R_sec2_GP_Bulk
7669_R_sec2_GP_DLX1
7669_R_sec2_Put_Bulk
7669_R_sec2_Thal_Bulk
7669_R_sec2_Thal_COUPTF2
7669_R_sec2_Thal_DLX1
7669_R_sec3_Cau_DLX1
7669_R_sec3_Hip_COUPTF2
7669_R_sec3_Hip_DLX1
7669_R_sec3_Hip_TBR1
7669_R_sec3_Put_DLX1
7669_R_Skin_Bulk
7669_R_T_1_Bulk
7669_R_T_1_COUPTF2
7669_R_T_1_DLX1
7669_R_T_1_TBR1
7669_RA9_Bulk
7669_RV10_Bulk
JGG

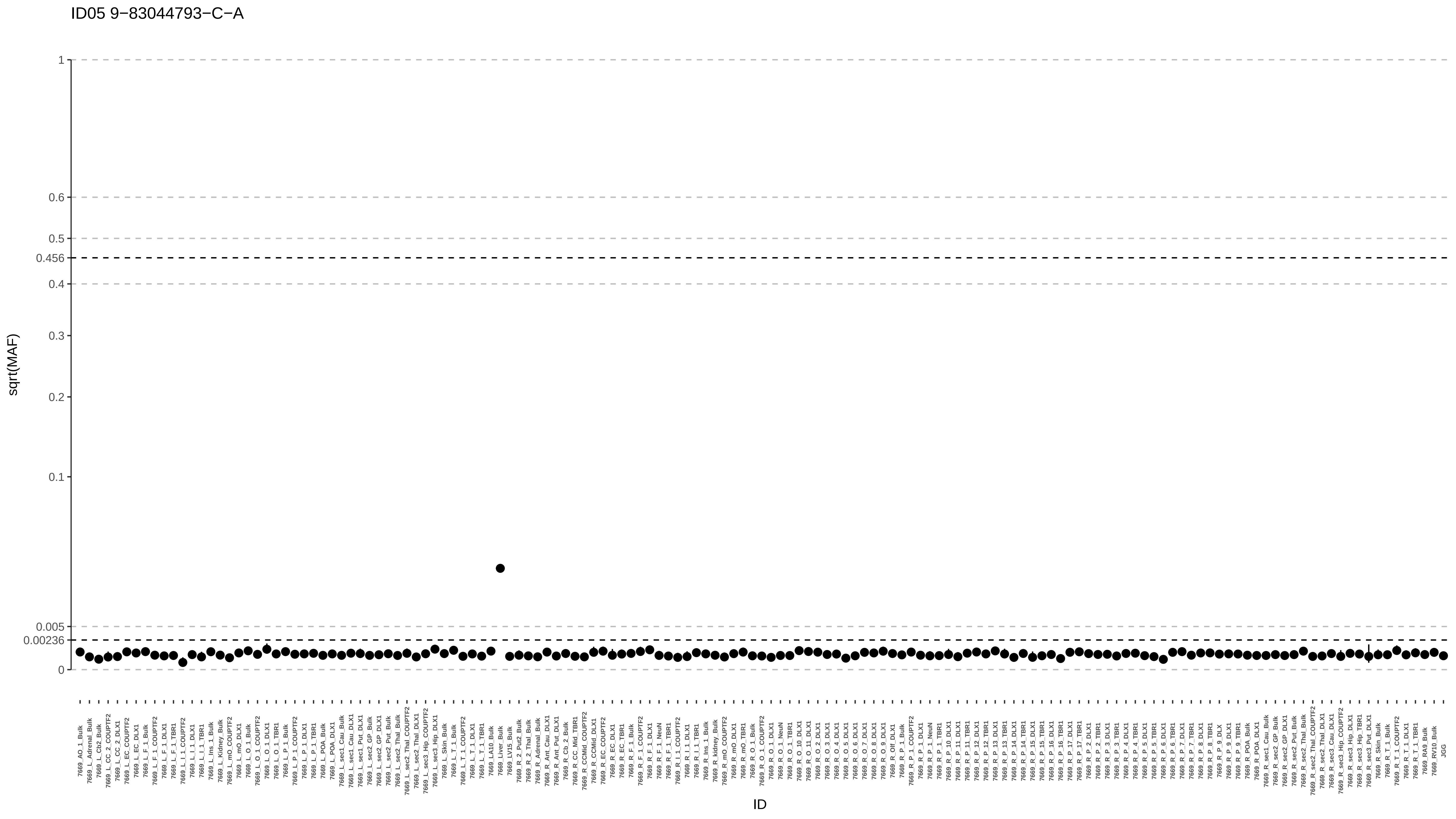
ID05 9−83044793−C−A
sqrt(MAF)
1
0.6
0.5
0.456
0.4
0.3
0.2
0.1
0.005
0.00236
0
ID
7669_AO_1_Bulk
7669_L_Adrenal_Bulk
7669_L_Cb2_Bulk
7669_L_CC_2_COUPTF2
7669_L_CC_2_DLX1
7669_L_EC_COUPTF2
7669_L_EC_DLX1
7669_L_F_1_Bulk
7669_L_F_1_COUPTF2
7669_L_F_1_DLX1
7669_L_F_1_TBR1
7669_L_I_1_COUPTF2
7669_L_I_1_DLX1
7669_L_I_1_TBR1
7669_L_Ins_1_Bulk
7669_L_Kidney_Bulk
7669_L_mO_COUPTF2
7669_L_mO_DLX1
7669_L_O_1_Bulk
7669_L_O_1_COUPTF2
7669_L_O_1_DLX1
7669_L_O_1_TBR1
7669_L_P_1_Bulk
7669_L_P_1_COUPTF2
7669_L_P_1_DLX1
7669_L_P_1_TBR1
7669_L_POA_Bulk
7669_L_POA_DLX1
7669_L_sec1_Cau_Bulk
7669_L_sec1_Cau_DLX1
7669_L_sec1_Put_DLX1
7669_L_sec2_GP_Bulk
7669_L_sec2_GP_DLX1
7669_L_sec2_Put_Bulk
7669_L_sec2_Thal_Bulk
7669_L_sec2_Thal_COUPTF2
7669_L_sec2_Thal_DLX1
7669_L_sec3_Hip_COUPTF2
7669_L_sec3_Hip_DLX1
7669_L_Skin_Bulk
7669_L_T_1_Bulk
7669_L_T_1_COUPTF2
7669_L_T_1_DLX1
7669_L_T_1_TBR1
7669_LA10_Bulk
7669_Liver_Bulk
7669_LV15_Bulk
7669_R_2_Put2_Bulk
7669_R_2_Thal_Bulk
7669_R_Adrenal_Bulk
7669_R_Ant_Cau_DLX1
7669_R_Ant_Put_DLX1
7669_R_Cb_2_Bulk
7669_R_CC_Mid_TBR1
7669_R_CCMid_COUPTF2
7669_R_CCMid_DLX1
7669_R_EC_COUPTF2
7669_R_EC_DLX1
7669_R_EC_TBR1
7669_R_F_1_Bulk
7669_R_F_1_COUPTF2
7669_R_F_1_DLX1
7669_R_F_1_NeuN
7669_R_F_1_TBR1
7669_R_I_1_COUPTF2
7669_R_I_1_DLX1
7669_R_I_1_TBR1
7669_R_Ins_1_Bulk
7669_R_kidney_Bulk
7669_R_mO_COUPTF2
7669_R_mO_DLX1
7669_R_mO_TBR1
7669_R_O_1_Bulk
7669_R_O_1_COUPTF2
7669_R_O_1_DLX1
7669_R_O_1_NeuN
7669_R_O_1_TBR1
7669_R_O_10_DLX1
7669_R_O_11_DLX1
7669_R_O_2_DLX1
7669_R_O_3_DLX1
7669_R_O_4_DLX1
7669_R_O_5_DLX1
7669_R_O_6_DLX1
7669_R_O_7_DLX1
7669_R_O_8_DLX1
7669_R_O_9_DLX1
7669_R_Olf_DLX1
7669_R_P_1_Bulk
7669_R_P_1_COUPTF2
7669_R_P_1_DLX1
7669_R_P_1_NeuN
7669_R_P_1_TBR1
7669_R_P_10_DLX1
7669_R_P_11_DLX1
7669_R_P_11_TBR1
7669_R_P_12_DLX1
7669_R_P_12_TBR1
7669_R_P_13_DLX1
7669_R_P_13_TBR1
7669_R_P_14_DLX1
7669_R_P_14_TBR1
7669_R_P_15_DLX1
7669_R_P_15_TBR1
7669_R_P_16_DLX1
7669_R_P_16_TBR1
7669_R_P_17_DLX1
7669_R_P_17_TBR1
7669_R_P_2_DLX1
7669_R_P_2_TBR1
7669_R_P_3_DLX1
7669_R_P_3_TBR1
7669_R_P_4_DLX1
7669_R_P_4_TBR1
7669_R_P_5_DLX1
7669_R_P_5_TBR1
7669_R_P_6_DLX1
7669_R_P_6_TBR1
7669_R_P_7_DLX1
7669_R_P_7_TBR1
7669_R_P_8_DLX1
7669_R_P_8_TBR1
7669_R_P_9_DLX
7669_R_P_9_DLX1
7669_R_P_9_TBR1
7669_R_POA_Bulk
7669_R_POA_DLX1
7669_R_sec1_Cau_Bulk
7669_R_sec2_GP_Bulk
7669_R_sec2_GP_DLX1
7669_R_sec2_Put_Bulk
7669_R_sec2_Thal_Bulk
7669_R_sec2_Thal_COUPTF2
7669_R_sec2_Thal_DLX1
7669_R_sec3_Cau_DLX1
7669_R_sec3_Hip_COUPTF2
7669_R_sec3_Hip_DLX1
7669_R_sec3_Hip_TBR1
7669_R_sec3_Put_DLX1
7669_R_Skin_Bulk
7669_R_T_1_Bulk
7669_R_T_1_COUPTF2
7669_R_T_1_DLX1
7669_R_T_1_TBR1
7669_RA9_Bulk
7669_RV10_Bulk
JGG

ID05 9−83538985−A−G

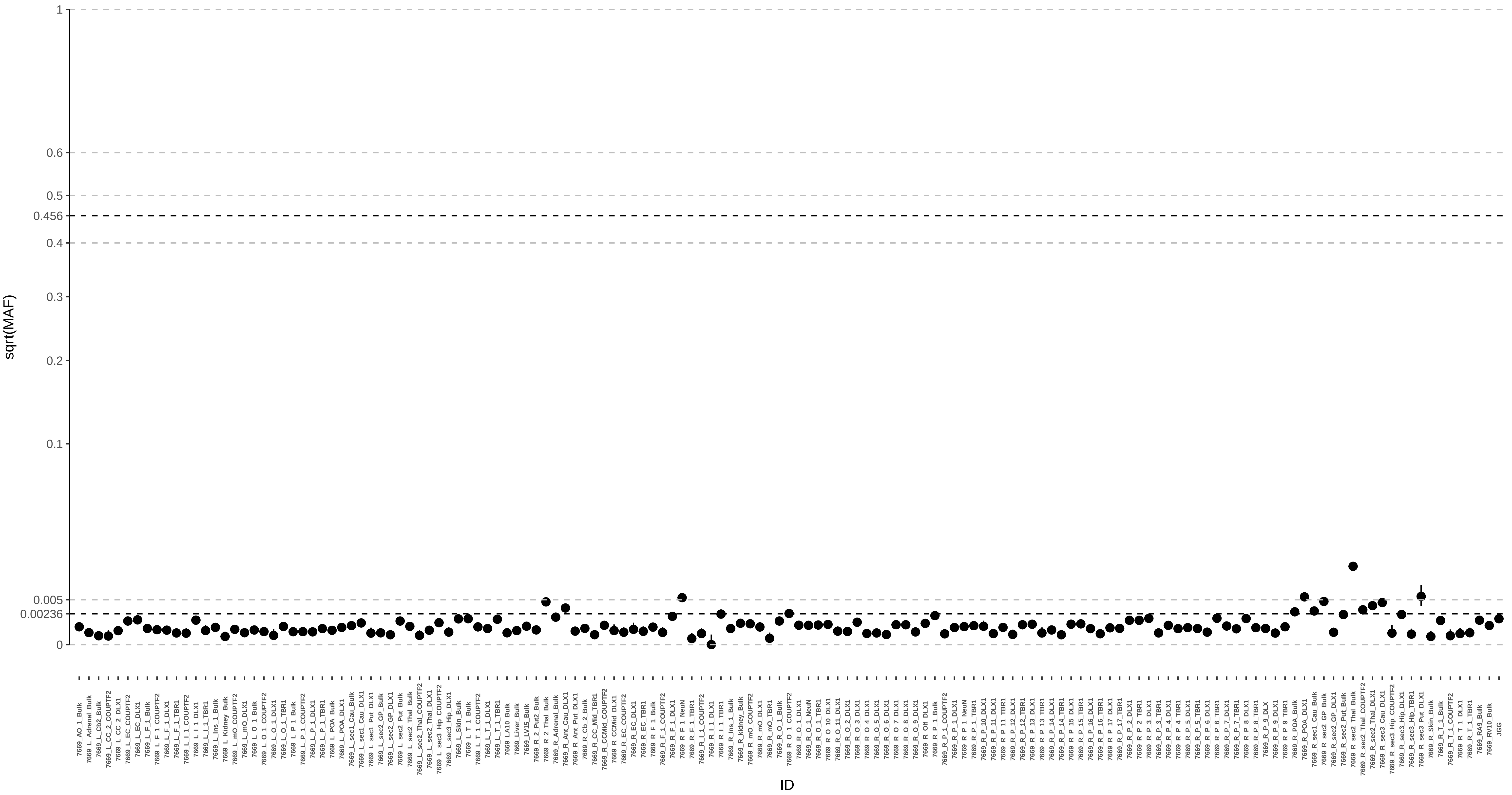

ID05 9−96677164−G−A

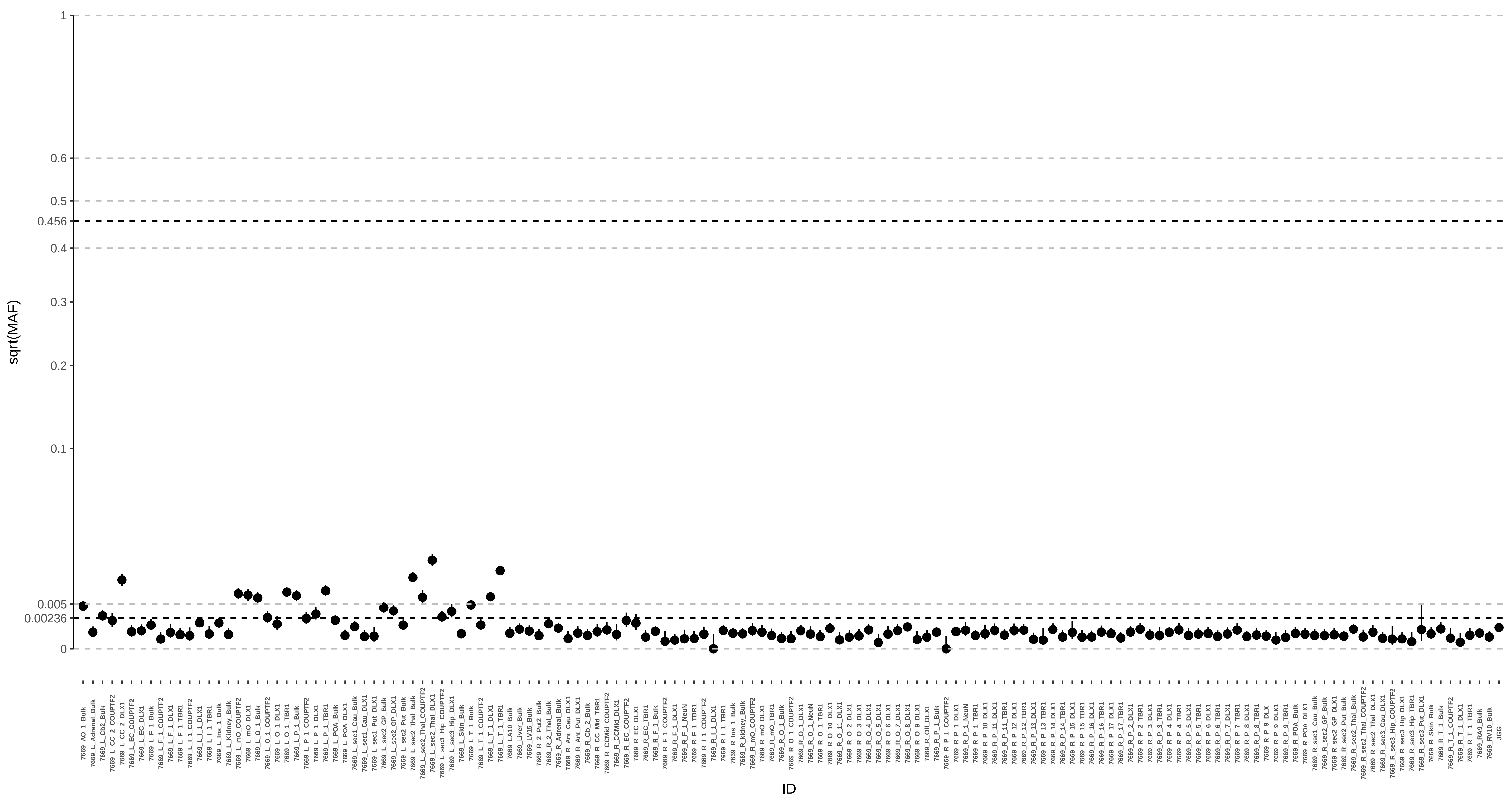

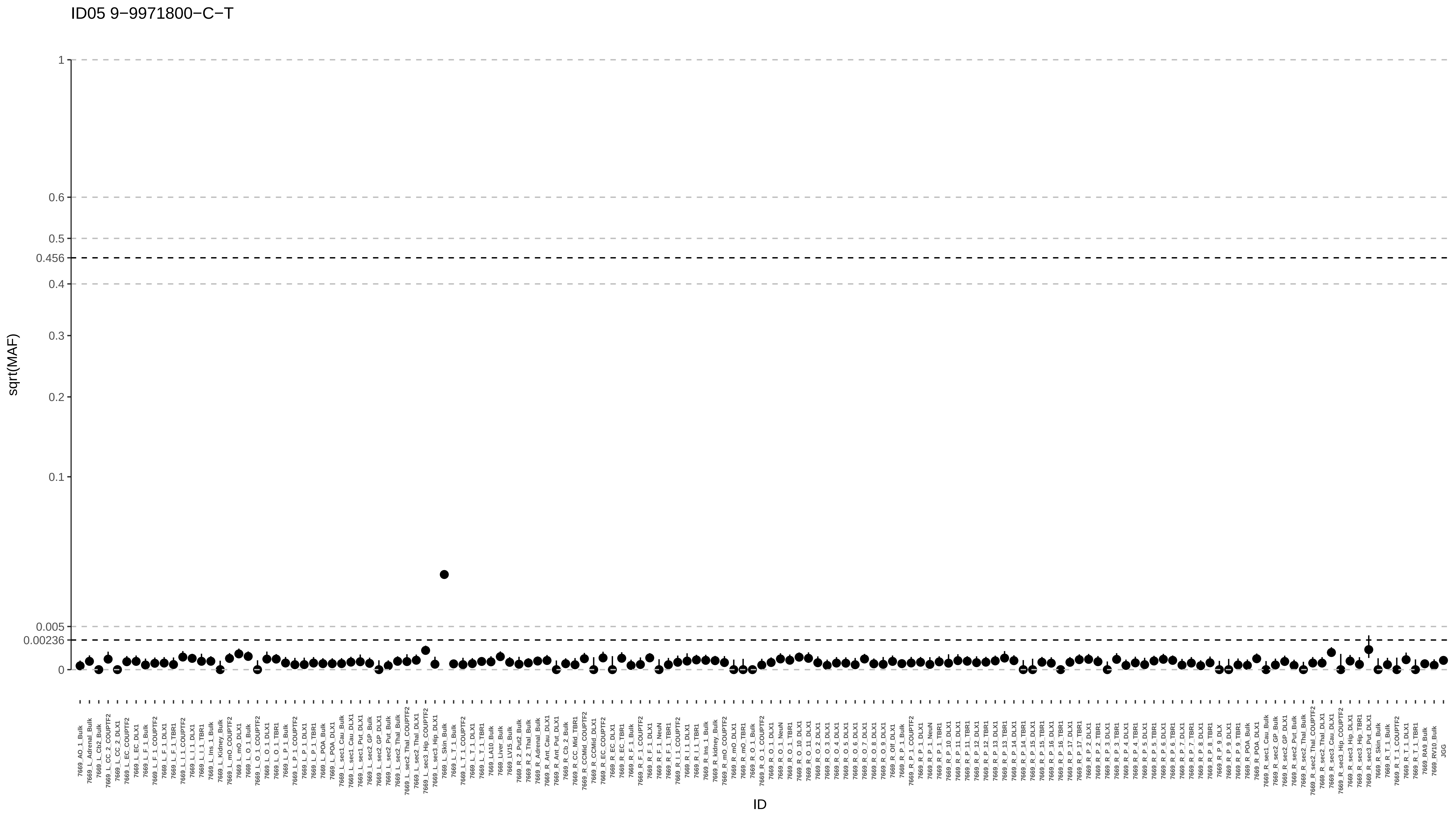
ID05 9−9971800−C−T
sqrt(MAF)
1
0.6
0.5
0.456
0.4
0.3
0.2
0.1
0.005
0.00236
0
ID

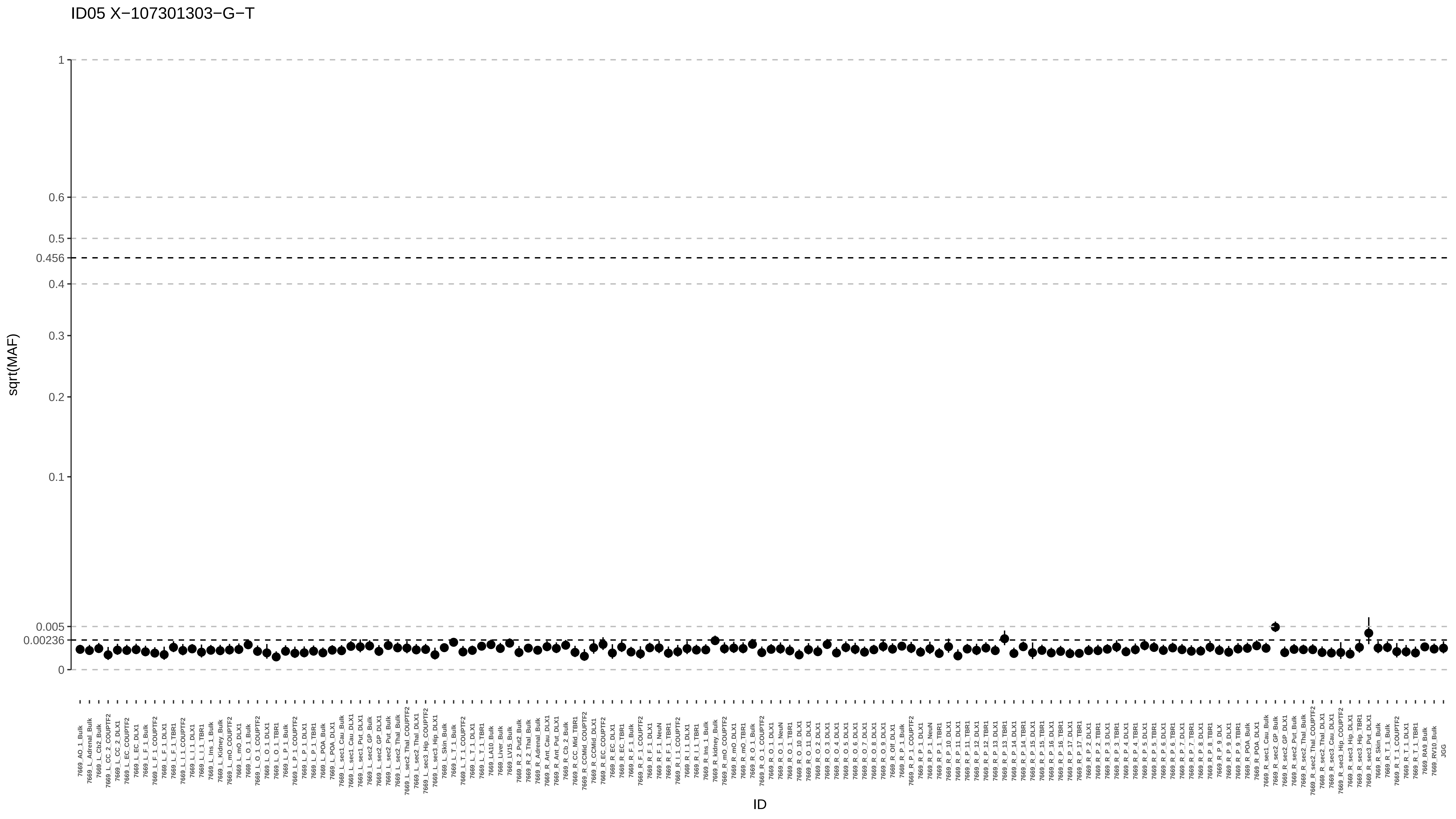
ID05 X−107301303−G−T
sqrt(MAF)
1
0.6
0.5
0.456
0.4
0.3
0.2
0.1
0.005
0.00236
0
ID

ID05 X−109753054−C−T

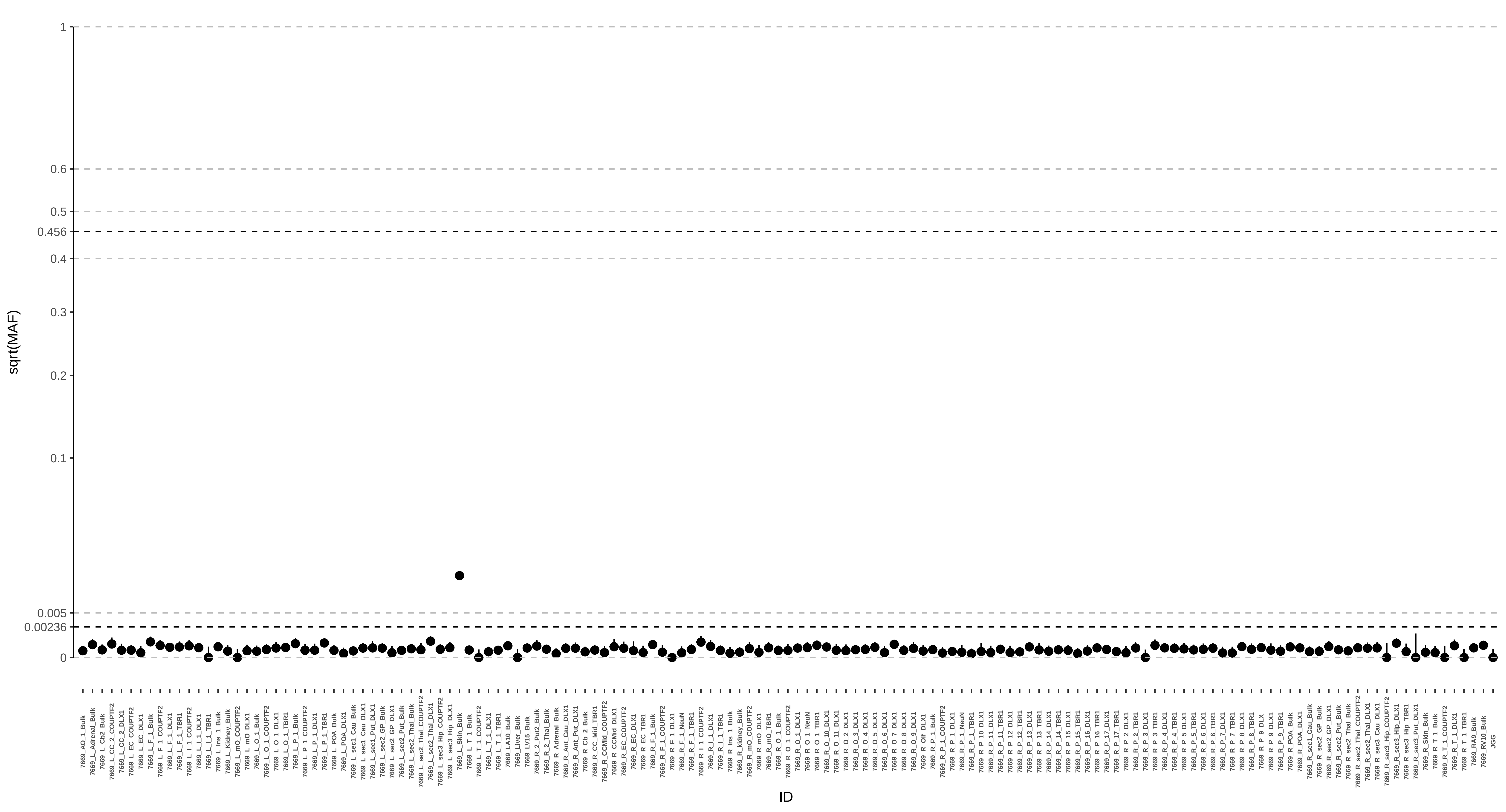

ID05 X−109939493−G−A

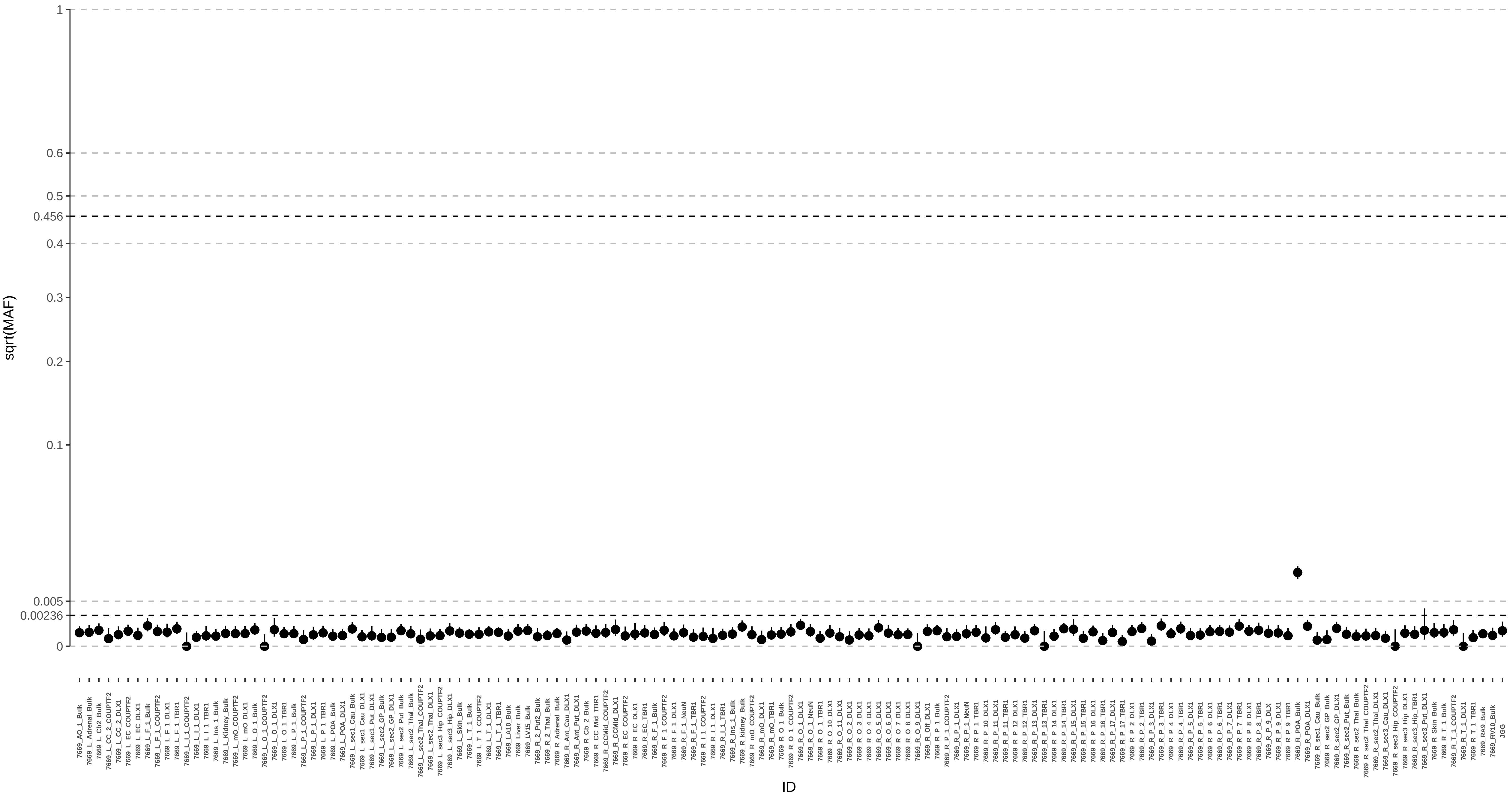

# ID05 X−117240525−A−C

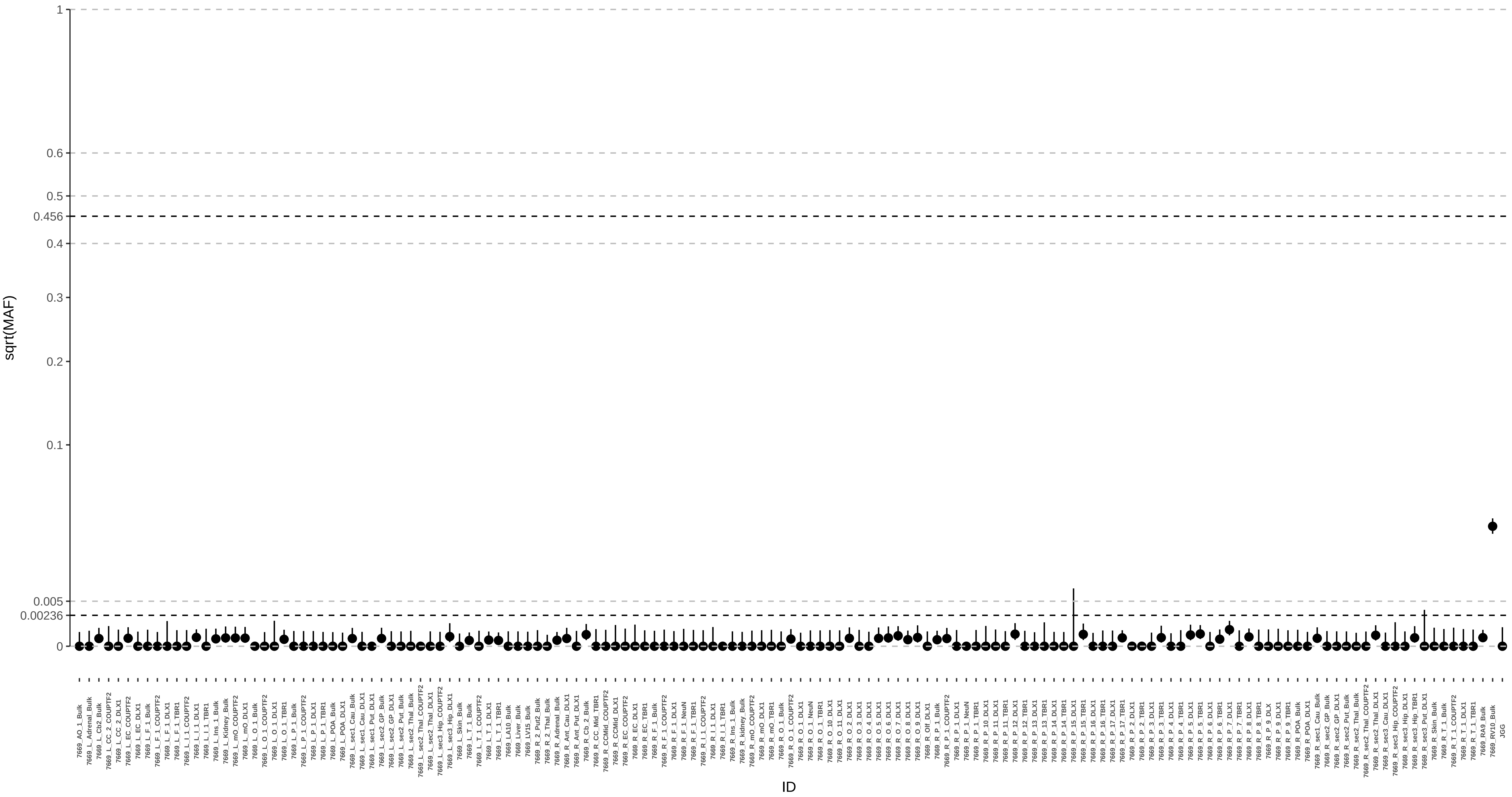

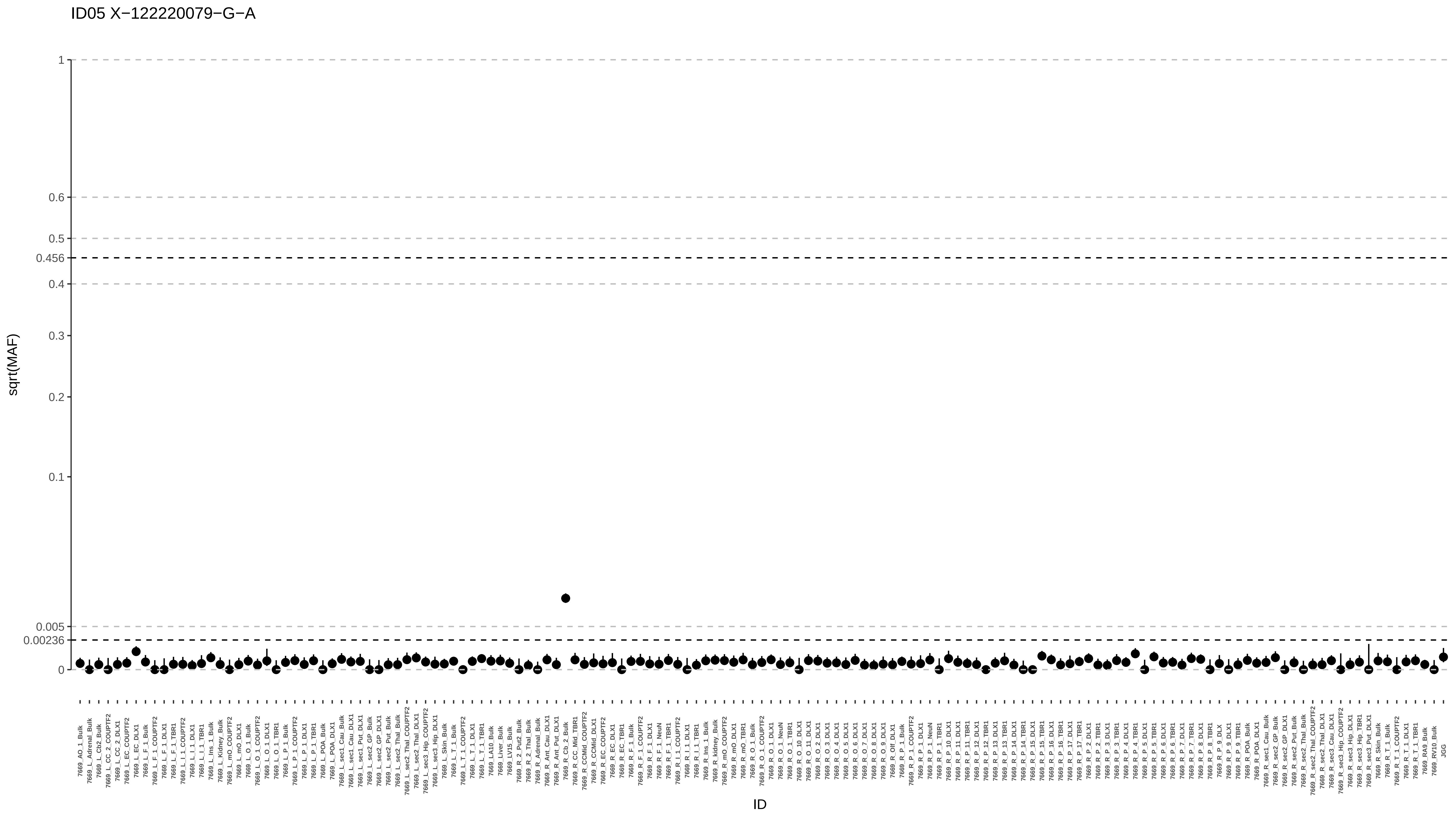
ID05 X−122220079−G−A
sqrt(MAF)
ID
1
0.6
0.5
0.456
0.4
0.3
0.2
0.1
0.005
0.00236
0

ID05 X−125464024−G−A

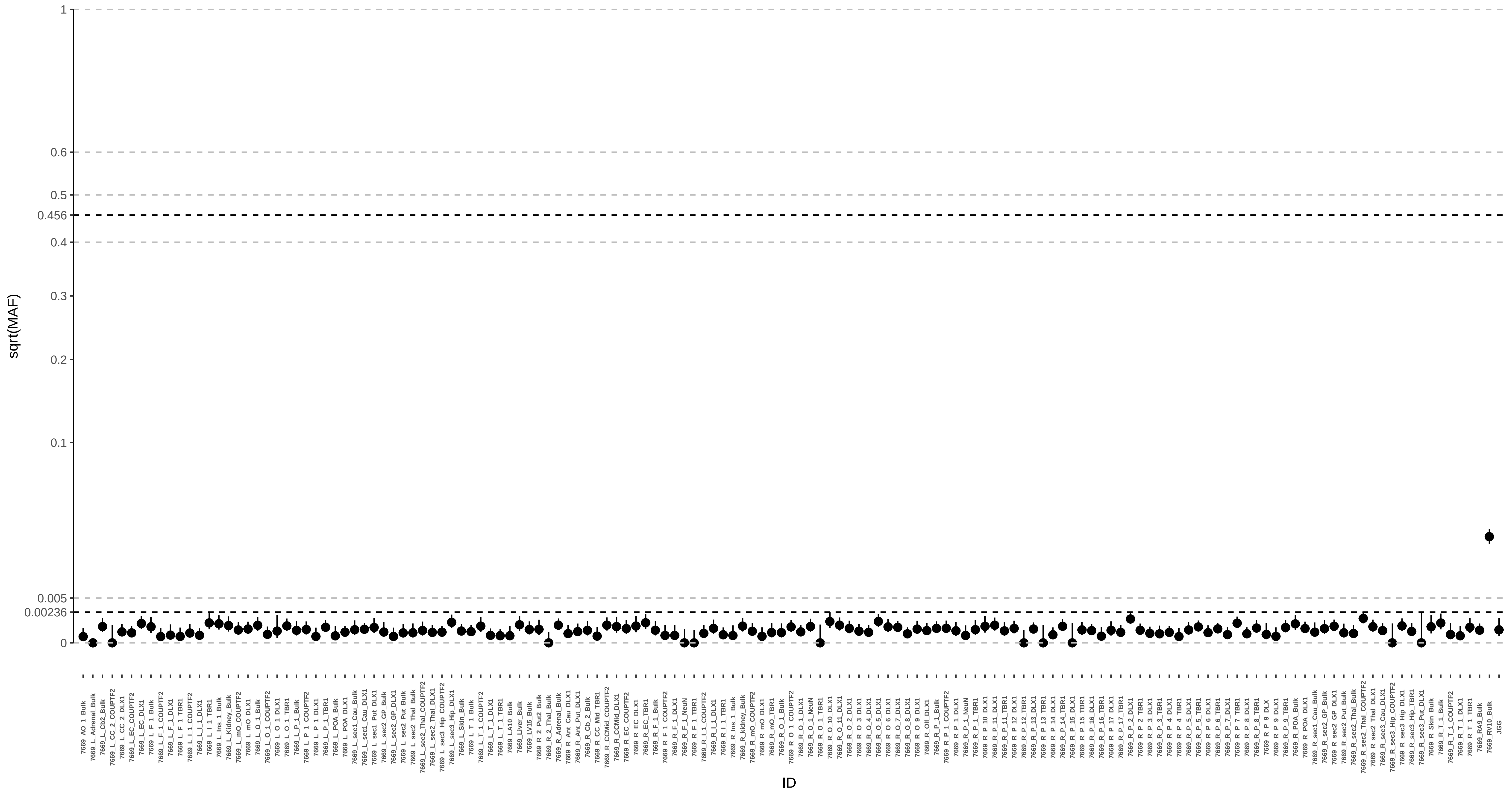

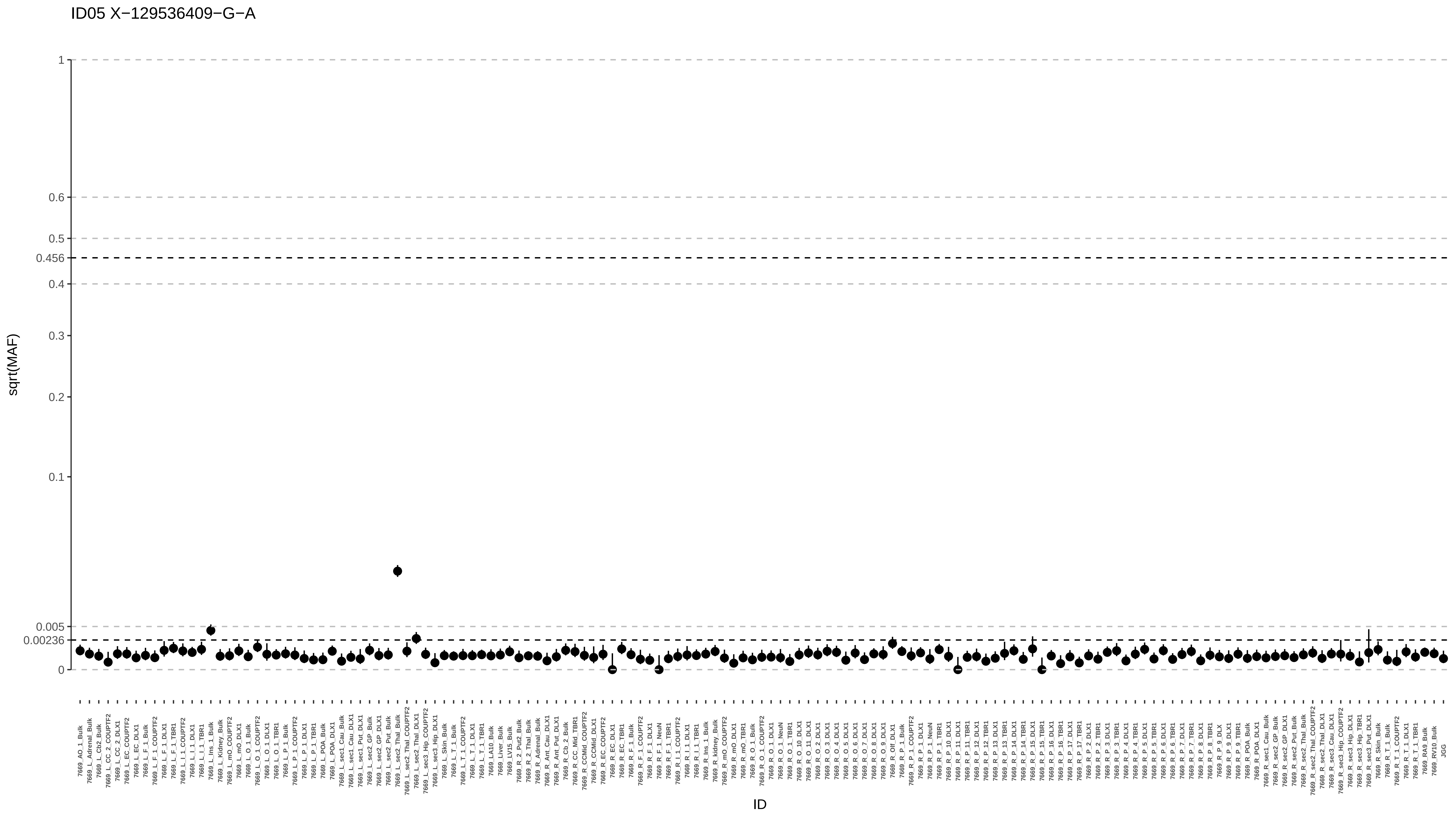

ID05 X−129536409−G−A
sqrt(MAF)
1
0.6
0.5
0.456
0.4
0.3
0.2
0.1
0.005
0.00236
0
ID

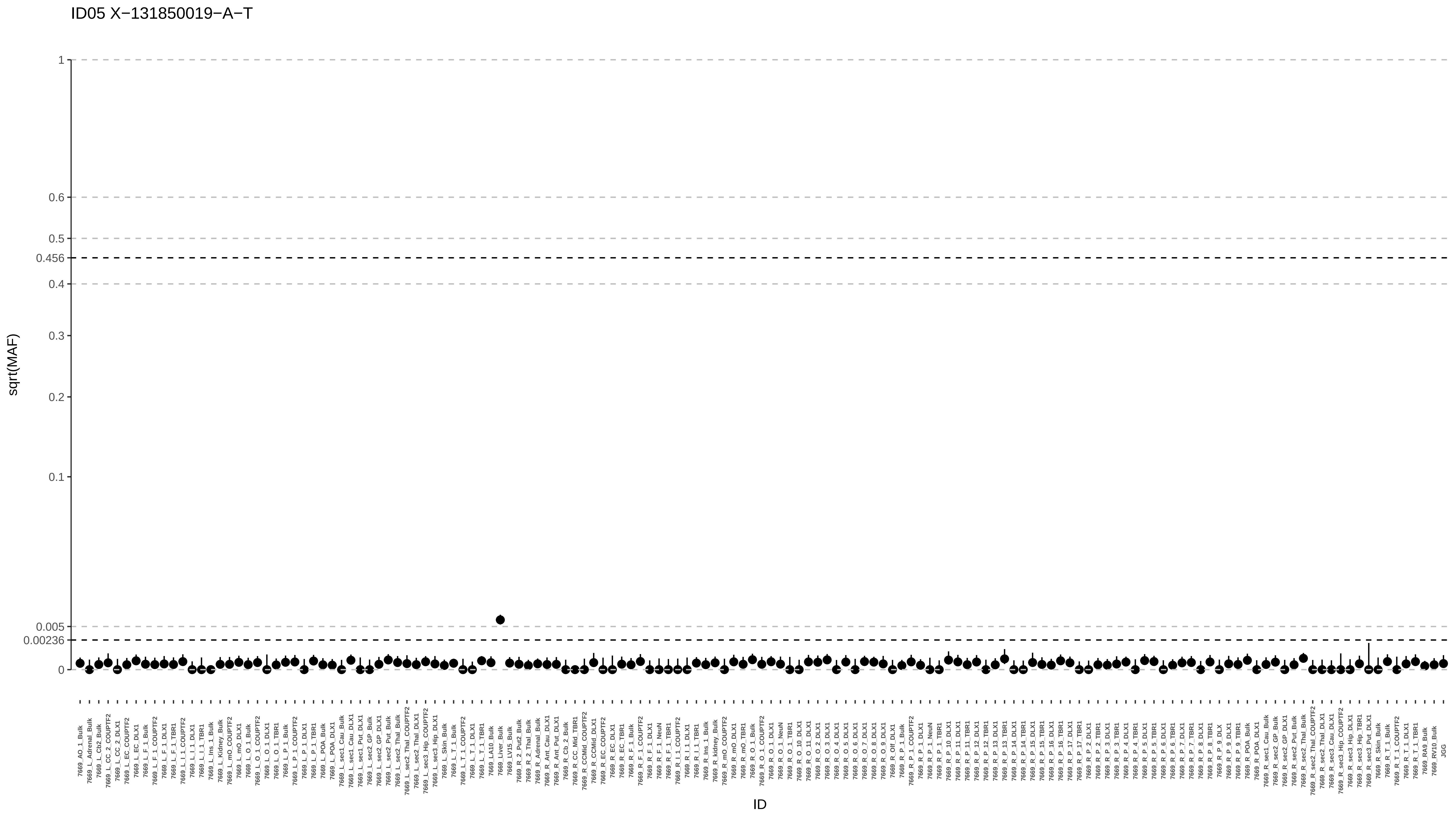
ID05 X−131850019−A−T
sqrt(MAF)
1
0.6
0.5
0.456
0.4
0.3
0.2
0.1
0.005
0.00236
0
ID
7669_AO_1_Bulk
7669_L_Adrenal_Bulk
7669_L_Cb2_Bulk
7669_L_CC_2_COUPTF2
7669_L_CC_2_DLX1
7669_L_EC_COUPTF2
7669_L_EC_DLX1
7669_L_F_1_Bulk
7669_L_F_1_COUPTF2
7669_L_F_1_DLX1
7669_L_F_1_TBR1
7669_L_I_1_COUPTF2
7669_L_I_1_DLX1
7669_L_I_1_TBR1
7669_L_Ins_1_Bulk
7669_L_Kidney_Bulk
7669_L_mO_COUPTF2
7669_L_mO_DLX1
7669_L_O_1_Bulk
7669_L_O_1_COUPTF2
7669_L_O_1_DLX1
7669_L_O_1_TBR1
7669_L_P_1_Bulk
7669_L_P_1_COUPTF2
7669_L_P_1_DLX1
7669_L_P_1_TBR1
7669_L_POA_Bulk
7669_L_POA_DLX1
7669_L_sec1_Cau_Bulk
7669_L_sec1_Cau_DLX1
7669_L_sec1_Put_DLX1
7669_L_sec2_GP_Bulk
7669_L_sec2_GP_DLX1
7669_L_sec2_Put_Bulk
7669_L_sec2_Thal_Bulk
7669_L_sec2_Thal_COUPTF2
7669_L_sec2_Thal_DLX1
7669_L_sec3_Hip_COUPTF2
7669_L_sec3_Hip_DLX1
7669_L_Skin_Bulk
7669_L_T_1_Bulk
7669_L_T_1_COUPTF2
7669_L_T_1_DLX1
7669_L_T_1_TBR1
7669_LA10_Bulk
7669_Liver_Bulk
7669_LV15_Bulk
7669_R_2_Put2_Bulk
7669_R_2_Thal_Bulk
7669_R_Adrenal_Bulk
7669_R_Ant_Cau_DLX1
7669_R_Ant_Put_DLX1
7669_R_Cb_2_Bulk
7669_R_CC_Mid_TBR1
7669_R_CCMid_COUPTF2
7669_R_CCMid_DLX1
7669_R_EC_COUPTF2
7669_R_EC_DLX1
7669_R_EC_TBR1
7669_R_F_1_Bulk
7669_R_F_1_COUPTF2
7669_R_F_1_DLX1
7669_R_F_1_NeuN
7669_R_F_1_TBR1
7669_R_I_1_COUPTF2
7669_R_I_1_DLX1
7669_R_I_1_TBR1
7669_R_Ins_1_Bulk
7669_R_kidney_Bulk
7669_R_mO_COUPTF2
7669_R_mO_DLX1
7669_R_mO_TBR1
7669_R_O_1_Bulk
7669_R_O_1_COUPTF2
7669_R_O_1_DLX1
7669_R_O_1_NeuN
7669_R_O_1_TBR1
7669_R_O_10_DLX1
7669_R_O_11_DLX1
7669_R_O_2_DLX1
7669_R_O_3_DLX1
7669_R_O_4_DLX1
7669_R_O_5_DLX1
7669_R_O_6_DLX1
7669_R_O_7_DLX1
7669_R_O_8_DLX1
7669_R_O_9_DLX1
7669_R_Olf_DLX1
7669_R_P_1_Bulk
7669_R_P_1_COUPTF2
7669_R_P_1_DLX1
7669_R_P_1_NeuN
7669_R_P_1_TBR1
7669_R_P_10_DLX1
7669_R_P_11_DLX1
7669_R_P_11_TBR1
7669_R_P_12_DLX1
7669_R_P_12_TBR1
7669_R_P_13_DLX1
7669_R_P_13_TBR1
7669_R_P_14_DLX1
7669_R_P_14_TBR1
7669_R_P_15_DLX1
7669_R_P_15_TBR1
7669_R_P_16_DLX1
7669_R_P_16_TBR1
7669_R_P_17_DLX1
7669_R_P_17_TBR1
7669_R_P_2_DLX1
7669_R_P_2_TBR1
7669_R_P_3_DLX1
7669_R_P_3_TBR1
7669_R_P_4_DLX1
7669_R_P_4_TBR1
7669_R_P_5_DLX1
7669_R_P_5_TBR1
7669_R_P_6_DLX1
7669_R_P_6_TBR1
7669_R_P_7_DLX1
7669_R_P_7_TBR1
7669_R_P_8_DLX1
7669_R_P_8_TBR1
7669_R_P_9_DLX
7669_R_P_9_DLX1
7669_R_P_9_TBR1
7669_R_POA_Bulk
7669_R_POA_DLX1
7669_R_sec1_Cau_Bulk
7669_R_sec2_GP_Bulk
7669_R_sec2_GP_DLX1
7669_R_sec2_Put_Bulk
7669_R_sec2_Thal_Bulk
7669_R_sec2_Thal_COUPTF2
7669_R_sec2_Thal_DLX1
7669_R_sec3_Cau_DLX1
7669_R_sec3_Hip_COUPTF2
7669_R_sec3_Hip_DLX1
7669_R_sec3_Hip_TBR1
7669_R_sec3_Put_DLX1
7669_R_Skin_Bulk
7669_R_T_1_Bulk
7669_R_T_1_COUPTF2
7669_R_T_1_DLX1
7669_R_T_1_TBR1
7669_RA9_Bulk
7669_RV10_Bulk
JGG

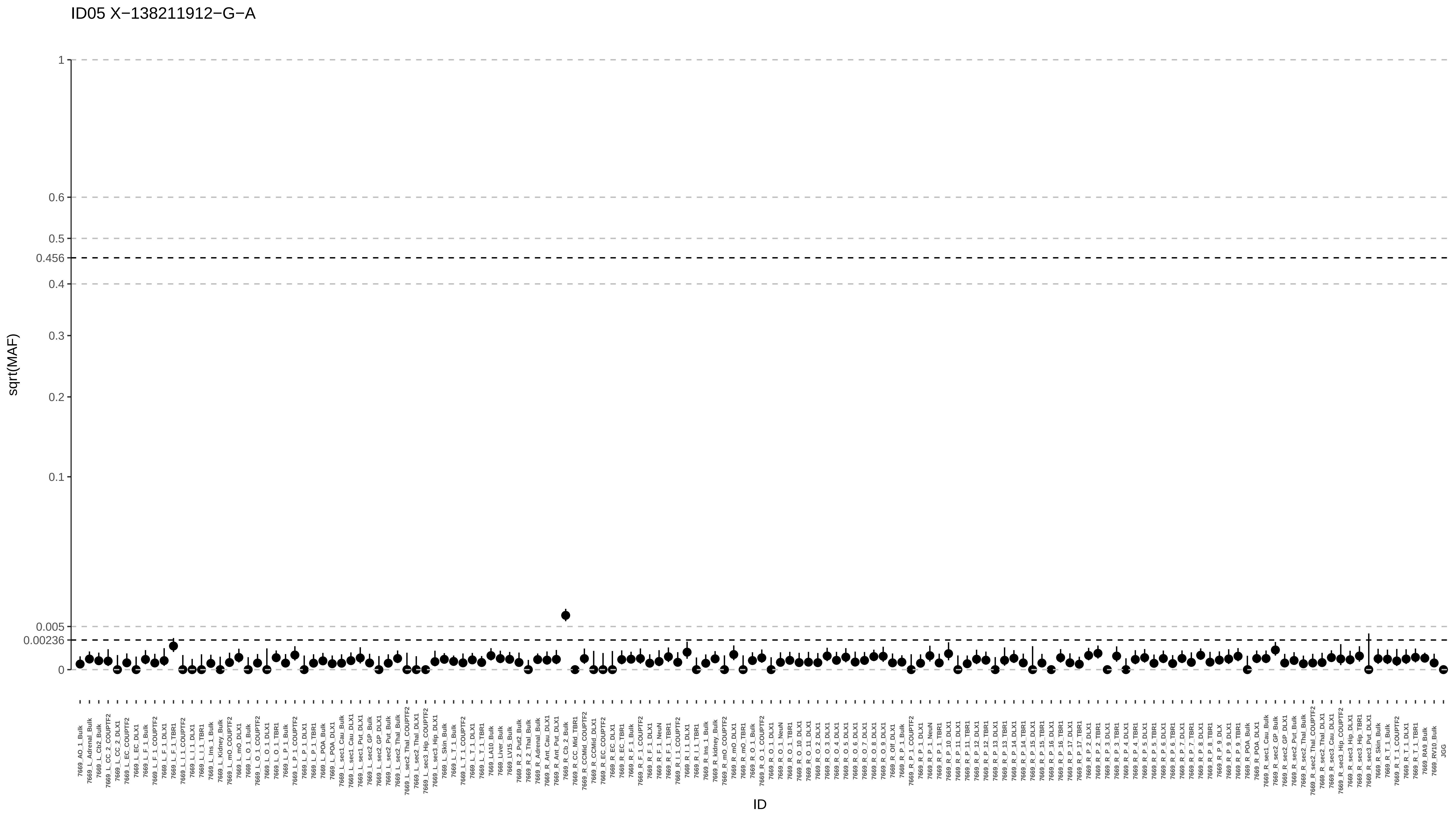
ID05 X−138211912−G−A
sqrt(MAF)
1
0.6
0.5
0.456
0.4
0.3
0.2
0.1
0.005
0.00236
0
ID

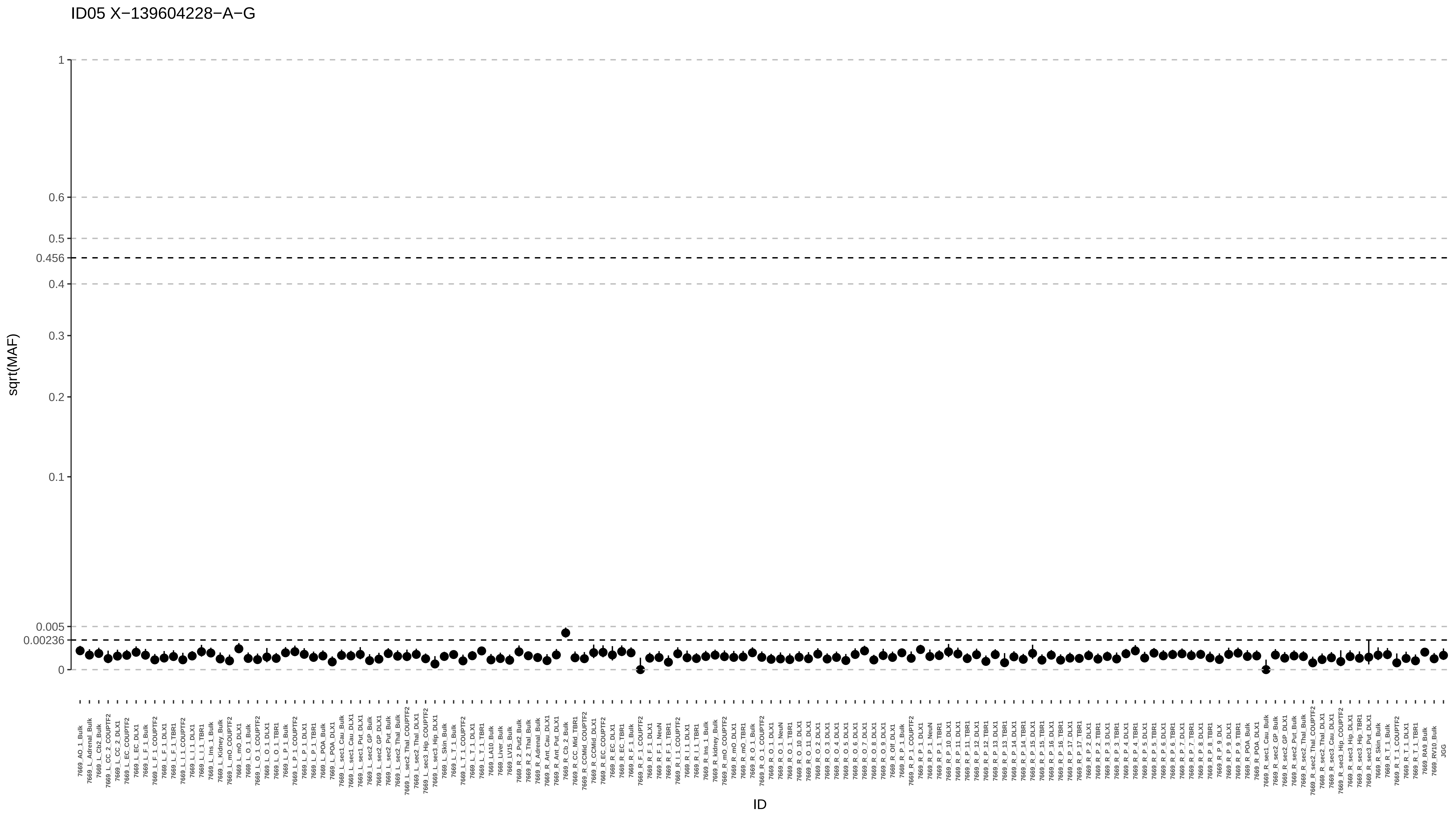
ID05 X−139604228−A−G
sqrt(MAF)
1
0.6
0.5
0.456
0.4
0.3
0.2
0.1
0.005
0.00236
0
ID

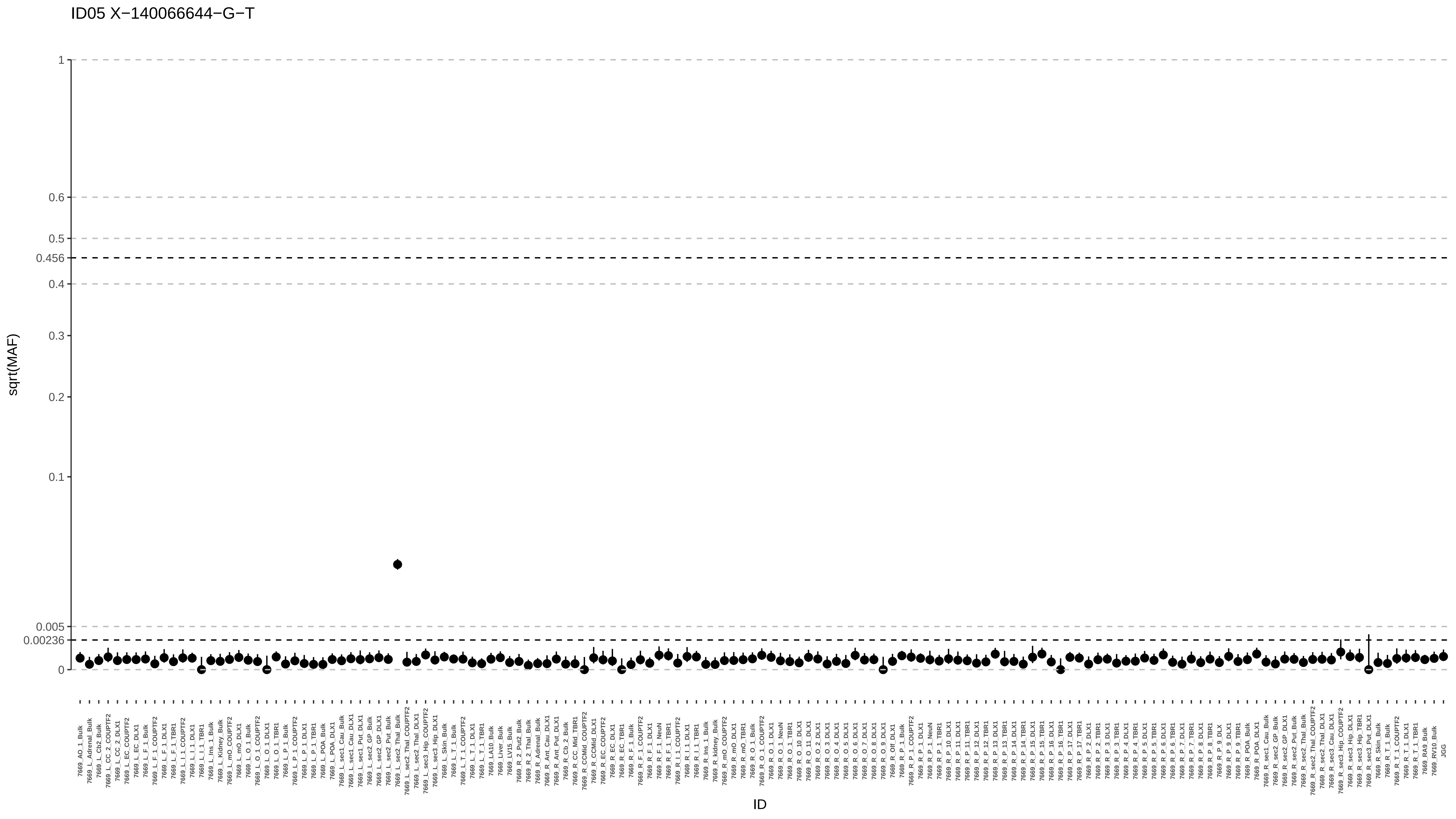
ID05 X−140066644−G−T
sqrt(MAF)
ID

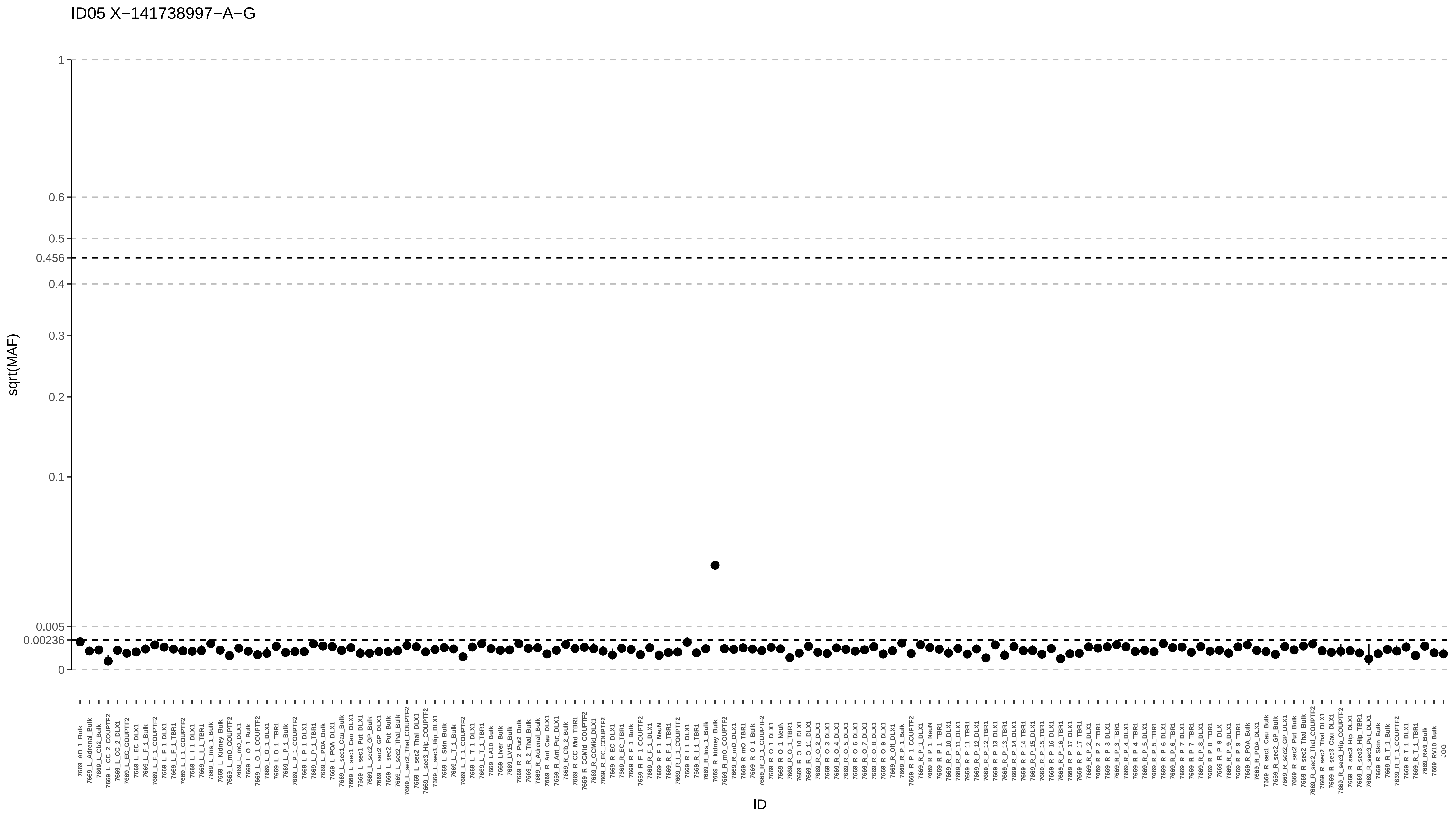
ID05 X−141738997−A−G
sqrt(MAF)
1
0.6
0.5
0.456
0.4
0.3
0.2
0.1
0.005
0.00236
0
ID

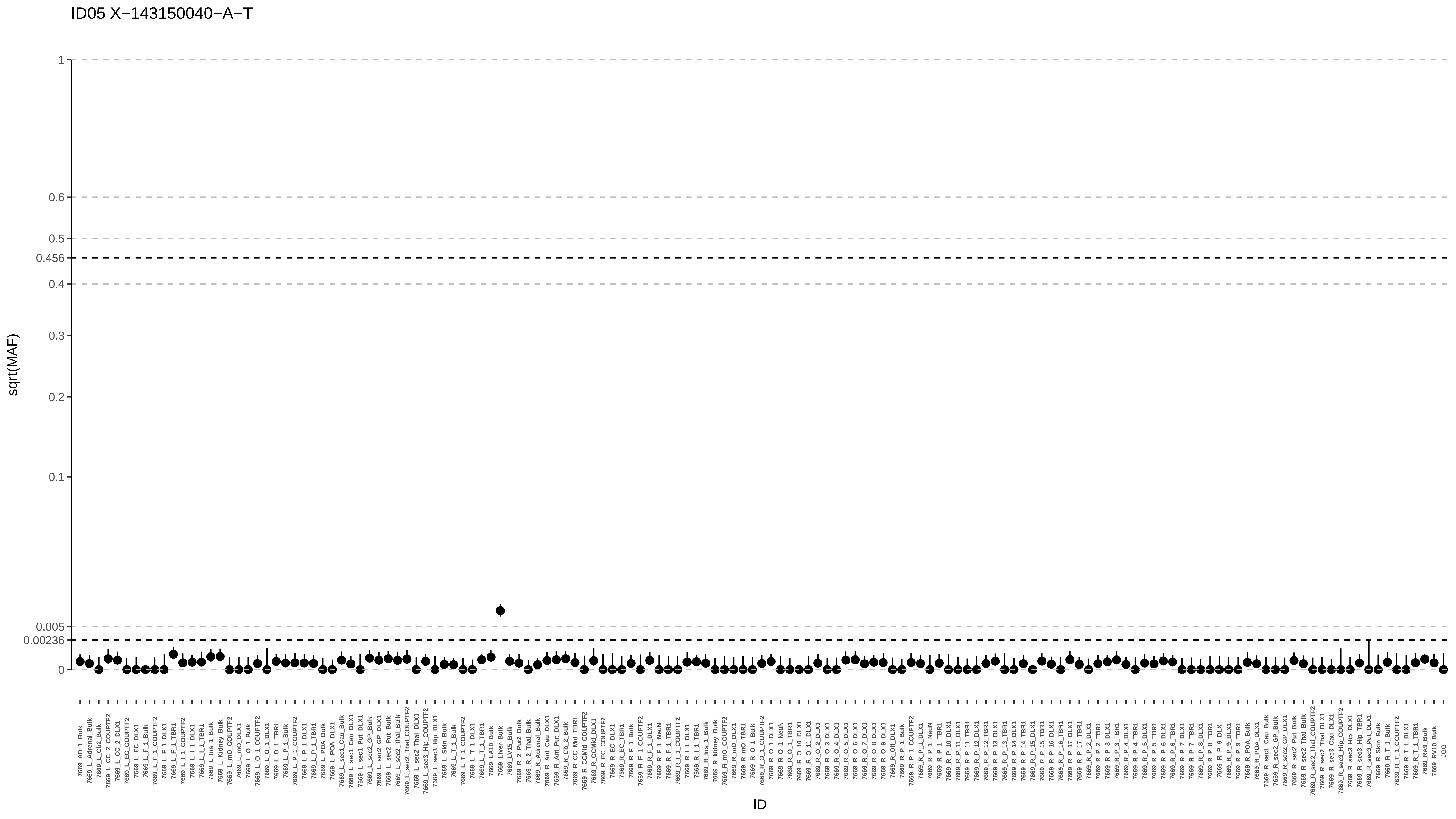
ID05 X−143150040−A−T
sqrt(MAF)
1
0.6
0.5
0.456
0.4
0.3
0.2
0.1
0.005
0.00236
0
ID
7669_AO_1_Bulk
7669_L_Adrenal_Bulk
7669_L_Cb2_Bulk
7669_L_CC_2_COUPTF2
7669_L_CC_2_DLX1
7669_L_EC_COUPTF2
7669_L_EC_DLX1
7669_L_F_1_Bulk
7669_L_F_1_COUPTF2
7669_L_F_1_DLX1
7669_L_F_1_TBR1
7669_L_I_1_COUPTF2
7669_L_I_1_DLX1
7669_L_I_1_TBR1
7669_L_Ins_1_Bulk
7669_L_Kidney_Bulk
7669_L_mO_COUPTF2
7669_L_mO_DLX1
7669_L_O_1_Bulk
7669_L_O_1_COUPTF2
7669_L_O_1_DLX1
7669_L_O_1_TBR1
7669_L_P_1_Bulk
7669_L_P_1_COUPTF2
7669_L_P_1_DLX1
7669_L_P_1_TBR1
7669_L_POA_Bulk
7669_L_POA_DLX1
7669_L_sec1_Cau_Bulk
7669_L_sec1_Cau_DLX1
7669_L_sec1_Put_DLX1
7669_L_sec2_GP_Bulk
7669_L_sec2_GP_DLX1
7669_L_sec2_Put_Bulk
7669_L_sec2_Thal_Bulk
7669_L_sec2_Thal_COUPTF2
7669_L_sec2_Thal_DLX1
7669_L_sec3_Hip_COUPTF2
7669_L_sec3_Hip_DLX1
7669_L_Skin_Bulk
7669_L_T_1_Bulk
7669_L_T_1_COUPTF2
7669_L_T_1_DLX1
7669_L_T_1_TBR1
7669_LA10_Bulk
7669_Liver_Bulk
7669_LV15_Bulk
7669_R_2_Put2_Bulk
7669_R_2_Thal_Bulk
7669_R_Adrenal_Bulk
7669_R_Ant_Cau_DLX1
7669_R_Ant_Put_DLX1
7669_R_Cb_2_Bulk
7669_R_CC_Mid_TBR1
7669_R_CCMid_COUPTF2
7669_R_CCMid_DLX1
7669_R_EC_COUPTF2
7669_R_EC_DLX1
7669_R_EC_TBR1
7669_R_F_1_Bulk
7669_R_F_1_COUPTF2
7669_R_F_1_DLX1
7669_R_F_1_NeuN
7669_R_F_1_TBR1
7669_R_I_1_COUPTF2
7669_R_I_1_DLX1
7669_R_I_1_TBR1
7669_R_Ins_1_Bulk
7669_R_kidney_Bulk
7669_R_mO_COUPTF2
7669_R_mO_DLX1
7669_R_mO_TBR1
7669_R_O_1_Bulk
7669_R_O_1_COUPTF2
7669_R_O_1_DLX1
7669_R_O_1_NeuN
7669_R_O_1_TBR1
7669_R_O_10_DLX1
7669_R_O_11_DLX1
7669_R_O_2_DLX1
7669_R_O_3_DLX1
7669_R_O_4_DLX1
7669_R_O_5_DLX1
7669_R_O_6_DLX1
7669_R_O_7_DLX1
7669_R_O_8_DLX1
7669_R_O_9_DLX1
7669_R_Olf_DLX1
7669_R_P_1_Bulk
7669_R_P_1_COUPTF2
7669_R_P_1_DLX1
7669_R_P_1_NeuN
7669_R_P_1_TBR1
7669_R_P_10_DLX1
7669_R_P_11_DLX1
7669_R_P_11_TBR1
7669_R_P_12_DLX1
7669_R_P_12_TBR1
7669_R_P_13_DLX1
7669_R_P_13_TBR1
7669_R_P_14_DLX1
7669_R_P_14_TBR1
7669_R_P_15_DLX1
7669_R_P_15_TBR1
7669_R_P_16_DLX1
7669_R_P_16_TBR1
7669_R_P_17_DLX1
7669_R_P_17_TBR1
7669_R_P_2_DLX1
7669_R_P_2_TBR1
7669_R_P_3_DLX1
7669_R_P_3_TBR1
7669_R_P_4_DLX1
7669_R_P_4_TBR1
7669_R_P_5_DLX1
7669_R_P_5_TBR1
7669_R_P_6_DLX1
7669_R_P_6_TBR1
7669_R_P_7_DLX1
7669_R_P_7_TBR1
7669_R_P_8_DLX1
7669_R_P_8_TBR1
7669_R_P_9_DLX
7669_R_P_9_DLX1
7669_R_P_9_TBR1
7669_R_POA_Bulk
7669_R_POA_DLX1
7669_R_sec1_Cau_Bulk
7669_R_sec2_GP_Bulk
7669_R_sec2_GP_DLX1
7669_R_sec2_Put_Bulk
7669_R_sec2_Thal_Bulk
7669_R_sec2_Thal_COUPTF2
7669_R_sec2_Thal_DLX1
7669_R_sec3_Cau_DLX1
7669_R_sec3_Hip_COUPTF2
7669_R_sec3_Hip_DLX1
7669_R_sec3_Hip_TBR1
7669_R_sec3_Put_DLX1
7669_R_Skin_Bulk
7669_R_T_1_Bulk
7669_R_T_1_COUPTF2
7669_R_T_1_DLX1
7669_R_T_1_TBR1
7669_RA9_Bulk
7669_RV10_Bulk
JGG

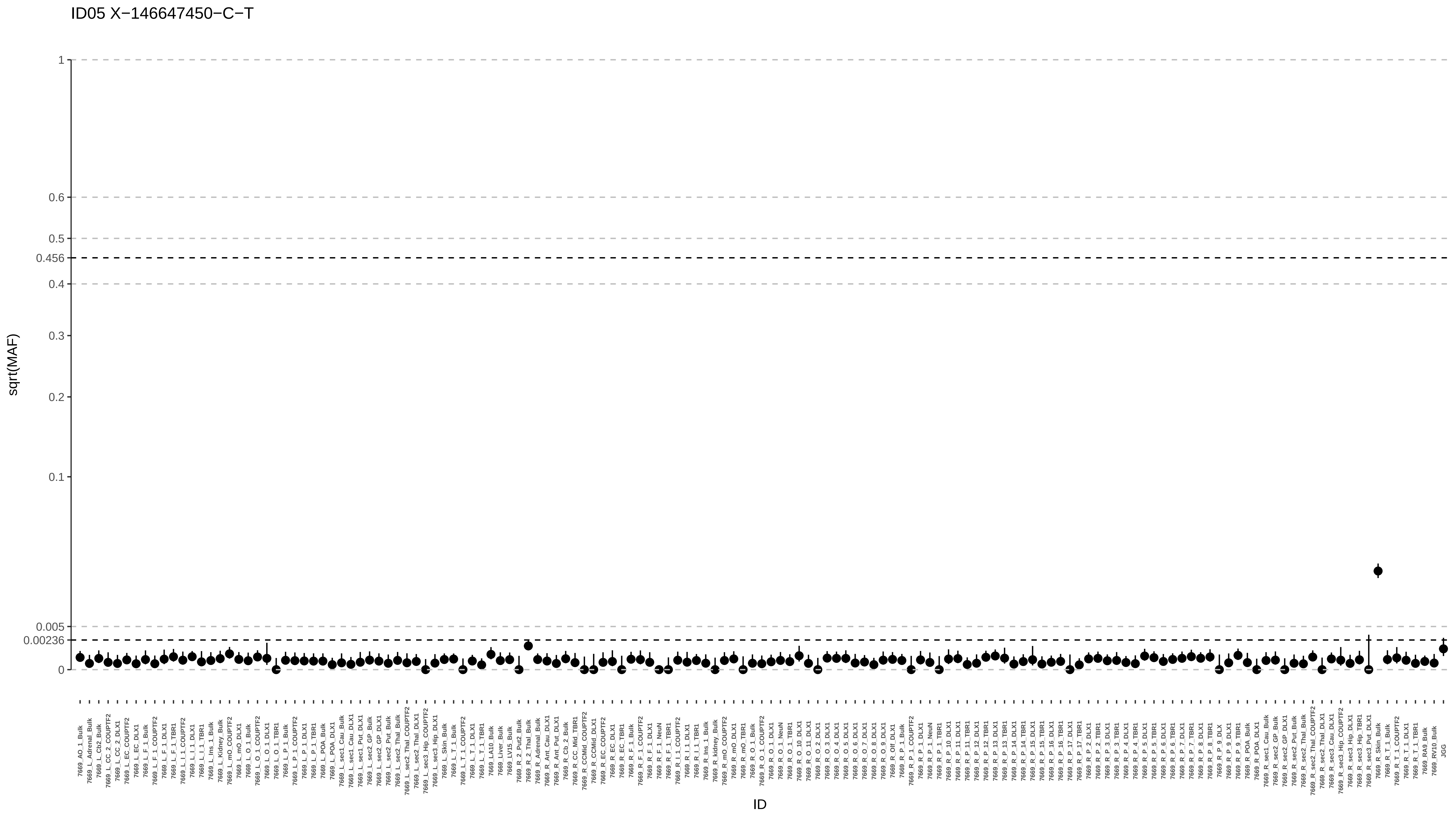

ID05 X−146647450−C−T
sqrt(MAF)
1
0.6
0.5
0.456
0.4
0.3
0.2
0.1
0.005
0.00236
0
7669_AO_1_Bulk
7669_L_Adrenal_Bulk
7669_L_Cb2_Bulk
7669_L_CC_2_COUPTF2
7669_L_CC_2_DLX1
7669_L_EC_COUPTF2
7669_L_EC_DLX1
7669_L_F_1_Bulk
7669_L_F_1_COUPTF2
7669_L_F_1_DLX1
7669_L_F_1_TBR1
7669_L_I_1_COUPTF2
7669_L_I_1_DLX1
7669_L_I_1_TBR1
7669_L_Ins_1_Bulk
7669_L_Kidney_Bulk
7669_L_mO_COUPTF2
7669_L_mO_DLX1
7669_L_O_1_Bulk
7669_L_O_1_COUPTF2
7669_L_O_1_DLX1
7669_L_O_1_TBR1
7669_L_P_1_Bulk
7669_L_P_1_COUPTF2
7669_L_P_1_DLX1
7669_L_P_1_TBR1
7669_L_POA_Bulk
7669_L_POA_DLX1
7669_L_sec1_Cau_Bulk
7669_L_sec1_Cau_DLX1
7669_L_sec1_Put_DLX1
7669_L_sec2_GP_Bulk
7669_L_sec2_GP_DLX1
7669_L_sec2_Put_Bulk
7669_L_sec2_Thal_Bulk
7669_L_sec2_Thal_COUPTF2
7669_L_sec2_Thal_DLX1
7669_L_sec3_Hip_COUPTF2
7669_L_sec3_Hip_DLX1
7669_L_Skin_Bulk
7669_L_T_1_Bulk
7669_L_T_1_COUPTF2
7669_L_T_1_DLX1
7669_L_T_1_TBR1
7669_LA10_Bulk
7669_Liver_Bulk
7669_LV15_Bulk
7669_R_2_Put2_Bulk
7669_R_2_Thal_Bulk
7669_R_Adrenal_Bulk
7669_R_Ant_Cau_DLX1
7669_R_Ant_Put_DLX1
7669_R_Cb_2_Bulk
7669_R_CC_Mid_TBR1
7669_R_CCMid_COUPTF2
7669_R_CCMid_DLX1
7669_R_EC_COUPTF2
7669_R_EC_DLX1
7669_R_EC_TBR1
7669_R_F_1_Bulk
7669_R_F_1_COUPTF2
7669_R_F_1_DLX1
7669_R_F_1_NeuN
7669_R_F_1_TBR1
7669_R_I_1_COUPTF2
7669_R_I_1_DLX1
7669_R_I_1_TBR1
7669_R_Ins_1_Bulk
7669_R_kidney_Bulk
7669_R_mO_COUPTF2
7669_R_mO_DLX1
7669_R_mO_TBR1
7669_R_O_1_Bulk
7669_R_O_1_COUPTF2
7669_R_O_1_DLX1
7669_R_O_1_NeuN
7669_R_O_1_TBR1
7669_R_O_10_DLX1
7669_R_O_11_DLX1
7669_R_O_2_DLX1
7669_R_O_3_DLX1
7669_R_O_4_DLX1
7669_R_O_5_DLX1
7669_R_O_6_DLX1
7669_R_O_7_DLX1
7669_R_O_8_DLX1
7669_R_O_9_DLX1
7669_R_Olf_DLX1
7669_R_P_1_Bulk
7669_R_P_1_COUPTF2
7669_R_P_1_DLX1
7669_R_P_1_NeuN
7669_R_P_1_TBR1
7669_R_P_10_DLX1
7669_R_P_11_DLX1
7669_R_P_11_TBR1
7669_R_P_12_DLX1
7669_R_P_12_TBR1
7669_R_P_13_DLX1
7669_R_P_13_TBR1
7669_R_P_14_DLX1
7669_R_P_14_TBR1
7669_R_P_15_DLX1
7669_R_P_15_TBR1
7669_R_P_16_DLX1
7669_R_P_16_TBR1
7669_R_P_17_DLX1
7669_R_P_17_TBR1
7669_R_P_2_DLX1
7669_R_P_2_TBR1
7669_R_P_3_DLX1
7669_R_P_3_TBR1
7669_R_P_4_DLX1
7669_R_P_4_TBR1
7669_R_P_5_DLX1
7669_R_P_5_TBR1
7669_R_P_6_DLX1
7669_R_P_6_TBR1
7669_R_P_7_DLX1
7669_R_P_7_TBR1
7669_R_P_8_DLX1
7669_R_P_8_TBR1
7669_R_P_9_DLX
7669_R_P_9_DLX1
7669_R_P_9_TBR1
7669_R_POA_Bulk
7669_R_POA_DLX1
7669_R_sec1_Cau_Bulk
7669_R_sec2_GP_Bulk
7669_R_sec2_GP_DLX1
7669_R_sec2_Put_Bulk
7669_R_sec2_Thal_Bulk
7669_R_sec2_Thal_COUPTF2
7669_R_sec2_Thal_DLX1
7669_R_sec3_Cau_DLX1
7669_R_sec3_Hip_COUPTF2
7669_R_sec3_Hip_DLX1
7669_R_sec3_Hip_TBR1
7669_R_sec3_Put_DLX1
7669_R_Skin_Bulk
7669_R_T_1_Bulk
7669_R_T_1_COUPTF2
7669_R_T_1_DLX1
7669_R_T_1_TBR1
7669_RA9_Bulk
7669_RV10_Bulk
JGG
ID

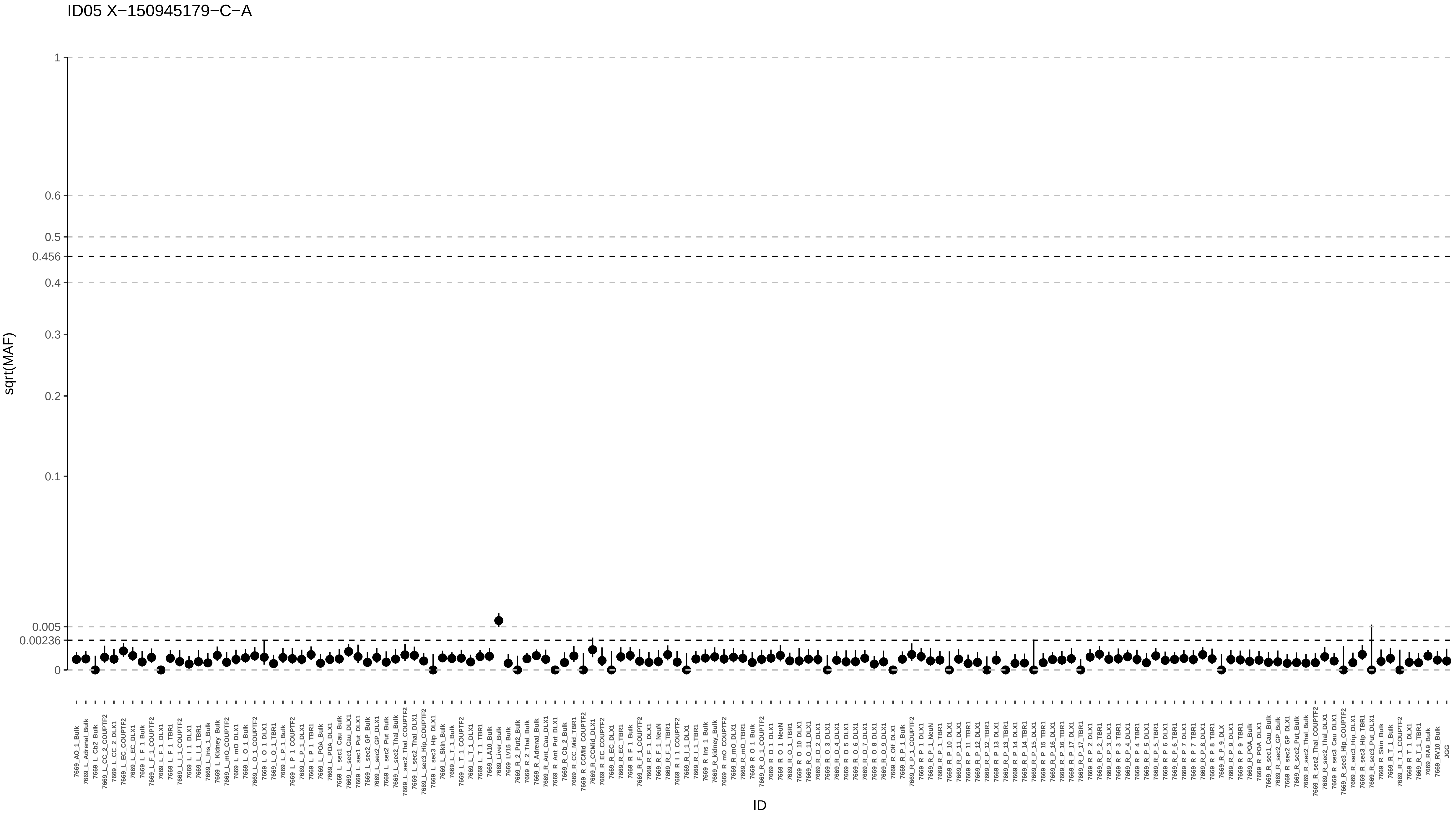
ID05 X−150945179−C−A
sqrt(MAF)
1
0.6
0.5
0.456
0.4
0.3
0.2
0.1
0.005
0.00236
0
ID
7669_AO_1_Bulk
7669_L_Adrenal_Bulk
7669_L_Cb2_Bulk
7669_L_CC_2_COUPTF2
7669_L_CC_2_DLX1
7669_L_EC_COUPTF2
7669_L_EC_DLX1
7669_L_F_1_Bulk
7669_L_F_1_COUPTF2
7669_L_F_1_DLX1
7669_L_F_1_TBR1
7669_L_I_1_COUPTF2
7669_L_I_1_DLX1
7669_L_I_1_TBR1
7669_L_Ins_1_Bulk
7669_L_Kidney_Bulk
7669_L_mO_COUPTF2
7669_L_mO_DLX1
7669_L_O_1_Bulk
7669_L_O_1_COUPTF2
7669_L_O_1_DLX1
7669_L_O_1_TBR1
7669_L_P_1_Bulk
7669_L_P_1_COUPTF2
7669_L_P_1_DLX1
7669_L_P_1_TBR1
7669_L_POA_Bulk
7669_L_POA_DLX1
7669_L_sec1_Cau_Bulk
7669_L_sec1_Cau_DLX1
7669_L_sec1_Put_DLX1
7669_L_sec2_GP_Bulk
7669_L_sec2_GP_DLX1
7669_L_sec2_Put_Bulk
7669_L_sec2_Thal_Bulk
7669_L_sec2_Thal_COUPTF2
7669_L_sec2_Thal_DLX1
7669_L_sec3_Hip_COUPTF2
7669_L_sec3_Hip_DLX1
7669_L_Skin_Bulk
7669_L_T_1_Bulk
7669_L_T_1_COUPTF2
7669_L_T_1_DLX1
7669_L_T_1_TBR1
7669_LA10_Bulk
7669_Liver_Bulk
7669_LV15_Bulk
7669_R_2_Put2_Bulk
7669_R_2_Thal_Bulk
7669_R_Adrenal_Bulk
7669_R_Ant_Cau_DLX1
7669_R_Ant_Put_DLX1
7669_R_Cb_2_Bulk
7669_R_CC_Mid_TBR1
7669_R_CCMid_COUPTF2
7669_R_CCMid_DLX1
7669_R_EC_COUPTF2
7669_R_EC_DLX1
7669_R_EC_TBR1
7669_R_F_1_Bulk
7669_R_F_1_COUPTF2
7669_R_F_1_DLX1
7669_R_F_1_NeuN
7669_R_F_1_TBR1
7669_R_I_1_COUPTF2
7669_R_I_1_DLX1
7669_R_I_1_TBR1
7669_R_Ins_1_Bulk
7669_R_kidney_Bulk
7669_R_mO_COUPTF2
7669_R_mO_DLX1
7669_R_mO_TBR1
7669_R_O_1_Bulk
7669_R_O_1_COUPTF2
7669_R_O_1_DLX1
7669_R_O_1_NeuN
7669_R_O_1_TBR1
7669_R_O_10_DLX1
7669_R_O_11_DLX1
7669_R_O_2_DLX1
7669_R_O_3_DLX1
7669_R_O_4_DLX1
7669_R_O_5_DLX1
7669_R_O_6_DLX1
7669_R_O_7_DLX1
7669_R_O_8_DLX1
7669_R_O_9_DLX1
7669_R_Olf_DLX1
7669_R_P_1_Bulk
7669_R_P_1_COUPTF2
7669_R_P_1_DLX1
7669_R_P_1_NeuN
7669_R_P_1_TBR1
7669_R_P_10_DLX1
7669_R_P_11_DLX1
7669_R_P_11_TBR1
7669_R_P_12_DLX1
7669_R_P_12_TBR1
7669_R_P_13_DLX1
7669_R_P_13_TBR1
7669_R_P_14_DLX1
7669_R_P_14_TBR1
7669_R_P_15_DLX1
7669_R_P_15_TBR1
7669_R_P_16_DLX1
7669_R_P_16_TBR1
7669_R_P_17_DLX1
7669_R_P_17_TBR1
7669_R_P_2_DLX1
7669_R_P_2_TBR1
7669_R_P_3_DLX1
7669_R_P_3_TBR1
7669_R_P_4_DLX1
7669_R_P_4_TBR1
7669_R_P_5_DLX1
7669_R_P_5_TBR1
7669_R_P_6_DLX1
7669_R_P_6_TBR1
7669_R_P_7_DLX1
7669_R_P_7_TBR1
7669_R_P_8_DLX1
7669_R_P_8_TBR1
7669_R_P_9_DLX
7669_R_P_9_DLX1
7669_R_P_9_TBR1
7669_R_POA_Bulk
7669_R_POA_DLX1
7669_R_sec1_Cau_Bulk
7669_R_sec2_GP_Bulk
7669_R_sec2_GP_DLX1
7669_R_sec2_Put_Bulk
7669_R_sec2_Thal_Bulk
7669_R_sec2_Thal_COUPTF2
7669_R_sec2_Thal_DLX1
7669_R_sec3_Cau_DLX1
7669_R_sec3_Hip_COUPTF2
7669_R_sec3_Hip_DLX1
7669_R_sec3_Hip_TBR1
7669_R_sec3_Put_DLX1
7669_R_Skin_Bulk
7669_R_T_1_Bulk
7669_R_T_1_COUPTF2
7669_R_T_1_DLX1
7669_R_T_1_TBR1
7669_RA9_Bulk
7669_RV10_Bulk
JGG

ID05 X−151132909−G−A

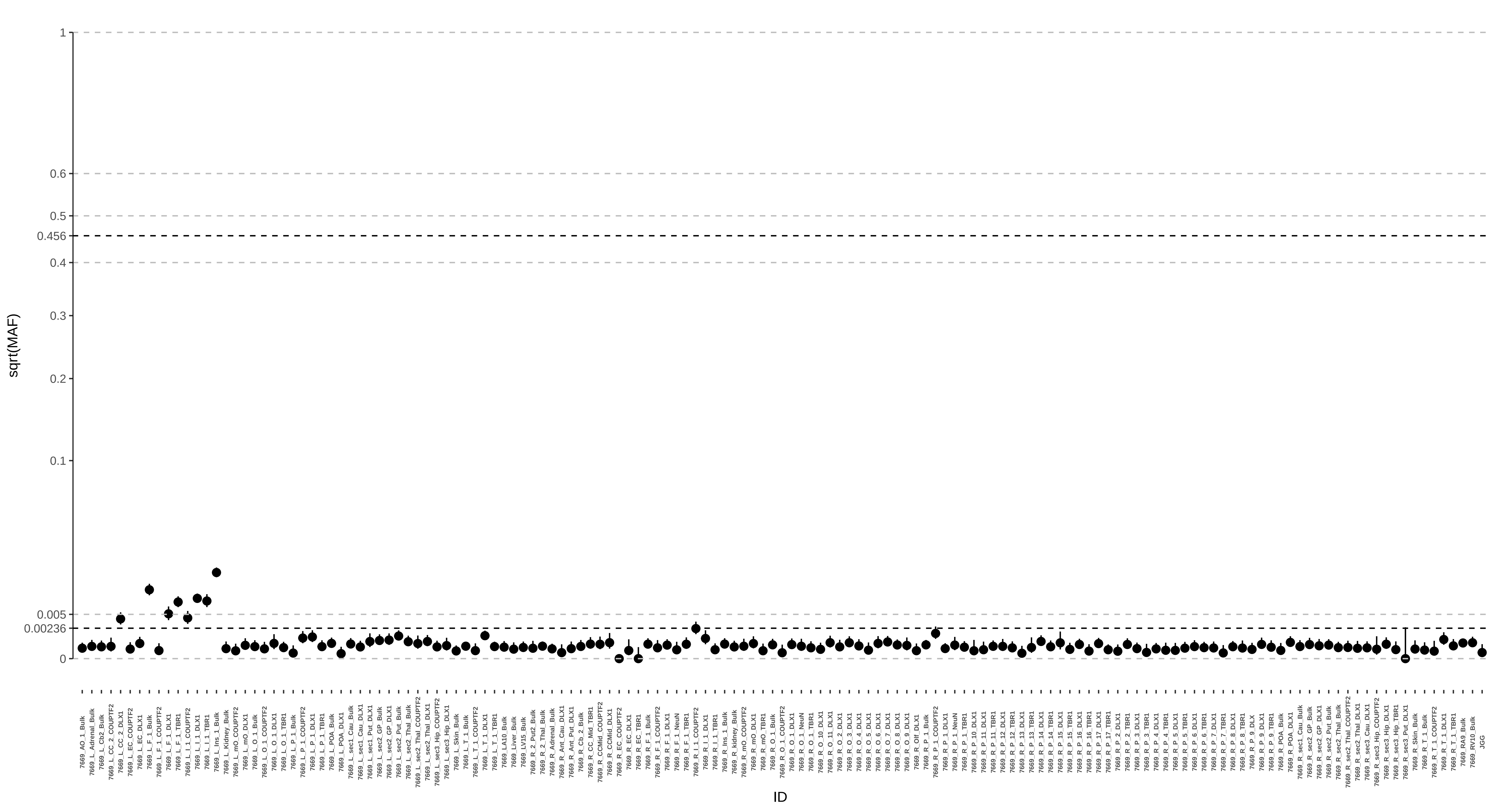

ID05 X−16444039−G−A

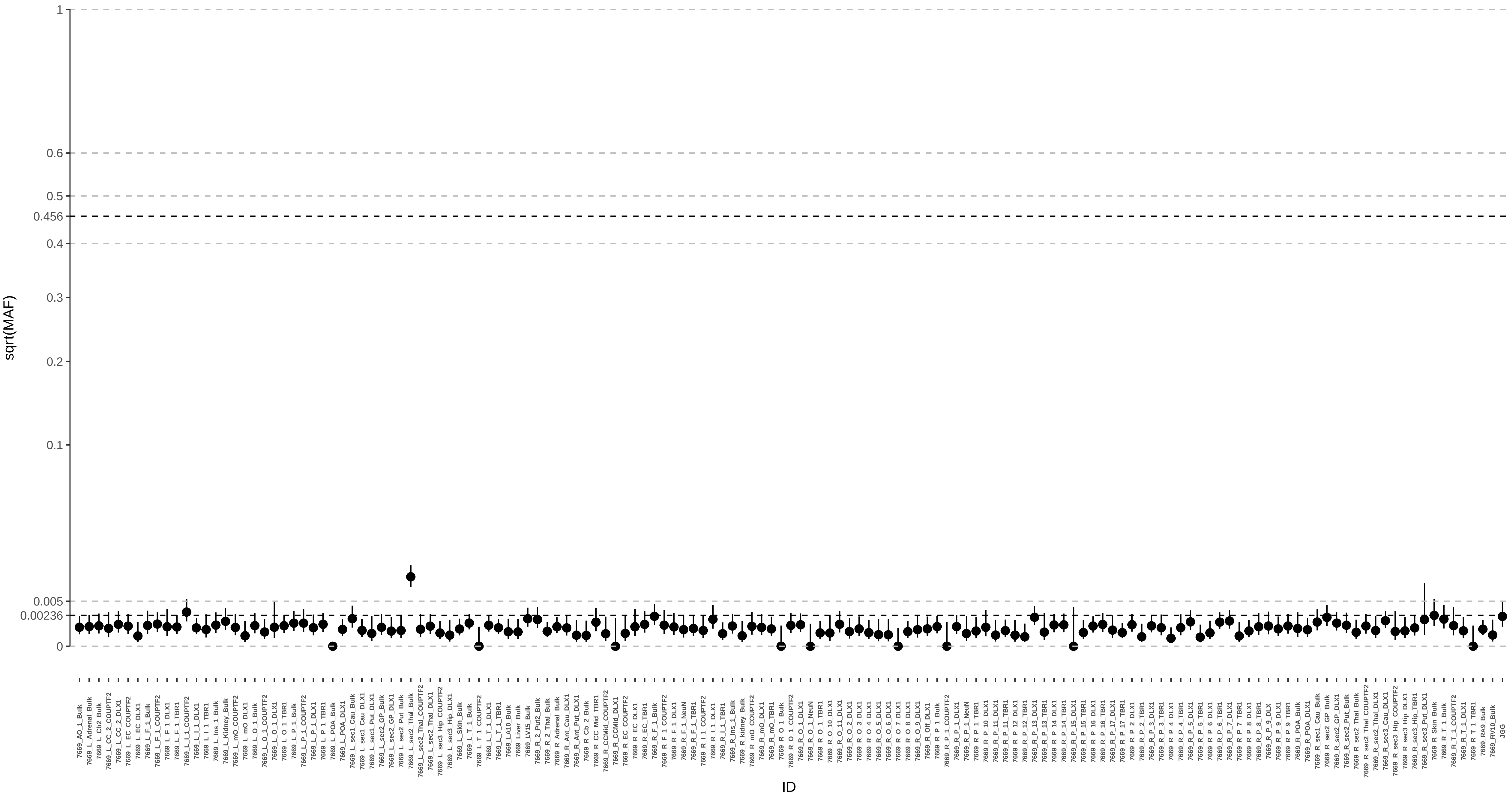

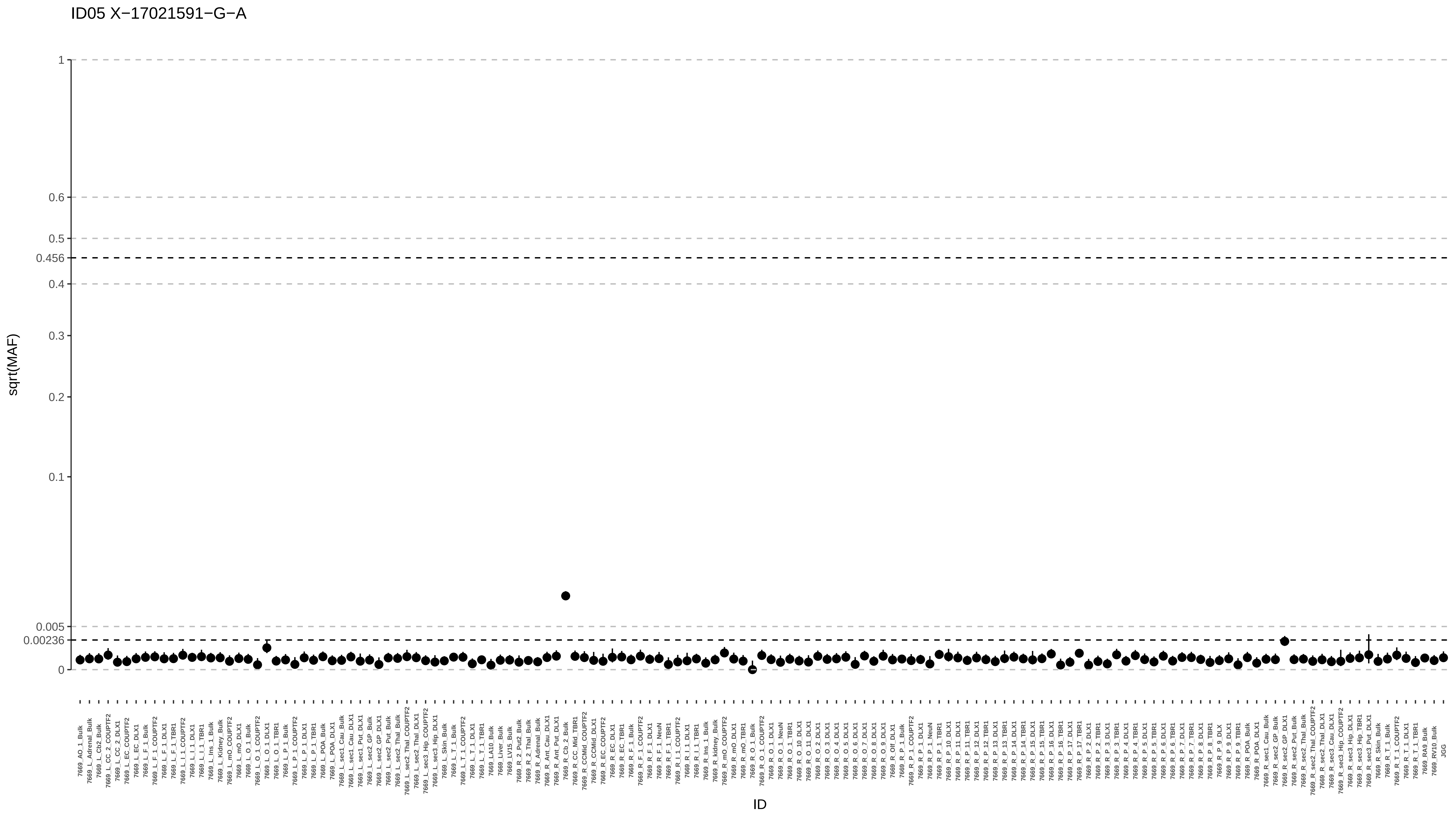
ID05 X−17021591−G−A
sqrt(MAF)
1
0.6
0.5
0.456
0.4
0.3
0.2
0.1
0.005
0.00236
0
ID

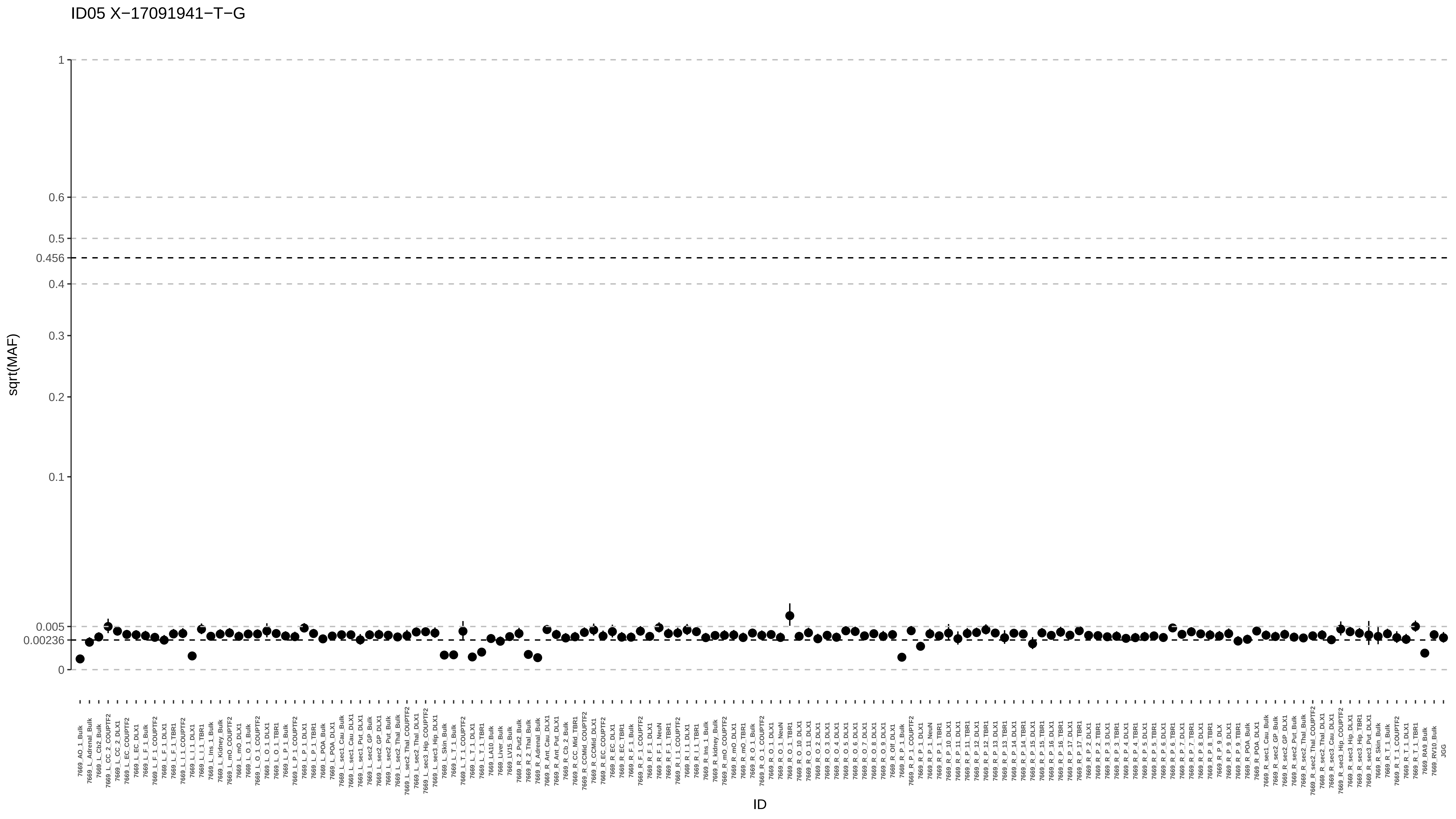
ID05 X−17091941−T−G
sqrt(MAF)
1
0.6
0.5
0.456
0.4
0.3
0.2
0.1
0.005
0.00236
0
ID

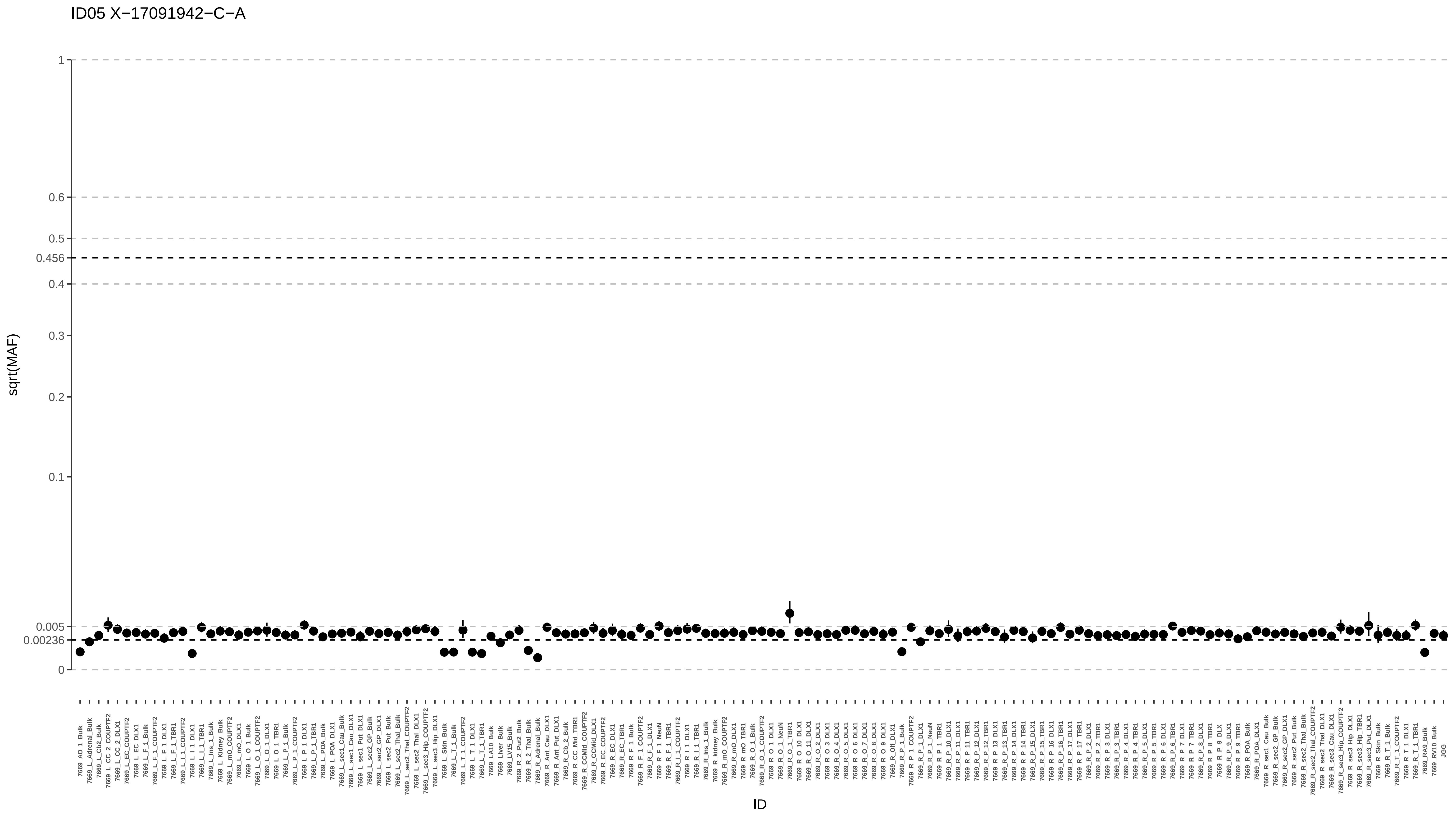
ID05 X−17091942−C−A
sqrt(MAF)
1
0.6
0.5
0.456
0.4
0.3
0.2
0.1
0.005
0.00236
0
ID

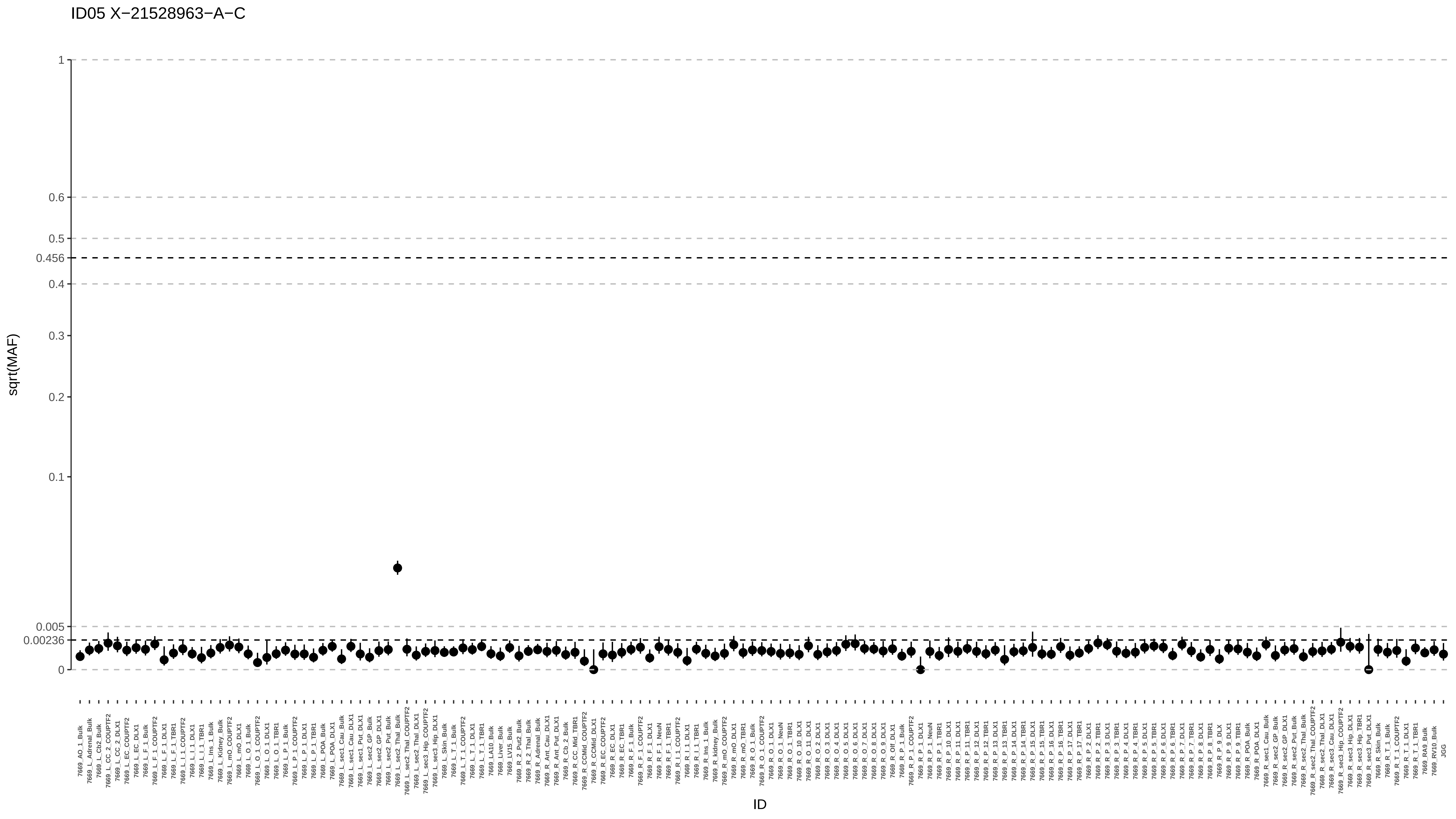

ID05 X−21528963−A−C
sqrt(MAF)
1
0.6
0.5
0.456
0.4
0.3
0.2
0.1
0.005
0.00236
0
ID

ID05 X−24541835−C−A

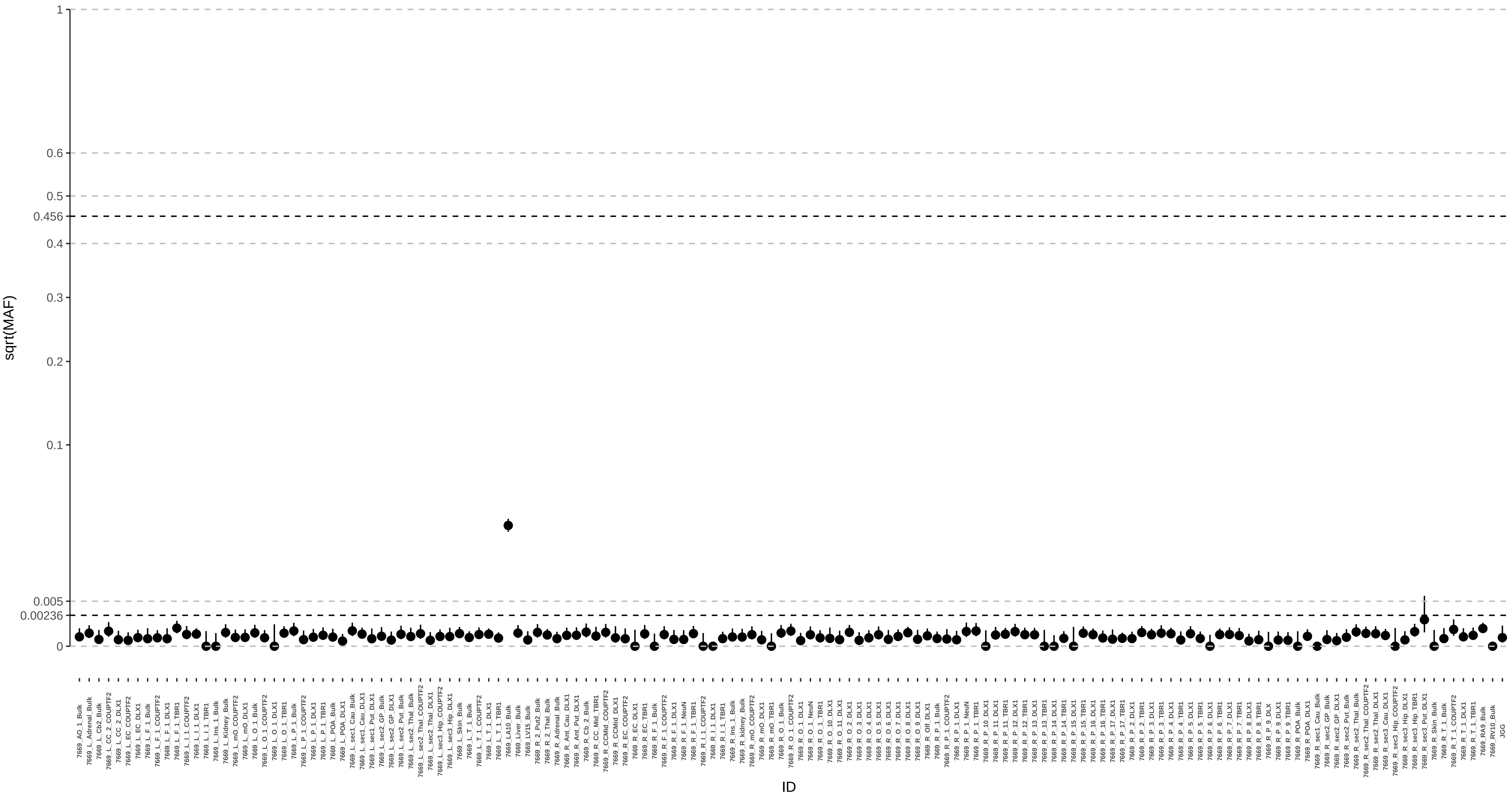

ID05 X−27063252−A−G

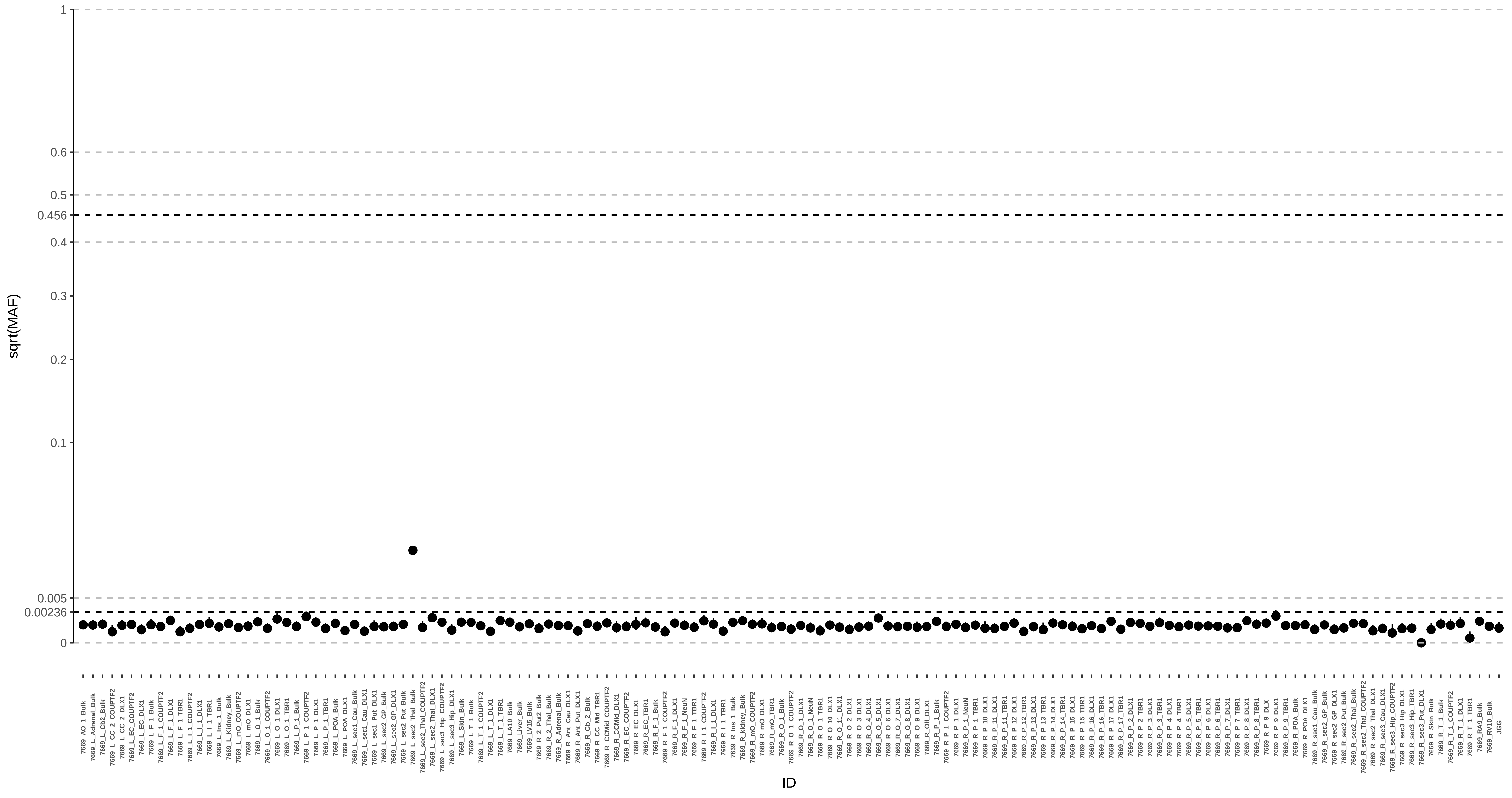

ID05 X−28357791−T−A

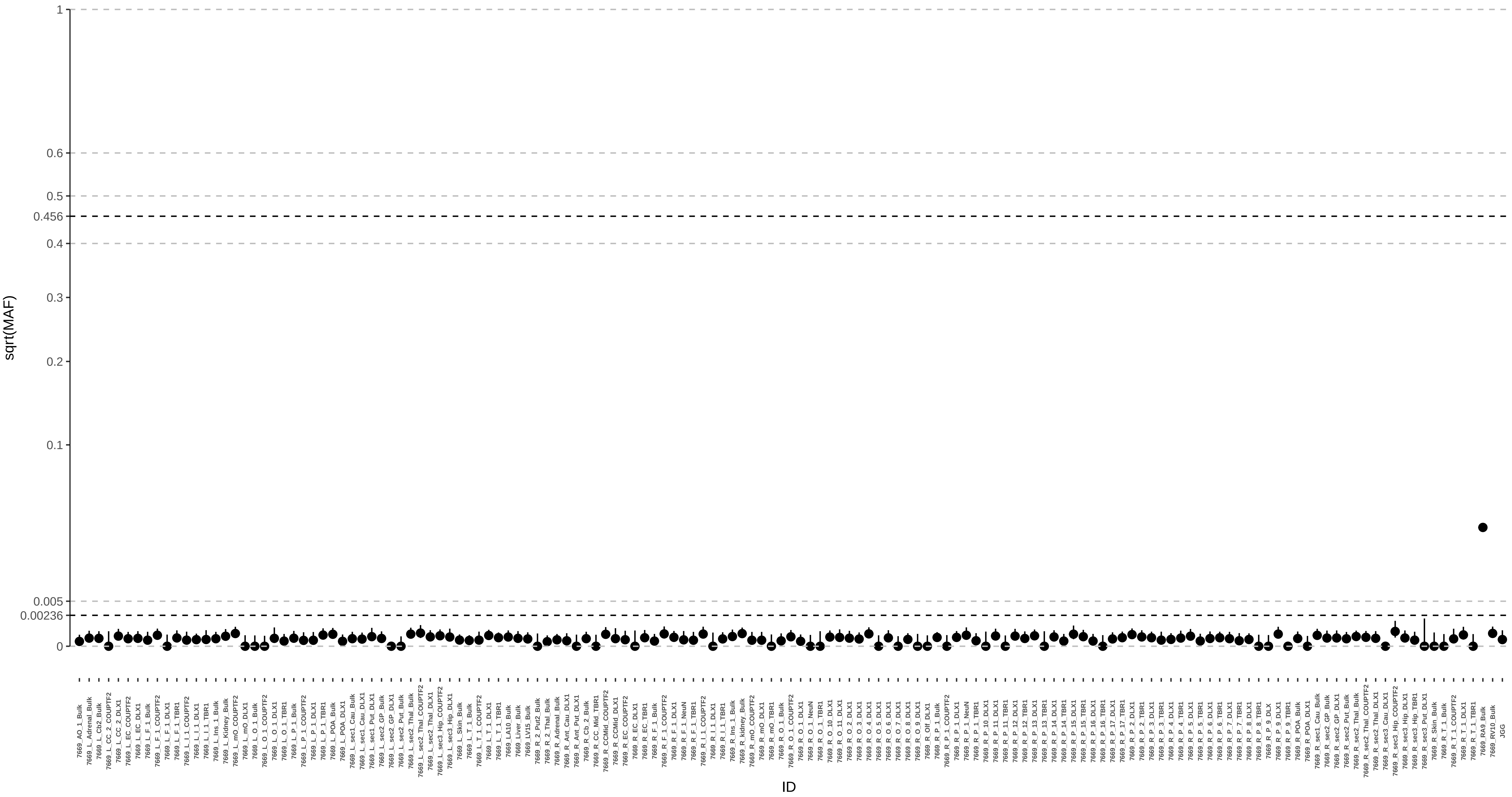

ID05 X−32465017−G−A

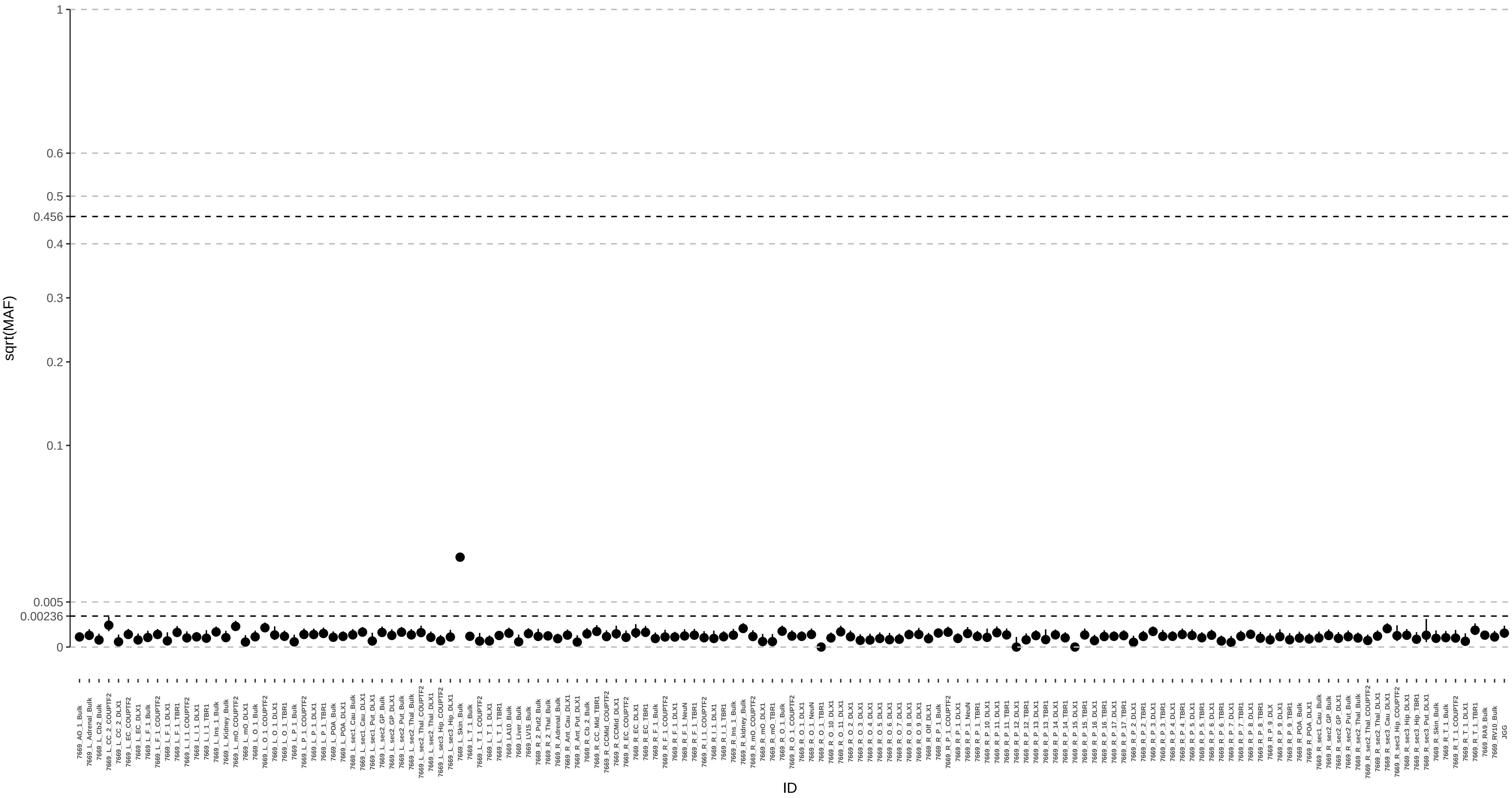

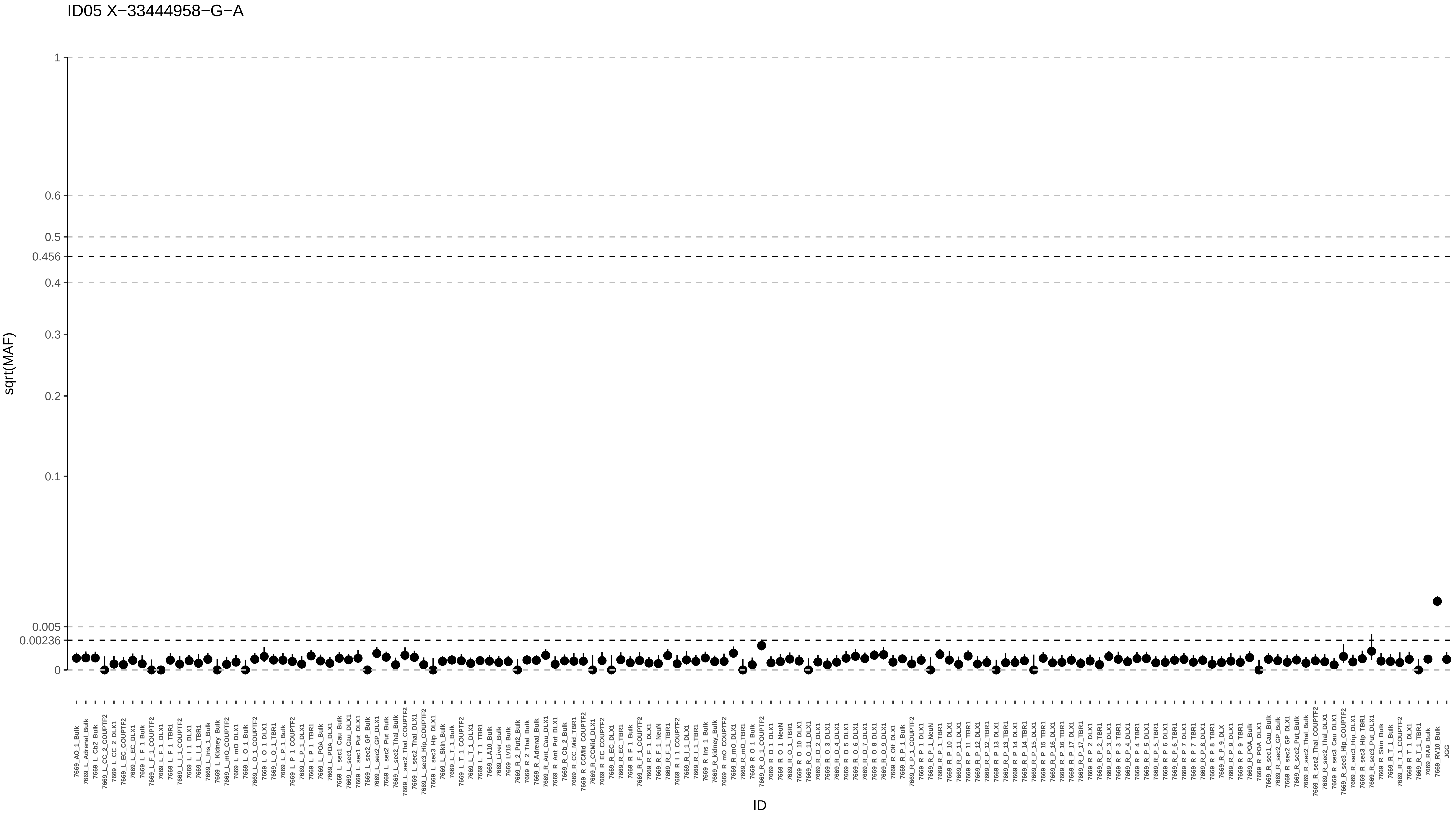
ID05 X−33444958−G−A
sqrt(MAF)
1
0.6
0.5
0.456
0.4
0.3
0.2
0.1
0.005
0.00236
0
ID
7669_AO_1_Bulk
7669_L_Adrenal_Bulk
7669_L_Cb2_Bulk
7669_L_CC_2_COUPTF2
7669_L_CC_2_DLX1
7669_L_EC_COUPTF2
7669_L_EC_DLX1
7669_L_F_1_Bulk
7669_L_F_1_COUPTF2
7669_L_F_1_DLX1
7669_L_F_1_TBR1
7669_L_I_1_COUPTF2
7669_L_I_1_DLX1
7669_L_I_1_TBR1
7669_L_Ins_1_Bulk
7669_L_Kidney_Bulk
7669_L_mO_COUPTF2
7669_L_mO_DLX1
7669_L_O_1_Bulk
7669_L_O_1_COUPTF2
7669_L_O_1_DLX1
7669_L_O_1_TBR1
7669_L_P_1_Bulk
7669_L_P_1_COUPTF2
7669_L_P_1_DLX1
7669_L_P_1_TBR1
7669_L_POA_Bulk
7669_L_POA_DLX1
7669_L_sec1_Cau_Bulk
7669_L_sec1_Cau_DLX1
7669_L_sec1_Put_DLX1
7669_L_sec2_GP_Bulk
7669_L_sec2_GP_DLX1
7669_L_sec2_Put_Bulk
7669_L_sec2_Thal_Bulk
7669_L_sec2_Thal_COUPTF2
7669_L_sec2_Thal_DLX1
7669_L_sec3_Hip_COUPTF2
7669_L_sec3_Hip_DLX1
7669_L_Skin_Bulk
7669_L_T_1_Bulk
7669_L_T_1_COUPTF2
7669_L_T_1_DLX1
7669_L_T_1_TBR1
7669_LA10_Bulk
7669_Liver_Bulk
7669_LV15_Bulk
7669_R_2_Put2_Bulk
7669_R_2_Thal_Bulk
7669_R_Adrenal_Bulk
7669_R_Ant_Cau_DLX1
7669_R_Ant_Put_DLX1
7669_R_Cb_2_Bulk
7669_R_CC_Mid_TBR1
7669_R_CCMid_COUPTF2
7669_R_CCMid_DLX1
7669_R_EC_COUPTF2
7669_R_EC_DLX1
7669_R_EC_TBR1
7669_R_F_1_Bulk
7669_R_F_1_COUPTF2
7669_R_F_1_DLX1
7669_R_F_1_NeuN
7669_R_F_1_TBR1
7669_R_I_1_COUPTF2
7669_R_I_1_DLX1
7669_R_I_1_TBR1
7669_R_Ins_1_Bulk
7669_R_kidney_Bulk
7669_R_mO_COUPTF2
7669_R_mO_DLX1
7669_R_mO_TBR1
7669_R_O_1_Bulk
7669_R_O_1_COUPTF2
7669_R_O_1_DLX1
7669_R_O_1_NeuN
7669_R_O_1_TBR1
7669_R_O_10_DLX1
7669_R_O_11_DLX1
7669_R_O_2_DLX1
7669_R_O_3_DLX1
7669_R_O_4_DLX1
7669_R_O_5_DLX1
7669_R_O_6_DLX1
7669_R_O_7_DLX1
7669_R_O_8_DLX1
7669_R_O_9_DLX1
7669_R_Olf_DLX1
7669_R_P_1_Bulk
7669_R_P_1_COUPTF2
7669_R_P_1_DLX1
7669_R_P_1_NeuN
7669_R_P_1_TBR1
7669_R_P_10_DLX1
7669_R_P_11_DLX1
7669_R_P_11_TBR1
7669_R_P_12_DLX1
7669_R_P_12_TBR1
7669_R_P_13_DLX1
7669_R_P_13_TBR1
7669_R_P_14_DLX1
7669_R_P_14_TBR1
7669_R_P_15_DLX1
7669_R_P_15_TBR1
7669_R_P_16_DLX1
7669_R_P_16_TBR1
7669_R_P_17_DLX1
7669_R_P_17_TBR1
7669_R_P_2_DLX1
7669_R_P_2_TBR1
7669_R_P_3_DLX1
7669_R_P_3_TBR1
7669_R_P_4_DLX1
7669_R_P_4_TBR1
7669_R_P_5_DLX1
7669_R_P_5_TBR1
7669_R_P_6_DLX1
7669_R_P_6_TBR1
7669_R_P_7_DLX1
7669_R_P_7_TBR1
7669_R_P_8_DLX1
7669_R_P_8_TBR1
7669_R_P_9_DLX
7669_R_P_9_DLX1
7669_R_P_9_TBR1
7669_R_POA_Bulk
7669_R_POA_DLX1
7669_R_sec1_Cau_Bulk
7669_R_sec2_GP_Bulk
7669_R_sec2_GP_DLX1
7669_R_sec2_Put_Bulk
7669_R_sec2_Thal_Bulk
7669_R_sec2_Thal_COUPTF2
7669_R_sec2_Thal_DLX1
7669_R_sec3_Cau_DLX1
7669_R_sec3_Hip_COUPTF2
7669_R_sec3_Hip_DLX1
7669_R_sec3_Hip_TBR1
7669_R_sec3_Put_DLX1
7669_R_Skin_Bulk
7669_R_T_1_Bulk
7669_R_T_1_COUPTF2
7669_R_T_1_DLX1
7669_R_T_1_TBR1
7669_RA9_Bulk
7669_RV10_Bulk
JGG

ID05 X−34424349−A−T

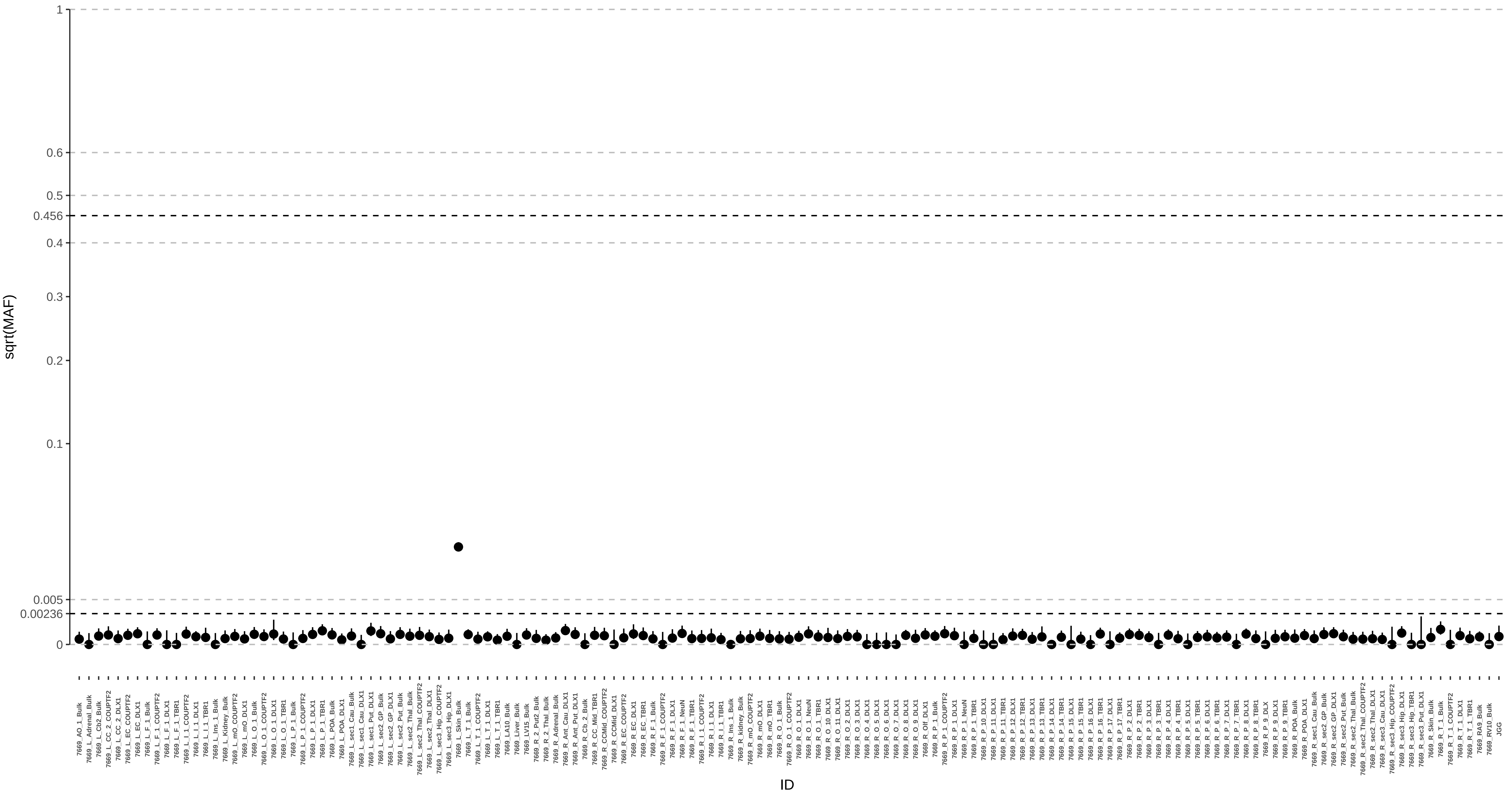

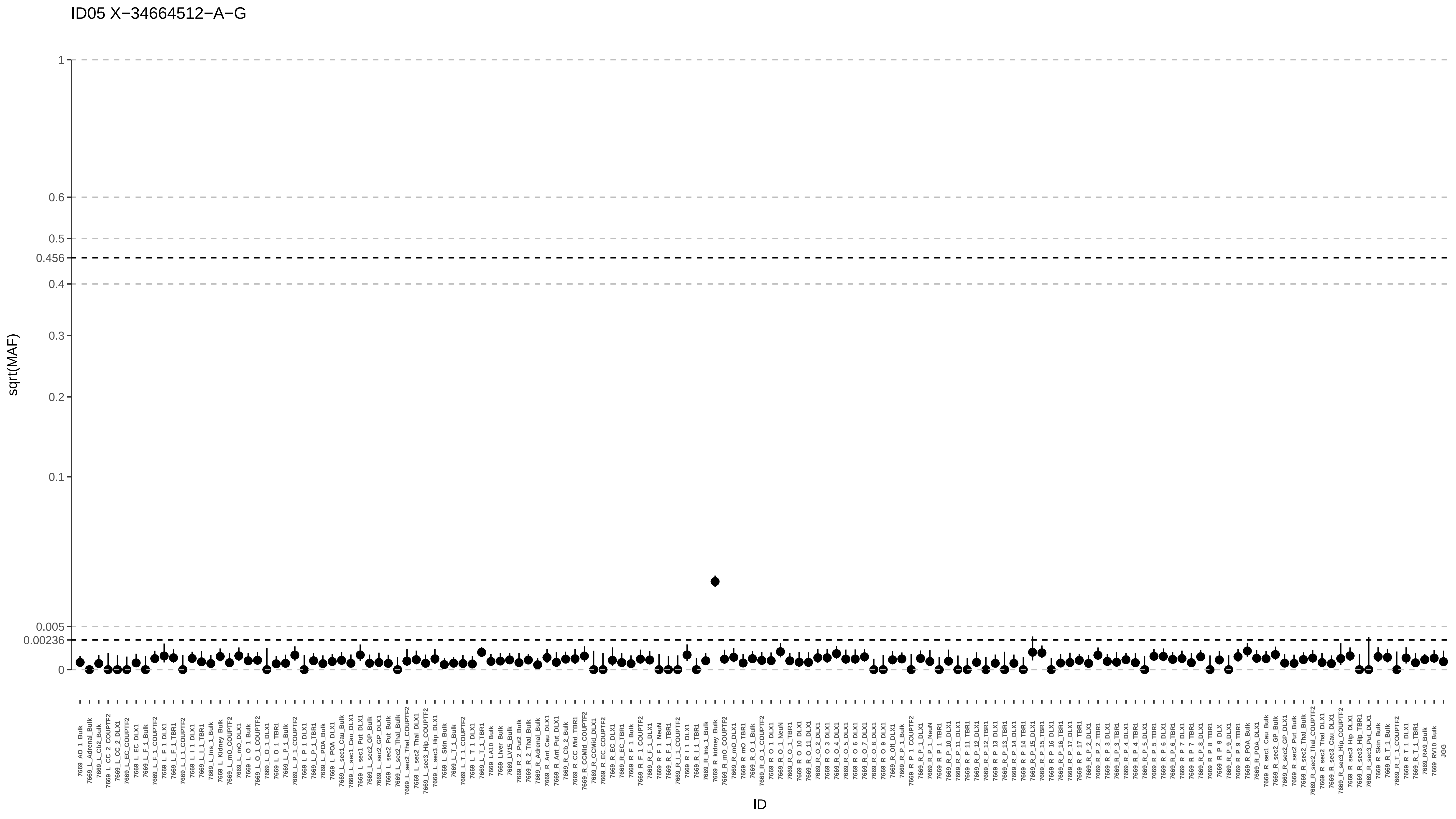
ID05 X−34664512−A−G
sqrt(MAF)
1
0.6
0.5
0.456
0.4
0.3
0.2
0.1
0.005
0.00236
0
ID
7669_AO_1_Bulk
7669_L_Adrenal_Bulk
7669_L_Cb2_Bulk
7669_L_CC_2_COUPTF2
7669_L_CC_2_DLX1
7669_L_EC_COUPTF2
7669_L_EC_DLX1
7669_L_F_1_Bulk
7669_L_F_1_COUPTF2
7669_L_F_1_DLX1
7669_L_F_1_TBR1
7669_L_I_1_COUPTF2
7669_L_I_1_DLX1
7669_L_I_1_TBR1
7669_L_Ins_1_Bulk
7669_L_Kidney_Bulk
7669_L_mO_COUPTF2
7669_L_mO_DLX1
7669_L_O_1_Bulk
7669_L_O_1_COUPTF2
7669_L_O_1_DLX1
7669_L_O_1_TBR1
7669_L_P_1_Bulk
7669_L_P_1_COUPTF2
7669_L_P_1_DLX1
7669_L_P_1_TBR1
7669_L_POA_Bulk
7669_L_POA_DLX1
7669_L_sec1_Cau_Bulk
7669_L_sec1_Cau_DLX1
7669_L_sec1_Put_DLX1
7669_L_sec2_GP_Bulk
7669_L_sec2_GP_DLX1
7669_L_sec2_Put_Bulk
7669_L_sec2_Thal_Bulk
7669_L_sec2_Thal_COUPTF2
7669_L_sec2_Thal_DLX1
7669_L_sec3_Hip_COUPTF2
7669_L_sec3_Hip_DLX1
7669_L_Skin_Bulk
7669_L_T_1_Bulk
7669_L_T_1_COUPTF2
7669_L_T_1_DLX1
7669_L_T_1_TBR1
7669_LA10_Bulk
7669_Liver_Bulk
7669_LV15_Bulk
7669_R_2_Put2_Bulk
7669_R_2_Thal_Bulk
7669_R_Adrenal_Bulk
7669_R_Ant_Cau_DLX1
7669_R_Ant_Put_DLX1
7669_R_Cb_2_Bulk
7669_R_CC_Mid_TBR1
7669_R_CCMid_COUPTF2
7669_R_CCMid_DLX1
7669_R_EC_COUPTF2
7669_R_EC_DLX1
7669_R_EC_TBR1
7669_R_F_1_Bulk
7669_R_F_1_COUPTF2
7669_R_F_1_DLX1
7669_R_F_1_NeuN
7669_R_F_1_TBR1
7669_R_I_1_COUPTF2
7669_R_I_1_DLX1
7669_R_I_1_TBR1
7669_R_Ins_1_Bulk
7669_R_kidney_Bulk
7669_R_mO_COUPTF2
7669_R_mO_DLX1
7669_R_mO_TBR1
7669_R_O_1_Bulk
7669_R_O_1_COUPTF2
7669_R_O_1_DLX1
7669_R_O_1_NeuN
7669_R_O_1_TBR1
7669_R_O_10_DLX1
7669_R_O_11_DLX1
7669_R_O_2_DLX1
7669_R_O_3_DLX1
7669_R_O_4_DLX1
7669_R_O_5_DLX1
7669_R_O_6_DLX1
7669_R_O_7_DLX1
7669_R_O_8_DLX1
7669_R_O_9_DLX1
7669_R_Olf_DLX1
7669_R_P_1_Bulk
7669_R_P_1_COUPTF2
7669_R_P_1_DLX1
7669_R_P_1_NeuN
7669_R_P_1_TBR1
7669_R_P_10_DLX1
7669_R_P_11_DLX1
7669_R_P_11_TBR1
7669_R_P_12_DLX1
7669_R_P_12_TBR1
7669_R_P_13_DLX1
7669_R_P_13_TBR1
7669_R_P_14_DLX1
7669_R_P_14_TBR1
7669_R_P_15_DLX1
7669_R_P_15_TBR1
7669_R_P_16_DLX1
7669_R_P_16_TBR1
7669_R_P_17_DLX1
7669_R_P_17_TBR1
7669_R_P_2_DLX1
7669_R_P_2_TBR1
7669_R_P_3_DLX1
7669_R_P_3_TBR1
7669_R_P_4_DLX1
7669_R_P_4_TBR1
7669_R_P_5_DLX1
7669_R_P_5_TBR1
7669_R_P_6_DLX1
7669_R_P_6_TBR1
7669_R_P_7_DLX1
7669_R_P_7_TBR1
7669_R_P_8_DLX1
7669_R_P_8_TBR1
7669_R_P_9_DLX
7669_R_P_9_DLX1
7669_R_P_9_TBR1
7669_R_POA_Bulk
7669_R_POA_DLX1
7669_R_sec1_Cau_Bulk
7669_R_sec2_GP_Bulk
7669_R_sec2_GP_DLX1
7669_R_sec2_Put_Bulk
7669_R_sec2_Thal_Bulk
7669_R_sec2_Thal_COUPTF2
7669_R_sec2_Thal_DLX1
7669_R_sec3_Cau_DLX1
7669_R_sec3_Hip_COUPTF2
7669_R_sec3_Hip_DLX1
7669_R_sec3_Hip_TBR1
7669_R_sec3_Put_DLX1
7669_R_Skin_Bulk
7669_R_T_1_Bulk
7669_R_T_1_COUPTF2
7669_R_T_1_DLX1
7669_R_T_1_TBR1
7669_RA9_Bulk
7669_RV10_Bulk
JGG

ID05 X−34801213−T−C

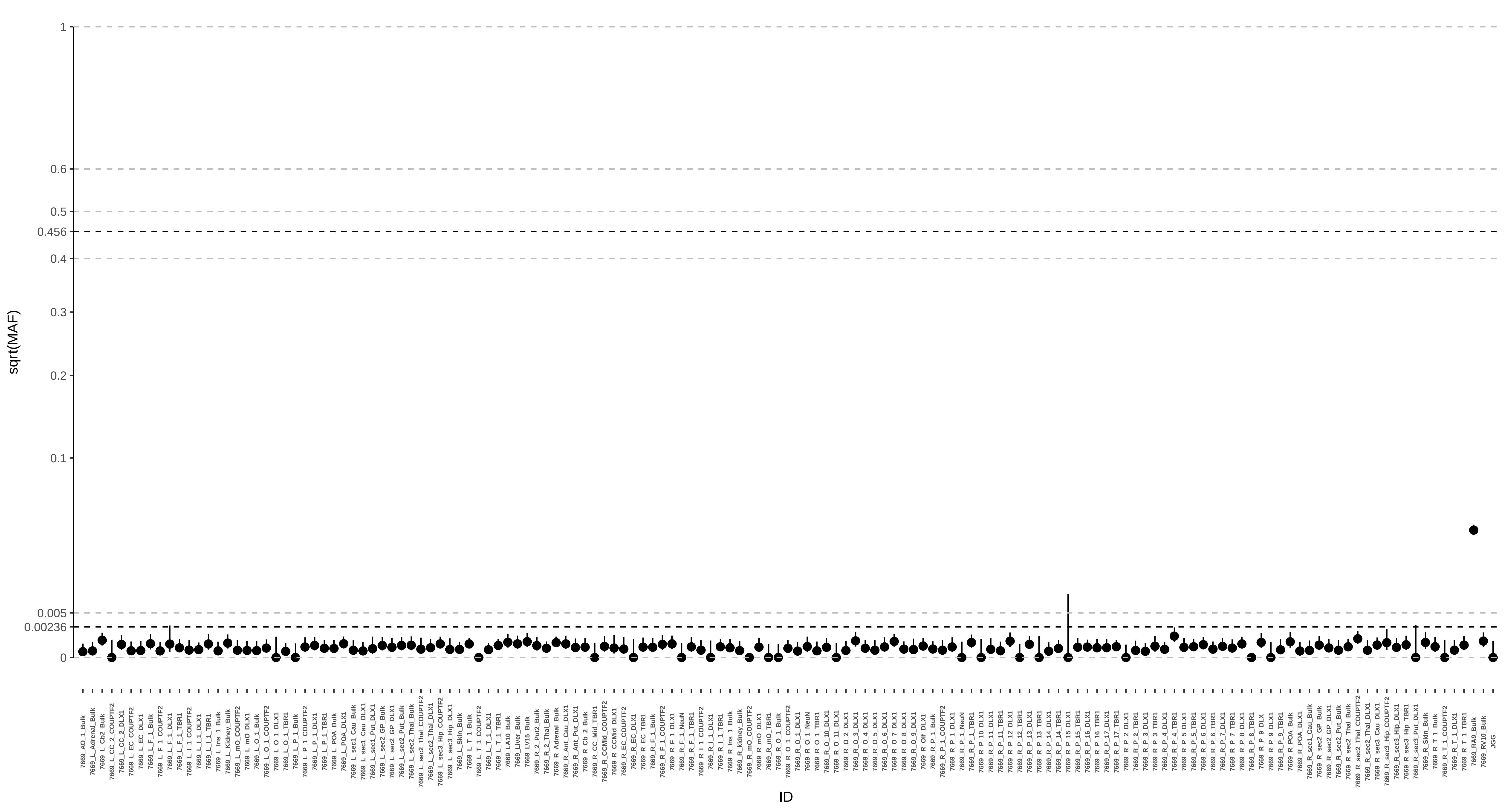

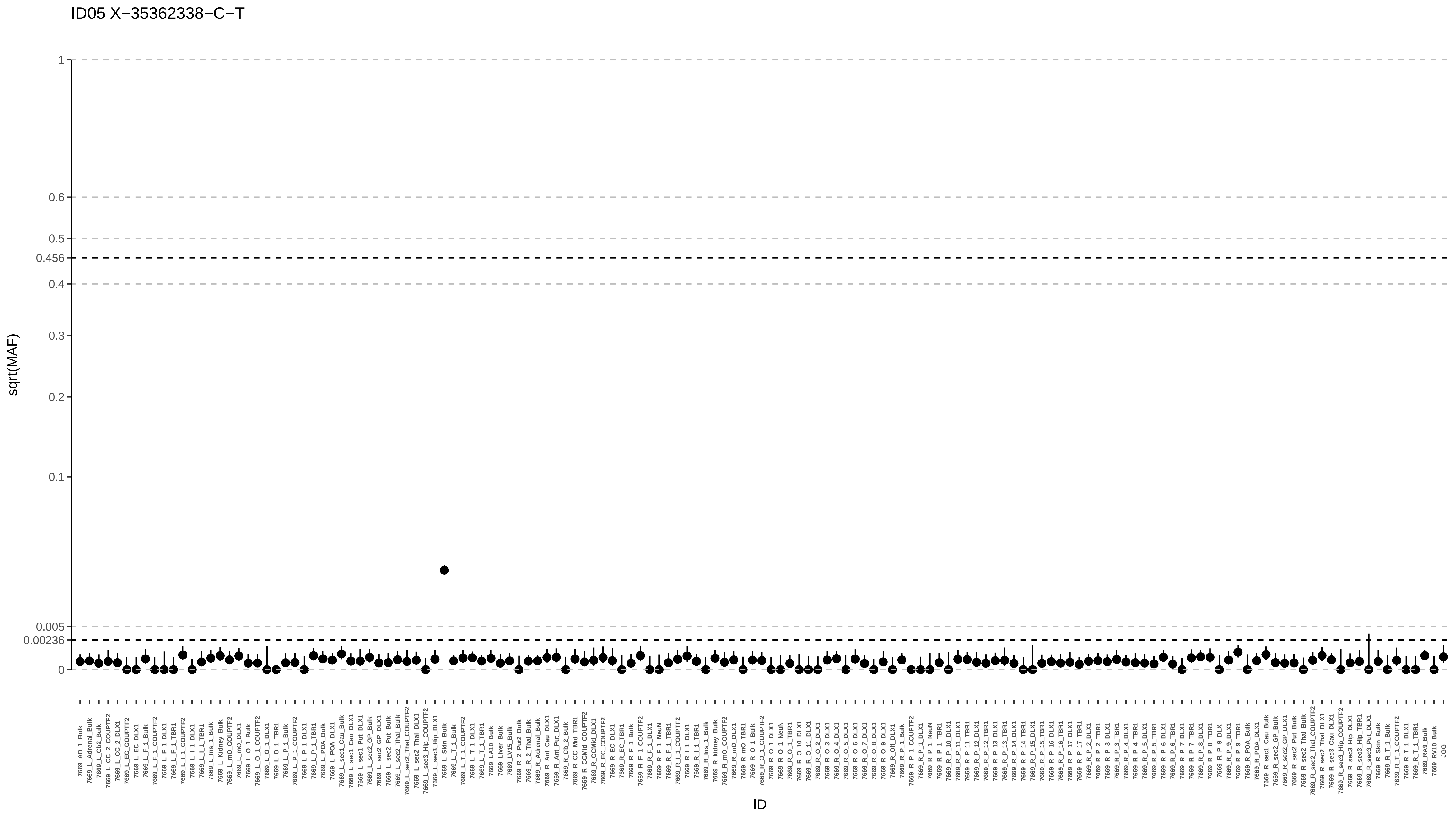
ID05 X−35362338−C−T
sqrt(MAF)
1
0.6
0.5
0.456
0.4
0.3
0.2
0.1
0.005
0.00236
0
ID
7669_AO_1_Bulk
7669_L_Adrenal_Bulk
7669_L_Cb2_Bulk
7669_L_CC_2_COUPTF2
7669_L_CC_2_DLX1
7669_L_EC_COUPTF2
7669_L_EC_DLX1
7669_L_F_1_Bulk
7669_L_F_1_COUPTF2
7669_L_F_1_DLX1
7669_L_F_1_TBR1
7669_L_I_1_COUPTF2
7669_L_I_1_DLX1
7669_L_I_1_TBR1
7669_L_Ins_1_Bulk
7669_L_Kidney_Bulk
7669_L_mO_COUPTF2
7669_L_mO_DLX1
7669_L_O_1_Bulk
7669_L_O_1_COUPTF2
7669_L_O_1_DLX1
7669_L_O_1_TBR1
7669_L_P_1_Bulk
7669_L_P_1_COUPTF2
7669_L_P_1_DLX1
7669_L_P_1_TBR1
7669_L_POA_Bulk
7669_L_POA_DLX1
7669_L_sec1_Cau_Bulk
7669_L_sec1_Cau_DLX1
7669_L_sec1_Put_DLX1
7669_L_sec2_GP_Bulk
7669_L_sec2_GP_DLX1
7669_L_sec2_Put_Bulk
7669_L_sec2_Thal_Bulk
7669_L_sec2_Thal_COUPTF2
7669_L_sec2_Thal_DLX1
7669_L_sec3_Hip_COUPTF2
7669_L_sec3_Hip_DLX1
7669_L_Skin_Bulk
7669_L_T_1_Bulk
7669_L_T_1_COUPTF2
7669_L_T_1_DLX1
7669_L_T_1_TBR1
7669_LA10_Bulk
7669_Liver_Bulk
7669_LV15_Bulk
7669_R_2_Put2_Bulk
7669_R_2_Thal_Bulk
7669_R_Adrenal_Bulk
7669_R_Ant_Cau_DLX1
7669_R_Ant_Put_DLX1
7669_R_Cb_2_Bulk
7669_R_CC_Mid_TBR1
7669_R_CCMid_COUPTF2
7669_R_CCMid_DLX1
7669_R_EC_COUPTF2
7669_R_EC_DLX1
7669_R_EC_TBR1
7669_R_F_1_Bulk
7669_R_F_1_COUPTF2
7669_R_F_1_DLX1
7669_R_F_1_NeuN
7669_R_F_1_TBR1
7669_R_I_1_COUPTF2
7669_R_I_1_DLX1
7669_R_I_1_TBR1
7669_R_Ins_1_Bulk
7669_R_kidney_Bulk
7669_R_mO_COUPTF2
7669_R_mO_DLX1
7669_R_mO_TBR1
7669_R_O_1_Bulk
7669_R_O_1_COUPTF2
7669_R_O_1_DLX1
7669_R_O_1_NeuN
7669_R_O_1_TBR1
7669_R_O_10_DLX1
7669_R_O_11_DLX1
7669_R_O_2_DLX1
7669_R_O_3_DLX1
7669_R_O_4_DLX1
7669_R_O_5_DLX1
7669_R_O_6_DLX1
7669_R_O_7_DLX1
7669_R_O_8_DLX1
7669_R_O_9_DLX1
7669_R_Olf_DLX1
7669_R_P_1_Bulk
7669_R_P_1_COUPTF2
7669_R_P_1_DLX1
7669_R_P_1_NeuN
7669_R_P_1_TBR1
7669_R_P_10_DLX1
7669_R_P_11_DLX1
7669_R_P_11_TBR1
7669_R_P_12_DLX1
7669_R_P_12_TBR1
7669_R_P_13_DLX1
7669_R_P_13_TBR1
7669_R_P_14_DLX1
7669_R_P_14_TBR1
7669_R_P_15_DLX1
7669_R_P_15_TBR1
7669_R_P_16_DLX1
7669_R_P_16_TBR1
7669_R_P_17_DLX1
7669_R_P_17_TBR1
7669_R_P_2_DLX1
7669_R_P_2_TBR1
7669_R_P_3_DLX1
7669_R_P_3_TBR1
7669_R_P_4_DLX1
7669_R_P_4_TBR1
7669_R_P_5_DLX1
7669_R_P_5_TBR1
7669_R_P_6_DLX1
7669_R_P_6_TBR1
7669_R_P_7_DLX1
7669_R_P_7_TBR1
7669_R_P_8_DLX1
7669_R_P_8_TBR1
7669_R_P_9_DLX
7669_R_P_9_DLX1
7669_R_P_9_TBR1
7669_R_POA_Bulk
7669_R_POA_DLX1
7669_R_sec1_Cau_Bulk
7669_R_sec2_GP_Bulk
7669_R_sec2_GP_DLX1
7669_R_sec2_Put_Bulk
7669_R_sec2_Thal_Bulk
7669_R_sec2_Thal_COUPTF2
7669_R_sec2_Thal_DLX1
7669_R_sec3_Cau_DLX1
7669_R_sec3_Hip_COUPTF2
7669_R_sec3_Hip_DLX1
7669_R_sec3_Hip_TBR1
7669_R_sec3_Put_DLX1
7669_R_Skin_Bulk
7669_R_T_1_Bulk
7669_R_T_1_COUPTF2
7669_R_T_1_DLX1
7669_R_T_1_TBR1
7669_RA9_Bulk
7669_RV10_Bulk
JGG

ID05 X−35471077−C−T

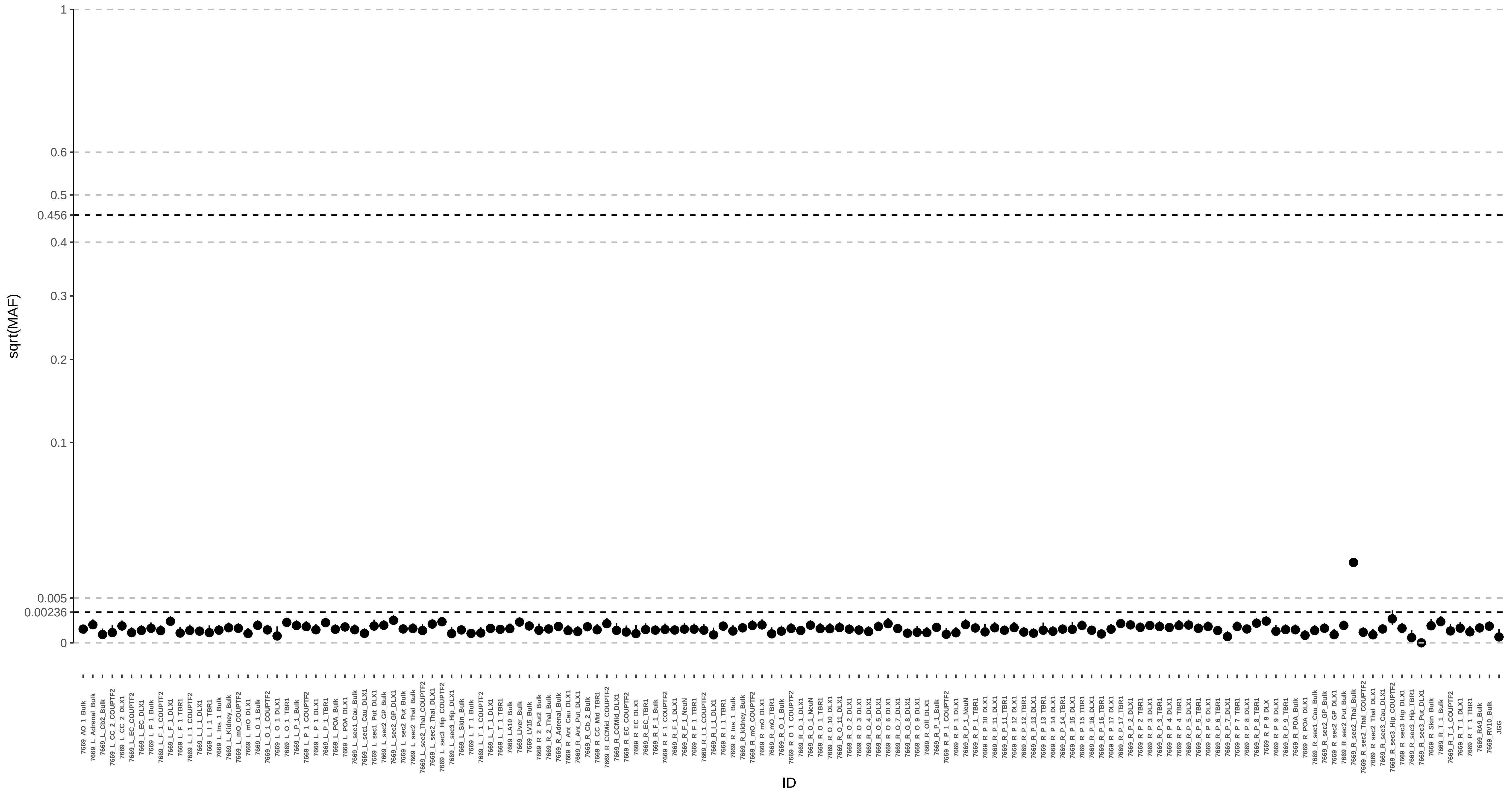

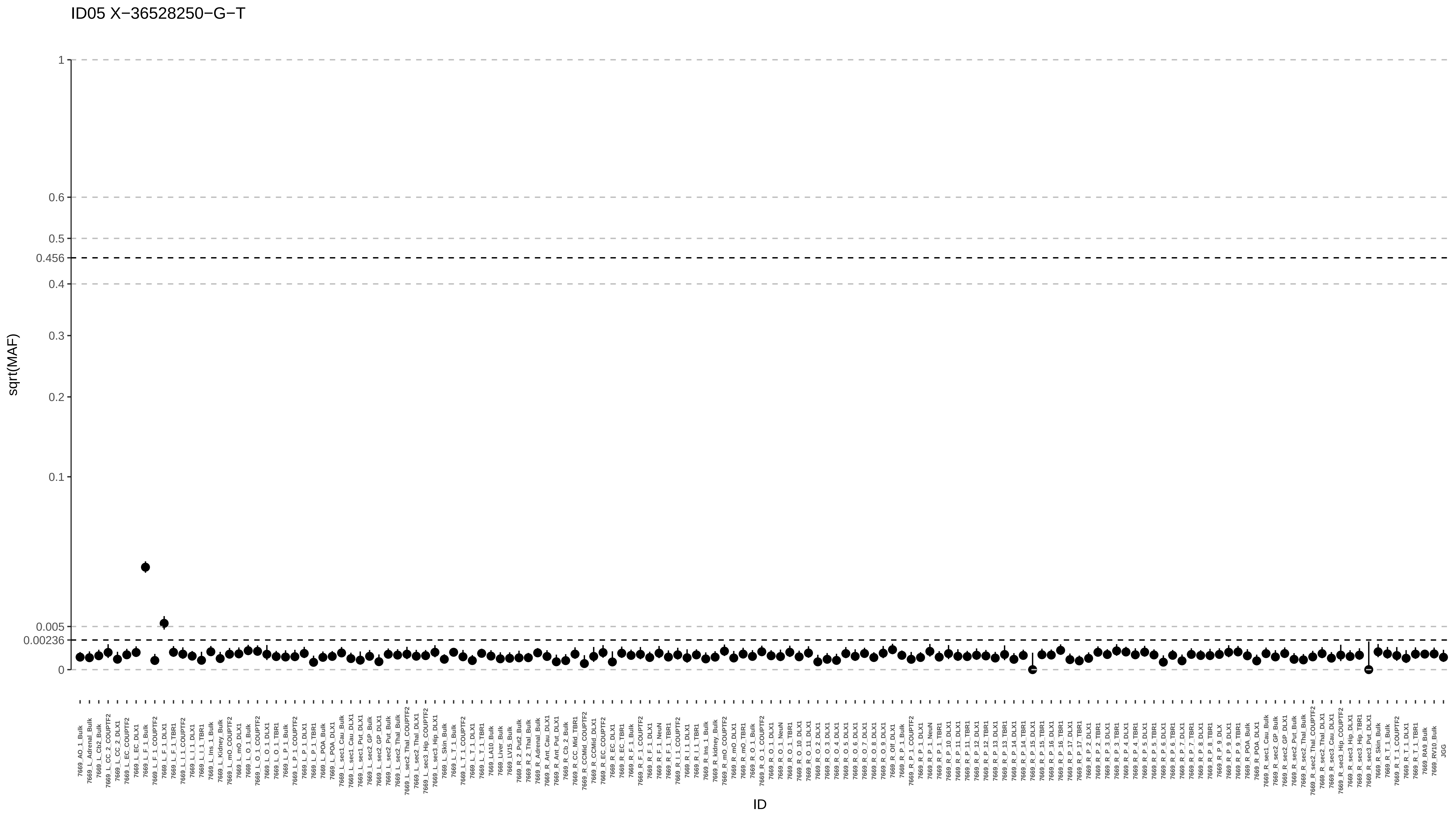

ID05 X−36528250−G−T
sqrt(MAF)
1
0.6
0.5
0.456
0.4
0.3
0.2
0.1
0.005
0.00236
0
ID

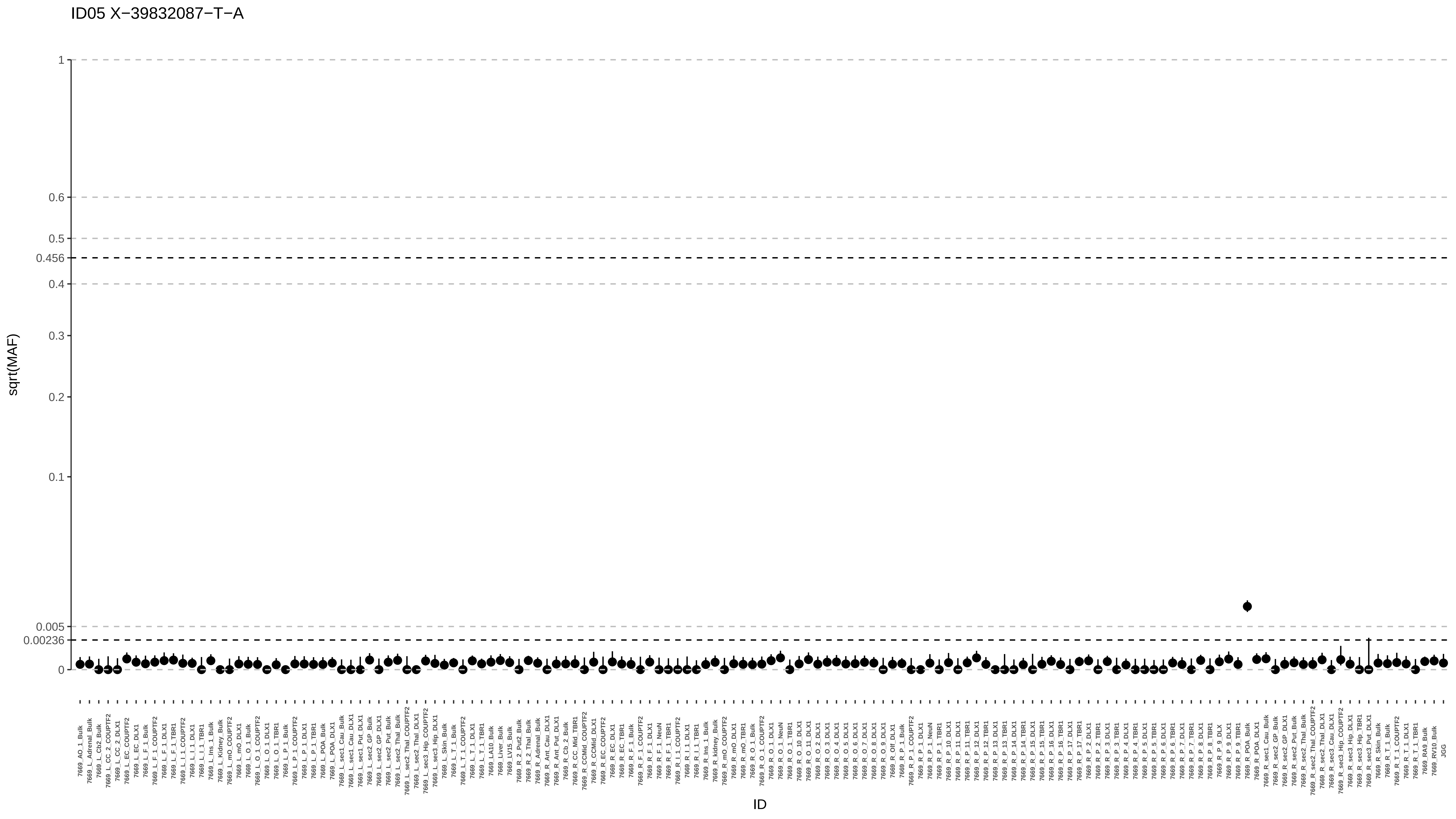
ID05 X−39832087−T−A
sqrt(MAF)
1
0.6
0.5
0.456
0.4
0.3
0.2
0.1
0.005
0.00236
0
ID
7669_AO_1_Bulk
7669_L_Adrenal_Bulk
7669_L_Cb2_Bulk
7669_L_CC_2_COUPTF2
7669_L_CC_2_DLX1
7669_L_EC_COUPTF2
7669_L_EC_DLX1
7669_L_F_1_Bulk
7669_L_F_1_COUPTF2
7669_L_F_1_DLX1
7669_L_F_1_TBR1
7669_L_I_1_COUPTF2
7669_L_I_1_DLX1
7669_L_I_1_TBR1
7669_L_Ins_1_Bulk
7669_L_Kidney_Bulk
7669_L_mO_COUPTF2
7669_L_mO_DLX1
7669_L_O_1_Bulk
7669_L_O_1_COUPTF2
7669_L_O_1_DLX1
7669_L_O_1_TBR1
7669_L_P_1_Bulk
7669_L_P_1_COUPTF2
7669_L_P_1_DLX1
7669_L_P_1_TBR1
7669_L_POA_Bulk
7669_L_POA_DLX1
7669_L_sec1_Cau_Bulk
7669_L_sec1_Cau_DLX1
7669_L_sec1_Put_DLX1
7669_L_sec2_GP_Bulk
7669_L_sec2_GP_DLX1
7669_L_sec2_Put_Bulk
7669_L_sec2_Thal_Bulk
7669_L_sec2_Thal_COUPTF2
7669_L_sec2_Thal_DLX1
7669_L_sec3_Hip_COUPTF2
7669_L_sec3_Hip_DLX1
7669_L_Skin_Bulk
7669_L_T_1_Bulk
7669_L_T_1_COUPTF2
7669_L_T_1_DLX1
7669_L_T_1_TBR1
7669_LA10_Bulk
7669_Liver_Bulk
7669_LV15_Bulk
7669_R_2_Put2_Bulk
7669_R_2_Thal_Bulk
7669_R_Adrenal_Bulk
7669_R_Ant_Cau_DLX1
7669_R_Ant_Put_DLX1
7669_R_Cb_2_Bulk
7669_R_CC_Mid_TBR1
7669_R_CCMid_COUPTF2
7669_R_CCMid_DLX1
7669_R_EC_COUPTF2
7669_R_EC_DLX1
7669_R_EC_TBR1
7669_R_F_1_Bulk
7669_R_F_1_COUPTF2
7669_R_F_1_DLX1
7669_R_F_1_NeuN
7669_R_F_1_TBR1
7669_R_I_1_COUPTF2
7669_R_I_1_DLX1
7669_R_I_1_TBR1
7669_R_Ins_1_Bulk
7669_R_kidney_Bulk
7669_R_mO_COUPTF2
7669_R_mO_DLX1
7669_R_mO_TBR1
7669_R_O_1_Bulk
7669_R_O_1_COUPTF2
7669_R_O_1_DLX1
7669_R_O_1_NeuN
7669_R_O_1_TBR1
7669_R_O_10_DLX1
7669_R_O_11_DLX1
7669_R_O_2_DLX1
7669_R_O_3_DLX1
7669_R_O_4_DLX1
7669_R_O_5_DLX1
7669_R_O_6_DLX1
7669_R_O_7_DLX1
7669_R_O_8_DLX1
7669_R_O_9_DLX1
7669_R_Olf_DLX1
7669_R_P_1_Bulk
7669_R_P_1_COUPTF2
7669_R_P_1_DLX1
7669_R_P_1_NeuN
7669_R_P_1_TBR1
7669_R_P_10_DLX1
7669_R_P_11_DLX1
7669_R_P_11_TBR1
7669_R_P_12_DLX1
7669_R_P_12_TBR1
7669_R_P_13_DLX1
7669_R_P_13_TBR1
7669_R_P_14_DLX1
7669_R_P_14_TBR1
7669_R_P_15_DLX1
7669_R_P_15_TBR1
7669_R_P_16_DLX1
7669_R_P_16_TBR1
7669_R_P_17_DLX1
7669_R_P_17_TBR1
7669_R_P_2_DLX1
7669_R_P_2_TBR1
7669_R_P_3_DLX1
7669_R_P_3_TBR1
7669_R_P_4_DLX1
7669_R_P_4_TBR1
7669_R_P_5_DLX1
7669_R_P_5_TBR1
7669_R_P_6_DLX1
7669_R_P_6_TBR1
7669_R_P_7_DLX1
7669_R_P_7_TBR1
7669_R_P_8_DLX1
7669_R_P_8_TBR1
7669_R_P_9_DLX
7669_R_P_9_DLX1
7669_R_P_9_TBR1
7669_R_POA_Bulk
7669_R_POA_DLX1
7669_R_sec1_Cau_Bulk
7669_R_sec2_GP_Bulk
7669_R_sec2_GP_DLX1
7669_R_sec2_Put_Bulk
7669_R_sec2_Thal_Bulk
7669_R_sec2_Thal_COUPTF2
7669_R_sec2_Thal_DLX1
7669_R_sec3_Cau_DLX1
7669_R_sec3_Hip_COUPTF2
7669_R_sec3_Hip_DLX1
7669_R_sec3_Hip_TBR1
7669_R_sec3_Put_DLX1
7669_R_Skin_Bulk
7669_R_T_1_Bulk
7669_R_T_1_COUPTF2
7669_R_T_1_DLX1
7669_R_T_1_TBR1
7669_RA9_Bulk
7669_RV10_Bulk
JGG

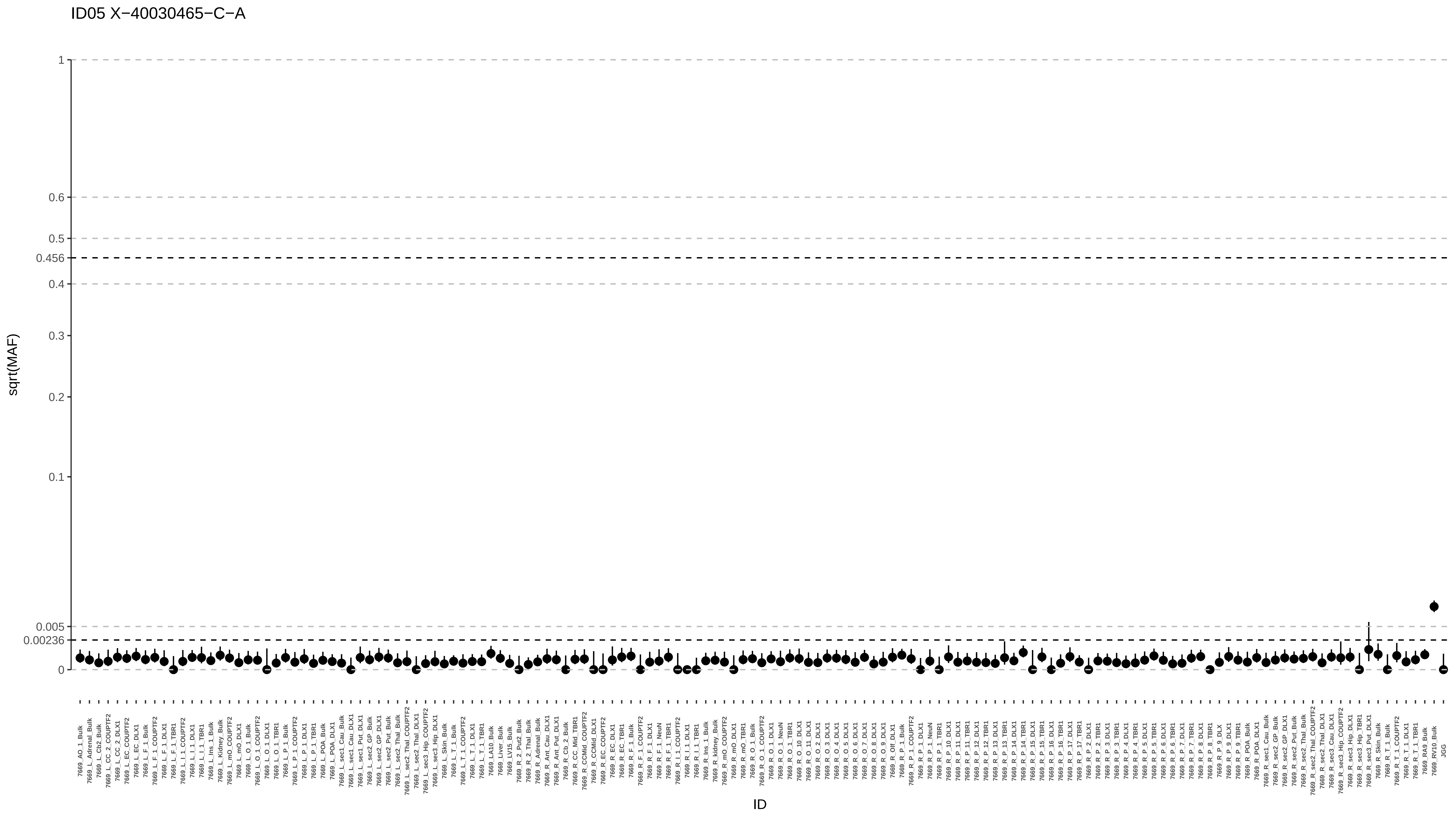
ID05 X−40030465−C−A
sqrt(MAF)
1
0.6
0.5
0.456
0.4
0.3
0.2
0.1
0.005
0.00236
0
ID
7669_AO_1_Bulk
7669_L_Adrenal_Bulk
7669_L_Cb2_Bulk
7669_L_CC_2_COUPTF2
7669_L_CC_2_DLX1
7669_L_EC_COUPTF2
7669_L_EC_DLX1
7669_L_F_1_Bulk
7669_L_F_1_COUPTF2
7669_L_F_1_DLX1
7669_L_F_1_TBR1
7669_L_I_1_COUPTF2
7669_L_I_1_DLX1
7669_L_I_1_TBR1
7669_L_Ins_1_Bulk
7669_L_Kidney_Bulk
7669_L_mO_COUPTF2
7669_L_mO_DLX1
7669_L_O_1_Bulk
7669_L_O_1_COUPTF2
7669_L_O_1_DLX1
7669_L_O_1_TBR1
7669_L_P_1_Bulk
7669_L_P_1_COUPTF2
7669_L_P_1_DLX1
7669_L_P_1_TBR1
7669_L_POA_Bulk
7669_L_POA_DLX1
7669_L_sec1_Cau_Bulk
7669_L_sec1_Cau_DLX1
7669_L_sec1_Put_DLX1
7669_L_sec2_GP_Bulk
7669_L_sec2_GP_DLX1
7669_L_sec2_Put_Bulk
7669_L_sec2_Thal_Bulk
7669_L_sec2_Thal_COUPTF2
7669_L_sec2_Thal_DLX1
7669_L_sec3_Hip_COUPTF2
7669_L_sec3_Hip_DLX1
7669_L_Skin_Bulk
7669_L_T_1_Bulk
7669_L_T_1_COUPTF2
7669_L_T_1_DLX1
7669_L_T_1_TBR1
7669_LA10_Bulk
7669_Liver_Bulk
7669_LV15_Bulk
7669_R_2_Put2_Bulk
7669_R_2_Thal_Bulk
7669_R_Adrenal_Bulk
7669_R_Ant_Cau_DLX1
7669_R_Ant_Put_DLX1
7669_R_Cb_2_Bulk
7669_R_CC_Mid_TBR1
7669_R_CCMid_COUPTF2
7669_R_CCMid_DLX1
7669_R_EC_COUPTF2
7669_R_EC_DLX1
7669_R_EC_TBR1
7669_R_F_1_Bulk
7669_R_F_1_COUPTF2
7669_R_F_1_DLX1
7669_R_F_1_NeuN
7669_R_F_1_TBR1
7669_R_I_1_COUPTF2
7669_R_I_1_DLX1
7669_R_I_1_TBR1
7669_R_Ins_1_Bulk
7669_R_kidney_Bulk
7669_R_mO_COUPTF2
7669_R_mO_DLX1
7669_R_mO_TBR1
7669_R_O_1_Bulk
7669_R_O_1_COUPTF2
7669_R_O_1_DLX1
7669_R_O_1_NeuN
7669_R_O_1_TBR1
7669_R_O_10_DLX1
7669_R_O_11_DLX1
7669_R_O_2_DLX1
7669_R_O_3_DLX1
7669_R_O_4_DLX1
7669_R_O_5_DLX1
7669_R_O_6_DLX1
7669_R_O_7_DLX1
7669_R_O_8_DLX1
7669_R_O_9_DLX1
7669_R_Olf_DLX1
7669_R_P_1_Bulk
7669_R_P_1_COUPTF2
7669_R_P_1_DLX1
7669_R_P_1_NeuN
7669_R_P_1_TBR1
7669_R_P_10_DLX1
7669_R_P_11_DLX1
7669_R_P_11_TBR1
7669_R_P_12_DLX1
7669_R_P_12_TBR1
7669_R_P_13_DLX1
7669_R_P_13_TBR1
7669_R_P_14_DLX1
7669_R_P_14_TBR1
7669_R_P_15_DLX1
7669_R_P_15_TBR1
7669_R_P_16_DLX1
7669_R_P_16_TBR1
7669_R_P_17_DLX1
7669_R_P_17_TBR1
7669_R_P_2_DLX1
7669_R_P_2_TBR1
7669_R_P_3_DLX1
7669_R_P_3_TBR1
7669_R_P_4_DLX1
7669_R_P_4_TBR1
7669_R_P_5_DLX1
7669_R_P_5_TBR1
7669_R_P_6_DLX1
7669_R_P_6_TBR1
7669_R_P_7_DLX1
7669_R_P_7_TBR1
7669_R_P_8_DLX1
7669_R_P_8_TBR1
7669_R_P_9_DLX
7669_R_P_9_DLX1
7669_R_P_9_TBR1
7669_R_POA_Bulk
7669_R_POA_DLX1
7669_R_sec1_Cau_Bulk
7669_R_sec2_GP_Bulk
7669_R_sec2_GP_DLX1
7669_R_sec2_Put_Bulk
7669_R_sec2_Thal_Bulk
7669_R_sec2_Thal_COUPTF2
7669_R_sec2_Thal_DLX1
7669_R_sec3_Cau_DLX1
7669_R_sec3_Hip_COUPTF2
7669_R_sec3_Hip_DLX1
7669_R_sec3_Hip_TBR1
7669_R_sec3_Put_DLX1
7669_R_Skin_Bulk
7669_R_T_1_Bulk
7669_R_T_1_COUPTF2
7669_R_T_1_DLX1
7669_R_T_1_TBR1
7669_RA9_Bulk
7669_RV10_Bulk
JGG

ID05 X−40790933−G−A

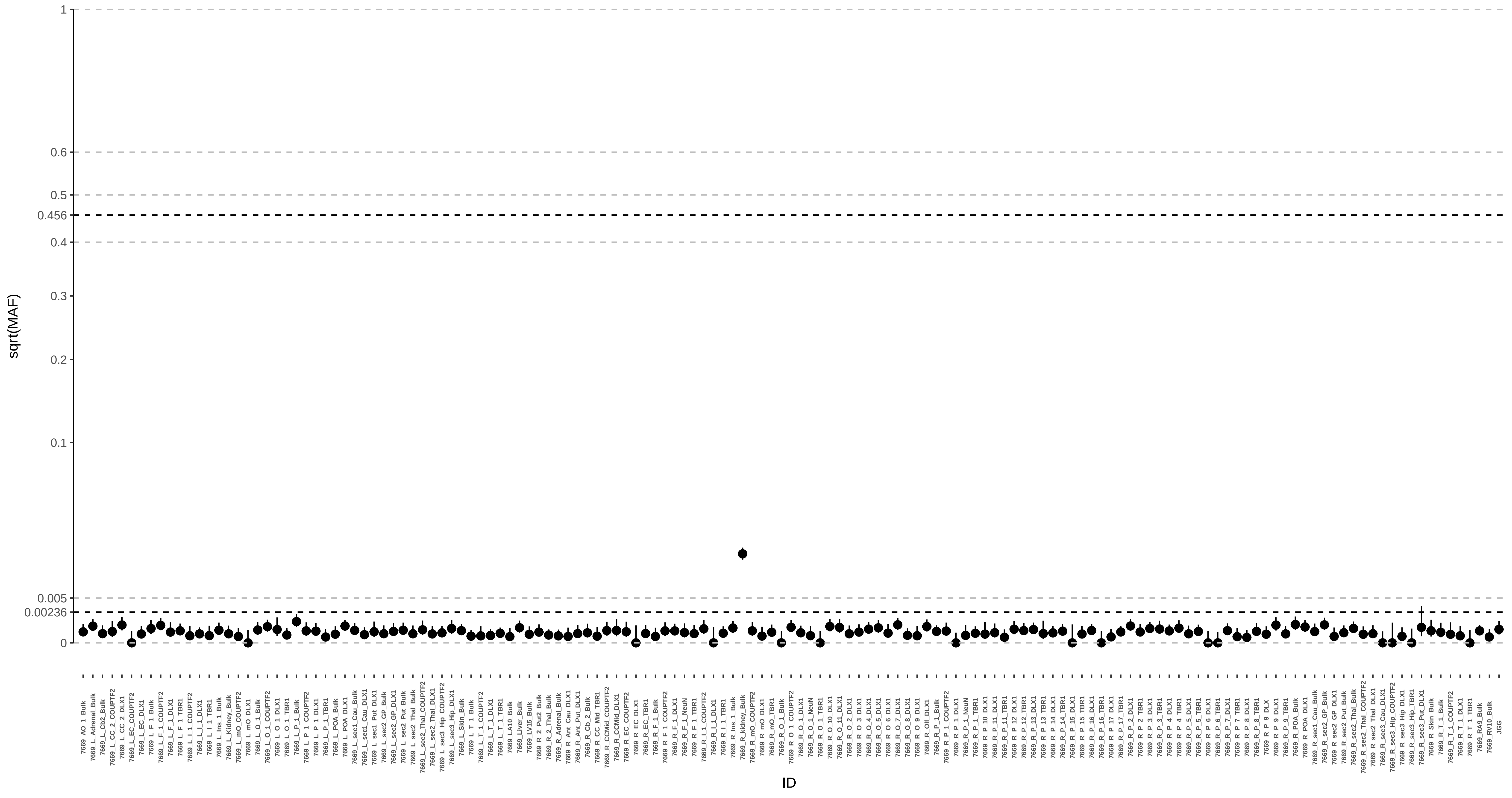

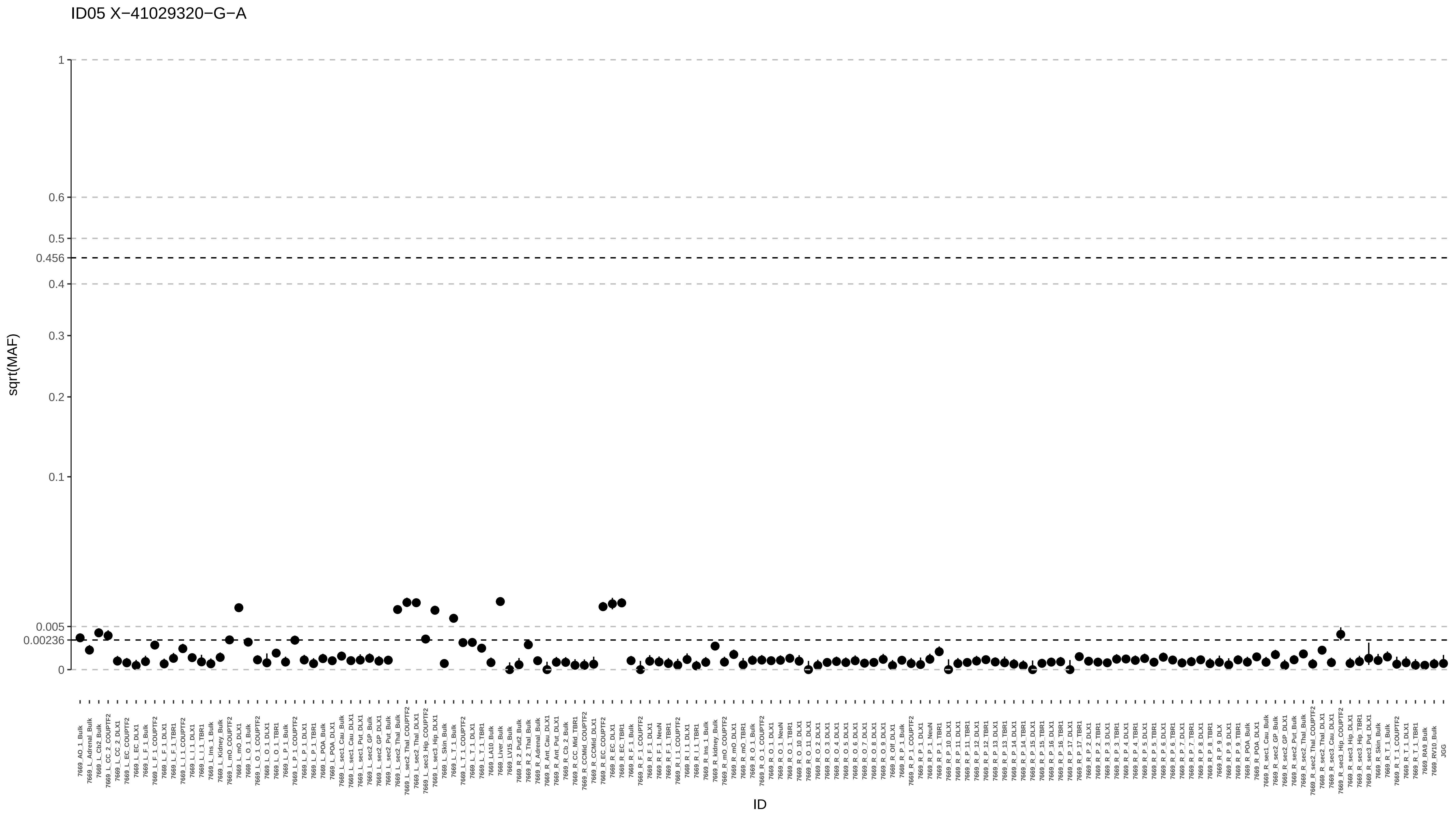

ID05 X−41029320−G−A
sqrt(MAF)
1
0.6
0.5
0.456
0.4
0.3
0.2
0.1
0.005
0.00236
0
ID
7669_AO_1_Bulk
7669_L_Adrenal_Bulk
7669_L_Cb2_Bulk
7669_L_CC_2_COUPTF2
7669_L_CC_2_DLX1
7669_L_EC_COUPTF2
7669_L_EC_DLX1
7669_L_F_1_Bulk
7669_L_F_1_COUPTF2
7669_L_F_1_DLX1
7669_L_F_1_TBR1
7669_L_I_1_COUPTF2
7669_L_I_1_DLX1
7669_L_I_1_TBR1
7669_L_Ins_1_Bulk
7669_L_Kidney_Bulk
7669_L_mO_COUPTF2
7669_L_mO_DLX1
7669_L_O_1_Bulk
7669_L_O_1_COUPTF2
7669_L_O_1_DLX1
7669_L_O_1_TBR1
7669_L_P_1_Bulk
7669_L_P_1_COUPTF2
7669_L_P_1_DLX1
7669_L_P_1_TBR1
7669_L_POA_Bulk
7669_L_POA_DLX1
7669_L_sec1_Cau_Bulk
7669_L_sec1_Cau_DLX1
7669_L_sec1_Put_DLX1
7669_L_sec2_GP_Bulk
7669_L_sec2_GP_DLX1
7669_L_sec2_Put_Bulk
7669_L_sec2_Thal_Bulk
7669_L_sec2_Thal_COUPTF2
7669_L_sec2_Thal_DLX1
7669_L_sec3_Hip_COUPTF2
7669_L_sec3_Hip_DLX1
7669_L_Skin_Bulk
7669_L_T_1_Bulk
7669_L_T_1_COUPTF2
7669_L_T_1_DLX1
7669_L_T_1_TBR1
7669_LA10_Bulk
7669_Liver_Bulk
7669_LV15_Bulk
7669_R_2_Put2_Bulk
7669_R_2_Thal_Bulk
7669_R_Adrenal_Bulk
7669_R_Ant_Cau_DLX1
7669_R_Ant_Put_DLX1
7669_R_Cb_2_Bulk
7669_R_CC_Mid_TBR1
7669_R_CCMid_COUPTF2
7669_R_CCMid_DLX1
7669_R_EC_COUPTF2
7669_R_EC_DLX1
7669_R_EC_TBR1
7669_R_F_1_Bulk
7669_R_F_1_COUPTF2
7669_R_F_1_DLX1
7669_R_F_1_NeuN
7669_R_F_1_TBR1
7669_R_I_1_COUPTF2
7669_R_I_1_DLX1
7669_R_I_1_TBR1
7669_R_Ins_1_Bulk
7669_R_kidney_Bulk
7669_R_mO_COUPTF2
7669_R_mO_DLX1
7669_R_mO_TBR1
7669_R_O_1_Bulk
7669_R_O_1_COUPTF2
7669_R_O_1_DLX1
7669_R_O_1_NeuN
7669_R_O_1_TBR1
7669_R_O_10_DLX1
7669_R_O_11_DLX1
7669_R_O_2_DLX1
7669_R_O_3_DLX1
7669_R_O_4_DLX1
7669_R_O_5_DLX1
7669_R_O_6_DLX1
7669_R_O_7_DLX1
7669_R_O_8_DLX1
7669_R_O_9_DLX1
7669_R_Olf_DLX1
7669_R_P_1_Bulk
7669_R_P_1_COUPTF2
7669_R_P_1_DLX1
7669_R_P_1_NeuN
7669_R_P_1_TBR1
7669_R_P_10_DLX1
7669_R_P_11_DLX1
7669_R_P_11_TBR1
7669_R_P_12_DLX1
7669_R_P_12_TBR1
7669_R_P_13_DLX1
7669_R_P_13_TBR1
7669_R_P_14_DLX1
7669_R_P_14_TBR1
7669_R_P_15_DLX1
7669_R_P_15_TBR1
7669_R_P_16_DLX1
7669_R_P_16_TBR1
7669_R_P_17_DLX1
7669_R_P_17_TBR1
7669_R_P_2_DLX1
7669_R_P_2_TBR1
7669_R_P_3_DLX1
7669_R_P_3_TBR1
7669_R_P_4_DLX1
7669_R_P_4_TBR1
7669_R_P_5_DLX1
7669_R_P_5_TBR1
7669_R_P_6_DLX1
7669_R_P_6_TBR1
7669_R_P_7_DLX1
7669_R_P_7_TBR1
7669_R_P_8_DLX1
7669_R_P_8_TBR1
7669_R_P_9_DLX
7669_R_P_9_DLX1
7669_R_P_9_TBR1
7669_R_POA_Bulk
7669_R_POA_DLX1
7669_R_sec1_Cau_Bulk
7669_R_sec2_GP_Bulk
7669_R_sec2_GP_DLX1
7669_R_sec2_Put_Bulk
7669_R_sec2_Thal_Bulk
7669_R_sec2_Thal_COUPTF2
7669_R_sec2_Thal_DLX1
7669_R_sec3_Cau_DLX1
7669_R_sec3_Hip_COUPTF2
7669_R_sec3_Hip_DLX1
7669_R_sec3_Hip_TBR1
7669_R_sec3_Put_DLX1
7669_R_Skin_Bulk
7669_R_T_1_Bulk
7669_R_T_1_COUPTF2
7669_R_T_1_DLX1
7669_R_T_1_TBR1
7669_RA9_Bulk
7669_RV10_Bulk
JGG

ID05 X−4557817−T−C

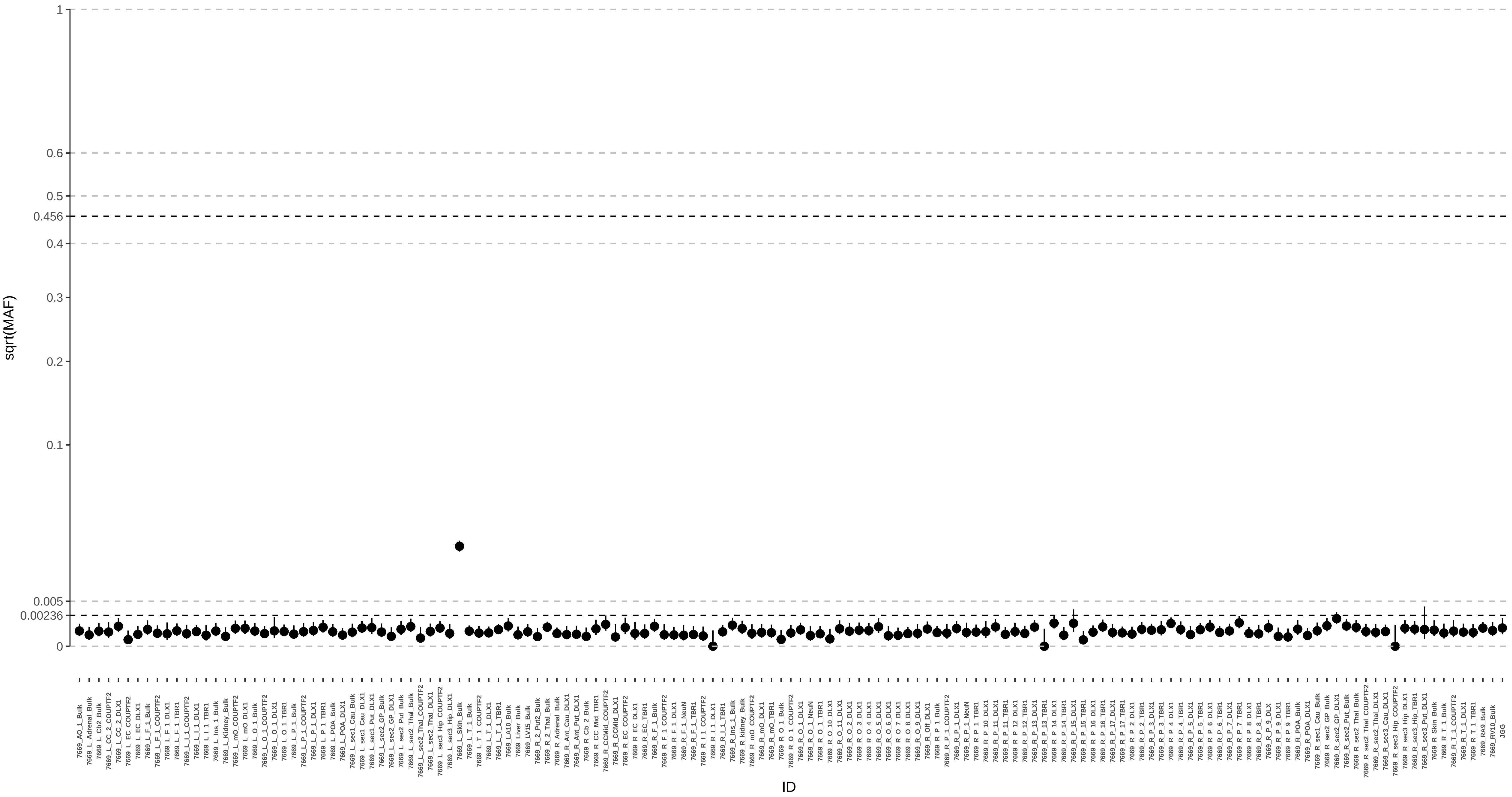

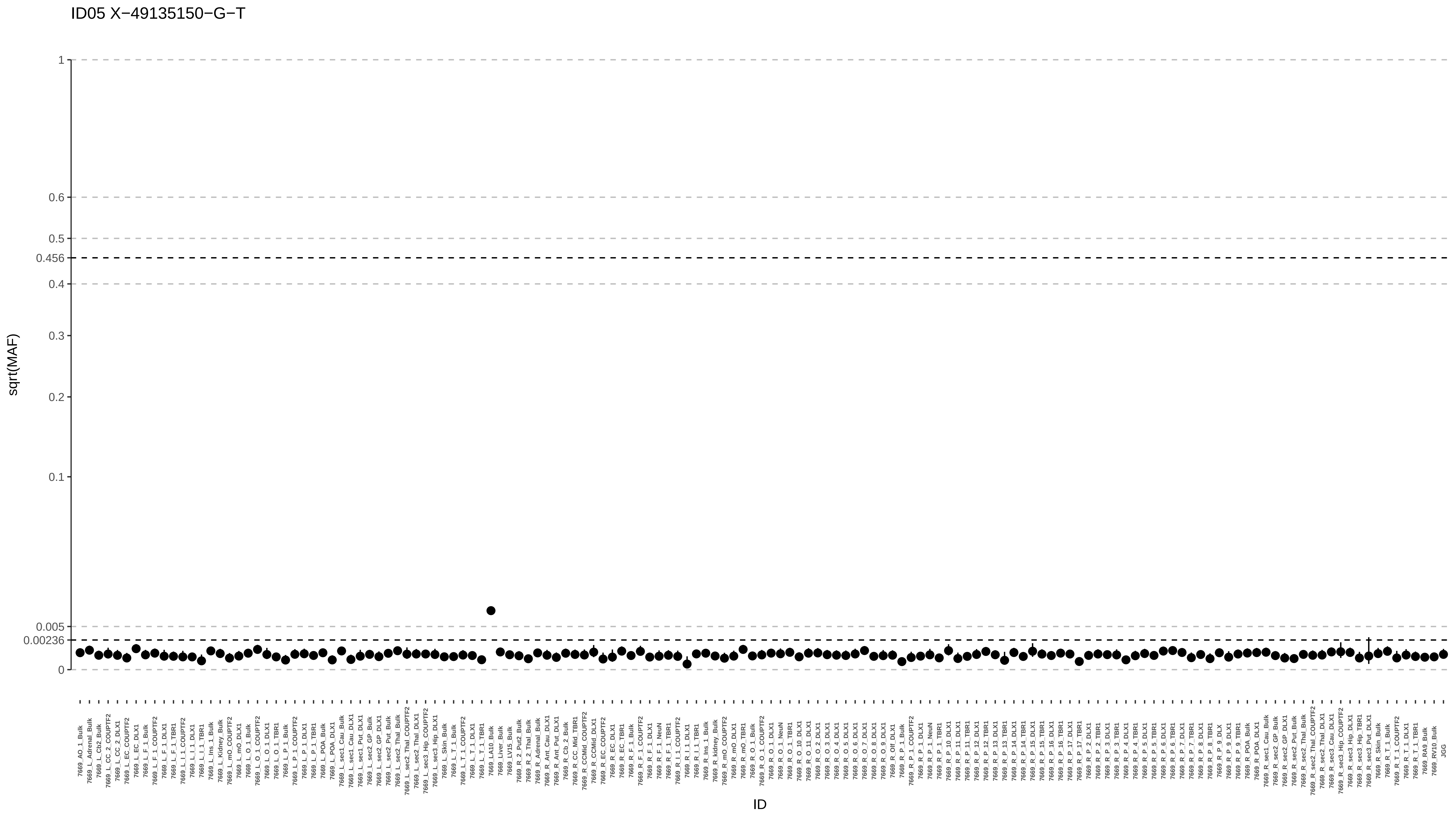
ID05 X−49135150−G−T
sqrt(MAF)
1
0.6
0.5
0.456
0.4
0.3
0.2
0.1
0.005
0.00236
0
ID
7669_AO_1_Bulk
7669_L_Adrenal_Bulk
7669_L_Cb2_Bulk
7669_L_CC_2_COUPTF2
7669_L_CC_2_DLX1
7669_L_EC_COUPTF2
7669_L_EC_DLX1
7669_L_F_1_Bulk
7669_L_F_1_COUPTF2
7669_L_F_1_DLX1
7669_L_F_1_TBR1
7669_L_I_1_COUPTF2
7669_L_I_1_DLX1
7669_L_I_1_TBR1
7669_L_Ins_1_Bulk
7669_L_Kidney_Bulk
7669_L_mO_COUPTF2
7669_L_mO_DLX1
7669_L_O_1_Bulk
7669_L_O_1_COUPTF2
7669_L_O_1_DLX1
7669_L_O_1_TBR1
7669_L_P_1_Bulk
7669_L_P_1_COUPTF2
7669_L_P_1_DLX1
7669_L_P_1_TBR1
7669_L_POA_Bulk
7669_L_POA_DLX1
7669_L_sec1_Cau_Bulk
7669_L_sec1_Cau_DLX1
7669_L_sec1_Put_DLX1
7669_L_sec2_GP_DLX1
7669_L_sec2_GP_Bulk
7669_L_sec2_Put_Bulk
7669_L_sec2_Thal_Bulk
7669_L_sec2_Thal_COUPTF2
7669_L_sec2_Thal_DLX1
7669_L_sec3_Hip_COUPTF2
7669_L_sec3_Hip_DLX1
7669_L_Skin_Bulk
7669_L_T_1_Bulk
7669_L_T_1_COUPTF2
7669_L_T_1_DLX1
7669_L_T_1_TBR1
7669_LA10_Bulk
7669_Liver_Bulk
7669_LV15_Bulk
7669_R_2_Put2_Bulk
7669_R_2_Thal_Bulk
7669_R_Adrenal_Bulk
7669_R_Ant_Cau_DLX1
7669_R_Ant_Put_DLX1
7669_R_Cb_2_Bulk
7669_R_CC_Mid_TBR1
7669_R_CCMid_COUPTF2
7669_R_CCMid_DLX1
7669_R_EC_COUPTF2
7669_R_EC_DLX1
7669_R_EC_TBR1
7669_R_F_1_Bulk
7669_R_F_1_COUPTF2
7669_R_F_1_DLX1
7669_R_F_1_NeuN
7669_R_F_1_TBR1
7669_R_I_1_COUPTF2
7669_R_I_1_DLX1
7669_R_I_1_TBR1
7669_R_Ins_1_Bulk
7669_R_kidney_Bulk
7669_R_mO_COUPTF2
7669_R_mO_DLX1
7669_R_mO_TBR1
7669_R_O_1_Bulk
7669_R_O_1_COUPTF2
7669_R_O_1_DLX1
7669_R_O_1_NeuN
7669_R_O_1_TBR1
7669_R_O_10_DLX1
7669_R_O_11_DLX1
7669_R_O_2_DLX1
7669_R_O_3_DLX1
7669_R_O_4_DLX1
7669_R_O_5_DLX1
7669_R_O_6_DLX1
7669_R_O_7_DLX1
7669_R_O_8_DLX1
7669_R_O_9_DLX1
7669_R_Olf_DLX1
7669_R_P_1_Bulk
7669_R_P_1_COUPTF2
7669_R_P_1_DLX1
7669_R_P_1_NeuN
7669_R_P_1_TBR1
7669_R_P_10_DLX1
7669_R_P_11_DLX1
7669_R_P_11_TBR1
7669_R_P_12_DLX1
7669_R_P_12_TBR1
7669_R_P_13_DLX1
7669_R_P_13_TBR1
7669_R_P_14_DLX1
7669_R_P_14_TBR1
7669_R_P_15_DLX1
7669_R_P_15_TBR1
7669_R_P_16_DLX1
7669_R_P_16_TBR1
7669_R_P_17_DLX1
7669_R_P_17_TBR1
7669_R_P_2_DLX1
7669_R_P_2_TBR1
7669_R_P_3_DLX1
7669_R_P_3_TBR1
7669_R_P_4_DLX1
7669_R_P_4_TBR1
7669_R_P_5_DLX1
7669_R_P_5_TBR1
7669_R_P_6_DLX1
7669_R_P_6_TBR1
7669_R_P_7_DLX1
7669_R_P_7_TBR1
7669_R_P_8_DLX1
7669_R_P_8_TBR1
7669_R_P_9_DLX
7669_R_P_9_DLX1
7669_R_P_9_TBR1
7669_R_POA_Bulk
7669_R_POA_DLX1
7669_R_sec1_Cau_Bulk
7669_R_sec2_GP_Bulk
7669_R_sec2_GP_DLX1
7669_R_sec2_Put_Bulk
7669_R_sec2_Thal_Bulk
7669_R_sec2_Thal_COUPTF2
7669_R_sec2_Thal_DLX1
7669_R_sec3_Cau_DLX1
7669_R_sec3_Hip_COUPTF2
7669_R_sec3_Hip_DLX1
7669_R_sec3_Hip_TBR1
7669_R_sec3_Put_DLX1
7669_R_Skin_Bulk
7669_R_T_1_Bulk
7669_R_T_1_COUPTF2
7669_R_T_1_DLX1
7669_R_T_1_TBR1
7669_RA9_Bulk
7669_RV10_Bulk
JGG

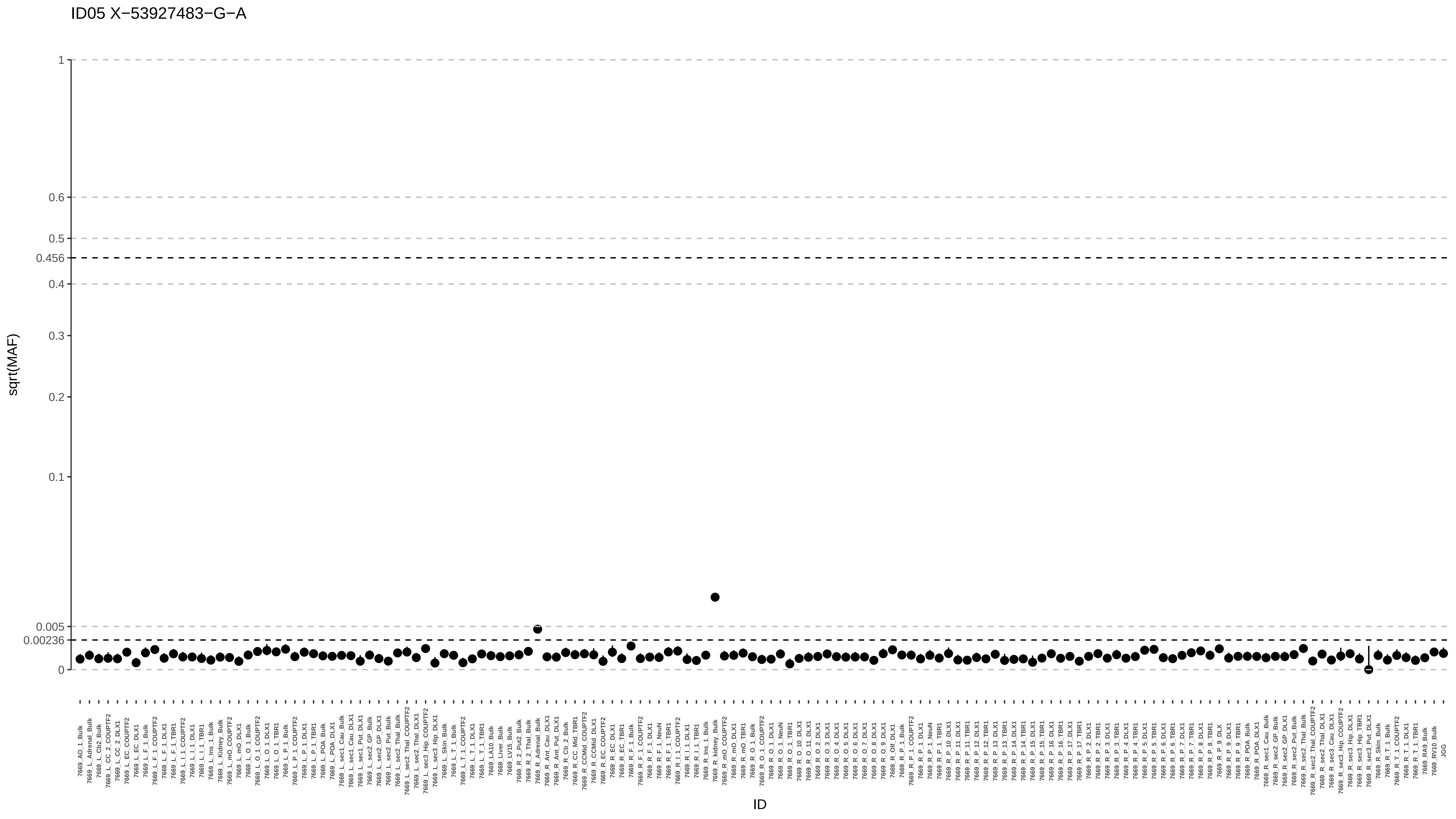
ID05 X−53927483−G−A
sqrt(MAF)
1
0.6
0.5
0.456
0.4
0.3
0.2
0.1
0.005
0.00236
0
ID

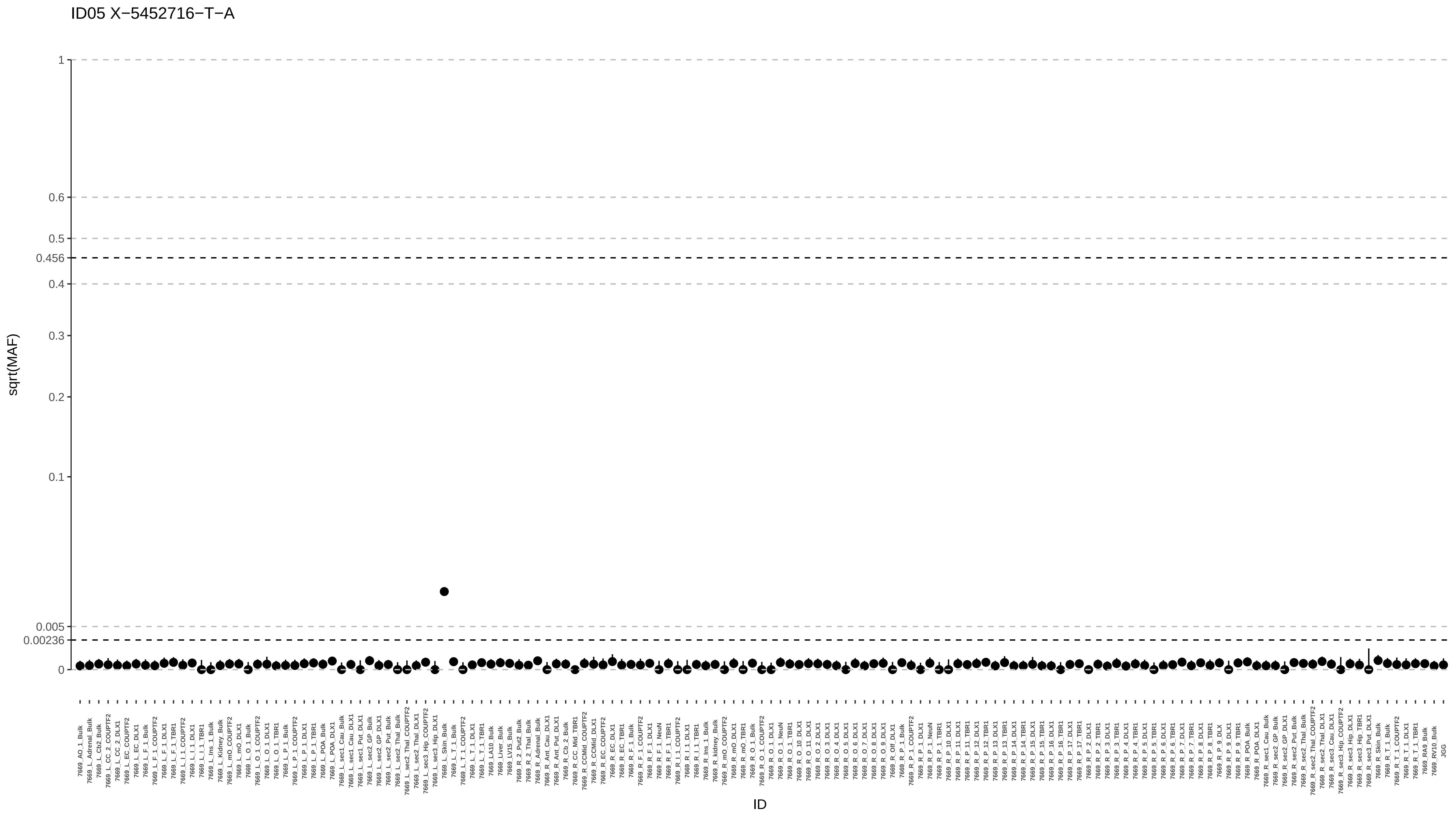
ID05 X−5452716−T−A
sqrt(MAF)
1
0.6
0.5
0.456
0.4
0.3
0.2
0.1
0.005
0.00236
0
ID
7669_AO_1_Bulk
7669_L_Adrenal_Bulk
7669_L_Cb2_Bulk
7669_L_CC_2_COUPTF2
7669_L_CC_2_DLX1
7669_L_EC_COUPTF2
7669_L_EC_DLX1
7669_L_F_1_Bulk
7669_L_F_1_COUPTF2
7669_L_F_1_DLX1
7669_L_F_1_TBR1
7669_L_I_1_COUPTF2
7669_L_I_1_DLX1
7669_L_I_1_TBR1
7669_L_Ins_1_Bulk
7669_L_Kidney_Bulk
7669_L_mO_COUPTF2
7669_L_mO_DLX1
7669_L_O_1_Bulk
7669_L_O_1_COUPTF2
7669_L_O_1_DLX1
7669_L_O_1_TBR1
7669_L_P_1_Bulk
7669_L_P_1_COUPTF2
7669_L_P_1_DLX1
7669_L_P_1_TBR1
7669_L_POA_Bulk
7669_L_POA_DLX1
7669_L_sec1_Cau_Bulk
7669_L_sec1_Cau_DLX1
7669_L_sec1_Put_DLX1
7669_L_sec2_GP_Bulk
7669_L_sec2_GP_DLX1
7669_L_sec2_Put_Bulk
7669_L_sec2_Thal_Bulk
7669_L_sec2_Thal_COUPTF2
7669_L_sec2_Thal_DLX1
7669_L_sec3_Hip_COUPTF2
7669_L_sec3_Hip_DLX1
7669_L_Skin_Bulk
7669_L_T_1_Bulk
7669_L_T_1_COUPTF2
7669_L_T_1_DLX1
7669_L_T_1_TBR1
7669_LA10_Bulk
7669_Liver_Bulk
7669_LV15_Bulk
7669_R_2_Put2_Bulk
7669_R_2_Thal_Bulk
7669_R_Adrenal_Bulk
7669_R_Ant_Cau_DLX1
7669_R_Ant_Put_DLX1
7669_R_Cb_2_Bulk
7669_R_CC_Mid_TBR1
7669_R_CCMid_COUPTF2
7669_R_CCMid_DLX1
7669_R_EC_COUPTF2
7669_R_EC_DLX1
7669_R_EC_TBR1
7669_R_F_1_Bulk
7669_R_F_1_COUPTF2
7669_R_F_1_DLX1
7669_R_F_1_NeuN
7669_R_F_1_TBR1
7669_R_I_1_COUPTF2
7669_R_I_1_DLX1
7669_R_I_1_TBR1
7669_R_Ins_1_Bulk
7669_R_kidney_Bulk
7669_R_mO_COUPTF2
7669_R_mO_DLX1
7669_R_mO_TBR1
7669_R_O_1_Bulk
7669_R_O_1_COUPTF2
7669_R_O_1_DLX1
7669_R_O_1_NeuN
7669_R_O_1_TBR1
7669_R_O_10_DLX1
7669_R_O_11_DLX1
7669_R_O_2_DLX1
7669_R_O_3_DLX1
7669_R_O_4_DLX1
7669_R_O_5_DLX1
7669_R_O_6_DLX1
7669_R_O_7_DLX1
7669_R_O_8_DLX1
7669_R_O_9_DLX1
7669_R_Olf_DLX1
7669_R_P_1_Bulk
7669_R_P_1_COUPTF2
7669_R_P_1_DLX1
7669_R_P_1_NeuN
7669_R_P_1_TBR1
7669_R_P_10_DLX1
7669_R_P_11_DLX1
7669_R_P_11_TBR1
7669_R_P_12_DLX1
7669_R_P_12_TBR1
7669_R_P_13_DLX1
7669_R_P_13_TBR1
7669_R_P_14_DLX1
7669_R_P_14_TBR1
7669_R_P_15_DLX1
7669_R_P_15_TBR1
7669_R_P_16_DLX1
7669_R_P_16_TBR1
7669_R_P_17_DLX1
7669_R_P_17_TBR1
7669_R_P_2_DLX1
7669_R_P_2_TBR1
7669_R_P_3_DLX1
7669_R_P_3_TBR1
7669_R_P_4_DLX1
7669_R_P_4_TBR1
7669_R_P_5_DLX1
7669_R_P_5_TBR1
7669_R_P_6_DLX1
7669_R_P_6_TBR1
7669_R_P_7_DLX1
7669_R_P_7_TBR1
7669_R_P_8_DLX1
7669_R_P_8_TBR1
7669_R_P_9_DLX
7669_R_P_9_DLX1
7669_R_P_9_TBR1
7669_R_POA_Bulk
7669_R_POA_DLX1
7669_R_sec1_Cau_Bulk
7669_R_sec2_GP_Bulk
7669_R_sec2_GP_DLX1
7669_R_sec2_Put_Bulk
7669_R_sec2_Thal_Bulk
7669_R_sec2_Thal_COUPTF2
7669_R_sec2_Thal_DLX1
7669_R_sec3_Cau_DLX1
7669_R_sec3_Hip_COUPTF2
7669_R_sec3_Hip_DLX1
7669_R_sec3_Hip_TBR1
7669_R_sec3_Put_DLX1
7669_R_Skin_Bulk
7669_R_T_1_Bulk
7669_R_T_1_COUPTF2
7669_R_T_1_DLX1
7669_R_T_1_TBR1
7669_RA9_Bulk
7669_RV10_Bulk
JGG

ID05 X−6657783−T−A

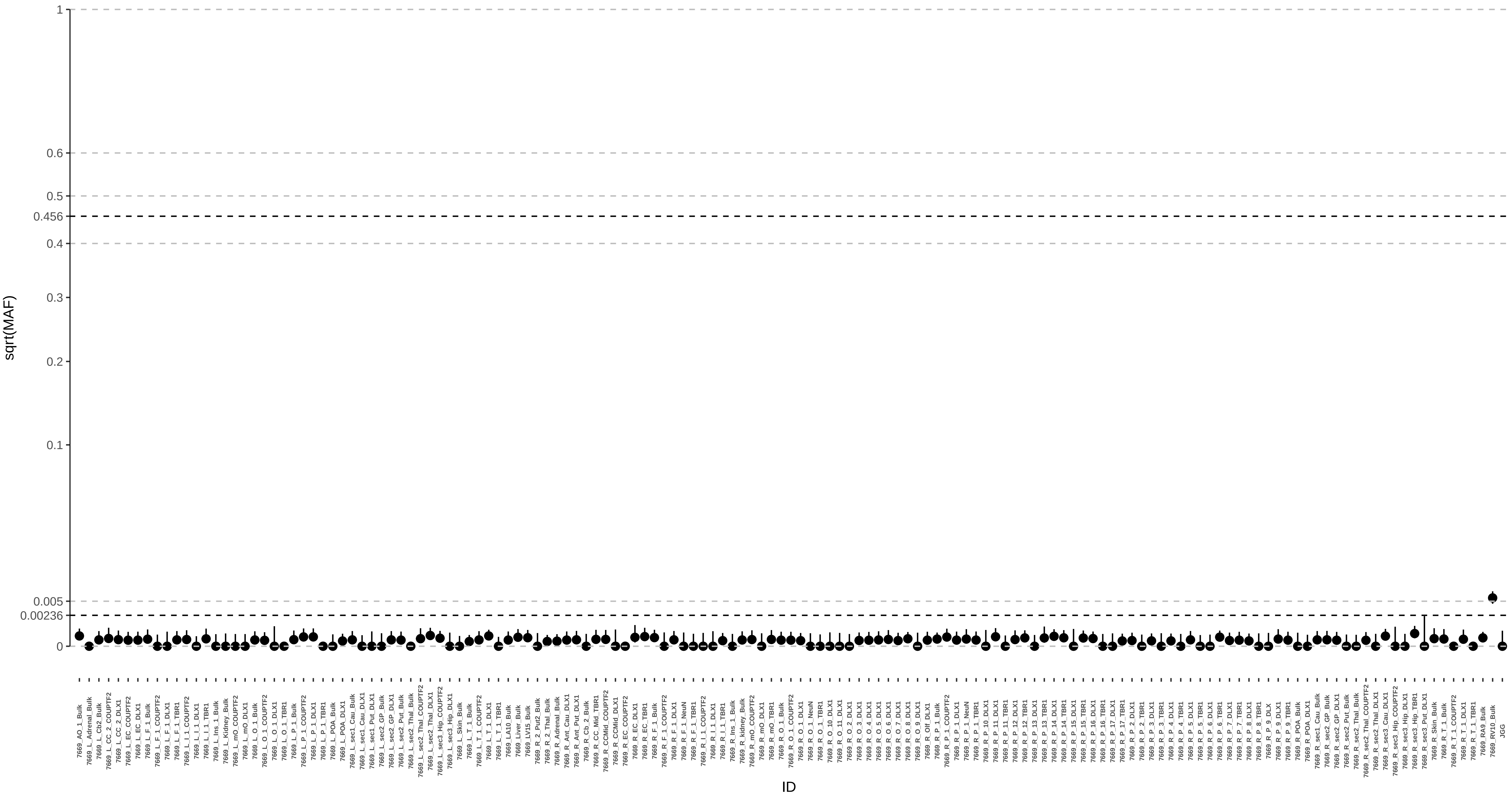

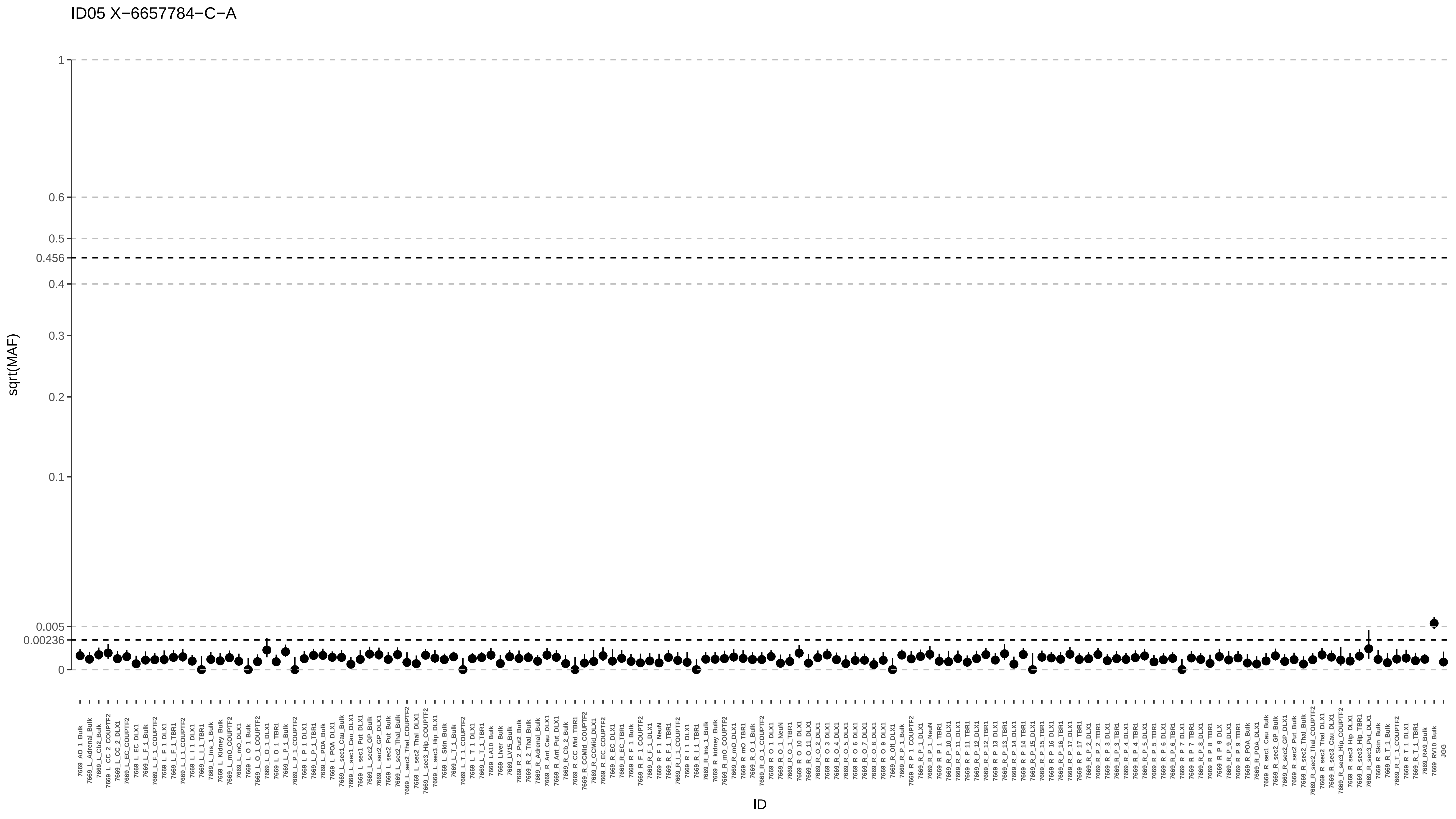

ID05 X−6657784−C−A
sqrt(MAF)
1
0.6
0.5
0.456
0.4
0.3
0.2
0.1
0.005
0.00236
0
ID
7669_AO_1_Bulk
7669_L_Adrenal_Bulk
7669_L_Cb2_Bulk
7669_L_CC_2_COUPTF2
7669_L_CC_2_DLX1
7669_L_EC_COUPTF2
7669_L_EC_DLX1
7669_L_F_1_Bulk
7669_L_F_1_COUPTF2
7669_L_F_1_DLX1
7669_L_F_1_TBR1
7669_L_I_1_COUPTF2
7669_L_I_1_DLX1
7669_L_I_1_TBR1
7669_L_Ins_1_Bulk
7669_L_Kidney_Bulk
7669_L_mO_COUPTF2
7669_L_mO_DLX1
7669_L_O_1_Bulk
7669_L_O_1_COUPTF2
7669_L_O_1_DLX1
7669_L_O_1_TBR1
7669_L_P_1_Bulk
7669_L_P_1_COUPTF2
7669_L_P_1_DLX1
7669_L_P_1_TBR1
7669_L_POA_Bulk
7669_L_POA_DLX1
7669_L_sec1_Cau_Bulk
7669_L_sec1_Cau_DLX1
7669_L_sec1_Put_DLX1
7669_L_sec2_GP_Bulk
7669_L_sec2_GP_DLX1
7669_L_sec2_Put_Bulk
7669_L_sec2_Thal_Bulk
7669_L_sec2_Thal_COUPTF2
7669_L_sec2_Thal_DLX1
7669_L_sec3_Hip_COUPTF2
7669_L_sec3_Hip_DLX1
7669_L_Skin_Bulk
7669_L_T_1_Bulk
7669_L_T_1_COUPTF2
7669_L_T_1_DLX1
7669_L_T_1_TBR1
7669_LA10_Bulk
7669_Liver_Bulk
7669_LV15_Bulk
7669_R_2_Put2_Bulk
7669_R_2_Thal_Bulk
7669_R_Adrenal_Bulk
7669_R_Ant_Cau_DLX1
7669_R_Ant_Put_DLX1
7669_R_Cb_2_Bulk
7669_R_CC_Mid_TBR1
7669_R_CCMid_COUPTF2
7669_R_CCMid_DLX1
7669_R_EC_COUPTF2
7669_R_EC_DLX1
7669_R_EC_TBR1
7669_R_F_1_Bulk
7669_R_F_1_COUPTF2
7669_R_F_1_DLX1
7669_R_F_1_NeuN
7669_R_F_1_TBR1
7669_R_I_1_COUPTF2
7669_R_I_1_DLX1
7669_R_I_1_TBR1
7669_R_Ins_1_Bulk
7669_R_kidney_Bulk
7669_R_mO_COUPTF2
7669_R_mO_DLX1
7669_R_mO_TBR1
7669_R_O_1_Bulk
7669_R_O_1_COUPTF2
7669_R_O_1_DLX1
7669_R_O_1_NeuN
7669_R_O_1_TBR1
7669_R_O_10_DLX1
7669_R_O_11_DLX1
7669_R_O_2_DLX1
7669_R_O_3_DLX1
7669_R_O_4_DLX1
7669_R_O_5_DLX1
7669_R_O_6_DLX1
7669_R_O_7_DLX1
7669_R_O_8_DLX1
7669_R_O_9_DLX1
7669_R_Olf_DLX1
7669_R_P_1_Bulk
7669_R_P_1_COUPTF2
7669_R_P_1_DLX1
7669_R_P_1_NeuN
7669_R_P_1_TBR1
7669_R_P_10_DLX1
7669_R_P_11_DLX1
7669_R_P_11_TBR1
7669_R_P_12_DLX1
7669_R_P_12_TBR1
7669_R_P_13_DLX1
7669_R_P_13_TBR1
7669_R_P_14_DLX1
7669_R_P_14_TBR1
7669_R_P_15_DLX1
7669_R_P_15_TBR1
7669_R_P_16_DLX1
7669_R_P_16_TBR1
7669_R_P_17_DLX1
7669_R_P_17_TBR1
7669_R_P_2_DLX1
7669_R_P_2_TBR1
7669_R_P_3_DLX1
7669_R_P_3_TBR1
7669_R_P_4_DLX1
7669_R_P_4_TBR1
7669_R_P_5_DLX1
7669_R_P_5_TBR1
7669_R_P_6_DLX1
7669_R_P_6_TBR1
7669_R_P_7_DLX1
7669_R_P_7_TBR1
7669_R_P_8_DLX1
7669_R_P_8_TBR1
7669_R_P_9_DLX
7669_R_P_9_DLX1
7669_R_P_9_TBR1
7669_R_POA_Bulk
7669_R_POA_DLX1
7669_R_sec1_Cau_Bulk
7669_R_sec2_GP_Bulk
7669_R_sec2_GP_DLX1
7669_R_sec2_Put_Bulk
7669_R_sec2_Thal_Bulk
7669_R_sec2_Thal_COUPTF2
7669_R_sec2_Thal_DLX1
7669_R_sec3_Cau_DLX1
7669_R_sec3_Hip_COUPTF2
7669_R_sec3_Hip_DLX1
7669_R_sec3_Hip_TBR1
7669_R_sec3_Put_DLX1
7669_R_Skin_Bulk
7669_R_T_1_Bulk
7669_R_T_1_COUPTF2
7669_R_T_1_DLX1
7669_R_T_1_TBR1
7669_RA9_Bulk
7669_RV10_Bulk
JGG

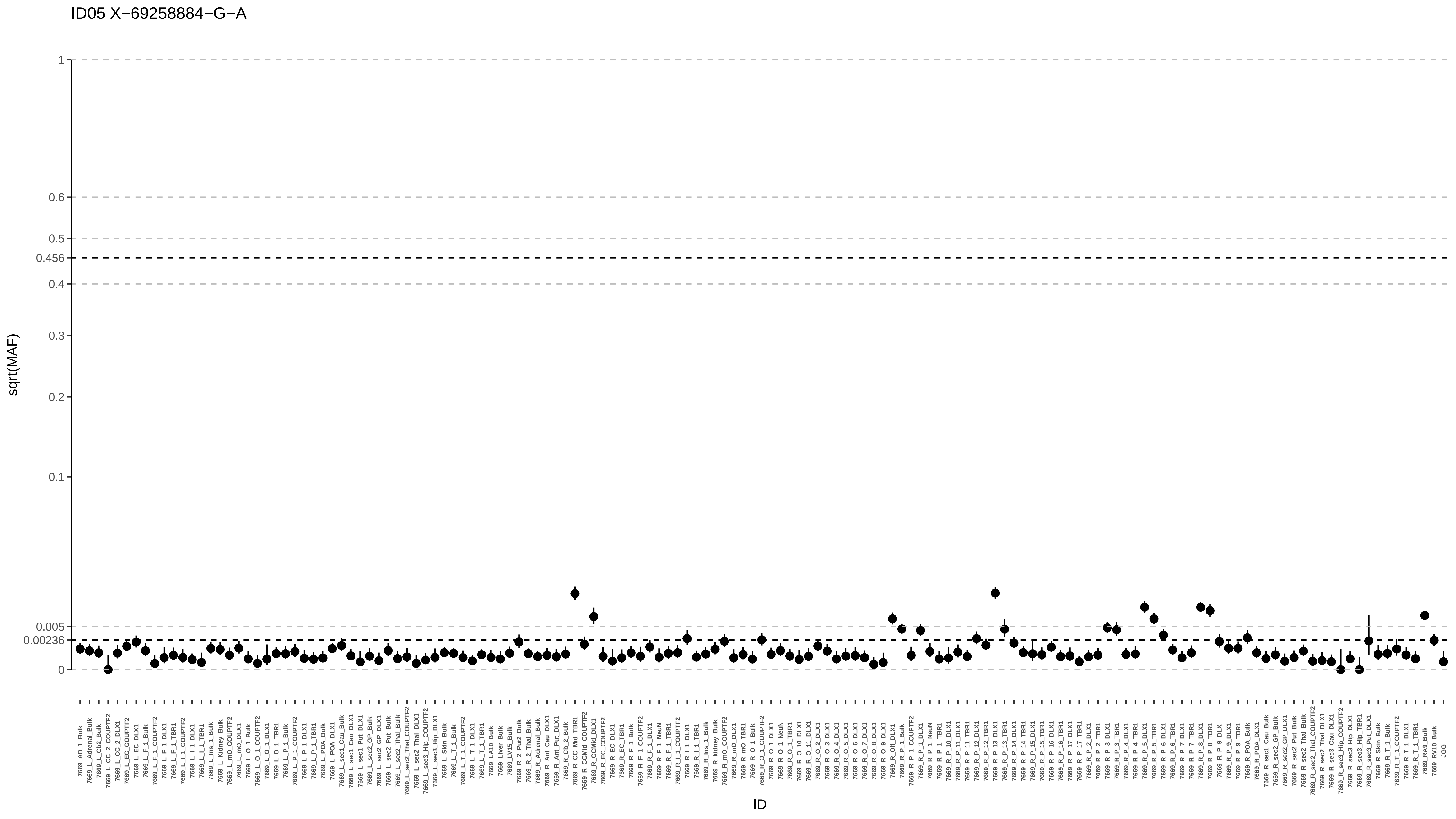
ID05 X−69258884−G−A
sqrt(MAF)
1
0.6
0.5
0.456
0.4
0.3
0.2
0.1
0.005
0.00236
0
ID
7669_AO_1_Bulk
7669_L_Adrenal_Bulk
7669_L_Cb2_Bulk
7669_L_CC_2_COUPTF2
7669_L_CC_2_DLX1
7669_L_EC_COUPTF2
7669_L_EC_DLX1
7669_L_F_1_Bulk
7669_L_F_1_COUPTF2
7669_L_F_1_DLX1
7669_L_F_1_TBR1
7669_L_I_1_COUPTF2
7669_L_I_1_DLX1
7669_L_I_1_TBR1
7669_L_Ins_1_Bulk
7669_L_Kidney_Bulk
7669_L_mO_COUPTF2
7669_L_mO_DLX1
7669_L_O_1_Bulk
7669_L_O_1_COUPTF2
7669_L_O_1_DLX1
7669_L_O_1_TBR1
7669_L_P_1_Bulk
7669_L_P_1_COUPTF2
7669_L_P_1_DLX1
7669_L_P_1_TBR1
7669_L_POA_Bulk
7669_L_POA_DLX1
7669_L_sec1_Cau_Bulk
7669_L_sec1_Cau_DLX1
7669_L_sec1_Put_DLX1
7669_L_sec2_GP_Bulk
7669_L_sec2_GP_DLX1
7669_L_sec2_Put_Bulk
7669_L_sec2_Thal_Bulk
7669_L_sec2_Thal_COUPTF2
7669_L_sec2_Thal_DLX1
7669_L_sec3_Hip_COUPTF2
7669_L_sec3_Hip_DLX1
7669_L_Skin_Bulk
7669_L_T_1_Bulk
7669_L_T_1_COUPTF2
7669_L_T_1_DLX1
7669_L_T_1_TBR1
7669_LA10_Bulk
7669_Liver_Bulk
7669_LV15_Bulk
7669_R_2_Put2_Bulk
7669_R_2_Thal_Bulk
7669_R_Adrenal_Bulk
7669_R_Ant_Cau_DLX1
7669_R_Ant_Put_DLX1
7669_R_Cb_2_Bulk
7669_R_CC_Mid_TBR1
7669_R_CCMid_COUPTF2
7669_R_CCMid_DLX1
7669_R_EC_COUPTF2
7669_R_EC_DLX1
7669_R_EC_TBR1
7669_R_F_1_Bulk
7669_R_F_1_COUPTF2
7669_R_F_1_DLX1
7669_R_F_1_NeuN
7669_R_F_1_TBR1
7669_R_I_1_COUPTF2
7669_R_I_1_DLX1
7669_R_I_1_TBR1
7669_R_Ins_1_Bulk
7669_R_kidney_Bulk
7669_R_mO_COUPTF2
7669_R_mO_DLX1
7669_R_mO_TBR1
7669_R_O_1_Bulk
7669_R_O_1_COUPTF2
7669_R_O_1_DLX1
7669_R_O_1_NeuN
7669_R_O_1_TBR1
7669_R_O_10_DLX1
7669_R_O_11_DLX1
7669_R_O_2_DLX1
7669_R_O_3_DLX1
7669_R_O_4_DLX1
7669_R_O_5_DLX1
7669_R_O_6_DLX1
7669_R_O_7_DLX1
7669_R_O_8_DLX1
7669_R_O_9_DLX1
7669_R_Olf_DLX1
7669_R_P_1_Bulk
7669_R_P_1_COUPTF2
7669_R_P_1_DLX1
7669_R_P_1_NeuN
7669_R_P_1_TBR1
7669_R_P_10_DLX1
7669_R_P_11_DLX1
7669_R_P_11_TBR1
7669_R_P_12_DLX1
7669_R_P_12_TBR1
7669_R_P_13_DLX1
7669_R_P_13_TBR1
7669_R_P_14_DLX1
7669_R_P_14_TBR1
7669_R_P_15_DLX1
7669_R_P_15_TBR1
7669_R_P_16_DLX1
7669_R_P_16_TBR1
7669_R_P_17_DLX1
7669_R_P_17_TBR1
7669_R_P_2_DLX1
7669_R_P_2_TBR1
7669_R_P_3_DLX1
7669_R_P_3_TBR1
7669_R_P_4_DLX1
7669_R_P_4_TBR1
7669_R_P_5_DLX1
7669_R_P_5_TBR1
7669_R_P_6_DLX1
7669_R_P_6_TBR1
7669_R_P_7_DLX1
7669_R_P_7_TBR1
7669_R_P_8_DLX1
7669_R_P_8_TBR1
7669_R_P_9_DLX
7669_R_P_9_DLX1
7669_R_P_9_TBR1
7669_R_POA_Bulk
7669_R_POA_DLX1
7669_R_sec1_Cau_Bulk
7669_R_sec2_GP_Bulk
7669_R_sec2_GP_DLX1
7669_R_sec2_Put_Bulk
7669_R_sec2_Thal_Bulk
7669_R_sec2_Thal_COUPTF2
7669_R_sec2_Thal_DLX1
7669_R_sec3_Cau_DLX1
7669_R_sec3_Hip_COUPTF2
7669_R_sec3_Hip_DLX1
7669_R_sec3_Hip_TBR1
7669_R_sec3_Put_DLX1
7669_R_Skin_Bulk
7669_R_T_1_Bulk
7669_R_T_1_COUPTF2
7669_R_T_1_DLX1
7669_R_T_1_TBR1
7669_RA9_Bulk
7669_RV10_Bulk
JGG

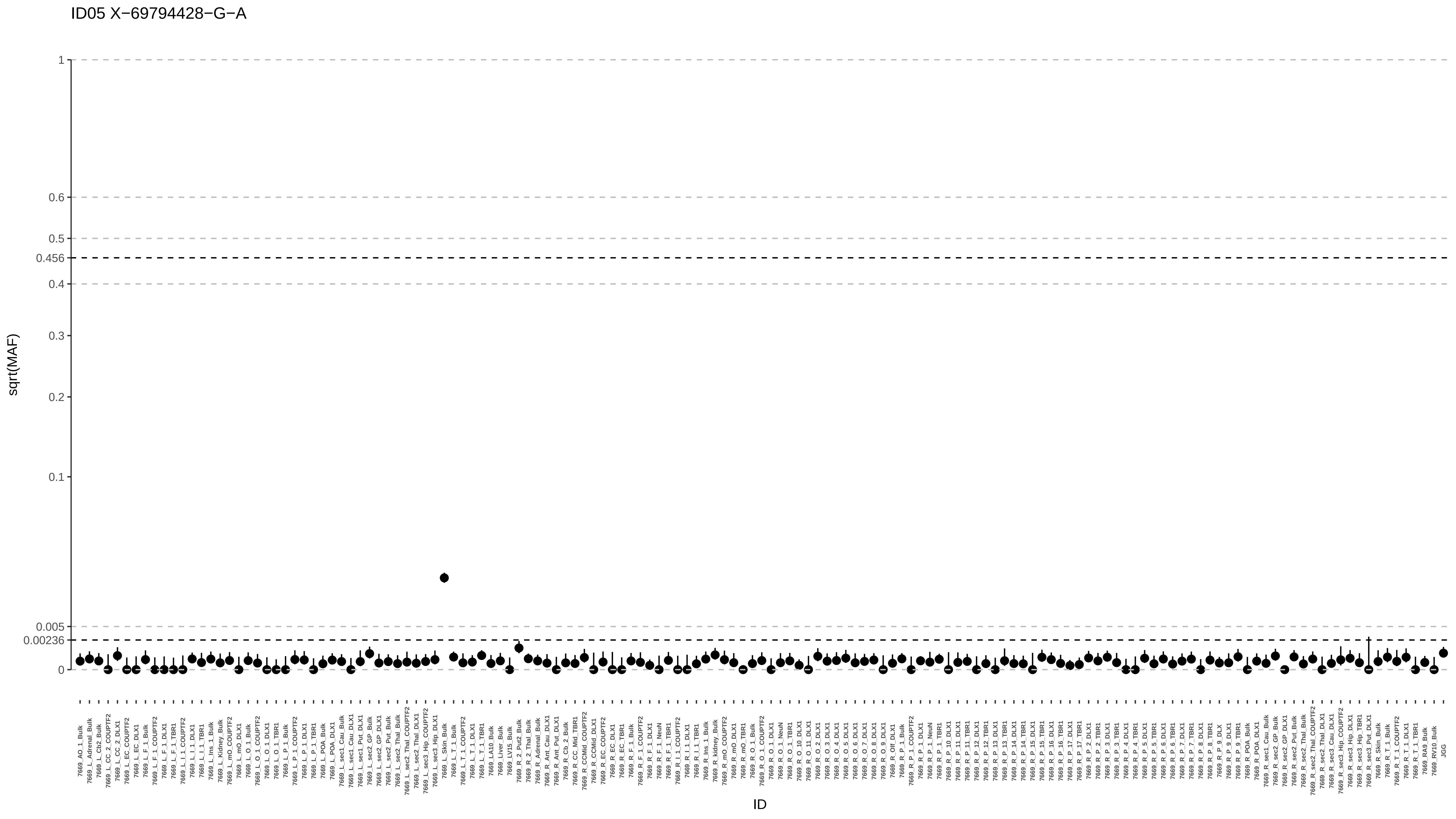
ID05 X−69794428−G−A
sqrt(MAF)
1
0.6
0.5
0.456
0.4
0.3
0.2
0.1
0.005
0.00236
0
ID
7669_AO_1_Bulk
7669_L_Adrenal_Bulk
7669_L_Cb2_Bulk
7669_L_CC_2_COUPTF2
7669_L_CC_2_DLX1
7669_L_EC_COUPTF2
7669_L_EC_DLX1
7669_L_F_1_Bulk
7669_L_F_1_COUPTF2
7669_L_F_1_DLX1
7669_L_F_1_TBR1
7669_L_I_1_COUPTF2
7669_L_I_1_DLX1
7669_L_I_1_TBR1
7669_L_Ins_1_Bulk
7669_L_Kidney_Bulk
7669_L_mO_COUPTF2
7669_L_mO_DLX1
7669_L_O_1_Bulk
7669_L_O_1_COUPTF2
7669_L_O_1_DLX1
7669_L_O_1_TBR1
7669_L_P_1_Bulk
7669_L_P_1_COUPTF2
7669_L_P_1_DLX1
7669_L_P_1_TBR1
7669_L_POA_Bulk
7669_L_POA_DLX1
7669_L_sec1_Cau_Bulk
7669_L_sec1_Cau_DLX1
7669_L_sec1_Put_DLX1
7669_L_sec2_GP_Bulk
7669_L_sec2_GP_DLX1
7669_L_sec2_Put_Bulk
7669_L_sec2_Thal_Bulk
7669_L_sec2_Thal_COUPTF2
7669_L_sec2_Thal_DLX1
7669_L_sec3_Hip_COUPTF2
7669_L_sec3_Hip_DLX1
7669_L_Skin_Bulk
7669_L_T_1_Bulk
7669_L_T_1_COUPTF2
7669_L_T_1_DLX1
7669_L_T_1_TBR1
7669_LA10_Bulk
7669_Liver_Bulk
7669_LV15_Bulk
7669_R_2_Put2_Bulk
7669_R_2_Thal_Bulk
7669_R_Adrenal_Bulk
7669_R_Ant_Cau_DLX1
7669_R_Ant_Put_DLX1
7669_R_Cb_2_Bulk
7669_R_CC_Mid_TBR1
7669_R_CCMid_COUPTF2
7669_R_CCMid_DLX1
7669_R_EC_COUPTF2
7669_R_EC_DLX1
7669_R_EC_TBR1
7669_R_F_1_Bulk
7669_R_F_1_COUPTF2
7669_R_F_1_DLX1
7669_R_F_1_NeuN
7669_R_F_1_TBR1
7669_R_I_1_COUPTF2
7669_R_I_1_DLX1
7669_R_I_1_TBR1
7669_R_Ins_1_Bulk
7669_R_kidney_Bulk
7669_R_mO_COUPTF2
7669_R_mO_DLX1
7669_R_mO_TBR1
7669_R_O_1_Bulk
7669_R_O_1_COUPTF2
7669_R_O_1_DLX1
7669_R_O_1_NeuN
7669_R_O_1_TBR1
7669_R_O_10_DLX1
7669_R_O_11_DLX1
7669_R_O_2_DLX1
7669_R_O_3_DLX1
7669_R_O_4_DLX1
7669_R_O_5_DLX1
7669_R_O_6_DLX1
7669_R_O_7_DLX1
7669_R_O_8_DLX1
7669_R_O_9_DLX1
7669_R_Olf_DLX1
7669_R_P_1_Bulk
7669_R_P_1_COUPTF2
7669_R_P_1_DLX1
7669_R_P_1_NeuN
7669_R_P_1_TBR1
7669_R_P_10_DLX1
7669_R_P_11_DLX1
7669_R_P_11_TBR1
7669_R_P_12_DLX1
7669_R_P_12_TBR1
7669_R_P_13_DLX1
7669_R_P_13_TBR1
7669_R_P_14_DLX1
7669_R_P_14_TBR1
7669_R_P_15_DLX1
7669_R_P_15_TBR1
7669_R_P_16_DLX1
7669_R_P_16_TBR1
7669_R_P_17_DLX1
7669_R_P_17_TBR1
7669_R_P_2_DLX1
7669_R_P_2_TBR1
7669_R_P_3_DLX1
7669_R_P_3_TBR1
7669_R_P_4_DLX1
7669_R_P_4_TBR1
7669_R_P_5_DLX1
7669_R_P_5_TBR1
7669_R_P_6_DLX1
7669_R_P_6_TBR1
7669_R_P_7_DLX1
7669_R_P_7_TBR1
7669_R_P_8_DLX1
7669_R_P_8_TBR1
7669_R_P_9_DLX
7669_R_P_9_DLX1
7669_R_P_9_TBR1
7669_R_POA_Bulk
7669_R_POA_DLX1
7669_R_sec1_Cau_Bulk
7669_R_sec2_GP_Bulk
7669_R_sec2_GP_DLX1
7669_R_sec2_Put_Bulk
7669_R_sec2_Thal_Bulk
7669_R_sec2_Thal_COUPTF2
7669_R_sec2_Thal_DLX1
7669_R_sec3_Cau_DLX1
7669_R_sec3_Hip_COUPTF2
7669_R_sec3_Hip_DLX1
7669_R_sec3_Hip_TBR1
7669_R_sec3_Put_DLX1
7669_R_Skin_Bulk
7669_R_T_1_Bulk
7669_R_T_1_COUPTF2
7669_R_T_1_DLX1
7669_R_T_1_TBR1
7669_RA9_Bulk
7669_RV10_Bulk
JGG

ID05 X−70346122−G−A

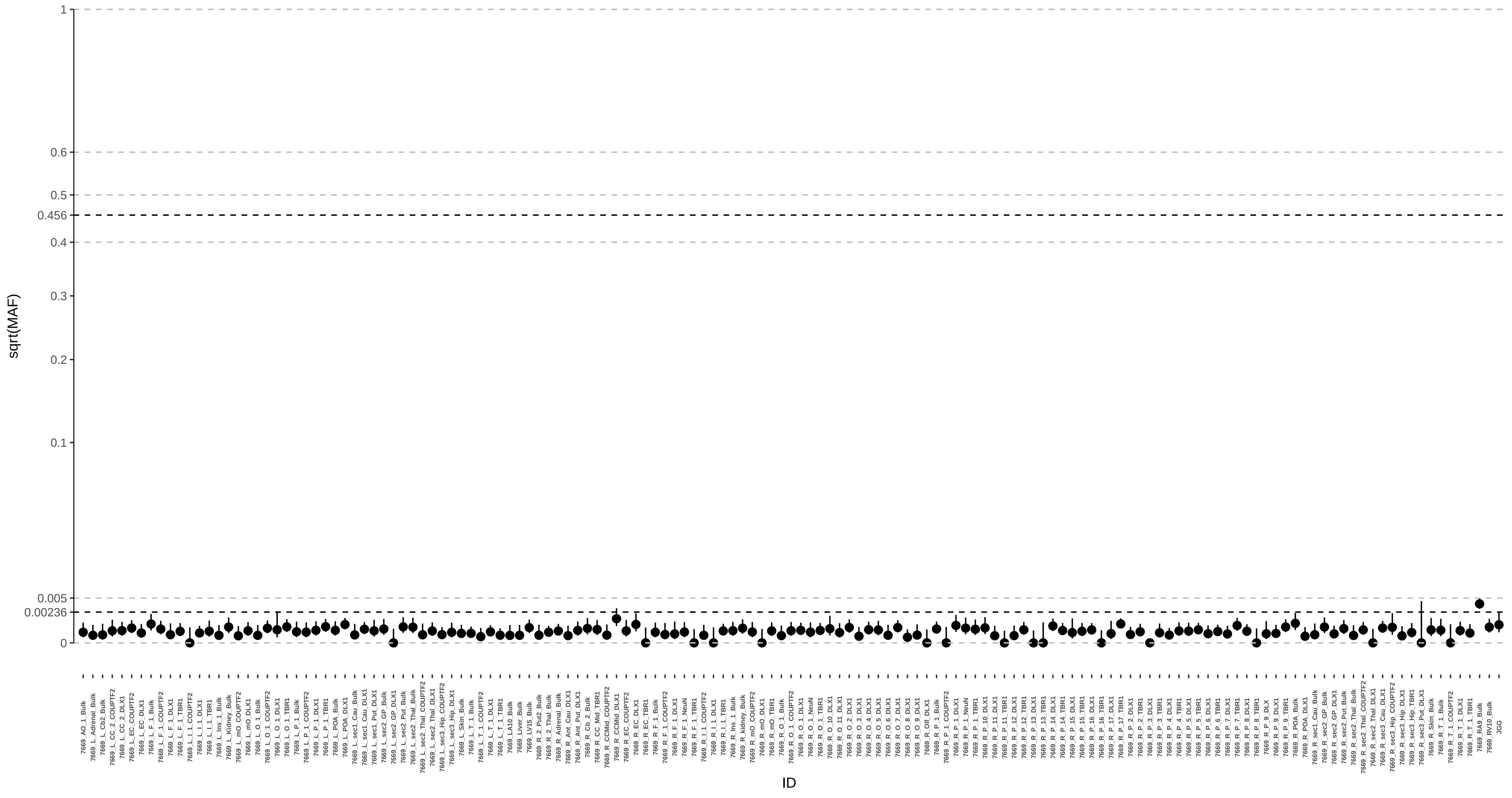

ID05 X−72495149−C−A

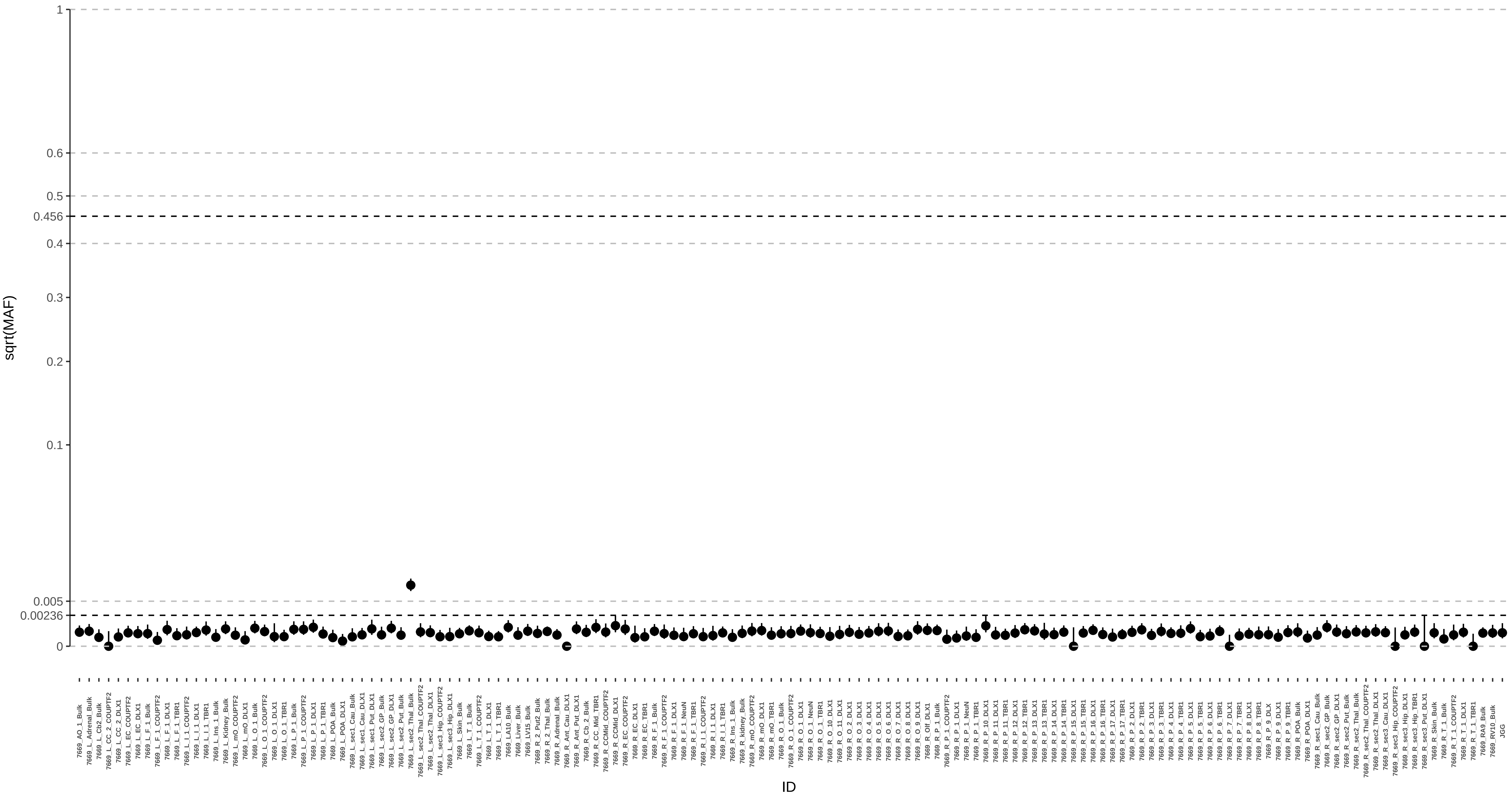

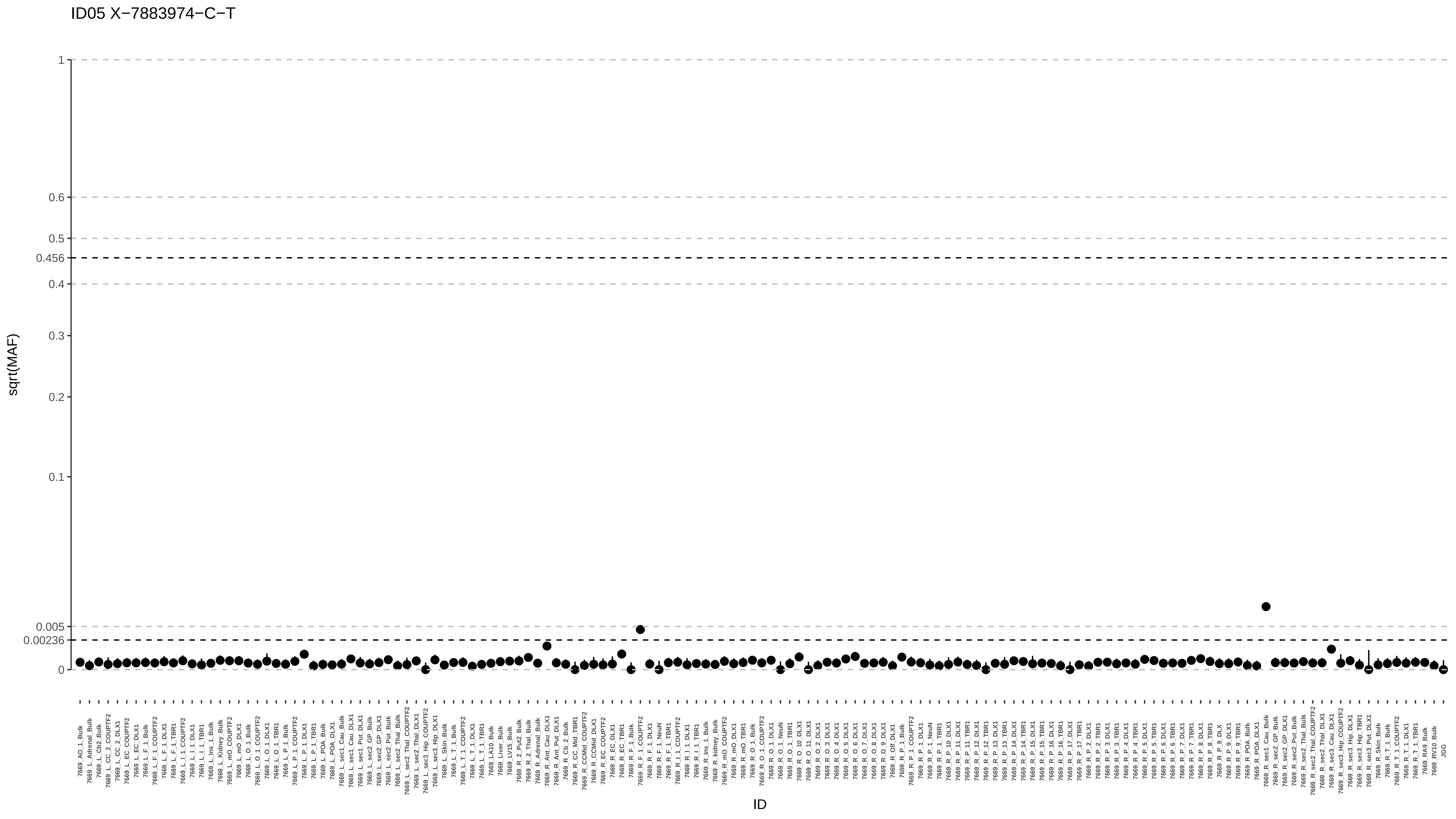

ID05 X−7883974−C−T
sqrt(MAF)
1
0.6
0.5
0.456
0.4
0.3
0.2
0.1
0.005
0.00236
0
ID

ID05 X−79691731−A−T

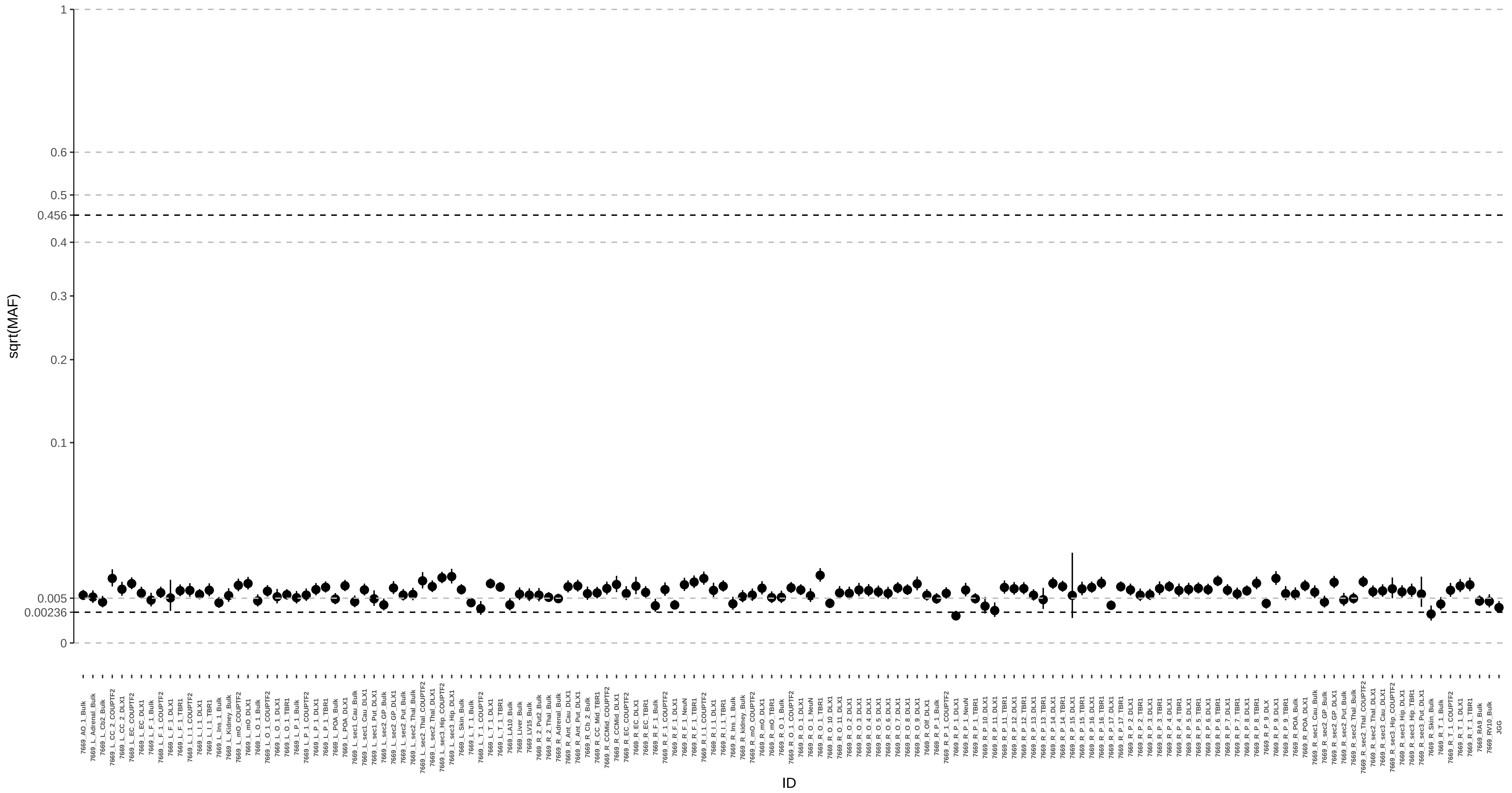

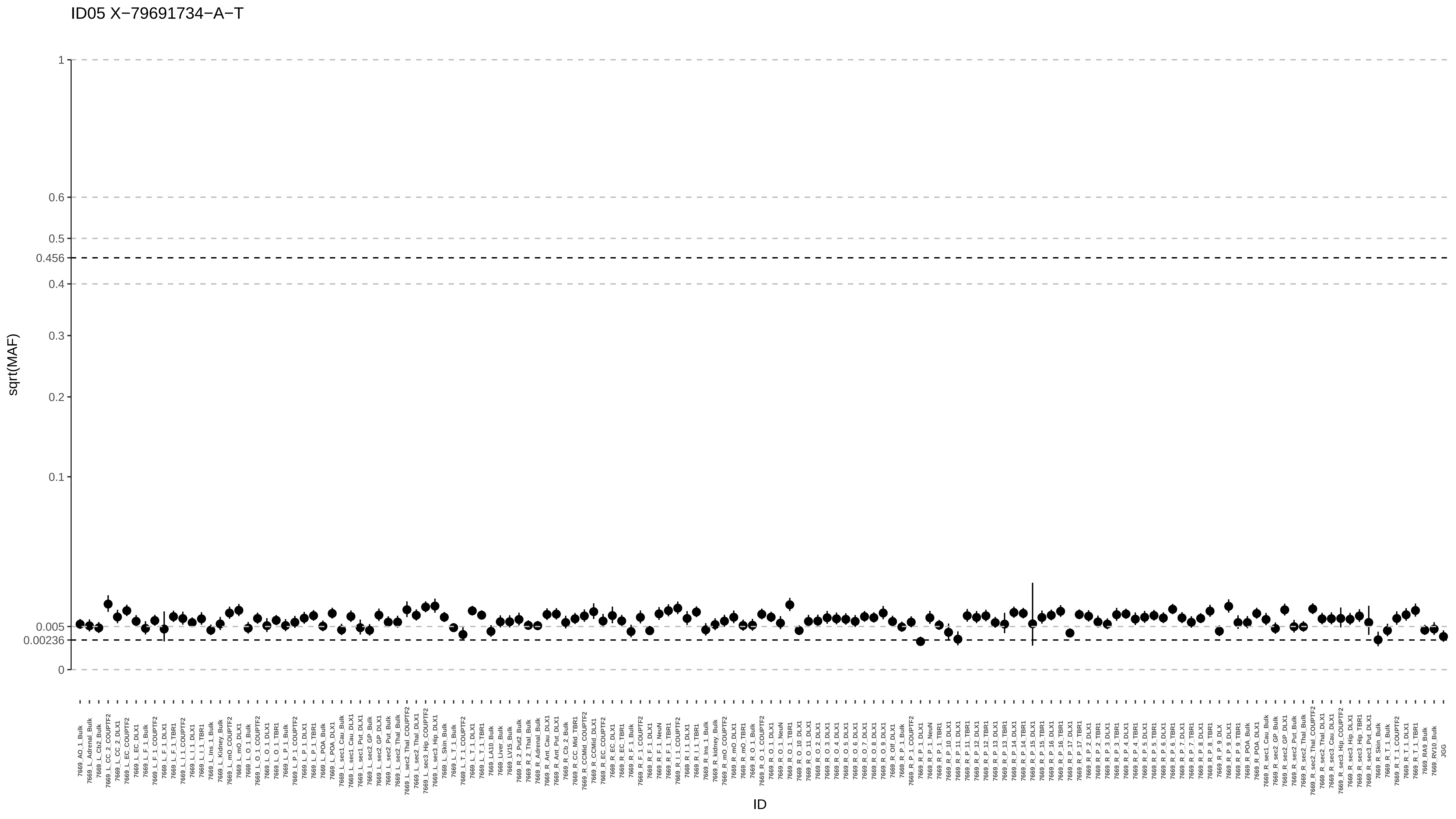
ID05 X−79691734−A−T
sqrt(MAF)
1
0.6
0.5
0.456
0.4
0.3
0.2
0.1
0.005
0.00236
0
ID
7669_AO_1_Bulk
7669_L_Adrenal_Bulk
7669_L_Cb2_Bulk
7669_L_CC_2_COUPTF2
7669_L_CC_2_DLX1
7669_L_EC_COUPTF2
7669_L_EC_DLX1
7669_L_F_1_Bulk
7669_L_F_1_COUPTF2
7669_L_F_1_DLX1
7669_L_F_1_TBR1
7669_L_I_1_COUPTF2
7669_L_I_1_DLX1
7669_L_I_1_TBR1
7669_L_Ins_1_Bulk
7669_L_Kidney_Bulk
7669_L_mO_COUPTF2
7669_L_mO_DLX1
7669_L_O_1_Bulk
7669_L_O_1_COUPTF2
7669_L_O_1_DLX1
7669_L_O_1_TBR1
7669_L_P_1_Bulk
7669_L_P_1_COUPTF2
7669_L_P_1_DLX1
7669_L_P_1_TBR1
7669_L_POA_Bulk
7669_L_POA_DLX1
7669_L_sec1_Cau_Bulk
7669_L_sec1_Cau_DLX1
7669_L_sec1_Put_DLX1
7669_L_sec2_GP_Bulk
7669_L_sec2_GP_DLX1
7669_L_sec2_Put_Bulk
7669_L_sec2_Thal_Bulk
7669_L_sec2_Thal_COUPTF2
7669_L_sec2_Thal_DLX1
7669_L_sec3_Hip_COUPTF2
7669_L_sec3_Hip_DLX1
7669_L_Skin_Bulk
7669_L_T_1_Bulk
7669_L_T_1_COUPTF2
7669_L_T_1_DLX1
7669_L_T_1_TBR1
7669_LA10_Bulk
7669_Liver_Bulk
7669_LV15_Bulk
7669_R_2_Put2_Bulk
7669_R_2_Thal_Bulk
7669_R_Adrenal_Bulk
7669_R_Ant_Cau_DLX1
7669_R_Ant_Put_DLX1
7669_R_Cb_2_Bulk
7669_R_CC_Mid_TBR1
7669_R_CCMid_COUPTF2
7669_R_CCMid_DLX1
7669_R_EC_COUPTF2
7669_R_EC_DLX1
7669_R_EC_TBR1
7669_R_F_1_Bulk
7669_R_F_1_COUPTF2
7669_R_F_1_DLX1
7669_R_F_1_NeuN
7669_R_F_1_TBR1
7669_R_I_1_COUPTF2
7669_R_I_1_DLX1
7669_R_I_1_TBR1
7669_R_Ins_1_Bulk
7669_R_kidney_Bulk
7669_R_mO_COUPTF2
7669_R_mO_DLX1
7669_R_mO_TBR1
7669_R_O_1_Bulk
7669_R_O_1_COUPTF2
7669_R_O_1_DLX1
7669_R_O_1_NeuN
7669_R_O_1_TBR1
7669_R_O_10_DLX1
7669_R_O_11_DLX1
7669_R_O_2_DLX1
7669_R_O_3_DLX1
7669_R_O_4_DLX1
7669_R_O_5_DLX1
7669_R_O_6_DLX1
7669_R_O_7_DLX1
7669_R_O_8_DLX1
7669_R_O_9_DLX1
7669_R_Olf_DLX1
7669_R_P_1_Bulk
7669_R_P_1_COUPTF2
7669_R_P_1_DLX1
7669_R_P_1_NeuN
7669_R_P_1_TBR1
7669_R_P_10_DLX1
7669_R_P_11_DLX1
7669_R_P_11_TBR1
7669_R_P_12_DLX1
7669_R_P_12_TBR1
7669_R_P_13_DLX1
7669_R_P_13_TBR1
7669_R_P_14_DLX1
7669_R_P_14_TBR1
7669_R_P_15_DLX1
7669_R_P_15_TBR1
7669_R_P_16_DLX1
7669_R_P_16_TBR1
7669_R_P_17_DLX1
7669_R_P_17_TBR1
7669_R_P_2_DLX1
7669_R_P_2_TBR1
7669_R_P_3_DLX1
7669_R_P_3_TBR1
7669_R_P_4_DLX1
7669_R_P_4_TBR1
7669_R_P_5_DLX1
7669_R_P_5_TBR1
7669_R_P_6_DLX1
7669_R_P_6_TBR1
7669_R_P_7_DLX1
7669_R_P_7_TBR1
7669_R_P_8_DLX1
7669_R_P_8_TBR1
7669_R_P_9_DLX
7669_R_P_9_DLX1
7669_R_P_9_TBR1
7669_R_POA_Bulk
7669_R_POA_DLX1
7669_R_sec1_Cau_Bulk
7669_R_sec2_GP_Bulk
7669_R_sec2_GP_DLX1
7669_R_sec2_Put_Bulk
7669_R_sec2_Thal_Bulk
7669_R_sec2_Thal_COUPTF2
7669_R_sec2_Thal_DLX1
7669_R_sec3_Cau_DLX1
7669_R_sec3_Hip_COUPTF2
7669_R_sec3_Hip_DLX1
7669_R_sec3_Hip_TBR1
7669_R_sec3_Put_DLX1
7669_R_Skin_Bulk
7669_R_T_1_Bulk
7669_R_T_1_COUPTF2
7669_R_T_1_DLX1
7669_R_T_1_TBR1
7669_RA9_Bulk
7669_RV10_Bulk
JGG

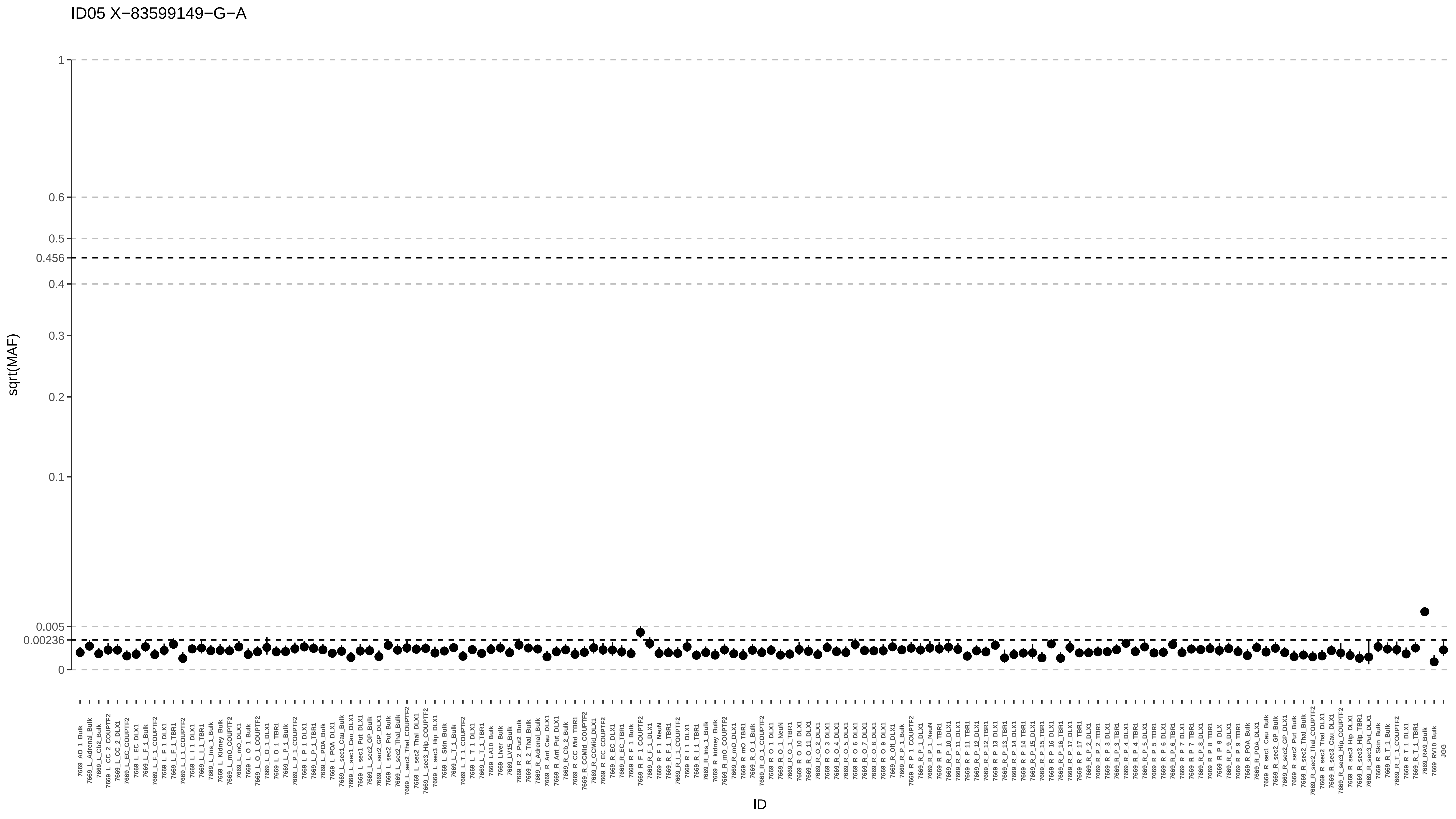
ID05 X−83599149−G−A
sqrt(MAF)
1
0.6
0.5
0.456
0.4
0.3
0.2
0.1
0.005
0.00236
0
ID
7669_AO_1_Bulk
7669_L_Adrenal_Bulk
7669_L_Cb2_Bulk
7669_L_CC_2_COUPTF2
7669_L_CC_2_DLX1
7669_L_EC_COUPTF2
7669_L_EC_DLX1
7669_L_F_1_Bulk
7669_L_F_1_COUPTF2
7669_L_F_1_DLX1
7669_L_F_1_TBR1
7669_L_I_1_COUPTF2
7669_L_I_1_DLX1
7669_L_I_1_TBR1
7669_L_Ins_1_Bulk
7669_L_Kidney_Bulk
7669_L_mO_COUPTF2
7669_L_mO_DLX1
7669_L_O_1_Bulk
7669_L_O_1_COUPTF2
7669_L_O_1_DLX1
7669_L_O_1_TBR1
7669_L_P_1_Bulk
7669_L_P_1_COUPTF2
7669_L_P_1_DLX1
7669_L_P_1_TBR1
7669_L_POA_Bulk
7669_L_POA_DLX1
7669_L_sec1_Cau_Bulk
7669_L_sec1_Cau_DLX1
7669_L_sec1_Put_DLX1
7669_L_sec2_GP_Bulk
7669_L_sec2_GP_DLX1
7669_L_sec2_Put_Bulk
7669_L_sec2_Thal_Bulk
7669_L_sec2_Thal_COUPTF2
7669_L_sec2_Thal_DLX1
7669_L_sec3_Hip_COUPTF2
7669_L_sec3_Hip_DLX1
7669_L_Skin_Bulk
7669_L_T_1_Bulk
7669_L_T_1_COUPTF2
7669_L_T_1_DLX1
7669_L_T_1_TBR1
7669_LA10_Bulk
7669_Liver_Bulk
7669_LV15_Bulk
7669_R_2_Put2_Bulk
7669_R_2_Thal_Bulk
7669_R_Adrenal_Bulk
7669_R_Ant_Cau_DLX1
7669_R_Ant_Put_DLX1
7669_R_Cb_2_Bulk
7669_R_CC_Mid_TBR1
7669_R_CCMid_COUPTF2
7669_R_CCMid_DLX1
7669_R_EC_COUPTF2
7669_R_EC_DLX1
7669_R_EC_TBR1
7669_R_F_1_Bulk
7669_R_F_1_COUPTF2
7669_R_F_1_DLX1
7669_R_F_1_NeuN
7669_R_F_1_TBR1
7669_R_I_1_COUPTF2
7669_R_I_1_DLX1
7669_R_I_1_TBR1
7669_R_Ins_1_Bulk
7669_R_kidney_Bulk
7669_R_mO_COUPTF2
7669_R_mO_DLX1
7669_R_mO_TBR1
7669_R_O_1_Bulk
7669_R_O_1_COUPTF2
7669_R_O_1_DLX1
7669_R_O_1_NeuN
7669_R_O_1_TBR1
7669_R_O_10_DLX1
7669_R_O_11_DLX1
7669_R_O_2_DLX1
7669_R_O_3_DLX1
7669_R_O_4_DLX1
7669_R_O_5_DLX1
7669_R_O_6_DLX1
7669_R_O_7_DLX1
7669_R_O_8_DLX1
7669_R_O_9_DLX1
7669_R_Olf_DLX1
7669_R_P_1_Bulk
7669_R_P_1_COUPTF2
7669_R_P_1_DLX1
7669_R_P_1_NeuN
7669_R_P_1_TBR1
7669_R_P_10_DLX1
7669_R_P_11_DLX1
7669_R_P_11_TBR1
7669_R_P_12_DLX1
7669_R_P_12_TBR1
7669_R_P_13_DLX1
7669_R_P_13_TBR1
7669_R_P_14_DLX1
7669_R_P_14_TBR1
7669_R_P_15_DLX1
7669_R_P_15_TBR1
7669_R_P_16_DLX1
7669_R_P_16_TBR1
7669_R_P_17_DLX1
7669_R_P_17_TBR1
7669_R_P_2_DLX1
7669_R_P_2_TBR1
7669_R_P_3_DLX1
7669_R_P_3_TBR1
7669_R_P_4_DLX1
7669_R_P_4_TBR1
7669_R_P_5_DLX1
7669_R_P_5_TBR1
7669_R_P_6_DLX1
7669_R_P_6_TBR1
7669_R_P_7_DLX1
7669_R_P_7_TBR1
7669_R_P_8_DLX1
7669_R_P_8_TBR1
7669_R_P_9_DLX
7669_R_P_9_DLX1
7669_R_P_9_TBR1
7669_R_POA_Bulk
7669_R_POA_DLX1
7669_R_sec1_Cau_Bulk
7669_R_sec2_GP_Bulk
7669_R_sec2_GP_DLX1
7669_R_sec2_Put_Bulk
7669_R_sec2_Thal_Bulk
7669_R_sec2_Thal_COUPTF2
7669_R_sec2_Thal_DLX1
7669_R_sec3_Cau_DLX1
7669_R_sec3_Hip_COUPTF2
7669_R_sec3_Hip_DLX1
7669_R_sec3_Hip_TBR1
7669_R_sec3_Put_DLX1
7669_R_Skin_Bulk
7669_R_T_1_Bulk
7669_R_T_1_COUPTF2
7669_R_T_1_DLX1
7669_R_T_1_TBR1
7669_RA9_Bulk
7669_RV10_Bulk
JGG

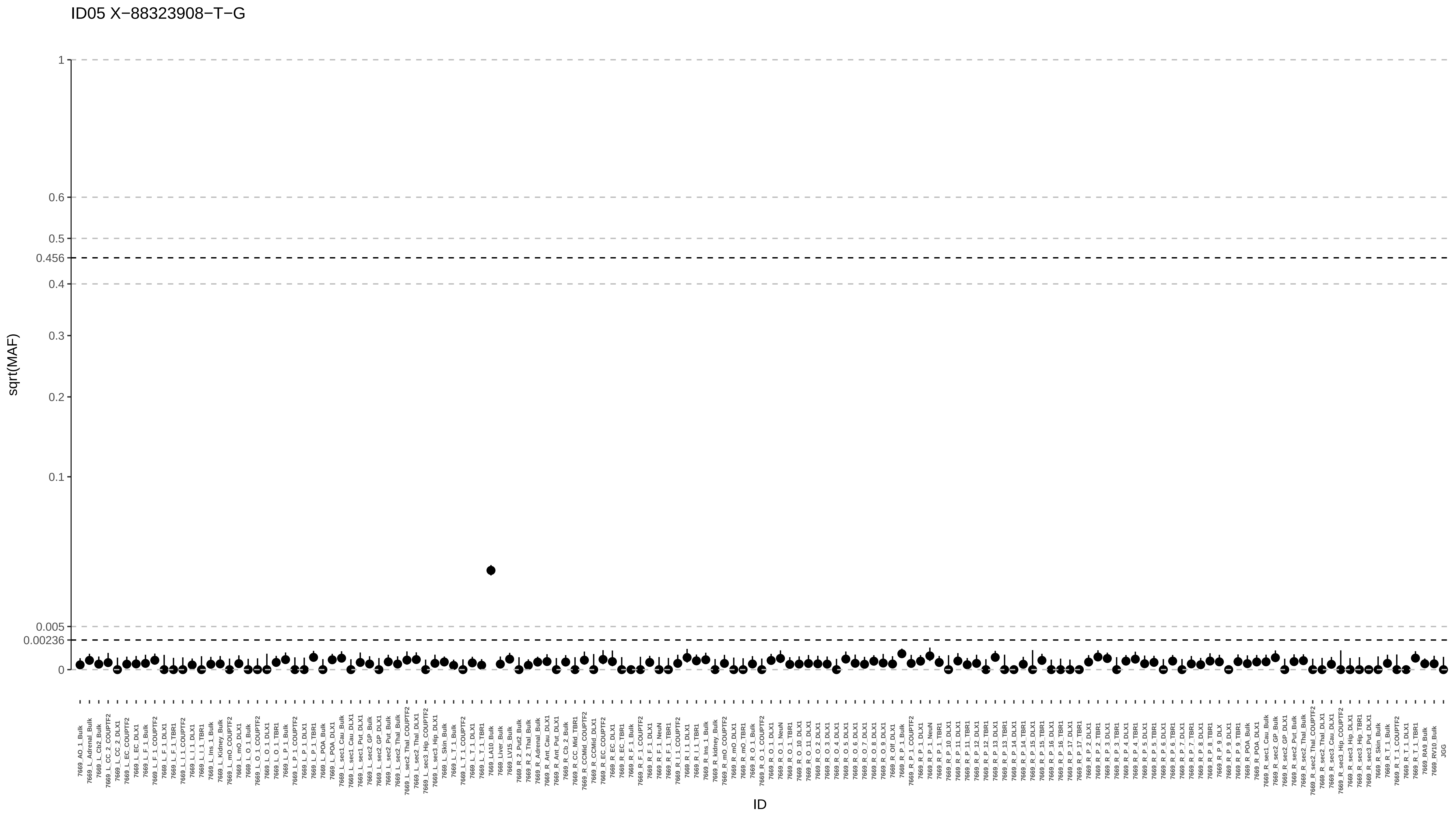
ID05 X−88323908−T−G
sqrt(MAF)
1
0.6
0.5
0.456
0.4
0.3
0.2
0.1
0.005
0.00236
0
ID
7669_AO_1_Bulk
7669_L_Adrenal_Bulk
7669_L_Cb2_Bulk
7669_L_CC_2_COUPTF2
7669_L_CC_2_DLX1
7669_L_EC_COUPTF2
7669_L_EC_DLX1
7669_L_F_1_Bulk
7669_L_F_1_COUPTF2
7669_L_F_1_DLX1
7669_L_F_1_TBR1
7669_L_I_1_COUPTF2
7669_L_I_1_DLX1
7669_L_I_1_TBR1
7669_L_Ins_1_Bulk
7669_L_Kidney_Bulk
7669_L_mO_COUPTF2
7669_L_mO_DLX1
7669_L_O_1_Bulk
7669_L_O_1_COUPTF2
7669_L_O_1_DLX1
7669_L_O_1_TBR1
7669_L_P_1_Bulk
7669_L_P_1_COUPTF2
7669_L_P_1_DLX1
7669_L_P_1_TBR1
7669_L_POA_Bulk
7669_L_POA_DLX1
7669_L_sec1_Cau_Bulk
7669_L_sec1_Cau_DLX1
7669_L_sec1_Put_DLX1
7669_L_sec2_GP_Bulk
7669_L_sec2_GP_DLX1
7669_L_sec2_Put_Bulk
7669_L_sec2_Thal_Bulk
7669_L_sec2_Thal_COUPTF2
7669_L_sec2_Thal_DLX1
7669_L_sec3_Hip_COUPTF2
7669_L_sec3_Hip_DLX1
7669_L_Skin_Bulk
7669_L_T_1_Bulk
7669_L_T_1_COUPTF2
7669_L_T_1_DLX1
7669_L_T_1_TBR1
7669_LA10_Bulk
7669_Liver_Bulk
7669_LV15_Bulk
7669_R_2_Put2_Bulk
7669_R_2_Thal_Bulk
7669_R_Adrenal_Bulk
7669_R_Ant_Cau_DLX1
7669_R_Ant_Put_DLX1
7669_R_Cb_2_Bulk
7669_R_CC_Mid_TBR1
7669_R_CCMid_COUPTF2
7669_R_CCMid_DLX1
7669_R_EC_COUPTF2
7669_R_EC_DLX1
7669_R_EC_TBR1
7669_R_F_1_Bulk
7669_R_F_1_COUPTF2
7669_R_F_1_DLX1
7669_R_F_1_NeuN
7669_R_F_1_TBR1
7669_R_I_1_COUPTF2
7669_R_I_1_DLX1
7669_R_I_1_TBR1
7669_R_Ins_1_Bulk
7669_R_kidney_Bulk
7669_R_mO_COUPTF2
7669_R_mO_DLX1
7669_R_mO_TBR1
7669_R_O_1_Bulk
7669_R_O_1_COUPTF2
7669_R_O_1_DLX1
7669_R_O_1_NeuN
7669_R_O_1_TBR1
7669_R_O_10_DLX1
7669_R_O_11_DLX1
7669_R_O_2_DLX1
7669_R_O_3_DLX1
7669_R_O_4_DLX1
7669_R_O_5_DLX1
7669_R_O_6_DLX1
7669_R_O_7_DLX1
7669_R_O_8_DLX1
7669_R_O_9_DLX1
7669_R_Olf_DLX1
7669_R_P_1_Bulk
7669_R_P_1_COUPTF2
7669_R_P_1_DLX1
7669_R_P_1_NeuN
7669_R_P_1_TBR1
7669_R_P_10_DLX1
7669_R_P_11_DLX1
7669_R_P_11_TBR1
7669_R_P_12_DLX1
7669_R_P_12_TBR1
7669_R_P_13_DLX1
7669_R_P_13_TBR1
7669_R_P_14_DLX1
7669_R_P_14_TBR1
7669_R_P_15_DLX1
7669_R_P_15_TBR1
7669_R_P_16_DLX1
7669_R_P_16_TBR1
7669_R_P_17_DLX1
7669_R_P_17_TBR1
7669_R_P_2_DLX1
7669_R_P_2_TBR1
7669_R_P_3_DLX1
7669_R_P_3_TBR1
7669_R_P_4_DLX1
7669_R_P_4_TBR1
7669_R_P_5_DLX1
7669_R_P_5_TBR1
7669_R_P_6_DLX1
7669_R_P_6_TBR1
7669_R_P_7_DLX1
7669_R_P_7_TBR1
7669_R_P_8_DLX1
7669_R_P_8_TBR1
7669_R_P_9_DLX
7669_R_P_9_DLX1
7669_R_P_9_TBR1
7669_R_POA_Bulk
7669_R_POA_DLX1
7669_R_sec1_Cau_Bulk
7669_R_sec2_GP_Bulk
7669_R_sec2_GP_DLX1
7669_R_sec2_Put_Bulk
7669_R_sec2_Thal_Bulk
7669_R_sec2_Thal_COUPTF2
7669_R_sec2_Thal_DLX1
7669_R_sec3_Cau_DLX1
7669_R_sec3_Hip_COUPTF2
7669_R_sec3_Hip_DLX1
7669_R_sec3_Hip_TBR1
7669_R_sec3_Put_DLX1
7669_R_Skin_Bulk
7669_R_T_1_Bulk
7669_R_T_1_COUPTF2
7669_R_T_1_DLX1
7669_R_T_1_TBR1
7669_RA9_Bulk
7669_RV10_Bulk
JGG

ID05 X−95198288−C−T

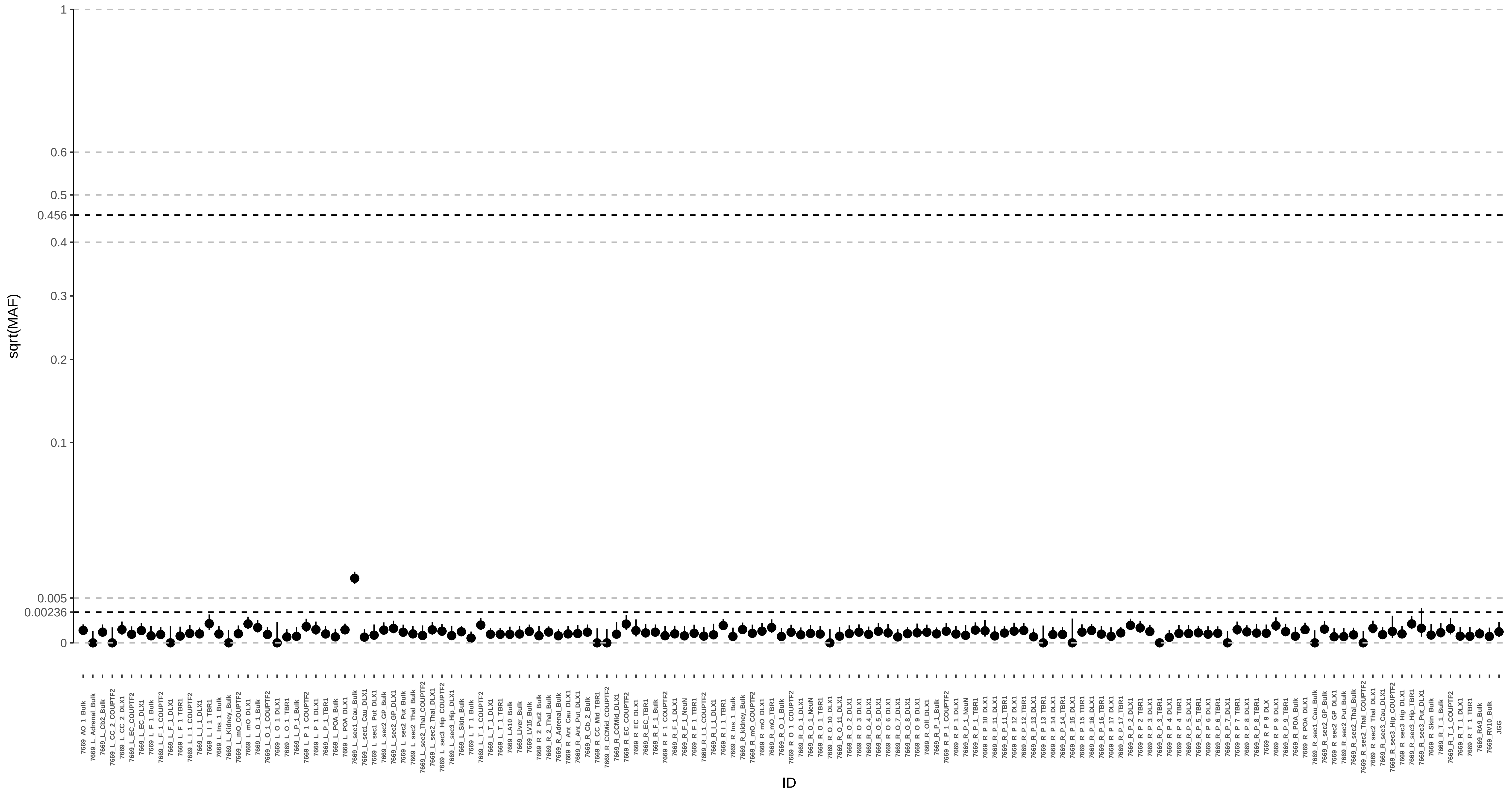

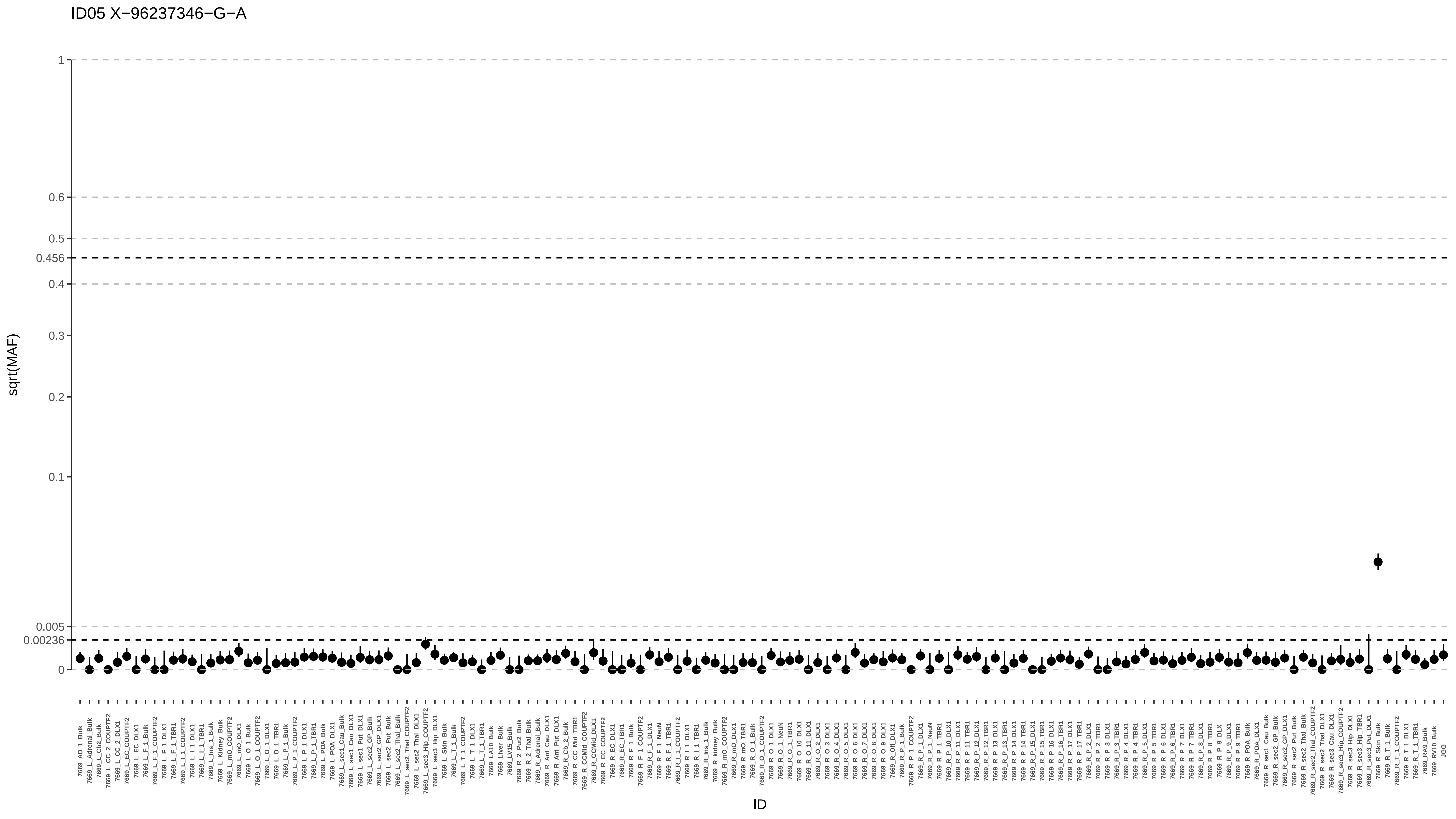
ID05 X−96237346−G−A
sqrt(MAF)
1
0.6
0.5
0.456
0.4
0.3
0.2
0.1
0.005
0.00236
0
ID

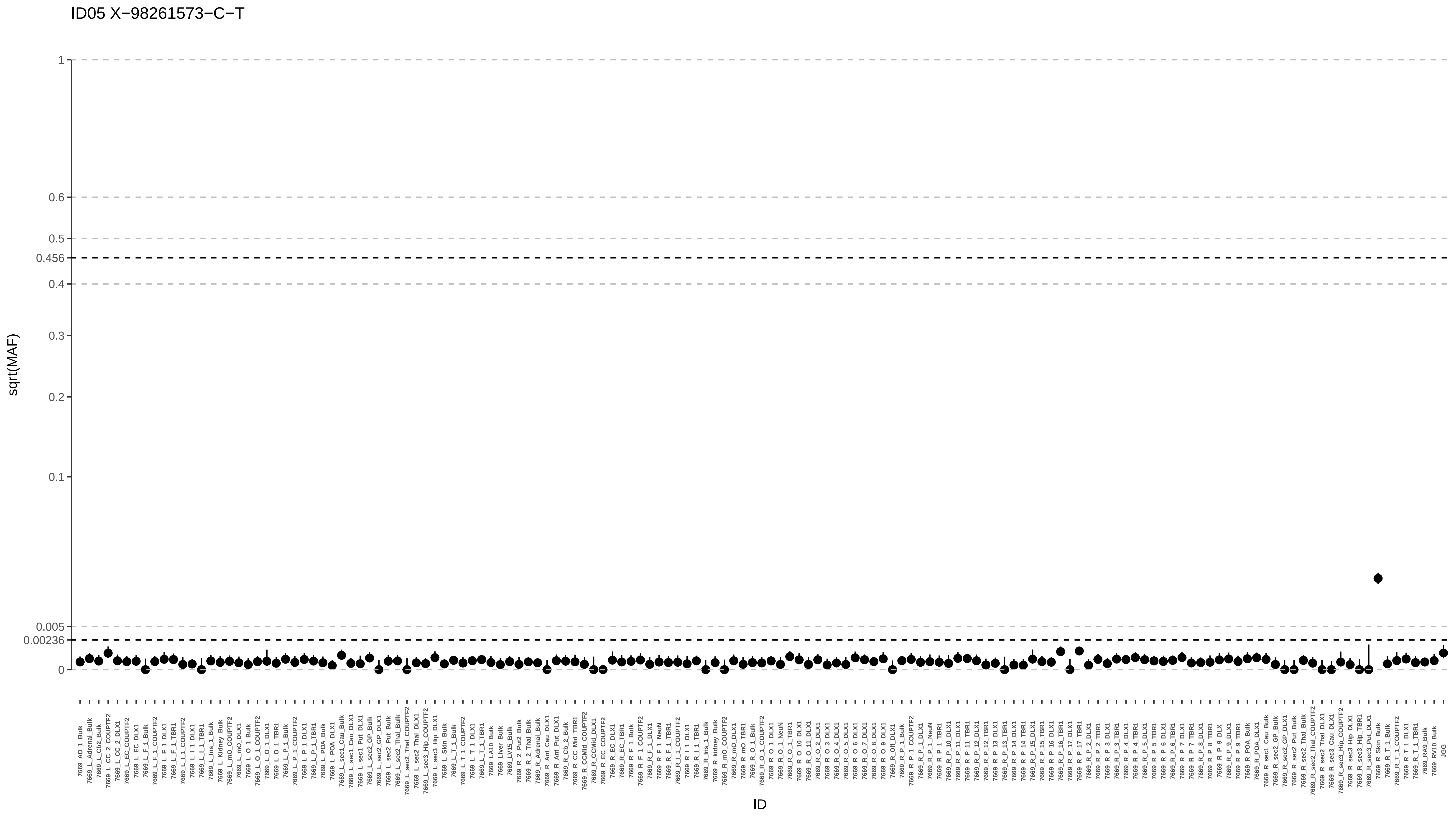

ID05 X−98261573−C−T
sqrt(MAF)
1
0.6
0.5
0.456
0.4
0.3
0.2
0.1
0.005
0.00236
0
ID

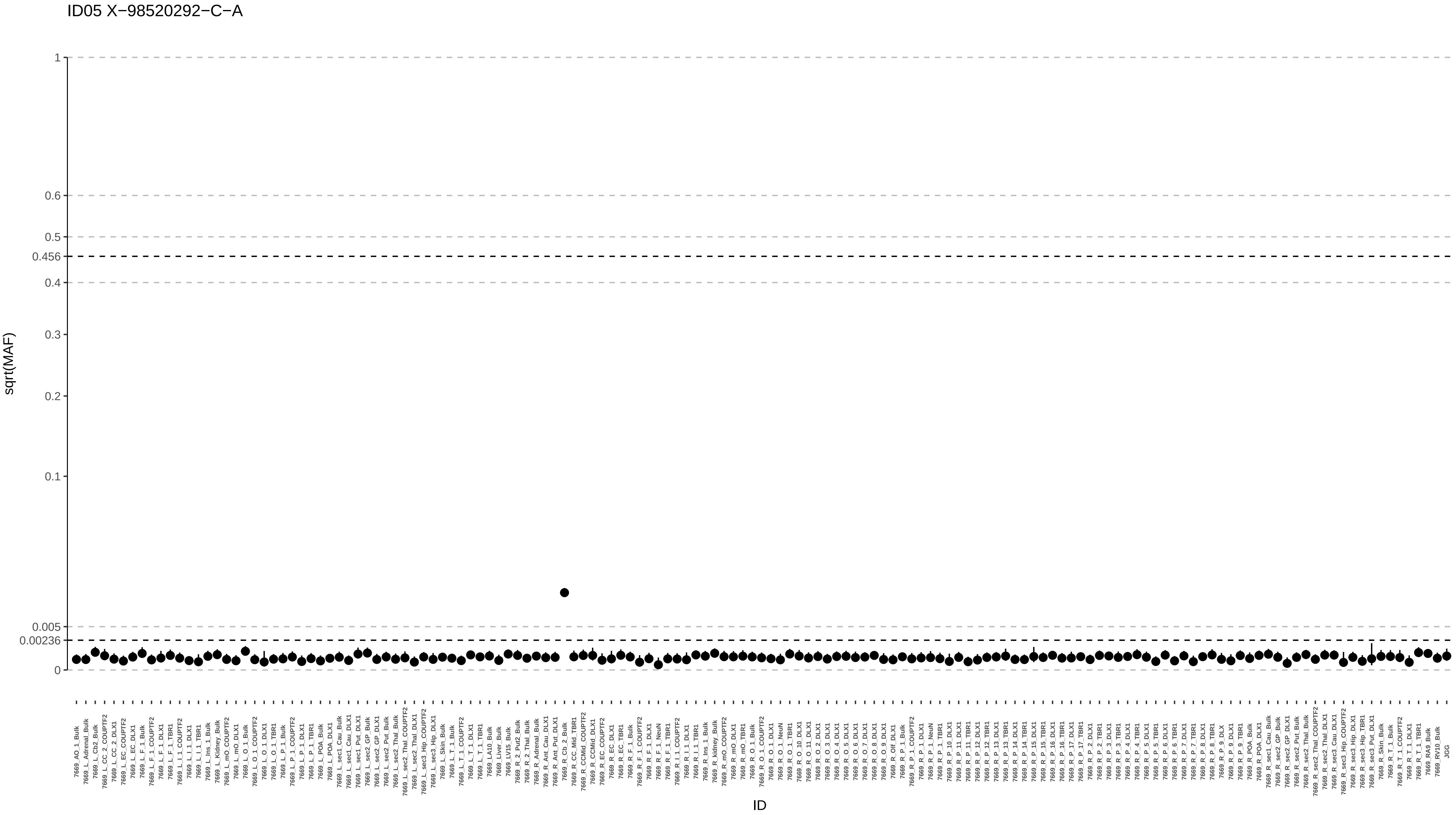
ID05 X−98520292−C−A
sqrt(MAF)
1
0.6
0.5
0.456
0.4
0.3
0.2
0.1
0.005
0.00236
0
ID
7669_AO_1_Bulk
7669_L_Adrenal_Bulk
7669_L_Cb2_Bulk
7669_L_CC_2_COUPTF2
7669_L_CC_2_DLX1
7669_L_EC_COUPTF2
7669_L_EC_DLX1
7669_L_F_1_Bulk
7669_L_F_1_COUPTF2
7669_L_F_1_DLX1
7669_L_F_1_TBR1
7669_L_I_1_COUPTF2
7669_L_I_1_DLX1
7669_L_I_1_TBR1
7669_L_Ins_1_Bulk
7669_L_Kidney_Bulk
7669_L_mO_COUPTF2
7669_L_mO_DLX1
7669_L_O_1_Bulk
7669_L_O_1_COUPTF2
7669_L_O_1_DLX1
7669_L_O_1_TBR1
7669_L_P_1_Bulk
7669_L_P_1_COUPTF2
7669_L_P_1_DLX1
7669_L_P_1_TBR1
7669_L_POA_Bulk
7669_L_POA_DLX1
7669_L_sec1_Cau_Bulk
7669_L_sec1_Cau_DLX1
7669_L_sec1_Put_DLX1
7669_L_sec2_GP_Bulk
7669_L_sec2_GP_DLX1
7669_L_sec2_Put_Bulk
7669_L_sec2_Thal_Bulk
7669_L_sec2_Thal_COUPTF2
7669_L_sec2_Thal_DLX1
7669_L_sec3_Hip_COUPTF2
7669_L_sec3_Hip_DLX1
7669_L_Skin_Bulk
7669_L_T_1_Bulk
7669_L_T_1_COUPTF2
7669_L_T_1_DLX1
7669_L_T_1_TBR1
7669_LA10_Bulk
7669_Liver_Bulk
7669_LV15_Bulk
7669_R_2_Put2_Bulk
7669_R_2_Thal_Bulk
7669_R_Adrenal_Bulk
7669_R_Ant_Cau_DLX1
7669_R_Ant_Put_DLX1
7669_R_Cb_2_Bulk
7669_R_CC_Mid_TBR1
7669_R_CCMid_COUPTF2
7669_R_CCMid_DLX1
7669_R_EC_COUPTF2
7669_R_EC_DLX1
7669_R_EC_TBR1
7669_R_F_1_Bulk
7669_R_F_1_COUPTF2
7669_R_F_1_DLX1
7669_R_F_1_NeuN
7669_R_F_1_TBR1
7669_R_I_1_COUPTF2
7669_R_I_1_DLX1
7669_R_I_1_TBR1
7669_R_Ins_1_Bulk
7669_R_kidney_Bulk
7669_R_mO_COUPTF2
7669_R_mO_DLX1
7669_R_mO_TBR1
7669_R_O_1_Bulk
7669_R_O_1_COUPTF2
7669_R_O_1_DLX1
7669_R_O_1_NeuN
7669_R_O_1_TBR1
7669_R_O_10_DLX1
7669_R_O_11_DLX1
7669_R_O_2_DLX1
7669_R_O_3_DLX1
7669_R_O_4_DLX1
7669_R_O_5_DLX1
7669_R_O_6_DLX1
7669_R_O_7_DLX1
7669_R_O_8_DLX1
7669_R_O_9_DLX1
7669_R_Olf_DLX1
7669_R_P_1_Bulk
7669_R_P_1_COUPTF2
7669_R_P_1_DLX1
7669_R_P_1_NeuN
7669_R_P_1_TBR1
7669_R_P_10_DLX1
7669_R_P_11_DLX1
7669_R_P_11_TBR1
7669_R_P_12_DLX1
7669_R_P_12_TBR1
7669_R_P_13_DLX1
7669_R_P_13_TBR1
7669_R_P_14_DLX1
7669_R_P_14_TBR1
7669_R_P_15_DLX1
7669_R_P_15_TBR1
7669_R_P_16_DLX1
7669_R_P_16_TBR1
7669_R_P_17_DLX1
7669_R_P_17_TBR1
7669_R_P_2_DLX1
7669_R_P_2_TBR1
7669_R_P_3_DLX1
7669_R_P_3_TBR1
7669_R_P_4_DLX1
7669_R_P_4_TBR1
7669_R_P_5_DLX1
7669_R_P_5_TBR1
7669_R_P_6_DLX1
7669_R_P_6_TBR1
7669_R_P_7_DLX1
7669_R_P_7_TBR1
7669_R_P_8_DLX1
7669_R_P_8_TBR1
7669_R_P_9_DLX
7669_R_P_9_DLX1
7669_R_P_9_TBR1
7669_R_POA_Bulk
7669_R_POA_DLX1
7669_R_sec1_Cau_Bulk
7669_R_sec2_GP_Bulk
7669_R_sec2_GP_DLX1
7669_R_sec2_Put_Bulk
7669_R_sec2_Thal_Bulk
7669_R_sec2_Thal_COUPTF2
7669_R_sec2_Thal_DLX1
7669_R_sec3_Cau_DLX1
7669_R_sec3_Hip_COUPTF2
7669_R_sec3_Hip_DLX1
7669_R_sec3_Hip_TBR1
7669_R_sec3_Put_DLX1
7669_R_Skin_Bulk
7669_R_T_1_Bulk
7669_R_T_1_COUPTF2
7669_R_T_1_DLX1
7669_R_T_1_TBR1
7669_RA9_Bulk
7669_RV10_Bulk
JGG

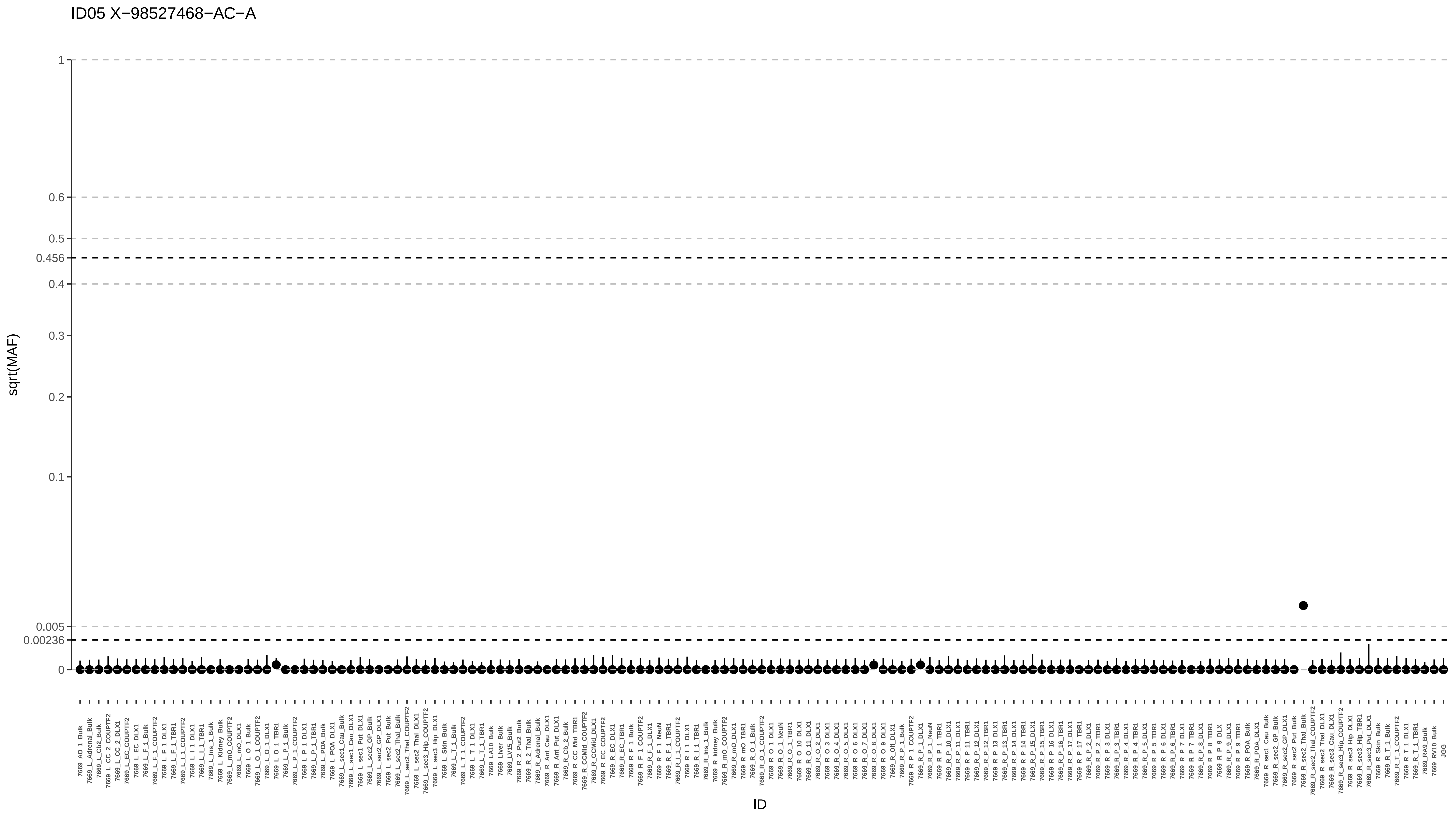
ID05 X−98527468−AC−A
sqrt(MAF)
1
0.6
0.5
0.456
0.4
0.3
0.2
0.1
0.005
0.00236
0
ID
7669_AO_1_Bulk
7669_L_Adrenal_Bulk
7669_L_Cb2_Bulk
7669_L_CC_2_COUPTF2
7669_L_CC_2_DLX1
7669_L_EC_COUPTF2
7669_L_EC_DLX1
7669_L_F_1_Bulk
7669_L_F_1_COUPTF2
7669_L_F_1_DLX1
7669_L_F_1_TBR1
7669_L_I_1_COUPTF2
7669_L_I_1_DLX1
7669_L_I_1_TBR1
7669_L_Ins_1_Bulk
7669_L_Kidney_Bulk
7669_L_mO_COUPTF2
7669_L_mO_DLX1
7669_L_O_1_Bulk
7669_L_O_1_COUPTF2
7669_L_O_1_DLX1
7669_L_O_1_TBR1
7669_L_P_1_Bulk
7669_L_P_1_COUPTF2
7669_L_P_1_DLX1
7669_L_P_1_TBR1
7669_L_POA_Bulk
7669_L_POA_DLX1
7669_L_sec1_Cau_Bulk
7669_L_sec1_Cau_DLX1
7669_L_sec1_Put_DLX1
7669_L_sec2_GP_Bulk
7669_L_sec2_GP_DLX1
7669_L_sec2_Put_Bulk
7669_L_sec2_Thal_Bulk
7669_L_sec2_Thal_COUPTF2
7669_L_sec2_Thal_DLX1
7669_L_sec3_Hip_COUPTF2
7669_L_sec3_Hip_DLX1
7669_L_Skin_Bulk
7669_L_T_1_Bulk
7669_L_T_1_COUPTF2
7669_L_T_1_DLX1
7669_L_T_1_TBR1
7669_LA10_Bulk
7669_Liver_Bulk
7669_LV15_Bulk
7669_R_2_Put2_Bulk
7669_R_2_Thal_Bulk
7669_R_Adrenal_Bulk
7669_R_Ant_Cau_DLX1
7669_R_Ant_Put_DLX1
7669_R_Cb_2_Bulk
7669_R_CC_Mid_TBR1
7669_R_CCMid_COUPTF2
7669_R_CCMid_DLX1
7669_R_EC_COUPTF2
7669_R_EC_DLX1
7669_R_EC_TBR1
7669_R_F_1_Bulk
7669_R_F_1_COUPTF2
7669_R_F_1_DLX1
7669_R_F_1_NeuN
7669_R_F_1_TBR1
7669_R_I_1_COUPTF2
7669_R_I_1_DLX1
7669_R_I_1_TBR1
7669_R_Ins_1_Bulk
7669_R_kidney_Bulk
7669_R_mO_COUPTF2
7669_R_mO_DLX1
7669_R_mO_TBR1
7669_R_O_1_Bulk
7669_R_O_1_COUPTF2
7669_R_O_1_DLX1
7669_R_O_1_NeuN
7669_R_O_1_TBR1
7669_R_O_10_DLX1
7669_R_O_11_DLX1
7669_R_O_2_DLX1
7669_R_O_3_DLX1
7669_R_O_4_DLX1
7669_R_O_5_DLX1
7669_R_O_6_DLX1
7669_R_O_7_DLX1
7669_R_O_8_DLX1
7669_R_O_9_DLX1
7669_R_Olf_DLX1
7669_R_P_1_Bulk
7669_R_P_1_COUPTF2
7669_R_P_1_DLX1
7669_R_P_1_NeuN
7669_R_P_1_TBR1
7669_R_P_10_DLX1
7669_R_P_11_DLX1
7669_R_P_11_TBR1
7669_R_P_12_DLX1
7669_R_P_12_TBR1
7669_R_P_13_DLX1
7669_R_P_13_TBR1
7669_R_P_14_DLX1
7669_R_P_14_TBR1
7669_R_P_15_DLX1
7669_R_P_15_TBR1
7669_R_P_16_DLX1
7669_R_P_16_TBR1
7669_R_P_17_DLX1
7669_R_P_17_TBR1
7669_R_P_2_DLX1
7669_R_P_2_TBR1
7669_R_P_3_DLX1
7669_R_P_3_TBR1
7669_R_P_4_DLX1
7669_R_P_4_TBR1
7669_R_P_5_DLX1
7669_R_P_5_TBR1
7669_R_P_6_DLX1
7669_R_P_6_TBR1
7669_R_P_7_DLX1
7669_R_P_7_TBR1
7669_R_P_8_DLX1
7669_R_P_8_TBR1
7669_R_P_9_DLX
7669_R_P_9_DLX1
7669_R_P_9_TBR1
7669_R_POA_Bulk
7669_R_POA_DLX1
7669_R_sec1_Cau_Bulk
7669_R_sec2_GP_Bulk
7669_R_sec2_GP_DLX1
7669_R_sec2_Put_Bulk
7669_R_sec2_Thal_Bulk
7669_R_sec2_Thal_COUPTF2
7669_R_sec2_Thal_DLX1
7669_R_sec3_Cau_DLX1
7669_R_sec3_Hip_COUPTF2
7669_R_sec3_Hip_DLX1
7669_R_sec3_Hip_TBR1
7669_R_sec3_Put_DLX1
7669_R_Skin_Bulk
7669_R_T_1_Bulk
7669_R_T_1_COUPTF2
7669_R_T_1_DLX1
7669_R_T_1_TBR1
7669_RA9_Bulk
7669_RV10_Bulk
JGG

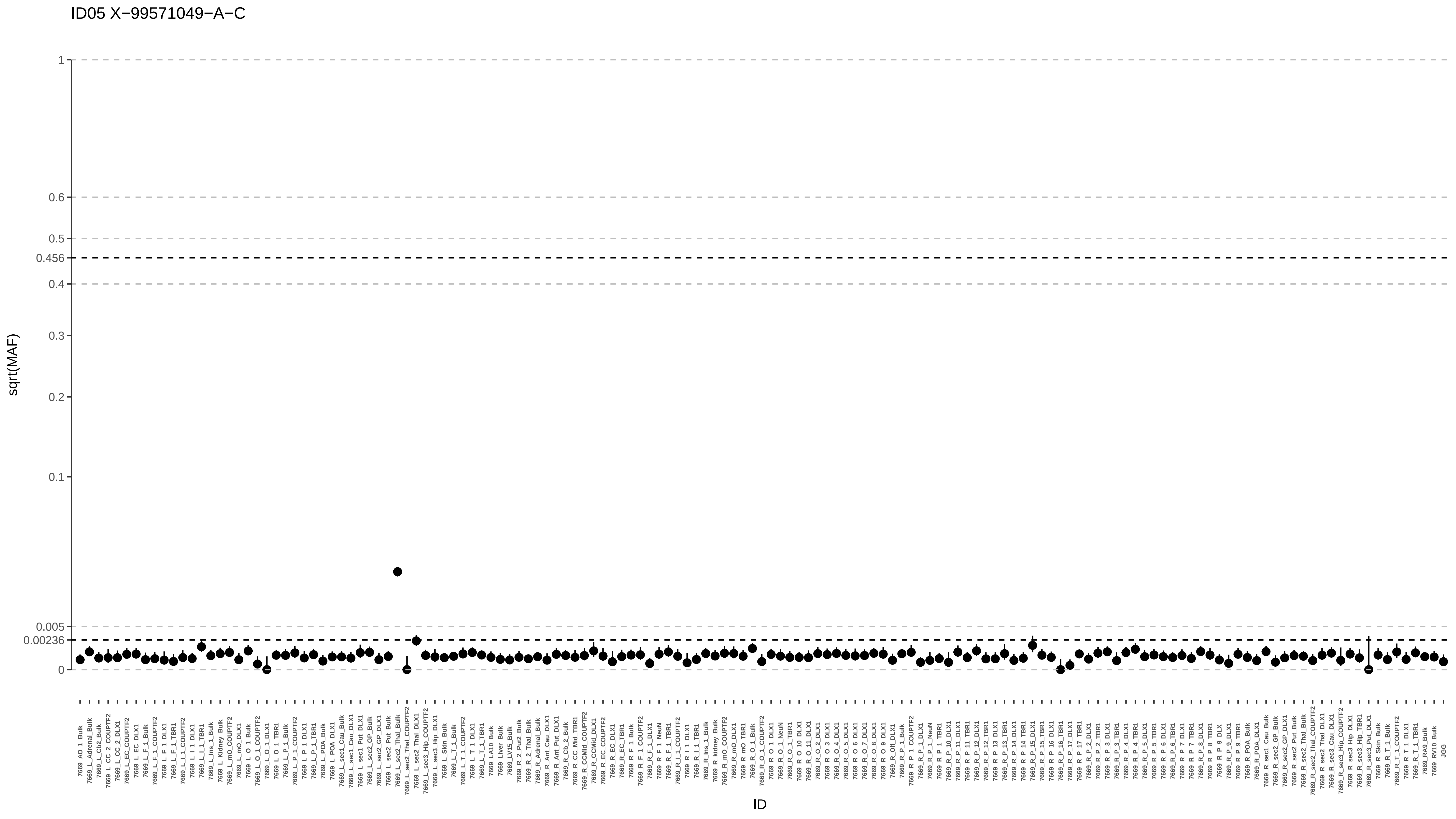

ID05 X−99571049−A−C
sqrt(MAF)
1
0.6
0.5
0.456
0.4
0.3
0.2
0.1
0.005
0.00236
0
ID

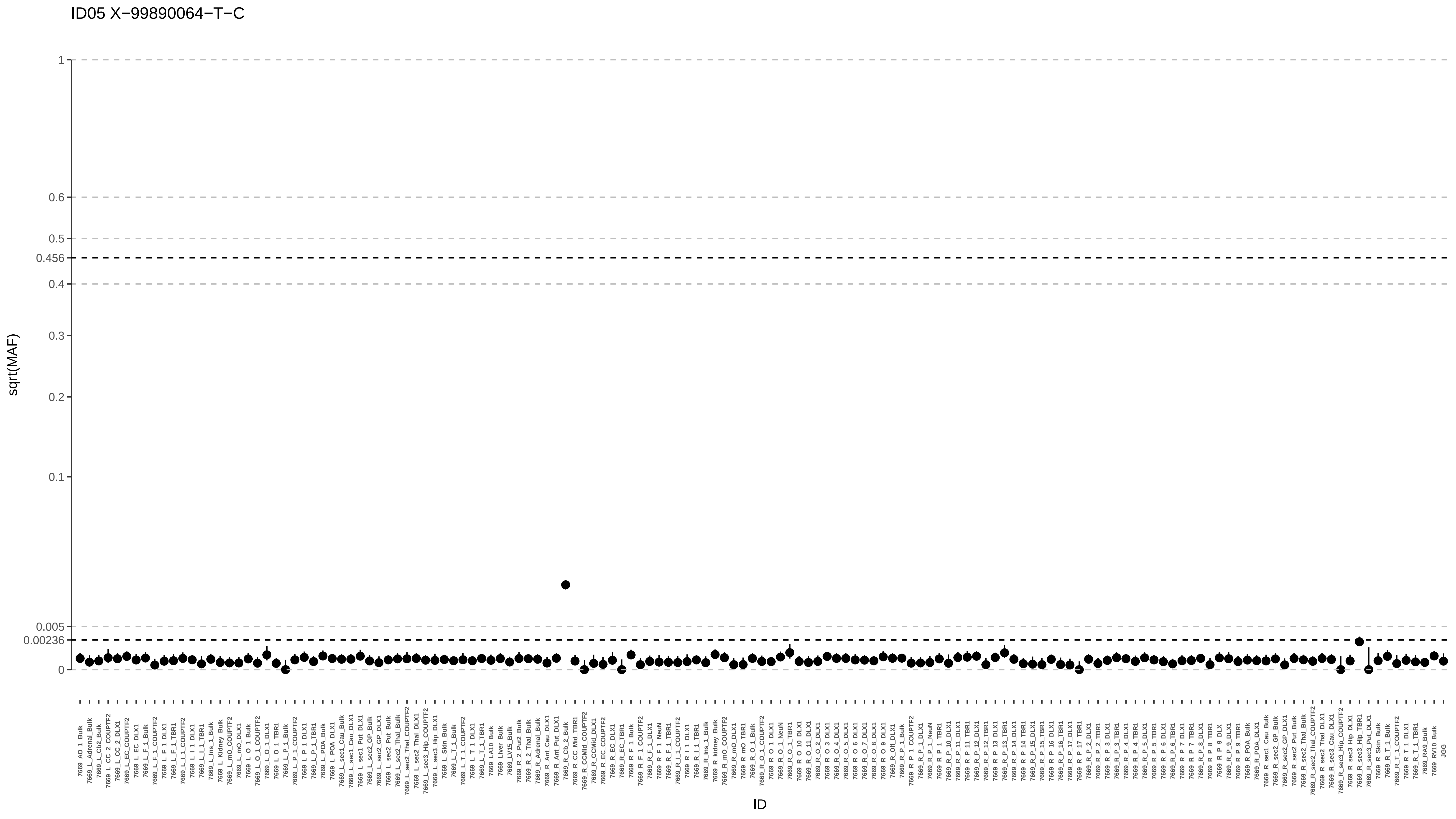
ID05 X−99890064−T−C
sqrt(MAF)
1
0.6
0.5
0.456
0.4
0.3
0.2
0.1
0.005
0.00236
0
ID
