## Supplementary figures and images for "Cell-type-resolved somatic mosaicism reveals clonal dynamics of the human forebrain"

### Supplementary Data 5

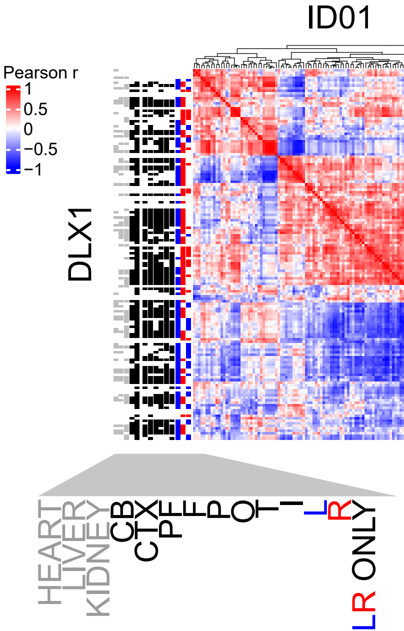

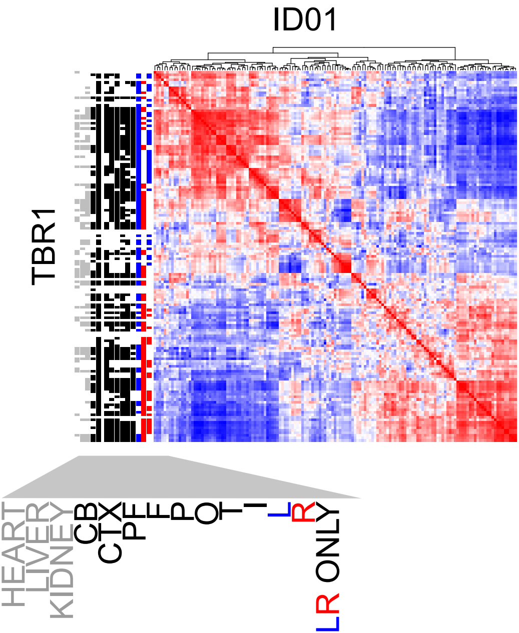

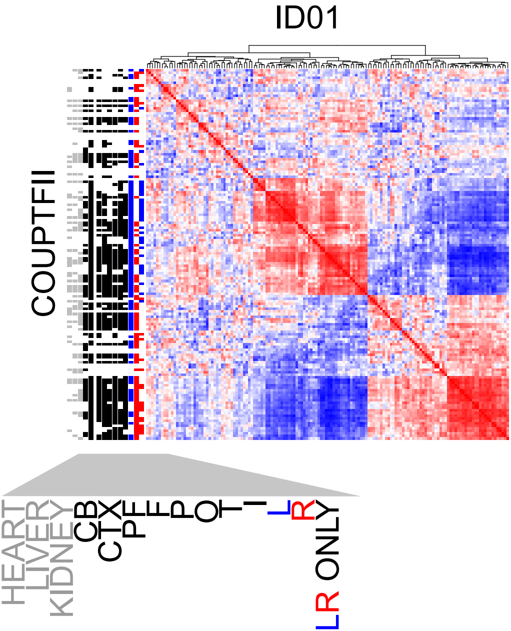

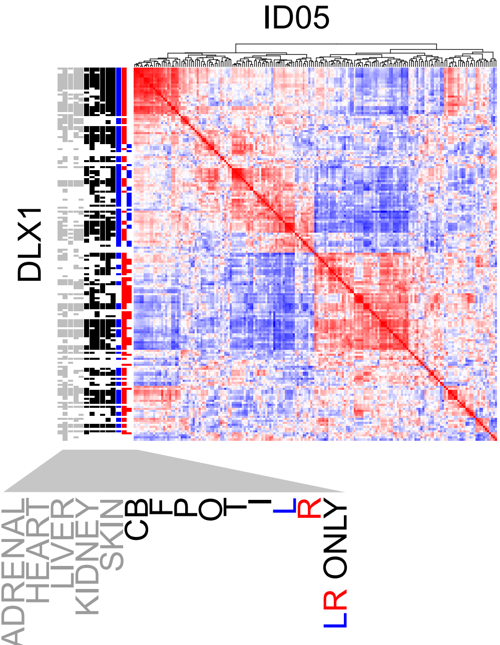

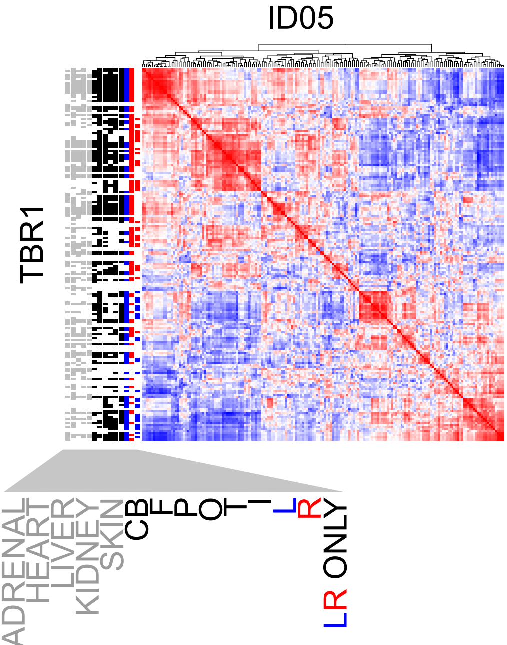

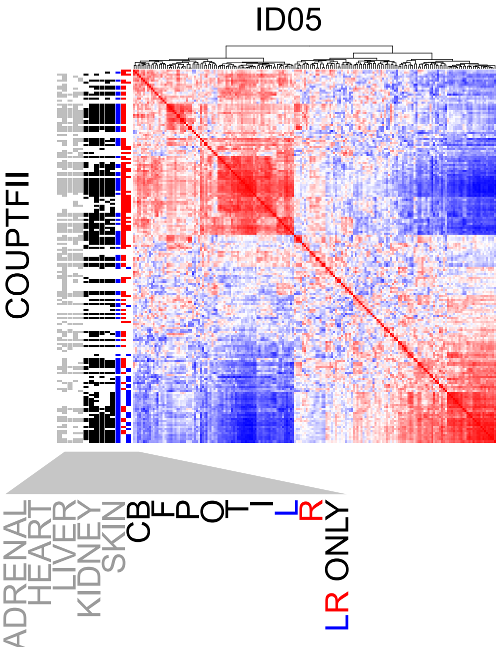
